# Role of channels in the oxygen permeability of red blood cells

**DOI:** 10.1101/2020.08.28.265066

**Authors:** Pan Zhao, R. Ryan Geyer, Ahlam I. Salameh, Amanda B. Wass, Sara Taki, Dale E. Huffman, Howard J Meyerson, Gerolf Gros, Rossana Occhipinti, Fraser J. Moss, Walter F. Boron

## Abstract

Many have believed that oxygen (O_2_) crosses red blood cell (RBC) membranes by dissolving in lipids that offer no resistance to diffusion. However, using stopped-flow (SF) analyses of hemoglobin (Hb) absorbance spectra during O_2_ off-loading from mouse RBCs, we now report that most O_2_ traverses membrane-protein channels. Two agents excluded from the RBC interior markedly slow O_2_ off-loading: p-chloromercuribenzenesulfonate (pCMBS) reduces inferred membrane O_2_ permeability (*P*_Membrane_) by ∼82%, and 4,4’-diisothiocyanatostilbene-2,2’-disulfonate (DIDS), by ∼56%. Because neither likely produces these effects via membrane lipids, we examined RBCs from mice genetically deficient in aquaporin-1 (AQP1), the Rh complex (i.e., rhesus proteins RhAG + mRh), or both. The double knockout (dKO) reduces *P*_Membrane_ by ∼55%, and pCMBS+dKO, by ∼91%. Proteomic analyses of RBC membranes, flow cytometry, hematology, and mathematical simulations rule out explanations involving other membrane proteins, RBC geometry, or extracellular unconvected fluid (EUF). By identifying the first two O_2_ channels and pointing to the existence of other O_2_ channel(s), all of which could be subject to physiological regulation and pharmacological intervention, our work represents a paradigm shift for O_2_ handling.

## Results & Discussion

Central to human life is the inhalation of O_2_, its diffusion across the alveolar wall to pulmonary capillary blood plasma, O_2_ diffusion across the RBC membrane and transfer to Hb (O_2_ + Hb[O_2_]_3_ → Hb[O_2_]_4_), the distribution of RBCs throughout the body, and the offloading of this O_2_ from Hb to support metabolism. A century ago, Krogh’s mathematical analysis of O_2_ diffusion—from Hb, through the intracellular fluid (ICF), across the RBC membrane, to the bulk extracellular fluid (bECF) of the capillary blood plasma, and then across several membranes to the ICF of O_2_-metabolizing cells^1^—required the implicit simplifying assumption^2^ that the membranes offer no resistance to O_2_ diffusion (i.e., *R*_Membrane_ = 0; see Supplementary Discussion). Although some argued otherwise^3,4^, Krogh’s assumption became engrained in the literature, and most have assumed that total resistance to O_2_ diffusion out of RBCs (*R*_Total_ = *R*_ICF_ + *R*_Membrane_ + *R*_EUF_) comprises exclusively *R*_ICF_ and *R*_EUF_^5–7^.

Calling the above prevailing view into question were the discoveries of the first membranes with no detectable CO_2_ permeability^8^ and the first membrane protein (i.e., AQP1^9,10^) with an additional role as a CO_2_ channel.^11^ pCMBS, which reacts with the –SH of Cys-189 in the extracellular vestibule of the AQP1 monomeric pore and thereby inhibits H_2_O movement through AQP1^12^, also blocks a component of CO_2_ traffic through AQP1.^13^ Moreover, various AQPs exhibit CO_2_ vs. NH_3_ selectivity.^14,15^ Other work showed that rhesus (Rh) proteins also can conduct CO_2_.^16^ These observations indicated that gas transport involves more than diffusion through lipids. Work on human RBCs genetically deficient in AQP1^17,18^ or RhAG^18,19^ showed that these two channels account for >90% of CO_2_ permeability of RBCs, and that DIDS, which binds to and can react with the –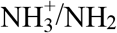 of Lys residues, blocks a component of CO_2_ traffic through both AQP1 and the Rh complex. Although neither pCMBS nor DIDS is specific, both are valuable tools because they do not interact with membrane lipids.^20,21^ Thus, we hypothesized that, like CO_2_, O_2_ crosses RBC membranes mainly via channels, and that pCMBS or DIDS would block these hypothetical O_2_ channels.

In the present study, we use SF absorbance spectroscopy to monitor Hb deoxygenation as we mix oxygenated RBCs with a solution containing the O_2_ scavenger sodium dithionite (NDT; final concentration, 25 mM). The 3D graphs in Figure 1*a–b* show examples of time courses of Hb absorbance spectra of RBCs—either untreated (panel *a*) or pCMBS-treated (panel *b*)—from wild-type (WT) mice. Figure 1*c* shows the time courses of relative HbO_2_ saturation (HbSat). An SF assay^22^ based on the release of carbonic anhydrase (CA) from RBC cytosol to bECF (Methods/Hemolysis) quantitates hemolysis within the SF chamber, and ImageStream flow cytometry (Methods/ImageStream) measures the frequency and diameter of spherocytes (Supplementary Table 8). We find that a 15-min incubation with 1 mM pCMBS increases hemolysis from ∼5.2% to ∼8.2% (Extended Data Figure 1*a*), and increases the prevalence of spherocytes from ∼1.4% to ∼8.7% (Extended Data Figure 1*b*). A novel protocol (Extended Data Figure 2) accounts for the effects of extracellular Hb on the Hb-deoxygenation rate constant 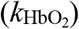, and mathematical simulations (below) account for the presence of spherocytes (Methods/Accommodation for Spherocytes). We conclude that that pCMBS decreases 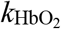 by ∼61% (Figure 1*d*).

**Figure 1.**
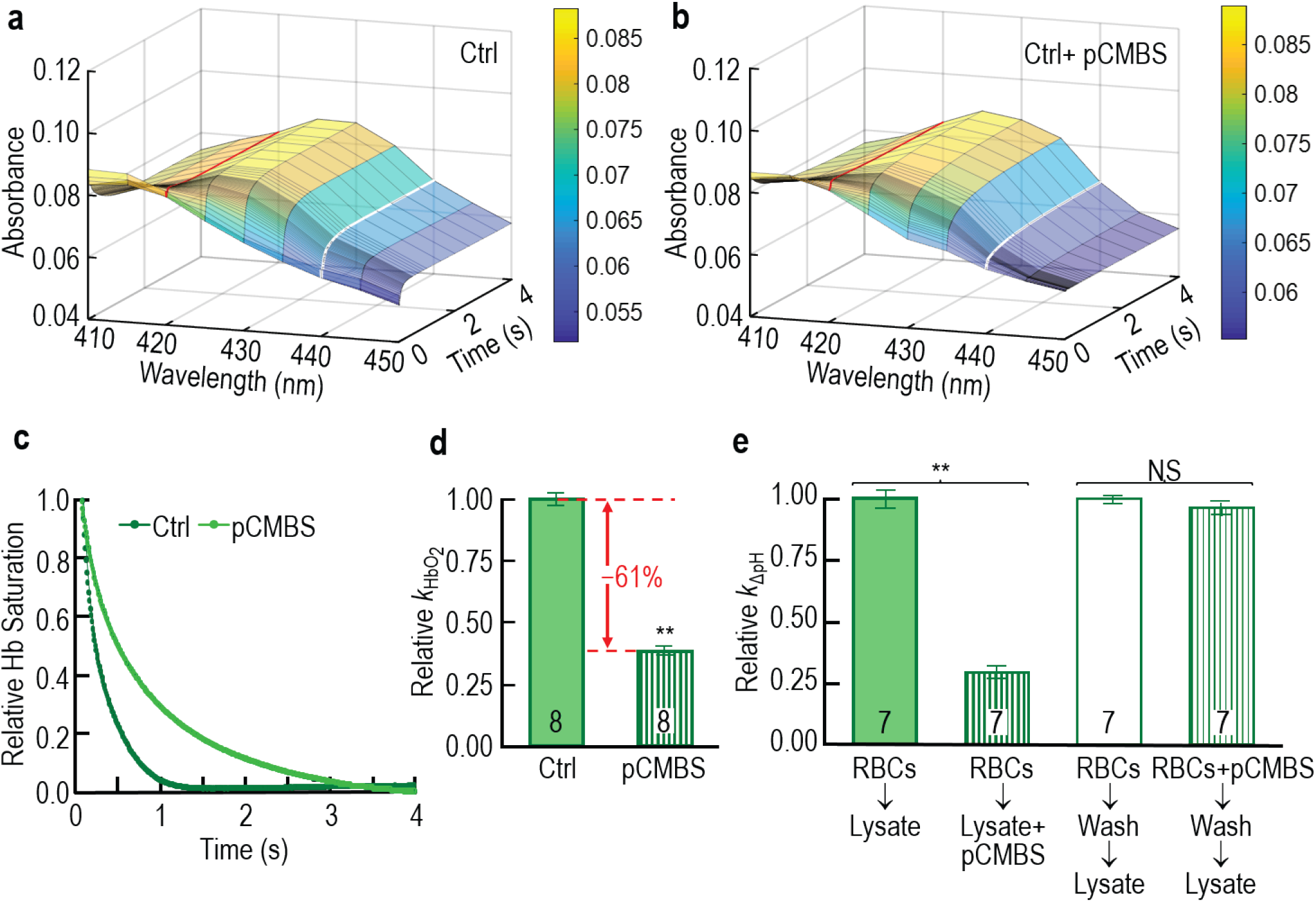
Effect of extracellular pCMBS on rate constant of hemoglobin deoxygenation 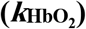 during O_2_ efflux from RBCs. **a & b**, Representative time courses of hemoglobin absorbance spectra during deoxygenation of intact RBCs, from which we compute 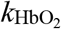. **c**, Time courses of relative HbSat (Supplementary Methods) for the same two data sets in the previous panels. **d**, Summary of effect of pCMBS on 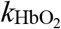 in intact RBCs, corrected for hemolysis, as described in Extended Data Figure 1. **e**, Exclusion of pCMBS from RBC cytosol. Wash solution contained 0.2% bovine serum albumin. Each “n” represents RBCs from 1 WT mouse. We performed paired t-tests (see Methods/Statistical Analysis). \*\**P*<0.01; NS, not significant.

To determine whether pCMBS accesses RBC cytosol, we used CA as a sentinel. Although incubating hemolysate with pCMBS for 15 minutes markedly reduces lysate CA activity, treating intact RBCs with pCMBS has no effect (Figure 1*e*). Thus, pCMBS does not significantly cross RBC membranes.

Figure 2*a-b* show representative time courses of Hb absorbance spectra of untreated and DIDS-treated RBCs from WT mice, and Figure 2*c* shows time courses of relative HbSat. For DIDS-treated RBCs, we used a variant of the aforementioned approach to correct raw 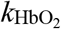 values for hemolysis (Extended Data Figure 3). A 60-min incubation with 200 μM DIDS increases mean hemolysis from ∼5.2% to ∼13.4% (Extended Data Figure 1*a*), and increases spherocytosis to ∼41% (Extended Data Figure 1*b*). After accounting for hemolysis and spherocytosis, we conclude that DIDS reduces 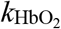 by ∼31% (Figure 2*d*). Although the degree of DIDS-induced spherocytosis is cause for pause, we will see below that protocols with substantially less spherocytosis reveal even greater decreases in 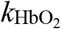.

**Figure 2.**
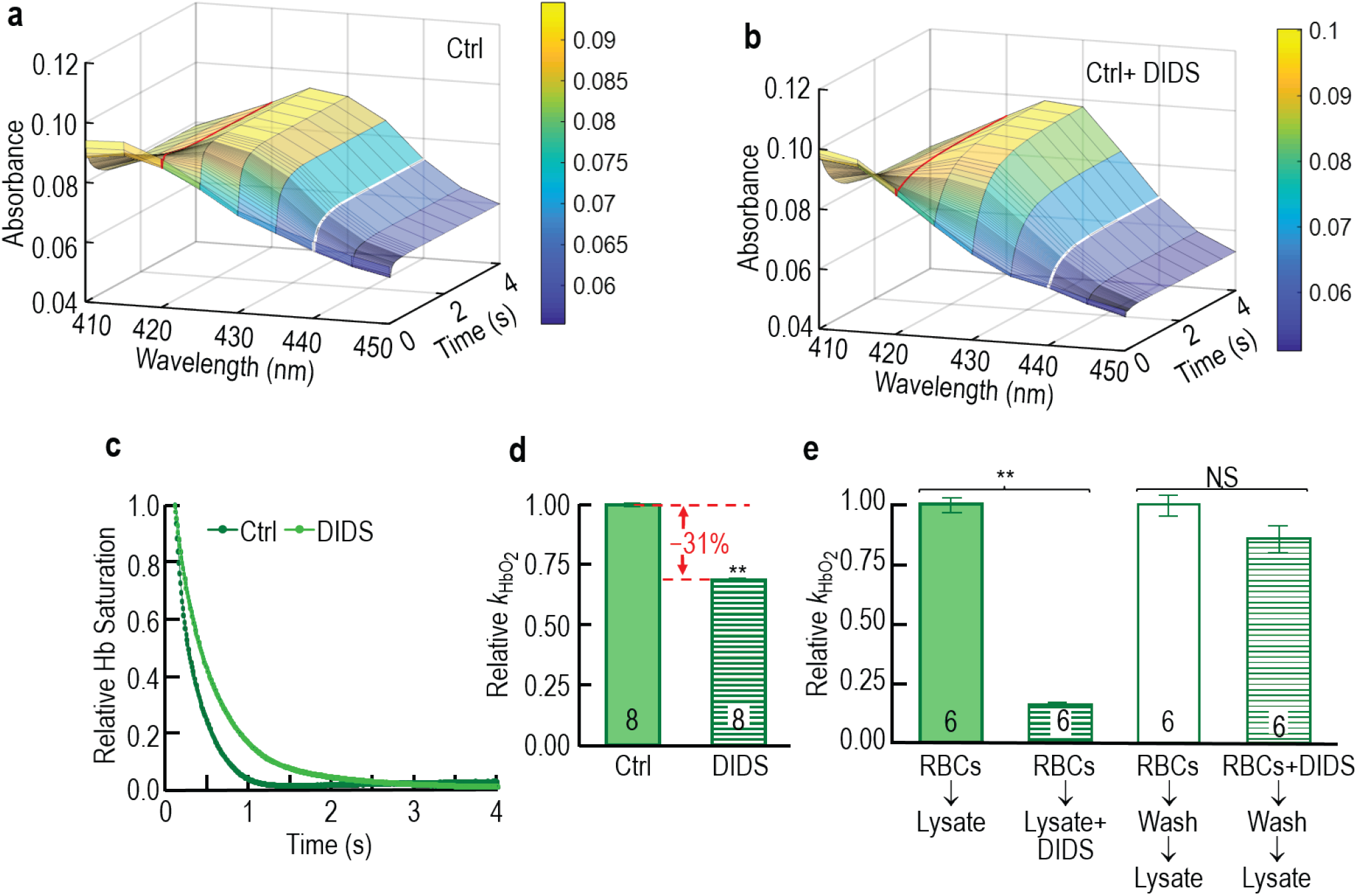
Effect of extracellular DIDS on rate constant of hemoglobin deoxygenation 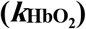 during O_2_ efflux from RBCs. **a & b**, Representative time courses of hemoglobin absorbance spectra during deoxygenation of intact RBCs, from which we compute 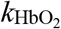. **c**, Time courses of relative hemoglobin saturation for the same two data sets in the previous panels. **d**, Summary of effect of DIDS on 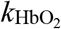 in intact RBCs, corrected for hemolysis as described in Extended Data Figure 3. **e**, Exclusion of DIDS from RBC cytosol. Each “n” represents RBCs from 1 WT mouse. We performed paired t-tests (see below). \*\**P*<0.01; NS, not significant.

To determine whether DIDS accesses RBC cytosol, we used Hb as a sentinel (DIDS has little effect on CA). Figure 2*e* shows that, although incubating hemolysate with DIDS substantially reduces lysate 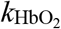, treating intact RBCs with DIDS does not. Thus, DIDS does not significantly cross RBC membranes.

Because pCMBS and DIDS each decrease 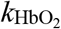 by substantial amounts, without crossing the membrane (Figure 1 and Figure 2) or interacting with membrane lipid,^20,21^ they must block O_2_-conducting proteins. We began our search for such proteins by examining RBCs from mice genetically deficient in AQP1 or RhAG, which have established roles as major CO_2_ channels in human RBCs. Figure 3*a* shows representative experiments comparing 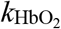 of RBCs from WT mice with those from *Aqp1*–/–, *RHag*–/–, or dKO mice. Our observation that the individual and double knockouts do not significantly affect hemolysis (Extended Data Figure 1*c*) is consistent with the report that deletion of *mRh* and *RHag* do not increase RBC fragility^23^. Spherocytosis of RBCs from dKO mice is low (Extended Data Figure 1*d*). The summary in Figure 3*d* shows that KOs decrease hemolysis-/shape-corrected 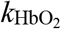 values by ∼9% for *Aqp1*–/–, ∼17% for *RHag*–/–, and ∼30% for the dKO. Compared to RBCs from WT mice, those from dKO mice reveal substantially less drug-induced spherocytosis (Extended Data Figure 1*d*), presumably reflecting the absence of two drug targets. Figure 3*e* reveal that the combination of dKO+pCMBS decreases 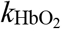 by 78%, and dKO+DIDS, by 53%. These results show that AQP1, RhAG, and at least one other protein make substantial contributions to RBC O_2_ permeability.

**Figure 3.**
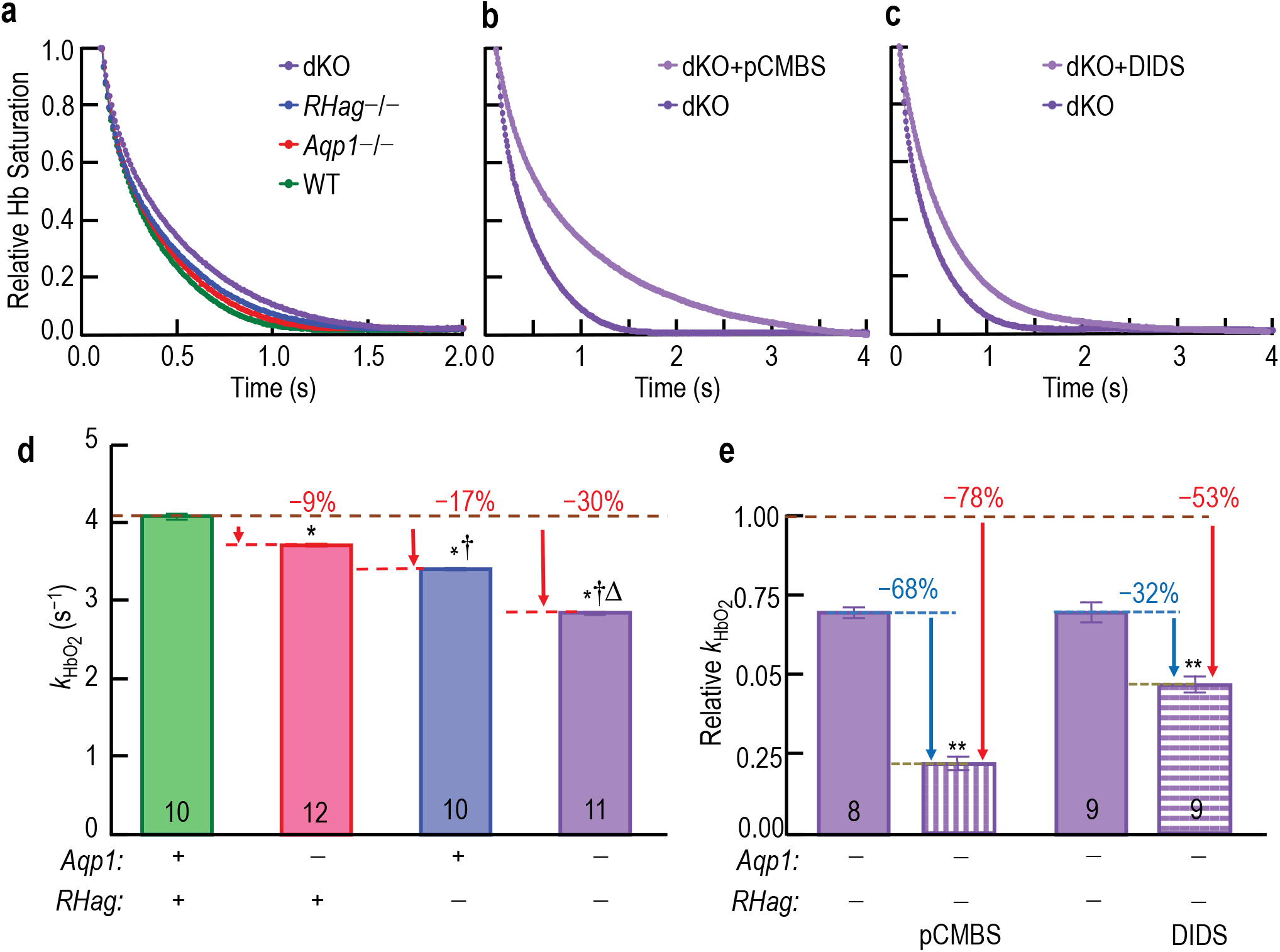
Effect of genetic deletions of *Aqp1, RHag*, or both on rate constant of hemoglobin deoxygenation 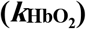 during O_2_ efflux from RBCs. **a**, Effect of genetic deletions on 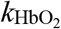, in the absence of inhibitors. For each mouse, we computed 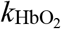 as in Figure 1*a*. **b & c**, Effects of pCMBS or DIDS on 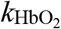, on a dKO background. **d & e**, Summary of 3 full data sets (accommodated for hemolysis and spherocytosis) of which panels *a–c* are representative. In *e*, we normalize each filled bar to the height of the violet bar in *d*, and appropriately scale the hatched bar. Each “n” represents RBCs from 1 mouse, with an independent hemolysis correction (Extended Data Figure 1 & Extended Data Figure 3). For *d*, we performed a one-way ANOVA, followed by a Holm-Bonferroni correction (see below). *Significant vs. WT, ^†^Also significant vs. *Aqp1*–/–, ^*Δ*^Also significant vs. *RHag*–/–. For *e*, we performed paired t-tests (see below). **P<0.01 vs. dKO.

Because the deletion of one protein could alter the expression of others, we assessed effects of the knockouts on RBC membrane-protein levels by purifying proteins from RBC ghosts, and quantitating proteins by mass spectrometry (label-free LC/MS/MS). We detected 211 plasma-membrane–associated (PMA) proteins. Figure 4*a* and *b* show that *Aqp1* or *RHag* deletions produce the expected elimination of the cognate proteins from RBC membranes, as well as mRh, which forms heterotrimers with RhAG^24^ and falls pari passu with RhAG. The deletions do not significantly affect levels of any of the other 47 PMA proteins with the greatest inferred abundance (Extended Data Figure 4*a*). Of the 27 proteins in the RBC-ghost preparation exhibiting a significant change with at least 1 genetic deletion (Extended Data Figure 4*b*), the most abundant is <1% as abundant as AE1. Thus, secondary changes in membrane proteins other than AQP1 and RhAG/mRh are unlikely to account for the observed decreases in 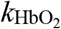.

**Figure 4.**
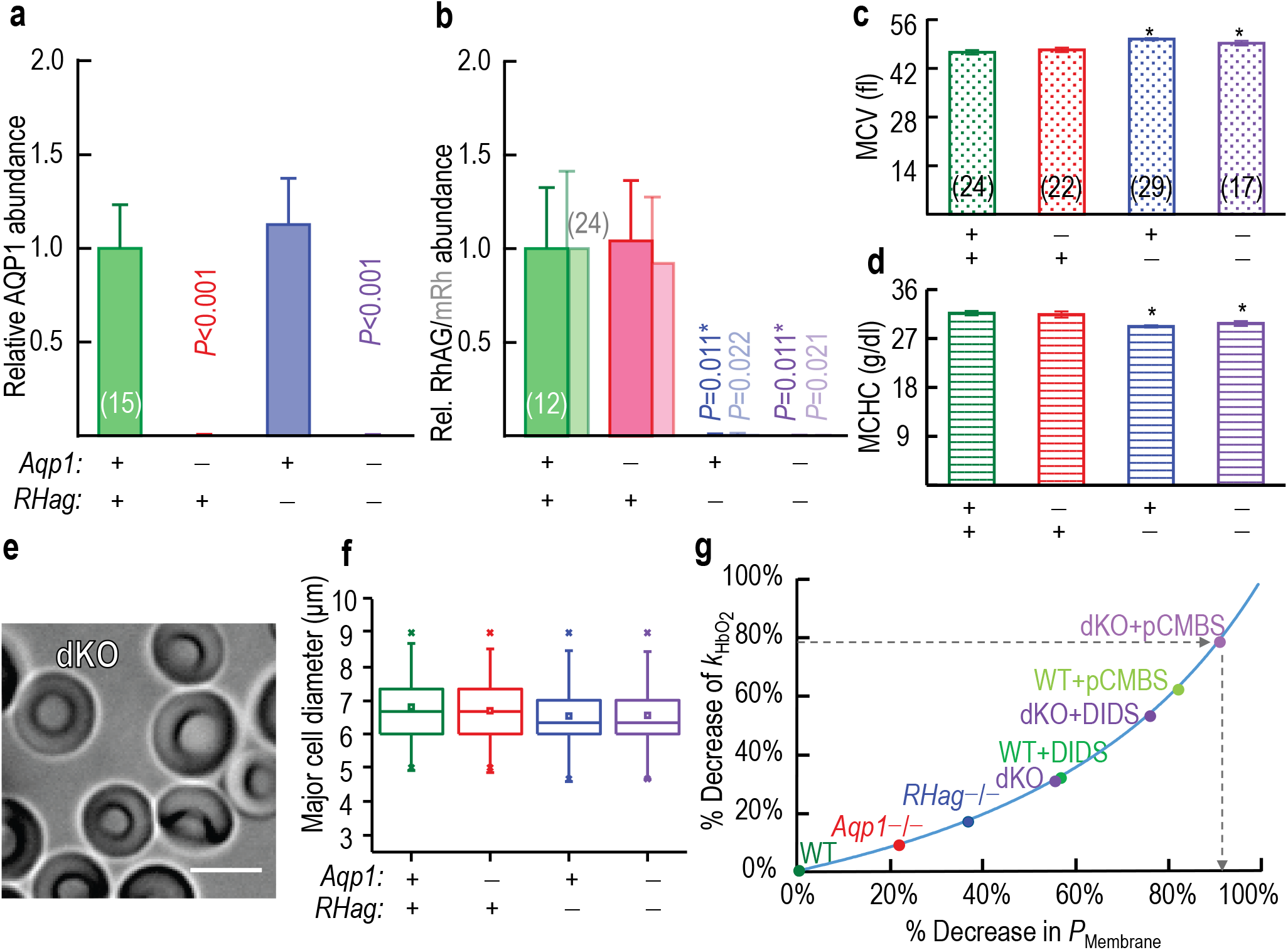
Effect of genetic deletions of *Aqp1, RHag*, or both on AQP1/RhAG protein expression, RBC morphology, and predicted effects of altered morphology on 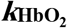. **a & b**, Proteomic analysis by LC/MS/MS of all peptides of AQP1, RhAG (intense colors), and mRh (muted colors) from RBC ghosts, normalized to 1 for WT. See Extended Data Figure 4*a* for 22 “membrane-associated” proteins among the 50 most abundant proteins with greatest inferred abundance in WTs; see Supplementary Data Figure 1 for how each genetic deletion affects each (none significantly affected). Extended Data Figure 4*b* shows inferred abundance in WTs of all 27 proteins (out of 1104 detected), for which levels did change significantly with ≥1 genetic deletion; see Supplementary Data Figure 2 for how genetic deletions affected each of these proteins of much lower inferred abundance. Bars represent mean peak intensity (AUC) ±s.e.m. We performed unpaired Welch’s t-tests, followed by Holm-Bonferroni corrections (see below). Each “n” represents number of peptides per protein, 3 mice/genotype. **c & d**, Mean corpuscular volume (MCV) and mean corpuscular hemoglobin concentration (MCHC) data from automated hematological analyses (below). Each “n” represents 1 mouse. Bars represent means ± s.e.m. We performed a one-way ANOVA, followed by the Holm-Bonferroni correction (see Methods/Statistical Analysis). *denotes statistical significance vs. WT. See Extended Data Figure 5 and (Supplementary Table 2) for additional hematology data. **e**, DIC micrograph of fresh RBCs from dKO mouse, tumbling through plane of focus and revealing normal biconcave disks. Bar represents 5 μm. See Extended Data Figure 6 for blood smears, Extended Data Figure 7 for additional DIC still micrographs, and Supplementary Videos of RBCs tumbling through the plane of focus. **f**, ImageStream flow cytometry, showing major cell diameters for all cell types. Blood samples from 4 mice analyzed for each group. Boxes, interquartile range (25-75%); horizontal line within box, median; open square within box, mean; whiskers, 2 s.d.; × symbols, 1% and 99%. See Extended Data Figure 8 and Supplementary Table 3 through Supplementary Table 5 for additional data. **g**, Predicted effects of decreasing the O_2_ permeability of the RBC membrane (*P*_Membrane_, expressed as % decrease vs. WT without inhibitors) on 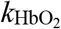. Using the reaction-diffusion model (see below), we simulated the time course of HbO_2_ desaturation, employing hematological and morphological data gathered for RBCs from WT mice. Because the time course of the simulated HbO_2_ desaturation is not precisely exponential, we determined a quasi-time constant, the simulated time to 37% of complete HbO_2_ desaturation (i.e., τ ≅ *t*_37_), from which we estimated the experimental rate constant of HbO_2_ desaturation from *t*_37_ (i.e., 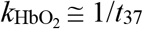). We then systematically decreased *P*_Membrane_ (x-axis), and computed the predicted % decrease in 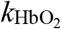 (y-axis; blue curve). For our 8 experimental conditions (labeled points), we used experimentally determined 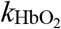 values to determine position on the y-axis, and then placed the points on the blue curve. Thus, the observed 79% decrease in 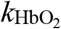 for dKO+pCMBS (horizontal arrow) corresponds to a 92% decrease in *P*_Membrane_ (downward arrow).

Because altered RBC geometry, by altering intracellular diffusion distances, could change 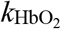, we performed hematological analyses on fresh blood from all genotypes (Supplementary Table 2). We observe no significant sex- or genotype-related differences (Extended Data Figure 5*a-c*) except—in *RHag*–/– and dKO mice—slightly higher mean corpuscular volume (MCV; see Figure 4*c*), slightly lower mean corpuscular hemoglobin concentration (MCHC; see Figure 4*d*), and slightly greater RBC distribution width. Review of blood smears reveals unremarkable RBC morphology, with no differences among genotypes for either control cells (Extended Data Figure 6) or drug-treated cells (Supplementary Data Figure 7). Previous authors noted normal RBC morphology for *RHag*–/– mice^23^. DIC microscopy, micrographs (Figure 4*e*, Extended Data Figure 7) and videos of live, tumbling RBCs (Supplementary Videos) confirm that RBCs of all genotypes are normal biconcave disks. DIC studies (Supplementary Data Figure 8) show that the misshapen cells identified/quantitated by ImageStream are, in fact, spherocytes. Forward light scattering during flow cytometry (Extended Data Figure 8) is consistent with the slightly greater MCVs in knockouts. ImageStream flow cytometry reveals that mean maximal diameters of RBCs+precursors are slightly less in knockouts than WTs (Figure 4*f* and Supplementary Tables 5 *–* 6), which implies, together with increased MCVs, that knockout cells are slightly thicker.

To assess effects of small changes in RBC dimensions and [Hb], we performed mathematical simulations using a reaction-diffusion model (see Methods and Extended Data Figure 9*a*) populated with measured parameter values and known constants. We assume a 1-μm EUF thickness (ℒ_EUF_), which—given our [NTD]_o_ of 25 mM—is probably an overestimate (Supplementary Discussion). For untreated WT RBCs, our model yields a simulated 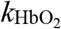 value within ∼1% of experimental values. In simulating 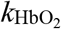 for KO mice, we use our measured genotype-specific values of mean corpuscular volume, mean corpuscular Hb, and major diameter, and find that gene deletions would either increase 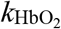 (*Aqp1*) or produce decreases (*RHag*, dKO) far smaller than those observed (Extended Data Figure 9*b*).

Recalling that *R*_Total_ = *R*_ICF_ + *R*_Membrane_ + *R*_EUF_, what do the present data tell us about the contribution of *R*_Membrane_ and O_2_ channels to *R*_Total_? *R*_ICF_ is doubtlessly important, as emphasized recently by Richardson and colleagues^25^. However, their low [NDT]_o_ of 1 mM (vs. our value of 25 mM) likely leads to a large ℒ_EUF_ and thus an overestimated *R*_ICF_. ^5,6,26^ Thus, in our simulations, we have assumed that the intracellular diffusion constant for O_2_ is the arithmetic mean of the generally accepted value and the Richardson value (Supplementary Methods/Parameter values). Our simulations argue that *R*_Membrane_—even for WT RBCs without inhibitors—is ∼33% of *R*_Total_ (Supplementary Discussion). To the extent that our ℒ_EUF_ estimate is low, the *R*_Membrane_ percentage is even higher. Thus, even for control RBCs from WT mice, the membrane offers substantial resistance to O_2_ diffusion. However, *R*_Total_ and *R*_Membrane_ are as low as they are only because of O_2_ channels. Figure 4*g* and Extended Data Figure 9*c-d* summarize the predicted dependence of 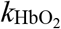 on *P*_Membrane_. Thus, the 30% decrease in 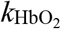 caused by dKO corresponds to a ∼55% decrease in *P*_Membrane_, so that *R*_Membrane_ now represents nearly half of *R*_Total_. The 78% decrease in 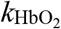 caused by dKO+pCMBS corresponds to a ∼91% decrease in *P*_Membrane_, so that *R*_Membrane_ now represents ∼83% of *R*_Total_. It is not clear how drugs confined to the outside of the membrane or deletion of membrane proteins could produce the observed decreases of 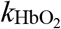 in any way other than decreasing O_2_ egress through channels. Moreover, an analysis of 8 other key parameters (e.g., ℒ_EUF_, cell thickness) shows that no reasonable change in any could explain our 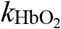 data (Extended Data Figure 10).

In summary, two channel proteins—AQP1 and the Rh complex—contribute ∼55% of the RBCs O_2_ permeability. Other proteins blocked by pCMBS likely contribute at least another 35%. Together with earlier CO_2_ studies, the present work shows that, of the O_2_ and CO_2_ traffic across RBC membranes, >90% occurs through channels. Our work raises questions that should trigger new research: What is the molecular mechanism of O_2_ diffusion through the channels? We suggest that the most likely routes are the hydrophobic central pores at the middle of the AQP1 tetramers and Rh trimers, or other pathways between monomers. Which other membrane proteins contribute to *P*_Membrane_ in RBCs? And to what extent do AQPs, Rh proteins, and other membrane proteins contribute to O_2_ permeability in other cell types? An important implication of O_2_ channels is that they provide a mechanism for cellular regulation of O_2_ permeability. They also offer the future physician the possibility of raising O_2_ permeability (e.g., to enhance performance or wound healing) or lowering O_2_ permeability (e.g., to treat oxygen toxicity) by altering the number of or intrinsic permeability of channels.

## Supporting information

Supplementary Tables (MassSpecData)

Supplementary Discussion

Supplementary Video 2 AQP1

Supplementary Video 3 RhAG

Supplementary Video 4 dKO

## End notes

**Supplementary Information** is available in the online version of the paper.

## Acknowledgements

We thank Jean-Pierre Cartron for the gift of the *RHag*–/– mice and for helpful discussions. We thank Thomas Radford for organizing the husbandry of the mouse colonies; Gerald Babcock for his role as laboratory manager; James W. Jacobberger and Philip G. Woost of the CWRU Flow Cytometry and Imaging Microscopy Core (FCIMC) for their assistance with flow cytometry; Daniela Schlatzer of the CWRU Center for Proteomics and Bioinformatics for their assistance with mass spectrometry. This work was supported by Office of Naval Research (ONR) grant N00014-11-1-0889, N00014-14-1-0716, and N00014-15-1-2060; a Multidisciplinary University Research Initiative (MURI) grant N00014-16-1-2535 from the DoD, NIH grant multi-scale modeling grant 5U01GM111251 (to WFB). R.O. and the modeling were supported in part by NIH grant K01-DK107787. R.R.G was supported by a fellowship grant from the ONR (N00014-12-1-0326). The authors gratefully acknowledge Daniela Calvetti and Erkki Somersalo for having developed the engine of an earlier version of the CO_2_/pH reaction-diffusion model of an oocyte, which in part served as the starting point for the RBC model.

## Author Contributions

P.Z., R.R.G., R.O., G. G., F.J.M. & W.F.B. designed the study; R.R.G., performed the initial stopped-flow experiments; P.Z., A.I.S., A.B.W., S.T., R.O., & F.J.M. performed experiments and collected data; P.Z., A.I.S., A.B.W., D.E.H., H.J.M., R.O., F.J.M., & W.F.B. analyzed data; P.Z., R.O., F.J.M., & W.F.B. wrote the manuscript.

## Author Information

The authors declare no competing financial interests. Readers are welcome to comment on the online version of the paper. Correspondence and requests for materials should be addressed to W.F.B., P.Z. or R.R.G.

## Methods

### Mice

WT C57BL/6 mice (originally obtained as *Aqp1*+/– mice on a C57BL/6 background, a generous gift of Alan Verkman) were backcrossed for >20 generations by our laboratory to define our lab-standard C57BL/6 strain. We backcrossed *Aqp1*–/– mice, with a targeted disruption of the *Aqp1* gene^27^, into our WT mice for >18 generations, and confirmed genotypes by real-time PCR (performed by TransnetYX, Inc.). We backcrossed *RHag*–/– mice (provided by Jean-Pierre Cartron as *RHag*+/– mice on a C57/BL6 background), with a targeted disruption of the *RHag* gene^23^, into our WT mice for at least 7 generations, and confirmed genotypes of *RHag*–/– mice were confirmed by standard PCR (Supplementary Methods). From *Aqp1*–/– and *RHag*–/– mice, we generated dKO (i.e., *Aqp1*–/–*RHag*–/–) mice, and confirmed genotypes as noted above. Mice were allowed access to food and water ad libitum. In experiments, we used mice of both sexes (8 to 19 weeks old). Not shown are data showing that no changes in kinetic or hematological parameters occur up to 6 months of age for all genotypes. All animal procedures were reviewed and approved by the Institutional Animal Care and Use Committee (IACUC) at Case Western Reserve University.

### Physiological solutions

Supplementary Table 12 shows the compositions of all solutions used in this study. We made pH measurements—either at room temperature (RT) or at 10°C—using a portable pH meter (model A121 Orion Star, Thermo Fischer Scientific (TFS), Waltham, MA) fitted with a pH electrode (Ross Sure-Flow combination pH Electrode, TFS). Beakers containing pH calibration buffers (pH at 6, 8 and 10; TFS), physiological solutions to be titrated, and the pH electrode in its storage solution (Beckman Coulter, Inc. Brea, CA) were equilibrated at either RT or 10°C, as appropriate. For work at 10°C, we used a refrigerated, constant-temperature, shaker water bath (model RWB 3220, TFS) and submersible Telesystem magnetic stirrers (TFS). We adjusted pH with NaOH (5 M) or HCl (5 M). Osmolality was measured using a vapor pressure osmometer (Vapro 5520; Wescor, Inc., Logan, UT) and, as necessary, adjusted upward by addition of NaCl.

### Inhibitors

Stock solutions of 2 mM 4-(chloromercuri)benzenesulfonic acid sodium salt (pCMBS; catalog no. C367750; Toronto Research Chemicals, Toronto, ON, Canada) and 2 mM 4,4’-diisothiocya-natostilbene-2,2’-disulfonate (DIDS; catalog no. 1226913; Invitrogen, Eugene, OR) were freshly prepared by dissolving directly into oxygenated solutions, preventing exposure to light. In some experiments, RBC samples were pre-incubated with 1 mM pCMBS for 15 min (Figure 1) or with 200 μM DIDS for 60 min (Figure 2 and Extended Data Figure 3*a*, right bar). For assessing the effect of millisecond exposures to DIDS on the Hb absorbance spectrum (Extended Data Figure 3*a*, stippled bar), we added 200 μM DIDS to the deoxygenated solution.

### Preparation of RBCs

We collected fresh blood from WT or KO mice for seven assays: SF, proteomics/mass spectrometry, hematology, blood smears, still microphotography, microvideography, and flow cytometry. We used the cardiac-puncture method^28^ for some SF studies, and we used the submandibular-bleed method^29^ for the remaining SF and all other studies. For cardiac puncture, a 1-ml syringe (Becton, Dickinson and Co., Franklin Lakes, New Jersey, USA) and attached 23-gauge PrecisionGlide needle (Becton, Dickinson and Co.) was used. For bleeds of the submandibular vein, a 3-, 4-, or 5-mm point-length sterile animal lancet (MEDIpoint, Inc., Mineola, NY) was used, and blood samples (∼250 μl) were taken from the same mouse no sooner than 72 h from a previous sample on the contralateral side.

We collected blood samples for mass spectrometry into 0.6-ml microcentrifuge tubes that we had previously rinsed with 0.1% sodium heparin (H4784, Sigma-Aldrich, St. Louis, MO), and processed the blood as described below. We collected blood for hematology studies into special micropipettes (see below), and for blood smears into 2 ml K2E K2EDTA VACUETTE® tubes (Monroe, NC 28110, USA) (see below). For the other 4 assays (SF, still microphotography, microvideography, flow cytometry), we collected blood into heparinized (see above) microcentrifuge tubes, which we centrifuged in a Microfuge 16 Microcentrifuge (Beckman Coulter, Inc.) at 600 × g for 10 min. We aspirated and discarded the resulting supernatant and buffy coat. To remove residual extracellular Hb, we resuspended the pelleted RBCs in our oxygenated solution (Supplementary Table 12) to a hematocrit (Hct) of 5% to 10%, centrifuged at 600 × g for 5 min. After 4 such washes, RBCs were resuspended in oxygenated solution to a final Hct of 25% to 30%, and maintained on ice (for up to ∼6 h) for experiments. At this point, the RBCs were directed to studies of SF (see below), still microphotography and microvideography (see below), or flow-cytometry (see below).

For SF studies, we computed [Hb] in a suspension of intact RBCs from the Hb absorbance measured on a UV/Vis spectrophotometer (model 730 Life Science, Beckman Coulter, Inc.), using an equation described previously^22^. As necessary, we produced an RBC lysate by osmotic lysis of 20 μl of freshly prepared, packed mouse RBCs in a 1:8 dilution of Milli-Q H_2_O (Milli-Q® Integral Water Purification System, EMD Millipore Corporation, Billerica, MA), followed by centrifugation at 15000 × g for 5 min in a Microfuge 16 Microcentrifuge, at RT. We transferred the supernatant to a clean 1.5-ml tube for a spectroscopic determination of free [Hb], as described above^22^ for intact RBCs. For all stopped-flow experiments—on intact RBCs or lysates or mixtures thereof—we use a final [Hb] of 2.5 μM in the SF reaction cell.

### Quantitation of hemolysis in SF reaction cell

We have developed a novel assay^22^, based on carbonic-anhydrase activity, for determining RBC hemolysis in the SF reaction cell in a time domain similar to that of our O_2_-efflux experiments. We use a SX-20 stopped-flow apparatus (Applied Photophysics, Leatherhead, UK; fluorescence spectroscopy mode) to mix solutions A and B (Supplementary Table 12) in the SF reaction cell, and create an out-of-equilibrium (OOE) 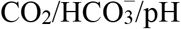 state that re-equilibrates according to the reaction 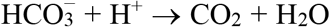, catalyzed by CA released from hemolyzed RBCs. Solution A contains HEPES/pH 7.03 + RBCs or hemolysate. Solution B contains ∼1% CO_2_/44 mM 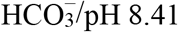 + the fluorescent pH-sensitive dye pyranine (H348, Invitrogen), which reports extracellular pH (pH_o_). We excite the dye at 460 nm or 415 nm, while monitoring total fluorescence emission using a 488-nm long-pass filter. Immediately upon mixing of A and B, the solution in the SF reaction cell (10°C) has the composition ∼0.5% 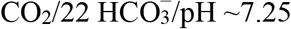, which is out of equilibrium. The pH_o_ rises exponentially (rate constant, *k*_ΔpH_, computed as described previously^22^) to ∼7.50.

We calculate the actual percent hemolysis (*Act%H*) of ostensibly 100% intact RBCs (i.e., apparent percent hemolysis, *App%H*, is 0%) during a SF experiment, as described previously^22^, using a procedure that requires three rate constants: *k*_uncat_, *k*_RBC,Lysate_, and *k*_RBC,OstInt_. (1) *k*_uncat_ is the uncatalyzed rate constant (without CA). (2) *k*_RBC,Lysate_ is *k*_ΔpH_ in the presence of fully lysed RBCs. And (3) *k*_RBC,OstInt_ is *k*_ΔpH_ in the presence of ostensibly 100% intact RBCs. From these, we compute *k*_cat,min_ = *k*_RBC,OstInt_ – *k*_uncat_ and *k*_cat,max_ = *k*_RBC,Lysate_ – *k*_uncat_. Finally:

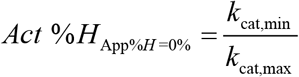

### Stopped-flow absorbance spectroscopy and calculation of 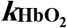

We used the SF apparatus (absorbance spectroscopy mode) to determine the rate constant 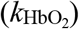 of hemoglobin deoxygenation. We load RBCs ([Hb] = 5 μM, Hct ≅ 0.3%) in the oxygenated solution (Supplementary Table 12) into one syringe, and load the deoxygenated solution containing the O_2_ scavenger NDT (50 mM) in the other syringe. We record absorbance (*A*) in the SF reaction cell (10°C) over the portion of the Hb spectrum with highest Signal/Noise (wavelength [λ]: 410–450 nm, at 5 nm intervals), sampling at 10-ms intervals for 4000 ms. For each new set of loaded samples, we began by performing ≥6 “shots” to ensure that the new solutions are loaded into the SF reaction cell, and then sequentially acquired *A*_λ_ vs. time during 9 separate shots at λ = 410, 415 … 450 nm. We compiled such data set into 3-D graphs (e.g., *A*_λ_ vs. λ vs. time plots in Figure 1*a*). Then, as described in Supplementary Methods, we use absorbances at 6 wavelengths (i.e., 410, 415, 425, 430, 435, 440 nm)—sufficiently distant from isosbestic wavelengths—to compute Hb saturation at each time point (e.g., Figure 1*c*) beginning at t = 100 ms, and finally computed the quasi-rate constant 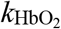 (e.g., Figure 1*d*). We accept a data set only if each of the 9 records of *A*_λ_ vs. time approaches an asymptote. We accept 1 to 5 such data sets for each set of conditions, and then average these 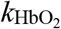 values to obtain the 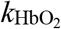 value used in further analyses.

### Correction of 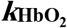 for hemolysis

Because Hb released into free solution deoxygenates faster than Hb inside RBCs, we correct raw 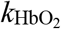 values, based on *Act*%*H*, determined as described above. For each mouse, we generate simulated hemolysis samples for apparent hemolysis levels of 0% (i.e., ostensibly 100% intact RBCs), 2.5%, 5 %, 10% and 25% by mixing ostensibly 100% intact RBCs (100%, 97.5%, 95%, 90%, and 75%) with hemolysate (0%, 2.5%, 5%, 10%, and 25%)—keeping total [Hb] constant at 5 μM. We determine 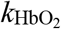, as described above, for each simulated hemolysis sample, and—using an algebraic translation of the x-axis—convert a plot 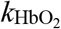 vs. apparent % hemolysis (e.g., blue elements in Extended Data Figure 1 and Extended Data Figure 3*b*) to a plot of 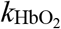 vs. actual % hemolysis (green elements). Extrapolating 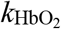 back to 0% actual hemolysis yields an estimate of the actual 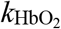 of 1 mouse. We used this approach for RBCs treated with pCMBS or no drug.

For RBCs treated with DIDS, we modified the above approach because DIDS decreases the 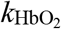 of free Hb, so that the actual slope of the 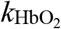 vs. *Act%H* regression line (*Slope* _DIDS,Act_) is less than the control value (*Slope* _Ctrl,Act_). Two factors are at work. (1) During the 1-h pretreatment with DIDS, the drug produces a substantial reduction in 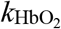 (compare hatched and open bars in Extended Data Figure 3*a*) for the minute fraction of Hb already released^22^ (∼0.4%) from RBCs at this stage. (2) During the actual SF experiment, DIDS produces a small reduction in 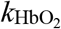 (compare stippled and open bars in Extended Data Figure 3*a*) for a larger fraction of Hb newly released in the SF reaction cell. Thus, in a DIDS experiment, the actual slope of the regression line becomes:

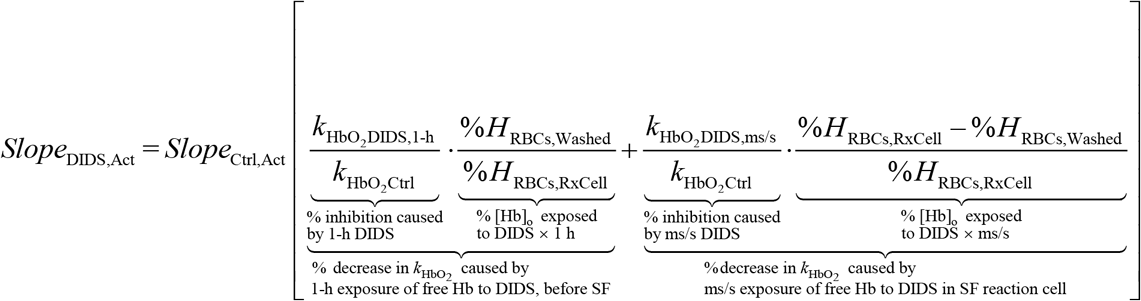

Extended Data Figure 3*b* shows the 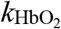 hemolysis correction for RBCs—from 1 mouse—studied under control conditions (i.e., no DIDS). Here, *Slope* _Ctrl,Act_ is 6.51 s^−1^/*Act%H*. For RBCs from the same mouse, but treated ×1 h with 200 μM DIDS, the *Act%H* was 12.7%. Inserting these and the other values for this mouse into the above equation yields:

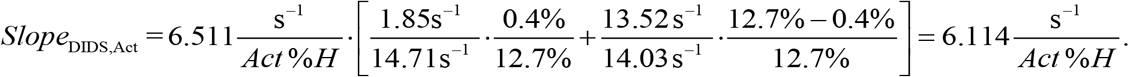

Thus, knowing the 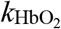 measured in ostensibly 100% intact RBCs 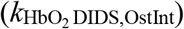, we can estimate the actual 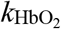 for DIDS-treated RBCs as:

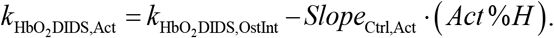

As summarized in Extended Data Figure 3*b*, for this particular mouse:

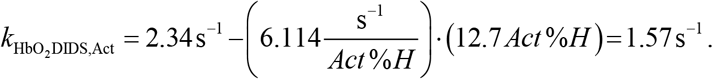

### Accommodation for spherocytes

As described in Supplementary Methods, we generated preparations of RBCs from WT and dKO mice, and then treated these with no drug, pCMBS or DIDS as in 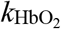 experiments. Here, however, after treatment ±drug, we performed hematology (Supplementary Table 7, Supplementary Table 9), blood-smears (Supplementary Data Figure 7), microphotography (Supplementary Data Figure 8), and ImageStream (Supplementary Table 8, Supplementary Data Figure 9) assays. For each of six conditions ([WT vs. dKO] × [control vs. pCMBS vs. DIDS]), ImageStream flow cytometry (see Methods/ImageStream) revealed spherocyte abundance and diameter, from which mathematical simulations yielded spherocyte 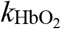 (Supplementary Methods). Linear combinations of HbSat vs. time for normal RBCs and spherocytes allowed us to reconstruct the 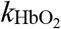 of normal RBCs, from which we computed *P*_Membrane_. See Methods/Accommodation for Spherocytes for an overview of this reconstruction procedure.

### RBC-ghosts preparation and Proteomic analysis

Blood samples (0.2 to 0.5 ml) were collected into heparinized microcentrifuge tubes from 3 mice for each genotype, and then placed the tubes on ice, pending immediate processing into ghosts. We generated erythrocyte ghosts as previously described^30^, with the following changes to buffer composition (Supplementary Table 12): RBC lysis and post-lysis wash buffers contained 5 mM Tris-HCl/pH 8.0 and complete Protease Inhibitor Cocktail tablets (cPIC; Roche, 04693116001), as opposed to sodium-phosphate buffer/pH 7.5 and PMSF and pepstatin A—a modification required to obtain good mass-spectrometry data. Ghosts were then flash-frozen and held at −80 °C prior to mass-spectrometry analysis.

Mass spectrometry experiments were performed by the CWRU Center for Proteomics and Bioinformatics. Briefly, RBC ghost samples were lysed with 2% SDS and cPIC, using pulse sonification. SDS was removed by filter-aided sample preparation^31^ detergent cleanup, and total protein concentration was determined by the Bio-Rad Protein assay kit (Bio-Rad, Hercules CA). A 10-µg sample was then digested with LysC/Trypsin, and 300 ng of the resulting product was analyzed via 4-hr LC/MS/MS. Data were processed and quantified using Elucidator, and analyzed statistically using one-way ANOVA. See Supplementary Methods for details.

### Automated hematological analyses

We collected fresh blood into a 20-μl plastic Boule MPA Micro pipettes (EDTA-K2, Boule Medical AB, Stockholm, Sweden), which we inserted into a Heska HemaTrue® Veterinary Hematology Analyzer (Heska corporation, Loveland, CO) according to the manufacturer’s protocol. See Supplementary Methods/Inhibitor Studies for details on preparation of RBCs for automated hematological analyses in inhibitor studies.

### Blood smears

We collected fresh blood from 3 mice of each genotype into 2-ml K2E K2EDTA VACUETTE® tubes (Monroe, NC 28110, USA). Blood smears were prepared using microscope slides (Fisher scientific, Pittsburgh, USA), and were stained using Wright’s stain on a Sysmex SP-10 autostainer (Kobe, Japan). Air-dried smears were placed in staining cassettes, and staining performed per manufacturer’s specifications. Blood smears images (1000× magnification) were taken on an Olympus BH-2 microscope (Tokyo, Japan) with a DP73 digital camera attachment (Olympus) and visualized with cellSens software (Olympus). See Supplementary Methods/Inhibitor studies for details on preparation of RBCs for blood smears in inhibitor studies.

### Still microphotography and microvideography of living RBCs

We performed experiments (still or video) on fresh blood collected as described above from one mouse of each of four genotypes, all on the same day. We repeated this on three different days, for a total of 3 mice of each genotype, for both still and video studies (i.e., a total of 3+3 mice/genotype). RBCs at an initial Hct of 25% to 30% (see above) were suspended 1:10 in oxygenated solution (Supplementary Table 12), for a final Hct of 0.5% to 1%, and stored on ice for 30 – 120 min before imaging. A droplet containing suspended RBCs from one mouse was placed on a glass coverslip that served as the bottom of a recording chamber. The chamber was then mounted on Olympus IX-81 inverted microscope equipped for differential interference contrast (DIC) studies, using either of two oil immersion objectives (60× objective, NA 1.42 for still micrographs or 40× objective, NA 1.35 for microvideos) with a 1.5× magnification selector. The light was detected with an intensified EM-CCD camera (C9100-13, Hamamatsu Corporation, Bridgewater, NJ) with 512 × 512 pixels, and data acquired using SlideBook 5.0 software (Intelligent Imaging Innovation, Denver, CO) for the Hamamatsu camera. We recorded still micrographs or microvideos (1 frame per 5 sec) of the RBC droplet as RBCs fell freely through the plane of focus, toward the coverslip surface. See Supplementary Methods/Inhibitor studies for details on preparation of RBCs for blood smears in inhibitor studies.

### Flow cytometry

On one day, fresh blood was collected (see above) from 2 mice of each genotype. Experiments were repeated on a second day, for a total of 4 mice per genotype. To permit gating of viable RBC precursors, 100-μl samples of cells were diluted to 1% Hct in oxygenated solution containing 1 μM Calcein Violet (CV; viability marker; ex/em 405/450 nm; TFS, C3099), 0.1 μg/ml Thiazol Orange (TO; to stain RNA; ex/em 490/530 nm; Sigma-Aldrich 390062), and 5 μM DRAQ-5 (to stain DNA; ex/em 647/683 nm; TFS, 62252), and then incubated for 20 min at RT in the dark. The dye-loaded RBC samples were then washed ×3 in 1 ml oxygenated solution, centrifuged at 600×g for 5 min between washes. Experiments were performed at either 0.06% Hct (∼2 million cells/ml) for light scattering on the LSRII or 1 % Hct for imaging on the ImageStream. Stained cells were maintained on ice for up to ∼2 h on ice for experiments performed that day.

Forward-scatter intensity area (FSC-A) is a measure of cell size; side-scatter intensity area (SSC-A) is a measure of size, granularity, surface projections, and (for asymmetric cells) orientation^32^. FSC-A, SSC-A, as well as the fluorescence of CV, TO, and DRAQ-5 were measured with an LSRII Flow cytometer (BD Biosciences; San Jose, CA). Data were gated on FSC-A vs. FSC-W (width), enabling separation of RBCs from very small events and aggregates (Supplementary Data Figure 3*a*). CV-positive cells (>99.9%) were analyzed for TO and DRAQ-5 fluorescence, yielding gated populations (Supplementary Data Figure 3*b*) of TO-negative/DRAQ-5–negative cells (>96%) and TO-Positive/DRAQ-5–positive cells (∼3%). TO-negative/DRAQ-5–negative cells were then analyzed for light-scatter characteristics in WT, *Aqp1*–/–, *RHag*–/–, and dKO mice (Extended Data Figure 8). To evaluate FSC (size) of RBC populations quantitatively, gates were set on the center 90% of “dim” or negative cells in CV vs. DRAQ5, TO vs. CV, and TO vs. DRAQ5 plots. These gates were combined (Boolean AND) with a center 90% gate on an FSC vs. SSC plot. See legend of Extended Data Figure 8 for additional details on analysis.

ImageStream flow cytometry (Amnis, EMD Millipore) analyzes individual cells, in flow, by bright-field and multiple fluorescence parameters. Data are collected as images—two-dimensional spatial grids, in which a third dimension of intensity is captured for each pixel. RBCs are loaded with CV (for viability), TO (for RNA), and DRAQ-5 (for DNA) as described above. Gating schemes (Supplementary Data Figure 4) were established to allow size analysis of individual RBCs (Supplementary Data Figure 5) and RBC precursor types separately from one another. See Supplementary Methods/Inhibitor studies for details on preparation of RBCs for ImageStream analyses in inhibitor studies. Supplementary Data Figure 9 shows images of abnormally shaped cells, and the legend of that figure describes how we identified them using ImageStream. DIC still microphotography (Supplementary Data Figure 8) and DIC microvideography (not shown) show that these misshapen cells are spheres.

### Mathematical modeling

We used a reaction-diffusion model—based on one originally developed for CO_2_ fluxes^33^—of O_2_ efflux from an RBC that we modeled as a sphere (Supplementary Methods: Model formulation), as has been done by others previously (Supplementary Methods: Sphere). The diameter of the sphere equals RBC thickness, which we computed from measured hematological and morphological parameters (Supplementary Methods: Thickness). As illustrated in Extended Data Figure 9*a*, the spherical RBC, a membrane encompassing ICF, is surrounded by a thin layer of EUF, which is in turn surrounded by an infinite reservoir of bECF. Reactions among O_2_, Hb and HbO_2_ occur only in the ICF. We model the reaction term as the simple one-step reversible reaction

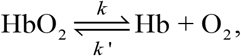

combined with a variation (Supplementary Methods: VRC) of the “variable rate coefficient” model proposed by Moll^34^, which guarantees that the hemoglobin-O_2_ saturation curve is sigmoidal. All solutes diffuse within the cytosol and bECF. O_2_ is the only solute that can move through the plasma membrane. Extended Data Figure 10*b* and *c* show examples of fluxes through selected reaction and diffusion events in two simulated HbO_2_ desaturation time courses. SI Methods contains details on the computational model, parameter values, and the simulation of the time course of deoxygenation of HbO_2_.

### Statistical Analysis

We report results as mean ± s.e.m. In each figure legend, we report which statistical tests we performed, from among the following, to generate unadjusted p-values: (1) paired two-tailed t-test, (2) unpaired two-tailed Welch’s t-test^35^ or (3) one-way analysis of variance (ANOVA). For comparisons of 2 means, we performed #1. For comparisons among >2 means, we performed #2 or #3 and then, to control for type I errors across multiple means comparisons, we applied the Holm-Bonferroni correction, setting the familywise error rate (FWER) to α = 0.05. Briefly, we order the unadjusted p-values for all 𝒩 comparisons in each dataset from lowest to highest. For the first test, we compare the lowest unadjusted p-value to the first adjusted α value, α/𝒩. If the null hypothesis is rejected, then we compare the second-lowest p-value to the second adjusted α value, α/(𝒩–1), and so on. If, at any point, the unadjusted p-value is ≥ the adjusted α, the null hypothesis is accepted and all subsequent hypotheses in the test group are considered null.

### Data availability

The data supporting the findings of this study are available within the paper and its Supplementary Information files. Any further relevant data are available from the corresponding author upon reasonable request.

## Extended Data

**Extended Data Figure 1.**
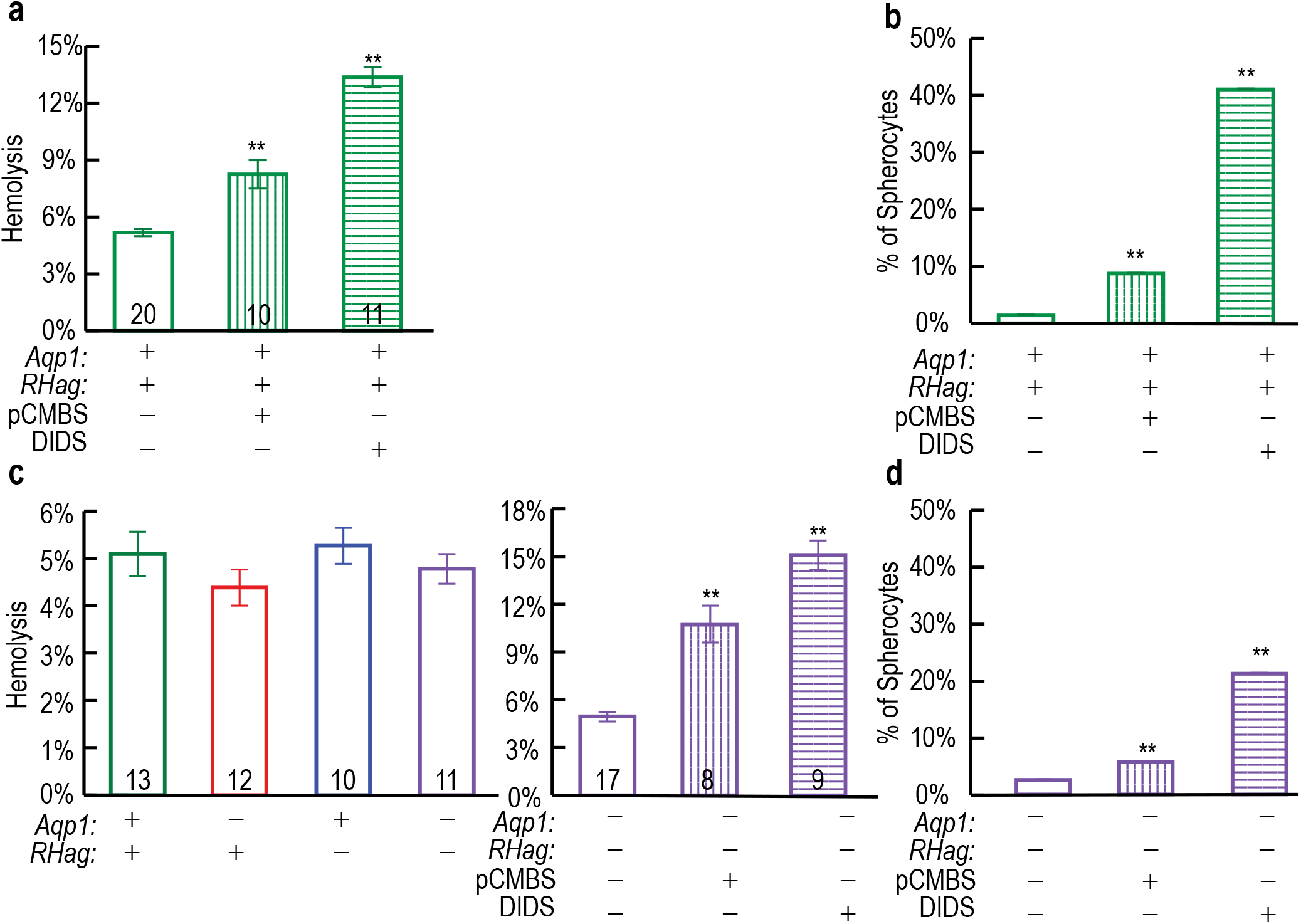
Degree of hemolysis and spherocyte formation in experiments on RBCs. **a**, Hemolysis in experiments on RBCs treated with pCMBS or DIDS. RBCs were treated with 1 mM pCMBS for 15 min or 200 μM DIDS for 60 min. **b**, Prevalence of spherocytes among RBCs, treated with pCMBS or DIDS, from WT mice. **c**, Hemolysis in experiments on mouse RBCs of different genotypes. **d**, Prevalence of spherocytes among RBCs, treated with pCMBS or DIDS, from dKO mice. Bars represent means ± s.e.m. We performed a one-way ANOVA, followed by the Holm-Bonferroni correction for four genotypes results, and we performed unpaired two-tailed t-test for drug treated WT or dKO results (see Methods/Statistical Analysis). \*\**P*<0.01 vs. WT or dKO.

**Extended Data Figure 2.**
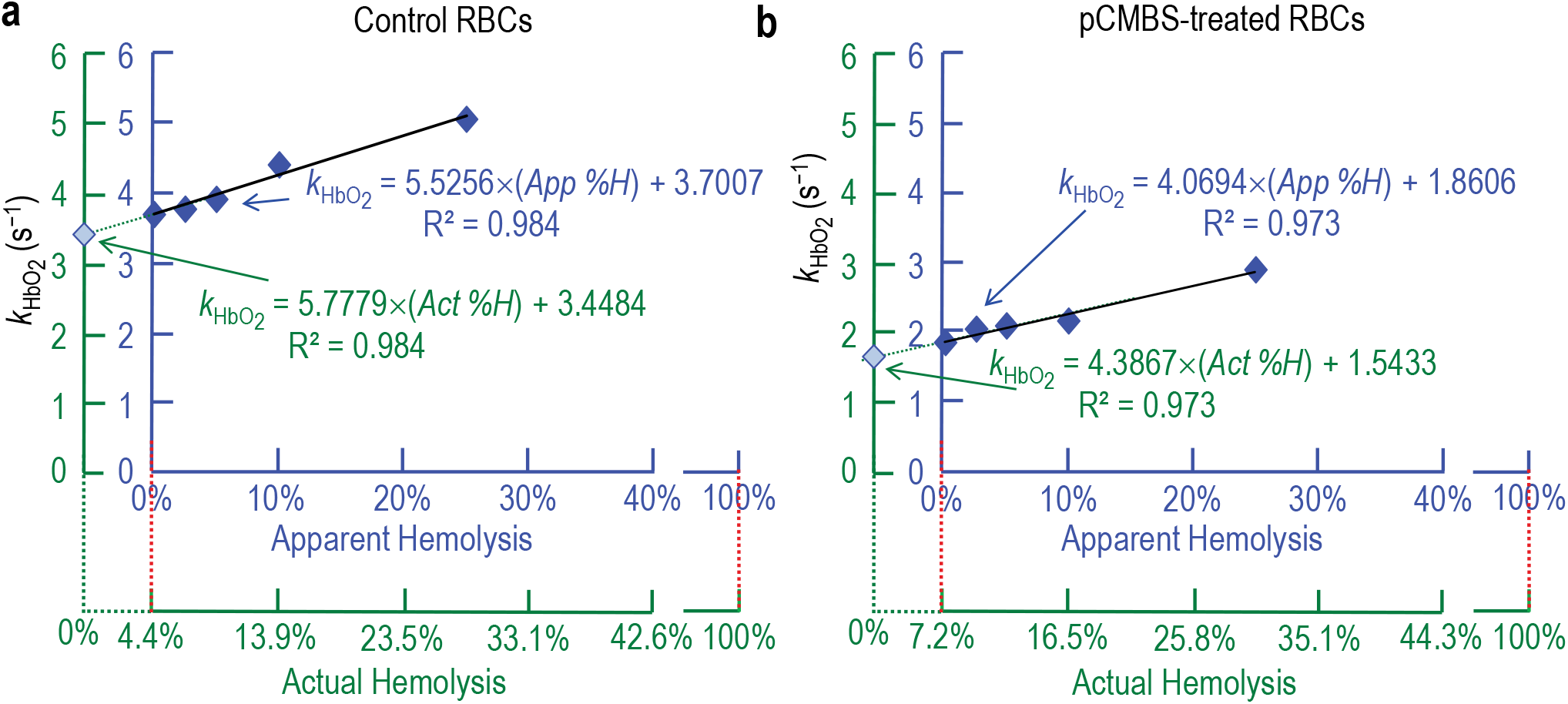
Correction of 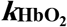 for hemolysis in experiments on RBCs without inhibitors (a), or with pCMBS (b) on blood from the same mouse. We used a novel approach to determine the rate constant 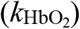 for the deoxygenation of hemoglobin for ostensibly 100% intact RBCs (i.e., apparent percent hemolysis [*App%H*] = 0%) as described above, as well as for mixtures of ostensibly intact RBCs (97.5%, 95%, 90%, and 75%) with fully hemolyzed RBCs (2.5%, 5%, 10%, and 25%) to keep total [Hb] = 2.5 μM in the SF reaction cell. We determined actual percent hemolysis (*Act%H*) of ostensibly 100% intact RBCs as described above for both control RBCs (4.4% for mouse in panel *a*) and pCMBS-treated RBCs (7.2% for mouse in panel *b*). For mixtures of ostensibly 100% intact RBCs and 100% hemolysate, we computed *Act%H* for translation of the x-axis as (*Act%H*) = (*App%H*) + (*Act%H*_App=0_)*(100% – *App%H*). Extrapolating 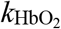 back to *Act%H* = 0% yields an actual 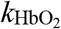, which for the sample in panel *a* was 3.45 s^−1^ for control RBCs and, for the sample in panel *b*, 1.54 s^−1^ for pCMBS-treated RBCs. We repeat this analysis for each mouse.

**Extended Data Figure 3.**
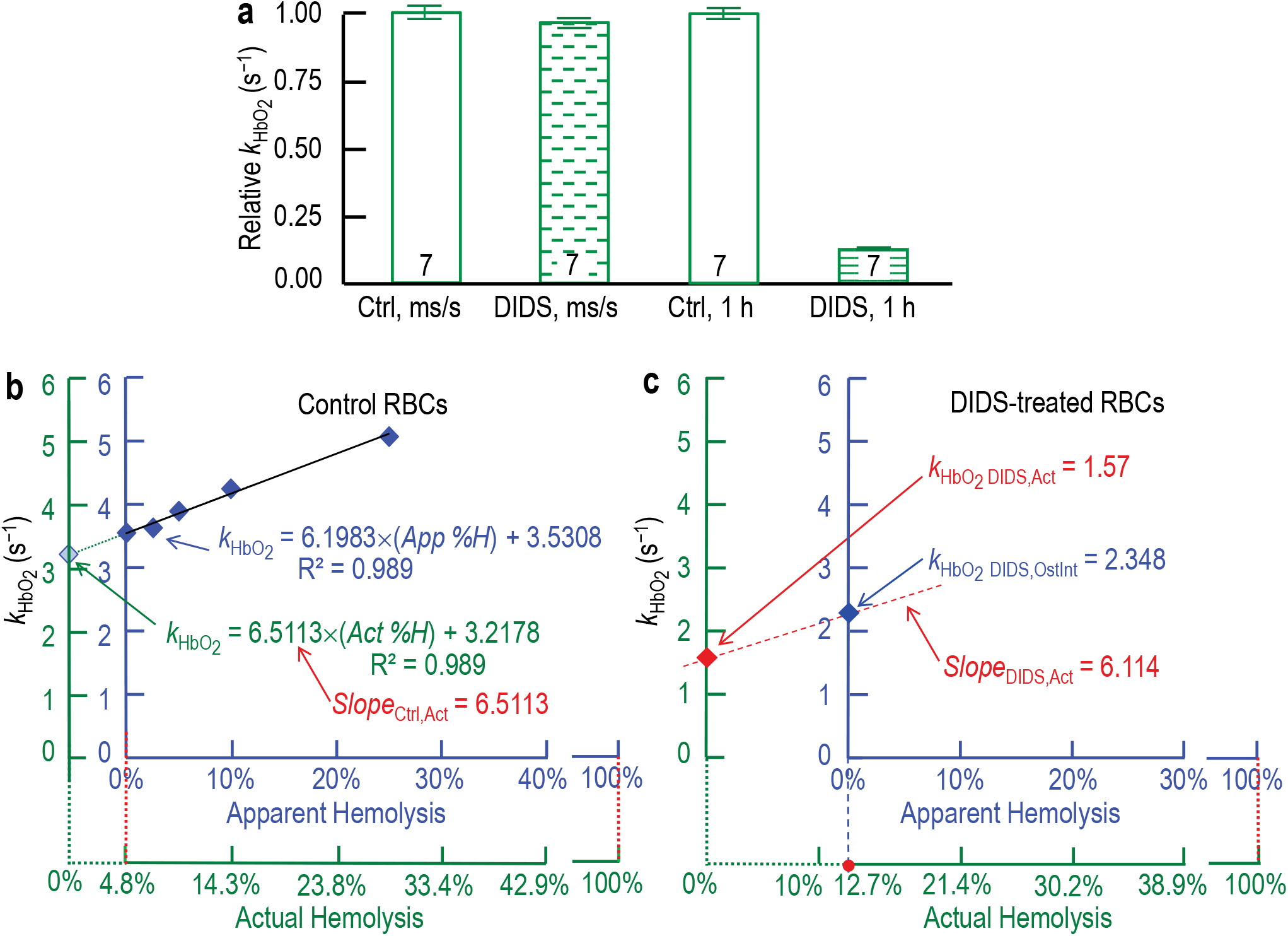
Correction of 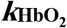 for hemolysis in experiments on RBCs treated with DIDS. **a**, Effect of DIDS on 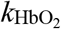 in 100% hemolysate. In SF experiments, we computed the rate constant 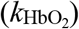 for the deoxygenation of hemoglobin (Hb, from hemolysate) exposed to DIDS for the first time in the SF reaction cell (i.e., for milliseconds to seconds [ms/s]), or for Hb pretreated with 200 μM DIDS for 60 min (1 h). For each of 7 mice, we normalized 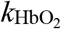 values for ms/s DIDS exposures and 1-h DIDS exposures to the 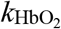 of the matched, untreated Hb. **b**, Determination of control slope (*Slope*_Ctrl,Act_) in plot of 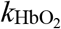 vs. actual percent hemolysis (*Act%H*, green) for RBCs without inhibitors—using the same approach as in Extended Data Figure 2*a*. Extrapolating 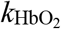 back to *Act%H* = 0% yields an actual 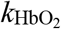, which for sample in panel *b* was 3.22 s^−1^ for control RBCs. **c**, Correction of 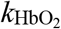 for hemolysis in experiments on RBCs with DIDS. 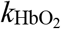 values were determined for ostensibly “100% intact” RBCs as described above. For the sample in panel *c* (from same mouse as in panel *b*), this uncorrected value (blue) was 2.35 s^−1^ for DIDS-treated RBCs. We calculated the slope (*Slope*_DIDS,Act_) of the regression line (red dashed line) using the data in panel *a*, as described above. Extrapolation of this line yielded a corrected 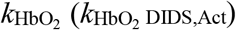 of 1.57 s^−1^.

**Extended Data Figure 4.**
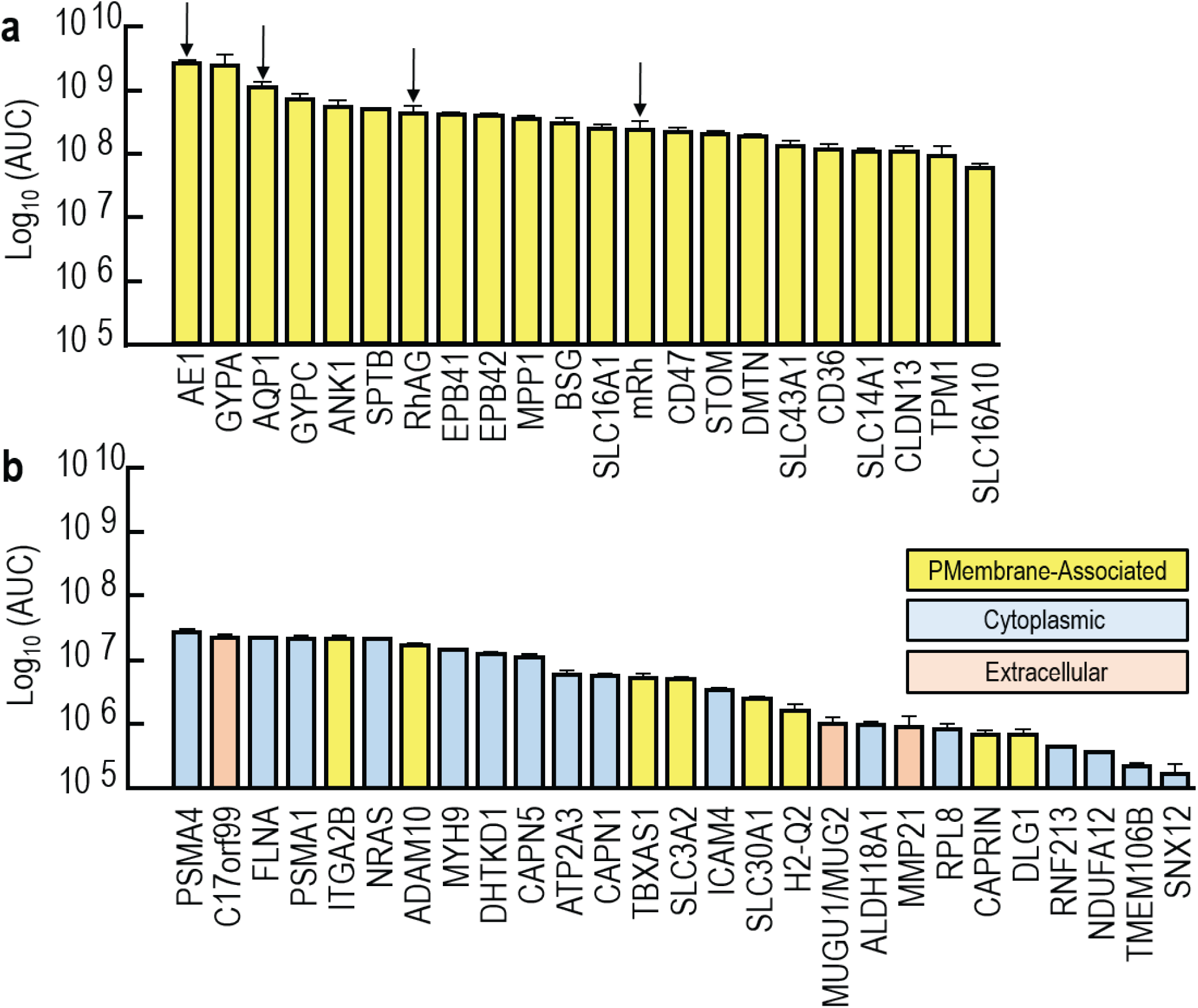
Proteomic analysis by LC/MS/MS of proteins in samples of RBC ghosts from WT mice. **a**, Plasma-membrane–associated proteins, ranked by abundance, as inferred from mass-spectrometry. AUC is area under curve. We detected a total of 7,188 unique peptides from 1,104 unique proteins, not all of RBC origin. Of these, 1,902 peptides and 211 proteins are “plasma-membrane– associated” (integral-membrane proteins + others) as defined by the data-analysis software Elucidator (see Methods/RBC ghosts). None of the 47 PMA proteins of greatest inferred abundance exhibited a significant change in response to any of the deletions. Of these 47, panel *a* shows the 22 PMA proteins, from among the 50 proteins with greatest inferred abundance (Supplementary Table 1*a* provides protein glossary and rank order of abundance). These 22 proteins—which spanning a ∼40-fold range in abundance—include AE1 (represented by 57 peptides), AQP1 (5), RhAG (4), and mRh (8). Arrows indicate AE1 (most abundant species), AQP1, RhAG, and mRh. Of these 22 most abundant PMA protein species, only the intended target proteins in each KO strain display a significant change in abundance. See Figure 4*a* and *b* for effects of gene deletions on abundance of AQP1, RhAG, and mRh. See Supplementary Data Figure 1, for effects of gene deletions on abundance of each of these 22 proteins. Others have shown that mRh expression requires *RHag* in mice^23^. Neither we nor others^36^ have been able to detect GLUT4 by mass spectrometry in RBC ghosts from adult mice. However, like others^37^, we do detect GLUT4 by western blot in RBC ghosts from newborn but not adult mice. See above and Supplementary Methods for details. **b**, The 27 proteins (out of all 1,104 detected by mass spectrometry) from RBC ghosts, ranked by inferred abundance, that demonstrated a significant difference from WT in one or more of the KO mouse strains. For protein glossary and rank order by inferred abundance, see Supplementary Table 1*b* for PMA proteins and Supplementary Table 1*c* for other proteins. Supplementary Data Figure 2 shows the effects of gene deletions on abundance of each of these 27 proteins. The PMA protein with the greatest inferred abundance was ITGA2B, which ranked 48^th^ among PMA proteins, and 136^th^ among all proteins. Consistently falling in parallel with AQP1 is DHTKD1; falling with RhAG are C17orf99, NRAS, and ICAM4; and rising with Rhag are ADAM10 and H2-Q2. Note that even PSMA4 (i.e., most abundant in panel *b*) is ∼4.2-fold less abundant than SLC16A10 (i.e., the least abundant in panel *a*), and 103-fold less than AE1 (i.e., most abundant in panel *a*). Not all of these proteins are from RBCs, let alone RBC integral membrane proteins. The colors of the bars indicate the location of the proteins, as assigned by the data-analysis software Elucidator (see above): yellow (plasma-membrane–associated), blue (cytoplasmic), and tan (extracellular). For both panels *a* and *b*, we purified proteins from RBC ghosts of 3 mice/genotype.

**Extended Data Figure 5.**
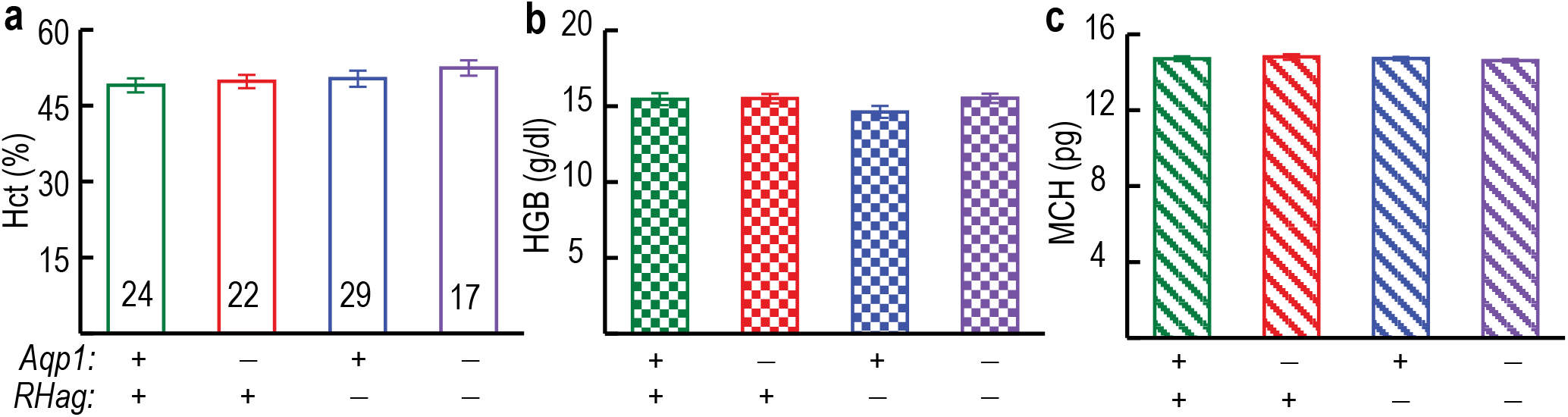
Summary of key hematological parameters. The bar graphs summarize key parameter values from automated hematological analyses (see above) for WT, *Aqp1*–/–, *RHag*–/–, and dKO mice. **a**, Hematocrit (Hct). **b**, Hemoglobin content of whole blood (HGB). **c**, Mean corpuscular hemoglobin content (MCH). Each “n” represents 1 mouse. Results are presented as means ± s.e.m. We performed a one-way ANOVA, followed by the Holm-Bonferroni correction (see below). *Significant vs. WT. See Figure 4*c* for data on mean corpuscular volume, and Supplementary Table 2 for the results of the full hematology screens on fresh blood from WT and KO mice.

**Extended Data Figure 6.**
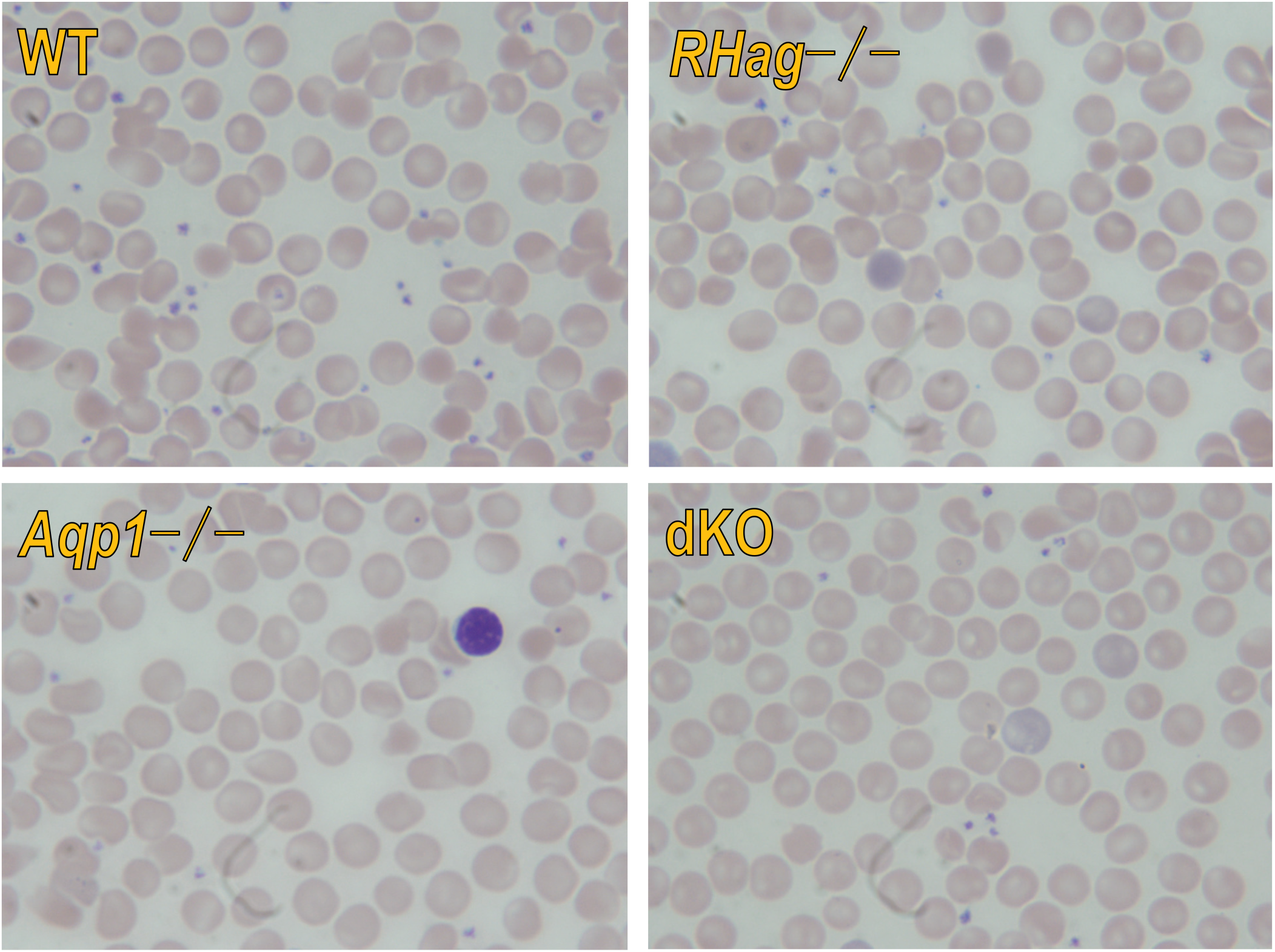
Representative blood smears from each of four genotypes. We reviewed 12 blood smears for each genotype (i.e., 4 smears from each of 3 separate mice), for a total of 48 smears. Red cell morphology was similar in all groups and was unremarkable, demonstrating only occasional target cells and stomatocytes, with no differences noted among knockout strains and WT mice. The dried cells are stained with Wright’s stain; photos taken at 1000× magnification.

**Extended Data Figure 7.**
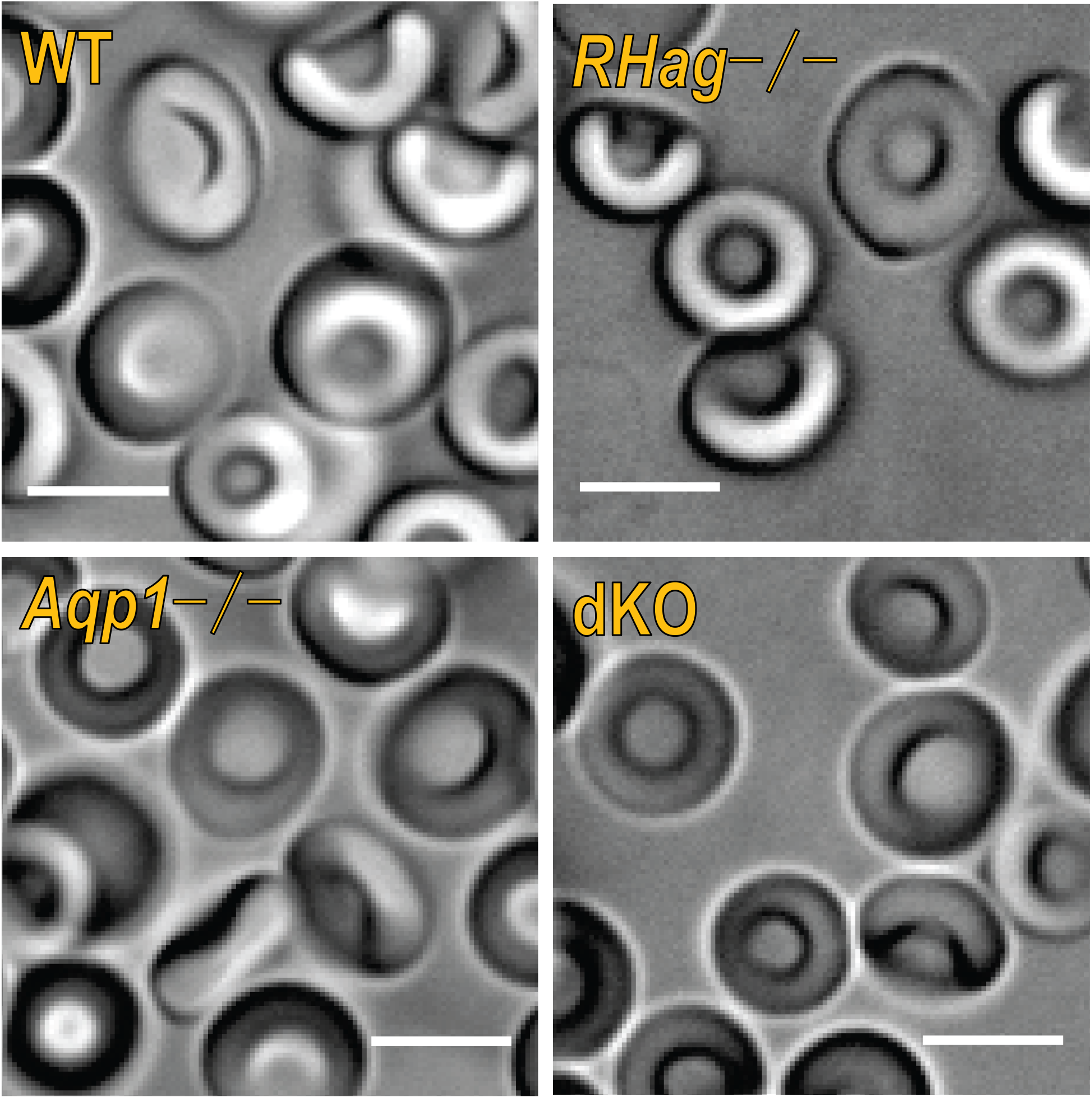
Representative DIC still micrographs of living RBCs. On a given day, we reviewed 4 to 5 samples (i.e., droplets) of RBCs suspended in saline for 1 mouse of each of genotype. We executed this protocol on 3 separate days, for a total of 3 separate mice/genotype (i.e., a total of 12 to 15 samples/genotype). Red cell morphology is similar in all groups and is unremarkable, with no differences noted among knockout strains and WT mice. The bar represents 5 μm. See Supplementary videos for a representative DIC video clip of each genotype (i.e., WT, *Aqp1–/–, RHag–/–*, dKO); each clip shows RBCs tumbling through the plane of focus.

**Extended Data Figure 8.**
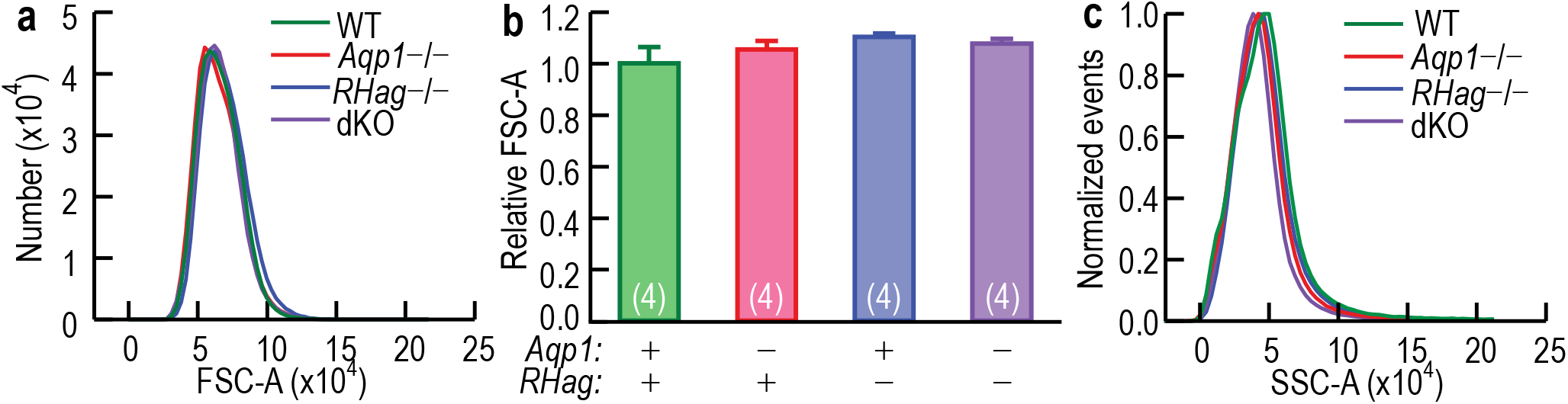
Light-scattering flow cytometry. **a**, Frequency distributions of forward-scatter intensity area (FSC-A; see above) of RBCs from WT, *Aqp1*–/–, *RHag*–/–, and dKO mice. Plots, obtained on the same day, are similar for all genotypes, and are representative data from 1 of the 4 mouse blood samples analyzed for each genotype. **b**, Summary of gated FSC-A data (from experiments like those in panel *a*). We normalized data to constant area (counts), integrated, and computed relative FSC-A values at ½ maximum frequency as the center measure of FSC-A. Within experiments, we normalized these values to the mean of WT samples (i.e., KO values are relative to WT value of unity), combined data from 2 independent experiment days (total of 4 mice/genotype), and performed a one-way ANOVA, followed by the Holm-Bonferroni correction (see below). The mean relative FSC-A values for the RBCs from *Aqp1*–/–, *RHag*–/–, and dKO mice are not significantly different from WT. The numbers in parentheses represent the number of mice from which blood samples were analyzed for each genotype. **c**, Frequency distributions of side-scatter intensity area (SSC-A; see above) for RBCs from WT, *Aqp1*–/–, *RHag*–/–, and dKO mice. Plots, which are similar for all genotypes, are representative data from one of the 4 mouse blood samples analyzed for each genotype.

**Extended Data Figure 9.**
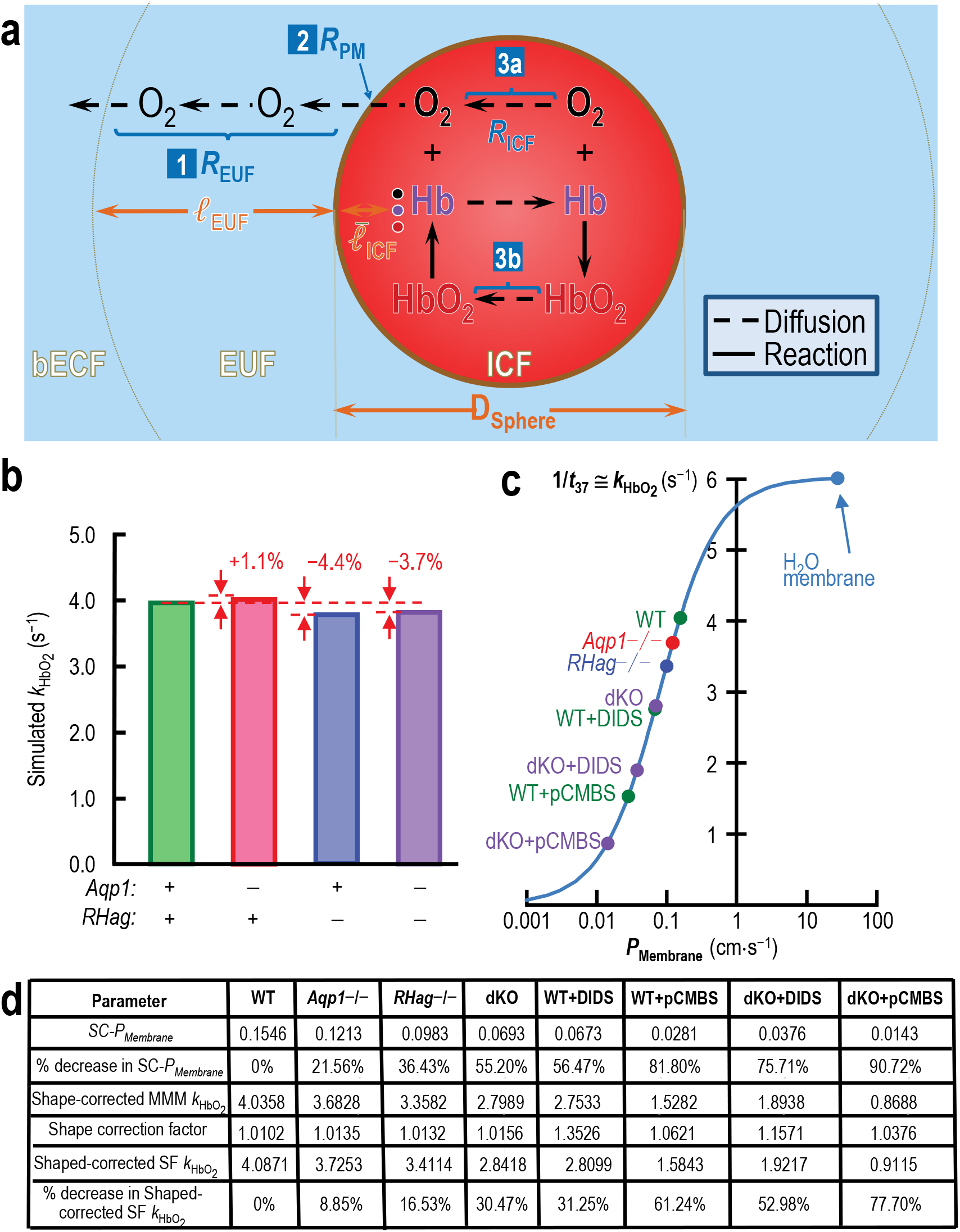
Mathematical simulations mimicking experimental conditions. **a**, Major components of mathematical model of spherical cell, the diameter of which matches the computed thickness of an average RBC from a WT or KO mouse. See Methods/Mathematical_Modeling and Supplementary Methods/Mathematical_modeling_and_simulations for details on the model. For panel ‘a’, the abbreviations are: bECF, bulk extracellular fluid; EUF, extracellular unconvected fluid; ICF, intracellular fluid; PM, plasma membrane; D_Sphere_, diameter of sphere (matches thickness of RBC); ℓ_EUF_, thickness of EUF; 3 values of 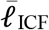 are average distance from PM to the following 3 molecules in ICF: (a) O_2_ (black dot and lettering), (b) Hb (violet), or HbO_2_ (red) in ICF; [1] *R*_EUF_, effective resistance offered by EUF to O_2_ diffusion from PM to bECF; [2] *R*_PM_, effective resistance offered by PM to O_2_ diffusion; [3] *R*_ICF_, effective resistance offered by ICF to combined diffusion of O_2_ and HbO_2_. 1/*R*_ICF_ = 1/*R*_ICFa_ + 1/*R*_ICFb_, where *R*_ICFa_ is effective resistance offered by ICF to O_2_ diffusion over the distance 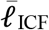, from the position of the average O_2_ to PM; *R*_ICFb_ is effective resistance offered by ICF to HbO_2_ diffusion over the distance 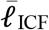, from the position of the average HbO_2_ to PM. **b**, Predicted 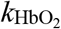 values from simulations of RBCs from WT and KO mice. Supplementary Methods explains how we use measured hematological and morphological values for each genotype to estimate parameters used in the simulations. Supplementary Methods also explains how we compute 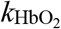 values from mathematical simulations. The model predicts that for (1) RBCs from *Aqp1*–/– mice, 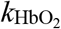 should have increased by 1.1%, rather than decreased by ∼9%, as observed; (2) RBCs from *RHag*–/– mice, 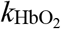 should have decreased 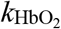 by 4.4%, not the observed ∼17%; and (3) RBCs from dKO mice, 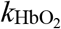 should have decreased 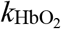 by only 3.7%, not the observed ∼30%. Thus, the hematological/morphological changes cannot explain our observed 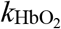 data. **c**, Predicted effects on hemolysis-corrected/shape-corrected 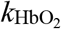 of varying *P*_Membrane_ according to our reaction-diffusion model. See Supplementary Methods/Accommodation_for_spherocytes for details on how we estimate *P*_Membrane_ and then use this value to simulate 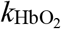 for normoshapen cells. To obtain the sigmoidal curve, we systematically vary *P*_Membrane_ (x-axis) and use our mathematical simulation to compute 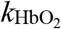 (y-axis). The point labeled WT has the coordinates (*P*_Membrane_ = 0.1546 cm s^−1^, 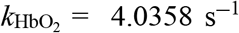. We obtained the other labeled points plotted on the sigmoidal curve from the experimentally determined fractional decrease in 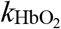, the measured prevalence of spherocytes, and spherocyte diameter, as detailed in Supplementary Methods/Accommodation_for_spherocytes. Note that the dKO and the WT+DIDS points nearly overlie each other. The point labeled “H_2_O membrane” is the result of a simulation in which we assumed that the diffusion constant of O_2_ in the plasma membrane is the same as in water. **d**, Numerical values associated with each of the 8 points to the left of the y-axis in panel *c*. All derived values in this table are the result of calculations based on numbers reported to 16 digits by the computer-simulation software. SC-*P*_Membrane_ is the estimate of the O_2_ permeability, after shape correction (i.e., correcting for the presence of spherocytes). MMM is the macroscopic mathematical model. The shape correction factor is the number by which one must multiple the simulated 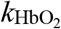 or normoshapen cells to obtain the simulated 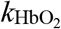 of a mixed population of normo- and misshapen cells. The shape-corrected SF 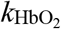 is the product of the shape correction factor and the hemolysis-corrected 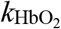 obtained in stopped-flow experiments, and is our best estimate of the 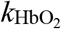—based on wet-laboratory experiments—of a mixed population of normo- and misshapen cells.

**Extended Data Figure 10.**
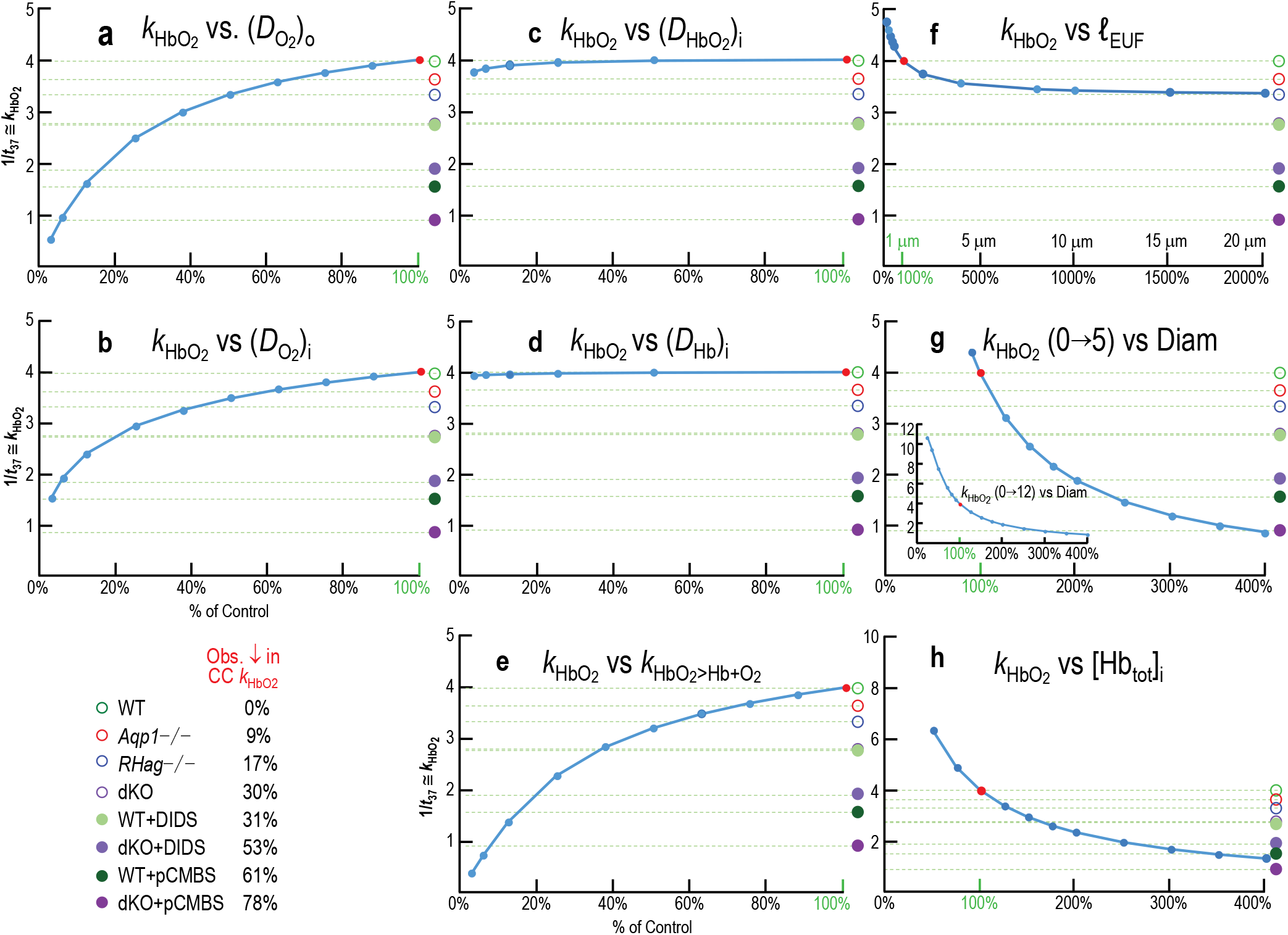
Mathematical simulations exploring the predicted sensitivity of 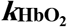 to 7 key kinetic and length parameters. Starting with the parameter values that generated the “WT” point in the first round of simulations, we systematically varied 8 parameters, one at a time, performed simulations, and computed 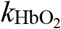 (see Methods/Mathematical modeling and Supplementary Methods). In each panel, the red dot—associated with 100% of the value of the varied parameter (green text with green tick mark)—indicates the conditions for control RBCs (from WT mice, no inhibitors), which we summarize in Supplementary Table 10. The horizontal dashed lines indicate the 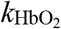 values, relative to the control, for each of the other 7 experimental conditions. As summarized in the legend at the lower left, for each condition, we multiplied the control simulated 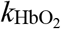 (i.e., 3.995 s^−1^) by experimentally observed decrease in 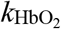 (e.g., 8.85% for RBCs from *Aqp1*–/– mice) to obtain the simulated 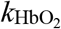 for that condition (e.g., 3.641 s^−1^ for *Aqp1*–/– RBCs). **a**, Dependence on diffusion constant for O_2_ 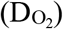 in 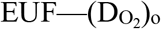. Although this is one of the parameters to which 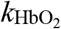 is most sensitive, it is also the one known with greatest confidence under control conditions, and the least subject to change with alterations in our experimental conditions. **b**, Dependence on diffusion constant for O_2_ 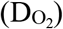 in 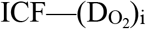. Only very large decreases in 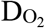 could explain even our smallest experimental effect on 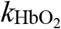 (i.e., *Aqp1*–/– RBCs). Note: Extended Data Figure 9*c* illustrates the dependence of 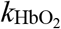 on the diffusion constant for O_2_ in the plasma membrane (*P*_Membrane_, shown as membrane permeability to O_2_). **c**, Dependence on diffusion constant for oxy-Hb 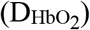 in 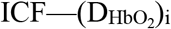. **d**, Dependence on diffusion constant for deoxy-Hb (D_Hb_) in ICF—(D_Hb_)_i_. **e**, Dependence on the rate constant 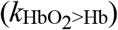 for deoxygenation of HbO_2_: HbO_2_ → Hb + O_2_. Although the 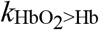 measured in hemolysates decreased by ∼9% in the dKO (Supplementary Figure 11), it would have had to have decreased by more than 60% to account for the observed 30% decrease in 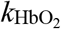 (Figure 3*d*). The measured 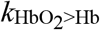 did not change for other genotypes. **f**, Dependence on thickness of the EUF (ℓ_EUF_). Note that a near-doubling of ℓ_EUF_ would be necessary to account for the *Aqp1*–/– RBC 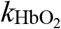 data, more than a doubling would be necessary to account for the *RHag*–/– RBC data, and no amount of increase could explain the other 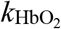 data. **g**, Dependence on the diameter of the sphere (D_Sphere_) that mimics the thickness of the RBC. Note that panel *g* shows the y-axis from 0 to 5 s^−1^, whereas the inset expands the y-axis to extend from 0 to 12 s^−1^. **h**, Dependence on total concentration of Hb in ICF—[Hb_tot_]_i_. Only very large increases in [Hb_tot_]_i_ could explain our experimental data. See Supplementary Figure 10 for simulated time courses—both for WT RBCs without inhibitors and dKO+pCMBS RBCs—of intracellular [O_2_]_Free_, [HbO_2_], [O_2_]_Free_ + [HbO_2_], HbSat, [Hb], and the transmembrane flux 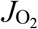.

