## Supplementary Methods for "Role of channels in the oxygen permeability of red blood cells"

### Supplementary Information: Methods

#### Genotyping

*RHag*–/– mice are genotyped by real-time PCR (performed by TransnetYX, Inc.) to determine the presence of a KO allele. If we detect a KO allele in a mouse, we must differentiate between a *RHag*–/– and *RHag*+/– mouse. We prepare highly purified genomic DNA from tail tissue using the DNeasy Blood and Tissue Kit (Qiagen), and perform *RHag*-WT genotyping. Primer "*RHag*-WT 3F" (5′-TGAAATGTTTAATGCTCATCGACCAC-3′) is designed to anneal to the end of Intron 1–2 of *RHag*, and Primer "*RHag*-WT 3R" (5′-GAGTGACTTCACCTTCTTATCTTCTAAG-3′), to the middle of intron 6–7. The *RHag*-WT PCR product is ~8 kb, using the Qiagen Long Range (LR) PCR Kit. Presence of the WT-PCR product confirms a *RHag*+/– genotype for any mouse reported to be positive for *RHag*–/– by TransnetYX. Absence of a WT-PCR product confirms the *RHag*–/– genotype.

#### Inhibitor studies

We collected fresh blood from WT or KO mice, centrifuged, removed the buffy coat, and suspended the pelleted RBCs in our oxygenated solution (Supplementary Table 12) to a hematocrit (Hct) of 5% to 10%, centrifuged at 600 × g for 5 min (see Methods). After 2 such washes, we resuspended RBCs in oxygenated solution without drugs or with either DIDS (200 µM for 1 hour) or pCMBS (1 mM for 15 min), to a final Hct of 25%. After centrifuging RBCs at 600 × g for 5 min, we aspirated and discarded the resulting supernatant to remove inhibitors. RBCs were then resuspended in oxygenated solution to a hematocrit (Hct) of 5% to 10%, centrifuged at 600 × g for 5 min for 1 more wash. Finally, we either resuspended packed RBCs at ostensibly 50% Hct for hematology assays, or directed the RBCs to studies of flow-cytometry (see Methods/Flow_cytometry).

For blood-smear studies with inhibitors, after 2 washes, we resuspended packed RBCs in oxygenated solution ([Hb] = 5 µM, Hct ≅ 0.3%) without drugs or with either DIDS (200 µM for 1 hour) or pCMBS (1 mM for 15 min). After centrifuging RBCs at 600 × g for 5 min, we aspirated and discarded the resulting supernatant to remove inhibitors. RBCs were then resuspended in oxygenated solution to make blood smears, following our standard procedure (see Methods/Blood_Smears). We repeated this on three different days, for a total of 3 mice of each genotype.

For still microphotography of living RBCs study with inhibitors, after 4 washes, we resuspended packed RBCs were in oxygenated solution without drugs or with either DIDS (200 µM for 1 hour) or pCMBS (1 mM for 15 min) with a final Hct of 1% to 2% and directed the RBCs for imaging studies, following our standard procedures (see Methods/Still_microphotography_and_microvideography_of_living_RBCs). We repeated this on three different days, for a total of 3 mice of each genotype.

#### Calculation of $k_{HbO_2}$

The time course of the decay $HbO_2 \rightarrow Hb + O_2$ does not follow a perfect exponential function—nor should it according to our mathematical modeling (see Supplementary Data Figure 10d). We therefore developed an unbiased approach* for computing the quasi-rate constant $k_{HbO_2}$, based on the time ($t_{37}$) required for Hb saturation (HbSat) to fall to 1/e of its initial value. We proceeded in the following steps:

**Preprocess data.** We eliminate noisy absorbance data—which we standardize as the first 9 points (0.010 – 0.090 s)—and translate the time axis so that raw data at times 0.100 – 4.000 s now covers the time range 0.000 – 3.900 s. We also eliminate data at 3 wavelengths (420, 445, 450 nm) that are close to isosbestic points, and thus have absorbance ($A$) values that do not change consistently during an experiment. The result is a data set describing $A_\lambda$ vs. time (from 0.000 to 3.900 s) at each of 6 wavelengths ($\lambda$ = 410, 415, 425, 430, 435, and 440 nm).

* The "Pro-KIV" software supplied by Applied Photophysics assumes that the absorbances at all wavelengths decay as a perfect exponential, and fits them all simultaneously, producing a single time constant/rate. Because this approach could introduce a bias into the analysis, we developed the method described here. Nevertheless, the two approaches produce numerical results that differ only slightly, and do not affect the conclusions that we draw.

**Obtain data parameters.** For each of 6 plots of $A_\lambda$ vs. time, obtain the minimum ($A_{\lambda,\mathrm{min}}$) and maximum values ($A_{\lambda,\mathrm{max}}$), as well as the difference $\Delta A_\lambda = A_{\lambda,\mathrm{max}} - A_{\lambda,\mathrm{min}}$ (i.e,. magnitude of the absorbance change). The 6 values of $\Delta A_\lambda$ are the weighting factors.

**Normalization.** Perform a vertical flip of plots of $A_\lambda$ vs. time that were originally concave downwards (these are $\lambda$ = 425, 430, 435, 440 nm), so that all plots are now concave upwards. Vertically translate each plot of $A_\lambda$ vs. time so that the minimum value is zero. Normalize each plot of $A_\lambda$ vs. time so that $A_{\lambda,\mathrm{max}}$ is now unity.

**Apply weighting factors.** Normalize the 6 weighting factors (i.e., $\Delta A_\lambda$) so that they sum to unity. Multiple each plot of $A_\lambda$ vs. time by its weighting factor. At each time point, sum the 6 weighted $A_\lambda$ values to obtain relative HbSat. The resulting plot of relative HbSat falls from unity to zero as time rises from 0.000 to 3.900 s. Note that the actual HbSat falls from a value of ~98% when the RBCs first enter the SF chamber at time 0.000, and has already fallen to some extent by the time of our first sample at 0.100 s, and then continues to fall to zero during the during to total 4.000 s of the "shot."

**Find $t_{37}$ and $k_{\mathrm{HbO_2}}$.** Fit the 5 HbSat points to the left[†] and the 5 HbSat points to the right of the estimated $t_{37}$ with a polynomial of degree 2. Solve the polynomial for $t_{37}$ (i.e., the time at which HbSat has fallen to 1/e). Obtain $k_{\mathrm{HbO_2}}$ as $1/t_{37}$.

### Mass Spectrometry

Relative abundance was determined by comparing the mean area under curve (AUC) for all detected peptides for each protein, from 3 mice of each genotype. For example, for AE1, we detected 57 different peptides. Therefore, in samples from 3 animals, the relative abundance of AE1 is the mean AUC of the 57 × 3 = 171 total peptides.

[†] In some cases, HbSat fell so rapidly that we were forced to use < 5 points to the left.

### Mathematical modeling and simulations

**Model formulation.** To examine the possibility that increases in intracellular diffusion distances can explain the observed decreases in $k_{HbO_2}$, we developed a distributed (i.e., space-dependent) and dynamic (i.e., time-dependent) reaction-diffusion mathematical model of an RBC—simplified as a perfectly symmetric sphere with diameter equal to the thickness of the RBC. Note that the thickness of the biconcave disk near its edge (~2 µm)—not the major diameter (~6.8 µm)—is the dimension of the RBC that is critically important for gas exchange, as pointed out by several authors[1–4]. Previous modelers have used a sphere in their simulations of an RBC[5–8]. As described below, we chose the diameter of the sphere to match the minor diameter of a torus having the volume of an RBC. In our model (see Extended Data Figure 9*a*), the spherical RBC is enveloped by a layer of extracellular unconvected fluid (EUF), which is in turn enveloped by the bulk extracellular fluid (bECF). The model accounts for diffusion of $O_2$, Hb and $HbO_2$ within the intracellular fluid (ICF). Only $O_2$ can move—via diffusion—between the ICF and EUF (i.e., across the plasma membrane, PM), within the EUF (where there is no Hb or $HbO_2$), and between the EUF and bECF. Reactions among $O_2$, Hb, and $HbO_2$ occur only in the ICF.

Assuming spherical radial symmetry, then for each solute *S* (i.e., $O_2$, $HbO_2$, and Hb), the concentration ($C_S$) changes in time (*t*) and space according to the reaction-diffusion equation

$$\frac{\partial}{\partial t} C_S(t,r) = \underbrace{\frac{1}{r^2}\frac{\partial}{\partial r}\left( D_S(r)\, r^2 \frac{\partial}{\partial r} C_S(t,r)\right)}_{\text{Diffusion term}} + \text{Reaction term}, \quad (1)$$

where *r* is the radial distance from the center of the RBC, and $D_S$ is the diffusion coefficient of solute *S*.

We modeled the kinetics of the oxygen-hemoglobin reaction—that is, the reaction term in Equation (1)—as a simple, one-step reaction in which we used the variable-rate-coefficient (VRC) approach proposed by Moll[9,10]. Thus, we assume that each hemoglobin tetramer ($Hb_4$) is replaced by four independent heme molecules, 4×Hb, and that each heme combines with $O_2$ via the one-step reaction

$$HbO_2 \underset{k'}{\overset{k}{\rightleftharpoons}} Hb + O_2, \quad (2)$$

where $k$ and $k'$ are the dissociation and association reaction rate constants. Rather than assuming that both $k$ and $k'$ are constant—an assumption that would lead to a non-realistic hyperbolic hemoglobin-oxygen saturation curve—we follow Moll's VRC approach, in which we assume that only one of the two reaction rates is constant, while the other one becomes a function of the partial pressure of $O_2$ ($P_{O_2}$) so that the mathematical representation of the hemoglobin-oxygen saturation curve corresponds to the measured sigmoidal curve. Yap and Hellums[11] showed that the VRC model is appropriate in the physiological range, and avoids the use of the more complex Adair model[12]. In his VRC approach, Moll assumed that $k'$ is constant and $k$ is a function of $P_{O_2}$. However, in 1985 Clark et al pointed out that the function used by Moll to describe $k$ as a function of $P_{O_2}$ approaches infinity as $P_{O_2}$ approaches zero.[10] These authors overcame this problem by assuming that $k$ is constant and $k'$ is a function of $P_{O_2}$, as described by

$$k' = k\frac{1}{\alpha \times P_{O_2}}\left(\frac{P_{O_2}}{P_{50}}\right)^n, \qquad (3)$$

where $\alpha$ is the solubility coefficient for $O_2$ in water, $P_{50}$ is $P_{O_2}$ when [Hb] = [$HbO_2$], and $n$ is the Hill coefficient. With this choice and $n > 1$, $k'$ approaches zero when $P_{O_2}$ approaches zero. Thus, in our model, we follow Clark's modification of the VRC approach, which we implement by using [$O_2$] rather than $P_{O_2}$:

$$k' = k\frac{[O_2]^{n-1}}{([O_2]_{50})^n}. \qquad (4)$$

**Calculation of RBC thickness based on the geometry of a torus.** For each mouse strain, we calculate the RBC thickness as follows. We start from the mean major diameter of all RBC-related cell types[‡] (e.g., 6.80 μm for WT cells; see Supplementary Table 5)—a measured value—and assume that the RBC is a torus, the outer diameter ($OD_{Torus}$) of which equals the aforementioned major RBC diameter in Supplementary Table 5 (i.e., 6.80 μm in our example). Next, we use the mean corpuscular volume (e.g., MCV = 47.9 pl for WT cells; see Supplementary Table 6)—also a measured value, which we assume to be equal

[‡] These include immature RBCs (i.e., reticulocytes and nucleated cells) as determined by ImageStream flow cytometry.

to the volume of the torus ($V_{\text{Torus}}$)—to calculate the minor radius ($r$) of the torus (equivalent to half the thickness of the RBC). Thus, $r$ is a root of the polynomial:

$$2\pi^2 r^3 - \pi^2 \left(OD_{\text{Torus}}\right) r^2 + V_{\text{Torus}} = 0.$$

The calculated value for the WT cells is $r$ = 1.01 μm. In Supplementary Table 11, we provide—for each of the 4 mouse strains—the computed values for $r$, the major radius of the torus ($R$), and the diameter ($D_{\text{Sphere}}$) of the equivalent sphere = 2$r$ (i.e., RBC thickness). The equivalent sphere diameters serve as inputs to our mathematical model.

**Calculation of Hb content.** For each mouse strain, we calculate the concentration (in mM) of total Hb tetramers (i.e., [THb] = [$HbO_2$] + [Hb] ) as follows. Based on the hematology data reported in Supplementary Table 6—specifically, the MCH (in pg) and MCV (in fl)—and using a value of 64316 g/mol for the molecular weight of mouse Hb, we calculate the MCHC (in mM). For WT mice, the concentration of total Hb tetramers was ~4.7 mM. However, because our model deals with monomers, we calculate the concentration of total of Hb monomers as 4 × MCHC. In the case of RBCs from WT mice, this figure is 18.73 mM. Supplementary Table 11 lists this value, as well as the comparable values for the other three genotypes. These values inform our mathematical model.

**Computational model.** In order to solve the system of the three reaction-diffusion equations (1)—one for $O_2$, one for $HbO_2$ monomers and one for Hb monomers—we set the following boundary conditions.

At the center of the RBC, we posit the no-flux Neumann boundary condition:

$$\frac{\partial C_S}{\partial r}(t,0) = 0.$$

At the interface between the ICF and the EUF (i.e., at the PM), we establish continuity of Fick's diffusive flux:

$$D_S(R+)\frac{\partial}{\partial r}C_S(t,R+) = P_{\text{PM,S}} \cdot (C_S(t,R+) - C_S(t,R-)).$$

$$D_S(R-)\frac{\partial}{\partial r}C_S(t,R-) = P_{\text{PM,S}} \cdot (C_S(t,R+) - C_S(t,R-)).$$

Here, $R$ is the radius of the sphere, $C_S(t, R–)$ is the time-dependent concentration of solute $S$ in the aqueous phase adjacent to the intracellular (−) side of the PM; $C_S(t, R+)$, the comparable value on the extracellular (+) side; and $P_{PM,S}$ is the true permeability of the membrane to solute $S$. Because $O_2$ is the only solute that can diffuse across the PM, $P_{PM,S} \neq 0$ only for $S = O_2$.

In the bECF, we assign Dirichlet boundary conditions and assume that the concentration of $O_2$ is constant and equal to zero.

Finally, we set the initial conditions by assuming that, at the beginning of the experiment, there is no $O_2$ in the EUF and that the RBC contains 21% $O_2$. The initial concentrations of $HbO_2$, Hb, and $O_2$ in the ICF are those reported in Supplementary Table 10 for RBCs from WT mice, and in Supplementary Table 11 for RBCs from mice of each of four genotypes.

We solve the system of the three reaction-diffusion equations (i.e., Eqn (1) for the solutes $O_2$, $HbO_2$, and Hb) using the method of lines and the approach described by Somersalo et al for the numerical implementation of the diffusion component[13]. We solve the resulting system of time-dependent ordinary differential equations in Matlab R2015a using the stiff solver ode15s with AbsTol=RelTol=1e–12. The Matlab code is based on the original implementation of Somersalo et al.

**Parameter values.** Supplementary Table 10 lists the parameter values used in the model for RBCs from WT mice. Supplementary Table 11 lists a subset of these values that depend on each of the 4 genotypes; we used these values for computing values for Extended Data Figure 9*b*. All parameter values correspond to a temperature of 10°C—the temperature at which we perform the stopped-flow experiments. For parameter values not available at the temperature of 10°C, we derive them using Arrhenius equation and values at 25°C and 37°C from references indicated in Supplementary Table 10. We used this approach for calculating the diffusion coefficient of $HbO_2$ and Hb in the Bulk ICF.

For the diffusion coefficient of $O_2$ in the bulk ICF, the commonly used value (adjusted for 10°C using the Arrhenius equation, as described above) is $5.09 \times 10^{-6}$ $cm^2$ $s^{-1}$.[10] Recently, Richardson et al[14] obtained the much lower value of $4.5907 \times 10^{-7}$ $cm^2$ $s^{-1}$ (see Supplementary Discussion/PaperByRichardson).

However, this latter value is likely to be an underestimate because Richardson et al used a very low extra-cellular concentration of NDT (i.e., 1 mM vs. the 25 mM in the present study). This low $[NDT]_o$ would have increased the thickness of the EUF ($\ell_{EUF}$) surrounding the RBC, and thus increased the resistance to the diffusion of $O_2$ away from the RBC.[15–17] Thus, to some extent, the authors attributed to the ICF a component of total resistance to $O_2$ diffusion that was actually due to the EUF. The result is an inappropriately low estimate for the intracellular diffusion constant for $O_2$. Because we do not know the magnitude of this error, we chose to compute the arithmetic mean of the commonly used value of $5.09 \times 10^{-6}$ cm$^2$ s$^{-1}$ and the more recent value of $4.5907 \times 10^{-7}$ cm$^2$ s$^{-1}$. The result is $2.7745 \times 10^{-6}$ cm$^2$ s$^{-1}$, which we report in Supplementary Table 10.

Because the molecular weight of $O_2$ is much less than the molecular weight of Hb, and because the diffusion coefficient mainly depends on molecular weight, we assume that the diffusion coefficients of monomeric $HbO_2$ and Hb are the same[18]. We assume that the permeability of the RBC plasma membrane to $O_2$ is 0.15 cm/sec, the value estimated by Endeward et al[19] for the permeability of the RBC plasma membrane to $CO_2$.

**Simulation of time course of $HbO_2$ deoxygenation, and estimation of $k_{HbO_2}$.** Using the reaction-diffusion model, we simulate the time course of the decline of $HbO_2$ saturation (i.e., HbSat = $[HbO_2]/([Hb]+[HbO_2])$ vs. time, employing hematological and morphological data that we gathered for RBCs from WT mice (see Supplementary Table 10). Because neither the actual nor the simulated time courses of $HbO_2$ desaturation are precisely exponential, we compute $k_{HbO_2}$ from simulations in a manner analogous to that for our biological data (see Methods/StoppedFlowAbsorbanceSpectroscopy and above): we determine the time for the simulated saturation to fall to 1/e ≅ 37% of its initial value (i.e., $\tau_{Quasi} \cong t_{37}$), and then compute $k_{HbO_2}$ as $1/t_{37}$.

### Accommodation for spherocytes

We computed the data in Extended Data Figure 9*c* and *d* using a combination of our MMM simulations, morphological data from hematology and ImageStream, and a mathematical reconstruction of the time courses of HbSat for normoshapen RBCs and misshapen RBCs (i.e., spherocytes).

For WT RBCs, we first compute the equivalent of a hemolysis-corrected $k_{HbO_2}$, using the parameters for WT RBCs in Supplementary Table 5, Supplementary Table 6, and Supplementary Table 10, and an initial $P_{Membrane}$ estsimate of 0.15 cm·s$^{-1}$, the value obtained by Endeward et al[19] for the permeability of the RBC plasma membrane to $CO_2$. In our SF experiments, the analogous hemolysis-corrected $k_{HbO_2}$ value pertains to a mixed population of normoshapen and misshapen RBCs, but we take it as our first estimate of the $k_{HbO_2}$ of normoshapen cells.

Second, we computed the $k_{HbO_2}$ of a misshapen cell (i.e., spherocyte), using the mean measured diameter from our ImageStream data.

Third, knowing the fraction of normoshapen vs. misshapen cells, we obtain the linear combination of the time courses of Hb saturation for normo- and misshapen cells, and estimate the $k_{HbO_2}$ of the mixed population.

Finally, we back-calculate $P_{Membrane}$, and obtain a value of 0.1546 cm·s$^{-1}$ for normoshapen, control, WT cells. This $P_{Membrane}$ yields a simulated $k_{HbO_2}$ of 4.0358 s$^{-1}$—this is the shape-corrected MMM $k_{HbO_2}$ in Extended Data Figure 9*d*—which is within ~1% of the experimentally observed value.
