## Supplementary Tables for "Role of channels in the oxygen permeability of red blood cells"

### Supplementary Information: Tables

#### Supplementary Table 1 | Protein IDs and Entrez Gene names for Mass Spec Figures

##### a. 22 "Plasma-membrane–associated proteins"—none of which exhibit statistically significant changes—from among the 50 proteins with highest inferred abundance in samples of RBC ghosts from WT mice

| Rank order in abundance of all proteins | Rank order in abundance of PMAPs$^{\Delta}$ | Protein ID | Entrez Gene Name |
|---|---|---|---|
| 1 | 1 | AE1* | solute carrier family 4 (anion exchanger), member 1 (Diego blood group) |
| 2 | 2 | GYPA* | glycophorin A |
| 4 | 3 | AQP1* | aquaporin 1 (Colton blood group) |
| 8 | 4 | GYPC* | glycophorin C (Gerbich blood group) |
| 9 | 5 | ANK1 | ankyrin 1, erythrocytic (Cytoskeletal protein) |
| 12 | 6 | SPTB | spectrin, beta, erythrocytic (Cytoskeletal protein) |
| 14 | 7 | RhAG* | Rh-associated glycoprotein |
| 15 | 8 | EPB41 | erythrocyte membrane protein band 4.1 (Cytoskeletal protein) |
| 16 | 9 | EPB42 | erythrocyte membrane protein band 4.2 (Cytoskeletal protein) |
| 17 | 10 | MPP1 | membrane protein, palmitoylated 1, 55 kDa (Membrane-associated protein) |
| 21 | 11 | BSG* | basigin (Ok blood group) |
| 23 | 12 | SLC16A1* | solute carrier family 16 (monocarboxylate transporter), member 1 (MCT1) |
| 24 | 13 | mRh* | Rh blood group, CE/D homolog |
| 25 | 14 | CD47* | CD47 molecule |
| 26 | 15 | STOM* | Stomatin |
| 28 | 16 | DMTN | dematin actin binding protein (Cytoskeletal protein) |
| 33 | 17 | SLC43A1* | solute carrier family 43 (amino acid system L transporter), member 1 |
| 36 | 18 | CD36† | CD36 molecule (thrombospondin receptor) |
| 37 | 19 | SLC14A1* | solute carrier family 14 (urea transporter), member 1 (Kidd blood group) (UT1) |
| 38 | 20 | CLDN13* | claudin 13 |
| 42 | 21 | TPM1 | tropomyosin 1, alpha (Cytoskeletal protein) |
| 50 | 22 | SLC16A10* | solute carrier family 16 (aromatic amino acid transporter), member 10 |

##### b. All plasma-membrane–associated proteins with significant changes

| Rank order in abundance of all proteins | Rank order in abundance of PMAPs$^{\Delta}$ | Protein ID | Entrez Gene Name |
|---|---|---|---|
| 136 | 48 | ITGA2B† | integrin, alpha 2b (platelet glycoprotein IIb of IIb/IIIa complex, antigen CD41) |
| 138 | 49 | NRAS | neuroblastoma RAS viral (v-ras) oncogene homolog (GTPase) |
| 177 | 63 | ADAM10 | ADAM metallopeptidase domain 10 (peptidase) |
| 469 | 117 | TBXAS1† | thromboxane A synthase 1 (platelet) |
| 490 | 122 | SLC3A2* | solute carrier family 3 (amino acid transporter heavy chain), member 2 |
| 655 | 143 | ICAM4* | intercellular adhesion molecule 4 (Landsteiner-Wiener blood group |

| | | | |
|---|---|---|---|
| **793** | **172** | **SLC30A1*** | solute carrier family 30 (zinc transporter), member 1 |
| **952** | **193** | **H2-Q2‡** | major histocompatibility complex |
| **1066** | **205** | **CAPRIN1‡** | cell cycle associated protein 1 |
| **1069** | **207** | **DLG1‡** | discs, large homolog 1 (Drosophila) |
| **1100** | **211** | **TMEM106B** | transmembrane protein 106B |

##### c. All non-membrane-associated proteins with significant changes

| Rank order in abundance of all proteins | Rank order in abundance of MAP[Δ] | Protein ID | Entrez Gene Name |
|---|---|---|---|
| **101** | **-** | **PSMA4** | proteasome (prosome, macropain) subunit, alpha type, 4 |
| **130** | **-** | **C17orf99** | chromosome 17 open reading frame 99 |
| **132** | **-** | **FLNA** | filamin A, alpha (Cytoskeletal protein) |
| **133** | **-** | **PSMA1** | proteasome (prosome, macropain) subunit, alpha type, 1 |
| **195** | **-** | **MYH9** | myosin, heavy chain 9, non-muscle (Cytoskeletal protein) |
| **214** | **-** | **DHTKD1** | dehydrogenase E1 and transketolase domain containing 1 (Mitochondrion) |
| **236** | **-** | **CAPN5** | calpain 5 (Protease) |
| **433** | **-** | **ATP2A3** | ATPase, $Ca^{++}$-transporting, ubiquitous (a SERCA) |
| **448** | **-** | **CAPN1** | calpain 1 (Protease) |
| **1045** | **-** | **MUG1/MUG2** | murinoglobulin 1 (Proteinase) |
| **1051** | **-** | **ALDH18A1** | aldehyde dehydrogenase 18 family, member A1 |
| **1052** | **-** | **MMP21** | matrix metallopeptidase 21 |
| **1059** | **-** | **RPL8** | ribosomal protein L18 |
| **1087** | **-** | **RNF213** | ring finger protein 213 |
| **1095** | **-** | **NDUFA12** | NADH dehydrogenase (ubiquinone) 1 alpha subcomplex, 12 |
| **1102** | **-** | **SNX12** | sorting nexin 12 |

[Δ]Plasma-membrane–associated proteins

*Integral membrane protein, known to be present in erythrocytes

†Integral membrane protein, known to be present in platelets

‡ Membrane-associated protein, known to be present in white cells

**Supplementary Table 2 | Summary of all hematological data obtained from four strains of mice†**

| | Strains | WT | | | *Aqp1–/–* mice | | | *RHag–/–* mice | | | dKO | | |
|---|---|---|---|---|---|---|---|---|---|---|---|---|---|
| | **Sex** | **M** | **F** | **M+F** | **M** | **F** | **M+F** | **M** | **F** | **M+F** | **M** | **F** | **M+F** |
| | **n** | 9 | 15 | 24 | 14 | 8 | 22 | 17 | 12 | 29 | 10 | 7 | 17 |
| hematological parameters | **Age (weeks)** | 12.2 ±0.8 | 13.8 ±0.8 | 13.2 ±0.6 | 12.1 ±0.7 | 14.0 ±1.0 | 12.8 ±0.6 | 14.1 ±0.7 | 14.4 ±1.2 | 14.2 ±0.6 | 12.6 ±0.9 | 15.0 ±0.2 | 13.6 ±0.6 |
| | **WBC ($10^3$/ul)** | 12.1 ±1.6 | 10.2 ±0.7 | 10.9 ±0.7 | 12.9 ±1.0 | 14.5 ±1.4 | 13.5 ±0.8 | 13.6 ±1.0 | 10.7 ±1.3 | 12.4 ±0.8 | 12.0 ±1.1 | 10.7 ±0.9 | 11.5 ±0.8 |
| | **LYM ($10^3$/ul)** | 9.8 ±1.3 | 8.5 ±0.6 | 9.0 ±0.6 | 11.0 ±0.9 | 12.3 ±1.0 | 11.5 ±0.7 | 11.4 ±0.9 | 9.0 ±1.1 | 10.4 ±0.7 | 10.1 ±1.0 | 9.1 ±0.9 | 9.7 ±0.7 |
| | **MONO ($10^3$/ul)** | 0.6 ±0.1 | 0.4 ±0.0 | 0.5 ±0.0 | 0.5 ±0.0 | 0.6 ±0.1 | 0.5 ±0.0 | 0.5 ±0.1 | 0.5 ±0.0 | 0.5 ±0.0 | 0.5 ±0.0 | 0.5 ±0.0 | 0.5 ±0.0 |
| | **GRAN ($10^3$/ul)** | 1.7 ±0.3 | 1.2 ±0.1 | 1.4 ±0.1 | 1.4 ±0.2 | 1.6 ±0.4 | 1.5 ±0.2 | 1.7 ±0.2 | 1.2 ±0.2 | 1.5 ±0.2 | 1.4 ±0.1 | 1.2 ±0.1 | 1.3 ±0.1 |
| | **LYM %** | 81.9 ±1.8 | 83.8 ±1.1 | 83.1 ±0.9 | 84.8 ±1.5 | 85.7 ±1.5 | 85.1 ±1.1 | 83.4 ±1.7 | 83.5 ±1.2 | 83.4 ±1.1 | 83.8 ±1.0 | 84.1 ±1.3 | 83.9 ±0.8 |
| | **MONO %** | 4.4 ±0.4 | 3.9 ±0.2 | 4.1 ±0.2 | 3.7 ±0.3 | 3.4 ±0.3 | 3.6 ±0.2 | 3.8 ±0.2 | 4.0 ±0.2 | 3.9 ±0.1 | 3.6 ±0.2 | 4.2 ±0.4 | 3.8 ±0.2 |
| | **GRAN %** | 13.7 ±1.6 | 12.3 ±1.0 | 12.8 ±0.8 | 11.6 ±1.3 | 10.9 ±1.3 | 11.3 ±0.9 | 12.8 ±1.6 | 12.6 ±1.0 | 12.7 ±1.0 | 12.6 ±0.9 | 11.7 ±1.0 | 12.2 ±0.7 |
| | **HCT (%)** | 49.0 ±2.0 | 49.0 ±1.9 | 49.0 ±1.4 | 48.8 ±1.8 | 51.6 ±1.5 | 49.8 ±1.3 | 50.2 ±1.9 | 50.6 ±2.8 | 50.3 ±1.6 | 54.4 ±2.2 | 49.6 ±1.6 | 52.4 ±1.5 |
| | **HGB (g/dl)** | 15.5 ±0.4 | 15.5 ±0.6 | 15.5 ±0.4 | 15.2 ±0.5 | 16.1 ±0.2 | 15.5 ±0.3 | 14.6 ±0.5 | 14.7 ±0.7 | 14.6 ±0.4 | 16.0 ±0.4 | 14.8 ±0.3 | 15.5 ±0.3 |
| | **RDWa (fl)** | 33.0 ±1.2 | 31.8 ±0.6 | 32.3 ±0.6 | 32.7 ±0.6 | 32.4 ±0.9 | 32.6 ±0.5 | 35.1 ±0.5 | 34.6 ±0.6 | 34.9 ±0.4* | 35.3 ±0.9 | 34.0 ±0.6 | 34.7 ±0.6* |
| | **RDW % (%)** | 18.4 ±0.5 | 17.5 ±0.3 | 17.9 ±0.3 | 17.6 ±0.1 | 17.4 ±0.2 | 17.6 ±0.1 | 17.5 ±0.1 | 17.2 ±0.1 | 17.4 ±0.1 | 18.2 ±0.2 | 17.5 ±0.2 | 17.9 ±0.2 |
| | **MCV (fl)** | 46.6 ±0.9 | 46.6 ±0.9 | 46.6 ±0.7 | 47.3 ±0.7 | 47.5 ±1.3 | 47.3 ±0.6 | 50.3 ±0.4 | 50.5 ±0.4 | 50.4 ±0.3* | 49.1 ±1.1 | 49.4 ±0.4 | 49.2 ±0.6* |
| | **MCH (pg)** | 14.8 ±0.3 | 14.7 ±0.1 | 14.7 ±0.1 | 14.8 ±0.2 | 14.9 ±0.1 | 14.8 ±0.1 | 14.7 ±0.1 | 14.8 ±0.1 | 14.7 ±0.1 | 14.5 ±0.1 | 14.8 ±0.3 | 14.6 ±0.1 |
| | **MCHC (g/dl)** | 31.7 ±0.7 | 31.6 ±0.5 | 31.7 ±0.4 | 31.4 ±0.8 | 31.5 ±0.8 | 31.4 ±0.6 | 29.2 ±0.3 | 29.3 ±0.4 | 29.2 ±0.2* | 29.6 ±0.7 | 30.0 ±0.4 | 29.8 ±0.4* |
| | **RBC ($10^6$/ul)** | 10.5 ±0.3 | 10.6 ±0.4 | 10.5 ±0.3 | 10.3 ±0.4 | 10.8 ±0.2 | 10.5 ±0.2 | 10.0 ±0.4 | 10.0 ±0.5 | 10.0 ±0.3 | 11.1 ±0.3 | 10.0 ±0.3 | 10.6 ±0.2 |
| | **PLT ($10^3$/ul)** | 294.1 ± 50.7 | 320.9 ± 46.7 | 310.8 ± 34.2 | 303.6 ± 39.2 | 325.9 ± 41.6 | 311.7 ± 28.6 | 377.6 ± 43.6 | 261.4 ± 40.7 | 329.6 ± 32.0 | 353.2 ± 50.3 | 255.0 ± 44.9 | 313.1 ± 35.9 |
| | **MPV (fl)** | 6.4 ±0.2 | 6.4 ±0.2 | 6.4 ±0.1 | 6.4 ±0.1 | 6.6 ±0.2 | 6.5 ±0.1 | 6.6 ±0.1 | 6.5 ±0.1 | 6.5 ±0.1 | 6.4 ±0.2 | 6.3 ±0.2 | 6.4 ±0.1 |

†Results are presented as means ± s.e.m. Each "n" represents an analysis of the blood from 1 mouse. We performed full hematology screens on fresh blood from WT and KO mice. We analyzed data using a one-way ANOVA, followed by the Holm-Bonferroni[1] correction (see Methods/Statistical_Analysis). For each parameter (e.g., MCV), we made all possible comparisons within a row. The only 6 significant differences (indicated by *) involved *RHag*–/– vs. WT or dKO vs. WT, and none were not sex related. M, male; F, female; WBC, white blood cells; LYM, lymphocytes; MONO, monocytes; GRAN, granulocytes; HCT, hematocrit; HGB, hemoglobin; RDW, RBC distribution width; MCV, mean corpuscular volume; MCH, mean corpuscular hemoglobin; MCHC, mean corpuscular hemoglobin concentration; RBC, red blood cells; PLT, platelet; MPV, mean platelet volume. In Supplementary Table 6, we summarizes a subset of the above data from mice that we used in ImageStream flow cytometry and mathematical simulations. dKO, double knockout (i.e., *Aqp1*–/–*RHag*–/–).

**Supplementary Table 3 | Percent Gated Cells by stain type in light-scattering, flow cytometry analyses**

| | CV positive | | CV negative | | TO positive | | TO negative | | DRAQ-5 positive | | DRAQ-5 negative | |
|---|---|---|---|---|---|---|---|---|---|---|---|---|
| | Mean | s.e.m. | Mean | s.e.m. | Mean | s.e.m. | Mean | s.e.m. | Mean | s.e.m. | Mean | s.e.m. |
| **WT** | 99.96 | 0.02 | 0.03 | 0.02 | 2.53 | 0.59 | 97.44 | 0.61 | 1.50 | 0.33 | 98.48 | 0.34 |
| ***Aqp1–/–*** | 99.93 | 0.03 | 0.06 | 0.03 | 2.74 | 0.36 | 97.24 | 0.36 | 1.85 | 0.16 | 98.11 | 0.18 |
| ***RHag–/–*** | 99.93 | 0.04 | 0.05 | 0.05 | 3.80 | 0.73 | 96.18 | 0.74 | 2.55 | 0.41 | 97.36 | 0.46 |
| **dKO** | 99.88 | 0.07 | 0.11 | 0.07 | 3.42 | 0.55 | 96.56 | 0.55 | 2.36 | 0.33 | 97.60 | 0.34 |

Each value is reflects a percentage of all cells. CV, calcein violet (reports cell viability); TO; thiazole orange (reports RNA); DRAQ5, Deep Red Anthraquinone 5 (reports DNA). CV staining was analyzed in cells from R1 on the FSC-A vs. FSC-W histogram (Supplementary Data Figure 3), that is, cells separated from very small events and aggregates. TO and DRAQ-5 staining were analyzed in CV-positive cells, which also had been gated through R1 (Supplementary Data Figure 3). Mature RBCs are CV positive and TO negative. Reticulocytes are CV positive, TO positive, but DRAQ5 negative. Nucleated cells are CV positive, TO positive, and DRAQ5 positive; they include both erythrocyte precursors and white blood cells. See Supplementary Data Figure 6 for additional data on gating of cell types. dKO, double knockout (i.e., *Aqp1–/–RHag–/–*).

**Supplementary Table 4 | Percent mature vs. immature erythrocytes present in mouse blood, determined using ImageStream flow cytometry**

| | WT | s.e.m. | *Aqp1–/–* | s.e.m. | *RHag–/–* | s.e.m. | dKO | s.e.m. |
|---|---|---|---|---|---|---|---|---|
| **Mature RBCs** | 96.06 | 0.29 | 95.25 | 0.34 | 92.90 | 0.33 | 93.94 | 0.73 |
| **Reticulocytes** | 2.40 | 0.37 | 3.06 | 0.44 | 4.60 | 0.70 | 3.53 | 0.55 |
| **Nucleated cells** | 1.54 | 0.08 | 1.69 | 0.27 | 2.50 | 0.48 | 2.53 | 0.21 |
| **All Immature cells** | 3.94 | 0.29 | 4.75 | 0.34 | 7.10 | 0.33 | 6.06 | 0.73 |

RBCs constitute ≥93% of all cells in the blood samples acquired from WT, *Aqp1–/–*, *RHag–/–*, and double KO mice. We performed a one-way ANOVA, followed by the Holm-Bonferroni[1] correction (see Methods/Statistical_Analysis), and observed no significant differences. dKO, double knockout (i.e., *Aqp1–/–RHag–/–*).

**Supplementary Table 5 | Quantification of major diameter, among various blood-cell types, from four mouse genotypes ***

| Strains | Cell types | n total | Mean (μm) | s.d. | s.e.m. | Mode (μm) | Median (μm) | IQR |
|---|---|---|---|---|---|---|---|---|
| WT | Mature RBCs | 1156720 | 6.79 | 0.93 | 0.00 | 6.67 | 6.67 | 1.33 |
| | Reticulocytes | 29317 | 7.06 | 1.15 | 0.01 | 6.67 | 7.00 | 1.33 |
| | Nucleated cells | 18492 | 7.36 | 1.05 | 0.01 | 7.00 | 7.33 | 1.33 |
| | All Immature cells | 47809 | 7.18 | 1.12 | 0.01 | 7.00 | 7.00 | 1.33 |
| | ALL CELL TYPES | 1204530 | 6.80 | 0.94 | 0.00 | 6.67 | 6.67 | 1.33 |
| *Aqp1–/–* | Mature RBCs | 1139890 | 6.68 | 0.90 | 0.00 | 6.67 | 6.67 | 1.33 |
| | Reticulocytes | 36806 | 6.77 | 1.15 | 0.01 | 6.67 | 6.67 | 1.33 |
| | Nucleated cells | 20044 | 7.14 | 1.01 | 0.01 | 7.00 | 7.00 | 1.00 |
| | All Immature cells | 56850 | 6.90 | 1.12 | 0.00 | 6.67 | 7.00 | 1.33 |
| | ALL CELL TYPES | 1196740 | 6.69 | 0.92 | 0.00 | 6.67 | 6.67 | 1.33 |
| *RHag–/–* | Mature RBCs | 1077350 | 6.47 | 0.92 | 0.00 | 6.00 | 6.33 | 1.33 |
| | Reticulocytes | 53283 | 7.18 | 1.22 | 0.01 | 7.00 | 7.00 | 1.67 |
| | Nucleated cells | 29162 | 7.46 | 1.13 | 0.01 | 7.33 | 7.33 | 1.33 |
| | All Immature cells | 82445 | 7.28 | 1.20 | 0.00 | 7.33 | 7.33 | 1.33 |
| | ALL CELL TYPES | 1159800 | 6.53 | 0.97 | 0.00 | 6.00 | 6.33 | 1.00 |
| dKO | Mature RBCs | 1166840 | 6.52 | 0.92 | 0.00 | 6.33 | 6.33 | 1.00 |
| | Reticulocytes | 44110 | 6.94 | 1.21 | 0.01 | 6.67 | 7.00 | 1.67 |
| | Nucleated cells | 31462 | 7.32 | 1.04 | 0.01 | 7.33 | 7.33 | 1.33 |
| | All Immature cells | 75572 | 7.10 | 1.16 | 0.00 | 7.00 | 7.00 | 1.33 |
| | ALL CELL TYPES | 1242420 | 6.55 | 0.95 | 0.00 | 6.33 | 6.33 | 1.00 |

*Blood samples were taken from four age-matched mice of each genotype. The major diameter of all RBCs and RBC precursor cells from blood of four strains of mice were analyzed after gating according to their fluorescence signature in an ImageStream flow cytometer. Due to the extremely high number of individual cells analyzed (n), one-way ANOVA with a Holm-Bonferroni means comparison[1] (see Methods) shows that differences between all pairs of means are statistically significant. However, even the greatest difference (i.e., between WT and *RHag–/–*) among only mature RBCs is ~5% (i.e., 6.79 vs. 7.18), and for "all cell types" (i.e., the sum of mature RBCs + reticulocytes + nucleated cells) is <5% (i.e., 6.80 vs. 6.53). In our mathematical modeling, we take as the diameter, the diameter of "all cell types." This approach reflects the reality of an SF experiment, which necessarily includes all cell types. Note that the inclusion of larger-diameter RBC precursor cells in the size analyses, does not substantially increase the mean major blood cell diameter because they represent only 4% (WT) to 7% (*RHag–/–*) of the total cells in the sample. Thus, even in the case of cells from *RHag–/–* mice, the inclusion of immature cells increases the mean major diameter by <1% (i.e., 6.53 vs. 6.47). These data, in a different format, are also presented in Figure 4*e*. dKO, double knockout (i.e., *Aqp1–/–RHag–/–*).

**Supplementary Table 6 | Summary of hematological data obtained from the subset of mice used in ImageStream analysis and as basis for mathematical simulations***

| | Strains | WT | *Aqp1–/–* | *RHag–/–* | dKO |
|---|---|---|---|---|---|
| hematological parameters | n | 4 | 4 | 4 | 4 |
| | Age (weeks) | 13.0 ±0.6 | 13 ±0.0 | 8.7 ±0.3 | 11.5 ±2.0 |
| | WBC ($10^3$/ul) | 14.5 ±1.4 | 14.0 ±1.6 | 11.9 ±1.2 | 9.9 ±1.1 |
| | LYM ($10^3$/ul) | 12 ±1.0 | 11.8 ±1.4 | 10.3 ±1.0 | 8.3 ±0.9 |
| | MONO ($10^3$/ul) | 0.7 ±0.1 | 0.6 ±0.1 | 0.1 ±0.0 | 0.4 ±0.1 |
| | GRAN ($10^3$/ul) | 1.9 ±0.4 | 1.7 ±0.2 | 1.2 ±0.2 | 1.3 ±0.3 |
| | LYM % | 83.1 ±2.6 | 83.8 ±0.5 | 86.7 ±1.0 | 83.8 ±1.5 |
| | MONO % | 4.0 ±0.4 | 3.7 ±0.3 | 3.3 ±0.2 | 3.6 ±0.5 |
| | GRAN % | 12.9 ±2.2 | 12.5 ±0.5 | 10.0 ±0.8 | 12.7 ±1.2 |
| | HCT (%) | 54.5 ±2.0 | 52.7 ±0.9 | 57.9 ±2.2 | 58.9 ±3.9 |
| | HGB (g/dl) | 16.4 ±0.1 | 15.1 ±0.2 | 16.6 ±0.4 | 17.1 ±0.7 |
| | RDWa (fl) | 32.9 ±0.9 | 34.5 ±0.5 | 36.6 ±0.4 | 36.4 ±1.6 |
| | RDW % (%) | 17.3 ±0.3 | 17.5 ±0.2 | 17.9 ±0.2 | 18.4 ±0.2 |
| | MCV (fl) | 47.9 ±1.30 | 49.5 ±0.7 | 51.0 ±0.4 | 49.8 ±1.8 |
| | MCH (pg) | 14.4 ±0.2 | 14.1 ±0.1 | 14.7 ±0.2 | 14.5 ±0.1 |
| | MCHC (g/dl) | 30.1 ±1.1 | 28.6 ±0.3 | 28.7 ±0.2 | 29.4 ±1.2 |
| | RBC ($10^6$/ul) | 11.4 ±0.2 | 10.6 ±0.1 | 11.3 ±0.2 | 11.8 ±0.5 |
| | PLT ($10^3$/ul) | 286.3 ± 35.1 | 417.3 ± 29.8 | 467.8 ± 68.1 | 443.0 ± 75.0 |
| | MPV (fl) | 6.5 ±0.2 | 6.5 ±0.2 | 6.5 ±0.2 | 6.2 ±0.1 |

*These data on 4×4 = 16 mice are a subset of the hematology data in Supplementary Table 2. We used these mice in the ImageStream flow-cytometry studies (see Supplementary Table 4 and Supplementary Table 5) and as the basis for the mathematical simulations. Because we observed no statistically significant differences between sexes in Supplementary Table 2, we grouped a males and females for this study. dKO, double knockout (i.e., *Aqp1*–/–*RHag*–/–).

**Supplementary Table 7 | Summary of hematological data obtained from mice used in Inhibitor study***

| | Strains | WT | | | dKO | | |
|---|---|---|---|---|---|---|---|
| | n | 13 | | | 13 | | |
| | Age (weeks) | 11.9 ±0.3 | | | 11.2 ±0.2 | | |
| | groups | Ctrl | pCMBS | DIDS | Ctrl | pCMBS | DIDS |
| hematological parameters | WBC ($10^3$/ul) | 3.3 ±0.8 | 1.6 ±0.2* | 2.2 ±1.1 | 6.0 ±1.3 | 2.3 ±0.5* | 3.3 ±0.5* |
| | LYM ($10^3$/ul) | 2.6 ±0.6 | 1.0 ±0.1* | 1.7 ±1.0 | 4.6 ±1.1 | 1.6 ±0.4* | 2.5 ±0.4* |
| | MONO ($10^3$/ul) | 0.2 ±0.0 | 0.2 ±0.0 | 0.2 ±0.1 | 0.3 ±0.1 | 0.2 ±0.1 | 0.1 ±0.0* |
| | GRAN ($10^3$/ul) | 0.5 ±0.1 | 0.4 ±0.0 | 0.3 ±0.1 | 1.0 ±0.2 | 0.5 ±0.1* | 0.6 ±0.2* |
| | LYM % | 77.0 ±1.1 | 62.4 ±1.8* | 70.4 ±2.3* | 78.0 ±3.2 | 65.9 ±3.3* | 77.1 ±3.3 |
| | MONO % | 4.0 ±0.3 | 9.7 ±0.6* | 4.4 ±0.2 | 3.9 ±0.3 | 8.2 ±0.7* | 3.5 ±0.3 |
| | GRAN % | 19.0 ±1.1 | 27.9 ±1.5* | 25.2 ±2.3* | 18.2 ±3.2 | 25.8 ±3.4* | 19.4 ±3.1 |
| | HCT (%) | 33.1 ±1.3 | 34.9 ±0.9 | 35.4 ±1.0* | 37.0 ±1.7 | 39.1 ±1.2 | 40.0 ±1.1* |
| | HGB (g/dl) | 11.3 ±0.5 | 11.8 ±0.2* | 11.1 ±0.2 | 11.3 ±0.5 | 12.5 ±0.4* | 12.1 ±0.3 |
| | RDWa (fl) | 29.7 ±0.6 | 28.7 ±0.5 | 31.7 ±0.7* | 32.6 ±0.4 | 31.8 ±0.4 | 35.2 ±0.5 |
| | RDW % (%) | 16.8 ±0.2 | 17.6 ±0.3* | 17.3 ±0.2* | 16.5 ±0.1 | 17.1 ±0.1 | 16.9 ±0.1 |
| | MCV (fl) | 46.4 ±0.9 | 44.3 ±0.8* | 47.5 ±0.8* | 50.2 ±0.2 | 48.2 ±0.3* | 51.8 ±0.4* |
| | MCH (pg) | 14.8 ±0.1 | 15.0 ±0.1* | 15.0 ±0.1* | 15.3 ±0.1 | 15.4 ±0.1 | 15.6 ±0.1* |
| | MCHC (g/dl) | 32.1 ±0.6 | 34.0 ±0.6* | 31.6 ±0.4* | 30.6 ±0.4 | 32.0 ±0.2* | 30.2 ±0.3 |
| | RBC ($10^6$/ul) | 7.1 ±0.2 | 7.9 ±0.2* | 7.4 ±0.2 | 7.4 ±0.3 | 8.1 ±0.3* | 7.7 ±0.2 |
| | PLT ($10^3$/ul) | 91.7 ±24.2 | 64.7 ±5.1 | 85.9 ±30.9 | 127.5 ±31.4 | 68.0 ±12.6 | 84.3 ±15.5 |
| | MPV (fl) (n) | 6.4 ±0.4 (9) | 6.1 ±0.1 (12) | 6.2 ±0.2 (11) | 6.2 ±0.3 (13) | 6.1 ±0.1 (11) | 7.7 ±0.4* (9) |

*These data on 13×2 = 26 mice are the hematology data obtained from mice used in inhibitor study. We included mice used in the ImageStream flow-cytometry with RBCs treated with inhibitor studies (see Supplementary Table 9). Because we observed no statistically significant differences between sexes in Supplementary Table 2, we grouped a males and females (same gender for one pair of WT and dKO mice on that day) for this study. dKO, double knockout (i.e., *Aqp1–/–RHag–/–*). Results are presented as means ± s.e.m. Each “n” represents an analysis of the blood from 1 mouse. We performed paired t-tests. *P<0.05 compared with Ctrl. Samples are tested at ostensibly 50% hematocrit (See Methods/Hematology).

**Supplementary Table 8 | Quantification of major diameter, among mature RBCs, from WT and dKO mice treated with inhibitors ***

| Strains | Drug | Cell shape | n total | % | Major Diameter (μm) | s.d. | s.e.m. |
|---|---|---|---|---|---|---|---|
| WT | No Drug | Normal | 224756 | 98.6 | 6.73 | 1.07 | 0.00225 |
| | | Misshapen | 3223 | 1.4 | 6.47 | 0.89 | 0.0157 |
| | pCMBS | Normal | 111534 | 91.3 | 6.67 | 0.87 | 0.00183 |
| | | Misshapen | 10646 | 8.7 | 5.93 | 0.57 | 0.0101 |
| | DIDS | Normal | 102170 | 59.0 | 6.43 | 0.79 | 0.0017 |
| | | Misshapen | 71075 | 41.0 | 5.77 | 0.70 | 0.0123 |
| dKO | No Drug | Normal | 183684 | 97.5 | 6.22 | 0.82 | 0.00174 |
| | | Misshapen | 4726 | 2.5 | 6.11 | 0.57 | 0.00996 |
| | pCMBS | Normal | 151278 | 94.3 | 6.59 | 0.92 | 0.00193 |
| | | Misshapen | 9184 | 5.7 | 5.90 | 0.59 | 0.0103 |
| | DIDS | Normal | 141579 | 78.7 | 6.47 | 0.87 | 0.00184 |
| | | Misshapen | 38259 | 21.3 | 5.85 | 0.59 | 0.0104 |

*We split blood samples from three age-matched pairs of WT and dKO mice into three equal parts and incubated each of these with a different inhibitor (see Methods/Statistical_Analysis). We sorted RBCs according to their fluorescence signature in an ImageStream flow cytometer and analyzed the number and major diameter of normal and misshapen RBCs following each drug treatment (see Supplementary Data Figure 9). We used the mean major diameters and percentage of normal vs. misshapen cells in each condition in mathematical modeling to determine the contribution that the inhibitor-induced changes in RBC morphology had on $k_{HbO_2}$.

**Supplementary Table 9 | Summary of hematological data obtained from the same samples used for ImageStream analysis and for mathematical simulations***

| | Strains | WT | | | dKO | | |
|---|---|---|---|---|---|---|---|
| | n | 3 | | | 3 | | |
| | Age (weeks) | 11.0 ±0.0 | | | 10.3 ±0.3 | | |
| | groups | Ctrl | pCMBS | DIDS | Ctrl | pCMBS | DIDS |
| hematological parameters | WBC ($10^3$/ul) | 6.1 ±3.0 | 2.1 ±0.4 | 6.8 ±4.4 | 4.2 ±1.1 | 2.1 ±0.9 | 4.9 ±1.8 |
| | LYM ($10^3$/ul) | 4.8 ±2.5 | 1.4 ±0.3 | 5.6 ±3.8 | 2.6 ±0.6 | 1.3 ±0.7 | 3.2 ±1.5 |
| | MONO ($10^3$/ul) | 0.4 ±0.2 | 0.2 ±0.0 | 0.4 ±0.2 | 0.2 ±0.0 | 0.2 ±0.1 | 0.2 ±0.1 |
| | GRAN ($10^3$/ul) | 0.9 ±0.4 | 0.5 ±0.1 | 0.8 ±0.5 | 1.4 ±0.9 | 0.6 ±0.1 | 1.5 ±0.9 |
| | LYM % | 77.8 ±1.8 | 67.4 ±3.0* | 77.3 ±6.0 | 66.9 ±12.2 | 55.8 ±11.7* | 66.8 ±12.7 |
| | MONO % | 4.6 ±1.1 | 8.9 ±1.5* | 3.9 ±0.3 | 3.1 ±0.3 | 6.6 ±1.2* | 4.1 ±0.6 |
| | GRAN % | 17.6 ±2.7 | 23.6 ±1.6 | 18.8 ±6.1 | 30.0 ±12.3 | 37.6 ±12.8* | 29.0 ±12.4 |
| | HCT (%) | 35.5 ±3.2 | 36.1 ±3.0 | 36.5 ±1.9 | 33.9 ±2.8 | 38.7 ±2.8 | 35.7 ±1.7 |
| | HGB (g/dl) | 11.3 ±0.5 | 11.9 ±0.5 | 11.5 ±0.3 | 10.3 ±0.7 | 12.2 ±0.7 | 11.3 ±0.7 |
| | RDWa (fl) | 29.7 ±1.7 | 27.9 ±1.4* | 31.1 ±1.5* | 32.1 ±1.5 | 31.4 ±1.7* | 34.1 ±1.1 |
| | RDW % (%) | 16.9 ±0.3 | 17.1 ±0.5 | 17.2 ±0.1 | 16.5 ±0.5 | 16.9 ±0.3 | 17.1 ±0.6 |
| | MCV (fl) | 46.0 ±2.2 | 44.2 ±2.1* | 47.2 ±1.7 | 49.7 ±0.8 | 48.1 ±1.1* | 50.7 ±0.2 |
| | MCH (pg) | 14.7 ±0.1 | 14.8 ±0.1 | 14.9 ±0.2 | 15.1 ±0.1 | 15.1 ±0.1 | 16.0* ±0.3 |
| | MCHC (g/dl) | 32.1 ±1.8 | 33.5 ±1.4* | 31.6 ±1.1 | 30.5 ±0.5 | 31.5 ±0.6* | 31.6 ±0.6 |
| | RBC ($10^6$/ul) | 7.7 ±0.4 | 8.1 ±0.3 | 7.7 ±0.3 | 6.8 ±0.5 | 8.0 ±0.5 | 7.1 ±0.3 |
| | PLT ($10^3$/ul) | 198.3 ±83.7 | 89.0 ±13.3 | 201.0 ±127.0 | 138.0 ±84.5 | 64.3 ±3.7 | 174.7 ±19.3 |
| | MPV (fl) | 5.7 ±0.1 | 5.8 ±0.2 | 6.0 ±0.2 | 5.8 ±0.2 | 6.4 ±0.5 | 7.5 ±0.5 |

*These data on 3×2 = 6 mice are a subset of the hematology data in Supplementary Table 7. We used these mice in the ImageStream flow-cytometry studies on RBCs treated with inhibitors (see Supplementary Table 8) and as the basis for the mathematical simulations.

Because we observed no statistically significant differences between sexes in Supplementary Table 2, we grouped a males and females (same gender for one pair of WT and dKO mice on that day) for this study. dKO, double knockout (i.e., *Aqp1–/–RHag–/–*). Results are presented as means ± s.e.m. Each "n" represents an analysis of the blood from 1 mouse. We performed paired t-tests. *P<0.05 compared with Ctrl. Samples are tested at ostensibly 50% hematocrit (See Methods/Hematology).

**Supplementary Table 10 | Parameter values used in mathematical simulations of RBCs from WT mice**

| | | Bulk ECF | Bulk ICF | Reference |
|---|---|---|---|---|
| **Temperature** | | 10 °C | 10 °C | |
| **Sphere diameters** | | $4.02 \times 10^{-4}$ cm ($\ell_{\mathrm{EUF}}$ = 1 μm) | $2.02 \times 10^{-4}$ cm (= 2.02 μm) | See Supplementary Methods/CalculationRBCthickness |
| **Shell thickness** | | $10^{-6}$ cm (= $10^{-2}$ μm) | $10^{-6}$ cm (= $10^{-2}$ μm) | |
| **Diffusion coefficient (*D*)** | $O_2$ | $1.3313 \times 10^{-5}$ cm²/sec | $2.7745 \times 10^{-6}$ cm²/sec | Computed using data from ref. 2<br>See Supplementary Discussion/OurCalculationOfD$_{O_2}$ |
| | Hb | 0 | $6.07 \times 10^{-8}$ cm²/sec | Computed using data from ref. 3 |
| | $HbO_2$ | 0 | $6.07 \times 10^{-8}$ cm²/sec | Computed using data from ref. 3 |
| **$Hb/HbO_2$ reaction**<br>**Rate constants** | *k* | 0 | 11.60 sec$^{-1}$ | Measured |
| | *k´* | 0 | $6.7481 \times 10^{4}$ mM$^{-1}$sec$^{-1}$ (*k′* varies with [$O_2$]; this is initial value) | Computed using Eqn 4 in Supplementary Methods |
| **Equilibrium constant** | K | | $1.1790 \times 10^{-4}$ mM (K varies with *k′*; this is initial value) | Calculated as *k/k´* |
| **$O_2$ permeability** | | | 0.150 cm/sec | Assumed same as $CO_2$ permeability in RBCs[4]. |
| **Partial pressure of $H_2O$ vapor ($P_{H_2O}$)** | | | 9.2 mmHg | From ref. [5] |
| **$P_{50}$** | | | 9.35 mmHg | From ref. [6] |
| **$O_2$ solubility** | | | $2.24 \times 10^{-3}$ mM/mmHg | From ref. [5] |
| **Hill coefficient** | | | 2.7 | From ref. [7] |
| **Concentration** | Total Hb | 0 mM | 18.73 mM | See Suppl Methods/HbContent |
| | $O_2$ | 0 mM | 0.3532 mM (21% $O_2$) | Computed from $O_2$ solubility (above), using Henry's law |
| | $HbO_2$ monomer | 0 mM | 18.6767 mM | Computed using the relation $[HbO_2] = [Hb]_{tot} ([O_2]^n/(K+[O_2]^n)$ |
| | Hb monomer | 0 mM | 0.0533 mM | Computed as $[Hb]_{tot} - [HbO_2]$ |

Parameter values correspond to the experimental temperature of 10°C.

**Supplementary Table 11 | Genotype-specific parameter values used in mathematical simulations of RBCs in Extended Data Figure 9*b* ***

| Parameter | WT | *Aqp1–/–* | *RHag–/–* | dKO | Reference |
|---|---|---|---|---|---|
| **Major diameter of RBC (µm)** | 6.80 | 6.69 | 6.53 | 6.55 | From Supplementary Table 5 (We will assume that $OD_{Torus}$= RBC major diameter) |
| **MCV (fl)** | 47.9 | 49.5 | 51.0 | 49.8 | From Supplementary Table 6 |
| ***r*, minor radius of torus (µm)** | 1.01 | 1.04 | 1.09 | 1.07 | R & r are computed such that torus volume = MCV |
| ***R*, major radius of torus (µm)** | 2.39 | 2.31 | 2.18 | 2.21 | R & r are computed such that torus volume = MCV |
| **$D_{Sphere}$, diameter of sphere (µm)** | 2.02 | 2.08 | 2.18 | 2.14 | Computed from r: D = 2×r. See SI Methods/RBC_Thickness |
| **MCH, mean corpuscular hemoglobin (pg)** | 14.4 | 14.1 | 14.7 | 14.5 | From Supplementary Table 6 |
| **[Total Hb monomers]$_i$ (mM)** | 18.73 | 17.71 | 17.87 | 18.16 | Computed from MCH |
| **Initial [$HbO_2$ monomers]$_i$ (mM)** | 18.6767 | 17.6596 | 17.8191 | 18.1083 | Computed from [Total Hb monomers]$_i$ and $P_{O_2}$ |
| **Initial [Hb monomers]$_i$ (mM)** | 0.0533 | 0.0504 | 0.0509 | 0.0517 | Computed from [Total Hb monomers]$_i$ and $P_{O_2}$ |

*Values listed here for WT mice are the same as those in Supplementary Table 10. Values not listed here, but necessary for mathematical simulations, also are the same as in Supplementary Table 10. dKO, double knockout (i.e., *Aqp1–/–RHag–/–*).

**Supplementary Table 12 | Physiological solutions**

| Component or parameter | Oxygenated solution (also, RBC washing) | De-oxygenated solution | Components for OOE Hemolysis assay | | | Solutions for preparation of mouse erythrocyte ghosts | | |
|---|---|---|---|---|---|---|---|---|
| | | | A | B | Mix¶ | Wash Buffer | Sedimentation Buffer | Lysis Buffer‖ |
| NaCl (mM) | 92.5 | 15 | 140 | 116 | 128 | 150 | 150 | 0 |
| KCl (mM) | 0 (ref. [8]) | 0 (ref. [8]) | 3 | 3 | 3 | 0 | 0 | 0 |
| $CaCl_2$ (mM) | 0.01 | 0.01 | 2 | 0 | 1 | 0 | 0 | 0 |
| $Na_2HPO_4$ / $NaH_2PO_4$ (mM)* | 46.98/11.02 | 46.98/11.02 | 0 | 0 | 0 | 0 | 4.06/0.94 | 0 |
| HEPES (mM) | 0 | 0 | 16 | 0 | 8 | 0 | 0 | 0 |
| $NaHCO_3$ (mM)‡ | 0 | 0 | ~0 | 44 | 22 | 0 | 0 | 0 |
| $CO_2$ (%) | 0 | 0 | ~0 | ~1 | 0.5 | 0 | 0 | 0 |
| pH | ~7.40* | ~7.40* | 7.03† | 8.41‡ | ~7.25 | ~7. 0 | ~7.50* | ~8.0 |
| Pyranine (μM)§ | 0 | 0 | 0 | 2 or 0 | 1 or 0 | 0 | 0 | 0 |
| RBCs, lysate | ++ or 0 | 0 | ++ or 0 | 0 | + or 0 | + or 0 | + or 0 | + or 0 |
| NDT$^{\Delta}$ | 0 | 50 | 0 | 0 | 0 | 0 | 0 | 0 |
| Tris-HCl (mM) | 0 | 0 | 0 | 0 | 0 | 0 | 0 | 5 |
| Dextran500(w/v) | 0 | 0 | 0 | 0 | 0 | 0 | 0.75% | 0 |
| Temperature (°C) | 10 (RT or ice) | 10 | 10 | 10 | 10 | RT | RT | RT |
| Osmolality (mOsm) | ~300 | ~300 | ~295 | ~300 | ~298 | ~290 | / | / |

*The ratio $[HPO_4^{=}]/[H_2PO_4^{-}]$ determines the pH at RT ( ~22°C).

†We titrated HEPES free acid (pK ~7.41) to pH 7.50 with NaOH, and then in some aliquots added either HCl or more NaOH to achieve pH values from 5.50 to 8.50 at 10°C. After the titration, we added pyranine to equal concentrations in each solution.

‡The addition of $HCO_3^{-}$ generates some $CO_2$ and $CO_3^{=}$; this mixture determined the final pH at 10°C.

§[Pyranine] in the reaction cell was either 1 μM (to obtain pH data) or 0 μM (to obtain background data).

¶The values in this pyranine column are those at the instant of mixing solutions A and B. The solution is out of equilibrium because the pH of 7.25 is far too low for the system to be in equilibrium, given $[HCO_3^{-}]$ = 22 mM and $CO_2$ = 0.5%.

ΔNDT is freshly added and used within ~2 hours. Final [NDT] in SF reaction cell is 25 mM.

‖Complete protease inhibitor tablet (Roche) – only for lysis and first wash following lysis step.
