## Supplementary Tables (MassSpecData) for "Role of channels in the oxygen permeability of red blood cells"

### Mass Spec. Data - All Data

| Charge Group ID | Charge State | Factor 1 P-value | Intensity [dKO_1] | Intensity [dKO_2] | Intensity [dKO_3] | Intensity [AQP1KO_1] | Intensity [AQP1KO_2] | Intensity [AQP1KO_3] | Intensity [RhAGKO_1] | Intensity [RhAGKO_2] | Intensity [RhAGKO_3] | Intensity [WT_1] | Intensity [WT_2] | Intensity [WT_3] | M + H (Feature) | Modified Peptide Sequence | Primary Protein Name | ID | Symbol | Entrez Gene Name | Primary Location | Type(s) |
|---|---|---|---|---|---|---|---|---|---|---|---|---|---|---|---|---|---|---|---|---|---|---|
| 15554680 | 2 | 0.786 | 1023527.188 | 979379.125 | 990138.5 | 1315149.25 | 641206.062 | 569730.875 | 1101591.125 | 872663.438 | 758731.375 | 1313602.625 | 862820.188 | 701254.125 | 1567.8044 | AQATPSQPLQSSQPK | Q3UHJ0 | Q3UHJ0 | AAK1 | AP2 associated kinase 1 | Cytoplasm | kinase |
| 15552647 | 3 | 0.232 | 1.16E+07 | 3.62E+07 | 1.47E+07 | 1.48E+07 | 1.28E+07 | 2.35E+07 | 1.07E+07 | 1.68E+07 | 1.81E+07 | 1.14E+07 | 7289731 | 9925025 | 2778.4136 | ILLLDEATSALDAENEHLVQEALDR | Q9JI39 | Q9JI39 | ABCB10 | ATP-binding cassette, sub-family B (MDR/TAP), member 10 | Cytoplasm | transporter |
| 15550136 | 3 | 0.139 | 1.32E+07 | 2.49E+07 | 7081912 | 1.21E+07 | 1.20E+07 | 1.63E+07 | 9621738 | 1.44E+07 | 1.55E+07 | 6332840 | 6041680.5 | 8413606 | 3839.869 | IGTVSQEPVLFSC[160.0307]SVAENIAYGADNLSSVTAQQVER | Q9JI39 | Q9JI39 | ABCB10 | ATP-binding cassette, sub-family B (MDR/TAP), member 10 | Cytoplasm | transporter |
| 15533437 | 2 | 0.172 | 5669490 | 1.34E+07 | 4231947.5 | 5134905 | 6097008.5 | 7486709 | 3691963.75 | 7599960.5 | 7532876 | 3209369 | 3364578.5 | 3541787.5 | 1032.5665 | LSSDTALLGR | Q9JI39 | Q9JI39 | ABCB10 | ATP-binding cassette, sub-family B (MDR/TAP), member 10 | Cytoplasm | transporter |
| 15559362 | 2 | 0.226 | 2837780.5 | 3538272 | 2299725 | 2767352.5 | 2076733.375 | 1790009.625 | 2215662.5 | 2250750.5 | 2641814.5 | 2353307.25 | 2162250 | 1908195.25 | 1732.8247 | ATQDSLAEATQLAEER | Q9JI39 | Q9JI39 | ABCB10 | ATP-binding cassette, sub-family B (MDR/TAP), member 10 | Cytoplasm | transporter |
| 15540728 | 2 | 0.043 | 4108594.75 | 6175137 | 2903497.75 | 2965125.25 | 3346896.75 | 4309698.5 | 2719897.25 | 5039134.5 | 5116280 | 1960281.625 | 2021167.125 | 2155089.5 | 1284.6661 | NANFVAVLDHGK | Q9JI39 | Q9JI39 | ABCB10 | ATP-binding cassette, sub-family B (MDR/TAP), member 10 | Cytoplasm | transporter |
| 15542750 | 2 | 0.139 | 3418209.25 | 7147102.5 | 3531161.25 | 3482082.5 | 3801826.75 | 5171538 | 2394133.25 | 5452794.5 | 5765157 | 1775389.125 | 2598952.75 | 2456438.25 | 983.4815 | QEVAFFDK | Q9JI39 | Q9JI39 | ABCB10 | ATP-binding cassette, sub-family B (MDR/TAP), member 10 | Cytoplasm | transporter |
| 15537481 | 2 | 0.158 | 3176291.75 | 7876071 | 3211201.5 | 3455503 | 2893680.75 | 6031781.5 | 2608369.75 | 4341638.5 | 5416581.5 | 1770016.25 | 1686377.75 | 2823162 | 987.6181 | STVVSLLLR | Q9JI39 | Q9JI39 | ABCB10 | ATP-binding cassette, sub-family B (MDR/TAP), member 10 | Cytoplasm | transporter |
| 15536116 | 2 | 0.178 | 3040666.5 | 6827866.5 | 2506444.25 | 2971433.25 | 3278868 | 4440326 | 1847704.125 | 4225914 | 4726169 | 1759276.375 | 1801890.625 | 2021915.375 | 958.53375 | LWELLER | Q9JI39 | Q9JI39 | ABCB10 | ATP-binding cassette, sub-family B (MDR/TAP), member 10 | Cytoplasm | transporter |
| 15550359 | 2 | 0.221 | 2065191.75 | 4554023.5 | 1554486 | 1887297.375 | 2486945.75 | 3856554.5 | 1909124.125 | 2744847.5 | 3237393 | 1398149.375 | 1578077.375 | 1401543.25 | 1581.8021 | VYLMQSSGQSIVNR | Q9JI39 | Q9JI39 | ABCB10 | ATP-binding cassette, sub-family B (MDR/TAP), member 10 | Cytoplasm | transporter |
| 15534077 | 2 | 0.158 | 3610339.75 | 8416091 | 2357357.75 | 2538405.5 | 3991995.5 | 6175970.5 | 2761271.5 | 5047171 | 5684115.5 | 1326483.875 | 1826603.75 | 2568060 | 1023.5813 | TSLFSSILR | Q9JI39 | Q9JI39 | ABCB10 | ATP-binding cassette, sub-family B (MDR/TAP), member 10 | Cytoplasm | transporter |
| 15544827 | 2 | 0.303 | 1871996.25 | 1991290.125 | 1742957.5 | 1238706.5 | 1098555.25 | 2793082.5 | 1345937.25 | 1636913.375 | 2260400.5 | 1138085.625 | 1278864.125 | 1113010 | 1410.686 | SFPQGFDTVVGEK | Q9JI39 | Q9JI39 | ABCB10 | ATP-binding cassette, sub-family B (MDR/TAP), member 10 | Cytoplasm | transporter |
| 15556410 | 2 | 0.071 | 1180126.25 | 2130069 | 1045574.312 | 945867.812 | 915423.375 | 1316312.125 | 725046.5 | 1078015.25 | 1292958.125 | 627558.125 | 855074.875 | 469807.688 | 1858.8906 | LYDPNSGTVSLDGHDIR | Q9JI39 | Q9JI39 | ABCB10 | ATP-binding cassette, sub-family B (MDR/TAP), member 10 | Cytoplasm | transporter |
| 15549769 | 2 | 0.496 | 2.71E+07 | 2.63E+07 | 3.63E+07 | 2.71E+07 | 2.07E+07 | 2.55E+07 | 2.90E+07 | 2.57E+07 | 2.42E+07 | 3.35E+07 | 2.49E+07 | 2.41E+07 | 1833.9282 | VLLLDEATSALDTESEK | P21440 | P21440 | ABCB4 | ATP-binding cassette, sub-family B (MDR/TAP), member 4 | Plasma Membrane | transporter |
| 15538710 | 2 | 0.161 | 1.60E+07 | 1.47E+07 | 1.55E+07 | 1.60E+07 | 1.15E+07 | 1.12E+07 | 1.57E+07 | 1.24E+07 | 1.07E+07 | 1.88E+07 | 1.50E+07 | 1.58E+07 | 1227.6663 | AGAVAEEALGAIR | P21440 | P21440 | ABCB4 | ATP-binding cassette, sub-family B (MDR/TAP), member 4 | Plasma Membrane | transporter |
| 15532894 | 3 | 0.617 | 1.28E+07 | 1.32E+07 | 2.01E+07 | 1.49E+07 | 1.49E+07 | 1.24E+07 | 1.55E+07 | 1.80E+07 | 1.45E+07 | 1.86E+07 | 1.56E+07 | 1.54E+07 | 1754.8098 | GNLEFSDVHFSYPSR | P21440 | P21440 | ABCB4 | ATP-binding cassette, sub-family B (MDR/TAP), member 4 | Plasma Membrane | transporter |
| 15549730 | 2, 3 | 0.123 | 1.31E+07 | 1.46E+07 | 1.62E+07 | 9780653 | 1.08E+07 | 1.18E+07 | 1.28E+07 | 1.40E+07 | 1.20E+07 | 1.77E+07 | 1.10E+07 | 1.40E+07 | 2415.307, 2415.3083 | EIIGVVSQEPVLFSTTIAENIR | P21440 | P21440 | ABCB4 | ATP-binding cassette, sub-family B (MDR/TAP), member 4 | Plasma Membrane | transporter |
| 15531693 | 2 | 0.46 | 1.67E+07 | 1.44E+07 | 1.63E+07 | 1.73E+07 | 1.34E+07 | 1.23E+07 | 1.47E+07 | 1.47E+07 | 1.16E+07 | 1.73E+07 | 1.53E+07 | 1.42E+07 | 1459.747 | LATDAAQVQGATGTR | P21440 | P21440 | ABCB4 | ATP-binding cassette, sub-family B (MDR/TAP), member 4 | Plasma Membrane | transporter |
| 15550143 | 3 | 0.174 | 1.29E+07 | 1.53E+07 | 1.49E+07 | 1.15E+07 | 1.11E+07 | 1.48E+07 | 1.19E+07 | 1.41E+07 | 1.25E+07 | 1.66E+07 | 1.39E+07 | 1.49E+07 | 3080.4922 | AQLGIVSQEPILFDC[160.0307]SIAENIAYGDNSR | P21440 | P21440 | ABCB4 | ATP-binding cassette, sub-family B (MDR/TAP), member 4 | Plasma Membrane | transporter |
| 15533643 | 3 | 0.18 | 1.38E+07 | 1.60E+07 | 1.70E+07 | 1.49E+07 | 1.30E+07 | 1.30E+07 | 1.87E+07 | 1.52E+07 | 1.54E+07 | 1.60E+07 | 1.67E+07 | 1.45E+07 | 1636.8663 | EANIHPFIETLPQK | P21440 | P21440 | ABCB4 | ATP-binding cassette, sub-family B (MDR/TAP), member 4 | Plasma Membrane | transporter |
| 15531732 | 2 | 0.716 | 1.28E+07 | 1.18E+07 | 1.55E+07 | 1.30E+07 | 1.06E+07 | 1.01E+07 | 1.26E+07 | 1.53E+07 | 6689358 | 1.47E+07 | 1.21E+07 | 1.15E+07 | 1129.6184 | IATEAIENIR | P21440 | P21440 | ABCB4 | ATP-binding cassette, sub-family B (MDR/TAP), member 4 | Plasma Membrane | transporter |
| 15534335 | 2 | 0.577 | 1.17E+07 | 1.15E+07 | 1.41E+07 | 1.28E+07 | 1.01E+07 | 1.04E+07 | 1.17E+07 | 1.09E+07 | 1.03E+07 | 1.38E+07 | 1.01E+07 | 1.17E+07 | 1466.8552 | ANVPVLQGLSLEVK | P21440 | P21440 | ABCB4 | ATP-binding cassette, sub-family B (MDR/TAP), member 4 | Plasma Membrane | transporter |
| 15549267 | 2 | 0.797 | 9305181 | 9843795 | 9044523 | 1.14E+07 | 8578539 | 1.06E+07 | 1.14E+07 | 8664280 | 7927802.5 | 1.15E+07 | 9274393 | 9403828 | 1840.8728 | FYDPMAGSVLLDGQEAK | P21440 | P21440 | ABCB4 | ATP-binding cassette, sub-family B (MDR/TAP), member 4 | Plasma Membrane | transporter |
| 15535583 | 2 | 0.804 | 9542266 | 8283619 | 1.25E+07 | 9552567 | 9319960 | 9306429 | 9868161 | 8502768 | 8821640 | 9855601 | 8606534 | 1.06E+07 | 1045.5989 | STTVQLLQR | P21440 | P21440 | ABCB4 | ATP-binding cassette, sub-family B (MDR/TAP), member 4 | Plasma Membrane | transporter |
| 15554867 | 2 | 0.733 | 8146750.5 | 8291964.5 | 1.17E+07 | 9177367 | 7542390 | 8882015 | 1.05E+07 | 7459775.5 | 7902644.5 | 9769593 | 9839147 | 8891646 | 1804.8727 | QPLIDSYSGEGLWPDK | P21440 | P21440 | ABCB4 | ATP-binding cassette, sub-family B (MDR/TAP), member 4 | Plasma Membrane | transporter |
| 15537972 | 2 | 0.088 | 7134308 | 6072062 | 6545002.5 | 7792765 | 6725211 | 5455553 | 6200758.5 | 6147119 | 5345316.5 | 8194694.5 | 8179264.5 | 7097662.5 | 1434.698 | SGQTVALVGNSGC[160.0307]GK | P21440 | P21440 | ABCB4 | ATP-binding cassette, sub-family B (MDR/TAP), member 4 | Plasma Membrane | transporter |
| 15532231 | 2 | 0.32 | 6987972 | 5877782 | 7808233.5 | 7419671.5 | 6667009 | 6861859 | 6999255.5 | 6248495 | 5500906 | 7888313.5 | 7074446.5 | 7082108 | 1034.5603 | TVIAFGGQNK | P21440 | P21440 | ABCB4 | ATP-binding cassette, sub-family B (MDR/TAP), member 4 | Plasma Membrane | transporter |
| 15533364 | 2 | 0.468 | 8284547.5 | 8338817.5 | 1.07E+07 | 8661057 | 7294413.5 | 7978149 | 6202433 | 9398491 | 7333127 | 7877104 | 8102595 | 8801876 | 1032.5648 | TVVSLTQER | P21440 | P21440 | ABCB4 | ATP-binding cassette, sub-family B (MDR/TAP), member 4 | Plasma Membrane | transporter |
| 15537383 | 2 | 0.328 | 6649497.5 | 8578936 | 8676622 | 8620775 | 6410018 | 8433190 | 7034590.5 | 6801267.5 | 6082633 | 7236325.5 | 7541278.5 | 7329213.5 | 1416.7502 | LSAAYLFSLFER | P21440 | P21440 | ABCB4 | ATP-binding cassette, sub-family B (MDR/TAP), member 4 | Plasma Membrane | transporter |
| 15534464 | 3 | 0.797 | 3979119.5 | 4856122.5 | 4435005 | 5508740.5 | 4483818.5 | 5447695.5 | 4690289.5 | 5250238 | 4308785.5 | 6195433 | 3270584.25 | 4918480 | 1649.8502 | GAAYVIFDIIDNNPK | P21440 | P21440 | ABCB4 | ATP-binding cassette, sub-family B (MDR/TAP), member 4 | Plasma Membrane | transporter |
| 15533551 | 2 | 0.29 | 4631437.5 | 5387041 | 7062705 | 4113016.25 | 4170613.5 | 5050830.5 | 5875553 | 4916612.5 | 4507142 | 5902773 | 5193599.5 | 5001338.5 | 928.53406 | LNVQWLR | P21440 | P21440 | ABCB4 | ATP-binding cassette, sub-family B (MDR/TAP), member 4 | Plasma Membrane | transporter |
| 15534292 | 2 | 0.85 | 3819291.75 | 2793517.25 | 3884543.5 | 4005628.25 | 2923840 | 2982802 | 4030943.75 | 3111476.75 | 3095747.25 | 5077668.5 | 3535773.5 | 2969077 | 1020.4996 | ILEEEMTR | P21440 | P21440 | ABCB4 | ATP-binding cassette, sub-family B (MDR/TAP), member 4 | Plasma Membrane | transporter |
| 15533375 | 2 | 0.379 | 5146337.5 | 2401103.75 | 1414049 | 3789498.5 | 2222289.75 | 2097144.75 | 3514818 | 2554964 | 3034778 | 5029358 | 3940184.5 | 3984126 | 1160.5626 | KFESMYVEK | P21440 | P21440 | ABCB4 | ATP-binding cassette, sub-family B (MDR/TAP), member 4 | Plasma Membrane | transporter |
| 15544007 | 2 | 0.862 | 3280503.75 | 3418023.25 | 4758375 | 3352497 | 3567431.25 | 3395247.75 | 3869112 | 3859194.5 | 2722052.25 | 4236595 | 3351697 | 3424961.75 | 860.4818 | AGEILTTR | P21440 | P21440 | ABCB4 | ATP-binding cassette, sub-family B (MDR/TAP), member 4 | Plasma Membrane | transporter |
| 15558327, 15560805, 15561087 | 2 | 0.294 | 3937934.5 | 4479635.5 | 4670759 | 3727024.75 | 3882297.75 | 2361420.25 | 4413075.5 | 4294058.5 | 3252051.5 | 4078936.25 | 3543186 | 4498200 | 1636.8661, 1636.8662, 1636.8668 | EANIHPFIETLPQK | P21440 | P21440 | ABCB4 | ATP-binding cassette, sub-family B (MDR/TAP), member 4 | Plasma Membrane | transporter |
| 15537579 | 3 | 0.095 | 2733030 | 3001182.75 | 2181185 | 2225233.25 | 1799774.5 | 1563902.75 | 3649639 | 2710339.25 | 2183691.25 | 2634747.25 | 2245413 | 2317968.75 | 1308.5277 | QDMSWFDDHK | P21440 | P21440 | ABCB4 | ATP-binding cassette, sub-family B (MDR/TAP), member 4 | Plasma Membrane | transporter |
| 15538630 | 2 | 0.708 | 1675143.875 | 1674961.375 | 2615903.25 | 450019.094 | 1579705.625 | 1967734 | 1371637 | 1954752.5 | 2024310.5 | 467062.344 | 2633680.75 | 1967988.625 | 969.5284 | TC[160.0307]IVIAHR | P21440 | P21440 | ABCB4 | ATP-binding cassette, sub-family B (MDR/TAP), member 4 | Plasma Membrane | transporter |
| 15548651 | 2, 3 | 0.171 | 1.08E+08 | 1.07E+08 | 1.28E+08 | 8.75E+07 | 9.55E+07 | 8.46E+07 | 8.39E+07 | 8.19E+07 | 1.04E+08 | 1.28E+08 | 8.45E+07 | 9.63E+07 | 2423.2642, 2423.2646 | SHIGVVPQDTVLFNDTIANNIR | Q9DC29 | Q9DC29 | ABCB6 | ATP-binding cassette, sub-family B (MDR/TAP), member 6 (Langereis blood group) | Cytoplasm | transporter |
| 15547805 | 3 | 0.673 | 9.33E+07 | 9.94E+07 | 1.26E+08 | 8.32E+07 | 9.58E+07 | 1.04E+08 | 8.33E+07 | 1.02E+08 | 1.03E+08 | 1.06E+08 | 8.40E+07 | 9.50E+07 | 3588.6987 | VTAGDSEVEAAAQAAGIHDAILSFPEGYETQVGER | Q9DC29 | Q9DC29 | ABCB6 | ATP-binding cassette, sub-family B (MDR/TAP), member 6 (Langereis blood group) | Cytoplasm | transporter |
| 15551810 | 3 | 0.294 | 3.51E+07 | 4.67E+07 | 4.76E+07 | 3.36E+07 | 4.18E+07 | 2.95E+07 | 4.06E+07 | 4.08E+07 | 3.83E+07 | 5.38E+07 | 3.72E+07 | 4.30E+07 | 3134.4324 | GGVYAEMWQLQQQGQETVPEESKPQDTA | Q9DC29 | Q9DC29 | ABCB6 | ATP-binding cassette, sub-family B (MDR/TAP), member 6 (Langereis blood group) | Cytoplasm | transporter |
| 15555472 | 2 | 0.922 | 3.51E+07 | 3.43E+07 | 5.41E+07 | 3.18E+07 | 4.29E+07 | 3.54E+07 | 3.26E+07 | 3.75E+07 | 4.21E+07 | 4.94E+07 | 3.12E+07 | 3.87E+07 | 1665.8203 | EVPAGPEELSWAAGPR | Q9DC29 | Q9DC29 | ABCB6 | ATP-binding cassette, sub-family B (MDR/TAP), member 6 (Langereis blood group) | Cytoplasm | transporter |
| 15547767 | 2 | 0.115 | 4.75E+07 | 5.33E+07 | 6.60E+07 | 4.28E+07 | 4.59E+07 | 4.06E+07 | 4.07E+07 | 4.86E+07 | 4.99E+07 | 4.87E+07 | 3.79E+07 | 4.32E+07 | 2144.0667 | APDIILLDEATSALDTSNER | Q9DC29 | Q9DC29 | ABCB6 | ATP-binding cassette, sub-family B (MDR/TAP), member 6 (Langereis blood group) | Cytoplasm | transporter |
| 15532663 | 2 | 0.531 | 3.65E+07 | 3.53E+07 | 5.00E+07 | 2.35E+07 | 4.07E+07 | 3.36E+07 | 3.41E+07 | 3.17E+07 | 3.97E+07 | 4.23E+07 | 3.10E+07 | 4.35E+07 | 1335.7135 | AVDSLLNFETVK | Q9DC29 | Q9DC29 | ABCB6 | ATP-binding cassette, sub-family B (MDR/TAP), member 6 (Langereis blood group) | Cytoplasm | transporter |
| 15549591 | 2 | 0.519 | 2.52E+07 | 2.48E+07 | 3.80E+07 | 2.09E+07 | 2.85E+07 | 2.43E+07 | 2.60E+07 | 2.76E+07 | 3.10E+07 | 3.40E+07 | 2.83E+07 | 2.67E+07 | 1672.8823 | IDGQDISQVTQISLR | Q9DC29 | Q9DC29 | ABCB6 | ATP-binding cassette, sub-family B (MDR/TAP), member 6 (Langereis blood group) | Cytoplasm | transporter |
| 15549131 | 2 | 0.129 | 1.49E+07 | 2.01E+07 | 1.66E+07 | 1.27E+07 | 1.71E+07 | 1.42E+07 | 1.76E+07 | 1.75E+07 | 2.09E+07 | 2.71E+07 | 1.74E+07 | 1.92E+07 | 2102.9907 | MIQTNFIDMENMFDLLK | Q9DC29 | Q9DC29 | ABCB6 | ATP-binding cassette, sub-family B (MDR/TAP), member 6 (Langereis blood group) | Cytoplasm | transporter |
| 15533860 | 2 | 0.335 | 1.85E+07 | 1.97E+07 | 2.26E+07 | 1.45E+07 | 2.37E+07 | 1.83E+07 | 1.98E+07 | 2.01E+07 | 2.18E+07 | 2.53E+07 | 2.28E+07 | 2.16E+07 | 1217.5592 | FYDISSGC[160.0307]IR | Q9DC29 | Q9DC29 | ABCB6 | ATP-binding cassette, sub-family B (MDR/TAP), member 6 (Langereis blood group) | Cytoplasm | transporter |
| 15541858 | 3 | 0.24 | 1.41E+07 | 1.58E+07 | 2.14E+07 | 1.51E+07 | 1.23E+07 | 1.24E+07 | 1.32E+07 | 1.79E+07 | 1.20E+07 | 2.26E+07 | 1.56E+07 | 1.63E+07 | 2144.072 | APDIILLDEATSALDTSNER | Q9DC29 | Q9DC29 | ABCB6 | ATP-binding cassette, sub-family B (MDR/TAP), member 6 (Langereis blood group) | Cytoplasm | transporter |
| 15538498 | 2 | 0.149 | 1.32E+07 | 1.32E+07 | 2.07E+07 | 1.01E+07 | 1.37E+07 | 9591124 | 1.33E+07 | 1.44E+07 | 1.63E+07 | 1.66E+07 | 1.39E+07 | 1.48E+07 | 907.4655 | FQGLEWK | Q9DC29 | Q9DC29 | ABCB6 | ATP-binding cassette, sub-family B (MDR/TAP), member 6 (Langereis blood group) | Cytoplasm | transporter |
| 15533436 | 2 | 0.495 | 1.10E+07 | 1.76E+07 | 2.50E+07 | 6505775.5 | 2.08E+07 | 1.57E+07 | 1.52E+07 | 1.82E+07 | 1.45E+07 | 1.28E+07 | 7401402 | 1.23E+07 | 1304.7699 | ALNVLVPIFYR | Q9DC29 | Q9DC29 | ABCB6 | ATP-binding cassette, sub-family B (MDR/TAP), member 6 (Langereis blood group) | Cytoplasm | transporter |
| 15538149 | 2 | 0.553 | 1.02E+07 | 1.14E+07 | 1.67E+07 | 7223106 | 1.25E+07 | 1.09E+07 | 1.08E+07 | 1.00E+07 | 1.15E+07 | 1.03E+07 | 8700079 | 1.14E+07 | 1134.6279 | LLSSYLWPR | Q9DC29 | Q9DC29 | ABCB6 | ATP-binding cassette, sub-family B (MDR/TAP), member 6 (Langereis blood group) | Cytoplasm | transporter |
| 15534349 | 2 | 0.608 | 8335962 | 9068953 | 1.18E+07 | 4488753 | 1.20E+07 | 7160934.5 | 8234651.5 | 1.06E+07 | 7678055.5 | 9897208 | 5276236.5 | 6783860 | 1321.5676 | YYGAEGYEVDR | Q9DC29 | Q9DC29 | ABCB6 | ATP-binding cassette, sub-family B (MDR/TAP), member 6 (Langereis blood group) | Cytoplasm | transporter |
| 15542340 | 2 | 0.527 | 7156440.5 | 1.02E+07 | 1.53E+07 | 3426374.5 | 1.22E+07 | 8845434 | 8100268.5 | 9513779 | 9555593 | 8426053 | 5063723 | 6640051 | 986.5857 | DIVNLLTAK | Q9DC29 | Q9DC29 | ABCB6 | ATP-binding cassette, sub-family B (MDR/TAP), member 6 (Langereis blood group) | Cytoplasm | transporter |
| 15550459 | 2 | 0.479 | 5310749 | 6215591 | 1.08E+07 | 4943896 | 5511619.5 | 5755294 | 3895307 | 5667289 | 9318729 | 5134540.5 | 3849089.75 | 5466966 | 1512.8961 | LSTVVNADQILVIK | Q9DC29 | Q9DC29 | ABCB6 | ATP-binding cassette, sub-family B (MDR/TAP), member 6 (Langereis blood group) | Cytoplasm | transporter |
| 15542091 | 3 | 0.026 | 3702328 | 3154973.5 | 3476106.5 | 3077742.75 | 3730957.5 | 2783491 | 4004374.5 | 3068810.5 | 2928204.75 | 4867352.5 | 4088024.5 | 5507182 | 1672.8828 | IDGQDISQVTQISLR | Q9DC29 | Q9DC29 | ABCB6 | ATP-binding cassette, sub-family B (MDR/TAP), member 6 (Langereis blood group) | Cytoplasm | transporter |
| 15549132 | 2 | 0.371 | 3459273.75 | 4081198.75 | 7818826.5 | 1602856.75 | 4574552.5 | 3682101.25 | 3048911.5 | 3893749.5 | 3980988.5 | 3572953.75 | 2119797.75 | 2889002.25 | 1697.8564 | FLQGGGTGSTGFVSNLR | Q9DC29 | Q9DC29 | ABCB6 | ATP-binding cassette, sub-family B (MDR/TAP), member 6 (Langereis blood group) | Cytoplasm | transporter |
| 15545328 | 2 | 0.252 | 1866624.875 | 1953659.625 | 371294.25 | 550334.688 | 1509370.875 | 213271.109 | 2429824.5 | 932137.125 | 363525.531 | 2299364.5 | 2729942.25 | 2456853.5 | 1351.745 | LFSHLHELSLR | Q9DC29 | Q9DC29 | ABCB6 | ATP-binding cassette, sub-family B (MDR/TAP), member 6 (Langereis blood group) | Cytoplasm | transporter |
| 15542237 | 3 | 0.074 | 2642590 | 3617678.75 | 2740117.25 | 257122.781 | 1038832.125 | 2279535.5 | 3185954.75 | 3251284.5 | 3514383.75 | 1792665.25 | 4620910.5 | 3089992.75 | 2102.9922 | MIQTNFIDMENMFDLLK | Q9DC29 | Q9DC29 | ABCB6 | ATP-binding cassette, sub-family B (MDR/TAP), member 6 (Langereis blood group) | Cytoplasm | transporter |
| 15551998 | 2 | 0.27 | 1392166 | 3213153.5 | 1211359.125 | 1824504.375 | 1791831.75 | 2696493 | 1285864.125 | 2088254.625 | 3198073.25 | 780524.688 | 1136027.25 | 1530991.375 | 1620.8999 | QALIDMNTLFTLLK | Q61102 | Q61102 | ABCB7 | ATP-binding cassette, sub-family B (MDR/TAP), member 7 | Cytoplasm | transporter |
| 15542172 | 2 | 0.815 | 2008977 | 1.02E+07 | 2799373.75 | 8725452 | 7470840 | 9011007 | 3971241.25 | 3865634.75 | 2.17E+07 | 1205371.5 | 9973444 | 4.74E+07 | 845.4713 | SLLEDIR | Q9JJ59 | Q9JJ59 | ABCB9 | ATP-binding cassette, sub-family B (MDR/TAP), member 9 | Cytoplasm | transporter |
| 15549308 | 3 | 0.416 | 1.57E+07 | 1.81E+07 | 2.04E+07 | 1.53E+07 | 1.36E+07 | 1.50E+07 | 1.48E+07 | 1.66E+07 | 1.50E+07 | 2.09E+07 | 1.30E+07 | 1.41E+07 | 2596.2522 | AVYSNSDIYLFDDPLSAVDAHVGK | O35379 | O35379 | ABCC1 | ATP-binding cassette, sub-family C (CFTR/MRP), member 1 | Plasma Membrane | transporter |
| 15549050 | 3 | 0.404 | 1.48E+07 | 1.69E+07 | 1.67E+07 | 1.44E+07 | 1.41E+07 | 1.48E+07 | 1.26E+07 | 1.58E+07 | 1.48E+07 | 1.62E+07 | 1.25E+07 | 1.42E+07 | 2827.4148 | AVMEAC[160.0307]ALLPDLEILPSGDRTEIGEK | O35379 | O35379 | ABCC1 | ATP-binding cassette, sub-family C (CFTR/MRP), member 1 | Plasma Membrane | transporter |
| 15551324 | 3 | 0.69 | 1.02E+07 | 1.16E+07 | 1.29E+07 | 1.01E+07 | 1.14E+07 | 1.12E+07 | 1.14E+07 | 1.26E+07 | 9956485 | 1.39E+07 | 7276206 | 8979338 | 2275.0852 | EAPWQIQETAPPSTWPHSGR | O35379 | O35379 | ABCC1 | ATP-binding cassette, sub-family C (CFTR/MRP), member 1 | Plasma Membrane | transporter |
| 15548909 | 3 | 0.286 | 1.20E+07 | 1.44E+07 | 1.26E+07 | 1.21E+07 | 1.22E+07 | 1.38E+07 | 1.16E+07 | 1.24E+07 | 8819103 | 1.24E+07 | 1.01E+07 | 1.18E+07 | 2547.2131 | ISEMGSYQELLDRDGAFAEFLR | O35379 | O35379 | ABCC1 | ATP-binding cassette, sub-family C (CFTR/MRP), member 1 | Plasma Membrane | transporter |
| 15548826 | 2 | 0.122 | 9628028 | 1.02E+07 | 1.11E+07 | 1.00E+07 | 9390528 | 1.01E+07 | 8441256 | 8980383 | 8961841 | 1.03E+07 | 6812133 | 7704773.5 | 1755.9966 | ITIIPQDPVLFSGSLR | O35379 | O35379 | ABCC1 | ATP-binding cassette, sub-family C (CFTR/MRP), member 1 | Plasma Membrane | transporter |
| 15534524 | 3 | 0.544 | 7152297 | 8057718.5 | 7771078.5 | 7110079.5 | 7067996.5 | 7574361 | 7546892.5 | 7178152 | 5602325 | 8061344 | 6377788 | 7164922.5 | 2018.0658 | SPVYSHFNETLLGVSVIR | O35379 | O35379 | ABCC1 | ATP-binding cassette, sub-family C (CFTR/MRP), member 1 | Plasma Membrane | transporter |
| 15551651 | 3 | 0.759 | 4960573.5 | 6894369.5 | 2396755 | 4972443 | 5429021.5 | 3036780.75 | 6975342.5 | 5205438 | 3544792.75 | 7733731 | 4570019 | 5488231 | 2866.338 | MNLDPFSQYSDEEVWMALELAHLK | O35379 | O35379 | ABCC1 | ATP-binding cassette, sub-family C (CFTR/MRP), member 1 | Plasma Membrane | transporter |
| 15553906 | 2 | 0.411 | 7109862.5 | 6558180 | 6477259.5 | 7044457 | 6453937.5 | 7363698 | 5973929 | 6399028 | 6390397.5 | 7321330 | 6073652.5 | 5261912 | 1857.9193 | EDTSEEVVPVLVNNWK | O35379 | O35379 | ABCC1 | ATP-binding cassette, sub-family C (CFTR/MRP), member 1 | Plasma Membrane | transporter |
| 15532844 | 2 | 0.791 | 6844025.5 | 5519387.5 | 4740703 | 5575399 | 5651611.5 | 5486385.5 | 5837191.5 | 5161817.5 | 4614613 | 5682589 | 5386241.5 | 4924024 | 1387.66 | EC[160.0307]GAPSELLQQR | O35379 | O35379 | ABCC1 | ATP-binding cassette, sub-family C (CFTR/MRP), member 1 | Plasma Membrane | transporter |
| 15538992 | 2 | 0.586 | 4482250.5 | 5428920.5 | 5067834.5 | 4647667 | 4982912 | 4820566 | 4716741 | 4705216 | 3855684 | 5274039 | 3735119 | 4698339.5 | 1383.692 | LYAWELAFQDK | O35379 | O35379 | ABCC1 | ATP-binding cassette, sub-family C (CFTR/MRP), member 1 | Plasma Membrane | transporter |
| 15552514 | 2 | 0.873 | 3861807 | 4298573 | 3907110.25 | 3434105.25 | 4516966.5 | 3701582.5 | 4366592 | 4634296 | 3392243.25 | 4661681.5 | 3081238.25 | 3589325.25 | 1805.901 | SSTVGEIVNLMSVDAQR | O35379 | O35379 | ABCC1 | ATP-binding cassette, sub-family C (CFTR/MRP), member 1 | Plasma Membrane | transporter |
| 15534864 | 2 | 0.809 | 4624044.5 | 4386111 | 4650264.5 | 3295895.25 | 4505546.5 | 4897783 | 3525126.25 | 5996264 | 5167687.5 | 4554708 | 4421247 | 4233541.5 | 1377.7274 | SSLLSALLAEMDK | O35379 | O35379 | ABCC1 | ATP-binding cassette, sub-family C (CFTR/MRP), member 1 | Plasma Membrane | transporter |
| 15554692 | 3 | 0.158 | 3302337.25 | 3742042 | 4662286.5 | 2519416.25 | 2989538.75 | 4368776 | 3975111.25 | 4470059 | 4651216.5 | 3838998.25 | 1945024.875 | 2648368.25 | 2613.3916 | ILVLDEATAAVDLETDNLIQSTIR | O35379 | O35379 | ABCC1 | ATP-binding cassette, sub-family C (CFTR/MRP), member 1 | Plasma Membrane | transporter |
| 15559645 | 3 | 0.685 | 2746128 | 3067809 | 2538383.75 | 3145939.25 | 2502879.75 | 2389394.25 | 3066203 | 3415052.5 | 2591467 | 3294396.5 | 2356518.5 | 2300696 | 2867.4905 | GEPPTLNGITFSIPEGALVAVVGQVGC[160.0307]GK | O35379 | O35379 | ABCC1 | ATP-binding cassette, sub-family C (CFTR/MRP), member 1 | Plasma Membrane | transporter |
| 15550049 | 2 | 0.355 | 2395267.75 | 3283179.5 | 3695683.5 | 2809037.5 | 3667410 | 3153335.75 | 2451720.25 | 3997100.75 | 2134533 | 3289552.5 | 1220762.5 | 2023858.5 | 2032.0284 | GSVAYVPQQAWIQNDSLR | O35379 | O35379 | ABCC1 | ATP-binding cassette, sub-family C (CFTR/MRP), member 1 | Plasma Membrane | transporter |
| 15555834 | 2 | 0.227 | 3574275.5 | 2712634.5 | 3233396.5 | 3166235.25 | 2999772.25 | 3335512.25 | 2459440.5 | 3615507.75 | 2617426.25 | 2826008.5 | 2480730.75 | 2290722 | 2143.9229 | TYANAEQDLASEDDSVSGSGK | O35379 | O35379 | ABCC1 | ATP-binding cassette, sub-family C (CFTR/MRP), member 1 | Plasma Membrane | transporter |
| 15537904 | 2 | 0.309 | 2544655.5 | 2632320.75 | 2827228 | 2241797.25 | 2268059.5 | 2476069.5 | 2510386.75 | 2063901.875 | 2604212.75 | 1951959.375 | 1924058.375 | 2735724.25 | 993.5706 | SSLTLGLFR | O35379 | O35379 | ABCC1 | ATP-binding cassette, sub-family C (CFTR/MRP), member 1 | Plasma Membrane | transporter |
| 15542216 | 2 | 0.307 | 1489198.625 | 1615236.875 | 1795276.25 | 1411711.375 | 2044942.875 | 2173169.25 | 1740471.375 | 2335973.75 | 2263636.5 | 1873533.875 | 1848649.625 | 1821641.625 | 859.48016 | QLVC[160.0307]LAR | O35379 | O35379 | ABCC1 | ATP-binding cassette, sub-family C (CFTR/MRP), member 1 | Plasma Membrane | transporter |
| 15534162 | 2 | 0.759 | 5126124.5 | 7426640.5 | 1.02E+07 | 2968568.25 | 7557686 | 8341698 | 4761171 | 8132295 | 7780860 | 3329372.75 | 6542358.5 | 6563059.5 | 842.4359 | QM[147.0354]HAALR | Q9R1S7 | Q9R1S7 | ABCC6 | ATP-binding cassette, sub-family C (CFTR/MRP), member 6 | Plasma Membrane | transporter |
| 15550584 | 2 | 0.16 | 5396297 | 6704207 | 3846566.5 | 4314818.5 | 5352311.5 | 2307816.75 | 3966254.5 | 4507437 | 4060850.75 | 3888062 | 2192450.5 | 2478496.75 | 1949.9907 | EGINIFLDGVYPTENLR | P61222 | P61222 | ABCE1 | ATP-binding cassette, sub-family E (OABP), member 1 | Cytoplasm | transporter |
| 15536975 | 2 | 0.176 | 2536084.5 | 3060709 | 2529491.75 | 2670108.5 | 3431863.75 | 1678209.25 | 2473629.5 | 2357840.75 | 2039743.25 | 2107911.25 | 1437464.875 | 1814865.625 | 1087.5759 | PQYVDQIPK | P61222 | P61222 | ABCE1 | ATP-binding cassette, sub-family E (OABP), member 1 | Cytoplasm | transporter |
| 15550149 | 2 | 0.025 | 4862459.5 | 4171471.5 | 3075554.75 | 3716667.5 | 4168675.5 | 2104121 | 2412858.25 | 2946005 | 2434740.5 | 2086094.75 | 1806652.625 | 1941820.75 | 1658.8502 | NTVANSPQTLLAGMNK | P61222 | P61222 | ABCE1 | ATP-binding cassette, sub-family E (OABP), member 1 | Cytoplasm | transporter |
| 15553623 | 2 | 0.092 | 2158236 | 2877913.75 | 1801100.125 | 1854091.875 | 1925215.125 | 949026.188 | 1212791 | 1405815 | 1659792.25 | 1704641.25 | 658151.688 | 660480.625 | 2480.2957 | PLQIENIIDQEVQTLSGGELQR | P61222 | P61222 | ABCE1 | ATP-binding cassette, sub-family E (OABP), member 1 | Cytoplasm | transporter |
| 15537335 | 3, 4 | 0.003 | 1.37E+08 | 1.36E+08 | 1.28E+08 | 7.97E+07 | 9.91E+07 | 7.29E+07 | 1.26E+08 | 1.29E+08 | 1.12E+08 | 1.12E+08 | 1.03E+08 | 8.91E+07 | 1858.9393 | TLAEGDVLSFHHITYR | Q7TMS5 | Q7TMS5 | ABCG2 | ATP-binding cassette, sub-family G (WHITE), member 2 (Junior blood group) | Plasma Membrane | transporter |
| 15532315 | 2 | 0.003 | 1.08E+08 | 1.02E+08 | 1.13E+08 | 6.53E+07 | 7.98E+07 | 6.13E+07 | 9.93E+07 | 9.40E+07 | 8.62E+07 | 8.81E+07 | 8.38E+07 | 7.00E+07 | 1298.657 | AELDQLPGAQEK | Q7TMS5 | Q7TMS5 | ABCG2 | ATP-binding cassette, sub-family G (WHITE), member 2 (Junior blood group) | Plasma Membrane | transporter |
| 15531387 | 2 | 0.013 | 8.92E+07 | 9.86E+07 | 1.04E+08 | 5.94E+07 | 7.37E+07 | 4.53E+07 | 9.48E+07 | 9.80E+07 | 8.48E+07 | 8.56E+07 | 6.34E+07 | 6.43E+07 | 1251.7023 | PGLNAILGPTGGGK | Q7TMS5 | Q7TMS5 | ABCG2 | ATP-binding cassette, sub-family G (WHITE), member 2 (Junior blood group) | Plasma Membrane | transporter |
| 15536177 | 2 | 0.004 | 7.74E+07 | 8.01E+07 | 6.73E+07 | 5.41E+07 | 5.96E+07 | 5.40E+07 | 7.34E+07 | 7.91E+07 | 7.15E+07 | 6.69E+07 | 5.60E+07 | 5.42E+07 | 1044.603 | SSLLDVLAAR | Q7TMS5 | Q7TMS5 | ABCG2 | ATP-binding cassette, sub-family G (WHITE), member 2 (Junior blood group) | Plasma Membrane | transporter |
| 15539354 | 2 | 0.008 | 5.23E+07 | 6.11E+07 | 6.67E+07 | 4.04E+07 | 4.56E+07 | 4.12E+07 | 5.67E+07 | 5.68E+07 | 5.65E+07 | 5.58E+07 | 4.38E+07 | 4.72E+07 | 1264.7119 | LFDSLTLLASGK | Q7TMS5 | Q7TMS5 | ABCG2 | ATP-binding cassette, sub-family G (WHITE), member 2 (Junior blood group) | Plasma Membrane | transporter |
| 15533320 | 2 | 0.011 | 4.91E+07 | 3.94E+07 | 4.64E+07 | 2.98E+07 | 2.72E+07 | 2.69E+07 | 3.55E+07 | 3.94E+07 | 3.93E+07 | 4.37E+07 | 3.22E+07 | 2.93E+07 | 820.4658 | VGTQFIR | Q7TMS5 | Q7TMS5 | ABCG2 | ATP-binding cassette, sub-family G (WHITE), member 2 (Junior blood group) | Plasma Membrane | transporter |
| 15532322 | 3 | 6.49E-04 | 5.02E+07 | 5.47E+07 | 5.28E+07 | 2.85E+07 | 3.51E+07 | 3.07E+07 | 4.36E+07 | 4.79E+07 | 3.85E+07 | 3.69E+07 | 4.18E+07 | 3.60E+07 | 1820.9618 | GLSGDVLINGAPQPAHFK | Q7TMS5 | Q7TMS5 | ABCG2 | ATP-binding cassette, sub-family G (WHITE), member 2 (Junior blood group) | Plasma Membrane | transporter |
| 15532624 | 3 | 0.068 | 2.89E+07 | 2.98E+07 | 3.52E+07 | 2.17E+07 | 2.67E+07 | 2.00E+07 | 2.95E+07 | 3.08E+07 | 2.88E+07 | 3.30E+07 | 2.47E+07 | 2.35E+07 | 1211.6865 | TIIFSIHQPR | Q7TMS5 | Q7TMS5 | ABCG2 | ATP-binding cassette, sub-family G (WHITE), member 2 (Junior blood group) | Plasma Membrane | transporter |
| 15551869 | 4 | 0.06 | 3.83E+07 | 5.63E+07 | 3.93E+07 | 2.65E+07 | 3.32E+07 | 2.39E+07 | 4.27E+07 | 4.81E+07 | 3.08E+07 | 3.05E+07 | 3.16E+07 | 3.66E+07 | 5225.307 | ALEYFASAGYHC[160.0307]EPYNNPADFFLDVINGDSSAVMLNREEQDNEANK | Q7TMS5 | Q7TMS5 | ABCG2 | ATP-binding cassette, sub-family G (WHITE), member 2 (Junior blood group) | Plasma Membrane | transporter |
| 15535369 | 2 | 9.31E-04 | 3.23E+07 | 3.14E+07 | 3.75E+07 | 1.99E+07 | 2.23E+07 | 2.03E+07 | 2.48E+07 | 2.86E+07 | 2.74E+07 | 2.70E+07 | 2.17E+07 | 2.32E+07 | 701.45435 | INTIIK | Q7TMS5 | Q7TMS5 | ABCG2 | ATP-binding cassette, sub-family G (WHITE), member 2 (Junior blood group) | Plasma Membrane | transporter |
| 15538881 | 2 | 0.013 | 2.71E+07 | 2.42E+07 | 2.75E+07 | 1.34E+07 | 1.98E+07 | 1.29E+07 | 2.46E+07 | 2.35E+07 | 2.12E+07 | 2.35E+07 | 1.68E+07 | 1.75E+07 | 1148.6034 | ENLQFSAALR | Q7TMS5 | Q7TMS5 | ABCG2 | ATP-binding cassette, sub-family G (WHITE), member 2 (Junior blood group) | Plasma Membrane | transporter |
| 15532693 | 2 | 0.018 | 1.96E+07 | 1.77E+07 | 2.13E+07 | 9569948 | 1.44E+07 | 1.18E+07 | 1.77E+07 | 1.78E+07 | 1.45E+07 | 1.63E+07 | 1.51E+07 | 1.19E+07 | 996.56146 | LVFHGPAQK | Q7TMS5 | Q7TMS5 | ABCG2 | ATP-binding cassette, sub-family G (WHITE), member 2 (Junior blood group) | Plasma Membrane | transporter |
| 15532526 | 3 | 0.007 | 1.48E+07 | 1.61E+07 | 1.67E+07 | 1.17E+07 | 1.08E+07 | 9239083 | 1.27E+07 | 1.49E+07 | 1.27E+07 | 1.49E+07 | 1.39E+07 | 1.19E+07 | 1635.7927 | EPVYVTSFC[160.0307]HQLR | Q7TMS5 | Q7TMS5 | ABCG2 | ATP-binding cassette, sub-family G (WHITE), member 2 (Junior blood group) | Plasma Membrane | transporter |
| 15540539 | 2 | 0.001 | 2.14E+07 | 2.10E+07 | 2.47E+07 | 1.36E+07 | 1.58E+07 | 1.58E+07 | 2.19E+07 | 2.04E+07 | 1.96E+07 | 1.49E+07 | 1.59E+07 | 1.18E+07 | 1232.6511 | EILSDINGIMK | Q7TMS5 | Q7TMS5 | ABCG2 | ATP-binding cassette, sub-family G (WHITE), member 2 (Junior blood group) | Plasma Membrane | transporter |
| 15534926 | 2 | 0.042 | 1.02E+07 | 1.19E+07 | 1.19E+07 | 5411348.5 | 9329692 | 7469204.5 | 1.30E+07 | 1.31E+07 | 9509156 | 1.37E+07 | 1.09E+07 | 9807314 | 1061.5454 | VMSDLLPMR | Q7TMS5 | Q7TMS5 | ABCG2 | ATP-binding cassette, sub-family G (WHITE), member 2 (Junior blood group) | Plasma Membrane | transporter |
| 15553996 | 2 | 0.01 | 1.21E+07 | 1.07E+07 | 1.20E+07 | 6022609 | 8341622.5 | 6983919.5 | 8927720 | 1.06E+07 | 9750185 | 1.00E+07 | 1.02E+07 | 7591999.5 | 1858.9417 | TLAEGDVLSFHHITYR | Q7TMS5 | Q7TMS5 | ABCG2 | ATP-binding cassette, sub-family G (WHITE), member 2 (Junior blood group) | Plasma Membrane | transporter |
| 15552806 | 2 | 0.073 | 5616932.5 | 5040974 | 5341339.5 | 3195923.25 | 4193627.75 | 1784041.125 | 3828686.5 | 4798760 | 4750544 | 4003344.75 | 4350097.5 | 3154034.25 | 1635.7915 | EPVYVTSFC[160.0307]HQLR | Q7TMS5 | Q7TMS5 | ABCG2 | ATP-binding cassette, sub-family G (WHITE), member 2 (Junior blood group) | Plasma Membrane | transporter |
| 15553994 | 4 | 0.045 | 9112585 | 8942083 | 1.34E+07 | 2581528 | 6011800.5 | 9196885 | 6076298.5 | 1.02E+07 | 9874654 | 3719093 | 3765471.75 | 4217887.5 | 5241.3037 | ALEYFASAGYHC[160.0307]EPYNNPADFFLDVINGDSSAVM[147.0354]LNREEQDNEANK | Q7TMS5 | Q7TMS5 | ABCG2 | ATP-binding cassette, sub-family G (WHITE), member 2 (Junior blood group) | Plasma Membrane | transporter |
| 15550005 | 2 | 0.032 | 7114677 | 6309670.5 | 7497577.5 | 4231501.5 | 4409261.5 | 4648851.5 | 6078470 | 6761047 | 6204644 | 2353872.5 | 5236518.5 | 4607198.5 | 1820.9591 | GLSGDVLINGAPQPAHFK | Q7TMS5 | Q7TMS5 | ABCG2 | ATP-binding cassette, sub-family G (WHITE), member 2 (Junior blood group) | Plasma Membrane | transporter |
| 15561607 | 2 | 0.168 | 1378525.375 | 1411005 | 808709.375 | 1021413.562 | 1119728.75 | 593855 | 1314429.125 | 1085062.25 | 814962 | 946253.438 | 543225.438 | 478101.594 | 2287.143 | PVIENLSEFYINSAIYGETK | Q7TMS5 | Q7TMS5 | ABCG2 | ATP-binding cassette, sub-family G (WHITE), member 2 (Junior blood group) | Plasma Membrane | transporter |
| 15554576 | 3 | 0.415 | 3278546 | 4031609.5 | 5327089.5 | 5239455.5 | 4237711 | 4334915 | 4284823 | 5624232 | 5688654.5 | 4735394.5 | 4193960 | 3964909 | 2826.316 | AASSVPETPTAVPAASSSSWDTYYQPR | Q9Z1Q2 | Q9Z1Q2 | ABHD16A | abhydrolase domain containing 16A | Cytoplasm | enzyme |
| 15537416 | 2 | 0.747 | 3164801.25 | 4393438.5 | 4875318 | 3024969.5 | 5315220.5 | 4630233.5 | 4219069 | 5597855 | 4651507.5 | 4210057 | 4296399 | 3873923 | 1028.6222 | FQGPVLLVR | Q9Z1Q2 | Q9Z1Q2 | ABHD16A | abhydrolase domain containing 16A | Cytoplasm | enzyme |
| 15543757 | 2 | 0.502 | 940071.875 | 970536.25 | 865880.812 | 710930.438 | 936081.125 | 1080353 | 788688.312 | 919572.375 | 1158311.75 | 811453.938 | 823464.25 | 736385.75 | 772.4544 | GNDLLLK | Q9Z1Q2 | Q9Z1Q2 | ABHD16A | abhydrolase domain containing 16A | Cytoplasm | enzyme |
| 15550980 | 3 | 0.243 | 451583.969 | 1635119.25 | 1786344.375 | 3403160.25 | 3570933.5 | 4651105.5 | 2893885.5 | 2988005 | 5172271 | 180827.562 | 3635310 | 2313762.75 | 2521.3386 | LKPEQLGDISVGNVLEPGAGAVMAR | Q921H8 | Q921H8 | ACAA1 | acetyl-CoA acyltransferase 1 | Cytoplasm | enzyme |
| 15550788 | 3 | 0.181 | 6759508.5 | 1.63E+07 | 7558998.5 | 8353512.5 | 8423863 | 1.33E+07 | 6196118.5 | 1.06E+07 | 1.43E+07 | 3940763.75 | 5071712 | 6953793 | 2719.3533 | VPPETIDSVIVGNVMQSSSDAAYLAR | Q8BWT1 | Q8BWT1 | ACAA2 | acetyl-CoA acyltransferase 2 | Cytoplasm | enzyme |
| 15538138 | 2 | 0.187 | 4978159.5 | 9757014 | 4545632.5 | 6468088.5 | 6025313 | 8508107 | 4197395.5 | 7099633.5 | 8645298 | 3215672.5 | 4549188.5 | 4123992.75 | 1427.7822 | VGVPTETGALTLNR | Q8BWT1 | Q8BWT1 | ACAA2 | acetyl-CoA acyltransferase 2 | Cytoplasm | enzyme |
| 15536927 | 3 | 0.358 | 3772808.75 | 7515826 | 3732188 | 4876221 | 3916401 | 7226459.5 | 3057260.25 | 5146860.5 | 5945297.5 | 3210928.75 | 2735370.25 | 3848553.25 | 1807.9342 | TNVSGGAIALGHPLGGSGSR | Q8BWT1 | Q8BWT1 | ACAA2 | acetyl-CoA acyltransferase 2 | Cytoplasm | enzyme |
| 15549663 | 3 | 0.19 | 3926109.25 | 7301543.5 | 3524835.5 | 5300833.5 | 4615069 | 5117999 | 2653808 | 5661116.5 | 6009948 | 2519394.5 | 3510918.75 | 2535776 | 2381.1829 | QTMQVDEHARPQTTLEQLQK | Q8BWT1 | Q8BWT1 | ACAA2 | acetyl-CoA acyltransferase 2 | Cytoplasm | enzyme |
| 15556739 | 2 | 0.517 | 2788119.25 | 6866519 | 2214532.75 | 3630944 | 3411631.5 | 4481592 | 1477558.125 | 3469064.75 | 5590259 | 1576919 | 2784727.25 | 2279460.75 | 1982.9131 | AANEAGYFNEEMAPIEVK | Q8BWT1 | Q8BWT1 | ACAA2 | acetyl-CoA acyltransferase 2 | Cytoplasm | enzyme |
| 15555224 | 2 | 0.115 | 3361545 | 7543653 | 2871782 | 3239688.5 | 4774247 | 5557515 | 1867031.875 | 4257934.5 | 5056492.5 | 1414916.625 | 1542763.375 | 2672238.25 | 2280.1182 | DMDLIDVNEAFAPQFLSVQK | Q8BWT1 | Q8BWT1 | ACAA2 | acetyl-CoA acyltransferase 2 | Cytoplasm | enzyme |
| 15553440 | 3 | 0.224 | 1602437.125 | 3013673.5 | 812292.25 | 2784238.25 | 2146972 | 2834082 | 1789085.5 | 2163761 | 3428151 | 803627.812 | 1765506.5 | 1464645.5 | 2776.3147 | LSSGEHIAAFC[160.0307]LTEPASGSDAASIQTR | Q8JZN5 | Q8JZN5 | ACAD9 | acyl-CoA dehydrogenase family, member 9 | Cytoplasm | enzyme |
| 15547892 | 3 | 0.344 | 2.50E+07 | 5.45E+07 | 2.14E+07 | 3.69E+07 | 2.81E+07 | 4.53E+07 | 1.97E+07 | 3.04E+07 | 4.29E+07 | 1.52E+07 | 2.36E+07 | 2.41E+07 | 2790.3892 | AQDTAELFFEDVRLPANALLGEENK | P51174 | P51174 | ACADL | acyl-CoA dehydrogenase, long chain | Cytoplasm | enzyme |
| 15533034 | 3 | 0.218 | 9025784 | 1.65E+07 | 7754814 | 1.21E+07 | 1.19E+07 | 1.37E+07 | 8102927.5 | 1.26E+07 | 1.41E+07 | 4478708 | 1.03E+07 | 7365655 | 1955.9244 | FFQEEVIPHHTEWEK | P51174 | P51174 | ACADL | acyl-CoA dehydrogenase, long chain | Cytoplasm | enzyme |
| 15534249 | 2 | 0.234 | 7533382.5 | 1.48E+07 | 5722670 | 1.02E+07 | 8795517 | 1.07E+07 | 6332003 | 1.09E+07 | 1.10E+07 | 4477133 | 7297601.5 | 5678019.5 | 1449.7374 | VQPIYGGTNEIMK | P51174 | P51174 | ACADL | acyl-CoA dehydrogenase, long chain | Cytoplasm | enzyme |
| 15551610 | 2 | 0.287 | 5010864.5 | 8190993.5 | 3808237 | 5382427 | 5062054 | 6047133.5 | 3765376.5 | 5706222.5 | 8118376.5 | 3691303.25 | 3878784.5 | 3558191.5 | 1673.7928 | GFYYLMQELPQER | P51174 | P51174 | ACADL | acyl-CoA dehydrogenase, long chain | Cytoplasm | enzyme |
| 15556278 | 2 | 0.653 | 3878344 | 8063578.5 | 1691779.625 | 8174849 | 3815104 | 4740339 | 3698395.25 | 4028103.75 | 3583493.75 | 2794245.75 | 3991923.5 | 3489407.75 | 2002.964 | C[160.0307]IGAIAMTEPGAGSDLQGVR | P51174 | P51174 | ACADL | acyl-CoA dehydrogenase, long chain | Cytoplasm | enzyme |
| 15552450 | 2 | 0.107 | 4679130.5 | 8835343 | 5589536.5 | 5440412 | 6683849 | 7718229.5 | 4175099.75 | 7330080.5 | 1.06E+07 | 1510848.625 | 4471136 | 3682018.5 | 1155.6713 | QGLLGINIAEK | P51174 | P51174 | ACADL | acyl-CoA dehydrogenase, long chain | Cytoplasm | enzyme |
| 15535198 | 2 | 0.099 | 2369883.5 | 5494702.5 | 2414424.25 | 1742383 | 6193355.5 | 4535748 | 2428315.75 | 5392485 | 4444865 | 1465261.75 | 1565990.375 | 1333748.375 | 1268.6813 | LPANALLGEENK | P51174 | P51174 | ACADL | acyl-CoA dehydrogenase, long chain | Cytoplasm | enzyme |
| 15544113 | 3 | 0.231 | 134632.188 | 525366.812 | 546418.938 | 1169462.375 | 1085542.75 | 1455036.375 | 821226.938 | 1070649.75 | 1775297.5 | 93058.625 | 1498694.375 | 720594.688 | 1467.8353 | TRPTVAAGAVGLAQR | P45952 | P45952 | ACADM | acyl-CoA dehydrogenase, C-4 to C-12 straight chain | Cytoplasm | enzyme |
| 15551394 | 3 | 0.477 | 2420891 | 4874939 | 2297463 | 3232987.5 | 3446462.5 | 4409895 | 2065991.75 | 4657697.5 | 5153030.5 | 2194312.75 | 2330774.5 | 2927524.5 | 2737.4062 | GQLTIDQVFPYPSVLSEEQAQFLK | P50544 | P50544 | ACADVL | acyl-CoA dehydrogenase, very long chain | Cytoplasm | enzyme |
| 15538088 | 3 | 0.121 | 2142302.75 | 4785995.5 | 2315763 | 2027679.5 | 4117153.75 | 3277306.5 | 1908065.5 | 3583086.5 | 3351230 | 1123271.25 | 1668165.75 | 1773534.5 | 2119.0457 | EATQAVLDKPETLSSDASTR | P50544 | P50544 | ACADVL | acyl-CoA dehydrogenase, very long chain | Cytoplasm | enzyme |
| 15541373 | 2 | 0.72 | 1107695.75 | 2784873.25 | 1068220.875 | 1547262.25 | 1428928.375 | 1775688.5 | 974240.375 | 1603273.75 | 2559655.75 | 835604.5 | 1211175.625 | 1482632.25 | 934.53357 | ITAFVVER | P50544 | P50544 | ACADVL | acyl-CoA dehydrogenase, very long chain | Cytoplasm | enzyme |
| 15537491 | 2 | 0.64 | 935734.562 | 2709101.25 | 549805.812 | 1125155.875 | 1500051.25 | 1850945.125 | 1050075.875 | 2251417.75 | 1643020.625 | 691454.938 | 1336254.125 | 1047583.812 | 1195.5593 | FFEEVNDPAK | P50544 | P50544 | ACADVL | acyl-CoA dehydrogenase, very long chain | Cytoplasm | enzyme |
| 15541573 | 2 | 0.224 | 555725.625 | 1171175.625 | 511302.781 | 772376.312 | 765357.5 | 873456.5 | 467639.812 | 772689.938 | 1026903.625 | 408683.312 | 567698.562 | 407535.969 | 1446.6836 | SLSEGYPTAQHEK | P50544 | P50544 | ACADVL | acyl-CoA dehydrogenase, very long chain | Cytoplasm | enzyme |
| 15548914 | 2, 3 | 0.309 | 8127389 | 1.83E+07 | 9320676 | 1.53E+07 | 1.76E+07 | 2.02E+07 | 9861399 | 2.03E+07 | 2.11E+07 | 4641630 | 1.39E+07 | 1.28E+07 | 2443.2866, 2443.288 | IAAFADAAVDPIDFPLAPAYAVPK | Q8QZT1 | Q8QZT1 | ACAT1 | acetyl-CoA acetyltransferase 1 | Cytoplasm | enzyme |
| 15533044 | 2 | 0.278 | 6260538 | 1.05E+07 | 8949111 | 1.02E+07 | 1.17E+07 | 1.45E+07 | 6764851 | 1.09E+07 | 1.50E+07 | 3108211 | 9686164 | 7991976 | 1171.6659 | LGTAAIQGAIEK | Q8QZT1 | Q8QZT1 | ACAT1 | acetyl-CoA acetyltransferase 1 | Cytoplasm | enzyme |
| 15549833 | 2 | 0.278 | 4026975.75 | 8858672 | 4882132 | 7180518.5 | 7993669 | 8978147 | 4687725.5 | 8124648.5 | 1.09E+07 | 2428965.25 | 6875972.5 | 4717857.5 | 1561.7103 | QEQDTYALSSYTR | Q8QZT1 | Q8QZT1 | ACAT1 | acetyl-CoA acetyltransferase 1 | Cytoplasm | enzyme |
| 15533069 | 2 | 0.228 | 5309213.5 | 1.46E+07 | 5690967 | 1.23E+07 | 1.20E+07 | 1.31E+07 | 8215136.5 | 9998823 | 1.37E+07 | 2151835.75 | 7946749 | 8273781.5 | 1405.7904 | FASEITPITISVK | Q8QZT1 | Q8QZT1 | ACAT1 | acetyl-CoA acetyltransferase 1 | Cytoplasm | enzyme |
| 15554212 | 2 | 0.12 | 3018626 | 7691887 | 4477385.5 | 6599672 | 7100870 | 9515958 | 4736350.5 | 6600695 | 8634755 | 1299048.125 | 4805533 | 3813740.5 | 1905.91 | EVYMGNVIQGGEGQAPTR | Q8QZT1 | Q8QZT1 | ACAT1 | acetyl-CoA acetyltransferase 1 | Cytoplasm | enzyme |
| 15557754 | 2 | 0.223 | 2191266.75 | 5362400.5 | 3120472.25 | 4644215.5 | 5333631 | 7915275.5 | 3403127.75 | 4650966 | 7610692 | 1073544.125 | 3686724 | 3895485.25 | 1674.9017 | TPIGSFLGSLASQPATK | Q8QZT1 | Q8QZT1 | ACAT1 | acetyl-CoA acetyltransferase 1 | Cytoplasm | enzyme |
| 15553604 | 2 | 0.454 | 2063514.875 | 4500338.5 | 1782278.875 | 3293615.25 | 5842858.5 | 6112809 | 2872545.75 | 4300661.5 | 7384713.5 | 1050215.125 | 5843628 | 3491918.25 | 1929.0063 | QATLGAGLPISTPC[160.0307]TTVNK | Q8QZT1 | Q8QZT1 | ACAT1 | acetyl-CoA acetyltransferase 1 | Cytoplasm | enzyme |
| 15531512 | 2 | 0.673 | 3.19E+07 | 3.13E+07 | 3.43E+07 | 2.45E+07 | 3.06E+07 | 2.88E+07 | 3.73E+07 | 3.74E+07 | 2.34E+07 | 3.89E+07 | 2.09E+07 | 2.47E+07 | 1119.5438 | QASQMDALAGK | Q9QUI6 | Q9QUI6 | ACKR1 | atypical chemokine receptor 1 (Duffy blood group) | Plasma Membrane | G-protein coupled receptor |
| 15532640 | 2 | 0.009 | 3.32E+07 | 3.77E+07 | 4.14E+07 | 2.77E+07 | 3.28E+07 | 3.23E+07 | 3.43E+07 | 3.64E+07 | 3.03E+07 | 2.85E+07 | 2.13E+07 | 1.97E+07 | 1060.5979 | SLSSLSLPTR | Q9QUI6 | Q9QUI6 | ACKR1 | atypical chemokine receptor 1 (Duffy blood group) | Plasma Membrane | G-protein coupled receptor |
| 15531663 | 2 | 0.209 | 1.01E+07 | 8555344 | 1.25E+07 | 6240862 | 8494639 | 8486843 | 9735634 | 1.11E+07 | 8596145 | 1.04E+07 | 6683118.5 | 6690175 | 813.4932 | LNIGQLR | Q9QUI6 | Q9QUI6 | ACKR1 | atypical chemokine receptor 1 (Duffy blood group) | Plasma Membrane | G-protein coupled receptor |
| 15542643 | 2 | 0.47 | 8112890.5 | 1222138.875 | 72376.461 | 5320406.5 | 480069.844 | 319805.531 | 1154238.875 | 925885.125 | 3087784 | 3621377.25 | 8534182 | 4350955 | 1206.5754 | QASQMDALAGKS | Q9QUI6 | Q9QUI6 | ACKR1 | atypical chemokine receptor 1 (Duffy blood group) | Plasma Membrane | G-protein coupled receptor |
| 15532657 | 3 | 0.108 | 1.49E+07 | 3.15E+07 | 1.30E+07 | 2.21E+07 | 2.67E+07 | 3.00E+07 | 1.67E+07 | 2.58E+07 | 2.48E+07 | 1.03E+07 | 1.60E+07 | 1.40E+07 | 1868.9094 | IVYGHLDDPANQEIER | Q99KI0 | Q99KI0 | ACO2 | aconitase 2, mitochondrial | Cytoplasm | enzyme |
| 15556417 | 2 | 0.052 | 8989799 | 1.80E+07 | 8036088 | 1.36E+07 | 1.35E+07 | 1.91E+07 | 7736579.5 | 1.38E+07 | 1.60E+07 | 5251240.5 | 7170701 | 6804730.5 | 1778.8922 | QGLLPLTFADPSDYNK | Q99KI0 | Q99KI0 | ACO2 | aconitase 2, mitochondrial | Cytoplasm | enzyme |
| 15537615 | 2 | 0.226 | 6994929.5 | 1.47E+07 | 6485314 | 1.07E+07 | 9399966 | 1.51E+07 | 6844038.5 | 1.08E+07 | 1.25E+07 | 4010804.25 | 7720188.5 | 7193413 | 1463.7467 | SQFTITPGSEQIR | Q99KI0 | Q99KI0 | ACO2 | aconitase 2, mitochondrial | Cytoplasm | enzyme |
| 15551472 | 3 | 0.114 | 4529160.5 | 1.79E+07 | 1.02E+07 | 5342364 | 1.29E+07 | 1.76E+07 | 7990550.5 | 1.43E+07 | 8283662.5 | 3908626.5 | 3301727.25 | 4505804.5 | 2709.3127 | DHLVPDPGC[160.0307]QYDQVIEINLNELK | Q99KI0 | Q99KI0 | ACO2 | aconitase 2, mitochondrial | Cytoplasm | enzyme |
| 15535779 | 3 | 0.112 | 7980269.5 | 1.38E+07 | 5714092.5 | 1.26E+07 | 8700822 | 9234056 | 7174218.5 | 9343600 | 1.35E+07 | 3804413.25 | 6268843.5 | 5791349.5 | 1565.7389 | VAMSHFEPSEYIR | Q99KI0 | Q99KI0 | ACO2 | aconitase 2, mitochondrial | Cytoplasm | enzyme |
| 15549920 | 3 | 0.281 | 7098956.5 | 1.15E+07 | 5469559 | 7700695 | 1.10E+07 | 1.12E+07 | 5222633.5 | 9600828 | 1.28E+07 | 3766848.5 | 8345527 | 4675131.5 | 2260.1345 | VAVPSTIHC[160.0307]DHLIEAQVGGEK | Q99KI0 | Q99KI0 | ACO2 | aconitase 2, mitochondrial | Cytoplasm | enzyme |
| 15549795 | 2 | 0.319 | 4492683 | 9554408 | 3643365.5 | 7570821 | 7957017 | 1.08E+07 | 4006125.5 | 7933632 | 1.04E+07 | 3621111.25 | 5806333.5 | 5355704 | 1845.8064 | VGLIGSC[160.0307]TNSSYEDMGR | Q99KI0 | Q99KI0 | ACO2 | aconitase 2, mitochondrial | Cytoplasm | enzyme |
| 15534919 | 2 | 0.099 | 4126492.75 | 1.22E+07 | 5325448 | 5627404.5 | 8332205 | 1.23E+07 | 5363646.5 | 1.11E+07 | 1.10E+07 | 3359214.5 | 3404942.25 | 4164021.25 | 1102.6129 | LQLLEPFDK | Q99KI0 | Q99KI0 | ACO2 | aconitase 2, mitochondrial | Cytoplasm | enzyme |
| 15532927 | 2 | 0.097 | 5513518 | 1.42E+07 | 5508364.5 | 8905280 | 8800583 | 1.22E+07 | 6234146 | 9677584 | 1.01E+07 | 3115078.75 | 6021115 | 4389224.5 | 1268.6139 | FNPETDFLTGK | Q99KI0 | Q99KI0 | ACO2 | aconitase 2, mitochondrial | Cytoplasm | enzyme |
| 15551267 | 3 | 0.889 | 6635038.5 | 9485228 | 3202646.75 | 6628208 | 5130331 | 7753342 | 4306839 | 6049857.5 | 1.06E+07 | 2956934.75 | 9047802 | 4634811.5 | 2286.1082 | DVGGIVLANAC[160.0307]GPC[160.0307]IGQWDRK | Q99KI0 | Q99KI0 | ACO2 | aconitase 2, mitochondrial | Cytoplasm | enzyme |
| 15535001 | 2 | 0.231 | 5818976 | 1.19E+07 | 4787151 | 5922236.5 | 7042861 | 1.09E+07 | 4386912.5 | 7470377 | 1.05E+07 | 2378627.25 | 4495395.5 | 5220057 | 1199.6863 | DLEDLQILIK | Q99KI0 | Q99KI0 | ACO2 | aconitase 2, mitochondrial | Cytoplasm | enzyme |
| 15559716 | 2 | 0.324 | 3090829.25 | 4782580.5 | 2920765.75 | 3038568.75 | 4419363.5 | 4964733.5 | 2233532.75 | 5021766.5 | 6068960 | 2064706.625 | 3311779.25 | 2223136 | 1667.7646 | WVVIGDENYGEGSSR | Q99KI0 | Q99KI0 | ACO2 | aconitase 2, mitochondrial | Cytoplasm | enzyme |
| 15537322 | 2 | 0.296 | 3764494.25 | 8432482 | 3562402 | 4112533.5 | 5305063.5 | 9532339 | 3971745.25 | 5845429.5 | 7735856.5 | 1949686 | 4698447.5 | 3206908.25 | 1233.6438 | LTGSLSGWTSPK | Q99KI0 | Q99KI0 | ACO2 | aconitase 2, mitochondrial | Cytoplasm | enzyme |
| 15537369 | 3 | 0.144 | 3602171.5 | 5239920 | 2482769.5 | 5671462.5 | 4327980.5 | 5954977.5 | 3498108.25 | 4816708 | 4859374.5 | 1835935.125 | 4163280 | 2813515.5 | 1556.7479 | C[160.0307]TTDHISAAGPWLK | Q99KI0 | Q99KI0 | ACO2 | aconitase 2, mitochondrial | Cytoplasm | enzyme |
| 15535626 | 2 | 0.26 | 2989468.5 | 5795459.5 | 2715832.25 | 3852182.25 | 3903481.5 | 5765372 | 2364834.25 | 4414387.5 | 5174033 | 1506207.375 | 3087834 | 2783830 | 772.491 | LTIQGLK | Q99KI0 | Q99KI0 | ACO2 | aconitase 2, mitochondrial | Cytoplasm | enzyme |
| 15545075 | 3 | 0.43 | 1536379.25 | 3705934 | 868933.5 | 2626181.5 | 2601220.5 | 3089010.5 | 1547509.25 | 2651457.5 | 2239256.5 | 1343663.875 | 2005365.125 | 1664663.625 | 1753.8728 | DINQEVYNFLATAGAK | Q99KI0 | Q99KI0 | ACO2 | aconitase 2, mitochondrial | Cytoplasm | enzyme |
| 15550986 | 2 | 0.2 | 1773258.75 | 3672473.5 | 1424841.625 | 1933348.75 | 2388569.75 | 2822915.5 | 1564816.625 | 2597304.5 | 2984468.5 | 925687.875 | 1679130.25 | 1378438.375 | 1868.9104 | IVYGHLDDPANQEIER | Q99KI0 | Q99KI0 | ACO2 | aconitase 2, mitochondrial | Cytoplasm | enzyme |
| 15541706 | 4 | 0.14 | 1800177.75 | 6941248 | 3783044 | 1730987.625 | 6137217.5 | 6248390.5 | 2905327.75 | 6054566.5 | 3729898.25 | 864702.688 | 1288070.875 | 2267207.25 | 2482.2688 | PHINGPFTPDLAHPVADVGTVAEK | Q99KI0 | Q99KI0 | ACO2 | aconitase 2, mitochondrial | Cytoplasm | enzyme |
| 15538554 | 2 | 0.1 | 1215279.125 | 4014162.75 | 2232768 | 1113869.875 | 4441517 | 4464518.5 | 1682276.5 | 4242059 | 2598629 | 781536.438 | 713391.312 | 1114344.5 | 1225.6033 | LEAPDADELPR | Q99KI0 | Q99KI0 | ACO2 | aconitase 2, mitochondrial | Cytoplasm | enzyme |
| 15554033 | 3 | 0.102 | 1458429 | 2498486.25 | 1931276.75 | 1661490.875 | 1824922.625 | 3742749.25 | 1585944.25 | 2219149.25 | 3713439 | 733141.125 | 1412602 | 1141242.375 | 2338.1575 | VAMQDATAQMAMLQFISSGLPK | Q99KI0 | Q99KI0 | ACO2 | aconitase 2, mitochondrial | Cytoplasm | enzyme |
| 15556755 | 3 | 0.096 | 688005.375 | 4848783.5 | 2528138.75 | 1146711.125 | 3850089 | 2239378.5 | 2107452.5 | 2905477.75 | 1667184.625 | 314656.75 | 533008.688 | 1029271.875 | 2482.2659 | PHINGPFTPDLAHPVADVGTVAEK | Q99KI0 | Q99KI0 | ACO2 | aconitase 2, mitochondrial | Cytoplasm | enzyme |
| 15533775 | 2 | 0.149 | 1.65E+07 | 1.96E+07 | 1.59E+07 | 2.12E+07 | 2.15E+07 | 1.85E+07 | 1.86E+07 | 2.03E+07 | 1.65E+07 | 1.87E+07 | 1.57E+07 | 1.57E+07 | 1218.6339 | LAQGEYIAPEK | P41216 | P41216 | ACSL1 | acyl-CoA synthetase long-chain family member 1 | Cytoplasm | enzyme |
| 15542112 | 2 | 0.152 | 1.16E+07 | 1.53E+07 | 1.55E+07 | 1.34E+07 | 1.90E+07 | 1.75E+07 | 1.79E+07 | 1.73E+07 | 1.61E+07 | 1.46E+07 | 1.25E+07 | 1.35E+07 | 1052.55 | IGFFQGDIR | P41216 | P41216 | ACSL1 | acyl-CoA synthetase long-chain family member 1 | Cytoplasm | enzyme |
| 15531654 | 3 | 0.271 | 1.44E+07 | 1.97E+07 | 1.34E+07 | 1.49E+07 | 1.89E+07 | 1.63E+07 | 1.84E+07 | 1.73E+07 | 1.84E+07 | 1.35E+07 | 1.45E+07 | 1.57E+07 | 2235.2349 | GIAVHPELFSIDNGLLTPTLK | P41216 | P41216 | ACSL1 | acyl-CoA synthetase long-chain family member 1 | Cytoplasm | enzyme |
| 15533550 | 3 | 0.4 | 9751175 | 1.35E+07 | 7928309 | 1.17E+07 | 1.35E+07 | 1.13E+07 | 1.31E+07 | 1.27E+07 | 1.24E+07 | 1.31E+07 | 1.17E+07 | 9686220 | 2018.9724 | GAMITHQNIINDC[160.0307]SGFIK | P41216 | P41216 | ACSL1 | acyl-CoA synthetase long-chain family member 1 | Cytoplasm | enzyme |
| 15549017 | 3 | 0.202 | 8082120.5 | 2.18E+07 | 2.48E+07 | 9455292 | 2.70E+07 | 2.08E+07 | 1.87E+07 | 2.25E+07 | 1.53E+07 | 1.18E+07 | 5756730.5 | 8776750 | 2469.2283 | PKPPEPEDLAIIC[160.0307]FTSGTTGNPK | P41216 | P41216 | ACSL1 | acyl-CoA synthetase long-chain family member 1 | Cytoplasm | enzyme |
| 15531974 | 2 | 0.284 | 1.33E+07 | 1.82E+07 | 8975064 | 1.49E+07 | 1.54E+07 | 1.78E+07 | 1.39E+07 | 1.53E+07 | 1.68E+07 | 1.09E+07 | 1.23E+07 | 1.27E+07 | 1365.8257 | VLQPTIFPVVPR | P41216 | P41216 | ACSL1 | acyl-CoA synthetase long-chain family member 1 | Cytoplasm | enzyme |
| 15552025 | 2 | 0.234 | 7886282 | 1.08E+07 | 9503651 | 9681499 | 1.15E+07 | 1.06E+07 | 9135525 | 1.10E+07 | 1.32E+07 | 1.09E+07 | 6955306 | 7016425.5 | 1510.7433 | LVDVEEMNYLASK | P41216 | P41216 | ACSL1 | acyl-CoA synthetase long-chain family member 1 | Cytoplasm | enzyme |
| 15532583 | 2 | 0.126 | 6518475.5 | 7798856.5 | 7585465 | 8184385 | 8652286 | 9024755 | 8547671 | 8931860 | 9913636 | 8576849 | 5059523.5 | 7183400 | 1280.6704 | SQIDELYATIK | P41216 | P41216 | ACSL1 | acyl-CoA synthetase long-chain family member 1 | Cytoplasm | enzyme |
| 15534244 | 2 | 0.346 | 5321556.5 | 5540848.5 | 5593167 | 5942039 | 6993791.5 | 4868333 | 6135434.5 | 6771773.5 | 6687060.5 | 6571424.5 | 5507931 | 5225931 | 1018.55786 | C[160.0307]GVEIISLK | P41216 | P41216 | ACSL1 | acyl-CoA synthetase long-chain family member 1 | Cytoplasm | enzyme |
| 15554729 | 2 | 0.504 | 6275690 | 7138894 | 3818883.75 | 5221691.5 | 4976304.5 | 2993884.25 | 5459532 | 5947277.5 | 5265624 | 6516312 | 6637441.5 | 4339017 | 1552.774 | KPNQPYEWISYK | P41216 | P41216 | ACSL1 | acyl-CoA synthetase long-chain family member 1 | Cytoplasm | enzyme |
| 15548822 | 2 | 0.284 | 5976722.5 | 7561833.5 | 7490760 | 6870411.5 | 1.01E+07 | 6923664 | 6897682.5 | 8767256 | 9665460 | 6330786 | 7447746.5 | 5306162.5 | 1716.8806 | EVAELAEC[160.0307]IGSGLIQK | P41216 | P41216 | ACSL1 | acyl-CoA synthetase long-chain family member 1 | Cytoplasm | enzyme |
| 15540090 | 2 | 0.319 | 5137861 | 6305430.5 | 5953156.5 | 5314878 | 6867937.5 | 4768271 | 5695799.5 | 6895201.5 | 6810443 | 5735175 | 5229208 | 4780474.5 | 986.5321 | MPELIDIR | P41216 | P41216 | ACSL1 | acyl-CoA synthetase long-chain family member 1 | Cytoplasm | enzyme |
| 15543372 | 2 | 0.685 | 3148930 | 4218548 | 2846861.25 | 3477832.5 | 3836652 | 3699547.75 | 3447805 | 4096292.5 | 3531936.5 | 3815801 | 3335259 | 2868196.5 | 847.45795 | LLMDDLK | P41216 | P41216 | ACSL1 | acyl-CoA synthetase long-chain family member 1 | Cytoplasm | enzyme |
| 15534094 | 2 | 0.854 | 3727631 | 3361280.25 | 3118009.5 | 4377544.5 | 3766194 | 2753496 | 3052027.25 | 3809931 | 4870096 | 3618039.75 | 3775586.25 | 3447739 | 979.5245 | WLLDFASK | P41216 | P41216 | ACSL1 | acyl-CoA synthetase long-chain family member 1 | Cytoplasm | enzyme |
| 15533149 | 3 | 0.058 | 3135098.25 | 3033277.75 | 3805241.25 | 3403043 | 4499561 | 3750281.75 | 3651635.75 | 4815878.5 | 4319239 | 3596686.5 | 2547358.75 | 2573327.5 | 1198.5825 | DGWLHTGDIGK | P41216 | P41216 | ACSL1 | acyl-CoA synthetase long-chain family member 1 | Cytoplasm | enzyme |
| 15539071 | 2 | 0.136 | 2967759 | 3913673.25 | 3226984.25 | 3587876 | 4460751.5 | 4055691.5 | 3381137 | 4534499.5 | 4629027 | 3437578.25 | 3331642.5 | 3295464.25 | 1458.7068 | GIQVSNNGPC[160.0307]LGSR | P41216 | P41216 | ACSL1 | acyl-CoA synthetase long-chain family member 1 | Cytoplasm | enzyme |
| 15537761 | 3 | 0.249 | 3785055.75 | 5165479 | 4092781.5 | 4307706 | 6238236.5 | 4475142.5 | 4750165.5 | 5790971 | 6591943.5 | 3394197.75 | 4336839 | 5050888.5 | 1716.8812 | EVAELAEC[160.0307]IGSGLIQK | P41216 | P41216 | ACSL1 | acyl-CoA synthetase long-chain family member 1 | Cytoplasm | enzyme |
| 15550148 | 3 | 0.448 | 1.44E+07 | 1.31E+07 | 1.20E+07 | 1.22E+07 | 1.43E+07 | 1.31E+07 | 1.26E+07 | 1.21E+07 | 1.30E+07 | 1.89E+07 | 1.48E+07 | 1.19E+07 | 2536.131 | WFC[160.0307]TGDIGEFHPDGC[160.0307]LQIIDR | Q9QUJ7 | Q9QUJ7 | ACSL4 | acyl-CoA synthetase long-chain family member 4 | Cytoplasm | enzyme |
| 15532539 | 3 | 0.543 | 1.43E+07 | 1.39E+07 | 1.13E+07 | 1.44E+07 | 1.45E+07 | 1.44E+07 | 1.27E+07 | 1.55E+07 | 1.36E+07 | 1.51E+07 | 1.33E+07 | 1.47E+07 | 2214.1257 | SVTHFDSLAVIDIPGADTLDK | Q9QUJ7 | Q9QUJ7 | ACSL4 | acyl-CoA synthetase long-chain family member 4 | Cytoplasm | enzyme |
| 15549532 | 3 | 0.21 | 1.08E+07 | 1.21E+07 | 1.39E+07 | 8199007 | 1.13E+07 | 1.24E+07 | 1.11E+07 | 1.35E+07 | 1.21E+07 | 1.06E+07 | 8710410 | 9955279 | 2717.3838 | EAVVHGLNESEASYLITSVELLESK | Q9QUJ7 | Q9QUJ7 | ACSL4 | acyl-CoA synthetase long-chain family member 4 | Cytoplasm | enzyme |
| 15550827 | 2 | 0.912 | 5275939.5 | 3985809.25 | 6701319.5 | 4193813.75 | 4992214.5 | 5518231.5 | 5505640 | 4844066 | 5626921 | 7631305 | 5039822 | 4316190 | 1582.7921 | IGYSSPLTLSDQSSK | Q9QUJ7 | Q9QUJ7 | ACSL4 | acyl-CoA synthetase long-chain family member 4 | Cytoplasm | enzyme |
| 15553873 | 2 | 0.296 | 2462831.75 | 1928496 | 2863131.25 | 1951520.625 | 1997247 | 2042157 | 2457823 | 2353661.25 | 2489198 | 3415101.5 | 4401632.5 | 1850504.25 | 1523.792 | GYDAPLC[160.0307]NLILFK | Q9QUJ7 | Q9QUJ7 | ACSL4 | acyl-CoA synthetase long-chain family member 4 | Cytoplasm | enzyme |
| 15552968 | 3 | 0.257 | 1543978.625 | 1910621.125 | 1352354.25 | 2653015 | 2104527 | 1999470.625 | 1707130.75 | 2240006.25 | 2216840.25 | 3212008 | 1981424.375 | 1698552 | 2332.0903 | GVEGSWVDIC[160.0307]NNPAMEAEILK | Q9QUJ7 | Q9QUJ7 | ACSL4 | acyl-CoA synthetase long-chain family member 4 | Cytoplasm | enzyme |
| 15551129 | 2 | 0.342 | 2931637.25 | 3183504.5 | 2848794.75 | 2650751.5 | 2364088.25 | 2822411.5 | 2705980 | 2834920.75 | 3955518.25 | 2989535.75 | 2724314.5 | 2373719.5 | 1712.8646 | GEIVIGGQNISMGYFK | Q9QUJ7 | Q9QUJ7 | ACSL4 | acyl-CoA synthetase long-chain family member 4 | Cytoplasm | enzyme |
| 15540602 | 2 | 0.866 | 2713761.25 | 2525870.25 | 2453291.5 | 2992508.25 | 2410333 | 1948890 | 2874234 | 2216901.5 | 2263299.25 | 2979645.75 | 3460170 | 1962310.5 | 1138.5532 | VQEMNYVQK | Q9QUJ7 | Q9QUJ7 | ACSL4 | acyl-CoA synthetase long-chain family member 4 | Cytoplasm | enzyme |
| 15552522 | 2 | 0.921 | 1999701.75 | 1669444.875 | 2139533.5 | 2316331.75 | 1861156.375 | 1916341.25 | 2418201.5 | 1643336.625 | 1632346 | 2304811.5 | 1882837 | 1415451 | 1556.6252 | TAEDYC[160.0307]VDENGQR | Q9QUJ7 | Q9QUJ7 | ACSL4 | acyl-CoA synthetase long-chain family member 4 | Cytoplasm | enzyme |
| 15554149 | 3 | 0.004 | 7886127.5 | 1.10E+07 | 8931511 | 9413270 | 1.08E+07 | 1.20E+07 | 1.29E+07 | 1.33E+07 | 1.32E+07 | 1.44E+07 | 1.38E+07 | 1.45E+07 | 3066.521 | FLEPIFQPTSDDVTISYLPLAHMFER | Q8JZR0 | Q8JZR0 | ACSL5 | acyl-CoA synthetase long-chain family member 5 | Cytoplasm | enzyme |
| 15552957 | 3 | 0.308 | 4227993.5 | 7471057.5 | 8581474 | 9620572 | 8704911 | 8729794 | 9699091 | 7374624 | 8153309 | 7879329 | 8948880 | 7510522 | 2353.2764 | SIFVHPEPFTIENGLLTPTLK | Q8JZR0 | Q8JZR0 | ACSL5 | acyl-CoA synthetase long-chain family member 5 | Cytoplasm | enzyme |
| 15550846 | 2 | 0.523 | 5161773.5 | 6063510.5 | 7014631.5 | 6594268 | 7894585 | 6002260 | 6226064.5 | 7916854 | 7051911.5 | 7113353.5 | 6043303.5 | 6427417.5 | 2059.106 | SFLIGVVVPDPDSLPSFAAK | Q8JZR0 | Q8JZR0 | ACSL5 | acyl-CoA synthetase long-chain family member 5 | Cytoplasm | enzyme |
| 15539135 | 2 | 0.096 | 4610396 | 5311745 | 4097450 | 5882927.5 | 6774613.5 | 4382935 | 6064719.5 | 5091953.5 | 4969020.5 | 6911968.5 | 6869392.5 | 6051373 | 1455.7488 | ADIPVVIC[160.0307]DTPQK | Q8JZR0 | Q8JZR0 | ACSL5 | acyl-CoA synthetase long-chain family member 5 | Cytoplasm | enzyme |
| 15537862 | 3 | 0.144 | 3429365 | 5065368.5 | 4228451.5 | 4759923.5 | 5555086.5 | 5298388 | 5739404 | 5572367 | 4692637.5 | 5966748 | 4713316.5 | 5605816.5 | 1772.8263 | C[160.0307]GVEMLSLHDAENIGK | Q8JZR0 | Q8JZR0 | ACSL5 | acyl-CoA synthetase long-chain family member 5 | Cytoplasm | enzyme |
| 15550125 | 2 | 0.205 | 3344693 | 3383849.25 | 4391552 | 3766090 | 3938395 | 4046987.75 | 4315034.5 | 4758280 | 4958370 | 4971323.5 | 3429108.25 | 4476678 | 1935.0731 | LMITGAAPISTPVLTFFR | Q8JZR0 | Q8JZR0 | ACSL5 | acyl-CoA synthetase long-chain family member 5 | Cytoplasm | enzyme |
| 15555858 | 2 | 0.136 | 3469606.25 | 4098774 | 5295944 | 4904918 | 5280137.5 | 4481269 | 5149123 | 5269190 | 5916086 | 4317570 | 4337125 | 4686498 | 1575.765 | NNDLILYYFSDAK | Q8JZR0 | Q8JZR0 | ACSL5 | acyl-CoA synthetase long-chain family member 5 | Cytoplasm | enzyme |
| 15536047 | 2 | 0.203 | 2684284.75 | 1765435.375 | 2223160.25 | 2848003 | 2526100.5 | 2359055.75 | 3896600.75 | 2637216.25 | 3159321 | 3553604.25 | 5068241 | 2238061.25 | 1278.6842 | LVQGILFSC[160.0307]GGK | Q8JZR0 | Q8JZR0 | ACSL5 | acyl-CoA synthetase long-chain family member 5 | Cytoplasm | enzyme |
| 15537280 | 2 | 0.696 | 2945362 | 2562031.25 | 3673241 | 2919851 | 3429340.5 | 3108521.25 | 3547422.5 | 3158977.25 | 3437151.75 | 3217172.25 | 2936129.75 | 3474739.75 | 1070.5239 | TLYENFQR | Q8JZR0 | Q8JZR0 | ACSL5 | acyl-CoA synthetase long-chain family member 5 | Cytoplasm | enzyme |
| 15535388 | 2 | 0.344 | 2787404.25 | 2266508 | 2062975.5 | 2169691.5 | 3249140.5 | 2611426.25 | 2617016.25 | 2636054.5 | 2741781.5 | 2921652.5 | 2991352.75 | 2764439.5 | 1478.7036 | GLAVSDNGPC[160.0307]LGYR | Q8JZR0 | Q8JZR0 | ACSL5 | acyl-CoA synthetase long-chain family member 5 | Cytoplasm | enzyme |
| 15534674 | 2 | 0.487 | 2068223.5 | 2574115.25 | 2638026 | 3382529.75 | 2580326.25 | 2275947.5 | 3223479 | 3084609.5 | 2549193.5 | 2915704.75 | 2663721 | 2265779.75 | 1133.5826 | ATMLVENVEK | Q8JZR0 | Q8JZR0 | ACSL5 | acyl-CoA synthetase long-chain family member 5 | Cytoplasm | enzyme |
| 15538151 | 3 | 0.173 | 2947933.25 | 5583638 | 1642782 | 4427494 | 2358082.75 | 3031669.5 | 1729559.375 | 3518974 | 5673662.5 | 1587054.375 | 1732576 | 992115.312 | 2198.1096 | SPETIALIWERDEPGTEVR | Q99NB1 | Q99NB1 | ACSS1 | acyl-CoA synthetase short-chain family member 1 | Cytoplasm | enzyme |
| 15552672 | 2 | 0.13 | 2113767 | 4446065.5 | 1444991 | 2136824.5 | 1902518.5 | 2890814.75 | 1348604.875 | 2305774.75 | 3448860 | 1228105.625 | 842429.438 | 1266518.125 | 1757.8773 | VPMGSLDIPLEQMAK | Q99NB1 | Q99NB1 | ACSS1 | acyl-CoA synthetase short-chain family member 1 | Cytoplasm | enzyme |
| 15537496 | 2 | 0.298 | 1771693.625 | 3559760.75 | 1022739.188 | 1137554.625 | 1343116.875 | 2311676.5 | 1043189.625 | 2179105 | 1900535.5 | 932819.312 | 870621.312 | 1073860.75 | 1336.6975 | INQFYGAPTAVR | Q99NB1 | Q99NB1 | ACSS1 | acyl-CoA synthetase short-chain family member 1 | Cytoplasm | enzyme |
| 15536921 | 3 | 0.04 | 4.49E+09 | 4.45E+09 | 4.71E+09 | 4.46E+09 | 4.57E+09 | 4.50E+09 | 4.93E+09 | 4.68E+09 | 4.33E+09 | 5.29E+09 | 5.18E+09 | 4.79E+09 | 1954.0632 | VAPEEHPVLLTEAPLNPK | P60710 | P60710 | ACTB | actin, beta | Cytoplasm | other |
| 15550510, 15550526, 15552360 | 3, 4 | 0.339 | 2.86E+09 | 2.98E+09 | 3.01E+09 | 3.05E+09 | 2.65E+09 | 2.95E+09 | 2.37E+09 | 2.78E+09 | 2.82E+09 | 3.06E+09 | 2.66E+09 | 3.11E+09 | 3183.607, 3183.6082, 3183.6147 | TTGIVMDSGDGVTHTVPIYEGYALPHAILR | P60710 | P60710 | ACTB | actin, beta | Cytoplasm | other |
| 15550774 | 2 | 0.813 | 2.56E+09 | 2.60E+09 | 3.01E+09 | 2.71E+09 | 2.47E+09 | 2.56E+09 | 2.82E+09 | 2.71E+09 | 2.53E+09 | 3.00E+09 | 2.41E+09 | 2.32E+09 | 1790.8878 | SYELPDGQVITIGNER | P60710 | P60710 | ACTB | actin, beta | Cytoplasm | other |
| 15537167, 15541768 | 2 | 0.352 | 1.25E+09 | 1.25E+09 | 1.43E+09 | 1.39E+09 | 1.29E+09 | 1.25E+09 | 1.37E+09 | 1.36E+09 | 1.30E+09 | 1.71E+09 | 1.40E+09 | 1.32E+09 | 1161.6124, 1161.6167 | EITALAPSTMK | P60710 | P60710 | ACTB | actin, beta | Cytoplasm | other |
| 15547671, 15547674 | 3, 4 | 0.052 | 1.59E+09 | 1.31E+09 | 1.17E+09 | 1.82E+09 | 1.05E+09 | 1.33E+09 | 1.31E+09 | 1.10E+09 | 1.03E+09 | 1.64E+09 | 1.93E+09 | 2.18E+09 | 3534.624, 3534.6265 | FRC[160.0307]PEALFQPSFLGMESC[160.0307]GIHETTFNSIMK | P60710 | P60710 | ACTB | actin, beta | Cytoplasm | other |
| 15547959, 15550572 | 3, 4 | 0.972 | 8.13E+08 | 1.31E+09 | 1.19E+09 | 8.30E+08 | 1.61E+09 | 1.19E+09 | 1.06E+09 | 1.43E+09 | 1.04E+09 | 1.48E+09 | 9.23E+08 | 1.22E+09 | 3231.4575, 3231.4604 | C[160.0307]PEALFQPSFLGMESC[160.0307]GIHETTFNSIMK | P60710 | P60710 | ACTB | actin, beta | Cytoplasm | other |
| 15534113, 15541593 | 2 | 0.88 | 1.20E+09 | 1.32E+09 | 1.46E+09 | 1.11E+09 | 1.44E+09 | 1.27E+09 | 1.38E+09 | 1.43E+09 | 1.25E+09 | 1.46E+09 | 1.24E+09 | 1.25E+09 | 1132.5244, 1132.526 | GYSFTTTAER | P60710 | P60710 | ACTB | actin, beta | Cytoplasm | other |
| 15535999 | 2 | 0.088 | 1.11E+09 | 1.09E+09 | 1.23E+09 | 9.78E+08 | 1.07E+09 | 1.07E+09 | 1.11E+09 | 1.06E+09 | 9.79E+08 | 1.38E+09 | 1.22E+09 | 1.10E+09 | 800.5338 | RGILTLK | P60710 | P60710 | ACTB | actin, beta | Cytoplasm | other |
| 15531331 | 2 | 0.983 | 1.07E+09 | 9.73E+08 | 1.19E+09 | 1.28E+09 | 9.53E+08 | 1.08E+09 | 1.24E+09 | 1.06E+09 | 9.73E+08 | 1.38E+09 | 1.04E+09 | 9.85E+08 | 976.4474 | AGFAGDDAPR | P60710 | P60710 | ACTB | actin, beta | Cytoplasm | other |
| 15531687 | 3 | 1 | 8.47E+08 | 9.24E+08 | 9.09E+08 | 7.97E+08 | 1.14E+09 | 7.44E+08 | 9.81E+08 | 1.01E+09 | 7.11E+08 | 1.14E+09 | 8.08E+08 | 7.72E+08 | 1516.7018 | QEYDESGPSIVHR | P60710 | P60710 | ACTB | actin, beta | Cytoplasm | other |
| 15533734 | 2, 3 | 0.853 | 1.02E+09 | 9.58E+08 | 1.13E+09 | 1.19E+09 | 9.49E+08 | 1.05E+09 | 1.13E+09 | 9.99E+08 | 1.03E+09 | 1.13E+09 | 1.05E+09 | 1.10E+09 | 1198.7036 | AVFPSIVGRPR | P60710 | P60710 | ACTB | actin, beta | Cytoplasm | other |
| 15553216, 15557291, 15560733 | 2 | 0.942 | 8.85E+08 | 9.16E+08 | 1.03E+09 | 8.75E+08 | 9.29E+08 | 9.95E+08 | 9.65E+08 | 9.64E+08 | 9.45E+08 | 1.03E+09 | 9.38E+08 | 8.98E+08 | 2215.0603, 2215.0608, 2215.0662 | DLYANTVLSGGTTMYPGIADR | P60710 | P60710 | ACTB | actin, beta | Cytoplasm | other |
| 15539033, 15546263 | 2 | 0.817 | 5.79E+08 | 6.23E+08 | 7.68E+08 | 5.51E+08 | 7.10E+08 | 6.75E+08 | 7.79E+08 | 7.21E+08 | 6.26E+08 | 7.85E+08 | 6.10E+08 | 6.45E+08 | 998.48486, 998.4849 | DLTDYLMK | P60710 | P60710 | ACTB | actin, beta | Cytoplasm | other |
| 15549858 | 2 | 0.26 | 6.19E+08 | 5.78E+08 | 6.59E+08 | 5.58E+08 | 5.69E+08 | 5.80E+08 | 6.49E+08 | 5.75E+08 | 5.13E+08 | 7.55E+08 | 6.91E+08 | 5.68E+08 | 1954.0593 | VAPEEHPVLLTEAPLNPK | P60710 | P60710 | ACTB | actin, beta | Cytoplasm | other |
| 15531249 | 3 | 0.934 | 3.79E+08 | 4.02E+08 | 4.64E+08 | 4.85E+08 | 3.12E+08 | 4.41E+08 | 5.37E+08 | 3.76E+08 | 3.76E+08 | 4.11E+08 | 4.48E+08 | 4.65E+08 | 1644.7942 | QEYDESGPSIVHRK | P60710 | P60710 | ACTB | actin, beta | Cytoplasm | other |
| 15547699, 15547940, 15555859, 15557142, 15558607, 15559211, 15562516 | 3, 4 | 0.016 | 4.79E+08 | 4.14E+08 | 5.48E+08 | 3.37E+08 | 3.26E+08 | 3.22E+08 | 3.70E+08 | 4.59E+08 | 3.11E+08 | 3.30E+08 | 2.29E+08 | 2.95E+08 | 3199.5955, 3199.6016, 3199.6025, 3199.6028, 3199.6052, 3199.613 | TTGIVM[147.0354]DSGDGVTHTVPIYEGYALPHAILR | P60710 | P60710 | ACTB | actin, beta | Cytoplasm | other |

| | | | | | | | | | | | | | | | | | | | | | | |
|---|---|---|---|---|---|---|---|---|---|---|---|---|---|---|---|---|---|---|---|---|---|---|
| 15554933, 15556341, 15556984, 15558293, 15558981, 15559872, 15560459, 15560958, 15561179, 15561464, 15561705, 15562320, 15562526, 15562729, 15562803, 15562829, 15562898, 15562939, 15562999, 15563118, 15563392, 15563405, 15563696, 15563718, 15563882, 15563883 | 2, 3 | 0.805 | 2.57E+08 | 2.92E+08 | 3.86E+08 | 2.41E+08 | 3.48E+08 | 3.32E+08 | 2.81E+08 | 3.35E+08 | 3.64E+08 | 2.85E+08 | 2.91E+08 | 2.82E+08 | 2550.1553, 2550.1565, 2550.1606, 2550.1653, 2550.1655, 2550.1658, 2550.1667, 2550.167, 2550.1682, 2550.169, 2550.1692, 2550.1702, 2550.1707, 2550.1711, 2550.1716, 2550.1719, 2550.1724, 2550.1726, 2550.1736, 2550.1743, 2550.1746, 2550.1748, 2550.1755, 2550.1758, 2550.1821, 2550.1829 | LC[160.0307]YVALDFEQEMATAASSSSLEK | P60710 | P60710 | ACTB | actin, beta | Cytoplasm | other |
| 15547768 | 2 | 0.947 | 2.12E+08 | 1.98E+08 | 2.28E+08 | 1.78E+08 | 2.50E+08 | 1.55E+08 | 2.05E+08 | 2.30E+08 | 1.72E+08 | 2.82E+08 | 1.97E+08 | 1.44E+08 | 1516.7007 | QEYDESGPSIVHR | P60710 | P60710 | ACTB | actin, beta | Cytoplasm | other |
| 15536376 | 2 | 0.964 | 1.52E+08 | 1.60E+08 | 1.96E+08 | 1.56E+08 | 1.90E+08 | 1.41E+08 | 1.84E+08 | 1.99E+08 | 1.45E+08 | 2.45E+08 | 1.41E+08 | 1.49E+08 | 795.4708 | IIAPPER | P60710 | P60710 | ACTB | actin, beta | Cytoplasm | other |
| 15539571, 15544549, 15545756 | 3 | 0.41 | 2.90E+08 | 3.18E+08 | 2.67E+08 | 2.05E+08 | 3.51E+08 | 2.12E+08 | 3.54E+08 | 4.18E+08 | 2.92E+08 | 2.16E+08 | 3.36E+08 | 3.90E+08 | 2215.0652, 2215.066, 2215.067 | DLYANTVLSGGTTMYPGIADR | P60710 | P60710 | ACTB | actin, beta | Cytoplasm | other |
| 15551435, 15555333, 15559098, 15561543 | 3, 4 | 0.668 | 2.00E+08 | 1.68E+08 | 2.19E+08 | 2.13E+08 | 1.26E+08 | 2.12E+08 | 1.82E+08 | 1.56E+08 | 1.84E+08 | 1.62E+08 | 1.54E+08 | 1.74E+08 | 3550.6155, 3550.6167, 3550.617, 3550.619, 3550.6191, 3550.6196 | FRC[160.0307]PEALFQPSFLGMESC[160.0307]GIHETTFNSIM[147.0354]K | P60710 | P60710 | ACTB | actin, beta | Cytoplasm | other |
| 15548368, 15549804 | 3, 4 | 0.623 | 1.72E+08 | 1.41E+08 | 1.85E+08 | 1.74E+08 | 9.14E+07 | 2.09E+08 | 1.34E+08 | 1.59E+08 | 1.79E+08 | 1.29E+08 | 1.12E+08 | 1.42E+08 | 3550.6165, 3550.6182 | FRC[160.0307]PEALFQPSFLGM[147.0354]ESC[160.0307]GIHETTFNSIMK | P60710 | P60710 | ACTB | actin, beta | Cytoplasm | other |
| 15531824, 15547348 | 3 | 0.496 | 2.26E+08 | 7.06E+07 | 3.06E+07 | 3.90E+08 | 1.80E+07 | 1.15E+08 | 1.17E+08 | 3.56E+07 | 1.58E+08 | 1.21E+08 | 5.25E+08 | 3.12E+08 | 1623.8386, 1623.8396 | LDLAGRDLTDYLMK | P60710 | P60710 | ACTB | actin, beta | Cytoplasm | other |
| 15541551, 15546126, 15546614, 15546660, 15546846 | 2 | 0.321 | 8.90E+07 | 1.03E+08 | 1.38E+08 | 6.52E+07 | 9.89E+07 | 9.57E+07 | 1.02E+08 | 1.05E+08 | 7.82E+07 | 1.10E+08 | 4.50E+07 | 6.35E+07 | 1014.4766, 1014.47815, 1014.4794, 1014.4799, 1014.4804 | DLTDYLM[147.0354]K | P60710 | P60710 | ACTB | actin, beta | Cytoplasm | other |
| 15552572 | 2 | 0.41 | 8.58E+07 | 7.83E+07 | 9.29E+07 | 8.82E+07 | 5.57E+07 | 8.40E+07 | 1.07E+08 | 6.70E+07 | 7.27E+07 | 1.09E+08 | 9.33E+07 | 9.09E+07 | 1644.7933 | QEYDESGPSIVHRK | P60710 | P60710 | ACTB | actin, beta | Cytoplasm | other |
| 15531536 | 3 | 0.55 | 1.88E+08 | 5.58E+07 | 4.98E+07 | 2.83E+08 | 1.39E+07 | 8.65E+07 | 9.17E+07 | 3.32E+07 | 1.29E+08 | 1.07E+08 | 3.09E+08 | 2.14E+08 | 1629.8196 | GYSFTTTAEREIVR | P60710 | P60710 | ACTB | actin, beta | Cytoplasm | other |
| 15552729, 15559263 | 3, 4 | 0.155 | 1.01E+08 | 1.22E+08 | 1.37E+08 | 8.14E+07 | 1.33E+08 | 1.03E+08 | 9.46E+07 | 1.30E+08 | 7.99E+07 | 1.04E+08 | 4.72E+07 | 6.37E+07 | 3247.4456, 3247.4495, 3247.4502 | C[160.0307]PEALFQPSFLGMESC[160.0307]GIHETTFNSIM[147.0354]K | P60710 | P60710 | ACTB | actin, beta | Cytoplasm | other |
| 15536161, 15540047, 15543335, 15544697, 15544702, 15545142, 15545483, 15546373, 15546687, 15547109 | 2 | 0.147 | 8.03E+07 | 1.02E+08 | 1.14E+08 | 7.94E+07 | 1.01E+08 | 8.53E+07 | 8.65E+07 | 8.98E+07 | 7.50E+07 | 8.16E+07 | 6.61E+07 | 7.51E+07 | 1177.6078, 1177.6079, 1177.6085, 1177.6107, 1177.6113, 1177.6118, 1177.6122, 1177.6133, 1177.6139, 1177.618 | EITALAPSTM[147.0354]K | P60710 | P60710 | ACTB | actin, beta | Cytoplasm | other |
| 15531278, 15543362 | 2 | 0.442 | 8.68E+07 | 7.40E+07 | 6.76E+07 | 7.35E+07 | 6.05E+07 | 6.25E+07 | 8.69E+07 | 6.29E+07 | 5.80E+07 | 7.01E+07 | 8.20E+07 | 8.02E+07 | 1171.5691, 1171.57 | HQGVMVGMGQK | P60710 | P60710 | ACTB | actin, beta | Cytoplasm | other |
| 15553763, 15556956, 15560551, 15561854, 15562484 | 2 | 0.102 | 7.19E+07 | 7.13E+07 | 9.55E+07 | 5.34E+07 | 6.92E+07 | 6.86E+07 | 7.49E+07 | 6.77E+07 | 6.05E+07 | 6.76E+07 | 4.44E+07 | 5.07E+07 | 2231.0574, 2231.0588, 2231.0593, 2231.0615 | DLYANTVLSGGTTM[147.0354]YPGIADR | P60710 | P60710 | ACTB | actin, beta | Cytoplasm | other |
| 15547703, 15562490 | 3 | 0.253 | 7.30E+07 | 7.05E+07 | 9.24E+07 | 4.49E+07 | 9.12E+07 | 6.94E+07 | 7.19E+07 | 9.84E+07 | 4.63E+07 | 6.07E+07 | 3.89E+07 | 4.36E+07 | 3247.4385, 3247.4517 | C[160.0307]PEALFQPSFLGM[147.0354]ESC[160.0307]GIHETTFNSIMK | P60710 | P60710 | ACTB | actin, beta | Cytoplasm | other |
| 15536396 | 3 | 0.427 | 1.23E+08 | 1.75E+08 | 1.75E+08 | 1.38E+08 | 1.45E+08 | 1.75E+08 | 1.57E+08 | 2.17E+08 | 1.51E+08 | 5.41E+07 | 1.32E+08 | 1.77E+08 | 1790.8883 | SYELPDGQVITIGNER | P60710 | P60710 | ACTB | actin, beta | Cytoplasm | other |
| 15540063, 15546612, 15546864, 15546888, 15547287 | 3 | 0.45 | 4.46E+07 | 5.53E+07 | 5.74E+07 | 3.40E+07 | 5.23E+07 | 5.09E+07 | 5.10E+07 | 5.47E+07 | 3.97E+07 | 3.88E+07 | 3.50E+07 | 4.94E+07 | 2231.0608, 2231.0632, 2231.065, 2231.0652, 2231.0735 | DLYANTVLSGGTTM[147.0354]YPGIADR | P60710 | P60710 | ACTB | actin, beta | Cytoplasm | other |
| 15537372 | 3 | 0.719 | 6.99E+07 | 2.07E+07 | 1.96E+07 | 8.32E+07 | 9261604 | 2.70E+07 | 3.62E+07 | 1.48E+07 | 3.20E+07 | 3.55E+07 | 6.51E+07 | 4.75E+07 | 1639.8314 | LDLAGRDLTDYLM[147.0354]K | P60710 | P60710 | ACTB | actin, beta | Cytoplasm | other |
| 15536560 | 5 | 0.145 | 2.15E+07 | 2.64E+07 | 2.00E+07 | 2.71E+07 | 1.61E+07 | 2.36E+07 | 1.92E+07 | 2.17E+07 | 1.79E+07 | 3.51E+07 | 2.32E+07 | 3.08E+07 | 3183.6045 | TTGIVMDSGDGVTHTVPIYEGYALPHAILR | P60710 | P60710 | ACTB | actin, beta | Cytoplasm | other |
| 15540348 | 3 | 0.35 | 3.54E+07 | 3.23E+07 | 2.73E+07 | 2.73E+07 | 2.98E+07 | 1.74E+07 | 2.59E+07 | 2.79E+07 | 2.33E+07 | 3.01E+07 | 2.39E+07 | 2.34E+07 | 1946.8918 | YPIEHGIVTNWDDMEK | P60710 | P60710 | ACTB | actin, beta | Cytoplasm | other |
| 15555268, 15557308 | 2 | 0.463 | 4.34E+07 | 1.30E+07 | 9487024 | 6.58E+07 | 2794779.25 | 1.10E+07 | 1.99E+07 | 5861383.5 | 2.47E+07 | 2.64E+07 | 7.06E+07 | 4.43E+07 | 1629.8188, 1629.8192 | GYSFTTTAEREIVR | P60710 | P60710 | ACTB | actin, beta | Cytoplasm | other |
| 15549497 | 4 | 0.413 | 2.49E+07 | 2.01E+07 | 4.30E+07 | 1.77E+07 | 1.73E+07 | 4.14E+07 | 1.62E+07 | 3.70E+07 | 2.54E+07 | 1.70E+07 | 1.33E+07 | 1.89E+07 | 3566.6123 | FRC[160.0307]PEALFQPSFLGM[147.0354]ESC[160.0307]GIHETTFNSIM[147.0354]K | P60710 | P60710 | ACTB | actin, beta | Cytoplasm | other |
| 15556606 | 2 | 0.604 | 4.42E+07 | 1.29E+07 | 3240734 | 5.73E+07 | 2989513 | 2.00E+07 | 2.19E+07 | 5400546.5 | 2.73E+07 | 1.53E+07 | 7.57E+07 | 5.90E+07 | 1623.8375 | LDLAGRDLTDYLMK | P60710 | P60710 | ACTB | actin, beta | Cytoplasm | other |
| 15542577, 15545690, 15546076, 15547174, 15547194, 15547394, 15547482 | 2 | 0.621 | 1.26E+07 | 5565249.5 | 3271442.75 | 1.22E+07 | 3742182.75 | 2199691.25 | 1.03E+07 | 5287849.5 | 1613013.625 | 1.52E+07 | 9080671 | 6127025 | 1187.5569, 1187.5593, 1187.5599, 1187.5626, 1187.5635, 1187.5659, 1187.5709 | HQGVMVGM[147.0354]GQK | P60710 | P60710 | ACTB | actin, beta | Cytoplasm | other |
| 15535598 | 5 | 0.077 | 9184216 | 9570591 | 8219672 | 1.44E+07 | 8248124.5 | 9482930 | 9563600 | 6729707.5 | 1763952.625 | 1.46E+07 | 1.55E+07 | 1.91E+07 | 3534.6282 | FRC[160.0307]PEALFQPSFLGMESC[160.0307]GIHETTFNSIMK | P60710 | P60710 | ACTB | actin, beta | Cytoplasm | other |
| 15532474, 15544481, 15546358 | 3 | 0.336 | 7566030 | 1.87E+08 | 5.36E+08 | 1.59E+07 | 3.74E+08 | 3.97E+08 | 1.83E+08 | 3.99E+08 | 5.57E+08 | 1.40E+07 | 1.37E+07 | 1.06E+08 | 1515.7446, 1515.7562, 1515.7573 | IWHHTFYNELR | P60710 | P60710 | ACTB | actin, beta | Cytoplasm | other |
| 15559139, 15559988, 15561729 | 3 | 0.358 | 7618167.5 | 1.45E+07 | 2.30E+07 | 3831635 | 1.01E+07 | 1.55E+07 | 6777736.5 | 1.16E+07 | 1.46E+07 | 8338241 | 5152283 | 5853039.5 | 3301.5847, 3301.5854, 3301.587 | M[147.0354]TQIM[147.0354]FETFNTPAM[147.0354]YVAIQAVLSLYASGR | P60710 | P60710 | ACTB | actin, beta | Cytoplasm | other |
| 15553151 | 3 | 0.205 | 9263654 | 7610002.5 | 1.25E+07 | 5105090 | 9403684 | 8724199 | 6686072 | 1.16E+07 | 5055884.5 | 8152939.5 | 2374233 | 3970828.25 | 3263.4434 | C[160.0307]PEALFQPSFLGM[147.0354]ESC[160.0307]GIHETTFNSIM[147.0354]K | P60710 | P60710 | ACTB | actin, beta | Cytoplasm | other |
| 15545021, 15546074, 15546457, 15546463, 15546527, 15546658 | 2, 3 | 0.158 | 4432620.5 | 5470412.5 | 7068134.5 | 3131245.75 | 6066276.5 | 5334903 | 4994997.5 | 5650425.5 | 6303084.5 | 3448422 | 3812777.5 | 3940716.75 | 1187.5582, 1187.5635, 1187.5637, 1187.5646, 1187.5663 | HQGVM[147.0354]VGMGQK | P60710 | P60710 | ACTB | actin, beta | Cytoplasm | other |
| 15552260 | 2 | 0.015 | 4084241.75 | 3811468 | 3268632 | 2586749 | 2461937 | 1720724.5 | 3038101.75 | 2898810.25 | 2492030.5 | 2841460.75 | 2215682.5 | 2438840.5 | 1946.89 | YPIEHGIVTNWDDMEK | P60710 | P60710 | ACTB | actin, beta | Cytoplasm | other |
| 15559462, 15559622 | 2 | 0.48 | 3015160 | 4.31E+07 | 1.04E+08 | 936325.062 | 6.91E+07 | 7.99E+07 | 3.99E+07 | 6.86E+07 | 8.24E+07 | 1757990.625 | 3179331.75 | 3.51E+07 | 1515.744, 1515.7446 | IWHHTFYNELR | P60710 | P60710 | ACTB | actin, beta | Cytoplasm | other |
| 15547418, 15547615, 15547642 | 2, 3 | 0.943 | 97009.805 | 184765.859 | 297432.5 | 141865.328 | 162757.547 | 341222.625 | 153940.797 | 165873.969 | 356620.5 | 176601.453 | 260447.406 | 106389.68 | 1203.558, 1203.5593, 1203.5614 | HQGVM[147.0354]VGM[147.0354]GQK | P60710 | P60710 | ACTB | actin, beta | Cytoplasm | other |
| 15531836 | 3 | 0.748 | 2.89E+09 | 2.73E+09 | 3.19E+09 | 2.93E+09 | 3.11E+09 | 2.81E+09 | 3.16E+09 | 3.06E+09 | 2.55E+09 | 3.37E+09 | 2.95E+09 | 3.03E+09 | 1960.9089 | YPIEHGIITNWDDMEK | P68033 | P68033 | ACTC1 | actin, alpha, cardiac muscle 1 | Cytoplasm | enzyme |
| 15555926 | 2 | 0.534 | 4.86E+08 | 4.58E+08 | 5.48E+08 | 4.09E+08 | 5.10E+08 | 3.80E+08 | 5.19E+08 | 4.88E+08 | 4.46E+08 | 4.02E+08 | 5.42E+08 | 4.63E+08 | 1960.9061 | YPIEHGIITNWDDMEK | P68033 | P68033 | ACTC1 | actin, alpha, cardiac muscle 1 | Cytoplasm | enzyme |
| 15531433, 15543309, 15543594, 15545805, 15545971 | 3 | 0.548 | 2.38E+08 | 1.94E+08 | 2.24E+08 | 1.92E+08 | 2.04E+08 | 1.60E+08 | 2.46E+08 | 2.09E+08 | 1.60E+08 | 2.30E+08 | 1.66E+08 | 1.55E+08 | 1976.9021, 1976.9061, 1976.9067, 1976.9089, 1976.9109 | YPIEHGIITNWDDM[147.0354]EK | P68033 | P68033 | ACTC1 | actin, alpha, cardiac muscle 1 | Cytoplasm | enzyme |
| 15553242 | 2 | 0.417 | 3.45E+07 | 2.62E+07 | 2.70E+07 | 2.35E+07 | 2.44E+07 | 8166488.5 | 2.88E+07 | 2.19E+07 | 1.76E+07 | 3.15E+07 | 2.06E+07 | 1.69E+07 | 1976.9017 | YPIEHGIITNWDDM[147.0354]EK | P68033 | P68033 | ACTC1 | actin, alpha, cardiac muscle 1 | Cytoplasm | enzyme |
| 15534502 | 2 | 0.126 | 5188532 | 4661830.5 | 4162914.5 | 8407380 | 6888169.5 | 5412120.5 | 6048916.5 | 4993587 | 7032741.5 | 3344871.75 | 5529105 | 5327661.5 | 1353.7407 | ILLTEPPMNPTK | P61161 | P61161 | ACTR2 | ARP2 actin-related protein 2 homolog (yeast) | Plasma Membrane | other |
| 15534738 | 2 | 0.507 | 2804784.25 | 2534045 | 2711249 | 2739122 | 3390076 | 2354100.25 | 2362344.75 | 3327558.5 | 3404515 | 1513266.125 | 3098958.25 | 2424161.5 | 1294.643 | VVVC[160.0307]DNGTGFVK | P61161 | P61161 | ACTR2 | ARP2 actin-related protein 2 homolog (yeast) | Plasma Membrane | other |
| 15551067 | 2 | 0.434 | 2458840.5 | 2062756.25 | 2722954.5 | 3303913.75 | 3682038 | 2660748.5 | 2734565 | 3266924.25 | 4108600.75 | 1350946 | 3888503.5 | 2596609 | 1643.7717 | LC[160.0307]YVGYNIEQEQK | P61161 | P61161 | ACTR2 | ARP2 actin-related protein 2 homolog (yeast) | Plasma Membrane | other |
| 15551454 | 3 | 0.452 | 9842705 | 9753599 | 8755115 | 1.56E+07 | 1.42E+07 | 1.10E+07 | 1.08E+07 | 1.02E+07 | 1.80E+07 | 7399365 | 1.66E+07 | 1.12E+07 | 3058.5698 | TLTGTVIDSGDGVTHVIPVAEGYVIGSC[160.0307]IK | Q99JY9 | Q99JY9 | ACTR3 | ARP3 actin-related protein 3 homolog (yeast) | Plasma Membrane | other |
| 15551786 | 2 | 0.138 | 2345687.75 | 2687812.75 | 4115494.25 | 1941864.375 | 4587637.5 | 2344265.75 | 2508569.5 | 3615419.75 | 2887375.25 | 1525828.125 | 2077454.5 | 1536111 | 1540.7101 | LPAC[160.0307]VVDC[160.0307]GTGYTK | Q99JY9 | Q99JY9 | ACTR3 | ARP3 actin-related protein 3 homolog (yeast) | Plasma Membrane | other |
| 15558038 | 2 | 0.077 | 1642877.375 | 2941752.75 | 4878932 | 2476832.25 | 5447100.5 | 4246502.5 | 3854013.75 | 3893645.25 | 4155091.25 | 1418809.375 | 1384665.375 | 2451029.75 | 1782.8391 | GVDDLDFFIGDEAIEK | Q99JY9 | Q99JY9 | ACTR3 | ARP3 actin-related protein 3 homolog (yeast) | Plasma Membrane | other |
| 15534938 | 2 | 0.097 | 2131875 | 1981760.125 | 1873902.625 | 2326682.25 | 2619361 | 1666422.875 | 2375804.25 | 2372478.5 | 2933500 | 1189395.25 | 2142234.25 | 1542695.125 | 1281.6599 | NIVLSGGSTMFR | Q99JY9 | Q99JY9 | ACTR3 | ARP3 actin-related protein 3 homolog (yeast) | Plasma Membrane | other |
| 15542479 | 2 | 0.345 | 2237112.5 | 1956977.625 | 2577281.75 | 2763004 | 3106105 | 2328722.75 | 2379104.25 | 2251113.5 | 3555259.75 | 1109860.625 | 2527601.25 | 2256036.5 | 1080.5654 | QYTGVNAISK | Q99JY9 | Q99JY9 | ACTR3 | ARP3 actin-related protein 3 homolog (yeast) | Plasma Membrane | other |
| 15535261 | 2 | 0.238 | 2887796.75 | 3838322 | 4123394.5 | 3382200.5 | 3839868.5 | 4981982.5 | 4018302.5 | 3800045.5 | 6817445 | 2243707.75 | 3960124.75 | 2959788.5 | 1409.7761 | DITYFIQQLLR | Q641P0 | Q641P0 | ACTR3B | ARP3 actin-related protein 3 homolog B (yeast) | Cytoplasm | other |
| 15537548 | 3 | 0.006 | 1.09E+08 | 1.03E+08 | 7.75E+07 | 6.42E+07 | 3.97E+07 | 4.46E+07 | 9.04E+07 | 7.22E+07 | 8.34E+07 | 4.52E+07 | 6.20E+07 | 5.78E+07 | 1730.8823 | DPTNPFRFPNIGVEK | O35598 | O35598 | ADAM10 | ADAM metallopeptidase domain 10 | Plasma Membrane | peptidase |
| 15553947 | 2 | 0.007 | 4.83E+07 | 3.97E+07 | 4.73E+07 | 2.17E+07 | 3.52E+07 | 2.52E+07 | 5.11E+07 | 5.31E+07 | 5.32E+07 | 4.28E+07 | 3.81E+07 | 2.83E+07 | 1605.765 | TITLQPGSPC[160.0307]NDFR | O35598 | O35598 | ADAM10 | ADAM metallopeptidase domain 10 | Plasma Membrane | peptidase |
| 15533215 | 2 | 3.12E-04 | 6.17E+07 | 5.68E+07 | 6.58E+07 | 3.28E+07 | 3.31E+07 | 2.57E+07 | 5.69E+07 | 5.81E+07 | 5.33E+07 | 4.26E+07 | 3.67E+07 | 2.94E+07 | 1026.5548 | LVDADGPLAR | O35598 | O35598 | ADAM10 | ADAM metallopeptidase domain 10 | Plasma Membrane | peptidase |
| 15553211 | 2 | 2.08E-05 | 6.68E+07 | 7.05E+07 | 7.99E+07 | 4.38E+07 | 4.29E+07 | 4.06E+07 | 6.59E+07 | 6.35E+07 | 6.03E+07 | 4.05E+07 | 3.13E+07 | 3.45E+07 | 1700.844 | AIDTIYQTTDFSGIR | O35598 | O35598 | ADAM10 | ADAM metallopeptidase domain 10 | Plasma Membrane | peptidase |
| 15549878 | 2 | 2.94E-04 | 6.57E+07 | 5.08E+07 | 6.16E+07 | 3.10E+07 | 3.47E+07 | 2.67E+07 | 5.69E+07 | 5.36E+07 | 5.05E+07 | 3.97E+07 | 3.51E+07 | 3.11E+07 | 1711.774 | MAPSTC[160.0307]ASTGSLQWSK | O35598 | O35598 | ADAM10 | ADAM metallopeptidase domain 10 | Plasma Membrane | peptidase |
| [illegible] | [illegible] | [illegible] | [illegible] | [illegible] | [illegible] | [illegible] | [illegible] | [illegible] | [illegible] | [illegible] | [illegible] | [illegible] | [illegible] | [illegible] | [illegible] | [illegible] | [illegible] | [illegible] | [illegible] | [illegible] | [illegible] | [illegible] |

| | | | | | | | | | | | | | | | | | | | | | | | |
|---|---|---|---|---|---|---|---|---|---|---|---|---|---|---|---|---|---|---|---|---|---|---|---|
| 15540146 | | 3 | 0.288 | 3.11E+07 | 2.70E+07 | 1.28E+07 | 3.73E+07 | 2.08E+07 | 2.17E+07 | 2.90E+07 | 1.90E+07 | 6816319.5 | 3.32E+07 | 3.71E+07 | 3.37E+07 | 1266.6305 | ASTKM[147.0354]LEDDLK | O55112 | O55112 | Aff2 | AF4/FMR2 family, member 2 | Nucleus | other |
| 15539549 | | 3 | 0.504 | 1633864 | 1908039 | 2173981 | 1785666.625 | 3426958.25 | 1802567.25 | 2337962.25 | 2528213.25 | 2265261.5 | 1625057 | 1819505.875 | 2264383.5 | 1360.683 | SSAIPDFRDPQK | Q8K2K6 | Q8K2K6 | AGFG1 | ArfGAP with FG repeats 1 | Nucleus | other |
| 15550681 | | 2 | 0.601 | 4035461.25 | 6129987 | 3407416.75 | 3788762.75 | 4612348 | 2664434.75 | 3478510 | 3907921.5 | 3224673.25 | 3801892.25 | 3644136 | 3766827 | 2086.0505 | SISMTTFTEPEVLFLQSR | Q80WC7 | Q80WC7 | AGFG2 | ArfGAP with FG repeats 2 | Cytoplasm | other |
| 15550228 | | 2 | 0.297 | 2014133.25 | 4308313 | 5156975.5 | 1888354.125 | 4009889.5 | 2706228.25 | 2798751.25 | 3298583.25 | 2239008.75 | 2916000 | 1395985.5 | 1582251.625 | 1499.7672 | GSVSATPVQGSVPEGK | Q80WC7 | Q80WC7 | AGFG2 | ArfGAP with FG repeats 2 | Cytoplasm | other |
| 15541661 | | 2 | 0.375 | 2070372.75 | 2809172.75 | 2298819 | 2266999.5 | 1905713 | 1626327.75 | 2161893.5 | 1808166.5 | 2367622.75 | 2129671 | 2405574.25 | 1993957.625 | 1090.6023 | IWLGLFDAR | Q80WC7 | Q80WC7 | AGFG2 | ArfGAP with FG repeats 2 | Cytoplasm | other |
| 15554554 | | 3 | 0.345 | 3516751.25 | 4815412 | 2945294.75 | 2070786.5 | 4297166 | 2126621.5 | 2364910.25 | 4165229.5 | 2806955.75 | 2023472.875 | 2142645.75 | 2907607.5 | 2683.3108 | TLLGDPVPSLSDPASTSSQPGSQSQAR | Q80WC7 | Q80WC7 | AGFG2 | ArfGAP with FG repeats 2 | Cytoplasm | other |
| 15552563 | | 3 | 0.25 | 5.49E+07 | 4.71E+07 | 4.93E+07 | 3.76E+07 | 5.14E+07 | 3.72E+07 | 4.63E+07 | 4.27E+07 | 3.92E+07 | 4.76E+07 | 4.50E+07 | 4.16E+07 | 2276.1748 | DGVSEGQFQQVLHHELLAIR | Q8CJG0 | Q8CJG0 | AGO2 | argonaute RISC catalytic component 2 | Cytoplasm | translation regulator |
| 15549594 | | 2 | 0.321 | 3.92E+07 | 3.56E+07 | 3.53E+07 | 3.30E+07 | 3.25E+07 | 2.55E+07 | 3.28E+07 | 3.17E+07 | 3.36E+07 | 4.41E+07 | 3.71E+07 | 2.86E+07 | 1743.7708 | SFFTASEGC[160.0307]SNPLGGGR | Q8CJG0 | Q8CJG0 | AGO2 | argonaute RISC catalytic component 2 | Cytoplasm | translation regulator |
| 15537697 | | 2 | 0.295 | 4.01E+07 | 3.64E+07 | 3.44E+07 | 3.42E+07 | 3.31E+07 | 2.68E+07 | 3.34E+07 | 3.09E+07 | 2.41E+07 | 4.08E+07 | 3.19E+07 | 2.92E+07 | 1260.6406 | SGNIPAGTTVDTK | Q8CJG0 | Q8CJG0 | AGO2 | argonaute RISC catalytic component 2 | Cytoplasm | translation regulator |
| 15531756 | | 2 | 0.051 | 2.81E+07 | 3.05E+07 | 2.17E+07 | 2.48E+07 | 2.04E+07 | 1.83E+07 | 2.22E+07 | 1.96E+07 | 1.86E+07 | 3.54E+07 | 2.90E+07 | 2.44E+07 | 1443.7395 | AIATPVQGVWDMR | Q8CJG0 | Q8CJG0 | AGO2 | argonaute RISC catalytic component 2 | Cytoplasm | translation regulator |
| 15531996 | | 2 | 0.167 | 2.53E+07 | 2.89E+07 | 2.76E+07 | 2.77E+07 | 2.00E+07 | 2.15E+07 | 2.69E+07 | 2.12E+07 | 1.76E+07 | 3.53E+07 | 2.91E+07 | 2.44E+07 | 1452.7168 | LQANFFEMDIPK | Q8CJG0 | Q8CJG0 | AGO2 | argonaute RISC catalytic component 2 | Cytoplasm | translation regulator |
| 15531856 | | 3 | 0.204 | 3.37E+07 | 3.35E+07 | 3.30E+07 | 2.98E+07 | 3.03E+07 | 2.15E+07 | 2.92E+07 | 2.75E+07 | 2.04E+07 | 3.14E+07 | 3.60E+07 | 2.63E+07 | 1775.9188 | IDIYHYELDIKPEK | Q8CJG0 | Q8CJG0 | AGO2 | argonaute RISC catalytic component 2 | Cytoplasm | translation regulator |
| 15540933 | | 2 | 0.187 | 3.11E+07 | 3.10E+07 | 3.15E+07 | 2.98E+07 | 2.23E+07 | 2.17E+07 | 3.11E+07 | 2.12E+07 | 1.79E+07 | 3.12E+07 | 2.85E+07 | 2.57E+07 | 1299.7396 | VLQPPSILYGGR | Q8CJG0 | Q8CJG0 | AGO2 | argonaute RISC catalytic component 2 | Cytoplasm | translation regulator |
| 15534976 | | 3 | 0.091 | 2.73E+07 | 2.25E+07 | 1.83E+07 | 2.17E+07 | 1.91E+07 | 1.69E+07 | 2.32E+07 | 2.14E+07 | 1.88E+07 | 3.12E+07 | 3.55E+07 | 2.22E+07 | 1696.8436 | YPHLPC[160.0307]LQVGQEQK | Q8CJG0 | Q8CJG0 | AGO2 | argonaute RISC catalytic component 2 | Cytoplasm | translation regulator |
| 15555977 | | 2 | 0.5 | 2.33E+07 | 2.69E+07 | 2.74E+07 | 2.18E+07 | 2.26E+07 | 2.34E+07 | 2.44E+07 | 2.21E+07 | 2.48E+07 | 3.04E+07 | 2.70E+07 | 2.03E+07 | 1793.9103 | AQPVIEFVC[160.0307]EVLDFK | Q8CJG0 | Q8CJG0 | AGO2 | argonaute RISC catalytic component 2 | Cytoplasm | translation regulator |
| 15532282 | | 2 | 0.75 | 2.79E+07 | 3.06E+07 | 2.78E+07 | 2.35E+07 | 2.38E+07 | 3.35E+07 | 2.78E+07 | 2.83E+07 | 2.09E+07 | 2.91E+07 | 2.60E+07 | 2.30E+07 | 1386.7384 | QEIIQDLAAMVR | Q8CJG0 | Q8CJG0 | AGO2 | argonaute RISC catalytic component 2 | Cytoplasm | translation regulator |
| 15531575 | | 2 | 0.086 | 2.74E+07 | 2.60E+07 | 2.56E+07 | 2.35E+07 | 2.32E+07 | 1.92E+07 | 2.26E+07 | 2.06E+07 | 1.71E+07 | 2.89E+07 | 2.22E+07 | 2.26E+07 | 1256.5891 | SASFNTDPYVR | Q8CJG0 | Q8CJG0 | AGO2 | argonaute RISC catalytic component 2 | Cytoplasm | translation regulator |
| 15548698 | | 2 | 0.807 | 1.53E+07 | 2.01E+07 | 2.15E+07 | 1.29E+07 | 1.99E+07 | 1.45E+07 | 1.58E+07 | 1.80E+07 | 1.46E+07 | 2.88E+07 | 1.34E+07 | 1.42E+07 | 1915.0315 | LPSVPFETIQALDVVMR | Q8CJG0 | Q8CJG0 | AGO2 | argonaute RISC catalytic component 2 | Cytoplasm | translation regulator |
| 15531605 | | 2 | 0.38 | 2.34E+07 | 2.16E+07 | 1.92E+07 | 2.15E+07 | 1.84E+07 | 1.72E+07 | 2.39E+07 | 2.10E+07 | 1.46E+07 | 2.84E+07 | 2.60E+07 | 1.93E+07 | 1470.6661 | YAQGADSVEPMFR | Q8CJG0 | Q8CJG0 | AGO2 | argonaute RISC catalytic component 2 | Cytoplasm | translation regulator |
| 15555110 | | 2 | 0.059 | 2.67E+07 | 2.86E+07 | 3.55E+07 | 2.33E+07 | 2.55E+07 | 2.20E+07 | 2.40E+07 | 2.19E+07 | 2.59E+07 | 2.71E+07 | 2.54E+07 | 2.35E+07 | 1507.8126 | DYQPGITFIVVQK | Q8CJG0 | Q8CJG0 | AGO2 | argonaute RISC catalytic component 2 | Cytoplasm | translation regulator |
| 15531921 | | 2 | 0.133 | 2.65E+07 | 2.42E+07 | 2.48E+07 | 2.32E+07 | 1.96E+07 | 1.74E+07 | 2.37E+07 | 2.00E+07 | 1.73E+07 | 2.71E+07 | 2.58E+07 | 2.04E+07 | 1008.5446 | SFTEQLRK | Q8CJG0 | Q8CJG0 | AGO2 | argonaute RISC catalytic component 2 | Cytoplasm | translation regulator |
| 15540412 | | 2 | 0.081 | 3.30E+07 | 3.26E+07 | 2.75E+07 | 2.46E+07 | 2.54E+07 | 2.09E+07 | 2.65E+07 | 2.50E+07 | 2.06E+07 | 2.69E+07 | 3.05E+07 | 2.29E+07 | 1375.7202 | TTPQTLSNLC[160.0307]LK | Q8CJG0 | Q8CJG0 | AGO2 | argonaute RISC catalytic component 2 | Cytoplasm | translation regulator |
| 15532075 | | 3 | 0.141 | 2.42E+07 | 1.83E+07 | 1.33E+07 | 1.74E+07 | 1.88E+07 | 1.40E+07 | 2.21E+07 | 1.52E+07 | 1.50E+07 | 2.32E+07 | 3.39E+07 | 2.24E+07 | 1869.9603 | HTYLPLEVC[160.0307]NIVAGQR | Q8CJG0 | Q8CJG0 | AGO2 | argonaute RISC catalytic component 2 | Cytoplasm | translation regulator |
| 15536327 | | 2 | 0.009 | 2.31E+07 | 2.27E+07 | 2.45E+07 | 1.69E+07 | 2.04E+07 | 1.68E+07 | 1.94E+07 | 1.74E+07 | 1.66E+07 | 2.27E+07 | 2.17E+07 | 1.92E+07 | 1053.5958 | ELLIQFYK | Q8CJG0 | Q8CJG0 | AGO2 | argonaute RISC catalytic component 2 | Cytoplasm | translation regulator |
| 15549421 | | 3 | 0.495 | 1.61E+07 | 1.35E+07 | 2.04E+07 | 8726380 | 2.09E+07 | 1.20E+07 | 1.34E+07 | 1.62E+07 | 1.25E+07 | 2.19E+07 | 1.71E+07 | 1.50E+07 | 2373.15 | FSSDELQILTYQLC[160.0307]HTYVR | Q8CJG0 | Q8CJG0 | AGO2 | argonaute RISC catalytic component 2 | Cytoplasm | translation regulator |
| 15548792 | | 2 | 0.142 | 1.82E+07 | 1.18E+07 | 9713735 | 1.19E+07 | 1.32E+07 | 9251144 | 1.48E+07 | 1.20E+07 | 1.23E+07 | 2.17E+07 | 2.76E+07 | 1.30E+07 | 1608.6941 | DAGMPIQGQPC[160.0307]FC[160.0307]K | Q8CJG0 | Q8CJG0 | AGO2 | argonaute RISC catalytic component 2 | Cytoplasm | translation regulator |
| 15533010 | | 2 | 0.125 | 2.53E+07 | 2.29E+07 | 2.12E+07 | 2.25E+07 | 2.05E+07 | 1.65E+07 | 2.10E+07 | 1.72E+07 | 1.56E+07 | 2.05E+07 | 2.26E+07 | 2.16E+07 | 1345.7283 | NLYTAMPLPIGR | Q8CJG0 | Q8CJG0 | AGO2 | argonaute RISC catalytic component 2 | Cytoplasm | translation regulator |
| 15536745 | | 2 | 0.274 | 1.53E+07 | 1.48E+07 | 1.42E+07 | 1.32E+07 | 1.13E+07 | 1.14E+07 | 1.39E+07 | 1.10E+07 | 1.10E+07 | 1.90E+07 | 1.41E+07 | 1.09E+07 | 906.4924 | TPVYAEVK | Q8CJG0 | Q8CJG0 | AGO2 | argonaute RISC catalytic component 2 | Cytoplasm | translation regulator |
| 15535592 | | 3 | 0.136 | 1.69E+07 | 1.21E+07 | 9627371 | 1.67E+07 | 9894008 | 1.07E+07 | 1.48E+07 | 1.28E+07 | 1.25E+07 | 1.79E+07 | 2.14E+07 | 1.67E+07 | 1893.9302 | RVGDTVLGMATQC[160.0307]VQMK | Q8CJG0 | Q8CJG0 | AGO2 | argonaute RISC catalytic component 2 | Cytoplasm | translation regulator |
| 15544070 | | 2 | 0.587 | 1.30E+07 | 1.27E+07 | 1.27E+07 | 1.27E+07 | 1.14E+07 | 1.05E+07 | 1.27E+07 | 1.11E+07 | 8915387 | 1.53E+07 | 1.03E+07 | 1.07E+07 | 1023.4389 | DEMTDVTGR | Q8CJG0 | Q8CJG0 | AGO2 | argonaute RISC catalytic component 2 | Cytoplasm | translation regulator |
| [illegible] | [illegible] | [illegible] | [illegible] | [illegible] | [illegible] | [illegible] | [illegible] | [illegible] | [illegible] | [illegible] | [illegible] | [illegible] | [illegible] | [illegible] | [illegible] | [illegible] | [illegible] | [illegible] | [illegible] | [illegible] | [illegible] | [illegible] | [illegible] |

| | | | | | | | | | | | | | | | | | | | | | |
|---|---|---|---|---|---|---|---|---|---|---|---|---|---|---|---|---|---|---|---|---|---|
| 15554922, 15558823 | 3 | 0.373 | 7.25E+08 | 7.36E+08 | 8.38E+08 | 6.83E+08 | 7.92E+08 | 8.23E+08 | 7.20E+08 | 7.22E+08 | 4.30E+08 | 7.74E+08 | 6.76E+08 | 5.81E+08 | 2705.4482, 2705.449 | TGFTPLHIAAHYENLNVAQLLLNR | Q02357 | Q02357 | ANK1 | ankyrin 1, erythrocytic | Plasma Membrane | other |
| 15548618 | 3, 4 | 0.783 | 5.36E+08 | 6.59E+08 | 6.55E+08 | 5.38E+08 | 6.03E+08 | 5.06E+08 | 5.18E+08 | 6.76E+08 | 5.49E+08 | 7.30E+08 | 5.10E+08 | 5.52E+08 | 3012.524, 3012.5242 | VENPNSLLDQSTALLTLWVDREGENAK | Q02357 | Q02357 | ANK1 | ankyrin 1, erythrocytic | Plasma Membrane | other |
| 15531819 | 2 | 0.014 | 5.31E+08 | 5.23E+08 | 5.41E+08 | 5.06E+08 | 5.87E+08 | 5.77E+08 | 5.68E+08 | 5.36E+08 | 4.71E+08 | 7.10E+08 | 6.70E+08 | 6.15E+08 | 1122.5862 | ELTAVPYMAK | Q02357 | Q02357 | ANK1 | ankyrin 1, erythrocytic | Plasma Membrane | other |
| 15534876 | 2 | 0.063 | 4.61E+08 | 4.49E+08 | 5.08E+08 | 4.93E+08 | 4.50E+08 | 4.62E+08 | 4.89E+08 | 4.43E+08 | 4.50E+08 | 6.98E+08 | 5.14E+08 | 5.34E+08 | 914.56506 | TLTPLTLR | Q02357 | Q02357 | ANK1 | ankyrin 1, erythrocytic | Plasma Membrane | other |
| 15552905 | 4 | 0.242 | 6.36E+08 | 8.30E+08 | 8.72E+08 | 5.84E+08 | 8.92E+08 | 6.67E+08 | 5.72E+08 | 7.73E+08 | 4.76E+08 | 6.89E+08 | 4.49E+08 | 5.04E+08 | 4804.235 | EPSLWAPESAFSQEVQGDELQNIPGEQVTEEQFTDEQGNIVTK | Q02357 | Q02357 | ANK1 | ankyrin 1, erythrocytic | Plasma Membrane | other |
| 15534520 | 3 | 0.653 | 3.94E+08 | 4.72E+08 | 5.55E+08 | 4.58E+08 | 5.18E+08 | 4.85E+08 | 5.02E+08 | 4.41E+08 | 3.76E+08 | 6.72E+08 | 4.26E+08 | 4.87E+08 | 1726.9725 | LVPLVQATFPENAVTK | Q02357 | Q02357 | ANK1 | ankyrin 1, erythrocytic | Plasma Membrane | other |
| 15531246 | 2 | 0.201 | 6.16E+08 | 5.16E+08 | 6.01E+08 | 4.66E+08 | 5.73E+08 | 5.21E+08 | 5.00E+08 | 5.18E+08 | 4.51E+08 | 6.60E+08 | 5.83E+08 | 5.06E+08 | 1040.5874 | GFTPLHVAAK | Q02357 | Q02357 | ANK1 | ankyrin 1, erythrocytic | Plasma Membrane | other |
| 15534080, 15542121 | 2 | 0.78 | 5.81E+08 | 6.01E+08 | 7.91E+08 | 6.14E+08 | 6.25E+08 | 6.41E+08 | 6.29E+08 | 6.51E+08 | 5.71E+08 | 6.35E+08 | 5.85E+08 | 5.87E+08 | 960.53735, 960.538 | ELLDFVPK | Q02357 | Q02357 | ANK1 | ankyrin 1, erythrocytic | Plasma Membrane | other |
| 15531559 | 2 | 0.735 | 4.89E+08 | 5.20E+08 | 6.74E+08 | 4.37E+08 | 5.39E+08 | 5.68E+08 | 5.11E+08 | 5.54E+08 | 4.43E+08 | 6.19E+08 | 5.10E+08 | 5.20E+08 | 951.4886 | ADAATSFLR | Q02357 | Q02357 | ANK1 | ankyrin 1, erythrocytic | Plasma Membrane | other |
| 15549283 | 3 | 0.872 | 4.62E+08 | 5.10E+08 | 5.19E+08 | 3.81E+08 | 5.65E+08 | 5.12E+08 | 4.40E+08 | 5.29E+08 | 3.82E+08 | 6.01E+08 | 3.42E+08 | 4.28E+08 | 2827.557 | IIALGPTGAQFLSPVIVEIPHFASHGR | Q02357 | Q02357 | ANK1 | ankyrin 1, erythrocytic | Plasma Membrane | other |
| 15555174, 15558022 | 3 | 0.967 | 3.25E+08 | 3.40E+08 | 4.22E+08 | 2.52E+08 | 4.68E+08 | 4.59E+08 | 3.56E+08 | 4.69E+08 | 3.65E+08 | 5.94E+08 | 2.53E+08 | 4.35E+08 | 2756.2307, 2756.2327 | MSYPETVDEILDVSEDEGDELVGSK | Q02357 | Q02357 | ANK1 | ankyrin 1, erythrocytic | Plasma Membrane | other |
| 15550236, 15554985 | 2, 3 | 0.104 | 5.68E+08 | 5.02E+08 | 4.92E+08 | 5.63E+08 | 4.68E+08 | 3.88E+08 | 4.83E+08 | 4.63E+08 | 4.35E+08 | 5.91E+08 | 5.98E+08 | 5.43E+08 | 2526.3381, 2526.3386 | LLLENGASPNLATTAGHTPLHTAAR | Q02357 | Q02357 | ANK1 | ankyrin 1, erythrocytic | Plasma Membrane | other |
| 15537668, 15542051, 15546954 | 2, 3 | 0.17 | 6.23E+08 | 5.52E+08 | 5.75E+08 | 6.54E+08 | 5.26E+08 | 5.78E+08 | 6.00E+08 | 5.05E+08 | 4.75E+08 | 5.73E+08 | 7.48E+08 | 6.68E+08 | 1363.7899, 1363.7905, 1363.7915, 1363.7927 | RLGYISVTDVLK | Q02357 | Q02357 | ANK1 | ankyrin 1, erythrocytic | Plasma Membrane | other |
| 15531527 | 3 | 0.488 | 6.36E+08 | 5.45E+08 | 6.11E+08 | 5.40E+08 | 6.36E+08 | 4.25E+08 | 5.62E+08 | 5.98E+08 | 5.47E+08 | 5.68E+08 | 6.77E+08 | 5.91E+08 | 2080.0896 | GASVNFTPQNGITPLHIASR | Q02357 | Q02357 | ANK1 | ankyrin 1, erythrocytic | Plasma Membrane | other |
| 15538435 | 2 | 0.01 | 3.89E+08 | 3.33E+08 | 3.46E+08 | 3.75E+08 | 3.31E+08 | 2.95E+08 | 4.07E+08 | 3.84E+08 | 3.73E+08 | 5.62E+08 | 7.29E+08 | 4.37E+08 | 1080.4912 | FWLSDC[160.0307]PR | Q02357 | Q02357 | ANK1 | ankyrin 1, erythrocytic | Plasma Membrane | other |
| 15532232 | 2, 3 | 0.993 | 3.14E+08 | 3.93E+08 | 5.63E+08 | 2.71E+08 | 4.71E+08 | 4.71E+08 | 4.48E+08 | 4.44E+08 | 2.85E+08 | 5.56E+08 | 3.49E+08 | 3.28E+08 | 1283.577, 1283.5775 | DDQTPLHC[160.0307]AAR | Q02357 | Q02357 | ANK1 | ankyrin 1, erythrocytic | Plasma Membrane | other |
| 15534558 | 2, 3 | 0.97 | 3.05E+08 | 5.04E+08 | 6.90E+08 | 3.15E+08 | 5.71E+08 | 5.42E+08 | 5.78E+08 | 5.20E+08 | 3.59E+08 | 5.55E+08 | 3.56E+08 | 3.93E+08 | 1282.6183, 1282.6189 | DELTPLHC[160.0307]AAR | Q02357 | Q02357 | ANK1 | ankyrin 1, erythrocytic | Plasma Membrane | other |
| 15535477, 15543763 | 3 | 0.049 | 3.87E+08 | 4.27E+08 | 4.43E+08 | 3.86E+08 | 3.03E+08 | 4.07E+08 | 3.48E+08 | 3.49E+08 | 3.24E+08 | 5.50E+08 | 3.97E+08 | 4.59E+08 | 1842.009, 1842.0106 | LLGNQATFSPIVTVEPR | Q02357 | Q02357 | ANK1 | ankyrin 1, erythrocytic | Plasma Membrane | other |
| 15547656 | 2 | 0.198 | 4.20E+08 | 3.98E+08 | 4.47E+08 | 4.38E+08 | 3.94E+08 | 3.46E+08 | 3.82E+08 | 3.92E+08 | 3.46E+08 | 5.41E+08 | 4.46E+08 | 3.92E+08 | 1571.7784 | TLEQHENFVEVAR | Q02357 | Q02357 | ANK1 | ankyrin 1, erythrocytic | Plasma Membrane | other |
| 15534124 | 2 | 0.2 | 3.58E+08 | 3.34E+08 | 3.76E+08 | 3.76E+08 | 4.21E+08 | 3.27E+08 | 3.24E+08 | 3.04E+08 | 3.60E+08 | 4.97E+08 | 4.03E+08 | 3.53E+08 | 724.43823 | FVIFAK | Q02357 | Q02357 | ANK1 | ankyrin 1, erythrocytic | Plasma Membrane | other |
| 15531637 | 3 | 0.993 | 2.98E+08 | 3.59E+08 | 3.35E+08 | 2.53E+08 | 3.95E+08 | 3.16E+08 | 2.81E+08 | 3.69E+08 | 3.03E+08 | 4.79E+08 | 1.96E+08 | 3.18E+08 | 1567.7842 | DSSREPGGFLSFLR | Q02357 | Q02357 | ANK1 | ankyrin 1, erythrocytic | Plasma Membrane | other |
| 15539138, 15545247 | 2 | 0.022 | 3.98E+08 | 3.65E+08 | 3.58E+08 | 4.12E+08 | 3.06E+08 | 4.23E+08 | 2.98E+08 | 3.30E+08 | 2.81E+08 | 4.61E+08 | 4.17E+08 | 4.16E+08 | 845.51434, 845.51556 | VMELLLK | Q02357 | Q02357 | ANK1 | ankyrin 1, erythrocytic | Plasma Membrane | other |
| 15553430 | 2 | 0.395 | 3.34E+08 | 3.10E+08 | 5.39E+07 | 3.66E+08 | 1.81E+08 | 7.57E+07 | 2.60E+08 | 1.10E+08 | 7.21E+07 | 4.35E+08 | 3.75E+08 | 3.01E+08 | 1604.8976 | ISEILLDHGAPIQAK | Q02357 | Q02357 | ANK1 | ankyrin 1, erythrocytic | Plasma Membrane | other |
| 15535790, 15542949, 15543289 | 2, 3 | 0.541 | 4.50E+08 | 3.76E+08 | 4.16E+08 | 5.42E+08 | 3.38E+08 | 4.02E+08 | 4.11E+08 | 3.58E+08 | 3.81E+08 | 4.33E+08 | 4.90E+08 | 4.39E+08 | 1216.7589, 1216.7595, 1216.7603, 1216.7606, 1216.7612 | VRLPALHIAAR | Q02357 | Q02357 | ANK1 | ankyrin 1, erythrocytic | Plasma Membrane | other |
| 15531646 | 2 | 0.817 | 2.59E+08 | 3.89E+08 | 4.25E+08 | 2.24E+08 | 4.92E+08 | 3.21E+08 | 3.65E+08 | 4.64E+08 | 2.66E+08 | 4.07E+08 | 2.15E+08 | 2.61E+08 | 1291.6302 | SEIVNMLEGSGR | Q02357 | Q02357 | ANK1 | ankyrin 1, erythrocytic | Plasma Membrane | other |
| 15534528 | 3 | 0.131 | 2.01E+08 | 2.14E+08 | 2.28E+08 | 1.16E+08 | 1.95E+08 | 2.31E+08 | 2.36E+08 | 2.43E+08 | 2.37E+08 | 3.62E+08 | 2.32E+08 | 2.58E+08 | 1794.9579 | TAAVLLQNDPNPDVLSK | Q02357 | Q02357 | ANK1 | ankyrin 1, erythrocytic | Plasma Membrane | other |
| 15533992 | 5 | 0.45 | 3.75E+08 | 2.17E+08 | 1.61E+08 | 5.15E+08 | 6.52E+07 | 2.79E+08 | 3.72E+08 | 1.33E+08 | 3.42E+08 | 3.41E+08 | 6.41E+08 | 5.29E+08 | 3155.7065 | IIALGPTGAQFLSPVIVEIPHFASHGRGDR | Q02357 | Q02357 | ANK1 | ankyrin 1, erythrocytic | Plasma Membrane | other |
| 15542888, 15543012 | 2 | 0.381 | 2.60E+08 | 2.48E+08 | 2.99E+08 | 2.71E+08 | 2.55E+08 | 2.45E+08 | 2.49E+08 | 2.66E+08 | 2.26E+08 | 3.40E+08 | 2.53E+08 | 2.74E+08 | 777.37665 | TLDWDK | Q02357 | Q02357 | ANK1 | ankyrin 1, erythrocytic | Plasma Membrane | other |
| 15553346 | 2 | 0.587 | 3.90E+08 | 2.52E+08 | 2.39E+08 | 4.46E+08 | 2.01E+08 | 2.29E+08 | 3.36E+08 | 2.11E+08 | 2.16E+08 | 3.34E+08 | 4.17E+08 | 3.01E+08 | 1789.8799 | NIDRSEIVNMLEGSGR | Q02357 | Q02357 | ANK1 | ankyrin 1, erythrocytic | Plasma Membrane | other |
| 15548357 | 2 | 0.372 | 2.40E+08 | 2.04E+08 | 2.20E+08 | 2.73E+08 | 2.07E+08 | 1.77E+08 | 2.31E+08 | 1.79E+08 | 1.63E+08 | 3.11E+08 | 2.42E+08 | 2.06E+08 | 1526.7241 | LGFTSDTDRVEMR | Q02357 | Q02357 | ANK1 | ankyrin 1, erythrocytic | Plasma Membrane | other |
| 15558084 | 3, 4 | 0.377 | 9.26E+07 | 1.04E+08 | 1.61E+08 | 1.24E+08 | 5.07E+07 | 2.53E+08 | 2.15E+08 | 1.18E+08 | 3.31E+08 | 3.02E+08 | 1.41E+08 | 2.23E+08 | 3155.7046, 3155.706 | IIALGPTGAQFLSPVIVEIPHFASHGRGDR | Q02357 | Q02357 | ANK1 | ankyrin 1, erythrocytic | Plasma Membrane | other |
| 15532787 | 2 | 0.555 | 2.01E+08 | 3.05E+08 | 3.66E+08 | 1.46E+08 | 3.81E+08 | 2.76E+08 | 2.47E+08 | 3.31E+08 | 1.94E+08 | 2.87E+08 | 9.35E+07 | 1.93E+08 | 1122.5927 | EPGGFLSFLR | Q02357 | Q02357 | ANK1 | ankyrin 1, erythrocytic | Plasma Membrane | other |
| 15537354 | 3 | 0.22 | 2.78E+08 | 1.98E+08 | 1.67E+08 | 2.59E+08 | 1.83E+08 | 1.02E+08 | 2.21E+08 | 1.57E+08 | 1.44E+08 | 2.73E+08 | 3.12E+08 | 2.51E+08 | 2091.1416 | NGLTPLHVAVHHNNLDIVK | Q02357 | Q02357 | ANK1 | ankyrin 1, erythrocytic | Plasma Membrane | other |
| 15538112 | 3, 4 | 0.375 | 3.28E+08 | 1.14E+08 | 2.26E+07 | 2.79E+08 | 9.92E+07 | 5.91E+07 | 1.03E+08 | 8.58E+07 | 2.13E+08 | 2.52E+08 | 3.98E+08 | 2.86E+08 | 1963.0681, 1963.0687 | KGNTALHIAALAGQDEVVR | Q02357 | Q02357 | ANK1 | ankyrin 1, erythrocytic | Plasma Membrane | other |
| 15547789 | 2 | 0.556 | 1.97E+08 | 2.49E+08 | 3.12E+08 | 1.57E+08 | 2.38E+08 | 2.61E+08 | 2.10E+08 | 2.17E+08 | 1.60E+08 | 2.51E+08 | 1.68E+08 | 1.86E+08 | 1834.9728 | GNTALHIAALAGQDEVVR | Q02357 | Q02357 | ANK1 | ankyrin 1, erythrocytic | Plasma Membrane | other |
| 15532371 | 3 | 0.215 | 2.88E+08 | 1.66E+08 | 1.73E+08 | 2.42E+08 | 1.91E+08 | 2.12E+08 | 2.38E+08 | 2.12E+08 | 2.17E+08 | 2.40E+08 | 4.24E+08 | 2.68E+08 | 2090.9805 | NGLSPIHMAAQGDHLDC[160.0307]VR | Q02357 | Q02357 | ANK1 | ankyrin 1, erythrocytic | Plasma Membrane | other |
| 15547658 | 2 | 0.199 | 2.12E+08 | 1.67E+08 | 2.14E+08 | 2.20E+08 | 1.65E+08 | 1.99E+08 | 1.99E+08 | 1.50E+08 | 1.84E+08 | 2.36E+08 | 2.32E+08 | 2.13E+08 | 1927.9679 | NGASPNEVSSNGTTPLAIAK | Q02357 | Q02357 | ANK1 | ankyrin 1, erythrocytic | Plasma Membrane | other |
| 15531635 | 2, 3 | 0.085 | 2.45E+08 | 2.03E+08 | 1.80E+08 | 1.91E+08 | 1.88E+08 | 2.26E+08 | 2.01E+08 | 1.76E+08 | 1.93E+08 | 2.29E+08 | 4.63E+08 | 2.68E+08 | 1441.7595, 1441.7599 | ALNGFTPLHIAC[160.0307]K | Q02357 | Q02357 | ANK1 | ankyrin 1, erythrocytic | Plasma Membrane | other |
| 15552815, 15559406 | 2 | 0.454 | 1.71E+08 | 1.10E+08 | 2.62E+07 | 2.12E+08 | 9.74E+07 | 5.30E+07 | 1.57E+08 | 6.35E+07 | 2.41E+07 | 2.23E+08 | 1.77E+08 | 1.31E+08 | 2047.027, 2047.0272 | GFTPLYMAAQENHLEVVK | Q02357 | Q02357 | ANK1 | ankyrin 1, erythrocytic | Plasma Membrane | other |
| 15534429 | 3 | 0.199 | 1.87E+08 | 2.50E+08 | 8.66E+07 | 2.43E+08 | 2.06E+08 | 2.38E+08 | 2.37E+08 | 2.55E+08 | 3.12E+08 | 2.16E+08 | 3.41E+08 | 2.81E+08 | 1815.9702 | IITTDFPLYFVIMSR | Q02357 | Q02357 | ANK1 | ankyrin 1, erythrocytic | Plasma Membrane | other |
| 15547839, 15548721, 15550377, 15556641, 15560213, 15561200, 15562614, 15563120 | 2, 3 | 0.405 | 1.88E+08 | 2.12E+08 | 3.84E+08 | 1.40E+08 | 1.83E+08 | 2.62E+08 | 1.60E+08 | 2.62E+08 | 1.61E+08 | 2.11E+08 | 1.01E+08 | 1.59E+08 | 2629.3745, 2629.3843, 2629.3848, 2629.3862, 2629.3867, 2629.398 | SRDIEVLEGM[147.0354]PLFAELSGNLVPVK | Q02357 | Q02357 | ANK1 | ankyrin 1, erythrocytic | Plasma Membrane | other |
| 15547785, 15548021, 15560986, 15562997 | 2 | 0.314 | 2.21E+08 | 1.84E+08 | 3.23E+08 | 1.98E+08 | 2.26E+08 | 2.76E+08 | 2.15E+08 | 2.49E+08 | 2.07E+08 | 2.07E+08 | 1.36E+08 | 1.88E+08 | 1831.9635, 1831.9637, 1831.9647, 1831.9669 | IITTDFPLYFVIM[147.0354]SR | Q02357 | Q02357 | ANK1 | ankyrin 1, erythrocytic | Plasma Membrane | other |
| 15551741 | 2, 3 | 0.773 | 1.18E+08 | 2.37E+08 | 2.23E+08 | 9.25E+07 | 2.46E+08 | 1.78E+08 | 1.92E+08 | 2.08E+08 | 1.12E+08 | 1.83E+08 | 9.54E+07 | 1.31E+08 | 2653.4265, 2653.4272 | IRVENPNSLLDQSTALLTLWVDR | Q02357 | Q02357 | ANK1 | ankyrin 1, erythrocytic | Plasma Membrane | other |
| 15554538, 15558879 | 2 | 0.365 | 1.22E+08 | 1.12E+08 | 7.16E+07 | 1.02E+08 | 1.01E+08 | 6.87E+07 | 1.32E+08 | 7.33E+07 | 5.98E+07 | 1.71E+08 | 1.30E+08 | 9.82E+07 | 1798.9414 | LAELLLEHDAHPNAAGK | Q02357 | Q02357 | ANK1 | ankyrin 1, erythrocytic | Plasma Membrane | other |
| 15531652 | 2 | 0.051 | 1.36E+08 | 1.11E+08 | 1.15E+08 | 1.13E+08 | 1.03E+08 | 1.02E+08 | 1.17E+08 | 1.09E+08 | 9.43E+07 | 1.58E+08 | 1.26E+08 | 1.27E+08 | 1056.6133 | GDRELVVLR | Q02357 | Q02357 | ANK1 | ankyrin 1, erythrocytic | Plasma Membrane | other |
| 15532344 | 3 | 0.333 | 1.17E+08 | 1.19E+08 | 1.06E+08 | 1.19E+08 | 8.47E+07 | 2.50E+07 | 1.10E+08 | 9.55E+07 | 1.06E+08 | 1.52E+08 | 9.46E+07 | 1.07E+08 | 1693.8727 | LNTPPPLAEEEGLASR | Q02357 | Q02357 | ANK1 | ankyrin 1, erythrocytic | Plasma Membrane | other |
| 15541884 | 3 | 0.06 | 1.23E+08 | 8.69E+07 | 7.06E+07 | 9.19E+07 | 9.56E+07 | 1.12E+08 | 1.05E+08 | 1.14E+08 | 1.16E+08 | 1.47E+08 | 1.51E+08 | 1.20E+08 | 1780.8143 | LC[160.0307]QDYDTIGPEGGSLR | Q02357 | Q02357 | ANK1 | ankyrin 1, erythrocytic | Plasma Membrane | other |
| 15534628 | 4 | 0.059 | 1.90E+08 | 2.39E+08 | 2.39E+08 | 1.71E+08 | 1.54E+08 | 2.03E+08 | 1.52E+08 | 1.82E+08 | 1.44E+08 | 1.42E+08 | 7.70E+07 | 1.58E+08 | 2964.5442 | TGASIDAVTESGLTPLHVASFM[147.0354]GHLPIVK | Q02357 | Q02357 | ANK1 | ankyrin 1, erythrocytic | Plasma Membrane | other |
| 15543495, 15543761, 15544130 | 2 | 0.749 | 8.92E+07 | 1.60E+08 | 1.33E+08 | 7.63E+07 | 1.77E+08 | 1.50E+08 | 1.44E+08 | 1.38E+08 | 1.08E+08 | 1.41E+08 | 8.29E+07 | 8.57E+07 | 1392.7214, 1392.7247, 1392.725 | IPLPPSWTDNPR | Q02357 | Q02357 | ANK1 | ankyrin 1, erythrocytic | Plasma Membrane | other |
| 15534261 | 2 | 0.29 | 1.28E+08 | 1.47E+08 | 1.55E+08 | 1.36E+08 | 1.23E+08 | 1.20E+08 | 1.15E+08 | 1.12E+08 | 8.77E+07 | 1.35E+08 | 4.10E+07 | 1.07E+08 | 1138.5787 | ELTAVPYM[147.0354]AK | Q02357 | Q02357 | ANK1 | ankyrin 1, erythrocytic | Plasma Membrane | other |
| 15550167 | 4 | 0.392 | 2.16E+08 | 4.77E+07 | 6957061 | 2.18E+08 | 3.29E+07 | 3.10E+07 | 4.14E+07 | 4.54E+07 | 1.33E+08 | 1.34E+08 | 2.85E+08 | 1.98E+08 | 4932.3276 | EPSLWAPESAFSQEVQGDELQNIPGEQVTEEQFTDEQGNIVTKK | Q02357 | Q02357 | ANK1 | ankyrin 1, erythrocytic | Plasma Membrane | other |
| 15547689 | 2 | 0.791 | 8.06E+07 | 8.34E+07 | 9.36E+07 | 7.31E+07 | 9.47E+07 | 6.30E+07 | 6.78E+07 | 7.11E+07 | 6.54E+07 | 1.33E+08 | 4.98E+07 | 6.74E+07 | 1567.7834 | DSSREPGGFLSFLR | Q02357 | Q02357 | ANK1 | ankyrin 1, erythrocytic | Plasma Membrane | other |
| 15547745 | 2 | 0.254 | 1.10E+08 | 4.78E+07 | 1.16E+07 | 7.20E+07 | 4.33E+07 | 3.02E+07 | 5.25E+07 | 4.30E+07 | 6.07E+07 | 1.18E+08 | 1.43E+08 | 8.31E+07 | 1511.7616 | TKDELTPLHC[160.0307]AAR | Q02357 | Q02357 | ANK1 | ankyrin 1, erythrocytic | Plasma Membrane | other |
| 15532623, 15533462, 15547395, 15547630, 15547641 | 3 | 0.615 | 1.23E+08 | 9.28E+07 | 1.78E+08 | 1.16E+08 | 1.18E+08 | 1.66E+08 | 1.39E+08 | 1.57E+08 | 1.11E+08 | 1.18E+08 | 8.17E+07 | 1.21E+08 | 1831.9609, 1831.9613, 1831.9635, 1831.9636 | IITTDFPLYFVIM[147.0354]SR | Q02357 | Q02357 | ANK1 | ankyrin 1, erythrocytic | Plasma Membrane | other |
| 15550045, 15556814, 15557340 | 3 | 0.724 | 1.08E+08 | 1.03E+08 | 9.28E+07 | 1.02E+08 | 1.07E+08 | 8.83E+07 | 1.02E+08 | 9.84E+07 | 8.14E+07 | 1.09E+08 | 9.53E+07 | 9.98E+07 | 3920.016, 3920.0164, 3920.019 | SLLQYGGSANAESVQGVTPLHLAAQEGHTEMVALLLSK | Q02357 | Q02357 | ANK1 | ankyrin 1, erythrocytic | Plasma Membrane | other |
| 15558710 | 2 | 0.066 | 8.39E+07 | 6.75E+07 | 5.08E+07 | 6.53E+07 | 5.28E+07 | 4.70E+07 | 6.66E+07 | 4.62E+07 | 3.95E+07 | 9.87E+07 | 9.43E+07 | 6.91E+07 | 1787.8856 | MGYTPLHVASHYGNIK | Q02357 | Q02357 | ANK1 | ankyrin 1, erythrocytic | Plasma Membrane | other |
| 15549328 | 4 | 0.051 | 1.03E+08 | 1.96E+08 | 1.37E+08 | 1.57E+08 | 1.85E+08 | 1.30E+08 | 9.51E+07 | 1.79E+08 | 1.33E+08 | 9.21E+07 | 7.20E+07 | 7.98E+07 | 4611.06 | LEGAHSEDTQGPELGSQDLVEDDTVDSDATNGLADLLGQEEGQR | Q02357 | Q02357 | ANK1 | ankyrin 1, erythrocytic | Plasma Membrane | other |
| 15542813 | 2, 3 | 0.828 | 1.50E+08 | 3.76E+08 | 6.35E+08 | 5.71E+07 | 4.75E+08 | 5.35E+08 | 2.17E+08 | 4.97E+08 | 4.92E+08 | 8.85E+07 | 2.71E+08 | 3.84E+08 | 1055.504, 1055.5046 | SFHFQSFR | Q02357 | Q02357 | ANK1 | ankyrin 1, erythrocytic | Plasma Membrane | other |
| 15540907 | 4 | 0.722 | 6.27E+07 | 7.27E+07 | 7.89E+07 | 6.02E+07 | 6.29E+07 | 7.25E+07 | 6.79E+07 | 6.91E+07 | 5.65E+07 | 8.27E+07 | 3.13E+07 | 6.50E+07 | 2487.1406 | YEDTQHILC[160.0307]HLNITM[147.0354]PPC[160.0307]TK | Q02357 | Q02357 | ANK1 | ankyrin 1, erythrocytic | Plasma Membrane | other |
| 15531743 | 3 | 0.168 | 7.45E+07 | 5.60E+07 | 6.01E+07 | 8.17E+07 | 4.78E+07 | 6.22E+07 | 7.36E+07 | 5.60E+07 | 6.19E+07 | 8.21E+07 | 1.17E+08 | 7.45E+07 | 2239.0852 | NGVDINTC[160.0307]NQNGLNGLHLASK | Q02357 | Q02357 | ANK1 | ankyrin 1, erythrocytic | Plasma Membrane | other |
| 15531656 | 3 | 0.32 | 9.24E+07 | 8.49E+07 | 7.64E+07 | 8.75E+07 | 6.86E+07 | 6.40E+07 | 7.28E+07 | 7.32E+07 | 4.33E+07 | 8.14E+07 | 5.95E+07 | 6.90E+07 | 1803.8811 | M[147.0354]GYTPLHVASHYGNIK | Q02357 | Q02357 | ANK1 | ankyrin 1, erythrocytic | Plasma Membrane | other |
| 15535683, 15544562, 15546108, 15546690, 15546962 | 2 | 0.202 | 6.91E+07 | 8.63E+07 | 1.01E+08 | 7.16E+07 | 7.32E+07 | 7.13E+07 | 5.91E+07 | 6.22E+07 | 5.28E+07 | 8.13E+07 | 3.17E+07 | 5.84E+07 | 984.55225, 984.5525, 984.5526, 984.553, 984.5531 | M[147.0354]VVELLHK | Q02357 | Q02357 | ANK1 | ankyrin 1, erythrocytic | Plasma Membrane | other |
| 15543229, 15543504, 15544863 | 4 | 0.045 | 5.51E+07 | 3.93E+07 | 3.12E+07 | 4.11E+07 | 5.84E+07 | 3.81E+07 | 7.58E+07 | 6.42E+07 | 5.75E+07 | 6.44E+07 | 1.62E+08 | 7.54E+07 | 2471.1426, 2471.143 | YEDTQHILC[160.0307]HLNITMPPC[160.0307]TK | Q02357 | Q02357 | ANK1 | ankyrin 1, erythrocytic | Plasma Membrane | other |
| 15548062 | 2, 3 | 0.783 | 2.97E+07 | 6.60E+07 | 9.27E+07 | 2.70E+07 | 1.17E+08 | 6.81E+07 | 4.59E+07 | 9.10E+07 | 4.63E+07 | 6.37E+07 | 2.91E+07 | 3.38E+07 | 2384.2407, 2384.2417 | VENPNSLLDQSTALLTLWVDR | Q02357 | Q02357 | ANK1 | ankyrin 1, erythrocytic | Plasma Membrane | other |
| 15540021, 15541242, 15544866, 15545976, 15546110, 15547321 | 2 | 0.695 | 6.60E+07 | 5.20E+07 | 6.48E+07 | 5.59E+07 | 5.95E+07 | 5.59E+07 | 5.87E+07 | 6.03E+07 | 4.74E+07 | 6.13E+07 | 5.29E+07 | 5.31E+07 | 1126.5527, 1126.553, 1126.5536, 1126.5537, 1126.5542, 1126.5544 | M[147.0354]ENLYTALR | Q02357 | Q02357 | ANK1 | ankyrin 1, erythrocytic | Plasma Membrane | other |
| 15537228 | 2, 3 | 0.321 | 7.90E+07 | 7.59E+07 | 7.46E+07 | 6.18E+07 | 4.61E+07 | 7.29E+07 | 7.41E+07 | 6.35E+07 | 7.54E+07 | 5.85E+07 | 7.98E+07 | 9.90E+07 | 1250.6864, 1250.6865 | EPGGFLSFLRK | Q02357 | Q02357 | ANK1 | ankyrin 1, erythrocytic | Plasma Membrane | other |
| 15538058 | 2 | 0.891 | 3.50E+07 | 7.12E+07 | 9.98E+07 | 2.98E+07 | 1.13E+08 | 6.83E+07 | 5.39E+07 | 7.41E+07 | 4.88E+07 | 5.65E+07 | 3.37E+07 | 5.80E+07 | 1207.6921 | LGYISVTDVLK | Q02357 | Q02357 | ANK1 | ankyrin 1, erythrocytic | Plasma Membrane | other |
| 15551197 | 3 | 0.1 | 7.04E+07 | 7.63E+07 | 8.49E+07 | 5.50E+07 | 6.08E+07 | 7.23E+07 | 5.43E+07 | 6.44E+07 | 5.03E+07 | 5.50E+07 | 2.87E+07 | 5.88E+07 | 2964.5432 | TGASIDAVTESGLTPLHVASFM[147.0354]GHLPIVK | Q02357 | Q02357 | ANK1 | ankyrin 1, erythrocytic | Plasma Membrane | other |
| 15534274 | 2 | 0.913 | 4.14E+07 | 4.25E+07 | 5.03E+07 | 3.24E+07 | 5.32E+07 | 4.42E+07 | 4.28E+07 | 4.87E+07 | 3.69E+07 | 5.21E+07 | 3.39E+07 | 3.59E+07 | 961.59216 | LPALHIAAR | Q02357 | Q02357 | ANK1 | ankyrin 1, erythrocytic | Plasma Membrane | other |
| 15543995 | 2 | 0.781 | 3.97E+07 | 7.28E+07 | 9.32E+07 | 9959154 | 8.94E+07 | 9.07E+07 | 5.90E+07 | 8.63E+07 | 8.54E+07 | 5.11E+07 | 3.33E+07 | 7.82E+07 | 995.5876 | FHRPIGLR | Q02357 | Q02357 | ANK1 | ankyrin 1, erythrocytic | Plasma Membrane | other |
| 15538730, 15543987, 15544993, 15545213 | 3, 4 | 0.36 | 9.46E+07 | 2.30E+07 | 7630253 | 1.46E+08 | 4755778.5 | 3.31E+07 | 2.97E+07 | 1.03E+07 | 3.28E+07 | 4.76E+07 | 2.28E+08 | 1.29E+08 | 2054.1091, 2054.1099, 2054.11, 2054.1125 | VRLAELLLEHDAHPNAAGK | Q02357 | Q02357 | ANK1 | ankyrin 1, erythrocytic | Plasma Membrane | other |
| 15534853 | 4 | 0.27 | 2.42E+07 | 2.78E+07 | 2.35E+07 | 2.97E+07 | 2.04E+07 | 2.67E+07 | 1.95E+07 | 2.06E+07 | 1.97E+07 | 4.35E+07 | 2.28E+07 | 2.38E+07 | 2685.4158 | RITTIQGPEPGALQEYEQVLVSTR | Q02357 | Q02357 | ANK1 | ankyrin 1, erythrocytic | Plasma Membrane | other |
| 15556214, 15558530 | 3 | 0.529 | 3.50E+07 | 3.68E+07 | 4.11E+07 | 2.71E+07 | 3.58E+07 | 3.27E+07 | 3.04E+07 | 3.38E+07 | 2.43E+07 | 4.25E+07 | 1.60E+07 | 2.91E+07 | 2487.1426, 2487.1455 | YEDTQHILC[160.0307]HLNITM[147.0354]PPC[160.0307]TK | Q02357 | Q02357 | ANK1 | ankyrin 1, erythrocytic | Plasma Membrane | other |
| 15540572, 15544449 | 2 | 0.296 | 3.31E+07 | 4.09E+07 | 5.18E+07 | 3.31E+07 | 3.48E+07 | 3.38E+07 | 2.99E+07 | 3.34E+07 | 2.39E+07 | 4.10E+07 | 1.53E+07 | 2.84E+07 | 1140.5792, 1140.5797 | VETPLHM[147.0354]AAR | Q02357 | Q02357 | ANK1 | ankyrin 1, erythrocytic | Plasma Membrane | other |
| 15537156 | 2 | 0.741 | 3.13E+07 | 3.03E+07 | 4.60E+07 | 2.96E+07 | 3.33E+07 | 3.06E+07 | 2.84E+07 | 2.56E+07 | 2.68E+07 | 4.06E+07 | 1.04E+07 | 3.76E+07 | 861.5097 | VM[147.0354]ELLLK | Q02357 | Q02357 | ANK1 | ankyrin 1, erythrocytic | Plasma Membrane | other |
| 15548128, 15548778 | 3 | 0.479 | 6.01E+07 | 5.99E+07 | 1.15E+08 | 4.64E+07 | 4.92E+07 | 1.25E+08 | 4.67E+07 | 8.11E+07 | 1.02E+08 | 3.99E+07 | 3.31E+07 | 6.75E+07 | 3133.445, 3133.4456 | SRYGESYLDQILNGM[147.0354]DEELGSLEELEK | Q02357 | Q02357 | ANK1 | ankyrin 1, erythrocytic | Plasma Membrane | other |
| 15553324 | 2 | 0.375 | 3.12E+07 | 1.72E+07 | 4952294 | 3.40E+07 | 1.05E+07 | 2353896 | 2.17E+07 | 8150715 | 7376175.5 | 3.56E+07 | 3.84E+07 | 2.48E+07 | 2061.026 | GGSPHSPAWNGYTPLHIAAK | Q02357 | Q02357 | ANK1 | ankyrin 1, erythrocytic | Plasma Membrane | other |
| 15557771 | 3 | 0.209 | 3.36E+07 | 1.93E+07 | 1.56E+07 | 2.81E+07 | 2.54E+07 | 1.44E+07 | 2.85E+07 | 2.56E+07 | 2.38E+07 | 3.49E+07 | 8.36E+07 | 2.67E+07 | 2471.1443 | YEDTQHILC[160.0307]HLNITMPPC[160.0307]TK | Q02357 | Q02357 | ANK1 | ankyrin 1, erythrocytic | Plasma Membrane | other |
| 15531817 | 3 | 0.106 | 3.30E+07 | 2.72E+07 | 2.44E+07 | 3.34E+07 | 2.29E+07 | 2.39E+07 | 2.67E+07 | 2.04E+07 | 2.62E+07 | 3.37E+07 | 3.46E+07 | 3.39E+07 | 1369.7402 | GSGLQPDLIEGRK | Q02357 | Q02357 | ANK1 | ankyrin 1, erythrocytic | Plasma Membrane | other |
| [illegible] | [illegible] | [illegible] | [illegible] | [illegible] | [illegible] | [illegible] | [illegible] | [illegible] | [illegible] | [illegible] | [illegible] | [illegible] | [illegible] | [illegible] | [illegible] | [illegible] | [illegible] | [illegible] | [illegible] | [illegible] | [illegible] | [illegible] |

| | | | | | | | | | | | | | | | | | | | | | | |
|---|---|---|---|---|---|---|---|---|---|---|---|---|---|---|---|---|---|---|---|---|---|---|
| 15543433 | 2 | 0.38 | 1.02E+07 | 7711392 | 6063773.5 | 7779071 | 1.25E+07 | 7938409.5 | 9563253 | 9763066 | 6292268.5 | 1.49E+07 | 1.23E+07 | 8308336 | 1319.5394 | EIC[160.0307]C[160.0307]YSISC[160.0307]K | Q9CQW2 | Q9CQW2 | ARL8B | ADP-ribosylation factor-like 8B | Plasma Membrane | enzyme |
| 15534480 | 2 | 0.658 | 7886910 | 2.17E+07 | 4.21E+07 | 1.06E+07 | 2.42E+07 | 3.18E+07 | 2.28E+07 | 1.97E+07 | 1.08E+07 | 1.47E+07 | 7981868.5 | 1.38E+07 | 1170.6085 | RDLPNALDEK | Q9CQW2 | Q9CQW2 | ARL8B | ADP-ribosylation factor-like 8B | Plasma Membrane | enzyme |
| 15531898 | 2 | 0.484 | 7798953 | 1.98E+07 | 2.42E+07 | 6185725 | 2.28E+07 | 1.69E+07 | 1.28E+07 | 1.95E+07 | 9576875 | 1.23E+07 | 5788943 | 7340182.5 | 1095.5781 | NELHNLLDK | Q9CQW2 | Q9CQW2 | ARL8B | ADP-ribosylation factor-like 8B | Plasma Membrane | enzyme |
| 15550056 | 2 | 0.135 | 8580291 | 8955772 | 9439133 | 1.01E+07 | 9748227 | 9712276 | 8477824 | 9505324 | 1.08E+07 | 1.05E+07 | 1.00E+07 | 1.09E+07 | 1764.8917 | GVNAIVYMIDAADREK | Q9CQW2 | Q9CQW2 | ARL8B | ADP-ribosylation factor-like 8B | Plasma Membrane | enzyme |
| 15550194 | 2 | 0.659 | 9852335 | 1.00E+07 | 8420994 | 1.04E+07 | 9786528 | 9700058 | 8984633 | 1.07E+07 | 8116839 | 1.03E+07 | 1.05E+07 | 9223816 | 1823.9607 | DNIDITLQWLIQHSK | Q9CQW2 | Q9CQW2 | ARL8B | ADP-ribosylation factor-like 8B | Plasma Membrane | enzyme |
| 15551234 | 3 | 0.267 | 7419721 | 1.06E+07 | 5579031 | 1.01E+07 | 1.25E+07 | 9190632 | 9479298 | 1.03E+07 | 1.18E+07 | 1.02E+07 | 6652010.5 | 9281009 | 2552.4502 | NELHNLLDKPQLQGIPVLVLGNK | Q9CQW2 | Q9CQW2 | ARL8B | ADP-ribosylation factor-like 8B | Plasma Membrane | enzyme |
| 15550963 | 3 | 0.815 | 3063814 | 2215812.75 | 3074631.5 | 2869529.25 | 2918672.5 | 2286601 | 2170913.75 | 3347403.25 | 4821753 | 2029635 | 4518540 | 3444140.75 | 2544.2507 | STVLSLDWHPNNVLLAAGSC[160.0307]DFK | Q9R0Q6 | Q9R0Q6 | ARPC1A | actin related protein 2/3 complex, subunit 1A, 41kDa | Extracellular Space | other |
| 15540109 | 2 | 0.777 | 1.84E+07 | 1.54E+07 | 2.33E+07 | 1.47E+07 | 1.60E+07 | 2.03E+07 | 1.73E+07 | 1.86E+07 | 2.34E+07 | 1.25E+07 | 1.87E+07 | 2.13E+07 | 1175.5309 | YFQFQEEGK | Q9CVB6 | Q9CVB6 | ARPC2 | actin related protein 2/3 complex, subunit 2, 34kDa | Cytoplasm | other |
| 15551744 | 3 | 0.009 | 7820744 | 7734852.5 | 7040257.5 | 1.02E+07 | 1.12E+07 | 8793437 | 9350217 | 9779645 | 1.14E+07 | 4602995 | 6933100.5 | 7139568 | 3254.6672 | RVYGSFLVNPEPGYNVSLLYDLENLPASK | Q9CVB6 | Q9CVB6 | ARPC2 | actin related protein 2/3 complex, subunit 2, 34kDa | Cytoplasm | other |
| 15552401 | 4 | 0.149 | 4261648.5 | 5282208.5 | 4947077 | 5324866 | 9055652 | 7449146.5 | 6343547 | 6323433 | 8193369.5 | 2393943.75 | 5677299 | 5527103.5 | 3636.721 | FENAAAGNKPEAVEVTFADFDGVLYHISNPNGDK | Q9CVB6 | Q9CVB6 | ARPC2 | actin related protein 2/3 complex, subunit 2, 34kDa | Cytoplasm | other |
| 15552120 | 2 | 0.082 | 3524492.25 | 3714823.5 | 4691464.5 | 3852507.75 | 4978972 | 4168160.5 | 2724609.75 | 3902422 | 4001578 | 2163676.25 | 2692697 | 3300035.25 | 1591.7513 | DTDIVDEAIYYFK | Q9JM76 | Q9JM76 | ARPC3 | actin related protein 2/3 complex, subunit 3, 21kDa | Cytoplasm | other |
| 15535240 | 2 | 0.543 | 2286443.5 | 1579721.5 | 2058459.5 | 3079382.75 | 2937548.25 | 2107207.75 | 1889520.375 | 2316590.25 | 3122318 | 1096367.375 | 2698600.75 | 2463252.5 | 1210.7324 | LIGNMALLPLR | Q9JM76 | Q9JM76 | ARPC3 | actin related protein 2/3 complex, subunit 3, 21kDa | Cytoplasm | other |
| 15537102 | 2 | 0.484 | 2198986.5 | 2397224 | 4150592.5 | 2960291.5 | 4076601 | 2922303.75 | 2508965.75 | 3466707.75 | 4538621 | 1978910.375 | 3145740.75 | 2413694.5 | 1009.545 | AENFFILR | P59999 | P59999 | ARPC4 | actin related protein 2/3 complex, subunit 4, 20kDa | Cytoplasm | other |
| 15540314 | 2 | 0.453 | 1146444.625 | 1153239.375 | 1521097.625 | 1675400 | 1936957.5 | 1803071 | 1440001.25 | 1533064 | 2363661.5 | 744103.875 | 2136788.5 | 1531078 | 1070.6293 | ALAAGGVGSIVR | Q9CPW4 | Q9CPW4 | ARPC5 | actin related protein 2/3 complex, subunit 5, 16kDa | Cytoplasm | other |
| 15551273 | 3 | 0.397 | 3.68E+07 | 4.67E+07 | 3.43E+07 | 2.74E+07 | 3.59E+07 | 3.11E+07 | 3.32E+07 | 3.85E+07 | 3.05E+07 | 4.68E+07 | 3.05E+07 | 3.64E+07 | 3884.6733 | VDVDLTPDSFDDQYQGC[160.0307]SEQMVEELNQGDYFIK | Q9CRA0 | Q9CRA0 | ART4 | ADP-ribosyltransferase 4 (Dombrock blood group) | Nucleus | enzyme |
| 15532433 | 3 | 0.004 | 3.32E+07 | 3.63E+07 | 3.50E+07 | 2.67E+07 | 2.72E+07 | 2.82E+07 | 3.38E+07 | 3.14E+07 | 2.86E+07 | 3.09E+07 | 2.85E+07 | 2.94E+07 | 1611.7506 | AAGSPGQYSQSFHFK | Q9CRA0 | Q9CRA0 | ART4 | ADP-ribosyltransferase 4 (Dombrock blood group) | Nucleus | enzyme |
| 15539165 | 2 | 0.069 | 1.43E+07 | 9317760 | 1.04E+07 | 9870896 | 8270045 | 6287461.5 | 9367681 | 8855587 | 7016137.5 | 1.20E+07 | 1.35E+07 | 1.06E+07 | 1181.64 | AHLTWLNQAK | Q9CRA0 | Q9CRA0 | ART4 | ADP-ribosyltransferase 4 (Dombrock blood group) | Nucleus | enzyme |
| 15531954 | 2 | 0.233 | 7382318.5 | 7360850 | 1.00E+07 | 6105257.5 | 6214657 | 7323918.5 | 7334498.5 | 6724710.5 | 7895462.5 | 8256485.5 | 6777426 | 7319884 | 800.4605 | GDLINLR | Q9CRA0 | Q9CRA0 | ART4 | ADP-ribosyltransferase 4 (Dombrock blood group) | Nucleus | enzyme |
| 15549299 | 2 | 0.445 | 6768201.5 | 5224146 | 4911119.5 | 4464644.5 | 4018885.75 | 5056130.5 | 6035187 | 4347425 | 4367436 | 5730085 | 4183508 | 4064367.5 | 1611.7512 | AAGSPGQYSQSFHFK | Q9CRA0 | Q9CRA0 | ART4 | ADP-ribosyltransferase 4 (Dombrock blood group) | Nucleus | enzyme |
| 15544486 | 3 | 0.09 | 3178180 | 3871622 | 2825130.25 | 2138841.5 | 2521924 | 2803852.5 | 3857009 | 3708361.5 | 3192380 | 1525266.5 | 2552068.5 | 3031961.75 | 1753.9597 | YLHYYLTSAIQLLR | Q9CRA0 | Q9CRA0 | ART4 | ADP-ribosyltransferase 4 (Dombrock blood group) | Nucleus | enzyme |
| 15544135, 15546171 | 3 | 0.708 | 2225341.75 | 784754.438 | 5220284 | 2130033 | 2161355.75 | 3746650.25 | 2719010.5 | 4468148 | 3333255.5 | 1132105.125 | 2548592.25 | 3248945.25 | 1761.9641 | EVLIPPYELFEVVSK | Q9CRA0 | Q9CRA0 | ART4 | ADP-ribosyltransferase 4 (Dombrock blood group) | Nucleus | enzyme |
| 15553788 | 3 | 0.973 | 3490032.25 | 2847425.5 | 3942353.25 | 3663400.75 | 3450699 | 2709231 | 3560855.25 | 2598876 | 3682820 | 4961593.5 | 2889137.25 | 2910980.75 | 2535.1697 | AIYNSGQDHVQNEENYAQVLDK | Q9QWY8 | Q9QWY8 | ASAP1 | ArfGAP with SH3 domain, ankyrin repeat and PH domain 1 | Plasma Membrane | other |
| 15543561 | 2 | 0.617 | 1696136.125 | 1566503 | 1397010.875 | 1810370.25 | 1381255.125 | 965191.812 | 1787224 | 987655.812 | 1360411.75 | 2528839.75 | 1646532.375 | 1283157.125 | 1440.6937 | FEGLSQQASTSSAK | Q9QWY8 | Q9QWY8 | ASAP1 | ArfGAP with SH3 domain, ankyrin repeat and PH domain 1 | Plasma Membrane | other |
| 15539934 | 2 | 0.144 | 1.20E+07 | 1.24E+07 | 4466174 | 1.54E+07 | 1.24E+07 | 3.07E+07 | 1.07E+07 | 5155600.5 | 1.19E+07 | 1.33E+07 | 1.90E+07 | 1.87E+07 | 973.5293 | EIISEVQR | Q7SIG6 | Q7SIG6 | ASAP2 | ArfGAP with SH3 domain, ankyrin repeat and PH domain 2 | Nucleus | other |
| 15532061 | 2 | 0.389 | 8482723 | 8788258 | 5472545 | 9070696 | 7111267 | 6158828.5 | 6185663 | 8235535.5 | 7079676 | 6690752.5 | 5823183.5 | 4845228.5 | 963.5479 | KDFLEALK | Q9WV71 | Q9WV71 | ASB4 | ankyrin repeat and SOCS box containing 4 | Nucleus | transcription regulator |
| 15534379 | 2 | 0.281 | 1.33E+07 | 1.36E+07 | 1.35E+07 | 1.82E+07 | 9606087 | 1.64E+07 | 1.57E+07 | 1.06E+07 | 1.29E+07 | 1.72E+07 | 1.80E+07 | 1.79E+07 | 1117.5933 | GVKNSSLEQR | Q4QY64 | Q4QY64 | ATAD5 | ATPase family, AAA domain containing 5 | Extracellular Space | other |
| 15541709 | 2 | 0.076 | 6.16E+07 | 6.52E+07 | 6.66E+07 | 7.36E+07 | 5.72E+07 | 4.58E+07 | 6.08E+07 | 6.67E+07 | 7.06E+07 | 1.18E+08 | 7.27E+07 | 8.36E+07 | 822.51776 | QLLLAHK | P16951 | P16951 | ATF2 | activating transcription factor 2 | Nucleus | transcription regulator |
| 15545473 | 2 | 0.889 | 1116163.125 | 680845.25 | 855979.375 | 978144.25 | 565415.938 | 1002484.188 | 912470 | 895156 | 850212.062 | 1414573.5 | 990145.438 | 667051.688 | 795.4375 | YSWVLK | Q811C2 | Q811C2 | ATG4C | autophagy related 4C, cysteine peptidase | Cytoplasm | peptidase |
| 15548769 | 3 | 0.699 | 1.16E+07 | 1.16E+07 | 1.19E+07 | 8026591 | 9383494 | 9990447 | 1.18E+07 | 7623802.5 | 5119889 | 1.83E+07 | 3839903.25 | 8233701.5 | 2578.2957 | RAEISNAIDQYVTGTIGEGEDLVK | Q9CWJ9 | Q9CWJ9 | ATIC | 5-aminoimidazole-4-carboxamide ribonucleotide formyltransferase/IMP cyclohydrolase | Cytoplasm | enzyme |
| 15550713 | 3 | 0.713 | 9395874 | 6943016.5 | 6922108.5 | 6012429 | 5260898 | 5961675 | 7308569 | 4986261 | 6418966 | 1.41E+07 | 4153529 | 6274783 | 2451.0752 | VTVVC[160.0307]EPEDYAGVAAEMHGSDSK | Q9CWJ9 | Q9CWJ9 | ATIC | 5-aminoimidazole-4-carboxamide ribonucleotide formyltransferase/IMP cyclohydrolase | Cytoplasm | enzyme |
| 15533885 | 3 | 0.302 | 1.04E+07 | 5636892.5 | 5235079.5 | 6979489.5 | 2309548.25 | 3855792.5 | 5263305.5 | 2306283 | 2988615.25 | 9375410 | 3600440.75 | 5059130 | 2215.0762 | LSGVSVSSDAFFPFRDNVDR | Q9CWJ9 | Q9CWJ9 | ATIC | 5-aminoimidazole-4-carboxamide ribonucleotide formyltransferase/IMP cyclohydrolase | Cytoplasm | enzyme |
| 15535524 | 3 | 0.584 | 5923025 | 4215332 | 3941694.75 | 4619380.5 | 3628494.5 | 3542098.75 | 4603485 | 3842946.25 | 2384423 | 9056291 | 3287543.25 | 4194730.5 | 1861.9216 | YGMNPHQTPAQLYTLK | Q9CWJ9 | Q9CWJ9 | ATIC | 5-aminoimidazole-4-carboxamide ribonucleotide formyltransferase/IMP cyclohydrolase | Cytoplasm | enzyme |
| 15550772 | 2 | 0.607 | 5759279.5 | 4042565.75 | 4518898 | 4252438 | 3546395.25 | 4062541 | 3926471 | 3621062.75 | 2375582.25 | 8395289 | 2474540.5 | 3953776.5 | 2159.9941 | MSSFGDFVALSDIC[160.0307]DVPTAK | Q9CWJ9 | Q9CWJ9 | ATIC | 5-aminoimidazole-4-carboxamide ribonucleotide formyltransferase/IMP cyclohydrolase | Cytoplasm | enzyme |
| 15550280 | 2 | 0.732 | 3544694.75 | 5760277 | 7053795.5 | 3038481.5 | 4223376 | 5523299.5 | 4985675.5 | 4257770 | 2717767.5 | 7872085.5 | 1391482.125 | 3152852.5 | 2033.0063 | WEALFEEVPELLTEAEK | Q9CWJ9 | Q9CWJ9 | ATIC | 5-aminoimidazole-4-carboxamide ribonucleotide formyltransferase/IMP cyclohydrolase | Cytoplasm | enzyme |
| 15536724 | 3 | 0.589 | 4422977.5 | 2871799.75 | 3841765.5 | 3266133.5 | 2073328.25 | 3017099 | 3486692.25 | 2817058.25 | 1225427.25 | 6835483 | 1862974.875 | 3314272.75 | 1861.9358 | HVSPAGAAVGVPLSEDEAR | Q9CWJ9 | Q9CWJ9 | ATIC | 5-aminoimidazole-4-carboxamide ribonucleotide formyltransferase/IMP cyclohydrolase | Cytoplasm | enzyme |
| 15543096 | 2 | 0.803 | 2397036.25 | 1333618.375 | 1552362.125 | 1676564.875 | 1276415.75 | 1601657.5 | 1991232.125 | 1273849.625 | 959946 | 3814508.5 | 1174212.75 | 1360032.125 | 1320.5865 | YTQSNSVC[160.0307]YAK | Q9CWJ9 | Q9CWJ9 | ATIC | 5-aminoimidazole-4-carboxamide ribonucleotide formyltransferase/IMP cyclohydrolase | Cytoplasm | enzyme |
| 15542891 | 3 | 0.972 | 516474.781 | 1670513.5 | 1743403 | 443575.406 | 1424389.875 | 1566960.375 | 1395428.75 | 886129.562 | 584746.25 | 2712634.25 | 292932.281 | 843935.938 | 2033.0112 | WEALFEEVPELLTEAEK | Q9CWJ9 | Q9CWJ9 | ATIC | 5-aminoimidazole-4-carboxamide ribonucleotide formyltransferase/IMP cyclohydrolase | Cytoplasm | enzyme |
| 15541926 | 2 | 0.557 | 2024127.75 | 2172585 | 2601013.25 | 648661.625 | 2049686.875 | 2158437.75 | 1838588 | 2018852.75 | 1982221 | 2176307.5 | 999395.562 | 1977253.375 | 1385.7109 | DGQVIGIGAGQQSR | Q9CWJ9 | Q9CWJ9 | ATIC | 5-aminoimidazole-4-carboxamide ribonucleotide formyltransferase/IMP cyclohydrolase | Cytoplasm | enzyme |
| 15548709 | 2 | 0.272 | 5779194 | 6554889.5 | 7576699 | 5345405.5 | 5858325 | 6735228.5 | 4578341 | 5335934.5 | 5530440 | 5746725.5 | 4557817 | 6868219.5 | 1661.9789 | NLVPLLLAPENLVEK | Q6PA06 | Q6PA06 | ATL2 | atlastin GTPase 2 | Cytoplasm | other |
| 15550497 | 2 | 0.101 | 1564572.125 | 2261680 | 2192280 | 2371042.75 | 2343803.75 | 1873253.25 | 1710224.375 | 1974426.5 | 2670339.5 | 1550851.625 | 1402108.625 | 1566978.75 | 1858.9414 | SMLQATAEANNLAAVAGAR | Q6PA06 | Q6PA06 | ATL2 | atlastin GTPase 2 | Cytoplasm | other |
| 15541574 | 2 | 0.738 | 1172948.625 | 1650280.875 | 2492965.5 | 1439278.125 | 1777795 | 1659390.625 | 1101638 | 1493251.625 | 2198995 | 1357422.25 | 1278183.875 | 1400569.125 | 1063.5143 | VATNPSFDGR | Q6PA06 | Q6PA06 | ATL2 | atlastin GTPase 2 | Cytoplasm | other |
| 15547902, 15554760 | 2, 3 | 0.538 | 5.31E+07 | 6.08E+07 | 5.88E+07 | 5.24E+07 | 5.81E+07 | 5.00E+07 | 6.52E+07 | 5.82E+07 | 5.52E+07 | 6.55E+07 | 5.34E+07 | 5.44E+07 | 2372.0515, 2372.0562 | EGGHSDWLGDPEEPLTGFSWR | Q91YH5 | Q91YH5 | ATL3 | atlastin GTPase 3 | Cytoplasm | other |
| 15549239 | 2 | 0.477 | 3.27E+07 | 3.62E+07 | 3.59E+07 | 3.35E+07 | 3.42E+07 | 3.05E+07 | 3.12E+07 | 3.24E+07 | 3.62E+07 | 3.34E+07 | 3.12E+07 | 3.27E+07 | 1712.9543 | EQLQALIPYVLNPSK | Q91YH5 | Q91YH5 | ATL3 | atlastin GTPase 3 | Cytoplasm | other |
| 15555195 | 4 | 0.004 | 2.13E+07 | 1.93E+07 | 1.53E+07 | 1.70E+07 | 1.98E+07 | 1.62E+07 | 2.48E+07 | 2.18E+07 | 2.41E+07 | 2.87E+07 | 3.59E+07 | 2.66E+07 | 3495.6492 | NHIHSC[160.0307]FSDVTC[160.0307]FLLPHPGLQVATSPNFDGK | Q91YH5 | Q91YH5 | ATL3 | atlastin GTPase 3 | Cytoplasm | other |
| 15534270 | 3 | 0.373 | 2.13E+07 | 1.79E+07 | 2.11E+07 | 1.75E+07 | 1.91E+07 | 1.87E+07 | 2.16E+07 | 1.80E+07 | 1.95E+07 | 2.71E+07 | 2.32E+07 | 1.78E+07 | 1296.655 | IYQGEDLPHPK | Q91YH5 | Q91YH5 | ATL3 | atlastin GTPase 3 | Cytoplasm | other |
| 15531445 | 2 | 0.227 | 2.53E+07 | 2.75E+07 | 2.98E+07 | 2.27E+07 | 2.78E+07 | 2.08E+07 | 2.71E+07 | 2.63E+07 | 2.84E+07 | 2.69E+07 | 2.76E+07 | 2.50E+07 | 1303.6753 | PYLSPDILEEK | Q91YH5 | Q91YH5 | ATL3 | atlastin GTPase 3 | Cytoplasm | other |
| 15531768 | 2 | 0.372 | 2.30E+07 | 2.26E+07 | 2.58E+07 | 1.94E+07 | 2.48E+07 | 2.16E+07 | 2.47E+07 | 2.26E+07 | 2.39E+07 | 2.49E+07 | 2.69E+07 | 2.29E+07 | 1308.6296 | YQQELEEEIK | Q91YH5 | Q91YH5 | ATL3 | atlastin GTPase 3 | Cytoplasm | other |
| 15547808 | 3 | 0.066 | 2.94E+07 | 3.60E+07 | 3.33E+07 | 2.80E+07 | 4.05E+07 | 2.55E+07 | 2.85E+07 | 3.42E+07 | 3.39E+07 | 2.32E+07 | 2.25E+07 | 2.44E+07 | 2972.4639 | ELGGAIDSGAAYVLEQASSHIGNSTQAAVR | Q91YH5 | Q91YH5 | ATL3 | atlastin GTPase 3 | Cytoplasm | other |
| 15548247 | 2 | 0.86 | 1.97E+07 | 2.10E+07 | 2.09E+07 | 2.15E+07 | 2.04E+07 | 1.82E+07 | 2.12E+07 | 1.92E+07 | 2.01E+07 | 2.10E+07 | 2.01E+07 | 1.77E+07 | 1832.9132 | SMLQATAEANNLAAAASAK | Q91YH5 | Q91YH5 | ATL3 | atlastin GTPase 3 | Cytoplasm | other |
| 15533192 | 2 | 0.829 | 1.66E+07 | 1.76E+07 | 1.88E+07 | 1.76E+07 | 2.03E+07 | 1.59E+07 | 2.08E+07 | 1.66E+07 | 1.93E+07 | 1.89E+07 | 1.75E+07 | 1.76E+07 | 1094.5532 | LAMDEIFQK | Q91YH5 | Q91YH5 | ATL3 | atlastin GTPase 3 | Cytoplasm | other |
| 15538024 | 2 | 0.303 | 1.14E+07 | 1.20E+07 | 9815630 | 8817311 | 1.13E+07 | 1.09E+07 | 1.01E+07 | 9950593 | 1.05E+07 | 1.05E+07 | 7871976.5 | 9318293 | 1265.7048 | PFQTLMFLIR | Q91YH5 | Q91YH5 | ATL3 | atlastin GTPase 3 | Cytoplasm | other |
| 15533229 | 2, 3 | 0.802 | 7412292 | 1.10E+07 | 1.19E+07 | 5972772 | 1.25E+07 | 1.03E+07 | 1.15E+07 | 1.16E+07 | 7859714.5 | 9258101 | 7542681.5 | 8438100 | 1335.7705, 1335.7706 | ALASVLLQDHIR | Q91YH5 | Q91YH5 | ATL3 | atlastin GTPase 3 | Cytoplasm | other |
| 15538276 | 2 | 0.77 | 4845286.5 | 5777435 | 6918269 | 3728524.25 | 6313440 | 6334940 | 6235776 | 6455172 | 5559989 | 7186214 | 6183320.5 | 5364611 | 870.4451 | NVFSTFR | Q91YH5 | Q91YH5 | ATL3 | atlastin GTPase 3 | Cytoplasm | other |
| 15532448 | 2 | 0.257 | 6060225 | 7014062.5 | 5516481 | 4692125 | 5724483.5 | 5165560 | 6020405.5 | 5661997 | 7371430.5 | 6424005.5 | 6460958 | 5217709 | 869.47424 | GLLEYFK | Q91YH5 | Q91YH5 | ATL3 | atlastin GTPase 3 | Cytoplasm | other |
| 15533454 | 2 | 0.258 | 4543284 | 4477150 | 5452693 | 4415335.5 | 5248775.5 | 4684785 | 5578581 | 5020796 | 4359095 | 6175788 | 5353624 | 5345477 | 1303.5885 | EQHSFELEER | Q91YH5 | Q91YH5 | ATL3 | atlastin GTPase 3 | Cytoplasm | other |
| 15539515 | 2 | 0.285 | 4133710.25 | 2566480.25 | 2754793.75 | 4641468 | 3381196.5 | 3110471.5 | 5322697 | 3019124.5 | 3940410.25 | 5878640.5 | 4225414 | 4029663 | 1141.6057 | SFILDFMLR | Q91YH5 | Q91YH5 | ATL3 | atlastin GTPase 3 | Cytoplasm | other |
| 15535989 | 2 | 0.114 | 4144815 | 6371800 | 6342582 | 3353097.25 | 5121145 | 4814366.5 | 5390285.5 | 5137613 | 4659071 | 4538636 | 2445662.75 | 3331203.75 | 1281.7003 | PFQTLM[147.0354]FLIR | Q91YH5 | Q91YH5 | ATL3 | atlastin GTPase 3 | Cytoplasm | other |
| 15533724 | 3 | 0.615 | 1.22E+07 | 9902421 | 1.01E+07 | 1.10E+07 | 1.41E+07 | 1.07E+07 | 1.11E+07 | 1.06E+07 | 9624194 | 1.19E+07 | 1.20E+07 | 9493398 | 2086.0676 | TVFVGNHPISGTEPYIAQR | Q9QZW0 | Q9QZW0 | ATP11C | ATPase, class VI, type 11C | Plasma Membrane | transporter |
| 15553488 | 3 | 0.146 | 1.20E+07 | 1.17E+07 | 1.22E+07 | 1.02E+07 | 1.23E+07 | 9861329 | 9520511 | 9908568 | 9989875 | 1.19E+07 | 7927912 | 8819225 | 2342.0054 | TGTLTENSMEFIEC[160.0307]C[160.0307]IDGHK | Q9QZW0 | Q9QZW0 | ATP11C | ATPase, class VI, type 11C | Plasma Membrane | transporter |
| 15548664 | 2 | 0.078 | 7061061 | 6543919.5 | 6119707 | 5648226 | 5848202 | 6064737 | 6791158.5 | 5632330.5 | 5268114 | 5748351.5 | 4995090.5 | 5062351 | 1650.8911 | LFQTNTELLELTTK | Q9QZW0 | Q9QZW0 | ATP11C | ATPase, class VI, type 11C | Plasma Membrane | transporter |
| 15554825 | 2 | 0.123 | 4894664 | 4372446.5 | 5298418.5 | 4291182.5 | 4880578.5 | 5016120 | 3538648.5 | 4722889 | 4103860.75 | 4637733.5 | 3231498.75 | 3176995.5 | 1719.7982 | ATIEC[160.0307]EQPQPDLYR | Q9QZW0 | Q9QZW0 | ATP11C | ATPase, class VI, type 11C | Plasma Membrane | transporter |
| 15536356 | 3 | 0.122 | 2945875.25 | 4937097.5 | 5485752 | 4388736.5 | 4388060 | 3984852.5 | 4339954.5 | 4799327.5 | 4128391.5 | 3297193.25 | 2181527.75 | 3406545 | 2107.0298 | EIEEYELLHTLNFDSVR | Q9QZW0 | Q9QZW0 | ATP11C | ATPase, class VI, type 11C | Plasma Membrane | transporter |
| 15540900 | 2 | 0.289 | 2457250 | 2997752.25 | 3078922.5 | 2362093.25 | 3255539.25 | 3132817 | 2873017.25 | 2630099.75 | 3015096.25 | 2556644 | 2155431.25 | 2516475.5 | 917.50757 | VWVLTGDK | Q9QZW0 | Q9QZW0 | ATP11C | ATPase, class VI, type 11C | Plasma Membrane | transporter |
| 15541941 | 2 | 0.982 | 1.35E+07 | 9282360 | 1.21E+07 | 1.10E+07 | 1.20E+07 | 1.25E+07 | 1.21E+07 | 1.19E+07 | 9807084 | 1.65E+07 | 7898092 | 1.32E+07 | 763.4704 | VLALGYK | Q9EPE9 | Q9EPE9 | ATP13A1 | ATPase type 13A1 | Extracellular Space | transporter |
| 15548380 | 2 | 0.319 | 3.36E+07 | 3.70E+07 | 3.94E+07 | 3.32E+07 | 3.71E+07 | 2.83E+07 | 3.04E+07 | 3.21E+07 | 3.37E+07 | 3.72E+07 | 2.71E+07 | 2.54E+07 | 1519.6976 | SPDFTNENPLETR | Q8VDN2 | Q8VDN2 | ATP1A1 | ATPase, Na+/K+ transporting, alpha 1 polypeptide | Plasma Membrane | transporter |
| 15534090 | 2 | 0.15 | 3.83E+07 | 3.47E+07 | 3.72E+07 | 3.09E+07 | 3.31E+07 | 2.37E+07 | 3.26E+07 | 4.06E+07 | 3.24E+07 | 3.65E+07 | 3.42E+07 | 3.20E+07 | 1261.6758 | IVEIPFNSTNK | Q8VDN2 | Q8VDN2 | ATP1A1 | ATPase, Na+/K+ transporting, alpha 1 polypeptide | Plasma Membrane | transporter |
| 15548727 | 2 | 0.403 | 3.20E+07 | 3.50E+07 | 3.20E+07 | 2.96E+07 | 3.21E+07 | 2.63E+07 | 2.83E+07 | 3.08E+07 | 2.88E+07 | 3.41E+07 | 2.99E+07 | 2.54E+07 | 1811.9457 | QGAIVAVTGDGVNDSPALK | Q8VDN2 | Q8VDN2 | ATP1A1 | ATPase, Na+/K+ transporting, alpha 1 polypeptide | Plasma Membrane | transporter |
| 15547804 | 2 | 0.064 | 3.25E+07 | 3.36E+07 | 4.32E+07 | 2.75E+07 | 3.02E+07 | 3.25E+07 | 2.96E+07 | 3.41E+07 | 3.91E+07 | 3.08E+07 | 2.25E+07 | 2.41E+07 | 1829.9205 | GVGIISEGNETVEDIAAR | Q8VDN2 | Q8VDN2 | ATP1A1 | ATPase, Na+/K+ transporting, alpha 1 polypeptide | Plasma Membrane | transporter |
| 15549438 | 2 | 0.184 | 3.12E+07 | 3.43E+07 | 2.89E+07 | 2.69E+07 | 3.10E+07 | 2.69E+07 | 2.30E+07 | 2.86E+07 | 2.97E+07 | 2.94E+07 | 2.19E+07 | 2.35E+07 | 1752.8512 | DAFQNAYLELGGLGER | Q8VDN2 | Q8VDN2 | ATP1A1 | ATPase, Na+/K+ transporting, alpha 1 polypeptide | Plasma Membrane | transporter |
| 15548389 | 2 | 0.034 | 3.13E+07 | 3.45E+07 | 3.41E+07 | 2.58E+07 | 2.63E+07 | 2.31E+07 | 2.19E+07 | 2.92E+07 | 2.95E+07 | 2.54E+07 | 2.40E+07 | 1.77E+07 | 1584.8728 | AVFQANQENLPILK | Q8VDN2 | Q8VDN2 | ATP1A1 | ATPase, Na+/K+ transporting, alpha 1 polypeptide | Plasma Membrane | transporter |
| 15555769 | 3 | 0.505 | 2.33E+07 | 2.37E+07 | 1.76E+07 | 1.98E+07 | 1.80E+07 | 1.83E+07 | 2.09E+07 | 1.78E+07 | 2.02E+07 | 2.13E+07 | 1.82E+07 | 1.78E+07 | 3105.3184 | ETWDDRWVNDVEDSYGQQWTYEQR | Q8VDN2 | Q8VDN2 | ATP1A1 | ATPase, Na+/K+ transporting, alpha 1 polypeptide | Plasma Membrane | transporter |
| 15535600 | 2 | 0.329 | 2.39E+07 | 2.42E+07 | 2.09E+07 | 2.18E+07 | 2.01E+07 | 1.27E+07 | 1.98E+07 | 2.18E+07 | 1.88E+07 | 2.09E+07 | 1.87E+07 | 1.89E+07 | 1268.712 | NMVPQQALVIR | Q8VDN2 | Q8VDN2 | ATP1A1 | ATPase, Na+/K+ transporting, alpha 1 polypeptide | Plasma Membrane | transporter |
| 15552797 | 2 | 0.296 | 1.97E+07 | 1.82E+07 | 2.34E+07 | 2.06E+07 | 1.86E+07 | 1.62E+07 | 1.77E+07 | 1.82E+07 | 1.87E+07 | 1.99E+07 | 1.53E+07 | 1.50E+07 | 1619.7466 | VDNSSLTGESEPQTR | Q8VDN2 | Q8VDN2 | ATP1A1 | ATPase, Na+/K+ transporting, alpha 1 polypeptide | Plasma Membrane | transporter |
| 15531590 | 2 | 0.204 | 2.23E+07 | 1.93E+07 | 1.88E+07 | 1.86E+07 | 1.66E+07 | 1.59E+07 | 1.80E+07 | 1.76E+07 | 2.00E+07 | 1.92E+07 | 2.11E+07 | 1.71E+07 | 1139.5818 | TSATWFALSR | Q8VDN2 | Q8VDN2 | ATP1A1 | ATPase, Na+/K+ transporting, alpha 1 polypeptide | Plasma Membrane | transporter |
| 15531949 | 2, 3 | 0.315 | 2.08E+07 | 2.04E+07 | 1.76E+07 | 2.00E+07 | 1.69E+07 | 1.49E+07 | 1.76E+07 | 1.65E+07 | 1.76E+07 | 1.87E+07 | 1.48E+07 | 1.67E+07 | 1110.6238, 1110.6244 | LSLDELHRK | Q8VDN2 | Q8VDN2 | ATP1A1 | ATPase, Na+/K+ transporting, alpha 1 polypeptide | Plasma Membrane | transporter |
| 15554384 | 4 | 0.752 | 1.34E+07 | 1.46E+07 | 9902374 | 1.26E+07 | 1.18E+07 | 9449345 | 1.10E+07 | 1.13E+07 | 1.08E+07 | 1.37E+07 | 9498249 | 1.04E+07 | 3238.4136 | MTVAHMWFDNQIHEADTTENQSGVSFDK | Q8VDN2 | Q8VDN2 | ATP1A1 | ATPase, Na+/K+ transporting, alpha 1 polypeptide | Plasma Membrane | transporter |
| 15535413 | 2 | 0.299 | 1.27E+07 | 1.32E+07 | 1.26E+07 | 1.08E+07 | 1.20E+07 | 1.06E+07 | 1.21E+07 | 1.33E+07 | 1.18E+07 | 1.36E+07 | 1.14E+07 | 1.00E+07 | 979.51917 | GIVVYTGDR | Q8VDN2 | Q8VDN2 | ATP1A1 | ATPase, Na+/K+ transporting, alpha 1 polypeptide | Plasma Membrane | transporter |
| 15535264 | 2 | 0.584 | 1.30E+07 | 1.12E+07 | 1.07E+07 | 1.30E+07 | 1.00E+07 | 7441013.5 | 1.05E+07 | 9599230 | 9196227 | 1.35E+07 | 1.10E+07 | 9134823 | 1100.5416 | EQPLDEELK | Q8VDN2 | Q8VDN2 | ATP1A1 | ATPase, Na+/K+ transporting, alpha 1 polypeptide | Plasma Membrane | transporter |
| 15556464 | 2 | 0.755 | 1.01E+07 | 1.39E+07 | 9383740 | 9457678 | 1.14E+07 | 1.03E+07 | 1.06E+07 | 1.22E+07 | 1.18E+07 | 1.10E+07 | 8935065 | 1.13E+07 | 1816.9316 | MSINAEDVVVGDLVEVK | Q8VDN2 | Q8VDN2 | ATP1A1 | ATPase, Na+/K+ transporting, alpha 1 polypeptide | Plasma Membrane | transporter |
| 15533460 | 2 | 0.109 | 1.08E+07 | 1.45E+07 | 1.56E+07 | 9448187 | 1.30E+07 | 1.16E+07 | 1.29E+07 | 1.44E+07 | 1.28E+07 | 1.08E+07 | 6808627 | 1.06E+07 | 862.5013 | LIFDNLK | Q8VDN2 | Q8VDN2 | ATP1A1 | ATPase, Na+/K+ transporting, alpha 1 polypeptide | Plasma Membrane | transporter |
| 15549821 | 2 | 0.193 | 9409468 | 1.40E+07 | 1.29E+07 | 9064645 | 1.21E+07 | 9470847 | 9397707 | 1.36E+07 | 1.25E+07 | 1.08E+07 | 6310456 | 8163742.5 | 2464.1982 | QAADMILLDDNFASIVTGVEEGR | Q8VDN2 | Q8VDN2 | ATP1A1 | ATPase, Na+/K+ transporting, alpha 1 polypeptide | Plasma Membrane | transporter |
| 15533452 | 3 | 0.105 | 6536166 | 8836742 | 5657808.5 | 4970028.5 | 5515599.5 | 4463771 | 5833054.5 | 6249016.5 | 6205455 | 7016080.5 | 5668538 | 5809364.5 | 1135.5873 | YHTEIVFAR | Q8VDN2 | Q8VDN2 | ATP1A1 | ATPase, Na+/K+ transporting, alpha 1 polypeptide | Plasma Membrane | transporter |
| 15540029 | 2 | 0.131 | 7473941.5 | 7291999 | 6589497.5 | 6477543.5 | 6743990 | 5345311 | 6659386 | 6761176 | 6578482 | 6509640.5 | 6124833 | 5628164 | 1038.5018 | NSVFQQGMK | Q8VDN2 | Q8VDN2 | ATP1A1 | ATPase, Na+/K+ transporting, alpha 1 polypeptide | Plasma Membrane | transporter |
| 15533190 | 2 | 0.317 | 4892341 | 8382456 | 8449559 | 3986630.75 | 9119403 | 5549081.5 | 5959317.5 | 8644353 | 6016756 | 5942002.5 | 3392989.5 | 4189935 | 1231.65 | AVAGDASESALLK | Q8VDN2 | Q8VDN2 | ATP1A1 | ATPase, Na+/K+ transporting, alpha 1 polypeptide | Plasma Membrane | transporter |
| 15559230 | 2 | 0.449 | 5936675.5 | 6880251.5 | 7795315.5 | 5104016 | 7171029.5 | 6449788.5 | 5808957 | 7019770 | 5326004.5 | 5521442 | 5576213 | 6000720 | 1924.9126 | NLEAVETLGSTSTIC[160.0307]SDK | Q8VDN2 | Q8VDN2 | ATP1A1 | ATPase, Na+/K+ transporting, alpha 1 polypeptide | Plasma Membrane | transporter |
| 15543749, 15545687 | 3 | 0.591 | 3449582.5 | 4440887.5 | 2964465.75 | 4173564 | 4532870.5 | 3591052.25 | 2714836.75 | 3667984 | 3683509 | 5144643.5 | 2995472.25 | 4025440.25 | 1752.8518, 1752.8528 | DAFQNAYLELGGLGER | Q8VDN2 | Q8VDN2 | ATP1A1 | ATPase, Na+/K+ transporting, alpha 1 polypeptide | Plasma Membrane | transporter |
| 15531639 | 2 | 0.065 | 6820444 | 6000135.5 | 5678493.5 | 4481470 | 5629759.5 | 3956826.5 | 5694625.5 | 5690236.5 | 5234962.5 | 5042261 | 5535017 | 5045738.5 | 1381.7484 | VIMVTGDHPITAK | Q8VDN2 | Q8VDN2 | ATP1A1 | ATPase, Na+/K+ transporting, alpha 1 polypeptide | Plasma Membrane | transporter |
| 15539331 | 2, 3 | 0.164 | 4970260.5 | 8556989 | 1.11E+07 | 3172186 | 7540850.5 | 7735720 | 6490176.5 | 7789256.5 | 5255165 | 3418501.25 | 3702842 | 4475377.5 | 1302.5931, 1302.5936 | YEPAAVSEHGDK | Q8VDN2 | Q8VDN2 | ATP1A1 | ATPase, Na+/K+ transporting, alpha 1 polypeptide | Plasma Membrane | transporter |
| 15536254 | 3 | 0.381 | 4707349.5 | 5856939 | 8382100 | 4535880 | 6724335.5 | 4420258 | 5662178 | 6928967.5 | 7750435.5 | 3132450.75 | 4571305.5 | 6696864.5 | 1829.9193 | GVGIISEGNETVEDIAAR | Q8VDN2 | Q8VDN2 | ATP1A1 | ATPase, Na+/K+ transporting, alpha 1 polypeptide | Plasma Membrane | transporter |
| 15539709 | 3 | 0.351 | 3489912.25 | 2492793 | 2664629 | 3298550 | 1972703.25 | 2445918.5 | 2790942 | 2266281.75 | 2950854.5 | 2930533 | 3955678.5 | 3187777 | 1387.7512 | RAVAGDASESALLK | Q8VDN2 | Q8VDN2 | ATP1A1 | ATPase, Na+/K+ transporting, alpha 1 polypeptide | Plasma Membrane | transporter |
| 15544581 | 3 | 0.308 | 2286002.5 | 3418997.25 | 2027691.125 | 2105940 | 2673004.75 | 2216575.5 | 2953167 | 3445490.5 | 2857204.25 | 2548940.25 | 1846766.25 | 2740533.25 | 1816.932 | MSINAEDVVVGDLVEVK | Q8VDN2 | Q8VDN2 | ATP1A1 | ATPase, Na+/K+ transporting, alpha 1 polypeptide | Plasma Membrane | transporter |
| 15546141 | 3 | 0.206 | 1912672.375 | 954795.375 | 1037909.875 | 1602253.75 | 1005601 | 750263.312 | 1195311.75 | 971405.312 | 1207517 | 2293840 | 1906027.875 | 1383651.375 | 1811.9464 | QGAIVAVTGDGVNDSPALK | Q8VDN2 | Q8VDN2 | ATP1A1 | ATPase, Na+/K+ transporting, alpha 1 polypeptide | Plasma Membrane | transporter |
| 15543172, 15546460 | 2 | 0.236 | 3269145.25 | 3544418.75 | 3156833.75 | 1723359 | 2809041.25 | 3010980.25 | 2626659.25 | 3410976.75 | 3104105.75 | 2183116.75 | 2912123 | 2742945.5 | 811.3926, 811.39276 | YGTDLSR | Q8VDN2 | Q8VDN2 | ATP1A1 | ATPase, Na+/K+ transporting, alpha 1 polypeptide | Plasma Membrane | transporter |
| 15547945 | 3 | 0.513 | 6.87E+07 | 6.20E+07 | 6.30E+07 | 6.71E+07 | 6.29E+07 | 5.30E+07 | 6.29E+07 | 5.18E+07 | 4.92E+07 | 7.31E+07 | 5.74E+07 | 4.95E+07 | 2958.3289 | TQLGDC[160.0307]SGIGDPTHYGYSTGQPC[160.0307]VFIK | P14231 | P14231 | ATP1B2 | ATPase, Na+/K+ transporting, beta 2 polypeptide | Plasma Membrane | transporter |
| 15535244 | 3 | 0.241 | 4.22E+07 | 4.04E+07 | 3.59E+07 | 3.59E+07 | 3.40E+07 | 2.74E+07 | 3.42E+07 | 3.18E+07 | 2.76E+07 | 4.13E+07 | 2.87E+07 | 3.03E+07 | 1545.7466 | INAANIATDDERDK | P14231 | P14231 | ATP1B2 | ATPase, Na+/K+ transporting, beta 2 polypeptide | Plasma Membrane | transporter |
| 15536458 | 3 | 0.014 | 1.00E+07 | 9094637 | 8480550 | 8036977.5 | 7820937.5 | 7362742.5 | 7695976 | 7117454 | 6610249.5 | 9633963 | 7917677.5 | 9278553 | 1196.7157 | LATPGLMIRPK | P14231 | P14231 | ATP1B2 | ATPase, Na+/K+ transporting, beta 2 polypeptide | Plasma Membrane | transporter |
| 15548414 | 2 | 0.704 | 7274703 | 6301067 | 5859468.5 | 6057817.5 | 6076965.5 | 4297367 | 5938627 | 5319653 | 5339910 | 8182511 | 5004554 | 4551032 | 1545.7477 | INAANIATDDERDK | P14231 | P14231 | ATP1B2 | ATPase, Na+/K+ transporting, beta 2 polypeptide | Plasma Membrane | transporter |
| 15533802 | 2 | 0.509 | 3800491.75 | 2590612 | 2582560.75 | 3094578.5 | 2898047.5 | 2085686 | 2528254.25 | 2707380.5 | 2731706 | 3927156.75 | 3220352.5 | 2694094.25 | 1221.5536 | SC[160.0307]GQVVEEWK | P14231 | P14231 | ATP1B2 | ATPase, Na+/K+ transporting, beta 2 polypeptide | Plasma Membrane | transporter |
| 15550778 | 2 | 0.25 | 3426221 | 5931593.5 | 1780797.25 | 4277534 | 2946381.75 | 4208573 | 3384697.25 | 5133068 | 7502350.5 | 1667524.75 | 2991771.5 | 2861556 | 1942.0309 | IIDLIPDGYPQISC[160.0307]LPK | P97370 | P97370 | ATP1B3 | ATPase, Na+/K+ transporting, beta 3 polypeptide | Plasma Membrane | transporter |
| 15549601 | 2 | 0.139 | 2613103.75 | 2804765.75 | 1572689.625 | 1719060 | 2129996.75 | 2843763.5 | 2833127 | 3050116.75 | 5138442 | 1094043.125 | 2107849 | 2296720.5 | 1599.8153 | LFIYNPSSGEFLGR | P97370 | P97370 | ATP1B3 | ATPase, Na+/K+ transporting, beta 3 polypeptide | Plasma Membrane | transporter |
| 15539544 | 3 | 0.167 | 2060846.625 | 3655878.25 | 1626001.375 | 1707237.25 | 2095547 | 1987249.625 | 2301252.75 | 2776378 | 3193913.25 | 892384.688 | 1936690.125 | 1798436.5 | 1747.9175 | YRDQIPSPGLMVFPK | P97370 | P97370 | ATP1B3 | ATPase, Na+/K+ transporting, beta 3 polypeptide | Plasma Membrane | transporter |
| 15538400 | 2 | 0.042 | 1622155.25 | 3556071 | 2024881 | 1504280.5 | 2621272.5 | 2648211.25 | 2464791.25 | 4383864 | 4485995 | 740454.562 | 1327486.625 | 1738637.5 | 1192.644 | LVEDLESFLK | P97370 | P97370 | ATP1B3 | ATPase, Na+/K+ transporting, beta 3 polypeptide | Plasma Membrane | transporter |
| 15548476 | 2 | 0.139 | 4145930.75 | 6905910.5 | 6473206.5 | 1.00E+07 | 1.06E+07 | 1.06E+07 | 5750271.5 | 9571304 | 1.11E+07 | 2364437.25 | 7904156.5 | 6155909 | 1539.824 | DIVPGDIVEIAVGDK | O55143 | O55143 | ATP2A2 | ATPase, Ca++ transporting, cardiac muscle, slow twitch 2 | Cytoplasm | transporter |
| 15548749 | 2 | 0.221 | 2366220.5 | 5570998.5 | 1.01E+07 | 2.05E+07 | 1.99E+07 | 1.81E+07 | 1.16E+07 | 1.40E+07 | 2.28E+07 | 1593060.875 | 2.16E+07 | 1.24E+07 | 1952.926 | SMSVETLGC[160.0307]TSVIC[160.0307]SDK | O55143 | O55143 | ATP2A2 | ATPase, Ca++ transporting, cardiac muscle, slow twitch 2 | Cytoplasm | transporter |
| 15549286 | 2 | 0.085 | 1270840.875 | 3256487.5 | 3449805 | 4182196.75 | 7927473.5 | 6941506.5 | 3230310 | 4516647.5 | 6243152.5 | 1010624.688 | 3651827.5 | 2515131.75 | 1574.8604 | VDQSILTGESVSVIK | O55143 | O55143 | ATP2A2 | ATPase, Ca++ transporting, cardiac muscle, slow twitch 2 | Cytoplasm | transporter |
| 15539232 | 2 | 0.176 | 698139.125 | 2108494 | 2029000.875 | 3661277 | 4636806.5 | 3266197.5 | 2564219.25 | 3286296 | 4018114.75 | 676065.375 | 3351031.75 | 2607387.5 | 1408.6907 | IGIFGQDEDVTSK | O55143 | O55143 | ATP2A2 | ATPase, Ca++ transporting, cardiac muscle, slow twitch 2 | Cytoplasm | transporter |
| 15538471 | 2 | 0.102 | 910225.625 | 1703594.75 | 1638206.625 | 1999590.625 | 4481465.5 | 2648200 | 1377423.75 | 2521755.5 | 2550339 | 533587.562 | 1423178 | 1640212.125 | 1278.593 | EFDELSPSAQR | O55143 | O55143 | ATP2A2 | ATPase, Ca++ transporting, cardiac muscle, slow twitch 2 | Cytoplasm | transporter |
| 15535432 | 2 | 0.239 | 748435.188 | 1478620.75 | 1242401 | 1567434.375 | 3279730.5 | 2212243.25 | 1079762.625 | 2190519.5 | 2212523.25 | 303837.188 | 1553781.25 | 1428492.875 | 1072.5608 | NAENAIEALK | O55143 | O55143 | ATP2A2 | ATPase, Ca++ transporting, cardiac muscle, slow twitch 2 | Cytoplasm | transporter |
| 15547828 | 2 | 0.162 | 3899190.25 | 9316973 | 1.75E+07 | 3.14E+07 | 3.79E+07 | 3.26E+07 | 1.84E+07 | 2.54E+07 | 4.02E+07 | 3022483 | 2.94E+07 | 2.04E+07 | 1620.8119 | VGEATETALTC[160.0307]LVEK | Q64518 | Q64518 | ATP2A3 | ATPase, Ca++ transporting, ubiquitous | Cytoplasm | transporter |
| 15535548 | 2 | 0.152 | 2331896.25 | 4478951.5 | 5458065.5 | 1.39E+07 | 1.38E+07 | 1.70E+07 | 9722097 | 1.02E+07 | 1.36E+07 | 1686538.125 | 1.82E+07 | 1.05E+07 | 1468.6848 | TGTLTTNQMSVC[160.0307]R | Q64518 | Q64518 | ATP2A3 | ATPase, Ca++ transporting, ubiquitous | Cytoplasm | transporter |
| 15535531 | 2 | 0.047 | 691564.25 | 2746950 | 2938158.5 | 8727691 | 9375571 | 1.02E+07 | 5609867.5 | 7471670.5 | 9426381 | 1358915.875 | 7959052 | 5294448.5 | 1363.7118 | LGIFGDTEDVLGK | Q64518 | Q64518 | ATP2A3 | ATPase, Ca++ transporting, ubiquitous | Cytoplasm | transporter |
| 15537768 | 2 | 0.178 | 863393.125 | 1921315.375 | 4685970 | 8665942 | 9833986 | 8259964 | 6240040.5 | 6450668 | 1.23E+07 | 835203.438 | 1.04E+07 | 7411475 | 1082.5167 | MNVFDTDLK | Q64518 | Q64518 | ATP2A3 | ATPase, Ca++ transporting, ubiquitous | Cytoplasm | transporter |
| 15532037 | 3 | 0.203 | 943548.062 | 3812837.75 | 9651418 | 2.07E+07 | 2.07E+07 | 2.13E+07 | 1.22E+07 | 1.72E+07 | 2.72E+07 | 791212.938 | 2.28E+07 | 1.39E+07 | 2036.989 | LHEFTISGTTYTPEGEVR | Q64518 | Q64518 | ATP2A3 | ATPase, Ca++ transporting, ubiquitous | Cytoplasm | transporter |
| 15554359 | 3 | 0.085 | 298167.969 | 1038489.5 | 1986521.125 | 5405627.5 | 5935061 | 5752634 | 3295521.5 | 5990451 | 6756423.5 | 513953.25 | 5974276.5 | 4693777.5 | 3147.3982 | C[160.0307]GQFDGLVELATIC[160.0307]ALC[160.0307]NDSALDYNEAK | Q64518 | Q64518 | ATP2A3 | ATPase, Ca++ transporting, ubiquitous | Cytoplasm | transporter |
| 15549880 | 2 | 0.088 | 197587.25 | 870960.438 | 3581178.75 | 7742346.5 | 1.02E+07 | 1.03E+07 | 5126516 | 6742275.5 | 1.24E+07 | 451844.25 | 6787288 | 5925467 | 1525.8091 | DIVPGDIVEVAVGDK | Q64518 | Q64518 | ATP2A3 | ATPase, Ca++ transporting, ubiquitous | Cytoplasm | transporter |
| 15533412 | 3 | 0.255 | 789539.062 | 1282254 | 2881264.75 | 1.36E+07 | 5233159.5 | 1.13E+07 | 5565116.5 | 4907088 | 1.14E+07 | 406390.25 | 1.79E+07 | 8670085 | 1911.8793 | AYTGREFDDLSPEQQR | Q64518 | Q64518 | ATP2A3 | ATPase, Ca++ transporting, ubiquitous | Cytoplasm | transporter |
| 15539681 | 3 | 0.201 | 200368.312 | 1711221.875 | 2715115.25 | 6840914.5 | 6783646.5 | 7247861 | 4357770 | 5145127 | 8298976.5 | 375688.719 | 9108343 | 5088089.5 | 1362.674 | IRDWGSGSDTLR | Q64518 | Q64518 | ATP2A3 | ATPase, Ca++ transporting, ubiquitous | Cytoplasm | transporter |
| 15538815 | 3 | 0.209 | 393799.469 | 1890121.5 | 4534231 | 1.19E+07 | 9220781 | 1.28E+07 | 5992374.5 | 7969165 | 1.29E+07 | 360334.25 | 1.17E+07 | 8189618 | 1712.8711 | SQMAAVEPERTPLQR | Q64518 | Q64518 | ATP2A3 | ATPase, Ca++ transporting, ubiquitous | Cytoplasm | transporter |
| 15554663 | 3 | 0.21 | 80995.766 | 77790.359 | 0 | 2321967 | 8706.535 | 951268.938 | 258071.062 | 73976.656 | 2303870.75 | 293732.938 | 3259874.5 | 2159574.25 | 2990.4849 | SRIVENLQSFNEITAMTGDGVNDAPALK | Q64518 | Q64518 | ATP2A3 | ATPase, Ca++ transporting, ubiquitous | Cytoplasm | transporter |
| 15540986 | 2 | 0.074 | 108844.375 | 269886.625 | 745066.875 | 1083913.875 | 2263243 | 1655546.375 | 984976.438 | 1790053.625 | 1466182 | 98958.156 | 1038232.125 | 647382.125 | 1117.5292 | SQMAAVEPER | Q64518 | Q64518 | ATP2A3 | ATPase, Ca++ transporting, ubiquitous | Cytoplasm | transporter |
| 15553972 | 2 | 0.217 | 0 | 223246.234 | 513107.75 | 1673188 | 1443952.875 | 1501839.75 | 638709.062 | 934806.312 | 1707628.375 | 42089.805 | 2228157.5 | 1022111.938 | 1712.8717 | SQMAAVEPERTPLQR | Q64518 | Q64518 | ATP2A3 | ATPase, Ca++ transporting, ubiquitous | Cytoplasm | transporter |
| 15543337 | 2 | 0.336 | 5255.968 | 734233.25 | 2261549.25 | 3487138.25 | 6457057 | 4452122.5 | 3108770.5 | 4128281.25 | 3831503 | 7771.243 | 3978208 | 2654645.25 | 1363.6085 | EFDDLSPEQQR | Q64518 | Q64518 | ATP2A3 | ATPase, Ca++ transporting, ubiquitous | Cytoplasm | transporter |
| 15545694 | 2 | 0.228 | 61439.359 | 107261.984 | 250507.828 | 1124603.5 | 1212971.875 | 1416023.5 | 452231.938 | 724507.25 | 1612270.875 | 5623.83 | 1356851.375 | 654877.25 | 957.49896 | GAPESVIER | Q64518 | Q64518 | ATP2A3 | ATPase, Ca++ transporting, ubiquitous | Cytoplasm | transporter |
| 15556074 | 2 | 0.371 | 78347.227 | 973220.438 | 3249582.75 | 5208655.5 | 7543926.5 | 9260906 | 3679664.75 | 6219331.5 | 8416866 | 624.626 | 5262080 | 4627066.5 | 1562.824 | VDQSILTGESVSVTK | Q64518 | Q64518 | ATP2A3 | ATPase, Ca++ transporting, ubiquitous | Cytoplasm | transporter |
| 15547737, 15552550 | 2 | 0.115 | 1.35E+08 | 1.52E+08 | 1.47E+08 | 1.01E+08 | 1.26E+08 | 1.01E+08 | 1.15E+08 | 1.22E+08 | 1.07E+08 | 1.61E+08 | 1.16E+08 | 1.08E+08 | 1641.8351, 1641.842 | QVVAVTGDGTNDGPALK | Q9R0K7 | Q9R0K7 | ATP2B2 | ATPase, Ca++ transporting, plasma membrane 2 | Plasma Membrane | transporter |
| 15532977 | 3 | 0.033 | 4.58E+07 | 5.29E+07 | 5.11E+07 | 3.35E+07 | 4.32E+07 | 3.68E+07 | 3.98E+07 | 4.74E+07 | 3.74E+07 | 4.86E+07 | 4.79E+07 | 4.45E+07 | 1993.838 | HLDAC[160.0307]ETMGNATAIC[160.0307]SDK | Q9R0K7 | Q9R0K7 | ATP2B2 | ATPase, Ca++ transporting, plasma membrane 2 | Plasma Membrane | transporter |
| 15531338 | 2 | 0.37 | 3.57E+07 | 4.09E+07 | 5.32E+07 | 2.49E+07 | 4.01E+07 | 3.23E+07 | 3.31E+07 | 4.06E+07 | 3.27E+07 | 4.36E+07 | 2.41E+07 | 2.90E+07 | 985.5087 | VYTFNSVR | Q9R0K7 | Q9R0K7 | ATP2B2 | ATPase, Ca++ transporting, plasma membrane 2 | Plasma Membrane | transporter |
| 15540745 | 3 | 0.587 | 2.49E+07 | 2.13E+07 | 1.92E+07 | 3.04E+07 | 2.15E+07 | 1.32E+07 | 2.23E+07 | 1.74E+07 | 1.47E+07 | 4.36E+07 | 2.19E+07 | 1.87E+07 | 1686.791 | DPMLLSGTHVMEGSGR | Q9R0K7 | Q9R0K7 | ATP2B2 | ATPase, Ca++ transporting, plasma membrane 2 | Plasma Membrane | transporter |
| 15549386 | 2 | 0.213 | 2.53E+07 | 2.48E+07 | 2.67E+07 | 1.90E+07 | 2.29E+07 | 1.49E+07 | 2.41E+07 | 2.09E+07 | 1.86E+07 | 2.75E+07 | 1.84E+07 | 2.01E+07 | 1686.7904 | DPMLLSGTHVMEGSGR | Q9R0K7 | Q9R0K7 | ATP2B2 | ATPase, Ca++ transporting, plasma membrane 2 | Plasma Membrane | transporter |
| 15550031 | 2 | 0.009 | 1.10E+07 | 1.14E+07 | 1.04E+07 | 7803906.5 | 8989889 | 7423683 | 9004919 | 9207908 | 8076875 | 1.19E+07 | 1.07E+07 | 9261237 | 1993.8379 | HLDAC[160.0307]ETMGNATAIC[160.0307]SDK | Q9R0K7 | Q9R0K7 | ATP2B2 | ATPase, Ca++ transporting, plasma membrane 2 | Plasma Membrane | transporter |
| 15534978 | 3 | 0.233 | 8227184 | 7137824.5 | 5840977.5 | 5906376 | 5220667.5 | 4372962 | 8178421.5 | 6159310.5 | 4935114.5 | 7889190.5 | 6759325 | 5967035.5 | 1844.8629 | C[160.0307]GIIHPGEDFLC[160.0307]LEGK | Q9R0K7 | Q9R0K7 | ATP2B2 | ATPase, Ca++ transporting, plasma membrane 2 | Plasma Membrane | transporter |
| 15532836 | 2 | 0.377 | 1.04E+07 | 5687791 | 4619347.5 | 9303539 | 2394148.25 | 4832871 | 6161362 | 3447566.25 | 5865756 | 7141959.5 | 1.11E+07 | 8410669 | 1113.602 | VYTFNSVRK | Q9R0K7 | Q9R0K7 | ATP2B2 | ATPase, Ca++ transporting, plasma membrane 2 | Plasma Membrane | transporter |
| 15534682 | 2 | 0.267 | 3921733 | 3202868.25 | 2777614.75 | 2590890.25 | 2081910.5 | 2697181.5 | 3775788 | 3075426.75 | 2350872.25 | 3915931.75 | 2755355.25 | 3118420.25 | 1191.5759 | MVTGDNINTAR | Q9R0K7 | Q9R0K7 | ATP2B2 | ATPase, Ca++ transporting, plasma membrane 2 | Plasma Membrane | transporter |
| 15537524 | 3 | 0.125 | 2108489 | 1922903.625 | 1854225.25 | 1405122.625 | 1661955.625 | 1406877 | 1869784.625 | 1584558.625 | 1352532.25 | 1723174.625 | 836460 | 1283896.375 | 2009.8314 | HLDAC[160.0307]ETM[147.0354]GNATAIC[160.0307]SDK | Q9R0K7 | Q9R0K7 | ATP2B2 | ATPase, Ca++ transporting, plasma membrane 2 | Plasma Membrane | transporter |
| 15531295 | 3 | 0.335 | 5.31E+07 | 1.02E+08 | 5.20E+07 | 6.75E+07 | 7.31E+07 | 9.71E+07 | 4.85E+07 | 7.82E+07 | 9.52E+07 | 2.53E+07 | 6.23E+07 | 5.68E+07 | 1780.989 | RTGAIVDVPVGEELLGR | Q03265 | Q03265 | ATP5A1 | ATP synthase, H+ transporting, mitochondrial F1 complex, alpha subunit 1, cardiac muscle | Cytoplasm | transporter |
| 15548118, 15548365 | 2, 3 | 0.22 | 4.49E+07 | 9.68E+07 | 5.20E+07 | 7.23E+07 | 6.24E+07 | 8.85E+07 | 4.20E+07 | 6.91E+07 | 1.05E+08 | 2.48E+07 | 4.95E+07 | 4.59E+07 | 2338.1619, 2338.1646 | EVAAFAQFGSDLDAATQQLLSR | Q03265 | Q03265 | ATP5A1 | ATP synthase, H+ transporting, mitochondrial F1 complex, alpha subunit 1, cardiac muscle | Cytoplasm | transporter |
| 15534669 | 4 | 0.293 | 3.22E+07 | 7.38E+07 | 2.84E+07 | 6.31E+07 | 5.62E+07 | 7.36E+07 | 3.35E+07 | 5.40E+07 | 8.10E+07 | 2.27E+07 | 4.31E+07 | 4.16E+07 | 2410.258 | FENAFLSHVISQHQSLLGNIR | Q03265 | Q03265 | ATP5A1 | ATP synthase, H+ transporting, mitochondrial F1 complex, alpha subunit 1, cardiac muscle | Cytoplasm | transporter |
| 15533227 | 2 | 0.2 | 3.02E+07 | 6.91E+07 | 3.62E+07 | 5.46E+07 | 5.35E+07 | 6.55E+07 | 3.71E+07 | 5.17E+07 | 6.02E+07 | 2.17E+07 | 4.04E+07 | 3.77E+07 | 1171.63 | VVDALGNAIDGK | Q03265 | Q03265 | ATP5A1 | ATP synthase, H+ transporting, mitochondrial F1 complex, alpha subunit 1, cardiac muscle | Cytoplasm | transporter |
| 15537977 | 2 | 0.332 | 3.24E+07 | 6.89E+07 | 3.16E+07 | 5.11E+07 | 5.43E+07 | 4.94E+07 | 3.33E+07 | 4.88E+07 | 5.77E+07 | 2.03E+07 | 3.94E+07 | 3.77E+07 | 1423.6699 | TGTAEMSSILEER | Q03265 | Q03265 | ATP5A1 | ATP synthase, H+ transporting, mitochondrial F1 complex, alpha subunit 1, cardiac muscle | Cytoplasm | transporter |
| 15547794 | 3 | 0.198 | 3.84E+07 | 6.73E+07 | 3.18E+07 | 5.70E+07 | 5.37E+07 | 6.43E+07 | 3.67E+07 | 4.93E+07 | 7.99E+07 | 1.97E+07 | 4.24E+07 | 3.52E+07 | 2410.2578 | FENAFLSHVISQHQSLLGNIR | Q03265 | Q03265 | ATP5A1 | ATP synthase, H+ transporting, mitochondrial F1 complex, alpha subunit 1, cardiac muscle | Cytoplasm | transporter |
| 15535167 | 2 | 0.409 | 2.96E+07 | 6.80E+07 | 3.21E+07 | 4.97E+07 | 4.82E+07 | 5.79E+07 | 3.01E+07 | 5.06E+07 | 6.34E+07 | 1.92E+07 | 4.26E+07 | 3.59E+07 | 1287.6917 | HALIIYDDLSK | Q03265 | Q03265 | ATP5A1 | ATP synthase, H+ transporting, mitochondrial F1 complex, alpha subunit 1, cardiac muscle | Cytoplasm | transporter |
| 15538170 | 2 | 0.239 | 3.12E+07 | 5.37E+07 | 2.88E+07 | 3.72E+07 | 4.20E+07 | 5.64E+07 | 2.85E+07 | 4.61E+07 | 6.29E+07 | 1.77E+07 | 3.42E+07 | 2.87E+07 | 1000.5773 | VLSIGDGIAR | Q03265 | Q03265 | ATP5A1 | ATP synthase, H+ transporting, mitochondrial F1 complex, alpha subunit 1, cardiac muscle | Cytoplasm | transporter |
| 15551123 | 2 | 0.232 | 2.49E+07 | 5.51E+07 | 2.11E+07 | 4.09E+07 | 3.70E+07 | 4.86E+07 | 2.28E+07 | 3.62E+07 | 5.02E+07 | 1.63E+07 | 2.51E+07 | 2.57E+07 | 2104.0364 | GMSLNLEPDNVGVVVFGNDK | Q03265 | Q03265 | ATP5A1 | ATP synthase, H+ transporting, mitochondrial F1 complex, alpha subunit 1, cardiac muscle | Cytoplasm | transporter |
| 15531375 | 2 | 0.3 | 2.32E+07 | 4.55E+07 | 2.32E+07 | 3.57E+07 | 3.46E+07 | 4.24E+07 | 2.38E+07 | 3.83E+07 | 4.27E+07 | 1.62E+07 | 2.74E+07 | 2.83E+07 | 1026.5919 | AVDSLVPIGR | Q03265 | Q03265 | ATP5A1 | ATP synthase, H+ transporting, mitochondrial F1 complex, alpha subunit 1, cardiac muscle | Cytoplasm | transporter |
| 15534324 | 2 | 0.26 | 2.34E+07 | 5.10E+07 | 2.57E+07 | 3.43E+07 | 3.85E+07 | 5.53E+07 | 2.35E+07 | 3.88E+07 | 4.74E+07 | 1.59E+07 | 2.74E+07 | 2.77E+07 | 1336.677 | EIVTNFLAGFEP | Q03265 | Q03265 | ATP5A1 | ATP synthase, H+ transporting, mitochondrial F1 complex, alpha subunit 1, cardiac muscle | Cytoplasm | transporter |
| 15533316 | 2 | 0.161 | 2.18E+07 | 4.97E+07 | 2.61E+07 | 3.64E+07 | 4.42E+07 | 5.16E+07 | 2.49E+07 | 3.81E+07 | 5.31E+07 | 1.26E+07 | 2.79E+07 | 2.52E+07 | 1316.7396 | TSIAIDTIINQK | Q03265 | Q03265 | ATP5A1 | ATP synthase, H+ transporting, mitochondrial F1 complex, alpha subunit 1, cardiac muscle | Cytoplasm | transporter |
| 15553064, 15554726 | 2, 3 | 0.468 | 1.64E+07 | 3.28E+07 | 1.49E+07 | 2.51E+07 | 2.61E+07 | 3.04E+07 | 1.67E+07 | 2.43E+07 | 3.75E+07 | 1.08E+07 | 2.38E+07 | 2.01E+07 | 2309.1545, 2309.159 | QGQYSPMAIEEQVAVIYAGVR | Q03265 | Q03265 | ATP5A1 | ATP synthase, H+ transporting, mitochondrial F1 complex, alpha subunit 1, cardiac muscle | Cytoplasm | transporter |
| 15534838 | 2, 3 | 0.738 | 1.93E+07 | 3.26E+07 | 1.20E+07 | 3.37E+07 | 1.72E+07 | 3.37E+07 | 1.59E+07 | 2.05E+07 | 3.67E+07 | 1.07E+07 | 2.81E+07 | 2.05E+07 | 1358.7434, 1358.7435 | ISVREPMQTGIK | Q03265 | Q03265 | ATP5A1 | ATP synthase, H+ transporting, mitochondrial F1 complex, alpha subunit 1, cardiac muscle | Cytoplasm | transporter |
| 15534186 | 2 | 0.277 | 1.31E+07 | 2.63E+07 | 1.37E+07 | 1.94E+07 | 2.03E+07 | 2.50E+07 | 1.28E+07 | 2.11E+07 | 2.82E+07 | 7950870.5 | 1.63E+07 | 1.41E+07 | 892.4872 | LELAQYR | Q03265 | Q03265 | ATP5A1 | ATP synthase, H+ transporting, mitochondrial F1 complex, alpha subunit 1, cardiac muscle | Cytoplasm | transporter |
| 15557856 | 2 | 0.895 | 1.07E+07 | 1.82E+07 | 9442879 | 1.37E+07 | 1.34E+07 | 1.62E+07 | 7526870 | 1.35E+07 | 2.24E+07 | 7940947.5 | 1.56E+07 | 1.24E+07 | 1575.7811 | ILGADTSVDLEETGR | Q03265 | Q03265 | ATP5A1 | ATP synthase, H+ transporting, mitochondrial F1 complex, alpha subunit 1, cardiac muscle | Cytoplasm | transporter |
| 15532255 | 2 | 0.215 | 1.13E+07 | 1.93E+07 | 1.25E+07 | 1.68E+07 | 1.66E+07 | 2.18E+07 | 1.03E+07 | 1.79E+07 | 2.55E+07 | 7092159.5 | 1.28E+07 | 1.19E+07 | 723.4498 | APGIIPR | Q03265 | Q03265 | ATP5A1 | ATP synthase, H+ transporting, mitochondrial F1 complex, alpha subunit 1, cardiac muscle | Cytoplasm | transporter |
| 15533520 | 2 | 0.289 | 7564590 | 1.54E+07 | 7507359 | 1.22E+07 | 1.19E+07 | 1.57E+07 | 8194565.5 | 1.21E+07 | 1.54E+07 | 5245350.5 | 1.06E+07 | 8448330 | 716.4541 | LTELLK | Q03265 | Q03265 | ATP5A1 | ATP synthase, H+ transporting, mitochondrial F1 complex, alpha subunit 1, cardiac muscle | Cytoplasm | transporter |
| 15540158 | 2 | 0.331 | 5609806 | 1.08E+07 | 3942146.25 | 9205461 | 8010911.5 | 8769464 | 4025152.25 | 8636753 | 9291033 | 4290945 | 5355014 | 5157822 | 845.5069 | STVAQLVK | Q03265 | Q03265 | ATP5A1 | ATP synthase, H+ transporting, mitochondrial F1 complex, alpha subunit 1, cardiac muscle | Cytoplasm | transporter |
| 15542137 | 3 | 0.308 | 3387733.75 | 1.12E+07 | 4313914.5 | 7293860 | 8168432.5 | 1.02E+07 | 3499070 | 5696044 | 9084145 | 3510241.5 | 4640443 | 4746524.5 | 2104.035 | GMSLNLEPDNVGVVVFGNDK | Q03265 | Q03265 | ATP5A1 | ATP synthase, H+ transporting, mitochondrial F1 complex, alpha subunit 1, cardiac muscle | Cytoplasm | transporter |
| 15556131 | 2 | 0.211 | 6699533 | 1.36E+07 | 7885766.5 | 7329115.5 | 1.07E+07 | 1.40E+07 | 6253794.5 | 1.10E+07 | 1.47E+07 | 2856056.25 | 6874706.5 | 6825639 | 1553.7356 | EAYPGDVFYLHSR | Q03265 | Q03265 | ATP5A1 | ATP synthase, H+ transporting, mitochondrial F1 complex, alpha subunit 1, cardiac muscle | Cytoplasm | transporter |
| 15556027 | 2 | 0.409 | 5085148 | 1.34E+07 | 6095746.5 | 5465716.5 | 1.04E+07 | 1.25E+07 | 4673699.5 | 1.07E+07 | 1.24E+07 | 2231530.75 | 5662424 | 7127865 | 1780.988 | RTGAIVDVPVGEELLGR | Q03265 | Q03265 | ATP5A1 | ATP synthase, H+ transporting, mitochondrial F1 complex, alpha subunit 1, cardiac muscle | Cytoplasm | transporter |
| 15539737 | 2 | 0.631 | 3183618.25 | 2.20E+07 | 1.52E+07 | 661429 | 1.90E+07 | 2.78E+07 | 3806138.25 | 1.69E+07 | 2.63E+07 | 83571.609 | 7598651.5 | 1.10E+07 | 1001.6073 | RSTVAQLVK | Q03265 | Q03265 | ATP5A1 | ATP synthase, H+ transporting, mitochondrial F1 complex, alpha subunit 1, cardiac muscle | Cytoplasm | transporter |
| 15549754 | 3 | 0.366 | 6.92E+07 | 1.75E+08 | 8.13E+07 | 1.16E+08 | 1.15E+08 | 1.39E+08 | 8.21E+07 | 1.38E+08 | 1.53E+08 | 5.91E+07 | 9.24E+07 | 8.82E+07 | 3350.6387 | GFQQILAGEYDHLPEQAFYMVGPIEEAVAK | P56480 | P56480 | ATP5B | ATP synthase, H+ transporting, mitochondrial F1 complex, beta polypeptide | Cytoplasm | transporter |
| 15541641 | 3 | 0.207 | 5.91E+07 | 1.01E+08 | 4.61E+07 | 9.43E+07 | 8.54E+07 | 1.03E+08 | 6.97E+07 | 9.16E+07 | 8.59E+07 | 3.85E+07 | 7.80E+07 | 6.17E+07 | 1842.8751 | IMDPNIVGNEHYDVAR | P56480 | P56480 | ATP5B | ATP synthase, H+ transporting, mitochondrial F1 complex, beta polypeptide | Cytoplasm | transporter |
| 15549748 | 2 | 0.211 | 5.35E+07 | 1.12E+08 | 5.74E+07 | 6.30E+07 | 8.66E+07 | 1.15E+08 | 5.02E+07 | 7.49E+07 | 1.18E+08 | 3.40E+07 | 4.93E+07 | 5.45E+07 | 1988.0278 | AIAELGIYPAVDPLDSTSR | P56480 | P56480 | ATP5B | ATP synthase, H+ transporting, mitochondrial F1 complex, beta polypeptide | Cytoplasm | transporter |
| 15531386 | 2 | 0.315 | 4.22E+07 | 7.74E+07 | 4.17E+07 | 6.07E+07 | 6.06E+07 | 8.71E+07 | 3.71E+07 | 6.35E+07 | 8.45E+07 | 2.75E+07 | 4.88E+07 | 4.73E+07 | 1038.5919 | IPVGPETLGR | P56480 | P56480 | ATP5B | ATP synthase, H+ transporting, mitochondrial F1 complex, beta polypeptide | Cytoplasm | transporter |
| 15533413 | 3 | 0.476 | 4.26E+07 | 8.41E+07 | 3.66E+07 | 5.81E+07 | 5.49E+07 | 7.17E+07 | 3.44E+07 | 5.70E+07 | 8.44E+07 | 2.05E+07 | 5.40E+07 | 4.28E+07 | 2023.0073 | FLSQPFQVAEVFTGHMGK | P56480 | P56480 | ATP5B | ATP synthase, H+ transporting, mitochondrial F1 complex, beta polypeptide | Cytoplasm | transporter |
| 15540560 | 2 | 0.363 | 2.67E+07 | 4.25E+07 | 2.31E+07 | 3.73E+07 | 3.05E+07 | 3.47E+07 | 2.24E+07 | 3.26E+07 | 4.07E+07 | 1.98E+07 | 2.79E+07 | 2.40E+07 | 899.5182 | VLDSGAPIK | P56480 | P56480 | ATP5B | ATP synthase, H+ transporting, mitochondrial F1 complex, beta polypeptide | Cytoplasm | transporter |
| 15531651 | 2 | 0.275 | 2.76E+07 | 5.66E+07 | 2.74E+07 | 3.88E+07 | 4.57E+07 | 5.77E+07 | 3.04E+07 | 4.39E+07 | 5.42E+07 | 1.87E+07 | 3.56E+07 | 3.12E+07 | 1262.6384 | TIAMDGTEGLVR | P56480 | P56480 | ATP5B | ATP synthase, H+ transporting, mitochondrial F1 complex, beta polypeptide | Cytoplasm | transporter |
| 15551890, 15561448 | 3, 4 | 0.262 | 3.64E+07 | 1.14E+08 | 5.53E+07 | 4.20E+07 | 8.43E+07 | 9.67E+07 | 3.55E+07 | 8.24E+07 | 8.93E+07 | 1.83E+07 | 3.84E+07 | 4.38E+07 | 3714.8792, 3714.8801, 3714.8804 | GSITSVQAIYVPADDLTDPAPATTFAHLDATTVLSR | P56480 | P56480 | ATP5B | ATP synthase, H+ transporting, mitochondrial F1 complex, beta polypeptide | Cytoplasm | transporter |
| 15547935 | 2 | 0.33 | 1.92E+07 | 5.19E+07 | 2.38E+07 | 3.46E+07 | 3.17E+07 | 4.72E+07 | 2.38E+07 | 4.09E+07 | 4.88E+07 | 1.80E+07 | 2.49E+07 | 2.54E+07 | 2119.0437 | SLQDIIAILGMDELSEEDK | P56480 | P56480 | ATP5B | ATP synthase, H+ transporting, mitochondrial F1 complex, beta polypeptide | Cytoplasm | transporter |
| 15531516 | 2 | 0.099 | 2.56E+07 | 5.90E+07 | 2.72E+07 | 3.64E+07 | 4.72E+07 | 5.69E+07 | 3.44E+07 | 4.95E+07 | 6.03E+07 | 1.69E+07 | 2.93E+07 | 2.59E+07 | 1088.6334 | VVDLLAPYAK | P56480 | P56480 | ATP5B | ATP synthase, H+ transporting, mitochondrial F1 complex, beta polypeptide | Cytoplasm | transporter |
| 15531611 | 3 | 0.693 | 2.90E+07 | 4.52E+07 | 1.93E+07 | 4.28E+07 | 2.24E+07 | 5.29E+07 | 3.35E+07 | 4.16E+07 | 5.91E+07 | 1.63E+07 | 4.58E+07 | 3.95E+07 | 1780.9607 | IMNVIGEPIDERGPIK | P56480 | P56480 | ATP5B | ATP synthase, H+ transporting, mitochondrial F1 complex, beta polypeptide | Cytoplasm | transporter |
| 15532621 | 2 | 0.194 | 2.91E+07 | 6.30E+07 | 3.69E+07 | 4.11E+07 | 5.09E+07 | 5.98E+07 | 3.16E+07 | 5.39E+07 | 7.21E+07 | 1.49E+07 | 3.50E+07 | 3.24E+07 | 1435.7512 | FTQAGSEVSALLGR | P56480 | P56480 | ATP5B | ATP synthase, H+ transporting, mitochondrial F1 complex, beta polypeptide | Cytoplasm | transporter |
| 15555437 | 3 | 0.286 | 1.66E+07 | 3.92E+07 | 2.37E+07 | 2.33E+07 | 3.07E+07 | 3.25E+07 | 1.85E+07 | 3.16E+07 | 2.94E+07 | 1.41E+07 | 1.94E+07 | 2.04E+07 | 3026.5479 | QFAPIHAEAPEFIEMSVEQEILVTGIK | P56480 | P56480 | ATP5B | ATP synthase, H+ transporting, mitochondrial F1 complex, beta polypeptide | Cytoplasm | transporter |
| 15531930 | 2 | 0.255 | 2.10E+07 | 4.53E+07 | 2.28E+07 | 2.93E+07 | 3.51E+07 | 3.26E+07 | 2.45E+07 | 3.88E+07 | 5.33E+07 | 1.37E+07 | 2.63E+07 | 2.41E+07 | 975.56055 | IGLFGGAGVGK | P56480 | P56480 | ATP5B | ATP synthase, H+ transporting, mitochondrial F1 complex, beta polypeptide | Cytoplasm | transporter |
| 15557112 | 3 | 0.281 | 1.87E+07 | 4.25E+07 | 2.18E+07 | 1.83E+07 | 3.48E+07 | 4.77E+07 | 2.24E+07 | 3.80E+07 | 5.11E+07 | 1.32E+07 | 1.65E+07 | 2.53E+07 | 2318.1436 | TREGNDLYHEMIESGVINLK | P56480 | P56480 | ATP5B | ATP synthase, H+ transporting, mitochondrial F1 complex, beta polypeptide | Cytoplasm | transporter |
| 15539025 | 3 | 0.137 | 2.14E+07 | 4.96E+07 | 2.46E+07 | 3.14E+07 | 4.59E+07 | 5.86E+07 | 2.89E+07 | 3.97E+07 | 5.28E+07 | 1.28E+07 | 2.78E+07 | 2.42E+07 | 1650.9144 | LVLEVAQHLGESTVR | P56480 | P56480 | ATP5B | ATP synthase, H+ transporting, mitochondrial F1 complex, beta polypeptide | Cytoplasm | transporter |
| 15533029 | 2, 3 | 0.152 | 1.80E+07 | 3.10E+07 | 2.11E+07 | 2.92E+07 | 2.85E+07 | 5.35E+07 | 2.51E+07 | 4.13E+07 | 4.26E+07 | 9951699 | 2.45E+07 | 2.49E+07 | 1457.8375, 1457.838 | TVLIMELINNVAK | P56480 | P56480 | ATP5B | ATP synthase, H+ transporting, mitochondrial F1 complex, beta polypeptide | Cytoplasm | transporter |
| 15547976 | 2 | 0.168 | 1.60E+07 | 2.97E+07 | 1.45E+07 | 2.53E+07 | 2.39E+07 | 3.65E+07 | 1.60E+07 | 2.33E+07 | 2.86E+07 | 9923226 | 2.01E+07 | 1.54E+07 | 2266.0784 | IPSAVGYQPTLATDMGTMQER | P56480 | P56480 | ATP5B | ATP synthase, H+ transporting, mitochondrial F1 complex, beta polypeptide | Cytoplasm | transporter |
| 15534337 | 2 | 0.242 | 9428117 | 2.72E+07 | 1.49E+07 | 1.06E+07 | 3.02E+07 | 2.02E+07 | 1.19E+07 | 2.67E+07 | 1.95E+07 | 8624374 | 8954405 | 1.05E+07 | 1385.7069 | IMNVIGEPIDER | P56480 | P56480 | ATP5B | ATP synthase, H+ transporting, mitochondrial F1 complex, beta polypeptide | Cytoplasm | transporter |
| 15532051 | 3 | 0.241 | 1.32E+07 | 3.56E+07 | 1.40E+07 | 2.56E+07 | 2.84E+07 | 3.20E+07 | 1.64E+07 | 2.86E+07 | 3.13E+07 | 7790438.5 | 1.79E+07 | 1.92E+07 | 2119.0442 | SLQDIIAILGMDELSEEDK | P56480 | P56480 | ATP5B | ATP synthase, H+ transporting, mitochondrial F1 complex, beta polypeptide | Cytoplasm | transporter |
| 15556009, 15560207 | 2 | 0.209 | 9892710 | 2.53E+07 | 1.28E+07 | 1.83E+07 | 1.91E+07 | 2.91E+07 | 1.30E+07 | 2.09E+07 | 2.82E+07 | 7010379.5 | 1.39E+07 | 1.36E+07 | 1601.7987, 1601.8071 | VALVYGQMNEPPGAR | P56480 | P56480 | ATP5B | ATP synthase, H+ transporting, mitochondrial F1 complex, beta polypeptide | Cytoplasm | transporter |
| 15554496 | 2 | 0.179 | 1.42E+07 | 2.67E+07 | 1.62E+07 | 1.68E+07 | 2.23E+07 | 2.41E+07 | 1.46E+07 | 2.13E+07 | 3.37E+07 | 7007126.5 | 1.69E+07 | 1.22E+07 | 1650.9133 | LVLEVAQHLGESTVR | P56480 | P56480 | ATP5B | ATP synthase, H+ transporting, mitochondrial F1 complex, beta polypeptide | Cytoplasm | transporter |
| 15539498 | 3 | 0.077 | 6875457.5 | 1.70E+07 | 8613347 | 1.16E+07 | 1.55E+07 | 1.63E+07 | 8961665 | 1.60E+07 | 1.58E+07 | 5694345.5 | 8195824 | 6869690.5 | 2060.9917 | EGNDLYHEMIESGVINLK | P56480 | P56480 | ATP5B | ATP synthase, H+ transporting, mitochondrial F1 complex, beta polypeptide | Cytoplasm | transporter |
| 15532993 | 3 | 0.647 | 9339719 | 1.77E+07 | 5507794.5 | 8817736 | 1.05E+07 | 1.69E+07 | 7024567 | 1.02E+07 | 1.65E+07 | 5592985.5 | 9347862 | 8246310.5 | 1988.0294 | AIAELGIYPAVDPLDSTSR | P56480 | P56480 | ATP5B | ATP synthase, H+ transporting, mitochondrial F1 complex, beta polypeptide | Cytoplasm | transporter |
| 15556316 | 2 | 0.373 | 8628209 | 1.04E+07 | 5062092.5 | 8511939 | 8576862 | 9072613 | 6096296 | 8108464.5 | 1.03E+07 | 5007482.5 | 7688263 | 5610585 | 1842.874 | IMDPNIVGNEHYDVAR | P56480 | P56480 | ATP5B | ATP synthase, H+ transporting, mitochondrial F1 complex, beta polypeptide | Cytoplasm | transporter |
| 15532041 | 2 | 0.178 | 5132776.5 | 2.08E+07 | 8860294 | 6214992 | 1.89E+07 | 1.52E+07 | 5957131.5 | 1.45E+07 | 1.37E+07 | 3191018 | 4991914.5 | 5470303 | 1439.7871 | VALTGLTVAEYFR | P56480 | P56480 | ATP5B | ATP synthase, H+ transporting, mitochondrial F1 complex, beta polypeptide | Cytoplasm | transporter |
| 15552460, 15552745 | 3 | 0.218 | 4542371.5 | 7417813.5 | 6051436.5 | 3771577.5 | 7197741.5 | 1.08E+07 | 3096393.25 | 7511991 | 7683251 | 2581953.25 | 2739505.25 | 4521016 | 3042.541, 3042.5425 | QFAPIHAEAPEFIEM[147.0354]SVEQEILVTGIK | P56480 | P56480 | ATP5B | ATP synthase, H+ transporting, mitochondrial F1 complex, beta polypeptide | Cytoplasm | transporter |
| 15555796 | 4 | 0.901 | 9266699 | 4977853 | 1358128.75 | 1.42E+07 | 2885088.25 | 3088123.25 | 2305192.25 | 4151266.25 | 1.30E+07 | 2419253.5 | 1.72E+07 | 8386847.5 | 3842.973 | KGSITSVQAIYVPADDLTDPAPATTFAHLDATTVLSR | P56480 | P56480 | ATP5B | ATP synthase, H+ transporting, mitochondrial F1 complex, beta polypeptide | Cytoplasm | transporter |
| 15543386 | 3 | 0.522 | 4099425.25 | 7342249 | 4435932 | 4895416 | 5284691.5 | 7009677.5 | 3890734.5 | 6137487 | 7673698 | 1846790.125 | 5384692.5 | 5034450.5 | 1406.6787 | AHGGYSVFAGVGER | P56480 | P56480 | ATP5B | ATP synthase, H+ transporting, mitochondrial F1 complex, beta polypeptide | Cytoplasm | transporter |
| 15548739 | 2 | 0.73 | 4711491 | 6949110.5 | 2716176 | 5234750 | 3217298.5 | 8038653 | 3797004 | 4690359 | 8175438 | 1363568 | 5914489 | 4569872.5 | 1780.963 | IMNVIGEPIDERGPIK | P56480 | P56480 | ATP5B | ATP synthase, H+ transporting, mitochondrial F1 complex, beta polypeptide | Cytoplasm | transporter |
| 15559519 | 2 | 0.012 | 945876.875 | 1197966.875 | 1239903.125 | 697452.25 | 1191464.625 | 2793513.5 | 630987.062 | 1911819.625 | 1391715 | 289277.844 | 261591.094 | 333630.688 | 2135.0383 | SLQDIIAILGM[147.0354]DELSEEDK | P56480 | P56480 | ATP5B | ATP synthase, H+ transporting, mitochondrial F1 complex, beta polypeptide | Cytoplasm | transporter |
| 15561386 | 3 | 0.2 | 554931.875 | 1253974.625 | 2122065.5 | 702502.312 | 1549485.875 | 3334920 | 282489.781 | 2138093.25 | 1648805.25 | 264072.062 | 360478.906 | 513134.312 | 3366.6233 | GFQQILAGEYDHLPEQAFYM[147.0354]VGPIEEAVAK | P56480 | P56480 | ATP5B | ATP synthase, H+ transporting, mitochondrial F1 complex, beta polypeptide | Cytoplasm | transporter |
| 15537970 | 2 | 0.261 | 6510883 | 1.48E+07 | 7049691.5 | 1.13E+07 | 1.17E+07 | 1.35E+07 | 7159643.5 | 1.23E+07 | 1.59E+07 | 3697722.75 | 9014425 | 7871819 | 1044.5665 | NEVAALTAAGK | Q91VR2 | Q91VR2 | ATP5C1 | ATP synthase, H+ transporting, mitochondrial F1 complex, gamma polypeptide 1 | Cytoplasm | transporter |
| 15535564 | 2 | 0.13 | 4406596.5 | 9652468 | 5510281 | 8338221 | 7302815 | 1.04E+07 | 4956149 | 7134107 | 1.03E+07 | 2513397 | 6070370 | 3995069.5 | 1300.676 | VYGTGSLALYEK | Q91VR2 | Q91VR2 | ATP5C1 | ATP synthase, H+ transporting, mitochondrial F1 complex, gamma polypeptide 1 | Cytoplasm | transporter |
| 15539697 | 2 | 0.94 | 3579827 | 6088817 | 1465485.125 | 5261766 | 3244014.5 | 2517785.5 | 2838847.75 | 2943328.25 | 2878718.5 | 1968314.875 | 4473069.5 | 2658031.25 | 1308.6553 | THSDQFLVSFK | Q91VR2 | Q91VR2 | ATP5C1 | ATP synthase, H+ transporting, mitochondrial F1 complex, gamma polypeptide 1 | Cytoplasm | transporter |
| 15542906 | 2 | 0.769 | 231095.484 | 1281749 | 1598350.625 | 415335.438 | 1728389.875 | 3193501.75 | 352249.062 | 1531177.5 | 3582867.75 | 208675.609 | 1222402.375 | 856700.688 | 1060.5682 | EVMIVGVGEK | Q91VR2 | Q91VR2 | ATP5C1 | ATP synthase, H+ transporting, mitochondrial F1 complex, gamma polypeptide 1 | Cytoplasm | transporter |
| 15539760 | 2 | 0.189 | 1.06E+07 | 1.82E+07 | 1.01E+07 | 1.29E+07 | 1.55E+07 | 1.80E+07 | 8887071 | 1.49E+07 | 1.75E+07 | 6729322.5 | 1.07E+07 | 9581481 | 986.5492 | IEANEALVK | Q9D3D9 | Q9D3D9 | ATP5D | ATP synthase, H+ transporting, mitochondrial F1 complex, delta subunit | Cytoplasm | transporter |
| 15532601 | 2 | 0.058 | 1.11E+07 | 2.40E+07 | 1.17E+07 | 1.56E+07 | 1.64E+07 | 1.92E+07 | 1.16E+07 | 1.83E+07 | 1.87E+07 | 6364967 | 1.02E+07 | 9086542 | 1375.6416 | AQSELSGAADEAAR | Q9D3D9 | Q9D3D9 | ATP5D | ATP synthase, H+ transporting, mitochondrial F1 complex, delta subunit | Cytoplasm | transporter |
| 15534196 | 2 | 0.06 | 4376866 | 7691237.5 | 3896572.75 | 6702678.5 | 7820262.5 | 9258949 | 3797618 | 6783287 | 8913346 | 2349075.75 | 4684083 | 2866636.25 | 1190.5682 | QIQDAIDMEK | Q9CQQ7 | Q9CQQ7 | ATP5F1 | ATP synthase, H+ transporting, mitochondrial Fo complex, subunit B1 | Cytoplasm | transporter |
| 15531904 | 3 | 0.504 | 8278266.5 | 1.84E+07 | 6394731 | 1.15E+07 | 1.09E+07 | 1.31E+07 | 7958790 | 1.23E+07 | 1.46E+07 | 5742954 | 1.05E+07 | 7741065 | 1980.0194 | LASLSEKPPAIDWAYYR | Q9DCX2 | Q9DCX2 | ATP5H | ATP synthase, H+ transporting, mitochondrial Fo complex, subunit d | Cytoplasm | enzyme |
| 15536206 | 2 | 0.336 | 5317767.5 | 1.27E+07 | 5239937.5 | 9166241 | 8709675 | 1.05E+07 | 4899435.5 | 9728889 | 1.19E+07 | 3881063.75 | 6337818 | 6110809 | 1340.6245 | SC[160.0307]AEFVSGSQLR | Q9DCX2 | Q9DCX2 | ATP5H | ATP synthase, H+ transporting, mitochondrial Fo complex, subunit d | Cytoplasm | enzyme |
| 15534580 | 2 | 0.229 | 4676965.5 | 9757186 | 4038125.25 | 7477090 | 7853804 | 8271966.5 | 5098576 | 7388198.5 | 9380151 | 3215293.5 | 5710261 | 4984125 | 1195.5814 | YTALVDQEEK | Q9DCX2 | Q9DCX2 | ATP5H | ATP synthase, H+ transporting, mitochondrial Fo complex, subunit d | Cytoplasm | enzyme |
| 15549729 | 2 | 0.106 | 2690774.5 | 6781669.5 | 3606880 | 4223950.5 | 5804677 | 5921282 | 3215875.25 | 4288057 | 5858411.5 | 2497807.75 | 2729235 | 2728766 | 2493.2178 | NIIPFDQMTIDDLNEIFPETK | Q9DCX2 | Q9DCX2 | ATP5H | ATP synthase, H+ transporting, mitochondrial Fo complex, subunit d | Cytoplasm | enzyme |
| 15556333 | 2 | 0.354 | 2960803.25 | 7263426 | 2994610.25 | 3428424.25 | 5424014 | 5589976.5 | 3163378.75 | 4471238.5 | 6868897.5 | 2479783.25 | 2848673.75 | 3284019.5 | 1920.9476 | TIDWVSFVEVMPQNQK | Q9DCX2 | Q9DCX2 | ATP5H | ATP synthase, H+ transporting, mitochondrial Fo complex, subunit d | Cytoplasm | enzyme |
| 15553249 | 2 | 0.056 | 2485209.25 | 4599440.5 | 3051918.25 | 2614275.5 | 3847692 | 5075958.5 | 2280767 | 3604175 | 4143521.75 | 1562960.5 | 1973557.375 | 1912362.125 | 1556.7506 | YPYWPHQPIENL | Q9DCX2 | Q9DCX2 | ATP5H | ATP synthase, H+ transporting, mitochondrial Fo complex, subunit d | Cytoplasm | enzyme |
| 15534534 | 2 | 0.229 | 3874501.75 | 1.00E+07 | 4002611.25 | 5685848.5 | 7318376 | 8154421 | 4015837.25 | 6741564.5 | 8641863 | 2811276.25 | 4731390 | 3561946.25 | 1172.5272 | GEMDTFPTFK | P97450 | P97450 | ATP5J | ATP synthase, H+ transporting, mitochondrial Fo complex, subunit F6 | Cytoplasm | transporter |
| 15533765 | 2 | 0.443 | 3876427 | 7288017 | 3010980.25 | 6774985.5 | 5457552.5 | 7264535 | 3814308.25 | 5741099 | 7566499 | 2279644.25 | 5674063 | 4787921 | 1332.6776 | LGELPSWIMMR | P56135 | P56135 | ATP5J2 | ATP synthase, H+ transporting, mitochondrial Fo complex, subunit F2 | Cytoplasm | transporter |
| 15533613 | 2 | 0.244 | 8369690 | 1.90E+07 | 9749608 | 1.46E+07 | 1.33E+07 | 1.72E+07 | 8852504 | 1.54E+07 | 1.92E+07 | 7253758.5 | 1.07E+07 | 8504727 | 1331.665 | ELAEAQDDSILK | Q06185 | Q06185 | Atp5k | ATP synthase, H+ transporting, mitochondrial F1F0 complex, subunit E | Cytoplasm | enzyme |
| 15538687 | 2 | 0.143 | 3818220.25 | 8303308.5 | 4196947 | 4346669.5 | 6715727.5 | 6783261.5 | 4331670.5 | 7314885.5 | 9020116 | 1796963.875 | 3925616.5 | 3896708.5 | 1176.7328 | VPPVQVSPLIK | Q06185 | Q06185 | Atp5k | ATP synthase, H+ transporting, mitochondrial F1F0 complex, subunit E | Cytoplasm | enzyme |
| 15534384 | 2 | 0.472 | 4870598 | 1.00E+07 | 4099875.25 | 6626712.5 | 6988308.5 | 9133412 | 5204802 | 9896165 | 1.02E+07 | 2681157.25 | 6473283.5 | 6857173 | 1295.6648 | APSMVAAAVTYSK | Q9CPQ8 | Q9CPQ8 | ATP5L | ATP synthase, H+ transporting, mitochondrial Fo complex, subunit G | Cytoplasm | enzyme |
| 15544643, 15546300 | 3 | 0.123 | 3210249.75 | 8083747.5 | 2707473.5 | 5218832 | 6274444.5 | 6116744.5 | 2706220.25 | 5109295.5 | 4981017.5 | 2051359.375 | 3119617.25 | 2605689 | 1989.1229, 1989.1235 | VELVPPTPAEIPTAIQSVK | Q9CPQ8 | Q9CPQ8 | ATP5L | ATP synthase, H+ transporting, mitochondrial Fo complex, subunit G | Cytoplasm | enzyme |
| 15533988 | 3 | 0.477 | 1.29E+07 | 2.38E+07 | 1.12E+07 | 1.77E+07 | 1.45E+07 | 1.90E+07 | 1.12E+07 | 1.82E+07 | 2.17E+07 | 8294045.5 | 1.52E+07 | 1.24E+07 | 1582.9045 | LVRPPVQVYGIEGR | Q9DB20 | Q9DB20 | ATP5O | ATP synthase, H+ transporting, mitochondrial F1 complex, O subunit | Cytoplasm | transporter |
| 15532989 | 3 | 0.2 | 8168845.5 | 1.90E+07 | 7928978.5 | 1.31E+07 | 1.61E+07 | 1.94E+07 | 1.18E+07 | 1.52E+07 | 2.32E+07 | 6528194.5 | 1.03E+07 | 1.16E+07 | 2032.0604 | LGNTQGIISAFSTIMSVHR | Q9DB20 | Q9DB20 | ATP5O | ATP synthase, H+ transporting, mitochondrial F1 complex, O subunit | Cytoplasm | transporter |
| 15538382 | 2 | 0.366 | 7035510 | 1.35E+07 | 5270238.5 | 1.11E+07 | 9561128 | 1.13E+07 | 6561519 | 9521329 | 1.19E+07 | 5763950 | 8279672 | 6460227.5 | 1145.5792 | YATALYSAASK | Q9DB20 | Q9DB20 | ATP5O | ATP synthase, H+ transporting, mitochondrial F1 complex, O subunit | Cytoplasm | transporter |
| 15531641 | 2 | 0.402 | 8502394 | 1.61E+07 | 8786062 | 1.16E+07 | 1.13E+07 | 1.46E+07 | 6702691.5 | 1.18E+07 | 2.02E+07 | 4200996.5 | 8564470 | 1.02E+07 | 1216.7278 | VSLAVLNPYIK | Q9DB20 | Q9DB20 | ATP5O | ATP synthase, H+ transporting, mitochondrial F1 complex, O subunit | Cytoplasm | transporter |
| 15555930 | 4 | 0.92 | 5926199.5 | 6028559.5 | 2045314.375 | 1.28E+07 | 1932616.625 | 5853816 | 3717004.75 | 3536840.25 | 9253112 | 3017887 | 1.19E+07 | 6801826 | 4315.185 | LGNTQGIISAFSTIMSVHRGEVPC[160.0307]TVTTASPLDDAVLSELK | Q9DB20 | Q9DB20 | ATP5O | ATP synthase, H+ transporting, mitochondrial F1 complex, O subunit | Cytoplasm | transporter |
| 15552018 | 3 | 0.338 | 2727015.75 | 7250060.5 | 3494227.25 | 3661200.25 | 8548997 | 6502058.5 | 2873246.5 | 6772365 | 8134529.5 | 2664701.75 | 3646937 | 3051310.75 | 2302.146 | GEVPC[160.0307]TVTTASPLDDAVLSELK | Q9DB20 | Q9DB20 | ATP5O | ATP synthase, H+ transporting, mitochondrial F1 complex, O subunit | Cytoplasm | transporter |
| 15534549 | 2 | 0.418 | 4034839 | 7494645.5 | 2176025 | 6626051.5 | 5425745.5 | 5759692 | 3266815 | 5210522 | 5505198.5 | 2393033.75 | 4375988.5 | 3956564 | 1276.6367 | TDPSIMGGMIVR | Q9DB20 | Q9DB20 | ATP5O | ATP synthase, H+ transporting, mitochondrial F1 complex, O subunit | Cytoplasm | transporter |
| 15551544 | 2 | 0.208 | 3368978.5 | 6204345 | 3311833 | 5460766.5 | 5941549 | 7358592.5 | 3666445.75 | 5681905.5 | 8854857 | 2230448.5 | 4334195 | 4179473.5 | 1860.9592 | FSPLTANLMNLLAENGR | Q9DB20 | Q9DB20 | ATP5O | ATP synthase, H+ transporting, mitochondrial F1 complex, O subunit | Cytoplasm | transporter |
| 15540433 | 2 | 0.699 | 1141359.75 | 2932373 | 2342948.5 | 1547313.25 | 2726216.25 | 1939960.875 | 1849522.25 | 2606769 | 1779266.375 | 1849545.25 | 1248186.75 | 1641505.625 | 1042.6135 | GPTITLTQIV | Q9R1Q9 | Q9R1Q9 | ATP6AP1 | ATPase, H+ transporting, lysosomal accessory protein 1 | Cytoplasm | transporter |

| | | | | | | | | | | | | | | | | | | | | | | |
|---|---|---|---|---|---|---|---|---|---|---|---|---|---|---|---|---|---|---|---|---|---|---|
| 15554216 | 2 | 0.21 | 1846802.75 | 3333938.25 | 3294536.5 | 2136095 | 3475192.75 | 3055483.5 | 2390660 | 2815771.75 | 2992928.25 | 2443582 | 1578674.625 | 1675674.125 | 1555.7357 | QENTQSPYNLAYK | Q9CYN9 | Q9CYN9 | ATP6AP2 | ATPase, H+ transporting, lysosomal accessory protein 2 | Cytoplasm | transporter |
| 15548485 | 2 | 0.289 | 7720586.5 | 1.30E+07 | 9466086 | 1.25E+07 | 1.19E+07 | 9793523 | 9313085 | 9090244 | 1.08E+07 | 9719886 | 6587901 | 8838134 | 1996.9745 | ANIPIMDTGENPEVPFPR | Q9Z1G4 | Q9Z1G4 | ATP6V0A1 | ATPase, H+ transporting, lysosomal V0 subunit a1 | Cytoplasm | transporter |
| 15553937 | 3 | 0.932 | 4652679.5 | 1.03E+07 | 6670658 | 5949383.5 | 8953013 | 8196872 | 5782677.5 | 7718800.5 | 8036214.5 | 9595206 | 4868707 | 5856930.5 | 3163.4553 | SEEMTLAQLFLQSEAAYC[160.0307]C[160.0307]VSELGELGK | Q9Z1G4 | Q9Z1G4 | ATP6V0A1 | ATPase, H+ transporting, lysosomal V0 subunit a1 | Cytoplasm | transporter |
| 15532445 | 3 | 0.389 | 5607333.5 | 8829571 | 7339906 | 6895317 | 9073271 | 7974089.5 | 8063640 | 8721598 | 8599820 | 7760668.5 | 6156176 | 7090258 | 2154.0305 | QAEIENPLEDPVTGDYVHK | Q9Z1G4 | Q9Z1G4 | ATP6V0A1 | ATPase, H+ transporting, lysosomal V0 subunit a1 | Cytoplasm | transporter |
| 15536633 | 3 | 0.615 | 7125573.5 | 1.35E+07 | 9859146 | 6846827.5 | 1.46E+07 | 7515037.5 | 8933594 | 1.14E+07 | 8080902.5 | 7679975 | 6015559 | 8381024.5 | 1826.8666 | IDDLQMVLNQTEDHR | Q9Z1G4 | Q9Z1G4 | ATP6V0A1 | ATPase, H+ transporting, lysosomal V0 subunit a1 | Cytoplasm | transporter |
| 15532901 | 3 | 0.633 | 8218397.5 | 1.20E+07 | 1.05E+07 | 8837177 | 1.14E+07 | 9176224 | 8347955.5 | 9562255 | 1.08E+07 | 7091966.5 | 8593046 | 1.02E+07 | 2059 | FTHGFQNIVDAYGIGTYR | Q9Z1G4 | Q9Z1G4 | ATP6V0A1 | ATPase, H+ transporting, lysosomal V0 subunit a1 | Cytoplasm | transporter |
| 15540226 | 2 | 0.677 | 3057628 | 4597481 | 3626581.5 | 4810213 | 4533357 | 4094873.75 | 3926232.75 | 4181889.25 | 3460845.5 | 5447661.5 | 2976825.25 | 3759521 | 1422.6653 | MQTNQTPPTYNK | Q9Z1G4 | Q9Z1G4 | ATP6V0A1 | ATPase, H+ transporting, lysosomal V0 subunit a1 | Cytoplasm | transporter |
| 15532930 | 2 | 0.626 | 3734261.75 | 5779852.5 | 5333541.5 | 5179810.5 | 6126078 | 4884147 | 4673108.5 | 5103436 | 4836354.5 | 5022845 | 3593598 | 5000100.5 | 1159.5925 | EINTNQEALK | Q9Z1G4 | Q9Z1G4 | ATP6V0A1 | ATPase, H+ transporting, lysosomal V0 subunit a1 | Cytoplasm | transporter |
| 15553851 | 2, 3 | 0.009 | 2577528 | 3080438.5 | 3141755.5 | 3077383 | 3295673.5 | 3531434.25 | 3080351.75 | 3995453 | 3816276.5 | 4853748.5 | 4108104.75 | 4215928 | 2506.2415, 2506.242 | C[160.0307]LIAEVWC[160.0307]PVTDLDSIQFALR | Q9Z1G4 | Q9Z1G4 | ATP6V0A1 | ATPase, H+ transporting, lysosomal V0 subunit a1 | Cytoplasm | transporter |
| 15554085 | 2 | 0.72 | 1988431.875 | 3628115.25 | 3327385.5 | 1982379.5 | 4011784.5 | 2961815.5 | 2376795 | 3530077.75 | 2889411.75 | 2921845 | 1647292 | 2471982.75 | 1547.7123 | ASLYPC[160.0307]PETPQER | Q9Z1G4 | Q9Z1G4 | ATP6V0A1 | ATPase, H+ transporting, lysosomal V0 subunit a1 | Cytoplasm | transporter |
| 15532374 | 3 | 0.96 | 1.72E+07 | 3.71E+07 | 1.67E+07 | 2.21E+07 | 2.40E+07 | 2.59E+07 | 2.07E+07 | 2.43E+07 | 2.30E+07 | 1.85E+07 | 2.12E+07 | 2.59E+07 | 1891.9767 | SGTGIAAMSVMRPELIMK | P63082 | P63082 | ATP6V0C | ATPase, H+ transporting, lysosomal 16kDa, V0 subunit c | Cytoplasm | transporter |
| 15549843 | 2 | 0.581 | 2381000 | 3819565.25 | 770547.375 | 3010375.5 | 3491831.75 | 3051727.5 | 2422707.5 | 2944693.25 | 2837473 | 1407564.875 | 2295389.5 | 3280682.5 | 1891.9757 | SGTGIAAMSVMRPELIMK | P63082 | P63082 | ATP6V0C | ATPase, H+ transporting, lysosomal 16kDa, V0 subunit c | Cytoplasm | transporter |
| 15550065 | 3 | 0.387 | 1.46E+07 | 2.58E+07 | 2.50E+07 | 2.27E+07 | 2.14E+07 | 2.04E+07 | 1.40E+07 | 1.95E+07 | 2.44E+07 | 1.64E+07 | 1.37E+07 | 1.79E+07 | 2947.4612 | LHLQSTDYGNFLANEASPLTVSVIDDK | P51863 | P51863 | ATP6V0D1 | ATPase, H+ transporting, lysosomal 38kDa, V0 subunit d1 | Cytoplasm | transporter |
| 15551210 | 3 | 0.502 | 9848618 | 1.47E+07 | 8468071 | 1.11E+07 | 1.35E+07 | 1.10E+07 | 8697125 | 1.15E+07 | 1.06E+07 | 1.20E+07 | 1.20E+07 | 1.37E+07 | 2744.2769 | FC[160.0307]TLLGGTTADAMC[160.0307]PILEFEADRR | P51863 | P51863 | ATP6V0D1 | ATPase, H+ transporting, lysosomal 38kDa, V0 subunit d1 | Cytoplasm | transporter |
| 15538132 | 2 | 0.282 | 6078003.5 | 9305887 | 8075865.5 | 8909907 | 1.17E+07 | 9421715 | 7122739 | 8843354 | 7509737 | 8645496 | 6122683 | 8833788 | 1304.6461 | LLFEGAGSNPGDK | P51863 | P51863 | ATP6V0D1 | ATPase, H+ transporting, lysosomal 38kDa, V0 subunit d1 | Cytoplasm | transporter |
| 15551896 | 3 | 0.643 | 3127153.5 | 9067513 | 8484629 | 7129225 | 9797055 | 8435178 | 5632513.5 | 9099011 | 6876717.5 | 7423739.5 | 4793219.5 | 5919615.5 | 2479.2307 | AGVLSQADYLNLVQC[160.0307]ETLEDLK | P51863 | P51863 | ATP6V0D1 | ATPase, H+ transporting, lysosomal 38kDa, V0 subunit d1 | Cytoplasm | transporter |
| 15555668 | 2 | 0.195 | 3804395.25 | 6220785.5 | 5220177 | 4621148.5 | 4836694 | 5229600 | 3915401.25 | 5727300 | 6702502.5 | 3399867.75 | 2799277.75 | 4603067.5 | 1640.8867 | AFIITINSFGTELSK | P51863 | P51863 | ATP6V0D1 | ATPase, H+ transporting, lysosomal 38kDa, V0 subunit d1 | Cytoplasm | transporter |
| 15535482 | 2 | 0.964 | 2277343.25 | 3663974 | 3898285.75 | 2690407 | 4039004 | 3289389 | 2487589.25 | 3808157 | 3237252.25 | 3395860.25 | 2558149.5 | 3131522.25 | 1020.50195 | AYLESFYK | P51863 | P51863 | ATP6V0D1 | ATPase, H+ transporting, lysosomal 38kDa, V0 subunit d1 | Cytoplasm | transporter |
| 15536652 | 3 | 0.876 | 5977438.5 | 5962212.5 | 3697601 | 6099126.5 | 4325870.5 | 5829045.5 | 4972815.5 | 4751806.5 | 4562477.5 | 5130404 | 4393935 | 5546874 | 2186.1401 | VGSHITGGDIYGIVNENSLIK | P50516 | P50516 | ATP6V1A | ATPase, H+ transporting, lysosomal 70kDa, V1 subunit A | Plasma Membrane | transporter |
| 15552207 | 2 | 0.594 | 2522357.75 | 2232469.75 | 2828724 | 2289328.75 | 2401698 | 2411103.25 | 1532059 | 1997988.25 | 2185196.75 | 3339848.5 | 981321.562 | 1977905.375 | 1784.8247 | ADYAQLLEDMQNAFR | P50516 | P50516 | ATP6V1A | ATPase, H+ transporting, lysosomal 70kDa, V1 subunit A | Plasma Membrane | transporter |
| 15543743 | 3 | 0.5 | 2676461.5 | 3734178 | 3945833.25 | 2734840.25 | 2569070.5 | 3334423.25 | 2982291 | 2677891.5 | 1947464.5 | 3144007 | 1644246.375 | 3365421.75 | 2055.0828 | EILQEEEDLAEIVQLVGK | P50516 | P50516 | ATP6V1A | ATPase, H+ transporting, lysosomal 70kDa, V1 subunit A | Plasma Membrane | transporter |
| 15536955 | 3 | 0.585 | 1851574.625 | 2435140.25 | 1731854.375 | 1914899.875 | 1851381 | 2162322 | 1689096.5 | 1385471.125 | 1522540.25 | 2480087.25 | 1088639 | 2348122.5 | 1671.8921 | RTALVANTSNMPVAAR | P50516 | P50516 | ATP6V1A | ATPase, H+ transporting, lysosomal 70kDa, V1 subunit A | Plasma Membrane | transporter |
| 15537616 | 2 | 0.551 | 1391899.375 | 937943.312 | 1029529.25 | 1067521.75 | 1307942.125 | 726095.312 | 775010 | 619480.125 | 838558.812 | 1671671.875 | 866154.75 | 549703.5 | 902.50336 | GVNVSALSR | P50516 | P50516 | ATP6V1A | ATPase, H+ transporting, lysosomal 70kDa, V1 subunit A | Plasma Membrane | transporter |
| 15532862 | 3 | 0.43 | 8709978 | 7807350.5 | 4812993 | 7114942 | 6045933.5 | 8374071 | 8427954 | 7508566 | 5486508.5 | 1.09E+07 | 8271646.5 | 8376620 | 2178.145 | IPIFSAAGLPHNEIAAQIC[160.0307]R | P62814 | P62814 | ATP6V1B2 | ATPase, H+ transporting, lysosomal 56/58kDa, V1 subunit B2 | Cytoplasm | transporter |
| 15554974 | 2 | 0.698 | 2090044.5 | 2758319.25 | 3005567 | 1962865.75 | 2353262.25 | 2613088.75 | 2021776.625 | 2413760.5 | 2762019.25 | 3390886.75 | 1352597.375 | 1676448.375 | 1520.7916 | AVVQVFEGTSGIDAK | P62814 | P62814 | ATP6V1B2 | ATPase, H+ transporting, lysosomal 56/58kDa, V1 subunit B2 | Cytoplasm | transporter |
| 15541375 | 2 | 0.749 | 1963591.125 | 2135881.5 | 1905974.25 | 2096123.5 | 1684367.375 | 1650488.25 | 1616064.375 | 1813197.625 | 1741096.75 | 2538100 | 1563677.125 | 1410293.375 | 1298.6029 | TSC[160.0307]EFTGDILR | P62814 | P62814 | ATP6V1B2 | ATPase, H+ transporting, lysosomal 56/58kDa, V1 subunit B2 | Cytoplasm | transporter |
| 15550278 | 2 | 0.287 | 2632676.5 | 2799122.75 | 1812721.75 | 1838479.375 | 2102869.25 | 2561311.25 | 2049214.375 | 1792960.125 | 2242533 | 1957698.625 | 1087643.625 | 1976568.25 | 1586.8346 | ARDDLITDLLNEAK | P50518 | P50518 | ATP6V1E1 | ATPase, H+ transporting, lysosomal 31kDa, V1 subunit E1 | Cytoplasm | transporter |
| 15542604 | 2 | 0.372 | 1335872.25 | 1147743.375 | 1067281.875 | 1302340.75 | 1078923.875 | 1228138 | 1138858.625 | 898819.75 | 669603.812 | 2144919.25 | 898343.75 | 1155494.625 | 1427.7212 | QLQSEQPQTAAAR | Q8BVE3 | Q8BVE3 | ATP6V1H | ATPase, H+ transporting, lysosomal 50/57kDa, V1 subunit H | Cytoplasm | transporter |
| 15557396 | 2 | 0.428 | 349429.656 | 200821.641 | 179808.422 | 432603.969 | 234117.359 | 170589.453 | 452165.219 | 249448.828 | 121440.844 | 814540.812 | 372027.688 | 289065.156 | 1509.6643 | QEMLQTEGSQC[160.0307]AK | Q8BVE3 | Q8BVE3 | ATP6V1H | ATPase, H+ transporting, lysosomal 50/57kDa, V1 subunit H | Cytoplasm | transporter |
| 15548202 | 3 | 0.003 | 2.17E+07 | 2.43E+07 | 2.35E+07 | 1.47E+07 | 1.85E+07 | 1.36E+07 | 2.02E+07 | 2.05E+07 | 2.12E+07 | 1.85E+07 | 1.62E+07 | 1.72E+07 | 3045.5317 | SDLTAHPVGVGNPPTGEGAGPQTFSVLIGNR | Q64446 | Q64446 | ATP7B | ATPase, Cu++ transporting, beta polypeptide | Cytoplasm | transporter |
| 15550295 | 3 | 0.261 | 3818180.25 | 3371608.5 | 4018042 | 2455054 | 3472440.75 | 2567442.5 | 3957478.25 | 3452193.25 | 3042909.75 | 3915294.75 | 2912384.5 | 2320024.25 | 2416.1445 | GLPTHQGPGHSSETPSSPGATASQK | Q64446 | Q64446 | ATP7B | ATPase, Cu++ transporting, beta polypeptide | Cytoplasm | transporter |
| 15539842 | 2 | 0.032 | 3327749 | 4347947.5 | 4298829 | 2807274.5 | 3214889.75 | 2553506.75 | 3282812.5 | 3794893.25 | 3709492.5 | 3431349.5 | 2761436.25 | 2607327.75 | 1146.609 | VSLEQGSATVR | Q64446 | Q64446 | ATP7B | ATPase, Cu++ transporting, beta polypeptide | Cytoplasm | transporter |
| 15536836 | 2 | 0.009 | 5136106 | 4136818.25 | 5414279.5 | 2769622.75 | 3224437.5 | 3390380 | 4384387 | 4385950 | 4507641 | 3383569.75 | 3246469.75 | 4447780 | 1014.59155 | AIATQVGINK | Q64446 | Q64446 | ATP7B | ATPase, Cu++ transporting, beta polypeptide | Cytoplasm | transporter |
| 15535738 | 2 | 0.005 | 4035214.5 | 3724412.5 | 4292507.5 | 2304769.5 | 2775208 | 2235583.25 | 2949543.5 | 2829939.5 | 3083183 | 3027189.75 | 2866064.25 | 2067537.125 | 1217.6135 | YDPEIIQSPR | Q64446 | Q64446 | ATP7B | ATPase, Cu++ transporting, beta polypeptide | Cytoplasm | transporter |
| 15551552 | 3 | 0.968 | 3371613.25 | 2698460.75 | 1722651.5 | 4755615 | 1186022.25 | 1921958.25 | 2071186 | 2030927.5 | 3531306.25 | 2545043.25 | 2955090.5 | 2365346.25 | 2805.4417 | TAVLTISGITC[160.0307]ASSVQPIEDMLSQRK | Q64446 | Q64446 | ATP7B | ATPase, Cu++ transporting, beta polypeptide | Cytoplasm | transporter |
| 15535278 | 2 | 0.056 | 3265575.5 | 2778071 | 3517958 | 1820784.75 | 2642831 | 2020396 | 2708648.75 | 2744577.5 | 2423926.75 | 2188402.25 | 2764671.75 | 1803696.375 | 1126.6234 | VFAEVLPSHK | Q64446 | Q64446 | ATP7B | ATPase, Cu++ transporting, beta polypeptide | Cytoplasm | transporter |
| 15556765 | 2 | 0.01 | 2228795 | 2567870.25 | 2231253.75 | 1600526.5 | 1695494 | 1354865.125 | 1545683 | 1795250 | 1952342.875 | 1906638.375 | 1693125.75 | 1468345.5 | 2306.1213 | VLEGNTMADESLITGEAMPVTK | Q64446 | Q64446 | ATP7B | ATPase, Cu++ transporting, beta polypeptide | Cytoplasm | transporter |
| 15549885, 15551761 | 3, 4 | 0.071 | 1.97E+07 | 2.26E+07 | 2.20E+07 | 2.20E+07 | 2.02E+07 | 2.42E+07 | 2.33E+07 | 2.34E+07 | 2.47E+07 | 3.01E+07 | 2.29E+07 | 2.78E+07 | 3194.6138, 3194.6165 | TPDSVIIDSLGQEERYELLNVLEFTSAR | P70704 | P70704 | ATP8A1 | ATPase, aminophospholipid transporter (APLT), class I, type 8A, member 1 | Cytoplasm | transporter |
| 15548598 | 2 | 0.406 | 1.13E+07 | 1.44E+07 | 1.55E+07 | 1.45E+07 | 1.71E+07 | 1.64E+07 | 1.40E+07 | 1.48E+07 | 1.37E+07 | 1.72E+07 | 1.34E+07 | 1.44E+07 | 1602.8453 | IIYQAASPDEGALVR | P70704 | P70704 | ATP8A1 | ATPase, aminophospholipid transporter (APLT), class I, type 8A, member 1 | Cytoplasm | transporter |
| 15531658 | 3 | 0.876 | 1.55E+07 | 2.00E+07 | 1.95E+07 | 1.67E+07 | 2.07E+07 | 2.16E+07 | 1.82E+07 | 1.85E+07 | 1.93E+07 | 1.65E+07 | 1.78E+07 | 2.10E+07 | 1774.9768 | LDGHGTVPLGADQILLR | P70704 | P70704 | ATP8A1 | ATPase, aminophospholipid transporter (APLT), class I, type 8A, member 1 | Cytoplasm | transporter |
| 15534436 | 3 | 0.039 | 1.09E+07 | 1.23E+07 | 1.10E+07 | 1.23E+07 | 1.20E+07 | 1.17E+07 | 1.19E+07 | 1.16E+07 | 1.16E+07 | 1.47E+07 | 1.35E+07 | 1.22E+07 | 1954.9712 | NTQWVHGIVVYTGHDTK | P70704 | P70704 | ATP8A1 | ATPase, aminophospholipid transporter (APLT), class I, type 8A, member 1 | Cytoplasm | transporter |
| 15535086 | 2 | 0.398 | 1.07E+07 | 1.05E+07 | 1.13E+07 | 1.18E+07 | 1.27E+07 | 8973433 | 1.19E+07 | 9586052 | 1.06E+07 | 1.28E+07 | 1.15E+07 | 1.28E+07 | 1302.7385 | TIFINQPQLTK | P70704 | P70704 | ATP8A1 | ATPase, aminophospholipid transporter (APLT), class I, type 8A, member 1 | Cytoplasm | transporter |
| 15531774 | 2 | 0.27 | 9373353 | 9864579 | 1.13E+07 | 1.12E+07 | 1.15E+07 | 9632686 | 1.02E+07 | 9837101 | 1.12E+07 | 1.20E+07 | 1.17E+07 | 1.10E+07 | 1331.6763 | TSNLNEELGQVK | P70704 | P70704 | ATP8A1 | ATPase, aminophospholipid transporter (APLT), class I, type 8A, member 1 | Cytoplasm | transporter |
| 15549316 | 2 | 0.388 | 7963358.5 | 9330658 | 8980571 | 7676151 | 1.11E+07 | 1.04E+07 | 1.07E+07 | 1.04E+07 | 8899365 | 1.06E+07 | 1.06E+07 | 9952060 | 1644.848 | LQDQVPETIETLMK | P70704 | P70704 | ATP8A1 | ATPase, aminophospholipid transporter (APLT), class I, type 8A, member 1 | Cytoplasm | transporter |
| 15550062 | 3 | 0.283 | 9699679 | 9415444 | 9701877 | 9959953 | 9314191 | 9806149 | 8228699 | 9360275 | 8146148 | 9973173 | 8315941 | 1.12E+07 | 3688.7188 | EYIPADTVLLSSSEPQAMC[160.0307]YIETSNLDGETNLK | P70704 | P70704 | ATP8A1 | ATPase, aminophospholipid transporter (APLT), class I, type 8A, member 1 | Cytoplasm | transporter |
| 15538506 | 2 | 0.171 | 6783869.5 | 6224116 | 7451040.5 | 7472743.5 | 7236726 | 8231199.5 | 6932965 | 7412744.5 | 8080116 | 8820811 | 7355402 | 7969876 | 1081.5774 | QLNFVFTGR | P70704 | P70704 | ATP8A1 | ATPase, aminophospholipid transporter (APLT), class I, type 8A, member 1 | Cytoplasm | transporter |
| 15536228 | 2 | 0.081 | 5319292.5 | 5621200.5 | 6646605 | 6859051 | 7881107 | 6561732 | 6756326.5 | 7135884 | 5927090 | 7887504.5 | 6876832 | 7177361 | 1252.6278 | LEESYELIEK | P70704 | P70704 | ATP8A1 | ATPase, aminophospholipid transporter (APLT), class I, type 8A, member 1 | Cytoplasm | transporter |
| 15540674 | 2 | 0.985 | 5377796 | 7329077 | 8040306 | 5540502 | 7395430.5 | 7864826.5 | 6397773 | 6558186 | 7762334.5 | 7526191.5 | 6970625.5 | 6863204.5 | 1070.5833 | SQDPGAVVLGK | P70704 | P70704 | ATP8A1 | ATPase, aminophospholipid transporter (APLT), class I, type 8A, member 1 | Cytoplasm | transporter |
| 15549484 | 3 | 0.278 | 5352219.5 | 6120197.5 | 4998180.5 | 5653532 | 4036601.5 | 4587129.5 | 4845753.5 | 4699671.5 | 3664206.75 | 7357988.5 | 5410522.5 | 4608254 | 3248.5757 | SESLQQNLLHGYAFSQDENGIVSQSEVIR | P70704 | P70704 | ATP8A1 | ATPase, aminophospholipid transporter (APLT), class I, type 8A, member 1 | Cytoplasm | transporter |
| 15534456 | 2 | 0.704 | 5957569 | 7900978.5 | 8035827.5 | 7563182.5 | 7425358 | 8473061 | 7009978.5 | 7500819 | 7042471 | 7197700.5 | 7641825.5 | 7450027.5 | 1069.66 | VNVGDIVIIK | P70704 | P70704 | ATP8A1 | ATPase, aminophospholipid transporter (APLT), class I, type 8A, member 1 | Cytoplasm | transporter |
| 15532839 | 2 | 0.166 | 6210478 | 5738623.5 | 5923748.5 | 5980679 | 6054506 | 5788164.5 | 6953521 | 6345875 | 4779240.5 | 6661979 | 7384245 | 7317475 | 1215.6382 | LMQNSTSPPLK | P70704 | P70704 | ATP8A1 | ATPase, aminophospholipid transporter (APLT), class I, type 8A, member 1 | Cytoplasm | transporter |
| 15535480 | 2 | 0.876 | 7485552 | 7349611.5 | 8709322 | 5520417 | 7262255 | 9201313 | 7029022 | 6894820 | 7809749.5 | 6097965.5 | 7692177 | 7801081.5 | 1122.629 | YNVITFLPR | P70704 | P70704 | ATP8A1 | ATPase, aminophospholipid transporter (APLT), class I, type 8A, member 1 | Cytoplasm | transporter |
| 15535798 | 2, 3 | 0.123 | 4627141 | 3252504.5 | 4800754.5 | 4448629 | 3737312.25 | 3478797 | 4698555 | 3252363.25 | 3023663 | 5938791.5 | 6232239.5 | 4399606 | 1385.6559, 1385.6562 | QETAINIGHSC[160.0307]R | P70704 | P70704 | ATP8A1 | ATPase, aminophospholipid transporter (APLT), class I, type 8A, member 1 | Cytoplasm | transporter |
| 15538126 | 2 | 0.183 | 4756960.5 | 6245619 | 5162485 | 4795697 | 4529683.5 | 4587035.5 | 5661767.5 | 5394031.5 | 5119329.5 | 5556128 | 5970925.5 | 4868258.5 | 1300.6617 | HLEQFATEGLR | P70704 | P70704 | ATP8A1 | ATPase, aminophospholipid transporter (APLT), class I, type 8A, member 1 | Cytoplasm | transporter |
| 15534913 | 2 | 0.757 | 4317349.5 | 3912276.5 | 4709643 | 4346222 | 4903380.5 | 4957017 | 5323268.5 | 3621050.5 | 4540733 | 4626159 | 4332113 | 4043690.5 | 1233.6355 | HLYDFVGNIR | P70704 | P70704 | ATP8A1 | ATPase, aminophospholipid transporter (APLT), class I, type 8A, member 1 | Cytoplasm | transporter |
| 15535066 | 2 | 0.822 | 3250904.25 | 3638071.25 | 5168306.5 | 3010777.5 | 5319368.5 | 4877033 | 4569436.5 | 4727107 | 4183865.25 | 4512726.5 | 4131109.25 | 5042487 | 1238.598 | TSQNALDFNTK | P70704 | P70704 | ATP8A1 | ATPase, aminophospholipid transporter (APLT), class I, type 8A, member 1 | Cytoplasm | transporter |
| 15541133 | 2 | 0.744 | 2931768.5 | 4265271.5 | 5495788 | 3401567 | 4849401 | 5132120.5 | 3951897.5 | 4720620 | 4760898 | 3964031 | 3320268.75 | 4047337.25 | 960.4916 | FLYSQFR | P70704 | P70704 | ATP8A1 | ATPase, aminophospholipid transporter (APLT), class I, type 8A, member 1 | Cytoplasm | transporter |
| 15533758 | 2 | 0.043 | 2780696.75 | 2870395 | 2417508.75 | 2459546 | 3292288.5 | 2617806 | 2691592.75 | 2441743.75 | 2396808.5 | 3900865.25 | 3157273.75 | 3232634 | 920.4737 | SEVVEMVK | P70704 | P70704 | ATP8A1 | ATPase, aminophospholipid transporter (APLT), class I, type 8A, member 1 | Cytoplasm | transporter |
| 15550172 | 2 | 0.876 | 1842807.625 | 2680949 | 4039746.25 | 1536271 | 5203407 | 2884602.25 | 2515644.5 | 3392528.25 | 1606980.375 | 2920926 | 1699427.75 | 2136523.5 | 1658.8203 | TPDSVIIDSLGQEER | P70704 | P70704 | ATP8A1 | ATPase, aminophospholipid transporter (APLT), class I, type 8A, member 1 | Cytoplasm | transporter |
| 15541992 | 2 | 0.32 | 5615129 | 1.35E+07 | 6487152 | 6282756.5 | 9186363 | 1.43E+07 | 6829398 | 1.07E+07 | 1.54E+07 | 2669393.5 | 7639777.5 | 6440296.5 | 1085.4988 | AEEDRYFR | O35143 | O35143 | ATPIF1 | ATPase inhibitory factor 1 | Cytoplasm | other |
| 15553235 | 4 | 0.72 | 3179703.5 | 3006208.25 | 2410110 | 3786944 | 4277891 | 3536136.5 | 2412859.5 | 3568655.75 | 4563396.5 | 2094437.75 | 6137419.5 | 3376555.5 | 4236.127 | VLISNHVTPFDHNIVNLLTTC[160.0307]STPLLNSPPSFVC[160.0307]WSR | P70295 | P70295 | AUP1 | ancient ubiquitous protein 1 | Cytoplasm | other |
| 15532388 | 2 | 0.009 | 6156096.5 | 8234439.5 | 7131402 | 9011798 | 8544231 | 1.06E+07 | 7290124 | 1.09E+07 | 1.21E+07 | 3706802.5 | 6289381.5 | 4267590.5 | 1091.5823 | TPQIQVYSR | P01887 | P01887 | B2M | beta-2-microglobulin | Plasma Membrane | transmembrane receptor |
| 15532438 | 2 | 0.497 | 2.10E+07 | 1.85E+07 | 1.85E+07 | 2.19E+07 | 1.29E+07 | 2.05E+07 | 2.02E+07 | 1.50E+07 | 2.03E+07 | 1.97E+07 | 2.28E+07 | 2.45E+07 | 1200.6921 | RVGFDPILQR | Q09199 | Q09199 | B4GALNT2 | beta-1,4-N-acetyl-galactosaminyl transferase 2 | Cytoplasm | enzyme |
| 15532840 | 2 | 0.64 | 1.75E+08 | 2.33E+08 | 3.13E+08 | 1.07E+08 | 2.91E+08 | 2.03E+08 | 2.10E+08 | 2.80E+08 | 1.86E+08 | 2.39E+08 | 1.28E+08 | 1.56E+08 | 884.5073 | VELHKMK | Q9JJ04 | Q9JJ04 | B4GALT4 | UDP-Gal:betaGlcNAc beta 1,4- galactosyltransferase, polypeptide 4 | Cytoplasm | enzyme |
| 15533115 | 3 | 0.211 | 4.07E+07 | 4.25E+07 | 2.44E+07 | 4.42E+07 | 3.44E+07 | 3.15E+07 | 3.76E+07 | 3.00E+07 | 2.70E+07 | 5.78E+07 | 3.99E+07 | 4.50E+07 | 2057.015 | EILLEMIHSIQNSQDMR | Q91YN9 | Q91YN9 | BAG2 | BCL2-associated athanogene 2 | Cytoplasm | other |
| 15536566 | 2 | 0.96 | 2.56E+07 | 3.31E+07 | 2.60E+07 | 2.50E+07 | 3.14E+07 | 2.21E+07 | 3.06E+07 | 2.72E+07 | 1.99E+07 | 3.72E+07 | 1.89E+07 | 2.68E+07 | 1328.7393 | LLESLDQLELR | Q91YN9 | Q91YN9 | BAG2 | BCL2-associated athanogene 2 | Cytoplasm | other |
| 15538803 | 3 | 0.062 | 1.59E+07 | 2.04E+07 | 1.70E+07 | 1.62E+07 | 1.48E+07 | 1.61E+07 | 2.00E+07 | 1.49E+07 | 1.46E+07 | 2.15E+07 | 1.92E+07 | 2.31E+07 | 1844.9066 | QISDGEREELNLTANR | Q91YN9 | Q91YN9 | BAG2 | BCL2-associated athanogene 2 | Cytoplasm | other |
| 15542256 | 2 | 0.81 | 1.28E+07 | 1.33E+07 | 1.24E+07 | 1.00E+07 | 1.31E+07 | 1.09E+07 | 1.28E+07 | 1.25E+07 | 7915713.5 | 2.04E+07 | 9242177 | 1.08E+07 | 1061.5084 | DAATAVEQEK | Q91YN9 | Q91YN9 | BAG2 | BCL2-associated athanogene 2 | Cytoplasm | other |
| 15539850 | 3 | 0.264 | 1.15E+07 | 8357837 | 7056329.5 | 1.47E+07 | 5380583.5 | 9018829 | 9458084 | 7514697 | 8461292 | 1.58E+07 | 1.26E+07 | 1.20E+07 | 1629.8419 | VEALRDAATAVEQEK | Q91YN9 | Q91YN9 | BAG2 | BCL2-associated athanogene 2 | Cytoplasm | other |
| 15557970 | 2 | 0.715 | 1.05E+07 | 1.44E+07 | 9783863 | 8122707 | 1.12E+07 | 1.12E+07 | 9409279 | 9991209 | 1.16E+07 | 1.36E+07 | 8918603 | 1.32E+07 | 2057.0117 | EILLEMIHSIQNSQDMR | Q91YN9 | Q91YN9 | BAG2 | BCL2-associated athanogene 2 | Cytoplasm | other |
| 15554780 | 2 | 0.22 | 9854012 | 1.09E+07 | 1.08E+07 | 8957734 | 1.27E+07 | 9609859 | 1.00E+07 | 9815522 | 1.03E+07 | 1.33E+07 | 1.38E+07 | 1.04E+07 | 1607.806 | FQSIVIGC[160.0307]ALEDQK | Q91YN9 | Q91YN9 | BAG2 | BCL2-associated athanogene 2 | Cytoplasm | other |
| 15533574 | 3 | 0.178 | 1.26E+07 | 7419123 | 2897122.25 | 8210220 | 6100741.5 | 6160680 | 6628562 | 6276432.5 | 7979850.5 | 1.12E+07 | 1.56E+07 | 1.26E+07 | 1735.9012 | FQSIVIGC[160.0307]ALEDQKK | Q91YN9 | Q91YN9 | BAG2 | BCL2-associated athanogene 2 | Cytoplasm | other |
| 15548661 | 2 | 0.374 | 7492947.5 | 5003045.5 | 4864657 | 7755321 | 1783173.875 | 5165686 | 5323931.5 | 3759532 | 4673599.5 | 9946498 | 6784258 | 6399696.5 | 1629.8408 | VEALRDAATAVEQEK | Q91YN9 | Q91YN9 | BAG2 | BCL2-associated athanogene 2 | Cytoplasm | other |
| 15536109 | 3 | 0.501 | 9471064 | 6608181 | 2687552 | 1.03E+07 | 2105431.25 | 5606383.5 | 6238582 | 3097472 | 5494825.5 | 8597400 | 8213538 | 1.03E+07 | 1897.071 | LLESLDQLELRVEALR | Q91YN9 | Q91YN9 | BAG2 | BCL2-associated athanogene 2 | Cytoplasm | other |
| 15555797 | 2 | 0.359 | 2064054 | 1036191.875 | 281907.312 | 1137820.125 | 782320.062 | 770849.625 | 848270.438 | 928148.25 | 1359816.75 | 1300084.25 | 2754581 | 1764170.375 | 1735.9012 | FQSIVIGC[160.0307]ALEDQKK | Q91YN9 | Q91YN9 | BAG2 | BCL2-associated athanogene 2 | Cytoplasm | other |
| 15555201 | 3 | 0.093 | 2068220.75 | 4532211 | 2568132.75 | 3099165.75 | 4017658.5 | 4868607 | 2477329 | 3953823.5 | 5093494 | 1213742.125 | 2371321.25 | 2112427 | 3089.5195 | RYDTEFQNLLEQLQPTAGNAYELFTK | O08734 | O08734 | BAK1 | BCL2-antagonist/killer 1 | Cytoplasm | other |
| 15536557 | 2 | 0.628 | 4.63E+07 | 1.05E+08 | 5.98E+07 | 1.20E+08 | 8.74E+07 | 1.13E+08 | 5.53E+07 | 4.27E+07 | 2.02E+08 | 5.39E+07 | 7.70E+07 | 7.20E+07 | 1312.7445 | PVGSLAGIGDVLSK | O54962 | O54962 | BANF1 | barrier to autointegration factor 1 | Nucleus | other |
| 15541667, 15545921 | 2, 3 | 0.493 | 3.07E+07 | 5.72E+07 | 2.96E+07 | 6.17E+07 | 5.26E+07 | 6.26E+07 | 3.34E+07 | 2.36E+07 | 8.47E+07 | 2.88E+07 | 3.96E+07 | 4.61E+07 | 1415.6704, 1415.6709 | HRDFVAEPMGEK | O54962 | O54962 | BANF1 | barrier to autointegration factor 1 | Nucleus | other |
| 15536197, 15544312 | 2 | 0.629 | 1.03E+07 | 1.99E+07 | 1.38E+07 | 2.66E+07 | 1.83E+07 | 2.59E+07 | 1.24E+07 | 9015960 | 4.52E+07 | 1.12E+07 | 1.88E+07 | 1.66E+07 | 927.37854, 927.3791 | DC[160.0307]FGC[160.0307]LR | O54962 | O54962 | BANF1 | barrier to autointegration factor 1 | Nucleus | other |
| 15540911 | 2 | 0.674 | 6806454.5 | 1.28E+07 | 9200400 | 1.39E+07 | 8149813 | 1.54E+07 | 5234358 | 5787018.5 | 2.08E+07 | 5481346 | 8731618 | 8943328 | 1349.8188 | AYVVLGQFLVLK | O54962 | O54962 | BANF1 | barrier to autointegration factor 1 | Nucleus | other |
| 15542366 | 2 | 0.07 | 647437.312 | 1120898.625 | 2175808.75 | 4149083.5 | 3978022 | 4115344.75 | 2818634.75 | 2994327 | 3981309.5 | 803633.938 | 3851847.5 | 2532297.5 | 1104.5327 | AENEALAMQK | Q61335 | Q61335 | BCAP31 | B-cell receptor-associated protein 31 | Cytoplasm | transporter |
| 15551690 | 3 | 0.964 | 2000261.5 | 3548577.75 | 1977365.125 | 2893576 | 3000787.5 | 2465533 | 1586062.5 | 2829608.75 | 5381199.5 | 1571558.375 | 5067547 | 2185010 | 2479.1702 | EVFGSGTAC[160.0307]QVC[160.0307]PVHQILYEGK | O35855 | O35855 | BCAT2 | branched chain amino-acid transaminase 2, mitochondrial | Cytoplasm | enzyme |
| 15553543 | 3 | 0.221 | 3848112.5 | 1.01E+07 | 4299514.5 | 4286657.5 | 5505868.5 | 7533824.5 | 3072372.25 | 7728332 | 8456970 | 1973846.75 | 3131872.25 | 3740281.5 | 2611.4075 | QQGPSPPGVQLDVAPQSLNPEVLLK | P59017 | P59017 | BCL2L13 | BCL2-like 13 (apoptosis facilitator) | Cytoplasm | other |
| 15537692 | 2 | 0.165 | 2646405.25 | 5458716.5 | 2084524 | 3460822.5 | 3077700 | 3939872.5 | 2232400.25 | 3258852.25 | 4037514.5 | 1778908.625 | 2278329.75 | 1744592.625 | 1160.5521 | AEGAAQLSEER | P59017 | P59017 | BCL2L13 | BCL2-like 13 (apoptosis facilitator) | Cytoplasm | other |
| 15535220 | 3 | 0.062 | 2833519 | 5774324 | 2500474 | 3778713.75 | 3207955.25 | 4603293.5 | 2336357.5 | 3455019.75 | 4252953.5 | 1623786.625 | 1786117 | 2155064.75 | 1658.7574 | SENNSSNSDIVHVEK | P59017 | P59017 | BCL2L13 | BCL2-like 13 (apoptosis facilitator) | Cytoplasm | other |
| 15532893 | 2 | 0.325 | 1.88E+08 | 1.69E+08 | 1.77E+08 | 1.95E+08 | 1.68E+08 | 1.39E+08 | 1.67E+08 | 1.71E+08 | 1.57E+08 | 2.38E+08 | 1.77E+08 | 1.82E+08 | 706.38715 | LEAAFR | Q8C6A8 | Q8C6A8 | BHLHE22 | basic helix-loop-helix family, member e22 | Nucleus | transcription regulator |
| 15540821 | 2 | 0.728 | 4.28E+07 | 3.15E+07 | 4.04E+07 | 4.16E+07 | 3.13E+07 | 3.37E+07 | 4.67E+07 | 3.04E+07 | 2.43E+07 | 6.17E+07 | 3.30E+07 | 3.46E+07 | 915.4876 | QELEQLR | Q8BR07 | Q8BR07 | BICD1 | bicaudal D homolog 1 (Drosophila) | Cytoplasm | other |
| 15532864 | 2 | 0.475 | 9094759 | 6779981 | 8508561 | 7158236.5 | 7067517.5 | 7753486 | 8658946 | 5464910.5 | 4767843 | 1.66E+07 | 5380424 | 9421275 | 1465.7651 | TLYNNQPIDFLK | Q8R016 | Q8R016 | BLMH | bleomycin hydrolase | Cytoplasm | peptidase |
| 15535185 | 4 | 0.403 | 1.07E+07 | 5176282.5 | 2213203.75 | 9629925 | 4574251 | 5202150.5 | 7172209.5 | 4856315 | 4456683.5 | 1.34E+07 | 7463331.5 | 8815439 | 2727.4258 | ATVQGAQHVFQHVVPQEGKPVTNQK | Q8R016 | Q8R016 | BLMH | bleomycin hydrolase | Cytoplasm | peptidase |
| 15551569 | 3 | 0.69 | 7710833 | 6379632.5 | 6396265.5 | 5394507.5 | 5854815.5 | 6117868 | 7398532 | 5661186.5 | 3754363 | 1.32E+07 | 3942430.5 | 7781424 | 2839.4346 | LNSDPQFVLAQNVGTTHDLLDIC[160.0307]LR | Q8R016 | Q8R016 | BLMH | bleomycin hydrolase | Cytoplasm | peptidase |
| 15538440 | 2 | 0.284 | 4985005 | 2872237 | 3232628 | 3470053 | 3471090 | 3855496.25 | 4346821 | 3376128.25 | 2026729.75 | 8179085 | 3742186.5 | 4704922 | 1439.6234 | DGEAVWFGC[160.0307]DVGK | Q8R016 | Q8R016 | BLMH | bleomycin hydrolase | Cytoplasm | peptidase |
| 15555331 | 3 | 0.874 | 3230862 | 2933238.5 | 2824235 | 3096904.75 | 3125498.5 | 2466127.5 | 4366612.5 | 3343566.25 | 1772472 | 7962530.5 | 2366368.5 | 2663837 | 2890.4429 | LVQYLLMNPTNDGGQWDMLVNIVEK | Q8R016 | Q8R016 | BLMH | bleomycin hydrolase | Cytoplasm | peptidase |
| 15535572 | 2 | 0.378 | 3958620.75 | 2140159.5 | 2522340.5 | 2570468 | 2623985.75 | 3244854 | 3435451 | 2697141.5 | 1744925.125 | 6364238.5 | 2862322 | 3473226 | 993.4679 | PLFNMEDK | Q8R016 | Q8R016 | BLMH | bleomycin hydrolase | Cytoplasm | peptidase |
| 15557912 | 2 | 0.093 | 2402350.5 | 1902179.5 | 1829417.375 | 1927727.875 | 1651694.75 | 1765262.875 | 1647373.875 | 1417687.625 | 1451466.625 | 3475974 | 1896203.625 | 2002361.375 | 1723.8094 | C[160.0307]YFFLNAFVDTAQK | Q8R016 | Q8R016 | BLMH | bleomycin hydrolase | Cytoplasm | peptidase |
| 15533825 | 3, 4 | 0.098 | 1.20E+07 | 6920345.5 | 7697139.5 | 1.42E+07 | 4815083.5 | 7427243 | 6045310 | 4394749 | 4881183.5 | 1.27E+07 | 9808062 | 1.07E+07 | 2202.1208, 2202.1218 | DSSRLPSEGPQPAHVVVGDVR | Q923D2 | Q923D2 | BLVRB | biliverdin reductase B | Cytoplasm | enzyme |
| 15548999 | 3 | 0.035 | 9926066 | 7731448 | 6280635 | 9806139 | 6123959.5 | 7781940 | 5485095 | 5085007.5 | 4815839.5 | 1.19E+07 | 9473946 | 7399072 | 2301.9307 | C[160.0307]LTTNEYDGHTTYPSHQYD | Q923D2 | Q923D2 | BLVRB | biliverdin reductase B | Cytoplasm | enzyme |
| 15538778 | 3 | 0.089 | 3236844.25 | 3541806.25 | 5288101.5 | 2711808.25 | 3474337 | 3965228.75 | 2022191.5 | 2371571.25 | 1691425.25 | 5510980 | 2211989.75 | 3568735.5 | 1756.9298 | LPSEGPQPAHVVVGDVR | Q923D2 | Q923D2 | BLVRB | biliverdin reductase B | Cytoplasm | enzyme |
| 15533420 | 2 | 0.016 | 1580581.375 | 1146034.125 | 1491273 | 1231675.125 | 849611.625 | 1210413.25 | 852409.688 | 741277.312 | 480903.719 | 1294493 | 1026507.812 | 1455409.75 | 1211.598 | LQDVTDDHIR | Q923D2 | Q923D2 | BLVRB | biliverdin reductase B | Cytoplasm | enzyme |
| 15536595 | 3 | 0.039 | 1.54E+07 | 1.40E+07 | 1.70E+07 | 1.48E+07 | 1.24E+07 | 1.84E+07 | 1.01E+07 | 8495713 | 5678006 | 1.33E+07 | 5017869.5 | 7624191.5 | 1992.0643 | AVGPHQFLGNQEAIQAAIK | P15327 | P15327 | BPGM | 2,3-bisphosphoglycerate mutase | Extracellular Space | phosphatase |
| 15538892 | 2 | 0.079 | 5367929.5 | 4379813.5 | 6195489 | 5567254.5 | 4035476.5 | 5701623 | 3151322.25 | 2428996.25 | 1687606.75 | 6145480 | 1806176 | 2497716.75 | 1130.5404 | LNNDGLEEAR | P15327 | P15327 | BPGM | 2,3-bisphosphoglycerate mutase | Extracellular Space | phosphatase |
| 15541313 | 2 | 0.025 | 3212943.25 | 2032352.375 | 3235513 | 2416245 | 2263811.5 | 2548512 | 1233087.875 | 1493156.75 | 838527.562 | 2179085 | 851445.438 | 1505582.75 | 1113.6124 | HYGALIGLNR | P15327 | P15327 | BPGM | 2,3-bisphosphoglycerate mutase | Extracellular Space | phosphatase |
| 15534735 | 2 | 0.238 | 908880.875 | 466639.906 | 890671.875 | 539011.5 | 566956.25 | 846123.5 | 425956.25 | 372369.531 | 49515.602 | 956905.25 | 171371.578 | 275738.406 | 1283.614 | MALNHGEEQVR | P15327 | P15327 | BPGM | 2,3-bisphosphoglycerate mutase | Extracellular Space | phosphatase |
| 15533386 | 2 | 0.511 | 1.14E+07 | 1.09E+07 | 1.40E+07 | 8050664.5 | 1.21E+07 | 1.05E+07 | 1.26E+07 | 1.20E+07 | 9975071 | 1.14E+07 | 1.22E+07 | 1.04E+07 | 1282.5526 | M[147.0354]SGNNLTSAEDK | A2AHJ4 | A2AHJ4 | BRWD3 | bromodomain and WD repeat domain containing 3 | Nucleus | other |
| 15532459 | 3 | 0.547 | 8.73E+08 | 7.42E+08 | 1.01E+09 | 6.97E+08 | 8.72E+08 | 7.52E+08 | 8.55E+08 | 8.29E+08 | 8.13E+08 | 1.10E+09 | 8.42E+08 | 7.84E+08 | 1508.7949 | VLQEDTLPDLHTK | P18572 | P18572 | BSG | basigin (Ok blood group) | Plasma Membrane | transporter |
| 15531266, 15546712 | 2 | 0.68 | 7.03E+08 | 6.48E+08 | 7.71E+08 | 6.90E+08 | 7.30E+08 | 6.48E+08 | 6.61E+08 | 6.85E+08 | 6.29E+08 | 8.48E+08 | 7.03E+08 | 6.30E+08 | 1097.5575, 1097.5576 | SEINVEGPPR | P18572 | P18572 | BSG | basigin (Ok blood group) | Plasma Membrane | transporter |
| 15550520, 15552802, 15559246, 15560163, 15563053 | 2, 3 | 0.809 | 8.78E+08 | 9.03E+08 | 7.70E+08 | 7.81E+08 | 7.48E+08 | 9.78E+08 | 8.94E+08 | 7.86E+08 | 1.04E+09 | 7.58E+08 | 8.92E+08 | 8.54E+08 | 2658.2307, 2658.2473, 2658.249, 2658.264 | YIVDADDRSGEYSC[160.0307]IFLPEPVGR | P18572 | P18572 | BSG | basigin (Ok blood group) | Plasma Membrane | transporter |
| 15532159 | 2 | 0.209 | 5.80E+08 | 5.74E+08 | 6.58E+08 | 5.98E+08 | 5.95E+08 | 5.81E+08 | 5.89E+08 | 5.64E+08 | 4.78E+08 | 7.37E+08 | 5.78E+08 | 6.31E+08 | 1021.5547 | YVVVSTPEK | P18572 | P18572 | BSG | basigin (Ok blood group) | Plasma Membrane | transporter |
| 15556657, 15558620, 15558776, 15559298 | 2 | 0.819 | 2.35E+08 | 2.50E+08 | 3.57E+08 | 2.35E+08 | 2.62E+08 | 2.45E+08 | 2.29E+08 | 2.72E+08 | 2.99E+08 | 3.73E+08 | 2.32E+08 | 2.62E+08 | 1859.8604, 1859.8608, 1859.8613, 1859.8618 | SDASYPPITDWFWFK | P18572 | P18572 | BSG | basigin (Ok blood group) | Plasma Membrane | transporter |
| 15547670 | 2 | 0.533 | 2.20E+08 | 1.86E+08 | 2.53E+08 | 1.89E+08 | 2.12E+08 | 1.75E+08 | 2.20E+08 | 1.95E+08 | 1.92E+08 | 2.92E+08 | 2.18E+08 | 1.86E+08 | 1508.7931 | VLQEDTLPDLHTK | P18572 | P18572 | BSG | basigin (Ok blood group) | Plasma Membrane | transporter |
| 15550685 | 2 | 0.69 | 1.53E+08 | 2.42E+08 | 3.91E+08 | 1.33E+08 | 4.06E+08 | 2.32E+08 | 2.27E+08 | 2.95E+08 | 2.10E+08 | 2.57E+08 | 1.21E+08 | 1.58E+08 | 1710.8115 | SGEYSC[160.0307]IFLPEPVGR | P18572 | P18572 | BSG | basigin (Ok blood group) | Plasma Membrane | transporter |
| 15534968 | 2 | 0.526 | 4.34E+07 | 7.74E+07 | 1.18E+08 | 3.50E+07 | 1.24E+08 | 7.36E+07 | 6.49E+07 | 1.01E+08 | 6.04E+07 | 6.41E+07 | 3.20E+07 | 4.34E+07 | 966.4514 | YIVDADDR | P18572 | P18572 | BSG | basigin (Ok blood group) | Plasma Membrane | transporter |
| 15531934 | 3 | 0.735 | 3.85E+07 | 4.84E+07 | 5.61E+07 | 3.28E+07 | 7.54E+07 | 4.25E+07 | 5.76E+07 | 7.35E+07 | 4.62E+07 | 6.02E+07 | 4.07E+07 | 4.02E+07 | 1809.8811 | KPDQTLDEDDPGAAPLK | P18572 | P18572 | BSG | basigin (Ok blood group) | Plasma Membrane | transporter |
| 15536179 | 2 | 0.407 | 4.82E+07 | 5.78E+07 | 7.87E+07 | 4.42E+07 | 5.50E+07 | 5.37E+07 | 4.33E+07 | 5.48E+07 | 5.80E+07 | 4.59E+07 | 5.01E+07 | 4.89E+07 | 718.4079 | ETISLR | P18572 | P18572 | BSG | basigin (Ok blood group) | Plasma Membrane | transporter |
| 15555975, 15558765 | 2 | 0.983 | 1.11E+07 | 1.09E+07 | 1.45E+07 | 9339973 | 1.91E+07 | 1.05E+07 | 1.50E+07 | 1.63E+07 | 9400585 | 2.01E+07 | 1.09E+07 | 9941418 | 1809.8817, 1809.8828 | KPDQTLDEDDPGAAPLK | P18572 | P18572 | BSG | basigin (Ok blood group) | Plasma Membrane | transporter |
| 15537529 | 3 | 0.911 | 5158075.5 | 8722651 | 1.33E+07 | 2113046.25 | 1.44E+07 | 9593504 | 8081916 | 9491021 | 7821352 | 1.02E+07 | 4457598 | 6062637 | 1710.8114 | SGEYSC[160.0307]IFLPEPVGR | P18572 | P18572 | BSG | basigin (Ok blood group) | Plasma Membrane | transporter |
| 15546222 | 3 | 0.025 | 2042919.375 | 1907705.5 | 1320302.75 | 1231934.75 | 1432357 | 1345001.625 | 2109684.5 | 1654580.875 | 1810117 | 4108788.75 | 2081852.875 | 2454507.5 | 1859.8607 | SDASYPPITDWFWFK | P18572 | P18572 | BSG | basigin (Ok blood group) | Plasma Membrane | transporter |
| 15556258 | 2 | 0.686 | 817711.312 | 5172031.5 | 1.37E+07 | 1791299.5 | 5270113 | 7544049 | 5729828 | 4983304.5 | 1684611.5 | 3844355.25 | 767312.125 | 1859835.875 | 1681.7885 | PDQTLDEDDPGAAPLK | P18572 | P18572 | BSG | basigin (Ok blood group) | Plasma Membrane | transporter |
| 15555500 | 3 | 0.101 | 1.22E+07 | 1.79E+07 | 1.13E+07 | 9940584 | 9651146 | 5081071 | 9342308 | 1.07E+07 | 8308237 | 1.01E+07 | 7193396 | 6248784 | 2910.2852 | MDSSNTQGIENPGFETTPPFQGMPEAK | Q9D659 | Q9D659 | C10orf54 | chromosome 10 open reading frame 54 | Plasma Membrane | other |
| 15537549 | 2 | 0.997 | 566741 | 1510920.125 | 1458963.875 | 764087.375 | 1912164.5 | 960904.938 | 899071.812 | 1411220.375 | 1176406.5 | 1162080.5 | 1120985.375 | 1168326.75 | 1490.7053 | APLC[160.0307]TEQFGSGAPR | Q91W34 | Q91W34 | C16orf58 | chromosome 16 open reading frame 58 | Cytoplasm | other |
| 15554155 | 3 | 8.96E-06 | 1324321.625 | 1159894.5 | 1909059.375 | 5.39E+07 | 6.71E+07 | 4.78E+07 | 393037.594 | 2175781 | 1229781.75 | 5.03E+07 | 3.19E+07 | 3.24E+07 | 2488.1746 | HGDSGPTVELSC[160.0307]LASSGPPIPTYR | Q9CX63 | Q9CX63 | C17orf99 | chromosome 17 open reading frame 99 | Extracellular Space | cytokine |
| 15550014 | 2, 3 | 2.05E-04 | 83912.844 | 171575.609 | 98313.445 | 3.80E+07 | 5.16E+07 | 3.25E+07 | 76100.117 | 0 | 373510.719 | 4.33E+07 | 2.07E+07 | 2.69E+07 | 2405.2332, 2405.234 | VLITC[160.0307]DAPEASQPITYSLLASR | Q9CX63 | Q9CX63 | C17orf99 | chromosome 17 open reading frame 99 | Extracellular Space | cytokine |
| 15548978 | 3 | 5.56E-05 | 99593.188 | 0 | 0 | 3.94E+07 | 3.58E+07 | 2.64E+07 | 83021.305 | 14481.472 | 0 | 2.86E+07 | 2.39E+07 | 2.18E+07 | 2561.3342 | RVLITC[160.0307]DAPEASQPITYSLLASR | Q9CX63 | Q9CX63 | C17orf99 | chromosome 17 open reading frame 99 | Extracellular Space | cytokine |
| 15531952 | 2 | 1.54E-05 | 28804.061 | 116814.828 | 158075.359 | 2.73E+07 | 3.47E+07 | 2.63E+07 | 51642.344 | 4.70E-06 | 49485.91 | 2.67E+07 | 1.70E+07 | 2.01E+07 | 1090.5875 | VLEVYPQSR | Q9CX63 | Q9CX63 | C17orf99 | chromosome 17 open reading frame 99 | Extracellular Space | cytokine |
| 15539320 | 2 | 2.51E-07 | 458344.094 | 656259.562 | 300408.031 | 1.95E+07 | 2.07E+07 | 1.56E+07 | 605911.438 | 354088.031 | 733894.375 | 1.69E+07 | 1.36E+07 | 1.29E+07 | 1309.6588 | LQMYQELWAK | Q9CX63 | Q9CX63 | C17orf99 | chromosome 17 open reading frame 99 | Extracellular Space | cytokine |
| 15542461 | 2 | 6.75E-05 | 192143.141 | 637091.688 | 510405.781 | 1.32E+07 | 1.24E+07 | 8526329 | 624940.75 | 485583.875 | 120158.977 | 1.07E+07 | 7021441.5 | 7822702.5 | 1372.7549 | PVSQLQADFVLR | Q9CX63 | Q9CX63 | C17orf99 | chromosome 17 open reading frame 99 | Extracellular Space | cytokine |
| 15555414 | 2 | 1.72E-04 | 90410.859 | 125396.188 | 565.98 | 1.20E+07 | 1.23E+07 | 8408848 | 46673.957 | 0 | 0 | 9628616 | 8337708 | 7081642 | 2536.1243 | SSPDLLTYSC[160.0307]QATSNSGTYGPSSR | Q9CX63 | Q9CX63 | C17orf99 | chromosome 17 open reading frame 99 | Extracellular Space | cytokine |
| 15552804 | 2 | 0.125 | 330236.844 | 72789.961 | 12287.765 | 414262.5 | 1081726.75 | 1409944 | 59353.238 | 1944655.125 | 355959.938 | 322046.281 | 1639049.375 | 1.03E+07 | 1575.8192 | QNPMTLGNVVIFDK | P98086 | P98086 | C1QA | complement component 1, q subcomponent, A chain | Extracellular Space | other |
| 15559274 | 2 | 0.535 | 170433.047 | 0 | 0 | 54010.32 | 150453.281 | 683602.438 | 4021.414 | 672844.125 | 177005.609 | 0 | 636995 | 5827700 | 1571.9254 | GLFQVLAGGTVLQLR | P98086 | P98086 | C1QA | complement component 1, q subcomponent, A chain | Extracellular Space | other |
| 15536490 | 2 | 0.082 | 84053.5 | 142381.109 | 4.70E-06 | 323430.312 | 773713.5 | 1417271.25 | 43892.996 | 2110693.25 | 394637.5 | 436133.469 | 1684360.875 | 1.27E+07 | 1419.6844 | VITNANENYEPR | P14106 | P14106 | C1QB | complement component 1, q subcomponent, B chain | Extracellular Space | other |
| 15535927 | 2 | 0.279 | 2.14E+07 | 4.43E+07 | 1.64E+07 | 2.46E+07 | 2.29E+07 | 3.76E+07 | 1.39E+07 | 2.73E+07 | 2.91E+07 | 1.08E+07 | 1.54E+07 | 1.93E+07 | 1311.6802 | AFVEFLTDEIK | O35658 | O35658 | C1QBP | complement component 1, q subcomponent binding protein | Cytoplasm | transcription regulator |
| 15549802, 15552130 | 3 | 0.326 | 3703595.25 | 2.06E+07 | 1.49E+07 | 905106 | 1.09E+07 | 3.71E+07 | 2935978.5 | 1.77E+07 | 1.97E+07 | 1520354.75 | 1051411.625 | 3809624 | 3134.3943, 3134.4006 | DTNYTLNTDSLDWALYDHLM[147.0354]DFLADR | O35658 | O35658 | C1QBP | complement component 1, q subcomponent binding protein | Cytoplasm | transcription regulator |
| 15550276 | 3 | 0.275 | 5634322 | 1.09E+07 | 3952314.5 | 7521694 | 7183143 | 9776968 | 6011795 | 7672625.5 | 1.08E+07 | 3089628.5 | 5585934.5 | 5910218.5 | 2455.2676 | ITSLAQLNAANHDAAIFPGGFGAAK | Q9D172 | Q9D172 | C21orf33/LOC102724023 | chromosome 21 open reading frame 33 | Cytoplasm | other |
| 15557342 | 2 | 0.39 | 128563.281 | 599606.25 | 299785.719 | 113251.297 | 438667.438 | 620786.75 | 255358.703 | 332039.094 | 304419.969 | 137636.438 | 102541.57 | 193466.609 | 1554.7374 | GVEVTVGHEQEEGGK | Q9D172 | Q9D172 | C21orf33/LOC102724023 | chromosome 21 open reading frame 33 | Cytoplasm | other |
| 15548140 | 3 | 0.075 | 2.17E+07 | 1.84E+07 | 1.73E+07 | 2.11E+07 | 1.74E+07 | 1.87E+07 | 1.96E+07 | 2.01E+07 | 1.80E+07 | 2.67E+07 | 2.19E+07 | 2.23E+07 | 2660.2712 | SSSC[160.0307]GDAELLGQATLPVGSPSRPMSR | Q80X80 | Q80X80 | C2CD2L | C2CD2-like | Cytoplasm | other |
| 15532617 | 3 | 0.272 | 2.66E+07 | 2.78E+07 | 3.39E+07 | 2.27E+07 | 2.76E+07 | 2.41E+07 | 2.95E+07 | 2.62E+07 | 2.44E+07 | 2.64E+07 | 2.70E+07 | 2.86E+07 | 2044.0034 | ERDEEQPELSTVEELIK | Q80X80 | Q80X80 | C2CD2L | C2CD2-like | Cytoplasm | other |
| 15532060 | 3 | 0.102 | 1.90E+07 | 1.83E+07 | 2.23E+07 | 1.73E+07 | 1.73E+07 | 1.52E+07 | 1.77E+07 | 1.80E+07 | 1.54E+07 | 2.50E+07 | 1.85E+07 | 1.92E+07 | 1588.7346 | ASISHVTC[160.0307]VDQSER | Q80X80 | Q80X80 | C2CD2L | C2CD2-like | Cytoplasm | other |
| 15536397 | 2, 3 | 0.315 | 8343676 | 6689756.5 | 8306826.5 | 7133998.5 | 6235041 | 6665984.5 | 9009714 | 7870677.5 | 5558116.5 | 1.10E+07 | 8323573 | 7559943.5 | 1309.7015, 1309.7024 | RQVC[160.0307]PLTPGPGK | Q80X80 | Q80X80 | C2CD2L | C2CD2-like | Cytoplasm | other |
| 15540698 | 2 | 0.265 | 7637715 | 8445424 | 8405872 | 6718467.5 | 6562658.5 | 8823049 | 8296577.5 | 6826323 | 6787168.5 | 1.10E+07 | 7692977 | 8575700 | 1066.6277 | GLLASLFAFK | Q80X80 | Q80X80 | C2CD2L | C2CD2-like | Cytoplasm | other |
| 15535039 | 3 | 0.735 | 9863577 | 1.03E+07 | 8760513 | 7872356 | 1.06E+07 | 6493273.5 | 8802103 | 1.06E+07 | 6617784.5 | 9881142 | 8437190 | 7609566.5 | 2186.026 | EAGLSQSHDDLSNTTATPSVR | Q80X80 | Q80X80 | C2CD2L | C2CD2-like | Cytoplasm | other |
| 15549435 | 3 | 0.813 | 8024286 | 1.36E+07 | 1.40E+07 | 7989754 | 1.05E+07 | 1.29E+07 | 1.02E+07 | 1.28E+07 | 1.21E+07 | 9865549 | 9175458 | 1.20E+07 | 2945.3015 | ASHLGSELGGTEELC[160.0307]C[160.0307]AAELDNPMQQK | Q80X80 | Q80X80 | C2CD2L | C2CD2-like | Cytoplasm | other |
| 15534699 | 2 | 0.269 | 1.23E+07 | 8163774 | 1.14E+07 | 8557295 | 1.03E+07 | 7814922 | 9853345 | 9513316 | 8379361.5 | 9753543 | 1.23E+07 | 1.14E+07 | 1160.697 | RSTLIISGVSK | Q80X80 | Q80X80 | C2CD2L | C2CD2-like | Cytoplasm | other |
| 15548775 | 2 | 0.614 | 7082115 | 8609387 | 7093249 | 7622689.5 | 6215276.5 | 6904527 | 8923897 | 7502010.5 | 5553242.5 | 9471285 | 7952418 | 7208407.5 | 1645.8374 | DAIVSTQPAMMVNLR | Q80X80 | Q80X80 | C2CD2L | C2CD2-like | Cytoplasm | other |
| 15534303 | 3 | 0.875 | 7633966.5 | 7655340 | 6787981 | 7772543 | 7175337.5 | 5760549 | 8061847.5 | 6808393 | 5088490 | 9110655 | 6084807 | 6413196 | 2021.9135 | MTTVLSESSGPSNASHSSSR | Q80X80 | Q80X80 | C2CD2L | C2CD2-like | Cytoplasm | other |
| 15548828 | 2 | 0.313 | 4242960 | 3672941.5 | 4973665 | 3541576.5 | 3672887.25 | 3578040.5 | 4678483.5 | 3649036.75 | 3498078.25 | 7352834 | 4216235.5 | 3960845 | 1588.7352 | ASISHVTC[160.0307]VDQSER | Q80X80 | Q80X80 | C2CD2L | C2CD2-like | Cytoplasm | other |
| 15550535 | 2 | 0.08 | 3010056.5 | 1747188.375 | 2598439.25 | 1975102.75 | 1442657.5 | 1777906.625 | 2846535.5 | 2187681.75 | 1614892.375 | 3698331.75 | 2721319.5 | 2868954 | 1854.9886 | NLGTPTSSTPRPSITPTK | Q80X80 | Q80X80 | C2CD2L | C2CD2-like | Cytoplasm | other |
| 15545283 | 3 | 0.565 | 1467269.125 | 611198 | 919536.75 | 988847.125 | 447967.75 | 818118.062 | 918340.812 | 367404.562 | 956070.125 | 650348.375 | 1590327.25 | 1336270.75 | 1122.5417 | SFRENWQR | Q80X80 | Q80X80 | C2CD2L | C2CD2-like | Cytoplasm | other |
| 15553970 | 3 | 0.781 | 2969547.75 | 1.21E+07 | 6525728.5 | 2068107.25 | 4490996 | 1.58E+07 | 3944685 | 5456631 | 4487911 | 4170201.75 | 3557308.5 | 3346540.5 | 2766.3403 | STQDTVVTLDALSAYWIASHTTEEK | P01029 | P01029 | C4A/C4B | complement component 4B (Chido blood group) | Extracellular Space | other |
| 15549207 | 4 | 0.717 | 3836406.75 | 1.27E+07 | 5666429 | 2935743 | 3685761.75 | 1.45E+07 | 3697200.5 | 4516296 | 6224707 | 4048991 | 4554438.5 | 3219420 | 3075.5042 | LQETASWLLAQQLGDGSFHDPC[160.0307]PVIHR | P01029 | P01029 | C4A/C4B | complement component 4B (Chido blood group) | Extracellular Space | other |
| 15542701 | 3 | 0.768 | 1928738.75 | 5110600 | 3211428.25 | 2214794.75 | 1638262.125 | 8070655 | 2162343.5 | 2584442.5 | 3127834 | 1867686.125 | 2914625 | 1753505.75 | 2085.1667 | ASAGLLGAHAAAITAYALTLTK | P01029 | P01029 | C4A/C4B | complement component 4B (Chido blood group) | Extracellular Space | other |
| 15553571 | 3 | 0.641 | 898662.25 | 5065779.5 | 3374698 | 1029642.125 | 871172.625 | 5751907 | 1338617.25 | 1513588.375 | 1250334.125 | 1826297.875 | 1098504.375 | 890858.438 | 2529.2803 | VTASEPLETMGSEGALSPGGVASLLR | P01029 | P01029 | C4A/C4B | complement component 4B (Chido blood group) | Extracellular Space | other |
| 15555475 | 2 | 0.884 | 573870.5 | 5147666 | 2043850.75 | 1230660.375 | 701028.438 | 5716454 | 1842305.125 | 2112292.75 | 1319799.125 | 1523534.125 | 1572649 | 598562.75 | 2415.1975 | TLEIPGSSDPNIVPDGDFSSLVR | P01029 | P01029 | C4A/C4B | complement component 4B (Chido blood group) | Extracellular Space | other |
| 15547988 | 3 | 0.002 | 2.83E+07 | 3.14E+07 | 3.58E+07 | 3.49E+07 | 3.53E+07 | 3.86E+07 | 2.18E+07 | 2.47E+07 | 2.26E+07 | 2.62E+07 | 1.83E+07 | 1.88E+07 | 2836.4297 | QSPVDIDTATAQHDPALQPLLISYDK | P00920 | P00920 | CA2 | carbonic anhydrase II | Cytoplasm | enzyme |
| 15550492 | 3 | 0.017 | 1.16E+07 | 1.10E+07 | 1.35E+07 | 1.19E+07 | 1.06E+07 | 1.50E+07 | 7959352 | 7917062 | 8420870 | 1.01E+07 | 1.17E+07 | 8330627.5 | 2874.3286 | TLNFNEEGDAEEAMVDNWRPAQPLK | P00920 | P00920 | CA2 | carbonic anhydrase II | Cytoplasm | enzyme |
| 15532788 | 2 | 0.081 | 7402147 | 6283251 | 9708779 | 8528692 | 6545714.5 | 8902086 | 5024997.5 | 5510415 | 5570634 | 8020493.5 | 7039539 | 5508575 | 998.5606 | IGPASQGLQK | P00920 | P00920 | CA2 | carbonic anhydrase II | Cytoplasm | enzyme |
| 15536081 | 2 | 0.162 | 3204917.5 | 2745110.75 | 1.29E+07 | 3566809.75 | 7509178.5 | 1.26E+07 | 2357153 | 2379432.5 | 2290430.75 | 7178987.5 | 4001631.5 | 1.08E+07 | 1009.6033 | VLEALHSIK | P00920 | P00920 | CA2 | carbonic anhydrase II | Cytoplasm | enzyme |
| 15552739 | 2 | 0.501 | 5789606 | 4715838.5 | 1.26E+07 | 7955085.5 | 6826499 | 1.14E+07 | 4039945 | 6173655.5 | 7820878 | 5943655.5 | 6268209.5 | 5453726.5 | 1668.9614 | AVQQPDGLAVLGIFLK | P00920 | P00920 | CA2 | carbonic anhydrase II | Cytoplasm | enzyme |
| 15545575 | 3 | 0.322 | 329728.969 | 82198.234 | 726669.312 | 260157.984 | 193091.328 | 359953.844 | 222670.188 | 195759.969 | 480888.969 | 87757.695 | 193432.875 | 76148.445 | 1668.9647 | AVQQPDGLAVLGIFLK | P00920 | P00920 | CA2 | carbonic anhydrase II | Cytoplasm | enzyme |
| 15536149 | 3 | 0.128 | 2.51E+07 | 3.12E+07 | 3.49E+07 | 2.50E+07 | 3.67E+07 | 3.48E+07 | 3.79E+07 | 3.31E+07 | 2.88E+07 | 2.89E+07 | 2.07E+07 | 1.96E+07 | 1596.7103 | DTDSEEEIREAFR | P62204 | P62204 | CALM1 | Calmodulin | Cytoplasm | |
| 15543114, 15544793 | 2 | 0.392 | 1.23E+07 | 9249110 | 9788587 | 1.22E+07 | 1.13E+07 | 1.02E+07 | 1.01E+07 | 1.16E+07 | 1.07E+07 | 9569761 | 1.07E+07 | 8576285 | 1265.609, 1265.61 | DGNGYISAAELR | P62204 | P62204 | CALM1 | Calmodulin | Cytoplasm | |
| 15548639 | 2 | 0.287 | 7439419.5 | 6690254.5 | 8298481.5 | 6079201 | 8225202.5 | 6917352 | 7524256.5 | 5956864.5 | 6634228 | 7838481 | 4912387 | 3840933.25 | 1596.7097 | DTDSEEEIREAFR | Q9D6P8 | Q9D6P8 | CALML3 | calmodulin-like 3 | Cytoplasm | other |
| 15547819 | 3 | 0.372 | 1.39E+08 | 1.95E+08 | 1.72E+08 | 1.62E+08 | 2.70E+08 | 2.03E+08 | 2.30E+08 | 2.19E+08 | 1.70E+08 | 2.00E+08 | 1.53E+08 | 1.52E+08 | 2960.2847 | KPEDWDEEMDGEWEPPVIQNPEYK | P14211 | P14211 | CALR | calreticulin | Cytoplasm | transcription regulator |
| 15548029 | 2, 3 | 0.635 | 9.05E+07 | 1.32E+08 | 1.29E+08 | 1.08E+08 | 1.78E+08 | 1.24E+08 | 1.38E+08 | 1.46E+08 | 1.17E+08 | 1.35E+08 | 9.93E+07 | 1.12E+08 | 2441.0166, 2441.0178 | DMHGDSEYNIMFGPDIC[160.0307]GPGTK | P14211 | P14211 | CALR | calreticulin | Cytoplasm | transcription regulator |
| 15548302, 15550824 | 2, 3 | 0.329 | 9.27E+07 | 1.75E+08 | 2.22E+08 | 1.13E+08 | 2.44E+08 | 2.09E+08 | 1.68E+08 | 1.90E+08 | 1.49E+08 | 1.18E+08 | 8.77E+07 | 1.28E+08 | 2568.256, 2568.259 | DDEFTHLYTLIVRPDNTYEVK | P14211 | P14211 | CALR | calreticulin | Cytoplasm | transcription regulator |
| 15531481 | 2, 3 | 0.528 | 7.72E+07 | 7.84E+07 | 8.79E+07 | 1.00E+08 | 1.06E+08 | 8.59E+07 | 1.21E+08 | 9.14E+07 | 7.55E+07 | 1.01E+08 | 9.44E+07 | 7.66E+07 | 1476.6509, 1476.6512 | HEQNIDC[160.0307]GGGYVK | P14211 | P14211 | CALR | calreticulin | Cytoplasm | transcription regulator |
| 15532148, 15542739 | 2 | 0.626 | 9.65E+07 | 9.74E+07 | 1.08E+08 | 9.15E+07 | 1.32E+08 | 1.13E+08 | 1.11E+08 | 1.05E+08 | 9.95E+07 | 9.90E+07 | 1.00E+08 | 1.03E+08 | 1451.6526, 1451.6528 | EQFLDGDAWTNR | P14211 | P14211 | CALR | calreticulin | Cytoplasm | transcription regulator |
| 15549664 | 3 | 0.756 | 5.25E+07 | 8.00E+07 | 6.94E+07 | 6.88E+07 | 1.12E+08 | 6.44E+07 | 7.44E+07 | 8.10E+07 | 7.80E+07 | 9.56E+07 | 5.61E+07 | 6.30E+07 | 3268.4902 | SGTIFDNFLITNDEAYAEEFGNETWGVTK | P14211 | P14211 | CALR | calreticulin | Cytoplasm | transcription regulator |
| 15531509 | 2 | 0.524 | 7.34E+07 | 6.93E+07 | 9.40E+07 | 8.39E+07 | 9.68E+07 | 8.23E+07 | 8.51E+07 | 8.07E+07 | 7.52E+07 | 8.07E+07 | 7.90E+07 | 7.75E+07 | 1004.5402 | LFPSGLDQK | P14211 | P14211 | CALR | calreticulin | Cytoplasm | transcription regulator |
| 15531413 | 4 | 0.528 | 4.34E+07 | 1.13E+08 | 1.23E+08 | 6.46E+07 | 1.35E+08 | 1.25E+08 | 9.72E+07 | 1.37E+08 | 8.28E+07 | 7.10E+07 | 5.02E+07 | 8.40E+07 | 2568.2576 | DDEFTHLYTLIVRPDNTYEVK | P14211 | P14211 | CALR | calreticulin | Cytoplasm | transcription regulator |
| 15541885 | 2 | 0.717 | 4.15E+07 | 3.92E+07 | 5.05E+07 | 4.85E+07 | 5.72E+07 | 4.13E+07 | 4.75E+07 | 4.60E+07 | 4.71E+07 | 5.36E+07 | 5.12E+07 | 4.08E+07 | 871.4174 | FYGDLEK | P14211 | P14211 | CALR | calreticulin | Cytoplasm | transcription regulator |
| 15535982 | 2 | 0.451 | 3.14E+07 | 3.08E+07 | 3.25E+07 | 3.28E+07 | 3.53E+07 | 2.54E+07 | 3.03E+07 | 2.95E+07 | 2.59E+07 | 3.35E+07 | 3.57E+07 | 3.00E+07 | 1019.565 | VHVIFNYK | P14211 | P14211 | CALR | calreticulin | Cytoplasm | transcription regulator |
| 15556405, 15559288 | 3 | 0.401 | 3.36E+07 | 2.80E+07 | 1.99E+07 | 3.92E+07 | 3.68E+07 | 2.68E+07 | 3.32E+07 | 2.71E+07 | 3.65E+07 | 3.22E+07 | 3.78E+07 | 3.20E+07 | 2519.1921, 2519.1934 | IDNSQVESGSLEDDWDFLPPKK | P14211 | P14211 | CALR | calreticulin | Cytoplasm | transcription regulator |
| 15554486 | 4 | 0.783 | 3910023.75 | 4483461.5 | 2568938.25 | 1928736.375 | 4391615 | 3192128.75 | 4320183 | 1915211.25 | 2058160 | 6864311.5 | 3151137.5 | 2106372 | 4136.9824 | GTWIHPEIDNPEYSPDANIYAYDSFAVLGLDLWQVK | P14211 | P14211 | CALR | calreticulin | Cytoplasm | transcription regulator |
| 15551873 | 3 | 0.466 | 6931946.5 | 5653065.5 | 7775984.5 | 5202158 | 9787947 | 6502502 | 8342685 | 6785640.5 | 5075832.5 | 6482116 | 4751788.5 | 4497777 | 2457.0107 | DMHGDSEYNIM[147.0354]FGPDIC[160.0307]GPGTK | P14211 | P14211 | CALR | calreticulin | Cytoplasm | transcription regulator |
| 15532010 | 2 | 0.726 | 1.05E+07 | 1.64E+07 | 2.64E+07 | 5792803 | 2.55E+07 | 1.90E+07 | 1.13E+07 | 2.00E+07 | 2.23E+07 | 4824956.5 | 1.56E+07 | 1.49E+07 | 737.4175 | FVLSSGK | P14211 | P14211 | CALR | calreticulin | Cytoplasm | transcription regulator |
| 15556964 | 3 | 0.105 | 5500418 | 4368244.5 | 5413295.5 | 3655941.25 | 6459472.5 | 4261329 | 7011945 | 4587399 | 4133262.5 | 3453964.75 | 3062445.5 | 3308087.25 | 2457.011 | DM[147.0354]HGDSEYNIMFGPDIC[160.0307]GPGTK | P14211 | P14211 | CALR | calreticulin | Cytoplasm | transcription regulator |
| 15561734, 15562193 | 2 | 0.768 | 1057792.875 | 1728041.75 | 1242051.25 | 1538676.125 | 1337567.75 | 1458091 | 1329874.25 | 1405293.75 | 1019678.25 | 1987998.875 | 832486 | 858204.125 | 2391.094, 2391.0947 | IDNSQVESGSLEDDWDFLPPK | P14211 | P14211 | CALR | calreticulin | Cytoplasm | transcription regulator |
| 15555817 | 2 | 0.454 | 1144002 | 1626077.875 | 5645556.5 | 780762.438 | 5319424.5 | 2700517.5 | 2636360.5 | 2841428 | 1371098.75 | 1513870.375 | 695973.875 | 927416.75 | 1543.6616 | DPDAAKPEDWDER | P14211 | P14211 | CALR | calreticulin | Cytoplasm | transcription regulator |
| 15547246 | 2 | 0.696 | 470887.75 | 446598.156 | 227616.828 | 607189.75 | 374931 | 214599.953 | 454257.312 | 263213.594 | 235623.984 | 386844.812 | 502872.719 | 448966.281 | 1219.7036 | GQTLVVQFTVK | P14211 | P14211 | CALR | calreticulin | Cytoplasm | transcription regulator |
| 15541471 | 3 | 0.702 | 918808.938 | 4083974.75 | 1.01E+07 | 229343.156 | 8436856 | 5238800 | 2058878.25 | 3566813.25 | 1371292.875 | 310433.656 | 1097266.875 | 2649334.25 | 1660.7286 | IDDPTDSKPEDWDK | P14211 | P14211 | CALR | calreticulin | Cytoplasm | transcription regulator |
| 15533803 | 3 | 0.218 | 7021485 | 7517189 | 7697764 | 8378694 | 1.07E+07 | 9021656 | 9552239 | 8518393 | 6945332 | 7864879 | 8944493 | 6816417 | 2048.8904 | EEFTAFLHPEEYDYMK | O35887 | O35887 | CALU | calumenin | Cytoplasm | other |
| 15537346 | 2 | 0.186 | 6365611 | 7095042.5 | 7159947.5 | 7071116 | 9528862 | 8196176 | 8130464 | 8425129 | 7354721.5 | 7077758 | 7975516 | 6481475.5 | 1280.5978 | SFDQLTPEESK | O35887 | O35887 | CALU | calumenin | Cytoplasm | other |
| 15552639 | 3 | 0.4 | 5928818 | 6505614 | 5171153 | 7607828 | 1.11E+07 | 5983144.5 | 8540123 | 8058725.5 | 5303793 | 6110564 | 6728853.5 | 7073464.5 | 3329.4207 | NADGFIDLEEYIGDMYSHDGNADEPEWVK | O35887 | O35887 | CALU | calumenin | Cytoplasm | other |
| 15551158 | 3 | 0.485 | 5858772.5 | 6761437.5 | 9168515 | 5466705 | 7364994.5 | 9575208 | 7712521.5 | 7451055 | 6894274.5 | 4283506.5 | 5960571.5 | 7176659 | 2516.1663 | YDLFVGSQATDFGEALVRHDEF | O35887 | O35887 | CALU | calumenin | Cytoplasm | other |
| 15553175 | 3 | 0.344 | 3550078.5 | 4060934 | 4134097 | 3456690.75 | 4238726.5 | 1391900.5 | 3869270.75 | 5015402 | 3348128.75 | 3245123.75 | 2783472.75 | 2244842 | 2506.082 | VHNDAQNFDYDHDAFLGAEEAK | O35887 | O35887 | CALU | calumenin | Cytoplasm | other |
| 15538441 | 4 | 0.878 | 3832717.25 | 3709503.25 | 2540300.25 | 3063193.75 | 5589861.5 | 2114292.75 | 3419731.75 | 4931209.5 | 2910091.5 | 3160751 | 3443877.5 | 2388996 | 2506.0845 | VHNDAQNFDYDHDAFLGAEEAK | O35887 | O35887 | CALU | calumenin | Cytoplasm | other |
| 15537532 | 3 | 0.211 | 2215013.75 | 2836803.5 | 3138585.25 | 2497752.5 | 4011568 | 3514200.75 | 3331882.75 | 3426363 | 2921935.5 | 2505173.75 | 2647478.25 | 2501253.75 | 1340.656 | TEREQFVEFR | O35887 | O35887 | CALU | calumenin | Cytoplasm | other |
| 15545120 | 3 | 0.71 | 817836.125 | 1254338 | 310806.031 | 92156.711 | 1315763.75 | 649373.125 | 1285984.75 | 712219 | 148170.297 | 857007 | 929895.25 | 1380231 | 1239.6216 | RWIHEDVER | O35887 | O35887 | CALU | calumenin | Cytoplasm | other |
| 15554169 | 2 | 0.288 | 469767.781 | 496627.062 | 473360.406 | 1764320.375 | 883163.188 | 3658241.5 | 1157775.25 | 635396.688 | 1.10E+07 | 570814.188 | 2478540.5 | 2469058.75 | 1993.921 | LLDLDPEPQGDEDPDTPK | P51437 | P51437 | CAMP | cathelicidin antimicrobial peptide | Cytoplasm | other |
| 15552039 | 2 | 0.351 | 800346.188 | 398071.938 | 602737.625 | 2135753.5 | 1407295.75 | 5223625 | 1376070.75 | 1061631.375 | 1.31E+07 | 248912.594 | 3475728.5 | 3419808.75 | 1898.8849 | AVDDFNQQSLDTNLYR | P51437 | P51437 | CAMP | cathelicidin antimicrobial peptide | Cytoplasm | other |
| 15532224 | 3 | 0.516 | 2.20E+07 | 1.83E+07 | 1.84E+07 | 1.78E+07 | 1.82E+07 | 1.89E+07 | 2.08E+07 | 1.48E+07 | 1.30E+07 | 2.83E+07 | 1.54E+07 | 1.90E+07 | 2047.0796 | VIRPLDQPSSFDATPYIK | Q6ZQ38 | Q6ZQ38 | CAND1 | cullin-associated and neddylation-dissociated 1 | Cytoplasm | transcription regulator |
| 15541568 | 3 | 0.88 | 1.38E+07 | 1.55E+07 | 1.18E+07 | 1.30E+07 | 9877410 | 1.38E+07 | 1.64E+07 | 1.15E+07 | 1.29E+07 | 2.25E+07 | 1.12E+07 | 1.13E+07 | 2065.2122 | IDLRPVLGEGVPILASFLR | Q6ZQ38 | Q6ZQ38 | CAND1 | cullin-associated and neddylation-dissociated 1 | Cytoplasm | transcription regulator |
| 15549427 | 3 | 0.415 | 1.55E+07 | 1.14E+07 | 1.06E+07 | 1.53E+07 | 9421603 | 1.13E+07 | 1.49E+07 | 1.16E+07 | 1.40E+07 | 2.13E+07 | 1.43E+07 | 1.29E+07 | 2661.4443 | QGGLLVNFHPSILTC[160.0307]LLPQLTSPR | Q6ZQ38 | Q6ZQ38 | CAND1 | cullin-associated and neddylation-dissociated 1 | Cytoplasm | transcription regulator |
| 15553215 | 2, 3 | 0.568 | 1.05E+07 | 9608864 | 9005563 | 7980330.5 | 7951508 | 9743936 | 1.01E+07 | 6601029 | 7837210 | 1.79E+07 | 7139800 | 9763174 | 2259.0457, 2259.0474 | EYQVETIVDTLC[160.0307]TNMLSDK | Q6ZQ38 | Q6ZQ38 | CAND1 | cullin-associated and neddylation-dissociated 1 | Cytoplasm | transcription regulator |
| 15550176 | 3 | 0.528 | 1.16E+07 | 8555124 | 8415164 | 9841480 | 8989481 | 1.05E+07 | 1.10E+07 | 9673047 | 5808012 | 1.72E+07 | 8611848 | 1.10E+07 | 3882.8716 | QTRPVQSWLC[160.0307]DPDAMEQGDTPLTMLQSQVPNIVK | Q6ZQ38 | Q6ZQ38 | CAND1 | cullin-associated and neddylation-dissociated 1 | Cytoplasm | transcription regulator |
| 15536147 | 2 | 0.869 | 1.44E+07 | 1.21E+07 | 1.13E+07 | 1.34E+07 | 1.09E+07 | 1.14E+07 | 1.59E+07 | 9392383 | 9008326 | 1.66E+07 | 1.03E+07 | 1.17E+07 | 1486.787 | EGPAVVGQFIQDVK | Q6ZQ38 | Q6ZQ38 | CAND1 | cullin-associated and neddylation-dissociated 1 | Cytoplasm | transcription regulator |
| 15541116 | 3 | 0.322 | 1.12E+07 | 8989726 | 8431316 | 7039834.5 | 7127256 | 6837221.5 | 8458519 | 7283929.5 | 6127820 | 1.51E+07 | 6648466.5 | 8154783 | 1733.8856 | MLTGPVYSQSTALTHK | Q6ZQ38 | Q6ZQ38 | CAND1 | cullin-associated and neddylation-dissociated 1 | Cytoplasm | transcription regulator |
| 15532335 | 2 | 0.946 | 9369924 | 1.04E+07 | 8263120.5 | 8396214 | 8268490.5 | 1.05E+07 | 1.07E+07 | 9287471 | 1.03E+07 | 1.39E+07 | 5577610 | 9879596 | 1483.8817 | ISGSILNELIGLVR | Q6ZQ38 | Q6ZQ38 | CAND1 | cullin-associated and neddylation-dissociated 1 | Cytoplasm | transcription regulator |
| 15541208 | 2 | 0.394 | 1.28E+07 | 9886479 | 9887337 | 1.10E+07 | 7440001.5 | 7747204.5 | 9675917 | 7769658.5 | 6568259 | 1.34E+07 | 7817457.5 | 8341431.5 | 1325.7656 | AVAALLTIPEAEK | Q6ZQ38 | Q6ZQ38 | CAND1 | cullin-associated and neddylation-dissociated 1 | Cytoplasm | transcription regulator |
| 15548684 | 2 | 0.527 | 1.09E+07 | 8762348 | 1.06E+07 | 8190098.5 | 7737118.5 | 7307659 | 1.01E+07 | 6563745 | 7499659.5 | 1.29E+07 | 6233251 | 7740758.5 | 1542.9072 | ITSEALLVTQQLVK | Q6ZQ38 | Q6ZQ38 | CAND1 | cullin-associated and neddylation-dissociated 1 | Cytoplasm | transcription regulator |
| 15532354 | 2 | 0.574 | 1.19E+07 | 9727808 | 9599363 | 9215377 | 7109340 | 8395257 | 1.23E+07 | 7952255 | 8238435.5 | 1.29E+07 | 7320719 | 1.01E+07 | 1256.7792 | LGTLSALDILIK | Q6ZQ38 | Q6ZQ38 | CAND1 | cullin-associated and neddylation-dissociated 1 | Cytoplasm | transcription regulator |
| 15535663 | 3 | 0.486 | 9298743 | 6910089 | 7708085.5 | 7138887 | 5473394 | 6944162.5 | 8611145 | 5270371.5 | 4987141 | 1.11E+07 | 5909397.5 | 7320778 | 1268.6572 | HTVDDGLDIRK | Q6ZQ38 | Q6ZQ38 | CAND1 | cullin-associated and neddylation-dissociated 1 | Cytoplasm | transcription regulator |
| 15534921 | 2 | 0.86 | 6745145 | 5786033 | 4810968 | 5027928 | 5319442 | 5578964.5 | 6202081 | 3760506.25 | 4034975.75 | 9592854 | 2984696.25 | 5134582 | 1193.5637 | HEMLPEFYK | Q6ZQ38 | Q6ZQ38 | CAND1 | cullin-associated and neddylation-dissociated 1 | Cytoplasm | transcription regulator |
| 15538509 | 2, 3 | 0.773 | 8678178 | 7048659 | 6943235 | 6221500.5 | 5324889 | 7501191 | 7699181.5 | 5512624 | 6454516 | 9406335 | 4691740 | 6695095 | 1376.7549, 1376.755 | ADVFHAYLSLLK | Q6ZQ38 | Q6ZQ38 | CAND1 | cullin-associated and neddylation-dissociated 1 | Cytoplasm | transcription regulator |
| 15552288 | 3 | 0.223 | 5097778 | 1363120.375 | 1260212.75 | 3910262.5 | 2276032.5 | 3144645 | 3525109.5 | 2701772 | 3426508.75 | 8849629 | 5759079 | 2987816.75 | 2812.2278 | FC[160.0307]NVDDDELREYC[160.0307]IQAFESFVR | Q6ZQ38 | Q6ZQ38 | CAND1 | cullin-associated and neddylation-dissociated 1 | Cytoplasm | transcription regulator |
| 15533020 | 2 | 0.578 | 7625999 | 5575875 | 6577201.5 | 5502756.5 | 4911627.5 | 5521431 | 6058580 | 4360863.5 | 5090875 | 8793516 | 4093723 | 5491447.5 | 1083.6758 | ALTLIAGSPLK | Q6ZQ38 | Q6ZQ38 | CAND1 | cullin-associated and neddylation-dissociated 1 | Cytoplasm | transcription regulator |
| 15532694 | 2 | 0.435 | 6669283.5 | 4925953.5 | 4293325 | 4581059.5 | 4340464 | 4516281 | 4587490.5 | 3330245.25 | 4079864.5 | 8372154 | 4006339.5 | 4586980.5 | 927.56 | TVSPALIAR | Q6ZQ38 | Q6ZQ38 | CAND1 | cullin-associated and neddylation-dissociated 1 | Cytoplasm | transcription regulator |
| 15542567 | 3 | 0.877 | 5683345.5 | 6530238 | 5817463.5 | 4009200.5 | 5132137.5 | 6657029.5 | 5000296 | 5965498.5 | 4672306 | 8371463 | 3593200.25 | 5480051.5 | 1756.9076 | DLLDSVLPHLYNETK | Q6ZQ38 | Q6ZQ38 | CAND1 | cullin-associated and neddylation-dissociated 1 | Cytoplasm | transcription regulator |
| 15536952 | 3 | 0.715 | 1838207.75 | 3739646 | 3891637 | 1455949.5 | 2635765 | 2834195.25 | 2822550.25 | 2399451.25 | 1841759.25 | 4236413 | 816880.375 | 1799296.625 | 1371.7354 | VALVTFNSAAHNK | Q6ZQ38 | Q6ZQ38 | CAND1 | cullin-associated and neddylation-dissociated 1 | Cytoplasm | transcription regulator |
| 15548246 | 2 | 0.35 | 3.45E+07 | 8.24E+07 | 1.18E+08 | 5.67E+07 | 1.34E+08 | 1.15E+08 | 9.50E+07 | 1.01E+08 | 9.19E+07 | 6.32E+07 | 3.92E+07 | 6.69E+07 | 1707.8229 | IPNPDFFEDLEPFK | P35564 | P35564 | CANX | calnexin | Cytoplasm | other |
| 15548065 | 3 | 0.066 | 5.54E+07 | 5.55E+07 | 5.81E+07 | 7.90E+07 | 7.74E+07 | 6.13E+07 | 6.83E+07 | 6.79E+07 | 6.33E+07 | 6.21E+07 | 7.37E+07 | 5.93E+07 | 2479.09 | C[160.0307]ESAPGC[160.0307]GVWQRPMIDNPNYK | P35564 | P35564 | CANX | calnexin | Cytoplasm | other |
| 15532331 | 2, 3 | 0.085 | 5.32E+07 | 5.91E+07 | 6.58E+07 | 6.95E+07 | 8.30E+07 | 6.78E+07 | 7.28E+07 | 7.52E+07 | 6.49E+07 | 6.20E+07 | 7.03E+07 | 6.49E+07 | 1403.6771, 1403.6775 | TAELSLDQFHDK | P35564 | P35564 | CANX | calnexin | Cytoplasm | other |
| 15554494 | 2 | 0.761 | 3.37E+07 | 5.77E+07 | 7.68E+07 | 4.36E+07 | 8.08E+07 | 5.54E+07 | 5.33E+07 | 6.38E+07 | 5.63E+07 | 5.89E+07 | 3.68E+07 | 4.58E+07 | 1770.8291 | APVPTGEVYFADSFDR | P35564 | P35564 | CANX | calnexin | Cytoplasm | other |
| 15532020, 15543642, 15543932 | 3 | 0.373 | 2.00E+07 | 1.68E+07 | 1.95E+07 | 2.27E+07 | 2.05E+07 | 2.23E+07 | 3.25E+07 | 2.10E+07 | 1.58E+07 | 3.76E+07 | 2.27E+07 | 2.30E+07 | 2122.8853, 2122.8855 | SDAEEDGVTGSQDEEDSKPK | P35564 | P35564 | CANX | calnexin | Cytoplasm | other |
| 15534089 | 3 | 0.415 | 3.38E+07 | 4.62E+07 | 5.03E+07 | 4.36E+07 | 5.92E+07 | 5.32E+07 | 5.36E+07 | 4.62E+07 | 4.94E+07 | 3.43E+07 | 4.57E+07 | 4.93E+07 | 1986.9862 | WKPPMIDNPNYQGIWK | P35564 | P35564 | CANX | calnexin | Cytoplasm | other |
| 15539501, 15543807 | 3 | 0.612 | 1.64E+07 | 3.98E+07 | 7.09E+07 | 2.02E+07 | 6.94E+07 | 4.82E+07 | 4.36E+07 | 4.98E+07 | 2.61E+07 | 3.11E+07 | 1.81E+07 | 2.37E+07 | 2083.9407, 2083.9417 | IADPDAVKPDDWDEDAPSK | P35564 | P35564 | CANX | calnexin | Cytoplasm | other |
| 15531373 | 2 | 0.283 | 2.46E+07 | 3.28E+07 | 3.47E+07 | 3.44E+07 | 4.70E+07 | 3.60E+07 | 3.81E+07 | 3.99E+07 | 3.05E+07 | 3.05E+07 | 3.55E+07 | 3.29E+07 | 1269.617 | TPYTIMFGPDK | P35564 | P35564 | CANX | calnexin | Cytoplasm | other |
| 15531405 | 2 | 0.13 | 2.47E+07 | 2.52E+07 | 3.25E+07 | 3.04E+07 | 4.25E+07 | 3.14E+07 | 3.06E+07 | 3.63E+07 | 3.29E+07 | 2.96E+07 | 2.77E+07 | 2.55E+07 | 1088.5204 | AEEDEILNR | P35564 | P35564 | CANX | calnexin | Cytoplasm | other |

| | | | | | | | | | | |
|---|---|---|---|---|---|---|---|---|---|---|
| 15532776 | 2 | 0.57 | 867.45935 | PFLFDTK | P35564 | P35564 | CANX | calnexin | Cytoplasm | other |
| 15533983 | 2, 3 | 0.261 | 1399.6179, 1399.619 | YDGKWEVDEMK | P35564 | P35564 | CANX | calnexin | Cytoplasm | other |
| 15540624 | 3 | 0.243 | 1835.9178 | KIPNPDFFEDLEPFK | P35564 | P35564 | CANX | calnexin | Cytoplasm | other |
| 15539917 | 2 | 0.054 | 1474.6937 | VVDDWANDGWGLK | P35564 | P35564 | CANX | calnexin | Cytoplasm | other |
| 15532289 | 2 | 0.386 | 1047.5814 | GSLSGWILSK | P35564 | P35564 | CANX | calnexin | Cytoplasm | other |
| 15540750 | 2 | 0.017 | 775.4481 | GLVLMSR | P35564 | P35564 | CANX | calnexin | Cytoplasm | other |
| 15540851 | 3 | 0.962 | 1770.8301 | APVPTGEVYFADSFDR | P35564 | P35564 | CANX | calnexin | Cytoplasm | other |
| 15536291 | 2 | 0.246 | 936.4115 | WEVDEMK | P35564 | P35564 | CANX | calnexin | Cytoplasm | other |
| 15554436 | 4 | 0.412 | 4453.224 | THLYTLILNPDNSFEILVDQSVVNSGNLLNDM[147.0354]TPPVNPSR | P35564 | P35564 | CANX | calnexin | Cytoplasm | other |
| 15552233 | 3 | 0.708 | 2799.3928 | APVPTGEVYFADSFDRGSLSGWILSK | P35564 | P35564 | CANX | calnexin | Cytoplasm | other |
| 15551985 | 2 | 0.648 | 2083.9412 | IADPDAVKPDDWDEDAPSK | P35564 | P35564 | CANX | calnexin | Cytoplasm | other |
| 15557767, 15561180, 15563452 | 2 | 0.183 | 2122.8855, 2122.8862, 2122.8867 | SDAEEDGVTGSQDEEDSKPK | P35564 | P35564 | CANX | calnexin | Cytoplasm | other |
| 15563192, 15563327 | 2 | 0.817 | 1897.7372, 1897.7374 | SDAEEDGVTGSQDEEDSK | P35564 | P35564 | CANX | calnexin | Cytoplasm | other |
| 15550878 | 2 | 0.699 | 2172.185 | LSDLLAPISEQIQEVITFR | P40124 | P40124 | CAP1 | CAP, adenylate cyclase-associated protein 1 (yeast) | Plasma Membrane | other |
| 15552395 | 2 | 0.715 | 2365.2378 | GAVPYVQAFDSLLANPVAEYLK | P40124 | P40124 | CAP1 | CAP, adenylate cyclase-associated protein 1 (yeast) | Plasma Membrane | other |
| 15544229 | 3 | 0.819 | 2172.1873 | LSDLLAPISEQIQEVITFR | P40124 | P40124 | CAP1 | CAP, adenylate cyclase-associated protein 1 (yeast) | Plasma Membrane | other |
| 15555841 | 3 | 0.08 | 2563.2585 | C[160.0307]LQSGVLFQDEAFPPVSHSLGFK | O35350 | O35350 | CAPN1 | calpain 1, (mu/I) large subunit | Cytoplasm | peptidase |
| 15539790 | 3 | 0.139 | 2032.023 | RPTELMSNPQFIVDGATR | O35350 | O35350 | CAPN1 | calpain 1, (mu/I) large subunit | Cytoplasm | peptidase |
| 15533033 | 2 | 0.208 | 1260.7175 | APSDLYQIILK | O35350 | O35350 | CAPN1 | calpain 1, (mu/I) large subunit | Cytoplasm | peptidase |
| 15538969 | 2 | 0.031 | 1437.6715 | VLSEEEIDDNFK | O35350 | O35350 | CAPN1 | calpain 1, (mu/I) large subunit | Cytoplasm | peptidase |
| 15552776 | 3 | 0.17 | 2422.2573 | GSLLGC[160.0307]SINISDIRDLEAITFK | O35350 | O35350 | CAPN1 | calpain 1, (mu/I) large subunit | Cytoplasm | peptidase |
| 15554689 | 2 | 0.716 | 1984.9421 | AGTQELDDQIQANLPDEK | O35350 | O35350 | CAPN1 | calpain 1, (mu/I) large subunit | Cytoplasm | peptidase |
| 15554723 | 2 | 0.017 | 2319.0364 | YSEPDLAVDFDNFVC[160.0307]C[160.0307]LVR | O35350 | O35350 | CAPN1 | calpain 1, (mu/I) large subunit | Cytoplasm | peptidase |
| 15552055 | 2 | 0.31 | 2133.0388 | VLFEDPLFPATDDSLYYK | O08688 | O08688 | CAPN5 | calpain 5 | Cytoplasm | peptidase |
| 15548080 | 2 | 0.278 | 1761.8447 | DTFFQNPQYVFEVK | O08688 | O08688 | CAPN5 | calpain 5 | Cytoplasm | peptidase |
| 15540065 | 2 | 0.005 | 1194.5552 | VFTDVPSNC[160.0307]R | O08688 | O08688 | CAPN5 | calpain 5 | Cytoplasm | peptidase |
| 15534843 | 2 | 0.038 | 1364.7015 | DSPTGANSYVIIK | O08688 | O08688 | CAPN5 | calpain 5 | Cytoplasm | peptidase |
| 15534034 | 3 | 0.124 | 1561.8567 | LAQPITVQVWNHR | O08688 | O08688 | CAPN5 | calpain 5 | Cytoplasm | peptidase |
| 15534800 | 2, 3 | 0.021 | 1288.7233, 1288.7235 | LINTSYLSIHK | O08688 | O08688 | CAPN5 | calpain 5 | Cytoplasm | peptidase |
| 15536512 | 2 | 0.426 | 1251.7037 | QPSDLPGIVAVR | O08688 | O08688 | CAPN5 | calpain 5 | Cytoplasm | peptidase |
| 15542406, 15544152 | 2 | 0.046 | 1095.5289 | GTSTPEYNVK | O08688 | O08688 | CAPN5 | calpain 5 | Cytoplasm | peptidase |
| 15533269 | 2 | 0.102 | 1301.6002 | AYEDQNYSALK | O08688 | O08688 | CAPN5 | calpain 5 | Cytoplasm | peptidase |
| 15533161 | 2 | 0.068 | 1081.5615 | AASSIYINSR | O08688 | O08688 | CAPN5 | calpain 5 | Cytoplasm | peptidase |
| 15534263 | 2 | 0.054 | 1034.4907 | AVTAADMEAR | O08688 | O08688 | CAPN5 | calpain 5 | Cytoplasm | peptidase |
| 15534488 | 2 | 3.91E-04 | 1295.6064 | NEFWC[160.0307]ALVEK | O08688 | O08688 | CAPN5 | calpain 5 | Cytoplasm | peptidase |
| 15537623 | 2 | 0.005 | 899.4851 | YFTDIIK | O08688 | O08688 | CAPN5 | calpain 5 | Cytoplasm | peptidase |
| 15549369 | 2 | 0.01 | 2190.1545 | YVIIPTTFEPGHTGEFLLR | O08688 | O08688 | CAPN5 | calpain 5 | Cytoplasm | peptidase |
| 15545047, 15546223 | 2 | 0.467 | 1102.6389, 1102.6394 | LGHGLLAFFK | O08688 | O08688 | CAPN5 | calpain 5 | Cytoplasm | peptidase |
| 15542179 | 2 | 0.438 | 1088.5464 | GHAYAVTDVR | O08688 | O08688 | CAPN5 | calpain 5 | Cytoplasm | peptidase |
| 15560222 | 2 | 0.421 | 1865.7896 | EWTGPWSDTSEEWQK | O08688 | O08688 | CAPN5 | calpain 5 | Cytoplasm | peptidase |
| 15561434 | 2 | 0.881 | 1561.8563 | LAQPITVQVWNHR | O08688 | O08688 | CAPN5 | calpain 5 | Cytoplasm | peptidase |
| 15560666 | 3 | 0.962 | 2605.132 | NPWGEREWTGPWSDTSEEWQK | O08688 | O08688 | CAPN5 | calpain 5 | Cytoplasm | peptidase |
| 15545958 | 3 | 0.991 | 1216.6405 | GHAYAVTDVRK | O08688 | O08688 | CAPN5 | calpain 5 | Cytoplasm | peptidase |
| 15542126 | 2 | 0.069 | 1141.4913 | TDGFGIDTC[160.0307]R | O88456 | O88456 | CAPNS1 | calpain, small subunit 1 | Cytoplasm | peptidase |
| 15556148 | 2 | 0.14 | 2466.2612 | QGLSGVPILSEEELSLLDEFYK | Q60865 | Q60865 | CAPRIN1 | cell cycle associated protein 1 | Plasma Membrane | other |
| 15553918 | 4 | 0.03 | 4877.431 | LAQSNQVPVQPEATQVPLVSSTSEGYTASQPLYQPSHATEQRPQK | Q60865 | Q60865 | CAPRIN1 | cell cycle associated protein 1 | Plasma Membrane | other |
| 15534557 | 3 | 0.012 | 2089.0098 | FITHAPPGEFNEVFNDVR | P47753 | P47753 | CAPZA1 | capping protein (actin filament) muscle Z-line, alpha 1 | Cytoplasm | other |
| 15552944 | 2 | 0.231 | 1556.7051 | EASDPQPEDVDGGLK | P47753 | P47753 | CAPZA1 | capping protein (actin filament) muscle Z-line, alpha 1 | Cytoplasm | other |
| 15542796 | 2 | 0.038 | 1171.5963 | STLNEIYFGK | P47757 | P47757 | CAPZB | capping protein (actin filament) muscle Z-line, beta | Cytoplasm | other |
| 15538157 | 2 | 0.133 | 1337.644 | SGSGTMNLGGSLTR | P47757 | P47757 | CAPZB | capping protein (actin filament) muscle Z-line, beta | Cytoplasm | other |
| 15537581 | 3 | 0.22 | 1675.9717 | LQAVEVVITHLAPGTK | Q9CR86 | Q9CR86 | CARHSP1 | calcium regulated heat stable protein 1, 24kDa | Cytoplasm | other |
| 15551175 | 3 | 0.691 | 3313.5452 | PSGNMFPTIGDVHLAPFTDEQLYMEQFTK | Q9WVG6 | Q9WVG6 | CARM1 | coactivator-associated arginine methyltransferase 1 | Nucleus | transcription regulator |
| 15537962 | 3 | 0.358 | 2198.0852 | GAAVDEYFRQPVVDTFDIR | Q9WVG6 | Q9WVG6 | CARM1 | coactivator-associated arginine methyltransferase 1 | Nucleus | transcription regulator |
| 15540864 | 3 | 0.737 | 2108.0305 | ANFWYQPSFHGVDLSALR | Q9WVG6 | Q9WVG6 | CARM1 | coactivator-associated arginine methyltransferase 1 | Nucleus | transcription regulator |
| 15550873 | 3 | 0.273 | 2463.377 | VLEVLQLSDALRDDILPELGVR | Q9ER72 | Q9ER72 | CARS | cysteinyl-tRNA synthetase | Cytoplasm | enzyme |
| 15541438 | 2 | 0.315 | 1226.6714 | LATEPLEQAVR | Q9ER72 | Q9ER72 | CARS | cysteinyl-tRNA synthetase | Cytoplasm | enzyme |
| 15547961 | 3 | 0.732 | 2858.4822 | LSAAISEVVSQTPAPSTHAAAPLPGTEQK | P51125 | P51125 | CAST | calpastatin | Cytoplasm | peptidase |
| 15539358 | 3 | 0.366 | 1870.9487 | PFTPASPVQSTPSKPSDK | P51125 | P51125 | CAST | calpastatin | Cytoplasm | peptidase |
| 15536051 | 2 | 0.959 | 1239.6439 | PLLPEPEETSK | P51125 | P51125 | CAST | calpastatin | Cytoplasm | peptidase |
| 15554138 | 2 | 0.598 | 1770.919 | GVVPEDAVETLAGSLGTR | P51125 | P51125 | CAST | calpastatin | Cytoplasm | peptidase |
| 15533224 | 3 | 0.609 | 1770.92 | GVVPEDAVETLAGSLGTR | P51125 | P51125 | CAST | calpastatin | Cytoplasm | peptidase |
| 15532846 | 2 | 0.792 | 1395.7081 | NEGITQPLPDSPK | P51125 | P51125 | CAST | calpastatin | Cytoplasm | peptidase |
| 15551230 | 4 | 0.916 | 3914.9268 | VEEEVINDQALQALSDSLGTRQPDPPSHVSQAEQVK | P51125 | P51125 | CAST | calpastatin | Cytoplasm | peptidase |
| 15554534 | 2 | 0.542 | 1867.8851 | SLSESELIGELSADFDR | P51125 | P51125 | CAST | calpastatin | Cytoplasm | peptidase |
| 15536710 | 2 | 0.565 | 1443.7443 | PFTPASPVQSTPSK | P51125 | P51125 | CAST | calpastatin | Cytoplasm | peptidase |
| 15551466 | 3 | 0.528 | 3762.7402 | NEGITQPLPDSPKPMGTDQAIDALSSDFTC[160.0307]SSPTGK | P51125 | P51125 | CAST | calpastatin | Cytoplasm | peptidase |
| 15553364 | 2 | 0.969 | 2234.0894 | FQDVPSAGGESVAGGGTVATALDK | P51125 | P51125 | CAST | calpastatin | Cytoplasm | peptidase |
| 15553032 | 2 | 0.904 | 2386.0503 | PMGTDQAIDALSSDFTC[160.0307]SSPTGK | P51125 | P51125 | CAST | calpastatin | Cytoplasm | peptidase |
| 15532013 | 3 | 0.035 | 1882.9585 | ASQRPDVLTTGGGNPIGDK | P24270 | P24270 | CAT | catalase | Cytoplasm | enzyme |
| 15533038 | 4 | 0.162 | 2474.209 | GPLLVQDVVFTDEMAHFDRER | P24270 | P24270 | CAT | catalase | Cytoplasm | enzyme |
| 15548361 | 2 | 0.053 | 1655.7925 | EAETFPFNPFDLTK | P24270 | P24270 | CAT | catalase | Cytoplasm | enzyme |
| 15549644, 15555369 | 2, 3 | 0.05 | 2518.1936, 2518.1985 | FYTEDGNWDLVGNNTPIFFIR | P24270 | P24270 | CAT | catalase | Cytoplasm | enzyme |
| 15551446 | 3 | 0.028 | 3168.2937 | DGPMC[160.0307]MHDNQGGAPNYYPNSFSAPEQQR | P24270 | P24270 | CAT | catalase | Cytoplasm | enzyme |
| 15542369 | 2 | 0.026 | 1338.6329 | LVNADGEAVYC[160.0307]K | P24270 | P24270 | CAT | catalase | Cytoplasm | enzyme |
| 15532341 | 3 | 0.08 | 1649.7966 | RFNSANEDNVTQVR | P24270 | P24270 | CAT | catalase | Cytoplasm | enzyme |
| 15541387 | 2 | 0.037 | 1276.6141 | LAQEDPDYGLR | P24270 | P24270 | CAT | catalase | Cytoplasm | enzyme |
| 15549180 | 2 | 0.181 | 1803.9166 | LGPNYLQIPVNC[160.0307]PYR | P24270 | P24270 | CAT | catalase | Cytoplasm | enzyme |
| 15533181 | 3 | 0.023 | 1542.7543 | SALEHSVQC[160.0307]AVDVK | P24270 | P24270 | CAT | catalase | Cytoplasm | enzyme |
| 15538370 | 3 | 0.013 | 1310.6973 | RLC[160.0307]ENIAGHLK | P24270 | P24270 | CAT | catalase | Cytoplasm | enzyme |
| 15555618 | 3 | 0.157 | 3220.4573 | NPVNYFAEVEQMAFDPSNMPPGIEPSPDK | P24270 | P24270 | CAT | catalase | Cytoplasm | enzyme |
| 15539981 | 2 | 0.028 | 962.5284 | DAQLFIQK | P24270 | P24270 | CAT | catalase | Cytoplasm | enzyme |
| 15540918 | 3 | 0.132 | 1119.5565 | LFAYPDTHR | P24270 | P24270 | CAT | catalase | Cytoplasm | enzyme |
| 15555227 | 3 | 0.051 | 2474.2139 | GPLLVQDVVFTDEMAHFDRER | P24270 | P24270 | CAT | catalase | Cytoplasm | enzyme |
| 15551878 | 2 | 0.021 | 1542.7551 | SALEHSVQC[160.0307]AVDVK | P24270 | P24270 | CAT | catalase | Cytoplasm | enzyme |
| 15551835 | 2 | 0.074 | 1649.7966 | RFNSANEDNVTQVR | P24270 | P24270 | CAT | catalase | Cytoplasm | enzyme |
| 15552634 | 2 | 0.012 | 1882.9587 | ASQRPDVLTTGGGNPIGDK | P24270 | P24270 | CAT | catalase | Cytoplasm | enzyme |
| 15541153 | 2 | 0.261 | 1493.6953 | FNSANEDNVTQVR | P24270 | P24270 | CAT | catalase | Cytoplasm | enzyme |
| 15543900 | 2 | 0.484 | 1391.6289 | NFTDVHPDYGAR | P24270 | P24270 | CAT | catalase | Cytoplasm | enzyme |
| 15536224 | 2 | 0.103 | 1038.613 | KTPVLPNGGR | Q8C9M2 | Q8C9M2 | CCDC15 | coiled-coil domain containing 15 | Cytoplasm | other |
| 15540542 | 3 | 0.259 | 1912.9056 | GLVHAEQGQGSGSPTGSSTR | Q3URS9 | Q3URS9 | CCDC51 | coiled-coil domain containing 51 | Cytoplasm | other |
| 15552592 | 2 | 0.344 | 1857.9675 | SLAEANNLSFPLEPLSR | Q8BGU5 | Q8BGU5 | CCNY | cyclin Y | Nucleus | other |
| 15539476 | 3 | 0.047 | 1784.9213 | MLLDIFDENLHPLSK | Q8BGU5 | Q8BGU5 | CCNY | cyclin Y | Nucleus | other |
| 15533533 | 3 | 0.511 | 2097.1191 | LALVTGGEIASTFDHPELVK | P80314 | P80314 | CCT2 | chaperonin containing TCP1, subunit 2 (beta) | Cytoplasm | kinase |
| 15548218 | 2 | 0.74 | 1912.9492 | LGGSLADSYLDEGFLLDK | P80314 | P80314 | CCT2 | chaperonin containing TCP1, subunit 2 (beta) | Cytoplasm | kinase |
| 15550146 | 3 | 0.852 | 2366.1013 | TVYGGGC[160.0307]SEMLMAHAVTQLANR | P80314 | P80314 | CCT2 | chaperonin containing TCP1, subunit 2 (beta) | Cytoplasm | kinase |
| 15542248 | 2 | 0.789 | 1330.6576 | GATQQILDEAER | P80314 | P80314 | CCT2 | chaperonin containing TCP1, subunit 2 (beta) | Cytoplasm | kinase |
| 15549895 | 3 | 0.162 | 2288.1614 | VQDDEVGDGTTSVTVLAAELLR | P80314 | P80314 | CCT2 | chaperonin containing TCP1, subunit 2 (beta) | Cytoplasm | kinase |
| 15550288 | 3 | 0.777 | 3209.5686 | QLIYNYPEQLFGAAGVMAIEHADFAGVER | P80314 | P80314 | CCT2 | chaperonin containing TCP1, subunit 2 (beta) | Cytoplasm | kinase |
| 15533720 | 3 | 0.718 | 1554.8275 | SLHDALC[160.0307]VLAQTVK | P80314 | P80314 | CCT2 | chaperonin containing TCP1, subunit 2 (beta) | Cytoplasm | kinase |
| 15548868 | 2 | 0.562 | 1532.7954 | DAALMVTNDGATILK | P80314 | P80314 | CCT2 | chaperonin containing TCP1, subunit 2 (beta) | Cytoplasm | kinase |
| 15536575 | 2 | 0.446 | 1275.648 | LIEEVMIGEDK | P80314 | P80314 | CCT2 | chaperonin containing TCP1, subunit 2 (beta) | Cytoplasm | kinase |
| 15532613 | 2 | 0.364 | 1082.5162 | EAVAMESFAK | P80314 | P80314 | CCT2 | chaperonin containing TCP1, subunit 2 (beta) | Cytoplasm | kinase |
| 15556712 | 3 | 0.515 | 2347.1912 | MLPTIIADNAGYDSADLVAQLR | P80314 | P80314 | CCT2 | chaperonin containing TCP1, subunit 2 (beta) | Cytoplasm | kinase |
| 15534500 | 2 | 0.642 | 1237.6865 | GSGNLEAIHVIK | P80314 | P80314 | CCT2 | chaperonin containing TCP1, subunit 2 (beta) | Cytoplasm | kinase |
| 15540964 | 2 | 0.841 | 1291.6534 | ILIANTGMDTDK | P80314 | P80314 | CCT2 | chaperonin containing TCP1, subunit 2 (beta) | Cytoplasm | kinase |
| 15556020 | 2 | 0.88 | 1554.8286 | SLHDALC[160.0307]VLAQTVK | P80314 | P80314 | CCT2 | chaperonin containing TCP1, subunit 2 (beta) | Cytoplasm | kinase |
| 15547969 | 3 | 0.168 | 2617.3582 | NVLLDPQLVPGGGASEMAVAHALTEK | P80318 | P80318 | CCT3 | chaperonin containing TCP1, subunit 3 (gamma) | Cytoplasm | other |
| 15533807 | 2 | 0.436 | 1185.717 | TAVETAVLLLR | P80318 | P80318 | CCT3 | chaperonin containing TCP1, subunit 3 (gamma) | Cytoplasm | other |
| 15531889 | 2 | 0.912 | 1428.749 | IPGGIIEDSC[160.0307]VLR | P80318 | P80318 | CCT3 | chaperonin containing TCP1, subunit 3 (gamma) | Cytoplasm | other |
| 15531837 | 2 | 0.676 | 1166.6866 | AVAQALEVIPR | P80318 | P80318 | CCT3 | chaperonin containing TCP1, subunit 3 (gamma) | Cytoplasm | other |
| 15532373 | 2 | 0.785 | 1254.7059 | ELGIWEPLAVK | P80318 | P80318 | CCT3 | chaperonin containing TCP1, subunit 3 (gamma) | Cytoplasm | other |
| 15541305 | 2 | 0.559 | 1333.6852 | TLIQNC[160.0307]GASTIR | P80318 | P80318 | CCT3 | chaperonin containing TCP1, subunit 3 (gamma) | Cytoplasm | other |
| 15535288 | 2 | 0.88 | 1237.6145 | MALDDMISTLK | P80318 | P80318 | CCT3 | chaperonin containing TCP1, subunit 3 (gamma) | Cytoplasm | other |
| 15537492 | 2 | 0.662 | 1114.582 | ISTPVDVNNR | P80318 | P80318 | CCT3 | chaperonin containing TCP1, subunit 3 (gamma) | Cytoplasm | other |
| 15539771 | 2 | 0.569 | 1234.5627 | NLQDAMQVC[160.0307]R | P80318 | P80318 | CCT3 | chaperonin containing TCP1, subunit 3 (gamma) | Cytoplasm | other |
| 15550479 | 2 | 0.729 | 1608.7213 | IGDEYFTFITDC[160.0307]K | P80318 | P80318 | CCT3 | chaperonin containing TCP1, subunit 3 (gamma) | Cytoplasm | other |
| 15542753 | 2 | 0.504 | 1337.6279 | AMTGVEQWPYR | P80318 | P80318 | CCT3 | chaperonin containing TCP1, subunit 3 (gamma) | Cytoplasm | other |
| 15532393 | 2 | 0.718 | 974.51306 | EILSEVER | P80318 | P80318 | CCT3 | chaperonin containing TCP1, subunit 3 (gamma) | Cytoplasm | other |
| 15539179 | 2 | 0.713 | 1279.7108 | IVLLDSSLEYK | P80318 | P80318 | CCT3 | chaperonin containing TCP1, subunit 3 (gamma) | Cytoplasm | other |
| 15534526 | 3 | 0.806 | 1403.7058 | GISDLAQHYLMR | P80318 | P80318 | CCT3 | chaperonin containing TCP1, subunit 3 (gamma) | Cytoplasm | other |
| 15550987 | 2 | 0.965 | 2131.0671 | MLLDPMGGIVMTNDGNAILR | P80318 | P80318 | CCT3 | chaperonin containing TCP1, subunit 3 (gamma) | Cytoplasm | other |
| 15538875 | 2 | 0.695 | 835.4322 | SMIEISR | P80318 | P80318 | CCT3 | chaperonin containing TCP1, subunit 3 (gamma) | Cytoplasm | other |
| 15554702, 15558591 | 3 | 0.535 | 3030.538, 3030.5383 | AFADAMEVIPSTLAENAGLNPISTVTELR | P80315 | P80315 | CCT4 | chaperonin containing TCP1, subunit 4 (delta) | Cytoplasm | other |
| 15534994 | 2 | 0.587 | 1357.7291 | VIDPATATSVDLR | P80315 | P80315 | CCT4 | chaperonin containing TCP1, subunit 4 (delta) | Cytoplasm | other |
| 15533028 | 2, 3 | 0.874 | 1456.7759, 1456.7761 | GIHPTIISESFQK | P80315 | P80315 | CCT4 | chaperonin containing TCP1, subunit 4 (delta) | Cytoplasm | other |
| 15541754 | 2, 3 | 0.801 | 1456.7766, 1456.7771 | DALSDLALHFLNK | P80315 | P80315 | CCT4 | chaperonin containing TCP1, subunit 4 (delta) | Cytoplasm | other |
| 15548987 | 2 | 0.751 | 1550.8883 | ALIAGGGAPEIELALR | P80315 | P80315 | CCT4 | chaperonin containing TCP1, subunit 4 (delta) | Cytoplasm | other |
| 15532301 | 2 | 0.831 | 958.5181 | LVIEEAER | P80315 | P80315 | CCT4 | chaperonin containing TCP1, subunit 4 (delta) | Cytoplasm | other |
| 15534581 | 2 | 0.426 | 1346.7477 | IGLIQFC[160.0307]LSAPK | P80315 | P80315 | CCT4 | chaperonin containing TCP1, subunit 4 (delta) | Cytoplasm | other |
| 15541226 | 2 | 0.826 | 931.482 | IDDVVNTR | P80315 | P80315 | CCT4 | chaperonin containing TCP1, subunit 4 (delta) | Cytoplasm | other |
| 15536666 | 2 | 0.527 | 1145.6317 | TGC[160.0307]NVLLIQK | P80315 | P80315 | CCT4 | chaperonin containing TCP1, subunit 4 (delta) | Cytoplasm | other |
| 15551440 | 2 | 0.484 | 2040.0449 | VVSQYSSLLSPMSVNAVMK | P80315 | P80315 | CCT4 | chaperonin containing TCP1, subunit 4 (delta) | Cytoplasm | other |
| 15532286 | 2 | 0.784 | 933.5747 | AYILNLVK | P80315 | P80315 | CCT4 | chaperonin containing TCP1, subunit 4 (delta) | Cytoplasm | other |
| 15549370 | 2 | 0.529 | 2060.0547 | LGGTIDDC[160.0307]ELVEGLVLTQK | P80315 | P80315 | CCT4 | chaperonin containing TCP1, subunit 4 (delta) | Cytoplasm | other |
| 15532734 | 2 | 0.354 | 1302.577 | TLSGMESYC[160.0307]VR | P80315 | P80315 | CCT4 | chaperonin containing TCP1, subunit 4 (delta) | Cytoplasm | other |
| 15555316 | 2 | 0.565 | 1552.7267 | DVEREDIEFIC[160.0307]K | P80315 | P80315 | CCT4 | chaperonin containing TCP1, subunit 4 (delta) | Cytoplasm | other |
| 15540467 | 3 | 0.523 | 2060.0557 | LGGTIDDC[160.0307]ELVEGLVLTQK | P80315 | P80315 | CCT4 | chaperonin containing TCP1, subunit 4 (delta) | Cytoplasm | other |
| 15540925 | 3 | 0.497 | 2188.1482 | KLGGTIDDC[160.0307]ELVEGLVLTQK | P80315 | P80315 | CCT4 | chaperonin containing TCP1, subunit 4 (delta) | Cytoplasm | other |
| 15537420 | 3 | 0.522 | 1150.6124 | QMQVLHPAAR | P80315 | P80315 | CCT4 | chaperonin containing TCP1, subunit 4 (delta) | Cytoplasm | other |
| 15544236 | 3 | 0.284 | 1706.988 | RALIAGGGAPEIELALR | P80315 | P80315 | CCT4 | chaperonin containing TCP1, subunit 4 (delta) | Cytoplasm | other |
| 15547994 | 2 | 0.875 | 1763.9862 | VLVDINNPEPLIQTAK | P80316 | P80316 | CCT5 | chaperonin containing TCP1, subunit 5 (epsilon) | Cytoplasm | other |
| 15557918 | 3 | 0.508 | 2660.2478 | DGDVTITNDGATILSMMDVDHQIAK | P80316 | P80316 | CCT5 | chaperonin containing TCP1, subunit 5 (epsilon) | Cytoplasm | other |
| 15552320 | 3 | 0.884 | 3135.509 | AFADALEVIPMALSENSGMNPIQTMTEVR | P80316 | P80316 | CCT5 | chaperonin containing TCP1, subunit 5 (epsilon) | Cytoplasm | other |
| 15556622 | 2 | 0.879 | 1738.9453 | WVGGPEIELIAIATGGR | P80316 | P80316 | CCT5 | chaperonin containing TCP1, subunit 5 (epsilon) | Cytoplasm | other |
| 15548811 | 2 | 0.493 | 1653.8806 | LGFAGVVQEISFGTTK | P80316 | P80316 | CCT5 | chaperonin containing TCP1, subunit 5 (epsilon) | Cytoplasm | other |
| 15538847 | 3 | 0.928 | 1453.7063 | ESNPALGIDC[160.0307]LHK | P80316 | P80316 | CCT5 | chaperonin containing TCP1, subunit 5 (epsilon) | Cytoplasm | other |
| 15537119 | 2 | 0.801 | 940.46014 | FSELTSEK | P80316 | P80316 | CCT5 | chaperonin containing TCP1, subunit 5 (epsilon) | Cytoplasm | other |
| 15532442 | 2 | 0.51 | 1274.6859 | QQISLATQMVR | P80316 | P80316 | CCT5 | chaperonin containing TCP1, subunit 5 (epsilon) | Cytoplasm | other |
| 15533484 | 2 | 0.802 | 1093.5255 | IADGYEQAAR | P80316 | P80316 | CCT5 | chaperonin containing TCP1, subunit 5 (epsilon) | Cytoplasm | other |
| 15541417 | 2 | 0.568 | 1268.5719 | C[160.0307]PTLEQYAMR | P80316 | P80316 | CCT5 | chaperonin containing TCP1, subunit 5 (epsilon) | Cytoplasm | other |
| 15537751 | 2 | 0.564 | 937.5454 | IAIQHLDK | P80316 | P80316 | CCT5 | chaperonin containing TCP1, subunit 5 (epsilon) | Cytoplasm | other |
| 15534783 | 3 | 0.568 | 1339.7235 | RSLHDALC[160.0307]VIR | P80316 | P80316 | CCT5 | chaperonin containing TCP1, subunit 5 (epsilon) | Cytoplasm | other |
| 15542808 | 3 | 0.401 | 1463.717 | HKLDVMSVEDYK | P80316 | P80316 | CCT5 | chaperonin containing TCP1, subunit 5 (epsilon) | Cytoplasm | other |
| 15549326 | 3 | 0.538 | 2315.2122 | DGNVLLHEMQIQHPTASLIAK | P80317 | P80317 | CCT6A | chaperonin containing TCP1, subunit 6A (zeta 1) | Cytoplasm | other |
| 15548761 | 2, 3 | 0.851 | 2452.2717, 2452.272 | NAIDDGC[160.0307]VVPGAGAVEVALAEALIK | P80317 | P80317 | CCT6A | chaperonin containing TCP1, subunit 6A (zeta 1) | Cytoplasm | other |
| 15549893 | 2, 3 | 0.904 | 2544.3342, 2544.3362 | VATAQDDITGDGTTSNVLIIGELLK | P80317 | P80317 | CCT6A | chaperonin containing TCP1, subunit 6A (zeta 1) | Cytoplasm | other |
| 15532616 | 2 | 0.746 | 1246.6646 | GIDPFSLDALAK | P80317 | P80317 | CCT6A | chaperonin containing TCP1, subunit 6A (zeta 1) | Cytoplasm | other |
| 15532803 | 2 | 0.647 | 1078.5753 | IITEGFEAAK | P80317 | P80317 | CCT6A | chaperonin containing TCP1, subunit 6A (zeta 1) | Cytoplasm | other |
| 15534929 | 2, 3 | 0.552 | 1447.7177, 1447.7183 | EMDRETLIDVAR | P80317 | P80317 | CCT6A | chaperonin containing TCP1, subunit 6A (zeta 1) | Cytoplasm | other |
| 15535715 | 2 | 0.758 | 1075.6113 | ALQFLEQVK | P80317 | P80317 | CCT6A | chaperonin containing TCP1, subunit 6A (zeta 1) | Cytoplasm | other |
| 15539488 | 2 | 0.599 | 1021.51385 | SETDTSLIR | P80317 | P80317 | CCT6A | chaperonin containing TCP1, subunit 6A (zeta 1) | Cytoplasm | other |
| 15540768 | 2 | 0.94 | 1191.567 | TEVNSGFFYK | P80317 | P80317 | CCT6A | chaperonin containing TCP1, subunit 6A (zeta 1) | Cytoplasm | other |
| 15541376 | 2 | 0.592 | 990.5265 | MLVSGAGDIK | P80317 | P80317 | CCT6A | chaperonin containing TCP1, subunit 6A (zeta 1) | Cytoplasm | other |
| 15555300 | 2 | 0.97 | 1839.8546 | DEPIDLFMVEIMEMK | P80317 | P80317 | CCT6A | chaperonin containing TCP1, subunit 6A (zeta 1) | Cytoplasm | other |
| 15538393 | 3 | 0.99 | 1967.949 | KDEPIDLFMVEIMEMK | P80317 | P80317 | CCT6A | chaperonin containing TCP1, subunit 6A (zeta 1) | Cytoplasm | other |
| 15551013 | 2, 3 | 0.481 | 2252.1233, 2252.1243 | SQDAEVGDGTTSVTLLAAEFLK | P80313 | P80313 | CCT7 | chaperonin containing TCP1, subunit 7 (eta) | Cytoplasm | other |
| 15548274 | 3 | 0.85 | 2403.1162 | LPIGDVATQYFADRDMFC[160.0307]AGR | P80313 | P80313 | CCT7 | chaperonin containing TCP1, subunit 7 (eta) | Cytoplasm | other |
| 15552711 | 3 | 0.541 | 2829.3992 | EGTDSSQGIPQLVSNISAC[160.0307]QVIAEAVR | P80313 | P80313 | CCT7 | chaperonin containing TCP1, subunit 7 (eta) | Cytoplasm | other |
| 15533689 | 2 | 0.662 | 1203.6542 | ATISNDGATILK | P80313 | P80313 | CCT7 | chaperonin containing TCP1, subunit 7 (eta) | Cytoplasm | other |
| 15549538 | 3 | 0.949 | 2289.1953 | INALTAASEAAC[160.0307]LIVSVDETIK | P80313 | P80313 | CCT7 | chaperonin containing TCP1, subunit 7 (eta) | Cytoplasm | other |
| 15550361 | 2 | 0.757 | 1793.8457 | QLC[160.0307]DNAGFDATNILNK | P80313 | P80313 | CCT7 | chaperonin containing TCP1, subunit 7 (eta) | Cytoplasm | other |
| 15532230 | 2 | 0.479 | 1383.582 | GGAEQFMEETER | P80313 | P80313 | CCT7 | chaperonin containing TCP1, subunit 7 (eta) | Cytoplasm | other |
| 15538055 | 2 | 0.795 | 1405.6411 | TFSYAGFEMQPK | P80313 | P80313 | CCT7 | chaperonin containing TCP1, subunit 7 (eta) | Cytoplasm | other |
| 15541601 | 2 | 0.571 | 1154.5765 | STVDPPAPSAGR | P80313 | P80313 | CCT7 | chaperonin containing TCP1, subunit 7 (eta) | Cytoplasm | other |
| 15537718 | 3 | 0.393 | 1663.9126 | PYVEEGLHPQIIIR | P80313 | P80313 | CCT7 | chaperonin containing TCP1, subunit 7 (eta) | Cytoplasm | other |
| 15533691 | 2 | 0.653 | 1104.638 | QQLLIGAYAK | P80313 | P80313 | CCT7 | chaperonin containing TCP1, subunit 7 (eta) | Cytoplasm | other |
| 15538213 | 2 | 0.195 | 876.4953 | TC[160.0307]TIILR | P80313 | P80313 | CCT7 | chaperonin containing TCP1, subunit 7 (eta) | Cytoplasm | other |
| 15536340 | 2 | 0.836 | 1062.6283 | LLDVVHPAAK | P80313 | P80313 | CCT7 | chaperonin containing TCP1, subunit 7 (eta) | Cytoplasm | other |
| 15551494 | 2 | 0.69 | 1565.7927 | LPIGDVATQYFADR | P80313 | P80313 | CCT7 | chaperonin containing TCP1, subunit 7 (eta) | Cytoplasm | other |
| 15534660 | 2 | 0.744 | 1154.6323 | SLHDAIMIVR | P80313 | P80313 | CCT7 | chaperonin containing TCP1, subunit 7 (eta) | Cytoplasm | other |
| 15537968 | 2 | 0.84 | 1333.7446 | LFVTNDAATILR | P42932 | P42932 | CCT8 | chaperonin containing TCP1, subunit 8 (theta) | Cytoplasm | enzyme |
| 15531737 | 2 | 0.63 | 1150.5859 | FAEAFEAIPR | P42932 | P42932 | CCT8 | chaperonin containing TCP1, subunit 8 (theta) | Cytoplasm | enzyme |
| 15548228 | 2 | 0.851 | 1831.8551 | IAVYSC[160.0307]PFDGMITETK | P42932 | P42932 | CCT8 | chaperonin containing TCP1, subunit 8 (theta) | Cytoplasm | enzyme |
| 15548581 | 2 | 0.87 | 2057.981 | QITSYGETC[160.0307]PGLEQYAIK | P42932 | P42932 | CCT8 | chaperonin containing TCP1, subunit 8 (theta) | Cytoplasm | enzyme |
| 15532584 | 2 | 0.541 | 1128.6693 | LATNAAVTVLR | P42932 | P42932 | CCT8 | chaperonin containing TCP1, subunit 8 (theta) | Cytoplasm | enzyme |
| 15533358 | 2 | 0.576 | 1173.6454 | EDGAISTIVLR | P42932 | P42932 | CCT8 | chaperonin containing TCP1, subunit 8 (theta) | Cytoplasm | enzyme |
| 15554697 | 2 | 0.752 | 1568.7832 | DMLEASILDTYLGK | P42932 | P42932 | CCT8 | chaperonin containing TCP1, subunit 8 (theta) | Cytoplasm | enzyme |
| 15534979 | 2 | 0.717 | 1233.575 | GEENLMDAQVK | P42932 | P42932 | CCT8 | chaperonin containing TCP1, subunit 8 (theta) | Cytoplasm | enzyme |
| 15536200 | 2 | 0.852 | 1365.5923 | GSTDNLMDDIER | P42932 | P42932 | CCT8 | chaperonin containing TCP1, subunit 8 (theta) | Cytoplasm | enzyme |
| 15537857 | 3 | 0.105 | 1895.0049 | ILGSGIYSSSVLHGMVFK | P42932 | P42932 | CCT8 | chaperonin containing TCP1, subunit 8 (theta) | Cytoplasm | enzyme |
| 15538200 | 2 | 0.938 | 962.51086 | APGFAQMLK | P42932 | P42932 | CCT8 | chaperonin containing TCP1, subunit 8 (theta) | Cytoplasm | enzyme |
| 15536241 | 2 | 0.735 | 1065.5198 | AVDDGVNTFK | P42932 | P42932 | CCT8 | chaperonin containing TCP1, subunit 8 (theta) | Cytoplasm | enzyme |
| 15536531 | 2 | 0.674 | 1307.6328 | HFSGLEEAVYR | P42932 | P42932 | CCT8 | chaperonin containing TCP1, subunit 8 (theta) | Cytoplasm | enzyme |
| 15532639 | 3 | 0.532 | 1510.8556 | RLVPGGGATEIELAK | P42932 | P42932 | CCT8 | chaperonin containing TCP1, subunit 8 (theta) | Cytoplasm | enzyme |
| 15555441 | 2 | 0.607 | 1510.8542 | RLVPGGGATEIELAK | P42932 | P42932 | CCT8 | chaperonin containing TCP1, subunit 8 (theta) | Cytoplasm | enzyme |
| 15552736 | 3 | 0.395 | 2412.094 | GSTDNLMDDIERAVDDGVNTFK | P42932 | P42932 | CCT8 | chaperonin containing TCP1, subunit 8 (theta) | Cytoplasm | enzyme |
| 15538493 | 2 | 0.24 | 1453.786 | NVGLDIEAEVPAVK | P42932 | P42932 | CCT8 | chaperonin containing TCP1, subunit 8 (theta) | Cytoplasm | enzyme |
| 15534252 | 3 | 0.38 | 1515.8092 | NLRDVDEVSSLLR | P42932 | P42932 | CCT8 | chaperonin containing TCP1, subunit 8 (theta) | Cytoplasm | enzyme |
| 15539617 | 3 | 0.106 | 1626.7942 | AHEILPELVC[160.0307]C[160.0307]SAK | P42932 | P42932 | CCT8 | chaperonin containing TCP1, subunit 8 (theta) | Cytoplasm | enzyme |
| 15541255 | 2 | 0.847 | 917.5105 | VDQIIMAK | P42932 | P42932 | CCT8 | chaperonin containing TCP1, subunit 8 (theta) | Cytoplasm | enzyme |
| 15540988 | 2 | 0.257 | 908.5001 | YNIMLVR | P42932 | P42932 | CCT8 | chaperonin containing TCP1, subunit 8 (theta) | Cytoplasm | enzyme |
| 15540018 | 2 | 0.413 | 1214.65 | VADIALHYANK | P42932 | P42932 | CCT8 | chaperonin containing TCP1, subunit 8 (theta) | Cytoplasm | enzyme |
| 15551987 | 2 | 0.347 | 1515.8093 | NLRDVDEVSSLLR | P42932 | P42932 | CCT8 | chaperonin containing TCP1, subunit 8 (theta) | Cytoplasm | enzyme |
| 15537109 | 2 | 0.368 | 1132.5831 | DVDEVSSLLR | P42932 | P42932 | CCT8 | chaperonin containing TCP1, subunit 8 (theta) | Cytoplasm | enzyme |
| 15543069 | 2 | 0.168 | 1404.6461 | SPTSPIQSTDDEK | Q8K4F0 | Q8K4F0 | CD226 | CD226 molecule | Plasma Membrane | other |
| 15548320 | 3 | 0.032 | 2777.2761 | FVLPANAFASPLQNPDNHC[160.0307]FC[160.0307]TEK | Q08857 | Q08857 | CD36 | CD36 molecule (thrombospondin receptor) | Plasma Membrane | transmembrane receptor |
| 15547773 | 2 | 0.22 | 1821.8773 | QFWIFDVQNPDDVAK | Q08857 | Q08857 | CD36 | CD36 molecule (thrombospondin receptor) | Plasma Membrane | transmembrane receptor |
| 15548835 | 2 | 0.819 | 1943.0123 | TYLDVEPITGFTLQFAK | Q08857 | Q08857 | CD36 | CD36 molecule (thrombospondin receptor) | Plasma Membrane | transmembrane receptor |
| 15531688 | 2 | 0.013 | 922.52325 | VAIIESYK | Q08857 | Q08857 | CD36 | CD36 molecule (thrombospondin receptor) | Plasma Membrane | transmembrane receptor |
| 15538982 | 2 | 0.078 | 1322.6808 | EVVLEEGTTAFK | Q08857 | Q08857 | CD36 | CD36 molecule (thrombospondin receptor) | Plasma Membrane | transmembrane receptor |
| 15532164 | 5 | 0.797 | 3531.7234 | PVYISLPHFLHASPDVSEPIEGLHPNEDEHR | Q08857 | Q08857 | CD36 | CD36 molecule (thrombospondin receptor) | Plasma Membrane | transmembrane receptor |
| 15550523 | 5 | 0.432 | 3845.876 | EGKPVYISLPHFLHASPDVSEPIEGLHPNEDEHR | Q08857 | Q08857 | CD36 | CD36 molecule (thrombospondin receptor) | Plasma Membrane | transmembrane receptor |
| 15540554 | 2 | 0.835 | 908.4856 | ELLWGYK | Q08857 | Q08857 | CD36 | CD36 molecule (thrombospondin receptor) | Plasma Membrane | transmembrane receptor |
| 15531480 | 2 | 0.002 | 1031.4613 | FFSSDIC[160.0307]R | Q08857 | Q08857 | CD36 | CD36 molecule (thrombospondin receptor) | Plasma Membrane | transmembrane receptor |
| 15532192 | 2, 3 | 0.504 | 1478.7822, 1478.7832 | REVVLEEGTTAFK | Q08857 | Q08857 | CD36 | CD36 molecule (thrombospondin receptor) | Plasma Membrane | transmembrane receptor |
| 15548248 | 4 | 0.793 | 3531.7195 | PVYISLPHFLHASPDVSEPIEGLHPNEDEHR | Q08857 | Q08857 | CD36 | CD36 molecule (thrombospondin receptor) | Plasma Membrane | transmembrane receptor |
| 15532381 | 3 | 0.98 | 1943.0122 | TYLDVEPITGFTLQFAK | Q08857 | Q08857 | CD36 | CD36 molecule (thrombospondin receptor) | Plasma Membrane | transmembrane receptor |

| | | | | | | | | | | | | | | | | | | | | | | |
|---|---|---|---|---|---|---|---|---|---|---|---|---|---|---|---|---|---|---|---|---|---|---|
| 15538428 | 4 | 0.107 | 1.71E+07 | 2.08E+07 | 1.56E+07 | 2.49E+07 | 2.10E+07 | 2.23E+07 | 2.18E+07 | 2.61E+07 | 1.58E+07 | 2.26E+07 | 3.02E+07 | 2.64E+07 | 2777.2751 | FVLPANAFASPLQNPDNHC[160.0307]FC[160.0307]TEK | Q08857 | Q08857 | CD36 | CD36 molecule (thrombospondin receptor) | Plasma Membrane | transmembrane receptor |
| 15533154 | 3 | 0.208 | 1.09E+07 | 1.23E+07 | 1.21E+07 | 1.11E+07 | 1.14E+07 | 1.54E+07 | 1.14E+07 | 1.35E+07 | 6387666 | 2.22E+07 | 1.21E+07 | 1.70E+07 | 1821.8766 | QFWIFDVQNPDDVAK | Q08857 | Q08857 | CD36 | CD36 molecule (thrombospondin receptor) | Plasma Membrane | transmembrane receptor |
| 15544984 | 2 | 0.049 | 3764255.25 | 2793972 | 1772084.75 | 3561800 | 2280475 | 2257873 | 1725860.25 | 2034529.625 | 2250784.75 | 7482531.5 | 4732767.5 | 3209600.25 | 926.6017 | LQVNILVK | Q08857 | Q08857 | CD36 | CD36 molecule (thrombospondin receptor) | Plasma Membrane | transmembrane receptor |
| 15542872 | 2 | 0.702 | 89459.055 | 1463662.625 | 4523801.5 | 4.70E-06 | 3294649.5 | 5944497.5 | 2706700.75 | 2500758.25 | 4814061 | 2107007.5 | 1756463.375 | 2774380 | 1082.7019 | RLQVNILVK | Q08857 | Q08857 | CD36 | CD36 molecule (thrombospondin receptor) | Plasma Membrane | transmembrane receptor |
| 15540403 | 2 | 0.625 | 1.59E+07 | 1.72E+07 | 2.06E+07 | 1.65E+07 | 1.49E+07 | 1.74E+07 | 1.34E+07 | 1.78E+07 | 2.16E+07 | 1.35E+07 | 1.75E+07 | 1.47E+07 | 1363.7346 | YGFIEGNVVIPR | P15379 | P15379 | CD44 | CD44 molecule (Indian blood group) | Plasma Membrane | enzyme |
| 15537458, 15543937 | 2, 3 | 0.178 | 5156774 | 5049507.5 | 8143719.5 | 5524198 | 8106175 | 6657902.5 | 3818282 | 6688802.5 | 9313152 | 2571301.25 | 4939246 | 3890110.75 | 1049.5388, 1049.5399 | YAGVFHVEK | P15379 | P15379 | CD44 | CD44 molecule (Indian blood group) | Plasma Membrane | enzyme |
| 15556255 | 2 | 0.041 | 2792644.25 | 4338609 | 6449034.5 | 2355496 | 4207019 | 3651010.25 | 3300277.75 | 4206760 | 4257892 | 2290726.25 | 1898552.5 | 1921812.25 | 1866.7463 | EPSETPDQC[160.0307]MTADETR | P15379 | P15379 | CD44 | CD44 molecule (Indian blood group) | Plasma Membrane | enzyme |
| 15541626 | 2 | 0.171 | 740926.688 | 1518439.875 | 1576001.125 | 966277 | 1432368 | 1011714.688 | 1331573.875 | 1722060.375 | 1369544.75 | 796863 | 936192.625 | 894955.562 | 1184.6068 | SQEMVHLVNK | P15379 | P15379 | CD44 | CD44 molecule (Indian blood group) | Plasma Membrane | enzyme |
| 15531772, 15534216 | 2 | 0.247 | 5.95E+08 | 5.65E+08 | 6.17E+08 | 5.18E+08 | 5.38E+08 | 5.76E+08 | 5.70E+08 | 5.59E+08 | 4.72E+08 | 5.57E+08 | 5.24E+08 | 5.38E+08 | 1219.598 | SYIFIYDGNK | Q61735 | Q61735 | CD47 | CD47 molecule | Plasma Membrane | transmembrane receptor |
| 15531647 | 2, 3 | 0.947 | 2.96E+08 | 2.70E+08 | 3.59E+08 | 3.40E+08 | 2.67E+08 | 3.28E+08 | 3.25E+08 | 3.48E+08 | 3.03E+08 | 3.00E+08 | 2.78E+08 | 3.92E+08 | 1429.8232, 1429.8247 | ISVSDLINGIASLK | Q61735 | Q61735 | CD47 | CD47 molecule | Plasma Membrane | transmembrane receptor |
| 15556695 | 2 | 0.499 | 1.93E+08 | 1.90E+08 | 2.56E+08 | 1.37E+08 | 2.03E+08 | 1.96E+08 | 1.63E+08 | 2.01E+08 | 1.92E+08 | 1.97E+08 | 1.76E+08 | 1.65E+08 | 1511.7014 | NVEAQSTEEMFVK | Q61735 | Q61735 | CD47 | CD47 molecule | Plasma Membrane | transmembrane receptor |
| 15531431 | 2 | 0.821 | 1.38E+08 | 1.21E+08 | 1.55E+08 | 1.29E+08 | 1.18E+08 | 1.25E+08 | 1.17E+08 | 1.37E+08 | 1.33E+08 | 1.81E+08 | 1.21E+08 | 1.14E+08 | 702.4382 | TVIELK | Q61735 | Q61735 | CD47 | CD47 molecule | Plasma Membrane | transmembrane receptor |
| 15531563 | 2 | 0.926 | 7.69E+07 | 9.81E+07 | 1.30E+08 | 7.10E+07 | 1.06E+08 | 9.50E+07 | 1.05E+08 | 1.12E+08 | 7.79E+07 | 1.13E+08 | 8.76E+07 | 9.02E+07 | 1194.5769 | TVSWFSPNEK | Q61735 | Q61735 | CD47 | CD47 molecule | Plasma Membrane | transmembrane receptor |
| 15553836 | 2 | 0.193 | 8866831 | 9769444 | 1.65E+07 | 7037595 | 8485070 | 9719181 | 7560798 | 8181114 | 7826991.5 | 9135320 | 3660716.25 | 7127191.5 | 1527.6969 | NVEAQSTEEM[147.0354]FVK | Q61735 | Q61735 | CD47 | CD47 molecule | Plasma Membrane | transmembrane receptor |
| 15538767 | 2 | 0.048 | 2663666.25 | 2956068 | 2447786.5 | 1945206.625 | 2560806 | 1965509.875 | 1258305.75 | 1607800.25 | 2344229.75 | 2316017.25 | 3188429.75 | 2941624.75 | 1257.7069 | ITLEVFDPVPK | P18181 | P18181 | CD48 | CD48 molecule | Plasma Membrane | other |
| 15551543 | 2 | 0.142 | 1.23E+07 | 1.45E+07 | 1.45E+07 | 1.13E+07 | 1.30E+07 | 1.22E+07 | 1.26E+07 | 1.34E+07 | 1.40E+07 | 1.18E+07 | 1.26E+07 | 1.25E+07 | 2023.9569 | ATC[160.0307]LEDLVWSPVAQFC[160.0307]K | Q61475 | Q61475 | CD55 | CD55 molecule, decay accelerating factor for complement (Cromer blood group) | Plasma Membrane | other |
| 15552759 | 3 | 0.177 | 1.03E+07 | 1.23E+07 | 1.05E+07 | 1.15E+07 | 1.19E+07 | 1.07E+07 | 8159055.5 | 1.21E+07 | 1.09E+07 | 9419374 | 7339787 | 9630807 | 3979.7776 | LVGVSSTFC[160.0307]SVTGNTVDWDDEFPVC[160.0307]TEIHC[160.0307]PEPPK | Q61475 | Q61475 | CD55 | CD55 molecule, decay accelerating factor for complement (Cromer blood group) | Plasma Membrane | other |
| 15554392 | 2 | 0.187 | 6034464.5 | 7321515 | 8528358 | 4254778.5 | 7672391.5 | 6772439 | 6360854 | 6458798 | 6267109 | 5784862.5 | 4251971.5 | 4831361 | 2088.8667 | GESDSYTYSQVVTYSC[160.0307]DK | Q61475 | Q61475 | CD55 | CD55 molecule, decay accelerating factor for complement (Cromer blood group) | Plasma Membrane | other |
| 15553303 | 3 | 0.731 | 5114195.5 | 7552266 | 6583420.5 | 4249687.5 | 6309682.5 | 6458394 | 4599210 | 6284195.5 | 5411397.5 | 4985323.5 | 6432816.5 | 6757013.5 | 3120.4775 | DLDNGHINIPTGILFGSEINFSC[160.0307]NPGYR | Q61475 | Q61475 | CD55 | CD55 molecule, decay accelerating factor for complement (Cromer blood group) | Plasma Membrane | other |
| 15534535 | 2 | 0.03 | 5141186 | 6634307 | 6047476.5 | 5695951.5 | 5088053 | 5900782.5 | 5671013 | 4801980 | 4940483.5 | 3547051.75 | 4351169 | 4757378.5 | 765.44855 | LSFASLK | Q61475 | Q61475 | CD55 | CD55 molecule, decay accelerating factor for complement (Cromer blood group) | Plasma Membrane | other |
| 15545662, 15546170 | 2 | 0.244 | 3403652.25 | 3062492 | 4964380.5 | 1978302.75 | 3995142 | 3553894.25 | 3525959.25 | 4179417.5 | 3905672.75 | 2501027 | 2923368.25 | 2550474.25 | 1312.624, 1312.6263 | SDVGQWSSPPPR | Q61475 | Q61475 | CD55 | CD55 molecule, decay accelerating factor for complement (Cromer blood group) | Plasma Membrane | other |
| 15533337 | 3 | 0.132 | 1.08E+08 | 9.54E+07 | 1.33E+08 | 9.90E+07 | 8.88E+07 | 1.12E+08 | 1.08E+08 | 9.89E+07 | 9.78E+07 | 1.27E+08 | 1.63E+08 | 1.12E+08 | 2032.8911 | QSDC[160.0307]HGEIIMDQLEETK | O55186 | O55186 | Cd59a | CD59a antigen | Plasma Membrane | other |
| 15548489 | 2 | 0.084 | 8759535 | 7487705.5 | 9372789 | 7721246 | 7681169.5 | 4977050.5 | 8022166 | 6560070 | 7654858.5 | 9722399 | 1.23E+07 | 8573033 | 2032.8912 | QSDC[160.0307]HGEIIMDQLEETK | O55186 | O55186 | Cd59a | CD59a antigen | Plasma Membrane | other |
| 15541400 | 2 | 0.069 | 4520506.5 | 1263368.5 | 1049274.125 | 2516504.25 | 1904868.5 | 1474524.875 | 2487252 | 2194727.5 | 3275597.25 | 4612024.5 | 1.62E+07 | 4347839.5 | 1243.4988 | C[160.0307]C[160.0307]QFNLC[160.0307]NK | O55186 | O55186 | Cd59a | CD59a antigen | Plasma Membrane | other |
| 15542040 | 2 | 0.136 | 6882867.5 | 1.11E+07 | 1.59E+07 | 2694257.75 | 9644478 | 7733070.5 | 8383307.5 | 1.17E+07 | 9943614 | 5474135.5 | 3150463.5 | 5887102.5 | 1278.6155 | EMGNTVMDIIR | P40237 | P40237 | CD82 | CD82 molecule | Plasma Membrane | other |
| 15538475 | 2 | 0.071 | 6861266.5 | 8941533 | 1.40E+07 | 3517701.25 | 8371678.5 | 3054009.25 | 6684652.5 | 1.05E+07 | 6930825 | 4383738.5 | 4870393.5 | 4681102.5 | 1465.6917 | EEAWDYVQAQVK | P40237 | P40237 | CD82 | CD82 molecule | Plasma Membrane | other |
| 15556835 | 3 | 0.146 | 1954708 | 6422758.5 | 1.26E+07 | 3.37E+07 | 5.12E+07 | 3.81E+07 | 2.75E+07 | 3.77E+07 | 4.79E+07 | 2123187.5 | 6.27E+07 | 3.83E+07 | 2804.2856 | AIHMALDC[160.0307]C[160.0307]GIAGPLEQFISDTC[160.0307]PK | P40240 | P40240 | CD9 | CD9 molecule | Plasma Membrane | other |
| 15534351 | 2 | 0.225 | 193291.203 | 1064804.625 | 4413105 | 1993519.625 | 6459962.5 | 7167189 | 4596347 | 4946335.5 | 4013369.75 | 141426.75 | 2369567.75 | 2169750.5 | 1091.6073 | QLLESFQVK | P40240 | P40240 | CD9 | CD9 molecule | Plasma Membrane | other |
| 15553193 | 3 | 0.241 | 11435.842 | 74475.766 | 203420.703 | 2341904.25 | 1090500.25 | 2541798.75 | 1116697 | 1229952.25 | 3747528.5 | 5.29 | 4257961 | 2483987.5 | 2577.295 | QLLESFQVKPC[160.0307]PEAISEVFNNK | P40240 | P40240 | CD9 | CD9 molecule | Plasma Membrane | other |
| 15533075 | 2 | 0.15 | 3885166.5 | 5013648 | 6531665 | 5244124.5 | 9347171 | 7731145.5 | 6530004.5 | 7033496.5 | 6583125.5 | 5369976 | 5401760 | 5335438 | 748.4587 | LIDFLK | Q8CII2 | Q8CII2 | CDC123 | cell division cycle 123 | Cytoplasm | other |
| 15533258 | 2 | 0.255 | 9937222 | 7487637.5 | 4616749.5 | 1.21E+07 | 7848287.5 | 6730854 | 9624855 | 8411592 | 9450332 | 1.02E+07 | 1.33E+07 | 1.03E+07 | 1198.575 | YVEC[160.0307]SALTQK | P60766 | P60766 | Cdc42 | cell division cycle 42 | Plasma Membrane | enzyme |
| 15560485 | 3 | 0.658 | 6767033 | 4499840 | 4717047 | 1.16E+07 | 4999144 | 4974296.5 | 5970282 | 4448608 | 6192605 | 9797046 | 6581568 | 5251984.5 | 2358.2522 | TPFLLVGTQIDLRDDPSTIEK | P60766 | P60766 | Cdc42 | cell division cycle 42 | Plasma Membrane | enzyme |
| 15542827 | 2 | 0.272 | 6140952 | 6001484 | 8140107.5 | 8035463.5 | 7772131 | 7072205.5 | 9402487 | 8023876.5 | 1.02E+07 | 5965740.5 | 1.05E+07 | 7715108 | 1313.6727 | TC[160.0307]LLISYTTNK | P60766 | P60766 | Cdc42 | cell division cycle 42 | Plasma Membrane | enzyme |
| 15558166, 15559465 | 2 | 1 | 3309733 | 3659739.75 | 6476793 | 3996927 | 4382540 | 4607143.5 | 3832307.75 | 5019483.5 | 4257839 | 5376683 | 3678116.25 | 4034223.25 | 1852.9669, 1852.9683 | NVFDEAILAALEPPEPK | P60766 | P60766 | Cdc42 | cell division cycle 42 | Plasma Membrane | enzyme |
| 15545558 | 3 | 0.408 | 2376935.5 | 2748276 | 3008163 | 2491397.25 | 2871424 | 2676925.25 | 3224693 | 4279700 | 2375484.5 | 2359156 | 3042659.75 | 1873144.75 | 1852.9664 | NVFDEAILAALEPPEPK | P60766 | P60766 | Cdc42 | cell division cycle 42 | Plasma Membrane | enzyme |
| 15541974 | 2 | 0.628 | 2.78E+07 | 2.37E+07 | 1.21E+07 | 3.06E+07 | 1.57E+07 | 2.34E+07 | 2.98E+07 | 1.85E+07 | 2.19E+07 | 2.24E+07 | 3.46E+07 | 2.93E+07 | 1460.7285 | AESQENSDALLKR | Q922M5 | Q922M5 | CDCA7L | cell division cycle associated 7-like | Nucleus | other |
| 15536130 | 2 | 0.009 | 1426089.875 | 1795205.875 | 1966618.75 | 2469102 | 2857943.5 | 2412590.75 | 2267210.5 | 2383142.5 | 2321506.5 | 2353717 | 2638305.5 | 2136336.5 | 1047.4867 | NMAALDAADR | Q8VDP6 | Q8VDP6 | CDIPT | CDP-diacylglycerol--inositol 3-phosphatidyltransferase | Cytoplasm | enzyme |
| 15555166 | 3 | 0.117 | 379439.375 | 1125093.5 | 1772610.875 | 5690125.5 | 3384527.75 | 4837291.5 | 3278182.75 | 2133408 | 4425885 | 700368.938 | 6073029.5 | 3672762.75 | 3272.5334 | LDGETASDSESRAETAPLPTSVDDTPEVLNR | Q99L43 | Q99L43 | CDS2 | CDP-diacylglycerol synthase (phosphatidate cytidylyltransferase) 2 | Cytoplasm | enzyme |
| 15551795 | 3 | 0.037 | 1933214.75 | 2422043.5 | 1540914.25 | 3448284 | 2526248 | 1510580.5 | 1739132.875 | 1469491 | 2836168.75 | 1037011.438 | 954311.562 | 1080000.875 | 2445.1418 | AANAASGAGGSSAAAGSRPGDGGSLGSGAR | Q4VAA2 | Q4VAA2 | CDV3 | CDV3 homolog (mouse) | Cytoplasm | other |
| 15534152 | 2 | 0.447 | 1.67E+07 | 1.61E+07 | 1.46E+07 | 1.42E+07 | 1.68E+07 | 1.57E+07 | 1.64E+07 | 1.65E+07 | 1.76E+07 | 1.73E+07 | 1.61E+07 | 1.55E+07 | 927.4447 | YWANFAR | Q6AW46 | Q6AW46 | CES5A | carboxylesterase 5A | Extracellular Space | enzyme |
| 15550196 | 2 | 0.167 | 3259561.75 | 4772908 | 8715398 | 1.19E+07 | 1.23E+07 | 1.36E+07 | 7376042 | 1.25E+07 | 1.76E+07 | 2649475 | 1.16E+07 | 9003909 | 2196.1028 | EILVGDVGQTVDDPYTTFVK | P18760 | P18760 | CFL1 | cofilin 1 (non-muscle) | Nucleus | other |
| 15554996 | 2 | 0.131 | 1055845 | 1175639.125 | 2981152.5 | 2630517 | 3873394.5 | 3686277 | 2288383.5 | 3455600.5 | 4346993.5 | 1548557.75 | 3282158 | 2415868.25 | 1888.978 | EDLVFIFWAPENAPLK | P18760 | P18760 | CFL1 | cofilin 1 (non-muscle) | Nucleus | other |
| 15541852 | 2 | 0.075 | 838394.688 | 1064673.75 | 2274093 | 3524848.75 | 4648995.5 | 5176880 | 2894903.5 | 4016078.75 | 5908648 | 946846.438 | 4603737.5 | 3672254.25 | 1337.623 | YALYDATYETK | P18760 | P18760 | CFL1 | cofilin 1 (non-muscle) | Nucleus | other |
| 15535147 | 3 | 0.436 | 1397794.875 | 2782733.75 | 3446543 | 4201902.5 | 5810939 | 5396878.5 | 3422559.75 | 4362922 | 6144152 | 804261.188 | 4901408.5 | 5964326 | 2017.0754 | KEDLVFIFWAPENAPLK | P18760 | P18760 | CFL1 | cofilin 1 (non-muscle) | Nucleus | other |
| 15554971 | 2 | 0.362 | 634877.188 | 690671.062 | 1370855 | 1366941.375 | 2073288 | 2289586.75 | 1216603.25 | 2056935.625 | 3439989.5 | 152180.938 | 2358792.5 | 1279584.375 | 1519.6823 | HELQANC[160.0307]YEEVK | P18760 | P18760 | CFL1 | cofilin 1 (non-muscle) | Nucleus | other |
| 15539947 | 2 | 0.181 | 2670061.25 | 7161604 | 3093124 | 4032050 | 4656372.5 | 5861668.5 | 2647526.25 | 5538504 | 6241862.5 | 1565400.125 | 2389231 | 3160568.25 | 1200.6444 | VAEELALEQAK | Q9CRB9 | Q9CRB9 | CHCHD3 | coiled-coil-helix-coiled-coil-helix domain containing 3 | Cytoplasm | other |
| 15552432 | 2 | 0.082 | 389558.938 | 262614.625 | 104281.391 | 1246010.125 | 562210.875 | 1641805.625 | 773427 | 546694.312 | 4597617 | 525609.812 | 1084381.625 | 1554353.125 | 1610.8033 | SADLNVDSIISYWK | O35744 | O35744 | Chil3/Chil4 | chitinase-like 3 | Cytoplasm | enzyme |
| 15541915 | 3 | 0.15 | 277283.625 | 510482.438 | 359354.094 | 3477832.5 | 1049261.125 | 4476415.5 | 1600697.5 | 1456287 | 1.10E+07 | 287673.875 | 3284999 | 6400453.5 | 2062.0981 | LIVGFPAYGHTFILSDPSK | O35744 | O35744 | Chil3/Chil4 | chitinase-like 3 | Cytoplasm | enzyme |

| | | | | | | | | | | | | | | | | | | | | | | |
|---|---|---|---|---|---|---|---|---|---|---|---|---|---|---|---|---|---|---|---|---|---|---|
| 15540495 | 4 | 0.175 | 712469.438 | 2526287.5 | 1108908.375 | 1894105 | 3179392.25 | 3789095.25 | 1110369.625 | 3035808 | 3555921.75 | 293736.938 | 1607813.25 | 1218238.75 | 2955.4573 | GHFYVFDVLDQDGNIVNPSEIQAHLK | P52825 | P52825 | CPT2 | carnitine palmitoyltransferase 2 | Cytoplasm | enzyme |
| 15556293 | 2 | 0.259 | 417467.938 | 514621.375 | 161233.875 | 850690.25 | 1046899.312 | 1247968.875 | 413933.219 | 1019424.188 | 891876.562 | 71752.266 | 928847.062 | 547189 | 1877.9896 | LSPDAVAQLAFQMAFLR | P52825 | P52825 | CPT2 | carnitine palmitoyltransferase 2 | Cytoplasm | enzyme |
| 15548160, 15554613 | 3, 4 | 0.205 | 6.66E+07 | 8.72E+07 | 6.25E+07 | 5.01E+07 | 6.29E+07 | 4.58E+07 | 5.75E+07 | 6.14E+07 | 5.50E+07 | 7.56E+07 | 5.01E+07 | 7.21E+07 | 3861.8372, 3861.8408 | ALFNLVGEPSLYC[160.0307]TSNDGEIGVWSGPPPQC[160.0307]IELNK | Q64735 | Q64735 | CR1L | complement component (3b/4b) receptor 1-like | Plasma Membrane | other |
| 15532733 | 2 | 0.345 | 3.59E+07 | 2.95E+07 | 3.11E+07 | 3.37E+07 | 3.26E+07 | 2.55E+07 | 3.27E+07 | 3.08E+07 | 3.32E+07 | 4.68E+07 | 3.31E+07 | 3.37E+07 | 1164.5704 | LPQEMSGFQK | Q64735 | Q64735 | CR1L | complement component (3b/4b) receptor 1-like | Plasma Membrane | other |
| 15541544 | 2 | 0.899 | 2.94E+07 | 2.37E+07 | 2.96E+07 | 2.84E+07 | 2.58E+07 | 2.20E+07 | 3.25E+07 | 2.60E+07 | 2.13E+07 | 3.25E+07 | 2.62E+07 | 2.40E+07 | 1267.5409 | QDSTWTSAEDK | Q64735 | Q64735 | CR1L | complement component (3b/4b) receptor 1-like | Plasma Membrane | other |
| 15537421 | 2 | 0.023 | 3.09E+07 | 2.60E+07 | 2.83E+07 | 2.14E+07 | 2.35E+07 | 2.08E+07 | 2.85E+07 | 2.35E+07 | 2.20E+07 | 2.68E+07 | 2.76E+07 | 2.97E+07 | 1039.5328 | RQFSITC[160.0307]K | Q64735 | Q64735 | CR1L | complement component (3b/4b) receptor 1-like | Plasma Membrane | other |
| 15549075 | 2 | 0.365 | 9455333 | 8387067.5 | 9758150 | 7115841.5 | 1.00E+07 | 6700578 | 8502313 | 8885566 | 7109462.5 | 1.09E+07 | 9991443 | 8047815.5 | 1919.963 | LQSLLTSQENSSTTSPAR | Q64735 | Q64735 | CR1L | complement component (3b/4b) receptor 1-like | Plasma Membrane | other |
| 15532481 | 2 | 0.102 | 7449228 | 7689807 | 9831585 | 5762688 | 7086900.5 | 5488267.5 | 7676883.5 | 8230406.5 | 8650178 | 9144118 | 6148227.5 | 7565986.5 | 722.41797 | SLFSLR | Q64735 | Q64735 | CR1L | complement component (3b/4b) receptor 1-like | Plasma Membrane | other |
| 15537923 | 4 | 0.375 | 9043078 | 8826847 | 7518723.5 | 9659487 | 5663534 | 5469700 | 6950473 | 7307317 | 6212214.5 | 8202852.5 | 7191306.5 | 7916601 | 2248.108 | TPSDPENGLVHVHTGIQFGSR | Q64735 | Q64735 | CR1L | complement component (3b/4b) receptor 1-like | Plasma Membrane | other |
| 15555203 | 3 | 0.396 | 6440364 | 4823840.5 | 3682398.25 | 5201632 | 3762518.75 | 2484131 | 5453528.5 | 3204204.75 | 2921790.75 | 5868418 | 5340098.5 | 4433655 | 2248.105 | TPSDPENGLVHVHTGIQFGSR | Q64735 | Q64735 | CR1L | complement component (3b/4b) receptor 1-like | Plasma Membrane | other |
| 15540853 | 2 | 0.201 | 1202025.125 | 1803936 | 1961397.875 | 1188715.875 | 1304358 | 1303521 | 1009608.5 | 1258331.625 | 693796.812 | 1297826.125 | 350059.625 | 1009928.688 | 1180.5642 | LPQEM[147.0354]SGFQK | Q64735 | Q64735 | CR1L | complement component (3b/4b) receptor 1-like | Plasma Membrane | other |
| 15536509 | 2 | 0.073 | 1950393.75 | 5129770 | 2293161 | 3431786.75 | 3776714.25 | 4974322.5 | 2930324.5 | 3947345.5 | 4552552.5 | 1230308.125 | 2065352.25 | 2284837 | 1458.822 | RTLIAEGYPIPTK | Q9Z125 | Q9Z125 | CREB3L1 | cAMP responsive element binding protein 3-like 1 | Nucleus | transcription regulator |
| 15536436 | 3 | 0.031 | 2637737.5 | 3700150 | 3573407 | 5101440 | 5696968 | 6499286 | 3508161.5 | 4277632 | 5580902.5 | 2915070.25 | 4063791.5 | 4209997 | 1962.0295 | FVTHVSDWGSLATISTIK | O88668 | O88668 | CREG1 | cellular repressor of E1A-stimulated genes 1 | Nucleus | transcription regulator |
| 15537997 | 2 | 0.05 | 1648635.5 | 3144224.75 | 2900660.5 | 4030366.75 | 4599445 | 5301513.5 | 3134799 | 3199022.75 | 3540700.75 | 1930638.25 | 2863574 | 3403127.5 | 1294.6808 | IWVLDYFGGPK | O88668 | O88668 | CREG1 | cellular repressor of E1A-stimulated genes 1 | Nucleus | transcription regulator |
| 15532910 | 2 | 0.4 | 2.85E+07 | 3.07E+07 | 3.68E+07 | 3.33E+07 | 3.85E+07 | 3.25E+07 | 4.77E+07 | 4.11E+07 | 2.84E+07 | 4.81E+07 | 3.45E+07 | 4.04E+07 | 1427.712 | GVNIGGAGSYIYEK | Q9DCT8 | Q9DCT8 | Crip2 | cysteine rich protein 2 | Plasma Membrane | other |
| 15552376 | 2 | 0.861 | 2.77E+07 | 3.20E+07 | 4.58E+07 | 2.47E+07 | 5.26E+07 | 3.33E+07 | 3.75E+07 | 4.58E+07 | 3.59E+07 | 4.28E+07 | 3.74E+07 | 3.63E+07 | 1917.041 | PQTEAPQVTGPIEVPVVR | Q9DCT8 | Q9DCT8 | Crip2 | cysteine rich protein 2 | Plasma Membrane | other |
| 15553039 | 2 | 0.099 | 2.69E+07 | 1.96E+07 | 2.09E+07 | 2.51E+07 | 3.40E+07 | 2.10E+07 | 3.61E+07 | 2.95E+07 | 2.78E+07 | 3.80E+07 | 4.49E+07 | 2.70E+07 | 1754.7803 | ASSVTTFTGEPNMC[160.0307]PR | Q9DCT8 | Q9DCT8 | Crip2 | cysteine rich protein 2 | Plasma Membrane | other |
| 15533968 | 2 | 0.884 | 2.18E+07 | 3.31E+07 | 4.12E+07 | 2.45E+07 | 4.50E+07 | 3.28E+07 | 4.50E+07 | 4.02E+07 | 2.55E+07 | 3.57E+07 | 2.25E+07 | 3.35E+07 | 1443.7086 | GVNTGAVGSYIYDK | Q9DCT8 | Q9DCT8 | Crip2 | cysteine rich protein 2 | Plasma Membrane | other |
| 15532156 | 2 | 0.161 | 1.59E+07 | 1.28E+07 | 1.51E+07 | 1.75E+07 | 2.03E+07 | 1.06E+07 | 2.21E+07 | 1.81E+07 | 1.63E+07 | 2.03E+07 | 2.67E+07 | 1.93E+07 | 1151.5901 | PC[160.0307]YGILFGPK | Q9DCT8 | Q9DCT8 | Crip2 | cysteine rich protein 2 | Plasma Membrane | other |
| 15538206 | 3 | 0.278 | 1.65E+07 | 1.92E+07 | 1.92E+07 | 1.26E+07 | 2.17E+07 | 8120762 | 2.18E+07 | 2.03E+07 | 1.60E+07 | 1.75E+07 | 1.97E+07 | 2.34E+07 | 1917.04 | PQTEAPQVTGPIEVPVVR | Q9DCT8 | Q9DCT8 | Crip2 | cysteine rich protein 2 | Plasma Membrane | other |
| 15532314 | 2 | 0.046 | 1.06E+07 | 8048337.5 | 7990032 | 1.01E+07 | 1.26E+07 | 6794574 | 1.46E+07 | 1.35E+07 | 1.16E+07 | 1.73E+07 | 2.30E+07 | 1.17E+07 | 1153.5692 | PC[160.0307]YATLFGPK | Q9DCT8 | Q9DCT8 | Crip2 | cysteine rich protein 2 | Plasma Membrane | other |
| 15541838 | 3 | 0.27 | 1.30E+07 | 9560654 | 7483742 | 9909173 | 1.49E+07 | 6521038.5 | 1.42E+07 | 1.18E+07 | 7460996 | 1.41E+07 | 2.53E+07 | 1.30E+07 | 2004.894 | TLTPGGHAEHDGQPYC[160.0307]HK | Q9DCT8 | Q9DCT8 | Crip2 | cysteine rich protein 2 | Plasma Membrane | other |
| 15549170 | 2 | 0.419 | 8804711 | 1049863.5 | 82387.344 | 1.19E+07 | 822657.062 | 0 | 3359280.5 | 1862207.375 | 5607004.5 | 6956829 | 1.41E+07 | 6131239 | 2267.0786 | GVNTGAVGSYIYDKDPEGTVQP | Q9DCT8 | Q9DCT8 | Crip2 | cysteine rich protein 2 | Plasma Membrane | other |
| 15550281 | 3 | 0.645 | 6280293.5 | 3977667.25 | 3516244.5 | 6335565.5 | 3310453.5 | 3569567 | 3686224.25 | 3255907.75 | 3869339.25 | 7517866.5 | 3751239.5 | 4271466 | 2828.4536 | LVTLLQEVDTIEQETIKPLDDC[160.0307]QK | Q9Z2L7 | Q9Z2L7 | CRLF3 | cytokine receptor-like factor 3 | Cytoplasm | transcription regulator |
| 15531418 | 2 | 0.362 | 2.37E+07 | 5.65E+07 | 2.28E+07 | 3.28E+07 | 3.10E+07 | 4.33E+07 | 2.16E+07 | 3.88E+07 | 4.69E+07 | 1.71E+07 | 2.35E+07 | 2.50E+07 | 1223.7329 | IVPNILLEQGK | Q9CZU6 | Q9CZU6 | CS | citrate synthase | Cytoplasm | enzyme |
| 15548790 | 4 | 0.81 | 1.05E+07 | 1.95E+07 | 4776886 | 1.26E+07 | 1.61E+07 | 1.15E+07 | 8430266 | 1.80E+07 | 1.58E+07 | 1.47E+07 | 1.58E+07 | 9380688 | 4166.0684 | AALPSHVVTMLDNFPTNLHPMSQLSAAITALNSESNFAR | Q9CZU6 | Q9CZU6 | CS | citrate synthase | Cytoplasm | enzyme |
| 15551892 | 2 | 0.108 | 1.42E+07 | 3.85E+07 | 1.83E+07 | 1.71E+07 | 2.90E+07 | 2.90E+07 | 1.49E+07 | 2.68E+07 | 2.98E+07 | 1.07E+07 | 1.16E+07 | 1.25E+07 | 1762.8807 | GLVYETSVLDPDEGIR | Q9CZU6 | Q9CZU6 | CS | citrate synthase | Cytoplasm | enzyme |
| 15538433 | 2 | 0.281 | 7296891.5 | 1.71E+07 | 8165211 | 8256615.5 | 9764743 | 1.37E+07 | 6772978.5 | 1.13E+07 | 1.51E+07 | 5487016 | 7065802 | 6598610.5 | 834.50653 | LVAQLYK | Q9CZU6 | Q9CZU6 | CS | citrate synthase | Cytoplasm | enzyme |
| 15540512 | 2 | 0.252 | 6518245.5 | 1.56E+07 | 7706273 | 7860374.5 | 1.05E+07 | 1.45E+07 | 6344367.5 | 1.05E+07 | 1.47E+07 | 4698243.5 | 6446589 | 6634918 | 998.5865 | DVLSNLIPK | Q9CZU6 | Q9CZU6 | CS | citrate synthase | Cytoplasm | enzyme |
| 15534857 | 3 | 0.218 | 6960447 | 1.57E+07 | 4780394.5 | 1.09E+07 | 8862336 | 1.32E+07 | 6038735 | 1.14E+07 | 1.52E+07 | 3761863.25 | 7672161 | 4526812 | 1607.8257 | LRDYIWNTLNSGR | Q9CZU6 | Q9CZU6 | CS | citrate synthase | Cytoplasm | enzyme |
| 15533141 | 3 | 0.991 | 9878635 | 8934072 | 3349037.75 | 1.29E+07 | 2871938.25 | 1.06E+07 | 4678458.5 | 3765700 | 1.50E+07 | 3456218.75 | 1.35E+07 | 9138746 | 2066.0508 | GLVYETSVLDPDEGIRFR | Q9CZU6 | Q9CZU6 | CS | citrate synthase | Cytoplasm | enzyme |
| 15554035, 15558564 | 2 | 0.807 | 4650799.5 | 6073617.5 | 2534069.5 | 6430969 | 3565052 | 4169557 | 3068762.5 | 3710778.25 | 4632231 | 3278909 | 5150017 | 2695978 | 1607.8253, 1607.826 | LRDYIWNTLNSGR | Q9CZU6 | Q9CZU6 | CS | citrate synthase | Cytoplasm | enzyme |
| 15535128 | 2 | 0.491 | 4109741.75 | 7626832.5 | 3040646 | 4957674.5 | 4624188 | 4946220 | 2838731.25 | 4454827.5 | 5312335.5 | 2575773.5 | 4476547.5 | 3046061.75 | 969.43616 | SMSTDGLMK | Q9CZU6 | Q9CZU6 | CS | citrate synthase | Cytoplasm | enzyme |
| 15538212 | 2 | 0.231 | 4160051 | 8211081.5 | 3922464.75 | 3753019.25 | 4149556.75 | 6418525 | 2986936.75 | 5531553.5 | 8032719.5 | 1304386.625 | 3184613.75 | 3700199.5 | 1326.787 | ALGVLAQLIWSR | Q9CZU6 | Q9CZU6 | CS | citrate synthase | Cytoplasm | enzyme |
| 15538957 | 2 | 0.526 | 1.08E+07 | 1.09E+07 | 7699027 | 9546768 | 7250752 | 9366224 | 1.28E+07 | 1.03E+07 | 8330814 | 1.24E+07 | 9118581 | 1.10E+07 | 1377.7208 | TVLMLADQMISR | Q8BK63 | Q8BK63 | CSNK1A1 | casein kinase 1, alpha 1 | Cytoplasm | kinase |
| 15543164 | 3 | 0.593 | 1.18E+07 | 4017219.25 | 110182.344 | 8895995 | 1371285 | 543719.188 | 9971751 | 725946.5 | 72585.742 | 1.04E+07 | 7999345 | 4074111.75 | 1341.7246 | ARHPQLLYESK | Q8BK63 | Q8BK63 | CSNK1A1 | casein kinase 1, alpha 1 | Cytoplasm | kinase |
| 15533779 | 2 | 0.228 | 1.01E+07 | 1.09E+07 | 1.13E+07 | 6221738 | 6011301.5 | 9973706 | 1.14E+07 | 1.09E+07 | 6515777 | 1.02E+07 | 9267167 | 8571855 | 1136.6698 | LFLIDFGLAK | Q8BK63 | Q8BK63 | CSNK1A1 | casein kinase 1, alpha 1 | Cytoplasm | kinase |
| 15535711 | 3 | 0.2 | 6746612.5 | 6241672 | 8813379 | 4813845 | 4139588.75 | 5679398.5 | 6218471 | 8157117.5 | 6228236 | 8683685 | 4507271 | 6515739.5 | 1259.7551 | ILQGGVGIPHIR | Q8BK63 | Q8BK63 | CSNK1A1 | casein kinase 1, alpha 1 | Cytoplasm | kinase |
| 15539573 | 2 | 0.062 | 6976354 | 4759013 | 5956480.5 | 4450981.5 | 5361100.5 | 5691733 | 7890447.5 | 6441279.5 | 6090839.5 | 7080030.5 | 8941319 | 6849261.5 | 1163.5769 | MSTPVEVLC[160.0307]K | Q8BK63 | Q8BK63 | CSNK1A1 | casein kinase 1, alpha 1 | Cytoplasm | kinase |
| 15537725 | 3 | 0.57 | 4324921 | 3334038.25 | 3528366.25 | 3170399.75 | 2990389 | 3417284.5 | 5096686 | 3326008.75 | 3047805.5 | 4554549 | 3290862 | 3903485.25 | 2075.9492 | TLNHQYDYTFDWTMLK | Q8BK63 | Q8BK63 | CSNK1A1 | casein kinase 1, alpha 1 | Cytoplasm | kinase |
| 15535549 | 2 | 0.373 | 4523134.5 | 5286169.5 | 5149641 | 3257123 | 4057838.25 | 3827402 | 7042108.5 | 4879030 | 3164154.75 | 4481100 | 3458723.75 | 4198531.5 | 1433.6373 | FEEAPDYMYLR | Q8BK63 | Q8BK63 | CSNK1A1 | casein kinase 1, alpha 1 | Cytoplasm | kinase |
| 15538425 | 3 | 0.603 | 2600986 | 2599122.75 | 1802084.25 | 1875700.75 | 2198996 | 1243093.125 | 4266407.5 | 1881663.375 | 925112.875 | 4302931.5 | 2075771.375 | 2567177.5 | 2008.9221 | RDDMESLGYVLMYFNR | Q8BK63 | Q8BK63 | CSNK1A1 | casein kinase 1, alpha 1 | Cytoplasm | kinase |
| 15534547 | 2 | 0.574 | 2985797.75 | 2803938.25 | 3131504.25 | 2369045.25 | 2216705.5 | 1994346.125 | 3221596.25 | 2070492.5 | 1987580.875 | 3781728.25 | 1420420.75 | 2151364.25 | 1310.7657 | GGPNIITLADIVK | Q60737 | Q60737 | CSNK2A1 | casein kinase 2, alpha 1 polypeptide | Nucleus | kinase |
| 15535804 | 3 | 0.54 | 3562640 | 3548343.75 | 3308734.5 | 2876129.5 | 3480966 | 2370355.5 | 2795629 | 2614764.5 | 2422632 | 5982439.5 | 2891535 | 2287644.75 | 1687.852 | FNLTGLNEQVPHYR | P67871 | P67871 | CSNK2B | casein kinase 2, beta polypeptide | Cytoplasm | kinase |
| 15539017 | 2 | 0.183 | 9390273 | 1.55E+07 | 1.66E+07 | 2.65E+07 | 3.15E+07 | 2.50E+07 | 1.77E+07 | 2.24E+07 | 2.69E+07 | 7522641.5 | 2.82E+07 | 1.87E+07 | 1474.6787 | GYGYGQGAGTLSTDK | P97315 | P97315 | CSRP1 | cysteine and glycine-rich protein 1 | Nucleus | other |
| 15532226 | 2 | 0.165 | 7447050 | 1.34E+07 | 1.78E+07 | 2.38E+07 | 3.10E+07 | 3.02E+07 | 1.77E+07 | 2.42E+07 | 2.98E+07 | 5621010 | 2.40E+07 | 1.93E+07 | 1433.6777 | GFGFGQGAGALVHSE | P97315 | P97315 | CSRP1 | cysteine and glycine-rich protein 1 | Nucleus | other |
| 15537969 | 2 | 0.183 | 6869418.5 | 1.26E+07 | 1.39E+07 | 2.22E+07 | 2.66E+07 | 2.22E+07 | 1.34E+07 | 2.14E+07 | 2.47E+07 | 5008381 | 2.17E+07 | 1.53E+07 | 1034.5334 | GLESTTLADK | P97315 | P97315 | CSRP1 | cysteine and glycine-rich protein 1 | Nucleus | other |
| 15542933 | 3 | 0.292 | 6441525.5 | 9065718 | 6521639 | 1.78E+07 | 2.14E+07 | 1.65E+07 | 1.06E+07 | 1.56E+07 | 2.30E+07 | 4261929 | 3.62E+07 | 1.22E+07 | 2044.9023 | TVYFAEEVQC[160.0307]EGNSFHK | P97315 | P97315 | CSRP1 | cysteine and glycine-rich protein 1 | Nucleus | other |
| 15534889 | 2 | 0.419 | 668717.75 | 1886954.5 | 463411.281 | 4791477 | 3545879 | 3708586.25 | 1209763 | 2137665.25 | 6027783 | 329082.469 | 1.60E+07 | 3390545.25 | 1204.4957 | SC[160.0307]FLC[160.0307]MVC[160.0307]K | P97315 | P97315 | CSRP1 | cysteine and glycine-rich protein 1 | Nucleus | other |
| 15548315 | 2 | 0.258 | 7292677.5 | 8666347 | 1.11E+07 | 1.12E+07 | 1.55E+07 | 1.30E+07 | 1.03E+07 | 1.01E+07 | 9450030 | 1.36E+07 | 6431021.5 | 1.02E+07 | 1794.8495 | VAFGEDIDLPETFDAR | P10605 | P10605 | CTSB | cathepsin B | Cytoplasm | peptidase |
| 15551272 | 2 | 0.088 | 7962743 | 1.20E+07 | 8982525 | 9615856 | 1.60E+07 | 1.25E+07 | 7905668.5 | 1.05E+07 | 9885771 | 9178435 | 5060890 | 7141441.5 | 1588.7506 | EQWSNC[160.0307]PTIGQIR | P10605 | P10605 | CTSB | cathepsin B | Cytoplasm | peptidase |
| 15553359 | 2 | 0.136 | 6132447 | 1.14E+07 | 7391156 | 9895461 | 1.15E+07 | 1.35E+07 | 6304137 | 9043716 | 1.15E+07 | 7288712.5 | 5111202.5 | 7609686.5 | 1991.9698 | NGPVEGAFTVFSDFLTYK | P10605 | P10605 | CTSB | cathepsin B | Cytoplasm | peptidase |
| 15535446 | 3 | 0.32 | 2145772.5 | 6536041.5 | 7785828 | 5366646.5 | 1.06E+07 | 8487001 | 5133825 | 7134209.5 | 3598625 | 5414235.5 | 1928043.375 | 3897091.75 | 1872.9443 | PSFHPLSDDLINYINK | P10605 | P10605 | CTSB | cathepsin B | Cytoplasm | peptidase |
| 15538146 | 3 | 0.178 | 4165082.5 | 4251488.5 | 2488923.25 | 4549577.5 | 4814929.5 | 4355186 | 2866161.25 | 3555412.5 | 2922009.5 | 4368428 | 2895488 | 3437648.75 | 1381.6096 | HEAGDMMGGHAIR | P10605 | P10605 | CTSB | cathepsin B | Cytoplasm | peptidase |
| 15551253 | 3 | 0.147 | 1.69E+07 | 3.23E+07 | 1.36E+07 | 2.83E+07 | 1.91E+07 | 2.42E+07 | 1.09E+07 | 1.94E+07 | 2.22E+07 | 1.31E+07 | 1.09E+07 | 1.25E+07 | 2942.393 | NIFSFYLNRDPEGQPGGELMLGGTDSK | P18242 | P18242 | CTSD | cathepsin D | Cytoplasm | peptidase |
| 15549628 | 3 | 0.744 | 8935913 | 2.47E+07 | 5631011.5 | 1.41E+07 | 1.12E+07 | 1.35E+07 | 7847771 | 1.34E+07 | 1.41E+07 | 8524043 | 8340246 | 9685370 | 3146.5715 | FDGILGMGYPHISVNNVLPVFDNLMQQK | P18242 | P18242 | CTSD | cathepsin D | Cytoplasm | peptidase |
| 15548892 | 2 | 0.179 | 9069520 | 2.07E+07 | 9845959 | 1.56E+07 | 1.57E+07 | 1.60E+07 | 7777731.5 | 1.28E+07 | 1.62E+07 | 8362487 | 8726742 | 8810479 | 1989.0139 | AIGAVPLIQGEYMIPC[160.0307]EK | P18242 | P18242 | CTSD | cathepsin D | Cytoplasm | peptidase |
| 15538745 | 3 | 0.19 | 9117246 | 1.94E+07 | 7373037.5 | 1.18E+07 | 1.22E+07 | 1.70E+07 | 6114518.5 | 1.30E+07 | 1.35E+07 | 6590934 | 6140946 | 7423275.5 | 2230.1233 | GGC[160.0307]EAIVDTGTSLLVGPVEEVK | P18242 | P18242 | CTSD | cathepsin D | Cytoplasm | peptidase |
| 15534151 | 2 | 0.18 | 8857442 | 1.69E+07 | 7942080 | 1.02E+07 | 9933251 | 1.16E+07 | 6911292.5 | 1.12E+07 | 1.35E+07 | 6571506 | 6530185 | 6869336.5 | 1106.6439 | VSSLPTVYLK | P18242 | P18242 | CTSD | cathepsin D | Cytoplasm | peptidase |
| 15537089 | 3 | 0.404 | 4088213.75 | 6542134 | 2020184.75 | 5068468 | 3891263.75 | 6010799.5 | 2387163.75 | 4100321.5 | 6319982.5 | 1520279.5 | 4058115.25 | 2514010.25 | 1391.7214 | ILDIAC[160.0307]WVHHK | P18242 | P18242 | CTSD | cathepsin D | Cytoplasm | peptidase |
| 15557271 | 3 | 0.028 | 2409514 | 4870519.5 | 3466569 | 3067513.25 | 3596340 | 5880688 | 1556228.75 | 3097198.75 | 3477675 | 1297339.25 | 1445231.375 | 1730651.5 | 3162.5667 | FDGILGM[147.0354]GYPHISVNNVLPVFDNLMQQK | P18242 | P18242 | CTSD | cathepsin D | Cytoplasm | peptidase |
| 15531372 | 2 | 0.851 | 9.88E+07 | 1.33E+08 | 1.33E+08 | 8.35E+07 | 1.25E+08 | 1.18E+08 | 1.29E+08 | 1.27E+08 | 1.01E+08 | 1.42E+08 | 1.04E+08 | 1.09E+08 | 1221.5989 | AQGQLSEFWR | P70269 | P70269 | CTSE | cathepsin E | Cytoplasm | peptidase |
| 15536399 | 2 | 0.407 | 8.71E+07 | 1.14E+08 | 1.20E+08 | 8.49E+07 | 1.21E+08 | 9.24E+07 | 1.21E+08 | 1.27E+08 | 1.07E+08 | 1.21E+08 | 6.83E+07 | 8.50E+07 | 1061.503 | QFYSVFDR | P70269 | P70269 | CTSE | cathepsin E | Cytoplasm | peptidase |
| 15553467 | 2 | 0.572 | 1.88E+07 | 6406016.5 | 4013875.5 | 2.86E+07 | 1021903.188 | 8966125 | 9858220 | 3738866.25 | 1.70E+07 | 1.26E+07 | 2.84E+07 | 2.22E+07 | 2179.0894 | QFYSVFDRGNNQVGLAPAVP | P70269 | P70269 | CTSE | cathepsin E | Cytoplasm | peptidase |
| 15541882 | 3 | 0.562 | 4487478.5 | 3346920.5 | 2371275.75 | 4146118.75 | 2438598 | 2742725.5 | 3645819.5 | 2203142.25 | 1872192.125 | 6000261.5 | 2697817.75 | 3143794 | 2070.0254 | HLEIFHTEFQNLLDADK | Q9WTX6 | Q9WTX6 | CUL1 | cullin 1 | Nucleus | enzyme |
| 15552112 | 2 | 0.804 | 4266346.5 | 3210104.5 | 4089622.75 | 3432104.75 | 3229452.75 | 3573004.75 | 3906709.5 | 3137576 | 2572474.25 | 5316420.5 | 1875983.25 | 2935355 | 1664.8602 | GQTPGGAQFVGLELYK | Q9WTX6 | Q9WTX6 | CUL1 | cullin 1 | Nucleus | enzyme |
| 15555216 | 2 | 0.877 | 3004757.75 | 2739732.75 | 2723809.75 | 2629931.75 | 2619131.25 | 2451114 | 2795708 | 2311640.25 | 3151555.75 | 4103238 | 2016064.75 | 2981353.25 | 2205.0693 | NPEEAELEDTLNQVMVVFK | Q9WTX6 | Q9WTX6 | CUL1 | cullin 1 | Nucleus | enzyme |
| 15539404 | 3 | 0.538 | 3150011 | 3000671.25 | 2377295.25 | 2005105.625 | 2549619.5 | 1996323.5 | 2283827 | 2788473 | 2444852.5 | 3845012 | 1842483.375 | 2526733.75 | 1708.932 | HQQLLGEVLTQLSSR | Q9WTX6 | Q9WTX6 | CUL1 | cullin 1 | Nucleus | enzyme |
| 15538811 | 3 | 0.24 | 2380403 | 2661046.75 | 2495835.75 | 2102078.75 | 1506404.5 | 1916968 | 2705420.5 | 2708276.25 | 2044567.875 | 3138792 | 1705522.125 | 2357974.75 | 2205.0715 | NPEEAELEDTLNQVMVVFK | Q9WTX6 | Q9WTX6 | CUL1 | cullin 1 | Nucleus | enzyme |
| 15538444 | 2 | 0.791 | 2404599.5 | 1645040 | 1427384.625 | 1337877.25 | 1469142.375 | 1530862.125 | 2448127.75 | 1497770.125 | 1576958.5 | 2429992.75 | 1017623 | 1873227 | 996.6545 | LLIQAAIVR | Q9WTX6 | Q9WTX6 | CUL1 | cullin 1 | Nucleus | enzyme |
| 15555675 | 3 | 0.515 | 1130429.375 | 597925.875 | 179014.922 | 1272488.75 | 29141.814 | 497727.562 | 819435.312 | 133374.422 | 795140.75 | 1521952.25 | 1212548.625 | 980204.5 | 2488.2888 | QIGLDQIWDDLRAGIQQVYTR | Q9WTX6 | Q9WTX6 | CUL1 | cullin 1 | Nucleus | enzyme |
| 15546202 | 3 | 0.707 | 1440832.75 | 562177.312 | 55956.891 | 1296734.875 | 355387.5 | 203469.125 | 1320136.75 | 114614.531 | 122363.25 | 1277193.875 | 1112816.375 | 415100.656 | 2058.9797 | RLVHQNSASDDAEASMISK | Q9WTX6 | Q9WTX6 | CUL1 | cullin 1 | Nucleus | enzyme |
| 15557880 | 2 | 0.251 | 1096220.625 | 1064793.5 | 1201361.625 | 1202067.625 | 1176646.75 | 784029 | 995783.062 | 1107071 | 565632.375 | 1111395.5 | 402979.094 | 573153.375 | 1943.8994 | ESTEFLQQNPVTEYMK | Q9WTX6 | Q9WTX6 | CUL1 | cullin 1 | Nucleus | enzyme |
| 15540462 | 2 | 0.861 | 4649665.5 | 3934589 | 3050276.75 | 3572892 | 2791793.5 | 3070787.25 | 4145877.25 | 2765071.75 | 2566559.75 | 6351584 | 2097023.5 | 2725681.5 | 1474.7611 | GLTEQEVETILDK | Q9JLV5 | Q9JLV5 | CUL3 | cullin 3 | Nucleus | enzyme |
| 15551302 | 2 | 0.669 | 3696565.5 | 2857642.25 | 2914477.75 | 2669824.5 | 2847742.25 | 2341640.25 | 2392106.5 | 2222200 | 3020079 | 4445454.5 | 2631092.25 | 2035393 | 1649.8508 | VLTTGYWPTQSATPK | Q9JLV5 | Q9JLV5 | CUL3 | cullin 3 | Nucleus | enzyme |
| 15543436 | 3 | 0.099 | 4830342 | 5117161 | 3708917.25 | 3760118.5 | 3910230.75 | 4131016.5 | 4768582.5 | 4311566 | 2846991.75 | 3533394.75 | 2544324 | 2782678 | 1288.6034 | SSAQHALNSMSR | P70658 | P70658 | CXCR4 | chemokine (C-X-C motif) receptor 4 | Plasma Membrane | G-protein coupled receptor |
| 15539427 | 3 | 0.117 | 5863612.5 | 9561538 | 1.27E+07 | 1.20E+07 | 2.64E+07 | 1.06E+07 | 1.16E+07 | 1.43E+07 | 1.03E+07 | 6737722.5 | 8148662.5 | 7314355 | 2205.9224 | EQAGGDATENFEDVGHSTDAR | P56395 | P56395 | CYB5A | cytochrome b5 type A (microsomal) | Cytoplasm | enzyme |
| 15538109 | 2 | 0.081 | 5351708.5 | 7870873 | 1.01E+07 | 1.14E+07 | 1.74E+07 | 1.08E+07 | 8136648.5 | 1.10E+07 | 1.21E+07 | 5886942.5 | 9163290 | 7626479.5 | 1186.5968 | YYTLEEIQK | P56395 | P56395 | CYB5A | cytochrome b5 type A (microsomal) | Cytoplasm | enzyme |
| 15535615 | 2 | 0.151 | 1320372.75 | 2465692.75 | 3087230 | 2240951.75 | 5468194.5 | 3632904.25 | 2836025 | 3510633.5 | 3041026.25 | 1576257.875 | 2328747.25 | 1909657.375 | 1428.7075 | TYIIGELHPDDR | P56395 | P56395 | CYB5A | cytochrome b5 type A (microsomal) | Cytoplasm | enzyme |
| 15549952 | 2 | 0.182 | 949414.438 | 2233363.25 | 3468468.75 | 2492779 | 6140474.5 | 2306018 | 2796169.75 | 3061299.5 | 3072060 | 1223085.625 | 1757471.75 | 1592351.625 | 1511.7451 | FLEEHPGGEEVLR | P56395 | P56395 | CYB5A | cytochrome b5 type A (microsomal) | Cytoplasm | enzyme |
| 15553257 | 4 | 0.786 | 3231875.25 | 2674213.75 | 2088974.875 | 5397509.5 | 3227022.75 | 3366382.75 | 2579994 | 2129098.75 | 4914147.5 | 1048102.125 | 5164298.5 | 3816968.75 | 3698.6511 | FLEEHPGGEEVLREQAGGDATENFEDVGHSTDAR | P56395 | P56395 | CYB5A | cytochrome b5 type A (microsomal) | Cytoplasm | enzyme |
| 15549144 | 4 | 0.005 | 9206906 | 1.31E+07 | 9849760 | 2.33E+07 | 2.09E+07 | 2.20E+07 | 1.29E+07 | 1.96E+07 | 1.79E+07 | 1.21E+07 | 1.73E+07 | 1.38E+07 | 3596.636 | FLSEHPGGEEVLLEQAGADATESFEDVGHSPDAR | Q9CQX2 | Q9CQX2 | CYB5B | cytochrome b5 type B (outer mitochondrial membrane) | Cytoplasm | enzyme |
| 15541398 | 3 | 0.588 | 1486270.375 | 1764674 | 50778.609 | 2062294.875 | 995262.375 | 71574.094 | 1985788.5 | 508315.875 | 18230.643 | 1952118.75 | 3610007.25 | 1265825.25 | 1685.8142 | RNSAEETWMVIHGR | Q9CQX2 | Q9CQX2 | CYB5B | cytochrome b5 type B (outer mitochondrial membrane) | Cytoplasm | enzyme |
| 15546097 | 2 | 0.587 | 804805.375 | 987536.75 | 15570.956 | 930944.938 | 1221699 | 612793.938 | 1392307.875 | 752591.562 | 85096.273 | 733704.312 | 1520607.125 | 859292.875 | 1386.65 | VEGSEPSVTYYR | Q9CQX2 | Q9CQX2 | CYB5B | cytochrome b5 type B (outer mitochondrial membrane) | Cytoplasm | enzyme |
| 15531678 | 3 | 0.316 | 2.80E+07 | 3.03E+07 | 3.11E+07 | 3.54E+07 | 4.56E+07 | 3.18E+07 | 2.81E+07 | 3.90E+07 | 3.62E+07 | 2.73E+07 | 3.37E+07 | 3.39E+07 | 2104.0518 | IDGNLVIRPYTPVSSDDDK | Q9DCN2 | Q9DCN2 | Cyb5r3 | cytochrome b5 reductase 3 | Cytoplasm | enzyme |
| 15531754, 15541118 | 2 | 0.774 | 1.42E+07 | 2.04E+07 | 2.24E+07 | 8376768 | 3.21E+07 | 1.37E+07 | 1.95E+07 | 2.83E+07 | 1.35E+07 | 2.08E+07 | 1.05E+07 | 1.26E+07 | 885.46515, 885.46655 | PELEELR | Q9DCN2 | Q9DCN2 | Cyb5r3 | cytochrome b5 reductase 3 | Cytoplasm | enzyme |
| 15534598 | 2 | 0.482 | 1.84E+07 | 2.15E+07 | 2.49E+07 | 2.29E+07 | 2.76E+07 | 2.20E+07 | 2.22E+07 | 2.95E+07 | 2.70E+07 | 1.99E+07 | 2.67E+07 | 2.60E+07 | 1398.7441 | STPAITLENPDIK | Q9DCN2 | Q9DCN2 | Cyb5r3 | cytochrome b5 reductase 3 | Cytoplasm | enzyme |
| 15542791 | 4 | 0.417 | 7927375 | 1.31E+07 | 1.44E+07 | 1.24E+07 | 1.40E+07 | 1.81E+07 | 1.32E+07 | 1.41E+07 | 2.46E+07 | 1.25E+07 | 1.53E+07 | 1.59E+07 | 2558.4187 | FALPSPQHILGLPIGQHIYLSTR | Q9DCN2 | Q9DCN2 | Cyb5r3 | cytochrome b5 reductase 3 | Cytoplasm | enzyme |
| 15536057 | 3 | 0.2 | 9672881 | 1.14E+07 | 1.45E+07 | 1.27E+07 | 1.73E+07 | 1.28E+07 | 1.28E+07 | 1.53E+07 | 1.76E+07 | 9358659 | 1.37E+07 | 1.06E+07 | 2150.9749 | DPNDHTVC[160.0307]YLLFANQSEK | Q9DCN2 | Q9DCN2 | Cyb5r3 | cytochrome b5 reductase 3 | Cytoplasm | enzyme |
| 15548650 | 2 | 0.282 | 7695307.5 | 1.19E+07 | 9465807 | 1.40E+07 | 1.37E+07 | 1.06E+07 | 8896831 | 1.10E+07 | 1.36E+07 | 9233875 | 1.06E+07 | 1.03E+07 | 1901.0311 | SVGMIAGGTGITPMLQVIR | Q9DCN2 | Q9DCN2 | Cyb5r3 | cytochrome b5 reductase 3 | Cytoplasm | enzyme |
| 15534024 | 2 | 0.181 | 5441632.5 | 7170670 | 6026748 | 7204791 | 1.05E+07 | 7371050.5 | 7283679.5 | 7687338.5 | 8851191 | 6732157.5 | 8611199 | 6805841 | 1143.515 | MSQYLENMK | Q9DCN2 | Q9DCN2 | Cyb5r3 | cytochrome b5 reductase 3 | Cytoplasm | enzyme |
| 15531739 | 2 | 0.123 | 6760269.5 | 7237675 | 8750606 | 6218165.5 | 1.07E+07 | 7932531 | 7632634 | 8708063 | 1.01E+07 | 6318005 | 5193052 | 6448714.5 | 950.4917 | IGDTIEFR | Q9DCN2 | Q9DCN2 | Cyb5r3 | cytochrome b5 reductase 3 | Cytoplasm | enzyme |
| 15535299 | 2 | 0.057 | 5886684 | 8005300 | 8141661 | 7398934.5 | 8136567 | 7367756.5 | 8327008 | 7877983.5 | 8639209 | 6187829.5 | 6712720 | 5933538 | 876.5166 | GFVDLVVK | Q9DCN2 | Q9DCN2 | Cyb5r3 | cytochrome b5 reductase 3 | Cytoplasm | enzyme |
| 15540957 | 2 | 0.511 | 4841838 | 5055011 | 6068285.5 | 5164653 | 5734619.5 | 5121971 | 5355555.5 | 5617382.5 | 5861719 | 4512703 | 5278341.5 | 5323530 | 1145.629 | GPNGLLVYQGK | Q9DCN2 | Q9DCN2 | Cyb5r3 | cytochrome b5 reductase 3 | Cytoplasm | enzyme |
| 15558547 | 3 | 0.154 | 1704515.875 | 3610796.75 | 5265995.5 | 3460829.25 | 5817843.5 | 6142900.5 | 5971387.5 | 5624787.5 | 9253929 | 4487136 | 4465185.5 | 4475217.5 | 2558.4138 | FALPSPQHILGLPIGQHIYLSTR | Q9DCN2 | Q9DCN2 | Cyb5r3 | cytochrome b5 reductase 3 | Cytoplasm | enzyme |
| 15550978 | 2 | 0.68 | 1601224.875 | 4601765.5 | 5366764.5 | 1453004.375 | 8774100 | 4082188 | 3769411.25 | 5269280.5 | 3239696.25 | 3145106 | 1592163 | 2412967.75 | 2127.9395 | APDAWDYSQGFVNEEMIR | Q9DCN2 | Q9DCN2 | Cyb5r3 | cytochrome b5 reductase 3 | Cytoplasm | enzyme |
| 15551018 | 3 | 0.09 | 1.18E+07 | 2.39E+07 | 9988560 | 1.83E+07 | 2.12E+07 | 1.97E+07 | 1.21E+07 | 1.62E+07 | 1.87E+07 | 9425621 | 1.18E+07 | 9501267 | 2634.1755 | ALAEEVEVQDGPNDDGEMFMRPGK | Q9D0M3 | Q9D0M3 | CYC1 | cytochrome c-1 | Cytoplasm | enzyme |
| 15537735 | 2 | 0.07 | 7063810 | 9727678 | 7792683 | 1.04E+07 | 1.11E+07 | 1.15E+07 | 7096501.5 | 1.20E+07 | 1.21E+07 | 5411839 | 8615587 | 6200149 | 1430.6527 | TGQAAGFSYTDANK | P62897 | P62897 | CYCS | cytochrome c, somatic | Cytoplasm | transporter |
| 15536320 | 2 | 0.532 | 3528344.5 | 6349881.5 | 3438837 | 4669116.5 | 4727594.5 | 4393803 | 3881615.75 | 6486994.5 | 7057602 | 3288825.5 | 5158277.5 | 4326708.5 | 779.44666 | MIFAGIK | P62897 | P62897 | CYCS | cytochrome c, somatic | Cytoplasm | transporter |
| 15538553 | 3 | 0.891 | 4423579 | 3232262.25 | 1822627.75 | 7495929 | 2112938.25 | 4262661 | 3527107 | 3298299.75 | 5902089 | 1575789.5 | 7664172 | 4271449.5 | 1296.7124 | TGPNLHGLFGRK | P62897 | P62897 | CYCS | cytochrome c, somatic | Cytoplasm | transporter |
| 15540121 | 3 | 0.118 | 1636319.875 | 4409341.5 | 2638009.5 | 3357050.75 | 4524184 | 4394696.5 | 3191921.25 | 5313658.5 | 4326663 | 1290945 | 2411451.75 | 2875385.5 | 1168.6195 | TGPNLHGLFGR | P62897 | P62897 | CYCS | cytochrome c, somatic | Cytoplasm | transporter |
| 15537843 | 2 | 0.192 | 1400539.75 | 1198114 | 1818593.625 | 1061836.375 | 1529867 | 1631860.625 | 1165917.75 | 2044009.5 | 1892296.625 | 413606.625 | 1224444.75 | 1118248.125 | 906.52655 | ADLIAYLK | P62897 | P62897 | CYCS | cytochrome c, somatic | Cytoplasm | transporter |
| 15532743 | 3 | 0.282 | 6861394.5 | 4863824 | 6667848.5 | 8639254 | 5044710.5 | 7681557.5 | 7405075 | 7217350.5 | 7082777 | 8061328.5 | 8639246 | 7821139 | 1795.988 | VLNLSEELRDIPQAAK | Q8BPM0 | Q8BPM0 | DAAM1 | dishevelled associated activator of morphogenesis 1 | Cytoplasm | other |
| 15539688 | 2 | 0.31 | 4977572.5 | 3833906.5 | 5721982 | 5598811 | 6133310 | 5098763 | 5117415 | 4198972 | 4029093.75 | 4830547.5 | 5410679.5 | 4182232.5 | 1264.5869 | EADAIDDTLSSK | Q8BPM0 | Q8BPM0 | DAAM1 | dishevelled associated activator of morphogenesis 1 | Cytoplasm | other |
| 15549158 | 2 | 0.04 | 4054662 | 4133270.5 | 5429340 | 7524181 | 6240811.5 | 6476802 | 4457428 | 4858781.5 | 5715878.5 | 4658813 | 6289051.5 | 4875967.5 | 1869.8314 | ESSEESGEFDDLVSALR | Q8BPM0 | Q8BPM0 | DAAM1 | dishevelled associated activator of morphogenesis 1 | Cytoplasm | other |
| 15532121 | 2 | 0.119 | 5517826 | 5454590 | 5562937 | 7348454.5 | 4793036.5 | 5532783.5 | 4592734 | 4548288 | 4034436.75 | 4301201 | 5614647 | 4862429.5 | 1115.5917 | ILDLEDLER | Q8BPM0 | Q8BPM0 | DAAM1 | dishevelled associated activator of morphogenesis 1 | Cytoplasm | other |
| 15534806 | 2, 3 | 0.604 | 2488095.25 | 3805197.5 | 7010563.5 | 4084929.75 | 5651835.5 | 6301975 | 4689141.5 | 4761900 | 3369071.5 | 4072896.75 | 3079384 | 4198802 | 1198.5925, 1198.5929 | SDIDLLEEHK | Q8BPM0 | Q8BPM0 | DAAM1 | dishevelled associated activator of morphogenesis 1 | Cytoplasm | other |
| 15551055 | 2 | 0.013 | 2008671.375 | 1615286.625 | 1874905.75 | 2965355 | 2328104 | 2205108.75 | 1351304.375 | 1673301.5 | 2208912.25 | 2981830 | 3342891.25 | 2587218.25 | 1564.8087 | SGNTVQYWLLLDR | Q8BPM0 | Q8BPM0 | DAAM1 | dishevelled associated activator of morphogenesis 1 | Cytoplasm | other |
| 15536757 | 2 | 0.367 | 3076155 | 4669228 | 6605874 | 4220285.5 | 6553310.5 | 6756549 | 4989978.5 | 5467795 | 5823208.5 | 2960204 | 4480268 | 4624155 | 1206.6367 | FQTLINDLDK | Q8BPM0 | Q8BPM0 | DAAM1 | dishevelled associated activator of morphogenesis 1 | Cytoplasm | other |
| 15541509 | 2 | 0.855 | 2289547.25 | 2569048.5 | 4185954 | 2752719.25 | 3068545.5 | 3848318 | 2114359.25 | 3590632.25 | 2641463.75 | 2882910 | 3058793 | 2851152 | 1209.5968 | AVETELEYQK | Q8BPM0 | Q8BPM0 | DAAM1 | dishevelled associated activator of morphogenesis 1 | Cytoplasm | other |
| 15538931 | 3 | 0.761 | 2970762.5 | 2713178.25 | 3947044.25 | 3517056.5 | 3362162.75 | 3820996.25 | 3623380.75 | 2982080.25 | 2856315 | 2682732.25 | 4097561.5 | 3393593.75 | 1658.838 | RAILTMDEQEDLPK | Q8BPM0 | Q8BPM0 | DAAM1 | dishevelled associated activator of morphogenesis 1 | Cytoplasm | other |
| 15543927 | 3 | 0.875 | 2334020.5 | 2644384.75 | 2162516 | 2490289 | 2254185.5 | 2883079.25 | 3089293.5 | 2812676.25 | 1851666.625 | 2427994.5 | 2807604.25 | 2714165.5 | 1784.0278 | NITLLHYLITIVENK | Q8BPM0 | Q8BPM0 | DAAM1 | dishevelled associated activator of morphogenesis 1 | Cytoplasm | other |
| 15544191 | 2 | 0.51 | 1082146.25 | 1453948.625 | 1616253.5 | 1323871.125 | 2085804.625 | 2006083.5 | 1861591.625 | 1767349 | 1167037.625 | 1460587.125 | 2053553 | 1734327.375 | 989.53204 | DMLEQLLK | Q8BPM0 | Q8BPM0 | DAAM1 | dishevelled associated activator of morphogenesis 1 | Cytoplasm | other |
| 15545192 | 3 | 0.438 | 2237612.5 | 278778.344 | 2783.96 | 2563478.75 | 148317.203 | 430878.625 | 240330.812 | 371180.906 | 760635.438 | 865231.312 | 4317057 | 1546506.875 | 1848.9044 | SDIDLLEEHKHELDR | Q8BPM0 | Q8BPM0 | DAAM1 | dishevelled associated activator of morphogenesis 1 | Cytoplasm | other |
| 15541171 | 2 | 0.095 | 231880.531 | 1467544.75 | 1377093.375 | 2929263 | 3246854 | 3217554.75 | 2360395.25 | 2366586.25 | 3887394 | 812363 | 2878856 | 2024320.125 | 1469.729 | FLEEYLSSTPQR | P61804 | P61804 | DAD1 | defender against cell death 1 | Cytoplasm | other |
| 15548872 | 2 | 0.852 | 7293348.5 | 7487745 | 9683212 | 6792364 | 9976432 | 6558921.5 | 7782616 | 7487445 | 7669903 | 9250279 | 8035731.5 | 7798406.5 | 1860.7922 | YSPGYC[160.0307]SSDSPLDSPTK | Q91WC9 | Q91WC9 | DAGLB | diacylglycerol lipase, beta | Plasma Membrane | enzyme |
| 15551901 | 3 | 0.889 | 4355221.5 | 5796388.5 | 7518860 | 3824705.5 | 7462523 | 4986748.5 | 4016473.75 | 6154035.5 | 4670122 | 6562082.5 | 4778554.5 | 4748953 | 2461.1042 | DIEYDAVEGDQHNC[160.0307]HFASILK | Q91WC9 | Q91WC9 | DAGLB | diacylglycerol lipase, beta | Plasma Membrane | enzyme |
| 15537788 | 2 | 0.508 | 3110480.25 | 3170799.25 | 3543214.5 | 2514495.25 | 3922088.75 | 2654056 | 3349424.75 | 3937466.5 | 2958894.25 | 5109595.5 | 2988673.75 | 3676529.5 | 876.47424 | PGMVLFGR | Q91WC9 | Q91WC9 | DAGLB | diacylglycerol lipase, beta | Plasma Membrane | enzyme |
| 15543424 | 3 | 0.603 | 2963887.25 | 5341232 | 7970362.5 | 2118313 | 8211638 | 3941407 | 5233444 | 7023391 | 3510082.5 | 4877642.5 | 1914622.75 | 3006614.25 | 1599.885 | VYELPFIVVLDHR | Q91WC9 | Q91WC9 | DAGLB | diacylglycerol lipase, beta | Plasma Membrane | enzyme |
| 15536446 | 3 | 0.217 | 3235888.25 | 3381130.5 | 2532066.5 | 3544058.5 | 3237836.25 | 2911531 | 3232263.75 | 3326429.5 | 2409424 | 3845131.25 | 3338198.75 | 4208974.5 | 1470.7235 | MLIDHMPDVMIR | Q91WC9 | Q91WC9 | DAGLB | diacylglycerol lipase, beta | Plasma Membrane | enzyme |
| 15536517 | 4 | 0.07 | 1970185.5 | 2255314 | 1945139.75 | 2036505 | 1327921.875 | 1952991.625 | 2536817 | 2071382.5 | 2109509.25 | 2776855.25 | 2189366 | 3039353.25 | 1727.9802 | VYELPFIVVLDHRK | Q91WC9 | Q91WC9 | DAGLB | diacylglycerol lipase, beta | Plasma Membrane | enzyme |
| 15537677 | 2 | 0.339 | 5639079.5 | 6775738.5 | 6493397.5 | 4724412.5 | 6683472.5 | 6621849 | 4562965.5 | 6790258 | 6862772.5 | 6081907.5 | 3723877.5 | 4484785.5 | 1059.541 | GEEILSGAQR | Q922B2 | Q922B2 | DARS | aspartyl-tRNA synthetase | Cytoplasm | enzyme |
| 15537531 | 3 | 0.404 | 3958543.25 | 3819239 | 3343713 | 5658694 | 5734165.5 | 3461623.75 | 2687626.5 | 2819061.25 | 4382274 | 3639441.25 | 1756101.75 | 4773110.5 | 1435.7411 | FGAPPHAGGGIGLER | Q922B2 | Q922B2 | DARS | aspartyl-tRNA synthetase | Cytoplasm | enzyme |
| 15540380 | 2 | 0.538 | 2258736.75 | 2502229.75 | 2393303.5 | 4376437.5 | 3512346.5 | 2680590.25 | 1653962.875 | 1658584.625 | 3157622 | 2942748.25 | 879775.312 | 3988549 | 1180.6173 | FQTEIQTVSK | Q922B2 | Q922B2 | DARS | aspartyl-tRNA synthetase | Cytoplasm | enzyme |
| 15541715 | 2 | 0.656 | 2584447.75 | 2770799.75 | 1889855.125 | 3810120.25 | 3725306.75 | 2641949.25 | 2301639.5 | 1928666.875 | 3241244.25 | 2825922 | 1004538.125 | 4062812.25 | 1368.6632 | LEYC[160.0307]EALAMLR | Q922B2 | Q922B2 | DARS | aspartyl-tRNA synthetase | Cytoplasm | enzyme |
| 15536479 | 3 | 0.652 | 2007868.25 | 1461577.75 | 1581656.375 | 2378001.5 | 2316683 | 1531146 | 1280898.875 | 983791.25 | 1623353 | 1881957.25 | 632419.062 | 3082185.5 | 1221.6565 | IHDPQLLTER | Q922B2 | Q922B2 | DARS | aspartyl-tRNA synthetase | Cytoplasm | enzyme |
| 15539486 | 2 | 0.408 | 1923096.5 | 1902133.75 | 1678484.25 | 2100875.5 | 1965309.875 | 1833242.25 | 1198542 | 1008074.438 | 1742233.25 | 1689513.75 | 773590.938 | 2418948.5 | 875.4963 | FLEPTLR | Q922B2 | Q922B2 | DARS | aspartyl-tRNA synthetase | Cytoplasm | enzyme |
| 15555513 | 2 | 0.647 | 2371217.25 | 1989580.375 | 2641147.5 | 2408204 | 3299109.25 | 1716973.75 | 1312966.75 | 1546461.5 | 2902080.5 | 1687348.375 | 863653.438 | 3065221 | 1509.7662 | NNAYLAQSPQLYK | Q922B2 | Q922B2 | DARS | aspartyl-tRNA synthetase | Cytoplasm | enzyme |
| 15546112 | 3 | 0.535 | 1526763.375 | 2016235.875 | 1953369.375 | 2417758 | 2554718.75 | 1557631.25 | 2005687.875 | 1203000.125 | 1906862.875 | 1154301.875 | 639483.438 | 2760649.5 | 1132.608 | ALHHGIDLEK | Q922B2 | Q922B2 | DARS | aspartyl-tRNA synthetase | Cytoplasm | enzyme |
| 15552410 | 2 | 0.007 | 1480889.5 | 2216266.75 | 1556394.5 | 2252456.75 | 1779254 | 945128.25 | 799815.312 | 1245831.375 | 945069.438 | 674517.812 | 621285.25 | 384976.562 | 1540.7224 | NQAPGQPGASQWGSR | Q9JII5 | Q9JII5 | Dazap1 | DAZ associated protein 1 | Cytoplasm | other |
| 15549846 | 2 | 0.132 | 3454810.5 | 2724339 | 2794003 | 2672045.75 | 3220432.75 | 2175506.75 | 1991872.625 | 1893198.25 | 2113595.5 | 2916710 | 1465672.5 | 2078844 | 2055.0137 | LDVATDNFFQNPELYIR | Q9QZ73 | Q9QZ73 | DCUN1D1 | DCN1, defective in cullin neddylation 1, domain containing 1 | Nucleus | other |
| 15551130 | 4 | 0.781 | 7257974.5 | 6322642.5 | 5796734.5 | 6006832 | 5052378 | 4965124.5 | 7656695 | 5624057.5 | 5211775 | 1.13E+07 | 4213168.5 | 6130335 | 3139.542 | ALC[160.0307]ELESGIPAAESQIVYAERPLTDNHR | A2ADY9 | A2ADY9 | DDI2 | DNA-damage inducible 1 homolog 2 | Plasma Membrane | transporter |
| 15553587 | 3 | 0.73 | 5451161.5 | 3927426.75 | 3972661 | 3148434.25 | 2936792.75 | 5020707.5 | 5643014 | 3016366.25 | 3143632 | 5905494.5 | 3208572.75 | 4805015 | 2592.2415 | TQAQHSSPGEMASSPQGLDNPALLR | A2ADY9 | A2ADY9 | DDI2 | DNA-damage inducible 1 homolog 2 | Plasma Membrane | transporter |
| 15549718 | 2 | 0.185 | 952786.5 | 3095313 | 3019319.25 | 3791752.75 | 6199886 | 5334717.5 | 2904372.75 | 4623760 | 6834143 | 1362321.125 | 3886259 | 3563992 | 1835.8922 | WVPFDGDDIQLEFVR | O54734 | O54734 | DDOST | dolichyl-diphosphooligosaccharide--protein glycosyltransferase subunit (non-catalytic) | Cytoplasm | enzyme |
| 15536542 | 2 | 0.286 | 647876.438 | 1813132.625 | 2488991 | 2860424.75 | 4382703 | 4412328 | 1794374.375 | 3305992.5 | 4346761 | 642513.688 | 3661502.25 | 2516175.25 | 1046.5963 | SSLNPILFR | O54734 | O54734 | DDOST | dolichyl-diphosphooligosaccharide--protein glycosyltransferase subunit (non-catalytic) | Cytoplasm | enzyme |
| 15543067 | 2 | 0.191 | 185634.75 | 1329659.5 | 2490087 | 2440902.25 | 4857059 | 4427378 | 2350908 | 2817658 | 4868871.5 | 321106.875 | 1880523.25 | 2299789.5 | 1159.6185 | TADDPSLSLIK | O54734 | O54734 | DDOST | dolichyl-diphosphooligosaccharide--protein glycosyltransferase subunit (non-catalytic) | Cytoplasm | enzyme |
| 15535236 | 2 | 0.006 | 2078808.875 | 2993762.5 | 1990222.5 | 2441461.5 | 2718762.5 | 3080972 | 2342002.25 | 3078540.5 | 2801746.75 | 1758484.125 | 1465145.5 | 1389225.875 | 876.529 | FLKGLGNK | Q8C4S8 | Q8C4S8 | DENND2A | DENN/MADD domain containing 2A | Cytoplasm | other |
| 15535177 | 2 | 0.506 | 2994934.5 | 3012525.5 | 2337905.25 | 1951351.625 | 2515591.5 | 2365102 | 4023468 | 2288961.25 | 2846425 | 3749578.5 | 2245251 | 2497676.25 | 1145.6543 | ESLAWLSLVK | Q91YP3 | Q91YP3 | DERA | deoxyribose-phosphate aldolase (putative) | Cytoplasm | enzyme |
| 15541747 | 2 | 0.345 | 1510374.375 | 1114844 | 1648429.125 | 1399423.75 | 1202934 | 1213010.75 | 1754867.375 | 1178331.125 | 1574823.125 | 2499267.5 | 1443510.125 | 1537862.875 | 1144.6129 | EWQAAWLLK | Q91YP3 | Q91YP3 | DERA | deoxyribose-phosphate aldolase (putative) | Cytoplasm | enzyme |
| 15538338 | 2 | 0.081 | 1700830.75 | 1187450.375 | 1195902.75 | 1395540.75 | 1483250.875 | 880472.375 | 2068565.25 | 1470503.5 | 1365110.875 | 2418871 | 2171370.5 | 1717742.875 | 1378.7124 | GITTAAVC[160.0307]VYPAR | Q91YP3 | Q91YP3 | DERA | deoxyribose-phosphate aldolase (putative) | Cytoplasm | enzyme |
| 15531851 | 2 | 0.146 | 7198370 | 1.02E+07 | 7790497.5 | 1.08E+07 | 1.59E+07 | 1.40E+07 | 1.02E+07 | 1.11E+07 | 1.40E+07 | 5077290.5 | 1.15E+07 | 9904221 | 1385.7188 | NFLSTPQFLYR | Q99J56 | Q99J56 | DERL1 | derlin 1 | Cytoplasm | other |
| 15553689 | 2 | 0.364 | 2658040.5 | 4816349.5 | 5496218 | 4445162 | 5755934 | 5858501 | 4103424 | 4689685.5 | 8798291 | 2755922.5 | 4872667 | 3424759.25 | 1535.8591 | YWFAATVAVPLIGK | Q99J56 | Q99J56 | DERL1 | derlin 1 | Cytoplasm | other |
| 15552256 | 3 | 0.561 | 1899515.75 | 4295981 | 2769824 | 4145268.25 | 4710614 | 2785606.25 | 2801483.5 | 2856196.75 | 3952840.75 | 1434796.375 | 3850588.75 | 2748579.25 | 3404.537 | TIFDTPDEDPNYNPLPEERPGGFAWGEGQR | Q8BNI4 | Q8BNI4 | DERL2 | derlin 2 | Cytoplasm | other |
| 15552595 | 3 | 0.03 | 4390066.5 | 3748891.5 | 4543375 | 5877451.5 | 2714917.25 | 5295416 | 2441957.25 | 2188129 | 2366125.5 | 6944219 | 3758062.75 | 5150894.5 | 2512.3452 | YAVTALTEGLRQELLEAQTHIR | Q3U0B3 | Q3U0B3 | DHRS11 | dehydrogenase/reductase (SDR family) member 11 | Cytoplasm | other |
| 15539477 | 2 | 6.17E-07 | 369929.312 | 147544.125 | 186130.625 | 512878.812 | 260300.266 | 204787.453 | 1.21E+07 | 8458144 | 1.12E+07 | 1.07E+07 | 1.28E+07 | 1.24E+07 | 980.5143 | QM[147.0354]IRNFR | A2ATU0 | A2ATU0 | DHTKD1 | dehydrogenase E1 and transketolase domain containing 1 | Cytoplasm | enzyme |
| 15534866 | 2 | 0.609 | 8536104 | 8454522 | 4306728 | 3971806 | 5848249.5 | 5750296 | 7030876.5 | 5875758 | 5735566 | 8761595 | 4704690.5 | 7282880 | 1131.6306 | EAISDSLLRK | Q80VY9 | Q80VY9 | DHX33 | DEAH (Asp-Glu-Ala-His) box polypeptide 33 | Nucleus | enzyme |
| 15551853 | 2 | 0.279 | 1329012 | 4910601.5 | 1909111.375 | 2972198 | 4079477.25 | 4051451.5 | 1669196.875 | 3498691.25 | 4138814.75 | 1592134.75 | 2120655.25 | 1663348.375 | 1838.0127 | VAPAPAGVFTDIPISNIR | Q8BMF4 | Q8BMF4 | DLAT | dihydrolipoamide S-acetyltransferase | Cytoplasm | enzyme |
| 15539733 | 2 | 0.248 | 1295411.625 | 3953287.5 | 1519694.125 | 2552736.25 | 2721324.5 | 3879354.5 | 1589597.25 | 4150578.75 | 3960457.5 | 806291.125 | 1827218 | 1891457.125 | 1489.808 | GLETIASDVVSLASK | Q8BMF4 | Q8BMF4 | DLAT | dihydrolipoamide S-acetyltransferase | Cytoplasm | enzyme |
| 15553510 | 3 | 0.191 | 1.53E+07 | 2.51E+07 | 1.18E+07 | 1.59E+07 | 2.03E+07 | 2.56E+07 | 1.20E+07 | 2.28E+07 | 2.27E+07 | 1.02E+07 | 1.26E+07 | 1.22E+07 | 3354.6392 | VLGAHILGPGAGEMVNEAALALEYGASC[160.0307]EDIAR | O08749 | O08749 | DLD | dihydrolipoamide dehydrogenase | Cytoplasm | enzyme |
| 15542484 | 3 | 0.251 | 9431054 | 1.78E+07 | 7226104 | 1.12E+07 | 1.21E+07 | 1.50E+07 | 8735992 | 1.34E+07 | 1.54E+07 | 4217257.5 | 7701650 | 1.05E+07 | 1979.0724 | IPNIYAIGDVVAGPMLAHK | O08749 | O08749 | DLD | dihydrolipoamide dehydrogenase | Cytoplasm | enzyme |
| 15554991 | 2 | 0.594 | 7076390.5 | 1.03E+07 | 3243853.75 | 7559651.5 | 7210495 | 7590468 | 4865966.5 | 7987284.5 | 1.09E+07 | 3678121.5 | 8015047.5 | 4115211.5 | 1819.864 | NETLGGTC[160.0307]LNVGC[160.0307]IPSK | O08749 | O08749 | DLD | dihydrolipoamide dehydrogenase | Cytoplasm | enzyme |
| 15542480 | 3 | 0.019 | 3105587.75 | 3434277.25 | 1819945.75 | 1817706.875 | 2487432.5 | 1219697.875 | 2169088.5 | 2491181 | 2545542.5 | 1135162.875 | 1402978.875 | 909930.875 | 2152.0735 | NAGQAVTIVAQYRPEEYSR | Q811D0 | Q811D0 | DLG1 | discs, large homolog 1 (Drosophila) | Plasma Membrane | kinase |
| 15553469 | 3 | 0.267 | 2036264.125 | 1771106.75 | 648598.312 | 1602549.625 | 967116.562 | 438444 | 1739806.75 | 932699.938 | 747906.938 | 486482.312 | 836101.375 | 350466.938 | 2272.0735 | FGDILHVINASDDEWWQAR | Q811D0 | Q811D0 | DLG1 | discs, large homolog 1 (Drosophila) | Plasma Membrane | kinase |
| 15547023 | 2 | 0.012 | 997075.5 | 1113409.25 | 725918 | 584208.812 | 787116.938 | 292578.406 | 736352.625 | 729185.438 | 722970.125 | 194389.594 | 422393.812 | 260740.422 | 1129.5933 | IITGGAAAQDGR | Q811D0 | Q811D0 | DLG1 | discs, large homolog 1 (Drosophila) | Plasma Membrane | kinase |
| 15549728 | 2 | 0.08 | 9508734 | 2.03E+07 | 8989880 | 1.43E+07 | 1.28E+07 | 1.76E+07 | 8770594 | 1.50E+07 | 1.67E+07 | 5215223 | 8396903 | 7512768.5 | 2187.1238 | ASAFALQEQPVVNAVIDDATK | Q9D2G2 | Q9D2G2 | DLST | dihydrolipoamide S-succinyltransferase (E2 component of 2-oxo-glutarate complex) | Cytoplasm | enzyme |
| 15539692 | 2 | 0.235 | 5470232.5 | 1.18E+07 | 5458076 | 7747031 | 8623312 | 1.10E+07 | 5907633 | 8609806 | 9428310 | 4571454 | 6024439.5 | 5849937.5 | 1454.6554 | NVETMNYADIER | Q9D2G2 | Q9D2G2 | DLST | dihydrolipoamide S-succinyltransferase (E2 component of 2-oxo-glutarate complex) | Cytoplasm | enzyme |
| 15539253 | 3 | 0.589 | 7145531 | 1.56E+07 | 5187257 | 8332336.5 | 7517615.5 | 1.35E+07 | 5158075.5 | 1.10E+07 | 1.16E+07 | 4552185.5 | 7386926 | 6512175.5 | 2187.1255 | ASAFALQEQPVVNAVIDDATK | Q9D2G2 | Q9D2G2 | DLST | dihydrolipoamide S-succinyltransferase (E2 component of 2-oxo-glutarate complex) | Cytoplasm | enzyme |
| 15549794 | 3 | 0.977 | 7511063 | 1.02E+07 | 3179996.5 | 1.07E+07 | 3274680 | 9413495 | 6336605.5 | 4971717 | 1.06E+07 | 3273383 | 9135643 | 6764744.5 | 2691.3193 | NDVITVQTPAFAESVTEGDVRWEK | Q9D2G2 | Q9D2G2 | DLST | dihydrolipoamide S-succinyltransferase (E2 component of 2-oxo-glutarate complex) | Cytoplasm | enzyme |
| 15535168 | 2 | 0.051 | 4398424.5 | 1.16E+07 | 6197653 | 4701282 | 1.07E+07 | 9947100 | 4887933 | 1.00E+07 | 9124032 | 3129844.25 | 2897463.75 | 3142670.25 | 1189.655 | VEGGTPLFTLR | Q9D2G2 | Q9D2G2 | DLST | dihydrolipoamide S-succinyltransferase (E2 component of 2-oxo-glutarate complex) | Cytoplasm | enzyme |
| 15539412 | 2 | 0.099 | 2500729.5 | 1.11E+07 | 7015351 | 2933641.5 | 8629439 | 9215421 | 5356938 | 8226630.5 | 6304841 | 2040430.125 | 2146933.75 | 2544423.5 | 1264.687 | PTAAPPLAEAGAAK | Q9D2G2 | Q9D2G2 | DLST | dihydrolipoamide S-succinyltransferase (E2 component of 2-oxo-glutarate complex) | Cytoplasm | enzyme |
| 15534695 | 2 | 0.532 | 3933432.5 | 7627141.5 | 3281070.75 | 3709165 | 4723281 | 6041340 | 3324245.5 | 5129902.5 | 7693509 | 1791127.125 | 4429277.5 | 4017564.5 | 999.5314 | LGFMSAFVK | Q9D2G2 | Q9D2G2 | DLST | dihydrolipoamide S-succinyltransferase (E2 component of 2-oxo-glutarate complex) | Cytoplasm | enzyme |
| 15534494 | 2 | 0.927 | 4655499.5 | 6020710 | 5533787 | 4519629 | 5749171.5 | 4702136.5 | 5161620.5 | 5171816.5 | 4990812 | 6360629.5 | 5213658 | 4014143.5 | 909.456 | WSVGSAFR | Q80WT2 | Q80WT2 | DMRT3 | doublesex and mab-3 related transcription factor 3 | Nucleus | transcription regulator |
| 15549168, 15552033 | 2, 3 | 0.805 | 4.73E+08 | 6.05E+08 | 6.61E+08 | 3.56E+08 | 6.36E+08 | 4.86E+08 | 5.11E+08 | 6.65E+08 | 4.30E+08 | 6.41E+08 | 3.59E+08 | 5.05E+08 | 2448.1482, 2448.151 | IETDYWPC[160.0307]PPSLAVVETEWR | Q9WV69 | Q9WV69 | DMTN | dematin actin binding protein | Plasma Membrane | other |
| 15538150, 15542905 | 2, 3 | 0.361 | 6.09E+08 | 7.03E+08 | 7.09E+08 | 4.49E+08 | 6.73E+08 | 6.01E+08 | 7.02E+08 | 7.21E+08 | 5.97E+08 | 5.74E+08 | 6.54E+08 | 6.14E+08 | 1477.7396, 1477.7397, 1477.7401 | FPAAQPPDPNQPAK | Q9WV69 | Q9WV69 | DMTN | dematin actin binding protein | Plasma Membrane | other |
| 15531289 | 2 | 0.464 | 4.24E+08 | 3.84E+08 | 4.21E+08 | 3.77E+08 | 3.56E+08 | 2.94E+08 | 3.93E+08 | 3.76E+08 | 3.07E+08 | 4.86E+08 | 3.52E+08 | 3.26E+08 | 1302.6621 | QPLTSPGSVSSSR | Q9WV69 | Q9WV69 | DMTN | dematin actin binding protein | Plasma Membrane | other |
| 15533921 | 2 | 0.381 | 3.87E+08 | 3.58E+08 | 3.58E+08 | 3.42E+08 | 2.91E+08 | 2.70E+08 | 3.52E+08 | 3.44E+08 | 2.85E+08 | 4.63E+08 | 3.10E+08 | 3.08E+08 | 1061.4996 | HLSAEDFSR | Q9WV69 | Q9WV69 | DMTN | dematin actin binding protein | Plasma Membrane | other |
| 15531293 | 2 | 0.121 | 4.73E+08 | 4.50E+08 | 4.05E+08 | 4.10E+08 | 3.66E+08 | 3.47E+08 | 4.18E+08 | 4.10E+08 | 3.80E+08 | 4.55E+08 | 4.08E+08 | 3.90E+08 | 1241.5844 | VFAMSPEEFGK | Q9WV69 | Q9WV69 | DMTN | dematin actin binding protein | Plasma Membrane | other |
| 15536917 | 2 | 0.369 | 3.45E+08 | 3.41E+08 | 4.09E+08 | 2.99E+08 | 3.10E+08 | 3.19E+08 | 3.18E+08 | 3.16E+08 | 2.85E+08 | 4.34E+08 | 3.00E+08 | 2.91E+08 | 1243.6981 | TLGIISQASTPR | Q9WV69 | Q9WV69 | DMTN | dematin actin binding protein | Plasma Membrane | other |
| 15533377, 15544924, 15545096 | 2, 3 | 0.861 | 3.81E+08 | 3.54E+08 | 3.80E+08 | 3.95E+08 | 3.23E+08 | 3.25E+08 | 3.80E+08 | 3.87E+08 | 3.09E+08 | 4.14E+08 | 3.33E+08 | 3.46E+08 | 1396.7634, 1396.7643, 1396.7645, 1396.7653 | HLIEDLIIESSK | Q9WV69 | Q9WV69 | DMTN | dematin actin binding protein | Plasma Membrane | other |
| 15548506 | 2 | 0.295 | 3.63E+08 | 3.39E+08 | 4.20E+08 | 3.25E+08 | 3.16E+08 | 3.12E+08 | 3.45E+08 | 3.44E+08 | 3.22E+08 | 4.13E+08 | 3.32E+08 | 3.09E+08 | 1626.7734 | STSPPPSPEVWAESR | Q9WV69 | Q9WV69 | DMTN | dematin actin binding protein | Plasma Membrane | other |
| 15548217, 15548532 | 3, 4 | 0.485 | 2.95E+08 | 2.68E+08 | 1.65E+08 | 3.06E+08 | 1.66E+08 | 1.55E+08 | 2.86E+08 | 2.62E+08 | 1.44E+08 | 3.43E+08 | 2.49E+08 | 3.16E+08 | 3509.806, 3509.8086 | AILDIERPDLMIYEPHFTYSLLEHVELPR | Q9WV69 | Q9WV69 | DMTN | dematin actin binding protein | Plasma Membrane | other |
| 15531583 | 2 | 0.603 | 2.41E+08 | 2.28E+08 | 2.28E+08 | 2.06E+08 | 2.04E+08 | 1.96E+08 | 2.23E+08 | 2.08E+08 | 1.92E+08 | 2.89E+08 | 2.00E+08 | 1.80E+08 | 1067.5183 | MDNQVLGYK | Q9WV69 | Q9WV69 | DMTN | dematin actin binding protein | Plasma Membrane | other |
| 15535169, 15543538 | 2 | 0.716 | 2.07E+08 | 1.83E+08 | 2.13E+08 | 2.07E+08 | 1.65E+08 | 1.73E+08 | 2.06E+08 | 1.99E+08 | 1.73E+08 | 2.25E+08 | 1.90E+08 | 1.73E+08 | 953.4679, 953.4681 | SSSLPSYGR | Q9WV69 | Q9WV69 | DMTN | dematin actin binding protein | Plasma Membrane | other |
| 15532022 | 2, 3 | 0.394 | 1.27E+08 | 1.54E+08 | 1.71E+08 | 8.64E+07 | 1.80E+08 | 1.14E+08 | 1.45E+08 | 1.76E+08 | 1.00E+08 | 1.42E+08 | 7.65E+07 | 8.74E+07 | 1399.6932, 1399.6937 | TPFHTSLHSGTSK | Q9WV69 | Q9WV69 | DMTN | dematin actin binding protein | Plasma Membrane | other |
| 15536106 | 3 | 0.49 | 1.33E+08 | 8.70E+07 | 6.32E+07 | 1.18E+08 | 6.05E+07 | 7.23E+07 | 9.07E+07 | 7.07E+07 | 7.11E+07 | 1.07E+08 | 1.26E+08 | 9.25E+07 | 1967.9911 | SLPDRTPFHTSLHSGTSK | Q9WV69 | Q9WV69 | DMTN | dematin actin binding protein | Plasma Membrane | other |
| 15532106 | 5 | 0.65 | 1.21E+08 | 1.05E+08 | 2.15E+08 | 6.22E+07 | 1.14E+08 | 2.16E+08 | 9.02E+07 | 1.88E+08 | 1.81E+08 | 9.64E+07 | 9.29E+07 | 1.05E+08 | 3525.8052 | AILDIERPDLM[147.0354]IYEPHFTYSLLEHVELPR | Q9WV69 | Q9WV69 | DMTN | dematin actin binding protein | Plasma Membrane | other |
| 15548566 | 3, 4 | 0.554 | 1.14E+08 | 9.95E+07 | 1.99E+08 | 5.65E+07 | 1.04E+08 | 1.96E+08 | 8.21E+07 | 1.63E+08 | 1.59E+08 | 9.02E+07 | 6.93E+07 | 9.45E+07 | 3525.7996, 3525.8013 | AILDIERPDLM[147.0354]IYEPHFTYSLLEHVELPR | Q9WV69 | Q9WV69 | DMTN | dematin actin binding protein | Plasma Membrane | other |
| 15537329 | 2 | 0.565 | 5.00E+07 | 5.71E+07 | 8.53E+07 | 3.47E+07 | 6.36E+07 | 6.93E+07 | 4.55E+07 | 7.53E+07 | 7.37E+07 | 4.25E+07 | 4.50E+07 | 5.50E+07 | 727.43365 | DLAAIPK | Q9WV69 | Q9WV69 | DMTN | dematin actin binding protein | Plasma Membrane | other |
| 15560421, 15560438 | 3 | 0.855 | 1.12E+07 | 5.84E+07 | 2.36E+07 | 3981590.75 | 4.18E+07 | 2.62E+07 | 5.18E+07 | 3.39E+07 | 1.54E+07 | 3.26E+07 | 1.05E+07 | 6.11E+07 | 2377.1636, 2377.1643 | TSLPHFHHPETTRPDSNIYK | Q9WV69 | Q9WV69 | DMTN | dematin actin binding protein | Plasma Membrane | other |
| 15538944 | 3 | 0.501 | 1.92E+07 | 2.08E+07 | 2.67E+07 | 1.82E+07 | 1.84E+07 | 2.13E+07 | 2.10E+07 | 2.29E+07 | 1.90E+07 | 2.93E+07 | 1.93E+07 | 2.24E+07 | 1626.7748 | STSPPPSPEVWAESR | Q9WV69 | Q9WV69 | DMTN | dematin actin binding protein | Plasma Membrane | other |
| 15532570 | 2 | 0.071 | 2.07E+07 | 2.25E+07 | 1.85E+07 | 1.51E+07 | 1.88E+07 | 1.38E+07 | 1.96E+07 | 1.74E+07 | 1.63E+07 | 1.77E+07 | 1.51E+07 | 1.46E+07 | 1469.7684 | IYPYEMLVVTNK | Q9WV69 | Q9WV69 | DMTN | dematin actin binding protein | Plasma Membrane | other |
| 15549791 | 3 | 0.351 | 1.47E+07 | 1.29E+07 | 1.38E+07 | 1.27E+07 | 1.19E+07 | 1.22E+07 | 1.84E+07 | 1.47E+07 | 1.30E+07 | 1.64E+07 | 1.23E+07 | 1.12E+07 | 2614.327 | QPLTSPGSVSSSRDSSVPGSPSSIVAK | Q9WV69 | Q9WV69 | DMTN | dematin actin binding protein | Plasma Membrane | other |
| 15553256 | 2 | 0.447 | 1.19E+07 | 6239635.5 | 2294548 | 1.04E+07 | 2639490 | 2871756 | 6623839.5 | 2533080 | 3179430.25 | 9282284 | 1.06E+07 | 6410458.5 | 1967.989 | SLPDRTPFHTSLHSGTSK | Q9WV69 | Q9WV69 | DMTN | dematin actin binding protein | Plasma Membrane | other |
| 15538833, 15546679 | 2 | 0.096 | 1.18E+07 | 8746124 | 1.12E+07 | 8029880.5 | 8481590 | 8204292.5 | 8958887 | 9628712 | 7206437.5 | 9169044 | 6705989 | 7070843 | 1257.5786, 1257.5792 | VFAM[147.0354]SPEEFGK | Q9WV69 | Q9WV69 | DMTN | dematin actin binding protein | Plasma Membrane | other |
| 15553042 | 2 | 0.09 | 4890891.5 | 4739054 | 5759480.5 | 5078337.5 | 5247606 | 5472428 | 4366713 | 4656064 | 4249144 | 6872027.5 | 4681623 | 6616446 | 2110.7903 | GAEEEEEEDDDSEEEIK | Q9WV69 | Q9WV69 | DMTN | dematin actin binding protein | Plasma Membrane | other |
| 15550717 | 2 | 0.627 | 7894513.5 | 2572210.25 | 1820622.5 | 1.05E+07 | 184766.594 | 3166357.75 | 3656688 | 1057676.5 | 4675981.5 | 5859832 | 8793303 | 5345722 | 2576.2441 | IETDYWPC[160.0307]PPSLAVVETEWRK | Q9WV69 | Q9WV69 | DMTN | dematin actin binding protein | Plasma Membrane | other |
| 15555396 | 3 | 0.165 | 1402356.5 | 1965206.125 | 1557831.875 | 1724589.75 | 2327390.25 | 723629.875 | 725960.312 | 1710968.875 | 1065295.25 | 754471.875 | 655716.188 | 895589.5 | 2520.1538 | LQSTEFSPSGSEAGSPGLQNGEGQR | Q9WV69 | Q9WV69 | DMTN | dematin actin binding protein | Plasma Membrane | other |
| 15551911 | 3 | 0.16 | 3366177 | 3299803.75 | 1662855.875 | 3190072.5 | 1620414.375 | 1979343.625 | 1794445 | 1488136.375 | 1058272.625 | 2198675 | 1786006.5 | 1984727 | 2776.3965 | ILFHGEGDQEPELDPGDVIIVLDQK | Q9JMC3 | Q9JMC3 | DNAJA4 | DnaJ (Hsp40) homolog, subfamily A, member 4 | Nucleus | other |
| 15551314 | 2 | 0.105 | 2.19E+07 | 1.88E+07 | 2.27E+07 | 1.50E+07 | 1.85E+07 | 1.65E+07 | 1.68E+07 | 1.72E+07 | 1.89E+07 | 2.23E+07 | 1.66E+07 | 2.04E+07 | 1959.9984 | EGDQTSNNIPADIVFVLK | Q9QYJ3 | Q9QYJ3 | DNAJB1 | DnaJ (Hsp40) homolog, subfamily B, member 1 | Nucleus | other |
| 15551811 | 2 | 0.038 | 1.42E+07 | 1.16E+07 | 1.04E+07 | 1.06E+07 | 9776371 | 1.08E+07 | 9497692 | 8266913.5 | 1.17E+07 | 1.60E+07 | 1.52E+07 | 1.22E+07 | 1761.8225 | EALC[160.0307]GC[160.0307]TVNVPTLDGR | Q9QYJ3 | Q9QYJ3 | DNAJB1 | DnaJ (Hsp40) homolog, subfamily B, member 1 | Nucleus | other |
| 15538442 | 2 | 0.343 | 1.56E+07 | 8764354 | 3882038 | 9731864 | 7139609 | 5267453 | 9355169 | 6168553 | 6104532.5 | 1.34E+07 | 1.13E+07 | 1.26E+07 | 1477.7169 | EIAEAYDVLSDPR | Q9QYJ3 | Q9QYJ3 | DNAJB1 | DnaJ (Hsp40) homolog, subfamily B, member 1 | Nucleus | other |
| 15535392 | 2 | 0.658 | 1.12E+07 | 1.48E+07 | 1.48E+07 | 8777566 | 1.48E+07 | 1.17E+07 | 1.24E+07 | 1.15E+07 | 1.19E+07 | 1.25E+07 | 9911792 | 1.29E+07 | 1006.59125 | VPGEGLPLPK | Q9QYJ3 | Q9QYJ3 | DNAJB1 | DnaJ (Hsp40) homolog, subfamily B, member 1 | Nucleus | other |
| 15536367 | 2 | 6.87E-04 | 5378250.5 | 5190924 | 5227252.5 | 4295023.5 | 3940321.75 | 3775667.25 | 4277048 | 4565542 | 3956377 | 6669328 | 5821450 | 5328477.5 | 815.5224 | ILTIEVK | Q9QYJ3 | Q9QYJ3 | DNAJB1 | DnaJ (Hsp40) homolog, subfamily B, member 1 | Nucleus | other |
| 15539287 | 2 | 0.559 | 5701661 | 6857261.5 | 5359377 | 4129323.5 | 6628533.5 | 4592604.5 | 4361458.5 | 6032343 | 4901542 | 6200912 | 3476149.75 | 4474423 | 1228.5724 | NPFDTFFGQR | Q9QYJ3 | Q9QYJ3 | DNAJB1 | DnaJ (Hsp40) homolog, subfamily B, member 1 | Nucleus | other |
| 15536740 | 2 | 0.068 | 4320616 | 4373940.5 | 5651393.5 | 3266367.5 | 3957051.25 | 3279129.5 | 4136474.75 | 3667112.5 | 3168384.75 | 4627097 | 4386416.5 | 3670588 | 1248.6295 | RDGSDVIYPAR | Q9QYJ3 | Q9QYJ3 | DNAJB1 | DnaJ (Hsp40) homolog, subfamily B, member 1 | Nucleus | other |
| 15543831 | 2 | 0.108 | 3545254.5 | 2832356 | 2917811.25 | 2778645.25 | 2032567 | 2843271.75 | 2977783 | 2366491.25 | 2578438.5 | 4132118.5 | 2908920.5 | 3511947.5 | 1025.622 | TILEQVLPI | Q9QYJ3 | Q9QYJ3 | DNAJB1 | DnaJ (Hsp40) homolog, subfamily B, member 1 | Nucleus | other |
| 15546200 | 3 | 0.643 | 424514.594 | 1479212.875 | 4849327 | 4.70E-06 | 2324982 | 2900253.5 | 336420.031 | 2331450.75 | 3499407.25 | 14947.229 | 369493.188 | 1205248.75 | 1305.688 | KQDPPVTHDLR | Q9QYJ3 | Q9QYJ3 | DNAJB1 | DnaJ (Hsp40) homolog, subfamily B, member 1 | Nucleus | other |
| 15535954 | 3 | 0.431 | 4362686.5 | 4897409 | 4666575.5 | 4991597.5 | 5720503 | 4761393.5 | 5909561 | 6354627 | 5084221.5 | 5081723.5 | 6556975.5 | 3909458.25 | 2185.0947 | LALQLHPDRNPDDPQAQEK | Q99KV1 | Q99KV1 | DNAJB11 | DnaJ (Hsp40) homolog, subfamily B, member 11 | Cytoplasm | other |
| 15558162 | 2 | 0.883 | 2669254.75 | 3619233.25 | 3573654.5 | 3293813.25 | 4059748.25 | 2654896.5 | 3316803.25 | 3401654.75 | 3261441.75 | 2992117 | 3202209.75 | 2992112.75 | 1983.8563 | FQMTQEVVC[160.0307]DEC[160.0307]PNVK | Q99KV1 | Q99KV1 | DNAJB11 | DnaJ (Hsp40) homolog, subfamily B, member 11 | Cytoplasm | other |
| 15543639 | 2 | 0.248 | 1328417.75 | 1866493 | 1906740 | 2054983.375 | 3073833 | 1830203.625 | 2231176.75 | 2425344.75 | 1964894.125 | 1966288.25 | 2124137.5 | 1897150.875 | 1302.5818 | QYDTYGEEGLK | Q99KV1 | Q99KV1 | DNAJB11 | DnaJ (Hsp40) homolog, subfamily B, member 11 | Cytoplasm | other |
| 15548121 | 2 | 0.145 | 2.17E+07 | 2.22E+07 | 2.47E+07 | 2.19E+07 | 2.11E+07 | 2.33E+07 | 1.71E+07 | 2.08E+07 | 1.83E+07 | 2.44E+07 | 1.82E+07 | 2.18E+07 | 1943.9917 | EGDETPNSIPADIVFVIK | Q9D832 | Q9D832 | DNAJB4 | DnaJ (Hsp40) homolog, subfamily B, member 4 | Nucleus | other |
| 15533563 | 2 | 0.054 | 2.42E+07 | 1.86E+07 | 1.58E+07 | 1.60E+07 | 1.67E+07 | 1.43E+07 | 1.79E+07 | 1.66E+07 | 1.93E+07 | 2.18E+07 | 2.23E+07 | 2.11E+07 | 1385.6595 | VSLEEIYSGC[160.0307]TK | Q9D832 | Q9D832 | DNAJB4 | DnaJ (Hsp40) homolog, subfamily B, member 4 | Nucleus | other |
| 15534170 | 2 | 0.083 | 1.35E+07 | 1.31E+07 | 1.36E+07 | 1.23E+07 | 1.38E+07 | 1.38E+07 | 1.26E+07 | 1.21E+07 | 1.03E+07 | 1.70E+07 | 1.50E+07 | 1.28E+07 | 1449.7073 | EVAEAYEVLSDPK | Q9D832 | Q9D832 | DNAJB4 | DnaJ (Hsp40) homolog, subfamily B, member 4 | Nucleus | other |
| 15550844 | 2 | 0.091 | 6851833.5 | 3865247.5 | 2426029.5 | 5738530.5 | 4330931 | 3810376.75 | 5293855 | 4238356.5 | 5415264 | 9025905 | 9845950 | 6267611.5 | 1779.7795 | EALC[160.0307]GC[160.0307]SLNVPTMDGR | Q9D832 | Q9D832 | DNAJB4 | DnaJ (Hsp40) homolog, subfamily B, member 4 | Nucleus | other |
| 15534142 | 2 | 0.24 | 8044058 | 7937595.5 | 9053013 | 8326420 | 7299986.5 | 8692086 | 5411895.5 | 7564070 | 7915374 | 8414192 | 7799530 | 8999899 | 1090.6284 | VIGYGLPFPK | Q9D832 | Q9D832 | DNAJB4 | DnaJ (Hsp40) homolog, subfamily B, member 4 | Nucleus | other |
| 15542293, 15544016 | 2 | 0.88 | 7389516.5 | 6493979.5 | 2959572.5 | 7787154 | 4362018.5 | 6823230 | 3814029.75 | 5418859 | 6671087.5 | 6705394 | 6125222 | 5158171.5 | 1067.5353, 1067.5453 | DGSNIVYTAK | Q9D832 | Q9D832 | DNAJB4 | DnaJ (Hsp40) homolog, subfamily B, member 4 | Nucleus | other |
| 15534388 | 3 | 0.542 | 5810487 | 5348783.5 | 6274061.5 | 6496087 | 5825170.5 | 5573912 | 5923709 | 5701830 | 4244499 | 6442318.5 | 5178295.5 | 6497019.5 | 1189.6555 | QDPPIIHELK | Q9D832 | Q9D832 | DNAJB4 | DnaJ (Hsp40) homolog, subfamily B, member 4 | Nucleus | other |
| 15542698 | 2 | 0.898 | 4418737.5 | 4397163 | 6228283.5 | 5045401.5 | 4535337 | 6102351.5 | 5298737 | 4550485 | 5238962.5 | 4698483 | 4872441.5 | 4782922.5 | 1236.622 | DYYHILGIDK | Q9D832 | Q9D832 | DNAJB4 | DnaJ (Hsp40) homolog, subfamily B, member 4 | Nucleus | other |
| 15535903 | 2 | 0.742 | 3830172.25 | 3667558.75 | 4286236 | 2694049.25 | 5449718 | 2972911.5 | 3686408.75 | 3731933.25 | 1999797.875 | 4332919 | 2796163 | 3241494.75 | 1427.666 | EIYDQFGEEGLK | Q9D832 | Q9D832 | DNAJB4 | DnaJ (Hsp40) homolog, subfamily B, member 4 | Nucleus | other |
| 15544206 | 2 | 0.775 | 1208704.5 | 3027240.25 | 5369617.5 | 920827.625 | 3839994.75 | 4785630 | 3185856 | 3063261 | 2100041 | 1430434.875 | 1757677.875 | 2128131.75 | 1223.6353 | RDGSNIVYTAK | Q9D832 | Q9D832 | DNAJB4 | DnaJ (Hsp40) homolog, subfamily B, member 4 | Nucleus | other |
| 15543935 | 2 | 0.614 | 480544.281 | 367514.812 | 246788.906 | 526934.625 | 506716.125 | 220092.984 | 471543.906 | 355508.719 | 304404.906 | 552244.5 | 575459.625 | 389373.969 | 1497.6432 | VDC[160.0307]QAYPQTC[160.0307]QK | Q9DC23 | Q9DC23 | DNAJC10 | DnaJ (Hsp40) homolog, subfamily C, member 10 | Cytoplasm | enzyme |
| 15550755 | 2 | 0.151 | 2354640.25 | 4312249.5 | 1673542.125 | 2674096.25 | 2491481 | 3357158.25 | 1265604.875 | 2579882 | 3014407.25 | 1404570.625 | 1329757.875 | 1541142.5 | 2091.0498 | GWGELEFGAGDLQGPLFGLK | Q5U458 | Q5U458 | DNAJC11 | DnaJ (Hsp40) homolog, subfamily C, member 11 | Cytoplasm | other |
| 15558079 | 2 | 0.007 | 1598790.125 | 3276327.75 | 1907300.5 | 1437341 | 1875623.125 | 2743525.75 | 1013225.312 | 2532323.25 | 2062891.125 | 654947.25 | 513709.719 | 630905.438 | 1516.7402 | AGFFGTIVEYGAER | Q5U458 | Q5U458 | DNAJC11 | DnaJ (Hsp40) homolog, subfamily C, member 11 | Cytoplasm | other |
| 15536763 | 3 | 0.997 | 2.80E+07 | 4.84E+07 | 4.58E+07 | 3.28E+07 | 4.68E+07 | 4.29E+07 | 4.38E+07 | 3.83E+07 | 3.55E+07 | 4.32E+07 | 3.50E+07 | 4.02E+07 | 1510.7844 | EINNAHAILTDATK | P60904 | P60904 | DNAJC5 | DnaJ (Hsp40) homolog, subfamily C, member 5 | Plasma Membrane | other |
| 15549056 | 3 | 0.955 | 2.65E+07 | 3.43E+07 | 4.43E+07 | 2.70E+07 | 3.74E+07 | 4.20E+07 | 2.78E+07 | 3.92E+07 | 3.09E+07 | 4.18E+07 | 3.09E+07 | 3.40E+07 | 3615.6626 | EATDTPIVIQPASATETTQLTADSHPSYHTDGFN | P60904 | P60904 | DNAJC5 | DnaJ (Hsp40) homolog, subfamily C, member 5 | Plasma Membrane | other |
| 15547985 | 2 | 0.903 | 1.14E+07 | 1.47E+07 | 1.95E+07 | 1.26E+07 | 1.66E+07 | 1.51E+07 | 1.46E+07 | 1.33E+07 | 1.26E+07 | 1.78E+07 | 1.15E+07 | 1.25E+07 | 1510.7833 | EINNAHAILTDATK | P60904 | P60904 | DNAJC5 | DnaJ (Hsp40) homolog, subfamily C, member 5 | Plasma Membrane | other |
| 15536045 | 3 | 0.49 | 1.48E+07 | 2302671.75 | 32408.523 | 7716076 | 2071698.125 | 321718.344 | 1645440.125 | 2472415.75 | 4749805 | 8255965.5 | 1.69E+07 | 7120041.5 | 1666.8824 | EINNAHAILTDATKR | P60904 | P60904 | DNAJC5 | DnaJ (Hsp40) homolog, subfamily C, member 5 | Plasma Membrane | other |

| | | | | | | | | | | | | |
|---|---|---|---|---|---|---|---|---|---|---|---|---|
| 15556038 | 2 | 0.312 | 1095196.25 | [illegible] | 1666.8838 | EINNAHAILTDATKR | P60904 | P60904 | DNAJC5 | DnaJ (Hsp40) homolog, subfamily C, member 5 | Plasma Membrane | other |
| 15545168 | 2 | 0.279 | 550801.25 | [illegible] | 1474.7916 | ESVMDTLVQILAR | Q9D7J6 | Q9D7J6 | DNASE1L1 | deoxyribonuclease I-like 1 | Cytoplasm | enzyme |
| 15548066 | 3 | 0.087 | 5.85E+07 | [illegible] | 2983.4875 | LQDAFSSIGQSC[160.0307]HLDLPQIAVVGGQSAGK | P39054 | P39054 | DNM2 | dynamin 2 | Plasma Membrane | enzyme |
| 15548339 | 2 | 0.057 | 4.22E+07 | [illegible] | 1697.8706 | VPVGDQPPDIEYQIK | P39054 | P39054 | DNM2 | dynamin 2 | Plasma Membrane | enzyme |
| 15531642 | 3 | 0.006 | 4.26E+07 | [illegible] | 2093.196 | VYSPHVLNLTLIDLPGITK | P39054 | P39054 | DNM2 | dynamin 2 | Plasma Membrane | enzyme |
| 15537612 | 3 | 0.115 | 3.21E+07 | [illegible] | 1933.9111 | LDLMDEGTDARDVLENK | P39054 | P39054 | DNM2 | dynamin 2 | Plasma Membrane | enzyme |
| 15531940 | 2 | 0.036 | 2.44E+07 | [illegible] | 1348.7461 | NLVDSYVAIINK | P39054 | P39054 | DNM2 | dynamin 2 | Plasma Membrane | enzyme |
| 15538300 | 2 | 0.288 | 2.33E+07 | [illegible] | 1158.6704 | LQSQLLSLEK | P39054 | P39054 | DNM2 | dynamin 2 | Plasma Membrane | enzyme |
| 15550093 | 2 | 0.099 | 1.69E+07 | [illegible] | 1621.8802 | TGLFTPDLAFEAIVK | P39054 | P39054 | DNM2 | dynamin 2 | Plasma Membrane | enzyme |
| 15549297 | 2 | 0.01 | 1.46E+07 | [illegible] | 1922.8401 | QIELAC[160.0307]DSQEDVDSWK | P39054 | P39054 | DNM2 | dynamin 2 | Plasma Membrane | enzyme |
| 15548345 | 2 | 0.01 | 1.66E+07 | [illegible] | 2273.1614 | ESSLILAVTPANMDLANSDALK | P39054 | P39054 | DNM2 | dynamin 2 | Plasma Membrane | enzyme |
| 15533153 | 2 | 0.009 | 1.40E+07 | [illegible] | 1065.6403 | GISPVPINLR | P39054 | P39054 | DNM2 | dynamin 2 | Plasma Membrane | enzyme |
| 15534869 | 2 | 0.035 | 1.29E+07 | [illegible] | 1122.5953 | DMILQFISR | P39054 | P39054 | DNM2 | dynamin 2 | Plasma Membrane | enzyme |
| 15535497 | 2 | 0.041 | 1.04E+07 | [illegible] | 1107.5769 | SSVLENFVGR | P39054 | P39054 | DNM2 | dynamin 2 | Plasma Membrane | enzyme |
| 15533060 | 2 | 0.044 | 7723695.5 | [illegible] | 865.5126 | IVTTYIR | P39054 | P39054 | DNM2 | dynamin 2 | Plasma Membrane | enzyme |
| 15553990 | 3 | 0.041 | 7153853 | [illegible] | 2524.0479 | DQAENEDGAQENTFSMDPQLER | P39054 | P39054 | DNM2 | dynamin 2 | Plasma Membrane | enzyme |
| 15558819 | 4 | 0.348 | 4393084 | [illegible] | 3424.6047 | AFIHHELLAYLYSSADQSSLMEESAEQAQR | P39054 | P39054 | DNM2 | dynamin 2 | Plasma Membrane | enzyme |
| 15542103 | 2 | 0.036 | 4980224 | [illegible] | 877.487 | GYIGVVNR | P39054 | P39054 | DNM2 | dynamin 2 | Plasma Membrane | enzyme |
| 15542874 | 2 | 0.052 | 6417493 | [illegible] | 913.47174 | EVDPQGLR | P39054 | P39054 | DNM2 | dynamin 2 | Plasma Membrane | enzyme |
| 15533434 | 2 | 0.313 | 5872070 | [illegible] | 1337.7263 | TLNQQLTNHIR | P39054 | P39054 | DNM2 | dynamin 2 | Plasma Membrane | enzyme |
| 15556860 | 2 | 0.067 | 3903017.75 | [illegible] | 1752.8956 | ALLQMVQQFGVDFEK | P39054 | P39054 | DNM2 | dynamin 2 | Plasma Membrane | enzyme |
| 15551360 | 2 | 0.415 | 3824847.75 | [illegible] | 1644.8962 | C[160.0307]VDLVIQELISTVR | P39054 | P39054 | DNM2 | dynamin 2 | Plasma Membrane | enzyme |
| 15551170 | 2 | 0.198 | 3925751.5 | [illegible] | 1933.9163 | LDLMDEGTDARDVLENK | P39054 | P39054 | DNM2 | dynamin 2 | Plasma Membrane | enzyme |
| 15546005 | 3 | 0.035 | 2141893.5 | [illegible] | 1327.843 | RPLILQLIFSK | P39054 | P39054 | DNM2 | dynamin 2 | Plasma Membrane | enzyme |
| 15543240 | 3 | 0.124 | 2043529.25 | [illegible] | 2099.9463 | FTDFDEVRQEIEAETDR | P39054 | P39054 | DNM2 | dynamin 2 | Plasma Membrane | enzyme |
| 15533920 | 2 | 0.081 | 1.21E+07 | [illegible] | 1361.6925 | HVFALFNTEQR | Q8BZ98 | Q8BZ98 | DNM3 | dynamin 3 | Cytoplasm | enzyme |
| 15543529 | 2 | 8.02E-04 | 1.41E+07 | [illegible] | 879.4951 | FPFEIVK | Q8BZ98 | Q8BZ98 | DNM3 | dynamin 3 | Cytoplasm | enzyme |
| 15557510 | 2 | 0.088 | 3778502.75 | [illegible] | 1639.8358 | TGLFTPDMAFEAIVK | Q8BZ98 | Q8BZ98 | DNM3 | dynamin 3 | Cytoplasm | enzyme |
| 15548515 | 3 | 0.295 | 2.74E+07 | [illegible] | 3132.6143 | NINENFGPNTEIHLVPILATAVQEELEK | Q9Z2W0 | Q9Z2W0 | DNPEP | aspartyl aminopeptidase | Cytoplasm | peptidase |
| 15548535 | 2 | 0.362 | 7347869 | [illegible] | 1710.9177 | EVAGQVGVPLQDLMVR | Q9Z2W0 | Q9Z2W0 | DNPEP | aspartyl aminopeptidase | Cytoplasm | peptidase |
| 15552621 | 3 | 0.689 | 5417900 | [illegible] | 3066.5227 | MVTLYDNEEVGSESAQGAQSLLTELILR | Q9Z2W0 | Q9Z2W0 | DNPEP | aspartyl aminopeptidase | Cytoplasm | peptidase |
| 15535440 | 2 | 0.276 | 7091219 | [illegible] | 1118.6064 | LTAFEEAIPK | Q9Z2W0 | Q9Z2W0 | DNPEP | aspartyl aminopeptidase | Cytoplasm | peptidase |
| 15538586 | 2 | 0.125 | 5829082 | [illegible] | 1416.6611 | ETEGWDIVPENK | Q9Z2W0 | Q9Z2W0 | DNPEP | aspartyl aminopeptidase | Cytoplasm | peptidase |
| 15553252 | 2 | 0.319 | 4200429 | [illegible] | 1658.8142 | NDSPC[160.0307]GTTIGPILASR | Q9Z2W0 | Q9Z2W0 | DNPEP | aspartyl aminopeptidase | Cytoplasm | peptidase |
| 15536534 | 3 | 0.12 | 5651808.5 | [illegible] | 2120.9463 | SFMISADMAHAVHPNYSDK | Q9Z2W0 | Q9Z2W0 | DNPEP | aspartyl aminopeptidase | Cytoplasm | peptidase |
| 15549349 | 2 | 0.186 | 5227125.5 | [illegible] | 1782.9264 | ETAC[160.0307]TTGVLQTLTLFK | Q9Z2W0 | Q9Z2W0 | DNPEP | aspartyl aminopeptidase | Cytoplasm | peptidase |
| 15542518 | 2 | 0.107 | 2709053.5 | [illegible] | 804.47125 | LLQAGFR | Q9Z2W0 | Q9Z2W0 | DNPEP | aspartyl aminopeptidase | Cytoplasm | peptidase |
| 15559289 | 2 | 0.469 | 181611.859 | [illegible] | 1564.6703 | SASSGAEGDVSSEREP | Q9DD18 | Q9DD18 | DTD1 | D-tyrosyl-tRNA deacylase 1 | Cytoplasm | enzyme |
| 15545452 | 2 | 0.162 | 1998786.75 | [illegible] | 787.4703 | LVEAISR | Q9JHU4 | Q9JHU4 | DYNC1H1 | dynein, cytoplasmic 1, heavy chain 1 | Cytoplasm | peptidase |
| 15536902 | 3 | 0.358 | 3072562.75 | [illegible] | 1735.8829 | VGVVDEVVPEDQVHSK | P42125 | P42125 | ECI1 | enoyl-CoA delta isomerase 1 | Cytoplasm | enzyme |
| 15534593 | 3 | 0.36 | 1673618.5 | [illegible] | 1783.8928 | QREADIQNFTSFISK | P42125 | P42125 | ECI1 | enoyl-CoA delta isomerase 1 | Cytoplasm | enzyme |
| 15540133 | 3 | 0.935 | 1715350.875 | [illegible] | 1823.8949 | QQHQEQQALQQSATAK | Q8BL66 | Q8BL66 | EEA1 | early endosome antigen 1 | Cytoplasm | other |
| 15549279 | 2, 3 | 0.02 | 2.35E+08 | [illegible] | 2628.313, 2628.3147 | DGSASGTTLLEALDC[160.0307]ILPPTRPTDK | P10126 | P10126 | EEF1A1 | eukaryotic translation elongation factor 1 alpha 1 | Cytoplasm | translation regulator |
| 15531337 | 2 | 0.154 | 9.25E+07 | [illegible] | 1025.6085 | IGGIGTVPVGR | P10126 | P10126 | EEF1A1 | eukaryotic translation elongation factor 1 alpha 1 | Cytoplasm | translation regulator |
| 15548934 | 3 | 0.843 | 8.86E+07 | [illegible] | 2995.3933 | SGDAAIVDMVPGKPMC[160.0307]VESFSDYPPLGR | P10126 | P10126 | EEF1A1 | eukaryotic translation elongation factor 1 alpha 1 | Cytoplasm | translation regulator |
| 15532705 | 2, 3 | 0.087 | 4.73E+07 | [illegible] | 1404.7242, 1404.7247 | YYVTIIDAPGHR | P10126 | P10126 | EEF1A1 | eukaryotic translation elongation factor 1 alpha 1 | Cytoplasm | translation regulator |
| 15549935 | 3 | 0.776 | 2.39E+07 | [illegible] | 3567.6655 | IGYNPDTVAFVPISGWNGDNMLEPSANMPWFK | P10126 | P10126 | EEF1A1 | eukaryotic translation elongation factor 1 alpha 1 | Cytoplasm | translation regulator |
| 15539642 | 2 | 0.071 | 4.55E+07 | [illegible] | 1282.5593 | MDSTEPPYSQK | P10126 | P10126 | EEF1A1 | eukaryotic translation elongation factor 1 alpha 1 | Cytoplasm | translation regulator |
| 15531629 | 2 | 0.108 | 7.05E+07 | [illegible] | 1120.597 | STTTGHLIYK | P10126 | P10126 | EEF1A1 | eukaryotic translation elongation factor 1 alpha 1 | Cytoplasm | translation regulator |
| 15548329 | 2, 3 | 0.886 | 8.68E+07 | [illegible] | 2515.3777, 2515.3809 | VETGVLKPGMVVTFAPVNVTTEVK | P10126 | P10126 | EEF1A1 | eukaryotic translation elongation factor 1 alpha 1 | Cytoplasm | translation regulator |
| 15531327 | 3 | 0.255 | 5.63E+07 | [illegible] | 1341.7854 | PLRLPLQDVYK | P10126 | P10126 | EEF1A1 | eukaryotic translation elongation factor 1 alpha 1 | Cytoplasm | translation regulator |
| 15555041, 15560700 | 2 | 0.101 | 3.27E+07 | [illegible] | 1788.9525, 1788.9529 | PGMVVTFAPVNVTTEVK | P10126 | P10126 | EEF1A1 | eukaryotic translation elongation factor 1 alpha 1 | Cytoplasm | translation regulator |
| 15540024 | 2 | 0.047 | 4.63E+07 | [illegible] | 870.5392 | QLIVGVNK | P10126 | P10126 | EEF1A1 | eukaryotic translation elongation factor 1 alpha 1 | Cytoplasm | translation regulator |
| 15531572 | 2 | 0.026 | 4.00E+07 | [illegible] | 914.56494 | QTVAVGVIK | P10126 | P10126 | EEF1A1 | eukaryotic translation elongation factor 1 alpha 1 | Cytoplasm | translation regulator |
| 15535835 | 2 | 0.219 | 2.62E+07 | [illegible] | 1259.6279 | SGDAAIVDMVPGK | P10126 | P10126 | EEF1A1 | eukaryotic translation elongation factor 1 alpha 1 | Cytoplasm | translation regulator |
| 15533232 | 2, 3 | 0.201 | 3.17E+07 | [illegible] | 1314.7385, 1314.7391 | EHALLAYTLGVK | P10126 | P10126 | EEF1A1 | eukaryotic translation elongation factor 1 alpha 1 | Cytoplasm | translation regulator |
| 15531566 | 2 | 0.131 | 2.19E+07 | [illegible] | 975.55005 | LPLQDVYK | P10126 | P10126 | EEF1A1 | eukaryotic translation elongation factor 1 alpha 1 | Cytoplasm | translation regulator |
| 15534178 | 3 | 0.978 | 4.59E+07 | [illegible] | 1907.9977 | YYVTIIDAPGHRDFIK | P10126 | P10126 | EEF1A1 | eukaryotic translation elongation factor 1 alpha 1 | Cytoplasm | translation regulator |
| 15543415 | 2 | 0.042 | 9998456 | [illegible] | 839.44965 | EVSTYIK | P10126 | P10126 | EEF1A1 | eukaryotic translation elongation factor 1 alpha 1 | Cytoplasm | translation regulator |
| 15541602 | 3 | 0.904 | 2763017.25 | [illegible] | 1788.9531 | PGMVVTFAPVNVTTEVK | P10126 | P10126 | EEF1A1 | eukaryotic translation elongation factor 1 alpha 1 | Cytoplasm | translation regulator |
| 15558092 | 3 | 0.163 | 3153784.25 | [illegible] | 3583.6567 | IGYNPDTVAFVPISGWNGDNMLEPSANM[147.0354]PWFK | P10126 | P10126 | EEF1A1 | eukaryotic translation elongation factor 1 alpha 1 | Cytoplasm | translation regulator |
| 15558453 | 3 | 0.041 | 2674635.25 | [illegible] | 2909.4329 | NMITGTSQADC[160.0307]AVLIVAAGVGEFEAGISK | P10126 | P10126 | EEF1A1 | eukaryotic translation elongation factor 1 alpha 1 | Cytoplasm | translation regulator |
| 15557680, 15560626 | 3 | 0.484 | 4353254 | [illegible] | 3011.3875, 3011.3918 | SGDAAIVDMVPGKPM[147.0354]C[160.0307]VESFSDYPPLGR | P10126 | P10126 | EEF1A1 | eukaryotic translation elongation factor 1 alpha 1 | Cytoplasm | translation regulator |
| 15554472 | 2 | 0.951 | 5261809 | [illegible] | 1907.9962 | YYVTIIDAPGHRDFIK | P10126 | P10126 | EEF1A1 | eukaryotic translation elongation factor 1 alpha 1 | Cytoplasm | translation regulator |
| 15547005, 15547367 | 2 | 0.003 | 579919.125 | [illegible] | 1298.5544, 1298.5548 | M[147.0354]DSTEPPYSQK | P10126 | P10126 | EEF1A1 | eukaryotic translation elongation factor 1 alpha 1 | Cytoplasm | translation regulator |
| 15561293 | 2 | 0.117 | 1133765 | [illegible] | 1712.7585 | YGPSSVEDTTGSGAADAK | O70251 | O70251 | EEF1B2 | eukaryotic translation elongation factor 1 beta 2 | Cytoplasm | translation regulator |
| 15536255 | 2 | 0.126 | 1380496.875 | [illegible] | 1415.7457 | ITSLEVENQNLR | P57776 | P57776 | EEF1D | eukaryotic translation elongation factor 1 delta (guanine nucleotide exchange protein) | Cytoplasm | translation regulator |
| 15552232 | 3 | 0.203 | 2142054.5 | [illegible] | 2461.1313 | YSNEDTLSVALPYFWEHFDK | Q9D8N0 | Q9D8N0 | EEF1G | eukaryotic translation elongation factor 1 gamma | Cytoplasm | translation regulator |
| 15537938 | 2 | 0.303 | 606387.312 | [illegible] | 1444.724 | LDPGSEETQTLVR | Q9D8N0 | Q9D8N0 | EEF1G | eukaryotic translation elongation factor 1 gamma | Cytoplasm | translation regulator |
| 15539689 | 2 | 0.04 | 1158554.875 | [illegible] | 1347.7346 | ALIAAQYSGAQVR | Q9D8N0 | Q9D8N0 | EEF1G | eukaryotic translation elongation factor 1 gamma | Cytoplasm | translation regulator |
| 15557423 | 2 | 0.196 | 2090381.875 | [illegible] | 2204.109 | STAISLFYELSENDLNFIK | P58252 | P58252 | EEF2 | eukaryotic translation elongation factor 2 | Cytoplasm | translation regulator |
| 15553251 | 2 | 0.045 | 3322380.5 | [illegible] | 2105.033 | IWC[160.0307]FGPDGTGPNILTDITK | P58252 | P58252 | EEF2 | eukaryotic translation elongation factor 2 | Cytoplasm | translation regulator |
| 15553405 | 3 | 0.052 | 4605535 | [illegible] | 2759.4287 | YVEPIEDVPC[160.0307]GNIVGLVGVDQFLVK | P58252 | P58252 | EEF2 | eukaryotic translation elongation factor 2 | Cytoplasm | translation regulator |
| 15551723 | 2 | 0.052 | 2897135 | [illegible] | 2220.1475 | ALLELQLEPEELYQTFQR | P58252 | P58252 | EEF2 | eukaryotic translation elongation factor 2 | Cytoplasm | translation regulator |
| 15540574 | 2 | 0.032 | 691994.375 | [illegible] | 1439.681 | DLEEDHAC[160.0307]IPIK | P58252 | P58252 | EEF2 | eukaryotic translation elongation factor 2 | Cytoplasm | translation regulator |
| 15552103 | 3 | 0.444 | 4990119 | [illegible] | 2877.4905 | TIVTALPGSFLDPLLSPSLMEDYELR | Q8BG67 | Q8BG67 | EFR3A | EFR3 homolog A | Plasma Membrane | other |
| 15549383 | 2 | 0.067 | 3844788.25 | [illegible] | 1687.9592 | GSIGPTVLEVFNTLLK | Q8BG67 | Q8BG67 | EFR3A | EFR3 homolog A | Plasma Membrane | other |
| 15538439 | 2 | 0.021 | 4847626.5 | [illegible] | 1424.6982 | IAESLGGSGYSVER | Q8BG67 | Q8BG67 | EFR3A | EFR3 homolog A | Plasma Membrane | other |
| 15537878 | 2 | 0.278 | 4744215 | [illegible] | 1205.6859 | LQVLGTNSFVK | Q8BG67 | Q8BG67 | EFR3A | EFR3 homolog A | Plasma Membrane | other |
| 15536670 | 3 | 0.094 | 3708306.5 | [illegible] | 1578.7156 | FANIEEDTPSYHR | Q8BG67 | Q8BG67 | EFR3A | EFR3 homolog A | Plasma Membrane | other |
| 15536194 | 2 | 0.016 | 4620136.5 | [illegible] | 1126.572 | LGPPSSPSAADK | Q8BG67 | Q8BG67 | EFR3A | EFR3 homolog A | Plasma Membrane | other |
| 15542139 | 3 | 0.516 | 2969147.75 | [illegible] | 1597.8154 | QLVLEVMHNLMDR | Q8BG67 | Q8BG67 | EFR3A | EFR3 homolog A | Plasma Membrane | other |
| 15535461, 15538385, 15543588 | 2 | 0.025 | 5760337.5 | [illegible] | 1394.6953, 1394.6959, 1394.7024 | LDISDEFSEVIK | Q9WVK4 | Q9WVK4 | EHD1 | EH-domain containing 1 | Cytoplasm | other |
| 15553088 | 3 | 0.187 | 8244411 | [illegible] | 3272.6533 | IGPEPTTDSFIAVMHGPTEGVVPGNALVVDPR | Q9WVK4 | Q9WVK4 | EHD1 | EH-domain containing 1 | Cytoplasm | other |
| 15536267 | 2 | 0.397 | 3375580.5 | [illegible] | 1467.7129 | MQELLQTQDFSK | Q9WVK4 | Q9WVK4 | EHD1 | EH-domain containing 1 | Cytoplasm | other |
| 15551365 | 2 | 0.003 | 2867821.75 | [illegible] | 1674.8344 | LLDTVDDMLANDIAR | Q9WVK4 | Q9WVK4 | EHD1 | EH-domain containing 1 | Cytoplasm | other |
| 15553745 | 2 | 0.372 | 539207.188 | [illegible] | 1616.7225 | AMQEQLENYDFTK | Q9EQP2 | Q9EQP2 | EHD4 | EH-domain containing 4 | Plasma Membrane | enzyme |
| 15544160 | 2 | 0.463 | 923514.188 | [illegible] | 1304.6244 | ADQVDTQQLMR | Q9EQP2 | Q9EQP2 | EHD4 | EH-domain containing 4 | Plasma Membrane | enzyme |
| 15533173 | 2 | 0.438 | 7183040 | [illegible] | 1348.655 | VVTDTDETELAR | Q6ZWX6 | Q6ZWX6 | EIF2S1 | eukaryotic translation initiation factor 2, subunit 1 alpha, 35kDa | Cytoplasm | translation regulator |
| 15549353 | 2 | 0.181 | 5393141 | [illegible] | 1573.8593 | TEGLSVLNQAMAVIK | Q6ZWX6 | Q6ZWX6 | EIF2S1 | eukaryotic translation initiation factor 2, subunit 1 alpha, 35kDa | Cytoplasm | translation regulator |
| 15535518 | 2 | 0.246 | 5406762.5 | [illegible] | 1421.6729 | AGLNC[160.0307]STETMPIK | Q6ZWX6 | Q6ZWX6 | EIF2S1 | eukaryotic translation initiation factor 2, subunit 1 alpha, 35kDa | Cytoplasm | translation regulator |
| 15541300 | 2 | 0.47 | 3253245 | [illegible] | 1264.6124 | DEQLESLFQR | Q6ZWX6 | Q6ZWX6 | EIF2S1 | eukaryotic translation initiation factor 2, subunit 1 alpha, 35kDa | Cytoplasm | translation regulator |
| 15551204 | 3 | 0.423 | 3627956 | [illegible] | 2264.942 | LERENAEVDGDDDAEEMEAK | Q6ZWX6 | Q6ZWX6 | EIF2S1 | eukaryotic translation initiation factor 2, subunit 1 alpha, 35kDa | Cytoplasm | translation regulator |
| 15551413 | 4 | 0.149 | 3178929.5 | [illegible] | 3190.5947 | HAVSDPSILDSLDLNEDEREVLINNINR | Q6ZWX6 | Q6ZWX6 | EIF2S1 | eukaryotic translation initiation factor 2, subunit 1 alpha, 35kDa | Cytoplasm | translation regulator |
| 15535359 | 2 | 0.175 | 7210158 | [illegible] | 1248.5952 | FPEEDEILEK | Q99L45 | Q99L45 | EIF2S2 | eukaryotic translation initiation factor 2, subunit 2 beta, 38kDa | Cytoplasm | translation regulator |
| 15542400 | 2 | 0.256 | 6198797.5 | [illegible] | 1392.6869 | IFDIDEAEEAIK | Q99L45 | Q99L45 | EIF2S2 | eukaryotic translation initiation factor 2, subunit 2 beta, 38kDa | Cytoplasm | translation regulator |
| 15550552 | 2 | 0.33 | 5319513.5 | [illegible] | 1756.765 | DASDDLDDLNFFNQK | Q99L45 | Q99L45 | EIF2S2 | eukaryotic translation initiation factor 2, subunit 2 beta, 38kDa | Cytoplasm | translation regulator |
| 15552558 | 2 | 0.712 | 1.05E+07 | [illegible] | 1558.8116 | IVLTNPVC[160.0307]TEVGEK | Q9Z0N1 | Q9Z0N1 | EIF2S3 | eukaryotic translation initiation factor 2, subunit 3 gamma, 52kDa | Cytoplasm | translation regulator |
| 15549382 | 3 | 0.378 | 1.30E+07 | [illegible] | 2570.3547 | IVSLFAEHNDLQYAAPGGLIGVGTK | Q9Z0N1 | Q9Z0N1 | EIF2S3 | eukaryotic translation initiation factor 2, subunit 3 gamma, 52kDa | Cytoplasm | translation regulator |
| 15535200 | 3 | 0.189 | 1.05E+07 | [illegible] | 1510.8561 | VGQEIEVRPGIVSK | Q9Z0N1 | Q9Z0N1 | EIF2S3 | eukaryotic translation initiation factor 2, subunit 3 gamma, 52kDa | Cytoplasm | translation regulator |
| 15549799 | 3 | 0.477 | 6701025.5 | [illegible] | 3013.617 | EQYEQILAFVQGTVAEGAPIIPISAQLK | Q9Z0N1 | Q9Z0N1 | EIF2S3 | eukaryotic translation initiation factor 2, subunit 3 gamma, 52kDa | Cytoplasm | translation regulator |
| 15549522 | 2 | 0.698 | 7760113.5 | [illegible] | 1895.8285 | SC[160.0307]GSSTPDEFPTDIPGTK | Q9Z0N1 | Q9Z0N1 | EIF2S3 | eukaryotic translation initiation factor 2, subunit 3 gamma, 52kDa | Cytoplasm | translation regulator |
| 15533799 | 3 | 0.875 | 5959479.5 | [illegible] | 1664.712 | LDDPSC[160.0307]PRPEC[160.0307]YR | Q9Z0N1 | Q9Z0N1 | EIF2S3 | eukaryotic translation initiation factor 2, subunit 3 gamma, 52kDa | Cytoplasm | translation regulator |
| 15540550 | 2 | 0.312 | 5395349.5 | [illegible] | 1103.5924 | QDLATLDVTK | Q9Z0N1 | Q9Z0N1 | EIF2S3 | eukaryotic translation initiation factor 2, subunit 3 gamma, 52kDa | Cytoplasm | translation regulator |
| 15533604 | 2 | 0.552 | 4702134.5 | [illegible] | 978.6071 | HILILQNK | Q9Z0N1 | Q9Z0N1 | EIF2S3 | eukaryotic translation initiation factor 2, subunit 3 gamma, 52kDa | Cytoplasm | translation regulator |
| 15538453 | 3 | 0.216 | 3176226.75 | [illegible] | 1616.8828 | QATINIGTIGHVAHGK | Q9Z0N1 | Q9Z0N1 | EIF2S3 | eukaryotic translation initiation factor 2, subunit 3 gamma, 52kDa | Cytoplasm | translation regulator |
| 15551812 | 2 | 0.092 | 2206270.5 | [illegible] | 1510.855 | VGQEIEVRPGIVSK | Q9Z0N1 | Q9Z0N1 | EIF2S3 | eukaryotic translation initiation factor 2, subunit 3 gamma, 52kDa | Cytoplasm | translation regulator |
| 15534805 | 2 | 0.082 | 1981718.375 | [illegible] | 1251.7025 | LTPLSHEVISR | Q9Z0N1 | Q9Z0N1 | EIF2S3 | eukaryotic translation initiation factor 2, subunit 3 gamma, 52kDa | Cytoplasm | translation regulator |
| 15540280 | 2 | 0.61 | 1.22E+07 | [illegible] | 1242.657 | VLTPEEQLADK | Q3UGC7 | Q3UGC7 | EIF3J | eukaryotic translation initiation factor 3, subunit J | Cytoplasm | translation regulator |
| 15554017 | 3 | 0.819 | 1.23E+07 | [illegible] | 2248.9531 | VAGGGTAGGDRWEGEDEDEDVK | Q3UGC7 | Q3UGC7 | EIF3J | eukaryotic translation initiation factor 3, subunit J | Cytoplasm | translation regulator |
| 15535143 | 2 | 0.843 | 6654410 | [illegible] | 1364.7059 | ITNSLTVLC[160.0307]SEK | Q3UGC7 | Q3UGC7 | EIF3J | eukaryotic translation initiation factor 3, subunit J | Cytoplasm | translation regulator |
| 15547519 | 2 | 0.955 | 51610.098 | [illegible] | 1380.4785 | DNWDDDDENK | Q3UGC7 | Q3UGC7 | EIF3J | eukaryotic translation initiation factor 3, subunit J | Cytoplasm | translation regulator |
| 15549964 | 3 | 0.016 | 1.02E+07 | [illegible] | 3111.3938 | ETFGVNNTVYGIDAMNPSSRDDFTEFGK | Q3UGC7 | Q3UGC7 | EIF3J | eukaryotic translation initiation factor 3, subunit J | Cytoplasm | translation regulator |
| 15551431 | 2 | 0.163 | 8122498 | [illegible] | 2286.1072 | TISLTDFLAEDGGTGGGSTYVPK | Q8BGD9 | Q8BGD9 | EIF4B | eukaryotic translation initiation factor 4B | Cytoplasm | translation regulator |
| 15549035 | 2 | 0.654 | 5688502.5 | [illegible] | 1591.747 | VDVVGATQGQAGSC[160.0307]SR | Q8BGD9 | Q8BGD9 | EIF4B | eukaryotic translation initiation factor 4B | Cytoplasm | translation regulator |
| 15533566 | 2, 3 | 0.808 | 7573396 | [illegible] | 1357.7035, 1357.7036 | IRVDVADQAQDK | Q8BGD9 | Q8BGD9 | EIF4B | eukaryotic translation initiation factor 4B | Cytoplasm | translation regulator |
| 15552979 | 3 | 0.281 | 3655737 | [illegible] | 3224.3489 | PVSWADETDDLEGDVSTTWHSNDDDVYR | Q8BGD9 | Q8BGD9 | EIF4B | eukaryotic translation initiation factor 4B | Cytoplasm | translation regulator |
| 15541717 | 2 | 0.292 | 3641074.25 | [illegible] | 1047.5295 | SLENETLNK | Q8BGD9 | Q8BGD9 | EIF4B | eukaryotic translation initiation factor 4B | Cytoplasm | translation regulator |
| 15559104 | 2 | 0.1 | 2441190.25 | [illegible] | 2327.1438 | SPPYTAFLGNLPYDVTEDSIK | Q8BGD9 | Q8BGD9 | EIF4B | eukaryotic translation initiation factor 4B | Cytoplasm | translation regulator |
| 15536478 | 3 | 0.281 | 3846733.5 | [illegible] | 1960.9802 | FSALQQTLPAENTDNRR | Q6NZJ6 | Q6NZJ6 | EIF4G1 | eukaryotic translation initiation factor 4 gamma, 1 | Cytoplasm | translation regulator |
| 15555014 | 3 | 0.034 | 2052180.875 | [illegible] | 2306.212 | EAVQC[160.0307]VQELASPSLLFIFVR | Q6NZJ6 | Q6NZJ6 | EIF4G1 | eukaryotic translation initiation factor 4 gamma, 1 | Cytoplasm | translation regulator |
| 15545711 | 2 | 0.079 | 778334.562 | [illegible] | 1251.5125 | EMDEAATAEER | Q6NZJ6 | Q6NZJ6 | EIF4G1 | eukaryotic translation initiation factor 4 gamma, 1 | Cytoplasm | translation regulator |
| 15534906 | 2 | 0.537 | 2766443.5 | [illegible] | 1094.6223 | VNILFDFVK | P59325 | P59325 | EIF5 | eukaryotic translation initiation factor 5 | Cytoplasm | translation regulator |
| 15552989 | 3 | 0.116 | 3340892.75 | [illegible] | 2736.3909 | RNDFQLIGIQDGYLSLLQDSGEVR | P63242 | P63242 | EIF5A | eukaryotic translation initiation factor 5A | Cytoplasm | translation regulator |
| 15550589 | 3 | 0.287 | 234640.703 | [illegible] | 3144.6619 | LQLVGSGLHEAEAAGEAQQAVLEGLQGLLSR | Q99K41 | Q99K41 | EMILIN1 | elastin microfibril interfacer 1 | Extracellular Space | other |
| 15555795 | 2 | 0.064 | 19605.104 | [illegible] | 1722.7131 | TC[160.0307]GQIC[160.0307]SGAPGEQDSR | Q99K41 | Q99K41 | EMILIN1 | elastin microfibril interfacer 1 | Extracellular Space | other |
| 15548237 | 2 | 0.148 | 1.22E+07 | [illegible] | 1570.8037 | ILEVVNQVQDEER | Q8C522 | Q8C522 | ENDOD1 | endonuclease domain containing 1 | Extracellular Space | enzyme |
| 15532660 | 3 | 0.118 | 1.07E+07 | [illegible] | 2169.0818 | HTQDIDVIEDVMLRDLEK | Q8C522 | Q8C522 | ENDOD1 | endonuclease domain containing 1 | Extracellular Space | enzyme |
| 15540590 | 4 | 0.085 | 8880537 | [illegible] | 2657.2268 | LLPHKPQLFQDNC[160.0307]GEMEQDTEK | Q8C522 | Q8C522 | ENDOD1 | endonuclease domain containing 1 | Extracellular Space | enzyme |
| 15537217 | 2, 3 | 0.144 | 4653192 | [illegible] | 1148.5818, 1148.5819 | FATLYSPGHR | Q8C522 | Q8C522 | ENDOD1 | endonuclease domain containing 1 | Extracellular Space | enzyme |
| 15535879 | 2 | 0.439 | 6966436.5 | [illegible] | 936.5279 | IPVFSAFR | Q8C522 | Q8C522 | ENDOD1 | endonuclease domain containing 1 | Extracellular Space | enzyme |
| 15538319 | 3 | 0.292 | 4959742.5 | [illegible] | 1570.8164 | DLYILTGAVPSEHR | Q8C522 | Q8C522 | ENDOD1 | endonuclease domain containing 1 | Extracellular Space | enzyme |
| 15549334 | 3 | 0.319 | 6878275.5 | [illegible] | 2657.2283 | LLPHKPQLFQDNC[160.0307]GEMEQDTEK | Q8C522 | Q8C522 | ENDOD1 | endonuclease domain containing 1 | Extracellular Space | enzyme |
| 15549419 | 2 | 0.877 | 5682491.5 | [illegible] | 1570.8182 | DLYILTGAVPSEHR | Q8C522 | Q8C522 | ENDOD1 | endonuclease domain containing 1 | Extracellular Space | enzyme |
| 15543998 | 2 | 0.013 | 840676.625 | [illegible] | 1166.6542 | LLLVSFDGFR | Q8BTJ4 | Q8BTJ4 | ENPP4 | ectonucleotide pyrophosphatase/phosphodiesterase 4 (putative) | Cytoplasm | enzyme |
| 15532638 | 3 | 1.10E-05 | 1.07E+08 | [illegible] | 2012.0405 | QFEQLQAAIHTDLGALEK | P10404 | P10404 | ENV1 | MLV-related proviral Env polyprotein | Other | |
| 15531381 | 3 | 1.64E-04 | 7.73E+07 | [illegible] | 1802.8457 | VTYHSPGYVYGQFER | P10404 | P10404 | ENV1 | MLV-related proviral Env polyprotein | Other | |
| 15532242 | 3 | 2.45E-04 | 9.62E+07 | [illegible] | 2224.9866 | TFDFYVC[160.0307]PGHTVPTGC[160.0307]GGPR | P10404 | P10404 | ENV1 | MLV-related proviral Env polyprotein | Other | |
| 15549953, 15551636 | 2, 3 | 1.56E-04 | 5.55E+07 | [illegible] | 2533.0972, 2533.1003 | LYFDLC[160.0307]DLIGDDWDETGLGC[160.0307]R | P10404 | P10404 | ENV1 | MLV-related proviral Env polyprotein | Other | |
| 15532089 | 2, 3 | 2.19E-04 | 4.72E+07 | [illegible] | 1445.7916, 1445.7926 | SLTSLSEVVLQNR | P10404 | P10404 | ENV1 | MLV-related proviral Env polyprotein | Other | |
| 15555016 | 2 | 2.00E-05 | 6.90E+07 | [illegible] | 1828.9802 | LTLSEVTGQGLC[160.0307]VGAVPK | P10404 | P10404 | ENV1 | MLV-related proviral Env polyprotein | Other | |
| 15535409 | 2 | 2.12E-05 | 6.60E+07 | [illegible] | 1463.7164 | C[160.0307]NPLVLEFTDAGK | P10404 | P10404 | ENV1 | MLV-related proviral Env polyprotein | Other | |
| 15531511 | 2 | 4.33E-05 | 5.64E+07 | [illegible] | 1232.65 | PSSSWDLISLK | P10404 | P10404 | ENV1 | MLV-related proviral Env polyprotein | Other | |
| 15537115 | 3 | 2.83E-04 | 3.85E+07 | [illegible] | 1742.7219 | EEC[160.0307]C[160.0307]FYADHTGVVR | P10404 | P10404 | ENV1 | MLV-related proviral Env polyprotein | Other | |
| 15533645 | 3 | 1.77E-05 | 4.37E+07 | [illegible] | 1601.8939 | SLTSLSEVVLQNRR | P10404 | P10404 | ENV1 | MLV-related proviral Env polyprotein | Other | |
| 15531571 | 2 | 8.70E-05 | 3.46E+07 | [illegible] | 896.5295 | QVLNIGPR | P10404 | P10404 | ENV1 | MLV-related proviral Env polyprotein | Other | |
| 15540888 | 2 | 5.06E-05 | 3.16E+07 | [illegible] | 1486.639 | WGC[160.0307]ETTGQAYWK | P10404 | P10404 | ENV1 | MLV-related proviral Env polyprotein | Other | |
| 15538707 | 2 | 4.38E-05 | 1.76E+07 | [illegible] | 918.4696 | EGGLC[160.0307]AALK | P10404 | P10404 | ENV1 | MLV-related proviral Env polyprotein | Other | |
| 15532173 | 2 | 1.60E-06 | 2.15E+07 | [illegible] | 918.5635 | GLDLLFLK | P10404 | P10404 | ENV1 | MLV-related proviral Env polyprotein | Other | |
| 15539371 | 3 | 0.004 | 2.21E+07 | [illegible] | 1591.8109 | C[160.0307]NPLVLEFTDAGKK | P10404 | P10404 | ENV1 | MLV-related proviral Env polyprotein | Other | |
| 15532443 | 2 | 9.25E-06 | 1.50E+07 | [illegible] | 1074.6644 | RGLDLLFLK | P10404 | P10404 | ENV1 | MLV-related proviral Env polyprotein | Other | |
| 15549462 | 2 | 9.42E-06 | 8374275 | [illegible] | 2012.0404 | QFEQLQAAIHTDLGALEK | P10404 | P10404 | ENV1 | MLV-related proviral Env polyprotein | Other | |
| 15555704 | 2 | 1.92E-04 | 9658096 | [illegible] | 1802.846 | VTYHSPGYVYGQFER | P10404 | P10404 | ENV1 | MLV-related proviral Env polyprotein | Other | |
| 15535459 | 3 | 4.91E-04 | 7101171.5 | [illegible] | 1828.9789 | LTLSEVTGQGLC[160.0307]VGAVPK | P10404 | P10404 | ENV1 | MLV-related proviral Env polyprotein | Other | |
| 15533897 | 3 | 8.30E-04 | 8355074.5 | [illegible] | 1968.1229 | ISVVQALVLTQQYHQLK | P10404 | P10404 | ENV1 | MLV-related proviral Env polyprotein | Other | |
| 15550321 | 2 | 0.012 | 5192645.5 | [illegible] | 1786.8502 | LFESGQGWFEGLFNR | P10404 | P10404 | ENV1 | MLV-related proviral Env polyprotein | Other | |
| 15553583 | 2 | 8.71E-04 | 6750205.5 | [illegible] | 1742.7234 | EEC[160.0307]C[160.0307]FYADHTGVVR | P10404 | P10404 | ENV1 | MLV-related proviral Env polyprotein | Other | |
| 15551804 | 2 | 0.14 | 4276154.5 | [illegible] | 2224.9863 | TFDFYVC[160.0307]PGHTVPTGC[160.0307]GGPR | P10404 | P10404 | ENV1 | MLV-related proviral Env polyprotein | Other | |
| 15539147 | 3 | 7.99E-04 | 2168495 | [illegible] | 2239.2515 | DRISVVQALVLTQQYHQLK | P10404 | P10404 | ENV1 | MLV-related proviral Env polyprotein | Other | |
| 15548188, 15548626, 15550388, 15558245, 15560909 | 2, 3 | 0.564 | 1.71E+09 | [illegible] | 2360.2498, 2360.2507, 2360.2527, 2360.2542, 2360.2563 | SPRPTSAPAIAQSQVTEGPGAPIK | P48193 | P48193 | EPB41 | erythrocyte membrane protein band 4.1 | Plasma Membrane | other |
| 15537100 | 3, 4 | 0.188 | 1.88E+09 | [illegible] | 1707.891, 1707.8911 | QASALIDRPAPHFER | P48193 | P48193 | EPB41 | erythrocyte membrane protein band 4.1 | Plasma Membrane | other |
| 15531268 | 3 | 0.489 | 1.83E+09 | [illegible] | 1947.8877 | NFMESVPEPRPSEWDK | P48193 | P48193 | EPB41 | erythrocyte membrane protein band 4.1 | Plasma Membrane | other |
| 15532369 | 3 | 0.606 | 1.51E+09 | [illegible] | 2022.9397 | RGEEPAEPAEPEPTEAWK | P48193 | P48193 | EPB41 | erythrocyte membrane protein band 4.1 | Plasma Membrane | other |
| 15550945 | 2 | 0.094 | 1.33E+09 | [illegible] | 1768.8646 | VSLLDDTVYEC[160.0307]VVEK | P48193 | P48193 | EPB41 | erythrocyte membrane protein band 4.1 | Plasma Membrane | other |
| 15547691 | 2 | 0.574 | 1.30E+09 | [illegible] | 1764.8456 | SMTPAQADLEFLENAK | P48193 | P48193 | EPB41 | erythrocyte membrane protein band 4.1 | Plasma Membrane | other |
| 15534820, 15544482 | 2, 3 | 0.137 | 1.45E+09 | [illegible] | 1448.7556, 1448.7572, 1448.7573 | TQTVTISDTANAVK | P48193 | P48193 | EPB41 | erythrocyte membrane protein band 4.1 | Plasma Membrane | other |
| 15538572, 15543806, 15544528, 15547259 | 3, 4 | 0.247 | 1.30E+09 | [illegible] | 1890.9238, 1890.9321, 1890.9337, 1890.9338, 1890.9342 | IRPGEQEHYESTIGFK | P48193 | P48193 | EPB41 | erythrocyte membrane protein band 4.1 | Plasma Membrane | other |
| 15537236, 15539695, 15545057 | 2, 3 | 0.426 | 1.22E+09 | [illegible] | 1314.6335, 1314.6345 | HSNLMLEDLDK | P48193 | P48193 | EPB41 | erythrocyte membrane protein band 4.1 | Plasma Membrane | other |
| 15536804 | 2 | 0.321 | 1.10E+09 | [illegible] | 1308.6722 | GVPWNFTFNVK | P48193 | P48193 | EPB41 | erythrocyte membrane protein band 4.1 | Plasma Membrane | other |
| 15531432, 15532464, 15537530 | 2 | 0.433 | 1.06E+09 | [illegible] | 1092.5659, 1092.5663, 1092.5664 | LDGENIYIR | P48193 | P48193 | EPB41 | erythrocyte membrane protein band 4.1 | Plasma Membrane | other |
| 15539153 | 2 | 0.101 | 9.81E+08 | [illegible] | 975.5351 | LTSTDTIPK | P48193 | P48193 | EPB41 | erythrocyte membrane protein band 4.1 | Plasma Membrane | other |
| 15534214 | 2, 3 | 0.121 | 9.96E+08 | [illegible] | 1162.5903, 1162.5907 | LSMYGVDLHK | P48193 | P48193 | EPB41 | erythrocyte membrane protein band 4.1 | Plasma Membrane | other |
| 15534143 | 2 | 0.13 | 1.07E+09 | [illegible] | 1138.6084 | DVPIVHTETK | P48193 | P48193 | EPB41 | erythrocyte membrane protein band 4.1 | Plasma Membrane | other |
| 15538130, 15543934 | 2, 3 | 0.056 | 9.68E+08 | [illegible] | 1331.6294, 1331.6295, 1331.6302, 1331.6306 | VC[160.0307]VEHHTFFR | P48193 | P48193 | EPB41 | erythrocyte membrane protein band 4.1 | Plasma Membrane | other |
| 15531302 | 2 | 0.526 | 5.59E+08 | [illegible] | 735.4391 | FLALGSK | P48193 | P48193 | EPB41 | erythrocyte membrane protein band 4.1 | Plasma Membrane | other |
| 15532221 | 3 | 0.578 | 4.45E+08 | [illegible] | 1764.8463 | SMTPAQADLEFLENAK | P48193 | P48193 | EPB41 | erythrocyte membrane protein band 4.1 | Plasma Membrane | other |
| 15555225 | 2 | 0.142 | 4.32E+08 | [illegible] | 1762.8624 | FYPPDPAQLTEDITR | P48193 | P48193 | EPB41 | erythrocyte membrane protein band 4.1 | Plasma Membrane | other |
| 15531346, 15536263, 15536336, 15543448 | 2 | 0.341 | 4.74E+08 | [illegible] | 1263.5775, 1263.5791, 1263.5792, 1263.5802 | SLDGAAAAESTDR | P48193 | P48193 | EPB41 | erythrocyte membrane protein band 4.1 | Plasma Membrane | other |
| 15552185 | 2 | 0.378 | 4.22E+08 | [illegible] | 1947.8856 | NFMESVPEPRPSEWDK | P48193 | P48193 | EPB41 | erythrocyte membrane protein band 4.1 | Plasma Membrane | other |
| 15539796, 15547423 | 3 | 0.535 | 2.38E+08 | [illegible] | 2163.1619 | IVITGDADIDHDQVLVQAIK | P48193 | P48193 | EPB41 | erythrocyte membrane protein band 4.1 | Plasma Membrane | other |
| 15548442 | 2 | 0.557 | 2.49E+08 | [illegible] | 2022.937 | RGEEPAEPAEPEPTEAWK | P48193 | P48193 | EPB41 | erythrocyte membrane protein band 4.1 | Plasma Membrane | other |
| 15532834, 15541958, 15543686, 15544583 | 2 | 0.468 | 3.17E+08 | [illegible] | 1128.5836, 1128.5847, 1128.5853 | YYLC[160.0307]LQLR | P48193 | P48193 | EPB41 | erythrocyte membrane protein band 4.1 | Plasma Membrane | other |
| 15539755 | 2 | 0.087 | 2.96E+08 | [illegible] | 728.3961 | SSFFIK | P48193 | P48193 | EPB41 | erythrocyte membrane protein band 4.1 | Plasma Membrane | other |
| 15531363 | 3 | 0.113 | 3.16E+08 | [illegible] | 1057.5919 | INRFPWPK | P48193 | P48193 | EPB41 | erythrocyte membrane protein band 4.1 | Plasma Membrane | other |
| 15532413, 15532525 | 2, 3 | 0.031 | 1.57E+08 | [illegible] | 1377.7094, 1377.7097, 1377.7108 | ERLDGENIYIR | P48193 | P48193 | EPB41 | erythrocyte membrane protein band 4.1 | Plasma Membrane | other |
| 15543964, 15544459 | 2 | 0.216 | 1.23E+08 | [illegible] | 1100.5931, 1100.5936 | RLSTHSPFR | P48193 | P48193 | EPB41 | erythrocyte membrane protein band 4.1 | Plasma Membrane | other |
| 15557751 | 2 | 0.283 | 1.04E+08 | [illegible] | 1707.8894 | QASALIDRPAPHFER | P48193 | P48193 | EPB41 | erythrocyte membrane protein band 4.1 | Plasma Membrane | other |
| 15548115 | 2 | 0.019 | 1.47E+08 | [illegible] | 1819.9835 | TLNINGQVPTGDGPPLVK | P48193 | P48193 | EPB41 | erythrocyte membrane protein band 4.1 | Plasma Membrane | other |
| 15531301 | 2 | 0.633 | 6.17E+07 | [illegible] | 1205.5598 | NFMESVPEPR | P48193 | P48193 | EPB41 | erythrocyte membrane protein band 4.1 | Plasma Membrane | other |
| 15555058, 15556688, 15557805, 15557821, 15560597, 15561966, 15561980, 15562737, 15563840 | 2 | 0.374 | 8.27E+07 | [illegible] | 1780.8318, 1780.8358, 1780.8363, 1780.8374, 1780.8386, 1780.8397, 1780.841, 1780.8425 | SM[147.0354]TPAQADLEFLENAK | P48193 | P48193 | EPB41 | erythrocyte membrane protein band 4.1 | Plasma Membrane | other |
| 15535353 | 4 | 0.613 | 5.30E+07 | [illegible] | 2360.2556 | SPRPTSAPAIAQSQVTEGPGAPIK | P48193 | P48193 | EPB41 | erythrocyte membrane protein band 4.1 | Plasma Membrane | other |
| 15542887, 15544568, 15544716, 15545072, 15545165, 15545528, 15545736, 15545998, 15546294, 15546551, 15546791, 15546999, 15547441 | 2 | 0.215 | 8.57E+07 | [illegible] | 1398.6638, 1398.666, 1398.6671, 1398.6722, 1398.6727, 1398.6732, 1398.6736, 1398.675, 1398.677, 1398.6775, 1398.6781 | VVVHQETEISEE | P48193 | P48193 | EPB41 | erythrocyte membrane protein band 4.1 | Plasma Membrane | other |
| 15531944, 15541775, 15541879, 15544272, 15544538, 15544711, 15545802, 15546078 | 2 | 0.078 | 7.49E+07 | [illegible] | 1330.625, 1330.6262, 1330.6271, 1330.6272, 1330.6273, 1330.6279, 1330.6284, 1330.6301 | HSNLM[147.0354]LEDLDK | P48193 | P48193 | EPB41 | erythrocyte membrane protein band 4.1 | Plasma Membrane | other |
| 15552110, 15557077, 15558039, 15558207, 15558679, 15558719, 15558971, 15558990, 15558999, 15559164, 15559223, 15559388, 15559633, 15560635, 15560742, 15561534, 15561676, 15561973, 15562511, 15562554, 15562983, 15563458, 15563726, 15563830, 15563890, 15563894, 15563962 | 3, 6 | 0.207 | 5.24E+07 | [illegible] | 2319.2532, 2319.2593, 2319.2605, 2319.2607, 2319.2622, 2319.2625, 2319.264, 2319.2642, 2319.2646, 2319.265, 2319.2654, 2319.2656, 2319.2664, 2319.2666, 2319.267, 2319.2686, 2319.269, 2319.2725, 2319.2732, 4635.5166 | RIVITGDADIDHDQVLVQAIK | P48193 | P48193 | EPB41 | erythrocyte membrane protein band 4.1 | Plasma Membrane | other |
| 15542970 | 3 | 0.879 | 3.54E+07 | [illegible] | 2020.0635 | PTSAPAIAQSQVTEGPGAPIK | P48193 | P48193 | EPB41 | erythrocyte membrane protein band 4.1 | Plasma Membrane | other |
| 15533447 | 2 | 0.01 | 4.37E+07 | [illegible] | 1135.6335 | APIAAPEPELK | P48193 | P48193 | EPB41 | erythrocyte membrane protein band 4.1 | Plasma Membrane | other |
| 15541467, 15543444, 15544777 | 3 | 0.273 | 2.82E+07 | [illegible] | 1762.8624, 1762.8641 | FYPPDPAQLTEDITR | P48193 | P48193 | EPB41 | erythrocyte membrane protein band 4.1 | Plasma Membrane | other |
| 15543214, 15544042, 15544240, 15544553, 15544567, 15544725, 15545041, 15545059, 15545128, 15545212, 15545779, 15546588, 15547106 | 4 | 0.122 | 5.92E+07 | [illegible] | 2319.254, 2319.257, 2319.2598, 2319.2605, 2319.2607, 2319.2627, 2319.263, 2319.2637, 2319.2678, 2319.2708, 2319.2737 | RIVITGDADIDHDQVLVQAIK | P48193 | P48193 | EPB41 | erythrocyte membrane protein band 4.1 | Plasma Membrane | other |
| 15531409, 15542654, 15545238, 15546125, 15546884, 15546950, 15547290, 15547324 | 2, 3 | 0.213 | 3.87E+07 | [illegible] | 1178.5819, 1178.5842, 1178.5846, 1178.5848, 1178.5851, 1178.5854, 1178.5864, 1178.587 | LSM[147.0354]YGVDLHK | P48193 | P48193 | EPB41 | erythrocyte membrane protein band 4.1 | Plasma Membrane | other |
| 15549512 | 2 | 0.553 | 1.94E+07 | [illegible] | 2020.0663 | PTSAPAIAQSQVTEGPGAPIK | P48193 | P48193 | EPB41 | erythrocyte membrane protein band 4.1 | Plasma Membrane | other |
| 15553426 | 2 | 0.436 | 1.89E+07 | [illegible] | 2163.1614 | IVITGDADIDHDQVLVQAIK | P48193 | P48193 | EPB41 | erythrocyte membrane protein band 4.1 | Plasma Membrane | other |
| 15548469 | 2 | 0.058 | 2.63E+07 | [illegible] | 1523.6503 | EGEGIEEC[160.0307]SGTEVK | P48193 | P48193 | EPB41 | erythrocyte membrane protein band 4.1 | Plasma Membrane | other |
| 15536978 | 3 | 0.239 | 2.26E+07 | [illegible] | 1668.7894 | SLAAEAENSQHQQQK | P48193 | P48193 | EPB41 | erythrocyte membrane protein band 4.1 | Plasma Membrane | other |
| 15534983 | 3 | 0.721 | 2.99E+07 | [illegible] | 2148.1602 | PTSAPAIAQSQVTEGPGAPIKK | P48193 | P48193 | EPB41 | erythrocyte membrane protein band 4.1 | Plasma Membrane | other |
| 15531734 | 3 | 0.313 | 1.59E+07 | [illegible] | 1603.7644 | SLDGAAAAESTDRSPR | P48193 | P48193 | EPB41 | erythrocyte membrane protein band 4.1 | Plasma Membrane | other |
| 15544639, 15545677 | 3 | 0.332 | 1.04E+07 | [illegible] | 1780.8395, 1780.8397 | SM[147.0354]TPAQADLEFLENAK | P48193 | P48193 | EPB41 | erythrocyte membrane protein band 4.1 | Plasma Membrane | other |
| 15558557 | 2 | 0.537 | 2.64E+07 | [illegible] | 1890.9313 | IRPGEQEHYESTIGFK | P48193 | P48193 | EPB41 | erythrocyte membrane protein band 4.1 | Plasma Membrane | other |

| | | | | | | | | | | | | | | | | | | | | |
|---|---|---|---|---|---|---|---|---|---|---|---|---|---|---|---|---|---|---|---|---|
| 15548935 | 2 | 0.231 | 2.03E+07 | 1.25E+07 | 1.28E+07 | 9924210 | 1.10E+07 | 1.03E+07 | 1.33E+07 | 9765265 | 8047228 | 1.34E+07 | 1.15E+07 | 7879250 | 1963.8805 | NFM[147.0354]ESVPEPRPSEWDK | P48193 | P48193 | EPB41 | erythrocyte membrane protein band 4.1 | Plasma Membrane | other |
| 15532574 | 3 | 0.541 | 6.81E+07 | 7.46E+07 | 9.16E+07 | 5.03E+07 | 7.26E+07 | 6.52E+07 | 6.97E+07 | 8.57E+07 | 8.76E+07 | 1.24E+07 | 7.70E+07 | 8.55E+07 | 1768.8646 | VSLLDDTVYEC[160.0307]VVEK | P48193 | P48193 | EPB41 | erythrocyte membrane protein band 4.1 | Plasma Membrane | other |
| 15533463 | 3 | 0.342 | 4.88E+07 | 2589064 | 758817 | 2.52E+07 | 898600 | 986245.625 | 2055439.75 | 1776146.5 | 1.41E+07 | 1.21E+07 | 7.51E+07 | 2.65E+07 | 1290.6852 | KLSMYGVDLHK | P48193 | P48193 | EPB41 | erythrocyte membrane protein band 4.1 | Plasma Membrane | other |
| 15535884 | 2 | 0.25 | 1.01E+07 | 9581549 | 1.25E+07 | 9360744 | 6556566 | 9118898 | 9087729 | 8664830 | 1.12E+07 | 9602065 | 8930680 | 9763094 | 807.4957 | LLSSFLK | P48193 | P48193 | EPB41 | erythrocyte membrane protein band 4.1 | Plasma Membrane | other |
| 15543688, 15544336, 15546904 | 4 | 0.487 | 7026474 | 5742351 | 5853290.5 | 5317280 | 5457115 | 5610539 | 4267767.5 | 5695468 | 3125826.75 | 8458447 | 3233043 | 5025093.5 | 2966.4255, 2966.427, 2966.4302 | RVC[160.0307]EHLNLLEEDYFGLALWDSATSK | P48193 | P48193 | EPB41 | erythrocyte membrane protein band 4.1 | Plasma Membrane | other |
| 15549702 | 2 | 0.169 | 7681305 | 6752228 | 7011778 | 6251769 | 4880038 | 3895424 | 5574013.5 | 5788949 | 5674285.5 | 6805896 | 6471980.5 | 4203316.5 | 1721.7787 | SQVSEEEGREVESEK | P48193 | P48193 | EPB41 | erythrocyte membrane protein band 4.1 | Plasma Membrane | other |
| 15552951 | 2 | 0.158 | 3927839 | 4281467 | 5423028.5 | 2495758.5 | 3573380 | 2561711.25 | 4938247 | 3733253.5 | 3115462.75 | 4471718.5 | 2196947.5 | 2553063.25 | 1603.7635 | SLDGAAAAESTDRSPR | P48193 | P48193 | EPB41 | erythrocyte membrane protein band 4.1 | Plasma Membrane | other |
| 15557135, 15561263 | 2 | 0.057 | 4314557 | 5061778.5 | 5774764.5 | 2993665.5 | 3428167.5 | 2825705 | 2856644 | 4268831 | 5714109 | 3483710.25 | 2741377.5 | 3319025 | 1700.8888, 1700.8923 | EIELGNSLDEDIILK | P48193 | P48193 | EPB41 | erythrocyte membrane protein band 4.1 | Plasma Membrane | other |
| 15558089 | 2 | 0.049 | 4559542 | 4757231 | 7744559 | 2886766.25 | 4303108.5 | 3117911.25 | 4373928.5 | 4741010.5 | 5039716.5 | 3357036.5 | 4204508 | 3296679 | 1668.7905 | SLAAEAENSQHQQQK | P48193 | P48193 | EPB41 | erythrocyte membrane protein band 4.1 | Plasma Membrane | other |
| 15553189 | 2 | 0.445 | 1.11E+07 | 1081147.5 | 83654.383 | 5623236.5 | 277727.562 | 314186.281 | 1194701.625 | 340311.812 | 1892888.125 | 3033120 | 1.30E+07 | 4398865.5 | 1892.9352 | SMTPAQADLEFLENAKK | P48193 | P48193 | EPB41 | erythrocyte membrane protein band 4.1 | Plasma Membrane | other |
| 15560917, 15562863, 15563317, 15563737, 15563753, 15563766, 15563780 | 2 | 0.175 | 1145389.375 | 1929271.625 | 1775774.5 | 925819.812 | 1453049.5 | 1259618.25 | 1943912.25 | 1740095.25 | 1719547.875 | 1694013.25 | 1257090.125 | 1075004.125 | 2150.1323, 2150.1353, 2150.1375, 2150.1387, 2150.139 | DLEGVDIILGVC[160.0307]SSGLLVYK | P48193 | P48193 | EPB41 | erythrocyte membrane protein band 4.1 | Plasma Membrane | other |
| 15547649, 15547650 | 3 | 0.066 | 29129.771 | 114095.102 | 118112.82 | 83717.43 | 262414.406 | 108034.273 | 427306.312 | 375895.969 | 205579.156 | 236730.203 | 118700.266 | 228880.469 | 2150.1382 | DLEGVDIILGVC[160.0307]SSGLLVYK | P48193 | P48193 | EPB41 | erythrocyte membrane protein band 4.1 | Plasma Membrane | other |
| 15543057 | 2, 3 | 0.931 | 3276926.5 | 2714103.25 | 4065521.5 | 564741.562 | 449439.281 | 9064386 | 651020.5 | 1719151.625 | 1.85E+07 | 96426.969 | 1.52E+07 | 1.70E+07 | 1089.5869, 1089.588 | GGISETRIEK | P48193 | P48193 | EPB41 | erythrocyte membrane protein band 4.1 | Plasma Membrane | other |
| 15546997 | 2 | 0.152 | 671114.812 | 240695.5 | 356141.344 | 4157.404 | 482019.812 | 130002.398 | 420015.531 | 669377.688 | 608365.438 | 4.70E-06 | 197335.922 | 50336.359 | 1021.5402 | HHASISELK | P48193 | P48193 | EPB41 | erythrocyte membrane protein band 4.1 | Plasma Membrane | other |
| 15548500, 15550597 | 2 | 0.51 | 1.77E+09 | 1.93E+09 | 2.22E+09 | 1.77E+09 | 1.98E+09 | 1.77E+09 | 1.80E+09 | 1.73E+09 | 1.78E+09 | 2.38E+09 | 1.81E+09 | 1.81E+09 | 2017.0912, 2017.0941 | SVLPISQTQAAQEGALLYK | P49222 | P49222 | EPB42 | erythrocyte membrane protein band 4.2 | Plasma Membrane | transporter |
| 15531316, 15547636 | 2 | 0.315 | 1.68E+09 | 1.67E+09 | 1.73E+09 | 1.81E+09 | 1.69E+09 | 1.56E+09 | 1.73E+09 | 1.65E+09 | 1.44E+09 | 2.15E+09 | 1.79E+09 | 1.66E+09 | 1341.7286, 1341.7352 | VALIAQTGEQPSK | P49222 | P49222 | EPB42 | erythrocyte membrane protein band 4.2 | Plasma Membrane | transporter |
| 15531265 | 3 | 0.188 | 1.63E+09 | 1.42E+09 | 1.41E+09 | 1.90E+09 | 1.36E+09 | 1.40E+09 | 1.49E+09 | 1.43E+09 | 1.40E+09 | 2.14E+09 | 1.77E+09 | 1.58E+09 | 1783.8281 | VVGSDRC[160.0307]EDITQNYK | P49222 | P49222 | EPB42 | erythrocyte membrane protein band 4.2 | Plasma Membrane | transporter |
| 15548242 | 2 | 0.826 | 1.54E+09 | 1.71E+09 | 1.98E+09 | 1.71E+09 | 1.78E+09 | 1.57E+09 | 1.63E+09 | 1.60E+09 | 1.66E+09 | 1.98E+09 | 1.61E+09 | 1.63E+09 | 1504.7987 | TQAIFPISSLGDQK | P49222 | P49222 | EPB42 | erythrocyte membrane protein band 4.2 | Plasma Membrane | transporter |
| 15547653 | 3 | 0.897 | 1.19E+09 | 1.69E+09 | 1.63E+09 | 1.29E+09 | 1.39E+09 | 1.28E+09 | 1.46E+09 | 1.59E+09 | 1.04E+09 | 1.97E+09 | 1.10E+09 | 1.34E+09 | 3381.6619 | QYPLGQFTLLFNPWNRDDAVFLQNEAER | P49222 | P49222 | EPB42 | erythrocyte membrane protein band 4.2 | Plasma Membrane | transporter |
| 15551489 | 2, 3 | 0.046 | 1.44E+09 | 1.49E+09 | 1.26E+09 | 1.31E+09 | 1.24E+09 | 1.25E+09 | 1.30E+09 | 1.32E+09 | 1.26E+09 | 1.71E+09 | 1.53E+09 | 1.37E+09 | 2474.1616, 2474.1626 | LSTNLSFSC[160.0307]FEQTPPENSFLR | P49222 | P49222 | EPB42 | erythrocyte membrane protein band 4.2 | Plasma Membrane | transporter |
| 15547765, 15548679, 15556262, 15557968 | 3, 4 | 0.049 | 1.25E+09 | 1.37E+09 | 9.72E+08 | 1.28E+09 | 1.08E+09 | 1.16E+09 | 1.32E+09 | 1.23E+09 | 1.26E+09 | 1.66E+09 | 1.42E+09 | 1.53E+09 | 3080.4797, 3080.4844, 3080.4893, 3080.4907 | YSHTSLSC[160.0307]FAQENMAIGKPDLIIEMPK | P49222 | P49222 | EPB42 | erythrocyte membrane protein band 4.2 | Plasma Membrane | transporter |
| 15541018 | 2 | 0.531 | 9.45E+08 | 9.19E+08 | 1.02E+09 | 1.00E+09 | 9.35E+08 | 9.60E+08 | 8.93E+08 | 9.04E+08 | 8.29E+08 | 1.20E+09 | 9.18E+08 | 8.47E+08 | 992.5031 | YPAGSLQEK | P49222 | P49222 | EPB42 | erythrocyte membrane protein band 4.2 | Plasma Membrane | transporter |
| 15548439, 15563217 | 3 | 0.844 | 8.05E+08 | 9.64E+08 | 1.04E+09 | 8.59E+08 | 9.29E+08 | 1.03E+09 | 8.32E+08 | 1.18E+09 | 9.52E+08 | 1.19E+09 | 9.65E+08 | 9.13E+08 | 2742.379, 2742.3816 | EGELQLDPAVPELFAAVNASC[160.0307]VVWK | P49222 | P49222 | EPB42 | erythrocyte membrane protein band 4.2 | Plasma Membrane | transporter |
| 15548164, 15548865 | 3, 4 | 0.505 | 9.94E+08 | 7.42E+08 | 7.43E+08 | 8.38E+08 | 1.06E+09 | 6.21E+08 | 8.40E+08 | 9.66E+08 | 9.07E+08 | 1.14E+09 | 1.07E+09 | 8.29E+08 | 3028.5352, 3028.543 | YGLGSLWPGSSLHTQFQFTPTHLGLQR | P49222 | P49222 | EPB42 | erythrocyte membrane protein band 4.2 | Plasma Membrane | transporter |
| 15536027, 15546702 | 3 | 0.265 | 3.76E+08 | 4.98E+08 | 6.21E+08 | 3.38E+08 | 4.83E+08 | 5.24E+08 | 4.42E+08 | 6.39E+08 | 3.92E+08 | 1.09E+09 | 4.61E+08 | 7.54E+08 | 2017.093, 2017.0935 | SVLPISQTQAAQEGALLYK | P49222 | P49222 | EPB42 | erythrocyte membrane protein band 4.2 | Plasma Membrane | transporter |
| 15552331 | 3, 4 | 0.07 | 9.65E+08 | 1.09E+09 | 1.17E+09 | 8.30E+08 | 9.80E+08 | 7.95E+08 | 8.87E+08 | 1.03E+09 | 8.85E+08 | 9.56E+08 | 7.12E+08 | 7.13E+08 | 3575.6729, 3575.6738 | VVTTFDSAQGTVGSLLVDEYYNEEGLQNGEGQR | P49222 | P49222 | EPB42 | erythrocyte membrane protein band 4.2 | Plasma Membrane | transporter |
| 15534421, 15536897 | 2, 3 | 0.218 | 7.10E+08 | 5.98E+08 | 7.10E+08 | 7.05E+08 | 5.91E+08 | 6.85E+08 | 7.20E+08 | 6.80E+08 | 5.84E+08 | 9.18E+08 | 7.92E+08 | 6.83E+08 | 1193.5786, 1193.5798 | DWNQPAHVAR | P49222 | P49222 | EPB42 | erythrocyte membrane protein band 4.2 | Plasma Membrane | transporter |
| 15531261 | 3 | 0.009 | 6.96E+08 | 6.05E+08 | 5.00E+08 | 7.46E+08 | 6.62E+08 | 6.05E+08 | 6.39E+08 | 6.87E+08 | 6.24E+08 | 9.18E+08 | 1.29E+09 | 8.79E+08 | 2218.054 | MHNSLEAPMQNC[160.0307]IISIFGR | P49222 | P49222 | EPB42 | erythrocyte membrane protein band 4.2 | Plasma Membrane | transporter |
| 15539297 | 2 | 0.096 | 5.80E+08 | 5.30E+08 | 6.06E+08 | 5.70E+08 | 5.51E+08 | 5.16E+08 | 5.31E+08 | 5.64E+08 | 4.92E+08 | 7.65E+08 | 6.12E+08 | 5.83E+08 | 945.4889 | QWLTGQGR | P49222 | P49222 | EPB42 | erythrocyte membrane protein band 4.2 | Plasma Membrane | transporter |
| 15540174, 15545533 | 2 | 0.09 | 5.71E+08 | 4.65E+08 | 4.97E+08 | 5.05E+08 | 4.51E+08 | 3.97E+08 | 4.70E+08 | 5.20E+08 | 4.91E+08 | 6.69E+08 | 5.87E+08 | 5.09E+08 | 1020.6543, 1020.65497 | VVGALLHALK | P49222 | P49222 | EPB42 | erythrocyte membrane protein band 4.2 | Plasma Membrane | transporter |

[illegible]

| | | | | | | | | | | | | | | | | | | | | | |
|---|---|---|---|---|---|---|---|---|---|---|---|---|---|---|---|---|---|---|---|---|---|
| 15549703 | 2 | 0.16 | 2944920.75 | 2120253.25 | 2706325 | 3536700 | 3005020.25 | 2761874.25 | 3241586.25 | 1928099.875 | 1640114.625 | 6254880.5 | 2950891.5 | 3234399.75 | 1520.8647 | QILENELQIPVPK | P54731 | P54731 | FAF1 | Fas (TNFRSF6) associated factor 1 | Nucleus | other |
| 15537423 | 2 | 0.301 | 2605160.25 | 1867886.875 | 2340934.5 | 2886942.25 | 2573590.5 | 2367038.25 | 3114123 | 1971746.25 | 1967640 | 5925961.5 | 2370935 | 2994797.25 | 1163.5764 | TGDVEDSTVLK | P54731 | P54731 | FAF1 | Fas (TNFRSF6) associated factor 1 | Nucleus | other |
| 15553471 | 2 | 0.191 | 2331505.75 | 2089662.25 | 2305986 | 2546588.25 | 2235147 | 2614592.5 | 3353566.25 | 2369058 | 2423303.25 | 5279918.5 | 2501992.25 | 3041926.75 | 1522.8314 | TDQFPLFLIIMGK | P54731 | P54731 | FAF1 | Fas (TNFRSF6) associated factor 1 | Nucleus | other |
| 15536182 | 2 | 0.82 | 1967182.375 | 4879025 | 1670588.5 | 2360784.25 | 2825126.25 | 2667014 | 1541488.875 | 2859317.75 | 3181116.5 | 2015540.75 | 1645676.75 | 2442168 | 1332.7753 | STVLSEPVLFLK | Q8R0F8 | Q8R0F8 | FAHD1 | fumarylacetoacetate hydrolase domain containing 1 | Cytoplasm | enzyme |
| 15550287 | 2 | 0.595 | 1807420.875 | 4803564 | 1836210.75 | 1834203.375 | 2753119 | 3368764.5 | 1384807 | 3449344 | 4368459 | 1505973.125 | 1631857.125 | 2078752.625 | 1712.9636 | IITLEEGDLILTGTPK | Q8R0F8 | Q8R0F8 | FAHD1 | fumarylacetoacetate hydrolase domain containing 1 | Cytoplasm | enzyme |
| 15540221 | 2 | 0.552 | 1995553.375 | 2658611.25 | 2237656.25 | 2645562.75 | 2973780.25 | 2615209 | 2662777.5 | 2579938.25 | 2164750.75 | 3005661.25 | 2376823.25 | 1793013.75 | 1376.6145 | QALDTEEEGAADK | Q6NS82 | Q6NS82 | FAM134A | family with sequence similarity 134, member A | Extracellular Space | other |
| 15535660 | 3 | 0.091 | 2717135.5 | 5090780 | 1970766.125 | 3357324.75 | 2626754.5 | 3134622.5 | 2408904 | 3407964.75 | 3299209.75 | 1771695.375 | 1890572.875 | 1576114.125 | 1883.9045 | QEPNRTPEPGGSVTGTEK | Q9D8B6 | Q9D8B6 | FAM210B | family with sequence similarity 210, member B | Extracellular Space | other |
| 15547030 | 2 | 0.184 | 243894.375 | 772773.562 | 273935.375 | 145395.984 | 501119.719 | 327647.531 | 205797.125 | 528804.75 | 323369.438 | 380025.344 | 43935.184 | 33399.945 | 1259.6084 | TPEPGGSVTGTEK | Q9D8B6 | Q9D8B6 | FAM210B | family with sequence similarity 210, member B | Extracellular Space | other |
| 15531370 | 2 | 0.529 | 1.42E+08 | 1.74E+08 | 1.93E+08 | 1.38E+08 | 2.25E+08 | 1.50E+08 | 1.57E+08 | 1.96E+08 | 1.36E+08 | 1.58E+08 | 9.81E+07 | 1.49E+08 | 1415.8123 | LDELGVPLYAVVK | Q9CYH2 | Q9CYH2 | FAM213A | family with sequence similarity 213, member A | Extracellular Space | other |
| 15536425 | 2 | 0.278 | 9.79E+07 | 1.27E+08 | 1.13E+08 | 9.02E+07 | 1.85E+08 | 1.34E+08 | 1.27E+08 | 1.31E+08 | 1.03E+08 | 9.58E+07 | 7.40E+07 | 1.04E+08 | 1392.7234 | AALEYLEDIDLK | Q9CYH2 | Q9CYH2 | FAM213A | family with sequence similarity 213, member A | Extracellular Space | other |
| 15532929 | 2 | 0.402 | 6.02E+07 | 1.05E+08 | 1.19E+08 | 5.69E+07 | 1.78E+08 | 1.10E+08 | 9.19E+07 | 1.19E+08 | 7.39E+07 | 8.83E+07 | 3.35E+07 | 6.31E+07 | 1168.6915 | VNPLSVLEAVK | Q9CYH2 | Q9CYH2 | FAM213A | family with sequence similarity 213, member A | Extracellular Space | other |
| 15536467 | 2, 3 | 0.343 | 4.95E+07 | 1.10E+08 | 1.34E+08 | 6.52E+07 | 1.46E+08 | 1.13E+08 | 1.12E+08 | 1.18E+08 | 7.06E+07 | 7.21E+07 | 2.97E+07 | 7.45E+07 | 1457.7025, 1457.7029 | REVEDFQPYFK | Q9CYH2 | Q9CYH2 | FAM213A | family with sequence similarity 213, member A | Extracellular Space | other |
| 15531384 | 3 | 0.932 | 6.79E+07 | 5.04E+07 | 4.72E+07 | 8.36E+07 | 3.23E+07 | 6.13E+07 | 6.90E+07 | 4.26E+07 | 5.09E+07 | 5.87E+07 | 5.11E+07 | 7.84E+07 | 1772.9508 | EFGDRVNPLSVLEAVK | Q9CYH2 | Q9CYH2 | FAM213A | family with sequence similarity 213, member A | Extracellular Space | other |
| 15540627 | 2 | 0.623 | 3.69E+07 | 4.27E+07 | 5.70E+07 | 3.93E+07 | 6.63E+07 | 4.89E+07 | 4.70E+07 | 5.18E+07 | 4.02E+07 | 5.04E+07 | 2.83E+07 | 4.14E+07 | 950.48004 | GEIFLDEK | Q9CYH2 | Q9CYH2 | FAM213A | family with sequence similarity 213, member A | Extracellular Space | other |
| 15538517, 15538698 | 2, 3 | 0.495 | 2.54E+07 | 1.30E+07 | 5820431 | 2.36E+07 | 1.69E+07 | 1.15E+07 | 1.63E+07 | 1.62E+07 | 1.88E+07 | 2.41E+07 | 2.09E+07 | 1.96E+07 | 1273.678, 1273.6797 | AEAADLMSLKPK | Q9CYH2 | Q9CYH2 | FAM213A | family with sequence similarity 213, member A | Extracellular Space | other |
| 15533435 | 2 | 0.548 | 1.05E+07 | 1.97E+07 | 3.00E+07 | 1.08E+07 | 3.74E+07 | 2.50E+07 | 2.43E+07 | 2.69E+07 | 1.36E+07 | 1.97E+07 | 7997017.5 | 1.21E+07 | 1048.5323 | AEAADLMSLK | Q9CYH2 | Q9CYH2 | FAM213A | family with sequence similarity 213, member A | Extracellular Space | other |
| 15539068 | 2 | 0.836 | 1.22E+07 | 1.44E+07 | 1.59E+07 | 9455955 | 1.86E+07 | 1.29E+07 | 1.48E+07 | 1.55E+07 | 1.02E+07 | 1.56E+07 | 8922594 | 1.15E+07 | 965.5517 | QGILLEHR | Q9CYH2 | Q9CYH2 | FAM213A | family with sequence similarity 213, member A | Extracellular Space | other |
| 15551974 | 2 | 0.978 | 1.53E+07 | 1.16E+07 | 1.23E+07 | 1.87E+07 | 7042330 | 1.36E+07 | 1.47E+07 | 8519458 | 1.44E+07 | 1.40E+07 | 1.11E+07 | 1.50E+07 | 1772.9498 | EFGDRVNPLSVLEAVK | Q9CYH2 | Q9CYH2 | FAM213A | family with sequence similarity 213, member A | Extracellular Space | other |
| 15536622 | 2 | 0.595 | 9362526 | 9338984 | 9280164 | 9847650 | 1.24E+07 | 9066777 | 1.09E+07 | 9668822 | 7768984 | 1.03E+07 | 7208785.5 | 8933991 | 930.5172 | NGAVIMAVR | Q9CYH2 | Q9CYH2 | FAM213A | family with sequence similarity 213, member A | Extracellular Space | other |
| 15537452 | 2 | 0.819 | 1060227.875 | 1639488.25 | 1868466.875 | 792973.938 | 2885447.5 | 1060182.25 | 1597062.125 | 1713113.75 | 1090729.625 | 1264065.625 | 640439.562 | 1537472.75 | 998.49664 | MMFMGLIR | Q9CYH2 | Q9CYH2 | FAM213A | family with sequence similarity 213, member A | Extracellular Space | other |
| 15537295 | 2 | 0.137 | 1342911.5 | 1625817.25 | 644209.188 | 868776.75 | 1262312.125 | 950108.938 | 442159.688 | 770174.375 | 620420.125 | 869051.562 | 748386.438 | 461463.438 | 1148.6539 | LFEISDVVVK | Q9WUA2 | Q9WUA2 | FARSB | phenylalanyl-tRNA synthetase, beta subunit | Cytoplasm | enzyme |
| 15549409 | 3 | 0.766 | 9204763 | 7836952 | 6644344.5 | 7655979.5 | 7653737.5 | 7637121 | 8857227 | 7116551 | 6613886.5 | 1.49E+07 | 7190521 | 6957124.5 | 2449.324 | TLLEGSGLESIINIIHSSLAEPR | P19096 | P19096 | FASN | fatty acid synthase | Cytoplasm | enzyme |
| 15552252 | 3 | 0.435 | 1.31E+07 | 8577529 | 7205863.5 | 1.03E+07 | 6094851 | 7565622 | 9197122 | 4690101.5 | 7458480.5 | 1.39E+07 | 8231270.5 | 9028225 | 2873.5933 | VLVQVREEEPEAVLPGAQPTLISAISK | P19096 | P19096 | FASN | fatty acid synthase | Cytoplasm | enzyme |
| 15551607 | 3 | 0.891 | 1.04E+07 | 6023910.5 | 9706482 | 6793245.5 | 7354767 | 8217931.5 | 8896551 | 6719253 | 6781857 | 1.22E+07 | 6769267 | 6369159 | 2499.346 | VHLTGINVNPNALFPPVEFPAPR | P19096 | P19096 | FASN | fatty acid synthase | Cytoplasm | enzyme |
| 15552997 | 3 | 0.329 | 6041744 | 4711749.5 | 3507398.25 | 4216690.5 | 3528668 | 4224384 | 4989352.5 | 3883277.75 | 3155416.5 | 9891502 | 4889242 | 4365769 | 3455.7156 | LSVPTYGLQC[160.0307]TQAAPLDSIPNLAAYYIDC[160.0307]IK | P19096 | P19096 | FASN | fatty acid synthase | Cytoplasm | enzyme |
| 15549361 | 3 | 0.299 | 6039965.5 | 5755165 | 5778460.5 | 4205126 | 4449465.5 | 6250015.5 | 6500380.5 | 5020645.5 | 4491926 | 9207947 | 5632258 | 5856553.5 | 2832.3735 | QHSQDLAFVSMLNDIAATPTAAMPFR | P19096 | P19096 | FASN | fatty acid synthase | Cytoplasm | enzyme |
| 15549887 | 3 | 0.443 | 6848036 | 5357500.5 | 4694360 | 5233571.5 | 5275224.5 | 3627180.5 | 6238394.5 | 3775429.5 | 4257229.5 | 8422895 | 4388135.5 | 6665619 | 2325.0818 | FVFTPHMEAEC[160.0307]LSESTALQK | P19096 | P19096 | FASN | fatty acid synthase | Cytoplasm | enzyme |
| 15532765 | 2 | 0.355 | 7049630.5 | 5423134.5 | 4527358.5 | 5010391 | 5138775 | 5490646 | 7168272.5 | 4823787 | 2988352.75 | 7868072.5 | 6396021.5 | 6549294.5 | 1423.7738 | LDPGSPELQQVLK | P19096 | P19096 | FASN | fatty acid synthase | Cytoplasm | enzyme |
| 15551278 | 2 | 0.115 | 6720673 | 4949390.5 | 6378442.5 | 5231616.5 | 3742458 | 4746687.5 | 5046054 | 4067941.75 | 4236213 | 7495178 | 5066473.5 | 5338608 | 2034.1798 | LLLPEDPLISGLLNSQALK | P19096 | P19096 | FASN | fatty acid synthase | Cytoplasm | enzyme |
| 15551457 | 3 | 0.376 | 5791067.5 | 4322371 | 3763025 | 5222780.5 | 2849228.5 | 4129589.75 | 3996871.75 | 3759740.25 | 2341941 | 6571211.5 | 4328024 | 3945185.75 | 2541.2893 | GGNVGINSFGFGGSNVHVILQPNTR | P19096 | P19096 | FASN | fatty acid synthase | Cytoplasm | enzyme |
| 15550725 | 2 | 0.23 | 4261728 | 2570176 | 2348645.5 | 2812386.5 | 2273708 | 1984131.75 | 2583530.25 | 1401632.25 | 1981094.25 | 5189619 | 2704754.5 | 2526156.25 | 1959.9884 | WLSTSIPEAQWQSSLAR | P19096 | P19096 | FASN | fatty acid synthase | Cytoplasm | enzyme |
| 15535029 | 2 | 0.454 | 3912254 | 3170381.75 | 3262587.5 | 3191225.25 | 2771212.5 | 2815074 | 3897541.75 | 2362324.5 | 2349639 | 4757691.5 | 2982323.75 | 3102606.75 | 1410.7241 | DNLEFFLTNLGK | P19096 | P19096 | FASN | fatty acid synthase | Cytoplasm | enzyme |
| 15539262 | 3 | 0.322 | 3970267.25 | 1846254.75 | 2739565.25 | 3011871.5 | 1743911 | 2478897.5 | 2655156.25 | 1802514.75 | 1599935.875 | 4698077.5 | 2837628.5 | 2576265.5 | 1356.7246 | VTAIYIDPATHR | P19096 | P19096 | FASN | fatty acid synthase | Cytoplasm | enzyme |
| 15556115 | 4 | 0.108 | 3961665.5 | 3884925.75 | 3539852.75 | 3422886.75 | 3286853 | 2689876.75 | 3678829 | 2716361.5 | 2128977.75 | 4600893.5 | 3337681.25 | 4094573 | 3355.7595 | VLLSLEHGVWAPNLHFHNPNPEIPALLDGR | P19096 | P19096 | FASN | fatty acid synthase | Cytoplasm | enzyme |
| 15542447 | 3 | 0.626 | 4924423.5 | 3211060.5 | 2852465.75 | 3623405 | 2382800 | 3271339.75 | 3702393.25 | 2966449 | 2100772.25 | 4578983.5 | 3251764 | 3084139.75 | 1680.8331 | SNMGHPEPASGLAALTK | P19096 | P19096 | FASN | fatty acid synthase | Cytoplasm | enzyme |
| 15554140 | 3 | 0.848 | 3903323.25 | 2473005 | 1955282.25 | 2811795.75 | 2870720 | 1733331.375 | 2498139.25 | 2298366 | 2023606.25 | 4253708.5 | 1970774 | 2509813 | 3638.7668 | GHALGETLAC[160.0307]LPSEVQPAPSLLSQEEWESLFSR | P19096 | P19096 | FASN | fatty acid synthase | Cytoplasm | enzyme |
| 15537426 | 2 | 0.179 | 2966607.75 | 2214953.5 | 2249760.75 | 1981257.5 | 2122621.5 | 1854110.125 | 2166402.25 | 1844624 | 1326576.375 | 3731947.5 | 2268921.5 | 2059339.5 | 1259.6561 | VQEVQQVSTNK | P19096 | P19096 | FASN | fatty acid synthase | Cytoplasm | enzyme |
| 15539146 | 2 | 0.373 | 2999186.25 | 2985897.5 | 3052295.25 | 2536140.25 | 2164701.25 | 2396225 | 3091586 | 2084775.875 | 1773009.25 | 3605159 | 1998332.375 | 2733472.25 | 1471.7717 | TLEAVQDLLEQGR | P19096 | P19096 | FASN | fatty acid synthase | Cytoplasm | enzyme |
| 15542903 | 2 | 0.304 | 2107179.5 | 1844459.25 | 1474283.125 | 1625369.125 | 1347710 | 1259265.625 | 1283106.25 | 1189951.5 | 291197.938 | 2450622.25 | 768971.75 | 1236115.5 | 1146.5703 | HDLVMNVYR | P19096 | P19096 | FASN | fatty acid synthase | Cytoplasm | enzyme |
| 15559235 | 3 | 0.805 | 1087072.625 | 830920.75 | 479066.531 | 996300.938 | 893416 | 314591.156 | 965870.562 | 567442.562 | 160882.859 | 1758770.125 | 395716.625 | 517607.844 | 2447.1948 | NVTFHGILLDALFEEANDSWR | P19096 | P19096 | FASN | fatty acid synthase | Cytoplasm | enzyme |
| 15537198 | 3 | 0.15 | 2534761.5 | 1847807.375 | 2251613.5 | 2450187 | 1538968.375 | 1417630.25 | 1916905.75 | 1177060.875 | 999884.25 | 1679448.375 | 1633373 | 1729580.875 | 2034.1809 | LLLPEDPLISGLLNSQALK | P19096 | P19096 | FASN | fatty acid synthase | Cytoplasm | enzyme |
| 15537858 | 2 | 0.077 | 2019423.625 | 1579548.625 | 2351399.75 | 1078480.25 | 1510853.75 | 1821528.5 | 999126.938 | 1410060.875 | 1125571.875 | 1513286.375 | 1031671 | 1369989.25 | 1386.744 | GVDLVLNSLAEEK | P19096 | P19096 | FASN | fatty acid synthase | Cytoplasm | enzyme |
| 15545973 | 2 | 0.321 | 648038.188 | 352083.188 | 351331.438 | 244464.094 | 347999.219 | 357231.531 | 490509.062 | 321549.688 | 304296.281 | 890550.812 | 401831.719 | 408131.906 | 1250.4713 | SFDDSGSGYC[160.0307]R | P19096 | P19096 | FASN | fatty acid synthase | Cytoplasm | enzyme |
| 15546175 | 2 | 0.177 | 749950.812 | 662367.312 | 925487 | 372400.344 | 748165.375 | 691649.375 | 588888.812 | 478640.688 | 405284.719 | 678784.188 | 597199.812 | 868261.312 | 937.4968 | LTQGEVYK | P19096 | P19096 | FASN | fatty acid synthase | Cytoplasm | enzyme |
| 15539482 | 2 | 0.064 | 5521088.5 | 9869738 | 3759157 | 5375533.5 | 6835382 | 6186480.5 | 3194959.25 | 5389002.5 | 6301449 | 3356014.25 | 2111934.5 | 3019254.25 | 1107.6173 | FVNVVPTFGK | P62862 | P62862 | FAU | Finkel-Biskis-Reilly murine sarcoma virus (FBR-MuSV) ubiquitously expressed | Cytoplasm | other |
| 15556271 | 2 | 0.786 | 2879511.75 | 1984317.5 | 1761018.875 | 1991970.5 | 1462085.375 | 2447470 | 2102671.5 | 1541185.875 | 1866399.75 | 2774863.75 | 1165441.25 | 1493204.25 | 2145.078 | SVQLQPGSYVAAGVEPGESAAK | Q3U7U3 | Q3U7U3 | FBXO7 | F-box protein 7 | Cytoplasm | enzyme |
| 15547766 | 2 | 0.102 | 6.45E+07 | 1.22E+08 | 5.37E+07 | 5.06E+07 | 5.57E+07 | 7.63E+07 | 3.87E+07 | 7.00E+07 | 9.96E+07 | 3.34E+07 | 3.57E+07 | 3.58E+07 | 1706.9088 | VGPVPWLGPQTDEAIK | P22315 | P22315 | FECH | ferrochelatase | Cytoplasm | enzyme |
| 15533108 | 2, 3 | 0.085 | 4.24E+07 | 7.80E+07 | 3.43E+07 | 3.43E+07 | 3.91E+07 | 4.49E+07 | 2.50E+07 | 5.33E+07 | 5.02E+07 | 2.49E+07 | 2.09E+07 | 2.36E+07 | 1391.7496, 1391.7501 | LLDELSPATAPHK | P22315 | P22315 | FECH | ferrochelatase | Cytoplasm | enzyme |
| 15535638 | 3 | 0.51 | 3.38E+07 | 5.78E+07 | 1.62E+07 | 2.54E+07 | 2.02E+07 | 3.69E+07 | 1.65E+07 | 3.32E+07 | 3.57E+07 | 1.64E+07 | 1.60E+07 | 2.42E+07 | 1816.9966 | LFLDRDLMTLPIQNK | P22315 | P22315 | FECH | ferrochelatase | Cytoplasm | enzyme |
| 15550430 | 2, 3 | 0.412 | 2.39E+07 | 4.84E+07 | 1.57E+07 | 2.12E+07 | 1.95E+07 | 3.20E+07 | 1.37E+07 | 2.85E+07 | 3.80E+07 | 1.44E+07 | 1.56E+07 | 1.59E+07 | 2519.2578, 2519.2593 | TGILMLNMGGPETLGEVQDFLQR | P22315 | P22315 | FECH | ferrochelatase | Cytoplasm | enzyme |
| 15548479 | 3 | 0.248 | 1.26E+07 | 2.23E+07 | 1.07E+07 | 1.16E+07 | 1.13E+07 | 1.76E+07 | 7295433 | 1.52E+07 | 2.78E+07 | 8156954 | 7970591 | 6961968.5 | 2699.2769 | AIAFTQYPQYSC[160.0307]STTGSSLNAIYR | P22315 | P22315 | FECH | ferrochelatase | Cytoplasm | enzyme |
| 15540371 | 2 | 0.233 | 1.01E+07 | 2.02E+07 | 8234117.5 | 7718520.5 | 9087027 | 1.43E+07 | 6063525 | 1.22E+07 | 1.46E+07 | 5505966 | 6118793.5 | 7008069 | 818.41797 | YYIGFR | P22315 | P22315 | FECH | ferrochelatase | Cytoplasm | enzyme |
| 15543326 | 3 | 0.414 | 6433691 | 1.50E+07 | 1.12E+07 | 2740537 | 1.24E+07 | 1.79E+07 | 7880847.5 | 2.01E+07 | 2.40E+07 | 4689293.5 | 5688380 | 9078947 | 1532.8131 | ALADLVHSHIQSNK | P22315 | P22315 | FECH | ferrochelatase | Cytoplasm | enzyme |
| 15556865 | 3 | 0.453 | 9730425 | 4.64E+07 | 2.60E+07 | 3932284 | 2.39E+07 | 4.10E+07 | 8227494 | 3.37E+07 | 5.14E+07 | 4336555.5 | 6434526 | 1.58E+07 | 2416.143 | YVHPLTEEAIEEMERDGLER | P22315 | P22315 | FECH | ferrochelatase | Cytoplasm | enzyme |
| 15538019 | 2, 3 | 0.135 | 7754514.5 | 1.39E+07 | 5284643.5 | 7996716 | 7469610.5 | 9070013 | 5635771 | 7810655 | 1.14E+07 | 3970905.75 | 5243749 | 4700658.5 | 1432.7502, 1432.7512 | RAESLNGNPLFSK | P22315 | P22315 | FECH | ferrochelatase | Cytoplasm | enzyme |
| 15554537 | 3 | 0.962 | 4540239 | 3377303.5 | 775341.875 | 4983112 | 1738377.125 | 1905578.625 | 1189052.375 | 2396997.5 | 4348450 | 3138363.75 | 7845285.5 | 1200312.75 | 2291.0723 | LC[160.0307]STQLSLNC[160.0307]PLC[160.0307]VNPVC[160.0307]R | P22315 | P22315 | FECH | ferrochelatase | Cytoplasm | enzyme |
| 15534174 | 2 | 0.199 | 3796610.5 | 8256654.5 | 2587182.25 | 2171674.5 | 3441871.25 | 3789252.5 | 2397378.5 | 4889298 | 3813158 | 2785111.25 | 1382286.625 | 1965411.25 | 1497.73 | GDPYPQEVGATVHK | P22315 | P22315 | FECH | ferrochelatase | Cytoplasm | enzyme |
| 15549854 | 2 | 0.269 | 5709691.5 | 7282095 | 2821165.5 | 3684330 | 2819758.75 | 5427936 | 2137009.25 | 3677426.5 | 6126832.5 | 2713883.25 | 2235612.5 | 2377321.75 | 1816.9938 | LFLDRDLMTLPIQNK | P22315 | P22315 | FECH | ferrochelatase | Cytoplasm | enzyme |
| 15556963, 15557539 | 3 | 0.044 | 4409392 | 6541519 | 3118874.5 | 3478173 | 2656599.25 | 5228465 | 2501384.25 | 5100683 | 3375601.75 | 2340733.75 | 891138.562 | 1733061.375 | 2535.254, 2535.2546 | TGILM[147.0354]LNMGGPETLGEVQDFLQR | P22315 | P22315 | FECH | ferrochelatase | Cytoplasm | enzyme |
| 15539502 | 3 | 0.212 | 3054146.25 | 9135795 | 4515167 | 1947074.875 | 7485091.5 | 6734811 | 3933227.75 | 6324166 | 4481350 | 2238553 | 1685657.625 | 2794273.5 | 1845.8657 | YVHPLTEEAIEEMER | P22315 | P22315 | FECH | ferrochelatase | Cytoplasm | enzyme |
| 15544379 | 2 | 0.287 | 2638658.25 | 6298903 | 2912656.75 | 3048668.25 | 3002429.25 | 3299676 | 1701572.75 | 3756343.25 | 5085218.5 | 1723303.875 | 2060545.5 | 2188503.75 | 957.4659 | SFFTSQQL | P22315 | P22315 | FECH | ferrochelatase | Cytoplasm | enzyme |
| 15537700 | 2 | 0.903 | 3413372.25 | 3967093 | 933938.812 | 2409667.75 | 1650530.875 | 2115246 | 1747086.125 | 2738175.25 | 2981313 | 1622761.375 | 2405850 | 1808148.375 | 1457.7068 | YYNEVGQKPTMK | P22315 | P22315 | FECH | ferrochelatase | Cytoplasm | enzyme |
| 15535981 | 2 | 0.088 | 2468967.75 | 5715573 | 1813150 | 2792644 | 2209900.5 | 3482900.25 | 1903953.5 | 3131387.75 | 4048297.25 | 1488379.75 | 1017349 | 1468840.5 | 1276.6498 | AESLNGNPLFSK | P22315 | P22315 | FECH | ferrochelatase | Cytoplasm | enzyme |
| 15550930 | 3 | 0.791 | 3138010.25 | 827373.188 | 181140.109 | 2448349 | 481592.219 | 904301.688 | 721887.125 | 412438.812 | 4321175 | 710480.062 | 1.17E+07 | 1203826.875 | 2419.1675 | LC[160.0307]STQLSLNC[160.0307]PLC[160.0307]VNPVC[160.0307]RK | P22315 | P22315 | FECH | ferrochelatase | Cytoplasm | enzyme |
| 15559316 | 2 | 0.397 | 1631812.75 | 4405646 | 1670071.875 | 471144.406 | 2310885.5 | 3282914.75 | 927030.812 | 3046290.25 | 4535876 | 488817.125 | 1072759.875 | 1236636.25 | 1532.8146 | ALADLVHSHIQSNK | P22315 | P22315 | FECH | ferrochelatase | Cytoplasm | enzyme |
| 15552084 | 2 | 0.344 | 1.00E+07 | 1.09E+07 | 1.44E+07 | 2.39E+07 | 2.34E+07 | 1.97E+07 | 1.70E+07 | 1.25E+07 | 4.32E+07 | 7427753 | 2.04E+07 | 2.29E+07 | 1775.879 | VFVGEEDPEAQSVTLR | Q8K1B8 | Q8K1B8 | FERMT3 | fermitin family member 3 | Cytoplasm | enzyme |
| 15535221 | 3 | 0.37 | 1.20E+07 | 1.63E+07 | 1.68E+07 | 2.89E+07 | 2.73E+07 | 2.54E+07 | 1.84E+07 | 1.45E+07 | 4.23E+07 | 7223967.5 | 2.07E+07 | 2.64E+07 | 2134.1572 | ILEAHQNVAQLSLTEAQLR | Q8K1B8 | Q8K1B8 | FERMT3 | fermitin family member 3 | Cytoplasm | enzyme |
| 15551463 | 3 | 0.35 | 6645749 | 8893568 | 9129000 | 1.71E+07 | 1.46E+07 | 1.49E+07 | 1.06E+07 | 9044290 | 2.50E+07 | 6050117 | 1.18E+07 | 2.10E+07 | 2347.117 | ARGEELDEDLFLQLTGGHEAF | Q8K1B8 | Q8K1B8 | FERMT3 | fermitin family member 3 | Cytoplasm | enzyme |
| 15552147 | 2 | 0.414 | 6965636 | 9704348 | 7852410.5 | 1.78E+07 | 1.54E+07 | 1.58E+07 | 1.08E+07 | 8676386 | 3.15E+07 | 5407405 | 1.43E+07 | 2.24E+07 | 1703.8999 | LLVPSPEGMSEIYLR | Q8K1B8 | Q8K1B8 | FERMT3 | fermitin family member 3 | Cytoplasm | enzyme |
| 15533252 | 2 | 0.694 | 7533745.5 | 8980498 | 8946375 | 1.66E+07 | 9334277 | 1.49E+07 | 1.18E+07 | 7435956.5 | 2.80E+07 | 4767208.5 | 1.26E+07 | 2.15E+07 | 1299.7755 | VVLAGGVAPTLFR | Q8K1B8 | Q8K1B8 | FERMT3 | fermitin family member 3 | Cytoplasm | enzyme |
| 15554602 | 2 | 0.416 | 7633897.5 | 8176994 | 9764375 | 1.67E+07 | 1.48E+07 | 1.33E+07 | 1.23E+07 | 9772007 | 2.72E+07 | 4691706.5 | 1.25E+07 | 1.98E+07 | 1740.8364 | SQDEAPGDPTQQLNLK | Q8K1B8 | Q8K1B8 | FERMT3 | fermitin family member 3 | Cytoplasm | enzyme |
| 15533641 | 2 | 0.402 | 6474151.5 | 6682532 | 7013427 | 1.36E+07 | 1.21E+07 | 1.21E+07 | 1.12E+07 | 7441560 | 2.02E+07 | 3942658 | 1.33E+07 | 1.71E+07 | 1487.7126 | GC[160.0307]EVVPDVNVSGQK | Q8K1B8 | Q8K1B8 | FERMT3 | fermitin family member 3 | Cytoplasm | enzyme |
| 15548567 | 2 | 0.35 | 6078563 | 8223825.5 | 8671702 | 1.48E+07 | 1.30E+07 | 1.37E+07 | 1.05E+07 | 9625580 | 2.64E+07 | 3927008.75 | 1.01E+07 | 1.73E+07 | 1874.9376 | GSAPSDMLDSLTTIPELK | Q8K1B8 | Q8K1B8 | FERMT3 | fermitin family member 3 | Cytoplasm | enzyme |
| 15533615 | 2 | 0.381 | 4104070 | 5286356 | 6096307.5 | 1.07E+07 | 8497557 | 1.07E+07 | 7811936.5 | 5678594 | 1.91E+07 | 3312055.75 | 9106180 | 1.31E+07 | 1294.5963 | YYSFFDLDPK | Q8K1B8 | Q8K1B8 | FERMT3 | fermitin family member 3 | Cytoplasm | enzyme |
| 15531810 | 2 | 0.289 | 4754580.5 | 5727625 | 4772186 | 1.29E+07 | 9560043 | 9893366 | 7473945 | 5955210.5 | 1.72E+07 | 3128183.5 | 7988016 | 1.30E+07 | 1121.5918 | LTQLYEQAR | Q8K1B8 | Q8K1B8 | FERMT3 | fermitin family member 3 | Cytoplasm | enzyme |
| 15532484 | 2 | 0.529 | 4497127 | 5511225 | 6190877 | 8837906 | 8530405 | 1.06E+07 | 6964021 | 5222718 | 1.88E+07 | 3080519 | 6970346 | 1.30E+07 | 1028.597 | IDLAVGDVVK | Q8K1B8 | Q8K1B8 | FERMT3 | fermitin family member 3 | Cytoplasm | enzyme |
| 15539819 | 2 | 0.229 | 3574200.25 | 4254719.5 | 3880573 | 4685458 | 5380604 | 5179721.5 | 4186183.75 | 2732029.5 | 5994968 | 2690958 | 3427437.5 | 3975448.75 | 1114.5865 | DEILGIANNR | Q8K1B8 | Q8K1B8 | FERMT3 | fermitin family member 3 | Cytoplasm | enzyme |
| 15533367 | 2 | 0.454 | 2283352.5 | 2870355.25 | 3636668.5 | 5396698.5 | 4330893.5 | 5701378.5 | 3885785.75 | 3042703.5 | 1.04E+07 | 1830048.875 | 4038398.75 | 6635760 | 878.47156 | YGILADAR | Q8K1B8 | Q8K1B8 | FERMT3 | fermitin family member 3 | Cytoplasm | enzyme |
| 15540002 | 2 | 0.484 | 2236600.75 | 3115647 | 2952370.5 | 4990288 | 4271559.5 | 4507247.5 | 3597467.75 | 2877594.75 | 8534218 | 1700691.875 | 4821817.5 | 5330166 | 1230.5992 | QWNVNWDIR | Q8K1B8 | Q8K1B8 | FERMT3 | fermitin family member 3 | Cytoplasm | enzyme |
| 15535569 | 2 | 0.452 | 2519297 | 2699865 | 3492491.5 | 5457932.5 | 4276801 | 5129545 | 4467587.5 | 3165884.5 | 7188857.5 | 1661441.25 | 4667268 | 6758822 | 1309.744 | VTGESHIGGVLLK | Q8K1B8 | Q8K1B8 | FERMT3 | fermitin family member 3 | Cytoplasm | enzyme |
| 15554183, 15556740 | 2, 3 | 0.468 | 1651697.875 | 3035680.75 | 1256616 | 7297761.5 | 4114824.5 | 3203496.75 | 3787255 | 1726257.25 | 9342816 | 794674.188 | 3861226.75 | 5674816 | 2257.1453, 2257.1472 | FIQAWQSLPDFGISYVMVR | Q8K1B8 | Q8K1B8 | FERMT3 | fermitin family member 3 | Cytoplasm | enzyme |
| 15551991 | 3 | 0.854 | 8362247.5 | 2.79E+07 | 1.57E+07 | 2.18E+07 | 2.03E+07 | 3.92E+07 | 1.31E+07 | 1.50E+07 | 5.06E+07 | 7941250.5 | 1.91E+07 | 1.38E+08 | 3072.569 | SSIAELNNNIQSVSDTSSVTFQYLTLLK | Q8K0E8 | Q8K0E8 | FGB | fibrinogen beta chain | Extracellular Space | other |
| 15536079 | 3 | 0.849 | 8924944 | 6.99E+07 | 2.62E+07 | 3.48E+07 | 6.20E+07 | 6.69E+07 | 3.03E+07 | 3.08E+07 | 8.47E+07 | 6123591.5 | 2.47E+07 | 2.08E+08 | 1977.0156 | EEPPSLRPAPPPISGGGYR | Q8K0E8 | Q8K0E8 | FGB | fibrinogen beta chain | Extracellular Space | other |
| 15548325 | 2 | 0.923 | 1.14E+07 | 5.29E+07 | 1.63E+07 | 3.63E+07 | 3.43E+07 | 3.90E+07 | 2.01E+07 | 1.84E+07 | 1.04E+08 | 5414073 | 3.98E+07 | 2.26E+08 | 2051.947 | ENENVINEYSSILEDQR | Q8K0E8 | Q8K0E8 | FGB | fibrinogen beta chain | Extracellular Space | other |
| 15548102 | 2 | 0.685 | 6452684.5 | 2.71E+07 | 6833572.5 | 2.02E+07 | 2.20E+07 | 2.24E+07 | 1.17E+07 | 1.24E+07 | 8.08E+07 | 4580159 | 4.78E+07 | 1.49E+08 | 1618.7899 | TPC[160.0307]TVSC[160.0307]NIPVVSGK | Q8K0E8 | Q8K0E8 | FGB | fibrinogen beta chain | Extracellular Space | other |
| 15551178 | 2 | 0.926 | 6410180 | 4.07E+07 | 1.46E+07 | 2.47E+07 | 2.72E+07 | 3.08E+07 | 1.45E+07 | 1.40E+07 | 8.01E+07 | 4133639.25 | 2.27E+07 | 1.56E+08 | 1671.7435 | YC[160.0307]GLPGEYWLGNDK | Q8K0E8 | Q8K0E8 | FGB | fibrinogen beta chain | Extracellular Space | other |
| 15531779 | 3 | 0.576 | 8840132 | 2.18E+07 | 595652 | 3.34E+07 | 9764030 | 1.08E+07 | 9596955 | 5741241.5 | 5.45E+07 | 3922837.5 | 3.37E+07 | 1.44E+08 | 1991.9431 | MGPTELLIEMEDWKGDK | Q8K0E8 | Q8K0E8 | FGB | fibrinogen beta chain | Extracellular Space | other |
| 15549195 | 2 | 0.925 | 6203812 | 3.42E+07 | 1.20E+07 | 2.31E+07 | 2.39E+07 | 2.58E+07 | 1.18E+07 | 1.22E+07 | 6.44E+07 | 3467629.5 | 2.26E+07 | 1.29E+08 | 1868.8906 | GGETSEMYLIQPDTSIK | Q8K0E8 | Q8K0E8 | FGB | fibrinogen beta chain | Extracellular Space | other |
| 15533997 | 3 | 0.91 | 7333532.5 | 2.94E+07 | 9028135 | 2.22E+07 | 2.03E+07 | 2.23E+07 | 1.10E+07 | 1.13E+07 | 4.98E+07 | 3378407.5 | 2.40E+07 | 1.23E+08 | 1508.7119 | AHYGGFTVQNEASK | Q8K0E8 | Q8K0E8 | FGB | fibrinogen beta chain | Extracellular Space | other |
| 15547761 | 2 | 0.893 | 5808605.5 | 3.56E+07 | 9939412 | 2.47E+07 | 2.36E+07 | 3.04E+07 | 1.21E+07 | 1.28E+07 | 7.37E+07 | 2692473.25 | 2.76E+07 | 1.48E+08 | 1901.9933 | LYIDETVNDNIPLNLR | Q8K0E8 | Q8K0E8 | FGB | fibrinogen beta chain | Extracellular Space | other |
| 15539784 | 2 | 0.902 | 4981288.5 | 3.56E+07 | 6882700 | 1.99E+07 | 2.27E+07 | 2.03E+07 | 1.06E+07 | 1.05E+07 | 5.05E+07 | 2554367.5 | 2.13E+07 | 1.33E+08 | 1236.5831 | GFGNIATNEDAK | Q8K0E8 | Q8K0E8 | FGB | fibrinogen beta chain | Extracellular Space | other |
| 15538609 | 2 | 0.821 | 4372217 | 1.48E+07 | 4032377.25 | 1.35E+07 | 1.17E+07 | 1.08E+07 | 6603334.5 | 6004560 | 3.47E+07 | 2096629 | 1.86E+07 | 7.89E+07 | 1239.5153 | EDGGGWWYNR | Q8K0E8 | Q8K0E8 | FGB | fibrinogen beta chain | Extracellular Space | other |
| 15550151 | 2 | 0.933 | 2454591 | 2.45E+07 | 6882260.5 | 8077256.5 | 1.62E+07 | 1.88E+07 | 6822670 | 7891125 | 2.64E+07 | 1701762.625 | 7025743.5 | 6.78E+07 | 1691.7998 | MGPTELLIEMEDWK | Q8K0E8 | Q8K0E8 | FGB | fibrinogen beta chain | Extracellular Space | other |
| 15551465 | 2 | 0.746 | 2466226.5 | 9023894 | 1983136.125 | 9277229 | 6226813 | 4977670 | 4076801.5 | 2952806.25 | 2.27E+07 | 1624958.125 | 1.10E+07 | 4.84E+07 | 1655.7168 | YYWGGLYSWDMSK | Q8K0E8 | Q8K0E8 | FGB | fibrinogen beta chain | Extracellular Space | other |
| 15531711 | 2 | 0.884 | 3554903.5 | 1.49E+07 | 4616295.5 | 1.37E+07 | 9520470 | 1.22E+07 | 5978801.5 | 5319660.5 | 2.86E+07 | 1597271.125 | 1.48E+07 | 7.99E+07 | 1374.6738 | TENGGWTVIQNR | Q8K0E8 | Q8K0E8 | FGB | fibrinogen beta chain | Extracellular Space | other |
| 15550007 | 2 | 0.876 | 2530157.5 | 1.11E+07 | 2791909.75 | 7255641 | 7110649.5 | 7072506 | 4074069 | 3352006 | 1.78E+07 | 1467967.375 | 9320680 | 4.33E+07 | 1508.7109 | AHYGGFTVQNEASK | Q8K0E8 | Q8K0E8 | FGB | fibrinogen beta chain | Extracellular Space | other |
| 15536859 | 3 | 0.661 | 1610644.125 | 1.25E+07 | 1172047.25 | 8375577 | 7159199.5 | 1.08E+07 | 3381078.75 | 4931014 | 1.91E+07 | 1315827.875 | 8639238 | 4.61E+07 | 1901.994 | LYIDETVNDNIPLNLR | Q8K0E8 | Q8K0E8 | FGB | fibrinogen beta chain | Extracellular Space | other |
| 15549076 | 2 | 0.968 | 647858.188 | 1.02E+07 | 3993552.5 | 3043518.5 | 4384970.5 | 8211383.5 | 1950645.625 | 3098477.75 | 1.77E+07 | 579548.125 | 2609623 | 3.13E+07 | 1671.7098 | YYWGGLYSWDM[147.0354]SK | Q8K0E8 | Q8K0E8 | FGB | fibrinogen beta chain | Extracellular Space | other |
| 15537130 | 3 | 0.793 | 444527.062 | 4806734.5 | 1501136.875 | 3096045.75 | 1932189.875 | 3365568 | 2079699.625 | 1180275.25 | 8209299.5 | 507333.75 | 5362300.5 | 2.86E+07 | 1544.6913 | HGTDDGVVWMNWK | Q8K0E8 | Q8K0E8 | FGB | fibrinogen beta chain | Extracellular Space | other |
| 15539659 | 3 | 0.95 | 394139.906 | 7573519 | 1603421.5 | 1686733.125 | 3082329.75 | 6022827 | 1758080.125 | 1208792.375 | 7379691.5 | 489172.469 | 1489711.75 | 1.95E+07 | 1691.7999 | MGPTELLIEMEDWK | Q8K0E8 | Q8K0E8 | FGB | fibrinogen beta chain | Extracellular Space | other |
| 15542183 | 3 | 0.74 | 717634 | 1309191.875 | 102611.695 | 2227769 | 695039.812 | 676055.062 | 326554.062 | 581600.812 | 3888787.5 | 248036.891 | 1498645 | 1.12E+07 | 2007.9381 | M[147.0354]GPTELLIEMEDWKGDK | Q8K0E8 | Q8K0E8 | FGB | fibrinogen beta chain | Extracellular Space | other |
| 15554811 | 2 | 0.73 | 49753.203 | 2504531.75 | 623380.688 | 1188139.25 | 1712182.25 | 1757164.5 | 885985.688 | 716944.438 | 2482782.75 | 113684.266 | 630717.125 | 5862494 | 1977.0167 | EEPPSLRPAPPPISGGGYR | Q8K0E8 | Q8K0E8 | FGB | fibrinogen beta chain | Extracellular Space | other |
| 15539822 | 3 | 0.922 | 590456.5 | 3465840.75 | 1143282.25 | 1577884 | 3393650.25 | 2457710 | 1248857 | 2030147.5 | 9780214 | 93941.633 | 2124824.25 | 2.16E+07 | 1868.8922 | GGETSEMYLIQPDTSIK | Q8K0E8 | Q8K0E8 | FGB | fibrinogen beta chain | Extracellular Space | other |
| 15537278 | 3 | 0.912 | 157144.828 | 3824906 | 849442.125 | 953951.188 | 2347827.75 | 2911405.75 | 1199022.875 | 1405113.875 | 8677528 | 31483.266 | 2768204.5 | 2.89E+07 | 2051.9473 | ENENVINEYSSILEDQR | Q8K0E8 | Q8K0E8 | FGB | fibrinogen beta chain | Extracellular Space | other |
| 15537765 | 2 | 0.854 | 196515.297 | 753452.938 | 5867.889 | 1198106 | 497776.875 | 244445.203 | 217146.516 | 190966.609 | 2365067 | 4.70E-06 | 1478020.25 | 5774544 | 1364.6783 | GFGNIATNEDAKK | Q8K0E8 | Q8K0E8 | FGB | fibrinogen beta chain | Extracellular Space | other |
| 15552633 | 3 | 0.717 | 1.88E+07 | 5.49E+07 | 1.37E+07 | 5.75E+07 | 3.40E+07 | 3.12E+07 | 2.10E+07 | 2.10E+07 | 1.21E+08 | 1.27E+07 | 7.13E+07 | 2.87E+08 | 2561.2654 | AIQVYYNPDQPPKPGMIDSATQK | Q8VCM7 | Q8VCM7 | FGG | fibrinogen gamma chain | Extracellular Space | other |
| 15532938 | 3 | 0.808 | 2.01E+07 | 8.32E+07 | 2.07E+07 | 7.07E+07 | 6.28E+07 | 5.74E+07 | 3.58E+07 | 3.82E+07 | 1.46E+08 | 1.14E+07 | 7.85E+07 | 3.98E+08 | 2207.0374 | EGFGHLSPTGTTEFWLGNEK | Q8VCM7 | Q8VCM7 | FGG | fibrinogen gamma chain | Extracellular Space | other |
| 15549555 | 2 | 0.866 | 8630670 | 4.29E+07 | 1.28E+07 | 3.48E+07 | 2.96E+07 | 2.71E+07 | 1.44E+07 | 1.23E+07 | 7.80E+07 | 5443341 | 4.04E+07 | 1.74E+08 | 1560.7129 | VAQLEAQC[160.0307]QEPC[160.0307]K | Q8VCM7 | Q8VCM7 | FGG | fibrinogen gamma chain | Extracellular Space | other |
| 15548161 | 3 | 0.876 | 3697174 | 3.36E+07 | 7161894.5 | 1.31E+07 | 2.05E+07 | 1.92E+07 | 8840271 | 9395694 | 3.85E+07 | 4842494.5 | 1.23E+07 | 1.02E+08 | 3086.3467 | FGSFC[160.0307]PTTC[160.0307]GIADFLSSYQTDVDNDLR | Q8VCM7 | Q8VCM7 | FGG | fibrinogen gamma chain | Extracellular Space | other |
| 15532735 | 2, 3 | 0.923 | 7970815 | 3.31E+07 | 9398904 | 2.03E+07 | 2.28E+07 | 1.71E+07 | 1.04E+07 | 1.18E+07 | 6.11E+07 | 3373127.75 | 3.37E+07 | 1.45E+08 | 1322.7439, 1322.7443 | ESGLYFIRPLK | Q8VCM7 | Q8VCM7 | FGG | fibrinogen gamma chain | Extracellular Space | other |
| 15547803 | 2 | 0.9 | 5747720.5 | 2.81E+07 | 7423321.5 | 1.94E+07 | 1.64E+07 | 1.73E+07 | 8705588 | 8996688 | 5.58E+07 | 3185512.5 | 2.50E+07 | 1.08E+08 | 1513.7258 | YLQEIYNSNNQK | Q8VCM7 | Q8VCM7 | FGG | fibrinogen gamma chain | Extracellular Space | other |
| 15537382 | 2 | 0.839 | 5120636.5 | 2.30E+07 | 6544940.5 | 1.50E+07 | 1.53E+07 | 1.77E+07 | 1.03E+07 | 8873723 | 4.33E+07 | 3052397.25 | 2.36E+07 | 1.06E+08 | 1162.5178 | TSTADYAMFR | Q8VCM7 | Q8VCM7 | FGG | fibrinogen gamma chain | Extracellular Space | other |
| 15534710 | 3 | 0.875 | 4022041.75 | 2.53E+07 | 7313971.5 | 1.45E+07 | 1.35E+07 | 1.85E+07 | 8832600 | 8903241 | 6.23E+07 | 2976528.25 | 1.98E+07 | 1.04E+08 | 1545.8247 | YEALLLTHETSIR | Q8VCM7 | Q8VCM7 | FGG | fibrinogen gamma chain | Extracellular Space | other |
| 15554828 | 2 | 0.894 | 1718757.75 | 1.61E+07 | 1.04E+07 | 7149292 | 1.54E+07 | 1.61E+07 | 7014668.5 | 6808952 | 2.18E+07 | 2601879.25 | 7967074.5 | 3.79E+07 | 1532.7704 | AIQVYYNPDQPPK | Q8VCM7 | Q8VCM7 | FGG | fibrinogen gamma chain | Extracellular Space | other |
| 15535700 | 3 | 0.797 | 2951113.75 | 2.07E+07 | 3364761.75 | 1.46E+07 | 1.06E+07 | 1.39E+07 | 7849174.5 | 5887847 | 3.72E+07 | 2000611.75 | 1.73E+07 | 8.83E+07 | 1742.9596 | IHLISMQSTIPYALR | Q8VCM7 | Q8VCM7 | FGG | fibrinogen gamma chain | Extracellular Space | other |
| 15540043 | 2 | 0.875 | 2693666 | 1.04E+07 | 3840871 | 6860746.5 | 6766016 | 7426084.5 | 4095503 | 3580818 | 1.91E+07 | 1897584 | 8979753 | 4.60E+07 | 851.4399 | NWIQYK | Q8VCM7 | Q8VCM7 | FGG | fibrinogen gamma chain | Extracellular Space | other |
| 15548632 | 2 | 0.823 | 2138889.75 | 1.45E+07 | 3807077.75 | 9908530 | 8838382 | 1.20E+07 | 5018782 | 5835326 | 3.29E+07 | 1522869.625 | 1.41E+07 | 5.95E+07 | 1898.8883 | SSTTNGFDDGIIWATWK | Q8VCM7 | Q8VCM7 | FGG | fibrinogen gamma chain | Extracellular Space | other |
| 15531978 | 2 | 0.886 | 2284681 | 1.89E+07 | 4261522 | 5628048.5 | 1.29E+07 | 1.31E+07 | 7176440 | 7134897 | 2.48E+07 | 1465901.875 | 6434808.5 | 4.33E+07 | 1006.55536 | TLEDILFR | Q8VCM7 | Q8VCM7 | FGG | fibrinogen gamma chain | Extracellular Space | other |
| 15541166 | 3 | 0.404 | 1882617.25 | 5048828.5 | 3621154.25 | 1.11E+07 | 1.12E+07 | 1.23E+07 | 6786747.5 | 6551391 | 1.59E+07 | 1255526.375 | 1.50E+07 | 3.13E+07 | 1565.745 | LSIGEGQQHHMGGSK | Q8VCM7 | Q8VCM7 | FGG | fibrinogen gamma chain | Extracellular Space | other |
| 15553856, 15554832 | 2, 3 | 0.932 | 3558568.5 | 1.99E+07 | 5649617 | 1.25E+07 | 1.25E+07 | 1.69E+07 | 8244863.5 | 6076565.5 | 3.77E+07 | 1131395.125 | 1.68E+07 | 8.24E+07 | 2290.938, 2290.9424 | FFTSHNGMQFSTWDNDNDK | Q8VCM7 | Q8VCM7 | FGG | fibrinogen gamma chain | Extracellular Space | other |
| 15549944 | 2 | 0.952 | 2936514 | 1.47E+07 | 3077522.5 | 7752828 | 7844088 | 8866916 | 4845605.5 | 4406910 | 2.13E+07 | 1042707.375 | 9393745 | 5.54E+07 | 1545.8242 | YEALLLTHETSIR | Q8VCM7 | Q8VCM7 | FGG | fibrinogen gamma chain | Extracellular Space | other |
| 15548063 | 2 | 0.831 | 3291853.5 | 1.73E+07 | 4171973 | 1.46E+07 | 1.55E+07 | 1.66E+07 | 8710693 | 7159377 | 3.25E+07 | 950358.75 | 2.13E+07 | 7.12E+07 | 2015.7972 | FEGNC[160.0307]AEQDGSGWWMNK | Q8VCM7 | Q8VCM7 | FGG | fibrinogen gamma chain | Extracellular Space | other |
| 15543163 | 2 | 0.79 | 202240.516 | 3377074.25 | 1304677.875 | 1689483.875 | 2721804 | 3143319.75 | 1360232.875 | 1140821.75 | 3193968 | 339068.5 | 875355.812 | 8443994 | 1047.5112 | PGMIDSATQK | Q8VCM7 | Q8VCM7 | FGG | fibrinogen gamma chain | Extracellular Space | other |
| 15536686 | 2 | 0.907 | 803007.875 | 1.15E+07 | 4767602 | 1327187.125 | 1.04E+07 | 1.52E+07 | 3030082.5 | 6871847.5 | 4.27E+07 | 189784.516 | 4931793 | 6.39E+07 | 1194.4845 | DNC[160.0307]C[160.0307]ILDER | Q8VCM7 | Q8VCM7 | FGG | fibrinogen gamma chain | Extracellular Space | other |
| 15549200 | 2 | 0.939 | 1679592.375 | 8039286 | 826774.688 | 5150983 | 4709998.5 | 3651385 | 2515057.25 | 2408048.75 | 1.35E+07 | 187244.312 | 6048232.5 | 3.36E+07 | 2207.038 | EGFGHLSPTGTTEFWLGNEK | Q8VCM7 | Q8VCM7 | FGG | fibrinogen gamma chain | Extracellular Space | other |
| 15551888 | 2 | 0.77 | 822528.312 | 4496843.5 | 90678.109 | 3545973.75 | 2417287.25 | 3146277 | 1626792 | 1277910.75 | 9072536 | 69635.594 | 5077409.5 | 2.22E+07 | 1742.9592 | IHLISMQSTIPYALR | Q8VCM7 | Q8VCM7 | FGG | fibrinogen gamma chain | Extracellular Space | other |
| 15555758 | 2 | 0.478 | 0 | 1166249.5 | 0 | 475589.281 | 482461.375 | 870113.938 | 244075.453 | 100212.656 | 1696869.875 | 0 | 433356.344 | 3398242.25 | 2031.7921 | FEGNC[160.0307]AEQDGSGWWM[147.0354]NK | Q8VCM7 | Q8VCM7 | FGG | fibrinogen gamma chain | Extracellular Space | other |
| 15559966 | 2 | 0.766 | 87994.156 | 436635.219 | 248464.703 | 468746.062 | 918233.062 | 1166126.25 | 268512.688 | 487602.594 | 1680834.125 | 0 | 1035681.562 | 2699354.75 | 1565.7462 | LSIGEGQQHHMGGSK | Q8VCM7 | Q8VCM7 | FGG | fibrinogen gamma chain | Extracellular Space | other |
| 15548585 | 3 | 0.275 | 8473457 | 1.89E+07 | 8188637.5 | 1.31E+07 | 1.16E+07 | 1.45E+07 | 8138205 | 1.26E+07 | 1.36E+07 | 5112919.5 | 9221991 | 8973921 | 3166.5583 | THTQDAVPLTLGQEFSGYVQQVQYAMVR | P97807 | P97807 | FH | fumarate hydratase | Cytoplasm | enzyme |
| 15550315 | 2 | 0.159 | 5416682.5 | 1.02E+07 | 5272484.5 | 6813218.5 | 6517636 | 7973911 | 4236023 | 6114130.5 | 8247396.5 | 2750331.5 | 4132639.25 | 5328927.5 | 2125.0796 | SGLGELILPENEPGSSIMPGK | P97807 | P97807 | FH | fumarate hydratase | Cytoplasm | enzyme |
| 15550571 | 3 | 0.386 | 2955178 | 5585759 | 2262535.5 | 3650026 | 3785881.5 | 4067724.25 | 1907064.5 | 3119575.75 | 5362968.5 | 1774300.875 | 2436678 | 2645372.25 | 2259.1467 | LMNESLMLVTALNPHIGYDK | P97807 | P97807 | FH | fumarate hydratase | Cytoplasm | enzyme |
| 15536832 | 3 | 0.042 | 2628967.5 | 5294125 | 2866458.25 | 4223345.5 | 3643714.75 | 4023854 | 2455282.25 | 4029470.5 | 4561663.5 | 1406756 | 1961654.5 | 2296791.5 | 2242.0872 | ETAIELGYLTAEQFDEWVK | P97807 | P97807 | FH | fumarate hydratase | Cytoplasm | enzyme |
| 15552167 | 2 | 0.098 | 1705301.75 | 3509684 | 2059112.625 | 2273235.75 | 2432419 | 3035863.5 | 1939246.125 | 3015288 | 3842429.5 | 1182109.375 | 1542934.375 | 1791908.75 | 2242.087 | ETAIELGYLTAEQFDEWVK | P97807 | P97807 | FH | fumarate hydratase | Cytoplasm | enzyme |
| 15539874 | 2 | 0.101 | 1504168.375 | 3193533.75 | 1838564.25 | 1159329.5 | 2766900.5 | 2707856.25 | 987962.938 | 2794052 | 2635052.75 | 962527.625 | 567424.562 | 1008908 | 1184.5812 | VEFDTFGELK | P97807 | P97807 | FH | fumarate hydratase | Cytoplasm | enzyme |
| 15555015 | 3 | 0.226 | 2481158.25 | 4769946 | 2879436.25 | 4072024.25 | 2711002 | 4120326.25 | 2183082.75 | 3323362 | 4966283 | 757772.562 | 2136909.25 | 2879193 | 3270.5918 | LNDHFPLVVWQTGSGTQTNMNVNEVISNR | P97807 | P97807 | FH | fumarate hydratase | Cytoplasm | enzyme |
| 15541844 | 2 | 0.242 | 4682755.5 | 6583323 | 3670442.5 | 6066951 | 4961726 | 5813609.5 | 4688497 | 6801274 | 6198454.5 | 3725287 | 4633931 | 4412968 | 1440.7413 | GLLQTEPQNNQAK | Q9CQ92 | Q9CQ92 | FIS1 | fission, mitochondrial 1 | Cytoplasm | other |
| 15538485 | 2 | 0.058 | 3955659.75 | 6256769.5 | 4694116 | 5555874 | 5763053.5 | 6729528 | 4565733.5 | 6290827.5 | 6656118 | 3445081 | 4360926 | 3968996 | 1238.598 | FQSEQAAGSVSK | Q9CQ92 | Q9CQ92 | FIS1 | fission, mitochondrial 1 | Cytoplasm | other |
| 15546192 | 3 | 0.007 | 1178087.75 | 1110411 | 695788.312 | 1264216.75 | 1299533.875 | 614303.312 | 330949.844 | 332883.469 | 92176.938 | 1002949.375 | 1057415.375 | 923614.938 | 2185.986 | VPRPTAQPEEEDEDEVSMK | Q6P9Q6 | Q6P9Q6 | FKBP15 | FK506 binding protein 15, 133kDa | Plasma Membrane | enzyme |
| 15544600 | 3 | 0.18 | 538711.5 | 441911.312 | 368427.562 | 503245.875 | 545696.188 | 209179.297 | 130802.492 | 222478.188 | 300123.156 | 480408.438 | 224029.875 | 323290.344 | 1703.8375 | RNNSLQTATENTQAR | Q6P9Q6 | Q6P9Q6 | FKBP15 | FK506 binding protein 15, 133kDa | Plasma Membrane | enzyme |
| 15539206 | 2 | 0.411 | 1.34E+07 | 1.70E+07 | 1.74E+07 | 1.65E+07 | 2.24E+07 | 1.76E+07 | 1.82E+07 | 2.14E+07 | 1.56E+07 | 1.75E+07 | 1.63E+07 | 1.48E+07 | 1332.7131 | LVIPSELGYGER | P45878 | P45878 | FKBP2 | FK506 binding protein 2, 13kDa | Cytoplasm | enzyme |
| 15551224 | 3 | 0.381 | 9612339 | 1.51E+07 | 1.73E+07 | 1.37E+07 | 1.92E+07 | 1.54E+07 | 1.43E+07 | 1.86E+07 | 1.81E+07 | 1.36E+07 | 1.23E+07 | 1.37E+07 | 3053.5037 | LEDGTEFDSSLPQNQPFVFSLGTGQVIK | P45878 | P45878 | FKBP2 | FK506 binding protein 2, 13kDa | Cytoplasm | enzyme |
| 15535662 | 2 | 0.129 | 3.13E+07 | 2.12E+07 | 2.96E+07 | 1.92E+07 | 2.47E+07 | 1.15E+07 | 2.49E+07 | 2.65E+07 | 2.28E+07 | 2.81E+07 | 2.71E+07 | 2.76E+07 | 1280.5936 | EKMWGELMEK | O35450 | O35450 | FKBPL | FK506 binding protein like | Nucleus | other |
| 15531492 | 2 | 0.466 | 4.14E+07 | 3.42E+07 | 3.62E+07 | 4.66E+07 | 2.72E+07 | 3.91E+07 | 4.29E+07 | 3.26E+07 | 3.83E+07 | 3.97E+07 | 4.39E+07 | 5.36E+07 | 1394.6384 | QDELSSSEESRK | Q2VIS4 | Q2VIS4 | Flg2 | filaggrin family member 2 | Extracellular Space | other |
| 15547696 | 3 | 0.126 | 1.20E+07 | 3.64E+07 | 7.18E+07 | 1.89E+08 | 1.79E+08 | 1.63E+08 | 9.07E+07 | 1.06E+08 | 1.95E+08 | 1.54E+07 | 1.72E+08 | 1.14E+08 | 2963.4956 | FGGEHVPNSPFQVTALAGDQPTVQTPLR | Q8BTM8 | Q8BTM8 | FLNA | filamin A, alpha | Cytoplasm | other |
| 15531803 | 3 | 0.125 | 9411645 | 2.87E+07 | 5.24E+07 | 1.33E+08 | 1.39E+08 | 1.22E+08 | 7.64E+07 | 1.01E+08 | 1.33E+08 | 1.01E+07 | 1.26E+08 | 8.80E+07 | 1818.9315 | VATVPQHATSGPGPADVSK | Q8BTM8 | Q8BTM8 | FLNA | filamin A, alpha | Cytoplasm | other |
| 15547741, 15549643 | 3, 4 | 0.179 | 8307888.5 | 2.77E+07 | 5.30E+07 | 1.64E+08 | 8.81E+07 | 1.18E+08 | 7.42E+07 | 8.97E+07 | 1.63E+08 | 9843644 | 1.41E+08 | 9.85E+07 | 3762.9758, 3762.9785 | VGSAADIPINISETDLSLLTATVVPPSGREEPC[160.0307]LLK | Q8BTM8 | Q8BTM8 | FLNA | filamin A, alpha | Cytoplasm | other |
| 15551592 | 3 | 0.176 | 7875699 | 3.03E+07 | 7.05E+07 | 1.16E+08 | 1.41E+08 | 1.55E+08 | 8.43E+07 | 1.03E+08 | 1.76E+08 | 9459023 | 1.30E+08 | 9.60E+07 | 2467.1843 | FNEEHIPDSPFVVPVASPSGDAR | Q8BTM8 | Q8BTM8 | FLNA | filamin A, alpha | Cytoplasm | other |
| 15549337, 15549689 | 2, 3 | 0.148 | 8188743 | 2.64E+07 | 5.07E+07 | 1.04E+08 | 1.18E+08 | 1.16E+08 | 6.54E+07 | 9.09E+07 | 1.66E+08 | 9211251 | 1.18E+08 | 7.63E+07 | 2338.2131, 2338.2156 | YIPVQQGPVGVNVTYGGDHIPK | Q8BTM8 | Q8BTM8 | FLNA | filamin A, alpha | Cytoplasm | other |
| 15553024 | 2 | 0.201 | 7417760.5 | 1.19E+07 | 2.35E+07 | 2.28E+07 | 3.66E+07 | 2.70E+07 | 1.87E+07 | 2.75E+07 | 4.27E+07 | 8637301 | 2.77E+07 | 2.11E+07 | 1500.7516 | DAGEGGLSLAIEGPSK | Q8BTM8 | Q8BTM8 | FLNA | filamin A, alpha | Cytoplasm | other |
| 15553315 | 2, 3 | 0.172 | 6696111 | 2.02E+07 | 4.26E+07 | 8.44E+07 | 1.05E+08 | 7.60E+07 | 4.84E+07 | 6.29E+07 | 1.04E+08 | 7586862 | 9.25E+07 | 6.07E+07 | 2313.0942, 2313.0955 | SAGQGEVLVYVEDPAGHQEEAK | Q8BTM8 | Q8BTM8 | FLNA | filamin A, alpha | Cytoplasm | other |
| 15553713, 15553769, 15558190 | 2, 3 | 0.164 | 6930818.5 | 2.13E+07 | 4.54E+07 | 7.71E+07 | 9.61E+07 | 9.83E+07 | 4.75E+07 | 6.89E+07 | 1.23E+08 | 7113750.5 | 8.02E+07 | 5.89E+07 | 2277.0288, 2277.03 | VHSPSGALEEC[160.0307]YVTEIDQDK | Q8BTM8 | Q8BTM8 | FLNA | filamin A, alpha | Cytoplasm | other |
| 15550178, 15554021 | 2, 3 | 0.107 | 5065983 | 1.30E+07 | 2.23E+07 | 5.97E+07 | 4.69E+07 | 6.05E+07 | 3.33E+07 | 4.66E+07 | 6.53E+07 | 6885863 | 6.65E+07 | 4.45E+07 | 2591.1475, 2591.1504 | ALGALVDSC[160.0307]APGLC[160.0307]PDWDSWDASK | Q8BTM8 | Q8BTM8 | FLNA | filamin A, alpha | Cytoplasm | other |
| 15551136 | 3 | 0.127 | 6057505 | 1.78E+07 | 2.34E+07 | 6.47E+07 | 7.04E+07 | 6.71E+07 | 3.56E+07 | 5.24E+07 | 8.66E+07 | 6293526.5 | 6.58E+07 | 5.56E+07 | 3579.6304 | DGSC[160.0307]SVEYIPYEAGTYSLNVTYGGHQVPGSPFK | Q8BTM8 | Q8BTM8 | FLNA | filamin A, alpha | Cytoplasm | other |
| 15550115 | 4 | 0.08 | 3412043.25 | 1.15E+07 | 2.19E+07 | 4.45E+07 | 5.04E+07 | 5.76E+07 | 2.38E+07 | 4.11E+07 | 4.67E+07 | 6039133 | 3.18E+07 | 3.40E+07 | 4282.931 | VEYTPYEEGVHSVDVTYDGSPVPSSPFQVPVTEGC[160.0307]DPSR | Q8BTM8 | Q8BTM8 | FLNA | filamin A, alpha | Cytoplasm | other |
| 15548560 | 2 | 0.131 | 6841688.5 | 1.41E+07 | 3.09E+07 | 6.25E+07 | 5.98E+07 | 6.48E+07 | 3.28E+07 | 4.33E+07 | 7.96E+07 | 6008070.5 | 5.05E+07 | 4.09E+07 | 1531.7872 | AEVGVPAEFGIWTR | Q8BTM8 | Q8BTM8 | FLNA | filamin A, alpha | Cytoplasm | other |
| 15539247 | 4 | 0.104 | 5921293 | 1.31E+07 | 1.60E+07 | 4.94E+07 | 4.20E+07 | 4.35E+07 | 3.79E+07 | 2.63E+07 | 3.96E+07 | 5860944.5 | 4.68E+07 | 3.68E+07 | 2304.149 | GQHVPGSPFQFTVGPLGEGGAHK | Q8BTM8 | Q8BTM8 | FLNA | filamin A, alpha | Cytoplasm | other |
| 15549999 | 4 | 0.083 | 3631760.25 | 1.11E+07 | 1.93E+07 | 4.67E+07 | 6.30E+07 | 6.00E+07 | 3.00E+07 | 4.64E+07 | 6.39E+07 | 5838531 | 5.39E+07 | 4.71E+07 | 4570.165 | EAMQQADDWLGIPQVITPEEIVDPNVDEHSVMTYLSQFPK | Q8BTM8 | Q8BTM8 | FLNA | filamin A, alpha | Cytoplasm | other |
| 15547736 | 3 | 0.12 | 6038695 | 1.71E+07 | 2.97E+07 | 7.55E+07 | 8.79E+07 | 8.45E+07 | 4.12E+07 | 5.60E+07 | 1.05E+08 | 5791217 | 6.98E+07 | 5.66E+07 | 2763.336 | LQVEPAVDTSGVQC[160.0307]YGPGIEGQGVFR | Q8BTM8 | Q8BTM8 | FLNA | filamin A, alpha | Cytoplasm | other |
| 15531466 | 3 | 0.147 | 4460898.5 | 1.41E+07 | 3.21E+07 | 6.77E+07 | 7.60E+07 | 6.89E+07 | 4.51E+07 | 4.97E+07 | 7.93E+07 | 5405163.5 | 7.11E+07 | 5.34E+07 | 1602.7623 | YNDQHIPGSPFTAR | Q8BTM8 | Q8BTM8 | FLNA | filamin A, alpha | Cytoplasm | other |
| 15537943 | 2 | 0.136 | 3840071.75 | 1.14E+07 | 2.42E+07 | 4.55E+07 | 5.34E+07 | 6.04E+07 | 3.40E+07 | 4.03E+07 | 6.28E+07 | 5028355 | 5.87E+07 | 3.92E+07 | 1400.6455 | YGGDEIPFSPYR | Q8BTM8 | Q8BTM8 | FLNA | filamin A, alpha | Cytoplasm | other |
| 15548041 | 2, 3 | 0.142 | 5269994.5 | 1.72E+07 | 3.13E+07 | 7.97E+07 | 8.18E+07 | 8.68E+07 | 3.78E+07 | 5.22E+07 | 8.86E+07 | 4969175 | 6.88E+07 | 5.20E+07 | 2342.218, 2342.2195 | ASGPGLNTTGVPASLPVEFTIDAK | Q8BTM8 | Q8BTM8 | FLNA | filamin A, alpha | Cytoplasm | other |
| 15531955 | 2 | 0.131 | 4616458.5 | 1.40E+07 | 2.38E+07 | 6.67E+07 | 6.11E+07 | 5.78E+07 | 3.72E+07 | 4.50E+07 | 6.16E+07 | 4883427 | 6.04E+07 | 4.57E+07 | 1185.6267 | IVSPSGAAVPC[160.0307]K | Q8BTM8 | Q8BTM8 | FLNA | filamin A, alpha | Cytoplasm | other |
| 15549798 | 2 | 0.192 | 4362324.5 | 1.19E+07 | 3.14E+07 | 4.55E+07 | 6.01E+07 | 5.98E+07 | 3.06E+07 | 4.23E+07 | 7.34E+07 | 4723682.5 | 5.73E+07 | 3.86E+07 | 1570.8043 | GAGTGGLGLAVEGPSEAK | Q8BTM8 | Q8BTM8 | FLNA | filamin A, alpha | Cytoplasm | other |
| 15550534 | 3 | 0.182 | 4543652.5 | 1.56E+07 | 2.82E+07 | 5.38E+07 | 6.82E+07 | 6.40E+07 | 2.96E+07 | 4.05E+07 | 6.45E+07 | 4710627.5 | 5.41E+07 | 4.32E+07 | 3301.554 | AEISC[160.0307]TDNQDGTC[160.0307]SVSYLPVLPGDYSILVK | Q8BTM8 | Q8BTM8 | FLNA | filamin A, alpha | Cytoplasm | other |
| 15531482 | 2 | 0.163 | 4462799.5 | 1.48E+07 | 2.53E+07 | 5.46E+07 | 6.22E+07 | 5.98E+07 | 3.55E+07 | 4.95E+07 | 7.21E+07 | 4625529.5 | 6.37E+07 | 4.48E+07 | 1290.6453 | YGGQPVPNFPSK | Q8BTM8 | Q8BTM8 | FLNA | filamin A, alpha | Cytoplasm | other |
| 15549394 | 2 | 0.141 | 5328236 | 1.29E+07 | 2.59E+07 | 5.96E+07 | 6.32E+07 | 5.95E+07 | 3.77E+07 | 4.48E+07 | 7.39E+07 | 4501861 | 5.99E+07 | 4.34E+07 | 1652.8574 | VTAQGPGLEPSGNIANK | Q8BTM8 | Q8BTM8 | FLNA | filamin A, alpha | Cytoplasm | other |
| 15548971 | 3, 4 | 0.193 | 2936745.25 | 1.95E+07 | 4.97E+07 | 6.83E+07 | 7.79E+07 | 1.12E+08 | 5.04E+07 | 6.49E+07 | 7.20E+07 | 4480270.5 | 5.65E+07 | 4.74E+07 | 3672.8428, 3672.844 | FVPREEGPYEVEVTYDGVPVPGSPFPLEAVAPTK | Q8BTM8 | Q8BTM8 | FLNA | filamin A, alpha | Cytoplasm | other |
| 15538052 | 2 | 0.128 | 4767006 | 1.13E+07 | 2.35E+07 | 6.47E+07 | 6.43E+07 | 6.24E+07 | 3.68E+07 | 4.10E+07 | 5.53E+07 | 4330596 | 6.56E+07 | 4.41E+07 | 1456.7264 | AFGPGLQGGNAGSPAR | Q8BTM8 | Q8BTM8 | FLNA | filamin A, alpha | Cytoplasm | other |
| 15550656 | 2 | 0.095 | 3338850.75 | 8885914 | 1.96E+07 | 2.51E+07 | 4.12E+07 | 3.56E+07 | 1.97E+07 | 3.41E+07 | 4.23E+07 | 4275081 | 2.09E+07 | 2.10E+07 | 1740.9214 | VTYTPMAPGSYLISIK | Q8BTM8 | Q8BTM8 | FLNA | filamin A, alpha | Cytoplasm | other |
| 15538813 | 4 | 0.368 | 7080270.5 | 5651132 | 5959309 | 3.99E+07 | 8385359.5 | 1.84E+07 | 1.13E+07 | 6485011 | 1.63E+07 | 4082188 | 4.26E+07 | 2.12E+07 | 2283.0942 | GDAVRDVDIIDHHDNTYTVK | Q8BTM8 | Q8BTM8 | FLNA | filamin A, alpha | Cytoplasm | other |
| 15547844 | 2 | 0.148 | 3493321.75 | 1.28E+07 | 3.07E+07 | 5.75E+07 | 6.38E+07 | 7.22E+07 | 3.34E+07 | 4.24E+07 | 7.59E+07 | 4043399.25 | 5.00E+07 | 3.80E+07 | 1519.7977 | SPFSVGVSPSLDLSK | Q8BTM8 | Q8BTM8 | FLNA | filamin A, alpha | Cytoplasm | other |
| 15552519 | 2, 3 | 0.113 | 3357643.75 | 1.19E+07 | 2.22E+07 | 5.26E+07 | 6.28E+07 | 5.62E+07 | 3.22E+07 | 3.67E+07 | 6.19E+07 | 3957325.25 | 4.87E+07 | 3.18E+07 | 2258.024, 2258.0247 | DGSC[160.0307]GVAYVVQEPGDYEVSVK | Q8BTM8 | Q8BTM8 | FLNA | filamin A, alpha | Cytoplasm | other |
| 15532906 | 2 | 0.168 | 4296774 | 1.14E+07 | 2.27E+07 | 5.65E+07 | 6.02E+07 | 5.45E+07 | 3.42E+07 | 4.21E+07 | 5.88E+07 | 3880023 | 6.62E+07 | 4.73E+07 | 1312.6833 | VEPGLGADNSVVR | Q8BTM8 | Q8BTM8 | FLNA | filamin A, alpha | Cytoplasm | other |
| 15533425 | 2, 3 | 0.159 | 3589309.5 | 1.04E+07 | 2.39E+07 | 5.12E+07 | 5.63E+07 | 4.86E+07 | 3.10E+07 | 3.87E+07 | 5.79E+07 | 3799392.75 | 5.71E+07 | 3.51E+07 | 1413.6512, 1413.6516 | WGDEHIPGSPYR | Q8BTM8 | Q8BTM8 | FLNA | filamin A, alpha | Cytoplasm | other |
| 15531612 | 2 | 0.14 | 3776535.75 | 1.06E+07 | 2.19E+07 | 4.86E+07 | 5.30E+07 | 5.80E+07 | 2.97E+07 | 3.78E+07 | 5.95E+07 | 3797328 | 5.13E+07 | 3.81E+07 | 1434.7196 | ANLPQSFQVDTSK | Q8BTM8 | Q8BTM8 | FLNA | filamin A, alpha | Cytoplasm | other |
| 15536803 | 2 | 0.096 | 2318531.25 | 5670139.5 | 1.11E+07 | 2.16E+07 | 2.61E+07 | 2.30E+07 | 1.57E+07 | 1.78E+07 | 2.50E+07 | 3777850 | 2.82E+07 | 1.68E+07 | 1202.5598 | AHVAPC[160.0307]FDASK | Q8BTM8 | Q8BTM8 | FLNA | filamin A, alpha | Cytoplasm | other |
| 15531744 | 3 | 0.151 | 4762313.5 | 1.38E+07 | 2.93E+07 | 3.95E+07 | 7.50E+07 | 6.24E+07 | 3.24E+07 | 4.70E+07 | 6.17E+07 | 3727083.25 | 4.41E+07 | 3.05E+07 | 1784.8429 | DVDIIDHHDNTYTVK | Q8BTM8 | Q8BTM8 | FLNA | filamin A, alpha | Cytoplasm | other |
| 15533411 | 4 | 0.166 | 3455706 | 8950494 | 1.64E+07 | 3.74E+07 | 3.19E+07 | 4.21E+07 | 2.68E+07 | 3.01E+07 | 5.08E+07 | 3522507.75 | 4.70E+07 | 3.57E+07 | 2683.3071 | C[160.0307]SYQPTMEGVHTVHVTFAGVPIPR | Q8BTM8 | Q8BTM8 | FLNA | filamin A, alpha | Cytoplasm | other |
| 15536303 | 3 | 0.071 | 3487504.5 | 7825031.5 | 9312081 | 4.75E+07 | 3.71E+07 | 3.15E+07 | 2.05E+07 | 2.54E+07 | 4.89E+07 | 3394627.25 | 3.88E+07 | 2.60E+07 | 1938.0145 | DAGEGLLAVQITDPEGKPK | Q8BTM8 | Q8BTM8 | FLNA | filamin A, alpha | Cytoplasm | other |
| 15548025 | 2, 3 | 0.122 | 2092046.625 | 1.02E+07 | 2.32E+07 | 3.23E+07 | 5.85E+07 | 5.60E+07 | 2.94E+07 | 4.06E+07 | 5.91E+07 | 3271094.75 | 3.30E+07 | 2.74E+07 | 2375.0403, 2375.0415 | VANPSGNLTDTYVQDC[160.0307]GDGTYK | Q8BTM8 | Q8BTM8 | FLNA | filamin A, alpha | Cytoplasm | other |
| 15539420 | 3 | 0.207 | 4141797.75 | 1.18E+07 | 2.50E+07 | 4.93E+07 | 5.72E+07 | 6.06E+07 | 3.05E+07 | 4.02E+07 | 6.95E+07 | 3253970.25 | 6.34E+07 | 4.07E+07 | 1750.8756 | TFSVWYVPEVTGTHK | Q8BTM8 | Q8BTM8 | FLNA | filamin A, alpha | Cytoplasm | other |
| 15531622 | 2 | 0.141 | 2411742 | 8886796 | 1.93E+07 | 3.48E+07 | 4.71E+07 | 5.02E+07 | 2.47E+07 | 3.15E+07 | 4.73E+07 | 3190125.75 | 3.87E+07 | 3.02E+07 | 1240.666 | AWGPGLEGGIVGK | Q8BTM8 | Q8BTM8 | FLNA | filamin A, alpha | Cytoplasm | other |
| 15534132 | 2 | 0.111 | 2984543.75 | 7978864.5 | 1.81E+07 | 3.73E+07 | 4.22E+07 | 4.97E+07 | 2.60E+07 | 3.07E+07 | 5.40E+07 | 3184270.25 | 3.37E+07 | 3.12E+07 | 1226.7695 | LIALLEVLSQK | Q8BTM8 | Q8BTM8 | FLNA | filamin A, alpha | Cytoplasm | other |
| 15552283 | 2 | 0.121 | 2713176 | 1.10E+07 | 2.01E+07 | 4.60E+07 | 4.59E+07 | 5.99E+07 | 2.91E+07 | 3.45E+07 | 5.55E+07 | 3089342.5 | 3.97E+07 | 2.79E+07 | 2002.0127 | IPEISIQDMTAQVTSPSGK | Q8BTM8 | Q8BTM8 | FLNA | filamin A, alpha | Cytoplasm | other |
| 15554948 | 3 | 0.173 | 2655396 | 4894799.5 | 9206002 | 1.53E+07 | 1.50E+07 | 1.41E+07 | 9887941 | 1.40E+07 | 1.73E+07 | 3006399 | 1.89E+07 | 1.25E+07 | 2349.2078 | QMQLENVSVALEFLDRESIK | Q8BTM8 | Q8BTM8 | FLNA | filamin A, alpha | Cytoplasm | other |
| 15531753 | 2 | 0.149 | 3134103.25 | 9934867 | 2.09E+07 | 4.13E+07 | 5.00E+07 | 5.90E+07 | 2.92E+07 | 3.33E+07 | 5.30E+07 | 3003828.25 | 4.40E+07 | 3.34E+07 | 1083.5337 | SPFEVYVDK | Q8BTM8 | Q8BTM8 | FLNA | filamin A, alpha | Cytoplasm | other |
| 15532483 | 2 | 0.143 | 2896429.5 | 7223908.5 | 1.69E+07 | 4.04E+07 | 3.74E+07 | 3.72E+07 | 2.00E+07 | 2.82E+07 | 4.25E+07 | 2950167.25 | 3.90E+07 | 2.62E+07 | 1099.5725 | GTVEPQLEAR | Q8BTM8 | Q8BTM8 | FLNA | filamin A, alpha | Cytoplasm | other |
| 15535021 | 2 | 0.124 | 2613597.75 | 6797673.5 | 1.26E+07 | 2.38E+07 | 3.06E+07 | 3.29E+07 | 1.55E+07 | 2.20E+07 | 3.32E+07 | 2862714.25 | 2.54E+07 | 2.11E+07 | 1426.7512 | EAGAGGLAIAVEGPSK | Q8BTM8 | Q8BTM8 | FLNA | filamin A, alpha | Cytoplasm | other |
| 15535920 | 2 | 0.182 | 3018598.5 | 1.10E+07 | 2.39E+07 | 4.68E+07 | 5.47E+07 | 6.19E+07 | 3.27E+07 | 3.93E+07 | 7.55E+07 | 2837896.75 | 5.35E+07 | 4.12E+07 | 1285.725 | LPQLPITNFSR | Q8BTM8 | Q8BTM8 | FLNA | filamin A, alpha | Cytoplasm | other |
| 15556309, 15557914 | 2, 3 | 0.197 | 2613322.25 | 1.45E+07 | 2.45E+07 | 3.73E+07 | 6.60E+07 | 6.46E+07 | 3.06E+07 | 4.23E+07 | 6.82E+07 | 2829787 | 4.46E+07 | 3.74E+07 | 2544.2488, 2544.255 | GLVEPVDVVDNADGTQTVNYVPSR | Q8BTM8 | Q8BTM8 | FLNA | filamin A, alpha | Cytoplasm | other |
| 15533606 | 2, 3 | 0.169 | 2940072 | 1.21E+07 | 2.07E+07 | 4.24E+07 | 5.72E+07 | 5.35E+07 | 2.99E+07 | 3.63E+07 | 5.65E+07 | 2771699.25 | 4.68E+07 | 3.28E+07 | 1283.7446 | VTVLFAGQHIAK | Q8BTM8 | Q8BTM8 | FLNA | filamin A, alpha | Cytoplasm | other |
| 15535556 | 3 | 0.273 | 3397710.25 | 4307256 | 6411813.5 | 9169626 | 1.15E+07 | 1.10E+07 | 8268388 | 7831432 | 1.34E+07 | 2710822 | 1.68E+07 | 9518686 | 1571.849 | RIANLQTDLSDGLR | Q8BTM8 | Q8BTM8 | FLNA | filamin A, alpha | Cytoplasm | other |
| 15536372 | 2 | 0.132 | 2701417.25 | 7864500 | 1.74E+07 | 4.37E+07 | 4.02E+07 | 4.42E+07 | 2.42E+07 | 3.06E+07 | 4.46E+07 | 2683673.75 | 4.03E+07 | 2.67E+07 | 1072.5609 | VAQPSITDNK | Q8BTM8 | Q8BTM8 | FLNA | filamin A, alpha | Cytoplasm | other |
| 15548474 | 2 | 0.163 | 2839608.5 | 5782636 | 1.23E+07 | 2.87E+07 | 3.02E+07 | 3.20E+07 | 1.59E+07 | 2.20E+07 | 4.06E+07 | 2639415.5 | 4.33E+07 | 2.19E+07 | 1851.843 | SPYTVTVGQAC[160.0307]NPAAC[160.0307]R | Q8BTM8 | Q8BTM8 | FLNA | filamin A, alpha | Cytoplasm | other |
| 15532711 | 2 | 0.103 | 2310479.25 | 7373804.5 | 1.35E+07 | 3.58E+07 | 4.53E+07 | 3.70E+07 | 2.56E+07 | 2.75E+07 | 3.45E+07 | 2625822.75 | 3.63E+07 | 2.58E+07 | 1433.7069 | AYGPGIEPTGNMVK | Q8BTM8 | Q8BTM8 | FLNA | filamin A, alpha | Cytoplasm | other |
| 15532495 | 3 | 0.17 | 3125414.75 | 9712438 | 1.63E+07 | 3.68E+07 | 4.36E+07 | 5.09E+07 | 2.76E+07 | 3.51E+07 | 5.03E+07 | 2592097.75 | 4.79E+07 | 3.55E+07 | 2200.1206 | LVSNHSLHETSSVFVDSLTK | Q8BTM8 | Q8BTM8 | FLNA | filamin A, alpha | Cytoplasm | other |
| 15531831 | 2 | 0.161 | 2387269.75 | 6818749 | 1.77E+07 | 2.94E+07 | 3.05E+07 | 3.09E+07 | 2.11E+07 | 2.47E+07 | 4.35E+07 | 2562692 | 2.74E+07 | 2.26E+07 | 1077.5918 | LDVQFSGLAK | Q8BTM8 | Q8BTM8 | FLNA | filamin A, alpha | Cytoplasm | other |
| 15532920 | 2 | 0.145 | 2127148.25 | 7879626 | 1.48E+07 | 3.89E+07 | 3.59E+07 | 3.79E+07 | 2.16E+07 | 3.09E+07 | 4.84E+07 | 2557403.5 | 4.00E+07 | 2.89E+07 | 1088.5636 | TPC[160.0307]EEILVK | Q8BTM8 | Q8BTM8 | FLNA | filamin A, alpha | Cytoplasm | other |
| 15536497 | 2, 3 | 0.231 | 3283776.25 | 1.04E+07 | 2.27E+07 | 3.67E+07 | 4.97E+07 | 5.11E+07 | 2.80E+07 | 3.68E+07 | 5.71E+07 | 2483189.25 | 5.24E+07 | 3.40E+07 | 1379.6714, 1379.6721 | YGGPYHIGGSPFK | Q8BTM8 | Q8BTM8 | FLNA | filamin A, alpha | Cytoplasm | other |
| 15531906 | 2, 3 | 0.142 | 2374178.25 | 7210127 | 1.21E+07 | 2.98E+07 | 3.36E+07 | 4.20E+07 | 2.27E+07 | 2.41E+07 | 3.86E+07 | 2471771.25 | 4.25E+07 | 2.69E+07 | 1434.7607, 1434.7611 | AGNNMLLVGVHGPR | Q8BTM8 | Q8BTM8 | FLNA | filamin A, alpha | Cytoplasm | other |
| 15550083 | 3 | 0.178 | 2610192.5 | 7405428.5 | 1.59E+07 | 3.02E+07 | 4.22E+07 | 3.86E+07 | 1.40E+07 | 2.27E+07 | 4.84E+07 | 2437103.75 | 3.21E+07 | 2.36E+07 | 3079.4612 | C[160.0307]APGVVGPTEADIDFDIIRNDNDTFTVK | Q8BTM8 | Q8BTM8 | FLNA | filamin A, alpha | Cytoplasm | other |
| 15538567 | 2 | 0.063 | 1632656 | 2509353.75 | 4306797.5 | 1.04E+07 | 1.23E+07 | 9579644 | 5819764.5 | 7428872.5 | 1.32E+07 | 2374957 | 1.32E+07 | 9059774 | 809.43195 | YVIC[160.0307]VR | Q8BTM8 | Q8BTM8 | FLNA | filamin A, alpha | Cytoplasm | other |
| 15543030 | 4 | 0.182 | 679959 | 1571297.5 | 1176149 | 6458924 | 1309278.25 | 6818596.5 | 3454357 | 1089382.625 | 8282275.5 | 2170188.75 | 1.06E+07 | 6454747 | 2623.2827 | FNEEHIPDSPFVVPVASPSGDARR | Q8BTM8 | Q8BTM8 | FLNA | filamin A, alpha | Cytoplasm | other |
| 15532133 | 2 | 0.085 | 1141304.625 | 5654573 | 1.12E+07 | 1.73E+07 | 2.67E+07 | 2.64E+07 | 1.56E+07 | 1.82E+07 | 2.35E+07 | 2158629.5 | 1.29E+07 | 1.21E+07 | 977.49207 | PAEFTVDAK | Q8BTM8 | Q8BTM8 | FLNA | filamin A, alpha | Cytoplasm | other |
| 15537540 | 3 | 0.188 | 2164274.25 | 7556172 | 1.27E+07 | 2.43E+07 | 3.05E+07 | 3.08E+07 | 1.89E+07 | 2.32E+07 | 3.35E+07 | 2048597.625 | 3.07E+07 | 2.27E+07 | 2057.929 | THEAEIVEGENHTYC[160.0307]IR | Q8BTM8 | Q8BTM8 | FLNA | filamin A, alpha | Cytoplasm | other |
| 15531694 | 2 | 0.106 | 1476106.125 | 4353524 | 6638347 | 1.96E+07 | 1.94E+07 | 1.81E+07 | 1.18E+07 | 1.37E+07 | 2.12E+07 | 2007240.25 | 2.29E+07 | 1.52E+07 | 1217.5736 | C[160.0307]SGPGLSPGMVR | Q8BTM8 | Q8BTM8 | FLNA | filamin A, alpha | Cytoplasm | other |
| 15534668 | 2 | 0.114 | 2269663.75 | 3260896.75 | 6482734 | 1.12E+07 | 1.32E+07 | 1.41E+07 | 8779911 | 9807295 | 1.56E+07 | 1989154.375 | 1.32E+07 | 1.01E+07 | 1094.5454 | AEISFEDRK | Q8BTM8 | Q8BTM8 | FLNA | filamin A, alpha | Cytoplasm | other |
| 15533873 | 2 | 0.07 | 1693465.875 | 4448323.5 | 8750857 | 2.98E+07 | 2.50E+07 | 2.69E+07 | 1.39E+07 | 2.03E+07 | 2.74E+07 | 1964204.5 | 1.85E+07 | 1.93E+07 | 1415.7467 | IANLQTDLSDGLR | Q8BTM8 | Q8BTM8 | FLNA | filamin A, alpha | Cytoplasm | other |
| 15532578 | 2 | 0.134 | 1756012 | 5189718 | 1.13E+07 | 2.98E+07 | 2.49E+07 | 2.72E+07 | 1.91E+07 | 1.77E+07 | 2.99E+07 | 1961437.25 | 2.73E+07 | 2.06E+07 | 1049.5421 | LSPFMADIR | Q8BTM8 | Q8BTM8 | FLNA | filamin A, alpha | Cytoplasm | other |
| 15533049 | 2 | 0.129 | 1794177.25 | 5161537 | 1.21E+07 | 2.68E+07 | 3.18E+07 | 3.23E+07 | 1.55E+07 | 2.28E+07 | 3.37E+07 | 1948855.5 | 2.96E+07 | 1.94E+07 | 882.53925 | AGVAPLQVK | Q8BTM8 | Q8BTM8 | FLNA | filamin A, alpha | Cytoplasm | other |
| 15550333 | 3 | 0.164 | 940415.75 | 6353878.5 | 1.51E+07 | 1.21E+07 | 4.48E+07 | 2.43E+07 | 1.31E+07 | 2.71E+07 | 3.13E+07 | 1886388 | 1.16E+07 | 1.36E+07 | 2893.5457 | VGSAADIPINISETDLSLLTATVVPPSGR | Q8BTM8 | Q8BTM8 | FLNA | filamin A, alpha | Cytoplasm | other |
| 15532592 | 2, 3 | 0.149 | 1294035.25 | 3460860.5 | 7440143.5 | 1.82E+07 | 1.33E+07 | 1.85E+07 | 8991030 | 1.11E+07 | 2.11E+07 | 1783800.5 | 1.91E+07 | 1.39E+07 | 1417.7966, 1417.7977 | RLTVSSLQESGLK | Q8BTM8 | Q8BTM8 | FLNA | filamin A, alpha | Cytoplasm | other |
| 15534045 | 3 | 0.121 | 1891102.125 | 5461874 | 8187942.5 | 2.96E+07 | 2.01E+07 | 2.48E+07 | 1.44E+07 | 1.79E+07 | 2.91E+07 | 1695998.375 | 2.58E+07 | 1.51E+07 | 1942.9586 | HTAMVSWGGVSIPNSPFR | Q8BTM8 | Q8BTM8 | FLNA | filamin A, alpha | Cytoplasm | other |
| 15556110 | 2 | 0.153 | 2265704.25 | 6127478.5 | 1.46E+07 | 2.15E+07 | 2.75E+07 | 2.28E+07 | 1.38E+07 | 1.85E+07 | 2.62E+07 | 1694507.75 | 1.60E+07 | 1.30E+07 | 1913.9072 | EGSYSISVLYGEEEVPR | Q8BTM8 | Q8BTM8 | FLNA | filamin A, alpha | Cytoplasm | other |
| 15543972 | 2 | 0.129 | 1269740.625 | 4014668.5 | 9431693 | 2.19E+07 | 2.25E+07 | 2.56E+07 | 1.39E+07 | 1.72E+07 | 2.80E+07 | 1607489.125 | 2.70E+07 | 1.47E+07 | 1007.5234 | IQQNTFTR | Q8BTM8 | Q8BTM8 | FLNA | filamin A, alpha | Cytoplasm | other |
| 15550657 | 2 | 0.111 | 1411038 | 4336924 | 1.09E+07 | 2.58E+07 | 2.75E+07 | 2.82E+07 | 1.46E+07 | 1.85E+07 | 3.47E+07 | 1606681 | 2.30E+07 | 1.57E+07 | 1619.7418 | C[160.0307]GQSAAVASPGGSIDSR | Q8BTM8 | Q8BTM8 | FLNA | filamin A, alpha | Cytoplasm | other |
| 15541563 | 2 | 0.178 | 1351322 | 5656286.5 | 9382313 | 2.18E+07 | 2.10E+07 | 2.55E+07 | 1.36E+07 | 1.53E+07 | 2.58E+07 | 1547657.875 | 2.25E+07 | 1.77E+07 | 1085.6213 | LYSVSYLLK | Q8BTM8 | Q8BTM8 | FLNA | filamin A, alpha | Cytoplasm | other |

| | | | | | | | | | | | | | | | | | | | | | | |
|---|---|---|---|---|---|---|---|---|---|---|---|---|---|---|---|---|---|---|---|---|---|---|
| 15534499 | 3 | 0.224 | 2203724.25 | 4153083 | 3169018.5 | 5582868 | 8500602 | 1.18E+07 | 5105681.5 | 6105187.5 | 1.28E+07 | 1505381.375 | 7597742 | 8024910 | 2002.0146 | IPEISIQDMTAQVTSPSGK | Q8BTM8 | Q8BTM8 | FLNA | filamin A, alpha | Cytoplasm | other |
| 15542313 | 2 | 0.126 | 1292963.75 | 3649876 | 5626601.5 | 1.88E+07 | 1.63E+07 | 1.53E+07 | 1.02E+07 | 1.04E+07 | 1.18E+07 | 1424131.875 | 1.87E+07 | 1.28E+07 | 1225.5667 | EATTEFSVDAR | Q8BTM8 | Q8BTM8 | FLNA | filamin A, alpha | Cytoplasm | other |
| 15531961 | 3 | 0.125 | 2097491.25 | 3266641.75 | 2981594.75 | 2.51E+07 | 1.54E+07 | 1.45E+07 | 1.00E+07 | 1.21E+07 | 2.69E+07 | 1334345.25 | 3.16E+07 | 1.84E+07 | 1698.9133 | TGVELGKPTHFTVNAK | Q8BTM8 | Q8BTM8 | FLNA | filamin A, alpha | Cytoplasm | other |
| 15535457 | 3 | 0.093 | 689147 | 2263972.75 | 3452730.25 | 8444284 | 8203471 | 1.07E+07 | 5241113 | 6971147.5 | 1.23E+07 | 1312421.625 | 1.07E+07 | 7318997.5 | 1301.6606 | FNGTHIPGSPFK | Q8BTM8 | Q8BTM8 | FLNA | filamin A, alpha | Cytoplasm | other |
| 15535444 | 2 | 0.114 | 980944.438 | 2556041.25 | 5669117 | 9945540 | 1.34E+07 | 1.36E+07 | 6724483 | 1.04E+07 | 1.38E+07 | 1230048.75 | 1.10E+07 | 8589365 | 1261.697 | LTVSSLQESGLK | Q8BTM8 | Q8BTM8 | FLNA | filamin A, alpha | Cytoplasm | other |
| 15541294, 15547158 | 2 | 0.192 | 1887065.375 | 4651034.5 | 8775988 | 2.15E+07 | 2.01E+07 | 2.33E+07 | 1.34E+07 | 1.31E+07 | 2.36E+07 | 1226967.75 | 2.32E+07 | 1.52E+07 | 1108.5703, 1108.5714 | RAEFTVETR | Q8BTM8 | Q8BTM8 | FLNA | filamin A, alpha | Cytoplasm | other |
| 15534518 | 2 | 0.197 | 1200856.875 | 3735347.25 | 8239650 | 1.73E+07 | 1.53E+07 | 1.66E+07 | 1.13E+07 | 1.35E+07 | 2.57E+07 | 1172162.625 | 1.94E+07 | 1.43E+07 | 971.56494 | LLGWIQNK | Q8BTM8 | Q8BTM8 | FLNA | filamin A, alpha | Cytoplasm | other |
| 15554174 | 3 | 0.134 | 877400.625 | 2770690.25 | 5455834.5 | 1.30E+07 | 1.19E+07 | 1.65E+07 | 7080858 | 9762810 | 1.96E+07 | 1149439.375 | 1.38E+07 | 1.06E+07 | 2683.3083 | C[160.0307]SYQPTMEGVHTVHVTFAGVPIPR | Q8BTM8 | Q8BTM8 | FLNA | filamin A, alpha | Cytoplasm | other |
| 15553818 | 3 | 0.204 | 1730431.625 | 3893946.5 | 8667091 | 2.76E+07 | 2.11E+07 | 1.76E+07 | 1.11E+07 | 1.40E+07 | 2.26E+07 | 1145488.5 | 2.68E+07 | 1.46E+07 | 2455.1592 | THIQDNHDGTYTVAYVPDVPGR | Q8BTM8 | Q8BTM8 | FLNA | filamin A, alpha | Cytoplasm | other |
| 15534428 | 2 | 0.154 | 1831257.375 | 3191817.75 | 4728370.5 | 1.75E+07 | 1.64E+07 | 1.66E+07 | 9475257 | 1.18E+07 | 2.01E+07 | 1144258.5 | 2.54E+07 | 1.31E+07 | 1172.592 | DLAEDAPWKK | Q8BTM8 | Q8BTM8 | FLNA | filamin A, alpha | Cytoplasm | other |
| 15533290 | 3 | 0.139 | 1936451.5 | 1488868.5 | 16576.775 | 2.62E+07 | 4396680.5 | 4509579.5 | 4773600 | 5447453.5 | 2.10E+07 | 1118515 | 3.24E+07 | 1.85E+07 | 1646.8744 | TGVAVNKPAEFTVDAK | Q8BTM8 | Q8BTM8 | FLNA | filamin A, alpha | Cytoplasm | other |
| 15531650 | 2 | 0.176 | 819910.812 | 3444416.75 | 7274290 | 1.37E+07 | 1.40E+07 | 1.75E+07 | 8417560 | 1.14E+07 | 1.97E+07 | 1113352.875 | 1.53E+07 | 1.05E+07 | 750.47437 | YTILIK | Q8BTM8 | Q8BTM8 | FLNA | filamin A, alpha | Cytoplasm | other |
| 15550751 | 3 | 0.023 | 557614 | 2076722.5 | 888693.812 | 8687348 | 8571790 | 5613251.5 | 3807670 | 7082790.5 | 1.21E+07 | 1110210.375 | 6170060 | 6474500.5 | 3485.726 | EEGPYEVEVTYDGVPVPGSPFPLEAVAPTKPSK | Q8BTM8 | Q8BTM8 | FLNA | filamin A, alpha | Cytoplasm | other |
| 15533155 | 2 | 0.026 | 461572 | 842036.875 | 1782396 | 2366147.75 | 5494977.5 | 4113187 | 2446745.5 | 3101628 | 3687513.75 | 1053441.5 | 2747633.25 | 2127383 | 1211.541 | EAPQDFHPDR | Q8BTM8 | Q8BTM8 | FLNA | filamin A, alpha | Cytoplasm | other |
| 15541855 | 3 | 0.11 | 764718.312 | 1900757 | 4553417 | 1.04E+07 | 9863384 | 1.00E+07 | 6781572.5 | 7206076.5 | 8770916 | 1030713.188 | 1.11E+07 | 6564857 | 1824.9343 | RAPSVANIGSHC[160.0307]DLSLK | Q8BTM8 | Q8BTM8 | FLNA | filamin A, alpha | Cytoplasm | other |
| 15532979 | 2, 3 | 0.456 | 1857948.75 | 9545821 | 2.36E+07 | 1.37E+07 | 5.22E+07 | 4.71E+07 | 1.45E+07 | 3.67E+07 | 5.65E+07 | 851893.812 | 3.75E+07 | 3.04E+07 | 1167.598, 1167.5985 | VPVHDVTDASK | Q8BTM8 | Q8BTM8 | FLNA | filamin A, alpha | Cytoplasm | other |
| 15549467 | 3 | 0.412 | 1009988.312 | 1497851.5 | 2407108.5 | 1.58E+07 | 2145966.5 | 8220601.5 | 5034928.5 | 2074704.25 | 1.39E+07 | 849375.812 | 2.20E+07 | 1.02E+07 | 2373.1584 | EGSYSISVLYGEEEVPRSPFK | Q8BTM8 | Q8BTM8 | FLNA | filamin A, alpha | Cytoplasm | other |
| 15544800, 15545459 | 2 | 0.155 | 681846.562 | 2420094.5 | 6033957.5 | 9838715 | 1.36E+07 | 1.43E+07 | 6524545.5 | 9346997 | 1.52E+07 | 821291.438 | 1.07E+07 | 7972971 | 866.4235, 866.42377 | DQEFTVK | Q8BTM8 | Q8BTM8 | FLNA | filamin A, alpha | Cytoplasm | other |
| 15535668 | 2 | 0.117 | 849832.062 | 3603248 | 7472041.5 | 1.36E+07 | 2.12E+07 | 1.70E+07 | 1.11E+07 | 1.49E+07 | 1.73E+07 | 820339.438 | 1.16E+07 | 8736992 | 1444.5266 | MDC[160.0307]QEC[160.0307]PEGYR | Q8BTM8 | Q8BTM8 | FLNA | filamin A, alpha | Cytoplasm | other |
| 15548804 | 2 | 0.156 | 851996.938 | 2063743.5 | 4902915 | 1.19E+07 | 1.04E+07 | 9852565 | 6417995.5 | 6884066.5 | 1.03E+07 | 807817.562 | 1.17E+07 | 6644630 | 1602.7627 | YNDQHIPGSPFTAR | Q8BTM8 | Q8BTM8 | FLNA | filamin A, alpha | Cytoplasm | other |
| 15548580 | 2 | 0.126 | 560566.438 | 1812471.5 | 4432254.5 | 9364723 | 1.17E+07 | 9512667 | 7035503.5 | 7231353 | 1.27E+07 | 796509.188 | 1.18E+07 | 6627699 | 1818.932 | VATVPQHATSGPGPADVSK | Q8BTM8 | Q8BTM8 | FLNA | filamin A, alpha | Cytoplasm | other |
| 15541933 | 2 | 0.155 | 767879.062 | 2564646.5 | 5910187 | 1.21E+07 | 1.38E+07 | 1.39E+07 | 7005274.5 | 1.03E+07 | 1.64E+07 | 792033.812 | 1.22E+07 | 8914647 | 724.3867 | FTIDTK | Q8BTM8 | Q8BTM8 | FLNA | filamin A, alpha | Cytoplasm | other |
| 15533612 | 2 | 0.101 | 687490.5 | 3045432.25 | 5532221.5 | 1.07E+07 | 1.46E+07 | 1.45E+07 | 8286227 | 1.15E+07 | 1.50E+07 | 789855.875 | 8475709 | 7322846 | 908.5082 | GEYTLVVK | Q8BTM8 | Q8BTM8 | FLNA | filamin A, alpha | Cytoplasm | other |
| 15540067 | 2 | 0.1 | 698529.812 | 1937378.5 | 3186911.5 | 1.14E+07 | 1.02E+07 | 1.07E+07 | 5494081 | 7641672.5 | 1.45E+07 | 743550.375 | 1.12E+07 | 7404414.5 | 1163.6283 | ENGIYLIDVK | Q8BTM8 | Q8BTM8 | FLNA | filamin A, alpha | Cytoplasm | other |
| 15542920 | 4 | 0.036 | 250405.531 | 1374857.5 | 937940.188 | 6855021 | 6570192.5 | 6347064.5 | 2905946.5 | 3797036.25 | 6071879.5 | 731578.188 | 7583403 | 4422258 | 2963.4946 | FGGEHVPNSPFQVTALAGDQPTVQTPLR | Q8BTM8 | Q8BTM8 | FLNA | filamin A, alpha | Cytoplasm | other |
| 15550767 | 2 | 0.099 | 704798.062 | 1416300.875 | 2904636 | 1.11E+07 | 8991531 | 8966249 | 5155728.5 | 5771355 | 1.20E+07 | 723647.188 | 1.13E+07 | 7219337 | 1938.0139 | DAGEGLLAVQITDPEGKPK | Q8BTM8 | Q8BTM8 | FLNA | filamin A, alpha | Cytoplasm | other |
| 15554358 | 3 | 0.373 | 1290898.625 | 2326673.75 | 3829527.25 | 2.57E+07 | 5279112 | 1.46E+07 | 8556511 | 5303193.5 | 2.16E+07 | 631688.375 | 2.76E+07 | 2.00E+07 | 3166.4282 | MDC[160.0307]QEC[160.0307]PEGYRVTYTPMAPGSYLISIK | Q8BTM8 | Q8BTM8 | FLNA | filamin A, alpha | Cytoplasm | other |
| 15553283 | 2 | 0.141 | 469273.094 | 1815936 | 3463413.5 | 5926830 | 1.01E+07 | 9103691 | 4768259.5 | 6667727.5 | 1.02E+07 | 498597.156 | 6670632.5 | 5386828 | 1604.7336 | YSPSEAGLHEMDIR | Q8BTM8 | Q8BTM8 | FLNA | filamin A, alpha | Cytoplasm | other |
| 15548718 | 2 | 0.192 | 362721.031 | 1300005.25 | 3452213 | 6595419.5 | 6833299 | 7337291 | 4587493 | 4818678 | 8016053 | 403488.312 | 7721404 | 5131846 | 1532.7567 | FVPAEMGMHTVSVK | Q8BTM8 | Q8BTM8 | FLNA | filamin A, alpha | Cytoplasm | other |
| 15552321 | 2 | 0.031 | 404653.125 | 511180.219 | 777598.375 | 3495952 | 1922689 | 2779197.25 | 1943058.375 | 1552906.25 | 3215172.25 | 365720.031 | 2188495.75 | 1561003 | 1515.7567 | FADQHVPGSPFSVK | Q8BTM8 | Q8BTM8 | FLNA | filamin A, alpha | Cytoplasm | other |
| 15556385 | 2 | 0.363 | 373911.25 | 2236975 | 4438797 | 3199244.25 | 9809297 | 1.11E+07 | 5247616 | 7557531 | 1.38E+07 | 300436.125 | 1.05E+07 | 6883790.5 | 1750.8751 | TFSVWYVPEVTGTHK | Q8BTM8 | Q8BTM8 | FLNA | filamin A, alpha | Cytoplasm | other |
| 15554401 | 2 | 0.342 | 485310.781 | 828543.062 | 2624145 | 4576484.5 | 4632403.5 | 3542966.25 | 3193960.75 | 3241321.75 | 6282628 | 300161.062 | 8439203 | 4622419 | 1571.8469 | RIANLQTDLSDGLR | Q8BTM8 | Q8BTM8 | FLNA | filamin A, alpha | Cytoplasm | other |
| 15549107 | 2 | 0.242 | 474188.844 | 3053113 | 1.14E+07 | 1.01E+07 | 1.85E+07 | 2.11E+07 | 8612854 | 1.41E+07 | 1.61E+07 | 228988.672 | 9323789 | 6952675 | 1712.8652 | DAGEGLLAVQITDPEGK | Q8BTM8 | Q8BTM8 | FLNA | filamin A, alpha | Cytoplasm | other |
| 15551549 | 2 | 0.135 | 264187.906 | 844448.25 | 1671962.25 | 2715728.75 | 5182097 | 4058103.75 | 2794419.25 | 3247530 | 4738050.5 | 183247.547 | 3342092 | 2004502.75 | 1784.8419 | DVDIIDHHDNTYTVK | Q8BTM8 | Q8BTM8 | FLNA | filamin A, alpha | Cytoplasm | other |
| 15542485 | 3 | 0.233 | 422385.281 | 1264802.375 | 1750990.5 | 6037983.5 | 5122051.5 | 5201656.5 | 4516761.5 | 4068048.25 | 9323144 | 151313.672 | 6568011.5 | 4898509.5 | 1652.8563 | VTAQGPGLEPSGNIANK | Q8BTM8 | Q8BTM8 | FLNA | filamin A, alpha | Cytoplasm | other |
| 15550928 | 2 | 0.201 | 304504.156 | 189500.828 | 70583.602 | 5680504.5 | 983862.5 | 661331.125 | 814450.312 | 641484.812 | 4610092.5 | 137004.922 | 8005372.5 | 3340352.25 | 1646.8722 | TGVAVNKPAEFTVDAK | Q8BTM8 | Q8BTM8 | FLNA | filamin A, alpha | Cytoplasm | other |
| 15551357 | 2 | 0.027 | 139755.75 | 173543.047 | 51387.816 | 2670514 | 1378881.875 | 1563249.5 | 970286.75 | 1114468.25 | 2763265.5 | 136050.391 | 3310451.75 | 2053297.375 | 1698.9133 | TGVELGKPTHFTVNAK | Q8BTM8 | Q8BTM8 | FLNA | filamin A, alpha | Cytoplasm | other |
| 15535952 | 3 | 0.464 | 302418.156 | 273781.75 | 412385.125 | 8142496 | 363274.375 | 3333562.5 | 1317808 | 583982 | 3930263.25 | 109188.602 | 1.04E+07 | 4534690 | 1495.8341 | VRAWGPGLEGGIVGK | Q8BTM8 | Q8BTM8 | FLNA | filamin A, alpha | Cytoplasm | other |
| 15553122 | 2 | 0.214 | 0 | 635451.812 | 1618121.375 | 2122227.75 | 5631922 | 4290337 | 1417262.25 | 2747709.5 | 3664993.75 | 102014.195 | 1844494.875 | 843297.562 | 2044.9967 | C[160.0307]APGVVGPTEADIDFDIIR | Q8BTM8 | Q8BTM8 | FLNA | filamin A, alpha | Cytoplasm | other |
| 15539719 | 3 | 0.102 | 112183.773 | 293373.625 | 1038248.5 | 3796576.25 | 2024127.75 | 3402911.5 | 5797085.5 | 2321405.75 | 4157917.5 | 101070.602 | 3474779.75 | 2461037.75 | 1154.6289 | NGHVGISFVPK | Q8BTM8 | Q8BTM8 | FLNA | filamin A, alpha | Cytoplasm | other |
| 15554990 | 2 | 0.122 | 138137.609 | 488400 | 318736.656 | 4554810 | 2202861.75 | 2276152 | 1438144.875 | 1289297.125 | 3084718.75 | 77218.297 | 3473050.5 | 1541746.375 | 1942.9548 | HTAMVSWGGVSIPNSPFR | Q8BTM8 | Q8BTM8 | FLNA | filamin A, alpha | Cytoplasm | other |
| 15556168 | 2 | 0.249 | 226909.984 | 306678.438 | 2989605.75 | 2431862.75 | 3108589.75 | 4698046 | 1570613.375 | 1806655.25 | 5926239 | 57680.559 | 1498596.75 | 1906109.25 | 1636.7081 | VDINTEDLEDGTC[160.0307]R | Q8BTM8 | Q8BTM8 | FLNA | filamin A, alpha | Cytoplasm | other |
| 15546565 | 3 | 0.364 | 4.70E-06 | 243657.016 | 1398398.875 | 733752.062 | 1533521.875 | 2850267.5 | 1028783.5 | 1174781.625 | 1475107 | 4.70E-06 | 757350.25 | 777595.625 | 1550.8358 | VRVHGPGIQSGTTNK | Q8BTM8 | Q8BTM8 | FLNA | filamin A, alpha | Cytoplasm | other |
| 15532686 | 2 | 0.482 | 141780.688 | 223230.797 | 192694.703 | 2556491.5 | 883214.438 | 1808486.125 | 1524430 | 767407.188 | 2025930.5 | 4.70E-06 | 3932205 | 1857409.25 | 1438.7053 | EAPQDFHPDRVK | Q8BTM8 | Q8BTM8 | FLNA | filamin A, alpha | Cytoplasm | other |
| 15535106 | 2 | 0.26 | 1.74E+07 | 1.97E+07 | 2.16E+07 | 1.41E+07 | 1.41E+07 | 1.71E+07 | 1.49E+07 | 1.79E+07 | 1.85E+07 | 2.24E+07 | 1.51E+07 | 1.60E+07 | 1201.6764 | VTGEVLDILSR | O08917 | O08917 | FLOT1 | flotillin 1 | Plasma Membrane | other |
| 15533986 | 3 | 0.547 | 2.04E+07 | 1.99E+07 | 2.11E+07 | 1.86E+07 | 2.08E+07 | 1.91E+07 | 2.14E+07 | 2.18E+07 | 1.88E+07 | 2.11E+07 | 2.11E+07 | 1.98E+07 | 1655.7618 | DIHDDQDYLHSLGK | O08917 | O08917 | FLOT1 | flotillin 1 | Plasma Membrane | other |
| 15538849 | 3 | 0.854 | 2.10E+07 | 1.33E+07 | 1.07E+07 | 1.89E+07 | 1.23E+07 | 1.01E+07 | 1.99E+07 | 1.42E+07 | 9581453 | 1.84E+07 | 2.02E+07 | 1.22E+07 | 2018.0259 | TEAEIAHIALETLEGHQR | O08917 | O08917 | FLOT1 | flotillin 1 | Plasma Membrane | other |
| 15557820, 15558037 | 2 | 0.877 | 1.20E+07 | 1.07E+07 | 1.29E+07 | 1.40E+07 | 1.13E+07 | 1.27E+07 | 1.22E+07 | 1.25E+07 | 1.14E+07 | 1.40E+07 | 1.16E+07 | 1.05E+07 | 1753.928, 1753.9282 | LPQVAEEISGPLTSANK | O08917 | O08917 | FLOT1 | flotillin 1 | Plasma Membrane | other |
| 15533175 | 2 | 0.282 | 1.14E+07 | 1.31E+07 | 1.23E+07 | 9996720 | 1.04E+07 | 1.05E+07 | 1.17E+07 | 1.26E+07 | 9786500 | 1.38E+07 | 1.01E+07 | 1.20E+07 | 1465.699 | VSAQC[160.0307]LSEIEMAK | O08917 | O08917 | FLOT1 | flotillin 1 | Plasma Membrane | other |
| 15533627 | 3 | 0.93 | 9814656 | 7739385 | 1.22E+07 | 1.02E+07 | 9919506 | 8500205 | 9779911 | 1.10E+07 | 1.04E+07 | 1.35E+07 | 1.00E+07 | 7953746 | 1983.0436 | VASSDLVNMGISVVSYTLK | O08917 | O08917 | FLOT1 | flotillin 1 | Plasma Membrane | other |
| 15554072 | 3 | 0.579 | 1.34E+07 | 1.28E+07 | 7946089.5 | 1.36E+07 | 1.02E+07 | 7668915 | 1.19E+07 | 1.04E+07 | 1.01E+07 | 1.25E+07 | 1.44E+07 | 1.20E+07 | 2357.1475 | KAEAFQMYQEAAQLDMLLEK | O08917 | O08917 | FLOT1 | flotillin 1 | Plasma Membrane | other |
| 15534835 | 3 | 0.105 | 1.17E+07 | 9172461 | 7232550 | 1.05E+07 | 5515052 | 7670119 | 8212766 | 7072836.5 | 8842371 | 1.17E+07 | 1.33E+07 | 1.15E+07 | 1625.869 | AQQVAVQEQEIARR | O08917 | O08917 | FLOT1 | flotillin 1 | Plasma Membrane | other |
| 15532167 | 2 | 0.264 | 9872422 | 1.18E+07 | 1.39E+07 | 1.03E+07 | 1.10E+07 | 1.05E+07 | 1.12E+07 | 1.01E+07 | 1.06E+07 | 9220069 | 1.02E+07 | 1.04E+07 | 1215.73 | ISLNTLTLNVK | O08917 | O08917 | FLOT1 | flotillin 1 | Plasma Membrane | other |
| 15548621 | 2 | 0.267 | 6578184 | 5931455 | 5844972 | 5575284.5 | 6061116 | 5628352 | 5887187 | 6321225 | 7463890.5 | 7438660.5 | 6668173 | 5950272 | 1500.8333 | VFVLPC[160.0307]IQQIQR | O08917 | O08917 | FLOT1 | flotillin 1 | Plasma Membrane | other |
| 15534209 | 2 | 0.33 | 6640149 | 6893268 | 4578861 | 5678699.5 | 6669003.5 | 5734712.5 | 7518849.5 | 6376173 | 6192043.5 | 6990486 | 9472139 | 6473166.5 | 1469.6921 | EMLAAAC[160.0307]QMFLGK | O08917 | O08917 | FLOT1 | flotillin 1 | Plasma Membrane | other |
| 15538680 | 2 | 0.753 | 3673944.25 | 7232791.5 | 8953351 | 3338006 | 7541102 | 6427733 | 5619216 | 8654317 | 5711432.5 | 6166481.5 | 3884063.5 | 4805225.5 | 1469.7676 | AQQVAVQEQEIAR | O08917 | O08917 | FLOT1 | flotillin 1 | Plasma Membrane | other |
| 15537488 | 2 | 0.98 | 3157245.25 | 3649430 | 4119252.75 | 3194886 | 3684352.75 | 3321693.75 | 3233084.5 | 4246259.5 | 3206669.5 | 4819165.5 | 2583634.75 | 3555758.75 | 1181.5775 | ATYDIEVNTR | O08917 | O08917 | FLOT1 | flotillin 1 | Plasma Membrane | other |
| 15540639 | 3 | 0.018 | 4867122.5 | 6379538 | 6085590.5 | 4740157.5 | 5513286.5 | 4575770.5 | 4698833 | 6649361 | 5768497.5 | 3949401.25 | 3129440.75 | 4080589 | 1396.7505 | LTGVSISQVNHNK | O08917 | O08917 | FLOT1 | flotillin 1 | Plasma Membrane | other |
| 15534401 | 2 | 0.764 | 3111493 | 3455968.25 | 5584983 | 2182096.5 | 3484667.75 | 4551225.5 | 3314239.5 | 4084791.25 | 4094076.5 | 3063553.5 | 3303935.75 | 3711079 | 829.4394 | LPESVER | O08917 | O08917 | FLOT1 | flotillin 1 | Plasma Membrane | other |
| 15537729 | 3 | 0.404 | 1667905.125 | 305978.031 | 211699.75 | 2907334 | 59832.258 | 244698.562 | 870251 | 442733.344 | 1473239.5 | 1035180.75 | 2948645.5 | 1976023.375 | 2012.0986 | VTGEVLDILSRLPESVER | O08917 | O08917 | FLOT1 | flotillin 1 | Plasma Membrane | other |
| 15548378 | 3 | 0.501 | 3.52E+07 | 2.80E+07 | 3.30E+07 | 3.28E+07 | 2.11E+07 | 2.86E+07 | 2.68E+07 | 2.66E+07 | 3.09E+07 | 3.34E+07 | 3.16E+07 | 2.74E+07 | 2325.2046 | SILGTLTVEQIYQDRDQFAK | Q60634 | Q60634 | FLOT2 | flotillin 2 | Plasma Membrane | other |
| 15532451 | 3 | 0.208 | 3.31E+07 | 2.56E+07 | 2.86E+07 | 2.22E+07 | 2.15E+07 | 2.70E+07 | 2.53E+07 | 2.26E+07 | 2.37E+07 | 2.98E+07 | 2.20E+07 | 2.76E+07 | 1934.0925 | LLAELPASVHALTGVDLSK | Q60634 | Q60634 | FLOT2 | flotillin 2 | Plasma Membrane | other |
| 15531521 | 2 | 0.185 | 1.88E+07 | 1.69E+07 | 2.03E+07 | 1.66E+07 | 1.57E+07 | 1.60E+07 | 1.83E+07 | 1.73E+07 | 1.50E+07 | 1.93E+07 | 1.66E+07 | 1.81E+07 | 1199.66 | QVLLAQAEAEK | Q60634 | Q60634 | FLOT2 | flotillin 2 | Plasma Membrane | other |
| 15532379 | 2, 3 | 0.35 | 1.86E+07 | 2.12E+07 | 2.27E+07 | 1.75E+07 | 1.75E+07 | 2.01E+07 | 2.16E+07 | 2.08E+07 | 1.87E+07 | 1.69E+07 | 2.12E+07 | 1.81E+07 | 1378.7771, 1378.7777 | NVVLQTLEGHLR | Q60634 | Q60634 | FLOT2 | flotillin 2 | Plasma Membrane | other |
| 15535170 | 2 | 0.404 | 1.56E+07 | 1.74E+07 | 1.63E+07 | 1.64E+07 | 1.49E+07 | 1.38E+07 | 1.59E+07 | 1.62E+07 | 1.32E+07 | 1.52E+07 | 1.53E+07 | 1.28E+07 | 1388.7062 | IGEAEAAVIEAMGK | Q60634 | Q60634 | FLOT2 | flotillin 2 | Plasma Membrane | other |
| 15531807 | 2 | 0.531 | 1.30E+07 | 1.42E+07 | 1.46E+07 | 1.49E+07 | 1.20E+07 | 1.18E+07 | 1.40E+07 | 1.29E+07 | 1.20E+07 | 1.37E+07 | 1.31E+07 | 1.55E+07 | 1467.8583 | MALVLEALPQIAAK | Q60634 | Q60634 | FLOT2 | flotillin 2 | Plasma Membrane | other |
| 15548530 | 2 | 0.178 | 1.37E+07 | 1.28E+07 | 1.64E+07 | 1.07E+07 | 1.28E+07 | 1.19E+07 | 1.14E+07 | 1.25E+07 | 1.35E+07 | 1.33E+07 | 1.21E+07 | 1.08E+07 | 1520.7662 | TAEAQLAYELQGAR | Q60634 | Q60634 | FLOT2 | flotillin 2 | Plasma Membrane | other |
| 15532957 | 2 | 0.023 | 1.11E+07 | 1.09E+07 | 1.16E+07 | 9704218 | 9876025 | 1.06E+07 | 1.02E+07 | 1.04E+07 | 1.17E+07 | 1.22E+07 | 1.19E+07 | 1.13E+07 | 1477.7688 | ELLAVAC[160.0307]EQFLGK | Q60634 | Q60634 | FLOT2 | flotillin 2 | Plasma Membrane | other |
| 15537326 | 2 | 0.44 | 8890060 | 8220460 | 1.13E+07 | 7490574.5 | 7721533 | 8548283 | 8572039 | 8874600 | 8131042 | 9613687 | 6443185 | 8192040.5 | 981.52295 | VDYLSSLGK | Q60634 | Q60634 | FLOT2 | flotillin 2 | Plasma Membrane | other |
| 15532625 | 2 | 0.975 | 8669638 | 8651812 | 1.06E+07 | 8270696 | 1.03E+07 | 9373429 | 9594936 | 1.05E+07 | 8277334.5 | 9385558 | 9931588 | 9400522 | 1374.7083 | VDEIVVLSGDNSK | Q60634 | Q60634 | FLOT2 | flotillin 2 | Plasma Membrane | other |
| 15539582 | 2 | 0.288 | 7316648 | 7275067.5 | 8600263 | 7700761 | 6412771 | 6189686.5 | 8005911 | 6294739.5 | 6559162.5 | 9253779 | 6765244.5 | 8759690 | 1123.5608 | SAFSEEVNIK | Q60634 | Q60634 | FLOT2 | flotillin 2 | Plasma Membrane | other |
| 15540367 | 2 | 0.436 | 4969148.5 | 6338102 | 7561584.5 | 3943632.75 | 5676397 | 5840548 | 5130514 | 5550220 | 4900155.5 | 5388457.5 | 4707259.5 | 5489524.5 | 1269.7126 | QIAVEAQEILR | Q60634 | Q60634 | FLOT2 | flotillin 2 | Plasma Membrane | other |
| 15537721 | 2 | 0.009 | 4260944.5 | 4240033 | 4344102.5 | 3172066.5 | 3042449.25 | 3020421.75 | 4767929.5 | 3956333.75 | 3667684.75 | 5166686.5 | 3963367.25 | 5841950 | 1251.6992 | MGIEILSFTIK | Q60634 | Q60634 | FLOT2 | flotillin 2 | Plasma Membrane | other |
| 15549988 | 2 | 0.601 | 3731725 | 4061445.75 | 3827658 | 3554432 | 3091235 | 3748990 | 3007643.5 | 3748045.25 | 4195657.5 | 3825738.75 | 3323739.75 | 3413211.25 | 1934.0938 | LLAELPASVHALTGVDLSK | Q60634 | Q60634 | FLOT2 | flotillin 2 | Plasma Membrane | other |
| 15534137 | 2 | 0.27 | 3950926.25 | 4563692.5 | 3071719.5 | 2520578.25 | 3868166.75 | 3867352.75 | 4797976.5 | 4466986 | 4258347 | 3184426.75 | 4025290.5 | 3704068.75 | 1300.7274 | ISLEIMTLQPR | Q60634 | Q60634 | FLOT2 | flotillin 2 | Plasma Membrane | other |
| 15542285 | 2 | 0.72 | 1800752.25 | 2633995 | 4185440.75 | 763160.5 | 4360351.5 | 2749959.75 | 2407176 | 3486321.25 | 2412047.25 | 2011119.625 | 1443230.125 | 2031368.5 | 1145.5409 | DADIGVAEAER | Q60634 | Q60634 | FLOT2 | flotillin 2 | Plasma Membrane | other |
| 15545922 | 3 | 0.942 | 1337417.625 | 683247.25 | 382507.344 | 1861339.125 | 442749.5 | 171868.578 | 730872.938 | 487504.375 | 529378.562 | 830125.312 | 745085.812 | 518923.781 | 1520.7673 | TAEAQLAYELQGAR | Q60634 | Q60634 | FLOT2 | flotillin 2 | Plasma Membrane | other |
| 15540544 | 3 | 0.425 | 296085.156 | 385820.781 | 112757.367 | 303568.125 | 349461.594 | 358762.219 | 336343.469 | 286694.719 | 456202.969 | 255702.484 | 393946.375 | 6187295.5 | 1690.8135 | HALQSASAGSGSFTDVR | P11276 | P11276 | FN1 | fibronectin 1 | Extracellular Space | enzyme |
| 15553698 | 2 | 0.425 | 344507.375 | 186476.156 | 247514.656 | 735051.75 | 0 | 300489.688 | 516128.312 | 376694.875 | 557366.75 | 119432.672 | 1219314.25 | 1.28E+07 | 1927.0089 | SSPVIIDASTAIDAPSNLR | P11276 | P11276 | FN1 | fibronectin 1 | Extracellular Space | enzyme |
| 15531950 | 3 | 0.215 | 1.61E+07 | 1.34E+07 | 1.67E+07 | 1.23E+07 | 1.23E+07 | 1.21E+07 | 1.61E+07 | 1.27E+07 | 1.28E+07 | 2.09E+07 | 1.24E+07 | 1.67E+07 | 1781.959 | VIDLPGGGAVFVMEHLK | Q9ER35 | Q9ER35 | FN3K | fructosamine 3 kinase | Cytoplasm | kinase |
| 15539630 | 3 | 0.051 | 1.67E+07 | 1.15E+07 | 1.55E+07 | 1.16E+07 | 1.11E+07 | 1.06E+07 | 1.56E+07 | 1.20E+07 | 1.13E+07 | 1.88E+07 | 1.56E+07 | 1.52E+07 | 1668.7572 | DYADRETQELWSR | Q9ER35 | Q9ER35 | FN3K | fructosamine 3 kinase | Cytoplasm | kinase |
| 15532062 | 3 | 0.409 | 1.43E+07 | 1.05E+07 | 1.30E+07 | 1.12E+07 | 1.41E+07 | 1.07E+07 | 1.48E+07 | 1.02E+07 | 1.06E+07 | 1.76E+07 | 1.38E+07 | 1.29E+07 | 1659.8131 | LGEQMADLHLYNQK | Q9ER35 | Q9ER35 | FN3K | fructosamine 3 kinase | Cytoplasm | kinase |
| 15549404 | 3 | 0.053 | 1.08E+07 | 6024114.5 | 5758568 | 7618483.5 | 6419934 | 4014542.25 | 9832396 | 6114364 | 7709049 | 1.70E+07 | 1.66E+07 | 9235558 | 3506.5647 | FGFHTVTC[160.0307]C[160.0307]GFIPQVNEWQEDWPTFFTR | Q9ER35 | Q9ER35 | FN3K | fructosamine 3 kinase | Cytoplasm | kinase |
| 15532505 | 3 | 0.62 | 1.53E+07 | 9921740 | 1.11E+07 | 1.31E+07 | 9754650 | 8184408.5 | 1.71E+07 | 1.05E+07 | 1.05E+07 | 1.33E+07 | 1.46E+07 | 1.10E+07 | 2059.9778 | TRQNTVGC[160.0307]GAEGAEPQGVTK | Q9ER35 | Q9ER35 | FN3K | fructosamine 3 kinase | Cytoplasm | kinase |
| 15552864 | 2 | 0.713 | 2729377 | 2269929.75 | 3589976.25 | 1918846.125 | 3469046.5 | 1482148.875 | 3182974.25 | 3005643 | 1890632 | 3731263.75 | 1268089.5 | 1766351 | 1802.8312 | QNTVGC[160.0307]GAEGAEPQGVTK | Q9ER35 | Q9ER35 | FN3K | fructosamine 3 kinase | Cytoplasm | kinase |
| 15558101 | 2 | 0.343 | 2960325.75 | 2286711.25 | 2696166 | 1952655.25 | 2540693.5 | 1470780.875 | 2404157.75 | 1848352.375 | 2586439 | 2998606.75 | 2117774.25 | 2256900 | 1659.8132 | LGEQMADLHLYNQK | Q9ER35 | Q9ER35 | FN3K | fructosamine 3 kinase | Cytoplasm | kinase |
| 15532274 | 2, 3 | 0.148 | 2.27E+07 | 2.40E+07 | 2.37E+07 | 2.28E+07 | 2.94E+07 | 2.09E+07 | 2.02E+07 | 2.17E+07 | 1.89E+07 | 3.41E+07 | 2.34E+07 | 2.53E+07 | 1451.782, 1451.7826 | LGTQLADLHLENK | Q8K274 | Q8K274 | FN3KRP | fructosamine 3 kinase related protein | Cytoplasm | kinase |
| 15534059 | 3 | 0.834 | 1.57E+07 | 1.20E+07 | 1.32E+07 | 1.62E+07 | 1.55E+07 | 1.18E+07 | 1.60E+07 | 1.33E+07 | 8626550 | 1.94E+07 | 1.28E+07 | 1.14E+07 | 2025.852 | ATGHSGGGC[160.0307]ISQGQSYDTDK | Q8K274 | Q8K274 | FN3KRP | fructosamine 3 kinase related protein | Cytoplasm | kinase |
| 15537801 | 3 | 0.101 | 9155662 | 8680592 | 9415974 | 9975366 | 1.07E+07 | 8110553.5 | 1.13E+07 | 8175675.5 | 7828720.5 | 1.31E+07 | 1.12E+07 | 1.14E+07 | 2013.9503 | VVDAPGGGSMLVMEHLDMR | Q8K274 | Q8K274 | FN3KRP | fructosamine 3 kinase related protein | Cytoplasm | kinase |
| 15535529 | 2 | 0.269 | 7824050 | 7128225.5 | 8453123 | 8222028.5 | 9102641 | 7603507 | 7445224.5 | 7183674 | 6552672.5 | 8992419 | 7215766.5 | 7400265.5 | 1084.5189 | NWVEFYAR | Q8K274 | Q8K274 | FN3KRP | fructosamine 3 kinase related protein | Cytoplasm | kinase |
| 15543696 | 3 | 0.6 | 5213026 | 6406869 | 5756697.5 | 6147588.5 | 5181165 | 7274182.5 | 5566954.5 | 5900321 | 4428155.5 | 5992617.5 | 5003494 | 6081835 | 1815.9564 | SGDREALELWSALQLK | Q8K274 | Q8K274 | FN3KRP | fructosamine 3 kinase related protein | Cytoplasm | kinase |
| 15533139 | 2 | 0.292 | 4821994.5 | 4800866.5 | 6835900.5 | 4544268 | 5116258 | 6804519.5 | 4297266.5 | 4250594 | 3796280.75 | 5594832.5 | 3566440.5 | 5005509.5 | 760.4336 | IPDLFR | Q8K274 | Q8K274 | FN3KRP | fructosamine 3 kinase related protein | Cytoplasm | kinase |
| 15552499 | 4 | 0.554 | 3467957.25 | 5162325 | 5143453 | 4527805.5 | 7208731 | 6142874 | 4236518.5 | 5779147 | 6007973 | 7033191.5 | 4981982.5 | 4657737.5 | 3317.7307 | AADVLVYLADDTVVPLAVENLSSISAHELHR | Q3UFK8 | Q3UFK8 | FRMD8 | FERM domain containing 8 | Cytoplasm | other |
| 15537275 | 2 | 0.089 | 1113640.375 | 1028693.938 | 853677.75 | 1836814.875 | 1873934 | 3172387.25 | 659294.062 | 1321630.5 | 1685636.75 | 894550.562 | 1264939.625 | 1840231.75 | 876.5172 | IFLQDIK | P09528 | P09528 | FTH1 | ferritin, heavy polypeptide 1 | Cytoplasm | enzyme |
| 15536095 | 3 | 0.111 | 2276661.5 | 3837243.25 | 1982807.125 | 5908476 | 4501758 | 1.06E+07 | 2121547.5 | 3901738.25 | 5121587 | 2047089.625 | 4150624.25 | 4177479.25 | 1705.9305 | NLNQALLDLHALGSAR | P29391 | P29391 | FTL | ferritin, light polypeptide | Cytoplasm | enzyme |
| 15538774 | 2 | 0.054 | 1056365.125 | 1465649 | 1348565.125 | 2480834.5 | 2902324 | 4733930.5 | 933229.062 | 2133887.5 | 2498690 | 1461663.75 | 2093418.5 | 2086099.375 | 1034.5245 | LLEFQNDR | P29391 | P29391 | FTL | ferritin, light polypeptide | Cytoplasm | enzyme |
| 15543883 | 2 | 0.193 | 747559.812 | 154991.078 | 560933.625 | 1203320 | 855761.438 | 1446902.5 | 345334.281 | 1131369.125 | 446090.625 | 859589.438 | 620395.5 | 840206.75 | 1480.7003 | QNYSTEVEAAVNR | P29391 | P29391 | FTL | ferritin, light polypeptide | Cytoplasm | enzyme |
| 15540951 | 2 | 0.12 | 2226226 | 5736130 | 3002496.5 | 6934546 | 8313268 | 9271200 | 4593676.5 | 6768398.5 | 8095069.5 | 2038589.25 | 7185178.5 | 4344026 | 1249.6053 | LFGQESGPSAEK | Q9D6K8 | Q9D6K8 | FUNDC2 | FUN14 domain containing 2 | Cytoplasm | other |
| 15533698 | 2 | 0.151 | 2261084.75 | 7149206.5 | 3986233.25 | 3071093.5 | 6055881 | 7709968.5 | 3186227 | 5537370.5 | 4503094 | 1780018.75 | 2342777.25 | 2714138.25 | 1249.6644 | FEALDLAELTK | Q9D6K8 | Q9D6K8 | FUNDC2 | FUN14 domain containing 2 | Cytoplasm | other |
| 15534774 | 2 | 0.451 | 2580039.25 | 5358328.5 | 1478907.625 | 3066425.25 | 2725103.75 | 3270231.5 | 1873778.25 | 4232872.5 | 3229788.5 | 1913078 | 2060805.375 | 1656141.25 | 1249.6771 | LGGDLGTYVINK | O35943 | O35943 | FXN | frataxin | Cytoplasm | kinase |
| 15538552 | 3 | 0.133 | 2113266.25 | 5931913 | 1684922.625 | 3271099.25 | 2708996.5 | 3035137.25 | 2494311 | 3061863.5 | 3349789.25 | 1098987.625 | 1697759.875 | 1734367.125 | 1996.0001 | NWVYSHDGVSLHELLAR | O35943 | O35943 | FXN | frataxin | Cytoplasm | kinase |
| 15540236 | 3 | 0.294 | 127347.414 | 834015.438 | 1467668.875 | 3531200 | 2704041.5 | 3127478.75 | 2098798.75 | 1553570.875 | 4875346 | 86972.867 | 2923260.75 | 2334456.25 | 1814.8109 | VSESDNNEGSSLPSQHK | O35601 | O35601 | FYB | FYN binding protein | Nucleus | other |
| 15556215 | 2 | 0.223 | 45161.117 | 318618.219 | 729568.5 | 1348019.75 | 1305440 | 1385614.625 | 996189.438 | 655609.688 | 1990548.875 | 54072.773 | 1199155.25 | 712914.438 | 1651.7267 | PSLSTEDSQEENTSK | O35601 | O35601 | FYB | FYN binding protein | Nucleus | other |
| 15545567 | 2 | 0.346 | 112323.102 | 531749.062 | 792063.75 | 1761404.625 | 1481920.375 | 1707286.25 | 1055873.625 | 805445.938 | 2538462.25 | 38222 | 1505669.125 | 1235415.5 | 1273.6467 | VAGQSSPSGIQSR | O35601 | O35601 | FYB | FYN binding protein | Nucleus | other |
| 15549513 | 2 | 0.053 | 5033478 | 1.03E+07 | 4111123 | 8508107 | 6950029 | 4481356.5 | 3361002.75 | 4927855.5 | 5428757 | 3882090.25 | 1764401.25 | 2204256 | 1937.9608 | LPNFGFVVFDDSEPVQK | P97855 | P97855 | G3BP1 | GTPase activating protein (SH3 domain) binding protein 1 | Nucleus | enzyme |
| 15556557 | 2 | 0.006 | 4970944 | 8501957 | 5734026 | 6649377 | 5728587 | 4688580 | 3222140.75 | 4561539.5 | 4721485.5 | 3147992.5 | 1810228 | 2608102.25 | 1584.7773 | DFFQNFGNVVELR | P97855 | P97855 | G3BP1 | GTPase activating protein (SH3 domain) binding protein 1 | Nucleus | enzyme |
| 15554111 | 2 | 0.017 | 776099.5 | 640935.438 | 1408151.5 | 2439961 | 2642534.75 | 3353619 | 1583802.125 | 2424460.5 | 2986413.75 | 1212083.25 | 2735211.25 | 2213066.5 | 1854.8591 | NLFQFC[160.0307]FPAAYYASR | Q6NSQ9 | Q6NSQ9 | G6PC3 | glucose 6 phosphatase, catalytic, 3 | Cytoplasm | phosphatase |
| 15536647 | 2 | 0.018 | 1661111 | 1352184.5 | 1707122.25 | 2343171.25 | 1039685.562 | 1899593.5 | 1167803.375 | 805018.5 | 826599.375 | 3078938.75 | 2347786.25 | 1868943.5 | 911.46063 | LEEFFAR | Q00612 | Q00612 | G6PD | glucose-6-phosphate dehydrogenase | Cytoplasm | enzyme |
| 15545451 | 2 | 0.052 | 1047866.438 | 542283 | 810146 | 1464955.125 | 619535.938 | 1177824.25 | 820053.688 | 984958.562 | 656376.812 | 1963779.625 | 1778456.625 | 1376507 | 1394.7595 | IYPTIWWLFR | Q00612 | Q00612 | G6PD | glucose-6-phosphate dehydrogenase | Cytoplasm | enzyme |
| 15542282 | 3 | 0.137 | 587102.562 | 1099168.5 | 1348266 | 1058909.875 | 690812.688 | 766609.562 | 635896.688 | 957910.125 | 691533.75 | 1747573.625 | 1457038.125 | 1040079.438 | 2173.064 | DVMQNHLLQMLC[160.0307]LVAMEK | Q00612 | Q00612 | G6PD | glucose-6-phosphate dehydrogenase | Cytoplasm | enzyme |
| 15554586 | 2 | 0.109 | 1403060.375 | 676055.062 | 457787.844 | 1327496 | 517922.312 | 696405.75 | 694437.438 | 432404.969 | 130379.289 | 1540861.25 | 1866900.75 | 920447.312 | 1724.7445 | NIQETC[160.0307]MSQTGWNR | Q00612 | Q00612 | G6PD | glucose-6-phosphate dehydrogenase | Cytoplasm | enzyme |
| 15538755 | 2 | 0.016 | 1.69E+07 | 2.26E+07 | 1.70E+07 | 1.71E+07 | 1.67E+07 | 1.22E+07 | 1.26E+07 | 1.75E+07 | 1.68E+07 | 1.24E+07 | 9151313 | 8524438 | 1160.6125 | VSGSQIVDIDK | P60521 | P60521 | GABARAPL2 | GABA(A) receptor-associated protein like 2 | Cytoplasm | other |
| 15533744 | 3 | 0.089 | 1.21E+07 | 1.54E+07 | 1.28E+07 | 1.33E+07 | 1.36E+07 | 9063504 | 9684649 | 1.10E+07 | 1.19E+07 | 1.10E+07 | 6932630.5 | 8366908 | 1314.7389 | YPDRVPVIVEK | P60521 | P60521 | GABARAPL2 | GABA(A) receptor-associated protein like 2 | Cytoplasm | other |
| 15548905 | 2 | 0.112 | 1.02E+07 | 1.46E+07 | 1.09E+07 | 1.14E+07 | 1.22E+07 | 7356434 | 9258075 | 9424201 | 9492963 | 9877923 | 5438261 | 6356037 | 1681.8435 | TVPQSSLTMGQLYEK | P60521 | P60521 | GABARAPL2 | GABA(A) receptor-associated protein like 2 | Cytoplasm | other |
| 15541444 | 2 | 0.056 | 6075298.5 | 1.14E+07 | 8378706 | 6398918 | 9020012 | 7058171 | 5614288 | 7661668 | 1.02E+07 | 5355831.5 | 3377796.5 | 4500538 | 952.5486 | AIFLFVDK | P60521 | P60521 | GABARAPL2 | GABA(A) receptor-associated protein like 2 | Cytoplasm | other |
| 15554374 | 2 | 0.347 | 1456264.5 | 2251175 | 1050209.125 | 1505132.875 | 1803430.625 | 699368.688 | 1152258.25 | 1340589.25 | 1255938 | 1435485.625 | 717129.188 | 476027.781 | 2052.0938 | YLVPSDITVAQFMWIIR | P60521 | P60521 | GABARAPL2 | GABA(A) receptor-associated protein like 2 | Cytoplasm | other |
| 15553805 | 4 | 0.533 | 1.58E+07 | 1.87E+07 | 1.57E+07 | 1.66E+07 | 1.49E+07 | 1.42E+07 | 1.62E+07 | 2.02E+07 | 1.57E+07 | 1.90E+07 | 1.47E+07 | 1.87E+07 | 3241.554 | PYGPTSVGLDFSLPGMEHVYGIPEHADSLR | Q8BHN3 | Q8BHN3 | GANAB | glucosidase, alpha; neutral AB | Cytoplasm | enzyme |
| 15550158 | 3 | 0.188 | 1.41E+07 | 1.47E+07 | 1.32E+07 | 1.60E+07 | 1.65E+07 | 1.62E+07 | 1.88E+07 | 1.52E+07 | 1.42E+07 | 1.72E+07 | 2.10E+07 | 1.51E+07 | 2921.2144 | TRDGSDYEGWC[160.0307]WPGSASYPDFTNPR | Q8BHN3 | Q8BHN3 | GANAB | glucosidase, alpha; neutral AB | Cytoplasm | enzyme |
| 15536703 | 4 | 0.016 | 1.01E+07 | 1.15E+07 | 1.36E+07 | 1.53E+07 | 1.52E+07 | 1.59E+07 | 1.66E+07 | 1.61E+07 | 1.31E+07 | 1.62E+07 | 1.62E+07 | 1.58E+07 | 2497.398 | ALLDTLQLGPDALTVHLIHEVTK | Q8BHN3 | Q8BHN3 | GANAB | glucosidase, alpha; neutral AB | Cytoplasm | enzyme |
| 15548884 | 4 | 0.396 | 8092371 | 7467637 | 4018772.5 | 9885804 | 8800784 | 5593895.5 | 1.06E+07 | 7249884 | 4939131 | 1.49E+07 | 8215270 | 9058620 | 3620.8284 | LYNLDVFQYELNNPMALYGSVPVLLAHSFHR | Q8BHN3 | Q8BHN3 | GANAB | glucosidase, alpha; neutral AB | Cytoplasm | enzyme |
| 15535617 | 3 | 0.233 | 1.04E+07 | 1.23E+07 | 1.19E+07 | 1.43E+07 | 1.38E+07 | 1.21E+07 | 1.57E+07 | 1.51E+07 | 1.12E+07 | 1.28E+07 | 1.43E+07 | 1.52E+07 | 1754.9398 | LSFQHDPETSVLILR | Q8BHN3 | Q8BHN3 | GANAB | glucosidase, alpha; neutral AB | Cytoplasm | enzyme |
| 15536314 | 3 | 0.137 | 7486316 | 8576088 | 9969620 | 9363919 | 1.10E+07 | 1.13E+07 | 1.02E+07 | 9700127 | 8543318 | 1.01E+07 | 1.13E+07 | 1.00E+07 | 1668.9027 | YRVPDVLVADPPTAR | Q8BHN3 | Q8BHN3 | GANAB | glucosidase, alpha; neutral AB | Cytoplasm | enzyme |
| 15550009 | 3 | 0.047 | 9218666 | 1.10E+07 | 1.31E+07 | 1.02E+07 | 1.31E+07 | 1.14E+07 | 1.23E+07 | 1.21E+07 | 1.37E+07 | 9918678 | 8785249 | 8568258 | 2535.292 | QYASLTGTQALPPLFSLGYHQSR | Q8BHN3 | Q8BHN3 | GANAB | glucosidase, alpha; neutral AB | Cytoplasm | enzyme |
| 15552783 | 2 | 0.061 | 7743957 | 6687679.5 | 7674057 | 7959747 | 8324732.5 | 9082837 | 7720321.5 | 7384164 | 7823516 | 9318827 | 8680029 | 7904001.5 | 1923.9084 | MLDYLQGSGETPQTDIR | Q8BHN3 | Q8BHN3 | GANAB | glucosidase, alpha; neutral AB | Cytoplasm | enzyme |
| 15534858 | 3 | 0.28 | 7692496 | 8249871.5 | 8049988.5 | 8027487 | 9611591 | 7993428.5 | 1.27E+07 | 1.07E+07 | 8069862 | 8877280 | 1.06E+07 | 7619988 | 1946.8972 | FGAVWTGDNTAEWDHLK | Q8BHN3 | Q8BHN3 | GANAB | glucosidase, alpha; neutral AB | Cytoplasm | enzyme |
| 15533783 | 2 | 0.53 | 5757370 | 7666391.5 | 8204621.5 | 5917396 | 9404079 | 8023180 | 1.09E+07 | 1.03E+07 | 6734682.5 | 8480449 | 8422268 | 7678462 | 1066.5869 | NPEPELLVR | Q8BHN3 | Q8BHN3 | GANAB | glucosidase, alpha; neutral AB | Cytoplasm | enzyme |
| 15532523 | 3 | 0.441 | 6039562 | 6751055.5 | 6711481 | 5680743 | 7659928.5 | 7181964 | 7348170.5 | 8034486 | 6380898.5 | 7066776 | 8629942 | 7117701.5 | 1317.7246 | SGGIERPFVLSR | Q8BHN3 | Q8BHN3 | GANAB | glucosidase, alpha; neutral AB | Cytoplasm | enzyme |
| 15533999 | 3 | 0.518 | 6294067 | 4141614 | 3049838.5 | 1.24E+07 | 1985620.625 | 6141558.5 | 6933080.5 | 2877909.25 | 7289818 | 6868488 | 1.07E+07 | 8943772 | 1913.082 | IILTAQPFRLDLLEDR | Q8BHN3 | Q8BHN3 | GANAB | glucosidase, alpha; neutral AB | Cytoplasm | enzyme |
| 15536225 | 3 | 0.834 | 3703259.25 | 4832292.5 | 5138245.5 | 3968895.75 | 7455456.5 | 4866101 | 6279412.5 | 6059407 | 3934812.25 | 6782941 | 4238771 | 5125890 | 1326.5947 | DAVHYGGWEHR | Q8BHN3 | Q8BHN3 | GANAB | glucosidase, alpha; neutral AB | Cytoplasm | enzyme |
| 15532503 | 2 | 0.753 | 3503458 | 5343299.5 | 6239653.5 | 3120135.25 | 6406685.5 | 5749909 | 5060136.5 | 6438038.5 | 4409011 | 4887658 | 3286831 | 4303408 | 1058.6334 | IILTAQPFR | Q8BHN3 | Q8BHN3 | GANAB | glucosidase, alpha; neutral AB | Cytoplasm | enzyme |
| 15540190 | 2 | 0.042 | 3601339.5 | 4135426.75 | 4163097.25 | 3908400.5 | 4090864 | 4829710.5 | 4562917.5 | 5702036.5 | 5848124 | 4003575.75 | 4697999 | 4485012.5 | 1352.8473 | VLLVLELQGLQK | Q8BHN3 | Q8BHN3 | GANAB | glucosidase, alpha; neutral AB | Cytoplasm | enzyme |
| 15533102 | 2 | 0.72 | 2894884.25 | 3572585.25 | 3724183.25 | 3491683.25 | 4169950.75 | 3131320.5 | 4058711.75 | 3964553.25 | 3361038 | 3476305.75 | 3874418.5 | 3293542.75 | 1104.6764 | LVAIVDPHIK | Q8BHN3 | Q8BHN3 | GANAB | glucosidase, alpha; neutral AB | Cytoplasm | enzyme |
| 15538071 | 4 | 0.729 | 2059559.75 | 3199407.5 | 2962249.25 | 2636197.75 | 3832730.25 | 2914938 | 3594522.75 | 3184504.25 | 2230492.75 | 3287398.75 | 1752163.5 | 2646564.5 | 2400.208 | DIHNIYGLYVHMATADGLIQR | Q8BHN3 | Q8BHN3 | GANAB | glucosidase, alpha; neutral AB | Cytoplasm | enzyme |
| 15553293 | 2 | 0.104 | 1891799.875 | 1865025.625 | 2781582.5 | 2712380.75 | 2760556 | 2794348.5 | 3235680 | 2763681 | 2284314.75 | 2774777.5 | 3360704.5 | 2974107 | 1593.7307 | WYQMGAYQPFFR | Q8BHN3 | Q8BHN3 | GANAB | glucosidase, alpha; neutral AB | Cytoplasm | enzyme |
| 15540779 | 2 | 0.17 | 1680168.25 | 1497847.375 | 1420813.875 | 1261655.75 | 1517669.5 | 1523661.75 | 1773294.75 | 1814253.625 | 1117105.5 | 1935941.125 | 1949678.5 | 1889861.5 | 1088.5281 | GLMAFEHQR | Q8BHN3 | Q8BHN3 | GANAB | glucosidase, alpha; neutral AB | Cytoplasm | enzyme |
| 15556413 | 3 | 0.231 | 322174.219 | 1096558.25 | 1028147.125 | 868007 | 1386274.375 | 1065581.125 | 971551 | 949040.25 | 738823.062 | 701999.5 | 347902.656 | 564773.562 | 2667.1616 | DPAEGNGAQPEATPGDGDKPEETQEK | Q8BHN3 | Q8BHN3 | GANAB | glucosidase, alpha; neutral AB | Cytoplasm | enzyme |
| 15540144 | 2 | 0.182 | 5.22E+08 | 3.59E+08 | 3.08E+08 | 2.57E+08 | 2.49E+08 | 2.26E+08 | 3.64E+08 | 2.39E+08 | 2.17E+08 | 3.65E+08 | 2.08E+08 | 2.88E+08 | 1369.7395 | GAAQNIIPASTGAAK | P16858 | P16858 | GAPDH | glyceraldehyde-3-phosphate dehydrogenase | Cytoplasm | enzyme |
| 15532281 | 4 | 0.333 | 4.14E+08 | 2.72E+08 | 1.94E+08 | 1.82E+08 | 1.13E+08 | 2.06E+08 | 3.48E+08 | 2.28E+08 | 1.62E+08 | 2.89E+08 | 1.68E+08 | 3.10E+08 | 2595.3586 | VIHDNFGIVEGLMTTVHAITATQK | P16858 | P16858 | GAPDH | glyceraldehyde-3-phosphate dehydrogenase | Cytoplasm | enzyme |
| 15547705 | 2 | 0.431 | 3.31E+08 | 2.17E+08 | 2.52E+08 | 1.88E+08 | 1.81E+08 | 1.98E+08 | 2.73E+08 | 1.68E+08 | 1.65E+08 | 2.70E+08 | 1.24E+08 | 2.09E+08 | 1779.7944 | LISWYDNEYGYSNR | P16858 | P16858 | GAPDH | glyceraldehyde-3-phosphate dehydrogenase | Cytoplasm | enzyme |
| 15531256 | 3 | 0.413 | 3.81E+08 | 1.60E+08 | 1.23E+08 | 2.06E+08 | 6.42E+07 | 1.40E+08 | 2.15E+08 | 9.64E+07 | 1.44E+08 | 2.63E+08 | 1.84E+08 | 2.48E+08 | 1927.0282 | VPTPNVSVVDLTC[160.0307]RLEK | P16858 | P16858 | GAPDH | glyceraldehyde-3-phosphate dehydrogenase | Cytoplasm | enzyme |
| 15547662 | 2 | 0.329 | 2.42E+08 | 2.51E+08 | 2.73E+08 | 1.06E+08 | 2.52E+08 | 1.78E+08 | 2.37E+08 | 1.84E+08 | 1.49E+08 | 2.49E+08 | 8.00E+07 | 1.36E+08 | 1556.8075 | VPTPNVSVVDLTC[160.0307]R | P16858 | P16858 | GAPDH | glyceraldehyde-3-phosphate dehydrogenase | Cytoplasm | enzyme |
| 15547716 | 2 | 0.176 | 2.38E+08 | 1.17E+08 | 1.08E+08 | 8.40E+07 | 1.15E+08 | 9.43E+07 | 1.57E+08 | 1.02E+08 | 1.18E+08 | 2.21E+08 | 1.85E+08 | 1.27E+08 | 1819.9005 | IVSNASC[160.0307]TTNC[160.0307]LAPLAK | P16858 | P16858 | GAPDH | glyceraldehyde-3-phosphate dehydrogenase | Cytoplasm | enzyme |
| 15548531 | 2, 3 | 0.242 | 2.99E+08 | 2.15E+08 | 1.83E+08 | 1.41E+08 | 1.36E+08 | 1.52E+08 | 2.44E+08 | 1.53E+08 | 1.47E+08 | 2.15E+08 | 1.17E+08 | 2.02E+08 | 2595.357 | VIHDNFGIVEGLMTTVHAITATQK | P16858 | P16858 | GAPDH | glyceraldehyde-3-phosphate dehydrogenase | Cytoplasm | enzyme |
| 15550637, 15555712 | 2, 3 | 0.192 | 2.43E+08 | 1.81E+08 | 1.36E+08 | 1.15E+08 | 1.04E+08 | 1.02E+08 | 1.89E+08 | 9.68E+07 | 1.05E+08 | 1.80E+08 | 1.04E+08 | 1.40E+08 | 2369.2053, 2369.2092 | RVIISAPSADAPMFVMGVNHEK | P16858 | P16858 | GAPDH | glyceraldehyde-3-phosphate dehydrogenase | Cytoplasm | enzyme |
| 15532593 | 2 | 0.412 | 1.47E+08 | 1.61E+08 | 1.38E+08 | 7.24E+07 | 8.79E+07 | 1.12E+08 | 1.54E+08 | 9.32E+07 | 6.97E+07 | 1.49E+08 | 4.45E+07 | 1.17E+08 | 1227.6102 | VVDLMAYMASK | P16858 | P16858 | GAPDH | glyceraldehyde-3-phosphate dehydrogenase | Cytoplasm | enzyme |
| 15532207 | 3 | 0.346 | 1.33E+08 | 1.98E+08 | 2.57E+08 | 6.41E+07 | 1.45E+08 | 1.69E+08 | 2.31E+08 | 1.40E+08 | 7.51E+07 | 1.45E+08 | 3.74E+07 | 9.59E+07 | 1671.9044 | PITIFQERDPTNIK | P16858 | P16858 | GAPDH | glyceraldehyde-3-phosphate dehydrogenase | Cytoplasm | enzyme |
| 15538210 | 4 | 0.358 | 1.11E+08 | 1.03E+08 | 8.05E+07 | 8.09E+07 | 6.34E+07 | 6.11E+07 | 1.02E+08 | 5.82E+07 | 5.55E+07 | 1.00E+08 | 5.47E+07 | 8.49E+07 | 2369.2063 | RVIISAPSADAPMFVMGVNHEK | P16858 | P16858 | GAPDH | glyceraldehyde-3-phosphate dehydrogenase | Cytoplasm | enzyme |
| 15531311 | 2 | 0.588 | 8.03E+07 | 4.31E+07 | 4.66E+07 | 4.45E+07 | 3.34E+07 | 2.68E+07 | 6.11E+07 | 2.65E+07 | 2.65E+07 | 7.81E+07 | 2.16E+07 | 4.89E+07 | 795.4167 | LTGMAFR | P16858 | P16858 | GAPDH | glyceraldehyde-3-phosphate dehydrogenase | Cytoplasm | enzyme |
| 15552361 | 2, 3 | 0.136 | 9.20E+07 | 8.17E+07 | 7.45E+07 | 5.71E+07 | 5.82E+07 | 6.89E+07 | 6.71E+07 | 5.31E+07 | 6.18E+07 | 7.80E+07 | 4.21E+07 | 5.78E+07 | 2291.0474, 2291.0483 | WGEAGAEYVVESTGVFTTMEK | P16858 | P16858 | GAPDH | glyceraldehyde-3-phosphate dehydrogenase | Cytoplasm | enzyme |
| 15534173 | 4 | 0.116 | 7.67E+07 | 6.48E+07 | 1.08E+08 | 3.61E+07 | 3.94E+07 | 7.35E+07 | 5.34E+07 | 5.37E+07 | 4.09E+07 | 4.87E+07 | 1.48E+07 | 4.32E+07 | 2611.3503 | VIHDNFGIVEGLM[147.0354]TTVHAITATQK | P16858 | P16858 | GAPDH | glyceraldehyde-3-phosphate dehydrogenase | Cytoplasm | enzyme |
| 15532666 | 2 | 0.657 | 8.48E+07 | 1.97E+07 | 2720732.25 | 5.16E+07 | 7956148 | 8122013.5 | 2.31E+07 | 1.05E+07 | 3.06E+07 | 3.75E+07 | 4.00E+07 | 4.56E+07 | 1356.6511 | VVDLMAYMASKE | P16858 | P16858 | GAPDH | glyceraldehyde-3-phosphate dehydrogenase | Cytoplasm | enzyme |
| 15533143 | 2 | 0.302 | 4.50E+07 | 3.18E+07 | 3.20E+07 | 2.64E+07 | 2.18E+07 | 2.68E+07 | 3.55E+07 | 2.15E+07 | 2.08E+07 | 3.29E+07 | 1.91E+07 | 3.27E+07 | 805.42993 | VGVNGFGR | P16858 | P16858 | GAPDH | glyceraldehyde-3-phosphate dehydrogenase | Cytoplasm | enzyme |
| 15538911 | 3 | 0.157 | 2.27E+07 | 2.03E+07 | 1.60E+07 | 1.23E+07 | 1.66E+07 | 1.76E+07 | 1.69E+07 | 1.60E+07 | 2.09E+07 | 2.76E+07 | 1.98E+07 | 1.98E+07 | 1819.8989 | IVSNASC[160.0307]TTNC[160.0307]LAPLAK | P16858 | P16858 | GAPDH | glyceraldehyde-3-phosphate dehydrogenase | Cytoplasm | enzyme |
| 15532755 | 3 | 0.409 | 1.63E+07 | 1.86E+07 | 1.60E+07 | 7805172.5 | 1.64E+07 | 1.23E+07 | 1.62E+07 | 1.39E+07 | 9740864 | 1.74E+07 | 5581983.5 | 1.05E+07 | 1556.807 | VPTPNVSVVDLTC[160.0307]R | P16858 | P16858 | GAPDH | glyceraldehyde-3-phosphate dehydrogenase | Cytoplasm | enzyme |
| 15554686 | 2 | 0.437 | 2.83E+07 | 9251991 | 5843681 | 1.33E+07 | 2800979.25 | 6738873.5 | 1.35E+07 | 4248296.5 | 7452082.5 | 1.63E+07 | 1.33E+07 | 1.28E+07 | 1927.0272 | VPTPNVSVVDLTC[160.0307]RLEK | P16858 | P16858 | GAPDH | glyceraldehyde-3-phosphate dehydrogenase | Cytoplasm | enzyme |
| 15550791, 15552574, 15555799, 15556389 | 3, 4 | 0.201 | 3.44E+07 | 4.78E+07 | 1.11E+08 | 8883443 | 3.27E+07 | 7.19E+07 | 2.31E+07 | 4.08E+07 | 3.67E+07 | 1.58E+07 | 6742356 | 2.61E+07 | 3354.5593, 3354.56, 3354.5625, 3354.5662 | VEIVAINDPFIDLNYM[147.0354]VYM[147.0354]FQYDSTHGK | P16858 | P16858 | GAPDH | glyceraldehyde-3-phosphate dehydrogenase | Cytoplasm | enzyme |
| 15534486 | 3 | 0.924 | 1.76E+07 | 1.05E+07 | 1.05E+07 | 1.04E+07 | 1.00E+07 | 1.15E+07 | 1.59E+07 | 1.07E+07 | 9768656 | 1.58E+07 | 7127140.5 | 1.46E+07 | 1779.7942 | LISWYDNEYGYSNR | P16858 | P16858 | GAPDH | glyceraldehyde-3-phosphate dehydrogenase | Cytoplasm | enzyme |
| 15548894 | 3 | 0.152 | 2.43E+07 | 1.41E+07 | 1.51E+07 | 1.04E+07 | 9158458 | 9531339 | 1.63E+07 | 8172607 | 6979313.5 | 1.30E+07 | 5949891.5 | 1.01E+07 | 2385.1997 | RVIISAPSADAPM[147.0354]FVMGVNHEK | P16858 | P16858 | GAPDH | glyceraldehyde-3-phosphate dehydrogenase | Cytoplasm | enzyme |
| 15553837 | 2 | 0.379 | 1.09E+07 | 1.33E+07 | 1.25E+07 | 5286719.5 | 7883254 | 5703879 | 1.75E+07 | 7360492 | 4274190 | 1.25E+07 | 4048085 | 7045361 | 1671.9039 | PITIFQERDPTNIK | P16858 | P16858 | GAPDH | glyceraldehyde-3-phosphate dehydrogenase | Cytoplasm | enzyme |
| 15541702 | 2 | 0.388 | 5782594.5 | 1.24E+07 | 2.36E+07 | 2612309 | 1.68E+07 | 8746592 | 1.12E+07 | 1.20E+07 | 3563614.25 | 9559272 | 939909.5 | 3559016.75 | 1003.55475 | PITIFQER | P16858 | P16858 | GAPDH | glyceraldehyde-3-phosphate dehydrogenase | Cytoplasm | enzyme |
| 15548772 | 3 | 0.088 | 1.62E+07 | 1.30E+07 | 1.25E+07 | 6808510 | 7613924 | 7587817.5 | 1.07E+07 | 7370825.5 | 4406336 | 8973523 | 2638188.5 | 6368749 | 2385.202 | RVIISAPSADAPMFVM[147.0354]GVNHEK | P16858 | P16858 | GAPDH | glyceraldehyde-3-phosphate dehydrogenase | Cytoplasm | enzyme |
| 15560629, 15560819 | 3 | 0.126 | 8707336 | 1.02E+07 | 1.09E+07 | 3622883 | 7621188 | 6900769.5 | 8325634 | 7370917 | 5762072 | 6704856.5 | 3346527.75 | 6468736.5 | 3338.5662, 3338.5793 | VEIVAINDPFIDLNYM[147.0354]VYMFQYDSTHGK | P16858 | P16858 | GAPDH | glyceraldehyde-3-phosphate dehydrogenase | Cytoplasm | enzyme |
| 15537512 | 3 | 0.82 | 9485359 | 4871562.5 | 2308100.75 | 2923327.75 | 4220734.5 | 4229450.5 | 6004374 | 4015973.5 | 5706203 | 6355048.5 | 3038816.5 | 5058273 | 1627.9486 | LVINGKPITIFQER | P16858 | P16858 | GAPDH | glyceraldehyde-3-phosphate dehydrogenase | Cytoplasm | enzyme |
| 15535463 | 4 | 0.314 | 5871448 | 4745589 | 4671147 | 2259685 | 3072199 | 3275403.5 | 5486403.5 | 2569676.5 | 1586897.375 | 4979394.5 | 919195.875 | 2068637.375 | 2385.1997 | RVIISAPSADAPMFVM[147.0354]GVNHEK | P16858 | P16858 | GAPDH | glyceraldehyde-3-phosphate dehydrogenase | Cytoplasm | enzyme |
| 15536896 | 2 | 0.319 | 2915532.25 | 841600.938 | 294537.688 | 1206913.375 | 221169.391 | 241503.859 | 655009.938 | 545805.312 | 1187984.75 | 1956307.625 | 1015983.562 | 1590448.875 | 1372.6449 | VVDLMAYM[147.0354]ASKE | P16858 | P16858 | GAPDH | glyceraldehyde-3-phosphate dehydrogenase | Cytoplasm | enzyme |
| 15537483 | 3 | 0.6 | 1.02E+07 | 8806584 | 7196001 | 9933166 | 6693638.5 | 6673529 | 1.03E+07 | 7587010.5 | 6061821.5 | 1.65E+07 | 6948205.5 | 9544437 | 1961.975 | LPNFGSHVLTAAEMEAFK | Q6PAR5 | Q6PAR5 | GAPVD1 | GTPase activating protein and VPS9 domains 1 | Cytoplasm | other |
| 15552462 | 2 | 0.835 | 9327040 | 6872427.5 | 8273107 | 8048963.5 | 8977145 | 8081130 | 9134157 | 6477043 | 8707718 | 1.51E+07 | 7010918 | 7901393 | 1500.7878 | LSAQAQVAEDILDK | Q6PAR5 | Q6PAR5 | GAPVD1 | GTPase activating protein and VPS9 domains 1 | Cytoplasm | other |
| 15534243 | 2 | 0.547 | 1.07E+07 | 7098350 | 5639846 | 8561378 | 6385101 | 5707227 | 8255528.5 | 5553547 | 5685135 | 1.42E+07 | 6782582 | 7570669 | 1427.7023 | ILEDTQFVDGYK | Q6PAR5 | Q6PAR5 | GAPVD1 | GTPase activating protein and VPS9 domains 1 | Cytoplasm | other |
| 15532948 | 2, 3 | 0.51 | 7684490 | 5751821.5 | 5204901.5 | 5437890 | 6669802 | 5170612.5 | 6714065 | 5905832.5 | 5556658.5 | 1.41E+07 | 6172389 | 5815695.5 | 1381.7148, 1381.716 | QGLQTTQAHLER | Q6PAR5 | Q6PAR5 | GAPVD1 | GTPase activating protein and VPS9 domains 1 | Cytoplasm | other |
| 15533659 | 2 | 0.425 | 9463165 | 6842439 | 6658595 | 6364065.5 | 5979848.5 | 6796652 | 9728996 | 5277294 | 6071960.5 | 1.29E+07 | 7201066 | 7615511.5 | 1201.6917 | ALQIPEVYLR | Q6PAR5 | Q6PAR5 | GAPVD1 | GTPase activating protein and VPS9 domains 1 | Cytoplasm | other |
| 15551803 | 2 | 0.359 | 7227276.5 | 6178375.5 | 4873725 | 5973353.5 | 5882767.5 | 6086726 | 6850326.5 | 5452038 | 6219310.5 | 1.18E+07 | 5779491.5 | 7383787 | 1899.9801 | AVETPPMSSVNLLEGLSR | Q6PAR5 | Q6PAR5 | GAPVD1 | GTPase activating protein and VPS9 domains 1 | Cytoplasm | other |
| 15548568 | 2 | 0.435 | 6504410 | 6894915.5 | 6529615 | 5904052 | 4976544.5 | 5539490 | 5973661 | 5660020 | 6295700.5 | 1.13E+07 | 5304944.5 | 6192029 | 1511.8207 | MALDNLLANLPQAK | Q6PAR5 | Q6PAR5 | GAPVD1 | GTPase activating protein and VPS9 domains 1 | Cytoplasm | other |
| 15539336 | 3 | 0.351 | 6329602 | 4091090.5 | 2630397.5 | 4200186.5 | 4645988.5 | 3282320.75 | 5512442.5 | 3628975.75 | 2983156.5 | 8701083 | 5152305 | 5002952 | 1297.7346 | LDIHTLAHHLK | Q6PAR5 | Q6PAR5 | GAPVD1 | GTPase activating protein and VPS9 domains 1 | Cytoplasm | other |
| 15538330 | 2 | 0.479 | 6128513 | 4239972.5 | 3776941.5 | 4703328 | 4219357 | 3789916.25 | 5920878 | 4457398.5 | 3417400.25 | 8608940 | 4532390.5 | 5103200 | 1370.6671 | EAPWPSAQSEIR | Q6PAR5 | Q6PAR5 | GAPVD1 | GTPase activating protein and VPS9 domains 1 | Cytoplasm | other |
| 15557509 | 2 | 0.563 | 4876735 | 3171073.5 | 3967028.75 | 2696421 | 3245509.25 | 3718133 | 3894085.75 | 2887673 | 3913654.75 | 8369902 | 3242304.25 | 3477832.5 | 1758.8656 | LLQQLAMTGTEEGDPR | Q6PAR5 | Q6PAR5 | GAPVD1 | GTPase activating protein and VPS9 domains 1 | Cytoplasm | other |
| 15538534 | 3 | 0.072 | 4156185.75 | 2423465 | 2517641.25 | 2869520.75 | 2537731.75 | 3754760.5 | 4873025 | 2992529.5 | 3085675.75 | 7064956 | 5121291.5 | 4274463 | 1632.8341 | TLSC[160.0307]VDRLEVGEVR | Q6PAR5 | Q6PAR5 | GAPVD1 | GTPase activating protein and VPS9 domains 1 | Cytoplasm | other |
| 15536020 | 2 | 0.592 | 3951390.25 | 2933440 | 3874501.75 | 3404094 | 2893604.75 | 2870504.5 | 4271777.5 | 2813216 | 3203087 | 6011367.5 | 3243921.75 | 3190660.25 | 1341.7346 | VQIAEAINLQDK | Q6PAR5 | Q6PAR5 | GAPVD1 | GTPase activating protein and VPS9 domains 1 | Cytoplasm | other |
| 15535493 | 3 | 0.952 | 3370205.25 | 2735988 | 3618036.25 | 2646034 | 3040281.75 | 2677486.25 | 4190341 | 3294525.25 | 2060389.125 | 6007586 | 2033822.5 | 2638475.5 | 1274.6577 | DQVLHEHIQR | Q6PAR5 | Q6PAR5 | GAPVD1 | GTPase activating protein and VPS9 domains 1 | Cytoplasm | other |
| 15544278 | 2 | 0.946 | 3165370.75 | 1991468 | 1162238.25 | 1994134.75 | 2140704.25 | 728771.75 | 3425910 | 1637856 | 762250.5 | 5124843 | 1115773 | 1241932.5 | 1249.6036 | FSTLTDEPSPR | Q6PAR5 | Q6PAR5 | GAPVD1 | GTPase activating protein and VPS9 domains 1 | Cytoplasm | other |
| 15540126 | 2 | 0.217 | 3019053.75 | 1951629.375 | 2194106 | 2532744.75 | 1788647.625 | 1836072.75 | 2634696.75 | 1873877.625 | 2036683 | 4518165 | 2609905.75 | 2519289.75 | 772.4552 | LNADVLK | Q6PAR5 | Q6PAR5 | GAPVD1 | GTPase activating protein and VPS9 domains 1 | Cytoplasm | other |
| 15541312 | 2 | 0.695 | 1891540.25 | 1185333 | 1389392.5 | 1339533.875 | 1796719.25 | 1540314.625 | 2179706.5 | 1222024.25 | 1158959.375 | 3424143 | 1472535.625 | 1445898.125 | 1249.57 | VQEMVDSNEAK | Q6PAR5 | Q6PAR5 | GAPVD1 | GTPase activating protein and VPS9 domains 1 | Cytoplasm | other |
| 15545565 | 2 | 0.067 | 1691783.25 | 1574276.25 | 1005810.312 | 912000.375 | 1228980.125 | 964737.125 | 1454806.25 | 1105437.875 | 1055203.75 | 3004082.25 | 1611504.625 | 1597966.5 | 700.41315 | APIPFR | Q6PAR5 | Q6PAR5 | GAPVD1 | GTPase activating protein and VPS9 domains 1 | Cytoplasm | other |
| 15537310 | 2 | 0.672 | 3028669.75 | 2233834.25 | 3076191.75 | 3704114.75 | 1418055.125 | 3732596 | 2243666 | 1690126.375 | 3183252.25 | 2639749.25 | 3541109.75 | 3767842.75 | 781.45935 | FGVLFAK | Q3V0G7 | Q3V0G7 | GARNL3 | GTPase activating Rap/RanGAP domain-like 3 | Cytoplasm | other |
| 15550342 | 2 | 0.018 | 4464148 | 5475643 | 5658805 | 7119186 | 4892882.5 | 6728850.5 | 4560906.5 | 4103056 | 3349535.25 | 8739606 | 6378689 | 6196322.5 | 1744.8817 | TLNEWSSQISPDLVR | O09172 | O09172 | GCLM | glutamate-cysteine ligase, modifier subunit | Cytoplasm | enzyme |
| 15535968 | 2 | 0.015 | 5095266 | 3294045.75 | 4370644 | 6781637 | 3501103 | 5081570.5 | 3431698.25 | 2340555.75 | 2539110.75 | 7363614 | 5518634.5 | 6049646.5 | 1131.6234 | IVAIGTSDLDK | O09172 | O09172 | GCLM | glutamate-cysteine ligase, modifier subunit | Cytoplasm | enzyme |
| 15536167 | 3 | 0.016 | 3876075.5 | 3011663 | 3154832 | 4477492 | 2865684 | 3531658.25 | 2953781.75 | 2035458.75 | 2902688.5 | 5385853.5 | 4894384 | 4330265 | 1780.9408 | ASTLHLQTGNLLNWGR | O09172 | O09172 | GCLM | glutamate-cysteine ligase, modifier subunit | Cytoplasm | enzyme |
| 15548099 | 3 | 0.194 | 1.88E+07 | 1.51E+07 | 2.00E+07 | 2.95E+07 | 2.00E+07 | 2.80E+07 | 2.15E+07 | 1.44E+07 | 9383405 | 3.54E+07 | 1.58E+07 | 2.11E+07 | 2693.4258 | GASIAHC[160.0307]PNSNLSLSSGLLNVLEVLK | Q9R111 | Q9R111 | GDA | guanine deaminase | Cytoplasm | enzyme |
| 15550184 | 2, 3 | 0.901 | 1.62E+07 | 1.28E+07 | 1.56E+07 | 1.68E+07 | 1.29E+07 | 2.18E+07 | 1.90E+07 | 1.32E+07 | 8625365 | 3.21E+07 | 8599787 | 1.18E+07 | 2586.2563, 2586.257 | ASDSPIDLFYGDFVGDISEAVIQK | Q9R111 | Q9R111 | GDA | guanine deaminase | Cytoplasm | enzyme |
| 15533504 | 3 | 0.513 | 1.68E+07 | 1.40E+07 | 1.30E+07 | 2.32E+07 | 1.49E+07 | 2.01E+07 | 1.79E+07 | 1.36E+07 | 7576193 | 2.95E+07 | 1.07E+07 | 1.53E+07 | 2087.0398 | FLYLGDDRNIEEVYVGGK | Q9R111 | Q9R111 | GDA | guanine deaminase | Cytoplasm | enzyme |
| 15549275 | 2 | 0.588 | 1.07E+07 | 7081432.5 | 9631868 | 1.45E+07 | 8052166 | 9109360 | 9353964 | 7307090.5 | 3337316.5 | 2.01E+07 | 5555338.5 | 7100341.5 | 1583.7043 | VC[160.0307]MDLNDTVPEYK | Q9R111 | Q9R111 | GDA | guanine deaminase | Cytoplasm | enzyme |
| 15541864 | 2 | 0.402 | 1.20E+07 | 8977466 | 9030017 | 1.60E+07 | 1.18E+07 | 1.34E+07 | 1.30E+07 | 8910332 | 4259568.5 | 1.91E+07 | 7802566.5 | 1.04E+07 | 1436.7241 | IVFLEESSQQEK | Q9R111 | Q9R111 | GDA | guanine deaminase | Cytoplasm | enzyme |
| 15550314 | 2 | 0.393 | 6253869 | 3535461.5 | 4385360 | 8690465 | 5011854.5 | 7531341.5 | 5873171 | 4654182.5 | 2290171.75 | 1.38E+07 | 4227453 | 5181793 | 2014.9747 | FTLSC[160.0307]TETLMSELGNIAK | Q9R111 | Q9R111 | GDA | guanine deaminase | Cytoplasm | enzyme |
| 15534091 | 2 | 0.458 | 1.07E+07 | 6864727.5 | 5466428.5 | 1.17E+07 | 7114979 | 1.20E+07 | 9365701 | 6631573 | 4000629 | 1.37E+07 | 6509519.5 | 8033118 | 1013.61206 | TPPLALVFR | Q9R111 | Q9R111 | GDA | guanine deaminase | Cytoplasm | enzyme |
| 15534772 | 2 | 0.49 | 5270547.5 | 3694214.25 | 4664861 | 7548862 | 5357142.5 | 6274301 | 6007253.5 | 4038876.75 | 2359447.75 | 1.24E+07 | 3245665.5 | 5210609 | 981.5059 | FVSEMLQK | Q9R111 | Q9R111 | GDA | guanine deaminase | Cytoplasm | enzyme |
| 15550262 | 2 | 0.694 | 6779030 | 5818504.5 | 6386475 | 9637286 | 5317186.5 | 9982816 | 7825673.5 | 5648450 | 4188573.75 | 1.03E+07 | 3480044.25 | 6369232 | 2276.172 | LATLGGSQALGLDSEIGNFEVGK | Q9R111 | Q9R111 | GDA | guanine deaminase | Cytoplasm | enzyme |
| 15538209 | 3 | 0.538 | 3370184.5 | 1404001 | 1593323.75 | 3359286.75 | 2599137.5 | 2630936.25 | 2485784.5 | 1855766.125 | 849706.375 | 7862515.5 | 1488806.625 | 2029327.375 | 2014.9818 | FTLSC[160.0307]TETLMSELGNIAK | Q9R111 | Q9R111 | GDA | guanine deaminase | Cytoplasm | enzyme |
| 15551297 | 3 | 0.674 | 2226183.75 | 1562834.25 | 1743950.625 | 3051915.25 | 1485276.875 | 2453104.25 | 2569705 | 2050725 | 1261230.625 | 4284504 | 1882422.375 | 1970587.625 | 2829.357 | NGTTTAC[160.0307]YFGTIHTDSSLILAEITDK | Q9R111 | Q9R111 | GDA | guanine deaminase | Cytoplasm | enzyme |
| 15550964 | 3 | 0.502 | 7334498.5 | 8465919 | 7068826 | 5943148 | 7971161 | 6112167.5 | 6517148 | 6027377.5 | 6780307.5 | 9837031 | 6074594.5 | 7003203 | 2836.182 | NYYESHLGSSYITDSMLEDC[160.0307]APHF | Q9JL56 | Q9JL56 | GDE1 | glycerophosphodiester phosphodiesterase 1 | Plasma Membrane | enzyme |
| 15534720 | 2 | 0.127 | 2812488.25 | 2684460.5 | 3255469.5 | 2882257.75 | 2393472.75 | 2239904.5 | 3060069.25 | 3250721.25 | 2953860.5 | 3021548 | 2470038.25 | 2632804.5 | 1508.7423 | GGSHDAPENTLAAIR | Q9JL56 | Q9JL56 | GDE1 | glycerophosphodiester phosphodiesterase 1 | Plasma Membrane | enzyme |
| 15552969 | 2 | 0.051 | 1250168.625 | 969658.938 | 1191673.125 | 671754.875 | 745082.375 | 365445.75 | 974157.125 | 622221.438 | 926717.688 | 1577454.75 | 1127491.125 | 843055.438 | 1508.7417 | GGSHDAPENTLAAIR | Q9JL56 | Q9JL56 | GDE1 | glycerophosphodiester phosphodiesterase 1 | Plasma Membrane | enzyme |
| 15551577 | 2 | 0.896 | 2023211 | 2758177 | 2786921.5 | 2441463 | 2690917 | 2721309.25 | 3532595.75 | 2684773.25 | 2127639.25 | 2905225.5 | 2218155.5 | 2346432.75 | 1861.9537 | EVFEAFPETPINIDIK | Q9CRY7 | Q9CRY7 | GDPD1 | glycerophosphodiester phosphodiesterase domain containing 1 | Plasma Membrane | enzyme |
| 15536117 | 2 | 0.4 | 1760308.625 | 3392044 | 1109000.125 | 2151141.25 | 1799230.5 | 3226479 | 1473610.5 | 2393240.75 | 2443287.75 | 1259025.375 | 1215613.25 | 1729386.625 | 1120.5763 | VATMLATGSNR | Q91VC9 | Q91VC9 | GHITM | growth hormone inducible transmembrane protein | Cytoplasm | other |
| 15537976 | 2 | 0.1 | 1267365.5 | 2714508.75 | 776477.375 | 1554343 | 1597478 | 1909693.75 | 1516920.625 | 1751984.375 | 1248074.5 | 581460.312 | 692109.688 | 1049236.5 | 1364.6969 | RAEITPMYGAQK | Q91VC9 | Q91VC9 | GHITM | growth hormone inducible transmembrane protein | Cytoplasm | other |
| 15544331 | 2 | 0.306 | 4251147.5 | 3754986.25 | 3324728 | 3620591 | 3195244.25 | 2838973.75 | 2634693 | 3462504.25 | 3240116.75 | 3539715.75 | 3541057.5 | 2963643.75 | 1453.6887 | EAQQYSEALASTR | Q9CYL5 | Q9CYL5 | GLIPR2 | GLI pathogenesis-related 2 | Cytoplasm | other |
| 15548724 | 2 | 0.076 | 2.15E+07 | 4.57E+07 | 2.10E+07 | 2.37E+07 | 2.33E+07 | 3.59E+07 | 1.42E+07 | 2.75E+07 | 2.75E+07 | 1.63E+07 | 1.03E+07 | 1.25E+07 | 1653.7715 | DYAAYNVLDDPELR | Q80Y14 | Q80Y14 | GLRX5 | glutaredoxin 5 | Cytoplasm | other |
| 15556939 | 2 | 0.076 | 1.41E+07 | 2.39E+07 | 9267809 | 1.58E+07 | 1.24E+07 | 2.03E+07 | 8741724 | 1.49E+07 | 1.54E+07 | 7592246 | 7221152.5 | 8021289.5 | 2101.044 | GTPEQPQC[160.0307]GFSNAVVQILR | Q80Y14 | Q80Y14 | GLRX5 | glutaredoxin 5 | Cytoplasm | other |
| 15541435 | 2 | 0.118 | 845038.938 | 1430108.875 | 1474469.875 | 2886873.5 | 3007232.75 | 2987508.5 | 2249323.5 | 2696496.5 | 3682746 | 544117.812 | 2942642.75 | 1918760.5 | 1241.5875 | YSTDVSVDEVK | P26443 | P26443 | GLUD1 | glutamate dehydrogenase 1 | Cytoplasm | enzyme |
| 15554719 | 2 | 0.192 | 184272.672 | 1355358 | 874624.25 | 2790341.5 | 2621868.5 | 2820065.5 | 1221264.5 | 1836312.5 | 3616601.75 | 296485 | 2329780.75 | 1924567 | 1583.7872 | IIAEGANGPTTPEADK | P26443 | P26443 | GLUD1 | glutamate dehydrogenase 1 | Cytoplasm | enzyme |
| 15540458 | 2 | 0.453 | 4.70E-06 | 193309.75 | 55953.293 | 325784.344 | 119385.023 | 742926.5 | 213980.375 | 51483.703 | 4454073 | 4005.559 | 180944.531 | 613953.625 | 1279.6616 | GEHPGLSIGDVAK | P63158 | P63158 | Gm21596/Hmgb1 | high mobility group box 1 | Nucleus | transcription regulator |
| 15547124 | 2 | 0.106 | 4.70E-06 | 68712.766 | 4.70E-06 | 222034.625 | 384033.844 | 682741.438 | 183234.5 | 20054.871 | 2604563.25 | 4.70E-06 | 62229.852 | 124468.305 | 1464.6401 | LGEMWNNTAADDK | P63158 | P63158 | Gm21596/Hmgb1 | high mobility group box 1 | Nucleus | transcription regulator |
| 15539594 | 2 | 0.169 | 683507.5 | 1297951.5 | 1568094.25 | 3321035.75 | 3796911.25 | 3071435 | 2162282.75 | 2764972.75 | 3760488.25 | 556117.188 | 4085799.25 | 2523728.25 | 1087.6328 | LLLLGTGESGK | P21278 | P21278 | GNA11 | guanine nucleotide binding protein (G protein), alpha 11 (Gq class) | Plasma Membrane | enzyme |
| 15531453 | 2 | 0.139 | 5.05E+07 | 6.35E+07 | 6.68E+07 | 5.60E+07 | 6.88E+07 | 3.68E+07 | 4.28E+07 | 5.24E+07 | 5.87E+07 | 3.73E+07 | 3.94E+07 | 4.11E+07 | 1057.6246 | ILLLGAGESGK | P27600 | P27600 | GNA12 | guanine nucleotide binding protein (G protein) alpha 12 | Plasma Membrane | enzyme |
| 15548923 | 2 | 0.045 | 9.08E+07 | 1.02E+08 | 1.08E+08 | 9.16E+07 | 1.14E+08 | 6.51E+07 | 7.43E+07 | 9.31E+07 | 1.07E+08 | 6.79E+07 | 6.10E+07 | 5.44E+07 | 1746.8973 | IAQSDYIPTQQDVLR | P08752 | P08752 | GNAI2 | guanine nucleotide binding protein (G protein), alpha inhibiting activity polypeptide 2 | Plasma Membrane | other |
| 15555248 | 2, 3 | 0.123 | 8.11E+07 | 9.28E+07 | 9.30E+07 | 8.90E+07 | 9.51E+07 | 6.20E+07 | 6.22E+07 | 7.63E+07 | 9.27E+07 | 6.70E+07 | 6.31E+07 | 5.60E+07 | 2320.078, 2320.0793 | SREYQLNDSAAYYLNDLER | P08752 | P08752 | GNAI2 | guanine nucleotide binding protein (G protein), alpha inhibiting activity polypeptide 2 | Plasma Membrane | other |
| 15531385 | 3 | 0.147 | 5.18E+07 | 6.29E+07 | 6.25E+07 | 5.32E+07 | 8.00E+07 | 3.94E+07 | 5.42E+07 | 6.12E+07 | 5.04E+07 | 4.92E+07 | 3.60E+07 | 3.03E+07 | 1636.6876 | IIHEDGYSEEEC[160.0307]R | P08752 | P08752 | GNAI2 | guanine nucleotide binding protein (G protein), alpha inhibiting activity polypeptide 2 | Plasma Membrane | other |
| 15531377 | 3 | 0.65 | 6.49E+07 | 5.14E+07 | 4.50E+07 | 7.96E+07 | 5.42E+07 | 4.17E+07 | 5.33E+07 | 4.79E+07 | 4.92E+07 | 4.30E+07 | 4.90E+07 | 4.75E+07 | 2103.983 | AMGNLQIDFADPQRADDAR | P08752 | P08752 | GNAI2 | guanine nucleotide binding protein (G protein), alpha inhibiting activity polypeptide 2 | Plasma Membrane | other |
| 15549915 | 2, 3 | 0.358 | 4.13E+07 | 7.55E+07 | 7.85E+07 | 4.57E+07 | 8.63E+07 | 5.13E+07 | 5.37E+07 | 6.15E+07 | 8.71E+07 | 4.18E+07 | 4.56E+07 | 4.53E+07 | 2634.2842, 2634.2856 | QLFALSC[160.0307]AAEEQGMLPEDLSGVIR | P08752 | P08752 | GNAI2 | guanine nucleotide binding protein (G protein), alpha inhibiting activity polypeptide 2 | Plasma Membrane | other |

| | | | | | | | | | | | | | | | | | | | |
|---|---|---|---|---|---|---|---|---|---|---|---|---|---|---|---|---|---|---|---|
| 15531811 | 2 | 0.062 | 5.45E+07 | 6.72E+07 | 6.05E+07 | 5.79E+07 | 7.37E+07 | 4.13E+07 | 4.78E+07 | 5.78E+07 | 5.47E+07 | 4.11E+07 | 4.11E+07 | 3.74E+07 | 1274.5875 | YDEAASYIQSK | P08752 | P08752 | GNAI2 | guanine nucleotide binding protein (G protein), alpha inhibiting activity polypeptide 2 | Plasma Membrane | other |
| 15547968 | 2 | 0.04 | 4.42E+07 | 5.67E+07 | 5.49E+07 | 5.31E+07 | 5.86E+07 | 3.70E+07 | 4.21E+07 | 5.05E+07 | 5.31E+07 | 3.64E+07 | 3.25E+07 | 3.36E+07 | 2029.9858 | ITQSSLTIC[160.0307]FPEYTGANK | P08752 | P08752 | GNAI2 | guanine nucleotide binding protein (G protein), alpha inhibiting activity polypeptide 2 | Plasma Membrane | other |
| 15531804 | 3 | 0.168 | 3.90E+07 | 4.35E+07 | 4.34E+07 | 3.46E+07 | 4.92E+07 | 2.78E+07 | 3.03E+07 | 4.75E+07 | 4.16E+07 | 2.99E+07 | 3.05E+07 | 2.60E+07 | 1586.7113 | EIYTHFTC[160.0307]ATDTK | P08752 | P08752 | GNAI2 | guanine nucleotide binding protein (G protein), alpha inhibiting activity polypeptide 2 | Plasma Membrane | other |
| 15532582 | 2 | 0.344 | 3.56E+07 | 3.87E+07 | 3.69E+07 | 3.68E+07 | 5.28E+07 | 3.08E+07 | 2.94E+07 | 3.78E+07 | 4.48E+07 | 2.75E+07 | 3.48E+07 | 2.71E+07 | 1110.5226 | LFDSIC[160.0307]NNK | P08752 | P08752 | GNAI2 | guanine nucleotide binding protein (G protein), alpha inhibiting activity polypeptide 2 | Plasma Membrane | other |
| 15547753 | 2 | 0.022 | 3.82E+07 | 4.98E+07 | 5.03E+07 | 3.61E+07 | 4.37E+07 | 3.13E+07 | 2.96E+07 | 3.90E+07 | 4.19E+07 | 2.51E+07 | 2.89E+07 | 2.85E+07 | 1736.9568 | AVVYSNTIQSIMAIVK | P08752 | P08752 | GNAI2 | guanine nucleotide binding protein (G protein), alpha inhibiting activity polypeptide 2 | Plasma Membrane | other |
| 15539269 | 2 | 0.234 | 1.97E+07 | 2.49E+07 | 2.42E+07 | 2.15E+07 | 2.77E+07 | 1.62E+07 | 1.62E+07 | 2.25E+07 | 2.28E+07 | 1.64E+07 | 1.83E+07 | 1.49E+07 | 909.4234 | MFDVGGQR | P08752 | P08752 | GNAI2 | guanine nucleotide binding protein (G protein), alpha inhibiting activity polypeptide 2 | Plasma Membrane | other |
| 15550154 | 2 | 0.088 | 1.37E+07 | 2.19E+07 | 2.00E+07 | 1.71E+07 | 2.52E+07 | 1.46E+07 | 1.62E+07 | 1.88E+07 | 1.62E+07 | 1.44E+07 | 1.02E+07 | 1.06E+07 | 2076.946 | EYQLNDSAAYYLNDLER | P08752 | P08752 | GNAI2 | guanine nucleotide binding protein (G protein), alpha inhibiting activity polypeptide 2 | Plasma Membrane | other |
| 15534559 | 3 | 0.49 | 1.33E+07 | 1.60E+07 | 1.24E+07 | 1.43E+07 | 1.40E+07 | 6971051 | 1.10E+07 | 1.08E+07 | 1.04E+07 | 1.20E+07 | 9460516 | 1.10E+07 | 1380.7136 | TTGIVETHFTFK | P08752 | P08752 | GNAI2 | guanine nucleotide binding protein (G protein), alpha inhibiting activity polypeptide 2 | Plasma Membrane | other |
| 15543799, 15544578 | 3 | 0.472 | 1.77E+07 | 2.16E+07 | 2.43E+07 | 1.22E+07 | 2.64E+07 | 1.91E+07 | 1.89E+07 | 1.99E+07 | 2.66E+07 | 1.18E+07 | 1.91E+07 | 1.72E+07 | 1672.8077, 1672.8091 | RLWADHGVQAC[160.0307]FGR | P08752 | P08752 | GNAI2 | guanine nucleotide binding protein (G protein), alpha inhibiting activity polypeptide 2 | Plasma Membrane | other |
| 15555782 | 2 | 0.207 | 1.50E+07 | 1.56E+07 | 1.67E+07 | 1.42E+07 | 1.68E+07 | 9277668 | 1.12E+07 | 1.49E+07 | 1.55E+07 | 1.17E+07 | 1.21E+07 | 8222354.5 | 1586.7096 | EIYTHFTC[160.0307]ATDTK | P08752 | P08752 | GNAI2 | guanine nucleotide binding protein (G protein), alpha inhibiting activity polypeptide 2 | Plasma Membrane | other |
| 15536798 | 3 | 0.119 | 1.41E+07 | 1.66E+07 | 1.49E+07 | 1.55E+07 | 2.29E+07 | 1.16E+07 | 1.38E+07 | 1.62E+07 | 1.81E+07 | 1.03E+07 | 1.17E+07 | 1.10E+07 | 1516.7097 | LWADHGVQAC[160.0307]FGR | P08752 | P08752 | GNAI2 | guanine nucleotide binding protein (G protein), alpha inhibiting activity polypeptide 2 | Plasma Membrane | other |
| 15538939 | 3 | 0.243 | 8309236 | 1.80E+07 | 1.44E+07 | 9239506 | 1.70E+07 | 1.17E+07 | 1.14E+07 | 1.87E+07 | 1.57E+07 | 1.02E+07 | 4567577 | 1.04E+07 | 2029.9852 | ITQSSLTIC[160.0307]FPEYTGANK | P08752 | P08752 | GNAI2 | guanine nucleotide binding protein (G protein), alpha inhibiting activity polypeptide 2 | Plasma Membrane | other |
| 15534023 | 2 | 0.17 | 1.18E+07 | 1.31E+07 | 1.31E+07 | 1.37E+07 | 1.52E+07 | 7943777.5 | 1.09E+07 | 1.08E+07 | 1.20E+07 | 9449670 | 9229309 | 7767087 | 780.3762 | DLFEEK | P08752 | P08752 | GNAI2 | guanine nucleotide binding protein (G protein), alpha inhibiting activity polypeptide 2 | Plasma Membrane | other |
| 15553653 | 2 | 0.008 | 1.20E+07 | 1.21E+07 | 1.45E+07 | 1.29E+07 | 1.54E+07 | 1.10E+07 | 1.09E+07 | 1.34E+07 | 1.15E+07 | 9441614 | 7991518.5 | 7507606 | 1597.8596 | WFTDTSIILFLNK | P08752 | P08752 | GNAI2 | guanine nucleotide binding protein (G protein), alpha inhibiting activity polypeptide 2 | Plasma Membrane | other |
| 15552849 | 3 | 0.63 | 1.66E+07 | 1.07E+07 | 7370815 | 1.95E+07 | 5115230.5 | 7859546 | 8230273 | 3677982.75 | 1.14E+07 | 9420435 | 1.50E+07 | 1.13E+07 | 2790.3796 | QLFALSC[160.0307]AAEEQGMLPEDLSGVIRR | P08752 | P08752 | GNAI2 | guanine nucleotide binding protein (G protein), alpha inhibiting activity polypeptide 2 | Plasma Membrane | other |
| 15549113 | 2 | 0.041 | 1.16E+07 | 1.87E+07 | 2.13E+07 | 1.17E+07 | 2.76E+07 | 1.18E+07 | 1.26E+07 | 1.63E+07 | 1.53E+07 | 8769519 | 6406731.5 | 7061000.5 | 1575.7529 | AMGNLQIDFADPQR | P08752 | P08752 | GNAI2 | guanine nucleotide binding protein (G protein), alpha inhibiting activity polypeptide 2 | Plasma Membrane | other |
| 15553738 | 2 | 0.284 | 8189458 | 9266752 | 1.02E+07 | 6314747.5 | 1.16E+07 | 5268458.5 | 8135232 | 8728665 | 7905649.5 | 8003666 | 5960301 | 4409770.5 | 1636.6876 | IIHEDGYSEEEC[160.0307]R | P08752 | P08752 | GNAI2 | guanine nucleotide binding protein (G protein), alpha inhibiting activity polypeptide 2 | Plasma Membrane | other |
| 15541087 | 3 | 0.04 | 7894558.5 | 1.07E+07 | 9742593 | 7378185 | 7191231 | 6947420 | 6579257 | 8501271 | 7029186.5 | 5602391.5 | 7711859 | 5860312.5 | 1736.9585 | AVVYSNTIQSIMAIVK | P08752 | P08752 | GNAI2 | guanine nucleotide binding protein (G protein), alpha inhibiting activity polypeptide 2 | Plasma Membrane | other |
| 15538924 | 3 | 0.791 | 7669511.5 | 2710578.75 | 1095657.75 | 1.11E+07 | 819462.75 | 1649673.125 | 3790162.25 | 1318588 | 3440933.5 | 3307013.75 | 8440064 | 4069510.5 | 2083.9102 | IIHEDGYSEEEC[160.0307]RQYR | P08752 | P08752 | GNAI2 | guanine nucleotide binding protein (G protein), alpha inhibiting activity polypeptide 2 | Plasma Membrane | other |
| 15539825 | 3 | 0.004 | 5745730 | 5202865.5 | 4475407 | 5515783.5 | 4308286 | 3229150.5 | 3595356.5 | 3341825.5 | 3731037.5 | 2784798.75 | 2529764.75 | 2052250.125 | 2119.9802 | AM[147.0354]GNLQIDFADPQRADDAR | P08752 | P08752 | GNAI2 | guanine nucleotide binding protein (G protein), alpha inhibiting activity polypeptide 2 | Plasma Membrane | other |
| 15549603 | 2 | 0.126 | 4140960 | 4022502.5 | 3429681.25 | 4200403 | 4944485.5 | 2885950.5 | 3106566.5 | 3735081.75 | 4192663.25 | 1437207.125 | 3066358.25 | 2855734.5 | 1516.7074 | LWADHGVQAC[160.0307]FGR | P08752 | P08752 | GNAI2 | guanine nucleotide binding protein (G protein), alpha inhibiting activity polypeptide 2 | Plasma Membrane | other |
| 15556493 | 3 | 0.19 | 1544578.5 | 2842940.75 | 1914029.75 | 3362029.75 | 2236774.25 | 3164621.75 | 1674573.75 | 2507897.75 | 2853512 | 811886.812 | 1809655.625 | 2091243.625 | 3959.7808 | RSPLTIC[160.0307]YPEYTGSNTYEEAAAYIQC[160.0307]QFEDLNR | Q9DC51 | Q9DC51 | GNAI3 | guanine nucleotide binding protein (G protein), alpha inhibiting activity polypeptide 3 | Cytoplasm | enzyme |
| 15550808 | 3 | 0.23 | 1992616.625 | 2204164.5 | 5006025.5 | 8623020 | 4944568.5 | 1.13E+07 | 5513429 | 5794693.5 | 1.17E+07 | 1347370.5 | 8183264.5 | 6665861 | 2450.3418 | VRVPTTGIIEYPFDLQSVIFR | P21279 | P21279 | GNAQ | guanine nucleotide binding protein (G protein), q polypeptide | Plasma Membrane | enzyme |
| 15534923 | 2 | 0.324 | 2315305.25 | 2734876.75 | 3788390 | 3417284.5 | 3605587.5 | 3166889.5 | 3135129.75 | 5258215 | 3646195.5 | 3009337.75 | 3366207.25 | 3365580.75 | 1094.6063 | VLTSGIFETK | Q6R0H7 | Q6R0H7 | GNAS | GNAS complex locus | Plasma Membrane | enzyme |
| 15550531 | 2 | 1.88E-04 | 3627753.5 | 3480781.75 | 3468722 | 4042151 | 4568885 | 3815825 | 4401681 | 5225173 | 5421931.5 | 2798957 | 2967702.75 | 2464848 | 1774.7732 | YTTPEDATPEPGEDPR | Q6R0H7 | Q6R0H7 | GNAS | GNAS complex locus | Plasma Membrane | enzyme |
| 15551385 | 2 | 0.092 | 183617.906 | 949914.688 | 1016222.812 | 2534133.75 | 3149255 | 3644871.75 | 1719548.625 | 2544087.25 | 3524559.5 | 331646.625 | 2705516.5 | 1872584.125 | 1780.907 | AYDAVQLFALTGPAESK | O70443 | O70443 | GNAZ | guanine nucleotide binding protein (G protein), alpha z polypeptide | Plasma Membrane | enzyme |
| 15543225 | 2 | 0.402 | 29750.379 | 446749.344 | 1518881 | 2504030.5 | 3143996.5 | 4397740 | 1589121.125 | 2453285.5 | 3864371.25 | 4.70E-06 | 3003634.75 | 2559056 | 1102.6439 | LLLLGTSNSGK | O70443 | O70443 | GNAZ | guanine nucleotide binding protein (G protein), alpha z polypeptide | Plasma Membrane | enzyme |
| 15538968 | 2 | 0.504 | 6.85E+07 | 8.17E+07 | 7.73E+07 | 8.22E+07 | 1.03E+08 | 4.98E+07 | 5.94E+07 | 7.81E+07 | 8.76E+07 | 5.32E+07 | 6.94E+07 | 5.51E+07 | 1353.7018 | LIIWDSYTTNK | P62874 | P62874 | GNB1 | guanine nucleotide binding protein (G protein), beta polypeptide 1 | Plasma Membrane | enzyme |
| 15532872 | 2 | 0.024 | 6.21E+07 | 6.32E+07 | 6.32E+07 | 6.59E+07 | 7.23E+07 | 4.53E+07 | 4.92E+07 | 5.27E+07 | 5.94E+07 | 4.40E+07 | 4.08E+07 | 4.05E+07 | 1017.5544 | LLVSASQDGK | P62874 | P62874 | GNB1 | guanine nucleotide binding protein (G protein), beta polypeptide 1 | Plasma Membrane | enzyme |
| 15536556 | 2, 3 | 0.838 | 2.79E+07 | 2.46E+07 | 2.17E+07 | 2.50E+07 | 2.71E+07 | 1.82E+07 | 2.44E+07 | 2.38E+07 | 2.45E+07 | 2.33E+07 | 2.59E+07 | 1.80E+07 | 1336.6068, 1336.6074 | IYAMHWGTDSR | P62874 | P62874 | GNB1 | guanine nucleotide binding protein (G protein), beta polypeptide 1 | Plasma Membrane | enzyme |
| [illegible] | | | | | | | | | | | | | | | | | | | | | | |

| | | | | | | | | | | | | | | | | | | | | | | |
|---|---|---|---|---|---|---|---|---|---|---|---|---|---|---|---|---|---|---|---|---|---|---|
| 15531258, 15531457 | 2, 3 | 0.134 | 2.10E+09 | 1.66E+09 | 1.85E+09 | 2.06E+09 | 1.02E+09 | 1.54E+09 | 1.16E+09 | 1.10E+09 | 1.02E+09 | 1.29E+09 | 1.55E+09 | 7.20E+08 | 1126.5614, 1126.5626 | LHVDPENFR | P02088 | P02088 | Hbb-b1 | hemoglobin, beta adult major chain | Extracellular Space | transporter |
| 15531244 | 2 | 0.492 | 1.54E+09 | 9.75E+08 | 6.07E+08 | 1.03E+09 | 4.03E+08 | 5.20E+08 | 6.72E+08 | 6.07E+08 | 6.55E+08 | 6.04E+08 | 9.99E+08 | 4.62E+07 | 991.5021 | AAVSC[160.0307]LWGK | P02088 | P02088 | Hbb-b1 | hemoglobin, beta adult major chain | Extracellular Space | transporter |
| 15548004, 15548565, 15551806 | 2 | 0.005 | 1.11E+09 | 8.28E+08 | 1.10E+09 | 9.56E+08 | 5.96E+08 | 8.26E+08 | 5.57E+08 | 4.79E+08 | 4.57E+08 | 5.07E+08 | 5.91E+08 | 3.63E+08 | 1756.9177, 1756.9178 | VITAFNDGLNHLDSLK | P02088 | P02088 | Hbb-b1 | hemoglobin, beta adult major chain | Extracellular Space | transporter |
| 15535423, 15547413 | 3 | 0.028 | 4.43E+08 | 2.89E+08 | 3.10E+08 | 3.86E+08 | 2.14E+08 | 3.06E+08 | 1.85E+08 | 1.55E+08 | 1.58E+08 | 2.89E+08 | 2.29E+08 | 1.62E+08 | 1980.8961, 1980.8989 | YFDSFGDLSSASAIMGNAK | P02088 | P02088 | Hbb-b1 | hemoglobin, beta adult major chain | Extracellular Space | transporter |
| 15539941, 15541079, 15544114, 15547569 | 3, 4 | 0.015 | 5.36E+08 | 4.87E+08 | 3.44E+08 | 4.82E+08 | 2.72E+08 | 5.28E+08 | 2.42E+08 | 2.58E+08 | 2.91E+08 | 2.12E+08 | 2.58E+08 | 2.05E+08 | 1730.0096, 1730.0117, 1730.0122, 1730.0123 | LLGNM[147.0354]IVIVLGHHLGK | P02088 | P02088 | Hbb-b1 | hemoglobin, beta adult major chain | Extracellular Space | transporter |
| 15549173, 15555319, 15557933, 15559229, 15559935, 15560808, 15562269, 15562294, 15562733 | 2 | 0.01 | 2.04E+08 | 2.07E+08 | 2.33E+08 | 2.16E+08 | 9.64E+07 | 1.63E+08 | 1.02E+08 | 8.24E+07 | 7.75E+07 | 1.24E+08 | 5.87E+07 | 6.96E+07 | 1996.8823, 1996.89, 1996.8903, 1996.8911, 1996.8912, 1996.8917, 1996.8921, 1996.8972 | YFDSFGDLSSASAIM[147.0354]GNAK | P02088 | P02088 | Hbb-b1 | hemoglobin, beta adult major chain | Extracellular Space | transporter |
| 15544275 | 3, 4 | 0.904 | 5.79E+07 | 1.25E+08 | 2.12E+08 | 5.56E+07 | 1.11E+08 | 2.15E+08 | 1.28E+08 | 1.37E+08 | 1.68E+08 | 8.59E+07 | 1.06E+08 | 1.54E+08 | 1714.015, 1714.0157 | LLGNMIVIVLGHHLGK | P02088 | P02088 | Hbb-b1 | hemoglobin, beta adult major chain | Extracellular Space | transporter |
| 15533432 | 3, 4 | 0.568 | 1.40E+08 | 1.63E+07 | 2428104.75 | 1.29E+08 | 7551379.5 | 7462437.5 | 1.01E+07 | 8667116 | 2.00E+07 | 3.67E+07 | 1.10E+08 | 4.48E+07 | 1885.013, 1885.0143 | KVITAFNDGLNHLDSLK | P02088 | P02088 | Hbb-b1 | hemoglobin, beta adult major chain | Extracellular Space | transporter |
| 15548622, 15560570, 15560610 | 2 | 0.428 | 2.56E+07 | 1.18E+07 | 1.64E+07 | 2.45E+07 | 9363816 | 1.52E+07 | 9705908 | 1.00E+07 | 1.24E+07 | 1.49E+07 | 2.41E+07 | 1.20E+07 | 1714.0139, 1714.0143, 1714.0145 | LLGNMIVIVLGHHLGK | P02088 | P02088 | Hbb-b1 | hemoglobin, beta adult major chain | Extracellular Space | transporter |
| 15535770 | 3 | 0.005 | 2.53E+07 | 2.89E+07 | 3.06E+07 | 2.65E+07 | 1.13E+07 | 2.36E+07 | 1.25E+07 | 1.38E+07 | 1.06E+07 | 9799074 | 6531341 | 1.07E+07 | 1996.8906 | YFDSFGDLSSASAIM[147.0354]GNAK | P02088 | P02088 | Hbb-b1 | hemoglobin, beta adult major chain | Extracellular Space | transporter |
| 15558671 | 2 | 0.333 | 1.51E+07 | 1.58E+07 | 3327276 | 9400307 | 1.27E+07 | 2.09E+07 | 9213896 | 1.09E+07 | 1.51E+07 | 5736652.5 | 6184548 | 6874867.5 | 1730.0121 | LLGNM[147.0354]IVIVLGHHLGK | P02088 | P02088 | Hbb-b1 | hemoglobin, beta adult major chain | Extracellular Space | transporter |
| 15553614 | 2 | 0.647 | 1.30E+07 | 1267424.75 | 57092.895 | 1.04E+07 | 110109.953 | 211313.188 | 923531.688 | 216121.078 | 1179325.25 | 2142712.25 | 8284761 | 2800137 | 1885.0127 | KVITAFNDGLNHLDSLK | P02088 | P02088 | Hbb-b1 | hemoglobin, beta adult major chain | Extracellular Space | transporter |
| 15531264 | 2, 3 | 0.329 | 1.05E+09 | 7.06E+08 | 5.49E+08 | 1.10E+09 | 3.36E+08 | 4.46E+08 | 5.31E+08 | 4.97E+08 | 4.19E+08 | 4.20E+08 | 5.50E+08 | 3.36E+07 | 1312.6467, 1312.6472 | VNPDEVGGEALGR | P02089 | P02089 | Hbb-b2 | hemoglobin, beta adult minor chain | Extracellular Space | other |
| 15539599 | 2 | 0.255 | 7.81E+08 | 6.39E+08 | 5.32E+08 | 7.77E+08 | 3.03E+08 | 4.33E+08 | 4.27E+08 | 4.26E+08 | 3.77E+08 | 3.30E+08 | 4.46E+08 | 2.67E+07 | 1091.6085 | VITAFNEGLK | P02089 | P02089 | Hbb-b2 | hemoglobin, beta adult minor chain | Extracellular Space | other |
| 15548767, 15556012 | 2 | 0.287 | 7.16E+08 | 5.52E+08 | 3.08E+08 | 7.30E+08 | 2.09E+08 | 3.14E+08 | 3.25E+08 | 3.13E+08 | 3.14E+08 | 2.76E+08 | 3.19E+08 | 2.64E+07 | 2006.9121, 2006.9133 | YFDSFGDLSSASAIMGNPK | P02089 | P02089 | Hbb-b2 | hemoglobin, beta adult minor chain | Extracellular Space | other |
| 15531392 | 2 | 0.406 | 3.15E+08 | 1.99E+08 | 1.10E+08 | 2.35E+08 | 8.51E+07 | 1.14E+08 | 1.48E+08 | 1.39E+08 | 1.37E+08 | 9.35E+07 | 1.97E+08 | 7414173 | 1021.5105 | SAVSC[160.0307]LWAK | P02089 | P02089 | Hbb-b2 | hemoglobin, beta adult minor chain | Extracellular Space | other |
| 15535653, 15543497, 15543905 | 3, 4 | 0.346 | 2.23E+08 | 1.91E+08 | 9.27E+07 | 2.40E+08 | 6.69E+07 | 1.30E+08 | 1.32E+08 | 1.45E+08 | 1.30E+08 | 6.87E+07 | 1.68E+08 | 8631263 | 1654.0123, 1654.0125, 1654.0128, 1654.0135 | LLGNAIVIVLGHHLGK | P02089 | P02089 | Hbb-b2 | hemoglobin, beta adult minor chain | Extracellular Space | other |
| 15533027 | 3 | 0.349 | 9.73E+07 | 9.39E+07 | 4.02E+07 | 1.06E+08 | 3.06E+07 | 4.63E+07 | 5.26E+07 | 4.67E+07 | 4.56E+07 | 4.50E+07 | 4.94E+07 | 5470733.5 | 2006.9143 | YFDSFGDLSSASAIMGNPK | P02089 | P02089 | Hbb-b2 | hemoglobin, beta adult minor chain | Extracellular Space | other |
| 15553827 | 2 | 0.122 | 5.86E+07 | 5.56E+07 | 4.00E+07 | 6.63E+07 | 1.72E+07 | 2.93E+07 | 2.97E+07 | 2.50E+07 | 1.95E+07 | 2.21E+07 | 1.37E+07 | 838473.312 | 2022.9049 | YFDSFGDLSSASAIM[147.0354]GNPK | P02089 | P02089 | Hbb-b2 | hemoglobin, beta adult minor chain | Extracellular Space | other |
| 15560237 | 2 | 0.079 | 2266482.25 | 3690181.25 | 7804677 | 717281.312 | 4927152.5 | 8840465 | 2879511.75 | 7730035.5 | 9794098 | 983132.375 | 1300754.875 | 187932.297 | 1654.0128 | LLGNAIVIVLGHHLGK | P02089 | P02089 | Hbb-b2 | hemoglobin, beta adult minor chain | Extracellular Space | other |
| [illegible] | [illegible] | [illegible] | [illegible] | [illegible] | [illegible] | [illegible] | [illegible] | [illegible] | [illegible] | [illegible] | [illegible] | [illegible] | [illegible] | [illegible] | [illegible] | [illegible] | [illegible] | [illegible] | [illegible] | [illegible] | [illegible] | [illegible] |

| | | | | | | | | | | | | | | | | | | | | | | |
|---|---|---|---|---|---|---|---|---|---|---|---|---|---|---|---|---|---|---|---|---|---|---|
| 15548124 | 3 | 0.127 | 2.11E+07 | 4.86E+07 | 2.00E+07 | 2.40E+07 | 3.00E+07 | 4.52E+07 | 1.79E+07 | 3.53E+07 | 3.71E+07 | 1.21E+07 | 1.68E+07 | 1.63E+07 | 2560.2437 | LVQDVANNTNEEAGDGTTTATVLAR | P63038 | P63038 | HSPD1 | heat shock 60kDa protein 1 (chaperonin) | Cytoplasm | enzyme |
| 15535814 | 2 | 0.27 | 2.11E+07 | 3.51E+07 | 1.52E+07 | 2.82E+07 | 2.01E+07 | 2.70E+07 | 1.57E+07 | 2.58E+07 | 2.57E+07 | 1.20E+07 | 1.86E+07 | 1.60E+07 | 1344.7125 | TVIIEQSWGSPK | P63038 | P63038 | HSPD1 | heat shock 60kDa protein 1 (chaperonin) | Cytoplasm | enzyme |
| 15538053 | 2 | 0.139 | 1.74E+07 | 3.73E+07 | 1.79E+07 | 2.33E+07 | 3.07E+07 | 4.31E+07 | 1.59E+07 | 2.70E+07 | 2.83E+07 | 1.14E+07 | 1.79E+07 | 1.67E+07 | 1233.5936 | VGGTSDVEVNEK | P63038 | P63038 | HSPD1 | heat shock 60kDa protein 1 (chaperonin) | Cytoplasm | enzyme |
| 15551594 | 2 | 0.13 | 1.48E+07 | 4.02E+07 | 1.93E+07 | 2.59E+07 | 2.76E+07 | 3.10E+07 | 1.43E+07 | 2.78E+07 | 3.49E+07 | 1.10E+07 | 1.30E+07 | 1.56E+07 | 1504.7535 | TLNDELEIIEGMK | P63038 | P63038 | HSPD1 | heat shock 60kDa protein 1 (chaperonin) | Cytoplasm | enzyme |
| 15550710, 15552049 | 2, 3 | 0.313 | 1.22E+07 | 3.69E+07 | 1.45E+07 | 2.02E+07 | 2.07E+07 | 3.02E+07 | 1.34E+07 | 2.59E+07 | 2.32E+07 | 9564700 | 1.36E+07 | 1.54E+07 | 2532.1956, 2532.1985 | ILQSSSEVGYDAMLGDFVNMVEK | P63038 | P63038 | HSPD1 | heat shock 60kDa protein 1 (chaperonin) | Cytoplasm | enzyme |
| 15536387 | 3 | 0.124 | 1.63E+07 | 3.16E+07 | 1.47E+07 | 1.83E+07 | 2.31E+07 | 2.49E+07 | 1.26E+07 | 2.43E+07 | 2.52E+07 | 9300651 | 1.33E+07 | 1.25E+07 | 1905.0521 | ISSVQSIVPALEIANAHR | P63038 | P63038 | HSPD1 | heat shock 60kDa protein 1 (chaperonin) | Cytoplasm | enzyme |
| 15533717 | 2 | 0.276 | 1.52E+07 | 3.27E+07 | 1.50E+07 | 1.66E+07 | 2.05E+07 | 2.76E+07 | 1.12E+07 | 2.18E+07 | 2.62E+07 | 7295339 | 1.46E+07 | 1.37E+07 | 912.5861 | VGLQVVAVK | P63038 | P63038 | HSPD1 | heat shock 60kDa protein 1 (chaperonin) | Cytoplasm | enzyme |
| 15550923 | 2 | 0.1 | 1.56E+07 | 3.23E+07 | 1.51E+07 | 2.35E+07 | 2.15E+07 | 2.33E+07 | 1.08E+07 | 2.01E+07 | 2.54E+07 | 7191341 | 1.51E+07 | 9362642 | 1601.7488 | C[160.0307]EFQDAYVLLSEK | P63038 | P63038 | HSPD1 | heat shock 60kDa protein 1 (chaperonin) | Cytoplasm | enzyme |
| 15535575 | 2 | 0.132 | 9385195 | 2.48E+07 | 1.29E+07 | 1.52E+07 | 1.82E+07 | 2.37E+07 | 1.04E+07 | 1.90E+07 | 2.21E+07 | 6452783 | 9882702 | 1.05E+07 | 901.5336 | LSDGVAVLK | P63038 | P63038 | HSPD1 | heat shock 60kDa protein 1 (chaperonin) | Cytoplasm | enzyme |
| 15539605 | 2 | 0.063 | 1.11E+07 | 2.29E+07 | 1.08E+07 | 1.62E+07 | 1.46E+07 | 2.00E+07 | 9017203 | 1.71E+07 | 1.72E+07 | 6191913.5 | 8574796 | 8574285 | 844.51276 | VGEVIVTK | P63038 | P63038 | HSPD1 | heat shock 60kDa protein 1 (chaperonin) | Cytoplasm | enzyme |
| 15532490 | 2 | 0.154 | 1.01E+07 | 2.23E+07 | 1.00E+07 | 1.44E+07 | 1.33E+07 | 1.84E+07 | 1.05E+07 | 1.59E+07 | 1.51E+07 | 5911611 | 1.06E+07 | 8967879 | 1231.6357 | DRVTDALNATR | P63038 | P63038 | HSPD1 | heat shock 60kDa protein 1 (chaperonin) | Cytoplasm | enzyme |
| 15532183 | 2 | 0.176 | 8276706.5 | 2.05E+07 | 8532318 | 1.36E+07 | 1.12E+07 | 1.63E+07 | 7527642 | 1.51E+07 | 1.50E+07 | 5231197 | 7851871 | 7784364.5 | 844.4945 | IPAMTIAK | P63038 | P63038 | HSPD1 | heat shock 60kDa protein 1 (chaperonin) | Cytoplasm | enzyme |
| 15531621 | 2 | 0.102 | 9655525 | 1.83E+07 | 9002930 | 1.29E+07 | 1.16E+07 | 1.53E+07 | 7124101.5 | 1.39E+07 | 1.48E+07 | 5066988.5 | 7791832 | 7297786 | 785.5117 | IGIEIIK | P63038 | P63038 | HSPD1 | heat shock 60kDa protein 1 (chaperonin) | Cytoplasm | enzyme |
| 15532251 | 2 | 0.073 | 9836379 | 2.49E+07 | 1.40E+07 | 1.37E+07 | 1.78E+07 | 2.15E+07 | 1.00E+07 | 1.92E+07 | 1.96E+07 | 4913015 | 7892676 | 1.01E+07 | 1176.5902 | DIGNIISDAMK | P63038 | P63038 | HSPD1 | heat shock 60kDa protein 1 (chaperonin) | Cytoplasm | enzyme |
| 15556818 | 2 | 0.12 | 6862072 | 1.43E+07 | 8706148 | 8615636 | 9725966 | 1.26E+07 | 6820636 | 1.01E+07 | 1.47E+07 | 4203513 | 6665120 | 6068853 | 1684.9032 | AAVEEGIVLGGGC[160.0307]ALLR | P63038 | P63038 | HSPD1 | heat shock 60kDa protein 1 (chaperonin) | Cytoplasm | enzyme |
| 15535935 | 3, 4 | 0.174 | 1.05E+07 | 3.38E+07 | 1.70E+07 | 1.44E+07 | 1.59E+07 | 3.24E+07 | 1.29E+07 | 2.09E+07 | 2.55E+07 | 4072581.75 | 9381749 | 1.34E+07 | 2033.146, 2033.1461 | ISSVQSIVPALEIANAHRK | P63038 | P63038 | HSPD1 | heat shock 60kDa protein 1 (chaperonin) | Cytoplasm | enzyme |
| 15537890 | 2 | 0.155 | 2736391.75 | 7178799 | 3270540.5 | 4242698.5 | 4110846 | 6757377.5 | 3047443.25 | 4971610.5 | 5010020.5 | 2485234 | 2771530 | 2343468.75 | 1428.811 | GVMLAVDAVIAELK | P63038 | P63038 | HSPD1 | heat shock 60kDa protein 1 (chaperonin) | Cytoplasm | enzyme |
| 15538045 | 2 | 0.321 | 2156240.5 | 7284099.5 | 3713070.25 | 2153025 | 3817941.5 | 5778496 | 2048179 | 4158102 | 5709710.5 | 1070608.75 | 2286894.25 | 2501996.75 | 1215.6555 | NAGVEGSLIVEK | P63038 | P63038 | HSPD1 | heat shock 60kDa protein 1 (chaperonin) | Cytoplasm | enzyme |
| 15543136 | 3 | 0.265 | 1534958 | 4451748.5 | 2350738.5 | 4232464.5 | 2431139.25 | 3223532.75 | 2285461 | 2344145.75 | 3465253.75 | 1063047.75 | 1889940 | 2289580 | 2196.0999 | RIQEITEQLDITTSEYEK | P63038 | P63038 | HSPD1 | heat shock 60kDa protein 1 (chaperonin) | Cytoplasm | enzyme |
| 15543302 | 3 | 0.272 | 2321605.25 | 6830069 | 2344995.5 | 2197084.75 | 4928020 | 6115504.5 | 1729018.25 | 4980704 | 7131762.5 | 841815.625 | 1789061.25 | 2535332.75 | 2041.0405 | PVTTPEEIAQVATISANGDK | P63038 | P63038 | HSPD1 | heat shock 60kDa protein 1 (chaperonin) | Cytoplasm | enzyme |
| 15544720 | 2 | 0.417 | 1272414.125 | 2650261.5 | 797512.188 | 1190399.875 | 1729058.75 | 1818260.875 | 789650.188 | 2037172.25 | 2147175.75 | 547434.812 | 1169715.125 | 964004.938 | 960.5104 | VTDALNATR | P63038 | P63038 | HSPD1 | heat shock 60kDa protein 1 (chaperonin) | Cytoplasm | enzyme |
| 15537405 | 3 | 0.015 | 1681040.125 | 4920940.5 | 2739383.5 | 2487381.25 | 2214867.75 | 3335937.25 | 1851754.125 | 3187042.25 | 2244398 | 520541.625 | 795837.188 | 1318951 | 2129.1304 | ALMLQGVDLLADAVAVTM[147.0354]GPK | P63038 | P63038 | HSPD1 | heat shock 60kDa protein 1 (chaperonin) | Cytoplasm | enzyme |
| 15553843 | 2 | 0.213 | 1733936.75 | 2840061.25 | 1227464.125 | 1922628.5 | 1868139.75 | 2080550.25 | 596412.75 | 1868688.625 | 2297803 | 171063.312 | 1412833.625 | 862089.625 | 1905.0513 | ISSVQSIVPALEIANAHR | P63038 | P63038 | HSPD1 | heat shock 60kDa protein 1 (chaperonin) | Cytoplasm | enzyme |
| 15535036 | 2 | 0.039 | 1.77E+07 | 3.12E+07 | 1.63E+07 | 2.47E+07 | 2.43E+07 | 3.24E+07 | 1.45E+07 | 2.41E+07 | 2.70E+07 | 1.13E+07 | 1.39E+07 | 1.26E+07 | 1076.5964 | VLLPEYGGTK | Q64433 | Q64433 | HSPE1 | heat shock 10kDa protein 1 | Cytoplasm | enzyme |
| 15535273 | 2 | 0.217 | 1.21E+07 | 2.28E+07 | 1.01E+07 | 1.63E+07 | 1.58E+07 | 2.05E+07 | 9717058 | 1.75E+07 | 1.97E+07 | 8880456 | 1.16E+07 | 9514886 | 1044.5543 | SGEIEPVSVK | Q64433 | Q64433 | HSPE1 | heat shock 10kDa protein 1 | Cytoplasm | enzyme |
| 15533379 | 2 | 0.097 | 1.10E+07 | 2.09E+07 | 1.05E+07 | 1.32E+07 | 1.51E+07 | 2.02E+07 | 9239453 | 1.57E+07 | 1.61E+07 | 5489705.5 | 1.02E+07 | 8320115.5 | 1285.745 | VLQATVVAVGSGGK | Q64433 | Q64433 | HSPE1 | heat shock 10kDa protein 1 | Cytoplasm | enzyme |
| 15539312 | 2 | 0.197 | 3975230.5 | 1.01E+07 | 5320526 | 5962272 | 6614189.5 | 9686783 | 3795380.5 | 7085274.5 | 8950917 | 3553638 | 4216599.5 | 3458733.75 | 860.4283 | DYFLFR | Q64433 | Q64433 | HSPE1 | heat shock 10kDa protein 1 | Cytoplasm | enzyme |
| 15535919 | 2 | 0.24 | 4395908.5 | 8712904 | 3454015.75 | 5819537 | 4856438 | 6353781.5 | 3328127 | 5493106 | 6229264 | 2712855.75 | 4247193.5 | 2937572.25 | 844.458 | GGIMLPEK | Q64433 | Q64433 | HSPE1 | heat shock 10kDa protein 1 | Cytoplasm | enzyme |
| 15551973 | 2 | 0.062 | 1125817.375 | 1764483.875 | 1176613.5 | 2424699 | 2897010.5 | 1728998.75 | 861454.5 | 2359234 | 464610.406 | 727778.688 | 844873.5 | 342860.094 | 2354.2104 | FFGSLPDSWASGYAVPVVTVVR | Q9Z2G9 | Q9Z2G9 | HTATIP2 | HIV-1 Tat interactive protein 2, 30kDa | Nucleus | transcription regulator |
| 15538301 | 2 | 0.068 | 610220.375 | 1232799.625 | 521410.812 | 1683613.5 | 917770.688 | 924633.625 | 601318.812 | 1072760.25 | 465039.875 | 620662.812 | 302391.781 | 249696.359 | 1320.638 | NVNQEVVDFEK | Q9Z2G9 | Q9Z2G9 | HTATIP2 | HIV-1 Tat interactive protein 2, 30kDa | Nucleus | transcription regulator |
| 15534985 | 2 | 0.354 | 2517158 | 2062539.125 | 2583363.5 | 2301852.75 | 2505018.25 | 2406527.75 | 2515339.25 | 2036116 | 2372839.75 | 3747639.25 | 2547869.75 | 2406845 | 700.4336 | TPLLTR | Q7TMY8 | Q7TMY8 | HUWE1 | HECT, UBA and WWE domain containing 1, E3 ubiquitin protein ligase | Nucleus | transcription regulator |
| 15545512 | 2 | 0.059 | 3848885.5 | 4040910.75 | 4605829 | 5463096.5 | 4963992 | 2748070 | 4353636 | 4373475.5 | 4395573 | 3065421.25 | 2793508 | 2139973.5 | 966.45416 | DRAQM[147.0354]AMK | Q91ZJ9 | Q91ZJ9 | HYAL1 | hyaluronoglucosaminidase 1 | Cytoplasm | enzyme |
| 15543330 | 2 | 0.144 | 3140902.25 | 4277562.5 | 4307589 | 4066113.25 | 5206332 | 4308557.5 | 5092016.5 | 4941721.5 | 4549477.5 | 4643533 | 5205090 | 4686312 | 1347.6759 | SLAEDFAEQPIK | Q9JKR6 | Q9JKR6 | HYOU1 | hypoxia up-regulated 1 | Cytoplasm | other |
| 15539403 | 2 | 0.408 | 3222066.75 | 3528674.5 | 3478491.5 | 4549985.5 | 5103611.5 | 2998100 | 3882054.75 | 4096191 | 3554116.5 | 4482860 | 3990058.75 | 3973334 | 1196.6681 | PGVPMEIVLNK | Q9JKR6 | Q9JKR6 | HYOU1 | hypoxia up-regulated 1 | Cytoplasm | other |
| 15555428 | 3 | 0.156 | 2591775.25 | 3127022.25 | 3438039.5 | 3079378.5 | 3199830.75 | 4624097.5 | 4310322 | 3504324.5 | 4224077.5 | 3921883 | 3787954.5 | 4172192 | 2253.0986 | LIPEMDQVFTEVEMTTLEK | Q9JKR6 | Q9JKR6 | HYOU1 | hypoxia up-regulated 1 | Cytoplasm | other |
| 15551955 | 2 | 0.196 | 2542886.25 | 2512300.75 | 3080620.75 | 2906988.5 | 3970726 | 2613316 | 3778647 | 3336709.75 | 3294568 | 3492840 | 3107297.5 | 3256489 | 1683.8163 | AANSLEAFIFETQDK | Q9JKR6 | Q9JKR6 | HYOU1 | hypoxia up-regulated 1 | Cytoplasm | other |
| 15541759 | 4 | 0.327 | 3288320 | 2860273.25 | 2135788.75 | 4066707.25 | 3476692.25 | 2136279.5 | 4062300.5 | 2643294.25 | 2536274.5 | 3400331.75 | 4302357 | 4484152 | 2726.2578 | YSHDFNFHINYGDLGFLGPEDLR | Q9JKR6 | Q9JKR6 | HYOU1 | hypoxia up-regulated 1 | Cytoplasm | other |
| 15551347 | 2 | 0.251 | 1431023.375 | 1409598.375 | 2248838.25 | 2842710.25 | 2882108.25 | 1986871.25 | 2918322.75 | 2318986.5 | 1834444.375 | 2564755.75 | 2321846 | 1535926.125 | 1523.6938 | GQAGPEGAAPAPEEDK | Q9JKR6 | Q9JKR6 | HYOU1 | hypoxia up-regulated 1 | Cytoplasm | other |
| 15542528 | 2 | 0.497 | 1213609.375 | 2135357.5 | 2011367.5 | 1454590.75 | 3588794 | 3114076.75 | 2144380.5 | 2280043.75 | 2755744.75 | 1576167.5 | 2355466.5 | 2273677.25 | 1040.5323 | LC[160.0307]QGLFFR | Q9JKR6 | Q9JKR6 | HYOU1 | hypoxia up-regulated 1 | Cytoplasm | other |
| 15551398 | 2 | 0.432 | 1366246.375 | 3500475 | 1242202 | 2239343.25 | 2352547.75 | 3519249.75 | 1157895.875 | 2680691.75 | 3650097.25 | 1385184.25 | 1447847.75 | 1622318.375 | 2048.066 | FLLGNLTGFNPETDSVPVK | Q8BIJ6 | Q8BIJ6 | IARS2 | isoleucyl-tRNA synthetase 2, mitochondrial | Cytoplasm | enzyme |
| 15543891 | 2 | 4.29E-05 | 37919.27 | 5419.727 | 60818.859 | 1960857.375 | 2722378.25 | 1568382 | 34660.59 | 22197.473 | 1880.717 | 3228329 | 2868337.5 | 2203116 | 1152.5107 | C[160.0307]VVTC[160.0307]AGETR | Q9ERM2 | Q9ERM2 | ICAM4 | intercellular adhesion molecule 4 (Landsteiner-Wiener blood group) | Plasma Membrane | other |
| 15544432 | 3 | 6.96E-07 | 235327.016 | 99386.531 | 163396.688 | 2981533.75 | 5425608.5 | 3496267.25 | 113493.664 | 147105.484 | 187040.5 | 3173480.75 | 4407467 | 3618243.5 | 1751.8602 | AGLNDLWQPLTC[160.0307]HAR | Q9ERM2 | Q9ERM2 | ICAM4 | intercellular adhesion molecule 4 (Landsteiner-Wiener blood group) | Plasma Membrane | other |
| 15556866 | 2 | 0.034 | 2907899 | 1919112.875 | 2104998.25 | 4046053.25 | 3980428 | 4199595.5 | 2564229.25 | 1508484.5 | 1707192 | 4973240.5 | 2933583 | 2193915.75 | 1793.9275 | NEFIPTNFEILSLEK | Q9JHR7 | Q9JHR7 | IDE | insulin-degrading enzyme | Extracellular Space | peptidase |
| 15542259 | 4 | 0.036 | 3295333.75 | 2364336.5 | 2094907.625 | 3668903.25 | 5142967 | 3346414.25 | 2937950.75 | 1996102.375 | 1220719.875 | 4552961 | 4188216 | 3291575.25 | 2322.153 | QAALGVMQMVEDTLIEHAHTK | Q9JHR7 | Q9JHR7 | IDE | insulin-degrading enzyme | Extracellular Space | peptidase |
| 15543024 | 3 | 0.162 | 2760645.5 | 7051087 | 3987926.75 | 7270838 | 1.11E+07 | 1.17E+07 | 3901503.75 | 9061801 | 1.04E+07 | 3144451.25 | 6268535.5 | 6535187 | 1920.9663 | LNEHFLNTTDFLDTIK | P54071 | P54071 | IDH2 | isocitrate dehydrogenase 2 (NADP+), mitochondrial | Cytoplasm | enzyme |
| 15538217 | 2 | 0.21 | 1077793 | 2133104 | 1523044.5 | 1693440 | 2063846.625 | 2410051 | 2049340.375 | 1806593.625 | 1761601.625 | 1051288.875 | 1672152 | 1462094.75 | 1355.6864 | TIEAEAAHGTVTR | P54071 | P54071 | IDH2 | isocitrate dehydrogenase 2 (NADP+), mitochondrial | Cytoplasm | enzyme |
| 15540485 | 2 | 0.258 | 1104624.125 | 2466850.25 | 1749741.25 | 2816638.5 | 3329069.25 | 3935486.5 | 1502038.75 | 3260822 | 3730419.75 | 765490.5 | 2546442 | 2310842.25 | 1244.69 | LIDDMVAQVLK | P54071 | P54071 | IDH2 | isocitrate dehydrogenase 2 (NADP+), mitochondrial | Cytoplasm | enzyme |
| 15539210 | 3 | 0.156 | 784379.562 | 1757591.375 | 782705.438 | 2126544 | 2641640.25 | 3363997.25 | 1341358.5 | 2123936.75 | 3282206.25 | 526056.75 | 1835558 | 2027515.875 | 1874.983 | GRPTSTNPIASIFAWTR | P54071 | P54071 | IDH2 | isocitrate dehydrogenase 2 (NADP+), mitochondrial | Cytoplasm | enzyme |
| 15540876 | 2 | 0.286 | 712440.75 | 2227595.5 | 1145961.25 | 1873739.75 | 2116863.75 | 3569601 | 1124359.375 | 2275571.5 | 3332940.5 | 454765.312 | 1648229.125 | 1549515.75 | 1154.5708 | DIFQEIFDK | P54071 | P54071 | IDH2 | isocitrate dehydrogenase 2 (NADP+), mitochondrial | Cytoplasm | enzyme |
| 15541921 | 2 | 0.809 | 566827.938 | 1245659.375 | 406674.656 | 1061076 | 1252094.5 | 889870.562 | 936319.875 | 612591 | 1834256.5 | 144624.578 | 2082088.5 | 947164 | 1383.7023 | DLAGC[160.0307]IHGLSNVK | P54071 | P54071 | IDH2 | isocitrate dehydrogenase 2 (NADP+), mitochondrial | Cytoplasm | enzyme |
| 15548261 | 3 | 0.059 | 9372785 | 2.60E+07 | 1.25E+07 | 1.24E+07 | 1.89E+07 | 2.19E+07 | 1.00E+07 | 2.18E+07 | 2.03E+07 | 6049421.5 | 5830681.5 | 8869420 | 2502.2307 | ENTEGEYSGIEHVIVDGVVQSIK | Q9D6R2 | Q9D6R2 | IDH3A | isocitrate dehydrogenase 3 (NAD+) alpha | Cytoplasm | enzyme |
| 15535255 | 2 | 0.056 | 4354550.5 | 8048345 | 4100658 | 6528680.5 | 6101443 | 7768954.5 | 3831614.5 | 6346280.5 | 7122358.5 | 2802443.5 | 3647050 | 3543097.75 | 1041.5664 | NVTAIQGPGGK | Q9D6R2 | Q9D6R2 | IDH3A | isocitrate dehydrogenase 3 (NAD+) alpha | Cytoplasm | enzyme |
| 15537136 | 2 | 0.069 | 3479172.25 | 8611934 | 4431611.5 | 4530409.5 | 6450803.5 | 8826493 | 2977808.75 | 6579637 | 6600607 | 2008503.125 | 2453347.5 | 3030324 | 1391.7479 | TPYTDVNIVTIR | Q9D6R2 | Q9D6R2 | IDH3A | isocitrate dehydrogenase 3 (NAD+) alpha | Cytoplasm | enzyme |
| 15536335 | 2 | 0.117 | 3656829.25 | 7147068.5 | 3232649.5 | 4986705.5 | 5660057.5 | 6257235.5 | 3048099.25 | 4965901.5 | 7094504.5 | 1938796.625 | 3326628 | 2953305 | 1028.5138 | APIQWEER | Q9D6R2 | Q9D6R2 | IDH3A | isocitrate dehydrogenase 3 (NAD+) alpha | Cytoplasm | enzyme |
| 15540993 | 3 | 0.05 | 4020921 | 8147488.5 | 3498589 | 4481949 | 5074982.5 | 6174954 | 4046145.75 | 6099889 | 6276342.5 | 1805548.875 | 2611541 | 3326688.25 | 1590.8757 | TPIAAGHPSMNLLLR | Q9D6R2 | Q9D6R2 | IDH3A | isocitrate dehydrogenase 3 (NAD+) alpha | Cytoplasm | enzyme |
| 15555490 | 3 | 0.379 | 4691068.5 | 1.06E+07 | 3880704.75 | 6315953 | 5762569.5 | 7779190 | 4212406.5 | 7694460 | 8180325.5 | 3060005.25 | 4776758 | 4611886 | 3283.562 | AVLASMDNENMHTPDIGGQGTTSQAIQDIIR | P70404 | P70404 | IDH3G | isocitrate dehydrogenase 3 (NAD+) gamma | Cytoplasm | enzyme |
| 15542554 | 2 | 0.051 | 1043961.688 | 2501200 | 1377268.375 | 1109845.125 | 1523808.125 | 2537206 | 1145929.5 | 1912141.625 | 2130943.5 | 520348.875 | 670527.625 | 897624.062 | 956.57574 | DIDILIVR | P70404 | P70404 | IDH3G | isocitrate dehydrogenase 3 (NAD+) gamma | Cytoplasm | enzyme |
| 15550829 | 2 | 0.553 | 7883047 | 1.07E+07 | 1.23E+07 | 1.13E+07 | 1.14E+07 | 1.26E+07 | 1.24E+07 | 1.03E+07 | 1.13E+07 | 1.13E+07 | 1.12E+07 | 1.15E+07 | 2031.9728 | NIGWGTDQGIGGFGEEPGIK | Q9CR20 | Q9CR20 | IER3IP1 | immediate early response 3 interacting protein 1 | Cytoplasm | other |
| 15535570 | 2 | 0.559 | 2433208.75 | 3435375.5 | 3101690 | 3217843.5 | 3729690 | 3583334.5 | 3239512 | 3603702 | 2891366.75 | 2748630.25 | 4076318.75 | 3509375.5 | 974.5432 | SQLMNLIR | Q9CR20 | Q9CR20 | IER3IP1 | immediate early response 3 interacting protein 1 | Cytoplasm | other |
| 15550995 | 5 | 0.579 | 2157699.25 | 1.13E+07 | 1584477.25 | 4953099.5 | 8468592 | 6180731 | 6124520.5 | 4460866.5 | 2936522.25 | 3856173.75 | 6072276 | 9402836 | 5706.661 | YVTSAPMPEPGAPGFYFTHSILTVTEEEWNSGETYTC[160.0307]VVGHEALPHLVTER | P01872 | P01872 | IGHM | immunoglobulin heavy constant mu | Plasma Membrane | transmembrane receptor |
| 15541015 | 2 | 0.52 | 841765.25 | 2654207 | 226169.938 | 1385706.875 | 2197517 | 1614153 | 1926157.5 | 687003.625 | 1597856.625 | 927818.562 | 2799667.75 | 2831824.5 | 1107.5698 | ESATVTC[160.0307]LVK | P01872 | P01872 | IGHM | immunoglobulin heavy constant mu | Plasma Membrane | transmembrane receptor |
| 15536847 | 2 | 0.505 | 552486.875 | 2841938 | 514083.406 | 1185242.875 | 3373852.25 | 2476574.25 | 2581957.75 | 1284220.5 | 1933100.875 | 732308.312 | 3158699.75 | 3294943.5 | 1293.6478 | LIC[160.0307]EATNFTPK | P01872 | P01872 | IGHM | immunoglobulin heavy constant mu | Plasma Membrane | transmembrane receptor |
| 15538544 | 2 | 0.595 | 488890.625 | 2487265 | 810153.375 | 1285393.5 | 2864807.25 | 2084157.75 | 2290530.25 | 1338306.375 | 2316778.25 | 543437.688 | 2470045.25 | 2849647.25 | 943.55945 | PITVSWLK | P01872 | P01872 | IGHM | immunoglobulin heavy constant mu | Plasma Membrane | transmembrane receptor |
| 15534392 | 2 | 0.185 | 3973199.25 | 9051826 | 4493197.5 | 5392863 | 5770958.5 | 7589131.5 | 4327573.5 | 7009787.5 | 9323869 | 2412461.75 | 4129327.5 | 4225369.5 | 998.45496 | FSFQC[160.0307]PGR | O55222 | O55222 | ILK | integrin-linked kinase | Plasma Membrane | kinase |
| 15552981 | 2 | 0.129 | 237813.844 | 816043.625 | 521217.031 | 2927401.5 | 2086775.25 | 2288501.5 | 1579860.75 | 1215503.875 | 6765277 | 285665.5 | 1498909.625 | 3299992.75 | 1680.7931 | EVPFADLSNMEIGMK | O55222 | O55222 | ILK | integrin-linked kinase | Plasma Membrane | kinase |
| 15535594 | 2 | 0.269 | 433386.469 | 675415.688 | 871092.25 | 3699708.5 | 2627609.75 | 3318926.75 | 2153092 | 1049947.75 | 6417971 | 119944.789 | 1775763.125 | 4208115 | 1204.6627 | FDMIVPILEK | O55222 | O55222 | ILK | integrin-linked kinase | Plasma Membrane | kinase |
| 15540243 | 3 | 0.215 | 69922.703 | 516412.594 | 446805.125 | 2494715.75 | 2054374 | 1969326.625 | 1303600.125 | 861037.875 | 5213324.5 | 64539.828 | 1418945.125 | 3224696.25 | 1594.8745 | GMAFLHTLEPLIPR | O55222 | O55222 | ILK | integrin-linked kinase | Plasma Membrane | kinase |
| 15555635 | 3 | 0.463 | 94710.141 | 343886.781 | 286683.031 | 1101525.25 | 1262510 | 1073257.125 | 1239628.75 | 239648.031 | 4379400 | 6199.497 | 785781.562 | 1950048.625 | 3110.3752 | LWLDNTENDLNQGDDHGFSPLHWAC[160.0307]R | O55222 | O55222 | ILK | integrin-linked kinase | Plasma Membrane | kinase |
| 15544417 | 2 | 0.465 | 21190.99 | 527027.625 | 417850.812 | 1258869.5 | 878933.312 | 1079750.75 | 895776.062 | 574790.375 | 3790185.75 | 4.70E-06 | 655702.312 | 2047266.625 | 1148.6152 | SAVVEMLIMR | O55222 | O55222 | ILK | integrin-linked kinase | Plasma Membrane | kinase |
| 15541341 | 3 | 0.157 | 204513.594 | 802940.25 | 588830.938 | 612862.625 | 1266760.375 | 1059112.375 | 825252.125 | 1612855.625 | 1031487.938 | 876660.375 | 762646.25 | 984080.688 | 2179.2388 | NAQVAQSPVLLLGGAASTLLQK | Q8BU33 | Q8BU33 | ILVBL | ilvB (bacterial acetolactate synthase)-like | Cytoplasm | enzyme |
| 15551874 | 2 | 0.323 | 3490383.75 | 7248836 | 3700097.25 | 4582109 | 5198645 | 6846504 | 3649041.75 | 5802334.5 | 7487509 | 2904217.25 | 4568096 | 3147118.25 | 1571.7942 | ELDSITPDITPGWK | Q8CAQ8 | Q8CAQ8 | IMMT | inner membrane protein, mitochondrial | Cytoplasm | other |
| 15550245 | 2 | 0.229 | 4041040 | 8151772.5 | 3970902 | 5864228 | 6042441.5 | 7128072.5 | 3992517.75 | 5733894.5 | 8164572 | 2458898 | 4209607.5 | 4431751.5 | 1929.9901 | TSSAEMPTIPLGSAVEAIR | Q8CAQ8 | Q8CAQ8 | IMMT | inner membrane protein, mitochondrial | Cytoplasm | other |
| 15537072 | 2 | 0.216 | 3882064 | 6482224 | 4017225 | 5202128.5 | 5053414.5 | 6574798.5 | 3106274.25 | 5689493 | 6286669.5 | 2328095.75 | 4465179 | 3193026 | 1115.592 | AVDEAADALLK | Q8CAQ8 | Q8CAQ8 | IMMT | inner membrane protein, mitochondrial | Cytoplasm | other |
| 15539597 | 2 | 0.192 | 3118378.5 | 7118231.5 | 4494089.5 | 3272253.25 | 5363452 | 5871330.5 | 3708418 | 6318918.5 | 7217643 | 2082687.125 | 3150585.25 | 3612932.25 | 1229.5663 | YEFEQGLSEK | Q8CAQ8 | Q8CAQ8 | IMMT | inner membrane protein, mitochondrial | Cytoplasm | other |
| 15536750 | 3 | 0.16 | 2725280.75 | 5781466 | 3110009.75 | 5060498.5 | 4167289.25 | 5278709 | 2735014 | 5268858 | 6279993.5 | 1765967.5 | 2812857.5 | 3140265 | 1527.8241 | VVSQYHELVVQAR | Q8CAQ8 | Q8CAQ8 | IMMT | inner membrane protein, mitochondrial | Cytoplasm | other |
| 15536125 | 2 | 0.243 | 1604139.5 | 3540822.5 | 1113378.375 | 2317940 | 2545216.75 | 2059716.125 | 1780623.375 | 2095921.25 | 2542375 | 978959.25 | 1609338.75 | 1365158.5 | 1053.5189 | GVYSEETLR | Q8CAQ8 | Q8CAQ8 | IMMT | inner membrane protein, mitochondrial | Cytoplasm | other |
| 15551625 | 2 | 0.242 | 1483996.375 | 4908118.5 | 2340243 | 2763007 | 2908934.75 | 3080095.75 | 1616222.25 | 3112739.5 | 4351472 | 928822.625 | 2058002.75 | 1515777 | 1789.9883 | LFGMVLGSAPYTVPLPK | Q8CAQ8 | Q8CAQ8 | IMMT | inner membrane protein, mitochondrial | Cytoplasm | other |
| 15541292 | 2 | 0.116 | 654071.688 | 1172125.125 | 551948.5 | 862609.688 | 877190.812 | 1316698 | 757739.062 | 1130740.125 | 1175396.875 | 484969.281 | 748418.688 | 499713.625 | 1245.5923 | YSTSSSSGLTAGK | Q8CAQ8 | Q8CAQ8 | IMMT | inner membrane protein, mitochondrial | Cytoplasm | other |
| 15539980 | 3 | 0.26 | 707815 | 2927423.5 | 1240995.5 | 1900481.875 | 2132150.75 | 2624421.75 | 1195922.25 | 1945868.125 | 2383804.75 | 462564.219 | 1300144 | 1317643.625 | 1509.806 | LHNMIVDLDNVVK | Q8CAQ8 | Q8CAQ8 | IMMT | inner membrane protein, mitochondrial | Cytoplasm | other |
| 15545924 | 4 | 0.507 | 4707064 | 1498009.875 | 84165.477 | 1459235 | 353662.938 | 296958.5 | 2341658 | 236163.672 | 31239.066 | 2736869.75 | 1509013.25 | 1630537.375 | 2072.0103 | RTSSAQVEGGVHSLHSYEK | P24547 | P24547 | IMPDH2 | IMP (inosine 5'-monophosphate) dehydrogenase 2 | Cytoplasm | enzyme |
| 15552835 | 2 | 0.551 | 6782375 | 6770610.5 | 6746706 | 5000372.5 | 7438032 | 5068357 | 5344923.5 | 6161206 | 6555521 | 7412012.5 | 5996307 | 5701318.5 | 1836.8978 | SFLREDTLYEPEPQI | Q8C5L6 | Q8C5L6 | INPP5K | inositol polyphosphate-5-phosphatase K | Cytoplasm | phosphatase |
| 15553001 | 3 | 0.103 | 5694351 | 7701569 | 7625312 | 7620359 | 7591970.5 | 7761065 | 6710520.5 | 6684407 | 7682356.5 | 6357432 | 4288642 | 6119295.5 | 2491.3481 | RQPSQASPLASSVPTSYFLLTLK | Q8C5L6 | Q8C5L6 | INPP5K | inositol polyphosphate-5-phosphatase K | Cytoplasm | phosphatase |
| 15537935 | 2 | 0.421 | 3.55E+07 | 4.05E+07 | 4.98E+07 | 2.91E+07 | 3.92E+07 | 3.30E+07 | 3.39E+07 | 3.51E+07 | 3.56E+07 | 4.46E+07 | 2.57E+07 | 3.09E+07 | 878.4716 | KLM[147.0354]SDLR | Q8CIM8 | Q8CIM8 | INTS4 | integrator complex subunit 4 | Nucleus | other |
| 15550092, 15551738 | 2, 3 | 0.607 | 1.18E+07 | 1.23E+07 | 1.05E+07 | 1.24E+07 | 1.27E+07 | 8415714 | 9079783 | 9453700 | 7747500.5 | 1.65E+07 | 7317842 | 7516647.5 | 2364.2275, 2364.23 | LLSSAFDEVYPALPSDVQTAIK | Q8BKC5 | Q8BKC5 | IPO5 | importin 5 | Nucleus | transporter |
| 15555484 | 4 | 0.583 | 7034671 | 5755585 | 2695315.25 | 5943029 | 5580365 | 2191288.75 | 4735467 | 3812868.75 | 1819942.625 | 1.32E+07 | 4856586.5 | 3611815.25 | 3087.5347 | FKPDC[160.0307]VNVEEVLPHWLSWLPLHEDK | Q8BKC5 | Q8BKC5 | IPO5 | importin 5 | Nucleus | transporter |
| 15537942 | 3 | 0.133 | 1.36E+07 | 1.39E+07 | 1.55E+07 | 1.17E+07 | 1.39E+07 | 1.09E+07 | 1.26E+07 | 1.41E+07 | 9886582 | 1.24E+07 | 9793145 | 9871310 | 2102.119 | LVLEQVVTSIASVADTAEEK | Q8BKC5 | Q8BKC5 | IPO5 | importin 5 | Nucleus | transporter |
| 15553486 | 3 | 0.229 | 1.78E+07 | 2.08E+07 | 1.63E+07 | 1.16E+07 | 1.93E+07 | 1.24E+07 | 1.64E+07 | 1.95E+07 | 1.05E+07 | 1.13E+07 | 1.11E+07 | 1.35E+07 | 3512.7283 | EEAVQTFSYLC[160.0307]DLIESNHPIVLGPNNTNLPK | Q8BKC5 | Q8BKC5 | IPO5 | importin 5 | Nucleus | transporter |
| 15533650 | 2 | 0.855 | 8050691.5 | 5382391 | 6401980.5 | 7434531 | 6901704.5 | 4990154.5 | 6498200.5 | 6071966 | 4428265.5 | 9439133 | 6045852.5 | 4781252.5 | 1427.7023 | EGFVEYTEQVVK | Q8BKC5 | Q8BKC5 | IPO5 | importin 5 | Nucleus | transporter |
| 15548808 | 2 | 0.849 | 9237902 | 6332704 | 6446819.5 | 7089757.5 | 7319842.5 | 5434992.5 | 7507173 | 6434667 | 4964674.5 | 9148708 | 6076849.5 | 3854744 | 1605.7511 | ENVNATENC[160.0307]ISAVGK | Q8BKC5 | Q8BKC5 | IPO5 | importin 5 | Nucleus | transporter |
| 15542389 | 3 | 0.36 | 6652356 | 6399534.5 | 5476620 | 5875096.5 | 5875887 | 4856220 | 6790384 | 6595347 | 5831329.5 | 8755796 | 8428413 | 5098162 | 1955.9592 | VQAHAAAALINFTEDC[160.0307]PK | Q8BKC5 | Q8BKC5 | IPO5 | importin 5 | Nucleus | transporter |
| 15535118 | 2 | 0.194 | 1.07E+07 | 7641373.5 | 7571561 | 7213849.5 | 7114096.5 | 6137955.5 | 7118836 | 6441270.5 | 5909070.5 | 8513527 | 6537260.5 | 6008526 | 1171.6543 | SLVEIADTVPK | Q8BKC5 | Q8BKC5 | IPO5 | importin 5 | Nucleus | transporter |
| 15536619 | 2 | 0.769 | 7397911 | 4782788 | 4780141.5 | 5982713.5 | 5466681.5 | 2690187.25 | 5974080 | 4796521 | 3304800 | 8392086 | 5490968 | 3896957.75 | 1406.6516 | QAEETYENIPGR | Q8BKC5 | Q8BKC5 | IPO5 | importin 5 | Nucleus | transporter |
| 15551504 | 3 | 0.827 | 6539035.5 | 7663360.5 | 4560734.5 | 5451148.5 | 6080674.5 | 5067682.5 | 5923587.5 | 5142695 | 4842640 | 8258522 | 4986991.5 | 4990119 | 3370.6482 | QVQTSGGLWTEC[160.0307]IAQLSPEQQAAIQELLNSA | Q8BKC5 | Q8BKC5 | IPO5 | importin 5 | Nucleus | transporter |
| 15557875, 15559404 | 2 | 0.725 | 7000858.5 | 4940606.5 | 4733470 | 4876079 | 5171893.5 | 4013199.5 | 5561520 | 4736545 | 4338114 | 7092992 | 6086714.5 | 3810982.25 | 1630.8259, 1630.8263 | VAAAESMPLLLEC[160.0307]AR | Q8BKC5 | Q8BKC5 | IPO5 | importin 5 | Nucleus | transporter |
| 15539311 | 2 | 0.466 | 7180652 | 6474543.5 | 5066466 | 5608331.5 | 5074408.5 | 4396825.5 | 5161081.5 | 5102542.5 | 5367460.5 | 6820786 | 4573186 | 3522813.75 | 1074.6653 | ITFLLQAIR | Q8BKC5 | Q8BKC5 | IPO5 | importin 5 | Nucleus | transporter |
| 15550075 | 2 | 0.93 | 6317408.5 | 4528333 | 3998073.5 | 5225567 | 4795775 | 2965519 | 5810160 | 4426850 | 3371383.25 | 5737244 | 4533162.5 | 3382016.75 | 1786.9078 | VIAALLQTMEDQGNQR | Q8BKC5 | Q8BKC5 | IPO5 | importin 5 | Nucleus | transporter |
| 15533262 | 2 | 0.464 | 5347571 | 3935608.5 | 3386503.5 | 3859793.25 | 4773149 | 2261876.5 | 4963265 | 4229892.5 | 3254417.75 | 5639631 | 6370268 | 3738079.25 | 1131.5804 | IC[160.0307]DIAAELAR | Q8BKC5 | Q8BKC5 | IPO5 | importin 5 | Nucleus | transporter |
| 15552157 | 3 | 0.349 | 4175361 | 3553070.25 | 2915035.5 | 4692118.5 | 4547294.5 | 2659219.5 | 3257157.5 | 3891204.5 | 2895043.25 | 5513794.5 | 5114498 | 3591645.5 | 2634.1929 | C[160.0307]IEVMGDGC[160.0307]LNNEHFEELGGILK | Q8BKC5 | Q8BKC5 | IPO5 | importin 5 | Nucleus | transporter |
| 15549763 | 2 | 0.767 | 3588042.5 | 2781702 | 4023182 | 2924790.75 | 2923171.75 | 2806827.25 | 2804063 | 3672804.75 | 3203573.25 | 5192650.5 | 2945891 | 2560365 | 1638.8198 | FMQDASDVMQLLLK | Q8BKC5 | Q8BKC5 | IPO5 | importin 5 | Nucleus | transporter |
| 15540411 | 2 | 0.458 | 3761276.25 | 3033167.5 | 1349416.125 | 3056729 | 2420054 | 1833617 | 3589229 | 2449790.5 | 1761578.875 | 5149073.5 | 5091360.5 | 2307169.75 | 1431.639 | STAC[160.0307]QMLVC[160.0307]YAK | Q8BKC5 | Q8BKC5 | IPO5 | importin 5 | Nucleus | transporter |
| 15551612 | 3 | 0.951 | 4645119.5 | 3824063 | 3564284.75 | 5086203.5 | 3909354 | 3015298 | 4562787 | 3925472.5 | 3190395.5 | 5055865 | 4252622.5 | 3457311.25 | 3371.6497 | QAAAYGLGVMAQFGGDNYRPFC[160.0307]TDALPLLVR | Q8BKC5 | Q8BKC5 | IPO5 | importin 5 | Nucleus | transporter |
| 15555086 | 2 | 0.673 | 2822100.5 | 2812535.5 | 2694700.5 | 2876356.25 | 2462810.5 | 2564689.75 | 2625923.5 | 2524777 | 2160356 | 3781683.25 | 1781466.875 | 1526512.875 | 2102.119 | LVLEQVVTSIASVADTAEEK | Q8BKC5 | Q8BKC5 | IPO5 | importin 5 | Nucleus | transporter |
| 15538038 | 3 | 0.418 | 2706200.75 | 2887140.75 | 1711069 | 2907302 | 3384453.25 | 2171796.75 | 4115582.5 | 3054831.5 | 2541893.25 | 3594316.75 | 3268802 | 2688642.75 | 1786.9082 | VIAALLQTMEDQGNQR | Q8BKC5 | Q8BKC5 | IPO5 | importin 5 | Nucleus | transporter |
| 15554674 | 3 | 0.487 | 8050557 | 6067700.5 | 2591384.25 | 3376429.75 | 4231865 | 2815822.75 | 3885832.25 | 3385403.25 | 2770563.25 | 4317617.5 | 4238099.5 | 2744965.5 | 2313.229 | VLQQTLNYINQGVSHALTWK | Q9EPL8 | Q9EPL8 | IPO7 | importin 7 | Nucleus | transporter |
| 15552471 | 3 | 0.871 | 4081670 | 3403364.75 | 1528490.875 | 3209510 | 2868366.5 | 1194013.5 | 2824935 | 2403393.25 | 2251412.75 | 3129323 | 2745932.75 | 2532552.75 | 2835.2852 | DQMEYMLQNHVFPLFSSELGYMR | Q9EPL8 | Q9EPL8 | IPO7 | importin 7 | Nucleus | transporter |
| 15538159 | 2 | 0.193 | 4448779.5 | 3041942.5 | 2047712.5 | 2353387.75 | 2258729.25 | 1383345.25 | 1864391.125 | 2254239.5 | 1555955.375 | 2390682.5 | 1955943.25 | 1648153.5 | 1090.5992 | AIFQTIQNR | Q9EPL8 | Q9EPL8 | IPO7 | importin 7 | Nucleus | transporter |
| 15534156 | 2 | 0.089 | 3872668 | 3549553 | 2390106 | 2298732.25 | 2416714.75 | 1578855.25 | 2789116 | 2449803.25 | 2178958.25 | 2185588 | 2400695.75 | 2241863.75 | 1272.7645 | AFAVGVQQVLLK | Q9EPL8 | Q9EPL8 | IPO7 | importin 7 | Nucleus | transporter |
| 15543678 | 3 | 0.65 | 2210128.25 | 2461419.75 | 1107949 | 1534051 | 1602273.625 | 917798.375 | 1718543.125 | 1199616.875 | 1295468.5 | 1937654.25 | 1445828.375 | 846123.938 | 1680.9315 | DGALHMIGSLAEILLK | Q9EPL8 | Q9EPL8 | IPO7 | importin 7 | Nucleus | transporter |
| 15557028 | 2 | 0.527 | 2484548.5 | 1681255 | 1313843.125 | 1613034 | 1800612.5 | 1130925.125 | 1411155.375 | 1488267.875 | 1360862.875 | 1685769.25 | 1379331.375 | 912762.438 | 1518.724 | ETENDDLTNVIQK | Q9EPL8 | Q9EPL8 | IPO7 | importin 7 | Nucleus | transporter |
| 15559663 | 2 | 0.717 | 981217.312 | 1211221.875 | 315708 | 591010.062 | 818914.562 | 329086.844 | 817903.875 | 671674.75 | 897909.188 | 558498.688 | 712298.75 | 523805.156 | 1680.9319 | DGALHMIGSLAEILLK | Q9EPL8 | Q9EPL8 | IPO7 | importin 7 | Nucleus | transporter |
| 15540503 | 3 | 0.662 | 1.70E+07 | 1.68E+07 | 1.31E+07 | 1.83E+07 | 1.75E+07 | 1.45E+07 | 1.71E+07 | 2.03E+07 | 1.37E+07 | 1.82E+07 | 1.33E+07 | 1.21E+07 | 1622.8098 | VIGHDEDASAPTGVVR | Q60766 | Q60766 | Irgm1 | immunity-related GTPase family M member 1 | Cytoplasm | other |
| 15544015 | 2 | 0.646 | 1.79E+07 | 1.12E+07 | 9935246 | 1.40E+07 | 1.31E+07 | 8031236.5 | 1.22E+07 | 1.55E+07 | 1.23E+07 | 1.60E+07 | 1.73E+07 | 1.17E+07 | 1165.6266 | MLFLVAQDTK | Q60766 | Q60766 | Irgm1 | immunity-related GTPase family M member 1 | Cytoplasm | other |
| 15550410 | 3 | 0.089 | 1.21E+07 | 1.39E+07 | 1.24E+07 | 1.31E+07 | 1.36E+07 | 1.57E+07 | 1.07E+07 | 1.65E+07 | 1.30E+07 | 1.13E+07 | 1.00E+07 | 9955497 | 2773.3389 | LIFGVDDESVQQVAQSMGTVVMEYK | Q60766 | Q60766 | Irgm1 | immunity-related GTPase family M member 1 | Cytoplasm | other |
| 15532585 | 3 | 0.259 | 1.00E+07 | 1.24E+07 | 1.01E+07 | 1.28E+07 | 1.04E+07 | 1.04E+07 | 1.32E+07 | 1.41E+07 | 8121761.5 | 1.13E+07 | 6720650.5 | 6686413 | 1589.8452 | LDRDLSTSVLSEVR | Q60766 | Q60766 | Irgm1 | immunity-related GTPase family M member 1 | Cytoplasm | other |
| 15553518 | 3 | 0.278 | 8382756 | 8495434 | 6640405.5 | 9705876 | 7714240 | 9003922 | 9372718 | 9533886 | 8902991 | 9679809 | 6553870 | 5394392 | 3211.6223 | ETVATLSQIPVSIFVTGDSGNGMSSFINALR | Q60766 | Q60766 | Irgm1 | immunity-related GTPase family M member 1 | Cytoplasm | other |
| 15536543 | 2 | 0.114 | 7988458 | 9664418 | 6629536 | 7765086.5 | 7970590.5 | 9560002 | 8793516 | 9572778 | 6864180.5 | 6851140.5 | 5036459 | 6469597.5 | 1047.6427 | LLELVYGIK | Q60766 | Q60766 | Irgm1 | immunity-related GTPase family M member 1 | Cytoplasm | other |
| 15540647 | 3 | 0.11 | 5061489 | 7249237 | 6468755 | 6581995 | 7648288 | 7760017 | 7399368.5 | 1.07E+07 | 6470715 | 5577870.5 | 5280233 | 5900321 | 1822.8025 | NSLGVRDDDNMGEC[160.0307]LK | Q60766 | Q60766 | Irgm1 | immunity-related GTPase family M member 1 | Cytoplasm | other |
| 15550376 | 2 | 0.182 | 4790158 | 3926860.75 | 3625276.75 | 5493310.5 | 4002281 | 3792199.5 | 4011108.75 | 4475433 | 3293443.75 | 4035024 | 2680686.5 | 2583562.25 | 1622.8102 | VIGHDEDASAPTGVVR | Q60766 | Q60766 | Irgm1 | immunity-related GTPase family M member 1 | Cytoplasm | other |
| 15554271 | 4 | 0.236 | 3532155.75 | 5512879 | 9256256 | 3869169.25 | 3878416.25 | 1.17E+07 | 3513070.75 | 9065847 | 8266191.5 | 2150337.5 | 2674550.75 | 3739929.75 | 4085.9346 | TRTEYSSSHFPNVVLWDLPGLGATAQTVEDYVEEM[147.0354]K | Q60766 | Q60766 | Irgm1 | immunity-related GTPase family M member 1 | Cytoplasm | other |
| 15559461 | 3 | 0.307 | 898379.75 | 1946122.25 | 2351644.75 | 962032.5 | 1976564.5 | 3040298 | 1459981.875 | 3832688.25 | 2506644.5 | 1285342.375 | 836303.438 | 1240961 | 3828.7896 | TEYSSSHFPNVVLWDLPGLGATAQTVEDYVEEM[147.0354]K | Q60766 | Q60766 | Irgm1 | immunity-related GTPase family M member 1 | Cytoplasm | other |
| 15549241 | 2 | 0.053 | 2676604.75 | 5011309 | 2412616.5 | 4259231 | 4379684.5 | 4889806.5 | 2063059.625 | 3945987.75 | 6495777 | 1769508.5 | 1356346.875 | 2070368.125 | 1508.7722 | LSSEFVFNNPNIK | Q9D924 | Q9D924 | ISCA1 | iron-sulfur cluster assembly 1 | Cytoplasm | other |
| 15535453 | 3 | 0.427 | 1806906 | 2289556 | 837177.75 | 2528644.25 | 2077860 | 1875859.25 | 980395.125 | 2020973.125 | 2897574.5 | 1107482.375 | 1493457.125 | 1159095.125 | 1647.7782 | TKGDSDEEVIQDGVR | Q9D924 | Q9D924 | ISCA1 | iron-sulfur cluster assembly 1 | Cytoplasm | other |
| 15540892 | 3 | 0.411 | 2048063.5 | 5396853 | 1178904.25 | 2142904.25 | 1911728.875 | 3364377.75 | 1764914.25 | 2729117.5 | 3871330 | 1534471.75 | 922902.438 | 1851431.375 | 2236.1008 | FSLDTVINPDDRVFEQGGAR | Q9DCB8 | Q9DCB8 | ISCA2 | iron-sulfur cluster assembly 2 | Cytoplasm | other |
| 15541092, 15543383 | 3 | 0.233 | 4.51E+07 | 5.82E+07 | 5.40E+07 | 4.88E+07 | 7.40E+07 | 7.64E+07 | 5.85E+07 | 5.97E+07 | 3.45E+07 | 5.31E+07 | 3.12E+07 | 4.16E+07 | 2065.0552, 2065.0574 | GHSNIYEVFLTQTVDTLK | Q64339 | Q64339 | ISG15 | ISG15 ubiquitin-like modifier | Extracellular Space | other |
| 15533509 | 2 | 0.129 | 4.44E+07 | 3.92E+07 | 3.23E+07 | 4.97E+07 | 5.34E+07 | 5.91E+07 | 4.83E+07 | 4.75E+07 | 2.97E+07 | 4.67E+07 | 2.72E+07 | 2.80E+07 | 1015.56555 | IGVPAFQQR | Q64339 | Q64339 | ISG15 | ISG15 ubiquitin-like modifier | Extracellular Space | other |
| 15537779 | 4 | 0.392 | 2.44E+07 | 3.21E+07 | 1.86E+07 | 3.42E+07 | 3.56E+07 | 2.96E+07 | 3.68E+07 | 2.58E+07 | 1.65E+07 | 2.57E+07 | 2.52E+07 | 1.88E+07 | 2507.126 | EQVHEDQFWLSFEGRPMEDK | Q64339 | Q64339 | ISG15 | ISG15 ubiquitin-like modifier | Extracellular Space | other |
| 15532052 | 3 | 0.426 | 2.28E+07 | 1.09E+07 | 1869897.125 | 2.62E+07 | 1.34E+07 | 1.25E+07 | 1.56E+07 | 1.07E+07 | 1.34E+07 | 2.18E+07 | 2.66E+07 | 1.40E+07 | 1847.9896 | ELLGEYGLKPQC[160.0307]TVIK | Q64339 | Q64339 | ISG15 | ISG15 ubiquitin-like modifier | Extracellular Space | other |
| 15556125 | 3 | 0.025 | 2.28E+07 | 2.62E+07 | 2.00E+07 | 2.42E+07 | 4.14E+07 | 3.19E+07 | 2.68E+07 | 2.74E+07 | 2.35E+07 | 2.16E+07 | 1.71E+07 | 1.65E+07 | 2507.1252 | EQVHEDQFWLSFEGRPMEDK | Q64339 | Q64339 | ISG15 | ISG15 ubiquitin-like modifier | Extracellular Space | other |
| 15535283 | 2 | 0.653 | 6973279.5 | 1.22E+07 | 1.40E+07 | 8493878 | 1.92E+07 | 1.33E+07 | 1.53E+07 | 1.42E+07 | 6378454.5 | 1.40E+07 | 5843803.5 | 6765654.5 | 1021.55475 | ELLGEYGLK | Q64339 | Q64339 | ISG15 | ISG15 ubiquitin-like modifier | Extracellular Space | other |
| 15549166 | 2 | 0.535 | 5263290.5 | 2245267.5 | 114942.672 | 4489121.5 | 2322165.5 | 2060433.625 | 3586660.75 | 2360906.25 | 2868704.75 | 2915585.25 | 6860664.5 | 2915217.75 | 1847.988 | ELLGEYGLKPQC[160.0307]TVIK | Q64339 | Q64339 | ISG15 | ISG15 ubiquitin-like modifier | Extracellular Space | other |
| 15556926 | 2 | 0.052 | 2412780 | 3456199.25 | 3547157 | 2621000 | 3761046.5 | 4927869.5 | 3820883.5 | 4553743 | 3380199.75 | 2333331 | 2198868.25 | 2311962.75 | 2242.1033 | MLGGNDFLVSVTNSMTVSELK | Q64339 | Q64339 | ISG15 | ISG15 ubiquitin-like modifier | Extracellular Space | other |
| 15534301 | 3 | 0.159 | 8485183 | 1.04E+07 | 8334860.5 | 7666691 | 9927449 | 5329461.5 | 5503358 | 6121339.5 | 5917974.5 | 9033550 | 5705922.5 | 7014156.5 | 1511.7461 | YIRPEGEITDYR | Q9JL16 | Q9JL16 | ISG20 | interferon stimulated exonuclease gene 20kDa | Nucleus | enzyme |
| 15533662 | 3 | 0.148 | 6957041 | 8056277 | 5189777.5 | 6671558.5 | 7443374 | 3551483.25 | 4333680 | 4547830 | 2723742 | 6773534.5 | 5270555 | 5815254.5 | 1439.739 | TQVSGVTPQHMVR | Q9JL16 | Q9JL16 | ISG20 | interferon stimulated exonuclease gene 20kDa | Nucleus | enzyme |
| 15537748 | 2 | 0.119 | 6161409 | 6505437 | 5071141 | 4479526.5 | 7060090.5 | 3259494.75 | 3845810.25 | 3973127.25 | 3243901.5 | 5712277.5 | 5665134 | 4805597 | 1234.5549 | YTIYDTSTDR | Q9JL16 | Q9JL16 | ISG20 | interferon stimulated exonuclease gene 20kDa | Nucleus | enzyme |
| 15533057 | 2 | 0.003 | 5959630 | 7194415 | 6169262.5 | 5180075 | 5468152.5 | 4824883 | 3655293.75 | 4571009 | 4174552 | 4234949.5 | 4657217 | 5136691.5 | 969.63196 | LEILQLLK | Q9JL16 | Q9JL16 | ISG20 | interferon stimulated exonuclease gene 20kDa | Nucleus | enzyme |
| 15535290 | 2 | 0.065 | 2623933.25 | 1540455.125 | 1618942.125 | 1728472.375 | 1910528.625 | 1581908.75 | 2000069.125 | 1734286.75 | 2099090.5 | 4667257 | 2718686 | 2372653 | 1017.5378 | IVADQLC[160.0307]AK | Q9CX00 | Q9CX00 | IST1 | increased sodium tolerance 1 homolog (yeast) | Cytoplasm | other |
| 15551912 | 3 | 0.127 | 582924.5 | 1518512.875 | 3684863 | 6568184 | 6820219.5 | 9007617 | 4044244.5 | 6340297.5 | 8240762 | 738286.938 | 6800307.5 | 5143662 | 2341.2368 | ADNSVSLTIPLLYDAELHLTR | Q62469 | Q62469 | ITGA2 | integrin, alpha 2 (CD49B, alpha 2 subunit of VLA-2 receptor) | Plasma Membrane | transmembrane receptor |
| 15552399 | 3 | 0.083 | 207570.547 | 821856.938 | 1546719.5 | 5437062.5 | 5759584.5 | 5553343 | 3223726.75 | 3650616.5 | 6720243.5 | 286018.25 | 5341265.5 | 3096993.25 | 3790.8567 | EC[160.0307]GSDGIC[160.0307]ISDLILDVQQLPAIQTQSFIVSNQNK | Q62469 | Q62469 | ITGA2 | integrin, alpha 2 (CD49B, alpha 2 subunit of VLA-2 receptor) | Plasma Membrane | transmembrane receptor |
| 15536726 | 2 | 0.141 | 45019.371 | 710094.375 | 2514610.5 | 5941747 | 4554486 | 6022922 | 3123252.5 | 4971750.5 | 6627275 | 279853.25 | 5291953.5 | 3070451.5 | 1353.7024 | IIFNLNDFETK | Q62469 | Q62469 | ITGA2 | integrin, alpha 2 (CD49B, alpha 2 subunit of VLA-2 receptor) | Plasma Membrane | transmembrane receptor |
| 15551124 | 2 | 0.15 | 367272.438 | 946341.688 | 1556309.75 | 3807219.5 | 3469540 | 4397553.5 | 1938685.75 | 2713822.25 | 5342170 | 266110.125 | 3209534.5 | 2589108.75 | 1629.8131 | YFFNVADEAALLEK | Q62469 | Q62469 | ITGA2 | integrin, alpha 2 (CD49B, alpha 2 subunit of VLA-2 receptor) | Plasma Membrane | transmembrane receptor |
| 15534138 | 3 | 0.109 | 313306.625 | 892025.625 | 2188686.5 | 1.02E+07 | 7590424 | 7699141 | 4379960 | 6128799.5 | 7772001.5 | 258169.062 | 7952467.5 | 5452576 | 2038.0055 | GILNQHQFLEGPEGTGNAR | Q62469 | Q62469 | ITGA2 | integrin, alpha 2 (CD49B, alpha 2 subunit of VLA-2 receptor) | Plasma Membrane | transmembrane receptor |
| 15538114 | 2 | 0.083 | 131483.906 | 571550.75 | 1024248.875 | 3160577.5 | 4598521 | 4577889 | 2211952.75 | 3045288.25 | 4858514.5 | 222840.172 | 4534670.5 | 2573695 | 1390.6332 | C[160.0307]PVDLPTATC[160.0307]EK | Q62469 | Q62469 | ITGA2 | integrin, alpha 2 (CD49B, alpha 2 subunit of VLA-2 receptor) | Plasma Membrane | transmembrane receptor |
| 15551195 | 2 | 0.17 | 36323.781 | 554063.188 | 883179.25 | 4978131 | 4156552.75 | 4839665 | 2381423 | 2444757 | 5069111.5 | 51219.523 | 5067622 | 2961867.75 | 1603.7722 | TVIQQC[160.0307]NDDEILR | Q62469 | Q62469 | ITGA2 | integrin, alpha 2 (CD49B, alpha 2 subunit of VLA-2 receptor) | Plasma Membrane | transmembrane receptor |
| 15552870 | 2 | 0.151 | 47693.641 | 427283.844 | 876001.5 | 1193846.875 | 2907796.25 | 3139526.5 | 1397577.5 | 2566306.5 | 2841085 | 34594.875 | 915637.125 | 1445332.125 | 2045.9442 | GDQIGSYFGSVLC[160.0307]SVDVDK | Q62469 | Q62469 | ITGA2 | integrin, alpha 2 (CD49B, alpha 2 subunit of VLA-2 receptor) | Plasma Membrane | transmembrane receptor |
| 15537408 | 2 | 0.408 | 66276.664 | 216620.469 | 1315665.625 | 2147050.75 | 2449122.75 | 2715241 | 1438021.375 | 1806868.875 | 3906899.5 | 4.70E-06 | 2781221 | 2082546.875 | 1000.54877 | VFSIPFYK | Q62469 | Q62469 | ITGA2 | integrin, alpha 2 (CD49B, alpha 2 subunit of VLA-2 receptor) | Plasma Membrane | transmembrane receptor |
| 15555362 | 2 | 0.277 | 0 | 170889.5 | 431137.281 | 1369697.75 | 1053646.625 | 1209547.25 | 1009687.375 | 835672.062 | 1501067.75 | 0 | 1194041.5 | 618135.688 | 1658.7739 | DYAYSQTSGGRPGATK | Q62469 | Q62469 | ITGA2 | integrin, alpha 2 (CD49B, alpha 2 subunit of VLA-2 receptor) | Plasma Membrane | transmembrane receptor |
| 15554924 | 2 | 0.408 | 161120.359 | 925319.875 | 1647910.625 | 4680939.5 | 4556127.5 | 5068545.5 | 2800163.75 | 3276623.25 | 5273762.5 | 0 | 4706388 | 2938590.5 | 2228.068 | STNINFYEISSDENAPSVIK | Q62469 | Q62469 | ITGA2 | integrin, alpha 2 (CD49B, alpha 2 subunit of VLA-2 receptor) | Plasma Membrane | transmembrane receptor |
| 15547742, 15548201 | 3, 4, 5 | 0.186 | 8693661 | 3.76E+07 | 8.68E+07 | 1.91E+08 | 1.89E+08 | 2.18E+08 | 9.38E+07 | 1.67E+08 | 2.57E+08 | 8218264.5 | 1.79E+08 | 1.34E+08 | 3813.0168, 3813.0198, 3813.0205 | LLIHIPGQSQPSDLLYILDVQPQGGLLC[160.0307]STQPSPK | Q9QUM0 | Q9QUM0 | ITGA2B | integrin, alpha 2b (platelet glycoprotein IIb of IIb/IIIa complex, antigen CD41) | Plasma Membrane | transmembrane receptor |
| 15548610 | 3, 4 | 0.126 | 4622922 | 1.47E+07 | 2.19E+07 | 5.63E+07 | 5.30E+07 | 5.40E+07 | 3.10E+07 | 3.82E+07 | 5.43E+07 | 6308755.5 | 6.00E+07 | 3.83E+07 | 3321.6409, 3321.6423 | TGQGLGASVVSSWNDVIVAC[160.0307]APWQHWNVLEK | Q9QUM0 | Q9QUM0 | ITGA2B | integrin, alpha 2b (platelet glycoprotein IIb of IIb/IIIa complex, antigen CD41) | Plasma Membrane | transmembrane receptor |
| 15548549 | 2 | 0.155 | 2398609.5 | 1.21E+07 | 2.78E+07 | 5.48E+07 | 6.63E+07 | 6.25E+07 | 3.67E+07 | 4.51E+07 | 7.29E+07 | 3557464.75 | 6.09E+07 | 3.52E+07 | 1841.8092 | FTYDNSNPVFFDGYR | Q9QUM0 | Q9QUM0 | ITGA2B | integrin, alpha 2b (platelet glycoprotein IIb of IIb/IIIa complex, antigen CD41) | Plasma Membrane | transmembrane receptor |
| 15532689 | 2 | 0.151 | 3146123.75 | 1.07E+07 | 2.01E+07 | 4.54E+07 | 4.96E+07 | 5.55E+07 | 2.80E+07 | 4.11E+07 | 5.73E+07 | 3138354.5 | 5.19E+07 | 3.32E+07 | 1135.5707 | ALSNIEGFER | Q9QUM0 | Q9QUM0 | ITGA2B | integrin, alpha 2b (platelet glycoprotein IIb of IIb/IIIa complex, antigen CD41) | Plasma Membrane | transmembrane receptor |
| 15550957 | 2 | 0.199 | 3207190 | 1.87E+07 | 3.80E+07 | 8.20E+07 | 8.71E+07 | 8.75E+07 | 4.67E+07 | 6.26E+07 | 1.00E+08 | 3126101.5 | 7.00E+07 | 5.59E+07 | 2111.1555 | LTATAGDSPLLIGADNVLELK | Q9QUM0 | Q9QUM0 | ITGA2B | integrin, alpha 2b (platelet glycoprotein IIb of IIb/IIIa complex, antigen CD41) | Plasma Membrane | transmembrane receptor |
| 15547749 | 2 | 0.177 | 2689421 | 1.38E+07 | 3.38E+07 | 6.81E+07 | 7.96E+07 | 7.79E+07 | 4.12E+07 | 5.39E+07 | 9.02E+07 | 3103637.75 | 6.94E+07 | 4.37E+07 | 1915.9216 | IILDC[160.0307]GEDDLC[160.0307]VPQLR | Q9QUM0 | Q9QUM0 | ITGA2B | integrin, alpha 2b (platelet glycoprotein IIb of IIb/IIIa complex, antigen CD41) | Plasma Membrane | transmembrane receptor |
| 15531868 | 2 | 0.145 | 1999689 | 6554655 | 1.57E+07 | 3.30E+07 | 3.94E+07 | 3.51E+07 | 2.33E+07 | 3.46E+07 | 4.44E+07 | 2205154.75 | 3.83E+07 | 2.77E+07 | 1231.6147 | VALC[160.0307]ELGNPMK | Q9QUM0 | Q9QUM0 | ITGA2B | integrin, alpha 2b (platelet glycoprotein IIb of IIb/IIIa complex, antigen CD41) | Plasma Membrane | transmembrane receptor |
| 15531623 | 2 | 0.189 | 2132092.75 | 8587325 | 1.91E+07 | 4.02E+07 | 3.89E+07 | 4.99E+07 | 2.50E+07 | 2.94E+07 | 5.48E+07 | 1960731 | 4.13E+07 | 2.86E+07 | 1342.7136 | NLGFQIFQTFK | Q9QUM0 | Q9QUM0 | ITGA2B | integrin, alpha 2b (platelet glycoprotein IIb of IIb/IIIa complex, antigen CD41) | Plasma Membrane | transmembrane receptor |
| 15551062 | 3 | 0.078 | 1176328.375 | 2442230.25 | 8169325 | 8894281 | 2.48E+07 | 2.00E+07 | 7681711.5 | 1.92E+07 | 2.00E+07 | 1864715.625 | 1.03E+07 | 9246194 | 2763.2253 | PGQQDPVLVSC[160.0307]DGSASC[160.0307]TVVEC[160.0307]ELR | Q9QUM0 | Q9QUM0 | ITGA2B | integrin, alpha 2b (platelet glycoprotein IIb of IIb/IIIa complex, antigen CD41) | Plasma Membrane | transmembrane receptor |
| 15552379 | 3 | 0.308 | 1836238 | 6871844 | 1.19E+07 | 1.38E+07 | 1.96E+07 | 2.29E+07 | 1.05E+07 | 1.38E+07 | 2.06E+07 | 1625246.125 | 1.79E+07 | 1.35E+07 | 2383.2698 | AQPVVMATVQLMVQDSLNPTLK | Q9QUM0 | Q9QUM0 | ITGA2B | integrin, alpha 2b (platelet glycoprotein IIb of IIb/IIIa complex, antigen CD41) | Plasma Membrane | transmembrane receptor |
| 15547835 | 2 | 0.184 | 1729550.625 | 8321746.5 | 1.88E+07 | 4.25E+07 | 3.80E+07 | 4.96E+07 | 2.30E+07 | 3.12E+07 | 5.37E+07 | 1571457.75 | 4.17E+07 | 2.22E+07 | 2087.1462 | GPQALSTPTLLLTGTQLYGR | Q9QUM0 | Q9QUM0 | ITGA2B | integrin, alpha 2b (platelet glycoprotein IIb of IIb/IIIa complex, antigen CD41) | Plasma Membrane | transmembrane receptor |
| 15531497 | 2 | 0.177 | 1232947.5 | 7330927 | 1.83E+07 | 2.36E+07 | 5.57E+07 | 3.85E+07 | 2.04E+07 | 3.19E+07 | 3.87E+07 | 1466583.625 | 2.70E+07 | 2.02E+07 | 1330.7085 | FGSAIAPLGDLNR | Q9QUM0 | Q9QUM0 | ITGA2B | integrin, alpha 2b (platelet glycoprotein IIb of IIb/IIIa complex, antigen CD41) | Plasma Membrane | transmembrane receptor |
| 15538297 | 2 | 0.268 | 1716879.875 | 6669404 | 1.67E+07 | 2.76E+07 | 3.42E+07 | 2.56E+07 | 1.95E+07 | 2.64E+07 | 4.57E+07 | 1413696.25 | 3.84E+07 | 2.40E+07 | 1007.59076 | VYLFLQPK | Q9QUM0 | Q9QUM0 | ITGA2B | integrin, alpha 2b (platelet glycoprotein IIb of IIb/IIIa complex, antigen CD41) | Plasma Membrane | transmembrane receptor |
| 15537758 | 2 | 0.193 | 1356688 | 6408177.5 | 1.30E+07 | 2.35E+07 | 3.21E+07 | 3.89E+07 | 1.75E+07 | 2.47E+07 | 3.93E+07 | 1366705 | 3.41E+07 | 1.96E+07 | 1057.5862 | AELQLDLQK | Q9QUM0 | Q9QUM0 | ITGA2B | integrin, alpha 2b (platelet glycoprotein IIb of IIb/IIIa complex, antigen CD41) | Plasma Membrane | transmembrane receptor |
| 15537901 | 3 | 0.18 | 1290032 | 5252794 | 1.25E+07 | 2.54E+07 | 2.55E+07 | 3.64E+07 | 1.66E+07 | 2.47E+07 | 3.19E+07 | 1284584.625 | 2.94E+07 | 1.99E+07 | 2087.1465 | GPQALSTPTLLLTGTQLYGR | Q9QUM0 | Q9QUM0 | ITGA2B | integrin, alpha 2b (platelet glycoprotein IIb of IIb/IIIa complex, antigen CD41) | Plasma Membrane | transmembrane receptor |
| 15548553 | 2 | 0.261 | 1087193.75 | 7776055 | 1.42E+07 | 3.17E+07 | 2.96E+07 | 3.77E+07 | 1.76E+07 | 2.22E+07 | 3.79E+07 | 1239702.25 | 3.64E+07 | 2.67E+07 | 2210.057 | GAVDIDDNGYPDLIVGAYGASK | Q9QUM0 | Q9QUM0 | ITGA2B | integrin, alpha 2b (platelet glycoprotein IIb of IIb/IIIa complex, antigen CD41) | Plasma Membrane | transmembrane receptor |
| 15536392 | 3 | 0.146 | 411804 | 3412624.5 | 6313295.5 | 1.22E+07 | 1.66E+07 | 1.53E+07 | 9550367 | 1.01E+07 | 1.49E+07 | 1035102.062 | 1.30E+07 | 8369135 | 1915.9225 | IILDC[160.0307]GEDDLC[160.0307]VPQLR | Q9QUM0 | Q9QUM0 | ITGA2B | integrin, alpha 2b (platelet glycoprotein IIb of IIb/IIIa complex, antigen CD41) | Plasma Membrane | transmembrane receptor |
| 15553764 | 3 | 0.198 | 1001098.562 | 4802342.5 | 1.29E+07 | 2.43E+07 | 2.94E+07 | 3.23E+07 | 1.54E+07 | 2.16E+07 | 4.05E+07 | 809862.25 | 2.72E+07 | 1.65E+07 | 2679.4297 | VPIENIISSYRPGTLLWHVSNQR | Q9QUM0 | Q9QUM0 | ITGA2B | integrin, alpha 2b (platelet glycoprotein IIb of IIb/IIIa complex, antigen CD41) | Plasma Membrane | transmembrane receptor |
| 15533849 | 3 | 0.221 | 670869.875 | 3827433.25 | 6224261.5 | 1.85E+07 | 1.84E+07 | 1.32E+07 | 1.11E+07 | 1.40E+07 | 2.07E+07 | 653780.312 | 2.34E+07 | 1.22E+07 | 2120.084 | LEHTYELHNIGPGTVNGLR | Q9QUM0 | Q9QUM0 | ITGA2B | integrin, alpha 2b (platelet glycoprotein IIb of IIb/IIIa complex, antigen CD41) | Plasma Membrane | transmembrane receptor |
| 15537191 | 2 | 0.182 | 829543 | 4300561 | 1.04E+07 | 2.39E+07 | 3.13E+07 | 2.69E+07 | 1.50E+07 | 1.93E+07 | 2.61E+07 | 580118.812 | 2.32E+07 | 1.39E+07 | 1485.6312 | NRPPLEEDEEEE | Q9QUM0 | Q9QUM0 | ITGA2B | integrin, alpha 2b (platelet glycoprotein IIb of IIb/IIIa complex, antigen CD41) | Plasma Membrane | transmembrane receptor |
| 15533250 | 3 | 0.211 | 332252.156 | 2952911.75 | 7859894.5 | 1.49E+07 | 2.28E+07 | 2.02E+07 | 8965201 | 1.49E+07 | 2.29E+07 | 543821.188 | 1.78E+07 | 1.56E+07 | 2111.156 | LTATAGDSPLLIGADNVLELK | Q9QUM0 | Q9QUM0 | ITGA2B | integrin, alpha 2b (platelet glycoprotein IIb of IIb/IIIa complex, antigen CD41) | Plasma Membrane | transmembrane receptor |
| 15552882 | 2 | 0.042 | 215811.625 | 593804.062 | 1278303 | 4905725.5 | 4535847.5 | 4224033 | 3053610 | 2709021.5 | 5014112.5 | 490191.094 | 5303606 | 2754162.75 | 1762.8027 | ANTMSSVYAESFRGDK | Q9QUM0 | Q9QUM0 | ITGA2B | integrin, alpha 2b (platelet glycoprotein IIb of IIb/IIIa complex, antigen CD41) | Plasma Membrane | transmembrane receptor |
| 15554618 | 3 | 0.311 | 474024.531 | 1663623.375 | 1850045.625 | 6012110.5 | 3223479.25 | 2943137.5 | 3676057.75 | 2432141.5 | 4000235.75 | 258821.312 | 4319330 | 2775155.25 | 2459.1597 | TPVSC[160.0307]FNIQMC[160.0307]VGATGHNIPQK | Q9QUM0 | Q9QUM0 | ITGA2B | integrin, alpha 2b (platelet glycoprotein IIb of IIb/IIIa complex, antigen CD41) | Plasma Membrane | transmembrane receptor |
| 15533085 | 2 | 0.222 | 555468.5 | 1523559.125 | 3013410 | 8100265 | 1.17E+07 | 8538616 | 4534889 | 7266857.5 | 8532480 | 133876.906 | 8206484.5 | 5165858.5 | 1462.6606 | ANTMSSVYAESFR | Q9QUM0 | Q9QUM0 | ITGA2B | integrin, alpha 2b (platelet glycoprotein IIb of IIb/IIIa complex, antigen CD41) | Plasma Membrane | transmembrane receptor |
| 15557205 | 2 | 0.283 | 15638.983 | 1628963.375 | 4371143 | 3462114.5 | 1.21E+07 | 8267619 | 4047719.75 | 8547003 | 7532840 | 70608.547 | 3722207.5 | 3227563.5 | 2096.0386 | PSQVLDSPFPTGSGFGFSLR | Q9QUM0 | Q9QUM0 | ITGA2B | integrin, alpha 2b (platelet glycoprotein IIb of IIb/IIIa complex, antigen CD41) | Plasma Membrane | transmembrane receptor |
| 15545227 | 2 | 0.297 | 88762.281 | 883872.75 | 1855876 | 3343784.75 | 4186066 | 4332989.5 | 2395463 | 3291996.25 | 4193701.5 | 60680.543 | 4530457 | 2358559 | 1178.661 | HGSVSIVVGAPR | Q9QUM0 | Q9QUM0 | ITGA2B | integrin, alpha 2b (platelet glycoprotein IIb of IIb/IIIa complex, antigen CD41) | Plasma Membrane | transmembrane receptor |
| 15533505 | 2 | 0.233 | 201854.641 | 964672.5 | 2508813.5 | 2456399.75 | 8816369 | 4634625.5 | 1928995.75 | 4765312 | 4603170.5 | 49820.078 | 2369098.25 | 2131926.75 | 1147.5908 | C[160.0307]NPLLFDLR | Q9QUM0 | Q9QUM0 | ITGA2B | integrin, alpha 2b (platelet glycoprotein IIb of IIb/IIIa complex, antigen CD41) | Plasma Membrane | transmembrane receptor |
| 15549545 | 3 | 0.108 | 8608459 | 1.00E+07 | 7201887 | 1.02E+07 | 1.00E+07 | 9750413 | 7062533 | 8854962 | 8210098.5 | 8364438 | 8049259 | 8410127 | 3051.5732 | VYIYLQRPAGIDPTPTLTLTGQDEFSR | P11688 | P11688 | ITGA5 | integrin, alpha 5 (fibronectin receptor, alpha polypeptide) | Plasma Membrane | transmembrane receptor |
| 15552544 | 2 | 0.063 | 5374551.5 | 5800068 | 4894143.5 | 4973710 | 5045558 | 4670783.5 | 4607770.5 | 4525196.5 | 4358604.5 | 4729253 | 4694239 | 3748131.25 | 1504.7976 | VTAPLEAEYSGLVR | P11688 | P11688 | ITGA5 | integrin, alpha 5 (fibronectin receptor, alpha polypeptide) | Plasma Membrane | transmembrane receptor |
| 15537987 | 2 | 0.129 | 4111513.25 | 4077515.25 | 3462674.5 | 4048336 | 3733946.5 | 2966272 | 3144527.75 | 3011368.25 | 3020387.25 | 2526112.75 | 3232993.5 | 3490760 | 1110.556 | TPDFFGSALR | P11688 | P11688 | ITGA5 | integrin, alpha 5 (fibronectin receptor, alpha polypeptide) | Plasma Membrane | transmembrane receptor |
| 15543316 | 3 | 0.158 | 1834677.125 | 2809097.5 | 2336629.75 | 2111668.25 | 2726765.5 | 2815375 | 1813302.375 | 1881624.75 | 2016385.125 | 1223086.25 | 2112071.5 | 1973305.375 | 1788.8821 | AHGSSILAC[160.0307]APLYSWR | P11688 | P11688 | ITGA5 | integrin, alpha 5 (fibronectin receptor, alpha polypeptide) | Plasma Membrane | transmembrane receptor |
| 15548939 | 2 | 0.126 | 171915.625 | 1369753.375 | 2140571.75 | 8763647 | 7270436 | 8823719 | 3888214.25 | 5645540 | 1.03E+07 | 314225.625 | 8326964.5 | 5042444 | 1731.8928 | LIATFPDTLTYSAYR | Q61739 | Q61739 | ITGA6 | integrin, alpha 6 | Plasma Membrane | transmembrane receptor |
| 15538179 | 3 | 0.099 | 79228.195 | 291045.562 | 1100422.75 | 2491602 | 2701744.75 | 3445758.5 | 1209990.875 | 1706407.25 | 2995389.25 | 221642.641 | 3104642.25 | 1998900.375 | 1187.6395 | TVQTDVHFLK | Q61739 | Q61739 | ITGA6 | integrin, alpha 6 | Plasma Membrane | transmembrane receptor |
| 15551587 | 2 | 0.184 | 108916.164 | 739032.562 | 1568317.25 | 4448816 | 4558323.5 | 5484841 | 2541169 | 3431431.5 | 6289582.5 | 130493.203 | 5735199.5 | 3309107.5 | 1501.7073 | FGSC[160.0307]QQGVAATFTK | Q61739 | Q61739 | ITGA6 | integrin, alpha 6 | Plasma Membrane | transmembrane receptor |
| 15559042 | 2 | 0.14 | 0 | 210312.203 | 720048.5 | 2002234.25 | 1981887.25 | 2479726.5 | 1232231.625 | 1535648 | 2205867 | 85107.633 | 2299828.25 | 978527.75 | 1581.6233 | EGC[160.0307]GDDNVC[160.0307]NSNLK | Q61739 | Q61739 | ITGA6 | integrin, alpha 6 | Plasma Membrane | transmembrane receptor |
| 15541848 | 2 | 0.284 | 4.70E-06 | 304683.375 | 151988.094 | 132117.094 | 1853277 | 74544.875 | 1913608.875 | 2202353.5 | 3718026.25 | 4.70E-06 | 1197913 | 1289473.5 | 1132.6705 | LNYLDILLR | Q61739 | Q61739 | ITGA6 | integrin, alpha 6 | Plasma Membrane | transmembrane receptor |
| 15539998 | 2 | 0.375 | 44715.48 | 441422 | 639462.312 | 2295372.75 | 2164582.25 | 2149717 | 989193.875 | 1771261.25 | 2970008.25 | 4.70E-06 | 2241546.75 | 1502889.5 | 994.65094 | RLLLVGAPR | Q61739 | Q61739 | ITGA6 | integrin, alpha 6 | Plasma Membrane | transmembrane receptor |
| 15549514 | 2 | 0.368 | 0 | 857679.438 | 1759660.875 | 5109291 | 5141789.5 | 5552617 | 2556011 | 3711931.25 | 6423448 | 0 | 5080319.5 | 3119895.75 | 1715.8779 | LETTSNQDNLAPITAK | Q61739 | Q61739 | ITGA6 | integrin, alpha 6 | Plasma Membrane | transmembrane receptor |
| 15541587 | 3 | 0.15 | 6860193 | 1.28E+07 | 1.54E+07 | 2.65E+07 | 2.37E+07 | 2.99E+07 | 2.06E+07 | 2.06E+07 | 3.23E+07 | 6371836.5 | 2.70E+07 | 2.07E+07 | [illegible] | [illegible] | [illegible] | [illegible] | [illegible] | [illegible] | [illegible] | [illegible] |
| 15551600 | 3 | 0.068 | 6486864.5 | 8599861 | 8938317 | 1.79E+07 | 1.78E+07 | 2.31E+07 | 1.15E+07 | 1.54E+07 | 2.61E+07 | 5569709.5 | 1.73E+07 | 1.27E+07 | [illegible] | [illegible] | [illegible] | [illegible] | [illegible] | [illegible] | [illegible] | [illegible] |
| 15537996 | 3 | 0.139 | 5826207 | 8620384 | 1.09E+07 | 1.80E+07 | 2.22E+07 | 2.39E+07 | 1.21E+07 | 1.92E+07 | 2.34E+07 | 4705972.5 | 2.22E+07 | 1.52E+07 | [illegible] | [illegible] | [illegible] | [illegible] | [illegible] | [illegible] | [illegible] | [illegible] |
| 15551586 | 2, 3 | 0.055 | 5235777 | 4790535.5 | 9737232 | 1.72E+07 | 1.94E+07 | 2.08E+07 | 9338975 | 1.62E+07 | 2.20E+07 | 3452025.25 | 1.47E+07 | 8304354 | [illegible] | [illegible] | [illegible] | [illegible] | [illegible] | [illegible] | [illegible] | [illegible] |
| 15534560 | 2 | 0.08 | 3057601.75 | 4736038.5 | 5574154 | 8589947 | 1.12E+07 | 1.39E+07 | 7800713 | 9550048 | 1.34E+07 | 2663949.75 | 1.02E+07 | 7385932.5 | [illegible] | [illegible] | [illegible] | [illegible] | [illegible] | [illegible] | [illegible] | [illegible] |
| 15541396 | 2 | 0.076 | 2309010.25 | 4028911 | 4602878.5 | 1.03E+07 | 9688770 | 9580617 | 5844004.5 | 6728028.5 | 9672175 | 2268335.5 | 8755750 | 6704079 | [illegible] | [illegible] | [illegible] | [illegible] | [illegible] | [illegible] | [illegible] | [illegible] |
| 15540447 | 2 | 0.143 | 3249111.25 | 4005831 | 5905838 | 1.05E+07 | 1.03E+07 | 1.06E+07 | 6145512 | 8042439 | 1.10E+07 | 2243173.25 | 9811010 | 7614999 | [illegible] | [illegible] | [illegible] | [illegible] | [illegible] | [illegible] | [illegible] | [illegible] |
| 15533097 | 2 | 0.027 | 1578215 | 3169678 | 4363444.5 | 6382390.5 | 9504313 | 8712920 | 6187682 | 7145885.5 | 7901559 | 1939862.625 | 5359983 | 4290672.5 | [illegible] | [illegible] | [illegible] | [illegible] | [illegible] | [illegible] | [illegible] | [illegible] |
| 15532832 | 2 | 0.077 | 2312334.5 | 3291762.25 | 4112998.75 | 8876776 | 8674333 | 9237730 | 5658149 | 6660326.5 | 8603176 | 1893342.625 | 8759067 | 5716129.5 | [illegible] | [illegible] | [illegible] | [illegible] | [illegible] | [illegible] | [illegible] | [illegible] |
| 15536895 | 3 | 0.054 | 1448843.875 | 1850458 | 3068104.5 | 6723581.5 | 6314548.5 | 6587435.5 | 4459145 | 4973646 | 7021287 | 1579137.375 | 7446702 | 4157389.75 | [illegible] | [illegible] | [illegible] | [illegible] | [illegible] | [illegible] | [illegible] | [illegible] |
| 15552388 | 2 | 0.168 | 1087009.75 | 1778115.5 | 2304589.25 | 4507120.5 | 5434173.5 | 4163526.5 | 2028332.625 | 3408677.25 | 6477065 | 1229182.25 | 5564599 | 3121842.75 | [illegible] | [illegible] | [illegible] | [illegible] | [illegible] | [illegible] | [illegible] | [illegible] |
| 15551097 | 2 | 0.13 | 820148.688 | 932662.938 | 1569847.375 | 2914721 | 2962562 | 3145368.75 | 1563716.375 | 2440173 | 3103857 | 755479.375 | 3929082 | 1873193.375 | [illegible] | [illegible] | [illegible] | [illegible] | [illegible] | [illegible] | [illegible] | [illegible] |
| 15556503 | 3 | 0.098 | 324831.469 | 671165.25 | 835816.875 | 2577891 | 1727600.25 | 2002875.875 | 1446485.375 | 1447447.125 | 2646641.75 | 484393.75 | 3361728 | 1578983.875 | [illegible] | [illegible] | [illegible] | [illegible] | [illegible] | [illegible] | [illegible] | [illegible] |
| 15553164 | 2 | 0.386 | 0 | 102721 | 1204179.75 | 857676.125 | 1515175.875 | 2308708.5 | 370733.875 | 1214443.375 | 2607503.5 | 0 | 1161715.5 | 1158911.125 | [illegible] | [illegible] | [illegible] | [illegible] | [illegible] | [illegible] | [illegible] | [illegible] |
| 15547717 | 3 | 0.168 | 7740913.5 | 2.89E+07 | 4.76E+07 | 1.52E+08 | 1.41E+08 | 1.55E+08 | 7.10E+07 | 9.38E+07 | 1.65E+08 | 6613565 | 1.61E+08 | 9.57E+07 | [illegible] | [illegible] | [illegible] | [illegible] | [illegible] | [illegible] | [illegible] | [illegible] |
| 15553917 | 2 | 0.244 | 8307781.5 | 9559299 | 1.37E+07 | 1.69E+07 | 1.85E+07 | 2.00E+07 | 1.05E+07 | 1.46E+07 | 2.03E+07 | 4998616.5 | 1.85E+07 | 1.08E+07 | [illegible] | [illegible] | [illegible] | [illegible] | [illegible] | [illegible] | [illegible] | [illegible] |
| 15547888 | 2 | 0.127 | 2973615.75 | 1.01E+07 | 2.35E+07 | 5.83E+07 | 4.75E+07 | 4.95E+07 | 2.74E+07 | 3.28E+07 | 5.70E+07 | 4080349.5 | 4.82E+07 | 2.65E+07 | [illegible] | [illegible] | [illegible] | [illegible] | [illegible] | [illegible] | [illegible] | [illegible] |
| 15536587 | 3 | 0.125 | 2198646.25 | 8692426 | 2.06E+07 | 5.02E+07 | 6.22E+07 | 5.09E+07 | 3.03E+07 | 4.14E+07 | 6.04E+07 | 3637579.75 | 6.76E+07 | 3.35E+07 | [illegible] | [illegible] | [illegible] | [illegible] | [illegible] | [illegible] | [illegible] | [illegible] |
| 15533209 | 3 | 0.167 | 2984147.75 | 1.07E+07 | 2.13E+07 | 5.07E+07 | 5.41E+07 | 6.50E+07 | 2.87E+07 | 3.91E+07 | 6.18E+07 | 2600505.5 | 5.30E+07 | 3.81E+07 | [illegible] | [illegible] | [illegible] | [illegible] | [illegible] | [illegible] | [illegible] | [illegible] |
| 15533582, 15538787 | 2, 3 | 0.166 | 2148106 | 1.04E+07 | 2.03E+07 | 5.60E+07 | 5.23E+07 | 4.79E+07 | 2.98E+07 | 3.59E+07 | 5.51E+07 | 2331472 | 5.17E+07 | 3.07E+07 | [illegible] | [illegible] | [illegible] | [illegible] | [illegible] | [illegible] | [illegible] | [illegible] |
| 15550812 | 4 | 0.142 | 1749434.875 | 7893542 | 8866252 | 3.11E+07 | 2.73E+07 | 2.93E+07 | 1.74E+07 | 1.94E+07 | 2.90E+07 | 2182758.5 | 3.42E+07 | 2.04E+07 | [illegible] | [illegible] | [illegible] | [illegible] | [illegible] | [illegible] | [illegible] | [illegible] |
| 15547917 | 3 | 0.171 | 1842798.75 | 1.16E+07 | 1.71E+07 | 6.21E+07 | 5.72E+07 | 5.44E+07 | 2.45E+07 | 3.65E+07 | 7.31E+07 | 2092230.5 | 5.99E+07 | 3.29E+07 | [illegible] | [illegible] | [illegible] | [illegible] | [illegible] | [illegible] | [illegible] | [illegible] |
| 15535502 | 2 | 0.229 | 1581664.625 | 6413217.5 | 1.33E+07 | 2.94E+07 | 3.25E+07 | 3.15E+07 | 1.69E+07 | 2.31E+07 | 3.55E+07 | 1514218 | 3.86E+07 | 2.59E+07 | [illegible] | [illegible] | [illegible] | [illegible] | [illegible] | [illegible] | [illegible] | [illegible] |
| 15532093 | 2 | 0.149 | 986916.25 | 3411212.75 | 8713984 | 2.06E+07 | 1.59E+07 | 1.99E+07 | 9870204 | 1.30E+07 | 2.48E+07 | 1438720.5 | 2.04E+07 | 1.16E+07 | [illegible] | [illegible] | [illegible] | [illegible] | [illegible] | [illegible] | [illegible] | [illegible] |
| 15532519 | 2 | 0.189 | 2104517.5 | 5313042.5 | 9990918 | 2.28E+07 | 2.49E+07 | 2.27E+07 | 1.45E+07 | 1.90E+07 | 2.54E+07 | 1427940 | 2.78E+07 | 1.59E+07 | [illegible] | [illegible] | [illegible] | [illegible] | [illegible] | [illegible] | [illegible] | [illegible] |
| 15548791 | 2 | 0.172 | 886463.188 | 4499584.5 | 1.15E+07 | 2.70E+07 | 2.94E+07 | 2.57E+07 | 1.47E+07 | 2.12E+07 | 3.67E+07 | 1379546.25 | 3.44E+07 | 1.88E+07 | [illegible] | [illegible] | [illegible] | [illegible] | [illegible] | [illegible] | [illegible] | [illegible] |
| 15542693 | 2 | 0.166 | 1532787.125 | 6573171 | 1.68E+07 | 2.39E+07 | 3.45E+07 | 4.17E+07 | 1.82E+07 | 2.53E+07 | 3.82E+07 | 1342123.5 | 2.20E+07 | 1.82E+07 | [illegible] | [illegible] | [illegible] | [illegible] | [illegible] | [illegible] | [illegible] | [illegible] |
| 15548502 | 3 | 0.16 | 922489.938 | 4576342.5 | 1.12E+07 | 3.54E+07 | 3.18E+07 | 3.40E+07 | 1.61E+07 | 2.12E+07 | 3.40E+07 | 1174390.25 | 3.88E+07 | 2.07E+07 | [illegible] | [illegible] | [illegible] | [illegible] | [illegible] | [illegible] | [illegible] | [illegible] |
| 15537208 | 2 | 0.203 | 1084003.625 | 5953670.5 | 1.50E+07 | 1.93E+07 | 3.70E+07 | 3.89E+07 | 1.56E+07 | 2.63E+07 | 4.19E+07 | 958925.688 | 2.53E+07 | 1.70E+07 | [illegible] | [illegible] | [illegible] | [illegible] | [illegible] | [illegible] | [illegible] | [illegible] |
| 15543750 | 2 | 0.066 | 671234.375 | 730374.375 | 1710403 | 7127314 | 2605901.5 | 4643639 | 1827829.125 | 3136216.25 | 4197824 | 858958.625 | 3819475.75 | 3649334.25 | [illegible] | [illegible] | [illegible] | [illegible] | [illegible] | [illegible] | [illegible] | [illegible] |
| 15538676 | 2 | 0.189 | 626810.5 | 4890126 | 1.40E+07 | 1.48E+07 | 2.68E+07 | 3.61E+07 | 1.73E+07 | 2.16E+07 | 2.80E+07 | 807008.938 | 1.77E+07 | 1.24E+07 | [illegible] | [illegible] | [illegible] | [illegible] | [illegible] | [illegible] | [illegible] | [illegible] |
| 15532806 | 2 | 0.14 | 448677.719 | 3694203.75 | 9646351 | 1.57E+07 | 2.41E+07 | 2.33E+07 | 1.16E+07 | 1.67E+07 | 1.86E+07 | 775514.375 | 1.23E+07 | 9516289 | [illegible] | [illegible] | [illegible] | [illegible] | [illegible] | [illegible] | [illegible] | [illegible] |

| | | | | | | | | | | | | | | | | | | | | | | |
|---|---|---|---|---|---|---|---|---|---|---|---|---|---|---|---|---|---|---|---|---|---|---|
| 15552553 | 2, 3 | 0.117 | 682326.875 | 3811920.75 | 7642332.5 | 8714321 | 3.28E+07 | 1.93E+07 | 7790877 | 1.64E+07 | 1.55E+07 | 746593.062 | 7782820 | 6782824.5 | 2375.1128, 2375.1135 | DNC[160.0307]APESIEFPVSEAQILEAR | O54890 | O54890 | ITGB3 | integrin, beta 3 (platelet glycoprotein IIIa, antigen CD61) | Plasma Membrane | transmembrane receptor |
| 15548951 | 2, 3 | 0.136 | 316891.938 | 2009652.875 | 6185188 | 2.20E+07 | 2.16E+07 | 2.07E+07 | 9830183 | 1.35E+07 | 2.22E+07 | 586541.938 | 2.66E+07 | 1.23E+07 | 2523.9263, 2523.927 | GNC[160.0307]EC[160.0307]GSC[160.0307]VC[160.0307]VQPGSYGDTC[160.0307]EK | O54890 | O54890 | ITGB3 | integrin, beta 3 (platelet glycoprotein IIIa, antigen CD61) | Plasma Membrane | transmembrane receptor |
| 15558131 | 2 | 0.318 | 561540.312 | 2816082.25 | 8964543 | 3328484.75 | 1.88E+07 | 2.15E+07 | 4607983.5 | 1.36E+07 | 2.24E+07 | 400180.688 | 6540658 | 7164632 | 1531.7722 | NDASHLLVFTTDAK | O54890 | O54890 | ITGB3 | integrin, beta 3 (platelet glycoprotein IIIa, antigen CD61) | Plasma Membrane | transmembrane receptor |
| 15557759 | 2 | 0.234 | 178227.203 | 2774734 | 6706817.5 | 8966690 | 1.44E+07 | 1.52E+07 | 6365359 | 1.00E+07 | 1.50E+07 | 378623.719 | 9268383 | 6993771.5 | 1690.8835 | PVSPYMYISPPQAIK | O54890 | O54890 | ITGB3 | integrin, beta 3 (platelet glycoprotein IIIa, antigen CD61) | Plasma Membrane | transmembrane receptor |
| 15537471 | 2 | 0.295 | 180638.047 | 4528365 | 1.14E+07 | 8730548 | 3.22E+07 | 3.60E+07 | 1.18E+07 | 2.35E+07 | 4.33E+07 | 149171.969 | 1.36E+07 | 1.63E+07 | 1403.6769 | EATSTFTNITYR | O54890 | O54890 | ITGB3 | integrin, beta 3 (platelet glycoprotein IIIa, antigen CD61) | Plasma Membrane | transmembrane receptor |
| 15537986 | 2 | 0.07 | 23106.236 | 4.70E-06 | 4.70E-06 | 1549454.375 | 396002.812 | 455963.188 | 568662.938 | 285953.25 | 1444237.375 | 4.70E-06 | 3038429.25 | 1101700.625 | 1470.6162 | C[160.0307]PTC[160.0307]PDAC[160.0307]SFKK | O54890 | O54890 | ITGB3 | integrin, beta 3 (platelet glycoprotein IIIa, antigen CD61) | Plasma Membrane | transmembrane receptor |
| 15534442 | 2 | 0.24 | 2261562 | 3281514 | 4446159 | 7580600 | 8385315.5 | 1.14E+07 | 5548433 | 6721471.5 | 1.01E+07 | 1006529.312 | 1.02E+07 | 5536826.5 | 983.5177 | LGFGSFVEK | Q9Z0T9 | Q9Z0T9 | ITGB6 | integrin, beta 6 | Plasma Membrane | other |
| 15539837 | 2 | 0.1 | 1099144.875 | 1710544.25 | 2279450.5 | 2418748.5 | 2185706.75 | 2051816 | 2014990.25 | 2387643.5 | 2353975.5 | 1281434.125 | 1788545.125 | 1471160.625 | 1454.7126 | LLDSWFTSTQEK | Q5XKN4 | Q5XKN4 | JAGN1 | jagunal homolog 1 | Cytoplasm | other |
| 15538877 | 2 | 0.154 | 181486.531 | 181221.109 | 707589.938 | 2827465.75 | 2042635.5 | 2288858.5 | 1230005.75 | 1824368 | 2145729.5 | 47587.082 | 2396046.5 | 1303037.5 | 1450.7588 | MFVLNELVQTEK | A2CG49 | A2CG49 | KALRN | kalirin, RhoGEF kinase | Cytoplasm | kinase |
| 15545833 | 2 | 0.313 | 23171.74 | 87780.914 | 202950.281 | 1040330.188 | 985055.125 | 851109.312 | 517539.531 | 490062.5 | 980580.375 | 4.70E-06 | 1720799.75 | 518481.094 | 1121.513 | AGLEC[160.0307]SDIEK | A2CG49 | A2CG49 | KALRN | kalirin, RhoGEF kinase | Cytoplasm | kinase |
| 15551580 | 2 | 0.598 | 2852305.25 | 2956803.5 | 2885403.75 | 1679760.125 | 2855132 | 2573378.25 | 3079216.75 | 2693609.75 | 2973567.5 | 3927423 | 1634932.5 | 2020418.5 | 1617.9543 | SLFVQELLLSTLVR | Q80UY2 | Q80UY2 | KCMF1 | potassium channel modulatory factor 1 | Extracellular Space | other |
| 15542455 | 3 | 0.168 | 1180150.5 | 1518179.375 | 1293202.875 | 574234 | 662730.75 | 1203447.75 | 1672099.25 | 1413959.75 | 1366678.25 | 1766349.75 | 628729.562 | 1126126.875 | 1617.9536 | SLFVQELLLSTLVR | Q80UY2 | Q80UY2 | KCMF1 | potassium channel modulatory factor 1 | Extracellular Space | other |
| 15540732 | 2 | 0.865 | 1022800.812 | 957647.125 | 768022.875 | 565790.125 | 1079955.25 | 760195.25 | 822833.875 | 1002378.188 | 536065.75 | 1400699.75 | 701337.812 | 690535.438 | 1275.5428 | HEGVSC[160.0307]DAC[160.0307]LK | Q80UY2 | Q80UY2 | KCMF1 | potassium channel modulatory factor 1 | Extracellular Space | other |
| 15536308 | 3 | 0.312 | 1.52E+08 | 1.50E+08 | 1.62E+08 | 1.39E+08 | 1.59E+08 | 1.23E+08 | 1.73E+08 | 1.58E+08 | 1.21E+08 | 2.02E+08 | 1.89E+08 | 1.44E+08 | 1817.8729 | GLSSQDPQDPHSPPSLR | Q9EQF2 | Q9EQF2 | KEL | Kell blood group, metallo-endopeptidase | Plasma Membrane | peptidase |
| 15550307 | 3, 4 | 0.754 | 1.90E+08 | 2.22E+08 | 1.55E+08 | 1.77E+08 | 1.73E+08 | 1.62E+08 | 1.75E+08 | 1.81E+08 | 1.61E+08 | 1.89E+08 | 1.57E+08 | 1.85E+08 | 3890.0159, 3890.0176 | AYLRPHPAPPHTPIIQIDQPEFDILLQQEQEQK | Q9EQF2 | Q9EQF2 | KEL | Kell blood group, metallo-endopeptidase | Plasma Membrane | peptidase |
| 15548318 | 2 | 0.989 | 1.28E+08 | 1.34E+08 | 1.42E+08 | 1.23E+08 | 1.51E+08 | 1.22E+08 | 1.29E+08 | 1.41E+08 | 1.31E+08 | 1.50E+08 | 1.32E+08 | 1.18E+08 | 1765.7551 | SVAPC[160.0307]TDFFSFAC[160.0307]EK | Q9EQF2 | Q9EQF2 | KEL | Kell blood group, metallo-endopeptidase | Plasma Membrane | peptidase |
| 15547797 | 3 | 0.888 | 9.39E+07 | 1.43E+08 | 1.49E+08 | 9.08E+07 | 1.59E+08 | 1.21E+08 | 1.14E+08 | 1.31E+08 | 9.77E+07 | 1.47E+08 | 1.33E+08 | 1.09E+08 | 2702.4146 | IVEHTGELTLPNLDLSPYQLFFR | Q9EQF2 | Q9EQF2 | KEL | Kell blood group, metallo-endopeptidase | Plasma Membrane | peptidase |
| 15531818 | 3 | 0.201 | 1.25E+08 | 1.24E+08 | 1.44E+08 | 1.23E+08 | 1.14E+08 | 1.23E+08 | 1.45E+08 | 1.43E+08 | 1.22E+08 | 1.38E+08 | 1.28E+08 | 1.39E+08 | 1709.8464 | LLEAPGSWHLGSGEEK | Q9EQF2 | Q9EQF2 | KEL | Kell blood group, metallo-endopeptidase | Plasma Membrane | peptidase |
| 15547923, 15547928 | 2, 3 | 0.806 | 7.68E+07 | 1.21E+08 | 1.01E+08 | 7.80E+07 | 1.05E+08 | 8.63E+07 | 9.40E+07 | 1.05E+08 | 1.09E+08 | 1.25E+08 | 8.00E+07 | 7.64E+07 | 2394.2415, 2394.2432 | ESIQSYMILGLVDTLSPALDTK | Q9EQF2 | Q9EQF2 | KEL | Kell blood group, metallo-endopeptidase | Plasma Membrane | peptidase |
| 15531504, 15537563 | 2, 3 | 0.16 | 9.44E+07 | 9.15E+07 | 1.05E+08 | 7.30E+07 | 8.00E+07 | 8.45E+07 | 9.27E+07 | 9.59E+07 | 7.31E+07 | 1.07E+08 | 9.64E+07 | 8.39E+07 | 1382.7036, 1382.7053 | VHGPLSNTPDFAK | Q9EQF2 | Q9EQF2 | KEL | Kell blood group, metallo-endopeptidase | Plasma Membrane | peptidase |
| 15532402 | 3 | 0.484 | 1.01E+08 | 1.10E+08 | 1.15E+08 | 8.25E+07 | 1.17E+08 | 9.72E+07 | 1.13E+08 | 1.08E+08 | 7.76E+07 | 1.03E+08 | 7.48E+07 | 8.69E+07 | 2003.0416 | LFQFLRPQQQQQAQDK | Q9EQF2 | Q9EQF2 | KEL | Kell blood group, metallo-endopeptidase | Plasma Membrane | peptidase |
| 15532095 | 3 | 0.329 | 7.92E+07 | 9.33E+07 | 7.50E+07 | 7.79E+07 | 6.32E+07 | 7.52E+07 | 9.37E+07 | 7.54E+07 | 7.33E+07 | 8.74E+07 | 8.06E+07 | 8.53E+07 | 1642.8167 | LLMSQYGHFPFFR | Q9EQF2 | Q9EQF2 | KEL | Kell blood group, metallo-endopeptidase | Plasma Membrane | peptidase |
| 15539532 | 3 | 0.146 | 8.35E+07 | 9.25E+07 | 8.72E+07 | 7.17E+07 | 8.34E+07 | 7.42E+07 | 1.01E+08 | 8.32E+07 | 8.12E+07 | 8.33E+07 | 9.29E+07 | 9.14E+07 | 1632.8246 | NVQSFLQPFPYHR | Q9EQF2 | Q9EQF2 | KEL | Kell blood group, metallo-endopeptidase | Plasma Membrane | peptidase |
| 15548273 | 2 | 0.01 | 4.63E+07 | 5.90E+07 | 4.88E+07 | 4.30E+07 | 4.89E+07 | 5.27E+07 | 6.07E+07 | 6.05E+07 | 6.10E+07 | 7.49E+07 | 6.36E+07 | 6.29E+07 | 2039.0111 | EAFGPSIQSAAMELFAEIK | Q9EQF2 | Q9EQF2 | KEL | Kell blood group, metallo-endopeptidase | Plasma Membrane | peptidase |
| 15534503 | 2 | 0.404 | 5.91E+07 | 6.03E+07 | 6.26E+07 | 5.99E+07 | 5.54E+07 | 5.83E+07 | 7.54E+07 | 6.50E+07 | 4.76E+07 | 7.26E+07 | 6.61E+07 | 6.82E+07 | 1284.6943 | LAQLQVEMGAPK | Q9EQF2 | Q9EQF2 | KEL | Kell blood group, metallo-endopeptidase | Plasma Membrane | peptidase |
| 15531589 | 2 | 0.45 | 5.63E+07 | 6.80E+07 | 7.29E+07 | 4.80E+07 | 6.48E+07 | 7.38E+07 | 8.02E+07 | 7.57E+07 | 6.58E+07 | 7.05E+07 | 5.54E+07 | 5.86E+07 | 1220.6135 | LSWISEETQK | Q9EQF2 | Q9EQF2 | KEL | Kell blood group, metallo-endopeptidase | Plasma Membrane | peptidase |
| 15548435 | 3 | 0.876 | 2.75E+07 | 4.74E+07 | 4.85E+07 | 2.67E+07 | 4.71E+07 | 3.44E+07 | 4.09E+07 | 5.39E+07 | 3.40E+07 | 6.03E+07 | 3.32E+07 | 3.45E+07 | 3296.6536 | AVKPDIATQEYNDIQLGPSFLQSFLSC[160.0307]VR | Q9EQF2 | Q9EQF2 | KEL | Kell blood group, metallo-endopeptidase | Plasma Membrane | peptidase |
| 15549631 | 2 | 0.007 | 4.35E+07 | 4.57E+07 | 5.07E+07 | 4.10E+07 | 3.91E+07 | 3.84E+07 | 4.85E+07 | 5.06E+07 | 4.65E+07 | 5.37E+07 | 6.23E+07 | 4.83E+07 | 2156.08 | C[160.0307]VEQTGAFFEPTLAALFVR | Q9EQF2 | Q9EQF2 | KEL | Kell blood group, metallo-endopeptidase | Plasma Membrane | peptidase |
| 15534908, 15542189 | 2 | 0.725 | 5.11E+07 | 4.25E+07 | 4.73E+07 | 6.33E+07 | 3.02E+07 | 5.24E+07 | 4.92E+07 | 3.67E+07 | 4.50E+07 | 5.28E+07 | 5.19E+07 | 5.37E+07 | 1300.6843, 1300.6919 | LAQLQVEM[147.0354]GAPK | Q9EQF2 | Q9EQF2 | KEL | Kell blood group, metallo-endopeptidase | Plasma Membrane | peptidase |
| 15535043 | 4 | 0.692 | 5.80E+07 | 4.75E+07 | 5.72E+07 | 5.98E+07 | 2.93E+07 | 5.23E+07 | 6.32E+07 | 4.83E+07 | 5.52E+07 | 5.03E+07 | 4.43E+07 | 5.99E+07 | 2858.5142 | RIVEHTGELTLPNLDLSPYQLFFR | Q9EQF2 | Q9EQF2 | KEL | Kell blood group, metallo-endopeptidase | Plasma Membrane | peptidase |
| 15531495 | 3 | 0.031 | 3.38E+07 | 3.30E+07 | 3.62E+07 | 2.86E+07 | 3.15E+07 | 2.64E+07 | 3.49E+07 | 3.58E+07 | 2.66E+07 | 4.21E+07 | 3.68E+07 | 3.95E+07 | 1195.656 | ERPPLPAYPR | Q9EQF2 | Q9EQF2 | KEL | Kell blood group, metallo-endopeptidase | Plasma Membrane | peptidase |
| 15532377 | 2 | 0.602 | 3.68E+07 | 3.88E+07 | 4.26E+07 | 3.35E+07 | 3.88E+07 | 3.67E+07 | 3.29E+07 | 3.88E+07 | 3.97E+07 | 4.03E+07 | 3.45E+07 | 3.26E+07 | 928.51935 | GTLLNPSAR | Q9EQF2 | Q9EQF2 | KEL | Kell blood group, metallo-endopeptidase | Plasma Membrane | peptidase |
| 15531425 | 3 | 0.026 | 4.33E+07 | 4.17E+07 | 4.55E+07 | 3.47E+07 | 3.52E+07 | 3.30E+07 | 5.02E+07 | 4.96E+07 | 4.18E+07 | 3.94E+07 | 5.59E+07 | 4.30E+07 | 2156.0808 | C[160.0307]VEQTGAFFEPTLAALFVR | Q9EQF2 | Q9EQF2 | KEL | Kell blood group, metallo-endopeptidase | Plasma Membrane | peptidase |
| 15532796 | 2 | 0.064 | 2.62E+07 | 2.04E+07 | 1.84E+07 | 2.19E+07 | 2.08E+07 | 1.71E+07 | 2.69E+07 | 2.25E+07 | 1.94E+07 | 3.27E+07 | 3.31E+07 | 2.46E+07 | 1014.4473 | SYAQVMC[160.0307]R | Q9EQF2 | Q9EQF2 | KEL | Kell blood group, metallo-endopeptidase | Plasma Membrane | peptidase |
| 15548268 | 3 | 0.845 | 3.64E+07 | 2.49E+07 | 3.48E+07 | 4.11E+07 | 1.21E+07 | 3.21E+07 | 3.73E+07 | 2.62E+07 | 3.13E+07 | 2.58E+07 | 3.15E+07 | 3.60E+07 | 2858.513 | RIVEHTGELTLPNLDLSPYQLFFR | Q9EQF2 | Q9EQF2 | KEL | Kell blood group, metallo-endopeptidase | Plasma Membrane | peptidase |
| 15532567 | 3 | 0.146 | 2.06E+07 | 2.67E+07 | 2.02E+07 | 1.17E+07 | 2.12E+07 | 2.29E+07 | 2.75E+07 | 2.67E+07 | 2.38E+07 | 2.30E+07 | 3.15E+07 | 2.93E+07 | 2039.0115 | EAFGPSIQSAAMELFAEIK | Q9EQF2 | Q9EQF2 | KEL | Kell blood group, metallo-endopeptidase | Plasma Membrane | peptidase |
| 15536313 | 2 | 0.245 | 2.18E+07 | 2.59E+07 | 3.27E+07 | 1.85E+07 | 2.74E+07 | 2.20E+07 | 2.55E+07 | 3.27E+07 | 1.96E+07 | 2.27E+07 | 1.57E+07 | 1.79E+07 | 1057.5294 | EYVTYLNR | Q9EQF2 | Q9EQF2 | KEL | Kell blood group, metallo-endopeptidase | Plasma Membrane | peptidase |
| 15548990 | 2 | 0.278 | 1.81E+07 | 1.65E+07 | 1.68E+07 | 1.32E+07 | 1.59E+07 | 1.14E+07 | 1.74E+07 | 1.58E+07 | 1.29E+07 | 2.14E+07 | 1.76E+07 | 1.34E+07 | 1817.8738 | GLSSQDPQDPHSPPSLR | Q9EQF2 | Q9EQF2 | KEL | Kell blood group, metallo-endopeptidase | Plasma Membrane | peptidase |
| 15554655 | 2 | 0.717 | 2.09E+07 | 2.70E+07 | 3.42E+07 | 2.02E+07 | 2.78E+07 | 2.52E+07 | 2.78E+07 | 2.75E+07 | 2.66E+07 | 1.87E+07 | 2.54E+07 | 2.79E+07 | 1632.8237 | NVQSFLQPFPYHR | Q9EQF2 | Q9EQF2 | KEL | Kell blood group, metallo-endopeptidase | Plasma Membrane | peptidase |
| 15553025 | 2 | 0.014 | 1.47E+07 | 1.62E+07 | 1.50E+07 | 1.23E+07 | 1.16E+07 | 1.25E+07 | 1.61E+07 | 1.33E+07 | 1.54E+07 | 1.76E+07 | 1.48E+07 | 1.48E+07 | 1642.8181 | LLMSQYGHFPFFR | Q9EQF2 | Q9EQF2 | KEL | Kell blood group, metallo-endopeptidase | Plasma Membrane | peptidase |
| 15539806 | 2 | 0.253 | 1.55E+07 | 1.91E+07 | 2.44E+07 | 1.37E+07 | 2.08E+07 | 1.76E+07 | 1.79E+07 | 2.27E+07 | 1.56E+07 | 1.70E+07 | 1.09E+07 | 1.34E+07 | 862.51276 | VYAQILR | Q9EQF2 | Q9EQF2 | KEL | Kell blood group, metallo-endopeptidase | Plasma Membrane | peptidase |
| 15555896, 15556454 | 3 | 0.078 | 1.62E+07 | 1.98E+07 | 1.72E+07 | 1.46E+07 | 1.58E+07 | 1.52E+07 | 1.62E+07 | 1.47E+07 | 1.49E+07 | 1.62E+07 | 9456068 | 1.15E+07 | 2727.3796, 2727.38 | LGTLLGSNPQEAQQHASWSIVFTSR | Q9EQF2 | Q9EQF2 | KEL | Kell blood group, metallo-endopeptidase | Plasma Membrane | peptidase |
| 15548461 | 2 | 0.233 | 2.18E+07 | 1.98E+07 | 2.50E+07 | 1.78E+07 | 1.78E+07 | 1.45E+07 | 2.18E+07 | 2.02E+07 | 1.78E+07 | 1.28E+07 | 2.17E+07 | 1.92E+07 | 1709.8463 | LLEAPGSWHLGSGEEK | Q9EQF2 | Q9EQF2 | KEL | Kell blood group, metallo-endopeptidase | Plasma Membrane | peptidase |
| 15533127 | 3 | 0.128 | 1.49E+07 | 1.79E+07 | 1.75E+07 | 8031516 | 1.51E+07 | 1.97E+07 | 1.53E+07 | 1.95E+07 | 1.70E+07 | 1.12E+07 | 9566789 | 1.08E+07 | 1658.8109 | LLM[147.0354]SQYGHFPFFR | Q9EQF2 | Q9EQF2 | KEL | Kell blood group, metallo-endopeptidase | Plasma Membrane | peptidase |
| 15536886 | 2 | 0.888 | 5154862.5 | 6516355 | 9729659 | 4099016.5 | 8637664 | 5200842.5 | 7225925 | 9004180 | 4787187.5 | 8826897 | 5061223 | 5529962.5 | 900.512 | ELIQELR | Q9EQF2 | Q9EQF2 | KEL | Kell blood group, metallo-endopeptidase | Plasma Membrane | peptidase |
| 15556387, 15559028 | 2 | 0.148 | 8356800.5 | 8349907 | 1.00E+07 | 5422231.5 | 8744719 | 7998727 | 9538011 | 7803419 | 6281270.5 | 6202084 | 5623875 | 6498535.5 | 2003.041, 2003.0422 | LFQFLRPQQQQQAQDK | Q9EQF2 | Q9EQF2 | KEL | Kell blood group, metallo-endopeptidase | Plasma Membrane | peptidase |
| 15545178, 15546273 | 2 | 0.77 | 3189060 | 4232794 | 6091013.5 | 1779120.25 | 5526702.5 | 3907222 | 5228544.5 | 5099665.5 | 3364795.5 | 4532133 | 3032409 | 3893526.25 | 1485.6561, 1485.6573 | GPPWSQESSPEER | Q9EQF2 | Q9EQF2 | KEL | Kell blood group, metallo-endopeptidase | Plasma Membrane | peptidase |
| 15557565 | 3 | 0.465 | 3169596.5 | 1313608.25 | 1725530.875 | 4142412.75 | 945452.062 | 2329634 | 3378069.25 | 1645854.75 | 2479366.5 | 2943672.75 | 3844896.5 | 3869158.25 | 2355.0928 | GPPWSQESSPEERLPTEWSR | Q9EQF2 | Q9EQF2 | KEL | Kell blood group, metallo-endopeptidase | Plasma Membrane | peptidase |
| 15536923 | 3 | 0.445 | 4848945 | 1068522.875 | 61503.285 | 3920539.25 | 647018.062 | 494754.281 | 865258.25 | 765490.188 | 2096175.5 | 2644623.25 | 6127390.5 | 3496035.25 | 1348.707 | KLSWISEETQK | Q9EQF2 | Q9EQF2 | KEL | Kell blood group, metallo-endopeptidase | Plasma Membrane | peptidase |
| 15553155 | 3 | 0.649 | 5877224.5 | 5911273 | 3738561.25 | 5369574 | 5643590.5 | 3392756.5 | 4924604.5 | 5534146 | 4537166 | 5716296 | 3636969 | 2962953.75 | 2666.3982 | FIEAEQVPELEAVLHLVIASSDTR | Q6PDI5 | Q6PDI5 | KIAA0368 | KIAA0368 | Cytoplasm | other |
| 15552194 | 3 | 0.762 | 5196480 | 5444107 | 2917389.5 | 2508246.25 | 5831961 | 2919094.5 | 5006211.5 | 3267117.5 | 3415836.5 | 4892173 | 2276346.5 | 2937322.75 | 2387.2078 | GLLIESLATMETDNRPELQEK | Q6PDI5 | Q6PDI5 | KIAA0368 | KIAA0368 | Cytoplasm | other |
| 15541518 | 3 | 0.674 | 4864197.5 | 3302718.5 | 3336637.75 | 4200515 | 4284279 | 2212069 | 3424301.5 | 3843342.5 | 3308925.75 | 3461511.25 | 2477341.75 | 3067193.75 | 2019.0714 | ERELQLESLLGAFESLGK | Q6PDI5 | Q6PDI5 | KIAA0368 | KIAA0368 | Cytoplasm | other |
| 15540596 | 2 | 0.085 | 2987564.25 | 2340082 | 2142516.5 | 2262887.5 | 2646883 | 2089405.5 | 2187800.5 | 2097964 | 1573242.375 | 2143001.5 | 1431155.625 | 1421981.5 | 1314.7855 | TEALSVVELLLK | Q6PDI5 | Q6PDI5 | KIAA0368 | KIAA0368 | Cytoplasm | other |
| 15552631 | 2 | 0.676 | 2163615.5 | 2054670.625 | 1471162.5 | 1451643.5 | 2349508 | 1024597.938 | 1907449.875 | 1564025 | 1717603.75 | 2015869.375 | 1105561.625 | 1341799.5 | 1937.9481 | ELC[160.0307]SLASDLSQPDLVYK | Q6PDI5 | Q6PDI5 | KIAA0368 | KIAA0368 | Cytoplasm | other |
| 15553214 | 2 | 0.277 | 1921342.375 | 1306354.5 | 1300076.75 | 1073350 | 1534537.5 | 1185727.5 | 1550240.5 | 1689417.25 | 1730725 | 1549961.25 | 1339832.125 | 1010680.312 | 1647.8744 | DIALVQQLFEALC[160.0307]K | Q6PDI5 | Q6PDI5 | KIAA0368 | KIAA0368 | Cytoplasm | other |
| 15554711 | 2 | 0.173 | 1525143 | 2121738.25 | 1706896 | 1220312 | 2054648.125 | 1266729.375 | 2035328.875 | 1377796.25 | 2030224.375 | 1473742 | 891590.25 | 1197320.375 | 1722.9756 | VNDVVPWVLDVILNK | Q6PDI5 | Q6PDI5 | KIAA0368 | KIAA0368 | Cytoplasm | other |
| 15542527 | 3 | 0.254 | 890018.75 | 964260.5 | 976549.5 | 521391.562 | 905211.812 | 443148.125 | 873347.875 | 828925.062 | 820235 | 1023624.938 | 477741 | 766258.375 | 1722.9753 | VNDVVPWVLDVILNK | Q6PDI5 | Q6PDI5 | KIAA0368 | KIAA0368 | Cytoplasm | other |
| 15543208 | 2 | 0.675 | 711996 | 533058.562 | 388219.188 | 629481.438 | 594804 | 327753.844 | 600527.562 | 479711 | 364426.719 | 630314.25 | 272323.75 | 312461.844 | 1469.6875 | ALMSSSQATASSSNK | Q6PDI5 | Q6PDI5 | KIAA0368 | KIAA0368 | Cytoplasm | other |
| 15550709 | 2 | 0.477 | 2766772.75 | 4039810 | 3974950 | 2736201.25 | 5242061 | 3381471.25 | 4587359.5 | 4164862 | 4972901 | 4280803 | 3227549.75 | 3428491.5 | 1814.8976 | ADSVPNLEPSEESLVTK | Q7TQ95 | Q7TQ95 | KIAA1715 | KIAA1715 | Cytoplasm | other |
| 15533751 | 2 | 0.172 | 2568825.25 | 4082324.25 | 4664983.5 | 2538626 | 5001952 | 4148206.5 | 5439859.5 | 5465197.5 | 3818774.25 | 3293849.25 | 1957446.875 | 2912138.25 | 1471.797 | IPVTEQTSQVIEK | Q7TQ95 | Q7TQ95 | KIAA1715 | KIAA1715 | Cytoplasm | other |
| 15542228 | 2 | 0.202 | 3436223.25 | 3207796 | 2532343.5 | 2401283.75 | 3967856 | 2821906.25 | 4414530.5 | 4111111 | 3238437.5 | 3116863.5 | 2616278 | 2725846.75 | 1459.7518 | IVEYLVGDGPQNR | Q7TQ95 | Q7TQ95 | KIAA1715 | KIAA1715 | Cytoplasm | other |
| 15561092, 15561661 | 3 | 0.372 | 2282761 | 2640905.5 | 2487305.5 | 2497670.5 | 2618175 | 2340797.5 | 2107291.5 | 3170076.25 | 2955023 | 2575665.75 | 2159483.5 | 1795897.625 | 2325.1812, 2325.1829 | NLSPAPASSSQGPPPQGPVSPGPAK | Q7TQ95 | Q7TQ95 | KIAA1715 | KIAA1715 | Cytoplasm | other |
| 15540967 | 2 | 0.529 | 1623377 | 2422746.5 | 1916154.125 | 1856539.125 | 2314950.5 | 1809993.25 | 3035064.75 | 2189954.75 | 1996704.875 | 2417260.5 | 2162969.5 | 1967434.25 | 990.51575 | ILEEVMEK | Q7TQ95 | Q7TQ95 | KIAA1715 | KIAA1715 | Cytoplasm | other |
| 15534622 | 2 | 0.753 | 2099571.75 | 3108856.5 | 6720391 | 912794.062 | 1898198.625 | 6491471 | 1564541 | 3865684.5 | 7614599.5 | 1000077.5 | 2438165 | 3803556.25 | 1074.613 | SPWFILVGR | Q60575 | Q60575 | KIF1B | kinesin family member 1B | Cytoplasm | transporter |
| 15545281 | 2 | 0.212 | 4.70E-06 | 575898.75 | 987848.625 | 2564679.75 | 2702397.5 | 2337230 | 1099895.625 | 1866875.5 | 3316736.75 | 185009.875 | 2122745.25 | 1334273.125 | 700.4591 | VVELLK | O88448 | O88448 | KLC2 | kinesin light chain 2 | Cytoplasm | other |
| 15532852 | 3 | 0.65 | 6252631 | 6407410 | 8776871 | 5048062.5 | 6782827.5 | 7176049.5 | 7317925.5 | 5845014 | 4624837 | 6822361 | 4867761.5 | 6329052.5 | 1166.5454 | SVSDSSRDTGR | Q60682 | Q60682 | Klra4 (includes others) | killer cell lectin-like receptor, subfamily A, member 4 | Plasma Membrane | transmembrane receptor |
| 15556335 | 3 | 0.281 | 3346416 | 2085437.375 | 2848479.25 | 2544795.5 | 2505181.25 | 1810937.125 | 2129632.5 | 1612393 | 2038688 | 2753752.5 | 1879679.5 | 1859007.25 | 3021.5916 | AQNVTLEAILQNATSDNPVVQLSAVQAAR | O35344 | O35344 | KPNA3 | karyopherin alpha 3 (importin alpha 4) | Nucleus | transporter |
| 15548904 | 2 | 0.185 | 1.71E+07 | 1.20E+07 | 1.26E+07 | 8925267 | 9423450 | 8364246.5 | 1.02E+07 | 8262092 | 8348322 | 1.89E+07 | 8413057 | 9023283 | 1658.9077 | AAVENLPTFLVELSR | P70168 | P70168 | KPNB1 | karyopherin (importin) beta 1 | Nucleus | transporter |
| 15547872 | 2 | 0.076 | 1.89E+07 | 1.54E+07 | 1.51E+07 | 1.18E+07 | 1.18E+07 | 9594327 | 1.15E+07 | 1.02E+07 | 1.21E+07 | 1.71E+07 | 9756209 | 1.19E+07 | 1605.8813 | LAATNALLNSLEFTK | P70168 | P70168 | KPNB1 | karyopherin (importin) beta 1 | Nucleus | transporter |
| 15532480 | 3 | 0.205 | 1.51E+07 | 1.24E+07 | 1.04E+07 | 1.64E+07 | 1.35E+07 | 1.34E+07 | 1.39E+07 | 1.22E+07 | 1.34E+07 | 1.43E+07 | 1.79E+07 | 1.53E+07 | 1833.8872 | GDQENVHPDVMLVQPR | P70168 | P70168 | KPNB1 | karyopherin (importin) beta 1 | Nucleus | transporter |
| 15534007 | 2 | 0.521 | 1.22E+07 | 9244814 | 7771460 | 1.01E+07 | 6300111 | 5349181.5 | 9562524 | 6208514 | 5195277.5 | 1.34E+07 | 6951557.5 | 6752237.5 | 1225.6627 | VLANPGNSQVAR | P70168 | P70168 | KPNB1 | karyopherin (importin) beta 1 | Nucleus | transporter |
| 15548722 | 3 | 0.278 | 1.69E+07 | 1.42E+07 | 7892085 | 1.02E+07 | 8169200.5 | 6949777 | 9352969 | 7005996.5 | 8406794 | 1.24E+07 | 8331223.5 | 7700783 | 2406.207 | YRDAAVMAFGSILEGPEPNQLK | P70168 | P70168 | KPNB1 | karyopherin (importin) beta 1 | Nucleus | transporter |
| 15533529 | 3 | 0.522 | 1.13E+07 | 7773533 | 5078640 | 7786477.5 | 5847097 | 4324993 | 6884697 | 4906509.5 | 3912614.25 | 1.13E+07 | 5815096.5 | 5304372.5 | 1878.9375 | LLETTDRPDGHQNNLR | P70168 | P70168 | KPNB1 | karyopherin (importin) beta 1 | Nucleus | transporter |
| 15539615 | 3 | 0.36 | 9510666 | 1.08E+07 | 7802358 | 5572021.5 | 7957882.5 | 6730157 | 9699415 | 7256863 | 5575369.5 | 1.07E+07 | 6388492 | 8256580 | 2123.124 | VQHQDALQISDVVMASLLR | P70168 | P70168 | KPNB1 | karyopherin (importin) beta 1 | Nucleus | transporter |
| 15539127 | 2 | 0.399 | 9003707 | 7640889 | 5398158.5 | 5547967 | 5609898.5 | 3729328.5 | 6766500 | 4505899 | 4218880 | 1.05E+07 | 4674993 | 5340791.5 | 1356.6692 | SSAYESLMEIVK | P70168 | P70168 | KPNB1 | karyopherin (importin) beta 1 | Nucleus | transporter |
| 15550353 | 3 | 0.514 | 8836173 | 9321503 | 6347446.5 | 8047589 | 6811369 | 5336385 | 8431662 | 7229866 | 6361761.5 | 9619471 | 7622809.5 | 7241678 | 2958.453 | AAGVC[160.0307]LMLLSTC[160.0307]C[160.0307]EDDIVPHVLPFIK | P70168 | P70168 | KPNB1 | karyopherin (importin) beta 1 | Nucleus | transporter |
| 15541821 | 3 | 0.21 | 9530316 | 8768649 | 6956381 | 7950343.5 | 5968818 | 5971267.5 | 7189622 | 5421072.5 | 6787715.5 | 9415767 | 7189584.5 | 6933199.5 | 1759.0331 | GALQYLVPILTQTLTK | P70168 | P70168 | KPNB1 | karyopherin (importin) beta 1 | Nucleus | transporter |
| 15532016 | 2 | 0.074 | 8651589 | 6530752 | 6479299 | 5636481 | 5866998.5 | 4739678 | 5465980 | 4291751.5 | 4500686 | 8911918 | 5475177.5 | 5869767.5 | 955.5916 | VAALQNLVK | P70168 | P70168 | KPNB1 | karyopherin (importin) beta 1 | Nucleus | transporter |
| 15548833 | 2 | 0.291 | 6775221.5 | 6972222 | 4808953 | 4774885 | 4671104.5 | 4421146 | 5242756 | 4447455.5 | 5220779.5 | 6933921 | 4405938.5 | 4708043.5 | 1696.9723 | PLVIQAMPTLIELMK | P70168 | P70168 | KPNB1 | karyopherin (importin) beta 1 | Nucleus | transporter |
| 15539744 | 3 | 0.557 | 7392570 | 5291693 | 4962625.5 | 3399079 | 5641477 | 4994280 | 5016086.5 | 4745227 | 3779339.5 | 6638495.5 | 4261994 | 3839120.75 | 1999.9823 | LQQVLQMESHIQSTSDR | P70168 | P70168 | KPNB1 | karyopherin (importin) beta 1 | Nucleus | transporter |
| 15549055 | 2 | 0.289 | 7285712.5 | 6282157.5 | 5040510 | 3420628 | 5429231 | 3617231.75 | 5345842 | 4074920.75 | 3707577.75 | 6636697 | 3153445.5 | 3937754.25 | 1616.7224 | SDFDMVDYLNELR | P70168 | P70168 | KPNB1 | karyopherin (importin) beta 1 | Nucleus | transporter |
| 15551231 | 3 | 0.533 | 6014486 | 2947020.5 | 1418263.625 | 5532979 | 1063247.625 | 2215361 | 3791432.25 | 1536137.5 | 2665067 | 6015169 | 4577679.5 | 3706374.75 | 3265.5293 | SDFDMVDYLNELRESC[160.0307]LEAYTGIVQGLK | P70168 | P70168 | KPNB1 | karyopherin (importin) beta 1 | Nucleus | transporter |
| 15536062 | 2 | 0.077 | 7120578.5 | 6620124 | 5138179.5 | 4001383.25 | 4321298.5 | 3085646.25 | 5400788.5 | 3848233.5 | 3334340.5 | 5817691 | 3817425.75 | 3989666.5 | 1445.7761 | SNEILTAIIQGMR | P70168 | P70168 | KPNB1 | karyopherin (importin) beta 1 | Nucleus | transporter |
| 15549140 | 2 | 0.056 | 6554720.5 | 5115457.5 | 5841989 | 4002466.25 | 3573825.25 | 3277477.25 | 4233895 | 3041309.75 | 3002545.5 | 5472211 | 3062153 | 2800325.5 | 2565.1968 | ESTLEAIGYIC[160.0307]QDIDPEQLQDK | P70168 | P70168 | KPNB1 | karyopherin (importin) beta 1 | Nucleus | transporter |
| 15536653 | 2 | 0.697 | 4791657.5 | 2900876.75 | 1895531.375 | 2809956 | 2913912.75 | 1818800.375 | 2898159.75 | 1806399.375 | 1882866.25 | 4545192 | 2340147.75 | 2111572.5 | 962.532 | TTLVIMER | P70168 | P70168 | KPNB1 | karyopherin (importin) beta 1 | Nucleus | transporter |
| 15557874 | 2 | 0.019 | 5151494 | 5407154.5 | 5298241.5 | 4132700.5 | 4207687.5 | 4075080 | 4148701.5 | 4028177 | 5289166.5 | 4412991.5 | 3766724.5 | 3822946.25 | 1759.0332 | GALQYLVPILTQTLTK | P70168 | P70168 | KPNB1 | karyopherin (importin) beta 1 | Nucleus | transporter |
| 15553109 | 2 | 0.51 | 4358612.5 | 2556965.5 | 2157639.5 | 2673352.25 | 2448450.75 | 1459042.375 | 2794099 | 1255879.875 | 1747754.375 | 3881323.25 | 2295580.25 | 1772633.625 | 1556.8247 | TVSPDRLELEAAQK | P70168 | P70168 | KPNB1 | karyopherin (importin) beta 1 | Nucleus | transporter |
| 15538900 | 2 | 0.174 | 5149761.5 | 3294344.5 | 2991564 | 2923008.25 | 2658459 | 2398387 | 3285025.75 | 2103934.75 | 1634836 | 3208595.5 | 2355440.5 | 2333626 | 844.52625 | PFLGIGLK | P70168 | P70168 | KPNB1 | karyopherin (importin) beta 1 | Nucleus | transporter |
| 15531584 | 2 | 0.262 | 1.90E+07 | 2.60E+07 | 6717932.5 | 1.86E+07 | 2.93E+07 | 5.43E+07 | 1.78E+07 | 9698517 | 1.76E+07 | 1.62E+07 | 3.15E+07 | 2.82E+07 | 1475.7822 | FLEQQNQVLQTK | P04104 | P04104 | KRT1 | keratin 1, type II | Cytoplasm | other |
| 15541633 | 3 | 0.07 | 771769 | 105586.625 | 193991.75 | 728554.375 | 5793029.5 | 8734984 | 152116.859 | 777976.188 | 2840527.25 | 4059023.75 | 6507250 | 1425600 | 1393.729 | TNAENEFVTIKK | P04104 | P04104 | KRT1 | keratin 1, type II | Cytoplasm | other |
| 15533426 | 3 | 0.236 | 3776790 | 7635125 | 1636820.375 | 5445471 | 7190517.5 | 1.62E+07 | 4984650.5 | 2470215.25 | 3901701 | 3983660.25 | 4847393 | 7723854 | 1421.7361 | RTNAENEFVTIK | P04104 | P04104 | KRT1 | keratin 1, type II | Cytoplasm | other |
| 15532886 | 2 | 0.078 | 4052569 | 1847812.75 | 534610.688 | 3074894 | 4260847 | 9554317 | 1333066.5 | 1435697.875 | 2178515 | 3163564.25 | 7043055.5 | 6700360.5 | 1265.6365 | TNAENEFVTIK | P04104 | P04104 | KRT1 | keratin 1, type II | Cytoplasm | other |
| 15540363 | 2 | 0.216 | 9353656 | 7943872 | 2721945.5 | 8510603 | 7786043 | 1.00E+07 | 6953222.5 | 2701601.5 | 6595765.5 | 7679788 | 1.43E+07 | 1.22E+07 | 1381.6445 | ALEESNYELEGK | P02535 | P02535 | Krt10 | keratin 10 | Cytoplasm | other |
| 15538858 | 3 | 0.158 | 1241959.25 | 880981.5 | 84476.617 | 1181707.125 | 2675373 | 6412235.5 | 1013572.188 | 312469.531 | 2252061.75 | 4194596.5 | 2886050.75 | 1535376.125 | 2104.0479 | TEELNREVATNSELVQSGK | Q61781 | Q61781 | KRT14 | keratin 14, type I | Cytoplasm | other |
| 15537660 | 3 | 0.176 | 4684447.5 | 3443305.75 | 321941.156 | 8006854.5 | 2966207 | 9919837 | 2731431 | 662889.5 | 3082014.75 | 4023986.5 | 9870816 | 1.19E+07 | 1523.7849 | LLRDYQELMNTK | Q3TTY5 | Q3TTY5 | Krt2 | keratin 2 | Cytoplasm | other |
| 15532800 | 2 | 0.075 | 1.40E+07 | 1.99E+07 | 6768121 | 1.84E+07 | 3.36E+07 | 6.28E+07 | 1.13E+07 | 1.38E+07 | 2.11E+07 | 2.49E+07 | 2.78E+07 | 2.52E+07 | 1302.713 | SLDLDSIIAEVK | Q922U2 | Q922U2 | KRT5 | keratin 5, type II | Cytoplasm | other |
| 15533041 | 2 | 0.411 | 6374868 | 6270841.5 | 7941905.5 | 4629462.5 | 6903183 | 6356000.5 | 6400310 | 7911655.5 | 7569929 | 8427649 | 7481453.5 | 5920733.5 | 1154.5593 | KDVDAAYMNK | Q922U2 | Q922U2 | KRT5 | keratin 5, type II | Cytoplasm | other |
| 15544201 | 2 | 0.115 | 928013.438 | 1072908.625 | 282465.656 | 1036743 | 1603281.375 | 2548817.5 | 904725.125 | 412650.906 | 1394854 | 3606912.75 | 1470051.125 | 1707843.375 | 1194.572 | YEELQQTAGR | Q922U2 | Q922U2 | KRT5 | keratin 5, type II | Cytoplasm | other |
| 15542101 | 2 | 0.779 | 3827791.25 | 6356446.5 | 2051211.5 | 2449629 | 6826824.5 | 1.78E+07 | 3978481.5 | 2455832 | 4592442.5 | 859591.062 | 7706479 | 7641060 | 1073.592 | LRSEIDNVK | Q922U2 | Q922U2 | KRT5 | keratin 5, type II | Cytoplasm | other |
| 15546001 | 2 | 0.106 | 1050164.25 | 1138245.125 | 138209.984 | 2295151.5 | 1423709.25 | 2009131.5 | 945537.75 | 579997 | 556948.5 | 1448962.25 | 1458266.875 | 1915136.75 | 1277.7086 | LALDIEIATYR | Q6NXH9 | Q6NXH9 | KRT73 | keratin 73, type II | Extracellular Space | other |
| 15541253 | 2 | 0.056 | 1.25E+07 | 1.26E+07 | 5030399.5 | 1.50E+07 | 2.32E+07 | 4.19E+07 | 1.14E+07 | 7976265.5 | 1.53E+07 | 2.57E+07 | 2.40E+07 | 1.61E+07 | 1065.5193 | AQYEDIAQK | Q3UV17 | Q3UV17 | KRT76 | keratin 76, type II | Cytoplasm | other |
| 15532887 | 2 | 0.141 | 3809798.25 | 5551965.5 | 4838101.5 | 4602531.5 | 9970124 | 1.49E+07 | 3529611 | 2731505.25 | 8904074 | 1.39E+07 | 7739244.5 | 9014537 | 1329.7235 | NLDLDSIIAEVK | Q8VED5 | Q8VED5 | KRT79 | keratin 79, type II | Extracellular Space | other |
| 15532279 | 2 | 0.519 | 1.55E+07 | 1.30E+07 | 1.99E+07 | 9697870 | 1.56E+07 | 1.47E+07 | 1.56E+07 | 1.75E+07 | 1.39E+07 | 1.92E+07 | 1.57E+07 | 1.43E+07 | 1067.5012 | QLYEEETR | Q6IMF0 | Q6IMF0 | Krt83 | keratin 83 | Extracellular Space | other |
| 15536463 | 2 | 0.253 | 9822041 | 1.38E+07 | 1.08E+07 | 1.29E+07 | 1.28E+07 | 1.46E+07 | 9595046 | 1.31E+07 | 1.28E+07 | 7592876 | 1.25E+07 | 9599633 | 1416.6545 | EIYTMDSTTDIK | P11438 | P11438 | LAMP1 | lysosomal-associated membrane protein 1 | Plasma Membrane | other |
| 15536568 | 2 | 0.103 | 4987174.5 | 7313336.5 | 5824120 | 9377829 | 8905004 | 8455812 | 6100686.5 | 7805440.5 | 8418845 | 3616334 | 7152215 | 6310950 | 1151.6029 | ALQATVGNSYK | P11438 | P11438 | LAMP1 | lysosomal-associated membrane protein 1 | Plasma Membrane | other |
| 15536549 | 2 | 0.387 | 1869536.375 | 2558558.75 | 1880708.25 | 2925377 | 2866953.75 | 2874710.25 | 2147185 | 2276442 | 2916466.25 | 695970.938 | 2973811.5 | 2056775.25 | 1363.6279 | C[160.0307]NTEEHIFVSK | P11438 | P11438 | LAMP1 | lysosomal-associated membrane protein 1 | Plasma Membrane | other |
| 15533178 | 2 | 0.248 | 3050687.25 | 7842984 | 5990662.5 | 6342400 | 7841323 | 1.02E+07 | 4094000.25 | 7381600 | 9977546 | 1855237 | 5322791 | 4822066 | 930.5638 | VPLDVIFK | P17047 | P17047 | LAMP2 | lysosomal-associated membrane protein 2 | Plasma Membrane | enzyme |
| 15536513 | 2 | 0.163 | 1940388 | 3021043.25 | 2366056.25 | 3671101.25 | 4220821 | 4949607.5 | 2365894.75 | 4039304.75 | 5436860 | 868242.625 | 3536050.25 | 2234236.5 | 957.5592 | TITIAVPDK | P17047 | P17047 | LAMP2 | lysosomal-associated membrane protein 2 | Plasma Membrane | enzyme |
| 15548648 | 4 | 0.057 | 2.14E+07 | 2.85E+07 | 2.57E+07 | 2.37E+07 | 2.40E+07 | 2.13E+07 | 2.05E+07 | 2.64E+07 | 2.61E+07 | 1.56E+07 | 1.52E+07 | 2.19E+07 | 3409.8394 | KLPPLPSLTSQPHQVLASEPIPFSDLQQVSR | Q9CQ22 | Q9CQ22 | LAMTOR1 | late endosomal/lysosomal adaptor, MAPK and MTOR activator 1 | Plasma Membrane | other |
| 15532936 | 2 | 0.518 | 6881042.5 | 8164311 | 8250060.5 | 8386723.5 | 7741674.5 | 6036022 | 5941463.5 | 7345598 | 7825509.5 | 7065188.5 | 5209060.5 | 7154153.5 | 1388.759 | TDEQALLSSILAK | Q9CQ22 | Q9CQ22 | LAMTOR1 | late endosomal/lysosomal adaptor, MAPK and MTOR activator 1 | Plasma Membrane | other |
| 15549520 | 3 | 0.068 | 6975931.5 | 8736272 | 8682869 | 6535530 | 7946768 | 6608594 | 5898182 | 7391970.5 | 7960229 | 5232415.5 | 4440966 | 6747288.5 | 2668.1562 | TASNIIDVSAADSQGMEQHEYMDR | Q9CQ22 | Q9CQ22 | LAMTOR1 | late endosomal/lysosomal adaptor, MAPK and MTOR activator 1 | Plasma Membrane | other |
| 15541945 | 2 | 0.124 | 4311897.5 | 4179993.75 | 3524251.5 | 3679884 | 3668323.75 | 3057155.25 | 2428308.75 | 3304542.75 | 3884708.5 | 2335861 | 2931183.5 | 3169283.25 | 1426.7656 | IAAYAYSALSQIR | Q9CQ22 | Q9CQ22 | LAMTOR1 | late endosomal/lysosomal adaptor, MAPK and MTOR activator 1 | Plasma Membrane | other |
| 15532934 | 4 | 0.029 | 1.38E+07 | 1.90E+07 | 1.48E+07 | 1.29E+07 | 1.72E+07 | 1.18E+07 | 1.02E+07 | 1.27E+07 | 1.29E+07 | 9705052 | 8516297 | 1.10E+07 | 2900.4487 | VANDSAPEHALRPGFLSTFALATDQGSK | O88653 | O88653 | LAMTOR3 | late endosomal/lysosomal adaptor, MAPK and MTOR activator 3 | Cytoplasm | other |
| 15537834 | 2 | 0.15 | 5148083.5 | 7059481 | 7506428 | 4882059.5 | 6845018 | 5180250.5 | 3879712 | 6271616.5 | 4966617 | 3960483.75 | 3252801 | 5337704 | 1301.7329 | ELAPLFEELIK | O88653 | O88653 | LAMTOR3 | late endosomal/lysosomal adaptor, MAPK and MTOR activator 3 | Cytoplasm | other |
| 15555724 | 3 | 0.499 | 4792661.5 | 6090921 | 2757595.25 | 3770611.25 | 5880766.5 | 3371452.5 | 2954408 | 3644104.5 | 5081801 | 3202432.75 | 2359832.25 | 3617050.5 | 2900.4468 | VANDSAPEHALRPGFLSTFALATDQGSK | O88653 | O88653 | LAMTOR3 | late endosomal/lysosomal adaptor, MAPK and MTOR activator 3 | Cytoplasm | other |
| 15532261 | 3 | 0.031 | 1.16E+07 | 1.49E+07 | 1.21E+07 | 9692616 | 1.23E+07 | 8945184 | 8495041 | 1.10E+07 | 9549870 | 6685934.5 | 7558508 | 9618549 | 2023.0527 | GTLSDEHAGVISVLAQQAAR | Q9D1L9 | Q9D1L9 | LAMTOR5 | late endosomal/lysosomal adaptor, MAPK and MTOR activator 5 | Cytoplasm | other |
| 15549947 | 2 | 0.073 | 4972291.5 | 6119403.5 | 7785768.5 | 4134335.75 | 5885013 | 5200453.5 | 3994353 | 5068703 | 5899327.5 | 4384521 | 3024633.25 | 3827502.75 | 2160.0486 | NPSIVGVLC[160.0307]TDSQGLNLGC[160.0307]R | Q9D1L9 | Q9D1L9 | LAMTOR5 | late endosomal/lysosomal adaptor, MAPK and MTOR activator 5 | Cytoplasm | other |
| 15540820 | 3 | 0.337 | 2726596.25 | 7283950.5 | 6009240 | 3849297 | 3866871 | 3569827.5 | 3108059.75 | 3918855.75 | 3897359.5 | 3695318.25 | 2314841 | 3375044 | 2160.0503 | NPSIVGVLC[160.0307]TDSQGLNLGC[160.0307]R | Q9D1L9 | Q9D1L9 | LAMTOR5 | late endosomal/lysosomal adaptor, MAPK and MTOR activator 5 | Cytoplasm | other |
| 15556997 | 3 | 0.566 | 1885209 | 1979779.5 | 2109886.5 | 1643422.25 | 3431020.5 | 2500488.75 | 2612747 | 2672468 | 2188691 | 4754973 | 1800464.5 | 2809680 | 2537.2766 | IPQSHIQQIC[160.0307]ENILTSGENLSR | O89112 | O89112 | LANCL1 | LanC lantibiotic synthetase component C-like 1 (bacterial) | Plasma Membrane | other |
| 15532360 | 2 | 0.522 | 9024841 | 8921849 | 8542152 | 7485180.5 | 8916319 | 8902757 | 8862518 | 7533921.5 | 8002314.5 | 1.50E+07 | 8069866 | 8510782 | 1131.6234 | EVVSAIIESGK | Q9JJK2 | Q9JJK2 | LANCL2 | LanC lantibiotic synthetase component C-like 2 (bacterial) | Plasma Membrane | other |
| 15541179 | 2 | 0.762 | 8663561 | 1.05E+07 | 1.08E+07 | 7639340 | 9645720 | 8666649 | 8974751 | 8335031.5 | 7903104.5 | 1.29E+07 | 5876198 | 8058987 | 1296.6968 | FPAFELGFLQK | Q9JJK2 | Q9JJK2 | LANCL2 | LanC lantibiotic synthetase component C-like 2 (bacterial) | Plasma Membrane | other |
| 15550648 | 2 | 0.529 | 8339418.5 | 6940371 | 5967978 | 7622882.5 | 7859366 | 8830077 | 8871772 | 7076677.5 | 8652175 | 1.21E+07 | 7625890 | 7257143.5 | 2021.9807 | TIVC[160.0307]QESELPDELLYGR | Q9JJK2 | Q9JJK2 | LANCL2 | LanC lantibiotic synthetase component C-like 2 (bacterial) | Plasma Membrane | other |
| 15537186 | 2 | 0.986 | 4535635.5 | 6586807 | 6212350.5 | 4828839.5 | 5687031.5 | 5677968 | 7108682.5 | 5042792.5 | 3918203 | 1.15E+07 | 3554089.25 | 4456817.5 | 1292.6382 | VDQETLTEMVK | Q9JJK2 | Q9JJK2 | LANCL2 | LanC lantibiotic synthetase component C-like 2 (bacterial) | Plasma Membrane | other |
| 15534275 | 2 | 0.764 | 7323712 | 6415301.5 | 7011292.5 | 6420868 | 6221776.5 | 6163858 | 6278445 | 5821730.5 | 5868446 | 1.00E+07 | 5080111 | 5984883 | 1303.6532 | DLLQQMEEGLK | Q9JJK2 | Q9JJK2 | LANCL2 | LanC lantibiotic synthetase component C-like 2 (bacterial) | Plasma Membrane | other |
| 15549913 | 2 | 0.976 | 4691944 | 4534434.5 | 5313473.5 | 3277388.25 | 6109625.5 | 4094788 | 4866503 | 4737211.5 | 3787365.25 | 8949738 | 3344895.75 | 3451847.25 | 1526.67 | SGNYPSSLSNETDR | Q9JJK2 | Q9JJK2 | LANCL2 | LanC lantibiotic synthetase component C-like 2 (bacterial) | Plasma Membrane | other |
| 15538835 | 2 | 0.408 | 4636027.5 | 4442905.5 | 4743827.5 | 3559154.25 | 4468664 | 3595945 | 4830118.5 | 4910685.5 | 4002307.5 | 7223929.5 | 4212736 | 4109655.5 | 1165.6185 | VTGDQTYLLR | Q9JJK2 | Q9JJK2 | LANCL2 | LanC lantibiotic synthetase component C-like 2 (bacterial) | Plasma Membrane | other |
| 15538791 | 2 | 0.987 | 2674961.75 | 3654913.5 | 3700648 | 2659601.5 | 3662475.75 | 3134860.25 | 4037830.5 | 3763671.5 | 2117524 | 5597943 | 1607883.875 | 2888171.75 | 849.445 | PSIDYVR | Q9JJK2 | Q9JJK2 | LANCL2 | LanC lantibiotic synthetase component C-like 2 (bacterial) | Plasma Membrane | other |
| 15550514 | 3 | 0.206 | 4183671.75 | 3159437.25 | 2462994 | 3755158.5 | 3454804 | 2915042.5 | 3647877 | 3331704.75 | 3657520.5 | 5437726.5 | 3861178.75 | 4043252.5 | 2429.1392 | QYVGAAHGMAGIYYMLMQPEAK | Q9JJK2 | Q9JJK2 | LANCL2 | LanC lantibiotic synthetase component C-like 2 (bacterial) | Plasma Membrane | other |
| 15558833 | 2 | 0.077 | 2606420.75 | 1051789.875 | 2440822 | 2059397.75 | 2914313.5 | 2306916.5 | 3077431.5 | 2737737.75 | 2516908.25 | 4859961.5 | 4323443.5 | 3048347 | 1523.6584 | EAMEC[160.0307]SDVIWQR | Q9JJK2 | Q9JJK2 | LANCL2 | LanC lantibiotic synthetase component C-like 2 (bacterial) | Plasma Membrane | other |
| 15543592 | 3 | 0.084 | 3131328 | 1220293.375 | 1648301.625 | 1707243.75 | 2294268.25 | 1765982.75 | 2452342.75 | 2603480.5 | 2406588.75 | 4766242 | 5720560 | 2346893 | 1741.7175 | FAEWC[160.0307]LDYGAHGC[160.0307]R | Q9JJK2 | Q9JJK2 | LANCL2 | LanC lantibiotic synthetase component C-like 2 (bacterial) | Plasma Membrane | other |
| 15543445 | 3 | 0.26 | 5608197.5 | 965712.75 | 565064.688 | 4679329.5 | 494387.375 | 338759.406 | 1107402.875 | 1156001.875 | 2284376.5 | 4614056 | 6873122.5 | 3524672 | 2123.063 | VDQETLTEMVKPSIDYVR | Q9JJK2 | Q9JJK2 | LANCL2 | LanC lantibiotic synthetase component C-like 2 (bacterial) | Plasma Membrane | other |
| 15532560 | 2 | 0.038 | 1.13E+07 | 8274164.5 | 1.02E+07 | 9009249 | 7232073.5 | 6691969 | 6445608 | 6020394 | 5181147.5 | 8442347 | 5071964 | 4759581 | 1144.5814 | VTLTPEEEAR | P06151 | P06151 | LDHA | lactate dehydrogenase A | Cytoplasm | enzyme |
| 15549360 | 2 | 0.088 | 7797026.5 | 7220398 | 8674015 | 7389710.5 | 6292556.5 | 6399678 | 5886982 | 6293553.5 | 6599379 | 6715481 | 6041222 | 4140436.5 | 1944.1161 | LLIVSNPVDILTYVAWK | P06151 | P06151 | LDHA | lactate dehydrogenase A | Cytoplasm | enzyme |
| 15537101 | 2 | 0.02 | 8681942 | 6523845.5 | 7342967 | 5726461 | 5803865.5 | 5608251 | 5154515.5 | 5235149.5 | 3774604.25 | 5677670 | 3308281.25 | 3514685.5 | 1250.6598 | QVVDSAYEVIK | P06151 | P06151 | LDHA | lactate dehydrogenase A | Cytoplasm | enzyme |
| 15534677 | 2 | 0.001 | 6111686.5 | 5143465.5 | 6268550.5 | 4290281.5 | 4388661 | 4455493 | 4149105.25 | 3486076.75 | 4055345 | 3855549.75 | 2886522 | 3511991 | 943.5947 | FIIPNIVK | P06151 | P06151 | LDHA | lactate dehydrogenase A | Cytoplasm | enzyme |
| 15539921 | 2 | 8.07E-05 | 6922769 | 6207649 | 6102980.5 | 5572537.5 | 5069426 | 5800704 | 4652886 | 5053887 | 4993370 | 3514880.25 | 2608742.25 | 3279173.25 | 1055.6437 | DQLIVNLLK | P06151 | P06151 | LDHA | lactate dehydrogenase A | Cytoplasm | enzyme |
| 15551797 | 2 | 0.016 | 4862582.5 | 3348039.25 | 6156543 | 3088030.25 | 4003516.75 | 2737011.25 | 2715680.75 | 2981783 | 2089782 | 2689568.5 | 1872234.5 | 2053028.75 | 1675.8043 | DLADELALVDVMEDK | P06151 | P06151 | LDHA | lactate dehydrogenase A | Cytoplasm | enzyme |
| 15540148 | 2 | 0.023 | 4069374.5 | 3010666.75 | 4378453.5 | 3155836.5 | 1983064.75 | 2184980.5 | 2474549.5 | 1512510.75 | 1905204.25 | 2483845 | 1815876.625 | 2229961.5 | 913.58075 | LVIITAGAR | P06151 | P06151 | LDHA | lactate dehydrogenase A | Cytoplasm | enzyme |
| 15553999 | 3 | 0.119 | 2635160 | 2858167.5 | 2454581.5 | 2803245.25 | 1507493.75 | 1676670.25 | 1818938 | 1956808.75 | 1930067.875 | 2139432.25 | 1513483.5 | 1641349.25 | 2906.4893 | GLYGINEDVFLSVPC[160.0307]ILGQNGISDVVK | P06151 | P06151 | LDHA | lactate dehydrogenase A | Cytoplasm | enzyme |
| 15539588 | 3 | 0.247 | 1782614.75 | 1921915.625 | 2398538.25 | 1739234.75 | 1282715.625 | 2003486 | 1442826.75 | 1362414.5 | 1498521.125 | 1667594.375 | 2024259.25 | 1211233.5 | 1944.1163 | LLIVSNPVDILTYVAWK | P06151 | P06151 | LDHA | lactate dehydrogenase A | Cytoplasm | enzyme |
| 15545054 | 2 | 0.388 | 840998.125 | 1949211.125 | 1908089.125 | 5963671.5 | 7931490.5 | 5358648 | 4663103 | 1790803.75 | 2.28E+07 | 161864.859 | 4686392.5 | 7117566 | 715.4087 | EVLQAR | Q8CBY3 | Q8CBY3 | LENG8 | leukocyte receptor cluster (LRC) member 8 | Nucleus | other |
| 15535026 | 2 | 0.881 | 3818222 | 3043182.75 | 3060237.25 | 3163981 | 3431416.5 | 2896319 | 2994608.75 | 3106142.25 | 2985363 | 4770287 | 3388244.75 | 2430432.5 | 1432.5983 | VTYDSDATSSAC[160.0307]R | O89013 | O89013 | LEPROT | leptin receptor overlapping transcript | Plasma Membrane | other |
| 15561954 | 2 | 0.055 | 400263.969 | 314347.312 | 238396.359 | 224755.25 | 254874.5 | 183906.016 | 248785.234 | 168006.609 | 208409.641 | 276130.281 | 354086.125 | 303577 | 1588.6992 | RVTYDSDATSSAC[160.0307]R | O89013 | O89013 | LEPROT | leptin receptor overlapping transcript | Plasma Membrane | other |
| 15552554 | 3 | 0.7 | 1493294.875 | 2669611.5 | 906362.812 | 2091175.625 | 1876937.875 | 3103020 | 1218856.75 | 2183263 | 3306562.5 | 440723.25 | 2772544 | 1953939 | 2488.3079 | LFEDELTLDNLTRPQLVALC[160.0307]K | Q9Z2I0 | Q9Z2I0 | LETM1 | leucine zipper-EF-hand containing transmembrane protein 1 | Cytoplasm | other |
| 15531916 | 2 | 0.002 | 1.86E+07 | 1.75E+07 | 1.38E+07 | 2.72E+07 | 2.31E+07 | 2.88E+07 | 1.86E+07 | 2.29E+07 | 2.08E+07 | 2.92E+07 | 2.58E+07 | 2.75E+07 | 1402.6273 | FEEGGYVVC[160.0307]NTK | O08573 | O08573 | LGALS9B | lectin, galactoside-binding, soluble, 9B | Cytoplasm | other |
| 15553274 | 2 | 0.073 | 1.19E+07 | 1.08E+07 | 1.15E+07 | 2.12E+07 | 1.44E+07 | 2.33E+07 | 1.12E+07 | 1.71E+07 | 1.86E+07 | 2.45E+07 | 1.58E+07 | 1.66E+07 | 1575.7119 | NTQINNSWGQEER | O08573 | O08573 | LGALS9B | lectin, galactoside-binding, soluble, 9B | Cytoplasm | other |
| 15532236 | 2 | 9.28E-04 | 1.21E+07 | 1.17E+07 | 1.03E+07 | 2.08E+07 | 1.67E+07 | 2.30E+07 | 1.59E+07 | 1.69E+07 | 1.51E+07 | 2.34E+07 | 1.78E+07 | 2.22E+07 | 1473.7708 | SIMISGNVLPDATR | O08573 | O08573 | LGALS9B | lectin, galactoside-binding, soluble, 9B | Cytoplasm | other |
| 15538142 | 2 | 0.029 | 1.29E+07 | 1.46E+07 | 1.09E+07 | 2.11E+07 | 1.74E+07 | 2.62E+07 | 1.48E+07 | 1.93E+07 | 1.98E+07 | 2.28E+07 | 1.61E+07 | 2.06E+07 | 948.4884 | FNENAVVR | O08573 | O08573 | LGALS9B | lectin, galactoside-binding, soluble, 9B | Cytoplasm | other |
| 15535303 | 2, 3 | 0.104 | 6609603 | 8619866 | 7134195.5 | 1.05E+07 | 9375106 | 1.74E+07 | 8637696 | 1.18E+07 | 9862547 | 1.25E+07 | 9010934 | 1.12E+07 | 1124.5615, 1124.5616 | FFVQYQHR | O08573 | O08573 | LGALS9B | lectin, galactoside-binding, soluble, 9B | Cytoplasm | other |
| 15542951 | 3 | 0.023 | 6274157 | 4916014.5 | 3549556.5 | 6925709 | 6061538 | 9658257 | 6574842.5 | 7519007 | 7030088.5 | 1.06E+07 | 1.01E+07 | 7832754.5 | 1356.6438 | C[160.0307]GGDIAFHLNPR | O08573 | O08573 | LGALS9B | lectin, galactoside-binding, soluble, 9B | Cytoplasm | other |
| 15533603 | 2 | 0.068 | 5956523 | 5623845.5 | 3592727 | 7134386 | 5916255.5 | 1.05E+07 | 7701620.5 | 7919196 | 6081083.5 | 9589569 | 8687899 | 7669614 | 1496.7365 | GMPFELC[160.0307]FLVQR | O08573 | O08573 | LGALS9B | lectin, galactoside-binding, soluble, 9B | Cytoplasm | other |
| 15543918 | 2 | 0.029 | 3542754.5 | 3692173.25 | 2465172.5 | 4243555 | 3703261 | 5673581.5 | 3679362.5 | 5319592 | 4332745.5 | 5407038.5 | 5685214 | 5973034 | 799.4559 | FHINLR | O08573 | O08573 | LGALS9B | lectin, galactoside-binding, soluble, 9B | Cytoplasm | other |
| 15536466 | 2 | 0.089 | 310348 | 721731.688 | 2537077.5 | 2202717.25 | 5385931 | 5063711 | 2087767.75 | 3725326.25 | 4489564.5 | 453413.812 | 2163059.25 | 1698677.375 | 716.42896 | ILDLSR | A2ARI4 | A2ARI4 | LGR4 | leucine-rich repeat containing G protein-coupled receptor 4 | Plasma Membrane | transmembrane receptor |
| 15540577 | 3 | 0.312 | 84391.266 | 415529.656 | 401405.188 | 1642042.125 | 1779912.5 | 1281245.5 | 1167797.875 | 856635.5 | 4879383.5 | 31175.572 | 1872589.125 | 2339744.5 | 1583.8215 | C[160.0307]HAIIDEQPLIFK | Q99JW4 | Q99JW4 | LIMS1 | LIM and senescent cell antigen-like domains 1 | Plasma Membrane | other |
| 15534327 | 2 | 0.364 | 1.02E+07 | 1.33E+07 | 1.88E+07 | 1.62E+07 | 1.81E+07 | 1.42E+07 | 1.85E+07 | 1.54E+07 | 1.04E+07 | 1.31E+07 | 9473643 | 1.13E+07 | 1073.5671 | RDIDSLAQR | Q9D0F3 | Q9D0F3 | LMAN1 | lectin, mannose-binding, 1 | Cytoplasm | other |
| 15535744 | 3 | 0.511 | 8489944 | 8517910 | 1.16E+07 | 9381315 | 1.03E+07 | 9884229 | 1.04E+07 | 9384649 | 6402884.5 | 1.07E+07 | 9625530 | 1.11E+07 | 1416.7235 | QLDMILDEQRR | Q9D0F3 | Q9D0F3 | LMAN1 | lectin, mannose-binding, 1 | Cytoplasm | other |
| 15538736 | 2 | 0.415 | 9985649 | 8455631 | 1.48E+07 | 1.15E+07 | 1.44E+07 | 1.26E+07 | 1.15E+07 | 1.28E+07 | 8595456 | 9571075 | 9748338 | 1.00E+07 | 1283.6448 | YVSSLTEEISR | Q9D0F3 | Q9D0F3 | LMAN1 | lectin, mannose-binding, 1 | Cytoplasm | other |
| 15548942 | 3 | 0.122 | 1.11E+07 | 1.17E+07 | 1.15E+07 | 1.32E+07 | 1.49E+07 | 9378116 | 1.21E+07 | 1.15E+07 | 8597506 | 8757028 | 8572992 | 9070208 | 2941.4634 | GPHLVQSDGTVPFWAHAGNAIPSADQIR | Q9D0F3 | Q9D0F3 | LMAN1 | lectin, mannose-binding, 1 | Cytoplasm | other |
| 15555620 | 3 | 0.458 | 8529249 | 6399476.5 | 7615315 | 1.03E+07 | 1.14E+07 | 8732609 | 1.19E+07 | 8757789 | 5473075.5 | 8046207 | 8857590 | 8087763 | 2615.2852 | LVSGIQHPGSAGVYETTQHFMDIK | Q9D0F3 | Q9D0F3 | LMAN1 | lectin, mannose-binding, 1 | Cytoplasm | other |
| 15551200 | 2 | 0.762 | 5558369 | 5030676 | 6167420 | 6079566.5 | 6006574.5 | 2684540 | 5071076 | 5910328 | 3856332 | 7576240 | 5283595 | 4730033.5 | 1869.8252 | YQEEFEHFQQELDK | Q9D0F3 | Q9D0F3 | LMAN1 | lectin, mannose-binding, 1 | Cytoplasm | other |
| 15550881 | 3 | 0.018 | 9210350 | 1.23E+07 | 1.01E+07 | 1.85E+07 | 1.59E+07 | 1.32E+07 | 1.26E+07 | 1.27E+07 | 1.39E+07 | 1.26E+07 | 1.22E+07 | 1.15E+07 | 2712.3096 | LPTGYYFGASAGTGDLSDNHDIISIK | Q9DBH5 | Q9DBH5 | LMAN2 | lectin, mannose-binding 2 | Cytoplasm | transporter |
| 15550789 | 3 | 0.02 | 1.31E+07 | 1.68E+07 | 1.77E+07 | 2.16E+07 | 2.32E+07 | 1.83E+07 | 1.79E+07 | 1.93E+07 | 1.69E+07 | 1.22E+07 | 1.45E+07 | 1.55E+07 | 2468.1304 | DNFHGLAIFLDTYPNDETTER | Q9DBH5 | Q9DBH5 | LMAN2 | lectin, mannose-binding 2 | Cytoplasm | transporter |
| 15536725 | 2 | 0.019 | 3159562.25 | 3995422.25 | 4307338.5 | 7534399.5 | 5432143.5 | 5695087.5 | 5174559 | 4999358.5 | 4273559.5 | 4487189.5 | 5510843.5 | 4739517.5 | 1412.6229 | WSELAGC[160.0307]TADFR | Q9DBH5 | Q9DBH5 | LMAN2 | lectin, mannose-binding 2 | Cytoplasm | transporter |
| 15533962 | 2 | 0.166 | 2542541.25 | 4008351.25 | 4310149 | 4062370.25 | 5472289.5 | 4552289.5 | 5192400.5 | 5195716 | 3677177 | 3719739 | 2944733.75 | 3561555.25 | 1005.5712 | LFQLTVER | Q9DBH5 | Q9DBH5 | LMAN2 | lectin, mannose-binding 2 | Cytoplasm | transporter |
| 15538759 | 2 | 0.009 | 2461936.75 | 3346695.25 | 3442730.5 | 4429013.5 | 5077106.5 | 4645017.5 | 4066949.75 | 3852430.5 | 3437023.75 | 3217332 | 3553458.25 | 3645640.5 | 1047.5234 | NC[160.0307]IDITGVR | Q9DBH5 | Q9DBH5 | LMAN2 | lectin, mannose-binding 2 | Cytoplasm | transporter |
| 15532717 | 2 | 0.688 | 4389932.5 | 1.09E+07 | 1.86E+07 | 1061291.125 | 1.40E+07 | 1.51E+07 | 3230655 | 1.34E+07 | 1.46E+07 | 560108.75 | 6680425 | 8661214 | 706.3787 | LLMDSK | Q8VEE1 | Q8VEE1 | LMCD1 | LIM and cysteine-rich domains 1 | Cytoplasm | transcription regulator |
| 15549861 | 3 | 0.335 | 4340539 | 9693675 | 3017449.25 | 5868709 | 4848781 | 7360516.5 | 4289565 | 6245943.5 | 8953586 | 3862734.25 | 3448868.25 | 3598735 | 3684.8252 | VVDNPIYLSDMGAALTGAESHELQDVLEETNILK | Q8CGK3 | Q8CGK3 | LONP1 | lon peptidase 1, mitochondrial | Cytoplasm | peptidase |
| 15551622 | 3 | 0.938 | 4366931.5 | 4711493 | 188288.219 | 4497764.5 | 1187963.125 | 2223371.75 | 1287190 | 2984344.5 | 5250418.5 | 1009053 | 3560226.5 | 2434974.25 | 3187.6543 | IVSGEAQTVQVTPENLQDFVGKPVFTVER | Q8CGK3 | Q8CGK3 | LONP1 | lon peptidase 1, mitochondrial | Cytoplasm | peptidase |
| 15548803 | 2, 3 | 0.541 | 1.60E+07 | 2.75E+07 | 2.82E+07 | 1.93E+07 | 2.29E+07 | 2.31E+07 | 2.23E+07 | 2.55E+07 | 2.80E+07 | 2.31E+07 | 1.58E+07 | 2.11E+07 | 2393.2036, 2393.2039 | TALGVSELTVTDLFQAIDQEDK | Q3TFD2 | Q3TFD2 | LPCAT1 | lysophosphatidylcholine acyltransferase 1 | Cytoplasm | enzyme |
| 15554952 | 2 | 0.403 | 1.04E+07 | 1.03E+07 | 8890107 | 8554295 | 9886663 | 8471776 | 1.23E+07 | 1.02E+07 | 1.12E+07 | 1.30E+07 | 8596600 | 7872716.5 | 2198.9902 | MYGSPEDGSIDEANLSC[160.0307]ILK | Q3TFD2 | Q3TFD2 | LPCAT1 | lysophosphatidylcholine acyltransferase 1 | Cytoplasm | enzyme |
| 15539712 | 2 | 0.154 | 9437665 | 9643567 | 1.01E+07 | 5959752.5 | 9401975 | 8633717 | 1.15E+07 | 9747883 | 1.04E+07 | 1.11E+07 | 9335809 | 8745861 | 1405.7125 | LPADTC[160.0307]LLEFAR | Q3TFD2 | Q3TFD2 | LPCAT1 | lysophosphatidylcholine acyltransferase 1 | Cytoplasm | enzyme |
| 15549448 | 3 | 0.549 | 6403621 | 9968736 | 5757356.5 | 5790161.5 | 7535722.5 | 6213438.5 | 8037104 | 1.01E+07 | 7442412 | 1.01E+07 | 6328705.5 | 6614631.5 | 3634.8494 | QALPTEAAILTLAPHSSYFDAIPVTMTMSSIVMK | Q3TFD2 | Q3TFD2 | LPCAT1 | lysophosphatidylcholine acyltransferase 1 | Cytoplasm | enzyme |
| 15549030 | 2 | 0.824 | 7625111.5 | 7939731.5 | 7130119.5 | 7342189.5 | 9302874 | 6477676.5 | 7234969 | 8846236 | 8944519 | 9521133 | 8401479 | 6638948.5 | 1686.8784 | LDTITWTWQGPGALK | Q3TFD2 | Q3TFD2 | LPCAT1 | lysophosphatidylcholine acyltransferase 1 | Cytoplasm | enzyme |
| 15550972 | 3 | 0.198 | 8891201 | 3685450.25 | 1266621 | 6390339.5 | 2736266.5 | 3305530.5 | 3150627.75 | 3353836.75 | 7943216.5 | 9062505 | 1.23E+07 | 9180725 | 2509.4312 | TC[160.0307]LITFKPGAFIPGVPVQPVVLR | Q3TFD2 | Q3TFD2 | LPCAT1 | lysophosphatidylcholine acyltransferase 1 | Cytoplasm | enzyme |
| 15533246 | 2 | 0.382 | 7906547 | 8974700 | 8592086 | 6851192.5 | 8073701 | 8183174.5 | 8668658 | 1.03E+07 | 7696258.5 | 8550265 | 8997225 | 8174852.5 | 1183.681 | DIPIWGTLIR | Q3TFD2 | Q3TFD2 | LPCAT1 | lysophosphatidylcholine acyltransferase 1 | Cytoplasm | enzyme |
| 15534122 | 2 | 0.823 | 4215141 | 4009068.5 | 4268967.5 | 2817473.75 | 4659177.5 | 3553338 | 4430930.5 | 3933595.75 | 3359277 | 5035740.5 | 3357448.25 | 3218054 | 1260.678 | RNPALYASNVR | Q3TFD2 | Q3TFD2 | LPCAT1 | lysophosphatidylcholine acyltransferase 1 | Cytoplasm | enzyme |
| 15535334 | 2 | 0.694 | 1.97E+07 | 2.38E+07 | 2.39E+07 | 2.44E+07 | 2.69E+07 | 2.16E+07 | 2.48E+07 | 2.33E+07 | 1.78E+07 | 2.53E+07 | 1.80E+07 | 2.06E+07 | 1427.7502 | GLWPGVEDLSLNK | Q91V01 | Q91V01 | LPCAT3 | lysophosphatidylcholine acyltransferase 3 | Plasma Membrane | enzyme |
| 15531938 | 2 | 0.471 | 7808743.5 | 9594552 | 1.11E+07 | 1.02E+07 | 1.21E+07 | 1.01E+07 | 1.04E+07 | 1.09E+07 | 9295315 | 9989870 | 9402244 | 9123771 | 928.50757 | GQLTDIPGK | Q91V01 | Q91V01 | LPCAT3 | lysophosphatidylcholine acyltransferase 3 | Plasma Membrane | enzyme |
| 15541515 | 2 | 0.743 | 5768819.5 | 7952907.5 | 9818758 | 7096677.5 | 9915513 | 8900116 | 9041270 | 8319520 | 8289180.5 | 9553547 | 7788041.5 | 9457908 | 1148.6078 | VWLFETTPR | Q91V01 | Q91V01 | LPCAT3 | lysophosphatidylcholine acyltransferase 3 | Plasma Membrane | enzyme |
| 15536865 | 2 | 0.711 | 1231090 | 1844575.5 | 2505617.5 | 1710440.5 | 2938819 | 2067479.875 | 2065881 | 2284708.25 | 1545023.875 | 2088286.25 | 992205.438 | 2019294.375 | 1316.7162 | LATSLGASEQALR | Q91V01 | Q91V01 | LPCAT3 | lysophosphatidylcholine acyltransferase 3 | Plasma Membrane | enzyme |
| 15542312 | 3 | 0.462 | 1665522.75 | 2313918.25 | 2039990.75 | 1043917.75 | 2521343.75 | 1363890.125 | 1934062.25 | 2214844.5 | 1477991.25 | 1737223.625 | 1299094.125 | 1301660.625 | 1823.8142 | VSHQGYGSTTEFEEPR | P55302 | P55302 | LRPAP1 | low density lipoprotein receptor-related protein associated protein 1 | Plasma Membrane | transmembrane receptor |
| 15536163 | 2 | 0.385 | 6896501 | 7849598 | 4636027.5 | 8193463 | 6768324 | 5276380 | 6989492 | 7078088.5 | 4466916 | 9909821 | 6740365 | 9036498 | 1414.7584 | LTVLPDELC[160.0307]NLK | Q9D1G5 | Q9D1G5 | LRRC57 | leucine rich repeat containing 57 | Cytoplasm | other |
| 15533480 | 2 | 0.071 | 6005385 | 7980391 | 6540888.5 | 7924278.5 | 6477896 | 5859708.5 | 6019868.5 | 5251112.5 | 4540269.5 | 9553697 | 6297193.5 | 9217553 | 1390.7543 | ELPSTFGQLSALK | Q9D1G5 | Q9D1G5 | LRRC57 | leucine rich repeat containing 57 | Cytoplasm | other |
| 15540111 | 2 | 0.031 | 8367522.5 | 9236391 | 7666153.5 | 7297156 | 6791524.5 | 5766736 | 6483340.5 | 5842660.5 | 4924442.5 | 8926421 | 6606148 | 9879549 | 1165.7167 | IDSLPPLIIGK | Q9D1G5 | Q9D1G5 | LRRC57 | leucine rich repeat containing 57 | Cytoplasm | other |
| 15536122 | 3 | 0.095 | 6636022.5 | 5321582 | 3282519.5 | 7226304 | 5326139 | 4093734.75 | 3993432.25 | 4558380 | 2906329.25 | 6695021.5 | 6569384 | 8195660 | 1551.8564 | KLETLSLNNNHLR | Q9D1G5 | Q9D1G5 | LRRC57 | leucine rich repeat containing 57 | Cytoplasm | other |
| 15533508 | 2 | 0.027 | 5396433 | 4445946.5 | 4431195 | 5431073 | 5291693 | 3391331.75 | 3712443 | 3784017.25 | 3248242.75 | 6160721 | 5329329 | 6403367 | 1025.5604 | HLDVVDLSK | Q9D1G5 | Q9D1G5 | LRRC57 | leucine rich repeat containing 57 | Cytoplasm | other |
| 15533569 | 2 | 0.22 | 2847727 | 2849858 | 1873338 | 3289186.5 | 2521898.75 | 1570399.125 | 2165470.5 | 1979526.75 | 1638205.75 | 3004086.5 | 2693110.5 | 3451632.75 | 792.45966 | TGVFQLK | Q9D1G5 | Q9D1G5 | LRRC57 | leucine rich repeat containing 57 | Cytoplasm | other |
| 15550747 | 2 | 0.091 | 706299.812 | 2601494 | 1991573.875 | 3773615.75 | 5418593 | 4838914.5 | 2651198.25 | 3773797.5 | 3852344.25 | 671108.688 | 2533629.25 | 2594695.75 | 2070.055 | LDGNELDLSLSDLNEVPVK | Q922Q8 | Q922Q8 | LRRC59 | leucine rich repeat containing 59 | Cytoplasm | other |
| 15538306 | 2 | 0.048 | 1329179.875 | 1813929.625 | 1610768 | 3003918.75 | 2445968.75 | 2602715 | 1679054.5 | 2219717 | 2690160.25 | 887613.188 | 1887676.5 | 1641350.375 | 930.4986 | VLNLSDNR | Q80TE7 | Q80TE7 | LRRC7 | leucine rich repeat containing 7 | Plasma Membrane | other |
| 15541035 | 3 | 0.421 | 489141.25 | 227060.953 | 21132.502 | 941995.625 | 478881.031 | 1965274.25 | 729337 | 106522.203 | 8370732 | 151456.359 | 2855186.25 | 1812229.625 | 1854.8187 | DRFPNLC[160.0307]SSC[160.0307]AGTGANK | P08071 | P08071 | LTF | lactotransferrin | Extracellular Space | other |
| 15540552 | 2 | 0.056 | 318948.469 | 4.70E-06 | 4.70E-06 | 616683.5 | 238704.547 | 1241443.5 | 512833.781 | 470072.031 | 3785402.75 | 77825.352 | 731560.312 | 876370.125 | 1303.735 | LRPVAAEVYGTK | P08071 | P08071 | LTF | lactotransferrin | Extracellular Space | other |
| 15545411 | 2 | 0.618 | 516097.094 | 274176.719 | 273987.812 | 1005929.125 | 456228.156 | 1831395.5 | 808559.125 | 368222.469 | 3474877.5 | 71204.836 | 1496353.75 | 1566208.125 | 1327.701 | C[160.0307]GLVPVLAENQK | P08071 | P08071 | LTF | lactotransferrin | Extracellular Space | other |
| 15560138 | 2 | 0.161 | 2557.816 | 38530.703 | 160673.422 | 610742.562 | 160087.125 | 801214.812 | 129202.398 | 215484.469 | 2904648.5 | 66585.469 | 860765.062 | 730541.438 | 1801.7812 | FNEFFSQSC[160.0307]APGADPK | P08071 | P08071 | LTF | lactotransferrin | Extracellular Space | other |
| 15553124 | 2 | 0.311 | 101116.234 | 6379.625 | 321024.062 | 1371890.5 | 406033.688 | 2238528.75 | 711110.875 | 689814.375 | 5902249.5 | 1332.102 | 1934571.625 | 1444696.125 | 1814.7982 | C[160.0307]ASSPEEPYSGYAGALR | P08071 | P08071 | LTF | lactotransferrin | Extracellular Space | other |
| 15551905 | 3 | 0.337 | 3350109.25 | 6303758 | 9634377 | 1.04E+07 | 1.68E+07 | 1.43E+07 | 6315190.5 | 1.26E+07 | 1.74E+07 | 1976459.75 | 1.22E+07 | 9887634 | 2932.3826 | DPEEQGDIVVALYPYDGIHPDDLSFK | P25911 | P25911 | LYN | LYN proto-oncogene, Src family tyrosine kinase | Cytoplasm | kinase |
| 15553085 | 2 | 0.235 | 3204457.25 | 3530226 | 5699485.5 | 8230163 | 8876759 | 8432579 | 4342329 | 6812659 | 1.01E+07 | 1965834.125 | 7835318.5 | 6184925.5 | 1527.86 | QLLAPGNSAGAFLIR | P25911 | P25911 | LYN | LYN proto-oncogene, Src family tyrosine kinase | Cytoplasm | kinase |
| 15549709 | 3 | 0.075 | 2367242 | 4108100.75 | 4518460.5 | 8933160 | 1.14E+07 | 1.22E+07 | 5387519 | 9004227 | 1.22E+07 | 1944894.75 | 8686469 | 6184610 | 3439.552 | AEERPTFDYLQSVLDDFYTATEGQYQQQP | P25911 | P25911 | LYN | LYN proto-oncogene, Src family tyrosine kinase | Cytoplasm | kinase |
| 15549596 | 2 | 0.066 | 1920265.375 | 3007844.5 | 4058839.75 | 8510766 | 7415217 | 6282882.5 | 4189391.5 | 5451408.5 | 8790581 | 1756887.5 | 5273521 | 4423087.5 | 1542.7452 | VNTLETEEWFFK | P25911 | P25911 | LYN | LYN proto-oncogene, Src family tyrosine kinase | Cytoplasm | kinase |
| 15559252 | 2 | 0.247 | 682277.188 | 1468865.5 | 2004433.75 | 3380017.25 | 3447566.5 | 3684968.5 | 1106129.125 | 2227753.25 | 3375132.75 | 519725.719 | 3161360.75 | 1805105.125 | 1512.7079 | TNADVMSALSQGYR | P25911 | P25911 | LYN | LYN proto-oncogene, Src family tyrosine kinase | Cytoplasm | kinase |
| 15552532 | 2 | 0.1 | 565855.875 | 668108.688 | 872929.125 | 2058268.125 | 3401465 | 2913506.75 | 2014827 | 2421297.75 | 3230390 | 322615.062 | 3005571.25 | 2402921.75 | 1865.9122 | PGTMSVQAFLEEANLMK | P25911 | P25911 | LYN | LYN proto-oncogene, Src family tyrosine kinase | Cytoplasm | kinase |
| 15542027 | 2 | 0.153 | 491019.875 | 867514.562 | 876270.75 | 2377713 | 3158020 | 3419746.5 | 1481487.875 | 2122151.25 | 3171273 | 191064.844 | 2149598 | 2243291 | 892.5119 | GSLLDFLK | P25911 | P25911 | LYN | LYN proto-oncogene, Src family tyrosine kinase | Cytoplasm | kinase |
| 15546916 | 2 | 0.234 | 154586.75 | 255160.109 | 633461.5 | 1216934.875 | 1145463.375 | 740916.75 | 763689.062 | 832077.25 | 926753.375 | 79078.414 | 1120194.5 | 600926.375 | 1263.5305 | DNLNDDEVDSK | P25911 | P25911 | LYN | LYN proto-oncogene, Src family tyrosine kinase | Cytoplasm | kinase |
| 15551429 | 3 | 0.268 | 2356275.5 | 1871387.875 | 6747353 | 1.46E+07 | 7035654.5 | 1.80E+07 | 1.06E+07 | 7285166.5 | 1.69E+07 | 1328900.125 | 1.85E+07 | 1.35E+07 | 2420.1162 | ATNYNRGDQSTDYGIFQINSR | P08905 | P08905 | LYZ | lysozyme | Extracellular Space | enzyme |
| 15555860 | 3 | 0.423 | 535075.25 | 605234.5 | 2156059.5 | 3073346.25 | 5086126 | 3646810 | 3601360.5 | 4485543.5 | 7162192 | 173001.969 | 1.80E+07 | 5012304 | 2677.2673 | AVNAC[160.0307]GINC[160.0307]SALLQDDITAAIQC[160.0307]AK | P08905 | P08905 | LYZ | lysozyme | Extracellular Space | enzyme |
| 15554196 | 2 | 0.056 | 4011089.5 | 5964135.5 | 4650637.5 | 4545170.5 | 5544477 | 5441084.5 | 3779345 | 4571931.5 | 6928590.5 | 2288483 | 2747899.75 | 3924339.25 | 2043.9254 | SFESTVGQGSDTYSYIFR | P24668 | P24668 | M6PR | mannose-6-phosphate receptor (cation dependent) | Cytoplasm | transporter |
| 15552917 | 2 | 0.915 | 1478078.625 | 1485018.625 | 973621.875 | 976703.938 | 1209125.75 | 1253245.875 | 665955.562 | 1532303.5 | 2421783.5 | 1000132.688 | 1156313.625 | 1300396.25 | 1584.7734 | HTLAANFNPVSEER | P24668 | P24668 | M6PR | mannose-6-phosphate receptor (cation dependent) | Cytoplasm | transporter |
| 15534939 | 3 | 0.114 | 1.34E+07 | 1.93E+07 | 2.05E+07 | 1.68E+07 | 2.15E+07 | 2.20E+07 | 1.77E+07 | 2.44E+07 | 2.13E+07 | 1.39E+07 | 1.27E+07 | 1.70E+07 | 1962.9974 | DRDVTFSPATIEEELIK | Q9CXI5 | Q9CXI5 | MANF | mesencephalic astrocyte-derived neurotrophic factor | Extracellular Space | other |
| 15537120 | 2 | 0.065 | 3664451.5 | 3575979.75 | 4564381 | 4354286.5 | 5705748.5 | 4865383.5 | 5213981 | 5303466.5 | 5138853.5 | 4092092 | 5051705.5 | 3842605.25 | 992.469 | DSQIC[160.0307]ELK | Q9CXI5 | Q9CXI5 | MANF | mesencephalic astrocyte-derived neurotrophic factor | Extracellular Space | other |
| 15550927 | 2 | 0.635 | 1558112.5 | 2366888.25 | 1930426.5 | 1607872.75 | 3688181 | 2406843 | 1868764.5 | 2562632.25 | 3252118.25 | 2533623.25 | 1711180.5 | 1939971.375 | 1561.7174 | LC[160.0307]YYIGATDDAATK | Q9CXI5 | Q9CXI5 | MANF | mesencephalic astrocyte-derived neurotrophic factor | Extracellular Space | other |
| 15557575 | 2 | 0.002 | 608957.812 | 716326.625 | 864726.25 | 735372.5 | 630731.625 | 1050573.75 | 708648 | 626281.625 | 565409.75 | 1583700.75 | 1780166.25 | 1246714.875 | 1766.9608 | QGQETAVAPSLVAPALSK | Q61166 | Q61166 | MAPRE1 | microtubule-associated protein, RP/EB family, member 1 | Cytoplasm | other |
| 15545618 | 2 | 0.14 | 732617.688 | 1261006.375 | 554903.312 | 994026.312 | 1424500.125 | 1335464.25 | 744978.75 | 1173694.25 | 1750946.125 | 371559.312 | 968976.625 | 515078.031 | 1087.6075 | QLQQVGTVSK | Q922Q1 | Q922Q1 | MARC_2 | mitochondrial amidoxime reducing component 2 | Cytoplasm | enzyme |
| 15549377 | 2 | 0.252 | 5079018 | 5451182 | 5512774 | 3715649.5 | 5521990 | 3971199.75 | 4662395.5 | 6014425.5 | 5157399 | 5322463.5 | 3737036 | 4098138.5 | 1619.7993 | AGSSPPSTTAPVSAFSR | Q9DBD2 | Q9DBD2 | MARH8 | membrane-associated ring finger (C3HC4) 8, E3 ubiquitin protein ligase | Cytoplasm | enzyme |
| 15537527 | 2 | 0.211 | 3283295.25 | 5014844.5 | 5007114.5 | 2823939.75 | 4597027 | 3492806.5 | 4426163.5 | 4645210.5 | 3766823.25 | 3940972.5 | 2123005.75 | 2920609.5 | 1437.701 | VIYVQNC[160.0307]PETSK | Q9DBD2 | Q9DBD2 | MARH8 | membrane-associated ring finger (C3HC4) 8, E3 ubiquitin protein ligase | Cytoplasm | enzyme |
| 15536890 | 3 | 0.458 | 2361924 | 3266565.5 | 2541881 | 2615019.25 | 3461860 | 2023495.75 | 2341315.75 | 2562654 | 2300770.25 | 2226624.75 | 2366204.25 | 2008676.75 | 2196.0364 | DQQNLPYGVTPASPSGHSQGR | Q05512 | Q05512 | MARK2 | MAP/microtubule affinity-regulating kinase 2 | Cytoplasm | kinase |
| 15552303 | 2 | 0.38 | 1408048.5 | 1784461.375 | 1151041.875 | 1271435.75 | 1217969.625 | 457986.781 | 1167903.375 | 1520030.75 | 886318.125 | 977098.812 | 1023682.875 | 855627.875 | 1733.8635 | TTPAPSTNSVLSTSTNR | Q05512 | Q05512 | MARK2 | MAP/microtubule affinity-regulating kinase 2 | Cytoplasm | kinase |
| 15549508 | 4 | 0.256 | 3289000 | 8282963 | 5300469 | 5891561.5 | 7108618 | 8467756 | 5077433.5 | 9077973 | 1.17E+07 | 2337254.75 | 5664388.5 | 5340916.5 | 3307.68 | SLEPLQLPDFPAAVSGPSAFAPGHNIPDHGLR | Q8VCF0 | Q8VCF0 | MAVS | mitochondrial antiviral signaling protein | Cytoplasm | other |
| 15538999 | 2 | 0.133 | 1369146.25 | 2644908.75 | 1712443.625 | 3907422.25 | 3303639.25 | 4105966.25 | 2295766.25 | 3143055.75 | 3599267.5 | 918494.188 | 3267819.25 | 2176320 | 1488.7373 | VTTGGNQTGPNSSIR | Q8VCF0 | Q8VCF0 | MAVS | mitochondrial antiviral signaling protein | Cytoplasm | other |

| | | | | | | | | | | | | | | | | | | | |
|---|---|---|---|---|---|---|---|---|---|---|---|---|---|---|---|---|---|---|---|
| 15535149 | 2 | 0.038 | 1.66E+07 | 1.43E+07 | 1.70E+07 | 1.38E+07 | 1.25E+07 | 1.21E+07 | 1.33E+07 | 1.31E+07 | 1.51E+07 | 1.74E+07 | 1.60E+07 | 1.42E+07 | 871.51306 | ANIWLVR | Q8BL86 | Q8BL86 | MBLAC2 | metallo-beta-lactamase domain containing 2 | Cytoplasm | other |
| 15553288 | 3 | 0.157 | 194161.656 | 1458971.25 | 2708667 | 4253361.5 | 6698066 | 7020343 | 3490547 | 4961503.5 | 9374158 | 314667.656 | 4398651 | 3528132.5 | 2640.413 | TYLDWLEQPFPEAVPSLRPLLR | Q8CHK3 | Q8CHK3 | MBOAT7 | membrane bound O-acyltransferase domain containing 7 | Plasma Membrane | other |
| 15549943 | 3 | 0.361 | 6660400 | 1.06E+07 | 9205273 | 9107340 | 1.18E+07 | 1.08E+07 | 1.17E+07 | 1.16E+07 | 9367337 | 1.07E+07 | 7329500 | 9023339 | 3075.4233 | EEGSEQAPVMSEDELVSIIDGVLRDDDK | Q8K5B2 | Q8K5B2 | MCFD2 | multiple coagulation factor deficiency 2 | Cytoplasm | other |
| 15536674 | 2 | 0.166 | 166134.156 | 747509 | 1050257.625 | 1706888.25 | 1915372.75 | 2477308.75 | 1168810 | 1657710.875 | 2274434 | 409436.875 | 2639371.25 | 1410931.75 | 1076.5718 | VAIYSPDGVR | Q3UMR5 | Q3UMR5 | MCU | mitochondrial calcium uniporter | Cytoplasm | ion channel |
| 15547882 | 2 | 0.131 | 3.40E+07 | 5.89E+07 | 4.53E+07 | 5.27E+07 | 5.05E+07 | 7.34E+07 | 3.41E+07 | 6.57E+07 | 7.75E+07 | 2.35E+07 | 3.11E+07 | 4.13E+07 | 1793.0862 | VAVLGASGGIGQPLSLLLK | P08249 | P08249 | MDH2 | malate dehydrogenase 2, NAD (mitochondrial) | Cytoplasm | enzyme |
| 15539471 | 2 | 0.191 | 2.89E+07 | 4.69E+07 | 2.42E+07 | 4.16E+07 | 4.46E+07 | 5.01E+07 | 2.78E+07 | 4.06E+07 | 5.17E+07 | 1.80E+07 | 3.32E+07 | 3.04E+07 | 1489.7334 | GYLGPEQLPDC[160.0307]LK | P08249 | P08249 | MDH2 | malate dehydrogenase 2, NAD (mitochondrial) | Cytoplasm | enzyme |
| 15557038, 15559722, 15560274 | 2 | 0.132 | 2.66E+07 | 5.59E+07 | 3.41E+07 | 4.10E+07 | 4.44E+07 | 5.51E+07 | 2.50E+07 | 4.22E+07 | 6.30E+07 | 1.78E+07 | 2.82E+07 | 2.47E+07 | 1560.7977, 1560.7986, 1560.7999 | VDFPQDQLATLTGR | P08249 | P08249 | MDH2 | malate dehydrogenase 2, NAD (mitochondrial) | Cytoplasm | enzyme |
| 15533370 | 2 | 0.079 | 2.41E+07 | 4.74E+07 | 2.34E+07 | 4.18E+07 | 3.82E+07 | 4.70E+07 | 2.69E+07 | 3.98E+07 | 4.64E+07 | 1.59E+07 | 2.57E+07 | 2.40E+07 | 1073.5819 | IQEAGTEVVK | P08249 | P08249 | MDH2 | malate dehydrogenase 2, NAD (mitochondrial) | Cytoplasm | enzyme |
| 15531376 | 2 | 0.214 | 2.61E+07 | 5.97E+07 | 2.53E+07 | 3.94E+07 | 3.82E+07 | 4.68E+07 | 2.44E+07 | 4.39E+07 | 5.67E+07 | 1.46E+07 | 2.57E+07 | 2.71E+07 | 1233.7179 | IFGVTTLDIVR | P08249 | P08249 | MDH2 | malate dehydrogenase 2, NAD (mitochondrial) | Cytoplasm | enzyme |
| 15531448 | 2 | 0.497 | 2.86E+07 | 4.96E+07 | 2.01E+07 | 3.61E+07 | 3.88E+07 | 3.57E+07 | 2.55E+07 | 3.72E+07 | 4.68E+07 | 1.28E+07 | 3.51E+07 | 2.85E+07 | 1370.7694 | TIIPLISQC[160.0307]TPK | P08249 | P08249 | MDH2 | malate dehydrogenase 2, NAD (mitochondrial) | Cytoplasm | enzyme |
| 15547724 | 3 | 0.22 | 2.40E+07 | 4.58E+07 | 2.32E+07 | 3.35E+07 | 3.01E+07 | 4.27E+07 | 1.84E+07 | 3.37E+07 | 5.12E+07 | 1.20E+07 | 2.24E+07 | 2.19E+07 | 2393.2444 | LTLYDIAHTPGVAADLSHIETR | P08249 | P08249 | MDH2 | malate dehydrogenase 2, NAD (mitochondrial) | Cytoplasm | enzyme |
| 15531394 | 2 | 0.177 | 1.99E+07 | 3.70E+07 | 1.81E+07 | 3.62E+07 | 3.00E+07 | 4.24E+07 | 2.25E+07 | 3.53E+07 | 3.90E+07 | 1.15E+07 | 2.57E+07 | 2.33E+07 | 1338.7168 | GC[160.0307]DVVVIPAGVPR | P08249 | P08249 | MDH2 | malate dehydrogenase 2, NAD (mitochondrial) | Cytoplasm | enzyme |
| 15540283, 15543560 | 2 | 0.207 | 1.72E+07 | 3.61E+07 | 1.56E+07 | 2.42E+07 | 2.88E+07 | 3.31E+07 | 1.54E+07 | 2.93E+07 | 3.66E+07 | 1.02E+07 | 1.74E+07 | 1.76E+07 | 1114.6145, 1114.6147 | MIAEAIPELK | P08249 | P08249 | MDH2 | malate dehydrogenase 2, NAD (mitochondrial) | Cytoplasm | enzyme |
| 15538379 | 2 | 0.476 | 1.35E+07 | 1.72E+07 | 9190119 | 1.61E+07 | 1.33E+07 | 1.70E+07 | 9339550 | 1.37E+07 | 1.71E+07 | 8108692.5 | 1.40E+07 | 1.09E+07 | 1454.7046 | AGAGSATLSMAYAGAR | P08249 | P08249 | MDH2 | malate dehydrogenase 2, NAD (mitochondrial) | Cytoplasm | enzyme |
| 15532558 | 2 | 0.558 | 1.34E+07 | 2.19E+07 | 8003116 | 2.01E+07 | 1.79E+07 | 1.93E+07 | 1.15E+07 | 1.77E+07 | 3.05E+07 | 7635212 | 1.95E+07 | 1.39E+07 | 1368.6437 | EGVVEC[160.0307]SFVQSK | P08249 | P08249 | MDH2 | malate dehydrogenase 2, NAD (mitochondrial) | Cytoplasm | enzyme |
| 15534711 | 2 | 0.197 | 7010744 | 1.42E+07 | 6609906 | 1.01E+07 | 1.10E+07 | 1.37E+07 | 6165690.5 | 1.07E+07 | 1.25E+07 | 3706547.75 | 8343783 | 6187561 | 1147.6556 | VNVPVIGGHAGK | P08249 | P08249 | MDH2 | malate dehydrogenase 2, NAD (mitochondrial) | Cytoplasm | enzyme |
| 15541004 | 3 | 0.219 | 2281296.25 | 263581.156 | 2721250.5 | 3378704.25 | 3642226.75 | 6197413.5 | 1916325.375 | 4457966 | 7083017 | 1637529.75 | 2331577.75 | 2826691 | 1793.0864 | VAVLGASGGIGQPLSLLLK | P08249 | P08249 | MDH2 | malate dehydrogenase 2, NAD (mitochondrial) | Cytoplasm | enzyme |
| 15555161 | 3 | 0.41 | 2629813.75 | 7438163.5 | 4058597.75 | 4279390 | 5969635.5 | 6039562 | 4536516.5 | 6151696 | 5411322.5 | 3549060.25 | 4043495.75 | 3662393.75 | 3811.8162 | AMASINERPIIFALSNPTAQAEC[160.0307]TAEDAYTLTEGR | Q99KE1 | Q99KE1 | ME2 | malic enzyme 2, NAD(+)-dependent, mitochondrial | Cytoplasm | enzyme |
| 15550089 | 3 | 0.105 | 4638517 | 1.21E+07 | 5274701 | 7200939 | 8675442 | 1.08E+07 | 4647254.5 | 9194504 | 1.10E+07 | 3485035.5 | 3988336 | 4333686.5 | 3073.4302 | TASIDSNQEPYAHAAPESIPATFEDAVNK | Q99KE1 | Q99KE1 | ME2 | malic enzyme 2, NAD(+)-dependent, mitochondrial | Cytoplasm | enzyme |
| 15535218 | 2 | 0.425 | 1675235.875 | 3961230.5 | 1465413.625 | 1995517.125 | 2064055.625 | 2538166.75 | 1315572.5 | 2528319.5 | 2946824.75 | 1008154.5 | 1829072.75 | 1409065.875 | 1186.6401 | IETQDIQALR | Q99KE1 | Q99KE1 | ME2 | malic enzyme 2, NAD(+)-dependent, mitochondrial | Cytoplasm | enzyme |
| 15550097 | 2 | 0.198 | 1419734.5 | 3636604.5 | 1377841.625 | 3116802.75 | 2453854 | 3525685 | 1867745.375 | 3069597.5 | 3669195.75 | 962973.188 | 2024962.5 | 1783609.125 | 1587.8308 | ALTTQLTDAELAQGR | Q99KE1 | Q99KE1 | ME2 | malic enzyme 2, NAD(+)-dependent, mitochondrial | Cytoplasm | enzyme |
| [illegible] | | | | | | | | | | | | | | | | | | | | | | |

| | | | | | | | | | | | | | | | | | | | | | |
|---|---|---|---|---|---|---|---|---|---|---|---|---|---|---|---|---|---|---|---|---|---|
| 15551184 | 2 | 0.115 | 2175524.5 | 5702967 | 9728290 | 2.76E+07 | 2.07E+07 | 1.86E+07 | 1.18E+07 | 1.43E+07 | 2.86E+07 | 3200391.25 | 2.12E+07 | 1.69E+07 | 1601.7997 | IMGIPEDEQMGLLR | Q8VDD5 | Q8VDD5 | MYH9 | myosin, heavy chain 9, non-muscle | Cytoplasm | enzyme |
| 15532170 | 2 | 0.098 | 2272418.25 | 5313131.5 | 9715121 | 1.93E+07 | 2.55E+07 | 1.86E+07 | 1.39E+07 | 1.60E+07 | 2.79E+07 | 3141915.75 | 1.92E+07 | 1.88E+07 | 908.4861 | FVSELWK | Q8VDD5 | Q8VDD5 | MYH9 | myosin, heavy chain 9, non-muscle | Cytoplasm | enzyme |
| 15537213 | 3 | 0.209 | 2316822.5 | 7169490 | 1.10E+07 | 1.46E+07 | 1.54E+07 | 2.45E+07 | 1.44E+07 | 1.90E+07 | 3.76E+07 | 3131051.5 | 2.53E+07 | 1.81E+07 | 2110.064 | PAGPPGILALLDEEC[160.0307]WFPK | Q8VDD5 | Q8VDD5 | MYH9 | myosin, heavy chain 9, non-muscle | Cytoplasm | enzyme |
| 15548801 | 2 | 0.14 | 2692631.25 | 6615076 | 1.03E+07 | 2.20E+07 | 2.50E+07 | 1.96E+07 | 1.20E+07 | 1.44E+07 | 2.27E+07 | 2983621 | 1.84E+07 | 1.88E+07 | 2048.9583 | SMEAEMIQLQEELAAAER | Q8VDD5 | Q8VDD5 | MYH9 | myosin, heavy chain 9, non-muscle | Cytoplasm | enzyme |
| 15533468 | 3 | 0.038 | 2185387.5 | 4313655 | 2942780.5 | 1.25E+07 | 1.31E+07 | 9765139 | 7607650 | 8714001 | 1.39E+07 | 2954444.5 | 1.29E+07 | 1.25E+07 | 1961.9146 | TQLEELEDELQATEDAK | Q8VDD5 | Q8VDD5 | MYH9 | myosin, heavy chain 9, non-muscle | Cytoplasm | enzyme |
| 15548103 | 3 | 0.068 | 1997778.375 | 5227434.5 | 8609235 | 1.86E+07 | 2.96E+07 | 2.02E+07 | 1.26E+07 | 2.19E+07 | 2.40E+07 | 2851028.5 | 1.65E+07 | 1.80E+07 | 2467.2034 | LQQLFNHTMFILEQEEYQR | Q8VDD5 | Q8VDD5 | MYH9 | myosin, heavy chain 9, non-muscle | Cytoplasm | enzyme |
| 15533687 | 2 | 0.122 | 1713517.25 | 4478197 | 8161321 | 2.02E+07 | 1.65E+07 | 1.48E+07 | 9610372 | 1.08E+07 | 1.98E+07 | 2764340.5 | 1.84E+07 | 1.29E+07 | 873.50244 | ASIAALEAK | Q8VDD5 | Q8VDD5 | MYH9 | myosin, heavy chain 9, non-muscle | Cytoplasm | enzyme |
| 15532629 | 2 | 0.086 | 1750085.625 | 4379853.5 | 6561096.5 | 1.88E+07 | 1.60E+07 | 1.56E+07 | 1.12E+07 | 1.18E+07 | 1.98E+07 | 2761913.5 | 1.84E+07 | 1.69E+07 | 1388.5615 | DMFQETMEAMR | Q8VDD5 | Q8VDD5 | MYH9 | myosin, heavy chain 9, non-muscle | Cytoplasm | enzyme |
| 15533289 | 3 | 0.096 | 2008540.5 | 5768676 | 6876461.5 | 1.88E+07 | 1.88E+07 | 1.73E+07 | 8662743 | 1.19E+07 | 1.59E+07 | 2745100.25 | 1.52E+07 | 1.46E+07 | 2048.9592 | SMEAEMIQLQEELAAAER | Q8VDD5 | Q8VDD5 | MYH9 | myosin, heavy chain 9, non-muscle | Cytoplasm | enzyme |
| 15541448 | 2 | 0.071 | 1780933.875 | 4068693 | 6899074.5 | 1.78E+07 | 1.71E+07 | 1.49E+07 | 9708811 | 1.11E+07 | 1.82E+07 | 2542464.75 | 1.47E+07 | 1.20E+07 | 1048.4952 | ALEEAMEQK | Q8VDD5 | Q8VDD5 | MYH9 | myosin, heavy chain 9, non-muscle | Cytoplasm | enzyme |
| 15533493 | 2 | 0.053 | 2167002.5 | 3799920.5 | 6586738 | 2.12E+07 | 1.60E+07 | 1.56E+07 | 1.09E+07 | 1.30E+07 | 1.81E+07 | 2532394.25 | 1.50E+07 | 1.33E+07 | 1034.5167 | LQEMESAVK | Q8VDD5 | Q8VDD5 | MYH9 | myosin, heavy chain 9, non-muscle | Cytoplasm | enzyme |
| 15539493 | 2 | 0.097 | 3312822.5 | 3641658.25 | 3864589 | 1.97E+07 | 1.53E+07 | 1.18E+07 | 8104427.5 | 1.17E+07 | 2.48E+07 | 2388483.75 | 2.06E+07 | 1.38E+07 | 1116.5758 | DLGEELEALK | Q8VDD5 | Q8VDD5 | MYH9 | myosin, heavy chain 9, non-muscle | Cytoplasm | enzyme |
| 15533557 | 2 | 0.1 | 1473940.125 | 4065768 | 9601703 | 1.73E+07 | 2.17E+07 | 2.13E+07 | 1.05E+07 | 1.53E+07 | 2.73E+07 | 2319219.75 | 1.69E+07 | 1.32E+07 | 924.492 | VVFQEFR | Q8VDD5 | Q8VDD5 | MYH9 | myosin, heavy chain 9, non-muscle | Cytoplasm | enzyme |
| 15538608 | 2 | 0.117 | 1990230 | 4621327 | 7380733.5 | 1.40E+07 | 1.76E+07 | 1.48E+07 | 8585822 | 1.22E+07 | 1.81E+07 | 2095247.75 | 1.28E+07 | 1.13E+07 | 1284.713 | VEAQLQELQVK | Q8VDD5 | Q8VDD5 | MYH9 | myosin, heavy chain 9, non-muscle | Cytoplasm | enzyme |
| 15533605 | 2 | 0.094 | 1795948.75 | 4333128 | 7090616.5 | 1.59E+07 | 1.85E+07 | 1.56E+07 | 1.18E+07 | 1.37E+07 | 1.98E+07 | 2021389.125 | 1.32E+07 | 1.56E+07 | 1170.499 | TEMEDLMSSK | Q8VDD5 | Q8VDD5 | MYH9 | myosin, heavy chain 9, non-muscle | Cytoplasm | enzyme |
| 15532375 | 2, 3 | 0.188 | 1760186.75 | 4833300 | 7224808 | 1.38E+07 | 1.56E+07 | 1.36E+07 | 1.08E+07 | 9857990 | 1.85E+07 | 1881755.125 | 1.53E+07 | 1.34E+07 | 1455.7589, 1455.7594 | QIATLHAQVTDMK | Q8VDD5 | Q8VDD5 | MYH9 | myosin, heavy chain 9, non-muscle | Cytoplasm | enzyme |
| 15533588 | 2, 3 | 0.185 | 871392.812 | 4379375 | 1.16E+07 | 7327521.5 | 1.73E+07 | 1.28E+07 | 8234483 | 9831627 | 1.08E+07 | 1869056 | 4538008 | 6058765 | 1272.6765, 1272.6771 | RDLGEELEALK | Q8VDD5 | Q8VDD5 | MYH9 | myosin, heavy chain 9, non-muscle | Cytoplasm | enzyme |
| 15539690 | 3 | 0.112 | 1320182.875 | 3777898.75 | 4167494 | 1.26E+07 | 1.30E+07 | 6278636.5 | 8683462 | 9735773 | 1.35E+07 | 1850678.125 | 1.13E+07 | 1.07E+07 | 1919.9154 | TELEDTLDSTAAQQELR | Q8VDD5 | Q8VDD5 | MYH9 | myosin, heavy chain 9, non-muscle | Cytoplasm | enzyme |
| 15533945 | 3 | 0.149 | 1659160.75 | 3758523.75 | 6126066.5 | 8231788.5 | 1.52E+07 | 1.18E+07 | 6540966.5 | 8167992.5 | 1.69E+07 | 1775117.75 | 8366149.5 | 9242742 | 2018.0613 | IIGLDQVAGMSETALPGAFK | Q8VDD5 | Q8VDD5 | MYH9 | myosin, heavy chain 9, non-muscle | Cytoplasm | enzyme |
| 15539861 | 3 | 0.171 | 1186772.75 | 2117729.25 | 3977934.75 | 1.86E+07 | 6304705.5 | 1.05E+07 | 6011990 | 4342341 | 1.57E+07 | 1683464.75 | 1.74E+07 | 1.28E+07 | 1737.79 | NSFREQLEEEEEAK | Q8VDD5 | Q8VDD5 | MYH9 | myosin, heavy chain 9, non-muscle | Cytoplasm | enzyme |
| 15553362 | 2 | 0.096 | 793302.062 | 1757903.25 | 4065931.25 | 8183245 | 7378963 | 7398652 | 5516631 | 5037093.5 | 1.11E+07 | 1357574.5 | 7976045 | 5933860.5 | 1571.8513 | VSHLLGINVTDFTR | Q8VDD5 | Q8VDD5 | MYH9 | myosin, heavy chain 9, non-muscle | Cytoplasm | enzyme |
| 15549396 | 2 | 0.106 | 1360879.125 | 2403847.75 | 3701640.75 | 1.11E+07 | 1.03E+07 | 7478595.5 | 5878974 | 5917842 | 1.05E+07 | 1349308.875 | 1.09E+07 | 6963812 | 1729.869 | QTLENERGELANEVK | Q8VDD5 | Q8VDD5 | MYH9 | myosin, heavy chain 9, non-muscle | Cytoplasm | enzyme |
| 15542316 | 3 | 0.464 | 677340.688 | 6686043 | 1.11E+07 | 2208347.5 | 2.36E+07 | 2.94E+07 | 1.12E+07 | 2.11E+07 | 4.10E+07 | 1303846.875 | 1.35E+07 | 2.19E+07 | 1579.8406 | AGVLAHLEEERDLK | Q8VDD5 | Q8VDD5 | MYH9 | myosin, heavy chain 9, non-muscle | Cytoplasm | enzyme |
| 15541625 | 2, 3 | 0.093 | 985893.312 | 2661980.5 | 4156887.75 | 9207714 | 1.16E+07 | 1.05E+07 | 6102779.5 | 6964911.5 | 1.20E+07 | 1281842.125 | 8931460 | 7788981.5 | 1242.5863, 1242.5869 | THEAQIQEMR | Q8VDD5 | Q8VDD5 | MYH9 | myosin, heavy chain 9, non-muscle | Cytoplasm | enzyme |
| 15537063 | 2 | 0.083 | 915182.188 | 2194116.5 | 3741125.75 | 9812334 | 1.05E+07 | 7191659.5 | 5424242 | 6575175 | 1.15E+07 | 1238330.5 | 7961607.5 | 6933272.5 | 800.4173 | YLYVDK | Q8VDD5 | Q8VDD5 | MYH9 | myosin, heavy chain 9, non-muscle | Cytoplasm | enzyme |
| 15551713 | 3 | 0.516 | 2286230.75 | 2686128.5 | 2413081.75 | 2.72E+07 | 2676994.5 | 9648104 | 4758198 | 2670405.75 | 1.64E+07 | 1207656.125 | 2.30E+07 | 1.50E+07 | 2503.2815 | DVDRIIGLDQVAGMSETALPGAFK | Q8VDD5 | Q8VDD5 | MYH9 | myosin, heavy chain 9, non-muscle | Cytoplasm | enzyme |
| 15538549 | 3 | 0.152 | 815399.375 | 2873014.5 | 5579632.5 | 1.84E+07 | 1.40E+07 | 1.29E+07 | 9436035 | 8776620 | 1.93E+07 | 1176544.875 | 1.63E+07 | 1.62E+07 | 1640.8347 | FLSNGHVTIPGQQDK | Q8VDD5 | Q8VDD5 | MYH9 | myosin, heavy chain 9, non-muscle | Cytoplasm | enzyme |
| 15550809 | 2 | 0.028 | 292249.75 | 740591.312 | 1439610.125 | 3112438 | 4019248.25 | 2731075.25 | 1888012.25 | 2385295.5 | 3098341.75 | 1137683 | 3617815.5 | 1961665.25 | 1591.6987 | NTDQASMPDNTAAQK | Q8VDD5 | Q8VDD5 | MYH9 | myosin, heavy chain 9, non-muscle | Cytoplasm | enzyme |
| 15534496 | 2 | 0.082 | 1100767 | 2014592.875 | 3632495 | 7049331.5 | 8732372 | 8515675 | 4741399.5 | 5687677 | 8811088 | 1114390.875 | 6195583.5 | 6031827.5 | 1274.6968 | YEILTPNSIPK | Q8VDD5 | Q8VDD5 | MYH9 | myosin, heavy chain 9, non-muscle | Cytoplasm | enzyme |
| 15532501 | 2 | 0.129 | 838005.562 | 2283650.25 | 3455788.25 | 7129626.5 | 9656912 | 7595959 | 4034954.5 | 5964522.5 | 8616733 | 941624.312 | 7400057 | 5522321.5 | 1487.7054 | RQLEEAEEEAQR | Q8VDD5 | Q8VDD5 | MYH9 | myosin, heavy chain 9, non-muscle | Cytoplasm | enzyme |
| 15543228 | 2 | 0.102 | 281912.062 | 2009437.625 | 2920134 | 6326854.5 | 8094853 | 5547324 | 4090465.75 | 4372947.5 | 5953415 | 887042.125 | 4065496.25 | 3876173.5 | 917.4669 | DLEGLSQR | Q8VDD5 | Q8VDD5 | MYH9 | myosin, heavy chain 9, non-muscle | Cytoplasm | enzyme |
| 15538492 | 2 | 0.19 | 402032.406 | 95070.977 | 57194.77 | 4000556.5 | 94775.758 | 175534.359 | 213366.484 | 278320.875 | 1984723.75 | 695129.375 | 5837717.5 | 2200005.25 | 945.5221 | EQEVSILK | Q8VDD5 | Q8VDD5 | MYH9 | myosin, heavy chain 9, non-muscle | Cytoplasm | enzyme |
| 15555764 | 2 | 0.167 | 243851.828 | 1213032.125 | 4413200 | 4446857 | 7763260.5 | 5930843.5 | 3796166.75 | 4291395.5 | 5753776 | 646711 | 3447309.75 | 3893004.25 | 1638.6954 | MEDGVGC[160.0307]LETAEEAK | Q8VDD5 | Q8VDD5 | MYH9 | myosin, heavy chain 9, non-muscle | Cytoplasm | enzyme |
| 15544277 | 2 | 0.121 | 310169.844 | 1439041.625 | 4447438.5 | 6626718.5 | 9527171 | 8898410 | 3437227.25 | 5100595.5 | 1.06E+07 | 637607.188 | 5178305.5 | 3383116 | 1159.656 | RGDLPFVVTR | Q8VDD5 | Q8VDD5 | MYH9 | myosin, heavy chain 9, non-muscle | Cytoplasm | enzyme |
| 15542173 | 2 | 0.106 | 507541.438 | 957602.688 | 1420483.25 | 5689227 | 4328026 | 3738958.75 | 3035640 | 3506827.25 | 7128120 | 581840.938 | 7960722.5 | 4471676 | 1017.5116 | C[160.0307]NGVLEGIR | Q8VDD5 | Q8VDD5 | MYH9 | myosin, heavy chain 9, non-muscle | Cytoplasm | enzyme |
| 15548548 | 2 | 0.24 | 1479350.5 | 2759919.75 | 4037320 | 1.32E+07 | 1.06E+07 | 9254688 | 7152048 | 5264059 | 1.01E+07 | 571154.188 | 1.10E+07 | 7806166.5 | 1825.8888 | HSQAVEELADQLEQTK | Q8VDD5 | Q8VDD5 | MYH9 | myosin, heavy chain 9, non-muscle | Cytoplasm | enzyme |
| 15553955 | 2 | 0.23 | 316331.938 | 1930402.625 | 4754161 | 5622080 | 6595265.5 | 7521353.5 | 3348066.25 | 4054194 | 8843059 | 556226.688 | 3974216 | 3980050.25 | 1743.8615 | EEVGEEAIVELVENGK | Q8VDD5 | Q8VDD5 | MYH9 | myosin, heavy chain 9, non-muscle | Cytoplasm | enzyme |
| 15555678 | 2 | 0.078 | 420925.625 | 1249609.875 | 649478.062 | 5599703.5 | 4575323 | 3269028.5 | 2246217.5 | 2861407 | 4609301 | 405707.406 | 4797896.5 | 3480168.75 | 2126.0146 | NMDPLNDNIATLLHQSSDK | Q8VDD5 | Q8VDD5 | MYH9 | myosin, heavy chain 9, non-muscle | Cytoplasm | enzyme |
| 15539201 | 3 | 0.209 | 257154.547 | 1088480.25 | 2419668 | 3352660.25 | 4014557.5 | 4348597 | 1828691.875 | 3167414 | 6733204.5 | 400142.125 | 2933262 | 3086553.5 | 1918.9952 | LQVELDSVTGLLSQSDSK | Q8VDD5 | Q8VDD5 | MYH9 | myosin, heavy chain 9, non-muscle | Cytoplasm | enzyme |
| 15544624 | 3 | 0.407 | 262904.656 | 2124389 | 5309267 | 667487.062 | 9685231 | 1.09E+07 | 5458346.5 | 6944683 | 1.11E+07 | 388734.969 | 1725131.5 | 6055497 | 1747.9675 | PLLNSIRHEDELLAK | Q8VDD5 | Q8VDD5 | MYH9 | myosin, heavy chain 9, non-muscle | Cytoplasm | enzyme |
| 15555189 | 2 | 0.481 | 708161.125 | 766221.812 | 756738.5 | 6476589 | 1167807.875 | 1972899.5 | 2038162.25 | 834856.938 | 3621453.5 | 344933.656 | 6518992 | 3250505.75 | 1737.7896 | NSFREQLEEEEEAK | Q8VDD5 | Q8VDD5 | MYH9 | myosin, heavy chain 9, non-muscle | Cytoplasm | enzyme |
| 15553018 | 3 | 0.662 | 1140256.75 | 1492112.375 | 1680346 | 1.73E+07 | 1296078.5 | 6820054 | 4765564.5 | 1222276.625 | 1.00E+07 | 334966.781 | 1.32E+07 | 9761697 | 2713.3467 | IAQLEEELEEEQGNTELINDRLK | Q8VDD5 | Q8VDD5 | MYH9 | myosin, heavy chain 9, non-muscle | Cytoplasm | enzyme |
| 15541369 | 2 | 0.223 | 222816.562 | 742173.062 | 2002832.5 | 1963494.5 | 5602017.5 | 4042810.5 | 2134569.25 | 3117708.75 | 2970429.25 | 297012.594 | 3465440.5 | 3175941.75 | 1223.635 | AGVLAHLEEER | Q8VDD5 | Q8VDD5 | MYH9 | myosin, heavy chain 9, non-muscle | Cytoplasm | enzyme |
| 15540416 | 2 | 0.154 | 6678.666 | 488141.844 | 673164.625 | 1692185 | 2318117.25 | 1604262.125 | 837604.562 | 1444776.5 | 1872965.75 | 295611.562 | 1197101.75 | 1622304.75 | 1215.6189 | DVLLQVEDER | Q8VDD5 | Q8VDD5 | MYH9 | myosin, heavy chain 9, non-muscle | Cytoplasm | enzyme |
| 15553844 | 2 | 0.14 | 348288 | 601030.625 | 1350103 | 3884868.25 | 2695579.25 | 2616264.5 | 1602249.5 | 2030806.25 | 3723115.5 | 286954.312 | 2813412.5 | 2283885 | 1640.8362 | FLSNGHVTIPGQQDK | Q8VDD5 | Q8VDD5 | MYH9 | myosin, heavy chain 9, non-muscle | Cytoplasm | enzyme |
| 15556034 | 2 | 0.239 | 526835 | 1289464.75 | 1702035.75 | 4036734.25 | 4433310 | 3963346.5 | 2170616.25 | 2810294.25 | 5568173.5 | 249349.234 | 3852610.75 | 3158736.25 | 1892.8693 | KLEEDQIIMEDQNC[160.0307]K | Q8VDD5 | Q8VDD5 | MYH9 | myosin, heavy chain 9, non-muscle | Cytoplasm | enzyme |
| 15541913 | 2 | 0.204 | 133832.188 | 920522.375 | 2979337 | 1555861 | 6165852 | 3664622.75 | 2279055.5 | 3477110.5 | 3583266 | 205106.547 | 1124753 | 1969238.25 | 812.4974 | PLLNSIR | Q8VDD5 | Q8VDD5 | MYH9 | myosin, heavy chain 9, non-muscle | Cytoplasm | enzyme |
| 15546521 | 2 | 0.108 | 110579.508 | 128009.953 | 300526.875 | 436683.406 | 862731.875 | 303895.594 | 262102.344 | 412152.562 | 317949.156 | 125065.867 | 274603.219 | 368151.812 | 1204.5842 | ALEQQVEEMK | Q8VDD5 | Q8VDD5 | MYH9 | myosin, heavy chain 9, non-muscle | Cytoplasm | enzyme |
| 15560464, 15562332 | 2 | 0.586 | 263508.219 | 424481.656 | 819325.062 | 808535.938 | 1823546 | 897498.938 | 535860.312 | 1183942 | 1997828 | 118993.273 | 1999102.125 | 1215606.25 | 1876.8423, 1876.8425 | ERNTDQASMPDNTAAQK | Q8VDD5 | Q8VDD5 | MYH9 | myosin, heavy chain 9, non-muscle | Cytoplasm | enzyme |
| 15540873 | 2 | 0.68 | 25316.838 | 236787.969 | 1068839.25 | 212778.719 | 2545760 | 2256717.5 | 400817.469 | 1701635.375 | 3329418.75 | 4.70E-06 | 3215270.5 | 2040119.375 | 1010.5054 | C[160.0307]IIPNHEK | Q8VDD5 | Q8VDD5 | MYH9 | myosin, heavy chain 9, non-muscle | Cytoplasm | enzyme |
| 15561270 | 2 | 0.337 | 0 | 2303.207 | 1147373.125 | 23609.678 | 2153095.25 | 1571074.125 | 198168.469 | 718117.938 | 1733155 | 0 | 40528.535 | 460327.344 | 1769.848 | KQELEEIC[160.0307]HDLEAR | Q8VDD5 | Q8VDD5 | MYH9 | myosin, heavy chain 9, non-muscle | Cytoplasm | enzyme |
| 15532881 | 2 | 0.161 | 414561.562 | 998749.5 | 2373897 | 7633071 | 5634120.5 | 6287818 | 3529228.75 | 3908296 | 7750683 | 294347.25 | 6456431.5 | 4620410 | 1233.5875 | EGNGTVMGAELR | P05977 | P05977 | MYL1 | myosin, light chain 1, alkali; skeletal, fast | Cytoplasm | other |
| 15550897 | 3 | 0.032 | 4379364.5 | 8817643 | 1.05E+07 | 1.94E+07 | 2.01E+07 | 1.95E+07 | 1.29E+07 | 1.63E+07 | 2.50E+07 | 7665141 | 1.61E+07 | 1.34E+07 | 2432.1223 | ELLTTMGDRFTDEEVDELYR | Q3THE2 | Q3THE2 | MYL12A | myosin, light chain 12A, regulatory, non-sarcomeric | Cytoplasm | other |
| 15553138 | 2 | 0.088 | 3212932.75 | 7979843 | 1.25E+07 | 3.18E+07 | 2.82E+07 | 3.06E+07 | 1.69E+07 | 2.07E+07 | 3.95E+07 | 4245570.5 | 2.30E+07 | 2.54E+07 | 2090.9812 | ATSNVFAMFDQSQIQEFK | Q3THE2 | Q3THE2 | MYL12A | myosin, light chain 12A, regulatory, non-sarcomeric | Cytoplasm | other |
| 15541391 | 2 | 0.191 | 763388.562 | 2254505.25 | 3382352.25 | 1867484.625 | 9126514 | 3470854.75 | 2944683 | 3972543.25 | 3564815.25 | 1577698.5 | 1055128.625 | 1969754.625 | 1415.6295 | FTDEEVDELYR | Q3THE2 | Q3THE2 | MYL12A | myosin, light chain 12A, regulatory, non-sarcomeric | Cytoplasm | other |
| 15538417 | 2 | 0.083 | 638384 | 1885539.125 | 2859459.75 | 4265107 | 1.02E+07 | 5445707.5 | 3233237.5 | 5649606 | 5307180 | 773469.812 | 3317610.5 | 3509859.5 | 1035.5116 | ELLTTMGDR | Q3THE2 | Q3THE2 | MYL12A | myosin, light chain 12A, regulatory, non-sarcomeric | Cytoplasm | other |
| 15551008 | 2 | 0.479 | 1377444 | 1676333.875 | 1699603.125 | 8483835 | 3278776 | 5635585.5 | 3468300 | 2313068 | 5966244.5 | 290136.031 | 7880182.5 | 5603474.5 | 1912.8823 | EAFNMIDQNRDGFIDK | Q3THE2 | Q3THE2 | MYL12A | myosin, light chain 12A, regulatory, non-sarcomeric | Cytoplasm | other |
| 15532254 | 3 | 0.103 | 9416315 | 1.96E+07 | 2.95E+07 | 6.78E+07 | 6.57E+07 | 5.86E+07 | 4.49E+07 | 4.78E+07 | 7.50E+07 | 1.11E+07 | 6.46E+07 | 5.76E+07 | 1888.0012 | VLDFEHFLPMLQTVAK | Q60605 | Q60605 | MYL6 | myosin, light chain 6, alkali, smooth muscle and non-muscle | Cytoplasm | other |
| 15532083 | 2 | 0.118 | 9322379 | 1.61E+07 | 3.06E+07 | 5.93E+07 | 6.00E+07 | 6.47E+07 | 3.41E+07 | 4.53E+07 | 8.01E+07 | 1.02E+07 | 5.73E+07 | 5.34E+07 | 1354.7296 | ALGQNPTNAEVLK | Q60605 | Q60605 | MYL6 | myosin, light chain 6, alkali, smooth muscle and non-muscle | Cytoplasm | other |
| 15552117 | 2 | 0.06 | 8421099 | 1.46E+07 | 2.56E+07 | 4.49E+07 | 5.04E+07 | 5.06E+07 | 2.85E+07 | 3.75E+07 | 5.86E+07 | 9180454 | 3.49E+07 | 3.00E+07 | 1544.6847 | DQGTYEDYVEGLR | Q60605 | Q60605 | MYL6 | myosin, light chain 6, alkali, smooth muscle and non-muscle | Cytoplasm | other |
| 15531826 | 2 | 0.088 | 5726087 | 1.01E+07 | 1.67E+07 | 4.03E+07 | 3.60E+07 | 3.40E+07 | 2.19E+07 | 2.69E+07 | 4.26E+07 | 6972108.5 | 3.86E+07 | 3.00E+07 | 995.5868 | HVLVTLGEK | Q60605 | Q60605 | MYL6 | myosin, light chain 6, alkali, smooth muscle and non-muscle | Cytoplasm | other |
| 15534605 | 2 | 0.099 | 4245029.5 | 9431104 | 1.70E+07 | 2.94E+07 | 4.05E+07 | 4.09E+07 | 2.00E+07 | 2.50E+07 | 4.57E+07 | 4607807.5 | 2.74E+07 | 2.46E+07 | 1025.5033 | EAFQLFDR | Q60605 | Q60605 | MYL6 | myosin, light chain 6, alkali, smooth muscle and non-muscle | Cytoplasm | other |
| 15531407 | 2 | 0.108 | 4304685.5 | 7901977 | 1.17E+07 | 2.98E+07 | 2.82E+07 | 2.79E+07 | 1.60E+07 | 1.65E+07 | 3.03E+07 | 4055974 | 2.58E+07 | 2.27E+07 | 1341.6268 | ILYSQC[160.0307]GDVMR | Q60605 | Q60605 | MYL6 | myosin, light chain 6, alkali, smooth muscle and non-muscle | Cytoplasm | other |
| 15552019 | 2 | 0.096 | 1449507.125 | 2393151.75 | 4988324 | 1.24E+07 | 1.01E+07 | 1.01E+07 | 5814874.5 | 7377840 | 1.34E+07 | 1711515.75 | 1.04E+07 | 8716229 | 1887.999 | VLDFEHFLPMLQTVAK | Q60605 | Q60605 | MYL6 | myosin, light chain 6, alkali, smooth muscle and non-muscle | Cytoplasm | other |
| 15532990 | 2 | 0.113 | 3591633.25 | 5985634 | 8580176 | 2.18E+07 | 1.95E+07 | 1.97E+07 | 1.25E+07 | 1.23E+07 | 2.15E+07 | 3197310 | 2.08E+07 | 1.57E+07 | 1228.614 | LNGTDPEDVIR | Q9CQ19 | Q9CQ19 | MYL9 | myosin, light chain 9, regulatory | Cytoplasm | other |
| 15532386 | 4 | 0.156 | 2841952.75 | 5865479 | 1.39E+07 | 3.31E+07 | 2.69E+07 | 2.88E+07 | 1.70E+07 | 1.90E+07 | 3.29E+07 | 2536024 | 2.83E+07 | 2.30E+07 | 2365.0283 | NAFAC[160.0307]FDEEASGFIHEDHLR | Q9CQ19 | Q9CQ19 | MYL9 | myosin, light chain 9, regulatory | Cytoplasm | other |
| 15533666 | 2 | 0.065 | 1313392.375 | 3925657.75 | 5381646 | 8293784 | 1.82E+07 | 9010122 | 7459565.5 | 1.08E+07 | 9514355 | 2384037 | 6774226 | 7436581 | 1237.5602 | EAFNMIDQNR | Q9CQ19 | Q9CQ19 | MYL9 | myosin, light chain 9, regulatory | Cytoplasm | other |
| 15533051 | 3 | 0.133 | 894927.312 | 3372937.75 | 4012094.5 | 1.34E+07 | 1.18E+07 | 1.06E+07 | 6153518.5 | 6050354 | 1.11E+07 | 1206364.125 | 9586313 | 1.08E+07 | 2090.9822 | ATSNVFAMFDQSQIQEFK | Q9CQ19 | Q9CQ19 | MYL9 | myosin, light chain 9, regulatory | Cytoplasm | other |
| 15533415 | 2, 3 | 0.175 | 879850.688 | 2218419 | 6441083 | 1.47E+07 | 1.61E+07 | 1.24E+07 | 8604087 | 1.01E+07 | 2.02E+07 | 916139.5 | 1.51E+07 | 1.56E+07 | 1328.6498, 1328.6499 | EDLHDMLASLGK | Q9CQ19 | Q9CQ19 | MYL9 | myosin, light chain 9, regulatory | Cytoplasm | other |
| 15551045 | 3 | 0.183 | 1161701 | 3487551.25 | 7100304.5 | 1.33E+07 | 1.90E+07 | 1.35E+07 | 8405026 | 1.15E+07 | 2.08E+07 | 868990.375 | 1.17E+07 | 1.18E+07 | 2450.0796 | ELLTTMGDRFTDEEVDEMYR | Q9CQ19 | Q9CQ19 | MYL9 | myosin, light chain 9, regulatory | Cytoplasm | other |
| 15542382 | 3 | 0.148 | 210510.422 | 965665.688 | 2095082.625 | 5571623 | 4915742.5 | 5643744 | 3154756.75 | 3737897.25 | 5902379 | 288763.062 | 4696156.5 | 4444023 | 2086.9243 | FTDEEVDEMYREAPIDK | Q9CQ19 | Q9CQ19 | MYL9 | myosin, light chain 9, regulatory | Cytoplasm | other |
| 15535990 | 2 | 0.236 | 724698.125 | 2370435.75 | 6209746 | 1.37E+07 | 1.59E+07 | 1.29E+07 | 8128374 | 1.14E+07 | 1.68E+07 | 268326.125 | 1.08E+07 | 1.13E+07 | 1246.5835 | GNFNYVEFTR | Q9CQ19 | Q9CQ19 | MYL9 | myosin, light chain 9, regulatory | Cytoplasm | other |
| 15546168 | 2 | 0.031 | 415848.188 | 272501.906 | 141929.547 | 231818.344 | 180854.453 | 256722 | 491182.719 | 500801.938 | 607389.438 | 332908.625 | 561938.312 | 383464.375 | 1314.7789 | MSLLQLVEILR | Q9WTI7 | Q9WTI7 | MYO1C | myosin IC | Cytoplasm | other |
| 15535473 | 2 | 0.054 | 2358987 | 3870776 | 1664061 | 2771127.75 | 2401289.5 | 2830669 | 1079455 | 2174298.5 | 2301601 | 1478718.25 | 964508.688 | 1199879.625 | 749.4176 | DM[147.0354]KVLK | Q64331 | Q64331 | MYO6 | myosin VI | Cytoplasm | other |
| 15534993 | 2 | 0.247 | 8267023.5 | 9265864 | 5075764.5 | 6607207 | 6594661 | 7195493 | 5230840 | 5091110.5 | 3703861.75 | 1.01E+07 | 5024350 | 5300838 | 1484.7233 | SPASDTYIVFGEAK | P70670 | P70670 | NACA | nascent polypeptide-associated complex alpha subunit | Cytoplasm | transcription regulator |
| 15548885 | 2 | 0.122 | 7265408.5 | 7477104.5 | 4437011.5 | 4505549 | 5718268 | 5125562 | 3352284.5 | 3852039 | 3744198 | 6690468.5 | 3389899.75 | 4071678.5 | 1614.8308 | IEDLSQQAQLAAAEK | P70670 | P70670 | NACA | nascent polypeptide-associated complex alpha subunit | Cytoplasm | transcription regulator |
| 15555995 | 2 | 0.528 | 1438997.375 | 1564469 | 748454.312 | 654833.375 | 891786.5 | 1173918.125 | 752830.312 | 941160.25 | 871582.5 | 1289032.75 | 554393.562 | 757586.812 | 1678.774 | NNSNDIVNAIMELTM | P70670 | P70670 | NACA | nascent polypeptide-associated complex alpha subunit | Cytoplasm | transcription regulator |
| 15535180 | 3 | 0.538 | 3.81E+07 | 3.05E+07 | 3.26E+07 | 5.28E+07 | 4.81E+07 | 2.76E+07 | 4.71E+07 | 4.17E+07 | 2.64E+07 | 7.04E+07 | 3.40E+07 | 4.66E+07 | 2095.034 | NVDLLSDMVQEHDEPILK | P28656 | P28656 | NAP1L1 | nucleosome assembly protein 1-like 1 | Nucleus | other |
| 15552219 | 2 | 0.997 | 2.34E+07 | 2.12E+07 | 2.86E+07 | 2.83E+07 | 3.26E+07 | 1.68E+07 | 3.28E+07 | 2.71E+07 | 1.82E+07 | 4.79E+07 | 1.69E+07 | 2.12E+07 | 1817.9497 | LDGLVDTPTGYIESLPK | P28656 | P28656 | NAP1L1 | nucleosome assembly protein 1-like 1 | Nucleus | other |
| 15532822 | 2 | 0.611 | 2.35E+07 | 1.98E+07 | 2.17E+07 | 2.76E+07 | 2.50E+07 | 1.63E+07 | 2.88E+07 | 2.11E+07 | 1.68E+07 | 4.39E+07 | 1.90E+07 | 2.77E+07 | 1356.7169 | YAVLYQPLFDK | P28656 | P28656 | NAP1L1 | nucleosome assembly protein 1-like 1 | Nucleus | other |
| 15554292 | 3 | 0.713 | 1.09E+07 | 9387644 | 4996041.5 | 1.73E+07 | 1.21E+07 | 3464157.5 | 1.27E+07 | 8553908 | 4080706 | 2.34E+07 | 8975261 | 1.04E+07 | 3409.562 | FSDAGQPMSFVLEFHFEPNDYFTNEVLTK | P28656 | P28656 | NAP1L1 | nucleosome assembly protein 1-like 1 | Nucleus | other |
| 15531762 | 2 | 0.503 | 1.04E+07 | 8018072 | 1.77E+07 | 1.65E+07 | 1.39E+07 | 1.12E+07 | 1.59E+07 | 1.47E+07 | 1.19E+07 | 1.86E+07 | 1.26E+07 | 1.70E+07 | 1336.728 | GIPEFWLTVFK | P28656 | P28656 | NAP1L1 | nucleosome assembly protein 1-like 1 | Nucleus | other |
| 15552617 | 3 | 0.475 | 5956040 | 6420705.5 | 4703054.5 | 8558797 | 8735805 | 4935105 | 8533535 | 7420319.5 | 4489989 | 1.47E+07 | 5987662 | 7838537 | 3377.3765 | MRSEPDDSDPFSFDGPEIMGC[160.0307]TGC[160.0307]QIDWK | P28656 | P28656 | NAP1L1 | nucleosome assembly protein 1-like 1 | Nucleus | other |
| 15533738 | 3 | 0.492 | 6528577.5 | 6155412.5 | 5317879 | 7189848.5 | 7835710.5 | 5284671.5 | 6313497.5 | 5314706 | 3460366.75 | 9754608 | 4581658.5 | 6520506.5 | 2212.1545 | QLTVQMMQNPQILAALQER | P28656 | P28656 | NAP1L1 | nucleosome assembly protein 1-like 1 | Nucleus | other |
| 15553407 | 2 | 0.653 | 3376600.75 | 2681508.25 | 2859382 | 5149415 | 4023149.5 | 2053130.75 | 4023374 | 3135830.25 | 1608236.25 | 6443283.5 | 2894189.25 | 3502278 | 2095.0327 | NVDLLSDMVQEHDEPILK | P28656 | P28656 | NAP1L1 | nucleosome assembly protein 1-like 1 | Nucleus | other |
| 15538838 | 2, 3 | 0.371 | 2.78E+07 | 1.94E+07 | 2.62E+07 | 3.52E+07 | 2.76E+07 | 1.75E+07 | 3.65E+07 | 2.22E+07 | 1.84E+07 | 5.05E+07 | 3.04E+07 | 3.12E+07 | 1464.7085, 1464.7086 | FYEEVHDLERK | Q78ZA7 | Q78ZA7 | NAP1L4 | nucleosome assembly protein 1-like 4 | Cytoplasm | other |
| 15532652 | 3 | 0.648 | 1.91E+07 | 1.47E+07 | 1.52E+07 | 2.17E+07 | 2.17E+07 | 1.20E+07 | 2.62E+07 | 1.84E+07 | 1.22E+07 | 3.30E+07 | 1.73E+07 | 1.98E+07 | 1900.9603 | LDNVSHTPSSYIETLPK | Q78ZA7 | Q78ZA7 | NAP1L4 | nucleosome assembly protein 1-like 4 | Cytoplasm | other |
| 15549974 | 3 | 0.584 | 1.15E+07 | 1.06E+07 | 4906631.5 | 1.74E+07 | 1.18E+07 | 5282932 | 1.48E+07 | 1.03E+07 | 5444847.5 | 2.63E+07 | 1.03E+07 | 1.24E+07 | 3403.5916 | FSDPGQPMSFVLEFHFEPNDYFTNPVLTK | Q78ZA7 | Q78ZA7 | NAP1L4 | nucleosome assembly protein 1-like 4 | Cytoplasm | other |
| 15551731 | 2 | 0.435 | 8293143.5 | 7267596 | 1.08E+07 | 1.19E+07 | 1.18E+07 | 9638284 | 1.28E+07 | 1.08E+07 | 7173024 | 1.85E+07 | 8605380 | 1.23E+07 | 1800.8934 | QVPNESFFNFFSPLK | Q78ZA7 | Q78ZA7 | NAP1L4 | nucleosome assembly protein 1-like 4 | Cytoplasm | other |
| 15534025 | 2 | 0.73 | 1.05E+07 | 9314005 | 8157930.5 | 1.23E+07 | 1.26E+07 | 7786689.5 | 1.44E+07 | 1.00E+07 | 5828398 | 1.80E+07 | 8957187 | 1.11E+07 | 1328.6855 | YAALYQPLFDK | Q78ZA7 | Q78ZA7 | NAP1L4 | nucleosome assembly protein 1-like 4 | Cytoplasm | other |
| 15536060 | 2 | 0.49 | 7134777.5 | 6537469.5 | 6858341 | 1.06E+07 | 9334393 | 6391528.5 | 1.11E+07 | 7901604 | 5136532.5 | 1.55E+07 | 7121108 | 9336494 | 1200.6079 | EAATVEELNPK | Q78ZA7 | Q78ZA7 | NAP1L4 | nucleosome assembly protein 1-like 4 | Cytoplasm | other |
| 15537504 | 2, 3 | 0.985 | 6294529 | 8074402 | 1.00E+07 | 5431306.5 | 1.45E+07 | 5518726.5 | 1.04E+07 | 1.03E+07 | 6074646.5 | 1.41E+07 | 4858081 | 7374328.5 | 1336.6143, 1336.6152 | FYEEVHDLER | Q78ZA7 | Q78ZA7 | NAP1L4 | nucleosome assembly protein 1-like 4 | Cytoplasm | other |
| 15554182 | 2 | 0.306 | 4213542.5 | 4985944.5 | 5092473.5 | 7163405 | 6439473 | 6264165 | 6308659 | 6059141 | 3687283.5 | 1.15E+07 | 4377595 | 7732222.5 | 2135.052 | NVDMLSELVQEYDEPILK | Q78ZA7 | Q78ZA7 | NAP1L4 | nucleosome assembly protein 1-like 4 | Cytoplasm | other |
| 15540245 | 2 | 0.201 | 4882830.5 | 3672107 | 5370655 | 5530714.5 | 5703361.5 | 4900302.5 | 7773718 | 5506533 | 4201880.5 | 1.00E+07 | 5296755.5 | 7578452 | 1412.734 | GIPEFWFTIFR | Q78ZA7 | Q78ZA7 | NAP1L4 | nucleosome assembly protein 1-like 4 | Cytoplasm | other |
| 15549987 | 3 | 0.594 | 1784359.875 | 2499431.25 | 2838186.75 | 2522517.75 | 4540566.5 | 3206672.75 | 4538452 | 4069860.25 | 973457 | 7647452.5 | 2560373.5 | 3686132.75 | 2864.197 | EFITGDVEPTDAESAWHSENEEEDK | Q78ZA7 | Q78ZA7 | NAP1L4 | nucleosome assembly protein 1-like 4 | Cytoplasm | other |
| 15556499 | 2 | 0.889 | 1774517.875 | 2383962.25 | 2990807.5 | 2826007.25 | 4498678.5 | 2502606.25 | 4052078 | 3794160 | 1019383.75 | 4415725.5 | 1404940.5 | 2326225 | 2558.0803 | ADPFSFEGPEIVDC[160.0307]DGC[160.0307]TIDWK | Q78ZA7 | Q78ZA7 | NAP1L4 | nucleosome assembly protein 1-like 4 | Cytoplasm | other |
| 15542726 | 3 | 0.056 | 3574505.5 | 3472294.5 | 2636954.25 | 4234248.5 | 3868877.5 | 2619433.75 | 3880934 | 3592358.25 | 3455539.25 | 1908906.875 | 2468848.5 | 2671454.75 | 2074.989 | LLEAHEEQNVDSYTEAVK | Q9DB05 | Q9DB05 | NAPA | N-ethylmaleimide-sensitive factor attachment protein, alpha | Cytoplasm | transporter |
| 15540161 | 3 | 0.454 | 2439121 | 2098742.75 | 1472065 | 1907278.25 | 4.70E-06 | 338506.5 | 4.70E-06 | 383381.594 | 1307004.75 | 872099.125 | 392237.656 | 199387.188 | 2056.9011 | GPEASTQDSHGDTDPGLTGSK | Q8K2Z4 | Q8K2Z4 | NCAPD2 | non-SMC condensin I complex, subunit D2 | Nucleus | other |
| 15554284 | 3 | 0.069 | 2060700.375 | 2194982.5 | 1799268.625 | 1577030.5 | 3281309 | 909884.562 | 1533286.375 | 1082914.625 | 897890.25 | 1113911 | 721813.125 | 605673 | 2543.06 | EADTEANHTESGQEGAPEEVADEK | Q8C156 | Q8C156 | NCAPH | non-SMC condensin I complex, subunit H | Nucleus | other |
| 15549037 | 2 | 0.958 | 1.13E+07 | 1.18E+07 | 1.38E+07 | 8869394 | 1.35E+07 | 1.30E+07 | 1.25E+07 | 1.16E+07 | 1.27E+07 | 1.30E+07 | 1.04E+07 | 1.35E+07 | 2029.0394 | WVLTDGPNPPYMVLLEGK | P57716 | P57716 | NCSTN | nicastrin | Plasma Membrane | peptidase |
| 15538431 | 2 | 0.761 | 1.23E+07 | 1.15E+07 | 1.31E+07 | 1.17E+07 | 1.18E+07 | 1.12E+07 | 1.22E+07 | 1.16E+07 | 1.00E+07 | 1.28E+07 | 9735355 | 1.30E+07 | 1372.7292 | NQVEDLLATLEK | P57716 | P57716 | NCSTN | nicastrin | Plasma Membrane | peptidase |
| 15533736 | 2 | 0.125 | 8073759 | 7508816.5 | 8243361 | 7912353.5 | 8314529 | 6922346.5 | 6629134.5 | 6714696 | 6454479 | 1.13E+07 | 7394520.5 | 7972910.5 | 959.51416 | APDVTTLSR | P57716 | P57716 | NCSTN | nicastrin | Plasma Membrane | peptidase |
| 15550048 | 3 | 0.877 | 4363024 | 6856466.5 | 3476536.5 | 4408447.5 | 6109503 | 4336030 | 4828487.5 | 4582336.5 | 3698958.25 | 8757012 | 3280598 | 5396042 | 3290.57 | SFIQSTFSINPEIVC[160.0307]DPLSDYNVWSMLK | P57716 | P57716 | NCSTN | nicastrin | Plasma Membrane | peptidase |
| 15531986 | 2 | 1 | 7654931 | 6937186.5 | 9258795 | 6568086.5 | 7693388 | 9606087 | 7787454.5 | 8244017.5 | 7560701 | 6324018 | 7295732.5 | 1.04E+07 | 1098.661 | ALANVATVLAR | P57716 | P57716 | NCSTN | nicastrin | Plasma Membrane | peptidase |
| 15532731 | 3 | 0.025 | 4160785.5 | 4446218 | 3841494.25 | 4032975.75 | 2853065 | 3657815.25 | 3428021.5 | 3506517.75 | 3958201.5 | 5084752.5 | 5041835 | 4424233.5 | 1239.7141 | SGAGVPEVVLRR | P57716 | P57716 | NCSTN | nicastrin | Plasma Membrane | peptidase |
| 15557288, 15559569 | 3 | 0.779 | 5112239 | 4502426 | 5364145.5 | 4299912.5 | 4635045.5 | 5491277.5 | 4447105.5 | 4485121.5 | 4840891 | 4469046 | 3815368 | 5479521 | 2302.2515, 2302.2522 | FPVRLENIDSFVELGQVALR | P57716 | P57716 | NCSTN | nicastrin | Plasma Membrane | peptidase |
| 15540311 | 2 | 0.392 | 5596824 | 6071540.5 | 5025920.5 | 4553617 | 5037648.5 | 6396000 | 5038160.5 | 5307403 | 6467804.5 | 4382926.5 | 3985579 | 5550951 | 980.59106 | LLYGFLVR | P57716 | P57716 | NCSTN | nicastrin | Plasma Membrane | peptidase |
| 15554188 | 2 | 0.81 | 737004.375 | 944122.25 | 2364124.5 | 101937.305 | 1691129.125 | 1558047.75 | 490731.469 | 1681071.875 | 1535380.875 | 1270834.75 | 311220.969 | 691536.688 | 1802.9601 | LENIDSFVELGQVALR | P57716 | P57716 | NCSTN | nicastrin | Plasma Membrane | peptidase |
| 15551071 | 3 | 0.129 | 3648091.25 | 7720689.5 | 4831421 | 6618094 | 5294729 | 9554007 | 4938157.5 | 7725729.5 | 8632828 | 2534977 | 4684783 | 4196793 | 2425.3042 | LLQYADALEHLLSTGQGVVLER | Q99LC3 | Q99LC3 | NDUFA10 | NADH dehydrogenase (ubiquinone) 1 alpha subcomplex, 10, 42kDa | Cytoplasm | transporter |
| 15550215 | 3 | 0.337 | 2864222.25 | 7466682.5 | 3127081.75 | 5317399.5 | 4482370.5 | 6649592.5 | 2664587.25 | 4261437 | 6061966 | 2130680.25 | 3397775.25 | 3522410.5 | 2772.4495 | TEVLNYTTIPVYLPEITIGAHQGSR | Q99LC3 | Q99LC3 | NDUFA10 | NADH dehydrogenase (ubiquinone) 1 alpha subcomplex, 10, 42kDa | Cytoplasm | transporter |
| 15546209 | 2 | 0.029 | 622227.562 | 899822.812 | 388347.625 | 724440.312 | 860207.938 | 767423 | 494517.875 | 660185.062 | 863663.438 | 359609.625 | 392722.281 | 311634.5 | 931.5073 | IGTLVGEDK | Q7TMF3 | Q7TMF3 | NDUFA12 | NADH dehydrogenase (ubiquinone) 1 alpha subcomplex, 12 | Cytoplasm | enzyme |
| 15549708 | 2 | 0.177 | 1942932.125 | 5072491.5 | 2088586.75 | 3644316.75 | 2807309.5 | 4074119.5 | 2209930.25 | 3173004 | 3827740.25 | 1511379.75 | 1882047.5 | 2131184 | 1518.7727 | TVSLNNLSADEVTR | Q9CQ75 | Q9CQ75 | NDUFA2 | NADH dehydrogenase (ubiquinone) 1 alpha subcomplex, 2, 8kDa | Cytoplasm | enzyme |
| 15543589 | 2 | 0.192 | 5199175.5 | 9699045 | 4479913.5 | 7611323.5 | 7063449.5 | 9537552 | 4453592.5 | 7810230 | 9126318 | 3163005.75 | 5533499 | 4545474 | 948.47644 | LGPNEQYK | Q62425 | Q62425 | NDUFA4 | NDUFA4, mitochondrial complex associated | Cytoplasm | enzyme |
| 15533526 | 2 | 0.244 | 3595991.5 | 7216871.5 | 3865033.5 | 3572912.5 | 6518457 | 7518831.5 | 3087104.75 | 8540380 | 6260334.5 | 3145937.5 | 3018255.5 | 2916134.75 | 1432.7196 | LALFNPDVSWDR | Q62425 | Q62425 | NDUFA4 | NDUFA4, mitochondrial complex associated | Cytoplasm | enzyme |
| 15540228 | 2 | 0.031 | 935522.812 | 2310398.75 | 1152861.75 | 1575730.875 | 1690581 | 2257635.75 | 1003694.688 | 1778212.875 | 1870020.375 | 543943.25 | 671431.25 | 876099.25 | 1125.5396 | YTEQITNEK | Q9CPP6 | Q9CPP6 | NDUFA5 | NADH dehydrogenase (ubiquinone) 1 alpha subcomplex, 5 | Cytoplasm | enzyme |
| 15535899 | 3 | 0.249 | 1234822.125 | 4133493.25 | 1953789.25 | 1418298.375 | 2544134.5 | 3909092.5 | 1707030.125 | 3010572 | 2852039.75 | 1039292.312 | 1152767.5 | 1532993.625 | 1534.8192 | ARPEPNPVIEGDLK | Q9DCJ5 | Q9DCJ5 | NDUFA8 | NADH dehydrogenase (ubiquinone) 1 alpha subcomplex, 8, 19kDa | Cytoplasm | enzyme |
| 15551120 | 3 | 0.366 | 6838471 | 1.62E+07 | 6727615 | 9699494 | 8295715 | 1.40E+07 | 6245371.5 | 1.02E+07 | 1.11E+07 | 5007905.5 | 6616828 | 7426049 | 2927.539 | IHISDVMPTDLPGLEDLGVQPTPLELK | Q9DC69 | Q9DC69 | NDUFA9 | NADH dehydrogenase (ubiquinone) 1 alpha subcomplex, 9, 39kDa | Cytoplasm | enzyme |
| 15555890 | 2 | 0.233 | 1423035.75 | 3794234.25 | 2559934.75 | 2598429.5 | 3386074 | 4846933 | 1920468 | 3254023.25 | 4721336.5 | 889203.438 | 2036263.125 | 2275771.75 | 1522.7916 | LFGLSPFEPWTTK | Q9DC69 | Q9DC69 | NDUFA9 | NADH dehydrogenase (ubiquinone) 1 alpha subcomplex, 9, 39kDa | Cytoplasm | enzyme |
| 15539485 | 2 | 0.145 | 830532.25 | 2521867.25 | 905725.625 | 1710041.875 | 1369869.5 | 2796172.25 | 1275682 | 2256877 | 3372382.5 | 480359.812 | 1191334.75 | 918064.5 | 1476.859 | WFLAVPLVSLGFK | Q9DC69 | Q9DC69 | NDUFA9 | NADH dehydrogenase (ubiquinone) 1 alpha subcomplex, 9, 39kDa | Cytoplasm | enzyme |
| 15549102 | 2 | 0.135 | 2705750.25 | 1.15E+07 | 6913116 | 4458726 | 8246444.5 | 1.17E+07 | 5227222.5 | 8785940 | 7683368 | 2626969.25 | 2721449.5 | 3673292.25 | 1694.8105 | LMC[160.0307]PQEIVDYIADK | Q9CR21 | Q9CR21 | NDUFAB1 | NADH dehydrogenase (ubiquinone) 1, alpha/beta subcomplex, 1, 8kDa | Cytoplasm | enzyme |
| 15537814 | 2 | 0.38 | 1616954 | 2802808.75 | 1227913.75 | 1776120.25 | 2013719.375 | 2304217 | 1339050.5 | 1956000.25 | 2370748.25 | 985675.938 | 1695944.125 | 1361631.375 | 1062.5377 | LSVNSHFMK | Q9CR21 | Q9CR21 | NDUFAB1 | NADH dehydrogenase (ubiquinone) 1, alpha/beta subcomplex, 1, 8kDa | Cytoplasm | enzyme |
| 15551279 | 2 | 0.097 | 1913262 | 4347087 | 2027583 | 2922718.25 | 3185013.75 | 3577122 | 2058877.625 | 3384274 | 4050052 | 1343870.5 | 1925737.5 | 1705564.125 | 1608.7832 | FASEIAGVDDLGTTGR | Q91VD9 | Q91VD9 | NDUFS1 | NADH dehydrogenase (ubiquinone) Fe-S protein 1, 75kDa (NADH-coenzyme Q reductase) | Cytoplasm | enzyme |
| 15535967 | 2 | 0.12 | 1473904.875 | 5550572 | 1998871.75 | 2023782.75 | 3686227.75 | 3976222.75 | 2110466.5 | 3832166.75 | 3515964 | 1131302.125 | 1474871.5 | 1345564.875 | 1403.7867 | VALIGSPVDLTYR | Q91VD9 | Q91VD9 | NDUFS1 | NADH dehydrogenase (ubiquinone) Fe-S protein 1, 75kDa (NADH-coenzyme Q reductase) | Cytoplasm | enzyme |
| 15551270 | 2 | 0.183 | 1335736.125 | 3229716.25 | 1565090.25 | 2209990.5 | 2029543.875 | 3003149 | 1474739.5 | 2515205 | 2949715.5 | 937118.25 | 1425460.75 | 1516223.125 | 2228.0447 | TQPYDVYDQVEFDVPIGSR | Q91WD5 | Q91WD5 | NDUFS2 | NADH dehydrogenase (ubiquinone) Fe-S protein 2, 49kDa (NADH-coenzyme Q reductase) | Cytoplasm | enzyme |
| 15542513 | 2 | 0.231 | 1170965.125 | 3377029.25 | 1338055.125 | 1307556.5 | 2837315.5 | 3707749.25 | 1454796.5 | 3841019.25 | 3640586 | 470151.906 | 1384430 | 1543059.25 | 1357.7291 | SLADLTAVDVPTR | Q9DCT2 | Q9DCT2 | NDUFS3 | NADH dehydrogenase (ubiquinone) Fe-S protein 3, 30kDa (NADH-coenzyme Q reductase) | Cytoplasm | enzyme |
| 15535565 | 3 | 0.348 | 2247920 | 7640973 | 3010267.5 | 3725406.25 | 4333426 | 6208508 | 2201290.5 | 5598683 | 5883118 | 1065595.75 | 2860330.5 | 3147359.75 | 1665.8982 | LDITTLTGVPEEHIK | Q9CXZ1 | Q9CXZ1 | NDUFS4 | NADH dehydrogenase (ubiquinone) Fe-S protein 4, 18kDa (NADH-coenzyme Q reductase) | Cytoplasm | enzyme |
| 15555464 | 3 | 0.515 | 871905.812 | 2664313.75 | 507167.375 | 2038823.25 | 1723850.125 | 1190102.875 | 1064314.5 | 1471295.375 | 1658791.125 | 849433.562 | 1081861.75 | 917109 | 3051.4631 | ERWENPLMGWASTADPLSNMVLTFSAK | Q9CXZ1 | Q9CXZ1 | NDUFS4 | NADH dehydrogenase (ubiquinone) Fe-S protein 4, 18kDa (NADH-coenzyme Q reductase) | Cytoplasm | enzyme |
| 15549053 | 3 | 0.649 | 3426844 | 9402101 | 3601917.25 | 5182977.5 | 4247929.5 | 8180825 | 2807524 | 5586108 | 7170916.5 | 2739606 | 4476642 | 3842466 | 2614.3943 | HAGGVTGGWDNLLAVIPGGSSTPLIPK | Q91YT0 | Q91YT0 | NDUFV1 | NADH dehydrogenase (ubiquinone) flavoprotein 1, 51kDa | Cytoplasm | enzyme |
| 15556665 | 2 | 0.116 | 1395409.875 | 3562737 | 1617564.875 | 2301337.5 | 2550539 | 2691871.75 | 1368440.875 | 2597475.75 | 3603683.25 | 737524.75 | 1385792.125 | 1302117.875 | 1651.7605 | YLVVNADEGEPGTC[160.0307]K | Q91YT0 | Q91YT0 | NDUFV1 | NADH dehydrogenase (ubiquinone) flavoprotein 1, 51kDa | Cytoplasm | enzyme |
| 15545093 | 2 | 0.262 | 860458.688 | 1657754.5 | 696706.688 | 1096766.375 | 1916080 | 2216742.75 | 1055803 | 1595876 | 2317176.5 | 371595.562 | 1519713.625 | 844097.938 | 1059.5587 | LVEGC[160.0307]LVGGR | Q91YT0 | Q91YT0 | NDUFV1 | NADH dehydrogenase (ubiquinone) flavoprotein 1, 51kDa | Cytoplasm | enzyme |
| 15539360 | 2 | 0.146 | 1347790.625 | 3042237.25 | 1411616.25 | 1887800 | 2185429.25 | 3748803.75 | 1284396.75 | 2278060 | 2840619.5 | 853393 | 1245930.375 | 1354211.375 | 1216.629 | DIEEIIDELK | Q9D6J6 | Q9D6J6 | NDUFV2 | NADH dehydrogenase (ubiquinone) flavoprotein 2, 24kDa | Cytoplasm | enzyme |
| 15549929 | 3 | 0.089 | 8022529.5 | 1.19E+07 | 5903277.5 | 8065948 | 9033231 | 5265769.5 | 5519324.5 | 7411220.5 | 6179728.5 | 5691915 | 4123127 | 3798600.75 | 2796.3262 | TSGELFAQAPVDQFPGTAVESVTDSSR | Q9D1J1 | Q9D1J1 | NECAP2 | NECAP endocytosis associated 2 | Cytoplasm | other |
| 15540901 | 3 | 0.162 | 1.83E+07 | 1.61E+07 | 1.65E+07 | 1.25E+07 | 1.48E+07 | 1.54E+07 | 1.88E+07 | 2.13E+07 | 1.37E+07 | 1.61E+07 | 1.28E+07 | 1.07E+07 | 2142.196 | IPALLYLPPTHTFLAFAEK | Q9JMH7 | Q9JMH7 | NEU3 | sialidase 3 (membrane sialidase) | Plasma Membrane | enzyme |
| 15554125 | 3 | 0.776 | 9770138 | 1.01E+07 | 5538245.5 | 7401035 | 7230245.5 | 5936213.5 | 1.02E+07 | 8341893 | 6729829.5 | 1.03E+07 | 1.04E+07 | 5481498 | 3123.4805 | FIEPQVTGEC[160.0307]QVAEVAGTAGNPVLYC[160.0307]SAR | Q9JMH7 | Q9JMH7 | NEU3 | sialidase 3 (membrane sialidase) | Plasma Membrane | enzyme |
| 15557932 | 2 | 0.286 | 3385799.25 | 2139196.25 | 1302803.875 | 1111462.375 | 1823856.875 | 1135567.875 | 1335108 | 1933367.125 | 1360245.125 | 1202581.5 | 1722249.125 | 959077.812 | 1535.7954 | TGC[160.0307]QGSVVSFRPLK | Q9JMH7 | Q9JMH7 | NEU3 | sialidase 3 (membrane sialidase) | Plasma Membrane | enzyme |
| 15539452 | 2 | 0.802 | 5023746 | 2201207.5 | 3213395.5 | 5605728.5 | 2115466.25 | 3213595 | 3276521.5 | 2595559 | 2842403.25 | 3299586.25 | 4476221 | 3701294.5 | 1313.7141 | KMNALHMLDIK | Q9EST8 | Q9EST8 | NFKBIZ | nuclear factor of kappa light polypeptide gene enhancer in B-cells inhibitor, zeta | Nucleus | transcription regulator |
| 15542647 | 3 | 0.325 | 8069.789 | 159068.906 | 1500031.25 | 309516.938 | 3466426.5 | 4681243.5 | 859026 | 2662050.5 | 4630761.5 | 39016.742 | 2142959.5 | 2047958.875 | 2012.057 | QFGPTFALDTVHVDPVIR | Q3TL44 | Q3TL44 | NLRX1 | NLR family member X1 | Cytoplasm | other |
| 15538757 | 3 | 0.995 | 1375167 | 2687118.25 | 1079878.875 | 1292491.625 | 1772156.625 | 1683546 | 1165099.125 | 1852561 | 1778786.25 | 1728207.375 | 1312389 | 1488404.75 | 1507.7203 | NVIHGSDSVESAHR | Q9WV85 | Q9WV85 | NME3 | NME/NM23 nucleoside diphosphate kinase 3 | Cytoplasm | kinase |
| 15531446 | 3 | 0.38 | 1.20E+08 | 1.03E+08 | 8.51E+07 | 8.97E+07 | 9.91E+07 | 7.99E+07 | 1.03E+08 | 9.44E+07 | 8.33E+07 | 1.22E+08 | 1.07E+08 | 9.36E+07 | 2095.9136 | AEDSGEYHC[160.0307]VYHFVSAPK | P97300 | P97300 | Nptn | neuroplastin | Plasma Membrane | other |
| 15533814 | 2 | 0.507 | 4.25E+07 | 3.73E+07 | 3.14E+07 | 3.88E+07 | 3.27E+07 | 2.62E+07 | 3.77E+07 | 3.23E+07 | 2.56E+07 | 4.56E+07 | 3.50E+07 | 3.42E+07 | 1346.5134 | NEGQDAMMYC[160.0307]K | P97300 | P97300 | Nptn | neuroplastin | Plasma Membrane | other |
| 15558812 | 2 | 0.436 | 1.49E+07 | 1.99E+07 | 2.24E+07 | 9947931 | 1.85E+07 | 1.63E+07 | 1.52E+07 | 1.60E+07 | 1.70E+07 | 1.50E+07 | 1.55E+07 | 1.51E+07 | 1526.7505 | SVGYPHPEWIWR | P97300 | P97300 | Nptn | neuroplastin | Plasma Membrane | other |
| 15552453 | 2 | 0.231 | 1.36E+07 | 9621270 | 6447139.5 | 9384497 | 8790438 | 7121624.5 | 9158881 | 6778939.5 | 6704757.5 | 1.48E+07 | 1.07E+07 | 9796420 | 2095.9133 | AEDSGEYHC[160.0307]VYHFVSAPK | P97300 | P97300 | Nptn | neuroplastin | Plasma Membrane | other |
| 15542041 | 2 | 0.629 | 8529864 | 1.21E+07 | 1.80E+07 | 7332658 | 9486830 | 1.20E+07 | 7783514 | 9512876 | 1.18E+07 | 1.11E+07 | 6466280.5 | 1.14E+07 | 1158.6715 | IVTSEEVIIR | P97300 | P97300 | Nptn | neuroplastin | Plasma Membrane | other |
| 15537052 | 3 | 0.785 | 5290257 | 8796537 | 1.46E+07 | 5203046.5 | 1.04E+07 | 9452302 | 9931048 | 1.12E+07 | 5173073 | 9619563 | 4545159.5 | 5729232 | 1767.781 | RPDEVPDDDEPAGPMK | P97300 | P97300 | Nptn | neuroplastin | Plasma Membrane | other |
| 15542343 | 2 | 0.226 | 3725164.5 | 5006326 | 5308813.5 | 2705595 | 4484841 | 3739953 | 3958124 | 4142780.25 | 2776535.75 | 4147555.25 | 1970788.75 | 2615353.75 | 960.509 | NGVELTATR | P97300 | P97300 | Nptn | neuroplastin | Plasma Membrane | other |
| 15543378 | 2 | 0.066 | 881180.062 | 573345.188 | 485964.344 | 615575.625 | 363307.25 | 285082.25 | 506013.281 | 279689.5 | 379318.562 | 959932.688 | 892007.312 | 563431.125 | 1088.6028 | NGVELTATRK | P97300 | P97300 | Nptn | neuroplastin | Plasma Membrane | other |
| 15547051 | 2 | 0.786 | 336115.938 | 126443.695 | 107272.961 | 72731.602 | 109746.852 | 182047.516 | 151164.672 | 346912.25 | 40915.688 | 240136.891 | 179255.219 | 155883.875 | 1362.5095 | NEGQDAMM[147.0354]YC[160.0307]K | P97300 | P97300 | Nptn | neuroplastin | Plasma Membrane | other |
| 15542004 | 2 | 3.21E-06 | 8975000 | 9576039 | 1.06E+07 | 1.96E+07 | 1.97E+07 | 1.87E+07 | 1.17E+07 | 1.27E+07 | 1.11E+07 | 2.10E+07 | 2.10E+07 | 1.79E+07 | 1312.6765 | SYGIPFIETSAK | P08556 | P08556 | NRAS | neuroblastoma RAS viral (v-ras) oncogene homolog | Plasma Membrane | enzyme |
| 15534907 | 2 | 0.645 | 1.02E+07 | 8551807 | 7798168 | 1.02E+07 | 9598291 | 7719846 | 9351586 | 7387856.5 | 6459676 | 1.14E+07 | 7683481.5 | 7749433.5 | 1457.7812 | VLDDGELLVQQTK | P46460 | P46460 | NSF | N-ethylmaleimide-sensitive factor | Cytoplasm | transporter |
| 15535553 | 2 | 0.39 | 7919789 | 7438848.5 | 7635405 | 6818524.5 | 7206427 | 5614673.5 | 7242096 | 7112556 | 5918423.5 | 9163282 | 6948182 | 6473416.5 | 1145.614 | AENSSLNLIGK | P46460 | P46460 | NSF | N-ethylmaleimide-sensitive factor | Cytoplasm | transporter |
| 15541797 | 2 | 0.651 | 6777180.5 | 6277717 | 7332091.5 | 6798627 | 9331174 | 6782811.5 | 7730843 | 7736801 | 7047156 | 7957305.5 | 6933606.5 | 6303054.5 | 1142.6544 | LLDYVPIGPR | P46460 | P46460 | NSF | N-ethylmaleimide-sensitive factor | Cytoplasm | transporter |
| 15534229 | 2 | 0.935 | 8229421 | 5886408.5 | 5132157 | 8122792.5 | 6185486 | 5701169 | 7905163 | 5139677.5 | 4961536.5 | 7712970 | 4779079 | 7356237.5 | 1291.6604 | NFSGAELEGLVR | P46460 | P46460 | NSF | N-ethylmaleimide-sensitive factor | Cytoplasm | transporter |
| 15541839 | 4 | 0.739 | 5598554.5 | 5368749 | 5480771 | 6511526 | 4969689 | 4246091 | 4866270.5 | 5022190.5 | 4608812.5 | 6167164 | 4447875.5 | 5121790 | 2103.1155 | NSDRTPLVSVLLEGPPHSGK | P46460 | P46460 | NSF | N-ethylmaleimide-sensitive factor | Cytoplasm | transporter |
| 15539381 | 2 | 0.166 | 7614966.5 | 5620624 | 6848846.5 | 5839718 | 6773851 | 6175036.5 | 7946692 | 6454812 | 5706080.5 | 5702863.5 | 5276254 | 4927730.5 | 1490.7235 | QSIINPDWNFEK | P46460 | P46460 | NSF | N-ethylmaleimide-sensitive factor | Cytoplasm | transporter |
| 15538776 | 2 | 0.868 | 4736088.5 | 3512155.25 | 4227594 | 3944462.75 | 3872250.25 | 4039194.75 | 4830655.5 | 4120459.75 | 3217814.25 | 5314292.5 | 3845008.25 | 4043229.25 | 1231.6223 | DIEAMDPSILK | P46460 | P46460 | NSF | N-ethylmaleimide-sensitive factor | Cytoplasm | transporter |
| 15539950 | 2 | 0.349 | 4280976.5 | 5656049 | 5134682.5 | 3795603.25 | 5494589.5 | 4490610 | 5210092 | 5254290 | 3485534.25 | 5095448.5 | 2609735 | 3189018.75 | 1321.6611 | GDFLASLENDIK | P46460 | P46460 | NSF | N-ethylmaleimide-sensitive factor | Cytoplasm | transporter |
| 15535973 | 2 | 0.954 | 3105739 | 3438487.25 | 3642066.75 | 3011226 | 4197700 | 2912783.5 | 3750210 | 3332910.25 | 2904789.75 | 4059112.75 | 3135945.25 | 3404480.75 | 1031.5056 | MEIGLPDEK | P46460 | P46460 | NSF | N-ethylmaleimide-sensitive factor | Cytoplasm | transporter |
| 15552904 | 2 | 0.471 | 3520096.25 | 2760894.75 | 2350829.75 | 5485082 | 1541277.25 | 3210490.75 | 2851736.25 | 2233955.25 | 3781963 | 3868735.5 | 4559357 | 4639183.5 | 1810.8615 | FLALMREEGASPLDFD | P46460 | P46460 | NSF | N-ethylmaleimide-sensitive factor | Cytoplasm | transporter |
| 15542024 | 2 | 0.264 | 3922839.25 | 3601063 | 4428901.5 | 3273679.5 | 3640587.75 | 3638736.25 | 3369864.25 | 3758419.5 | 3289514.5 | 3736648.75 | 2677864.75 | 3478978 | 1082.6179 | VVNGPEILNK | P46460 | P46460 | NSF | N-ethylmaleimide-sensitive factor | Cytoplasm | transporter |
| 15533761 | 2 | 0.266 | 3327942.75 | 2459167.75 | 2414813.75 | 2185531.5 | 2423217.25 | 2032099.125 | 2806328.5 | 2084528 | 1813113.625 | 3689041.75 | 2869833.25 | 2277427 | 1265.6453 | YVGESEANIRK | P46460 | P46460 | NSF | N-ethylmaleimide-sensitive factor | Cytoplasm | transporter |
| 15537619 | 2 | 0.936 | 1384998.375 | 1065554 | 1508224.625 | 1342496.25 | 1205277 | 1348802.25 | 1154037.5 | 1504278.25 | 991454.938 | 1641370.75 | 1073417.625 | 1127250.625 | 715.4487 | VWIGIK | P46460 | P46460 | NSF | N-ethylmaleimide-sensitive factor | Cytoplasm | transporter |
| 15543038 | 2 | 0.408 | 281432.125 | 78670.367 | 270096.438 | 161564.328 | 499636.188 | 161667.359 | 489277.656 | 441172.781 | 170363.109 | 594309.812 | 210155.469 | 722534.688 | 1303.6163 | DFQSGQHVMVR | P46460 | P46460 | NSF | N-ethylmaleimide-sensitive factor | Cytoplasm | transporter |
| 15549476 | 2 | 0.573 | 4993415.5 | 2397077.75 | 2872273 | 3752260 | 2727509 | 3258620.75 | 3691190.5 | 2547029.5 | 1819012.625 | 6631159 | 2722341 | 3394199.75 | 1648.7789 | LGAAPEEESAYVAGER | Q9CZ44 | Q9CZ44 | NSFL1C | NSFL1 (p97) cofactor (p47) | Cytoplasm | other |
| 15551934 | 2 | 0.487 | 3978622 | 2568037 | 2066642.125 | 1515657.5 | 1616920.5 | 2614172 | 3647490.5 | 1195204.875 | 884990.375 | 4855307.5 | 1771562.875 | 2211649 | 2057.0833 | ASSSILINEAEPTTNIQIR | Q9CZ44 | Q9CZ44 | NSFL1C | NSFL1 (p97) cofactor (p47) | Cytoplasm | other |
| 15551619 | 3 | 0.944 | 2914707.25 | 1877731.375 | 1729605.125 | 2660986 | 2135025.25 | 1658199.25 | 2276093.25 | 1552143 | 1634428 | 4573027 | 1145025.25 | 1841923.25 | 2341.1582 | LGSTAPQVLNTSSPAQQAENEAK | Q9CZ44 | Q9CZ44 | NSFL1C | NSFL1 (p97) cofactor (p47) | Cytoplasm | other |
| 15540165 | 2 | 0.821 | 2318589.75 | 1525839.375 | 1384564.25 | 1253205.25 | 764508.562 | 1773512.5 | 1475685 | 987916.188 | 979277.75 | 3037179.25 | 422688.719 | 1564338.5 | 1038.566 | SGQQIVGPPR | Q9CZ44 | Q9CZ44 | NSFL1C | NSFL1 (p97) cofactor (p47) | Cytoplasm | other |
| 15531738 | 2 | 0.002 | 1.77E+07 | 1.50E+07 | 1.62E+07 | 1.22E+07 | 1.14E+07 | 1.10E+07 | 1.37E+07 | 1.15E+07 | 1.16E+07 | 1.49E+07 | 1.51E+07 | 1.37E+07 | 1027.6128 | FPPLLWVR | A6H603 | A6H603 | NWD1 | NACHT and WD repeat domain containing 1 | Cytoplasm | other |
| 15535487 | 2 | 0.341 | 9.25E+07 | 9.23E+07 | 9.79E+07 | 5.98E+07 | 9.32E+07 | 7.98E+07 | 7.87E+07 | 1.07E+08 | 8.27E+07 | 1.07E+08 | 8.45E+07 | 9.33E+07 | 1225.6145 | ELTAFDNLFR | P83503 | P83503 | NYX | nyctalopin | Extracellular Space | other |
| 15554817 | 2 | 0.083 | 1844302.75 | 2897361.75 | 1541339.125 | 2315021.5 | 2210867.5 | 3578515.25 | 1316869.625 | 2559017 | 3106231.25 | 820448.25 | 1112651.375 | 1604581.375 | 1952.9229 | AFYNNVLGEYEEYITK | P29758 | P29758 | OAT | ornithine aminotransferase | Cytoplasm | enzyme |
| 15535016 | 3 | 0.431 | 1992854.25 | 3143812 | 2636936.75 | 2261898 | 2660884 | 1905338 | 1725108.5 | 2349601.75 | 3320617.25 | 1757516.625 | 1456221.625 | 2353952 | 1771.9023 | ELQAQDVDEGATARLR | A2AAJ9 | A2AAJ9 | OBSCN | obscurin, cytoskeletal calmodulin and titin-interacting RhoGEF | Cytoplasm | kinase |
| 15554197 | 3 | 0.217 | 1342696.25 | 3377517.75 | 1396434.625 | 2649512.75 | 2416597.25 | 3236625.25 | 1002143.438 | 2622467 | 3115721.5 | 699345.25 | 1161641.625 | 1769092.75 | 3783.7654 | YEPIPFSASMNESTPTGITDHIAQGPEPNLEESPK | Q9CRD0 | Q9CRD0 | OCIAD1 | OCIA domain containing 1 | Cytoplasm | other |
| 15548795 | 3 | 0.205 | 8527512 | 1.61E+07 | 6968188.5 | 1.27E+07 | 1.20E+07 | 1.43E+07 | 1.05E+07 | 1.37E+07 | 1.81E+07 | 6007293.5 | 1.13E+07 | 7913947 | 2504.1917 | VVNAPIFHVNSDDPEAVMYVC[160.0307]K | Q60597 | Q60597 | OGDH | oxoglutarate (alpha-ketoglutarate) dehydrogenase (lipoamide) | Cytoplasm | enzyme |
| 15548784 | 3, 4 | 0.076 | 9743244 | 1.61E+07 | 9704279 | 1.84E+07 | 1.45E+07 | 2.55E+07 | 1.34E+07 | 2.15E+07 | 1.93E+07 | 4973316.5 | 1.16E+07 | 1.19E+07 | 3199.6052, 3199.606 | GHHVAQLDPLGILDADLDSSVPADIISSTDK | Q60597 | Q60597 | OGDH | oxoglutarate (alpha-ketoglutarate) dehydrogenase (lipoamide) | Cytoplasm | enzyme |
| 15550305 | 3 | 0.43 | 5047101.5 | 1.23E+07 | 6113404 | 6108346 | 7103309.5 | 1.05E+07 | 3878529.25 | 8378178.5 | 9552066 | 4008244.25 | 5219522 | 5059424.5 | 2524.2397 | YAELLVSQGVVNQPEYEEEISK | Q60597 | Q60597 | OGDH | oxoglutarate (alpha-ketoglutarate) dehydrogenase (lipoamide) | Cytoplasm | enzyme |
| 15532605 | 3 | 0.249 | 4780839.5 | 9891545 | 5822092 | 7277865 | 8569492 | 1.17E+07 | 5100546.5 | 9168480 | 1.04E+07 | 3255471 | 5533839.5 | 6702224.5 | 1492.8456 | LVEDHLAVQSLIR | Q60597 | Q60597 | OGDH | oxoglutarate (alpha-ketoglutarate) dehydrogenase (lipoamide) | Cytoplasm | enzyme |
| 15535319 | 3 | 0.134 | 5313494 | 1.57E+07 | 6885153.5 | 6267952.5 | 1.10E+07 | 1.93E+07 | 8540832 | 1.60E+07 | 1.26E+07 | 3155304.75 | 6019044 | 5473712.5 | 2194.1987 | VFHLPTTTFIGGQEPALPLR | Q60597 | Q60597 | OGDH | oxoglutarate (alpha-ketoglutarate) dehydrogenase (lipoamide) | Cytoplasm | enzyme |
| 15533330 | 3 | 0.199 | 5343600 | 1.24E+07 | 5296189 | 8123986 | 7853800.5 | 9567944 | 6581090.5 | 1.10E+07 | 1.11E+07 | 3078706.5 | 5934078.5 | 6354860.5 | 2197.0872 | SSLATMAHAQSLVEAQPNVDK | Q60597 | Q60597 | OGDH | oxoglutarate (alpha-ketoglutarate) dehydrogenase (lipoamide) | Cytoplasm | enzyme |
| 15548642 | 3 | 0.263 | 4891020 | 1.31E+07 | 4965854 | 5610665 | 8725165 | 9427060 | 4804627.5 | 9708006 | 1.09E+07 | 2716809 | 5055392.5 | 4975898.5 | 2376.2542 | NITLSLVANPSHLEAADPVVMGK | Q60597 | Q60597 | OGDH | oxoglutarate (alpha-ketoglutarate) dehydrogenase (lipoamide) | Cytoplasm | enzyme |
| 15552887 | 2 | 0.301 | 4590869.5 | 7521927.5 | 4257148.5 | 5069426 | 5395537.5 | 8343178.5 | 4734897.5 | 6623672 | 9670228 | 2587989.25 | 5515594 | 4328075.5 | 1543.7076 | ELEQIFC[160.0307]QFDSK | Q60597 | Q60597 | OGDH | oxoglutarate (alpha-ketoglutarate) dehydrogenase (lipoamide) | Cytoplasm | enzyme |
| 15538637 | 3 | 0.351 | 4463713.5 | 9256189 | 3219158.5 | 6797699.5 | 5257530 | 7254810.5 | 4221550.5 | 6515307.5 | 8864298 | 2537397.25 | 5060308 | 4065799 | 2120.9944 | SMTC[160.0307]PSTGLEEDVLFHIGK | Q60597 | Q60597 | OGDH | oxoglutarate (alpha-ketoglutarate) dehydrogenase (lipoamide) | Cytoplasm | enzyme |
| 15537772 | 2 | 0.182 | 3534632.75 | 1.00E+07 | 4645254.5 | 6124582 | 6048250 | 9139369 | 3936166.5 | 7994797.5 | 7472508 | 2140798.25 | 3994391 | 4120009.5 | 1402.6864 | FLDTAFDLDAFK | Q60597 | Q60597 | OGDH | oxoglutarate (alpha-ketoglutarate) dehydrogenase (lipoamide) | Cytoplasm | enzyme |
| 15532677 | 2 | 0.134 | 4281250.5 | 7493912.5 | 3598171.75 | 5500774.5 | 5053783.5 | 6407835 | 3697242.75 | 5673080.5 | 6216557.5 | 2130368.75 | 4335126 | 3008968.75 | 1025.4698 | IC[160.0307]EEAFTR | Q60597 | Q60597 | OGDH | oxoglutarate (alpha-ketoglutarate) dehydrogenase (lipoamide) | Cytoplasm | enzyme |
| 15560095 | 2 | 0.305 | 3175249.25 | 7548819 | 4063692.75 | 3747807.5 | 4177547.25 | 7760254 | 3243558 | 6447155 | 8490017 | 1896731.625 | 3528577 | 3519505 | 1659.9325 | IEQLSPFPFDLLLK | Q60597 | Q60597 | OGDH | oxoglutarate (alpha-ketoglutarate) dehydrogenase (lipoamide) | Cytoplasm | enzyme |
| 15538827 | 2 | 0.191 | 2139985.75 | 4621545.5 | 1931854.875 | 3141116.5 | 2930742.75 | 4697429.5 | 2308574.75 | 3829631.25 | 4707158.5 | 1404495.625 | 2039682.875 | 2371999.25 | 970.47626 | SWDIFFR | Q60597 | Q60597 | OGDH | oxoglutarate (alpha-ketoglutarate) dehydrogenase (lipoamide) | Cytoplasm | enzyme |
| 15534515 | 2 | 0.198 | 1575437.25 | 4598992.5 | 3078430.25 | 2275073.75 | 2884717.5 | 5177097.5 | 1982415.125 | 3373944 | 4138739.5 | 904403.938 | 1716739.625 | 2069652.125 | 914.56885 | PLIVFTPK | Q60597 | Q60597 | OGDH | oxoglutarate (alpha-ketoglutarate) dehydrogenase (lipoamide) | Cytoplasm | enzyme |
| 15545352 | 2 | 0.27 | 1488396.875 | 2825667 | 1279779.625 | 1798930.5 | 2104545.75 | 2644113.25 | 1686410 | 2290159.5 | 2314363.5 | 780671.5 | 1879829.125 | 1389703.25 | 1222.5226 | LEAADEGSGDMK | Q60597 | Q60597 | OGDH | oxoglutarate (alpha-ketoglutarate) dehydrogenase (lipoamide) | Cytoplasm | enzyme |
| 15543575 | 2 | 0.028 | 1149494.875 | 2758870.75 | 1543222.25 | 1554447 | 1939626.375 | 2257260.75 | 1477158.125 | 1922268.125 | 1850531.375 | 691930.062 | 1105274.625 | 835656.875 | 1404.5708 | AEQFYC[160.0307]GDTEGK | Q60597 | Q60597 | OGDH | oxoglutarate (alpha-ketoglutarate) dehydrogenase (lipoamide) | Cytoplasm | enzyme |
| 15556007 | 2 | 0.162 | 2248357.25 | 5531440 | 1897729.375 | 3549533 | 3162391.5 | 3956687 | 2175807 | 3833445.5 | 5769970 | 659325.688 | 2452960.25 | 1750149.625 | 2001.994 | NTNAGAPPGTAYQSPLSLSR | Q60597 | Q60597 | OGDH | oxoglutarate (alpha-ketoglutarate) dehydrogenase (lipoamide) | Cytoplasm | enzyme |
| 15552353 | 2 | 0.205 | 1628919.875 | 2238647.25 | 2749802.5 | 4407166.5 | 1808724.125 | 3187786.5 | 1688510.375 | 1766274.125 | 2482882.5 | 1833071 | 1075093.125 | 1960115.25 | 1568.9381 | IPAFLNVVDIAGLVK | Q9CZ30 | Q9CZ30 | OLA1 | Obg-like ATPase 1 | Cytoplasm | enzyme |
| 15544940 | 2 | 0.019 | 831373.875 | 1312022.75 | 727280.312 | 707289.75 | 992712 | 710934.562 | 528001.312 | 820266.438 | 707347 | 465486.344 | 385042.781 | 491286.938 | 1049.5024 | C[160.0307]GGSLLTADR | Q8BGT0 | Q8BGT0 | OSTM1 | osteopetrosis associated transmembrane protein 1 | Cytoplasm | other |
| 15544116 | 2 | 0.153 | 1141196 | 2204288.25 | 1443212.875 | 3549663.25 | 3186742.5 | 3526755.25 | 2398076.75 | 2593522.5 | 3096056 | 634377.062 | 3183776.5 | 1952916.125 | 1231.577 | DEADADLINAGK | Q9D0K2 | Q9D0K2 | OXCT1 | 3-oxoacid CoA transferase 1 | Cytoplasm | enzyme |
| 15535193 | 3 | 0.317 | 1.63E+07 | 2.06E+07 | 2.37E+07 | 2.34E+07 | 2.48E+07 | 2.09E+07 | 2.46E+07 | 2.76E+07 | 2.14E+07 | 2.37E+07 | 2.23E+07 | 2.16E+07 | 1965.0394 | HNQLPLVIEFTEQTAPK | P09103 | P09103 | P4HB | prolyl 4-hydroxylase, beta polypeptide | Cytoplasm | enzyme |
| 15532237 | 2, 3 | 0.046 | 1.51E+07 | 1.43E+07 | 1.66E+07 | 1.88E+07 | 2.13E+07 | 1.55E+07 | 2.18E+07 | 1.88E+07 | 1.81E+07 | 2.26E+07 | 2.10E+07 | 1.89E+07 | 1409.6753, 1409.6765 | YKPESDELTAEK | P09103 | P09103 | P4HB | prolyl 4-hydroxylase, beta polypeptide | Cytoplasm | enzyme |
| 15550366, 15553799 | 2, 3 | 0.473 | 1.55E+07 | 2.27E+07 | 2.41E+07 | 2.15E+07 | 2.54E+07 | 2.02E+07 | 2.26E+07 | 2.70E+07 | 2.50E+07 | 2.19E+07 | 2.40E+07 | 2.16E+07 | 2640.3455, 2640.3525 | QFLLAAEAIDDIPFGITSNSGVFSK | P09103 | P09103 | P4HB | prolyl 4-hydroxylase, beta polypeptide | Cytoplasm | enzyme |
| 15534054 | 2, 3 | 0.241 | 1.53E+07 | 1.53E+07 | 2.04E+07 | 1.94E+07 | 2.45E+07 | 1.88E+07 | 1.74E+07 | 2.03E+07 | 2.05E+07 | 2.11E+07 | 2.30E+07 | 1.85E+07 | 1216.593, 1216.5938 | SNFEEALAAHK | P09103 | P09103 | P4HB | prolyl 4-hydroxylase, beta polypeptide | Cytoplasm | enzyme |
| 15548950 | 3 | 0.705 | 1.53E+07 | 2.21E+07 | 2.97E+07 | 2.17E+07 | 2.29E+07 | 2.28E+07 | 2.17E+07 | 2.22E+07 | 2.27E+07 | 1.96E+07 | 1.88E+07 | 1.98E+07 | 3141.5872 | RTGPAATTLSDTAAAESLVDSSEVTVIGFFK | P09103 | P09103 | P4HB | prolyl 4-hydroxylase, beta polypeptide | Cytoplasm | enzyme |
| 15549425 | 2 | 0.757 | 1.37E+07 | 1.81E+07 | 2.62E+07 | 1.45E+07 | 2.41E+07 | 1.89E+07 | 1.77E+07 | 1.98E+07 | 2.45E+07 | 1.80E+07 | 1.53E+07 | 1.71E+07 | 1666.8301 | VLVGANFEEVAFDEK | P09103 | P09103 | P4HB | prolyl 4-hydroxylase, beta polypeptide | Cytoplasm | enzyme |
| 15540563 | 2 | 0.173 | 1.21E+07 | 1.06E+07 | 1.25E+07 | 1.43E+07 | 1.79E+07 | 1.16E+07 | 1.36E+07 | 1.34E+07 | 1.26E+07 | 1.59E+07 | 1.50E+07 | 1.36E+07 | 1293.5968 | MDSTANEVEAVK | P09103 | P09103 | P4HB | prolyl 4-hydroxylase, beta polypeptide | Cytoplasm | enzyme |
| 15532498 | 2 | 0.414 | 1.19E+07 | 1.17E+07 | 1.51E+07 | 1.39E+07 | 1.80E+07 | 1.25E+07 | 1.49E+07 | 1.58E+07 | 1.38E+07 | 1.43E+07 | 1.70E+07 | 1.48E+07 | 1052.5708 | DHENIIIAK | P09103 | P09103 | P4HB | prolyl 4-hydroxylase, beta polypeptide | Cytoplasm | enzyme |
| 15549871 | 3 | 0.386 | 9352179 | 1.13E+07 | 9299839 | 9316413 | 1.63E+07 | 1.02E+07 | 1.06E+07 | 1.49E+07 | 1.48E+07 | 1.38E+07 | 1.22E+07 | 1.10E+07 | 2985.4866 | TGPAATTLSDTAAAESLVDSSEVTVIGFFK | P09103 | P09103 | P4HB | prolyl 4-hydroxylase, beta polypeptide | Cytoplasm | enzyme |
| 15533068 | 2 | 0.647 | 1.20E+07 | 1.49E+07 | 1.97E+07 | 1.35E+07 | 2.19E+07 | 1.61E+07 | 1.73E+07 | 1.90E+07 | 1.84E+07 | 1.34E+07 | 1.60E+07 | 1.73E+07 | 1202.6029 | EADDIVNWLK | P09103 | P09103 | P4HB | prolyl 4-hydroxylase, beta polypeptide | Cytoplasm | enzyme |
| 15533837 | 3 | 0.262 | 1.35E+07 | 1.08E+07 | 5478012.5 | 1.74E+07 | 1.34E+07 | 8555103 | 1.19E+07 | 1.17E+07 | 1.49E+07 | 1.33E+07 | 2.20E+07 | 1.59E+07 | 1951.9532 | IKPHLMSQEVPEDWDK | P09103 | P09103 | P4HB | prolyl 4-hydroxylase, beta polypeptide | Cytoplasm | enzyme |
| 15533226 | 2, 3 | 0.393 | 8374990 | 9064823 | 1.17E+07 | 8529660 | 1.28E+07 | 1.01E+07 | 9391303 | 1.11E+07 | 1.25E+07 | 1.19E+07 | 1.35E+07 | 1.11E+07 | 1081.6744, 1081.6755 | THILLFLPK | P09103 | P09103 | P4HB | prolyl 4-hydroxylase, beta polypeptide | Cytoplasm | enzyme |
| 15532144 | 2 | 0.17 | 7475810 | 7913210.5 | 1.03E+07 | 8986690 | 1.34E+07 | 9937876 | 1.10E+07 | 1.13E+07 | 1.04E+07 | 1.12E+07 | 1.09E+07 | 1.00E+07 | 970.53345 | QLAPIWDK | P09103 | P09103 | P4HB | prolyl 4-hydroxylase, beta polypeptide | Cytoplasm | enzyme |
| 15557070 | 2 | 0.102 | 8047527.5 | 9160651 | 1.16E+07 | 1.16E+07 | 1.65E+07 | 1.19E+07 | 1.21E+07 | 1.13E+07 | 1.26E+07 | 1.10E+07 | 1.21E+07 | 1.12E+07 | 1780.8323 | VDATEESDLAQQYGVR | P09103 | P09103 | P4HB | prolyl 4-hydroxylase, beta polypeptide | Cytoplasm | enzyme |
| 15532398 | 2 | 0.785 | 7901657 | 8498206 | 1.16E+07 | 1.06E+07 | 1.13E+07 | 1.02E+07 | 7681198.5 | 8809701 | 1.32E+07 | 1.09E+07 | 8803631 | 9871404 | 966.5641 | ILEFFGLK | P09103 | P09103 | P4HB | prolyl 4-hydroxylase, beta polypeptide | Cytoplasm | enzyme |
| 15554685 | 3 | 0.604 | 4831349.5 | 9050509 | 9439862 | 7129735.5 | 1.31E+07 | 7845285.5 | 8452026 | 1.14E+07 | 1.04E+07 | 8396262 | 7844814 | 9344973 | 3626.4543 | FLESGGQDGAGDDEDLDLEEALEPDMEEDDDQK | P09103 | P09103 | P4HB | prolyl 4-hydroxylase, beta polypeptide | Cytoplasm | enzyme |
| 15535231 | 2 | 0.636 | 3963851.25 | 6098413.5 | 6558730.5 | 4784201 | 1.04E+07 | 5151263 | 6912595 | 7549578.5 | 5771798 | 5861632.5 | 4647485 | 5301303.5 | 1051.5038 | NNFEGEITK | P09103 | P09103 | P4HB | prolyl 4-hydroxylase, beta polypeptide | Cytoplasm | enzyme |
| 15543563 | 2 | 0.07 | 3915532 | 3574181.5 | 4375512.5 | 4871191 | 5855402 | 4720604.5 | 5217978 | 4813105.5 | 3726869.75 | 5141784.5 | 5858110 | 4954257.5 | 777.4488 | DGVVLFK | P09103 | P09103 | P4HB | prolyl 4-hydroxylase, beta polypeptide | Cytoplasm | enzyme |
| 15541674 | 3 | 0.656 | 2458322.75 | 3364988 | 3599832.5 | 2253117 | 6054254.5 | 5722184.5 | 3496584.25 | 3999565.75 | 4082466.5 | 3273070.25 | 3369644 | 4440091 | 1780.8315 | VDATEESDLAQQYGVR | P09103 | P09103 | P4HB | prolyl 4-hydroxylase, beta polypeptide | Cytoplasm | enzyme |
| 15537386 | 2 | 0.195 | 8110093 | 1.16E+07 | 4476550 | 8434244 | 8475883 | 5629211.5 | 4635698 | 5847753 | 5769290 | 5558607.5 | 3868665 | 4709881.5 | 1285.6716 | TIIQNPTDQQK | P50580 | P50580 | PA2G4 | proliferation-associated 2G4, 38kDa | Nucleus | transcription regulator |
| 15533403 | 2 | 0.119 | 6303998.5 | 9823311 | 3886995 | 6241401 | 5794168.5 | 5502712 | 3400950 | 4407681.5 | 5295246.5 | 4276062.5 | 2598435.5 | 3759354.25 | 1366.6864 | ITSGPFEPDLYK | P50580 | P50580 | PA2G4 | proliferation-associated 2G4, 38kDa | Nucleus | transcription regulator |
| 15535865 | 2 | 0.215 | 5468390 | 7164956.5 | 2919175.25 | 5650781.5 | 5902587.5 | 3965588.25 | 2761120 | 4053290.25 | 3513548.5 | 4102991.25 | 2557973.75 | 3276865.5 | 1278.5852 | SEMEVQDAELK | P50580 | P50580 | PA2G4 | proliferation-associated 2G4, 38kDa | Nucleus | transcription regulator |
| 15549287 | 2 | 0.152 | 4336471 | 5963771.5 | 2884479.5 | 3913604.25 | 3666225.25 | 3010295 | 2468313.75 | 2841193.5 | 3768010 | 3215390.25 | 2457927.5 | 1978584.75 | 1586.7523 | PGNQNTQVTEAWNK | P50580 | P50580 | PA2G4 | proliferation-associated 2G4, 38kDa | Nucleus | transcription regulator |
| 15552808 | 2 | 0.019 | 3772544.25 | 6231282.5 | 2838972.75 | 3802986.5 | 3978136 | 3235582.5 | 1834508.375 | 2847711 | 3421702.5 | 2002222.75 | 1336703 | 1764733.375 | 1629.8596 | HELLQPFNVLYEK | P50580 | P50580 | PA2G4 | proliferation-associated 2G4, 38kDa | Nucleus | transcription regulator |
| 15554037 | 3 | 0.189 | 4008405 | 7346288.5 | 1734201.75 | 3564099.25 | 2766117.5 | 2349239.75 | 1782897 | 2750708.5 | 2105647.75 | 2707767.25 | 1219102.125 | 1019493.938 | 2740.3823 | ITGMLLEIDNSELLHMLESPESLR | P29341 | P29341 | PABPC1 | poly(A) binding protein, cytoplasmic 1 | Cytoplasm | translation regulator |
| 15551700 | 2 | 0.007 | 5478941 | 7636910 | 3678556.75 | 5427255 | 5200609.5 | 3351771 | 3410345.25 | 3331919.5 | 3401679.25 | 2636947 | 1548713.875 | 1750791.75 | 1662.7439 | GFGFVC[160.0307]FSSPEEATK | P29341 | P29341 | PABPC1 | poly(A) binding protein, cytoplasmic 1 | Cytoplasm | translation regulator |

| | | | | | | | | | | | | | | | | | | | | |
|---|---|---|---|---|---|---|---|---|---|---|---|---|---|---|---|---|---|---|---|---|
| 15542424 | 2 | 0.073 | 3100218.75 | 5813831.5 | 2502886.75 | 3467339.75 | 3796467.5 | 1947742.625 | 1871385.75 | 3140133.25 | 3222616 | 2261363.5 | 1155894.625 | 923934.5 | 1284.6444 | EFSPFGTITSAK | P29341 | P29341 | PABPC1 | poly(A) binding protein, cytoplasmic 1 | Cytoplasm | translation regulator |
| 15552907 | 2 | 0.036 | 3856365 | 5804090 | 2362741 | 3575261.5 | 2383962.25 | 2356724.5 | 2399615.25 | 2154642.25 | 3352335.75 | 2238248.5 | 870084.375 | 680220.938 | 1928.9092 | SLGYAYVNFQQPADAER | P29341 | P29341 | PABPC1 | poly(A) binding protein, cytoplasmic 1 | Cytoplasm | translation regulator |
| 15541408 | 2 | 0.175 | 2289769.25 | 3640888.5 | 1828785.375 | 1900421.125 | 1853871 | 1150170.5 | 1110165.25 | 1743478.75 | 1804143.625 | 1841495.375 | 419548.281 | 1147742.875 | 934.5334 | SGVGNIFIK | P29341 | P29341 | PABPC1 | poly(A) binding protein, cytoplasmic 1 | Cytoplasm | translation regulator |
| 15533755 | 2 | 0.26 | 1.16E+07 | 7002234.5 | 7271545.5 | 9939706 | 6943738.5 | 7673156.5 | 5908163 | 5071342 | 6873857 | 1.21E+07 | 6913788.5 | 7527541.5 | 1246.6406 | HLDLSNVASYK | Q9WVE8 | Q9WVE8 | PACSIN2 | protein kinase C and casein kinase substrate in neurons 2 | Cytoplasm | transporter |
| 15548800 | 2 | 0.44 | 1.04E+07 | 9270209 | 7773547.5 | 9081191 | 7162240 | 5635096.5 | 6762266.5 | 5413414.5 | 6940092.5 | 1.07E+07 | 4499239 | 6946263.5 | 2150.0828 | GRLDSGQVGLYPANYVEAIQ | Q9WVE8 | Q9WVE8 | PACSIN2 | protein kinase C and casein kinase substrate in neurons 2 | Cytoplasm | transporter |
| 15534945 | 2 | 0.262 | 6047072.5 | 4312173 | 4024651 | 4404471 | 4206870 | 3521500.75 | 5209147.5 | 4074450 | 4667838 | 9574715 | 4568928.5 | 5276921 | 1336.6604 | AYAQQLTEWAR | Q9WVE8 | Q9WVE8 | PACSIN2 | protein kinase C and casein kinase substrate in neurons 2 | Cytoplasm | transporter |
| 15532983 | 2 | 0.434 | 5453856 | 4463059.5 | 4297933 | 5748225 | 3531833.5 | 3860778.75 | 4303603 | 3564240.25 | 4725321 | 9326569 | 4188142 | 5115762.5 | 1183.5266 | ASLMNEDFEK | Q9WVE8 | Q9WVE8 | PACSIN2 | protein kinase C and casein kinase substrate in neurons 2 | Cytoplasm | transporter |
| 15537055 | 2 | 0.777 | 5418600.5 | 4549905 | 3685036.75 | 5145009 | 3703165.5 | 3963783.25 | 3523784 | 3641081.25 | 4175311 | 6649998.5 | 3018218.25 | 4609406 | 959.4771 | AADAVEDLR | Q9WVE8 | Q9WVE8 | PACSIN2 | protein kinase C and casein kinase substrate in neurons 2 | Cytoplasm | transporter |
| 15538839 | 2 | 0.627 | 7035634.5 | 5190498 | 3987222.5 | 5510499 | 4132718 | 3613206.5 | 4729370 | 3731913.75 | 3785285.25 | 6242976.5 | 3465209 | 3679071 | 1408.565 | IEDEDEQGWC[160.0307]K | Q9WVE8 | Q9WVE8 | PACSIN2 | protein kinase C and casein kinase substrate in neurons 2 | Cytoplasm | transporter |
| 15533071 | 3 | 0.51 | 1604627 | 1864965.5 | 1763632.375 | 1481168.625 | 1206507.75 | 1666930.875 | 1306407.625 | 1215062.375 | 1643113.25 | 3687123.25 | 1249676.875 | 1601569.5 | 1053.5919 | VSELHLEVK | Q9WVE8 | Q9WVE8 | PACSIN2 | protein kinase C and casein kinase substrate in neurons 2 | Cytoplasm | transporter |
| 15555254 | 2 | 0.433 | 681295.438 | 1837583 | 2323121.5 | 657401.312 | 2698919.25 | 1991565.375 | 1650600.625 | 1487406.875 | 407628.844 | 1127115.375 | 281503.938 | 709377.75 | 2103.0283 | AVDGVTLTGINQTGDQSGQNK | Q9WVE8 | Q9WVE8 | PACSIN2 | protein kinase C and casein kinase substrate in neurons 2 | Cytoplasm | transporter |
| 15550488 | 2 | 0.274 | 692658.188 | 1511482.375 | 2032524.5 | 3371652 | 2967253.75 | 2330634.25 | 2926227.75 | 1582477.625 | 7554673 | 1079500.5 | 2678270.25 | 3305306.5 | 1677.8346 | QLEEDLYDGQVLQK | Q9ES46 | Q9ES46 | PARVB | parvin, beta | Cytoplasm | other |
| 15550086 | 2 | 0.221 | 678735.375 | 1616371.25 | 2947474.75 | 6546995 | 5780288.5 | 6913953.5 | 4865430 | 2647927.25 | 1.69E+07 | 776674.5 | 4315308.5 | 9378706 | 1684.8882 | VLLDWINDVLAEER | Q9ES46 | Q9ES46 | PARVB | parvin, beta | Cytoplasm | other |
| 15550667 | 2 | 0.312 | 397960.219 | 1192122.125 | 1940048.375 | 4343021 | 4576128.5 | 4549859.5 | 3220870.75 | 1810657.75 | 1.08E+07 | 126296.883 | 2932230.5 | 4813432 | 1515.7981 | LNVAEVTQSEIGQK | Q9ES46 | Q9ES46 | PARVB | parvin, beta | Cytoplasm | other |
| 15541585 | 2, 3 | 0.326 | 143826.719 | 727747.812 | 1152667.125 | 2862643.25 | 3766607.5 | 2658162 | 3296091.75 | 1189815 | 9591339 | 64823.355 | 2287491.75 | 5399513.5 | 1368.7451, 1368.7456 | ARPEDVVNLDLK | Q9ES46 | Q9ES46 | PARVB | parvin, beta | Cytoplasm | other |
| 15545444 | 4 | 0.338 | 340087.25 | 1046719.438 | 553302.125 | 2270484.5 | 2840362 | 2588627.75 | 3166851.75 | 1503285.25 | 7512833 | 39041.285 | 2049926.625 | 4158536.5 | 2341.3083 | LQTVLEAVQDLLRPHGWPLR | Q9ES46 | Q9ES46 | PARVB | parvin, beta | Cytoplasm | other |
| 15540140 | 2 | 0.459 | 148417.406 | 965770.5 | 625407.812 | 2264292.75 | 1980331.375 | 2751661.75 | 1742855.875 | 1052498.25 | 6043981 | 4.70E-06 | 1729070.25 | 2838048.5 | 921.5388 | SLITFVNK | Q9ES46 | Q9ES46 | PARVB | parvin, beta | Cytoplasm | other |
| 15556823 | 4 | 0.349 | 2104838.5 | 5478912 | 1555888.875 | 2334675 | 2342261 | 3545963.75 | 1984392.375 | 3528853.25 | 3494824.25 | 1888640 | 1189606.75 | 1882832.5 | 4643.353 | TNIPFLQNVLNNQQFLAGTVDTQFIDENPELFQLRPAQNR | Q05920 | Q05920 | PC | pyruvate carboxylase | Cytoplasm | enzyme |
| 15550253 | 3 | 0.183 | 2778327.25 | 6312765.5 | 2812119.25 | 2750092.5 | 3516524.75 | 3368219.5 | 1786731.875 | 3558690.75 | 5161108.5 | 1701950.375 | 2375047 | 1402967.875 | 2345.206 | LDNASAFQGAVISPHYDSLLVK | Q05920 | Q05920 | PC | pyruvate carboxylase | Cytoplasm | enzyme |
| 15554932 | 3 | 0.242 | 2565854.5 | 4071443.25 | 1474717 | 2699439.75 | 2088577.75 | 3877121 | 1837748 | 2667768 | 3235632 | 1491031.875 | 1331651.375 | 1918594.125 | 2446.2507 | SVVEFLQGYIGIPHGGFPEPFR | Q05920 | Q05920 | PC | pyruvate carboxylase | Cytoplasm | enzyme |
| 15545674 | 2 | 0.573 | 1049931.75 | 1761973.5 | 485453.156 | 1414082.75 | 920321.125 | 1244074.625 | 863768.812 | 1069563.75 | 1085088.625 | 713762.812 | 687417.125 | 929360.688 | 1171.6047 | SSSAPVASPNVR | Q05920 | Q05920 | PC | pyruvate carboxylase | Cytoplasm | enzyme |
| 15553525 | 2 | 0.158 | 1128218.625 | 2305372.75 | 716748.375 | 1275707.5 | 1173607.75 | 1718078.125 | 728740.562 | 1544527.625 | 1828466 | 532138.125 | 693019.25 | 718123.812 | 1564.8009 | AYVEANQMLGDLIK | Q05920 | Q05920 | PC | pyruvate carboxylase | Cytoplasm | enzyme |
| 15540631 | 2 | 0.363 | 4152472.5 | 6020756 | 1970087.625 | 3335072.75 | 2938706.75 | 1725767 | 1805633.375 | 2622912.5 | 2501640.75 | 3200195.5 | 1785305 | 1385207 | 1442.8005 | LVVPATQC[160.0307]GSLIGK | P60335 | P60335 | PCBP1 | poly(rC) binding protein 1 | Nucleus | translation regulator |
| 15554205 | 3 | 0.068 | 4126725.25 | 6841499 | 2364895.5 | 3217412.25 | 3633929.25 | 3012669.5 | 2269483.25 | 3446311.25 | 3153793.25 | 2490461.75 | 1593494 | 1229018.25 | 2489.224 | LEEDINSSMTNSTAASRPPVTLR | P60335 | P60335 | PCBP1 | poly(rC) binding protein 1 | Nucleus | translation regulator |
| 15534128 | 2 | 0.037 | 3730501.5 | 7317042.5 | 3616782.75 | 5383025 | 3369922.25 | 2290150.75 | 2586466 | 2521479.75 | 3708290.75 | 2307923.25 | 934689.312 | 1553723 | 1388.8125 | IITLTGPTNAIFK | P60335 | P60335 | PCBP1 | poly(rC) binding protein 1 | Nucleus | translation regulator |
| 15537499 | 2 | 0.127 | 2661187.75 | 4062064 | 2051756.25 | 2555061.5 | 1962175.5 | 1296088.5 | 1201213.5 | 1842430.5 | 1632452.5 | 1979243.125 | 524131.406 | 1229239.25 | 1086.5515 | IANPVEGSSGR | P60335 | P60335 | PCBP1 | poly(rC) binding protein 1 | Nucleus | translation regulator |
| 15539467 | 2 | 0.048 | 2081920.5 | 3015935.5 | 1714667.625 | 2349609.75 | 1545995 | 1244755.125 | 1243943.875 | 1948573.5 | 1292693.75 | 1425187.125 | 586620.625 | 733570.062 | 1014.5299 | QGANINEIR | P60335 | P60335 | PCBP1 | poly(rC) binding protein 1 | Nucleus | translation regulator |
| 15549408 | 4 | 0.603 | 1.41E+07 | 2.01E+07 | 7512715 | 1.07E+07 | 1.42E+07 | 1.50E+07 | 7138972 | 1.94E+07 | 1.82E+07 | 9264772 | 8996354 | 9480782 | 3758.8418 | SAHLC[160.0307]QPEGIHIC[160.0307]DGTEAENTAILALLEEQGLIR | Q8BH04 | Q8BH04 | PCK2 | phosphoenolpyruvate carboxykinase 2 (mitochondrial) | Cytoplasm | kinase |
| 15548453 | 2 | 0.082 | 1.44E+07 | 2.52E+07 | 1.30E+07 | 1.59E+07 | 1.33E+07 | 2.15E+07 | 1.00E+07 | 1.88E+07 | 2.35E+07 | 6785180.5 | 8977928 | 9998513 | 1643.8967 | LGTPVLQALGDGDFIK | Q8BH04 | Q8BH04 | PCK2 | phosphoenolpyruvate carboxykinase 2 (mitochondrial) | Cytoplasm | kinase |
| 15550071 | 3 | 0.39 | 4940835 | 1.29E+07 | 4105768.5 | 6447016.5 | 7928150.5 | 8186107 | 4637393 | 9632309 | 1.13E+07 | 4767254 | 4260690.5 | 4831979 | 2739.323 | QC[160.0307]PIMDPAWEAPEGVPIDAIIFGGR | Q8BH04 | Q8BH04 | PCK2 | phosphoenolpyruvate carboxykinase 2 (mitochondrial) | Cytoplasm | kinase |
| 15535246 | 3 | 0.103 | 8539541 | 1.61E+07 | 6571183 | 1.07E+07 | 1.18E+07 | 1.38E+07 | 7362145.5 | 1.14E+07 | 1.42E+07 | 4211675 | 7615678.5 | 6022867.5 | 1939.0084 | RLEGEDSAQETPIGLVPK | Q8BH04 | Q8BH04 | PCK2 | phosphoenolpyruvate carboxykinase 2 (mitochondrial) | Cytoplasm | kinase |
| 15542632 | 4 | 0.521 | 7060427.5 | 9531939 | 4154886.75 | 7228090 | 6293895.5 | 7856483 | 5403051.5 | 9376572 | 8862497 | 3666900.75 | 8452639 | 4142047 | 2770.2778 | C[160.0307]LHSVGQPLTGHGDPVGQWPC[160.0307]NPEK | Q8BH04 | Q8BH04 | PCK2 | phosphoenolpyruvate carboxykinase 2 (mitochondrial) | Cytoplasm | kinase |
| 15538283 | 2 | 0.144 | 5924593.5 | 1.46E+07 | 4157955.25 | 7096843.5 | 7764427 | 1.19E+07 | 5135103.5 | 9334762 | 1.04E+07 | 2892317.75 | 3720855.75 | 4988324 | 1328.7042 | EVLAELEALEGR | Q8BH04 | Q8BH04 | PCK2 | phosphoenolpyruvate carboxykinase 2 (mitochondrial) | Cytoplasm | kinase |
| 15536469 | 2 | 0.152 | 4868398 | 1.05E+07 | 4079239.75 | 4434262 | 5605025 | 8857336 | 4097113.75 | 6332133 | 8889695 | 2148582 | 3555892.75 | 3632505.5 | 1069.5979 | DTVPLLAGGAR | Q8BH04 | Q8BH04 | PCK2 | phosphoenolpyruvate carboxykinase 2 (mitochondrial) | Cytoplasm | kinase |
| 15542068 | 2 | 0.19 | 3731609.25 | 7657659.5 | 2799257.5 | 4435601.5 | 3885758 | 5952864 | 2745155.25 | 4979694 | 6643485.5 | 2079954.875 | 2706252.5 | 2665283 | 1450.7457 | GVPLVYEAFNWR | Q8BH04 | Q8BH04 | PCK2 | phosphoenolpyruvate carboxykinase 2 (mitochondrial) | Cytoplasm | kinase |
| 15534610 | 2 | 0.189 | 2869451 | 7081557.5 | 2050964.625 | 4545853.5 | 3292684.5 | 5283199 | 3386358.25 | 4777606 | 5240910 | 2048084.125 | 2261631.25 | 2610024 | 1450.6421 | VEC[160.0307]VGDDIAWMR | Q8BH04 | Q8BH04 | PCK2 | phosphoenolpyruvate carboxykinase 2 (mitochondrial) | Cytoplasm | kinase |
| 15549096 | 2 | 0.039 | 4374189 | 8040509.5 | 4159751.25 | 4693719.5 | 4703952.5 | 6539428.5 | 3561965.25 | 5768594 | 5466726 | 2042939.25 | 2715368.25 | 3282527.75 | 1664.7429 | GQLGNWMSPDEFQR | Q8BH04 | Q8BH04 | PCK2 | phosphoenolpyruvate carboxykinase 2 (mitochondrial) | Cytoplasm | kinase |
| 15533593 | 2 | 0.086 | 4089749.5 | 8720999 | 3014023 | 5922499.5 | 4926809 | 7342014 | 3034130.5 | 5866674 | 7427005.5 | 1841099.5 | 2897616 | 2619888.75 | 1000.5762 | TVIVTPSQR | Q8BH04 | Q8BH04 | PCK2 | phosphoenolpyruvate carboxykinase 2 (mitochondrial) | Cytoplasm | kinase |
| 15537111 | 3 | 0.321 | 2121963.75 | 6818557 | 1920470.5 | 1939304.125 | 3456682.5 | 3629722.25 | 2330362.5 | 4989357 | 2641224.25 | 1797953.75 | 1355406.75 | 1834185.5 | 2165.0999 | DEGWLAEHMLILGITNPAGK | Q8BH04 | Q8BH04 | PCK2 | phosphoenolpyruvate carboxykinase 2 (mitochondrial) | Cytoplasm | kinase |
| 15554190 | 2 | 0.073 | 6598475.5 | 9595801 | 2749528.25 | 5099466.5 | 4813372 | 7877393.5 | 4568582 | 7016807 | 7153986.5 | 1779396 | 2805932.25 | 3099261.75 | 2248.1707 | EGALDLSGLSAVDTSQLFSIPK | Q8BH04 | Q8BH04 | PCK2 | phosphoenolpyruvate carboxykinase 2 (mitochondrial) | Cytoplasm | kinase |
| 15552506 | 2 | 0.042 | 3656037.25 | 9355887 | 3670919.25 | 4306391 | 5073244 | 8469663 | 3788227.5 | 5585524.5 | 7729427 | 1706637.5 | 2798647 | 1940109.5 | 1684.8483 | EIVSFGSGYGGNSLLGK | Q8BH04 | Q8BH04 | PCK2 | phosphoenolpyruvate carboxykinase 2 (mitochondrial) | Cytoplasm | kinase |
| 15543608 | 2 | 0.027 | 2578244 | 6074095.5 | 2410772.5 | 4101356.5 | 3470137.5 | 5433162 | 2916971.5 | 4263723 | 4794750 | 1591140.5 | 1956391.625 | 1572064.875 | 951.4513 | FDSEGQLR | Q8BH04 | Q8BH04 | PCK2 | phosphoenolpyruvate carboxykinase 2 (mitochondrial) | Cytoplasm | kinase |
| 15540475 | 2 | 0.16 | 2376325 | 3622576.75 | 1704693.75 | 3105178.75 | 2358556.75 | 3707043.25 | 2498353.5 | 3645602 | 3903291.5 | 1475595.5 | 1931705.375 | 2424313.25 | 961.4901 | VLDWIC[160.0307]R | Q8BH04 | Q8BH04 | PCK2 | phosphoenolpyruvate carboxykinase 2 (mitochondrial) | Cytoplasm | kinase |
| 15540581 | 2 | 0.463 | 2726747.25 | 4314155 | 791291.75 | 2314298.25 | 1804108.25 | 2225950.75 | 1721255.25 | 2472459.25 | 2475203 | 762292.812 | 1633976.375 | 1451806 | 1060.5323 | HGVFVGSAMR | Q8BH04 | Q8BH04 | PCK2 | phosphoenolpyruvate carboxykinase 2 (mitochondrial) | Cytoplasm | kinase |
| 15533868 | 2 | 0.093 | 9735365 | 1.17E+07 | 1.18E+07 | 6911731 | 8784289 | 8224500 | 8786581 | 1.01E+07 | 9336762 | 1.19E+07 | 8394316 | 8504163 | 778.4079 | IDVFER | Q9CQF9 | Q9CQF9 | PCYOX1 | prenylcysteine oxidase 1 | Cytoplasm | enzyme |
| 15540737 | 2 | 0.228 | 1.49E+07 | 1.12E+07 | 1.31E+07 | 1.52E+07 | 8734271 | 1.12E+07 | 1.43E+07 | 9887469 | 1.28E+07 | 1.49E+07 | 1.71E+07 | 1.58E+07 | 1015.5765 | RDEEILLK | P27664 | P27664 | PDE6A | phosphodiesterase 6A, cGMP-specific, rod, alpha | Plasma Membrane | enzyme |
| 15542599 | 2 | 0.401 | 949672.25 | 2271248.75 | 952583 | 1148707.5 | 2020520.5 | 1760111.125 | 911735.125 | 1986885.125 | 1270318.625 | 1014873.75 | 980904.125 | 861971.688 | 1433.7263 | MVNSNLASVEELK | P35486 | P35486 | PDHA1 | pyruvate dehydrogenase (lipoamide) alpha 1 | Cytoplasm | enzyme |
| 15536768 | 2 | 0.112 | 1318668.625 | 2451330.25 | 1034299.875 | 1859658.75 | 1804157.375 | 1982430.25 | 1323088.625 | 1534406.5 | 2209444.5 | 631532.375 | 1091990.5 | 1214376 | 943.5548 | AILAELTGR | P35486 | P35486 | PDHA1 | pyruvate dehydrogenase (lipoamide) alpha 1 | Cytoplasm | enzyme |
| 15542279 | 2 | 0.476 | 1281485.625 | 2937063.25 | 1151727.25 | 1617933.875 | 2221313.75 | 2109603 | 757588.125 | 2181687 | 3374625.25 | 274702.844 | 1662304.25 | 1363426 | 1320.6931 | GPILMELQTYR | P35486 | P35486 | PDHA1 | pyruvate dehydrogenase (lipoamide) alpha 1 | Cytoplasm | enzyme |
| 15543260 | 2 | 0.176 | 1433039.625 | 3650322 | 2068962.375 | 3465574.5 | 3017368.25 | 3583440.5 | 1619890.75 | 3092157 | 4825827 | 256243.281 | 2145890.25 | 1507736 | 1227.6543 | VLEDNSVPQVK | Q9D051 | Q9D051 | PDHB | pyruvate dehydrogenase (lipoamide) beta | Cytoplasm | enzyme |
| 15548668 | 2, 3 | 0.105 | 6.57E+07 | 1.02E+08 | 9.53E+07 | 1.27E+08 | 1.34E+08 | 1.17E+08 | 1.11E+08 | 1.25E+08 | 1.13E+08 | 1.04E+08 | 7.89E+07 | 1.23E+08 | 2591.2932, 2591.2952 | TFSHELSDFGLESTTGEVPVVAIR | P27773 | P27773 | PDIA3 | protein disulfide isomerase family A, member 3 | Cytoplasm | peptidase |
| 15547807 | 2 | 0.055 | 4.03E+07 | 4.82E+07 | 5.12E+07 | 5.51E+07 | 6.99E+07 | 5.43E+07 | 5.44E+07 | 5.61E+07 | 5.47E+07 | 4.75E+07 | 4.12E+07 | 5.11E+07 | 1834.8672 | VVVAENFDDIVNEEDK | P27773 | P27773 | PDIA3 | protein disulfide isomerase family A, member 3 | Cytoplasm | peptidase |
| 15533286 | 2 | 0.146 | 2.96E+07 | 3.57E+07 | 4.18E+07 | 4.31E+07 | 5.84E+07 | 4.21E+07 | 3.95E+07 | 4.80E+07 | 4.18E+07 | 4.14E+07 | 3.40E+07 | 3.92E+07 | 1382.6766 | SEPIPESNEGPVK | P27773 | P27773 | PDIA3 | protein disulfide isomerase family A, member 3 | Cytoplasm | peptidase |
| 15548903 | 2 | 0.135 | 1.98E+07 | 2.17E+07 | 2.98E+07 | 2.77E+07 | 3.53E+07 | 3.00E+07 | 2.64E+07 | 2.85E+07 | 2.81E+07 | 2.77E+07 | 2.21E+07 | 2.37E+07 | 1636.748 | MDATANDVPSPYEVK | P27773 | P27773 | PDIA3 | protein disulfide isomerase family A, member 3 | Cytoplasm | peptidase |
| 15539517 | 2 | 0.246 | 1.64E+07 | 2.18E+07 | 2.45E+07 | 2.22E+07 | 3.09E+07 | 2.93E+07 | 2.73E+07 | 2.79E+07 | 2.48E+07 | 2.77E+07 | 1.82E+07 | 2.38E+07 | 1172.5381 | FVMQEEFSR | P27773 | P27773 | PDIA3 | protein disulfide isomerase family A, member 3 | Cytoplasm | peptidase |
| 15534689 | 3 | 0.005 | 2.30E+07 | 2.18E+07 | 2.33E+07 | 3.71E+07 | 2.98E+07 | 3.07E+07 | 3.15E+07 | 2.80E+07 | 2.72E+07 | 2.72E+07 | 2.67E+07 | 3.07E+07 | 1652.7639 | IFRDGEEAGAYDGPR | P27773 | P27773 | PDIA3 | protein disulfide isomerase family A, member 3 | Cytoplasm | peptidase |
| 15534257 | 2 | 0.405 | 1.66E+07 | 2.11E+07 | 2.66E+07 | 2.35E+07 | 2.58E+07 | 2.45E+07 | 2.54E+07 | 2.63E+07 | 2.53E+07 | 2.42E+07 | 1.54E+07 | 2.41E+07 | 1397.7025 | ELNDFISYLQR | P27773 | P27773 | PDIA3 | protein disulfide isomerase family A, member 3 | Cytoplasm | peptidase |
| 15536626 | 2 | 0.434 | 1.37E+07 | 2.43E+07 | 3.18E+07 | 2.26E+07 | 3.52E+07 | 2.84E+07 | 2.75E+07 | 2.98E+07 | 2.28E+07 | 2.41E+07 | 1.42E+07 | 2.21E+07 | 1368.697 | EATNPPIIQEEK | P27773 | P27773 | PDIA3 | protein disulfide isomerase family A, member 3 | Cytoplasm | peptidase |
| 15548508 | 2 | 0.231 | 2.23E+07 | 2.21E+07 | 3.18E+07 | 2.47E+07 | 3.72E+07 | 3.60E+07 | 2.93E+07 | 3.12E+07 | 3.21E+07 | 2.35E+07 | 1.84E+07 | 3.03E+07 | 1814.8938 | ALEQFLQEYFDGNLK | P27773 | P27773 | PDIA3 | protein disulfide isomerase family A, member 3 | Cytoplasm | peptidase |
| 15532063 | 3 | 0.476 | 1.47E+07 | 1.82E+07 | 2.52E+07 | 1.78E+07 | 3.02E+07 | 2.30E+07 | 2.50E+07 | 2.26E+07 | 2.12E+07 | 2.05E+07 | 1.64E+07 | 2.01E+07 | 1758.8976 | QAGPASVPLRTEEEFK | P27773 | P27773 | PDIA3 | protein disulfide isomerase family A, member 3 | Cytoplasm | peptidase |
| 15532172 | 3 | 0.637 | 1.43E+07 | 1.49E+07 | 2.20E+07 | 1.87E+07 | 2.00E+07 | 1.96E+07 | 1.82E+07 | 2.28E+07 | 1.66E+07 | 2.00E+07 | 1.74E+07 | 1.94E+07 | 1258.6769 | FAHTNIESLVK | P27773 | P27773 | PDIA3 | protein disulfide isomerase family A, member 3 | Cytoplasm | peptidase |
| 15542908 | 2 | 0.147 | 1.41E+07 | 1.66E+07 | 1.95E+07 | 2.03E+07 | 2.25E+07 | 2.02E+07 | 1.96E+07 | 2.05E+07 | 2.05E+07 | 1.89E+07 | 1.63E+07 | 2.20E+07 | 1341.681 | GFPTIYFSPANK | P27773 | P27773 | PDIA3 | protein disulfide isomerase family A, member 3 | Cytoplasm | peptidase |
| 15534416 | 3 | 0.145 | 1.12E+07 | 1.37E+07 | 1.91E+07 | 1.72E+07 | 2.37E+07 | 2.27E+07 | 1.93E+07 | 2.17E+07 | 2.22E+07 | 1.82E+07 | 1.43E+07 | 2.28E+07 | 1814.893 | ALEQFLQEYFDGNLK | P27773 | P27773 | PDIA3 | protein disulfide isomerase family A, member 3 | Cytoplasm | peptidase |
| 15533257 | 2 | 0.105 | 1.17E+07 | 1.30E+07 | 1.24E+07 | 1.54E+07 | 1.75E+07 | 1.30E+07 | 1.58E+07 | 1.51E+07 | 1.32E+07 | 1.41E+07 | 1.35E+07 | 1.40E+07 | 1347.6993 | RLAPEYEAAATR | P27773 | P27773 | PDIA3 | protein disulfide isomerase family A, member 3 | Cytoplasm | peptidase |
| 15535272 | 2 | 0.184 | 9948543 | 1.24E+07 | 1.67E+07 | 1.49E+07 | 1.91E+07 | 1.61E+07 | 1.44E+07 | 1.55E+07 | 1.57E+07 | 1.40E+07 | 1.02E+07 | 1.33E+07 | 1040.5709 | TADGIVSHLK | P27773 | P27773 | PDIA3 | protein disulfide isomerase family A, member 3 | Cytoplasm | peptidase |
| 15531733 | 3 | 0.012 | 1.11E+07 | 1.11E+07 | 1.14E+07 | 1.52E+07 | 1.75E+07 | 1.27E+07 | 1.52E+07 | 1.72E+07 | 1.75E+07 | 1.38E+07 | 1.87E+07 | 1.67E+07 | 2151.9822 | FIQDSIFGLC[160.0307]PHMTEDNK | P27773 | P27773 | PDIA3 | protein disulfide isomerase family A, member 3 | Cytoplasm | peptidase |
| 15542247 | 2 | 0.11 | 9770972 | 1.40E+07 | 1.64E+07 | 1.96E+07 | 1.79E+07 | 1.89E+07 | 1.49E+07 | 1.63E+07 | 1.80E+07 | 1.32E+07 | 1.16E+07 | 1.69E+07 | 997.5076 | DASVVGFFR | P27773 | P27773 | PDIA3 | protein disulfide isomerase family A, member 3 | Cytoplasm | peptidase |
| 15541799 | 2 | 0.462 | 7221893 | 8699390 | 1.17E+07 | 1.02E+07 | 1.26E+07 | 1.11E+07 | 1.10E+07 | 1.18E+07 | 1.14E+07 | 1.22E+07 | 6980985 | 1.11E+07 | 877.4872 | LNFAVASR | P27773 | P27773 | PDIA3 | protein disulfide isomerase family A, member 3 | Cytoplasm | peptidase |
| 15554658 | 2 | 0.743 | 7783733.5 | 9204522 | 1.41E+07 | 9481311 | 1.15E+07 | 1.13E+07 | 1.10E+07 | 9836008 | 1.25E+07 | 1.05E+07 | 6518289 | 1.11E+07 | 1607.7446 | DLLTAYYDVDYEK | P27773 | P27773 | PDIA3 | protein disulfide isomerase family A, member 3 | Cytoplasm | peptidase |
| 15557393, 15559261 | 3 | 0.072 | 5983099 | 1.30E+07 | 1.28E+07 | 8451800 | 1.75E+07 | 1.29E+07 | 1.08E+07 | 1.54E+07 | 1.34E+07 | 5219584 | 4589183 | 8433058 | 2302.1423 | EYDDNGEGITIFRPLHLANK | P27773 | P27773 | PDIA3 | protein disulfide isomerase family A, member 3 | Cytoplasm | peptidase |
| 15553265 | 2 | 0.743 | 3280305.25 | 5327110 | 5356289 | 3750391 | 7305459.5 | 5211367.5 | 5092454 | 5312138 | 4127462 | 5118775.5 | 3464734.5 | 4157229.25 | 1758.8983 | QAGPASVPLRTEEEFK | P27773 | P27773 | PDIA3 | protein disulfide isomerase family A, member 3 | Cytoplasm | peptidase |
| 15553163 | 2 | 0.361 | 2400056.5 | 2052223 | 1000370.5 | 2964163.25 | 2358107.75 | 1551000.625 | 2712821 | 1645921.625 | 1330230.25 | 3352215.75 | 2801329.75 | 2333585 | 1652.7645 | IFRDGEEAGAYDGPR | P27773 | P27773 | PDIA3 | protein disulfide isomerase family A, member 3 | Cytoplasm | peptidase |
| 15545742 | 2 | 0.302 | 1067860.875 | 1469471 | 2979668.5 | 1957151.5 | 4445136.5 | 2513350 | 2304899.25 | 3128591 | 1710379.25 | 2144656 | 626591.25 | 1640612.625 | 1236.5115 | DGEEAGAYDGPR | P27773 | P27773 | PDIA3 | protein disulfide isomerase family A, member 3 | Cytoplasm | peptidase |
| 15551380 | 3 | 0.049 | 5789457 | 7500064 | 7589045 | 1.17E+07 | 8489337 | 8949751 | 8385551.5 | 6695031 | 8926298 | 8996315 | 1.15E+07 | 1.00E+07 | 2379.251 | VREVSQPDWTPPPEVTLSLTK | P08003 | P08003 | PDIA4 | protein disulfide isomerase family A, member 4 | Cytoplasm | enzyme |
| 15549576 | 3 | 0.066 | 2630915 | 3273159.5 | 4967442 | 6094086 | 4622253.5 | 6612993.5 | 6271170.5 | 6019432 | 4472352.5 | 4309579.5 | 4261178.5 | 4576227 | 2509.154 | FAMEPEEFDSDTLREFVTAFK | P08003 | P08003 | PDIA4 | protein disulfide isomerase family A, member 4 | Cytoplasm | enzyme |
| 15555547 | 2 | 0.777 | 2120604.75 | 3137074.25 | 4407467 | 2962935.5 | 4732286.5 | 3474610.75 | 3434570.75 | 4185113.5 | 3472788.75 | 4017307.75 | 3060385 | 2996516.25 | 1599.6936 | MDATANDITNDQYK | P08003 | P08003 | PDIA4 | protein disulfide isomerase family A, member 4 | Cytoplasm | enzyme |
| 15533240 | 2 | 0.059 | 2659852.75 | 3077415.25 | 3266958.25 | 4216674 | 4333744.5 | 3172290.5 | 4627351 | 3717034.5 | 3513528.25 | 3940981.75 | 4209448.5 | 4045443 | 1248.6793 | QLEPIYTSLGK | P08003 | P08003 | PDIA4 | protein disulfide isomerase family A, member 4 | Cytoplasm | enzyme |
| 15535126 | 2 | 0.041 | 1966720.25 | 2608952.75 | 3396878.25 | 3546026.25 | 4619792.5 | 4222181 | 3541941.5 | 3460991 | 3515487.5 | 3393114.25 | 3961788.25 | 3365168 | 1194.6008 | IDATSASMLASK | P08003 | P08003 | PDIA4 | protein disulfide isomerase family A, member 4 | Cytoplasm | enzyme |
| 15535692 | 2 | 0.089 | 1859530 | 2629531 | 2665037.5 | 3076404.75 | 3135879.5 | 2908890 | 2602419.25 | 3282776.5 | 2511203.25 | 3381279 | 3741727.5 | 2870026.25 | 978.6067 | RSPPIPLAK | P08003 | P08003 | PDIA4 | protein disulfide isomerase family A, member 4 | Cytoplasm | enzyme |
| 15538720 | 3 | 0.132 | 1403319.5 | 1948796.875 | 2530618.75 | 2407984 | 3437261.75 | 3007703.5 | 2406137.25 | 2813647.25 | 2384039.25 | 2183269 | 3144347.75 | 2812164.75 | 1719.8333 | FHVMDVQGSTEASAIK | P08003 | P08003 | PDIA4 | protein disulfide isomerase family A, member 4 | Cytoplasm | enzyme |
| 15542407 | 2 | 0.305 | 1113261.75 | 1444044.125 | 1985686.125 | 1650012.875 | 2190207.75 | 1934451.125 | 1721300.5 | 1788949.25 | 1763383.875 | 1723291.5 | 2259982.75 | 1781268.25 | 1039.5394 | DNDPPIAVAK | P08003 | P08003 | PDIA4 | protein disulfide isomerase family A, member 4 | Cytoplasm | enzyme |
| 15554566 | 3 | 0.082 | 1.06E+07 | 1.84E+07 | 2.21E+07 | 2.00E+07 | 2.53E+07 | 2.82E+07 | 2.58E+07 | 2.75E+07 | 2.23E+07 | 1.84E+07 | 1.63E+07 | 1.64E+07 | 2637.3606 | ALDLFSDNAPPPELLEIINEDIAK | Q922R8 | Q922R8 | PDIA6 | protein disulfide isomerase family A, member 6 | Cytoplasm | enzyme |
| 15548347 | 2 | 0.071 | 8995234 | 1.14E+07 | 1.57E+07 | 1.48E+07 | 1.86E+07 | 1.62E+07 | 1.66E+07 | 1.73E+07 | 1.53E+07 | 1.34E+07 | 1.36E+07 | 1.23E+07 | 1528.7612 | NLEPEWAAAATEVK | Q922R8 | Q922R8 | PDIA6 | protein disulfide isomerase family A, member 6 | Cytoplasm | enzyme |
| 15548163 | 2 | 0.031 | 6477170 | 9337911 | 1.24E+07 | 1.28E+07 | 1.60E+07 | 1.37E+07 | 1.34E+07 | 1.30E+07 | 1.33E+07 | 9328958 | 9599711 | 1.00E+07 | 1601.8246 | GSTAPVGGGSFPTITPR | Q922R8 | Q922R8 | PDIA6 | protein disulfide isomerase family A, member 6 | Cytoplasm | enzyme |
| 15533549 | 3 | 0.067 | 6433384 | 9216801 | 9894989 | 1.37E+07 | 1.32E+07 | 1.13E+07 | 1.19E+07 | 1.14E+07 | 9118533 | 8734592 | 1.11E+07 | 1.01E+07 | 1816.9298 | HQSLGGQYGVQGFPTIK | Q922R8 | Q922R8 | PDIA6 | protein disulfide isomerase family A, member 6 | Cytoplasm | enzyme |
| 15536515 | 2 | 0.025 | 5938777 | 7305658 | 1.02E+07 | 1.19E+07 | 1.36E+07 | 1.22E+07 | 1.08E+07 | 1.19E+07 | 1.10E+07 | 8380281 | 9862406 | 1.14E+07 | 1483.7164 | GSFSEQGINEFLR | Q922R8 | Q922R8 | PDIA6 | protein disulfide isomerase family A, member 6 | Cytoplasm | enzyme |
| 15550223 | 2 | 0.482 | 6893692 | 1.08E+07 | 4292446 | 1.57E+07 | 7074409 | 1.25E+07 | 4881400.5 | 5812503 | 1.43E+07 | 8230128 | 1.24E+07 | 1.04E+07 | 1527.8485 | LAAVDATVNQVLASR | Q922R8 | Q922R8 | PDIA6 | protein disulfide isomerase family A, member 6 | Cytoplasm | enzyme |
| 15533706 | 2 | 0.098 | 6715932.5 | 7445878 | 1.05E+07 | 9470693 | 1.52E+07 | 1.36E+07 | 1.20E+07 | 1.19E+07 | 1.05E+07 | 8158716.5 | 1.08E+07 | 1.09E+07 | 1396.6449 | DVVELTDDTFDK | Q922R8 | Q922R8 | PDIA6 | protein disulfide isomerase family A, member 6 | Cytoplasm | enzyme |
| 15533521 | 2 | 0.12 | 6674844 | 7596245 | 1.15E+07 | 1.18E+07 | 1.25E+07 | 1.42E+07 | 1.08E+07 | 1.28E+07 | 1.07E+07 | 6725224 | 1.06E+07 | 1.09E+07 | 1386.7562 | TGEAIVDAALSALR | Q922R8 | Q922R8 | PDIA6 | protein disulfide isomerase family A, member 6 | Cytoplasm | enzyme |
| 15555677 | 2 | 0.306 | 3453201.75 | 6698609.5 | 9644262 | 6058089 | 1.07E+07 | 1.04E+07 | 6295664.5 | 1.03E+07 | 9859184 | 4557269 | 4265135.5 | 7562176 | 2473.1304 | DGELPVEDDIDLSDVELDDLEK | Q922R8 | Q922R8 | PDIA6 | protein disulfide isomerase family A, member 6 | Cytoplasm | enzyme |
| 15537905 | 4 | 0.138 | 3696826.25 | 4097454 | 3451972.25 | 5114922.5 | 7032101 | 5317511 | 5745650.5 | 6639275 | 6356167.5 | 3849117 | 1.00E+07 | 5554326 | 2646.3071 | TC[160.0307]EEHQLC[160.0307]VVAVLPHILDTGAAGR | Q922R8 | Q922R8 | PDIA6 | protein disulfide isomerase family A, member 6 | Cytoplasm | enzyme |
| 15542341 | 2 | 0.144 | 2582237.75 | 3363661.75 | 4687703 | 4897488.5 | 5796385.5 | 4915449 | 4509112.5 | 4691144.5 | 4929405 | 3096696 | 4812942.5 | 4611819.5 | 1078.6122 | NSYLEVLLK | Q922R8 | Q922R8 | PDIA6 | protein disulfide isomerase family A, member 6 | Cytoplasm | enzyme |
| 15534630 | 2 | 0.032 | 3.52E+07 | 4.19E+07 | 6.28E+07 | 1.14E+08 | 1.28E+08 | 1.26E+08 | 1.07E+08 | 9.13E+07 | 1.61E+08 | 3.14E+07 | 1.07E+08 | 7.33E+07 | 1039.6125 | HITSLEVIK | Q9Z126 | Q9Z126 | PF4 | platelet factor 4 | Extracellular Space | cytokine |
| 15533135 | 2, 3 | 0.079 | 3.23E+07 | 2.91E+07 | 3.11E+07 | 9.49E+07 | 7.89E+07 | 9.69E+07 | 7.70E+07 | 7.05E+07 | 1.20E+08 | 2.49E+07 | 1.50E+08 | 5.87E+07 | 1350.753, 1350.7539 | HC[160.0307]AVPQLIATLK | Q9Z126 | Q9Z126 | PF4 | platelet factor 4 | Extracellular Space | cytokine |
| 15531557 | 2 | 0.033 | 1.70E+07 | 1.90E+07 | 2.83E+07 | 4.50E+07 | 4.95E+07 | 5.65E+07 | 4.43E+07 | 4.23E+07 | 6.31E+07 | 1.31E+07 | 4.34E+07 | 2.82E+07 | 955.5552 | TISSGIHLK | Q9Z126 | Q9Z126 | PF4 | platelet factor 4 | Extracellular Space | cytokine |
| 15533857 | 2 | 0.159 | 3166736.75 | 4202366 | 3405485 | 2471148.25 | 2809863.5 | 2352370 | 2278612.5 | 2614568.5 | 1953714.625 | 3098677.25 | 751746.312 | 1821286.625 | 1189.6798 | VVAFSPVTELK | P12382 | P12382 | PFKL | phosphofructokinase, liver | Cytoplasm | kinase |
| 15540833 | 3 | 0.658 | 7543725 | 5800610 | 4884823 | 3423432.25 | 5242638.5 | 3466259.25 | 6437381.5 | 6033813.5 | 3563112.75 | 9579114 | 2520473.5 | 3718070.75 | 1840.8716 | NVLGHMQQGGSPTPFDR | P47857 | P47857 | PFKM | phosphofructokinase, muscle | Cytoplasm | kinase |
| 15552008 | 2 | 0.377 | 4865829.5 | 2569273.25 | 2818387.25 | 2667754 | 2726985.75 | 2483338.75 | 3278172 | 2631848.5 | 3603084.75 | 5783861 | 3174971.75 | 3202775.25 | 1637.7761 | IFANTPDSGC[160.0307]VLGMR | P47857 | P47857 | PFKM | phosphofructokinase, muscle | Cytoplasm | kinase |
| 15536777 | 2 | 0.42 | 4781795.5 | 3606542.5 | 3344086.25 | 2355797 | 2886184.5 | 2176624.5 | 4085391.75 | 2641326.25 | 2837512 | 5533937.5 | 2042467.75 | 2469847.25 | 1421.7717 | DLQVNVEHLVQK | P47857 | P47857 | PFKM | phosphofructokinase, muscle | Cytoplasm | kinase |
| 15553226 | 3 | 0.572 | 4994111 | 3165717 | 3281293.25 | 2608076.25 | 2853994.25 | 2846134.5 | 3226895 | 3655084.5 | 3072300.5 | 5398756 | 2408501.75 | 2842258.25 | 2280.135 | RGITNLC[160.0307]VIGGDGSLTGADTFR | P47857 | P47857 | PFKM | phosphofructokinase, muscle | Cytoplasm | kinase |
| 15552082 | 2 | 0.332 | 3304506.75 | 2482973.75 | 2317774.25 | 1975478.25 | 2205549 | 1396223.75 | 1802519.375 | 1617652.875 | 2008540.875 | 4031149.75 | 1711294.125 | 1820310.375 | 2153.983 | GQIEEAGWSYVGGWTGQGGSK | P47857 | P47857 | PFKM | phosphofructokinase, muscle | Cytoplasm | kinase |
| 15537683 | 2 | 0.259 | 3702426.75 | 3226402 | 3542895 | 2177655.75 | 2622653.25 | 2613736.25 | 4409354 | 3088378 | 2886987.75 | 4016696 | 1682038.625 | 2352571 | 1447.703 | SEWSDLLNDLQK | P47857 | P47857 | PFKM | phosphofructokinase, muscle | Cytoplasm | kinase |
| 15554866 | 3 | 0.348 | 3783725 | 2560753.75 | 3925882.75 | 4228926 | 3084118 | 3917102 | 3069281 | 2741838 | 2310292 | 3947451.25 | 1581356.75 | 2578395.75 | 2324.1528 | DAFERNPELQNLLLDDFFK | Q9DCD0 | Q9DCD0 | PGD | phosphogluconate dehydrogenase | Cytoplasm | enzyme |
| 15556404, 15559320 | 3 | 0.723 | 3768220.5 | 3075458 | 4134103 | 6001659 | 4494147.5 | 4851743.5 | 3118293.75 | 3661429.25 | 7178121 | 2351876.25 | 7615170 | 2919543.25 | 2417.1245, 2417.1282 | IVRGDQPGASGDNDDDEPPPLPR | O55022 | O55022 | PGRMC1 | progesterone receptor membrane component 1 | Plasma Membrane | transmembrane receptor |
| 15541085 | 2 | 0.794 | 1711968.375 | 2707831.75 | 3185168.75 | 2441065.25 | 2704707.75 | 2624746.25 | 2345972 | 2309242.75 | 3679984.25 | 2049852.25 | 2135462.5 | 2688610.75 | 1472.734 | FYGPAGPYGIFAGR | Q80UU9 | Q80UU9 | PGRMC2 | progesterone receptor membrane component 2 | Nucleus | other |
| 15539339 | 2 | 0.012 | 8220538.5 | 1.37E+07 | 9122335 | 9280761 | 9813682 | 1.06E+07 | 7894355.5 | 1.04E+07 | 1.17E+07 | 6080216 | 5732278 | 5712997 | 1061.545 | AAIISAEGDSK | P67778 | P67778 | PHB | prohibitin | Nucleus | transcription regulator |
| 15537294 | 2 | 0.092 | 8217268.5 | 2.15E+07 | 8929999 | 7285524.5 | 1.09E+07 | 1.59E+07 | 7074676 | 1.40E+07 | 1.43E+07 | 3975310.25 | 4898056.5 | 6350039 | 1213.7385 | VLPSITTEILK | P67778 | P67778 | PHB | prohibitin | Nucleus | transcription regulator |
| 15533353 | 3 | 0.062 | 6590305.5 | 1.19E+07 | 5800795.5 | 9011183 | 7824277.5 | 1.01E+07 | 5898061 | 8706809 | 1.18E+07 | 3701790.75 | 4961818.5 | 5081187.5 | 1396.8395 | ILFRPVASQLPR | P67778 | P67778 | PHB | prohibitin | Nucleus | transcription regulator |
| 15534255 | 3 | 0.257 | 5089009.5 | 1.38E+07 | 6669019 | 5434619 | 1.19E+07 | 1.10E+07 | 5945135.5 | 1.26E+07 | 1.13E+07 | 3552802 | 5432167 | 5791219.5 | 2222.1917 | GVQDIVVGEGTHFLIPWVQK | P67778 | P67778 | PHB | prohibitin | Nucleus | transcription regulator |
| 15535438 | 2 | 0.091 | 6422084 | 1.45E+07 | 6150707 | 5678759 | 8668670 | 9752101 | 4954872.5 | 9559974 | 9773322 | 2896089.5 | 4271575.5 | 4555479.5 | 1460.6511 | IYTSIGEDYDER | P67778 | P67778 | PHB | prohibitin | Nucleus | transcription regulator |
| 15534962 | 2 | 0.023 | 4602408.5 | 8419866 | 3937960.75 | 5375849 | 4676041.5 | 6073429 | 4264506.5 | 5579505 | 6069910 | 2597952.75 | 3173549 | 2567072 | 1062.504 | QVSDDLTER | P67778 | P67778 | PHB | prohibitin | Nucleus | transcription regulator |
| 15556450 | 2 | 0.16 | 2976403.5 | 1.11E+07 | 6565555.5 | 3484837.5 | 5551292.5 | 8236329.5 | 3272972 | 7612653 | 8716803 | 2113315 | 2010707 | 3493205.25 | 1855.0292 | NITYLPAGQSVLLQLPQ | P67778 | P67778 | PHB | prohibitin | Nucleus | transcription regulator |
| 15534380 | 3 | 0.093 | 4865390.5 | 1.62E+07 | 6201788 | 6341096 | 6900615 | 1.29E+07 | 5346788 | 8203522 | 1.17E+07 | 2014381.125 | 2774740.25 | 5096695.5 | 2126.1777 | AAELIANSLATAGDGLIELRK | P67778 | P67778 | PHB | prohibitin | Nucleus | transcription regulator |
| 15545240 | 2 | 0.454 | 2186191.25 | 3666113.25 | 2240863 | 1246986.75 | 3545848.5 | 4194608.5 | 1489570.125 | 3913757.25 | 4688620 | 1240822.25 | 1778059.875 | 1881697.5 | 920.46405 | PIIFDC[160.0307]R | P67778 | P67778 | PHB | prohibitin | Nucleus | transcription regulator |
| 15542262 | 3 | 0.03 | 2310095.25 | 4427739.5 | 2504934.5 | 2505568.5 | 2435627.5 | 3193919.25 | 3493428.25 | 4307015.5 | 3763725.25 | 1235570.25 | 1667490 | 2406879.5 | 2119.1401 | AATFGLILDDVSLTHLTFGK | P67778 | P67778 | PHB | prohibitin | Nucleus | transcription regulator |
| 15536339 | 2 | 0.458 | 1.09E+07 | 1.45E+07 | 7680477 | 1.22E+07 | 9200707 | 1.13E+07 | 8595505 | 9600558 | 8340349 | 1.15E+07 | 7984477 | 6392178 | 1188.6396 | DLQMVNISLR | O35129 | O35129 | PHB2 | prohibitin 2 | Cytoplasm | transcription regulator |
| 15531985 | 3 | 0.402 | 1.47E+07 | 2.65E+07 | 9020012 | 1.23E+07 | 1.35E+07 | 1.49E+07 | 1.02E+07 | 1.38E+07 | 1.83E+07 | 7516773 | 1.28E+07 | 8931890 | 1888.9653 | VLSRPNAQELPSMYQR | O35129 | O35129 | PHB2 | prohibitin 2 | Cytoplasm | transcription regulator |
| 15534003 | 2 | 0.62 | 9428315 | 1.70E+07 | 6190091.5 | 8147310 | 9290834 | 1.14E+07 | 7089676.5 | 1.13E+07 | 1.11E+07 | 7041262 | 8256647 | 6939189 | 1177.6316 | FNASQLITQR | O35129 | O35129 | PHB2 | prohibitin 2 | Cytoplasm | transcription regulator |
| 15538695 | 3 | 0.159 | 1.56E+07 | 3.63E+07 | 1.50E+07 | 1.29E+07 | 2.29E+07 | 2.14E+07 | 1.45E+07 | 2.30E+07 | 2.72E+07 | 6156346.5 | 1.26E+07 | 1.31E+07 | 1853.9835 | IGGVQQDTILAEGLHFR | O35129 | O35129 | PHB2 | prohibitin 2 | Cytoplasm | transcription regulator |
| 15548588 | 2 | 0.155 | 8838222 | 2.10E+07 | 1.03E+07 | 1.04E+07 | 1.34E+07 | 1.36E+07 | 7852553 | 1.46E+07 | 1.83E+07 | 5639015 | 8161118.5 | 7182043 | 1723.9186 | IPWFQYPIIYDIR | O35129 | O35129 | PHB2 | prohibitin 2 | Cytoplasm | transcription regulator |
| 15534666 | 2 | 0.043 | 9136771 | 1.76E+07 | 8520580 | 1.18E+07 | 1.15E+07 | 1.40E+07 | 7993043.5 | 1.18E+07 | 1.38E+07 | 5535529.5 | 6900374.5 | 6429711.5 | 1215.6185 | IVQAEGEAEAAK | O35129 | O35129 | PHB2 | prohibitin 2 | Cytoplasm | transcription regulator |
| 15540709 | 3 | 0.341 | 702379.25 | 2639784.5 | 1106974.625 | 1331151.375 | 1856165 | 1849117.625 | 1428428 | 2134906.75 | 2379611 | 1354902.75 | 846296.875 | 1132034.25 | 1723.9177 | IPWFQYPIIYDIR | O35129 | O35129 | PHB2 | prohibitin 2 | Cytoplasm | transcription regulator |
| 15553506 | 2 | 0.184 | 3751420.25 | 1.02E+07 | 3703103.75 | 5000329.5 | 3981167 | 6032294 | 3343919 | 5119833 | 7132936.5 | 1124377.125 | 2614103.25 | 3938030.5 | 2225.1409 | IYLTADNLVLNLQDESFTR | O35129 | O35129 | PHB2 | prohibitin 2 | Cytoplasm | transcription regulator |
| 15541788 | 3 | 0.006 | 1345875.5 | 2229718.75 | 1059431.625 | 1143903.5 | 1343053.25 | 1295531.75 | 885256.062 | 1461194.375 | 1441168 | 320740.062 | 533713 | 617662.875 | 1904.9607 | VLSRPNAQELPSM[147.0354]YQR | O35129 | O35129 | PHB2 | prohibitin 2 | Cytoplasm | transcription regulator |
| 15537629 | 2 | 0.173 | 3857154.5 | 2534085 | 3048487.75 | 4523063.5 | 2827418.5 | 2451302.25 | 3268781.75 | 2045599.25 | 1590345.5 | 6351678 | 3694286.75 | 3264050 | 1331.7507 | GTIQVVTQGTSLK | Q61753 | Q61753 | PHGDH | phosphoglycerate dehydrogenase | Cytoplasm | enzyme |
| 15542508 | 2 | 0.123 | 4249545 | 2315441.25 | 2718530.25 | 3980397.5 | 3056180.5 | 2572733 | 3020312.25 | 1688470.375 | 1104044.875 | 5090146 | 3661345 | 2918300.25 | 1130.6034 | VTADVINAAEK | Q61753 | Q61753 | PHGDH | phosphoglycerate dehydrogenase | Cytoplasm | enzyme |
| 15536806 | 3 | 0.091 | 3007548.5 | 1895621 | 2169660.5 | 2734376.75 | 1990769 | 1760303.625 | 1955282 | 1370573.5 | 1315890.125 | 4228223.5 | 2453659.25 | 2272070 | 2048.017 | ALVDHENVISC[160.0307]PHLGASTK | Q61753 | Q61753 | PHGDH | phosphoglycerate dehydrogenase | Cytoplasm | enzyme |
| 15545904 | 2 | 0.487 | 1987738.375 | 311656.281 | 43983.551 | 3215448.5 | 352609 | 126976.492 | 1753066 | 440311.562 | 69856.93 | 3823066.5 | 1687760.75 | 1126314.75 | 1298.729 | ILQDGGLQVVEK | Q61753 | Q61753 | PHGDH | phosphoglycerate dehydrogenase | Cytoplasm | enzyme |
| 15544134 | 2 | 0.075 | 2039672.125 | 1458103.75 | 1106030.125 | 2061868.5 | 1832423.625 | 1500842.875 | 1416636.75 | 1084511.75 | 1125911.5 | 2554668.75 | 1882294.375 | 1723005.875 | 1071.5774 | QADVNLVNAK | Q61753 | Q61753 | PHGDH | phosphoglycerate dehydrogenase | Cytoplasm | enzyme |
| 15535643 | 2 | 0.649 | 2194691.75 | 2783948.25 | 3012716.75 | 2004866.125 | 2196781.5 | 3465399.25 | 2618754.75 | 2029774.25 | 2301200 | 2866010.75 | 2266542.75 | 3678382.75 | 1376.755 | IAAIDNGLAFPFK | Q8CBQ5 | Q8CBQ5 | PI4K2B | phosphatidylinositol 4-kinase type 2 beta | Cytoplasm | kinase |
| 15535408 | 2 | 0.076 | 2289010.5 | 2938007 | 2928077.5 | 1914376.75 | 2395704.5 | 1928998.625 | 1859311.5 | 1902024.75 | 1921090.625 | 2285436 | 1816280.875 | 2774452.75 | 1458.7568 | AEQAIEIGVFPER | Q8CBQ5 | Q8CBQ5 | PI4K2B | phosphatidylinositol 4-kinase type 2 beta | Cytoplasm | kinase |
| 15541283 | 2 | 0.552 | 1862509.5 | 2561089.75 | 3450703.75 | 2083987.5 | 3009632.25 | 2119269.25 | 2008829.25 | 2351499.75 | 2410299.5 | 1878765.375 | 1548193.375 | 2521058.5 | 1259.6233 | ISQGSSGSYFVK | Q8CBQ5 | Q8CBQ5 | PI4K2B | phosphatidylinositol 4-kinase type 2 beta | Cytoplasm | kinase |
| 15540705 | 2 | 0.287 | 1282238 | 1285247.25 | 1465291.875 | 1120159.25 | 1096358.5 | 730650.375 | 849162.062 | 1044697.938 | 1091383.875 | 1546207 | 833316.375 | 1033432.562 | 1041.497 | QFQSQFEK | Q8CBQ5 | Q8CBQ5 | PI4K2B | phosphatidylinositol 4-kinase type 2 beta | Cytoplasm | kinase |
| 15548554 | 3, 4 | 0.141 | 1.86E+07 | 3.67E+07 | 2.20E+07 | 2.56E+07 | 2.71E+07 | 2.29E+07 | 2.07E+07 | 2.28E+07 | 2.29E+07 | 1.94E+07 | 1.73E+07 | 1.51E+07 | 3771.9312, 3771.9338 | VAEQVGIDRGDIPDLSQAPSSLLDALEQHLASLEGK | Q7M6Y3 | Q7M6Y3 | PICALM | phosphatidylinositol binding clathrin assembly protein | Cytoplasm | other |
| 15538404 | 2, 3 | 0.103 | 8917560 | 1.39E+07 | 8793663 | 7749740.5 | 9977498 | 6987532 | 7594999 | 8845632 | 8225127.5 | 8143793.5 | 4661635 | 6129475 | 1456.7728, 1456.7738 | ITAAQHSVTGSAVSK | Q7M6Y3 | Q7M6Y3 | PICALM | phosphatidylinositol binding clathrin assembly protein | Cytoplasm | other |
| 15537113 | 2 | 0.124 | 3912121.5 | 4361515 | 3142290.75 | 3450606.75 | 3426981.25 | 2744270.75 | 2876717 | 3121887.25 | 4147410.75 | 3098461.25 | 2025451.625 | 2592232.75 | 1474.6975 | SGLQGYDMSTFIR | Q7M6Y3 | Q7M6Y3 | PICALM | phosphatidylinositol binding clathrin assembly protein | Cytoplasm | other |
| 15544247 | 2 | 0.13 | 668532.25 | 3283944 | 3023520.25 | 791693 | 3148580 | 2187214.5 | 1474738.375 | 2506196.5 | 1675251.125 | 739613.312 | 235778.422 | 809120.562 | 1439.7816 | PTVASQNQSLPVAK | Q7M6Y3 | Q7M6Y3 | PICALM | phosphatidylinositol binding clathrin assembly protein | Cytoplasm | other |
| 15538817 | 4 | 0.63 | 2.69E+07 | 2.66E+07 | 2.63E+07 | 2.62E+07 | 2.36E+07 | 2.05E+07 | 2.79E+07 | 2.84E+07 | 2.09E+07 | 2.89E+07 | 2.41E+07 | 2.26E+07 | 2798.28 | ETPREDEEHELELDHLEPEPQAR | E2JF22 | E2JF22 | PIEZO1 | piezo-type mechanosensitive ion channel component 1 | Cytoplasm | ion channel |
| 15551414 | 2, 3 | 0.118 | 2.38E+07 | 2.58E+07 | 2.42E+07 | 2.27E+07 | 2.05E+07 | 2.49E+07 | 1.90E+07 | 2.14E+07 | 1.99E+07 | 2.55E+07 | 1.98E+07 | 2.18E+07 | 2466.34, 2466.341 | VPAALLEVTVADTEPTQTQTLLR | E2JF22 | E2JF22 | PIEZO1 | piezo-type mechanosensitive ion channel component 1 | Cytoplasm | ion channel |
| 15535387 | 3 | 0.491 | 1.89E+07 | 1.47E+07 | 1.42E+07 | 1.54E+07 | 1.56E+07 | 1.55E+07 | 1.81E+07 | 1.79E+07 | 1.11E+07 | 2.14E+07 | 1.94E+07 | 1.60E+07 | 1947.9409 | QHQQAPLPAQAVC[160.0307]ADGTR | E2JF22 | E2JF22 | PIEZO1 | piezo-type mechanosensitive ion channel component 1 | Cytoplasm | ion channel |
| 15549103 | 2 | 0.312 | 1.34E+07 | 1.42E+07 | 1.42E+07 | 1.33E+07 | 1.23E+07 | 1.21E+07 | 1.21E+07 | 1.26E+07 | 1.21E+07 | 1.88E+07 | 1.33E+07 | 1.23E+07 | 1690.7871 | QLQPDEEEDYLGVR | E2JF22 | E2JF22 | PIEZO1 | piezo-type mechanosensitive ion channel component 1 | Cytoplasm | ion channel |
| 15539043 | 2 | 0.777 | 1.23E+07 | 1.27E+07 | 9660031 | 8866911 | 1.25E+07 | 1.21E+07 | 1.29E+07 | 1.27E+07 | 1.09E+07 | 1.65E+07 | 1.10E+07 | 1.11E+07 | 1314.6565 | GPVDPANWFGVR | E2JF22 | E2JF22 | PIEZO1 | piezo-type mechanosensitive ion channel component 1 | Cytoplasm | ion channel |
| 15553228 | 3 | 0.705 | 1.23E+07 | 1.48E+07 | 1.15E+07 | 1.11E+07 | 1.21E+07 | 9744405 | 1.38E+07 | 1.38E+07 | 9592410 | 1.47E+07 | 1.05E+07 | 1.11E+07 | 3329.502 | DATQGEMPMTTEPDLDNC[160.0307]TVHVLTSQSPVR | E2JF22 | E2JF22 | PIEZO1 | piezo-type mechanosensitive ion channel component 1 | Cytoplasm | ion channel |
| 15549135 | 3 | 0.212 | 1.33E+07 | 1.25E+07 | 1.31E+07 | 1.30E+07 | 1.28E+07 | 1.08E+07 | 1.28E+07 | 1.18E+07 | 1.23E+07 | 1.42E+07 | 1.54E+07 | 1.24E+07 | 2810.2737 | AEQLASGGDLNPDVEPVDVPEDEMAGR | E2JF22 | E2JF22 | PIEZO1 | piezo-type mechanosensitive ion channel component 1 | Cytoplasm | ion channel |
| [illegible] | [illegible] | [illegible] | [illegible] | [illegible] | [illegible] | [illegible] | [illegible] | [illegible] | [illegible] | [illegible] | [illegible] | [illegible] | [illegible] | [illegible] | [illegible] | [illegible] | [illegible] | [illegible] | [illegible] | [illegible] | [illegible] | [illegible] |

| | | | | | | | | | | | | | | | | | | | | |
|---|---|---|---|---|---|---|---|---|---|---|---|---|---|---|---|---|---|---|---|---|
| 15532606 | 4 | 0.02 | 3.49E+07 | 3.45E+07 | 3.34E+07 | 2.72E+07 | 3.36E+07 | 3.93E+07 | 4.32E+07 | 3.20E+07 | 3.63E+07 | 5.66E+07 | 5.11E+07 | 4.38E+07 | 2375.192 | YTIGSLRPHFLAIC[160.0307]NPDWSK | Q61469 | Q61469 | Ppap2a | phosphatidic acid phosphatase type 2A | Plasma Membrane | phosphatase |
| 15532132 | 3 | 0.495 | 4.37E+07 | 7.02E+07 | 8.33E+07 | 2.79E+07 | 7.30E+07 | 5.88E+07 | 6.20E+07 | 6.56E+07 | 3.90E+07 | 5.10E+07 | 2.82E+07 | 4.29E+07 | 1709.769 | EEDPHTTLHETASSR | Q61469 | Q61469 | Ppap2a | phosphatidic acid phosphatase type 2A | Plasma Membrane | phosphatase |
| 15531347 | 2 | 0.744 | 3.25E+07 | 3.34E+07 | 4.36E+07 | 2.09E+07 | 3.76E+07 | 3.36E+07 | 3.82E+07 | 3.95E+07 | 2.68E+07 | 4.41E+07 | 2.79E+07 | 3.63E+07 | 1168.5281 | GIFC[160.0307]NDDSIK | Q61469 | Q61469 | Ppap2a | phosphatidic acid phosphatase type 2A | Plasma Membrane | phosphatase |
| 15549058 | 2 | 0.339 | 5613075 | 8538303 | 1.05E+07 | 1912553.875 | 8889195 | 4068403.75 | 4293835 | 7768799 | 5012318.5 | 3386180.5 | 4604170.5 | 5245215.5 | 1709.7694 | EEDPHTTLHETASSR | Q61469 | Q61469 | Ppap2a | phosphatidic acid phosphatase type 2A | Plasma Membrane | phosphatase |
| 15533283 | 2 | 0.093 | 4530042 | 7874376.5 | 1.49E+07 | 1.94E+07 | 2.56E+07 | 3.02E+07 | 1.62E+07 | 2.30E+07 | 3.28E+07 | 4620299.5 | 1.83E+07 | 1.82E+07 | 1154.571 | FEDENFILK | P17742 | P17742 | PPIA | peptidylprolyl isomerase A (cyclophilin A) | Cytoplasm | enzyme |
| 15549270 | 2 | 0.134 | 1656342.625 | 4117050 | 6665097.5 | 1.60E+07 | 1.54E+07 | 1.70E+07 | 6260266 | 1.08E+07 | 2.64E+07 | 2581346.75 | 1.70E+07 | 1.29E+07 | 1598.7418 | IIPGFMC[160.0307]QGGDFTR | P17742 | P17742 | PPIA | peptidylprolyl isomerase A (cyclophilin A) | Cytoplasm | enzyme |
| 15531980 | 2 | 0.109 | 2133076.5 | 6034369 | 1.03E+07 | 2.05E+07 | 2.02E+07 | 2.32E+07 | 1.08E+07 | 1.69E+07 | 2.52E+07 | 2274293 | 1.32E+07 | 1.44E+07 | 1055.5387 | VSFELFADK | P17742 | P17742 | PPIA | peptidylprolyl isomerase A (cyclophilin A) | Cytoplasm | enzyme |
| 15555266 | 3 | 0.121 | 1195396.375 | 2210366 | 1738714.25 | 1.46E+07 | 5308132.5 | 6771483 | 4971643.5 | 4351273 | 9547333 | 1061969.625 | 1.25E+07 | 5342050.5 | 2791.321 | HTGPGILSMANAGPNTNGSQFFIC[160.0307]TAK | P17742 | P17742 | PPIA | peptidylprolyl isomerase A (cyclophilin A) | Cytoplasm | enzyme |
| 15548437 | 2 | 0.261 | 1188912.5 | 5023734 | 8207870.5 | 1.65E+07 | 1.37E+07 | 1.52E+07 | 7972199.5 | 1.38E+07 | 2.12E+07 | 771341 | 1.19E+07 | 1.22E+07 | 2005.9813 | VNPTVFFDITADDEPLGR | P17742 | P17742 | PPIA | peptidylprolyl isomerase A (cyclophilin A) | Cytoplasm | enzyme |
| 15543205 | 2 | 0.119 | 340174.969 | 494826.219 | 202399.828 | 3905361.5 | 1403989.625 | 778208.25 | 674374.688 | 1070588.875 | 4715930.5 | 554436.25 | 3773185 | 2582045.25 | 1134.5803 | KITISDC[160.0307]GQL | P17742 | P17742 | PPIA | peptidylprolyl isomerase A (cyclophilin A) | Cytoplasm | enzyme |
| 15545214 | 2 | 0.433 | 386903.75 | 1302617.25 | 1694794.75 | 1040245.125 | 4978204.5 | 4469293 | 1665230.75 | 3055482 | 6303734 | 260682.547 | 3902883.5 | 2604756 | 848.41327 | TEWLDGK | P17742 | P17742 | PPIA | peptidylprolyl isomerase A (cyclophilin A) | Cytoplasm | enzyme |
| 15532985 | 2 | 0.392 | 6.58E+07 | 6.50E+07 | 8.99E+07 | 8.61E+07 | 9.10E+07 | 6.81E+07 | 8.56E+07 | 8.58E+07 | 8.34E+07 | 8.39E+07 | 8.85E+07 | 8.35E+07 | 1364.703 | TVDNFVALATGEK | P24369 | P24369 | PPIB | peptidylprolyl isomerase B (cyclophilin B) | Cytoplasm | enzyme |
| 15534793 | 3 | 0.16 | 3.49E+07 | 3.83E+07 | 3.78E+07 | 5.03E+07 | 3.75E+07 | 4.76E+07 | 4.87E+07 | 3.56E+07 | 4.53E+07 | 4.64E+07 | 5.08E+07 | 4.45E+07 | 1601.7573 | SIYGERFPDENFK | P24369 | P24369 | PPIB | peptidylprolyl isomerase B (cyclophilin B) | Cytoplasm | enzyme |
| 15532271 | 2 | 0.756 | 1.50E+07 | 2.36E+07 | 2.91E+07 | 1.63E+07 | 3.54E+07 | 2.50E+07 | 3.14E+07 | 3.21E+07 | 2.20E+07 | 2.99E+07 | 2.12E+07 | 1.90E+07 | 1286.5808 | DFMIQGGDFTR | P24369 | P24369 | PPIB | peptidylprolyl isomerase B (cyclophilin B) | Cytoplasm | enzyme |
| 15537151 | 2 | 0.314 | 2.15E+07 | 2.48E+07 | 2.56E+07 | 2.28E+07 | 3.01E+07 | 2.85E+07 | 3.10E+07 | 2.81E+07 | 2.47E+07 | 2.51E+07 | 2.42E+07 | 2.48E+07 | 945.52325 | DVIIVDSGK | P24369 | P24369 | PPIB | peptidylprolyl isomerase B (cyclophilin B) | Cytoplasm | enzyme |
| 15538363 | 2 | 0.045 | 2.14E+07 | 2.00E+07 | 2.31E+07 | 2.67E+07 | 2.93E+07 | 2.56E+07 | 2.78E+07 | 2.52E+07 | 2.68E+07 | 2.05E+07 | 2.73E+07 | 2.45E+07 | 1457.7252 | DTNGSQFFITTVK | P24369 | P24369 | PPIB | peptidylprolyl isomerase B (cyclophilin B) | Cytoplasm | enzyme |
| 15534247 | 2, 3 | 0.18 | 1.62E+07 | 2.17E+07 | 2.18E+07 | 2.08E+07 | 2.20E+07 | 2.82E+07 | 3.03E+07 | 2.43E+07 | 2.36E+07 | 2.00E+07 | 2.17E+07 | 2.30E+07 | 1145.6317, 1145.6324 | VLEGMDVVRK | P24369 | P24369 | PPIB | peptidylprolyl isomerase B (cyclophilin B) | Cytoplasm | enzyme |
| 15532547 | 2 | 0.38 | 1.27E+07 | 1.43E+07 | 1.94E+07 | 1.52E+07 | 1.91E+07 | 2.02E+07 | 1.85E+07 | 1.66E+07 | 2.19E+07 | 1.76E+07 | 1.63E+07 | 1.71E+07 | 866.5128 | VVFGLFGK | P24369 | P24369 | PPIB | peptidylprolyl isomerase B (cyclophilin B) | Cytoplasm | enzyme |
| 15532847 | 2 | 0.31 | 1.72E+07 | 1.22E+07 | 1.20E+07 | 1.31E+07 | 1.52E+07 | 1.15E+07 | 1.75E+07 | 1.37E+07 | 1.16E+07 | 1.62E+07 | 2.12E+07 | 1.52E+07 | 1474.687 | HYGPGWVSMANAGK | P24369 | P24369 | PPIB | peptidylprolyl isomerase B (cyclophilin B) | Cytoplasm | enzyme |
| 15554444 | 2 | 0.314 | 1.29E+07 | 9308533 | 9148428 | 1.33E+07 | 9921896 | 1.11E+07 | 1.30E+07 | 8496500 | 1.16E+07 | 1.48E+07 | 1.67E+07 | 1.10E+07 | 1601.7556 | SIYGERFPDENFK | P24369 | P24369 | PPIB | peptidylprolyl isomerase B (cyclophilin B) | Cytoplasm | enzyme |
| 15533854 | 2 | 0.071 | 1.08E+07 | 9508158 | 1.22E+07 | 1.13E+07 | 1.45E+07 | 1.22E+07 | 1.53E+07 | 1.34E+07 | 1.46E+07 | 1.42E+07 | 1.40E+07 | 1.15E+07 | 806.4029 | TSWLDGK | P24369 | P24369 | PPIB | peptidylprolyl isomerase B (cyclophilin B) | Cytoplasm | enzyme |
| 15551859 | 2 | 0.786 | 9648348 | 9661796 | 1.31E+07 | 1.28E+07 | 1.19E+07 | 9592740 | 9470616 | 1.12E+07 | 1.48E+07 | 1.18E+07 | 8867410 | 1.00E+07 | 1597.7816 | VYFDLQIGDESVGR | P24369 | P24369 | PPIB | peptidylprolyl isomerase B (cyclophilin B) | Cytoplasm | enzyme |
| 15550670 | 2 | 0.377 | 3943800.25 | 2744685 | 1843008.25 | 5620618.5 | 981702.312 | 1276272.875 | 3586190 | 1325989.875 | 3585996.75 | 3543652.75 | 6978621 | 4403660 | 1858.8353 | DFMIQGGDFTRGDGTGGK | P24369 | P24369 | PPIB | peptidylprolyl isomerase B (cyclophilin B) | Cytoplasm | enzyme |
| 15550380 | 3 | 0.108 | 1177891.75 | 2641407.25 | 560974.062 | 9056425 | 7205942.5 | 9766434 | 4540588 | 5376775 | 1.23E+07 | 638831.5 | 1.27E+07 | 5463636 | 2817.3762 | HVGPGVLSMANAGPNTNGSQFFIC[160.0307]TIK | Q99KR7 | Q99KR7 | PPIF | peptidylprolyl isomerase F | Cytoplasm | enzyme |
| 15550692 | 2 | 0.303 | 455276.219 | 2111235 | 3617061 | 7431932 | 7692969.5 | 8135632 | 4181111 | 6364539 | 8617605 | 96668.367 | 5718273.5 | 5339143 | 2344.1487 | GANSSSGNPLVYLDVGADGQPLGR | Q99KR7 | Q99KR7 | PPIF | peptidylprolyl isomerase F | Cytoplasm | enzyme |
| 15548069 | 3 | 0.174 | 2.66E+07 | 5.34E+07 | 2.36E+07 | 2.61E+07 | 2.47E+07 | 3.27E+07 | 1.71E+07 | 3.37E+07 | 3.69E+07 | 1.93E+07 | 1.76E+07 | 1.55E+07 | 2735.4038 | AAGHQLSPELFQQQAQEAAATQLGLK | P51175 | P51175 | PPOX | protoporphyrinogen oxidase | Cytoplasm | enzyme |
| 15552029 | 3 | 0.149 | 2.74E+07 | 5.62E+07 | 2.45E+07 | 2.42E+07 | 2.94E+07 | 4.50E+07 | 1.77E+07 | 3.63E+07 | 4.90E+07 | 1.68E+07 | 1.55E+07 | 1.72E+07 | 2439.2546 | VSLGDSSLEADHIISAIPASELSK | P51175 | P51175 | PPOX | protoporphyrinogen oxidase | Cytoplasm | enzyme |
| 15533904 | 2 | 0.242 | 1.15E+07 | 2.30E+07 | 7765923.5 | 8341769.5 | 9379310 | 1.37E+07 | 6765599.5 | 1.38E+07 | 1.47E+07 | 6726892.5 | 6210371 | 6942427 | 1121.6654 | LLPAEAAPLAR | P51175 | P51175 | PPOX | protoporphyrinogen oxidase | Cytoplasm | enzyme |
| 15553035 | 4 | 0.387 | 7457971.5 | 1.98E+07 | 6490585 | 1.01E+07 | 9570450 | 9806935 | 6894547 | 1.30E+07 | 1.66E+07 | 6227639 | 6610401 | 6967832.5 | 4422.161 | GAC[160.0307]LPVQGFGHLVPSSEDPTVLGIVYDSVAFPEQDGNPPSLR | P51175 | P51175 | PPOX | protoporphyrinogen oxidase | Cytoplasm | enzyme |
| 15534259 | 3 | 0.811 | 4858452 | 1.73E+07 | 5556934 | 9192157 | 6088050.5 | 1.03E+07 | 5288238 | 1.12E+07 | 1.37E+07 | 6087097.5 | 6598516.5 | 7026745.5 | 2235.2678 | SILLGLLLGAGQSPQPDSSLIR | P51175 | P51175 | PPOX | protoporphyrinogen oxidase | Cytoplasm | enzyme |
| 15548787 | 2 | 0.265 | 7814609 | 1.47E+07 | 5342338.5 | 7337245.5 | 5903179 | 6520435 | 4373223 | 7910153.5 | 9035584 | 4919424 | 5194487 | 4154863.25 | 2054.0637 | GVTVLSGQPVC[160.0307]GLSLQPEGR | P51175 | P51175 | PPOX | protoporphyrinogen oxidase | Cytoplasm | enzyme |
| 15534540 | 2 | 0.213 | 6577390 | 1.44E+07 | 5746278.5 | 6246510 | 7518881.5 | 1.03E+07 | 4446700 | 1.00E+07 | 9518427 | 4074251.5 | 4440514.5 | 4846919 | 1380.6909 | LDSAMQFLTAQR | P51175 | P51175 | PPOX | protoporphyrinogen oxidase | Cytoplasm | enzyme |
| 15549164 | 2 | 0.275 | 7692955 | 1.33E+07 | 4042756.75 | 7028596.5 | 6511529 | 8685110 | 3674116.25 | 8018818 | 1.01E+07 | 3949938.75 | 3729944.5 | 4391573 | 2235.2654 | SILLGLLLGAGQSPQPDSSLIR | P51175 | P51175 | PPOX | protoporphyrinogen oxidase | Cytoplasm | enzyme |
| 15532329 | 3 | 0.279 | 1.02E+07 | 1.95E+07 | 4376045 | 7824759 | 5942665.5 | 1.05E+07 | 5715450.5 | 7562490 | 1.09E+07 | 3858931.25 | 4869517.5 | 5177413.5 | 1726.9602 | FLYVGGTLHPLPSGLR | P51175 | P51175 | PPOX | protoporphyrinogen oxidase | Cytoplasm | enzyme |
| 15556377, 15558622 | 2 | 0.216 | 4996027.5 | 7873978.5 | 2206654 | 4272554.5 | 3096060.75 | 4042417.25 | 2747329.25 | 4194434.5 | 6316718 | 2224063 | 2633613 | 1857557.75 | 1618.7894, 1618.7896 | LGPEVASLAMDSLC[160.0307]R | P51175 | P51175 | PPOX | protoporphyrinogen oxidase | Cytoplasm | enzyme |
| 15541101 | 2 | 0.356 | 4407437.5 | 7655245.5 | 2309097.25 | 3392175 | 3868535.75 | 4257085.5 | 2240380.5 | 3818453.5 | 6539597 | 2221721.25 | 2524505.5 | 2379600.75 | 962.4825 | WSQWSLR | P51175 | P51175 | PPOX | protoporphyrinogen oxidase | Cytoplasm | enzyme |
| 15534571 | 2 | 0.089 | 3843214 | 8520962 | 2275682.25 | 2231129.75 | 3676619 | 5770283.5 | 2083885 | 4717810 | 5366298 | 1239045.25 | 1258840.625 | 1832693.25 | 959.54425 | PLFWAGLR | P51175 | P51175 | PPOX | protoporphyrinogen oxidase | Cytoplasm | enzyme |
| 15554071 | 3 | 0.038 | 3808119 | 6629893.5 | 3121900.75 | 4150202.5 | 2597489.25 | 3960918.75 | 1933961.875 | 2976745.75 | 3518791.25 | 1224059.5 | 1870915 | 2207029 | 2913.5718 | TLLLVSELGLESEVLPVRGDHPAAQNR | P51175 | P51175 | PPOX | protoporphyrinogen oxidase | Cytoplasm | enzyme |
| 15535481 | 3 | 0.117 | 2704472.75 | 4075247.25 | 3034938.75 | 6674726 | 4859755 | 6049465 | 3996930.75 | 4495477 | 7213226.5 | 3059607.75 | 6040449.5 | 4504338.5 | 2196.2612 | SREIFLSQPILLELEAPLK | P62137 | P62137 | PPP1CA | protein phosphatase 1, catalytic subunit, alpha isozyme | Cytoplasm | phosphatase |
| 15549696 | 3 | 0.137 | 2450598 | 4499372 | 4770403.5 | 6797826 | 8170354 | 8429217 | 5583987 | 6073461 | 1.08E+07 | 2990844.75 | 7684710 | 6480223.5 | 2684.3008 | IMRPTDVPDQGLLC[160.0307]DLLWSDPDK | P62137 | P62137 | PPP1CA | protein phosphatase 1, catalytic subunit, alpha isozyme | Cytoplasm | phosphatase |
| 15535684 | 2 | 0.129 | 2288276 | 2719127.25 | 2695481 | 2457921.25 | 2759237 | 2330823 | 2209498.5 | 2928681.5 | 3133917.75 | 2091906.5 | 2329420.25 | 1817639.125 | 1198.6233 | YPENFFLLR | P62137 | P62137 | PPP1CA | protein phosphatase 1, catalytic subunit, alpha isozyme | Cytoplasm | phosphatase |
| 15539322 | 3 | 0.16 | 1934305.25 | 2486210.75 | 2665209.25 | 3145301 | 4025778.5 | 3998978 | 2676177.75 | 3388747.75 | 3938643 | 1487836.5 | 3477171.75 | 2710191.75 | 1639.7722 | AHQVVEDGYEFFAK | P62137 | P62137 | PPP1CA | protein phosphatase 1, catalytic subunit, alpha isozyme | Cytoplasm | phosphatase |
| 15553345 | 2 | 0.031 | 1353190.25 | 2358035.75 | 2485990.25 | 2543595.5 | 4064297.75 | 2725487.75 | 1634635 | 3377411.5 | 3250574 | 1158544.875 | 1205480.375 | 1449879.125 | 1953.1266 | EIFLSQPILLELEAPLK | P62137 | P62137 | PPP1CA | protein phosphatase 1, catalytic subunit, alpha isozyme | Cytoplasm | phosphatase |
| 15556741 | 2 | 0.167 | 1103354.125 | 909869.75 | 930574.688 | 1655064.375 | 1477246.875 | 1510883 | 1222137 | 1274629.875 | 1256108 | 759168.188 | 1742152.875 | 1307738.875 | 1639.7711 | AHQVVEDGYEFFAK | P62137 | P62137 | PPP1CA | protein phosphatase 1, catalytic subunit, alpha isozyme | Cytoplasm | phosphatase |
| 15532939 | 2 | 0.294 | 3776049.25 | 4072407 | 5595056.5 | 2195394 | 4882094.5 | 2240164.25 | 3473405 | 2932948.5 | 2720658.25 | 4246512.5 | 1461211.875 | 2100976 | 729.44885 | LGIEIGK | Q62415 | Q62415 | PPP1R13B | protein phosphatase 1, regulatory subunit 13B | Cytoplasm | phosphatase |
| 15541465 | 2 | 0.18 | 9290315 | 1.19E+07 | 1.66E+07 | 5202436 | 1.64E+07 | 1.15E+07 | 1.22E+07 | 1.62E+07 | 1.04E+07 | 2908225 | 7182241.5 | 8519368 | 1302.5792 | TPDSEAPDVESR | Q3TDD9 | Q3TDD9 | PPP1R21 | protein phosphatase 1, regulatory subunit 21 | Cytoplasm | other |
| 15551093 | 3 | 0.337 | 4043775.5 | 3002035.5 | 3807099.5 | 5474830.5 | 4301646.5 | 3964303.75 | 4468159 | 3250988.25 | 2633904.75 | 9398877 | 4099507.75 | 3714144.5 | 2655.2732 | GAGYTFGQDISETFNHANGLTLVSR | P63330 | P63330 | PPP2CA | protein phosphatase 2, catalytic subunit, alpha isozyme | Cytoplasm | phosphatase |
| 15553121 | 4 | 0.109 | 4801135 | 3076575.25 | 1963158.25 | 6652956 | 2482689.25 | 4404763.5 | 4070813.75 | 1909186.125 | 3037808.75 | 9156457 | 5682358.5 | 5992915 | 3319.5176 | LQEVPHEGPMC[160.0307]DLLWSDPDDRGGWGISPR | P63330 | P63330 | PPP2CA | protein phosphatase 2, catalytic subunit, alpha isozyme | Cytoplasm | phosphatase |
| 15536657 | 2 | 0.015 | 3685186.25 | 2576637 | 2447944.25 | 4303116.5 | 3188531.75 | 4216847 | 3301492.25 | 2768392.25 | 1985130.5 | 6032990.5 | 4453633 | 4938902 | 1340.6625 | YSFLQFDPAPR | P63330 | P63330 | PPP2CA | protein phosphatase 2, catalytic subunit, alpha isozyme | Cytoplasm | phosphatase |
| 15551925 | 2 | 0.057 | 2019480.875 | 1266601.625 | 1650609.25 | 2821342 | 1737945.25 | 2272787.5 | 1903314.625 | 1167744.75 | 674396.312 | 4838261 | 2236387.5 | 2575200.75 | 1704.7866 | ELDQWIEQLNEC[160.0307]K | P63330 | P63330 | PPP2CA | protein phosphatase 2, catalytic subunit, alpha isozyme | Cytoplasm | phosphatase |
| 15556134 | 2 | 0.038 | 1483186.875 | 1970992.375 | 1259155.375 | 2277861.25 | 2453966.5 | 2567628.75 | 2291794 | 1760533.625 | 1550835.375 | 4388952 | 2397817.25 | 2415132.25 | 1703.8177 | NVVTIFSAPNYC[160.0307]YR | P63330 | P63330 | PPP2CA | protein phosphatase 2, catalytic subunit, alpha isozyme | Cytoplasm | phosphatase |
| 15536826 | 3 | 0.192 | 9574879 | 6122385 | 3946420.75 | 7323656.5 | 6997105.5 | 7799388.5 | 8321207 | 4792437.5 | 5722864 | 1.32E+07 | 7790672.5 | 1.04E+07 | 2194.2288 | IGPILDNSTLQSEVKPILEK | Q76MZ3 | Q76MZ3 | PPP2R1A | protein phosphatase 2, regulatory subunit A, alpha | Cytoplasm | phosphatase |
| 15556150 | 2 | 0.409 | 2359181 | 1728656.875 | 2591975.5 | 2495863.25 | 2464713.75 | 2723815 | 2690185 | 2218629 | 2105396.5 | 9282895 | 2206743.5 | 2669423 | 1925.0554 | QLSQSLLPAIVELAEDAK | Q76MZ3 | Q76MZ3 | PPP2R1A | protein phosphatase 2, regulatory subunit A, alpha | Cytoplasm | phosphatase |
| 15540665 | 3 | 0.269 | 2181168.25 | 2123751 | 2295815.5 | 2411287.25 | 1876159.5 | 2695625.25 | 3037057.5 | 2086234.875 | 1828991.5 | 7494091.5 | 2274587.25 | 3067422.25 | 1925.057 | QLSQSLLPAIVELAEDAK | Q76MZ3 | Q76MZ3 | PPP2R1A | protein phosphatase 2, regulatory subunit A, alpha | Cytoplasm | phosphatase |
| 15541113 | 3 | 0.454 | 4165114.25 | 3916789.75 | 3642962 | 4045784.75 | 3845146 | 5671078 | 6021548.5 | 4174352.75 | 3675741.75 | 7370656.5 | 3910781.25 | 5397105 | 1865.0493 | DNTIEHLLPLFLAQLK | Q76MZ3 | Q76MZ3 | PPP2R1A | protein phosphatase 2, regulatory subunit A, alpha | Cytoplasm | phosphatase |
| 15551164 | 2 | 0.233 | 3221393.5 | 2282365.25 | 1446521.25 | 3724732.75 | 3086577.25 | 3222579.25 | 3328867.5 | 1888188.125 | 1726150.875 | 5265910.5 | 2304608 | 3860854.5 | 2138.0017 | SEIIPMFSNLASDEQDSVR | Q76MZ3 | Q76MZ3 | PPP2R1A | protein phosphatase 2, regulatory subunit A, alpha | Cytoplasm | phosphatase |
| 15535311 | 2 | 0.542 | 2665062.75 | 1584392 | 1824160.625 | 2915050.75 | 1952239 | 2408255.5 | 2754989.75 | 1595577.875 | 1450122.5 | 4664187 | 1893206.875 | 2220740 | 1160.5764 | LTQDQDVDVK | Q76MZ3 | Q76MZ3 | PPP2R1A | protein phosphatase 2, regulatory subunit A, alpha | Cytoplasm | phosphatase |
| 15551117 | 2 | 0.266 | 1529233.75 | 1009962.188 | 1400335.375 | 1411055.625 | 1346413.25 | 1870990.125 | 1656815.875 | 1231772.75 | 1253682.125 | 2961889 | 1386332.625 | 1828899.25 | 1556.906 | SALASVIMGLSPILGK | Q76MZ3 | Q76MZ3 | PPP2R1A | protein phosphatase 2, regulatory subunit A, alpha | Cytoplasm | phosphatase |
| 15553143 | 3 | 0.903 | 2386374.25 | 876833.562 | 1181249 | 3170106.5 | 1870585.375 | 838424.75 | 2695864.75 | 1585220 | 1144624.625 | 7614890 | 1187725.125 | 1040626.938 | 3088.4512 | GAVDDDVAEADIISTVEFNHSGELLATGDK | Q6P1F6 | Q6P1F6 | PPP2R2A | protein phosphatase 2, regulatory subunit B, alpha | Cytoplasm | phosphatase |
| 15551616 | 2, 3 | 0.424 | 6232103 | 5945198 | 5774764.5 | 4737376.5 | 7165172 | 5258285.5 | 6910877 | 5368362.5 | 5647824.5 | 1.20E+07 | 5667421.5 | 6656531 | 2536.14 | GAFC[160.0307]DLVWSDPEDVDTWAISPR | Q9CQR6 | Q9CQR6 | PPP6C | protein phosphatase 6, catalytic subunit | Nucleus | phosphatase |
| 15553263 | 2 | 0.707 | 5243476.5 | 3914635.25 | 2480697.5 | 3489206.25 | 2914968.75 | 2342288.75 | 3881979 | 2690143.75 | 3198270.25 | 5824406 | 3164461.5 | 2747827.5 | 1879.8495 | QITQVYGFYDEC[160.0307]QTK | Q9CQR6 | Q9CQR6 | PPP6C | protein phosphatase 6, catalytic subunit | Nucleus | phosphatase |
| 15549652 | 4 | 0.411 | 1.54E+07 | 1.10E+07 | 8651564 | 1.01E+07 | 1.24E+07 | 6888912 | 1.38E+07 | 8449132 | 8641220 | 2.15E+07 | 1.25E+07 | 1.10E+07 | 3032.4783 | LVEIVHPSQEEDRHSNASQSLC[160.0307]EIVR | Q922D4 | Q922D4 | PPP6R3 | protein phosphatase 6, regulatory subunit 3 | Cytoplasm | other |
| 15537631 | 3 | 0.491 | 8986690 | 9247213 | 5042367 | 6363194 | 7196522.5 | 8106051.5 | 1.19E+07 | 7825490.5 | 8209703 | 1.14E+07 | 7342417 | 9352657 | 1711.9342 | QEIIEQLLSNIFHK | Q922D4 | Q922D4 | PPP6R3 | protein phosphatase 6, regulatory subunit 3 | Cytoplasm | other |
| 15549217 | 2 | 0.734 | 7036168 | 5369876 | 4993148.5 | 6056505.5 | 5449819.5 | 3835950.75 | 5543794.5 | 3944054.5 | 4464261 | 9478525 | 3917558.25 | 5121707 | 1598.7053 | FADQDDIGNVSFDR | Q922D4 | Q922D4 | PPP6R3 | protein phosphatase 6, regulatory subunit 3 | Cytoplasm | other |
| 15532634 | 2 | 0.906 | 7104787 | 5547125.5 | 5166462 | 5307691.5 | 5928167.5 | 5329052 | 6684011.5 | 5649838 | 4022895.75 | 9391523 | 4245013.5 | 5778201 | 1267.7346 | GPNSALVQQLIK | Q922D4 | Q922D4 | PPP6R3 | protein phosphatase 6, regulatory subunit 3 | Cytoplasm | other |
| 15554479 | 2 | 0.597 | 5118697 | 3478461.75 | 3718798.5 | 4359103.5 | 4097286 | 3296022.75 | 4506882 | 3463610.25 | 3890712.5 | 8846557 | 3976904.25 | 3814909 | 1637.7004 | ELMDEEDVLQEC[160.0307]K | Q922D4 | Q922D4 | PPP6R3 | protein phosphatase 6, regulatory subunit 3 | Cytoplasm | other |
| 15548953 | 2 | 0.801 | 6302552.5 | 4528646 | 3987776.5 | 5234555.5 | 4501141.5 | 3049530 | 5536459.5 | 3815144 | 3568781.75 | 8767766 | 4069572.75 | 3857397.5 | 1530.7191 | C[160.0307]TAPLTPSSSPEQR | Q922D4 | Q922D4 | PPP6R3 | protein phosphatase 6, regulatory subunit 3 | Cytoplasm | other |
| 15539184 | 2 | 0.293 | 5566784.5 | 4007613 | 5928430.5 | 4778921.5 | 4743576.5 | 4451323.5 | 5582074 | 4048677.75 | 4049659.5 | 7743192.5 | 5203848.5 | 5303664 | 1415.6876 | QDVLNWLNEER | Q922D4 | Q922D4 | PPP6R3 | protein phosphatase 6, regulatory subunit 3 | Cytoplasm | other |
| 15555034 | 2 | 0.819 | 4908884.5 | 3704588.5 | 3477595.25 | 3455595.5 | 5137525 | 2964696.75 | 5067293 | 3506280.25 | 3362915.5 | 6437565.5 | 3619780.75 | 3992295 | 1686.7395 | WETFC[160.0307]TNSLGETNK | Q922D4 | Q922D4 | PPP6R3 | protein phosphatase 6, regulatory subunit 3 | Cytoplasm | other |
| 15535358 | 2 | 0.711 | 5029094 | 3224528.5 | 3579792.75 | 3793700 | 3879693.5 | 2620262.75 | 4505351 | 2821926.5 | 2401825 | 6299008 | 3188652 | 3267970.5 | 1218.5968 | ILEAWDTNEK | Q922D4 | Q922D4 | PPP6R3 | protein phosphatase 6, regulatory subunit 3 | Cytoplasm | other |
| 15556257, 15558919 | 2 | 0.738 | 3916518.75 | 3197115 | 3140857.5 | 2414281.5 | 3195282.75 | 2178085.75 | 4482244 | 2991281.5 | 2067339 | 5339499.5 | 2287187.25 | 2639049 | 1570.6841, 1570.6843 | QDLFESSSANTEDK | Q922D4 | Q922D4 | PPP6R3 | protein phosphatase 6, regulatory subunit 3 | Cytoplasm | other |
| 15540893 | 3 | 0.722 | 2780512.25 | 1816626.25 | 2116525.25 | 2827226.75 | 1953777.125 | 1881191.625 | 3304667.5 | 1999199.125 | 1798201.125 | 4951365 | 2145855.5 | 2196931.5 | 1257.6243 | IANC[160.0307]IVHSTDK | Q922D4 | Q922D4 | PPP6R3 | protein phosphatase 6, regulatory subunit 3 | Cytoplasm | other |
| 15554231 | 2 | 0.637 | 1310908 | 1303103.5 | 682675.375 | 747797.375 | 1034199.562 | 1288733.375 | 1478910.125 | 875096.5 | 1142961.5 | 1493845.125 | 1023324.062 | 1590277.5 | 1711.9343 | QEIIEQLLSNIFHK | Q922D4 | Q922D4 | PPP6R3 | protein phosphatase 6, regulatory subunit 3 | Cytoplasm | other |
| 15553094 | 3 | 0.122 | 2704089.5 | 7391970.5 | 3736292.25 | 4788476.5 | 5068364 | 8311376.5 | 3048165 | 5172221.5 | 5897917.5 | 1670567.375 | 2729709.25 | 3310859.5 | 2518.252 | DSQHLNFLTSEQALADFAELIR | Q7TMR0 | Q7TMR0 | PRCP | prolylcarboxypeptidase (angiotensinase C) | Cytoplasm | peptidase |
| 15547964 | 2 | 0.913 | 2.45E+07 | 2.49E+07 | 2.52E+07 | 2.41E+07 | 2.41E+07 | 2.65E+07 | 2.47E+07 | 2.09E+07 | 1.91E+07 | 5.29E+07 | 1.31E+07 | 2.56E+07 | 1636.8679 | QGGLGPMNIPLISDPK | P35700 | P35700 | PRDX1 | peroxiredoxin 1 | Cytoplasm | enzyme |
| 15536942 | 2 | 0.734 | 3.30E+07 | 2.30E+07 | 2.49E+07 | 3.14E+07 | 1.92E+07 | 2.23E+07 | 2.66E+07 | 1.91E+07 | 1.97E+07 | 5.23E+07 | 1.85E+07 | 2.41E+07 | 1196.629 | LVQAFQFTDK | P35700 | P35700 | PRDX1 | peroxiredoxin 1 | Cytoplasm | enzyme |
| 15537487 | 2 | 0.457 | 2.07E+07 | 1.56E+07 | 1.54E+07 | 1.59E+07 | 1.50E+07 | 1.89E+07 | 1.74E+07 | 1.17E+07 | 1.07E+07 | 3.75E+07 | 1.27E+07 | 1.75E+07 | 1164.57 | ATAVMPDGQFK | P35700 | P35700 | PRDX1 | peroxiredoxin 1 | Cytoplasm | enzyme |
| 15538178 | 2, 3 | 0.376 | 3.24E+07 | 2.10E+07 | 2.31E+07 | 2.22E+07 | 1.96E+07 | 2.42E+07 | 2.28E+07 | 1.52E+07 | 1.44E+07 | 3.59E+07 | 1.57E+07 | 2.44E+07 | 1263.7032 | RTIAQDYGVLK | P35700 | P35700 | PRDX1 | peroxiredoxin 1 | Cytoplasm | enzyme |
| 15531507 | 2 | 0.551 | 2.30E+07 | 1.79E+07 | 2.04E+07 | 1.62E+07 | 1.70E+07 | 2.43E+07 | 2.09E+07 | 1.52E+07 | 1.35E+07 | 3.46E+07 | 1.44E+07 | 2.32E+07 | 1225.6881 | QITINDLPVGR | P35700 | P35700 | PRDX1 | peroxiredoxin 1 | Cytoplasm | enzyme |
| 15532476 | 2 | 0.774 | 1.48E+07 | 1.24E+07 | 1.57E+07 | 1.57E+07 | 1.48E+07 | 1.64E+07 | 1.27E+07 | 1.36E+07 | 1.28E+07 | 2.67E+07 | 9739484 | 1.54E+07 | 920.5071 | GLFIIDDK | P35700 | P35700 | PRDX1 | peroxiredoxin 1 | Cytoplasm | enzyme |
| 15540040 | 4 | 0.406 | 1.40E+07 | 8561464 | 6585204.5 | 8181855.5 | 1.02E+07 | 1.18E+07 | 1.36E+07 | 9337684 | 9672166 | 1.80E+07 | 1.24E+07 | 1.10E+07 | 2767.3384 | LNC[160.0307]QVIGASVDSHFC[160.0307]HLAWINTPK | P35700 | P35700 | PRDX1 | peroxiredoxin 1 | Cytoplasm | enzyme |
| 15540920 | 2 | 0.384 | 9633793 | 5995394.5 | 8066521.5 | 7867783.5 | 6063163.5 | 8201006 | 6710837.5 | 5054061 | 3744466 | 1.47E+07 | 4861626.5 | 7210209.5 | 954.475 | DISLSEYK | P35700 | P35700 | PRDX1 | peroxiredoxin 1 | Cytoplasm | enzyme |
| 15541900, 15545241 | 2 | 0.845 | 4338631.5 | 6805016 | 1.41E+07 | 4250009.5 | 9449534 | 1.02E+07 | 9028467 | 8142608.5 | 4156641.75 | 1.18E+07 | 2335077.25 | 4790418.5 | 1140.5222, 1140.5228 | HGEVC[160.0307]PAGWK | P35700 | P35700 | PRDX1 | peroxiredoxin 1 | Cytoplasm | enzyme |
| 15544806 | 2 | 0.334 | 3649240.25 | 3427727.25 | 1382144.625 | 4708434.5 | 2318089.5 | 1833555.75 | 3273521.75 | 1613397.375 | 665124.25 | 9924490 | 3578119 | 2388910.5 | 894.4298 | ADEGISFR | P35700 | P35700 | PRDX1 | peroxiredoxin 1 | Cytoplasm | enzyme |
| 15534871 | 2 | 0.707 | 994441 | 1551495.25 | 2550580 | 377857.125 | 1819660.625 | 1489583.625 | 918108.438 | 1377260.5 | 653717.938 | 1872093.75 | 393501.844 | 975510.188 | 1270.6616 | PGSDTIKPDVNK | P35700 | P35700 | PRDX1 | peroxiredoxin 1 | Cytoplasm | enzyme |
| 15547667 | 2 | 0.839 | 5.10E+08 | 3.37E+08 | 4.43E+08 | 3.67E+08 | 3.78E+08 | 4.51E+08 | 3.84E+08 | 2.82E+08 | 2.95E+08 | 8.85E+08 | 2.07E+08 | 2.95E+08 | 1706.966 | EGGLGPLNIPLLADVTK | Q61171 | Q61171 | PRDX2 | peroxiredoxin 2 | Cytoplasm | enzyme |
| 15537291 | 2 | 0.427 | 2.91E+08 | 1.97E+08 | 2.41E+08 | 2.39E+08 | 1.82E+08 | 2.51E+08 | 2.21E+08 | 1.51E+08 | 1.34E+08 | 5.02E+08 | 1.63E+08 | 2.21E+08 | 1211.6721 | QITVNDLPVGR | Q61171 | Q61171 | PRDX2 | peroxiredoxin 2 | Cytoplasm | enzyme |
| 15531747 | 2 | 0.442 | 2.69E+08 | 2.11E+08 | 2.24E+08 | 1.92E+08 | 1.80E+08 | 1.99E+08 | 2.15E+08 | 1.53E+08 | 1.42E+08 | 4.56E+08 | 1.49E+08 | 2.20E+08 | 1108.598 | SLSQNYGVLK | Q61171 | Q61171 | PRDX2 | peroxiredoxin 2 | Cytoplasm | enzyme |
| 15548617, 15549046 | 2, 3 | 0.638 | 1.38E+08 | 1.92E+08 | 3.52E+08 | 9.52E+07 | 1.71E+08 | 2.68E+08 | 1.78E+08 | 1.46E+08 | 9.06E+07 | 2.89E+08 | 5.48E+07 | 1.03E+08 | 2335.0715, 2335.075 | LVQAFQYTDEHGEVC[160.0307]PAGWK | Q61171 | Q61171 | PRDX2 | peroxiredoxin 2 | Cytoplasm | enzyme |
| 15554791, 15554972, 15557072 | 2 | 0.802 | 1.32E+08 | 8.78E+07 | 1.10E+08 | 1.07E+08 | 9.04E+07 | 1.02E+08 | 9.68E+07 | 7.01E+07 | 8.34E+07 | 2.27E+08 | 5.84E+07 | 9.16E+07 | 1596.7877, 1596.7888 | SAPDFTATAVVDGAFK | Q61171 | Q61171 | PRDX2 | peroxiredoxin 2 | Cytoplasm | enzyme |
| 15535864 | 2 | 0.734 | 1.20E+08 | 8.58E+07 | 1.15E+08 | 9.28E+07 | 8.36E+07 | 1.07E+08 | 1.05E+08 | 7.24E+07 | 6.52E+07 | 2.00E+08 | 5.93E+07 | 9.47E+07 | 876.5167 | GLFIIDAK | Q61171 | Q61171 | PRDX2 | peroxiredoxin 2 | Cytoplasm | enzyme |
| 15557428, 15562282 | 3 | 0.969 | 1.11E+08 | 1.00E+08 | 1.57E+08 | 9.01E+07 | 1.29E+08 | 1.65E+08 | 1.33E+08 | 1.12E+08 | 1.24E+08 | 1.96E+08 | 6.35E+07 | 1.06E+08 | 2699.349, 2699.3533 | LGC[160.0307]EVLGVSVDSQFTHLAWINTPR | Q61171 | Q61171 | PRDX2 | peroxiredoxin 2 | Cytoplasm | enzyme |
| 15532196 | 2, 3 | 0.813 | 5.95E+07 | 6.85E+07 | 9.91E+07 | 5.17E+07 | 6.98E+07 | 8.74E+07 | 8.20E+07 | 5.67E+07 | 2.77E+07 | 1.41E+08 | 2.66E+07 | 4.65E+07 | 1472.7192, 1472.7197 | PGSDTIKPNVDDSK | Q61171 | Q61171 | PRDX2 | peroxiredoxin 2 | Cytoplasm | enzyme |
| 15543478, 15544518 | 2 | 0.643 | 5.03E+07 | 3.57E+07 | 5.22E+07 | 4.46E+07 | 4.06E+07 | 4.81E+07 | 3.76E+07 | 3.21E+07 | 2.85E+07 | 1.03E+08 | 2.66E+07 | 3.61E+07 | 937.43616, 937.43634 | NDEGIAYR | Q61171 | Q61171 | PRDX2 | peroxiredoxin 2 | Cytoplasm | enzyme |
| 15534909 | 3 | 0.815 | 1.45E+07 | 1.37E+07 | 9375423 | 1.04E+07 | 1.18E+07 | 1.52E+07 | 1.39E+07 | 3245152.75 | 7937357 | 4.08E+07 | 9659971 | 2499952.5 | 1706.9652 | EGGLGPLNIPLLADVTK | Q61171 | Q61171 | PRDX2 | peroxiredoxin 2 | Cytoplasm | enzyme |
| 15548388 | 3 | 0.806 | 2.08E+07 | 2.35E+07 | 3.08E+07 | 1.48E+07 | 2.57E+07 | 2.75E+07 | 2.61E+07 | 1.90E+07 | 1.17E+07 | 4.01E+07 | 9910558 | 1.58E+07 | 3033.4302 | LVQAFQYTDEHGEVC[160.0307]PAGWKPGSDTIK | Q61171 | Q61171 | PRDX2 | peroxiredoxin 2 | Cytoplasm | enzyme |
| 15534741 | 4 | 0.415 | 4.31E+07 | 9601556 | 8893479 | 4.79E+07 | 6360842 | 1.58E+07 | 2.05E+07 | 6324274.5 | 1.63E+07 | 3.41E+07 | 3.86E+07 | 3.33E+07 | 2827.449 | LGC[160.0307]EVLGVSVDSQFTHLAWINTPRK | Q61171 | Q61171 | PRDX2 | peroxiredoxin 2 | Cytoplasm | enzyme |
| 15537114 | 3 | 0.195 | 1.40E+07 | 2054810.625 | 476983.062 | 6503333.5 | 2982649.75 | 1035687.875 | 2510543.75 | 2333408.75 | 3765452 | 2.68E+07 | 1.10E+07 | 8684427 | 1835.0602 | KEGGLGPLNIPLLADVTK | Q61171 | Q61171 | PRDX2 | peroxiredoxin 2 | Cytoplasm | enzyme |
| 15552945 | 3 | 0.416 | 1.10E+07 | 2826219 | 1985292 | 1.35E+07 | 1781295.75 | 5218921.5 | 5218237 | 2066630.25 | 5077836 | 8623255 | 1.21E+07 | 8898291 | 2827.4485 | LGC[160.0307]EVLGVSVDSQFTHLAWINTPRK | Q61171 | Q61171 | PRDX2 | peroxiredoxin 2 | Cytoplasm | enzyme |
| 15551304 | 3 | 0.102 | 7.18E+07 | 1.99E+08 | 7.85E+07 | 7.69E+07 | 1.14E+08 | 1.21E+08 | 6.98E+07 | 1.15E+08 | 1.17E+08 | 4.61E+07 | 5.01E+07 | 5.85E+07 | 2745.2947 | AFQFVETHGEVC[160.0307]PANWTPESPTIK | P20108 | P20108 | PRDX3 | peroxiredoxin 3 | Cytoplasm | enzyme |
| 15531443 | 2, 3 | 0.106 | 6.49E+07 | 9.44E+07 | 5.50E+07 | 6.74E+07 | 5.51E+07 | 8.11E+07 | 3.75E+07 | 6.62E+07 | 7.25E+07 | 3.39E+07 | 4.16E+07 | 4.60E+07 | 1206.6565, 1206.6566 | HLSVNDLPVGR | P20108 | P20108 | PRDX3 | peroxiredoxin 3 | Cytoplasm | enzyme |
| 15541111 | 2 | 0.134 | 3.74E+07 | 6.32E+07 | 3.86E+07 | 4.14E+07 | 3.37E+07 | 5.36E+07 | 2.39E+07 | 4.86E+07 | 4.92E+07 | 2.10E+07 | 2.52E+07 | 2.93E+07 | 1271.7336 | GLFIIDPNGVVK | P20108 | P20108 | PRDX3 | peroxiredoxin 3 | Cytoplasm | enzyme |
| 15533417 | 3 | 0.273 | 1.93E+07 | 3.68E+07 | 1.54E+07 | 2.75E+07 | 1.96E+07 | 2.87E+07 | 1.34E+07 | 2.18E+07 | 2.73E+07 | 1.25E+07 | 1.55E+07 | 1.62E+07 | 1783.9343 | NGGLGHMNITLLSDITK | P20108 | P20108 | PRDX3 | peroxiredoxin 3 | Cytoplasm | enzyme |
| 15532791 | 2 | 0.079 | 2.27E+07 | 4.17E+07 | 2.54E+07 | 2.77E+07 | 2.95E+07 | 4.58E+07 | 1.84E+07 | 3.47E+07 | 3.34E+07 | 1.23E+07 | 1.70E+07 | 2.02E+07 | 966.47595 | ELSLDDFK | P20108 | P20108 | PRDX3 | peroxiredoxin 3 | Cytoplasm | enzyme |
| 15549945 | 3, 4 | 0.704 | 2.52E+07 | 6207214 | 285143.781 | 2.67E+07 | 2151590.5 | 1948040.625 | 2180135.5 | 1880794.875 | 2.23E+07 | 6988498.5 | 2.61E+07 | 1.69E+07 | 3413.6392, 3413.641 | AFQFVETHGEVC[160.0307]PANWTPESPTIKPSPTASK | P20108 | P20108 | PRDX3 | peroxiredoxin 3 | Cytoplasm | enzyme |
| 15549352 | 2 | 0.577 | 4077772.5 | 6842465 | 2727468.75 | 5604672 | 3309819.75 | 5463803 | 2412488.5 | 3902198.5 | 5857914.5 | 3089876.25 | 2711118.5 | 3583199.5 | 1783.9364 | NGGLGHMNITLLSDITK | P20108 | P20108 | PRDX3 | peroxiredoxin 3 | Cytoplasm | enzyme |
| 15549447 | 3 | 0.68 | 7675957 | 1.12E+07 | 7012721 | 1.04E+07 | 1.10E+07 | 7877912 | 8182878 | 9846908 | 9849280 | 8968067 | 8264780 | 8129405.5 | 2845.3872 | SINTEVVAC[160.0307]SVDSQFTHLAWINTPR | O08807 | O08807 | PRDX4 | peroxiredoxin 4 | Cytoplasm | enzyme |
| 15550615 | 3 | 0.696 | 5800812 | 4703155.5 | 1953506.875 | 4994104 | 4797290 | 4329228 | 3990447.25 | 4167052.75 | 6212961.5 | 4495310 | 5662518.5 | 5208531 | 2403.171 | HGEVC[160.0307]PAGWKPGSETIIPDPAGK | O08807 | O08807 | PRDX4 | peroxiredoxin 4 | Cytoplasm | enzyme |
| 15545548 | 2 | 0.493 | 938004.812 | 1645037.625 | 3154741.5 | 805148.75 | 2901754.75 | 2017127.625 | 1918724.875 | 2275792 | 892722.125 | 1535138.375 | 428795.281 | 962034.125 | 1281.6665 | PGSETIIPDPAGK | O08807 | O08807 | PRDX4 | peroxiredoxin 4 | Cytoplasm | enzyme |
| 15539337 | 2 | 0.035 | 1492612.5 | 2847337 | 3093711.75 | 6296442.5 | 6275124 | 5905649 | 3032419.25 | 4571868 | 5613385.5 | 2326102.5 | 3819527 | 4913880.5 | 1089.591 | LLADPTGAFGK | P99029 | P99029 | PRDX5 | peroxiredoxin 5 | Cytoplasm | enzyme |
| 15537147 | 2 | 0.612 | 95078.961 | 348702.469 | 222347.828 | 1047241.188 | 577979.875 | 2662797.75 | 610044.062 | 239482.25 | 1.46E+07 | 21925.25 | 3400959.75 | 4272558.5 | 1284.6791 | GINQGQVWIGGR | Q61878 | Q61878 | PRG2 | proteoglycan 2, bone marrow (natural killer cell activator, eosinophil granule major basic protein) | Extracellular Space | other |
| 15541716 | 3 | 0.006 | 5089048.5 | 4253619 | 6088936.5 | 4749551 | 6027553 | 5898198.5 | 5526523 | 5078092.5 | 4651809 | 3667030.25 | 3353601.25 | 3176094.75 | 2063.974 | KWETPSQNTAQLDQFDR | P05132 | P05132 | PRKACA | protein kinase, cAMP-dependent, catalytic, alpha | Cytoplasm | kinase |
| 15537137 | 2 | 0.156 | 84385.445 | 229969 | 547002.312 | 2548198.25 | 1677726.75 | 1686389 | 1026311.938 | 1187758.625 | 1880619.375 | 51849.727 | 2075945.75 | 1307320.25 | 1397.7485 | LTDFNFLMVLGK | P20444 | P20444 | PRKCA | protein kinase C, alpha | Cytoplasm | kinase |
| 15534965 | 3 | 0.702 | 1.09E+07 | 1.16E+07 | 1.17E+07 | 1.18E+07 | 1.40E+07 | 1.24E+07 | 1.50E+07 | 1.34E+07 | 9282466 | 1.21E+07 | 1.28E+07 | 1.39E+07 | 2149.9626 | HGGSPTSLGTWGSWAGPDHDK | O08795 | O08795 | PRKCSH | protein kinase C substrate 80K-H | Cytoplasm | enzyme |
| 15535701 | 2 | 0.258 | 8780058 | 8817790 | 8695084 | 8444385 | 1.11E+07 | 1.16E+07 | 1.18E+07 | 1.05E+07 | 9110225 | 1.01E+07 | 1.08E+07 | 9787079 | 1189.6034 | SLEDQVETLR | O08795 | O08795 | PRKCSH | protein kinase C substrate 80K-H | Cytoplasm | enzyme |
| 15543022 | 3 | 0.016 | 6746191 | 7426559 | 7493051 | 8571569 | 1.17E+07 | 9712350 | 9388652 | 8274871 | 8453816 | 9224265 | 8732851 | 9042629 | 2147.9895 | MPPYDEETQAIIDAAQEAR | O08795 | O08795 | PRKCSH | protein kinase C substrate 80K-H | Cytoplasm | enzyme |
| 15534099 | 2 | 0.333 | 6404626 | 6240061.5 | 7265051.5 | 6412826 | 8002196 | 7190521 | 7899135.5 | 7100243.5 | 5604177.5 | 8231156.5 | 7870223 | 7496519 | 1173.5868 | LWEEQQAAAK | O08795 | O08795 | PRKCSH | protein kinase C substrate 80K-H | Cytoplasm | enzyme |
| 15535076 | 2 | 0.607 | 4864285.5 | 5297490.5 | 6000708 | 4982049 | 7101073 | 3727389 | 5582218 | 7912810.5 | 5070161 | 7553271 | 5544860.5 | 6065405.5 | 948.5488 | PLYILSSR | O08795 | O08795 | PRKCSH | protein kinase C substrate 80K-H | Cytoplasm | enzyme |
| 15552793 | 2 | 0.201 | 5493874 | 4710830 | 5989762 | 6139311.5 | 6443815.5 | 5354659 | 6115373 | 5588776.5 | 5608893.5 | 6852039.5 | 6176407 | 6092170.5 | 1552.6571 | YEQGTGC[160.0307]WQGPNR | O08795 | O08795 | PRKCSH | protein kinase C substrate 80K-H | Cytoplasm | enzyme |
| 15532384 | 2 | 0.087 | 4564614 | 4703833 | 6246995.5 | 5364025 | 6075089.5 | 5405031.5 | 7222399.5 | 6396439.5 | 6015059.5 | 6627142.5 | 5943533.5 | 6280497 | 930.5284 | ILIEEWK | O08795 | O08795 | PRKCSH | protein kinase C substrate 80K-H | Cytoplasm | enzyme |
| 15542314 | 2 | 0.788 | 3992254.75 | 4553619 | 5547181 | 4137916 | 7172877 | 4739515.5 | 5868745.5 | 5492802 | 3558485.25 | 6257360.5 | 3184841.75 | 3547197.5 | 1306.6456 | ETVVTSTTEPSR | O08795 | O08795 | PRKCSH | protein kinase C substrate 80K-H | Cytoplasm | enzyme |
| 15557273 | 3 | 0.527 | 2796542.25 | 2773767 | 2770128.25 | 3111438.25 | 4893980 | 2821006.75 | 2915180.25 | 3828551.25 | 3436074.25 | 2387720.25 | 4028928.5 | 2596305.25 | 3057.0962 | VNDGVC[160.0307]DC[160.0307]C[160.0307]DGTDEYNSGTVC[160.0307]ENTC[160.0307]R | O08795 | O08795 | PRKCSH | protein kinase C substrate 80K-H | Cytoplasm | enzyme |
| 15550260 | 3 | 0.199 | 3694258.5 | 2883867 | 2599633 | 3949804.75 | 4134760 | 2372085.75 | 4532877 | 4434522.5 | 3455219.5 | 2340806.25 | 3332051.5 | 2675423 | 2543.0132 | DGSDEPGTAAC[160.0307]PNGSFHC[160.0307]TNTGYK | O08795 | O08795 | PRKCSH | protein kinase C substrate 80K-H | Cytoplasm | enzyme |
| 15544352 | 2 | 0.274 | 1308183.5 | 495399.25 | 188616.391 | 1515423.875 | 970914.25 | 407888.781 | 796526.625 | 827428.25 | 1229530.25 | 1350851.25 | 2048984.125 | 1144770.125 | 999.61816 | LLELQAGKK | O08795 | O08795 | PRKCSH | protein kinase C substrate 80K-H | Cytoplasm | enzyme |
| 15551022 | 3 | 0.133 | 6757400 | 8059759.5 | 6696008.5 | 3903779.75 | 5475551.5 | 2921040.5 | 6184441.5 | 6226979.5 | 4983385.5 | 7362398.5 | 3139734.75 | 4598885.5 | 3288.6028 | ADIIVSELLGSFADNELSPEC[160.0307]LDGAQHFLK | Q8CIG8 | Q8CIG8 | PRMT5 | protein arginine methyltransferase 5 | Cytoplasm | enzyme |
| 15538451 | 2 | 0.165 | 2202575.5 | 1324688.875 | 1693852.625 | 1090564.25 | 1167929.625 | 498324.844 | 1430815.75 | 1034146 | 710846.625 | 1373456 | 667128.688 | 824839.062 | 1291.6293 | YSQYQQAIYK | Q8CIG8 | Q8CIG8 | PRMT5 | protein arginine methyltransferase 5 | Cytoplasm | enzyme |
| 15549543 | 2 | 0.004 | 8166718.5 | 9446132 | 8718196 | 1.47E+07 | 1.44E+07 | 1.82E+07 | 7945009 | 8698013 | 5863268.5 | 1.09E+07 | 1.05E+07 | 6976946.5 | 1512.7153 | VVEQMC[160.0307]VTQYQK | P04925 | P04925 | PRNP | prion protein | Plasma Membrane | other |
| 15539761 | 2 | 0.312 | 1.31E+07 | 8954965 | 9279562 | 7318023 | 8479850 | 4753001.5 | 8438571 | 7340673 | 7134315 | 1.29E+07 | 6549175.5 | 6824323.5 | 1393.7258 | VTAVIPC[160.0307]FPYAR | Q9D7G0 | Q9D7G0 | PRPS1 | phosphoribosyl pyrophosphate synthetase 1 | Cytoplasm | kinase |
| 15534294 | 2 | 0.822 | 1.10E+07 | 6719285 | 5770729.5 | 7628619 | 6882611.5 | 4921057.5 | 7555300 | 6324863 | 5026486.5 | 9897321 | 6156220 | 5532343 | 1289.6125 | NC[160.0307]TIVSPDAGGAK | Q9D7G0 | Q9D7G0 | PRPS1 | phosphoribosyl pyrophosphate synthetase 1 | Cytoplasm | kinase |
| 15537785 | 3 | 0.494 | 6984446 | 5193010 | 3589116 | 3792092.75 | 4723626 | 3014143.75 | 3490985 | 3323741 | 3608391.5 | 5907413 | 2150926.75 | 3544283.5 | 1786.8723 | THNGESVSYLFSHVPL | Q9D7G0 | Q9D7G0 | PRPS1 | phosphoribosyl pyrophosphate synthetase 1 | Cytoplasm | kinase |
| 15536318 | 2 | 0.323 | 7209016 | 5266749.5 | 4378549.5 | 5017204 | 4776063 | 3525430.75 | 4333529 | 3968465.75 | 3928414.5 | 5669409 | 2840207.25 | 3325500.75 | 860.4878 | MVLVGDVK | Q9D7G0 | Q9D7G0 | PRPS1 | phosphoribosyl pyrophosphate synthetase 1 | Cytoplasm | kinase |
| 15532980 | 2 | 0.35 | 3984457 | 3161440.5 | 3245455 | 2668094 | 2938203.25 | 2028765.375 | 2887129.75 | 2662214.5 | 2137642.75 | 3930707.75 | 1906873.875 | 2206896 | 1433.6991 | IFSGSSHQDLSQK | Q9D7G0 | Q9D7G0 | PRPS1 | phosphoribosyl pyrophosphate synthetase 1 | Cytoplasm | kinase |
| 15538128 | 3 | 0.137 | 3198309.75 | 3199908.5 | 4149311.25 | 2260736.25 | 3664456.75 | 2316220.75 | 2588491.75 | 2513673 | 2232079.25 | 2821385 | 756363.062 | 1770813.75 | 1169.6648 | LNVDFALIHK | Q9D7G0 | Q9D7G0 | PRPS1 | phosphoribosyl pyrophosphate synthetase 1 | Cytoplasm | kinase |
| 15554049 | 3 | 0.692 | 7851605 | 6950075 | 7268019 | 5894989 | 5053140 | 5514191.5 | 7781550.5 | 5496315 | 4382142.5 | 1.13E+07 | 3760221 | 5674166 | 2489.3083 | LIEESPIDEVVVTNTVPHELQK | Q9D0M1 | Q9D0M1 | PRPSAP1 | phosphoribosyl pyrophosphate synthetase-associated protein 1 | Extracellular Space | other |
| 15553456 | 2 | 0.438 | 4189851.25 | 3451733.5 | 3344107.25 | 3557149 | 2787629.5 | 2164900 | 3828730.5 | 2918176.5 | 2679785.75 | 4218425 | 2787913 | 3159657.5 | 1668.8358 | EIQGFFSFPVDNLR | Q9D0M1 | Q9D0M1 | PRPSAP1 | phosphoribosyl pyrophosphate synthetase-associated protein 1 | Extracellular Space | other |
| 15554587 | 2 | 0.695 | 3312471.75 | 1850718.25 | 2770454.75 | 2417834.5 | 2585287.25 | 1680484.125 | 2351894 | 1952898.75 | 1714061.25 | 3932122.5 | 1603227.375 | 2611036.25 | 2081.0657 | ASPFLLQYIQEEIPNYR | Q9D0M1 | Q9D0M1 | PRPSAP1 | phosphoribosyl pyrophosphate synthetase-associated protein 1 | Extracellular Space | other |
| 15542036 | 2 | 0.762 | 3646002.25 | 2664509.5 | 1853455.75 | 2656913.5 | 2647564 | 1959121.875 | 2636127.5 | 2031297.625 | 1699923.5 | 3358509.25 | 1578780.625 | 1784086 | 1400.7863 | GQDIFIIQTIPR | Q9D0M1 | Q9D0M1 | PRPSAP1 | phosphoribosyl pyrophosphate synthetase-associated protein 1 | Extracellular Space | other |
| 15552939 | 2 | 0.734 | 2523176.75 | 1049240.125 | 1528566 | 1604872.75 | 1751611.625 | 1154599.375 | 1286150.625 | 1355704.5 | 1826577.875 | 3115181.75 | 1323137.125 | 1781543.375 | 1569.7545 | VFSANSTAAC[160.0307]TELAK | Q9D0M1 | Q9D0M1 | PRPSAP1 | phosphoribosyl pyrophosphate synthetase-associated protein 1 | Extracellular Space | other |
| 15550316 | 2 | 0.808 | 2368691.5 | 2220704.25 | 2831766.75 | 2461619.75 | 1774981.625 | 1920905.625 | 2774308.75 | 1718446.625 | 1651574 | 4700916.5 | 1404911.625 | 2238337.25 | 1661.8608 | EIQGFFNIPVDNLR | Q8R574 | Q8R574 | PRPSAP2 | phosphoribosyl pyrophosphate synthetase-associated protein 2 | Cytoplasm | other |
| 15552915 | 3 | 0.9 | 3530713 | 2993918.5 | 2489095.75 | 3477231.75 | 3583333 | 2832749.5 | 3473080.5 | 2619860.25 | 2709123.5 | 4188525.75 | 2073076.125 | 2913364.75 | 2477.3083 | LIEESAIDEVVVTNTIPHEIQK | Q8R574 | Q8R574 | PRPSAP2 | phosphoribosyl pyrophosphate synthetase-associated protein 2 | Cytoplasm | other |
| 15537557 | 2 | 0.038 | 2178099 | 5558671 | 2320005 | 4561437 | 4560932 | 5852375 | 2270406.25 | 3520329.5 | 4989666.5 | 1502420 | 1844618.75 | 2022984.5 | 1059.6019 | LVSDVQTAVK | Q61207 | Q61207 | PSAP | prosaposin | Extracellular Space | other |
| 15557352 | 2 | 0.392 | 4807088 | 4368378 | 5013856.5 | 3316067.25 | 3683090.75 | 5149223.5 | 5246239.5 | 4913111.5 | 5668935.5 | 6330520 | 3769265.5 | 4058826.25 | 1627.9061 | LGLGDFIFYSVLVGK | P49769 | P49769 | PSEN1 | presenilin 1 | Plasma Membrane | peptidase |
| 15542290 | 2 | 0.749 | 1239452.75 | 1828284.875 | 2834759.5 | 1307447.75 | 2602542.25 | 1443844 | 1951237 | 2477260.25 | 1750041.375 | 1545284.5 | 1467642.125 | 1749391.125 | 1076.5377 | MLVETAQER | P49769 | P49769 | PSEN1 | presenilin 1 | Plasma Membrane | peptidase |
| 15543651 | 3 | 0.315 | 1920675 | 1896864.5 | 1300244 | 1596527.25 | 1680302.25 | 1010088.625 | 1128021.125 | 1262355 | 1256911.75 | 1615870.5 | 1268783.875 | 1174579.5 | 2146.9185 | SC[160.0307]QEGRPGPEDGESTAQWR | Q61144 | Q61144 | PSEN2 | presenilin 2 | Cytoplasm | peptidase |
| 15533228 | 2 | 0.265 | 3362422.75 | 2869223.5 | 3439930.25 | 5211059 | 2266722.25 | 3199305.25 | 2723934.5 | 3211400.75 | 2912402.5 | 3893297.75 | 5360648 | 4042158.5 | 902.5361 | SISQNLLK | Q91YA2 | Q91YA2 | PSKH1 | protein serine kinase H1 | Nucleus | kinase |
| 15532633 | 2, 3 | 0.675 | 5.05E+07 | 4.13E+07 | 4.32E+07 | 3.64E+07 | 3.59E+07 | 4.41E+07 | 4.17E+07 | 3.49E+07 | 3.73E+07 | 5.75E+07 | 2.89E+07 | 4.97E+07 | 1332.7395, 1332.7396 | FVFDRPLPVSR | Q9R1P4 | Q9R1P4 | PSMA1 | proteasome (prosome, macropain) subunit, alpha type, 1 | Cytoplasm | peptidase |
| 15554104, 15560984 | 4 | 0.35 | 2.86E+07 | 1.64E+07 | 7825151 | 2.54E+07 | 2.07E+07 | 7374712 | 2.66E+07 | 1.30E+07 | 9091276 | 5.63E+07 | 2.42E+07 | 2.56E+07 | 3690.6797, 3690.687 | RPYGVGLLIAGYDDMGPHIFQTC[160.0307]PSANYFDC[160.0307]R | Q9R1P4 | Q9R1P4 | PSMA1 | proteasome (prosome, macropain) subunit, alpha type, 1 | Cytoplasm | peptidase |
| 15533084 | 2, 3 | 0.42 | 3.33E+07 | 2.39E+07 | 1.84E+07 | 2.08E+07 | 2.16E+07 | 1.93E+07 | 2.94E+07 | 2.12E+07 | 1.86E+07 | 4.85E+07 | 2.07E+07 | 2.88E+07 | 1431.7268 | IHQIEYAMEAVK | Q9R1P4 | Q9R1P4 | PSMA1 | proteasome (prosome, macropain) subunit, alpha type, 1 | Cytoplasm | peptidase |
| 15533866 | 2 | 0.919 | 2.53E+07 | 2.79E+07 | 2.60E+07 | 1.69E+07 | 2.96E+07 | 2.07E+07 | 2.88E+07 | 2.60E+07 | 1.71E+07 | 4.35E+07 | 1.43E+07 | 2.35E+07 | 1345.682 | ETLPAEQDLTTK | Q9R1P4 | Q9R1P4 | PSMA1 | proteasome (prosome, macropain) subunit, alpha type, 1 | Cytoplasm | peptidase |
| 15548095 | 2 | 0.691 | 2.45E+07 | 1.86E+07 | 2.13E+07 | 2.05E+07 | 1.81E+07 | 1.77E+07 | 2.24E+07 | 1.81E+07 | 1.73E+07 | 3.93E+07 | 1.54E+07 | 2.16E+07 | 1778.8052 | NQYDNDVTVWSPQGR | Q9R1P4 | Q9R1P4 | PSMA1 | proteasome (prosome, macropain) subunit, alpha type, 1 | Cytoplasm | peptidase |
| 15532812 | 3 | 0.329 | 3.30E+07 | 1.59E+07 | 1.26E+07 | 3.24E+07 | 8304655 | 1.57E+07 | 2.36E+07 | 1.10E+07 | 1.64E+07 | 3.90E+07 | 2.61E+07 | 3.09E+07 | 1685.9026 | ALRETLPAEQDLTTK | Q9R1P4 | Q9R1P4 | PSMA1 | proteasome (prosome, macropain) subunit, alpha type, 1 | Cytoplasm | peptidase |
| 15548809 | 3 | 0.957 | 1.38E+07 | 1.88E+07 | 1.74E+07 | 8809058 | 2.57E+07 | 1.48E+07 | 2.01E+07 | 2.06E+07 | 9609047 | 3.19E+07 | 5723462 | 1.22E+07 | 2984.3723 | DLEFTIYDDDDVSPFLDGLEERPQR | Q9R1P4 | Q9R1P4 | PSMA1 | proteasome (prosome, macropain) subunit, alpha type, 1 | Cytoplasm | peptidase |
| 15540203 | 2 | 0.512 | 1.96E+07 | 1.86E+07 | 1.57E+07 | 1.40E+07 | 1.39E+07 | 1.49E+07 | 1.85E+07 | 1.50E+07 | 1.28E+07 | 3.00E+07 | 1.26E+07 | 1.75E+07 | 886.5338 | NVSIGIVGK | Q9R1P4 | Q9R1P4 | PSMA1 | proteasome (prosome, macropain) subunit, alpha type, 1 | Cytoplasm | peptidase |
| 15533083 | 2 | 0.377 | 1.91E+07 | 1.46E+07 | 1.39E+07 | 1.30E+07 | 1.44E+07 | 1.24E+07 | 1.95E+07 | 1.43E+07 | 1.25E+07 | 2.90E+07 | 1.41E+07 | 1.69E+07 | 951.5974 | THAVLVALK | Q9R1P4 | Q9R1P4 | PSMA1 | proteasome (prosome, macropain) subunit, alpha type, 1 | Cytoplasm | peptidase |
| 15531560 | 2 | 0.505 | 1.74E+07 | 1.30E+07 | 1.47E+07 | 1.41E+07 | 1.40E+07 | 1.46E+07 | 1.58E+07 | 1.34E+07 | 1.38E+07 | 2.88E+07 | 1.18E+07 | 1.90E+07 | 816.51794 | LVSLIGSK | Q9R1P4 | Q9R1P4 | PSMA1 | proteasome (prosome, macropain) subunit, alpha type, 1 | Cytoplasm | peptidase |
| 15536284 | 3 | 0.31 | 1.60E+07 | 1.24E+07 | 8422444 | 1.21E+07 | 1.07E+07 | 9621609 | 1.74E+07 | 1.07E+07 | 9823297 | 2.81E+07 | 1.14E+07 | 1.77E+07 | 1881.8148 | HMSEFMEC[160.0307]NLDELVK | Q9R1P4 | Q9R1P4 | PSMA1 | proteasome (prosome, macropain) subunit, alpha type, 1 | Cytoplasm | peptidase |
| 15552916 | 2 | 0.26 | 9156496 | 6502790.5 | 4300854.5 | 5520356.5 | 5825765 | 4859018 | 7941102.5 | 5220837 | 4294903 | 1.44E+07 | 7056266 | 7311173.5 | 1881.8146 | HMSEFMEC[160.0307]NLDELVK | Q9R1P4 | Q9R1P4 | PSMA1 | proteasome (prosome, macropain) subunit, alpha type, 1 | Cytoplasm | peptidase |
| 15535816 | 2 | 0.024 | 6909070.5 | 3612684 | 2744343 | 4157086.25 | 3950909.25 | 2754880.5 | 5084680 | 4712160.5 | 3961052.75 | 1.24E+07 | 8896428 | 6847732.5 | 953.46704 | LLC[160.0307]NFMR | Q9R1P4 | Q9R1P4 | PSMA1 | proteasome (prosome, macropain) subunit, alpha type, 1 | Cytoplasm | peptidase |
| 15539868 | 3 | 0.495 | 3489863.75 | 2796913.25 | 2260341 | 3767391.25 | 2867870.75 | 2430373.25 | 4046416 | 3048306 | 2203799 | 8351313 | 2382034.5 | 4268395 | 1778.8048 | NQYDNDVTVWSPQGR | Q9R1P4 | Q9R1P4 | PSMA1 | proteasome (prosome, macropain) subunit, alpha type, 1 | Cytoplasm | peptidase |
| 15550983 | 2 | 0.215 | 5740124 | 1941356.125 | 1536153.625 | 4519437 | 805471.812 | 1974512.375 | 2870810.25 | 1118994.875 | 1447575.875 | 6472292.5 | 4331780.5 | 4449911.5 | 1685.9028 | ALRETLPAEQDLTTK | Q9R1P4 | Q9R1P4 | PSMA1 | proteasome (prosome, macropain) subunit, alpha type, 1 | Cytoplasm | peptidase |
| 15554598 | 4 | 0.663 | 7973568.5 | 9009099 | 1.14E+07 | 2866042.5 | 9095783 | 1.21E+07 | 6473348.5 | 1.04E+07 | 1.07E+07 | 6201379.5 | 4032125.25 | 9642183 | 3706.678 | RPYGVGLLIAGYDDM[147.0354]GPHIFQTC[160.0307]PSANYFDC[160.0307]R | Q9R1P4 | Q9R1P4 | PSMA1 | proteasome (prosome, macropain) subunit, alpha type, 1 | Cytoplasm | peptidase |
| 15547866 | 2, 3 | 0.529 | 8.58E+07 | 7.46E+07 | 6.84E+07 | 6.56E+07 | 6.79E+07 | 5.15E+07 | 6.56E+07 | 6.66E+07 | 6.30E+07 | 1.16E+08 | 5.23E+07 | 7.55E+07 | 2521.3406, 2521.342 | LAQQYYLVYQEPIPTAQLVQR | P49722 | P49722 | PSMA2 | proteasome (prosome, macropain) subunit, alpha type, 2 | Cytoplasm | peptidase |
| 15551851 | 2, 3 | 0.635 | 6.41E+07 | 3.35E+07 | 1.87E+07 | 4.28E+07 | 2.17E+07 | 1.86E+07 | 5.13E+07 | 1.98E+07 | 1.23E+07 | 9.17E+07 | 2.61E+07 | 3.28E+07 | 2357.2312 | RYNEDLELEDAIHTAILTLK | P49722 | P49722 | PSMA2 | proteasome (prosome, macropain) subunit, alpha type, 2 | Cytoplasm | peptidase |
| 15538436 | 4 | 0.425 | 2.09E+07 | 2.37E+07 | 1.47E+07 | 2.05E+07 | 1.29E+07 | 1.29E+07 | 2.68E+07 | 1.47E+07 | 1.01E+07 | 4.28E+07 | 1.47E+07 | 2.63E+07 | 2357.231 | RYNEDLELEDAIHTAILTLK | P49722 | P49722 | PSMA2 | proteasome (prosome, macropain) subunit, alpha type, 2 | Cytoplasm | peptidase |
| 15536262 | 2 | 0.451 | 3.27E+07 | 2.56E+07 | 2.37E+07 | 2.14E+07 | 2.13E+07 | 2.19E+07 | 3.14E+07 | 2.03E+07 | 2.09E+07 | 4.18E+07 | 2.03E+07 | 2.89E+07 | 1478.7142 | GYSFSLTTFSPSGK | P49722 | P49722 | PSMA2 | proteasome (prosome, macropain) subunit, alpha type, 2 | Cytoplasm | peptidase |
| 15535717 | 3 | 0.894 | 1.10E+07 | 1.22E+07 | 1.29E+07 | 9584571 | 1.17E+07 | 1.35E+07 | 1.17E+07 | 1.25E+07 | 1.22E+07 | 1.94E+07 | 8305368.5 | 1.44E+07 | 2027.1494 | LVQIEYALAAVAGGAPSVGIK | P49722 | P49722 | PSMA2 | proteasome (prosome, macropain) subunit, alpha type, 2 | Cytoplasm | peptidase |
| 15548930 | 2 | 0.382 | 9460773 | 4422661 | 3466421.75 | 1.09E+07 | 1135729.625 | 4663014 | 5412613.5 | 2109723.25 | 6117839 | 1.27E+07 | 8485131 | 9360132 | 1532.8287 | LTPTEVRDYLAAIA | P49722 | P49722 | PSMA2 | proteasome (prosome, macropain) subunit, alpha type, 2 | Cytoplasm | peptidase |
| 15549458 | 2 | 0.404 | 5926920.5 | 4099535 | 3152124.25 | 4526637 | 4252484.5 | 1961705.5 | 4391485 | 2933985 | 3165931.5 | 9464894 | 4088194.5 | 4349525 | 1564.754 | HIGLVYSGMGPDYR | P49722 | P49722 | PSMA2 | proteasome (prosome, macropain) subunit, alpha type, 2 | Cytoplasm | peptidase |
| 15537467 | 3 | 0.944 | 3693916.5 | 3887199.25 | 3648489.75 | 2892154.75 | 6372809.5 | 4981022.5 | 4486226 | 6135610.5 | 2603408.5 | 8099449.5 | 1930695.5 | 3187109.25 | 2201.1304 | YNEDLELEDAIHTAILTLK | P49722 | P49722 | PSMA2 | proteasome (prosome, macropain) subunit, alpha type, 2 | Cytoplasm | peptidase |
| 15551576 | 2 | 0.991 | 4070293.25 | 4435293 | 4582537.5 | 3300374 | 3780916.25 | 5322273 | 4802287.5 | 4405104 | 3445556.25 | 7104912.5 | 2832891.5 | 3684266.75 | 2027.1495 | LVQIEYALAAVAGGAPSVGIK | P49722 | P49722 | PSMA2 | proteasome (prosome, macropain) subunit, alpha type, 2 | Cytoplasm | peptidase |
| 15537746 | 3 | 0.559 | 3380565.75 | 2755701.25 | 3594792 | 2281631.75 | 2164740.5 | 2325357.25 | 3356251.25 | 2537720.75 | 1673273.875 | 3739760.25 | 1314894.875 | 2652098 | 1580.749 | HIGLVYSGM[147.0354]GPDYR | P49722 | P49722 | PSMA2 | proteasome (prosome, macropain) subunit, alpha type, 2 | Cytoplasm | peptidase |
| 15536208 | 2 | 0.677 | 3.57E+07 | 2.74E+07 | 2.83E+07 | 3.02E+07 | 2.65E+07 | 2.25E+07 | 3.65E+07 | 2.80E+07 | 2.43E+07 | 6.39E+07 | 2.27E+07 | 3.10E+07 | 1152.6025 | SNFGYNIPLK | O70435 | O70435 | PSMA3 | proteasome (prosome, macropain) subunit, alpha type, 3 | Cytoplasm | peptidase |
| 15533427 | 2 | 0.43 | 3.42E+07 | 2.33E+07 | 2.49E+07 | 2.08E+07 | 2.45E+07 | 2.23E+07 | 2.67E+07 | 2.17E+07 | 2.21E+07 | 4.52E+07 | 2.06E+07 | 2.97E+07 | 1217.6448 | AVENSSTAIGIR | O70435 | O70435 | PSMA3 | proteasome (prosome, macropain) subunit, alpha type, 3 | Cytoplasm | peptidase |
| 15533003, 15534730 | 2, 3 | 0.638 | 2.81E+07 | 2.10E+07 | 2.20E+07 | 2.32E+07 | 2.15E+07 | 1.98E+07 | 2.91E+07 | 2.11E+07 | 1.87E+07 | 4.10E+07 | 1.83E+07 | 2.83E+07 | 1228.6898, 1228.6943 | IIYIVHDEVK | O70435 | O70435 | PSMA3 | proteasome (prosome, macropain) subunit, alpha type, 3 | Cytoplasm | peptidase |
| 15535223 | 2 | 0.542 | 2.33E+07 | 1.64E+07 | 1.58E+07 | 1.39E+07 | 1.37E+07 | 1.32E+07 | 2.14E+07 | 1.35E+07 | 1.26E+07 | 3.32E+07 | 1.16E+07 | 1.87E+07 | 1114.5565 | VFQVEYAMK | O70435 | O70435 | PSMA3 | proteasome (prosome, macropain) subunit, alpha type, 3 | Cytoplasm | peptidase |
| 15533254 | 2 | 0.697 | 1.91E+07 | 1.46E+07 | 1.45E+07 | 1.50E+07 | 1.47E+07 | 1.21E+07 | 1.77E+07 | 1.54E+07 | 1.12E+07 | 3.06E+07 | 1.12E+07 | 1.75E+07 | 949.4971 | DGVVFGVEK | O70435 | O70435 | PSMA3 | proteasome (prosome, macropain) subunit, alpha type, 3 | Cytoplasm | peptidase |
| 15537352 | 2 | 0.674 | 1.88E+07 | 1.54E+07 | 1.64E+07 | 1.38E+07 | 1.53E+07 | 1.49E+07 | 1.69E+07 | 1.41E+07 | 1.27E+07 | 2.65E+07 | 1.16E+07 | 1.73E+07 | 919.5096 | RLFNVDR | O70435 | O70435 | PSMA3 | proteasome (prosome, macropain) subunit, alpha type, 3 | Cytoplasm | peptidase |
| 15535195 | 2, 3 | 0.362 | 1.48E+07 | 1.07E+07 | 8301728 | 1.18E+07 | 1.15E+07 | 6103385 | 1.09E+07 | 9170951 | 8144224.5 | 2.25E+07 | 1.01E+07 | 1.33E+07 | 1380.739 | HVGMAVAGLLADAR | O70435 | O70435 | PSMA3 | proteasome (prosome, macropain) subunit, alpha type, 3 | Cytoplasm | peptidase |
| 15535458 | 3 | 0.41 | 1.31E+07 | 8381796 | 5446949.5 | 1.50E+07 | 2746369 | 5939080.5 | 1.04E+07 | 3573434.5 | 8761336 | 1.43E+07 | 1.10E+07 | 1.54E+07 | 1578.7837 | SLADIAREEASNFR | O70435 | O70435 | PSMA3 | proteasome (prosome, macropain) subunit, alpha type, 3 | Cytoplasm | peptidase |
| 15554712 | 2 | 0.574 | 9513253 | 4144208 | 3848867.5 | 9219431 | 1754711 | 4576323 | 6163681.5 | 2024142.625 | 6375360.5 | 1.11E+07 | 5993715.5 | 7485630.5 | 1578.7843 | SLADIAREEASNFR | O70435 | O70435 | PSMA3 | proteasome (prosome, macropain) subunit, alpha type, 3 | Cytoplasm | peptidase |
| 15547824 | 2, 3 | 0.025 | 3.86E+07 | 2.60E+07 | 2.52E+07 | 2.61E+07 | 2.97E+07 | 2.53E+07 | 3.25E+07 | 2.77E+07 | 3.14E+07 | 9.03E+07 | 6.14E+07 | 3.62E+07 | 2694.3472, 2694.3477 | YLLQYQEPIPC[160.0307]EQLVTALC[160.0307]DIK | Q9R1P0 | Q9R1P0 | PSMA4 | proteasome (prosome, macropain) subunit, alpha type, 4 | Cytoplasm | peptidase |
| 15548417, 15550651 | 2, 3 | 0.123 | 4.16E+07 | 3.33E+07 | 2.68E+07 | 3.51E+07 | 3.32E+07 | 3.09E+07 | 4.07E+07 | 3.93E+07 | 3.19E+07 | 8.05E+07 | 3.88E+07 | 4.63E+07 | 2593.2166, 2593.2195 | LNEDMAC[160.0307]SVAGITSDANVLTNELR | Q9R1P0 | Q9R1P0 | PSMA4 | proteasome (prosome, macropain) subunit, alpha type, 4 | Cytoplasm | peptidase |
| 15533061 | 2 | 0.737 | 4.47E+07 | 3.49E+07 | 3.78E+07 | 3.59E+07 | 3.60E+07 | 3.32E+07 | 4.86E+07 | 3.27E+07 | 3.04E+07 | 6.36E+07 | 2.92E+07 | 4.33E+07 | 1226.6279 | LLDEVFFSEK | Q9R1P0 | Q9R1P0 | PSMA4 | proteasome (prosome, macropain) subunit, alpha type, 4 | Cytoplasm | peptidase |
| 15540766 | 3 | 0.185 | 3.18E+07 | 2.28E+07 | 2.28E+07 | 2.14E+07 | 2.44E+07 | 2.49E+07 | 3.55E+07 | 2.36E+07 | 2.57E+07 | 4.60E+07 | 2.82E+07 | 3.13E+07 | 2203.9785 | HYGFQLYQSDPSGNYGGWK | Q9R1P0 | Q9R1P0 | PSMA4 | proteasome (prosome, macropain) subunit, alpha type, 4 | Cytoplasm | peptidase |
| 15532311 | 2 | 0.548 | 2.77E+07 | 2.19E+07 | 2.25E+07 | 2.14E+07 | 1.93E+07 | 2.09E+07 | 2.50E+07 | 2.15E+07 | 2.01E+07 | 4.48E+07 | 1.75E+07 | 2.67E+07 | 1007.5141 | TTIFSPEGR | Q9R1P0 | Q9R1P0 | PSMA4 | proteasome (prosome, macropain) subunit, alpha type, 4 | Cytoplasm | peptidase |
| 15537582 | 2 | 0.724 | 2.18E+07 | 1.91E+07 | 1.83E+07 | 2.07E+07 | 1.58E+07 | 1.62E+07 | 2.18E+07 | 1.80E+07 | 1.56E+07 | 3.63E+07 | 1.40E+07 | 2.12E+07 | 902.52985 | VEIATLTR | Q9R1P0 | Q9R1P0 | PSMA4 | proteasome (prosome, macropain) subunit, alpha type, 4 | Cytoplasm | peptidase |
| 15548899 | 2 | 0.268 | 2.13E+07 | 1.52E+07 | 1.43E+07 | 1.46E+07 | 1.54E+07 | 1.31E+07 | 1.86E+07 | 1.40E+07 | 1.53E+07 | 3.59E+07 | 1.84E+07 | 1.62E+07 | 1607.7842 | ATC[160.0307]IGNNSAAAVSMLK | Q9R1P0 | Q9R1P0 | PSMA4 | proteasome (prosome, macropain) subunit, alpha type, 4 | Cytoplasm | peptidase |
| 15535842 | 2 | 0.373 | 1.44E+07 | 1.14E+07 | 1.01E+07 | 1.15E+07 | 1.14E+07 | 9476473 | 1.31E+07 | 9501027 | 9119146 | 2.58E+07 | 1.06E+07 | 1.31E+07 | 858.49097 | EVEQLIK | Q9R1P0 | Q9R1P0 | PSMA4 | proteasome (prosome, macropain) subunit, alpha type, 4 | Cytoplasm | peptidase |
| 15535590 | 2 | 0.427 | 1.07E+07 | 7789015 | 9485260 | 8722942 | 7491961 | 7062812.5 | 1.25E+07 | 8011366.5 | 7726525.5 | 2.13E+07 | 7976471 | 1.00E+07 | 999.4872 | QAYTQFGGK | Q9R1P0 | Q9R1P0 | PSMA4 | proteasome (prosome, macropain) subunit, alpha type, 4 | Cytoplasm | peptidase |
| 15539920 | 2 | 0.7 | 1.05E+07 | 9963483 | 1.01E+07 | 9177402 | 9644552 | 7744987.5 | 1.09E+07 | 8085521.5 | 8155471.5 | 1.78E+07 | 6836742 | 1.09E+07 | 772.4914 | SALALAVK | Q9R1P0 | Q9R1P0 | PSMA4 | proteasome (prosome, macropain) subunit, alpha type, 4 | Cytoplasm | peptidase |
| 15540729 | 3 | 0.282 | 6383100 | 4036751.5 | 4644246.5 | 3904495 | 3778859.75 | 4796731.5 | 6105091.5 | 5191082.5 | 5181783 | 9182442 | 4186169.75 | 6260164.5 | 1650.8973 | RPFGVSLLYIGWDK | Q9R1P0 | Q9R1P0 | PSMA4 | proteasome (prosome, macropain) subunit, alpha type, 4 | Cytoplasm | peptidase |
| 15555924 | 2 | 0.275 | 5772131.5 | 2718801.75 | 1436563 | 3124220.25 | 2621375.5 | 1100480.75 | 4172244 | 2286947 | 2043583.25 | 7127525 | 3978872.75 | 4339141 | 2203.9792 | HYGFQLYQSDPSGNYGGWK | Q9R1P0 | Q9R1P0 | PSMA4 | proteasome (prosome, macropain) subunit, alpha type, 4 | Cytoplasm | peptidase |
| 15555470 | 2 | 0.528 | 1053177.625 | 660103.562 | 1169621.625 | 763023.25 | 617588.438 | 696977 | 706790.125 | 604045.438 | 149761.547 | 1544026.625 | 216929.781 | 875868.188 | 1623.7803 | ATC[160.0307]IGNNSAAAVSM[147.0354]LK | Q9R1P0 | Q9R1P0 | PSMA4 | proteasome (prosome, macropain) subunit, alpha type, 4 | Cytoplasm | peptidase |
| 15548385 | 2, 3 | 0.473 | 1.02E+08 | 8.68E+07 | 6.40E+07 | 9.22E+07 | 8.24E+07 | 7.21E+07 | 8.96E+07 | 7.41E+07 | 6.70E+07 | 1.40E+08 | 7.75E+07 | 9.38E+07 | 2474.245, 2474.2466 | LNATNIELATVQPGQNFHMFTK | Q9Z2U1 | Q9Z2U1 | PSMA5 | proteasome (prosome, macropain) subunit, alpha type, 5 | Cytoplasm | peptidase |
| 15535114 | 3 | 0.287 | 9.22E+07 | 6.21E+07 | 6.67E+07 | 6.13E+07 | 6.60E+07 | 5.55E+07 | 7.88E+07 | 5.89E+07 | 4.68E+07 | 1.11E+08 | 6.04E+07 | 9.21E+07 | 1961.9519 | AIGSASEGAQSSLQEVYHK | Q9Z2U1 | Q9Z2U1 | PSMA5 | proteasome (prosome, macropain) subunit, alpha type, 5 | Cytoplasm | peptidase |
| 15535795 | 3 | 0.547 | 8.10E+07 | 6.72E+07 | 5.08E+07 | 6.13E+07 | 6.31E+07 | 5.42E+07 | 7.83E+07 | 4.81E+07 | 4.16E+07 | 1.02E+08 | 5.40E+07 | 7.16E+07 | 2162.9685 | GPQLFHMDPSGTFVQC[160.0307]DAR | Q9Z2U1 | Q9Z2U1 | PSMA5 | proteasome (prosome, macropain) subunit, alpha type, 5 | Cytoplasm | peptidase |
| 15535696 | 2 | 0.624 | 3.73E+07 | 3.06E+07 | 2.37E+07 | 3.37E+07 | 2.56E+07 | 2.68E+07 | 3.44E+07 | 2.70E+07 | 2.02E+07 | 4.77E+07 | 2.43E+07 | 3.64E+07 | 1423.7822 | LFQVEYAIEAIK | Q9Z2U1 | Q9Z2U1 | PSMA5 | proteasome (prosome, macropain) subunit, alpha type, 5 | Cytoplasm | peptidase |
| 15533211 | 2 | 0.203 | 2.69E+07 | 2.15E+07 | 2.21E+07 | 1.75E+07 | 1.86E+07 | 1.70E+07 | 2.25E+07 | 1.92E+07 | 1.61E+07 | 3.87E+07 | 1.79E+07 | 2.50E+07 | 988.51776 | EELEEVIK | Q9Z2U1 | Q9Z2U1 | PSMA5 | proteasome (prosome, macropain) subunit, alpha type, 5 | Cytoplasm | peptidase |
| 15536972 | 3 | 0.627 | 2.10E+07 | 1.69E+07 | 1.30E+07 | 1.83E+07 | 1.76E+07 | 1.53E+07 | 2.38E+07 | 1.33E+07 | 1.13E+07 | 3.05E+07 | 1.50E+07 | 2.01E+07 | 1587.8386 | RITSPLMEPSSIEK | Q9Z2U1 | Q9Z2U1 | PSMA5 | proteasome (prosome, macropain) subunit, alpha type, 5 | Cytoplasm | peptidase |
| 15534177 | 2 | 0.071 | 1.41E+07 | 1.22E+07 | 1.33E+07 | 1.04E+07 | 1.16E+07 | 1.11E+07 | 1.32E+07 | 1.06E+07 | 9888870 | 2.17E+07 | 1.25E+07 | 1.53E+07 | 773.5129 | SSLIILK | Q9Z2U1 | Q9Z2U1 | PSMA5 | proteasome (prosome, macropain) subunit, alpha type, 5 | Cytoplasm | peptidase |
| 15548925 | 2 | 0.465 | 1.24E+07 | 9933938 | 7618818 | 9667567 | 9039809 | 7436641.5 | 1.23E+07 | 6769887.5 | 6178576 | 1.93E+07 | 8633281 | 1.04E+07 | 1587.8379 | RITSPLMEPSSIEK | Q9Z2U1 | Q9Z2U1 | PSMA5 | proteasome (prosome, macropain) subunit, alpha type, 5 | Cytoplasm | peptidase |
| 15537610 | 3 | 0.406 | 1.57E+07 | 3832545.5 | 2603695.5 | 1.56E+07 | 821227.375 | 4963982.5 | 7088682 | 2736250.5 | 6293142 | 1.20E+07 | 1.65E+07 | 1.39E+07 | 1713.7788 | SEYDRGVNTFSPEGR | Q9Z2U1 | Q9Z2U1 | PSMA5 | proteasome (prosome, macropain) subunit, alpha type, 5 | Cytoplasm | peptidase |
| 15536341 | 2 | 0.806 | 5438491.5 | 5499071.5 | 5562900 | 3373451.25 | 7520858 | 2949328 | 5247355.5 | 4632719.5 | 2455635.25 | 9836777 | 2619092.25 | 4612519.5 | 1431.7367 | ITSPLMEPSSIEK | Q9Z2U1 | Q9Z2U1 | PSMA5 | proteasome (prosome, macropain) subunit, alpha type, 5 | Cytoplasm | peptidase |
| 15549025 | 2 | 0.316 | 6675901.5 | 4191957 | 4808751 | 3653047.75 | 4203044 | 3914809 | 4487900.5 | 3724467.75 | 3399486.5 | 8287116.5 | 3672109 | 4886350 | 1961.9524 | AIGSASEGAQSSLQEVYHK | Q9Z2U1 | Q9Z2U1 | PSMA5 | proteasome (prosome, macropain) subunit, alpha type, 5 | Cytoplasm | peptidase |
| 15550499 | 2 | 0.759 | 7289522.5 | 5102172.5 | 4253430.5 | 5760725 | 4726052 | 3114502.25 | 6182076 | 2914829.75 | 5287337.5 | 7958319 | 4335064 | 5032313 | 2162.966 | GPQLFHMDPSGTFVQC[160.0307]DAR | Q9Z2U1 | Q9Z2U1 | PSMA5 | proteasome (prosome, macropain) subunit, alpha type, 5 | Cytoplasm | peptidase |
| 15539729 | 3 | 0.78 | 4970524 | 2630707.5 | 3270902.5 | 2775364.25 | 2936337 | 4011738.25 | 3218273.5 | 2183915.5 | 2566352.75 | 6330333 | 2177074.75 | 2592408.5 | 2084.0469 | IVEIDAHIGC[160.0307]AMSGLIADAK | Q9Z2U1 | Q9Z2U1 | PSMA5 | proteasome (prosome, macropain) subunit, alpha type, 5 | Cytoplasm | peptidase |
| 15538494 | 3 | 0.692 | 4856317.5 | 2860004.25 | 2082769.75 | 2509397 | 2989961 | 1867394.5 | 3181374.75 | 2087351 | 1511964.375 | 3804411.25 | 2077820.375 | 1891303.5 | 2178.965 | GPQLFHM[147.0354]DPSGTFVQC[160.0307]DAR | Q9Z2U1 | Q9Z2U1 | PSMA5 | proteasome (prosome, macropain) subunit, alpha type, 5 | Cytoplasm | peptidase |
| 15533713 | 2 | 0.4 | 5.69E+07 | 4.07E+07 | 4.32E+07 | 4.26E+07 | 4.08E+07 | 4.25E+07 | 4.68E+07 | 3.70E+07 | 3.34E+07 | 8.48E+07 | 3.70E+07 | 5.16E+07 | 1285.7197 | AINQGGLTSVAVR | Q9QUM9 | Q9QUM9 | PSMA6 | proteasome (prosome, macropain) subunit, alpha type, 6 | Cytoplasm | peptidase |
| 15532408 | 2, 3 | 0.538 | 5.54E+07 | 4.30E+07 | 4.07E+07 | 3.63E+07 | 3.67E+07 | 3.99E+07 | 5.54E+07 | 3.33E+07 | 3.13E+07 | 7.28E+07 | 3.35E+07 | 5.03E+07 | 1360.7429, 1360.7433 | LLDSSTVTHLFK | Q9QUM9 | Q9QUM9 | PSMA6 | proteasome (prosome, macropain) subunit, alpha type, 6 | Cytoplasm | peptidase |
| 15538955 | 2 | 0.289 | 3.41E+07 | 2.83E+07 | 3.05E+07 | 2.72E+07 | 2.78E+07 | 2.74E+07 | 3.53E+07 | 2.59E+07 | 2.21E+07 | 5.71E+07 | 2.89E+07 | 3.33E+07 | 1169.568 | QTESTSFLEK | Q9QUM9 | Q9QUM9 | PSMA6 | proteasome (prosome, macropain) subunit, alpha type, 6 | Cytoplasm | peptidase |
| 15539555 | 2 | 0.438 | 3.15E+07 | 2.40E+07 | 2.26E+07 | 2.79E+07 | 2.26E+07 | 1.88E+07 | 3.24E+07 | 2.14E+07 | 1.83E+07 | 5.15E+07 | 2.40E+07 | 2.80E+07 | 1156.61 | HITIFSPEGR | Q9QUM9 | Q9QUM9 | PSMA6 | proteasome (prosome, macropain) subunit, alpha type, 6 | Cytoplasm | peptidase |
| 15532947 | 2 | 0.379 | 2.93E+07 | 2.57E+07 | 2.35E+07 | 2.50E+07 | 2.21E+07 | 1.68E+07 | 3.33E+07 | 2.27E+07 | 2.21E+07 | 5.04E+07 | 2.18E+07 | 2.90E+07 | 1160.5969 | LYQVEYAFK | Q9QUM9 | Q9QUM9 | PSMA6 | proteasome (prosome, macropain) subunit, alpha type, 6 | Cytoplasm | peptidase |
| 15532859 | 2 | 0.266 | 2.59E+07 | 2.03E+07 | 1.97E+07 | 1.77E+07 | 2.25E+07 | 1.80E+07 | 2.85E+07 | 1.91E+07 | 1.68E+07 | 4.46E+07 | 2.17E+07 | 2.63E+07 | 1487.6873 | YGYEIPVDMLC[160.0307]K | Q9QUM9 | Q9QUM9 | PSMA6 | proteasome (prosome, macropain) subunit, alpha type, 6 | Cytoplasm | peptidase |
| 15542159 | 2 | 0.489 | 2.12E+07 | 1.35E+07 | 1.28E+07 | 1.52E+07 | 1.45E+07 | 1.18E+07 | 2.08E+07 | 1.33E+07 | 1.09E+07 | 3.69E+07 | 1.46E+07 | 1.66E+07 | 1033.5322 | DC[160.0307]AVIVTQK | Q9QUM9 | Q9QUM9 | PSMA6 | proteasome (prosome, macropain) subunit, alpha type, 6 | Cytoplasm | peptidase |
| 15535869 | 2 | 0.163 | 2.50E+07 | 1.40E+07 | 1.48E+07 | 1.55E+07 | 1.56E+07 | 1.27E+07 | 1.95E+07 | 1.54E+07 | 1.28E+07 | 3.68E+07 | 1.93E+07 | 2.06E+07 | 1337.5272 | C[160.0307]DPAGYYC[160.0307]GFK | Q9QUM9 | Q9QUM9 | PSMA6 | proteasome (prosome, macropain) subunit, alpha type, 6 | Cytoplasm | peptidase |
| 15548184 | 2 | 0.241 | 1.96E+07 | 1.37E+07 | 1.37E+07 | 1.48E+07 | 1.55E+07 | 1.28E+07 | 1.86E+07 | 1.25E+07 | 1.34E+07 | 3.08E+07 | 1.70E+07 | 1.73E+07 | 1828.8182 | ITESIGC[160.0307]VMTGMTADSR | Q9QUM9 | Q9QUM9 | PSMA6 | proteasome (prosome, macropain) subunit, alpha type, 6 | Cytoplasm | peptidase |
| 15540128 | 3 | 0.83 | 2.06E+07 | 1.54E+07 | 1.54E+07 | 1.22E+07 | 1.72E+07 | 1.49E+07 | 1.75E+07 | 1.45E+07 | 1.19E+07 | 2.34E+07 | 1.14E+07 | 1.51E+07 | 1979.0432 | ILTEAEIDAHLVALAERD | Q9QUM9 | Q9QUM9 | PSMA6 | proteasome (prosome, macropain) subunit, alpha type, 6 | Cytoplasm | peptidase |

| | | | | | | | | |
|---|---|---|---|---|---|---|---|---|
| 15544747 | 2 | 0.566 | 2530999.5 | 515785.219 | 35584.461 | 1959672.125 | 338115.906 | 134968.844 |
| 15548046 | 2, 3 | 0.825 | 1.06E+08 | 8.52E+07 | 9.86E+07 | 8.89E+07 | 7.83E+07 | 7.68E+07 |
| 15532181 | 2 | 0.406 | 5.51E+07 | 4.28E+07 | 4.48E+07 | 4.54E+07 | 4.17E+07 | 4.33E+07 |
| 15552335 | 2 | 0.531 | 5.38E+07 | 4.33E+07 | 4.70E+07 | 3.87E+07 | 4.42E+07 | 3.72E+07 |
| 15533471 | 3 | 0.46 | 4.04E+07 | 2.61E+07 | 2.70E+07 | 2.94E+07 | 2.94E+07 | 2.36E+07 |
| 15547987 | 2 | 0.734 | 4.14E+07 | 3.15E+07 | 3.44E+07 | 3.21E+07 | 3.27E+07 | 3.07E+07 |
| 15540279 | 2 | 0.345 | 2.25E+07 | 2.01E+07 | 2.04E+07 | 1.87E+07 | 1.77E+07 | 1.20E+07 |
| 15531591 | 2 | 0.312 | 1.29E+07 | 1.16E+07 | 1.16E+07 | 9814885 | 8680476 | 6073528 |
| 15548993 | 2 | 0.036 | 1.09E+07 | 6196586 | 4790688.5 | 7682826 | 6489981.5 | 6370742.5 |
| 15557363 | 2 | 0.503 | 9824066 | 4984282 | 6016859.5 | 6180433 | 5845164.5 | 4811553.5 |
| 15536317 | 3 | 0.023 | 3866496.5 | 2369634.5 | 2171603.75 | 4374176.5 | 3353273.25 | 3036896.5 |
| 15545858 | 3 | 0.805 | 7044484 | 5602063 | 4376657 | 4663125.5 | 6046772.5 | 5658799.5 |
| 15550183 | 3 | 0.481 | 7535852 | 1062635.625 | 54923.918 | 5946670.5 | 623584.812 | 478667.312 |
| 15548579 | 2 | 0.82 | 4.42E+07 | 3.97E+07 | 4.01E+07 | 4.31E+07 | 3.65E+07 | 3.32E+07 |
| 15532029 | 3 | 0.382 | 3.64E+07 | 3.12E+07 | 2.71E+07 | 2.87E+07 | 2.94E+07 | 2.83E+07 |
| 15548444 | 2 | 0.786 | 3.48E+07 | 2.67E+07 | 2.24E+07 | 2.83E+07 | 2.51E+07 | 2.31E+07 |
| 15539131 | 3 | 0.206 | 2.56E+07 | 2.09E+07 | 1.89E+07 | 2.29E+07 | 1.40E+07 | 1.97E+07 |
| 15537957 | 3 | 0.53 | 1.69E+07 | 1.69E+07 | 1.19E+07 | 1.64E+07 | 1.46E+07 | 1.28E+07 |
| 15552665 | 2 | 0.334 | 1.19E+07 | 7462203 | 6765454.5 | 7839670 | 6219355 | 5437788.5 |
| 15548856 | 2 | 0.555 | 7570319 | 4841434 | 4117741.5 | 6191459 | 4959544 | 5311958 |
| 15534133 | 2 | 0.971 | 7099796 | 5936947 | 5453414 | 3934491.25 | 8065251.5 | 4824590.5 |
| 15536662 | 2 | 0.827 | 6439418 | 6826131 | 9269776 | 3660148.25 | 8350831 | 6090473 |
| 15549722 | 2 | 0.527 | 7378076 | 3625242.25 | 3651751 | 5691708.5 | 2573065 | 4072799.75 |
| 15552079 | 2 | 0.999 | 4625644.5 | 3202507.75 | 4512746 | 4386774.5 | 3873621.75 | 3996130 |

| | | | | | | | |
|---|---|---|---|---|---|---|---|
| 1108.5508 | ARYEAANWK | Q9QUM9 | Q9QUM9 | PSMA6 | proteasome (prosome, macropain) subunit, alpha type, 6 | Cytoplasm | peptidase |
| 2449.2332, 2449.2344 | AITVFSPDGHLFQVEYAQEAVK | Q9Z2U0 | Q9Z2U0 | PSMA7 | proteasome (prosome, macropain) subunit, alpha type, 7 | Cytoplasm | peptidase |
| 1084.5868 | ILNPEEIEK | Q9Z2U0 | Q9Z2U0 | PSMA7 | proteasome (prosome, macropain) subunit, alpha type, 7 | Cytoplasm | peptidase |
| 1634.8583 | LTVEDPVTVEYITR | Q9Z2U0 | Q9Z2U0 | PSMA7 | proteasome (prosome, macropain) subunit, alpha type, 7 | Cytoplasm | peptidase |
| 1666.7833 | LYQTDPSGTYHAWK | Q9Z2U0 | Q9Z2U0 | PSMA7 | proteasome (prosome, macropain) subunit, alpha type, 7 | Cytoplasm | peptidase |
| 1726.7983 | NYTDDAIETDDLTIK | Q9Z2U0 | Q9Z2U0 | PSMA7 | proteasome (prosome, macropain) subunit, alpha type, 7 | Cytoplasm | peptidase |
| 971.57526 | DIVVLGVEK | Q9Z2U0 | Q9Z2U0 | PSMA7 | proteasome (prosome, macropain) subunit, alpha type, 7 | Cytoplasm | peptidase |
| 945.5163 | NIELAVMR | Q9Z2U0 | Q9Z2U0 | PSMA7 | proteasome (prosome, macropain) subunit, alpha type, 7 | Cytoplasm | peptidase |
| 2183.985 | IC[160.0307]ALDDNVC[160.0307]MAFAGLTADAR | Q9Z2U0 | Q9Z2U0 | PSMA7 | proteasome (prosome, macropain) subunit, alpha type, 7 | Cytoplasm | peptidase |
| 1666.7827 | LYQTDPSGTYHAWK | Q9Z2U0 | Q9Z2U0 | PSMA7 | proteasome (prosome, macropain) subunit, alpha type, 7 | Cytoplasm | peptidase |
| 2183.9856 | IC[160.0307]ALDDNVC[160.0307]MAFAGLTADAR | Q9Z2U0 | Q9Z2U0 | PSMA7 | proteasome (prosome, macropain) subunit, alpha type, 7 | Cytoplasm | peptidase |
| 2065.0835 | RPFGISALIVGFDFDGTPR | Q9Z2U0 | Q9Z2U0 | PSMA7 | proteasome (prosome, macropain) subunit, alpha type, 7 | Cytoplasm | peptidase |
| 2577.3293 | AITVFSPDGHLFQVEYAQEAVKK | Q9Z2U0 | Q9Z2U0 | PSMA7 | proteasome (prosome, macropain) subunit, alpha type, 7 | Cytoplasm | peptidase |
| 1545.7308 | GAVYSFDPVGSYQR | O09061 | O09061 | PSMB1 | proteasome (prosome, macropain) subunit, beta type, 1 | Cytoplasm | peptidase |
| 1665.8326 | NMQNVEHVPLTLDR | O09061 | O09061 | PSMB1 | proteasome (prosome, macropain) subunit, beta type, 1 | Cytoplasm | peptidase |
| 2004.0162 | AGGSASAMLQPLLDNQVGFK | O09061 | O09061 | PSMB1 | proteasome (prosome, macropain) subunit, beta type, 1 | Cytoplasm | peptidase |
| 1997.99 | DVFISAAERDVYTGDALR | O09061 | O09061 | PSMB1 | proteasome (prosome, macropain) subunit, beta type, 1 | Cytoplasm | peptidase |
| 2004.0171 | AGGSASAMLQPLLDNQVGFK | O09061 | O09061 | PSMB1 | proteasome (prosome, macropain) subunit, beta type, 1 | Cytoplasm | peptidase |
| 1665.8335 | NMQNVEHVPLTLDR | O09061 | O09061 | PSMB1 | proteasome (prosome, macropain) subunit, beta type, 1 | Cytoplasm | peptidase |
| 1870.9728 | AMTTGAIAAMLSTILYSR | O09061 | O09061 | PSMB1 | proteasome (prosome, macropain) subunit, beta type, 1 | Cytoplasm | peptidase |
| 1009.494 | DVYTGDALR | O09061 | O09061 | PSMB1 | proteasome (prosome, macropain) subunit, beta type, 1 | Cytoplasm | peptidase |
| 1007.51385 | DVFISAAER | O09061 | O09061 | PSMB1 | proteasome (prosome, macropain) subunit, beta type, 1 | Cytoplasm | peptidase |
| 1997.989 | DVFISAAERDVYTGDALR | O09061 | O09061 | PSMB1 | proteasome (prosome, macropain) subunit, beta type, 1 | Cytoplasm | peptidase |
| 2124.0286 | FFPYYVYNIIGGLDEEGK | O09061 | O09061 | PSMB1 | proteasome (prosome, macropain) subunit, beta type, 1 | Cytoplasm | peptidase |
| 1870.9728 | AMTTGAIAAMLSTILYSR | O09061 | O09061 | PSMB1 | proteasome (prosome, macropain) subunit, beta type, 1 | Cytoplasm | peptidase |
| 1886.9673 | AM[147.0354]TTGAIAAMLSTILYSR | O09061 | O09061 | PSMB1 | proteasome (prosome, macropain) subunit, beta type, 1 | Cytoplasm | peptidase |
| 2124.0278 | FFPYYVYNIIGGLDEEGK | O09061 | O09061 | PSMB1 | proteasome (prosome, macropain) subunit, beta type, 1 | Cytoplasm | peptidase |
| 1886.9674 | AM[147.0354]TTGAIAAMLSTILYSR | O09061 | O09061 | PSMB1 | proteasome (prosome, macropain) subunit, beta type, 1 | Cytoplasm | peptidase |
| 1681.8242 | NM[147.0354]QNVEHVPLTLDR | O09061 | O09061 | PSMB1 | proteasome (prosome, macropain) subunit, beta type, 1 | Cytoplasm | peptidase |
| 1865.8839 | TVIGC[160.0307]SGFHGDC[160.0307]LTLTK | O09061 | O09061 | PSMB1 | proteasome (prosome, macropain) subunit, beta type, 1 | Cytoplasm | peptidase |

[illegible]

| | | | | | | | | | | | | | | | | | | | | | |
|---|---|---|---|---|---|---|---|---|---|---|---|---|---|---|---|---|---|---|---|---|---|
| 15551821 | 2 | 0.303 | 4729442.5 | 5212703 | 5023324.5 | 3379189.75 | 4787165 | 3215025 | 3868800 | 4134679 | 4084904.5 | 5126533 | 2233717.25 | 3417747.75 | 2126.028 | EAASILQELQVETYGSMEK | Q9D8W5 | Q9D8W5 | PSMD12 | proteasome (prosome, macropain) 26S subunit, non-ATPase, 12 | Cytoplasm | other |
| 15554717 | 2 | 0.683 | 3101299.5 | 2490948 | 2567707.25 | 1774317.875 | 3536251.25 | 1488533.875 | 1805256.875 | 2098831.25 | 1822791.75 | 4208601 | 1577105 | 1795383 | 1566.777 | WSTLVEDYGVELR | Q9D8W5 | Q9D8W5 | PSMD12 | proteasome (prosome, macropain) 26S subunit, non-ATPase, 12 | Cytoplasm | other |
| 15533317 | 2 | 0.187 | 4526334.5 | 3585604.5 | 3356168 | 2478870.5 | 3232004.5 | 2085544.625 | 3672527.75 | 3248444.5 | 2619954 | 3980485 | 2541870.25 | 3455758.75 | 1242.7284 | LAGVINFQRPK | Q9D8W5 | Q9D8W5 | PSMD12 | proteasome (prosome, macropain) 26S subunit, non-ATPase, 12 | Cytoplasm | other |
| 15539226 | 2 | 0.563 | 2207699.25 | 1442832.25 | 901106.812 | 1151189.375 | 1187361.25 | 749970.938 | 1094317.25 | 1283616 | 1160526.875 | 2581272.5 | 1267921.125 | 991202.625 | 703.4154 | LC[160.0307]LAVK | Q9D8W5 | Q9D8W5 | PSMD12 | proteasome (prosome, macropain) 26S subunit, non-ATPase, 12 | Cytoplasm | other |
| 15548142 | 3 | 0.328 | 2.56E+07 | 2.88E+07 | 2.67E+07 | 1.65E+07 | 2.40E+07 | 1.73E+07 | 2.70E+07 | 2.08E+07 | 2.35E+07 | 3.44E+07 | 1.70E+07 | 2.22E+07 | 2469.1873 | ETIEDVEEMLNNLPGVTSVHSR | Q9WVJ2 | Q9WVJ2 | PSMD13 | proteasome (prosome, macropain) 26S subunit, non-ATPase, 13 | Cytoplasm | peptidase |
| 15550255 | 3 | 0.465 | 3.20E+07 | 2.77E+07 | 2.59E+07 | 1.69E+07 | 3.13E+07 | 1.57E+07 | 2.58E+07 | 2.22E+07 | 1.93E+07 | 3.13E+07 | 1.32E+07 | 1.79E+07 | 2366.1592 | DVPAFLQQSQSSGPGQAAVWHR | Q9WVJ2 | Q9WVJ2 | PSMD13 | proteasome (prosome, macropain) 26S subunit, non-ATPase, 13 | Cytoplasm | peptidase |
| 15550590 | 3 | 0.652 | 1.96E+07 | 1.89E+07 | 1.16E+07 | 1.31E+07 | 1.38E+07 | 8328867.5 | 1.88E+07 | 1.34E+07 | 9156134 | 2.90E+07 | 9358141 | 1.58E+07 | 3042.531 | LWHQLTLEVLDFVQDPC[160.0307]FAQGDGLIK | Q9WVJ2 | Q9WVJ2 | PSMD13 | proteasome (prosome, macropain) 26S subunit, non-ATPase, 13 | Cytoplasm | peptidase |
| 15536809 | 3 | 0.605 | 1.93E+07 | 1.71E+07 | 1.86E+07 | 1.50E+07 | 1.83E+07 | 1.38E+07 | 1.64E+07 | 1.58E+07 | 1.61E+07 | 1.99E+07 | 9785921 | 1.63E+07 | 1583.7452 | LYENFISEFEHR | Q9WVJ2 | Q9WVJ2 | PSMD13 | proteasome (prosome, macropain) 26S subunit, non-ATPase, 13 | Cytoplasm | peptidase |
| 15549812 | 2 | 0.489 | 1.64E+07 | 1.59E+07 | 1.43E+07 | 1.16E+07 | 1.48E+07 | 8947741 | 1.36E+07 | 1.17E+07 | 1.37E+07 | 1.85E+07 | 8982958 | 1.22E+07 | 1744.8209 | SMEMLVEHQAQDILT | Q9WVJ2 | Q9WVJ2 | PSMD13 | proteasome (prosome, macropain) 26S subunit, non-ATPase, 13 | Cytoplasm | peptidase |
| 15554776 | 2 | 0.725 | 1.03E+07 | 1.14E+07 | 1.09E+07 | 8078813.5 | 1.02E+07 | 7276667.5 | 1.02E+07 | 8217688 | 9312454 | 1.56E+07 | 5086575 | 7960115 | 1606.8109 | QMTDPNVALTFLEK | Q9WVJ2 | Q9WVJ2 | PSMD13 | proteasome (prosome, macropain) 26S subunit, non-ATPase, 13 | Cytoplasm | peptidase |
| 15538716 | 3 | 0.5 | 1.38E+07 | 1.14E+07 | 9165787 | 7917870 | 1.08E+07 | 6523123.5 | 1.21E+07 | 8570817 | 8576518 | 1.38E+07 | 8512483 | 8048022.5 | 1607.7462 | YYQTIGNHASYYK | Q9WVJ2 | Q9WVJ2 | PSMD13 | proteasome (prosome, macropain) 26S subunit, non-ATPase, 13 | Cytoplasm | peptidase |
| 15533345 | 2, 3 | 0.405 | 1.44E+07 | 1.20E+07 | 1.14E+07 | 8458144 | 1.18E+07 | 9029255 | 1.30E+07 | 9556992 | 9830291 | 1.28E+07 | 7772090.5 | 1.05E+07 | 1434.7013, 1434.702 | DRLELWC[160.0307]TDVK | Q9WVJ2 | Q9WVJ2 | PSMD13 | proteasome (prosome, macropain) 26S subunit, non-ATPase, 13 | Cytoplasm | peptidase |
| 15535401 | 2 | 0.213 | 1.10E+07 | 1.05E+07 | 1.06E+07 | 7957582.5 | 8664400 | 5035152 | 9541078 | 7979020.5 | 7828380.5 | 1.28E+07 | 6926264 | 8394060 | 1078.5759 | QLTFEEIAK | Q9WVJ2 | Q9WVJ2 | PSMD13 | proteasome (prosome, macropain) 26S subunit, non-ATPase, 13 | Cytoplasm | peptidase |
| 15537882 | 2 | 0.463 | 1.35E+07 | 1.02E+07 | 1.02E+07 | 8152231.5 | 1.12E+07 | 6611555 | 1.18E+07 | 1.03E+07 | 9187778 | 1.26E+07 | 7500618.5 | 9801940 | 1072.5974 | LNIGDLQATK | Q9WVJ2 | Q9WVJ2 | PSMD13 | proteasome (prosome, macropain) 26S subunit, non-ATPase, 13 | Cytoplasm | peptidase |
| 15538945 | 3 | 0.49 | 8117230 | 5423799.5 | 4585057 | 6506123 | 6919599.5 | 4099045.75 | 8537423 | 5678320.5 | 7004125 | 8834298 | 7619352 | 6238314 | 2169.0828 | C[160.0307]AWGQQPDLAANEAQLLRK | Q9WVJ2 | Q9WVJ2 | PSMD13 | proteasome (prosome, macropain) 26S subunit, non-ATPase, 13 | Cytoplasm | peptidase |
| 15540549 | 2 | 0.351 | 5851590 | 4479462.5 | 5427270.5 | 3382518.5 | 5557234 | 3299742 | 5217888.5 | 4749340.5 | 4235879.5 | 7765531 | 4259475 | 5204489.5 | 951.49347 | FLGC[160.0307]VDIK | Q9WVJ2 | Q9WVJ2 | PSMD13 | proteasome (prosome, macropain) 26S subunit, non-ATPase, 13 | Cytoplasm | peptidase |
| 15543253 | 4 | 0.76 | 3901848.25 | 1655990.125 | 967238.062 | 1771322.125 | 1515922.125 | 1217996.5 | 1616408 | 1401996.25 | 1807897.125 | 1499190.75 | 2220427.25 | 2575334.5 | 2231.1199 | IQLLC[160.0307]LMEMTFTRPANHR | Q9WVJ2 | Q9WVJ2 | PSMD13 | proteasome (prosome, macropain) 26S subunit, non-ATPase, 13 | Cytoplasm | peptidase |
| 15548138 | 3 | 0.499 | 2.14E+07 | 2.33E+07 | 1.88E+07 | 1.27E+07 | 1.80E+07 | 1.24E+07 | 2.12E+07 | 2.08E+07 | 1.68E+07 | 3.27E+07 | 1.02E+07 | 1.65E+07 | 3323.714 | LGGGMPGLGQGPPTDAPAVDTAEQVYISSLALLK | O35593 | O35593 | PSMD14 | proteasome (prosome, macropain) 26S subunit, non-ATPase, 14 | Cytoplasm | peptidase |
| 15548785 | 3 | 0.774 | 1.53E+07 | 1.88E+07 | 1.63E+07 | 9669818 | 1.99E+07 | 1.12E+07 | 1.47E+07 | 1.40E+07 | 1.34E+07 | 2.03E+07 | 8406177 | 1.69E+07 | 2691.3616 | VIDVFAMPQSGTGVSVEAVDPVFQAK | O35593 | O35593 | PSMD14 | proteasome (prosome, macropain) 26S subunit, non-ATPase, 14 | Cytoplasm | peptidase |
| 15538028 | 2 | 0.712 | 7138931.5 | 4136009 | 5488456 | 3582166.75 | 5643038.5 | 3503995 | 5108735 | 3841567.5 | 3683029 | 7276973 | 2701946 | 4537577 | 819.4704 | VVIDAFR | O35593 | O35593 | PSMD14 | proteasome (prosome, macropain) 26S subunit, non-ATPase, 14 | Cytoplasm | peptidase |
| 15543204 | 3 | 0.594 | 2973990.25 | 4061951.5 | 2572174.25 | 1357548 | 4055968.25 | 2056987.375 | 3760761 | 3583917.75 | 2304154.25 | 3019500.75 | 1253549.75 | 2955087.5 | 1318.7562 | PSIQALIHGLNR | O35593 | O35593 | PSMD14 | proteasome (prosome, macropain) 26S subunit, non-ATPase, 14 | Cytoplasm | peptidase |
| 15549593 | 3, 4 | 0.666 | 2.34E+07 | 2.87E+07 | 6885377 | 2.14E+07 | 2.13E+07 | 8946558 | 2.60E+07 | 1.47E+07 | 8731980 | 4.62E+07 | 1.71E+07 | 2.24E+07 | 3571.6143, 3571.6145 | EIVPYNMAHNAEHEAC[160.0307]DLLMEIEQVDMLEK | Q8VDM4 | Q8VDM4 | PSMD2 | proteasome (prosome, macropain) 26S subunit, non-ATPase, 2 | Cytoplasm | other |
| 15533138 | 3 | 0.713 | 2.33E+07 | 1.96E+07 | 1.87E+07 | 1.72E+07 | 1.94E+07 | 1.30E+07 | 2.39E+07 | 1.68E+07 | 1.42E+07 | 3.05E+07 | 1.32E+07 | 2.07E+07 | 1959.9496 | LVGSQEELASWGHEYVR | Q8VDM4 | Q8VDM4 | PSMD2 | proteasome (prosome, macropain) 26S subunit, non-ATPase, 2 | Cytoplasm | other |
| 15548110 | 2 | 0.967 | 1.56E+07 | 1.22E+07 | 1.37E+07 | 1.53E+07 | 1.36E+07 | 1.02E+07 | 1.94E+07 | 1.13E+07 | 1.06E+07 | 2.60E+07 | 1.00E+07 | 1.18E+07 | 1928.9166 | TPVQSQQPSATTPSGADEK | Q8VDM4 | Q8VDM4 | PSMD2 | proteasome (prosome, macropain) 26S subunit, non-ATPase, 2 | Cytoplasm | other |
| 15534204 | 2, 3 | 0.46 | 2.12E+07 | 1.04E+07 | 3546547.5 | 1.37E+07 | 8731764 | 5611990 | 9150205 | 7292684 | 1.06E+07 | 2.20E+07 | 1.28E+07 | 1.52E+07 | 1452.8141, 1452.8142 | VGQAVDVVGQAGKPK | Q8VDM4 | Q8VDM4 | PSMD2 | proteasome (prosome, macropain) 26S subunit, non-ATPase, 2 | Cytoplasm | other |
| 15549313 | 2 | 0.624 | 1.34E+07 | 1.09E+07 | 1.00E+07 | 1.02E+07 | 1.04E+07 | 8561987 | 1.11E+07 | 8128463 | 8901602 | 1.80E+07 | 7901604 | 1.07E+07 | 1502.749 | QLQDELEMLVER | Q8VDM4 | Q8VDM4 | PSMD2 | proteasome (prosome, macropain) 26S subunit, non-ATPase, 2 | Cytoplasm | other |
| 15550330 | 3 | 0.487 | 1.26E+07 | 9540623 | 6590506.5 | 9687980 | 6807029.5 | 8463294 | 1.26E+07 | 7164632 | 1.10E+07 | 1.60E+07 | 9805438 | 1.03E+07 | 2585.4536 | AELATEEFLPVTPILEGFVILRK | Q8VDM4 | Q8VDM4 | PSMD2 | proteasome (prosome, macropain) 26S subunit, non-ATPase, 2 | Cytoplasm | other |
| 15533777 | 3 | 0.473 | 1.63E+07 | 1.17E+07 | 1.50E+07 | 9237113 | 1.37E+07 | 1.16E+07 | 1.30E+07 | 1.21E+07 | 1.15E+07 | 1.60E+07 | 1.18E+07 | 1.10E+07 | 1840.8955 | VQQLLHIC[160.0307]SEHFDSK | Q8VDM4 | Q8VDM4 | PSMD2 | proteasome (prosome, macropain) 26S subunit, non-ATPase, 2 | Cytoplasm | other |
| 15541090 | 2 | 0.903 | 1.09E+07 | 9475397 | 8975338 | 8738510 | 1.02E+07 | 6881685.5 | 9785585 | 8561966 | 7963719.5 | 1.39E+07 | 6099723 | 8717331 | 1262.5508 | EWQELDDAEK | Q8VDM4 | Q8VDM4 | PSMD2 | proteasome (prosome, macropain) 26S subunit, non-ATPase, 2 | Cytoplasm | other |
| 15532021 | 2 | 0.402 | 1.23E+07 | 9808694 | 9703330 | 6888780.5 | 9717744 | 7037608 | 1.00E+07 | 9045213 | 7807220.5 | 1.30E+07 | 6962280.5 | 8913941 | 1280.6034 | YLYSSEDYIK | Q8VDM4 | Q8VDM4 | PSMD2 | proteasome (prosome, macropain) 26S subunit, non-ATPase, 2 | Cytoplasm | other |
| 15535212 | 3 | 0.601 | 1.10E+07 | 1.03E+07 | 7849729 | 4773308.5 | 1.14E+07 | 5088169.5 | 8577803 | 8421605 | 5924073.5 | 1.21E+07 | 5678802.5 | 7042253 | 2079.0942 | TITGFQTHTTPVLLAHGER | Q8VDM4 | Q8VDM4 | PSMD2 | proteasome (prosome, macropain) 26S subunit, non-ATPase, 2 | Cytoplasm | other |
| 15535534 | 3 | 0.667 | 9108982 | 1.05E+07 | 6348851.5 | 6164649 | 9317257 | 7080186 | 7135240.5 | 6815549 | 7643328 | 1.20E+07 | 6322992.5 | 8991034 | 2001.0793 | MLVTFDEELRPLPVSVR | Q8VDM4 | Q8VDM4 | PSMD2 | proteasome (prosome, macropain) 26S subunit, non-ATPase, 2 | Cytoplasm | other |
| 15533488 | 2 | 0.27 | 1.24E+07 | 1.12E+07 | 1.10E+07 | 7169141.5 | 1.12E+07 | 8556740 | 1.13E+07 | 8672935 | 8174938.5 | 1.12E+07 | 8685798 | 1.07E+07 | 1016.5957 | LNILDTLSK | Q8VDM4 | Q8VDM4 | PSMD2 | proteasome (prosome, macropain) 26S subunit, non-ATPase, 2 | Cytoplasm | other |
| 15553332 | 3 | 0.283 | 7627312.5 | 6914442 | 5284318.5 | 7335691.5 | 5972270.5 | 4407818.5 | 7064147 | 4911855 | 4469502 | 1.12E+07 | 6710434 | 6877596.5 | 2974.302 | FSHDADPEVSYNSIFAMGMVGSGTNNAR | Q8VDM4 | Q8VDM4 | PSMD2 | proteasome (prosome, macropain) 26S subunit, non-ATPase, 2 | Cytoplasm | other |
| 15536916 | 2 | 0.734 | 8224100 | 9232334 | 7346695 | 6461146 | 7846775 | 6071094.5 | 7216951 | 8256347.5 | 5656143.5 | 1.08E+07 | 4550854 | 6137314 | 1473.7809 | SGALLAC[160.0307]GIVNSGVR | Q8VDM4 | Q8VDM4 | PSMD2 | proteasome (prosome, macropain) 26S subunit, non-ATPase, 2 | Cytoplasm | other |
| 15535419 | 2 | 0.447 | 9913991 | 8297860 | 7885115.5 | 6314792.5 | 8041449 | 5124652.5 | 7068394.5 | 6960845.5 | 5771602.5 | 1.05E+07 | 4852155 | 6128299.5 | 1167.5366 | FGGSGSQVDSAR | Q8VDM4 | Q8VDM4 | PSMD2 | proteasome (prosome, macropain) 26S subunit, non-ATPase, 2 | Cytoplasm | other |
| 15534506 | 2 | 0.515 | 6653197 | 1.09E+07 | 1.40E+07 | 5340042 | 1.16E+07 | 7879363 | 1.06E+07 | 8869919 | 4420215.5 | 1.05E+07 | 3085273.5 | 5188131 | 1227.6661 | VGQAVDVVGQAGK | Q8VDM4 | Q8VDM4 | PSMD2 | proteasome (prosome, macropain) 26S subunit, non-ATPase, 2 | Cytoplasm | other |
| 15531947 | 2 | 0.488 | 9650198 | 6239370.5 | 5844238.5 | 5437062 | 5928685 | 4391350.5 | 7343054.5 | 5015440 | 5702975 | 9526617 | 4904466.5 | 6437019 | 1037.6068 | LAQGLTHLGK | Q8VDM4 | Q8VDM4 | PSMD2 | proteasome (prosome, macropain) 26S subunit, non-ATPase, 2 | Cytoplasm | other |
| 15555006 | 3 | 0.887 | 3437202.75 | 4079966 | 2838226 | 3283284.5 | 4250198.5 | 2241704.5 | 4473940.5 | 5002577.5 | 3456011.25 | 9508634 | 1843975.125 | 2442036.5 | 2533.1707 | NEC[160.0307]DPALALLSDYVLHNSNTMR | Q8VDM4 | Q8VDM4 | PSMD2 | proteasome (prosome, macropain) 26S subunit, non-ATPase, 2 | Cytoplasm | other |
| 15549231 | 2 | 0.669 | 6999586 | 6813920.5 | 7287318 | 6019549.5 | 6708792 | 4711306.5 | 6244349.5 | 4821946.5 | 6372830.5 | 8482435 | 3888212.25 | 6076330.5 | 1520.9153 | AVPLALALISVSNPR | Q8VDM4 | Q8VDM4 | PSMD2 | proteasome (prosome, macropain) 26S subunit, non-ATPase, 2 | Cytoplasm | other |
| 15533830 | 3 | 0.79 | 5808295 | 5149076 | 4224926 | 3891731.75 | 5003931 | 4580100 | 3541623.75 | 4620546.5 | 4831036 | 7621079 | 3356655 | 5099911.5 | 1417.8282 | WLPLGLGLNHLGK | Q8VDM4 | Q8VDM4 | PSMD2 | proteasome (prosome, macropain) 26S subunit, non-ATPase, 2 | Cytoplasm | other |
| 15537726 | 3 | 0.641 | 6758415.5 | 5313544.5 | 4937785.5 | 5409722 | 4313356.5 | 3274075 | 6022063 | 3820950.75 | 4100550.25 | 7216854.5 | 3453712.5 | 4417508 | 1267.7722 | AQREPLLTLVK | Q8VDM4 | Q8VDM4 | PSMD2 | proteasome (prosome, macropain) 26S subunit, non-ATPase, 2 | Cytoplasm | other |
| 15551872 | 2 | 0.2 | 6329859 | 5322247.5 | 4907331.5 | 4197742 | 4533247 | 4256527 | 4464131 | 3747557 | 5543493 | 6920141 | 4505488.5 | 5517636.5 | 2086.023 | SFANTLVDVC[160.0307]AYAGSGNVLK | Q8VDM4 | Q8VDM4 | PSMD2 | proteasome (prosome, macropain) 26S subunit, non-ATPase, 2 | Cytoplasm | other |
| 15541528 | 2 | 0.25 | 6741117.5 | 4968193 | 3639300.5 | 4483228 | 4664531.5 | 2979226.25 | 4529065.5 | 3611482.5 | 3108828.25 | 6619682 | 4615496 | 5095594 | 1105.5426 | DPNNLFMVR | Q8VDM4 | Q8VDM4 | PSMD2 | proteasome (prosome, macropain) 26S subunit, non-ATPase, 2 | Cytoplasm | other |
| 15538345 | 3 | 0.276 | 2679442.75 | 2864816.75 | 2548204.25 | 2357948.75 | 2169082.25 | 1471924.375 | 2303883.75 | 2064598.875 | 2826147.25 | 5533908 | 2124126.5 | 2809638 | 2117.0085 | MNLASSFVNGFVNAAFGQDK | Q8VDM4 | Q8VDM4 | PSMD2 | proteasome (prosome, macropain) 26S subunit, non-ATPase, 2 | Cytoplasm | other |
| 15540308 | 2 | 0.467 | 4429330.5 | 3069288 | 2183389 | 3617226.5 | 2240001.75 | 1898168.625 | 3137045.75 | 2015589.875 | 2138513.75 | 5086271.5 | 2598292.75 | 3182169 | 953.46747 | QMAFMLGR | Q8VDM4 | Q8VDM4 | PSMD2 | proteasome (prosome, macropain) 26S subunit, non-ATPase, 2 | Cytoplasm | other |
| 15536221 | 2 | 0.878 | 2448720.5 | 2996297.25 | 2936557.25 | 1592364.125 | 3613308.25 | 2170092.5 | 2335905.75 | 2548930.5 | 1751827.125 | 4406073 | 1338470.25 | 2203281.25 | 822.42725 | C[160.0307]ALGVFR | Q8VDM4 | Q8VDM4 | PSMD2 | proteasome (prosome, macropain) 26S subunit, non-ATPase, 2 | Cytoplasm | other |
| 15536260 | 3 | 0.795 | 3829316.75 | 3259295.5 | 2144792.25 | 2172506.25 | 3080301.5 | 2221188.75 | 3182854 | 2230761.25 | 2947740.5 | 4361806 | 1919758.75 | 3413615 | 1677.0134 | RAVPLALALISVSNPR | Q8VDM4 | Q8VDM4 | PSMD2 | proteasome (prosome, macropain) 26S subunit, non-ATPase, 2 | Cytoplasm | other |
| 15551796 | 2 | 0.511 | 3319535.25 | 2912422.25 | 3211608 | 3132459.75 | 2507620 | 2435127.75 | 2377347.75 | 2403252.25 | 3157228.75 | 4198100.5 | 2257581.25 | 3357578.25 | 2117.0076 | MNLASSFVNGFVNAAFGQDK | Q8VDM4 | Q8VDM4 | PSMD2 | proteasome (prosome, macropain) 26S subunit, non-ATPase, 2 | Cytoplasm | other |
| 15540446 | 3 | 0.329 | 3440949.75 | 3866551.75 | 2257284.5 | 1937417.375 | 2682313.5 | 2309687.25 | 2837019 | 3040518.25 | 1630709.5 | 2594818 | 1198572.375 | 2137805 | 2017.0743 | M[147.0354]LVTFDEELRPLPVSVR | Q8VDM4 | Q8VDM4 | PSMD2 | proteasome (prosome, macropain) 26S subunit, non-ATPase, 2 | Cytoplasm | other |
| 15550859 | 3 | 0.628 | 2.64E+07 | 3.18E+07 | 2.23E+07 | 2.14E+07 | 2.67E+07 | 1.80E+07 | 2.37E+07 | 2.04E+07 | 1.78E+07 | 3.96E+07 | 1.62E+07 | 2.50E+07 | 2679.3083 | DFLLPFLEEPMDTEADLQFRPR | P14685 | P14685 | PSMD3 | proteasome (prosome, macropain) 26S subunit, non-ATPase, 3 | Cytoplasm | other |
| 15556763 | 3 | 0.613 | 3.28E+07 | 2.85E+07 | 2.44E+07 | 2.36E+07 | 2.91E+07 | 1.54E+07 | 2.62E+07 | 2.09E+07 | 2.09E+07 | 3.95E+07 | 1.93E+07 | 2.36E+07 | 2296.1233 | AKPPPGGEQEPPPPAPQDVEMK | P14685 | P14685 | PSMD3 | proteasome (prosome, macropain) 26S subunit, non-ATPase, 3 | Cytoplasm | other |
| 15549203 | 2 | 0.452 | 2.36E+07 | 2.34E+07 | 2.52E+07 | 1.89E+07 | 2.21E+07 | 1.52E+07 | 1.92E+07 | 1.65E+07 | 1.98E+07 | 3.01E+07 | 1.44E+07 | 1.74E+07 | 1674.8541 | LQLDSPEDAEFIVAK | P14685 | P14685 | PSMD3 | proteasome (prosome, macropain) 26S subunit, non-ATPase, 3 | Cytoplasm | other |
| 15550239 | 2 | 0.744 | 1.46E+07 | 1.20E+07 | 1.43E+07 | 1.15E+07 | 1.32E+07 | 1.00E+07 | 1.26E+07 | 9693606 | 1.21E+07 | 1.83E+07 | 9191925 | 1.06E+07 | 1661.7633 | SVFPEQANNNEWAR | P14685 | P14685 | PSMD3 | proteasome (prosome, macropain) 26S subunit, non-ATPase, 3 | Cytoplasm | other |
| 15538584 | 2, 3 | 0.53 | 1.53E+07 | 1.02E+07 | 1.03E+07 | 7852358 | 1.11E+07 | 6823474 | 1.22E+07 | 8913818 | 7873047 | 1.65E+07 | 7952255 | 9361950 | 1421.6873, 1421.6876 | AVHGFFTSNNATR | P14685 | P14685 | PSMD3 | proteasome (prosome, macropain) 26S subunit, non-ATPase, 3 | Cytoplasm | other |
| 15532615 | 3, 4 | 0.464 | 1.92E+07 | 9238202 | 7445270.5 | 1.29E+07 | 8871565 | 6762353.5 | 1.05E+07 | 7656377.5 | 9429808 | 1.53E+07 | 1.19E+07 | 1.27E+07 | 1651.8723, 1651.8729 | AIRDGVIEASINHEK | P14685 | P14685 | PSMD3 | proteasome (prosome, macropain) 26S subunit, non-ATPase, 3 | Cytoplasm | other |
| 15533236 | 3 | 0.394 | 1.22E+07 | 9107809 | 1.04E+07 | 7841603.5 | 1.05E+07 | 6860412.5 | 1.05E+07 | 8731847 | 8046978.5 | 1.42E+07 | 8756381 | 9550134 | 1643.7667 | NYLHYSLYDQAEK | P14685 | P14685 | PSMD3 | proteasome (prosome, macropain) 26S subunit, non-ATPase, 3 | Cytoplasm | other |
| 15537759 | 3 | 0.582 | 9654503 | 9208760 | 1.09E+07 | 5376849.5 | 9598355 | 7194415 | 9607154 | 7404432 | 9108504 | 1.23E+07 | 5304937 | 9595028 | 1649.8938 | HDADGQATLLNLLLR | P14685 | P14685 | PSMD3 | proteasome (prosome, macropain) 26S subunit, non-ATPase, 3 | Cytoplasm | other |
| 15549623 | 2 | 0.486 | 8639337 | 4251482.5 | 1824388.375 | 5386488.5 | 3943926.5 | 2258972.25 | 3184846.5 | 3442980.5 | 4638962 | 8977552 | 5439927 | 5269466.5 | 1538.8368 | SLMPYFLLTQAVR | P14685 | P14685 | PSMD3 | proteasome (prosome, macropain) 26S subunit, non-ATPase, 3 | Cytoplasm | other |
| 15532739 | 2 | 0.565 | 6505623.5 | 4479934.5 | 5696253 | 3552912.25 | 5167626 | 3739330 | 5349932 | 4521488 | 4802610.5 | 7500371.5 | 3457786.25 | 4356454 | 913.4647 | VYEFLDK | P14685 | P14685 | PSMD3 | proteasome (prosome, macropain) 26S subunit, non-ATPase, 3 | Cytoplasm | other |
| 15539932 | 2 | 0.715 | 4403120 | 4766292 | 6945150 | 2812146 | 6530845.5 | 4651372 | 5563393.5 | 5323329 | 5395292.5 | 5843856.5 | 2686647.5 | 4825438 | 958.55475 | ISLADIAQK | P14685 | P14685 | PSMD3 | proteasome (prosome, macropain) 26S subunit, non-ATPase, 3 | Cytoplasm | other |
| 15554714 | 2 | 0.703 | 3607126.25 | 1760888.5 | 1298787.375 | 2991675.75 | 2024195.75 | 1178438.125 | 2459812.75 | 1462298.5 | 1055291.25 | 4786413.5 | 1975544.25 | 1721934.125 | 1643.7673 | NYLHYSLYDQAEK | P14685 | P14685 | PSMD3 | proteasome (prosome, macropain) 26S subunit, non-ATPase, 3 | Cytoplasm | other |
| 15539808 | 2 | 0.687 | 3282609 | 3208235.5 | 3068970.25 | 2505555.25 | 3732651 | 1380753.5 | 2839503.75 | 2800186.25 | 2342212.5 | 3651733.25 | 1398990.375 | 2505297 | 1179.5989 | AIQLEYSEAR | P14685 | P14685 | PSMD3 | proteasome (prosome, macropain) 26S subunit, non-ATPase, 3 | Cytoplasm | other |
| 15533939 | 2 | 0.658 | 2646826 | 1615345.625 | 2133735.75 | 1645909.375 | 2538852.5 | 1201006.75 | 2385118.5 | 1885519.375 | 1934572.25 | 3157252.75 | 1893736 | 1946628.5 | 800.48645 | ALDLVAAK | P14685 | P14685 | PSMD3 | proteasome (prosome, macropain) 26S subunit, non-ATPase, 3 | Cytoplasm | other |
| 15540393 | 3 | 0.415 | 6396351 | 1493952.5 | 1092722.875 | 4384031 | 299001 | 1082890.625 | 2959928.5 | 861514.375 | 2737396.5 | 2966590.75 | 5023916 | 4855805.5 | 2227.1162 | AAATEHSQRELDTVTLEDIK | P14685 | P14685 | PSMD3 | proteasome (prosome, macropain) 26S subunit, non-ATPase, 3 | Cytoplasm | other |
| 15535468 | 3 | 0.434 | 3073651.25 | 3438951.75 | 2845547.5 | 3201023.5 | 3326216.75 | 2467891.75 | 3416495 | 3300098 | 2960228 | 2807921.75 | 1742298.125 | 3219940.75 | 1694.9387 | RSLMPYFLLTQAVR | P14685 | P14685 | PSMD3 | proteasome (prosome, macropain) 26S subunit, non-ATPase, 3 | Cytoplasm | other |
| 15532685 | 3 | 0.449 | 3.76E+07 | 3.01E+07 | 2.41E+07 | 2.18E+07 | 2.92E+07 | 1.81E+07 | 2.85E+07 | 2.54E+07 | 2.22E+07 | 3.22E+07 | 1.95E+07 | 2.05E+07 | 1710.8546 | LQAQQDAVNIVC[160.0307]HSK | O35226 | O35226 | PSMD4 | proteasome (prosome, macropain) 26S subunit, non-ATPase, 4 | Cytoplasm | other |
| 15547937 | 2 | 0.274 | 3.13E+07 | 2.18E+07 | 2.41E+07 | 1.96E+07 | 2.15E+07 | 1.61E+07 | 1.90E+07 | 1.79E+07 | 2.23E+07 | 2.61E+07 | 1.18E+07 | 1.54E+07 | 1517.7805 | IIAFVGSPVEDNEK | O35226 | O35226 | PSMD4 | proteasome (prosome, macropain) 26S subunit, non-ATPase, 4 | Cytoplasm | other |
| 15548893 | 2 | 0.226 | 1.53E+07 | 1.45E+07 | 1.52E+07 | 9003510 | 1.63E+07 | 8348688.5 | 1.00E+07 | 1.08E+07 | 1.33E+07 | 1.28E+07 | 7710829 | 9741808 | 1948.9453 | VNVDIINFGEEEVNTEK | O35226 | O35226 | PSMD4 | proteasome (prosome, macropain) 26S subunit, non-ATPase, 4 | Cytoplasm | other |
| 15535094 | 2 | 0.223 | 1.42E+07 | 1.07E+07 | 1.13E+07 | 8696984 | 1.03E+07 | 5565695.5 | 1.12E+07 | 7427937 | 7751822 | 1.09E+07 | 5003792.5 | 6642718.5 | 1163.6068 | SVMGALASQATK | O35226 | O35226 | PSMD4 | proteasome (prosome, macropain) 26S subunit, non-ATPase, 4 | Cytoplasm | other |
| 15556637, 15558060 | 3 | 0.288 | 1.36E+07 | 1.51E+07 | 9188966 | 1.11E+07 | 1.36E+07 | 7333319.5 | 1.28E+07 | 1.11E+07 | 7675408 | 9099155 | 5959519.5 | 8726059 | 3632.6606, 3632.661 | EEDDYDVMQDPEFLQSVLENLPGVDPNNAAIR | O35226 | O35226 | PSMD4 | proteasome (prosome, macropain) 26S subunit, non-ATPase, 4 | Cytoplasm | other |
| 15550038 | 3 | 0.73 | 9195543 | 4499295 | 2849363.25 | 9833025 | 1086284.375 | 3782587.5 | 5099646.5 | 2362481.5 | 4850961 | 5928623 | 6205547 | 6115895.5 | 3271.6182 | TRSNPENNVGLITLANDC[160.0307]EVLTTLTPDTGR | O35226 | O35226 | PSMD4 | proteasome (prosome, macropain) 26S subunit, non-ATPase, 4 | Cytoplasm | other |
| 15541654 | 2 | 0.619 | 4560690.5 | 3721200.25 | 3081366 | 3858093.25 | 3251934.75 | 2226975.5 | 3812601 | 2725604.5 | 2553482.25 | 4240067.5 | 2273613.25 | 2368465 | 1177.655 | LTAFVNTLNGK | O35226 | O35226 | PSMD4 | proteasome (prosome, macropain) 26S subunit, non-ATPase, 4 | Cytoplasm | other |
| 15552209 | 2 | 0.383 | 3826911 | 2838686.75 | 2593644.25 | 2254339.75 | 3424249.5 | 1822953.75 | 3311291 | 2006488.25 | 1755327.25 | 2790342.75 | 1816652.625 | 1378623.5 | 2316.1172 | AAAASAAEAGIATPGTEDSDDALLK | O35226 | O35226 | PSMD4 | proteasome (prosome, macropain) 26S subunit, non-ATPase, 4 | Cytoplasm | other |
| 15551342 | 2 | 0.3 | 2759054.25 | 2369326 | 2058791.125 | 1604897.25 | 2108978.75 | 1342676.375 | 2160906 | 1444249.625 | 1739113.75 | 2532051.25 | 1310988.875 | 1370798 | 1710.8542 | LQAQQDAVNIVC[160.0307]HSK | O35226 | O35226 | PSMD4 | proteasome (prosome, macropain) 26S subunit, non-ATPase, 4 | Cytoplasm | other |
| 15551339 | 2, 3 | 0.102 | 3964960.25 | 3908454.75 | 3136501 | 3528060 | 2307534.75 | 3646261.75 | 3196492.25 | 1805807.125 | 2207654.25 | 6263101 | 3717063.25 | 3365682.25 | 2290.214, 2290.2163 | ISLTQAGLEALFESNLLDDLK | Q8BJY1 | Q8BJY1 | PSMD5 | proteasome (prosome, macropain) 26S subunit, non-ATPase, 5 | Cytoplasm | other |
| 15554854 | 3 | 0.467 | 3507071.75 | 2652561.75 | 1886454.75 | 6805496.5 | 2264184.75 | 2238975.5 | 2901529.25 | 2140291.75 | 2950031 | 4374784 | 3502077.5 | 4292223 | 2888.4065 | FFGNLAVMDSPQQIC[160.0307]ERYPVFLEK | Q8BJY1 | Q8BJY1 | PSMD5 | proteasome (prosome, macropain) 26S subunit, non-ATPase, 5 | Cytoplasm | other |
| 15549714 | 2 | 0.003 | 3490180.5 | 2575320 | 3139781.25 | 2962420.75 | 2528675.75 | 2461569.25 | 1957108.75 | 1231100.5 | 1733061.375 | 3973959.75 | 2992488.25 | 3357437.25 | 1502.7717 | VFTAIADQPWAQR | Q8BJY1 | Q8BJY1 | PSMD5 | proteasome (prosome, macropain) 26S subunit, non-ATPase, 5 | Cytoplasm | other |
| 15538121 | 3 | 0.033 | 3103813.75 | 2724328.5 | 3574283.75 | 2997051.25 | 3757680.75 | 3284961.5 | 3009849.25 | 2122030.75 | 2434554.25 | 3761118.25 | 3412745 | 3879523 | 1659.9502 | ALQSVVQAVPLHELR | Q8BJY1 | Q8BJY1 | PSMD5 | proteasome (prosome, macropain) 26S subunit, non-ATPase, 5 | Cytoplasm | other |
| 15550787 | 2 | 0.026 | 2234010.75 | 1557297.5 | 2214248.5 | 2617561.25 | 2257018.5 | 1997355.75 | 1735811.375 | 1184212.625 | 1819086.5 | 3280917.5 | 2931225.75 | 2369173.25 | 1650.8496 | QIVYC[160.0307]IGGENLSVAK | Q8BJY1 | Q8BJY1 | PSMD5 | proteasome (prosome, macropain) 26S subunit, non-ATPase, 5 | Cytoplasm | other |
| 15551685 | 2 | 0.053 | 2511669.75 | 1653829.75 | 1385243.125 | 2519059 | 872242.188 | 1486788.75 | 1080263.75 | 745328.938 | 1055457.75 | 3278015.5 | 2346532.5 | 1956382.625 | 2099.1182 | IVENSEAVTEILNNAELLK | Q8BJY1 | Q8BJY1 | PSMD5 | proteasome (prosome, macropain) 26S subunit, non-ATPase, 5 | Cytoplasm | other |
| 15544895 | 2 | 0.02 | 2211164 | 1998396.375 | 1240223.25 | 2250362.25 | 1418606.125 | 2026101.75 | 1208270 | 1076995.375 | 1079534.5 | 3072952.75 | 1933596.75 | 2424477.5 | 1483.7518 | TVAEIFGNSNYLR | Q8BJY1 | Q8BJY1 | PSMD5 | proteasome (prosome, macropain) 26S subunit, non-ATPase, 5 | Cytoplasm | other |
| 15553838 | 3 | 0.154 | 1512570.375 | 1766707.875 | 1337521.625 | 2183927.25 | 1723832.875 | 1892318.625 | 1619140.625 | 1260639.25 | 1787680.25 | 2740406.5 | 1572975.875 | 2276245.5 | 3052.4792 | VFEMADSQDPTMIGVAVDTVGILGSSVEGK | Q8BJY1 | Q8BJY1 | PSMD5 | proteasome (prosome, macropain) 26S subunit, non-ATPase, 5 | Cytoplasm | other |
| 15553894 | 2 | 0.004 | 1756601.375 | 1323347.75 | 1297328 | 975730.688 | 1005900.938 | 1742058 | 990683.312 | 482469.938 | 634100.625 | 2709471.5 | 2127704.25 | 1946224.5 | 1788.8397 | LMFNSPGFVEFVMDR | Q8BJY1 | Q8BJY1 | PSMD5 | proteasome (prosome, macropain) 26S subunit, non-ATPase, 5 | Cytoplasm | other |
| 15532348 | 3 | 0.778 | 2.05E+07 | 1.72E+07 | 1.65E+07 | 1.60E+07 | 1.57E+07 | 1.14E+07 | 2.06E+07 | 1.51E+07 | 1.18E+07 | 2.53E+07 | 1.12E+07 | 1.30E+07 | 1500.7617 | VNEIVETNRPDSK | Q99JI4 | Q99JI4 | PSMD6 | proteasome (prosome, macropain) 26S subunit, non-ATPase, 6 | Cytoplasm | enzyme |
| 15532728 | 2 | 0.604 | 1.85E+07 | 1.78E+07 | 1.66E+07 | 1.37E+07 | 1.74E+07 | 1.16E+07 | 1.71E+07 | 1.47E+07 | 1.29E+07 | 2.15E+07 | 8166882 | 1.27E+07 | 1367.7018 | PLENLEEEGLPK | Q99JI4 | Q99JI4 | PSMD6 | proteasome (prosome, macropain) 26S subunit, non-ATPase, 6 | Cytoplasm | enzyme |
| 15531827 | 2 | 0.664 | 1.81E+07 | 1.26E+07 | 1.15E+07 | 1.14E+07 | 1.25E+07 | 7680803 | 1.56E+07 | 1.02E+07 | 8546860 | 1.72E+07 | 9195653 | 9619150 | 1346.6395 | RLDEELEDAEK | Q99JI4 | Q99JI4 | PSMD6 | proteasome (prosome, macropain) 26S subunit, non-ATPase, 6 | Cytoplasm | enzyme |
| 15534479 | 2, 3 | 0.605 | 1.48E+07 | 9925692 | 1.08E+07 | 9085094 | 1.12E+07 | 6600676 | 1.23E+07 | 9605024 | 7620126.5 | 1.56E+07 | 8231572.5 | 9561379 | 1432.6782, 1432.6787 | SLIEEGGDWDRR | Q99JI4 | Q99JI4 | PSMD6 | proteasome (prosome, macropain) 26S subunit, non-ATPase, 6 | Cytoplasm | enzyme |
| 15532543 | 2 | 0.695 | 1.25E+07 | 1.05E+07 | 1.02E+07 | 8823942 | 1.12E+07 | 7139895 | 1.06E+07 | 8751412 | 8575205 | 1.43E+07 | 6884020.5 | 8379741 | 1196.5586 | AEYLC[160.0307]QIGDK | Q99JI4 | Q99JI4 | PSMD6 | proteasome (prosome, macropain) 26S subunit, non-ATPase, 6 | Cytoplasm | enzyme |
| 15550349 | 2 | 0.686 | 1.08E+07 | 8101536.5 | 7526109 | 6985085.5 | 8568789 | 5461181 | 8960793 | 6010940.5 | 7022406 | 1.07E+07 | 5517147 | 7065037 | 1602.6892 | ENNMAPYYEALC[160.0307]K | Q99JI4 | Q99JI4 | PSMD6 | proteasome (prosome, macropain) 26S subunit, non-ATPase, 6 | Cytoplasm | enzyme |
| 15534970 | 3 | 0.471 | 1.09E+07 | 9007866 | 7109014.5 | 6398768.5 | 7837681 | 6126128 | 7171789 | 7929061.5 | 7444052.5 | 9386117 | 5038117.5 | 8199480.5 | 1603.9131 | GAEILEVLHSLPAVR | Q99JI4 | Q99JI4 | PSMD6 | proteasome (prosome, macropain) 26S subunit, non-ATPase, 6 | Cytoplasm | enzyme |
| 15548913 | 2 | 0.389 | 5520954.5 | 4260359.5 | 4710693 | 3036535.75 | 4395806 | 2596642.5 | 4789742.5 | 3558811.5 | 3195595.25 | 5968949 | 3184009 | 3168170.25 | 1500.7623 | VNEIVETNRPDSK | Q99JI4 | Q99JI4 | PSMD6 | proteasome (prosome, macropain) 26S subunit, non-ATPase, 6 | Cytoplasm | enzyme |
| 15534183 | 2 | 0.131 | 4699752.5 | 4208926.5 | 3893520.75 | 3111727.75 | 4113962.5 | 2624925.5 | 4696649.5 | 4168514.25 | 3431921 | 5844068.5 | 4176187.75 | 4269450 | 1378.6439 | QYLFSLYEC[160.0307]R | Q99JI4 | Q99JI4 | PSMD6 | proteasome (prosome, macropain) 26S subunit, non-ATPase, 6 | Cytoplasm | enzyme |
| 15536835 | 2 | 0.4 | 4848336.5 | 3512564.25 | 3698312.25 | 2401810 | 3992346.25 | 1977937.125 | 4292223 | 2787425.5 | 4011650.25 | 5120140.5 | 2751305.75 | 3137036.75 | 1341.6953 | VYQGLYC[160.0307]VAIR | Q99JI4 | Q99JI4 | PSMD6 | proteasome (prosome, macropain) 26S subunit, non-ATPase, 6 | Cytoplasm | enzyme |
| 15549212 | 2 | 0.296 | 6487780.5 | 4653563 | 5442585.5 | 3692779.75 | 5555603.5 | 3128675 | 4761182.5 | 4275875 | 4825152.5 | 5093676.5 | 3243307.25 | 4121663.5 | 1603.9133 | GAEILEVLHSLPAVR | Q99JI4 | Q99JI4 | PSMD6 | proteasome (prosome, macropain) 26S subunit, non-ATPase, 6 | Cytoplasm | enzyme |
| 15555152, 15561592 | 2 | 0.555 | 3822302.25 | 3231397 | 2649607.75 | 2982415 | 2268539.5 | 1992779.375 | 2480046.25 | 1848070 | 2416337.75 | 4871893 | 1682495.5 | 2287365.25 | 1549.754, 1549.7545 | SLDWQMDVDLLSK | Q99JI4 | Q99JI4 | PSMD6 | proteasome (prosome, macropain) 26S subunit, non-ATPase, 6 | Cytoplasm | enzyme |
| 15543602 | 2 | 0.599 | 4339876 | 5736050.5 | 6603388 | 2492141 | 6007118.5 | 4706774.5 | 5078582 | 5292175.5 | 5777016 | 4572027.5 | 2968678.75 | 6005371 | 928.5551 | KGDLLLNR | Q99JI4 | Q99JI4 | PSMD6 | proteasome (prosome, macropain) 26S subunit, non-ATPase, 6 | Cytoplasm | enzyme |
| 15540138 | 2 | 0.153 | 1933377 | 2323766.75 | 1775581.625 | 1046484.688 | 2599730.75 | 768958.125 | 1833850 | 1836526.25 | 1518164.125 | 1538129.625 | 378283.375 | 779191 | 1151.6802 | LDIVFYLLR | Q99JI4 | Q99JI4 | PSMD6 | proteasome (prosome, macropain) 26S subunit, non-ATPase, 6 | Cytoplasm | enzyme |
| 15552790 | 3 | 0.605 | 2.80E+07 | 3.14E+07 | 3.04E+07 | 2.22E+07 | 3.30E+07 | 1.97E+07 | 2.81E+07 | 2.48E+07 | 2.42E+07 | 3.50E+07 | 1.61E+07 | 2.14E+07 | 2573.2239 | DLGLPTEAYISVEEVHDDGTPTSK | P26516 | P26516 | PSMD7 | proteasome (prosome, macropain) 26S subunit, non-ATPase, 7 | Cytoplasm | other |
| 15548470 | 2 | 0.846 | 2.24E+07 | 1.81E+07 | 1.54E+07 | 1.71E+07 | 1.89E+07 | 1.24E+07 | 1.94E+07 | 1.46E+07 | 1.38E+07 | 2.44E+07 | 1.28E+07 | 1.47E+07 | 1896.8821 | VLDVSNSFAVPFDEDDK | P26516 | P26516 | PSMD7 | proteasome (prosome, macropain) 26S subunit, non-ATPase, 7 | Cytoplasm | other |
| 15535542 | 3, 4 | 0.873 | 2.63E+07 | 1.79E+07 | 1.48E+07 | 1.96E+07 | 1.85E+07 | 1.15E+07 | 1.99E+07 | 1.47E+07 | 1.72E+07 | 2.38E+07 | 1.50E+07 | 1.50E+07 | 1943.1188, 1943.1189 | VVVHPLVLLSVVDHFNR | P26516 | P26516 | PSMD7 | proteasome (prosome, macropain) 26S subunit, non-ATPase, 7 | Cytoplasm | other |
| 15548358 | 3 | 0.693 | 1.63E+07 | 1.27E+07 | 7395878.5 | 1.37E+07 | 1.41E+07 | 4438701.5 | 1.37E+07 | 1.01E+07 | 7546316.5 | 2.33E+07 | 1.05E+07 | 1.27E+07 | 2525.074 | DDSVWFLDHDYLENMYGMFK | P26516 | P26516 | PSMD7 | proteasome (prosome, macropain) 26S subunit, non-ATPase, 7 | Cytoplasm | other |
| 15537585 | 2 | 0.762 | 9716775 | 8321715 | 7921119 | 7725704 | 7557726 | 5159350.5 | 9728156 | 7014414 | 6453158.5 | 1.26E+07 | 4954094.5 | 7607966 | 1077.551 | DTTVGTLSQR | P26516 | P26516 | PSMD7 | proteasome (prosome, macropain) 26S subunit, non-ATPase, 7 | Cytoplasm | other |
| 15542694 | 2 | 0.675 | 6049104 | 4490820 | 3818808.75 | 3870242 | 5277047 | 3315158.75 | 5737452 | 3449969.5 | 3925506 | 8575548 | 3901366 | 4698138 | 1160.5952 | NDIAINELMK | P26516 | P26516 | PSMD7 | proteasome (prosome, macropain) 26S subunit, non-ATPase, 7 | Cytoplasm | other |
| 15541181 | 4 | 0.133 | 8007733 | 1.08E+07 | 6088149 | 6615256 | 6565643 | 5452407 | 6803779.5 | 7791048 | 6654797 | 6407734 | 5334178 | 5429754.5 | 2681.3032 | TFEHVTSEIGAEEAEEVGVEHLLR | P26516 | P26516 | PSMD7 | proteasome (prosome, macropain) 26S subunit, non-ATPase, 7 | Cytoplasm | other |
| 15550139 | 2 | 0.334 | 4152762 | 4491990.5 | 3785305.25 | 3860729.25 | 3398660.75 | 2834788.25 | 4373417 | 3742084.75 | 4152199.25 | 4187620.5 | 2689121.75 | 4046717.5 | 1721.9221 | TNDQMVVVYLASLIR | P26516 | P26516 | PSMD7 | proteasome (prosome, macropain) 26S subunit, non-ATPase, 7 | Cytoplasm | other |
| 15538873 | 3 | 0.348 | 2732829 | 3209928 | 2298573.25 | 2475482 | 2246336.25 | 1932845.125 | 3612494.25 | 2339119 | 2704909.25 | 2734165.25 | 2210621.25 | 2953815.5 | 1721.9219 | TNDQMVVVYLASLIR | P26516 | P26516 | PSMD7 | proteasome (prosome, macropain) 26S subunit, non-ATPase, 7 | Cytoplasm | other |
| 15545712 | 2 | 0.619 | 2225947.5 | 1151190.5 | 177850.078 | 1066068.25 | 1062820 | 93168.68 | 1540486.125 | 537810.5 | 24553.502 | 1899956.375 | 1124022.375 | 1010133.562 | 1157.6075 | IVGWYHTGPK | P26516 | P26516 | PSMD7 | proteasome (prosome, macropain) 26S subunit, non-ATPase, 7 | Cytoplasm | other |
| 15551721 | 2 | 0.845 | 1.60E+07 | 1.35E+07 | 1.27E+07 | 1.23E+07 | 1.33E+07 | 1.14E+07 | 1.38E+07 | 1.16E+07 | 1.38E+07 | 1.96E+07 | 9021820 | 9735671 | 1658.9724 | LVLLELNFLPTTGTK | Q9CX56 | Q9CX56 | PSMD8 | proteasome (prosome, macropain) 26S subunit, non-ATPase, 8 | Cytoplasm | other |
| 15536728 | 2 | 0.18 | 8456655 | 8898329 | 9180655 | 5552376 | 6823386 | 5726756 | 7476230.5 | 5836211 | 6329502.5 | 1.01E+07 | 4948557 | 6593915.5 | 952.5483 | ILFFSTPK | Q9CX56 | Q9CX56 | PSMD8 | proteasome (prosome, macropain) 26S subunit, non-ATPase, 8 | Cytoplasm | other |
| 15534870 | 2 | 0.544 | 8872047 | 6793391 | 6750128 | 5346017.5 | 6556606.5 | 4211170.5 | 7191097 | 5524024 | 6184155.5 | 8891583 | 3444653.5 | 6330849 | 920.5186 | ILFAEATR | Q9CX56 | Q9CX56 | PSMD8 | proteasome (prosome, macropain) 26S subunit, non-ATPase, 8 | Cytoplasm | other |
| 15534109 | 3 | 0.607 | 1.67E+07 | 1.30E+07 | 1.19E+07 | 1.50E+07 | 1.58E+07 | 1.22E+07 | 1.79E+07 | 1.51E+07 | 1.03E+07 | 3.09E+07 | 1.23E+07 | 1.64E+07 | 1871.9083 | QLVHELDEAEYQEIR | P97371 | P97371 | PSME1 | proteasome (prosome, macropain) activator subunit 1 (PA28 alpha) | Cytoplasm | other |
| 15532594 | 2 | 0.838 | 1.30E+07 | 1.03E+07 | 1.38E+07 | 1.11E+07 | 1.31E+07 | 1.26E+07 | 1.37E+07 | 1.08E+07 | 9569131 | 2.46E+07 | 8237567 | 1.37E+07 | 1268.6493 | TENLLGSYFPK | P97371 | P97371 | PSME1 | proteasome (prosome, macropain) activator subunit 1 (PA28 alpha) | Cytoplasm | other |
| 15532171 | 2 | 0.874 | 1.40E+07 | 1.27E+07 | 1.32E+07 | 1.36E+07 | 1.09E+07 | 1.15E+07 | 1.67E+07 | 1.17E+07 | 1.12E+07 | 2.38E+07 | 8036138 | 1.65E+07 | 1395.7855 | NAYAVLYDIILK | P97371 | P97371 | PSME1 | proteasome (prosome, macropain) activator subunit 1 (PA28 alpha) | Cytoplasm | other |
| 15531740 | 3 | 0.836 | 1.28E+07 | 8875247 | 1.03E+07 | 1.07E+07 | 9733568 | 9550435 | 1.23E+07 | 8979813 | 7500235.5 | 2.07E+07 | 7287805 | 1.08E+07 | 1367.6605 | VDVFREDLC[160.0307]SK | P97371 | P97371 | PSME1 | proteasome (prosome, macropain) activator subunit 1 (PA28 alpha) | Cytoplasm | other |
| 15532618 | 2 | 0.981 | 9807131 | 1.09E+07 | 1.03E+07 | 1.04E+07 | 8260318.5 | 1.06E+07 | 1.32E+07 | 9536314 | 7432403.5 | 1.76E+07 | 4813925.5 | 9055026 | 1035.5693 | ISELDAFLK | P97371 | P97371 | PSME1 | proteasome (prosome, macropain) activator subunit 1 (PA28 alpha) | Cytoplasm | other |
| 15533495 | 2 | 0.218 | 7675536 | 4413385 | 4327889.5 | 5532570 | 5373297.5 | 4542729 | 6946932.5 | 5036468.5 | 4552146 | 1.39E+07 | 7452509 | 5831518.5 | 1328.5697 | GPPC[160.0307]GPVNC[160.0307]NEK | P97371 | P97371 | PSME1 | proteasome (prosome, macropain) activator subunit 1 (PA28 alpha) | Cytoplasm | other |
| 15549810 | 2 | 0.68 | 6034810 | 4772678 | 4839431.5 | 6503584.5 | 5750707.5 | 2441130.75 | 6882710 | 4967364 | 4109677.25 | 1.26E+07 | 4195629.5 | 5963546.5 | 1871.9081 | QLVHELDEAEYQEIR | P97371 | P97371 | PSME1 | proteasome (prosome, macropain) activator subunit 1 (PA28 alpha) | Cytoplasm | other |
| 15532300 | 2 | 0.783 | 6821964 | 6179811 | 7364569.5 | 5987359.5 | 5652944 | 7134355.5 | 7441411 | 5401634 | 5169714.5 | 1.22E+07 | 4543457.5 | 7227428 | 840.56525 | IVVLLQR | P97371 | P97371 | PSME1 | proteasome (prosome, macropain) activator subunit 1 (PA28 alpha) | Cytoplasm | other |
| 15533688 | 3 | 0.663 | 5764211 | 4811978 | 4901996 | 4710953.5 | 5317074.5 | 6657299 | 8049408.5 | 5433844 | 4379826 | 9694632 | 4607765.5 | 6722484.5 | 1332.6948 | VFELMTNLHTK | P97371 | P97371 | PSME1 | proteasome (prosome, macropain) activator subunit 1 (PA28 alpha) | Cytoplasm | other |
| 15534615 | 2 | 0.501 | 5563537 | 4199080.5 | 4276480.5 | 4261841.5 | 4111838.75 | 4747063.5 | 5842245.5 | 4157455.25 | 3225870.75 | 9397281 | 4385755 | 4864594.5 | 1159.6097 | LEGFHTQISK | P97371 | P97371 | PSME1 | proteasome (prosome, macropain) activator subunit 1 (PA28 alpha) | Cytoplasm | other |
| 15552441 | 2 | 0.623 | 7099014 | 5637403.5 | 7911131 | 5326998 | 6378296.5 | 5836247.5 | 6221563 | 5427053 | 5009148 | 8629320 | 4605919.5 | 5625208.5 | 1519.7352 | IEDGNNFGVAVQEK | P97371 | P97371 | PSME1 | proteasome (prosome, macropain) activator subunit 1 (PA28 alpha) | Cytoplasm | other |
| 15533230 | 2 | 0.298 | 4019228.75 | 1669829 | 1202399.875 | 3287038 | 729185.438 | 1670090.625 | 2772350.5 | 919109.188 | 1426273.875 | 4687396.5 | 3256627.25 | 3161685 | 1242.6085 | YFSERGDAVAK | P97371 | P97371 | PSME1 | proteasome (prosome, macropain) activator subunit 1 (PA28 alpha) | Cytoplasm | other |
| 15548404, 15549028 | 2, 3 | 0.986 | 1.81E+07 | 2.25E+07 | 2.59E+07 | 1.87E+07 | 2.00E+07 | 2.62E+07 | 2.62E+07 | 2.50E+07 | 1.97E+07 | 3.93E+07 | 1.10E+07 | 2.28E+07 | 2414.3088, 2414.312 | IISLSQLLQEDSLNVADLSSLR | P97372 | P97372 | PSME2 | proteasome (prosome, macropain) activator subunit 2 (PA28 beta) | Cytoplasm | peptidase |
| 15532943 | 2 | 0.647 | 1.43E+07 | 1.14E+07 | 1.29E+07 | 1.41E+07 | 1.43E+07 | 1.16E+07 | 1.74E+07 | 1.19E+07 | 8677698 | 2.77E+07 | 1.02E+07 | 1.63E+07 | 1123.5969 | VEAFQTTISK | P97372 | P97372 | PSME2 | proteasome (prosome, macropain) activator subunit 2 (PA28 beta) | Cytoplasm | peptidase |
| 15535275 | 3 | 0.454 | 1.50E+07 | 8988538 | 8444989 | 1.32E+07 | 9935919 | 1.07E+07 | 1.42E+07 | 9430663 | 8105889 | 2.26E+07 | 1.13E+07 | 1.24E+07 | 1670.8593 | ALVHERDEAAYGALR | P97372 | P97372 | PSME2 | proteasome (prosome, macropain) activator subunit 2 (PA28 beta) | Cytoplasm | peptidase |
| 15549310 | 2 | 0.858 | 4545005.5 | 3959764 | 5711127.5 | 2978634.75 | 4597733.5 | 5004497 | 5785290.5 | 4096345.5 | 4060207.5 | 8444027 | 3625716.25 | 4188993.5 | 1534.7341 | IEDGNDFGVAIQEK | P97372 | P97372 | PSME2 | proteasome (prosome, macropain) activator subunit 2 (PA28 beta) | Cytoplasm | peptidase |
| 15543071 | 2 | 0.2 | 2312787.25 | 1809754.75 | 2072431.375 | 2484012.25 | 2028853.5 | 2475815.25 | 2267869.75 | 1382488.75 | 1559257.5 | 5780241.5 | 2058444.75 | 2518038 | 817.4575 | AMVLDLR | P97372 | P97372 | PSME2 | proteasome (prosome, macropain) activator subunit 2 (PA28 beta) | Cytoplasm | peptidase |
| 15541688 | 3 | 0.206 | 3347991 | 2937196.5 | 2224083 | 3140470.5 | 1821275.375 | 2341671.25 | 3467326.5 | 2648583 | 1650906.25 | 4778445 | 4374867.5 | 2911632.25 | 1807.1426 | LLALLALVKPEVWTLK | P97372 | P97372 | PSME2 | proteasome (prosome, macropain) activator subunit 2 (PA28 beta) | Cytoplasm | peptidase |
| 15550511 | 3 | 0.575 | 5257292 | 2921186.75 | 2361615.5 | 4862394.5 | 936667.375 | 4240080 | 4546964 | 1509023.75 | 2961534.25 | 3482978.75 | 5583116 | 6396348 | 3765.0627 | IISLSQLLQEDSLNVADLSSLRAPLDIPIPDPPPK | P97372 | P97372 | PSME2 | proteasome (prosome, macropain) activator subunit 2 (PA28 beta) | Cytoplasm | peptidase |
| 15535047 | 2 | 0.032 | 2758270.25 | 3170801 | 4793018 | 2480456.75 | 2964617.25 | 2654817.75 | 2706366.25 | 2771458.5 | 2998600.75 | 2326270.25 | 1904969.5 | 2112359 | 917.5286 | LNTLESLK | Q8R3K3 | Q8R3K3 | PTCD2 | pentatricopeptide repeat domain 2 | Cytoplasm | other |
| 15532023 | 2 | 0.797 | 2.78E+07 | 2.94E+07 | 3.15E+07 | 2.40E+07 | 2.67E+07 | 3.35E+07 | 3.28E+07 | 3.01E+07 | 2.56E+07 | 2.93E+07 | 2.97E+07 | 3.25E+07 | 761.43616 | QNKMLK | O08586 | O08586 | PTEN | phosphatase and tensin homolog | Cytoplasm | phosphatase |
| 15535908 | 2 | 0.124 | 1912251.25 | 2351276.25 | 2377217 | 1692810 | 2166138.5 | 1806317.5 | 1912708 | 2272255.5 | 1974969.5 | 1803146 | 1818368.625 | 1721830.875 | 908.5183 | YGVTTIVR | Q63739 | Q63739 | PTP4A1 | protein tyrosine phosphatase type IVA, member 1 | Cytoplasm | phosphatase |
| 15552569 | 3 | 0.129 | 461199.531 | 1532042.125 | 2295777.5 | 4248675.5 | 4892303.5 | 5359384.5 | 3248160.75 | 3347452.75 | 4242251.5 | 629611.562 | 4684242 | 2915153.75 | 2830.2996 | MVGGGLQSAQASVPTEEELSSTEEEHK | P35821 | P35821 | PTPN1 | protein tyrosine phosphatase, non-receptor type 1 | Cytoplasm | phosphatase |
| 15532727 | 2 | 0.037 | 2.59E+07 | 2.81E+07 | 3.05E+07 | 2.01E+07 | 2.28E+07 | 2.54E+07 | 2.18E+07 | 2.14E+07 | 2.47E+07 | 2.52E+07 | 2.60E+07 | 2.93E+07 | 1219.515 | QEGTEEQEGGR | Q62136 | Q62136 | PTPN21 | protein tyrosine phosphatase, non-receptor type 21 | Cytoplasm | phosphatase |
| 15549423 | 2 | 0.679 | 7220281 | 9313756 | 7652854 | 9371909 | 6921999.5 | 7843246 | 7485452 | 7071959.5 | 9678669 | 9538780 | 1.02E+07 | 7681209.5 | 2623.258 | LFREEFNALPAC[160.0307]PIQATC[160.0307]EAASK | P18052 | P18052 | PTPRA | protein tyrosine phosphatase, receptor type, A | Plasma Membrane | phosphatase |
| 15551036 | 3 | 0.203 | 3540743.25 | 4212167.5 | 3968625 | 3899082.25 | 3315872.5 | 4502851.5 | 3812381.25 | 5290587.5 | 3822461 | 5135390.5 | 4511669.5 | 5002310 | 2248.099 | DSYIASQGPLLHTIEDFWR | P18052 | P18052 | PTPRA | protein tyrosine phosphatase, receptor type, A | Plasma Membrane | phosphatase |
| 15550298 | 3 | 0.19 | 3081100.25 | 3142402 | 3327469.75 | 4158890 | 2933444.25 | 3075189.25 | 2973252.5 | 3111866 | 2919579.5 | 3773567 | 3551708.75 | 3592530 | 2892.399 | VHLTPVEGVPDSDYINASFINGYQEK | P18052 | P18052 | PTPRA | protein tyrosine phosphatase, receptor type, A | Plasma Membrane | phosphatase |
| 15543192 | 3 | 0.348 | 571604.75 | 619892.625 | 743892.25 | 270460.188 | 625240.875 | 242013.578 | 275440.438 | 650665.312 | 684913.188 | 388253.156 | 863641.125 | 603009.312 | 2132.9956 | QQQQSGNHPITVHC[160.0307]SAGAGR | P18052 | P18052 | PTPRA | protein tyrosine phosphatase, receptor type, A | Plasma Membrane | phosphatase |
| 15553741 | 3 | 0.387 | 1638559.25 | 2140633 | 3925082.75 | 8178031.5 | 1.10E+07 | 8593435 | 5385900 | 6960553.5 | 7261877.5 | 115111.141 | 8788019 | 6563169 | 4014.955 | ILTQVGDGTTSLVPGWNLFC[160.0307]TEPEPVTSFHC[160.0307]EVVPK | Q64455 | Q64455 | PTPRJ | protein tyrosine phosphatase, receptor type, J | Plasma Membrane | phosphatase |
| 15544110 | 2 | 0.474 | 114561.227 | 150605.781 | 313855.719 | 77416.773 | 581853.438 | 831952.25 | 538985.938 | 532723.25 | 1209420.625 | 53274.434 | 615742.938 | 775275.312 | 1165.5864 | NNEVSFSQIK | Q64455 | Q64455 | PTPRJ | protein tyrosine phosphatase, receptor type, J | Plasma Membrane | phosphatase |
| 15546384 | 2 | 0.256 | 212366.25 | 370154.75 | 609669.625 | 1227887.25 | 927593.562 | 917555.125 | 851782.938 | 838991.188 | 1184631 | 33700.512 | 984229.438 | 505156.75 | 1066.5127 | YTAEIAENR | Q64455 | Q64455 | PTPRJ | protein tyrosine phosphatase, receptor type, J | Plasma Membrane | phosphatase |
| 15532678 | 3 | 0.263 | 8003421.5 | 1.28E+07 | 6678229.5 | 8283445 | 8649934 | 1.10E+07 | 5258049.5 | 1.06E+07 | 8957165 | 5221813.5 | 6609265.5 | 6189651.5 | 1664.8823 | APDEDTLIQLLTHAK | Q8R2Y8 | Q8R2Y8 | PTRH2 | peptidyl-tRNA hydrolase 2 | Cytoplasm | enzyme |
| 15540423 | 2 | 0.435 | 241477.953 | 333048.875 | 166497.984 | 258894.875 | 220090.859 | 324614.812 | 173171.781 | 271731.594 | 252732.547 | 43739.188 | 266787.531 | 195587.906 | 1462.7074 | VAAQC[160.0307]SHAAVSAYK | Q8R2Y8 | Q8R2Y8 | PTRH2 | peptidyl-tRNA hydrolase 2 | Cytoplasm | enzyme |
| 15553245 | 3 | 0.961 | 3192043.5 | 5170950.5 | 2887393.25 | 4518533.5 | 3830503.25 | 4081720.5 | 4774978.5 | 4452424 | 2422487.75 | 6927430.5 | 2024609.625 | 3099241 | 2733.2913 | LIDDYGVEEEPAELPEGTSLTVDNK | P42669 | P42669 | PURA | purine-rich element binding protein A | Nucleus | transcription regulator |
| 15556153 | 3 | 0.348 | 1851673.125 | 399724.969 | 87850.188 | 1182787.25 | 293266.469 | 654.293 | 321694.812 | 538548.375 | 493613.781 | 4434945.5 | 1109671.125 | 1056446 | 2889.3938 | LIDDYGVEEEPAELPEGTSLTVDNKR | P42669 | P42669 | PURA | purine-rich element binding protein A | Nucleus | transcription regulator |
| 15552718 | 2 | 0.077 | 3156187.75 | 5615282 | 2215576.75 | 3728353.5 | 2760037 | 3669306 | 1861998.375 | 4757415 | 4489693.5 | 1411655.25 | 1825103.125 | 1470433.75 | 1570.8286 | LESPTVSTLAPPSSGK | Q922Q4 | Q922Q4 | PYCR2 | pyrroline-5-carboxylate reductase family, member 2 | Cytoplasm | enzyme |
| 15537334 | 2 | 0.683 | 4649621 | 3161131 | 4385816 | 3600107.5 | 3996604.5 | 3812377.25 | 5266076 | 3366959.25 | 2056721.125 | 8575434 | 3085659.5 | 4166416.25 | 1086.6495 | IAAQTLLGTAK | Q9DCC4 | Q9DCC4 | PYCRL | pyrroline-5-carboxylate reductase-like | Cytoplasm | enzyme |
| 15538481 | 2 | 0.331 | 3323869.5 | 2524528.5 | 2475935.75 | 2901583 | 2444525.75 | 2271283.5 | 2854371.5 | 2298212 | 1961869.125 | 6890430.5 | 2367439.5 | 3210340.25 | 1338.6118 | AATMSAVEAATC[160.0307]R | Q9DCC4 | Q9DCC4 | PYCRL | pyrroline-5-carboxylate reductase-like | Cytoplasm | enzyme |
| 15537737 | 3 | 0.219 | 2942116.25 | 1739899.5 | 1444882.125 | 2176803.25 | 1942002.125 | 1935251.5 | 2580171 | 1965594.375 | 1571030.25 | 4294927 | 2990295 | 2239465 | 1842.8884 | QVLASAPTDNNLC[160.0307]HFR | Q9DCC4 | Q9DCC4 | PYCRL | pyrroline-5-carboxylate reductase-like | Cytoplasm | enzyme |
| 15552690 | 3 | 0.056 | 2601545 | 3469367.75 | 5945992.5 | 1349723 | 2257681.5 | 2129047 | 2784258 | 2161144.5 | 5717119.5 | 1489379.875 | 1341739.125 | 1981028 | 2346.2422 | VNTNYRPGLPFSGQVLLVDEK | Q61838 | Q61838 | Pzp | pregnancy zone protein | Extracellular Space | other |
| 15542657 | 3 | 0.273 | 978666.125 | 579340.562 | 230167.281 | 687325.875 | 637392.75 | 1015503.562 | 1443077 | 887214.438 | 898254.812 | 1739074.375 | 667210.5 | 989324 | 1671.8978 | RPNSGSLIQVVTTDGK | Q8BVI4 | Q8BVI4 | QDPR | quinoid dihydropteridine reductase | Cytoplasm | enzyme |
| 15531690 | 2 | 0.569 | 2.52E+08 | 2.97E+08 | 3.16E+08 | 2.26E+08 | 2.64E+08 | 2.95E+08 | 2.78E+08 | 2.85E+08 | 2.71E+08 | 2.78E+08 | 2.91E+08 | 2.86E+08 | 1071.6395 | LLLIGDSGVGK | P61027 | P61027 | RAB10 | RAB10, member RAS oncogene family | Cytoplasm | enzyme |
| 15531475, 15547646 | 2, 3 | 0.571 | 1.45E+08 | 1.32E+08 | 1.68E+08 | 1.51E+08 | 1.44E+08 | 1.43E+08 | 1.50E+08 | 1.47E+08 | 1.19E+08 | 1.96E+08 | 1.45E+08 | 1.44E+08 | 1440.6322, 1440.6323, 1440.6333 | NIDEHANEDVER | P61027 | P61027 | RAB10 | RAB10, member RAS oncogene family | Cytoplasm | enzyme |
| 15531496 | 2, 3 | 0.421 | 1.26E+08 | 1.50E+08 | 1.53E+08 | 1.71E+08 | 1.13E+08 | 1.63E+08 | 1.39E+08 | 1.56E+08 | 1.34E+08 | 1.04E+08 | 1.00E+08 | 1.53E+08 | 1389.8069, 1389.8073 | AFLTLAEDILRK | P61027 | P61027 | RAB10 | RAB10, member RAS oncogene family | Cytoplasm | enzyme |
| 15541030 | 2 | 0.421 | 3.17E+07 | 4.02E+07 | 4.17E+07 | 2.90E+07 | 4.51E+07 | 3.20E+07 | 3.43E+07 | 4.87E+07 | 2.66E+07 | 8.17E+07 | 4.89E+07 | 3.34E+07 | 1261.7135 | AFLTLAEDILR | P61027 | P61027 | RAB10 | RAB10, member RAS oncogene family | Cytoplasm | enzyme |
| 15532198 | 2 | 0.827 | 3.49E+07 | 3.72E+07 | 5.05E+07 | 3.37E+07 | 4.13E+07 | 4.23E+07 | 4.59E+07 | 4.34E+07 | 3.46E+07 | 4.36E+07 | 3.40E+07 | 3.32E+07 | 899.4856 | TYDLLFK | P61027 | P61027 | RAB10 | RAB10, member RAS oncogene family | Cytoplasm | enzyme |
| 15548522 | 2 | 0.636 | 3.04E+07 | 3.89E+07 | 5.76E+07 | 2.97E+07 | 4.95E+07 | 4.61E+07 | 4.55E+07 | 5.13E+07 | 3.60E+07 | 4.12E+07 | 2.57E+07 | 3.54E+07 | 1932.8895 | EPNSENVDISSGGGVTGWK | P61027 | P61027 | RAB10 | RAB10, member RAS oncogene family | Cytoplasm | enzyme |
| 15532304 | 2 | 0.477 | 3.25E+07 | 3.04E+07 | 4.14E+07 | 3.15E+07 | 2.89E+07 | 3.00E+07 | 3.50E+07 | 3.38E+07 | 2.87E+07 | 3.51E+07 | 3.11E+07 | 3.43E+07 | 829.4081 | FFETSAK | P61027 | P61027 | RAB10 | RAB10, member RAS oncogene family | Cytoplasm | enzyme |
| 15533690 | 2 | 0.603 | 2.31E+07 | 2.04E+07 | 2.07E+07 | 1.93E+07 | 2.14E+07 | 1.87E+07 | 2.29E+07 | 2.32E+07 | 1.79E+07 | 2.20E+07 | 2.18E+07 | 2.14E+07 | 1288.6305 | FHTITTSYYR | P61027 | P61027 | RAB10 | RAB10, member RAS oncogene family | Cytoplasm | enzyme |
| 15542749 | 2 | 0.736 | 1.47E+07 | 1.80E+07 | 3.38E+07 | 8905072 | 2.84E+07 | 3.08E+07 | 1.76E+07 | 2.97E+07 | 3.34E+07 | 1.05E+07 | 2.26E+07 | 1.90E+07 | 795.4165 | TC[160.0307]VLFR | P61027 | P61027 | RAB10 | RAB10, member RAS oncogene family | Cytoplasm | enzyme |
| 15542892 | 2 | 0.691 | 1.09E+07 | 1.41E+07 | 2.55E+07 | 9810654 | 1.99E+07 | 1.97E+07 | 1.52E+07 | 1.77E+07 | 1.58E+07 | 8057676 | 1.48E+07 | 1.45E+07 | 824.4132 | SFENISK | P61027 | P61027 | RAB10 | RAB10, member RAS oncogene family | Cytoplasm | enzyme |
| 15554634 | 2 | 0.588 | 4797672.5 | 4391122.5 | 3768655.75 | 5865434 | 4072381.75 | 4722341.5 | 4401996 | 4603608 | 4197872.5 | 4973945 | 4478122.5 | 4671728.5 | 1582.7943 | GAMGIMLVYDITNGK | P61027 | P61027 | RAB10 | RAB10, member RAS oncogene family | Cytoplasm | enzyme |
| 15532871 | 2 | 0.497 | 2.44E+07 | 3.27E+07 | 3.78E+07 | 3.72E+07 | 4.14E+07 | 4.44E+07 | 3.15E+07 | 3.87E+07 | 4.42E+07 | 2.30E+07 | 4.17E+07 | 4.05E+07 | 1043.6074 | VVLIGDSGVGK | P46638 | P46638 | RAB11B | RAB11B, member RAS oncogene family | Cytoplasm | enzyme |
| 15534959 | 2 | 0.296 | 1.67E+07 | 2.14E+07 | 2.35E+07 | 2.25E+07 | 2.78E+07 | 2.72E+07 | 2.22E+07 | 2.61E+07 | 2.58E+07 | 1.37E+07 | 2.59E+07 | 2.07E+07 | 1160.5665 | HLTYENVER | P46638 | P46638 | RAB11B | RAB11B, member RAS oncogene family | Cytoplasm | enzyme |
| 15531667 | 2 | 0.388 | 1.59E+07 | 2.41E+07 | 2.42E+07 | 2.34E+07 | 2.73E+07 | 3.19E+07 | 2.03E+07 | 2.56E+07 | 2.99E+07 | 1.36E+07 | 2.62E+07 | 2.17E+07 | 1289.744 | GAVGALLVYDIAK | P46638 | P46638 | RAB11B | RAB11B, member RAS oncogene family | Cytoplasm | enzyme |
| 15550585 | 2, 3 | 0.248 | 1.51E+07 | 2.42E+07 | 2.23E+07 | 1.98E+07 | 1.89E+07 | 2.29E+07 | 1.41E+07 | 2.02E+07 | 2.41E+07 | 1.07E+07 | 1.35E+07 | 1.88E+07 | 2329.1135, 2329.114 | NNLSFIETSALDSTNVEEAFK | P46638 | P46638 | RAB11B | RAB11B, member RAS oncogene family | Cytoplasm | enzyme |
| 15535247 | 2 | 0.667 | 7562565.5 | 8940274 | 1.22E+07 | 6931873.5 | 8633495 | 8387027.5 | 7324627.5 | 9983049 | 1.12E+07 | 5423998.5 | 9400173 | 9347724 | 1021.5656 | NILTEIYR | P46638 | P46638 | RAB11B | RAB11B, member RAS oncogene family | Cytoplasm | enzyme |
| 15533017 | 2 | 0.323 | 5997448.5 | 8233744.5 | 8326126.5 | 9574623 | 9120264 | 1.05E+07 | 7053062 | 9562323 | 1.06E+07 | 4939484 | 9497977 | 7620755.5 | 944.48175 | AITSAYYR | P46638 | P46638 | RAB11B | RAB11B, member RAS oncogene family | Cytoplasm | enzyme |
| 15538163 | 2 | 0.692 | 6215647.5 | 6141658 | 5837160.5 | 6730780 | 6948378 | 6505096 | 4641087.5 | 6192903.5 | 7445512 | 4059587.25 | 7242386 | 5873224 | 1080.5671 | STIGVEFATR | P46638 | P46638 | RAB11B | RAB11B, member RAS oncogene family | Cytoplasm | enzyme |
| 15536274 | 3 | 0.91 | 4052336.75 | 4273775.5 | 2812120.5 | 5601255.5 | 3213188.5 | 3463758.75 | 3248382.25 | 4401712.5 | 3109779.75 | 2921971.75 | 5393058.5 | 4180011.75 | 2024.0538 | ELRDHADSNIVIMLVGNK | P46638 | P46638 | RAB11B | RAB11B, member RAS oncogene family | Cytoplasm | enzyme |
| 15536143 | 3 | 0.71 | 3307154.25 | 2375591 | 1992760 | 9270842 | 1120997.25 | 3872984 | 3762949.5 | 2221010.5 | 5084267.5 | 2082270.5 | 9578332 | 6105222.5 | 1384.6826 | FTRNEFNLESK | P46638 | P46638 | RAB11B | RAB11B, member RAS oncogene family | Cytoplasm | enzyme |
| 15542278 | 3 | 0.378 | 584195.125 | 1647054.75 | 1542306.625 | 515430.5 | 2231549.75 | 1393456.125 | 690904.375 | 1185962.75 | 1299534.5 | 659208.812 | 488547.031 | 643739.562 | 1625.832 | DHADSNIVIMLVGNK | P46638 | P46638 | RAB11B | RAB11B, member RAS oncogene family | Cytoplasm | enzyme |
| 15548584 | 3 | 0.194 | 2.06E+07 | 3.12E+07 | 3.78E+07 | 2.20E+07 | 3.65E+07 | 3.89E+07 | 3.52E+07 | 3.21E+07 | 3.14E+07 | 2.68E+07 | 1.53E+07 | 2.09E+07 | 2400.1726 | IYQNIQDGSLDLNAAESGVQHK | Q91V41 | Q91V41 | RAB14 | RAB14, member RAS oncogene family | Cytoplasm | enzyme |
| 15534208 | 2 | 0.058 | 1.73E+07 | 1.83E+07 | 2.49E+07 | 1.56E+07 | 2.39E+07 | 2.06E+07 | 2.05E+07 | 2.27E+07 | 2.15E+07 | 1.40E+07 | 1.31E+07 | 1.62E+07 | 1493.7097 | TGENVEDAFLEAAK | Q91V41 | Q91V41 | RAB14 | RAB14, member RAS oncogene family | Cytoplasm | enzyme |
| 15541888 | 3 | 0.28 | 7734236.5 | 6989269 | 6280041.5 | 1.31E+07 | 5557788 | 7884491 | 7736982 | 6939225 | 8417978 | 9300677 | 1.11E+07 | 1.06E+07 | 1737.824 | ADLEAQRDVTYEEAK | Q91V41 | Q91V41 | RAB14 | RAB14, member RAS oncogene family | Cytoplasm | enzyme |
| 15538047 | 3 | 0.012 | 9406102 | 1.04E+07 | 1.16E+07 | 9699586 | 1.14E+07 | 1.16E+07 | 1.15E+07 | 1.14E+07 | 9684935 | 8063336 | 8514225 | 7926732 | 1650.7833 | STYNHLSSWLTDAR | Q91V41 | Q91V41 | RAB14 | RAB14, member RAS oncogene family | Cytoplasm | enzyme |
| 15538172 | 3 | 0.106 | 5915561 | 6217456.5 | 5760876 | 6823282 | 6468514 | 8912381 | 8827954 | 8442762 | 7190394 | 7516066.5 | 6717381 | 5116887.5 | 1837.8315 | FMADC[160.0307]PHTIGVEFGTR | Q91V41 | Q91V41 | RAB14 | RAB14, member RAS oncogene family | Cytoplasm | enzyme |
| 15535437 | 2 | 0.554 | 5273508.5 | 4169722 | 5042068.5 | 4254616 | 5884592 | 5149926 | 4917053.5 | 5390288 | 5915979 | 5033540 | 7784565 | 5062327 | 1262.6158 | SC[160.0307]LLHQFTEK | Q91V41 | Q91V41 | RAB14 | RAB14, member RAS oncogene family | Cytoplasm | enzyme |
| 15541782 | 3 | 0.47 | 5863400 | 1069067.25 | 159843.375 | 5082838.5 | 624718.25 | 321976.188 | 898749.188 | 1122974.25 | 3362801.5 | 3313755.5 | 6476854.5 | 3530637 | 1621.8032 | TGENVEDAFLEAAKK | Q91V41 | Q91V41 | RAB14 | RAB14, member RAS oncogene family | Cytoplasm | enzyme |
| 15551106 | 2 | 0.667 | 3011887.5 | 1716616.125 | 1444841.125 | 3903772.25 | 660764.812 | 2847636.25 | 2226338 | 1992132.25 | 3402729.75 | 3034781 | 4111236.75 | 2537781.5 | 1737.8242 | ADLEAQRDVTYEEAK | Q91V41 | Q91V41 | RAB14 | RAB14, member RAS oncogene family | Cytoplasm | enzyme |
| 15554745 | 3 | 0.461 | 1825844.25 | 410250.406 | 0 | 1763299.25 | 139271.406 | 0 | 287472.5 | 296036.312 | 1078657.75 | 1062527.375 | 2936654 | 1584958 | 3150.5483 | IYQNIQDGSLDLNAAESGVQHKPSAPQGGR | Q91V41 | Q91V41 | RAB14 | RAB14, member RAS oncogene family | Cytoplasm | enzyme |
| 15532399, 15535863 | 2 | 0.492 | 3.84E+08 | 3.92E+08 | 4.59E+08 | 4.25E+08 | 4.26E+08 | 4.21E+08 | 4.42E+08 | 4.33E+08 | 3.84E+08 | 5.12E+08 | 4.53E+08 | 4.08E+08 | 1316.657, 1316.6572 | IQIWDTAGQER | Q8K386 | Q8K386 | Rab15 | RAB15, member RAS oncogene family | Cytoplasm | enzyme |
| 15548863 | 2 | 0.254 | 1.24E+07 | 1.11E+07 | 1.55E+07 | 1.49E+07 | 1.74E+07 | 1.22E+07 | 1.51E+07 | 1.63E+07 | 1.13E+07 | 2.24E+07 | 2.02E+07 | 1.37E+07 | 1784.7787 | TC[160.0307]DGVQC[160.0307]AFEELVEK | P35293 | P35293 | RAB18 | RAB18, member RAS oncogene family | Cytoplasm | enzyme |
| 15550191 | 2 | 0.401 | 2.09E+07 | 1.85E+07 | 2.30E+07 | 2.15E+07 | 2.51E+07 | 1.84E+07 | 2.05E+07 | 1.86E+07 | 1.53E+07 | 2.12E+07 | 1.87E+07 | 1.61E+07 | 1756.8817 | IIQTPGLWESENQNK | P35293 | P35293 | RAB18 | RAB18, member RAS oncogene family | Cytoplasm | enzyme |
| 15540538 | 2 | 0.173 | 1.49E+07 | 1.30E+07 | 1.86E+07 | 1.93E+07 | 2.08E+07 | 1.47E+07 | 1.34E+07 | 1.31E+07 | 1.16E+07 | 2.02E+07 | 1.51E+07 | 1.31E+07 | 1085.6539 | ILIIGESGVGK | P35293 | P35293 | RAB18 | RAB18, member RAS oncogene family | Cytoplasm | enzyme |
| 15554735 | 2 | 0.245 | 1.21E+07 | 1.23E+07 | 1.55E+07 | 1.51E+07 | 1.60E+07 | 1.56E+07 | 1.47E+07 | 1.38E+07 | 1.07E+07 | 1.51E+07 | 1.39E+07 | 1.39E+07 | 1826.8341 | LDNWLNELETYC[160.0307]TR | P35293 | P35293 | RAB18 | RAB18, member RAS oncogene family | Cytoplasm | enzyme |
| 15538547 | 2 | 0.358 | 1.01E+07 | 8995980 | 1.12E+07 | 1.10E+07 | 1.23E+07 | 1.15E+07 | 1.17E+07 | 1.13E+07 | 7502332.5 | 1.50E+07 | 1.10E+07 | 1.12E+07 | 1216.634 | NDIVNMLVGNK | P35293 | P35293 | RAB18 | RAB18, member RAS oncogene family | Cytoplasm | enzyme |
| 15553654 | 2 | 0.974 | 5.76E+07 | 6.31E+07 | 9.64E+07 | 6.34E+07 | 7.90E+07 | 8.47E+07 | 7.12E+07 | 7.73E+07 | 7.30E+07 | 8.37E+07 | 6.29E+07 | 7.39E+07 | 1724.8696 | EFADSLGIPFLETSAK | P62821 | P62821 | RAB1A | RAB1A, member RAS oncogene family | Cytoplasm | enzyme |
| 15556538 | 2 | 0.874 | 2.16E+07 | 2.72E+07 | 4.14E+07 | 2.93E+07 | 3.38E+07 | 3.64E+07 | 2.77E+07 | 3.28E+07 | 3.15E+07 | 3.67E+07 | 2.38E+07 | 2.95E+07 | 1884.8789 | NATNVEQSFMTMAAEIK | P62821 | P62821 | RAB1A | RAB1A, member RAS oncogene family | Cytoplasm | enzyme |
| 15534258 | 3 | 0.307 | 8179963 | 9112367 | 1.19E+07 | 1.40E+07 | 1.34E+07 | 1.47E+07 | 1.12E+07 | 1.35E+07 | 1.09E+07 | 1.89E+07 | 8497736 | 1.18E+07 | 1884.8795 | NATNVEQSFMTMAAEIK | P62821 | P62821 | RAB1A | RAB1A, member RAS oncogene family | Cytoplasm | enzyme |
| 15551111 | 2 | 0.606 | 6692259.5 | 3785415.5 | 5146571 | 5705776 | 3187351.25 | 4978603.5 | 5648557.5 | 2831041 | 5592292 | 4274697.5 | 8188134 | 6878492.5 | 1992.9719 | QWLQEIDRYASENVNK | P62821 | P62821 | RAB1A | RAB1A, member RAS oncogene family | Cytoplasm | enzyme |
| 15554872 | 2 | 0.438 | 1804450 | 2923735.5 | 4879386 | 2690138.75 | 2607049.25 | 3047866.75 | 2892336 | 3054661 | 2027784.5 | 2963471.5 | 662171.25 | 2114167.75 | 1900.8727 | NATNVEQSFM[147.0354]TMAAEIK | P62821 | P62821 | RAB1A | RAB1A, member RAS oncogene family | Cytoplasm | enzyme |
| 15561609 | 2 | 0.066 | 1762765.25 | 1572515.25 | 1797701.625 | 2693837.25 | 2683092.25 | 2184895.25 | 1909386.75 | 2426831.25 | 1974903.75 | 2678188.5 | 1742111.25 | 2436533.75 | 2171.9983 | FADDTYTESYISTIGVDFK | P62821 | P62821 | RAB1A | RAB1A, member RAS oncogene family | Cytoplasm | enzyme |
| 15534974 | 3 | 0.444 | 5.02E+07 | 3.56E+07 | 4.70E+07 | 5.62E+07 | 3.01E+07 | 5.21E+07 | 5.45E+07 | 3.96E+07 | 5.57E+07 | 4.91E+07 | 6.80E+07 | 5.98E+07 | 1992.9718 | QWLQEIDRYASENVNK | Q9D1G1 | Q9D1G1 | RAB1B | RAB1B, member RAS oncogene family | Cytoplasm | other |
| 15548120 | 2 | 0.538 | 2.08E+07 | 2.38E+07 | 3.12E+07 | 2.76E+07 | 2.53E+07 | 3.17E+07 | 2.63E+07 | 2.73E+07 | 3.22E+07 | 2.71E+07 | 2.31E+07 | 2.59E+07 | 1710.8545 | EFADSLGVPFLETSAK | Q9D1G1 | Q9D1G1 | RAB1B | RAB1B, member RAS oncogene family | Cytoplasm | other |
| 15531669 | 2 | 0.888 | 8951950 | 1.46E+07 | 2.32E+07 | 8142927 | 2.52E+07 | 1.83E+07 | 1.70E+07 | 2.03E+07 | 1.44E+07 | 2.19E+07 | 8269796.5 | 1.16E+07 | 1087.5513 | QWLQEIDR | Q9D1G1 | Q9D1G1 | RAB1B | RAB1B, member RAS oncogene family | Cytoplasm | other |
| 15534262 | 2 | 0.12 | 1.28E+07 | 1.04E+07 | 1.75E+07 | 1.31E+07 | 1.31E+07 | 1.49E+07 | 1.32E+07 | 1.52E+07 | 1.71E+07 | 1.94E+07 | 1.73E+07 | 1.82E+07 | 761.4323 | SC[160.0307]LLLR | Q9D1G1 | Q9D1G1 | RAB1B | RAB1B, member RAS oncogene family | Cytoplasm | other |
| 15538993 | 3 | 0.204 | 3788883.5 | 4015207 | 6766054.5 | 4407160 | 7539440.5 | 8083755.5 | 7457672.5 | 6716951.5 | 8731959 | 6134489.5 | 5475279.5 | 7307990 | 1868.8849 | NATNVEQAFMTMAAEIK | Q9D1G1 | Q9D1G1 | RAB1B | RAB1B, member RAS oncogene family | Cytoplasm | other |
| 15535447 | 2 | 0.303 | 7954212.5 | 8744297 | 1.30E+07 | 1.08E+07 | 1.31E+07 | 1.37E+07 | 8973467 | 1.16E+07 | 8805529 | 1.03E+07 | 9154243 | 1.14E+07 | 1336.6798 | GIEELFLDLC[160.0307]K | P35282 | P35282 | RAB21 | RAB21, member RAS oncogene family | Cytoplasm | enzyme |
| 15540234 | 3 | 0.775 | 6806983.5 | 2906705.25 | 1797613.75 | 9862067 | 3382809.25 | 727536.688 | 1.07E+07 | 1993078.875 | 742059.312 | 6661982 | 7346681 | 3757302.25 | 2090.008 | ERHVSIQEAESYAESVGAK | P35282 | P35282 | RAB21 | RAB21, member RAS oncogene family | Cytoplasm | enzyme |
| 15537940 | 3 | 0.609 | 2944130.75 | 2799367 | 4681994 | 4497174 | 3895541 | 3710477 | 3755588.5 | 2828705 | 3534707 | 4262913.5 | 3608317.5 | 3166418 | 1236.6498 | FHALGPIYYR | P35282 | P35282 | RAB21 | RAB21, member RAS oncogene family | Cytoplasm | enzyme |
| 15538087 | 3 | 0.603 | 2508711.75 | 2675946.5 | 4246373 | 2073404.5 | 5778887.5 | 4668281.5 | 597785.5 | 4661608 | 2527745 | 4250383 | 2336391 | 3139872.5 | 1804.8679 | HVSIQEAESYAESVGAK | P35282 | P35282 | RAB21 | RAB21, member RAS oncogene family | Cytoplasm | enzyme |
| 15540633 | 3 | 0.803 | 2632587 | 1389825.5 | 398417.469 | 3989272.25 | 1299042 | 1147947.625 | 4914886 | 1297782.125 | 301817.875 | 4040469.25 | 2288452 | 1220812.75 | 1628.8462 | RVNLAIWDTAGQER | P35282 | P35282 | RAB21 | RAB21, member RAS oncogene family | Cytoplasm | enzyme |
| 15539400 | 2 | 0.979 | 3490606.75 | 2925405 | 4155364.75 | 4012309 | 3469854.5 | 2974260 | 3715202.5 | 3396033.75 | 3161137.25 | 3042157.25 | 3191862 | 3824981 | 1359.76 | HITTLQASFLTK | P35282 | P35282 | RAB21 | RAB21, member RAS oncogene family | Cytoplasm | enzyme |
| 15557795 | 2 | 0.506 | 1578635.375 | 2042287.625 | 2994071.5 | 3174987 | 2287976 | 2926955.75 | 1695936.375 | 2083274.375 | 2762571.75 | 2449231.75 | 2005967 | 2712678.5 | 2115.9675 | DSNGAILVYDVTDEDSFQK | P35282 | P35282 | RAB21 | RAB21, member RAS oncogene family | Cytoplasm | enzyme |
| 15539871 | 2 | 0.323 | 469295.125 | 447813.125 | 623547.562 | 948608.812 | 794333.688 | 569843.625 | 837594.125 | 868314.5 | 346179.594 | 1210569.5 | 764784.25 | 687349.25 | 1191.565 | MIETAQVDER | P35282 | P35282 | RAB21 | RAB21, member RAS oncogene family | Cytoplasm | enzyme |
| 15537062 | 2 | 0.513 | 3484953.75 | 3593603.25 | 4238680 | 3382791.5 | 3282616.75 | 3550781.5 | 3451059.75 | 3977230.5 | 3708972 | 3953155.5 | 3168894.5 | 3513862 | 1118.5853 | VC[160.0307]LLGDTGVGK | P35285 | P35285 | RAB22A | RAB22A, member RAS oncogene family | Cytoplasm | enzyme |
| 15541151 | 2 | 0.207 | 3274662.5 | 3433126.75 | 4940686.5 | 3543209.5 | 3309653.75 | 2879749.5 | 2746898.75 | 3509742.25 | 3587985.75 | 2695873.5 | 3051470.75 | 2848749.5 | 1321.7352 | GSAAAIIVYDITK | P35285 | P35285 | RAB22A | RAB22A, member RAS oncogene family | Cytoplasm | enzyme |
| 15544406 | 2 | 0.096 | 410868 | 589170 | 1386806.625 | 2046176.75 | 3213713 | 2846172.5 | 2047174.625 | 2529750.25 | 2370241 | 394758.719 | 2514064.25 | 2149946 | 1274.6229 | TSVLYQYTDGK | Q9ERI2 | Q9ERI2 | RAB27A | RAB27A, member RAS oncogene family | Cytoplasm | enzyme |
| 15549266 | 2 | 0.152 | 1429881.75 | 3805150.5 | 1.09E+07 | 2.20E+07 | 2.62E+07 | 2.61E+07 | 1.34E+07 | 1.78E+07 | 2.66E+07 | 1243734.5 | 2.34E+07 | 1.54E+07 | 1974.9393 | YGIPYFETSAATGQNVEK | Q99P58 | Q99P58 | RAB27B | RAB27B, member RAS oncogene family | Cytoplasm | enzyme |
| 15549691 | 2, 3 | 0.085 | 805306.438 | 2765884 | 5005194 | 1.79E+07 | 1.74E+07 | 1.94E+07 | 9937602 | 1.35E+07 | 2.00E+07 | 921927.375 | 1.58E+07 | 1.17E+07 | 2678.264, 2678.2642 | NWMSQLQANAYC[160.0307]ENPDIVLIGNK | Q99P58 | Q99P58 | RAB27B | RAB27B, member RAS oncogene family | Cytoplasm | enzyme |
| 15531855 | 2 | 0.163 | 568656.125 | 2799404.5 | 6581304 | 1.64E+07 | 1.66E+07 | 1.97E+07 | 1.02E+07 | 1.46E+07 | 2.36E+07 | 711255 | 2.00E+07 | 1.34E+07 | 1261.7052 | SVETLLDLIMK | Q99P58 | Q99P58 | RAB27B | RAB27B, member RAS oncogene family | Cytoplasm | enzyme |
| 15536901 | 2 | 0.176 | 687372.312 | 3037304 | 6952851 | 1.47E+07 | 1.72E+07 | 2.11E+07 | 9119172 | 1.26E+07 | 2.01E+07 | 655330.125 | 1.54E+07 | 1.20E+07 | 1029.5913 | LLALGDSGVGK | Q99P58 | Q99P58 | RAB27B | RAB27B, member RAS oncogene family | Cytoplasm | enzyme |
| 15540595 | 3 | 0.1 | 250485.688 | 1498707.25 | 3414340 | 1.03E+07 | 8770853 | 1.23E+07 | 6889733.5 | 6773738 | 8216034 | 439377.719 | 7712113 | 5498877.5 | 1523.7408 | RVVYDTQGADGASGK | Q99P58 | Q99P58 | RAB27B | RAB27B, member RAS oncogene family | Cytoplasm | enzyme |
| 15545605 | 2 | 0.106 | 55430.348 | 460671.469 | 1147034.625 | 2178335 | 2520457.75 | 2821056.25 | 950459.438 | 1906640.75 | 1981585 | 149036.828 | 881193.125 | 1209260 | 1277.6992 | SVETLLDLIM[147.0354]K | Q99P58 | Q99P58 | RAB27B | RAB27B, member RAS oncogene family | Cytoplasm | enzyme |
| 15544362 | 2 | 0.045 | 22797.357 | 63429.32 | 115167.258 | 1925625.875 | 1565709.125 | 1329514 | 688124.062 | 1183365.875 | 1201152.625 | 28343.635 | 2514163.75 | 919418.438 | 1367.6407 | VVYDTQGADGASGK | Q99P58 | Q99P58 | RAB27B | RAB27B, member RAS oncogene family | Cytoplasm | enzyme |

| | | | | | | | | | | | | | | | |
|---|---|---|---|---|---|---|---|---|---|---|---|---|---|---|---|
| 15560850 | 2 | 0.484 | 76027.219 | 379080.719 | 1187489 | 782989.25 | 2219995 | 2071190 | 783876.25 | 1726663.75 | 2413825 | 3976.828 | 1892160.5 | 1264780.625 | [illegible] |
| 15532949 | 2 | 0.032 | 8328764 | 1.11E+07 | 1.09E+07 | 1.40E+07 | 1.27E+07 | 1.37E+07 | 1.31E+07 | 1.19E+07 | 1.09E+07 | 1.24E+07 | 1.20E+07 | 1.20E+07 | [illegible] |
| 15535880 | 3 | 0.221 | 6041752.5 | 6070222.5 | 8333171 | 7305316.5 | 8542894 | 8077214 | 8514473 | 8994955 | 7457352.5 | 9575304 | 7332375 | 8351532 | [illegible] |
| 15533349 | 2 | 0.394 | 2231591.25 | 3553828 | 4882218 | 3208690.5 | 4964984.5 | 4856841 | 4697237 | 4593803.5 | 5177819 | 4398691 | 4849873 | 3786530 | [illegible] |
| 15553475 | 2 | 0.007 | 2696321.75 | 3507624.25 | 3201784.75 | 4132327.75 | 4608579.5 | 4326271 | 3087094.5 | 2926239 | 3594752.75 | 3353529.5 | 3345595.25 | 3045890.25 | [illegible] |
| 15543904 | 2 | 0.415 | 1360988.5 | 2307478.25 | 2703593.75 | 2012277.375 | 3279600.5 | 2883924.5 | 2505227.5 | 2435963.5 | 2290776.25 | 1888142.75 | 1815031.125 | 2336509.25 | [illegible] |
| 15531989 | 2 | 0.164 | 9.51E+07 | 8.88E+07 | 1.05E+08 | 8.52E+07 | 7.87E+07 | 6.05E+07 | 8.62E+07 | 9.06E+07 | 8.02E+07 | 9.94E+07 | 7.81E+07 | 7.90E+07 | [illegible] |
| 15550602, 15556998 | 2 | 0.753 | 5.07E+07 | 5.21E+07 | 4.48E+07 | 4.91E+07 | 4.59E+07 | 4.64E+07 | 5.11E+07 | 4.61E+07 | 5.15E+07 | 5.59E+07 | 5.32E+07 | 4.39E+07 | [illegible] |
| 15539951 | 2 | 0.317 | 4.95E+07 | 4.55E+07 | 5.83E+07 | 4.25E+07 | 4.28E+07 | 4.26E+07 | 4.96E+07 | 4.86E+07 | 4.26E+07 | 5.45E+07 | 4.44E+07 | 4.09E+07 | [illegible] |
| 15531519 | 3 | 0.15 | 3.70E+07 | 1.77E+07 | 1.28E+07 | 2.99E+07 | 2.34E+07 | 1.30E+07 | 2.01E+07 | 2.35E+07 | 2.10E+07 | 5.28E+07 | 4.99E+07 | 2.75E+07 | [illegible] |
| 15540409 | 2 | 0.039 | 2.91E+07 | 2.83E+07 | 3.10E+07 | 2.04E+07 | 2.60E+07 | 2.37E+07 | 2.48E+07 | 2.64E+07 | 2.24E+07 | 2.44E+07 | 2.47E+07 | 2.09E+07 | [illegible] |
| 15543396 | 3 | 0.497 | 2.06E+07 | 1.63E+07 | 402892.062 | 1.60E+07 | 2129463.5 | 1856147.25 | 1.95E+07 | 2941909.75 | 579931.625 | 2.17E+07 | 2.53E+07 | 1.50E+07 | [illegible] |
| 15555020 | 2 | 0.035 | 5299933 | 4867269 | 1.22E+07 | 3417433 | 5845661 | 5550736.5 | 2983977 | 3627931.75 | 4343817 | 2743428.75 | 1149010.75 | 2815138.75 | [illegible] |
| 15557123 | 2 | 0.163 | 0 | 66767.188 | 280040.25 | 523765.969 | 788694.062 | 735762.062 | 405528.5 | 532021.5 | 734267.562 | 1708.659 | 297281.125 | 273965.094 | [illegible] |
| 15535495 | 3 | 0.794 | 5905942 | 4522219.5 | 4921565 | 5325445.5 | 5063981.5 | 3886234.5 | 7426754 | 5248515 | 3840119.5 | 7083831 | 5724901.5 | 4489926.5 | [illegible] |
| 15541918 | 3 | 0.606 | 7261843 | 1711801.5 | 1273039.5 | 5769989 | 1752462.375 | 1139442 | 3016639.5 | 1813886.25 | 720741.75 | 4181131 | 6030861 | 2130693.25 | [illegible] |
| 15547374 | 3 | 0.352 | 221795.531 | 244348.188 | 4.70E-06 | 145736.328 | 82181.453 | 33317.652 | 269110.375 | 191839.156 | 335449.781 | 376031.188 | 293674.375 | 219877 | [illegible] |
| 15538959 | 3 | 0.098 | 2076368.125 | 1334981.25 | 1129139.25 | 1257669.625 | 1556937 | 1171103.375 | 1810280.375 | 2008127.125 | 942939.312 | 2792122.75 | 2886219 | 1952221.375 | [illegible] |
| 15538677 | 3 | 0.538 | 1279627.875 | 474067.062 | 462316.031 | 1582070.5 | 781675.625 | 553398.438 | 1532721.25 | 925283.312 | 710231.438 | 1610560.25 | 1105431.625 | 845067.625 | [illegible] |
| 15555720 | 3 | 0.24 | 1389778.125 | 2837999.75 | 1660617.875 | 1644742.5 | 2424754.5 | 2365647.75 | 1492717.125 | 1585136 | 2236927 | 1775526 | 1165854.25 | 1014856.438 | [illegible] |
| 15548614 | 3 | 0.832 | 2.75E+07 | 3.16E+07 | 2.90E+07 | 2.99E+07 | 3.14E+07 | 2.67E+07 | 3.29E+07 | 2.77E+07 | 2.45E+07 | 3.04E+07 | 2.55E+07 | 2.66E+07 | [illegible] |
| 15532541 | 2 | 0.156 | 2.96E+07 | 3.21E+07 | 3.47E+07 | 3.14E+07 | 2.79E+07 | 3.08E+07 | 2.90E+07 | 3.15E+07 | 2.58E+07 | 2.67E+07 | 2.68E+07 | 2.91E+07 | [illegible] |
| 15552211 | 2 | 0.51 | 9569961 | 1.16E+07 | 1.70E+07 | 9164165 | 1.15E+07 | 1.10E+07 | 9886186 | 1.11E+07 | 1.06E+07 | 1.27E+07 | 7488284.5 | 8931937 | [illegible] |
| 15536489 | 2 | 0.676 | 6426476 | 1.14E+07 | 1.51E+07 | 5609444.5 | 1.64E+07 | 1.07E+07 | 1.09E+07 | 1.49E+07 | 8425920 | 9553114 | 6174279.5 | 7544956 | [illegible] |
| 15540164 | 3 | 0.807 | 6167508.5 | 9376250 | 9346418 | 6802958 | 8133629.5 | 7765708.5 | 7303372 | 1.02E+07 | 8018072 | 7109727 | 8307143.5 | 7527591.5 | [illegible] |
| 15539953 | 3 | 0.374 | 2.83E+07 | 2.80E+07 | 2.67E+07 | 2.55E+07 | 2.61E+07 | 2.30E+07 | 3.13E+07 | 2.66E+07 | 2.14E+07 | 2.98E+07 | 2.67E+07 | 3.10E+07 | [illegible] |
| 15536882 | 2 | 0.45 | 1.55E+07 | 1.71E+07 | 1.94E+07 | 2.02E+07 | 1.79E+07 | 1.80E+07 | 2.06E+07 | 1.92E+07 | 1.83E+07 | 2.00E+07 | 1.66E+07 | 1.71E+07 | [illegible] |
| 15548960 | 2 | 0.551 | 1.40E+07 | 1.51E+07 | 1.65E+07 | 1.41E+07 | 1.41E+07 | 1.56E+07 | 1.40E+07 | 1.46E+07 | 1.12E+07 | 1.56E+07 | 1.21E+07 | 1.65E+07 | [illegible] |
| 15533089 | 2 | 0.986 | 8174856.5 | 8427275 | 1.07E+07 | 8402729 | 8653974 | 9329964 | 9787494 | 9683674 | 7923836 | 1.05E+07 | 8317191 | 7987223.5 | [illegible] |
| 15538520 | 3 | 0.358 | 5694259 | 7792032.5 | 5104532 | 6209930 | 7719154 | 8897213 | 8707403 | 8809361 | 6482183.5 | 9641217 | 7594455.5 | 6624850.5 | [illegible] |
| [illegible] | 2 | 0.716 | 3990310.25 | 5215631 | 6638844 | 4813046 | 5255223 | 6352726.5 | 6171879.5 | 5763463.5 | 5586515.5 | 5128894 | 5148658.5 | 5181511 | [illegible] |
| [illegible] | 2 | 0.326 | 1.19E+08 | 1.25E+08 | 1.50E+08 | 1.20E+08 | 1.10E+08 | 1.13E+08 | 1.31E+08 | 1.27E+08 | 1.06E+08 | 1.22E+08 | 1.13E+08 | 1.15E+08 | [illegible] |
| [illegible] | 2 | 0.662 | 9.37E+07 | 1.08E+08 | 1.37E+08 | 8.35E+07 | 1.15E+08 | 1.12E+08 | 1.26E+08 | 1.25E+08 | 9.53E+07 | 1.17E+08 | 8.61E+07 | 9.32E+07 | [illegible] |
| [illegible] | 2 | 0.088 | 6.74E+07 | 7.07E+07 | 9.94E+07 | 6.26E+07 | 7.62E+07 | 5.70E+07 | 1.05E+08 | 8.43E+07 | 9.00E+07 | 9.56E+07 | 8.29E+07 | 7.80E+07 | [illegible] |
| [illegible] | 2 | 0.76 | 6.17E+07 | 7.40E+07 | 8.79E+07 | 5.55E+07 | 8.82E+07 | 6.61E+07 | 7.75E+07 | 8.89E+07 | 5.71E+07 | 8.16E+07 | 5.00E+07 | 5.82E+07 | [illegible] |
| [illegible] | 2 | 0.114 | 7.04E+07 | 7.80E+07 | 8.08E+07 | 6.17E+07 | 6.94E+07 | 6.59E+07 | 6.28E+07 | 6.90E+07 | 5.49E+07 | 6.88E+07 | 5.73E+07 | 7.09E+07 | [illegible] |
| [illegible] | 3 | 0.732 | 2.36E+07 | 2.42E+07 | 3.05E+07 | 2.44E+07 | 2.44E+07 | 2.38E+07 | 2.59E+07 | 2.77E+07 | 2.96E+07 | 3.19E+07 | 1.93E+07 | 2.63E+07 | [illegible] |
| [illegible] | 2 | 0.148 | 2.93E+07 | 3.14E+07 | 4.34E+07 | 2.74E+07 | 3.13E+07 | 3.05E+07 | 3.12E+07 | 3.38E+07 | 3.49E+07 | 3.01E+07 | 2.35E+07 | 2.64E+07 | [illegible] |
| [illegible] | 3 | 0.325 | 2.35E+07 | 2.68E+07 | 3.25E+07 | 2.84E+07 | 2.44E+07 | 2.55E+07 | 2.55E+07 | 2.68E+07 | 2.81E+07 | 2.64E+07 | 1.98E+07 | 2.30E+07 | [illegible] |
| [illegible] | 3 | 0.955 | 1.55E+07 | 1.68E+07 | 9979207 | 1.93E+07 | 1.44E+07 | 1.02E+07 | 1.66E+07 | 1.40E+07 | 1.03E+07 | 2.17E+07 | 1.37E+07 | 1.17E+07 | [illegible] |
| [illegible] | 2 | 0.433 | 1.78E+07 | 1.79E+07 | 1.56E+07 | 1.48E+07 | 1.69E+07 | 1.48E+07 | 1.85E+07 | 1.62E+07 | 1.31E+07 | 1.73E+07 | 1.69E+07 | 1.86E+07 | [illegible] |
| [illegible] | 2 | 0.133 | 1.03E+07 | 1.08E+07 | 1.40E+07 | 7455104.5 | 7983654.5 | 9640403 | 8910940 | 9758443 | 7334897 | 1.03E+07 | 4598488 | 6699977 | [illegible] |
| [illegible] | 3 | 0.244 | 3501659.5 | 3424221.5 | 2896887.25 | 2629173.25 | 2715691 | 2201089.75 | 2625391.75 | 2471022 | 626838.438 | 3831715.25 | 2234779.75 | 3685073.75 | [illegible] |
| [illegible] | 2 | 0.148 | 2790160.25 | 2818122.25 | 3637031.5 | 1771571.375 | 2276051 | 2250279.75 | 2416220 | 2037683.375 | 1727733.875 | 2754412.5 | 688376.562 | 1347408.5 | [illegible] |
| [illegible] | 2 | 0.467 | 2945052.75 | 3282663.75 | 5499730.5 | 4946973 | 5670561.5 | 5473754 | 4569033 | 4933807.5 | 7168461 | 2727863 | 6243721 | 5673080.5 | [illegible] |
| [illegible] | 2 | 0.099 | 1148824.5 | 1510668.125 | 2028876.75 | 2254453.5 | 2530349.25 | 3054475.75 | 2671022.5 | 2023497.625 | 2818326.5 | 1376999.25 | 2398189 | 2438682 | [illegible] |
| [illegible] | 2 | 0.357 | 1.47E+07 | 1.63E+07 | 2.23E+07 | 1.99E+07 | 2.26E+07 | 2.35E+07 | 1.70E+07 | 2.46E+07 | 1.97E+07 | 1.88E+07 | 1.75E+07 | 1.98E+07 | [illegible] |
| [illegible] | 2 | 0.713 | 1.60E+07 | 1.70E+07 | 2.56E+07 | 1.97E+07 | 2.03E+07 | 1.96E+07 | 1.51E+07 | 1.97E+07 | 2.17E+07 | 1.80E+07 | 1.67E+07 | 1.66E+07 | [illegible] |
| [illegible] | 2 | 0.299 | 1.24E+07 | 1.83E+07 | 2.39E+07 | 1.51E+07 | 2.72E+07 | 1.99E+07 | 1.70E+07 | 2.18E+07 | 1.72E+07 | 1.60E+07 | 1.02E+07 | 1.43E+07 | [illegible] |
| [illegible] | 2 | 0.485 | 1.12E+07 | 1.40E+07 | 1.67E+07 | 1.57E+07 | 1.53E+07 | 1.38E+07 | 1.48E+07 | 1.74E+07 | 1.60E+07 | 1.49E+07 | 1.25E+07 | 1.49E+07 | [illegible] |
| [illegible] | 2 | 0.937 | 1.16E+07 | 1.17E+07 | 1.70E+07 | 1.48E+07 | 1.39E+07 | 1.08E+07 | 1.25E+07 | 1.33E+07 | 1.59E+07 | 1.31E+07 | 1.55E+07 | 1.34E+07 | [illegible] |
| [illegible] | 2 | 0.855 | 7816309 | 1.16E+07 | 1.27E+07 | 1.00E+07 | 1.25E+07 | 1.25E+07 | 1.11E+07 | 1.20E+07 | 1.12E+07 | 1.24E+07 | 1.01E+07 | 1.04E+07 | [illegible] |
| [illegible] | [illegible] | [illegible] | [illegible] | [illegible] | [illegible] | [illegible] | [illegible] | [illegible] | [illegible] | [illegible] | [illegible] | [illegible] | [illegible] | [illegible] | [illegible] |

| | | | | | | | | | | | | | | | | | | | | | | |
|---|---|---|---|---|---|---|---|---|---|---|---|---|---|---|---|---|---|---|---|---|---|---|
| 15539523 | 3 | 0.109 | 2025269.75 | 5091088.5 | 1628025.5 | 3293506.5 | 2749631.5 | 3148593.5 | 1706580 | 1909670.125 | 2874428.75 | 1974580.875 | 1110287.375 | 998563.188 | 1685.8574 | HSGNITFDEIVNIAR | P35979 | P35979 | RPL12 | ribosomal protein L12 | Nucleus | other |
| 15542582 | 2 | 0.181 | 6221809.5 | 9984563 | 3506288.5 | 6947410 | 5923952 | 6049121.5 | 3439577.25 | 5570535.5 | 5738656.5 | 4329649.5 | 3149442.75 | 3289813 | 1232.62 | STESLQANVQR | P47963 | P47963 | RPL13 | ribosomal protein L13 | Nucleus | other |
| 15533026 | 2 | 0.137 | 3717302.5 | 6992117 | 2333227 | 2973456.5 | 3110925.75 | 3682623 | 1478918.875 | 3662503.75 | 3327614 | 1967085.625 | 1750673.5 | 1585483.25 | 957.5343 | TIGISVDPR | P47963 | P47963 | RPL13 | ribosomal protein L13 | Nucleus | other |
| 15534293 | 2 | 0.073 | 3040620 | 4173572 | 2071699.125 | 2771666.25 | 2701321.75 | 4201547.5 | 1457157.875 | 3030073.75 | 2921649.5 | 1570458.375 | 1515531.75 | 1605324.75 | 950.4921 | GFSLEELR | P47963 | P47963 | RPL13 | ribosomal protein L13 | Nucleus | other |
| 15540959 | 3 | 0.09 | 3352377 | 4931694 | 1885783.25 | 3085537.25 | 2455222.5 | 3625989.5 | 1610958.125 | 2160802.5 | 3137255.25 | 1447676.375 | 1789956.25 | 1532232.75 | 1361.6931 | VDTWFNQPARK | P47963 | P47963 | RPL13 | ribosomal protein L13 | Nucleus | other |
| 15537287 | 2 | 0.072 | 3873033.75 | 6464463.5 | 2961925.5 | 3244016.25 | 4477838 | 4787745 | 1718581.75 | 3945441.75 | 4029295.75 | 2052785.25 | 1691266.875 | 2111869 | 1252.64 | YQAVTATLEEK | P19253 | P19253 | RPL13A | ribosomal protein L13a | Cytoplasm | other |
| 15533101 | 2 | 0.078 | 8927865 | 1.36E+07 | 6235362.5 | 8977736 | 1.04E+07 | 9884404 | 6368861 | 8760057 | 1.16E+07 | 6096668.5 | 4941587 | 5258998 | 1354.7656 | LVAIVDVIDQNR | Q9CR57 | Q9CR57 | RPL14 | ribosomal protein L14 | Cytoplasm | other |
| 15539542 | 3 | 0.346 | 4006762 | 5920855 | 2484895.75 | 2341542.75 | 3799193.25 | 4051405 | 1727585.875 | 3128429.75 | 4189995 | 2725094.5 | 1938036.5 | 2145949.75 | 1246.656 | VAYISFGPHAGK | Q9CR57 | Q9CR57 | RPL14 | ribosomal protein L14 | Cytoplasm | other |
| 15534018 | 2 | 0.043 | 4489397.5 | 8232860.5 | 3241153 | 4730713 | 5707232.5 | 5496333.5 | 3121194.5 | 4345216 | 5734165.5 | 2384943.25 | 2382288 | 2637431.5 | 1041.6023 | AAIAAAAAAAAAK | Q9CR57 | Q9CR57 | RPL14 | ribosomal protein L14 | Cytoplasm | other |
| 15549261 | 2 | 0.111 | 5218508.5 | 6949193.5 | 3712099.5 | 4851127.5 | 4758815.5 | 5627168 | 3071175.75 | 4590585 | 6033169 | 3768939.75 | 2717916.75 | 2750624.25 | 1660.782 | VLNSYWVGEDSTYK | Q9CZM2 | Q9CZM2 | RPL15 | ribosomal protein L15 | Cytoplasm | other |
| 15539006 | 3 | 0.285 | 4368646.5 | 6230055 | 1994597.625 | 3638656.25 | 3821806 | 3270768.25 | 2084153.75 | 2973336 | 3437350.25 | 2687317.25 | 1136243.5 | 2677019.75 | 1504.817 | FFEVILIDPFHK | Q9CZM2 | Q9CZM2 | RPL15 | ribosomal protein L15 | Cytoplasm | other |
| 15546094 | 2 | 0.015 | 3690896 | 5625568 | 2346961.25 | 4343207.5 | 3740672.25 | 3170793.25 | 2189806.25 | 2769780.25 | 3133419.75 | 1669693.375 | 1708252.125 | 1696902.75 | 1258.6506 | RNPDTQWITK | Q9CZM2 | Q9CZM2 | RPL15 | ribosomal protein L15 | Cytoplasm | other |
| 15535203 | 2 | 0.048 | 3674819.25 | 7086011 | 2916254.5 | 3481732 | 5622112.5 | 4544749.5 | 2663541.5 | 4111372 | 4163439 | 2490455.5 | 1392214.75 | 2106943 | 1163.5553 | YSLDPENPTK | Q9CPR4 | Q9CPR4 | RPL17 | ribosomal protein L17 | Cytoplasm | other |
| 15537407 | 2 | 0.096 | 9533267 | 1.75E+07 | 5340037 | 1.19E+07 | 1.25E+07 | 1.43E+07 | 7108404 | 9955554 | 1.03E+07 | 6394480.5 | 4599100 | 6530023 | 1474.8123 | ILTFDQLALESPK | P35980 | P35980 | RPL18 | ribosomal protein L18 | Cytoplasm | other |
| 15538751 | 2 | 0.071 | 6479219 | 1.02E+07 | 4433031 | 7173961.5 | 7682869.5 | 7809064 | 3632836.5 | 6393218 | 6842537 | 4206936.5 | 2765605 | 4145647.75 | 1331.7136 | TAVVVGTVTDDVR | P35980 | P35980 | RPL18 | ribosomal protein L18 | Cytoplasm | other |
| 15535932 | 2 | 0.032 | 5400665 | 8377379 | 3756431 | 5744422.5 | 5217562 | 7023109.5 | 3017218.5 | 4536289 | 5114068.5 | 2974449 | 2829401.75 | 2896650.75 | 1250.6708 | TNSTFNQVVLK | P35980 | P35980 | RPL18 | ribosomal protein L18 | Cytoplasm | other |
| 15537118 | 3 | 0.065 | 3275847.5 | 4126914.25 | 1787579.125 | 3764454.75 | 3370162 | 3252237.5 | 1799957.625 | 2843972.25 | 3791942.5 | 1955193.25 | 1451747.125 | 1446836.75 | 1140.6461 | TNRPPLSLSR | P35980 | P35980 | RPL18 | ribosomal protein L18 | Cytoplasm | other |
| 15542496 | 2 | 0.278 | 3737952.5 | 5319246.5 | 2466395.75 | 3271303.75 | 3981119.5 | 4304248 | 1708978.625 | 4096419.75 | 4901271 | 2266390 | 2231967.5 | 2130804.25 | 1455.6875 | DLTTAGAVTQC[160.0307]YR | P62717 | P62717 | RPL18A | ribosomal protein L18a | Cytoplasm | other |
| 15551660 | 2 | 0.216 | 9680492 | 1.24E+07 | 6423132 | 9529166 | 8777190 | 1.11E+07 | 3246301 | 7979222.5 | 1.02E+07 | 6838673 | 4027540.75 | 5347683.5 | 1942.922 | VWLDPNETNEIANANSR | P84099 | P84099 | RPL19 | ribosomal protein L19 | Cytoplasm | other |
| 15533334 | 3 | 0.106 | 8731739 | 1.36E+07 | 6735871.5 | 9674164 | 8385851.5 | 8027935 | 4136038.5 | 8181020.5 | 9031615 | 4953449 | 4660567 | 5544090.5 | 1654.9259 | VYNVTQHAVGIIVNK | O09167 | O09167 | RPL21 | ribosomal protein L21 | Cytoplasm | other |
| 15539193 | 2 | 0.66 | 3459458.5 | 5820850.5 | 1502869.375 | 2942602 | 2739106.25 | 3902084.75 | 1762466 | 2786803 | 2989458.5 | 2157780.25 | 2253139.5 | 2468770.75 | 1243.6589 | HGVVPLATYMR | O09167 | O09167 | RPL21 | ribosomal protein L21 | Cytoplasm | other |
| 15549595 | 3 | 0.216 | 1.35E+07 | 2.42E+07 | 8452816 | 1.65E+07 | 1.63E+07 | 1.36E+07 | 7592626 | 1.18E+07 | 1.41E+07 | 1.07E+07 | 7779089.5 | 8201878.5 | 3083.3782 | FTLDC[160.0307]THPVEDGIMDAANFEQFLQER | P67984 | P67984 | RPL22 | ribosomal protein L22 | Nucleus | other |
| 15538907 | 2 | 0.103 | 6021982.5 | 1.02E+07 | 3800502.5 | 6436460 | 6348055 | 6204182 | 3671351.75 | 5492105 | 6341486.5 | 4002699 | 2519956 | 3519214.25 | 1242.6776 | AGNLGGGVVTIER | P67984 | P67984 | RPL22 | ribosomal protein L22 | Nucleus | other |
| 15549154 | 2 | 0.184 | 6241746.5 | 1.11E+07 | 4451481 | 7868211.5 | 7522885.5 | 8087331 | 2901537.5 | 5315773.5 | 8828771 | 5221544.5 | 2650713.5 | 3604225 | 1970.0331 | ISLGLPVGAVINC[160.0307]ADNTGAK | P62830 | P62830 | RPL23 | ribosomal protein L23 | Cytoplasm | other |
| 15534238 | 2 | 0.032 | 1.03E+07 | 1.68E+07 | 7591355 | 1.24E+07 | 1.05E+07 | 1.19E+07 | 6121310 | 9952392 | 1.09E+07 | 6631658.5 | 4445106.5 | 5132480.5 | 1404.6973 | LAPDYDALDVANK | P62751 | P62751 | Rpl23a | ribosomal protein L23A | Nucleus | other |
| 15535987 | 2 | 0.066 | 2347724.5 | 3287810 | 1367666.25 | 2121482.5 | 2031962.25 | 2273332 | 1382155.75 | 1709502 | 2113661.75 | 1111257.75 | 1389985.625 | 1121016 | 1241.6819 | VNTLIRPDGEK | P62751 | P62751 | Rpl23a | ribosomal protein L23A | Nucleus | other |
| 15534408 | 2 | 0.087 | 3872448 | 4914949.5 | 2160954.25 | 2887315.75 | 3981305.5 | 4265508 | 1893788.5 | 3183647.25 | 4044777 | 1891279.125 | 1870091.875 | 1748757 | 966.5383 | VFQFLNAK | Q8BP67 | Q8BP67 | RPL24 | ribosomal protein L24 | Cytoplasm | other |
| 15537103 | 2 | 0.436 | 2135526.75 | 3578882.25 | 1630992.375 | 2144789 | 3100201 | 3573431 | 1341480.875 | 2500060 | 2678533.5 | 1861678.75 | 2111950.5 | 1718263.25 | 1189.5532 | VELC[160.0307]SFSGYK | Q8BP67 | Q8BP67 | RPL24 | ribosomal protein L24 | Cytoplasm | other |
| 15536128 | 2 | 0.139 | 3110119.5 | 5066176 | 2018902.875 | 2743956.5 | 3091288 | 4036291 | 2259352.75 | 2860706.25 | 3706146.25 | 1726355.75 | 1662316.125 | 2216306.5 | 1192.6448 | QINWTVLYR | Q8BP67 | Q8BP67 | RPL24 | ribosomal protein L24 | Cytoplasm | other |
| 15541433 | 2 | 0.035 | 2223662.25 | 3439887.5 | 1324780.75 | 2439459.75 | 2217154.5 | 2101608 | 1374454.125 | 2101280 | 1859184.5 | 1324738.875 | 889983.812 | 926044.625 | 959.47845 | DDEVQVVR | P61255 | P61255 | RPL26 | ribosomal protein L26 | Cytoplasm | other |
| 15534050 | 2 | 0.157 | 3733834 | 5885007.5 | 2335124 | 3992592 | 4223087.5 | 4182064.75 | 1727243.375 | 3791821.25 | 4478588 | 2497628.5 | 1770078.875 | 1938072.625 | 1232.6598 | LWTLVSEQTR | P14115 | P14115 | RPL27A | ribosomal protein L27a | Cytoplasm | other |
| 15536126 | 2 | 0.051 | 4670003.5 | 6522924 | 2876899.75 | 5595847 | 4714549 | 4607308.5 | 3018294.5 | 4029916.75 | 4001030 | 3325123 | 2184636.25 | 2158204.75 | 1294.6245 | QTYSTEPNNLK | P41105 | P41105 | RPL28 | ribosomal protein L28 | Cytoplasm | other |
| 15546910 | 3 | 0.796 | 949068.188 | 3192447 | 356388.969 | 122581.828 | 1565912.625 | 1127949.375 | 843466.312 | 972340.25 | 755440.125 | 373370.156 | 418369.438 | 916547.25 | 1052.6339 | RLAFIAHPK | P47915 | P47915 | Rpl29 (includes others) | ribosomal protein L29 | Cytoplasm | other |
| 15549503 | 3 | 0.216 | 1.02E+07 | 2.16E+07 | 6985432 | 1.52E+07 | 1.19E+07 | 1.28E+07 | 6624493.5 | 1.17E+07 | 1.13E+07 | 8558810 | 5251285.5 | 7372668.5 | 2971.5125 | ERLEQQVPVNQVFGQDEMIDVIGVTK | P27659 | P27659 | RPL3 | ribosomal protein L3 | Nucleus | other |
| 15550128 | 3 | 0.887 | 4305435.5 | 6900490 | 2760370.25 | 3781690.5 | 5866240 | 7448791 | 3808569.75 | 3982546.75 | 9221089 | 3090547.5 | 5073295 | 5953364 | 2451.2341 | SINPLGGFVHYGEVTNDFIMLK | P27659 | P27659 | RPL3 | ribosomal protein L3 | Nucleus | other |
| 15536457 | 2 | 0.095 | 3404360.5 | 6268873.5 | 2356815.25 | 3879710 | 4014304.5 | 3697020.5 | 2042860.875 | 3147643.75 | 4177880 | 2056620 | 2012628.125 | 1838033 | 891.5286 | IGQGYLIK | P27659 | P27659 | RPL3 | ribosomal protein L3 | Nucleus | other |
| 15548711 | 2 | 0.514 | 8008963 | 1.04E+07 | 2818675.25 | 6810384.5 | 8346940 | 7056670 | 3551497 | 6501069 | 7815951.5 | 4895117 | 4428548.5 | 3908440 | 1757.9094 | VC[160.0307]TLAIIDPGDSDIIR | P62889 | P62889 | RPL30 | ribosomal protein L30 | Cytoplasm | other |
| 15534378 | 2 | 0.189 | 7103597.5 | 9890521 | 4080538.25 | 6968451 | 6262333 | 7862365.5 | 3210524.25 | 5572311.5 | 6661562.5 | 5337217.5 | 3526393.25 | 3187238.5 | 988.54193 | SAINEVVTR | P62900 | P62900 | RPL31 | ribosomal protein L31 | Cytoplasm | other |
| 15548757 | 2 | 0.181 | 6934549.5 | 1.34E+07 | 5100045.5 | 9530439 | 8021527 | 8280687.5 | 3989362 | 8017682 | 1.06E+07 | 4402473 | 4031438.5 | 4736260 | 1644.922 | LYTLVTYVPVTTFK | P62900 | P62900 | RPL31 | ribosomal protein L31 | Cytoplasm | other |
| 15532646 | 2 | 0.146 | 6360329 | 9993716 | 4721275.5 | 7717195 | 6338226 | 8515773 | 4004976.5 | 6381191 | 6463825 | 4874908.5 | 4736908.5 | 3953429 | 1143.671 | VLTVINQTQK | Q6ZWV7 | Q6ZWV7 | RPL35 | ribosomal protein L35 | Cytoplasm | other |
| 15537355 | 2 | 0.067 | 2322026.5 | 3855317.75 | 2223463.75 | 2699497.5 | 3335916.5 | 3748821.5 | 1531139.375 | 2960483.75 | 3453158.75 | 1747486.875 | 1272048 | 1649956.125 | 844.4876 | VELSQLR | Q6ZWV7 | Q6ZWV7 | RPL35 | ribosomal protein L35 | Cytoplasm | other |
| 15534096 | 2 | 0.093 | 2861546 | 4299916.5 | 1977303.375 | 2519982.5 | 3049821 | 3291994.75 | 1809137.625 | 2909503.75 | 3295739.5 | 1805941.375 | 1760513 | 1422836 | 1227.5436 | EVC[160.0307]GFAPYER | P47964 | P47964 | Rpl36 | ribosomal protein L36 | Nucleus | other |
| 15544592 | 2 | 0.016 | 2450845.75 | 3604513.75 | 1588875.125 | 2019662.875 | 2201015 | 2148927.75 | 1564069.375 | 2034331.5 | 1593752.375 | 1233600.25 | 1158458.5 | 1067566.75 | 992.5204 | YPMAVGLNK | P47964 | P47964 | Rpl36 | ribosomal protein L36 | Nucleus | other |
| 15534856 | 2 | 0.166 | 7491750 | 1.08E+07 | 4024117 | 5622847.5 | 5176724.5 | 6946121 | 3715516.5 | 4428635 | 7323097.5 | 4108749.5 | 3009495.75 | 3759365 | 1268.7549 | NIPGITLLNVSK | Q9D8E6 | Q9D8E6 | RPL4 | ribosomal protein L4 | Cytoplasm | enzyme |
| 15545248 | 2 | 0.305 | 4707111 | 6823868 | 2318038.5 | 5304775 | 4502956.5 | 4896941.5 | 2251863 | 4462544 | 4860372 | 2889567.5 | 2553644.5 | 2885582.5 | 1187.6964 | KLEAAATALATK | Q9D8E6 | Q9D8E6 | RPL4 | ribosomal protein L4 | Cytoplasm | enzyme |
| 15536089 | 4 | 0.532 | 4688573.5 | 5576183.5 | 1607462.25 | 4996640 | 2575687.5 | 5509965.5 | 2170865.75 | 3038963 | 4394470 | 2146040 | 2626146.25 | 2542011 | 2716.2756 | NNRQPYAVSELAGHQTSAESWGTGR | Q9D8E6 | Q9D8E6 | RPL4 | ribosomal protein L4 | Cytoplasm | enzyme |
| 15540994 | 2 | 0.048 | 1864690.375 | 3785232.75 | 2965109.75 | 1351749 | 3880086 | 2659081 | 1338192 | 3191079 | 2481136.75 | 1249145.375 | 375161.812 | 941872.562 | 1035.5692 | PLISVYSEK | Q9D8E6 | Q9D8E6 | RPL4 | ribosomal protein L4 | Cytoplasm | enzyme |
| 15542718 | 2 | 0.487 | 1501351.375 | 2388859 | 980008.75 | 1945173.125 | 1615141.125 | 1882075 | 858456.438 | 1176604.5 | 3981923.25 | 504075.969 | 1197973.125 | 1329017.25 | 1338.7009 | GAVDGGLSIPHSTK | P47962 | P47962 | RPL5 | ribosomal protein L5 | Cytoplasm | other |
| 15556350 | 2 | 0.034 | 4748812.5 | 6475470.5 | 4700869.5 | 4609309.5 | 5790277.5 | 6601413 | 2777193.5 | 5579656.5 | 6151745.5 | 3027717.25 | 2378726.25 | 2786809.75 | 1527.9084 | SSITPGTVLIILTGR | P47911 | P47911 | RPL6 | ribosomal protein L6 | Nucleus | other |
| 15540373 | 2 | 0.364 | 4890520.5 | 6711298.5 | 2513963.25 | 4133231 | 4820317.5 | 3881199 | 487017.75 | 4011489.5 | 3694450.5 | 2714354.5 | 2409482.75 | 1849443.625 | 1332.6034 | HQEGEIFDTEK | P47911 | P47911 | RPL6 | ribosomal protein L6 | Nucleus | other |
| 15550629 | 2 | 0.119 | 2577301.5 | 7234962 | 2275208.75 | 2185864.75 | 4299124.5 | 4033884.25 | 1102774.375 | 3656227.25 | 2906784.5 | 1688661.25 | 573615.5 | 1398971.5 | 1808.0576 | QLDSGLLLVTGPLVINR | P47911 | P47911 | RPL6 | ribosomal protein L6 | Nucleus | other |
| 15547001 | 3 | 0.638 | 1360284.25 | 2391245.25 | 320523.438 | 1247558.625 | 1027211.625 | 522894.625 | 2716304.25 | 316673.719 | 19613.844 | 854123.188 | 417701.281 | 633223 | 1585.7573 | PRHQEGEIFDTEK | P47911 | P47911 | RPL6 | ribosomal protein L6 | Nucleus | other |
| 15533414 | 3, 4 | 0.089 | 1.41E+07 | 2.38E+07 | 8231777 | 2.02E+07 | 1.37E+07 | 1.50E+07 | 7726153.5 | 1.28E+07 | 1.24E+07 | 9681199 | 6286726.5 | 6474262.5 | 2115.9736, 2115.975 | TTHFVEGGDAGNREDQINR | P14148 | P14148 | RPL7 | ribosomal protein L7 | Nucleus | transcription regulator |
| 15536203 | 3 | 0.393 | 6772529.5 | 1.58E+07 | 4980777.5 | 6331036.5 | 9845236 | 1.08E+07 | 6376312.5 | 9366439 | 8156105.5 | 6029296 | 4129380.75 | 6078136.5 | 2138.062 | FGIIC[160.0307]MEDLIHEIYTVGK | P14148 | P14148 | RPL7 | ribosomal protein L7 | Nucleus | transcription regulator |
| 15536591 | 2 | 0.148 | 5947958.5 | 1.00E+07 | 4523860 | 7162950.5 | 6988442 | 6592689 | 3249277 | 7713478 | 8012081 | 3833601 | 3205133.25 | 3852075.5 | 1192.5974 | AGNFYVPAEPK | P14148 | P14148 | RPL7 | ribosomal protein L7 | Nucleus | transcription regulator |
| 15549823 | 2 | 0.163 | 5429239 | 1.09E+07 | 4188468 | 6174097 | 5320011 | 6524188 | 2746319.5 | 5144825 | 7799504 | 3580989 | 2655584.75 | 3167186 | 1662.8862 | IVEPYIAWGYPNLK | P14148 | P14148 | RPL7 | ribosomal protein L7 | Nucleus | transcription regulator |
| 15535923 | 3 | 0.05 | 1025489.562 | 2311562.75 | 1345176.5 | 944332.062 | 1185415.75 | 2188046 | 757285.688 | 938091 | 1169875 | 499337.781 | 480902.562 | 853508.188 | 1342.7772 | RIALTDNSLIAR | P14148 | P14148 | RPL7 | ribosomal protein L7 | Nucleus | transcription regulator |
| 15552426 | 2 | 0.02 | 8005399.5 | 1.32E+07 | 6132862.5 | 9265541 | 8159989.5 | 8501632 | 4548924 | 7259858 | 8032003 | 4698297.5 | 3671238.25 | 3736832.75 | 1569.8354 | VPPAINQFTQALDR | P12970 | P12970 | RPL7A | ribosomal protein L7a | Cytoplasm | other |
| 15537083 | 2 | 0.103 | 5865221.5 | 9783564 | 4342094.5 | 6684104 | 7520126.5 | 7062149 | 2897563.5 | 6154505 | 6274423.5 | 3662280 | 3325754.75 | 3880356.25 | 1216.6295 | NFGIGQDIQPK | P12970 | P12970 | RPL7A | ribosomal protein L7a | Cytoplasm | other |
| 15550826 | 2 | 0.196 | 4301745 | 4555721 | 2501452.25 | 3449209 | 3980494.5 | 3781865.5 | 1311591.875 | 3550166.5 | 4476266 | 1776224.125 | 2301173.75 | 1813598.125 | 1700.777 | TC[160.0307]TTVAFTQVNSEDK | P12970 | P12970 | RPL7A | ribosomal protein L7a | Cytoplasm | other |
| 15544741 | 2 | 0.105 | 2019902 | 3584090.5 | 1509781.25 | 2044224.25 | 1712569.625 | 2454808.75 | 1197353 | 1750346.5 | 2219811.25 | 1258992.875 | 883759.812 | 1333925.625 | 699.4499 | VGLIAAR | P62918 | P62918 | RPL8 | ribosomal protein L8 | Cytoplasm | other |
| 15554905 | 2 | 0.006 | 1306108.5 | 2321386 | 770802.375 | 1423086.75 | 1137511.75 | 1411297 | 694143.5 | 924698.5 | 784321.438 | 484713.062 | 425225.312 | 440957.188 | 1688.8208 | ASGNYATVISHNPETK | P62918 | P62918 | RPL8 | ribosomal protein L8 | Cytoplasm | other |
| 15549149 | 2 | 0.058 | 8357554 | 1.26E+07 | 5648339.5 | 8497193 | 8631625 | 7836338.5 | 3925789 | 6057149.5 | 7301853.5 | 5504545 | 3239212.25 | 4472036.5 | 2127.1497 | TILSNQTVDIPENVEITLK | P51410 | P51410 | RPL9 | ribosomal protein L9 | Nucleus | other |
| 15537750 | 2 | 0.211 | 4787199 | 7256880.5 | 2570561.25 | 3568899 | 4496571 | 4824986.5 | 2280661.25 | 4354955.5 | 4346387.5 | 2659814.75 | 2323074.5 | 2542638.5 | 1235.602 | TGVAC[160.0307]SVSQAQK | P51410 | P51410 | RPL9 | ribosomal protein L9 | Nucleus | other |
| 15534212 | 2 | 0.018 | 5075946 | 8366034 | 4072383.5 | 4755202 | 5778725 | 6672507 | 3020165.25 | 6009036.5 | 5843700.5 | 2201406.25 | 1889303.75 | 2881766.75 | 1170.6014 | FLDGIYVSEK | P51410 | P51410 | RPL9 | ribosomal protein L9 | Nucleus | other |
| 15548890 | 3 | 0.035 | 1.63E+07 | 2.67E+07 | 1.26E+07 | 1.50E+07 | 2.07E+07 | 1.78E+07 | 9931114 | 1.43E+07 | 1.63E+07 | 9781133 | 7602796 | 9798406 | 2696.3982 | AFLADPSAFAAAAPAAAATTAAPAAAAAPAK | P14869 | P14869 | RPLP0 | ribosomal protein, large, P0 | Cytoplasm | other |
| 15543336 | 3 | 0.495 | 5585644 | 1.12E+07 | 4104203.25 | 9380375 | 7862669.5 | 3251975.25 | 3343930 | 5963495.5 | 5276161 | 4673878 | 2974980 | 4343792 | 2180.0967 | AGAIAPC[160.0307]EVTVPAQNTGLGPEK | P14869 | P14869 | RPLP0 | ribosomal protein, large, P0 | Cytoplasm | other |
| 15551490 | 2 | 0.046 | 8110534 | 1.17E+07 | 4094405 | 7294963.5 | 8962901 | 7887820.5 | 4080770.25 | 5731630 | 6158441 | 3905579.75 | 3763273 | 3527607.25 | 2180.0972 | AGAIAPC[160.0307]EVTVPAQNTGLGPEK | P14869 | P14869 | RPLP0 | ribosomal protein, large, P0 | Cytoplasm | other |
| 15532732 | 3 | 0.126 | 6129808.5 | 9493647 | 3785341.25 | 8416910 | 5396672.5 | 5390527.5 | 3463947.25 | 4596621 | 5978710.5 | 3257569.25 | 2933479 | 4055842.5 | 1660.8535 | EDLTEIRDMLLANK | P14869 | P14869 | RPLP0 | ribosomal protein, large, P0 | Cytoplasm | other |
| 15540177 | 2 | 0.028 | 5023489.5 | 8028869.5 | 3727501.25 | 8065198 | 6896583.5 | 6922155 | 2281030.5 | 5287670.5 | 5036132 | 3012962.75 | 2788714 | 2677703.75 | 1217.6748 | IIQLLDDYPK | P14869 | P14869 | RPLP0 | ribosomal protein, large, P0 | Cytoplasm | other |
| 15541120 | 2 | 0.104 | 4523596.5 | 6123536 | 3038205.75 | 4184328.5 | 4267102 | 3776845.75 | 2323533.75 | 4171584.75 | 4600588.5 | 2106581.75 | 3064545 | 2073702.5 | 1266.6108 | C[160.0307]FIVGADNVGSK | P14869 | P14869 | RPLP0 | ribosomal protein, large, P0 | Cytoplasm | other |
| 15534951 | 2 | 0.032 | 5734529 | 9078025 | 4870284.5 | 4128464.5 | 7199266.5 | 6965533 | 3453152 | 5424350.5 | 6111295.5 | 1833401.625 | 3140671.25 | 3165499.25 | 1221.6189 | GHLENNPALEK | P14869 | P14869 | RPLP0 | ribosomal protein, large, P0 | Cytoplasm | other |
| 15553711 | 2 | 0.107 | 2553986 | 3581394.25 | 1125640.25 | 2704504 | 1663201.375 | 2209777.25 | 1028261.688 | 1061584.625 | 1890048.75 | 968450.375 | 874031.375 | 1357032.875 | 1660.8528 | EDLTEIRDMLLANK | P14869 | P14869 | RPLP0 | ribosomal protein, large, P0 | Cytoplasm | other |
| 15552435 | 3, 4 | 0.098 | 3.40E+07 | 5.85E+07 | 2.53E+07 | 3.71E+07 | 3.73E+07 | 3.28E+07 | 1.76E+07 | 3.01E+07 | 3.20E+07 | 2.32E+07 | 1.37E+07 | 2.25E+07 | 3782.9075, 3782.909 | ALANVNIGSLIC[160.0307]NVGAGGPAPAAGAAPAGGAAPSTAAAPAEEK | P47955 | P47955 | Rplp1 (includes others) | ribosomal protein, large, P1 | Nucleus | other |
| 15551668 | 2 | 0.058 | 2.73E+07 | 5.26E+07 | 2.80E+07 | 3.05E+07 | 3.34E+07 | 3.57E+07 | 1.94E+07 | 2.40E+07 | 3.86E+07 | 1.97E+07 | 1.41E+07 | 1.92E+07 | 1675.8806 | AAGVSVEPFWPGLFAK | P47955 | P47955 | Rplp1 (includes others) | ribosomal protein, large, P1 | Nucleus | other |
| 15548517 | 3 | 0.014 | 3.36E+07 | 5.27E+07 | 2.51E+07 | 3.79E+07 | 4.72E+07 | 3.83E+07 | 1.67E+07 | 2.65E+07 | 2.53E+07 | 2.29E+07 | 1.39E+07 | 1.62E+07 | 2774.4207 | LASVPAGGAVAVSAAPGSAAPAAGSAPAAAEEK | P99027 | P99027 | RPLP2 | ribosomal protein, large, P2 | Cytoplasm | other |
| 15533130 | 3 | 0.022 | 3.60E+07 | 5.60E+07 | 2.70E+07 | 3.64E+07 | 4.67E+07 | 4.15E+07 | 2.23E+07 | 3.34E+07 | 3.13E+07 | 2.28E+07 | 1.63E+07 | 2.14E+07 | 1772.8988 | ILDSVGIEADDDRLNK | P99027 | P99027 | RPLP2 | ribosomal protein, large, P2 | Cytoplasm | other |
| 15548533 | 2 | 0.086 | 8182983.5 | 1.42E+07 | 6358833.5 | 9638546 | 9100444 | 8643459 | 3979015.25 | 6613139 | 9508925 | 6150129 | 3725242.75 | 3889275.75 | 1868.9712 | YVASYLLAALGGNSSPSAK | P99027 | P99027 | RPLP2 | ribosomal protein, large, P2 | Cytoplasm | other |
| 15549717 | 2 | 0.01 | 5329698 | 6925164 | 3902857.75 | 5007948.5 | 5306537 | 4590309 | 2715412.5 | 4192549.25 | 4455312.5 | 3001790.25 | 2164612.75 | 2180422 | 1772.899 | ILDSVGIEADDDRLNK | P99027 | P99027 | RPLP2 | ribosomal protein, large, P2 | Cytoplasm | other |
| 15538590 | 3 | 0.24 | 3613442.75 | 5847736 | 2113314.5 | 3625977.5 | 3994749.25 | 3960737 | 2050366.25 | 4008538.75 | 3708860.5 | 2861913.5 | 1670527.125 | 2061421.5 | 1868.9718 | YVASYLLAALGGNSSPSAK | P99027 | P99027 | RPLP2 | ribosomal protein, large, P2 | Cytoplasm | other |
| 15538290 | 3 | 0.028 | 1900961.25 | 2722079.5 | 3217585.5 | 5327427.5 | 5555895 | 6728359.5 | 4089318 | 3769860.75 | 6058493.5 | 2540542.75 | 5132164.5 | 4160499.5 | 1991.0089 | FVDHVFDEQVIDSLTVK | Q91YQ5 | Q91YQ5 | RPN1 | ribophorin I | Cytoplasm | enzyme |
| 15554810 | 3 | 0.248 | 1959634.125 | 4582841.5 | 5752041 | 7760461.5 | 8579112 | 1.04E+07 | 5384088.5 | 6510352 | 1.17E+07 | 2392452.75 | 8000925 | 8678252 | 2997.4607 | THYIVGYNLPSYEYLYNLGDQYALK | Q91YQ5 | Q91YQ5 | RPN1 | ribophorin I | Cytoplasm | enzyme |
| 15536331 | 3 | 0.117 | 2461395 | 5614416.5 | 5641035.5 | 9141091 | 9982273 | 9927894 | 5337861.5 | 9744480 | 1.12E+07 | 2389927 | 6707006 | 6618728.5 | 2197.1282 | NLVEQHIQDIVVHYTFNK | Q91YQ5 | Q91YQ5 | RPN1 | ribophorin I | Cytoplasm | enzyme |
| 15536829 | 2 | 0.019 | 1575253.625 | 2187183.25 | 2375860 | 4779382 | 4578249.5 | 4925840.5 | 2967293 | 4009124 | 5239420 | 1908482.875 | 3994751.25 | 3414752.5 | 1273.7075 | AVTSEIAVLQSR | Q91YQ5 | Q91YQ5 | RPN1 | ribophorin I | Cytoplasm | enzyme |
| 15534431 | 2 | 0.069 | 1108494.25 | 2106683.5 | 2154131.75 | 4009283 | 4419368 | 4496637.5 | 2850968.5 | 3793276.5 | 5029960.5 | 1486475.75 | 4601649 | 3429311 | 996.5691 | LPVALDPGSK | Q91YQ5 | Q91YQ5 | RPN1 | ribophorin I | Cytoplasm | enzyme |
| 15535837 | 3 | 0.254 | 1198322 | 3221121.25 | 2562305.5 | 4442613 | 4488506.5 | 5024849 | 3040367.5 | 4754489.5 | 5488655 | 1315596.5 | 5345885 | 3971766.25 | 1665.8884 | VTAEVVLVHPGGGSTSR | Q91YQ5 | Q91YQ5 | RPN1 | ribophorin I | Cytoplasm | enzyme |
| 15553602 | 2 | 0.295 | 284536.031 | 1036214.188 | 1518634.75 | 1620677.625 | 2852212.75 | 2889825.5 | 1513894.25 | 2046234.75 | 3053922 | 386684.031 | 2917279.5 | 1945089 | 1587.8749 | VAC[160.0307]ITEQVLTLVNK | Q91YQ5 | Q91YQ5 | RPN1 | ribophorin I | Cytoplasm | enzyme |
| 15544476 | 2 | 0.086 | 368905.344 | 1016564.938 | 1101231.875 | 1680535.875 | 2187104.75 | 2555043.25 | 1565928.25 | 1709186.75 | 1995997.5 | 234845.844 | 1161832.125 | 1158835.75 | 1418.6002 | GEDEEDNNLEVR | Q91YQ5 | Q91YQ5 | RPN1 | ribophorin I | Cytoplasm | enzyme |
| 15548723 | 3 | 0.11 | 4088061.75 | 7288389 | 7326487 | 1.05E+07 | 1.26E+07 | 1.38E+07 | 7486205.5 | 1.04E+07 | 1.44E+07 | 4027784.75 | 1.06E+07 | 7765383 | 2609.3457 | EETVLATVQALQTASHLSQQADLR | Q9DBG6 | Q9DBG6 | RPN2 | ribophorin II | Cytoplasm | enzyme |
| 15549499 | 3 | 0.091 | 1510802.875 | 4496801 | 7912206.5 | 1.08E+07 | 1.11E+07 | 1.58E+07 | 8774603 | 1.07E+07 | 1.53E+07 | 3222937.5 | 9018016 | 9081650 | 2524.297 | YHVPVVVVPEGSTSDTQEQAILR | Q9DBG6 | Q9DBG6 | RPN2 | ribophorin II | Cytoplasm | enzyme |
| 15549925 | 2 | 0.17 | 938579.188 | 2923523.5 | 3737335.5 | 5235477 | 4859458.5 | 7349990 | 4101732.5 | 4741375 | 7420188.5 | 1519849.875 | 4901759.5 | 4623755.5 | 1711.9031 | TPFSLVGNVFELNFK | Q9DBG6 | Q9DBG6 | RPN2 | ribophorin II | Cytoplasm | enzyme |
| 15533978 | 2 | 0.088 | 905822.75 | 2841727.75 | 3216688.75 | 5650107.5 | 5271679.5 | 7315904.5 | 3763299.75 | 5442208.5 | 5711547 | 1400848.25 | 4072583.75 | 4107422.5 | 1480.8511 | NPILWNVADVVIK | Q9DBG6 | Q9DBG6 | RPN2 | ribophorin II | Cytoplasm | enzyme |
| 15555638 | 2 | 0.133 | 935582.125 | 2286727.5 | 3345769.25 | 3456247 | 5585500.5 | 6567573 | 3248178 | 4206756 | 5758153 | 1301792.375 | 3368237.25 | 3722081.5 | 1665.8492 | EDQVIQLMNTIFSK | Q9DBG6 | Q9DBG6 | RPN2 | ribophorin II | Cytoplasm | enzyme |
| 15544773 | 2 | 0.079 | 709031.375 | 1680920.5 | 2578198.5 | 3502468.5 | 3794415.25 | 4014410 | 2627729 | 2931464.75 | 4154639 | 1238110.25 | 2725215.5 | 2492988.25 | 1399.7407 | NIVEEIEDLVAR | Q9DBG6 | Q9DBG6 | RPN2 | ribophorin II | Cytoplasm | enzyme |
| 15550306 | 2 | 0.129 | 796045.25 | 2217896 | 2775301 | 3842183.75 | 4579918.5 | 4676468 | 3143680.25 | 3516524.75 | 5493630.5 | 1018680.875 | 3478888.5 | 2892595.5 | 1766.0138 | LQVSNVLSQPLAQAAVK | Q9DBG6 | Q9DBG6 | RPN2 | ribophorin II | Cytoplasm | enzyme |
| 15539212 | 2 | 0.039 | 775203.688 | 1185448.625 | 1453099.75 | 2115894.75 | 2509261.5 | 3348803 | 1867489.875 | 2852366.5 | 3447910.25 | 928619.75 | 2236110.75 | 1909068.25 | 1078.5875 | YIANTVELR | Q9DBG6 | Q9DBG6 | RPN2 | ribophorin II | Cytoplasm | enzyme |
| 15553620 | 2 | 0.132 | 640264.188 | 1805519.875 | 2236146.25 | 3370842.5 | 4352502 | 4218491.5 | 1994716.125 | 2576894.25 | 4988621.5 | 636167.75 | 2600919 | 2079429.75 | 1790.9354 | ISTEVGITNVDLSTVDK | Q9DBG6 | Q9DBG6 | RPN2 | ribophorin II | Cytoplasm | enzyme |
| 15543280 | 2 | 0.559 | 212918.375 | 745519.062 | 1146889.25 | 671168.188 | 1387096.25 | 1629335.875 | 1032170.438 | 1589285.125 | 2642609 | 8017.086 | 1460686.125 | 1625716.375 | 996.4619 | FELDTSER | Q9DBG6 | Q9DBG6 | RPN2 | ribophorin II | Cytoplasm | enzyme |
| 15540261 | 2 | 0.045 | 6072302 | 1.21E+07 | 5391816 | 6315160 | 5685892 | 8140162 | 3623062.75 | 5521776.5 | 7375697 | 3677242 | 2857758 | 3294171.5 | 1109.6583 | IAIYELLFK | P63325 | P63325 | RPS10 | ribosomal protein S10 | Cytoplasm | other |
| 15534119 | 3 | 0.151 | 4114914.75 | 1.02E+07 | 3423500.75 | 3626410 | 5774792 | 6442573.5 | 2315234.5 | 5238335.5 | 6286789.5 | 2225664.75 | 2486292.5 | 2352748.5 | 1986.0282 | DVQIGDIVTVGEC[160.0307]RPLSK | P62281 | P62281 | RPS11 | ribosomal protein S11 | Cytoplasm | other |
| 15544918 | 2 | 0.07 | 984941.125 | 2187922.75 | 838254.688 | 425802.812 | 1387723.5 | 928184.875 | 290773.406 | 1201564.375 | 970425.75 | 495268.312 | 129016.344 | 210012.719 | 979.49866 | DYLHYIR | P62281 | P62281 | RPS11 | ribosomal protein S11 | Cytoplasm | other |
| 15539984 | 2 | 0.069 | 2249080.75 | 4510326 | 1898626.5 | 2005376.625 | 2649089 | 2651782.75 | 1073442.875 | 2303256.75 | 2138453.5 | 1683823.5 | 637288.625 | 979086.625 | 843.5284 | LILIESR | P62301 | P62301 | RPS13 | ribosomal protein S13 | Cytoplasm | other |
| 15552237 | 2 | 0.045 | 1996689 | 4570254.5 | 1600944.875 | 2199053 | 2681025.75 | 2335263.75 | 990028.125 | 2173923 | 2596769 | 957682.188 | 807453 | 995283.062 | 1693.911 | GLAPDLPEDLYHLIK | P62301 | P62301 | RPS13 | ribosomal protein S13 | Cytoplasm | other |
| 15548309 | 3 | 0.061 | 8532452 | 1.51E+07 | 6390458.5 | 1.17E+07 | 1.34E+07 | 1.08E+07 | 5042586 | 9677639 | 1.05E+07 | 6586160 | 4572211 | 4376302 | 2265.051 | ADRDESSPYAAMLAAQDVAQR | P62264 | P62264 | RPS14 | ribosomal protein S14 | Cytoplasm | translation regulator |
| 15540605 | 2 | 0.079 | 6762305 | 1.12E+07 | 4269666 | 7994271.5 | 6388385.5 | 7045939 | 3763738 | 6300487 | 6321071 | 4620425.5 | 2948824.25 | 3104042 | 1054.5615 | TPGPGAQSALR | P62264 | P62264 | RPS14 | ribosomal protein S14 | Cytoplasm | translation regulator |
| 15533910 | 2 | 0.097 | 3578248.75 | 1.15E+07 | 4054747 | 3570158 | 7782734.5 | 5990862.5 | 3099316.25 | 6768169 | 4634694 | 2927900 | 1193524.375 | 2307644.5 | 1429.7135 | IEDVTPIPSDSTR | P62264 | P62264 | RPS14 | ribosomal protein S14 | Cytoplasm | translation regulator |
| 15539324 | 2 | 0.084 | 3085631.5 | 4873760 | 2014948.875 | 2520437.5 | 2945630.75 | 2968847.25 | 1581128.25 | 2429526.5 | 2377600.75 | 2271613.25 | 1050682.125 | 1378786.5 | 871.5339 | IVVNLTGR | P62245 | P62245 | RPS15A | ribosomal protein S15a | Cytoplasm | other |
| 15535090 | 2 | 0.032 | 2865168.25 | 5045377.5 | 2685662.75 | 3159274.25 | 3629679 | 4840611.5 | 2956512.5 | 3436328.25 | 3805735 | 2106915.75 | 1334470.25 | 2233145.5 | 974.53217 | MNVLADALK | P62245 | P62245 | RPS15A | ribosomal protein S15a | Cytoplasm | other |
| 15534447 | 2 | 0.083 | 7633071 | 1.36E+07 | 5110458.5 | 8643001 | 8774732 | 9062743 | 4692116 | 9058793 | 8718957 | 4616897 | 3359011 | 4746060 | 1094.7163 | LLEPVLLLGK | P14131 | P14131 | RPS16 | ribosomal protein S16 | Cytoplasm | other |
| 15543901 | 2 | 0.173 | 5322674 | 8242834 | 2399017 | 4387335.5 | 4765234.5 | 4921689.5 | 1675151.875 | 4768851 | 5557045.5 | 1813196.75 | 1939114.25 | 2484486.5 | 1187.6506 | GPLQSVQVFGR | P14131 | P14131 | RPS16 | ribosomal protein S16 | Cytoplasm | other |
| 15553280 | 2 | 0.056 | 2839448.5 | 4350066.5 | 2220408.5 | 2920988.75 | 2687229 | 2737328.75 | 1387655 | 2653349.5 | 3279802.75 | 1532752.375 | 1310520.25 | 1551356.375 | 2489.1875 | DNYVPEVSALDQEIIEVDPDTK | P63276 | P63276 | RPS17 | ribosomal protein S17 | Cytoplasm | other |
| 15534819 | 2, 3 | 0.018 | 4571929.5 | 9313947 | 3700150 | 5895118 | 5695343 | 5734094 | 2772845.5 | 5326812.5 | 5200629.5 | 2144836.25 | 2147197.25 | 2373333.75 | 1403.6736, 1403.6738 | RAGELTEDEVER | P62270 | P62270 | RPS18 | ribosomal protein S18 | Cytoplasm | other |
| 15538596 | 2 | 0.013 | 4275191 | 6537591.5 | 2694628 | 3737815.25 | 3965968.5 | 3979930.25 | 1584777.375 | 2825684.5 | 2850557.5 | 2011250.25 | 1478958.5 | 1489474.75 | 1071.5952 | VITIMQNPR | P62270 | P62270 | RPS18 | ribosomal protein S18 | Cytoplasm | other |
| 15533832 | 2 | 0.011 | 2775887.5 | 5528187 | 2473861 | 3141127 | 4232212.5 | 5198525.5 | 1644815.25 | 2180901.75 | 3244643.25 | 1346723 | 1457846.625 | 1357167.875 | 947.5898 | IAFAITAIK | P62270 | P62270 | RPS18 | ribosomal protein S18 | Cytoplasm | other |
| 15549829 | 2 | 0.082 | 6004405.5 | 1.08E+07 | 5589568.5 | 5882127.5 | 6082319.5 | 6839443 | 3174702 | 6308057 | 8037330.5 | 4162565 | 2623701.75 | 3281163.25 | 1703.7661 | ELAPYDENWFYTR | Q9CZX8 | Q9CZX8 | RPS19 | ribosomal protein S19 | Cytoplasm | other |
| 15535514 | 2 | 0.147 | 5058328.5 | 8696204 | 3132343 | 5845608 | 4643716.5 | 5383456.5 | 2739788.75 | 5605701.5 | 4920569.5 | 3321324.25 | 2391354.75 | 2697213.75 | 1134.5509 | DVNQQEFVR | Q9CZX8 | Q9CZX8 | RPS19 | ribosomal protein S19 | Cytoplasm | other |
| 15533094 | 2 | 0.035 | 5229527.5 | 9261957 | 3774471 | 5288220.5 | 6269017 | 6703983 | 3170035.25 | 4815947 | 5378296.5 | 3273793.75 | 2466842 | 2609272.75 | 1072.5648 | VPEWVDTVK | Q9CZX8 | Q9CZX8 | RPS19 | ribosomal protein S19 | Cytoplasm | other |
| 15538491 | 3 | 0.112 | 9130859 | 1.65E+07 | 6587435.5 | 9333378 | 9840537 | 8857184 | 6066427 | 1.01E+07 | 9056823 | 7408788.5 | 3319175.5 | 4787146.5 | 1463.713 | SPYQEFTDHLVK | P25444 | P25444 | RPS2 | ribosomal protein S2 | Cytoplasm | other |
| 15548623 | 2 | 0.064 | 6262306.5 | 1.21E+07 | 5024748 | 6582711 | 7755939 | 8691887 | 3567961 | 9204592 | 7606932 | 3846496.75 | 2336495 | 3492146.75 | 1551.8643 | SLEEIYLFSLPIK | P25444 | P25444 | RPS2 | ribosomal protein S2 | Cytoplasm | other |
| 15532572 | 2 | 0.048 | 5999946.5 | 9378107 | 4190713 | 5398539.5 | 6761953.5 | 6082639 | 2849184.75 | 5849622.5 | 5565600 | 3231318.25 | 2375882.5 | 3121240.5 | 1025.5963 | GTGIVSAPVPK | P25444 | P25444 | RPS2 | ribosomal protein S2 | Cytoplasm | other |
| 15536632 | 2 | 0.083 | 4934551.5 | 7409753.5 | 3535906.75 | 4207850 | 5775742.5 | 4609663.5 | 2534005.25 | 5017419.5 | 5508927 | 2468945.25 | 2004435.375 | 3118240 | 1386.6913 | TYSYLTPDLWK | P25444 | P25444 | RPS2 | ribosomal protein S2 | Cytoplasm | other |
| 15536349 | 3 | 0.182 | 6332749.5 | 1.33E+07 | 4155061.25 | 7591449 | 6911249.5 | 7656406.5 | 4454651 | 7553746.5 | 7431559.5 | 4574746.5 | 3284977.25 | 4139237.25 | 1247.6714 | TPVEPEVAIHR | P60867 | P60867 | RPS20 | ribosomal protein S20 | Cytoplasm | other |
| 15542806 | 3 | 0.067 | 5635846.5 | 8060117 | 3253855.25 | 6213752.5 | 4817062 | 5764709 | 2791961.5 | 5391677 | 6112185 | 2483575.75 | 3009067.75 | 2278358 | 1506.8604 | RLIDLHSPSEIVK | P60867 | P60867 | RPS20 | ribosomal protein S20 | Cytoplasm | other |
| 15539070 | 2 | 0.108 | 5015971.5 | 7756945 | 3474892.5 | 4146498.5 | 4723091.5 | 5171010 | 2395434.5 | 4226668 | 4967581.5 | 2956677.75 | 2206375.5 | 3007555.75 | 1190.7114 | VANVSLLALYK | P62267 | P62267 | RPS23 | ribosomal protein S23 | Cytoplasm | translation regulator |
| 15540643 | 3 | 0.024 | 2719413 | 4128243.75 | 1500201.125 | 2979128 | 2766393.25 | 2675179 | 1111784.5 | 1830181.75 | 2048230.75 | 1257550.5 | 1019934.688 | 1185957.125 | 1236.6748 | QMVIDVLHPGK | P62849 | P62849 | RPS24 | ribosomal protein S24 | Cytoplasm | other |
| 15536231 | 2 | 0.024 | 9129592 | 1.17E+07 | 6046677.5 | 8506127 | 8971297 | 9264352 | 3923324.25 | 8001318 | 8121754 | 4723878.5 | 3507892 | 3779716.5 | 1084.6338 | LITPAVVSER | P62852 | P62852 | RPS25 | ribosomal protein S25 | Cytoplasm | other |
| 15537086 | 2 | 0.045 | 5308299.5 | 1.01E+07 | 4467988 | 7056208.5 | 7442955 | 7136745 | 3588962 | 5752754.5 | 7782712.5 | 3665584.75 | 2755243.5 | 3202279.75 | 1075.6124 | LNNLVLFDK | P62852 | P62852 | RPS25 | ribosomal protein S25 | Cytoplasm | other |
| 15533976 | 2 | 0.045 | 5231918.5 | 9350453 | 4279794 | 5231329.5 | 5831485 | 5048101 | 2763421 | 5541924.5 | 5351706 | 3232740.5 | 1986130.875 | 2706274.5 | 972.5706 | AALQELLSK | P62852 | P62852 | RPS25 | ribosomal protein S25 | Cytoplasm | other |
| 15541380 | 2 | 0.115 | 4221357 | 6439743.5 | 2476345 | 4220052 | 3895005.5 | 4288775 | 2131146 | 2873987 | 3685881 | 3022712.25 | 1742232.375 | 2175494 | 942.53424 | NIVEAAAVR | P62855 | P62855 | RPS26 | ribosomal protein S26 | Cytoplasm | other |
| 15543582 | 2 | 0.031 | 1845833.25 | 2743829.5 | 1430025.5 | 1371564.25 | 2003789.25 | 1962893.5 | 1168203.25 | 1961442.875 | 1784173.75 | 1190044.75 | 777335.812 | 760675.5 | 969.4429 | LTEGC[160.0307]SFR | Q6ZWY3 | Q6ZWY3 | RPS27L | ribosomal protein S27-like | Cytoplasm | translation regulator |
| 15545358 | 3 | 0.222 | 761032.125 | 1043748.562 | 239795.844 | 537130.5 | 363973.219 | 480864.031 | 440794.156 | 603191.438 | 444727.562 | 497122.844 | 189878.609 | 93298.406 | 1683.8458 | RLVQSPNSYFMDVK | Q6ZWY3 | Q6ZWY3 | RPS27L | ribosomal protein S27-like | Cytoplasm | translation regulator |
| 15541999 | 2 | 0.343 | 856012.938 | 3333887 | 757806.938 | 564345.5 | 2441046.75 | 1279476.75 | 697472.688 | 1850285.625 | 1465111.875 | 1131259.375 | 340309.406 | 313607.188 | 1360.6938 | EGDVLTLLESER | P62858 | P62858 | RPS28 | ribosomal protein S28 | Cytoplasm | other |
| 15552408 | 3 | 0.146 | 7962485 | 1.78E+07 | 5322948.5 | 9344693 | 7573859 | 8763033 | 4667001 | 7263991 | 7954987 | 6223344.5 | 3786604.25 | 3510154.75 | 2468.187 | FVDGLMIHSGDPVNYYVDTAVR | P62908 | P62908 | RPS3 | ribosomal protein S3 | Cytoplasm | enzyme |
| 15550283 | 2 | 0.106 | 7126192 | 1.36E+07 | 3755984.75 | 7495171 | 7657681.5 | 6106504.5 | 3785020 | 7077406.5 | 6474503.5 | 4376300 | 2597402.5 | 2714055.5 | 1573.8004 | GGKPEPPAMPQPVPTA | P62908 | P62908 | RPS3 | ribosomal protein S3 | Cytoplasm | enzyme |
| 15543551 | 2 | 0.296 | 3778757 | 6564669 | 2776934.75 | 3470319.75 | 4043636.5 | 3570831.25 | 2299506.25 | 3253214 | 3498263.5 | 3475406.5 | 2286298.25 | 2531281 | 1470.7611 | DEILPTTPISEQK | P62908 | P62908 | RPS3 | ribosomal protein S3 | Cytoplasm | enzyme |
| 15536493 | 2 | 0.084 | 5303618.5 | 1.00E+07 | 3874943.5 | 6956696 | 5358883.5 | 5475851.5 | 2725736 | 6198114.5 | 5632147.5 | 3325245.25 | 2076587.375 | 2817420 | 1092.5654 | AELNEFLTR | P62908 | P62908 | RPS3 | ribosomal protein S3 | Cytoplasm | enzyme |
| 15534382 | 2 | 0.051 | 3385298 | 6171638 | 2618631 | 4029132.25 | 4107679.25 | 3874682.75 | 2146494 | 3942010.75 | 3332030.5 | 2116357.5 | 1391199.75 | 2215735.25 | 1423.667 | ELAEDGYSGVEVR | P62908 | P62908 | RPS3 | ribosomal protein S3 | Cytoplasm | enzyme |
| 15536137 | 3 | 0.83 | 3900682.75 | 3255090.25 | 635418.5 | 4125329.25 | 1158232.5 | 2996071.25 | 1305686.625 | 1192319.875 | 2565158.5 | 1333372.375 | 1900496.5 | 1690682 | 1583.945 | VTPTRTEIIILATR | P62908 | P62908 | RPS3 | ribosomal protein S3 | Cytoplasm | enzyme |
| 15534100 | 2 | 0.195 | 1965016.125 | 3449897 | 952146.062 | 1503306.875 | 1880615.75 | 2659522.75 | 1136573.875 | 2049875.75 | 2091053.5 | 1326321.5 | 869044.938 | 687489.188 | 798.5174 | LLGGLAVR | P62908 | P62908 | RPS3 | ribosomal protein S3 | Cytoplasm | enzyme |
| 15540932 | 2 | 0.12 | 1378867.5 | 3054242.5 | 1147686.875 | 1055300.125 | 1607006.125 | 2015977.75 | 793652.188 | 1585622.625 | 1315865 | 1015936.438 | 751907.562 | 504081.281 | 1029.6293 | TEIIILATR | P62908 | P62908 | RPS3 | ribosomal protein S3 | Cytoplasm | enzyme |
| 15544988 | 2 | 0.031 | 2535711.5 | 3704629 | 1868452.625 | 2309796.5 | 2459178.5 | 2092304.125 | 1200540.625 | 2041206.875 | 2723222 | 1414804.625 | 1021711.438 | 903056.5 | 919.4801 | APAMFNIR | P97351 | P97351 | Rps3a1 | ribosomal protein S3A1 | Cytoplasm | other |
| 15542917 | 2 | 0.287 | 356898.656 | 2008861.875 | 500215.969 | 85298.562 | 643210.812 | 1024591.375 | 292042.094 | 805601.75 | 1092540.5 | 74937.336 | 249220.953 | 241842.625 | 1002.5448 | LITEDVQGK | P97351 | P97351 | Rps3a1 | ribosomal protein S3A1 | Cytoplasm | other |
| 15536024 | 3 | 0.149 | 7625370 | 1.30E+07 | 3524690.5 | 7990398 | 5863548.5 | 7000177 | 3749904 | 6546923.5 | 7633395 | 3321758.75 | 3197174.25 | 3633022 | 2141.0916 | VRTDITYPAGFMDVISIDK | P62702 | P62702 | RPS4Y1 | ribosomal protein S4, Y-linked 1 | Cytoplasm | other |
| 15533306 | 2 | 0.037 | 6152696.5 | 7598430.5 | 3987846.75 | 4714335.5 | 5294567 | 3258679.5 | 2567382.25 | 5028636 | 5940965 | 2824821.5 | 1917068.625 | 2031586.75 | 990.59625 | LSNIFVIGK | P62702 | P62702 | RPS4Y1 | ribosomal protein S4, Y-linked 1 | Cytoplasm | other |
| 15544198 | 4 | 0.222 | 2804058.75 | 1712952.5 | 664967.5 | 4305263 | 1472187.375 | 1362561 | 955269.125 | 1075186 | 147472.359 | 2075388.75 | 1438737.625 | 1153506.25 | 1506.775 | ERHPGSFDVVHVK | P62702 | P62702 | RPS4Y1 | ribosomal protein S4, Y-linked 1 | Cytoplasm | other |
| 15552183 | 2 | 0.171 | 2406084.75 | 5087781.5 | 2315091 | 2112140.5 | 3756454.5 | 2454356.5 | 1255148.875 | 3005452 | 2494126.5 | 2042302.25 | 1057678 | 1379773.75 | 1885.921 | TDITYPAGFMDVISIDK | P62702 | P62702 | RPS4Y1 | ribosomal protein S4, Y-linked 1 | Cytoplasm | other |
| 15536488 | 2 | 0.054 | 2132362.5 | 4191474.75 | 2000986.375 | 2158161.5 | 3171071.75 | 2545457.5 | 1411701.25 | 2617407.25 | 2596209.5 | 1817565.375 | 753151.062 | 876953.688 | 995.5027 | YALTGDEVK | P62702 | P62702 | RPS4Y1 | ribosomal protein S4, Y-linked 1 | Cytoplasm | other |
| 15536739 | 3 | 0.849 | 2966827.25 | 3231403 | 1022986.75 | 1968425.75 | 2213589.5 | 1756202.5 | 1370639.625 | 1903263.375 | 2088619.375 | 1757253.125 | 1844929.75 | 1555767.375 | 1542.9156 | LREC[160.0307]LPLIIFLR | P62702 | P62702 | RPS4Y1 | ribosomal protein S4, Y-linked 1 | Cytoplasm | other |
| 15545516 | 3 | 0.024 | 2237468.25 | 3793687.25 | 1972602.375 | 2300697.75 | 2627158.25 | 2509077.25 | 1214319.75 | 2783844.5 | 2745172.25 | 1020284.062 | 907393.062 | 1256649.75 | 1215.6567 | GIPHLVTHDAR | P62702 | P62702 | RPS4Y1 | ribosomal protein S4, Y-linked 1 | Cytoplasm | other |
| 15541316 | 3 | 0.016 | 2111768 | 3274395.25 | 1454345.375 | 2002193.75 | 2494810 | 1707738.125 | 1173602.75 | 1915749.75 | 1833189.625 | 970473.125 | 816319.438 | 972594.75 | 1221.6357 | HPGSFDVVHVK | P62702 | P62702 | RPS4Y1 | ribosomal protein S4, Y-linked 1 | Cytoplasm | other |
| 15536862 | 2 | 0.269 | 5242523.5 | 8958242 | 2969357.5 | 5623660 | 5597652 | 4087581.75 | 3181610 | 4756275.5 | 5989793.5 | 2900118 | 3154162 | 3172364.75 | 1335.6692 | LNISFPATGC[160.0307]QK | P62754 | P62754 | RPS6 | ribosomal protein S6 | Cytoplasm | other |
| 15540099 | 3 | 0.5 | 2330881 | 1367721.75 | 455916.125 | 2926484.75 | 825408.938 | 1088745.125 | 710959.375 | 860212.875 | 1671898.75 | 497713.812 | 699899.562 | 801094.688 | 1776.8546 | RMATEVAADALGEEWK | P62754 | P62754 | RPS6 | ribosomal protein S6 | Cytoplasm | other |
| 15549114 | 3 | 0.047 | 9639079 | 1.74E+07 | 8541985 | 1.28E+07 | 1.29E+07 | 1.32E+07 | 6581166 | 1.19E+07 | 1.27E+07 | 8688807 | 4085739 | 3638340.5 | 2367.2776 | TLTAVHDAILEDLVFPSEIVGK | P62082 | P62082 | RPS7 | ribosomal protein S7 | Cytoplasm | other |
| 15533378 | 2 | 0.049 | 6897116.5 | 1.16E+07 | 4784705.5 | 7640149 | 6674506.5 | 8352110 | 3265638 | 7308248 | 5429290.5 | 3731924.5 | 3369677.5 | 3360040.25 | 1337.8546 | AIIIFVPVPQLK | P62082 | P62082 | RPS7 | ribosomal protein S7 | Cytoplasm | other |
| 15544873 | 2 | 0.033 | 3172747.75 | 4181230.75 | 1942655.25 | 2497235 | 2449321.5 | 3236646.5 | 1579277 | 2958699.5 | 2938706.75 | 1217111.125 | 1100213.375 | 1621679.375 | 1384.6407 | DVNFEFPEFQL | P62082 | P62082 | RPS7 | ribosomal protein S7 | Cytoplasm | other |
| 15548394 | 2 | 0.047 | 1.00E+07 | 1.39E+07 | 7207733.5 | 7685853.5 | 1.06E+07 | 8849224 | 4813436.5 | 8555079 | 9953057 | 5395195 | 4294694 | 4896009.5 | 1506.7755 | ISSLLEEQFQQGK | P62242 | P62242 | RPS8 | ribosomal protein S8 | Cytoplasm | other |
| 15534410 | 2 | 0.056 | 8019652 | 1.34E+07 | 5223752 | 7751396.5 | 9163492 | 8135713 | 4416758 | 7796814.5 | 6993361 | 4844049.5 | 3146548.75 | 4158062.25 | 1314.6763 | LTPEEEEILNK | P62242 | P62242 | RPS8 | ribosomal protein S8 | Cytoplasm | other |
| 15548984 | 2 | 0.019 | 6493939.5 | 8041691 | 5114376 | 5736776 | 6129378.5 | 7162226 | 3317968.75 | 6742182 | 6380375 | 3679375 | 2573074.75 | 2828346 | 1718.9027 | IIDVVYNASNNELVR | P62242 | P62242 | RPS8 | ribosomal protein S8 | Cytoplasm | other |
| 15538203 | 2 | 0.137 | 3256456.25 | 5542937.5 | 1726661.75 | 3000655.5 | 3205656.25 | 3569948.5 | 2073389.625 | 3068935 | 3115754 | 1993324.5 | 1679720.375 | 1513851.875 | 1450.7306 | NC[160.0307]IVLIDSTPYR | P62242 | P62242 | RPS8 | ribosomal protein S8 | Cytoplasm | other |
| 15535068 | 2 | 0.039 | 2434635.75 | 3900187.5 | 1339373.125 | 2445617.25 | 2001427 | 2513155.75 | 1616867.5 | 2347440 | 2309284.75 | 1168399.375 | 947514.812 | 1126307.75 | 1219.6403 | YELGRPAANTK | P62242 | P62242 | RPS8 | ribosomal protein S8 | Cytoplasm | other |
| 15539591 | 2 | 0.163 | 7369271.5 | 1.16E+07 | 3248415 | 7638436.5 | 6759360.5 | 7949960.5 | 4490659.5 | 6344873 | 6669273.5 | 4392447 | 3199677.5 | 3602907.5 | 1271.745 | QVVNIPSFIVR | Q6ZWN5 | Q6ZWN5 | RPS9 | ribosomal protein S9 | Cytoplasm | translation regulator |
| 15538886 | 2 | 0.026 | 4133057.25 | 6161423.5 | 3118886 | 4667221 | 4816892 | 4314141 | 2180479 | 4098003.25 | 3935133.5 | 2572944.25 | 1927506.625 | 1952574.75 | 920.51855 | LIGEYGLR | Q6ZWN5 | Q6ZWN5 | RPS9 | ribosomal protein S9 | Cytoplasm | translation regulator |
| 15540106 | 2 | 0.05 | 2747157.25 | 4444233 | 642289 | 2731007.25 | 2651957.5 | 2262186.5 | 1480021.75 | 3120122 | 2809239.75 | 641135.625 | 391436.281 | 859791.438 | 934.5587 | LDYILGLK | Q6ZWN5 | Q6ZWN5 | RPS9 | ribosomal protein S9 | Cytoplasm | translation regulator |
| 15554858 | 3 | 0.093 | 1.65E+07 | 2.97E+07 | 1.12E+07 | 1.89E+07 | 1.92E+07 | 1.95E+07 | 9278539 | 1.66E+07 | 1.88E+07 | 1.10E+07 | 8166999 | 9136963 | 2996.4695 | ADHQPLTEASYVNLPTIALC[160.0307]NTDSPLR | P14206 | P14206 | RPSA | ribosomal protein SA | Cytoplasm | translation regulator |
| 15549074 | 2 | 0.085 | 8152309.5 | 1.54E+07 | 6538599 | 9827595 | 9676845 | 8843464 | 4554788.5 | 8788665 | 9482125 | 5858934.5 | 4592223.5 | 3777361.5 | 1740.9453 | AIVAIENPADVSVISSR | P14206 | P14206 | RPSA | ribosomal protein SA | Cytoplasm | translation regulator |
| 15537124 | 2 | 0.036 | 7052570.5 | 1.28E+07 | 6095426.5 | 6016896.5 | 7672548.5 | 7152986.5 | 4113265.5 | 8077788.5 | 7780685.5 | 4251892 | 2957008 | 3448912.5 | 1203.6449 | FAAATGATPIAGR | P14206 | P14206 | RPSA | ribosomal protein SA | Cytoplasm | translation regulator |
| 15541506 | 2 | 0.029 | 3748930.75 | 6675697.5 | 2990727.5 | 4836437.5 | 3671169.75 | 4217455 | 1833838.625 | 3695898.5 | 3389595.5 | 2299340.5 | 1216160.125 | 1749342.625 | 912.54987 | LLVVTDPR | P14206 | P14206 | RPSA | ribosomal protein SA | Cytoplasm | translation regulator |
| 15536176 | 3 | 0.738 | 1.42E+07 | 2.53E+07 | 7108668.5 | 1.12E+07 | 5734912 | 1.48E+07 | 7429294.5 | 8752827 | 2.08E+07 | 7855074 | 8244883 | 1.07E+07 | 2127.0464 | INFSGGEPFLQDRGEYLGK | Q8CBB9 | Q8CBB9 | RSAD2 | radical S-adenosyl methionine domain containing 2 | Cytoplasm | enzyme |
| 15552366 | 2 | 0.193 | 1.53E+07 | 2.89E+07 | 1.09E+07 | 7628633.5 | 1.06E+07 | 2.06E+07 | 5899065.5 | 1.27E+07 | 2.28E+07 | 6765735 | 6038530.5 | 6509373.5 | 1773.8678 | FLISNEEFETFLER | Q8CBB9 | Q8CBB9 | RSAD2 | radical S-adenosyl methionine domain containing 2 | Cytoplasm | enzyme |
| 15542547 | 2 | 0.69 | 8435077 | 1.69E+07 | 3781105.5 | 5977738 | 5125498.5 | 7956007.5 | 3978718.75 | 6068045 | 1.10E+07 | 6060543 | 4975034.5 | 4292286.5 | 1389.6655 | EVSC[160.0307]LVPESNQK | Q8CBB9 | Q8CBB9 | RSAD2 | radical S-adenosyl methionine domain containing 2 | Cytoplasm | enzyme |
| 15558124 | 3 | 0.678 | 1.22E+07 | 2.56E+07 | 8200016.5 | 6570424.5 | 7321833 | 1.91E+07 | 4246875.5 | 1.08E+07 | 2.85E+07 | 5055339 | 7569596.5 | 9490193 | 2535.209 | VFQC[160.0307]LLIEGENSGEDALREAER | Q8CBB9 | Q8CBB9 | RSAD2 | radical S-adenosyl methionine domain containing 2 | Cytoplasm | enzyme |
| 15537614 | 2, 3 | 0.462 | 6789966.5 | 1.59E+07 | 3812335.5 | 3127774.25 | 4546387 | 9145352 | 2529234.5 | 5955818.5 | 9958679 | 3479594 | 3326632.75 | 3428348.75 | 1490.6538, 1490.6553 | FNVDEDMNEHIK | Q8CBB9 | Q8CBB9 | RSAD2 | radical S-adenosyl methionine domain containing 2 | Cytoplasm | enzyme |
| 15540871 | 2 | 0.322 | 3651322 | 7706604 | 2509415 | 2321619.75 | 2212410.25 | 4696587 | 1901504.25 | 3610476 | 5893847 | 2114151.25 | 1678144.375 | 2236948 | 1417.6646 | DSYLILDEYMR | Q8CBB9 | Q8CBB9 | RSAD2 | radical S-adenosyl methionine domain containing 2 | Cytoplasm | enzyme |
| 15543010, 15546128 | 2 | 0.144 | 4835374 | 9464240 | 5186339 | 4478998.5 | 2064752.625 | 6602436.5 | 1938794.75 | 3345896.75 | 6503079 | 2042690.125 | 1479314 | 3073404.75 | 1272.702, 1272.7024 | SILDVGVEEAIK | Q8CBB9 | Q8CBB9 | RSAD2 | radical S-adenosyl methionine domain containing 2 | Cytoplasm | enzyme |
| 15555639 | 2 | 0.302 | 3258776.25 | 8356103 | 2410316 | 2950840.75 | 2038789.25 | 4634167.5 | 1556315.75 | 2537627.75 | 5531543 | 2026026.625 | 1468848 | 1753117.5 | 1884.0396 | EELALPSVSIVSNGSLIR | Q8CBB9 | Q8CBB9 | RSAD2 | radical S-adenosyl methionine domain containing 2 | Cytoplasm | enzyme |
| 15549354 | 3 | 0.24 | 1464681.125 | 4149949 | 3362557.75 | 1.02E+07 | 9665898 | 1.07E+07 | 9689436 | 4675914.5 | 2.86E+07 | 835526.125 | 7603209.5 | 1.23E+07 | 3030.503 | AENNPWVTPIADQFQLGVSHVFEYIR | Q01730 | Q01730 | RSU1 | Ras suppressor protein 1 | Cytoplasm | other |
| 15549679 | 3 | 0.295 | 1139153.25 | 2941045 | 3258837 | 9802048 | 6589966 | 7673270 | 5927087 | 3920951 | 2.10E+07 | 560545.188 | 4440443 | 1.13E+07 | 2818.4585 | NLEVLNFFNNQIEELPTQISSLQK | Q01730 | Q01730 | RSU1 | Ras suppressor protein 1 | Cytoplasm | other |
| 15538152 | 3 | 0.717 | 209292.406 | 475136.812 | 529050.812 | 3963509 | 700149 | 1719471.875 | 1427508.125 | 328425.875 | 5348645.5 | 4.70E-06 | 2458774.5 | 4267454 | 1838.0704 | LQILSLRDNDLISLPK | Q01730 | Q01730 | RSU1 | Ras suppressor protein 1 | Cytoplasm | other |
| 15539078 | 2 | 0.516 | 6537161 | 1.12E+07 | 1.70E+07 | 6433126.5 | 1.65E+07 | 1.56E+07 | 8623448 | 1.27E+07 | 1.03E+07 | 5093824.5 | 8053845 | 9476559 | 872.5181 | SVLQAVQK | Q8K0T0 | Q8K0T0 | RTN1 | reticulon 1 | Cytoplasm | other |
| 15531834 | 2, 3 | 0.438 | 5.33E+07 | 6.99E+07 | 8.66E+07 | 7.41E+07 | 9.03E+07 | 8.36E+07 | 6.38E+07 | 6.95E+07 | 7.22E+07 | 6.13E+07 | 8.22E+07 | 6.84E+07 | 1272.6675, 1272.6683 | TQIDHYVGIAR | Q9ES97 | Q9ES97 | RTN3 | reticulon 3 | Cytoplasm | other |
| 15543356 | 2 | 0.789 | 138885.031 | 704826.5 | 1791760.375 | 424607.156 | 1153544.875 | 1279254.75 | 781623.625 | 1018793.312 | 579428.625 | 853444.5 | 245609.484 | 521294.781 | 1390.7809 | LFLVEDLVDSLK | Q9ES97 | Q9ES97 | RTN3 | reticulon 3 | Cytoplasm | other |
| 15557692, 15558502 | 3 | 0.16 | 3170961.5 | 1.41E+07 | 1.60E+07 | 1.20E+07 | 2.39E+07 | 2.61E+07 | 1.57E+07 | 1.79E+07 | 1.28E+07 | 4973468 | 9286726 | 8372285.5 | 3282.6575, 3282.6584 | LPEDDEPPARPPAPAGASPLAEPAAPPSTPAAPK | Q99P72 | Q99P72 | RTN4 | reticulon 4 | Cytoplasm | other |
| 15538906 | 3 | 0.216 | 5605945 | 1.07E+07 | 8015533 | 1.42E+07 | 1.75E+07 | 1.29E+07 | 9268905 | 1.07E+07 | 1.46E+07 | 4580257.5 | 1.40E+07 | 1.03E+07 | 1607.8248 | HQAQIDHYLGLANK | Q99P72 | Q99P72 | RTN4 | reticulon 4 | Cytoplasm | other |
| 15541753 | 2 | 0.054 | 2857172.75 | 6390406.5 | 6705886.5 | 7913878.5 | 1.73E+07 | 9714843 | 7607004 | 1.02E+07 | 8505534 | 2486797.25 | 5925029 | 5066758.5 | 1273.6869 | GPLPAAPPTAPER | Q99P72 | Q99P72 | RTN4 | reticulon 4 | Cytoplasm | other |
| 15559899 | 2 | 0.215 | 333054.531 | 2677202.75 | 3461833.75 | 4276852.5 | 4758359.5 | 3768407.5 | 2854565 | 3686421.25 | 2738447.5 | 1622195.125 | 1543362.75 | 1732280.875 | 1731.9596 | SPAASAPSLPPAAAVLPSK | Q99P72 | Q99P72 | RTN4 | reticulon 4 | Cytoplasm | other |
| 15536729 | 2 | 0.11 | 1768202.75 | 2670901.5 | 1801182.375 | 4398865 | 4745254 | 4475939 | 3818679.5 | 3376771.25 | 3974762 | 1345171.375 | 5204246 | 3021340.25 | 1490.7555 | YSNSALGHVNSTIK | Q99P72 | Q99P72 | RTN4 | reticulon 4 | Cytoplasm | other |
| 15538308 | 2 | 0.04 | 2328865.25 | 3205095 | 3370668.75 | 4837035 | 5819831 | 7875154 | 2403123.5 | 5844609.5 | 2971692.75 | 2690843.75 | 2440840.5 | 3285264 | 1021.5291 | GSLNINEFK | P97352 | P97352 | S100A13 | S100 calcium binding protein A13 | Cytoplasm | other |
| 15541930 | 2 | 0.055 | 149088.797 | 265380.688 | 130912.617 | 1005405.25 | 732410 | 665457 | 730405.5 | 392219.562 | 2277821.75 | 334020.906 | 2711087.25 | 2885823.5 | 1396.6942 | QMVEAQLATFMK | P31725 | P31725 | S100A9 | S100 calcium binding protein A9 | Cytoplasm | other |
| 15559287 | 2 | 0.091 | 123725.742 | 185853.5 | 110067.648 | 985756.75 | 582468.312 | 283005.875 | 464014.188 | 312309.594 | 2356633.25 | 264457.5 | 1790793 | 1916995.75 | 1681.8503 | SITTIIDTFHQYSR | P31725 | P31725 | S100A9 | S100 calcium binding protein A9 | Cytoplasm | other |
| 15537568 | 2 | 0.567 | 8785177 | 9303521 | 1.16E+07 | 1.12E+07 | 1.05E+07 | 1.12E+07 | 1.20E+07 | 1.22E+07 | 9498625 | 1.40E+07 | 1.08E+07 | 9915830 | 1134.5869 | VVTNQDGVFR | Q9EP69 | Q9EP69 | SACM1L | SAC1 suppressor of actin mutations 1-like (yeast) | Cytoplasm | phosphatase |
| 15542220 | 4 | 0.837 | 9471091 | 8370117.5 | 1.11E+07 | 8953491 | 1.10E+07 | 8908258 | 1.24E+07 | 9157684 | 9752524 | 1.11E+07 | 9737984 | 9543877 | 2630.217 | GIDSEGHAANFVETEQIVHYSGNR | Q9EP69 | Q9EP69 | SACM1L | SAC1 suppressor of actin mutations 1-like (yeast) | Cytoplasm | phosphatase |
| 15551975 | 2 | 0.075 | 9176198 | 8226242 | 8646688 | 1.14E+07 | 1.09E+07 | 8934025 | 1.18E+07 | 1.06E+07 | 9612655 | 1.02E+07 | 1.01E+07 | 1.07E+07 | 1880.9031 | LSNTSPEFQEMSLLER | Q9EP69 | Q9EP69 | SACM1L | SAC1 suppressor of actin mutations 1-like (yeast) | Cytoplasm | phosphatase |
| 15532607 | 3 | 0.809 | 8847945 | 8166165 | 9034506 | 8383459.5 | 9391290 | 1.00E+07 | 9979902 | 9435018 | 6775577 | 8503830 | 9537829 | 9911447 | 1669.8534 | TMLHLTDIQLQDNK | Q9EP69 | Q9EP69 | SACM1L | SAC1 suppressor of actin mutations 1-like (yeast) | Cytoplasm | phosphatase |
| 15550022 | 2 | 0.08 | 4956371.5 | 3700883 | 4097213.5 | 5269326 | 5445250 | 4041369.75 | 4302948.5 | 5732510.5 | 6317568 | 7933863 | 6325717 | 5522490.5 | 1677.896 | TQLGLLMDGFNSLLR | Q9EP69 | Q9EP69 | SACM1L | SAC1 suppressor of actin mutations 1-like (yeast) | Cytoplasm | phosphatase |
| 15549470 | 3 | 0.48 | 5479045.5 | 5372067 | 7325393 | 4852021 | 7091412 | 3649893.5 | 7637540 | 5882568.5 | 6320088 | 6766897.5 | 5564190 | 5664650.5 | 2630.2166 | GIDSEGHAANFVETEQIVHYSGNR | Q9EP69 | Q9EP69 | SACM1L | SAC1 suppressor of actin mutations 1-like (yeast) | Cytoplasm | phosphatase |
| 15535628 | 2 | 0.556 | 3369761.5 | 5558477.5 | 8432334 | 3766490.75 | 7253443.5 | 5497602.5 | 7695080 | 7919672 | 4961378 | 6529621 | 2896818 | 4104599 | 1158.565 | ATDFDVLSYK | Q9EP69 | Q9EP69 | SACM1L | SAC1 suppressor of actin mutations 1-like (yeast) | Cytoplasm | phosphatase |
| 15533361 | 3 | 0.006 | 4267045 | 3104695.5 | 5026705 | 4201170.5 | 4726115 | 4352101 | 6885528 | 6547048 | 6031433.5 | 6205615 | 6405087.5 | 5775974 | 2038.0066 | PLEQTFANMVSSLGSGMIR | Q9EP69 | Q9EP69 | SACM1L | SAC1 suppressor of actin mutations 1-like (yeast) | Cytoplasm | phosphatase |
| 15537575 | 3 | 0.289 | 3723630.5 | 3202003 | 4335117.5 | 3146090.75 | 5670750.5 | 3641175.25 | 5285070 | 4085869.5 | 4013653.25 | 5596316.5 | 4793045.5 | 4944127 | 1374.6351 | VGEC[160.0307]FNHAVWR | Q9EP69 | Q9EP69 | SACM1L | SAC1 suppressor of actin mutations 1-like (yeast) | Cytoplasm | phosphatase |
| 15535829 | 2 | 0.113 | 4581864 | 4211940 | 4373636 | 4933654.5 | 4552554.5 | 3844841.25 | 5748038.5 | 5335565 | 4404862 | 5338554.5 | 5328546 | 4998538 | 1306.6758 | HFDSQVIIYGK | Q9EP69 | Q9EP69 | SACM1L | SAC1 suppressor of actin mutations 1-like (yeast) | Cytoplasm | phosphatase |
| 15536816 | 2 | 0.949 | 4255028 | 3774775.5 | 4808390.5 | 5501420.5 | 4673795.5 | 3637277.75 | 5576638.5 | 4322981.5 | 3668742 | 5317704 | 4411587 | 4148842 | 1305.5604 | NAWADNANAC[160.0307]AK | Q9EP69 | Q9EP69 | SACM1L | SAC1 suppressor of actin mutations 1-like (yeast) | Cytoplasm | phosphatase |
| 15539031 | 2 | 0.791 | 3492139.75 | 3656363.25 | 3666456.25 | 3633939.5 | 3539156.75 | 3751108.75 | 3685214.5 | 3550590 | 3155594 | 4176444.75 | 3455973 | 3367898 | 943.53 | SLQAQLQR | Q9EP69 | Q9EP69 | SACM1L | SAC1 suppressor of actin mutations 1-like (yeast) | Cytoplasm | phosphatase |
| 15536018 | 2 | 0.287 | 2706377.75 | 2550295.25 | 3542580.5 | 2375966.5 | 2362638 | 2730204.25 | 3055534.5 | 2924236.5 | 3028882.25 | 3594879.5 | 2950147.5 | 2570152.5 | 950.57605 | LGVLHVGQK | Q9EP69 | Q9EP69 | SACM1L | SAC1 suppressor of actin mutations 1-like (yeast) | Cytoplasm | phosphatase |
| 15542465 | 2 | 0.445 | 2515927.75 | 2272883.75 | 3016707.25 | 1723596.125 | 2608028.5 | 2444937.75 | 2735024.25 | 2916062.5 | 2476519.75 | 3044045.75 | 2291871.5 | 2701722.75 | 1212.6385 | YFDWILISR | Q9EP69 | Q9EP69 | SACM1L | SAC1 suppressor of actin mutations 1-like (yeast) | Cytoplasm | phosphatase |
| 15544145 | 2 | 0.294 | 2005754.875 | 1780492.375 | 2012941.5 | 1961883.125 | 1957387 | 2043749.875 | 2985424.5 | 2193359.25 | 1631730.5 | 2817713.25 | 2650208.25 | 2136184.5 | 1099.6307 | RSLQAQLQR | Q9EP69 | Q9EP69 | SACM1L | SAC1 suppressor of actin mutations 1-like (yeast) | Cytoplasm | phosphatase |
| 15540000 | 2 | 0.15 | 2134725.25 | 1454687.375 | 1160776.625 | 1853391.125 | 2274905.5 | 2015381 | 1931206.375 | 2194187.75 | 1728848.125 | 2648360.5 | 2545387 | 1871480.5 | 1170.4309 | SNC[160.0307]MDC[160.0307]LDR | Q9EP69 | Q9EP69 | SACM1L | SAC1 suppressor of actin mutations 1-like (yeast) | Cytoplasm | phosphatase |
| 15533776 | 2 | 0.554 | 2794374.75 | 2104463.25 | 2785733.5 | 2221976.75 | 2978439.75 | 2313458.75 | 2370159.75 | 2442642.75 | 2120026.25 | 2545700.5 | 2870268.75 | 2623776.5 | 1165.5931 | ELSAQPEVHR | Q9EP69 | Q9EP69 | SACM1L | SAC1 suppressor of actin mutations 1-like (yeast) | Cytoplasm | phosphatase |
| 15543540 | 2 | 0.323 | 2006730.375 | 2372790 | 2322800.5 | 1436883.125 | 2409421.75 | 1660849.25 | 2401606 | 2619103.5 | 1757256.375 | 2252514.5 | 1259431.125 | 1575797.125 | 956.42004 | NNFSDGFR | Q9EP69 | Q9EP69 | SACM1L | SAC1 suppressor of actin mutations 1-like (yeast) | Cytoplasm | phosphatase |
| 15545866 | 3 | 0.647 | 204914.5 | 1005622.188 | 2021257.875 | 356223.938 | 985312.938 | 2135591 | 2066586.5 | 1193707 | 1666294 | 496036.031 | 455862.719 | 1707262.5 | 1833.9983 | RTQLGLLMDGFNSLLR | Q9EP69 | Q9EP69 | SACM1L | SAC1 suppressor of actin mutations 1-like (yeast) | Cytoplasm | phosphatase |
| 15536369 | 2 | 0.276 | 1231128.75 | 2807694 | 1391466.375 | 1386888.125 | 2127904.25 | 3229070.25 | 1694474.25 | 2207687.5 | 3023696.25 | 1318309.375 | 1292035.75 | 1335643 | 1096.5763 | FYLGGPTSVR | Q8BGH2 | Q8BGH2 | SAMM50 | SAMM50 sorting and assembly machinery component | Cytoplasm | other |
| 15544301 | 3 | 0.841 | 1994086.125 | 2344034 | 329531.719 | 3099982 | 1117195.75 | 1890879 | 1349682.5 | 1188022.75 | 3880052.5 | 907609.312 | 2910374.5 | 1856901 | 2104.009 | ETSYGLSFFKPQPGNFER | Q8BGH2 | Q8BGH2 | SAMM50 | SAMM50 sorting and assembly machinery component | Cytoplasm | other |
| 15543908 | 2 | 0.207 | 1.87E+07 | 1.35E+07 | 1.26E+07 | 1.39E+07 | 1.32E+07 | 1.07E+07 | 1.73E+07 | 1.24E+07 | 8287543.5 | 2.51E+07 | 1.78E+07 | 1.52E+07 | 1410.6777 | AQQEFATGVMSNK | Q8K021 | Q8K021 | SCAMP1 | secretory carrier membrane protein 1 | Cytoplasm | transporter |
| 15549451 | 2 | 0.039 | 2494007.5 | 3857819.25 | 4712066.5 | 8217590 | 7121869.5 | 7510368 | 5122369.5 | 6606403.5 | 7976300 | 2310915.25 | 5818387.5 | 4151448.5 | 2205.0164 | TGASFQQAQEEFSQGIFSSR | Q9ERN0 | Q9ERN0 | SCAMP2 | secretory carrier membrane protein 2 | Cytoplasm | transporter |
| 15537613 | 3 | 0.095 | 2152617 | 3440857.75 | 3703851.25 | 5449707.5 | 7378670.5 | 6581866 | 4187476.5 | 5653399.5 | 6727024 | 1923008.25 | 5315142 | 4820948 | 2205.0159 | TGASFQQAQEEFSQGIFSSR | Q9ERN0 | Q9ERN0 | SCAMP2 | secretory carrier membrane protein 2 | Cytoplasm | transporter |
| 15532799 | 2 | 0.097 | 1.10E+07 | 1.06E+07 | 1.12E+07 | 8875946 | 8480040 | 9138619 | 1.06E+07 | 1.04E+07 | 8269603 | 1.07E+07 | 8938398 | 9108230 | 1292.5984 | AQTEWSAGTWR | Q9JKV5 | Q9JKV5 | SCAMP4 | secretory carrier membrane protein 4 | Cytoplasm | other |
| 15551960 | 3 | 0.743 | 5573460 | 8840052 | 1.02E+07 | 4635136.5 | 1.08E+07 | 8246047 | 9258022 | 1.10E+07 | 5923805 | 9396277 | 4264584 | 5770647.5 | 2450.1619 | PC[160.0307]FYQDFSDEIPVEHQVLVK | Q9JKV5 | Q9JKV5 | SCAMP4 | secretory carrier membrane protein 4 | Cytoplasm | other |
| 15538605 | 3 | 0.179 | 2015221.25 | 5606405 | 2295889 | 1270428.5 | 2600042.25 | 4445639 | 1136375 | 3583750 | 2191144 | 671360.812 | 1510171.625 | 1091739.875 | 2082.0803 | EEHESFVDINPLTGIILR | O35114 | O35114 | SCARB2 | scavenger receptor class B, member 2 | Plasma Membrane | other |
| 15549109 | 3 | 0.316 | 8938484 | 1.82E+07 | 8316877.5 | 1.10E+07 | 1.42E+07 | 1.66E+07 | 8989923 | 1.62E+07 | 1.46E+07 | 8429917 | 7911210.5 | 1.01E+07 | 3813.7053 | GSDWLGDQDAIHYMTEQAPASVVELENYGMPFSR | Q8K2B3 | Q8K2B3 | SDHA | succinate dehydrogenase complex, subunit A, flavoprotein (Fp) | Cytoplasm | enzyme |

| | | | | | | | | | | | | | | | | | | | | | |
|---|---|---|---|---|---|---|---|---|---|---|---|---|---|---|---|---|---|---|---|---|---|
| 15553325 | 3 | 0.171 | 5081743 | 1.50E+07 | 8036364.5 | 5605102.5 | 1.22E+07 | 1.30E+07 | 5887150.5 | 1.32E+07 | 1.11E+07 | 4112828 | 4729945.5 | 4872753 | 3366.5645 | AGLPC[160.0307]QDLEFVQFHPTGIYGAGC[160.0307]LITEGC[160.0307]R | Q8K2B3 | Q8K2B3 | SDHA | succinate dehydrogenase complex, subunit A, flavoprotein (Fp) | Cytoplasm | enzyme |
| 15532748 | 2 | 0.367 | 4327427 | 9170396 | 4385847.5 | 6120840 | 6878308.5 | 9338156 | 4544732.5 | 8205447.5 | 8478058 | 2404010.5 | 6278583 | 4969988 | 1473.8402 | LGANSLLDLVVFGR | Q8K2B3 | Q8K2B3 | SDHA | succinate dehydrogenase complex, subunit A, flavoprotein (Fp) | Cytoplasm | enzyme |
| 15535658 | 2 | 0.215 | 3815974 | 8508955 | 3249714.5 | 4857469 | 5463276 | 6034409.5 | 3606714.75 | 5349046 | 6074072.5 | 1879509.5 | 3990136.75 | 3260405 | 1377.6283 | ANAGEESVMNLDK | Q8K2B3 | Q8K2B3 | SDHA | succinate dehydrogenase complex, subunit A, flavoprotein (Fp) | Cytoplasm | enzyme |
| 15534211 | 2 | 0.333 | 3276202.5 | 5186240 | 2074188.875 | 3707837.75 | 4481517 | 4931750.5 | 3253784 | 4855759 | 5258882.5 | 1856170.375 | 3940053 | 2965282.5 | 1205.5793 | VGSVLQEGC[160.0307]EK | Q8K2B3 | Q8K2B3 | SDHA | succinate dehydrogenase complex, subunit A, flavoprotein (Fp) | Cytoplasm | enzyme |
| 15552148 | 3 | 0.313 | 3671056 | 6184580.5 | 2582558.5 | 4573035.5 | 4218983 | 4946409 | 2270964 | 4726717 | 7199953.5 | 1629261.5 | 3552805.75 | 2491720 | 2250.9707 | TYFSC[160.0307]TSAHTSTGDGTAMVTR | Q8K2B3 | Q8K2B3 | SDHA | succinate dehydrogenase complex, subunit A, flavoprotein (Fp) | Cytoplasm | enzyme |
| 15549996 | 2 | 0.181 | 3271193.25 | 6778373.5 | 3809647.5 | 3962331 | 4817512.5 | 7159254 | 2444314.75 | 5118741 | 8247211.5 | 1592518.25 | 2589086.5 | 3129024.25 | 1757.8491 | AAFGLSEAGFNTAC[160.0307]LTK | Q8K2B3 | Q8K2B3 | SDHA | succinate dehydrogenase complex, subunit A, flavoprotein (Fp) | Cytoplasm | enzyme |
| 15535867 | 2 | 0.355 | 1337322.375 | 2871816.25 | 927255.438 | 2125842.25 | 1677645.75 | 2385862.25 | 812178.312 | 2008181.875 | 2174269.5 | 565367.938 | 1528495.5 | 1048145.625 | 1273.6019 | VRVDEYDYSK | Q8K2B3 | Q8K2B3 | SDHA | succinate dehydrogenase complex, subunit A, flavoprotein (Fp) | Cytoplasm | enzyme |
| 15533131 | 2 | 0.04 | 3390192.5 | 4236659.5 | 3296822 | 4052483.75 | 4582918 | 5821650 | 3320652.25 | 3713389.25 | 3531457.75 | 6449459 | 4797674.5 | 4358725 | 1279.622 | QQYLQSIEDR | Q9CQA3 | Q9CQA3 | SDHB | succinate dehydrogenase complex, subunit B, iron sulfur (Ip) | Cytoplasm | enzyme |
| 15535207 | 2 | 0.282 | 2850911.25 | 5333821.5 | 2082795.625 | 3438047.5 | 3085569.5 | 5236711 | 2715545.75 | 4469248.5 | 5924675.5 | 1685644 | 3007637.5 | 2382146 | 1124.571 | LQDPFSVYR | Q9CQA3 | Q9CQA3 | SDHB | succinate dehydrogenase complex, subunit B, iron sulfur (Ip) | Cytoplasm | enzyme |
| 15535390 | 2 | 0.368 | 2129128 | 3706804.25 | 1594605.5 | 2729262 | 2642012.25 | 3596961.25 | 1378679 | 3038317.5 | 3600030.25 | 934268.625 | 2165674.25 | 1967525.375 | 1240.5853 | MQTYEVDLNK | Q9CQA3 | Q9CQA3 | SDHB | succinate dehydrogenase complex, subunit B, iron sulfur (Ip) | Cytoplasm | enzyme |
| 15536578 | 2 | 0.246 | 1707936.375 | 4312412 | 1709156.5 | 2051191.875 | 2907629.5 | 4299752.5 | 1491520.25 | 3366406.5 | 3906549 | 1032864.75 | 1603757.875 | 1701245.25 | 981.5499 | HLLWDLGK | Q9CZB0 | Q9CZB0 | SDHC | succinate dehydrogenase complex, subunit C, integral membrane protein, 15kDa | Cytoplasm | enzyme |
| 15553902 | 3 | 0.531 | 5494713 | 4632162.5 | 3775817 | 1.07E+07 | 3639120.25 | 6417511.5 | 6769412.5 | 4693813.5 | 7177987.5 | 4840632.5 | 1.06E+07 | 7109689.5 | 3163.492 | VADGLPLAASMQEDEQSGRDLQQYQSQAK | O08547 | O08547 | SEC22B | SEC22 vesicle trafficking protein homolog B (S. cerevisiae) (gene/pseudogene) | Cytoplasm | other |
| 15549350 | 3 | 0.56 | 4371627 | 3260976.25 | 2878666.5 | 9339982 | 1604281 | 4740748 | 4115623.75 | 2126965.75 | 6085460 | 4593113.5 | 8532688 | 6617194 | 2386.2598 | IMVANIEEVLQRGEALSALDSK | O08547 | O08547 | SEC22B | SEC22 vesicle trafficking protein homolog B (S. cerevisiae) (gene/pseudogene) | Cytoplasm | other |
| 15533839 | 2 | 0.449 | 2980655.75 | 5324828 | 4995464.5 | 4590503.5 | 6723267 | 4739253 | 5227232.5 | 6050521 | 4373991 | 4213450 | 4113114.5 | 4303459.5 | 1414.7693 | IMVANIEEVLQR | O08547 | O08547 | SEC22B | SEC22 vesicle trafficking protein homolog B (S. cerevisiae) (gene/pseudogene) | Cytoplasm | other |
| 15556301 | 3 | 0.144 | 3528132.5 | 4965136.5 | 5680694 | 5051006.5 | 5439680.5 | 5390638 | 4620086 | 5177194 | 4398477 | 3690635.5 | 4491759 | 3402208.5 | 2249.0464 | LAFAYLEDLHSEFDEQHGK | O08547 | O08547 | SEC22B | SEC22 vesicle trafficking protein homolog B (S. cerevisiae) (gene/pseudogene) | Cytoplasm | other |
| 15537774 | 2 | 0.997 | 2044749.5 | 2987813.75 | 2513735.25 | 2633198 | 2603387.25 | 2184493.25 | 2074072.875 | 2899461.75 | 2497834.75 | 3023710.75 | 2278862.75 | 2051578.875 | 990.5094 | GEALSALDSK | O08547 | O08547 | SEC22B | SEC22 vesicle trafficking protein homolog B (S. cerevisiae) (gene/pseudogene) | Cytoplasm | other |
| 15550625 | 3 | 0.085 | 1249768.25 | 2948343 | 2768454.5 | 2827070.5 | 3707162 | 3620492.5 | 2641069.25 | 3139493.5 | 2327773.75 | 1873715 | 1767193.25 | 1752619.25 | 2274.1755 | VPTVSRPYSFIEFDTFIQK | O08547 | O08547 | SEC22B | SEC22 vesicle trafficking protein homolog B (S. cerevisiae) (gene/pseudogene) | Cytoplasm | other |
| 15538685 | 2 | 0.206 | 1942787 | 2042818.875 | 1413181.5 | 2508006.5 | 2016801.125 | 2625901.75 | 2027924.5 | 1870474.25 | 2262826 | 1865090.25 | 2833661 | 2422054.75 | 916.56323 | VLLTMIAR | O08547 | O08547 | SEC22B | SEC22 vesicle trafficking protein homolog B (S. cerevisiae) (gene/pseudogene) | Cytoplasm | other |
| 15560173 | 2 | 0.79 | 1168631.75 | 1923560.75 | 1692128 | 839339 | 2648187.5 | 1776998.125 | 1504695 | 1719166.75 | 1192078.25 | 1169714 | 1284662.5 | 1250621.625 | 1973.9183 | VADGLPLAASMQEDEQSGR | O08547 | O08547 | SEC22B | SEC22 vesicle trafficking protein homolog B (S. cerevisiae) (gene/pseudogene) | Cytoplasm | other |
| 15531564 | 2 | 0.139 | 2580390.5 | 6370183 | 1.11E+07 | 2.85E+07 | 2.78E+07 | 2.92E+07 | 1.55E+07 | 2.00E+07 | 2.87E+07 | 2170252 | 3.02E+07 | 1.72E+07 | 1156.6041 | VSPASQGNLQR | Q3TZ89 | Q3TZ89 | SEC31B | SEC31 homolog B, COPII coating complex component | Cytoplasm | other |
| 15535523 | 2 | 0.059 | 1219127.25 | 1452672.875 | 1923271 | 3385534 | 3710562 | 3458428.25 | 1398913.5 | 2845068 | 3699629 | 1336641.25 | 2253100.75 | 2427252.75 | 971.5384 | IIEVGDTPK | P61620 | P61620 | SEC61A1 | Sec61 alpha 1 subunit (S. cerevisiae) | Cytoplasm | transporter |
| 15533796 | 2 | 0.036 | 1982029.875 | 3722720.75 | 3859749 | 6388044 | 8535371 | 6621065.5 | 4226295 | 5851328 | 6482125 | 2610136.25 | 5299796.5 | 4403483.5 | 1370.6627 | PGPTPSGTNVGSSGR | Q9CQS8 | Q9CQS8 | SEC61B | Sec61 beta subunit | Cytoplasm | transporter |
| 15539878 | 2 | 0.027 | 971767.875 | 1553953.875 | 2080427.625 | 2895015.5 | 4414535 | 3347356.75 | 2238666.25 | 2420477 | 3036961.75 | 1216587.625 | 2125141 | 2276430 | 1156.5493 | FYTEDSPGLK | Q9CQS8 | Q9CQS8 | SEC61B | Sec61 beta subunit | Cytoplasm | transporter |
| 15554153 | 3 | 0.021 | 552309.25 | 316153.469 | 270189.438 | 1845548 | 489758.688 | 967821.438 | 170060.016 | 221247.328 | 373456.125 | 799590.25 | 3169003.75 | 909399.25 | 2405.1619 | FLHDPSATQGFVGC[160.0307]ALSSNIQR | P17563 | P17563 | SELENBP1 | selenium binding protein 1 | Cytoplasm | other |
| 15540472 | 2 | 0.015 | 414138.719 | 655733.75 | 1612331.75 | 4290287.5 | 3831943.75 | 5108532.5 | 3311956.25 | 3121441.75 | 4054834.25 | 924869.938 | 4190201 | 2551689.75 | 1056.4552 | FEC[160.0307]QPGYR | Q01102 | Q01102 | SELP | selectin P (granule membrane protein 140kDa, antigen CD62) | Plasma Membrane | transmembrane receptor |
| 15533040 | 3 | 0.191 | 1271370.125 | 2818180 | 5939242 | 1.41E+07 | 1.90E+07 | 2.09E+07 | 1.26E+07 | 1.19E+07 | 1.46E+07 | 445077.062 | 1.73E+07 | 1.09E+07 | 2213.9856 | LDC[160.0307]SPSGHWTGTPPTC[160.0307]EAIK | Q01102 | Q01102 | SELP | selectin P (granule membrane protein 140kDa, antigen CD62) | Plasma Membrane | transmembrane receptor |
| 15550126 | 3 | 0.165 | 336238.75 | 1758944 | 3768833.75 | 1.12E+07 | 1.21E+07 | 1.26E+07 | 6997192 | 9831946 | 8910430 | 290464.156 | 1.06E+07 | 8097630 | 3648.6116 | GNDAIQC[160.0307]ADSGQWTAPAPFC[160.0307]EALQC[160.0307]PEFPVPSK | Q01102 | Q01102 | SELP | selectin P (granule membrane protein 140kDa, antigen CD62) | Plasma Membrane | transmembrane receptor |
| 15546559 | 2 | 0.059 | 80319.367 | 262200.125 | 702325.062 | 1914582.5 | 1834187.25 | 2160133 | 1216340.375 | 1361860.5 | 1646485.125 | 181035.578 | 1851835.5 | 1047256.75 | 1043.495 | WSAPPPTC[160.0307]K | Q01102 | Q01102 | SELP | selectin P (granule membrane protein 140kDa, antigen CD62) | Plasma Membrane | transmembrane receptor |
| 15554515 | 3 | 0.211 | 229344.672 | 1348290.375 | 1736589.25 | 3253821.5 | 4231435 | 5299419.5 | 2900269 | 3862231 | 4077898.75 | 133712.844 | 2901387.75 | 2941485.75 | 2681.3442 | NEIAHLNDVIPFFNSYYWIGIR | Q01102 | Q01102 | SELP | selectin P (granule membrane protein 140kDa, antigen CD62) | Plasma Membrane | transmembrane receptor |
| 15545848 | 2 | 0.106 | 4.70E-06 | 20517.076 | 87772.484 | 580034 | 1483228.875 | 1253307.625 | 681425 | 654780.562 | 939368.688 | 4.70E-06 | 1009688.188 | 1132056.375 | 1464.6655 | ASGQWTAVTPMC[160.0307]R | Q01102 | Q01102 | SELP | selectin P (granule membrane protein 140kDa, antigen CD62) | Plasma Membrane | transmembrane receptor |
| 15551006 | 2 | 0.947 | 2121722 | 3067416.5 | 4178145.25 | 2510359.5 | 4114979.75 | 3303024.5 | 3101478.75 | 3359492 | 2834916.75 | 2577790.25 | 3808008 | 2457886.25 | 1591.806 | IEGENYLPQPIYR | P62342 | P62342 | SELT | selenoprotein T | Cytoplasm | other |
| 15537589 | 2 | 0.163 | 1141999.5 | 2640711 | 852510.062 | 1308044.125 | 1241915.875 | 1472231 | 636843.812 | 1068359.875 | 1151568.75 | 1117366.125 | 503605.781 | 479980.688 | 1239.6917 | KEETQPPVALK | Q9CY58 | Q9CY58 | SERBP1 | SERPINE1 mRNA binding protein 1 | Cytoplasm | other |
| 15540089 | 3 | 0.067 | 1180073.25 | 3966091.75 | 869430.938 | 1038839.562 | 3125274.75 | 2119011 | 1161560.5 | 2734101.5 | 1707738.125 | 675587.562 | 101691.32 | 588871 | 1670.8016 | PGHLQEGFGC[160.0307]VVTNR | Q9CY58 | Q9CY58 | SERBP1 | SERPINE1 mRNA binding protein 1 | Cytoplasm | other |
| 15558393 | 2 | 0.092 | 565352 | 3004775 | 1139745 | 702951.625 | 2254752 | 1531411.25 | 790015.875 | 1989956.625 | 1052898.125 | 633420.375 | 94677.414 | 362578.875 | 1943.885 | FDQLFDDESDPFEVLK | Q9CY58 | Q9CY58 | SERBP1 | SERPINE1 mRNA binding protein 1 | Cytoplasm | other |
| 15536877, 15536925 | 2, 3 | 0.059 | 3.44E+08 | 3.00E+08 | 2.19E+08 | 2.78E+08 | 2.49E+08 | 2.67E+08 | 2.52E+08 | 2.16E+08 | 2.25E+08 | 3.38E+08 | 3.08E+08 | 3.31E+08 | 1480.7596, 1480.7628 | KIPGFC[160.0307]EGGFQIK | Q9QZI9 | Q9QZI9 | SERINC3 | serine incorporator 3 | Cytoplasm | transporter |
| 15538366 | 2 | 0.2 | 1.47E+08 | 1.65E+08 | 1.78E+08 | 1.21E+08 | 1.48E+08 | 1.19E+08 | 1.32E+08 | 1.39E+08 | 1.09E+08 | 1.63E+08 | 1.11E+08 | 1.53E+08 | 1052.5061 | DC[160.0307]DVLVGFK | Q9QZI9 | Q9QZI9 | SERINC3 | serine incorporator 3 | Cytoplasm | transporter |
| 15533543 | 2 | 0.224 | 1.23E+08 | 1.79E+08 | 2.21E+08 | 1.09E+08 | 1.67E+08 | 1.58E+08 | 1.22E+08 | 1.39E+08 | 8.49E+07 | 1.38E+08 | 1.16E+08 | 1.11E+08 | 1352.6632 | IPGFC[160.0307]EGGFQIK | Q9QZI9 | Q9QZI9 | SERINC3 | serine incorporator 3 | Cytoplasm | transporter |
| 15532417 | 2 | 0.665 | 2.87E+07 | 4.57E+07 | 7.05E+07 | 1.98E+07 | 4.54E+07 | 4.60E+07 | 3.81E+07 | 3.75E+07 | 2.81E+07 | 3.94E+07 | 2.24E+07 | 3.48E+07 | 949.52747 | WLAVWFK | Q9QZI9 | Q9QZI9 | SERINC3 | serine incorporator 3 | Cytoplasm | transporter |
| 15548958 | 3 | 0.047 | 1.26E+07 | 1.47E+07 | 2.67E+07 | 6082473.5 | 6539182 | 8455595 | 9331446 | 1.01E+07 | 2.02E+07 | 1.18E+07 | 1.20E+07 | 1.76E+07 | 2405.1697 | DQSPASHEIATNLGDFAISLYR | P07758 | P07758 | SERPINA1 | serpin peptidase inhibitor, clade A (alpha-1 antiproteinase, antitrypsin), member 1 | Extracellular Space | other |
| 15544006 | 3 | 0.072 | 3521465.5 | 5998307 | 6830163.5 | 1898297.375 | 1776959.5 | 3749687.5 | 4749422 | 3549751.25 | 7202540 | 3885670.75 | 3420469.5 | 5499326 | 1999.0859 | KLDQDTVFALANYILFK | P07758 | P07758 | SERPINA1 | serpin peptidase inhibitor, clade A (alpha-1 antiproteinase, antitrypsin), member 1 | Extracellular Space | other |
| 15554964 | 4 | 0.295 | 2772409 | 3433269.5 | 6563119 | 2205641.5 | 2305887 | 2562615.25 | 2877501.5 | 2926279.5 | 5557472.5 | 1849933.375 | 2725789.75 | 3902896.75 | 3498.7952 | SFQHLLQTLNRPDSELQLSTGNGLFVNNDLK | P07758 | P07758 | SERPINA1 | serpin peptidase inhibitor, clade A (alpha-1 antiproteinase, antitrypsin), member 1 | Extracellular Space | other |
| 15538916 | 2 | 0.036 | 2685937.25 | 3285105.75 | 6390001 | 1666126.875 | 1215576 | 1932670.75 | 2903226 | 2363235 | 4045918 | 1788388.75 | 2255784.5 | 2745287.75 | 1088.6108 | TLMSPLGITR | P07758 | P07758 | SERPINA1 | serpin peptidase inhibitor, clade A (alpha-1 antiproteinase, antitrypsin), member 1 | Extracellular Space | other |
| 15551294 | 2 | 0.31 | 2430443.25 | 2698388.5 | 4831543 | 1752973.5 | 1176314.875 | 2447959.75 | 2245516.25 | 1819203 | 4589743.5 | 1536066.25 | 2626626.75 | 2501110.75 | 2003.0043 | IFNNGADLSGITEENAPLK | P07758 | P07758 | SERPINA1 | serpin peptidase inhibitor, clade A (alpha-1 antiproteinase, antitrypsin), member 1 | Extracellular Space | other |
| 15548257 | 3 | 0.73 | 2.11E+07 | 1.92E+07 | 2.53E+07 | 2.03E+07 | 2.05E+07 | 1.95E+07 | 2.69E+07 | 2.31E+07 | 2.55E+07 | 3.67E+07 | 1.68E+07 | 1.65E+07 | 2270.2295 | AVLDVAETGTEAAAATGVIGGIRK | P07759 | P07759 | SERPINA3 | serpin peptidase inhibitor, clade A (alpha-1 antiproteinase, antitrypsin), member 3 | Extracellular Space | other |
| 15540020, 15545121 | 3 | 0.781 | 1.13E+07 | 1.08E+07 | 1.54E+07 | 1.63E+07 | 1.50E+07 | 1.29E+07 | 1.72E+07 | 1.94E+07 | 1.51E+07 | 3.35E+07 | 1.03E+07 | 8893996 | 2142.1353, 2142.1357 | AVLDVAETGTEAAAATGVIGGIR | P07759 | P07759 | SERPINA3 | serpin peptidase inhibitor, clade A (alpha-1 antiproteinase, antitrypsin), member 3 | Extracellular Space | other |
| 15556779 | 2 | 0.963 | 7809600.5 | 4831831.5 | 7936456 | 1.21E+07 | 6817383 | 6653267 | 1.02E+07 | 7322867 | 6987122 | 2.28E+07 | 5303570.5 | 3943760.75 | 2142.1355 | AVLDVAETGTEAAAATGVIGGIR | P07759 | P07759 | SERPINA3 | serpin peptidase inhibitor, clade A (alpha-1 antiproteinase, antitrypsin), member 3 | Extracellular Space | other |
| 15548599 | 2 | 0.106 | 2936958.25 | 6583920 | 3684288 | 5784730 | 5258456 | 7787154 | 4074154.5 | 5712111 | 7730208.5 | 2362287.5 | 3818222.25 | 3607956.25 | 1682.8823 | QGIVPAGLTENELWR | Q99JR1 | Q99JR1 | SFXN1 | sideroflexin 1 | Cytoplasm | transporter |
| 15539090 | 2 | 0.325 | 3466535.5 | 6401641.5 | 3642000.75 | 6075591 | 5457623 | 8173566 | 3467197.5 | 5161288 | 6942731.5 | 2177686 | 5418732 | 4264934 | 1114.6555 | QAITQVVISR | Q99JR1 | Q99JR1 | SFXN1 | sideroflexin 1 | Cytoplasm | transporter |
| 15535855 | 3 | 0.186 | 3007372 | 7029210 | 3548696 | 4897163.5 | 7696445.5 | 8178090 | 3788711.75 | 5411302 | 7083314 | 1702342.75 | 5265043.5 | 2993133.75 | 1600.6881 | YAYDSAFHPDTGEK | Q99JR1 | Q99JR1 | SFXN1 | sideroflexin 1 | Cytoplasm | transporter |
| 15538419 | 2 | 0.252 | 2717230.25 | 5036416 | 2073720.375 | 4420846.5 | 3822552 | 5203014.5 | 3093519.75 | 4364429.5 | 4312130 | 1617688.875 | 3743342 | 2711996 | 1257.6405 | VGIPVTDENGTR | Q99JR1 | Q99JR1 | SFXN1 | sideroflexin 1 | Cytoplasm | transporter |
| 15542630 | 2 | 0.229 | 1874009.125 | 3995841.75 | 1948850.875 | 3557490 | 3063372 | 4260060.5 | 2355424.75 | 3867559.25 | 4484922.5 | 988693.375 | 2940055.25 | 2239088.5 | 1109.5353 | WDQSTFIGR | Q99JR1 | Q99JR1 | SFXN1 | sideroflexin 1 | Cytoplasm | transporter |
| 15543276 | 3 | 0.142 | 546081.812 | 1164749.875 | 378700.656 | 1254969.5 | 1345987.5 | 1708456.125 | 1158472.375 | 1761461.875 | 2627835.75 | 358713.031 | 1909490.375 | 520885.562 | 2075.0903 | FVPFAAVAAANC[160.0307]INIPLMR | Q99JR1 | Q99JR1 | SFXN1 | sideroflexin 1 | Cytoplasm | transporter |
| 15539905 | 2 | 0.093 | 1.45E+07 | 1.30E+07 | 1.77E+07 | 1.58E+07 | 1.74E+07 | 1.57E+07 | 1.45E+07 | 1.23E+07 | 1.05E+07 | 1.50E+07 | 9882598 | 9546172 | 1464.7119 | QQDVLC[160.0307]FLEANK | Q9JJU8 | Q9JJU8 | SH3BGRL | SH3 domain binding glutamate-rich protein like | Cytoplasm | other |
| 15548269 | 2 | 0.045 | 1.08E+07 | 1.01E+07 | 1.06E+07 | 1.23E+07 | 7911682 | 8656154 | 7078737 | 8337242.5 | 8040206.5 | 8509621 | 6580083 | 6437538 | 1848.9452 | ENNAVYAFLGLTAPPGSK | Q9JJU8 | Q9JJU8 | SH3BGRL | SH3 domain binding glutamate-rich protein like | Cytoplasm | other |
| 15540848 | 2 | 0.114 | 8077734.5 | 1.22E+07 | 1.49E+07 | 1.15E+07 | 1.05E+07 | 1.07E+07 | 6794538.5 | 1.10E+07 | 1.07E+07 | 7465450 | 6076608.5 | 8026499 | 1283.6815 | VYIASSSGSTAIK | Q9JJU8 | Q9JJU8 | SH3BGRL | SH3 domain binding glutamate-rich protein like | Cytoplasm | other |
| 15535298 | 2 | 0.474 | 1627307 | 2333257.25 | 4362589 | 2345647.25 | 2768708 | 4296632.5 | 3941489.75 | 3718727.25 | 3714084 | 1004238.312 | 2926818.5 | 3329671.5 | 1341.7023 | ESNTVFSFLGLK | Q8BG73 | Q8BG73 | SH3BGRL2 | SH3 domain binding glutamate-rich protein like 2 | Nucleus | other |
| 15535056 | 2 | 0.376 | 5216044 | 5735919 | 6324295.5 | 3427575 | 5269841 | 5441905 | 5146995.5 | 5294559.5 | 4245420.5 | 4962292 | 4877868.5 | 4808083.5 | 1199.6389 | DSIVATNNPIR | Q91Y57 | Q91Y57 | SIGLEC9 | sialic acid binding Ig-like lectin 9 | Plasma Membrane | other |
| 15542639 | 2 | 0.284 | 155679.656 | 626118.812 | 1018823 | 1971408.25 | 1861083.125 | 1631692.25 | 1317143.625 | 1075448.875 | 3125053.75 | 95088.523 | 1630158.875 | 1409242.375 | 1241.577 | GEFAIDGYDVR | Q3UND0 | Q3UND0 | SKAP2 | src kinase associated phosphoprotein 2 | Cytoplasm | other |
| 15543072 | 2 | 0.484 | 97331.039 | 416332.531 | 950704.562 | 1201027.375 | 1700947.75 | 2149601 | 1441862.375 | 863661.75 | 3497400.5 | 4.70E-06 | 1734815.375 | 1727130.375 | 1414.5409 | DC[160.0307]C[160.0307]FEIC[160.0307]APDK | Q3UND0 | Q3UND0 | SKAP2 | src kinase associated phosphoprotein 2 | Cytoplasm | other |
| 15539160 | 3 | 0.172 | 70970.273 | 1088238.25 | 2012305.25 | 4283933.5 | 5782313 | 4727357.5 | 2569642.25 | 3832148.5 | 4563246 | 323918.062 | 5012791.5 | 2922227 | 1879.8207 | GSYPDHLEDGYHFQSK | Q9QUM4 | Q9QUM4 | SLAMF1 | signaling lymphocytic activation molecule family member 1 | Plasma Membrane | transmembrane receptor |
| 15534399 | 2 | 0.17 | 317099.156 | 977972.062 | 1401461.625 | 4343980.5 | 3523134.75 | 4575755 | 2140733 | 2570048.25 | 4045109 | 257002.922 | 4478447 | 2803544.75 | 1204.6587 | ILVTMATSPGSK | Q9QUM4 | Q9QUM4 | SLAMF1 | signaling lymphocytic activation molecule family member 1 | Plasma Membrane | transmembrane receptor |
| 15535012 | 3 | 0.192 | 263813.75 | 1261663.625 | 3539986.25 | 8222409 | 8071798.5 | 8847645 | 5258345.5 | 6028361 | 6199445 | 242449.75 | 8082699 | 5019680 | 2007.0238 | LGQDTWLPLTNEHQINK | Q9QUM4 | Q9QUM4 | SLAMF1 | signaling lymphocytic activation molecule family member 1 | Plasma Membrane | transmembrane receptor |
| 15549706 | 4 | 0.089 | 1.02E+07 | 1.51E+07 | 1.10E+07 | 7576084.5 | 9894036 | 8974246 | 9331650 | 1.14E+07 | 1.11E+07 | 9149633 | 7580611 | 9609171 | 3002.5918 | TPQVEQAVGIVGAVIMPHNMYLHSALVK | P49282 | P49282 | SLC11A2 | solute carrier family 11 (proton-coupled divalent metal ion transporter), member 2 | Plasma Membrane | transporter |
| 15549977 | 2 | 0.11 | 5507955 | 7478174.5 | 6832490.5 | 3820476.75 | 6031779 | 4329674.5 | 3914093.5 | 5970712 | 7041618 | 4957790.5 | 2772389 | 3671420.25 | 1773.8109 | IPIPEEEYSC[160.0307]FSFR | P49282 | P49282 | SLC11A2 | solute carrier family 11 (proton-coupled divalent metal ion transporter), member 2 | Plasma Membrane | transporter |
| 15552250 | 4 | 0.106 | 3773523.75 | 1.11E+07 | 3422502.75 | 7914909.5 | 7176875 | 7291759 | 3750292.5 | 6847216.5 | 6635320 | 3655913.25 | 2369435.75 | 3210637.75 | 4774.0205 | MPDDGASGDHGDSASLGAINPAYSNSSLPHSTGDSEEPFTTYFDEK | P49282 | P49282 | SLC11A2 | solute carrier family 11 (proton-coupled divalent metal ion transporter), member 2 | Plasma Membrane | transporter |
| 15550804 | 2 | 0.101 | 2656850.5 | 5024283 | 2847128 | 1879278.25 | 4204985.5 | 3039513 | 1315429.125 | 3853752.75 | 3878823.5 | 1371235.625 | 872912.375 | 1689967 | 2474.3118 | LGLTAQPELYLLNTVDADSVVSR | P49282 | P49282 | SLC11A2 | solute carrier family 11 (proton-coupled divalent metal ion transporter), member 2 | Plasma Membrane | transporter |
| 15531784, 15547201 | 2 | 0.048 | 3.24E+08 | 4.16E+08 | 4.92E+08 | 2.01E+08 | 3.07E+08 | 2.93E+08 | 4.04E+08 | 4.18E+08 | 3.39E+08 | 3.26E+08 | 1.97E+08 | 2.58E+08 | 925.5121, 925.5141 | IFYLQNK | Q8VHL0 | Q8VHL0 | SLC14A1 | solute carrier family 14 (urea transporter), member 1 (Kidd blood group) | Plasma Membrane | transporter |
| 15547727 | 2, 3 | 0.045 | 1.92E+08 | 2.68E+08 | 2.39E+08 | 1.29E+08 | 1.70E+08 | 1.57E+08 | 1.80E+08 | 2.36E+08 | 2.83E+08 | 1.97E+08 | 1.52E+08 | 1.24E+08 | 2544.3914, 2544.3926 | LFTPVSSVPNITWSELSALELLK | Q8VHL0 | Q8VHL0 | SLC14A1 | solute carrier family 14 (urea transporter), member 1 (Kidd blood group) | Plasma Membrane | transporter |
| 15531277 | 2, 3 | 0.008 | 2.36E+08 | 1.87E+08 | 1.94E+08 | 1.38E+08 | 1.40E+08 | 1.09E+08 | 2.28E+08 | 2.18E+08 | 1.83E+08 | 1.76E+08 | 1.88E+08 | 1.39E+08 | 1446.7087, 1446.7102 | VDRGENQILSC[160.0307]R | Q8VHL0 | Q8VHL0 | SLC14A1 | solute carrier family 14 (urea transporter), member 1 (Kidd blood group) | Plasma Membrane | transporter |
| 15538962 | 2 | 0.137 | 1.44E+08 | 1.93E+08 | 2.78E+08 | 9.14E+07 | 1.49E+08 | 1.73E+08 | 2.07E+08 | 2.17E+08 | 1.74E+08 | 1.52E+08 | 1.15E+08 | 1.37E+08 | 907.4657 | EFANWLK | Q8VHL0 | Q8VHL0 | SLC14A1 | solute carrier family 14 (urea transporter), member 1 (Kidd blood group) | Plasma Membrane | transporter |
| 15531422 | 2 | 0.002 | 1.67E+08 | 1.70E+08 | 1.87E+08 | 1.19E+08 | 1.22E+08 | 1.14E+08 | 1.55E+08 | 1.56E+08 | 1.45E+08 | 1.49E+08 | 1.13E+08 | 1.37E+08 | 831.4387 | RMVESPL | Q8VHL0 | Q8VHL0 | SLC14A1 | solute carrier family 14 (urea transporter), member 1 (Kidd blood group) | Plasma Membrane | transporter |
| 15534289 | 2 | 0.085 | 1.12E+08 | 1.36E+08 | 1.94E+08 | 8.93E+07 | 1.01E+08 | 1.15E+08 | 1.67E+08 | 1.52E+08 | 1.26E+08 | 1.29E+08 | 1.08E+08 | 1.08E+08 | 1082.5537 | VLGYVTGDMK | Q8VHL0 | Q8VHL0 | SLC14A1 | solute carrier family 14 (urea transporter), member 1 (Kidd blood group) | Plasma Membrane | transporter |
| 15531529 | 3 | 0.532 | 2.37E+08 | 1.00E+08 | 7.94E+07 | 2.22E+08 | 2.34E+07 | 8.91E+07 | 1.47E+08 | 5.68E+07 | 1.56E+08 | 1.24E+08 | 2.79E+08 | 2.00E+08 | 1903.9507 | VTYSEENRIFYLQNK | Q8VHL0 | Q8VHL0 | SLC14A1 | solute carrier family 14 (urea transporter), member 1 (Kidd blood group) | Plasma Membrane | transporter |
| 15538805, 15546213 | 2 | 0.009 | 4.81E+07 | 4.28E+07 | 6.05E+07 | 2.70E+07 | 3.37E+07 | 3.42E+07 | 4.17E+07 | 4.37E+07 | 3.89E+07 | 3.65E+07 | 2.46E+07 | 2.59E+07 | 1098.5464, 1098.5493 | VLGYVTGDM[147.0354]K | Q8VHL0 | Q8VHL0 | SLC14A1 | solute carrier family 14 (urea transporter), member 1 (Kidd blood group) | Plasma Membrane | transporter |
| 15554953, 15558491 | 2 | 0.324 | 4.40E+07 | 1.55E+07 | 1.28E+07 | 3.81E+07 | 2679587.5 | 6066928 | 2.20E+07 | 7099545.5 | 2.31E+07 | 2.52E+07 | 4.66E+07 | 3.04E+07 | 1903.9504, 1903.9507 | VTYSEENRIFYLQNK | Q8VHL0 | Q8VHL0 | SLC14A1 | solute carrier family 14 (urea transporter), member 1 (Kidd blood group) | Plasma Membrane | transporter |
| 15541354 | 2 | 0.137 | 2.57E+07 | 2.27E+07 | 2.84E+07 | 1.14E+07 | 2.58E+07 | 1.37E+07 | 2.32E+07 | 3.10E+07 | 2.01E+07 | 2.22E+07 | 1.38E+07 | 1.31E+07 | 1076.513 | GENQILSC[160.0307]R | Q8VHL0 | Q8VHL0 | SLC14A1 | solute carrier family 14 (urea transporter), member 1 (Kidd blood group) | Plasma Membrane | transporter |
| 15540064, 15540778, 15544446 | 3 | 0.15 | 2.47E+07 | 3.36E+07 | 5.33E+07 | 1.09E+07 | 3.20E+07 | 3.07E+07 | 3.41E+07 | 4.53E+07 | 2.59E+07 | 1.97E+07 | 1.04E+07 | 1.99E+07 | 1775.9468, 1775.9475, 1775.948 | DKPVVLQFM[147.0354]DWILR | Q8VHL0 | Q8VHL0 | SLC14A1 | solute carrier family 14 (urea transporter), member 1 (Kidd blood group) | Plasma Membrane | transporter |
| 15535611 | 2 | 0.492 | 1.26E+07 | 1397042.875 | 53888.582 | 6540539 | 369657.219 | 409503.969 | 1713555.625 | 1321509.875 | 6535732.5 | 4701699 | 1.02E+07 | 5758068 | 1053.6068 | IFYLQNKK | Q8VHL0 | Q8VHL0 | SLC14A1 | solute carrier family 14 (urea transporter), member 1 (Kidd blood group) | Plasma Membrane | transporter |
| 15549087 | 2 | 0.832 | 5.31E+08 | 4.86E+08 | 5.89E+08 | 6.41E+08 | 5.59E+08 | 4.65E+08 | 5.90E+08 | 4.90E+08 | 4.69E+08 | 7.78E+08 | 5.66E+08 | 4.54E+08 | 2080.9026 | AAQSPQQHSSGDPTEEESPV | P53986 | P53986 | SLC16A1 | solute carrier family 16 (monocarboxylate transporter), member 1 | Plasma Membrane | transporter |
| 15534086, 15537037 | 2 | 0.452 | 5.03E+08 | 4.53E+08 | 4.75E+08 | 5.28E+08 | 5.48E+08 | 4.09E+08 | 4.84E+08 | 5.01E+08 | 4.14E+08 | 6.53E+08 | 4.88E+08 | 5.13E+08 | 1274.6201, 1274.6218 | SDANTDLIGGSPK | P53986 | P53986 | SLC16A1 | solute carrier family 16 (monocarboxylate transporter), member 1 | Plasma Membrane | transporter |
| 15534736 | 2 | 0.576 | 4.00E+08 | 5.18E+08 | 6.88E+08 | 3.85E+08 | 4.76E+08 | 3.66E+08 | 5.01E+08 | 5.59E+08 | 3.89E+08 | 6.08E+08 | 3.78E+08 | 4.65E+08 | 1049.5967 | LSVFQTINK | P53986 | P53986 | SLC16A1 | solute carrier family 16 (monocarboxylate transporter), member 1 | Plasma Membrane | transporter |
| 15549676, 15552306 | 2, 3 | 0.002 | 1.75E+08 | 1.80E+08 | 1.90E+08 | 1.68E+08 | 1.97E+08 | 2.15E+08 | 2.32E+08 | 2.30E+08 | 2.32E+08 | 3.03E+08 | 2.75E+08 | 2.42E+08 | 2654.4363, 2654.4392 | FSSAVGLVTIVEC[160.0307]C[160.0307]PVLLGPPLLGR | P53986 | P53986 | SLC16A1 | solute carrier family 16 (monocarboxylate transporter), member 1 | Plasma Membrane | transporter |
| 15536046 | 3 | 0.351 | 1.36E+08 | 1.44E+08 | 1.50E+08 | 1.46E+08 | 1.43E+08 | 1.24E+08 | 1.41E+08 | 1.31E+08 | 1.50E+08 | 2.34E+08 | 1.28E+08 | 1.71E+08 | 2080.9023 | AAQSPQQHSSGDPTEEESPV | P53986 | P53986 | SLC16A1 | solute carrier family 16 (monocarboxylate transporter), member 1 | Plasma Membrane | transporter |
| 15531383 | 2, 3 | 0.808 | 7.17E+07 | 8.66E+07 | 1.03E+08 | 5.88E+07 | 9.71E+07 | 8.58E+07 | 8.33E+07 | 9.04E+07 | 8.29E+07 | 9.48E+07 | 6.22E+07 | 7.35E+07 | 1248.6713, 1248.6714 | FLDLSLFTHR | P53986 | P53986 | SLC16A1 | solute carrier family 16 (monocarboxylate transporter), member 1 | Plasma Membrane | transporter |
| 15531813 | 2 | 0.981 | 2.33E+07 | 2.93E+07 | 3.89E+07 | 2.18E+07 | 3.22E+07 | 3.14E+07 | 3.54E+07 | 2.87E+07 | 2.15E+07 | 3.62E+07 | 2.74E+07 | 2.52E+07 | 1118.4801 | LNDMYGDYK | P53986 | P53986 | SLC16A1 | solute carrier family 16 (monocarboxylate transporter), member 1 | Plasma Membrane | transporter |
| 15531600 | 2 | 0.223 | 3.19E+07 | 5.15E+07 | 8.36E+07 | 3.85E+07 | 5.13E+07 | 6.12E+07 | 5.20E+07 | 4.66E+07 | 3.80E+07 | 3.35E+07 | 2.72E+07 | 3.46E+07 | 841.4804 | SITVFFK | P53986 | P53986 | SLC16A1 | solute carrier family 16 (monocarboxylate transporter), member 1 | Plasma Membrane | transporter |
| 15539399 | 4 | 0.243 | 6406471.5 | 4497225.5 | 7562933.5 | 2741919 | 8019418.5 | 7998795.5 | 9808871 | 9902232 | 8696731 | 7172651 | 1.03E+07 | 9292629 | 2654.4365 | FSSAVGLVTIVEC[160.0307]C[160.0307]PVLLGPPLLGR | P53986 | P53986 | SLC16A1 | solute carrier family 16 (monocarboxylate transporter), member 1 | Plasma Membrane | transporter |
| 15547688 | 2 | 0.978 | 1.53E+08 | 1.78E+08 | 1.82E+08 | 1.71E+08 | 1.81E+08 | 1.82E+08 | 1.99E+08 | 1.76E+08 | 1.40E+08 | 2.20E+08 | 1.41E+08 | 1.59E+08 | 1797.9016 | MLENQSSLLSGSSGIFK | Q3U9N9 | Q3U9N9 | SLC16A10 | solute carrier family 16 (aromatic amino acid transporter), member 10 | Plasma Membrane | transporter |
| 15531313 | 2 | 0.704 | 1.21E+08 | 1.17E+08 | 1.42E+08 | 1.35E+08 | 1.39E+08 | 1.35E+08 | 1.30E+08 | 1.35E+08 | 1.09E+08 | 1.76E+08 | 1.26E+08 | 1.18E+08 | 845.4716 | VEVELTR | Q3U9N9 | Q3U9N9 | SLC16A10 | solute carrier family 16 (aromatic amino acid transporter), member 10 | Plasma Membrane | transporter |
| 15531285 | 2 | 0.883 | 7.11E+07 | 9.27E+07 | 1.45E+08 | 9.03E+07 | 1.06E+08 | 1.23E+08 | 8.76E+07 | 9.02E+07 | 9.37E+07 | 1.30E+08 | 6.70E+07 | 9.04E+07 | 985.5492 | VFNFALFK | Q3U9N9 | Q3U9N9 | SLC16A10 | solute carrier family 16 (aromatic amino acid transporter), member 10 | Plasma Membrane | transporter |
| 15531537 | 2 | 0.889 | 4.24E+07 | 5.99E+07 | 7.48E+07 | 4.29E+07 | 8.16E+07 | 7.11E+07 | 7.53E+07 | 6.44E+07 | 5.22E+07 | 7.04E+07 | 4.53E+07 | 5.19E+07 | 975.5496 | IADYLPGVK | Q3U9N9 | Q3U9N9 | SLC16A10 | solute carrier family 16 (aromatic amino acid transporter), member 10 | Plasma Membrane | transporter |
| 15538851 | 2 | 0.435 | 4.74E+07 | 4.35E+07 | 5.40E+07 | 4.91E+07 | 4.93E+07 | 4.58E+07 | 4.93E+07 | 4.48E+07 | 4.04E+07 | 5.95E+07 | 5.00E+07 | 4.62E+07 | 730.35034 | SSFFSR | Q3U9N9 | Q3U9N9 | SLC16A10 | solute carrier family 16 (aromatic amino acid transporter), member 10 | Plasma Membrane | transporter |
| 15550853 | 3 | 0.708 | 2.75E+07 | 5.17E+07 | 5.98E+07 | 2.18E+07 | 9.82E+07 | 6.01E+07 | 4.41E+07 | 7.03E+07 | 3.78E+07 | 4.86E+07 | 2.17E+07 | 3.25E+07 | 3212.4822 | ETNEAQPPGPAPSDDAPLPGPGPSDVSDVAAEK | Q3U9N9 | Q3U9N9 | SLC16A10 | solute carrier family 16 (aromatic amino acid transporter), member 10 | Plasma Membrane | transporter |
| 15550961 | 4 | 0.738 | 1.32E+07 | 1.58E+07 | 1.55E+07 | 1.80E+07 | 1.86E+07 | 1.58E+07 | 1.80E+07 | 1.55E+07 | 1.37E+07 | 2.32E+07 | 1.17E+07 | 1.07E+07 | 4537.0864 | MVPSQEEPAAERETNEAQPPGPAPSDDAPLPGPGPSDVSDVAAEK | Q3U9N9 | Q3U9N9 | SLC16A10 | solute carrier family 16 (aromatic amino acid transporter), member 10 | Plasma Membrane | transporter |
| 15537439 | 3 | 0.584 | 8018022.5 | 1.17E+07 | 8004864.5 | 6988398.5 | 8558846 | 1.06E+07 | 1.02E+07 | 8158125 | 7565048 | 1.34E+07 | 7877416 | 1.16E+07 | 1797.9017 | MLENQSSLLSGSSGIFK | Q3U9N9 | Q3U9N9 | SLC16A10 | solute carrier family 16 (aromatic amino acid transporter), member 10 | Plasma Membrane | transporter |
| 15552772 | 2 | 0.484 | 8704828 | 8796940 | 1.24E+07 | 6209284 | 8456841 | 8514644 | 9722979 | 8528367 | 5557536 | 1.20E+07 | 4872520.5 | 3809369.5 | 1813.8948 | M[147.0354]LENQSSLLSGSSGIFK | Q3U9N9 | Q3U9N9 | SLC16A10 | solute carrier family 16 (aromatic amino acid transporter), member 10 | Plasma Membrane | transporter |
| 15540947 | 3 | 0.335 | 1.99E+07 | 984663.188 | 196653.391 | 1.72E+07 | 255857.188 | 251714.719 | 1575778.25 | 1495925.5 | 1.07E+07 | 8832718 | 3.21E+07 | 1.38E+07 | 1925.994 | MLENQSSLLSGSSGIFKK | Q3U9N9 | Q3U9N9 | SLC16A10 | solute carrier family 16 (aromatic amino acid transporter), member 10 | Plasma Membrane | transporter |
| 15556719 | 2 | 0.078 | 36288.133 | 46.485 | 352707.906 | 901223.125 | 933646 | 1251318.875 | 634636.562 | 686259.938 | 1265266.125 | 28011.83 | 771169.625 | 431905.562 | 1871.0718 | ATVQLLTNPFIGLLTNR | Q8BRU6 | Q8BRU6 | SLC18A2 | solute carrier family 18 (vesicular monoamine transporter), member 2 | Plasma Membrane | transporter |
| 15544729 | 2 | 0.342 | 1996115.75 | 1800034.25 | 1723848.125 | 1105503 | 1767621.75 | 1433079.375 | 1594346.5 | 1866612.375 | 1514453.125 | 1927803.125 | 1436612 | 1257453.75 | 1290.6885 | HQPLAQAQELR | P41438 | P41438 | SLC19A1 | solute carrier family 19 (folate transporter), member 1 | Plasma Membrane | transporter |
| 15537122 | 2 | 0.024 | 2196385 | 1926574.25 | 1960555 | 1606564 | 1592139 | 1463613.75 | 2254844.5 | 2185773.75 | 1860231.75 | 1986577.75 | 1604835.5 | 1422874.125 | 1148.5217 | NC[160.0307]VSPESDIK | Q3UHH2 | Q3UHH2 | SLC22A23 | solute carrier family 22, member 23 | Plasma Membrane | transporter |
| 15549787 | 3 | 0.213 | 5261866.5 | 1.25E+07 | 4583231 | 1.08E+07 | 7582708.5 | 1.09E+07 | 5338740.5 | 7400385.5 | 9696908 | 4350388.5 | 5699455.5 | 5278644 | 2989.3591 | NIFLQYASTEVDGEHYMTPEDFVQR | Q8BH59 | Q8BH59 | SLC25A12 | solute carrier family 25 (aspartate/glutamate carrier), member 12 | Cytoplasm | transporter |
| 15555118 | 3 | 0.162 | 2421375.25 | 5977196 | 3193695.5 | 3318938 | 3331779.5 | 5416284.5 | 2559288.25 | 4981317 | 6659523 | 1602166.125 | 2287548.75 | 2468042.75 | 3279.7344 | RDGSIPLPAEILAGGC[160.0307]AGGSQVIFTNPLEIVK | Q8BH59 | Q8BH59 | SLC25A12 | solute carrier family 25 (aspartate/glutamate carrier), member 12 | Cytoplasm | transporter |
| 15551498 | 2 | 0.293 | 2035142.25 | 4787432.5 | 1456321.625 | 3740997.25 | 2540084.75 | 4990278.5 | 1604396.625 | 3189109 | 4260186.5 | 1237736.25 | 2014226.625 | 1948921 | 1939.0604 | IAPLAEGALPYNLAELQR | Q8BH59 | Q8BH59 | SLC25A12 | solute carrier family 25 (aspartate/glutamate carrier), member 12 | Cytoplasm | transporter |
| 15541951 | 2 | 0.339 | 1001974.562 | 2582882.5 | 1370152.375 | 1876163.125 | 1765578.75 | 2267573.25 | 802322.812 | 2032628.125 | 2759387.25 | 824249.375 | 1181206.125 | 1097041.75 | 963.524 | LTVNDFVR | Q8BH59 | Q8BH59 | SLC25A12 | solute carrier family 25 (aspartate/glutamate carrier), member 12 | Cytoplasm | transporter |
| 15534550 | 2 | 0.175 | 4058750.5 | 7643211 | 3093280.5 | 3554116.5 | 4369966.5 | 5848645.5 | 2791264.5 | 5618562 | 5722108 | 2502447 | 2057766.5 | 3075503.25 | 1262.708 | IQGPQPVPGEIK | Q8BZ09 | Q8BZ09 | SLC25A21 | solute carrier family 25 (mitochondrial oxoadipate carrier), member 21 | Cytoplasm | transporter |
| 15540041 | 2 | 0.038 | 3155582 | 5620908 | 2100155.25 | 2295811.25 | 2872590.75 | 3565298.5 | 2230128.75 | 3722292.75 | 3924747.25 | 1607433.875 | 1438651.75 | 1297570.625 | 1062.5548 | EGLGFQGLNK | Q8BZ09 | Q8BZ09 | SLC25A21 | solute carrier family 25 (mitochondrial oxoadipate carrier), member 21 | Cytoplasm | transporter |
| 15555583 | 2 | 0.465 | 1515372.875 | 4379379 | 1234184.125 | 1399925.25 | 1484239.625 | 2480951 | 1467843.25 | 2364562.5 | 3289172.5 | 1380009.125 | 1169856 | 1346040 | 2290.1519 | FGIGFVSGTMGSVFNIPFDVAK | Q8BZ09 | Q8BZ09 | SLC25A21 | solute carrier family 25 (mitochondrial oxoadipate carrier), member 21 | Cytoplasm | transporter |
| 15544267 | 2 | 0.224 | 1549750.5 | 2751037.75 | 855846.25 | 1434708.875 | 1418551.875 | 2081177.875 | 1235938.875 | 1826260 | 1837950.5 | 949343.812 | 931296.438 | 970828.312 | 785.46106 | VGLQVNR | Q8BZ09 | Q8BZ09 | SLC25A21 | solute carrier family 25 (mitochondrial oxoadipate carrier), member 21 | Cytoplasm | transporter |
| 15539491 | 2 | 0.198 | 1356933.125 | 3599533.75 | 1343475.75 | 1257612.5 | 1648811.125 | 1794802.125 | 833768.375 | 2030535.375 | 1542284.625 | 676167.438 | 1118147.625 | 903698.438 | 1169.5554 | EQPSTFAYAR | Q8BZ09 | Q8BZ09 | SLC25A21 | solute carrier family 25 (mitochondrial oxoadipate carrier), member 21 | Cytoplasm | transporter |
| 15536402 | 2 | 0.313 | 2.33E+07 | 4.50E+07 | 2.03E+07 | 3.71E+07 | 3.33E+07 | 4.32E+07 | 2.39E+07 | 3.58E+07 | 4.03E+07 | 1.39E+07 | 3.27E+07 | 2.34E+07 | 1361.7141 | IQTQPGYANTLR | Q8VEM8 | Q8VEM8 | SLC25A3 | solute carrier family 25 (mitochondrial carrier; phosphate carrier), member 3 | Cytoplasm | transporter |
| 15534749 | 2 | 0.135 | 1.50E+07 | 2.91E+07 | 1.57E+07 | 2.58E+07 | 2.31E+07 | 3.04E+07 | 1.80E+07 | 2.58E+07 | 2.91E+07 | 1.08E+07 | 1.96E+07 | 1.56E+07 | 1046.5565 | GSTASQVLQR | Q8VEM8 | Q8VEM8 | SLC25A3 | solute carrier family 25 (mitochondrial carrier; phosphate carrier), member 3 | Cytoplasm | transporter |
| 15548333 | 3 | 0.721 | 1.05E+07 | 1.68E+07 | 1.00E+07 | 1.35E+07 | 1.40E+07 | 1.91E+07 | 1.14E+07 | 2.17E+07 | 2.43E+07 | 7334383 | 2.80E+07 | 1.59E+07 | 2910.4832 | YYALC[160.0307]GFGGVLSC[160.0307]GLTHTAVVPLDLVK | Q8VEM8 | Q8VEM8 | SLC25A3 | solute carrier family 25 (mitochondrial carrier; phosphate carrier), member 3 | Cytoplasm | transporter |
| 15549917 | 2 | 0.312 | 1.02E+07 | 2.31E+07 | 1.31E+07 | 1.57E+07 | 1.84E+07 | 2.53E+07 | 1.27E+07 | 2.08E+07 | 3.08E+07 | 6074374 | 1.73E+07 | 1.21E+07 | 1941.9662 | ALYSNILGEENTYLWR | Q8VEM8 | Q8VEM8 | SLC25A3 | solute carrier family 25 (mitochondrial carrier; phosphate carrier), member 3 | Cytoplasm | transporter |
| 15531854 | 2 | 0.326 | 7989079 | 1.70E+07 | 7063058.5 | 1.09E+07 | 1.57E+07 | 1.98E+07 | 1.11E+07 | 1.58E+07 | 1.91E+07 | 4196712.5 | 1.38E+07 | 1.02E+07 | 1036.5117 | FGFYEVFK | Q8VEM8 | Q8VEM8 | SLC25A3 | solute carrier family 25 (mitochondrial carrier; phosphate carrier), member 3 | Cytoplasm | transporter |
| 15532225 | 2 | 0.525 | 5519011 | 1.20E+07 | 4809754 | 8050276.5 | 7846633 | 1.03E+07 | 5647197 | 7842572.5 | 9307268 | 3308205.5 | 7274713.5 | 7002959 | 929.5006 | GVAPLWMR | Q8VEM8 | Q8VEM8 | SLC25A3 | solute carrier family 25 (mitochondrial carrier; phosphate carrier), member 3 | Cytoplasm | transporter |
| 15551677 | 2 | 0.517 | 4804347.5 | 9294371 | 3610624.25 | 7313866.5 | 7770289 | 9619852 | 6293935 | 8601215 | 1.29E+07 | 3145571.25 | 1.09E+07 | 6852866.5 | 1781.8661 | GWAPTLIGYSMQGLC[160.0307]K | Q8VEM8 | Q8VEM8 | SLC25A3 | solute carrier family 25 (mitochondrial carrier; phosphate carrier), member 3 | Cytoplasm | transporter |
| 15542561 | 2 | 0.211 | 2012704.125 | 3500446.25 | 1923546.625 | 4972410 | 2817976.75 | 4718715 | 2244798.75 | 3257368.75 | 3802560 | 2152799 | 3423567.75 | 2407864.5 | 823.4553 | TVEALYK | Q8VEM8 | Q8VEM8 | SLC25A3 | solute carrier family 25 (mitochondrial carrier; phosphate carrier), member 3 | Cytoplasm | transporter |
| 15535993 | 2 | 0.534 | 2823271 | 4720307 | 2095089.625 | 4511602.5 | 2642138.5 | 4476043.5 | 2320994.5 | 3692895.75 | 5635290 | 1418827.25 | 3799550.5 | 2430642.5 | 1196.6653 | GIFNGFSITLK | Q8VEM8 | Q8VEM8 | SLC25A3 | solute carrier family 25 (mitochondrial carrier; phosphate carrier), member 3 | Cytoplasm | transporter |
| 15548524 | 4 | 0.327 | 9643682 | 2.03E+07 | 6703967.5 | 7590297.5 | 1.10E+07 | 1.25E+07 | 6081707.5 | 1.71E+07 | 1.50E+07 | 6166673 | 4059878 | 8440741 | 4281.9375 | DAGSEDYENLPTSASVSTHMTAGAMAGILEHSIMYPVDSVK | Q920G8 | Q920G8 | SLC25A37 | solute carrier family 25 (mitochondrial iron transporter), member 37 | Cytoplasm | other |
| 15541758 | 2 | 0.327 | 5255030 | 1.21E+07 | 4777043 | 4511958 | 5113266 | 8627785 | 4089640 | 7719868.5 | 9244796 | 4122916.5 | 3681753.25 | 3969015 | 1013.50305 | TEGLGAFYR | Q920G8 | Q920G8 | SLC25A37 | solute carrier family 25 (mitochondrial iron transporter), member 37 | Cytoplasm | other |
| 15553229 | 3 | 0.566 | 5131442 | 9598942 | 1971541.25 | 6799010 | 4258219 | 4476620.5 | 4142770.5 | 5452404.5 | 4633774 | 4017102.5 | 2795668 | 2752625 | 3645.7952 | SYTTQLTMNIPFQSIHFITYEFLQEQVNPR | Q920G8 | Q920G8 | SLC25A37 | solute carrier family 25 (mitochondrial iron transporter), member 37 | Cytoplasm | other |
| 15551238 | 4 | 0.601 | 5502817 | 8127649 | 4047515 | 4418798.5 | 4944693.5 | 7530249.5 | 3372239.25 | 7392697 | 1.09E+07 | 3592639.75 | 3997627.25 | 5190302.5 | 3067.5574 | RDYNPQSHIISGGLAGALAAAATTPLDVC[160.0307]K | Q920G8 | Q920G8 | SLC25A37 | solute carrier family 25 (mitochondrial iron transporter), member 37 | Cytoplasm | other |
| 15551595 | 2 | 0.113 | 4479144 | 8687182 | 3764483.5 | 4739002 | 3966544.25 | 5520098.5 | 3027048.25 | 4929915.5 | 6837949 | 3033140.25 | 2507302.75 | 2545992.5 | 2032.0448 | TLLNTQENMALSLANVSGR | Q920G8 | Q920G8 | SLC25A37 | solute carrier family 25 (mitochondrial iron transporter), member 37 | Cytoplasm | other |
| 15539082 | 3 | 0.477 | 3948044.5 | 8069292.5 | 2759578.25 | 4101117.75 | 4277150 | 4693363.5 | 2467929.25 | 4809939.5 | 5076697 | 2937694.25 | 2514143.5 | 3396398.5 | 2032.0438 | TLLNTQENMALSLANVSGR | Q920G8 | Q920G8 | SLC25A37 | solute carrier family 25 (mitochondrial iron transporter), member 37 | Cytoplasm | other |
| 15533354 | 2 | 0.072 | 3714908.25 | 7184620 | 3091851.5 | 3758401.75 | 2997278.5 | 4325059 | 2314835.5 | 4847733 | 4261943 | 2461033 | 1860513.375 | 1629750 | 1029.5001 | MQSLNPDPK | Q920G8 | Q920G8 | SLC25A37 | solute carrier family 25 (mitochondrial iron transporter), member 37 | Cytoplasm | other |
| 15538253 | 2 | 0.25 | 4388996 | 7888437.5 | 2661626 | 3589174.25 | 3563833.75 | 4686994 | 2452718.75 | 4222565.5 | 5109795.5 | 2164426.25 | 2481307.25 | 2813753.25 | 966.4804 | LSGMANAFR | Q920G8 | Q920G8 | SLC25A37 | solute carrier family 25 (mitochondrial iron transporter), member 37 | Cytoplasm | other |
| 15540903 | 2 | 0.112 | 2056833.125 | 3718324.5 | 1779747.5 | 2095302.75 | 1903008 | 2441629.5 | 1943237.125 | 3240328.5 | 3612625.25 | 1613269.5 | 1468265.75 | 1579270.25 | 1473.7712 | TVYQLNGLAGYFK | Q920G8 | Q920G8 | SLC25A37 | solute carrier family 25 (mitochondrial iron transporter), member 37 | Cytoplasm | other |
| 15554453 | 2 | 0.699 | 976052.375 | 3047042 | 788674.062 | 1326926.875 | 1164306.125 | 1502040 | 930125.125 | 2034553.25 | 1948832.625 | 1191357.5 | 786844.625 | 1083963.75 | 2296.1301 | VIYQMPSTAISWSVYEFFK | Q920G8 | Q920G8 | SLC25A37 | solute carrier family 25 (mitochondrial iron transporter), member 37 | Cytoplasm | other |
| 15551717 | 3 | 0.119 | 2523773 | 8922394 | 4638714 | 1539980.25 | 2692465.5 | 7338670 | 1715387.375 | 5104907 | 5688860.5 | 1010028.375 | 475881.344 | 2218605.75 | 3661.7922 | SYTTQLTM[147.0354]NIPFQSIHFITYEFLQEQVNPR | Q920G8 | Q920G8 | SLC25A37 | solute carrier family 25 (mitochondrial iron transporter), member 37 | Cytoplasm | other |
| 15536672 | 2 | 0.211 | 1775071.875 | 3455841 | 3190093.75 | 5007334.5 | 6322313.5 | 7463320.5 | 2717540.25 | 4650016 | 6194283 | 910990.75 | 5362308 | 3057652.75 | 1169.5715 | EQGFLSFWR | P48962 | P48962 | SLC25A4 | solute carrier family 25 (mitochondrial carrier; adenine nucleotide translocator), member 4 | Cytoplasm | transporter |
| 15539879 | 2 | 0.199 | 1285109.625 | 2517130.5 | 2192066 | 3746705.75 | 4640886 | 5288104 | 2476181.75 | 3607511.75 | 4668764.5 | 675727.375 | 4083961 | 2600228.5 | 998.4573 | DEGANAFFK | P48962 | P48962 | SLC25A4 | solute carrier family 25 (mitochondrial carrier; adenine nucleotide translocator), member 4 | Cytoplasm | transporter |
| 15543292 | 2 | 0.496 | 1015629.375 | 1427978.875 | 1600455.75 | 3300989.75 | 2931397.75 | 3872782.5 | 1717506.625 | 2728102.75 | 3554737.25 | 50471.242 | 3732701 | 2587454 | 1004.5491 | QIFLGGVDR | P48962 | P48962 | SLC25A4 | solute carrier family 25 (mitochondrial carrier; adenine nucleotide translocator), member 4 | Cytoplasm | transporter |
| 15533282 | 2 | 0.277 | 6.43E+07 | 1.53E+08 | 7.63E+07 | 1.23E+08 | 1.11E+08 | 1.63E+08 | 7.92E+07 | 1.21E+08 | 1.64E+08 | 4.04E+07 | 9.23E+07 | 9.06E+07 | 1446.7397 | YFPTQALNFAFK | P51881 | P51881 | SLC25A5 | solute carrier family 25 (mitochondrial carrier; adenine nucleotide translocator), member 5 | Cytoplasm | transporter |
| 15535034 | 2, 3 | 0.152 | 5.34E+07 | 9.13E+07 | 5.31E+07 | 7.98E+07 | 7.42E+07 | 9.29E+07 | 5.25E+07 | 7.84E+07 | 1.11E+08 | 2.94E+07 | 5.90E+07 | 5.13E+07 | 1136.6749, 1136.6758 | LLLQVQHASK | P51881 | P51881 | SLC25A5 | solute carrier family 25 (mitochondrial carrier; adenine nucleotide translocator), member 5 | Cytoplasm | transporter |
| 15531573 | 2 | 0.138 | 4.22E+07 | 7.78E+07 | 4.30E+07 | 6.90E+07 | 7.31E+07 | 7.75E+07 | 4.96E+07 | 6.34E+07 | 8.14E+07 | 2.42E+07 | 5.48E+07 | 4.43E+07 | 1121.5726 | EQGVLSFWR | P51881 | P51881 | SLC25A5 | solute carrier family 25 (mitochondrial carrier; adenine nucleotide translocator), member 5 | Cytoplasm | transporter |
| 15532595 | 2 | 0.283 | 2.29E+07 | 4.40E+07 | 2.35E+07 | 3.09E+07 | 3.83E+07 | 4.19E+07 | 2.24E+07 | 3.57E+07 | 5.12E+07 | 1.33E+07 | 2.90E+07 | 2.40E+07 | 856.4986 | GNLANVIR | P51881 | P51881 | SLC25A5 | solute carrier family 25 (mitochondrial carrier; adenine nucleotide translocator), member 5 | Cytoplasm | transporter |
| 15551634 | 2 | 0.351 | 1.16E+07 | 3.15E+07 | 1.76E+07 | 1.32E+07 | 3.09E+07 | 2.55E+07 | 1.51E+07 | 3.02E+07 | 3.40E+07 | 1.16E+07 | 1.66E+07 | 1.26E+07 | 1688.7367 | GTDIMYTGTLDC[160.0307]WR | P51881 | P51881 | SLC25A5 | solute carrier family 25 (mitochondrial carrier; adenine nucleotide translocator), member 5 | Cytoplasm | transporter |
| 15533872 | 2 | 0.087 | 1.55E+07 | 4.65E+07 | 3.11E+07 | 2.11E+07 | 4.64E+07 | 5.31E+07 | 3.06E+07 | 4.23E+07 | 4.07E+07 | 1.15E+07 | 1.56E+07 | 2.04E+07 | 976.5454 | QIFLGGVDK | P51881 | P51881 | SLC25A5 | solute carrier family 25 (mitochondrial carrier; adenine nucleotide translocator), member 5 | Cytoplasm | transporter |
| 15532262 | 2, 3 | 0.902 | 2.14E+07 | 2.01E+07 | 3290844.75 | 2.48E+07 | 1.20E+07 | 1.05E+07 | 7926021.5 | 1.51E+07 | 3.08E+07 | 1.03E+07 | 2.80E+07 | 1.72E+07 | 1132.6448, 1132.6453 | QIFLGGVDKR | P51881 | P51881 | SLC25A5 | solute carrier family 25 (mitochondrial carrier; adenine nucleotide translocator), member 5 | Cytoplasm | transporter |
| 15533906 | 2 | 0.382 | 1.50E+07 | 3.02E+07 | 1.64E+07 | 1.95E+07 | 2.38E+07 | 3.19E+07 | 1.47E+07 | 2.26E+07 | 3.11E+07 | 9394171 | 1.96E+07 | 1.72E+07 | 902.4827 | GAWSNVLR | P51881 | P51881 | SLC25A5 | solute carrier family 25 (mitochondrial carrier; adenine nucleotide translocator), member 5 | Cytoplasm | transporter |
| 15539351 | 3 | 0.911 | 1.41E+07 | 2.06E+07 | 8875391 | 1.88E+07 | 8867508 | 2.11E+07 | 1.56E+07 | 1.36E+07 | 2.68E+07 | 8546387 | 2.35E+07 | 2.01E+07 | 1816.8308 | GTDIMYTGTLDC[160.0307]WRK | P51881 | P51881 | SLC25A5 | solute carrier family 25 (mitochondrial carrier; adenine nucleotide translocator), member 5 | Cytoplasm | transporter |
| 15535125 | 2 | 0.384 | 9055829 | 2.97E+07 | 2.06E+07 | 9271080 | 2.01E+07 | 3.03E+07 | 1.06E+07 | 2.81E+07 | 3.52E+07 | 7521726.5 | 1.14E+07 | 1.15E+07 | 1219.6661 | DFLAGGVAAAISK | P51881 | P51881 | SLC25A5 | solute carrier family 25 (mitochondrial carrier; adenine nucleotide translocator), member 5 | Cytoplasm | transporter |
| 15541510, 15542659 | 2 | 0.078 | 1.45E+07 | 3.42E+07 | 1.37E+07 | 2.22E+07 | 2.40E+07 | 2.40E+07 | 1.22E+07 | 2.50E+07 | 3.52E+07 | 4394505.5 | 1.17E+07 | 1.09E+07 | 1219.5964, 1219.5973 | AAYFGIYDTAK | P51881 | P51881 | SLC25A5 | solute carrier family 25 (mitochondrial carrier; adenine nucleotide translocator), member 5 | Cytoplasm | transporter |
| 15540430 | 2 | 0.324 | 4786297 | 1.05E+07 | 4922006.5 | 7948204.5 | 7689223.5 | 9699780 | 5567209.5 | 7769922 | 9128325 | 3298000.75 | 7092020.5 | 5456230 | 757.3893 | GMLPDPK | P51881 | P51881 | SLC25A5 | solute carrier family 25 (mitochondrial carrier; adenine nucleotide translocator), member 5 | Cytoplasm | transporter |
| 15557463 | 2 | 0.354 | 1714383.875 | 6327425.5 | 6286921.5 | 1791973 | 4050776.75 | 7219557.5 | 2585125.5 | 4633314.5 | 6491660 | 1468549 | 1509143.375 | 2966055.75 | 1627.8365 | GM[147.0354]GGAFVLVLYDEIK | P51881 | P51881 | SLC25A5 | solute carrier family 25 (mitochondrial carrier; adenine nucleotide translocator), member 5 | Cytoplasm | transporter |
| 15554954 | 2 | 0.121 | 346994.094 | 2730396 | 1573342.75 | 1197386.75 | 2970051 | 2839810.25 | 537974 | 2947773 | 1336909.375 | 205595.453 | 792001.25 | 277168.188 | 1704.7307 | GTDIM[147.0354]YTGTLDC[160.0307]WR | P51881 | P51881 | SLC25A5 | solute carrier family 25 (mitochondrial carrier; adenine nucleotide translocator), member 5 | Cytoplasm | transporter |
| 15537556 | 2 | 0.582 | 876641.5 | 5625852.5 | 5456755.5 | 251195.734 | 7570680 | 1.06E+07 | 777049.688 | 6920963 | 1.09E+07 | 8065.055 | 2517893.75 | 2921990 | 856.4873 | TAVAPIER | P51881 | P51881 | SLC25A5 | solute carrier family 25 (mitochondrial carrier; adenine nucleotide translocator), member 5 | Cytoplasm | transporter |
| 15545506 | 3 | 0.666 | 549078.875 | 3443203.75 | 1149310.75 | 868693.562 | 1148930.5 | 2351979.25 | 1307824.625 | 909399.812 | 2280701.5 | 766489.188 | 1029574.062 | 687377.938 | 1960.0708 | SVAALLAGTTEAILTPFER | Q5HZI9 | Q5HZI9 | SLC25A51 | solute carrier family 25, member 51 | Cytoplasm | other |
| 15548586 | 3 | 0.02 | 1.11E+07 | 1.36E+07 | 1.19E+07 | 9132061 | 8247003 | 8092143 | 9937488 | 1.02E+07 | 1.20E+07 | 1.20E+07 | 9056866 | 1.00E+07 | 3045.4302 | GGFLGEEQGAENELLFPSVHSAVEAAC[160.0307]AR | P58735 | P58735 | SLC26A1 | solute carrier family 26 (anion exchanger), member 1 | Plasma Membrane | transporter |
| 15550817 | 2 | 0.054 | 4097160.75 | 4930800 | 5799193.5 | 3622243.25 | 3173335.5 | 3205756 | 3797966.25 | 3336216 | 4678809.5 | 4954867.5 | 4554023.5 | 3778025 | 1806.9714 | VAGNIPTGFVAPQVPDPK | P58735 | P58735 | SLC26A1 | solute carrier family 26 (anion exchanger), member 1 | Plasma Membrane | transporter |
| 15536168 | 2 | 0.734 | 2173211.75 | 1653294.5 | 1852183.25 | 1252418.875 | 2425488.5 | 1521566.5 | 1987176.5 | 1599809 | 1621706.5 | 2538028.5 | 1974314 | 1624019.375 | 1019.58636 | TILQEFIR | Q9WVC8 | Q9WVC8 | SLC26A3 | solute carrier family 26 (anion exchanger), member 3 | Plasma Membrane | transporter |
| 15531929 | 2 | 0.048 | 1.53E+08 | 1.71E+08 | 1.62E+08 | 1.30E+08 | 1.27E+08 | 1.27E+08 | 1.60E+08 | 1.53E+08 | 1.28E+08 | 1.62E+08 | 1.33E+08 | 1.50E+08 | 1194.6832 | ALADPTVALPAR | Q9JIM1 | Q9JIM1 | SLC29A1 | solute carrier family 29 (equilibrative nucleoside transporter), member 1 | Plasma Membrane | transporter |
| 15539520 | 3 | 0.009 | 8.30E+07 | 9.23E+07 | 9.33E+07 | 6.17E+07 | 6.67E+07 | 5.53E+07 | 7.89E+07 | 7.96E+07 | 7.85E+07 | 8.54E+07 | 6.25E+07 | 6.81E+07 | 1811.9222 | HYLQLNLAGPAEQETK | Q9JIM1 | Q9JIM1 | SLC29A1 | solute carrier family 29 (equilibrative nucleoside transporter), member 1 | Plasma Membrane | transporter |
| 15549367 | 2 | 0.129 | 4.45E+07 | 5.07E+07 | 6.40E+07 | 2.97E+07 | 5.00E+07 | 3.01E+07 | 4.60E+07 | 5.02E+07 | 3.95E+07 | 4.63E+07 | 2.55E+07 | 2.67E+07 | 1537.7218 | EESGVPGPNSPPTNR | Q9JIM1 | Q9JIM1 | SLC29A1 | solute carrier family 29 (equilibrative nucleoside transporter), member 1 | Plasma Membrane | transporter |
| 15532903 | 2 | 0.038 | 5.15E+07 | 5.28E+07 | 6.01E+07 | 2.50E+07 | 4.32E+07 | 3.82E+07 | 4.84E+07 | 4.39E+07 | 4.75E+07 | 4.36E+07 | 4.00E+07 | 4.49E+07 | 1040.6234 | WLPVLVASR | Q9JIM1 | Q9JIM1 | SLC29A1 | solute carrier family 29 (equilibrative nucleoside transporter), member 1 | Plasma Membrane | transporter |
| 15551662 | 2 | 0.03 | 3.77E+07 | 2.60E+07 | 3.38E+07 | 1.38E+07 | 2.01E+07 | 2.89E+07 | 3.70E+07 | 3.68E+07 | 5.09E+07 | 3.75E+07 | 4.88E+07 | 3.19E+07 | 1559.9028 | IVFIPLLMLC[160.0307]NVK | Q9JIM1 | Q9JIM1 | SLC29A1 | solute carrier family 29 (equilibrative nucleoside transporter), member 1 | Plasma Membrane | transporter |
| 15550256 | 2 | 0.277 | 2.99E+07 | 2.63E+07 | 1.86E+07 | 2.24E+07 | 1.65E+07 | 6421879 | 2.29E+07 | 1.64E+07 | 1.70E+07 | 3.18E+07 | 2.39E+07 | 1.72E+07 | 1811.9238 | HYLQLNLAGPAEQETK | Q9JIM1 | Q9JIM1 | SLC29A1 | solute carrier family 29 (equilibrative nucleoside transporter), member 1 | Plasma Membrane | transporter |
| 15548206 | 2 | 0.307 | 1.75E+07 | 1.38E+07 | 1.46E+07 | 1.06E+07 | 1.46E+07 | 8246192.5 | 1.24E+07 | 1.62E+07 | 1.26E+07 | 1.44E+07 | 1.60E+07 | 1.05E+07 | 1607.7284 | SLTAVC[160.0307]MWPGQDSR | Q9JIM1 | Q9JIM1 | SLC29A1 | solute carrier family 29 (equilibrative nucleoside transporter), member 1 | Plasma Membrane | transporter |
| 15548619 | 2 | 0.237 | 6658890.5 | 4307196.5 | 8712509 | 3204379.25 | 4482071 | 5950181 | 4768198 | 7046702.5 | 5653467.5 | 4617060 | 3636833.5 | 4352685 | 1575.8977 | IVFIPLLM[147.0354]LC[160.0307]NVK | Q9JIM1 | Q9JIM1 | SLC29A1 | solute carrier family 29 (equilibrative nucleoside transporter), member 1 | Plasma Membrane | transporter |
| 15536711 | 3 | 0.042 | 3217547 | 1946862.875 | 2930617 | 672873.438 | 1393674 | 2669691.75 | 3067928.75 | 3714206.25 | 4299284.5 | 3653377 | 4350789 | 3129514 | 1559.9015 | IVFIPLLMLC[160.0307]NVK | Q9JIM1 | Q9JIM1 | SLC29A1 | solute carrier family 29 (equilibrative nucleoside transporter), member 1 | Plasma Membrane | transporter |
| 15553627 | 2 | 0.022 | 5356049 | 5749256.5 | 6377563 | 3034197.5 | 3779554.25 | 3186029.5 | 3611534.25 | 4637331 | 2983978.5 | 3343207.25 | 1605208.625 | 3237337.5 | 1623.7223 | SLTAVC[160.0307]M[147.0354]WPGQDSR | Q9JIM1 | Q9JIM1 | SLC29A1 | solute carrier family 29 (equilibrative nucleoside transporter), member 1 | Plasma Membrane | transporter |
| 15558520 | 2 | 0.129 | 2114493 | 2171867.5 | 2377295.5 | 931077.5 | 1680854.25 | 1433172.25 | 1800541 | 2010730.125 | 1379314.125 | 2212157 | 1527193.625 | 1230393.875 | 1693.821 | REESGVPGPNSPPTNR | Q9JIM1 | Q9JIM1 | SLC29A1 | solute carrier family 29 (equilibrative nucleoside transporter), member 1 | Plasma Membrane | transporter |
| 15535817 | 2 | 0.635 | 2.12E+07 | 2.39E+07 | 3.22E+07 | 1.71E+07 | 2.80E+07 | 2.68E+07 | 2.88E+07 | 2.69E+07 | 2.88E+07 | 2.49E+07 | 2.49E+07 | 2.41E+07 | 758.4906 | LPIIFR | Q61672 | Q61672 | SLC29A2 | solute carrier family 29 (equilibrative nucleoside transporter), member 2 | Plasma Membrane | transporter |
| 15548151 | 2 | 0.077 | 4.28E+07 | 8400353 | 3.32E+07 | 3.55E+07 | 4.40E+07 | 4.04E+07 | 2.26E+07 | 2.82E+07 | 4.28E+07 | 8400353 | 3.32E+07 | 2.78E+07 | 1762.864 | LWGTSDVVQEIQEMK | P32037 | P32037 | SLC2A3 | solute carrier family 2 (facilitated glucose transporter), member 3 | Plasma Membrane | transporter |
| 15532650 | 2, 3 | 0.062 | 2.66E+07 | 7233695.5 | 3.23E+07 | 3.03E+07 | 3.12E+07 | 2.95E+07 | 1.98E+07 | 2.38E+07 | 2.66E+07 | 7233695.5 | 3.23E+07 | 1.92E+07 | 1459.7354, 1459.7361 | KEEDQATEILQR | P32037 | P32037 | SLC2A3 | solute carrier family 2 (facilitated glucose transporter), member 3 | Plasma Membrane | transporter |
| 15531709 | 2 | 0.07 | 2.68E+07 | 5277017 | 2.29E+07 | 2.60E+07 | 2.65E+07 | 2.97E+07 | 1.66E+07 | 2.06E+07 | 2.68E+07 | 5277017 | 2.29E+07 | 2.02E+07 | 1104.6389 | QVTVLELFR | P32037 | P32037 | SLC2A3 | solute carrier family 2 (facilitated glucose transporter), member 3 | Plasma Membrane | transporter |
| 15534172 | 2 | 0.073 | 1.77E+07 | 5051474 | 2.31E+07 | 2.12E+07 | 2.59E+07 | 1.92E+07 | 1.33E+07 | 1.75E+07 | 1.77E+07 | 5051474 | 2.31E+07 | 1.51E+07 | 1426.7335 | GPAGVELNSMQPVK | P32037 | P32037 | SLC2A3 | solute carrier family 2 (facilitated glucose transporter), member 3 | Plasma Membrane | transporter |
| 15537177 | 2 | 0.121 | 1.14E+07 | 1883593.125 | 9129679 | 1.12E+07 | 1.17E+07 | 1.19E+07 | 6628195 | 9060832 | 1.14E+07 | 1883593.125 | 9129679 | 7588056.5 | 747.475 | FLLINK | P32037 | P32037 | SLC2A3 | solute carrier family 2 (facilitated glucose transporter), member 3 | Plasma Membrane | transporter |
| 15540037 | 2 | 0.041 | 3326474 | 997509.125 | 4614210 | 4704138.5 | 5163751.5 | 3639842.5 | 2878830 | 3046982.25 | 3326474 | 997509.125 | 4614210 | 2741831.25 | 1331.6437 | EEDQATEILQR | P32037 | P32037 | SLC2A3 | solute carrier family 2 (facilitated glucose transporter), member 3 | Plasma Membrane | transporter |
| 15543493 | 3 | 0.029 | 1919837.625 | 513119.75 | 1416032.375 | 1429119.625 | 1880948.125 | 2479465.75 | 1139371.75 | 1603800.75 | 1919837.625 | 513119.75 | 1416032.375 | 1618724.875 | 1762.865 | LWGTSDVVQEIQEMK | P32037 | P32037 | SLC2A3 | solute carrier family 2 (facilitated glucose transporter), member 3 | Plasma Membrane | transporter |
| 15533120 | 2 | 8.79E-04 | 4449013.5 | 5197298 | 4077968.75 | 3925352.25 | 4462327 | 4331885.5 | 6454987.5 | 5999228.5 | 5120399.5 | 3595088.75 | 3331273.75 | 3304246.25 | 1311.7861 | ESALILLQTVPK | Q60738 | Q60738 | SLC30A1 | solute carrier family 30 (zinc transporter), member 1 | Plasma Membrane | transporter |
| 15550270 | 3 | 0.211 | 3388602.5 | 3367375.75 | 2843430.75 | 3065347 | 3123949 | 1534371.75 | 4176427 | 3841927 | 2861296.25 | 2592899 | 2701443.25 | 2165986.75 | 2492.227 | DVFHNHGIHATTIQPEFASVGSK | Q60738 | Q60738 | SLC30A1 | solute carrier family 30 (zinc transporter), member 1 | Plasma Membrane | transporter |
| 15540961 | 2 | 3.36E-04 | 3511697.75 | 3825287.5 | 3691796.25 | 2895839.25 | 2888526.25 | 2405123 | 4330513 | 4124075.5 | 3331496.25 | 2424198.75 | 2232355.5 | 2186857.25 | 1325.7651 | AEGSLPAVVIEIK | Q60738 | Q60738 | SLC30A1 | solute carrier family 30 (zinc transporter), member 1 | Plasma Membrane | transporter |
| 15550304 | 2 | 0.002 | 2346715 | 3217544.25 | 2340967.25 | 2185743.75 | 1897604.25 | 1946351.875 | 2672057 | 2920270.75 | 3129487.25 | 1770302.375 | 1475499.375 | 1212846.625 | 1889.9496 | APTVSISC[160.0307]LELSENLEK | Q60738 | Q60738 | SLC30A1 | solute carrier family 30 (zinc transporter), member 1 | Plasma Membrane | transporter |
| 15548696 | 2 | 0.008 | 1.21E+07 | 1.83E+07 | 1.32E+07 | 1.50E+07 | 1.45E+07 | 1.30E+07 | 9085246 | 1.25E+07 | 1.07E+07 | 9407040 | 8295683.5 | 8661784 | 1566.8816 | LGASNLPAGISLPASAK | P10852 | P10852 | SLC3A2 | solute carrier family 3 (amino acid transporter heavy chain), member 2 | Plasma Membrane | transporter |
| 15553183 | 2 | 0.072 | 7914653 | 1.36E+07 | 8481735 | 1.06E+07 | 1.11E+07 | 1.36E+07 | 7138557 | 1.08E+07 | 9944226 | 8393475 | 5783866.5 | 5986034 | 1954.9253 | EALSSWLQDGVDGFQFR | P10852 | P10852 | SLC3A2 | solute carrier family 3 (amino acid transporter heavy chain), member 2 | Plasma Membrane | transporter |
| 15533046 | 2 | 0.006 | 9980435 | 1.16E+07 | 7731082.5 | 1.10E+07 | 9283746 | 1.02E+07 | 7570629.5 | 8099044 | 7523994.5 | 6940880.5 | 5995477 | 6427681 | 1476.7303 | QINPTLGSQEDFK | P10852 | P10852 | SLC3A2 | solute carrier family 3 (amino acid transporter heavy chain), member 2 | Plasma Membrane | transporter |
| 15539438 | 2 | 0.048 | 9058961 | 1.10E+07 | 6314124 | 7669475 | 8157576.5 | 9093709 | 5860821.5 | 7401275 | 6330103.5 | 6297527 | 5490742.5 | 5531076 | 1075.5874 | IGDLQAFVGR | P10852 | P10852 | SLC3A2 | solute carrier family 3 (amino acid transporter heavy chain), member 2 | Plasma Membrane | transporter |
| 15554707, 15560410 | 3 | 0.022 | 9030388 | 1.17E+07 | 6414555 | 8953282 | 7265772.5 | 9469830 | 7187366 | 7319158 | 7138410.5 | 5622987 | 5094240.5 | 5137817 | 2362.1777, 2362.1782 | SLLHGDFHALSSSPDLFSYIR | P10852 | P10852 | SLC3A2 | solute carrier family 3 (amino acid transporter heavy chain), member 2 | Plasma Membrane | transporter |
| 15535585 | 2 | 0.118 | 4818595 | 6162720 | 3636859.5 | 6436438.5 | 4606851 | 6056930 | 3837748.75 | 4126776.5 | 4858111 | 4458240.5 | 3022410.5 | 3560367.5 | 1076.5083 | DEINETDLK | P10852 | P10852 | SLC3A2 | solute carrier family 3 (amino acid transporter heavy chain), member 2 | Plasma Membrane | transporter |
| 15532316 | 2 | 0.026 | 5159796 | 6554883 | 4940127.5 | 4784415.5 | 5356258 | 5553594.5 | 3422225 | 4926999.5 | 3979269.5 | 4086579.25 | 3772916.75 | 4004441.5 | 933.58575 | GLVLGPIHK | P10852 | P10852 | SLC3A2 | solute carrier family 3 (amino acid transporter heavy chain), member 2 | Plasma Membrane | transporter |
| 15548403 | 2 | 0.011 | 6025460 | 1.12E+07 | 6469900 | 6884762.5 | 7392312.5 | 8399743 | 4700044 | 8415364 | 5726193.5 | 3895254.75 | 3454454.5 | 2829718.25 | 1561.7583 | GQNEDPGSLLTQFR | P10852 | P10852 | SLC3A2 | solute carrier family 3 (amino acid transporter heavy chain), member 2 | Plasma Membrane | transporter |
| 15537201 | 2 | 0.014 | 3028208.5 | 4243978.5 | 2241203.25 | 3825413.5 | 3246889.75 | 3787610.75 | 2513976.5 | 3019996.5 | 2457386.25 | 2219523 | 1600469.625 | 1685802.625 | 1330.6033 | QPMNAADGAAAGEK | P10852 | P10852 | SLC3A2 | solute carrier family 3 (amino acid transporter heavy chain), member 2 | Plasma Membrane | transporter |
| 15542879 | 2 | 0.219 | 1931506.125 | 2940406 | 1339239.375 | 3355481 | 1522746.125 | 1975618.5 | 1651368.5 | 965014.062 | 1180450.75 | 1693886.625 | 1026765.125 | 1598479.625 | 1023.596 | YLVVLNFR | P10852 | P10852 | SLC3A2 | solute carrier family 3 (amino acid transporter heavy chain), member 2 | Plasma Membrane | transporter |
| 15531908 | 2 | 0.163 | 3.22E+07 | 2.99E+07 | 2.90E+07 | 2.58E+07 | 3.05E+07 | 2.65E+07 | 3.08E+07 | 2.72E+07 | 2.55E+07 | 3.68E+07 | 3.23E+07 | 2.90E+07 | 1004.51807 | LADMNATIR | Q9JHI9 | Q9JHI9 | SLC40A1 | solute carrier family 40 (iron-regulated transporter), member 1 | Plasma Membrane | transporter |
| 15540541 | 2, 3 | 0.889 | 2.06E+07 | 2.57E+07 | 2.84E+07 | 1.84E+07 | 2.80E+07 | 2.32E+07 | 3.00E+07 | 2.42E+07 | 1.56E+07 | 3.41E+07 | 1.57E+07 | 1.54E+07 | 1211.6051, 1211.6052 | PLEGTHLMGEK | Q9JHI9 | Q9JHI9 | SLC40A1 | solute carrier family 40 (iron-regulated transporter), member 1 | Plasma Membrane | transporter |
| 15552593 | 3 | 0.213 | 2.61E+07 | 2.09E+07 | 1.56E+07 | 2.72E+07 | 1.29E+07 | 1.99E+07 | 2.18E+07 | 1.48E+07 | 2.04E+07 | 2.81E+07 | 3.10E+07 | 2.73E+07 | 2897.2432 | DSNIRELEC[160.0307]EQEPTC[160.0307]ASQMAEPFR | Q9JHI9 | Q9JHI9 | SLC40A1 | solute carrier family 40 (iron-regulated transporter), member 1 | Plasma Membrane | transporter |
| 15549124 | 2 | 0.113 | 1.73E+07 | 1.79E+07 | 1.90E+07 | 1.58E+07 | 1.74E+07 | 1.46E+07 | 1.49E+07 | 1.52E+07 | 1.51E+07 | 2.58E+07 | 1.61E+07 | 1.90E+07 | 1577.76 | TLGNQIFVC[160.0307]GPDEK | Q9JHI9 | Q9JHI9 | SLC40A1 | solute carrier family 40 (iron-regulated transporter), member 1 | Plasma Membrane | transporter |
| 15531665 | 3 | 0.486 | 3.71E+07 | 8189231.5 | 756772.062 | 2.52E+07 | 4936577 | 4404528 | 6934526 | 5469979 | 1.67E+07 | 1.81E+07 | 3.83E+07 | 2.40E+07 | 1781.8702 | DTEPKPLEGTHLMGEK | Q9JHI9 | Q9JHI9 | SLC40A1 | solute carrier family 40 (iron-regulated transporter), member 1 | Plasma Membrane | transporter |
| 15555025, 15555816 | 2, 3 | 0.902 | 1.15E+07 | 1.28E+07 | 1.74E+07 | 1.01E+07 | 1.93E+07 | 1.19E+07 | 1.03E+07 | 1.58E+07 | 1.05E+07 | 1.71E+07 | 8868785 | 1.10E+07 | 2311.9536, 2311.9587 | ELEC[160.0307]EQEPTC[160.0307]ASQMAEPFR | Q9JHI9 | Q9JHI9 | SLC40A1 | solute carrier family 40 (iron-regulated transporter), member 1 | Plasma Membrane | transporter |
| 15544803 | 2 | 0.769 | 1.30E+07 | 1.44E+07 | 1.43E+07 | 6125262.5 | 1.70E+07 | 1.37E+07 | 1.50E+07 | 1.55E+07 | 1.00E+07 | 1.03E+07 | 9638647 | 1.40E+07 | 1431.6555 | EVTDENQPNTSVV | Q9JHI9 | Q9JHI9 | SLC40A1 | solute carrier family 40 (iron-regulated transporter), member 1 | Plasma Membrane | transporter |
| 15552523 | 2 | 0.397 | 2819328 | 615584.375 | 74172.32 | 2319755 | 230935.078 | 63381.707 | 445663.031 | 289938.656 | 1189807 | 1891902.875 | 2910512.5 | 1822286 | 1781.8706 | DTEPKPLEGTHLMGEK | Q9JHI9 | Q9JHI9 | SLC40A1 | solute carrier family 40 (iron-regulated transporter), member 1 | Plasma Membrane | transporter |
| 15536157 | 2 | 0.747 | 6.98E+08 | 7.03E+08 | 6.86E+08 | 5.88E+08 | 7.02E+08 | 6.33E+08 | 6.80E+08 | 6.80E+08 | 5.75E+08 | 7.89E+08 | 6.39E+08 | 5.77E+08 | 1277.6356 | EYATNLVDPQK | Q8BSM7 | Q8BSM7 | SLC43A1 | solute carrier family 43 (amino acid system L transporter), member 1 | Plasma Membrane | transporter |
| 15537338 | 2 | 0.429 | 4.56E+08 | 4.22E+08 | 4.81E+08 | 3.27E+08 | 4.25E+08 | 4.07E+08 | 4.62E+08 | 4.20E+08 | 3.67E+08 | 4.85E+08 | 4.03E+08 | 3.82E+08 | 979.5931 | LIGLALDHK | Q8BSM7 | Q8BSM7 | SLC43A1 | solute carrier family 43 (amino acid system L transporter), member 1 | Plasma Membrane | transporter |
| 15537197 | 2 | 0.447 | 2.09E+08 | 2.80E+08 | 2.91E+08 | 1.57E+08 | 2.19E+08 | 2.34E+08 | 2.84E+08 | 2.70E+08 | 1.82E+08 | 2.37E+08 | 1.63E+08 | 2.24E+08 | 1076.6344 | ILEFIVTGGK | Q8BSM7 | Q8BSM7 | SLC43A1 | solute carrier family 43 (amino acid system L transporter), member 1 | Plasma Membrane | transporter |
| 15548747 | 2 | 0.236 | 1.27E+08 | 1.29E+08 | 1.23E+08 | 9.58E+07 | 1.27E+08 | 1.09E+08 | 1.17E+08 | 1.21E+08 | 1.13E+08 | 1.24E+08 | 1.02E+08 | 9.25E+07 | 2238.9534 | DC[160.0307]VDAPTEGTLNENASFGDAR | Q8BSM7 | Q8BSM7 | SLC43A1 | solute carrier family 43 (amino acid system L transporter), member 1 | Plasma Membrane | transporter |
| 15536604, 15542155 | 2, 3 | 0.68 | 8.63E+07 | 9.00E+07 | 6.54E+07 | 6.43E+07 | 6.98E+07 | 5.20E+07 | 8.84E+07 | 7.01E+07 | 4.37E+07 | 9.85E+07 | 5.66E+07 | 5.45E+07 | 1320.7023, 1320.703, 1320.7031 | FYTHVTIVGQR | Q8BSM7 | Q8BSM7 | SLC43A1 | solute carrier family 43 (amino acid system L transporter), member 1 | Plasma Membrane | transporter |
| 15551166 | 3 | 0.173 | 8.75E+07 | 5.92E+07 | 7.42E+07 | 6.59E+07 | 5.43E+07 | 8.78E+07 | 8.24E+07 | 7.03E+07 | 7.60E+07 | 9.60E+07 | 1.08E+08 | 8.34E+07 | 2798.211 | DC[160.0307]VDAPTEGTLNENASFGDARDGASTK | Q8BSM7 | Q8BSM7 | SLC43A1 | solute carrier family 43 (amino acid system L transporter), member 1 | Plasma Membrane | transporter |
| 15532945 | 3 | 0.692 | 6.58E+07 | 7.55E+07 | 7.38E+07 | 5.38E+07 | 7.52E+07 | 6.66E+07 | 6.60E+07 | 9.57E+07 | 6.61E+07 | 8.09E+07 | 5.72E+07 | 6.20E+07 | 2238.9534 | DC[160.0307]VDAPTEGTLNENASFGDAR | Q8BSM7 | Q8BSM7 | SLC43A1 | solute carrier family 43 (amino acid system L transporter), member 1 | Plasma Membrane | transporter |
| 15536808, 15546148 | 3, 4 | 0.351 | 9.58E+07 | 6.23E+07 | 3.13E+07 | 9.39E+07 | 2.29E+07 | 4.84E+07 | 7.29E+07 | 3.39E+07 | 5.27E+07 | 7.98E+07 | 1.37E+08 | 8.63E+07 | 1848.9652, 1848.9669, 1848.968 | VTGDRFYTHVTIVGQR | Q8BSM7 | Q8BSM7 | SLC43A1 | solute carrier family 43 (amino acid system L transporter), member 1 | Plasma Membrane | transporter |
| 15549671 | 3 | 0.068 | 3.33E+07 | 3.43E+07 | 2.97E+07 | 2.42E+07 | 2.28E+07 | 2.97E+07 | 2.98E+07 | 2.75E+07 | 3.48E+07 | 3.77E+07 | 3.34E+07 | 3.01E+07 | 2790.2795 | SPSLEEGADAFISSPDIPGTSEETPEK | Q8BSM7 | Q8BSM7 | SLC43A1 | solute carrier family 43 (amino acid system L transporter), member 1 | Plasma Membrane | transporter |
| 15551597 | 2 | 0.301 | 1.26E+07 | 6286423.5 | 2194634.5 | 1.21E+07 | 1535684.625 | 2973955 | 6821147 | 1641285.625 | 3332924.75 | 1.14E+07 | 1.58E+07 | 8099009 | 1848.9677 | VTGDRFYTHVTIVGQR | Q8BSM7 | Q8BSM7 | SLC43A1 | solute carrier family 43 (amino acid system L transporter), member 1 | Plasma Membrane | transporter |
| 15537447 | 2 | 0.831 | 5934778 | 9293453 | 9214256 | 4576279 | 1.29E+07 | 8032823 | 1.07E+07 | 1.07E+07 | 6576332.5 | 9741798 | 5691244 | 6077699 | 1173.5776 | VIFYMGAMNK | Q8BSM7 | Q8BSM7 | SLC43A1 | solute carrier family 43 (amino acid system L transporter), member 1 | Plasma Membrane | transporter |
| 15550490 | 3 | 0.793 | 3995760 | 3443006.75 | 4195037 | 4103322.25 | 4299851 | 2580633.25 | 5441269 | 4435091.5 | 3252660 | 4864703.5 | 3582961.75 | 2660574.5 | 2363.9746 | TNTTQDEQHQWTSC[160.0307]DQQEK | Q8BSM7 | Q8BSM7 | SLC43A1 | solute carrier family 43 (amino acid system L transporter), member 1 | Plasma Membrane | transporter |
| 15541895 | 3 | 0.461 | 4591936.5 | 1335052.625 | 1478511.875 | 1.05E+07 | 725504.438 | 1894236.25 | 2308070 | 1895370.25 | 2490669.5 | 3287061.5 | 8216759 | 5987862 | 1674.8773 | LQREYATNLVDPQK | Q8BSM7 | Q8BSM7 | SLC43A1 | solute carrier family 43 (amino acid system L transporter), member 1 | Plasma Membrane | transporter |
| 15544434 | 2 | 0.223 | 2110028.25 | 3517254.75 | 3978304.75 | 805099.812 | 3759549.75 | 2066641.75 | 2657441.25 | 3767425.5 | 969400.375 | 1880323.875 | 321272.219 | 967332.938 | 1205.5669 | VIFYM[147.0354]GAM[147.0354]NK | Q8BSM7 | Q8BSM7 | SLC43A1 | solute carrier family 43 (amino acid system L transporter), member 1 | Plasma Membrane | transporter |
| 15548538 | 4 | 0.641 | 3.68E+07 | 5.05E+07 | 6.24E+07 | 4.50E+07 | 9.51E+07 | 4.73E+07 | 8.15E+07 | 8.45E+07 | 3.85E+07 | 6.69E+07 | 3.86E+07 | 3.43E+07 | 3861.673 | EENSGPEQQQQQEQQQQQQQQQEQHEQHSFR | A2AVZ9 | A2AVZ9 | SLC43A3 | solute carrier family 43, member 3 | Extracellular Space | other |
| 15534316 | 3 | 0.134 | 2.87E+07 | 1.98E+07 | 1.89E+07 | 1.71E+07 | 1.70E+07 | 1.34E+07 | 2.16E+07 | 2.25E+07 | 1.92E+07 | 1.72E+07 | 2.68E+07 | 2.51E+07 | 2091.0083 | GHIPYPLPPNYSYGLC[160.0307]SR | A2AVZ9 | A2AVZ9 | SLC43A3 | solute carrier family 43, member 3 | Extracellular Space | other |
| 15532302 | 2 | 0.247 | 1.75E+07 | 1.95E+07 | 2.35E+07 | 1.01E+07 | 1.91E+07 | 1.85E+07 | 1.81E+07 | 1.98E+07 | 1.65E+07 | 1.55E+07 | 1.34E+07 | 1.42E+07 | 1191.6708 | LLYEQGISLR | A2AVZ9 | A2AVZ9 | SLC43A3 | solute carrier family 43, member 3 | Extracellular Space | other |
| 15538250 | 2 | 0.536 | 7394859 | 8169668 | 8917777 | 6707812.5 | 7616579.5 | 7202925 | 8926259 | 9760622 | 6704143 | 9029134 | 7045499 | 6376297.5 | 849.48846 | TFLLMPK | A2AVZ9 | A2AVZ9 | SLC43A3 | solute carrier family 43, member 3 | Extracellular Space | other |
| 15538925 | 3 | 0.099 | 2765098.25 | 3665222.75 | 4400799 | 3985816.75 | 3993045.5 | 3502486.75 | 4962436 | 5307279 | 5528374.5 | 2728212 | 4645957.5 | 3837047.5 | 1548.8647 | LPVPASAPIPFFHR | Q6X893 | Q6X893 | SLC44A1 | solute carrier family 44 (choline transporter), member 1 | Plasma Membrane | transporter |
| 15540080 | 3 | 0.235 | 2027495.625 | 1871970.375 | 2517852.75 | 1330585.875 | 2300267.25 | 2055873.75 | 2269383.25 | 2982948.75 | 3187815 | 1959755.5 | 3127938.5 | 2074385 | 1918.9973 | FAEALITFVSDNSVLHR | Q6X893 | Q6X893 | SLC44A1 | solute carrier family 44 (choline transporter), member 1 | Plasma Membrane | transporter |
| 15556878 | 2 | 0.838 | 2500142.5 | 1623887.125 | 2321375 | 1866481.75 | 2541546.5 | 1663946.875 | 2162502 | 2162205.5 | 2638261.75 | 1416195.25 | 2592288.75 | 2152986.25 | 1688.769 | LVSGYDSYGNIC[160.0307]GQR | Q6X893 | Q6X893 | SLC44A1 | solute carrier family 44 (choline transporter), member 1 | Plasma Membrane | transporter |
| 15549459, 15552799, 15553841, 15558344, 15559291, 15559659, 15559681, 15559699, 15560507, 15560613, 15561396, 15562718, 15562753, 15563288, 15563544 | 3, 4, 5 | 0.997 | 2.63E+10 | 2.66E+10 | 3.18E+10 | 2.45E+10 | 2.98E+10 | 3.00E+10 | 2.97E+10 | 2.82E+10 | 2.54E+10 | 3.15E+10 | 2.58E+10 | 2.67E+10 | 3827.7913, 3827.7922, 3827.793, 3827.7932, 3827.7961, 3827.7969, 3827.797, 3827.7976, 3827.802, 3827.8035, 3827.8037, 3827.8042, 3827.8093 | SGGASEPLLPHQPSLETQLYC[160.0307]GQAEGGSEGPSTSGTLK | P04919 | P04919 | SLC4A1 | solute carrier family 4 (anion exchanger), member 1 (Diego blood group) | Plasma Membrane | transporter |

| | | | | | | | | | | | | | | | | | | | | | |
|---|---|---|---|---|---|---|---|---|---|---|---|---|---|---|---|---|---|---|---|---|---|
| 15536770, 15539711, 15544536, 15545696, 15546857, 15546877, 15547258, 15547405, 15547447, 15547457, 15547459, 15547515, 15547547, 15547589 | 2, 3 | 0.67 | 1.52E+10 | 1.52E+10 | 1.74E+10 | 1.46E+10 | 1.42E+10 | 1.58E+10 | 1.72E+10 | 1.53E+10 | 1.33E+10 | 1.83E+10 | 1.57E+10 | 1.49E+10 | 1496.8213, 1496.8235, 1496.8236, 1496.8243, 1496.8252, 1496.8259, 1496.8282, 1496.8292, 1496.8296, 1496.8303, 1496.8304, 1496.8319, 1496.8331, 1496.8379 | IPPDSETTLVLVGR | P04919 | P04919 | SLC4A1 | solute carrier family 4 (anion exchanger), member 1 (Diego blood group) | Plasma Membrane | transporter |
| 15537825, 15540675, 15546482 | 2, 3 | 0.563 | 9.76E+09 | 8.92E+09 | 1.01E+10 | 9.26E+09 | 8.47E+09 | 8.33E+09 | 1.03E+10 | 8.67E+09 | 7.50E+09 | 1.16E+10 | 9.16E+09 | 8.64E+09 | 1283.5948, 1283.5951, 1283.5967 | GGPGDEDDPLRR | P04919 | P04919 | SLC4A1 | solute carrier family 4 (anion exchanger), member 1 (Diego blood group) | Plasma Membrane | transporter |
| 15536480, 15544797, 15546396 | 2, 3 | 0.556 | 7.80E+09 | 7.41E+09 | 8.79E+09 | 7.78E+09 | 8.35E+09 | 7.08E+09 | 8.19E+09 | 8.35E+09 | 6.50E+09 | 1.11E+10 | 7.88E+09 | 7.91E+09 | 1326.6923, 1326.6975, 1326.6979, 1326.6992 | ASGPGAAAQIQEVK | P04919 | P04919 | SLC4A1 | solute carrier family 4 (anion exchanger), member 1 (Diego blood group) | Plasma Membrane | transporter |
| 15547801, 15551410, 15552448, 15559528, 15559557, 15563273 | 2 | 0.537 | 7.23E+09 | 7.39E+09 | 9.80E+09 | 6.46E+09 | 7.95E+09 | 5.66E+09 | 7.55E+09 | 9.09E+09 | 6.85E+09 | 1.08E+10 | 7.22E+09 | 6.90E+09 | 1505.6705, 1505.6743, 1505.6744, 1505.6764, 1505.6777 | ELELQC[160.0307]LDGDDAK | P04919 | P04919 | SLC4A1 | solute carrier family 4 (anion exchanger), member 1 (Diego blood group) | Plasma Membrane | transporter |
| 15554280, 15556720, 15559304, 15561597 | 2, 3 | 0.341 | 9.04E+09 | 8.26E+09 | 9.42E+09 | 7.57E+09 | 8.14E+09 | 8.42E+09 | 8.18E+09 | 8.17E+09 | 7.07E+09 | 1.01E+10 | 7.79E+09 | 8.10E+09 | 2305.1245, 2305.1252, 2305.127, 2305.1277 | SLESFLDC[160.0307]SLVLPPTDAPSEK | P04919 | P04919 | SLC4A1 | solute carrier family 4 (anion exchanger), member 1 (Diego blood group) | Plasma Membrane | transporter |
| 15533950, 15537202, 15541392, 15542624, 15547434, 15547622 | 2 | 0.466 | 9.79E+09 | 9.87E+09 | 1.15E+10 | 8.80E+09 | 9.19E+09 | 9.99E+09 | 1.03E+10 | 9.99E+09 | 8.43E+09 | 9.78E+09 | 9.28E+09 | 1.02E+10 | 1094.6904, 1094.6914, 1094.6917, 1094.6919, 1094.6925, 1094.6964 | ALLNLVPVQK | P04919 | P04919 | SLC4A1 | solute carrier family 4 (anion exchanger), member 1 (Diego blood group) | Plasma Membrane | transporter |
| 15533407, 15534299, 15540431, 15541219 | 2, 3 | 0.202 | 7.34E+09 | 6.31E+09 | 6.39E+09 | 6.31E+09 | 6.25E+09 | 5.73E+09 | 7.13E+09 | 6.20E+09 | 5.07E+09 | 9.45E+09 | 7.41E+09 | 6.49E+09 | 1328.6959, 1328.6964, 1328.6981, 1328.6987 | SVTHANALTVMGK | P04919 | P04919 | SLC4A1 | solute carrier family 4 (anion exchanger), member 1 (Diego blood group) | Plasma Membrane | transporter |
| 15532073, 15539240, 15546775 | 3 | 0.036 | 8.11E+09 | 7.07E+09 | 7.82E+09 | 5.70E+09 | 6.85E+09 | 6.53E+09 | 7.97E+09 | 7.27E+09 | 7.09E+09 | 8.09E+09 | 8.00E+09 | 7.18E+09 | 1640.8496, 1640.8508, 1640.853 | ITASMAHNREELLR | P04919 | P04919 | SLC4A1 | solute carrier family 4 (anion exchanger), member 1 (Diego blood group) | Plasma Membrane | transporter |
| 15536869 | 2, 3 | 0.596 | 5.65E+09 | 5.92E+09 | 7.26E+09 | 4.12E+09 | 6.17E+09 | 5.74E+09 | 6.39E+09 | 6.22E+09 | 5.13E+09 | 6.54E+09 | 5.02E+09 | 5.05E+09 | 1101.571, 1101.5713 | YHPDVPFVK | P04919 | P04919 | SLC4A1 | solute carrier family 4 (anion exchanger), member 1 (Diego blood group) | Plasma Membrane | transporter |
| 15532909, 15536849, 15539122 | 2 | 0.648 | 4.44E+09 | 4.35E+09 | 5.55E+09 | 4.32E+09 | 4.26E+09 | 4.44E+09 | 4.85E+09 | 4.81E+09 | 4.28E+09 | 5.52E+09 | 4.22E+09 | 4.71E+09 | 828.4809, 828.48145, 828.48175 | LSVPDGLK | P04919 | P04919 | SLC4A1 | solute carrier family 4 (anion exchanger), member 1 (Diego blood group) | Plasma Membrane | transporter |
| 15533042, 15534400, 15542192, 15543542 | 2 | 0.568 | 4.11E+09 | 3.76E+09 | 4.13E+09 | 4.42E+09 | 3.35E+09 | 3.31E+09 | 3.96E+09 | 3.39E+09 | 3.19E+09 | 5.50E+09 | 3.71E+09 | 3.50E+09 | 963.48975, 963.49054, 963.4911, 963.4919 | AAATLMTER | P04919 | P04919 | SLC4A1 | solute carrier family 4 (anion exchanger), member 1 (Diego blood group) | Plasma Membrane | transporter |
| 15532530, 15541530 | 2, 3 | 0.388 | 3.75E+09 | 3.69E+09 | 4.40E+09 | 2.65E+09 | 3.98E+09 | 3.47E+09 | 4.43E+09 | 3.86E+09 | 3.57E+09 | 4.71E+09 | 3.57E+09 | 3.89E+09 | 1310.7347, 1310.7362 | GWVIHPLGLYR | P04919 | P04919 | SLC4A1 | solute carrier family 4 (anion exchanger), member 1 (Diego blood group) | Plasma Membrane | transporter |
| 15541804, 15542131 | 3, 4 | 0.585 | 5.00E+09 | 6.84E+09 | 8.51E+09 | 2.71E+09 | 7.18E+09 | 7.33E+09 | 6.24E+09 | 7.41E+09 | 7.20E+09 | 3.97E+09 | 4.62E+09 | 6.79E+09 | 1524.7744, 1524.7755 | RSHAEDLGNLEGVK | P04919 | P04919 | SLC4A1 | solute carrier family 4 (anion exchanger), member 1 (Diego blood group) | Plasma Membrane | transporter |
| 15531718, 15546534, 15546753, 15547550 | 2 | 0.817 | 3.23E+09 | 3.97E+09 | 5.29E+09 | 2.88E+09 | 4.23E+09 | 4.55E+09 | 4.09E+09 | 4.48E+09 | 4.57E+09 | 3.72E+09 | 3.64E+09 | 4.45E+09 | 984.65814, 984.66016, 984.6604, 984.66156 | LILPLIFR | P04919 | P04919 | SLC4A1 | solute carrier family 4 (anion exchanger), member 1 (Diego blood group) | Plasma Membrane | transporter |
| 15548512 | 2 | 0.904 | 2.09E+09 | 2.90E+09 | 4.68E+09 | 2.07E+09 | 3.09E+09 | 2.95E+09 | 3.01E+09 | 2.96E+09 | 2.34E+09 | 3.32E+09 | 2.17E+09 | 2.61E+09 | 2041.045 | IFQDYPLQQTYAPVVMK | P04919 | P04919 | SLC4A1 | solute carrier family 4 (anion exchanger), member 1 (Diego blood group) | Plasma Membrane | transporter |
| 15548475, 15550107 | 2, 3 | 0.838 | 1.43E+09 | 2.44E+09 | 2.83E+09 | 1.16E+09 | 3.54E+09 | 1.90E+09 | 2.09E+09 | 2.87E+09 | 1.59E+09 | 2.67E+09 | 1.06E+09 | 1.47E+09 | 2435.2053, 2435.2095 | NQELQWVEAAHWIGLEENLR | P04919 | P04919 | SLC4A1 | solute carrier family 4 (anion exchanger), member 1 (Diego blood group) | Plasma Membrane | transporter |
| 15531471, 15533416 | 2, 3 | 0.569 | 1.01E+09 | 1.39E+09 | 2.09E+09 | 7.84E+08 | 1.90E+09 | 1.21E+09 | 2.03E+09 | 1.73E+09 | 9.18E+08 | 2.12E+09 | 9.54E+08 | 9.18E+08 | 1368.6742, 1368.676 | SHAEDLGNLEGVK | P04919 | P04919 | SLC4A1 | solute carrier family 4 (anion exchanger), member 1 (Diego blood group) | Plasma Membrane | transporter |
| 15540545 | 2 | 0.683 | 1.55E+09 | 1.59E+09 | 2.57E+09 | 1.52E+09 | 1.87E+09 | 1.82E+09 | 1.86E+09 | 1.89E+09 | 1.78E+09 | 1.94E+09 | 1.30E+09 | 1.54E+09 | 913.4408 | NSTYFPGK | P04919 | P04919 | SLC4A1 | solute carrier family 4 (anion exchanger), member 1 (Diego blood group) | Plasma Membrane | transporter |
| 15558100, 15558847 | 2 | 0.546 | 1.20E+09 | 2.03E+09 | 2.16E+09 | 4.72E+08 | 1.82E+09 | 1.76E+09 | 1.78E+09 | 1.81E+09 | 1.69E+09 | 1.05E+09 | 1.15E+09 | 1.67E+09 | 1524.773, 1524.775 | RSHAEDLGNLEGVK | P04919 | P04919 | SLC4A1 | solute carrier family 4 (anion exchanger), member 1 (Diego blood group) | Plasma Membrane | transporter |
| 15533518 | 2 | 0.85 | 7.14E+08 | 7.26E+08 | 7.65E+08 | 6.58E+08 | 7.76E+08 | 6.11E+08 | 7.86E+08 | 7.42E+08 | 5.91E+08 | 9.07E+08 | 6.51E+08 | 6.77E+08 | 1127.4946 | GGPGDEDDPLR | P04919 | P04919 | SLC4A1 | solute carrier family 4 (anion exchanger), member 1 (Diego blood group) | Plasma Membrane | transporter |
| 15531550, 15537074, 15542825, 15544052, 15544802, 15544842, 15544974, 15546307 | 3 | 0.104 | 1.01E+09 | 1.03E+09 | 1.20E+09 | 9.55E+08 | 8.04E+08 | 8.52E+08 | 8.70E+08 | 8.25E+08 | 6.59E+08 | 9.05E+08 | 4.27E+08 | 7.08E+08 | 1656.8359, 1656.8376, 1656.8431, 1656.8453, 1656.8458, 1656.8474, 1656.85, 1656.852 | ITASM[147.0354]AHNREELLR | P04919 | P04919 | SLC4A1 | solute carrier family 4 (anion exchanger), member 1 (Diego blood group) | Plasma Membrane | transporter |
| 15536012 | 2, 3 | 0.424 | 7.61E+08 | 9.00E+08 | 1.13E+09 | 5.80E+08 | 1.07E+09 | 8.49E+08 | 9.76E+08 | 9.50E+08 | 6.42E+08 | 6.80E+08 | 6.20E+08 | 7.24E+08 | 1068.7151, 1068.7152 | ILLLFKPPK | P04919 | P04919 | SLC4A1 | solute carrier family 4 (anion exchanger), member 1 (Diego blood group) | Plasma Membrane | transporter |
| 15531260, 15532206, 15540092, 15541330, 15541658, 15542619, 15542762, 15543369, 15543482, 15543724, 15544057, 15544271, 15545100, 15545365, 15545398, 15545433, 15545492, 15545651, 15546007, 15546103, 15546104, 15546118, 15546303, 15546394, 15546562, 15546673, 15546722, 15546943, 15547099, 15547119 | 2, 3 | 0.232 | 5.82E+08 | 6.49E+08 | 8.25E+08 | 5.88E+08 | 5.94E+08 | 5.91E+08 | 6.36E+08 | 5.71E+08 | 4.71E+08 | 6.57E+08 | 3.21E+08 | 4.52E+08 | 1344.6848, 1344.6859, 1344.6865, 1344.6882, 1344.6886, 1344.6893, 1344.6896, 1344.6904, 1344.6909, 1344.6914, 1344.6915, 1344.6925, 1344.6926, 1344.6927, 1344.6931, 1344.6943, 1344.6946, 1344.6948, 1344.696, 1344.6964, 1344.6968, 1344.6969, 1344.6973, 1344.6976, 1344.6987, 1344.7 | SVTHANALTVM[147.0354]GK | P04919 | P04919 | SLC4A1 | solute carrier family 4 (anion exchanger), member 1 (Diego blood group) | Plasma Membrane | transporter |
| 15536820 | 3 | 0.859 | 4.30E+08 | 5.46E+08 | 8.22E+08 | 3.73E+08 | 7.42E+08 | 6.16E+08 | 5.90E+08 | 6.97E+08 | 6.91E+08 | 6.32E+08 | 4.28E+08 | 6.49E+08 | 2041.0454 | IFQDYPLQQTYAPVVMK | P04919 | P04919 | SLC4A1 | solute carrier family 4 (anion exchanger), member 1 (Diego blood group) | Plasma Membrane | transporter |
| 15548870, 15556703, 15557982, 15558235, 15558654, 15559556, 15561743, 15561810, 15562877 | 2 | 0.349 | 4.26E+08 | 5.97E+08 | 9.57E+08 | 3.33E+08 | 6.40E+08 | 6.27E+08 | 6.76E+08 | 5.72E+08 | 3.49E+08 | 5.61E+08 | 2.33E+08 | 2.97E+08 | 2057.0283, 2057.029, 2057.0322, 2057.0352, 2057.0369, 2057.0374, 2057.0376, 2057.039, 2057.0403 | IFQDYPLQQTYAPVVM[147.0354]K | P04919 | P04919 | SLC4A1 | solute carrier family 4 (anion exchanger), member 1 (Diego blood group) | Plasma Membrane | transporter |
| 15547802 | 3, 4 | 0.96 | 2.23E+08 | 3.35E+08 | 4.11E+08 | 1.68E+08 | 5.96E+08 | 3.85E+08 | 3.37E+08 | 3.77E+08 | 2.18E+08 | 5.02E+08 | 1.58E+08 | 2.61E+08 | 3211.5315, 3211.535 | DHEEVLEIPDRDSEEELENIIGQIAYR | P04919 | P04919 | SLC4A1 | solute carrier family 4 (anion exchanger), member 1 (Diego blood group) | Plasma Membrane | transporter |
| 15558639, 15558775 | 2 | 0.441 | 5.33E+08 | 6.06E+08 | 8.06E+08 | 3.03E+08 | 6.08E+08 | 6.63E+08 | 6.03E+08 | 6.80E+08 | 6.95E+08 | 4.61E+08 | 4.93E+08 | 5.99E+08 | 1640.8497, 1640.8503 | ITASMAHNREELLR | P04919 | P04919 | SLC4A1 | solute carrier family 4 (anion exchanger), member 1 (Diego blood group) | Plasma Membrane | transporter |
| 15533636 | 3, 4 | 0.337 | 4.37E+08 | 4.27E+07 | 2434495 | 2.96E+08 | 4.07E+07 | 1.31E+07 | 1.45E+07 | 4.23E+07 | 1.87E+08 | 3.90E+08 | 6.61E+08 | 2.49E+08 | 2006.0623, 2006.0627 | SHAEDLGNLEGVKPAVLTR | P04919 | P04919 | SLC4A1 | solute carrier family 4 (anion exchanger), member 1 (Diego blood group) | Plasma Membrane | transporter |
| 15555727 | 3 | 0.88 | 2.49E+08 | 3.80E+08 | 2.78E+08 | 1.46E+08 | 5.11E+08 | 1.86E+08 | 2.85E+08 | 3.88E+08 | 1.14E+08 | 3.72E+08 | 1.27E+08 | 1.93E+08 | 3234.5637 | NQELQWVEAAHWIGLEENLREDGVWGR | P04919 | P04919 | SLC4A1 | solute carrier family 4 (anion exchanger), member 1 (Diego blood group) | Plasma Membrane | transporter |
| 15531763 | 2, 3 | 0.039 | 3.44E+08 | 3.32E+08 | 3.55E+08 | 2.14E+08 | 3.11E+08 | 3.01E+08 | 3.88E+08 | 3.60E+08 | 3.89E+08 | 3.50E+08 | 3.29E+08 | 3.03E+08 | 1140.7589, 1140.7593 | RLILPLIFR | P04919 | P04919 | SLC4A1 | solute carrier family 4 (anion exchanger), member 1 (Diego blood group) | Plasma Membrane | transporter |
| 15548013, 15553822 | 2, 3 | 0.087 | 6.76E+08 | 7.62E+08 | 4.68E+08 | 6.65E+08 | 6.72E+08 | 3.56E+08 | 3.07E+08 | 5.01E+08 | 5.87E+08 | 3.21E+08 | 3.65E+08 | 2.62E+08 | 2387.2192, 2387.2212 | YLPSPAKPDPNLYNTLDLNGGK | P04919 | P04919 | SLC4A1 | solute carrier family 4 (anion exchanger), member 1 (Diego blood group) | Plasma Membrane | transporter |
| 15552623 | 2 | 0.194 | 2.73E+08 | 5.99E+08 | 4.71E+08 | 3.24E+08 | 5.90E+08 | 2.53E+08 | 3.29E+08 | 5.11E+08 | 4.44E+08 | 2.44E+08 | 1.84E+08 | 2.94E+08 | 2197.9453 | VTFDEENGLDEYDEVPMPV | P04919 | P04919 | SLC4A1 | solute carrier family 4 (anion exchanger), member 1 (Diego blood group) | Plasma Membrane | transporter |
| 15535061, 15546357, 15546554, 15546574, 15546802, 15547049 | 3 | 0.321 | 1.55E+08 | 2.39E+08 | 3.87E+08 | 9.93E+07 | 2.63E+08 | 2.43E+08 | 2.28E+08 | 2.21E+08 | 1.45E+08 | 1.73E+08 | 7.39E+07 | 1.36E+08 | 2057.0334, 2057.0337, 2057.0352, 2057.0374, 2057.0415 | IFQDYPLQQTYAPVVM[147.0354]K | P04919 | P04919 | SLC4A1 | solute carrier family 4 (anion exchanger), member 1 (Diego blood group) | Plasma Membrane | transporter |
| 15539899 | 4 | 0.763 | 9.82E+07 | 1.23E+08 | 1.87E+08 | 7.47E+07 | 1.31E+08 | 7.72E+07 | 1.24E+08 | 1.56E+08 | 3.76E+07 | 1.57E+08 | 6.09E+07 | 8.21E+07 | 2774.4226 | EDGVWGRPHLSYLTFWSLLELQK | P04919 | P04919 | SLC4A1 | solute carrier family 4 (anion exchanger), member 1 (Diego blood group) | Plasma Membrane | transporter |
| 15532535 | 3 | 0.301 | 1.42E+08 | 1.74E+08 | 3.45E+08 | 7.39E+07 | 1.86E+08 | 1.60E+08 | 1.92E+08 | 2.81E+08 | 1.17E+08 | 1.47E+08 | 7.92E+07 | 1.07E+08 | 1975.0659 | PHLSYLTFWSLLELQK | P04919 | P04919 | SLC4A1 | solute carrier family 4 (anion exchanger), member 1 (Diego blood group) | Plasma Membrane | transporter |
| 15554290, 15558609 | 3 | 0.435 | 9.77E+07 | 8.29E+07 | 5.73E+07 | 1.92E+08 | 3.34E+07 | 7.67E+07 | 7.83E+07 | 5.99E+07 | 8.93E+07 | 1.19E+08 | 1.65E+08 | 1.23E+08 | 2471.317, 2471.3179 | LILPLIFRELELQC[160.0307]LDGDDAK | P04919 | P04919 | SLC4A1 | solute carrier family 4 (anion exchanger), member 1 (Diego blood group) | Plasma Membrane | transporter |
| 15548101 | 2 | 0.127 | 8.00E+07 | 2.30E+08 | 4.28E+08 | 1.02E+08 | 3.78E+08 | 1.79E+08 | 1.53E+08 | 1.99E+08 | 1.13E+08 | 9.33E+07 | 4.29E+07 | 5.95E+07 | 1630.8047 | PDPNLYNTLDLNGGK | P04919 | P04919 | SLC4A1 | solute carrier family 4 (anion exchanger), member 1 (Diego blood group) | Plasma Membrane | transporter |
| 15540104, 15542242, 15542350, 15543037, 15543505, 15543550, 15543701, 15543814, 15543884, 15544049, 15544719, 15544900, 15544930, 15545287 | 2 | 0.862 | 6.07E+07 | 7.74E+07 | 1.35E+08 | 6.81E+07 | 9.54E+07 | 9.76E+07 | 8.45E+07 | 1.13E+08 | 9.76E+07 | 9.19E+07 | 7.60E+07 | 8.42E+07 | 979.48065, 979.4812, 979.4827, 979.4839, 979.4845, 979.4856, 979.48584, 979.4862, 979.48755, 979.4894, 979.49066, 979.4911, 979.4919, 979.4931 | AAATLM[147.0354]TER | P04919 | P04919 | SLC4A1 | solute carrier family 4 (anion exchanger), member 1 (Diego blood group) | Plasma Membrane | transporter |
| 15550716, 15556825, 15561858 | 2 | 0.252 | 7.77E+07 | 9.03E+07 | 1.12E+08 | 5.11E+07 | 6.40E+07 | 7.06E+07 | 6.68E+07 | 6.47E+07 | 5.28E+07 | 8.74E+07 | 2.77E+07 | 6.02E+07 | 1656.8456, 1656.8458, 1656.8463 | ITASM[147.0354]AHNREELLR | P04919 | P04919 | SLC4A1 | solute carrier family 4 (anion exchanger), member 1 (Diego blood group) | Plasma Membrane | transporter |
| 15538273 | 3 | 0.295 | 1.34E+08 | 2.21E+07 | 1.06E+07 | 1.01E+08 | 1.95E+07 | 1.40E+07 | 2.56E+07 | 2.47E+07 | 6.63E+07 | 6.71E+07 | 2.20E+08 | 1.11E+08 | 1257.6719 | YHPDVPFVKR | P04919 | P04919 | SLC4A1 | solute carrier family 4 (anion exchanger), member 1 (Diego blood group) | Plasma Membrane | transporter |
| 15534320 | 2 | 0.481 | 9.67E+07 | 3.00E+08 | 5.64E+08 | 4.55E+07 | 3.26E+08 | 3.32E+08 | 2.09E+08 | 3.22E+08 | 1.41E+08 | 6.28E+07 | 9.32E+07 | 1.45E+08 | 931.534 | RYLPSPAK | P04919 | P04919 | SLC4A1 | solute carrier family 4 (anion exchanger), member 1 (Diego blood group) | Plasma Membrane | transporter |
| 15554167, 15559555 | 3 | 0.408 | 5.73E+07 | 6.02E+07 | 9.77E+07 | 4.08E+07 | 4.77E+07 | 3.37E+07 | 5.51E+07 | 7.42E+07 | 1.98E+07 | 6.22E+07 | 2.96E+07 | 3.83E+07 | 2774.424, 2774.4253 | EDGVWGRPHLSYLTFWSLLELQK | P04919 | P04919 | SLC4A1 | solute carrier family 4 (anion exchanger), member 1 (Diego blood group) | Plasma Membrane | transporter |
| 15553778 | 1 | 0.567 | 5.90E+07 | 7.43E+07 | 1.17E+08 | 4.78E+07 | 7.87E+07 | 8.07E+07 | 7.18E+07 | 7.66E+07 | 9.73E+07 | 5.78E+07 | 5.71E+07 | 7.65E+07 | 984.65814 | LILPLIFR | P04919 | P04919 | SLC4A1 | solute carrier family 4 (anion exchanger), member 1 (Diego blood group) | Plasma Membrane | transporter |
| 15538598 | 4 | 0.993 | 1.24E+07 | 2.90E+07 | 3.01E+07 | 1.33E+07 | 3.62E+07 | 2.48E+07 | 2.48E+07 | 3.70E+07 | 1.47E+07 | 3.76E+07 | 1.37E+07 | 1.88E+07 | 2435.205 | NQELQWVEAAHWIGLEENLR | P04919 | P04919 | SLC4A1 | solute carrier family 4 (anion exchanger), member 1 (Diego blood group) | Plasma Membrane | transporter |
| 15534782, 15543247, 15544606, 15545250 | 2 | 0.637 | 2.23E+07 | 2.61E+07 | 3.15E+07 | 1.64E+07 | 2.78E+07 | 2.70E+07 | 2.69E+07 | 3.30E+07 | 2.36E+07 | 2.80E+07 | 2.87E+07 | 2.67E+07 | 787.4788, 787.48016, 787.48035, 787.4808 | PVLGFVR | P04919 | P04919 | SLC4A1 | solute carrier family 4 (anion exchanger), member 1 (Diego blood group) | Plasma Membrane | transporter |
| 15548379 | 2 | 0.419 | 2.41E+07 | 1713397.25 | 0 | 1.71E+07 | 1686859.125 | 128122.227 | 539540.375 | 1898416.5 | 8007966 | 2.49E+07 | 3.97E+07 | 1.20E+07 | 2006.0626 | SHAEDLGNLEGVKPAVLTR | P04919 | P04919 | SLC4A1 | solute carrier family 4 (anion exchanger), member 1 (Diego blood group) | Plasma Membrane | transporter |
| 15549670 | 3 | 0.651 | 4.18E+07 | 1.28E+07 | 6147972.5 | 3.24E+07 | 1.14E+07 | 9733201 | 1.15E+07 | 1.49E+07 | 2.95E+07 | 2.35E+07 | 2.85E+07 | 2.94E+07 | 2282.1838 | IFQDYPLQQTYAPVVM[147.0354]KPK | P04919 | P04919 | SLC4A1 | solute carrier family 4 (anion exchanger), member 1 (Diego blood group) | Plasma Membrane | transporter |
| 15542829 | 3 | 0.219 | 1.02E+07 | 1.37E+07 | 1.91E+07 | 1.04E+07 | 1.58E+07 | 1786807.375 | 1.52E+07 | 2.49E+07 | 1.76E+07 | 2.27E+07 | 1.36E+07 | 2.19E+07 | 1505.6754 | ELELQC[160.0307]LDGDDAK | P04919 | P04919 | SLC4A1 | solute carrier family 4 (anion exchanger), member 1 (Diego blood group) | Plasma Membrane | transporter |
| 15540508 | 3 | 0.413 | 2.58E+07 | 3.68E+07 | 3.63E+07 | 9337038 | 5.28E+07 | 2.55E+07 | 3.15E+07 | 3.23E+07 | 3.08E+07 | 1.68E+07 | 1.96E+07 | 1.92E+07 | 2197.9443 | VTFDEENGLDEYDEVPMPV | P04919 | P04919 | SLC4A1 | solute carrier family 4 (anion exchanger), member 1 (Diego blood group) | Plasma Membrane | transporter |
| 15540003 | 2, 3 | 0.287 | 8799773 | 4255767 | 3974564.75 | 8347099.5 | 4357202.5 | 3166129.25 | 5515562.5 | 4266428 | 6490350 | 1.60E+07 | 6751294 | 7445888.5 | 1489.8496, 1489.85 | ANFLEKPVLGFVR | P04919 | P04919 | SLC4A1 | solute carrier family 4 (anion exchanger), member 1 (Diego blood group) | Plasma Membrane | transporter |
| 15548712, 15548776 | 2 | 0.049 | 1.99E+07 | 1.69E+07 | 1.80E+07 | 1.64E+07 | 1.82E+07 | 8165351 | 1.42E+07 | 1.26E+07 | 1.32E+07 | 1.19E+07 | 6994432 | 7545442 | 2213.9382, 2213.9387 | VTFDEENGLDEYDEVPM[147.0354]PV | P04919 | P04919 | SLC4A1 | solute carrier family 4 (anion exchanger), member 1 (Diego blood group) | Plasma Membrane | transporter |
| 15556088 | 2 | 0.088 | 1.61E+07 | 1.85E+07 | 4.59E+07 | 7945289.5 | 2.43E+07 | 1.69E+07 | 1.96E+07 | 3.10E+07 | 1.45E+07 | 7461587 | 6685615.5 | 1.11E+07 | 1975.0647 | PHLSYLTFWSLLELQK | P04919 | P04919 | SLC4A1 | solute carrier family 4 (anion exchanger), member 1 (Diego blood group) | Plasma Membrane | transporter |
| 15551021 | 3 | 0.257 | 4705400 | 986143 | 103117.867 | 6354545.5 | 1369266.5 | 957939.25 | 863289.375 | 864395.625 | 2030366.875 | 2912850.25 | 1.16E+07 | 5256331.5 | 2266.1885 | IFQDYPLQQTYAPVVMKPK | P04919 | P04919 | SLC4A1 | solute carrier family 4 (anion exchanger), member 1 (Diego blood group) | Plasma Membrane | transporter |
| 15533834 | 2 | 0.611 | 7825774.5 | 6.10E+07 | 2.33E+08 | 1917388.25 | 9.72E+07 | 1.19E+08 | 1.57E+07 | 9.54E+07 | 5.36E+07 | 2578158 | 1.30E+07 | 2.43E+07 | 775.4336 | YLPSPAK | P04919 | P04919 | SLC4A1 | solute carrier family 4 (anion exchanger), member 1 (Diego blood group) | Plasma Membrane | transporter |
| 15542590 | 2 | 0.324 | 1069349 | 2734712.25 | 7597600.5 | 579565.312 | 4147171.25 | 2850403.75 | 3335341.75 | 2876349 | 1193218 | 1107845.125 | 594502.188 | 896019.938 | 746.51575 | ILLLFK | P04919 | P04919 | SLC4A1 | solute carrier family 4 (anion exchanger), member 1 (Diego blood group) | Plasma Membrane | transporter |
| 15539254 | 3 | 0.025 | 2743986.5 | 7591134 | 1.18E+07 | 3762708.75 | 1.41E+07 | 6504326 | 6006611.5 | 7090167 | 3563619.5 | 190761.062 | 949166.688 | 2042982.875 | 1630.8044 | PDPNLYNTLDLNGGK | P04919 | P04919 | SLC4A1 | solute carrier family 4 (anion exchanger), member 1 (Diego blood group) | Plasma Membrane | transporter |
| 15533954 | 2 | 0.49 | 1.40E+07 | 1.10E+07 | 9286372 | 1.66E+07 | 7966550 | 1.19E+07 | 1.44E+07 | 8058994.5 | 1.29E+07 | 1.20E+07 | 1.82E+07 | 1.73E+07 | 843.53906 | LLLM[147.0354]PLK | O88343 | O88343 | SLC4A4 | solute carrier family 4 (sodium bicarbonate cotransporter), member 4 | Plasma Membrane | transporter |
| 15551379 | 2 | 0.206 | 99.788 | 775534.562 | 1709990 | 4986422.5 | 5089500.5 | 5997992 | 3199162 | 4112086.25 | 5815848.5 | 128707.109 | 6085983 | 3374649.75 | 1785.8646 | SITPETPTEIPC[160.0307]GDIR | Q60857 | Q60857 | SLC6A4 | solute carrier family 6 (neurotransmitter transporter), member 4 | Plasma Membrane | transporter |
| 15548687 | 3 | 0.002 | 9803132 | 1.47E+07 | 9439907 | 6397144.5 | 6817679 | 9777826 | 8456809 | 1.05E+07 | 1.16E+07 | 4210455 | 4481128 | 4840681 | 2426.1458 | YAPTTTPSPEDGFEVQPLHPDK | P28571 | P28571 | SLC6A9 | solute carrier family 6 (neurotransmitter transporter, glycine), member 9 | Plasma Membrane | transporter |
| 15538414 | 2 | 9.72E-04 | 2418587 | 3193914.75 | 2541207.5 | 1358096.375 | 1973295.875 | 2113745.5 | 2412600 | 3425795.5 | 2545543.75 | 1030832 | 1277554.75 | 832345.75 | 1321.6353 | LSDDIGNFGEVR | P28571 | P28571 | SLC6A9 | solute carrier family 6 (neurotransmitter transporter, glycine), member 9 | Plasma Membrane | transporter |
| 15552490 | 3 | 0.182 | 6628230 | 1.42E+07 | 5107948 | 6701965.5 | 5731017.5 | 6593752 | 4207713.5 | 6520880 | 9241503 | 3526371.75 | 4510210 | 4045111 | 3192.5486 | TTEELDRVDQNELVSASESQTGFLPVAEK | Q09143 | Q09143 | SLC7A1 | solute carrier family 7 (cationic amino acid transporter, y+ system), member 1 | Plasma Membrane | transporter |
| 15539217 | 2 | 0.075 | 1.16E+07 | 1.80E+07 | 8535171 | 1.39E+07 | 1.17E+07 | 1.34E+07 | 1.02E+07 | 1.19E+07 | 8853210 | 8885757 | 7118544 | 7434527.5 | 1356.6722 | AVATPAAAAAEEER | Q9Z127 | Q9Z127 | SLC7A5 | solute carrier family 7 (amino acid transporter light chain, L system), member 5 | Plasma Membrane | transporter |
| 15533544 | 3 | 0.103 | 6150721.5 | 8440543 | 3732521 | 8806105 | 6168444 | 6269301 | 4970054.5 | 5785492 | 4898893 | 4281211.5 | 3651837.75 | 4096341.5 | 1656.7921 | RGDGADPEGEGVTLQR | Q9Z127 | Q9Z127 | SLC7A5 | solute carrier family 7 (amino acid transporter light chain, L system), member 5 | Plasma Membrane | transporter |
| 15539100 | 2 | 0.016 | 2304675.75 | 3838820.25 | 2272580 | 2663014 | 2605462.5 | 2423130.5 | 2460477.25 | 2372015.25 | 3071668.25 | 1579649 | 1565904.375 | 1700470.875 | 825.4598 | LFFVGSR | Q9Z127 | Q9Z127 | SLC7A5 | solute carrier family 7 (amino acid transporter light chain, L system), member 5 | Plasma Membrane | transporter |
| 15541555 | 2 | 0.109 | 1639510.25 | 2964378.25 | 1515343.5 | 2115276.25 | 1765948.125 | 2203359 | 1875221.5 | 2188735.75 | 1589873.75 | 1495952.625 | 1204131.375 | 1157581.5 | 1391.6189 | DMGQGDASNLQQK | Q9Z127 | Q9Z127 | SLC7A5 | solute carrier family 7 (amino acid transporter light chain, L system), member 5 | Plasma Membrane | transporter |
| 15532641 | 3 | 0.55 | 1.60E+07 | 1.44E+07 | 1.39E+07 | 1.52E+07 | 1.68E+07 | 1.04E+07 | 1.39E+07 | 1.32E+07 | 1.21E+07 | 1.93E+07 | 1.50E+07 | 1.37E+07 | 2160.0022 | C[160.0307]LSDPGPHPEPGEGEPFIPK | Q61165 | Q61165 | SLC9A1 | solute carrier family 9, subfamily A (NHE1, cation proton antiporter 1), member 1 | Plasma Membrane | ion channel |
| 15540379 | 3 | 0.596 | 9726365 | 1.05E+07 | 6871965.5 | 9006895 | 1.07E+07 | 8392630 | 1.15E+07 | 7759721 | 7852583 | 1.23E+07 | 9506748 | 1.04E+07 | 1708.9005 | IPSAVSTVSMQNIHPK | Q61165 | Q61165 | SLC9A1 | solute carrier family 9, subfamily A (NHE1, cation proton antiporter 1), member 1 | Plasma Membrane | ion channel |
| 15534824 | 3 | 0.887 | 6374226.5 | 9019156 | 7449796.5 | 5994141.5 | 7454301 | 7716260 | 6910109 | 8068388 | 6310898.5 | 8952970 | 6316287 | 7639792 | 2186.0688 | HTLVADPYEEAWNQMLLR | Q61165 | Q61165 | SLC9A1 | solute carrier family 9, subfamily A (NHE1, cation proton antiporter 1), member 1 | Plasma Membrane | ion channel |
| 15533401 | 3 | 0.976 | 7615279 | 8745820 | 4885907 | 6732363 | 6722061.5 | 7062897 | 6192368.5 | 7229473 | 6614303 | 7117769.5 | 6326507.5 | 6171452.5 | 1541.8076 | AFPVLDIDYPHVR | Q61165 | Q61165 | SLC9A1 | solute carrier family 9, subfamily A (NHE1, cation proton antiporter 1), member 1 | Plasma Membrane | ion channel |
| 15534397 | 3 | 0.229 | 5756472 | 6138828 | 5003463 | 6004603 | 5323611 | 4679959 | 4675524 | 4953274.5 | 3868057.75 | 6871188.5 | 5002494 | 5047409.5 | 1256.697 | ITNYLTVPAHK | Q61165 | Q61165 | SLC9A1 | solute carrier family 9, subfamily A (NHE1, cation proton antiporter 1), member 1 | Plasma Membrane | ion channel |
| 15533519 | 2 | 0.121 | 6639902 | 5344979.5 | 6542118.5 | 4032627.25 | 5070826.5 | 4913505 | 5293718.5 | 4535166 | 5358569 | 6682930.5 | 5081117 | 4983349.5 | 1189.6085 | IGSDPLAYEPK | Q61165 | Q61165 | SLC9A1 | solute carrier family 9, subfamily A (NHE1, cation proton antiporter 1), member 1 | Plasma Membrane | ion channel |
| 15556206 | 2 | 0.215 | 3081322.25 | 3315921.75 | 3810889.5 | 2246253 | 2886074.5 | 2796933.5 | 3315035.5 | 2748600.25 | 2345483 | 4493960.5 | 2871887.75 | 3085600.75 | 2337.0073 | EPSSPGTDDVFTPGSSDSPSSQR | Q61165 | Q61165 | SLC9A1 | solute carrier family 9, subfamily A (NHE1, cation proton antiporter 1), member 1 | Plasma Membrane | ion channel |
| 15554659 | 3 | 0.247 | 4199527.5 | 86818.805 | 17351.52 | 2240945.75 | 89692.188 | 65633.062 | 116999.219 | 286684.969 | 663509.75 | 4038528.25 | 5250719 | 1659630.5 | 2552.133 | SKEPSSPGTDDVFTPGSSDSPSSQR | Q61165 | Q61165 | SLC9A1 | solute carrier family 9, subfamily A (NHE1, cation proton antiporter 1), member 1 | Plasma Membrane | ion channel |
| 15531812 | 2 | 0.682 | 3.48E+07 | 5.33E+07 | 8.11E+07 | 2.72E+07 | 6.57E+07 | 4.76E+07 | 2.92E+07 | 5.91E+07 | 4.23E+07 | 5.14E+07 | 2.64E+07 | 3.40E+07 | 927.4303 | PHLEM[147.0354]ER | Q6UJY2 | Q6UJY2 | SLC9C1 | solute carrier family 9, subfamily C (Na+-transporting carboxylic acid decarboxylase), member 1 | Extracellular Space | other |
| 15552762 | 2 | 0.033 | 2306535.5 | 3029135.5 | 2416738 | 1974976.375 | 2353706.25 | 1400498.75 | 1105809.625 | 1282572 | 1754694.25 | 1928116.5 | 1691469.25 | 1356761.75 | 1514.829 | TILEEEITPTIQK | Q8CG48 | Q8CG48 | SMC2 | structural maintenance of chromosomes 2 | Nucleus | transporter |
| 15542378 | 1 | 0.839 | 1.18E+07 | 1.20E+07 | 1.53E+07 | 1.15E+07 | 1.47E+07 | 1.07E+07 | 1.17E+07 | 1.31E+07 | 1.01E+07 | 1.83E+07 | 1.06E+07 | 1.19E+07 | 704.36 | NNSLEK | Q8CG47 | Q8CG47 | SMC4 | structural maintenance of chromosomes 4 | Nucleus | transporter |
| 15537637 | 2 | 0.824 | 1205323.625 | 1401759.125 | 421605.438 | 1015288.875 | 1176904.375 | 423362.969 | 815558.625 | 600136.812 | 516061.688 | 934168.062 | 733275.438 | 626047.125 | 1235.5914 | QLDEC[160.0307]ASAITK | Q8CG47 | Q8CG47 | SMC4 | structural maintenance of chromosomes 4 | Nucleus | transporter |
| 15553478 | 3 | 0.161 | 3.30E+07 | 3.61E+07 | 3.51E+07 | 4.12E+07 | 3.13E+07 | 2.99E+07 | 3.92E+07 | 3.80E+07 | 3.54E+07 | 3.03E+07 | 3.33E+07 | 2.41E+07 | 2805.3293 | TITMLDEQGEQLNRIEEGMDQINK | O09044 | O09044 | SNAP23 | synaptosomal-associated protein, 23kDa | Plasma Membrane | transporter |
| 15537753 | 3 | 0.171 | 3.58E+07 | 3.24E+07 | 3.21E+07 | 2.95E+07 | 2.94E+07 | 2.67E+07 | 4.16E+07 | 3.67E+07 | 2.71E+07 | 2.74E+07 | 3.13E+07 | 2.25E+07 | 1628.7949 | AHQVTDESLESTRR | O09044 | O09044 | SNAP23 | synaptosomal-associated protein, 23kDa | Plasma Membrane | transporter |
| 15547931 | 2 | 0.068 | 2.70E+07 | 2.18E+07 | 2.85E+07 | 2.14E+07 | 2.56E+07 | 1.86E+07 | 2.46E+07 | 2.66E+07 | 2.67E+07 | 1.94E+07 | 2.22E+07 | 1.74E+07 | 1690.7632 | ATWGDGGDNSPSNVVSK | O09044 | O09044 | SNAP23 | synaptosomal-associated protein, 23kDa | Plasma Membrane | transporter |
| 15532780 | 2 | 0.018 | 2.39E+07 | 3.57E+07 | 3.41E+07 | 2.50E+07 | 2.65E+07 | 2.77E+07 | 3.17E+07 | 3.42E+07 | 3.17E+07 | 1.86E+07 | 2.38E+07 | 2.22E+07 | 1427.808 | ILGLAIESQDAGIK | O09044 | O09044 | SNAP23 | synaptosomal-associated protein, 23kDa | Plasma Membrane | transporter |
| 15540995 | 3 | 0.522 | 1.04E+07 | 1.20E+07 | 8680434 | 1.28E+07 | 1.03E+07 | 5681951.5 | 1.32E+07 | 1.24E+07 | 8465155 | 7678758.5 | 8472948 | 8530938 | 2160.9988 | NMALDMGNEIDAQNQQIQK | O09044 | O09044 | SNAP23 | synaptosomal-associated protein, 23kDa | Plasma Membrane | transporter |
| 15552684 | 3 | 0.084 | 7898770 | 1.64E+07 | 2.11E+07 | 6956168.5 | 2.17E+07 | 1.18E+07 | 1.27E+07 | 2.12E+07 | 1.01E+07 | 7372809 | 3759067.25 | 5107424 | 2789.3552 | ITNDAREDEMEENLTQVGSILGNLK | O09044 | O09044 | SNAP23 | synaptosomal-associated protein, 23kDa | Plasma Membrane | transporter |
| 15548546 | 2 | 0.006 | 9868382 | 9301631 | 1.17E+07 | 1.05E+07 | 1.09E+07 | 1.10E+07 | 1.23E+07 | 1.15E+07 | 9991548 | 7101334 | 8460234 | 5827274.5 | 2160.9973 | NMALDMGNEIDAQNQQIQK | O09044 | O09044 | SNAP23 | synaptosomal-associated protein, 23kDa | Plasma Membrane | transporter |
| 15549265 | 2 | 0.196 | 3316501 | 4896621.5 | 6538405.5 | 2386154.25 | 7511138 | 4600786 | 5346732 | 6679322.5 | 3739260.5 | 3413788 | 2272695.25 | 2159218.5 | 1647.7972 | TITMLDEQGEQLNR | O09044 | O09044 | SNAP23 | synaptosomal-associated protein, 23kDa | Plasma Membrane | transporter |
| 15539728 | 2 | 0.191 | 1871299.875 | 2873334 | 3877785 | 1739485.625 | 3446391.75 | 1791182.75 | 3177065.25 | 3943433.25 | 2003813.25 | 2315954.25 | 1069938.25 | 1318848.875 | 802.4399 | IDIANTR | O09044 | O09044 | SNAP23 | synaptosomal-associated protein, 23kDa | Plasma Membrane | transporter |
| 15536153 | 2 | 0.204 | 2005495.375 | 2575124.5 | 3351945 | 1814134.125 | 4311424.5 | 1842950.75 | 2420694.25 | 3815249.5 | 1722619.875 | 1930070.375 | 860216.625 | 1407577.75 | 1176.5532 | IEEGMDQINK | O09044 | O09044 | SNAP23 | synaptosomal-associated protein, 23kDa | Plasma Membrane | transporter |
| 15544441 | 2 | 0.771 | 3366941.5 | 2077857 | 1290505.125 | 3106796 | 880120.625 | 1207051.25 | 2300562.75 | 1643258.5 | 1172855.625 | 1884945.75 | 2679509.25 | 1642124.5 | 1072.5833 | NRIDIANTR | O09044 | O09044 | SNAP23 | synaptosomal-associated protein, 23kDa | Plasma Membrane | transporter |
| 15557432 | 2 | 0.299 | 1274472.625 | 1416621.625 | 1653071.375 | 413335.656 | 1202225.125 | 950026.438 | 789621.875 | 1465905.75 | 1380522.375 | 479856.75 | 1307489.5 | 896610.875 | 1628.7954 | AHQVTDESLESTRR | O09044 | O09044 | SNAP23 | synaptosomal-associated protein, 23kDa | Plasma Membrane | transporter |
| 15548174 | 2 | 0.023 | 1.24E+07 | 1.29E+07 | 1.85E+07 | 1.41E+07 | 1.39E+07 | 2.09E+07 | 1.13E+07 | 1.42E+07 | 1.02E+07 | 1.04E+07 | 6083756 | 7778422 | 1505.7926 | TVEGAGNIAAATGFVK | O55042 | O55042 | SNCA | synuclein, alpha (non A4 component of amyloid precursor) | Cytoplasm | other |
| 15549296 | 2 | 0.042 | 9916895 | 9143939 | 1.41E+07 | 1.04E+07 | 9817531 | 1.45E+07 | 9645854 | 1.02E+07 | 5781394.5 | 7775157 | 4981845 | 5208098.5 | 1928.0394 | EQVTNVGGAVVTGVTAVAQK | O55042 | O55042 | SNCA | synuclein, alpha (non A4 component of amyloid precursor) | Cytoplasm | other |
| 15541925 | 3 | 0.056 | 8948884 | 7719875.5 | 9943120 | 8755391 | 8483924 | 1.27E+07 | 7345882 | 9580846 | 5627997.5 | 6881085 | 4157534.5 | 5733675.5 | 1928.04 | EQVTNVGGAVVTGVTAVAQK | O55042 | O55042 | SNCA | synuclein, alpha (non A4 component of amyloid precursor) | Cytoplasm | other |
| 15532330 | 2 | 0.023 | 7921942.5 | 6664935.5 | 7137266 | 7443899.5 | 6124959 | 8793831 | 5380237 | 6981328 | 4397354 | 5371162.5 | 3633062 | 4277619.5 | 951.5132 | EGVLYVGSK | O55042 | O55042 | SNCA | synuclein, alpha (non A4 component of amyloid precursor) | Cytoplasm | other |
| 15545374 | 2 | 0.016 | 770178.062 | 1370933.75 | 585627.938 | 603701.062 | 777064.562 | 547303.562 | 373096.125 | 1064836.75 | 687827.188 | 163284.047 | 45852.895 | 291246.781 | 832.4906 | TNLPIFK | O70493 | O70493 | SNX12 | sorting nexin 12 | Cytoplasm | transporter |
| 15545266 | 2 | 0.501 | 545023.25 | 1043846.375 | 1341737.375 | 76708.055 | 795946.125 | 979048.188 | 474929.344 | 697743.312 | 1005310.562 | 582725.5 | 435797.812 | 225828.719 | 1318.6637 | KLMGEEALAGGDK | Q0P5V2 | Q0P5V2 | SOBP | sine oculis binding protein homolog | Nucleus | other |
| 15539489 | 2 | 0.043 | 4028941.75 | 8988765 | 4090838.5 | 6850911.5 | 6313931 | 8655015 | 3915434.75 | 6898561 | 6532188.5 | 1857837 | 3292348.25 | 3502371.5 | 1440.8026 | GDVTTQVALQPALK | P09671 | P09671 | SOD2 | superoxide dismutase 2, mitochondrial | Cytoplasm | enzyme |
| 15556617 | 4 | 0.961 | 3134039.25 | 3397734.75 | 181224.297 | 5942966.5 | 1085916.125 | 904752.688 | 2346719.5 | 1210794.625 | 605663.688 | 1829399 | 2990004 | 739679.375 | 3224.588 | HSLPDLPYDYGALEPHINAQIMQLHHSK | P09671 | P09671 | SOD2 | superoxide dismutase 2, mitochondrial | Cytoplasm | enzyme |
| 15536537 | 2 | 0.025 | 1399161.625 | 2976202.75 | 1539828.25 | 1389597.75 | 2077333.25 | 2275822.5 | 1323632.625 | 2279333 | 2117248 | 694819.812 | 893814.625 | 953357.125 | 872.50714 | GELLEAIK | P09671 | P09671 | SOD2 | superoxide dismutase 2, mitochondrial | Cytoplasm | enzyme |
| 15533481 | 3 | 0.09 | 4329096 | 5551713.5 | 5719932.5 | 7844769 | 1.13E+07 | 7891151 | 7185998 | 8760688 | 9378787 | 4317866.5 | 9286873 | 7248815.5 | 2214.0132 | FFDHSGTLVMDAYEPEISR | Q9CYN2 | Q9CYN2 | SPCS2 | signal peptidase complex subunit 2 | Cytoplasm | other |
| 15535339 | 2 | 0.135 | 1.49E+07 | 1.87E+07 | 1.81E+07 | 1.18E+07 | 1.49E+07 | 1.35E+07 | 1.87E+07 | 1.39E+07 | 1.56E+07 | 1.95E+07 | 1.50E+07 | 1.74E+07 | 1090.5609 | ETLGDDITVK | Q9JJF9 | Q9JJF9 | SPPL2A | signal peptide peptidase like 2A | Plasma Membrane | peptidase |
| 15538340 | 3 | 0.151 | 2.25E+07 | 1.82E+07 | 1.42E+07 | 1.20E+07 | 1.55E+07 | 1.46E+07 | 1.72E+07 | 1.57E+07 | 1.59E+07 | 1.64E+07 | 2.11E+07 | 1.91E+07 | 1678.8485 | AVVVHWGPC[160.0307]HFLEK | Q9JJF9 | Q9JJF9 | SPPL2A | signal peptide peptidase like 2A | Plasma Membrane | peptidase |
| 15549911, 15552609 | 3, 4 | 0.765 | 3.55E+09 | 3.58E+09 | 3.60E+09 | 3.62E+09 | 3.86E+09 | 3.61E+09 | 3.45E+09 | 3.93E+09 | 3.34E+09 | 4.26E+09 | 3.53E+09 | 3.54E+09 | 3290.4421, 3290.4497 | ASAFNNWC[160.0307]ENAEEDLSEPVHC[160.0307]VSLNEIR | P08032 | P08032 | SPTA1 | spectrin, alpha, erythrocytic 1 | Cytoplasm | other |
| 15547740, 15550841 | 2, 3 | 0.091 | 3.21E+09 | 3.00E+09 | 3.81E+09 | 2.40E+09 | 2.65E+09 | 2.76E+09 | 2.92E+09 | 2.63E+09 | 2.64E+09 | 3.66E+09 | 2.61E+09 | 3.34E+09 | 2340.1924, 2340.1973 | ANVQLQQYLADLHEAEAWIK | P08032 | P08032 | SPTA1 | spectrin, alpha, erythrocytic 1 | Cytoplasm | other |
| 15538256, 15539026 | 3, 4 | 0.649 | 2.94E+09 | 2.91E+09 | 3.37E+09 | 2.14E+09 | 2.90E+09 | 2.76E+09 | 3.12E+09 | 3.03E+09 | 2.29E+09 | 3.29E+09 | 2.23E+09 | 3.07E+09 | 2152.1238, 2152.1282 | VADELLFEELLTPEGAHIR | P08032 | P08032 | SPTA1 | spectrin, alpha, erythrocytic 1 | Cytoplasm | other |
| 15550845, 15552430, 15562711 | 2, 3 | 0.96 | 2.13E+09 | 2.30E+09 | 2.93E+09 | 2.05E+09 | 2.55E+09 | 2.56E+09 | 2.41E+09 | 2.83E+09 | 2.31E+09 | 2.95E+09 | 2.09E+09 | 2.56E+09 | 2513.327, 2513.3313, 2513.3335 | TIGGVISSPELAEDLTGTEILLER | P08032 | P08032 | SPTA1 | spectrin, alpha, erythrocytic 1 | Cytoplasm | other |
| 15547668, 15553161 | 2, 3 | 0.703 | 2.04E+09 | 2.54E+09 | 2.97E+09 | 1.54E+09 | 3.03E+09 | 2.13E+09 | 2.27E+09 | 2.77E+09 | 2.18E+09 | 2.82E+09 | 1.60E+09 | 1.73E+09 | 2530.1167, 2530.1182, 2530.1196 | DSEDEEAWIQETEPSAASTHLGK | P08032 | P08032 | SPTA1 | spectrin, alpha, erythrocytic 1 | Cytoplasm | other |
| 15547657, 15547663, 15561960 | 2, 3 | 0.854 | 2.08E+09 | 2.47E+09 | 3.09E+09 | 2.39E+09 | 2.31E+09 | 2.20E+09 | 2.39E+09 | 2.68E+09 | 2.18E+09 | 2.65E+09 | 2.17E+09 | 2.35E+09 | 2592.2625, 2592.2651, 2592.27 | DLNDLAQELISSGTFNIDQIEEK | P08032 | P08032 | SPTA1 | spectrin, alpha, erythrocytic 1 | Cytoplasm | other |
| 15550340, 15550433 | 2 | 0.739 | 2.22E+09 | 2.31E+09 | 2.83E+09 | 2.36E+09 | 2.14E+09 | 2.48E+09 | 2.05E+09 | 2.48E+09 | 1.94E+09 | 2.62E+09 | 1.75E+09 | 2.41E+09 | 2234.1501, 2234.152 | LLVDSQLLQQLYQDSDDLK | P08032 | P08032 | SPTA1 | spectrin, alpha, erythrocytic 1 | Cytoplasm | other |
| 15531556, 15536910, 15539644 | 3, 4 | 0.496 | 2.32E+09 | 2.06E+09 | 2.31E+09 | 2.13E+09 | 2.22E+09 | 1.83E+09 | 2.28E+09 | 2.02E+09 | 1.78E+09 | 2.59E+09 | 2.03E+09 | 2.21E+09 | 2034.912, 2034.9128, 2034.9177 | TGQEMIEDGHYASEAVAAR | P08032 | P08032 | SPTA1 | spectrin, alpha, erythrocytic 1 | Cytoplasm | other |
| 15547652, 15553797, 15562368 | 2, 3 | 0.274 | 1.93E+09 | 1.65E+09 | 1.77E+09 | 1.74E+09 | 1.58E+09 | 1.41E+09 | 1.72E+09 | 1.79E+09 | 1.68E+09 | 2.57E+09 | 1.74E+09 | 1.73E+09 | 2440.1543, 2440.1584, 2440.1592, 2440.1606 | ALRDQAEVC[160.0307]QQQQAAPVDEAGR | P08032 | P08032 | SPTA1 | spectrin, alpha, erythrocytic 1 | Cytoplasm | other |
| 15547953, 15549605 | 2, 3 | 0.507 | 2.27E+09 | 2.23E+09 | 1.61E+09 | 2.20E+09 | 1.85E+09 | 2.09E+09 | 2.44E+09 | 2.10E+09 | 1.86E+09 | 2.49E+09 | 2.43E+09 | 2.16E+09 | 2680.228, 2680.2305 | QALTPEQVSFC[160.0307]TIHMQQYMDPR | P08032 | P08032 | SPTA1 | spectrin, alpha, erythrocytic 1 | Cytoplasm | other |
| 15538806 | 3 | 0.245 | 1.76E+09 | 1.88E+09 | 2.16E+09 | 1.60E+09 | 1.87E+09 | 1.71E+09 | 1.82E+09 | 1.81E+09 | 1.62E+09 | 2.39E+09 | 1.77E+09 | 1.98E+09 | 1862.8816 | SQLDDVHAFQQFNWK | P08032 | P08032 | SPTA1 | spectrin, alpha, erythrocytic 1 | Cytoplasm | other |
| 15548190, 15549779 | 2, 3 | 0.441 | 2.11E+09 | 2.12E+09 | 2.40E+09 | 2.05E+09 | 2.16E+09 | 1.86E+09 | 1.94E+09 | 1.93E+09 | 2.01E+09 | 2.34E+09 | 1.68E+09 | 1.89E+09 | 2438.1802, 2438.184 | GTQLYEANQLLQFENNAEDLK | P08032 | P08032 | SPTA1 | spectrin, alpha, erythrocytic 1 | Cytoplasm | other |
| 15548216 | 2 | 0.59 | 1.93E+09 | 1.85E+09 | 2.20E+09 | 1.80E+09 | 1.72E+09 | 1.83E+09 | 1.74E+09 | 1.97E+09 | 1.70E+09 | 2.30E+09 | 1.71E+09 | 1.74E+09 | 1681.7896 | TSDDIESAFQALAEGK | P08032 | P08032 | SPTA1 | spectrin, alpha, erythrocytic 1 | Cytoplasm | other |
| 15548283, 15549583, 15555078, 15557632 | 2, 3 | 0.43 | 2.06E+09 | 2.73E+09 | 3.18E+09 | 1.58E+09 | 2.27E+09 | 2.42E+09 | 2.44E+09 | 2.30E+09 | 1.78E+09 | 2.26E+09 | 1.71E+09 | 2.26E+09 | 2533.1885, 2533.19, 2533.1921, 2533.1943 | REDPTFQALEDFGTELIDSGHR | P08032 | P08032 | SPTA1 | spectrin, alpha, erythrocytic 1 | Cytoplasm | other |
| 15550224, 15550394 | 3, 4 | 0.45 | 1.99E+09 | 2.18E+09 | 2.07E+09 | 2.13E+09 | 1.25E+09 | 2.02E+09 | 1.96E+09 | 1.87E+09 | 2.00E+09 | 2.25E+09 | 1.93E+09 | 2.28E+09 | 3362.6924, 3362.7039 | QEEPVNIPQLQQQVETLYHSLLDRAEER | P08032 | P08032 | SPTA1 | spectrin, alpha, erythrocytic 1 | Cytoplasm | other |
| 15532150, 15535171 | 3 | 0.328 | 1.76E+09 | 1.88E+09 | 2.03E+09 | 1.73E+09 | 1.77E+09 | 1.52E+09 | 2.09E+09 | 1.81E+09 | 1.84E+09 | 2.24E+09 | 1.68E+09 | 1.86E+09 | 1801.8835, 1801.886 | VEADDHQGFVPAVYVR | P08032 | P08032 | SPTA1 | spectrin, alpha, erythrocytic 1 | Cytoplasm | other |
| 15532958 | 3 | 0.227 | 1.03E+09 | 4.74E+08 | 1.27E+09 | 1.23E+09 | 1.13E+09 | 1.29E+09 | 1.07E+09 | 1.40E+09 | 1.07E+09 | 2.23E+09 | 1.14E+09 | 1.48E+09 | 2234.155 | LLVDSQLLQQLYQDSDDLK | P08032 | P08032 | SPTA1 | spectrin, alpha, erythrocytic 1 | Cytoplasm | other |
| 15552174, 15562242, 15562841 | 3 | 0.194 | 1.94E+09 | 2.33E+09 | 2.16E+09 | 1.87E+09 | 1.96E+09 | 1.83E+09 | 2.06E+09 | 2.29E+09 | 1.91E+09 | 2.07E+09 | 1.68E+09 | 1.87E+09 | 2723.3228, 2723.3232, 2723.3242 | EHEAFLASLAGAQEDFNYLLELDK | P08032 | P08032 | SPTA1 | spectrin, alpha, erythrocytic 1 | Cytoplasm | other |
| 15547836 | 2, 3 | 0.448 | 1.67E+09 | 1.89E+09 | 2.19E+09 | 1.38E+09 | 1.81E+09 | 1.64E+09 | 1.71E+09 | 1.77E+09 | 1.70E+09 | 2.01E+09 | 1.49E+09 | 1.86E+09 | 2490.232, 2490.2322 | QEAFLENEDLGNSVGSVEALLQK | P08032 | P08032 | SPTA1 | spectrin, alpha, erythrocytic 1 | Cytoplasm | other |
| 15536242, 15540368, 15544143, 15544506 | 3, 4 | 0.274 | 1.75E+09 | 1.46E+09 | 1.37E+09 | 1.53E+09 | 1.46E+09 | 1.32E+09 | 1.71E+09 | 1.38E+09 | 1.18E+09 | 1.95E+09 | 1.68E+09 | 1.59E+09 | 1537.7732, 1537.7738, 1537.7743, 1537.7744, 1537.7754 | KHEALENDFAVHK | P08032 | P08032 | SPTA1 | spectrin, alpha, erythrocytic 1 | Cytoplasm | other |
| 15550040, 15561987 | 2 | 0.784 | 1.57E+09 | 1.62E+09 | 1.97E+09 | 1.78E+09 | 1.49E+09 | 1.52E+09 | 1.72E+09 | 1.50E+09 | 1.44E+09 | 1.95E+09 | 1.41E+09 | 1.55E+09 | 2240.1716, 2240.1736 | TALINADELPTDVASGEALLAR | P08032 | P08032 | SPTA1 | spectrin, alpha, erythrocytic 1 | Cytoplasm | other |
| 15532392 | 3 | 0.931 | 1.60E+09 | 1.45E+09 | 1.55E+09 | 1.49E+09 | 1.46E+09 | 1.43E+09 | 1.63E+09 | 1.70E+09 | 1.23E+09 | 1.92E+09 | 1.37E+09 | 1.45E+09 | 1642.7374 | LC[160.0307]ESHPDATEDLQK | P08032 | P08032 | SPTA1 | spectrin, alpha, erythrocytic 1 | Cytoplasm | other |
| 15534574, 15536281 | 2, 3 | 0.266 | 1.46E+09 | 1.41E+09 | 1.66E+09 | 1.45E+09 | 1.46E+09 | 1.33E+09 | 1.45E+09 | 1.43E+09 | 1.25E+09 | 1.89E+09 | 1.45E+09 | 1.51E+09 | 1421.687, 1421.6885 | NYGADEEAAGALLK | P08032 | P08032 | SPTA1 | spectrin, alpha, erythrocytic 1 | Cytoplasm | other |
| 15531728, 15532229 | 2, 3 | 0.406 | 1.61E+09 | 1.47E+09 | 1.64E+09 | 1.54E+09 | 1.36E+09 | 1.24E+09 | 1.44E+09 | 1.50E+09 | 1.29E+09 | 1.84E+09 | 1.45E+09 | 1.36E+09 | 1324.6832, 1324.6848 | VLETAEEIQHR | P08032 | P08032 | SPTA1 | spectrin, alpha, erythrocytic 1 | Cytoplasm | other |
| 15534104, 15543716 | 2 | 0.807 | 1.34E+09 | 1.27E+09 | 1.49E+09 | 1.25E+09 | 1.31E+09 | 1.38E+09 | 1.27E+09 | 1.26E+09 | 1.24E+09 | 1.80E+09 | 1.23E+09 | 1.18E+09 | 1117.5808, 1117.5815 | ISDLENVATR | P08032 | P08032 | SPTA1 | spectrin, alpha, erythrocytic 1 | Cytoplasm | other |
| 15531838, 15532925 | 2, 3 | 0.113 | 1.52E+09 | 1.34E+09 | 1.58E+09 | 1.18E+09 | 1.29E+09 | 1.32E+09 | 1.48E+09 | 1.35E+09 | 1.24E+09 | 1.79E+09 | 1.45E+09 | 1.41E+09 | 1285.6661, 1285.6678 | LAQFVQHWEK | P08032 | P08032 | SPTA1 | spectrin, alpha, erythrocytic 1 | Cytoplasm | other |
| 15550682 | 2 | 0.27 | 1.68E+09 | 1.72E+09 | 1.87E+09 | 1.54E+09 | 1.56E+09 | 1.55E+09 | 1.80E+09 | 1.62E+09 | 1.49E+09 | 1.76E+09 | 1.45E+09 | 1.55E+09 | 1939.0216 | ALNAADPGSDLLSVQALQR | P08032 | P08032 | SPTA1 | spectrin, alpha, erythrocytic 1 | Cytoplasm | other |
| 15537283 | 4, 5 | 0.444 | 1.85E+09 | 2.00E+09 | 1.63E+09 | 1.66E+09 | 1.55E+09 | 1.63E+09 | 1.95E+09 | 1.57E+09 | 1.41E+09 | 1.76E+09 | 1.77E+09 | 1.73E+09 | 2501.2295, 2501.231 | RLADEQYQLLSSAHAVEMFHR | P08032 | P08032 | SPTA1 | spectrin, alpha, erythrocytic 1 | Cytoplasm | other |
| 15547849 | 2 | 0.738 | 1.33E+09 | 1.30E+09 | 1.83E+09 | 1.12E+09 | 1.46E+09 | 1.21E+09 | 1.41E+09 | 1.40E+09 | 1.23E+09 | 1.74E+09 | 1.08E+09 | 1.13E+09 | 1588.7617 | VQDVC[160.0307]AQGEDILNK | P08032 | P08032 | SPTA1 | spectrin, alpha, erythrocytic 1 | Cytoplasm | other |
| 15531464 | 2, 3 | 0.356 | 1.49E+09 | 1.45E+09 | 1.67E+09 | 1.36E+09 | 1.34E+09 | 1.26E+09 | 1.48E+09 | 1.45E+09 | 1.28E+09 | 1.71E+09 | 1.27E+09 | 1.37E+09 | 1476.7573, 1476.7583 | LAPDELPGFPQHR | P08032 | P08032 | SPTA1 | spectrin, alpha, erythrocytic 1 | Cytoplasm | other |
| 15535671, 15537513 | 2, 3 | 0.21 | 1.34E+09 | 1.22E+09 | 1.20E+09 | 1.21E+09 | 1.09E+09 | 1.14E+09 | 1.29E+09 | 1.11E+09 | 9.68E+08 | 1.71E+09 | 1.20E+09 | 1.29E+09 | 1467.7351, 1467.7363 | MQHNLEQQIQAK | P08032 | P08032 | SPTA1 | spectrin, alpha, erythrocytic 1 | Cytoplasm | other |
| 15532209, 15543293 | 2, 3 | 0.2 | 1.42E+09 | 1.36E+09 | 1.49E+09 | 1.41E+09 | 1.27E+09 | 1.26E+09 | 1.45E+09 | 1.30E+09 | 1.12E+09 | 1.70E+09 | 1.50E+09 | 1.35E+09 | 1197.6179, 1197.6201, 1197.6208 | HGLLESDVTAR | P08032 | P08032 | SPTA1 | spectrin, alpha, erythrocytic 1 | Cytoplasm | other |
| 15535845, 15538098, 15543836 | 2, 3 | 0.484 | 1.38E+09 | 1.23E+09 | 1.34E+09 | 1.25E+09 | 1.22E+09 | 1.08E+09 | 1.44E+09 | 1.23E+09 | 1.00E+09 | 1.67E+09 | 1.30E+09 | 1.22E+09 | 1402.6362, 1402.6375, 1402.6378 | FAEDHFAHEATK | P08032 | P08032 | SPTA1 | spectrin, alpha, erythrocytic 1 | Cytoplasm | other |
| 15532034, 15541703, 15543849, 15546328 | 2, 3 | 0.381 | 1.31E+09 | 1.08E+09 | 1.12E+09 | 1.14E+09 | 1.16E+09 | 9.67E+08 | 1.17E+09 | 1.23E+09 | 1.01E+09 | 1.67E+09 | 1.19E+09 | 1.15E+09 | 1260.6118, 1260.6166, 1260.6193, 1260.62, 1260.6201 | HESFVSEVQAK | P08032 | P08032 | SPTA1 | spectrin, alpha, erythrocytic 1 | Cytoplasm | other |
| 15547694, 15548225, 15557358 | 3, 4 | 0.447 | 1.64E+09 | 9.86E+08 | 8.43E+08 | 1.85E+09 | 4.00E+08 | 1.12E+09 | 1.40E+09 | 6.79E+08 | 1.43E+09 | 1.66E+09 | 2.14E+09 | 1.60E+09 | 3328.533, 3328.5378, 3328.5413 | LQQIC[160.0307]RDSEDEEAWIQETEPSAASTHLGK | P08032 | P08032 | SPTA1 | spectrin, alpha, erythrocytic 1 | Cytoplasm | other |
| 15533585 | 2 | 0.832 | 1.45E+09 | 1.54E+09 | 1.56E+09 | 1.52E+09 | 1.42E+09 | 1.40E+09 | 1.54E+09 | 1.55E+09 | 1.35E+09 | 1.62E+09 | 1.39E+09 | 1.36E+09 | 1104.614 | IITLDETATK | P08032 | P08032 | SPTA1 | spectrin, alpha, erythrocytic 1 | Cytoplasm | other |
| 15539237 | 3, 4 | 0.823 | 1.16E+09 | 1.28E+09 | 1.53E+09 | 9.70E+08 | 1.53E+09 | 1.13E+09 | 1.28E+09 | 1.52E+09 | 1.07E+09 | 1.55E+09 | 9.34E+08 | 9.96E+08 | 1580.7783, 1580.7789 | FENVQSLAAAHHEK | P08032 | P08032 | SPTA1 | spectrin, alpha, erythrocytic 1 | Cytoplasm | other |
| 15534437, 15544464 | 3, 4 | 0.175 | 1.44E+09 | 1.32E+09 | 1.38E+09 | 1.24E+09 | 1.28E+09 | 1.00E+09 | 1.38E+09 | 1.30E+09 | 1.11E+09 | 1.54E+09 | 1.37E+09 | 1.30E+09 | 1742.8458, 1742.8468, 1742.8484 | SDDRDHLNSLQALMK | P08032 | P08032 | SPTA1 | spectrin, alpha, erythrocytic 1 | Cytoplasm | other |
| 15533342 | 2 | 0.854 | 1.22E+09 | 1.07E+09 | 1.35E+09 | 1.15E+09 | 1.11E+09 | 1.13E+09 | 1.26E+09 | 1.16E+09 | 9.71E+08 | 1.53E+09 | 1.06E+09 | 1.09E+09 | 1076.557 | VTTLGETAER | P08032 | P08032 | SPTA1 | spectrin, alpha, erythrocytic 1 | Cytoplasm | other |
| 15538740 | 2, 3 | 0.601 | 1.23E+09 | 1.28E+09 | 1.45E+09 | 1.13E+09 | 1.33E+09 | 1.27E+09 | 1.25E+09 | 1.28E+09 | 1.20E+09 | 1.52E+09 | 1.26E+09 | 1.25E+09 | 1419.6941, 1419.6942 | ETGTLESQLEANK | P08032 | P08032 | SPTA1 | spectrin, alpha, erythrocytic 1 | Cytoplasm | other |
| 15536539, 15536554 | 2, 3 | 0.664 | 1.27E+09 | 1.20E+09 | 1.43E+09 | 1.01E+09 | 1.18E+09 | 1.09E+09 | 1.35E+09 | 1.27E+09 | 9.24E+08 | 1.52E+09 | 9.70E+08 | 9.95E+08 | 1470.6975, 1470.6997 | LEESYHYQVFR | P08032 | P08032 | SPTA1 | spectrin, alpha, erythrocytic 1 | Cytoplasm | other |
| 15531576 | 3 | 0.486 | 1.22E+09 | 1.19E+09 | 1.27E+09 | 1.27E+09 | 1.16E+09 | 1.14E+09 | 1.24E+09 | 1.16E+09 | 1.10E+09 | 1.51E+09 | 1.10E+09 | 1.34E+09 | 2245.1187 | YSTIGLAQQWDQLHQLGMR | P08032 | P08032 | SPTA1 | spectrin, alpha, erythrocytic 1 | Cytoplasm | other |
| 15536781 | 2 | 0.31 | 1.21E+09 | 1.21E+09 | 1.46E+09 | 1.03E+09 | 1.23E+09 | 1.23E+09 | 1.40E+09 | 1.42E+09 | 1.18E+09 | 1.50E+09 | 1.25E+09 | 1.34E+09 | 1232.6143 | ADVVESWIGEK | P08032 | P08032 | SPTA1 | spectrin, alpha, erythrocytic 1 | Cytoplasm | other |
| 15533381 | 3 | 0.45 | 1.25E+09 | 1.17E+09 | 1.31E+09 | 1.30E+09 | 1.11E+09 | 1.20E+09 | 1.38E+09 | 1.19E+09 | 9.32E+08 | 1.48E+09 | 1.34E+09 | 1.25E+09 | 1797.8971 | LMISHSADAPQIQQMK | P08032 | P08032 | SPTA1 | spectrin, alpha, erythrocytic 1 | Cytoplasm | other |
| 15532040, 15547008 | 2 | 0.521 | 1.29E+09 | 1.25E+09 | 1.36E+09 | 1.22E+09 | 1.20E+09 | 1.22E+09 | 1.45E+09 | 1.25E+09 | 1.09E+09 | 1.47E+09 | 1.37E+09 | 1.22E+09 | 1388.6992, 1388.7004 | EIDNTLQNINSK | P08032 | P08032 | SPTA1 | spectrin, alpha, erythrocytic 1 | Cytoplasm | other |
| 15534108 | 2 | 0.301 | 1.15E+09 | 1.13E+09 | 1.27E+09 | 1.10E+09 | 1.21E+09 | 1.01E+09 | 1.14E+09 | 1.06E+09 | 1.08E+09 | 1.45E+09 | 1.16E+09 | 1.15E+09 | 1191.6085 | DTIGVSEETLK | P08032 | P08032 | SPTA1 | spectrin, alpha, erythrocytic 1 | Cytoplasm | other |
| 15547755, 15562688 | 2 | 0.436 | 1.15E+09 | 1.10E+09 | 1.14E+09 | 1.06E+09 | 1.10E+09 | 9.67E+08 | 1.13E+09 | 1.12E+09 | 9.58E+08 | 1.45E+09 | 1.06E+09 | 1.09E+09 | 1964.9276, 1964.9327 | GLNYYLPMVEEGEPEPK | P08032 | P08032 | SPTA1 | spectrin, alpha, erythrocytic 1 | Cytoplasm | other |
| 15533653, 15539747 | 2, 3 | 0.461 | 1.31E+09 | 1.29E+09 | 1.40E+09 | 1.22E+09 | 1.29E+09 | 1.06E+09 | 1.48E+09 | 1.45E+09 | 1.15E+09 | 1.42E+09 | 1.16E+09 | 1.20E+09 | 1476.7512, 1476.7529 | GDSGDTLAATQSLLK | P08032 | P08032 | SPTA1 | spectrin, alpha, erythrocytic 1 | Cytoplasm | other |
| 15547759 | 3 | 0.699 | 9.13E+08 | 1.65E+09 | 2.08E+09 | 6.09E+08 | 1.80E+09 | 1.38E+09 | 1.23E+09 | 1.85E+09 | 1.18E+09 | 1.42E+09 | 6.46E+08 | 1.12E+09 | 2877.4746 | QEEPVNIPQLQQQVETLYHSLLDR | P08032 | P08032 | SPTA1 | spectrin, alpha, erythrocytic 1 | Cytoplasm | other |
| 15533110 | 2 | 0.271 | 1.41E+09 | 1.31E+09 | 1.72E+09 | 1.20E+09 | 1.24E+09 | 1.28E+09 | 1.43E+09 | 1.52E+09 | 1.20E+09 | 1.40E+09 | 1.33E+09 | 1.22E+09 | 1196.6273 | VIALYDFEAR | P08032 | P08032 | SPTA1 | spectrin, alpha, erythrocytic 1 | Cytoplasm | other |
| 15539417 | 3, 4 | 0.292 | 1.27E+09 | 1.11E+09 | 6.50E+08 | 1.44E+09 | 9.08E+08 | 9.23E+08 | 1.11E+09 | 9.90E+08 | 8.88E+08 | 1.39E+09 | 1.31E+09 | 1.44E+09 | 1975.0255, 1975.0266 | KHEAFLVDLNAFENSIK | P08032 | P08032 | SPTA1 | spectrin, alpha, erythrocytic 1 | Cytoplasm | other |
| 15532556, 15546338 | 3 | 0.66 | 8.60E+08 | 9.40E+08 | 1.06E+09 | 1.08E+09 | 9.40E+08 | 9.46E+08 | 1.15E+09 | 9.69E+08 | 8.50E+08 | 1.37E+09 | 8.69E+08 | 1.13E+09 | 2240.1692, 2240.1765 | TALINADELPTDVASGEALLAR | P08032 | P08032 | SPTA1 | spectrin, alpha, erythrocytic 1 | Cytoplasm | other |
| 15532139 | 2 | 0.315 | 1.14E+09 | 1.06E+09 | 1.23E+09 | 1.09E+09 | 1.02E+09 | 9.75E+08 | 1.06E+09 | 9.93E+08 | 9.12E+08 | 1.35E+09 | 1.07E+09 | 9.81E+08 | 1088.5103 | HFDENLTGR | P08032 | P08032 | SPTA1 | spectrin, alpha, erythrocytic 1 | Cytoplasm | other |
| 15532512 | 3 | 0.563 | 1.17E+09 | 1.27E+09 | 1.34E+09 | 8.43E+08 | 1.39E+09 | 1.14E+09 | 1.31E+09 | 1.56E+09 | 1.02E+09 | 1.35E+09 | 8.73E+08 | 9.56E+08 | 1858.8895 | LIDNDHYDSENIAAIR | P08032 | P08032 | SPTA1 | spectrin, alpha, erythrocytic 1 | Cytoplasm | other |
| 15548805, 15551468, 15558927 | 3, 4 | 0.934 | 7.90E+08 | 1.18E+09 | 9.28E+08 | 7.19E+08 | 1.01E+09 | 7.91E+08 | 8.74E+08 | 1.11E+09 | 6.94E+08 | 1.33E+09 | 6.49E+08 | 7.74E+08 | 3186.4363, 3186.4407, 3186.4438 | GVNLEESLEYLQFMENAEEEEAWLGEK | P08032 | P08032 | SPTA1 | spectrin, alpha, erythrocytic 1 | Cytoplasm | other |
| 15534040, 15546282 | 3 | 0.702 | 9.96E+08 | 1.01E+09 | 1.20E+09 | 9.07E+08 | 1.08E+09 | 1.07E+09 | 1.09E+09 | 1.12E+09 | 9.88E+08 | 1.27E+09 | 1.06E+09 | 1.04E+09 | 1774.8275, 1774.8336 | HDTLDASLQSFQQER | P08032 | P08032 | SPTA1 | spectrin, alpha, erythrocytic 1 | Cytoplasm | other |
| 15531248, 15536315, 15545874 | 2, 3 | 0.604 | 1.10E+09 | 9.46E+08 | 1.05E+09 | 1.10E+09 | 8.62E+08 | 8.75E+08 | 1.18E+09 | 9.31E+08 | 7.88E+08 | 1.26E+09 | 1.03E+09 | 9.79E+08 | 1276.6727, 1276.6731, 1276.6753 | RAEVLNQYQR | P08032 | P08032 | SPTA1 | spectrin, alpha, erythrocytic 1 | Cytoplasm | other |
| 15551056, 15554014 | 2 | 0.608 | 9.66E+08 | 1.30E+09 | 1.39E+09 | 7.64E+08 | 1.38E+09 | 1.21E+09 | 1.27E+09 | 1.29E+09 | 1.05E+09 | 1.25E+09 | 7.62E+08 | 9.34E+08 | 1826.9575, 1826.9581 | AEQVDGVINLGNSLIER | P08032 | P08032 | SPTA1 | spectrin, alpha, erythrocytic 1 | Cytoplasm | other |
| 15531569, 15536366, 15546720, 15547308 | 2, 3 | 0.071 | 1.29E+09 | 1.05E+09 | 1.35E+09 | 8.71E+08 | 1.14E+09 | 7.79E+08 | 1.17E+09 | 1.18E+09 | 1.03E+09 | 1.22E+09 | 1.22E+09 | 1.21E+09 | 1351.6318, 1351.6323, 1351.6338 | AEELFMEFAHK | P08032 | P08032 | SPTA1 | spectrin, alpha, erythrocytic 1 | Cytoplasm | other |
| 15535811, 15542655 | 3, 4 | 0.437 | 1.05E+09 | 1.33E+09 | 1.42E+09 | 1.03E+09 | 1.47E+09 | 1.22E+09 | 1.29E+09 | 1.45E+09 | 9.56E+08 | 1.19E+09 | 8.29E+08 | 9.97E+08 | 1586.8248, 1586.8252, 1586.8303 | HEVILADIASHEPR | P08032 | P08032 | SPTA1 | spectrin, alpha, erythrocytic 1 | Cytoplasm | other |
| 15534377 | 4 | 0.737 | 8.49E+08 | 1.29E+09 | 1.44E+09 | 7.57E+08 | 9.98E+08 | 1.12E+09 | 1.23E+09 | 1.05E+09 | 7.94E+08 | 1.18E+09 | 8.03E+08 | 1.13E+09 | 2533.1956 | REDPTFQALEDFGTELIDSGHR | P08032 | P08032 | SPTA1 | spectrin, alpha, erythrocytic 1 | Cytoplasm | other |
| 15547841 | 2 | 0.776 | 9.84E+08 | 1.12E+09 | 1.13E+09 | 1.10E+09 | 1.04E+09 | 1.04E+09 | 9.34E+08 | 1.08E+09 | 1.03E+09 | 1.16E+09 | 8.78E+08 | 1.01E+09 | 2136.9548 | FYQYSQEC[160.0307]EDILEWVK | P08032 | P08032 | SPTA1 | spectrin, alpha, erythrocytic 1 | Cytoplasm | other |
| 15539812 | 2 | 0.515 | 9.65E+08 | 9.42E+08 | 1.03E+09 | 9.63E+08 | 9.57E+08 | 8.46E+08 | 9.19E+08 | 9.60E+08 | 7.93E+08 | 1.16E+09 | 9.07E+08 | 9.15E+08 | 1266.5831 | EPQLTSEDYGK | P08032 | P08032 | SPTA1 | spectrin, alpha, erythrocytic 1 | Cytoplasm | other |
| 15550495 | 2 | 0.914 | 8.98E+08 | 8.35E+08 | 1.23E+09 | 1.16E+09 | 8.60E+08 | 1.11E+09 | 9.35E+08 | 1.13E+09 | 1.11E+09 | 1.15E+09 | 9.49E+08 | 1.02E+09 | 2088.1106 | ALNVPSSPYTWLTVDVLGR | P08032 | P08032 | SPTA1 | spectrin, alpha, erythrocytic 1 | Cytoplasm | other |
| 15534722 | 2 | 0.475 | 9.77E+08 | 8.34E+08 | 9.59E+08 | 8.46E+08 | 8.33E+08 | 8.13E+08 | 9.49E+08 | 8.98E+08 | 7.59E+08 | 1.13E+09 | 8.96E+08 | 8.37E+08 | 969.478 | KFDEFQR | P08032 | P08032 | SPTA1 | spectrin, alpha, erythrocytic 1 | Cytoplasm | other |
| 15531374 | 2, 3 | 0.133 | 1.01E+09 | 9.68E+08 | 1.25E+09 | 1.01E+09 | 8.17E+08 | 9.23E+08 | 9.78E+08 | 9.53E+08 | 1.00E+09 | 1.11E+09 | 1.04E+09 | 1.12E+09 | 1444.8348, 1444.8354 | KNDVLTLLSSINK | P08032 | P08032 | SPTA1 | spectrin, alpha, erythrocytic 1 | Cytoplasm | other |
| 15532518 | 2 | 0.365 | 8.17E+08 | 7.82E+08 | 8.98E+08 | 7.65E+08 | 8.26E+08 | 7.88E+08 | 8.69E+08 | 8.24E+08 | 7.53E+08 | 1.11E+09 | 8.37E+08 | 8.28E+08 | 1031.5714 | DLTSAGNLLK | P08032 | P08032 | SPTA1 | spectrin, alpha, erythrocytic 1 | Cytoplasm | other |
| 15540736 | 2 | 0.351 | 9.10E+08 | 8.07E+08 | 9.35E+08 | 8.01E+08 | 7.99E+08 | 7.83E+08 | 8.33E+08 | 8.06E+08 | 7.52E+08 | 1.09E+09 | 8.22E+08 | 7.97E+08 | 1002.5455 | ELLEATAQK | P08032 | P08032 | SPTA1 | spectrin, alpha, erythrocytic 1 | Cytoplasm | other |
| 15533159, 15546093, 15546782 | 3 | 0.673 | 9.55E+08 | 1.06E+09 | 1.24E+09 | 6.89E+08 | 1.13E+09 | 8.79E+08 | 9.99E+08 | 1.21E+09 | 7.69E+08 | 1.09E+09 | 7.54E+08 | 9.84E+08 | 1661.756, 1661.7578 | MVEEGHFAAEDIASR | P08032 | P08032 | SPTA1 | spectrin, alpha, erythrocytic 1 | Cytoplasm | other |
| 15549927 | 2, 3 | 0.972 | 8.24E+08 | 8.39E+08 | 1.10E+09 | 7.08E+08 | 1.00E+09 | 9.22E+08 | 9.16E+08 | 9.40E+08 | 8.13E+08 | 1.07E+09 | 7.65E+08 | 9.29E+08 | 2406.1414, 2406.142 | QQDFEEELAVNEIMLNNLEK | P08032 | P08032 | SPTA1 | spectrin, alpha, erythrocytic 1 | Cytoplasm | other |
| 15538613 | 2, 3 | 0.18 | 8.18E+08 | 7.61E+08 | 7.19E+08 | 7.70E+08 | 7.45E+08 | 5.33E+08 | 8.06E+08 | 6.87E+08 | 5.67E+08 | 1.06E+09 | 8.28E+08 | 8.08E+08 | 1380.6011, 1380.6012 | C[160.0307]DQVESWMVAR | P08032 | P08032 | SPTA1 | spectrin, alpha, erythrocytic 1 | Cytoplasm | other |
| 15534911, 15536704 | 3, 4 | 0.192 | 1.08E+09 | 7.78E+08 | 7.05E+08 | 1.03E+09 | 5.19E+08 | 6.40E+08 | 9.61E+08 | 7.08E+08 | 8.51E+08 | 1.06E+09 | 1.16E+09 | 1.04E+09 | 1809.9475, 1809.9492 | QHEVFERDIIPLGEK | P08032 | P08032 | SPTA1 | spectrin, alpha, erythrocytic 1 | Cytoplasm | other |
| 15532773, 15540603 | 2 | 0.846 | 7.77E+08 | 8.96E+08 | 9.75E+08 | 6.63E+08 | 9.48E+08 | 7.79E+08 | 8.66E+08 | 9.03E+08 | 6.73E+08 | 1.06E+09 | 6.61E+08 | 6.52E+08 | 1203.5941, 1203.5947 | ESLSNAADLQR | P08032 | P08032 | SPTA1 | spectrin, alpha, erythrocytic 1 | Cytoplasm | other |
| 15552066 | 2 | 0.421 | 8.15E+08 | 9.15E+08 | 1.02E+09 | 9.22E+08 | 8.08E+08 | 8.95E+08 | 7.34E+08 | 8.15E+08 | 7.98E+08 | 1.04E+09 | 7.37E+08 | 8.75E+08 | 1590.7151 | FLSDYDELSGWMK | P08032 | P08032 | SPTA1 | spectrin, alpha, erythrocytic 1 | Cytoplasm | other |
| 15550044, 15559244, 15561516 | 2 | 0.601 | 7.49E+08 | 9.57E+08 | 7.68E+08 | 8.03E+08 | 7.28E+08 | 6.70E+08 | 7.70E+08 | 7.71E+08 | 7.16E+08 | 1.04E+09 | 7.10E+08 | 7.68E+08 | 1565.7722, 1565.7794, 1565.782 | MLDQC[160.0307]LELQLFR | P08032 | P08032 | SPTA1 | spectrin, alpha, erythrocytic 1 | Cytoplasm | other |
| 15531875 | 2 | 0.424 | 1.03E+09 | 1.05E+09 | 1.22E+09 | 1.01E+09 | 1.04E+09 | 1.04E+09 | 1.17E+09 | 1.08E+09 | 9.06E+08 | 1.03E+09 | 9.09E+08 | 9.74E+08 | 1237.5841 | ENWDYLLER | P08032 | P08032 | SPTA1 | spectrin, alpha, erythrocytic 1 | Cytoplasm | other |
| 15533899, 15534880, 15539077, 15545910 | 2 | 0.096 | 8.65E+08 | 9.56E+08 | 1.02E+09 | 7.74E+08 | 8.23E+08 | 7.82E+08 | 9.04E+08 | 8.83E+08 | 8.37E+08 | 1.03E+09 | 8.12E+08 | 9.12E+08 | 1126.6124, 1126.6133, 1126.6134, 1126.6141 | AYFLDGSLLK | P08032 | P08032 | SPTA1 | spectrin, alpha, erythrocytic 1 | Cytoplasm | other |
| 15537339 | 2 | 0.6 | 7.54E+08 | 7.50E+08 | 9.87E+08 | 7.40E+08 | 7.08E+08 | 9.03E+08 | 8.86E+08 | 8.28E+08 | 7.65E+08 | 1.01E+09 | 8.93E+08 | 7.98E+08 | 1131.5768 | LDLVSNWER | P08032 | P08032 | SPTA1 | spectrin, alpha, erythrocytic 1 | Cytoplasm | other |
| 15535087 | 2 | 0.079 | 9.08E+08 | 8.89E+08 | 9.66E+08 | 7.96E+08 | 8.77E+08 | 9.41E+08 | 8.89E+08 | 8.17E+08 | 8.21E+08 | 9.88E+08 | 9.92E+08 | 9.15E+08 | 1075.5404 | ASDLENWIK | P08032 | P08032 | SPTA1 | spectrin, alpha, erythrocytic 1 | Cytoplasm | other |
| 15531369, 15536351, 15539037, 15541744, 15544681 | 3 | 0.483 | 1.00E+09 | 9.12E+08 | 5.26E+08 | 9.46E+08 | 6.81E+08 | 6.21E+08 | 8.24E+08 | 8.53E+08 | 8.22E+08 | 9.72E+08 | 1.11E+09 | 8.50E+08 | 2003.9857, 2003.9872, 2003.9874, 2003.9885, 2003.9911 | KTENTGVELDDVWELQK | P08032 | P08032 | SPTA1 | spectrin, alpha, erythrocytic 1 | Cytoplasm | other |
| 15532210, 15545017, 15546590 | 2 | 0.662 | 7.16E+08 | 6.43E+08 | 8.06E+08 | 6.89E+08 | 7.30E+08 | 6.31E+08 | 6.76E+08 | 6.95E+08 | 7.32E+08 | 9.64E+08 | 6.68E+08 | 7.08E+08 | 902.5151, 902.5165, 902.51807 | LSEIAELK | P08032 | P08032 | SPTA1 | spectrin, alpha, erythrocytic 1 | Cytoplasm | other |
| 15531941, 15544995, 15545938 | 2, 3 | 0.741 | 6.49E+08 | 7.52E+08 | 7.97E+08 | 5.49E+08 | 7.43E+08 | 5.73E+08 | 7.05E+08 | 7.21E+08 | 5.19E+08 | 9.58E+08 | 5.43E+08 | 6.23E+08 | 1310.6854, 1310.6859, 1310.6863, 1310.6871 | HQLLEAEMLAR | P08032 | P08032 | SPTA1 | spectrin, alpha, erythrocytic 1 | Cytoplasm | other |
| 15536969 | 2 | 0.978 | 7.53E+08 | 8.43E+08 | 1.01E+09 | 8.00E+08 | 8.30E+08 | 8.85E+08 | 9.09E+08 | 9.03E+08 | 7.86E+08 | 9.16E+08 | 7.61E+08 | 9.22E+08 | 1030.576 | EQLLTELGK | P08032 | P08032 | SPTA1 | spectrin, alpha, erythrocytic 1 | Cytoplasm | other |
| 15537500 | 2 | 0.331 | 6.84E+08 | 6.35E+08 | 7.31E+08 | 6.31E+08 | 6.03E+08 | 5.94E+08 | 6.62E+08 | 6.08E+08 | 5.70E+08 | 8.94E+08 | 6.28E+08 | 6.32E+08 | 843.4928 | IQVLNEK | P08032 | P08032 | SPTA1 | spectrin, alpha, erythrocytic 1 | Cytoplasm | other |
| 15533196 | 2 | 0.319 | 8.27E+08 | 9.44E+08 | 1.03E+09 | 8.80E+08 | 7.92E+08 | 8.48E+08 | 8.33E+08 | 9.02E+08 | 7.39E+08 | 8.86E+08 | 8.59E+08 | 8.41E+08 | 1059.5814 | LSEVANLWK | P08032 | P08032 | SPTA1 | spectrin, alpha, erythrocytic 1 | Cytoplasm | other |
| 15548149, 15555194 | 2 | 0.919 | 6.18E+08 | 5.82E+08 | 6.61E+08 | 5.42E+08 | 6.71E+08 | 5.15E+08 | 5.98E+08 | 5.87E+08 | 5.10E+08 | 8.72E+08 | 5.53E+08 | 4.54E+08 | 1809.7228 | VC[160.0307]DGDEENMQEQLDK | P08032 | P08032 | SPTA1 | spectrin, alpha, erythrocytic 1 | Cytoplasm | other |
| 15533678 | 3 | 0.241 | 7.10E+08 | 6.25E+08 | 6.61E+08 | 6.02E+08 | 6.29E+08 | 5.77E+08 | 6.33E+08 | 7.12E+08 | 6.10E+08 | 8.70E+08 | 6.83E+08 | 6.36E+08 | 1965.8273 | RVC[160.0307]DGDEENMQEQLDK | P08032 | P08032 | SPTA1 | spectrin, alpha, erythrocytic 1 | Cytoplasm | other |
| 15534348, 15543675 | 2, 3 | 0.262 | 7.45E+08 | 7.55E+08 | 8.75E+08 | 5.69E+08 | 7.03E+08 | 7.39E+08 | 8.30E+08 | 6.77E+08 | 7.22E+08 | 8.50E+08 | 7.29E+08 | 7.63E+08 | 1398.6587, 1398.6599, 1398.6609 | QC[160.0307]LDFHLFYR | P08032 | P08032 | SPTA1 | spectrin, alpha, erythrocytic 1 | Cytoplasm | other |
| 15548432 | 3 | 0.977 | 8.04E+08 | 7.57E+08 | 7.47E+08 | 7.82E+08 | 7.78E+08 | 7.30E+08 | 7.70E+08 | 8.64E+08 | 6.19E+08 | 8.44E+08 | 6.87E+08 | 7.77E+08 | 2896.3118 | QNQVDTLTDMAAHFEEIGHPDSGDIR | P08032 | P08032 | SPTA1 | spectrin, alpha, erythrocytic 1 | Cytoplasm | other |
| 15531296 | 2 | 0.321 | 6.99E+08 | 6.05E+08 | 7.57E+08 | 6.65E+08 | 5.64E+08 | 5.72E+08 | 6.52E+08 | 6.30E+08 | 5.77E+08 | 8.39E+08 | 6.17E+08 | 6.62E+08 | 858.5035 | IQVITER | P08032 | P08032 | SPTA1 | spectrin, alpha, erythrocytic 1 | Cytoplasm | other |
| 15536160 | 2 | 0.652 | 7.01E+08 | 6.56E+08 | 8.31E+08 | 6.44E+08 | 6.68E+08 | 6.53E+08 | 7.03E+08 | 7.07E+08 | 6.68E+08 | 8.37E+08 | 6.44E+08 | 6.44E+08 | 894.4551 | LGDYADLK | P08032 | P08032 | SPTA1 | spectrin, alpha, erythrocytic 1 | Cytoplasm | other |
| 15536428, 15546351 | 2, 3 | 0.612 | 7.78E+08 | 7.38E+08 | 7.74E+08 | 8.27E+08 | 6.75E+08 | 7.06E+08 | 8.25E+08 | 7.50E+08 | 5.80E+08 | 8.26E+08 | 7.64E+08 | 8.04E+08 | 1268.6346, 1268.6353, 1268.6356 | HWEQLLEASR | P08032 | P08032 | SPTA1 | spectrin, alpha, erythrocytic 1 | Cytoplasm | other |
| 15535679 | 2 | 0.29 | 8.05E+08 | 8.90E+08 | 1.08E+09 | 4.94E+08 | 9.40E+08 | 6.12E+08 | 8.48E+08 | 9.06E+08 | 6.72E+08 | 7.76E+08 | 6.32E+08 | 7.58E+08 | 1157.6512 | DLQSVQNLLK | P08032 | P08032 | SPTA1 | spectrin, alpha, erythrocytic 1 | Cytoplasm | other |
| 15537110 | 2 | 0.685 | 6.53E+08 | 6.51E+08 | 6.68E+08 | 5.60E+08 | 6.47E+08 | 5.75E+08 | 7.26E+08 | 6.54E+08 | 5.11E+08 | 7.68E+08 | 6.00E+08 | 6.13E+08 | 1339.5554 | DSEQVDSWMSR | P08032 | P08032 | SPTA1 | spectrin, alpha, erythrocytic 1 | Cytoplasm | other |
| 15537941 | 2 | 0.764 | 5.02E+08 | 6.96E+08 | 7.69E+08 | 4.38E+08 | 6.91E+08 | 6.22E+08 | 6.98E+08 | 7.29E+08 | 4.70E+08 | 7.42E+08 | 4.06E+08 | 4.78E+08 | 1203.5367 | QDEVNAAWDR | P08032 | P08032 | SPTA1 | spectrin, alpha, erythrocytic 1 | Cytoplasm | other |
| 15531367, 15535186 | 4 | 0.562 | 7.95E+08 | 5.25E+08 | 5.64E+08 | 9.15E+08 | 2.73E+08 | 6.01E+08 | 7.08E+08 | 5.01E+08 | 6.59E+08 | 7.17E+08 | 9.31E+08 | 7.70E+08 | 2484.2393, 2484.2422 | LIDNDHYDSENIAAIRDGLLAR | P08032 | P08032 | SPTA1 | spectrin, alpha, erythrocytic 1 | Cytoplasm | other |
| 15531999 | 2 | 0.381 | 6.58E+08 | 8.70E+08 | 9.17E+08 | 4.83E+08 | 8.96E+08 | 7.55E+08 | 6.38E+08 | 7.73E+08 | 5.80E+08 | 6.99E+08 | 4.47E+08 | 5.98E+08 | 1098.6251 | GLADVQNLLR | P08032 | P08032 | SPTA1 | spectrin, alpha, erythrocytic 1 | Cytoplasm | other |
| 15531534, 15546567 | 3 | 0.995 | 4.80E+08 | 6.34E+08 | 8.34E+08 | 7.09E+08 | 5.17E+08 | 7.45E+08 | 9.02E+08 | 5.45E+08 | 4.74E+08 | 6.94E+08 | 5.60E+08 | 6.34E+08 | 1823.8275, 1823.8336 | VDRVNQYANEC[160.0307]AQEK | P08032 | P08032 | SPTA1 | spectrin, alpha, erythrocytic 1 | Cytoplasm | other |
| 15555509, 15557633 | 2, 3 | 0.404 | 5.98E+08 | 4.63E+08 | 4.44E+08 | 4.99E+08 | 7.03E+08 | 4.26E+08 | 5.53E+08 | 8.08E+08 | 6.49E+08 | 6.65E+08 | 4.52E+08 | 5.45E+08 | 2377.0872, 2377.092 | EDPTFQALEDFGTELIDSGHR | P08032 | P08032 | SPTA1 | spectrin, alpha, erythrocytic 1 | Cytoplasm | other |
| 15531292, 15531615 | 2, 3 | 0.247 | 5.92E+08 | 4.97E+08 | 5.05E+08 | 5.25E+08 | 3.75E+08 | 4.63E+08 | 5.67E+08 | 4.75E+08 | 5.27E+08 | 6.64E+08 | 5.34E+08 | 5.11E+08 | 1453.7977, 1453.7993 | VLPELEEIREAR | P08032 | P08032 | SPTA1 | spectrin, alpha, erythrocytic 1 | Cytoplasm | other |
| 15535340 | 3 | 0.76 | 4.65E+08 | 6.02E+08 | 6.63E+08 | 3.77E+08 | 7.06E+08 | 5.14E+08 | 5.17E+08 | 6.02E+08 | 4.31E+08 | 6.46E+08 | 3.46E+08 | 4.20E+08 | 1526.7466 | KFEEFQEELTAR | P08032 | P08032 | SPTA1 | spectrin, alpha, erythrocytic 1 | Cytoplasm | other |
| 15533718 | 3 | 0.652 | 5.42E+08 | 6.71E+08 | 5.68E+08 | 4.59E+08 | 5.02E+08 | 6.37E+08 | 6.33E+08 | 5.86E+08 | 4.93E+08 | 6.34E+08 | 5.30E+08 | 6.39E+08 | 1939.0233 | ALNAADPGSDLLSVQALQR | P08032 | P08032 | SPTA1 | spectrin, alpha, erythrocytic 1 | Cytoplasm | other |
| 15549847 | 2 | 0.067 | 5.78E+08 | 5.90E+08 | 6.57E+08 | 4.98E+08 | 4.93E+08 | 4.89E+08 | 4.68E+08 | 4.93E+08 | 5.73E+08 | 6.33E+08 | 5.05E+08 | 5.36E+08 | 1970.8508 | SQPAGYDYVGFTNSFFGN | P08032 | P08032 | SPTA1 | spectrin, alpha, erythrocytic 1 | Cytoplasm | other |
| 15540248 | 3 | 0.389 | 7.63E+08 | 4.30E+08 | 3.19E+08 | 9.44E+08 | 2.46E+08 | 5.20E+08 | 6.22E+08 | 3.68E+08 | 5.49E+08 | 6.24E+08 | 1.10E+09 | 8.90E+08 | 2050.0435 | VSSQDYGRDLQSVQNLLK | P08032 | P08032 | SPTA1 | spectrin, alpha, erythrocytic 1 | Cytoplasm | other |

| | | | | | | | | | | | | | | | | | | | | | |
|---|---|---|---|---|---|---|---|---|---|---|---|---|---|---|---|---|---|---|---|---|---|
| 15552873, 15557075, 15557399, 15558204, 15559437, 15562255, 15562727, 15563399, 15563688, 15563717, 15563919 | 2, 6 | 0.388 | 5.86E+08 | 5.83E+08 | 6.57E+08 | 5.39E+08 | 5.63E+08 | 5.70E+08 | 6.07E+08 | 5.34E+08 | 5.79E+08 | 6.18E+08 | 5.13E+08 | 5.31E+08 | 1749.848, 1749.8512, 1749.8535, 1749.8552, 1749.8557, 1749.8568, 1749.8695, 5244.529, 5244.5547, 5244.5557 | GYVSLEDYTSFLIDK | P08032 | P08032 | SPTA1 | spectrin, alpha, erythrocytic 1 | Cytoplasm | other |
| 15534339, 15546686 | 3 | 0.733 | 5.81E+08 | 5.78E+08 | 6.83E+08 | 5.65E+08 | 6.62E+08 | 5.58E+08 | 5.44E+08 | 6.26E+08 | 5.62E+08 | 6.13E+08 | 6.18E+08 | 6.07E+08 | 1695.7108, 1695.7162 | HDDFEEAFTAQEEK | P08032 | P08032 | SPTA1 | spectrin, alpha, erythrocytic 1 | Cytoplasm | other |
| 15554076, 15558337, 15559987 | 2 | 0.273 | 5.24E+08 | 4.45E+08 | 4.82E+08 | 4.78E+08 | 4.43E+08 | 3.66E+08 | 4.53E+08 | 3.82E+08 | 3.33E+08 | 6.03E+08 | 4.99E+08 | 3.91E+08 | 1774.8301, 1774.8314, 1774.832 | HDTLDASLQSFQQER | P08032 | P08032 | SPTA1 | spectrin, alpha, erythrocytic 1 | Cytoplasm | other |
| 15549222 | 2 | 0.536 | 4.15E+08 | 3.67E+08 | 3.89E+08 | 3.27E+08 | 3.85E+08 | 3.08E+08 | 4.25E+08 | 3.47E+08 | 3.17E+08 | 5.74E+08 | 3.86E+08 | 3.25E+08 | 1642.7356 | LC[160.0307]ESHPDATEDLQK | P08032 | P08032 | SPTA1 | spectrin, alpha, erythrocytic 1 | Cytoplasm | other |
| 15532671, 15543989 | 2 | 0.933 | 5.45E+08 | 5.33E+08 | 5.42E+08 | 6.18E+08 | 4.41E+08 | 5.64E+08 | 5.21E+08 | 6.10E+08 | 5.24E+08 | 5.73E+08 | 4.97E+08 | 6.40E+08 | 901.5165, 901.5167 | FFLFLSK | P08032 | P08032 | SPTA1 | spectrin, alpha, erythrocytic 1 | Cytoplasm | other |
| 15532411 | 2 | 0.46 | 5.16E+08 | 4.03E+08 | 4.72E+08 | 4.45E+08 | 4.09E+08 | 3.79E+08 | 4.81E+08 | 3.96E+08 | 3.73E+08 | 5.49E+08 | 4.90E+08 | 3.95E+08 | 1116.6129 | FLNAVDPGRK | P08032 | P08032 | SPTA1 | spectrin, alpha, erythrocytic 1 | Cytoplasm | other |
| 15531585 | 2 | 0.173 | 5.61E+08 | 4.71E+08 | 4.91E+08 | 3.84E+08 | 4.61E+08 | 3.73E+08 | 4.94E+08 | 4.73E+08 | 3.58E+08 | 5.40E+08 | 5.23E+08 | 4.38E+08 | 1250.6819 | AIEEQHAALLR | P08032 | P08032 | SPTA1 | spectrin, alpha, erythrocytic 1 | Cytoplasm | other |
| 15552669, 15558532 | 2 | 0.228 | 3.94E+08 | 3.48E+08 | 3.92E+08 | 3.21E+08 | 3.40E+08 | 2.96E+08 | 3.76E+08 | 3.04E+08 | 3.04E+08 | 5.26E+08 | 3.62E+08 | 3.36E+08 | 1862.8789, 1862.8793 | SQLDDVHAFQQFNWK | P08032 | P08032 | SPTA1 | spectrin, alpha, erythrocytic 1 | Cytoplasm | other |
| 15541172 | 2, 3 | 0.073 | 4.07E+08 | 3.04E+08 | 1.92E+08 | 3.13E+08 | 3.11E+08 | 2.12E+08 | 3.54E+08 | 2.81E+08 | 2.43E+08 | 5.22E+08 | 5.63E+08 | 3.97E+08 | 1491.676, 1491.6763 | DVNEAIQWMEEK | P08032 | P08032 | SPTA1 | spectrin, alpha, erythrocytic 1 | Cytoplasm | other |
| 15531288 | 2, 3 | 0.542 | 5.40E+08 | 6.58E+08 | 6.96E+08 | 4.94E+08 | 5.40E+08 | 6.38E+08 | 5.73E+08 | 5.78E+08 | 6.34E+08 | 5.21E+08 | 5.99E+08 | 6.02E+08 | 1248.7075, 1248.7081 | IWNHLPDIIK | P08032 | P08032 | SPTA1 | spectrin, alpha, erythrocytic 1 | Cytoplasm | other |
| 15549183 | 2 | 0.644 | 3.74E+08 | 4.13E+08 | 5.05E+08 | 2.51E+08 | 4.96E+08 | 3.32E+08 | 3.68E+08 | 4.15E+08 | 3.30E+08 | 4.80E+08 | 2.72E+08 | 2.77E+08 | 1526.7451 | KFEEFQEELTAR | P08032 | P08032 | SPTA1 | spectrin, alpha, erythrocytic 1 | Cytoplasm | other |
| 15535089 | 2 | 0.329 | 3.71E+08 | 3.38E+08 | 3.69E+08 | 4.02E+08 | 3.13E+08 | 3.11E+08 | 3.39E+08 | 3.19E+08 | 2.74E+08 | 4.73E+08 | 3.41E+08 | 3.44E+08 | 905.4384 | NLAVMDDK | P08032 | P08032 | SPTA1 | spectrin, alpha, erythrocytic 1 | Cytoplasm | other |
| 15550329, 15555878 | 2, 3 | 0.065 | 5.89E+08 | 5.96E+08 | 6.28E+08 | 5.77E+08 | 4.86E+08 | 5.11E+08 | 4.99E+08 | 5.33E+08 | 6.06E+08 | 4.64E+08 | 5.33E+08 | 4.91E+08 | 2452.2673, 2452.2705 | EAIVTLVELGDDWERTEVLHK | P08032 | P08032 | SPTA1 | spectrin, alpha, erythrocytic 1 | Cytoplasm | other |
| 15531245 | 2 | 0.712 | 2.64E+08 | 4.30E+08 | 5.79E+08 | 1.67E+08 | 6.56E+08 | 3.77E+08 | 3.98E+08 | 5.54E+08 | 3.02E+08 | 4.44E+08 | 1.76E+08 | 2.50E+08 | 1097.6179 | VLPELEEIR | P08032 | P08032 | SPTA1 | spectrin, alpha, erythrocytic 1 | Cytoplasm | other |
| 15532012, 15543045 | 4 | 0.819 | 2.56E+08 | 3.87E+08 | 4.41E+08 | 2.28E+08 | 4.90E+08 | 2.94E+08 | 3.04E+08 | 4.48E+08 | 2.43E+08 | 4.33E+08 | 1.76E+08 | 2.48E+08 | 2646.3823, 2646.3833 | RLEGELVAHEPAVQNVLDTAESLR | P08032 | P08032 | SPTA1 | spectrin, alpha, erythrocytic 1 | Cytoplasm | other |
| 15552222 | 2 | 0.091 | 4.50E+08 | 3.94E+08 | 4.84E+08 | 3.57E+08 | 4.20E+08 | 3.66E+08 | 3.76E+08 | 3.85E+08 | 3.61E+08 | 4.33E+08 | 4.48E+08 | 3.88E+08 | 1695.7104 | HDDFEEAFTAQEEK | P08032 | P08032 | SPTA1 | spectrin, alpha, erythrocytic 1 | Cytoplasm | other |
| 15553833, 15557419, 15561315, 15562006, 15562247, 15562251, 15562380, 15563334, 15563554, 15563588, 15563811, 15563842, 15563951 | 2 | 0.422 | 3.36E+08 | 4.23E+08 | 4.68E+08 | 2.92E+08 | 3.44E+08 | 3.69E+08 | 3.61E+08 | 3.75E+08 | 4.19E+08 | 4.32E+08 | 3.05E+08 | 3.59E+08 | 1903.9423, 1903.9468, 1903.9481, 1903.9508, 1903.9529, 1903.9531, 1903.9537, 1903.954, 1903.9542, 1903.9547, 1903.9558, 1903.9686 | IEDLGDSMEEALILDIK | P08032 | P08032 | SPTA1 | spectrin, alpha, erythrocytic 1 | Cytoplasm | other |
| 15531821 | 2 | 0.667 | 4.01E+08 | 4.14E+08 | 5.42E+08 | 3.87E+08 | 3.92E+08 | 5.02E+08 | 3.81E+08 | 4.13E+08 | 4.21E+08 | 4.07E+08 | 3.30E+08 | 4.47E+08 | 714.4751 | LIDLLK | P08032 | P08032 | SPTA1 | spectrin, alpha, erythrocytic 1 | Cytoplasm | other |
| 15536574 | 3 | 0.842 | 2.56E+08 | 3.28E+08 | 4.41E+08 | 2.15E+08 | 4.14E+08 | 2.93E+08 | 2.71E+08 | 3.70E+08 | 2.52E+08 | 3.96E+08 | 1.85E+08 | 2.57E+08 | 2077.0164 | TELNEAWDTLQGLTSDRK | P08032 | P08032 | SPTA1 | spectrin, alpha, erythrocytic 1 | Cytoplasm | other |
| 15534845 | 3 | 0.854 | 2.25E+08 | 3.75E+08 | 3.35E+08 | 1.83E+08 | 4.45E+08 | 2.46E+08 | 3.27E+08 | 4.57E+08 | 2.12E+08 | 3.95E+08 | 1.44E+08 | 2.50E+08 | 2099.939 | DQAEVC[160.0307]QQQQAAPVDEAGR | P08032 | P08032 | SPTA1 | spectrin, alpha, erythrocytic 1 | Cytoplasm | other |
| 15547911 | 2 | 0.467 | 3.38E+08 | 3.54E+08 | 3.82E+08 | 2.59E+08 | 3.44E+08 | 2.94E+08 | 3.38E+08 | 3.37E+08 | 2.64E+08 | 3.83E+08 | 2.55E+08 | 2.94E+08 | 2152.1235 | VADELLFEELLTPEGAHIR | P08032 | P08032 | SPTA1 | spectrin, alpha, erythrocytic 1 | Cytoplasm | other |
| 15547986 | 2 | 0.532 | 2.58E+08 | 3.34E+08 | 4.39E+08 | 1.96E+08 | 4.45E+08 | 2.53E+08 | 3.50E+08 | 3.78E+08 | 2.48E+08 | 3.67E+08 | 1.83E+08 | 1.72E+08 | 2099.9387 | DQAEVC[160.0307]QQQQAAPVDEAGR | P08032 | P08032 | SPTA1 | spectrin, alpha, erythrocytic 1 | Cytoplasm | other |
| 15549198 | 2, 3 | 0.932 | 2.09E+08 | 1.87E+08 | 9.59E+07 | 1.74E+08 | 2.13E+08 | 1.05E+08 | 2.39E+08 | 1.61E+08 | 1.05E+08 | 3.65E+08 | 1.45E+08 | 1.31E+08 | 2462.154, 2462.1553 | YNEFLLAYEAGDMLEWIQEK | P08032 | P08032 | SPTA1 | spectrin, alpha, erythrocytic 1 | Cytoplasm | other |
| 15550494, 15558628 | 2 | 0.539 | 2.82E+08 | 4.15E+08 | 5.32E+08 | 2.07E+08 | 5.11E+08 | 3.19E+08 | 3.07E+08 | 4.74E+08 | 2.64E+08 | 3.59E+08 | 1.91E+08 | 2.47E+08 | 1744.8711, 1744.874 | EAIVTLVELGDDWER | P08032 | P08032 | SPTA1 | spectrin, alpha, erythrocytic 1 | Cytoplasm | other |
| 15544454, 15544575, 15544577, 15544642, 15547233 | 3, 4, 5 | 0.524 | 2.87E+08 | 4.97E+08 | 4.87E+08 | 7.48E+07 | 3.12E+08 | 6.90E+08 | 5.37E+08 | 5.02E+08 | 7.27E+08 | 3.57E+08 | 2.17E+08 | 6.93E+08 | 2197.1763, 2197.178, 2197.179, 2197.1802, 2197.181 | NLLNRHEVILADIASHEPR | P08032 | P08032 | SPTA1 | spectrin, alpha, erythrocytic 1 | Cytoplasm | other |
| 15531992 | 3 | 0.205 | 3.47E+08 | 4.32E+08 | 5.31E+08 | 4.18E+08 | 4.52E+08 | 4.73E+08 | 4.72E+08 | 4.14E+08 | 3.51E+08 | 3.55E+08 | 2.96E+08 | 3.80E+08 | 1647.7759 | RDVNEAIQWMEEK | P08032 | P08032 | SPTA1 | spectrin, alpha, erythrocytic 1 | Cytoplasm | other |
| 15548483, 15554344 | 2, 3 | 0.472 | 4.51E+08 | 2.16E+08 | 1.38E+08 | 6.01E+08 | 5.59E+07 | 2.27E+08 | 2.84E+08 | 9.68E+07 | 3.16E+08 | 3.50E+08 | 6.97E+08 | 4.78E+08 | 2332.115, 2332.1182 | MVEEGHFAAEDIASRVESLNK | P08032 | P08032 | SPTA1 | spectrin, alpha, erythrocytic 1 | Cytoplasm | other |
| 15548675, 15550153 | 2, 3 | 0.594 | 4.26E+08 | 2.44E+08 | 2.63E+08 | 4.55E+08 | 1.26E+08 | 2.71E+08 | 3.32E+08 | 2.11E+08 | 3.16E+08 | 3.36E+08 | 4.74E+08 | 3.59E+08 | 2484.2393, 2484.2422 | LIDNDHYDSENIAAIRDGLLAR | P08032 | P08032 | SPTA1 | spectrin, alpha, erythrocytic 1 | Cytoplasm | other |
| 15554141 | 3 | 0.617 | 2.40E+08 | 3.08E+08 | 3.75E+08 | 1.86E+08 | 4.69E+08 | 2.40E+08 | 2.77E+08 | 3.65E+08 | 2.13E+08 | 3.33E+08 | 1.47E+08 | 1.80E+08 | 2646.381 | RLEGELVAHEPAVQNVLDTAESLR | P08032 | P08032 | SPTA1 | spectrin, alpha, erythrocytic 1 | Cytoplasm | other |
| 15531345, 15536138 | 2, 3 | 0.926 | 2.48E+08 | 3.38E+08 | 3.57E+08 | 3.12E+08 | 2.81E+08 | 3.08E+08 | 3.26E+08 | 3.05E+08 | 2.38E+08 | 3.32E+08 | 2.48E+08 | 3.02E+08 | 1316.739, 1316.7407 | NDVLTLLSSINK | P08032 | P08032 | SPTA1 | spectrin, alpha, erythrocytic 1 | Cytoplasm | other |
| 15531671 | 2, 3 | 0.361 | 4.08E+08 | 2.25E+08 | 1.84E+08 | 3.87E+08 | 8.57E+07 | 2.42E+08 | 2.73E+08 | 1.73E+08 | 3.27E+08 | 3.29E+08 | 5.28E+08 | 4.15E+08 | 1226.7185, 1226.7191 | GLADVQNLLRK | P08032 | P08032 | SPTA1 | spectrin, alpha, erythrocytic 1 | Cytoplasm | other |
| 15550538, 15552067 | 3, 4 | 0.929 | 2.89E+08 | 4.27E+08 | 5.67E+08 | 1.79E+08 | 6.15E+08 | 4.10E+08 | 3.84E+08 | 4.37E+08 | 2.20E+08 | 3.18E+08 | 3.14E+08 | 4.37E+08 | 3647.601, 3647.603 | HEIDSYDDRFQSADATGQELLDGNHEASEEIR | P08032 | P08032 | SPTA1 | spectrin, alpha, erythrocytic 1 | Cytoplasm | other |
| 15547714 | 2 | 0.469 | 1.90E+08 | 4.25E+08 | 7.36E+08 | 1.97E+08 | 4.86E+08 | 3.88E+08 | 3.96E+08 | 3.96E+08 | 2.39E+08 | 3.11E+08 | 1.49E+08 | 2.04E+08 | 1875.894 | TENTGVELDDVWELQK | P08032 | P08032 | SPTA1 | spectrin, alpha, erythrocytic 1 | Cytoplasm | other |
| 15547830, 15550994 | 3, 4 | 0.295 | 3.67E+08 | 1.20E+08 | 8.97E+07 | 4.67E+08 | 5.71E+07 | 1.40E+08 | 2.17E+08 | 9.59E+07 | 1.37E+08 | 3.09E+08 | 5.04E+08 | 3.96E+08 | 3443.5894, 3443.5918 | TRGVNLEESLEYLQFMENAEEEEAWLGEK | P08032 | P08032 | SPTA1 | spectrin, alpha, erythrocytic 1 | Cytoplasm | other |
| 15531849 | 2, 3 | 0.08 | 2.77E+08 | 2.90E+08 | 2.86E+08 | 2.57E+08 | 2.06E+08 | 1.96E+08 | 2.79E+08 | 2.43E+08 | 2.30E+08 | 3.07E+08 | 2.35E+08 | 2.79E+08 | 1340.7512, 1340.7516 | SRVLPELEEIR | P08032 | P08032 | SPTA1 | spectrin, alpha, erythrocytic 1 | Cytoplasm | other |
| 15548045, 15551819 | 2 | 0.604 | 2.87E+08 | 2.09E+08 | 2.50E+08 | 2.27E+08 | 2.26E+08 | 1.78E+08 | 2.44E+08 | 2.07E+08 | 1.81E+08 | 3.03E+08 | 2.45E+08 | 1.81E+08 | 2034.9144, 2034.9146 | TGQEMIEDGHYASEAVAAR | P08032 | P08032 | SPTA1 | spectrin, alpha, erythrocytic 1 | Cytoplasm | other |
| 15537042 | 3 | 0.403 | 2.73E+08 | 1.18E+08 | 1.08E+08 | 4.73E+08 | 3.72E+07 | 1.91E+08 | 2.13E+08 | 9.40E+07 | 1.93E+08 | 2.97E+08 | 4.11E+08 | 4.22E+08 | 2015.0333 | QDEVNAAWDRLWSLALK | P08032 | P08032 | SPTA1 | spectrin, alpha, erythrocytic 1 | Cytoplasm | other |
| 15549997, 15551411 | 2 | 0.44 | 2.39E+08 | 2.08E+08 | 1.20E+08 | 2.55E+08 | 1.45E+08 | 6.20E+07 | 2.08E+08 | 1.35E+08 | 1.26E+08 | 2.91E+08 | 2.50E+08 | 1.84E+08 | 1801.8824, 1801.8839 | VEADDHQGFVPAVYVR | P08032 | P08032 | SPTA1 | spectrin, alpha, erythrocytic 1 | Cytoplasm | other |
| 15531619 | 4 | 0.435 | 2.41E+08 | 3.19E+08 | 3.62E+08 | 2.20E+08 | 4.25E+08 | 2.80E+08 | 2.76E+08 | 3.48E+08 | 2.08E+08 | 2.88E+08 | 1.73E+08 | 2.00E+08 | 2774.2817 | FQSADATGQELLDGNHEASEEIREK | P08032 | P08032 | SPTA1 | spectrin, alpha, erythrocytic 1 | Cytoplasm | other |
| 15534947 | 3 | 0.666 | 1.68E+08 | 2.35E+08 | 1.97E+08 | 1.83E+08 | 2.21E+08 | 2.13E+08 | 1.96E+08 | 2.51E+08 | 1.79E+08 | 2.82E+08 | 1.98E+08 | 2.22E+08 | 2136.9556 | FYQYSQEC[160.0307]EDILEWVK | P08032 | P08032 | SPTA1 | spectrin, alpha, erythrocytic 1 | Cytoplasm | other |
| 15532921, 15537946 | 2, 4 | 0.689 | 2.64E+08 | 2.56E+08 | 1.60E+08 | 2.38E+08 | 2.10E+08 | 1.97E+08 | 2.43E+08 | 2.01E+08 | 1.73E+08 | 2.64E+08 | 2.38E+08 | 2.25E+08 | 1496.7457 | DLVSSEALFHNHK | P08032 | P08032 | SPTA1 | spectrin, alpha, erythrocytic 1 | Cytoplasm | other |
| 15548832 | 2 | 0.434 | 2.15E+08 | 2.20E+08 | 2.75E+08 | 1.40E+08 | 2.33E+08 | 1.75E+08 | 1.98E+08 | 2.24E+08 | 1.73E+08 | 2.52E+08 | 1.58E+08 | 1.72E+08 | 1661.7562 | MVEEGHFAAEDIASR | P08032 | P08032 | SPTA1 | spectrin, alpha, erythrocytic 1 | Cytoplasm | other |
| 15550387 | 2 | 0.739 | 1.87E+08 | 2.11E+08 | 2.47E+08 | 1.43E+08 | 2.46E+08 | 1.58E+08 | 2.06E+08 | 2.28E+08 | 1.45E+08 | 2.48E+08 | 1.44E+08 | 1.42E+08 | 1858.8883 | LIDNDHYDSENIAAIR | P08032 | P08032 | SPTA1 | spectrin, alpha, erythrocytic 1 | Cytoplasm | other |
| 15553813 | 2 | 0.117 | 3.24E+08 | 4.03E+08 | 4.65E+08 | 3.79E+08 | 3.76E+08 | 3.83E+08 | 4.02E+08 | 3.26E+08 | 2.68E+08 | 2.45E+08 | 2.87E+08 | 3.30E+08 | 1647.7759 | RDVNEAIQWMEEK | P08032 | P08032 | SPTA1 | spectrin, alpha, erythrocytic 1 | Cytoplasm | other |
| 15533315, 15540303, 15543315, 15544571, 15544721, 15545585, 15546283, 15546715 | 4, 5 | 0.263 | 2.98E+08 | 2.55E+08 | 2.48E+08 | 2.42E+08 | 2.56E+08 | 2.41E+08 | 2.92E+08 | 2.54E+08 | 2.03E+08 | 2.34E+08 | 2.23E+08 | 1.98E+08 | 2517.2144, 2517.2185, 2517.2222, 2517.2244, 2517.2275, 2517.2278, 2517.2288, 2517.229 | RLADEQYQLLSSAHAVEM[147.0354]FHR | P08032 | P08032 | SPTA1 | spectrin, alpha, erythrocytic 1 | Cytoplasm | other |
| 15533098 | 3 | 0.501 | 2.42E+08 | 1.43E+08 | 3.16E+08 | 3.43E+08 | 2.13E+08 | 3.13E+08 | 2.91E+08 | 3.08E+08 | 3.01E+08 | 2.34E+08 | 2.51E+08 | 2.95E+08 | 2088.111 | ALNVPSSPYTWLTVDVLGR | P08032 | P08032 | SPTA1 | spectrin, alpha, erythrocytic 1 | Cytoplasm | other |
| 15532197 | 4 | 0.954 | 1.31E+08 | 2.17E+08 | 2.26E+08 | 7.99E+07 | 3.53E+08 | 1.64E+08 | 1.82E+08 | 2.46E+08 | 1.29E+08 | 2.26E+08 | 1.10E+08 | 1.43E+08 | 2345.1316 | LADEQYQLLSSAHAVEMFHR | P08032 | P08032 | SPTA1 | spectrin, alpha, erythrocytic 1 | Cytoplasm | other |
| 15551480, 15558909, 15559463 | 2 | 0.391 | 1.73E+08 | 1.39E+08 | 1.43E+08 | 1.57E+08 | 1.29E+08 | 4.90E+07 | 1.45E+08 | 1.19E+08 | 1.08E+08 | 2.11E+08 | 1.59E+08 | 1.25E+08 | 1742.8448, 1742.8451, 1742.8452 | SDDRDHLNSLQALMK | P08032 | P08032 | SPTA1 | spectrin, alpha, erythrocytic 1 | Cytoplasm | other |
| 15532496 | 2 | 0.608 | 1.65E+08 | 1.37E+08 | 1.22E+08 | 1.62E+08 | 8.80E+07 | 8.81E+07 | 1.56E+08 | 8.93E+07 | 5.21E+07 | 2.08E+08 | 8.75E+07 | 9.86E+07 | 830.511 | LWSLALK | P08032 | P08032 | SPTA1 | spectrin, alpha, erythrocytic 1 | Cytoplasm | other |
| 15547739 | 2 | 0.35 | 1.81E+08 | 1.35E+08 | 1.26E+08 | 1.65E+08 | 1.26E+08 | 1.06E+08 | 1.41E+08 | 1.10E+08 | 1.01E+08 | 2.06E+08 | 1.62E+08 | 1.23E+08 | 1965.824 | RVC[160.0307]DGDEENMQEQLDK | P08032 | P08032 | SPTA1 | spectrin, alpha, erythrocytic 1 | Cytoplasm | other |
| 15548457 | 3 | 0.474 | 2.17E+08 | 9.36E+07 | 8.37E+07 | 2.89E+08 | 3.16E+07 | 1.20E+08 | 1.22E+08 | 5.68E+07 | 1.50E+08 | 2.02E+08 | 2.82E+08 | 1.97E+08 | 2456.1187 | DQAEVC[160.0307]QQQQAAPVDEAGREAR | P08032 | P08032 | SPTA1 | spectrin, alpha, erythrocytic 1 | Cytoplasm | other |
| 15543487 | 2 | 0.936 | 1.76E+08 | 1.67E+08 | 2.23E+08 | 1.82E+08 | 1.81E+08 | 1.77E+08 | 2.00E+08 | 1.78E+08 | 1.58E+08 | 2.01E+08 | 1.76E+08 | 1.63E+08 | 875.4129 | EFSTTYK | P08032 | P08032 | SPTA1 | spectrin, alpha, erythrocytic 1 | Cytoplasm | other |
| 15532692 | 3 | 0.423 | 2.17E+08 | 1.08E+08 | 9.20E+07 | 2.20E+08 | 4.27E+07 | 1.29E+08 | 1.51E+08 | 6.31E+07 | 1.23E+08 | 1.92E+08 | 2.40E+08 | 1.91E+08 | 1983.0583 | AEQVDGVINLGNSLIERR | P08032 | P08032 | SPTA1 | spectrin, alpha, erythrocytic 1 | Cytoplasm | other |
| 15531535 | 3 | 0.653 | 1.18E+08 | 2.33E+08 | 4.75E+08 | 1.03E+08 | 2.73E+08 | 2.93E+08 | 2.12E+08 | 2.67E+08 | 1.34E+08 | 1.90E+08 | 9.00E+07 | 1.52E+08 | 1846.9314 | HEAFLVDLNAFENSIK | P08032 | P08032 | SPTA1 | spectrin, alpha, erythrocytic 1 | Cytoplasm | other |
| 15547708, 15554549, 15557169, 15559969, 15560616 | 2, 3 | 0.183 | 2.23E+08 | 1.72E+08 | 3.09E+08 | 1.21E+08 | 1.86E+08 | 1.77E+08 | 2.05E+08 | 1.95E+08 | 1.42E+08 | 1.88E+08 | 1.24E+08 | 1.21E+08 | 2261.1082, 2261.1086, 2261.1091, 2261.111, 2261.1118 | YSTIGLAQQWDQLHQLGM[147.0354]R | P08032 | P08032 | SPTA1 | spectrin, alpha, erythrocytic 1 | Cytoplasm | other |
| 15549827 | 2 | 0.321 | 1.64E+08 | 1.47E+08 | 1.43E+08 | 1.57E+08 | 1.36E+08 | 1.27E+08 | 1.58E+08 | 1.28E+08 | 1.09E+08 | 1.85E+08 | 1.64E+08 | 1.38E+08 | 1797.895 | LMISHSADAPQIQQMK | P08032 | P08032 | SPTA1 | spectrin, alpha, erythrocytic 1 | Cytoplasm | other |
| 15531558 | 3 | 0.76 | 1.79E+08 | 2.76E+08 | 1.31E+08 | 1.22E+08 | 3.51E+08 | 2.73E+08 | 2.77E+08 | 2.65E+08 | 1.97E+08 | 1.85E+08 | 1.75E+08 | 2.25E+08 | 1826.9576 | AEQVDGVINLGNSLIER | P08032 | P08032 | SPTA1 | spectrin, alpha, erythrocytic 1 | Cytoplasm | other |
| 15537678 | 3 | 0.263 | 9.67E+07 | 1.06E+08 | 1.07E+08 | 6.89E+07 | 8.99E+07 | 1.08E+08 | 9.08E+07 | 1.03E+08 | 6.61E+07 | 1.78E+08 | 9.69E+07 | 1.07E+08 | 1681.7893 | TSDDIESAFQALAEGK | P08032 | P08032 | SPTA1 | spectrin, alpha, erythrocytic 1 | Cytoplasm | other |
| 15538972 | 3 | 0.437 | 2.56E+08 | 8.87E+07 | 4.34E+07 | 3.17E+08 | 2.54E+07 | 9.79E+07 | 1.40E+08 | 4.46E+07 | 9.19E+07 | 1.75E+08 | 3.47E+08 | 2.29E+08 | 1626.7993 | LEESYHYQVFRR | P08032 | P08032 | SPTA1 | spectrin, alpha, erythrocytic 1 | Cytoplasm | other |
| 15533093 | 5 | 0.899 | 1.66E+08 | 2.80E+08 | 2.87E+08 | 1.03E+08 | 3.11E+08 | 2.27E+08 | 2.31E+08 | 2.50E+08 | 1.17E+08 | 1.67E+08 | 1.89E+08 | 2.58E+08 | 3647.6104 | HEIDSYDDRFQSADATGQELLDGNHEASEEIR | P08032 | P08032 | SPTA1 | spectrin, alpha, erythrocytic 1 | Cytoplasm | other |
| 15548109 | 3 | 0.283 | 8.25E+07 | 2.77E+07 | 1.09E+07 | 5.93E+07 | 2.73E+07 | 1.46E+07 | 2.07E+07 | 2.93E+07 | 3.97E+07 | 1.65E+08 | 1.10E+08 | 3.39E+07 | 2594.2798 | GTQLYEANQLLQFENNAEDLKR | P08032 | P08032 | SPTA1 | spectrin, alpha, erythrocytic 1 | Cytoplasm | other |
| 15547812 | 2 | 0.274 | 1.29E+08 | 1.09E+08 | 1.12E+08 | 1.42E+08 | 1.02E+08 | 9.55E+07 | 1.23E+08 | 9.12E+07 | 9.09E+07 | 1.61E+08 | 1.20E+08 | 1.25E+08 | 2245.1135 | YSTIGLAQQWDQLHQLGMR | P08032 | P08032 | SPTA1 | spectrin, alpha, erythrocytic 1 | Cytoplasm | other |
| 15533795, 15538437 | 4, 5 | 0.428 | 2.64E+08 | 8.77E+07 | 6.51E+07 | 2.70E+08 | 3.90E+07 | 1.08E+08 | 1.22E+08 | 5.87E+07 | 1.08E+08 | 1.59E+08 | 2.70E+08 | 2.40E+08 | 2381.1355, 2381.1375 | MNGVNERFENVQSLAAAHHEK | P08032 | P08032 | SPTA1 | spectrin, alpha, erythrocytic 1 | Cytoplasm | other |
| 15534850 | 3 | 0.922 | 8.21E+07 | 8.96E+07 | 1.60E+08 | 1.03E+08 | 1.10E+08 | 1.04E+08 | 9.41E+07 | 1.06E+08 | 9.42E+07 | 1.56E+08 | 8.21E+07 | 1.11E+08 | 1588.7606 | VQDVC[160.0307]AQGEDILNK | P08032 | P08032 | SPTA1 | spectrin, alpha, erythrocytic 1 | Cytoplasm | other |
| 15532571 | 3 | 0.462 | 1.95E+08 | 1.75E+08 | 1.40E+08 | 1.23E+08 | 1.91E+08 | 1.09E+08 | 1.97E+08 | 2.03E+08 | 1.39E+08 | 1.53E+08 | 1.69E+08 | 2.21E+08 | 1964.9283 | GLNYYLPMVEEGEPEPK | P08032 | P08032 | SPTA1 | spectrin, alpha, erythrocytic 1 | Cytoplasm | other |
| 15548988 | 2 | 0.281 | 1.66E+08 | 7.33E+07 | 7.37E+07 | 1.64E+08 | 3.29E+07 | 7.91E+07 | 9.29E+07 | 4.50E+07 | 8.25E+07 | 1.52E+08 | 2.00E+08 | 1.30E+08 | 1858.905 | NRVQDVC[160.0307]AQGEDILNK | P08032 | P08032 | SPTA1 | spectrin, alpha, erythrocytic 1 | Cytoplasm | other |
| 15552369, 15562789, 15562815 | 3, 4 | 0.488 | 2.00E+08 | 3.45E+08 | 3.19E+08 | 1.07E+08 | 1.44E+08 | 3.07E+08 | 1.93E+08 | 2.20E+08 | 1.83E+08 | 1.51E+08 | 1.37E+08 | 3.44E+08 | 3322.4766, 3322.4858, 3322.4868, 3322.4888 | QFYRDLEDLEEWINEM[147.0354]LPIAC[160.0307]DESYK | P08032 | P08032 | SPTA1 | spectrin, alpha, erythrocytic 1 | Cytoplasm | other |
| 15556890 | 2 | 0.677 | 1.06E+08 | 1.30E+08 | 1.33E+08 | 7.49E+07 | 1.37E+08 | 9.43E+07 | 1.18E+08 | 1.21E+08 | 8.89E+07 | 1.42E+08 | 9.67E+07 | 1.05E+08 | 1580.7782 | FENVQSLAAAHHEK | P08032 | P08032 | SPTA1 | spectrin, alpha, erythrocytic 1 | Cytoplasm | other |
| 15547673, 15552116, 15553977, 15558211, 15562546 | 3 | 0.605 | 2.17E+08 | 4.04E+08 | 6.75E+08 | 1.17E+08 | 3.29E+08 | 7.09E+08 | 1.96E+08 | 5.01E+08 | 4.61E+08 | 1.41E+08 | 1.56E+08 | 3.52E+08 | 2922.274, 2922.2754, 2922.2769, 2922.278, 2922.2788 | NFEM[147.0354]C[160.0307]QEFEQNASAFLQWIQETR | P08032 | P08032 | SPTA1 | spectrin, alpha, erythrocytic 1 | Cytoplasm | other |
| 15537389, 15543769 | 4 | 0.355 | 2.39E+08 | 8.19E+07 | 5.17E+07 | 3.52E+08 | 3.25E+07 | 1.14E+08 | 9.93E+07 | 3.70E+07 | 1.23E+08 | 1.39E+08 | 4.47E+08 | 3.05E+08 | 2893.4993, 2893.5022 | VADELLFEELLTPEGAHIRQELNTR | P08032 | P08032 | SPTA1 | spectrin, alpha, erythrocytic 1 | Cytoplasm | other |
| 15532038, 15538841, 15544207 | 3 | 0.475 | 1.55E+08 | 1.36E+08 | 1.48E+08 | 1.20E+08 | 1.45E+08 | 1.22E+08 | 1.26E+08 | 1.54E+08 | 8.85E+07 | 1.39E+08 | 9.28E+07 | 1.22E+08 | 1813.8888, 1813.8906, 1813.8966 | LM[147.0354]ISHSADAPQIQQMK | P08032 | P08032 | SPTA1 | spectrin, alpha, erythrocytic 1 | Cytoplasm | other |
| 15550202, 15554180, 15558675, 15559022, 15560296 | 3 | 0.399 | 1.47E+08 | 1.07E+08 | 1.19E+08 | 9.10E+07 | 9.59E+07 | 1.09E+08 | 1.25E+08 | 1.06E+08 | 8.77E+07 | 1.29E+08 | 9.77E+07 | 9.14E+07 | 2696.2197, 2696.22, 2696.2207, 2696.222, 2696.2263 | QALTPEQVSFC[160.0307]TIHMQQYM[147.0354]DPR | P08032 | P08032 | SPTA1 | spectrin, alpha, erythrocytic 1 | Cytoplasm | other |
| 15554053 | 3 | 0.494 | 8.44E+07 | 3.06E+08 | 7.56E+08 | 5.55E+07 | 3.46E+08 | 5.05E+08 | 1.72E+08 | 5.45E+08 | 2.61E+08 | 1.28E+08 | 4.87E+07 | 1.54E+08 | 2728.1943 | DLEDLEEWINEM[147.0354]LPIAC[160.0307]DESYK | P08032 | P08032 | SPTA1 | spectrin, alpha, erythrocytic 1 | Cytoplasm | other |
| 15554887, 15559641 | 2 | 0.308 | 1.10E+08 | 7.35E+07 | 5.11E+07 | 1.28E+08 | 4.16E+07 | 2.02E+07 | 9.92E+07 | 5.15E+07 | 6.20E+07 | 1.27E+08 | 1.22E+08 | 1.02E+08 | 1809.9459 | QHEVFERDIIPLGEK | P08032 | P08032 | SPTA1 | spectrin, alpha, erythrocytic 1 | Cytoplasm | other |
| 15535343 | 2, 3 | 0.522 | 2.08E+08 | 4.31E+07 | 3033749.75 | 1.45E+08 | 3.19E+07 | 1.50E+07 | 4.03E+07 | 3.93E+07 | 9.41E+07 | 1.26E+08 | 1.74E+08 | 1.12E+08 | 1402.6682, 1402.6686 | AKQDEVNAAWDR | P08032 | P08032 | SPTA1 | spectrin, alpha, erythrocytic 1 | Cytoplasm | other |
| 15547816 | 2 | 0.771 | 6.57E+07 | 8.96E+07 | 1.22E+08 | 9.02E+07 | 6.74E+07 | 9.86E+07 | 1.12E+08 | 6.42E+07 | 5.71E+07 | 1.25E+08 | 8.89E+07 | 7.57E+07 | 1823.8315 | VDRVNQYANEC[160.0307]AQEK | P08032 | P08032 | SPTA1 | spectrin, alpha, erythrocytic 1 | Cytoplasm | other |
| 15549739, 15554580, 15555576 | 2 | 0.799 | 2.29E+08 | 1.58E+08 | 9.62E+07 | 1.81E+08 | 1.24E+08 | 1.11E+08 | 1.57E+08 | 1.39E+08 | 1.36E+08 | 1.24E+08 | 2.43E+08 | 1.70E+08 | 2003.985, 2003.9852, 2003.9883 | KTENTGVELDDVWELQK | P08032 | P08032 | SPTA1 | spectrin, alpha, erythrocytic 1 | Cytoplasm | other |
| 15550704 | 3 | 0.515 | 9.95E+07 | 1.22E+08 | 1.43E+08 | 8.06E+07 | 1.55E+08 | 1.05E+08 | 1.10E+08 | 1.27E+08 | 9.34E+07 | 1.19E+08 | 7.37E+07 | 8.00E+07 | 2774.2773 | FQSADATGQELLDGNHEASEEIREK | P08032 | P08032 | SPTA1 | spectrin, alpha, erythrocytic 1 | Cytoplasm | other |
| 15548018, 15555160, 15555658, 15558616, 15558786, 15559173, 15562190 | 2 | 0.095 | 1.49E+08 | 1.08E+08 | 1.48E+08 | 8.86E+07 | 9.37E+07 | 1.16E+08 | 1.09E+08 | 1.07E+08 | 1.00E+08 | 1.18E+08 | 8.78E+07 | 8.47E+07 | 1581.7676, 1581.7706, 1581.7712, 1581.7728, 1581.7745, 1581.7761, 1581.7766 | M[147.0354]LDQC[160.0307]LELQLFR | P08032 | P08032 | SPTA1 | spectrin, alpha, erythrocytic 1 | Cytoplasm | other |
| 15554608, 15559136, 15560249 | 2 | 0.149 | 9.75E+07 | 7.72E+07 | 6.89E+07 | 3.51E+07 | 7.17E+07 | 6.37E+07 | 9.18E+07 | 6.07E+07 | 5.45E+07 | 1.18E+08 | 8.82E+07 | 8.16E+07 | 1537.7728, 1537.7733, 1537.7737 | KHEALENDFAVHK | P08032 | P08032 | SPTA1 | spectrin, alpha, erythrocytic 1 | Cytoplasm | other |
| 15550946 | 3 | 0.355 | 4.26E+07 | 2368627 | 2185923.25 | 6.15E+07 | 1664601.25 | 1572178 | 9563600 | 5729218.5 | 3.45E+07 | 1.18E+08 | 6.20E+07 | 1.37E+07 | 2739.239 | DVNEAIQWMEEKEPQLTSEDYGK | P08032 | P08032 | SPTA1 | spectrin, alpha, erythrocytic 1 | Cytoplasm | other |
| 15538065 | 2 | 0.682 | 7.55E+07 | 9.94E+07 | 1.38E+08 | 5.25E+07 | 1.30E+08 | 8.76E+07 | 9.78E+07 | 1.24E+08 | 7.04E+07 | 1.14E+08 | 4.81E+07 | 6.50E+07 | 737.3927 | FNTSIR | P08032 | P08032 | SPTA1 | spectrin, alpha, erythrocytic 1 | Cytoplasm | other |
| 15533025 | 2, 3 | 0.497 | 7.13E+07 | 1.08E+08 | 2.06E+08 | 6.20E+07 | 1.16E+08 | 1.07E+08 | 5.96E+07 | 1.02E+08 | 6.44E+07 | 1.11E+08 | 6.63E+07 | 7.33E+07 | 1409.6775, 1409.6776 | HEALENDFAVHK | P08032 | P08032 | SPTA1 | spectrin, alpha, erythrocytic 1 | Cytoplasm | other |
| 15551060, 15551083, 15557890, 15558867, 15559724, 15560288, 15561861 | 2 | 0.331 | 9.30E+07 | 8.34E+07 | 1.28E+08 | 6.96E+07 | 7.29E+07 | 8.70E+07 | 7.96E+07 | 7.76E+07 | 6.66E+07 | 1.09E+08 | 5.75E+07 | 5.59E+07 | 1606.7013, 1606.7047, 1606.7051, 1606.7057, 1606.7064, 1606.7097, 1606.7102 | FLSDYDELSGWM[147.0354]K | P08032 | P08032 | SPTA1 | spectrin, alpha, erythrocytic 1 | Cytoplasm | other |
| 15557461 | 2 | 0.2 | 9.34E+07 | 9.12E+07 | 5.47E+07 | 4.96E+07 | 7.56E+07 | 7.84E+07 | 1.02E+08 | 7.85E+07 | 7.17E+07 | 1.08E+08 | 8.71E+07 | 1.22E+08 | 1975.0234 | KHEAFLVDLNAFENSIK | P08032 | P08032 | SPTA1 | spectrin, alpha, erythrocytic 1 | Cytoplasm | other |
| 15533185 | 3 | 0.731 | 7.28E+07 | 1.15E+08 | 1.15E+08 | 6.17E+07 | 1.59E+08 | 9.06E+07 | 9.46E+07 | 1.26E+08 | 7.06E+07 | 1.06E+08 | 5.20E+07 | 7.24E+07 | 2233.08 | QRTELNEAWDTLQGLTSDR | P08032 | P08032 | SPTA1 | spectrin, alpha, erythrocytic 1 | Cytoplasm | other |
| 15543283, 15543887 | 3 | 0.411 | 6.58E+07 | 9.37E+07 | 9.44E+07 | 6.38E+07 | 6.00E+07 | 8.73E+07 | 8.86E+07 | 7.47E+07 | 8.23E+07 | 1.05E+08 | 7.84E+07 | 8.20E+07 | 1903.9536, 1903.9541 | IEDLGDSMEEALILDIK | P08032 | P08032 | SPTA1 | spectrin, alpha, erythrocytic 1 | Cytoplasm | other |
| 15532722 | 2 | 0.392 | 7.59E+07 | 6.71E+07 | 7.97E+07 | 7.64E+07 | 6.67E+07 | 5.99E+07 | 7.63E+07 | 6.89E+07 | 6.64E+07 | 1.01E+08 | 7.30E+07 | 7.17E+07 | 1120.5714 | AEVLNQYQR | P08032 | P08032 | SPTA1 | spectrin, alpha, erythrocytic 1 | Cytoplasm | other |
| 15531354, 15543093, 15543537, 15544564, 15547514 | 3 | 0.185 | 1.02E+08 | 1.02E+08 | 1.17E+08 | 9.21E+07 | 9.33E+07 | 8.11E+07 | 9.64E+07 | 9.53E+07 | 6.85E+07 | 9.98E+07 | 5.53E+07 | 7.23E+07 | 1813.8871, 1813.8884, 1813.89, 1813.8906, 1813.8956 | LMISHSADAPQIQQM[147.0354]K | P08032 | P08032 | SPTA1 | spectrin, alpha, erythrocytic 1 | Cytoplasm | other |
| 15552615, 15552774 | 3, 4 | 0.477 | 1.49E+08 | 2.91E+07 | 1.77E+07 | 2.80E+08 | 4109204.75 | 3.27E+07 | 5.56E+07 | 9453335 | 4.66E+07 | 9.78E+07 | 2.58E+08 | 1.15E+08 | 3108.539, 3108.5415 | HQAFENEVNGRAEQVDGVINLGNSLIER | P08032 | P08032 | SPTA1 | spectrin, alpha, erythrocytic 1 | Cytoplasm | other |
| 15549260, 15552249, 15552502, 15556584 | 3 | 0.617 | 9.76E+07 | 1.65E+08 | 2.74E+08 | 5.69E+07 | 1.53E+08 | 2.93E+08 | 8.96E+07 | 2.20E+08 | 1.56E+08 | 9.65E+07 | 5.96E+07 | 1.27E+08 | 3202.433, 3202.4338, 3202.4346 | GVNLEESLEYLQFM[147.0354]ENAEEEEAWLGEK | P08032 | P08032 | SPTA1 | spectrin, alpha, erythrocytic 1 | Cytoplasm | other |
| 15537468 | 3 | 0.503 | 1.86E+08 | 5.28E+07 | 3.31E+07 | 2.60E+08 | 1.34E+07 | 6.32E+07 | 7.64E+07 | 2.52E+07 | 9.93E+07 | 8.86E+07 | 3.02E+08 | 2.06E+08 | 1892.9863 | HQLLEAEMLAREDPLK | P08032 | P08032 | SPTA1 | spectrin, alpha, erythrocytic 1 | Cytoplasm | other |
| 15553655, 15554752 | 2, 3 | 0.209 | 9.26E+07 | 6.88E+07 | 9.61E+07 | 6.27E+07 | 6.43E+07 | 7.47E+07 | 6.74E+07 | 6.14E+07 | 5.93E+07 | 8.73E+07 | 5.00E+07 | 5.40E+07 | 2422.1404, 2422.1433 | QQDFEEELAVNEIM[147.0354]LNNLEK | P08032 | P08032 | SPTA1 | spectrin, alpha, erythrocytic 1 | Cytoplasm | other |
| 15532333, 15544153, 15544696, 15546520 | 3 | 0.254 | 8.15E+07 | 9.33E+07 | 1.21E+08 | 6.85E+07 | 1.02E+08 | 9.81E+07 | 9.58E+07 | 9.31E+07 | 8.04E+07 | 8.43E+07 | 6.86E+07 | 6.54E+07 | 1677.7458, 1677.748, 1677.7495, 1677.7552 | M[147.0354]VEEGHFAAEDIASR | P08032 | P08032 | SPTA1 | spectrin, alpha, erythrocytic 1 | Cytoplasm | other |
| 15532185 | 4 | 0.624 | 1.47E+08 | 8.32E+07 | 5.11E+07 | 1.88E+08 | 1.74E+07 | 1.04E+08 | 1.06E+08 | 3.68E+07 | 1.25E+08 | 8.17E+07 | 2.19E+08 | 2.20E+08 | 2332.1191 | MVEEGHFAAEDIASRVESLNK | P08032 | P08032 | SPTA1 | spectrin, alpha, erythrocytic 1 | Cytoplasm | other |
| 15538101 | 2 | 0.533 | 1.27E+08 | 1.55E+08 | 2.02E+08 | 7.63E+07 | 1.64E+08 | 1.50E+08 | 1.18E+08 | 1.61E+08 | 1.50E+08 | 7.91E+07 | 1.27E+08 | 1.41E+08 | 832.4516 | QESLLSR | P08032 | P08032 | SPTA1 | spectrin, alpha, erythrocytic 1 | Cytoplasm | other |
| 15531767 | 2, 3 | 0.425 | 8.19E+07 | 4.44E+07 | 3.88E+07 | 9.46E+07 | 2.29E+07 | 4.48E+07 | 6.58E+07 | 3.47E+07 | 4.95E+07 | 7.72E+07 | 9.68E+07 | 7.23E+07 | 1480.7831, 1480.7839 | VLETAEEIQHRR | P08032 | P08032 | SPTA1 | spectrin, alpha, erythrocytic 1 | Cytoplasm | other |
| 15532937, 15538352, 15544981, 15546026, 15546742, 15547202 | 2, 3 | 0.459 | 6.99E+07 | 6.08E+07 | 6.75E+07 | 6.21E+07 | 5.78E+07 | 6.14E+07 | 6.21E+07 | 5.73E+07 | 4.57E+07 | 7.67E+07 | 5.31E+07 | 5.40E+07 | 1483.7202, 1483.7263, 1483.7279, 1483.7284, 1483.7289, 1483.7303, 1483.7311, 1483.7333, 1483.734 | M[147.0354]QHNLEQQIQAK | P08032 | P08032 | SPTA1 | spectrin, alpha, erythrocytic 1 | Cytoplasm | other |
| 15552416 | 2 | 0.285 | 8.28E+07 | 4.38E+07 | 3.77E+07 | 1.01E+08 | 3.21E+07 | 4.67E+07 | 5.91E+07 | 3.61E+07 | 4.92E+07 | 7.45E+07 | 1.11E+08 | 8.14E+07 | 2050.0427 | VSSQDYGRDLQSVQNLLK | P08032 | P08032 | SPTA1 | spectrin, alpha, erythrocytic 1 | Cytoplasm | other |
| 15554056, 15558354, 15560648, 15561375 | 3 | 0.607 | 9.85E+07 | 4.65E+07 | 1.33E+07 | 4.43E+07 | 5.04E+07 | 4.56E+07 | 2.89E+07 | 8.10E+07 | 1.50E+08 | 7.17E+07 | 5.58E+07 | 1.04E+08 | 3552.6042, 3552.607, 3552.6072, 3552.6108 | DLEDLEEWINEM[147.0354]LPIAC[160.0307]DESYKDPTNIQR | P08032 | P08032 | SPTA1 | spectrin, alpha, erythrocytic 1 | Cytoplasm | other |
| 15547782 | 2, 3 | 0.846 | 4.30E+07 | 7.23E+07 | 8.07E+07 | 2.32E+07 | 1.16E+08 | 5.20E+07 | 5.62E+07 | 7.53E+07 | 4.25E+07 | 6.99E+07 | 3.12E+07 | 3.93E+07 | 2345.1304, 2345.1309 | LADEQYQLLSSAHAVEMFHR | P08032 | P08032 | SPTA1 | spectrin, alpha, erythrocytic 1 | Cytoplasm | other |
| 15532860 | 4 | 0.973 | 4.18E+07 | 5.18E+07 | 4.11E+07 | 3.04E+07 | 5.51E+07 | 3.88E+07 | 3.88E+07 | 5.01E+07 | 4.00E+07 | 6.68E+07 | 2.94E+07 | 3.82E+07 | 2530.1155 | DSEDEEAWIQETEPSAASTHLGK | P08032 | P08032 | SPTA1 | spectrin, alpha, erythrocytic 1 | Cytoplasm | other |
| 15547715 | 2 | 0.441 | 5.80E+07 | 1.24E+08 | 2.25E+08 | 2.47E+07 | 1.63E+08 | 1.76E+08 | 7.44E+07 | 1.85E+08 | 1.26E+08 | 5.89E+07 | 2.96E+07 | 6.70E+07 | 1901.895 | DFEFWLSEAEGLLAM[147.0354]K | P08032 | P08032 | SPTA1 | spectrin, alpha, erythrocytic 1 | Cytoplasm | other |
| 15541847, 15544332, 15545159, 15545842, 15546413, 15546584, 15547168, 15547613 | 3 | 0.241 | 4.83E+07 | 3.11E+07 | 2.72E+07 | 4.85E+07 | 2.34E+07 | 2.52E+07 | 3.56E+07 | 2.12E+07 | 2.42E+07 | 5.82E+07 | 4.79E+07 | 3.49E+07 | 2154.0076, 2154.0093, 2154.0095, 2154.0107, 2154.0117, 2154.0203, 2154.026 | RWLEEVEWQVTSEDYGK | P08032 | P08032 | SPTA1 | spectrin, alpha, erythrocytic 1 | Cytoplasm | other |
| 15531723, 15533571 | 2, 3 | 0.125 | 6.78E+07 | 5.52E+07 | 6.75E+07 | 5.25E+07 | 5.09E+07 | 4.23E+07 | 6.01E+07 | 5.07E+07 | 4.30E+07 | 5.50E+07 | 3.24E+07 | 4.32E+07 | 1367.6271, 1367.6277, 1367.6278 | AEELFM[147.0354]EFAHK | P08032 | P08032 | SPTA1 | spectrin, alpha, erythrocytic 1 | Cytoplasm | other |
| 15534145, 15543259, 15545007, 15545110, 15545156, 15545210, 15545327, 15546297 | 3 | 0.229 | 6.46E+07 | 5.53E+07 | 5.34E+07 | 5.48E+07 | 4.60E+07 | 5.27E+07 | 6.09E+07 | 5.66E+07 | 4.11E+07 | 4.99E+07 | 4.32E+07 | 4.20E+07 | 2050.9077, 2050.909, 2050.9092, 2050.91, 2050.9102, 2050.912, 2050.9153 | TGQEM[147.0354]IEDGHYASEAVAAR | P08032 | P08032 | SPTA1 | spectrin, alpha, erythrocytic 1 | Cytoplasm | other |
| 15534912 | 3 | 0.451 | 9.77E+07 | 2.84E+07 | 1.79E+07 | 1.60E+08 | 8648845 | 4.19E+07 | 4.03E+07 | 1.27E+07 | 4.26E+07 | 4.74E+07 | 1.98E+08 | 1.11E+08 | 2163.1042 | C[160.0307]ALVSRGDSGDTLAATQSLLK | P08032 | P08032 | SPTA1 | spectrin, alpha, erythrocytic 1 | Cytoplasm | other |
| 15548012, 15555052, 15557200, 15558611, 15559352, 15560305, 15560481, 15560573, 15561289, 15561421, 15562307, 15562350, 15562507, 15562531, 15562974 | 2 | 0.362 | 5.30E+07 | 3.48E+07 | 5.30E+07 | 3.65E+07 | 3.85E+07 | 3.89E+07 | 4.27E+07 | 4.17E+07 | 3.57E+07 | 4.74E+07 | 3.08E+07 | 2.67E+07 | 1980.9133, 1980.916, 1980.9183, 1980.9187, 1980.919, 1980.9203, 1980.9204, 1980.9232, 1980.9247, 1980.925, 1980.9253, 1980.9254, 1980.926, 1980.9293 | GLNYYLPM[147.0354]VEEGEPEPK | P08032 | P08032 | SPTA1 | spectrin, alpha, erythrocytic 1 | Cytoplasm | other |
| 15537005 | 3 | 0.496 | 3.94E+07 | 3.97E+07 | 3.62E+07 | 3.40E+07 | 3.99E+07 | 4.06E+07 | 3.86E+07 | 3.68E+07 | 3.80E+07 | 4.64E+07 | 4.04E+07 | 3.77E+07 | 1749.8557 | GYVSLEDYTSFLIDK | P08032 | P08032 | SPTA1 | spectrin, alpha, erythrocytic 1 | Cytoplasm | other |
| 15547820 | 2 | 0.52 | 2.79E+07 | 5.47E+07 | 1.19E+08 | 2.85E+07 | 6.25E+07 | 6.31E+07 | 4.38E+07 | 5.53E+07 | 3.17E+07 | 4.51E+07 | 2.15E+07 | 3.09E+07 | 1846.9294 | HEAFLVDLNAFENSIK | P08032 | P08032 | SPTA1 | spectrin, alpha, erythrocytic 1 | Cytoplasm | other |
| 15547756 | 3 | 0.835 | 3.13E+07 | 7.14E+07 | 1.13E+08 | 1.53E+07 | 1.47E+08 | 4.78E+07 | 4.21E+07 | 7.46E+07 | 2.52E+07 | 4.50E+07 | 2.34E+07 | 4.83E+07 | 2517.1436 | FQSADATGQELLDGNHEASEEIR | P08032 | P08032 | SPTA1 | spectrin, alpha, erythrocytic 1 | Cytoplasm | other |
| 15531580 | 2 | 0.679 | 2.93E+07 | 4.11E+07 | 5.16E+07 | 2.43E+07 | 5.47E+07 | 3.63E+07 | 3.47E+07 | 5.00E+07 | 3.16E+07 | 4.25E+07 | 2.15E+07 | 2.73E+07 | 988.5191 | FLNAVDPGR | P08032 | P08032 | SPTA1 | spectrin, alpha, erythrocytic 1 | Cytoplasm | other |
| 15555986 | 3 | 0.281 | 7.51E+07 | 2.29E+07 | 1.30E+07 | 1.00E+08 | 9914274 | 3.12E+07 | 2.97E+07 | 1.01E+07 | 2.16E+07 | 4.02E+07 | 1.24E+08 | 9.05E+07 | 2893.499 | VADELLFEELLTPEGAHIRQELNTR | P08032 | P08032 | SPTA1 | spectrin, alpha, erythrocytic 1 | Cytoplasm | other |
| 15552625 | 2 | 0.655 | 2.97E+07 | 3.59E+07 | 4.09E+07 | 2.18E+07 | 4.16E+07 | 2.94E+07 | 2.90E+07 | 3.46E+07 | 2.59E+07 | 4.00E+07 | 2.07E+07 | 2.31E+07 | 2077.0156 | TELNEAWDTLQGLTSDRK | P08032 | P08032 | SPTA1 | spectrin, alpha, erythrocytic 1 | Cytoplasm | other |
| 15531391 | 3 | 0.537 | 4.14E+07 | 6.94E+07 | 1.50E+08 | 1.41E+07 | 1.27E+08 | 1.24E+08 | 5.61E+07 | 1.44E+08 | 8.01E+07 | 3.73E+07 | 2.40E+07 | 4.95E+07 | 1901.8953 | DFEFWLSEAEGLLAM[147.0354]K | P08032 | P08032 | SPTA1 | spectrin, alpha, erythrocytic 1 | Cytoplasm | other |
| 15531601 | 2 | 0.404 | 6.74E+07 | 1.09E+07 | 1195283.75 | 5.38E+07 | 6199820.5 | 2755513.25 | 1.20E+07 | 9671188 | 3.76E+07 | 3.68E+07 | 8.47E+07 | 4.95E+07 | 842.5693 | KLIDLLK | P08032 | P08032 | SPTA1 | spectrin, alpha, erythrocytic 1 | Cytoplasm | other |
| 15540828 | 3 | 0.403 | 2.53E+07 | 3.05E+07 | 2.50E+07 | 1.58E+07 | 2.68E+07 | 2.77E+07 | 3.18E+07 | 2.97E+07 | 2.34E+07 | 3.56E+07 | 2.94E+07 | 2.65E+07 | 1565.7783 | MLDQC[160.0307]LELQLFR | P08032 | P08032 | SPTA1 | spectrin, alpha, erythrocytic 1 | Cytoplasm | other |
| 15549807 | 2 | 0.071 | 8.82E+07 | 1.01E+08 | 1.16E+08 | 7.06E+07 | 1.10E+08 | 8.58E+07 | 9.96E+07 | 1.03E+08 | 6.97E+07 | 3.55E+07 | 6.52E+07 | 7.04E+07 | 1586.8248 | HEVILADIASHEPR | P08032 | P08032 | SPTA1 | spectrin, alpha, erythrocytic 1 | Cytoplasm | other |
| 15551994 | 2 | 0.411 | 3.07E+07 | 1.36E+07 | 1.28E+07 | 6.01E+07 | 3812748.75 | 2.17E+07 | 2.35E+07 | 9895405 | 2.28E+07 | 3.52E+07 | 5.04E+07 | 4.93E+07 | 2015.0295 | QDEVNAAWDRLWSLALK | P08032 | P08032 | SPTA1 | spectrin, alpha, erythrocytic 1 | Cytoplasm | other |
| 15548944 | 3 | 0.428 | 5.11E+07 | 1.68E+07 | 8100690 | 5.24E+07 | 8762556 | 1.80E+07 | 2.80E+07 | 1.23E+07 | 2.34E+07 | 3.52E+07 | 5.96E+07 | 4.25E+07 | 2381.1377 | MNGVNERFENVQSLAAAHHEK | P08032 | P08032 | SPTA1 | spectrin, alpha, erythrocytic 1 | Cytoplasm | other |
| 15533022, 15546440, 15547026, 15547427 | 3, 4 | 0.806 | 3.66E+07 | 3.08E+07 | 3.58E+07 | 3.08E+07 | 3.55E+07 | 3.15E+07 | 3.91E+07 | 3.54E+07 | 2.81E+07 | 3.41E+07 | 3.27E+07 | 2.81E+07 | 1758.8354, 1758.8391, 1758.8402, 1758.8406, 1758.8436 | SDDRDHLNSLQALM[147.0354]K | P08032 | P08032 | SPTA1 | spectrin, alpha, erythrocytic 1 | Cytoplasm | other |
| 15534602 | 3 | 0.039 | 2.41E+07 | 2.42E+07 | 2.72E+07 | 2.01E+07 | 2.43E+07 | 2.38E+07 | 2.40E+07 | 2.43E+07 | 1.97E+07 | 3.35E+07 | 2.58E+07 | 3.14E+07 | 1970.8519 | SQPAGYDYVGFTNSFFGN | P08032 | P08032 | SPTA1 | spectrin, alpha, erythrocytic 1 | Cytoplasm | other |
| 15539022, 15539213, 15545849 | 2, 3 | 0.219 | 3.65E+07 | 3.74E+07 | 5.16E+07 | 2.42E+07 | 4.21E+07 | 3.97E+07 | 4.06E+07 | 4.11E+07 | 2.62E+07 | 3.26E+07 | 2.05E+07 | 2.57E+07 | 1326.6803, 1326.6805, 1326.6809 | HQLLEAEM[147.0354]LAR | P08032 | P08032 | SPTA1 | spectrin, alpha, erythrocytic 1 | Cytoplasm | other |
| 15534732 | 3 | 0.096 | 1.23E+07 | 3361050.75 | 2052740.25 | 9324562 | 3181473.5 | 2823992.25 | 3029119.25 | 4625589.5 | 4055859.75 | 3.26E+07 | 2.01E+07 | 7141663 | 1575.7943 | ETGTLESQLEANKR | P08032 | P08032 | SPTA1 | spectrin, alpha, erythrocytic 1 | Cytoplasm | other |
| 15547754 | 2 | 0.058 | 5.55E+07 | 5.58E+07 | 8.68E+07 | 6.44E+07 | 6.77E+07 | 5.03E+07 | 4.46E+07 | 6.06E+07 | 7.42E+07 | 3.22E+07 | 4.10E+07 | 4.22E+07 | 1534.8458 | SNGADLTAFLTLLAK | P08032 | P08032 | SPTA1 | spectrin, alpha, erythrocytic 1 | Cytoplasm | other |
| 15534311 | 3 | 0.489 | 4.06E+07 | 5.87E+07 | 5.04E+07 | 2.97E+07 | 6.91E+07 | 4.30E+07 | 4.20E+07 | 3.93E+07 | 2.33E+07 | 3.06E+07 | 4.21E+07 | 4.67E+07 | 2003.9861 | TENTGVELDDVWELQKK | P08032 | P08032 | SPTA1 | spectrin, alpha, erythrocytic 1 | Cytoplasm | other |
| 15535685 | 3 | 0.148 | 1.34E+07 | 2444511.75 | 716214.25 | 8855858 | 1833242.25 | 1487010.625 | 2316695.25 | 2313812.25 | 4889930 | 2.98E+07 | 2.07E+07 | 7225373.5 | 1604.8512 | KLAPDELPGFPQHR | P08032 | P08032 | SPTA1 | spectrin, alpha, erythrocytic 1 | Cytoplasm | other |
| 15532827 | 5 | 0.225 | 2.78E+07 | 1.99E+07 | 1.43E+07 | 2.02E+07 | 7315335.5 | 2.31E+07 | 2.71E+07 | 1.15E+07 | 2.95E+07 | 2.90E+07 | 4.11E+07 | 3.53E+07 | 3328.5361 | LQQIC[160.0307]RDSEDEEAWIQETEPSAASTHLGK | P08032 | P08032 | SPTA1 | spectrin, alpha, erythrocytic 1 | Cytoplasm | other |
| 15539633, 15540676 | 2, 3 | 0.344 | 7.06E+07 | 4806748.5 | 901667.5 | 4.82E+07 | 1725184.25 | 1283034.5 | 9809957 | 2662025 | 5507773.5 | 2.80E+07 | 8.40E+07 | 3.01E+07 | 1438.7805, 1438.7827 | KHQLLEAEMLAR | P08032 | P08032 | SPTA1 | spectrin, alpha, erythrocytic 1 | Cytoplasm | other |
| 15532178, 15546481, 15546930, 15546964, 15547507 | 3 | 0.93 | 2.44E+07 | 2.16E+07 | 2.37E+07 | 2.83E+07 | 2.02E+07 | 2.26E+07 | 2.74E+07 | 2.20E+07 | 1.75E+07 | 2.77E+07 | 1.85E+07 | 1.97E+07 | 1981.8167, 1981.8175, 1981.8186, 1981.8192 | RVC[160.0307]DGDEENM[147.0354]QEQLDK | P08032 | P08032 | SPTA1 | spectrin, alpha, erythrocytic 1 | Cytoplasm | other |
| 15547894 | 3 | 0.599 | 5.50E+07 | 1.23E+07 | 1.70E+07 | 6.07E+07 | 4501483 | 2.13E+07 | 2.11E+07 | 8520031 | 2.69E+07 | 2.77E+07 | 6.75E+07 | 3.61E+07 | 2348.1133 | M[147.0354]VEEGHFAAEDIASRVESLNK | P08032 | P08032 | SPTA1 | spectrin, alpha, erythrocytic 1 | Cytoplasm | other |
| 15556078, 15558489, 15559662, 15560319, 15560324, 15561957 | 2 | 0.365 | 2.20E+07 | 1.40E+07 | 8731551 | 2.99E+07 | 8974486 | 8981814 | 1.64E+07 | 7155508.5 | 5417489 | 2.48E+07 | 2.32E+07 | 1.33E+07 | 2154.004, 2154.0054, 2154.009, 2154.0093, 2154.01, 2154.0107 | RWLEEVEWQVTSEDYGK | P08032 | P08032 | SPTA1 | spectrin, alpha, erythrocytic 1 | Cytoplasm | other |
| 15531948, 15545977, 15546089, 15546445, 15547050, 15547271 | 2 | 0.284 | 2.37E+07 | 2.40E+07 | 3.15E+07 | 2.04E+07 | 2.65E+07 | 2.33E+07 | 2.50E+07 | 2.54E+07 | 1.90E+07 | 2.48E+07 | 1.41E+07 | 1.91E+07 | 1355.5482, 1355.5497, 1355.5498, 1355.5509, 1355.5526 | DSEQVDSWM[147.0354]SR | P08032 | P08032 | SPTA1 | spectrin, alpha, erythrocytic 1 | Cytoplasm | other |
| 15540652 | 3 | 0.857 | 7561003.5 | 2.45E+07 | 4.22E+07 | 1.08E+07 | 2.61E+07 | 2.47E+07 | 2.69E+07 | 2.27E+07 | 1.47E+07 | 2.46E+07 | 8730706 | 1.34E+07 | 1875.894 | TENTGVELDDVWELQK | P08032 | P08032 | SPTA1 | spectrin, alpha, erythrocytic 1 | Cytoplasm | other |
| 15557120, 15557625 | 3 | 0.993 | 1.19E+07 | 1.34E+07 | 1.32E+07 | 9356664 | 2.23E+07 | 9967633 | 1.10E+07 | 1.74E+07 | 9350525 | 2.43E+07 | 7209181.5 | 9182784 | 2490.2773, 2490.278 | LEGELVAHEPAVQNVLDTAESLR | P08032 | P08032 | SPTA1 | spectrin, alpha, erythrocytic 1 | Cytoplasm | other |
| 15554027 | 2 | 0.406 | 3.40E+07 | 1.04E+07 | 4506191.5 | 4.71E+07 | 2949384.75 | 9295249 | 1.69E+07 | 4774440.5 | 8770652 | 2.41E+07 | 4.70E+07 | 2.76E+07 | 1626.7993 | LEESYHYQVFRR | P08032 | P08032 | SPTA1 | spectrin, alpha, erythrocytic 1 | Cytoplasm | other |
| 15548331 | 3 | 0.499 | 6.12E+07 | 5038271.5 | 2956858.5 | 9.69E+07 | 152036.078 | 9596044 | 1.29E+07 | 2184150.5 | 1.99E+07 | 2.35E+07 | 1.44E+08 | 5.63E+07 | 2700.2722 | VTTLGETAERLC[160.0307]ESHPDATEDLQK | P08032 | P08032 | SPTA1 | spectrin, alpha, erythrocytic 1 | Cytoplasm | other |
| 15550432 | 1 | 0.085 | 1.94E+07 | 1.68E+07 | 2.02E+07 | 1.56E+07 | 1.45E+07 | 1.82E+07 | 1.92E+07 | 1.53E+07 | 1.38E+07 | 2.16E+07 | 2.17E+07 | 1.86E+07 | 1131.5763 | LDLVSNWER | P08032 | P08032 | SPTA1 | spectrin, alpha, erythrocytic 1 | Cytoplasm | other |
| 15553298, 15561033, 15561827, 15561922, 15562479 | 2 | 0.803 | 1.70E+07 | 1.33E+07 | 1.83E+07 | 1.76E+07 | 1.54E+07 | 1.44E+07 | 2.00E+07 | 1.37E+07 | 1.08E+07 | 2.07E+07 | 1.05E+07 | 1.07E+07 | 1825.7177, 1825.7185, 1825.7188, 1825.719, 1825.7202 | VC[160.0307]DGDEENM[147.0354]QEQLDK | P08032 | P08032 | SPTA1 | spectrin, alpha, erythrocytic 1 | Cytoplasm | other |
| 15543820 | 2 | 0.339 | 1.37E+07 | 1.28E+07 | 1.84E+07 | 9702409 | 1.41E+07 | 8747964 | 1.01E+07 | 1.10E+07 | 7350691.5 | 1.89E+07 | 1.04E+07 | 7840668.5 | 1398.6483 | FEEFQEELTAR | P08032 | P08032 | SPTA1 | spectrin, alpha, erythrocytic 1 | Cytoplasm | other |
| 15540296, 15543518, 15545903, 15546579, 15546823, 15547006 | 2 | 0.197 | 1.98E+07 | 1.68E+07 | 2.13E+07 | 1.55E+07 | 1.66E+07 | 1.72E+07 | 1.74E+07 | 1.61E+07 | 1.39E+07 | 1.88E+07 | 1.42E+07 | 1.53E+07 | 1396.5917, 1396.5935, 1396.5947, 1396.5969 | C[160.0307]DQVESWM[147.0354]VAR | P08032 | P08032 | SPTA1 | spectrin, alpha, erythrocytic 1 | Cytoplasm | other |
| 15537760 | 3 | 0.982 | 1.07E+07 | 1.18E+07 | 1.94E+07 | 8540535 | 1.83E+07 | 1.38E+07 | 1.40E+07 | 1.63E+07 | 7586844 | 1.78E+07 | 8919054 | 1.41E+07 | 1744.869 | EAIVTLVELGDDWER | P08032 | P08032 | SPTA1 | spectrin, alpha, erythrocytic 1 | Cytoplasm | other |
| 15549234, 15561401, 15561823, 15562430 | 2 | 0.116 | 1.92E+07 | 1.76E+07 | 2.32E+07 | 1.37E+07 | 1.95E+07 | 1.27E+07 | 1.72E+07 | 1.60E+07 | 1.50E+07 | 1.72E+07 | 1.30E+07 | 1.27E+07 | 1677.7485, 1677.7488, 1677.7502, 1677.7516 | M[147.0354]VEEGHFAAEDIASR | P08032 | P08032 | SPTA1 | spectrin, alpha, erythrocytic 1 | Cytoplasm | other |
| 15548180 | 3 | 0.479 | 2.01E+07 | 8611552 | 9499531 | 4.94E+07 | 3618033 | 9308307 | 1.10E+07 | 8373883.5 | 1.15E+07 | 1.68E+07 | 4.04E+07 | 2.33E+07 | 3810.7727 | DSEQVDSWMSRQEAFLENEDLGNSVGSVEALLQK | P08032 | P08032 | SPTA1 | spectrin, alpha, erythrocytic 1 | Cytoplasm | other |
| 15550210 | 3 | 0.907 | 3.72E+07 | 2.73E+07 | 3.46E+07 | 2.92E+07 | 8640045 | 5.89E+07 | 1.79E+07 | 1.70E+07 | 4.98E+07 | 1.63E+07 | 3.69E+07 | 5.68E+07 | 3459.583 | TRGVNLEESLEYLQFM[147.0354]ENAEEEEAWLGEK | P08032 | P08032 | SPTA1 | spectrin, alpha, erythrocytic 1 | Cytoplasm | other |
| 15554092, 15555036, 15558107, 15559896, 15561036 | 2, 3 | 0.118 | 1.53E+07 | 1.76E+07 | 1.98E+07 | 1.04E+07 | 1.22E+07 | 1.48E+07 | 1.38E+07 | 1.51E+07 | 1.24E+07 | 1.62E+07 | 8251882.5 | 9543873 | 2478.147, 2478.1477, 2478.1492, 2478.1497, 2478.1501, 2478.1516 | YNEFLLAYEAGDM[147.0354]LEWIQEK | P08032 | P08032 | SPTA1 | spectrin, alpha, erythrocytic 1 | Cytoplasm | other |
| 15548023 | 2 | 0.489 | 2.00E+07 | 8606427 | 8483811 | 2.08E+07 | 3015230.5 | 1.04E+07 | 1.32E+07 | 5577812 | 1.16E+07 | 1.53E+07 | 2.24E+07 | 1.72E+07 | 1983.057 | AEQVDGVINLGNSLIERR | P08032 | P08032 | SPTA1 | spectrin, alpha, erythrocytic 1 | Cytoplasm | other |
| 15553008 | 2 | 0.46 | 1.05E+07 | 1.83E+07 | 2.50E+07 | 7378058.5 | 3.23E+07 | 1.48E+07 | 1.35E+07 | 2.61E+07 | 1.28E+07 | 1.48E+07 | 5341309 | 8837666 | 1948.9199 | TELNEAWDTLQGLTSDR | P08032 | P08032 | SPTA1 | spectrin, alpha, erythrocytic 1 | Cytoplasm | other |
| 15537883 | 4 | 0.366 | 9740822 | 1.12E+07 | 6563923.5 | 1.05E+07 | 7264594 | 9301085 | 7455037 | 9099328 | 8606041 | 1.47E+07 | 8714949 | 1.18E+07 | 2440.1584 | ALRDQAEVC[160.0307]QQQQAAPVDEAGR | P08032 | P08032 | SPTA1 | spectrin, alpha, erythrocytic 1 | Cytoplasm | other |
| 15537258 | 3 | 0.985 | 8692877 | 9544510 | 1.38E+07 | 9649011 | 1.16E+07 | 1.09E+07 | 7528152 | 1.46E+07 | 1.27E+07 | 1.46E+07 | 8267019.5 | 9142273 | 1544.7981 | REIDNTLQNINSK | P08032 | P08032 | SPTA1 | spectrin, alpha, erythrocytic 1 | Cytoplasm | other |
| 15550955, 15552919 | 2, 3 | 0.451 | 4.25E+07 | 4542785.5 | 2454150.5 | 6.20E+07 | 385444.312 | 7828851 | 6342675.5 | 4969063.5 | 1.48E+07 | 1.41E+07 | 1.03E+08 | 5.14E+07 | 2255.147, 2255.151 | C[160.0307]RALNAADPGSDLLSVQALQR | P08032 | P08032 | SPTA1 | spectrin, alpha, erythrocytic 1 | Cytoplasm | other |
| 15539187 | 4 | 0.503 | 2.15E+07 | 2.52E+07 | 2924737.5 | 1.89E+07 | 2.00E+07 | 2.26E+07 | 2.87E+07 | 2.50E+07 | 2.24E+07 | 1.35E+07 | 2.95E+07 | 2.95E+07 | 2680.2305 | QALTPEQVSFC[160.0307]TIHMQQYMDPR | P08032 | P08032 | SPTA1 | spectrin, alpha, erythrocytic 1 | Cytoplasm | other |
| 15533171 | 4 | 0.542 | 1.10E+07 | 1.35E+07 | 1.92E+07 | 8264263.5 | 2.39E+07 | 1.34E+07 | 1.50E+07 | 1.87E+07 | 6873939 | 1.30E+07 | 6981031.5 | 7475257 | 2361.1226 | LADEQYQLLSSAHAVEM[147.0354]FHR | P08032 | P08032 | SPTA1 | spectrin, alpha, erythrocytic 1 | Cytoplasm | other |
| 15532499 | 4 | 0.394 | 1.48E+07 | 4560416.5 | 6429889 | 1.70E+07 | 1101402.5 | 4847233.5 | 8204660.5 | 2837632.5 | 9314009 | 1.28E+07 | 1.92E+07 | 1.31E+07 | 2348.1143 | M[147.0354]VEEGHFAAEDIASRVESLNK | P08032 | P08032 | SPTA1 | spectrin, alpha, erythrocytic 1 | Cytoplasm | other |
| 15548226 | 2 | 0.307 | 1.37E+07 | 1.10E+07 | 1.22E+07 | 1.02E+07 | 1.08E+07 | 8019721 | 1.14E+07 | 9431185 | 6764896 | 1.24E+07 | 7726806 | 8257560.5 | 1813.8906 | LM[147.0354]ISHSADAPQIQQMK | P08032 | P08032 | SPTA1 | spectrin, alpha, erythrocytic 1 | Cytoplasm | other |
| 15542114 | 4 | 0.462 | 7217670.5 | 3011047.75 | 138596.203 | 4098023 | 1368507.75 | 851750.312 | 1772833.125 | 2258027.25 | 2848305 | 1.16E+07 | 7864897.5 | 2738687.75 | 2594.279 | GTQLYEANQLLQFENNAEDLKR | P08032 | P08032 | SPTA1 | spectrin, alpha, erythrocytic 1 | Cytoplasm | other |
| 15550375 | 2 | 0.415 | 9401733 | 6847438.5 | 9283569 | 7862527 | 7996983.5 | 6017103.5 | 7750587 | 7068232.5 | 6719169.5 | 1.14E+07 | 8732205 | 6907004.5 | 1544.7988 | REIDNTLQNINSK | P08032 | P08032 | SPTA1 | spectrin, alpha, erythrocytic 1 | Cytoplasm | other |
| 15550876 | 2 | 0.265 | 1.03E+07 | 8939959 | 9390726 | 7253935 | 6824258.5 | 5806325 | 7762942 | 7914199.5 | 5710590.5 | 1.05E+07 | 4622450 | 6354530 | 1813.8894 | LMISHSADAPQIQQM[147.0354]K | P08032 | P08032 | SPTA1 | spectrin, alpha, erythrocytic 1 | Cytoplasm | other |
| 15551711 | 1 | 0.831 | 1.69E+07 | 1.27E+07 | 1.21E+07 | 2.08E+07 | 1.48E+07 | 1.25E+07 | 1.69E+07 | 1.22E+07 | 1.18E+07 | 1.02E+07 | 1.70E+07 | 1.48E+07 | 1237.5847 | ENWDYLLER | P08032 | P08032 | SPTA1 | spectrin, alpha, erythrocytic 1 | Cytoplasm | other |
| 15541096 | 3 | 0.61 | 1.74E+07 | 5365696.5 | 5175618 | 1.55E+07 | 1386108.25 | 8485248 | 8545710 | 2620246.5 | 7220911 | 9088375 | 1.50E+07 | 1.28E+07 | 1908.9786 | HQLLEAEM[147.0354]LAREDPLK | P08032 | P08032 | SPTA1 | spectrin, alpha, erythrocytic 1 | Cytoplasm | other |
| 15534427 | 4 | 0.402 | 1.71E+07 | 3.42E+07 | 3.65E+07 | 4154315.75 | 3.05E+07 | 2.33E+07 | 2.31E+07 | 2.89E+07 | 2.01E+07 | 9023739 | 1.29E+07 | 1.86E+07 | 2877.4697 | QEEPVNIPQLQQQVETLYHSLLDR | P08032 | P08032 | SPTA1 | spectrin, alpha, erythrocytic 1 | Cytoplasm | other |
| 15559600, 15559739, 15561692 | 2 | 0.823 | 3736342.25 | 5838467 | 8443318 | 2920841 | 6798834.5 | 4247753 | 4781526.5 | 5060856 | 2981817 | 8841532 | 2546613.75 | 3329434.75 | 1997.9073, 1997.9087, 1997.91 | WLEEVEWQVTSEDYGK | P08032 | P08032 | SPTA1 | spectrin, alpha, erythrocytic 1 | Cytoplasm | other |
| 15533884 | 3 | 0.344 | 2.49E+07 | 117124.695 | 644.733 | 1.43E+07 | 16813.092 | 6074.841 | 258589.547 | 181728.312 | 6448628 | 8461054 | 3.74E+07 | 8896941 | 1544.7963 | EIDNTLQNINSKR | P08032 | P08032 | SPTA1 | spectrin, alpha, erythrocytic 1 | Cytoplasm | other |
| 15542602 | 3 | 0.156 | 5433473 | 5403843 | 5537220 | 4786706 | 4435007 | 5158230.5 | 5281042 | 4461179.5 | 4712963.5 | 8328092.5 | 4868574.5 | 6076855 | 1590.7112 | FLSDYDELSGWMK | P08032 | P08032 | SPTA1 | spectrin, alpha, erythrocytic 1 | Cytoplasm | other |
| 15550242 | 3 | 0.953 | 9370353 | 3195221.75 | 1508483 | 7751711 | 1953186.875 | 2070397.875 | 3327700 | 2558774 | 3024886 | 8303696.5 | 3985183.5 | 1930863.625 | 3007.417 | LKETYALFQFFQDLDDEEAWIEEK | P08032 | P08032 | SPTA1 | spectrin, alpha, erythrocytic 1 | Cytoplasm | other |
| 15550877 | 2 | 0.129 | 2896127 | 691253.875 | 266529.75 | 1895319 | 537406.625 | 278149.219 | 375618.969 | 818243.375 | 673387.125 | 7851583 | 423085 | 1379188.25 | 1575.7943 | ETGTLESQLEANKR | P08032 | P08032 | SPTA1 | spectrin, alpha, erythrocytic 1 | Cytoplasm | other |
| 15552955, 15556208 | 2 | 0.402 | 9896826 | 7199149.5 | 6962187.5 | 5793126.5 | 6935912.5 | 5942402 | 8085583.5 | 6217524.5 | 4312823.5 | 7421346 | 6044860.5 | 4769502 | 2050.9092, 2050.9102 | TGQEM[147.0354]IEDGHYASEAVAAR | P08032 | P08032 | SPTA1 | spectrin, alpha, erythrocytic 1 | Cytoplasm | other |
| 15536146 | 4 | 0.694 | 1.48E+07 | 4374575 | 2789754 | 1.40E+07 | 1813033.125 | 5947439.5 | 8022594.5 | 3774982.5 | 6553560.5 | 6291776 | 1.60E+07 | 1.04E+07 | 2397.1292 | M[147.0354]NGVNERFENVQSLAAAHHEK | P08032 | P08032 | SPTA1 | spectrin, alpha, erythrocytic 1 | Cytoplasm | other |
| 15533697 | 2, 3 | 0.338 | 1.87E+07 | 4.70E-06 | 4.70E-06 | 1.55E+07 | 4.70E-06 | 4.70E-06 | 446162.812 | 58811.836 | 6098716.5 | 6200211 | 1.52E+07 | 4682939 | 1229.6453, 1229.6456 | AAVGKEEIQER | P08032 | P08032 | SPTA1 | spectrin, alpha, erythrocytic 1 | Cytoplasm | other |
| 15540122 | 3 | 0.835 | 2385488.75 | 5828114 | 5317189 | 2617689.75 | 9998465 | 4522854 | 4036162 | 7069912 | 3385592.25 | 6124368.5 | 2192564.25 | 2851054 | 1948.9203 | TELNEAWDTLQGLTSDR | P08032 | P08032 | SPTA1 | spectrin, alpha, erythrocytic 1 | Cytoplasm | other |
| 15540307 | 4 | 0.664 | 2176497.75 | 3261461.75 | 6225583 | 3448459.75 | 4504923 | 3140520 | 4281857 | 3003564.5 | 4766828.5 | 6105301.5 | 4620456 | 3964608 | 2513.3303 | TIGGVISSPELAEDLTGTEILLER | P08032 | P08032 | SPTA1 | spectrin, alpha, erythrocytic 1 | Cytoplasm | other |
| 15538718 | 3 | 0.553 | 8541513 | 3401008.5 | 5989610.5 | 4096232 | 3438249.25 | 4719960 | 5848687.5 | 4501173.5 | 3555402.25 | 5621353 | 3364675 | 3287020.75 | 1581.773 | M[147.0354]LDQC[160.0307]LELQLFR | P08032 | P08032 | SPTA1 | spectrin, alpha, erythrocytic 1 | Cytoplasm | other |
| 15549078 | 2 | 0.066 | 5549701 | 7260994 | 1.10E+07 | 6656435.5 | 6747163 | 7341867 | 7239885.5 | 6441940.5 | 5374264.5 | 5032289 | 2790141.75 | 4850836 | 1663.7712 | RDVNEAIQWM[147.0354]EEK | P08032 | P08032 | SPTA1 | spectrin, alpha, erythrocytic 1 | Cytoplasm | other |
| 15542790 | 3 | 0.062 | 7535427.5 | 8035797 | 1.16E+07 | 4611412.5 | 8270384 | 7263779.5 | 5302806 | 9060435 | 5672377 | 4910489 | 2089769.75 | 4593529.5 | 1829.8823 | LM[147.0354]ISHSADAPQIQQM[147.0354]K | P08032 | P08032 | SPTA1 | spectrin, alpha, erythrocytic 1 | Cytoplasm | other |
| 15553904, 15557421 | 2 | 0.23 | 5750134.5 | 3457359 | 4575607 | 4420521.5 | 4063644.25 | 3208018.25 | 3808755 | 2809465.25 | 2227291.75 | 4552081 | 2669836 | 2203972.75 | 1758.843, 1758.8433 | SDDRDHLNSLQALM[147.0354]K | P08032 | P08032 | SPTA1 | spectrin, alpha, erythrocytic 1 | Cytoplasm | other |
| 15534046 | 3 | 0.5 | 3843080.25 | 5139086.5 | 8371874.5 | 2120206.5 | 4922203.5 | 7060458 | 4681516 | 5522511.5 | 5637839 | 3968943 | 2865158.75 | 3931912.25 | 1919.9475 | IEDLGDSM[147.0354]EEALILDIK | P08032 | P08032 | SPTA1 | spectrin, alpha, erythrocytic 1 | Cytoplasm | other |
| 15549375 | 2 | 0.657 | 6438564 | 7638502 | 5413641.5 | 3799378.5 | 9196574 | 5313580 | 6609997.5 | 4254035.5 | 4056555 | 3920343 | 5830995.5 | 5646173 | 2003.9841 | TENTGVELDDVWELQKK | P08032 | P08032 | SPTA1 | spectrin, alpha, erythrocytic 1 | Cytoplasm | other |
| 15548393 | 2 | 0.558 | 1.15E+07 | 2132174.5 | 1186177.625 | 1.38E+07 | 348206.594 | 2987176.5 | 4598426.5 | 3363132.75 | 4933085 | 2906368.25 | 2.80E+07 | 1.26E+07 | 2163.103 | C[160.0307]ALVSRGDSGDTLAATQSLLK | P08032 | P08032 | SPTA1 | spectrin, alpha, erythrocytic 1 | Cytoplasm | other |
| 15541344 | 3 | 0.654 | 1616828.875 | 9937857 | 1.27E+07 | 703025.438 | 1.27E+07 | 1.32E+07 | 4750061 | 1.40E+07 | 1.43E+07 | 2704364.25 | 1951505.375 | 5776820.5 | 1809.7219 | VC[160.0307]DGDEENMQEQLDK | P08032 | P08032 | SPTA1 | spectrin, alpha, erythrocytic 1 | Cytoplasm | other |
| 15555904, 15561188 | 2 | 0.371 | 6377809.5 | 1667089.75 | 875228 | 8437689 | 21760.238 | 398956 | 2428219.75 | 451366.125 | 2834670.5 | 2329428.25 | 1.16E+07 | 6779321 | 1892.985, 1892.9863 | HQLLEAEMLAREDPLK | P08032 | P08032 | SPTA1 | spectrin, alpha, erythrocytic 1 | Cytoplasm | other |
| 15536288 | 3 | 0.399 | 3341930.75 | 2049100.625 | 2106114 | 2554799.25 | 765292.25 | 1790400.875 | 1610116.25 | 1312031.5 | 2043558.5 | 1667181.75 | 1759742.875 | 1210660.375 | 1293.6881 | LLRVSSQDYGR | P08032 | P08032 | SPTA1 | spectrin, alpha, erythrocytic 1 | Cytoplasm | other |
| 15554611 | 2 | 0.028 | 2310938.5 | 2218014.5 | 2582468.5 | 1486555 | 2050586 | 1954723 | 1934358.25 | 1848545.75 | 1884421.25 | 1655466 | 1877500.875 | 1779020.25 | 1981.8191 | RVC[160.0307]DGDEENM[147.0354]QEQLDK | P08032 | P08032 | SPTA1 | spectrin, alpha, erythrocytic 1 | Cytoplasm | other |
| 15556812 | 3 | 0.509 | 3750119 | 960140.062 | 888174.312 | 2937276.5 | 275685.062 | 1134278.5 | 1765209.75 | 282158.188 | 1168677.5 | 1594036.125 | 3731643.25 | 2297154.75 | 2397.131 | M[147.0354]NGVNERFENVQSLAAAHHEK | P08032 | P08032 | SPTA1 | spectrin, alpha, erythrocytic 1 | Cytoplasm | other |
| 15539428 | 3 | 0.673 | 1037911.375 | 1052761.5 | 1196114.375 | 844488.688 | 1040174.938 | 804740.688 | 993257.312 | 1130803.5 | 750991.125 | 1579861.125 | 823264.875 | 880268.875 | 1359.6952 | RESLSNAADLQR | P08032 | P08032 | SPTA1 | spectrin, alpha, erythrocytic 1 | Cytoplasm | other |
| 15557056 | 2 | 0.427 | 6809598.5 | 8136299.5 | 5592177 | 3183611 | 6960248 | 6592626 | 7222389 | 5155728.5 | 5333081 | 1562770.125 | 4858633 | 6332000 | 2501.231 | RLADEQYQLLSSAHAVEMFHR | P08032 | P08032 | SPTA1 | spectrin, alpha, erythrocytic 1 | Cytoplasm | other |
| 15554998 | 2 | 0.218 | 1374921.625 | 476834.531 | 547841.5 | 3519757 | 466104.25 | 931391.125 | 508095.781 | 609396.938 | 1337163.75 | 1547656.125 | 3546033 | 2020272.875 | 2183.9722 | GRSQPAGYDYVGFTNSFFGN | P08032 | P08032 | SPTA1 | spectrin, alpha, erythrocytic 1 | Cytoplasm | other |
| 15541121 | 3 | 0.337 | 1515305.125 | 1860654.75 | 1590150.25 | 1266498.875 | 1536788.875 | 1241274.375 | 1257865 | 1468801.25 | 1115126.5 | 1493215.25 | 1307117.875 | 499851.75 | 2019.0215 | M[147.0354]TILANDWAALLELWDK | P08032 | P08032 | SPTA1 | spectrin, alpha, erythrocytic 1 | Cytoplasm | other |
| 15559523 | 3 | 0.578 | 308803.094 | 873924.312 | 917246.812 | 400462.906 | 1676655.875 | 2069122.75 | 967304 | 351815.156 | 1555477.125 | 1331895.75 | 1206305.875 | 1296583.5 | 2696.2212 | QALTPEQVSFC[160.0307]TIHM[147.0354]QQYMDPR | P08032 | P08032 | SPTA1 | spectrin, alpha, erythrocytic 1 | Cytoplasm | other |
| 15540392, 15546039, 15546553 | 2, 3 | 0.727 | 5269781 | 2.54E+07 | 9.86E+07 | 974162.062 | 4.08E+07 | 5.88E+07 | 4211232.5 | 4.24E+07 | 7.09E+07 | 1159140.5 | 8070374 | 2.10E+07 | 1059.5881, 1059.5883, 1059.589, 1059.5902 | ARQESLLSR | P08032 | P08032 | SPTA1 | spectrin, alpha, erythrocytic 1 | Cytoplasm | other |
| 15534513 | 2 | 0.47 | 3106649.25 | 4.70E-06 | 68737.125 | 7216989 | 4.70E-06 | 396491.469 | 382895.719 | 125588.984 | 893483.188 | 779801.5 | 1.02E+07 | 3757607.25 | 1420.7765 | QESLLSRFEALK | P08032 | P08032 | SPTA1 | spectrin, alpha, erythrocytic 1 | Cytoplasm | other |
| 15557532 | 2 | 0.409 | 6494190.5 | 123589.711 | 0 | 3272147.25 | 0 | 0 | 120010.539 | 0 | 1780274.25 | 619085.562 | 1.14E+07 | 4173012.5 | 1544.7979 | EIDNTLQNINSKR | P08032 | P08032 | SPTA1 | spectrin, alpha, erythrocytic 1 | Cytoplasm | other |
| 15540734 | 3 | 0.298 | 1399870.5 | 955245.812 | 3915455.25 | 2861398.5 | 1686447 | 1929042.75 | 2095704.125 | 2573059.75 | 3528336 | 524773.938 | 1469237.875 | 1817527.125 | 1534.8441 | SNGADLTAFLTLLAK | P08032 | P08032 | SPTA1 | spectrin, alpha, erythrocytic 1 | Cytoplasm | other |
| 15547062, 15547496 | 2 | 0.702 | 263557.625 | 96853.555 | 748607.25 | 100901.539 | 340460.25 | 797790.5 | 287272.688 | 514750.719 | 686904.625 | 399244.906 | 610789.812 | 503928.219 | 921.43414, 921.4361 | NLAVM[147.0354]DDK | P08032 | P08032 | SPTA1 | spectrin, alpha, erythrocytic 1 | Cytoplasm | other |
| 15546434 | 4 | 0.316 | 1022075.125 | 1352618.25 | 2273293 | 109567.188 | 1417408.75 | 1800574.25 | 752513.75 | 1994267.125 | 1901949.375 | 4.70E-06 | 437670.219 | 1024351.75 | 2922.2754 | NFEM[147.0354]C[160.0307]QEFEQNASAFLQWIQETR | P08032 | P08032 | SPTA1 | spectrin, alpha, erythrocytic 1 | Cytoplasm | other |
| 15537272 | 2 | 0.847 | 8600127 | 8464998 | 1.06E+07 | 5150154.5 | 1.09E+07 | 7431198 | 9566639 | 1.04E+07 | 6598359 | 1.41E+07 | 4628638.5 | 5985146 | 1173.6083 | DVTGAEALLER | P16546 | P16546 | SPTAN1 | spectrin, alpha, non-erythrocytic 1 | Plasma Membrane | other |
| 15547665 | 3 | 0.276 | 3.34E+09 | 3.69E+09 | 4.21E+09 | 2.77E+09 | 3.49E+09 | 2.83E+09 | 3.02E+09 | 3.08E+09 | 3.09E+09 | 4.48E+09 | 2.88E+09 | 3.26E+09 | 2705.3472 | AVASEDMPESLPEAEQLLQQHAAIK | P15508 | P15508 | SPTB | spectrin, beta, erythrocytic | Plasma Membrane | other |
| 15552469, 15556632, 15559861 | 3, 4 | 0.273 | 2.97E+09 | 2.64E+09 | 1.91E+09 | 2.58E+09 | 2.30E+09 | 2.07E+09 | 2.06E+09 | 2.57E+09 | 2.35E+09 | 3.09E+09 | 2.88E+09 | 2.68E+09 | 3178.5962, 3178.5981, 3178.601 | QAEAILSNQEYTLAHLEPPDSLAAAEAGIR | P15508 | P15508 | SPTB | spectrin, beta, erythrocytic | Plasma Membrane | other |
| 15547840, 15547863, 15551961, 15556514 | 2, 3 | 0.317 | 2.52E+09 | 2.52E+09 | 3.32E+09 | 2.57E+09 | 2.17E+09 | 2.14E+09 | 2.33E+09 | 2.57E+09 | 2.47E+09 | 2.90E+09 | 2.29E+09 | 2.40E+09 | 2420.23, 2420.232, 2420.2346 | TLMAQIDGVNLAANNLVESGHPR | P15508 | P15508 | SPTB | spectrin, beta, erythrocytic | Plasma Membrane | other |
| 15535008 | 3 | 0.753 | 2.24E+09 | 2.29E+09 | 2.24E+09 | 2.42E+09 | 2.18E+09 | 2.32E+09 | 2.39E+09 | 2.30E+09 | 2.02E+09 | 2.47E+09 | 2.19E+09 | 2.39E+09 | 1805.9491 | AQSLLSAGHPEGEQIIR | P15508 | P15508 | SPTB | spectrin, beta, erythrocytic | Plasma Membrane | other |
| 15549380 | 2 | 0.577 | 1.98E+09 | 1.82E+09 | 2.19E+09 | 1.87E+09 | 1.92E+09 | 1.55E+09 | 1.79E+09 | 1.75E+09 | 1.83E+09 | 2.45E+09 | 1.88E+09 | 1.65E+09 | 1546.7319 | LLSGEDVGQDEGATR | P15508 | P15508 | SPTB | spectrin, beta, erythrocytic | Plasma Membrane | other |
| 15549084, 15551301 | 2, 3 | 0.301 | 2.14E+09 | 2.10E+09 | 2.59E+09 | 1.92E+09 | 2.19E+09 | 2.19E+09 | 2.18E+09 | 2.70E+09 | 2.53E+09 | 2.39E+09 | 2.15E+09 | 2.02E+09 | 2382.2058, 2382.2085 | QLGIIPLLDPEDVFTENPDEK | P15508 | P15508 | SPTB | spectrin, beta, erythrocytic | Plasma Membrane | other |
| 15531547, 15542570 | 2 | 0.425 | 2.02E+09 | 1.67E+09 | 2.01E+09 | 1.67E+09 | 1.75E+09 | 1.82E+09 | 1.94E+09 | 1.69E+09 | 1.54E+09 | 2.26E+09 | 1.83E+09 | 1.79E+09 | 1465.7135, 1465.7147 | LLTSPDVSYDEAR | P15508 | P15508 | SPTB | spectrin, beta, erythrocytic | Plasma Membrane | other |
| 15536278, 15542637 | 2, 3 | 0.269 | 1.84E+09 | 1.52E+09 | 1.70E+09 | 1.74E+09 | 1.51E+09 | 1.32E+09 | 1.74E+09 | 1.68E+09 | 1.21E+09 | 2.16E+09 | 1.76E+09 | 1.77E+09 | 1255.64, 1255.6427 | HRPDLIDFDK | P15508 | P15508 | SPTB | spectrin, beta, erythrocytic | Plasma Membrane | other |
| 15532889 | 3 | 0.631 | 1.72E+09 | 1.65E+09 | 1.98E+09 | 1.59E+09 | 1.51E+09 | 1.62E+09 | 1.77E+09 | 1.70E+09 | 1.49E+09 | 2.05E+09 | 1.48E+09 | 1.52E+09 | 1590.8326 | NQTLQNEILGHAPR | P15508 | P15508 | SPTB | spectrin, beta, erythrocytic | Plasma Membrane | other |

| | | | | | | | | | | | | | | | | | | | | |
|---|---|---|---|---|---|---|---|---|---|---|---|---|---|---|---|---|---|---|---|---|
| 15551920 | 2 | 0.959 | 1.69E+09 | 1.50E+09 | 1.94E+09 | 1.53E+09 | 1.65E+09 | 1.66E+09 | 1.89E+09 | 1.61E+09 | 1.40E+09 | 2.04E+09 | 1.59E+09 | 1.42E+09 | 1843.9365 | GLQDALQGQELSLGEASK | P15508 | P15508 | SPTB | spectrin, beta, erythrocytic | Plasma Membrane | other |
| 15548157, 15553136 | 2, 3 | 0.694 | 1.38E+09 | 2.09E+09 | 2.67E+09 | 9.94E+08 | 2.58E+09 | 1.70E+09 | 1.67E+09 | 2.11E+09 | 1.64E+09 | 2.03E+09 | 9.47E+08 | 1.45E+09 | 2451.1938, 2451.1975 | WLDQMDIPNTLEDLEVVQHR | P15508 | P15508 | SPTB | spectrin, beta, erythrocytic | Plasma Membrane | other |
| 15535680 | 3 | 0.849 | 1.68E+09 | 1.74E+09 | 1.65E+09 | 1.90E+09 | 1.53E+09 | 1.50E+09 | 1.97E+09 | 1.74E+09 | 1.42E+09 | 2.03E+09 | 1.64E+09 | 1.70E+09 | 1711.864 | LIDAGHSEAATIAEWK | P15508 | P15508 | SPTB | spectrin, beta, erythrocytic | Plasma Membrane | other |
| 15537021, 15538920, 15544351, 15546169 | 2, 3 | 0.075 | 1.78E+09 | 1.56E+09 | 1.69E+09 | 1.60E+09 | 1.39E+09 | 1.32E+09 | 1.63E+09 | 1.47E+09 | 1.40E+09 | 2.02E+09 | 1.71E+09 | 1.61E+09 | 1437.6826, 1437.6855, 1437.6859, 1437.6864 | HNLEHAFDVAER | P15508 | P15508 | SPTB | spectrin, beta, erythrocytic | Plasma Membrane | other |
| 15537159, 15545135, 15546526, 15547277 | 3 | 0.999 | 2.32E+09 | 2.72E+09 | 1.72E+09 | 2.09E+09 | 2.07E+09 | 2.42E+09 | 2.34E+09 | 2.23E+09 | 2.10E+09 | 2.01E+09 | 2.38E+09 | 2.30E+09 | 2081.1106, 2081.1135, 2081.1143 | ELHLLGVQVQQFQDVATR | P15508 | P15508 | SPTB | spectrin, beta, erythrocytic | Plasma Membrane | other |
| 15532182, 15539338 | 2, 3 | 0.422 | 1.57E+09 | 1.73E+09 | 1.78E+09 | 1.39E+09 | 1.56E+09 | 1.52E+09 | 1.62E+09 | 1.57E+09 | 1.45E+09 | 2.00E+09 | 1.57E+09 | 1.45E+09 | 1412.7256, 1412.7258, 1412.7267 | LGHLQSSWDTLR | P15508 | P15508 | SPTB | spectrin, beta, erythrocytic | Plasma Membrane | other |
| 15535316, 15539876, 15540482, 15544982 | 2, 3 | 0.317 | 1.62E+09 | 1.66E+09 | 1.81E+09 | 1.59E+09 | 1.70E+09 | 1.64E+09 | 1.62E+09 | 1.60E+09 | 1.47E+09 | 1.94E+09 | 1.55E+09 | 1.78E+09 | 1452.734, 1452.7341, 1452.7354, 1452.7357 | LQAFLQDLDDFK | P15508 | P15508 | SPTB | spectrin, beta, erythrocytic | Plasma Membrane | other |
| 15535393, 15540241 | 2, 3 | 0.262 | 1.56E+09 | 1.53E+09 | 1.44E+09 | 1.45E+09 | 1.46E+09 | 1.36E+09 | 1.57E+09 | 1.47E+09 | 1.20E+09 | 1.91E+09 | 1.60E+09 | 1.48E+09 | 1448.6295, 1448.6301, 1448.6307 | AAEIDC[160.0307]QDIEER | P15508 | P15508 | SPTB | spectrin, beta, erythrocytic | Plasma Membrane | other |
| 15548948, 15561440, 15561758 | 2 | 0.73 | 1.42E+09 | 1.36E+09 | 1.67E+09 | 1.18E+09 | 1.57E+09 | 1.11E+09 | 1.36E+09 | 1.32E+09 | 1.29E+09 | 1.80E+09 | 1.18E+09 | 1.17E+09 | 1831.9133, 1831.9166, 1831.9286 | VIEGQTDPDYQLLGQR | P15508 | P15508 | SPTB | spectrin, beta, erythrocytic | Plasma Membrane | other |
| 15534725, 15539365, 15544839 | 2, 3 | 0.353 | 1.43E+09 | 1.35E+09 | 1.66E+09 | 1.30E+09 | 1.31E+09 | 1.16E+09 | 1.56E+09 | 1.46E+09 | 1.41E+09 | 1.77E+09 | 1.24E+09 | 1.32E+09 | 1320.7135, 1320.7145 | AITAATLQFAEGK | P15508 | P15508 | SPTB | spectrin, beta, erythrocytic | Plasma Membrane | other |
| 15531864, 15540798, 15546340 | 2, 3 | 0.423 | 1.40E+09 | 1.36E+09 | 1.56E+09 | 1.19E+09 | 1.20E+09 | 1.40E+09 | 1.51E+09 | 1.34E+09 | 1.17E+09 | 1.75E+09 | 1.30E+09 | 1.38E+09 | 1468.8093, 1468.8094, 1468.8097 | DLGQDLAGVIAIQR | P15508 | P15508 | SPTB | spectrin, beta, erythrocytic | Plasma Membrane | other |
| 15547862, 15548526, 15553840, 15560450 | 2, 3 | 0.845 | 1.38E+09 | 1.68E+09 | 1.58E+09 | 1.40E+09 | 1.64E+09 | 1.29E+09 | 1.36E+09 | 1.50E+09 | 1.44E+09 | 1.74E+09 | 1.35E+09 | 1.32E+09 | 2311.009, 2311.0098, 2311.0107, 2311.0142 | EMVASSQEMGQDFDHVTMLR | P15508 | P15508 | SPTB | spectrin, beta, erythrocytic | Plasma Membrane | other |
| 15535946, 15539696 | 3, 4 | 0.235 | 1.45E+09 | 1.31E+09 | 1.30E+09 | 1.51E+09 | 1.30E+09 | 1.06E+09 | 1.18E+09 | 1.23E+09 | 1.06E+09 | 1.73E+09 | 1.45E+09 | 1.24E+09 | 1762.8209, 1762.822 | KHEAIETDTAAYEER | P15508 | P15508 | SPTB | spectrin, beta, erythrocytic | Plasma Membrane | other |
| 15531280, 15547279 | 2, 3 | 0.284 | 1.46E+09 | 1.30E+09 | 1.59E+09 | 1.32E+09 | 1.30E+09 | 1.20E+09 | 1.42E+09 | 1.34E+09 | 1.15E+09 | 1.73E+09 | 1.32E+09 | 1.41E+09 | 1336.7445, 1336.7448, 1336.7457 | HLLEVEDLLQK | P15508 | P15508 | SPTB | spectrin, beta, erythrocytic | Plasma Membrane | other |
| 15548222, 15556826 | 2 | 0.875 | 1.37E+09 | 1.34E+09 | 1.90E+09 | 1.31E+09 | 1.49E+09 | 1.42E+09 | 1.44E+09 | 1.39E+09 | 1.57E+09 | 1.72E+09 | 1.37E+09 | 1.43E+09 | 1635.7776, 1635.7778 | DNLELQNFLQNC[160.0307]K | P15508 | P15508 | SPTB | spectrin, beta, erythrocytic | Plasma Membrane | other |
| 15535395 | 3 | 0.057 | 1.04E+09 | 1.01E+09 | 8.52E+08 | 1.05E+09 | 1.07E+09 | 8.05E+08 | 1.24E+09 | 1.07E+09 | 8.97E+08 | 1.71E+09 | 1.18E+09 | 1.37E+09 | 2020.9497 | LFQDMLHSIDWMDEIK | P15508 | P15508 | SPTB | spectrin, beta, erythrocytic | Plasma Membrane | other |
| 15547660 | 2 | 0.328 | 1.36E+09 | 1.23E+09 | 1.42E+09 | 1.10E+09 | 1.14E+09 | 1.06E+09 | 1.16E+09 | 1.29E+09 | 1.21E+09 | 1.69E+09 | 1.24E+09 | 1.06E+09 | 1575.7206 | GYQPC[160.0307]DPQVIQDR | P15508 | P15508 | SPTB | spectrin, beta, erythrocytic | Plasma Membrane | other |
| 15537065 | 2 | 0.125 | 1.36E+09 | 1.45E+09 | 1.48E+09 | 1.39E+09 | 1.35E+09 | 1.22E+09 | 1.49E+09 | 1.34E+09 | 1.26E+09 | 1.68E+09 | 1.49E+09 | 1.45E+09 | 1207.6306 | LVAEGNLYSNK | P15508 | P15508 | SPTB | spectrin, beta, erythrocytic | Plasma Membrane | other |
| 15536113 | 2 | 0.66 | 1.25E+09 | 1.21E+09 | 1.41E+09 | 1.31E+09 | 1.15E+09 | 1.09E+09 | 1.19E+09 | 1.14E+09 | 1.10E+09 | 1.63E+09 | 1.14E+09 | 1.07E+09 | 1029.5548 | ILSPVDSGNK | P15508 | P15508 | SPTB | spectrin, beta, erythrocytic | Plasma Membrane | other |
| 15548026 | 2 | 0.39 | 1.32E+09 | 1.43E+09 | 1.54E+09 | 1.32E+09 | 1.33E+09 | 1.25E+09 | 1.42E+09 | 1.32E+09 | 1.42E+09 | 1.61E+09 | 1.37E+09 | 1.32E+09 | 1770.8445 | GNTLTQC[160.0307]LGFQEFQK | P15508 | P15508 | SPTB | spectrin, beta, erythrocytic | Plasma Membrane | other |
| 15534441, 15535030, 15544976 | 2, 3 | 0.063 | 1.31E+09 | 1.13E+09 | 1.29E+09 | 1.05E+09 | 1.02E+09 | 8.90E+08 | 1.34E+09 | 1.20E+09 | 1.10E+09 | 1.59E+09 | 1.17E+09 | 1.21E+09 | 1417.7224, 1417.7246 | MQLLAASYDLHR | P15508 | P15508 | SPTB | spectrin, beta, erythrocytic | Plasma Membrane | other |
| 15551363, 15557233, 15561953 | 2 | 0.778 | 1.39E+09 | 1.54E+09 | 1.53E+09 | 1.48E+09 | 1.52E+09 | 1.40E+09 | 1.50E+09 | 1.42E+09 | 1.40E+09 | 1.58E+09 | 1.32E+09 | 1.35E+09 | 2215.973, 2215.9736, 2215.976 | DLQDAENFFQFQGDADDLK | P15508 | P15508 | SPTB | spectrin, beta, erythrocytic | Plasma Membrane | other |
| 15531252 | 2 | 0.368 | 1.36E+09 | 1.20E+09 | 1.62E+09 | 1.23E+09 | 1.41E+09 | 9.66E+08 | 1.37E+09 | 1.52E+09 | 1.28E+09 | 1.58E+09 | 1.36E+09 | 1.37E+09 | 1258.6505 | ALEDLAQELEK | P15508 | P15508 | SPTB | spectrin, beta, erythrocytic | Plasma Membrane | other |
| 15553691, 15555170, 15558273, 15559820 | 3, 4 | 0.65 | 1.29E+09 | 1.49E+09 | 1.59E+09 | 9.94E+08 | 1.62E+09 | 1.23E+09 | 1.49E+09 | 1.49E+09 | 1.22E+09 | 1.54E+09 | 1.13E+09 | 1.14E+09 | 3025.3767, 3025.3806, 3025.3818, 3025.3865 | GVMEHLEHQAQGFPEEFRDSPDVTNR | P15508 | P15508 | SPTB | spectrin, beta, erythrocytic | Plasma Membrane | other |
| 15547993, 15548067, 15562000 | 2, 3 | 0.694 | 1.11E+09 | 1.54E+09 | 1.62E+09 | 1.06E+09 | 1.62E+09 | 1.25E+09 | 1.24E+09 | 1.64E+09 | 1.22E+09 | 1.48E+09 | 8.73E+08 | 1.18E+09 | 2338.1982, 2338.2056, 2338.21 | LPLAQSADYGTNLQTVQLFMK | P15508 | P15508 | SPTB | spectrin, beta, erythrocytic | Plasma Membrane | other |
| 15548027, 15549980 | 3, 4 | 0.218 | 1.09E+09 | 8.70E+08 | 3.71E+08 | 1.28E+09 | 3.43E+08 | 5.35E+08 | 1.06E+09 | 5.50E+08 | 6.20E+08 | 1.47E+09 | 1.45E+09 | 1.30E+09 | 3570.717, 3570.728 | WLDQMDIPNTLEDLEVVQHRFDILDQEMK | P15508 | P15508 | SPTB | spectrin, beta, erythrocytic | Plasma Membrane | other |
| 15531397, 15531553 | 2, 3 | 0.811 | 1.22E+09 | 1.12E+09 | 1.34E+09 | 1.20E+09 | 9.90E+08 | 1.10E+09 | 1.40E+09 | 1.09E+09 | 9.68E+08 | 1.46E+09 | 1.02E+09 | 1.10E+09 | 1257.6775, 1257.6787 | ALADEREVVQK | P15508 | P15508 | SPTB | spectrin, beta, erythrocytic | Plasma Membrane | other |
| 15533382 | 3 | 0.336 | 1.49E+09 | 1.42E+09 | 1.49E+09 | 1.41E+09 | 1.36E+09 | 1.43E+09 | 1.60E+09 | 1.44E+09 | 1.40E+09 | 1.41E+09 | 1.72E+09 | 1.53E+09 | 1908.9203 | TAGYPHVNVTNFTSSWK | P15508 | P15508 | SPTB | spectrin, beta, erythrocytic | Plasma Membrane | other |
| 15548231 | 2 | 0.758 | 1.09E+09 | 1.10E+09 | 1.31E+09 | 1.10E+09 | 1.06E+09 | 1.06E+09 | 1.12E+09 | 1.02E+09 | 1.04E+09 | 1.39E+09 | 9.96E+08 | 9.64E+08 | 1522.7179 | QIFQEADDMVAQK | P15508 | P15508 | SPTB | spectrin, beta, erythrocytic | Plasma Membrane | other |
| 15532868 | 2 | 0.164 | 1.20E+09 | 1.10E+09 | 1.17E+09 | 9.25E+08 | 1.09E+09 | 1.04E+09 | 1.14E+09 | 1.15E+09 | 1.06E+09 | 1.37E+09 | 1.16E+09 | 1.07E+09 | 1204.6195 | LWEELQSTAK | P15508 | P15508 | SPTB | spectrin, beta, erythrocytic | Plasma Membrane | other |
| 15537859 | 2, 3 | 0.143 | 1.24E+09 | 1.23E+09 | 1.40E+09 | 1.08E+09 | 1.20E+09 | 1.06E+09 | 1.15E+09 | 1.31E+09 | 1.09E+09 | 1.35E+09 | 1.17E+09 | 1.32E+09 | 1461.8048, 1461.8049 | LYQQVLTQAELR | P15508 | P15508 | SPTB | spectrin, beta, erythrocytic | Plasma Membrane | other |
| 15532670 | 2, 3 | 0.285 | 1.05E+09 | 9.81E+08 | 1.10E+09 | 1.07E+09 | 9.40E+08 | 8.80E+08 | 9.95E+08 | 1.03E+09 | 8.74E+08 | 1.35E+09 | 1.05E+09 | 1.01E+09 | 1233.5719, 1233.572 | DFGHTVDSVEK | P15508 | P15508 | SPTB | spectrin, beta, erythrocytic | Plasma Membrane | other |
| 15548135, 15549747, 15563855 | 2 | 0.118 | 1.29E+09 | 1.54E+09 | 1.89E+09 | 1.22E+09 | 1.36E+09 | 1.29E+09 | 1.50E+09 | 1.39E+09 | 1.35E+09 | 1.34E+09 | 1.16E+09 | 1.18E+09 | 1924.9576, 1924.9609, 1924.9625 | LVTQDNFGYDLAAVEAAK | P15508 | P15508 | SPTB | spectrin, beta, erythrocytic | Plasma Membrane | other |
| 15534463 | 3 | 0.4 | 1.15E+09 | 1.12E+09 | 1.27E+09 | 9.18E+08 | 1.27E+09 | 8.09E+08 | 1.09E+09 | 1.30E+09 | 1.10E+09 | 1.33E+09 | 1.09E+09 | 1.07E+09 | 1559.7798 | LEGLDTDWDALRR | P15508 | P15508 | SPTB | spectrin, beta, erythrocytic | Plasma Membrane | other |
| 15553933 | 2 | 0.794 | 8.95E+08 | 1.04E+09 | 1.36E+09 | 7.82E+08 | 1.09E+09 | 1.02E+09 | 9.83E+08 | 1.03E+09 | 1.03E+09 | 1.31E+09 | 7.30E+08 | 8.11E+08 | 1510.6782 | AWESLEEAEYQR | P15508 | P15508 | SPTB | spectrin, beta, erythrocytic | Plasma Membrane | other |
| 15533850, 15536868, 15540317, 15544232 | 2, 3 | 0.625 | 1.14E+09 | 1.16E+09 | 1.37E+09 | 1.06E+09 | 1.16E+09 | 1.12E+09 | 1.25E+09 | 1.28E+09 | 1.05E+09 | 1.30E+09 | 1.16E+09 | 1.13E+09 | 1198.6554, 1198.6556, 1198.6561, 1198.6562, 1198.657 | DGLAFNALIHK | P15508 | P15508 | SPTB | spectrin, beta, erythrocytic | Plasma Membrane | other |
| 15531324 | 2 | 0.399 | 9.76E+08 | 9.15E+08 | 1.08E+09 | 9.20E+08 | 9.47E+08 | 8.08E+08 | 9.47E+08 | 9.43E+08 | 8.97E+08 | 1.28E+09 | 9.35E+08 | 9.29E+08 | 1302.6198 | EQIYSSLDYGK | P15508 | P15508 | SPTB | spectrin, beta, erythrocytic | Plasma Membrane | other |
| 15547883 | 2 | 0.194 | 1.35E+09 | 1.18E+09 | 1.25E+09 | 1.23E+09 | 1.08E+09 | 1.10E+09 | 1.16E+09 | 1.04E+09 | 1.14E+09 | 1.26E+09 | 1.28E+09 | 1.10E+09 | 2145.0688 | FANSLSGVQQQLQAFSTYR | P15508 | P15508 | SPTB | spectrin, beta, erythrocytic | Plasma Membrane | other |
| 15549282, 15556452, 15557253, 15563130 | 2 | 0.367 | 1.09E+09 | 1.13E+09 | 1.57E+09 | 9.48E+08 | 1.14E+09 | 1.24E+09 | 9.34E+08 | 1.08E+09 | 9.64E+08 | 1.23E+09 | 9.44E+08 | 1.15E+09 | 1761.8823, 1761.8824, 1761.8839, 1761.8928 | YFYTGTEILGLIDEK | P15508 | P15508 | SPTB | spectrin, beta, erythrocytic | Plasma Membrane | other |
| 15547654, 15549048, 15555807 | 2 | 0.263 | 1.15E+09 | 1.04E+09 | 1.19E+09 | 9.47E+08 | 9.98E+08 | 8.10E+08 | 1.03E+09 | 1.05E+09 | 9.28E+08 | 1.21E+09 | 9.49E+08 | 8.54E+08 | 1819.7325, 1819.7336, 1819.7346 | WDAPDDELDNDNSSAR | P15508 | P15508 | SPTB | spectrin, beta, erythrocytic | Plasma Membrane | other |
| 15538829 | 2 | 0.452 | 8.99E+08 | 1.02E+09 | 1.22E+09 | 8.59E+08 | 9.31E+08 | 9.36E+08 | 9.16E+08 | 9.54E+08 | 1.01E+09 | 1.19E+09 | 9.52E+08 | 9.40E+08 | 1102.6343 | DVSSVELLLK | P15508 | P15508 | SPTB | spectrin, beta, erythrocytic | Plasma Membrane | other |
| 15532754, 15539196 | 2, 3 | 0.159 | 1.02E+09 | 9.07E+08 | 9.70E+08 | 8.66E+08 | 9.36E+08 | 8.24E+08 | 9.51E+08 | 9.00E+08 | 7.66E+08 | 1.17E+09 | 9.95E+08 | 9.27E+08 | 1212.5986, 1212.5996 | EVSAQWDHLK | P15508 | P15508 | SPTB | spectrin, beta, erythrocytic | Plasma Membrane | other |
| 15550933 | 2 | 0.498 | 9.72E+08 | 1.05E+09 | 2.06E+09 | 7.79E+08 | 1.11E+09 | 9.91E+08 | 1.23E+09 | 1.07E+09 | 6.52E+08 | 1.16E+09 | 7.98E+08 | 7.91E+08 | 1632.7849 | READDLEQWITEK | P15508 | P15508 | SPTB | spectrin, beta, erythrocytic | Plasma Membrane | other |
| 15531959, 15536086 | 2, 3 | 0.03 | 1.19E+09 | 1.10E+09 | 1.35E+09 | 9.52E+08 | 9.42E+08 | 1.05E+09 | 1.09E+09 | 1.03E+09 | 1.05E+09 | 1.12E+09 | 1.02E+09 | 1.11E+09 | 1157.6865, 1157.6881 | DLTSVLILQR | P15508 | P15508 | SPTB | spectrin, beta, erythrocytic | Plasma Membrane | other |
| 15547659 | 2 | 0.785 | 9.02E+08 | 8.03E+08 | 9.73E+08 | 1.00E+09 | 7.55E+08 | 7.31E+08 | 9.44E+08 | 8.14E+08 | 7.76E+08 | 1.11E+09 | 8.87E+08 | 7.76E+08 | 1617.65 | VNNYC[160.0307]VDC[160.0307]EETSK | P15508 | P15508 | SPTB | spectrin, beta, erythrocytic | Plasma Membrane | other |
| 15531648, 15543491 | 2 | 0.338 | 9.65E+08 | 9.02E+08 | 9.54E+08 | 9.07E+08 | 8.34E+08 | 8.11E+08 | 9.31E+08 | 9.02E+08 | 8.60E+08 | 1.10E+09 | 8.83E+08 | 8.76E+08 | 972.5689, 972.5714 | AQEATVLLK | P15508 | P15508 | SPTB | spectrin, beta, erythrocytic | Plasma Membrane | other |
| 15549405, 15562129 | 2 | 0.374 | 7.80E+08 | 7.00E+08 | 5.68E+08 | 6.45E+08 | 6.89E+08 | 4.50E+08 | 7.94E+08 | 5.90E+08 | 5.15E+08 | 1.10E+09 | 6.59E+08 | 7.32E+08 | 1717.7527, 1717.7546 | FFWEMDEAESWIK | P15508 | P15508 | SPTB | spectrin, beta, erythrocytic | Plasma Membrane | other |
| 15531439 | 2 | 0.611 | 7.05E+08 | 8.66E+08 | 9.09E+08 | 5.86E+08 | 8.17E+08 | 6.27E+08 | 9.15E+08 | 8.37E+08 | 7.07E+08 | 1.07E+09 | 5.67E+08 | 6.89E+08 | 1218.5919 | WIGAMEDQLR | P15508 | P15508 | SPTB | spectrin, beta, erythrocytic | Plasma Membrane | other |
| 15531684, 15547278 | 3 | 0.701 | 9.41E+08 | 8.95E+08 | 1.18E+09 | 8.58E+08 | 9.26E+08 | 9.15E+08 | 9.87E+08 | 9.81E+08 | 7.85E+08 | 1.06E+09 | 8.51E+08 | 9.29E+08 | 1632.7844, 1632.7849 | READDLEQWITEK | P15508 | P15508 | SPTB | spectrin, beta, erythrocytic | Plasma Membrane | other |
| 15535533, 15536577 | 3, 4 | 0.423 | 1.25E+09 | 6.82E+08 | 7.28E+08 | 1.22E+09 | 4.00E+08 | 7.83E+08 | 1.09E+09 | 5.80E+08 | 9.26E+08 | 1.05E+09 | 1.44E+09 | 1.20E+09 | 1613.8248, 1613.8259 | AFEDELRGLDAHLK | P15508 | P15508 | SPTB | spectrin, beta, erythrocytic | Plasma Membrane | other |
| 15531251 | 3 | 0.019 | 9.08E+08 | 9.60E+08 | 9.63E+08 | 8.17E+08 | 8.33E+08 | 8.48E+08 | 7.95E+08 | 7.26E+08 | 7.87E+08 | 1.04E+09 | 8.21E+08 | 9.26E+08 | 2090.0535 | DASVAEAWLIAQEPYLASR | P15508 | P15508 | SPTB | spectrin, beta, erythrocytic | Plasma Membrane | other |
| 15537994 | 2 | 0.51 | 8.57E+08 | 8.22E+08 | 9.50E+08 | 6.99E+08 | 8.21E+08 | 7.74E+08 | 9.55E+08 | 8.80E+08 | 7.64E+08 | 1.02E+09 | 7.50E+08 | 7.74E+08 | 1189.5571 | ETWLNENQR | P15508 | P15508 | SPTB | spectrin, beta, erythrocytic | Plasma Membrane | other |
| 15548113 | 2 | 0.558 | 6.87E+08 | 9.07E+08 | 1.14E+09 | 5.58E+08 | 9.57E+08 | 7.52E+08 | 9.17E+08 | 9.46E+08 | 7.80E+08 | 1.01E+09 | 4.95E+08 | 6.45E+08 | 1533.7042 | DLEDETLWVEER | P15508 | P15508 | SPTB | spectrin, beta, erythrocytic | Plasma Membrane | other |
| 15549578, 15549762 | 3 | 0.273 | 8.47E+08 | 1.03E+09 | 1.31E+09 | 7.20E+08 | 9.37E+08 | 8.87E+08 | 7.89E+08 | 8.29E+08 | 8.76E+08 | 9.97E+08 | 7.79E+08 | 8.84E+08 | 2767.244, 2767.247 | EELEELFADAPSLGAEAGDTDMSIEK | P15508 | P15508 | SPTB | spectrin, beta, erythrocytic | Plasma Membrane | other |
| 15547977, 15557585 | 3 | 0.7 | 6.33E+08 | 6.39E+08 | 2.68E+08 | 5.77E+08 | 6.87E+08 | 3.61E+08 | 5.47E+08 | 6.75E+08 | 6.09E+08 | 9.27E+08 | 5.14E+08 | 5.64E+08 | 2895.3384, 2895.3418 | KEELEELFADAPSLGAEAGDTDMSIEK | P15508 | P15508 | SPTB | spectrin, beta, erythrocytic | Plasma Membrane | other |
| 15531574, 15541870 | 2, 3 | 0.164 | 8.55E+08 | 7.83E+08 | 9.70E+08 | 7.05E+08 | 8.23E+08 | 7.45E+08 | 7.99E+08 | 8.04E+08 | 7.16E+08 | 9.24E+08 | 8.47E+08 | 8.06E+08 | 1072.5764, 1072.5773, 1072.5775 | AHILSAEFGK | P15508 | P15508 | SPTB | spectrin, beta, erythrocytic | Plasma Membrane | other |
| 15534052 | 3 | 0.414 | 8.03E+08 | 6.63E+08 | 6.70E+08 | 7.15E+08 | 5.85E+08 | 6.16E+08 | 6.83E+08 | 6.66E+08 | 5.83E+08 | 9.09E+08 | 7.06E+08 | 6.36E+08 | 1605.7202 | ESQYLMESHPEQK | P15508 | P15508 | SPTB | spectrin, beta, erythrocytic | Plasma Membrane | other |
| 15532880, 15542000, 15544694, 15547322 | 2 | 0.154 | 7.28E+08 | 6.49E+08 | 7.50E+08 | 5.54E+08 | 6.32E+08 | 5.96E+08 | 6.76E+08 | 6.88E+08 | 5.87E+08 | 8.94E+08 | 6.97E+08 | 6.39E+08 | 1303.6482, 1303.6534, 1303.6545 | LMEADIAIQGDK | P15508 | P15508 | SPTB | spectrin, beta, erythrocytic | Plasma Membrane | other |
| 15532349, 15545893 | 3 | 0.452 | 6.56E+08 | 7.41E+08 | 4.95E+08 | 6.16E+08 | 5.85E+08 | 5.74E+08 | 5.13E+08 | 6.55E+08 | 5.64E+08 | 8.85E+08 | 6.42E+08 | 6.14E+08 | 2145.0679, 2145.0718 | FANSLSGVQQQLQAFSTYR | P15508 | P15508 | SPTB | spectrin, beta, erythrocytic | Plasma Membrane | other |
| 15550631, 15551867 | 3, 4 | 0.656 | 4.52E+08 | 1.19E+09 | 1.85E+09 | 5.03E+08 | 9.06E+08 | 1.46E+09 | 9.97E+08 | 8.27E+08 | 7.08E+08 | 8.56E+08 | 3.60E+08 | 6.82E+08 | 3306.6897, 3306.6934 | QAEAILSNQEYTLAHLEPPDSLAAAEAGIRK | P15508 | P15508 | SPTB | spectrin, beta, erythrocytic | Plasma Membrane | other |
| 15547655 | 2 | 0.074 | 7.70E+08 | 7.87E+08 | 9.12E+08 | 6.78E+08 | 7.20E+08 | 7.10E+08 | 6.70E+08 | 6.49E+08 | 6.93E+08 | 8.41E+08 | 6.83E+08 | 6.68E+08 | 2090.0513 | DASVAEAWLIAQEPYLASR | P15508 | P15508 | SPTB | spectrin, beta, erythrocytic | Plasma Membrane | other |
| 15547897, 15548956, 15561177 | 3, 4 | 0.426 | 8.95E+08 | 5.07E+08 | 3.02E+08 | 1.19E+09 | 1.43E+09 | 5.16E+08 | 5.87E+08 | 3.28E+08 | 5.76E+08 | 8.35E+08 | 1.15E+09 | 1.01E+09 | 3852.8936, 3852.8962 | DLEDETLWVEERLPLAQSADYGTNLQTVQLFMK | P15508 | P15508 | SPTB | spectrin, beta, erythrocytic | Plasma Membrane | other |
| 15531710, 15536485 | 2, 3 | 0.439 | 1.10E+09 | 5.40E+08 | 6.22E+08 | 1.04E+09 | 2.95E+08 | 6.38E+08 | 6.58E+08 | 4.30E+08 | 7.03E+08 | 8.28E+08 | 1.04E+09 | 9.84E+08 | 1395.7556, 1395.7584 | DVVSERLEALHK | P15508 | P15508 | SPTB | spectrin, beta, erythrocytic | Plasma Membrane | other |
| 15531841, 15537647 | 2 | 0.735 | 6.36E+08 | 6.98E+08 | 7.32E+08 | 5.86E+08 | 6.51E+08 | 5.90E+08 | 6.87E+08 | 6.76E+08 | 5.50E+08 | 8.27E+08 | 5.28E+08 | 6.48E+08 | 801.47076, 801.4709 | PTTLELK | P15508 | P15508 | SPTB | spectrin, beta, erythrocytic | Plasma Membrane | other |
| 15539387 | 3 | 0.006 | 7.37E+08 | 7.47E+08 | 7.84E+08 | 6.40E+08 | 7.18E+08 | 6.75E+08 | 7.36E+08 | 6.71E+08 | 7.43E+08 | 8.23E+08 | 8.03E+08 | 8.01E+08 | 1604.827 | RWQAFQAVVSEQR | P15508 | P15508 | SPTB | spectrin, beta, erythrocytic | Plasma Membrane | other |
| 15532424 | 3 | 0.312 | 7.51E+08 | 7.80E+08 | 7.33E+08 | 7.83E+08 | 8.47E+08 | 6.97E+08 | 9.95E+08 | 7.14E+08 | 6.40E+08 | 8.21E+08 | 1.09E+09 | 9.05E+08 | 1682.9491 | LLEVLSGEMLPRPTK | P15508 | P15508 | SPTB | spectrin, beta, erythrocytic | Plasma Membrane | other |
| 15531426 | 5 | 0.74 | 5.07E+08 | 7.85E+08 | 8.93E+08 | 3.26E+08 | 1.26E+09 | 5.26E+08 | 8.05E+08 | 1.16E+09 | 4.31E+08 | 8.20E+08 | 3.42E+08 | 4.08E+08 | 3594.632 | QTPERPTEEPGPQEEEGETAGEAPQVHHAATER | P15508 | P15508 | SPTB | spectrin, beta, erythrocytic | Plasma Membrane | other |
| 15532861 | 2 | 0.976 | 7.05E+08 | 4.75E+08 | 7.56E+08 | 5.01E+08 | 6.13E+08 | 7.39E+08 | 6.26E+08 | 6.67E+08 | 6.36E+08 | 7.88E+08 | 5.39E+08 | 6.45E+08 | 1320.7296 | LWSYLQELLR | P15508 | P15508 | SPTB | spectrin, beta, erythrocytic | Plasma Membrane | other |
| 15531325 | 3 | 0.907 | 6.21E+08 | 8.36E+08 | 7.96E+08 | 5.44E+08 | 9.52E+08 | 6.17E+08 | 8.82E+08 | 7.86E+08 | 5.59E+08 | 7.69E+08 | 6.43E+08 | 5.98E+08 | 2035.9332 | VSHLEQC[160.0307]FSELSNMAAGR | P15508 | P15508 | SPTB | spectrin, beta, erythrocytic | Plasma Membrane | other |
| 15532327 | 3 | 0.828 | 4.28E+08 | 4.33E+08 | 5.89E+08 | 4.41E+08 | 6.06E+08 | 5.22E+08 | 4.80E+08 | 4.39E+08 | 4.60E+08 | 7.29E+08 | 4.42E+08 | 4.32E+08 | 1831.9166 | VIEGQTDPDYQLLGQR | P15508 | P15508 | SPTB | spectrin, beta, erythrocytic | Plasma Membrane | other |
| 15533373 | 2 | 0.162 | 7.73E+08 | 8.00E+08 | 8.32E+08 | 5.84E+08 | 7.56E+08 | 7.48E+08 | 6.95E+08 | 7.18E+08 | 6.64E+08 | 7.10E+08 | 6.98E+08 | 7.55E+08 | 1131.6019 | ELTLWINDK | P15508 | P15508 | SPTB | spectrin, beta, erythrocytic | Plasma Membrane | other |
| 15537098, 15538183, 15546397, 15547572 | 2, 3 | 0.299 | 7.02E+08 | 6.52E+08 | 6.66E+08 | 5.87E+08 | 5.67E+08 | 6.25E+08 | 6.77E+08 | 6.16E+08 | 5.41E+08 | 7.02E+08 | 5.86E+08 | 6.17E+08 | 1489.8326, 1489.834, 1489.8344, 1489.8358 | GNLEVLLFTIQSR | P15508 | P15508 | SPTB | spectrin, beta, erythrocytic | Plasma Membrane | other |
| 15538391, 15545430, 15546608 | 2 | 0.094 | 5.82E+08 | 5.35E+08 | 6.57E+08 | 5.12E+08 | 5.48E+08 | 5.08E+08 | 5.44E+08 | 5.53E+08 | 5.19E+08 | 6.85E+08 | 5.62E+08 | 6.08E+08 | 986.5854, 986.587, 986.5914 | LEATLALQK | P15508 | P15508 | SPTB | spectrin, beta, erythrocytic | Plasma Membrane | other |
| 15531250 | 2 | 0.469 | 5.45E+08 | 7.77E+08 | 9.52E+08 | 4.92E+08 | 1.08E+09 | 7.45E+08 | 8.18E+08 | 8.81E+08 | 6.09E+08 | 6.80E+08 | 4.43E+08 | 5.35E+08 | 1280.5989 | EFLEELEESR | P15508 | P15508 | SPTB | spectrin, beta, erythrocytic | Plasma Membrane | other |
| 15532805 | 2 | 0.02 | 5.22E+08 | 4.32E+08 | 4.47E+08 | 4.52E+08 | 4.56E+08 | 3.89E+08 | 5.77E+08 | 5.71E+08 | 4.58E+08 | 6.69E+08 | 7.56E+08 | 5.39E+08 | 1277.6366 | DALLLWC[160.0307]QMK | P15508 | P15508 | SPTB | spectrin, beta, erythrocytic | Plasma Membrane | other |
| 15548752, 15559771, 15562219, 15563058 | 2 | 0.385 | 6.89E+08 | 7.57E+08 | 6.39E+08 | 5.84E+08 | 6.60E+08 | 6.34E+08 | 5.85E+08 | 6.15E+08 | 7.00E+08 | 6.68E+08 | 6.43E+08 | 5.76E+08 | 1766.8711, 1766.8727, 1766.8763, 1766.8773 | NFSTC[160.0307]LELGESLLQR | P15508 | P15508 | SPTB | spectrin, beta, erythrocytic | Plasma Membrane | other |
| 15532220 | 2, 3 | 0.115 | 5.64E+08 | 4.27E+08 | 4.48E+08 | 4.01E+08 | 4.88E+08 | 4.21E+08 | 5.49E+08 | 5.42E+08 | 4.34E+08 | 6.41E+08 | 9.44E+08 | 4.97E+08 | 1127.5483, 1127.5494 | IHC[160.0307]LENVDK | P15508 | P15508 | SPTB | spectrin, beta, erythrocytic | Plasma Membrane | other |
| 15536518 | 3 | 0.583 | 4.61E+08 | 6.94E+08 | 4.69E+08 | 3.22E+08 | 5.05E+08 | 4.57E+08 | 4.84E+08 | 5.42E+08 | 4.46E+08 | 6.37E+08 | 4.04E+08 | 4.83E+08 | 2215.975 | DLQDAENFFQFQGDADDLK | P15508 | P15508 | SPTB | spectrin, beta, erythrocytic | Plasma Membrane | other |
| 15550847, 15551082 | 3, 4 | 0.503 | 4.53E+08 | 6.84E+08 | 8.25E+08 | 3.02E+08 | 1.08E+09 | 4.72E+08 | 6.47E+08 | 9.39E+08 | 4.36E+08 | 6.35E+08 | 2.68E+08 | 3.15E+08 | 3594.6282, 3594.6304 | QTPERPTEEPGPQEEEGETAGEAPQVHHAATER | P15508 | P15508 | SPTB | spectrin, beta, erythrocytic | Plasma Membrane | other |
| 15536202 | 2, 3 | 0.665 | 8.37E+08 | 8.27E+08 | 1.03E+09 | 5.14E+08 | 8.80E+08 | 8.73E+08 | 6.85E+08 | 8.96E+08 | 8.20E+08 | 6.32E+08 | 8.27E+08 | 8.61E+08 | 1154.6025, 1154.6033 | VIDHAIETEK | P15508 | P15508 | SPTB | spectrin, beta, erythrocytic | Plasma Membrane | other |
| 15533739, 15546436 | 2, 3 | 0.224 | 6.58E+08 | 7.05E+08 | 7.82E+08 | 6.48E+08 | 6.22E+08 | 7.49E+08 | 6.46E+08 | 6.73E+08 | 6.28E+08 | 6.09E+08 | 5.98E+08 | 6.66E+08 | 1328.7661, 1328.7662, 1328.7667 | RFLDLLEPLGR | P15508 | P15508 | SPTB | spectrin, beta, erythrocytic | Plasma Membrane | other |
| 15532218, 15542964 | 3 | 0.312 | 4.54E+08 | 6.46E+08 | 6.50E+08 | 4.33E+08 | 5.14E+08 | 5.08E+08 | 5.98E+08 | 6.08E+08 | 5.56E+08 | 5.70E+08 | 4.74E+08 | 5.81E+08 | 1924.9624, 1924.9636 | LVTQDNFGYDLAAVEAAK | P15508 | P15508 | SPTB | spectrin, beta, erythrocytic | Plasma Membrane | other |
| 15540034, 15540514, 15546933 | 3, 4 | 0.488 | 5.35E+08 | 5.75E+08 | 5.28E+08 | 4.17E+08 | 6.05E+08 | 4.72E+08 | 4.99E+08 | 5.09E+08 | 3.36E+08 | 5.53E+08 | 4.18E+08 | 4.15E+08 | 1929.894, 1929.8943, 1929.8972 | VHLENMGSHDIVDGNHR | P15508 | P15508 | SPTB | spectrin, beta, erythrocytic | Plasma Membrane | other |
| 15531305 | 2 | 0.076 | 4.68E+08 | 4.10E+08 | 4.10E+08 | 4.31E+08 | 4.34E+08 | 3.66E+08 | 4.31E+08 | 4.02E+08 | 3.45E+08 | 5.45E+08 | 4.73E+08 | 4.64E+08 | 934.48016 | AWLSMAQK | P15508 | P15508 | SPTB | spectrin, beta, erythrocytic | Plasma Membrane | other |
| 15539186, 15540225 | 3, 4 | 0.345 | 1.04E+09 | 1.07E+09 | 7.08E+08 | 5.90E+08 | 5.63E+08 | 8.09E+08 | 1.26E+09 | 8.50E+08 | 1.01E+09 | 5.41E+08 | 1.57E+09 | 1.21E+09 | 2164.0232, 2164.0254 | VSHLEQC[160.0307]FSELSNMAAGRK | P15508 | P15508 | SPTB | spectrin, beta, erythrocytic | Plasma Membrane | other |
| 15533914, 15539904 | 3 | 0.247 | 5.58E+08 | 3.98E+08 | 4.70E+08 | 5.57E+08 | 3.58E+08 | 3.95E+08 | 4.83E+08 | 4.05E+08 | 4.28E+08 | 5.38E+08 | 6.06E+08 | 5.21E+08 | 1591.8379, 1591.8381 | RLENMYHLFQLK | P15508 | P15508 | SPTB | spectrin, beta, erythrocytic | Plasma Membrane | other |
| 15538188, 15544484, 15544778, 15546354, 15546495, 15546786, 15547084, 15547354, 15547378, 15547464, 15547561, 15547563, 15547627 | 4 | 0.044 | 4.95E+08 | 5.47E+08 | 4.86E+08 | 4.82E+08 | 4.31E+08 | 4.01E+08 | 4.93E+08 | 4.53E+08 | 3.78E+08 | 5.10E+08 | 5.69E+08 | 5.62E+08 | 2266.0667, 2266.0684, 2266.0718, 2266.0742, 2266.075, 2266.0767, 2266.0771, 2266.0776, 2266.079, 2266.0806, 2266.0813, 2266.0862 | HRELPEDVGLDASTAESFHR | P15508 | P15508 | SPTB | spectrin, beta, erythrocytic | Plasma Membrane | other |
| 15547726, 15561072 | 2 | 0.397 | 4.24E+08 | 3.91E+08 | 3.68E+08 | 4.05E+08 | 3.42E+08 | 2.71E+08 | 4.03E+08 | 3.64E+08 | 3.43E+08 | 5.02E+08 | 3.93E+08 | 3.53E+08 | 1711.8575, 1711.8611 | LIDAGHSEAATIAEWK | P15508 | P15508 | SPTB | spectrin, beta, erythrocytic | Plasma Membrane | other |
| 15540068, 15544436 | 3, 4 | 0.136 | 4.64E+08 | 3.48E+08 | 3.24E+08 | 4.56E+08 | 2.58E+08 | 4.01E+08 | 3.98E+08 | 2.88E+08 | 3.63E+08 | 4.82E+08 | 6.18E+08 | 4.83E+08 | 2131.0305, 2131.0308, 2131.032 | EQEVSAAWQALLDAC[160.0307]AGRR | P15508 | P15508 | SPTB | spectrin, beta, erythrocytic | Plasma Membrane | other |
| 15556682, 15563055 | 3 | 0.044 | 5.77E+08 | 6.43E+08 | 6.36E+08 | 4.61E+08 | 5.45E+08 | 5.29E+08 | 5.59E+08 | 5.83E+08 | 5.37E+08 | 4.76E+08 | 5.17E+08 | 5.70E+08 | 2610.199, 2610.2107 | HQAFMAELASHQGWLENIDAEGR | P15508 | P15508 | SPTB | spectrin, beta, erythrocytic | Plasma Membrane | other |
| 15533916 | 2, 3 | 0.868 | 3.25E+08 | 3.48E+08 | 3.34E+08 | 2.40E+08 | 4.35E+08 | 2.29E+08 | 3.42E+08 | 3.90E+08 | 2.68E+08 | 4.54E+08 | 2.33E+08 | 2.40E+08 | 1142.6251 | RVEEQVNLR | P15508 | P15508 | SPTB | spectrin, beta, erythrocytic | Plasma Membrane | other |
| 15550133 | 2 | 0.013 | 5.51E+08 | 5.89E+08 | 7.52E+08 | 5.40E+08 | 5.24E+08 | 4.96E+08 | 3.52E+08 | 4.94E+08 | 4.54E+08 | 4.52E+08 | 3.70E+08 | 3.75E+08 | 1660.8613 | FQIQDIVVQTQEGR | P15508 | P15508 | SPTB | spectrin, beta, erythrocytic | Plasma Membrane | other |
| 15531624, 15535287, 15537924, 15542771, 15542865, 15544572, 15545482, 15545494, 15545790, 15546083, 15546719, 15547296 | 3 | 0.219 | 4.44E+08 | 5.01E+08 | 5.72E+08 | 4.29E+08 | 3.97E+08 | 5.04E+08 | 3.91E+08 | 5.55E+08 | 3.63E+08 | 4.19E+08 | 2.35E+08 | 3.90E+08 | 2036.9381, 2036.9391, 2036.9393, 2036.9396, 2036.9402, 2036.9404, 2036.9414, 2036.9415, 2036.9427, 2036.9431, 2036.9437, 2036.946 | LFQDMLHSIDWM[147.0354]DEIK | P15508 | P15508 | SPTB | spectrin, beta, erythrocytic | Plasma Membrane | other |
| 15543196 | 4 | 0.66 | 3.70E+08 | 5.09E+08 | 5.74E+08 | 3.56E+08 | 4.52E+08 | 5.25E+08 | 4.27E+08 | 6.12E+08 | 5.17E+08 | 4.09E+08 | 4.07E+08 | 4.90E+08 | 2610.213 | HQAFMAELASHQGWLENIDAEGR | P15508 | P15508 | SPTB | spectrin, beta, erythrocytic | Plasma Membrane | other |
| 15531253 | 2 | 0.729 | 2.53E+08 | 3.42E+08 | 4.34E+08 | 1.90E+08 | 4.02E+08 | 2.92E+08 | 3.12E+08 | 3.72E+08 | 2.62E+08 | 4.06E+08 | 1.85E+08 | 2.11E+08 | 879.41876 | AFEDELR | P15508 | P15508 | SPTB | spectrin, beta, erythrocytic | Plasma Membrane | other |
| 15548708 | 2 | 0.294 | 3.48E+08 | 3.14E+08 | 3.36E+08 | 3.22E+08 | 2.51E+08 | 2.46E+08 | 3.00E+08 | 2.53E+08 | 2.45E+08 | 4.06E+08 | 2.91E+08 | 2.53E+08 | 1590.8308 | NQTLQNEILGHAPR | P15508 | P15508 | SPTB | spectrin, beta, erythrocytic | Plasma Membrane | other |
| 15532528 | 2 | 0.18 | 3.63E+08 | 3.01E+08 | 3.04E+08 | 2.58E+08 | 2.74E+08 | 2.68E+08 | 3.24E+08 | 2.92E+08 | 2.44E+08 | 4.06E+08 | 3.38E+08 | 2.79E+08 | 1212.6097 | QFGHPQIETR | P15508 | P15508 | SPTB | spectrin, beta, erythrocytic | Plasma Membrane | other |
| 15533249 | 2 | 0.048 | 3.48E+08 | 3.11E+08 | 3.30E+08 | 2.52E+08 | 2.64E+08 | 2.47E+08 | 3.29E+08 | 2.95E+08 | 2.52E+08 | 3.88E+08 | 3.15E+08 | 2.91E+08 | 996.5 | AWLQDAHR | P15508 | P15508 | SPTB | spectrin, beta, erythrocytic | Plasma Membrane | other |
| 15531828 | 3 | 0.499 | 4.76E+08 | 1.99E+08 | 1.36E+08 | 6.15E+08 | 4.78E+07 | 2.28E+08 | 3.20E+08 | 9.57E+07 | 2.85E+08 | 3.67E+08 | 6.34E+08 | 4.92E+08 | 1833.8762 | WIGAMEDQLRSDDLGK | P15508 | P15508 | SPTB | spectrin, beta, erythrocytic | Plasma Membrane | other |
| 15547896 | 2 | 0.315 | 2.99E+08 | 2.47E+08 | 2.50E+08 | 2.41E+08 | 2.08E+08 | 1.89E+08 | 2.70E+08 | 2.22E+08 | 1.89E+08 | 3.65E+08 | 2.71E+08 | 2.10E+08 | 1605.7195 | ESQYLMESHPEQK | P15508 | P15508 | SPTB | spectrin, beta, erythrocytic | Plasma Membrane | other |
| 15531393 | 2 | 0.814 | 2.79E+08 | 3.28E+08 | 3.66E+08 | 2.41E+08 | 3.30E+08 | 2.96E+08 | 3.12E+08 | 3.30E+08 | 2.80E+08 | 3.63E+08 | 2.35E+08 | 3.02E+08 | 1403.6775 | LEGLDTDWDALR | P15508 | P15508 | SPTB | spectrin, beta, erythrocytic | Plasma Membrane | other |
| 15556837, 15558312 | 2 | 0.164 | 3.27E+08 | 2.54E+08 | 2.73E+08 | 2.60E+08 | 2.33E+08 | 1.15E+08 | 2.62E+08 | 2.15E+08 | 2.46E+08 | 3.52E+08 | 3.21E+08 | 2.62E+08 | 1604.8253, 1604.8265 | RWQAFQAVVSEQR | P15508 | P15508 | SPTB | spectrin, beta, erythrocytic | Plasma Membrane | other |
| 15531467 | 2 | 0.361 | 3.84E+08 | 1.65E+08 | 1.52E+08 | 3.61E+08 | 6.95E+07 | 1.90E+08 | 2.35E+08 | 1.34E+08 | 2.60E+08 | 3.16E+08 | 4.42E+08 | 3.56E+08 | 957.5454 | DVLAIRDR | P15508 | P15508 | SPTB | spectrin, beta, erythrocytic | Plasma Membrane | other |
| 15550533 | 2 | 0.142 | 1.92E+08 | 1.50E+08 | 1.19E+08 | 1.64E+08 | 1.37E+08 | 1.00E+08 | 1.42E+08 | 1.15E+08 | 8.81E+07 | 2.86E+08 | 1.96E+08 | 1.44E+08 | 1762.8209 | KHEAIETDTAAYEER | P15508 | P15508 | SPTB | spectrin, beta, erythrocytic | Plasma Membrane | other |
| 15531254 | 2 | 0.141 | 2.21E+08 | 1.94E+08 | 2.12E+08 | 1.60E+08 | 1.92E+08 | 1.81E+08 | 2.25E+08 | 2.08E+08 | 1.75E+08 | 2.85E+08 | 2.30E+08 | 1.96E+08 | 982.51965 | WVNSHLAR | P15508 | P15508 | SPTB | spectrin, beta, erythrocytic | Plasma Membrane | other |
| 15549192 | 2 | 0.06 | 1.75E+08 | 1.14E+08 | 1.51E+08 | 1.70E+08 | 1.74E+08 | 1.34E+08 | 1.92E+08 | 1.63E+08 | 1.56E+08 | 2.74E+08 | 1.98E+08 | 2.05E+08 | 2020.9441 | LFQDMLHSIDWMDEIK | P15508 | P15508 | SPTB | spectrin, beta, erythrocytic | Plasma Membrane | other |
| 15550199 | 2 | 0.125 | 2.88E+08 | 2.72E+08 | 3.39E+08 | 1.70E+08 | 2.84E+08 | 1.20E+08 | 2.50E+08 | 2.63E+08 | 2.40E+08 | 2.72E+08 | 2.48E+08 | 2.34E+08 | 1559.7778 | LEGLDTDWDALRR | P15508 | P15508 | SPTB | spectrin, beta, erythrocytic | Plasma Membrane | other |
| 15534893, 15545491 | 3 | 0.378 | 2.80E+08 | 3.16E+08 | 2.44E+08 | 2.23E+08 | 2.54E+08 | 2.64E+08 | 2.54E+08 | 2.58E+08 | 2.70E+08 | 2.69E+08 | 2.64E+08 | 2.56E+08 | 1766.8713, 1766.873 | NFSTC[160.0307]LELGESLLQR | P15508 | P15508 | SPTB | spectrin, beta, erythrocytic | Plasma Membrane | other |
| 15555521, 15558652, 15558747 | 4 | 0.364 | 4.03E+08 | 4.05E+08 | 2.48E+08 | 2.00E+08 | 3.93E+08 | 1.84E+08 | 2.89E+08 | 2.34E+08 | 7.90E+07 | 2.66E+08 | 2.17E+08 | 2.57E+08 | 4097.8916, 4097.894, 4097.914 | LRDAHEAQQYYLDAGEAEAWISEQELYVFSDEPPK | P15508 | P15508 | SPTB | spectrin, beta, erythrocytic | Plasma Membrane | other |
| 15535250 | 3 | 0.577 | 1.55E+08 | 3.84E+08 | 6.39E+08 | 1.92E+08 | 3.55E+08 | 4.73E+08 | 4.31E+08 | 3.00E+08 | 2.09E+08 | 2.66E+08 | 1.29E+08 | 2.25E+08 | 1743.7947 | FEDFLVSMENNRDK | P15508 | P15508 | SPTB | spectrin, beta, erythrocytic | Plasma Membrane | other |
| 15532306 | 3 | 0.461 | 1.96E+08 | 1.78E+08 | 1.82E+08 | 1.75E+08 | 1.89E+08 | 1.16E+08 | 2.17E+08 | 1.67E+08 | 1.48E+08 | 2.57E+08 | 1.70E+08 | 1.95E+08 | 1843.9365 | GLQDALQGQELSLGEASK | P15508 | P15508 | SPTB | spectrin, beta, erythrocytic | Plasma Membrane | other |
| 15535620 | 2 | 0.532 | 4.48E+08 | 5.53E+08 | 5.84E+08 | 2.59E+08 | 5.53E+08 | 4.55E+08 | 4.90E+08 | 4.75E+08 | 2.88E+08 | 2.53E+08 | 4.10E+08 | 4.67E+08 | 1104.558 | DEEGAIVMLK | P15508 | P15508 | SPTB | spectrin, beta, erythrocytic | Plasma Membrane | other |
| 15548491 | 2 | 0.467 | 4.71E+08 | 3.98E+08 | 4.85E+08 | 4.60E+08 | 3.72E+08 | 3.85E+08 | 4.48E+08 | 3.91E+08 | 3.61E+08 | 2.43E+08 | 4.37E+08 | 4.02E+08 | 1805.9464 | AQSLLSAGHPEGEQIIR | P15508 | P15508 | SPTB | spectrin, beta, erythrocytic | Plasma Membrane | other |
| 15536762 | 3 | 0.378 | 1.33E+08 | 1.76E+08 | 1.55E+08 | 2.42E+08 | 1.55E+08 | 1.57E+08 | 1.92E+08 | 1.58E+08 | 1.60E+08 | 2.39E+08 | 1.64E+08 | 2.10E+08 | 1522.7177 | QIFQEADDMVAQK | P15508 | P15508 | SPTB | spectrin, beta, erythrocytic | Plasma Membrane | other |
| 15550406 | 2 | 0.228 | 2.24E+08 | 2.08E+08 | 2.35E+08 | 2.06E+08 | 2.15E+08 | 1.89E+08 | 2.41E+08 | 2.01E+08 | 1.88E+08 | 2.35E+08 | 2.66E+08 | 2.17E+08 | 1682.9473 | LLEVLSGEMLPRPTK | P15508 | P15508 | SPTB | spectrin, beta, erythrocytic | Plasma Membrane | other |
| 15537757, 15542972, 15545529, 15545638, 15546450 | 3 | 0.092 | 2.72E+08 | 2.72E+08 | 3.34E+08 | 2.30E+08 | 2.91E+08 | 2.74E+08 | 2.50E+08 | 2.94E+08 | 2.77E+08 | 2.32E+08 | 1.54E+08 | 2.38E+08 | 1698.9417, 1698.9419, 1698.9426, 1698.9432 | LLEVLSGEM[147.0354]LPRPTK | P15508 | P15508 | SPTB | spectrin, beta, erythrocytic | Plasma Membrane | other |
| 15538768 | 2, 3 | 0.348 | 1.84E+08 | 1.87E+08 | 1.88E+08 | 2.20E+08 | 1.80E+08 | 1.75E+08 | 1.87E+08 | 1.71E+08 | 1.44E+08 | 2.21E+08 | 1.83E+08 | 1.81E+08 | 1435.7374, 1435.7379 | LENMYHLFQLK | P15508 | P15508 | SPTB | spectrin, beta, erythrocytic | Plasma Membrane | other |
| 15535057 | 4 | 0.55 | 1.86E+08 | 2.16E+08 | 2.53E+08 | 2.22E+08 | 1.59E+08 | 1.81E+08 | 1.92E+08 | 2.40E+08 | 1.89E+08 | 2.15E+08 | 1.95E+08 | 2.39E+08 | 2420.2322 | TLMAQIDGVNLAANNLVESGHPR | P15508 | P15508 | SPTB | spectrin, beta, erythrocytic | Plasma Membrane | other |
| 15548539, 15556660, 15559361 | 2 | 0.495 | 2.68E+08 | 4.10E+08 | 8.12E+08 | 1.31E+08 | 3.81E+08 | 8.57E+08 | 2.73E+08 | 6.33E+08 | 6.42E+08 | 2.00E+08 | 1.90E+08 | 3.21E+08 | 2005.9498, 2005.95, 2005.9507 | DGLNDM[147.0354]WADLLELIDTR | P15508 | P15508 | SPTB | spectrin, beta, erythrocytic | Plasma Membrane | other |
| 15538975 | 2 | 0.399 | 1.65E+08 | 2.14E+08 | 3.35E+08 | 8.44E+07 | 2.26E+08 | 2.19E+08 | 2.32E+08 | 2.69E+08 | 1.83E+08 | 1.94E+08 | 1.06E+08 | 1.49E+08 | 1138.5413 | FDILDQEMK | P15508 | P15508 | SPTB | spectrin, beta, erythrocytic | Plasma Membrane | other |
| 15547687, 15551848, 15553900, 15558960, 15559088, 15559891, 15561112, 15561665, 15562148 | 2 | 0.382 | 2.37E+08 | 3.33E+08 | 5.27E+08 | 1.63E+08 | 2.58E+08 | 5.31E+08 | 1.81E+08 | 3.61E+08 | 3.23E+08 | 1.74E+08 | 1.46E+08 | 2.45E+08 | 1733.7446, 1733.7457, 1733.7462, 1733.748, 1733.7501, 1733.7511, 1733.7523, 1733.7529 | FFWEM[147.0354]DEAESWIK | P15508 | P15508 | SPTB | spectrin, beta, erythrocytic | Plasma Membrane | other |
| 15548335, 15550825, 15551692, 15556671, 15557039, 15558912, 15560188, 15560376, 15560531, 15561781, 15562460, 15562884 | 3 | 0.296 | 1.71E+08 | 1.80E+08 | 2.32E+08 | 1.02E+08 | 2.36E+08 | 1.58E+08 | 1.54E+08 | 2.19E+08 | 1.14E+08 | 1.70E+08 | 8.67E+07 | 8.75E+07 | 2467.1797, 2467.1802, 2467.1863, 2467.187, 2467.1895, 2467.19, 2467.191, 2467.1926, 2467.1929, 2467.1963, 2467.1987 | WLDQM[147.0354]DIPNTLEDLEVVQHR | P15508 | P15508 | SPTB | spectrin, beta, erythrocytic | Plasma Membrane | other |
| 15551833, 15552155 | 2, 3 | 0.472 | 2.16E+08 | 3.68E+07 | 2.76E+07 | 3.50E+08 | 6224695.5 | 7.22E+07 | 7.32E+07 | 2.15E+07 | 1.19E+08 | 1.59E+08 | 3.29E+08 | 2.28E+08 | 2308.1155, 2308.118 | LVSDINRAWESLEEAEYQR | P15508 | P15508 | SPTB | spectrin, beta, erythrocytic | Plasma Membrane | other |
| 15547731 | 2 | 0.009 | 5.91E+07 | 5.78E+07 | 8.17E+07 | 8.70E+07 | 7.20E+07 | 1.06E+08 | 7.38E+07 | 6.42E+07 | 6.80E+07 | 1.59E+08 | 9.80E+07 | 1.31E+08 | 1796.881 | LSSSWESLLPEPAHPF | P15508 | P15508 | SPTB | spectrin, beta, erythrocytic | Plasma Membrane | other |
| 15550118, 15559878 | 2 | 0.387 | 1.37E+08 | 1.76E+08 | 2.24E+08 | 1.05E+08 | 1.91E+08 | 1.87E+08 | 1.53E+08 | 1.97E+08 | 1.66E+08 | 1.58E+08 | 1.13E+08 | 1.13E+08 | 1974.9288, 1974.9292 | EQEVSAAWQALLDAC[160.0307]AGR | P15508 | P15508 | SPTB | spectrin, beta, erythrocytic | Plasma Membrane | other |
| 15548817, 15560550, 15560582 | 3 | 0.294 | 1.72E+08 | 1.43E+08 | 1.92E+08 | 1.30E+08 | 1.60E+08 | 1.39E+08 | 1.71E+08 | 1.36E+08 | 1.23E+08 | 1.57E+08 | 1.23E+08 | 1.18E+08 | 2721.3337, 2721.3342, 2721.3457 | AVASEDM[147.0354]PESLPEAEQLLQQHAAIK | P15508 | P15508 | SPTB | spectrin, beta, erythrocytic | Plasma Membrane | other |
| 15547772, 15560820 | 2, 3 | 0.661 | 1.68E+08 | 1.09E+08 | 1.22E+08 | 8.50E+07 | 1.15E+08 | 9.75E+07 | 1.55E+08 | 1.00E+08 | 6.38E+07 | 1.54E+08 | 8.38E+07 | 6.96E+07 | 2327.004, 2327.0051, 2327.0095 | EMVASSQEMGQDFDHVTM[147.0354]LR | P15508 | P15508 | SPTB | spectrin, beta, erythrocytic | Plasma Membrane | other |
| 15555556, 15558836 | 2 | 0.977 | 1.12E+08 | 1.25E+08 | 1.13E+08 | 1.06E+08 | 1.38E+08 | 8.79E+07 | 1.40E+08 | 1.14E+08 | 7.88E+07 | 1.53E+08 | 1.03E+08 | 8.10E+07 | 2035.9286, 2035.9294 | VSHLEQC[160.0307]FSELSNMAAGR | P15508 | P15508 | SPTB | spectrin, beta, erythrocytic | Plasma Membrane | other |
| 15552857 | 3 | 0.07 | 1.40E+08 | 1.34E+08 | 1.21E+08 | 1.37E+08 | 8.98E+07 | 9.37E+07 | 1.16E+08 | 8.38E+07 | 9.21E+07 | 1.53E+08 | 1.45E+08 | 1.19E+08 | 2266.0803 | HRELPEDVGLDASTAESFHR | P15508 | P15508 | SPTB | spectrin, beta, erythrocytic | Plasma Membrane | other |
| 15548019, 15553237 | 3, 4 | 0.875 | 2.19E+08 | 1.57E+08 | 2.33E+08 | 1.62E+08 | 7.08E+07 | 2.96E+08 | 1.45E+08 | 1.39E+08 | 3.42E+08 | 1.52E+08 | 2.15E+08 | 1.97E+08 | 3586.7085, 3586.7146 | WLDQMDIPNTLEDLEVVQHRFDILDQEM[147.0354]K | P15508 | P15508 | SPTB | spectrin, beta, erythrocytic | Plasma Membrane | other |
| 15547870 | 2 | 0.585 | 2.50E+08 | 2.32E+08 | 2.32E+08 | 1.79E+08 | 2.13E+08 | 2.03E+08 | 2.20E+08 | 2.02E+08 | 2.00E+08 | 1.46E+08 | 2.73E+08 | 2.19E+08 | 1908.9233 | TAGYPHVNVTNFTSSWK | P15508 | P15508 | SPTB | spectrin, beta, erythrocytic | Plasma Membrane | other |
| 15535627 | 2 | 0.11 | 6.85E+07 | 4.36E+07 | 3.89E+07 | 7.66E+07 | 4.68E+07 | 4.04E+07 | 3.79E+07 | 4.65E+07 | 4.35E+07 | 1.39E+08 | 7.99E+07 | 5.77E+07 | 1476.6831 | EADDLEQWITEK | P15508 | P15508 | SPTB | spectrin, beta, erythrocytic | Plasma Membrane | other |
| 15534904 | 3 | 0.46 | 2.45E+08 | 6.54E+07 | 1.54E+07 | 2.03E+08 | 4.55E+07 | 5.01E+07 | 8.14E+07 | 6.88E+07 | 1.48E+08 | 1.34E+08 | 2.71E+08 | 1.88E+08 | 2053.0552 | LVTQDNFGYDLAAVEAAKK | P15508 | P15508 | SPTB | spectrin, beta, erythrocytic | Plasma Membrane | other |
| 15547700, 15553278, 15555868, 15558551, 15561997 | 3, 4 | 0.9 | 2.08E+08 | 1.35E+08 | 2.37E+08 | 1.69E+08 | 5.93E+07 | 2.76E+08 | 1.33E+08 | 1.32E+08 | 3.30E+08 | 1.29E+08 | 1.75E+08 | 2.07E+08 | 3586.6958, 3586.7065, 3586.707, 3586.7136, 3586.716 | WLDQM[147.0354]DIPNTLEDLEVVQHRFDILDQEMK | P15508 | P15508 | SPTB | spectrin, beta, erythrocytic | Plasma Membrane | other |
| 15532005 | 3 | 0.984 | 9.47E+07 | 1.32E+08 | 9.96E+07 | 1.13E+08 | 1.15E+08 | 9.74E+07 | 9.89E+07 | 1.44E+08 | 9.20E+07 | 1.25E+08 | 9.08E+07 | 9.78E+07 | 1819.7335 | WDAPDDELDNDNSSAR | P15508 | P15508 | SPTB | spectrin, beta, erythrocytic | Plasma Membrane | other |
| 15534816 | 3 | 0.87 | 8.84E+07 | 9.07E+07 | 6.00E+07 | 7.31E+07 | 1.28E+08 | 6.47E+07 | 8.53E+07 | 1.20E+08 | 8.47E+07 | 1.24E+08 | 6.36E+07 | 8.34E+07 | 1628.7721 | KFEDFLVSMENNR | P15508 | P15508 | SPTB | spectrin, beta, erythrocytic | Plasma Membrane | other |
| 15555631, 15558099 | 2, 3 | 0.411 | 1.37E+08 | 1.13E+08 | 1.21E+08 | 9.17E+07 | 1.13E+08 | 1.01E+08 | 1.24E+08 | 1.09E+08 | 6.89E+07 | 1.22E+08 | 8.28E+07 | 7.60E+07 | 2327.0056, 2327.0059 | EM[147.0354]VASSQEMGQDFDHVTMLR | P15508 | P15508 | SPTB | spectrin, beta, erythrocytic | Plasma Membrane | other |
| 15531436 | 3 | 0.233 | 1.00E+08 | 1.03E+08 | 1.39E+08 | 9.29E+07 | 1.01E+08 | 9.42E+07 | 7.80E+07 | 1.05E+08 | 7.44E+07 | 1.21E+08 | 8.96E+07 | 9.50E+07 | 1634.7263 | HEAIETDTAAYEER | P15508 | P15508 | SPTB | spectrin, beta, erythrocytic | Plasma Membrane | other |
| 15551282, 15556409 | 2 | 0.067 | 1.31E+08 | 1.17E+08 | 1.45E+08 | 1.12E+08 | 1.05E+08 | 1.01E+08 | 1.06E+08 | 9.83E+07 | 9.14E+07 | 1.20E+08 | 7.43E+07 | 8.04E+07 | 1698.9404, 1698.9409 | LLEVLSGEM[147.0354]LPRPTK | P15508 | P15508 | SPTB | spectrin, beta, erythrocytic | Plasma Membrane | other |
| 15532698 | 2, 3 | 0.556 | 9.94E+07 | 1.06E+08 | 1.03E+08 | 8.24E+07 | 1.06E+08 | 9.91E+07 | 9.62E+07 | 9.89E+07 | 7.93E+07 | 1.19E+08 | 1.00E+08 | 8.88E+07 | 1448.7245, 1448.7251 | WQAFQAVVSEQR | P15508 | P15508 | SPTB | spectrin, beta, erythrocytic | Plasma Membrane | other |
| 15534651, 15540187, 15541415, 15546525 | 3 | 0.64 | 9.09E+07 | 1.11E+08 | 8.17E+07 | 7.38E+07 | 8.83E+07 | 7.73E+07 | 8.62E+07 | 8.66E+07 | 7.22E+07 | 1.18E+08 | 7.31E+07 | 7.64E+07 | 1532.7677, 1532.768, 1532.7683, 1532.7687 | LHMLLEVC[160.0307]QFSR | P15508 | P15508 | SPTB | spectrin, beta, erythrocytic | Plasma Membrane | other |
| 15542969 | 3 | 0.694 | 1.19E+08 | 6.66E+07 | 1.87E+08 | 6.57E+07 | 1.02E+08 | 7.61E+07 | 1.31E+08 | 1.50E+08 | 6.02E+07 | 1.16E+08 | 1.37E+08 | 8.17E+07 | 1837.949 | SIITYVVAFYHYFSK | P15508 | P15508 | SPTB | spectrin, beta, erythrocytic | Plasma Membrane | other |
| 15537183, 15546867 | 3 | 0.503 | 7.09E+07 | 8.05E+07 | 1.31E+08 | 7.12E+07 | 8.91E+07 | 9.83E+07 | 6.94E+07 | 8.27E+07 | 5.95E+07 | 1.15E+08 | 6.57E+07 | 1.17E+08 | 1761.8917, 1761.8918 | YFYTGTEILGLIDEK | P15508 | P15508 | SPTB | spectrin, beta, erythrocytic | Plasma Membrane | other |
| 15547893, 15548653, 15555177, 15561463, 15561527 | 2, 3 | 0.187 | 1.15E+08 | 1.02E+08 | 1.27E+08 | 6.27E+07 | 8.80E+07 | 7.22E+07 | 1.01E+08 | 8.19E+07 | 6.03E+07 | 1.14E+08 | 4.63E+07 | 4.92E+07 | 2354.202, 2354.2039, 2354.2046, 2354.2083, 2354.2144 | LPLAQSADYGTNLQTVQLFM[147.0354]K | P15508 | P15508 | SPTB | spectrin, beta, erythrocytic | Plasma Membrane | other |
| 15534341, 15544906, 15546951 | 2, 3 | 0.016 | 1.21E+08 | 1.02E+08 | 1.03E+08 | 1.08E+08 | 7.64E+07 | 9.02E+07 | 9.10E+07 | 6.86E+07 | 6.99E+07 | 1.13E+08 | 1.22E+08 | 1.11E+08 | 1270.7451, 1270.7452, 1270.7457, 1270.7472 | QRLEATLALQK | P15508 | P15508 | SPTB | spectrin, beta, erythrocytic | Plasma Membrane | other |
| 15547901, 15557449, 15558558, 15559660, 15559996, 15561476 | 3 | 0.471 | 1.05E+08 | 8.90E+07 | 1.07E+08 | 8.15E+07 | 8.45E+07 | 6.79E+07 | 1.09E+08 | 7.74E+07 | 5.76E+07 | 1.06E+08 | 6.96E+07 | 5.85E+07 | 2436.2222, 2436.2268, 2436.228, 2436.2288, 2436.229, 2436.2368 | TLM[147.0354]AQIDGVNLAANNLVESGHPR | P15508 | P15508 | SPTB | spectrin, beta, erythrocytic | Plasma Membrane | other |
| 15531845 | 3 | 0.771 | 9.49E+07 | 1.45E+08 | 1.01E+08 | 1.29E+08 | 1.26E+08 | 1.20E+08 | 1.23E+08 | 1.40E+08 | 1.04E+08 | 9.56E+07 | 1.46E+08 | 1.67E+08 | 1770.8436 | GNTLTQC[160.0307]LGFQEFQK | P15508 | P15508 | SPTB | spectrin, beta, erythrocytic | Plasma Membrane | other |
| 15547728, 15559132, 15560516, 15560605, 15561354, 15561740, 15563132 | 3 | 0.191 | 9.40E+07 | 8.73E+07 | 1.04E+08 | 7.50E+07 | 8.79E+07 | 9.07E+07 | 8.53E+07 | 7.66E+07 | 7.10E+07 | 9.47E+07 | 6.47E+07 | 6.31E+07 | 2783.2322, 2783.2385, 2783.2388, 2783.24, 2783.2402, 2783.2432, 2783.2454 | EELEELFADAPSLGAEAGDTDM[147.0354]SIEK | P15508 | P15508 | SPTB | spectrin, beta, erythrocytic | Plasma Membrane | other |
| 15536036, 15546736, 15547204 | 3 | 0.607 | 6.19E+07 | 1.77E+08 | 3.56E+08 | 6.23E+07 | 1.64E+08 | 3.70E+08 | 1.28E+08 | 2.70E+08 | 2.40E+08 | 9.37E+07 | 8.39E+07 | 1.16E+08 | 2005.9508, 2005.9514, 2005.9529 | DGLNDM[147.0354]WADLLELIDTR | P15508 | P15508 | SPTB | spectrin, beta, erythrocytic | Plasma Membrane | other |
| 15553107 | 2 | 0.317 | 1.06E+08 | 5.72E+07 | 5.30E+07 | 1.11E+08 | 2.99E+07 | 3.46E+07 | 8.40E+07 | 4.13E+07 | 6.43E+07 | 9.36E+07 | 1.22E+08 | 9.00E+07 | 1613.8259 | AFEDELRGLDAHLK | P15508 | P15508 | SPTB | spectrin, beta, erythrocytic | Plasma Membrane | other |
| 15548464 | 2 | 0.847 | 2.19E+08 | 1.99E+08 | 4.33E+07 | 1.93E+08 | 1.66E+08 | 1.70E+08 | 1.78E+08 | 1.56E+08 | 1.45E+08 | 9.29E+07 | 1.98E+08 | 1.69E+08 | 2081.1096 | ELHLLGVQVQQFQDVATR | P15508 | P15508 | SPTB | spectrin, beta, erythrocytic | Plasma Membrane | other |
| 15531424 | 2 | 0.669 | 1.29E+08 | 1.86E+08 | 2.79E+08 | 8.41E+07 | 1.92E+08 | 2.21E+08 | 1.42E+08 | 2.02E+08 | 2.06E+08 | 9.15E+07 | 1.64E+08 | 1.58E+08 | 860.446 | EAVDSALR | P15508 | P15508 | SPTB | spectrin, beta, erythrocytic | Plasma Membrane | other |
| 15541662, 15545032, 15545820, 15546315, 15546407, 15547081, 15547240 | 2 | 0.157 | 8.33E+07 | 1.06E+08 | 1.28E+08 | 9.34E+07 | 9.12E+07 | 9.54E+07 | 8.42E+07 | 8.80E+07 | 7.03E+07 | 9.04E+07 | 4.55E+07 | 7.49E+07 | 1319.6462, 1319.6473, 1319.6477, 1319.6482, 1319.6489, 1319.6494, 1319.6503 | LM[147.0354]EADIAIQGDK | P15508 | P15508 | SPTB | spectrin, beta, erythrocytic | Plasma Membrane | other |
| 15537403, 15540069 | 3, 4 | 0.285 | 1.09E+08 | 1.19E+08 | 7.55E+07 | 9.60E+07 | 9.09E+07 | 5.25E+07 | 8.18E+07 | 5.99E+07 | 2.98E+07 | 9.01E+07 | 2.59E+07 | 5.51E+07 | 1945.889, 1945.8903 | VHLENM[147.0354]GSHDIVDGNHR | P15508 | P15508 | SPTB | spectrin, beta, erythrocytic | Plasma Membrane | other |
| 15543392 | 2 | 0.697 | 2.05E+08 | 3.09E+08 | 4.25E+08 | 8.72E+07 | 3.43E+08 | 3.13E+08 | 1.74E+08 | 3.45E+08 | 3.24E+08 | 8.65E+07 | 2.68E+08 | 2.57E+08 | 980.5025 | LQTAYAGEK | P15508 | P15508 | SPTB | spectrin, beta, erythrocytic | Plasma Membrane | other |
| 15532762 | 3, 4 | 0.583 | 1.33E+08 | 8.26E+07 | 8.51E+07 | 1.12E+08 | 4.41E+07 | 6.69E+07 | 1.07E+08 | 7.04E+07 | 6.95E+07 | 8.58E+07 | 7.41E+07 | 8.13E+07 | 2180.018, 2180.019 | VSHLEQC[160.0307]FSELSNM[147.0354]AAGRK | P15508 | P15508 | SPTB | spectrin, beta, erythrocytic | Plasma Membrane | other |
| 15532367, 15536561, 15537003, 15544303, 15546009, 15546291, 15547365, 15547430 | 3, 4 | 0.706 | 1.01E+08 | 8.03E+07 | 6.74E+07 | 1.33E+08 | 4.14E+07 | 7.55E+07 | 6.99E+07 | 4.97E+07 | 8.89E+07 | 8.29E+07 | 1.27E+08 | 8.22E+07 | 1890.9216, 1890.9248, 1890.9258, 1890.928, 1890.9281, 1890.9291, 1890.9302, 1890.9309, 1890.9331 | SDRLHMLLEVC[160.0307]QFSR | P15508 | P15508 | SPTB | spectrin, beta, erythrocytic | Plasma Membrane | other |
| 15548465 | 3 | 0.515 | 1.02E+08 | 2.25E+07 | 1.79E+07 | 1.96E+08 | 2074150.5 | 5.05E+07 | 4.61E+07 | 1.91E+07 | 6.68E+07 | 7.83E+07 | 1.88E+08 | 1.46E+08 | 3216.5747 | VIEGQTDPDYQLLGQRLEGLDTDWDALR | P15508 | P15508 | SPTB | spectrin, beta, erythrocytic | Plasma Membrane | other |
| 15550423 | 3, 4 | 0.828 | 1.16E+08 | 3.93E+07 | 5.91E+07 | 1.07E+08 | 1.82E+07 | 8.87E+07 | 6.24E+07 | 3.96E+07 | 6.57E+07 | 7.56E+07 | 9.09E+07 | 7.66E+07 | 3868.888, 3868.8896 | DLEDETLWVEERLPLAQSADYGTNLQTVQLFM[147.0354]K | P15508 | P15508 | SPTB | spectrin, beta, erythrocytic | Plasma Membrane | other |
| 15535352 | 3 | 0.631 | 7.68E+07 | 7.01E+07 | 5.57E+07 | 1.08E+08 | 2.90E+07 | 5.14E+07 | 5.33E+07 | 3.43E+07 | 3.34E+07 | 7.53E+07 | 2.20E+07 | 6.61E+07 | 1430.686 | M[147.0354]RIHC[160.0307]LENVDK | P15508 | P15508 | SPTB | spectrin, beta, erythrocytic | Plasma Membrane | other |
| 15534657 | 3 | 0.364 | 1.77E+08 | 1.29E+07 | 2516756.75 | 1.28E+08 | 7177984 | 4173398.75 | 1.12E+07 | 1.31E+07 | 8.28E+07 | 7.44E+07 | 2.40E+08 | 1.24E+08 | 1460.8336 | FAALEKPTTLELK | P15508 | P15508 | SPTB | spectrin, beta, erythrocytic | Plasma Membrane | other |
| 15533134, 15543294, 15544014, 15546850, 15547245 | 2 | 0.05 | 8.70E+07 | 9.67E+07 | 1.01E+08 | 6.39E+07 | 7.71E+07 | 7.80E+07 | 7.20E+07 | 7.12E+07 | 5.63E+07 | 7.17E+07 | 4.05E+07 | 6.51E+07 | 1293.6268, 1293.6296, 1293.6307, 1293.6328 | DALLLWC[160.0307]QM[147.0354]K | P15508 | P15508 | SPTB | spectrin, beta, erythrocytic | Plasma Membrane | other |
| 15550032, 15555534, 15556811, 15561943 | 3 | 0.984 | 6.86E+07 | 4.49E+07 | 2.01E+07 | 3.51E+07 | 4.84E+07 | 2.60E+07 | 3.95E+07 | 3.96E+07 | 4.08E+07 | 6.85E+07 | 2.69E+07 | 2.97E+07 | 2911.3342, 2911.3347, 2911.3357, 2911.337 | KEELEELFADAPSLGAEAGDTDM[147.0354]SIEK | P15508 | P15508 | SPTB | spectrin, beta, erythrocytic | Plasma Membrane | other |
| 15534948 | 3 | 0.399 | 6.11E+07 | 7.66E+07 | 1.01E+08 | 3.87E+07 | 8.27E+07 | 6.43E+07 | 7.36E+07 | 9.16E+07 | 4.10E+07 | 6.69E+07 | 2.98E+07 | 4.67E+07 | 2051.923 | VSHLEQC[160.0307]FSELSNM[147.0354]AAGR | P15508 | P15508 | SPTB | spectrin, beta, erythrocytic | Plasma Membrane | other |
| 15532328, 15537926, 15544963, 15545543, 15545611, 15545804, 15546400, 15546892, 15547516 | 3 | 0.883 | 5.49E+07 | 4.53E+07 | 5.71E+07 | 6.68E+07 | 5.13E+07 | 4.87E+07 | 6.65E+07 | 4.93E+07 | 4.49E+07 | 6.66E+07 | 4.07E+07 | 4.18E+07 | 1621.7048, 1621.7083, 1621.7119, 1621.7124, 1621.7157, 1621.7174, 1621.7197, 1621.7229, 1621.7233 | ESQYLM[147.0354]ESHPEQK | P15508 | P15508 | SPTB | spectrin, beta, erythrocytic | Plasma Membrane | other |
| 15533167 | 3 | 0.531 | 3.71E+07 | 4.29E+07 | 5.42E+07 | 2.43E+07 | 4.46E+07 | 5.45E+07 | 4.61E+07 | 4.56E+07 | 4.97E+07 | 6.64E+07 | 4.39E+07 | 5.21E+07 | 1635.7761 | DNLELQNFLQNC[160.0307]K | P15508 | P15508 | SPTB | spectrin, beta, erythrocytic | Plasma Membrane | other |
| 15537068 | 3 | 0.532 | 6.25E+07 | 7.15E+07 | 8.71E+07 | 4.89E+07 | 7.36E+07 | 8.89E+07 | 6.18E+07 | 8.23E+07 | 6.91E+07 | 6.63E+07 | 4.17E+07 | 6.43E+07 | 2036.942 | LFQDM[147.0354]LHSIDWMDEIK | P15508 | P15508 | SPTB | spectrin, beta, erythrocytic | Plasma Membrane | other |
| 15553204, 15558869 | 2 | 0.48 | 8.41E+07 | 2.85E+07 | 2.13E+07 | 1.06E+08 | 6388117.5 | 3.38E+07 | 4.65E+07 | 1.61E+07 | 4.53E+07 | 6.58E+07 | 1.20E+08 | 6.97E+07 | 1833.8745, 1833.8751 | WIGAMEDQLRSDDLGK | P15508 | P15508 | SPTB | spectrin, beta, erythrocytic | Plasma Membrane | other |
| 15536477, 15545570 | 3 | 0.519 | 5.53E+07 | 9.49E+07 | 6.53E+07 | 3.00E+07 | 6.62E+07 | 6.37E+07 | 5.15E+07 | 1.19E+08 | 6.26E+07 | 6.52E+07 | 4.69E+07 | 5.19E+07 | 1974.9297, 1974.931 | EQEVSAAWQALLDAC[160.0307]AGR | P15508 | P15508 | SPTB | spectrin, beta, erythrocytic | Plasma Membrane | other |
| 15548353 | 2 | 0.231 | 4.56E+07 | 4.73E+07 | 6.71E+07 | 4.39E+07 | 4.49E+07 | 4.19E+07 | 3.20E+07 | 4.60E+07 | 3.68E+07 | 6.15E+07 | 4.53E+07 | 3.94E+07 | 1634.7269 | HEAIETDTAAYEER | P15508 | P15508 | SPTB | spectrin, beta, erythrocytic | Plasma Membrane | other |
| 15538919, 15538922, 15545423, 15545984, 15546642, 15547163, 15547421 | 2, 3 | 0.195 | 6.53E+07 | 6.13E+07 | 7.48E+07 | 5.32E+07 | 6.23E+07 | 6.05E+07 | 5.68E+07 | 6.79E+07 | 4.78E+07 | 5.94E+07 | 4.54E+07 | 5.19E+07 | 1433.7157, 1433.716, 1433.7174, 1433.7175, 1433.7179, 1433.718 | M[147.0354]QLLAASYDLHR | P15508 | P15508 | SPTB | spectrin, beta, erythrocytic | Plasma Membrane | other |
| 15533942 | 3 | 0.421 | 3.25E+07 | 3.64E+07 | 2.78E+07 | 3.76E+07 | 3.21E+07 | 2.15E+07 | 4.48E+07 | 2.83E+07 | 2.44E+07 | 5.88E+07 | 3.62E+07 | 3.54E+07 | 1717.7527 | FFWEMDEAESWIK | P15508 | P15508 | SPTB | spectrin, beta, erythrocytic | Plasma Membrane | other |
| 15535963 | 3 | 0.717 | 3.77E+07 | 8.25E+07 | 1.30E+08 | 2.04E+07 | 1.62E+08 | 6.79E+07 | 6.53E+07 | 1.19E+08 | 4.27E+07 | 5.80E+07 | 2.53E+07 | 4.47E+07 | 2140.9817 | GVMEHLEHQAQGFPEEFR | P15508 | P15508 | SPTB | spectrin, beta, erythrocytic | Plasma Membrane | other |
| 15549376, 15554387 | 2 | 0.107 | 6.19E+07 | 6.67E+07 | 8.75E+07 | 5.86E+07 | 5.64E+07 | 7.08E+07 | 5.11E+07 | 6.81E+07 | 5.37E+07 | 5.75E+07 | 2.97E+07 | 4.73E+07 | 2036.9414, 2036.9425 | LFQDMLHSIDWM[147.0354]DEIK | P15508 | P15508 | SPTB | spectrin, beta, erythrocytic | Plasma Membrane | other |
| 15544181 | 5 | 0.555 | 6.44E+07 | 9.98E+07 | 1.84E+08 | 2.65E+07 | 1.69E+08 | 1.44E+08 | 1.33E+08 | 1.47E+08 | 1.14E+08 | 5.71E+07 | 4.04E+07 | 1.06E+08 | 3025.379 | GVMEHLEHQAQGFPEEFRDSPDVTNR | P15508 | P15508 | SPTB | spectrin, beta, erythrocytic | Plasma Membrane | other |
| 15554341, 15555823, 15558230, 15559262 | 3, 4 | 0.377 | 5.17E+07 | 5.96E+07 | 9.25E+07 | 3.23E+07 | 8.04E+07 | 5.38E+07 | 5.88E+07 | 7.84E+07 | 4.47E+07 | 5.70E+07 | 2.96E+07 | 3.66E+07 | 3041.364, 3041.3723, 3041.3733, 3041.3752 | GVM[147.0354]EHLEHQAQGFPEEFRDSPDVTNR | P15508 | P15508 | SPTB | spectrin, beta, erythrocytic | Plasma Membrane | other |
| 15549294, 15558226 | 3 | 0.481 | 5.38E+07 | 5.20E+07 | 8.63E+07 | 3.32E+07 | 6.27E+07 | 7.08E+07 | 4.61E+07 | 7.41E+07 | 6.53E+07 | 5.28E+07 | 3.98E+07 | 4.10E+07 | 2864.2966, 2864.3037 | LQEALDLYTVFGESDAC[160.0307]ELWM[147.0354]TEK | P15508 | P15508 | SPTB | spectrin, beta, erythrocytic | Plasma Membrane | other |
| 15534847 | 3 | 0.796 | 1.97E+07 | 3.97E+07 | 4.21E+07 | 3.26E+07 | 3.42E+07 | 3.39E+07 | 3.93E+07 | 3.76E+07 | 3.81E+07 | 5.02E+07 | 1.85E+07 | 2.73E+07 | 1510.6786 | AWESLEEAEYQR | P15508 | P15508 | SPTB | spectrin, beta, erythrocytic | Plasma Membrane | other |
| 15556305 | 2 | 0.562 | 3.05E+07 | 6.87E+07 | 1.17E+08 | 4.07E+07 | 6.14E+07 | 8.81E+07 | 8.84E+07 | 5.53E+07 | 3.25E+07 | 4.99E+07 | 2.45E+07 | 4.01E+07 | 1743.795 | FEDFLVSMENNRDK | P15508 | P15508 | SPTB | spectrin, beta, erythrocytic | Plasma Membrane | other |
| 15532896, 15540608 | 2 | 0.851 | 4.90E+07 | 4.08E+07 | 6.26E+07 | 5.19E+07 | 4.62E+07 | 4.89E+07 | 4.81E+07 | 4.62E+07 | 4.73E+07 | 4.87E+07 | 4.56E+07 | 4.54E+07 | 950.4748, 950.4811 | AWLSM[147.0354]AQK | P15508 | P15508 | SPTB | spectrin, beta, erythrocytic | Plasma Membrane | other |
| 15552893 | 2 | 0.536 | 1.76E+07 | 8.16E+07 | 1.80E+08 | 1.89E+07 | 1.36E+08 | 7.87E+07 | 7.28E+07 | 8.59E+07 | 3.66E+07 | 4.84E+07 | 1.31E+07 | 2.49E+07 | 1500.6765 | FEDFLVSMENNR | P15508 | P15508 | SPTB | spectrin, beta, erythrocytic | Plasma Membrane | other |
| 15533676 | 2 | 0.591 | 3.12E+07 | 4.03E+07 | 4.22E+07 | 2.82E+07 | 4.16E+07 | 5.90E+07 | 2.67E+07 | 3.96E+07 | 2.69E+07 | 4.40E+07 | 2.37E+07 | 3.46E+07 | 1172.6654 | FLDLLEPLGR | P15508 | P15508 | SPTB | spectrin, beta, erythrocytic | Plasma Membrane | other |
| 15551571, 15560082 | 3 | 0.232 | 3.84E+07 | 3.22E+07 | 5.64E+07 | 2.71E+07 | 3.94E+07 | 3.82E+07 | 4.05E+07 | 4.97E+07 | 5.11E+07 | 4.14E+07 | 2.27E+07 | 3.14E+07 | 2848.3013, 2848.3018 | LQEALDLYTVFGESDAC[160.0307]ELWMTEK | P15508 | P15508 | SPTB | spectrin, beta, erythrocytic | Plasma Membrane | other |
| 15537209 | 2 | 0.485 | 2.97E+07 | 2.56E+07 | 3.90E+07 | 2.18E+07 | 3.60E+07 | 2.34E+07 | 2.45E+07 | 4.13E+07 | 2.79E+07 | 4.07E+07 | 9440227 | 1.70E+07 | 802.3897 | FFSM[147.0354]VR | P15508 | P15508 | SPTB | spectrin, beta, erythrocytic | Plasma Membrane | other |
| 15534372 | 3 | 0.236 | 7.06E+07 | 8.23E+07 | 1.20E+08 | 3.05E+07 | 7.22E+07 | 9.50E+07 | 5.67E+07 | 1.10E+08 | 9.74E+07 | 3.98E+07 | 3.82E+07 | 6.62E+07 | 1607.8331 | RLENM[147.0354]YHLFQLK | P15508 | P15508 | SPTB | spectrin, beta, erythrocytic | Plasma Membrane | other |
| 15551663 | 2 | 0.163 | 3.55E+07 | 2.48E+07 | 2.25E+07 | 3.46E+07 | 1.67E+07 | 2.45E+07 | 2.74E+07 | 1.71E+07 | 2.24E+07 | 3.51E+07 | 4.57E+07 | 3.17E+07 | 2131.03 | EQEVSAAWQALLDAC[160.0307]AGRR | P15508 | P15508 | SPTB | spectrin, beta, erythrocytic | Plasma Membrane | other |
| 15532790, 15546957 | 2 | 0.479 | 2.85E+07 | 3.12E+07 | 3.81E+07 | 1.49E+07 | 3.80E+07 | 3.06E+07 | 3.52E+07 | 3.46E+07 | 1.90E+07 | 3.28E+07 | 1.42E+07 | 1.64E+07 | 1154.5361, 1154.5375 | FDILDQEM[147.0354]K | P15508 | P15508 | SPTB | spectrin, beta, erythrocytic | Plasma Membrane | other |
| 15535326 | 3 | 0.48 | 6.05E+07 | 1.10E+07 | 6434268 | 1.07E+08 | 1171325.75 | 1.73E+07 | 2.22E+07 | 5619385 | 2.79E+07 | 3.28E+07 | 1.38E+08 | 6.78E+07 | 1618.7739 | AAMRETWLNENQR | P15508 | P15508 | SPTB | spectrin, beta, erythrocytic | Plasma Membrane | other |
| 15533394 | 1 | 0.121 | 2.58E+07 | 2.33E+07 | 1.94E+07 | 2.22E+07 | 1.78E+07 | 1.83E+07 | 2.46E+07 | 1.82E+07 | 1.48E+07 | 3.21E+07 | 2.63E+07 | 2.33E+07 | 692.34235 | WIMDK | P15508 | P15508 | SPTB | spectrin, beta, erythrocytic | Plasma Membrane | other |
| 15531809, 15545425, 15545501, 15545867, 15546508, 15546523, 15546610 | 2 | 0.288 | 2.96E+07 | 3.35E+07 | 4.17E+07 | 1.72E+07 | 3.61E+07 | 3.21E+07 | 3.40E+07 | 3.81E+07 | 2.45E+07 | 3.08E+07 | 1.49E+07 | 2.05E+07 | 1234.5814, 1234.5826, 1234.5847, 1234.5853, 1234.5864, 1234.5897, 1234.5907 | WIGAM[147.0354]EDQLR | P15508 | P15508 | SPTB | spectrin, beta, erythrocytic | Plasma Membrane | other |
| 15540942 | 3 | 0.372 | 2.11E+07 | 1.90E+07 | 1.40E+07 | 2.72E+07 | 1.57E+07 | 1.29E+07 | 1.84E+07 | 1.68E+07 | 1.42E+07 | 2.92E+07 | 2.19E+07 | 2.02E+07 | 1546.7289 | LLSGEDVGQDEGATR | P15508 | P15508 | SPTB | spectrin, beta, erythrocytic | Plasma Membrane | other |
| 15533965 | 2 | 0.818 | 1.31E+07 | 2.48E+07 | 3.13E+07 | 1.17E+07 | 4.07E+07 | 2.17E+07 | 2.39E+07 | 3.99E+07 | 1.51E+07 | 2.85E+07 | 1.18E+07 | 1.34E+07 | 1356.7527 | LLEVLSGEMLPR | P15508 | P15508 | SPTB | spectrin, beta, erythrocytic | Plasma Membrane | other |
| 15538836 | 3 | 0.063 | 6.08E+07 | 9.52E+07 | 7.89E+07 | 7.19E+07 | 6.48E+07 | 6.66E+07 | 4.61E+07 | 6.13E+07 | 5.07E+07 | 2.73E+07 | 4.96E+07 | 5.71E+07 | 1660.8613 | FQIQDIVVQTQEGR | P15508 | P15508 | SPTB | spectrin, beta, erythrocytic | Plasma Membrane | other |
| 15533927 | 3 | 0.03 | 3.84E+07 | 3.25E+07 | 5.07E+07 | 1.68E+07 | 2.52E+07 | 3.21E+07 | 2.86E+07 | 3.09E+07 | 2.23E+07 | 2.50E+07 | 1.27E+07 | 1.38E+07 | 2052.9365 | LFQDM[147.0354]LHSIDWM[147.0354]DEIK | P15508 | P15508 | SPTB | spectrin, beta, erythrocytic | Plasma Membrane | other |
| 15554099, 15562764, 15563799 | 2 | 0.399 | 2.43E+07 | 1.93E+07 | 2.77E+07 | 1.84E+07 | 2.11E+07 | 1.92E+07 | 2.22E+07 | 1.97E+07 | 1.66E+07 | 2.44E+07 | 1.43E+07 | 1.73E+07 | 1538.7119, 1538.7123, 1538.713 | QIFQEADDM[147.0354]VAQK | P15508 | P15508 | SPTB | spectrin, beta, erythrocytic | Plasma Membrane | other |
| 15533719 | 3 | 0.586 | 3.45E+07 | 1.64E+07 | 1.75E+07 | 3.20E+07 | 9638141 | 1.91E+07 | 2.44E+07 | 1.20E+07 | 1.64E+07 | 2.39E+07 | 2.83E+07 | 2.63E+07 | 1849.8712 | WIGAM[147.0354]EDQLRSDDLGK | P15508 | P15508 | SPTB | spectrin, beta, erythrocytic | Plasma Membrane | other |
| 15534256, 15535629, 15541266, 15544035, 15545517 | 2 | 0.083 | 2.46E+07 | 2.92E+07 | 3.77E+07 | 1.65E+07 | 2.78E+07 | 2.42E+07 | 2.55E+07 | 3.02E+07 | 2.20E+07 | 2.31E+07 | 9202722 | 1.48E+07 | 1378.7007, 1378.7008, 1378.7009, 1378.701, 1378.7015 | DLLSWM[147.0354]ESIIR | P15508 | P15508 | SPTB | spectrin, beta, erythrocytic | Plasma Membrane | other |
| 15534643 | 3 | 0.442 | 4.91E+07 | 7969564 | 5960710.5 | 7.93E+07 | 2073742.125 | 1.39E+07 | 1.25E+07 | 5622477 | 2.75E+07 | 2.23E+07 | 1.09E+08 | 5.93E+07 | 1596.9031 | DLGQDLAGVIAIQRK | P15508 | P15508 | SPTB | spectrin, beta, erythrocytic | Plasma Membrane | other |
| 15536178 | 2 | 0.339 | 2.12E+07 | 2.73E+07 | 2.48E+07 | 1.68E+07 | 3.71E+07 | 2.28E+07 | 1.91E+07 | 2.82E+07 | 1.98E+07 | 2.12E+07 | 1.06E+07 | 1.84E+07 | 786.395 | FFSMVR | P15508 | P15508 | SPTB | spectrin, beta, erythrocytic | Plasma Membrane | other |
| 15559495 | 2 | 0.578 | 2.67E+07 | 1.97E+07 | 6.27E+07 | 1.04E+07 | 3.03E+07 | 2.50E+07 | 3.00E+07 | 4.30E+07 | 2.20E+07 | 2.11E+07 | 3.03E+07 | 2.21E+07 | 1837.949 | SIITYVVAFYHYFSK | P15508 | P15508 | SPTB | spectrin, beta, erythrocytic | Plasma Membrane | other |
| 15549981 | 3 | 0.48 | 4.51E+07 | 6893577 | 5020188 | 6.78E+07 | 1163641.75 | 1.16E+07 | 1.37E+07 | 3297365 | 2.09E+07 | 2.08E+07 | 9.27E+07 | 4.45E+07 | 2554.1292 | EMVASSQEMGQDFDHVTMLRDK | P15508 | P15508 | SPTB | spectrin, beta, erythrocytic | Plasma Membrane | other |
| 15538438 | 2, 3 | 0.538 | 3.85E+07 | 7483720.5 | 4906521.5 | 5.39E+07 | 703003.25 | 1.44E+07 | 1.83E+07 | 6196952.5 | 2.08E+07 | 1.96E+07 | 7.21E+07 | 4.12E+07 | 1341.7822 | LSGLERDVLAIR | P15508 | P15508 | SPTB | spectrin, beta, erythrocytic | Plasma Membrane | other |
| 15553696 | 3 | 0.352 | 2.03E+07 | 1.79E+07 | 1.91E+07 | 1.74E+07 | 1.72E+07 | 1.24E+07 | 1.59E+07 | 1.73E+07 | 1.20E+07 | 1.95E+07 | 1.18E+07 | 1.32E+07 | 2327.0042 | EMVASSQEM[147.0354]GQDFDHVTMLR | P15508 | P15508 | SPTB | spectrin, beta, erythrocytic | Plasma Membrane | other |

| | | | | | | | | | | | | | | | |
|---|---|---|---|---|---|---|---|---|---|---|---|---|---|---|---|
| 15548561 | 1 | 0.302 | 1.18E+07 | 9516720 | 9446276 | 1.28E+07 | 7786871.5 | 7060444.5 | 1.02E+07 | 7659402 | 5588981.5 | 1.93E+07 | 1.07E+07 | 9284441 | [illegible] |
| 15533168 | 3 | 0.499 | 1.08E+07 | 2.03E+07 | 4.92E+07 | 1.38E+07 | 2.07E+07 | 3.26E+07 | 2.78E+07 | 1.92E+07 | 1.29E+07 | 1.91E+07 | 6584300 | 1.26E+07 | [illegible] |
| 15549098 | 3 | 0.345 | 1.58E+07 | 989135.875 | 1269622.875 | 4.33E+07 | 206597.328 | 5050628 | 5497894 | 1.20E+07 | 1.81E+07 | 1.69E+07 | 5.02E+07 | 2.16E+07 | [illegible] |
| 15548214, 15561246 | 4 | 0.833 | 3.03E+07 | 1.20E+07 | 7.25E+07 | 1.63E+07 | 1.14E+07 | 1.24E+08 | 1.64E+07 | 5.30E+07 | 5.92E+07 | 1.66E+07 | 1.91E+07 | 2.33E+07 | [illegible] |
| 15532191 | 2, 3 | 0.351 | 4.12E+07 | 1319712.125 | 1445248.875 | 3.58E+07 | 1546864.625 | 579689 | 1964314.25 | 1405911.875 | 1.31E+07 | 1.62E+07 | 7.30E+07 | 2.03E+07 | [illegible] |
| 15554285 | 2 | 0.41 | 1.31E+07 | 1.14E+07 | 5547212.5 | 1.04E+07 | 8983163 | 6171193.5 | 9961615 | 5776195 | 2846076.25 | 1.45E+07 | 7583573 | 8726900 | [illegible] |
| 15531840 | 2 | 0.266 | 4.71E+07 | 5.58E+07 | 7.32E+07 | 2.04E+07 | 5.41E+07 | 6.80E+07 | 4.16E+07 | 7.71E+07 | 5.31E+07 | 1.42E+07 | 2.57E+07 | 5.10E+07 | [illegible] |
| 15553678 | 3 | 0.334 | 1.23E+07 | 9764594 | 6799055.5 | 8473344 | 6861417.5 | 3708524.5 | 1.18E+07 | 6421159 | 2822790.25 | 1.36E+07 | 9306575 | 9777560 | [illegible] |
| 15537547 | 3 | 0.692 | 1.71E+07 | 1.20E+07 | 1.20E+07 | 2.08E+07 | 7616525 | 1.12E+07 | 1.22E+07 | 8697191 | 1.14E+07 | 1.30E+07 | 1.40E+07 | 1.44E+07 | [illegible] |
| 15548410, 15562084, 15562682, 15563040, 15563312, 15563567 | 2 | 0.574 | 1.26E+07 | 9609634 | 1.42E+07 | 1.14E+07 | 1.07E+07 | 1.07E+07 | 1.27E+07 | 1.03E+07 | 9684792 | 1.23E+07 | 9727163 | 7936244 | [illegible] |
| 15540802 | 4 | 0.025 | 1.81E+07 | 2.16E+07 | 2.26E+07 | 1.21E+07 | 1.67E+07 | 2.18E+07 | 1.64E+07 | 1.97E+07 | 2.09E+07 | 1.13E+07 | 1.33E+07 | 1.16E+07 | [illegible] |
| 15549690 | 2 | 0.191 | 1.07E+07 | 1.12E+07 | 1.35E+07 | 9062929 | 1.02E+07 | 1.21E+07 | 9344198 | 1.14E+07 | 9989608 | 1.01E+07 | 7092247.5 | 9556386 | [illegible] |
| 15558590, 15559159, 15560471 | 2 | 0.504 | 1.01E+07 | 1.86E+07 | 1.39E+07 | 4573587.5 | 1.65E+07 | 1.26E+07 | 1.62E+07 | 1.57E+07 | 1.35E+07 | 9851263 | 1.02E+07 | 1.23E+07 | [illegible] |
| 15556218, 15558670 | 2 | 0.633 | 7714166 | 7527441 | 8164338.5 | 4244391.5 | 7861874.5 | 6148043 | 7386500 | 7208204.5 | 3386820.75 | 9670547 | 2919813.75 | 4702089.5 | [illegible] |
| 15535082 | 3 | 0.42 | 3.55E+07 | 1591096.375 | 1304696.625 | 2.11E+07 | 2511547.25 | 1108441.875 | 2345640.5 | 1968274.5 | 6140623.5 | 9181023 | 4.63E+07 | 1.16E+07 | [illegible] |
| 15552501, 15560651 | 3 | 0.053 | 1.29E+07 | 1.27E+07 | 1.49E+07 | 7494699 | 8781767 | 1.37E+07 | 1.15E+07 | 1.27E+07 | 1.12E+07 | 8631547 | 9438255 | 8251516.5 | [illegible] |
| 15541367 | 4 | 0.656 | 5570690 | 6462650.5 | 5241433 | 3070151 | 5550212 | 3971070.5 | 7529926 | 5431830 | 2962041.5 | 8176951 | 4065597.25 | 4575587 | [illegible] |
| 15555013 | 2 | 0.479 | 1.49E+07 | 1871381.625 | 701194.688 | 2.33E+07 | 67561.172 | 2578132.25 | 4427517.5 | 594594.188 | 4907355 | 7683298.5 | 3.28E+07 | 1.35E+07 | [illegible] |
| 15551635 | 2 | 0.656 | 3.19E+07 | 2.26E+07 | 1.37E+07 | 1.20E+07 | 1.16E+07 | 1.54E+07 | 3.24E+07 | 1.48E+07 | 1.86E+07 | 7044581.5 | 4.42E+07 | 2.95E+07 | [illegible] |
| 15549831, 15557395, 15560816 | 2 | 0.514 | 1.28E+07 | 5488307 | 3985166.5 | 1.82E+07 | 3253161.5 | 4466223.5 | 5055329.5 | 2552329.5 | 4455973.5 | 6408492.5 | 1.52E+07 | 6346535 | [illegible] |
| 15541148 | 3 | 0.704 | 4685382 | 1.11E+07 | 1.54E+07 | 4918893.5 | 1.16E+07 | 1.19E+07 | 7001359 | 1.19E+07 | 8741244 | 5986933.5 | 5350355.5 | 8173324 | [illegible] |
| 15548881 | 2 | 0.527 | 1.39E+07 | 2199672.75 | 1964264.875 | 2.72E+07 | 156048.875 | 3496414.25 | 3153495.25 | 1335503.75 | 7103316 | 5961743 | 3.19E+07 | 1.57E+07 | [illegible] |
| 15537180 | 3 | 0.217 | 6055029 | 9030814 | 1.20E+07 | 2776509.25 | 8926485 | 8369299 | 9003540 | 1.11E+07 | 5317151 | 5159904.5 | 2354522 | 4793718 | [illegible] |
| 15559999 | 2 | 0.132 | 7044215 | 8228208.5 | 1.01E+07 | 5255576.5 | 9177174 | 5199612 | 6551675.5 | 8946985 | 6813514 | 5147344.5 | 5565584 | 5562855 | [illegible] |
| 15548516 | 2 | 0.641 | 2.09E+07 | 4244797 | 600482.625 | 1.29E+07 | 3209476.25 | 2043057.375 | 5136231 | 4815674 | 1.11E+07 | 5097712.5 | 2.21E+07 | 1.24E+07 | [illegible] |
| 15545083 | 3 | 0.47 | 4314075 | 705250.312 | 4.70E-06 | 3930139.5 | 607433.688 | 170547.953 | 1259331.125 | 993466.375 | 2368573.5 | 4718127.5 | 4205900.5 | 1713789.75 | [illegible] |
| 15540097 | 3 | 0.032 | 4783892.5 | 3873606.75 | 3838523.5 | 4079163.75 | 2346150.25 | 2891699.25 | 3173061.25 | 3363586.25 | 3410254 | 4497307 | 4857263 | 4624020.5 | [illegible] |
| 15557763, 15557868, 15557979, 15558141 | 3, 4 | 0.275 | 7334008.5 | 1.68E+07 | 1.19E+07 | 3694193.25 | 2.07E+07 | 9280328 | 1.00E+07 | 1.18E+07 | 4250555.5 | 4090346.75 | 3956568 | 5338284.5 | [illegible] |
| 15537041 | 3 | 0.303 | 4728955 | 2743096 | 3543644 | 4477795.5 | 4040870.25 | 3249351.5 | 3217448.75 | 2645117 | 3033053.25 | 3972297 | 2384643.75 | 2304146.5 | [illegible] |
| 15545317 | 3 | 0.585 | 2209491 | 3477196.75 | 5514489 | 539273.688 | 2580806.75 | 4396857 | 3788337.5 | 3146222.75 | 4164094.75 | 3467179.25 | 1088291.75 | 3673008 | [illegible] |
| 15553845 | 2 | 0.302 | 3377804.75 | 2469324 | 2108341.25 | 3934057.75 | 1101482.125 | 1786445.875 | 2262225.5 | 1110010.25 | 1401355.625 | 2840383.75 | 2740179 | 2575154.25 | [illegible] |
| 15540905 | 2 | 0.117 | 2651595.5 | 5890794 | 6941056 | 1554854 | 5509450 | 3066435.5 | 3173264.25 | 5849022.5 | 3746181.75 | 2763934.25 | 778263.062 | 1596068.125 | [illegible] |
| 15550655 | 2 | 0.451 | 1552209 | 2989126.25 | 7773807.5 | 1643542.375 | 3480715.25 | 5477487.5 | 4726410.5 | 2519083 | 1794042.125 | 2596079.75 | 656939.438 | 1809512.125 | [illegible] |
| 15533762 | 2 | 0.56 | 5149496 | 2001553.125 | 2432834.5 | 8087033.5 | 1145815.5 | 2081918.5 | 2834080.25 | 3933975 | 6088971 | 2423395.5 | 1.34E+07 | 6475844.5 | [illegible] |
| 15555360 | 2 | 0.601 | 3969549.25 | 1658527.25 | 1905250.125 | 2974868 | 981905.188 | 1948057.375 | 2402089 | 955242.438 | 1752069.375 | 2379127.25 | 2505003 | 2367243.25 | [illegible] |
| 15554746 | 3 | 0.267 | 5576346 | 1165847.5 | 727857.812 | 3947247.75 | 905548.688 | 793417.5 | 633407.312 | 867789.812 | 1265131.875 | 2332556.75 | 1.12E+07 | 2264005.25 | [illegible] |
| [illegible] | [illegible] | [illegible] | [illegible] | [illegible] | [illegible] | [illegible] | [illegible] | [illegible] | [illegible] | [illegible] | [illegible] | [illegible] | [illegible] | [illegible] | [illegible] |

| | | | | | | | | | | | | | | | | | | | | |
|---|---|---|---|---|---|---|---|---|---|---|---|---|---|---|---|---|---|---|---|---|
| 15552836 | 2 | 0.803 | 2610809.5 | 2174500.25 | 2389294 | 2990667.75 | 2157102.75 | 1424139.875 | 2437014.25 | 1848398.375 | 2124510.25 | 2975272.75 | 2108414.75 | 2231063 | 2179.1648 | ILATGANVILTTGGIDDMYLK | P11983 | P11983 | TCP1 | t-complex 1 | Cytoplasm | other |
| 15543146 | 2, 3 | 0.931 | 911863.812 | 1190357.25 | 2112994.75 | 774263.062 | 2493456 | 595838.75 | 1416036.375 | 1581396.375 | 596029 | 2109878 | 535778.125 | 762251.25 | 1411.667 | AFHNEAQVNPER | P11983 | P11983 | TCP1 | t-complex 1 | Cytoplasm | other |
| 15563486, 15563566, 15563715 | 2 | 0.299 | 498293.406 | 372273.188 | 235243.734 | 24404.488 | 498894.125 | 21253.744 | 75266.25 | 338622 | 106213.742 | 115347.695 | 265237.219 | 347060.219 | 1539.7627, 1539.7635, 1539.7639 | AFHNEAQVNPERK | P11983 | P11983 | TCP1 | t-complex 1 | Cytoplasm | other |
| 15533511, 15543822, 15545594 | 2, 3 | 0.147 | 5.74E+07 | 9.22E+07 | 2.98E+07 | 5.59E+07 | 5.27E+07 | 6.71E+07 | 3.53E+07 | 5.32E+07 | 4.85E+07 | 2.68E+07 | 3.52E+07 | 3.14E+07 | 1313.7053, 1313.7062, 1313.7069 | HTTIFEVLPEK | Q921I1 | Q921I1 | TF | transferrin | Extracellular Space | transporter |
| 15535007 | 3 | 0.145 | 2.84E+07 | 4.67E+07 | 2.17E+07 | 2.34E+07 | 4.31E+07 | 3.96E+07 | 2.08E+07 | 4.08E+07 | 3.20E+07 | 2.39E+07 | 1.34E+07 | 1.63E+07 | 1656.77 | KPVDQYEDC[160.0307]YLAR | Q921I1 | Q921I1 | TF | transferrin | Extracellular Space | transporter |
| 15533287 | 3 | 0.048 | 4.89E+07 | 9.17E+07 | 3.52E+07 | 4.54E+07 | 6.53E+07 | 5.68E+07 | 3.50E+07 | 8.06E+07 | 6.26E+07 | 2.36E+07 | 2.47E+07 | 2.67E+07 | 2007.9796 | DFASC[160.0307]HLAQAPNHVVVSR | Q921I1 | Q921I1 | TF | transferrin | Extracellular Space | transporter |
| 15549185 | 2 | 0.054 | 3.85E+07 | 6.15E+07 | 2.84E+07 | 3.53E+07 | 3.89E+07 | 5.03E+07 | 2.26E+07 | 5.11E+07 | 4.65E+07 | 2.15E+07 | 1.86E+07 | 1.88E+07 | 1957.8893 | NQQEGVC[160.0307]PEGSIDNSPVK | Q921I1 | Q921I1 | TF | transferrin | Extracellular Space | transporter |
| 15534253 | 3 | 0.037 | 2.84E+07 | 6.57E+07 | 2.68E+07 | 2.99E+07 | 4.78E+07 | 4.90E+07 | 2.70E+07 | 6.11E+07 | 4.36E+07 | 1.92E+07 | 1.29E+07 | 1.67E+07 | 1881.8729 | ADRDQYELLC[160.0307]LDNTR | Q921I1 | Q921I1 | TF | transferrin | Extracellular Space | transporter |
| 15551182 | 2 | 0.056 | 3.41E+07 | 5.87E+07 | 2.68E+07 | 3.53E+07 | 4.57E+07 | 5.55E+07 | 2.90E+07 | 5.61E+07 | 5.72E+07 | 1.91E+07 | 2.33E+07 | 2.28E+07 | 1781.7468 | FDEFFSQGC[160.0307]APGYEK | Q921I1 | Q921I1 | TF | transferrin | Extracellular Space | transporter |
| 15548158 | 2 | 0.079 | 3.14E+07 | 6.50E+07 | 2.94E+07 | 3.24E+07 | 4.54E+07 | 5.17E+07 | 2.34E+07 | 5.34E+07 | 4.66E+07 | 1.88E+07 | 1.97E+07 | 2.13E+07 | 1813.7427 | IEC[160.0307]ESAETTEDC[160.0307]IEK | Q921I1 | Q921I1 | TF | transferrin | Extracellular Space | transporter |
| 15540616 | 2 | 0.065 | 3.07E+07 | 6.40E+07 | 2.75E+07 | 3.98E+07 | 3.98E+07 | 5.29E+07 | 2.32E+07 | 5.16E+07 | 4.61E+07 | 1.84E+07 | 1.89E+07 | 2.17E+07 | 1238.6396 | DFQLFSSPLGK | Q921I1 | Q921I1 | TF | transferrin | Extracellular Space | transporter |
| 15531987 | 2 | 0.04 | 2.92E+07 | 5.08E+07 | 2.43E+07 | 3.52E+07 | 3.61E+07 | 4.49E+07 | 2.22E+07 | 4.06E+07 | 4.07E+07 | 1.68E+07 | 1.64E+07 | 2.11E+07 | 1349.6666 | ASDTSITWNNLK | Q921I1 | Q921I1 | TF | transferrin | Extracellular Space | transporter |
| 15532007 | 2 | 0.069 | 2.98E+07 | 5.08E+07 | 2.24E+07 | 2.64E+07 | 3.36E+07 | 4.73E+07 | 2.05E+07 | 3.96E+07 | 3.73E+07 | 1.47E+07 | 1.55E+07 | 1.94E+07 | 1323.5858 | C[160.0307]DEWSIISEGK | Q921I1 | Q921I1 | TF | transferrin | Extracellular Space | transporter |
| 15541470 | 3 | 0.234 | 1.72E+07 | 4.58E+07 | 2.33E+07 | 1.70E+07 | 2.65E+07 | 3.32E+07 | 1.58E+07 | 3.95E+07 | 3.77E+07 | 1.46E+07 | 1.13E+07 | 1.76E+07 | 1419.7711 | LYLGHNYVTAIR | Q921I1 | Q921I1 | TF | transferrin | Extracellular Space | transporter |
| 15536135 | 2 | 0.126 | 2.16E+07 | 5.12E+07 | 2.85E+07 | 1.96E+07 | 3.34E+07 | 5.45E+07 | 2.20E+07 | 4.46E+07 | 4.49E+07 | 1.41E+07 | 1.36E+07 | 1.99E+07 | 1158.6384 | EDLIWEILK | Q921I1 | Q921I1 | TF | transferrin | Extracellular Space | transporter |
| 15537588 | 2 | 0.073 | 2.03E+07 | 3.27E+07 | 1.40E+07 | 2.14E+07 | 2.12E+07 | 2.58E+07 | 1.28E+07 | 2.30E+07 | 2.31E+07 | 1.11E+07 | 1.25E+07 | 1.15E+07 | 964.4716 | DGGGDVAFVK | Q921I1 | Q921I1 | TF | transferrin | Extracellular Space | transporter |
| 15531703 | 2 | 0.085 | 2.74E+07 | 3.66E+07 | 1.68E+07 | 2.36E+07 | 2.08E+07 | 3.39E+07 | 1.50E+07 | 2.69E+07 | 3.02E+07 | 1.08E+07 | 1.42E+07 | 1.46E+07 | 1122.5768 | DLLFRDDTK | Q921I1 | Q921I1 | TF | transferrin | Extracellular Space | transporter |
| 15548058 | 2 | 0.529 | 1.64E+07 | 2.14E+07 | 9939735 | 1.51E+07 | 2.19E+07 | 2.07E+07 | 1.17E+07 | 2.10E+07 | 3.07E+07 | 1.07E+07 | 1.87E+07 | 1.18E+07 | 2230.0261 | AVSSFFSGSC[160.0307]VPC[160.0307]ADPVAFPK | Q921I1 | Q921I1 | TF | transferrin | Extracellular Space | transporter |
| 15531682 | 2 | 0.148 | 1.60E+07 | 3.56E+07 | 1.23E+07 | 1.46E+07 | 1.99E+07 | 2.46E+07 | 9876326 | 2.38E+07 | 2.36E+07 | 9023847 | 8701955 | 1.01E+07 | 969.5385 | GYYAVAVVK | Q921I1 | Q921I1 | TF | transferrin | Extracellular Space | transporter |
| 15549734, 15552259 | 2, 3 | 0.433 | 1.10E+07 | 2.09E+07 | 9759314 | 1.27E+07 | 1.46E+07 | 1.99E+07 | 8807748 | 2.09E+07 | 2.13E+07 | 8006892.5 | 1.30E+07 | 9428643 | 2552.174, 2552.177 | AVLTSQETLFGGSDC[160.0307]TGNFC[160.0307]LFK | Q921I1 | Q921I1 | TF | transferrin | Extracellular Space | transporter |
| 15551000 | 2 | 0.129 | 7992475.5 | 1.24E+07 | 6409461.5 | 5975294.5 | 9883626 | 7864012.5 | 5790051 | 1.06E+07 | 7667814 | 6760157 | 2998782.75 | 4232032.5 | 1656.7699 | KPVDQYEDC[160.0307]YLAR | Q921I1 | Q921I1 | TF | transferrin | Extracellular Space | transporter |
| 15548207 | 2 | 0.121 | 7321669 | 2.00E+07 | 8744193 | 6327685 | 1.84E+07 | 1.66E+07 | 7012149 | 1.83E+07 | 1.40E+07 | 6075298.5 | 3654165.5 | 4926863 | 1579.7013 | QEDFELLC[160.0307]PDGTR | Q921I1 | Q921I1 | TF | transferrin | Extracellular Space | transporter |
| 15533579 | 2 | 0.046 | 9926999 | 1.89E+07 | 1.02E+07 | 1.09E+07 | 1.13E+07 | 1.82E+07 | 6986492 | 1.71E+07 | 1.64E+07 | 6001432.5 | 4551972.5 | 6079024 | 878.47076 | DSAFGLLR | Q921I1 | Q921I1 | TF | transferrin | Extracellular Space | transporter |
| 15538964 | 2 | 0.317 | 1.18E+07 | 1.40E+07 | 4972478.5 | 8741724 | 1.20E+07 | 1.19E+07 | 7937164 | 1.41E+07 | 1.37E+07 | 5870899 | 8736892 | 5974881 | 1118.5637 | LLEAC[160.0307]TFHK | Q921I1 | Q921I1 | TF | transferrin | Extracellular Space | transporter |
| 15537017 | 2 | 0.053 | 6520170.5 | 2.11E+07 | 1.49E+07 | 7048077 | 1.53E+07 | 1.63E+07 | 8945675 | 1.92E+07 | 1.04E+07 | 5404500.5 | 3224413 | 4745795 | 1349.667 | GTDFQLNQLEGK | Q921I1 | Q921I1 | TF | transferrin | Extracellular Space | transporter |
| 15556974 | 2 | 0.095 | 8010957.5 | 1.40E+07 | 5558857 | 9538234 | 1.14E+07 | 1.36E+07 | 5910649.5 | 1.16E+07 | 1.40E+07 | 5238090.5 | 4778458.5 | 5685588.5 | 1623.7738 | TAGWNIPMGMLYNR | Q921I1 | Q921I1 | TF | transferrin | Extracellular Space | transporter |
| 15538165 | 3 | 0.763 | 5272451.5 | 1.22E+07 | 3463139 | 6438149 | 8126059.5 | 7991373.5 | 4160682.25 | 1.48E+07 | 1.16E+07 | 4678880.5 | 7629354 | 7261534.5 | 2230.0269 | AVSSFFSGSC[160.0307]VPC[160.0307]ADPVAFPK | Q921I1 | Q921I1 | TF | transferrin | Extracellular Space | transporter |
| 15553049 | 4 | 0.139 | 9301494 | 2.61E+07 | 8210823.5 | 9191526 | 1.25E+07 | 1.59E+07 | 7891783.5 | 1.46E+07 | 1.64E+07 | 4433117.5 | 6585625.5 | 6862327.5 | 5129.373 | IVNGEADAMTLDGGHAYIAGQC[160.0307]GLVPVMAEYYESSNC[160.0307]AIPSQQGIFPK | Q921I1 | Q921I1 | TF | transferrin | Extracellular Space | transporter |
| 15532775 | 2 | 0.068 | 9960418 | 1.71E+07 | 6498712 | 1.25E+07 | 1.10E+07 | 1.43E+07 | 6887449.5 | 1.36E+07 | 1.32E+07 | 4320020 | 6163546.5 | 6329427 | 1306.5677 | EEYNGYTGAFR | Q921I1 | Q921I1 | TF | transferrin | Extracellular Space | transporter |
| 15538796 | 3 | 0.069 | 5694215.5 | 1.15E+07 | 5865232.5 | 7966014 | 7796650.5 | 1.09E+07 | 4206360.5 | 1.12E+07 | 9272137 | 3690952.5 | 3489222.75 | 4252935 | 1957.8884 | NQQEGVC[160.0307]PEGSIDNSPVK | Q921I1 | Q921I1 | TF | transferrin | Extracellular Space | transporter |
| 15535174 | 3 | 0.078 | 4460377 | 1.48E+07 | 1.30E+07 | 4156856 | 1.18E+07 | 1.52E+07 | 6751767.5 | 1.39E+07 | 6776327 | 3414130 | 2376015.25 | 3284591.5 | 1707.7965 | QEDFELLC[160.0307]PDGTRK | Q921I1 | Q921I1 | TF | transferrin | Extracellular Space | transporter |
| 15550399 | 3 | 0.546 | 4706179.5 | 3745181 | 1209803.5 | 2943450.75 | 3268984.75 | 3188475.75 | 2920176 | 5791089.5 | 7718833.5 | 3309004.75 | 1.92E+07 | 2792458.75 | 2741.1907 | LC[160.0307]QLC[160.0307]PGC[160.0307]GC[160.0307]SSTQPFFGYVGAFK | Q921I1 | Q921I1 | TF | transferrin | Extracellular Space | transporter |
| 15532544 | 2 | 0.37 | 6622060.5 | 8129758 | 3348027.75 | 4981388.5 | 6768414.5 | 8076547.5 | 2729625.5 | 8098541.5 | 7381420.5 | 2619226 | 4736924.5 | 3401378.75 | 1360.5918 | WC[160.0307]AVSEHENTK | Q921I1 | Q921I1 | TF | transferrin | Extracellular Space | transporter |
| 15552686 | 2 | 0.025 | 5748148.5 | 8964530 | 3615730.25 | 5257796 | 6919890 | 3561416.25 | 4020610.25 | 7852092 | 6566150.5 | 2433430.75 | 2271919.5 | 2216344 | 1881.8723 | ADRDQYELLC[160.0307]LDNTR | Q921I1 | Q921I1 | TF | transferrin | Extracellular Space | transporter |
| 15538097 | 2, 3 | 0.382 | 1.07E+07 | 3.22E+07 | 1.91E+07 | 3531444.25 | 2.39E+07 | 3.37E+07 | 1.00E+07 | 3.13E+07 | 3.44E+07 | 2182385.25 | 8216547.5 | 1.41E+07 | 1171.5638, 1171.5654 | WC[160.0307]ALSHLER | Q921I1 | Q921I1 | TF | transferrin | Extracellular Space | transporter |
| 15533311 | 2 | 0.646 | 4338861.5 | 5551160 | 411433.375 | 4897439.5 | 4204799 | 4281794 | 1605658.5 | 4893440.5 | 4614210 | 1956895.375 | 2362472.25 | 2431159 | 1477.7616 | KGTDFQLNQLEGK | Q921I1 | Q921I1 | TF | transferrin | Extracellular Space | transporter |
| 15551366 | 2 | 0.047 | 2445827.75 | 3895645.25 | 1567340.375 | 2049729.25 | 2675921.25 | 2286854.25 | 1274902.875 | 2932282.25 | 2636781 | 965556.375 | 1133066.375 | 1099618 | 2007.9785 | DFASC[160.0307]HLAQAPNHVVVSR | Q921I1 | Q921I1 | TF | transferrin | Extracellular Space | transporter |
| 15540954 | 2 | 0.461 | 476077.344 | 3183726.25 | 2729843.25 | 44208.539 | 3289968.5 | 3817770 | 179030.141 | 3995775 | 4531363 | 4.70E-06 | 572623.25 | 684514.062 | 971.50275 | LPEGTTPEK | Q921I1 | Q921I1 | TF | transferrin | Extracellular Space | transporter |
| 15534193 | 3 | 0.051 | 6.42E+07 | 1.30E+08 | 5.00E+07 | 7.25E+07 | 7.44E+07 | 9.12E+07 | 4.56E+07 | 8.68E+07 | 8.34E+07 | 4.18E+07 | 3.50E+07 | 3.02E+07 | 1593.8026 | VEYHFLSPYVSPR | Q62351 | Q62351 | TFRC | transferrin receptor | Plasma Membrane | transporter |
| 15534131 | 3 | 0.073 | 7.71E+07 | 1.66E+08 | 6.49E+07 | 8.46E+07 | 7.64E+07 | 1.01E+08 | 5.76E+07 | 1.23E+08 | 1.10E+08 | 4.15E+07 | 4.65E+07 | 4.70E+07 | 1737.8528 | GYEEPDRYVVVGAQR | Q62351 | Q62351 | TFRC | transferrin receptor | Plasma Membrane | transporter |
| 15531308 | 2 | 0.103 | 4.78E+07 | 9.52E+07 | 4.22E+07 | 6.62E+07 | 5.28E+07 | 6.93E+07 | 4.19E+07 | 7.27E+07 | 6.35E+07 | 3.94E+07 | 3.30E+07 | 3.17E+07 | 1207.6025 | QLSQNTYTPR | Q62351 | Q62351 | TFRC | transferrin receptor | Plasma Membrane | transporter |
| 15531336 | 3 | 0.102 | 5.94E+07 | 1.55E+08 | 7.37E+07 | 6.45E+07 | 8.78E+07 | 1.10E+08 | 4.83E+07 | 1.01E+08 | 1.07E+08 | 3.74E+07 | 3.43E+07 | 5.16E+07 | 2032.9254 | DESLAYYIENQFHEFK | Q62351 | Q62351 | TFRC | transferrin receptor | Plasma Membrane | transporter |
| 15531276 | 2 | 0.119 | 5.52E+07 | 1.42E+08 | 5.28E+07 | 8.98E+07 | 5.92E+07 | 1.01E+08 | 5.26E+07 | 9.34E+07 | 9.10E+07 | 3.54E+07 | 3.87E+07 | 4.54E+07 | 1379.7584 | VSASPLLYTLMGK | Q62351 | Q62351 | TFRC | transferrin receptor | Plasma Membrane | transporter |
| 15547711 | 2 | 0.071 | 6.54E+07 | 1.36E+08 | 5.67E+07 | 7.67E+07 | 6.05E+07 | 9.01E+07 | 3.92E+07 | 9.16E+07 | 8.97E+07 | 3.50E+07 | 3.22E+07 | 3.79E+07 | 1745.8452 | SAFSNLFGGEPLSYTR | Q62351 | Q62351 | TFRC | transferrin receptor | Plasma Membrane | transporter |
| 15531451 | 2 | 0.057 | 5.31E+07 | 1.06E+08 | 4.78E+07 | 7.13E+07 | 5.50E+07 | 7.47E+07 | 3.89E+07 | 7.75E+07 | 7.54E+07 | 3.49E+07 | 3.26E+07 | 2.93E+07 | 1172.625 | LELSQNQNVK | Q62351 | Q62351 | TFRC | transferrin receptor | Plasma Membrane | transporter |
| 15537158 | 2 | 0.08 | 5.60E+07 | 1.21E+08 | 5.12E+07 | 6.28E+07 | 6.60E+07 | 9.71E+07 | 4.32E+07 | 8.22E+07 | 8.80E+07 | 3.35E+07 | 3.71E+07 | 3.60E+07 | 1250.6606 | LNSIEFADTIK | Q62351 | Q62351 | TFRC | transferrin receptor | Plasma Membrane | transporter |
| 15547701 | 2 | 0.022 | 5.79E+07 | 1.22E+08 | 5.23E+07 | 7.85E+07 | 6.58E+07 | 6.84E+07 | 4.39E+07 | 8.30E+07 | 8.38E+07 | 3.32E+07 | 2.99E+07 | 2.86E+07 | 1568.672 | SETMETEDVPTSSR | Q62351 | Q62351 | TFRC | transferrin receptor | Plasma Membrane | transporter |
| 15538723 | 2 | 0.159 | 5.13E+07 | 1.05E+08 | 4.30E+07 | 5.96E+07 | 6.66E+07 | 5.16E+07 | 3.60E+07 | 8.58E+07 | 7.43E+07 | 3.26E+07 | 3.66E+07 | 3.18E+07 | 1238.6448 | LAQVFSDMISK | Q62351 | Q62351 | TFRC | transferrin receptor | Plasma Membrane | transporter |
| 15532168 | 2 | 0.047 | 5.20E+07 | 1.06E+08 | 4.90E+07 | 6.66E+07 | 5.60E+07 | 7.56E+07 | 4.03E+07 | 8.46E+07 | 7.02E+07 | 3.13E+07 | 2.88E+07 | 3.45E+07 | 1181.6019 | LDTYEALTQK | Q62351 | Q62351 | TFRC | transferrin receptor | Plasma Membrane | transporter |
| 15547867 | 2 | 0.009 | 4.34E+07 | 8.94E+07 | 4.19E+07 | 6.13E+07 | 4.80E+07 | 6.01E+07 | 4.18E+07 | 6.67E+07 | 6.62E+07 | 2.66E+07 | 2.21E+07 | 2.16E+07 | 1563.6566 | LAADEEENADNNMK | Q62351 | Q62351 | TFRC | transferrin receptor | Plasma Membrane | transporter |
| 15548060, 15548287 | 3, 4 | 0.558 | 5.27E+07 | 8.69E+07 | 1.86E+07 | 7.23E+07 | 4.08E+07 | 5.19E+07 | 2.96E+07 | 6.27E+07 | 7.27E+07 | 2.56E+07 | 3.50E+07 | 3.42E+07 | 3965.9343, 3965.9375 | SSIGQNMVTIVQSNGNLDPVESPEGYVAFSKPTEVSGK | Q62351 | Q62351 | TFRC | transferrin receptor | Plasma Membrane | transporter |
| 15533973 | 2 | 0.083 | 4.06E+07 | 7.29E+07 | 3.34E+07 | 5.09E+07 | 3.49E+07 | 5.72E+07 | 2.92E+07 | 5.70E+07 | 5.25E+07 | 2.41E+07 | 2.46E+07 | 2.50E+07 | 1084.5908 | VPQLNQMVR | Q62351 | Q62351 | TFRC | transferrin receptor | Plasma Membrane | transporter |
| 15538331 | 2 | 0.263 | 2.66E+07 | 5.41E+07 | 2.12E+07 | 3.00E+07 | 2.53E+07 | 3.89E+07 | 2.16E+07 | 3.20E+07 | 3.51E+07 | 2.28E+07 | 1.88E+07 | 1.73E+07 | 908.48584 | LYWADLK | Q62351 | Q62351 | TFRC | transferrin receptor | Plasma Membrane | transporter |
| 15535478 | 2 | 0.042 | 3.82E+07 | 8.46E+07 | 3.66E+07 | 5.06E+07 | 4.23E+07 | 4.91E+07 | 2.97E+07 | 6.08E+07 | 6.51E+07 | 2.22E+07 | 2.09E+07 | 2.32E+07 | 1084.5654 | AFTYINLDK | Q62351 | Q62351 | TFRC | transferrin receptor | Plasma Membrane | transporter |
| 15532767 | 2 | 0.046 | 3.69E+07 | 9.10E+07 | 3.86E+07 | 5.14E+07 | 4.32E+07 | 5.83E+07 | 3.01E+07 | 6.40E+07 | 6.39E+07 | 2.15E+07 | 2.33E+07 | 2.21E+07 | 964.5443 | VVLGTSNFK | Q62351 | Q62351 | TFRC | transferrin receptor | Plasma Membrane | transporter |
| 15531343 | 2 | 0.078 | 3.63E+07 | 8.49E+07 | 3.52E+07 | 5.24E+07 | 4.15E+07 | 6.25E+07 | 2.47E+07 | 5.99E+07 | 6.92E+07 | 1.91E+07 | 2.01E+07 | 2.50E+07 | 1016.648 | ILNIFGVIK | Q62351 | Q62351 | TFRC | transferrin receptor | Plasma Membrane | transporter |
| 15547942 | 2 | 0.018 | 3.39E+07 | 6.93E+07 | 3.42E+07 | 3.61E+07 | 3.80E+07 | 4.48E+07 | 2.55E+07 | 4.47E+07 | 4.89E+07 | 1.71E+07 | 1.69E+07 | 2.03E+07 | 2032.9238 | DESLAYYIENQFHEFK | Q62351 | Q62351 | TFRC | transferrin receptor | Plasma Membrane | transporter |
| 15534021 | 2 | 0.134 | 2.63E+07 | 4.83E+07 | 1.90E+07 | 2.99E+07 | 2.81E+07 | 3.34E+07 | 1.94E+07 | 3.91E+07 | 3.81E+07 | 1.71E+07 | 1.83E+07 | 1.55E+07 | 1009.439 | WNIDSSC[160.0307]K | Q62351 | Q62351 | TFRC | transferrin receptor | Plasma Membrane | transporter |
| 15542375, 15544885, 15545038 | 3 | 0.04 | 2.68E+07 | 4.88E+07 | 2.30E+07 | 3.45E+07 | 2.61E+07 | 3.40E+07 | 2.11E+07 | 3.67E+07 | 3.77E+07 | 1.64E+07 | 1.70E+07 | 1.47E+07 | 1231.619, 1231.6191, 1231.6193 | VWRDEHYVK | Q62351 | Q62351 | TFRC | transferrin receptor | Plasma Membrane | transporter |
| 15534453 | 2 | 0.043 | 3.22E+07 | 7.56E+07 | 3.51E+07 | 4.82E+07 | 3.89E+07 | 6.15E+07 | 2.63E+07 | 5.12E+07 | 5.82E+07 | 1.55E+07 | 1.92E+07 | 2.42E+07 | 974.5867 | SSVGTGLLLK | Q62351 | Q62351 | TFRC | transferrin receptor | Plasma Membrane | transporter |
| 15538286 | 3, 4 | 0.325 | 2.05E+07 | 6.34E+07 | 2.39E+07 | 1.22E+07 | 4.13E+07 | 5.27E+07 | 2.37E+07 | 5.27E+07 | 5.47E+07 | 1.31E+07 | 1.22E+07 | 2.59E+07 | 2096.0876, 2096.089 | HIFWGSGSHTLSALVENLK | Q62351 | Q62351 | TFRC | transferrin receptor | Plasma Membrane | transporter |
| 15531701 | 2 | 0.108 | 1.62E+07 | 3.16E+07 | 1.30E+07 | 1.69E+07 | 1.44E+07 | 2.13E+07 | 1.10E+07 | 2.16E+07 | 2.13E+07 | 1.04E+07 | 9526707 | 9001165 | 1175.5664 | LTTDFHNAEK | Q62351 | Q62351 | TFRC | transferrin receptor | Plasma Membrane | transporter |
| 15540640 | 3 | 0.21 | 1.05E+07 | 2.00E+07 | 1.50E+07 | 9757810 | 1.48E+07 | 1.33E+07 | 1.04E+07 | 1.65E+07 | 1.49E+07 | 1.00E+07 | 8943960 | 1.01E+07 | 2099.955 | LTHDVELNLDYEM[147.0354]YNSK | Q62351 | Q62351 | TFRC | transferrin receptor | Plasma Membrane | transporter |
| 15557052 | 2 | 0.175 | 1.06E+07 | 2.63E+07 | 1.36E+07 | 1.39E+07 | 1.15E+07 | 1.85E+07 | 8075319 | 1.68E+07 | 2.09E+07 | 8889149 | 6843728.5 | 8238176 | 1954.0051 | VANAQSFNAIGVLIYMDK | Q62351 | Q62351 | TFRC | transferrin receptor | Plasma Membrane | transporter |
| 15553196 | 2 | 0.043 | 1.47E+07 | 2.84E+07 | 1.17E+07 | 1.65E+07 | 1.48E+07 | 1.64E+07 | 1.01E+07 | 1.68E+07 | 1.74E+07 | 8621770 | 7929391 | 7897225.5 | 1737.853 | GYEEPDRYVVVGAQR | Q62351 | Q62351 | TFRC | transferrin receptor | Plasma Membrane | transporter |
| 15551378 | 2 | 0.045 | 1.61E+07 | 2.81E+07 | 1.18E+07 | 1.47E+07 | 1.58E+07 | 1.65E+07 | 1.27E+07 | 1.96E+07 | 1.99E+07 | 7533637.5 | 1.06E+07 | 8531642 | 1593.8041 | VEYHFLSPYVSPR | Q62351 | Q62351 | TFRC | transferrin receptor | Plasma Membrane | transporter |
| 15549263 | 3 | 0.046 | 6567488 | 3.32E+07 | 2.28E+07 | 7753453 | 1.20E+07 | 2.01E+07 | 7971492 | 2.25E+07 | 1.23E+07 | 5560156.5 | 2205688 | 3435905.5 | 3267.5835 | SSIGQNMVTIVQSNGNLDPVESPEGYVAFSK | Q62351 | Q62351 | TFRC | transferrin receptor | Plasma Membrane | transporter |
| 15545196, 15545526 | 2 | 0.056 | 4203581 | 8361968.5 | 3813969.5 | 5094707 | 4789362.5 | 6106396.5 | 3133192.25 | 6073325 | 6061619 | 2893415.5 | 2529319 | 2404025.5 | 849.4088, 849.4107 | DSNWISK | Q62351 | Q62351 | TFRC | transferrin receptor | Plasma Membrane | transporter |
| 15545042 | 2 | 0.032 | 3722461.5 | 7339699.5 | 3063723 | 4280272 | 3158053 | 3647535.5 | 2682244.25 | 4076799.25 | 4963400 | 1837945.25 | 2225625.5 | 1524803.75 | 1213.712 | TAAEVAGQLIIK | Q62351 | Q62351 | TFRC | transferrin receptor | Plasma Membrane | transporter |
| 15550672 | 2 | 0.002 | 2310438.75 | 4594391 | 2247024.25 | 3420251 | 2595682 | 3839703.75 | 1999949.125 | 2640126.5 | 2792806.75 | 1196992.5 | 978427.125 | 847927.875 | 1584.666 | SETM[147.0354]ETEDVPTSSR | Q62351 | Q62351 | TFRC | transferrin receptor | Plasma Membrane | transporter |
| 15539356 | 2 | 0.053 | 1803300.125 | 3010421.25 | 1884830.5 | 2530696 | 1055523.75 | 2127760.75 | 1041713.812 | 2314153.5 | 2016153 | 1167238.5 | 444170.094 | 742086 | 1395.7522 | VSASPLLYTLM[147.0354]GK | Q62351 | Q62351 | TFRC | transferrin receptor | Plasma Membrane | transporter |
| 15541102 | 3 | 0.124 | 2318542.25 | 5720937 | 1203352.75 | 3028731.75 | 2884374.75 | 3521845.5 | 1773038.125 | 5086538.5 | 4274714 | 1050504.875 | 946386.312 | 1707782.125 | 1745.846 | SAFSNLFGGEPLSYTR | Q62351 | Q62351 | TFRC | transferrin receptor | Plasma Membrane | transporter |
| 15543805 | 3 | 0.162 | 1319961.375 | 4426362 | 2388383.5 | 2347165.25 | 2099093.5 | 3543224.75 | 1897624.75 | 3248709.5 | 4066404.5 | 918363.688 | 1256595.125 | 1858449.375 | 1954.0067 | VANAQSFNAIGVLIYMDK | Q62351 | Q62351 | TFRC | transferrin receptor | Plasma Membrane | transporter |
| 15556961 | 2 | 0.148 | 1347771.75 | 5656664.5 | 3240797.5 | 1195174.625 | 2518594.75 | 5535255 | 1246013 | 4580589.5 | 3189273.25 | 877151.688 | 655965.375 | 1147582.625 | 1555.7551 | DM[147.0354]GLSLQWLYSAR | Q62351 | Q62351 | TFRC | transferrin receptor | Plasma Membrane | transporter |
| 15563627 | 2 | 0.044 | 103382.984 | 143858.688 | 112595.812 | 111171.867 | 133937.984 | 258336.609 | 70753.289 | 245624.609 | 205472.281 | 67594.93 | 7431.859 | 41028.578 | 1579.6512 | LAADEEENADNNM[147.0354]K | Q62351 | Q62351 | TFRC | transferrin receptor | Plasma Membrane | transporter |
| 15541287 | 3 | 0.968 | 1.39E+07 | 1.98E+07 | 2.06E+07 | 1.63E+07 | 1.85E+07 | 2.04E+07 | 2.25E+07 | 1.67E+07 | 1.55E+07 | 2.30E+07 | 1.62E+07 | 1.86E+07 | 1377.7352 | QYWLITAFHAK | Q62312 | Q62312 | TGFBR2 | transforming growth factor, beta receptor II (70/80kDa) | Plasma Membrane | kinase |
| 15552368 | 3 | 0.348 | 7175283 | 5579473 | 3869522 | 6425449 | 5569592.5 | 6379221.5 | 5305987.5 | 4090280.5 | 3837344.25 | 1.19E+07 | 4311795 | 6836487.5 | 2497.208 | VVTNYNSAHDQNSNLLIEYFR | P21981 | P21981 | TGM2 | transglutaminase 2 | Cytoplasm | enzyme |
| 15550291 | 2 | 0.705 | 1779123.125 | 2690501 | 2280378.25 | 1367709.875 | 1411902.75 | 2356146 | 1869109.875 | 1340907.625 | 794479.812 | 4738098 | 631841.875 | 1540671.25 | 1729.9463 | GLLIEPAANSYLLAER | P21981 | P21981 | TGM2 | transglutaminase 2 | Cytoplasm | enzyme |
| 15532434 | 2 | 0.252 | 2034084.75 | 8710481 | 1.45E+07 | 3.27E+07 | 3.42E+07 | 2.78E+07 | 1.70E+07 | 2.05E+07 | 5.32E+07 | 982663.375 | 2.13E+07 | 2.56E+07 | 1394.729 | FVFGTTPEDILR | P35441 | P35441 | THBS1 | thrombospondin 1 | Extracellular Space | other |
| 15532531 | 2 | 0.253 | 1078262.125 | 5571216 | 9012155 | 1.69E+07 | 1.73E+07 | 1.68E+07 | 1.09E+07 | 1.21E+07 | 2.79E+07 | 770665.375 | 1.34E+07 | 1.32E+07 | 1271.7181 | AGTLDLSLSLPGK | P35441 | P35441 | THBS1 | thrombospondin 1 | Extracellular Space | other |
| 15534527 | 2 | 0.234 | 619057.188 | 3417923.75 | 6745747 | 1.13E+07 | 1.22E+07 | 1.22E+07 | 5901140 | 9007582 | 1.94E+07 | 527183.875 | 8808306 | 7365265 | 1207.6288 | SITLFVQEDR | P35441 | P35441 | THBS1 | thrombospondin 1 | Extracellular Space | other |
| 15550293 | 2 | 0.291 | 1155258.75 | 6582299.5 | 1.10E+07 | 2.44E+07 | 2.34E+07 | 2.11E+07 | 1.12E+07 | 1.44E+07 | 3.86E+07 | 520716.688 | 1.69E+07 | 1.64E+07 | 1835.9167 | MESAELDVPIQSIFTR | P35441 | P35441 | THBS1 | thrombospondin 1 | Extracellular Space | other |
| 15539134 | 2 | 0.225 | 570881.688 | 3121499.75 | 4874971.5 | 1.10E+07 | 1.14E+07 | 1.07E+07 | 5684744.5 | 7543966 | 2.01E+07 | 473251.562 | 8751512 | 8176256.5 | 1125.5223 | AQLYIDC[160.0307]DK | P35441 | P35441 | THBS1 | thrombospondin 1 | Extracellular Space | other |
| 15542820, 15546234 | 2 | 0.257 | 956770.188 | 2312039.75 | 3429119.5 | 7902116.5 | 6391507 | 6088648.5 | 3943905.5 | 4330713.5 | 1.11E+07 | 452276.906 | 5598659 | 5244507.5 | 1465.7246, 1465.7253 | DNTGQIFSVVSNGK | P35441 | P35441 | THBS1 | thrombospondin 1 | Extracellular Space | other |
| 15536797 | 2 | 0.195 | 664712.688 | 2471151.75 | 3543245 | 8923198 | 1.09E+07 | 1.06E+07 | 4274667 | 6077957 | 1.72E+07 | 398249 | 8447560 | 5863098 | 996.48865 | FTGSQPFGR | P35441 | P35441 | THBS1 | thrombospondin 1 | Extracellular Space | other |
| 15537025 | 3 | 0.233 | 864164.938 | 4091584.25 | 7068927 | 1.69E+07 | 1.98E+07 | 1.72E+07 | 9031624 | 9137486 | 2.74E+07 | 362813.938 | 1.31E+07 | 1.03E+07 | 1545.8087 | VTEENRELVSELK | P35441 | P35441 | THBS1 | thrombospondin 1 | Extracellular Space | other |
| 15533907 | 2 | 0.213 | 506656.062 | 4138354.25 | 6507703 | 1.36E+07 | 1.69E+07 | 1.41E+07 | 6810261 | 1.04E+07 | 2.44E+07 | 321261.688 | 8294722 | 8348640.5 | 1077.531 | GQDLSSPAFR | P35441 | P35441 | THBS1 | thrombospondin 1 | Extracellular Space | other |
| 15532353 | 2 | 0.203 | 738306.5 | 3273720.25 | 5127807.5 | 1.33E+07 | 1.69E+07 | 1.47E+07 | 7560048 | 8985524 | 2.10E+07 | 277398.094 | 1.13E+07 | 7826356.5 | 1372.6583 | GTSQNDPNWVVR | P35441 | P35441 | THBS1 | thrombospondin 1 | Extracellular Space | other |
| 15557186, 15558714 | 2 | 0.156 | 436881.688 | 2143945.5 | 3462684.75 | 1.05E+07 | 1.09E+07 | 1.05E+07 | 5773795 | 6959517.5 | 1.55E+07 | 266678.562 | 8359468 | 5394008.5 | 1660.7174, 1660.719 | NTDPGYNC[160.0307]LPC[160.0307]PPR | P35441 | P35441 | THBS1 | thrombospondin 1 | Extracellular Space | other |
| 15558127 | 2 | 0.126 | 472329.312 | 885511.438 | 1250502.625 | 2908940.25 | 3384894.25 | 2958061.5 | 1492723.5 | 2294802 | 4709832 | 264726.75 | 2545412.75 | 1613439.5 | 1550.7804 | NALWHTGNTPGQVR | P35441 | P35441 | THBS1 | thrombospondin 1 | Extracellular Space | other |
| 15549069 | 3 | 0.137 | 1992844.125 | 1.13E+07 | 1.23E+07 | 3.24E+07 | 4.98E+07 | 3.82E+07 | 1.89E+07 | 2.59E+07 | 5.29E+07 | 260687.938 | 2.05E+07 | 8135643.5 | 3255.326 | DDFDHDNVPDIDDIC[160.0307]PENFDISETDFR | P35441 | P35441 | THBS1 | thrombospondin 1 | Extracellular Space | other |
| 15548742 | 2 | 0.246 | 515371.219 | 3782706.75 | 6146049 | 1.56E+07 | 1.65E+07 | 1.48E+07 | 7521500.5 | 8751742 | 2.20E+07 | 239455.359 | 1.23E+07 | 7598608.5 | 1637.687 | DC[160.0307]VGDVTENQVC[160.0307]NK | P35441 | P35441 | THBS1 | thrombospondin 1 | Extracellular Space | other |
| 15533867 | 2 | 0.198 | 286450.875 | 1548209.625 | 3357546 | 7791850.5 | 8708553 | 9086602 | 4369309.5 | 5272175 | 1.31E+07 | 191506.594 | 6329810.5 | 5371580 | 1009.5304 | AQGYSGLSVK | P35441 | P35441 | THBS1 | thrombospondin 1 | Extracellular Space | other |
| 15541175 | 2 | 0.197 | 255136.594 | 2375196.5 | 2090906.75 | 1.04E+07 | 1.05E+07 | 9094815 | 5645112 | 5202349 | 1.06E+07 | 122902.953 | 7689858.5 | 5335626 | 1209.579 | IMADSGPIYDK | P35441 | P35441 | THBS1 | thrombospondin 1 | Extracellular Space | other |
| 15551261 | 2 | 0.16 | 136952.047 | 804983.062 | 1099911 | 3848896.75 | 3106719.25 | 3577613.75 | 1705002 | 1850765.125 | 5154289.5 | 113668.367 | 2804286.5 | 1646001.875 | 1545.8093 | VTEENRELVSELK | P35441 | P35441 | THBS1 | thrombospondin 1 | Extracellular Space | other |
| 15553004 | 2 | 0.3 | 373267.281 | 2996705.25 | 3894903.5 | 8413181 | 9993812 | 1.01E+07 | 5167261 | 6336636 | 1.69E+07 | 90801.422 | 7707769.5 | 5376754.5 | 1595.742 | QVTQSYWDTNPTR | P35441 | P35441 | THBS1 | thrombospondin 1 | Extracellular Space | other |
| 15552937 | 3 | 0.268 | 0 | 1127952.125 | 1761694.5 | 2044198.25 | 5590814 | 4822922 | 1876688.625 | 2631388.75 | 4057615.75 | 85446.32 | 1363446.125 | 1243983 | 2446.8809 | GDAC[160.0307]DHDDDNDGIPDDRDNC[160.0307]R | P35441 | P35441 | THBS1 | thrombospondin 1 | Extracellular Space | other |
| 15545179 | 3 | 0.266 | 21945.441 | 1116411.25 | 1252268.625 | 2292098 | 4813452.5 | 3352481 | 1524343.375 | 2609774.75 | 7471620.5 | 83529.766 | 2645171.25 | 2263023.75 | 1615.8126 | RPPLC[160.0307]FHNGVQYK | P35441 | P35441 | THBS1 | thrombospondin 1 | Extracellular Space | other |
| 15543674 | 2 | 0.263 | 175192.703 | 1020153.75 | 1317719.125 | 3243496 | 3942697.5 | 3466449.5 | 1961063.5 | 2407670.25 | 5899884.5 | 78581.586 | 3228931.5 | 2455091.25 | 972.49945 | FYVVMWK | P35441 | P35441 | THBS1 | thrombospondin 1 | Extracellular Space | other |
| 15559751 | 2 | 0.187 | 96470.055 | 765956.625 | 1245902.875 | 3360802.25 | 4118604.25 | 3310012.5 | 1923739 | 2052356.25 | 4392715 | 77112.742 | 2887473.25 | 1658897.25 | 1865.7582 | DNC[160.0307]PNLPNSGQEDYDK | P35441 | P35441 | THBS1 | thrombospondin 1 | Extracellular Space | other |
| 15535222 | 4 | 0.228 | 10569.437 | 2139643.5 | 3920024.75 | 6781805.5 | 1.25E+07 | 1.05E+07 | 5989405 | 6068803.5 | 8921458 | 67878.492 | 3550517.25 | 2918719.5 | 2446.8782 | GDAC[160.0307]DHDDDNDGIPDDRDNC[160.0307]R | P35441 | P35441 | THBS1 | thrombospondin 1 | Extracellular Space | other |
| 15543663 | 2 | 0.227 | 93548.961 | 740004.062 | 1855620.75 | 5456633.5 | 6272586 | 5794798.5 | 2944269.75 | 3408957 | 8486818 | 32715.977 | 5209940 | 2652541.25 | 1200.4966 | C[160.0307]TSYPDGSWK | P35441 | P35441 | THBS1 | thrombospondin 1 | Extracellular Space | other |
| 15543880 | 3 | 0.353 | 16188.354 | 82723.406 | 78046.969 | 6323149 | 72534.305 | 1809721.875 | 827978.938 | 172234.969 | 3459163 | 13352.175 | 5060617 | 2370575 | 1969.8406 | SC[160.0307]DSLNNRC[160.0307]EGSSVQTR | P35441 | P35441 | THBS1 | thrombospondin 1 | Extracellular Space | other |
| 15543843 | 2 | 0.418 | 76078.758 | 927152.5 | 1642584.25 | 2759771.75 | 2903842.75 | 3696718.75 | 2278489.75 | 1585928 | 5244257 | 5899.621 | 2289006 | 2768100 | 1115.6505 | TRGTLLAVER | P35441 | P35441 | THBS1 | thrombospondin 1 | Extracellular Space | other |
| 15543837 | 3 | 0.224 | 37672.055 | 305836.875 | 42335.871 | 3304210 | 1781958.375 | 1345062.75 | 1103346.625 | 1125014.375 | 3985187.25 | 4.70E-06 | 2606691.75 | 1732189.5 | 2131.9285 | LC[160.0307]NSPSPQMNGKPC[160.0307]EGEAR | P35441 | P35441 | THBS1 | thrombospondin 1 | Extracellular Space | other |
| 15543885 | 3 | 0.273 | 4.70E-06 | 913477.375 | 1125406.125 | 2387528.5 | 6449271 | 3599571.5 | 2371348.25 | 3065582.25 | 3834949.5 | 4.70E-06 | 1579085.375 | 1253850 | 2063.777 | C[160.0307]DNC[160.0307]PYNHNPDQADTDK | P35441 | P35441 | THBS1 | thrombospondin 1 | Extracellular Space | other |
| 15540137 | 2 | 0.34 | 132866.219 | 2212046.75 | 3613027 | 8825019 | 1.20E+07 | 1.02E+07 | 6057014 | 6780048.5 | 1.48E+07 | 4.70E-06 | 7322084.5 | 4272748 | 1332.6329 | LC[160.0307]NNPTPQFGGK | P35441 | P35441 | THBS1 | thrombospondin 1 | Extracellular Space | other |
| 15538620 | 2 | 0.423 | 198240.266 | 842562.562 | 1495935.5 | 4287860.5 | 3366278 | 4128998.25 | 1826529.5 | 2351233.5 | 8548491 | 4.70E-06 | 3294855.5 | 3058728.75 | 1023.5964 | GFIFLASLR | P35441 | P35441 | THBS1 | thrombospondin 1 | Extracellular Space | other |
| 15550357 | 2 | 0.332 | 135056.203 | 1173014.25 | 1755733.125 | 3512661.75 | 7632510 | 3945310 | 2084077.375 | 3573380 | 6400905.5 | 0 | 2957069 | 2065109 | 1573.6671 | DNC[160.0307]QYVYNVDQR | P35441 | P35441 | THBS1 | thrombospondin 1 | Extracellular Space | other |
| 15554585 | 2 | 0.34 | 111252.703 | 366248.562 | 556326.688 | 2337302.75 | 2240969.25 | 2296277.5 | 882813.75 | 996797.188 | 3217599.5 | 0 | 2219998 | 894012.562 | 1725.7118 | C[160.0307]GAC[160.0307]PPGYSGNGIQC[160.0307]K | P35441 | P35441 | THBS1 | thrombospondin 1 | Extracellular Space | other |
| 15534448 | 3 | 0.99 | 3440644.5 | 5167382 | 5608698 | 4341800.5 | 5671297 | 4594895.5 | 5559554 | 5446588 | 3725946.75 | 5072912.5 | 4839591 | 4572826 | 1274.6498 | LFEPFEVHTR | Q80ZW2 | Q80ZW2 | THEM6 | thioesterase superfamily member 6 | Extracellular Space | other |
| 15542096 | 2, 3 | 0.192 | 3133575.25 | 2769901.75 | 4111158 | 3393477 | 3267947 | 3170962.75 | 4571903 | 3795674 | 2723804.5 | 5012311 | 4172974.5 | 4196632.5 | 1099.6193, 1099.6201 | VRDLLAEQR | Q80ZW2 | Q80ZW2 | THEM6 | thioesterase superfamily member 6 | Extracellular Space | other |
| 15536067 | 3 | 0.007 | 1838259.5 | 1446415.125 | 2077749.875 | 1893075.875 | 1653417 | 1666739.875 | 2773658.25 | 1848074.875 | 1604980.5 | 3430897.5 | 3600183 | 2960011.75 | 1412.7365 | FRQHVLGTSPDR | Q80ZW2 | Q80ZW2 | THEM6 | thioesterase superfamily member 6 | Extracellular Space | other |
| 15542210 | 2 | 0.003 | 1892547.875 | 1826250.375 | 2258430.5 | 1869081.875 | 2148915.5 | 2241359.5 | 2820046.5 | 2542955.25 | 2533339.75 | 2839376.25 | 2857546.5 | 3603211.75 | 1050.5381 | DGFVC[160.0307]ALLR | Q80ZW2 | Q80ZW2 | THEM6 | thioesterase superfamily member 6 | Extracellular Space | other |
| 15545818 | 2 | 0.473 | 1027419.938 | 1125478.25 | 1615010.75 | 900302.188 | 1835840.375 | 1356763.25 | 1921822 | 1465804.5 | 1443179.125 | 2101212.75 | 1843425.25 | 1216958.75 | 814.4588 | VPC[160.0307]AVLR | Q80ZW2 | Q80ZW2 | THEM6 | thioesterase superfamily member 6 | Extracellular Space | other |
| 15550162 | 3 | 0.051 | 8028256.5 | 6230393.5 | 5601304 | 7055491.5 | 5317270 | 5286825.5 | 7778533 | 6595020 | 6180097 | 1.23E+07 | 1.11E+07 | 7164550 | 3613.7905 | QADC[160.0307]HINNLYNTVFWALIQQSGLTPVQAQQR | Q9CY52 | Q9CY52 | THG1L | tRNA-histidine guanylyltransferase 1-like (S. cerevisiae) | Cytoplasm | enzyme |
| 15534039 | 2 | 0.415 | 1.09E+07 | 7796003.5 | 7548433.5 | 8271804.5 | 6833628 | 6394401.5 | 9528689 | 7651182 | 6471196.5 | 1.17E+07 | 8537956 | 7795431 | 1261.7136 | VVLYPSNQTLK | Q9CY52 | Q9CY52 | THG1L | tRNA-histidine guanylyltransferase 1-like (S. cerevisiae) | Cytoplasm | enzyme |
| 15538693 | 3 | 0.051 | 7199929 | 3580626.5 | 3885765.5 | 4819998 | 4908952 | 2970989.25 | 7770700 | 5274006.5 | 5050924.5 | 8963568 | 1.19E+07 | 6924410.5 | 2159.013 | NFEVQDTC[160.0307]LPHC[160.0307]WVVVR | Q9CY52 | Q9CY52 | THG1L | tRNA-histidine guanylyltransferase 1-like (S. cerevisiae) | Cytoplasm | enzyme |
| 15538271 | 2, 3 | 0.461 | 5419872.5 | 3909288.5 | 4122658.75 | 4528544.5 | 3388354.75 | 2580014.75 | 5887046 | 4207546.5 | 2744736 | 6698408 | 4411467 | 4188929.5 | 1092.5084 | FAEEHNFAK | Q9CY52 | Q9CY52 | THG1L | tRNA-histidine guanylyltransferase 1-like (S. cerevisiae) | Cytoplasm | enzyme |
| 15539809 | 2 | 0.066 | 5721193.5 | 4193583.5 | 4599855 | 4071167.25 | 3783880 | 2465346.75 | 4664091 | 3974957.25 | 3123671.5 | 4808329 | 6278936 | 5407121 | 1180.5455 | EHPEILAEEN | Q9CY52 | Q9CY52 | THG1L | tRNA-histidine guanylyltransferase 1-like (S. cerevisiae) | Cytoplasm | enzyme |
| 15538016 | 3 | 0.265 | 2388208 | 5273450.5 | 1933044.625 | 3207750.25 | 3314587.5 | 3998376.75 | 1596579.875 | 4054048.75 | 4496590.5 | 1192362.625 | 2039182.25 | 2114462.25 | 1989.9767 | TVLEHYALEDDPLEAFK | Q9D880 | Q9D880 | TIMM50 | translocase of inner mitochondrial membrane 50 homolog (S. cerevisiae) | Cytoplasm | phosphatase |
| 15538852 | 3 | 0.14 | 1768835.5 | 3337190.5 | 1362093.5 | 2499502.5 | 2071856.375 | 2974771.25 | 1567916.75 | 2517882.75 | 2644074.25 | 1077006.75 | 1653567.875 | 1279119.125 | 1738.8104 | AQGPQHQPGSEGPSYAK | Q9D880 | Q9D880 | TIMM50 | translocase of inner mitochondrial membrane 50 homolog (S. cerevisiae) | Cytoplasm | phosphatase |
| 15538540 | 2 | 0.172 | 510921.25 | 2964447.5 | 4433202 | 1.04E+07 | 1.34E+07 | 1.49E+07 | 8575217 | 8601518 | 1.77E+07 | 557175 | 1.31E+07 | 8897128 | 1324.713 | EGPFGTLVYTIK | P39876 | P39876 | TIMP3 | TIMP metallopeptidase inhibitor 3 | Extracellular Space | other |
| 15537149 | 3 | 0.201 | 98920.305 | 887067.562 | 1412369.375 | 2350507.5 | 3927389 | 3124995.75 | 3229320 | 2307398.75 | 5340801.5 | 200831.328 | 4223208.5 | 2620236.5 | 1155.5883 | WDHLTLSQR | P39876 | P39876 | TIMP3 | TIMP metallopeptidase inhibitor 3 | Extracellular Space | other |
| 15534914 | 2 | 0.416 | 668532.625 | 1240114.25 | 3101673.75 | 3899309.25 | 5793220 | 6264442.5 | 4464957.5 | 4120719.5 | 7913818.5 | 112306.836 | 7243450.5 | 4938760.5 | 1013.54126 | YQYLLTGR | P39876 | P39876 | TIMP3 | TIMP metallopeptidase inhibitor 3 | Extracellular Space | other |
| 15556987 | 2 | 0.108 | 77538.812 | 119093.242 | 253255.531 | 1973564.875 | 1365426.5 | 1141776.75 | 1067646 | 1070652.875 | 2569670.5 | 84047.609 | 6438146 | 1289374.75 | 1524.6833 | SC[160.0307]YYLPC[160.0307]FVTSK | P39876 | P39876 | TIMP3 | TIMP metallopeptidase inhibitor 3 | Extracellular Space | other |
| 15545084 | 3 | 0.602 | 3096141.75 | 2837907.5 | 2732951.75 | 2875734 | 1820130.875 | 2568667 | 1902791.625 | 2013378 | 3492579.75 | 2644995.75 | 2917606.5 | 2619685 | 2008.9977 | NMAEQIIQEIYSQVQSK | P40142 | P40142 | TKT | transketolase | Cytoplasm | enzyme |
| 15558758 | 2 | 0.171 | 101089.836 | 187002.719 | 1149831.25 | 14892.106 | 1512709.125 | 852383.562 | 290159.969 | 507285.156 | 1563596.125 | 22214.49 | 42291.922 | 38373.375 | 1914.9127 | SVPMSTVFYPSDGVATEK | P40142 | P40142 | TKT | transketolase | Cytoplasm | enzyme |
| 15558740 | 4 | 0.591 | 423258.781 | 1268870.5 | 1495137 | 1224424.75 | 1569906.625 | 2522188 | 1271148.25 | 1051674.25 | 3870933 | 362514 | 1736256.625 | 1692891.25 | 4164.054 | QNLLQAAGNVGQASGELLQQIGESDTDPHFQDVLMQLAK | P26039 | P26039 | TLN1 | talin 1 | Plasma Membrane | other |
| 15543108 | 3 | 0.524 | 1696259.25 | 2849501.75 | 2228244.5 | 2195235.75 | 2064338.5 | 4920182 | 3664171.75 | 2827085 | 9856715 | 360567.25 | 3566532.25 | 4195435 | 2091.152 | GVGAAATAVTQALNELLQHVK | P26039 | P26039 | TLN1 | talin 1 | Plasma Membrane | other |
| 15538898 | 2 | 0.472 | 752031.062 | 926731.625 | 930095.062 | 1576123.125 | 1729677.5 | 1975995 | 1228327.625 | 1314571.25 | 3195137.5 | 170159.953 | 1546744.125 | 2200277.5 | 1243.6611 | VLVQNAAGSQEK | P26039 | P26039 | TLN1 | talin 1 | Plasma Membrane | other |
| 15554619 | 3 | 0.867 | 641827.938 | 813694 | 587512.75 | 979475.625 | 1065965.875 | 1768497.125 | 592530.25 | 736976.438 | 3546579.75 | 117604.102 | 1960786.625 | 2543142.5 | 2556.188 | MVAAATNNLC[160.0307]EAANAAVQGHASQEK | P26039 | P26039 | TLN1 | talin 1 | Plasma Membrane | other |
| 15552367 | 2 | 0.459 | 482455.594 | 1336040.625 | 1336347.25 | 1969462.375 | 1947835.125 | 2526858.25 | 1658852.875 | 1702238.25 | 5214446.5 | 108591.578 | 1605592.125 | 3065611.75 | 1521.8496 | TLAESALQLLYTAK | P26039 | P26039 | TLN1 | talin 1 | Plasma Membrane | other |
| 15539345 | 2 | 0.609 | 786023 | 1245935.75 | 1208445.125 | 1685262.375 | 1526845.125 | 2356522.75 | 1684137.25 | 1087878.625 | 3721497 | 28619.428 | 1532546.125 | 3101635.25 | 1456.8344 | ILAQATSDLVNAIK | P26039 | P26039 | TLN1 | talin 1 | Plasma Membrane | other |
| 15558402 | 2 | 0.778 | 8.569 | 61215.742 | 424958.031 | 1.05 | 223916.812 | 726402.312 | 131983.734 | 169430.625 | 1297177.375 | 725.884 | 182776.766 | 549944.938 | 1935.9143 | EADESLNFEEQILEAAK | P26039 | P26039 | TLN1 | talin 1 | Plasma Membrane | other |
| 15545760 | 2 | 0.532 | 239840.562 | 328756.312 | 290871.969 | 885502.625 | 582788.938 | 745643.625 | 991673.562 | 559600.875 | 2049623.25 | 4.70E-06 | 954387.062 | 1156498.875 | 1236.6305 | STVLQQQYNR | P26039 | P26039 | TLN1 | talin 1 | Plasma Membrane | other |
| 15537479 | 2 | 0.047 | 653859.25 | 984798.375 | 1475516.5 | 2147728.5 | 2258495.25 | 2285141.25 | 1685092.5 | 1822327.375 | 2489524.75 | 1039325.125 | 2332531 | 1690056.125 | 1289.6028 | NLSGQPNFPC[160.0307]R | Q9ET30 | Q9ET30 | TM9SF3 | transmembrane 9 superfamily member 3 | Cytoplasm | transporter |
| 15551311 | 2 | 0.621 | 2192993 | 2720883.25 | 3583899 | 1789201.75 | 3453556 | 2073191.875 | 2130586.75 | 2707302 | 2211894 | 2885811.25 | 1746122.125 | 1768989.625 | 1954.02 | GDQTLPGLEGAPALSSATLR | Q7TN60 | Q7TN60 | TMC6 | transmembrane channel-like 6 | Cytoplasm | transporter |
| 15536070 | 2 | 0.873 | 1597248.25 | 1287223.875 | 1640542.875 | 1298353.625 | 1544165.625 | 1251048 | 1393856.5 | 1753389.25 | 1209964.875 | 1662687.875 | 1367402.125 | 1201397.25 | 747.47485 | IFLINK | Q7TN60 | Q7TN60 | TMC6 | transmembrane channel-like 6 | Cytoplasm | transporter |
| 15546661 | 3 | 0.461 | 660254.375 | 681095.125 | 814348.188 | 460177.656 | 753664.062 | 797456.438 | 894929.875 | 613833.312 | 627193.625 | 536872.688 | 546204.125 | 602721.5 | 1218.5714 | LHSVYEEEGR | Q7TN60 | Q7TN60 | TMC6 | transmembrane channel-like 6 | Cytoplasm | transporter |
| 15539797 | 3 | 0.817 | 3128492.5 | 2914104.5 | 3012663.75 | 2986038.75 | 3603801.75 | 2310555.25 | 2854954.5 | 3873126.25 | 1959949.25 | 3371927 | 3600178 | 2899594.75 | 2173.0122 | RPESQAASEELWEQEVER | Q7TN58 | Q7TN58 | TMC8 | transmembrane channel-like 8 | Cytoplasm | other |
| 15544692 | 2 | 0.686 | 1562776.375 | 1538769.875 | 2030885.375 | 1522181 | 1777466.375 | 1354635.375 | 2245809.75 | 1897350.125 | 1129489.75 | 1582965.75 | 960053.938 | 1636410.625 | 1282.5892 | AGPESSSEYSIR | Q7TN58 | Q7TN58 | TMC8 | transmembrane channel-like 8 | Cytoplasm | other |
| 15551773 | 2 | 0.198 | 1305387 | 1849070.375 | 1504651.875 | 1834327.875 | 1642879 | 933971.25 | 1785071.75 | 1735349.5 | 1462298.5 | 1198108.875 | 1072234.625 | 1086884 | 1573.8158 | LSSSSLGAPSASVPASR | Q7TN58 | Q7TN58 | TMC8 | transmembrane channel-like 8 | Cytoplasm | other |
| 15550209 | 3 | 0.827 | 8838930 | 1.14E+07 | 1.30E+07 | 1.05E+07 | 1.16E+07 | 9892158 | 8349265.5 | 1.04E+07 | 1.22E+07 | 1.13E+07 | 7833330 | 1.02E+07 | 2412.1658 | LEEQLNDLTELHQNEMTNLK | Q80W04 | Q80W04 | TMCC2 | transmembrane and coiled-coil domain family 2 | Extracellular Space | other |
| 15549364 | 3 | 0.387 | 9581143 | 2.03E+07 | 1.28E+07 | 1.16E+07 | 1.67E+07 | 1.63E+07 | 9979402 | 1.44E+07 | 1.49E+07 | 1.11E+07 | 7553757.5 | 1.18E+07 | 3308.5957 | GDLVALSLPSGPGHGDSDGPISLDVPDGAPDPQR | Q80W04 | Q80W04 | TMCC2 | transmembrane and coiled-coil domain family 2 | Extracellular Space | other |
| 15532968 | 3 | 0.865 | 5405240.5 | 5287541.5 | 3724620.5 | 4342312 | 4351164.5 | 5692319.5 | 3772708 | 4412048 | 5137074 | 5368131.5 | 3568400 | 4120540.5 | 1792.8677 | IEQEARDDNVAEYLK | Q80W04 | Q80W04 | TMCC2 | transmembrane and coiled-coil domain family 2 | Extracellular Space | other |
| 15534713 | 3 | 0.835 | 3276083.5 | 5198401.5 | 3327193.25 | 3327868.25 | 4147139.75 | 3302157.25 | 3229175 | 3438835.25 | 3629732.75 | 4192921.25 | 3088257.25 | 4244649 | 1617.7035 | EGQSHLEDSMEDLK | Q80W04 | Q80W04 | TMCC2 | transmembrane and coiled-coil domain family 2 | Extracellular Space | other |
| 15553703 | 2 | 0.431 | 3076733.75 | 4208057 | 3400288 | 2732438 | 4018997 | 3273104.75 | 2278446.5 | 2758850 | 3649183 | 3305704 | 2314257.75 | 2869678.5 | 1736.7224 | DYTYMTQC[160.0307]LQEER | Q80W04 | Q80W04 | TMCC2 | transmembrane and coiled-coil domain family 2 | Extracellular Space | other |
| 15551517 | 2 | 0.218 | 1338950.625 | 2434841.75 | 1507565.75 | 873368.25 | 1837862.125 | 1385056.5 | 1056517.75 | 1240685.75 | 1766533.25 | 1090553.875 | 767054.688 | 1074834 | 1642.8729 | GSLSGLSQATHTAVVSK | Q80W04 | Q80W04 | TMCC2 | transmembrane and coiled-coil domain family 2 | Extracellular Space | other |
| 15534529 | 3 | 0.091 | 1931722 | 3143496.75 | 3950864 | 5853522.5 | 7905966.5 | 6842830.5 | 4933322.5 | 4275483 | 6898989 | 2008954 | 6269546.5 | 4318214.5 | 1728.9403 | LPFTPLSYIQGLSHR | Q921L3 | Q921L3 | TMCO1 | transmembrane and coiled-coil domains 1 | Cytoplasm | other |
| 15533431 | 2 | 0.081 | 1.61E+07 | 1.95E+07 | 2.10E+07 | 2.60E+07 | 3.05E+07 | 2.31E+07 | 2.29E+07 | 2.15E+07 | 1.86E+07 | 2.38E+07 | 1.70E+07 | 2.07E+07 | 1284.7208 | IPDQLVILDMK | Q9D1D4 | Q9D1D4 | TMED10 | transmembrane emp24-like trafficking protein 10 (yeast) | Cytoplasm | transporter |
| 15532186 | 3 | 0.01 | 1.35E+07 | 1.57E+07 | 1.72E+07 | 2.46E+07 | 2.12E+07 | 2.15E+07 | 1.95E+07 | 1.97E+07 | 1.71E+07 | 2.02E+07 | 1.68E+07 | 1.91E+07 | 2029.9865 | RLEDLSESIVNDFAYMK | Q9D1D4 | Q9D1D4 | TMED10 | transmembrane emp24-like trafficking protein 10 (yeast) | Cytoplasm | transporter |
| 15535604 | 2, 3 | 0.591 | 1.50E+07 | 1.50E+07 | 2.00E+07 | 2.08E+07 | 1.91E+07 | 1.71E+07 | 1.87E+07 | 1.77E+07 | 1.80E+07 | 1.67E+07 | 2.04E+07 | 1.26E+07 | 1288.6848, 1288.6858 | ITDSAGHILYAK | Q9D1D4 | Q9D1D4 | TMED10 | transmembrane emp24-like trafficking protein 10 (yeast) | Cytoplasm | transporter |
| 15532460 | 3 | 0.298 | 5161987.5 | 3324245.5 | 2097534.25 | 6652400.5 | 5333226 | 3170524 | 4245366 | 4102803 | 6032354.5 | 5137650 | 6467024 | 5215800 | 1096.67 | LKPLEVELR | Q9D1D4 | Q9D1D4 | TMED10 | transmembrane emp24-like trafficking protein 10 (yeast) | Cytoplasm | transporter |
| 15533070 | 2 | 0.332 | 2766223 | 5306696.5 | 1.13E+07 | 5058364.5 | 7699361.5 | 7804561.5 | 6898340.5 | 7004746.5 | 4569781 | 4586068 | 2676392.5 | 3752555.5 | 855.49054 | PLEVELR | Q9D1D4 | Q9D1D4 | TMED10 | transmembrane emp24-like trafficking protein 10 (yeast) | Cytoplasm | transporter |
| 15549526 | 2 | 0.066 | 4163306 | 4845300.5 | 5294145.5 | 7001119 | 6641590.5 | 5099999 | 5726688 | 5294244 | 5524786 | 4359139 | 4999611.5 | 5109313 | 2029.9852 | RLEDLSESIVNDFAYMK | Q9D1D4 | Q9D1D4 | TMED10 | transmembrane emp24-like trafficking protein 10 (yeast) | Cytoplasm | transporter |
| 15543604 | 2 | 0.306 | 7584000 | 7291181.5 | 9172636 | 8806088 | 1.05E+07 | 8824388 | 9663704 | 9669269 | 7748291.5 | 8411628 | 8685371 | 7510536 | 1220.5334 | HEQEYMEVR | Q9R0Q3 | Q9R0Q3 | TMED2 | transmembrane emp24 domain trafficking protein 2 | Cytoplasm | transporter |
| 15549262 | 2 | 0.438 | 6206512 | 1.17E+07 | 1.25E+07 | 6437974 | 1.67E+07 | 1.28E+07 | 1.14E+07 | 1.43E+07 | 9964857 | 6588114.5 | 6376535 | 9914383 | 1761.9122 | LEEMINELAVAMTAVK | Q9R0Q3 | Q9R0Q3 | TMED2 | transmembrane emp24 domain trafficking protein 2 | Cytoplasm | transporter |
| 15539171 | 3 | 0.862 | 2925525 | 5227217.5 | 5173791 | 3566144.25 | 5769692 | 5215924.5 | 4884329 | 5997497 | 3492271.5 | 3755796.5 | 3253380.25 | 5254175 | 1761.9099 | LEEMINELAVAMTAVK | Q9R0Q3 | Q9R0Q3 | TMED2 | transmembrane emp24 domain trafficking protein 2 | Cytoplasm | transporter |
| 15548278 | 2 | 0.032 | 9625231 | 1.40E+07 | 1.21E+07 | 1.83E+07 | 1.92E+07 | 1.51E+07 | 1.56E+07 | 1.67E+07 | 1.34E+07 | 1.38E+07 | 1.23E+07 | 1.42E+07 | 2085.9573 | C[160.0307]FIEEIPDETMVIGNYR | Q8R1V4 | Q8R1V4 | TMED4 | transmembrane emp24 protein transport domain containing 4 | Cytoplasm | transporter |
| 15539943 | 2 | 0.704 | 2222701.75 | 3357400 | 3351975.5 | 2211473.25 | 4284784 | 2829041.25 | 3052621.75 | 3012527 | 2665138 | 2070540.875 | 2341339.25 | 3033205.25 | 1326.6583 | YITNTDVLEMK | Q9CXE7 | Q9CXE7 | TMED5 | transmembrane emp24 protein transport domain containing 5 | Cytoplasm | other |
| 15534217 | 2 | 0.269 | 1.78E+07 | 1.87E+07 | 2.59E+07 | 2.50E+07 | 2.72E+07 | 2.29E+07 | 2.64E+07 | 2.61E+07 | 2.06E+07 | 2.07E+07 | 1.97E+07 | 2.29E+07 | 1341.7339 | QLVEQVEQIQK | Q99KF1 | Q99KF1 | TMED9 | transmembrane emp24 protein transport domain containing 9 | Cytoplasm | transporter |
| 15537756 | 3 | 0.323 | 4189875 | 8808365 | 1.09E+07 | 6861872.5 | 1.56E+07 | 9642330 | 1.12E+07 | 8838711 | 3953178.25 | 5298603 | 4128245.75 | 5600459.5 | 2179.08 | QREEYQPATPGLGMFVEVK | Q99KF1 | Q99KF1 | TMED9 | transmembrane emp24 protein transport domain containing 9 | Cytoplasm | transporter |
| 15536641 | 3 | 0.633 | 1584795.5 | 335533.906 | 432011.312 | 2881021.25 | 74258.406 | 952470.438 | 1147403.25 | 524157.312 | 1008893.562 | 920568.188 | 2868400.5 | 1794479.75 | 1596.9033 | VRQLVEQVEQIQK | Q99KF1 | Q99KF1 | TMED9 | transmembrane emp24 protein transport domain containing 9 | Cytoplasm | transporter |
| 15547017 | 2 | 7.63E-05 | 602615.188 | 767849.688 | 671895.5 | 308053.094 | 371788.781 | 339937.844 | 712850 | 615427.812 | 537054.25 | 267791.938 | 159069.25 | 213300.625 | 1438.6019 | DSVTC[160.0307]PTC[160.0307]QGTGR | Q80X71 | Q80X71 | TMEM106B | transmembrane protein 106B | Plasma Membrane | other |
| 15536009 | 2 | 0.177 | 9131730 | 1.53E+07 | 1.15E+07 | 8918875 | 1.47E+07 | 1.62E+07 | 1.14E+07 | 1.60E+07 | 1.26E+07 | 8549156 | 9234849 | 8430934 | 1106.618 | VGISLLSSPHP | Q9CQN6 | Q9CQN6 | TMEM14C | transmembrane protein 14C | Plasma Membrane | other |
| 15553700 | 3 | 0.335 | 2734300 | 5637812.5 | 4194358 | 2628118.25 | 5335203.5 | 5542215.5 | 3867050.25 | 4998819.5 | 6408688 | 3655883.5 | 3027970.25 | 2426021.5 | 2820.461 | AGSVPSLAAGLFFGGLAGLGAYQLSQDPR | Q9CQN6 | Q9CQN6 | TMEM14C | transmembrane protein 14C | Plasma Membrane | other |
| 15534839 | 2 | 0.483 | 2374265.5 | 2564739.75 | 3490323.5 | 2668662 | 3751461.25 | 2899603 | 3171111.25 | 3183849.25 | 2267777.75 | 2968664.5 | 3972132 | 3498903 | 1034.6013 | TGLLGIFWK | Q9CR64 | Q9CR64 | TMEM167A | transmembrane protein 167A | Extracellular Space | other |
| 15539205 | 3 | 0.657 | 2854652 | 3184063.5 | 3121817.25 | 2441768.25 | 3675917.25 | 2305936.25 | 2981098.25 | 3974638.75 | 2490248.5 | 3009648.25 | 2112273.25 | 2720389.25 | 2206.9348 | GDQEWSSETDALVGHHQSHS | Q9CY24 | Q9CY24 | TMEM179B | transmembrane protein 179B | Extracellular Space | other |
| 15551846 | 3 | 0.032 | 7884521.5 | 7358304.5 | 1.54E+07 | 1.01E+07 | 1.21E+07 | 1.20E+07 | 8052635 | 8309608.5 | 1.03E+07 | 1.99E+07 | 1.44E+07 | 1.88E+07 | 2598.2598 | EVDRYDAPGIAFYPGQAQLLSC[160.0307]K | Q9D771 | Q9D771 | TMEM206 | transmembrane protein 206 | Plasma Membrane | other |
| 15537640 | 3 | 0.307 | 2170448 | 2553434.75 | 4653678.5 | 2151361.25 | 4844190.5 | 3696798 | 3441013.75 | 3563939.25 | 2771363.25 | 7139340 | 3449538 | 5301981.5 | 1975.9807 | QETSVVNYIDQRPAAER | Q9D771 | Q9D771 | TMEM206 | transmembrane protein 206 | Plasma Membrane | other |
| 15541906 | 2 | 0.728 | 1563085.5 | 1512137.875 | 3525801 | 1207869.25 | 2490610 | 2528297.75 | 1849795.25 | 1999703.375 | 1619373 | 3746076.25 | 1678647 | 2652218.25 | 1051.5914 | ELVFLQFR | Q9D771 | Q9D771 | TMEM206 | transmembrane protein 206 | Plasma Membrane | other |
| 15549041 | 3 | 0.017 | 1.83E+07 | 3.18E+07 | 1.45E+07 | 2.39E+07 | 1.94E+07 | 2.67E+07 | 1.44E+07 | 1.96E+07 | 2.18E+07 | 8816878 | 1.12E+07 | 1.07E+07 | 2620.3113 | LGALSGAGALGLASYGAHGAQFPDAYGK | Q5F285 | Q5F285 | TMEM256 | transmembrane protein 256 | Cytoplasm | other |
| 15535227 | 2 | 0.939 | 1.80E+07 | 1.69E+07 | 1.71E+07 | 2.18E+07 | 1.86E+07 | 1.54E+07 | 2.16E+07 | 1.78E+07 | 1.34E+07 | 1.92E+07 | 1.70E+07 | 1.55E+07 | 1394.6517 | DEVDGGPAGPPGGAAK | Q8VEK0 | Q8VEK0 | TMEM30A | transmembrane protein 30A | Cytoplasm | other |
| 15533620 | 3 | 0.562 | 1.35E+07 | 1.52E+07 | 1.79E+07 | 1.63E+07 | 1.81E+07 | 1.81E+07 | 1.76E+07 | 1.54E+07 | 1.53E+07 | 1.44E+07 | 1.60E+07 | 1.85E+07 | 2013.9789 | SRDDSQLNGDPSALLNPSK | Q8VEK0 | Q8VEK0 | TMEM30A | transmembrane protein 30A | Cytoplasm | other |
| 15553601 | 2 | 0.868 | 9169902 | 7949843 | 1.00E+07 | 9533499 | 1.18E+07 | 8429297 | 9495119 | 1.01E+07 | 9409338 | 1.10E+07 | 9638619 | 7924422 | 1939.8545 | EIEIDYTGTEPSSPC[160.0307]NK | Q8VEK0 | Q8VEK0 | TMEM30A | transmembrane protein 30A | Cytoplasm | other |
| 15538279 | 2 | 0.578 | 8552831 | 7615289.5 | 9366144 | 8628896 | 9876458 | 9548818 | 9322997 | 9770991 | 8371858.5 | 8890064 | 8441280 | 1.00E+07 | 976.48724 | GIAWWTDK | Q8VEK0 | Q8VEK0 | TMEM30A | transmembrane protein 30A | Cytoplasm | other |
| 15542444 | 2 | 0.875 | 5163414 | 7051773.5 | 7424699.5 | 3704125.25 | 8145087 | 5771313.5 | 5372897.5 | 6735171 | 5135684.5 | 6502610.5 | 5644565 | 5199830.5 | 876.49207 | TAALPTFR | Q8VEK0 | Q8VEK0 | TMEM30A | transmembrane protein 30A | Cytoplasm | other |
| 15552750 | 3 | 0.22 | 2229310.25 | 3021856.75 | 3506492.75 | 2673326.75 | 4136471 | 3523589 | 4596191.5 | 4418971.5 | 3227907 | 3205413.25 | 1573197.625 | 2989564.5 | 2693.3462 | SPLLSASHSGNVTPTAPPYLQESSPR | Q9CZX7 | Q9CZX7 | TMEM55A | transmembrane protein 55A | Cytoplasm | phosphatase |
| 15540166 | 2 | 0.32 | 6790280.5 | 6289732.5 | 7021548.5 | 5243036 | 6635516 | 4999652 | 6392025.5 | 6576834.5 | 4803348 | 7863836 | 6032055 | 6356355.5 | 1230.6117 | VC[160.0307]QSPINVEGK | Q3TWL2 | Q3TWL2 | TMEM55B | transmembrane protein 55B | Cytoplasm | phosphatase |
| 15550909 | 3 | 0.611 | 4974284.5 | 5125765 | 4206489 | 3747133.25 | 4398091 | 2977742 | 3556417.5 | 4017583.75 | 3410625.25 | 7446244 | 2629729.75 | 4312583 | 2305.2449 | IINLGPVHPGPLSPEPQPMGVR | Q3TWL2 | Q3TWL2 | TMEM55B | transmembrane protein 55B | Cytoplasm | phosphatase |
| 15536031 | 2 | 0.093 | 4524635 | 3616938.25 | 3865944.75 | 2982086 | 3852005.75 | 2759990.5 | 4200110.5 | 3555136 | 3924882.25 | 5380254.5 | 4232323.5 | 3866177.25 | 1248.5681 | C[160.0307]GVC[160.0307]NEATPIK | Q3TWL2 | Q3TWL2 | TMEM55B | transmembrane protein 55B | Cytoplasm | phosphatase |
| 15539963 | 2 | 0.375 | 4708686 | 3936837 | 4451313 | 4206689.5 | 3495349.5 | 3325583.25 | 4299063 | 4531447.5 | 3707770.5 | 4716963.5 | 3517716.25 | 4753146.5 | 1429.6716 | NTFLWTEFTDR | Q3TWL2 | Q3TWL2 | TMEM55B | transmembrane protein 55B | Cytoplasm | phosphatase |
| 15536398 | 2 | 0.23 | 366063.656 | 1115744.125 | 445999.812 | 1621454 | 1511327.375 | 2485857 | 952208.5 | 1318788.375 | 5508874.5 | 373656.094 | 1922842.125 | 1823909.75 | 1202.6307 | IALVSEAGSEAR | Q91YT8 | Q91YT8 | TMEM63A | transmembrane protein 63A | Cytoplasm | other |
| 15552254 | 4 | 0.583 | 5496171 | 6802760.5 | 6028436 | 5639510 | 6859784 | 4161216.5 | 5011395.5 | 6695446.5 | 6600188 | 6952002 | 6126610 | 6679262 | 3429.4573 | DC[160.0307]NC[160.0307]LHVVEPMPVPGHDVEAYC[160.0307]LLC[160.0307]EC[160.0307]R | Q9CR23 | Q9CR23 | TMEM9 | transmembrane protein 9 | Cytoplasm | other |
| 15549008, 15551098 | 2, 3 | 0.684 | 2.13E+08 | 3.28E+08 | 3.49E+08 | 2.31E+08 | 3.24E+08 | 2.58E+08 | 2.74E+08 | 3.19E+08 | 2.47E+08 | 3.11E+08 | 1.82E+08 | 2.30E+08 | 2279.0981, 2279.1006 | YRDLDEDEILGALTEEELR | P49813 | P49813 | TMOD1 | tropomodulin 1 | Cytoplasm | enzyme |
| 15548061 | 2, 3 | 0.556 | 1.73E+08 | 2.01E+08 | 3.04E+08 | 1.36E+08 | 2.38E+08 | 2.17E+08 | 2.01E+08 | 2.31E+08 | 1.60E+08 | 2.28E+08 | 1.36E+08 | 1.24E+08 | 2308.1638, 2308.1648 | TLENELDELDPDNALLPAGLR | P49813 | P49813 | TMOD1 | tropomodulin 1 | Cytoplasm | enzyme |
| 15531880 | 3 | 0.447 | 2.27E+08 | 1.94E+08 | 2.26E+08 | 2.12E+08 | 1.69E+08 | 2.30E+08 | 2.05E+08 | 1.99E+08 | 1.64E+08 | 2.16E+08 | 2.13E+08 | 2.16E+08 | 1662.9398 | LVEALPHNTSLVELK | P49813 | P49813 | TMOD1 | tropomodulin 1 | Cytoplasm | enzyme |
| 15533113 | 2, 3 | 0.382 | 1.86E+08 | 1.96E+08 | 2.40E+08 | 1.57E+08 | 1.90E+08 | 1.98E+08 | 1.73E+08 | 1.84E+08 | 1.57E+08 | 2.15E+08 | 1.64E+08 | 1.63E+08 | 1281.6771, 1281.6776 | REELLDHLEK | P49813 | P49813 | TMOD1 | tropomodulin 1 | Cytoplasm | enzyme |
| 15551681 | 3 | 0.636 | 1.13E+08 | 2.25E+08 | 3.83E+08 | 9.59E+07 | 2.46E+08 | 1.73E+08 | 2.12E+08 | 2.77E+08 | 1.07E+08 | 2.13E+08 | 7.92E+07 | 1.12E+08 | 2673.2466 | PTQYKPVPDEEPNSTDVEETLER | P49813 | P49813 | TMOD1 | tropomodulin 1 | Cytoplasm | enzyme |
| 15553948 | 2 | 0.21 | 1.81E+08 | 1.77E+08 | 1.95E+08 | 1.93E+08 | 1.75E+08 | 1.62E+08 | 1.56E+08 | 1.63E+08 | 1.62E+08 | 1.90E+08 | 1.61E+08 | 1.55E+08 | 1782.8595 | NNDPELEEVNLNNIR | P49813 | P49813 | TMOD1 | tropomodulin 1 | Cytoplasm | enzyme |
| 15547946 | 2 | 0.476 | 1.15E+08 | 1.35E+08 | 1.63E+08 | 1.28E+08 | 1.15E+08 | 1.26E+08 | 1.20E+08 | 1.14E+08 | 1.29E+08 | 1.56E+08 | 1.24E+08 | 1.29E+08 | 1690.9077 | TLNVESNFISGAGILR | P49813 | P49813 | TMOD1 | tropomodulin 1 | Cytoplasm | enzyme |
| 15533888 | 2, 3 | 0.113 | 1.33E+08 | 1.24E+08 | 1.38E+08 | 1.31E+08 | 1.27E+08 | 1.13E+08 | 1.23E+08 | 1.16E+08 | 1.07E+08 | 1.46E+08 | 1.25E+08 | 1.32E+08 | 1337.6362, 1337.6366 | FGYHFTQQGPR | P49813 | P49813 | TMOD1 | tropomodulin 1 | Cytoplasm | enzyme |
| 15548055 | 2 | 0.985 | 9.46E+07 | 8.78E+07 | 1.35E+08 | 9.65E+07 | 1.08E+08 | 1.02E+08 | 9.31E+07 | 1.10E+08 | 9.53E+07 | 1.42E+08 | 8.74E+07 | 9.20E+07 | 1505.7645 | SNDPVAFALAEMLK | P49813 | P49813 | TMOD1 | tropomodulin 1 | Cytoplasm | enzyme |
| 15533292 | 2 | 0.74 | 8.51E+07 | 9.91E+07 | 9.68E+07 | 8.04E+07 | 1.02E+08 | 8.97E+07 | 9.52E+07 | 8.44E+07 | 6.64E+07 | 1.16E+08 | 7.53E+07 | 8.36E+07 | 1334.6672 | VEMEIVNMLEK | P49813 | P49813 | TMOD1 | tropomodulin 1 | Cytoplasm | enzyme |
| 15531498 | 2 | 0.386 | 6.30E+07 | 6.02E+07 | 6.10E+07 | 5.71E+07 | 6.02E+07 | 5.00E+07 | 6.29E+07 | 5.42E+07 | 4.41E+07 | 6.90E+07 | 5.77E+07 | 5.73E+07 | 1421.6887 | DREDLVPYTGEK | P49813 | P49813 | TMOD1 | tropomodulin 1 | Cytoplasm | enzyme |
| 15547693 | 2 | 0.138 | 4.61E+07 | 4.20E+07 | 5.39E+07 | 3.60E+07 | 3.84E+07 | 4.01E+07 | 4.26E+07 | 3.94E+07 | 3.74E+07 | 5.05E+07 | 4.45E+07 | 3.77E+07 | 1662.9396 | LVEALPHNTSLVELK | P49813 | P49813 | TMOD1 | tropomodulin 1 | Cytoplasm | enzyme |
| 15533870 | 2 | 0.764 | 3.31E+07 | 4.57E+07 | 5.97E+07 | 3.24E+07 | 4.78E+07 | 5.05E+07 | 3.99E+07 | 4.23E+07 | 3.64E+07 | 4.60E+07 | 3.15E+07 | 3.70E+07 | 779.4395 | FSIVGTR | P49813 | P49813 | TMOD1 | tropomodulin 1 | Cytoplasm | enzyme |
| 15549553 | 3 | 0.414 | 3.78E+07 | 2884402.5 | 113337.68 | 4.40E+07 | 1495476.375 | 916586.125 | 5635298 | 4387145 | 2.26E+07 | 2.82E+07 | 5.17E+07 | 1.90E+07 | 2529.2632 | IDNQSQPLGNKVEMEIVNMLEK | P49813 | P49813 | TMOD1 | tropomodulin 1 | Cytoplasm | enzyme |
| 15534053 | 3 | 0.745 | 1.35E+07 | 1.43E+07 | 2.22E+07 | 1.60E+07 | 1.94E+07 | 1.83E+07 | 1.56E+07 | 1.93E+07 | 1.66E+07 | 2.35E+07 | 1.54E+07 | 1.95E+07 | 1505.764 | SNDPVAFALAEMLK | P49813 | P49813 | TMOD1 | tropomodulin 1 | Cytoplasm | enzyme |
| 15540630 | 3 | 0.355 | 2.40E+07 | 1.32E+07 | 9565078 | 2.87E+07 | 7087924.5 | 1.56E+07 | 1.66E+07 | 1.01E+07 | 1.26E+07 | 1.99E+07 | 2.92E+07 | 2.57E+07 | 1550.7365 | ASNAMMSNNDLVRK | P49813 | P49813 | TMOD1 | tropomodulin 1 | Cytoplasm | enzyme |
| 15535732 | 2 | 0.486 | 1.16E+07 | 2.35E+07 | 4.53E+07 | 1.03E+07 | 2.50E+07 | 2.44E+07 | 2.33E+07 | 2.49E+07 | 1.34E+07 | 1.92E+07 | 7664943.5 | 1.19E+07 | 859.487 | EGLNSVIK | P49813 | P49813 | TMOD1 | tropomodulin 1 | Cytoplasm | enzyme |
| 15537887 | 3 | 0.275 | 7847774.5 | 1.31E+07 | 1.32E+07 | 1.12E+07 | 8368895.5 | 1.03E+07 | 1.07E+07 | 1.05E+07 | 9231722 | 1.89E+07 | 1.09E+07 | 1.34E+07 | 1690.9089 | TLNVESNFISGAGILR | P49813 | P49813 | TMOD1 | tropomodulin 1 | Cytoplasm | enzyme |
| 15541200 | 3 | 0.167 | 1.10E+07 | 1.18E+07 | 1.28E+07 | 1.24E+07 | 1.13E+07 | 1.22E+07 | 7384903.5 | 1.18E+07 | 9713318 | 1.12E+07 | 1.11E+07 | 1.24E+07 | 1782.8591 | NNDPELEEVNLNNIR | P49813 | P49813 | TMOD1 | tropomodulin 1 | Cytoplasm | enzyme |
| 15537314 | 2 | 0.71 | 1.92E+07 | 3.66E+07 | 5.28E+07 | 7464075.5 | 4.29E+07 | 3.49E+07 | 1.91E+07 | 4.04E+07 | 3.44E+07 | 1.12E+07 | 2.56E+07 | 2.60E+07 | 1213.6135 | IDNQSQPLGNK | P49813 | P49813 | TMOD1 | tropomodulin 1 | Cytoplasm | enzyme |
| 15550567 | 2 | 0.479 | 3613655 | 1.02E+07 | 2.58E+07 | 3432073.5 | 1.39E+07 | 1.16E+07 | 9458978 | 1.09E+07 | 4721882 | 7531298.5 | 2179655.75 | 3727798.5 | 2055.9312 | PVPDEEPNSTDVEETLER | P49813 | P49813 | TMOD1 | tropomodulin 1 | Cytoplasm | enzyme |
| 15551964, 15554032 | 2 | 0.247 | 5371385.5 | 6899031.5 | 1.07E+07 | 6367682.5 | 6250728 | 5787424.5 | 6459522 | 6389108 | 5722938 | 6481760 | 3207729 | 4851148.5 | 1521.7589, 1521.759 | SNDPVAFALAEM[147.0354]LK | P49813 | P49813 | TMOD1 | tropomodulin 1 | Cytoplasm | enzyme |
| 15549877 | 2 | 0.37 | 6974946.5 | 3094995.25 | 1909482.5 | 7377604.5 | 1121515.75 | 2558096 | 3945300.5 | 1912717.375 | 2417057.75 | 5956478 | 7181858 | 5283721 | 1550.7368 | ASNAMMSNNDLVRK | P49813 | P49813 | TMOD1 | tropomodulin 1 | Cytoplasm | enzyme |
| 15546156 | 2 | 0.791 | 932156.75 | 1547536.625 | 1651936.625 | 744878.188 | 1370763.375 | 1021570.25 | 1228789.25 | 1348899.625 | 598152.625 | 1719689.625 | 429815.781 | 1075103.75 | 1438.6384 | ASNAM[147.0354]MSNNDLVR | P49813 | P49813 | TMOD1 | tropomodulin 1 | Cytoplasm | enzyme |
| 15546535 | 3 | 0.568 | 332716.094 | 570963.312 | 700463.312 | 309907.344 | 515765.906 | 665560.938 | 323955.188 | 571804.875 | 685636.688 | 448601.688 | 368679.25 | 255888.922 | 1581.9685 | AVLAPLIALVYSVPR | Q9D710 | Q9D710 | TMX2 | thioredoxin-related transmembrane protein 2 | Cytoplasm | enzyme |
| 15537831 | 3 | 0.084 | 2499741.25 | 3264802.5 | 3785903 | 3567266 | 4436865.5 | 3388417.75 | 3966839.25 | 3737588.75 | 3775609.5 | 3487025.25 | 2141022.25 | 2089425.5 | 1937.8777 | EDGAHPADTQDVVEDALR | Q8C0L0 | Q8C0L0 | TMX4 | thioredoxin-related transmembrane protein 4 | Cytoplasm | enzyme |
| 15552438 | 3 | 0.752 | 273653.719 | 1642218.25 | 1159373.375 | 898543.562 | 1524180.625 | 1215987.5 | 1039118.5 | 1258128.25 | 1284082.25 | 1962653.875 | 768095.062 | 1264920.25 | 3134.63 | GDLSQHATPLPTPAVLPGDSPITPTPEQIGK | O88746 | O88746 | TOM1 | target of myb1 membrane trafficking protein | Cytoplasm | transporter |
| 15549768 | 3 | 0.668 | 3772972.75 | 4377826.5 | 4491812.5 | 3588080 | 3668070 | 4132984.5 | 4752745.5 | 3897298.25 | 2864908.5 | 8228039.5 | 2818325.5 | 5254165 | 2730.4722 | QEAGQISPNRPTSVPTAPALSSIIAPK | Q923U0 | Q923U0 | TOM1L1 | target of myb1 like 1 membrane trafficking protein | Cytoplasm | other |
| 15554442 | 3 | 0.125 | 1777562.625 | 1437767 | 974369.688 | 1359818.75 | 1394647.125 | 1401444.75 | 1628830.375 | 949563.875 | 1119608.75 | 3985153 | 2046856.25 | 1560495.125 | 3247.5637 | ATFAGVLTEDWGQFLHIC[160.0307]DIINTTQDGPK | Q923U0 | Q923U0 | TOM1L1 | target of myb1 like 1 membrane trafficking protein | Cytoplasm | other |
| 15550532 | 2 | 0.241 | 2133226.5 | 3791515.5 | 1371937.75 | 2564950.5 | 2545382.5 | 2947512.75 | 1754588.625 | 2138755 | 3097584.75 | 1310288.5 | 1812920 | 1554349.375 | 1557.7772 | LQMEQQQLQQR | Q9CPQ3 | Q9CPQ3 | TOMM22 | translocase of outer mitochondrial membrane 22 homolog (yeast) | Cytoplasm | transporter |
| 15542289 | 2 | 0.363 | 1935957.75 | 4915144 | 1672686.5 | 2846582.75 | 2632378.5 | 3600007.75 | 1657224.75 | 2981041.25 | 3845226.75 | 1278316.5 | 1892479.125 | 1983869.875 | 1378.6798 | LWGLTEMFPER | Q9CPQ3 | Q9CPQ3 | TOMM22 | translocase of outer mitochondrial membrane 22 homolog (yeast) | Cytoplasm | transporter |
| 15550241 | 3 | 0.216 | 5250125.5 | 9840067 | 3518102.75 | 6306005 | 5950976 | 6383581 | 3692853.75 | 6816866 | 6795858 | 2307647 | 4026339.5 | 4070130 | 3590.821 | QLSPTEAFPVLVGDMDNSGSLNAQVIHQLSPGLR | Q9QYA2 | Q9QYA2 | TOMM40 | translocase of outer mitochondrial membrane 40 homolog (yeast) | Cytoplasm | ion channel |

| | | | | | | | | | | | | | | | | | | | | | | |
|---|---|---|---|---|---|---|---|---|---|---|---|---|---|---|---|---|---|---|---|---|---|---|
| 15555697 | 2 | 0.366 | 1514596.875 | 2538754.75 | 1032999.812 | 2027571.375 | 2956885.5 | 2901484.75 | [illegible] | [illegible] | [illegible] | [illegible] | [illegible] | [illegible] | 1978.9419 | SYFSSFTDDIISQPMLK | Q9CZW5 | Q9CZW5 | TOMM70A | translocase of outer mitochondrial membrane 70 homolog A (S. cerevisiae) | Cytoplasm | transporter |
| 15555746 | 3 | 0.536 | 4587779.5 | 4132024.25 | 3159719.25 | 3124535 | 4401557 | 3046043 | [illegible] | [illegible] | [illegible] | [illegible] | [illegible] | [illegible] | 2299.16 | TPVVEGLTEGEEEELRAELAK | Q9CYZ2 | Q9CYZ2 | TPD52L2 | tumor protein D52-like 2 | Cytoplasm | other |
| 15535418, 15539325 | 2 | 0.217 | 3.62E+08 | 2.95E+08 | 3.82E+08 | 2.73E+08 | 2.70E+08 | 2.61E+08 | [illegible] | [illegible] | [illegible] | [illegible] | [illegible] | [illegible] | 1131.6057, 1131.6063 | MEIQEIQLK | P58771 | P58771 | Tpm1 | tropomyosin 1, alpha | Plasma Membrane | other |
| 15532614 | 3 | 0.934 | 5.61E+07 | 6.30E+07 | 7.34E+07 | 5.39E+07 | 5.92E+07 | 6.84E+07 | [illegible] | [illegible] | [illegible] | [illegible] | [illegible] | [illegible] | 1671.8875 | LVIIESDLERAEER | P58771 | P58771 | Tpm1 | tropomyosin 1, alpha | Plasma Membrane | other |
| 15534064 | 2 | 0.78 | 1.67E+07 | 1.83E+07 | 2.89E+07 | 8624579 | 2.22E+07 | 1.72E+07 | 2.20E+07 | 2.03E+07 | 1.17E+07 | 2.38E+07 | 1.05E+07 | 1.79E+07 | 1186.6654 | LVIIESDLER | P58771 | P58771 | Tpm1 | tropomyosin 1, alpha | Plasma Membrane | other |
| 15549477 | 2 | 0.17 | 1.29E+07 | 1.24E+07 | 1.76E+07 | 9670722 | 1.27E+07 | 1.44E+07 | 1.37E+07 | 1.42E+07 | 9876548 | 9804287 | 8709692 | 1.09E+07 | 1671.8872 | LVIIESDLERAEER | P58771 | P58771 | Tpm1 | tropomyosin 1, alpha | Plasma Membrane | other |
| 15539055 | 2 | 0.617 | 4861844.5 | 2846616.75 | 2717383.25 | 4361061 | 2616755 | 3048012 | 4438695 | 2777005 | 2134603 | 5570567.5 | 3886687.25 | 3311918.25 | 1120.5166 | C[160.0307]AELEEELK | P58771 | P58771 | Tpm1 | tropomyosin 1, alpha | Plasma Membrane | other |
| 15538458 | 3 | 0.297 | 6.14E+08 | 4.62E+08 | 5.83E+08 | 5.75E+08 | 4.34E+08 | 4.80E+08 | 6.25E+08 | 4.77E+08 | 4.84E+08 | 6.54E+08 | 6.87E+08 | 5.41E+08 | 1727.8903 | IQLVEEELDRAQER | P21107 | P21107 | TPM3 | tropomyosin 3 | Cytoplasm | other |
| 15548002 | 2 | 0.889 | 2.00E+08 | 1.54E+08 | 2.20E+08 | 1.77E+08 | 1.75E+08 | 1.87E+08 | 2.07E+08 | 1.81E+08 | 1.70E+08 | 2.59E+08 | 1.69E+08 | 1.77E+08 | 1552.7346 | TIDDLEDELYAQK | P21107 | P21107 | TPM3 | tropomyosin 3 | Cytoplasm | other |
| 15531282 | 2, 3 | 0.336 | 2.28E+08 | 1.77E+08 | 1.95E+08 | 2.02E+08 | 1.45E+08 | 1.74E+08 | 2.07E+08 | 1.75E+08 | 1.56E+08 | 2.50E+08 | 2.00E+08 | 1.91E+08 | 1472.7424, 1472.7434 | AADAEAEVASLNRR | P21107 | P21107 | TPM3 | tropomyosin 3 | Cytoplasm | other |
| 15531465 | 3 | 0.746 | 1.80E+08 | 2.04E+08 | 2.65E+08 | 1.46E+08 | 2.17E+08 | 1.87E+08 | 2.31E+08 | 2.18E+08 | 1.65E+08 | 2.45E+08 | 1.73E+08 | 1.95E+08 | 1671.8894 | LVIIEGDLERTEER | P21107 | P21107 | TPM3 | tropomyosin 3 | Cytoplasm | other |
| 15549027 | 2 | 0.279 | 1.07E+08 | 7.34E+07 | 9.58E+07 | 9.75E+07 | 6.67E+07 | 3.88E+07 | 9.83E+07 | 6.81E+07 | 7.45E+07 | 1.22E+08 | 1.10E+08 | 8.03E+07 | 1727.8893 | IQLVEEELDRAQER | P21107 | P21107 | TPM3 | tropomyosin 3 | Cytoplasm | other |
| 15554563 | 2 | 0.403 | 7.87E+07 | 9.98E+07 | 1.65E+08 | 5.64E+07 | 8.38E+07 | 9.84E+07 | 9.71E+07 | 1.20E+08 | 1.15E+08 | 1.19E+08 | 5.59E+07 | 8.24E+07 | 1744.8033 | AISDELDHALNDMTSI | P21107 | P21107 | TPM3 | tropomyosin 3 | Cytoplasm | other |
| 15535915 | 2 | 0.62 | 7.82E+07 | 1.04E+08 | 1.44E+08 | 4.78E+07 | 1.11E+08 | 8.71E+07 | 1.14E+08 | 1.14E+08 | 7.66E+07 | 1.17E+08 | 5.21E+07 | 7.83E+07 | 1156.6549 | LVIIEGDLER | P21107 | P21107 | TPM3 | tropomyosin 3 | Cytoplasm | other |
| 15538320 | 2 | 0.504 | 1.08E+08 | 4.78E+07 | 1.90E+07 | 8.80E+07 | 4.06E+07 | 3.60E+07 | 8.53E+07 | 4.96E+07 | 4.77E+07 | 1.14E+08 | 9.16E+07 | 6.63E+07 | 1182.5493 | EDKYEEEIK | P21107 | P21107 | TPM3 | tropomyosin 3 | Cytoplasm | other |
| 15538867 | 2, 3 | 0.894 | 6.43E+07 | 7.46E+07 | 8.79E+07 | 5.23E+07 | 8.29E+07 | 6.58E+07 | 8.05E+07 | 7.96E+07 | 4.92E+07 | 8.95E+07 | 5.67E+07 | 5.65E+07 | 1316.6412, 1316.6415 | AADAEAEVASLNR | P21107 | P21107 | TPM3 | tropomyosin 3 | Cytoplasm | other |
| 15532854 | 2 | 0.802 | 4.84E+07 | 6.42E+07 | 8.14E+07 | 3.48E+07 | 7.66E+07 | 4.10E+07 | 6.90E+07 | 8.83E+07 | 3.59E+07 | 8.08E+07 | 3.30E+07 | 4.80E+07 | 1243.6495 | IQLVEEELDR | P21107 | P21107 | TPM3 | tropomyosin 3 | Cytoplasm | other |
| 15548467 | 2 | 0.273 | 3.80E+07 | 3.95E+07 | 5.64E+07 | 2.62E+07 | 4.28E+07 | 2.40E+07 | 3.99E+07 | 4.06E+07 | 3.48E+07 | 4.76E+07 | 3.27E+07 | 3.45E+07 | 1671.888 | LVIIEGDLERTEER | P21107 | P21107 | TPM3 | tropomyosin 3 | Cytoplasm | other |
| 15533340 | 2, 3 | 0.882 | 9513502 | 1.24E+07 | 1.50E+07 | 5401969 | 1.68E+07 | 1.13E+07 | 1.32E+07 | 1.36E+07 | 7554254.5 | 1.46E+07 | 5887928 | 9426080 | 1399.7506, 1399.751 | RIQLVEEELDR | P21107 | P21107 | TPM3 | tropomyosin 3 | Cytoplasm | other |
| 15539480 | 3 | 0.688 | 9233580 | 7003173 | 1.28E+07 | 5784222.5 | 1.05E+07 | 9682547 | 1.19E+07 | 1.15E+07 | 1.01E+07 | 1.34E+07 | 6610543 | 1.19E+07 | 1552.7328 | TIDDLEDELYAQK | P21107 | P21107 | TPM3 | tropomyosin 3 | Cytoplasm | other |
| 15537182 | 2 | 0.268 | 1.12E+07 | 7410559.5 | 8802220 | 5540415 | 6368348 | 8666392 | 9869498 | 6352075 | 6084801 | 6977802 | 6069157 | 6137794.5 | 1147.6 | M[147.0354]ELQEIQLK | P21107 | P21107 | TPM3 | tropomyosin 3 | Cytoplasm | other |
| 15535595 | 3 | 0.16 | 4311109.5 | 9298717 | 4495517.5 | 6968973 | 8278886.5 | 8313573 | 4128679.25 | 8018661.5 | 8434180 | 2284530.5 | 4413206 | 5082897 | 2233.1252 | FRPSFPASSPYVTTVGGTSFK | O89023 | O89023 | TPP1 | tripeptidyl peptidase I | Cytoplasm | peptidase |
| 15548204 | 4 | 0.471 | 2.51E+07 | 1.41E+07 | 1.11E+07 | 1.75E+07 | 1.34E+07 | 1.59E+07 | 2.10E+07 | 1.28E+07 | 1.33E+07 | 3.78E+07 | 1.55E+07 | 2.03E+07 | 3564.7566 | GIYLRDPVQVAAPSDHGVGIEPVFPENTENSEK | Q64514 | Q64514 | TPP2 | tripeptidyl peptidase II | Cytoplasm | peptidase |
| 15548193 | 3 | 0.573 | 1.50E+07 | 1.01E+07 | 6241919.5 | 1.00E+07 | 1.09E+07 | 1.14E+07 | 1.36E+07 | 1.12E+07 | 9896934 | 2.50E+07 | 1.05E+07 | 1.14E+07 | 2566.1985 | EGLHYTEVC[160.0307]GYDIASPNAGPLFR | Q64514 | Q64514 | TPP2 | tripeptidyl peptidase II | Cytoplasm | peptidase |
| 15551462 | 2 | 0.401 | 1.44E+07 | 8898949 | 9928547 | 8763183 | 9512522 | 7952456 | 1.07E+07 | 7864320 | 8097703.5 | 2.39E+07 | 8472722 | 1.09E+07 | 1670.8027 | YNQPFFVTSLPDDK | Q64514 | Q64514 | TPP2 | tripeptidyl peptidase II | Cytoplasm | peptidase |
| 15548973 | 2 | 0.429 | 1.14E+07 | 6907492 | 6922795.5 | 7477989 | 7586221.5 | 7853377 | 1.06E+07 | 6970815 | 7854437.5 | 2.22E+07 | 7731330 | 9566329 | 1555.8126 | ETYPAYLPLYVAR | Q64514 | Q64514 | TPP2 | tripeptidyl peptidase II | Cytoplasm | peptidase |
| 15548807 | 3 | 0.746 | 1.44E+07 | 9687823 | 4283752 | 9982140 | 6920028.5 | 6498966.5 | 9591367 | 6255429.5 | 8462317 | 2.08E+07 | 6748183.5 | 9766811 | 2636.391 | GVLIAVLDTGVDPGAPGMQVTTDGKPK | Q64514 | Q64514 | TPP2 | tripeptidyl peptidase II | Cytoplasm | peptidase |
| 15531984 | 2 | 0.441 | 1.09E+07 | 8078732.5 | 7485477 | 8372996.5 | 7757663 | 8526459 | 1.01E+07 | 6311684.5 | 6787055 | 1.63E+07 | 7413130 | 1.02E+07 | 982.6281 | VPITAVIAAK | Q64514 | Q64514 | TPP2 | tripeptidyl peptidase II | Cytoplasm | peptidase |
| 15537127 | 3 | 0.227 | 8441042 | 6359076 | 4582992 | 6058250.5 | 5158604.5 | 4761998 | 9089411 | 5278341.5 | 5360834.5 | 1.55E+07 | 7182341 | 7432155 | 2225.967 | C[160.0307]DLVNYSYGEATHWPNSGR | Q64514 | Q64514 | TPP2 | tripeptidyl peptidase II | Cytoplasm | peptidase |
| 15536459 | 2 | 0.388 | 8988804 | 5806942.5 | 4865562.5 | 5538208 | 5066111 | 5859885 | 7668384.5 | 4242091.5 | 4771555 | 1.45E+07 | 5566670 | 7356630.5 | 1210.6555 | IPANWTNPLGK | Q64514 | Q64514 | TPP2 | tripeptidyl peptidase II | Cytoplasm | peptidase |
| 15550715 | 2 | 0.748 | 9563441 | 6247299.5 | 6789905 | 6459001 | 5574494.5 | 7027962.5 | 7640932.5 | 5173830.5 | 7208335.5 | 1.36E+07 | 5323060 | 7095733 | 1792.8928 | EELQSQVELLNSFEK | Q64514 | Q64514 | TPP2 | tripeptidyl peptidase II | Cytoplasm | peptidase |
| 15543312 | 2 | 0.407 | 7479637.5 | 5467717 | 4119795.25 | 4894960.5 | 4883166 | 4994461.5 | 6714472 | 4936296.5 | 5104600 | 1.35E+07 | 4583908.5 | 7384914 | 1301.7451 | GTLIEAFPVLGGK | Q64514 | Q64514 | TPP2 | tripeptidyl peptidase II | Cytoplasm | peptidase |
| 15533878 | 2 | 0.296 | 5605787 | 4012500.5 | 3146656.75 | 3876532.5 | 2942818.5 | 3571833.25 | 4594294.5 | 2650462.75 | 3230386.75 | 1.02E+07 | 3724849.75 | 4732636.5 | 1278.6704 | LSTMETGTGLIR | Q64514 | Q64514 | TPP2 | tripeptidyl peptidase II | Cytoplasm | peptidase |
| 15535934 | 2 | 0.302 | 4264779 | 2465376.25 | 2562898.75 | 3121216.75 | 2618204.75 | 2281068.5 | 3810251 | 2295976.75 | 2394506 | 8539036 | 3012813.25 | 3615797.25 | 1016.53424 | DGEIIGLSGR | Q64514 | Q64514 | TPP2 | tripeptidyl peptidase II | Cytoplasm | peptidase |
| 15556489 | 3 | 0.992 | 3294942.25 | 2343319 | 2312129.25 | 2484147.5 | 3852919.5 | 1278844.5 | 2976828.25 | 3271308.75 | 1225967.5 | 8475236 | 1207371 | 1566689.75 | 2962.4001 | DPVQVAAPSDHGVGIEPVFPENTENSEK | Q64514 | Q64514 | TPP2 | tripeptidyl peptidase II | Cytoplasm | peptidase |
| 15543893 | 2 | 0.341 | 6002280 | 4024756.5 | 3307937.25 | 4088428.25 | 3645819.5 | 2993149.5 | 5022933.5 | 3920400.75 | 2974324 | 8440797 | 3868428.75 | 5183647.5 | 1322.6461 | LPANQYTWSSR | Q64514 | Q64514 | TPP2 | tripeptidyl peptidase II | Cytoplasm | peptidase |
| 15551584 | 2 | 0.974 | 3622895.25 | 2058965.125 | 3527632.5 | 3885424.5 | 3114133.5 | 2644989.75 | 2933751 | 2865548.75 | 3195833.25 | 8415918 | 1820215.625 | 2659214.5 | 2288.146 | IIDIIDTTGSGDVNTATEVEPK | Q64514 | Q64514 | TPP2 | tripeptidyl peptidase II | Cytoplasm | peptidase |
| 15538569 | 3 | 0.3 | 3364678.25 | 2595391.75 | 1815274 | 3438037.75 | 1029471.625 | 2298011.25 | 3054197.25 | 2019587.625 | 2052901 | 7244141.5 | 2849201 | 3295841.75 | 1353.7081 | HEQISDLDRLK | Q64514 | Q64514 | TPP2 | tripeptidyl peptidase II | Cytoplasm | peptidase |
| 15536715 | 2 | 0.727 | 3249615.25 | 2785192.25 | 2418706 | 2465787 | 3072791.5 | 2715759.75 | 3389351 | 2251856.5 | 2588711.75 | 6804568 | 2180320.5 | 2954082 | 994.482 | EEFTEALR | Q64514 | Q64514 | TPP2 | tripeptidyl peptidase II | Cytoplasm | peptidase |
| 15534239 | 2 | 0.317 | 4634621 | 3176818 | 2908511 | 2891368 | 2412340.75 | 2002588.625 | 4748092 | 3047962.5 | 2523689.75 | 6536225.5 | 2631216.25 | 3797775.75 | 1083.5923 | DLPFIVSHR | Q64514 | Q64514 | TPP2 | tripeptidyl peptidase II | Cytoplasm | peptidase |
| 15542466 | 2 | 0.335 | 2514313.75 | 1379655.25 | 103034.938 | 1352144.375 | 343474.438 | 453417.75 | 1562474 | 241286.047 | 128655.859 | 4155005.75 | 1441231.375 | 1863025.5 | 853.5158 | VLIFAYK | Q64514 | Q64514 | TPP2 | tripeptidyl peptidase II | Cytoplasm | peptidase |
| 15543949 | 2 | 0.944 | 1421565 | 880770.75 | 609707.312 | 706293.812 | 919195 | 880716.812 | 1370787.625 | 962674.062 | 483988.094 | 3029340.5 | 521364.281 | 794951.5 | 1112.5289 | HEQISDLDR | Q64514 | Q64514 | TPP2 | tripeptidyl peptidase II | Cytoplasm | peptidase |
| 15544844 | 2 | 0.58 | 1660761.125 | 1077150.5 | 1095088.375 | 1218426.875 | 1054154.625 | 1133551.375 | 1440890.5 | 1045259.188 | 910386.812 | 2773610.75 | 1043126.25 | 1268291.5 | 1408.5984 | AC[160.0307]VDSNENGDLSK | Q64514 | Q64514 | TPP2 | tripeptidyl peptidase II | Cytoplasm | peptidase |
| 15539851 | 2 | 0.168 | 1519039.25 | 816480.812 | 611072.625 | 688382.562 | 574397.875 | 817390.75 | 1294468.25 | 731713.812 | 736106.188 | 1994703.375 | 1106708.875 | 1187764.375 | 1153.7064 | FVLHAVQLVK | Q64514 | Q64514 | TPP2 | tripeptidyl peptidase II | Cytoplasm | peptidase |
| 15548968 | 2 | 0.113 | 1.09E+07 | 1.22E+07 | 9279580 | 1.57E+07 | 1.56E+07 | 1.18E+07 | 1.28E+07 | 1.15E+07 | 9182832 | 1.29E+07 | 5490221.5 | 6457602.5 | 1513.7822 | GVVDSEDIPLNLSR | Q9CQN1 | Q9CQN1 | TRAP1 | TNF receptor-associated protein 1 | Cytoplasm | enzyme |
| 15537988 | 2 | 0.084 | 1355908.125 | 3290210.25 | 1254161.125 | 1525429 | 1698989.875 | 2424710.25 | 1431894.375 | 1949683.125 | 1802275.375 | 597055.562 | 996490.438 | 1110862.5 | 1289.6566 | ELISNASDALEK | Q9CQN1 | Q9CQN1 | TRAP1 | TNF receptor-associated protein 1 | Cytoplasm | enzyme |
| 15554895 | 3 | 0.176 | 3760328.75 | 1.09E+07 | 2.36E+07 | 4.14E+07 | 6.78E+07 | 5.50E+07 | 3.08E+07 | 3.97E+07 | 4.81E+07 | 3999043 | 6.67E+07 | 3.52E+07 | 2320.1116 | LVVC[160.0307]GPSQSTGVPGMDPPSAAHR | Q8K558 | Q8K558 | TREML1 | triggering receptor expressed on myeloid cells-like 1 | Plasma Membrane | other |
| 15538329 | 2 | 0.152 | 1483021.25 | 6251023 | 1.61E+07 | 3.48E+07 | 4.19E+07 | 4.31E+07 | 2.46E+07 | 3.14E+07 | 3.65E+07 | 2033835.125 | 4.68E+07 | 2.60E+07 | 1440.733 | SVTYATVVFPGGDK | Q8K558 | Q8K558 | TREML1 | triggering receptor expressed on myeloid cells-like 1 | Plasma Membrane | other |
| 15540841 | 3 | 0.133 | 1030017.812 | 4214061 | 1.14E+07 | 2.45E+07 | 3.01E+07 | 2.64E+07 | 1.60E+07 | 2.03E+07 | 2.46E+07 | 1666998.625 | 2.80E+07 | 1.73E+07 | 2051.9668 | IASC[160.0307]EPVQDPPNSQTPPSK | Q8K558 | Q8K558 | TREML1 | triggering receptor expressed on myeloid cells-like 1 | Plasma Membrane | other |
| 15531901, 15543398 | 2, 3 | 0.25 | 2153154.75 | 7921259 | 2.22E+07 | 3.07E+07 | 4.91E+07 | 4.67E+07 | 2.72E+07 | 3.00E+07 | 3.65E+07 | 1635676.625 | 4.35E+07 | 2.84E+07 | 1466.7214, 1466.7241 | SSDSGLPSDIPHVR | Q8K558 | Q8K558 | TREML1 | triggering receptor expressed on myeloid cells-like 1 | Plasma Membrane | other |
| 15549928 | 3 | 0.268 | 2181502 | 5982151.5 | 1.06E+07 | 2.12E+07 | 2.90E+07 | 2.85E+07 | 1.55E+07 | 1.46E+07 | 3.00E+07 | 1112229.625 | 3.05E+07 | 1.97E+07 | 2402.1355 | VWC[160.0307]QFLQEGC[160.0307]HPLVTSAVDR | Q8K558 | Q8K558 | TREML1 | triggering receptor expressed on myeloid cells-like 1 | Plasma Membrane | other |
| 15533066 | 2 | 0.183 | 1188938.75 | 4633778.5 | 7412412 | 1.65E+07 | 2.42E+07 | 2.24E+07 | 9003003 | 1.24E+07 | 2.29E+07 | 848164.5 | 1.58E+07 | 1.36E+07 | 1343.8381 | VSLLVLPPVPGPR | Q8K558 | Q8K558 | TREML1 | triggering receptor expressed on myeloid cells-like 1 | Plasma Membrane | other |
| 15549413 | 2 | 0.188 | 677123.938 | 3488805 | 8810500 | 1.99E+07 | 2.38E+07 | 1.95E+07 | 1.22E+07 | 1.43E+07 | 1.85E+07 | 724682.062 | 2.32E+07 | 1.07E+07 | 2051.967 | IASC[160.0307]EPVQDPPNSQTPPSK | Q8K558 | Q8K558 | TREML1 | triggering receptor expressed on myeloid cells-like 1 | Plasma Membrane | other |
| 15557983, 15561124, 15561504 | 3 | 0.194 | 106829.727 | 1269358.25 | 3236554.25 | 3719061.25 | 6463622 | 6984899 | 2461751.25 | 4191251 | 5348352 | 580427.312 | 4871165.5 | 2564015.25 | 3601.7522, 3601.7527, 3601.7568 | LDSPPSFDSIYTGSSLDPPSSEPPAPPSQPPLPPK | Q8K558 | Q8K558 | TREML1 | triggering receptor expressed on myeloid cells-like 1 | Plasma Membrane | other |
| 15550147 | 3 | 0.285 | 2.09E+07 | 2.04E+07 | 1.89E+07 | 1.53E+07 | 1.94E+07 | 1.49E+07 | 2.36E+07 | 2.05E+07 | 1.59E+07 | 2.24E+07 | 2.18E+07 | 1.75E+07 | 2336.1711 | LC[160.0307]PTVQLEDIC[160.0307]NHQGLTPLK | Q9WTR1 | Q9WTR1 | TRPV2 | transient receptor potential cation channel, subfamily V, member 2 | Plasma Membrane | ion channel |
| 15549498 | 2 | 0.824 | 1.30E+07 | 1.40E+07 | 1.86E+07 | 1.20E+07 | 1.63E+07 | 1.28E+07 | 1.32E+07 | 1.27E+07 | 1.36E+07 | 1.84E+07 | 1.18E+07 | 1.19E+07 | 1806.9076 | TLPTLSEDPSGAGITGYK | Q9WTR1 | Q9WTR1 | TRPV2 | transient receptor potential cation channel, subfamily V, member 2 | Plasma Membrane | ion channel |
| 15554665 | 2 | 0.877 | 5883738.5 | 7155764.5 | 1.19E+07 | 5106310.5 | 7787763 | 8513121 | 6442659.5 | 7755894.5 | 8839390 | 9559792 | 7186210.5 | 7890553 | 1627.7816 | VSLYDLSSVDSWEK | Q9WTR1 | Q9WTR1 | TRPV2 | transient receptor potential cation channel, subfamily V, member 2 | Plasma Membrane | ion channel |
| 15534171 | 2 | 0.206 | 9142003 | 7924581 | 9995604 | 5937216 | 9373804 | 5345558.5 | 8088959 | 1.01E+07 | 9132175 | 9113398 | 7627512.5 | 7449693.5 | 1360.65 | VEEVNWAAWEK | Q9WTR1 | Q9WTR1 | TRPV2 | transient receptor potential cation channel, subfamily V, member 2 | Plasma Membrane | ion channel |
| 15553590 | 3 | 0.084 | 7525523.5 | 6024066 | 8400513 | 4731650 | 7506929.5 | 5976420.5 | 7957715.5 | 7252516.5 | 8802577 | 8720546 | 1.05E+07 | 7905574 | 2300.08 | HQGTC[160.0307]FYFGELPLSLAAC[160.0307]TK | Q9WTR1 | Q9WTR1 | TRPV2 | transient receptor potential cation channel, subfamily V, member 2 | Plasma Membrane | ion channel |
| 15540748 | 2 | 0.1 | 9133638 | 8996401 | 8972799 | 8122277 | 8744940 | 6895902.5 | 9694771 | 9737301 | 8375197.5 | 8507518 | 7807172 | 8261520.5 | 1296.6555 | EFSGLYQPLSR | Q9WTR1 | Q9WTR1 | TRPV2 | transient receptor potential cation channel, subfamily V, member 2 | Plasma Membrane | ion channel |
| 15542429 | 2 | 0.208 | 6986345.5 | 6100226.5 | 6133254.5 | 5186458 | 6462663 | 4728948 | 6209624.5 | 5901190.5 | 5459795 | 7980029.5 | 8152402.5 | 5484286.5 | 1492.6777 | YLTDSAYTEGSTGK | Q9WTR1 | Q9WTR1 | TRPV2 | transient receptor potential cation channel, subfamily V, member 2 | Plasma Membrane | ion channel |
| 15540714 | 2, 3 | 0.007 | 6221382 | 5134792.5 | 4170947.75 | 3511451.25 | 4249381 | 4067574.75 | 5567326 | 5453505 | 4808214 | 6924007.5 | 7835852.5 | 6065570.5 | 1430.743, 1430.7433 | NSVLEIIAFHC[160.0307]K | Q9WTR1 | Q9WTR1 | TRPV2 | transient receptor potential cation channel, subfamily V, member 2 | Plasma Membrane | ion channel |
| 15550970 | 2, 3 | 0.707 | 3833635.75 | 4234347.5 | 6690839 | 4454513 | 6985465.5 | 4562669.5 | 5229493 | 7011533.5 | 5466707.5 | 5936194 | 4236221 | 4680421.5 | 2335.2756, 2335.2778 | AVLNLQDGVNAC[160.0307]ILPLLQIDR | Q9WTR1 | Q9WTR1 | TRPV2 | transient receptor potential cation channel, subfamily V, member 2 | Plasma Membrane | ion channel |
| 15551447 | 2 | 0.455 | 2851890 | 3908178.75 | 5472602.5 | 2951370.5 | 5554933 | 2561984 | 4452645 | 4171734.25 | 2737628 | 4223797 | 1925650.125 | 1781103.5 | 1679.7206 | LETSDGDEEGSAEVNK | Q9WTR1 | Q9WTR1 | TRPV2 | transient receptor potential cation channel, subfamily V, member 2 | Plasma Membrane | ion channel |
| 15535852 | 2 | 0.042 | 4055732 | 4066555.75 | 3929822.5 | 2686818.25 | 3418292.5 | 2943596.75 | 3132247.25 | 3631236.75 | 3619119 | 3858399 | 2983994 | 3474766.5 | 807.47046 | LFSVVSR | Q9WTR1 | Q9WTR1 | TRPV2 | transient receptor potential cation channel, subfamily V, member 2 | Plasma Membrane | ion channel |
| 15542422 | 2 | 0.45 | 2176921.75 | 3020295 | 4518589.5 | 1515240.625 | 3328973.75 | 2617431 | 4186913 | 3550183.25 | 2782031 | 3161141.75 | 2278322.25 | 2319912 | 1314.5906 | FTEWC[160.0307]YGPVR | Q9WTR1 | Q9WTR1 | TRPV2 | transient receptor potential cation channel, subfamily V, member 2 | Plasma Membrane | ion channel |
| 15541142 | 3 | 0.013 | 8243817 | 5060672.5 | 4613609 | 2763525.25 | 3371436 | 2593688.5 | 4537103 | 3810956.75 | 4086128.75 | 4092619.25 | 2894244.5 | 2952246.5 | 2194.1833 | EILTLLQGVHQGTGFQDIPK | Q62348 | Q62348 | TSN | translin | Nucleus | other |
| 15538418 | 2 | 0.296 | 2930296.5 | 1931354.875 | 1395822.25 | 1431522.875 | 1422447.75 | 634138.188 | 1629954.75 | 1380896.75 | 1412187.75 | 2016762.625 | 1115747.875 | 866788.75 | 1130.6152 | VVQSLEQTAR | Q62348 | Q62348 | TSN | translin | Nucleus | other |
| 15544174 | 3 | 0.151 | 420159.594 | 1486212.875 | 1932295 | 5838634 | 5513484 | 6005494 | 3337571.25 | 4416012 | 5880236.5 | 387457.812 | 6565571 | 3825592.5 | 2036.0276 | SYRDDIDLQNLIDSLQK | Q8QZY6 | Q8QZY6 | TSPAN14 | tetraspanin 14 | Plasma Membrane | other |
| 15550507, 15551644 | 3, 4 | 1.13E-04 | 2.17E+08 | 2.09E+08 | 2.32E+08 | 8.77E+07 | 1.19E+08 | 1.03E+08 | 1.91E+08 | 1.89E+08 | 1.64E+08 | 1.38E+08 | 1.07E+08 | 1.23E+08 | 3142.6016, 3142.6082 | VSEIINNAIVHYRDDLDLQNLIDFGQK | Q8R3S2 | Q8R3S2 | TSPAN33 | tetraspanin 33 | Plasma Membrane | other |
| 15532719 | 2, 3 | 6.92E-05 | 7.11E+07 | 7.31E+07 | 7.77E+07 | 3.68E+07 | 3.89E+07 | 3.44E+07 | 7.33E+07 | 6.55E+07 | 5.51E+07 | 4.43E+07 | 3.38E+07 | 3.81E+07 | 1199.6244, 1199.6249 | LQLYNQQHR | Q8R3S2 | Q8R3S2 | TSPAN33 | tetraspanin 33 | Plasma Membrane | other |
| 15550073 | 3 | 0.049 | 3.39E+07 | 1.90E+07 | 1.23E+07 | 1.71E+07 | 1.61E+07 | 1.12E+07 | 2.97E+07 | 3.10E+07 | 3.08E+07 | 2.67E+07 | 3.48E+07 | 2.49E+07 | 3963.7456 | C[160.0307]SVPYSC[160.0307]C[160.0307]LPTPNQAVINTMC[160.0307]GQGMQALDYLEASK | Q8R3S2 | Q8R3S2 | TSPAN33 | tetraspanin 33 | Plasma Membrane | other |
| 15542057, 15544300 | 2 | 0.021 | 2.68E+07 | 1.88E+07 | 1.80E+07 | 1.13E+07 | 1.39E+07 | 9562086 | 2.79E+07 | 2.20E+07 | 1.92E+07 | 2.62E+07 | 2.20E+07 | 1.47E+07 | 1206.4993, 1206.4995 | FSC[160.0307]C[160.0307]GGISYR | Q8R3S2 | Q8R3S2 | TSPAN33 | tetraspanin 33 | Plasma Membrane | other |
| 15532651 | 3 | 0.064 | 4557366.5 | 7593658.5 | 1.02E+07 | 2211281 | 6493797 | 4010207 | 6157260 | 8545258 | 5224632 | 4406335.5 | 1968992.75 | 1977303.375 | 1527.8242 | VSEIINNAIVHYR | Q8R3S2 | Q8R3S2 | TSPAN33 | tetraspanin 33 | Plasma Membrane | other |
| 15555074 | 2 | 0.064 | 2585493.25 | 5961857 | 8585454 | 1631449.5 | 4079787 | 3175681.25 | 4629908.5 | 8189641.5 | 4013894.75 | 2392170.5 | 1676769.625 | 1907299.5 | 1633.8054 | DDLDLQNLIDFGQK | Q8R3S2 | Q8R3S2 | TSPAN33 | tetraspanin 33 | Plasma Membrane | other |
| 15538275 | 2 | 0.021 | 2207507.25 | 1759359.625 | 3295511.5 | 1127262.5 | 1284261.625 | 1283956 | 3382642.75 | 2979079.75 | 2301025.25 | 2079618.125 | 2819225.5 | 1399616.375 | 1182.5793 | VIYTNGC[160.0307]IDK | Q8R3S2 | Q8R3S2 | TSPAN33 | tetraspanin 33 | Plasma Membrane | other |
| 15540601 | 2 | 0.382 | 5.67E+07 | 4.95E+07 | 5.32E+07 | 4.40E+07 | 4.35E+07 | 4.49E+07 | 5.45E+07 | 4.93E+07 | 3.67E+07 | 6.18E+07 | 4.62E+07 | 4.64E+07 | 1105.5353 | LLSDNTDEAK | Q8R3G9 | Q8R3G9 | TSPAN8 | tetraspanin 8 | Plasma Membrane | other |
| 15548513 | 2 | 0.114 | 3.00E+07 | 2.49E+07 | 2.54E+07 | 3.22E+07 | 2.52E+07 | 2.89E+07 | 3.37E+07 | 2.93E+07 | 2.72E+07 | 3.99E+07 | 3.84E+07 | 2.93E+07 | 2111.8867 | C[160.0307]C[160.0307]GLENGAADWGNNFVEAK | Q8R3G9 | Q8R3G9 | TSPAN8 | tetraspanin 8 | Plasma Membrane | other |
| 15539714 | 2 | 0.535 | 2.26E+07 | 2.71E+07 | 2.07E+07 | 2.42E+07 | 1.75E+07 | 2.14E+07 | 2.85E+07 | 2.32E+07 | 2.04E+07 | 2.47E+07 | 2.40E+07 | 2.44E+07 | 1199.6102 | AMIVFQSEFK | Q8R3G9 | Q8R3G9 | TSPAN8 | tetraspanin 8 | Plasma Membrane | other |
| 15548550 | 3 | 0.915 | 1.03E+07 | 9808853 | 1.41E+07 | 1.10E+07 | 1.11E+07 | 1.19E+07 | 1.13E+07 | 1.34E+07 | 1.20E+07 | 1.90E+07 | 1.07E+07 | 9404191 | 2398.9539 | ESC[160.0307]QC[160.0307]TGTDC[160.0307]ATYQGSSVYPK | Q8R3G9 | Q8R3G9 | TSPAN8 | tetraspanin 8 | Plasma Membrane | other |
| 15536189 | 3 | 0.143 | 6033868 | 5466913.5 | 5184661.5 | 7079662.5 | 5395444.5 | 4405635.5 | 7343643 | 5946437.5 | 5336033.5 | 7548808 | 9445249 | 6497456.5 | 2111.8862 | C[160.0307]C[160.0307]GLENGAADWGNNFVEAK | Q8R3G9 | Q8R3G9 | TSPAN8 | tetraspanin 8 | Plasma Membrane | other |
| 15531907 | 2 | 0.191 | 6670355.5 | 1.44E+07 | 8244824 | 9668490 | 1.36E+07 | 1.50E+07 | 7615082.5 | 1.40E+07 | 1.63E+07 | 4449875.5 | 8206684.5 | 8231730 | 895.4663 | WYASLQK | P50637 | P50637 | TSPO | translocator protein (18kDa) | Cytoplasm | transmembrane receptor |
| 15538299 | 2 | 0.368 | 4269340 | 4115474.5 | 4579545 | 3600968.25 | 4059754 | 3375587 | 3634069.5 | 3473388.5 | 3100921.75 | 5224931 | 3082237 | 3326251.5 | 844.4762 | KITPEEK | A2ASS6 | A2ASS6 | TTN | titin | Cytoplasm | kinase |
| 15552195 | 2 | 0.218 | 2832537.25 | 4255446.5 | 1989395 | 5078839 | 4446952.5 | 3168852.25 | 3108239.5 | 3049278.25 | 5067653.5 | 1036830.938 | 2601392.25 | 3122908.25 | 1701.9016 | AVFVDLEPTVIDEVR | P68369 | P68369 | TUBA1A | tubulin, alpha 1a | Cytoplasm | other |
| 15538464 | 2 | 0.263 | 2000416.875 | 2705121 | 2415974 | 2930894 | 3481713.75 | 1878840.75 | 2102566.75 | 2060680.375 | 2460166 | 954109.5 | 2213196.25 | 1993829 | 1446.6875 | EVDEQMLNVQNK | Q7TMM9 | Q7TMM9 | TUBB2A | tubulin, beta 2A class IIa | Cytoplasm | other |
| 15534787 | 2 | 0.321 | 1365722.375 | 2165716.5 | 2115666.25 | 3609109.75 | 4019234.5 | 2895792.25 | 2616905 | 2210232.75 | 2633019.5 | 349547.844 | 2471050.25 | 2781472 | 1143.6318 | LAVNMVPFPR | Q7TMM9 | Q7TMM9 | TUBB2A | tubulin, beta 2A class IIa | Cytoplasm | other |
| 15549340 | 3 | 0.369 | 4628716 | 6836898.5 | 2725741.5 | 7532430.5 | 6708404.5 | 3598812.5 | 3699087.25 | 4564174 | 5051255 | 1833632.25 | 4007379.75 | 4093326.5 | 3311.5283 | EAESC[160.0307]DC[160.0307]LQGFQLTHSLGGGTGSGMGTLLISK | Q9D6F9 | Q9D6F9 | TUBB4A | tubulin, beta 4A class IVa | Cytoplasm | other |
| 15552133 | 3 | 0.405 | 6657957 | 7671945 | 6017359 | 8237755.5 | 7867284 | 5081743 | 5582671 | 5907560 | 7591108.5 | 868243 | 6103105.5 | 6472481 | 2798.3354 | SGPFGQIFRPDNFVFGQSGAGNNWAK | Q9D6F9 | Q9D6F9 | TUBB4A | tubulin, beta 4A class IVa | Cytoplasm | other |
| 15551626 | 2 | 0.314 | 3505907 | 9910081 | 4815517.5 | 4791779 | 8735634 | 8114659.5 | 3929370.75 | 7370657 | 9847162 | 2777287.5 | 4320814 | 4375444 | 1572.8596 | LLDAVDTYIPVPTR | Q8BFR5 | Q8BFR5 | TUFM | Tu translation elongation factor, mitochondrial | Cytoplasm | translation regulator |
| 15550750 | 2 | 0.073 | 3528093.75 | 8745499 | 5848534 | 7311397 | 7818129 | 9582153 | 5179071.5 | 9586857 | 1.21E+07 | 2303558.25 | 4824241 | 4074959.5 | 1876.9187 | TIGTGLVTDVPAMTEEDK | Q8BFR5 | Q8BFR5 | TUFM | Tu translation elongation factor, mitochondrial | Cytoplasm | translation regulator |
| 15549003 | 2 | 0.179 | 4134805.25 | 8396038 | 4544992.5 | 6465019 | 6304636.5 | 9599587 | 3959002.5 | 7976281 | 9276477 | 2155946.25 | 4560447 | 4511831 | 1673.9238 | PFLLPVESVYSIPGR | Q8BFR5 | Q8BFR5 | TUFM | Tu translation elongation factor, mitochondrial | Cytoplasm | translation regulator |
| 15534328 | 2, 3 | 0.108 | 3032824.5 | 5755063.5 | 2739761.5 | 5664234 | 4684774 | 5301339 | 3932805.5 | 5267616.5 | 5824186.5 | 1971583.375 | 4065335.25 | 2842106.25 | 1492.6877, 1492.6887 | KYEEIDNAPEER | Q8BFR5 | Q8BFR5 | TUFM | Tu translation elongation factor, mitochondrial | Cytoplasm | translation regulator |
| 15552381 | 3 | 0.302 | 2360811.5 | 5275486.5 | 3018241.5 | 4111856.75 | 3850881 | 4726110.5 | 2224350.75 | 4488999 | 6297602 | 1058785.875 | 3709873.25 | 2276280.25 | 2812.3718 | NMITGTAPLDGC[160.0307]ILVVAANDGPMPQTR | Q8BFR5 | Q8BFR5 | TUFM | Tu translation elongation factor, mitochondrial | Cytoplasm | translation regulator |
| 15537826 | 2 | 0.309 | 1041724.312 | 2886130.75 | 1152463.75 | 1773899.625 | 2107498.5 | 3732455.25 | 1504621.875 | 2185419.75 | 2888484.75 | 444216.375 | 1812075.5 | 1478699.125 | 1149.6487 | VEAQVYILSK | Q8BFR5 | Q8BFR5 | TUFM | Tu translation elongation factor, mitochondrial | Cytoplasm | translation regulator |
| 15554203 | 2 | 0.932 | 1044250 | 1100683.375 | 1064585.875 | 1365890.625 | 1495870.875 | 680904.562 | 1044650.125 | 1148721.25 | 878584.625 | 1302285.25 | 722610.688 | 985996.75 | 1739.7296 | VDC[160.0307]TADSDVC[160.0307]SAQGVR | Q91W90 | Q91W90 | TXNDC5 | thioredoxin domain containing 5 (endoplasmic reticulum) | Cytoplasm | enzyme |
| 15542730 | 3 | 0.71 | 1.45E+07 | 7291922.5 | 1538972.75 | 1.04E+07 | 6416060.5 | 2316546 | 6510591 | 5268725 | 1.49E+07 | 8964218 | 8786208 | 1.21E+07 | 2199.0989 | VGVKPVGSDPDFQPELSGAGSR | Q8CDN6 | Q8CDN6 | TXNL1 | thioredoxin-like 1 | Cytoplasm | enzyme |
| 15551879 | 3 | 0.025 | 1.35E+07 | 9991776 | 7070401 | 7159855.5 | 8504772 | 4274708 | 8010369 | 5889189.5 | 6808935.5 | 5001140.5 | 3852816.75 | 3537635.25 | 2783.2498 | SMDFEEAERSEPTQALELTEDDIK | Q8CDN6 | Q8CDN6 | TXNL1 | thioredoxin-like 1 | Cytoplasm | enzyme |
| 15536541 | 4 | 0.054 | 9734719 | 7612820 | 6007649 | 6552691.5 | 7126732.5 | 3520938 | 5326566 | 3986037.25 | 3745463.25 | 4574709.5 | 3177182 | 4026771.75 | 2410.9768 | AGC[160.0307]EC[160.0307]LNESDEHGFDNC[160.0307]LRK | Q8CDN6 | Q8CDN6 | TXNL1 | thioredoxin-like 1 | Cytoplasm | enzyme |
| 15534322 | 3 | 0.069 | 8008653.5 | 6741011.5 | 5145888 | 4878371 | 5362044.5 | 2416599.25 | 3872145 | 4248401.5 | 3672440 | 4464934 | 1963537.625 | 2379960.75 | 1878.8452 | QHLENDPGSNEDADIPK | Q8CDN6 | Q8CDN6 | TXNL1 | thioredoxin-like 1 | Cytoplasm | enzyme |
| 15551701 | 2 | 0.082 | 2100534.75 | 4556216.5 | 6522131 | 1278237 | 4687909 | 2537953.75 | 2885457.25 | 3086413.75 | 1437355.5 | 2022193.75 | 448698.344 | 700605.188 | 1815.8474 | PVGSDPDFQPELSGAGSR | Q8CDN6 | Q8CDN6 | TXNL1 | thioredoxin-like 1 | Cytoplasm | enzyme |
| 15556099 | 3 | 0.196 | 4195583.5 | 2279215.75 | 1238967.625 | 2062093.375 | 1589547.5 | 602057.625 | 1540519.625 | 1095989 | 870764.562 | 1429594.875 | 495850.312 | 705001.312 | 2410.9744 | AGC[160.0307]EC[160.0307]LNESDEHGFDNC[160.0307]LRK | Q8CDN6 | Q8CDN6 | TXNL1 | thioredoxin-like 1 | Cytoplasm | enzyme |
| 15543759 | 3 | 0.115 | 2902555.5 | 3879550.75 | 1806932.75 | 2540192.25 | 1498665 | 672535.75 | 1268457 | 1452794.25 | 1366810.875 | 1342751 | 86455.617 | 666629.688 | 1890.9542 | VRIDQYQGADAVGLEEK | Q8CDN6 | Q8CDN6 | TXNL1 | thioredoxin-like 1 | Cytoplasm | enzyme |
| 15546986 | 2 | 0.027 | 733932.688 | 598190.812 | 503945.625 | 248965.734 | 444096.719 | 256320.953 | 312679.094 | 388781.688 | 353032.125 | 395769.75 | 177428.547 | 167476.453 | 1144.5354 | FQGPDNGQGPK | Q8CDN6 | Q8CDN6 | TXNL1 | thioredoxin-like 1 | Cytoplasm | enzyme |
| 15548514 | 4 | 0.896 | 1.72E+07 | 1.85E+07 | 1.36E+07 | 1.81E+07 | 1.27E+07 | 1.90E+07 | 1.12E+07 | 1.82E+07 | 1.50E+07 | 2.26E+07 | 9768311 | 1.33E+07 | 3474.8416 | IIVDAQEATSVPHIYAIGDVAEGRPELTPTAIK | Q9JLT4 | Q9JLT4 | TXNRD2 | thioredoxin reductase 2 | Cytoplasm | enzyme |
| 15539752 | 2 | 0.085 | 7408071 | 6336705 | 5501830 | 6490396.5 | 4988459.5 | 6707688 | 4422675.5 | 5070146.5 | 4625205.5 | 5965456 | 4761105 | 5646041 | 1302.6661 | VAVADYVEPSPR | Q9JLT4 | Q9JLT4 | TXNRD2 | thioredoxin reductase 2 | Cytoplasm | enzyme |
| 15536526 | 2 | 0.485 | 1166815.625 | 1204309.875 | 746677.625 | 1064957.375 | 846287.938 | 1052451 | 819227 | 1027123.188 | 511164.969 | 1270095.75 | 957268.312 | 815271.375 | 1227.6123 | TMAEAVQNHVK | Q9JLT4 | Q9JLT4 | TXNRD2 | thioredoxin reductase 2 | Cytoplasm | enzyme |
| 15552027, 15556585 | 2, 3 | 0.307 | 7877683 | 7320219.5 | 6255068.5 | 8415605 | 5426955 | 8033087.5 | 6327114.5 | 6374457.5 | 4896811 | 1.28E+07 | 7712013.5 | 6511349 | 2614.2007, 2614.2036 | IYDDDFFQNLDGVANALDNIDAR | Q02053 | Q02053 | UBA1 | ubiquitin-like modifier activating enzyme 1 | Cytoplasm | enzyme |
| 15553030 | 2 | 0.256 | 6784905.5 | 4349340 | 5538158 | 6330985 | 4385933.5 | 3900640 | 4734427.5 | 3887316.25 | 3084539 | 1.12E+07 | 5590955.5 | 4585753 | 1606.7676 | QPAENVNQYLTDSK | Q02053 | Q02053 | UBA1 | ubiquitin-like modifier activating enzyme 1 | Cytoplasm | enzyme |
| 15551950 | 2 | 0.417 | 7130266 | 5331455.5 | 6323167.5 | 7118985.5 | 4434607 | 6248623 | 6033888.5 | 4812162 | 3478531.5 | 1.11E+07 | 5875861.5 | 5182661 | 1885.0388 | AAVASLLQSVQVPEFTPK | Q02053 | Q02053 | UBA1 | ubiquitin-like modifier activating enzyme 1 | Cytoplasm | enzyme |
| 15542100 | 2 | 0.124 | 6611161 | 4361531.5 | 4665135 | 6451425.5 | 5580713.5 | 4975758.5 | 5249073.5 | 3641517.5 | 2412095.5 | 1.03E+07 | 6095688 | 5219776 | 1445.6663 | DNPGVVTC[160.0307]LDEAR | Q02053 | Q02053 | UBA1 | ubiquitin-like modifier activating enzyme 1 | Cytoplasm | enzyme |
| 15550331 | 2 | 0.195 | 4652821.5 | 4551117 | 4976974.5 | 5298727 | 3332143.75 | 4621764 | 3761324.75 | 3162451.75 | 2438413.25 | 9191075 | 3874681.25 | 4192567.25 | 1966.9785 | SLPASLVEPDFVMTDFAK | Q02053 | Q02053 | UBA1 | ubiquitin-like modifier activating enzyme 1 | Cytoplasm | enzyme |
| 15549544 | 2 | 0.241 | 5245731 | 3686062.5 | 4025525 | 5986988 | 3595360 | 4201507.5 | 4116138.5 | 2757653 | 2316526.25 | 7784981 | 5414261.5 | 3235215 | 1899.9365 | NEEDATELVGLAQAVNAR | Q02053 | Q02053 | UBA1 | ubiquitin-like modifier activating enzyme 1 | Cytoplasm | enzyme |
| 15535011 | 2 | 0.265 | 4579735 | 3224470 | 3239071.5 | 3730567.5 | 3218585.5 | 3057492.25 | 3192497.25 | 2575606.25 | 1628971.625 | 7662501 | 3365023.5 | 2900720.25 | 1250.5621 | AENYDISPADR | Q02053 | Q02053 | UBA1 | ubiquitin-like modifier activating enzyme 1 | Cytoplasm | enzyme |
| 15533823 | 3 | 0.052 | 5201113 | 3844439.5 | 3892337.25 | 4925626.5 | 3737865.25 | 4315874.5 | 3714564.5 | 2905212.75 | 1913089.875 | 7362872.5 | 5195431 | 4165768.25 | 1573.7976 | ERLDQPMTEIVSR | Q02053 | Q02053 | UBA1 | ubiquitin-like modifier activating enzyme 1 | Cytoplasm | enzyme |
| 15543201 | 2, 3 | 0.307 | 3404037.25 | 1909167.625 | 2419722.25 | 4017558.75 | 1676633.75 | 2594216 | 3000460.75 | 1276591.5 | 1128939.5 | 5836698.5 | 3601850 | 2036671.25 | 1401.7096, 1401.7097 | VVQGHQQLDSYK | Q02053 | Q02053 | UBA1 | ubiquitin-like modifier activating enzyme 1 | Cytoplasm | enzyme |
| 15532520 | 2 | 0.815 | 2278917.25 | 3181318.5 | 5392284.5 | 2660761 | 2972830.75 | 3967928 | 3727111.75 | 2583380.75 | 1592180.75 | 4691941.5 | 1637410.875 | 3054591 | 1115.6268 | PLLESGTLGTK | Q02053 | Q02053 | UBA1 | ubiquitin-like modifier activating enzyme 1 | Cytoplasm | enzyme |
| 15551573 | 2 | 0.073 | 3660129.25 | 2373087 | 2748675 | 2836840.5 | 2317399.25 | 2541434.75 | 2058163.875 | 1720980.25 | 2622493.25 | 4333632.5 | 3542673.75 | 2741504.25 | 1610.8584 | YFLVGAGAIGC[160.0307]ELLK | Q02053 | Q02053 | UBA1 | ubiquitin-like modifier activating enzyme 1 | Cytoplasm | enzyme |
| 15535538 | 3 | 0.184 | 2747809 | 2362836.5 | 1840504.25 | 1769465.875 | 2461784 | 3081157.25 | 2103269.75 | 1914142.75 | 1363788.375 | 4044267.5 | 3158356 | 2121965 | 1623.9033 | LAGTQPLEVLEAVQR | Q02053 | Q02053 | UBA1 | ubiquitin-like modifier activating enzyme 1 | Cytoplasm | enzyme |
| 15549516 | 2 | 0.198 | 939718.688 | 729071.125 | 283627.219 | 1277355 | 531756.438 | 1257833.625 | 1003532.125 | 581969.25 | 393134.125 | 1770721.875 | 1136826.625 | 1158404.75 | 2603.3855 | LAYVAAGDLAPINAFIGGLAAQEVMK | Q02053 | Q02053 | UBA1 | ubiquitin-like modifier activating enzyme 1 | Cytoplasm | enzyme |
| 15544078 | 2 | 0.528 | 2347392.75 | 363180.75 | 4.70E-06 | 1333265.875 | 261194.797 | 166.49 | 395455.062 | 426960.406 | 297382.344 | 1756163 | 2326392.25 | 799175.75 | 1243.7233 | KPLLESGTLGTK | Q02053 | Q02053 | UBA1 | ubiquitin-like modifier activating enzyme 1 | Cytoplasm | enzyme |
| 15548799 | 2 | 0.144 | 7.77E+08 | 8.54E+08 | 1.20E+09 | 1.06E+09 | 1.05E+09 | 1.07E+09 | 8.41E+08 | 8.53E+08 | 9.14E+08 | 1.56E+09 | 1.05E+09 | 1.06E+09 | 1787.9238 | TITLEVEPSDTIENVK | P62984 | P62984 | Uba52 | ubiquitin A-52 residue ribosomal protein fusion product 1 | Cytoplasm | other |
| 15536569 | 2 | 0.067 | 2.46E+08 | 3.17E+08 | 3.64E+08 | 3.73E+08 | 3.70E+08 | 4.31E+08 | 3.16E+08 | 3.19E+08 | 2.72E+08 | 5.78E+08 | 3.71E+08 | 3.88E+08 | 1081.5513 | TLSDYNIQK | P62984 | P62984 | Uba52 | ubiquitin A-52 residue ribosomal protein fusion product 1 | Cytoplasm | other |
| 15534664 | 2 | 0.012 | 1.42E+08 | 1.56E+08 | 1.72E+08 | 2.03E+08 | 1.58E+08 | 2.05E+08 | 1.74E+08 | 1.52E+08 | 1.67E+08 | 2.77E+08 | 2.10E+08 | 2.28E+08 | 765.4314 | MQIFVK | P62984 | P62984 | Uba52 | ubiquitin A-52 residue ribosomal protein fusion product 1 | Cytoplasm | other |
| 15531610, 15544917 | 2 | 0.002 | 1.60E+08 | 1.90E+08 | 1.84E+08 | 2.16E+08 | 1.99E+08 | 2.32E+08 | 1.77E+08 | 1.49E+08 | 1.43E+08 | 2.64E+08 | 2.54E+08 | 2.18E+08 | 1067.6189, 1067.6191 | ESTLHLVLR | P62984 | P62984 | Uba52 | ubiquitin A-52 residue ribosomal protein fusion product 1 | Cytoplasm | other |
| 15531463 | 3 | 0.26 | 6.18E+07 | 2.31E+07 | 1.27E+07 | 1.25E+08 | 1.15E+07 | 3.19E+07 | 3.03E+07 | 1.84E+07 | 4.77E+07 | 7.71E+07 | 1.15E+08 | 8.44E+07 | 1779.8828 | QLEDGRTLSDYNIQK | P62984 | P62984 | Uba52 | ubiquitin A-52 residue ribosomal protein fusion product 1 | Cytoplasm | other |
| 15549712 | 2 | 0.245 | 8541476 | 3403243 | 1625213.5 | 1.63E+07 | 1513898.625 | 3364339.5 | 4199671.5 | 2495848 | 7002378 | 1.26E+07 | 1.67E+07 | 1.05E+07 | 1779.8839 | QLEDGRTLSDYNIQK | P62984 | P62984 | Uba52 | ubiquitin A-52 residue ribosomal protein fusion product 1 | Cytoplasm | other |
| 15533114 | 3 | 0.911 | 2.37E+07 | 3.10E+07 | 3.43E+07 | 1.71E+07 | 3.84E+07 | 1.80E+07 | 2.55E+07 | 2.70E+07 | 2.70E+07 | 1.22E+07 | 3.20E+07 | 5.00E+07 | 1787.9244 | TITLEVEPSDTIENVK | P62984 | P62984 | Uba52 | ubiquitin A-52 residue ribosomal protein fusion product 1 | Cytoplasm | other |
| 15552747 | 3 | 0.775 | 1178952.625 | 128143.789 | 347051.031 | 974435.875 | 354904.844 | 1583835.375 | 15972.815 | 872144.938 | 7929315 | 611426.75 | 2113624.75 | 2635262 | 2288.219 | TLTGKTITLEVEPSDTIENVK | P62984 | P62984 | Uba52 | ubiquitin A-52 residue ribosomal protein fusion product 1 | Cytoplasm | other |
| 15551996 | 3, 4 | 0.906 | 7489434.5 | 1.04E+07 | 8419151 | 4887490 | 1.13E+07 | 7915472 | 9079480 | 1.21E+07 | 6261927 | 1.50E+07 | 4593902.5 | 5168215.5 | 3783.9722, 3783.9739 | KPSPEELDQGIDPNSPLFQAILDNPVVQLGLTNPK | Q8VDI7 | Q8VDI7 | UBAC1 | UBA domain containing 1 | Nucleus | other |
| 15532684 | 2 | 0.625 | 7122402.5 | 7016967.5 | 7863768.5 | 6244901 | 6059702 | 6871680 | 7533490.5 | 6266337.5 | 4601225 | 1.26E+07 | 5023111 | 7499781 | 1442.8231 | LLALNPDAVELFK | Q8VDI7 | Q8VDI7 | UBAC1 | UBA domain containing 1 | Nucleus | other |
| 15555723 | 3 | 0.376 | 5237018.5 | 5389643 | 6903792.5 | 4000026 | 5258870 | 5614312 | 4974555 | 4576323 | 4947801.5 | 6924308 | 5066024 | 5106203 | 2574.1606 | LHIC[160.0307]AADGAEWLEEATEDTSVEK | Q8VDI7 | Q8VDI7 | UBAC1 | UBA domain containing 1 | Nucleus | other |
| 15538946 | 3 | 0.559 | 5429992.5 | 4153984.75 | 3465568 | 4074737.75 | 4097602.5 | 3355734 | 4662053 | 4356269 | 2107710.75 | 6152470.5 | 4159693.75 | 4310027.5 | 2191.9744 | ANAMLDEDEDERVDETALR | Q8VDI7 | Q8VDI7 | UBAC1 | UBA domain containing 1 | Nucleus | other |
| 15537004 | 2 | 0.408 | 4875143.5 | 3009027.5 | 3802944.75 | 4492901.5 | 3120850.25 | 3459385.75 | 3784317 | 2932161.75 | 2171033.5 | 6022252.5 | 3446579.25 | 3499758 | 1212.754 | ILVSLIEVAQK | Q8VDI7 | Q8VDI7 | UBAC1 | UBA domain containing 1 | Nucleus | other |
| 15543863 | 2 | 0.386 | 3216289.5 | 2419716.5 | 2856529.25 | 2964674 | 2738610.75 | 2886979.5 | 3378322 | 2219739.25 | 1913895.875 | 5684926.5 | 2988106 | 2720184.25 | 1294.6085 | QLTEMGFPESR | Q8VDI7 | Q8VDI7 | UBAC1 | UBA domain containing 1 | Nucleus | other |
| 15554940 | 2 | 0.899 | 2673887 | 2589562.5 | 2218590.75 | 2842217.5 | 2485774.25 | 2605521.25 | 3256731.5 | 1816038.375 | 1595511.25 | 4914896 | 1491105.875 | 1933452.125 | 1997.1111 | TILEENIQDQDVLLLIK | Q8VDI7 | Q8VDI7 | UBAC1 | UBA domain containing 1 | Nucleus | other |
| 15535335 | 2 | 0.402 | 2094475.5 | 1124005.125 | 1678724 | 1964183.875 | 974297.688 | 1325395 | 1945548.5 | 1237922.125 | 513772.5 | 3556675.25 | 1587708.875 | 1692493 | 992.5506 | IFQTLNRT | Q8VDI7 | Q8VDI7 | UBAC1 | UBA domain containing 1 | Nucleus | other |
| 15536501 | 2 | 0.102 | 1613304.875 | 1537782 | 977696.562 | 2079818.75 | 1421406.125 | 1619045 | 1456851.5 | 1212840.25 | 644897.312 | 2302196.75 | 1839509.375 | 1792051.375 | 1469.6981 | AVISLMEMGFDEK | Q8VDI7 | Q8VDI7 | UBAC1 | UBA domain containing 1 | Nucleus | other |
| 15539686 | 2 | 0.74 | 841598.125 | 2070812.875 | 2793610.5 | 263599.469 | 2308478.75 | 2891275.5 | 2188868.5 | 2406391.25 | 2255908.5 | 609109.438 | 1142454.375 | 2532300 | 1276.5914 | ATANLPAC[160.0307]STDR | Q8VDI7 | Q8VDI7 | UBAC1 | UBA domain containing 1 | Nucleus | other |
| 15551866 | 4 | 0.375 | 9136070 | 6593028.5 | 8799525 | 8471877 | 7663174 | 1.02E+07 | 6700977.5 | 7355753 | 7081256.5 | 1.54E+07 | 6503460.5 | 1.06E+07 | 3489.7783 | TDQVIQSLIALVNDPQPEHPLRADLAEEYSK | P68037 | P68037 | UBE2L3 | ubiquitin-conjugating enzyme E2L 3 | Nucleus | enzyme |
| 15548697 | 3 | 0.751 | 2.22E+07 | 2.49E+07 | 8835701 | 1.63E+07 | 2.30E+07 | 2.20E+07 | 2.83E+07 | 2.57E+07 | 1.75E+07 | 3.22E+07 | 1.42E+07 | 9334313 | 2965.543 | QLSSDDANVLVWHMLLLPDQLPYGLK | Q9QZU9 | Q9QZU9 | UBE2L6 | ubiquitin-conjugating enzyme E2L 6 | Cytoplasm | enzyme |
| 15534458 | 2 | 0.567 | 5445427 | 5490554 | 2495026.75 | 4074189.5 | 6504546.5 | 4929913 | 5745099.5 | 5691219.5 | 4346074.5 | 7036889.5 | 2435420.5 | 2173713 | 777.3873 | FGVDRPS | Q9QZU9 | Q9QZU9 | UBE2L6 | ubiquitin-conjugating enzyme E2L 6 | Cytoplasm | enzyme |
| 15552809 | 2 | 0.329 | 2386245.5 | 3925234.5 | 3288467.75 | 1203137 | 6619382 | 4412823 | 3979761.5 | 5752085 | 2308361.25 | 4235990.5 | 457303.25 | 951598.938 | 1572.8976 | PYQVLEALNVLVSK | Q9QZU9 | Q9QZU9 | UBE2L6 | ubiquitin-conjugating enzyme E2L 6 | Cytoplasm | enzyme |
| 15555983 | 3 | 0.682 | 3787088.75 | 1935944.25 | 450714.688 | 1939635.125 | 1517554.625 | 1674431.375 | 3078279.25 | 2228839.75 | 2776863.25 | 3091884.25 | 1575229.125 | 1206919.125 | 2766.3965 | IYHPNVREDGLVC[160.0307]LPLISNENWK | Q9QZU9 | Q9QZU9 | UBE2L6 | ubiquitin-conjugating enzyme E2L 6 | Cytoplasm | enzyme |
| 15548123 | 3 | 0.575 | 2.27E+07 | 1.76E+07 | 2.60E+07 | 2.99E+07 | 4.00E+07 | 2.44E+07 | 3.71E+07 | 3.57E+07 | 1.82E+07 | 5.81E+07 | 1.96E+07 | 3.14E+07 | 3563.6746 | LYDVC[160.0307]PHVSDSGLFFDDSYGFYPGQVLIGPAK | Q6ZPJ3 | Q6ZPJ3 | UBE2O | ubiquitin-conjugating enzyme E2O | Nucleus | enzyme |
| 15548028 | 3 | 0.811 | 2.38E+07 | 1.74E+07 | 2.48E+07 | 2.71E+07 | 3.89E+07 | 1.60E+07 | 2.96E+07 | 2.99E+07 | 2.28E+07 | 4.89E+07 | 1.93E+07 | 2.37E+07 | 2863.3438 | AEWPSETPVLC[160.0307]QQC[160.0307]GGRPGVTFTSAK | Q6ZPJ3 | Q6ZPJ3 | UBE2O | ubiquitin-conjugating enzyme E2O | Nucleus | enzyme |
| 15552676 | 3 | 0.198 | 1.83E+07 | 1.14E+07 | 1.78E+07 | 2.05E+07 | 2.72E+07 | 1.60E+07 | 3.03E+07 | 2.12E+07 | 1.83E+07 | 4.78E+07 | 1.88E+07 | 2.83E+07 | 2612.3186 | NMTVEQLLTGSPTSPTVEPEKPTR | Q6ZPJ3 | Q6ZPJ3 | UBE2O | ubiquitin-conjugating enzyme E2O | Nucleus | enzyme |
| 15551306 | 3, 4 | 0.889 | 2.15E+07 | 1.55E+07 | 2.59E+07 | 2.58E+07 | 2.71E+07 | 2.11E+07 | 2.68E+07 | 2.88E+07 | 2.08E+07 | 4.65E+07 | 1.40E+07 | 2.09E+07 | 3287.5254, 3287.526 | LRPSGDDVELIGEEEDVSVYDIADHPDFR | Q6ZPJ3 | Q6ZPJ3 | UBE2O | ubiquitin-conjugating enzyme E2O | Nucleus | enzyme |
| 15532648 | 4 | 0.35 | 1.72E+07 | 1.19E+07 | 1.82E+07 | 1.84E+07 | 2.28E+07 | 1.64E+07 | 3.24E+07 | 2.12E+07 | 1.41E+07 | 4.53E+07 | 1.81E+07 | 2.29E+07 | 2815.2732 | SNDLFPVHHLDNNEFC[160.0307]PGDFVVDK | Q6ZPJ3 | Q6ZPJ3 | UBE2O | ubiquitin-conjugating enzyme E2O | Nucleus | enzyme |
| 15532961 | 3, 4 | 0.811 | 1.37E+07 | 1.02E+07 | 1.80E+07 | 1.86E+07 | 2.25E+07 | 1.18E+07 | 2.57E+07 | 1.93E+07 | 1.15E+07 | 3.59E+07 | 1.13E+07 | 1.52E+07 | 2234.0625, 2234.0652 | DHTEQTETAPDASAPPSVRPK | Q6ZPJ3 | Q6ZPJ3 | UBE2O | ubiquitin-conjugating enzyme E2O | Nucleus | enzyme |
| 15532533 | 3 | 0.823 | 1.42E+07 | 1.19E+07 | 1.64E+07 | 1.73E+07 | 2.10E+07 | 1.40E+07 | 2.01E+07 | 1.65E+07 | 1.43E+07 | 2.89E+07 | 1.01E+07 | 1.75E+07 | 1894.9303 | GEVFSVLEFAPSNHSFK | Q6ZPJ3 | Q6ZPJ3 | UBE2O | ubiquitin-conjugating enzyme E2O | Nucleus | enzyme |
| 15548299 | 3 | 0.344 | 1.32E+07 | 9469645 | 9879615 | 1.26E+07 | 1.81E+07 | 1.21E+07 | 2.12E+07 | 1.67E+07 | 1.13E+07 | 2.89E+07 | 1.18E+07 | 1.50E+07 | 2815.272 | SNDLFPVHHLDNNEFC[160.0307]PGDFVVDK | Q6ZPJ3 | Q6ZPJ3 | UBE2O | ubiquitin-conjugating enzyme E2O | Nucleus | enzyme |
| 15548233 | 2 | 0.884 | 1.27E+07 | 9651409 | 2.03E+07 | 1.35E+07 | 1.75E+07 | 1.25E+07 | 1.63E+07 | 1.70E+07 | 1.67E+07 | 2.88E+07 | 8557099 | 1.71E+07 | 1637.855 | SGYPDIGFPLFPLSK | Q6ZPJ3 | Q6ZPJ3 | UBE2O | ubiquitin-conjugating enzyme E2O | Nucleus | enzyme |
| 15533363 | 2 | 0.427 | 8380572.5 | 6005359.5 | 9119172 | 1.20E+07 | 1.31E+07 | 8318095.5 | 1.47E+07 | 1.03E+07 | 7541181 | 2.29E+07 | 8248014 | 1.07E+07 | 1186.5671 | EDEPSVGQVAR | Q6ZPJ3 | Q6ZPJ3 | UBE2O | ubiquitin-conjugating enzyme E2O | Nucleus | enzyme |
| 15536812 | 2, 3 | 0.53 | 5947802.5 | 5107711.5 | 7908904.5 | 1.02E+07 | 8465741 | 6156149.5 | 1.25E+07 | 8840372 | 6387736.5 | 1.87E+07 | 5474318.5 | 9554135 | 1243.6754, 1243.676 | LLFSHDLVSGR | Q6ZPJ3 | Q6ZPJ3 | UBE2O | ubiquitin-conjugating enzyme E2O | Nucleus | enzyme |
| 15555593 | 2 | 0.428 | 7493619.5 | 5253383 | 1.13E+07 | 1.00E+07 | 1.06E+07 | 7336724 | 1.09E+07 | 1.01E+07 | 7938701.5 | 1.84E+07 | 8771711 | 1.01E+07 | 1561.7396 | AALLEAGMPESTEDK | Q6ZPJ3 | Q6ZPJ3 | UBE2O | ubiquitin-conjugating enzyme E2O | Nucleus | enzyme |
| 15533186 | 2 | 0.614 | 8095849.5 | 5373566.5 | 1.07E+07 | 8390949 | 1.10E+07 | 7902501 | 1.22E+07 | 9709084 | 7245047 | 1.73E+07 | 7262854.5 | 1.07E+07 | 1105.5652 | VQWYPEGVK | Q6ZPJ3 | Q6ZPJ3 | UBE2O | ubiquitin-conjugating enzyme E2O | Nucleus | enzyme |
| 15550108 | 2 | 0.633 | 8729977 | 6037367 | 1.17E+07 | 1.23E+07 | 1.32E+07 | 6840598.5 | 1.13E+07 | 1.13E+07 | 9839645 | 1.69E+07 | 8854189 | 1.07E+07 | 1591.848 | LIGTNC[160.0307]IIYPVNSK | Q6ZPJ3 | Q6ZPJ3 | UBE2O | ubiquitin-conjugating enzyme E2O | Nucleus | enzyme |
| 15537577 | 2 | 0.676 | 7394104 | 5437529 | 8358028.5 | 9221885 | 9881557 | 5915727.5 | 1.13E+07 | 9708867 | 6421845 | 1.68E+07 | 6420966 | 8435270 | 1114.5709 | IGNTEDGALPK | Q6ZPJ3 | Q6ZPJ3 | UBE2O | ubiquitin-conjugating enzyme E2O | Nucleus | enzyme |
| 15540246 | 2 | 0.471 | 6339429.5 | 4852491 | 8137269.5 | 9830473 | 1.06E+07 | 6360235 | 1.20E+07 | 1.10E+07 | 7330654 | 1.65E+07 | 5498177 | 8702412 | 1331.6646 | TLDNVAIAEEEK | Q6ZPJ3 | Q6ZPJ3 | UBE2O | ubiquitin-conjugating enzyme E2O | Nucleus | enzyme |
| 15532479 | 2 | 0.521 | 7260620 | 5386797 | 8619882 | 8365762.5 | 1.05E+07 | 6763944 | 1.18E+07 | 9758490 | 6859849.5 | 1.63E+07 | 6803026.5 | 9294109 | 1146.576 | VEVVWADNSK | Q6ZPJ3 | Q6ZPJ3 | UBE2O | ubiquitin-conjugating enzyme E2O | Nucleus | enzyme |
| 15550529 | 2 | 0.893 | 7595821.5 | 5824714.5 | 1.25E+07 | 9284401 | 1.43E+07 | 7310591.5 | 1.32E+07 | 1.13E+07 | 7410510 | 1.52E+07 | 5710539 | 9648744 | 2164.0981 | IEELAAILPAEQPTAPEEDK | Q6ZPJ3 | Q6ZPJ3 | UBE2O | ubiquitin-conjugating enzyme E2O | Nucleus | enzyme |
| 15552573 | 2 | 0.686 | 4064195 | 4943690.5 | 7652521.5 | 8077665 | 1.20E+07 | 5631051.5 | 8616738 | 7993185 | 6022344.5 | 1.45E+07 | 3529842 | 7163032.5 | 1688.893 | EMALLATSLPDGIMVK | Q6ZPJ3 | Q6ZPJ3 | UBE2O | ubiquitin-conjugating enzyme E2O | Nucleus | enzyme |
| 15532808 | 2 | 0.475 | 4553919 | 4067617.5 | 6508324 | 6367424 | 8226599 | 6651442 | 1.03E+07 | 6814492.5 | 5281155 | 1.39E+07 | 4628634 | 6833986.5 | 1350.7393 | IFSSVQWLSGVK | Q6ZPJ3 | Q6ZPJ3 | UBE2O | ubiquitin-conjugating enzyme E2O | Nucleus | enzyme |
| 15538883 | 2 | 0.378 | 5274729 | 3639342.25 | 6482892 | 6325403 | 9591664 | 5835373 | 9247773 | 6775457.5 | 5501588.5 | 1.29E+07 | 5372513 | 7437709.5 | 1160.6432 | VVQSMTQLVR | Q6ZPJ3 | Q6ZPJ3 | UBE2O | ubiquitin-conjugating enzyme E2O | Nucleus | enzyme |
| 15551511 | 3 | 0.431 | 3799984 | 2512300.75 | 1439477.375 | 4591417 | 4045302.25 | 1059831.5 | 6301142.5 | 3510695.75 | 1951752.75 | 1.28E+07 | 3476179.75 | 4194652.5 | 2791.2625 | DLQHIWPFMYGDYIAYDC[160.0307]WLGK | Q6ZPJ3 | Q6ZPJ3 | UBE2O | ubiquitin-conjugating enzyme E2O | Nucleus | enzyme |
| 15536388 | 2 | 0.516 | 3431881.75 | 2211721.5 | 5093880.5 | 5529289.5 | 6182235 | 2975799.5 | 5472832.5 | 4768755.5 | 3268589.75 | 1.04E+07 | 3802104.5 | 4723617 | 997.5158 | C[160.0307]LYVFPAK | Q6ZPJ3 | Q6ZPJ3 | UBE2O | ubiquitin-conjugating enzyme E2O | Nucleus | enzyme |
| 15532745 | 2 | 0.838 | 2564692 | 3197394 | 6609612.5 | 3028120.25 | 7205086.5 | 4711048 | 6357128.5 | 6528609 | 4106448.5 | 9537897 | 1830091.75 | 3986769.75 | 1037.568 | MDLFSALIK | Q6ZPJ3 | Q6ZPJ3 | UBE2O | ubiquitin-conjugating enzyme E2O | Nucleus | enzyme |
| 15537967 | 2 | 0.602 | 2586750 | 1915202.625 | 3402294.5 | 3165881.5 | 3916397.25 | 2655852 | 4626660 | 3565892.5 | 2614910.5 | 7808289.5 | 2344164.75 | 3264556.25 | 1152.5265 | GLQEGYENSR | Q6ZPJ3 | Q6ZPJ3 | UBE2O | ubiquitin-conjugating enzyme E2O | Nucleus | enzyme |
| 15541698 | 2 | 0.569 | 2760967 | 2028537.75 | 3530512.25 | 3755905.75 | 4394314.5 | 2659136.75 | 4273900 | 3360340.25 | 2921267.5 | 7060980 | 2778575 | 3177585.25 | 817.4757 | TTDIVIR | Q6ZPJ3 | Q6ZPJ3 | UBE2O | ubiquitin-conjugating enzyme E2O | Nucleus | enzyme |
| 15543097 | 3 | 0.331 | 3772756.5 | 1905388 | 1407394.125 | 4759133.5 | 1001470 | 2032972.75 | 4768883 | 1568454.75 | 2206524 | 5527240.5 | 4057706.75 | 4478005 | 1685.8545 | TFEDRMDLFSALIK | Q6ZPJ3 | Q6ZPJ3 | UBE2O | ubiquitin-conjugating enzyme E2O | Nucleus | enzyme |
| 15542125 | 2 | 0.126 | 2882966.5 | 1979007.75 | 2673678.75 | 3214790 | 4657779.5 | 2246965 | 4706907 | 4256776.5 | 3658087.75 | 5293163 | 3473093.5 | 3280720 | 1332.7324 | VC[160.0307]VSLLGTWIGK | Q6ZPJ3 | Q6ZPJ3 | UBE2O | ubiquitin-conjugating enzyme E2O | Nucleus | enzyme |
| 15536475 | 2 | 0.337 | 1687459.25 | 941473 | 1462833.375 | 1700971.375 | 1786594.875 | 1311232.125 | 3035041.75 | 1796538.125 | 1179601.125 | 4591266 | 1495275.125 | 2272969.5 | 1269.6659 | EEKPEVQSPVK | Q6ZPJ3 | Q6ZPJ3 | UBE2O | ubiquitin-conjugating enzyme E2O | Nucleus | enzyme |
| 15543157 | 3 | 0.435 | 2599987.75 | 508914.719 | 4.70E-06 | 3154505.25 | 264032.656 | 92282.484 | 2478132 | 81940.438 | 105258.648 | 4513079.5 | 3088917.5 | 1968542.375 | 2112.0608 | LVNRIESWLETHAMQER | Q6ZPJ3 | Q6ZPJ3 | UBE2O | ubiquitin-conjugating enzyme E2O | Nucleus | enzyme |
| 15545413 | 2 | 0.437 | 1628703.875 | 1039175.5 | 1942279.5 | 1514393.125 | 2185008.75 | 1912898 | 2847360 | 2702228.5 | 1574960.125 | 3364655.75 | 1395942.375 | 2103209.5 | 1186.6437 | KIEFQPPEAK | Q6ZPJ3 | Q6ZPJ3 | UBE2O | ubiquitin-conjugating enzyme E2O | Nucleus | enzyme |
| 15543623 | 2, 3 | 0.378 | 1782730.625 | 1173862 | 2317457.5 | 1849378.25 | 2930751.25 | 1980219.875 | 2629376.75 | 2795367.5 | 1985541 | 3270231.5 | 1942047.625 | 2139381 | 1175.5693, 1175.5703 | MEAVPDTERK | Q6ZPJ3 | Q6ZPJ3 | UBE2O | ubiquitin-conjugating enzyme E2O | Nucleus | enzyme |
| 15561532 | 3 | 0.01 | 713561.375 | 479698.344 | 513311.344 | 1327686.25 | 1112686 | 732652.625 | 1334502 | 1239928 | 1084880.625 | 2796914.75 | 1378302.625 | 1253528.125 | 4030.9048 | TPYEDGLYLFDIQLPNIYPAVPPHFC[160.0307]YLSQC[160.0307]SGR | Q6ZPJ3 | Q6ZPJ3 | UBE2O | ubiquitin-conjugating enzyme E2O | Nucleus | enzyme |
| 15546810 | 3 | 0.748 | 209735.641 | 686955.625 | 497360.25 | 4.70E-06 | 1111038.875 | 885795.875 | 732535.938 | 1228523.75 | 385182.938 | 86223.445 | 167868.234 | 950071.188 | 1259.6033 | RLGC[160.0307]FDHAQR | Q6ZPJ3 | Q6ZPJ3 | UBE2O | ubiquitin-conjugating enzyme E2O | Nucleus | enzyme |
| 15541029 | 2 | 0.342 | 1820091.75 | 903608.062 | 745639.688 | 1886794.125 | 1832225.875 | 1612992 | 1805003.875 | 1669814.625 | 652352.812 | 4617791.5 | 1422824.375 | 1577062.375 | 1294.6681 | VPTTLAEYC[160.0307]IK | Q6ZWZ2 | Q6ZWZ2 | UBE2R2 | ubiquitin-conjugating enzyme E2R 2 | Cytoplasm | enzyme |
| 15551264 | 3 | 0.674 | 1.47E+07 | 1.17E+07 | 1.16E+07 | 1.16E+07 | 1.30E+07 | 9298406 | 1.20E+07 | 9967271 | 8713066 | 2.16E+07 | 8711179 | 1.15E+07 | 3311.5022 | FYC[160.0307]VLSTEASEEDLNRLDSEAC[160.0307]EVLFSK | A2AN08 | A2AN08 | UBR4 | ubiquitin protein ligase E3 component n-recognin 4 | Nucleus | enzyme |
| 15551754 | 3 | 0.502 | 9937763 | 6822003 | 6824542 | 5732885 | 7693098 | 5063501 | 6531353.5 | 7356283 | 5952268 | 1.18E+07 | 7625348 | 5669658 | 2759.3025 | VNEAAPEKPQEDSGTAGGISSTSASVNR | A2AN08 | A2AN08 | UBR4 | ubiquitin protein ligase E3 component n-recognin 4 | Nucleus | enzyme |
| 15553146 | 3 | 0.459 | 9130951 | 8602524 | 1487224.25 | 8237704.5 | 3139803.75 | 1574075.125 | 4263851.5 | 2417202 | 1473923 | 1.11E+07 | 5531532.5 | 4630280 | 2501.368 | HAAQELATLLLSLPAPASVQQQSK | A2AN08 | A2AN08 | UBR4 | ubiquitin protein ligase E3 component n-recognin 4 | Nucleus | enzyme |
| 15550424 | 2, 3 | 0.96 | 5751522.5 | 5871297 | 7306658.5 | 4056938.25 | 9586477 | 5096092.5 | 6804945 | 6739777 | 5888119.5 | 1.07E+07 | 3513964.5 | 4660863 | 2357.2126, 2357.2139 | LEQVSSDEGIGTLAENLLEALR | A2AN08 | A2AN08 | UBR4 | ubiquitin protein ligase E3 component n-recognin 4 | Nucleus | enzyme |
| 15534360 | 3 | 0.623 | 6808351 | 4876125.5 | 5072249.5 | 4591274.5 | 5060016 | 4463622 | 6491409 | 4533489 | 4584969.5 | 1.02E+07 | 5360903.5 | 4390181.5 | 1439.7446 | AQQALSELHTVDK | A2AN08 | A2AN08 | UBR4 | ubiquitin protein ligase E3 component n-recognin 4 | Nucleus | enzyme |
| 15549669 | 3 | 0.492 | 7312457.5 | 6694047.5 | 4491388 | 5593244.5 | 6183432.5 | 4278260.5 | 7020691 | 5681252.5 | 5805333.5 | 9551360 | 5435864 | 6880333.5 | 3815.9216 | NAFANDTIPSESYISAVQAAHLGTLC[160.0307]GQSLPLAASLK | A2AN08 | A2AN08 | UBR4 | ubiquitin protein ligase E3 component n-recognin 4 | Nucleus | enzyme |
| 15539979 | 2 | 0.579 | 7140767 | 5884839 | 6229044.5 | 5897619 | 4888019.5 | 4120845.25 | 5858546 | 6013203 | 4419330 | 9208158 | 4260170 | 5358656 | 1394.754 | IIPYLSFGEVEK | A2AN08 | A2AN08 | UBR4 | ubiquitin protein ligase E3 component n-recognin 4 | Nucleus | enzyme |
| 15551227 | 2, 3 | 0.535 | 6100797 | 5693615 | 4991950.5 | 3850831.5 | 4387884.5 | 4096210.75 | 6640843 | 4385236.5 | 4823037 | 9100526 | 4012856.75 | 4060190 | 2390.2483, 2390.2502 | IYDLSIDALSPTFYFLLPSSK | A2AN08 | A2AN08 | UBR4 | ubiquitin protein ligase E3 component n-recognin 4 | Nucleus | enzyme |
| 15540612 | 3 | 0.266 | 6393831 | 5253187.5 | 4145414.25 | 5098770.5 | 4065940.75 | 3653108.5 | 6261455 | 4144455 | 3559506 | 8937264 | 5475512 | 5295097.5 | 1816.8668 | PHTTSSPPDMSPFFLR | A2AN08 | A2AN08 | UBR4 | ubiquitin protein ligase E3 component n-recognin 4 | Nucleus | enzyme |
| 15533510 | 2 | 0.749 | 6509597 | 4777663 | 4050034.25 | 5503799 | 4133359 | 3276499.5 | 5611754.5 | 4807163.5 | 3343874 | 8481808 | 4005349.25 | 4597312 | 1485.7306 | TSVQPTFTASQYR | A2AN08 | A2AN08 | UBR4 | ubiquitin protein ligase E3 component n-recognin 4 | Nucleus | enzyme |
| 15538958 | 2 | 0.592 | 5943627 | 3771912.25 | 2999207.75 | 4264592 | 3605282.75 | 2873058.5 | 5345648 | 3676770 | 2958845 | 8418665 | 3576869 | 4475984 | 1302.7115 | QVLFTPATQAAR | A2AN08 | A2AN08 | UBR4 | ubiquitin protein ligase E3 component n-recognin 4 | Nucleus | enzyme |
| 15538655 | 3 | 0.495 | 8695357 | 4972400.5 | 5006586.5 | 5997259.5 | 5469141.5 | 4414897.5 | 5652005.5 | 4476887.5 | 4458179 | 8192361 | 5176262.5 | 5867872 | 1913.0184 | SIAVTRPNNLVHFTESK | A2AN08 | A2AN08 | UBR4 | ubiquitin protein ligase E3 component n-recognin 4 | Nucleus | enzyme |
| 15536831 | 3 | 0.781 | 5743559.5 | 6975948 | 5440241 | 4572933 | 6583376.5 | 4899311.5 | 6040982.5 | 5062558.5 | 5295782.5 | 8105742 | 4357088 | 6331103 | 1825.0189 | LLASLFQDLQVEALHK | A2AN08 | A2AN08 | UBR4 | ubiquitin protein ligase E3 component n-recognin 4 | Nucleus | enzyme |
| 15552787 | 2 | 0.658 | 4056009 | 3107546.75 | 3927790.25 | 3932188.25 | 3464084.75 | 2981538.25 | 4106687.75 | 2829745.25 | 3182629.5 | 7874598.5 | 2952013 | 3696556.5 | 2037.0967 | NLLPATLQLIDTYASFTR | A2AN08 | A2AN08 | UBR4 | ubiquitin protein ligase E3 component n-recognin 4 | Nucleus | enzyme |
| 15551575 | 2 | 0.64 | 3936653.25 | 4850796.5 | 5601344 | 3896385.25 | 5353966.5 | 2871442.25 | 5313296.5 | 3846612.5 | 3032575.75 | 7794014 | 4139865.5 | 4040087.5 | 1729.8562 | SQQSDEEPVAASQLLK | A2AN08 | A2AN08 | UBR4 | ubiquitin protein ligase E3 component n-recognin 4 | Nucleus | enzyme |
| 15540576 | 2 | 0.535 | 5805917.5 | 4770908.5 | 4081523.75 | 4626975.5 | 5337388 | 3944236.75 | 5536612.5 | 4141189.25 | 3936675.5 | 7750631.5 | 4570108 | 5110588 | 1187.661 | AVSNINTLLDK | A2AN08 | A2AN08 | UBR4 | ubiquitin protein ligase E3 component n-recognin 4 | Nucleus | enzyme |
| 15548952 | 2 | 0.554 | 6065721 | 5384898 | 6810992 | 5155696.5 | 5553844 | 4264256.5 | 5523650 | 4592162 | 4616542.5 | 7646066.5 | 3996911.75 | 4895192 | 1977.0265 | SVFIAQNVASLQELGGSEK | A2AN08 | A2AN08 | UBR4 | ubiquitin protein ligase E3 component n-recognin 4 | Nucleus | enzyme |
| 15542963 | 2 | 0.874 | 4152870.75 | 4941686 | 5040683 | 5316839 | 4028399.75 | 3462299.5 | 4522744 | 4182208.5 | 3176075 | 7592836 | 3383382.25 | 3374820.25 | 1199.6605 | VQLTPGQTEVK | A2AN08 | A2AN08 | UBR4 | ubiquitin protein ligase E3 component n-recognin 4 | Nucleus | enzyme |
| 15538156 | 4 | 0.641 | 7359512 | 5344485 | 4415274.5 | 4814837.5 | 6118565.5 | 3340401.5 | 5354635.5 | 4421407.5 | 3228164.25 | 7571611.5 | 4277587 | 4592425 | 2840.261 | TSPADHGGSVGSESGGSAVDSVAGEHSVSGR | A2AN08 | A2AN08 | UBR4 | ubiquitin protein ligase E3 component n-recognin 4 | Nucleus | enzyme |
| 15549934 | 3 | 0.031 | 4698196.5 | 3197000.5 | 2938193.75 | 3967852.5 | 4262262.5 | 3054900.25 | 3886817.25 | 3516393.75 | 4100300 | 7508652 | 5537162 | 5085820.5 | 2518.2832 | SPNPSLLHLC[160.0307]GSLAQLAC[160.0307]VEPVR | A2AN08 | A2AN08 | UBR4 | ubiquitin protein ligase E3 component n-recognin 4 | Nucleus | enzyme |
| 15550603 | 2 | 0.855 | 4232871 | 4205585.5 | 4865674 | 3851287.25 | 3756065.5 | 3751182.25 | 3989253.5 | 4146841 | 4693271.5 | 7338001.5 | 2773207 | 4222161 | 1891.9902 | DQLVMLLDQINSTFVR | A2AN08 | A2AN08 | UBR4 | ubiquitin protein ligase E3 component n-recognin 4 | Nucleus | enzyme |
| 15534553 | 2 | 0.616 | 6336303 | 5552757.5 | 5023626 | 5118585 | 5544178 | 2947616.75 | 5705680.5 | 4560610 | 4004479.75 | 7145203.5 | 4832288 | 4230609 | 1145.6544 | PGTFLIQEIK | A2AN08 | A2AN08 | UBR4 | ubiquitin protein ligase E3 component n-recognin 4 | Nucleus | enzyme |
| 15551887 | 2 | 0.663 | 5076246.5 | 4970735 | 5672588 | 4780203.5 | 4659873.5 | 4082813.5 | 4116439 | 4620624 | 5417370 | 6956042.5 | 3907753.5 | 4992970 | 1972.0532 | LASAPVPFTVLSLTGNPC[160.0307]K | A2AN08 | A2AN08 | UBR4 | ubiquitin protein ligase E3 component n-recognin 4 | Nucleus | enzyme |
| 15551698 | 3 | 0.455 | 4866681.5 | 4804375 | 2740648.5 | 4172524.5 | 4892014 | 3302431.75 | 3959238.75 | 4296206.5 | 5699542.5 | 6602320 | 4909861 | 4667205.5 | 2552.249 | C[160.0307]NGLLPVWGPHVPESAFATC[160.0307]LAR | A2AN08 | A2AN08 | UBR4 | ubiquitin protein ligase E3 component n-recognin 4 | Nucleus | enzyme |
| 15533501 | 2 | 0.332 | 5497647.5 | 4316185.5 | 4479701.5 | 3570832.75 | 4342345 | 2965790.75 | 4983523 | 3437734.25 | 3051076.25 | 6513120 | 3899247.75 | 4105764.5 | 1128.6344 | IQIGTQAIER | A2AN08 | A2AN08 | UBR4 | ubiquitin protein ligase E3 component n-recognin 4 | Nucleus | enzyme |
| 15537360 | 3 | 0.668 | 4308084.5 | 3594558.75 | 2924709.75 | 3519874.25 | 3733160.5 | 2898170.75 | 3693357.5 | 2617223.5 | 2631257.75 | 6410951 | 2741152 | 3286976.75 | 1192.6046 | DLHTLDSHVR | A2AN08 | A2AN08 | UBR4 | ubiquitin protein ligase E3 component n-recognin 4 | Nucleus | enzyme |
| 15540335 | 2 | 0.76 | 4328621 | 3174773.25 | 3203611.25 | 4297570 | 3544114.25 | 2831239.25 | 3329477.5 | 3579234.25 | 3076225.5 | 6357850.5 | 3052107.5 | 3493546.75 | 1294.6953 | VC[160.0307]LNLPYFLR | A2AN08 | A2AN08 | UBR4 | ubiquitin protein ligase E3 component n-recognin 4 | Nucleus | enzyme |
| 15536811 | 2 | 0.828 | 4151254 | 2688309 | 3476372 | 3050840.25 | 3542360.75 | 3239572.25 | 4094559 | 3172578.25 | 2547324.75 | 6294871.5 | 2887234.5 | 3190357 | 1460.8816 | QLLQLLTTYIVR | A2AN08 | A2AN08 | UBR4 | ubiquitin protein ligase E3 component n-recognin 4 | Nucleus | enzyme |
| 15552074 | 2 | 0.978 | 4759197 | 4132223.25 | 3959448.5 | 4087193.75 | 4655728.5 | 4114581 | 4839990.5 | 4121142.25 | 3189477.25 | 6282822.5 | 2785346.75 | 3888403.5 | 2145.1548 | AVWLPGSQTELAIVTADFVK | A2AN08 | A2AN08 | UBR4 | ubiquitin protein ligase E3 component n-recognin 4 | Nucleus | enzyme |
| 15552024 | 3 | 0.266 | 3923339.25 | 2515261.5 | 1772317.875 | 2947194.75 | 3759914.25 | 2049455.125 | 3142378 | 2395111 | 2969485.25 | 6273977.5 | 3944882.75 | 3207115 | 2754.2288 | TYMNQLAMNSPEMSEC[160.0307]DILHTLR | A2AN08 | A2AN08 | UBR4 | ubiquitin protein ligase E3 component n-recognin 4 | Nucleus | enzyme |
| 15536823 | 2 | 0.575 | 5394366 | 3987576.5 | 4115450.75 | 4485796 | 4397329 | 2775836 | 4655337.5 | 3257337.5 | 3040836.25 | 6231591.5 | 3961274 | 3769686.25 | 1445.6825 | DSDQLEVIQENR | A2AN08 | A2AN08 | UBR4 | ubiquitin protein ligase E3 component n-recognin 4 | Nucleus | enzyme |
| 15550379 | 3 | 0.644 | 3214722.75 | 4868591 | 4857342 | 2852264.5 | 4192617.25 | 2560846.75 | 3435398.75 | 4275699.5 | 3163448.25 | 6186162 | 2509802.25 | 3922835.75 | 2560.2485 | LC[160.0307]SIIYPEENEVTEFFVTLEK | A2AN08 | A2AN08 | UBR4 | ubiquitin protein ligase E3 component n-recognin 4 | Nucleus | enzyme |
| 15550973 | 2 | 0.603 | 4564337.5 | 5167715 | 5425896 | 5468557 | 5048609 | 4233054.5 | 4606510.5 | 4289068 | 3941335.5 | 6159155 | 3739787 | 4408472.5 | 1614.9437 | STLEVLQLFPINIK | A2AN08 | A2AN08 | UBR4 | ubiquitin protein ligase E3 component n-recognin 4 | Nucleus | enzyme |
| 15535210 | 2 | 0.858 | 3217527 | 4234749.5 | 4843127.5 | 2645579.25 | 4298527.5 | 4041427.5 | 4214099.5 | 3332678 | 2517628 | 5935706.5 | 1569839.375 | 3232067.75 | 1459.8353 | IISLDLPVAEVYK | A2AN08 | A2AN08 | UBR4 | ubiquitin protein ligase E3 component n-recognin 4 | Nucleus | enzyme |
| 15534180 | 2 | 0.756 | 5276976.5 | 2386002.5 | 2771993 | 3619775.5 | 2781963.5 | 1673258.75 | 4127558.25 | 2155478.5 | 2586600.5 | 5765410.5 | 2472416.75 | 3164105 | 1173.6246 | NLQGFLEQPK | A2AN08 | A2AN08 | UBR4 | ubiquitin protein ligase E3 component n-recognin 4 | Nucleus | enzyme |
| 15553730 | 2 | 0.884 | 4224315 | 4108910.5 | 4018237.5 | 3606751 | 3548374.5 | 3567550.5 | 4410103 | 3598601.25 | 3347192.25 | 5698221 | 2350869 | 3629199.5 | 2037.0172 | EVQMNFLNQLTSVFNPR | A2AN08 | A2AN08 | UBR4 | ubiquitin protein ligase E3 component n-recognin 4 | Nucleus | enzyme |
| 15552958 | 3 | 0.355 | 2583133 | 3232763.75 | 2101147.75 | 1962124.375 | 2673963.5 | 2114536.25 | 5016972 | 2063871.625 | 2083533 | 5509447.5 | 3000986.75 | 3138939 | 2789.3743 | GHWANPDLASSLQYEMLLLTDSISK | A2AN08 | A2AN08 | UBR4 | ubiquitin protein ligase E3 component n-recognin 4 | Nucleus | enzyme |
| 15534361 | 3 | 0.737 | 4552387.5 | 3470644.5 | 3748617.5 | 3192709 | 4257648 | 2825061.5 | 4323330 | 3071897 | 2901426.5 | 5500777.5 | 3026110.5 | 3648881.5 | 1803.8575 | GREEWESAALQNANTK | A2AN08 | A2AN08 | UBR4 | ubiquitin protein ligase E3 component n-recognin 4 | Nucleus | enzyme |
| 15536250 | 2 | 0.785 | 3825882.75 | 4252066.5 | 3122552 | 3297851.25 | 3916184.25 | 2572245.5 | 4543221 | 2708434 | 2827440 | 5362446 | 2776738.5 | 3768873.25 | 1152.602 | APSYIEIFGR | A2AN08 | A2AN08 | UBR4 | ubiquitin protein ligase E3 component n-recognin 4 | Nucleus | enzyme |

| | | | | | | | | | | | | | | | | | | | | | |
|---|---|---|---|---|---|---|---|---|---|---|---|---|---|---|---|---|---|---|---|---|---|
| 15535490 | 2 | 0.861 | 3468756.5 | 3529606 | 2078908.125 | 3020388.5 | 3130458.25 | 1889184.625 | 4091033.75 | 2306486.75 | 2012111.125 | 5269730.5 | 2244732.25 | 2932576 | 1253.6157 | GITQNALDYMK | A2AN08 | A2AN08 | UBR4 | ubiquitin protein ligase E3 component n-recognin 4 | Nucleus | enzyme |
| 15539120 | 2 | 0.361 | 4102131.75 | 2751622.25 | 3008747.5 | 2562851 | 3113436.25 | 2547656.75 | 3486863.75 | 2769521 | 2214288.25 | 5222615.5 | 3301963.5 | 2940108.5 | 1449.6212 | VWC[160.0307]ATNEGEPMR | A2AN08 | A2AN08 | UBR4 | ubiquitin protein ligase E3 component n-recognin 4 | Nucleus | enzyme |
| 15554267 | 3 | 0.461 | 4742239 | 1587821.5 | 337917.281 | 3329905.25 | 1462327.125 | 1066946.125 | 1811776.375 | 1296258.75 | 3015175.75 | 4638470.5 | 3225958.75 | 3440327.75 | 2278.1147 | AYLLQNLNEEGSTEKPSQEK | A2AN08 | A2AN08 | UBR4 | ubiquitin protein ligase E3 component n-recognin 4 | Nucleus | enzyme |
| 15551047 | 2 | 0.838 | 2213543.25 | 4065050.25 | 4404767.5 | 2345331.75 | 3903016 | 2486307 | 3102524 | 2916644.5 | 2517505.75 | 4453097.5 | 1833397.625 | 2472089 | 1708.9341 | SLTGLLSSFVEVESIK | A2AN08 | A2AN08 | UBR4 | ubiquitin protein ligase E3 component n-recognin 4 | Nucleus | enzyme |
| 15553642 | 2 | 0.953 | 3336063.25 | 3212058.75 | 2676158.75 | 2997577.75 | 2861194 | 2405963 | 2589410.5 | 2322619 | 3483475.75 | 4393593.5 | 1734902.75 | 2923869.5 | 1571.8445 | AAQFVDLLGYFSLK | A2AN08 | A2AN08 | UBR4 | ubiquitin protein ligase E3 component n-recognin 4 | Nucleus | enzyme |
| 15536092 | 2 | 0.342 | 3583206.25 | 3200146.75 | 2380589.25 | 2106501.25 | 2452099.25 | 1417882.375 | 3071492.25 | 2613182.25 | 2033567 | 4312146.5 | 1906707.125 | 2602279 | 1235.6284 | ALGTLGMTTNEK | A2AN08 | A2AN08 | UBR4 | ubiquitin protein ligase E3 component n-recognin 4 | Nucleus | enzyme |
| 15535808 | 2 | 0.449 | 3162607 | 1909723.875 | 1702399.625 | 1685346.125 | 2243435.75 | 1280065.875 | 2604652.75 | 2180415.25 | 1656176 | 4285786 | 2334835.25 | 1884201.375 | 1403.6223 | HNDMPIYEAADK | A2AN08 | A2AN08 | UBR4 | ubiquitin protein ligase E3 component n-recognin 4 | Nucleus | enzyme |
| 15542427 | 2 | 0.865 | 3065494.5 | 2512466 | 2143589 | 2660031.75 | 2790201.5 | 2189602.25 | 3178462 | 2431263.5 | 2209158.75 | 4151030.25 | 2366022.5 | 2431314.5 | 1369.6146 | MNYSGDQGQTIR | A2AN08 | A2AN08 | UBR4 | ubiquitin protein ligase E3 component n-recognin 4 | Nucleus | enzyme |
| 15558008 | 3 | 0.445 | 2665667.5 | 1946062.75 | 1292405 | 1602723.5 | 2222117.25 | 1664159.125 | 1997000.875 | 1976027.375 | 1570978.125 | 3519348.75 | 1923742.625 | 2287092.25 | 2541.2756 | FQDAVVANSFFIMPATVADATAVR | A2AN08 | A2AN08 | UBR4 | ubiquitin protein ligase E3 component n-recognin 4 | Nucleus | enzyme |
| 15554894 | 2 | 0.425 | 2080766.875 | 2180403.25 | 3394669.5 | 1564703.625 | 2682007.5 | 1589209.625 | 3275366.25 | 2667098.25 | 2361862.25 | 3391616.75 | 1769225.375 | 2484904.5 | 1786.864 | DNPEATQQMNDLIIGK | A2AN08 | A2AN08 | UBR4 | ubiquitin protein ligase E3 component n-recognin 4 | Nucleus | enzyme |
| 15541581 | 2 | 0.35 | 3200617.25 | 1142051.375 | 1264656.75 | 2351091 | 557784.5 | 823154.562 | 2195596.5 | 995543.312 | 584371.562 | 3078858 | 2951260.25 | 1490617.5 | 1460.6616 | LENPDEAC[160.0307]AVSQK | A2AN08 | A2AN08 | UBR4 | ubiquitin protein ligase E3 component n-recognin 4 | Nucleus | enzyme |
| 15535853 | 2 | 0.33 | 2490436.5 | 2439414.5 | 2447688.5 | 1579845.25 | 3476709 | 1683852.375 | 2169124.75 | 3204091.5 | 2520255.75 | 2247037 | 1472984.625 | 1633942.75 | 1008.54895 | TPVFIFER | A2AN08 | A2AN08 | UBR4 | ubiquitin protein ligase E3 component n-recognin 4 | Nucleus | enzyme |
| 15552468 | 2 | 0.735 | 1505160.25 | 1120920.625 | 1666693.25 | 1072772.25 | 1491574.125 | 955417.438 | 1383614.25 | 1671418.375 | 1206905.875 | 1868836.75 | 850700 | 1190504 | 2418.1973 | YDELYSSLTTLLAAGSQLDTTR | A2AN08 | A2AN08 | UBR4 | ubiquitin protein ligase E3 component n-recognin 4 | Nucleus | enzyme |
| 15539620 | 2 | 0.222 | 1739110.125 | 1345309.625 | 1309036.625 | 768976.562 | 1400285.375 | 856501.812 | 1340249.75 | 1175924.625 | 867295.812 | 1412495.5 | 1112491.75 | 1225416.75 | 1212.5729 | HFSSDAAPQPR | A2AN08 | A2AN08 | UBR4 | ubiquitin protein ligase E3 component n-recognin 4 | Nucleus | enzyme |
| 15552635 | 4 | 0.668 | 9920504 | 1.19E+07 | 7854782 | 8560937 | 1.03E+07 | 4762598 | 5971242 | 8401955 | 1.08E+07 | 1.18E+07 | 4783831 | 5513026.5 | 3809.8848 | QVINNAC[160.0307]ATQAIVSVLLNC[160.0307]THQDVHLGETLSEFK | Q9WUP7 | Q9WUP7 | UCHL5 | ubiquitin carboxyl-terminal hydrolase L5 | Cytoplasm | peptidase |
| 15549268 | 3 | 0.288 | 7112885.5 | 1.06E+07 | 6915138 | 5077591 | 1.17E+07 | 4270332.5 | 7031336.5 | 7600229 | 6392614 | 7199534 | 2560403 | 4051325.75 | 2696.3284 | EGPIDLGAC[160.0307]NQDDWITAVRPVIEK | Q9WUP7 | Q9WUP7 | UCHL5 | ubiquitin carboxyl-terminal hydrolase L5 | Cytoplasm | peptidase |
| 15553771 | 2 | 0.263 | 3130119.25 | 3426461.25 | 3757456.75 | 2033094.375 | 4029749.5 | 1170888.375 | 2002879.75 | 2679412.25 | 1892273.375 | 3312454.25 | 713652.625 | 1072961 | 1741.8099 | WQPGEEPAGSVVQDSR | Q9WUP7 | Q9WUP7 | UCHL5 | ubiquitin carboxyl-terminal hydrolase L5 | Cytoplasm | peptidase |
| 15543826 | 2 | 0.191 | 1075261.625 | 1583739.625 | 1983917.25 | 372082.625 | 1067419.125 | 1875277.875 | 718524 | 1479768.625 | 4533530 | 580276.5 | 291276.406 | 725446.75 | 1144.6255 | GLALSNSDVIR | Q9WUP7 | Q9WUP7 | UCHL5 | ubiquitin carboxyl-terminal hydrolase L5 | Cytoplasm | peptidase |
| 15550989 | 3 | 0.059 | 9099450 | 8719306 | 8104114.5 | 9471913 | 1.21E+07 | 1.06E+07 | 1.19E+07 | 1.11E+07 | 1.03E+07 | 1.33E+07 | 1.07E+07 | 9824042 | 2952.531 | IIGPLEDNELFNQDDFHLLENIILK | Q6P5E4 | Q6P5E4 | UGGT1 | UDP-glucose glycoprotein glucosyltransferase 1 | Cytoplasm | enzyme |
| 15552364 | 3 | 0.711 | 3414821 | 3663829 | 3707809.5 | 4664683 | 3938641.25 | 3869210 | 4386680.5 | 3915195.75 | 3780061.25 | 4718512.5 | 4114019.5 | 2603411.25 | 3463.685 | GQYQGLSQDPNSLSNLDQDLPNNMIHQVPIK | Q6P5E4 | Q6P5E4 | UGGT1 | UDP-glucose glycoprotein glucosyltransferase 1 | Cytoplasm | enzyme |
| 15553065 | 3 | 0.105 | 2057144.875 | 2619145 | 1829746 | 2964766 | 2732263.25 | 2256360 | 2738275 | 2696911.25 | 1741414.875 | 3743256.5 | 3319937.75 | 2808768 | 3051.489 | FLDMPQSPLFTLNLNTPESWMVESVR | Q6P5E4 | Q6P5E4 | UGGT1 | UDP-glucose glycoprotein glucosyltransferase 1 | Cytoplasm | enzyme |
| 15555353 | 3 | 0.022 | 2837794 | 4514346 | 3516508 | 4944901 | 5336881.5 | 5276418 | 4633089 | 5364808.5 | 4788869.5 | 3518227 | 4056067 | 4122637.25 | 2630.2666 | IYSHDGTDSPPDANDVVVILNNFK | Q6P5E4 | Q6P5E4 | UGGT1 | UDP-glucose glycoprotein glucosyltransferase 1 | Cytoplasm | enzyme |
| 15537989 | 2 | 0.906 | 2203245.5 | 2614984.25 | 2631462.5 | 2378827 | 2335144 | 2511770.25 | 3228674.25 | 2503319.5 | 2176955 | 3069949 | 2071321.25 | 2137856.25 | 1322.7079 | FLFVDADQIVR | Q6P5E4 | Q6P5E4 | UGGT1 | UDP-glucose glycoprotein glucosyltransferase 1 | Cytoplasm | enzyme |
| 15550640 | 2 | 0.115 | 1982332.125 | 2531491.5 | 1938570 | 2342580.75 | 2140275.25 | 2369859 | 2579022.25 | 2485554.5 | 2539282 | 2506517.5 | 2466332.5 | 2387308.75 | 1733.8668 | LNIQPSETDYAVDIR | Q6P5E4 | Q6P5E4 | UGGT1 | UDP-glucose glycoprotein glucosyltransferase 1 | Cytoplasm | enzyme |
| 15557140 | 3 | 0.889 | 1788082.5 | 3443642.75 | 4226813.5 | 2844579.25 | 4079853 | 3254595 | 3034933.25 | 3415055.75 | 3719769.5 | 2460689 | 2574755.75 | 4072819 | 2836.3508 | AYNYVAQEVDGYHAFQTLTQIYNK | Q6P5E4 | Q6P5E4 | UGGT1 | UDP-glucose glycoprotein glucosyltransferase 1 | Cytoplasm | enzyme |
| 15536292 | 2 | 0.644 | 1431623.5 | 1738059.5 | 1763144.75 | 2017373.375 | 2518546.5 | 1482351.75 | 1489482.625 | 2017522.75 | 1582488.625 | 1677157.75 | 1639453 | 2219993.75 | 1041.5913 | VDALLSAQPK | Q6P5E4 | Q6P5E4 | UGGT1 | UDP-glucose glycoprotein glucosyltransferase 1 | Cytoplasm | enzyme |
| 15536884 | 2 | 0.17 | 934456.812 | 3859353 | 2084573.75 | 1299579.25 | 2313134 | 2990624.75 | 1642158.25 | 2890762.25 | 3189133 | 905965.75 | 1057003.375 | 1115722.875 | 911.5331 | LYSLLFR | Q8R1I1 | Q8R1I1 | UQCR10 | ubiquinol-cytochrome c reductase, complex III subunit X | Cytoplasm | enzyme |
| 15533727 | 3 | 0.539 | 1.40E+07 | 2.42E+07 | 9648619 | 2.02E+07 | 1.54E+07 | 1.64E+07 | 1.07E+07 | 1.66E+07 | 1.95E+07 | 8703649 | 1.53E+07 | 1.19E+07 | 2053.9937 | NALVSHLDGTTPVC[160.0307]EDIGR | Q9CZ13 | Q9CZ13 | UQCRC1 | ubiquinol-cytochrome c reductase core protein I | Cytoplasm | enzyme |
| 15550906 | 2 | 0.169 | 4295626.5 | 8778517 | 4625894 | 6504072 | 6233756.5 | 9970381 | 4096896.75 | 7500300 | 8166991 | 3404963.5 | 4676488 | 3896773.75 | 1605.7545 | VYEEDAVPGLTPC[160.0307]R | Q9CZ13 | Q9CZ13 | UQCRC1 | ubiquinol-cytochrome c reductase core protein I | Cytoplasm | enzyme |
| 15537662 | 2 | 0.185 | 5801960.5 | 9546158 | 4511841.5 | 7964578 | 6514568 | 1.00E+07 | 4974144.5 | 8807290 | 8650351 | 3099674.25 | 6000530.5 | 4542620.5 | 1353.6285 | LC[160.0307]TSATESEVTR | Q9CZ13 | Q9CZ13 | UQCRC1 | ubiquinol-cytochrome c reductase core protein I | Cytoplasm | enzyme |
| 15534840 | 2 | 0.357 | 3779678.5 | 6203634.5 | 2742355 | 5200696.5 | 4248639 | 6622863 | 3281294.75 | 4498728 | 4868226 | 1964462.5 | 4300967 | 3670181.5 | 1256.6719 | RIPLAEWESR | Q9CZ13 | Q9CZ13 | UQCRC1 | ubiquinol-cytochrome c reductase core protein I | Cytoplasm | enzyme |
| 15535410 | 3 | 0.082 | 5214974 | 1.21E+07 | 6010415.5 | 6340757.5 | 6727637 | 1.12E+07 | 5231666.5 | 9436994 | 9173668 | 1392568.75 | 4713645 | 3849420.25 | 2092.1145 | MVLAAAGGVEHQQLLDLAQK | Q9CZ13 | Q9CZ13 | UQCRC1 | ubiquinol-cytochrome c reductase core protein I | Cytoplasm | enzyme |
| 15552706 | 2 | 0.214 | 1.02E+07 | 2.56E+07 | 1.09E+07 | 1.52E+07 | 1.96E+07 | 2.39E+07 | 1.01E+07 | 1.79E+07 | 2.31E+07 | 7035181.5 | 1.11E+07 | 1.12E+07 | 1956.0027 | TSAAPGGVPLQPQDLEFTK | Q9DB77 | Q9DB77 | UQCRC2 | ubiquinol-cytochrome c reductase core protein II | Cytoplasm | enzyme |
| 15548303 | 2 | 0.153 | 9350346 | 1.77E+07 | 8754063 | 1.32E+07 | 1.34E+07 | 1.50E+07 | 8032060.5 | 1.48E+07 | 1.60E+07 | 6716862 | 9262501 | 7581395.5 | 1529.7892 | AVAQGNLSSADVQAAK | Q9DB77 | Q9DB77 | UQCRC2 | ubiquinol-cytochrome c reductase core protein II | Cytoplasm | enzyme |
| 15550555 | 2 | 0.153 | 8073173.5 | 1.71E+07 | 9241908 | 1.23E+07 | 1.19E+07 | 1.56E+07 | 7440960 | 1.29E+07 | 1.71E+07 | 6512303 | 7244245.5 | 7729958 | 1759.828 | SMAASGNLGHTPFLDEL | Q9DB77 | Q9DB77 | UQCRC2 | ubiquinol-cytochrome c reductase core protein II | Cytoplasm | enzyme |
| 15539667 | 3 | 0.222 | 7769017.5 | 1.29E+07 | 5532464 | 9619604 | 9678687 | 9852034 | 5403987.5 | 9086147 | 1.10E+07 | 3601370.5 | 7001957 | 5970754.5 | 1575.8408 | RGNNTTSLLSQSVAK | Q9DB77 | Q9DB77 | UQCRC2 | ubiquinol-cytochrome c reductase core protein II | Cytoplasm | enzyme |
| 15550279 | 2 | 0.372 | 3847787.5 | 8058391 | 3287035 | 4872444 | 5888434 | 6852127.5 | 4096648.5 | 6779622 | 9501308 | 2814043.75 | 5658176 | 3335744.75 | 1566.7343 | NALANPLYC[160.0307]PDYR | Q9DB77 | Q9DB77 | UQCRC2 | ubiquinol-cytochrome c reductase core protein II | Cytoplasm | enzyme |
| 15537399 | 3 | 0.549 | 6295715.5 | 9947888 | 4529357 | 5795465 | 7197807 | 7183057 | 3952046.5 | 7803754 | 8397880 | 2679097.5 | 7359873.5 | 4455027.5 | 2179.0525 | ITSEELHYFVQNHFTSAR | Q9DB77 | Q9DB77 | UQCRC2 | ubiquinol-cytochrome c reductase core protein II | Cytoplasm | enzyme |
| 15535073 | 3 | 0.689 | 2630120 | 5710637 | 2608647.5 | 3470435.75 | 3145544 | 5054410.5 | 2391142.5 | 4567579 | 4405251 | 2400491.75 | 2830912.25 | 3273054.75 | 1410.8098 | MALVGLGVSHSVLK | Q9DB77 | Q9DB77 | UQCRC2 | ubiquinol-cytochrome c reductase core protein II | Cytoplasm | enzyme |
| 15536312 | 2 | 0.271 | 3839793.25 | 7461412.5 | 3463496 | 4604908.5 | 5240970.5 | 6421781 | 3930473.5 | 5488123.5 | 6027791.5 | 2163822.75 | 4244035 | 3781046 | 1235.6206 | QVAEQFLNMR | Q9DB77 | Q9DB77 | UQCRC2 | ubiquinol-cytochrome c reductase core protein II | Cytoplasm | enzyme |
| 15549452 | 2 | 0.21 | 3074696 | 7304888 | 3598809 | 5434881 | 4032881.5 | 6257050.5 | 3466157 | 5213282.5 | 7525735.5 | 1462442.5 | 3374651.25 | 3570804 | 1927.0599 | LPNGLVIASLENYAPLSR | Q9DB77 | Q9DB77 | UQCRC2 | ubiquinol-cytochrome c reductase core protein II | Cytoplasm | enzyme |
| 15534370 | 2 | 0.135 | 2730609.5 | 5177137 | 2533654.25 | 3525964.25 | 3451904.75 | 4926160.5 | 2519199.5 | 3962402.75 | 4272984.5 | 1457774.25 | 2970151.25 | 1937113 | 1314.7023 | IIENLHDVAYK | Q9DB77 | Q9DB77 | UQCRC2 | ubiquinol-cytochrome c reductase core protein II | Cytoplasm | enzyme |
| 15555044 | 2 | 0.279 | 2108890.25 | 4385674 | 2175699 | 2346315 | 2448241.5 | 3519132 | 1315043.125 | 2690953 | 3957519.75 | 799093.938 | 2043540.375 | 1695955.75 | 1575.8414 | RGNNTTSLLSQSVAK | Q9DB77 | Q9DB77 | UQCRC2 | ubiquinol-cytochrome c reductase core protein II | Cytoplasm | enzyme |
| 15542318 | 2 | 0.612 | 972022.625 | 3754631.5 | 2338586.25 | 558999.562 | 2636983.25 | 4469630 | 803934.812 | 3440601.75 | 4245850 | 212758.328 | 1770641.125 | 1577307.375 | 1121.5673 | AVAFQNSQTR | Q9DB77 | Q9DB77 | UQCRC2 | ubiquinol-cytochrome c reductase core protein II | Cytoplasm | enzyme |
| 15532259 | 3 | 0.19 | 2.33E+07 | 4.70E+07 | 1.85E+07 | 3.40E+07 | 2.86E+07 | 3.80E+07 | 2.09E+07 | 3.72E+07 | 3.52E+07 | 1.60E+07 | 2.19E+07 | 1.84E+07 | 2226.957 | SQTEEDC[160.0307]TEELFDFLHAR | P99028 | P99028 | UQCRHL | ubiquinol-cytochrome c reductase hinge protein like | Cytoplasm | other |
| 15548300 | 2 | 0.199 | 5506953.5 | 2.06E+07 | 1.04E+07 | 5225769 | 1.33E+07 | 1.75E+07 | 6793394.5 | 1.19E+07 | 1.36E+07 | 2511540.25 | 5391353 | 6048518.5 | 1787.8512 | EEEEEELVDPLTTVR | P99028 | P99028 | UQCRHL | ubiquinol-cytochrome c reductase hinge protein like | Cytoplasm | other |
| 15551604 | 3 | 0.281 | 2744621 | 2837150.5 | 2982474.75 | 3921286 | 2523676.75 | 3432986 | 2209394.25 | 2507923 | 1468558.875 | 4235815 | 1879567.25 | 2155612.75 | 2363.2688 | LRDPAAAASELGYVFQAITLTR | P70697 | P70697 | UROD | uroporphyrinogen decarboxylase | Cytoplasm | enzyme |
| 15553667 | 2 | 0.111 | 4168846.75 | 1.09E+07 | 8005865 | 8308740.5 | 9941793 | 1.27E+07 | 6080558.5 | 9174207 | 1.30E+07 | 3673416.5 | 5522369 | 5348324 | 1555.7352 | AGAESDGQFQFTGIK | Q78IK2 | Q78IK2 | USMG5 | up-regulated during skeletal muscle growth 5 homolog (mouse) | Cytoplasm | other |
| 15534185 | 2 | 0.431 | 4602327.5 | 8001978.5 | 4219828.5 | 4772359 | 5994779.5 | 7522720.5 | 3838624.25 | 6615720 | 7724631.5 | 2307618 | 5599206.5 | 4252754.5 | 1221.5875 | YFNSYTLTGR | Q78IK2 | Q78IK2 | USMG5 | up-regulated during skeletal muscle growth 5 homolog (mouse) | Cytoplasm | other |
| 15548836 | 3 | 0.152 | 7067912.5 | 5842504.5 | 8037913 | 6184188 | 6962699 | 5345964 | 4802741 | 4544819 | 4845171 | 8250871 | 5051807 | 4875069.5 | 2462.15 | LEAIEDDSGRETDSSSAPAVTPSK | Q9JMA1 | Q9JMA1 | USP14 | ubiquitin specific peptidase 14 (tRNA-guanine transglycosylase) | Cytoplasm | peptidase |
| 15540235 | 2 | 0.156 | 4440423.5 | 4093797.25 | 4124589 | 3825236.5 | 3209012 | 3010815 | 2768302.25 | 2624939.25 | 2336164.5 | 5148769 | 2196619 | 2814192.5 | 1241.7081 | VSIVTPEDILR | Q9JMA1 | Q9JMA1 | USP14 | ubiquitin specific peptidase 14 (tRNA-guanine transglycosylase) | Cytoplasm | peptidase |
| 15552869 | 2 | 0.183 | 3493938.75 | 2218681 | 2106543 | 2748986 | 2662557.75 | 2554234.75 | 3225258.25 | 2216846.75 | 2577661 | 1.37E+07 | 3279010.75 | 3496587.75 | 2128.1316 | IGNILDLC[160.0307]TALSALSGVPADK | Q8R5H1 | Q8R5H1 | USP15 | ubiquitin specific peptidase 15 | Cytoplasm | peptidase |
| 15548889 | 3 | 0.835 | 8132186.5 | 7506410 | 7222010 | 8344615 | 6684939.5 | 6673277.5 | 8593029 | 6828427 | 5016050.5 | 1.34E+07 | 5131413 | 7957404.5 | 2298.161 | EHLIDELDYILLPTEGWNK | Q8R5H1 | Q8R5H1 | USP15 | ubiquitin specific peptidase 15 | Cytoplasm | peptidase |
| 15551767 | 3 | 0.362 | 5680870.5 | 3634271 | 2409737.25 | 4768412 | 4066439.5 | 3923178.25 | 5227617 | 2977382.5 | 2646355 | 1.05E+07 | 3919220.5 | 5165521 | 3552.6775 | FAPQFSGYQQQDC[160.0307]QELLAFLLDGLHEDLNR | Q8R5H1 | Q8R5H1 | USP15 | ubiquitin specific peptidase 15 | Cytoplasm | peptidase |
| 15553144 | 3 | 0.987 | 3528154.5 | 6496610.5 | 4293030 | 3504476.75 | 4034610 | 5005435.5 | 6518883 | 4856832 | 3135125.25 | 9295528 | 2190342 | 4114999.25 | 3049.4565 | LDTLVDFPISDLDMSEFLINPNAGPC[160.0307]R | Q8R5H1 | Q8R5H1 | USP15 | ubiquitin specific peptidase 15 | Cytoplasm | peptidase |
| 15536282 | 2 | 0.502 | 4295392.5 | 2920017 | 3399635.5 | 4032030.75 | 2735117 | 1953347 | 3980897.25 | 2272542 | 2337711 | 7554391.5 | 2621345.25 | 3713013.5 | 1126.5422 | DC[160.0307]IELFTTK | Q8R5H1 | Q8R5H1 | USP15 | ubiquitin specific peptidase 15 | Cytoplasm | peptidase |
| 15551707 | 3 | 0.651 | 5309029 | 5232378 | 3123524 | 4019175.25 | 4015918 | 3494696 | 5281122.5 | 3395933.5 | 2469873.5 | 6790867 | 3306239 | 4739192 | 3626.7979 | YMSNTFEPLNKPDSTIQDAGLYQGQVLVIEQK | Q8R5H1 | Q8R5H1 | USP15 | ubiquitin specific peptidase 15 | Cytoplasm | peptidase |
| 15541704 | 3 | 0.655 | 3651698.5 | 2497547.5 | 1642840.875 | 3307223.5 | 1690709.5 | 2282222.25 | 2703453 | 2353313.75 | 2131365 | 6081518.5 | 1909574.375 | 3268569.5 | 1857.1323 | KLDLWSLPPVLVVHLK | Q8R5H1 | Q8R5H1 | USP15 | ubiquitin specific peptidase 15 | Cytoplasm | peptidase |
| 15539084 | 3 | 0.669 | 2521169 | 2259564.25 | 2425869.5 | 2815380.5 | 2670952.5 | 2771621.25 | 2425061 | 2620638.25 | 1863571.125 | 5430106.5 | 1878300.625 | 2575900.25 | 1637.8274 | AEDTEHVVIPVC[160.0307]LR | Q8R5H1 | Q8R5H1 | USP15 | ubiquitin specific peptidase 15 | Cytoplasm | peptidase |
| 15551749 | 2 | 0.656 | 3159203.5 | 3033134.5 | 2243574.25 | 2750333 | 2273954.5 | 2438278.25 | 2887320.25 | 1798838.125 | 2005656.625 | 4806212 | 1820737.25 | 2767044.75 | 1892.9535 | LVSWYTLMEGQEPIAR | Q8R5H1 | Q8R5H1 | USP15 | ubiquitin specific peptidase 15 | Cytoplasm | peptidase |
| 15540372 | 3 | 0.292 | 3198386 | 1745880.625 | 1885950.75 | 3142841.25 | 1452648.5 | 1962418.25 | 2923309.75 | 2121958.75 | 1408643 | 4610851.5 | 2726606.75 | 3119870.25 | 1773.9613 | RNDSIIVDIFHGLFK | Q8R5H1 | Q8R5H1 | USP15 | ubiquitin specific peptidase 15 | Cytoplasm | peptidase |
| 15537654 | 3 | 0.942 | 3015441.75 | 1874702.125 | 1913182.125 | 2273386.25 | 2255640.25 | 1649582.5 | 2673450.25 | 1817429 | 1218859.75 | 4068388 | 1259302.875 | 1572492.375 | 1505.6505 | MSTETEETDGHLR | Q8R5H1 | Q8R5H1 | USP15 | ubiquitin specific peptidase 15 | Cytoplasm | peptidase |
| 15542263 | 2 | 0.472 | 3242652.25 | 2239717.25 | 2586866 | 2265482.25 | 2329440.25 | 2645314 | 2971783.75 | 2232269 | 1669977.375 | 4044035.75 | 2756717 | 2357109 | 1093.6118 | VEVYLTELK | Q8R5H1 | Q8R5H1 | USP15 | ubiquitin specific peptidase 15 | Cytoplasm | peptidase |
| 15535995 | 2 | 0.324 | 3207107.25 | 2540171.75 | 1922572.125 | 2565356.75 | 2058710.125 | 1988124.875 | 2258795 | 1513449.75 | 1702534.625 | 3435574 | 2056458 | 2143991.25 | 1130.5974 | AAYVLFYQR | Q8R5H1 | Q8R5H1 | USP15 | ubiquitin specific peptidase 15 | Cytoplasm | peptidase |
| 15534141 | 2 | 0.703 | 2606680.5 | 2057433.375 | 2091684.75 | 1649066.375 | 1622117.25 | 2151340.75 | 2488574 | 1990648.25 | 1598802.75 | 3414667.75 | 1539038.25 | 2000595.5 | 978.56 | SLEVYLVR | Q8R5H1 | Q8R5H1 | USP15 | ubiquitin specific peptidase 15 | Cytoplasm | peptidase |
| 15542931 | 3 | 0.528 | 989048.125 | 1246152.5 | 574385.438 | 1018519 | 982577.375 | 948416.812 | 1023717.5 | 625135.125 | 496379.375 | 2359949 | 566893.188 | 1263124.625 | 1892.9581 | LVSWYTLMEGQEPIAR | Q8R5H1 | Q8R5H1 | USP15 | ubiquitin specific peptidase 15 | Cytoplasm | peptidase |
| 15532429 | 2 | 0.22 | 2162054.5 | 1330614.5 | 1378520.25 | 1327133.875 | 1230845.75 | 1329111.875 | 1434108.375 | 1145181.875 | 640250.875 | 2295611.75 | 1442983.25 | 1380797 | 1424.7137 | VVAEEAWENHLK | Q8R5H1 | Q8R5H1 | USP15 | ubiquitin specific peptidase 15 | Cytoplasm | peptidase |
| 15561546 | 2 | 0.651 | 93038 | 117299.773 | 63695.516 | 196195.484 | 109112.617 | 135099.297 | 109835.578 | 104959.273 | 79363.609 | 265051.219 | 90935.836 | 52217.777 | 1527.7375 | GEVGSGAGPGTQAGPSAK | Q3UJD6 | Q3UJD6 | USP19 | ubiquitin specific peptidase 19 | Cytoplasm | peptidase |
| 15532387 | 2 | 0.723 | 1.13E+09 | 1.13E+09 | 1.39E+09 | 9.62E+08 | 1.16E+09 | 1.10E+09 | 1.17E+09 | 1.23E+09 | 1.12E+09 | 1.47E+09 | 9.64E+08 | 1.06E+09 | 1172.6256 | VDNWKQNLR | Q5DU02 | Q5DU02 | USP22 | ubiquitin specific peptidase 22 | Nucleus | peptidase |
| 15550012 | 4 | 0.242 | 7689928 | 6604765 | 5590675 | 7800813.5 | 7030165.5 | 6121202 | 9148607 | 6050045 | 4686489 | 2.99E+07 | 6948762.5 | 9696667 | 4179.936 | LGTITPDGADVYSYDEDDMVLDPSLAEHLSHFGIDMLK | P56399 | P56399 | USP5 | ubiquitin specific peptidase 5 (isopeptidase T) | Cytoplasm | peptidase |
| 15554438 | 3 | 0.445 | 6493964.5 | 5288508 | 6198957 | 5157618 | 4535255 | 4496005 | 5138648 | 4246103.5 | 3580821.5 | 1.12E+07 | 3508314 | 5973208 | 2599.3098 | GTGLQPGEEELPDIAPPLVTPDEPK | P56399 | P56399 | USP5 | ubiquitin specific peptidase 5 (isopeptidase T) | Cytoplasm | peptidase |
| 15548861 | 2 | 0.669 | 3601704.25 | 2585532.75 | 4790222 | 4622743 | 3680990.75 | 3896556.25 | 4342874 | 3220790.75 | 2869562.25 | 1.03E+07 | 3065174.25 | 3707545.75 | 1697.9053 | IGEWELIQESGVPLK | P56399 | P56399 | USP5 | ubiquitin specific peptidase 5 (isopeptidase T) | Cytoplasm | peptidase |
| 15553057 | 2 | 0.762 | 4820975.5 | 3285863.25 | 4036403 | 4039491.75 | 3716052.25 | 2977952.5 | 4865072.5 | 3448779.25 | 2791977.75 | 9419316 | 2825834.5 | 3948297.25 | 1583.7517 | PALESGDGEQVPEQK | P56399 | P56399 | USP5 | ubiquitin specific peptidase 5 (isopeptidase T) | Cytoplasm | peptidase |
| 15537820 | 2 | 0.407 | 5254240 | 2957217 | 3499751.5 | 3895191.5 | 2684816.75 | 3767163 | 4156449.5 | 3182564 | 2057765.5 | 9352179 | 3052377 | 4736260 | 1457.7456 | SAAESISESVPVGPK | P56399 | P56399 | USP5 | ubiquitin specific peptidase 5 (isopeptidase T) | Cytoplasm | peptidase |
| 15552090 | 2 | 0.527 | 5965433.5 | 3516202.5 | 4016468 | 4637076.5 | 4198784 | 3881471.5 | 4693370 | 3195310.25 | 2564649.25 | 9289088 | 3516858.75 | 4142428.5 | 2108.0264 | IFQNAPTDPTQDFSTQVAK | P56399 | P56399 | USP5 | ubiquitin specific peptidase 5 (isopeptidase T) | Cytoplasm | peptidase |
| 15540885 | 3 | 0.34 | 5203784 | 3852215.25 | 3298077.75 | 4670977.5 | 3178853.5 | 3555799.5 | 5589600.5 | 2994808.75 | 3036374.75 | 8849405 | 3854094.5 | 5555229.5 | 2115.097 | KLDVSIEMPEELDISQLR | P56399 | P56399 | USP5 | ubiquitin specific peptidase 5 (isopeptidase T) | Cytoplasm | peptidase |
| 15534114 | 2 | 0.528 | 4097786.25 | 2610439 | 3790534.75 | 4073610 | 2925652.25 | 4255720.5 | 3511476.5 | 2918629 | 2620102.75 | 7336272.5 | 2536482.75 | 4717107.5 | 1414.8285 | IVILPDYLEIAR | P56399 | P56399 | USP5 | ubiquitin specific peptidase 5 (isopeptidase T) | Cytoplasm | peptidase |
| 15543717 | 3 | 0.393 | 4048917.5 | 3434088.75 | 1768941.125 | 2205477 | 1984081.25 | 1396079.75 | 4317254.5 | 2487173.5 | 1432803.375 | 7127297 | 2153128 | 3280464.75 | 2118.04 | QQDAQEFFLHLINMVER | P56399 | P56399 | USP5 | ubiquitin specific peptidase 5 (isopeptidase T) | Cytoplasm | peptidase |
| 15543321 | 3 | 0.353 | 1275021.125 | 395750.531 | 624910.188 | 1418381.125 | 796029.938 | 153999.906 | 1408999.25 | 717465.562 | 108930.328 | 3903479.75 | 1295113.125 | 1424069.25 | 2108.025 | IFQNAPTDPTQDFSTQVAK | P56399 | P56399 | USP5 | ubiquitin specific peptidase 5 (isopeptidase T) | Cytoplasm | peptidase |
| 15538171 | 2 | 0.974 | 990388.438 | 1245578.375 | 1623100.25 | 1201139.375 | 858960.25 | 1557024.125 | 1455275.5 | 971737.625 | 852223.312 | 2416830 | 436848.5 | 1235125.5 | 1209.628 | FTFGLDWVPK | P56399 | P56399 | USP5 | ubiquitin specific peptidase 5 (isopeptidase T) | Cytoplasm | peptidase |
| 15559685 | 2 | 0.989 | 758500 | 776956.062 | 1362155.125 | 664102.875 | 1235707.25 | 1386466.5 | 1056247 | 1365659.5 | 850463.062 | 2232834.25 | 362266.156 | 1224512.5 | 1860.9249 | ENLWLNLTDGSILC[160.0307]GR | P56399 | P56399 | USP5 | ubiquitin specific peptidase 5 (isopeptidase T) | Cytoplasm | peptidase |
| 15544985 | 3 | 0.852 | 1235717.375 | 945713.688 | 2477773.5 | 228383.266 | 1071494.875 | 2756844.75 | 946887.188 | 1568705 | 1292651.625 | 1374874.5 | 374738.688 | 1526643 | 2134.0352 | QQDAQEFFLHLINM[147.0354]VER | P56399 | P56399 | USP5 | ubiquitin specific peptidase 5 (isopeptidase T) | Cytoplasm | peptidase |
| 15542032 | 2 | 0.092 | 780573.562 | 684404.875 | 821910.812 | 1127051.25 | 793845.25 | 888376.312 | 861159 | 625269.062 | 528667.562 | 1015498.688 | 815959.25 | 1067404.375 | 1495.8286 | EVFGTFGIPFLLR | Q6A4J8 | Q6A4J8 | USP7 | ubiquitin specific peptidase 7 (herpes virus-associated) | Nucleus | peptidase |
| 15551817 | 3 | 0.773 | 6633978 | 4252188.5 | 3354510.75 | 5924161 | 2941099.75 | 3507540.5 | 4167993.25 | 2797745.25 | 3603463 | 6147368.5 | 3210754.25 | 3678154.75 | 2738.2737 | FNDYFEFPRELDMEPYTVAGVAK | P70398 | P70398 | USP9X | ubiquitin specific peptidase 9, X-linked | Plasma Membrane | peptidase |
| 15551358 | 2 | 0.478 | 3282947.5 | 2598886 | 3271219.75 | 2220426.5 | 2876279.25 | 1936353.875 | 2864454.5 | 2403617.25 | 2245713.5 | 3672221.25 | 2019265.875 | 2301933.25 | 1827.9568 | ALTLQDLDNIWAAQAGK | P70398 | P70398 | USP9X | ubiquitin specific peptidase 9, X-linked | Plasma Membrane | peptidase |
| 15539855 | 2 | 0.542 | 3041807.5 | 2291832 | 2014040.625 | 2241759.75 | 1991076.125 | 851730.75 | 2192364.5 | 1409129.875 | 1645944.875 | 2846538 | 1258053.25 | 1528451.75 | 1175.6295 | ELLAFQTPEK | P70398 | P70398 | USP9X | ubiquitin specific peptidase 9, X-linked | Plasma Membrane | peptidase |
| 15553043 | 2 | 0.89 | 1497372 | 1089230.625 | 1221812.75 | 956649.125 | 1488572.25 | 765768.5 | 1801168.125 | 1289231.875 | 883719.562 | 2138299.5 | 794729.375 | 897114.688 | 1509.6708 | IIEDC[160.0307]SNSDETVK | P70398 | P70398 | USP9X | ubiquitin specific peptidase 9, X-linked | Plasma Membrane | peptidase |
| 15538221 | 2 | 0.748 | 2669566.75 | 2992959.5 | 2843019.25 | 2131856.5 | 2642373.25 | 2596140.25 | 3183960.25 | 2352652 | 2227787 | 3370897 | 2189571 | 2203788.5 | 1319.7706 | FDLVSFIPLLR | Q80WQ2 | Q80WQ2 | VAC14 | Vac14 homolog (S. cerevisiae) | Cytoplasm | other |
| 15550626, 15554779 | 2, 3 | 0.348 | 6.94E+07 | 6.62E+07 | 9.87E+07 | 5.72E+07 | 8.03E+07 | 8.06E+07 | 9.41E+07 | 8.86E+07 | 9.82E+07 | 7.77E+07 | 9.70E+07 | 6.89E+07 | 2494.1982, 2494.201 | LSELDDRADALQAGASQFETSAAK | P63024 | P63024 | VAMP3 | vesicle-associated membrane protein 3 | Plasma Membrane | other |
| 15550911 | 2 | 0.203 | 1.27E+07 | 1.42E+07 | 1.76E+07 | 7398147.5 | 1.31E+07 | 1.07E+07 | 1.26E+07 | 1.45E+07 | 1.28E+07 | 1.39E+07 | 1.34E+07 | 9704473 | 1915.9503 | LQQTQNQVDEVVDIMR | P63024 | P63024 | VAMP3 | vesicle-associated membrane protein 3 | Plasma Membrane | other |
| 15541841, 15544947 | 3 | 0.395 | 5259379.5 | 8041936 | 7566668.5 | 4435115 | 7373867.5 | 6722218.5 | 6337352 | 7887335 | 1.01E+07 | 6810446 | 5986848 | 4987410 | 1915.9485, 1915.9498 | LQQTQNQVDEVVDIMR | P63024 | P63024 | VAMP3 | vesicle-associated membrane protein 3 | Plasma Membrane | other |
| 15537034 | 3 | 0.041 | 6717073.5 | 7705563.5 | 7206602.5 | 4383845 | 6309872 | 4334073 | 7716146 | 8385475.5 | 6993261 | 5906573.5 | 8841654 | 6289297.5 | 2072.052 | RLQQTQNQVDEVVDIMR | P63024 | P63024 | VAMP3 | vesicle-associated membrane protein 3 | Plasma Membrane | other |
| 15550129 | 2 | 0.87 | 1861203.5 | 2535172 | 4561862 | 1519038.25 | 6221055.5 | 3037563.75 | 2808651.25 | 5768093 | 2245587 | 2540502 | 2633679.75 | 2092830.125 | 1665.8071 | ADALQAGASQFETSAAK | P63024 | P63024 | VAMP3 | vesicle-associated membrane protein 3 | Plasma Membrane | other |
| 15548364 | 2, 3 | 0.427 | 2.15E+07 | 2.71E+07 | 2.65E+07 | 2.76E+07 | 3.13E+07 | 1.97E+07 | 2.33E+07 | 2.67E+07 | 2.28E+07 | 2.48E+07 | 1.80E+07 | 1.98E+07 | 2311.0283, 2311.0298 | QC[160.0307]QQQADEVTEIMLNNFDK | Q9Z2P8 | Q9Z2P8 | VAMP5 | vesicle-associated membrane protein 5 | Plasma Membrane | transporter |
| 15536718 | 2 | 0.213 | 1.02E+07 | 1.34E+07 | 2.17E+07 | 6170534 | 1.31E+07 | 1.58E+07 | 1.03E+07 | 1.93E+07 | 1.84E+07 | 8188872 | 7055023.5 | 9439187 | 1428.6654 | SDQLLDMSSAFSK | Q9Z2P8 | Q9Z2P8 | VAMP5 | vesicle-associated membrane protein 5 | Plasma Membrane | transporter |
| 15532265 | 2 | 0.313 | 1.00E+07 | 1.25E+07 | 1.75E+07 | 7807775.5 | 1.26E+07 | 1.48E+07 | 1.03E+07 | 1.47E+07 | 1.32E+07 | 6831828.5 | 1.05E+07 | 9389297 | 858.46686 | LAELEQR | Q9Z2P8 | Q9Z2P8 | VAMP5 | vesicle-associated membrane protein 5 | Plasma Membrane | transporter |
| 15534865 | 2, 3 | 0.046 | 1.08E+07 | 2.48E+07 | 2.05E+07 | 2.06E+07 | 2.78E+07 | 2.97E+07 | 2.05E+07 | 2.82E+07 | 2.51E+07 | 9798140 | 1.53E+07 | 1.34E+07 | 1406.6401, 1406.6407 | TEDLEATSEHFK | O70404 | O70404 | VAMP8 | vesicle-associated membrane protein 8 | Plasma Membrane | transporter |
| 15538181 | 2 | 0.104 | 1.08E+07 | 1.93E+07 | 1.34E+07 | 1.41E+07 | 1.76E+07 | 1.78E+07 | 1.28E+07 | 1.87E+07 | 2.07E+07 | 6223299.5 | 1.37E+07 | 9374801 | 1102.571 | NLQSEVEGVK | O70404 | O70404 | VAMP8 | vesicle-associated membrane protein 8 | Plasma Membrane | transporter |
| 15534989 | 3 | 0.495 | 4459238.5 | 560762.312 | 631.86 | 5740452.5 | 486836.406 | 155596.562 | 710237.562 | 698025.438 | 3288795.5 | 1751968.5 | 7815951 | 2726874.75 | 1648.7771 | NKTEDLEATSEHFK | O70404 | O70404 | VAMP8 | vesicle-associated membrane protein 8 | Plasma Membrane | transporter |
| 15531724 | 2 | 0.744 | 4.57E+07 | 4.59E+07 | 5.21E+07 | 4.85E+07 | 5.03E+07 | 5.25E+07 | 5.39E+07 | 5.19E+07 | 4.36E+07 | 5.14E+07 | 5.00E+07 | 5.08E+07 | 1291.687 | GPFTDVVTTNLK | Q9WV55 | Q9WV55 | VAPA | VAMP (vesicle-associated membrane protein)-associated protein A, 33kDa | Plasma Membrane | other |
| 15548235 | 2, 3 | 0.387 | 2.68E+07 | 3.22E+07 | 3.26E+07 | 3.56E+07 | 2.91E+07 | 2.62E+07 | 3.58E+07 | 3.42E+07 | 3.28E+07 | 3.16E+07 | 3.28E+07 | 3.31E+07 | 2324.1401, 2324.1414 | FMVQTIFAPPNISDMEAVWK | Q9WV55 | Q9WV55 | VAPA | VAMP (vesicle-associated membrane protein)-associated protein A, 33kDa | Plasma Membrane | other |
| 15538565 | 3 | 0.374 | 1.77E+07 | 1.79E+07 | 2.32E+07 | 2.37E+07 | 2.59E+07 | 2.01E+07 | 2.24E+07 | 2.04E+07 | 2.12E+07 | 2.35E+07 | 2.11E+07 | 1.99E+07 | 1603.8663 | HEQILVLDPPSDLK | Q9WV55 | Q9WV55 | VAPA | VAMP (vesicle-associated membrane protein)-associated protein A, 33kDa | Plasma Membrane | other |
| 15532115 | 3 | 0.336 | 1.93E+07 | 1.24E+07 | 6068187 | 1.93E+07 | 1.23E+07 | 8408812 | 1.59E+07 | 1.28E+07 | 9571920 | 2.06E+07 | 2.19E+07 | 1.76E+07 | 1645.8263 | VAHSDKPGSTSAVSFR | Q9WV55 | Q9WV55 | VAPA | VAMP (vesicle-associated membrane protein)-associated protein A, 33kDa | Plasma Membrane | other |
| 15532468 | 2 | 0.44 | 8593414 | 8620910 | 8290026.5 | 9219136 | 7773788.5 | 6271269.5 | 9841255 | 7486005.5 | 6121485.5 | 1.03E+07 | 9759346 | 8128284.5 | 1262.5914 | EAKPDELMDSK | Q9WV55 | Q9WV55 | VAPA | VAMP (vesicle-associated membrane protein)-associated protein A, 33kDa | Plasma Membrane | other |
| 15532414 | 2 | 0.017 | 1.14E+07 | 9939133 | 9668522 | 1.04E+07 | 1.13E+07 | 9890620 | 1.23E+07 | 1.22E+07 | 1.11E+07 | 9661740 | 9572098 | 8765391 | 1382.5677 | C[160.0307]VFEMPNENDK | Q9WV55 | Q9WV55 | VAPA | VAMP (vesicle-associated membrane protein)-associated protein A, 33kDa | Plasma Membrane | other |
| 15556361 | 2 | 0.997 | 8918390 | 7327228 | 1.14E+07 | 8912581 | 1.00E+07 | 7875785.5 | 9774161 | 9307978 | 8254178 | 8567506 | 9952677 | 8474108 | 1603.8662 | HEQILVLDPPSDLK | Q9WV55 | Q9WV55 | VAPA | VAMP (vesicle-associated membrane protein)-associated protein A, 33kDa | Plasma Membrane | other |
| 15542310 | 3 | 0.559 | 7789279 | 4495444.5 | 1398151 | 6146902 | 4369893.5 | 1618806 | 5704295 | 3546439 | 1399898.625 | 6791725.5 | 8485163 | 4833510 | 1483.7428 | PHSVSLNDTETRK | Q9WV55 | Q9WV55 | VAPA | VAMP (vesicle-associated membrane protein)-associated protein A, 33kDa | Plasma Membrane | other |
| 15555985 | 2 | 0.524 | 1791895.875 | 917808.312 | 55978.32 | 1816115.625 | 541541.938 | 426441.75 | 1510901 | 324157.531 | 155762.672 | 2176985.5 | 2107920.5 | 885508.438 | 1645.8275 | VAHSDKPGSTSAVSFR | Q9WV55 | Q9WV55 | VAPA | VAMP (vesicle-associated membrane protein)-associated protein A, 33kDa | Plasma Membrane | other |
| 15534254 | 3 | 0.036 | 3036443 | 4341906 | 4903065 | 5904795.5 | 7369567 | 6052565.5 | 5368142 | 5154346 | 6421618.5 | 4711760.5 | 5608154.5 | 4682829.5 | 1648.8867 | VEQVLSLEPQHELK | Q9QY76 | Q9QY76 | VAPB | VAMP (vesicle-associated membrane protein)-associated protein B and C | Plasma Membrane | other |
| 15554022 | 2 | 0.48 | 717735.625 | 1033518.188 | 1244593.875 | 1029484.562 | 2094076.75 | 1387463.375 | 1240026.375 | 1190174 | 1441522.5 | 806857.188 | 1586710.5 | 1544517.25 | 1648.8872 | VEQVLSLEPQHELK | Q9QY76 | Q9QY76 | VAPB | VAMP (vesicle-associated membrane protein)-associated protein B and C | Plasma Membrane | other |
| 15531970 | 2 | 0.003 | 2.12E+07 | 2.79E+07 | 3.00E+07 | 7537085 | 1.55E+07 | 1.27E+07 | 1.92E+07 | 2.51E+07 | 2.39E+07 | 8609572 | 1.00E+07 | 1.33E+07 | 725.4546 | IPAVPTK | Q3U2A8 | Q3U2A8 | VARS2 | valyl-tRNA synthetase 2, mitochondrial | Cytoplasm | enzyme |
| 15550371 | 3 | 0.482 | 2758705.25 | 3167851.25 | 4138706 | 5146602.5 | 7074794 | 3232512.25 | 3496139 | 4510186.5 | 7552849.5 | 2101634.5 | 5209145 | 4188807.75 | 2639.0967 | SSSSVTTSEAHPSTPC[160.0307]SSDDSDLER | P70460 | P70460 | VASP | vasodilator-stimulated phosphoprotein | Plasma Membrane | other |
| 15548166, 15550616 | 3, 4 | 0.975 | 2.05E+08 | 2.16E+08 | 1.69E+08 | 1.62E+08 | 2.34E+08 | 1.46E+08 | 2.25E+08 | 2.02E+08 | 1.14E+08 | 3.59E+08 | 1.21E+08 | 1.51E+08 | 3672.7834, 3672.7869 | LADDVDLEQVANETHGHVGADLAALC[160.0307]SEAALQAIR | Q01853 | Q01853 | VCP | valosin containing protein | Cytoplasm | enzyme |
| 15548731, 15550952 | 3, 4, 5 | 0.228 | 2.85E+08 | 1.78E+08 | 1.14E+08 | 2.20E+08 | 8.61E+07 | 1.84E+08 | 1.54E+08 | 9.67E+07 | 1.58E+08 | 3.31E+08 | 2.33E+08 | 2.31E+08 | 3800.8772, 3800.8782, 3800.8806 | LADDVDLEQVANETHGHVGADLAALC[160.0307]SEAALQAIRK | Q01853 | Q01853 | VCP | valosin containing protein | Cytoplasm | enzyme |
| 15549616, 15550015 | 2, 3 | 0.447 | 2.39E+08 | 1.18E+08 | 5.15E+07 | 1.82E+08 | 1.02E+08 | 9.13E+07 | 1.22E+08 | 1.18E+08 | 1.41E+08 | 3.15E+08 | 1.52E+08 | 1.75E+08 | 2498.2732, 2498.2776 | ETVVEVPQVTWEDIGGLEDVKR | Q01853 | Q01853 | VCP | valosin containing protein | Cytoplasm | enzyme |
| 15548006 | 3 | 0.878 | 1.65E+08 | 2.83E+08 | 3.69E+08 | 1.24E+08 | 3.31E+08 | 2.65E+08 | 2.45E+08 | 2.29E+08 | 1.58E+08 | 2.86E+08 | 1.19E+08 | 2.35E+08 | 2925.3955 | VVETDPSPYC[160.0307]IVAPDTVIHC[160.0307]EGEPIK | Q01853 | Q01853 | VCP | valosin containing protein | Cytoplasm | enzyme |
| 15547680 | 2 | 0.965 | 1.50E+08 | 1.73E+08 | 2.00E+08 | 1.47E+08 | 1.52E+08 | 1.78E+08 | 1.82E+08 | 1.68E+08 | 1.68E+08 | 2.67E+08 | 1.07E+08 | 1.77E+08 | 1810.9918 | NAPAIIFIDELDAIAPK | Q01853 | Q01853 | VCP | valosin containing protein | Cytoplasm | enzyme |
| 15533622 | 3 | 0.282 | 1.49E+08 | 8.18E+07 | 5.06E+07 | 1.25E+08 | 1.05E+08 | 6.53E+07 | 1.04E+08 | 8.08E+07 | 9.00E+07 | 2.20E+08 | 1.16E+08 | 1.35E+08 | 1823.9177 | ELQELVQYPVEHPDK | Q01853 | Q01853 | VCP | valosin containing protein | Cytoplasm | enzyme |
| 15552264 | 2 | 0.873 | 1.41E+08 | 8.98E+07 | 1.47E+08 | 1.10E+08 | 1.20E+08 | 1.21E+08 | 1.28E+08 | 1.17E+08 | 1.23E+08 | 1.96E+08 | 8.87E+07 | 1.48E+08 | 1923.948 | QAAPC[160.0307]VLFFDELDSIAK | Q01853 | Q01853 | VCP | valosin containing protein | Cytoplasm | enzyme |
| 15536866 | 2 | 0.677 | 1.21E+08 | 9.89E+07 | 8.70E+07 | 1.05E+08 | 1.04E+08 | 9.81E+07 | 1.33E+08 | 8.29E+07 | 7.23E+07 | 1.94E+08 | 8.38E+07 | 1.13E+08 | 1329.6882 | WALSQSNPSALR | Q01853 | Q01853 | VCP | valosin containing protein | Cytoplasm | enzyme |
| 15547713 | 2 | 0.635 | 1.08E+08 | 9.63E+07 | 9.32E+07 | 8.82E+07 | 9.79E+07 | 9.47E+07 | 1.06E+08 | 8.04E+07 | 8.28E+07 | 1.73E+08 | 8.15E+07 | 9.79E+07 | 2170.1204 | LIVDEAINEDNSVVSLSQPK | Q01853 | Q01853 | VCP | valosin containing protein | Cytoplasm | enzyme |
| 15533277 | 2 | 0.438 | 1.07E+08 | 8.11E+07 | 7.67E+07 | 9.27E+07 | 9.71E+07 | 7.30E+07 | 9.49E+07 | 6.55E+07 | 7.84E+07 | 1.69E+08 | 7.97E+07 | 9.89E+07 | 1251.6161 | GVLFYGPPGC[160.0307]GK | Q01853 | Q01853 | VCP | valosin containing protein | Cytoplasm | enzyme |
| 15548298 | 2 | 0.422 | 8.69E+07 | 8.09E+07 | 7.34E+07 | 7.75E+07 | 8.29E+07 | 7.71E+07 | 7.93E+07 | 7.46E+07 | 7.04E+07 | 1.63E+08 | 8.34E+07 | 7.42E+07 | 1540.8019 | LGDVISIQPC[160.0307]PDVK | Q01853 | Q01853 | VCP | valosin containing protein | Cytoplasm | enzyme |
| 15549079, 15550762 | 2, 3 | 0.846 | 7.64E+07 | 1.17E+08 | 1.06E+08 | 7.19E+07 | 9.69E+07 | 1.02E+08 | 5.63E+07 | 1.04E+08 | 7.62E+07 | 1.57E+08 | 5.12E+07 | 9.34E+07 | 2585.3787, 2585.3828 | IHVLPIDDTVEGITGNLFEVYLK | Q01853 | Q01853 | VCP | valosin containing protein | Cytoplasm | enzyme |
| 15547757 | 2 | 0.461 | 1.11E+08 | 8.22E+07 | 8.08E+07 | 7.77E+07 | 8.30E+07 | 7.98E+07 | 1.01E+08 | 7.48E+07 | 9.40E+07 | 1.54E+08 | 7.75E+07 | 1.02E+08 | 1578.7916 | AIANEC[160.0307]QANFISIK | Q01853 | Q01853 | VCP | valosin containing protein | Cytoplasm | enzyme |
| 15540442 | 3 | 0.365 | 7.96E+07 | 7.21E+07 | 5.70E+07 | 6.18E+07 | 6.32E+07 | 6.17E+07 | 6.61E+07 | 4.88E+07 | 4.40E+07 | 1.47E+08 | 5.73E+07 | 6.38E+07 | 2170.1204 | LIVDEAINEDNSVVSLSQPK | Q01853 | Q01853 | VCP | valosin containing protein | Cytoplasm | enzyme |
| 15547948 | 2 | 0.967 | 6.89E+07 | 7.97E+07 | 9.83E+07 | 5.30E+07 | 9.90E+07 | 7.87E+07 | 7.77E+07 | 7.58E+07 | 6.40E+07 | 1.35E+08 | 3.78E+07 | 6.92E+07 | 1556.8535 | LDQLIYIPLPDEK | Q01853 | Q01853 | VCP | valosin containing protein | Cytoplasm | enzyme |
| 15547847 | 3 | 0.36 | 8.12E+07 | 5.65E+07 | 6.19E+07 | 7.54E+07 | 5.13E+07 | 5.64E+07 | 6.87E+07 | 5.41E+07 | 5.80E+07 | 1.29E+08 | 6.31E+07 | 7.16E+07 | 2312.0776 | QTNPSAMEVEEDDPVPEIRR | Q01853 | Q01853 | VCP | valosin containing protein | Cytoplasm | enzyme |
| 15547747 | 2 | 0.974 | 7.44E+07 | 8.51E+07 | 7.82E+07 | 6.48E+07 | 8.71E+07 | 6.61E+07 | 9.09E+07 | 6.70E+07 | 6.31E+07 | 1.26E+08 | 4.46E+07 | 6.70E+07 | 1679.8314 | VINQILTEMDGMSTK | Q01853 | Q01853 | VCP | valosin containing protein | Cytoplasm | enzyme |
| 15532447 | 2, 3 | 0.49 | 6.77E+07 | 5.32E+07 | 5.80E+07 | 4.46E+07 | 5.05E+07 | 5.73E+07 | 6.36E+07 | 5.54E+07 | 5.14E+07 | 1.16E+08 | 4.76E+07 | 6.24E+07 | 1430.8263, 1430.827 | IVSQLLTLMDGLK | Q01853 | Q01853 | VCP | valosin containing protein | Cytoplasm | enzyme |
| 15548386 | 2, 3 | 0.827 | 3.86E+07 | 9.15E+07 | 1.27E+08 | 5.70E+07 | 9.84E+07 | 9.92E+07 | 9.28E+07 | 7.86E+07 | 5.02E+07 | 1.15E+08 | 2.39E+07 | 6.17E+07 | 2342.171, 2342.1711 | ETVVEVPQVTWEDIGGLEDVK | Q01853 | Q01853 | VCP | valosin containing protein | Cytoplasm | enzyme |
| 15531286 | 2 | 0.365 | 7.05E+07 | 5.85E+07 | 5.72E+07 | 6.35E+07 | 5.31E+07 | 4.77E+07 | 6.53E+07 | 5.25E+07 | 5.39E+07 | 1.12E+08 | 5.37E+07 | 6.85E+07 | 1172.6649 | GILLYGPPGTGK | Q01853 | Q01853 | VCP | valosin containing protein | Cytoplasm | enzyme |
| 15531963 | 2 | 0.992 | 5.07E+07 | 5.86E+07 | 6.66E+07 | 3.56E+07 | 6.32E+07 | 7.22E+07 | 7.37E+07 | 5.51E+07 | 4.45E+07 | 9.26E+07 | 3.67E+07 | 5.97E+07 | 1032.5552 | GDDLSTAILK | Q01853 | Q01853 | VCP | valosin containing protein | Cytoplasm | enzyme |
| 15532356 | 2 | 0.908 | 4.63E+07 | 4.32E+07 | 5.12E+07 | 3.23E+07 | 4.66E+07 | 4.72E+07 | 5.35E+07 | 4.73E+07 | 3.93E+07 | 8.29E+07 | 2.89E+07 | 4.97E+07 | 1051.5221 | MDELQLFR | Q01853 | Q01853 | VCP | valosin containing protein | Cytoplasm | enzyme |
| 15550467 | 2, 3 | 0.353 | 5.36E+07 | 3.80E+07 | 3.35E+07 | 4.31E+07 | 2.55E+07 | 3.85E+07 | 4.41E+07 | 2.67E+07 | 4.21E+07 | 7.35E+07 | 3.95E+07 | 4.94E+07 | 2318.2676, 2318.2678 | EVDIGIPDATGRLEILQIHTK | Q01853 | Q01853 | VCP | valosin containing protein | Cytoplasm | enzyme |
| 15551087 | 2 | 0.972 | 2.93E+07 | 5.19E+07 | 6.54E+07 | 3.39E+07 | 5.15E+07 | 4.60E+07 | 5.85E+07 | 3.88E+07 | 3.24E+07 | 6.94E+07 | 2.19E+07 | 4.16E+07 | 1501.707 | YEMFAQTLQQSR | Q01853 | Q01853 | VCP | valosin containing protein | Cytoplasm | enzyme |
| 15531474, 15531953 | 3, 4 | 0.888 | 2.06E+07 | 5.80E+07 | 1.02E+08 | 2.68E+07 | 6.83E+07 | 7.95E+07 | 7.44E+07 | 5.33E+07 | 2.70E+07 | 6.51E+07 | 1.83E+07 | 3.86E+07 | 1980.013, 1980.0156 | RELQELVQYPVEHPDK | Q01853 | Q01853 | VCP | valosin containing protein | Cytoplasm | enzyme |
| 15542612 | 2 | 0.899 | 4.80E+07 | 4.27E+07 | 4.45E+07 | 4.44E+07 | 4.00E+07 | 4.27E+07 | 5.27E+07 | 4.06E+07 | 3.99E+07 | 6.21E+07 | 3.39E+07 | 4.96E+07 | 1049.5488 | DVDLEFLAK | Q01853 | Q01853 | VCP | valosin containing protein | Cytoplasm | enzyme |
| 15547954 | 2 | 0.432 | 4.24E+07 | 2.53E+07 | 2.44E+07 | 3.39E+07 | 2.78E+07 | 1.17E+07 | 3.03E+07 | 2.27E+07 | 2.48E+07 | 5.97E+07 | 2.87E+07 | 3.16E+07 | 1823.9181 | ELQELVQYPVEHPDK | Q01853 | Q01853 | VCP | valosin containing protein | Cytoplasm | enzyme |
| 15531428 | 2 | 0.324 | 3.87E+07 | 2.87E+07 | 2.81E+07 | 2.91E+07 | 2.63E+07 | 2.58E+07 | 3.64E+07 | 2.55E+07 | 2.35E+07 | 5.73E+07 | 2.86E+07 | 3.43E+07 | 1118.577 | ESIESEIRR | Q01853 | Q01853 | VCP | valosin containing protein | Cytoplasm | enzyme |
| 15548008 | 2 | 0.903 | 3.17E+07 | 3.75E+07 | 3.58E+07 | 2.64E+07 | 3.60E+07 | 3.47E+07 | 3.19E+07 | 3.17E+07 | 3.06E+07 | 5.44E+07 | 2.15E+07 | 3.89E+07 | 1951.9176 | GPELLTM[147.0354]WFGESEANVR | Q01853 | Q01853 | VCP | valosin containing protein | Cytoplasm | enzyme |
| 15533285 | 2, 3 | 0.497 | 2.29E+07 | 1.52E+07 | 1.94E+07 | 1.83E+07 | 2.58E+07 | 1.41E+07 | 1.80E+07 | 1.92E+07 | 1.28E+07 | 5.04E+07 | 1.75E+07 | 1.97E+07 | 1424.7656, 1424.7657 | PYFLEAYRPIR | Q01853 | Q01853 | VCP | valosin containing protein | Cytoplasm | enzyme |
| 15548208 | 3 | 0.65 | 2.97E+07 | 2.70E+07 | 2.59E+07 | 2.36E+07 | 2.47E+07 | 2.19E+07 | 3.24E+07 | 2.27E+07 | 2.30E+07 | 4.87E+07 | 2.16E+07 | 2.57E+07 | 2441.1235 | ERQTNPSAMEVEEDDPVPEIR | Q01853 | Q01853 | VCP | valosin containing protein | Cytoplasm | enzyme |
| 15532122 | 3 | 0.742 | 4.26E+07 | 2.61E+07 | 4.24E+07 | 3.59E+07 | 3.34E+07 | 3.81E+07 | 4.57E+07 | 3.81E+07 | 3.79E+07 | 4.66E+07 | 3.08E+07 | 4.94E+07 | 1923.9493 | QAAPC[160.0307]VLFFDELDSIAK | Q01853 | Q01853 | VCP | valosin containing protein | Cytoplasm | enzyme |
| 15532018 | 2 | 0.959 | 2.07E+07 | 2.78E+07 | 3.53E+07 | 1.63E+07 | 4.15E+07 | 2.71E+07 | 3.03E+07 | 3.07E+07 | 1.94E+07 | 4.51E+07 | 1.21E+07 | 2.15E+07 | 1242.6296 | EVDIGIPDATGR | Q01853 | Q01853 | VCP | valosin containing protein | Cytoplasm | enzyme |
| 15535431 | 3 | 0.101 | 4.28E+07 | 2.97E+07 | 2.40E+07 | 2.91E+07 | 2.42E+07 | 2.94E+07 | 3.61E+07 | 2.25E+07 | 2.77E+07 | 4.40E+07 | 3.83E+07 | 4.49E+07 | 1679.9263 | EMVELPLRHPALFK | Q01853 | Q01853 | VCP | valosin containing protein | Cytoplasm | enzyme |
| 15548593 | 2, 3 | 0.434 | 2.67E+07 | 2.32E+07 | 2.29E+07 | 2.78E+07 | 1.89E+07 | 2.24E+07 | 2.40E+07 | 2.32E+07 | 2.38E+07 | 4.24E+07 | 2.16E+07 | 2.82E+07 | 2256.1313, 2256.1338 | AVANETGAFFFLINGPEIMSK | Q01853 | Q01853 | VCP | valosin containing protein | Cytoplasm | enzyme |
| 15547960 | 2 | 0.989 | 2.41E+07 | 2.54E+07 | 2.34E+07 | 2.14E+07 | 3.19E+07 | 2.18E+07 | 2.89E+07 | 2.40E+07 | 1.68E+07 | 4.23E+07 | 1.47E+07 | 2.14E+07 | 2155.9775 | QTNPSAMEVEEDDPVPEIR | Q01853 | Q01853 | VCP | valosin containing protein | Cytoplasm | enzyme |
| 15539598 | 3 | 0.254 | 3.87E+07 | 9610404 | 2495617.75 | 2.33E+07 | 9311499 | 5587899 | 1.36E+07 | 1.15E+07 | 1.50E+07 | 4.06E+07 | 3.51E+07 | 2.87E+07 | 1629.8037 | KYEMFAQTLQQSR | Q01853 | Q01853 | VCP | valosin containing protein | Cytoplasm | enzyme |
| 15541162 | 2 | 0.962 | 2.49E+07 | 2.54E+07 | 2.62E+07 | 2.73E+07 | 2.27E+07 | 2.86E+07 | 3.23E+07 | 2.30E+07 | 2.30E+07 | 4.06E+07 | 1.84E+07 | 2.91E+07 | 819.4704 | GDIFLVR | Q01853 | Q01853 | VCP | valosin containing protein | Cytoplasm | enzyme |
| 15552243 | 2 | 0.494 | 3.03E+07 | 2.35E+07 | 2.44E+07 | 2.59E+07 | 2.25E+07 | 2.34E+07 | 2.78E+07 | 2.01E+07 | 2.62E+07 | 4.00E+07 | 2.35E+07 | 2.77E+07 | 1799.8003 | MTNGFSGADLTEIC[160.0307]QR | Q01853 | Q01853 | VCP | valosin containing protein | Cytoplasm | enzyme |
| 15538780 | 3 | 0.611 | 3.56E+07 | 4.38E+07 | 4.53E+07 | 3.62E+07 | 3.53E+07 | 4.20E+07 | 4.77E+07 | 4.11E+07 | 4.16E+07 | 3.97E+07 | 2.87E+07 | 4.66E+07 | 1810.9918 | NAPAIIFIDELDAIAPK | Q01853 | Q01853 | VCP | valosin containing protein | Cytoplasm | enzyme |
| 15541994 | 2 | 0.662 | 2.19E+07 | 1.64E+07 | 1.67E+07 | 1.97E+07 | 1.44E+07 | 1.48E+07 | 2.38E+07 | 1.45E+07 | 1.28E+07 | 3.29E+07 | 1.56E+07 | 1.80E+07 | 1034.4337 | DHFEEAMR | Q01853 | Q01853 | VCP | valosin containing protein | Cytoplasm | enzyme |
| 15531897 | 2 | 0.145 | 1.45E+07 | 3723623.5 | 1812427.875 | 8610500 | 4501745 | 1995439 | 5664453 | 4795230.5 | 3596245.5 | 3.11E+07 | 1.35E+07 | 9824923 | 947.5655 | KGDIFLVR | Q01853 | Q01853 | VCP | valosin containing protein | Cytoplasm | enzyme |
| 15540182 | 3 | 0.706 | 1.36E+07 | 1.33E+07 | 1.15E+07 | 9088874 | 1.86E+07 | 1.30E+07 | 1.19E+07 | 1.40E+07 | 8631564 | 2.41E+07 | 1.16E+07 | 1.24E+07 | 2146.1514 | AHVIVMAATNRPNSIDPALR | Q01853 | Q01853 | VCP | valosin containing protein | Cytoplasm | enzyme |
| 15533400 | 2 | 0.988 | 6067923.5 | 8884172 | 9988822 | 5396296.5 | 1.26E+07 | 7118452 | 8531479 | 9004193 | 5462705.5 | 1.93E+07 | 3767098.25 | 4859022.5 | 1075.535 | LAGESESNLR | Q01853 | Q01853 | VCP | valosin containing protein | Cytoplasm | enzyme |
| 15540315 | 3 | 0.21 | 1.14E+07 | 286869.438 | 13050.796 | 7362785 | 295686.219 | 336850.188 | 997953.438 | 580794.75 | 1928233.125 | 1.93E+07 | 1.52E+07 | 7389767 | 1967.093 | NAPAIIFIDELDAIAPKR | Q01853 | Q01853 | VCP | valosin containing protein | Cytoplasm | enzyme |
| 15533006 | 3 | 0.367 | 2.15E+07 | 6275510.5 | 3002962.5 | 2.77E+07 | 1313963.125 | 1.03E+07 | 9121630 | 2776539.75 | 1.10E+07 | 1.84E+07 | 2.72E+07 | 3.16E+07 | 1777.9481 | MDELQLFRGDTVLLK | Q01853 | Q01853 | VCP | valosin containing protein | Cytoplasm | enzyme |
| 15531988 | 3 | 0.862 | 6547232.5 | 8398557 | 7134563 | 6574344 | 7351961 | 7655417 | 1.07E+07 | 6862903.5 | 5592361 | 1.75E+07 | 5405044.5 | 7540713.5 | 1679.8333 | VINQILTEMDGMSTK | Q01853 | Q01853 | VCP | valosin containing protein | Cytoplasm | enzyme |
| 15531994 | 2 | 0.876 | 9297182 | 1.23E+07 | 1.36E+07 | 6006955.5 | 1.43E+07 | 1.10E+07 | 1.30E+07 | 1.27E+07 | 7735911.5 | 1.71E+07 | 5503121.5 | 7722089.5 | 986.5322 | EMVELPLR | Q01853 | Q01853 | VCP | valosin containing protein | Cytoplasm | enzyme |
| 15532680 | 3 | 0.961 | 1.11E+07 | 1.21E+07 | 1.15E+07 | 9465608 | 1.45E+07 | 1.01E+07 | 1.10E+07 | 1.26E+07 | 7847048.5 | 1.71E+07 | 6954456.5 | 1.16E+07 | 2155.9778 | QTNPSAMEVEEDDPVPEIR | Q01853 | Q01853 | VCP | valosin containing protein | Cytoplasm | enzyme |
| 15548518 | 3 | 0.844 | 8529286 | 1.64E+07 | 2.21E+07 | 3776384.75 | 2.77E+07 | 1.62E+07 | 1.33E+07 | 1.64E+07 | 8401362 | 1.37E+07 | 5147943.5 | 1.06E+07 | 2341.9795 | REDEEESLNEVGYDDIGGC[160.0307]R | Q01853 | Q01853 | VCP | valosin containing protein | Cytoplasm | enzyme |
| 15554678, 15558115 | 2 | 0.739 | 7033312 | 4818712.5 | 6467795.5 | 6411835 | 5679518 | 4839540 | 7636545.5 | 4801458 | 4741540 | 1.22E+07 | 6379364.5 | 4679626.5 | 1840.7893, 1840.7899 | EAVC[160.0307]IVLSDDTC[160.0307]SDEK | Q01853 | Q01853 | VCP | valosin containing protein | Cytoplasm | enzyme |
| 15558061 | 3 | 0.449 | 1.15E+07 | 5669279 | 5580655 | 1.05E+07 | 5707913 | 8988795 | 1.52E+07 | 7103041.5 | 7841424.5 | 1.21E+07 | 9569765 | 1.21E+07 | 2741.4856 | RIHVLPIDDTVEGITGNLFEVYLK | Q01853 | Q01853 | VCP | valosin containing protein | Cytoplasm | enzyme |
| 15539944 | 3 | 0.832 | 6111535 | 8024860.5 | 6996281 | 6235445.5 | 7653459.5 | 7286758.5 | 6491365.5 | 7090092.5 | 4738604 | 1.19E+07 | 4575030.5 | 7175142.5 | 1951.9183 | GPELLTM[147.0354]WFGESEANVR | Q01853 | Q01853 | VCP | valosin containing protein | Cytoplasm | enzyme |
| 15535752 | 2 | 0.766 | 8259912.5 | 1.01E+07 | 1.12E+07 | 4995929.5 | 1.25E+07 | 9056257 | 1.16E+07 | 1.13E+07 | 6157380 | 1.02E+07 | 4671314 | 8138046 | 1094.6539 | LEILQIHTK | Q01853 | Q01853 | VCP | valosin containing protein | Cytoplasm | enzyme |
| 15537526 | 3 | 0.341 | 1.30E+07 | 2770249.5 | 1804834.875 | 9481953 | 1636527.875 | 1870421.5 | 3085599 | 1974977.875 | 6331991 | 7667627.5 | 1.28E+07 | 8704454 | 1807.9287 | VINQILTEMDGMSTKK | Q01853 | Q01853 | VCP | valosin containing protein | Cytoplasm | enzyme |
| 15556737, 15558708 | 2 | 0.8 | 3955814.75 | 4787848 | 5714812.5 | 3668917.25 | 5125811.5 | 4030345.75 | 6121567.5 | 4600518.5 | 3519704.75 | 6308800.5 | 2455331.75 | 3560066.75 | 1935.9204, 1935.9221 | GPELLTMWFGESEANVR | Q01853 | Q01853 | VCP | valosin containing protein | Cytoplasm | enzyme |
| 15555642 | 3 | 0.905 | 3062543.25 | 2872285.25 | 2455653 | 1985749 | 3876262.5 | 2280329.25 | 2533917 | 3242046 | 2218137.5 | 5127807.5 | 2100115.5 | 2701028 | 2313.9731 | EDEEESLNEVGYDDIGGC[160.0307]RK | Q01853 | Q01853 | VCP | valosin containing protein | Cytoplasm | enzyme |
| 15551384 | 3 | 0.31 | 6130007.5 | 161471.375 | 391444.531 | 9827885 | 369601.938 | 1127649.625 | 1985205.75 | 209840.672 | 2170039 | 4659787 | 9343810 | 6795294 | 2584.236 | GPELLTM[147.0354]WFGESEANVREIFDK | Q01853 | Q01853 | VCP | valosin containing protein | Cytoplasm | enzyme |
| 15550429 | 2 | 0.923 | 1216791.125 | 3276434 | 5449325.5 | 1415847.375 | 3257207.25 | 3953631.25 | 4374744.5 | 2175098.5 | 1598445.625 | 4583773 | 799576.812 | 2019905.25 | 1980.0146 | RELQELVQYPVEHPDK | Q01853 | Q01853 | VCP | valosin containing protein | Cytoplasm | enzyme |
| 15550720 | 2 | 0.542 | 6905713 | 5411033.5 | 3250408 | 2056871.75 | 3883250 | 5031962.5 | 6690315.5 | 3457230.25 | 4395036 | 3601116.25 | 5687547 | 7860775.5 | 1679.9264 | EMVELPLRHPALFK | Q01853 | Q01853 | VCP | valosin containing protein | Cytoplasm | enzyme |
| 15556434 | 3 | 0.689 | 2472098.5 | 2514502.5 | 3855023.75 | 2457288 | 2089192.25 | 3971899 | 2806117 | 3363780.75 | 3974291.75 | 3389736.25 | 1673662.875 | 2820368.5 | 3245.4832 | KMDLIDLEDETIDAEVMNSLAVTM[147.0354]DDFR | Q01853 | Q01853 | VCP | valosin containing protein | Cytoplasm | enzyme |
| 15560868 | 3 | 0.22 | 1585588.875 | 1655404.375 | 1048170.312 | 1364690 | 1272494.875 | 1182293.625 | 1095745.875 | 1153510.125 | 481955 | 2960290.25 | 1050003.125 | 1635230.125 | 3220.311 | FPSGNQGGAGPSQGSGGGTGGSVYTEDNDDDLYG | Q01853 | Q01853 | VCP | valosin containing protein | Cytoplasm | enzyme |
| 15540619 | 2 | 0.506 | 2073118.375 | 2163260.75 | 2713045 | 1267669 | 1434735.625 | 1865216.75 | 1580631.25 | 1952057.25 | 1037661.25 | 2450707.75 | 625690.812 | 1982508.875 | 1446.8203 | IVSQLLTLM[147.0354]DGLK | Q01853 | Q01853 | VCP | valosin containing protein | Cytoplasm | enzyme |
| 15552737 | 2 | 0.349 | 3020553.25 | 240081.375 | 112744.086 | 5462218 | 96626.641 | 1097085.375 | 895320.812 | 326367.469 | 1406255.125 | 2275093.75 | 4156755 | 5012108 | 1777.9475 | MDELQLFRGDTVLLK | Q01853 | Q01853 | VCP | valosin containing protein | Cytoplasm | enzyme |
| 15545220 | 4 | 0.257 | 3466380 | 2434077.75 | 1269275.875 | 2297836.75 | 1284696.375 | 2828764.5 | 2244346.5 | 1156236.25 | 905739.062 | 1854467.125 | 3290296.5 | 4753573 | 2741.4812 | RIHVLPIDDTVEGITGNLFEVYLK | Q01853 | Q01853 | VCP | valosin containing protein | Cytoplasm | enzyme |
| 15536034 | 2 | 0.282 | 2.03E+07 | 3.53E+07 | 1.74E+07 | 3.28E+07 | 3.13E+07 | 3.54E+07 | 1.92E+07 | 3.17E+07 | 4.17E+07 | 1.23E+07 | 2.96E+07 | 1.86E+07 | 1400.6122 | YQVDPDAC[160.0307]FSAK | Q60932 | Q60932 | VDAC1 | voltage-dependent anion channel 1 | Cytoplasm | ion channel |
| 15535191 | 2 | 0.2 | 1.05E+07 | 1.95E+07 | 9816702 | 1.35E+07 | 2.06E+07 | 1.60E+07 | 9557813 | 1.75E+07 | 1.64E+07 | 1.10E+07 | 9088783 | 8787898 | 1213.6198 | VTQSNFAVGYK | Q60932 | Q60932 | VDAC1 | voltage-dependent anion channel 1 | Cytoplasm | ion channel |
| 15539703 | 2 | 0.083 | 9806350 | 2.94E+07 | 1.92E+07 | 1.59E+07 | 2.95E+07 | 2.99E+07 | 1.48E+07 | 2.77E+07 | 2.12E+07 | 6900763 | 1.04E+07 | 1.20E+07 | 1400.6655 | LTFDSSFSPNTGK | Q60932 | Q60932 | VDAC1 | voltage-dependent anion channel 1 | Cytoplasm | ion channel |
| 15535504 | 2, 3 | 0.583 | 1.28E+07 | 1.78E+07 | 7251246.5 | 1.76E+07 | 1.17E+07 | 1.88E+07 | 1.16E+07 | 1.15E+07 | 1.82E+07 | 6052721 | 1.59E+07 | 1.10E+07 | 1456.7511, 1456.7513 | SRVTQSNFAVGYK | Q60932 | Q60932 | VDAC1 | voltage-dependent anion channel 1 | Cytoplasm | ion channel |
| 15531608 | 2 | 0.172 | 1.36E+07 | 3.10E+07 | 1.17E+07 | 2.01E+07 | 2.34E+07 | 3.13E+07 | 1.69E+07 | 2.47E+07 | 3.19E+07 | 6047257.5 | 1.71E+07 | 1.38E+07 | 1030.6122 | LTLSALLDGK | Q60932 | Q60932 | VDAC1 | voltage-dependent anion channel 1 | Cytoplasm | ion channel |
| 15536360 | 2 | 0.069 | 1.15E+07 | 1.98E+07 | 1.38E+07 | 1.30E+07 | 1.99E+07 | 1.65E+07 | 1.20E+07 | 2.01E+07 | 2.08E+07 | 6030771.5 | 1.08E+07 | 1.03E+07 | 1374.6549 | WTEYGLTFTEK | Q60932 | Q60932 | VDAC1 | voltage-dependent anion channel 1 | Cytoplasm | ion channel |
| 15548068 | 2 | 0.065 | 1.07E+07 | 2.60E+07 | 1.70E+07 | 1.53E+07 | 2.04E+07 | 2.38E+07 | 1.17E+07 | 1.85E+07 | 2.41E+07 | 4479381 | 1.05E+07 | 9756926 | 1707.9243 | VNNSSLIGLGYTQTLK | Q60932 | Q60932 | VDAC1 | voltage-dependent anion channel 1 | Cytoplasm | ion channel |
| 15539989 | 3 | 0.609 | 6279116 | 1170155.25 | 450218.156 | 1.35E+07 | 1791976.875 | 2112458 | 2167311.75 | 1924171.375 | 7936728.5 | 1670113 | 1.13E+07 | 5462020 | 2103.174 | VNNSSLIGLGYTQTLKPGIK | Q60932 | Q60932 | VDAC1 | voltage-dependent anion channel 1 | Cytoplasm | ion channel |
| 15536889 | 2 | 0.099 | 1.78E+07 | 3.72E+07 | 1.94E+07 | 2.63E+07 | 3.36E+07 | 3.93E+07 | 1.80E+07 | 2.97E+07 | 3.54E+07 | 1.19E+07 | 1.81E+07 | 1.81E+07 | 1293.6648 | YQLDPTASISAK | Q60930 | Q60930 | VDAC2 | voltage-dependent anion channel 2 | Cytoplasm | ion channel |
| 15533924 | 3 | 0.24 | 1.61E+07 | 4.46E+07 | 1.89E+07 | 3.05E+07 | 2.55E+07 | 3.88E+07 | 1.68E+07 | 2.84E+07 | 3.33E+07 | 1.07E+07 | 1.93E+07 | 1.84E+07 | 2103.1497 | VNNSSLIGVGYTQTLRPGVK | Q60930 | Q60930 | VDAC2 | voltage-dependent anion channel 2 | Cytoplasm | ion channel |
| 15533421 | 2 | 0.044 | 1.17E+07 | 3.17E+07 | 2.10E+07 | 2.03E+07 | 2.81E+07 | 3.03E+07 | 1.88E+07 | 3.08E+07 | 2.59E+07 | 7186625 | 1.24E+07 | 1.32E+07 | 1428.6987 | LTFDTTFSPNTGK | Q60930 | Q60930 | VDAC2 | voltage-dependent anion channel 2 | Cytoplasm | ion channel |
| 15535670 | 2 | 0.227 | 1.36E+07 | 2.69E+07 | 1.34E+07 | 1.91E+07 | 2.40E+07 | 1.81E+07 | 1.25E+07 | 2.26E+07 | 2.69E+07 | 6534015 | 1.53E+07 | 1.29E+07 | 1016.5973 | LTLSALVDGK | Q60930 | Q60930 | VDAC2 | voltage-dependent anion channel 2 | Cytoplasm | ion channel |
| 15538705 | 2 | 0.455 | 1.23E+07 | 1.93E+07 | 1.09E+07 | 1.62E+07 | 2.01E+07 | 1.40E+07 | 9717637 | 1.69E+07 | 2.11E+07 | 6481973.5 | 1.61E+07 | 1.10E+07 | 1433.6371 | WC[160.0307]EYGLTFTEK | Q60930 | Q60930 | VDAC2 | voltage-dependent anion channel 2 | Cytoplasm | ion channel |
| 15534617 | 2 | 0.402 | 4496129 | 8348907 | 4482573.5 | 5569595 | 8719531 | 6572396.5 | 3926986.25 | 8089001.5 | 1.06E+07 | 2354483.75 | 5923451.5 | 4829710.5 | 824.46484 | GFGFGLVK | Q60930 | Q60930 | VDAC2 | voltage-dependent anion channel 2 | Cytoplasm | ion channel |
| 15555961 | 3 | 0.209 | 3328252.75 | 6652388 | 3951597.75 | 3090408.75 | 6736401.5 | 9576551 | 3962938 | 8173425.5 | 9164973 | 1601600.5 | 3288569.75 | 4217352.5 | 2519.2012 | WNTDNTLGTEIAIEDQIC[160.0307]QGLK | Q60930 | Q60930 | VDAC2 | voltage-dependent anion channel 2 | Cytoplasm | ion channel |
| 15559097, 15563180 | 2 | 0.191 | 862624.062 | 1936827.375 | 1284948.625 | 1827876.625 | 1728839.375 | 1826136.5 | 947525 | 1554716.25 | 1818389.75 | 611147.812 | 1248077.5 | 1085020.125 | 1907.7894, 1907.7905 | SC[160.0307]SGVEFSTSGSSNTDTGK | Q60930 | Q60930 | VDAC2 | voltage-dependent anion channel 2 | Cytoplasm | ion channel |
| 15552156 | 2 | 0.189 | 1605470.125 | 3417393.75 | 2209495.5 | 3071977.75 | 2433012.5 | 3938987 | 1885934.875 | 2245761.75 | 3793553.5 | 142809.969 | 2312157 | 1221813.375 | 2103.1511 | VNNSSLIGVGYTQTLRPGVK | Q60930 | Q60930 | VDAC2 | voltage-dependent anion channel 2 | Cytoplasm | ion channel |
| 15533749 | 2 | 0.107 | 2.76E+07 | 6.09E+07 | 2.72E+07 | 3.59E+07 | 3.78E+07 | 4.67E+07 | 2.64E+07 | 3.90E+07 | 4.40E+07 | 1.40E+07 | 2.59E+07 | 2.24E+07 | 1254.6433 | LSQNNFALGYK | Q60931 | Q60931 | VDAC3 | voltage-dependent anion channel 3 | Cytoplasm | ion channel |
| 15531552 | 2 | 0.083 | 1.45E+07 | 4.39E+07 | 1.63E+07 | 2.38E+07 | 3.29E+07 | 4.25E+07 | 2.03E+07 | 3.04E+07 | 3.00E+07 | 8230736.5 | 1.51E+07 | 1.53E+07 | 1418.7844 | LTLDTIFVPNTGK | Q60931 | Q60931 | VDAC3 | voltage-dependent anion channel 3 | Cytoplasm | ion channel |
| 15535042 | 2 | 0.232 | 1.30E+07 | 3.06E+07 | 1.58E+07 | 2.06E+07 | 2.38E+07 | 2.88E+07 | 1.61E+07 | 2.18E+07 | 3.10E+07 | 7379339.5 | 1.64E+07 | 1.69E+07 | 1030.6119 | LTLSALIDGK | Q60931 | Q60931 | VDAC3 | voltage-dependent anion channel 3 | Cytoplasm | ion channel |
| 15532425 | 2 | 0.449 | 1.12E+07 | 1.96E+07 | 6905541.5 | 1.55E+07 | 1.95E+07 | 1.88E+07 | 1.03E+07 | 1.77E+07 | 2.10E+07 | 6033180.5 | 1.91E+07 | 1.01E+07 | 1330.6438 | VC[160.0307]NYGLTFTQK | Q60931 | Q60931 | VDAC3 | voltage-dependent anion channel 3 | Cytoplasm | ion channel |
| 15532626 | 3 | 0.786 | 8578093 | 9636477 | 2435147.25 | 1.37E+07 | 6484018 | 8523336 | 5148297 | 6580145.5 | 1.33E+07 | 3324626.25 | 1.06E+07 | 7283022.5 | 1546.8801 | LTLDTIFVPNTGKK | Q60931 | Q60931 | VDAC3 | voltage-dependent anion channel 3 | Cytoplasm | ion channel |
| 15549123 | 2 | 0.19 | 3058239.75 | 4875388 | 4220738.5 | 4045302.25 | 3795755.75 | 5327977 | 2772452.5 | 3990194 | 5192502 | 831716.125 | 3762646.25 | 2432972 | 2101.1697 | VNNASLIGLGYTQTLRPGVK | Q60931 | Q60931 | VDAC3 | voltage-dependent anion channel 3 | Cytoplasm | ion channel |
| 15539748 | 2 | 0.117 | 49799.281 | 19887.469 | 106254.797 | 1859551.25 | 762751 | 710586 | 823730.375 | 90621.352 | 6324920 | 37215.66 | 621407.188 | 889828.375 | 1169.7124 | ILLAELEQLK | P20152 | P20152 | VIM | vimentin | Cytoplasm | other |
| 15533461 | 2 | 0.142 | 1.45E+07 | 1.65E+07 | 2.12E+07 | 2.81E+07 | 2.03E+07 | 2.25E+07 | 2.13E+07 | 1.93E+07 | 2.08E+07 | 3.53E+07 | 2.13E+07 | 2.20E+07 | 837.5181 | VVEILHK | Q8BGQ1 | Q8BGQ1 | VIPAS39 | VPS33B interacting protein, apical-basolateral polarity regulator, spe-39 homolog | Plasma Membrane | other |
| 15548169 | 3 | 0.358 | 1.32E+07 | 1.79E+07 | 1.83E+07 | 1.94E+07 | 2.82E+07 | 1.92E+07 | 1.76E+07 | 2.29E+07 | 1.66E+07 | 2.13E+07 | 1.48E+07 | 1.89E+07 | 3443.443 | LSGAEPDDEEYQEFEEMLEHAEAAQDFASR | Q99KU0 | Q99KU0 | VMP1 | vacuole membrane protein 1 | Plasma Membrane | other |
| 15548406 | 3 | 0.274 | 1.71E+07 | 2.11E+07 | 1.64E+07 | 1.83E+07 | 1.65E+07 | 1.68E+07 | 2.13E+07 | 2.07E+07 | 1.90E+07 | 2.11E+07 | 1.71E+07 | 1.77E+07 | 3647.828 | NHFSIPISVFEGDTLLGIASPENEFNIPLASYR | Q5H8C4 | Q5H8C4 | VPS13A | vacuolar protein sorting 13 homolog A (S. cerevisiae) | Cytoplasm | transporter |
| 15548576 | 3 | 0.976 | 1.17E+07 | 1.27E+07 | 1.82E+07 | 1.26E+07 | 1.77E+07 | 1.29E+07 | 1.38E+07 | 1.76E+07 | 1.31E+07 | 1.80E+07 | 1.16E+07 | 1.24E+07 | 2959.5876 | ASYVIVPQYGNFSPTSNLLLLDLGHLK | Q5H8C4 | Q5H8C4 | VPS13A | vacuolar protein sorting 13 homolog A (S. cerevisiae) | Cytoplasm | transporter |
| 15548596 | 3 | 0.535 | 1.32E+07 | 1.29E+07 | 1.33E+07 | 1.45E+07 | 1.18E+07 | 1.34E+07 | 1.58E+07 | 1.11E+07 | 1.60E+07 | 1.30E+07 | 1.10E+07 | 1.24E+07 | 2476.3162 | GLVGAVTRPTGGIIDMASSTFQGIK | Q5H8C4 | Q5H8C4 | VPS13A | vacuolar protein sorting 13 homolog A (S. cerevisiae) | Cytoplasm | transporter |
| 15549465 | 3 | 0.791 | 6899950 | 1.24E+07 | 1.59E+07 | 8766821 | 1.37E+07 | 1.45E+07 | 1.25E+07 | 1.30E+07 | 9881147 | 1.25E+07 | 6450834.5 | 1.05E+07 | 2618.4153 | LQGIMELLGSIPKPEPVTDVSAPAR | Q5H8C4 | Q5H8C4 | VPS13A | vacuolar protein sorting 13 homolog A (S. cerevisiae) | Cytoplasm | transporter |
| 15533494 | 3 | 0.672 | 1.15E+07 | 1.19E+07 | 1.05E+07 | 1.09E+07 | 9176119 | 1.26E+07 | 1.02E+07 | 1.14E+07 | 9661533 | 1.22E+07 | 1.12E+07 | 1.09E+07 | 1840.9862 | SFQIQASALPVSHISQK | Q5H8C4 | Q5H8C4 | VPS13A | vacuolar protein sorting 13 homolog A (S. cerevisiae) | Cytoplasm | transporter |
| 15535914 | 3 | 0.021 | 9246688 | 1.01E+07 | 9849421 | 9617925 | 9585526 | 9897921 | 8883443 | 1.00E+07 | 8908543 | 8710360 | 6869428.5 | 8011978 | 1785.9868 | LFFILLTPANHSVADK | Q5H8C4 | Q5H8C4 | VPS13A | vacuolar protein sorting 13 homolog A (S. cerevisiae) | Cytoplasm | transporter |
| 15549146 | 3 | 0.436 | 5704910 | 8806777 | 5587395.5 | 5596375.5 | 7052661.5 | 7170201.5 | 7917802 | 7752702 | 8397945 | 7951416.5 | 5520401.5 | 6383352.5 | 2770.514 | APVVLIPQSPVSQNVFVADFGLITMK | Q5H8C4 | Q5H8C4 | VPS13A | vacuolar protein sorting 13 homolog A (S. cerevisiae) | Cytoplasm | transporter |
| 15533668 | 2 | 0.866 | 7268920 | 6761114.5 | 7563540 | 6932293.5 | 8079549.5 | 5755338 | 8007320.5 | 7767157.5 | 6359170 | 7898031.5 | 7401660 | 6808162.5 | 1449.7225 | SIIC[160.0307]QIDTVEGSK | Q5H8C4 | Q5H8C4 | VPS13A | vacuolar protein sorting 13 homolog A (S. cerevisiae) | Cytoplasm | transporter |
| 15535860 | 3 | 0.63 | 6016199 | 7218311 | 7954694.5 | 6411636 | 9010831 | 7264892 | 6291233 | 8887499 | 6960998.5 | 7557549 | 5008041.5 | 6480038 | 1742.921 | KPSPEILMSLEELEK | Q5H8C4 | Q5H8C4 | VPS13A | vacuolar protein sorting 13 homolog A (S. cerevisiae) | Cytoplasm | transporter |
| 15543511 | 3 | 0.462 | 6362138 | 6594381 | 6681391 | 7643641.5 | 8795245 | 5896184 | 8009666 | 7291498 | 7131511 | 7457480.5 | 6274815.5 | 6456256 | 1512.7568 | FFNEDGVIRPYR | Q5H8C4 | Q5H8C4 | VPS13A | vacuolar protein sorting 13 homolog A (S. cerevisiae) | Cytoplasm | transporter |
| 15537598 | 2 | 0.143 | 7643302.5 | 8443898 | 6754828 | 8393431 | 9156771 | 8128851 | 8079580.5 | 7919891 | 7634658.5 | 7302933 | 7599061.5 | 7591525 | 1298.7385 | ALVGGAVGGLAGAASK | Q5H8C4 | Q5H8C4 | VPS13A | vacuolar protein sorting 13 homolog A (S. cerevisiae) | Cytoplasm | transporter |
| 15538239 | 2 | 0.804 | 5966392.5 | 5099729 | 6371891 | 5267244.5 | 6460841 | 4709281.5 | 5896086 | 6669487 | 4966790 | 7027788 | 4768205 | 3862142.25 | 1288.6698 | GEPTSNTVTAAIK | Q5H8C4 | Q5H8C4 | VPS13A | vacuolar protein sorting 13 homolog A (S. cerevisiae) | Cytoplasm | transporter |
| 15554380 | 3 | 0.155 | 6878328 | 7749160 | 7422457.5 | 6969731 | 6390089.5 | 7255862.5 | 7151519.5 | 5959371.5 | 7007253.5 | 6838213 | 5992229 | 5863033.5 | 2736.3918 | NGIVNENIVPEGYDFVFRPISASAK | Q5H8C4 | Q5H8C4 | VPS13A | vacuolar protein sorting 13 homolog A (S. cerevisiae) | Cytoplasm | transporter |
| 15532492 | 2 | 0.489 | 6145075.5 | 7409944.5 | 8154962 | 7219857 | 7918259 | 6618615 | 8810668 | 7700011.5 | 6877540.5 | 6696580.5 | 5760260.5 | 7420227 | 1145.6539 | SFVINIVPEK | Q5H8C4 | Q5H8C4 | VPS13A | vacuolar protein sorting 13 homolog A (S. cerevisiae) | Cytoplasm | transporter |

| | | | | | | | | | | | | | | | | | | | | |
|---|---|---|---|---|---|---|---|---|---|---|---|---|---|---|---|---|---|---|---|---|
| 15534771 | 3 | 0.019 | 6926654 | 7580480.5 | 7138332 | 6414255 | 6240960.5 | 6175033.5 | 7697205.5 | 7169370.5 | 6428015.5 | 6399666 | 5612842 | 6151916 | 2070.1296 | DSEQHGGLIPVHSLNLLLK | Q5H8C4 | Q5H8C4 | VPS13A | vacuolar protein sorting 13 homolog A (S. cerevisiae) | Cytoplasm | transporter |
| 15535948 | 3 | 0.459 | 7092122 | 5338641 | 7607374.5 | 5751130 | 5448043.5 | 5735705.5 | 4596454.5 | 5582093 | 7617444 | 6079586.5 | 4913517 | 4796081.5 | 1949.9653 | PVLFSFQPNHFFNNNK | Q5H8C4 | Q5H8C4 | VPS13A | vacuolar protein sorting 13 homolog A (S. cerevisiae) | Cytoplasm | transporter |
| 15538511 | 2 | 0.185 | 6173858.5 | 6649871.5 | 7821206.5 | 7296073.5 | 5626958.5 | 6341465.5 | 4862480.5 | 6171320 | 5700967.5 | 5562364 | 5030678 | 6359422 | 1262.7064 | TLDVFNITIAR | Q5H8C4 | Q5H8C4 | VPS13A | vacuolar protein sorting 13 homolog A (S. cerevisiae) | Cytoplasm | transporter |
| 15536683 | 3 | 0.024 | 5754753 | 7276573.5 | 7328682 | 5588232.5 | 6523051.5 | 5898657.5 | 7077980.5 | 6564662.5 | 5754660 | 4899800 | 5219046 | 4895255 | 1604.7927 | IDSFHITGLPDDFK | Q5H8C4 | Q5H8C4 | VPS13A | vacuolar protein sorting 13 homolog A (S. cerevisiae) | Cytoplasm | transporter |
| 15553897 | 2 | 0.342 | 4274155 | 4009051.5 | 4652297.5 | 4331642 | 3948054.25 | 3087461.5 | 4609349 | 4358800 | 3563856 | 4344733 | 3256496.75 | 2747047.25 | 1583.7024 | EAIVESDSEAENYK | Q5H8C4 | Q5H8C4 | VPS13A | vacuolar protein sorting 13 homolog A (S. cerevisiae) | Cytoplasm | transporter |
| 15534718 | 2 | 0.353 | 3638632 | 4097772.5 | 4589167.5 | 2633042 | 5178498.5 | 3886577.75 | 4080709.75 | 5444637 | 3578457 | 4183945 | 1927707.375 | 2855937.5 | 1281.582 | EYTEASPLEDK | Q5H8C4 | Q5H8C4 | VPS13A | vacuolar protein sorting 13 homolog A (S. cerevisiae) | Cytoplasm | transporter |
| 15551376 | 2 | 0.003 | 4424242 | 5001145.5 | 5366667 | 4437164 | 5558687 | 5106101 | 5576117 | 5604252.5 | 5632088.5 | 4063566.5 | 3319197.75 | 3701441.25 | 1860.8956 | DALAEATVQAAEMAADGVK | Q5H8C4 | Q5H8C4 | VPS13A | vacuolar protein sorting 13 homolog A (S. cerevisiae) | Cytoplasm | transporter |
| 15533903 | 2 | 0.317 | 2839892.75 | 2187652.25 | 3789408 | 3825234.5 | 2413540.25 | 3870223.5 | 4174271 | 4027342.5 | 3830300 | 4054656.25 | 3527182.75 | 3063633.75 | 1371.7726 | IVTFIPFYMIK | Q5H8C4 | Q5H8C4 | VPS13A | vacuolar protein sorting 13 homolog A (S. cerevisiae) | Cytoplasm | transporter |
| 15532308 | 2 | 0.077 | 4358329.5 | 5513484 | 6688190 | 4537280.5 | 5155175 | 5364406.5 | 5388717 | 5848844 | 4797905.5 | 4023080.25 | 4076702 | 4096482.25 | 1016.59705 | LENIISSLK | Q5H8C4 | Q5H8C4 | VPS13A | vacuolar protein sorting 13 homolog A (S. cerevisiae) | Cytoplasm | transporter |
| 15549245 | 2 | 0.019 | 4217586 | 5033953.5 | 5427897.5 | 4346838 | 4305723 | 5300886.5 | 4759826 | 5968428 | 5324830.5 | 3469626 | 3774952.25 | 3901472.25 | 1916.0194 | NAPNLNSTYNNVLQLIK | Q5H8C4 | Q5H8C4 | VPS13A | vacuolar protein sorting 13 homolog A (S. cerevisiae) | Cytoplasm | transporter |
| 15537752 | 2 | 0.184 | 3221688.5 | 3191488.75 | 3816553.25 | 3088190.75 | 3285209.25 | 2914310.5 | 3475604 | 3724967 | 3292824.5 | 3203424.75 | 3172440.25 | 3248591.75 | 1031.5704 | DQLNITLSK | Q5H8C4 | Q5H8C4 | VPS13A | vacuolar protein sorting 13 homolog A (S. cerevisiae) | Cytoplasm | transporter |
| 15540266 | 2 | 0.681 | 2847213.25 | 2953654.75 | 2081942.375 | 2979815 | 3047347.25 | 2250325.75 | 3181021 | 2901891.75 | 2153998 | 2758936.75 | 2405027.75 | 1814669.75 | 1164.6067 | PFTDVSIVMR | Q5H8C4 | Q5H8C4 | VPS13A | vacuolar protein sorting 13 homolog A (S. cerevisiae) | Cytoplasm | transporter |
| 15536025 | 3 | 0.047 | 5543979.5 | 4755483.5 | 5481649.5 | 5619101 | 3829455.75 | 4621955.5 | 4902363 | 6218830 | 6314205.5 | 2675969.5 | 4225736.5 | 3983966.5 | 2203.2876 | LVTQIIQNLQVQISSIHIR | Q5H8C4 | Q5H8C4 | VPS13A | vacuolar protein sorting 13 homolog A (S. cerevisiae) | Cytoplasm | transporter |
| 15542064 | 2 | 0.156 | 2724006 | 2395722.75 | 2809317.5 | 2523652.25 | 2607127.25 | 2367848.75 | 2754146.75 | 2271522.25 | 2577466.25 | 2349008.5 | 2104513.5 | 2326178 | 836.4507 | YDGQLLK | Q5H8C4 | Q5H8C4 | VPS13A | vacuolar protein sorting 13 homolog A (S. cerevisiae) | Cytoplasm | transporter |
| 15536987 | 2 | 0.316 | 2020396 | 3028133.25 | 3126401 | 1424660.75 | 2413760.25 | 1680869.125 | 2516365 | 2942546 | 767539.812 | 2091773.625 | 858797.062 | 1001917.062 | 1484.7162 | AVYYTWADPVGSR | Q5H8C4 | Q5H8C4 | VPS13A | vacuolar protein sorting 13 homolog A (S. cerevisiae) | Cytoplasm | transporter |
| 15552036 | 2 | 0.02 | 2425181.75 | 1998695.5 | 2911249 | 1915172.75 | 2111961 | 2333621.75 | 2626733.5 | 3277621.25 | 2824544 | 2081943.625 | 1666733 | 1914986.25 | 1572.8397 | TIYLVSFFEGLQR | Q5H8C4 | Q5H8C4 | VPS13A | vacuolar protein sorting 13 homolog A (S. cerevisiae) | Cytoplasm | transporter |
| 15556789 | 2 | 0.515 | 1661843.25 | 1340495 | 1724428.25 | 1420702.375 | 2148810 | 1699467.5 | 1665046.625 | 1975433.875 | 1870317.375 | 1747583.25 | 2254966.25 | 1682285.75 | 2358.033 | GTFGQLTC[160.0307]EWQYTFDEFTK | Q5H8C4 | Q5H8C4 | VPS13A | vacuolar protein sorting 13 homolog A (S. cerevisiae) | Cytoplasm | transporter |
| 15557543 | 2 | 0.249 | 2286553 | 1458016.375 | 1667384.5 | 1654327.625 | 1980393.25 | 1579004.25 | 1594589.875 | 1803839 | 1428260.5 | 1279509.75 | 1503196.625 | 1383302.125 | 1586.6793 | GVAAMTMDEDYQQK | Q5H8C4 | Q5H8C4 | VPS13A | vacuolar protein sorting 13 homolog A (S. cerevisiae) | Cytoplasm | transporter |
| 15552202 | 2 | 0.387 | 989005.75 | 1451321.625 | 1300402.375 | 1315632.75 | 1180566.375 | 1239527.375 | 1035169.5 | 1387545 | 1248511.875 | 999617.625 | 988616 | 1151466.875 | 2296.18 | FLGDYVVNLDESQLSLGIWK | Q5H8C4 | Q5H8C4 | VPS13A | vacuolar protein sorting 13 homolog A (S. cerevisiae) | Cytoplasm | transporter |
| 15539705 | 3 | 0.06 | 1382467.625 | 4389203.5 | 5703070.5 | 2926314.25 | 4673815.5 | 4188301.75 | 4011748 | 6063947 | 3964432 | 787854.312 | 377793.562 | 2707659.75 | 1538.666 | WSC[160.0307]GQSYGEVTHK | Q5H8C4 | Q5H8C4 | VPS13A | vacuolar protein sorting 13 homolog A (S. cerevisiae) | Cytoplasm | transporter |
| 15542387 | 3 | 0.27 | 1821738.25 | 3498517.25 | 2368564.5 | 2673860 | 2245489.25 | 1747553.625 | 2113323.75 | 2436350.75 | 1520751.75 | 1893101.125 | 1444157.375 | 1606485.625 | 2064.189 | VQIIGLGTQVSQRPGAQALK | Q8BX70 | Q8BX70 | VPS13C | vacuolar protein sorting 13 homolog C (S. cerevisiae) | Cytoplasm | other |
| 15551072 | 2 | 0.322 | 2617560 | 2803269.25 | 961967.625 | 2705862.5 | 2076721.5 | 1600244.375 | 1559556.75 | 1521729 | 1541824.5 | 1814432.625 | 1200541.125 | 552803.688 | 1630.8152 | ILAENLGEATEDLDK | Q8BX70 | Q8BX70 | VPS13C | vacuolar protein sorting 13 homolog C (S. cerevisiae) | Cytoplasm | other |
| 15555941 | 3 | 0.009 | 1963027 | 4011905 | 2180587.5 | 2003213.625 | 3012537.25 | 2811630.75 | 2722914 | 2326419.25 | 2444324 | 1313201.5 | 720802.688 | 1171527.375 | 2591.3284 | VDQGFLGAVISLFTPTTDPEAERK | Q8BX70 | Q8BX70 | VPS13C | vacuolar protein sorting 13 homolog C (S. cerevisiae) | Cytoplasm | other |
| 15538726 | 2 | 0.077 | 1957390.75 | 1814331.625 | 1155163.125 | 1756132 | 1712218.625 | 1306574.375 | 1468566.125 | 1274251.625 | 1048040.375 | 1156654.25 | 1001559.625 | 1018774.312 | 1260.6388 | EALSAATAQAAEK | Q8BX70 | Q8BX70 | VPS13C | vacuolar protein sorting 13 homolog C (S. cerevisiae) | Cytoplasm | other |
| 15542774 | 3 | 0.697 | 1340357.375 | 1891590.875 | 554007.312 | 1350604 | 1278752.25 | 831528.625 | 1160494.125 | 1276377.375 | 566158.938 | 1082455.75 | 685897.812 | 633206 | 2241.245 | VEETRPVDIINVLLNFEIK | Q8BX70 | Q8BX70 | VPS13C | vacuolar protein sorting 13 homolog C (S. cerevisiae) | Cytoplasm | other |
| 15535877 | 2 | 0.866 | 2630651 | 1.13E+07 | 3439189.75 | 7095614.5 | 8643999 | 9131434 | 4331211.5 | 4430482.5 | 2.11E+07 | 1365972.375 | 1.15E+07 | 6.12E+07 | 799.51306 | LRLEIR | Q9D1C8 | Q9D1C8 | VPS28 | vacuolar protein sorting 28 homolog (S. cerevisiae) | Cytoplasm | transporter |
| 15534271 | 3 | 0.302 | 315222.375 | 2738039.5 | 2489380.75 | 9427182 | 6690513 | 6418020 | 5303972.5 | 4862322.5 | 9327725 | 443614.812 | 1.00E+07 | 1.26E+07 | 1991.0292 | VAVFPVGVGDRYDEAQLR | Q8CIZ8 | Q8CIZ8 | VWF | von Willebrand factor | Extracellular Space | other |
| 15544115 | 3 | 0.131 | 283979.281 | 812078.75 | 626554.062 | 2371522.5 | 2259730.25 | 2609104.5 | 1649969.5 | 1443682.625 | 2508275.75 | 330451.781 | 2653578.5 | 3670106.25 | 1867.7295 | QDETFQDGC[160.0307]DSHLC[160.0307]R | Q8CIZ8 | Q8CIZ8 | VWF | von Willebrand factor | Extracellular Space | other |
| 15554258 | 3 | 0.07 | 194370.984 | 549134.562 | 141874.359 | 3745615.25 | 2424603 | 3376919.75 | 1411626.875 | 1077160.375 | 3634659.5 | 182511.891 | 2950047.75 | 4988640.5 | 2445.005 | TTDFC[160.0307]AMSC[160.0307]PPSLVYNHC[160.0307]ER | Q8CIZ8 | Q8CIZ8 | VWF | von Willebrand factor | Extracellular Space | other |
| 15551075 | 3 | 0.249 | 321697.281 | 848442.562 | 1189060.375 | 4274430.5 | 3764153 | 3471479.5 | 3428376.75 | 2161914.5 | 4607293 | 168430.188 | 3743776.25 | 6188379 | 2951.4985 | AHLQSLVDLMQQEGGPSQIGDALAFAVR | Q8CIZ8 | Q8CIZ8 | VWF | von Willebrand factor | Extracellular Space | other |
| 15537691 | 3 | 0.347 | 3180.929 | 532514.25 | 1159218.125 | 2160840.75 | 1028414.062 | 1886386.75 | 2173162 | 731456.312 | 2174336 | 154886.109 | 2120266.25 | 4160811.25 | 1499.9272 | VIVIPVGIGPHASLK | Q8CIZ8 | Q8CIZ8 | VWF | von Willebrand factor | Extracellular Space | other |
| 15542276 | 2 | 0.126 | 62568.566 | 202767 | 489423.5 | 2613705 | 1122567.75 | 1685314.625 | 1034331.5 | 858453.562 | 2103788 | 130058.031 | 1954476.25 | 3546024.5 | 1148.5908 | EFVEEVIQR | Q8CIZ8 | Q8CIZ8 | VWF | von Willebrand factor | Extracellular Space | other |
| 15539807 | 2 | 0.423 | 394069.281 | 1065593.25 | 1247019 | 2725630.75 | 3053832.75 | 3393580.75 | 2014939.25 | 1870374.125 | 3783367.25 | 121052.305 | 2959720.75 | 4413031 | 1469.7455 | YTGSQVASTSEVLK | Q8CIZ8 | Q8CIZ8 | VWF | von Willebrand factor | Extracellular Space | other |
| 15539265 | 3 | 0.196 | 80178 | 799957.312 | 637720.25 | 3979563.75 | 2740490 | 3420587 | 2025947.875 | 1761470.25 | 3755732 | 107526.336 | 3360604.75 | 4566921 | 1936.8269 | C[160.0307]PC[160.0307]FHQGAEYAPGDTVK | Q8CIZ8 | Q8CIZ8 | VWF | von Willebrand factor | Extracellular Space | other |
| 15550325 | 4 | 0.366 | 163175.172 | 1762968.25 | 1683023.625 | 7395765.5 | 5016735 | 5854056 | 4041065.25 | 3157396.25 | 7310873.5 | 105893.945 | 7192840 | 9314574 | 3501.4436 | VVQC[160.0307]SQRPC[160.0307]EDSC[160.0307]QPGFSYVLHEGEC[160.0307]C[160.0307]GR | Q8CIZ8 | Q8CIZ8 | VWF | von Willebrand factor | Extracellular Space | other |
| 15540451 | 2 | 0.306 | 4.70E-06 | 1181734.75 | 946329.688 | 3449235.25 | 3448321.5 | 2886721.75 | 2089536.25 | 1889212.625 | 4178709.5 | 94770.016 | 3829592.75 | 4930426 | 1212.691 | ILAGPGASSNVVK | Q8CIZ8 | Q8CIZ8 | VWF | von Willebrand factor | Extracellular Space | other |
| 15538582 | 2 | 0.252 | 142620.672 | 741519.75 | 1035860.75 | 4180237.25 | 3629294.5 | 3673055.25 | 2716855.25 | 2173583.25 | 3972801.25 | 73244.594 | 3385034.75 | 4807422.5 | 1409.6813 | ILTSDVFQGC[160.0307]NR | Q8CIZ8 | Q8CIZ8 | VWF | von Willebrand factor | Extracellular Space | other |
| 15554457 | 2 | 0.381 | 218293.297 | 352993.75 | 450835.25 | 2056399.375 | 1643389.25 | 1494115 | 776881.312 | 978942.125 | 2133620.75 | 49588.555 | 2022805.125 | 2826350 | 1520.7053 | EAPDLVLQTC[160.0307]C[160.0307]SK | Q8CIZ8 | Q8CIZ8 | VWF | von Willebrand factor | Extracellular Space | other |
| 15538166 | 2 | 0.307 | 4.70E-06 | 461085.656 | 707993.688 | 2063656.5 | 2013802.125 | 2040501.25 | 767288.125 | 1072853.625 | 2832900.75 | 41865.191 | 1638839.875 | 3338603.5 | 1116.6068 | YTLFQIFGK | Q8CIZ8 | Q8CIZ8 | VWF | von Willebrand factor | Extracellular Space | other |
| 15537311 | 2 | 0.464 | 50209.148 | 444454.281 | 759512.75 | 2120355.25 | 1835231.875 | 2187728.5 | 1234463.25 | 1047305.812 | 2293469.75 | 6851.076 | 1944394.75 | 3280522.75 | 1041.5121 | LVC[160.0307]PADNPR | Q8CIZ8 | Q8CIZ8 | VWF | von Willebrand factor | Extracellular Space | other |
| 15545774 | 2 | 0.368 | 4.70E-06 | 171618.109 | 234105.641 | 1045746.188 | 938092.75 | 907883.375 | 581710.938 | 572069.75 | 1303645 | 4.70E-06 | 1302297.25 | 1737251 | 1139.4728 | IGC[160.0307]NTC[160.0307]VC[160.0307]R | Q8CIZ8 | Q8CIZ8 | VWF | von Willebrand factor | Extracellular Space | other |
| 15536944 | 2 | 0.704 | 3449283 | 2725305.25 | 2253723.75 | 3675106.75 | 2076966.125 | 2259966.5 | 3132827.5 | 2117593.75 | 1995457.125 | 4146130.5 | 2393424.75 | 2829058.75 | 1458.7401 | LAGEGQAIDGAELSK | Q8CBE3 | Q8CBE3 | WDR37 | WD repeat domain 37 | Nucleus | other |
| 15544034 | 2 | 0.192 | 1056794.125 | 645500.438 | 872618.438 | 1142177 | 891901.875 | 845442.25 | 819819.75 | 709667 | 712994.062 | 1706643.625 | 915208.5 | 989501.938 | 989.487 | EIDTLNER | Q8CBE3 | Q8CBE3 | WDR37 | WD repeat domain 37 | Nucleus | other |
| 15553957 | 3 | 0.501 | 2248664.5 | 1468794.625 | 1251216 | 2029056.375 | 1736782.625 | 556300.812 | 928099.438 | 1255452.5 | 1336819.25 | 974612.5 | 1091039 | 813363.375 | 2605.2166 | IINLDNESQTTSSSNNEKPEQEK | Q8BH57 | Q8BH57 | WDR48 | WD repeat domain 48 | Cytoplasm | other |
| 15546818 | 2 | 0.733 | 738523.625 | 592160 | 739242.25 | 923766.062 | 880746.312 | 434907.344 | 879866.312 | 623571.188 | 313397.469 | 1142846.75 | 652230.625 | 649802.625 | 973.4924 | QKVGEC[160.0307]PR | Q4KUS1 | Q4KUS1 | WFDC8 | WAP four-disulfide core domain 8 | Extracellular Space | other |
| 15551910 | 2 | 0.648 | 1443647.125 | 1713153.375 | 2004186.875 | 1285073.375 | 1874160.75 | 1117084.75 | 1483903.5 | 1413248.25 | 976988 | 2197521.5 | 687628.5 | 1155009.625 | 1562.7157 | SQQQDDIEELETK | P83741 | P83741 | WNK1 | WNK lysine deficient protein kinase 1 | Cytoplasm | kinase |
| 15534396 | 2 | 0.76 | 3.05E+07 | 2.59E+07 | 2.62E+07 | 2.91E+07 | 2.61E+07 | 2.33E+07 | 3.01E+07 | 2.58E+07 | 2.00E+07 | 3.53E+07 | 2.58E+07 | 2.51E+07 | 1467.6437 | AC[160.0307]TEQDVMPTTSK | Q9QXY7 | Q9QXY7 | XK | X-linked Kx blood group | Plasma Membrane | transporter |
| 15553321 | 2 | 0.882 | 1.80E+07 | 2.39E+07 | 2.81E+07 | 1.72E+07 | 2.43E+07 | 2.68E+07 | 2.05E+07 | 2.55E+07 | 2.36E+07 | 3.20E+07 | 2.09E+07 | 2.43E+07 | 1860.9015 | VIPEATDIWTAVDLC[160.0307]SA | Q9QXY7 | Q9QXY7 | XK | X-linked Kx blood group | Plasma Membrane | transporter |
| 15533532 | 2 | 0.53 | 2.60E+07 | 2.74E+07 | 3.30E+07 | 1.87E+07 | 2.67E+07 | 2.96E+07 | 3.51E+07 | 2.90E+07 | 2.57E+07 | 2.85E+07 | 3.03E+07 | 2.57E+07 | 1028.5591 | IDNPELISK | Q9QXY7 | Q9QXY7 | XK | X-linked Kx blood group | Plasma Membrane | transporter |
| 15535161 | 2 | 0.43 | 1.74E+07 | 2.00E+07 | 1.97E+07 | 1.81E+07 | 1.68E+07 | 1.55E+07 | 1.81E+07 | 1.66E+07 | 1.44E+07 | 2.17E+07 | 1.66E+07 | 1.58E+07 | 1005.47046 | DGLSEEVEK | Q9QXY7 | Q9QXY7 | XK | X-linked Kx blood group | Plasma Membrane | transporter |
| 15555159 | 3 | 0.546 | 2229619 | 1531278.875 | 1035422.062 | 2664938.5 | 1175208.625 | 1769888.125 | 1783219.875 | 1194289.25 | 1142006.625 | 2714303 | 1569724.375 | 1830287.875 | 2541.246 | LDINLLDNVVNC[160.0307]LYHGEGAQQR | Q6P5F9 | Q6P5F9 | XPO1 | exportin 1 | Nucleus | transporter |
| 15553038 | 2 | 0.173 | 2238021 | 2430344.5 | 1918399.875 | 2766321 | 2531358.5 | 1469543.75 | 2402555.75 | 2423599.5 | 1475000.25 | 1961839.5 | 1186313.25 | 919731.938 | 1810.919 | LYGDDALENALQTFIK | Q9EPK7 | Q9EPK7 | XPO7 | exportin 7 | Nucleus | transporter |
| 15536500 | 2 | 0.659 | 2387958.25 | 1805629.125 | 1312153.125 | 2591624 | 1764742.25 | 863732.875 | 1965359.625 | 1361873.5 | 793719.438 | 1823191.125 | 1084843.125 | 829876.25 | 1212.6195 | NSIVNSQPPEK | Q9EPK7 | Q9EPK7 | XPO7 | exportin 7 | Nucleus | transporter |
| 15549838 | 2 | 0.087 | 4187372.75 | 5686733 | 2592787.5 | 5283270 | 4504480.5 | 2708131.75 | 2301860.75 | 2467923.75 | 3714351.75 | 2662313.75 | 1617272.75 | 1918014.125 | 1695.8629 | GAEAANVTGPGGVPVQGSK | P62960 | P62960 | YBX1 | Y box binding protein 1 | Nucleus | transcription regulator |
| 15562422, 15563092 | 2 | 0.208 | 382000 | 405751.875 | 220719.828 | 597584.062 | 329062.625 | 262983.25 | 279654.875 | 209022.109 | 193464.641 | 356734.719 | 135046.484 | 135871.484 | 1688.7112, 1688.7119 | NYQQNYQNSESGEK | P62960 | P62960 | YBX1 | Y box binding protein 1 | Nucleus | transcription regulator |
| 15550689 | 2 | 0.732 | 3276867.25 | 2980034.25 | 1976773.125 | 2192187.75 | 3081788.5 | 1994655.125 | 2483617.5 | 1666866.875 | 2148740.25 | 3526171.25 | 1490498.25 | 1914401.875 | 1782.8585 | GAEAANVTGPDGVPVEGSR | Q9JKB3 | Q9JKB3 | YBX3 | Y box binding protein 3 | Nucleus | transcription regulator |
| 15541689 | 2 | 0.09 | 2919454 | 3910833.5 | 1941317.75 | 2431539.75 | 3028406.5 | 1883388 | 1824482.125 | 1566116.875 | 1833124 | 1993480.625 | 1233126.75 | 1836365.75 | 1286.6714 | EDVFVHQTAIK | Q9JKB3 | Q9JKB3 | YBX3 | Y box binding protein 3 | Nucleus | transcription regulator |
| 15557846 | 3 | 0.816 | 2068027.25 | 3887524 | 3770326.75 | 2851247.5 | 3619108.75 | 3540371.5 | 2698630.75 | 3338272.25 | 2793022.75 | 3379546.25 | 2502876.25 | 2713448 | 3112.5825 | LGLLVFPYLHQDWEVQYQQDTPVAPR | Q9CX30 | Q9CX30 | YIF1B | Yip1 interacting factor homolog B (S. cerevisiae) | Cytoplasm | other |
| 15536184 | 2 | 0.387 | 2306315 | 2601336 | 3872418.5 | 2616878.75 | 3872693.75 | 2917321 | 2760712.75 | 3067069.25 | 2794236.25 | 2658540.25 | 2095573 | 2402172.5 | 1294.7457 | ILAQAAAEGVPVR | Q9CX30 | Q9CX30 | YIF1B | Yip1 interacting factor homolog B (S. cerevisiae) | Cytoplasm | other |
| 15550931 | 2 | 0.033 | 3129744.25 | 4077718 | 5763309.5 | 6221014 | 8336678 | 7970062.5 | 5708749.5 | 6613227 | 9285451 | 5171631.5 | 5341714 | 4696224 | 2130.0588 | IEAELQDIC[160.0307]NDVLELLDK | Q9CQV8 | Q9CQV8 | YWHAB | tyrosine 3-monooxygenase/tryptophan 5-monooxygenase activation protein, beta | Cytoplasm | transcription regulator |
| 15533021 | 3 | 0.011 | 2754653 | 3729213 | 4910076.5 | 6648454 | 7570416.5 | 7080696 | 5254759 | 5658057 | 6471045 | 3982280.5 | 5594538.5 | 4809017 | 1598.7357 | AVTEQGHELSNEER | Q9CQV8 | Q9CQV8 | YWHAB | tyrosine 3-monooxygenase/tryptophan 5-monooxygenase activation protein, beta | Cytoplasm | transcription regulator |
| 15533657 | 2 | 0.137 | 1883001.25 | 2768285.25 | 3807244.75 | 3492179.75 | 4875441.5 | 5386640 | 3591593.75 | 4203661.5 | 4976183.5 | 2481308.5 | 4040403.5 | 3605208.75 | 1350.7229 | YLILNATQAESK | Q9CQV8 | Q9CQV8 | YWHAB | tyrosine 3-monooxygenase/tryptophan 5-monooxygenase activation protein, beta | Cytoplasm | transcription regulator |
| 15536912 | 2 | 0.067 | 1106256 | 1818037.75 | 2582262.25 | 3539425.25 | 3562383.25 | 3620160.75 | 2705192 | 2772739.5 | 3504918.25 | 1563105.75 | 2940354.25 | 2852519 | 1196.5768 | YLSEVASGENK | Q9CQV8 | Q9CQV8 | YWHAB | tyrosine 3-monooxygenase/tryptophan 5-monooxygenase activation protein, beta | Cytoplasm | transcription regulator |
| 15554629 | 2 | 0.565 | 385001.938 | 644330.312 | 918146.5 | 623969.812 | 1232228.625 | 736570.375 | 707172.75 | 922579.5 | 1408162.25 | 446530.562 | 1157733.75 | 679671.125 | 1598.7374 | AVTEQGHELSNEER | Q9CQV8 | Q9CQV8 | YWHAB | tyrosine 3-monooxygenase/tryptophan 5-monooxygenase activation protein, beta | Cytoplasm | transcription regulator |
| 15546657, 15547433 | 3 | 0.247 | 354953.875 | 438097.312 | 820610.812 | 4.70E-06 | 52864.539 | 1191035.75 | 883251.188 | 1078667 | 1168482.5 | 95819.727 | 1137289.25 | 720437.688 | 2130.0593, 2130.0603 | IEAELQDIC[160.0307]NDVLELLDK | Q9CQV8 | Q9CQV8 | YWHAB | tyrosine 3-monooxygenase/tryptophan 5-monooxygenase activation protein, beta | Cytoplasm | transcription regulator |
| 15532757 | 2 | 0.013 | 6420852.5 | 8370213 | 1.23E+07 | 1.89E+07 | 1.60E+07 | 1.77E+07 | 1.47E+07 | 1.36E+07 | 1.75E+07 | 1.16E+07 | 1.61E+07 | 1.44E+07 | 1476.7428 | LIC[160.0307]C[160.0307]DILDVLDK | P62259 | P62259 | YWHAE | tyrosine 3-monooxygenase/tryptophan 5-monooxygenase activation protein, epsilon | Cytoplasm | other |
| 15555999 | 2 | 0.027 | 4847601 | 5682510.5 | 8996315 | 1.05E+07 | 1.25E+07 | 1.27E+07 | 8227816 | 1.06E+07 | 1.31E+07 | 9167160 | 1.05E+07 | 9978127 | 2087.9622 | AAFDDAIAELDTLSEESYK | P62259 | P62259 | YWHAE | tyrosine 3-monooxygenase/tryptophan 5-monooxygenase activation protein, epsilon | Cytoplasm | other |
| 15533347 | 2 | 0.039 | 5216103.5 | 5159161 | 8471356 | 8962127 | 1.10E+07 | 9405761 | 9314893 | 8856010 | 8593009 | 6874644 | 8904180 | 7550680.5 | 1237.6515 | HLIPAANTGESK | P62259 | P62259 | YWHAE | tyrosine 3-monooxygenase/tryptophan 5-monooxygenase activation protein, epsilon | Cytoplasm | other |
| 15534413 | 3 | 0.01 | 2804137.75 | 5111397.5 | 5918567.5 | 8875671 | 9102966 | 9534713 | 9462439 | 7742398 | 8538702 | 6486524 | 6990309 | 7713308.5 | 1928.8679 | LAEQAERYDEMVESMK | P62259 | P62259 | YWHAE | tyrosine 3-monooxygenase/tryptophan 5-monooxygenase activation protein, epsilon | Cytoplasm | other |
| 15533385 | 2 | 0.053 | 2740446.75 | 3769177 | 7188963.5 | 8297393 | 9691992 | 8606862 | 6175470 | 7803251.5 | 7850875 | 4663239 | 6808812 | 6236975 | 1194.5969 | EAAENSLVAYK | P62259 | P62259 | YWHAE | tyrosine 3-monooxygenase/tryptophan 5-monooxygenase activation protein, epsilon | Cytoplasm | other |
| 15533902 | 2 | 0.059 | 2603203.25 | 2970788 | 5091156.5 | 5002355.5 | 5323812 | 5976546 | 4812646.5 | 5441435 | 5410997 | 4390967.5 | 4799183.5 | 4645330 | 917.5293 | IISSIEQK | P62259 | P62259 | YWHAE | tyrosine 3-monooxygenase/tryptophan 5-monooxygenase activation protein, epsilon | Cytoplasm | other |
| 15542495, 15545770 | 3 | 0.011 | 2324133 | 2810922 | 3878690 | 5759994 | 7746587.5 | 8001478.5 | 5096552 | 6547994.5 | 3927805.25 | 4321038.5 | 6171149 | 7135298.5 | 2087.9622 | AAFDDAIAELDTLSEESYK | P62259 | P62259 | YWHAE | tyrosine 3-monooxygenase/tryptophan 5-monooxygenase activation protein, epsilon | Cytoplasm | other |
| 15554425 | 2 | 0.013 | 2735997.5 | 3250540 | 4213721.5 | 8160180 | 6295526.5 | 5882770.5 | 5312670 | 5400832.5 | 5709599 | 4013073 | 6788405.5 | 5447768.5 | 2180.936 | DNLTLWTSDMQGDGEEQNK | P62259 | P62259 | YWHAE | tyrosine 3-monooxygenase/tryptophan 5-monooxygenase activation protein, epsilon | Cytoplasm | other |
| 15544347 | 2 | 0.628 | 1061966.375 | 1930005.875 | 2965981.75 | 1362944.625 | 4363717.5 | 3864790 | 2582452.25 | 3225472 | 3853182.5 | 775641.375 | 2922554.75 | 3584218.75 | 1289.546 | EALQDVEDENQ | P62259 | P62259 | YWHAE | tyrosine 3-monooxygenase/tryptophan 5-monooxygenase activation protein, epsilon | Cytoplasm | other |
| 15549449 | 2 | 0.173 | 1488744.375 | 3269662 | 3925451.75 | 4243353 | 6259881 | 5317536.5 | 3697124.75 | 4815143 | 7007755 | 2004465.375 | 4853818 | 3436415.5 | 1643.7838 | NVTELNEPLSNEER | P61982 | P61982 | YWHAG | tyrosine 3-monooxygenase/tryptophan 5-monooxygenase activation protein, gamma | Cytoplasm | other |
| 15537827 | 2 | 0.044 | 482023.281 | 1161420.625 | 1494635.875 | 2704403 | 3184908.75 | 2992044 | 1949744.625 | 2287010 | 2897421 | 885537.375 | 2533060.5 | 1928856.75 | 1080.5547 | YLAEVATGEK | P61982 | P61982 | YWHAG | tyrosine 3-monooxygenase/tryptophan 5-monooxygenase activation protein, gamma | Cytoplasm | other |
| 15548100 | 3 | 0.159 | 5680065.5 | 1.28E+07 | 2.07E+07 | 6.23E+07 | 4.49E+07 | 5.54E+07 | 3.07E+07 | 3.24E+07 | 5.97E+07 | 4670984 | 6.34E+07 | 3.55E+07 | 2475.2664 | AVTELNEPLSNEDRNLLSVAYK | P68510 | P68510 | YWHAH | tyrosine 3-monooxygenase/tryptophan 5-monooxygenase activation protein, eta | Cytoplasm | transcription regulator |
| 15548527 | 2 | 0.117 | 1601483.5 | 5187948 | 8112790 | 2.11E+07 | 1.92E+07 | 2.06E+07 | 1.06E+07 | 1.18E+07 | 2.08E+07 | 1937206.75 | 1.71E+07 | 1.24E+07 | 2157.9775 | QAFDDAIAELDTLNEDSYK | P68510 | P68510 | YWHAH | tyrosine 3-monooxygenase/tryptophan 5-monooxygenase activation protein, eta | Cytoplasm | transcription regulator |
| 15548837 | 2 | 0.218 | 2110091 | 6765199.5 | 1.05E+07 | 1.97E+07 | 2.44E+07 | 2.69E+07 | 1.29E+07 | 1.74E+07 | 3.19E+07 | 1706495.625 | 2.73E+07 | 1.58E+07 | 1731.8793 | ELETVC[160.0307]NDVLALLDK | P68510 | P68510 | YWHAH | tyrosine 3-monooxygenase/tryptophan 5-monooxygenase activation protein, eta | Cytoplasm | transcription regulator |
| 15542441 | 2 | 0.18 | 2235564 | 4396189.5 | 7456765.5 | 1.49E+07 | 1.69E+07 | 1.61E+07 | 8134813 | 1.15E+07 | 1.71E+07 | 1319439.75 | 1.46E+07 | 1.01E+07 | 1066.5392 | YLAEVASGEK | P68510 | P68510 | YWHAH | tyrosine 3-monooxygenase/tryptophan 5-monooxygenase activation protein, eta | Cytoplasm | transcription regulator |
| 15535901 | 3 | 0.213 | 980655.938 | 2055878.625 | 3100432 | 1.32E+07 | 5151929 | 9129705 | 4569314.5 | 5466371 | 1.19E+07 | 926292.312 | 1.45E+07 | 7694122 | 1828.8163 | LAEQAERYDDMASAMK | P68510 | P68510 | YWHAH | tyrosine 3-monooxygenase/tryptophan 5-monooxygenase activation protein, eta | Cytoplasm | transcription regulator |
| 15550018 | 2 | 0.154 | 356790.656 | 1601463.625 | 3319790.25 | 3262553.5 | 1.06E+07 | 6557144.5 | 2902594.5 | 6303175 | 6068795 | 408307.688 | 3351119.75 | 3070525 | 1586.7625 | AVTELNEPLSNEDR | P68510 | P68510 | YWHAH | tyrosine 3-monooxygenase/tryptophan 5-monooxygenase activation protein, eta | Cytoplasm | transcription regulator |
| 15540712 | 3 | 0.2 | 259691.484 | 2140945.5 | 3246485.25 | 9043160 | 7344018 | 1.02E+07 | 5418249 | 6550003.5 | 7426073.5 | 378693.562 | 8538763 | 6471672 | 2157.9802 | QAFDDAIAELDTLNEDSYK | P68510 | P68510 | YWHAH | tyrosine 3-monooxygenase/tryptophan 5-monooxygenase activation protein, eta | Cytoplasm | transcription regulator |
| 15542866 | 2 | 0.182 | 413722.094 | 1116591.25 | 2445522.75 | 6352308.5 | 6566896 | 6462912.5 | 3637225.75 | 4021899.5 | 7142337.5 | 316119.875 | 7492050 | 4695957.5 | 1304.5183 | NC[160.0307]NDFQYESK | P68510 | P68510 | YWHAH | tyrosine 3-monooxygenase/tryptophan 5-monooxygenase activation protein, eta | Cytoplasm | transcription regulator |
| 15555341 | 2 | 0.242 | 42328.465 | 87874.836 | 157723.891 | 2638784.5 | 1140812 | 1425427.375 | 917976.5 | 946540.938 | 2161429.25 | 0 | 2497416.5 | 1146593.25 | 1828.8165 | LAEQAERYDDMASAMK | P68510 | P68510 | YWHAH | tyrosine 3-monooxygenase/tryptophan 5-monooxygenase activation protein, eta | Cytoplasm | transcription regulator |
| 15537754 | 2 | 0.444 | 1768811 | 3038020.25 | 3860229.75 | 4486239 | 5746264.5 | 6099251.5 | 2728549.5 | 3271080.5 | 6224199.5 | 1216692.25 | 5801789 | 4230914 | 1391.7422 | SIC[160.0307]TTVLELLDK | P68254 | P68254 | YWHAQ | tyrosine 3-monooxygenase/tryptophan 5-monooxygenase activation protein, theta | Cytoplasm | other |
| 15549636 | 2 | 0.182 | 831673.938 | 2331047 | 3752901.25 | 5583782 | 5826557.5 | 4413490.5 | 2402108.5 | 3214808.75 | 5339224.5 | 673675.625 | 3575237.75 | 2247100.5 | 1532.7158 | AVTEQGAELSNEER | P68254 | P68254 | YWHAQ | tyrosine 3-monooxygenase/tryptophan 5-monooxygenase activation protein, theta | Cytoplasm | other |
| 15537327 | 2 | 0.277 | 837645.438 | 1675737.75 | 1840415.5 | 3152432.5 | 3706236.5 | 3284773.25 | 2491825.5 | 2674337.5 | 3813083.25 | 371173.75 | 3921849 | 2030625.5 | 1320.6776 | YLIANATNPESK | P68254 | P68254 | YWHAQ | tyrosine 3-monooxygenase/tryptophan 5-monooxygenase activation protein, theta | Cytoplasm | other |
| 15531294 | 2 | 0.079 | 1.76E+07 | 2.87E+07 | 4.62E+07 | 6.92E+07 | 7.83E+07 | 8.95E+07 | 5.03E+07 | 6.48E+07 | 9.19E+07 | 2.26E+07 | 7.04E+07 | 6.31E+07 | 1189.6594 | DSTLIMQLLR | P63101 | P63101 | YWHAZ | tyrosine 3-monooxygenase/tryptophan 5-monooxygenase activation protein, zeta | Cytoplasm | enzyme |
| 15551755 | 2 | 0.054 | 1.13E+07 | 1.95E+07 | 3.96E+07 | 4.39E+07 | 6.70E+07 | 6.43E+07 | 3.74E+07 | 4.47E+07 | 5.46E+07 | 1.63E+07 | 3.62E+07 | 3.26E+07 | 2040.9823 | GIVDQSQQAYQEAFEISK | P63101 | P63101 | YWHAZ | tyrosine 3-monooxygenase/tryptophan 5-monooxygenase activation protein, zeta | Cytoplasm | enzyme |
| 15547981 | 2 | 0.031 | 1.18E+07 | 1.42E+07 | 2.68E+07 | 5.19E+07 | 5.73E+07 | 5.11E+07 | 3.04E+07 | 3.69E+07 | 5.45E+07 | 1.41E+07 | 4.39E+07 | 3.12E+07 | 1548.7114 | SVTEQGAELSNEER | P63101 | P63101 | YWHAZ | tyrosine 3-monooxygenase/tryptophan 5-monooxygenase activation protein, zeta | Cytoplasm | enzyme |
| 15531614 | 3 | 0.092 | 8992441 | 1.59E+07 | 2.52E+07 | 6.42E+07 | 4.02E+07 | 5.83E+07 | 3.76E+07 | 3.50E+07 | 6.76E+07 | 1.32E+07 | 6.36E+07 | 5.09E+07 | 2160.1187 | IETELRDIC[160.0307]NDVLSLLEK | P63101 | P63101 | YWHAZ | tyrosine 3-monooxygenase/tryptophan 5-monooxygenase activation protein, zeta | Cytoplasm | enzyme |
| 15538305 | 2 | 0.089 | 9059061 | 1.67E+07 | 3.05E+07 | 4.99E+07 | 6.16E+07 | 6.43E+07 | 3.69E+07 | 4.77E+07 | 6.11E+07 | 1.22E+07 | 6.12E+07 | 4.39E+07 | 1330.6973 | FLIPNASQPESK | P63101 | P63101 | YWHAZ | tyrosine 3-monooxygenase/tryptophan 5-monooxygenase activation protein, zeta | Cytoplasm | enzyme |
| 15535323 | 3 | 0.03 | 7149752 | 1.05E+07 | 1.67E+07 | 3.96E+07 | 3.82E+07 | 4.56E+07 | 2.50E+07 | 1.79E+07 | 4.18E+07 | 9525490 | 3.35E+07 | 1.21E+07 | 2131.9885 | TAFDEAIAELDTLSEESYK | P63101 | P63101 | YWHAZ | tyrosine 3-monooxygenase/tryptophan 5-monooxygenase activation protein, zeta | Cytoplasm | enzyme |
| 15547979 | 2 | 0.075 | 1.08E+07 | 1.15E+07 | 3.00E+07 | 5.30E+07 | 6.17E+07 | 5.85E+07 | 3.07E+07 | 4.84E+07 | 6.36E+07 | 8994783 | 4.12E+07 | 4.21E+07 | 2131.9875 | TAFDEAIAELDTLSEESYK | P63101 | P63101 | YWHAZ | tyrosine 3-monooxygenase/tryptophan 5-monooxygenase activation protein, zeta | Cytoplasm | enzyme |
| 15531716 | 2 | 0.081 | 7698689.5 | 1.29E+07 | 1.98E+07 | 3.36E+07 | 3.67E+07 | 4.33E+07 | 2.38E+07 | 2.55E+07 | 3.94E+07 | 7550299 | 2.95E+07 | 2.63E+07 | 907.52295 | NLLSVAYK | P63101 | P63101 | YWHAZ | tyrosine 3-monooxygenase/tryptophan 5-monooxygenase activation protein, zeta | Cytoplasm | enzyme |
| 15534135 | 3 | 0.148 | 4560096.5 | 4843800 | 6807656 | 2.57E+07 | 1.22E+07 | 2.23E+07 | 1.23E+07 | 1.21E+07 | 2.37E+07 | 3949322.25 | 3.37E+07 | 2.15E+07 | 1901.8136 | LAEQAERYDDMAAC[160.0307]MK | P63101 | P63101 | YWHAZ | tyrosine 3-monooxygenase/tryptophan 5-monooxygenase activation protein, zeta | Cytoplasm | enzyme |
| 15543558 | 3 | 0.036 | 1307708.375 | 2353848.5 | 4254823 | 4603827.5 | 7333357.5 | 1.00E+07 | 5916114.5 | 7373998 | 6381105.5 | 2693096.5 | 4449136.5 | 5550694 | 2040.9828 | GIVDQSQQAYQEAFEISK | P63101 | P63101 | YWHAZ | tyrosine 3-monooxygenase/tryptophan 5-monooxygenase activation protein, zeta | Cytoplasm | enzyme |
| 15533183 | 3 | 0.202 | 2927682 | 1766482.25 | 344791.188 | 2.03E+07 | 2980735.25 | 4326192.5 | 3361821 | 3201975.5 | 2.02E+07 | 2186142.25 | 2.54E+07 | 1.39E+07 | 2169.0776 | GIVDQSQQAYQEAFEISKK | P63101 | P63101 | YWHAZ | tyrosine 3-monooxygenase/tryptophan 5-monooxygenase activation protein, zeta | Cytoplasm | enzyme |
| 15539522 | 2 | 0.128 | 1059926.25 | 2892895 | 8735513 | 9139888 | 1.37E+07 | 1.41E+07 | 6655956 | 8711175 | 8884922 | 1826668.25 | 7082020 | 6847663.5 | 1151.5552 | YLAEVAAGDDK | P63101 | P63101 | YWHAZ | tyrosine 3-monooxygenase/tryptophan 5-monooxygenase activation protein, zeta | Cytoplasm | enzyme |
| 15549621 | 2 | 0.09 | 1056775 | 1732055 | 2514409 | 6861663 | 4826110.5 | 6810950 | 2970041 | 4175759.5 | 8255379 | 1578915.25 | 8076008.5 | 5760871 | 2160.118 | IETELRDIC[160.0307]NDVLSLLEK | P63101 | P63101 | YWHAZ | tyrosine 3-monooxygenase/tryptophan 5-monooxygenase activation protein, zeta | Cytoplasm | enzyme |
| 15533741 | 2 | 0.172 | 1212377.625 | 2920864.5 | 9229846 | 6305529.5 | 1.88E+07 | 1.35E+07 | 9296623 | 1.22E+07 | 1.21E+07 | 1245976.25 | 7318054.5 | 8062355 | 1418.7197 | DIC[160.0307]NDVLSLLEK | P63101 | P63101 | YWHAZ | tyrosine 3-monooxygenase/tryptophan 5-monooxygenase activation protein, zeta | Cytoplasm | enzyme |
| 15551988 | 2 | 0.083 | 497053.062 | 515028.406 | 1044775.5 | 5740513 | 2414505 | 2950649.25 | 2503815.75 | 1201049.75 | 3734213.75 | 681270 | 6204969.5 | 2946985.25 | 1901.8142 | LAEQAERYDDMAAC[160.0307]MK | P63101 | P63101 | YWHAZ | tyrosine 3-monooxygenase/tryptophan 5-monooxygenase activation protein, zeta | Cytoplasm | enzyme |
| 15538905 | 3 | 0.23 | 391910.312 | 246382.25 | 107599.109 | 3503581.75 | 648070.125 | 471976.875 | 610385.062 | 385647.219 | 2457828.5 | 295811.062 | 4546467.5 | 1947301.75 | 2169.0789 | KGIVDQSQQAYQEAFEISK | P63101 | P63101 | YWHAZ | tyrosine 3-monooxygenase/tryptophan 5-monooxygenase activation protein, zeta | Cytoplasm | enzyme |
| 15539823 | 2 | 0.673 | 3857612.75 | 7834473.5 | 1.04E+07 | 2337472.25 | 9636721 | 6622503 | 9754315 | 8050691.5 | 4632702 | 4883685.5 | 3928988 | 5100172 | 1037.5349 | HSISHSELK | Q9D0B1 | Q9D0B1 | ZNF524 | zinc finger protein 524 | Nucleus | other |
| 15533204 | 2 | 0.277 | 2.28E+08 | 2.70E+08 | 2.60E+08 | 3.01E+08 | 2.73E+08 | 2.57E+08 | 2.57E+08 | 2.56E+08 | 2.07E+08 | 4.41E+08 | 2.67E+08 | 2.65E+08 | 941.5142 | SSLTIHQR | Q02525 | Q02525 | ZNF717 | zinc finger protein 717 | Nucleus | other |
| 15552077 | 3 | 0.453 | 2775721 | 2586307 | 2298430.75 | 1348374.75 | 2750872.5 | 1241030.375 | 2055350.75 | 2164368.75 | 2687013.5 | 3550270 | 1918355.875 | 1551248.875 | 2539.3706 | AQELGLLQEYLLALTTEDHLLR | Q5SSH7 | Q5SSH7 | ZZEF1 | zinc finger, ZZ-type with EF-hand domain 1 | Cytoplasm | other |

Wild Type

| Entrez Gene Name | Primary Location | Type(s) | Intensity [WT_1] | Intensity [WT_2] | Intensity [WT_3] | ID | Symbol | Mean Intensity | St. Dev | Std. Err | n |
|---|---|---|---|---|---|---|---|---|---|---|---|
| AP2 associated kinase 1 | Cytoplasm | kinase | 1313602.625 | 862820.188 | 701254.125 | Q3UHJ0 | AAK1 | 1313603 | 317353 | 183224 | 3 |
| ATP-binding cassette, sub-family B (MDR/TAP), member 10 | Cytoplasm | transporter | 1.14E+07 | 7289731 | 9925025 | | | | | | |
| ATP-binding cassette, sub-family B (MDR/TAP), member 10 | Cytoplasm | transporter | 6332840 | 6041680.5 | 8413606 | | | | | | |
| ATP-binding cassette, sub-family B (MDR/TAP), member 10 | Cytoplasm | transporter | 3209369 | 3364578.5 | 3541787.5 | | | | | | |
| ATP-binding cassette, sub-family B (MDR/TAP), member 10 | Cytoplasm | transporter | 2353307.25 | 2162250 | 1908195.25 | | | | | | |
| ATP-binding cassette, sub-family B (MDR/TAP), member 10 | Cytoplasm | transporter | 1960281.625 | 2021167.125 | 2155089.5 | | | | | | |
| ATP-binding cassette, sub-family B (MDR/TAP), member 10 | Cytoplasm | transporter | 1775389.125 | 2598952.75 | 2456438.25 | | | | | | |
| ATP-binding cassette, sub-family B (MDR/TAP), member 10 | Cytoplasm | transporter | 1770016.25 | 1686377.75 | 2823162 | | | | | | |
| ATP-binding cassette, sub-family B (MDR/TAP), member 10 | Cytoplasm | transporter | 1759276.375 | 1801890.625 | 2021915.375 | | | | | | |
| ATP-binding cassette, sub-family B (MDR/TAP), member 10 | Cytoplasm | transporter | 1398149.375 | 1578077.375 | 1401543.25 | | | | | | |
| ATP-binding cassette, sub-family B (MDR/TAP), member 10 | Cytoplasm | transporter | 1326483.875 | 1826603.75 | 2568060 | | | | | | |
| ATP-binding cassette, sub-family B (MDR/TAP), member 10 | Cytoplasm | transporter | 1138085.625 | 1278864.125 | 1113010 | | | | | | |
| ATP-binding cassette, sub-family B (MDR/TAP), member 10 | Cytoplasm | transporter | 627558.125 | 855074.875 | 469807.688 | Q9JI39 | ABCB10 | 2917563 | 2611269 | 435212 | 36 |
| ATP-binding cassette, sub-family B (MDR/TAP), member 4 | Plasma Membrane | transporter | 3.35E+07 | 2.49E+07 | 2.41E+07 | | | | | | |
| ATP-binding cassette, sub-family B (MDR/TAP), member 4 | Plasma Membrane | transporter | 1.88E+07 | 1.50E+07 | 1.58E+07 | | | | | | |
| ATP-binding cassette, sub-family B (MDR/TAP), member 4 | Plasma Membrane | transporter | 1.86E+07 | 1.56E+07 | 1.54E+07 | | | | | | |
| ATP-binding cassette, sub-family B (MDR/TAP), member 4 | Plasma Membrane | transporter | 1.77E+07 | 1.10E+07 | 1.40E+07 | | | | | | |
| ATP-binding cassette, sub-family B (MDR/TAP), member 4 | Plasma Membrane | transporter | 1.73E+07 | 1.53E+07 | 1.42E+07 | | | | | | |
| ATP-binding cassette, sub-family B (MDR/TAP), member 4 | Plasma Membrane | transporter | 1.66E+07 | 1.39E+07 | 1.49E+07 | | | | | | |
| ATP-binding cassette, sub-family B (MDR/TAP), member 4 | Plasma Membrane | transporter | 1.60E+07 | 1.67E+07 | 1.45E+07 | | | | | | |
| ATP-binding cassette, sub-family B (MDR/TAP), member 4 | Plasma Membrane | transporter | 1.47E+07 | 1.21E+07 | 1.15E+07 | | | | | | |
| ATP-binding cassette, sub-family B (MDR/TAP), member 4 | Plasma Membrane | transporter | 1.38E+07 | 1.01E+07 | 1.17E+07 | | | | | | |
| ATP-binding cassette, sub-family B (MDR/TAP), member 4 | Plasma Membrane | transporter | 1.15E+07 | 9274393 | 9403828 | | | | | | |
| ATP-binding cassette, sub-family B (MDR/TAP), member 4 | Plasma Membrane | transporter | 9855601 | 8606534 | 1.06E+07 | | | | | | |
| ATP-binding cassette, sub-family B (MDR/TAP), member 4 | Plasma Membrane | transporter | 9769593 | 9839147 | 8891646 | | | | | | |
| ATP-binding cassette, sub-family B (MDR/TAP), member 4 | Plasma Membrane | transporter | 8194694.5 | 8179264.5 | 7097662.5 | | | | | | |
| ATP-binding cassette, sub-family B (MDR/TAP), member 4 | Plasma Membrane | transporter | 7888313.5 | 7074446.5 | 7082108 | | | | | | |
| ATP-binding cassette, sub-family B (MDR/TAP), member 4 | Plasma Membrane | transporter | 7877104 | 8102595 | 8801876 | | | | | | |
| ATP-binding cassette, sub-family B (MDR/TAP), member 4 | Plasma Membrane | transporter | 7236325.5 | 7541278.5 | 7329213.5 | | | | | | |
| ATP-binding cassette, sub-family B (MDR/TAP), member 4 | Plasma Membrane | transporter | 6195433 | 3270584.25 | 4918480 | | | | | | |
| ATP-binding cassette, sub-family B (MDR/TAP), member 4 | Plasma Membrane | transporter | 5902773 | 5193599.5 | 5001338.5 | | | | | | |
| ATP-binding cassette, sub-family B (MDR/TAP), member 4 | Plasma Membrane | transporter | 5077668.5 | 3535773.5 | 2969077 | | | | | | |
| ATP-binding cassette, sub-family B (MDR/TAP), member 4 | Plasma Membrane | transporter | 5029358 | 3940184.5 | 3984126 | | | | | | |
| ATP-binding cassette, sub-family B (MDR/TAP), member 4 | Plasma Membrane | transporter | 4236595 | 3351697 | 3424961.75 | | | | | | |
| ATP-binding cassette, sub-family B (MDR/TAP), member 4 | Plasma Membrane | transporter | 4078936.25 | 3543186 | 4498200 | | | | | | |
| ATP-binding cassette, sub-family B (MDR/TAP), member 4 | Plasma Membrane | transporter | 2634747.25 | 2245413 | 2317968.75 | | | | | | |
| ATP-binding cassette, sub-family B (MDR/TAP), member 4 | Plasma Membrane | transporter | 467062.344 | 2633680.75 | 1967988.625 | P21440 | ABCB4 | 10950175 | 6154848 | 725356 | 72 |
| ATP-binding cassette, sub-family B (MDR/TAP), member 6 (Langereis blood group) | Cytoplasm | transporter | 1.28E+08 | 8.45E+07 | 9.63E+07 | | | | | | |
| ATP-binding cassette, sub-family B (MDR/TAP), member 6 (Langereis blood group) | Cytoplasm | transporter | 1.06E+08 | 8.40E+07 | 9.50E+07 | | | | | | |
| ATP-binding cassette, sub-family B (MDR/TAP), member 6 (Langereis blood group) | Cytoplasm | transporter | 5.38E+07 | 3.72E+07 | 4.30E+07 | | | | | | |
| ATP-binding cassette, sub-family B (MDR/TAP), member 6 (Langereis blood group) | Cytoplasm | transporter | 4.94E+07 | 3.12E+07 | 3.87E+07 | | | | | | |
| ATP-binding cassette, sub-family B (MDR/TAP), member 6 (Langereis blood group) | Cytoplasm | transporter | 4.87E+07 | 3.79E+07 | 4.32E+07 | | | | | | |
| ATP-binding cassette, sub-family B (MDR/TAP), member 6 (Langereis blood group) | Cytoplasm | transporter | 4.23E+07 | 3.10E+07 | 4.35E+07 | | | | | | |
| ATP-binding cassette, sub-family B (MDR/TAP), member 6 (Langereis blood group) | Cytoplasm | transporter | 3.40E+07 | 2.83E+07 | 2.67E+07 | | | | | | |
| ATP-binding cassette, sub-family B (MDR/TAP), member 6 (Langereis blood group) | Cytoplasm | transporter | 2.71E+07 | 1.74E+07 | 1.92E+07 | | | | | | |
| ATP-binding cassette, sub-family B (MDR/TAP), member 6 (Langereis blood group) | Cytoplasm | transporter | 2.53E+07 | 2.28E+07 | 2.16E+07 | | | | | | |
| ATP-binding cassette, sub-family B (MDR/TAP), member 6 (Langereis blood group) | Cytoplasm | transporter | 2.26E+07 | 1.56E+07 | 1.63E+07 | | | | | | |
| ATP-binding cassette, sub-family B (MDR/TAP), member 6 (Langereis blood group) | Cytoplasm | transporter | 1.66E+07 | 1.39E+07 | 1.48E+07 | | | | | | |
| ATP-binding cassette, sub-family B (MDR/TAP), member 6 (Langereis blood group) | Cytoplasm | transporter | 1.28E+07 | 7401402 | 1.23E+07 | | | | | | |
| ATP-binding cassette, sub-family B (MDR/TAP), member 6 (Langereis blood group) | Cytoplasm | transporter | 1.03E+07 | 8700079 | 1.14E+07 | | | | | | |
| ATP-binding cassette, sub-family B (MDR/TAP), member 6 (Langereis blood group) | Cytoplasm | transporter | 9897208 | 5276236.5 | 6783860 | | | | | | |
| ATP-binding cassette, sub-family B (MDR/TAP), member 6 (Langereis blood group) | Cytoplasm | transporter | 8426053 | 5063723 | 6640051 | | | | | | |
| ATP-binding cassette, sub-family B (MDR/TAP), member 6 (Langereis blood group) | Cytoplasm | transporter | 5134540.5 | 3849089.75 | 5466966 | | | | | | |
| ATP-binding cassette, sub-family B (MDR/TAP), member 6 (Langereis blood group) | Cytoplasm | transporter | 4867352.5 | 4088024.5 | 5507182 | | | | | | |
| ATP-binding cassette, sub-family B (MDR/TAP), member 6 (Langereis blood group) | Cytoplasm | transporter | 3572953.75 | 2119797.75 | 2889002.25 | | | | | | |
| ATP-binding cassette, sub-family B (MDR/TAP), member 6 (Langereis blood group) | Cytoplasm | transporter | 2299364.5 | 2729942.25 | 2456853.5 | | | | | | |
| ATP-binding cassette, sub-family B (MDR/TAP), member 6 (Langereis blood group) | Cytoplasm | transporter | 1792665.25 | 4620910.5 | 3089992.75 | Q9DC29 | ABCB6 | 30612507 | 28706249 | 3705961 | 60 |
| ATP-binding cassette, sub-family B (MDR/TAP), member 7 | Cytoplasm | transporter | 780524.688 | 1136027.25 | 1530991.375 | Q61102 | ABCB7 | 780525 | 375406 | 216741 | 3 |
| ATP-binding cassette, sub-family B (MDR/TAP), member 9 | Cytoplasm | transporter | 1205371.5 | 9973444 | 4.74E+07 | Q9JJ59 | ABCB9 | 1205372 | 24545589 | 14171402 | 3 |
| ATP-binding cassette, sub-family C (CFTR/MRP), member 1 | Plasma Membrane | transporter | 2.09E+07 | 1.30E+07 | 1.41E+07 | | | | | | |
| ATP-binding cassette, sub-family C (CFTR/MRP), member 1 | Plasma Membrane | transporter | 1.62E+07 | 1.25E+07 | 1.42E+07 | | | | | | |
| ATP-binding cassette, sub-family C (CFTR/MRP), member 1 | Plasma Membrane | transporter | 1.39E+07 | 7276206 | 8979338 | | | | | | |
| ATP-binding cassette, sub-family C (CFTR/MRP), member 1 | Plasma Membrane | transporter | 1.24E+07 | 1.01E+07 | 1.18E+07 | | | | | | |
| ATP-binding cassette, sub-family C (CFTR/MRP), member 1 | Plasma Membrane | transporter | 1.03E+07 | 6812133 | 7704773.5 | | | | | | |
| ATP-binding cassette, sub-family C (CFTR/MRP), member 1 | Plasma Membrane | transporter | 8061344 | 6377788 | 7164922.5 | | | | | | |
| ATP-binding cassette, sub-family C (CFTR/MRP), member 1 | Plasma Membrane | transporter | 7733731 | 4570019 | 5488231 | | | | | | |
| ATP-binding cassette, sub-family C (CFTR/MRP), member 1 | Plasma Membrane | transporter | 7321330 | 6073652.5 | 5261912 | | | | | | |
| ATP-binding cassette, sub-family C (CFTR/MRP), member 1 | Plasma Membrane | transporter | 5682589 | 5386241.5 | 4924024 | | | | | | |
| ATP-binding cassette, sub-family C (CFTR/MRP), member 1 | Plasma Membrane | transporter | 5274039 | 3735119 | 4698339.5 | | | | | | |
| ATP-binding cassette, sub-family C (CFTR/MRP), member 1 | Plasma Membrane | transporter | 4661681.5 | 3081238.25 | 3589325.25 | | | | | | |
| ATP-binding cassette, sub-family C (CFTR/MRP), member 1 | Plasma Membrane | transporter | 4554708 | 4421247 | 4233541.5 | | | | | | |
| ATP-binding cassette, sub-family C (CFTR/MRP), member 1 | Plasma Membrane | transporter | 3838998.25 | 1945024.875 | 2648368.25 | | | | | | |
| ATP-binding cassette, sub-family C (CFTR/MRP), member 1 | Plasma Membrane | transporter | 3294396.5 | 2356518.5 | 2300696 | | | | | | |
| ATP-binding cassette, sub-family C (CFTR/MRP), member 1 | Plasma Membrane | transporter | 3289552.5 | 1220762.5 | 2023858.5 | | | | | | |
| ATP-binding cassette, sub-family C (CFTR/MRP), member 1 | Plasma Membrane | transporter | 2826008.5 | 2480730.75 | 2290722 | | | | | | |
| ATP-binding cassette, sub-family C (CFTR/MRP), member 1 | Plasma Membrane | transporter | 1951959.375 | 1924058.375 | 2735724.25 | | | | | | |
| ATP-binding cassette, sub-family C (CFTR/MRP), member 1 | Plasma Membrane | transporter | 1873533.875 | 1848649.625 | 1821641.625 | O35379 | ABCC1 | 7444104 | 4390498 | 597471 | 54 |
| ATP-binding cassette, sub-family C (CFTR/MRP), member 6 | Plasma Membrane | transporter | 3329372.75 | 6542358.5 | 6563059.5 | Q9R1S7 | ABCC6 | 3329373 | 1861023 | 1074462 | 3 |
| ATP-binding cassette, sub-family E (OABP), member 1 | Cytoplasm | transporter | 3888062 | 2192450.5 | 2478496.75 | | | | | | |
| ATP-binding cassette, sub-family E (OABP), member 1 | Cytoplasm | transporter | 2107911.25 | 1437464.875 | 1814865.625 | | | | | | |
| ATP-binding cassette, sub-family E (OABP), member 1 | Cytoplasm | transporter | 2086094.75 | 1806652.625 | 1941820.75 | | | | | | |
| ATP-binding cassette, sub-family E (OABP), member 1 | Cytoplasm | transporter | 1704641.25 | 658151.688 | 660480.625 | P61222 | ABCE1 | 2446677 | 842011 | 243068 | 12 |
| ATP-binding cassette, sub-family G (WHITE), member 2 (Junior blood group) | Plasma Membrane | transporter | 1.12E+08 | 1.03E+08 | 8.91E+07 | | | | | | |
| ATP-binding cassette, sub-family G (WHITE), member 2 (Junior blood group) | Plasma Membrane | transporter | 8.81E+07 | 8.38E+07 | 7.00E+07 | | | | | | |
| ATP-binding cassette, sub-family G (WHITE), member 2 (Junior blood group) | Plasma Membrane | transporter | 8.56E+07 | 6.34E+07 | 6.43E+07 | | | | | | |
| ATP-binding cassette, sub-family G (WHITE), member 2 (Junior blood group) | Plasma Membrane | transporter | 6.69E+07 | 5.60E+07 | 5.42E+07 | | | | | | |
| ATP-binding cassette, sub-family G (WHITE), member 2 (Junior blood group) | Plasma Membrane | transporter | 5.58E+07 | 4.38E+07 | 4.72E+07 | | | | | | |
| ATP-binding cassette, sub-family G (WHITE), member 2 (Junior blood group) | Plasma Membrane | transporter | 4.37E+07 | 3.22E+07 | 2.93E+07 | | | | | | |
| ATP-binding cassette, sub-family G (WHITE), member 2 (Junior blood group) | Plasma Membrane | transporter | 3.69E+07 | 4.18E+07 | 3.60E+07 | | | | | | |
| ATP-binding cassette, sub-family G (WHITE), member 2 (Junior blood group) | Plasma Membrane | transporter | 3.30E+07 | 2.47E+07 | 2.35E+07 | | | | | | |
| ATP-binding cassette, sub-family G (WHITE), member 2 (Junior blood group) | Plasma Membrane | transporter | 3.05E+07 | 3.16E+07 | 3.66E+07 | | | | | | |
| ATP-binding cassette, sub-family G (WHITE), member 2 (Junior blood group) | Plasma Membrane | transporter | 2.70E+07 | 2.17E+07 | 2.32E+07 | | | | | | |
| ATP-binding cassette, sub-family G (WHITE), member 2 (Junior blood group) | Plasma Membrane | transporter | 2.35E+07 | 1.68E+07 | 1.75E+07 | | | | | | |
| ATP-binding cassette, sub-family G (WHITE), member 2 (Junior blood group) | Plasma Membrane | transporter | 1.63E+07 | 1.51E+07 | 1.19E+07 | | | | | | |

| | | | | | | | | | | | |
|---|---|---|---|---|---|---|---|---|---|---|---|
| ATP-binding cassette, sub-family G (WHITE), member 2 (Junior blood group) | Plasma Membrane | transporter | 1.49E+07 | 1.39E+07 | 1.19E+07 | | | | | | |
| ATP-binding cassette, sub-family G (WHITE), member 2 (Junior blood group) | Plasma Membrane | transporter | 1.49E+07 | 1.59E+07 | 1.18E+07 | | | | | | |
| ATP-binding cassette, sub-family G (WHITE), member 2 (Junior blood group) | Plasma Membrane | transporter | 1.37E+07 | 1.09E+07 | 9807314 | | | | | | |
| ATP-binding cassette, sub-family G (WHITE), member 2 (Junior blood group) | Plasma Membrane | transporter | 1.00E+07 | 1.02E+07 | 7591999.5 | | | | | | |
| ATP-binding cassette, sub-family G (WHITE), member 2 (Junior blood group) | Plasma Membrane | transporter | 4003344.75 | 4350097.5 | 3154034.25 | | | | | | |
| ATP-binding cassette, sub-family G (WHITE), member 2 (Junior blood group) | Plasma Membrane | transporter | 3719093 | 3765471.75 | 4217887.5 | | | | | | |
| ATP-binding cassette, sub-family G (WHITE), member 2 (Junior blood group) | Plasma Membrane | transporter | 2353872.5 | 5236518.5 | 4607198.5 | | | | | | |
| ATP-binding cassette, sub-family G (WHITE), member 2 (Junior blood group) | Plasma Membrane | transporter | 946253.438 | 543225.438 | 478101.594 | Q7TMS5 | ABCG2 | 34164128 | 28162130 | 3635715 | 60 |
| abhydrolase domain containing 16A | Cytoplasm | enzyme | 4735394.5 | 4193960 | 3964909 | | | | | | |
| abhydrolase domain containing 16A | Cytoplasm | enzyme | 4210057 | 4296399 | 3873923 | | | | | | |
| abhydrolase domain containing 16A | Cytoplasm | enzyme | 811453.938 | 823464.25 | 736385.75 | Q9Z1Q2 | ABHD16A | 3252302 | 1727767 | 575922 | 9 |
| acetyl-CoA acyltransferase 1 | Cytoplasm | enzyme | 180827.562 | 3635310 | 2313762.75 | Q921H8 | ACAA1 | 180828 | 1743050 | 1006351 | 3 |
| acetyl-CoA acyltransferase 2 | Cytoplasm | enzyme | 3940763.75 | 5071712 | 6953793 | | | | | | |
| acetyl-CoA acyltransferase 2 | Cytoplasm | enzyme | 3215672.5 | 4549188.5 | 4123992.75 | | | | | | |
| acetyl-CoA acyltransferase 2 | Cytoplasm | enzyme | 3210928.75 | 2735370.25 | 3848553.25 | | | | | | |
| acetyl-CoA acyltransferase 2 | Cytoplasm | enzyme | 2519394.5 | 3510918.75 | 2535776 | | | | | | |
| acetyl-CoA acyltransferase 2 | Cytoplasm | enzyme | 1576919 | 2784727.25 | 2279460.75 | | | | | | |
| acetyl-CoA acyltransferase 2 | Cytoplasm | enzyme | 1414916.625 | 1542763.375 | 2672238.25 | Q8BWT1 | ACAA2 | 2646433 | 1377915 | 324778 | 18 |
| acyl-CoA dehydrogenase family, member 9 | Cytoplasm | enzyme | 803627.812 | 1765506.5 | 1464645.5 | Q8JZN5 | ACAD9 | 803628 | 492049 | 284085 | 3 |
| acyl-CoA dehydrogenase, long chain | Cytoplasm | enzyme | 1.52E+07 | 2.36E+07 | 2.41E+07 | | | | | | |
| acyl-CoA dehydrogenase, long chain | Cytoplasm | enzyme | 4478708 | 1.03E+07 | 7365655 | | | | | | |
| acyl-CoA dehydrogenase, long chain | Cytoplasm | enzyme | 4477133 | 7297601.5 | 5678019.5 | | | | | | |
| acyl-CoA dehydrogenase, long chain | Cytoplasm | enzyme | 3691303.25 | 3878784.5 | 3558191.5 | | | | | | |
| acyl-CoA dehydrogenase, long chain | Cytoplasm | enzyme | 2794245.75 | 3991923.5 | 3489407.75 | | | | | | |
| acyl-CoA dehydrogenase, long chain | Cytoplasm | enzyme | 1510848.625 | 4471136 | 3682018.5 | | | | | | |
| acyl-CoA dehydrogenase, long chain | Cytoplasm | enzyme | 1465261.75 | 1565990.375 | 1333748.375 | P51174 | ACADL | 4806786 | 6595073 | 1439163 | 21 |
| acyl-CoA dehydrogenase, C-4 to C-12 straight chain | Cytoplasm | enzyme | 93058.625 | 1498694.375 | 720594.688 | P45952 | ACADM | 93059 | 704161 | 406547 | 3 |
| acyl-CoA dehydrogenase, very long chain | Cytoplasm | enzyme | 2194312.75 | 2330774.5 | 2927524.5 | | | | | | |
| acyl-CoA dehydrogenase, very long chain | Cytoplasm | enzyme | 1123271.25 | 1668165.75 | 1773534.5 | | | | | | |
| acyl-CoA dehydrogenase, very long chain | Cytoplasm | enzyme | 835604.5 | 1211175.625 | 1482632.25 | | | | | | |
| acyl-CoA dehydrogenase, very long chain | Cytoplasm | enzyme | 691454.938 | 1336254.125 | 1047583.812 | | | | | | |
| acyl-CoA dehydrogenase, very long chain | Cytoplasm | enzyme | 408683.312 | 567698.562 | 407535.969 | P50544 | ACADVL | 1050665 | 741838 | 191542 | 15 |
| acetyl-CoA acetyltransferase 1 | Cytoplasm | enzyme | 4641630 | 1.39E+07 | 1.28E+07 | | | | | | |
| acetyl-CoA acetyltransferase 1 | Cytoplasm | enzyme | 3108211 | 9686164 | 7991976 | | | | | | |
| acetyl-CoA acetyltransferase 1 | Cytoplasm | enzyme | 2428965.25 | 6875972.5 | 4717857.5 | | | | | | |
| acetyl-CoA acetyltransferase 1 | Cytoplasm | enzyme | 2151835.75 | 7946749 | 8273781.5 | | | | | | |
| acetyl-CoA acetyltransferase 1 | Cytoplasm | enzyme | 1299048.125 | 4805533 | 3813740.5 | | | | | | |
| acetyl-CoA acetyltransferase 1 | Cytoplasm | enzyme | 1073544.125 | 3686724 | 3895485.25 | | | | | | |
| acetyl-CoA acetyltransferase 1 | Cytoplasm | enzyme | 1050215.125 | 5843628 | 3491918.25 | Q8QZT1 | ACAT1 | 2250493 | 3606579 | 787020 | 21 |
| atypical chemokine receptor 1 (Duffy blood group) | Plasma Membrane | G-protein coupled receptor | 3.89E+07 | 2.09E+07 | 2.47E+07 | | | | | | |
| atypical chemokine receptor 1 (Duffy blood group) | Plasma Membrane | G-protein coupled receptor | 2.85E+07 | 2.13E+07 | 1.97E+07 | | | | | | |
| atypical chemokine receptor 1 (Duffy blood group) | Plasma Membrane | G-protein coupled receptor | 1.04E+07 | 6683118.5 | 6690175 | | | | | | |
| atypical chemokine receptor 1 (Duffy blood group) | Plasma Membrane | G-protein coupled receptor | 3621377.25 | 8534182 | 4350955 | Q9QUI6 | ACKR1 | 20327844 | 11155511 | 3220319 | 12 |
| aconitase 2, mitochondrial | Cytoplasm | enzyme | 1.03E+07 | 1.60E+07 | 1.40E+07 | | | | | | |
| aconitase 2, mitochondrial | Cytoplasm | enzyme | 5251240.5 | 7170701 | 6804730.5 | | | | | | |
| aconitase 2, mitochondrial | Cytoplasm | enzyme | 4010804.25 | 7720188.5 | 7193413 | | | | | | |
| aconitase 2, mitochondrial | Cytoplasm | enzyme | 3908626.5 | 3301727.25 | 4505804.5 | | | | | | |
| aconitase 2, mitochondrial | Cytoplasm | enzyme | 3804413.25 | 6268843.5 | 5791349.5 | | | | | | |
| aconitase 2, mitochondrial | Cytoplasm | enzyme | 3766848.5 | 8345527 | 4675131.5 | | | | | | |
| aconitase 2, mitochondrial | Cytoplasm | enzyme | 3621111.25 | 5806333.5 | 5355704 | | | | | | |
| aconitase 2, mitochondrial | Cytoplasm | enzyme | 3359214.5 | 3404942.25 | 4164021.25 | | | | | | |
| aconitase 2, mitochondrial | Cytoplasm | enzyme | 3115078.75 | 6021115 | 4389224.5 | | | | | | |
| aconitase 2, mitochondrial | Cytoplasm | enzyme | 2956934.75 | 9047802 | 4634811.5 | | | | | | |
| aconitase 2, mitochondrial | Cytoplasm | enzyme | 2378627.25 | 4495395.5 | 5220057 | | | | | | |
| aconitase 2, mitochondrial | Cytoplasm | enzyme | 2064706.625 | 3311779.25 | 2223136 | | | | | | |
| aconitase 2, mitochondrial | Cytoplasm | enzyme | 1949686 | 4698447.5 | 3206908.25 | | | | | | |
| aconitase 2, mitochondrial | Cytoplasm | enzyme | 1835935.125 | 4163280 | 2813515.5 | | | | | | |
| aconitase 2, mitochondrial | Cytoplasm | enzyme | 1506207.375 | 3087834 | 2783830 | | | | | | |
| aconitase 2, mitochondrial | Cytoplasm | enzyme | 1343663.875 | 2005365.125 | 1664663.625 | | | | | | |
| aconitase 2, mitochondrial | Cytoplasm | enzyme | 925687.875 | 1679130.25 | 1378438.375 | | | | | | |
| aconitase 2, mitochondrial | Cytoplasm | enzyme | 864702.688 | 1288070.875 | 2267207.25 | | | | | | |
| aconitase 2, mitochondrial | Cytoplasm | enzyme | 781536.438 | 713391.312 | 1114344.5 | | | | | | |
| aconitase 2, mitochondrial | Cytoplasm | enzyme | 733141.125 | 1412602 | 1141242.375 | | | | | | |
| aconitase 2, mitochondrial | Cytoplasm | enzyme | 314656.75 | 533008.688 | 1029271.875 | Q99KI0 | ACO2 | 2800134 | 3041910 | 383245 | 63 |
| acyl-CoA synthetase long-chain family member 1 | Cytoplasm | enzyme | 1.87E+07 | 1.57E+07 | 1.57E+07 | | | | | | |
| acyl-CoA synthetase long-chain family member 1 | Cytoplasm | enzyme | 1.46E+07 | 1.25E+07 | 1.35E+07 | | | | | | |
| acyl-CoA synthetase long-chain family member 1 | Cytoplasm | enzyme | 1.35E+07 | 1.45E+07 | 1.57E+07 | | | | | | |
| acyl-CoA synthetase long-chain family member 1 | Cytoplasm | enzyme | 1.31E+07 | 1.17E+07 | 9686220 | | | | | | |
| acyl-CoA synthetase long-chain family member 1 | Cytoplasm | enzyme | 1.18E+07 | 5756730.5 | 8776750 | | | | | | |
| acyl-CoA synthetase long-chain family member 1 | Cytoplasm | enzyme | 1.09E+07 | 1.23E+07 | 1.27E+07 | | | | | | |
| acyl-CoA synthetase long-chain family member 1 | Cytoplasm | enzyme | 1.09E+07 | 6955306 | 7016425.5 | | | | | | |
| acyl-CoA synthetase long-chain family member 1 | Cytoplasm | enzyme | 8576849 | 5059523.5 | 7183400 | | | | | | |
| acyl-CoA synthetase long-chain family member 1 | Cytoplasm | enzyme | 6571424.5 | 5507931 | 5225931 | | | | | | |
| acyl-CoA synthetase long-chain family member 1 | Cytoplasm | enzyme | 6516312 | 6637441.5 | 4339017 | | | | | | |
| acyl-CoA synthetase long-chain family member 1 | Cytoplasm | enzyme | 6330786 | 7447746.5 | 5306162.5 | | | | | | |
| acyl-CoA synthetase long-chain family member 1 | Cytoplasm | enzyme | 5735175 | 5229208 | 4780474.5 | | | | | | |
| acyl-CoA synthetase long-chain family member 1 | Cytoplasm | enzyme | 3815801 | 3335259 | 2868196.5 | | | | | | |
| acyl-CoA synthetase long-chain family member 1 | Cytoplasm | enzyme | 3618039.75 | 3775586.25 | 3447739 | | | | | | |
| acyl-CoA synthetase long-chain family member 1 | Cytoplasm | enzyme | 3596686.5 | 2547358.75 | 2573327.5 | | | | | | |
| acyl-CoA synthetase long-chain family member 1 | Cytoplasm | enzyme | 3437578.25 | 3331642.5 | 3295464.25 | | | | | | |
| acyl-CoA synthetase long-chain family member 1 | Cytoplasm | enzyme | 3394197.75 | 4336839 | 5050888.5 | P41216 | ACSL1 | 8529579 | 4392116 | 615019 | 51 |
| acyl-CoA synthetase long-chain family member 4 | Cytoplasm | enzyme | 1.89E+07 | 1.48E+07 | 1.19E+07 | | | | | | |
| acyl-CoA synthetase long-chain family member 4 | Cytoplasm | enzyme | 1.51E+07 | 1.33E+07 | 1.47E+07 | | | | | | |
| acyl-CoA synthetase long-chain family member 4 | Cytoplasm | enzyme | 1.06E+07 | 8710410 | 9955279 | | | | | | |
| acyl-CoA synthetase long-chain family member 4 | Cytoplasm | enzyme | 7631305 | 5039822 | 4316190 | | | | | | |
| acyl-CoA synthetase long-chain family member 4 | Cytoplasm | enzyme | 3415101.5 | 4401632.5 | 1850504.25 | | | | | | |
| acyl-CoA synthetase long-chain family member 4 | Cytoplasm | enzyme | 3212008 | 1981424.375 | 1698552 | | | | | | |
| acyl-CoA synthetase long-chain family member 4 | Cytoplasm | enzyme | 2989535.75 | 2724314.5 | 2373719.5 | | | | | | |
| acyl-CoA synthetase long-chain family member 4 | Cytoplasm | enzyme | 2979645.75 | 3460170 | 1962310.5 | | | | | | |
| acyl-CoA synthetase long-chain family member 4 | Cytoplasm | enzyme | 2304811.5 | 1882837 | 1415451 | Q9QUJ7 | ACSL4 | 7446934 | 5260813 | 1012444 | 27 |
| acyl-CoA synthetase long-chain family member 5 | Cytoplasm | enzyme | 1.44E+07 | 1.38E+07 | 1.45E+07 | | | | | | |
| acyl-CoA synthetase long-chain family member 5 | Cytoplasm | enzyme | 7879329 | 8948880 | 7510522 | | | | | | |
| acyl-CoA synthetase long-chain family member 5 | Cytoplasm | enzyme | 7113353.5 | 6043303.5 | 6427417.5 | | | | | | |
| acyl-CoA synthetase long-chain family member 5 | Cytoplasm | enzyme | 6911968.5 | 6869392.5 | 6051373 | | | | | | |
| acyl-CoA synthetase long-chain family member 5 | Cytoplasm | enzyme | 5966748 | 4713316.5 | 5605816.5 | | | | | | |

| | | | | | | | | | | | |
|---|---|---|---|---|---|---|---|---|---|---|---|
| acyl-CoA synthetase long-chain family member 5 | Cytoplasm | enzyme | 4971323.5 | 3429108.25 | 4476678 | | | | | | |
| acyl-CoA synthetase long-chain family member 5 | Cytoplasm | enzyme | 4317570 | 4337125 | 4686498 | | | | | | |
| acyl-CoA synthetase long-chain family member 5 | Cytoplasm | enzyme | 3553604.25 | 5068241 | 2238061.25 | | | | | | |
| acyl-CoA synthetase long-chain family member 5 | Cytoplasm | enzyme | 3217172.25 | 2936129.75 | 3474739.75 | | | | | | |
| acyl-CoA synthetase long-chain family member 5 | Cytoplasm | enzyme | 2921652.5 | 2991352.75 | 2764439.5 | | | | | | |
| acyl-CoA synthetase long-chain family member 5 | Cytoplasm | enzyme | 2915704.75 | 2663721 | 2265779.75 | Q8JZR0 | ACSL5 | 5834402 | 3277843 | 570599 | 33 |
| acyl-CoA synthetase short-chain family member 1 | Cytoplasm | enzyme | 1587054.375 | 1732576 | 992115.312 | | | | | | |
| acyl-CoA synthetase short-chain family member 1 | Cytoplasm | enzyme | 1228105.625 | 842429.438 | 1266518.125 | | | | | | |
| acyl-CoA synthetase short-chain family member 1 | Cytoplasm | enzyme | 932819.312 | 870621.312 | 1073860.75 | Q99NB1 | ACSS1 | 1249326 | 315671 | 105224 | 9 |
| actin, beta | Cytoplasm | other | 5.29E+09 | 5.18E+09 | 4.79E+09 | | | | | | |
| actin, beta | Cytoplasm | other | 3.06E+09 | 2.66E+09 | 3.11E+09 | | | | | | |
| actin, beta | Cytoplasm | other | 3.00E+09 | 2.41E+09 | 2.32E+09 | | | | | | |
| actin, beta | Cytoplasm | other | 1.71E+09 | 1.40E+09 | 1.32E+09 | | | | | | |
| actin, beta | Cytoplasm | other | 1.64E+09 | 1.93E+09 | 2.18E+09 | | | | | | |
| actin, beta | Cytoplasm | other | 1.48E+09 | 9.23E+08 | 1.22E+09 | | | | | | |
| actin, beta | Cytoplasm | other | 1.46E+09 | 1.24E+09 | 1.25E+09 | | | | | | |
| actin, beta | Cytoplasm | other | 1.38E+09 | 1.22E+09 | 1.10E+09 | | | | | | |
| actin, beta | Cytoplasm | other | 1.38E+09 | 1.04E+09 | 9.85E+08 | | | | | | |
| actin, beta | Cytoplasm | other | 1.14E+09 | 8.08E+08 | 7.72E+08 | | | | | | |
| actin, beta | Cytoplasm | other | 1.13E+09 | 1.05E+09 | 1.10E+09 | | | | | | |
| actin, beta | Cytoplasm | other | 1.03E+09 | 9.38E+08 | 8.98E+08 | | | | | | |
| actin, beta | Cytoplasm | other | 7.85E+08 | 6.10E+08 | 6.45E+08 | | | | | | |
| actin, beta | Cytoplasm | other | 7.55E+08 | 6.91E+08 | 5.68E+08 | | | | | | |
| actin, beta | Cytoplasm | other | 4.11E+08 | 4.48E+08 | 4.65E+08 | | | | | | |
| actin, beta | Cytoplasm | other | 3.30E+08 | 2.29E+08 | 2.95E+08 | | | | | | |
| actin, beta | Cytoplasm | other | 2.85E+08 | 2.91E+08 | 2.82E+08 | | | | | | |
| actin, beta | Cytoplasm | other | 2.82E+08 | 1.97E+08 | 1.44E+08 | | | | | | |
| actin, beta | Cytoplasm | other | 2.45E+08 | 1.41E+08 | 1.49E+08 | | | | | | |
| actin, beta | Cytoplasm | other | 2.16E+08 | 3.36E+08 | 3.90E+08 | | | | | | |
| actin, beta | Cytoplasm | other | 1.62E+08 | 1.54E+08 | 1.74E+08 | | | | | | |
| actin, beta | Cytoplasm | other | 1.29E+08 | 1.12E+08 | 1.42E+08 | | | | | | |
| actin, beta | Cytoplasm | other | 1.21E+08 | 5.25E+08 | 3.12E+08 | | | | | | |
| actin, beta | Cytoplasm | other | 1.10E+08 | 4.50E+07 | 6.35E+07 | | | | | | |
| actin, beta | Cytoplasm | other | 1.09E+08 | 9.33E+07 | 9.09E+07 | | | | | | |
| actin, beta | Cytoplasm | other | 1.07E+08 | 3.09E+08 | 2.14E+08 | | | | | | |
| actin, beta | Cytoplasm | other | 1.04E+08 | 4.72E+07 | 6.37E+07 | | | | | | |
| actin, beta | Cytoplasm | other | 8.16E+07 | 6.61E+07 | 7.51E+07 | | | | | | |
| actin, beta | Cytoplasm | other | 7.01E+07 | 8.20E+07 | 8.02E+07 | | | | | | |
| actin, beta | Cytoplasm | other | 6.76E+07 | 4.44E+07 | 5.07E+07 | | | | | | |
| actin, beta | Cytoplasm | other | 6.07E+07 | 3.89E+07 | 4.36E+07 | | | | | | |
| actin, beta | Cytoplasm | other | 5.41E+07 | 1.32E+08 | 1.77E+08 | | | | | | |
| actin, beta | Cytoplasm | other | 3.88E+07 | 3.50E+07 | 4.94E+07 | | | | | | |
| actin, beta | Cytoplasm | other | 3.55E+07 | 6.51E+07 | 4.75E+07 | | | | | | |
| actin, beta | Cytoplasm | other | 3.51E+07 | 2.32E+07 | 3.08E+07 | | | | | | |
| actin, beta | Cytoplasm | other | 3.01E+07 | 2.39E+07 | 2.34E+07 | | | | | | |
| actin, beta | Cytoplasm | other | 2.64E+07 | 7.06E+07 | 4.43E+07 | | | | | | |
| actin, beta | Cytoplasm | other | 1.70E+07 | 1.33E+07 | 1.89E+07 | | | | | | |
| actin, beta | Cytoplasm | other | 1.53E+07 | 7.57E+07 | 5.90E+07 | | | | | | |
| actin, beta | Cytoplasm | other | 1.52E+07 | 9080671 | 6127025 | | | | | | |
| actin, beta | Cytoplasm | other | 1.46E+07 | 1.55E+07 | 1.91E+07 | | | | | | |
| actin, beta | Cytoplasm | other | 1.40E+07 | 1.37E+07 | 1.06E+08 | | | | | | |
| actin, beta | Cytoplasm | other | 8338241 | 5152283 | 5853039.5 | | | | | | |
| actin, beta | Cytoplasm | other | 8152939.5 | 2374233 | 3970828.25 | | | | | | |
| actin, beta | Cytoplasm | other | 3448422 | 3812777.5 | 3940716.75 | | | | | | |
| actin, beta | Cytoplasm | other | 2841460.75 | 2215682.5 | 2438840.5 | | | | | | |
| actin, beta | Cytoplasm | other | 1757990.625 | 3179331.75 | 3.51E+07 | | | | | | |
| actin, beta | Cytoplasm | other | 176601.453 | 260447.406 | 106389.68 | P60710 | ACTB | 592322618 | 952229689 | 79352474 | 144 |
| actin, alpha, cardiac muscle 1 | Cytoplasm | enzyme | 3.37E+09 | 2.95E+09 | 3.03E+09 | | | | | | |
| actin, alpha, cardiac muscle 1 | Cytoplasm | enzyme | 4.02E+08 | 5.42E+08 | 4.63E+08 | | | | | | |
| actin, alpha, cardiac muscle 1 | Cytoplasm | enzyme | 2.30E+08 | 1.66E+08 | 1.55E+08 | | | | | | |
| actin, alpha, cardiac muscle 1 | Cytoplasm | enzyme | 3.15E+07 | 2.06E+07 | 1.69E+07 | P68033 | ACTC1 | 1008300000 | 1321631201 | 381522065 | 12 |
| ARP2 actin-related protein 2 homolog (yeast) | Plasma Membrane | other | 3344871.75 | 5529105 | 5327661.5 | | | | | | |
| ARP2 actin-related protein 2 homolog (yeast) | Plasma Membrane | other | 1513266.125 | 3098958.25 | 2424161.5 | | | | | | |
| ARP2 actin-related protein 2 homolog (yeast) | Plasma Membrane | other | 1350946 | 3888503.5 | 2596609 | P61161 | ACTR2 | 2069695 | 1486550 | 495517 | 9 |
| ARP3 actin-related protein 3 homolog (yeast) | Plasma Membrane | other | 7399365 | 1.66E+07 | 1.12E+07 | | | | | | |
| ARP3 actin-related protein 3 homolog (yeast) | Plasma Membrane | other | 1525828.125 | 2077454.5 | 1536111 | | | | | | |
| ARP3 actin-related protein 3 homolog (yeast) | Plasma Membrane | other | 1418809.375 | 1384665.375 | 2451029.75 | | | | | | |
| ARP3 actin-related protein 3 homolog (yeast) | Plasma Membrane | other | 1189395.25 | 2142234.25 | 1542695.125 | | | | | | |
| ARP3 actin-related protein 3 homolog (yeast) | Plasma Membrane | other | 1109860.625 | 2527601.25 | 2256036.5 | Q99JY9 | ACTR3 | 2528652 | 4494117 | 1160376 | 15 |
| ARP3 actin-related protein 3 homolog B (yeast) | Cytoplasm | other | 2243707.75 | 3960124.75 | 2959788.5 | Q641P0 | ACTR3B | 2243708 | 862123 | 497747 | 3 |
| ADAM metallopeptidase domain 10 | Plasma Membrane | peptidase | 4.52E+07 | 6.20E+07 | 5.78E+07 | | | | | | |
| ADAM metallopeptidase domain 10 | Plasma Membrane | peptidase | 4.28E+07 | 3.81E+07 | 2.83E+07 | | | | | | |
| ADAM metallopeptidase domain 10 | Plasma Membrane | peptidase | 4.26E+07 | 3.67E+07 | 2.94E+07 | | | | | | |
| ADAM metallopeptidase domain 10 | Plasma Membrane | peptidase | 4.05E+07 | 3.13E+07 | 3.45E+07 | | | | | | |
| ADAM metallopeptidase domain 10 | Plasma Membrane | peptidase | 3.97E+07 | 3.51E+07 | 3.11E+07 | | | | | | |
| ADAM metallopeptidase domain 10 | Plasma Membrane | peptidase | 3.29E+07 | 3.36E+07 | 2.57E+07 | | | | | | |
| ADAM metallopeptidase domain 10 | Plasma Membrane | peptidase | 3.27E+07 | 4.92E+07 | 3.49E+07 | | | | | | |
| ADAM metallopeptidase domain 10 | Plasma Membrane | peptidase | 2.14E+07 | 2.51E+07 | 2.34E+07 | | | | | | |
| ADAM metallopeptidase domain 10 | Plasma Membrane | peptidase | 1.72E+07 | 1.73E+07 | 1.41E+07 | | | | | | |
| ADAM metallopeptidase domain 10 | Plasma Membrane | peptidase | 1.69E+07 | 1.74E+07 | 1.39E+07 | | | | | | |
| ADAM metallopeptidase domain 10 | Plasma Membrane | peptidase | 1.55E+07 | 1.19E+07 | 1.09E+07 | | | | | | |
| ADAM metallopeptidase domain 10 | Plasma Membrane | peptidase | 1.51E+07 | 1.26E+07 | 9028169 | | | | | | |
| ADAM metallopeptidase domain 10 | Plasma Membrane | peptidase | 1.18E+07 | 5936120 | 9042608 | | | | | | |
| ADAM metallopeptidase domain 10 | Plasma Membrane | peptidase | 9855667 | 1.21E+07 | 9381623 | | | | | | |
| ADAM metallopeptidase domain 10 | Plasma Membrane | peptidase | 9162740 | 1.02E+07 | 9527662 | | | | | | |
| ADAM metallopeptidase domain 10 | Plasma Membrane | peptidase | 6809959 | 4671316 | 6789846.5 | | | | | | |
| ADAM metallopeptidase domain 10 | Plasma Membrane | peptidase | 6637894 | 7509862.5 | 7629496 | | | | | | |
| ADAM metallopeptidase domain 10 | Plasma Membrane | peptidase | 4158332.25 | 5429386.5 | 3304439 | | | | | | |
| ADAM metallopeptidase domain 10 | Plasma Membrane | peptidase | 3345994.25 | 4099042 | 2644577.75 | | | | | | |
| ADAM metallopeptidase domain 10 | Plasma Membrane | peptidase | 3310864.25 | 1400531.5 | 1316731.25 | | | | | | |
| ADAM metallopeptidase domain 10 | Plasma Membrane | peptidase | 3002908 | 1400119.5 | 234223.094 | | | | | | |
| ADAM metallopeptidase domain 10 | Plasma Membrane | peptidase | 2790663.5 | 1150253 | 2250576.25 | | | | | | |
| ADAM metallopeptidase domain 10 | Plasma Membrane | peptidase | 2531037 | 948246.062 | 1650399.5 | | | | | | |
| ADAM metallopeptidase domain 10 | Plasma Membrane | peptidase | 2482403.5 | 7729058.5 | 4782866 | | | | | | |
| ADAM metallopeptidase domain 10 | Plasma Membrane | peptidase | 2304488 | 2725428 | 3796067.25 | | | | | | |

| | | | | | | | | | | | |
|---|---|---|---|---|---|---|---|---|---|---|---|
| ADAM metallopeptidase domain 10 | Plasma Membrane | peptidase | 1532434.625 | 2064132.875 | 2389327 | O35598 | ADAM10 | 16620592 | 15347026 | 1737709 | 78 |
| ADAM metallopeptidase domain 17 | Plasma Membrane | peptidase | 7254658.5 | 6398081.5 | 5599455 | | | | | | |
| ADAM metallopeptidase domain 17 | Plasma Membrane | peptidase | 4.70E-06 | 3478616 | 2854150.75 | Q9Z0F8 | ADAM17 | 3627329 | 2685814 | 1096479 | 6 |
| adducin 1 (alpha) | Cytoplasm | other | 1.29E+09 | 8.16E+08 | 9.85E+08 | | | | | | |
| adducin 1 (alpha) | Cytoplasm | other | 7.40E+08 | 5.07E+08 | 4.74E+08 | | | | | | |
| adducin 1 (alpha) | Cytoplasm | other | 5.30E+08 | 3.38E+08 | 3.23E+08 | | | | | | |
| adducin 1 (alpha) | Cytoplasm | other | 5.08E+08 | 4.81E+08 | 3.96E+08 | | | | | | |
| adducin 1 (alpha) | Cytoplasm | other | 4.37E+08 | 3.20E+08 | 3.46E+08 | | | | | | |
| adducin 1 (alpha) | Cytoplasm | other | 3.12E+08 | 2.18E+08 | 2.48E+08 | | | | | | |
| adducin 1 (alpha) | Cytoplasm | other | 3.04E+08 | 2.17E+08 | 2.12E+08 | | | | | | |
| adducin 1 (alpha) | Cytoplasm | other | 3.02E+08 | 3.64E+08 | 2.80E+08 | | | | | | |
| adducin 1 (alpha) | Cytoplasm | other | 2.96E+08 | 1.46E+08 | 1.64E+08 | | | | | | |
| adducin 1 (alpha) | Cytoplasm | other | 2.90E+08 | 2.15E+08 | 2.03E+08 | | | | | | |
| adducin 1 (alpha) | Cytoplasm | other | 2.82E+08 | 2.07E+08 | 1.87E+08 | | | | | | |
| adducin 1 (alpha) | Cytoplasm | other | 2.81E+08 | 2.27E+08 | 2.72E+08 | | | | | | |
| adducin 1 (alpha) | Cytoplasm | other | 2.79E+08 | 3.43E+08 | 2.31E+08 | | | | | | |
| adducin 1 (alpha) | Cytoplasm | other | 2.79E+08 | 1.99E+08 | 2.04E+08 | | | | | | |
| adducin 1 (alpha) | Cytoplasm | other | 2.77E+08 | 2.20E+08 | 2.54E+08 | | | | | | |
| adducin 1 (alpha) | Cytoplasm | other | 1.89E+08 | 1.13E+08 | 1.21E+08 | | | | | | |
| adducin 1 (alpha) | Cytoplasm | other | 1.71E+08 | 1.61E+08 | 1.34E+08 | | | | | | |
| adducin 1 (alpha) | Cytoplasm | other | 1.19E+08 | 1.49E+08 | 1.45E+08 | | | | | | |
| adducin 1 (alpha) | Cytoplasm | other | 1.06E+08 | 1.03E+08 | 9.78E+07 | | | | | | |
| adducin 1 (alpha) | Cytoplasm | other | 8.13E+07 | 4.91E+07 | 4.71E+07 | | | | | | |
| adducin 1 (alpha) | Cytoplasm | other | 7.24E+07 | 6.31E+07 | 4.62E+07 | | | | | | |
| adducin 1 (alpha) | Cytoplasm | other | 7.15E+07 | 3.07E+07 | 3.83E+07 | | | | | | |
| adducin 1 (alpha) | Cytoplasm | other | 5.90E+07 | 4.84E+07 | 4.39E+07 | | | | | | |
| adducin 1 (alpha) | Cytoplasm | other | 5.49E+07 | 2.00E+07 | 2.98E+07 | | | | | | |
| adducin 1 (alpha) | Cytoplasm | other | 4.50E+07 | 3.31E+07 | 3.47E+07 | | | | | | |
| adducin 1 (alpha) | Cytoplasm | other | 3.46E+07 | 1.51E+07 | 2.58E+07 | | | | | | |
| adducin 1 (alpha) | Cytoplasm | other | 2.77E+07 | 2.27E+07 | 1.76E+07 | | | | | | |
| adducin 1 (alpha) | Cytoplasm | other | 2.48E+07 | 1.77E+07 | 2.47E+07 | | | | | | |
| adducin 1 (alpha) | Cytoplasm | other | 2.33E+07 | 1.39E+07 | 1.64E+07 | | | | | | |
| adducin 1 (alpha) | Cytoplasm | other | 1.89E+07 | 1.01E+07 | 1.24E+07 | | | | | | |
| adducin 1 (alpha) | Cytoplasm | other | 1.64E+07 | 7070846.5 | 7757456 | | | | | | |
| adducin 1 (alpha) | Cytoplasm | other | 1.51E+07 | 4247262.5 | 6222729.5 | | | | | | |
| adducin 1 (alpha) | Cytoplasm | other | 1.39E+07 | 9267694 | 9913244 | | | | | | |
| adducin 1 (alpha) | Cytoplasm | other | 1.08E+07 | 5794030 | 9764426 | | | | | | |
| adducin 1 (alpha) | Cytoplasm | other | 1.03E+07 | 4575124.5 | 4915852.5 | | | | | | |
| adducin 1 (alpha) | Cytoplasm | other | 1.03E+07 | 2.21E+07 | 1.75E+07 | | | | | | |
| adducin 1 (alpha) | Cytoplasm | other | 9782085 | 6856424 | 6061951.5 | | | | | | |
| adducin 1 (alpha) | Cytoplasm | other | 8827183 | 4989335.5 | 5963299 | | | | | | |
| adducin 1 (alpha) | Cytoplasm | other | 6195093 | 1.84E+07 | 1.41E+07 | | | | | | |
| adducin 1 (alpha) | Cytoplasm | other | 2506401.5 | 475304.031 | 1678323.5 | | | | | | |
| adducin 1 (alpha) | Cytoplasm | other | 450182.062 | 330053.375 | 379252.719 | Q9QYC0 | ADD1 | 185462950 | 205786193 | 18555116 | 123 |
| adducin 2 (beta) | Cytoplasm | other | 5.24E+08 | 4.24E+08 | 3.74E+08 | | | | | | |
| adducin 2 (beta) | Cytoplasm | other | 4.67E+08 | 3.27E+08 | 3.78E+08 | | | | | | |
| adducin 2 (beta) | Cytoplasm | other | 4.45E+08 | 3.62E+08 | 3.40E+08 | | | | | | |
| adducin 2 (beta) | Cytoplasm | other | 4.26E+08 | 4.17E+08 | 4.66E+08 | | | | | | |
| adducin 2 (beta) | Cytoplasm | other | 3.44E+08 | 2.31E+08 | 2.39E+08 | | | | | | |
| adducin 2 (beta) | Cytoplasm | other | 3.06E+08 | 2.51E+08 | 2.49E+08 | | | | | | |
| adducin 2 (beta) | Cytoplasm | other | 2.86E+08 | 2.93E+08 | 2.04E+08 | | | | | | |
| adducin 2 (beta) | Cytoplasm | other | 2.77E+08 | 3.87E+08 | 3.12E+08 | | | | | | |
| adducin 2 (beta) | Cytoplasm | other | 2.70E+08 | 2.47E+08 | 2.35E+08 | | | | | | |
| adducin 2 (beta) | Cytoplasm | other | 2.66E+08 | 2.46E+08 | 2.66E+08 | | | | | | |
| adducin 2 (beta) | Cytoplasm | other | 2.45E+08 | 1.79E+08 | 1.78E+08 | | | | | | |
| adducin 2 (beta) | Cytoplasm | other | 2.45E+08 | 1.40E+08 | 1.62E+08 | | | | | | |
| adducin 2 (beta) | Cytoplasm | other | 2.28E+08 | 1.79E+08 | 1.68E+08 | | | | | | |
| adducin 2 (beta) | Cytoplasm | other | 2.24E+08 | 2.34E+08 | 2.02E+08 | | | | | | |
| adducin 2 (beta) | Cytoplasm | other | 2.14E+08 | 1.55E+08 | 1.51E+08 | | | | | | |
| adducin 2 (beta) | Cytoplasm | other | 1.70E+08 | 2.12E+08 | 1.89E+08 | | | | | | |
| adducin 2 (beta) | Cytoplasm | other | 1.69E+08 | 1.78E+08 | 1.72E+08 | | | | | | |
| adducin 2 (beta) | Cytoplasm | other | 1.52E+08 | 1.05E+08 | 9.73E+07 | | | | | | |
| adducin 2 (beta) | Cytoplasm | other | 9.29E+07 | 4.65E+07 | 5.48E+07 | | | | | | |
| adducin 2 (beta) | Cytoplasm | other | 8.66E+07 | 7.86E+07 | 6.99E+07 | | | | | | |
| adducin 2 (beta) | Cytoplasm | other | 8.29E+07 | 4.11E+07 | 4.95E+07 | | | | | | |
| adducin 2 (beta) | Cytoplasm | other | 8.12E+07 | 1.23E+08 | 9.09E+07 | | | | | | |
| adducin 2 (beta) | Cytoplasm | other | 7.59E+07 | 4.55E+07 | 5.45E+07 | | | | | | |
| adducin 2 (beta) | Cytoplasm | other | 7.23E+07 | 4.62E+07 | 5.01E+07 | | | | | | |
| adducin 2 (beta) | Cytoplasm | other | 7.10E+07 | 1.38E+08 | 1.60E+08 | | | | | | |
| adducin 2 (beta) | Cytoplasm | other | 6.35E+07 | 3.24E+07 | 2.55E+07 | | | | | | |
| adducin 2 (beta) | Cytoplasm | other | 6.25E+07 | 5.19E+07 | 4.24E+07 | | | | | | |
| adducin 2 (beta) | Cytoplasm | other | 5.89E+07 | 2.74E+07 | 4.45E+07 | | | | | | |
| adducin 2 (beta) | Cytoplasm | other | 5.64E+07 | 4.07E+07 | 4.05E+07 | | | | | | |
| adducin 2 (beta) | Cytoplasm | other | 5.22E+07 | 3.74E+07 | 3.37E+07 | | | | | | |
| adducin 2 (beta) | Cytoplasm | other | 4.53E+07 | 3.77E+07 | 2.93E+07 | | | | | | |
| adducin 2 (beta) | Cytoplasm | other | 4.39E+07 | 4.18E+07 | 2.58E+07 | | | | | | |
| adducin 2 (beta) | Cytoplasm | other | 4.36E+07 | 2.50E+07 | 2.34E+07 | | | | | | |
| adducin 2 (beta) | Cytoplasm | other | 2.56E+07 | 1.10E+07 | 1.81E+07 | | | | | | |
| adducin 2 (beta) | Cytoplasm | other | 2.43E+07 | 1.18E+07 | 1.53E+07 | | | | | | |
| adducin 2 (beta) | Cytoplasm | other | 2.31E+07 | 5.01E+07 | 3.86E+07 | | | | | | |
| adducin 2 (beta) | Cytoplasm | other | 1.94E+07 | 1.99E+07 | 1.47E+07 | | | | | | |
| adducin 2 (beta) | Cytoplasm | other | 1.81E+07 | 8984054 | 1.44E+07 | | | | | | |
| adducin 2 (beta) | Cytoplasm | other | 1.70E+07 | 8400384 | 1.36E+07 | | | | | | |
| adducin 2 (beta) | Cytoplasm | other | 1.65E+07 | 1.93E+07 | 1.31E+07 | | | | | | |
| adducin 2 (beta) | Cytoplasm | other | 1.41E+07 | 3.63E+07 | 1.91E+07 | | | | | | |
| adducin 2 (beta) | Cytoplasm | other | 1.37E+07 | 1.47E+07 | 1.88E+07 | | | | | | |
| adducin 2 (beta) | Cytoplasm | other | 1.19E+07 | 7019777 | 1.81E+07 | | | | | | |
| adducin 2 (beta) | Cytoplasm | other | 1.17E+07 | 3166261 | 3062084.25 | | | | | | |
| adducin 2 (beta) | Cytoplasm | other | 8317667 | 4748305 | 6950055.5 | | | | | | |
| adducin 2 (beta) | Cytoplasm | other | 7984389.5 | 3632477.75 | 3759388.25 | | | | | | |
| adducin 2 (beta) | Cytoplasm | other | 7975272.5 | 4957180 | 5916106 | | | | | | |
| adducin 2 (beta) | Cytoplasm | other | 7216321 | 5942762.5 | 9571605 | | | | | | |
| adducin 2 (beta) | Cytoplasm | other | 6758989.5 | 3850219.5 | 5086567.5 | | | | | | |
| adducin 2 (beta) | Cytoplasm | other | 6713293.5 | 3952921.75 | 4645687 | | | | | | |
| adducin 2 (beta) | Cytoplasm | other | 6036215 | 3259440.25 | 3167054.25 | | | | | | |

| | | | | | | | | | | | |
|---|---|---|---|---|---|---|---|---|---|---|---|
| adducin 2 (beta) | Cytoplasm | other | 5898609.5 | 2.65E+07 | 1.11E+07 | | | | | | |
| adducin 2 (beta) | Cytoplasm | other | 5430309 | 3208290.75 | 4387788 | | | | | | |
| adducin 2 (beta) | Cytoplasm | other | 4876467.5 | 2660897 | 6402035.5 | | | | | | |
| adducin 2 (beta) | Cytoplasm | other | 4127209.75 | 2133150.25 | 2735792.5 | | | | | | |
| adducin 2 (beta) | Cytoplasm | other | 2659328.5 | 745082.188 | 2032200 | | | | | | |
| adducin 2 (beta) | Cytoplasm | other | 2478657.75 | 6998641 | 3067526.25 | | | | | | |
| adducin 2 (beta) | Cytoplasm | other | 1765985.625 | 311202.531 | 703864 | | | | | | |
| adducin 2 (beta) | Cytoplasm | other | 1733642.625 | 417926.062 | 3327573 | | | | | | |
| adducin 2 (beta) | Cytoplasm | other | 1685168.5 | 1487890.25 | 3729873.5 | | | | | | |
| adducin 2 (beta) | Cytoplasm | other | 991777.625 | 743449.062 | 676735.5 | | | | | | |
| adducin 2 (beta) | Cytoplasm | other | 957468.25 | 2988592.75 | 946930.438 | | | | | | |
| adducin 2 (beta) | Cytoplasm | other | 507488.656 | 359379.594 | 542831.438 | | | | | | |
| adducin 2 (beta) | Cytoplasm | other | 260716.766 | 3042231.5 | 3559786.25 | Q9QYB8 | ADD2 | 102009609 | 122359875 | 8830563 | 192 |
| adducin 3 (gamma) | Cytoplasm | other | 3.54E+07 | 2.71E+07 | 2.66E+07 | | | | | | |
| adducin 3 (gamma) | Cytoplasm | other | 1.80E+07 | 1.49E+07 | 1.34E+07 | | | | | | |
| adducin 3 (gamma) | Cytoplasm | other | 1.39E+07 | 1.26E+07 | 1.06E+07 | | | | | | |
| adducin 3 (gamma) | Cytoplasm | other | 1.15E+07 | 8988071 | 9511769 | | | | | | |
| adducin 3 (gamma) | Cytoplasm | other | 8632264 | 7275487.5 | 9351952 | | | | | | |
| adducin 3 (gamma) | Cytoplasm | other | 8527703 | 2911510.25 | 3404513.5 | | | | | | |
| adducin 3 (gamma) | Cytoplasm | other | 6598790 | 6447428.5 | 6301695.5 | | | | | | |
| adducin 3 (gamma) | Cytoplasm | other | 6507442.5 | 6694731 | 7217508.5 | | | | | | |
| adducin 3 (gamma) | Cytoplasm | other | 6366094 | 5275063.5 | 5530015 | | | | | | |
| adducin 3 (gamma) | Cytoplasm | other | 4585243 | 4317940.5 | 3866623.75 | | | | | | |
| adducin 3 (gamma) | Cytoplasm | other | 3848343.75 | 4610270.5 | 5591070 | | | | | | |
| adducin 3 (gamma) | Cytoplasm | other | 3703648 | 1892654.5 | 2206395.5 | | | | | | |
| adducin 3 (gamma) | Cytoplasm | other | 2179204.75 | 1025902.125 | 1287372.875 | | | | | | |
| adducin 3 (gamma) | Cytoplasm | other | 2130323.25 | 3971002.5 | 4030784.25 | | | | | | |
| adducin 3 (gamma) | Cytoplasm | other | 1796781.875 | 3127425 | 2554823.75 | Q9QYB5 | ADD3 | 8911056 | 7142873 | 1064797 | 45 |
| ADP-dependent glucokinase | Plasma Membrane | kinase | 3746707.75 | 3600499.5 | 2576452.75 | Q8VDL4 | ADPGK | 3746708 | 637645 | 368144 | 3 |
| adenylosuccinate lyase | Cytoplasm | enzyme | 2.69E+07 | 8982071 | 1.26E+07 | | | | | | |
| adenylosuccinate lyase | Cytoplasm | enzyme | 1.94E+07 | 6020457 | 9157055 | | | | | | |
| adenylosuccinate lyase | Cytoplasm | enzyme | 1.60E+07 | 7223946.5 | 7899124 | | | | | | |
| adenylosuccinate lyase | Cytoplasm | enzyme | 1.60E+07 | 8422015 | 8769335 | | | | | | |
| adenylosuccinate lyase | Cytoplasm | enzyme | 1.52E+07 | 6611804.5 | 9913820 | | | | | | |
| adenylosuccinate lyase | Cytoplasm | enzyme | 1.44E+07 | 5028895 | 6638204.5 | | | | | | |
| adenylosuccinate lyase | Cytoplasm | enzyme | 1.27E+07 | 3742343.5 | 4904946 | | | | | | |
| adenylosuccinate lyase | Cytoplasm | enzyme | 1.15E+07 | 4524961 | 6360563 | | | | | | |
| adenylosuccinate lyase | Cytoplasm | enzyme | 7937622 | 2074342.625 | 3535171.25 | | | | | | |
| adenylosuccinate lyase | Cytoplasm | enzyme | 4504508.5 | 2963624 | 2752853.25 | | | | | | |
| adenylosuccinate lyase | Cytoplasm | enzyme | 3762836.5 | 1062846.5 | 2406781 | | | | | | |
| adenylosuccinate lyase | Cytoplasm | enzyme | 1759404.5 | 622475.875 | 585427.25 | | | | | | |
| adenylosuccinate lyase | Cytoplasm | enzyme | 1567349.75 | 496321.75 | 1176138.25 | P54822 | ADSL | 11667825 | 5898748 | 944556 | 39 |
| AE binding protein 1 | Nucleus | peptidase | 1.46E+07 | 9319409 | 9096581 | Q640N1 | AEBP1 | 14610000 | 3120838 | 1801817 | 3 |
| AF4/FMR2 family, member 2 | Nucleus | other | 3.32E+07 | 3.71E+07 | 3.37E+07 | O55112 | Aff2 | 33240000 | 2095718 | 1209963 | 3 |
| ArfGAP with FG repeats 1 | Nucleus | other | 1625057 | 1819505.875 | 2264383.5 | Q8K2K6 | AGFG1 | 1625057 | 327736 | 189218 | 3 |
| ArfGAP with FG repeats 2 | Cytoplasm | other | 3801892.25 | 3644136 | 3766827 | | | | | | |
| ArfGAP with FG repeats 2 | Cytoplasm | other | 2916000 | 1395985.5 | 1582251.625 | | | | | | |
| ArfGAP with FG repeats 2 | Cytoplasm | other | 2129671 | 2405574.25 | 1993957.625 | | | | | | |
| ArfGAP with FG repeats 2 | Cytoplasm | other | 2023472.875 | 2142645.75 | 2907607.5 | Q80WC7 | AGFG2 | 2717759 | 837965 | 241900 | 12 |
| argonaute RISC catalytic component 2 | Cytoplasm | translation regulator | 4.76E+07 | 4.50E+07 | 4.16E+07 | | | | | | |
| argonaute RISC catalytic component 2 | Cytoplasm | translation regulator | 4.41E+07 | 3.71E+07 | 2.86E+07 | | | | | | |
| argonaute RISC catalytic component 2 | Cytoplasm | translation regulator | 4.08E+07 | 3.19E+07 | 2.92E+07 | | | | | | |
| argonaute RISC catalytic component 2 | Cytoplasm | translation regulator | 3.54E+07 | 2.90E+07 | 2.44E+07 | | | | | | |
| argonaute RISC catalytic component 2 | Cytoplasm | translation regulator | 3.53E+07 | 2.91E+07 | 2.44E+07 | | | | | | |
| argonaute RISC catalytic component 2 | Cytoplasm | translation regulator | 3.14E+07 | 3.60E+07 | 2.63E+07 | | | | | | |
| argonaute RISC catalytic component 2 | Cytoplasm | translation regulator | 3.12E+07 | 2.85E+07 | 2.57E+07 | | | | | | |
| argonaute RISC catalytic component 2 | Cytoplasm | translation regulator | 3.12E+07 | 3.55E+07 | 2.22E+07 | | | | | | |
| argonaute RISC catalytic component 2 | Cytoplasm | translation regulator | 3.04E+07 | 2.70E+07 | 2.03E+07 | | | | | | |
| argonaute RISC catalytic component 2 | Cytoplasm | translation regulator | 2.91E+07 | 2.60E+07 | 2.30E+07 | | | | | | |
| argonaute RISC catalytic component 2 | Cytoplasm | translation regulator | 2.89E+07 | 2.22E+07 | 2.26E+07 | | | | | | |
| argonaute RISC catalytic component 2 | Cytoplasm | translation regulator | 2.88E+07 | 1.34E+07 | 1.42E+07 | | | | | | |
| argonaute RISC catalytic component 2 | Cytoplasm | translation regulator | 2.84E+07 | 2.60E+07 | 1.93E+07 | | | | | | |
| argonaute RISC catalytic component 2 | Cytoplasm | translation regulator | 2.71E+07 | 2.54E+07 | 2.35E+07 | | | | | | |
| argonaute RISC catalytic component 2 | Cytoplasm | translation regulator | 2.71E+07 | 2.58E+07 | 2.04E+07 | | | | | | |
| argonaute RISC catalytic component 2 | Cytoplasm | translation regulator | 2.69E+07 | 3.05E+07 | 2.29E+07 | | | | | | |
| argonaute RISC catalytic component 2 | Cytoplasm | translation regulator | 2.32E+07 | 3.39E+07 | 2.24E+07 | | | | | | |
| argonaute RISC catalytic component 2 | Cytoplasm | translation regulator | 2.27E+07 | 2.17E+07 | 1.92E+07 | | | | | | |
| argonaute RISC catalytic component 2 | Cytoplasm | translation regulator | 2.19E+07 | 1.71E+07 | 1.50E+07 | | | | | | |
| argonaute RISC catalytic component 2 | Cytoplasm | translation regulator | 2.17E+07 | 2.76E+07 | 1.30E+07 | | | | | | |
| argonaute RISC catalytic component 2 | Cytoplasm | translation regulator | 2.05E+07 | 2.26E+07 | 2.16E+07 | | | | | | |
| argonaute RISC catalytic component 2 | Cytoplasm | translation regulator | 1.90E+07 | 1.41E+07 | 1.09E+07 | | | | | | |
| argonaute RISC catalytic component 2 | Cytoplasm | translation regulator | 1.79E+07 | 2.14E+07 | 1.67E+07 | | | | | | |
| argonaute RISC catalytic component 2 | Cytoplasm | translation regulator | 1.53E+07 | 1.03E+07 | 1.07E+07 | | | | | | |
| argonaute RISC catalytic component 2 | Cytoplasm | translation regulator | 1.35E+07 | 1.29E+07 | 8201479.5 | | | | | | |
| argonaute RISC catalytic component 2 | Cytoplasm | translation regulator | 1.17E+07 | 8040969.5 | 7986389 | | | | | | |
| argonaute RISC catalytic component 2 | Cytoplasm | translation regulator | 1.11E+07 | 1.23E+07 | 8196825 | | | | | | |
| argonaute RISC catalytic component 2 | Cytoplasm | translation regulator | 9520230 | 1.20E+07 | 8473320 | | | | | | |
| argonaute RISC catalytic component 2 | Cytoplasm | translation regulator | 8853915 | 7622937 | 7446595.5 | | | | | | |
| argonaute RISC catalytic component 2 | Cytoplasm | translation regulator | 8229370 | 1.38E+07 | 7571037.5 | | | | | | |
| argonaute RISC catalytic component 2 | Cytoplasm | translation regulator | 7299680 | 1.05E+07 | 5858867.5 | | | | | | |
| argonaute RISC catalytic component 2 | Cytoplasm | translation regulator | 6135868 | 4372480 | 4621622.5 | | | | | | |
| argonaute RISC catalytic component 2 | Cytoplasm | translation regulator | 6072311 | 5600660 | 4974287 | | | | | | |
| argonaute RISC catalytic component 2 | Cytoplasm | translation regulator | 5513668 | 2240988.25 | 1846807.75 | | | | | | |
| argonaute RISC catalytic component 2 | Cytoplasm | translation regulator | 5479651.5 | 3169363.5 | 5268996.5 | | | | | | |
| argonaute RISC catalytic component 2 | Cytoplasm | translation regulator | 5460079 | 4799094 | 2488786.75 | | | | | | |
| argonaute RISC catalytic component 2 | Cytoplasm | translation regulator | 3561325.75 | 1233593.75 | 1751315 | | | | | | |
| argonaute RISC catalytic component 2 | Cytoplasm | translation regulator | 1326818.625 | 1.07E+07 | 1.37E+07 | | | | | | |
| argonaute RISC catalytic component 2 | Cytoplasm | translation regulator | 1189507.875 | 429996.062 | 703394.938 | | | | | | |
| argonaute RISC catalytic component 2 | Cytoplasm | translation regulator | 772105.312 | 3023016.5 | 1321138.5 | | | | | | |
| argonaute RISC catalytic component 2 | Cytoplasm | translation regulator | 232148.516 | 32254.23 | 107250.141 | Q8CJG0 | AGO2 | 19310407 | 11738916 | 1058462 | 123 |
| 1-acylglycerol-3-phosphate O-acyltransferase 1 | Cytoplasm | enzyme | 1239139.375 | 1066633.875 | 765393.438 | O35083 | AGPAT1 | 1239139 | 239770 | 138432 | 3 |
| adenosylhomocysteinase | Cytoplasm | enzyme | 5960483 | 2533452.25 | 2782526.5 | | | | | | |
| adenosylhomocysteinase | Cytoplasm | enzyme | 4111328.75 | 1982442.625 | 2168503.5 | | | | | | |
| adenosylhomocysteinase | Cytoplasm | enzyme | 3509362.25 | 1328159.25 | 1927579.375 | P50247 | AHCY | 4527058 | 1420476 | 473492 | 9 |
| AHA1, activator of heat shock protein ATPase 2 | Cytoplasm | other | 2.96E+08 | 2.30E+08 | 2.61E+08 | Q8N9S3 | Ahsa2 | 295800000 | 32929166 | 19011663 | 3 |

| | | | | | | | | | | | |
|---|---|---|---|---|---|---|---|---|---|---|---|
| apoptosis-inducing factor, mitochondrion-associated, 1 | Cytoplasm | enzyme | 906296.125 | 1925884.25 | 1396864 | | | | | | |
| apoptosis-inducing factor, mitochondrion-associated, 1 | Cytoplasm | enzyme | 717068.125 | 1622229.625 | 1165800.625 | Q9Z0X1 | AIFM1 | 811682 | 451009 | 184124 | 6 |
| aminoacyl tRNA synthetase complex-interacting multifunctional protein 1 | Extracellular Space | cytokine | 2697913 | 2742310.5 | 2051936.5 | | | | | | |
| aminoacyl tRNA synthetase complex-interacting multifunctional protein 1 | Extracellular Space | cytokine | 615168.625 | 142600.031 | 1246771.625 | P31230 | AIMP1 | 1656541 | 1089168 | 444651 | 6 |
| adenylate kinase 2 | Cytoplasm | kinase | 4.70E-06 | 705672.312 | 605875.438 | Q9WTP6 | AK2 | 0 | 381885 | 220482 | 3 |
| aminolevulinate dehydratase | Cytoplasm | enzyme | 1.18E+08 | 4.81E+07 | 5.60E+07 | | | | | | |
| aminolevulinate dehydratase | Cytoplasm | enzyme | 5.93E+07 | 2.64E+07 | 2.73E+07 | | | | | | |
| aminolevulinate dehydratase | Cytoplasm | enzyme | 5.45E+07 | 4.13E+07 | 3.96E+07 | | | | | | |
| aminolevulinate dehydratase | Cytoplasm | enzyme | 3.72E+07 | 1.60E+07 | 1.60E+07 | | | | | | |
| aminolevulinate dehydratase | Cytoplasm | enzyme | 3.67E+07 | 1.83E+07 | 2.09E+07 | | | | | | |
| aminolevulinate dehydratase | Cytoplasm | enzyme | 2.42E+07 | 1.02E+07 | 1.24E+07 | | | | | | |
| aminolevulinate dehydratase | Cytoplasm | enzyme | 1.90E+07 | 1.00E+07 | 1.06E+07 | | | | | | |
| aminolevulinate dehydratase | Cytoplasm | enzyme | 8719543 | 6795203 | 7855070.5 | | | | | | |
| aminolevulinate dehydratase | Cytoplasm | enzyme | 7533070 | 4210392.5 | 4336833 | | | | | | |
| aminolevulinate dehydratase | Cytoplasm | enzyme | 7241022 | 1757627.875 | 2140046.5 | | | | | | |
| aminolevulinate dehydratase | Cytoplasm | enzyme | 6716628 | 2238078.75 | 5300203.5 | | | | | | |
| aminolevulinate dehydratase | Cytoplasm | enzyme | 6433252 | 3363960.5 | 3806385.5 | | | | | | |
| aminolevulinate dehydratase | Cytoplasm | enzyme | 5202240 | 2125215 | 2173539.75 | | | | | | |
| aminolevulinate dehydratase | Cytoplasm | enzyme | 4498152.5 | 1203986.375 | 3551074.75 | | | | | | |
| aminolevulinate dehydratase | Cytoplasm | enzyme | 3624640 | 1719615.625 | 2427653.75 | | | | | | |
| aminolevulinate dehydratase | Cytoplasm | enzyme | 1313853.875 | 348171.031 | 515427.062 | P10518 | ALAD | 24999525 | 22082069 | 3187272 | 48 |
| 5'-aminolevulinate synthase 2 | Cytoplasm | enzyme | 2.13E+07 | 1.24E+07 | 1.75E+07 | | | | | | |
| 5'-aminolevulinate synthase 2 | Cytoplasm | enzyme | 1.41E+07 | 1.74E+07 | 1.16E+07 | | | | | | |
| 5'-aminolevulinate synthase 2 | Cytoplasm | enzyme | 7552990 | 7560856 | 8683060 | | | | | | |
| 5'-aminolevulinate synthase 2 | Cytoplasm | enzyme | 6721779 | 6287320.5 | 5908309.5 | | | | | | |
| 5'-aminolevulinate synthase 2 | Cytoplasm | enzyme | 5792800 | 4763679.5 | 5404841 | | | | | | |
| 5'-aminolevulinate synthase 2 | Cytoplasm | enzyme | 4334162 | 2823158 | 5293036.5 | | | | | | |
| 5'-aminolevulinate synthase 2 | Cytoplasm | enzyme | 4061077.25 | 5181741 | 5240742.5 | | | | | | |
| 5'-aminolevulinate synthase 2 | Cytoplasm | enzyme | 3984554 | 4152950.25 | 5193166 | | | | | | |
| 5'-aminolevulinate synthase 2 | Cytoplasm | enzyme | 3460576.5 | 3504414.75 | 2750017.5 | | | | | | |
| 5'-aminolevulinate synthase 2 | Cytoplasm | enzyme | 2707139 | 2666190.5 | 3366704 | | | | | | |
| 5'-aminolevulinate synthase 2 | Cytoplasm | enzyme | 1600389.25 | 2571206.75 | 1716730.625 | | | | | | |
| 5'-aminolevulinate synthase 2 | Cytoplasm | enzyme | 263841.281 | 636090.625 | 855069.562 | P08680 | ALAS2 | 6324109 | 4975874 | 829312 | 36 |
| albumin | Extracellular Space | transporter | 4.89E+07 | 4.99E+07 | 6.40E+07 | | | | | | |
| albumin | Extracellular Space | transporter | 1.37E+07 | 2.00E+07 | 2.84E+07 | | | | | | |
| albumin | Extracellular Space | transporter | 1.31E+07 | 1.38E+07 | 1.65E+07 | | | | | | |
| albumin | Extracellular Space | transporter | 1.28E+07 | 1.39E+07 | 1.77E+07 | | | | | | |
| albumin | Extracellular Space | transporter | 1.20E+07 | 1.40E+07 | 2.11E+07 | | | | | | |
| albumin | Extracellular Space | transporter | 1.20E+07 | 1.48E+07 | 2.29E+07 | | | | | | |
| albumin | Extracellular Space | transporter | 1.16E+07 | 1.47E+07 | 2.03E+07 | | | | | | |
| albumin | Extracellular Space | transporter | 1.15E+07 | 1.57E+07 | 1.81E+07 | | | | | | |
| albumin | Extracellular Space | transporter | 1.15E+07 | 1.37E+07 | 1.86E+07 | | | | | | |
| albumin | Extracellular Space | transporter | 1.01E+07 | 1.38E+07 | 1.81E+07 | | | | | | |
| albumin | Extracellular Space | transporter | 7335768.5 | 9598319 | 1.42E+07 | | | | | | |
| albumin | Extracellular Space | transporter | 6938179.5 | 2.00E+07 | 1.60E+07 | | | | | | |
| albumin | Extracellular Space | transporter | 6776666.5 | 5522440 | 9498417 | | | | | | |
| albumin | Extracellular Space | transporter | 6456616.5 | 5517713 | 8984633 | | | | | | |
| albumin | Extracellular Space | transporter | 6023057 | 8512073 | 1.20E+07 | | | | | | |
| albumin | Extracellular Space | transporter | 5531714.5 | 6172495 | 8105053.5 | | | | | | |
| albumin | Extracellular Space | transporter | 4462478.5 | 6780935 | 6800894.5 | | | | | | |
| albumin | Extracellular Space | transporter | 3733531.25 | 2567962 | 5161758.5 | | | | | | |
| albumin | Extracellular Space | transporter | 3067810.25 | 2960444.25 | 4372473.5 | | | | | | |
| albumin | Extracellular Space | transporter | 1942190.875 | 1772486.375 | 2626302 | | | | | | |
| albumin | Extracellular Space | transporter | 1566063 | 3017969 | 4784011.5 | | | | | | |
| albumin | Extracellular Space | transporter | 1228943.125 | 2222567.25 | 3563810 | | | | | | |
| albumin | Extracellular Space | transporter | 1019089.125 | 1126841.625 | 2104571.75 | | | | | | |
| albumin | Extracellular Space | transporter | 974516.312 | 1029162.938 | 1305587.375 | P07724 | ALB | 8926943 | 11124638 | 1311051 | 72 |
| aldehyde dehydrogenase 16 family, member A1 | Cytoplasm | enzyme | 9407830 | 5269685 | 6095464 | | | | | | |
| aldehyde dehydrogenase 16 family, member A1 | Cytoplasm | enzyme | 8774699 | 3781892.5 | 4752956.5 | | | | | | |
| aldehyde dehydrogenase 16 family, member A1 | Cytoplasm | enzyme | 5376249 | 2684014.5 | 3622319.25 | | | | | | |
| aldehyde dehydrogenase 16 family, member A1 | Cytoplasm | enzyme | 3556653.25 | 898396.938 | 1029037.062 | | | | | | |
| aldehyde dehydrogenase 16 family, member A1 | Cytoplasm | enzyme | 3008388.5 | 2166104.5 | 1720827.75 | | | | | | |
| aldehyde dehydrogenase 16 family, member A1 | Cytoplasm | enzyme | 1446168.375 | 874939.562 | 1016785.188 | Q571I9 | ALDH16A1 | 5261665 | 2583208 | 608868 | 18 |
| aldehyde dehydrogenase 18 family, member A1 | Cytoplasm | kinase | 1932871 | 2134524.5 | 2047020.25 | | | | | | |
| aldehyde dehydrogenase 18 family, member A1 | Cytoplasm | kinase | 1350791.25 | 1393137.25 | 1777416.25 | | | | | | |
| aldehyde dehydrogenase 18 family, member A1 | Cytoplasm | kinase | 597806.375 | 614185.562 | 974219.812 | | | | | | |
| aldehyde dehydrogenase 18 family, member A1 | Cytoplasm | kinase | 593734.938 | 795714.125 | 1238661.5 | | | | | | |
| aldehyde dehydrogenase 18 family, member A1 | Cytoplasm | kinase | 592074.312 | 903818.188 | 717198 | | | | | | |
| aldehyde dehydrogenase 18 family, member A1 | Cytoplasm | kinase | 500495.906 | 517038.281 | 886087.562 | | | | | | |
| aldehyde dehydrogenase 18 family, member A1 | Cytoplasm | kinase | 253642.75 | 531851.375 | 590211.125 | | | | | | |
| aldehyde dehydrogenase 18 family, member A1 | Cytoplasm | kinase | 243240.547 | 500540.812 | 673051.062 | Q9Z110 | ALDH18A1 | 758082 | 560397 | 114391 | 24 |
| aldehyde dehydrogenase 1 family, member A1 | Cytoplasm | enzyme | 1529391.875 | 2153495 | 1359030.125 | P24549 | ALDH1A1 | 1529392 | 418271 | 241489 | 3 |
| aldehyde dehydrogenase 2 family (mitochondrial) | Cytoplasm | enzyme | 5501315.5 | 4.08E+07 | 3.46E+07 | | | | | | |
| aldehyde dehydrogenase 2 family (mitochondrial) | Cytoplasm | enzyme | 3337608 | 3.87E+07 | 3.09E+07 | | | | | | |
| aldehyde dehydrogenase 2 family (mitochondrial) | Cytoplasm | enzyme | 3025424.75 | 3.26E+07 | 1.42E+07 | | | | | | |
| aldehyde dehydrogenase 2 family (mitochondrial) | Cytoplasm | enzyme | 2654956 | 3.43E+07 | 2.56E+07 | | | | | | |
| aldehyde dehydrogenase 2 family (mitochondrial) | Cytoplasm | enzyme | 2635756.5 | 2.70E+07 | 2.08E+07 | | | | | | |
| aldehyde dehydrogenase 2 family (mitochondrial) | Cytoplasm | enzyme | 2575757.5 | 1.89E+07 | 1.30E+07 | | | | | | |
| aldehyde dehydrogenase 2 family (mitochondrial) | Cytoplasm | enzyme | 2517476.5 | 2.86E+07 | 2.24E+07 | | | | | | |
| aldehyde dehydrogenase 2 family (mitochondrial) | Cytoplasm | enzyme | 2325613.75 | 2.78E+07 | 1.93E+07 | | | | | | |
| aldehyde dehydrogenase 2 family (mitochondrial) | Cytoplasm | enzyme | 1989173.75 | 1.75E+07 | 1.26E+07 | | | | | | |
| aldehyde dehydrogenase 2 family (mitochondrial) | Cytoplasm | enzyme | 1804288.375 | 1.84E+07 | 1.19E+07 | | | | | | |
| aldehyde dehydrogenase 2 family (mitochondrial) | Cytoplasm | enzyme | 1561160.625 | 1.73E+07 | 1.21E+07 | | | | | | |
| aldehyde dehydrogenase 2 family (mitochondrial) | Cytoplasm | enzyme | 1514682.25 | 1.71E+07 | 9030663 | | | | | | |
| aldehyde dehydrogenase 2 family (mitochondrial) | Cytoplasm | enzyme | 1493524.25 | 6641435.5 | 6752009 | | | | | | |
| aldehyde dehydrogenase 2 family (mitochondrial) | Cytoplasm | enzyme | 1379940.75 | 2.21E+07 | 1.69E+07 | | | | | | |
| aldehyde dehydrogenase 2 family (mitochondrial) | Cytoplasm | enzyme | 1357681.5 | 1.58E+07 | 1.40E+07 | | | | | | |
| aldehyde dehydrogenase 2 family (mitochondrial) | Cytoplasm | enzyme | 1228324.625 | 2.35E+07 | 1.74E+07 | | | | | | |
| aldehyde dehydrogenase 2 family (mitochondrial) | Cytoplasm | enzyme | 1095043.25 | 1.23E+07 | 9999472 | | | | | | |
| aldehyde dehydrogenase 2 family (mitochondrial) | Cytoplasm | enzyme | 1055774.625 | 1.27E+07 | 1.10E+07 | | | | | | |
| aldehyde dehydrogenase 2 family (mitochondrial) | Cytoplasm | enzyme | 499835.188 | 1940686.5 | 1901733.125 | | | | | | |
| aldehyde dehydrogenase 2 family (mitochondrial) | Cytoplasm | enzyme | 467107.688 | 5194839 | 6716243.5 | | | | | | |
| aldehyde dehydrogenase 2 family (mitochondrial) | Cytoplasm | enzyme | 460092.188 | 9065078 | 7182766 | | | | | | |
| aldehyde dehydrogenase 2 family (mitochondrial) | Cytoplasm | enzyme | 302654.125 | 6010203 | 4416328 | | | | | | |
| aldehyde dehydrogenase 2 family (mitochondrial) | Cytoplasm | enzyme | 234433.359 | 7903440 | 5279837.5 | | | | | | |

| | | | | | | | | | | | |
|---|---|---|---|---|---|---|---|---|---|---|---|
| aldehyde dehydrogenase 2 family (mitochondrial) | Cytoplasm | enzyme | 185165.953 | 3263537.5 | 2114388.75 | P47738 | ALDH2 | 1716783 | 10772207 | 1269517 | 72 |
| aldolase A, fructose-bisphosphate | Cytoplasm | enzyme | 8.16E+07 | 4.29E+07 | 7.65E+07 | | | | | | |
| aldolase A, fructose-bisphosphate | Cytoplasm | enzyme | 5.25E+07 | 2.84E+07 | 4.62E+07 | | | | | | |
| aldolase A, fructose-bisphosphate | Cytoplasm | enzyme | 4.14E+07 | 1.98E+07 | 4.19E+07 | | | | | | |
| aldolase A, fructose-bisphosphate | Cytoplasm | enzyme | 3.04E+07 | 2.70E+07 | 3.66E+07 | | | | | | |
| aldolase A, fructose-bisphosphate | Cytoplasm | enzyme | 2.96E+07 | 1.21E+07 | 2.54E+07 | | | | | | |
| aldolase A, fructose-bisphosphate | Cytoplasm | enzyme | 1.56E+07 | 1.04E+07 | 1.39E+07 | | | | | | |
| aldolase A, fructose-bisphosphate | Cytoplasm | enzyme | 7084264 | 1956073.125 | 4889470.5 | | | | | | |
| aldolase A, fructose-bisphosphate | Cytoplasm | enzyme | 6711615.5 | 2583487 | 4890201 | | | | | | |
| aldolase A, fructose-bisphosphate | Cytoplasm | enzyme | 6010974.5 | 2899048.25 | 4200706 | | | | | | |
| aldolase A, fructose-bisphosphate | Cytoplasm | enzyme | 3898916.5 | 4049383 | 4836714.5 | | | | | | |
| aldolase A, fructose-bisphosphate | Cytoplasm | enzyme | 3408516.25 | 1676495.625 | 2562654.25 | | | | | | |
| aldolase A, fructose-bisphosphate | Cytoplasm | enzyme | 1078425.125 | 45367.453 | 642281.312 | P05064 | ALDOA | 23281059 | 21363857 | 3560643 | 36 |
| adenosine monophosphate deaminase 3 | Cytoplasm | enzyme | 3786270 | 3341985 | 3212842 | | | | | | |
| adenosine monophosphate deaminase 3 | Cytoplasm | enzyme | 3689355 | 1865984.875 | 4074323.5 | | | | | | |
| adenosine monophosphate deaminase 3 | Cytoplasm | enzyme | 3660077 | 4247696.5 | 8255828 | | | | | | |
| adenosine monophosphate deaminase 3 | Cytoplasm | enzyme | 2472610.5 | 1882085.875 | 2772738.25 | | | | | | |
| adenosine monophosphate deaminase 3 | Cytoplasm | enzyme | 2069213.375 | 1417521.25 | 2356432.75 | | | | | | |
| adenosine monophosphate deaminase 3 | Cytoplasm | enzyme | 1628326.125 | 1660922.625 | 1241268.875 | | | | | | |
| adenosine monophosphate deaminase 3 | Cytoplasm | enzyme | 1128198.125 | 1806284.625 | 1968152.25 | | | | | | |
| adenosine monophosphate deaminase 3 | Cytoplasm | enzyme | 933096 | 898049.375 | 936792.562 | O08739 | AMPD3 | 2420893 | 1610016 | 328643 | 24 |
| ankyrin 1, erythrocytic | Plasma Membrane | other | 4.45E+09 | 3.69E+09 | 3.35E+09 | | | | | | |
| ankyrin 1, erythrocytic | Plasma Membrane | other | 4.19E+09 | 3.59E+09 | 3.52E+09 | | | | | | |
| ankyrin 1, erythrocytic | Plasma Membrane | other | 3.12E+09 | 2.26E+09 | 2.22E+09 | | | | | | |
| ankyrin 1, erythrocytic | Plasma Membrane | other | 3.10E+09 | 2.31E+09 | 2.03E+09 | | | | | | |
| ankyrin 1, erythrocytic | Plasma Membrane | other | 3.06E+09 | 1.84E+09 | 2.30E+09 | | | | | | |
| ankyrin 1, erythrocytic | Plasma Membrane | other | 2.87E+09 | 2.37E+09 | 2.42E+09 | | | | | | |
| ankyrin 1, erythrocytic | Plasma Membrane | other | 2.74E+09 | 2.21E+09 | 2.45E+09 | | | | | | |
| ankyrin 1, erythrocytic | Plasma Membrane | other | 2.71E+09 | 2.07E+09 | 2.07E+09 | | | | | | |
| ankyrin 1, erythrocytic | Plasma Membrane | other | 2.65E+09 | 1.97E+09 | 2.47E+09 | | | | | | |
| ankyrin 1, erythrocytic | Plasma Membrane | other | 2.65E+09 | 2.05E+09 | 2.34E+09 | | | | | | |
| ankyrin 1, erythrocytic | Plasma Membrane | other | 2.53E+09 | 1.77E+09 | 1.84E+09 | | | | | | |
| ankyrin 1, erythrocytic | Plasma Membrane | other | 2.30E+09 | 1.56E+09 | 1.69E+09 | | | | | | |
| ankyrin 1, erythrocytic | Plasma Membrane | other | 2.19E+09 | 1.76E+09 | 1.54E+09 | | | | | | |
| ankyrin 1, erythrocytic | Plasma Membrane | other | 2.10E+09 | 1.69E+09 | 1.55E+09 | | | | | | |
| ankyrin 1, erythrocytic | Plasma Membrane | other | 2.09E+09 | 1.37E+09 | 1.33E+09 | | | | | | |
| ankyrin 1, erythrocytic | Plasma Membrane | other | 2.02E+09 | 1.50E+09 | 1.55E+09 | | | | | | |
| ankyrin 1, erythrocytic | Plasma Membrane | other | 2.00E+09 | 1.76E+09 | 1.55E+09 | | | | | | |
| ankyrin 1, erythrocytic | Plasma Membrane | other | 1.85E+09 | 1.48E+09 | 1.45E+09 | | | | | | |
| ankyrin 1, erythrocytic | Plasma Membrane | other | 1.83E+09 | 1.34E+09 | 1.51E+09 | | | | | | |
| ankyrin 1, erythrocytic | Plasma Membrane | other | 1.73E+09 | 1.80E+09 | 1.36E+09 | | | | | | |
| ankyrin 1, erythrocytic | Plasma Membrane | other | 1.72E+09 | 1.72E+09 | 1.82E+09 | | | | | | |
| ankyrin 1, erythrocytic | Plasma Membrane | other | 1.69E+09 | 1.38E+09 | 1.82E+09 | | | | | | |
| ankyrin 1, erythrocytic | Plasma Membrane | other | 1.68E+09 | 1.41E+09 | 1.55E+09 | | | | | | |
| ankyrin 1, erythrocytic | Plasma Membrane | other | 1.60E+09 | 1.17E+09 | 1.24E+09 | | | | | | |
| ankyrin 1, erythrocytic | Plasma Membrane | other | 1.45E+09 | 1.39E+09 | 1.35E+09 | | | | | | |
| ankyrin 1, erythrocytic | Plasma Membrane | other | 1.42E+09 | 1.52E+09 | 9.82E+08 | | | | | | |
| ankyrin 1, erythrocytic | Plasma Membrane | other | 1.40E+09 | 1.33E+09 | 1.19E+09 | | | | | | |
| ankyrin 1, erythrocytic | Plasma Membrane | other | 1.33E+09 | 9.32E+08 | 1.03E+09 | | | | | | |
| ankyrin 1, erythrocytic | Plasma Membrane | other | 1.25E+09 | 1.27E+09 | 1.24E+09 | | | | | | |
| ankyrin 1, erythrocytic | Plasma Membrane | other | 1.25E+09 | 1.01E+09 | 9.85E+08 | | | | | | |
| ankyrin 1, erythrocytic | Plasma Membrane | other | 1.24E+09 | 1.05E+09 | 1.04E+09 | | | | | | |
| ankyrin 1, erythrocytic | Plasma Membrane | other | 1.19E+09 | 8.93E+08 | 1.04E+09 | | | | | | |
| ankyrin 1, erythrocytic | Plasma Membrane | other | 1.18E+09 | 1.10E+09 | 1.12E+09 | | | | | | |
| ankyrin 1, erythrocytic | Plasma Membrane | other | 1.13E+09 | 8.51E+08 | 8.67E+08 | | | | | | |
| ankyrin 1, erythrocytic | Plasma Membrane | other | 1.12E+09 | 1.50E+09 | 1.34E+09 | | | | | | |
| ankyrin 1, erythrocytic | Plasma Membrane | other | 1.07E+09 | 9.21E+08 | 1.01E+09 | | | | | | |
| ankyrin 1, erythrocytic | Plasma Membrane | other | 1.04E+09 | 8.11E+08 | 8.81E+08 | | | | | | |
| ankyrin 1, erythrocytic | Plasma Membrane | other | 9.99E+08 | 1.05E+09 | 8.29E+08 | | | | | | |
| ankyrin 1, erythrocytic | Plasma Membrane | other | 9.82E+08 | 1.04E+09 | 9.48E+08 | | | | | | |
| ankyrin 1, erythrocytic | Plasma Membrane | other | 9.77E+08 | 6.82E+08 | 6.64E+08 | | | | | | |
| ankyrin 1, erythrocytic | Plasma Membrane | other | 9.54E+08 | 7.45E+08 | 7.95E+08 | | | | | | |
| ankyrin 1, erythrocytic | Plasma Membrane | other | 9.31E+08 | 8.95E+08 | 9.03E+08 | | | | | | |
| ankyrin 1, erythrocytic | Plasma Membrane | other | 9.13E+08 | 8.27E+08 | 7.50E+08 | | | | | | |
| ankyrin 1, erythrocytic | Plasma Membrane | other | 8.68E+08 | 6.81E+08 | 7.10E+08 | | | | | | |
| ankyrin 1, erythrocytic | Plasma Membrane | other | 8.60E+08 | 7.51E+08 | 6.07E+08 | | | | | | |
| ankyrin 1, erythrocytic | Plasma Membrane | other | 8.52E+08 | 7.45E+08 | 6.65E+08 | | | | | | |
| ankyrin 1, erythrocytic | Plasma Membrane | other | 8.48E+08 | 9.38E+08 | 9.02E+08 | | | | | | |
| ankyrin 1, erythrocytic | Plasma Membrane | other | 8.37E+08 | 9.83E+08 | 7.04E+08 | | | | | | |
| ankyrin 1, erythrocytic | Plasma Membrane | other | 7.74E+08 | 6.76E+08 | 5.81E+08 | | | | | | |
| ankyrin 1, erythrocytic | Plasma Membrane | other | 7.30E+08 | 5.10E+08 | 5.52E+08 | | | | | | |
| ankyrin 1, erythrocytic | Plasma Membrane | other | 7.10E+08 | 6.70E+08 | 6.15E+08 | | | | | | |
| ankyrin 1, erythrocytic | Plasma Membrane | other | 6.98E+08 | 5.14E+08 | 5.34E+08 | | | | | | |
| ankyrin 1, erythrocytic | Plasma Membrane | other | 6.89E+08 | 4.49E+08 | 5.04E+08 | | | | | | |
| ankyrin 1, erythrocytic | Plasma Membrane | other | 6.72E+08 | 4.26E+08 | 4.87E+08 | | | | | | |
| ankyrin 1, erythrocytic | Plasma Membrane | other | 6.60E+08 | 5.83E+08 | 5.06E+08 | | | | | | |
| ankyrin 1, erythrocytic | Plasma Membrane | other | 6.35E+08 | 5.85E+08 | 5.87E+08 | | | | | | |
| ankyrin 1, erythrocytic | Plasma Membrane | other | 6.19E+08 | 5.10E+08 | 5.20E+08 | | | | | | |
| ankyrin 1, erythrocytic | Plasma Membrane | other | 6.01E+08 | 3.42E+08 | 4.28E+08 | | | | | | |
| ankyrin 1, erythrocytic | Plasma Membrane | other | 5.94E+08 | 2.53E+08 | 4.35E+08 | | | | | | |
| ankyrin 1, erythrocytic | Plasma Membrane | other | 5.91E+08 | 5.98E+08 | 5.43E+08 | | | | | | |
| ankyrin 1, erythrocytic | Plasma Membrane | other | 5.73E+08 | 7.48E+08 | 6.68E+08 | | | | | | |
| ankyrin 1, erythrocytic | Plasma Membrane | other | 5.68E+08 | 6.77E+08 | 5.91E+08 | | | | | | |
| ankyrin 1, erythrocytic | Plasma Membrane | other | 5.62E+08 | 7.29E+08 | 4.37E+08 | | | | | | |
| ankyrin 1, erythrocytic | Plasma Membrane | other | 5.56E+08 | 3.49E+08 | 3.28E+08 | | | | | | |
| ankyrin 1, erythrocytic | Plasma Membrane | other | 5.55E+08 | 3.56E+08 | 3.93E+08 | | | | | | |
| ankyrin 1, erythrocytic | Plasma Membrane | other | 5.50E+08 | 3.97E+08 | 4.59E+08 | | | | | | |
| ankyrin 1, erythrocytic | Plasma Membrane | other | 5.41E+08 | 4.46E+08 | 3.92E+08 | | | | | | |
| ankyrin 1, erythrocytic | Plasma Membrane | other | 4.97E+08 | 4.03E+08 | 3.53E+08 | | | | | | |
| ankyrin 1, erythrocytic | Plasma Membrane | other | 4.79E+08 | 1.96E+08 | 3.18E+08 | | | | | | |
| ankyrin 1, erythrocytic | Plasma Membrane | other | 4.61E+08 | 4.17E+08 | 4.16E+08 | | | | | | |
| ankyrin 1, erythrocytic | Plasma Membrane | other | 4.35E+08 | 3.75E+08 | 3.01E+08 | | | | | | |
| ankyrin 1, erythrocytic | Plasma Membrane | other | 4.33E+08 | 4.90E+08 | 4.39E+08 | | | | | | |
| ankyrin 1, erythrocytic | Plasma Membrane | other | 4.07E+08 | 2.15E+08 | 2.61E+08 | | | | | | |
| ankyrin 1, erythrocytic | Plasma Membrane | other | 3.62E+08 | 2.32E+08 | 2.58E+08 | | | | | | |

| | | | | | | | | | | | |
|---|---|---|---|---|---|---|---|---|---|---|---|
| ankyrin 1, erythrocytic | Plasma Membrane | other | 3.41E+08 | 6.41E+08 | 5.29E+08 | | | | | | |
| ankyrin 1, erythrocytic | Plasma Membrane | other | 3.40E+08 | 2.53E+08 | 2.74E+08 | | | | | | |
| ankyrin 1, erythrocytic | Plasma Membrane | other | 3.34E+08 | 4.17E+08 | 3.01E+08 | | | | | | |
| ankyrin 1, erythrocytic | Plasma Membrane | other | 3.11E+08 | 2.42E+08 | 2.06E+08 | | | | | | |
| ankyrin 1, erythrocytic | Plasma Membrane | other | 3.02E+08 | 1.41E+08 | 2.23E+08 | | | | | | |
| ankyrin 1, erythrocytic | Plasma Membrane | other | 2.87E+08 | 9.35E+07 | 1.93E+08 | | | | | | |
| ankyrin 1, erythrocytic | Plasma Membrane | other | 2.73E+08 | 3.12E+08 | 2.51E+08 | | | | | | |
| ankyrin 1, erythrocytic | Plasma Membrane | other | 2.52E+08 | 3.98E+08 | 2.86E+08 | | | | | | |
| ankyrin 1, erythrocytic | Plasma Membrane | other | 2.51E+08 | 1.68E+08 | 1.86E+08 | | | | | | |
| ankyrin 1, erythrocytic | Plasma Membrane | other | 2.40E+08 | 4.24E+08 | 2.68E+08 | | | | | | |
| ankyrin 1, erythrocytic | Plasma Membrane | other | 2.36E+08 | 2.32E+08 | 2.13E+08 | | | | | | |
| ankyrin 1, erythrocytic | Plasma Membrane | other | 2.29E+08 | 4.63E+08 | 2.68E+08 | | | | | | |
| ankyrin 1, erythrocytic | Plasma Membrane | other | 2.23E+08 | 1.77E+08 | 1.31E+08 | | | | | | |
| ankyrin 1, erythrocytic | Plasma Membrane | other | 2.16E+08 | 3.41E+08 | 2.81E+08 | | | | | | |
| ankyrin 1, erythrocytic | Plasma Membrane | other | 2.11E+08 | 1.01E+08 | 1.59E+08 | | | | | | |
| ankyrin 1, erythrocytic | Plasma Membrane | other | 2.07E+08 | 1.36E+08 | 1.88E+08 | | | | | | |
| ankyrin 1, erythrocytic | Plasma Membrane | other | 1.83E+08 | 9.54E+07 | 1.31E+08 | | | | | | |
| ankyrin 1, erythrocytic | Plasma Membrane | other | 1.71E+08 | 1.30E+08 | 9.82E+07 | | | | | | |
| ankyrin 1, erythrocytic | Plasma Membrane | other | 1.58E+08 | 1.26E+08 | 1.27E+08 | | | | | | |
| ankyrin 1, erythrocytic | Plasma Membrane | other | 1.52E+08 | 9.46E+07 | 1.07E+08 | | | | | | |
| ankyrin 1, erythrocytic | Plasma Membrane | other | 1.47E+08 | 1.51E+08 | 1.20E+08 | | | | | | |
| ankyrin 1, erythrocytic | Plasma Membrane | other | 1.42E+08 | 7.70E+07 | 1.58E+08 | | | | | | |
| ankyrin 1, erythrocytic | Plasma Membrane | other | 1.41E+08 | 8.29E+07 | 8.57E+07 | | | | | | |
| ankyrin 1, erythrocytic | Plasma Membrane | other | 1.35E+08 | 4.10E+07 | 1.07E+08 | | | | | | |
| ankyrin 1, erythrocytic | Plasma Membrane | other | 1.34E+08 | 2.85E+08 | 1.98E+08 | | | | | | |
| ankyrin 1, erythrocytic | Plasma Membrane | other | 1.33E+08 | 4.98E+07 | 6.74E+07 | | | | | | |
| ankyrin 1, erythrocytic | Plasma Membrane | other | 1.18E+08 | 1.43E+08 | 8.31E+07 | | | | | | |
| ankyrin 1, erythrocytic | Plasma Membrane | other | 1.18E+08 | 8.17E+07 | 1.21E+08 | | | | | | |
| ankyrin 1, erythrocytic | Plasma Membrane | other | 1.09E+08 | 9.53E+07 | 9.98E+07 | | | | | | |
| ankyrin 1, erythrocytic | Plasma Membrane | other | 9.87E+07 | 9.43E+07 | 6.91E+07 | | | | | | |
| ankyrin 1, erythrocytic | Plasma Membrane | other | 9.21E+07 | 7.20E+07 | 7.98E+07 | | | | | | |
| ankyrin 1, erythrocytic | Plasma Membrane | other | 8.85E+07 | 2.71E+08 | 3.84E+08 | | | | | | |
| ankyrin 1, erythrocytic | Plasma Membrane | other | 8.27E+07 | 3.13E+07 | 6.50E+07 | | | | | | |
| ankyrin 1, erythrocytic | Plasma Membrane | other | 8.21E+07 | 1.17E+08 | 7.45E+07 | | | | | | |
| ankyrin 1, erythrocytic | Plasma Membrane | other | 8.14E+07 | 5.95E+07 | 6.90E+07 | | | | | | |
| ankyrin 1, erythrocytic | Plasma Membrane | other | 8.13E+07 | 3.17E+07 | 5.84E+07 | | | | | | |
| ankyrin 1, erythrocytic | Plasma Membrane | other | 6.44E+07 | 1.62E+08 | 7.54E+07 | | | | | | |
| ankyrin 1, erythrocytic | Plasma Membrane | other | 6.37E+07 | 2.91E+07 | 3.38E+07 | | | | | | |
| ankyrin 1, erythrocytic | Plasma Membrane | other | 6.13E+07 | 5.29E+07 | 5.31E+07 | | | | | | |
| ankyrin 1, erythrocytic | Plasma Membrane | other | 5.85E+07 | 7.98E+07 | 9.90E+07 | | | | | | |
| ankyrin 1, erythrocytic | Plasma Membrane | other | 5.65E+07 | 3.37E+07 | 5.80E+07 | | | | | | |
| ankyrin 1, erythrocytic | Plasma Membrane | other | 5.50E+07 | 2.87E+07 | 5.88E+07 | | | | | | |
| ankyrin 1, erythrocytic | Plasma Membrane | other | 5.21E+07 | 3.39E+07 | 3.59E+07 | | | | | | |
| ankyrin 1, erythrocytic | Plasma Membrane | other | 5.11E+07 | 3.33E+07 | 7.82E+07 | | | | | | |
| ankyrin 1, erythrocytic | Plasma Membrane | other | 4.76E+07 | 2.28E+08 | 1.29E+08 | | | | | | |
| ankyrin 1, erythrocytic | Plasma Membrane | other | 4.35E+07 | 2.28E+07 | 2.38E+07 | | | | | | |
| ankyrin 1, erythrocytic | Plasma Membrane | other | 4.25E+07 | 1.60E+07 | 2.91E+07 | | | | | | |
| ankyrin 1, erythrocytic | Plasma Membrane | other | 4.10E+07 | 1.53E+07 | 2.84E+07 | | | | | | |
| ankyrin 1, erythrocytic | Plasma Membrane | other | 4.06E+07 | 1.04E+07 | 3.76E+07 | | | | | | |
| ankyrin 1, erythrocytic | Plasma Membrane | other | 3.99E+07 | 3.31E+07 | 6.75E+07 | | | | | | |
| ankyrin 1, erythrocytic | Plasma Membrane | other | 3.56E+07 | 3.84E+07 | 2.48E+07 | | | | | | |
| ankyrin 1, erythrocytic | Plasma Membrane | other | 3.49E+07 | 8.36E+07 | 2.67E+07 | | | | | | |
| ankyrin 1, erythrocytic | Plasma Membrane | other | 3.37E+07 | 3.46E+07 | 3.39E+07 | | | | | | |
| ankyrin 1, erythrocytic | Plasma Membrane | other | 3.36E+07 | 1.39E+08 | 8.16E+07 | | | | | | |
| ankyrin 1, erythrocytic | Plasma Membrane | other | 3.34E+07 | 2.11E+07 | 2.94E+07 | | | | | | |
| ankyrin 1, erythrocytic | Plasma Membrane | other | 3.34E+07 | 1.63E+07 | 1.96E+07 | | | | | | |
| ankyrin 1, erythrocytic | Plasma Membrane | other | 3.02E+07 | 1.39E+07 | 1.78E+07 | | | | | | |
| ankyrin 1, erythrocytic | Plasma Membrane | other | 3.00E+07 | 7.25E+07 | 3.91E+07 | | | | | | |
| ankyrin 1, erythrocytic | Plasma Membrane | other | 2.38E+07 | 2.88E+07 | 2.77E+07 | | | | | | |
| ankyrin 1, erythrocytic | Plasma Membrane | other | 2.29E+07 | 2.78E+07 | 2.03E+07 | | | | | | |
| ankyrin 1, erythrocytic | Plasma Membrane | other | 2.24E+07 | 4.84E+07 | 4.06E+07 | | | | | | |
| ankyrin 1, erythrocytic | Plasma Membrane | other | 2.10E+07 | 1.16E+07 | 1.39E+07 | | | | | | |
| ankyrin 1, erythrocytic | Plasma Membrane | other | 1.84E+07 | 2.80E+07 | 1.88E+07 | | | | | | |
| ankyrin 1, erythrocytic | Plasma Membrane | other | 1.72E+07 | 3.58E+07 | 2.40E+07 | | | | | | |
| ankyrin 1, erythrocytic | Plasma Membrane | other | 1.59E+07 | 5202622 | 9865497 | | | | | | |
| ankyrin 1, erythrocytic | Plasma Membrane | other | 1.52E+07 | 1.65E+07 | 907907.688 | | | | | | |
| ankyrin 1, erythrocytic | Plasma Membrane | other | 1.44E+07 | 8221452.5 | 1.04E+07 | | | | | | |
| ankyrin 1, erythrocytic | Plasma Membrane | other | 1.43E+07 | 1.34E+07 | 1.36E+07 | | | | | | |
| ankyrin 1, erythrocytic | Plasma Membrane | other | 1.41E+07 | 5.55E+07 | 2.36E+07 | | | | | | |
| ankyrin 1, erythrocytic | Plasma Membrane | other | 1.40E+07 | 9625644 | 8720712 | | | | | | |
| ankyrin 1, erythrocytic | Plasma Membrane | other | 1.17E+07 | 7228897 | 9410608 | | | | | | |
| ankyrin 1, erythrocytic | Plasma Membrane | other | 1.16E+07 | 4615465.5 | 7727668.5 | | | | | | |
| ankyrin 1, erythrocytic | Plasma Membrane | other | 1.12E+07 | 1.25E+07 | 8360560.5 | | | | | | |
| ankyrin 1, erythrocytic | Plasma Membrane | other | 1.07E+07 | 3557527.75 | 2318531.5 | | | | | | |
| ankyrin 1, erythrocytic | Plasma Membrane | other | 1.07E+07 | 7846707.5 | 7581956 | | | | | | |
| ankyrin 1, erythrocytic | Plasma Membrane | other | 1.01E+07 | 5946032 | 8606763 | | | | | | |
| ankyrin 1, erythrocytic | Plasma Membrane | other | 9572568 | 1.97E+07 | 2.88E+07 | | | | | | |
| ankyrin 1, erythrocytic | Plasma Membrane | other | 9128412 | 9185982 | 4080168.5 | | | | | | |
| ankyrin 1, erythrocytic | Plasma Membrane | other | 7993718.5 | 5184167 | 4268925 | | | | | | |
| ankyrin 1, erythrocytic | Plasma Membrane | other | 7464481.5 | 3.11E+07 | 1.94E+07 | | | | | | |
| ankyrin 1, erythrocytic | Plasma Membrane | other | 7018139.5 | 6414818 | 4456807 | | | | | | |
| ankyrin 1, erythrocytic | Plasma Membrane | other | 6848588 | 1.49E+07 | 1.31E+07 | | | | | | |
| ankyrin 1, erythrocytic | Plasma Membrane | other | 4877342.5 | 2005447.5 | 3356269 | | | | | | |
| ankyrin 1, erythrocytic | Plasma Membrane | other | 3913271.75 | 2349076 | 2783492.5 | | | | | | |
| ankyrin 1, erythrocytic | Plasma Membrane | other | 3883012.75 | 2.09E+07 | 4860091 | | | | | | |
| ankyrin 1, erythrocytic | Plasma Membrane | other | 3753822 | 2145025 | 2257138 | | | | | | |
| ankyrin 1, erythrocytic | Plasma Membrane | other | 3399618 | 5238083 | 3442336.25 | | | | | | |
| ankyrin 1, erythrocytic | Plasma Membrane | other | 3063549.25 | 519016.156 | 2020551.75 | | | | | | |
| ankyrin 1, erythrocytic | Plasma Membrane | other | 2383019.5 | 8089036 | 2.03E+07 | | | | | | |
| ankyrin 1, erythrocytic | Plasma Membrane | other | 2135659.5 | 6549457 | 4534395.5 | | | | | | |
| ankyrin 1, erythrocytic | Plasma Membrane | other | 1623177.75 | 1243835 | 1095150.125 | | | | | | |
| ankyrin 1, erythrocytic | Plasma Membrane | other | 1448283.25 | 682868.688 | 1277366.75 | | | | | | |
| ankyrin 1, erythrocytic | Plasma Membrane | other | 1392849.375 | 2.33E+07 | 5577854.5 | | | | | | |
| ankyrin 1, erythrocytic | Plasma Membrane | other | 1346970.75 | 1642882.5 | 2486039.75 | | | | | | |
| ankyrin 1, erythrocytic | Plasma Membrane | other | 146735.5 | 1028530.625 | 3303189.75 | Q02357 | ANK1 | 638599546 | 762807356 | 33877464 | 507 |

| | | | | | | | | | | | |
|---|---|---|---|---|---|---|---|---|---|---|---|
| ANKH inorganic pyrophosphate transport regulator | Plasma Membrane | transporter | 2350574 | 1258139.125 | 1562437.125 | Q9JHZ2 | ANKH | 2350574 | 563792 | 325506 | 3 |
| ankyrin repeat domain 28 | Cytoplasm | other | 3003060.25 | 1106948.125 | 988998.188 | Q505D1 | ANKRD28 | 3003060 | 1130310 | 652585 | 3 |
| anoctamin 6 | Plasma Membrane | ion channel | 1100232.875 | 2939022.75 | 2130282.5 | Q6P9J9 | ANO6 | 1100233 | 921612 | 532093 | 3 |
| acidic (leucine-rich) nuclear phosphoprotein 32 family, member A | Nucleus | other | 2.17E+07 | 1.52E+07 | 1.62E+07 | | | | | | |
| acidic (leucine-rich) nuclear phosphoprotein 32 family, member A | Nucleus | other | 1.37E+07 | 5079534.5 | 7917617 | | | | | | |
| acidic (leucine-rich) nuclear phosphoprotein 32 family, member A | Nucleus | other | 1.26E+07 | 2920431.25 | 5162111 | O35381 | Anp32a | 15986667 | 6257015 | 2085672 | 9 |
| adaptor-related protein complex 1, beta 1 subunit | Cytoplasm | transporter | 1.38E+07 | 1.26E+07 | 9505374 | | | | | | |
| adaptor-related protein complex 1, beta 1 subunit | Cytoplasm | transporter | 1.33E+07 | 9837392 | 8516329 | | | | | | |
| adaptor-related protein complex 1, beta 1 subunit | Cytoplasm | transporter | 1.22E+07 | 9061044 | 9141868 | | | | | | |
| adaptor-related protein complex 1, beta 1 subunit | Cytoplasm | transporter | 1.15E+07 | 8783154 | 6574573 | | | | | | |
| adaptor-related protein complex 1, beta 1 subunit | Cytoplasm | transporter | 6591868 | 6972681 | 5847175.5 | | | | | | |
| adaptor-related protein complex 1, beta 1 subunit | Cytoplasm | transporter | 6220895 | 4806510 | 3557384.75 | | | | | | |
| adaptor-related protein complex 1, beta 1 subunit | Cytoplasm | transporter | 6092121 | 2542000.25 | 2358018.75 | | | | | | |
| adaptor-related protein complex 1, beta 1 subunit | Cytoplasm | transporter | 5636244.5 | 6051341 | 3805591.75 | | | | | | |
| adaptor-related protein complex 1, beta 1 subunit | Cytoplasm | transporter | 5485817.5 | 2294771.5 | 1375914.875 | | | | | | |
| adaptor-related protein complex 1, beta 1 subunit | Cytoplasm | transporter | 4599104.5 | 4414632 | 3102303 | | | | | | |
| adaptor-related protein complex 1, beta 1 subunit | Cytoplasm | transporter | 3344325.75 | 1776346.75 | 1485228.875 | | | | | | |
| adaptor-related protein complex 1, beta 1 subunit | Cytoplasm | transporter | 1332312.875 | 1382690.75 | 1139515 | O35643 | AP1B1 | 7509391 | 3727168 | 621195 | 36 |
| adaptor-related protein complex 2, alpha 1 subunit | Cytoplasm | transporter | 3329365 | 3454050.5 | 2734288.25 | | | | | | |
| adaptor-related protein complex 2, alpha 1 subunit | Cytoplasm | transporter | 2163564.75 | 2249783.5 | 2352358 | | | | | | |
| adaptor-related protein complex 2, alpha 1 subunit | Cytoplasm | transporter | 762957.188 | 772550.625 | 333650.094 | P17426 | AP2A1 | 2085296 | 1142639 | 380880 | 9 |
| adaptor-related protein complex 2, alpha 2 subunit | Cytoplasm | transporter | 1.06E+07 | 7172165.5 | 8191599 | | | | | | |
| adaptor-related protein complex 2, alpha 2 subunit | Cytoplasm | transporter | 5727813.5 | 4532892 | 3633915.25 | | | | | | |
| adaptor-related protein complex 2, alpha 2 subunit | Cytoplasm | transporter | 4794681 | 1.58E+07 | 1.40E+07 | | | | | | |
| adaptor-related protein complex 2, alpha 2 subunit | Cytoplasm | transporter | 3169770.5 | 3377764.25 | 2494040.5 | | | | | | |
| adaptor-related protein complex 2, alpha 2 subunit | Cytoplasm | transporter | 3104074.5 | 2289903.5 | 2328875.25 | | | | | | |
| adaptor-related protein complex 2, alpha 2 subunit | Cytoplasm | transporter | 2866158.75 | 2035431.875 | 1726192 | | | | | | |
| adaptor-related protein complex 2, alpha 2 subunit | Cytoplasm | transporter | 1847072.375 | 1486031.875 | 1104932.625 | | | | | | |
| adaptor-related protein complex 2, alpha 2 subunit | Cytoplasm | transporter | 1567525.75 | 1295772.625 | 878552.312 | P17427 | AP2A2 | 4209637 | 4018658 | 820305 | 24 |
| adaptor-related protein complex 2, beta 1 subunit | Plasma Membrane | other | 973984.75 | 828775.5 | 922348.688 | | | | | | |
| adaptor-related protein complex 2, beta 1 subunit | Plasma Membrane | other | 40027.32 | 168082.828 | 84860.711 | Q9DBG3 | Ap2b1 | 507006 | 448367 | 183045 | 6 |
| adaptor-related protein complex 2, mu 1 subunit | Cytoplasm | transporter | 8009326 | 8077726.5 | 5791819 | | | | | | |
| adaptor-related protein complex 2, mu 1 subunit | Cytoplasm | transporter | 5325237 | 5345798.5 | 3689006.5 | | | | | | |
| adaptor-related protein complex 2, mu 1 subunit | Cytoplasm | transporter | 3587316.5 | 4176215.75 | 1797979.875 | | | | | | |
| adaptor-related protein complex 2, mu 1 subunit | Cytoplasm | transporter | 3439137.25 | 3952084.25 | 2155182.25 | | | | | | |
| adaptor-related protein complex 2, mu 1 subunit | Cytoplasm | transporter | 2312126.75 | 2104587 | 1710235.5 | P84091 | AP2M1 | 4534629 | 2068255 | 534021 | 15 |
| acylaminoacyl-peptide hydrolase | Cytoplasm | peptidase | 1.34E+08 | 6.30E+07 | 8.06E+07 | | | | | | |
| acylaminoacyl-peptide hydrolase | Cytoplasm | peptidase | 1.01E+08 | 3.88E+07 | 5.04E+07 | | | | | | |
| acylaminoacyl-peptide hydrolase | Cytoplasm | peptidase | 8.47E+07 | 2.24E+07 | 4.05E+07 | | | | | | |
| acylaminoacyl-peptide hydrolase | Cytoplasm | peptidase | 7.74E+07 | 2.91E+07 | 3.61E+07 | | | | | | |
| acylaminoacyl-peptide hydrolase | Cytoplasm | peptidase | 7.04E+07 | 2.68E+07 | 3.90E+07 | | | | | | |
| acylaminoacyl-peptide hydrolase | Cytoplasm | peptidase | 6.23E+07 | 2.36E+07 | 3.28E+07 | | | | | | |
| acylaminoacyl-peptide hydrolase | Cytoplasm | peptidase | 5.76E+07 | 2.22E+07 | 3.05E+07 | | | | | | |
| acylaminoacyl-peptide hydrolase | Cytoplasm | peptidase | 5.64E+07 | 1.55E+07 | 2.80E+07 | | | | | | |
| acylaminoacyl-peptide hydrolase | Cytoplasm | peptidase | 5.27E+07 | 2.60E+07 | 3.16E+07 | | | | | | |
| acylaminoacyl-peptide hydrolase | Cytoplasm | peptidase | 5.13E+07 | 1.83E+07 | 2.86E+07 | | | | | | |
| acylaminoacyl-peptide hydrolase | Cytoplasm | peptidase | 4.90E+07 | 1.87E+07 | 2.47E+07 | | | | | | |
| acylaminoacyl-peptide hydrolase | Cytoplasm | peptidase | 4.88E+07 | 1.97E+07 | 2.99E+07 | | | | | | |
| acylaminoacyl-peptide hydrolase | Cytoplasm | peptidase | 4.84E+07 | 2.25E+07 | 2.63E+07 | | | | | | |
| acylaminoacyl-peptide hydrolase | Cytoplasm | peptidase | 4.82E+07 | 1.78E+07 | 2.67E+07 | | | | | | |
| acylaminoacyl-peptide hydrolase | Cytoplasm | peptidase | 4.73E+07 | 2.90E+07 | 3.84E+07 | | | | | | |
| acylaminoacyl-peptide hydrolase | Cytoplasm | peptidase | 4.21E+07 | 1.65E+07 | 2.40E+07 | | | | | | |
| acylaminoacyl-peptide hydrolase | Cytoplasm | peptidase | 3.40E+07 | 1.44E+07 | 1.97E+07 | | | | | | |
| acylaminoacyl-peptide hydrolase | Cytoplasm | peptidase | 3.30E+07 | 1.42E+07 | 1.59E+07 | | | | | | |
| acylaminoacyl-peptide hydrolase | Cytoplasm | peptidase | 2.34E+07 | 1.04E+07 | 1.11E+07 | | | | | | |
| acylaminoacyl-peptide hydrolase | Cytoplasm | peptidase | 1.34E+07 | 4979207 | 6436494 | | | | | | |
| acylaminoacyl-peptide hydrolase | Cytoplasm | peptidase | 1.16E+07 | 4888530.5 | 6031260.5 | | | | | | |
| acylaminoacyl-peptide hydrolase | Cytoplasm | peptidase | 1.13E+07 | 3452966 | 5011565 | | | | | | |
| acylaminoacyl-peptide hydrolase | Cytoplasm | peptidase | 9264684 | 2661031.75 | 3901973 | | | | | | |
| acylaminoacyl-peptide hydrolase | Cytoplasm | peptidase | 8864602 | 9750185 | 1.23E+07 | | | | | | |
| acylaminoacyl-peptide hydrolase | Cytoplasm | peptidase | 2943880.75 | 983911.438 | 1070761.5 | Q8R146 | APEH | 47137327 | 24733351 | 2855961 | 75 |
| APH1A gamma secretase subunit | Cytoplasm | peptidase | 2077638.75 | 759744.062 | 1403755.5 | Q8BVF7 | APH1A | 2077639 | 659004 | 380476 | 3 |
| apolipoprotein A-I | Extracellular Space | transporter | 3953480.25 | 1923246.5 | 2484260.25 | | | | | | |
| apolipoprotein A-I | Extracellular Space | transporter | 1907427.625 | 2332847.25 | 2416910.75 | | | | | | |
| apolipoprotein A-I | Extracellular Space | transporter | 612637.125 | 652331.25 | 683877.875 | Q00623 | APOA1 | 2157848 | 1101377 | 367126 | 9 |
| apolipoprotein H (beta-2-glycoprotein I) | Extracellular Space | transporter | 90291.609 | 330065 | 611046.25 | Q01339 | APOH | 90292 | 260649 | 150486 | 3 |
| aquaporin 1 (Colton blood group) | Plasma Membrane | transporter | 2.73E+09 | 2.31E+09 | 2.52E+09 | | | | | | |
| aquaporin 1 (Colton blood group) | Plasma Membrane | transporter | 2.56E+09 | 1.30E+09 | 1.40E+09 | | | | | | |
| aquaporin 1 (Colton blood group) | Plasma Membrane | transporter | 8.84E+08 | 7.44E+08 | 9.28E+08 | | | | | | |
| aquaporin 1 (Colton blood group) | Plasma Membrane | transporter | 3.96E+08 | 2.70E+08 | 3.14E+08 | | | | | | |
| aquaporin 1 (Colton blood group) | Plasma Membrane | transporter | 6.35E+07 | 4.92E+07 | 4.51E+07 | Q02013 | AQP1 | 1325990000 | 988767078 | 255298562 | 15 |
| ADP-ribosylation factor 5 | Cytoplasm | enzyme | 4250768.5 | 3207061.5 | 2824415.75 | | | | | | |
| ADP-ribosylation factor 5 | Cytoplasm | enzyme | 2747528.25 | 2499476.25 | 1561576.125 | P84084 | ARF5 | 3499148 | 881327 | 359800 | 6 |
| ADP-ribosylation factor 6 | Plasma Membrane | transporter | 1.72E+07 | 1.33E+07 | 1.62E+07 | | | | | | |
| ADP-ribosylation factor 6 | Plasma Membrane | transporter | 1.27E+07 | 1.23E+07 | 1.16E+07 | | | | | | |
| ADP-ribosylation factor 6 | Plasma Membrane | transporter | 8097000.5 | 4824600 | 6702806.5 | | | | | | |
| ADP-ribosylation factor 6 | Plasma Membrane | transporter | 7400890 | 8070162.5 | 7331399 | | | | | | |
| ADP-ribosylation factor 6 | Plasma Membrane | transporter | 4555814 | 3985141.5 | 5222907 | | | | | | |
| ADP-ribosylation factor 6 | Plasma Membrane | transporter | 3742909.75 | 4335241.5 | 5169273 | P62331 | ARF6 | 8941102 | 4315297 | 1017125 | 18 |
| Rho GTPase activating protein 33 | Plasma Membrane | transporter | 1.92E+07 | 1.11E+07 | 1.27E+07 | Q80YF9 | ARHGAP33 | 19240000 | 4324037 | 2496484 | 3 |
| AT rich interactive domain 3B (BRIGHT-like) | Nucleus | other | 9949065 | 1.43E+07 | 1.31E+07 | Q9Z1N7 | ARID3B | 9949065 | 2252749 | 1300625 | 3 |
| ADP-ribosylation factor-like 6 interacting protein 1 | Cytoplasm | other | 9676199 | 1.41E+07 | 1.03E+07 | Q9JKW0 | ARL6IP1 | 9676199 | 2416037 | 1394900 | 3 |
| ADP-ribosylation factor-like 6 interacting protein 5 | Cytoplasm | other | 2183505.75 | 3141837.75 | 3186400.25 | Q8R5J9 | ARL6IP5 | 2183506 | 566596 | 327124 | 3 |
| ADP-ribosylation factor-like 8A | Cytoplasm | enzyme | 3144759 | 2499926.25 | 2220844 | Q8VEH3 | ARL8A | 3144759 | 473870 | 273589 | 3 |
| ADP-ribosylation factor-like 8B | Plasma Membrane | enzyme | 5.89E+07 | 5.28E+07 | 5.41E+07 | | | | | | |
| ADP-ribosylation factor-like 8B | Plasma Membrane | enzyme | 3.20E+07 | 3.17E+07 | 2.60E+07 | | | | | | |
| ADP-ribosylation factor-like 8B | Plasma Membrane | enzyme | 3.08E+07 | 3.23E+07 | 2.83E+07 | | | | | | |
| ADP-ribosylation factor-like 8B | Plasma Membrane | enzyme | 2.80E+07 | 2.66E+07 | 2.44E+07 | | | | | | |
| ADP-ribosylation factor-like 8B | Plasma Membrane | enzyme | 2.41E+07 | 1.73E+07 | 1.69E+07 | | | | | | |
| ADP-ribosylation factor-like 8B | Plasma Membrane | enzyme | 1.94E+07 | 1.45E+07 | 1.61E+07 | | | | | | |
| ADP-ribosylation factor-like 8B | Plasma Membrane | enzyme | 1.90E+07 | 6959624 | 1.29E+07 | | | | | | |
| ADP-ribosylation factor-like 8B | Plasma Membrane | enzyme | 1.54E+07 | 3.55E+07 | 1.38E+07 | | | | | | |
| ADP-ribosylation factor-like 8B | Plasma Membrane | enzyme | 1.49E+07 | 1.23E+07 | 8308336 | | | | | | |
| ADP-ribosylation factor-like 8B | Plasma Membrane | enzyme | 1.47E+07 | 7981868.5 | 1.38E+07 | | | | | | |
| ADP-ribosylation factor-like 8B | Plasma Membrane | enzyme | 1.23E+07 | 5788943 | 7340182.5 | | | | | | |

| | | | | | | | | | | | |
|---|---|---|---|---|---|---|---|---|---|---|---|
| ADP-ribosylation factor-like 8B | Plasma Membrane | enzyme | 1.05E+07 | 1.00E+07 | 1.09E+07 | | | | | | |
| ADP-ribosylation factor-like 8B | Plasma Membrane | enzyme | 1.03E+07 | 1.05E+07 | 9223816 | | | | | | |
| ADP-ribosylation factor-like 8B | Plasma Membrane | enzyme | 1.02E+07 | 6652010.5 | 9281009 | Q9CQW2 | ARL8B | 21444286 | 13019634 | 2008973 | 42 |
| actin related protein 2/3 complex, subunit 1A, 41kDa | Extracellular Space | other | 2029635 | 4518540 | 3444140.75 | Q9R0Q6 | ARPC1A | 2029635 | 1248319 | 720718 | 3 |
| actin related protein 2/3 complex, subunit 2, 34kDa | Cytoplasm | other | 1.25E+07 | 1.87E+07 | 2.13E+07 | | | | | | |
| actin related protein 2/3 complex, subunit 2, 34kDa | Cytoplasm | other | 4602995 | 6933100.5 | 7139568 | | | | | | |
| actin related protein 2/3 complex, subunit 2, 34kDa | Cytoplasm | other | 2393943.75 | 5677299 | 5527103.5 | Q9CVB6 | ARPC2 | 6495646 | 6619946 | 2206649 | 9 |
| actin related protein 2/3 complex, subunit 3, 21kDa | Cytoplasm | other | 2163676.25 | 2992697 | 3300035.25 | | | | | | |
| actin related protein 2/3 complex, subunit 3, 21kDa | Cytoplasm | other | 1096367.375 | 2698600.75 | 2463252.5 | Q9JM76 | ARPC3 | 1630022 | 773734 | 315876 | 6 |
| actin related protein 2/3 complex, subunit 4, 20kDa | Cytoplasm | other | 1978910.375 | 3145740.75 | 2413694.5 | P59999 | ARPC4 | 1978910 | 589692 | 340459 | 3 |
| actin related protein 2/3 complex, subunit 5, 16kDa | Cytoplasm | other | 744103.875 | 2136788.5 | 1531078 | Q9CPW4 | ARPC5 | 744104 | 698306 | 403167 | 3 |
| ADP-ribosyltransferase 4 (Dombrock blood group) | Nucleus | enzyme | 4.68E+07 | 3.05E+07 | 3.64E+07 | | | | | | |
| ADP-ribosyltransferase 4 (Dombrock blood group) | Nucleus | enzyme | 3.09E+07 | 2.85E+07 | 2.94E+07 | | | | | | |
| ADP-ribosyltransferase 4 (Dombrock blood group) | Nucleus | enzyme | 1.20E+07 | 1.35E+07 | 1.06E+07 | | | | | | |
| ADP-ribosyltransferase 4 (Dombrock blood group) | Nucleus | enzyme | 8256485.5 | 6777426 | 7319884 | | | | | | |
| ADP-ribosyltransferase 4 (Dombrock blood group) | Nucleus | enzyme | 5730085 | 4183508 | 4064367.5 | | | | | | |
| ADP-ribosyltransferase 4 (Dombrock blood group) | Nucleus | enzyme | 1525266.5 | 2552068.5 | 3031961.75 | | | | | | |
| ADP-ribosyltransferase 4 (Dombrock blood group) | Nucleus | enzyme | 1132105.125 | 2548592.25 | 3248945.25 | Q9CRA0 | ART4 | 15184849 | 13805868 | 3012687 | 21 |
| ArfGAP with SH3 domain, ankyrin repeat and PH domain 1 | Plasma Membrane | other | 4961593.5 | 2889137.25 | 2910980.75 | | | | | | |
| ArfGAP with SH3 domain, ankyrin repeat and PH domain 1 | Plasma Membrane | other | 2528839.75 | 1646532.375 | 1283157.125 | Q9QWY8 | ASAP1 | 3745217 | 1291631 | 527306 | 6 |
| ArfGAP with SH3 domain, ankyrin repeat and PH domain 2 | Nucleus | other | 1.33E+07 | 1.90E+07 | 1.87E+07 | Q7SIG6 | ASAP2 | 13320000 | 3189457 | 1841434 | 3 |
| ankyrin repeat and SOCS box containing 4 | Nucleus | transcription regulator | 6690752.5 | 5823183.5 | 4845228.5 | Q9WV71 | ASB4 | 6690753 | 923312 | 533074 | 3 |
| ATPase family, AAA domain containing 5 | Extracellular Space | other | 1.72E+07 | 1.80E+07 | 1.79E+07 | Q4QY64 | ATAD5 | 17240000 | 408575 | 235891 | 3 |
| activating transcription factor 2 | Nucleus | transcription regulator | 1.18E+08 | 7.27E+07 | 8.36E+07 | P16951 | ATF2 | 118100000 | 23709511 | 13688693 | 3 |
| autophagy related 4C, cysteine peptidase | Cytoplasm | peptidase | 1414573.5 | 990145.438 | 667051.688 | Q811C2 | ATG4C | 1414574 | 374904 | 216451 | 3 |
| 5-aminoimidazole-4-carboxamide ribonucleotide formyltransferase/IMP cyclohydrolase | Cytoplasm | enzyme | 1.83E+07 | 3839903.25 | 8233701.5 | | | | | | |
| 5-aminoimidazole-4-carboxamide ribonucleotide formyltransferase/IMP cyclohydrolase | Cytoplasm | enzyme | 1.41E+07 | 4153529 | 6274783 | | | | | | |
| 5-aminoimidazole-4-carboxamide ribonucleotide formyltransferase/IMP cyclohydrolase | Cytoplasm | enzyme | 9375410 | 3600440.75 | 5059130 | | | | | | |
| 5-aminoimidazole-4-carboxamide ribonucleotide formyltransferase/IMP cyclohydrolase | Cytoplasm | enzyme | 9056291 | 3287543.25 | 4194730.5 | | | | | | |
| 5-aminoimidazole-4-carboxamide ribonucleotide formyltransferase/IMP cyclohydrolase | Cytoplasm | enzyme | 8395289 | 2474540.5 | 3953776.5 | | | | | | |
| 5-aminoimidazole-4-carboxamide ribonucleotide formyltransferase/IMP cyclohydrolase | Cytoplasm | enzyme | 7872085.5 | 1391482.125 | 3152852.5 | | | | | | |
| 5-aminoimidazole-4-carboxamide ribonucleotide formyltransferase/IMP cyclohydrolase | Cytoplasm | enzyme | 6835483 | 1862974.875 | 3314272.75 | | | | | | |
| 5-aminoimidazole-4-carboxamide ribonucleotide formyltransferase/IMP cyclohydrolase | Cytoplasm | enzyme | 3814508.5 | 1174212.75 | 1360032.125 | | | | | | |
| 5-aminoimidazole-4-carboxamide ribonucleotide formyltransferase/IMP cyclohydrolase | Cytoplasm | enzyme | 2712634.25 | 292932.281 | 843935.938 | | | | | | |
| 5-aminoimidazole-4-carboxamide ribonucleotide formyltransferase/IMP cyclohydrolase | Cytoplasm | enzyme | 2176307.5 | 999395.562 | 1977253.375 | Q9CWJ9 | ATIC | 8259801 | 4068219 | 742752 | 30 |
| atlastin GTPase 2 | Cytoplasm | other | 5746725.5 | 4557817 | 6868219.5 | | | | | | |
| atlastin GTPase 2 | Cytoplasm | other | 1550851.625 | 1402108.625 | 1566978.75 | | | | | | |
| atlastin GTPase 2 | Cytoplasm | other | 1357422.25 | 1278183.875 | 1400569.125 | Q6PA06 | ATL2 | 2885000 | 2227179 | 742393 | 9 |
| atlastin GTPase 3 | Cytoplasm | other | 6.55E+07 | 5.34E+07 | 5.44E+07 | | | | | | |
| atlastin GTPase 3 | Cytoplasm | other | 3.34E+07 | 3.12E+07 | 3.27E+07 | | | | | | |
| atlastin GTPase 3 | Cytoplasm | other | 2.87E+07 | 3.59E+07 | 2.66E+07 | | | | | | |
| atlastin GTPase 3 | Cytoplasm | other | 2.71E+07 | 2.32E+07 | 1.78E+07 | | | | | | |
| atlastin GTPase 3 | Cytoplasm | other | 2.69E+07 | 2.76E+07 | 2.50E+07 | | | | | | |
| atlastin GTPase 3 | Cytoplasm | other | 2.49E+07 | 2.69E+07 | 2.29E+07 | | | | | | |
| atlastin GTPase 3 | Cytoplasm | other | 2.32E+07 | 2.25E+07 | 2.44E+07 | | | | | | |
| atlastin GTPase 3 | Cytoplasm | other | 2.10E+07 | 2.01E+07 | 1.77E+07 | | | | | | |
| atlastin GTPase 3 | Cytoplasm | other | 1.89E+07 | 1.75E+07 | 1.76E+07 | | | | | | |
| atlastin GTPase 3 | Cytoplasm | other | 1.05E+07 | 7871976.5 | 9318293 | | | | | | |
| atlastin GTPase 3 | Cytoplasm | other | 9258101 | 7542681.5 | 8438100 | | | | | | |
| atlastin GTPase 3 | Cytoplasm | other | 7186214 | 6183320.5 | 5364611 | | | | | | |
| atlastin GTPase 3 | Cytoplasm | other | 6424005.5 | 6460958 | 5217709 | | | | | | |
| atlastin GTPase 3 | Cytoplasm | other | 6175788 | 5353624 | 5345477 | | | | | | |
| atlastin GTPase 3 | Cytoplasm | other | 5878640.5 | 4225414 | 4029663 | | | | | | |
| atlastin GTPase 3 | Cytoplasm | other | 4538636 | 2445662.75 | 3331203.75 | Q91YH5 | ATL3 | 19956962 | 14224386 | 2053113 | 48 |
| ATPase, class VI, type 11C | Plasma Membrane | transporter | 1.19E+07 | 1.20E+07 | 9493398 | | | | | | |
| ATPase, class VI, type 11C | Plasma Membrane | transporter | 1.19E+07 | 7927912 | 8819225 | | | | | | |
| ATPase, class VI, type 11C | Plasma Membrane | transporter | 5748351.5 | 4995090.5 | 5062351 | | | | | | |
| ATPase, class VI, type 11C | Plasma Membrane | transporter | 4637733.5 | 3231498.75 | 3176995.5 | | | | | | |
| ATPase, class VI, type 11C | Plasma Membrane | transporter | 3297193.25 | 2181527.75 | 3406545 | | | | | | |
| ATPase, class VI, type 11C | Plasma Membrane | transporter | 2556644 | 2155431.25 | 2516475.5 | Q9QZW0 | ATP11C | 6673320 | 3564056 | 840056 | 18 |
| ATPase type 13A1 | Extracellular Space | transporter | 1.65E+07 | 7898092 | 1.32E+07 | Q9EPE9 | ATP13A1 | 16460000 | 4322934 | 2495847 | 3 |
| ATPase, Na+/K+ transporting, alpha 1 polypeptide | Plasma Membrane | transporter | 3.72E+07 | 2.71E+07 | 2.54E+07 | | | | | | |
| ATPase, Na+/K+ transporting, alpha 1 polypeptide | Plasma Membrane | transporter | 3.65E+07 | 3.42E+07 | 3.20E+07 | | | | | | |
| ATPase, Na+/K+ transporting, alpha 1 polypeptide | Plasma Membrane | transporter | 3.41E+07 | 2.99E+07 | 2.54E+07 | | | | | | |
| ATPase, Na+/K+ transporting, alpha 1 polypeptide | Plasma Membrane | transporter | 3.08E+07 | 2.25E+07 | 2.41E+07 | | | | | | |
| ATPase, Na+/K+ transporting, alpha 1 polypeptide | Plasma Membrane | transporter | 2.94E+07 | 2.19E+07 | 2.35E+07 | | | | | | |
| ATPase, Na+/K+ transporting, alpha 1 polypeptide | Plasma Membrane | transporter | 2.54E+07 | 2.40E+07 | 1.77E+07 | | | | | | |
| ATPase, Na+/K+ transporting, alpha 1 polypeptide | Plasma Membrane | transporter | 2.13E+07 | 1.82E+07 | 1.78E+07 | | | | | | |
| ATPase, Na+/K+ transporting, alpha 1 polypeptide | Plasma Membrane | transporter | 2.09E+07 | 1.87E+07 | 1.89E+07 | | | | | | |
| ATPase, Na+/K+ transporting, alpha 1 polypeptide | Plasma Membrane | transporter | 1.99E+07 | 1.53E+07 | 1.50E+07 | | | | | | |
| ATPase, Na+/K+ transporting, alpha 1 polypeptide | Plasma Membrane | transporter | 1.92E+07 | 2.11E+07 | 1.71E+07 | | | | | | |
| ATPase, Na+/K+ transporting, alpha 1 polypeptide | Plasma Membrane | transporter | 1.87E+07 | 1.48E+07 | 1.67E+07 | | | | | | |
| ATPase, Na+/K+ transporting, alpha 1 polypeptide | Plasma Membrane | transporter | 1.37E+07 | 9498249 | 1.04E+07 | | | | | | |
| ATPase, Na+/K+ transporting, alpha 1 polypeptide | Plasma Membrane | transporter | 1.36E+07 | 1.14E+07 | 1.00E+07 | | | | | | |
| ATPase, Na+/K+ transporting, alpha 1 polypeptide | Plasma Membrane | transporter | 1.35E+07 | 1.10E+07 | 9134823 | | | | | | |
| ATPase, Na+/K+ transporting, alpha 1 polypeptide | Plasma Membrane | transporter | 1.10E+07 | 8935065 | 1.13E+07 | | | | | | |
| ATPase, Na+/K+ transporting, alpha 1 polypeptide | Plasma Membrane | transporter | 1.08E+07 | 6808627 | 1.06E+07 | | | | | | |
| ATPase, Na+/K+ transporting, alpha 1 polypeptide | Plasma Membrane | transporter | 1.08E+07 | 6310456 | 8163742.5 | | | | | | |
| ATPase, Na+/K+ transporting, alpha 1 polypeptide | Plasma Membrane | transporter | 7016080.5 | 5668538 | 5809364.5 | | | | | | |
| ATPase, Na+/K+ transporting, alpha 1 polypeptide | Plasma Membrane | transporter | 6509640.5 | 6124833 | 5628164 | | | | | | |
| ATPase, Na+/K+ transporting, alpha 1 polypeptide | Plasma Membrane | transporter | 5942002.5 | 3392989.5 | 4189935 | | | | | | |
| ATPase, Na+/K+ transporting, alpha 1 polypeptide | Plasma Membrane | transporter | 5521442 | 5576213 | 6000720 | | | | | | |
| ATPase, Na+/K+ transporting, alpha 1 polypeptide | Plasma Membrane | transporter | 5144643.5 | 2995472.25 | 4025440.25 | | | | | | |
| ATPase, Na+/K+ transporting, alpha 1 polypeptide | Plasma Membrane | transporter | 5042261 | 5535017 | 5045738.5 | | | | | | |
| ATPase, Na+/K+ transporting, alpha 1 polypeptide | Plasma Membrane | transporter | 3418501.25 | 3702842 | 4475377.5 | | | | | | |
| ATPase, Na+/K+ transporting, alpha 1 polypeptide | Plasma Membrane | transporter | 3132450.75 | 4571305.5 | 6696864.5 | | | | | | |
| ATPase, Na+/K+ transporting, alpha 1 polypeptide | Plasma Membrane | transporter | 2930533 | 3955678.5 | 3187777 | | | | | | |
| ATPase, Na+/K+ transporting, alpha 1 polypeptide | Plasma Membrane | transporter | 2548940.25 | 1846766.25 | 2740533.25 | | | | | | |
| ATPase, Na+/K+ transporting, alpha 1 polypeptide | Plasma Membrane | transporter | 2293840 | 1906027.875 | 1383651.375 | | | | | | |
| ATPase, Na+/K+ transporting, alpha 1 polypeptide | Plasma Membrane | transporter | 2183116.75 | 2912123 | 2742945.5 | Q8VDN2 | ATP1A1 | 14428395 | 9643633 | 1033906 | 87 |
| ATPase, Na+/K+ transporting, beta 2 polypeptide | Plasma Membrane | transporter | 7.31E+07 | 5.74E+07 | 4.95E+07 | | | | | | |
| ATPase, Na+/K+ transporting, beta 2 polypeptide | Plasma Membrane | transporter | 4.13E+07 | 2.87E+07 | 3.03E+07 | | | | | | |
| ATPase, Na+/K+ transporting, beta 2 polypeptide | Plasma Membrane | transporter | 9633963 | 7917677.5 | 9278553 | | | | | | |
| ATPase, Na+/K+ transporting, beta 2 polypeptide | Plasma Membrane | transporter | 8182511 | 5004554 | 4551032 | | | | | | |
| ATPase, Na+/K+ transporting, beta 2 polypeptide | Plasma Membrane | transporter | 3927156.75 | 3220352.5 | 2694094.25 | P14231 | ATP1B2 | 27226726 | 23068164 | 5956174 | 15 |

| | | | | | | | | | | | |
|---|---|---|---|---|---|---|---|---|---|---|---|
| ATPase, Na+/K+ transporting, beta 3 polypeptide | Plasma Membrane | transporter | 1667524.75 | 2991771.5 | 2861556 | | | | | | |
| ATPase, Na+/K+ transporting, beta 3 polypeptide | Plasma Membrane | transporter | 1094043.125 | 2107849 | 2296720.5 | | | | | | |
| ATPase, Na+/K+ transporting, beta 3 polypeptide | Plasma Membrane | transporter | 892384.688 | 1936690.125 | 1798436.5 | | | | | | |
| ATPase, Na+/K+ transporting, beta 3 polypeptide | Plasma Membrane | transporter | 740454.562 | 1327486.625 | 1738637.5 | P97370 | ATP1B3 | 1098602 | 713186 | 205879 | 12 |
| ATPase, Ca++ transporting, cardiac muscle, slow twitch 2 | Cytoplasm | transporter | 2364437.25 | 7904156.5 | 6155909 | | | | | | |
| ATPase, Ca++ transporting, cardiac muscle, slow twitch 2 | Cytoplasm | transporter | 1593060.875 | 2.16E+07 | 1.24E+07 | | | | | | |
| ATPase, Ca++ transporting, cardiac muscle, slow twitch 2 | Cytoplasm | transporter | 1010624.688 | 3651827.5 | 2515131.75 | | | | | | |
| ATPase, Ca++ transporting, cardiac muscle, slow twitch 2 | Cytoplasm | transporter | 676065.375 | 3351031.75 | 2607387.5 | | | | | | |
| ATPase, Ca++ transporting, cardiac muscle, slow twitch 2 | Cytoplasm | transporter | 533587.562 | 1423178 | 1640212.125 | | | | | | |
| ATPase, Ca++ transporting, cardiac muscle, slow twitch 2 | Cytoplasm | transporter | 303837.188 | 1537881.25 | 1428492.875 | O55143 | ATP2A2 | 1080269 | 5336259 | 1257768 | 18 |
| ATPase, Ca++ transporting, ubiquitous | Cytoplasm | transporter | 3022483 | 2.94E+07 | 2.04E+07 | | | | | | |
| ATPase, Ca++ transporting, ubiquitous | Cytoplasm | transporter | 1686538.125 | 1.82E+07 | 1.05E+07 | | | | | | |
| ATPase, Ca++ transporting, ubiquitous | Cytoplasm | transporter | 1358915.875 | 7959052 | 5294448.5 | | | | | | |
| ATPase, Ca++ transporting, ubiquitous | Cytoplasm | transporter | 835203.438 | 1.04E+07 | 7411475 | | | | | | |
| ATPase, Ca++ transporting, ubiquitous | Cytoplasm | transporter | 791212.938 | 2.28E+07 | 1.39E+07 | | | | | | |
| ATPase, Ca++ transporting, ubiquitous | Cytoplasm | transporter | 513953.25 | 5974276.5 | 4693777.5 | | | | | | |
| ATPase, Ca++ transporting, ubiquitous | Cytoplasm | transporter | 451844.25 | 6787288 | 5925467 | | | | | | |
| ATPase, Ca++ transporting, ubiquitous | Cytoplasm | transporter | 406390.25 | 1.79E+07 | 8670085 | | | | | | |
| ATPase, Ca++ transporting, ubiquitous | Cytoplasm | transporter | 375688.719 | 9108343 | 5088089.5 | | | | | | |
| ATPase, Ca++ transporting, ubiquitous | Cytoplasm | transporter | 360334.25 | 1.17E+07 | 8189618 | | | | | | |
| ATPase, Ca++ transporting, ubiquitous | Cytoplasm | transporter | 293732.938 | 3259874.5 | 2159574.25 | | | | | | |
| ATPase, Ca++ transporting, ubiquitous | Cytoplasm | transporter | 98958.156 | 1038232.125 | 647382.125 | | | | | | |
| ATPase, Ca++ transporting, ubiquitous | Cytoplasm | transporter | 42089.805 | 2228157.5 | 1022111.938 | | | | | | |
| ATPase, Ca++ transporting, ubiquitous | Cytoplasm | transporter | 7771.243 | 3978208 | 2654645.25 | | | | | | |
| ATPase, Ca++ transporting, ubiquitous | Cytoplasm | transporter | 5623.83 | 1356851.375 | 654877.25 | | | | | | |
| ATPase, Ca++ transporting, ubiquitous | Cytoplasm | transporter | 624.626 | 5262080 | 4627066.5 | Q64518 | ATP2A3 | 640710 | 6746505 | 973774 | 48 |
| ATPase, Ca++ transporting, plasma membrane 2 | Plasma Membrane | transporter | 1.61E+08 | 1.16E+08 | 1.08E+08 | | | | | | |
| ATPase, Ca++ transporting, plasma membrane 2 | Plasma Membrane | transporter | 4.86E+07 | 4.79E+07 | 4.45E+07 | | | | | | |
| ATPase, Ca++ transporting, plasma membrane 2 | Plasma Membrane | transporter | 4.36E+07 | 2.41E+07 | 2.90E+07 | | | | | | |
| ATPase, Ca++ transporting, plasma membrane 2 | Plasma Membrane | transporter | 4.36E+07 | 2.19E+07 | 1.87E+07 | | | | | | |
| ATPase, Ca++ transporting, plasma membrane 2 | Plasma Membrane | transporter | 2.75E+07 | 1.84E+07 | 2.01E+07 | | | | | | |
| ATPase, Ca++ transporting, plasma membrane 2 | Plasma Membrane | transporter | 1.19E+07 | 1.07E+07 | 9261237 | | | | | | |
| ATPase, Ca++ transporting, plasma membrane 2 | Plasma Membrane | transporter | 7889190.5 | 6759325 | 5967035.5 | | | | | | |
| ATPase, Ca++ transporting, plasma membrane 2 | Plasma Membrane | transporter | 7141959.5 | 1.11E+07 | 8410669 | | | | | | |
| ATPase, Ca++ transporting, plasma membrane 2 | Plasma Membrane | transporter | 3915931.75 | 2755355.25 | 3118420.25 | | | | | | |
| ATPase, Ca++ transporting, plasma membrane 2 | Plasma Membrane | transporter | 1723174.625 | 836460 | 1283896.375 | Q9R0K7 | ATP2B2 | 35639026 | 37544897 | 6854729 | 30 |
| ATP synthase, H+ transporting, mitochondrial F1 complex, alpha subunit 1, cardiac muscle | Cytoplasm | transporter | 2.53E+07 | 6.23E+07 | 5.68E+07 | | | | | | |
| ATP synthase, H+ transporting, mitochondrial F1 complex, alpha subunit 1, cardiac muscle | Cytoplasm | transporter | 2.48E+07 | 4.95E+07 | 4.59E+07 | | | | | | |
| ATP synthase, H+ transporting, mitochondrial F1 complex, alpha subunit 1, cardiac muscle | Cytoplasm | transporter | 2.27E+07 | 4.31E+07 | 4.16E+07 | | | | | | |
| ATP synthase, H+ transporting, mitochondrial F1 complex, alpha subunit 1, cardiac muscle | Cytoplasm | transporter | 2.17E+07 | 4.04E+07 | 3.77E+07 | | | | | | |
| ATP synthase, H+ transporting, mitochondrial F1 complex, alpha subunit 1, cardiac muscle | Cytoplasm | transporter | 2.03E+07 | 3.94E+07 | 3.77E+07 | | | | | | |
| ATP synthase, H+ transporting, mitochondrial F1 complex, alpha subunit 1, cardiac muscle | Cytoplasm | transporter | 1.97E+07 | 4.24E+07 | 3.52E+07 | | | | | | |
| ATP synthase, H+ transporting, mitochondrial F1 complex, alpha subunit 1, cardiac muscle | Cytoplasm | transporter | 1.92E+07 | 4.26E+07 | 3.59E+07 | | | | | | |
| ATP synthase, H+ transporting, mitochondrial F1 complex, alpha subunit 1, cardiac muscle | Cytoplasm | transporter | 1.77E+07 | 3.42E+07 | 2.87E+07 | | | | | | |
| ATP synthase, H+ transporting, mitochondrial F1 complex, alpha subunit 1, cardiac muscle | Cytoplasm | transporter | 1.63E+07 | 2.51E+07 | 2.57E+07 | | | | | | |
| ATP synthase, H+ transporting, mitochondrial F1 complex, alpha subunit 1, cardiac muscle | Cytoplasm | transporter | 1.62E+07 | 2.74E+07 | 2.83E+07 | | | | | | |
| ATP synthase, H+ transporting, mitochondrial F1 complex, alpha subunit 1, cardiac muscle | Cytoplasm | transporter | 1.59E+07 | 2.74E+07 | 2.77E+07 | | | | | | |
| ATP synthase, H+ transporting, mitochondrial F1 complex, alpha subunit 1, cardiac muscle | Cytoplasm | transporter | 1.26E+07 | 2.79E+07 | 2.52E+07 | | | | | | |
| ATP synthase, H+ transporting, mitochondrial F1 complex, alpha subunit 1, cardiac muscle | Cytoplasm | transporter | 1.08E+07 | 2.38E+07 | 2.01E+07 | | | | | | |
| ATP synthase, H+ transporting, mitochondrial F1 complex, alpha subunit 1, cardiac muscle | Cytoplasm | transporter | 1.07E+07 | 2.81E+07 | 2.05E+07 | | | | | | |
| ATP synthase, H+ transporting, mitochondrial F1 complex, alpha subunit 1, cardiac muscle | Cytoplasm | transporter | 7950870.5 | 1.63E+07 | 1.41E+07 | | | | | | |
| ATP synthase, H+ transporting, mitochondrial F1 complex, alpha subunit 1, cardiac muscle | Cytoplasm | transporter | 7940947.5 | 1.56E+07 | 1.24E+07 | | | | | | |
| ATP synthase, H+ transporting, mitochondrial F1 complex, alpha subunit 1, cardiac muscle | Cytoplasm | transporter | 7092159.5 | 1.28E+07 | 1.19E+07 | | | | | | |
| ATP synthase, H+ transporting, mitochondrial F1 complex, alpha subunit 1, cardiac muscle | Cytoplasm | transporter | 5245350.5 | 1.06E+07 | 8448330 | | | | | | |
| ATP synthase, H+ transporting, mitochondrial F1 complex, alpha subunit 1, cardiac muscle | Cytoplasm | transporter | 4290945 | 5355014 | 5157822 | | | | | | |
| ATP synthase, H+ transporting, mitochondrial F1 complex, alpha subunit 1, cardiac muscle | Cytoplasm | transporter | 3510241.5 | 4640443 | 4746524.5 | | | | | | |
| ATP synthase, H+ transporting, mitochondrial F1 complex, alpha subunit 1, cardiac muscle | Cytoplasm | transporter | 2856056.25 | 6874706.5 | 6825639 | | | | | | |
| ATP synthase, H+ transporting, mitochondrial F1 complex, alpha subunit 1, cardiac muscle | Cytoplasm | transporter | 2231530.75 | 5662424 | 7127865 | | | | | | |
| ATP synthase, H+ transporting, mitochondrial F1 complex, alpha subunit 1, cardiac muscle | Cytoplasm | transporter | 83571.609 | 7598651.5 | 1.10E+07 | Q03265 | ATP5A1 | 12827899 | 14422122 | 1736220 | 69 |
| ATP synthase, H+ transporting, mitochondrial F1 complex, beta polypeptide | Cytoplasm | transporter | 5.91E+07 | 9.24E+07 | 8.82E+07 | | | | | | |
| ATP synthase, H+ transporting, mitochondrial F1 complex, beta polypeptide | Cytoplasm | transporter | 3.85E+07 | 7.80E+07 | 6.17E+07 | | | | | | |
| ATP synthase, H+ transporting, mitochondrial F1 complex, beta polypeptide | Cytoplasm | transporter | 3.40E+07 | 4.93E+07 | 5.45E+07 | | | | | | |
| ATP synthase, H+ transporting, mitochondrial F1 complex, beta polypeptide | Cytoplasm | transporter | 2.75E+07 | 4.88E+07 | 4.73E+07 | | | | | | |
| ATP synthase, H+ transporting, mitochondrial F1 complex, beta polypeptide | Cytoplasm | transporter | 2.05E+07 | 5.40E+07 | 4.28E+07 | | | | | | |
| ATP synthase, H+ transporting, mitochondrial F1 complex, beta polypeptide | Cytoplasm | transporter | 1.98E+07 | 2.79E+07 | 2.40E+07 | | | | | | |
| ATP synthase, H+ transporting, mitochondrial F1 complex, beta polypeptide | Cytoplasm | transporter | 1.87E+07 | 3.56E+07 | 3.12E+07 | | | | | | |
| ATP synthase, H+ transporting, mitochondrial F1 complex, beta polypeptide | Cytoplasm | transporter | 1.83E+07 | 3.84E+07 | 4.38E+07 | | | | | | |
| ATP synthase, H+ transporting, mitochondrial F1 complex, beta polypeptide | Cytoplasm | transporter | 1.80E+07 | 2.49E+07 | 2.54E+07 | | | | | | |
| ATP synthase, H+ transporting, mitochondrial F1 complex, beta polypeptide | Cytoplasm | transporter | 1.69E+07 | 2.93E+07 | 2.59E+07 | | | | | | |
| ATP synthase, H+ transporting, mitochondrial F1 complex, beta polypeptide | Cytoplasm | transporter | 1.63E+07 | 4.58E+07 | 3.95E+07 | | | | | | |
| ATP synthase, H+ transporting, mitochondrial F1 complex, beta polypeptide | Cytoplasm | transporter | 1.49E+07 | 3.50E+07 | 3.24E+07 | | | | | | |
| ATP synthase, H+ transporting, mitochondrial F1 complex, beta polypeptide | Cytoplasm | transporter | 1.41E+07 | 1.94E+07 | 2.04E+07 | | | | | | |
| ATP synthase, H+ transporting, mitochondrial F1 complex, beta polypeptide | Cytoplasm | transporter | 1.37E+07 | 2.63E+07 | 2.41E+07 | | | | | | |
| ATP synthase, H+ transporting, mitochondrial F1 complex, beta polypeptide | Cytoplasm | transporter | 1.32E+07 | 1.65E+07 | 2.53E+07 | | | | | | |
| ATP synthase, H+ transporting, mitochondrial F1 complex, beta polypeptide | Cytoplasm | transporter | 1.28E+07 | 2.78E+07 | 2.42E+07 | | | | | | |
| ATP synthase, H+ transporting, mitochondrial F1 complex, beta polypeptide | Cytoplasm | transporter | 9951699 | 2.45E+07 | 2.49E+07 | | | | | | |
| ATP synthase, H+ transporting, mitochondrial F1 complex, beta polypeptide | Cytoplasm | transporter | 9923226 | 2.01E+07 | 1.54E+07 | | | | | | |
| ATP synthase, H+ transporting, mitochondrial F1 complex, beta polypeptide | Cytoplasm | transporter | 8624374 | 8954405 | 1.05E+07 | | | | | | |
| ATP synthase, H+ transporting, mitochondrial F1 complex, beta polypeptide | Cytoplasm | transporter | 7790438.5 | 1.79E+07 | 1.92E+07 | | | | | | |
| ATP synthase, H+ transporting, mitochondrial F1 complex, beta polypeptide | Cytoplasm | transporter | 7010379.5 | 1.39E+07 | 1.36E+07 | | | | | | |
| ATP synthase, H+ transporting, mitochondrial F1 complex, beta polypeptide | Cytoplasm | transporter | 7007126.5 | 1.69E+07 | 1.22E+07 | | | | | | |
| ATP synthase, H+ transporting, mitochondrial F1 complex, beta polypeptide | Cytoplasm | transporter | 5694345.5 | 8195824 | 6869690.5 | | | | | | |
| ATP synthase, H+ transporting, mitochondrial F1 complex, beta polypeptide | Cytoplasm | transporter | 5592985.5 | 9347862 | 8246310.5 | | | | | | |
| ATP synthase, H+ transporting, mitochondrial F1 complex, beta polypeptide | Cytoplasm | transporter | 5007482.5 | 7688263 | 5610585 | | | | | | |
| ATP synthase, H+ transporting, mitochondrial F1 complex, beta polypeptide | Cytoplasm | transporter | 3191018 | 4991914.5 | 5470303 | | | | | | |
| ATP synthase, H+ transporting, mitochondrial F1 complex, beta polypeptide | Cytoplasm | transporter | 2581953.25 | 2739505.25 | 4521016 | | | | | | |
| ATP synthase, H+ transporting, mitochondrial F1 complex, beta polypeptide | Cytoplasm | transporter | 2419253.5 | 1.72E+07 | 8386847.5 | | | | | | |
| ATP synthase, H+ transporting, mitochondrial F1 complex, beta polypeptide | Cytoplasm | transporter | 1846790.125 | 5384692.5 | 5034450.5 | | | | | | |
| ATP synthase, H+ transporting, mitochondrial F1 complex, beta polypeptide | Cytoplasm | transporter | 1363568 | 5914489 | 4569872.5 | | | | | | |
| ATP synthase, H+ transporting, mitochondrial F1 complex, beta polypeptide | Cytoplasm | transporter | 289277.844 | 261591.094 | 333630.688 | | | | | | |
| ATP synthase, H+ transporting, mitochondrial F1 complex, beta polypeptide | Cytoplasm | transporter | 264072.062 | 360478.906 | 513134.312 | P56480 | ATP5B | 13588062 | 19024547 | 1941685 | 96 |
| ATP synthase, H+ transporting, mitochondrial F1 complex, gamma polypeptide 1 | Cytoplasm | transporter | 3697722.75 | 9014425 | 7871819 | | | | | | |
| ATP synthase, H+ transporting, mitochondrial F1 complex, gamma polypeptide 1 | Cytoplasm | transporter | 2513397 | 6070370 | 3995069.5 | | | | | | |
| ATP synthase, H+ transporting, mitochondrial F1 complex, gamma polypeptide 1 | Cytoplasm | transporter | 1968314.875 | 4473069.5 | 2658031.25 | | | | | | |
| ATP synthase, H+ transporting, mitochondrial F1 complex, gamma polypeptide 1 | Cytoplasm | transporter | 208675.609 | 1222402.375 | 856700.688 | Q91VR2 | ATP5C1 | 2097028 | 2760983 | 797027 | 12 |

| | | | | | | | | | | | |
|---|---|---|---|---|---|---|---|---|---|---|---|
| ATP synthase, H+ transporting, mitochondrial F1 complex, delta subunit | Cytoplasm | transporter | 6729322.5 | 1.07E+07 | 9581481 | | | | | | |
| ATP synthase, H+ transporting, mitochondrial F1 complex, delta subunit | Cytoplasm | transporter | 6364967 | 1.02E+07 | 9086542 | Q9D3D9 | ATP5D | 6547145 | 1818004 | 742197 | 6 |
| ATP synthase, H+ transporting, mitochondrial Fo complex, subunit B1 | Cytoplasm | transporter | 2349075.75 | 4684083 | 2866636.25 | Q9CQQ7 | ATP5F1 | 2349076 | 1226325 | 708019 | 3 |
| ATP synthase, H+ transporting, mitochondrial Fo complex, subunit d | Cytoplasm | enzyme | 5742954 | 1.05E+07 | 7741065 | | | | | | |
| ATP synthase, H+ transporting, mitochondrial Fo complex, subunit d | Cytoplasm | enzyme | 3881063.75 | 6337818 | 6110809 | | | | | | |
| ATP synthase, H+ transporting, mitochondrial Fo complex, subunit d | Cytoplasm | enzyme | 3215293.5 | 5710261 | 4984125 | | | | | | |
| ATP synthase, H+ transporting, mitochondrial Fo complex, subunit d | Cytoplasm | enzyme | 2497807.75 | 2729235 | 2728766 | | | | | | |
| ATP synthase, H+ transporting, mitochondrial Fo complex, subunit d | Cytoplasm | enzyme | 2479783.25 | 2848673.75 | 3284019.5 | | | | | | |
| ATP synthase, H+ transporting, mitochondrial Fo complex, subunit d | Cytoplasm | enzyme | 1562960.5 | 1973557.375 | 1912362.125 | Q9DCX2 | ATP5H | 3229977 | 2382770 | 561624 | 18 |
| ATP synthase, H+ transporting, mitochondrial Fo complex, subunit F6 | Cytoplasm | transporter | 2811276.25 | 4731390 | 3561946.25 | P97450 | ATP5J | 2811276 | 967638 | 558666 | 3 |
| ATP synthase, H+ transporting, mitochondrial Fo complex, subunit F2 | Cytoplasm | transporter | 2279644.25 | 5674063 | 4787921 | P56135 | ATP5J2 | 2279644 | 1760624 | 1016497 | 3 |
| ATP synthase, H+ transporting, mitochondrial F1F0 complex, subunit E | Cytoplasm | enzyme | 7253758.5 | 1.07E+07 | 8504727 | | | | | | |
| ATP synthase, H+ transporting, mitochondrial F1F0 complex, subunit E | Cytoplasm | enzyme | 1796963.875 | 3925616.5 | 3896708.5 | Q06185 | Atp5k | 4525361 | 3364787 | 1373668 | 6 |
| ATP synthase, H+ transporting, mitochondrial Fo complex, subunit G | Cytoplasm | enzyme | 2681157.25 | 6473283.5 | 6857173 | | | | | | |
| ATP synthase, H+ transporting, mitochondrial Fo complex, subunit G | Cytoplasm | enzyme | 2051359.375 | 3119617.25 | 2605689 | Q9CPQ8 | ATP5L | 2366258 | 2122679 | 866580 | 6 |
| ATP synthase, H+ transporting, mitochondrial F1 complex, O subunit | Cytoplasm | transporter | 8294045.5 | 1.52E+07 | 1.24E+07 | | | | | | |
| ATP synthase, H+ transporting, mitochondrial F1 complex, O subunit | Cytoplasm | transporter | 6528194.5 | 1.03E+07 | 1.16E+07 | | | | | | |
| ATP synthase, H+ transporting, mitochondrial F1 complex, O subunit | Cytoplasm | transporter | 5763950 | 8279672 | 6460227.5 | | | | | | |
| ATP synthase, H+ transporting, mitochondrial F1 complex, O subunit | Cytoplasm | transporter | 4200996.5 | 8564470 | 1.02E+07 | | | | | | |
| ATP synthase, H+ transporting, mitochondrial F1 complex, O subunit | Cytoplasm | transporter | 3017887 | 1.19E+07 | 6801826 | | | | | | |
| ATP synthase, H+ transporting, mitochondrial F1 complex, O subunit | Cytoplasm | transporter | 2664701.75 | 3646937 | 3051310.75 | | | | | | |
| ATP synthase, H+ transporting, mitochondrial F1 complex, O subunit | Cytoplasm | transporter | 2393033.75 | 4375988.5 | 3956564 | | | | | | |
| ATP synthase, H+ transporting, mitochondrial F1 complex, O subunit | Cytoplasm | transporter | 2230448.5 | 4334195 | 4179473.5 | Q9DB20 | ATP5O | 4386657 | 3679583 | 751092 | 24 |
| ATPase, H+ transporting, lysosomal accessory protein 1 | Cytoplasm | transporter | 1849545.25 | 1248186.75 | 1641505.625 | Q9R1Q9 | ATP6AP1 | 1849545 | 305399 | 176322 | 3 |
| ATPase, H+ transporting, lysosomal accessory protein 2 | Cytoplasm | transporter | 2443582 | 1578674.625 | 1675674.125 | Q9CYN9 | ATP6AP2 | 2443582 | 473842 | 273573 | 3 |
| ATPase, H+ transporting, lysosomal V0 subunit a1 | Cytoplasm | transporter | 9719886 | 6587901 | 8838134 | | | | | | |
| ATPase, H+ transporting, lysosomal V0 subunit a1 | Cytoplasm | transporter | 9595206 | 4868707 | 5856930.5 | | | | | | |
| ATPase, H+ transporting, lysosomal V0 subunit a1 | Cytoplasm | transporter | 7760668.5 | 6156176 | 7090258 | | | | | | |
| ATPase, H+ transporting, lysosomal V0 subunit a1 | Cytoplasm | transporter | 7679975 | 6015559 | 8381024.5 | | | | | | |
| ATPase, H+ transporting, lysosomal V0 subunit a1 | Cytoplasm | transporter | 7091966.5 | 8593046 | 1.02E+07 | | | | | | |
| ATPase, H+ transporting, lysosomal V0 subunit a1 | Cytoplasm | transporter | 5447661.5 | 2976825.25 | 3759521 | | | | | | |
| ATPase, H+ transporting, lysosomal V0 subunit a1 | Cytoplasm | transporter | 5022845 | 3593598 | 5000100.5 | | | | | | |
| ATPase, H+ transporting, lysosomal V0 subunit a1 | Cytoplasm | transporter | 4853748.5 | 4108104.75 | 4215928 | | | | | | |
| ATPase, H+ transporting, lysosomal V0 subunit a1 | Cytoplasm | transporter | 2921845 | 1647292 | 2471982.75 | Q9Z1G4 | ATP6V0A1 | 6677089 | 2369484 | 456007 | 27 |
| ATPase, H+ transporting, lysosomal 16kDa, V0 subunit c | Cytoplasm | transporter | 1.85E+07 | 2.12E+07 | 2.59E+07 | | | | | | |
| ATPase, H+ transporting, lysosomal 16kDa, V0 subunit c | Cytoplasm | transporter | 1407564.875 | 2295389.5 | 3280682.5 | P63082 | ATP6V0C | 9968782 | 10986974 | 4485413 | 6 |
| ATPase, H+ transporting, lysosomal 38kDa, V0 subunit d1 | Cytoplasm | transporter | 1.64E+07 | 1.37E+07 | 1.79E+07 | | | | | | |
| ATPase, H+ transporting, lysosomal 38kDa, V0 subunit d1 | Cytoplasm | transporter | 1.20E+07 | 1.20E+07 | 1.37E+07 | | | | | | |
| ATPase, H+ transporting, lysosomal 38kDa, V0 subunit d1 | Cytoplasm | transporter | 8645496 | 6122683 | 8833788 | | | | | | |
| ATPase, H+ transporting, lysosomal 38kDa, V0 subunit d1 | Cytoplasm | transporter | 7423739.5 | 4793219.5 | 5919615.5 | | | | | | |
| ATPase, H+ transporting, lysosomal 38kDa, V0 subunit d1 | Cytoplasm | transporter | 3399867.75 | 2799277.75 | 4603067.5 | | | | | | |
| ATPase, H+ transporting, lysosomal 38kDa, V0 subunit d1 | Cytoplasm | transporter | 3395860.25 | 2558149.5 | 3131522.25 | P51863 | ATP6V0D1 | 8539161 | 4965665 | 1170419 | 18 |
| ATPase, H+ transporting, lysosomal 70kDa, V1 subunit A | Plasma Membrane | transporter | 5130404 | 4393935 | 5546874 | | | | | | |
| ATPase, H+ transporting, lysosomal 70kDa, V1 subunit A | Plasma Membrane | transporter | 3339848.5 | 981321.562 | 1977905.375 | | | | | | |
| ATPase, H+ transporting, lysosomal 70kDa, V1 subunit A | Plasma Membrane | transporter | 3144007 | 1644246.375 | 3365421.75 | | | | | | |
| ATPase, H+ transporting, lysosomal 70kDa, V1 subunit A | Plasma Membrane | transporter | 2480087.25 | 1088639 | 2348122.5 | | | | | | |
| ATPase, H+ transporting, lysosomal 70kDa, V1 subunit A | Plasma Membrane | transporter | 1671671.875 | 866154.75 | 549703.5 | P50516 | ATP6V1A | 3153204 | 1560081 | 402811 | 15 |
| ATPase, H+ transporting, lysosomal 56/58kDa, V1 subunit B2 | Cytoplasm | transporter | 1.09E+07 | 8271646.5 | 8376620 | | | | | | |
| ATPase, H+ transporting, lysosomal 56/58kDa, V1 subunit B2 | Cytoplasm | transporter | 3390886.75 | 1352597.375 | 1676448.375 | | | | | | |
| ATPase, H+ transporting, lysosomal 56/58kDa, V1 subunit B2 | Cytoplasm | transporter | 2538100 | 1563677.125 | 1410293.375 | P62814 | ATP6V1B2 | 5592996 | 3717777 | 1239259 | 9 |
| ATPase, H+ transporting, lysosomal 31kDa, V1 subunit E1 | Cytoplasm | transporter | 1957698.625 | 1087643.625 | 1976568.25 | P50518 | ATP6V1E1 | 1957699 | 507861 | 293214 | 3 |
| ATPase, H+ transporting, lysosomal 50/57kDa, V1 subunit H | Cytoplasm | transporter | 2144919.25 | 898343.75 | 1155494.625 | | | | | | |
| ATPase, H+ transporting, lysosomal 50/57kDa, V1 subunit H | Cytoplasm | transporter | 814540.812 | 372027.688 | 289065.156 | Q8BVE3 | ATP6V1H | 1479730 | 672585 | 274582 | 6 |
| ATPase, Cu++ transporting, beta polypeptide | Cytoplasm | transporter | 1.85E+07 | 1.62E+07 | 1.72E+07 | | | | | | |
| ATPase, Cu++ transporting, beta polypeptide | Cytoplasm | transporter | 3915294.75 | 2912384.5 | 2320024.25 | | | | | | |
| ATPase, Cu++ transporting, beta polypeptide | Cytoplasm | transporter | 3431349.5 | 2761436.25 | 2607327.75 | | | | | | |
| ATPase, Cu++ transporting, beta polypeptide | Cytoplasm | transporter | 3383569.75 | 3246469.75 | 4447780 | | | | | | |
| ATPase, Cu++ transporting, beta polypeptide | Cytoplasm | transporter | 3027189.75 | 2866064.25 | 2067537.125 | | | | | | |
| ATPase, Cu++ transporting, beta polypeptide | Cytoplasm | transporter | 2545043.25 | 2955090.5 | 2365346.25 | | | | | | |
| ATPase, Cu++ transporting, beta polypeptide | Cytoplasm | transporter | 2188402.25 | 2764671.75 | 1803696.375 | | | | | | |
| ATPase, Cu++ transporting, beta polypeptide | Cytoplasm | transporter | 1906638.375 | 1693125.75 | 1468345.5 | Q64446 | ATP7B | 4860936 | 4990502 | 1018682 | 24 |
| ATPase, aminophospholipid transporter (APLT), class I, type 8A, member 1 | Cytoplasm | transporter | 3.01E+07 | 2.29E+07 | 2.78E+07 | | | | | | |
| ATPase, aminophospholipid transporter (APLT), class I, type 8A, member 1 | Cytoplasm | transporter | 1.72E+07 | 1.34E+07 | 1.44E+07 | | | | | | |
| ATPase, aminophospholipid transporter (APLT), class I, type 8A, member 1 | Cytoplasm | transporter | 1.65E+07 | 1.78E+07 | 2.10E+07 | | | | | | |
| ATPase, aminophospholipid transporter (APLT), class I, type 8A, member 1 | Cytoplasm | transporter | 1.47E+07 | 1.35E+07 | 1.22E+07 | | | | | | |
| ATPase, aminophospholipid transporter (APLT), class I, type 8A, member 1 | Cytoplasm | transporter | 1.28E+07 | 1.15E+07 | 1.28E+07 | | | | | | |
| ATPase, aminophospholipid transporter (APLT), class I, type 8A, member 1 | Cytoplasm | transporter | 1.20E+07 | 1.17E+07 | 1.10E+07 | | | | | | |
| ATPase, aminophospholipid transporter (APLT), class I, type 8A, member 1 | Cytoplasm | transporter | 1.06E+07 | 1.06E+07 | 9952060 | | | | | | |
| ATPase, aminophospholipid transporter (APLT), class I, type 8A, member 1 | Cytoplasm | transporter | 9973173 | 8315941 | 1.12E+07 | | | | | | |
| ATPase, aminophospholipid transporter (APLT), class I, type 8A, member 1 | Cytoplasm | transporter | 8820811 | 7355402 | 7969876 | | | | | | |
| ATPase, aminophospholipid transporter (APLT), class I, type 8A, member 1 | Cytoplasm | transporter | 7887504.5 | 6876832 | 7177361 | | | | | | |
| ATPase, aminophospholipid transporter (APLT), class I, type 8A, member 1 | Cytoplasm | transporter | 7526191.5 | 6970625.5 | 6863204.5 | | | | | | |
| ATPase, aminophospholipid transporter (APLT), class I, type 8A, member 1 | Cytoplasm | transporter | 7357988.5 | 5410522.5 | 4608254 | | | | | | |
| ATPase, aminophospholipid transporter (APLT), class I, type 8A, member 1 | Cytoplasm | transporter | 7197700.5 | 7641825.5 | 7450027.5 | | | | | | |
| ATPase, aminophospholipid transporter (APLT), class I, type 8A, member 1 | Cytoplasm | transporter | 6661979 | 7384245 | 7317475 | | | | | | |
| ATPase, aminophospholipid transporter (APLT), class I, type 8A, member 1 | Cytoplasm | transporter | 6097965.5 | 7692177 | 7801081.5 | | | | | | |
| ATPase, aminophospholipid transporter (APLT), class I, type 8A, member 1 | Cytoplasm | transporter | 5938791.5 | 6232239.5 | 4399606 | | | | | | |
| ATPase, aminophospholipid transporter (APLT), class I, type 8A, member 1 | Cytoplasm | transporter | 5556128 | 5970925.5 | 4868258.5 | | | | | | |
| ATPase, aminophospholipid transporter (APLT), class I, type 8A, member 1 | Cytoplasm | transporter | 4626159 | 4332113 | 4043690.5 | | | | | | |
| ATPase, aminophospholipid transporter (APLT), class I, type 8A, member 1 | Cytoplasm | transporter | 4512726.5 | 4131109.25 | 5042487 | | | | | | |
| ATPase, aminophospholipid transporter (APLT), class I, type 8A, member 1 | Cytoplasm | transporter | 3964031 | 3320268.75 | 4047337.25 | | | | | | |
| ATPase, aminophospholipid transporter (APLT), class I, type 8A, member 1 | Cytoplasm | transporter | 3900865.25 | 3157273.75 | 3232634 | | | | | | |
| ATPase, aminophospholipid transporter (APLT), class I, type 8A, member 1 | Cytoplasm | transporter | 2920926 | 1699427.75 | 2136523.5 | P70704 | ATP8A1 | 9401952 | 5707003 | 702484 | 66 |
| ATPase inhibitory factor 1 | Cytoplasm | other | 2669393.5 | 7639777.5 | 6440296.5 | O35143 | ATPIF1 | 2669394 | 2593684 | 1497464 | 3 |
| ancient ubiquitous protein 1 | Cytoplasm | other | 2094437.75 | 6137419.5 | 3376555.5 | P70295 | AUP1 | 2094438 | 2066071 | 1192847 | 3 |
| beta-2-microglobulin | Plasma Membrane | transmembrane receptor | 3706802.5 | 6289381.5 | 4267590.5 | P01887 | B2M | 3706803 | 1358420 | 784284 | 3 |
| beta-1,4-N-acetyl-galactosaminyl transferase 2 | Cytoplasm | enzyme | 1.97E+07 | 2.28E+07 | 2.45E+07 | Q09199 | B4GALNT2 | 19740000 | 2411231 | 1392125 | 3 |
| UDP-Gal:betaGlcNAc beta 1,4- galactosyltransferase, polypeptide 4 | Cytoplasm | enzyme | 2.39E+08 | 1.28E+08 | 1.56E+08 | Q9JJ04 | B4GALT4 | 239200000 | 57952078 | 33458648 | 3 |
| BCL2-associated athanogene 2 | Cytoplasm | other | 5.78E+07 | 3.99E+07 | 4.50E+07 | | | | | | |
| BCL2-associated athanogene 2 | Cytoplasm | other | 3.72E+07 | 1.89E+07 | 2.68E+07 | | | | | | |
| BCL2-associated athanogene 2 | Cytoplasm | other | 2.15E+07 | 1.92E+07 | 2.31E+07 | | | | | | |
| BCL2-associated athanogene 2 | Cytoplasm | other | 2.04E+07 | 9242177 | 1.08E+07 | | | | | | |
| BCL2-associated athanogene 2 | Cytoplasm | other | 1.58E+07 | 1.26E+07 | 1.20E+07 | | | | | | |
| BCL2-associated athanogene 2 | Cytoplasm | other | 1.36E+07 | 8918603 | 1.32E+07 | | | | | | |
| BCL2-associated athanogene 2 | Cytoplasm | other | 1.33E+07 | 1.38E+07 | 1.04E+07 | | | | | | |

| | | | | | | | | | | | |
|---|---|---|---|---|---|---|---|---|---|---|---|
| BCL2-associated athanogene 2 | Cytoplasm | other | 1.12E+07 | 1.56E+07 | 1.26E+07 | | | | | | |
| BCL2-associated athanogene 2 | Cytoplasm | other | 9946498 | 6784258 | 6399696.5 | | | | | | |
| BCL2-associated athanogene 2 | Cytoplasm | other | 8597400 | 8213538 | 1.03E+07 | | | | | | |
| BCL2-associated athanogene 2 | Cytoplasm | other | 1300084.25 | 2754581 | 1764170.375 | Q91YN9 | BAG2 | 19143998 | 12539378 | 2182826 | 33 |
| BCL2-antagonist/killer 1 | Cytoplasm | other | 1213742.125 | 2371321.25 | 2112427 | O08734 | BAK1 | 1213742 | 607543 | 350765 | 3 |
| barrier to autointegration factor 1 | Nucleus | other | 5.39E+07 | 7.70E+07 | 7.20E+07 | | | | | | |
| barrier to autointegration factor 1 | Nucleus | other | 2.88E+07 | 3.96E+07 | 4.61E+07 | | | | | | |
| barrier to autointegration factor 1 | Nucleus | other | 1.12E+07 | 1.88E+07 | 1.66E+07 | | | | | | |
| barrier to autointegration factor 1 | Nucleus | other | 5481346 | 8731618 | 8943328 | O54962 | BANF1 | 24835337 | 25210116 | 7277533 | 12 |
| B-cell receptor-associated protein 31 | Cytoplasm | transporter | 803633.938 | 3851847.5 | 2532297.5 | Q61335 | BCAP31 | 803634 | 1528676 | 882581 | 3 |
| branched chain amino-acid transaminase 2, mitochondrial | Cytoplasm | enzyme | 1571558.375 | 5067547 | 2185010 | O35855 | BCAT2 | 1571558 | 1866694 | 1077736 | 3 |
| BCL2-like 13 (apoptosis facilitator) | Cytoplasm | other | 1973846.75 | 3131872.25 | 3740281.5 | | | | | | |
| BCL2-like 13 (apoptosis facilitator) | Cytoplasm | other | 1778908.625 | 2278329.75 | 1744592.625 | | | | | | |
| BCL2-like 13 (apoptosis facilitator) | Cytoplasm | other | 1623786.625 | 1786117 | 2155064.75 | P59017 | BCL2L13 | 1792181 | 721912 | 240637 | 9 |
| basic helix-loop-helix family, member e22 | Nucleus | transcription regulator | 2.38E+08 | 1.77E+08 | 1.82E+08 | Q8C6A8 | BHLHE22 | 238400000 | 34310494 | 19809173 | 3 |
| bicaudal D homolog 1 (Drosophila) | Cytoplasm | other | 6.17E+07 | 3.30E+07 | 3.46E+07 | Q8BR07 | BICD1 | 61650000 | 16101239 | 9296055 | 3 |
| bleomycin hydrolase | Cytoplasm | peptidase | 1.66E+07 | 5380424 | 9421275 | | | | | | |
| bleomycin hydrolase | Cytoplasm | peptidase | 1.34E+07 | 7463331.5 | 8815439 | | | | | | |
| bleomycin hydrolase | Cytoplasm | peptidase | 1.32E+07 | 3942430.5 | 7781424 | | | | | | |
| bleomycin hydrolase | Cytoplasm | peptidase | 8179085 | 3742186.5 | 4704922 | | | | | | |
| bleomycin hydrolase | Cytoplasm | peptidase | 7962530.5 | 2366368.5 | 2663837 | | | | | | |
| bleomycin hydrolase | Cytoplasm | peptidase | 6364238.5 | 2862322 | 3473226 | | | | | | |
| bleomycin hydrolase | Cytoplasm | peptidase | 3475974 | 1896203.625 | 2002361.375 | Q8R016 | BLMH | 9887404 | 4122174 | 899532 | 21 |
| biliverdin reductase B | Cytoplasm | enzyme | 1.27E+07 | 9808062 | 1.07E+07 | | | | | | |
| biliverdin reductase B | Cytoplasm | enzyme | 1.19E+07 | 9473946 | 7399072 | | | | | | |
| biliverdin reductase B | Cytoplasm | enzyme | 5510980 | 2211989.75 | 3568735.5 | | | | | | |
| biliverdin reductase B | Cytoplasm | enzyme | 1294493 | 1026507.812 | 1455409.75 | Q923D2 | BLVRB | 7853868 | 4431760 | 1279339 | 12 |
| 2,3-bisphosphoglycerate mutase | Extracellular Space | phosphatase | 1.33E+07 | 5017869.5 | 7624191.5 | | | | | | |
| 2,3-bisphosphoglycerate mutase | Extracellular Space | phosphatase | 6145480 | 1806176 | 2497716.75 | | | | | | |
| 2,3-bisphosphoglycerate mutase | Extracellular Space | phosphatase | 2179085 | 851445.438 | 1505582.75 | | | | | | |
| 2,3-bisphosphoglycerate mutase | Extracellular Space | phosphatase | 956905.25 | 171371.578 | 275738.406 | P15327 | BPGM | 5650368 | 3899738 | 1125757 | 12 |
| bromodomain and WD repeat domain containing 3 | Nucleus | other | 1.14E+07 | 1.22E+07 | 1.04E+07 | A2AHJ4 | BRWD3 | 11410000 | 873232 | 504160 | 3 |
| basigin (Ok blood group) | Plasma Membrane | transporter | 1.10E+09 | 8.42E+08 | 7.84E+08 | | | | | | |
| basigin (Ok blood group) | Plasma Membrane | transporter | 8.48E+08 | 7.03E+08 | 6.30E+08 | | | | | | |
| basigin (Ok blood group) | Plasma Membrane | transporter | 7.58E+08 | 8.92E+08 | 8.54E+08 | | | | | | |
| basigin (Ok blood group) | Plasma Membrane | transporter | 7.37E+08 | 5.78E+08 | 6.31E+08 | | | | | | |
| basigin (Ok blood group) | Plasma Membrane | transporter | 3.73E+08 | 2.32E+08 | 2.62E+08 | | | | | | |
| basigin (Ok blood group) | Plasma Membrane | transporter | 2.92E+08 | 2.18E+08 | 1.86E+08 | | | | | | |
| basigin (Ok blood group) | Plasma Membrane | transporter | 2.57E+08 | 1.21E+08 | 1.58E+08 | | | | | | |
| basigin (Ok blood group) | Plasma Membrane | transporter | 6.41E+07 | 3.20E+07 | 4.34E+07 | | | | | | |
| basigin (Ok blood group) | Plasma Membrane | transporter | 6.02E+07 | 4.07E+07 | 4.02E+07 | | | | | | |
| basigin (Ok blood group) | Plasma Membrane | transporter | 4.59E+07 | 5.01E+07 | 4.89E+07 | | | | | | |
| basigin (Ok blood group) | Plasma Membrane | transporter | 2.01E+07 | 1.09E+07 | 9941418 | | | | | | |
| basigin (Ok blood group) | Plasma Membrane | transporter | 1.02E+07 | 4457598 | 6062637 | | | | | | |
| basigin (Ok blood group) | Plasma Membrane | transporter | 4108788.75 | 2081852.875 | 2454507.5 | | | | | | |
| basigin (Ok blood group) | Plasma Membrane | transporter | 3844355.25 | 767312.125 | 1859835.875 | P18572 | BSG | 326675939 | 337511176 | 52079105 | 42 |
| chromosome 10 open reading frame 54 | Plasma Membrane | other | 1.01E+07 | 7193396 | 6248784 | Q9D659 | C10orf54 | 10080000 | 1995954 | 1152364 | 3 |
| chromosome 16 open reading frame 58 | Cytoplasm | other | 1162080.5 | 1120985.375 | 1168326.75 | Q91W34 | C16orf58 | 1162081 | 25720 | 14849 | 3 |
| chromosome 17 open reading frame 99 | Extracellular Space | cytokine | 5.03E+07 | 3.19E+07 | 3.24E+07 | | | | | | |
| chromosome 17 open reading frame 99 | Extracellular Space | cytokine | 4.33E+07 | 2.07E+07 | 2.69E+07 | | | | | | |
| chromosome 17 open reading frame 99 | Extracellular Space | cytokine | 2.86E+07 | 2.39E+07 | 2.18E+07 | | | | | | |
| chromosome 17 open reading frame 99 | Extracellular Space | cytokine | 2.67E+07 | 1.70E+07 | 2.01E+07 | | | | | | |
| chromosome 17 open reading frame 99 | Extracellular Space | cytokine | 1.69E+07 | 1.36E+07 | 1.29E+07 | | | | | | |
| chromosome 17 open reading frame 99 | Extracellular Space | cytokine | 1.07E+07 | 7021441.5 | 7822702.5 | | | | | | |
| chromosome 17 open reading frame 99 | Extracellular Space | cytokine | 9628616 | 8337708 | 7081642 | Q9CX63 | C17orf99 | 26592659 | 11936941 | 2604854 | 21 |
| complement component 1, q subcomponent, A chain | Extracellular Space | other | 322046.281 | 1639049.375 | 1.03E+07 | | | | | | |
| complement component 1, q subcomponent, A chain | Extracellular Space | other | 0 | 636995 | 5827700 | P98086 | C1QA | 161023 | 4104449 | 1675634 | 6 |
| complement component 1, q subcomponent, B chain | Extracellular Space | other | 436133.469 | 1684360.875 | 1.27E+07 | P14106 | C1QB | 436133 | 6743385 | 3893295 | 3 |
| complement component 1, q subcomponent binding protein | Cytoplasm | transcription regulator | 1.08E+07 | 1.54E+07 | 1.93E+07 | | | | | | |
| complement component 1, q subcomponent binding protein | Cytoplasm | transcription regulator | 1520354.75 | 1051411.625 | 3809624 | O35658 | C1QBP | 6145177 | 7697454 | 3142473 | 6 |
| chromosome 21 open reading frame 33 | Cytoplasm | other | 3089628.5 | 5585934.5 | 5910218.5 | | | | | | |
| chromosome 21 open reading frame 33 | Cytoplasm | other | 137636.438 | 102541.57 | 193466.609 | Q9D172 | C21orf33/LOC102724023 | 1613632 | 2762204 | 1127665 | 6 |
| C2CD2-like | Cytoplasm | other | 2.67E+07 | 2.19E+07 | 2.23E+07 | | | | | | |
| C2CD2-like | Cytoplasm | other | 2.64E+07 | 2.70E+07 | 2.86E+07 | | | | | | |
| C2CD2-like | Cytoplasm | other | 2.50E+07 | 1.85E+07 | 1.92E+07 | | | | | | |
| C2CD2-like | Cytoplasm | other | 1.10E+07 | 8323573 | 7559943.5 | | | | | | |
| C2CD2-like | Cytoplasm | other | 1.10E+07 | 7692977 | 8575700 | | | | | | |
| C2CD2-like | Cytoplasm | other | 9881142 | 8437190 | 7609566.5 | | | | | | |
| C2CD2-like | Cytoplasm | other | 9865549 | 9175458 | 1.20E+07 | | | | | | |
| C2CD2-like | Cytoplasm | other | 9753543 | 1.23E+07 | 1.14E+07 | | | | | | |
| C2CD2-like | Cytoplasm | other | 9471285 | 7952418 | 7208407.5 | | | | | | |
| C2CD2-like | Cytoplasm | other | 9110655 | 6084807 | 6413196 | | | | | | |
| C2CD2-like | Cytoplasm | other | 7352834 | 4216235.5 | 3960845 | | | | | | |
| C2CD2-like | Cytoplasm | other | 3698331.75 | 2721319.5 | 2868954 | | | | | | |
| C2CD2-like | Cytoplasm | other | 650348.375 | 1590327.25 | 1336270.75 | Q80X80 | C2CD2L | 12291053 | 7840465 | 1255479 | 39 |
| complement component 4B (Chido blood group) | Extracellular Space | other | 4170201.75 | 3557308.5 | 3346540.5 | | | | | | |
| complement component 4B (Chido blood group) | Extracellular Space | other | 4048991 | 4554438.5 | 3219420 | | | | | | |
| complement component 4B (Chido blood group) | Extracellular Space | other | 1867686.125 | 2914625 | 1753505.75 | | | | | | |
| complement component 4B (Chido blood group) | Extracellular Space | other | 1826297.875 | 1098504.375 | 890858.438 | | | | | | |
| complement component 4B (Chido blood group) | Extracellular Space | other | 1523534.125 | 1572649 | 598562.75 | P01029 | C4A/C4B | 2687342 | 1290249 | 333141 | 15 |
| carbonic anhydrase II | Cytoplasm | enzyme | 2.62E+07 | 1.83E+07 | 1.88E+07 | | | | | | |
| carbonic anhydrase II | Cytoplasm | enzyme | 1.01E+07 | 1.17E+07 | 8330627.5 | | | | | | |
| carbonic anhydrase II | Cytoplasm | enzyme | 8020493.5 | 7039539 | 5508575 | | | | | | |
| carbonic anhydrase II | Cytoplasm | enzyme | 7178987.5 | 4001631.5 | 1.08E+07 | | | | | | |
| carbonic anhydrase II | Cytoplasm | enzyme | 5943655.5 | 6268209.5 | 5453726.5 | | | | | | |
| carbonic anhydrase II | Cytoplasm | enzyme | 87757.695 | 193432.875 | 76148.445 | P00920 | CA2 | 9586816 | 6856093 | 1615997 | 18 |
| Calmodulin | Cytoplasm | | 2.89E+07 | 2.07E+07 | 1.96E+07 | | | | | | |
| Calmodulin | Cytoplasm | | 9569761 | 1.07E+07 | 8576285 | P62204 | CALM1 | 19229881 | 8076535 | 3297231 | 6 |
| calmodulin-like 3 | Cytoplasm | other | 7838481 | 4912387 | 3840933.25 | Q9D6P8 | CALML3 | 7838481 | 2069236 | 1194674 | 3 |
| calreticulin | Cytoplasm | transcription regulator | 2.00E+08 | 1.53E+08 | 1.52E+08 | | | | | | |
| calreticulin | Cytoplasm | transcription regulator | 1.35E+08 | 9.93E+07 | 1.12E+08 | | | | | | |
| calreticulin | Cytoplasm | transcription regulator | 1.18E+08 | 8.77E+07 | 1.28E+08 | | | | | | |
| calreticulin | Cytoplasm | transcription regulator | 1.01E+08 | 9.44E+07 | 7.66E+07 | | | | | | |
| calreticulin | Cytoplasm | transcription regulator | 9.90E+07 | 1.00E+08 | 1.03E+08 | | | | | | |
| calreticulin | Cytoplasm | transcription regulator | 9.56E+07 | 5.61E+07 | 6.30E+07 | | | | | | |

| | | | | | | | | | | | |
|---|---|---|---|---|---|---|---|---|---|---|---|
| calreticulin | Cytoplasm | transcription regulator | 8.07E+07 | 7.90E+07 | 7.75E+07 | | | | | | |
| calreticulin | Cytoplasm | transcription regulator | 7.10E+07 | 5.02E+07 | 8.40E+07 | | | | | | |
| calreticulin | Cytoplasm | transcription regulator | 5.36E+07 | 5.12E+07 | 4.08E+07 | | | | | | |
| calreticulin | Cytoplasm | transcription regulator | 3.35E+07 | 3.57E+07 | 3.00E+07 | | | | | | |
| calreticulin | Cytoplasm | transcription regulator | 3.22E+07 | 3.78E+07 | 3.20E+07 | | | | | | |
| calreticulin | Cytoplasm | transcription regulator | 6864311.5 | 3151137.5 | 2106372 | | | | | | |
| calreticulin | Cytoplasm | transcription regulator | 6482116 | 4751788.5 | 4497777 | | | | | | |
| calreticulin | Cytoplasm | transcription regulator | 4824956.5 | 1.56E+07 | 1.49E+07 | | | | | | |
| calreticulin | Cytoplasm | transcription regulator | 3453964.75 | 3062445.5 | 3308087.25 | | | | | | |
| calreticulin | Cytoplasm | transcription regulator | 1987998.875 | 832486 | 858204.125 | | | | | | |
| calreticulin | Cytoplasm | transcription regulator | 1513870.375 | 695973.875 | 927416.75 | | | | | | |
| calreticulin | Cytoplasm | transcription regulator | 386844.812 | 502872.719 | 448966.281 | | | | | | |
| calreticulin | Cytoplasm | transcription regulator | 310433.656 | 1097266.875 | 2649334.25 | P14211 | CALR | 54945500 | 50410330 | 6677011 | 57 |
| calumenin | Cytoplasm | other | 7864879 | 8944493 | 6816417 | | | | | | |
| calumenin | Cytoplasm | other | 7077758 | 7975516 | 6481475.5 | | | | | | |
| calumenin | Cytoplasm | other | 6110564 | 6728853.5 | 7073464.5 | | | | | | |
| calumenin | Cytoplasm | other | 4283506.5 | 5960571.5 | 7176659 | | | | | | |
| calumenin | Cytoplasm | other | 3245123.75 | 2783472.75 | 2244842 | | | | | | |
| calumenin | Cytoplasm | other | 3160751 | 3443877.5 | 2388996 | | | | | | |
| calumenin | Cytoplasm | other | 2505173.75 | 2647478.25 | 2501253.75 | | | | | | |
| calumenin | Cytoplasm | other | 857007 | 929895.75 | 1380231 | O35887 | CALU | 4388095 | 2523555 | 515119 | 24 |
| cathelicidin antimicrobial peptide | Cytoplasm | other | 570814.188 | 2478540.5 | 2469058.75 | | | | | | |
| cathelicidin antimicrobial peptide | Cytoplasm | other | 248912.594 | 3475728.5 | 3419808.75 | P51437 | CAMP | 409863 | 1391280 | 567988 | 6 |
| cullin-associated and neddylation-dissociated 1 | Cytoplasm | transcription regulator | 2.83E+07 | 1.54E+07 | 1.90E+07 | | | | | | |
| cullin-associated and neddylation-dissociated 1 | Cytoplasm | transcription regulator | 2.25E+07 | 1.12E+07 | 1.13E+07 | | | | | | |
| cullin-associated and neddylation-dissociated 1 | Cytoplasm | transcription regulator | 2.13E+07 | 1.43E+07 | 1.29E+07 | | | | | | |
| cullin-associated and neddylation-dissociated 1 | Cytoplasm | transcription regulator | 1.79E+07 | 7139800 | 9763174 | | | | | | |
| cullin-associated and neddylation-dissociated 1 | Cytoplasm | transcription regulator | 1.72E+07 | 8611848 | 1.10E+07 | | | | | | |
| cullin-associated and neddylation-dissociated 1 | Cytoplasm | transcription regulator | 1.66E+07 | 1.03E+07 | 1.17E+07 | | | | | | |
| cullin-associated and neddylation-dissociated 1 | Cytoplasm | transcription regulator | 1.51E+07 | 6648466.5 | 8154783 | | | | | | |
| cullin-associated and neddylation-dissociated 1 | Cytoplasm | transcription regulator | 1.39E+07 | 5577610 | 9879596 | | | | | | |
| cullin-associated and neddylation-dissociated 1 | Cytoplasm | transcription regulator | 1.34E+07 | 7817457.5 | 8341431.5 | | | | | | |
| cullin-associated and neddylation-dissociated 1 | Cytoplasm | transcription regulator | 1.29E+07 | 6233251 | 7740758.5 | | | | | | |
| cullin-associated and neddylation-dissociated 1 | Cytoplasm | transcription regulator | 1.29E+07 | 7320719 | 1.01E+07 | | | | | | |
| cullin-associated and neddylation-dissociated 1 | Cytoplasm | transcription regulator | 1.11E+07 | 5909397.5 | 7320778 | | | | | | |
| cullin-associated and neddylation-dissociated 1 | Cytoplasm | transcription regulator | 9592854 | 2984696.25 | 5134582 | | | | | | |
| cullin-associated and neddylation-dissociated 1 | Cytoplasm | transcription regulator | 9406335 | 4691740 | 6695095 | | | | | | |
| cullin-associated and neddylation-dissociated 1 | Cytoplasm | transcription regulator | 8849629 | 5759079 | 2987816.75 | | | | | | |
| cullin-associated and neddylation-dissociated 1 | Cytoplasm | transcription regulator | 8793516 | 4093723 | 5491447.5 | | | | | | |
| cullin-associated and neddylation-dissociated 1 | Cytoplasm | transcription regulator | 8372154 | 4006339.5 | 4586980.5 | | | | | | |
| cullin-associated and neddylation-dissociated 1 | Cytoplasm | transcription regulator | 8371463 | 3593200.25 | 5480051.5 | | | | | | |
| cullin-associated and neddylation-dissociated 1 | Cytoplasm | transcription regulator | 4236413 | 816880.375 | 1799296.625 | Q6ZQ38 | CAND1 | 13717493 | 5413853 | 717082 | 57 |
| calnexin | Cytoplasm | other | 6.32E+07 | 3.92E+07 | 6.69E+07 | | | | | | |
| calnexin | Cytoplasm | other | 6.21E+07 | 7.37E+07 | 5.93E+07 | | | | | | |
| calnexin | Cytoplasm | other | 6.20E+07 | 7.03E+07 | 6.49E+07 | | | | | | |
| calnexin | Cytoplasm | other | 5.89E+07 | 3.68E+07 | 4.58E+07 | | | | | | |
| calnexin | Cytoplasm | other | 3.76E+07 | 2.27E+07 | 2.30E+07 | | | | | | |
| calnexin | Cytoplasm | other | 3.43E+07 | 4.57E+07 | 4.93E+07 | | | | | | |
| calnexin | Cytoplasm | other | 3.11E+07 | 1.81E+07 | 2.37E+07 | | | | | | |
| calnexin | Cytoplasm | other | 3.05E+07 | 3.55E+07 | 3.29E+07 | | | | | | |
| calnexin | Cytoplasm | other | 2.96E+07 | 2.77E+07 | 2.55E+07 | | | | | | |
| calnexin | Cytoplasm | other | 2.46E+07 | 2.89E+07 | 3.15E+07 | | | | | | |
| calnexin | Cytoplasm | other | 2.32E+07 | 2.80E+07 | 2.09E+07 | | | | | | |
| calnexin | Cytoplasm | other | 2.30E+07 | 3.14E+07 | 1.84E+07 | | | | | | |
| calnexin | Cytoplasm | other | 2.19E+07 | 2.76E+07 | 2.93E+07 | | | | | | |
| calnexin | Cytoplasm | other | 1.39E+07 | 1.43E+07 | 1.81E+07 | | | | | | |
| calnexin | Cytoplasm | other | 7599782.5 | 9130180 | 9296854 | | | | | | |
| calnexin | Cytoplasm | other | 3518131 | 3020579 | 4020786.5 | | | | | | |
| calnexin | Cytoplasm | other | 3409147.25 | 2682907.5 | 4030472.5 | | | | | | |
| calnexin | Cytoplasm | other | 2453544.75 | 2666564.75 | 7777468.5 | | | | | | |
| calnexin | Cytoplasm | other | 2072954.25 | 4331484.5 | 4499112.5 | | | | | | |
| calnexin | Cytoplasm | other | 2033481.875 | 1123050 | 1407856.75 | | | | | | |
| calnexin | Cytoplasm | other | 1683782.875 | 1754108.5 | 1498889.625 | | | | | | |
| calnexin | Cytoplasm | other | 555742 | 122041.078 | 168011.188 | P35564 | CANX | 24506208 | 20949334 | 2578685 | 66 |
| CAP, adenylate cyclase-associated protein 1 (yeast) | Plasma Membrane | other | 4252245.5 | 2617415 | 2710088.5 | | | | | | |
| CAP, adenylate cyclase-associated protein 1 (yeast) | Plasma Membrane | other | 2887946 | 1525932.375 | 2257980 | | | | | | |
| CAP, adenylate cyclase-associated protein 1 (yeast) | Plasma Membrane | other | 1415966 | 1149258.5 | 1276237.625 | P40124 | CAP1 | 2852053 | 1008693 | 336231 | 9 |
| calpain 1, (mu/I) large subunit | Cytoplasm | peptidase | 9937218 | 8459912 | 7074190 | | | | | | |
| calpain 1, (mu/I) large subunit | Cytoplasm | peptidase | 9047008 | 7072556.5 | 6247624.5 | | | | | | |
| calpain 1, (mu/I) large subunit | Cytoplasm | peptidase | 6408834.5 | 5649681.5 | 5510252 | | | | | | |
| calpain 1, (mu/I) large subunit | Cytoplasm | peptidase | 6151860 | 5057795 | 5355908 | | | | | | |
| calpain 1, (mu/I) large subunit | Cytoplasm | peptidase | 5948183 | 6243912 | 4700008 | | | | | | |
| calpain 1, (mu/I) large subunit | Cytoplasm | peptidase | 4468510.5 | 2421525.5 | 3317867.25 | | | | | | |
| calpain 1, (mu/I) large subunit | Cytoplasm | peptidase | 1127525.125 | 1722969.25 | 780474.625 | O35350 | CAPN1 | 6155591 | 2449632 | 534554 | 21 |
| calpain 5 | Cytoplasm | peptidase | 3.75E+07 | 1.80E+07 | 3.96E+07 | | | | | | |
| calpain 5 | Cytoplasm | peptidase | 3.69E+07 | 1.92E+07 | 3.73E+07 | | | | | | |
| calpain 5 | Cytoplasm | peptidase | 2.26E+07 | 1.47E+07 | 2.17E+07 | | | | | | |
| calpain 5 | Cytoplasm | peptidase | 2.11E+07 | 1.32E+07 | 2.08E+07 | | | | | | |
| calpain 5 | Cytoplasm | peptidase | 2.09E+07 | 1.36E+07 | 2.23E+07 | | | | | | |
| calpain 5 | Cytoplasm | peptidase | 1.59E+07 | 1.20E+07 | 1.98E+07 | | | | | | |
| calpain 5 | Cytoplasm | peptidase | 1.57E+07 | 1.03E+07 | 1.72E+07 | | | | | | |
| calpain 5 | Cytoplasm | peptidase | 1.20E+07 | 6528992 | 1.05E+07 | | | | | | |
| calpain 5 | Cytoplasm | peptidase | 1.11E+07 | 5854089.5 | 1.14E+07 | | | | | | |
| calpain 5 | Cytoplasm | peptidase | 9356423 | 6610845.5 | 1.16E+07 | | | | | | |
| calpain 5 | Cytoplasm | peptidase | 9045683 | 4809873 | 7327004 | | | | | | |
| calpain 5 | Cytoplasm | peptidase | 8242075 | 8991707 | 9380957 | | | | | | |
| calpain 5 | Cytoplasm | peptidase | 8151224.5 | 6001991 | 9564823 | | | | | | |
| calpain 5 | Cytoplasm | peptidase | 3967774.75 | 2895820 | 4564213.5 | | | | | | |
| calpain 5 | Cytoplasm | peptidase | 3520067.5 | 1593717.375 | 3461863.25 | | | | | | |
| calpain 5 | Cytoplasm | peptidase | 2603435.75 | 846963 | 1449618.125 | | | | | | |
| calpain 5 | Cytoplasm | peptidase | 2573185.25 | 1200744.375 | 2358996.25 | | | | | | |
| calpain 5 | Cytoplasm | peptidase | 1125103.75 | 1114543.375 | 2498807.25 | | | | | | |
| calpain 5 | Cytoplasm | peptidase | 793874.188 | 632322.25 | 3658933 | | | | | | |
| calpain 5 | Cytoplasm | peptidase | 529915.625 | 1365750.375 | 4832064 | O08688 | CAPN5 | 12175938 | 9806778 | 1266050 | 60 |
| calpain, small subunit 1 | Cytoplasm | peptidase | 3130624 | 2408771.75 | 2281276.5 | O88456 | CAPNS1 | 3130624 | 458024 | 264440 | 3 |

| | | | | | | | | | | | |
|---|---|---|---|---|---|---|---|---|---|---|---|
| cell cycle associated protein 1 | Plasma Membrane | other | 1063310.75 | 261573.156 | 457417.656 | | | | | | |
| cell cycle associated protein 1 | Plasma Membrane | other | 813162.125 | 834745.438 | 614762.375 | Q60865 | CAPRIN1 | 938236 | 288888 | 117938 | 6 |
| capping protein (actin filament) muscle Z-line, alpha 1 | Cytoplasm | other | 5903694 | 5813590 | 6193045 | | | | | | |
| capping protein (actin filament) muscle Z-line, alpha 1 | Cytoplasm | other | 3186561.75 | 3173729.5 | 2221953.75 | P47753 | CAPZA1 | 4545128 | 1743156 | 711641 | 6 |
| capping protein (actin filament) muscle Z-line, beta | Cytoplasm | other | 3527108.75 | 3557008 | 3493835.25 | | | | | | |
| capping protein (actin filament) muscle Z-line, beta | Cytoplasm | other | 1393450.125 | 2474843.75 | 1911786.375 | P47757 | CAPZB | 2460279 | 940599 | 383998 | 6 |
| calcium regulated heat stable protein 1, 24kDa | Cytoplasm | other | 9698735 | 7480379.5 | 7194538.5 | Q9CR86 | CARHSP1 | 9698735 | 1370754 | 791405 | 3 |
| coactivator-associated arginine methyltransferase 1 | Nucleus | transcription regulator | 5081437.5 | 2248045 | 2813211 | | | | | | |
| coactivator-associated arginine methyltransferase 1 | Nucleus | transcription regulator | 4741377 | 2803180.75 | 2505475.25 | | | | | | |
| coactivator-associated arginine methyltransferase 1 | Nucleus | transcription regulator | 4653980.5 | 3630034.25 | 3512587.75 | Q9WVG6 | CARM1 | 4825598 | 1053435 | 351145 | 9 |
| cysteinyl-tRNA synthetase | Cytoplasm | enzyme | 4883926 | 2082659.625 | 2579245.25 | | | | | | |
| cysteinyl-tRNA synthetase | Cytoplasm | enzyme | 2989207.5 | 1007768.812 | 1551734 | Q9ER72 | CARS | 3936567 | 1358481 | 554598 | 6 |
| calpastatin | Cytoplasm | peptidase | 3.70E+07 | 2.16E+07 | 2.46E+07 | | | | | | |
| calpastatin | Cytoplasm | peptidase | 2.07E+07 | 1.42E+07 | 1.40E+07 | | | | | | |
| calpastatin | Cytoplasm | peptidase | 1.93E+07 | 7663557.5 | 1.22E+07 | | | | | | |
| calpastatin | Cytoplasm | peptidase | 1.46E+07 | 6693805 | 7711097.5 | | | | | | |
| calpastatin | Cytoplasm | peptidase | 1.27E+07 | 5456289.5 | 7038561.5 | | | | | | |
| calpastatin | Cytoplasm | peptidase | 1.08E+07 | 3744044.25 | 5731651.5 | | | | | | |
| calpastatin | Cytoplasm | peptidase | 9162819 | 5463513.5 | 7842131.5 | | | | | | |
| calpastatin | Cytoplasm | peptidase | 6199427.5 | 2591649 | 3243339.75 | | | | | | |
| calpastatin | Cytoplasm | peptidase | 5603904.5 | 2009769.75 | 3073498.5 | | | | | | |
| calpastatin | Cytoplasm | peptidase | 5191102.5 | 5092138 | 4270785 | | | | | | |
| calpastatin | Cytoplasm | peptidase | 5164421.5 | 2832468.25 | 3807546.5 | | | | | | |
| calpastatin | Cytoplasm | peptidase | 3758710.25 | 840392.188 | 1495031.25 | P51125 | CAST | 12512532 | 7702661 | 1283777 | 36 |
| catalase | Cytoplasm | enzyme | 1.90E+07 | 9970529 | 1.30E+07 | | | | | | |
| catalase | Cytoplasm | enzyme | 1.81E+07 | 1.05E+07 | 1.03E+07 | | | | | | |
| catalase | Cytoplasm | enzyme | 1.44E+07 | 6301614.5 | 7308216.5 | | | | | | |
| catalase | Cytoplasm | enzyme | 1.24E+07 | 6417655.5 | 7302721 | | | | | | |
| catalase | Cytoplasm | enzyme | 9708862 | 5139136 | 5509056 | | | | | | |
| catalase | Cytoplasm | enzyme | 8824763 | 6503007.5 | 4753763.5 | | | | | | |
| catalase | Cytoplasm | enzyme | 8412278 | 4908298.5 | 4657386 | | | | | | |
| catalase | Cytoplasm | enzyme | 7127916 | 2920127 | 3732617.25 | | | | | | |
| catalase | Cytoplasm | enzyme | 6236356 | 3966773.25 | 3742000.75 | | | | | | |
| catalase | Cytoplasm | enzyme | 5846673 | 4509900 | 4990909.5 | | | | | | |
| catalase | Cytoplasm | enzyme | 5773811.5 | 5414228 | 3421486.75 | | | | | | |
| catalase | Cytoplasm | enzyme | 5383428.5 | 2009891 | 4152585.75 | | | | | | |
| catalase | Cytoplasm | enzyme | 4856257 | 3379741.25 | 3209975.5 | | | | | | |
| catalase | Cytoplasm | enzyme | 4257695 | 1946242.75 | 3411180.25 | | | | | | |
| catalase | Cytoplasm | enzyme | 3629842 | 1668898.625 | 1923787.625 | | | | | | |
| catalase | Cytoplasm | enzyme | 3064121 | 1853976.75 | 1733025.875 | | | | | | |
| catalase | Cytoplasm | enzyme | 2748840.5 | 1436275.375 | 1391204.875 | | | | | | |
| catalase | Cytoplasm | enzyme | 2023126.875 | 1173853.625 | 1448561.875 | | | | | | |
| catalase | Cytoplasm | enzyme | 1280668.75 | 187252.141 | 379400.094 | | | | | | |
| catalase | Cytoplasm | enzyme | 159064.688 | 260649.844 | 1038737.25 | P24270 | CAT | 7162685 | 4118516 | 531698 | 60 |
| coiled-coil domain containing 15 | Cytoplasm | other | 1608951.25 | 966704.688 | 1757152.375 | Q8C9M2 | CCDC15 | 1608951 | 420169 | 242585 | 3 |
| coiled-coil domain containing 51 | Cytoplasm | other | 536402.938 | 578681.312 | 612591.875 | Q3URS9 | CCDC51 | 536403 | 38171 | 22038 | 3 |
| cyclin Y | Nucleus | other | 3980981 | 2301206.75 | 2688473.5 | | | | | | |
| cyclin Y | Nucleus | other | 2730995.5 | 2344580.25 | 2452358 | Q8BGU5 | CCNY | 3355988 | 628304 | 256504 | 6 |
| chaperonin containing TCP1, subunit 2 (beta) | Cytoplasm | kinase | 2.10E+07 | 1.14E+07 | 1.63E+07 | | | | | | |
| chaperonin containing TCP1, subunit 2 (beta) | Cytoplasm | kinase | 2.09E+07 | 6957954 | 1.32E+07 | | | | | | |
| chaperonin containing TCP1, subunit 2 (beta) | Cytoplasm | kinase | 1.54E+07 | 8792359 | 9263623 | | | | | | |
| chaperonin containing TCP1, subunit 2 (beta) | Cytoplasm | kinase | 1.43E+07 | 6792185.5 | 8916034 | | | | | | |
| chaperonin containing TCP1, subunit 2 (beta) | Cytoplasm | kinase | 1.39E+07 | 7664201 | 9267216 | | | | | | |
| chaperonin containing TCP1, subunit 2 (beta) | Cytoplasm | kinase | 1.35E+07 | 6520681 | 1.16E+07 | | | | | | |
| chaperonin containing TCP1, subunit 2 (beta) | Cytoplasm | kinase | 9935227 | 6721955.5 | 6894340 | | | | | | |
| chaperonin containing TCP1, subunit 2 (beta) | Cytoplasm | kinase | 9521733 | 4504411.5 | 6833859.5 | | | | | | |
| chaperonin containing TCP1, subunit 2 (beta) | Cytoplasm | kinase | 8448555 | 4251117 | 6148934.5 | | | | | | |
| chaperonin containing TCP1, subunit 2 (beta) | Cytoplasm | kinase | 8207870.5 | 4349468.5 | 6306712 | | | | | | |
| chaperonin containing TCP1, subunit 2 (beta) | Cytoplasm | kinase | 7644276 | 2807780 | 6379587 | | | | | | |
| chaperonin containing TCP1, subunit 2 (beta) | Cytoplasm | kinase | 6880533.5 | 3220941.25 | 4368617.5 | | | | | | |
| chaperonin containing TCP1, subunit 2 (beta) | Cytoplasm | kinase | 3928433.25 | 2604834.5 | 5788601 | | | | | | |
| chaperonin containing TCP1, subunit 2 (beta) | Cytoplasm | kinase | 3753573 | 2127840.5 | 3290840.25 | P80314 | CCT2 | 11236443 | 4600307 | 709843 | 42 |
| chaperonin containing TCP1, subunit 3 (gamma) | Cytoplasm | other | 2.83E+07 | 1.49E+07 | 2.29E+07 | | | | | | |
| chaperonin containing TCP1, subunit 3 (gamma) | Cytoplasm | other | 1.81E+07 | 1.06E+07 | 1.41E+07 | | | | | | |
| chaperonin containing TCP1, subunit 3 (gamma) | Cytoplasm | other | 1.68E+07 | 1.05E+07 | 1.21E+07 | | | | | | |
| chaperonin containing TCP1, subunit 3 (gamma) | Cytoplasm | other | 1.65E+07 | 8607457 | 1.40E+07 | | | | | | |
| chaperonin containing TCP1, subunit 3 (gamma) | Cytoplasm | other | 1.29E+07 | 6744360 | 1.21E+07 | | | | | | |
| chaperonin containing TCP1, subunit 3 (gamma) | Cytoplasm | other | 1.26E+07 | 6329916 | 7282470 | | | | | | |
| chaperonin containing TCP1, subunit 3 (gamma) | Cytoplasm | other | 1.17E+07 | 5608866.5 | 8978387 | | | | | | |
| chaperonin containing TCP1, subunit 3 (gamma) | Cytoplasm | other | 1.15E+07 | 4911656 | 7289234 | | | | | | |
| chaperonin containing TCP1, subunit 3 (gamma) | Cytoplasm | other | 9985821 | 4483902 | 5897563 | | | | | | |
| chaperonin containing TCP1, subunit 3 (gamma) | Cytoplasm | other | 9688290 | 3558213.75 | 4877379.5 | | | | | | |
| chaperonin containing TCP1, subunit 3 (gamma) | Cytoplasm | other | 9482640 | 4779988.5 | 5268174.5 | | | | | | |
| chaperonin containing TCP1, subunit 3 (gamma) | Cytoplasm | other | 9195179 | 3570248.25 | 6049453.5 | | | | | | |
| chaperonin containing TCP1, subunit 3 (gamma) | Cytoplasm | other | 8099221.5 | 3290323.25 | 5758455.5 | | | | | | |
| chaperonin containing TCP1, subunit 3 (gamma) | Cytoplasm | other | 4722093.5 | 2834203.5 | 3218570 | | | | | | |
| chaperonin containing TCP1, subunit 3 (gamma) | Cytoplasm | other | 3583473.25 | 1655909.75 | 3035718.5 | | | | | | |
| chaperonin containing TCP1, subunit 3 (gamma) | Cytoplasm | other | 3415590.25 | 1589936.75 | 2122714.75 | P80318 | CCT3 | 11652019 | 5628009 | 812333 | 48 |
| chaperonin containing TCP1, subunit 4 (delta) | Cytoplasm | other | 3.81E+07 | 1.60E+07 | 2.60E+07 | | | | | | |
| chaperonin containing TCP1, subunit 4 (delta) | Cytoplasm | other | 2.84E+07 | 1.27E+07 | 1.94E+07 | | | | | | |
| chaperonin containing TCP1, subunit 4 (delta) | Cytoplasm | other | 1.69E+07 | 8747359 | 1.52E+07 | | | | | | |
| chaperonin containing TCP1, subunit 4 (delta) | Cytoplasm | other | 1.66E+07 | 9591591 | 1.42E+07 | | | | | | |
| chaperonin containing TCP1, subunit 4 (delta) | Cytoplasm | other | 1.50E+07 | 5626075.5 | 9043785 | | | | | | |
| chaperonin containing TCP1, subunit 4 (delta) | Cytoplasm | other | 1.25E+07 | 5370688.5 | 7562490 | | | | | | |
| chaperonin containing TCP1, subunit 4 (delta) | Cytoplasm | other | 1.22E+07 | 8463847 | 8660181 | | | | | | |
| chaperonin containing TCP1, subunit 4 (delta) | Cytoplasm | other | 1.14E+07 | 4622643.5 | 6612760 | | | | | | |
| chaperonin containing TCP1, subunit 4 (delta) | Cytoplasm | other | 9732960 | 6041006 | 6588042 | | | | | | |
| chaperonin containing TCP1, subunit 4 (delta) | Cytoplasm | other | 8338447.5 | 4180786 | 6333761.5 | | | | | | |
| chaperonin containing TCP1, subunit 4 (delta) | Cytoplasm | other | 7810703.5 | 4585276 | 6308490.5 | | | | | | |
| chaperonin containing TCP1, subunit 4 (delta) | Cytoplasm | other | 6720198.5 | 2325205.5 | 4413509.5 | | | | | | |
| chaperonin containing TCP1, subunit 4 (delta) | Cytoplasm | other | 6717217.5 | 3655410.5 | 4855254 | | | | | | |
| chaperonin containing TCP1, subunit 4 (delta) | Cytoplasm | other | 5937479.5 | 2771282.75 | 2376402.5 | | | | | | |
| chaperonin containing TCP1, subunit 4 (delta) | Cytoplasm | other | 4827231.5 | 1821563.375 | 3779693 | | | | | | |
| chaperonin containing TCP1, subunit 4 (delta) | Cytoplasm | other | 3915683.25 | 3496901.5 | 3127086.25 | | | | | | |
| chaperonin containing TCP1, subunit 4 (delta) | Cytoplasm | other | 3041855.25 | 1390115.75 | 1549912.625 | | | | | | |

| | | | | | | | | | | | |
|---|---|---|---|---|---|---|---|---|---|---|---|
| chaperonin containing TCP1, subunit 4 (delta) | Cytoplasm | other | 2500994.75 | 2023895.75 | 2558887.25 | P80315 | CCT4 | 11701265 | 7174634 | 976344 | 54 |
| chaperonin containing TCP1, subunit 5 (epsilon) | Cytoplasm | other | 3.16E+07 | 1.25E+07 | 1.99E+07 | | | | | | |
| chaperonin containing TCP1, subunit 5 (epsilon) | Cytoplasm | other | 3.00E+07 | 1.29E+07 | 1.93E+07 | | | | | | |
| chaperonin containing TCP1, subunit 5 (epsilon) | Cytoplasm | other | 2.21E+07 | 7943008.5 | 1.14E+07 | | | | | | |
| chaperonin containing TCP1, subunit 5 (epsilon) | Cytoplasm | other | 1.61E+07 | 7220222.5 | 9235483 | | | | | | |
| chaperonin containing TCP1, subunit 5 (epsilon) | Cytoplasm | other | 1.41E+07 | 7418571 | 1.28E+07 | | | | | | |
| chaperonin containing TCP1, subunit 5 (epsilon) | Cytoplasm | other | 1.17E+07 | 4905238.5 | 9102193 | | | | | | |
| chaperonin containing TCP1, subunit 5 (epsilon) | Cytoplasm | other | 1.02E+07 | 4361017.5 | 7163839 | | | | | | |
| chaperonin containing TCP1, subunit 5 (epsilon) | Cytoplasm | other | 8325641.5 | 4600421.5 | 7059053.5 | | | | | | |
| chaperonin containing TCP1, subunit 5 (epsilon) | Cytoplasm | other | 8191849 | 2811623.75 | 4594384.5 | | | | | | |
| chaperonin containing TCP1, subunit 5 (epsilon) | Cytoplasm | other | 7701804 | 3990626.25 | 4734497.5 | | | | | | |
| chaperonin containing TCP1, subunit 5 (epsilon) | Cytoplasm | other | 4442329 | 1729737.75 | 2889422.5 | | | | | | |
| chaperonin containing TCP1, subunit 5 (epsilon) | Cytoplasm | other | 2928551.5 | 1713896.25 | 2807355 | | | | | | |
| chaperonin containing TCP1, subunit 5 (epsilon) | Cytoplasm | other | 1888826.375 | 1768675 | 1812580.25 | P80316 | CCT5 | 13019154 | 7335706 | 1174653 | 39 |
| chaperonin containing TCP1, subunit 6A (zeta 1) | Cytoplasm | other | 2.82E+07 | 1.32E+07 | 1.79E+07 | | | | | | |
| chaperonin containing TCP1, subunit 6A (zeta 1) | Cytoplasm | other | 2.24E+07 | 1.50E+07 | 1.71E+07 | | | | | | |
| chaperonin containing TCP1, subunit 6A (zeta 1) | Cytoplasm | other | 2.23E+07 | 9416257 | 1.96E+07 | | | | | | |
| chaperonin containing TCP1, subunit 6A (zeta 1) | Cytoplasm | other | 1.99E+07 | 9666428 | 1.41E+07 | | | | | | |
| chaperonin containing TCP1, subunit 6A (zeta 1) | Cytoplasm | other | 1.75E+07 | 8408800 | 1.25E+07 | | | | | | |
| chaperonin containing TCP1, subunit 6A (zeta 1) | Cytoplasm | other | 1.43E+07 | 7257676.5 | 9697564 | | | | | | |
| chaperonin containing TCP1, subunit 6A (zeta 1) | Cytoplasm | other | 1.22E+07 | 7220267 | 8864995 | | | | | | |
| chaperonin containing TCP1, subunit 6A (zeta 1) | Cytoplasm | other | 1.11E+07 | 5445411 | 7574411.5 | | | | | | |
| chaperonin containing TCP1, subunit 6A (zeta 1) | Cytoplasm | other | 8307361.5 | 3534432 | 5877956 | | | | | | |
| chaperonin containing TCP1, subunit 6A (zeta 1) | Cytoplasm | other | 7789613.5 | 3978304.75 | 5543688.5 | | | | | | |
| chaperonin containing TCP1, subunit 6A (zeta 1) | Cytoplasm | other | 2846165.75 | 781902.938 | 1117317.625 | | | | | | |
| chaperonin containing TCP1, subunit 6A (zeta 1) | Cytoplasm | other | 2508383.75 | 653089.938 | 1661849.25 | P80317 | CCT6A | 14114294 | 6908120 | 1151353 | 36 |
| chaperonin containing TCP1, subunit 7 (eta) | Cytoplasm | other | 2.16E+07 | 1.05E+07 | 1.59E+07 | | | | | | |
| chaperonin containing TCP1, subunit 7 (eta) | Cytoplasm | other | 1.83E+07 | 7115916 | 1.11E+07 | | | | | | |
| chaperonin containing TCP1, subunit 7 (eta) | Cytoplasm | other | 1.78E+07 | 9868961 | 1.19E+07 | | | | | | |
| chaperonin containing TCP1, subunit 7 (eta) | Cytoplasm | other | 1.61E+07 | 9706718 | 1.16E+07 | | | | | | |
| chaperonin containing TCP1, subunit 7 (eta) | Cytoplasm | other | 1.59E+07 | 1.08E+07 | 1.34E+07 | | | | | | |
| chaperonin containing TCP1, subunit 7 (eta) | Cytoplasm | other | 1.46E+07 | 8459714 | 9296845 | | | | | | |
| chaperonin containing TCP1, subunit 7 (eta) | Cytoplasm | other | 1.35E+07 | 6957346.5 | 8102823 | | | | | | |
| chaperonin containing TCP1, subunit 7 (eta) | Cytoplasm | other | 1.04E+07 | 5058176.5 | 6385424 | | | | | | |
| chaperonin containing TCP1, subunit 7 (eta) | Cytoplasm | other | 7407152.5 | 3659353.75 | 3692085.25 | | | | | | |
| chaperonin containing TCP1, subunit 7 (eta) | Cytoplasm | other | 7340512 | 5381690 | 5796380 | | | | | | |
| chaperonin containing TCP1, subunit 7 (eta) | Cytoplasm | other | 7008667.5 | 4381958.5 | 7018152.5 | | | | | | |
| chaperonin containing TCP1, subunit 7 (eta) | Cytoplasm | other | 6169830.5 | 4263314 | 3687550.75 | | | | | | |
| chaperonin containing TCP1, subunit 7 (eta) | Cytoplasm | other | 4734838.5 | 2702130.5 | 3366051.5 | | | | | | |
| chaperonin containing TCP1, subunit 7 (eta) | Cytoplasm | other | 4279829 | 1052047 | 2586683.5 | | | | | | |
| chaperonin containing TCP1, subunit 7 (eta) | Cytoplasm | other | 3793859.25 | 1905153.25 | 2421664 | P80313 | CCT7 | 11256979 | 4976047 | 741785 | 45 |
| chaperonin containing TCP1, subunit 8 (theta) | Cytoplasm | enzyme | 2.03E+07 | 8677371 | 1.50E+07 | | | | | | |
| chaperonin containing TCP1, subunit 8 (theta) | Cytoplasm | enzyme | 1.60E+07 | 7267803.5 | 1.06E+07 | | | | | | |
| chaperonin containing TCP1, subunit 8 (theta) | Cytoplasm | enzyme | 1.49E+07 | 7577111 | 1.01E+07 | | | | | | |
| chaperonin containing TCP1, subunit 8 (theta) | Cytoplasm | enzyme | 1.38E+07 | 7511367.5 | 9525590 | | | | | | |
| chaperonin containing TCP1, subunit 8 (theta) | Cytoplasm | enzyme | 1.33E+07 | 6313470.5 | 1.02E+07 | | | | | | |
| chaperonin containing TCP1, subunit 8 (theta) | Cytoplasm | enzyme | 1.16E+07 | 5982822 | 8649526 | | | | | | |
| chaperonin containing TCP1, subunit 8 (theta) | Cytoplasm | enzyme | 1.14E+07 | 5729746 | 7621446 | | | | | | |
| chaperonin containing TCP1, subunit 8 (theta) | Cytoplasm | enzyme | 1.14E+07 | 6450622 | 8513884 | | | | | | |
| chaperonin containing TCP1, subunit 8 (theta) | Cytoplasm | enzyme | 1.13E+07 | 3671376.25 | 6252115 | | | | | | |
| chaperonin containing TCP1, subunit 8 (theta) | Cytoplasm | enzyme | 1.12E+07 | 7754281.5 | 9041779 | | | | | | |
| chaperonin containing TCP1, subunit 8 (theta) | Cytoplasm | enzyme | 1.08E+07 | 3602340 | 6555433.5 | | | | | | |
| chaperonin containing TCP1, subunit 8 (theta) | Cytoplasm | enzyme | 9952416 | 3806370.75 | 6092318.5 | | | | | | |
| chaperonin containing TCP1, subunit 8 (theta) | Cytoplasm | enzyme | 9037937 | 4879216 | 6270168.5 | | | | | | |
| chaperonin containing TCP1, subunit 8 (theta) | Cytoplasm | enzyme | 6833791 | 4275266.5 | 4595910.5 | | | | | | |
| chaperonin containing TCP1, subunit 8 (theta) | Cytoplasm | enzyme | 6222887 | 2955291 | 3534435.75 | | | | | | |
| chaperonin containing TCP1, subunit 8 (theta) | Cytoplasm | enzyme | 6176398 | 5028038.5 | 6827655 | | | | | | |
| chaperonin containing TCP1, subunit 8 (theta) | Cytoplasm | enzyme | 5985757 | 4218678.5 | 6462709 | | | | | | |
| chaperonin containing TCP1, subunit 8 (theta) | Cytoplasm | enzyme | 5175047.5 | 4211940 | 5476915 | | | | | | |
| chaperonin containing TCP1, subunit 8 (theta) | Cytoplasm | enzyme | 4712700.5 | 5445507.5 | 3492688.25 | | | | | | |
| chaperonin containing TCP1, subunit 8 (theta) | Cytoplasm | enzyme | 4075350.25 | 1859995.375 | 3110991.25 | | | | | | |
| chaperonin containing TCP1, subunit 8 (theta) | Cytoplasm | enzyme | 3861444 | 1839846 | 2381552.25 | | | | | | |
| chaperonin containing TCP1, subunit 8 (theta) | Cytoplasm | enzyme | 3569538 | 2848241 | 3261829.25 | | | | | | |
| chaperonin containing TCP1, subunit 8 (theta) | Cytoplasm | enzyme | 2705844.5 | 3312111.25 | 2784712.75 | | | | | | |
| chaperonin containing TCP1, subunit 8 (theta) | Cytoplasm | enzyme | 779497.688 | 328114.406 | 1370426.125 | P42932 | CCT8 | 8963692 | 3922937 | 462323 | 72 |
| CD226 molecule | Plasma Membrane | other | 269696.75 | 3422520.5 | 2238747.5 | Q8K4F0 | CD226 | 269697 | 1592628 | 919504 | 3 |
| CD36 molecule (thrombospondin receptor) | Plasma Membrane | transmembrane receptor | 7.88E+08 | 6.60E+08 | 5.82E+08 | | | | | | |
| CD36 molecule (thrombospondin receptor) | Plasma Membrane | transmembrane receptor | 3.04E+08 | 2.22E+08 | 2.59E+08 | | | | | | |
| CD36 molecule (thrombospondin receptor) | Plasma Membrane | transmembrane receptor | 1.63E+08 | 7.87E+07 | 1.36E+08 | | | | | | |
| CD36 molecule (thrombospondin receptor) | Plasma Membrane | transmembrane receptor | 1.50E+08 | 1.32E+08 | 1.40E+08 | | | | | | |
| CD36 molecule (thrombospondin receptor) | Plasma Membrane | transmembrane receptor | 1.22E+08 | 1.17E+08 | 6.34E+07 | | | | | | |
| CD36 molecule (thrombospondin receptor) | Plasma Membrane | transmembrane receptor | 1.07E+08 | 3.10E+07 | 6.29E+07 | | | | | | |
| CD36 molecule (thrombospondin receptor) | Plasma Membrane | transmembrane receptor | 1.05E+08 | 9.46E+07 | 1.07E+08 | | | | | | |
| CD36 molecule (thrombospondin receptor) | Plasma Membrane | transmembrane receptor | 1.02E+08 | 8.15E+07 | 9.83E+07 | | | | | | |
| CD36 molecule (thrombospondin receptor) | Plasma Membrane | transmembrane receptor | 1.01E+08 | 9.36E+07 | 9.38E+07 | | | | | | |
| CD36 molecule (thrombospondin receptor) | Plasma Membrane | transmembrane receptor | 9.78E+07 | 7.31E+07 | 1.34E+08 | | | | | | |
| CD36 molecule (thrombospondin receptor) | Plasma Membrane | transmembrane receptor | 2.92E+07 | 8169882.5 | 1.68E+07 | | | | | | |
| CD36 molecule (thrombospondin receptor) | Plasma Membrane | transmembrane receptor | 2.51E+07 | 1.11E+07 | 2.85E+07 | | | | | | |
| CD36 molecule (thrombospondin receptor) | Plasma Membrane | transmembrane receptor | 2.26E+07 | 3.02E+07 | 2.64E+07 | | | | | | |
| CD36 molecule (thrombospondin receptor) | Plasma Membrane | transmembrane receptor | 2.22E+07 | 1.21E+07 | 1.70E+07 | | | | | | |
| CD36 molecule (thrombospondin receptor) | Plasma Membrane | transmembrane receptor | 7482531.5 | 4732767.5 | 3209600.25 | | | | | | |
| CD36 molecule (thrombospondin receptor) | Plasma Membrane | transmembrane receptor | 2107007.5 | 1756463.375 | 2774380 | Q08857 | CD36 | 134276846 | 162583649 | 23466928 | 48 |
| CD44 molecule (Indian blood group) | Plasma Membrane | enzyme | 1.35E+07 | 1.75E+07 | 1.47E+07 | | | | | | |
| CD44 molecule (Indian blood group) | Plasma Membrane | enzyme | 2571301.25 | 4939246 | 3890110.75 | | | | | | |
| CD44 molecule (Indian blood group) | Plasma Membrane | enzyme | 2290726.25 | 1898552.5 | 1921812.25 | | | | | | |
| CD44 molecule (Indian blood group) | Plasma Membrane | enzyme | 796863 | 936192.625 | 894955.562 | P15379 | CD44 | 4797223 | 6070981 | 1752541 | 12 |
| CD47 molecule | Plasma Membrane | transmembrane receptor | 5.57E+08 | 5.24E+08 | 5.38E+08 | | | | | | |
| CD47 molecule | Plasma Membrane | transmembrane receptor | 3.00E+08 | 2.78E+08 | 3.92E+08 | | | | | | |
| CD47 molecule | Plasma Membrane | transmembrane receptor | 1.97E+08 | 1.76E+08 | 1.65E+08 | | | | | | |
| CD47 molecule | Plasma Membrane | transmembrane receptor | 1.81E+08 | 1.21E+08 | 1.14E+08 | | | | | | |
| CD47 molecule | Plasma Membrane | transmembrane receptor | 1.13E+08 | 8.76E+07 | 9.02E+07 | | | | | | |
| CD47 molecule | Plasma Membrane | transmembrane receptor | 9135320 | 3660716.25 | 7127191.5 | Q61735 | CD47 | 226305887 | 180791264 | 42612909 | 18 |
| CD48 molecule | Plasma Membrane | other | 2316017.25 | 3188429.75 | 2941624.75 | P18181 | CD48 | 2316017 | 449704 | 259637 | 3 |
| CD55 molecule, decay accelerating factor for complement (Cromer blood group) | Plasma Membrane | other | 1.18E+07 | 1.26E+07 | 1.25E+07 | | | | | | |
| CD55 molecule, decay accelerating factor for complement (Cromer blood group) | Plasma Membrane | other | 9419374 | 7339787 | 9630807 | | | | | | |

| | | | | | | | | | | | |
|---|---|---|---|---|---|---|---|---|---|---|---|
| CD55 molecule, decay accelerating factor for complement (Cromer blood group) | Plasma Membrane | other | 5784862.5 | 4251971.5 | 4831361 | | | | | | |
| CD55 molecule, decay accelerating factor for complement (Cromer blood group) | Plasma Membrane | other | 4985323.5 | 6432816.5 | 6757013.5 | | | | | | |
| CD55 molecule, decay accelerating factor for complement (Cromer blood group) | Plasma Membrane | other | 3547051.75 | 4351169 | 4757378.5 | | | | | | |
| CD55 molecule, decay accelerating factor for complement (Cromer blood group) | Plasma Membrane | other | 2501027 | 2923368.25 | 2550474.25 | Q61475 | CD55 | 6344606 | 3359648 | 791877 | 18 |
| CD59a antigen | Plasma Membrane | other | 1.27E+08 | 1.63E+08 | 1.12E+08 | | | | | | |
| CD59a antigen | Plasma Membrane | other | 9722399 | 1.23E+07 | 8573033 | | | | | | |
| CD59a antigen | Plasma Membrane | other | 4612024.5 | 1.62E+07 | 4347839.5 | O55186 | Cd59a | 46944808 | 63749243 | 21249748 | 9 |
| CD82 molecule | Plasma Membrane | other | 5474135.5 | 3150463.5 | 5887102.5 | | | | | | |
| CD82 molecule | Plasma Membrane | other | 4383738.5 | 4870393.5 | 4681102.5 | P40237 | CD82 | 4928937 | 951715 | 388536 | 6 |
| CD9 molecule | Plasma Membrane | other | 2123187.5 | 6.27E+07 | 3.83E+07 | | | | | | |
| CD9 molecule | Plasma Membrane | other | 141426.75 | 2369567.75 | 2169750.5 | | | | | | |
| CD9 molecule | Plasma Membrane | other | 5.29 | 4257961 | 2483987.5 | P40240 | CD9 | 754873 | 22294887 | 7431629 | 9 |
| cell division cycle 123 | Cytoplasm | other | 5369976 | 5401760 | 5335438 | Q8CII2 | CDC123 | 5369976 | 33171 | 19151 | 3 |
| cell division cycle 42 | Plasma Membrane | enzyme | 1.02E+07 | 1.33E+07 | 1.03E+07 | | | | | | |
| cell division cycle 42 | Plasma Membrane | enzyme | 9797046 | 6581568 | 5251984.5 | | | | | | |
| cell division cycle 42 | Plasma Membrane | enzyme | 5965740.5 | 1.05E+07 | 7715108 | | | | | | |
| cell division cycle 42 | Plasma Membrane | enzyme | 5376683 | 3678116.25 | 4034223.25 | | | | | | |
| cell division cycle 42 | Plasma Membrane | enzyme | 2359156 | 3042659.75 | 1873144.75 | P60766 | Cdc42 | 6747725 | 3477308 | 897837 | 15 |
| cell division cycle associated 7-like | Nucleus | other | 2.24E+07 | 3.46E+07 | 2.93E+07 | Q922M5 | CDCA7L | 22360000 | 6127433 | 3537675 | 3 |
| CDP-diacylglycerol--inositol 3-phosphatidyltransferase | Cytoplasm | enzyme | 2353717 | 2638305.5 | 2136336.5 | Q8VDP6 | CDIPT | 2353717 | 251733 | 145338 | 3 |
| CDP-diacylglycerol synthase (phosphatidate cytidylyltransferase) 2 | Cytoplasm | enzyme | 700368.938 | 6073029.5 | 3672762.75 | Q99L43 | CDS2 | 700369 | 2691403 | 1553882 | 3 |
| CDV3 homolog (mouse) | Cytoplasm | other | 1037011.438 | 954311.562 | 1080000.875 | Q4VAA2 | CDV3 | 1037011 | 63882 | 36882 | 3 |
| carboxylesterase 5A | Extracellular Space | enzyme | 1.73E+07 | 1.61E+07 | 1.55E+07 | Q6AW46 | CES5A | 17260000 | 908974 | 524796 | 3 |
| cofilin 1 (non-muscle) | Nucleus | other | 2649475 | 1.16E+07 | 9003909 | | | | | | |
| cofilin 1 (non-muscle) | Nucleus | other | 1548557.75 | 3282158 | 2415868.25 | | | | | | |
| cofilin 1 (non-muscle) | Nucleus | other | 946846.438 | 4603737.5 | 3672254.25 | | | | | | |
| cofilin 1 (non-muscle) | Nucleus | other | 804261.188 | 4901408.5 | 5964326 | | | | | | |
| cofilin 1 (non-muscle) | Nucleus | other | 152180.938 | 2358792.5 | 1279584.375 | P18760 | CFL1 | 1220264 | 3174181 | 819570 | 15 |
| coiled-coil-helix-coiled-coil-helix domain containing 3 | Cytoplasm | other | 1565400.125 | 2389231 | 3160568.25 | Q9CRB9 | CHCHD3 | 1565400 | 797728 | 460568 | 3 |
| chitinase-like 3 | Cytoplasm | enzyme | 525609.812 | 1084381.625 | 1554353.125 | | | | | | |
| chitinase-like 3 | Cytoplasm | enzyme | 287673.875 | 3284999 | 6400453.5 | | | | | | |
| chitinase-like 3 | Cytoplasm | enzyme | 191082.891 | 2211302.25 | 3046999.5 | | | | | | |
| chitinase-like 3 | Cytoplasm | enzyme | 147188.891 | 1672688.5 | 2543483.75 | | | | | | |
| chitinase-like 3 | Cytoplasm | enzyme | 97800.086 | 742780.312 | 1061162.25 | | | | | | |
| chitinase-like 3 | Cytoplasm | enzyme | 5221.034 | 848343.562 | 1650780 | O35744 | Chil3/Chil4 | 209096 | 1586034 | 373832 | 18 |
| charged multivesicular body protein 2A | Cytoplasm | other | 2409814 | 1311991.375 | 1819598 | Q9DB34 | CHMP2A | 2409814 | 549429 | 317213 | 3 |
| charged multivesicular body protein 6 | Cytoplasm | other | 4848852.5 | 3725740.75 | 3590274.25 | P0C0A3 | CHMP6 | 4848853 | 690863 | 398870 | 3 |
| calcineurin-like EF-hand protein 1 | Cytoplasm | transporter | 8725651 | 8215003.5 | 7198730.5 | | | | | | |
| calcineurin-like EF-hand protein 1 | Cytoplasm | transporter | 5775539 | 4863125.5 | 4855754.5 | | | | | | |
| calcineurin-like EF-hand protein 1 | Cytoplasm | transporter | 4359267.5 | 2605735 | 2424811 | | | | | | |
| calcineurin-like EF-hand protein 1 | Cytoplasm | transporter | 4290828 | 2737665.5 | 3772589.25 | | | | | | |
| calcineurin-like EF-hand protein 1 | Cytoplasm | transporter | 4240015 | 4091481 | 3979450 | | | | | | |
| calcineurin-like EF-hand protein 1 | Cytoplasm | transporter | 1773113.5 | 4369601.5 | 3055044.25 | P61022 | CHP1 | 4860736 | 1916786 | 451791 | 18 |
| calcium and integrin binding 1 (calmyrin) | Nucleus | other | 9149066 | 4338240.5 | 6281069.5 | | | | | | |
| calcium and integrin binding 1 (calmyrin) | Nucleus | other | 8105773 | 6249708.5 | 7270578 | | | | | | |
| calcium and integrin binding 1 (calmyrin) | Nucleus | other | 5403340.5 | 3357775.25 | 4026970 | | | | | | |
| calcium and integrin binding 1 (calmyrin) | Nucleus | other | 996578.625 | 346188.406 | 732546.125 | Q9Z0F4 | CIB1 | 5913690 | 2926258 | 844738 | 12 |
| CDGSH iron sulfur domain 1 | Cytoplasm | other | 1382143.25 | 3303984.5 | 1698696 | | | | | | |
| CDGSH iron sulfur domain 1 | Cytoplasm | other | 1297509.25 | 3003398.25 | 2488164 | Q91WS0 | CISD1 | 1339826 | 858119 | 350326 | 6 |
| CDGSH iron sulfur domain 2 | Cytoplasm | other | 7414509 | 4942700 | 5828915 | | | | | | |
| CDGSH iron sulfur domain 2 | Cytoplasm | other | 5877319.5 | 8532191 | 4736242 | | | | | | |
| CDGSH iron sulfur domain 2 | Cytoplasm | other | 4939922.5 | 5180418.5 | 4866858 | | | | | | |
| CDGSH iron sulfur domain 2 | Cytoplasm | other | 4431229 | 4106615 | 3542277.75 | | | | | | |
| CDGSH iron sulfur domain 2 | Cytoplasm | other | 3773271.75 | 4273998 | 3150337.5 | Q9CQB5 | CISD2 | 5287250 | 1428431 | 368819 | 15 |
| claudin 13 | Plasma Membrane | other | 2.78E+08 | 2.17E+08 | 2.02E+08 | | | | | | |
| claudin 13 | Plasma Membrane | other | 9.31E+07 | 8.60E+07 | 7.15E+07 | | | | | | |
| claudin 13 | Plasma Membrane | other | 7.08E+07 | 6.08E+07 | 5.88E+07 | | | | | | |
| claudin 13 | Plasma Membrane | other | 4.32E+07 | 4.70E+07 | 3.59E+07 | Q9Z0S4 | Cldn13 | 121317500 | 80192897 | 23149695 | 12 |
| C-type lectin domain family 1, member B | Plasma Membrane | transmembrane receptor | 101775.469 | 1.09E+07 | 6501013 | Q9JL99 | CLEC1B | 101775 | 5429901 | 3134955 | 3 |
| ceroid-lipofuscinosis, neuronal 8 | Cytoplasm | other | 6625258.5 | 4522599.5 | 6327603.5 | Q9QUK3 | CLN8 | 6625259 | 1137820 | 656921 | 3 |
| chloride channel, nucleotide-sensitive, 1A | Plasma Membrane | ion channel | 8492958 | 2348399.5 | 3336131.5 | Q61189 | CLNS1A | 8492958 | 3299598 | 1905024 | 3 |
| cleft lip and palate associated transmembrane protein 1 | Plasma Membrane | other | 896680.062 | 1179958.125 | 856983.188 | Q8VBZ3 | CLPTM1 | 896680 | 176132 | 101690 | 3 |
| clathrin, heavy chain (Hc) | Plasma Membrane | other | 2.41E+07 | 9969972 | 9135970 | | | | | | |
| clathrin, heavy chain (Hc) | Plasma Membrane | other | 2.40E+07 | 1.12E+07 | 1.47E+07 | | | | | | |
| clathrin, heavy chain (Hc) | Plasma Membrane | other | 2.24E+07 | 1.23E+07 | 1.25E+07 | | | | | | |
| clathrin, heavy chain (Hc) | Plasma Membrane | other | 2.23E+07 | 9914880 | 1.35E+07 | | | | | | |
| clathrin, heavy chain (Hc) | Plasma Membrane | other | 2.01E+07 | 8281533 | 7867839.5 | | | | | | |
| clathrin, heavy chain (Hc) | Plasma Membrane | other | 1.98E+07 | 8300988 | 9819240 | | | | | | |
| clathrin, heavy chain (Hc) | Plasma Membrane | other | 1.92E+07 | 9987936 | 1.09E+07 | | | | | | |
| clathrin, heavy chain (Hc) | Plasma Membrane | other | 1.45E+07 | 4919726.5 | 6852068.5 | | | | | | |
| clathrin, heavy chain (Hc) | Plasma Membrane | other | 1.39E+07 | 7434850 | 7895225 | | | | | | |
| clathrin, heavy chain (Hc) | Plasma Membrane | other | 1.28E+07 | 6304979.5 | 7461761.5 | | | | | | |
| clathrin, heavy chain (Hc) | Plasma Membrane | other | 1.20E+07 | 6484117 | 9789553 | | | | | | |
| clathrin, heavy chain (Hc) | Plasma Membrane | other | 1.14E+07 | 6141499.5 | 7610841 | | | | | | |
| clathrin, heavy chain (Hc) | Plasma Membrane | other | 1.11E+07 | 3830391.5 | 5204181.5 | | | | | | |
| clathrin, heavy chain (Hc) | Plasma Membrane | other | 1.04E+07 | 4423870.5 | 4055823 | | | | | | |
| clathrin, heavy chain (Hc) | Plasma Membrane | other | 1.03E+07 | 5202543 | 7691259.5 | | | | | | |
| clathrin, heavy chain (Hc) | Plasma Membrane | other | 1.03E+07 | 5603303.5 | 5736116.5 | | | | | | |
| clathrin, heavy chain (Hc) | Plasma Membrane | other | 9446641 | 3379626.75 | 5908450.5 | | | | | | |
| clathrin, heavy chain (Hc) | Plasma Membrane | other | 9276216 | 2431818.75 | 4464303.5 | | | | | | |
| clathrin, heavy chain (Hc) | Plasma Membrane | other | 8957678 | 5126817 | 5527575 | | | | | | |
| clathrin, heavy chain (Hc) | Plasma Membrane | other | 8703483 | 5113949 | 5062604.5 | | | | | | |
| clathrin, heavy chain (Hc) | Plasma Membrane | other | 8574342 | 4691229.5 | 3308310 | | | | | | |
| clathrin, heavy chain (Hc) | Plasma Membrane | other | 8536503 | 3276523.25 | 3679627.75 | | | | | | |
| clathrin, heavy chain (Hc) | Plasma Membrane | other | 8533872 | 5410512 | 5149911.5 | | | | | | |
| clathrin, heavy chain (Hc) | Plasma Membrane | other | 8048986.5 | 7275557 | 7602255.5 | | | | | | |
| clathrin, heavy chain (Hc) | Plasma Membrane | other | 7943455.5 | 5324129.5 | 3793014 | | | | | | |
| clathrin, heavy chain (Hc) | Plasma Membrane | other | 7864560 | 3939865 | 4580863 | | | | | | |
| clathrin, heavy chain (Hc) | Plasma Membrane | other | 7710535 | 5353803 | 4935180 | | | | | | |
| clathrin, heavy chain (Hc) | Plasma Membrane | other | 7225963 | 2710411.75 | 3764794.25 | | | | | | |
| clathrin, heavy chain (Hc) | Plasma Membrane | other | 7138165 | 5135973.5 | 4221617 | | | | | | |
| clathrin, heavy chain (Hc) | Plasma Membrane | other | 6187844.5 | 2466543 | 4002916.75 | | | | | | |
| clathrin, heavy chain (Hc) | Plasma Membrane | other | 5693213 | 2454302.5 | 4173791 | | | | | | |
| clathrin, heavy chain (Hc) | Plasma Membrane | other | 5416866 | 1917499.375 | 2717023 | | | | | | |
| clathrin, heavy chain (Hc) | Plasma Membrane | other | 5340870.5 | 2582933 | 1922465.625 | | | | | | |

| | | | | | | | | | | | |
|---|---|---|---|---|---|---|---|---|---|---|---|
| clathrin, heavy chain (Hc) | Plasma Membrane | other | 5039173 | 1540957.25 | 2933634.5 | | | | | | |
| clathrin, heavy chain (Hc) | Plasma Membrane | other | 4380125 | 3558505.5 | 2568670.75 | | | | | | |
| clathrin, heavy chain (Hc) | Plasma Membrane | other | 4124717 | 1798134.125 | 2300378.25 | | | | | | |
| clathrin, heavy chain (Hc) | Plasma Membrane | other | 3864852.75 | 1061416 | 1739272.125 | | | | | | |
| clathrin, heavy chain (Hc) | Plasma Membrane | other | 3329255 | 1974661 | 2855887 | | | | | | |
| clathrin, heavy chain (Hc) | Plasma Membrane | other | 3211824 | 1024140.812 | 1799528.25 | | | | | | |
| clathrin, heavy chain (Hc) | Plasma Membrane | other | 3138028 | 1335348.75 | 1526228 | | | | | | |
| clathrin, heavy chain (Hc) | Plasma Membrane | other | 2590555.5 | 1135169.375 | 1271443 | | | | | | |
| clathrin, heavy chain (Hc) | Plasma Membrane | other | 1876427.5 | 2228219 | 1837593.25 | | | | | | |
| clathrin, heavy chain (Hc) | Plasma Membrane | other | 1862757.75 | 3047254.25 | 1455567.625 | | | | | | |
| clathrin, heavy chain (Hc) | Plasma Membrane | other | 1666534 | 1055235.625 | 1697020.75 | | | | | | |
| clathrin, heavy chain (Hc) | Plasma Membrane | other | 229604.266 | 678154.938 | 464103.656 | | | | | | |
| clathrin, heavy chain (Hc) | Plasma Membrane | other | 172573.859 | 60567.363 | 200778.266 | Q68FD5 | CLTC | 9229470 | 4966679 | 422792 | 138 |
| clusterin | Cytoplasm | other | 9714968 | 1.14E+07 | 9971323 | | | | | | |
| clusterin | Cytoplasm | other | 2591940.75 | 3055740 | 2864133 | | | | | | |
| clusterin | Cytoplasm | other | 1310189.5 | 4301293.5 | 2042416.375 | Q06890 | CLU | 4539033 | 3946182 | 1315394 | 9 |
| citrate lyase beta like | Cytoplasm | enzyme | 882020.75 | 1111949.875 | 1043156.812 | Q8R4N0 | CLYBL | 882021 | 118015 | 68136 | 3 |
| cornichon family AMPA receptor auxiliary protein 1 | Plasma Membrane | other | 4515070 | 7573223 | 5715668.5 | O35372 | CNIH1 | 4515070 | 1540792 | 889577 | 3 |
| calponin 2 | Cytoplasm | other | 297340.844 | 3770974.75 | 6793832 | Q08093 | CNN2 | 297341 | 3250851 | 1876880 | 3 |
| cyclin and CBS domain divalent metal cation transport mediator 2 | Plasma Membrane | other | 4867929 | 3816446 | 3347786.75 | | | | | | |
| cyclin and CBS domain divalent metal cation transport mediator 2 | Plasma Membrane | other | 4182645.5 | 3284516.25 | 4036401 | Q3TWN3 | CNNM2 | 4525287 | 587035 | 239656 | 6 |
| 2',3'-cyclic nucleotide 3' phosphodiesterase | Cytoplasm | enzyme | 2.86E+07 | 3.64E+07 | 3.42E+07 | | | | | | |
| 2',3'-cyclic nucleotide 3' phosphodiesterase | Cytoplasm | enzyme | 1.35E+07 | 1.91E+07 | 1.38E+07 | | | | | | |
| 2',3'-cyclic nucleotide 3' phosphodiesterase | Cytoplasm | enzyme | 8318063.5 | 9810205 | 7734030 | | | | | | |
| 2',3'-cyclic nucleotide 3' phosphodiesterase | Cytoplasm | enzyme | 5809450.5 | 8458725 | 7414520 | | | | | | |
| 2',3'-cyclic nucleotide 3' phosphodiesterase | Cytoplasm | enzyme | 4721179 | 4703348.5 | 4325435 | | | | | | |
| 2',3'-cyclic nucleotide 3' phosphodiesterase | Cytoplasm | enzyme | 3657516.75 | 4355190.5 | 3531921.25 | | | | | | |
| 2',3'-cyclic nucleotide 3' phosphodiesterase | Cytoplasm | enzyme | 3315277.5 | 4941344.5 | 4546695.5 | | | | | | |
| 2',3'-cyclic nucleotide 3' phosphodiesterase | Cytoplasm | enzyme | 659576.812 | 3925944.25 | 4147733.5 | P16330 | CNP | 8570133 | 9855222 | 2011689 | 24 |
| canopy FGF signaling regulator 2 | Plasma Membrane | other | 7927953.5 | 5266410.5 | 5426224.5 | | | | | | |
| canopy FGF signaling regulator 2 | Plasma Membrane | other | 6033816.5 | 5565698 | 6104937 | | | | | | |
| canopy FGF signaling regulator 2 | Plasma Membrane | other | 2319793.5 | 2397599.5 | 3111935.75 | Q9QXT0 | CNPY2 | 5427188 | 1899887 | 633296 | 9 |
| canopy FGF signaling regulator 4 | Extracellular Space | other | 3037772.5 | 1751827.5 | 2427436.25 | Q8BQ47 | CNPY4 | 3037773 | 643249 | 371380 | 3 |
| catechol-O-methyltransferase | Cytoplasm | enzyme | 4706174.5 | 4680071 | 4225748.5 | | | | | | |
| catechol-O-methyltransferase | Cytoplasm | enzyme | 4556201 | 5518157.5 | 5264146.5 | | | | | | |
| catechol-O-methyltransferase | Cytoplasm | enzyme | 3313836.5 | 2776842 | 3158458.75 | | | | | | |
| catechol-O-methyltransferase | Cytoplasm | enzyme | 3199300.75 | 4194932.5 | 3795413 | O88587 | COMT | 3943878 | 876959 | 253156 | 12 |
| catechol-O-methyltransferase domain containing 1 | Plasma Membrane | enzyme | 781073.812 | 3950826.5 | 3472074.5 | Q8BIG7 | COMTD1 | 781074 | 1708704 | 986521 | 3 |
| COP9 (constitutive photomorphogenic) homolog, subunit 2 (Arabidopsis thaliana) | Nucleus | transcription regulator | 9185846 | 4113575.75 | 6027558.5 | | | | | | |
| COP9 (constitutive photomorphogenic) homolog, subunit 2 (Arabidopsis thaliana) | Nucleus | transcription regulator | 6797382 | 3214894.5 | 3771790 | | | | | | |
| COP9 (constitutive photomorphogenic) homolog, subunit 2 (Arabidopsis thaliana) | Nucleus | transcription regulator | 4513056 | 1494211 | 1503185.375 | | | | | | |
| COP9 (constitutive photomorphogenic) homolog, subunit 2 (Arabidopsis thaliana) | Nucleus | transcription regulator | 2506780.5 | 1577466.375 | 1946643.375 | | | | | | |
| COP9 (constitutive photomorphogenic) homolog, subunit 2 (Arabidopsis thaliana) | Nucleus | transcription regulator | 876677.188 | 597296 | 472304.344 | P61202 | Cops2 | 4775948 | 2531428 | 653612 | 15 |
| COP9 signalosome subunit 3 | Cytoplasm | other | 6907801.5 | 3660475 | 3482784 | | | | | | |
| COP9 signalosome subunit 3 | Cytoplasm | other | 6408156 | 2856638.25 | 3709692.5 | | | | | | |
| COP9 signalosome subunit 3 | Cytoplasm | other | 6208069.5 | 2703570.75 | 3171084 | | | | | | |
| COP9 signalosome subunit 3 | Cytoplasm | other | 3537409 | 1500853.625 | 1971002.375 | | | | | | |
| COP9 signalosome subunit 3 | Cytoplasm | other | 3006670.25 | 952664.375 | 1019308.188 | | | | | | |
| COP9 signalosome subunit 3 | Cytoplasm | other | 2821674.75 | 1898077.5 | 2575515.5 | O88543 | COPS3 | 4814964 | 1724452 | 406457 | 18 |
| COP9 signalosome subunit 4 | Cytoplasm | other | 6438640.5 | 2314012.5 | 2710058.75 | | | | | | |
| COP9 signalosome subunit 4 | Cytoplasm | other | 4367375.5 | 2079542 | 3114729.75 | | | | | | |
| COP9 signalosome subunit 4 | Cytoplasm | other | 3668094.25 | 1783656.5 | 1980483.75 | | | | | | |
| COP9 signalosome subunit 4 | Cytoplasm | other | 3580714 | 1459161.625 | 1217051.75 | | | | | | |
| COP9 signalosome subunit 4 | Cytoplasm | other | 3428483 | 2286494.75 | 2319152.25 | | | | | | |
| COP9 signalosome subunit 4 | Cytoplasm | other | 2626042.25 | 1333609.5 | 2282662.5 | O88544 | COPS4 | 4018225 | 1263926 | 297910 | 18 |
| COP9 signalosome subunit 5 | Nucleus | transcription regulator | 1.53E+07 | 8731880 | 9876006 | | | | | | |
| COP9 signalosome subunit 5 | Nucleus | transcription regulator | 3826879.75 | 1672251 | 2326864.5 | O35864 | COPS5 | 9548440 | 5290386 | 2159791 | 6 |
| COP9 signalosome subunit 6 | Nucleus | other | 9748059 | 3841670.25 | 6817510 | | | | | | |
| COP9 signalosome subunit 6 | Nucleus | other | 8232821.5 | 3609837 | 6184683.5 | | | | | | |
| COP9 signalosome subunit 6 | Nucleus | other | 7905623 | 3135501 | 4475172.5 | | | | | | |
| COP9 signalosome subunit 6 | Nucleus | other | 3272155 | 1910512.125 | 1685275.25 | | | | | | |
| COP9 signalosome subunit 6 | Nucleus | other | 2789897.75 | 1216546.875 | 1673999.875 | O88545 | COPS6 | 6389711 | 2700957 | 697384 | 15 |
| COP9 signalosome subunit 7A | Cytoplasm | other | 1603109.5 | 960051.438 | 1107970.25 | Q9CZ04 | COPS7A | 1603110 | 336790 | 194446 | 3 |
| COP9 signalosome subunit 7B | Cytoplasm | other | 3283044.75 | 1408826.625 | 1977155.5 | Q8BV13 | COPS7B | 3283045 | 960992 | 554829 | 3 |
| COP9 signalosome subunit 8 | Nucleus | other | 4591310 | 1647076.375 | 2357216.25 | Q8VBV7 | COPS8 | 4591310 | 1536445 | 887067 | 3 |
| coronin, actin binding protein, 1A | Cytoplasm | other | 4427741.5 | 5.40E+07 | 3.64E+07 | | | | | | |
| coronin, actin binding protein, 1A | Cytoplasm | other | 1839199.25 | 6656607 | 5122222.5 | | | | | | |
| coronin, actin binding protein, 1A | Cytoplasm | other | 1214168.75 | 1.80E+07 | 1.86E+07 | | | | | | |
| coronin, actin binding protein, 1A | Cytoplasm | other | 1152365.125 | 2.00E+07 | 1.05E+07 | | | | | | |
| coronin, actin binding protein, 1A | Cytoplasm | other | 856001.875 | 8654028 | 4558484.5 | | | | | | |
| coronin, actin binding protein, 1A | Cytoplasm | other | 456414.281 | 1.53E+07 | 9362544 | | | | | | |
| coronin, actin binding protein, 1A | Cytoplasm | other | 454952.625 | 1.06E+07 | 4694044.5 | | | | | | |
| coronin, actin binding protein, 1A | Cytoplasm | other | 299458.844 | 4571962 | 3788968.5 | | | | | | |
| coronin, actin binding protein, 1A | Cytoplasm | other | 263760.094 | 4380177.5 | 3577296 | | | | | | |
| coronin, actin binding protein, 1A | Cytoplasm | other | 222634.016 | 9654162 | 5696748 | | | | | | |
| coronin, actin binding protein, 1A | Cytoplasm | other | 186777.062 | 1.32E+07 | 1.20E+07 | | | | | | |
| coronin, actin binding protein, 1A | Cytoplasm | other | 171800.766 | 4804274.5 | 6814938 | | | | | | |
| coronin, actin binding protein, 1A | Cytoplasm | other | 49477.574 | 6970722 | 4647434 | O89053 | CORO1A | 891904 | 10508445 | 1682698 | 39 |
| coronin, actin binding protein, 1C | Cytoplasm | other | 850139.188 | 7461690.5 | 7201643.5 | | | | | | |
| coronin, actin binding protein, 1C | Cytoplasm | other | 772952.812 | 1893536.5 | 1946255.125 | Q9WUM4 | CORO1C | 811546 | 3121644 | 1274406 | 6 |
| cytochrome c oxidase subunit IV isoform 1 | Cytoplasm | enzyme | 8673721 | 1.80E+07 | 1.51E+07 | | | | | | |
| cytochrome c oxidase subunit IV isoform 1 | Cytoplasm | enzyme | 4107283.25 | 9071354 | 8660363 | | | | | | |
| cytochrome c oxidase subunit IV isoform 1 | Cytoplasm | enzyme | 1887877 | 3883750.5 | 3685181.25 | P19783 | COX4I1 | 4889627 | 5485555 | 1828518 | 9 |
| cytochrome c oxidase subunit Va | Cytoplasm | enzyme | 4994385 | 8472916 | 8073797.5 | | | | | | |
| cytochrome c oxidase subunit Va | Cytoplasm | enzyme | 2973625.75 | 3804030.25 | 4599778.5 | | | | | | |
| cytochrome c oxidase subunit Va | Cytoplasm | enzyme | 2939794.25 | 4764275 | 3965864.5 | P12787 | COX5A | 3635935 | 2017324 | 672441 | 9 |
| cytochrome c oxidase subunit Vb | Cytoplasm | other | 5961387.5 | 1.05E+07 | 7283227.5 | P19536 | Cox5b | 5961388 | 2317699 | 1338124 | 3 |
| cytochrome c oxidase subunit VIb polypeptide 1 (ubiquitous) | Cytoplasm | enzyme | 3213345 | 6347785.5 | 4909357.5 | P56391 | COX6B1 | 3213345 | 1568983 | 905853 | 3 |
| cytochrome c oxidase subunit VIc | Cytoplasm | enzyme | 807133.375 | 1236455.375 | 953228.062 | Q9CPQ1 | Cox6c | 807133 | 218281 | 126024 | 3 |
| cytochrome c oxidase subunit VIIa polypeptide 2 (liver) | Cytoplasm | enzyme | 2917922.75 | 6707428.5 | 4490460 | P48771 | COX7A2 | 2917923 | 1903863 | 1099196 | 3 |
| copine II | Cytoplasm | other | 788190.5 | 3388500.5 | 1773534.875 | | | | | | |
| copine II | Cytoplasm | other | 493252.938 | 1648121.625 | 1521764 | P59108 | CPNE2 | 640722 | 1012045 | 413166 | 6 |
| coproporphyrinogen oxidase | Cytoplasm | enzyme | 7022895 | 7271403.5 | 6322467.5 | | | | | | |
| coproporphyrinogen oxidase | Cytoplasm | enzyme | 5739332.5 | 7427030 | 5977142 | | | | | | |
| coproporphyrinogen oxidase | Cytoplasm | enzyme | 5626988.5 | 6049823 | 5173423.5 | | | | | | |

| | | | | | | | | | | | |
|---|---|---|---|---|---|---|---|---|---|---|---|
| coproporphyrinogen oxidase | Cytoplasm | enzyme | 5259467 | 4678758 | 7143840.5 | | | | | | |
| coproporphyrinogen oxidase | Cytoplasm | enzyme | 4487358.5 | 3307321.5 | 4606743.5 | | | | | | |
| coproporphyrinogen oxidase | Cytoplasm | enzyme | 4421534 | 4230752.5 | 4536633.5 | | | | | | |
| coproporphyrinogen oxidase | Cytoplasm | enzyme | 3803834.25 | 4147636.5 | 3288190 | | | | | | |
| coproporphyrinogen oxidase | Cytoplasm | enzyme | 3328214.5 | 3733089.25 | 4360998.5 | | | | | | |
| coproporphyrinogen oxidase | Cytoplasm | enzyme | 2987796.5 | 3861403.25 | 4622937 | | | | | | |
| coproporphyrinogen oxidase | Cytoplasm | enzyme | 2623730.25 | 3205993 | 3122809.75 | | | | | | |
| coproporphyrinogen oxidase | Cytoplasm | enzyme | 1774732 | 1850930.875 | 997304.438 | | | | | | |
| coproporphyrinogen oxidase | Cytoplasm | enzyme | 1258192 | 1313814.875 | 792829 | P36552 | CPOX | 4027840 | 1805564 | 300927 | 36 |
| carnitine palmitoyltransferase 2 | Cytoplasm | enzyme | 895356.375 | 3915565.75 | 3435013.5 | | | | | | |
| carnitine palmitoyltransferase 2 | Cytoplasm | enzyme | 293736.938 | 1607813.25 | 1218238.75 | | | | | | |
| carnitine palmitoyltransferase 2 | Cytoplasm | enzyme | 71752.266 | 928847.062 | 547189 | P52825 | CPT2 | 420282 | 1356772 | 452257 | 9 |
| complement component (3b/4b) receptor 1-like | Plasma Membrane | other | 7.56E+07 | 5.01E+07 | 7.21E+07 | | | | | | |
| complement component (3b/4b) receptor 1-like | Plasma Membrane | other | 4.68E+07 | 3.31E+07 | 3.37E+07 | | | | | | |
| complement component (3b/4b) receptor 1-like | Plasma Membrane | other | 3.25E+07 | 2.62E+07 | 2.40E+07 | | | | | | |
| complement component (3b/4b) receptor 1-like | Plasma Membrane | other | 2.68E+07 | 2.76E+07 | 2.97E+07 | | | | | | |
| complement component (3b/4b) receptor 1-like | Plasma Membrane | other | 1.09E+07 | 9991443 | 8047815.5 | | | | | | |
| complement component (3b/4b) receptor 1-like | Plasma Membrane | other | 9144118 | 6148227.5 | 7565986.5 | | | | | | |
| complement component (3b/4b) receptor 1-like | Plasma Membrane | other | 8202852.5 | 7191306.5 | 7916601 | | | | | | |
| complement component (3b/4b) receptor 1-like | Plasma Membrane | other | 5868418 | 5340098.5 | 4433655 | | | | | | |
| complement component (3b/4b) receptor 1-like | Plasma Membrane | other | 1297826.125 | 350059.625 | 1009928.688 | Q64735 | CR1L | 24120357 | 20700859 | 3983882 | 27 |
| cAMP responsive element binding protein 3-like 1 | Nucleus | transcription regulator | 1230308.125 | 2065352.25 | 2284837 | Q9Z125 | CREB3L1 | 1230308 | 556403 | 321239 | 3 |
| cellular repressor of E1A-stimulated genes 1 | Nucleus | transcription regulator | 2915070.25 | 4063791.5 | 4209997 | | | | | | |
| cellular repressor of E1A-stimulated genes 1 | Nucleus | transcription regulator | 1930638.25 | 2863574 | 3403127.5 | O88668 | CREG1 | 2422854 | 849395 | 346764 | 6 |
| cysteine rich protein 2 | Plasma Membrane | other | 4.81E+07 | 3.45E+07 | 4.04E+07 | | | | | | |
| cysteine rich protein 2 | Plasma Membrane | other | 4.28E+07 | 3.74E+07 | 3.63E+07 | | | | | | |
| cysteine rich protein 2 | Plasma Membrane | other | 3.80E+07 | 4.49E+07 | 2.70E+07 | | | | | | |
| cysteine rich protein 2 | Plasma Membrane | other | 3.57E+07 | 2.25E+07 | 3.35E+07 | | | | | | |
| cysteine rich protein 2 | Plasma Membrane | other | 2.03E+07 | 2.67E+07 | 1.93E+07 | | | | | | |
| cysteine rich protein 2 | Plasma Membrane | other | 1.75E+07 | 1.97E+07 | 2.34E+07 | | | | | | |
| cysteine rich protein 2 | Plasma Membrane | other | 1.73E+07 | 2.30E+07 | 1.17E+07 | | | | | | |
| cysteine rich protein 2 | Plasma Membrane | other | 1.41E+07 | 2.53E+07 | 1.30E+07 | | | | | | |
| cysteine rich protein 2 | Plasma Membrane | other | 6956829 | 1.41E+07 | 6131239 | Q9DCT8 | Crip2 | 26757425 | 11838056 | 2278235 | 27 |
| cytokine receptor-like factor 3 | Cytoplasm | transcription regulator | 7517866.5 | 3751239.5 | 4271466 | Q9Z2L7 | CRLF3 | 7517867 | 2041128 | 1178446 | 3 |
| citrate synthase | Cytoplasm | enzyme | 1.71E+07 | 2.35E+07 | 2.50E+07 | | | | | | |
| citrate synthase | Cytoplasm | enzyme | 1.47E+07 | 1.58E+07 | 9380688 | | | | | | |
| citrate synthase | Cytoplasm | enzyme | 1.07E+07 | 1.16E+07 | 1.25E+07 | | | | | | |
| citrate synthase | Cytoplasm | enzyme | 5487016 | 7065802 | 6598610.5 | | | | | | |
| citrate synthase | Cytoplasm | enzyme | 4698243.5 | 6446589 | 6634918 | | | | | | |
| citrate synthase | Cytoplasm | enzyme | 3761863.25 | 7672161 | 4526812 | | | | | | |
| citrate synthase | Cytoplasm | enzyme | 3456218.75 | 1.35E+07 | 9138746 | | | | | | |
| citrate synthase | Cytoplasm | enzyme | 3278909 | 5150017 | 2695978 | | | | | | |
| citrate synthase | Cytoplasm | enzyme | 2575773.5 | 4476547.5 | 3046061.75 | | | | | | |
| citrate synthase | Cytoplasm | enzyme | 1304386.625 | 3184613.75 | 3700199.5 | Q9CZU6 | CS | 6702241 | 6086142 | 1111172 | 30 |
| casein kinase 1, alpha 1 | Cytoplasm | kinase | 1.24E+07 | 9118581 | 1.10E+07 | | | | | | |
| casein kinase 1, alpha 1 | Cytoplasm | kinase | 1.04E+07 | 7999345 | 4074111.75 | | | | | | |
| casein kinase 1, alpha 1 | Cytoplasm | kinase | 1.02E+07 | 9267167 | 8571855 | | | | | | |
| casein kinase 1, alpha 1 | Cytoplasm | kinase | 8683685 | 4507271 | 6515739.5 | | | | | | |
| casein kinase 1, alpha 1 | Cytoplasm | kinase | 7080030.5 | 8941319 | 6849261.5 | | | | | | |
| casein kinase 1, alpha 1 | Cytoplasm | kinase | 4554549 | 3290862 | 3903485.25 | | | | | | |
| casein kinase 1, alpha 1 | Cytoplasm | kinase | 4481100 | 3458723.75 | 4198531.5 | | | | | | |
| casein kinase 1, alpha 1 | Cytoplasm | kinase | 4302931.5 | 2075771.375 | 2567177.5 | Q8BK63 | CSNK1A1 | 7771537 | 2990116 | 610355 | 24 |
| casein kinase 2, alpha 1 polypeptide | Nucleus | kinase | 3781728.25 | 1420420.75 | 2151364.25 | Q60737 | CSNK2A1 | 3781728 | 1208866 | 697939 | 3 |
| casein kinase 2, beta polypeptide | Cytoplasm | kinase | 5982439.5 | 2891535 | 2287644.75 | P67871 | CSNK2B | 5982440 | 1981997 | 1144307 | 3 |
| cysteine and glycine-rich protein 1 | Nucleus | other | 7522641.5 | 2.82E+07 | 1.87E+07 | | | | | | |
| cysteine and glycine-rich protein 1 | Nucleus | other | 5621010 | 2.40E+07 | 1.93E+07 | | | | | | |
| cysteine and glycine-rich protein 1 | Nucleus | other | 5008381 | 2.17E+07 | 1.53E+07 | | | | | | |
| cysteine and glycine-rich protein 1 | Nucleus | other | 4261929 | 3.62E+07 | 1.22E+07 | | | | | | |
| cysteine and glycine-rich protein 1 | Nucleus | other | 329082.469 | 1.60E+07 | 3390545.25 | P97315 | CSRP1 | 4548609 | 10323920 | 2665625 | 15 |
| cathepsin B | Cytoplasm | peptidase | 1.36E+07 | 6431021.5 | 1.02E+07 | | | | | | |
| cathepsin B | Cytoplasm | peptidase | 9178435 | 5060890 | 7141441.5 | | | | | | |
| cathepsin B | Cytoplasm | peptidase | 7288712.5 | 5111202.5 | 7609686.5 | | | | | | |
| cathepsin B | Cytoplasm | peptidase | 5414235.5 | 1928043.375 | 3897091.75 | | | | | | |
| cathepsin B | Cytoplasm | peptidase | 4368428 | 2895488 | 3437648.75 | P10605 | CTSB | 7965962 | 3068601 | 792309 | 15 |
| cathepsin D | Cytoplasm | peptidase | 1.31E+07 | 1.09E+07 | 1.25E+07 | | | | | | |
| cathepsin D | Cytoplasm | peptidase | 8524043 | 8340246 | 9685370 | | | | | | |
| cathepsin D | Cytoplasm | peptidase | 8362487 | 8726742 | 8810479 | | | | | | |
| cathepsin D | Cytoplasm | peptidase | 6590934 | 6140946 | 7423275.5 | | | | | | |
| cathepsin D | Cytoplasm | peptidase | 6571506 | 6530185 | 6869336.5 | | | | | | |
| cathepsin D | Cytoplasm | peptidase | 1520279.5 | 4058115.25 | 2514010.25 | | | | | | |
| cathepsin D | Cytoplasm | peptidase | 1297339.25 | 1445231.375 | 1730651.5 | P18242 | CTSD | 6570941 | 3555613 | 775898 | 21 |
| cathepsin E | Cytoplasm | peptidase | 1.42E+08 | 1.04E+08 | 1.09E+08 | | | | | | |
| cathepsin E | Cytoplasm | peptidase | 1.21E+08 | 6.83E+07 | 8.50E+07 | | | | | | |
| cathepsin E | Cytoplasm | peptidase | 1.26E+07 | 2.84E+07 | 2.22E+07 | P70269 | CTSE | 92076667 | 46891913 | 15630638 | 9 |
| cullin 1 | Nucleus | enzyme | 6000261.5 | 2697817.75 | 3143794 | | | | | | |
| cullin 1 | Nucleus | enzyme | 5316420.5 | 1875983.25 | 2935355 | | | | | | |
| cullin 1 | Nucleus | enzyme | 4103238 | 2016064.75 | 2981353.25 | | | | | | |
| cullin 1 | Nucleus | enzyme | 3845012 | 1842483.375 | 2526733.75 | | | | | | |
| cullin 1 | Nucleus | enzyme | 3138792 | 1705522.125 | 2357974.75 | | | | | | |
| cullin 1 | Nucleus | enzyme | 2429992.75 | 1017623 | 1873227 | | | | | | |
| cullin 1 | Nucleus | enzyme | 1521952.25 | 1212548.625 | 980204.5 | | | | | | |
| cullin 1 | Nucleus | enzyme | 1277193.875 | 1112816.375 | 415100.656 | | | | | | |
| cullin 1 | Nucleus | enzyme | 1111395.5 | 402979.094 | 573153.375 | Q9WTX6 | CUL1 | 3193806 | 1400811 | 269586 | 27 |
| cullin 3 | Nucleus | enzyme | 6351584 | 2097023.5 | 2725681.5 | | | | | | |
| cullin 3 | Nucleus | enzyme | 4445454.5 | 2631092.25 | 2035393 | Q9JLV5 | CUL3 | 5398519 | 1697557 | 693025 | 6 |
| chemokine (C-X-C motif) receptor 4 | Plasma Membrane | G-protein coupled receptor | 3533394.75 | 2544324 | 2782678 | P70658 | CXCR4 | 3533395 | 516180 | 298017 | 3 |
| cytochrome b5 type A (microsomal) | Cytoplasm | enzyme | 6737722.5 | 8148662.5 | 7314355 | | | | | | |
| cytochrome b5 type A (microsomal) | Cytoplasm | enzyme | 5886942.5 | 9163290 | 7626479.5 | | | | | | |
| cytochrome b5 type A (microsomal) | Cytoplasm | enzyme | 1576257.875 | 2328747.25 | 1909657.375 | | | | | | |
| cytochrome b5 type A (microsomal) | Cytoplasm | enzyme | 1223085.625 | 1757471.75 | 1592351.625 | | | | | | |
| cytochrome b5 type A (microsomal) | Cytoplasm | enzyme | 1048102.125 | 5164298.5 | 3816968.75 | P56395 | CYB5A | 3294422 | 2913897 | 752365 | 15 |
| cytochrome b5 type B (outer mitochondrial membrane) | Cytoplasm | enzyme | 1.21E+07 | 1.73E+07 | 1.38E+07 | | | | | | |
| cytochrome b5 type B (outer mitochondrial membrane) | Cytoplasm | enzyme | 1952118.75 | 3610007.25 | 1265825.25 | | | | | | |
| cytochrome b5 type B (outer mitochondrial membrane) | Cytoplasm | enzyme | 733704.312 | 1520607.125 | 859292.875 | Q9CQX2 | CYB5B | 4941941 | 6576105 | 2192035 | 9 |
| cytochrome b5 reductase 3 | Cytoplasm | enzyme | 2.73E+07 | 3.37E+07 | 3.39E+07 | | | | | | |

| | | | | | | | | | | | |
|---|---|---|---|---|---|---|---|---|---|---|---|
| cytochrome b5 reductase 3 | Cytoplasm | enzyme | 2.08E+07 | 1.05E+07 | 1.26E+07 | | | | | | |
| cytochrome b5 reductase 3 | Cytoplasm | enzyme | 1.99E+07 | 2.67E+07 | 2.60E+07 | | | | | | |
| cytochrome b5 reductase 3 | Cytoplasm | enzyme | 1.25E+07 | 1.53E+07 | 1.59E+07 | | | | | | |
| cytochrome b5 reductase 3 | Cytoplasm | enzyme | 9358659 | 1.37E+07 | 1.06E+07 | | | | | | |
| cytochrome b5 reductase 3 | Cytoplasm | enzyme | 9233875 | 1.06E+07 | 1.03E+07 | | | | | | |
| cytochrome b5 reductase 3 | Cytoplasm | enzyme | 6732157.5 | 8611199 | 6805841 | | | | | | |
| cytochrome b5 reductase 3 | Cytoplasm | enzyme | 6318005 | 5193052 | 6448714.5 | | | | | | |
| cytochrome b5 reductase 3 | Cytoplasm | enzyme | 6187829.5 | 6712720 | 5933538 | | | | | | |
| cytochrome b5 reductase 3 | Cytoplasm | enzyme | 4512703 | 5278341.5 | 5323530 | | | | | | |
| cytochrome b5 reductase 3 | Cytoplasm | enzyme | 4487136 | 4465185.5 | 4475217.5 | | | | | | |
| cytochrome b5 reductase 3 | Cytoplasm | enzyme | 3145106 | 1592163 | 2412967.75 | Q9DCN2 | Cyb5r3 | 10870456 | 8691199 | 1448533 | 36 |
| cytochrome c-1 | Cytoplasm | enzyme | 9425621 | 1.18E+07 | 9501267 | Q9D0M3 | CYC1 | 9425621 | 1372626 | 792486 | 3 |
| cytochrome c, somatic | Cytoplasm | transporter | 5411839 | 8615587 | 6200149 | | | | | | |
| cytochrome c, somatic | Cytoplasm | transporter | 3288825.5 | 5158277.5 | 4326708.5 | | | | | | |
| cytochrome c, somatic | Cytoplasm | transporter | 1575789.5 | 7664172 | 4271449.5 | | | | | | |
| cytochrome c, somatic | Cytoplasm | transporter | 1290945 | 2411451.75 | 2875385.5 | | | | | | |
| cytochrome c, somatic | Cytoplasm | transporter | 413606.625 | 1224444.75 | 1118248.125 | P62897 | CYCS | 2396201 | 2511631 | 648500 | 15 |
| dishevelled associated activator of morphogenesis 1 | Cytoplasm | other | 8061328.5 | 8639246 | 7821139 | | | | | | |
| dishevelled associated activator of morphogenesis 1 | Cytoplasm | other | 4830547.5 | 5410679.5 | 4182232.5 | | | | | | |
| dishevelled associated activator of morphogenesis 1 | Cytoplasm | other | 4658813 | 6289051.5 | 4875967.5 | | | | | | |
| dishevelled associated activator of morphogenesis 1 | Cytoplasm | other | 4301201 | 5614647 | 4862429.5 | | | | | | |
| dishevelled associated activator of morphogenesis 1 | Cytoplasm | other | 4072896.75 | 3079384 | 4198802 | | | | | | |
| dishevelled associated activator of morphogenesis 1 | Cytoplasm | other | 2981830 | 3342891.25 | 2587218.25 | | | | | | |
| dishevelled associated activator of morphogenesis 1 | Cytoplasm | other | 2960204 | 4480268 | 4624155 | | | | | | |
| dishevelled associated activator of morphogenesis 1 | Cytoplasm | other | 2882910 | 3058793 | 2851152 | | | | | | |
| dishevelled associated activator of morphogenesis 1 | Cytoplasm | other | 2682732.25 | 4097561.5 | 3393593.75 | | | | | | |
| dishevelled associated activator of morphogenesis 1 | Cytoplasm | other | 2427994.5 | 2807604.25 | 2714165.5 | | | | | | |
| dishevelled associated activator of morphogenesis 1 | Cytoplasm | other | 1460587.125 | 2053553 | 1734327.375 | | | | | | |
| dishevelled associated activator of morphogenesis 1 | Cytoplasm | other | 865231.312 | 4317057 | 1546506.875 | Q8BPM0 | DAAM1 | 3515523 | 1800094 | 300016 | 36 |
| defender against cell death 1 | Cytoplasm | other | 812363 | 2878856 | 2024320.125 | P61804 | DAD1 | 812363 | 1038385 | 599512 | 3 |
| diacylglycerol lipase, beta | Plasma Membrane | enzyme | 9250279 | 8035731.5 | 7798406.5 | | | | | | |
| diacylglycerol lipase, beta | Plasma Membrane | enzyme | 6562082.5 | 4778554.5 | 4748953 | | | | | | |
| diacylglycerol lipase, beta | Plasma Membrane | enzyme | 5109595.5 | 2988673.75 | 3676529.5 | | | | | | |
| diacylglycerol lipase, beta | Plasma Membrane | enzyme | 4877642.5 | 1914622.75 | 3006614.25 | | | | | | |
| diacylglycerol lipase, beta | Plasma Membrane | enzyme | 3845131.25 | 3338198.75 | 4208974.5 | | | | | | |
| diacylglycerol lipase, beta | Plasma Membrane | enzyme | 2776855.25 | 2189366 | 3039353.25 | Q91WC9 | DAGLB | 5403598 | 2099871 | 494944 | 18 |
| aspartyl-tRNA synthetase | Cytoplasm | enzyme | 6081907.5 | 3723877.5 | 4484785.5 | | | | | | |
| aspartyl-tRNA synthetase | Cytoplasm | enzyme | 3639441.25 | 1756101.75 | 4773110.5 | | | | | | |
| aspartyl-tRNA synthetase | Cytoplasm | enzyme | 2942748.25 | 879775.312 | 3988549 | | | | | | |
| aspartyl-tRNA synthetase | Cytoplasm | enzyme | 2825922 | 1004538.125 | 4062812.25 | | | | | | |
| aspartyl-tRNA synthetase | Cytoplasm | enzyme | 1881957.25 | 632419.062 | 3082185.5 | | | | | | |
| aspartyl-tRNA synthetase | Cytoplasm | enzyme | 1689513.75 | 773590.938 | 2418948.5 | | | | | | |
| aspartyl-tRNA synthetase | Cytoplasm | enzyme | 1687348.375 | 863653.438 | 3065221 | | | | | | |
| aspartyl-tRNA synthetase | Cytoplasm | enzyme | 1154301.875 | 639483.438 | 2760649.5 | Q922B2 | DARS | 2737893 | 1502622 | 306721 | 24 |
| DAZ associated protein 1 | Cytoplasm | other | 674517.812 | 621285.25 | 384976.562 | Q9JII5 | Dazap1 | 674518 | 154116 | 88979 | 3 |
| DCN1, defective in cullin neddylation 1, domain containing 1 | Nucleus | other | 2916710 | 1465672.5 | 2078844 | Q9QZ73 | DCUN1D1 | 2916710 | 728412 | 420549 | 3 |
| DNA-damage inducible 1 homolog 2 | Plasma Membrane | transporter | 1.13E+07 | 4213168.5 | 6130335 | | | | | | |
| DNA-damage inducible 1 homolog 2 | Plasma Membrane | transporter | 5905494.5 | 3208572.75 | 4805015 | A2ADY9 | DDI2 | 8607747 | 2849716 | 1163391 | 6 |
| dolichyl-diphosphooligosaccharide--protein glycosyltransferase subunit (non-catalytic) | Cytoplasm | enzyme | 1362321.125 | 3886259 | 3563992 | | | | | | |
| dolichyl-diphosphooligosaccharide--protein glycosyltransferase subunit (non-catalytic) | Cytoplasm | enzyme | 642513.688 | 3661502.25 | 2516175.25 | | | | | | |
| dolichyl-diphosphooligosaccharide--protein glycosyltransferase subunit (non-catalytic) | Cytoplasm | enzyme | 321106.875 | 1880523.25 | 2299789.5 | O54734 | DDOST | 775314 | 1308493 | 436164 | 9 |
| DENN/MADD domain containing 2A | Cytoplasm | other | 1758484.125 | 1465145.5 | 1389225.875 | Q8C4S8 | DENND2A | 1758484 | 195006 | 112587 | 3 |
| deoxyribose-phosphate aldolase (putative) | Cytoplasm | enzyme | 3749578.5 | 2245251 | 2497676.25 | | | | | | |
| deoxyribose-phosphate aldolase (putative) | Cytoplasm | enzyme | 2499267.5 | 1443510.125 | 1537862.875 | | | | | | |
| deoxyribose-phosphate aldolase (putative) | Cytoplasm | enzyme | 2418871 | 2171370.5 | 1717742.875 | Q91YP3 | DERA | 2889239 | 693514 | 231171 | 9 |
| derlin 1 | Cytoplasm | other | 5077290.5 | 1.15E+07 | 9904221 | | | | | | |
| derlin 1 | Cytoplasm | other | 2755922.5 | 4872667 | 3424759.25 | Q99J56 | DERL1 | 3916607 | 3597051 | 1468490 | 6 |
| derlin 2 | Cytoplasm | other | 1434796.375 | 3850588.75 | 2748579.25 | Q8BNI4 | DERL2 | 1434796 | 1209442 | 698272 | 3 |
| dehydrogenase/reductase (SDR family) member 11 | Cytoplasm | other | 6944219 | 3758062.75 | 5150894.5 | Q3U0B3 | DHRS11 | 6944219 | 1597268 | 922183 | 3 |
| dehydrogenase E1 and transketolase domain containing 1 | Cytoplasm | enzyme | 1.07E+07 | 1.28E+07 | 1.24E+07 | A2ATU0 | DHTKD1 | 10680000 | 1137673 | 656836 | 3 |
| DEAH (Asp-Glu-Ala-His) box polypeptide 33 | Nucleus | enzyme | 8761595 | 4704690.5 | 7282880 | Q80VY9 | DHX33 | 8761595 | 2053133 | 1185377 | 3 |
| dihydrolipoamide S-acetyltransferase | Cytoplasm | enzyme | 1592134.75 | 2120655.25 | 1663348.375 | | | | | | |
| dihydrolipoamide S-acetyltransferase | Cytoplasm | enzyme | 806291.125 | 1827218 | 1891457.125 | Q8BMF4 | DLAT | 1199213 | 453121 | 184986 | 6 |
| dihydrolipoamide dehydrogenase | Cytoplasm | enzyme | 1.02E+07 | 1.26E+07 | 1.22E+07 | | | | | | |
| dihydrolipoamide dehydrogenase | Cytoplasm | enzyme | 4217257.5 | 7701650 | 1.05E+07 | | | | | | |
| dihydrolipoamide dehydrogenase | Cytoplasm | enzyme | 3678121.5 | 8015047.5 | 4115211.5 | O08749 | DLD | 6038460 | 3484487 | 1161496 | 9 |
| discs, large homolog 1 (Drosophila) | Plasma Membrane | kinase | 1135162.875 | 1402978.875 | 909930.875 | | | | | | |
| discs, large homolog 1 (Drosophila) | Plasma Membrane | kinase | 486482.312 | 836101.375 | 350466.938 | | | | | | |
| discs, large homolog 1 (Drosophila) | Plasma Membrane | kinase | 194389.594 | 422393.812 | 260740.422 | Q811D0 | DLG1 | 605345 | 422710 | 140903 | 9 |
| dihydrolipoamide S-succinyltransferase (E2 component of 2-oxo-glutarate complex) | Cytoplasm | enzyme | 5215223 | 8396903 | 7512768.5 | | | | | | |
| dihydrolipoamide S-succinyltransferase (E2 component of 2-oxo-glutarate complex) | Cytoplasm | enzyme | 4571454 | 6024439.5 | 5849937.5 | | | | | | |
| dihydrolipoamide S-succinyltransferase (E2 component of 2-oxo-glutarate complex) | Cytoplasm | enzyme | 4552185.5 | 7386926 | 6512175.5 | | | | | | |
| dihydrolipoamide S-succinyltransferase (E2 component of 2-oxo-glutarate complex) | Cytoplasm | enzyme | 3273383 | 9135643 | 6764744.5 | | | | | | |
| dihydrolipoamide S-succinyltransferase (E2 component of 2-oxo-glutarate complex) | Cytoplasm | enzyme | 3129844.25 | 2897463.75 | 3142670.25 | | | | | | |
| dihydrolipoamide S-succinyltransferase (E2 component of 2-oxo-glutarate complex) | Cytoplasm | enzyme | 2040430.125 | 2146933.75 | 2544423.5 | | | | | | |
| dihydrolipoamide S-succinyltransferase (E2 component of 2-oxo-glutarate complex) | Cytoplasm | enzyme | 1791127.125 | 4429277.5 | 4017564.5 | Q9D2G2 | DLST | 3510521 | 2196401 | 479294 | 21 |
| doublesex and mab-3 related transcription factor 3 | Nucleus | transcription regulator | 6360629.5 | 5213658 | 4014143.5 | Q80WT2 | DMRT3 | 6360630 | 1173341 | 677429 | 3 |
| dematin actin binding protein | Plasma Membrane | other | 6.41E+08 | 3.59E+08 | 5.05E+08 | | | | | | |
| dematin actin binding protein | Plasma Membrane | other | 5.74E+08 | 6.54E+08 | 6.14E+08 | | | | | | |
| dematin actin binding protein | Plasma Membrane | other | 4.86E+08 | 3.52E+08 | 3.26E+08 | | | | | | |
| dematin actin binding protein | Plasma Membrane | other | 4.63E+08 | 3.10E+08 | 3.08E+08 | | | | | | |
| dematin actin binding protein | Plasma Membrane | other | 4.55E+08 | 4.08E+08 | 3.90E+08 | | | | | | |
| dematin actin binding protein | Plasma Membrane | other | 4.34E+08 | 3.00E+08 | 2.91E+08 | | | | | | |
| dematin actin binding protein | Plasma Membrane | other | 4.14E+08 | 3.33E+08 | 3.46E+08 | | | | | | |
| dematin actin binding protein | Plasma Membrane | other | 4.13E+08 | 3.32E+08 | 3.09E+08 | | | | | | |
| dematin actin binding protein | Plasma Membrane | other | 3.43E+08 | 2.49E+08 | 3.16E+08 | | | | | | |
| dematin actin binding protein | Plasma Membrane | other | 2.89E+08 | 2.00E+08 | 1.80E+08 | | | | | | |
| dematin actin binding protein | Plasma Membrane | other | 2.25E+08 | 1.90E+08 | 1.73E+08 | | | | | | |
| dematin actin binding protein | Plasma Membrane | other | 1.42E+08 | 7.65E+07 | 8.74E+07 | | | | | | |
| dematin actin binding protein | Plasma Membrane | other | 1.07E+08 | 1.26E+08 | 9.25E+07 | | | | | | |
| dematin actin binding protein | Plasma Membrane | other | 9.64E+07 | 9.29E+07 | 1.05E+08 | | | | | | |
| dematin actin binding protein | Plasma Membrane | other | 9.02E+07 | 6.93E+07 | 9.45E+07 | | | | | | |
| dematin actin binding protein | Plasma Membrane | other | 4.25E+07 | 4.50E+07 | 5.50E+07 | | | | | | |
| dematin actin binding protein | Plasma Membrane | other | 3.26E+07 | 1.05E+07 | 6.11E+07 | | | | | | |
| dematin actin binding protein | Plasma Membrane | other | 2.93E+07 | 1.93E+07 | 2.24E+07 | | | | | | |

| | | | | | | | | | | | |
|---|---|---|---|---|---|---|---|---|---|---|---|
| dematin actin binding protein | Plasma Membrane | other | 1.77E+07 | 1.51E+07 | 1.46E+07 | | | | | | |
| dematin actin binding protein | Plasma Membrane | other | 1.64E+07 | 1.23E+07 | 1.12E+07 | | | | | | |
| dematin actin binding protein | Plasma Membrane | other | 9282284 | 1.06E+07 | 6410458.5 | | | | | | |
| dematin actin binding protein | Plasma Membrane | other | 9169044 | 6705989 | 7070843 | | | | | | |
| dematin actin binding protein | Plasma Membrane | other | 6872027.5 | 4681623 | 6616446 | | | | | | |
| dematin actin binding protein | Plasma Membrane | other | 5859832 | 8793303 | 5345722 | | | | | | |
| dematin actin binding protein | Plasma Membrane | other | 754471.875 | 655716.188 | 895589.5 | Q9WV69 | DMTN | 213698706 | 187247416 | 21621469 | 75 |
| DnaJ (Hsp40) homolog, subfamily A, member 4 | Nucleus | other | 2198675 | 1786006.5 | 1984727 | Q9JMC3 | DNAJA4 | 2198675 | 206381 | 119154 | 3 |
| DnaJ (Hsp40) homolog, subfamily B, member 1 | Nucleus | other | 2.23E+07 | 1.66E+07 | 2.04E+07 | | | | | | |
| DnaJ (Hsp40) homolog, subfamily B, member 1 | Nucleus | other | 1.60E+07 | 1.52E+07 | 1.22E+07 | | | | | | |
| DnaJ (Hsp40) homolog, subfamily B, member 1 | Nucleus | other | 1.34E+07 | 1.13E+07 | 1.26E+07 | | | | | | |
| DnaJ (Hsp40) homolog, subfamily B, member 1 | Nucleus | other | 1.25E+07 | 9911792 | 1.29E+07 | | | | | | |
| DnaJ (Hsp40) homolog, subfamily B, member 1 | Nucleus | other | 6669328 | 5821450 | 5328477.5 | | | | | | |
| DnaJ (Hsp40) homolog, subfamily B, member 1 | Nucleus | other | 6200912 | 3476149.75 | 4474423 | | | | | | |
| DnaJ (Hsp40) homolog, subfamily B, member 1 | Nucleus | other | 4627097 | 4386416.5 | 3670588 | | | | | | |
| DnaJ (Hsp40) homolog, subfamily B, member 1 | Nucleus | other | 4132118.5 | 2908920.5 | 3511947.5 | | | | | | |
| DnaJ (Hsp40) homolog, subfamily B, member 1 | Nucleus | other | 14947.229 | 369493.188 | 1205248.75 | Q9QYJ3 | DNAJB1 | 9541600 | 6160602 | 1185608 | 27 |
| DnaJ (Hsp40) homolog, subfamily B, member 11 | Cytoplasm | other | 5081723.5 | 6556975.5 | 3909458.25 | | | | | | |
| DnaJ (Hsp40) homolog, subfamily B, member 11 | Cytoplasm | other | 2992117 | 3202209.75 | 2992112.75 | | | | | | |
| DnaJ (Hsp40) homolog, subfamily B, member 11 | Cytoplasm | other | 1966288.25 | 2124137.5 | 1897150.875 | Q99KV1 | DNAJB11 | 3346710 | 1555890 | 518630 | 9 |
| DnaJ (Hsp40) homolog, subfamily B, member 4 | Nucleus | other | 2.44E+07 | 1.82E+07 | 2.18E+07 | | | | | | |
| DnaJ (Hsp40) homolog, subfamily B, member 4 | Nucleus | other | 2.18E+07 | 2.23E+07 | 2.11E+07 | | | | | | |
| DnaJ (Hsp40) homolog, subfamily B, member 4 | Nucleus | other | 1.70E+07 | 1.50E+07 | 1.28E+07 | | | | | | |
| DnaJ (Hsp40) homolog, subfamily B, member 4 | Nucleus | other | 9025905 | 9845950 | 6267611.5 | | | | | | |
| DnaJ (Hsp40) homolog, subfamily B, member 4 | Nucleus | other | 8414192 | 7799530 | 8999899 | | | | | | |
| DnaJ (Hsp40) homolog, subfamily B, member 4 | Nucleus | other | 6705394 | 6125222 | 5158171.5 | | | | | | |
| DnaJ (Hsp40) homolog, subfamily B, member 4 | Nucleus | other | 6442318.5 | 5178295.5 | 6497019.5 | | | | | | |
| DnaJ (Hsp40) homolog, subfamily B, member 4 | Nucleus | other | 4698483 | 4872441.5 | 4782922.5 | | | | | | |
| DnaJ (Hsp40) homolog, subfamily B, member 4 | Nucleus | other | 4332919 | 2796163 | 3241494.75 | | | | | | |
| DnaJ (Hsp40) homolog, subfamily B, member 4 | Nucleus | other | 1430434.875 | 1757677.875 | 2128131.75 | Q9D832 | DNAJB4 | 10421965 | 7047839 | 1286753 | 30 |
| DnaJ (Hsp40) homolog, subfamily C, member 10 | Cytoplasm | enzyme | 552244.5 | 575459.625 | 389373.969 | Q9DC23 | DNAJC10 | 552245 | 101402 | 58544 | 3 |
| DnaJ (Hsp40) homolog, subfamily C, member 11 | Cytoplasm | other | 1404570.625 | 1329757.875 | 1541142.5 | | | | | | |
| DnaJ (Hsp40) homolog, subfamily C, member 11 | Cytoplasm | other | 654947.25 | 513709.719 | 630905.438 | Q5U458 | DNAJC11 | 1029759 | 459583 | 187624 | 6 |
| DnaJ (Hsp40) homolog, subfamily C, member 5 | Plasma Membrane | other | 4.32E+07 | 3.50E+07 | 4.02E+07 | | | | | | |
| DnaJ (Hsp40) homolog, subfamily C, member 5 | Plasma Membrane | other | 4.18E+07 | 3.09E+07 | 3.40E+07 | | | | | | |
| DnaJ (Hsp40) homolog, subfamily C, member 5 | Plasma Membrane | other | 1.78E+07 | 1.15E+07 | 1.25E+07 | | | | | | |
| DnaJ (Hsp40) homolog, subfamily C, member 5 | Plasma Membrane | other | 8255965.5 | 1.69E+07 | 7120041.5 | | | | | | |
| DnaJ (Hsp40) homolog, subfamily C, member 5 | Plasma Membrane | other | 490463.094 | 1502944.5 | 467483.875 | P60904 | DNAJC5 | 22297286 | 15827009 | 4086516 | 15 |
| deoxyribonuclease I-like 1 | Cytoplasm | enzyme | 235750.391 | 549482.625 | 520685.969 | Q9D7J6 | DNASE1L1 | 235750 | 173419 | 100124 | 3 |
| dynamin 2 | Plasma Membrane | enzyme | 4.40E+07 | 3.47E+07 | 3.63E+07 | | | | | | |
| dynamin 2 | Plasma Membrane | enzyme | 2.76E+07 | 2.03E+07 | 2.04E+07 | | | | | | |
| dynamin 2 | Plasma Membrane | enzyme | 2.73E+07 | 2.39E+07 | 2.91E+07 | | | | | | |
| dynamin 2 | Plasma Membrane | enzyme | 2.40E+07 | 1.98E+07 | 1.98E+07 | | | | | | |
| dynamin 2 | Plasma Membrane | enzyme | 1.62E+07 | 1.21E+07 | 1.47E+07 | | | | | | |
| dynamin 2 | Plasma Membrane | enzyme | 1.57E+07 | 1.37E+07 | 1.31E+07 | | | | | | |
| dynamin 2 | Plasma Membrane | enzyme | 1.39E+07 | 9471524 | 1.18E+07 | | | | | | |
| dynamin 2 | Plasma Membrane | enzyme | 1.03E+07 | 7883130 | 8144057.5 | | | | | | |
| dynamin 2 | Plasma Membrane | enzyme | 1.01E+07 | 8267816.5 | 7419859 | | | | | | |
| dynamin 2 | Plasma Membrane | enzyme | 9075466 | 7292503 | 7332416.5 | | | | | | |
| dynamin 2 | Plasma Membrane | enzyme | 8735292 | 6181533 | 7509339.5 | | | | | | |
| dynamin 2 | Plasma Membrane | enzyme | 8284223.5 | 4254362 | 6367640 | | | | | | |
| dynamin 2 | Plasma Membrane | enzyme | 6268063 | 5091533 | 4999642.5 | | | | | | |
| dynamin 2 | Plasma Membrane | enzyme | 5963620.5 | 3839921.75 | 3918975.5 | | | | | | |
| dynamin 2 | Plasma Membrane | enzyme | 5225013.5 | 2673229.75 | 1226638.75 | | | | | | |
| dynamin 2 | Plasma Membrane | enzyme | 4835173 | 5833218.5 | 5008053.5 | | | | | | |
| dynamin 2 | Plasma Membrane | enzyme | 4489151 | 2244603.75 | 3132222 | | | | | | |
| dynamin 2 | Plasma Membrane | enzyme | 4357992.5 | 3443580 | 2809371.25 | | | | | | |
| dynamin 2 | Plasma Membrane | enzyme | 2972961.5 | 2020736 | 2650002.5 | | | | | | |
| dynamin 2 | Plasma Membrane | enzyme | 2802141.5 | 4301322.5 | 1802403.125 | | | | | | |
| dynamin 2 | Plasma Membrane | enzyme | 2055113.25 | 1410119.875 | 1423029.625 | | | | | | |
| dynamin 2 | Plasma Membrane | enzyme | 1501806.25 | 1501476.25 | 1687164.25 | | | | | | |
| dynamin 2 | Plasma Membrane | enzyme | 1321694.875 | 358133.406 | 939570.5 | P39054 | DNM2 | 11171640 | 9466933 | 1139685 | 69 |
| dynamin 3 | Cytoplasm | enzyme | 8162294 | 7245541.5 | 6622566 | | | | | | |
| dynamin 3 | Cytoplasm | enzyme | 7769380.5 | 5588720.5 | 7722365.5 | | | | | | |
| dynamin 3 | Cytoplasm | enzyme | 2988069.25 | 2755725.25 | 2851036.75 | Q8BZ98 | DNM3 | 6306581 | 2286073 | 762024 | 9 |
| aspartyl aminopeptidase | Cytoplasm | peptidase | 5.05E+07 | 2.29E+07 | 3.25E+07 | | | | | | |
| aspartyl aminopeptidase | Cytoplasm | peptidase | 1.55E+07 | 5094374 | 7757537.5 | | | | | | |
| aspartyl aminopeptidase | Cytoplasm | peptidase | 1.21E+07 | 4504517 | 6794172 | | | | | | |
| aspartyl aminopeptidase | Cytoplasm | peptidase | 1.19E+07 | 4949782.5 | 7531808.5 | | | | | | |
| aspartyl aminopeptidase | Cytoplasm | peptidase | 1.04E+07 | 5332577 | 5818387.5 | | | | | | |
| aspartyl aminopeptidase | Cytoplasm | peptidase | 8207921.5 | 4003396.5 | 4811165.5 | | | | | | |
| aspartyl aminopeptidase | Cytoplasm | peptidase | 7160306 | 4985642.5 | 5877353 | | | | | | |
| aspartyl aminopeptidase | Cytoplasm | peptidase | 6155973.5 | 4041024.75 | 4546309.5 | | | | | | |
| aspartyl aminopeptidase | Cytoplasm | peptidase | 4427055 | 1877650.375 | 2391373.25 | Q9Z2W0 | DNPEP | 14040140 | 10458424 | 2012725 | 27 |
| D-tyrosyl-tRNA deacylase 1 | Cytoplasm | enzyme | 375495.094 | 100507.953 | 209203.469 | Q9DD18 | DTD1 | 375495 | 138495 | 79960 | 3 |
| dynein, cytoplasmic 1, heavy chain 1 | Cytoplasm | peptidase | 1504662.875 | 1297798.875 | 1381943.125 | Q9JHU4 | DYNC1H1 | 1504663 | 104030 | 60062 | 3 |
| enoyl-CoA delta isomerase 1 | Cytoplasm | enzyme | 5632760.5 | 7765094 | 8549437 | | | | | | |
| enoyl-CoA delta isomerase 1 | Cytoplasm | enzyme | 2025293.875 | 4654020.5 | 4765332 | P42125 | ECI1 | 3829027 | 2355875 | 961782 | 6 |
| early endosome antigen 1 | Cytoplasm | other | 1748332.625 | 978624.938 | 807121 | Q8BL66 | EEA1 | 1748333 | 501289 | 289419 | 3 |
| eukaryotic translation elongation factor 1 alpha 1 | Cytoplasm | translation regulator | 1.86E+08 | 1.51E+08 | 1.50E+08 | | | | | | |
| eukaryotic translation elongation factor 1 alpha 1 | Cytoplasm | translation regulator | 7.39E+07 | 6.33E+07 | 8.76E+07 | | | | | | |
| eukaryotic translation elongation factor 1 alpha 1 | Cytoplasm | translation regulator | 5.03E+07 | 7.50E+07 | 6.68E+07 | | | | | | |
| eukaryotic translation elongation factor 1 alpha 1 | Cytoplasm | translation regulator | 4.90E+07 | 2.91E+07 | 4.32E+07 | | | | | | |
| eukaryotic translation elongation factor 1 alpha 1 | Cytoplasm | translation regulator | 4.26E+07 | 2.84E+07 | 3.60E+07 | | | | | | |
| eukaryotic translation elongation factor 1 alpha 1 | Cytoplasm | translation regulator | 4.17E+07 | 3.32E+07 | 3.47E+07 | | | | | | |
| eukaryotic translation elongation factor 1 alpha 1 | Cytoplasm | translation regulator | 4.03E+07 | 5.89E+07 | 5.85E+07 | | | | | | |
| eukaryotic translation elongation factor 1 alpha 1 | Cytoplasm | translation regulator | 3.92E+07 | 7.00E+07 | 5.81E+07 | | | | | | |
| eukaryotic translation elongation factor 1 alpha 1 | Cytoplasm | translation regulator | 3.82E+07 | 3.90E+07 | 4.64E+07 | | | | | | |
| eukaryotic translation elongation factor 1 alpha 1 | Cytoplasm | translation regulator | 3.77E+07 | 1.89E+07 | 2.97E+07 | | | | | | |
| eukaryotic translation elongation factor 1 alpha 1 | Cytoplasm | translation regulator | 3.69E+07 | 3.46E+07 | 3.66E+07 | | | | | | |
| eukaryotic translation elongation factor 1 alpha 1 | Cytoplasm | translation regulator | 3.39E+07 | 2.99E+07 | 3.09E+07 | | | | | | |
| eukaryotic translation elongation factor 1 alpha 1 | Cytoplasm | translation regulator | 3.33E+07 | 1.57E+07 | 2.40E+07 | | | | | | |
| eukaryotic translation elongation factor 1 alpha 1 | Cytoplasm | translation regulator | 2.75E+07 | 2.56E+07 | 2.78E+07 | | | | | | |
| eukaryotic translation elongation factor 1 alpha 1 | Cytoplasm | translation regulator | 2.40E+07 | 1.11E+07 | 2.10E+07 | | | | | | |
| eukaryotic translation elongation factor 1 alpha 1 | Cytoplasm | translation regulator | 2.18E+07 | 3.96E+07 | 3.03E+07 | | | | | | |

| | | | | | | | | | | | |
|---|---|---|---|---|---|---|---|---|---|---|---|
| eukaryotic translation elongation factor 1 alpha 1 | Cytoplasm | translation regulator | 6947098.5 | 6231874 | 6754409 | | | | | | |
| eukaryotic translation elongation factor 1 alpha 1 | Cytoplasm | translation regulator | 4554058 | 1485103.375 | 3122957.25 | | | | | | |
| eukaryotic translation elongation factor 1 alpha 1 | Cytoplasm | translation regulator | 3352687.75 | 1363364 | 2551694.75 | | | | | | |
| eukaryotic translation elongation factor 1 alpha 1 | Cytoplasm | translation regulator | 2862713 | 1540295.875 | 1375398.125 | | | | | | |
| eukaryotic translation elongation factor 1 alpha 1 | Cytoplasm | translation regulator | 2739208.5 | 1751067.75 | 2478547.5 | | | | | | |
| eukaryotic translation elongation factor 1 alpha 1 | Cytoplasm | translation regulator | 1220575.125 | 5238005.5 | 2888141.75 | | | | | | |
| eukaryotic translation elongation factor 1 alpha 1 | Cytoplasm | translation regulator | 227783.469 | 74906.672 | 260658.016 | P10126 | EEF1A1 | 34722353 | 35351625 | 4255836 | 69 |
| eukaryotic translation elongation factor 1 beta 2 | Cytoplasm | translation regulator | 830583.938 | 607328.5 | 815714.438 | O70251 | EEF1B2 | 830584 | 124826 | 72068 | 3 |
| eukaryotic translation elongation factor 1 delta (guanine nucleotide exchange protein) | Cytoplasm | translation regulator | 2129895.75 | 918248.125 | 2429997.5 | P57776 | EEF1D | 2129896 | 800368 | 462093 | 3 |
| eukaryotic translation elongation factor 1 gamma | Cytoplasm | translation regulator | 6348315.5 | 1860137.875 | 6132657.5 | | | | | | |
| eukaryotic translation elongation factor 1 gamma | Cytoplasm | translation regulator | 1944862.875 | 247948.875 | 1972707.875 | | | | | | |
| eukaryotic translation elongation factor 1 gamma | Cytoplasm | translation regulator | 1925674.625 | 676564.688 | 2404957.75 | Q9D8N0 | EEF1G | 3406284 | 2168603 | 722868 | 9 |
| eukaryotic translation elongation factor 2 | Cytoplasm | translation regulator | 1522938.75 | 587497.25 | 930760.375 | | | | | | |
| eukaryotic translation elongation factor 2 | Cytoplasm | translation regulator | 1511356 | 1363302.75 | 1077450.875 | | | | | | |
| eukaryotic translation elongation factor 2 | Cytoplasm | translation regulator | 1467149 | 1266789.375 | 1887378.5 | | | | | | |
| eukaryotic translation elongation factor 2 | Cytoplasm | translation regulator | 1007146.375 | 411961.344 | 899475.75 | | | | | | |
| eukaryotic translation elongation factor 2 | Cytoplasm | translation regulator | 111793.773 | 299710.156 | 382787.5 | P58252 | EEF2 | 1124077 | 530229 | 136905 | 15 |
| EFR3 homolog A | Plasma Membrane | other | 4328964 | 4079321.5 | 3002324.75 | | | | | | |
| EFR3 homolog A | Plasma Membrane | other | 4010919.25 | 3318228.5 | 2371056.25 | | | | | | |
| EFR3 homolog A | Plasma Membrane | other | 3551954.25 | 4082026.25 | 3299458.5 | | | | | | |
| EFR3 homolog A | Plasma Membrane | other | 3297903 | 4405492.5 | 3367881.75 | | | | | | |
| EFR3 homolog A | Plasma Membrane | other | 3095273 | 3135620.5 | 2772526.5 | | | | | | |
| EFR3 homolog A | Plasma Membrane | other | 2885842.75 | 3287264 | 2871170.5 | | | | | | |
| EFR3 homolog A | Plasma Membrane | other | 2383885.5 | 2527497.5 | 2142186 | Q8BG67 | EFR3A | 3364963 | 649946 | 141830 | 21 |
| EH-domain containing 1 | Cytoplasm | other | 5266061 | 2384202 | 3319839.25 | | | | | | |
| EH-domain containing 1 | Cytoplasm | other | 4803197 | 4314458 | 7237616.5 | | | | | | |
| EH-domain containing 1 | Cytoplasm | other | 2875660 | 1444653.625 | 1831366.625 | | | | | | |
| EH-domain containing 1 | Cytoplasm | other | 585678.5 | 341900.906 | 768363.188 | Q9WVK4 | EHD1 | 3382649 | 2135653 | 616510 | 12 |
| EH-domain containing 4 | Plasma Membrane | enzyme | 1523450.375 | 770473.625 | 1708351.25 | | | | | | |
| EH-domain containing 4 | Plasma Membrane | enzyme | 859094.25 | 1174326.25 | 1702884.375 | Q9EQP2 | EHD4 | 1191272 | 416883 | 170192 | 6 |
| eukaryotic translation initiation factor 2, subunit 1 alpha, 35kDa | Cytoplasm | translation regulator | 7042145.5 | 2887325.5 | 3539297 | | | | | | |
| eukaryotic translation initiation factor 2, subunit 1 alpha, 35kDa | Cytoplasm | translation regulator | 4267601 | 2297007.75 | 3089920.5 | | | | | | |
| eukaryotic translation initiation factor 2, subunit 1 alpha, 35kDa | Cytoplasm | translation regulator | 4231035 | 2495333 | 2610106.25 | | | | | | |
| eukaryotic translation initiation factor 2, subunit 1 alpha, 35kDa | Cytoplasm | translation regulator | 3101189.75 | 1541138.375 | 1756050.625 | | | | | | |
| eukaryotic translation initiation factor 2, subunit 1 alpha, 35kDa | Cytoplasm | translation regulator | 2976510 | 1157708.25 | 1342988.375 | | | | | | |
| eukaryotic translation initiation factor 2, subunit 1 alpha, 35kDa | Cytoplasm | translation regulator | 2653719.25 | 1210467.625 | 2891571.25 | Q6ZWX6 | EIF2S1 | 4045367 | 1400923 | 330201 | 18 |
| eukaryotic translation initiation factor 2, subunit 2 beta, 38kDa | Cytoplasm | translation regulator | 5870571.5 | 2231822.5 | 2804037.25 | | | | | | |
| eukaryotic translation initiation factor 2, subunit 2 beta, 38kDa | Cytoplasm | translation regulator | 5406391 | 1917830.625 | 3179264.75 | | | | | | |
| eukaryotic translation initiation factor 2, subunit 2 beta, 38kDa | Cytoplasm | translation regulator | 4601605 | 1806348 | 2213296 | Q99L45 | EIF2S2 | 5292856 | 1559270 | 519757 | 9 |
| eukaryotic translation initiation factor 2, subunit 3 gamma, 52kDa | Cytoplasm | translation regulator | 1.20E+07 | 6432276.5 | 6452804.5 | | | | | | |
| eukaryotic translation initiation factor 2, subunit 3 gamma, 52kDa | Cytoplasm | translation regulator | 1.16E+07 | 4711219 | 8611852 | | | | | | |
| eukaryotic translation initiation factor 2, subunit 3 gamma, 52kDa | Cytoplasm | translation regulator | 8335819 | 4759322 | 6538127.5 | | | | | | |
| eukaryotic translation initiation factor 2, subunit 3 gamma, 52kDa | Cytoplasm | translation regulator | 7909942.5 | 3481725.5 | 3986286.75 | | | | | | |
| eukaryotic translation initiation factor 2, subunit 3 gamma, 52kDa | Cytoplasm | translation regulator | 7283988.5 | 4002336.25 | 4295792.5 | | | | | | |
| eukaryotic translation initiation factor 2, subunit 3 gamma, 52kDa | Cytoplasm | translation regulator | 6814944.5 | 3991921.25 | 2927771.5 | | | | | | |
| eukaryotic translation initiation factor 2, subunit 3 gamma, 52kDa | Cytoplasm | translation regulator | 5558628.5 | 2754323 | 2837429.25 | | | | | | |
| eukaryotic translation initiation factor 2, subunit 3 gamma, 52kDa | Cytoplasm | translation regulator | 4679430 | 2738059 | 2072634.25 | | | | | | |
| eukaryotic translation initiation factor 2, subunit 3 gamma, 52kDa | Cytoplasm | translation regulator | 3012354.5 | 1473219.125 | 1775044 | | | | | | |
| eukaryotic translation initiation factor 2, subunit 3 gamma, 52kDa | Cytoplasm | translation regulator | 1771062.875 | 1041713.438 | 1029398.875 | | | | | | |
| eukaryotic translation initiation factor 2, subunit 3 gamma, 52kDa | Cytoplasm | translation regulator | 1233180.375 | 909151.438 | 906021.688 | Q9Z0N1 | EIF2S3 | 6376305 | 2963110 | 515811 | 33 |
| eukaryotic translation initiation factor 3, subunit J | Cytoplasm | translation regulator | 1.86E+07 | 6772856 | 9903928 | | | | | | |
| eukaryotic translation initiation factor 3, subunit J | Cytoplasm | translation regulator | 1.36E+07 | 6746670.5 | 6817907 | | | | | | |
| eukaryotic translation initiation factor 3, subunit J | Cytoplasm | translation regulator | 9273292 | 3646510.5 | 6022755.5 | | | | | | |
| eukaryotic translation initiation factor 3, subunit J | Cytoplasm | translation regulator | 90092.445 | 33162.621 | 39923.855 | | | | | | |
| eukaryotic translation initiation factor 3, subunit J | Cytoplasm | translation regulator | 10303.499 | 1547117.125 | 368.988 | Q3UGC7 | EIF3J | 8316738 | 5635240 | 1455013 | 15 |
| eukaryotic translation initiation factor 4B | Cytoplasm | translation regulator | 8102394.5 | 4780349.5 | 5493164 | | | | | | |
| eukaryotic translation initiation factor 4B | Cytoplasm | translation regulator | 8018172 | 3014601.5 | 3950562.75 | | | | | | |
| eukaryotic translation initiation factor 4B | Cytoplasm | translation regulator | 7451730 | 4078531.25 | 5251323 | | | | | | |
| eukaryotic translation initiation factor 4B | Cytoplasm | translation regulator | 4544103.5 | 1025679.562 | 1668880.375 | | | | | | |
| eukaryotic translation initiation factor 4B | Cytoplasm | translation regulator | 3900373.75 | 1886539 | 2578066.75 | | | | | | |
| eukaryotic translation initiation factor 4B | Cytoplasm | translation regulator | 2462909.25 | 1327090.625 | 1953197.25 | Q8BGD9 | EIF4B | 5746614 | 2236474 | 527142 | 18 |
| eukaryotic translation initiation factor 4 gamma, 1 | Cytoplasm | translation regulator | 2799959 | 1816621.875 | 1994684.375 | | | | | | |
| eukaryotic translation initiation factor 4 gamma, 1 | Cytoplasm | translation regulator | 1273712 | 857775.875 | 668071.875 | | | | | | |
| eukaryotic translation initiation factor 4 gamma, 1 | Cytoplasm | translation regulator | 540189.812 | 229809.109 | 230759.984 | Q6NZJ6 | EIF4G1 | 1537954 | 885748 | 295249 | 9 |
| eukaryotic translation initiation factor 5 | Cytoplasm | translation regulator | 3754414.75 | 604301.188 | 1983118.75 | P59325 | EIF5 | 3754415 | 1579126 | 911709 | 3 |
| eukaryotic translation initiation factor 5A | Cytoplasm | translation regulator | 2327407 | 1538573.25 | 2845712 | P63242 | EIF5A | 2327407 | 658219 | 380023 | 3 |
| elastin microfibril interfacer 1 | Extracellular Space | other | 64582.645 | 6164243.5 | 4583602.5 | | | | | | |
| elastin microfibril interfacer 1 | Extracellular Space | other | 47196.441 | 960585.25 | 576847.688 | Q99K41 | EMILIN1 | 55890 | 2632667 | 1074782 | 6 |
| endonuclease domain containing 1 | Extracellular Space | enzyme | 1.80E+07 | 1.62E+07 | 1.69E+07 | | | | | | |
| endonuclease domain containing 1 | Extracellular Space | enzyme | 1.47E+07 | 1.41E+07 | 1.64E+07 | | | | | | |
| endonuclease domain containing 1 | Extracellular Space | enzyme | 1.15E+07 | 1.36E+07 | 1.48E+07 | | | | | | |
| endonuclease domain containing 1 | Extracellular Space | enzyme | 7698652.5 | 6687411.5 | 6385920.5 | | | | | | |
| endonuclease domain containing 1 | Extracellular Space | enzyme | 7555671 | 8215215 | 8156218.5 | | | | | | |
| endonuclease domain containing 1 | Extracellular Space | enzyme | 6899946.5 | 7049261 | 6942718.5 | | | | | | |
| endonuclease domain containing 1 | Extracellular Space | enzyme | 5954756 | 1.05E+07 | 7432297 | | | | | | |
| endonuclease domain containing 1 | Extracellular Space | enzyme | 5421160.5 | 7002291 | 5934908 | Q8C522 | ENDOD1 | 9707523 | 4205853 | 858516 | 24 |
| ectonucleotide pyrophosphatase/phosphodiesterase 4 (putative) | Cytoplasm | enzyme | 187245.375 | 470609.375 | 423752 | Q8BTJ4 | ENPP4 | 187245 | 151891 | 87695 | 3 |
| MLV-related proviral Env polyprotein | other | | 2.67E+07 | 1.83E+07 | 2.73E+07 | | | | | | |
| MLV-related proviral Env polyprotein | other | | 1.88E+07 | 1.18E+07 | 1.60E+07 | | | | | | |
| MLV-related proviral Env polyprotein | other | | 1.54E+07 | 2.07E+07 | 1.73E+07 | | | | | | |
| MLV-related proviral Env polyprotein | other | | 1.24E+07 | 1.04E+07 | 9545867 | | | | | | |
| MLV-related proviral Env polyprotein | other | | 1.13E+07 | 7022660.5 | 8841342 | | | | | | |
| MLV-related proviral Env polyprotein | other | | 9772180 | 8830148 | 1.16E+07 | | | | | | |
| MLV-related proviral Env polyprotein | other | | 9142177 | 9671146 | 1.15E+07 | | | | | | |
| MLV-related proviral Env polyprotein | other | | 8211081.5 | 9217259 | 9446515 | | | | | | |
| MLV-related proviral Env polyprotein | other | | 8185884 | 1.23E+07 | 8568340 | | | | | | |
| MLV-related proviral Env polyprotein | other | | 7447593.5 | 8559610 | 8371371 | | | | | | |
| MLV-related proviral Env polyprotein | other | | 7279007.5 | 5630737 | 6869425 | | | | | | |
| MLV-related proviral Env polyprotein | other | | 5593826 | 6715266.5 | 5128686 | | | | | | |
| MLV-related proviral Env polyprotein | other | | 4707634.5 | 5482086 | 4170315 | | | | | | |
| MLV-related proviral Env polyprotein | other | | 3762425.25 | 3011502 | 3902637.75 | | | | | | |
| MLV-related proviral Env polyprotein | other | | 2542104.25 | 2834717.5 | 3101531.75 | | | | | | |
| MLV-related proviral Env polyprotein | other | | 2097367 | 1977198.625 | 2064186.75 | | | | | | |
| MLV-related proviral Env polyprotein | other | | 1764866.625 | 1395121.875 | 1903571.875 | | | | | | |
| MLV-related proviral Env polyprotein | other | | 1638065.375 | 1099165.375 | 1721550 | | | | | | |

| | | | | | | | | | | | |
|---|---|---|---|---|---|---|---|---|---|---|---|
| MLV-related proviral Env polyprotein | other | | 1577792.75 | 992152.312 | 1457104.25 | | | | | | |
| MLV-related proviral Env polyprotein | other | | 1509398.25 | 891906.562 | 1534635.25 | | | | | | |
| MLV-related proviral Env polyprotein | other | | 1291343 | 225512.812 | 1095358.625 | | | | | | |
| MLV-related proviral Env polyprotein | other | | 1094786.25 | 2235653 | 832193.188 | | | | | | |
| MLV-related proviral Env polyprotein | other | | 285261.75 | 748519.938 | 627933.25 | | | | | | |
| MLV-related proviral Env polyprotein | other | | 252013.344 | 323080.438 | 704096.062 | P10404 | ENV1 | 6786450 | 6201270 | 730827 | 72 |
| erythrocyte membrane protein band 4.1 | Plasma Membrane | other | 2.61E+09 | 1.23E+09 | 1.43E+09 | | | | | | |
| erythrocyte membrane protein band 4.1 | Plasma Membrane | other | 2.21E+09 | 1.85E+09 | 1.70E+09 | | | | | | |
| erythrocyte membrane protein band 4.1 | Plasma Membrane | other | 2.13E+09 | 1.71E+09 | 1.80E+09 | | | | | | |
| erythrocyte membrane protein band 4.1 | Plasma Membrane | other | 1.72E+09 | 1.32E+09 | 1.50E+09 | | | | | | |
| erythrocyte membrane protein band 4.1 | Plasma Membrane | other | 1.64E+09 | 1.75E+09 | 1.25E+09 | | | | | | |
| erythrocyte membrane protein band 4.1 | Plasma Membrane | other | 1.64E+09 | 1.26E+09 | 1.33E+09 | | | | | | |
| erythrocyte membrane protein band 4.1 | Plasma Membrane | other | 1.46E+09 | 1.37E+09 | 1.29E+09 | | | | | | |
| erythrocyte membrane protein band 4.1 | Plasma Membrane | other | 1.37E+09 | 1.20E+09 | 1.45E+09 | | | | | | |
| erythrocyte membrane protein band 4.1 | Plasma Membrane | other | 1.33E+09 | 1.20E+09 | 1.06E+09 | | | | | | |
| erythrocyte membrane protein band 4.1 | Plasma Membrane | other | 1.31E+09 | 9.14E+08 | 1.00E+09 | | | | | | |
| erythrocyte membrane protein band 4.1 | Plasma Membrane | other | 1.31E+09 | 1.02E+09 | 1.07E+09 | | | | | | |
| erythrocyte membrane protein band 4.1 | Plasma Membrane | other | 1.19E+09 | 9.16E+08 | 9.17E+08 | | | | | | |
| erythrocyte membrane protein band 4.1 | Plasma Membrane | other | 1.10E+09 | 9.35E+08 | 9.41E+08 | | | | | | |
| erythrocyte membrane protein band 4.1 | Plasma Membrane | other | 1.04E+09 | 9.30E+08 | 7.70E+08 | | | | | | |
| erythrocyte membrane protein band 4.1 | Plasma Membrane | other | 1.03E+09 | 1.20E+09 | 8.58E+08 | | | | | | |
| erythrocyte membrane protein band 4.1 | Plasma Membrane | other | 6.40E+08 | 5.06E+08 | 5.90E+08 | | | | | | |
| erythrocyte membrane protein band 4.1 | Plasma Membrane | other | 6.37E+08 | 4.61E+08 | 5.53E+08 | | | | | | |
| erythrocyte membrane protein band 4.1 | Plasma Membrane | other | 6.35E+08 | 4.57E+08 | 4.22E+08 | | | | | | |
| erythrocyte membrane protein band 4.1 | Plasma Membrane | other | 5.84E+08 | 2.92E+08 | 3.54E+08 | | | | | | |
| erythrocyte membrane protein band 4.1 | Plasma Membrane | other | 4.46E+08 | 3.98E+08 | 3.46E+08 | | | | | | |
| erythrocyte membrane protein band 4.1 | Plasma Membrane | other | 4.27E+08 | 1.60E+08 | 2.39E+08 | | | | | | |
| erythrocyte membrane protein band 4.1 | Plasma Membrane | other | 3.27E+08 | 2.09E+08 | 1.88E+08 | | | | | | |
| erythrocyte membrane protein band 4.1 | Plasma Membrane | other | 2.97E+08 | 3.75E+08 | 2.57E+08 | | | | | | |
| erythrocyte membrane protein band 4.1 | Plasma Membrane | other | 2.92E+08 | 2.64E+08 | 2.38E+08 | | | | | | |
| erythrocyte membrane protein band 4.1 | Plasma Membrane | other | 2.85E+08 | 3.28E+08 | 3.12E+08 | | | | | | |
| erythrocyte membrane protein band 4.1 | Plasma Membrane | other | 1.76E+08 | 1.66E+08 | 1.41E+08 | | | | | | |
| erythrocyte membrane protein band 4.1 | Plasma Membrane | other | 1.34E+08 | 1.24E+08 | 1.59E+08 | | | | | | |
| erythrocyte membrane protein band 4.1 | Plasma Membrane | other | 1.08E+08 | 8.70E+07 | 9.29E+07 | | | | | | |
| erythrocyte membrane protein band 4.1 | Plasma Membrane | other | 9.82E+07 | 9.68E+07 | 1.10E+08 | | | | | | |
| erythrocyte membrane protein band 4.1 | Plasma Membrane | other | 9.51E+07 | 4.04E+07 | 5.26E+07 | | | | | | |
| erythrocyte membrane protein band 4.1 | Plasma Membrane | other | 9.43E+07 | 4.24E+07 | 4.38E+07 | | | | | | |
| erythrocyte membrane protein band 4.1 | Plasma Membrane | other | 8.78E+07 | 3.67E+07 | 4.69E+07 | | | | | | |
| erythrocyte membrane protein band 4.1 | Plasma Membrane | other | 7.04E+07 | 5.71E+07 | 6.86E+07 | | | | | | |
| erythrocyte membrane protein band 4.1 | Plasma Membrane | other | 5.95E+07 | 4.10E+07 | 4.52E+07 | | | | | | |
| erythrocyte membrane protein band 4.1 | Plasma Membrane | other | 5.56E+07 | 5.88E+07 | 4.51E+07 | | | | | | |
| erythrocyte membrane protein band 4.1 | Plasma Membrane | other | 5.54E+07 | 2.15E+07 | 2.66E+07 | | | | | | |
| erythrocyte membrane protein band 4.1 | Plasma Membrane | other | 4.85E+07 | 3.99E+07 | 3.94E+07 | | | | | | |
| erythrocyte membrane protein band 4.1 | Plasma Membrane | other | 4.74E+07 | 2.86E+07 | 2.80E+07 | | | | | | |
| erythrocyte membrane protein band 4.1 | Plasma Membrane | other | 4.39E+07 | 5.23E+07 | 5.12E+07 | | | | | | |
| erythrocyte membrane protein band 4.1 | Plasma Membrane | other | 4.25E+07 | 1.95E+07 | 2.88E+07 | | | | | | |
| erythrocyte membrane protein band 4.1 | Plasma Membrane | other | 3.54E+07 | 1.39E+07 | 1.80E+07 | | | | | | |
| erythrocyte membrane protein band 4.1 | Plasma Membrane | other | 2.98E+07 | 1.20E+07 | 1.43E+07 | | | | | | |
| erythrocyte membrane protein band 4.1 | Plasma Membrane | other | 2.53E+07 | 2.23E+07 | 1.84E+07 | | | | | | |
| erythrocyte membrane protein band 4.1 | Plasma Membrane | other | 2.38E+07 | 1.79E+07 | 1.67E+07 | | | | | | |
| erythrocyte membrane protein band 4.1 | Plasma Membrane | other | 2.25E+07 | 2.03E+07 | 1.66E+07 | | | | | | |
| erythrocyte membrane protein band 4.1 | Plasma Membrane | other | 1.94E+07 | 8871180 | 1.03E+07 | | | | | | |
| erythrocyte membrane protein band 4.1 | Plasma Membrane | other | 1.64E+07 | 8752514 | 8227164.5 | | | | | | |
| erythrocyte membrane protein band 4.1 | Plasma Membrane | other | 1.48E+07 | 1.85E+07 | 5.10E+07 | | | | | | |
| erythrocyte membrane protein band 4.1 | Plasma Membrane | other | 1.34E+07 | 1.15E+07 | 7879250 | | | | | | |
| erythrocyte membrane protein band 4.1 | Plasma Membrane | other | 1.24E+07 | 7.70E+07 | 8.55E+07 | | | | | | |
| erythrocyte membrane protein band 4.1 | Plasma Membrane | other | 1.21E+07 | 7.51E+07 | 2.65E+07 | | | | | | |
| erythrocyte membrane protein band 4.1 | Plasma Membrane | other | 9602065 | 8930680 | 9763094 | | | | | | |
| erythrocyte membrane protein band 4.1 | Plasma Membrane | other | 8458447 | 3233043 | 5025093.5 | | | | | | |
| erythrocyte membrane protein band 4.1 | Plasma Membrane | other | 6805896 | 6471980.5 | 4203316.5 | | | | | | |
| erythrocyte membrane protein band 4.1 | Plasma Membrane | other | 4471718.5 | 2196947.5 | 2553063.25 | | | | | | |
| erythrocyte membrane protein band 4.1 | Plasma Membrane | other | 3483710.25 | 2741377.5 | 3319025 | | | | | | |
| erythrocyte membrane protein band 4.1 | Plasma Membrane | other | 3357036.5 | 4204508 | 3296679 | | | | | | |
| erythrocyte membrane protein band 4.1 | Plasma Membrane | other | 3033120 | 1.30E+07 | 4398865.5 | | | | | | |
| erythrocyte membrane protein band 4.1 | Plasma Membrane | other | 1694013.25 | 1257090.125 | 1075004.125 | | | | | | |
| erythrocyte membrane protein band 4.1 | Plasma Membrane | other | 236730.203 | 118700.266 | 228880.469 | | | | | | |
| erythrocyte membrane protein band 4.1 | Plasma Membrane | other | 96426.969 | 1.52E+07 | 1.70E+07 | | | | | | |
| erythrocyte membrane protein band 4.1 | Plasma Membrane | other | 4.70E-06 | 197335.922 | 50336.359 | P48193 | EPB41 | 470048696 | 577329892 | 42331882 | 186 |
| erythrocyte membrane protein band 4.2 | Plasma Membrane | transporter | 2.38E+09 | 1.81E+09 | 1.81E+09 | | | | | | |
| erythrocyte membrane protein band 4.2 | Plasma Membrane | transporter | 2.15E+09 | 1.79E+09 | 1.66E+09 | | | | | | |
| erythrocyte membrane protein band 4.2 | Plasma Membrane | transporter | 2.14E+09 | 1.77E+09 | 1.58E+09 | | | | | | |
| erythrocyte membrane protein band 4.2 | Plasma Membrane | transporter | 1.98E+09 | 1.61E+09 | 1.63E+09 | | | | | | |
| erythrocyte membrane protein band 4.2 | Plasma Membrane | transporter | 1.97E+09 | 1.10E+09 | 1.34E+09 | | | | | | |
| erythrocyte membrane protein band 4.2 | Plasma Membrane | transporter | 1.71E+09 | 1.53E+09 | 1.37E+09 | | | | | | |
| erythrocyte membrane protein band 4.2 | Plasma Membrane | transporter | 1.66E+09 | 1.42E+09 | 1.53E+09 | | | | | | |
| erythrocyte membrane protein band 4.2 | Plasma Membrane | transporter | 1.20E+09 | 9.18E+08 | 8.47E+08 | | | | | | |
| erythrocyte membrane protein band 4.2 | Plasma Membrane | transporter | 1.19E+09 | 9.65E+08 | 9.13E+08 | | | | | | |
| erythrocyte membrane protein band 4.2 | Plasma Membrane | transporter | 1.14E+09 | 1.07E+09 | 8.29E+08 | | | | | | |
| erythrocyte membrane protein band 4.2 | Plasma Membrane | transporter | 1.09E+09 | 4.61E+08 | 7.54E+08 | | | | | | |
| erythrocyte membrane protein band 4.2 | Plasma Membrane | transporter | 9.56E+08 | 7.12E+08 | 7.13E+08 | | | | | | |
| erythrocyte membrane protein band 4.2 | Plasma Membrane | transporter | 9.18E+08 | 7.92E+08 | 6.83E+08 | | | | | | |
| erythrocyte membrane protein band 4.2 | Plasma Membrane | transporter | 9.18E+08 | 1.29E+09 | 8.79E+08 | | | | | | |
| erythrocyte membrane protein band 4.2 | Plasma Membrane | transporter | 7.65E+08 | 6.12E+08 | 5.83E+08 | | | | | | |
| erythrocyte membrane protein band 4.2 | Plasma Membrane | transporter | 6.69E+08 | 5.87E+08 | 5.09E+08 | | | | | | |
| erythrocyte membrane protein band 4.2 | Plasma Membrane | transporter | 6.20E+08 | 4.24E+08 | 5.24E+08 | | | | | | |
| erythrocyte membrane protein band 4.2 | Plasma Membrane | transporter | 6.14E+08 | 5.83E+08 | 5.01E+08 | | | | | | |
| erythrocyte membrane protein band 4.2 | Plasma Membrane | transporter | 6.09E+08 | 4.61E+08 | 4.51E+08 | | | | | | |
| erythrocyte membrane protein band 4.2 | Plasma Membrane | transporter | 4.94E+08 | 3.87E+08 | 3.82E+08 | | | | | | |
| erythrocyte membrane protein band 4.2 | Plasma Membrane | transporter | 4.84E+08 | 2.89E+08 | 2.63E+08 | | | | | | |
| erythrocyte membrane protein band 4.2 | Plasma Membrane | transporter | 4.64E+08 | 3.61E+08 | 2.76E+08 | | | | | | |
| erythrocyte membrane protein band 4.2 | Plasma Membrane | transporter | 4.19E+08 | 4.04E+08 | 2.94E+08 | | | | | | |
| erythrocyte membrane protein band 4.2 | Plasma Membrane | transporter | 4.18E+08 | 2.86E+08 | 3.71E+08 | | | | | | |
| erythrocyte membrane protein band 4.2 | Plasma Membrane | transporter | 4.13E+08 | 2.84E+08 | 2.61E+08 | | | | | | |
| erythrocyte membrane protein band 4.2 | Plasma Membrane | transporter | 2.18E+08 | 1.02E+08 | 1.29E+08 | | | | | | |
| erythrocyte membrane protein band 4.2 | Plasma Membrane | transporter | 2.02E+08 | 2.05E+08 | 1.46E+08 | | | | | | |

| | | | | | | | | | | | |
|---|---|---|---|---|---|---|---|---|---|---|---|
| erythrocyte membrane protein band 4.2 | Plasma Membrane | transporter | 1.82E+08 | 1.02E+08 | 1.24E+08 | | | | | | |
| erythrocyte membrane protein band 4.2 | Plasma Membrane | transporter | 1.81E+08 | 1.53E+08 | 1.71E+08 | | | | | | |
| erythrocyte membrane protein band 4.2 | Plasma Membrane | transporter | 1.65E+08 | 6.62E+07 | 9.54E+07 | | | | | | |
| erythrocyte membrane protein band 4.2 | Plasma Membrane | transporter | 1.64E+08 | 1.09E+08 | 1.27E+08 | | | | | | |
| erythrocyte membrane protein band 4.2 | Plasma Membrane | transporter | 1.40E+08 | 9.33E+07 | 1.04E+08 | | | | | | |
| erythrocyte membrane protein band 4.2 | Plasma Membrane | transporter | 1.33E+08 | 8.72E+07 | 1.08E+08 | | | | | | |
| erythrocyte membrane protein band 4.2 | Plasma Membrane | transporter | 1.08E+08 | 1.66E+08 | 1.15E+08 | | | | | | |
| erythrocyte membrane protein band 4.2 | Plasma Membrane | transporter | 8.50E+07 | 9.84E+07 | 6.29E+07 | | | | | | |
| erythrocyte membrane protein band 4.2 | Plasma Membrane | transporter | 6.25E+07 | 6.67E+07 | 5.36E+07 | | | | | | |
| erythrocyte membrane protein band 4.2 | Plasma Membrane | transporter | 5.38E+07 | 4.38E+07 | 7.48E+07 | | | | | | |
| erythrocyte membrane protein band 4.2 | Plasma Membrane | transporter | 3.94E+07 | 1.47E+07 | 2.27E+07 | | | | | | |
| erythrocyte membrane protein band 4.2 | Plasma Membrane | transporter | 3.85E+07 | 3.57E+07 | 2.52E+07 | | | | | | |
| erythrocyte membrane protein band 4.2 | Plasma Membrane | transporter | 3.44E+07 | 2.83E+07 | 2.09E+07 | | | | | | |
| erythrocyte membrane protein band 4.2 | Plasma Membrane | transporter | 3.10E+07 | 1.51E+07 | 2.75E+07 | | | | | | |
| erythrocyte membrane protein band 4.2 | Plasma Membrane | transporter | 3.09E+07 | 9629484 | 3.32E+07 | | | | | | |
| erythrocyte membrane protein band 4.2 | Plasma Membrane | transporter | 2.88E+07 | 2.86E+07 | 3.05E+07 | | | | | | |
| erythrocyte membrane protein band 4.2 | Plasma Membrane | transporter | 2.22E+07 | 6635133 | 1.74E+07 | | | | | | |
| erythrocyte membrane protein band 4.2 | Plasma Membrane | transporter | 1.36E+07 | 1.43E+07 | 1.64E+07 | | | | | | |
| erythrocyte membrane protein band 4.2 | Plasma Membrane | transporter | 1.28E+07 | 1.97E+07 | 9394511 | | | | | | |
| erythrocyte membrane protein band 4.2 | Plasma Membrane | transporter | 1.09E+07 | 6.54E+08 | 1.99E+07 | | | | | | |
| erythrocyte membrane protein band 4.2 | Plasma Membrane | transporter | 9295253 | 8873563 | 1.03E+07 | | | | | | |
| erythrocyte membrane protein band 4.2 | Plasma Membrane | transporter | 8892546 | 5343954.5 | 4823649 | | | | | | |
| erythrocyte membrane protein band 4.2 | Plasma Membrane | transporter | 6967503.5 | 4285289 | 5588587 | | | | | | |
| erythrocyte membrane protein band 4.2 | Plasma Membrane | transporter | 5707695 | 1.93E+07 | 8248155.5 | | | | | | |
| erythrocyte membrane protein band 4.2 | Plasma Membrane | transporter | 4893228 | 2339149.25 | 3456585 | | | | | | |
| erythrocyte membrane protein band 4.2 | Plasma Membrane | transporter | 4687837.5 | 1266567.75 | 2998741.25 | | | | | | |
| erythrocyte membrane protein band 4.2 | Plasma Membrane | transporter | 4026633.5 | 1.54E+07 | 6545034 | | | | | | |
| erythrocyte membrane protein band 4.2 | Plasma Membrane | transporter | 3948378.25 | 5222601 | 4651738 | | | | | | |
| erythrocyte membrane protein band 4.2 | Plasma Membrane | transporter | 3797293.75 | 2125625 | 3423531.75 | | | | | | |
| erythrocyte membrane protein band 4.2 | Plasma Membrane | transporter | 2282415.5 | 2894232.25 | 1462025.25 | | | | | | |
| erythrocyte membrane protein band 4.2 | Plasma Membrane | transporter | 1700230.5 | 7351701.5 | 6104721.5 | | | | | | |
| erythrocyte membrane protein band 4.2 | Plasma Membrane | transporter | 1214680.25 | 499081.5 | 982453.312 | | | | | | |
| erythrocyte membrane protein band 4.2 | Plasma Membrane | transporter | 1135502.125 | 799457.812 | 944602.062 | | | | | | |
| erythrocyte membrane protein band 4.2 | Plasma Membrane | transporter | 448207.875 | 356691.844 | 900215.562 | | | | | | |
| erythrocyte membrane protein band 4.2 | Plasma Membrane | transporter | 222154.734 | 3.77E+07 | 507056.906 | | | | | | |
| erythrocyte membrane protein band 4.2 | Plasma Membrane | transporter | 198122.953 | 696555.875 | 447288.969 | | | | | | |
| erythrocyte membrane protein band 4.2 | Plasma Membrane | transporter | 125454.977 | 275413.438 | 1380559.25 | P49222 | EPB42 | 459205518 | 560909686 | 40480170 | 192 |
| epsin 1 | Plasma Membrane | other | 5757230 | 3324299.5 | 3515529.5 | | | | | | |
| epsin 1 | Plasma Membrane | other | 5564636 | 2048535.375 | 2795301 | | | | | | |
| epsin 1 | Plasma Membrane | other | 2241522.25 | 1045983.312 | 1572668.125 | | | | | | |
| epsin 1 | Plasma Membrane | other | 2086031.625 | 1371760.25 | 1346522.75 | Q80VP1 | EPN1 | 3912355 | 1573213 | 454147 | 12 |
| glutamyl-prolyl-tRNA synthetase | Cytoplasm | enzyme | 1522838.25 | 678424.625 | 3693722.75 | | | | | | |
| glutamyl-prolyl-tRNA synthetase | Cytoplasm | enzyme | 1124624.5 | 167249.562 | 4352942.5 | Q8CGC7 | EPRS | 1323731 | 1701131 | 694484 | 6 |
| epidermal growth factor receptor pathway substrate 15 | Cytoplasm | other | 7307512.5 | 3541841.75 | 4167979.5 | | | | | | |
| epidermal growth factor receptor pathway substrate 15 | Cytoplasm | other | 5956426.5 | 1724972.875 | 3030743.75 | | | | | | |
| epidermal growth factor receptor pathway substrate 15 | Cytoplasm | other | 4427931.5 | 2036523.5 | 2935744.25 | | | | | | |
| epidermal growth factor receptor pathway substrate 15 | Cytoplasm | other | 4373993 | 2156212.5 | 3468566.25 | | | | | | |
| epidermal growth factor receptor pathway substrate 15 | Cytoplasm | other | 3539146.75 | 1987867.875 | 1459508.875 | P42567 | EPS15 | 5121002 | 1618722 | 417952 | 15 |
| epidermal growth factor receptor pathway substrate 15-like 1 | Plasma Membrane | other | 385390.375 | 169461.031 | 203289.703 | Q60902 | EPS15L1 | 385390 | 116140 | 67053 | 3 |
| endoplasmic reticulum aminopeptidase 1 | Extracellular Space | peptidase | 4405343.5 | 3142988.25 | 1779758.5 | Q9EQH2 | ERAP1 | 4405344 | 1313115 | 758128 | 3 |
| erbb2 interacting protein | Plasma Membrane | other | 2409800.5 | 6082227 | 3922277.75 | | | | | | |
| erbb2 interacting protein | Plasma Membrane | other | 2320565 | 4129438 | 3255482 | | | | | | |
| erbb2 interacting protein | Plasma Membrane | other | 1783520.5 | 2936756.5 | 3557463 | | | | | | |
| erbb2 interacting protein | Plasma Membrane | other | 1507673.125 | 4394432.5 | 3648404.5 | | | | | | |
| erbb2 interacting protein | Plasma Membrane | other | 1199642.125 | 4063694.5 | 3385012 | | | | | | |
| erbb2 interacting protein | Plasma Membrane | other | 1028100.75 | 2239832.75 | 1734570.875 | | | | | | |
| erbb2 interacting protein | Plasma Membrane | other | 943108.312 | 3021970.75 | 2360900.5 | | | | | | |
| erbb2 interacting protein | Plasma Membrane | other | 655669.312 | 2532760.5 | 1806925.875 | Q80TH2 | ERBB2IP | 1481010 | 1304932 | 266368 | 24 |
| ER lipid raft associated 2 | Plasma Membrane | other | 2646321 | 2336153.75 | 2044631.75 | Q8BFZ9 | ERLIN2 | 2646321 | 300893 | 173721 | 3 |
| erythroblast membrane-associated protein (Scianna blood group) | Cytoplasm | other | 9.91E+07 | 9.31E+07 | 1.15E+08 | | | | | | |
| erythroblast membrane-associated protein (Scianna blood group) | Cytoplasm | other | 6.44E+07 | 5.58E+07 | 6.16E+07 | | | | | | |
| erythroblast membrane-associated protein (Scianna blood group) | Cytoplasm | other | 5.38E+07 | 4.78E+07 | 5.82E+07 | | | | | | |
| erythroblast membrane-associated protein (Scianna blood group) | Cytoplasm | other | 5.13E+07 | 4.49E+07 | 5.04E+07 | | | | | | |
| erythroblast membrane-associated protein (Scianna blood group) | Cytoplasm | other | 4.97E+07 | 3.83E+07 | 4.25E+07 | | | | | | |
| erythroblast membrane-associated protein (Scianna blood group) | Cytoplasm | other | 4.84E+07 | 3.83E+07 | 4.41E+07 | | | | | | |
| erythroblast membrane-associated protein (Scianna blood group) | Cytoplasm | other | 3.71E+07 | 2.61E+07 | 3.25E+07 | | | | | | |
| erythroblast membrane-associated protein (Scianna blood group) | Cytoplasm | other | 2.51E+07 | 1.95E+07 | 2.66E+07 | | | | | | |
| erythroblast membrane-associated protein (Scianna blood group) | Cytoplasm | other | 1.85E+07 | 1.72E+07 | 2.06E+07 | | | | | | |
| erythroblast membrane-associated protein (Scianna blood group) | Cytoplasm | other | 1.75E+07 | 1.66E+07 | 1.49E+07 | | | | | | |
| erythroblast membrane-associated protein (Scianna blood group) | Cytoplasm | other | 1.59E+07 | 1.21E+07 | 1.27E+07 | | | | | | |
| erythroblast membrane-associated protein (Scianna blood group) | Cytoplasm | other | 9637065 | 7655873.5 | 7497535 | | | | | | |
| erythroblast membrane-associated protein (Scianna blood group) | Cytoplasm | other | 7412507.5 | 6813556.5 | 6634953 | | | | | | |
| erythroblast membrane-associated protein (Scianna blood group) | Cytoplasm | other | 6850000 | 3967077.75 | 5268511 | | | | | | |
| erythroblast membrane-associated protein (Scianna blood group) | Cytoplasm | other | 6022557 | 6337007.5 | 6628299.5 | | | | | | |
| erythroblast membrane-associated protein (Scianna blood group) | Cytoplasm | other | 4591250.5 | 2757355.25 | 3910393 | | | | | | |
| erythroblast membrane-associated protein (Scianna blood group) | Cytoplasm | other | 3218176.75 | 2364254 | 3150086.25 | | | | | | |
| erythroblast membrane-associated protein (Scianna blood group) | Cytoplasm | other | 2845877.75 | 1977824.125 | 3231677.75 | | | | | | |
| erythroblast membrane-associated protein (Scianna blood group) | Cytoplasm | other | 1508231 | 3111683.25 | 5694636.5 | | | | | | |
| erythroblast membrane-associated protein (Scianna blood group) | Cytoplasm | other | 1116728 | 3270512.5 | 3586378.25 | Q9JLN5 | ERMAP | 26193120 | 26263218 | 3390567 | 60 |
| endoplasmic reticulum oxidoreductase alpha | Cytoplasm | enzyme | 2068036.125 | 1359000.125 | 1653151.25 | Q8R180 | ERO1A | 2068036 | 356227 | 205668 | 3 |
| endoplasmic reticulum protein 29 | Cytoplasm | transporter | 9095522 | 1.01E+07 | 9997497 | | | | | | |
| endoplasmic reticulum protein 29 | Cytoplasm | transporter | 7459334 | 5817410.5 | 6795644 | | | | | | |
| endoplasmic reticulum protein 29 | Cytoplasm | transporter | 7051248.5 | 3859782.25 | 5176887.5 | | | | | | |
| endoplasmic reticulum protein 29 | Cytoplasm | transporter | 3794140 | 2716446.75 | 3499758 | P57759 | ERP29 | 6850061 | 2560321 | 739101 | 12 |
| endoplasmic reticulum protein 44 | Cytoplasm | enzyme | 1.33E+07 | 1.64E+07 | 1.56E+07 | | | | | | |
| endoplasmic reticulum protein 44 | Cytoplasm | enzyme | 1.07E+07 | 1.39E+07 | 1.61E+07 | | | | | | |
| endoplasmic reticulum protein 44 | Cytoplasm | enzyme | 1.07E+07 | 8931396 | 8689437 | | | | | | |
| endoplasmic reticulum protein 44 | Cytoplasm | enzyme | 8564572 | 9466890 | 7765616 | | | | | | |
| endoplasmic reticulum protein 44 | Cytoplasm | enzyme | 4091506.25 | 1848429.75 | 3340347.25 | | | | | | |
| endoplasmic reticulum protein 44 | Cytoplasm | enzyme | 3230556.5 | 3228905.5 | 2676571.5 | Q9D1Q6 | ERP44 | 8437772 | 4925179 | 1160876 | 18 |
| extended synaptotagmin-like protein 1 | Cytoplasm | other | 2.75E+07 | 1.87E+07 | 2.10E+07 | | | | | | |
| extended synaptotagmin-like protein 1 | Cytoplasm | other | 2.67E+07 | 2.61E+07 | 2.82E+07 | | | | | | |
| extended synaptotagmin-like protein 1 | Cytoplasm | other | 2.44E+07 | 2.33E+07 | 2.39E+07 | | | | | | |
| extended synaptotagmin-like protein 1 | Cytoplasm | other | 1.85E+07 | 1.64E+07 | 1.57E+07 | | | | | | |
| extended synaptotagmin-like protein 1 | Cytoplasm | other | 1.75E+07 | 1.26E+07 | 1.21E+07 | | | | | | |

| | | | | | | | | | | | |
|---|---|---|---|---|---|---|---|---|---|---|---|
| extended synaptotagmin-like protein 1 | Cytoplasm | other | 1.47E+07 | 1.28E+07 | 1.27E+07 | | | | | | |
| extended synaptotagmin-like protein 1 | Cytoplasm | other | 1.45E+07 | 9363267 | 1.22E+07 | | | | | | |
| extended synaptotagmin-like protein 1 | Cytoplasm | other | 1.40E+07 | 1.40E+07 | 1.35E+07 | | | | | | |
| extended synaptotagmin-like protein 1 | Cytoplasm | other | 1.31E+07 | 1.00E+07 | 1.10E+07 | | | | | | |
| extended synaptotagmin-like protein 1 | Cytoplasm | other | 1.29E+07 | 1.09E+07 | 9762559 | | | | | | |
| extended synaptotagmin-like protein 1 | Cytoplasm | other | 1.10E+07 | 8389492 | 8292407.5 | | | | | | |
| extended synaptotagmin-like protein 1 | Cytoplasm | other | 1.05E+07 | 9492642 | 1.17E+07 | | | | | | |
| extended synaptotagmin-like protein 1 | Cytoplasm | other | 9626898 | 7594230.5 | 7994069.5 | | | | | | |
| extended synaptotagmin-like protein 1 | Cytoplasm | other | 9317875 | 8173741.5 | 7716584 | | | | | | |
| extended synaptotagmin-like protein 1 | Cytoplasm | other | 8481286 | 5139283 | 5923016.5 | | | | | | |
| extended synaptotagmin-like protein 1 | Cytoplasm | other | 8056911.5 | 7278337.5 | 7413869 | | | | | | |
| extended synaptotagmin-like protein 1 | Cytoplasm | other | 8045900 | 9320311 | 8300188 | | | | | | |
| extended synaptotagmin-like protein 1 | Cytoplasm | other | 8039746 | 7194065 | 7246948 | | | | | | |
| extended synaptotagmin-like protein 1 | Cytoplasm | other | 7958539.5 | 5286739.5 | 5894162 | | | | | | |
| extended synaptotagmin-like protein 1 | Cytoplasm | other | 7454045 | 7852965 | 8482739 | | | | | | |
| extended synaptotagmin-like protein 1 | Cytoplasm | other | 6860301.5 | 5178199.5 | 4748955.5 | | | | | | |
| extended synaptotagmin-like protein 1 | Cytoplasm | other | 6711641 | 6344679 | 5601683.5 | | | | | | |
| extended synaptotagmin-like protein 1 | Cytoplasm | other | 6398393 | 4071882.5 | 5106673.5 | | | | | | |
| extended synaptotagmin-like protein 1 | Cytoplasm | other | 5239315 | 5404196.5 | 6853795 | | | | | | |
| extended synaptotagmin-like protein 1 | Cytoplasm | other | 5132083.5 | 3455931.75 | 5294329.5 | | | | | | |
| extended synaptotagmin-like protein 1 | Cytoplasm | other | 4255237 | 3670647.5 | 4533175.5 | | | | | | |
| extended synaptotagmin-like protein 1 | Cytoplasm | other | 3518211.75 | 2148849 | 2731729.75 | | | | | | |
| extended synaptotagmin-like protein 1 | Cytoplasm | other | 3038236.25 | 2372254.25 | 3029138 | | | | | | |
| extended synaptotagmin-like protein 1 | Cytoplasm | other | 628605.125 | 499477.5 | 493822.844 | Q3U7R1 | ESYT1 | 10827697 | 6338468 | 679555 | 87 |
| extended synaptotagmin-like protein 2 | Plasma Membrane | other | 1.28E+07 | 1.01E+07 | 8247495 | | | | | | |
| extended synaptotagmin-like protein 2 | Plasma Membrane | other | 1.15E+07 | 1.02E+07 | 9264193 | | | | | | |
| extended synaptotagmin-like protein 2 | Plasma Membrane | other | 9891870 | 8402440 | 9858563 | | | | | | |
| extended synaptotagmin-like protein 2 | Plasma Membrane | other | 9877565 | 9782701 | 7678249.5 | | | | | | |
| extended synaptotagmin-like protein 2 | Plasma Membrane | other | 9369088 | 7778622 | 7202433.5 | | | | | | |
| extended synaptotagmin-like protein 2 | Plasma Membrane | other | 9187529 | 6989455.5 | 7745704.5 | | | | | | |
| extended synaptotagmin-like protein 2 | Plasma Membrane | other | 8980961 | 6209319.5 | 6127136.5 | | | | | | |
| extended synaptotagmin-like protein 2 | Plasma Membrane | other | 8557471 | 7368603.5 | 9676347 | | | | | | |
| extended synaptotagmin-like protein 2 | Plasma Membrane | other | 8222805.5 | 8457503 | 7300655 | | | | | | |
| extended synaptotagmin-like protein 2 | Plasma Membrane | other | 7579356 | 3916154.5 | 5099909 | | | | | | |
| extended synaptotagmin-like protein 2 | Plasma Membrane | other | 7116591.5 | 5640417 | 7227011 | | | | | | |
| extended synaptotagmin-like protein 2 | Plasma Membrane | other | 6308957 | 5541512 | 5572207.5 | | | | | | |
| extended synaptotagmin-like protein 2 | Plasma Membrane | other | 5995406 | 5117251.5 | 5312492.5 | | | | | | |
| extended synaptotagmin-like protein 2 | Plasma Membrane | other | 5789622.5 | 5244675 | 5155969.5 | | | | | | |
| extended synaptotagmin-like protein 2 | Plasma Membrane | other | 5430977.5 | 6532201 | 7985254 | | | | | | |
| extended synaptotagmin-like protein 2 | Plasma Membrane | other | 5408024 | 5834666 | 5931494.5 | | | | | | |
| extended synaptotagmin-like protein 2 | Plasma Membrane | other | 3479863 | 3171330.75 | 3103221.25 | | | | | | |
| extended synaptotagmin-like protein 2 | Plasma Membrane | other | 3192432 | 1437540.875 | 2215708 | | | | | | |
| extended synaptotagmin-like protein 2 | Plasma Membrane | other | 1393009.375 | 1647703.25 | 1823868.75 | | | | | | |
| extended synaptotagmin-like protein 2 | Plasma Membrane | other | 193294.938 | 378732.062 | 283297.188 | Q3TZZ7 | ESYT2 | 7014741 | 2910103 | 375693 | 60 |
| electron-transfer-flavoprotein, alpha polypeptide | Cytoplasm | transporter | 4087008.5 | 3992032 | 4949069.5 | | | | | | |
| electron-transfer-flavoprotein, alpha polypeptide | Cytoplasm | transporter | 2716884.25 | 3453231.25 | 5754448.5 | | | | | | |
| electron-transfer-flavoprotein, alpha polypeptide | Cytoplasm | transporter | 2209142.75 | 3499350.5 | 5031857 | | | | | | |
| electron-transfer-flavoprotein, alpha polypeptide | Cytoplasm | transporter | 2151758 | 2779385.75 | 4165096.5 | | | | | | |
| electron-transfer-flavoprotein, alpha polypeptide | Cytoplasm | transporter | 1154178.75 | 1836815.75 | 1448488.75 | | | | | | |
| electron-transfer-flavoprotein, alpha polypeptide | Cytoplasm | transporter | 928164.438 | 1791503.625 | 2635771.25 | | | | | | |
| electron-transfer-flavoprotein, alpha polypeptide | Cytoplasm | transporter | 891294.75 | 2266857.5 | 1925615.75 | | | | | | |
| electron-transfer-flavoprotein, alpha polypeptide | Cytoplasm | transporter | 388113.594 | 1061975.5 | 842793.25 | | | | | | |
| electron-transfer-flavoprotein, alpha polypeptide | Cytoplasm | transporter | 37997.195 | 258810.797 | 515771.594 | Q99LC5 | ETFA | 1618282 | 1587738 | 305560 | 27 |
| electron-transfer-flavoprotein, beta polypeptide | Cytoplasm | transporter | 7063749.5 | 8135042 | 8769498 | | | | | | |
| electron-transfer-flavoprotein, beta polypeptide | Cytoplasm | transporter | 1507081.25 | 2861114.5 | 3049040.75 | | | | | | |
| electron-transfer-flavoprotein, beta polypeptide | Cytoplasm | transporter | 981384.688 | 1741709.875 | 2389107.75 | | | | | | |
| electron-transfer-flavoprotein, beta polypeptide | Cytoplasm | transporter | 625053.5 | 2228617.25 | 1072837.375 | Q9DCW4 | ETFB | 2544317 | 2904070 | 838333 | 12 |
| exocyst complex component 3-like 4 | Cytoplasm | other | 269306.312 | 1744498 | 1078238.5 | Q6DIA2 | EXOC3L4 | 269306 | 738745 | 426515 | 3 |
| ezrin | Plasma Membrane | other | 2.65E+07 | 1.82E+07 | 1.34E+07 | | | | | | |
| ezrin | Plasma Membrane | other | 1.60E+07 | 1.46E+07 | 1.16E+07 | | | | | | |
| ezrin | Plasma Membrane | other | 8184802.5 | 8970955 | 6546751.5 | | | | | | |
| ezrin | Plasma Membrane | other | 7586232.5 | 5237940.5 | 5118543.5 | | | | | | |
| ezrin | Plasma Membrane | other | 6926591 | 5177922.5 | 5735897.5 | | | | | | |
| ezrin | Plasma Membrane | other | 4060937.75 | 1887611.625 | 563675.625 | | | | | | |
| ezrin | Plasma Membrane | other | 3817883 | 2021242.375 | 1877405.5 | | | | | | |
| ezrin | Plasma Membrane | other | 3786904.25 | 2445209.75 | 1817203.375 | | | | | | |
| ezrin | Plasma Membrane | other | 3364670.25 | 2309064.25 | 1325314 | | | | | | |
| ezrin | Plasma Membrane | other | 1727779.75 | 1112422.125 | 344055.688 | P26040 | EZR | 8196580 | 6114277 | 1116309 | 30 |
| F11 receptor | Plasma Membrane | other | 1716184.875 | 1.79E+07 | 8402958 | | | | | | |
| F11 receptor | Plasma Membrane | other | 445952.25 | 2170962 | 1948540 | | | | | | |
| F11 receptor | Plasma Membrane | other | 93551.297 | 3222396.25 | 2096900.75 | | | | | | |
| F11 receptor | Plasma Membrane | other | 86877.164 | 1738534 | 1578389.5 | | | | | | |
| F11 receptor | Plasma Membrane | other | 50246.09 | 4176092 | 2983420 | O88792 | F11R | 478562 | 4553875 | 1175805 | 15 |
| coagulation factor II (thrombin) | Extracellular Space | peptidase | 5378663.5 | 4495571.5 | 4492052.5 | P19221 | F2 | 5378664 | 510872 | 294952 | 3 |
| coagulation factor V (proaccelerin, labile factor) | Plasma Membrane | enzyme | 315108.625 | 5778389 | 3899830.5 | | | | | | |
| coagulation factor V (proaccelerin, labile factor) | Plasma Membrane | enzyme | 71853.75 | 1802647.75 | 1429116 | | | | | | |
| coagulation factor V (proaccelerin, labile factor) | Plasma Membrane | enzyme | 4.70E-06 | 1300713.375 | 721495.625 | O88783 | F5 | 128987 | 1940750 | 646917 | 9 |
| fatty acid desaturase 1 | Plasma Membrane | enzyme | 2915675.5 | 2823999 | 3505808.25 | Q920L1 | FADS1 | 2915676 | 370028 | 213636 | 3 |
| Fas (TNFRSF6) associated factor 1 | Nucleus | other | 6254880.5 | 2950891.5 | 3234399.75 | | | | | | |
| Fas (TNFRSF6) associated factor 1 | Nucleus | other | 5925961.5 | 2370935 | 2994797.25 | | | | | | |
| Fas (TNFRSF6) associated factor 1 | Nucleus | other | 5279918.5 | 2501992.25 | 3041926.75 | P54731 | FAF1 | 5820254 | 1529429 | 509810 | 9 |
| fumarylacetoacetate hydrolase domain containing 1 | Cytoplasm | enzyme | 2015540.75 | 1645676.75 | 2442168 | | | | | | |
| fumarylacetoacetate hydrolase domain containing 1 | Cytoplasm | enzyme | 1505973.125 | 1631857.125 | 2078752.625 | Q8R0F8 | FAHD1 | 1760757 | 354970 | 144916 | 6 |
| family with sequence similarity 134, member A | Extracellular Space | other | 3005661.25 | 2376823.25 | 1793013.75 | Q6NS82 | FAM134A | 3005661 | 606463 | 350142 | 3 |
| family with sequence similarity 210, member B | Extracellular Space | other | 1771695.375 | 1890572.875 | 1576114.125 | | | | | | |
| family with sequence similarity 210, member B | Extracellular Space | other | 380025.344 | 43935.184 | 33399.945 | Q9D8B6 | FAM210B | 1075860 | 887452 | 362301 | 6 |
| family with sequence similarity 213, member A | Extracellular Space | other | 1.58E+08 | 9.81E+07 | 1.49E+08 | | | | | | |
| family with sequence similarity 213, member A | Extracellular Space | other | 9.58E+07 | 7.40E+07 | 1.04E+08 | | | | | | |
| family with sequence similarity 213, member A | Extracellular Space | other | 8.83E+07 | 3.35E+07 | 6.31E+07 | | | | | | |
| family with sequence similarity 213, member A | Extracellular Space | other | 7.21E+07 | 2.97E+07 | 7.45E+07 | | | | | | |
| family with sequence similarity 213, member A | Extracellular Space | other | 5.87E+07 | 5.11E+07 | 7.84E+07 | | | | | | |
| family with sequence similarity 213, member A | Extracellular Space | other | 5.04E+07 | 2.83E+07 | 4.14E+07 | | | | | | |
| family with sequence similarity 213, member A | Extracellular Space | other | 2.41E+07 | 2.09E+07 | 1.96E+07 | | | | | | |
| family with sequence similarity 213, member A | Extracellular Space | other | 1.97E+07 | 7997017.5 | 1.21E+07 | | | | | | |
| family with sequence similarity 213, member A | Extracellular Space | other | 1.56E+07 | 8922594 | 1.15E+07 | | | | | | |

| | | | | | | | | | | | |
|---|---|---|---|---|---|---|---|---|---|---|---|
| family with sequence similarity 213, member A | Extracellular Space | other | 1.40E+07 | 1.11E+07 | 1.50E+07 | | | | | | |
| family with sequence similarity 213, member A | Extracellular Space | other | 1.03E+07 | 7208785.5 | 8933991 | | | | | | |
| family with sequence similarity 213, member A | Extracellular Space | other | 1264065.625 | 640439.562 | 1537472.75 | Q9CYH2 | FAM213A | 50685339 | 41311526 | 6885254 | 36 |
| phenylalanyl-tRNA synthetase, beta subunit | Cytoplasm | enzyme | 869051.562 | 748386.438 | 461463.438 | Q9WUA2 | FARSB | 869052 | 209369 | 120879 | 3 |
| fatty acid synthase | Cytoplasm | enzyme | 1.49E+07 | 7190521 | 6957124.5 | | | | | | |
| fatty acid synthase | Cytoplasm | enzyme | 1.39E+07 | 8231270.5 | 9028225 | | | | | | |
| fatty acid synthase | Cytoplasm | enzyme | 1.22E+07 | 6769267 | 6369159 | | | | | | |
| fatty acid synthase | Cytoplasm | enzyme | 9891502 | 4889242 | 4365769 | | | | | | |
| fatty acid synthase | Cytoplasm | enzyme | 9207947 | 5632258 | 5856553.5 | | | | | | |
| fatty acid synthase | Cytoplasm | enzyme | 8422895 | 4388135.5 | 6665619 | | | | | | |
| fatty acid synthase | Cytoplasm | enzyme | 7868072.5 | 6396021.5 | 6549294.5 | | | | | | |
| fatty acid synthase | Cytoplasm | enzyme | 7495178 | 5066473.5 | 5338608 | | | | | | |
| fatty acid synthase | Cytoplasm | enzyme | 6571211.5 | 4328024 | 3945185.75 | | | | | | |
| fatty acid synthase | Cytoplasm | enzyme | 5189619 | 2704754.5 | 2526156.25 | | | | | | |
| fatty acid synthase | Cytoplasm | enzyme | 4757691.5 | 2982323.75 | 3102606.75 | | | | | | |
| fatty acid synthase | Cytoplasm | enzyme | 4698077.5 | 2837628.5 | 2576265.5 | | | | | | |
| fatty acid synthase | Cytoplasm | enzyme | 4600893.5 | 3337681.25 | 4094573 | | | | | | |
| fatty acid synthase | Cytoplasm | enzyme | 4578983.5 | 3251764 | 3084139.75 | | | | | | |
| fatty acid synthase | Cytoplasm | enzyme | 4253708.5 | 1970774 | 2509813 | | | | | | |
| fatty acid synthase | Cytoplasm | enzyme | 3731947.5 | 2268921.5 | 2059339.5 | | | | | | |
| fatty acid synthase | Cytoplasm | enzyme | 3605159 | 1998332.375 | 2733472.25 | | | | | | |
| fatty acid synthase | Cytoplasm | enzyme | 2450622.25 | 768971.75 | 1236115.5 | | | | | | |
| fatty acid synthase | Cytoplasm | enzyme | 1758770.125 | 395716.625 | 517607.844 | | | | | | |
| fatty acid synthase | Cytoplasm | enzyme | 1679448.375 | 1633373 | 1729580.875 | | | | | | |
| fatty acid synthase | Cytoplasm | enzyme | 1513286.375 | 1031671 | 1369989.25 | | | | | | |
| fatty acid synthase | Cytoplasm | enzyme | 890550.812 | 401831.719 | 408131.906 | | | | | | |
| fatty acid synthase | Cytoplasm | enzyme | 678784.188 | 597199.812 | 868261.312 | P19096 | FASN | 5861059 | 3176669 | 382426 | 69 |
| Finkel-Biskis-Reilly murine sarcoma virus (FBR-MuSV) ubiquitously expressed | Cytoplasm | other | 3356014.25 | 2111934.5 | 3019254.25 | P62862 | FAU | 3356014 | 643476 | 371511 | 3 |
| F-box protein 7 | Cytoplasm | enzyme | 2774863.75 | 1165441.25 | 1493204.25 | Q3U7U3 | FBXO7 | 2774864 | 850521 | 491049 | 3 |
| ferrochelatase | Cytoplasm | enzyme | 3.34E+07 | 3.57E+07 | 3.58E+07 | | | | | | |
| ferrochelatase | Cytoplasm | enzyme | 2.49E+07 | 2.09E+07 | 2.36E+07 | | | | | | |
| ferrochelatase | Cytoplasm | enzyme | 1.64E+07 | 1.60E+07 | 2.42E+07 | | | | | | |
| ferrochelatase | Cytoplasm | enzyme | 1.44E+07 | 1.56E+07 | 1.59E+07 | | | | | | |
| ferrochelatase | Cytoplasm | enzyme | 8156954 | 7970591 | 6961968.5 | | | | | | |
| ferrochelatase | Cytoplasm | enzyme | 5505966 | 6118793.5 | 7008069 | | | | | | |
| ferrochelatase | Cytoplasm | enzyme | 4689293.5 | 5688380 | 9078947 | | | | | | |
| ferrochelatase | Cytoplasm | enzyme | 4336555.5 | 6434526 | 1.58E+07 | | | | | | |
| ferrochelatase | Cytoplasm | enzyme | 3970905.75 | 5243749 | 4700658.5 | | | | | | |
| ferrochelatase | Cytoplasm | enzyme | 3138363.75 | 7845285.5 | 1200312.75 | | | | | | |
| ferrochelatase | Cytoplasm | enzyme | 2785111.25 | 1382286.625 | 1965411.25 | | | | | | |
| ferrochelatase | Cytoplasm | enzyme | 2713883.25 | 2235612.5 | 2377321.75 | | | | | | |
| ferrochelatase | Cytoplasm | enzyme | 2340733.75 | 891138.562 | 1733061.375 | | | | | | |
| ferrochelatase | Cytoplasm | enzyme | 2238553 | 1685657.625 | 2794273.5 | | | | | | |
| ferrochelatase | Cytoplasm | enzyme | 1723303.875 | 2060545.5 | 2188503.75 | | | | | | |
| ferrochelatase | Cytoplasm | enzyme | 1622761.375 | 2405850 | 1808148.375 | | | | | | |
| ferrochelatase | Cytoplasm | enzyme | 1488379.75 | 1017349 | 1468840.5 | | | | | | |
| ferrochelatase | Cytoplasm | enzyme | 710480.062 | 1.17E+07 | 1203826.875 | | | | | | |
| ferrochelatase | Cytoplasm | enzyme | 488817.125 | 1072759.875 | 1236636.25 | P22315 | FECH | 7101056 | 9146618 | 1211499 | 57 |
| fermitin family member 3 | Cytoplasm | enzyme | 7427753 | 2.04E+07 | 2.29E+07 | | | | | | |
| fermitin family member 3 | Cytoplasm | enzyme | 7223967.5 | 2.07E+07 | 2.64E+07 | | | | | | |
| fermitin family member 3 | Cytoplasm | enzyme | 6050117 | 1.18E+07 | 2.10E+07 | | | | | | |
| fermitin family member 3 | Cytoplasm | enzyme | 5407405 | 1.43E+07 | 2.24E+07 | | | | | | |
| fermitin family member 3 | Cytoplasm | enzyme | 4767208.5 | 1.26E+07 | 2.15E+07 | | | | | | |
| fermitin family member 3 | Cytoplasm | enzyme | 4691706.5 | 1.25E+07 | 1.98E+07 | | | | | | |
| fermitin family member 3 | Cytoplasm | enzyme | 3942658 | 1.33E+07 | 1.71E+07 | | | | | | |
| fermitin family member 3 | Cytoplasm | enzyme | 3927008.75 | 1.01E+07 | 1.73E+07 | | | | | | |
| fermitin family member 3 | Cytoplasm | enzyme | 3312055.75 | 9106180 | 1.31E+07 | | | | | | |
| fermitin family member 3 | Cytoplasm | enzyme | 3128183.5 | 7988016 | 1.30E+07 | | | | | | |
| fermitin family member 3 | Cytoplasm | enzyme | 3080519 | 6970346 | 1.30E+07 | | | | | | |
| fermitin family member 3 | Cytoplasm | enzyme | 2690958 | 3427437.5 | 3975448.75 | | | | | | |
| fermitin family member 3 | Cytoplasm | enzyme | 1830048.875 | 4038398.75 | 6635760 | | | | | | |
| fermitin family member 3 | Cytoplasm | enzyme | 1700691.875 | 4821817.5 | 5330166 | | | | | | |
| fermitin family member 3 | Cytoplasm | enzyme | 1661441.25 | 4667268 | 6758822 | | | | | | |
| fermitin family member 3 | Cytoplasm | enzyme | 794674.188 | 3861226.75 | 5674816 | Q8K1B8 | FERMT3 | 3852275 | 6952112 | 1003451 | 48 |
| fibrinogen beta chain | Extracellular Space | other | 7941250.5 | 1.91E+07 | 1.38E+08 | | | | | | |
| fibrinogen beta chain | Extracellular Space | other | 6123591.5 | 2.47E+07 | 2.08E+08 | | | | | | |
| fibrinogen beta chain | Extracellular Space | other | 5414073 | 3.98E+07 | 2.26E+08 | | | | | | |
| fibrinogen beta chain | Extracellular Space | other | 4580159 | 4.78E+07 | 1.49E+08 | | | | | | |
| fibrinogen beta chain | Extracellular Space | other | 4133639.25 | 2.27E+07 | 1.56E+08 | | | | | | |
| fibrinogen beta chain | Extracellular Space | other | 3922837.5 | 3.37E+07 | 1.44E+08 | | | | | | |
| fibrinogen beta chain | Extracellular Space | other | 3467629.5 | 2.26E+07 | 1.29E+08 | | | | | | |
| fibrinogen beta chain | Extracellular Space | other | 3378407.5 | 2.40E+07 | 1.23E+08 | | | | | | |
| fibrinogen beta chain | Extracellular Space | other | 2692473.25 | 2.76E+07 | 1.48E+08 | | | | | | |
| fibrinogen beta chain | Extracellular Space | other | 2554367.5 | 2.13E+07 | 1.33E+08 | | | | | | |
| fibrinogen beta chain | Extracellular Space | other | 2096629 | 1.86E+07 | 7.89E+07 | | | | | | |
| fibrinogen beta chain | Extracellular Space | other | 1701762.625 | 7025743.5 | 6.78E+07 | | | | | | |
| fibrinogen beta chain | Extracellular Space | other | 1624958.125 | 1.10E+07 | 4.84E+07 | | | | | | |
| fibrinogen beta chain | Extracellular Space | other | 1597271.125 | 1.48E+07 | 7.99E+07 | | | | | | |
| fibrinogen beta chain | Extracellular Space | other | 1467967.375 | 9320680 | 4.33E+07 | | | | | | |
| fibrinogen beta chain | Extracellular Space | other | 1315827.875 | 8639238 | 4.61E+07 | | | | | | |
| fibrinogen beta chain | Extracellular Space | other | 579548.125 | 2609623 | 3.13E+07 | | | | | | |
| fibrinogen beta chain | Extracellular Space | other | 507333.75 | 5362300.5 | 2.86E+07 | | | | | | |
| fibrinogen beta chain | Extracellular Space | other | 489172.469 | 1489711.75 | 1.95E+07 | | | | | | |
| fibrinogen beta chain | Extracellular Space | other | 248036.891 | 1498645 | 1.12E+07 | | | | | | |
| fibrinogen beta chain | Extracellular Space | other | 113684.266 | 630717.125 | 5862494 | | | | | | |
| fibrinogen beta chain | Extracellular Space | other | 93941.633 | 2124824.25 | 2.16E+07 | | | | | | |
| fibrinogen beta chain | Extracellular Space | other | 31483.266 | 2768204.5 | 2.89E+07 | | | | | | |
| fibrinogen beta chain | Extracellular Space | other | 4.70E-06 | 1478020.25 | 5774544 | Q8K0E8 | FGB | 2336502 | 53395291 | 6292695 | 72 |
| fibrinogen gamma chain | Extracellular Space | other | 1.27E+07 | 7.13E+07 | 2.87E+08 | | | | | | |
| fibrinogen gamma chain | Extracellular Space | other | 1.14E+07 | 7.85E+07 | 3.98E+08 | | | | | | |
| fibrinogen gamma chain | Extracellular Space | other | 5443341 | 4.04E+07 | 1.74E+08 | | | | | | |
| fibrinogen gamma chain | Extracellular Space | other | 4842494.5 | 1.23E+07 | 1.02E+08 | | | | | | |
| fibrinogen gamma chain | Extracellular Space | other | 3373127.75 | 3.37E+07 | 1.45E+08 | | | | | | |
| fibrinogen gamma chain | Extracellular Space | other | 3185512.5 | 2.50E+07 | 1.08E+08 | | | | | | |
| fibrinogen gamma chain | Extracellular Space | other | 3052397.25 | 2.36E+07 | 1.06E+08 | | | | | | |

| | | | | | | | | | | | |
|---|---|---|---|---|---|---|---|---|---|---|---|
| fibrinogen gamma chain | Extracellular Space | other | 2976528.25 | 1.98E+07 | 1.04E+08 | | | | | | |
| fibrinogen gamma chain | Extracellular Space | other | 2601879.25 | 7967074.5 | 3.79E+07 | | | | | | |
| fibrinogen gamma chain | Extracellular Space | other | 2000611.75 | 1.73E+07 | 8.83E+07 | | | | | | |
| fibrinogen gamma chain | Extracellular Space | other | 1897584 | 8979753 | 4.60E+07 | | | | | | |
| fibrinogen gamma chain | Extracellular Space | other | 1522869.625 | 1.41E+07 | 5.95E+07 | | | | | | |
| fibrinogen gamma chain | Extracellular Space | other | 1465901.875 | 6434808.5 | 4.33E+07 | | | | | | |
| fibrinogen gamma chain | Extracellular Space | other | 1255526.375 | 1.50E+07 | 3.13E+07 | | | | | | |
| fibrinogen gamma chain | Extracellular Space | other | 1131395.125 | 1.68E+07 | 8.24E+07 | | | | | | |
| fibrinogen gamma chain | Extracellular Space | other | 1042707.375 | 9393745 | 5.54E+07 | | | | | | |
| fibrinogen gamma chain | Extracellular Space | other | 950358.75 | 2.13E+07 | 7.12E+07 | | | | | | |
| fibrinogen gamma chain | Extracellular Space | other | 339068.5 | 875355.812 | 8443994 | | | | | | |
| fibrinogen gamma chain | Extracellular Space | other | 189784.516 | 4931793 | 6.39E+07 | | | | | | |
| fibrinogen gamma chain | Extracellular Space | other | 187244.312 | 6048232.5 | 3.36E+07 | | | | | | |
| fibrinogen gamma chain | Extracellular Space | other | 69635.594 | 5077409.5 | 2.22E+07 | | | | | | |
| fibrinogen gamma chain | Extracellular Space | other | 0 | 433356.344 | 3398242.25 | | | | | | |
| fibrinogen gamma chain | Extracellular Space | other | 0 | 1035681.562 | 2699354.75 | Q8VCM7 | FGG | 2682086 | 66038085 | 7950051 | 69 |
| fumarate hydratase | Cytoplasm | enzyme | 5112919.5 | 9221991 | 8973921 | | | | | | |
| fumarate hydratase | Cytoplasm | enzyme | 2750331.5 | 4132639.25 | 5328927.5 | | | | | | |
| fumarate hydratase | Cytoplasm | enzyme | 1774300.875 | 2436678 | 2645372.25 | | | | | | |
| fumarate hydratase | Cytoplasm | enzyme | 1406756 | 1961654.5 | 2296791.5 | | | | | | |
| fumarate hydratase | Cytoplasm | enzyme | 1182109.375 | 1542934.375 | 1791908.75 | | | | | | |
| fumarate hydratase | Cytoplasm | enzyme | 962527.625 | 567424.562 | 1008908 | | | | | | |
| fumarate hydratase | Cytoplasm | enzyme | 757772.562 | 2136909.25 | 2879193 | P97807 | FH | 1992388 | 2430873 | 530460 | 21 |
| fission, mitochondrial 1 | Cytoplasm | other | 3725287 | 4633931 | 4412968 | | | | | | |
| fission, mitochondrial 1 | Cytoplasm | other | 3445081 | 4360926 | 3968996 | Q9CQ92 | FIS1 | 3585184 | 455443 | 185934 | 6 |
| FK506 binding protein 15, 133kDa | Plasma Membrane | enzyme | 1002949.375 | 1057415.375 | 923614.938 | | | | | | |
| FK506 binding protein 15, 133kDa | Plasma Membrane | enzyme | 480408.438 | 224029.875 | 323290.344 | Q6P9Q6 | FKBP15 | 741679 | 368862 | 150587 | 6 |
| FK506 binding protein 2, 13kDa | Cytoplasm | enzyme | 1.75E+07 | 1.63E+07 | 1.48E+07 | | | | | | |
| FK506 binding protein 2, 13kDa | Cytoplasm | enzyme | 1.36E+07 | 1.23E+07 | 1.37E+07 | P45878 | FKBP2 | 15570000 | 1926473 | 786479 | 6 |
| FK506 binding protein like | Nucleus | other | 2.81E+07 | 2.71E+07 | 2.76E+07 | O35450 | FKBPL | 28140000 | 545619 | 315013 | 3 |
| filaggrin family member 2 | Extracellular Space | other | 3.97E+07 | 4.39E+07 | 5.36E+07 | Q2VIS4 | Flg2 | 39700000 | 7130009 | 4116513 | 3 |
| filamin A, alpha | Cytoplasm | other | 1.54E+07 | 1.72E+08 | 1.14E+08 | | | | | | |
| filamin A, alpha | Cytoplasm | other | 1.01E+07 | 1.26E+08 | 8.80E+07 | | | | | | |
| filamin A, alpha | Cytoplasm | other | 9843644 | 1.41E+08 | 9.85E+07 | | | | | | |
| filamin A, alpha | Cytoplasm | other | 9459023 | 1.30E+08 | 9.60E+07 | | | | | | |
| filamin A, alpha | Cytoplasm | other | 9211251 | 1.18E+08 | 7.63E+07 | | | | | | |
| filamin A, alpha | Cytoplasm | other | 8637301 | 2.77E+07 | 2.11E+07 | | | | | | |
| filamin A, alpha | Cytoplasm | other | 7586862 | 9.25E+07 | 6.07E+07 | | | | | | |
| filamin A, alpha | Cytoplasm | other | 7113750.5 | 8.02E+07 | 5.89E+07 | | | | | | |
| filamin A, alpha | Cytoplasm | other | 6885863 | 6.65E+07 | 4.45E+07 | | | | | | |
| filamin A, alpha | Cytoplasm | other | 6293526.5 | 6.58E+07 | 5.56E+07 | | | | | | |
| filamin A, alpha | Cytoplasm | other | 6039133 | 3.18E+07 | 3.40E+07 | | | | | | |
| filamin A, alpha | Cytoplasm | other | 6008070.5 | 5.05E+07 | 4.09E+07 | | | | | | |
| filamin A, alpha | Cytoplasm | other | 5860944.5 | 4.68E+07 | 3.68E+07 | | | | | | |
| filamin A, alpha | Cytoplasm | other | 5838531 | 5.39E+07 | 4.71E+07 | | | | | | |
| filamin A, alpha | Cytoplasm | other | 5791217 | 6.98E+07 | 5.66E+07 | | | | | | |
| filamin A, alpha | Cytoplasm | other | 5405163.5 | 7.11E+07 | 5.34E+07 | | | | | | |
| filamin A, alpha | Cytoplasm | other | 5028355 | 5.87E+07 | 3.92E+07 | | | | | | |
| filamin A, alpha | Cytoplasm | other | 4969175 | 6.88E+07 | 5.20E+07 | | | | | | |
| filamin A, alpha | Cytoplasm | other | 4883427 | 6.04E+07 | 4.57E+07 | | | | | | |
| filamin A, alpha | Cytoplasm | other | 4723682.5 | 5.73E+07 | 3.86E+07 | | | | | | |
| filamin A, alpha | Cytoplasm | other | 4710627.5 | 5.41E+07 | 4.32E+07 | | | | | | |
| filamin A, alpha | Cytoplasm | other | 4625529.5 | 6.37E+07 | 4.48E+07 | | | | | | |
| filamin A, alpha | Cytoplasm | other | 4501861 | 5.99E+07 | 4.34E+07 | | | | | | |
| filamin A, alpha | Cytoplasm | other | 4480270.5 | 5.65E+07 | 4.74E+07 | | | | | | |
| filamin A, alpha | Cytoplasm | other | 4330596 | 6.56E+07 | 4.41E+07 | | | | | | |
| filamin A, alpha | Cytoplasm | other | 4275081 | 2.09E+07 | 2.10E+07 | | | | | | |
| filamin A, alpha | Cytoplasm | other | 4082188 | 4.26E+07 | 2.12E+07 | | | | | | |
| filamin A, alpha | Cytoplasm | other | 4043399.25 | 5.00E+07 | 3.80E+07 | | | | | | |
| filamin A, alpha | Cytoplasm | other | 3957325.25 | 4.87E+07 | 3.18E+07 | | | | | | |
| filamin A, alpha | Cytoplasm | other | 3880023 | 6.62E+07 | 4.73E+07 | | | | | | |
| filamin A, alpha | Cytoplasm | other | 3799392.75 | 5.71E+07 | 3.51E+07 | | | | | | |
| filamin A, alpha | Cytoplasm | other | 3797328 | 5.13E+07 | 3.81E+07 | | | | | | |
| filamin A, alpha | Cytoplasm | other | 3777850 | 2.82E+07 | 1.68E+07 | | | | | | |
| filamin A, alpha | Cytoplasm | other | 3727083.25 | 4.41E+07 | 3.05E+07 | | | | | | |
| filamin A, alpha | Cytoplasm | other | 3522507.75 | 4.70E+07 | 3.57E+07 | | | | | | |
| filamin A, alpha | Cytoplasm | other | 3394627.25 | 3.88E+07 | 2.60E+07 | | | | | | |
| filamin A, alpha | Cytoplasm | other | 3271094.75 | 3.30E+07 | 2.74E+07 | | | | | | |
| filamin A, alpha | Cytoplasm | other | 3253970.25 | 6.34E+07 | 4.07E+07 | | | | | | |
| filamin A, alpha | Cytoplasm | other | 3190125.75 | 3.87E+07 | 3.02E+07 | | | | | | |
| filamin A, alpha | Cytoplasm | other | 3184270.25 | 3.37E+07 | 3.12E+07 | | | | | | |
| filamin A, alpha | Cytoplasm | other | 3089342.5 | 3.97E+07 | 2.79E+07 | | | | | | |
| filamin A, alpha | Cytoplasm | other | 3006399 | 1.89E+07 | 1.25E+07 | | | | | | |
| filamin A, alpha | Cytoplasm | other | 3003828.25 | 4.40E+07 | 3.34E+07 | | | | | | |
| filamin A, alpha | Cytoplasm | other | 2950167.25 | 3.90E+07 | 2.62E+07 | | | | | | |
| filamin A, alpha | Cytoplasm | other | 2862714.25 | 2.54E+07 | 2.11E+07 | | | | | | |
| filamin A, alpha | Cytoplasm | other | 2837896.75 | 5.35E+07 | 4.12E+07 | | | | | | |
| filamin A, alpha | Cytoplasm | other | 2829787 | 4.46E+07 | 3.74E+07 | | | | | | |
| filamin A, alpha | Cytoplasm | other | 2771699.25 | 4.68E+07 | 3.28E+07 | | | | | | |
| filamin A, alpha | Cytoplasm | other | 2710822 | 1.68E+07 | 9518686 | | | | | | |
| filamin A, alpha | Cytoplasm | other | 2683673.75 | 4.03E+07 | 2.67E+07 | | | | | | |
| filamin A, alpha | Cytoplasm | other | 2639415.5 | 4.33E+07 | 2.19E+07 | | | | | | |
| filamin A, alpha | Cytoplasm | other | 2625822.75 | 3.63E+07 | 2.58E+07 | | | | | | |
| filamin A, alpha | Cytoplasm | other | 2592097.75 | 4.79E+07 | 3.55E+07 | | | | | | |
| filamin A, alpha | Cytoplasm | other | 2562692 | 2.74E+07 | 2.26E+07 | | | | | | |
| filamin A, alpha | Cytoplasm | other | 2557403.5 | 4.00E+07 | 2.89E+07 | | | | | | |
| filamin A, alpha | Cytoplasm | other | 2483189.25 | 5.24E+07 | 3.40E+07 | | | | | | |
| filamin A, alpha | Cytoplasm | other | 2471771.25 | 4.25E+07 | 2.69E+07 | | | | | | |
| filamin A, alpha | Cytoplasm | other | 2437103.75 | 3.21E+07 | 2.36E+07 | | | | | | |
| filamin A, alpha | Cytoplasm | other | 2374957 | 1.32E+07 | 9059774 | | | | | | |
| filamin A, alpha | Cytoplasm | other | 2170188.75 | 1.06E+07 | 6454747 | | | | | | |
| filamin A, alpha | Cytoplasm | other | 2158629.5 | 1.29E+07 | 1.21E+07 | | | | | | |
| filamin A, alpha | Cytoplasm | other | 2048597.625 | 3.07E+07 | 2.27E+07 | | | | | | |
| filamin A, alpha | Cytoplasm | other | 2007240.25 | 2.29E+07 | 1.52E+07 | | | | | | |
| filamin A, alpha | Cytoplasm | other | 1989154.375 | 1.32E+07 | 1.01E+07 | | | | | | |

| | | | | | | | | | | | |
|---|---|---|---|---|---|---|---|---|---|---|---|
| filamin A, alpha | Cytoplasm | other | 1964204.5 | 1.85E+07 | 1.93E+07 | | | | | | |
| filamin A, alpha | Cytoplasm | other | 1961437.25 | 2.73E+07 | 2.06E+07 | | | | | | |
| filamin A, alpha | Cytoplasm | other | 1948855.5 | 2.96E+07 | 1.94E+07 | | | | | | |
| filamin A, alpha | Cytoplasm | other | 1886388 | 1.16E+07 | 1.36E+07 | | | | | | |
| filamin A, alpha | Cytoplasm | other | 1783800.5 | 1.91E+07 | 1.39E+07 | | | | | | |
| filamin A, alpha | Cytoplasm | other | 1695998.375 | 2.58E+07 | 1.51E+07 | | | | | | |
| filamin A, alpha | Cytoplasm | other | 1694507.75 | 1.60E+07 | 1.30E+07 | | | | | | |
| filamin A, alpha | Cytoplasm | other | 1607489.125 | 2.70E+07 | 1.47E+07 | | | | | | |
| filamin A, alpha | Cytoplasm | other | 1606681 | 2.30E+07 | 1.57E+07 | | | | | | |
| filamin A, alpha | Cytoplasm | other | 1547657.875 | 2.25E+07 | 1.77E+07 | | | | | | |
| filamin A, alpha | Cytoplasm | other | 1505381.375 | 7597742 | 8024910 | | | | | | |
| filamin A, alpha | Cytoplasm | other | 1424131.875 | 1.87E+07 | 1.28E+07 | | | | | | |
| filamin A, alpha | Cytoplasm | other | 1334345.25 | 3.16E+07 | 1.84E+07 | | | | | | |
| filamin A, alpha | Cytoplasm | other | 1312421.625 | 1.07E+07 | 7318997.5 | | | | | | |
| filamin A, alpha | Cytoplasm | other | 1230048.75 | 1.10E+07 | 8589365 | | | | | | |
| filamin A, alpha | Cytoplasm | other | 1226967.75 | 2.32E+07 | 1.52E+07 | | | | | | |
| filamin A, alpha | Cytoplasm | other | 1172162.625 | 1.94E+07 | 1.43E+07 | | | | | | |
| filamin A, alpha | Cytoplasm | other | 1149439.375 | 1.38E+07 | 1.06E+07 | | | | | | |
| filamin A, alpha | Cytoplasm | other | 1145488.5 | 2.68E+07 | 1.46E+07 | | | | | | |
| filamin A, alpha | Cytoplasm | other | 1144258.5 | 2.54E+07 | 1.31E+07 | | | | | | |
| filamin A, alpha | Cytoplasm | other | 1118515 | 3.24E+07 | 1.85E+07 | | | | | | |
| filamin A, alpha | Cytoplasm | other | 1113352.875 | 1.53E+07 | 1.05E+07 | | | | | | |
| filamin A, alpha | Cytoplasm | other | 1110210.375 | 6170060 | 6474500.5 | | | | | | |
| filamin A, alpha | Cytoplasm | other | 1053441.5 | 2747633.25 | 2127383 | | | | | | |
| filamin A, alpha | Cytoplasm | other | 1030713.188 | 1.11E+07 | 6564857 | | | | | | |
| filamin A, alpha | Cytoplasm | other | 851893.812 | 3.75E+07 | 3.04E+07 | | | | | | |
| filamin A, alpha | Cytoplasm | other | 849375.812 | 2.20E+07 | 1.02E+07 | | | | | | |
| filamin A, alpha | Cytoplasm | other | 821291.438 | 1.07E+07 | 7972971 | | | | | | |
| filamin A, alpha | Cytoplasm | other | 820339.438 | 1.16E+07 | 8736992 | | | | | | |
| filamin A, alpha | Cytoplasm | other | 807817.562 | 1.17E+07 | 6644630 | | | | | | |
| filamin A, alpha | Cytoplasm | other | 796509.188 | 1.18E+07 | 6627699 | | | | | | |
| filamin A, alpha | Cytoplasm | other | 792033.812 | 1.22E+07 | 8914647 | | | | | | |
| filamin A, alpha | Cytoplasm | other | 789855.875 | 8475709 | 7322846 | | | | | | |
| filamin A, alpha | Cytoplasm | other | 743550.375 | 1.12E+07 | 7404414.5 | | | | | | |
| filamin A, alpha | Cytoplasm | other | 731578.188 | 7583403 | 4422258 | | | | | | |
| filamin A, alpha | Cytoplasm | other | 723647.188 | 1.13E+07 | 7219337 | | | | | | |
| filamin A, alpha | Cytoplasm | other | 631688.375 | 2.76E+07 | 2.00E+07 | | | | | | |
| filamin A, alpha | Cytoplasm | other | 498597.156 | 6670632.5 | 5386828 | | | | | | |
| filamin A, alpha | Cytoplasm | other | 403488.312 | 7721404 | 5131846 | | | | | | |
| filamin A, alpha | Cytoplasm | other | 365720.031 | 2188495.75 | 1561003 | | | | | | |
| filamin A, alpha | Cytoplasm | other | 300436.125 | 1.05E+07 | 6883790.5 | | | | | | |
| filamin A, alpha | Cytoplasm | other | 300161.062 | 8439203 | 4622419 | | | | | | |
| filamin A, alpha | Cytoplasm | other | 228988.672 | 9323789 | 6952675 | | | | | | |
| filamin A, alpha | Cytoplasm | other | 183247.547 | 3342092 | 2004502.75 | | | | | | |
| filamin A, alpha | Cytoplasm | other | 151313.672 | 6568011.5 | 4898509.5 | | | | | | |
| filamin A, alpha | Cytoplasm | other | 137004.922 | 8005372.5 | 3340352.25 | | | | | | |
| filamin A, alpha | Cytoplasm | other | 136050.391 | 3310451.75 | 2053297.375 | | | | | | |
| filamin A, alpha | Cytoplasm | other | 109188.602 | 1.04E+07 | 4534690 | | | | | | |
| filamin A, alpha | Cytoplasm | other | 102014.195 | 1844494.875 | 843297.562 | | | | | | |
| filamin A, alpha | Cytoplasm | other | 101070.602 | 3474779.75 | 2461037.75 | | | | | | |
| filamin A, alpha | Cytoplasm | other | 77218.297 | 3473050.5 | 1541746.375 | | | | | | |
| filamin A, alpha | Cytoplasm | other | 57680.559 | 1498596.75 | 1906109.25 | | | | | | |
| filamin A, alpha | Cytoplasm | other | 4.70E-06 | 757350.25 | 777595.625 | | | | | | |
| filamin A, alpha | Cytoplasm | other | 4.70E-06 | 3932205 | 1857409.25 | Q8BTM8 | FLNA | 2822698 | 25242727 | 1341636 | 354 |
| flotillin 1 | Plasma Membrane | other | 2.24E+07 | 1.51E+07 | 1.60E+07 | | | | | | |
| flotillin 1 | Plasma Membrane | other | 2.11E+07 | 2.11E+07 | 1.98E+07 | | | | | | |
| flotillin 1 | Plasma Membrane | other | 1.84E+07 | 2.02E+07 | 1.22E+07 | | | | | | |
| flotillin 1 | Plasma Membrane | other | 1.40E+07 | 1.16E+07 | 1.05E+07 | | | | | | |
| flotillin 1 | Plasma Membrane | other | 1.38E+07 | 1.01E+07 | 1.20E+07 | | | | | | |
| flotillin 1 | Plasma Membrane | other | 1.35E+07 | 1.00E+07 | 7953746 | | | | | | |
| flotillin 1 | Plasma Membrane | other | 1.25E+07 | 1.44E+07 | 1.20E+07 | | | | | | |
| flotillin 1 | Plasma Membrane | other | 1.17E+07 | 1.33E+07 | 1.15E+07 | | | | | | |
| flotillin 1 | Plasma Membrane | other | 9220069 | 1.02E+07 | 1.04E+07 | | | | | | |
| flotillin 1 | Plasma Membrane | other | 7438660.5 | 6668173 | 5950272 | | | | | | |
| flotillin 1 | Plasma Membrane | other | 6990486 | 9472139 | 6473166.5 | | | | | | |
| flotillin 1 | Plasma Membrane | other | 6166481.5 | 3884063.5 | 4805225.5 | | | | | | |
| flotillin 1 | Plasma Membrane | other | 4819165.5 | 2583634.75 | 3555758.75 | | | | | | |
| flotillin 1 | Plasma Membrane | other | 3949401.25 | 3129440.75 | 4080589 | | | | | | |
| flotillin 1 | Plasma Membrane | other | 3063553.5 | 3303935.75 | 3711079 | | | | | | |
| flotillin 1 | Plasma Membrane | other | 1035180.75 | 2948645.5 | 1976023.375 | O08917 | FLOT1 | 10630187 | 5721693 | 825855 | 48 |
| flotillin 2 | Plasma Membrane | other | 3.34E+07 | 3.16E+07 | 2.74E+07 | | | | | | |
| flotillin 2 | Plasma Membrane | other | 2.98E+07 | 2.20E+07 | 2.76E+07 | | | | | | |
| flotillin 2 | Plasma Membrane | other | 1.93E+07 | 1.66E+07 | 1.81E+07 | | | | | | |
| flotillin 2 | Plasma Membrane | other | 1.69E+07 | 2.12E+07 | 1.81E+07 | | | | | | |
| flotillin 2 | Plasma Membrane | other | 1.52E+07 | 1.53E+07 | 1.28E+07 | | | | | | |
| flotillin 2 | Plasma Membrane | other | 1.37E+07 | 1.31E+07 | 1.55E+07 | | | | | | |
| flotillin 2 | Plasma Membrane | other | 1.33E+07 | 1.21E+07 | 1.08E+07 | | | | | | |
| flotillin 2 | Plasma Membrane | other | 1.22E+07 | 1.19E+07 | 1.13E+07 | | | | | | |
| flotillin 2 | Plasma Membrane | other | 9613687 | 6443185 | 8192040.5 | | | | | | |
| flotillin 2 | Plasma Membrane | other | 9385558 | 9931588 | 9400522 | | | | | | |
| flotillin 2 | Plasma Membrane | other | 9253779 | 6765244.5 | 8759690 | | | | | | |
| flotillin 2 | Plasma Membrane | other | 5388457.5 | 4707259.5 | 5489524.5 | | | | | | |
| flotillin 2 | Plasma Membrane | other | 5166686.5 | 3963367.25 | 5841950 | | | | | | |
| flotillin 2 | Plasma Membrane | other | 3825738.75 | 3323739.75 | 3413211.25 | | | | | | |
| flotillin 2 | Plasma Membrane | other | 3184426.75 | 4025290.5 | 3704068.75 | | | | | | |
| flotillin 2 | Plasma Membrane | other | 2011119.625 | 1443230.125 | 2031368.5 | | | | | | |
| flotillin 2 | Plasma Membrane | other | 830125.312 | 745085.812 | 518923.781 | Q60634 | FLOT2 | 11911152 | 8406282 | 1177115 | 51 |
| fibronectin 1 | Extracellular Space | enzyme | 437431.75 | 112139.141 | 4111221 | | | | | | |
| fibronectin 1 | Extracellular Space | enzyme | 255702.484 | 393946.375 | 6187295.5 | | | | | | |
| fibronectin 1 | Extracellular Space | enzyme | 119432.672 | 1219314.25 | 1.28E+07 | P11276 | FN1 | 270856 | 4285613 | 1428538 | 9 |
| fructosamine 3 kinase | Cytoplasm | kinase | 2.09E+07 | 1.24E+07 | 1.67E+07 | | | | | | |
| fructosamine 3 kinase | Cytoplasm | kinase | 1.88E+07 | 1.56E+07 | 1.52E+07 | | | | | | |
| fructosamine 3 kinase | Cytoplasm | kinase | 1.76E+07 | 1.38E+07 | 1.29E+07 | | | | | | |
| fructosamine 3 kinase | Cytoplasm | kinase | 1.70E+07 | 1.66E+07 | 9235558 | | | | | | |
| fructosamine 3 kinase | Cytoplasm | kinase | 1.33E+07 | 1.46E+07 | 1.10E+07 | | | | | | |

| | | | | | | | | | | | |
|---|---|---|---|---|---|---|---|---|---|---|---|
| fructosamine 3 kinase | Cytoplasm | kinase | 3731263.75 | 1268089.5 | 1766351 | | | | | | |
| fructosamine 3 kinase | Cytoplasm | kinase | 2998606.75 | 2117774.25 | 2256900 | Q9ER35 | FN3K | 13467124 | 6412762 | 1399379 | 21 |
| fructosamine 3 kinase related protein | Cytoplasm | kinase | 3.41E+07 | 2.34E+07 | 2.53E+07 | | | | | | |
| fructosamine 3 kinase related protein | Cytoplasm | kinase | 1.94E+07 | 1.28E+07 | 1.14E+07 | | | | | | |
| fructosamine 3 kinase related protein | Cytoplasm | kinase | 1.31E+07 | 1.12E+07 | 1.14E+07 | | | | | | |
| fructosamine 3 kinase related protein | Cytoplasm | kinase | 8992419 | 7215766.5 | 7400265.5 | | | | | | |
| fructosamine 3 kinase related protein | Cytoplasm | kinase | 5992617.5 | 5003494 | 6081835 | | | | | | |
| fructosamine 3 kinase related protein | Cytoplasm | kinase | 5594832.5 | 3566440.5 | 5005509.5 | Q8K274 | FN3KRP | 14523312 | 8351114 | 1968376 | 18 |
| FERM domain containing 8 | Cytoplasm | other | 7033191.5 | 4981982.5 | 4657737.5 | Q3UFK8 | FRMD8 | 7033192 | 1288111 | 743691 | 3 |
| ferritin, heavy polypeptide 1 | Cytoplasm | enzyme | 894550.562 | 1264939.625 | 1840231.75 | P09528 | FTH1 | 894551 | 476526 | 275122 | 3 |
| ferritin, light polypeptide | Cytoplasm | enzyme | 2047089.625 | 4150624.25 | 4177479.25 | | | | | | |
| ferritin, light polypeptide | Cytoplasm | enzyme | 1461663.75 | 2093418.5 | 2086099.375 | | | | | | |
| ferritin, light polypeptide | Cytoplasm | enzyme | 859589.438 | 620395.5 | 840206.75 | P29391 | FTL | 1456114 | 1332814 | 444271 | 9 |
| FUN14 domain containing 2 | Cytoplasm | other | 2038589.25 | 7185178.5 | 4344026 | | | | | | |
| FUN14 domain containing 2 | Cytoplasm | other | 1780018.75 | 2342777.25 | 2714138.25 | Q9D6K8 | FUNDC2 | 1909304 | 2063234 | 842312 | 6 |
| frataxin | Cytoplasm | kinase | 1913078 | 2060805.375 | 1656141.25 | | | | | | |
| frataxin | Cytoplasm | kinase | 1098987.625 | 1697759.875 | 1734367.125 | O35943 | FXN | 1506033 | 328529 | 134121 | 6 |
| FYN binding protein | Nucleus | other | 86972.867 | 2923260.75 | 2334456.25 | | | | | | |
| FYN binding protein | Nucleus | other | 54072.773 | 1199155.25 | 712914.438 | | | | | | |
| FYN binding protein | Nucleus | other | 38222 | 1505669.125 | 1235415.5 | O35601 | FYB | 59756 | 1026784 | 342261 | 9 |
| GTPase activating protein (SH3 domain) binding protein 1 | Nucleus | enzyme | 3882090.25 | 1764401.25 | 2204256 | | | | | | |
| GTPase activating protein (SH3 domain) binding protein 1 | Nucleus | enzyme | 3147992.5 | 1810228 | 2608102.25 | P97855 | G3BP1 | 3515041 | 826694 | 337497 | 6 |
| glucose 6 phosphatase, catalytic, 3 | Cytoplasm | phosphatase | 1212083.25 | 2735211.25 | 2213066.5 | Q6NSQ9 | G6PC3 | 1212083 | 774007 | 446873 | 3 |
| glucose-6-phosphate dehydrogenase | Cytoplasm | enzyme | 3078938.75 | 2347786.25 | 1868943.5 | | | | | | |
| glucose-6-phosphate dehydrogenase | Cytoplasm | enzyme | 1963779.625 | 1778456.625 | 1376507 | | | | | | |
| glucose-6-phosphate dehydrogenase | Cytoplasm | enzyme | 1747573.625 | 1457038.125 | 1040079.438 | | | | | | |
| glucose-6-phosphate dehydrogenase | Cytoplasm | enzyme | 1540861.25 | 1866900.75 | 920447.312 | Q00612 | G6PD | 2082788 | 576743 | 166491 | 12 |
| GABA(A) receptor-associated protein like 2 | Cytoplasm | other | 1.24E+07 | 9151313 | 8524438 | | | | | | |
| GABA(A) receptor-associated protein like 2 | Cytoplasm | other | 1.10E+07 | 6932630.5 | 8366908 | | | | | | |
| GABA(A) receptor-associated protein like 2 | Cytoplasm | other | 9877923 | 5438261 | 6356037 | | | | | | |
| GABA(A) receptor-associated protein like 2 | Cytoplasm | other | 5355831.5 | 3377796.5 | 4500538 | | | | | | |
| GABA(A) receptor-associated protein like 2 | Cytoplasm | other | 1435485.625 | 717129.188 | 476027.781 | P60521 | GABARAPL2 | 8019848 | 3705971 | 956878 | 15 |
| glucosidase, alpha; neutral AB | Cytoplasm | enzyme | 1.90E+07 | 1.47E+07 | 1.87E+07 | | | | | | |
| glucosidase, alpha; neutral AB | Cytoplasm | enzyme | 1.72E+07 | 2.10E+07 | 1.51E+07 | | | | | | |
| glucosidase, alpha; neutral AB | Cytoplasm | enzyme | 1.62E+07 | 1.62E+07 | 1.58E+07 | | | | | | |
| glucosidase, alpha; neutral AB | Cytoplasm | enzyme | 1.49E+07 | 8215270 | 9058620 | | | | | | |
| glucosidase, alpha; neutral AB | Cytoplasm | enzyme | 1.28E+07 | 1.43E+07 | 1.52E+07 | | | | | | |
| glucosidase, alpha; neutral AB | Cytoplasm | enzyme | 1.01E+07 | 1.13E+07 | 1.00E+07 | | | | | | |
| glucosidase, alpha; neutral AB | Cytoplasm | enzyme | 9918678 | 8785249 | 8568258 | | | | | | |
| glucosidase, alpha; neutral AB | Cytoplasm | enzyme | 9318827 | 8680029 | 7904001.5 | | | | | | |
| glucosidase, alpha; neutral AB | Cytoplasm | enzyme | 8877280 | 1.06E+07 | 7619988 | | | | | | |
| glucosidase, alpha; neutral AB | Cytoplasm | enzyme | 8480449 | 8422268 | 7678462 | | | | | | |
| glucosidase, alpha; neutral AB | Cytoplasm | enzyme | 7066776 | 8629942 | 7117701.5 | | | | | | |
| glucosidase, alpha; neutral AB | Cytoplasm | enzyme | 6868488 | 1.07E+07 | 8943772 | | | | | | |
| glucosidase, alpha; neutral AB | Cytoplasm | enzyme | 6782941 | 4238771 | 5125890 | | | | | | |
| glucosidase, alpha; neutral AB | Cytoplasm | enzyme | 4887658 | 3286831 | 4303408 | | | | | | |
| glucosidase, alpha; neutral AB | Cytoplasm | enzyme | 4003575.75 | 4697999 | 4485012.5 | | | | | | |
| glucosidase, alpha; neutral AB | Cytoplasm | enzyme | 3476305.75 | 3874418.5 | 3293542.75 | | | | | | |
| glucosidase, alpha; neutral AB | Cytoplasm | enzyme | 3287398.75 | 1752163.5 | 2646564.5 | | | | | | |
| glucosidase, alpha; neutral AB | Cytoplasm | enzyme | 2774777.5 | 3360704.5 | 2974107 | | | | | | |
| glucosidase, alpha; neutral AB | Cytoplasm | enzyme | 1935941.125 | 1949678.5 | 1889861.5 | | | | | | |
| glucosidase, alpha; neutral AB | Cytoplasm | enzyme | 701999.5 | 347902.656 | 564773.562 | Q8BHN3 | GANAB | 8426555 | 5194783 | 670644 | 60 |
| glyceraldehyde-3-phosphate dehydrogenase | Cytoplasm | enzyme | 3.65E+08 | 2.08E+08 | 2.88E+08 | | | | | | |
| glyceraldehyde-3-phosphate dehydrogenase | Cytoplasm | enzyme | 2.89E+08 | 1.68E+08 | 3.10E+08 | | | | | | |
| glyceraldehyde-3-phosphate dehydrogenase | Cytoplasm | enzyme | 2.70E+08 | 1.24E+08 | 2.09E+08 | | | | | | |
| glyceraldehyde-3-phosphate dehydrogenase | Cytoplasm | enzyme | 2.63E+08 | 1.84E+08 | 2.48E+08 | | | | | | |
| glyceraldehyde-3-phosphate dehydrogenase | Cytoplasm | enzyme | 2.49E+08 | 8.00E+07 | 1.36E+08 | | | | | | |
| glyceraldehyde-3-phosphate dehydrogenase | Cytoplasm | enzyme | 2.21E+08 | 1.85E+08 | 1.27E+08 | | | | | | |
| glyceraldehyde-3-phosphate dehydrogenase | Cytoplasm | enzyme | 2.15E+08 | 1.17E+08 | 2.02E+08 | | | | | | |
| glyceraldehyde-3-phosphate dehydrogenase | Cytoplasm | enzyme | 1.80E+08 | 1.04E+08 | 1.40E+08 | | | | | | |
| glyceraldehyde-3-phosphate dehydrogenase | Cytoplasm | enzyme | 1.49E+08 | 4.45E+07 | 1.17E+08 | | | | | | |
| glyceraldehyde-3-phosphate dehydrogenase | Cytoplasm | enzyme | 1.45E+08 | 3.74E+07 | 9.59E+07 | | | | | | |
| glyceraldehyde-3-phosphate dehydrogenase | Cytoplasm | enzyme | 1.00E+08 | 5.47E+07 | 8.49E+07 | | | | | | |
| glyceraldehyde-3-phosphate dehydrogenase | Cytoplasm | enzyme | 7.81E+07 | 2.16E+07 | 4.89E+07 | | | | | | |
| glyceraldehyde-3-phosphate dehydrogenase | Cytoplasm | enzyme | 7.80E+07 | 4.21E+07 | 5.78E+07 | | | | | | |
| glyceraldehyde-3-phosphate dehydrogenase | Cytoplasm | enzyme | 4.87E+07 | 1.48E+07 | 4.32E+07 | | | | | | |
| glyceraldehyde-3-phosphate dehydrogenase | Cytoplasm | enzyme | 3.75E+07 | 4.00E+07 | 4.56E+07 | | | | | | |
| glyceraldehyde-3-phosphate dehydrogenase | Cytoplasm | enzyme | 3.29E+07 | 1.91E+07 | 3.27E+07 | | | | | | |
| glyceraldehyde-3-phosphate dehydrogenase | Cytoplasm | enzyme | 2.76E+07 | 1.98E+07 | 1.98E+07 | | | | | | |
| glyceraldehyde-3-phosphate dehydrogenase | Cytoplasm | enzyme | 1.74E+07 | 5581983.5 | 1.05E+07 | | | | | | |
| glyceraldehyde-3-phosphate dehydrogenase | Cytoplasm | enzyme | 1.63E+07 | 1.33E+07 | 1.28E+07 | | | | | | |
| glyceraldehyde-3-phosphate dehydrogenase | Cytoplasm | enzyme | 1.58E+07 | 6742356 | 2.61E+07 | | | | | | |
| glyceraldehyde-3-phosphate dehydrogenase | Cytoplasm | enzyme | 1.58E+07 | 7127140.5 | 1.46E+07 | | | | | | |
| glyceraldehyde-3-phosphate dehydrogenase | Cytoplasm | enzyme | 1.30E+07 | 5949891.5 | 1.01E+07 | | | | | | |
| glyceraldehyde-3-phosphate dehydrogenase | Cytoplasm | enzyme | 1.25E+07 | 4048085 | 7045361 | | | | | | |
| glyceraldehyde-3-phosphate dehydrogenase | Cytoplasm | enzyme | 9559272 | 939909.5 | 3559016.75 | | | | | | |
| glyceraldehyde-3-phosphate dehydrogenase | Cytoplasm | enzyme | 8973523 | 2638188.5 | 6368749 | | | | | | |
| glyceraldehyde-3-phosphate dehydrogenase | Cytoplasm | enzyme | 6704856.5 | 3346527.75 | 6468736.5 | | | | | | |
| glyceraldehyde-3-phosphate dehydrogenase | Cytoplasm | enzyme | 6355048.5 | 3038816.5 | 5058273 | | | | | | |
| glyceraldehyde-3-phosphate dehydrogenase | Cytoplasm | enzyme | 4979394.5 | 919195.875 | 2068637.375 | | | | | | |
| glyceraldehyde-3-phosphate dehydrogenase | Cytoplasm | enzyme | 1956307.625 | 1015983.562 | 1590448.875 | P16858 | GAPDH | 99250290 | 91661706 | 9827166 | 87 |
| GTPase activating protein and VPS9 domains 1 | Cytoplasm | other | 1.65E+07 | 6948205.5 | 9544437 | | | | | | |
| GTPase activating protein and VPS9 domains 1 | Cytoplasm | other | 1.51E+07 | 7010918 | 7901393 | | | | | | |
| GTPase activating protein and VPS9 domains 1 | Cytoplasm | other | 1.42E+07 | 6782582 | 7570669 | | | | | | |
| GTPase activating protein and VPS9 domains 1 | Cytoplasm | other | 1.41E+07 | 6172389 | 5815695.5 | | | | | | |
| GTPase activating protein and VPS9 domains 1 | Cytoplasm | other | 1.29E+07 | 7201066 | 7615511.5 | | | | | | |
| GTPase activating protein and VPS9 domains 1 | Cytoplasm | other | 1.18E+07 | 5779491.5 | 7383787 | | | | | | |
| GTPase activating protein and VPS9 domains 1 | Cytoplasm | other | 1.13E+07 | 5304944.5 | 6192029 | | | | | | |
| GTPase activating protein and VPS9 domains 1 | Cytoplasm | other | 8701083 | 5152305 | 5002952 | | | | | | |
| GTPase activating protein and VPS9 domains 1 | Cytoplasm | other | 8608940 | 4532390.5 | 5103200 | | | | | | |
| GTPase activating protein and VPS9 domains 1 | Cytoplasm | other | 8369902 | 3242304.25 | 3477832.5 | | | | | | |
| GTPase activating protein and VPS9 domains 1 | Cytoplasm | other | 7064956 | 5121291.5 | 4274463 | | | | | | |
| GTPase activating protein and VPS9 domains 1 | Cytoplasm | other | 6011367.5 | 3243921.75 | 3190660.25 | | | | | | |
| GTPase activating protein and VPS9 domains 1 | Cytoplasm | other | 6007586 | 2033822.5 | 2638475.5 | | | | | | |
| GTPase activating protein and VPS9 domains 1 | Cytoplasm | other | 5124843 | 1115773 | 1241932.5 | | | | | | |

| | | | | | | | | | | | |
|---|---|---|---|---|---|---|---|---|---|---|---|
| GTPase activating protein and VPS9 domains 1 | Cytoplasm | other | 4518165 | 2609905.75 | 2519289.75 | | | | | | |
| GTPase activating protein and VPS9 domains 1 | Cytoplasm | other | 3424143 | 1472535.625 | 1445898.125 | | | | | | |
| GTPase activating protein and VPS9 domains 1 | Cytoplasm | other | 3004082.25 | 1611504.625 | 1597966.5 | Q6PAR5 | GAPVD1 | 9213828 | 3779848 | 529285 | 51 |
| GTPase activating Rap/RanGAP domain-like 3 | Cytoplasm | other | 2639749.25 | 3541109.75 | 3767842.75 | Q3V0G7 | GARNL3 | 2639749 | 596721 | 344517 | 3 |
| glutamate-cysteine ligase, modifier subunit | Cytoplasm | enzyme | 8739606 | 6378689 | 6196322.5 | | | | | | |
| glutamate-cysteine ligase, modifier subunit | Cytoplasm | enzyme | 7363614 | 5518634.5 | 6049646.5 | | | | | | |
| glutamate-cysteine ligase, modifier subunit | Cytoplasm | enzyme | 5385853.5 | 4894384 | 4330265 | O09172 | GCLM | 7163025 | 1326979 | 442326 | 9 |
| guanine deaminase | Cytoplasm | enzyme | 3.54E+07 | 1.58E+07 | 2.11E+07 | | | | | | |
| guanine deaminase | Cytoplasm | enzyme | 3.21E+07 | 8599787 | 1.18E+07 | | | | | | |
| guanine deaminase | Cytoplasm | enzyme | 2.95E+07 | 1.07E+07 | 1.53E+07 | | | | | | |
| guanine deaminase | Cytoplasm | enzyme | 2.01E+07 | 5555338.5 | 7100341.5 | | | | | | |
| guanine deaminase | Cytoplasm | enzyme | 1.91E+07 | 7802566.5 | 1.04E+07 | | | | | | |
| guanine deaminase | Cytoplasm | enzyme | 1.38E+07 | 4227453 | 5181793 | | | | | | |
| guanine deaminase | Cytoplasm | enzyme | 1.37E+07 | 6509519.5 | 8033118 | | | | | | |
| guanine deaminase | Cytoplasm | enzyme | 1.24E+07 | 3245665.5 | 5210609 | | | | | | |
| guanine deaminase | Cytoplasm | enzyme | 1.03E+07 | 3480044.25 | 6369232 | | | | | | |
| guanine deaminase | Cytoplasm | enzyme | 7862515.5 | 1488806.625 | 2029327.375 | | | | | | |
| guanine deaminase | Cytoplasm | enzyme | 4284504 | 1882422.375 | 1970587.625 | Q9R111 | GDA | 18030638 | 8701552 | 1514746 | 33 |
| glycerophosphodiester phosphodiesterase 1 | Plasma Membrane | enzyme | 9837031 | 6074594.5 | 7003203 | | | | | | |
| glycerophosphodiester phosphodiesterase 1 | Plasma Membrane | enzyme | 3021548 | 2470038.25 | 2632804.5 | | | | | | |
| glycerophosphodiester phosphodiesterase 1 | Plasma Membrane | enzyme | 1577454.75 | 1127491.125 | 843055.438 | Q9JL56 | GDE1 | 4812011 | 3090834 | 1030278 | 9 |
| glycerophosphodiester phosphodiesterase domain containing 1 | Plasma Membrane | enzyme | 2905225.5 | 2218155.5 | 2346432.75 | Q9CRY7 | GDPD1 | 2905226 | 365324 | 210920 | 3 |
| growth hormone inducible transmembrane protein | Cytoplasm | other | 1259025.375 | 1215613.25 | 1729386.625 | | | | | | |
| growth hormone inducible transmembrane protein | Cytoplasm | other | 581460.312 | 692109.688 | 1049236.5 | Q91VC9 | GHITM | 920243 | 417547 | 170463 | 6 |
| GLI pathogenesis-related 2 | Cytoplasm | other | 3539715.75 | 3541057.5 | 2963643.75 | Q9CYL5 | GLIPR2 | 3539716 | 332983 | 192248 | 3 |
| glutaredoxin 5 | Cytoplasm | other | 1.63E+07 | 1.03E+07 | 1.25E+07 | | | | | | |
| glutaredoxin 5 | Cytoplasm | other | 7592246 | 7221152.5 | 8021289.5 | Q80Y14 | GLRX5 | 11951123 | 3549798 | 1449199 | 6 |
| glutamate dehydrogenase 1 | Cytoplasm | enzyme | 544117.812 | 2942642.75 | 1918760.5 | | | | | | |
| glutamate dehydrogenase 1 | Cytoplasm | enzyme | 296485 | 2329780.75 | 1924567 | P26443 | GLUD1 | 420301 | 1032966 | 421707 | 6 |
| high mobility group box 1 | Nucleus | transcription regulator | 4005.559 | 180944.531 | 613953.625 | | | | | | |
| high mobility group box 1 | Nucleus | transcription regulator | 4.70E-06 | 62229.852 | 124468.305 | P63158 | Gm21596/Hmgb1 | 2003 | 231153 | 94368 | 6 |
| guanine nucleotide binding protein (G protein), alpha 11 (Gq class) | Plasma Membrane | enzyme | 556117.188 | 4085799.25 | 2523728.25 | P21278 | GNA11 | 556117 | 1768720 | 1021171 | 3 |
| guanine nucleotide binding protein (G protein) alpha 12 | Plasma Membrane | enzyme | 3.73E+07 | 3.94E+07 | 4.11E+07 | P27600 | GNA12 | 37250000 | 1939356 | 1119687 | 3 |
| guanine nucleotide binding protein (G protein), alpha inhibiting activity polypeptide 2 | Plasma Membrane | other | 6.79E+07 | 6.10E+07 | 5.44E+07 | | | | | | |
| guanine nucleotide binding protein (G protein), alpha inhibiting activity polypeptide 2 | Plasma Membrane | other | 6.70E+07 | 6.31E+07 | 5.60E+07 | | | | | | |
| guanine nucleotide binding protein (G protein), alpha inhibiting activity polypeptide 2 | Plasma Membrane | other | 4.92E+07 | 3.60E+07 | 3.03E+07 | | | | | | |
| guanine nucleotide binding protein (G protein), alpha inhibiting activity polypeptide 2 | Plasma Membrane | other | 4.30E+07 | 4.90E+07 | 4.75E+07 | | | | | | |
| guanine nucleotide binding protein (G protein), alpha inhibiting activity polypeptide 2 | Plasma Membrane | other | 4.18E+07 | 4.56E+07 | 4.53E+07 | | | | | | |
| guanine nucleotide binding protein (G protein), alpha inhibiting activity polypeptide 2 | Plasma Membrane | other | 4.11E+07 | 4.11E+07 | 3.74E+07 | | | | | | |
| guanine nucleotide binding protein (G protein), alpha inhibiting activity polypeptide 2 | Plasma Membrane | other | 3.64E+07 | 3.25E+07 | 3.36E+07 | | | | | | |
| guanine nucleotide binding protein (G protein), alpha inhibiting activity polypeptide 2 | Plasma Membrane | other | 2.99E+07 | 3.05E+07 | 2.60E+07 | | | | | | |
| guanine nucleotide binding protein (G protein), alpha inhibiting activity polypeptide 2 | Plasma Membrane | other | 2.75E+07 | 3.48E+07 | 2.71E+07 | | | | | | |
| guanine nucleotide binding protein (G protein), alpha inhibiting activity polypeptide 2 | Plasma Membrane | other | 2.51E+07 | 2.89E+07 | 2.85E+07 | | | | | | |
| guanine nucleotide binding protein (G protein), alpha inhibiting activity polypeptide 2 | Plasma Membrane | other | 1.64E+07 | 1.83E+07 | 1.49E+07 | | | | | | |
| guanine nucleotide binding protein (G protein), alpha inhibiting activity polypeptide 2 | Plasma Membrane | other | 1.44E+07 | 1.02E+07 | 1.06E+07 | | | | | | |
| guanine nucleotide binding protein (G protein), alpha inhibiting activity polypeptide 2 | Plasma Membrane | other | 1.20E+07 | 9460516 | 1.10E+07 | | | | | | |
| guanine nucleotide binding protein (G protein), alpha inhibiting activity polypeptide 2 | Plasma Membrane | other | 1.18E+07 | 1.91E+07 | 1.72E+07 | | | | | | |
| guanine nucleotide binding protein (G protein), alpha inhibiting activity polypeptide 2 | Plasma Membrane | other | 1.17E+07 | 1.21E+07 | 8222354.5 | | | | | | |
| guanine nucleotide binding protein (G protein), alpha inhibiting activity polypeptide 2 | Plasma Membrane | other | 1.03E+07 | 1.17E+07 | 1.10E+07 | | | | | | |
| guanine nucleotide binding protein (G protein), alpha inhibiting activity polypeptide 2 | Plasma Membrane | other | 1.02E+07 | 4567577 | 1.04E+07 | | | | | | |
| guanine nucleotide binding protein (G protein), alpha inhibiting activity polypeptide 2 | Plasma Membrane | other | 9449670 | 9229309 | 7767087 | | | | | | |
| guanine nucleotide binding protein (G protein), alpha inhibiting activity polypeptide 2 | Plasma Membrane | other | 9441614 | 7991518.5 | 7507606 | | | | | | |
| guanine nucleotide binding protein (G protein), alpha inhibiting activity polypeptide 2 | Plasma Membrane | other | 9420435 | 1.50E+07 | 1.13E+07 | | | | | | |
| guanine nucleotide binding protein (G protein), alpha inhibiting activity polypeptide 2 | Plasma Membrane | other | 8769519 | 6406731.5 | 7061000.5 | | | | | | |
| guanine nucleotide binding protein (G protein), alpha inhibiting activity polypeptide 2 | Plasma Membrane | other | 8003666 | 5960301 | 4409770.5 | | | | | | |
| guanine nucleotide binding protein (G protein), alpha inhibiting activity polypeptide 2 | Plasma Membrane | other | 5602391.5 | 7711859 | 5860312.5 | | | | | | |
| guanine nucleotide binding protein (G protein), alpha inhibiting activity polypeptide 2 | Plasma Membrane | other | 3307013.75 | 8440064 | 4069510.5 | | | | | | |
| guanine nucleotide binding protein (G protein), alpha inhibiting activity polypeptide 2 | Plasma Membrane | other | 2784798.75 | 2529764.75 | 2052250.125 | | | | | | |
| guanine nucleotide binding protein (G protein), alpha inhibiting activity polypeptide 2 | Plasma Membrane | other | 1437207.125 | 3066358.25 | 2855734.5 | P08752 | GNAI2 | 22077551 | 17903553 | 2027178 | 78 |
| guanine nucleotide binding protein (G protein), alpha inhibiting activity polypeptide 3 | Cytoplasm | enzyme | 811886.812 | 1809655.625 | 2091243.625 | Q9DC51 | GNAI3 | 811887 | 672258 | 388129 | 3 |
| guanine nucleotide binding protein (G protein), q polypeptide | Plasma Membrane | enzyme | 1347370.5 | 8183264.5 | 6665861 | P21279 | GNAQ | 1347371 | 3589761 | 2072549 | 3 |
| GNAS complex locus | Plasma Membrane | enzyme | 3009337.75 | 3366207.25 | 3365580.75 | | | | | | |
| GNAS complex locus | Plasma Membrane | enzyme | 2798957 | 2967702.75 | 2464848 | Q6R0H7 | GNAS | 2904147 | 345129 | 140898 | 6 |
| guanine nucleotide binding protein (G protein), alpha z polypeptide | Plasma Membrane | enzyme | 331646.625 | 2705516.5 | 1872584.125 | | | | | | |
| guanine nucleotide binding protein (G protein), alpha z polypeptide | Plasma Membrane | enzyme | 4.70E-06 | 3003634.75 | 2559056 | O70443 | GNAZ | 165823 | 1282834 | 523715 | 6 |
| guanine nucleotide binding protein (G protein), beta polypeptide 1 | Plasma Membrane | enzyme | 5.32E+07 | 6.94E+07 | 5.51E+07 | | | | | | |
| guanine nucleotide binding protein (G protein), beta polypeptide 1 | Plasma Membrane | enzyme | 4.40E+07 | 4.08E+07 | 4.05E+07 | | | | | | |
| guanine nucleotide binding protein (G protein), beta polypeptide 1 | Plasma Membrane | enzyme | 2.33E+07 | 2.59E+07 | 1.80E+07 | | | | | | |
| guanine nucleotide binding protein (G protein), beta polypeptide 1 | Plasma Membrane | enzyme | 2.04E+07 | 2.76E+07 | 2.43E+07 | | | | | | |
| guanine nucleotide binding protein (G protein), beta polypeptide 1 | Plasma Membrane | enzyme | 1.47E+07 | 2.28E+07 | 1.46E+07 | | | | | | |
| guanine nucleotide binding protein (G protein), beta polypeptide 1 | Plasma Membrane | enzyme | 1.30E+07 | 1.46E+07 | 1.39E+07 | | | | | | |
| guanine nucleotide binding protein (G protein), beta polypeptide 1 | Plasma Membrane | enzyme | 1.22E+07 | 1.57E+07 | 1.08E+07 | | | | | | |
| guanine nucleotide binding protein (G protein), beta polypeptide 1 | Plasma Membrane | enzyme | 1.09E+07 | 1.33E+07 | 1.39E+07 | | | | | | |
| guanine nucleotide binding protein (G protein), beta polypeptide 1 | Plasma Membrane | enzyme | 8777910 | 1.23E+07 | 1.00E+07 | | | | | | |
| guanine nucleotide binding protein (G protein), beta polypeptide 1 | Plasma Membrane | enzyme | 7691058 | 9365907 | 7928350.5 | | | | | | |
| guanine nucleotide binding protein (G protein), beta polypeptide 1 | Plasma Membrane | enzyme | 4652577.5 | 1.13E+07 | 5888386 | | | | | | |
| guanine nucleotide binding protein (G protein), beta polypeptide 1 | Plasma Membrane | enzyme | 3416314.25 | 4213044 | 2590695.25 | | | | | | |
| guanine nucleotide binding protein (G protein), beta polypeptide 1 | Plasma Membrane | enzyme | 2182682.25 | 2476924.25 | 1956915.875 | P62874 | GNB1 | 16799272 | 15965815 | 2556576 | 39 |
| guanine nucleotide binding protein (G protein), beta polypeptide 2 | Plasma Membrane | enzyme | 3.88E+07 | 4.05E+07 | 4.61E+07 | | | | | | |
| guanine nucleotide binding protein (G protein), beta polypeptide 2 | Plasma Membrane | enzyme | 3.58E+07 | 3.76E+07 | 2.96E+07 | | | | | | |
| guanine nucleotide binding protein (G protein), beta polypeptide 2 | Plasma Membrane | enzyme | 3.33E+07 | 2.68E+07 | 3.73E+07 | | | | | | |
| guanine nucleotide binding protein (G protein), beta polypeptide 2 | Plasma Membrane | enzyme | 2.73E+07 | 3.27E+07 | 3.07E+07 | | | | | | |
| guanine nucleotide binding protein (G protein), beta polypeptide 2 | Plasma Membrane | enzyme | 2.19E+07 | 2.74E+07 | 2.25E+07 | | | | | | |
| guanine nucleotide binding protein (G protein), beta polypeptide 2 | Plasma Membrane | enzyme | 1.70E+07 | 2.19E+07 | 2.23E+07 | | | | | | |
| guanine nucleotide binding protein (G protein), beta polypeptide 2 | Plasma Membrane | enzyme | 9510771 | 9639778 | 1.05E+07 | | | | | | |
| guanine nucleotide binding protein (G protein), beta polypeptide 2 | Plasma Membrane | enzyme | 8517166 | 1.18E+07 | 1.03E+07 | | | | | | |
| guanine nucleotide binding protein (G protein), beta polypeptide 2 | Plasma Membrane | enzyme | 7858376 | 8735884 | 6692451 | | | | | | |
| guanine nucleotide binding protein (G protein), beta polypeptide 2 | Plasma Membrane | enzyme | 4616051 | 2984391.5 | 5479808.5 | | | | | | |
| guanine nucleotide binding protein (G protein), beta polypeptide 2 | Plasma Membrane | enzyme | 2565992.75 | 1933908.375 | 2430163.5 | | | | | | |
| guanine nucleotide binding protein (G protein), beta polypeptide 2 | Plasma Membrane | enzyme | 1610402.125 | 1682432.75 | 1675453.125 | P62880 | GNB2 | 17398230 | 13802409 | 2300402 | 36 |
| guanine nucleotide binding protein (G protein), beta polypeptide 2-like 1 | Cytoplasm | enzyme | 4836022 | 4085887 | 2870817 | | | | | | |
| guanine nucleotide binding protein (G protein), beta polypeptide 2-like 1 | Cytoplasm | enzyme | 3818384.25 | 2142283.5 | 3266049.5 | | | | | | |
| guanine nucleotide binding protein (G protein), beta polypeptide 2-like 1 | Cytoplasm | enzyme | 2693523.5 | 2701414.75 | 2624884.25 | | | | | | |
| guanine nucleotide binding protein (G protein), beta polypeptide 2-like 1 | Cytoplasm | enzyme | 2389175 | 2349702.75 | 2043192.625 | | | | | | |
| guanine nucleotide binding protein (G protein), beta polypeptide 2-like 1 | Cytoplasm | enzyme | 1695696.125 | 1946661.125 | 1292361.875 | P68040 | GNB2L1 | 3086560 | 948846 | 244991 | 15 |

| | | | | | | | | | | | |
|---|---|---|---|---|---|---|---|---|---|---|---|
| guanine nucleotide binding protein (G protein), gamma 12 | Plasma Membrane | enzyme | 6504702 | 5786938.5 | 6064685 | Q9DAS9 | GNG12 | 6504702 | 361926 | 208958 | 3 |
| golgin A7 | Cytoplasm | other | 4901208 | 4390590.5 | 3277616.5 | | | | | | |
| golgin A7 | Cytoplasm | other | 3866620 | 3620729.25 | 2997647.75 | Q91W53 | GOLGA7 | 4383914 | 707658 | 288900 | 6 |
| golgi transport 1B | Cytoplasm | other | 3371519.75 | 3223788.25 | 3306262.5 | Q9CR60 | GOLT1B | 3371520 | 74033 | 42743 | 3 |
| glutamic-oxaloacetic transaminase 2, mitochondrial | Cytoplasm | enzyme | 9908029 | 1.72E+07 | 1.66E+07 | | | | | | |
| glutamic-oxaloacetic transaminase 2, mitochondrial | Cytoplasm | enzyme | 8192478 | 1.12E+07 | 1.00E+07 | | | | | | |
| glutamic-oxaloacetic transaminase 2, mitochondrial | Cytoplasm | enzyme | 6297962.5 | 1.05E+07 | 8074028.5 | | | | | | |
| glutamic-oxaloacetic transaminase 2, mitochondrial | Cytoplasm | enzyme | 5643041 | 8647393 | 7105580 | | | | | | |
| glutamic-oxaloacetic transaminase 2, mitochondrial | Cytoplasm | enzyme | 4658204 | 5878856.5 | 5357406 | | | | | | |
| glutamic-oxaloacetic transaminase 2, mitochondrial | Cytoplasm | enzyme | 4101716.75 | 5464433.5 | 5440869 | | | | | | |
| glutamic-oxaloacetic transaminase 2, mitochondrial | Cytoplasm | enzyme | 2651984 | 4144983.25 | 4338037.5 | | | | | | |
| glutamic-oxaloacetic transaminase 2, mitochondrial | Cytoplasm | enzyme | 2044601.125 | 1900835 | 1501584.5 | | | | | | |
| glutamic-oxaloacetic transaminase 2, mitochondrial | Cytoplasm | enzyme | 1944807.5 | 1574925.875 | 1272864.75 | | | | | | |
| glutamic-oxaloacetic transaminase 2, mitochondrial | Cytoplasm | enzyme | 724007.375 | 454121.156 | 1096210.125 | | | | | | |
| glutamic-oxaloacetic transaminase 2, mitochondrial | Cytoplasm | enzyme | 504282.062 | 955753.25 | 997045.438 | P05202 | GOT2 | 4242828 | 4392931 | 764711 | 33 |
| glycoprotein Ib (platelet), alpha polypeptide | Plasma Membrane | other | 4155015.75 | 5.98E+07 | 3.99E+07 | | | | | | |
| glycoprotein Ib (platelet), alpha polypeptide | Plasma Membrane | other | 2646691.25 | 8.04E+07 | 4.88E+07 | | | | | | |
| glycoprotein Ib (platelet), alpha polypeptide | Plasma Membrane | other | 1784950.25 | 4.59E+07 | 2.57E+07 | | | | | | |
| glycoprotein Ib (platelet), alpha polypeptide | Plasma Membrane | other | 1715605.375 | 3.52E+07 | 1.98E+07 | | | | | | |
| glycoprotein Ib (platelet), alpha polypeptide | Plasma Membrane | other | 1414518.5 | 3.06E+07 | 1.69E+07 | | | | | | |
| glycoprotein Ib (platelet), alpha polypeptide | Plasma Membrane | other | 1080772.625 | 3.29E+07 | 1.81E+07 | | | | | | |
| glycoprotein Ib (platelet), alpha polypeptide | Plasma Membrane | other | 871186.438 | 1.82E+07 | 1.14E+07 | | | | | | |
| glycoprotein Ib (platelet), alpha polypeptide | Plasma Membrane | other | 681599.375 | 8561611 | 6503274.5 | | | | | | |
| glycoprotein Ib (platelet), alpha polypeptide | Plasma Membrane | other | 264890.906 | 6007563 | 4221284.5 | | | | | | |
| glycoprotein Ib (platelet), alpha polypeptide | Plasma Membrane | other | 182231.188 | 9387756 | 5261417.5 | | | | | | |
| glycoprotein Ib (platelet), alpha polypeptide | Plasma Membrane | other | 170889.391 | 5801996.5 | 4070539.75 | | | | | | |
| glycoprotein Ib (platelet), alpha polypeptide | Plasma Membrane | other | 91254.516 | 1.14E+07 | 6079183.5 | | | | | | |
| glycoprotein Ib (platelet), alpha polypeptide | Plasma Membrane | other | 4.70E-06 | 560866.625 | 11732.813 | O35930 | GP1BA | 1158431 | 19062408 | 3052428 | 39 |
| glycoprotein Ib (platelet), beta polypeptide | Plasma Membrane | other | 7593426.5 | 1.08E+08 | 7.54E+07 | | | | | | |
| glycoprotein Ib (platelet), beta polypeptide | Plasma Membrane | other | 2800151.75 | 5.85E+07 | 3.24E+07 | | | | | | |
| glycoprotein Ib (platelet), beta polypeptide | Plasma Membrane | other | 1944754.25 | 3.77E+07 | 1.87E+07 | | | | | | |
| glycoprotein Ib (platelet), beta polypeptide | Plasma Membrane | other | 1715008.5 | 4.79E+07 | 2.58E+07 | | | | | | |
| glycoprotein Ib (platelet), beta polypeptide | Plasma Membrane | other | 703166.562 | 1.28E+07 | 1.05E+07 | | | | | | |
| glycoprotein Ib (platelet), beta polypeptide | Plasma Membrane | other | 538006.562 | 1.92E+07 | 9553893 | P56400 | GP1BB | 2549086 | 29748226 | 7011724 | 18 |
| glycoprotein V (platelet) | Plasma Membrane | other | 3550158 | 5.00E+07 | 3.77E+07 | | | | | | |
| glycoprotein V (platelet) | Plasma Membrane | other | 2669800.25 | 4.96E+07 | 3.38E+07 | | | | | | |
| glycoprotein V (platelet) | Plasma Membrane | other | 2621001.25 | 1.95E+07 | 1.17E+07 | | | | | | |
| glycoprotein V (platelet) | Plasma Membrane | other | 2574104.25 | 3.64E+07 | 2.88E+07 | | | | | | |
| glycoprotein V (platelet) | Plasma Membrane | other | 2531511.75 | 2.85E+07 | 2.04E+07 | | | | | | |
| glycoprotein V (platelet) | Plasma Membrane | other | 1888041.125 | 3.83E+07 | 2.42E+07 | | | | | | |
| glycoprotein V (platelet) | Plasma Membrane | other | 665662.062 | 1.21E+07 | 8698100 | | | | | | |
| glycoprotein V (platelet) | Plasma Membrane | other | 388846.156 | 1.13E+07 | 7030608.5 | | | | | | |
| glycoprotein V (platelet) | Plasma Membrane | other | 206316.234 | 1.03E+07 | 7534518 | | | | | | |
| glycoprotein V (platelet) | Plasma Membrane | other | 131851.938 | 5393177 | 4392051 | O08742 | GP5 | 1722729 | 15520554 | 2833653 | 30 |
| glycoprotein IX (platelet) | Plasma Membrane | other | 4147555.25 | 1.01E+08 | 6.44E+07 | | | | | | |
| glycoprotein IX (platelet) | Plasma Membrane | other | 3214461.75 | 4.14E+07 | 1.86E+07 | | | | | | |
| glycoprotein IX (platelet) | Plasma Membrane | other | 138723.484 | 8444764 | 4360707.5 | | | | | | |
| glycoprotein IX (platelet) | Plasma Membrane | other | 74459.758 | 8123571 | 3884951.75 | O88186 | GP9 | 1893800 | 31678674 | 9144845 | 12 |
| glypican 4 | Plasma Membrane | transmembrane receptor | 3658246.75 | 3712595.5 | 4919034 | | | | | | |
| glypican 4 | Plasma Membrane | transmembrane receptor | 2755510.5 | 2416485 | 3517976.5 | | | | | | |
| glypican 4 | Plasma Membrane | transmembrane receptor | 2000883.125 | 2133433 | 2520007.75 | P51655 | GPC4 | 2804880 | 950475 | 316825 | 9 |
| glycerol-3-phosphate dehydrogenase 2 (mitochondrial) | Cytoplasm | enzyme | 931805.25 | 1.03E+07 | 6629865 | | | | | | |
| glycerol-3-phosphate dehydrogenase 2 (mitochondrial) | Cytoplasm | enzyme | 841204.25 | 8348254 | 7237123 | | | | | | |
| glycerol-3-phosphate dehydrogenase 2 (mitochondrial) | Cytoplasm | enzyme | 579757.938 | 3970214.25 | 3018026.5 | | | | | | |
| glycerol-3-phosphate dehydrogenase 2 (mitochondrial) | Cytoplasm | enzyme | 522007.688 | 7815090 | 5675479.5 | | | | | | |
| glycerol-3-phosphate dehydrogenase 2 (mitochondrial) | Cytoplasm | enzyme | 514841.906 | 894145.625 | 857269.875 | | | | | | |
| glycerol-3-phosphate dehydrogenase 2 (mitochondrial) | Cytoplasm | enzyme | 468800.062 | 5315251 | 3958007 | | | | | | |
| glycerol-3-phosphate dehydrogenase 2 (mitochondrial) | Cytoplasm | enzyme | 396316.969 | 4149933 | 3258671.75 | | | | | | |
| glycerol-3-phosphate dehydrogenase 2 (mitochondrial) | Cytoplasm | enzyme | 160490.938 | 3212734.75 | 2205379.25 | | | | | | |
| glycerol-3-phosphate dehydrogenase 2 (mitochondrial) | Cytoplasm | enzyme | 85352.125 | 3553365.25 | 3163717 | | | | | | |
| glycerol-3-phosphate dehydrogenase 2 (mitochondrial) | Cytoplasm | enzyme | 4.70E-06 | 4086799.5 | 3935774 | Q64521 | GPD2 | 450058 | 2803827 | 511906 | 30 |
| G protein pathway suppressor 1 | Cytoplasm | other | 9901736 | 3244834 | 4803841 | | | | | | |
| G protein pathway suppressor 1 | Cytoplasm | other | 7813949.5 | 5217383 | 4522282.5 | | | | | | |
| G protein pathway suppressor 1 | Cytoplasm | other | 4098971.5 | 2336550.5 | 2018417 | | | | | | |
| G protein pathway suppressor 1 | Cytoplasm | other | 2588005 | 1035248.562 | 1442561.25 | | | | | | |
| G protein pathway suppressor 1 | Cytoplasm | other | 1337471.125 | 322727.531 | 379102.25 | Q99LD4 | GPS1 | 5148027 | 2735653 | 706343 | 15 |
| glutathione peroxidase 1 | Cytoplasm | enzyme | 3.98E+07 | 2.43E+07 | 3.01E+07 | | | | | | |
| glutathione peroxidase 1 | Cytoplasm | enzyme | 2.40E+07 | 1.53E+07 | 2.20E+07 | | | | | | |
| glutathione peroxidase 1 | Cytoplasm | enzyme | 2.33E+07 | 1.12E+07 | 1.56E+07 | | | | | | |
| glutathione peroxidase 1 | Cytoplasm | enzyme | 1.44E+07 | 8344894 | 1.02E+07 | | | | | | |
| glutathione peroxidase 1 | Cytoplasm | enzyme | 9057004 | 4969978.5 | 6301725.5 | | | | | | |
| glutathione peroxidase 1 | Cytoplasm | enzyme | 8471247 | 5408339 | 7844776.5 | | | | | | |
| glutathione peroxidase 1 | Cytoplasm | enzyme | 6440149 | 3159372.25 | 4954974 | | | | | | |
| glutathione peroxidase 1 | Cytoplasm | enzyme | 5415558.5 | 1243564.5 | 3484794.25 | | | | | | |
| glutathione peroxidase 1 | Cytoplasm | enzyme | 4011001.5 | 1119299.875 | 2600788.75 | | | | | | |
| glutathione peroxidase 1 | Cytoplasm | enzyme | 3623874 | 2196052.75 | 3664850.5 | P11352 | GPX1 | 13843883 | 9611987 | 1754901 | 30 |
| glutathione peroxidase 4 | Cytoplasm | enzyme | 3.75E+07 | 3.93E+07 | 3.77E+07 | | | | | | |
| glutathione peroxidase 4 | Cytoplasm | enzyme | 3.35E+07 | 3.36E+07 | 2.98E+07 | | | | | | |
| glutathione peroxidase 4 | Cytoplasm | enzyme | 2.20E+07 | 1.63E+07 | 1.66E+07 | | | | | | |
| glutathione peroxidase 4 | Cytoplasm | enzyme | 1.97E+07 | 1.79E+07 | 1.50E+07 | | | | | | |
| glutathione peroxidase 4 | Cytoplasm | enzyme | 1.73E+07 | 1.77E+07 | 1.26E+07 | | | | | | |
| glutathione peroxidase 4 | Cytoplasm | enzyme | 1.05E+07 | 2362074 | 4929494.5 | | | | | | |
| glutathione peroxidase 4 | Cytoplasm | enzyme | 3630550.75 | 3014266.25 | 2418034.25 | O70325 | GPX4 | 20597222 | 12325609 | 2689668 | 21 |
| G1 to S phase transition 1 | Cytoplasm | translation regulator | 4293038 | 1305649.75 | 2005961.875 | Q8R050 | GSPT1 | 4293038 | 1562351 | 902024 | 3 |
| glutathione S-transferase pi 1 | Cytoplasm | enzyme | 1704730 | 1100140.875 | 1714233.125 | P19157 | GSTP1 | 1704730 | 351835 | 203132 | 3 |
| guanylyl cyclase domain containing 1 | Nucleus | other | 2072686 | 2254874.75 | 2176326.5 | Q8BZI6 | GUCD1 | 2072686 | 91382 | 52759 | 3 |
| glycophorin A | Plasma Membrane | other | 7.01E+09 | 3.75E+09 | 4.04E+09 | | | | | | |
| glycophorin A | Plasma Membrane | other | 3.69E+07 | 4.53E+07 | 3.11E+07 | P14220 | Gypa | 3521970000 | 2912563827 | 1189049203 | 6 |
| glycophorin C (Gerbich blood group) | Plasma Membrane | other | 1.19E+09 | 1.61E+09 | 1.30E+09 | | | | | | |
| glycophorin C (Gerbich blood group) | Plasma Membrane | other | 9.92E+08 | 5.21E+08 | 7.74E+08 | | | | | | |
| glycophorin C (Gerbich blood group) | Plasma Membrane | other | 5.36E+07 | 6.37E+07 | 6.72E+07 | Q78HU7 | GYPC | 743700000 | 587439847 | 195813282 | 9 |
| H2A histone family, member V | Nucleus | other | 9137.851 | 1900550.875 | 2326950 | Q3THW5 | H2AFV | 9138 | 1233661 | 712254 | 3 |
| H2A histone family, member X | Nucleus | transcription regulator | 1002475.688 | 5055619 | 6859908.5 | | | | | | |
| H2A histone family, member X | Nucleus | transcription regulator | 0 | 970586.625 | 1040412.938 | P27661 | H2AFX | 501238 | 2774932 | 1132861 | 6 |
| H2A histone family, member Y | Nucleus | other | 19064.715 | 1318064.375 | 2120048.5 | Q9QZQ8 | H2AFY | 19065 | 1060245 | 612132 | 3 |

| | | | | | | | | | | | |
|---|---|---|---|---|---|---|---|---|---|---|---|
| Histone H3.1 | Nucleus | | 1.08E+07 | 5.22E+07 | 7.33E+07 | | | | | | |
| Histone H3.1 | Nucleus | | 7638472.5 | 5.24E+07 | 6.04E+07 | | | | | | |
| Histone H3.1 | Nucleus | | 5965928.5 | 3.65E+07 | 2.35E+07 | | | | | | |
| Histone H3.1 | Nucleus | | 3604324.5 | 2.11E+07 | 3.26E+07 | | | | | | |
| Histone H3.1 | Nucleus | | 1577411.75 | 5377563 | 6790167.5 | | | | | | |
| Histone H3.1 | Nucleus | | 542923.438 | 1.20E+07 | 2.12E+07 | P68433 | H31 | 5013177 | 22527915 | 5309880 | 18 |
| Histone H4 | Nucleus | | 2.14E+07 | 1.02E+08 | 1.63E+08 | | | | | | |
| Histone H4 | Nucleus | | 1.96E+07 | 1.35E+08 | 1.95E+08 | | | | | | |
| Histone H4 | Nucleus | | 9019698 | 4.78E+07 | 4.74E+07 | | | | | | |
| Histone H4 | Nucleus | | 5699958.5 | 2.98E+07 | 4.95E+07 | | | | | | |
| Histone H4 | Nucleus | | 2580206.75 | 5991088 | 1.54E+07 | | | | | | |
| Histone H4 | Nucleus | | 913125.125 | 1.32E+07 | 3.37E+07 | | | | | | |
| Histone H4 | Nucleus | | 4.70E-06 | 181482.484 | 155744.641 | | | | | | |
| Histone H4 | Nucleus | | 4.70E-06 | 672203.312 | 1240838.25 | P62806 | H4 | 7401624 | 55183955 | 11264378 | 24 |
| HECT domain and ankyrin repeat containing E3 ubiquitin protein ligase 1 | Cytoplasm | enzyme | 1575575 | 703215.375 | 648861.5 | Q3U0D9 | HACE1 | 1575575 | 520058 | 300256 | 3 |
| hydroxyacyl-CoA dehydrogenase | Cytoplasm | enzyme | 413584.562 | 4014078.75 | 2546149.25 | | | | | | |
| hydroxyacyl-CoA dehydrogenase | Cytoplasm | enzyme | 49946.273 | 649326.562 | 622225.312 | Q61425 | HADH | 231765 | 1556235 | 635330 | 6 |
| hydroxyacyl-CoA dehydrogenase/3-ketoacyl-CoA thiolase/enoyl-CoA hydratase (trifunctional protein), alpha subunit | Cytoplasm | enzyme | 4188537.75 | 5164313.5 | 5612911.5 | | | | | | |
| hydroxyacyl-CoA dehydrogenase/3-ketoacyl-CoA thiolase/enoyl-CoA hydratase (trifunctional protein), alpha subunit | Cytoplasm | enzyme | 3276329 | 4899367.5 | 3885296.25 | | | | | | |
| hydroxyacyl-CoA dehydrogenase/3-ketoacyl-CoA thiolase/enoyl-CoA hydratase (trifunctional protein), alpha subunit | Cytoplasm | enzyme | 3154671 | 4808833.5 | 3860758.5 | | | | | | |
| hydroxyacyl-CoA dehydrogenase/3-ketoacyl-CoA thiolase/enoyl-CoA hydratase (trifunctional protein), alpha subunit | Cytoplasm | enzyme | 2292146.5 | 3061828.5 | 2925345.25 | | | | | | |
| hydroxyacyl-CoA dehydrogenase/3-ketoacyl-CoA thiolase/enoyl-CoA hydratase (trifunctional protein), alpha subunit | Cytoplasm | enzyme | 2138535.5 | 3746934.75 | 4250115.5 | | | | | | |
| hydroxyacyl-CoA dehydrogenase/3-ketoacyl-CoA thiolase/enoyl-CoA hydratase (trifunctional protein), alpha subunit | Cytoplasm | enzyme | 1999206.25 | 2686418.25 | 3096416.75 | | | | | | |
| hydroxyacyl-CoA dehydrogenase/3-ketoacyl-CoA thiolase/enoyl-CoA hydratase (trifunctional protein), alpha subunit | Cytoplasm | enzyme | 1741813 | 2979394 | 2926416.25 | | | | | | |
| hydroxyacyl-CoA dehydrogenase/3-ketoacyl-CoA thiolase/enoyl-CoA hydratase (trifunctional protein), alpha subunit | Cytoplasm | enzyme | 1680700.5 | 2399652.75 | 2463049 | | | | | | |
| hydroxyacyl-CoA dehydrogenase/3-ketoacyl-CoA thiolase/enoyl-CoA hydratase (trifunctional protein), alpha subunit | Cytoplasm | enzyme | 1579844.5 | 2307565.25 | 3120522.5 | | | | | | |
| hydroxyacyl-CoA dehydrogenase/3-ketoacyl-CoA thiolase/enoyl-CoA hydratase (trifunctional protein), alpha subunit | Cytoplasm | enzyme | 1403263.875 | 2320200 | 1936851.125 | | | | | | |
| hydroxyacyl-CoA dehydrogenase/3-ketoacyl-CoA thiolase/enoyl-CoA hydratase (trifunctional protein), alpha subunit | Cytoplasm | enzyme | 927043.188 | 1279304.875 | 1454873.375 | | | | | | |
| hydroxyacyl-CoA dehydrogenase/3-ketoacyl-CoA thiolase/enoyl-CoA hydratase (trifunctional protein), alpha subunit | Cytoplasm | enzyme | 705899.5 | 1404168.75 | 876292.562 | | | | | | |
| hydroxyacyl-CoA dehydrogenase/3-ketoacyl-CoA thiolase/enoyl-CoA hydratase (trifunctional protein), alpha subunit | Cytoplasm | enzyme | 573522.25 | 984044.938 | 1223675.25 | | | | | | |
| hydroxyacyl-CoA dehydrogenase/3-ketoacyl-CoA thiolase/enoyl-CoA hydratase (trifunctional protein), alpha subunit | Cytoplasm | enzyme | 241061 | 187815.531 | 252241.484 | Q8BMS1 | HADHA | 1850184 | 1405477 | 216870 | 42 |
| hydroxyacyl-CoA dehydrogenase/3-ketoacyl-CoA thiolase/enoyl-CoA hydratase (trifunctional protein), beta subunit | Cytoplasm | enzyme | 5476782 | 4629221.5 | 5436382.5 | | | | | | |
| hydroxyacyl-CoA dehydrogenase/3-ketoacyl-CoA thiolase/enoyl-CoA hydratase (trifunctional protein), beta subunit | Cytoplasm | enzyme | 4082829 | 7031192 | 5903114 | | | | | | |
| hydroxyacyl-CoA dehydrogenase/3-ketoacyl-CoA thiolase/enoyl-CoA hydratase (trifunctional protein), beta subunit | Cytoplasm | enzyme | 3606348 | 5050522 | 4603847 | | | | | | |
| hydroxyacyl-CoA dehydrogenase/3-ketoacyl-CoA thiolase/enoyl-CoA hydratase (trifunctional protein), beta subunit | Cytoplasm | enzyme | 2050701.125 | 8120630.5 | 2820124.5 | | | | | | |
| hydroxyacyl-CoA dehydrogenase/3-ketoacyl-CoA thiolase/enoyl-CoA hydratase (trifunctional protein), beta subunit | Cytoplasm | enzyme | 956529 | 1302231.875 | 1215549.25 | | | | | | |
| hydroxyacyl-CoA dehydrogenase/3-ketoacyl-CoA thiolase/enoyl-CoA hydratase (trifunctional protein), beta subunit | Cytoplasm | enzyme | 868024.938 | 2524612.75 | 1657081.375 | | | | | | |
| hydroxyacyl-CoA dehydrogenase/3-ketoacyl-CoA thiolase/enoyl-CoA hydratase (trifunctional protein), beta subunit | Cytoplasm | enzyme | 823385.062 | 1436140.375 | 1825578.875 | Q99JY0 | HADHB | 2552086 | 2200886 | 480273 | 21 |
| hydroxyacylglutathione hydrolase | Cytoplasm | enzyme | 3703110.75 | 2436869.75 | 2750989.25 | | | | | | |
| hydroxyacylglutathione hydrolase | Cytoplasm | enzyme | 795572.062 | 1091779.5 | 1387955.625 | | | | | | |
| hydroxyacylglutathione hydrolase | Cytoplasm | enzyme | 115903.922 | 149371.547 | 127553.711 | Q99KB8 | HAGH | 1538196 | 1298893 | 432964 | 9 |
| hemoglobin, alpha 1 | Extracellular Space | transporter | 2.47E+09 | 1.64E+09 | 9.71E+08 | | | | | | |
| hemoglobin, alpha 1 | Extracellular Space | transporter | 2.15E+09 | 2.16E+09 | 1.15E+09 | | | | | | |
| hemoglobin, alpha 1 | Extracellular Space | transporter | 1.45E+09 | 1.72E+09 | 8.88E+08 | | | | | | |
| hemoglobin, alpha 1 | Extracellular Space | transporter | 1.34E+09 | 3.82E+09 | 1.07E+09 | | | | | | |
| hemoglobin, alpha 1 | Extracellular Space | transporter | 1.31E+09 | 1.55E+09 | 1.04E+09 | | | | | | |
| hemoglobin, alpha 1 | Extracellular Space | transporter | 1.15E+09 | 1.42E+09 | 8.00E+08 | | | | | | |
| hemoglobin, alpha 1 | Extracellular Space | transporter | 4.00E+08 | 1.01E+09 | 2.75E+08 | | | | | | |
| hemoglobin, alpha 1 | Extracellular Space | transporter | 2.22E+08 | 1.68E+08 | 1.54E+08 | | | | | | |
| hemoglobin, alpha 1 | Extracellular Space | transporter | 2.21E+08 | 3.16E+08 | 2.92E+08 | | | | | | |
| hemoglobin, alpha 1 | Extracellular Space | transporter | 1.82E+08 | 1.19E+08 | 1.07E+08 | | | | | | |
| hemoglobin, alpha 1 | Extracellular Space | transporter | 1.69E+08 | 1.13E+08 | 7.49E+07 | P01942 | HBA1/HBA2 | 1005672727 | 860790764 | 149844438 | 33 |
| hemoglobin, beta adult major chain | Extracellular Space | transporter | 3.13E+09 | 3.23E+09 | 2.24E+09 | | | | | | |
| hemoglobin, beta adult major chain | Extracellular Space | transporter | 2.68E+09 | 2.76E+09 | 1.55E+09 | | | | | | |
| hemoglobin, beta adult major chain | Extracellular Space | transporter | 2.36E+09 | 2.66E+09 | 1.93E+09 | | | | | | |
| hemoglobin, beta adult major chain | Extracellular Space | transporter | 2.25E+09 | 1.81E+09 | 1.34E+09 | | | | | | |
| hemoglobin, beta adult major chain | Extracellular Space | transporter | 1.78E+09 | 2.83E+09 | 1.37E+08 | | | | | | |
| hemoglobin, beta adult major chain | Extracellular Space | transporter | 1.76E+09 | 2.86E+09 | 9.25E+08 | | | | | | |
| hemoglobin, beta adult major chain | Extracellular Space | transporter | 1.54E+09 | 2.05E+09 | 1.25E+08 | | | | | | |
| hemoglobin, beta adult major chain | Extracellular Space | transporter | 1.29E+09 | 1.55E+09 | 7.20E+08 | | | | | | |
| hemoglobin, beta adult major chain | Extracellular Space | transporter | 6.04E+08 | 9.99E+08 | 4.62E+07 | | | | | | |
| hemoglobin, beta adult major chain | Extracellular Space | transporter | 5.07E+08 | 5.91E+08 | 3.63E+08 | | | | | | |
| hemoglobin, beta adult major chain | Extracellular Space | transporter | 2.89E+08 | 2.29E+08 | 1.62E+08 | | | | | | |
| hemoglobin, beta adult major chain | Extracellular Space | transporter | 2.12E+08 | 2.58E+08 | 2.05E+08 | | | | | | |
| hemoglobin, beta adult major chain | Extracellular Space | transporter | 1.24E+08 | 5.87E+07 | 6.96E+07 | | | | | | |
| hemoglobin, beta adult major chain | Extracellular Space | transporter | 8.59E+07 | 1.06E+08 | 1.54E+08 | | | | | | |
| hemoglobin, beta adult major chain | Extracellular Space | transporter | 3.67E+07 | 1.10E+08 | 4.48E+07 | | | | | | |
| hemoglobin, beta adult major chain | Extracellular Space | transporter | 1.49E+07 | 2.41E+07 | 1.20E+07 | | | | | | |
| hemoglobin, beta adult major chain | Extracellular Space | transporter | 9799074 | 6531341 | 1.07E+07 | | | | | | |
| hemoglobin, beta adult major chain | Extracellular Space | transporter | 5736652.5 | 6184548 | 6874867.5 | | | | | | |
| hemoglobin, beta adult major chain | Extracellular Space | transporter | 2142712.25 | 8284761 | 2800137 | P02088 | Hbb-b1 | 983157813 | 1039775856 | 137721677 | 57 |
| hemoglobin, beta adult minor chain | Extracellular Space | other | 4.20E+08 | 5.50E+08 | 3.36E+07 | | | | | | |
| hemoglobin, beta adult minor chain | Extracellular Space | other | 3.30E+08 | 4.46E+08 | 2.67E+07 | | | | | | |
| hemoglobin, beta adult minor chain | Extracellular Space | other | 2.76E+08 | 3.19E+08 | 2.64E+07 | | | | | | |
| hemoglobin, beta adult minor chain | Extracellular Space | other | 9.35E+07 | 1.97E+08 | 7414173 | | | | | | |
| hemoglobin, beta adult minor chain | Extracellular Space | other | 6.87E+07 | 1.68E+08 | 8631263 | | | | | | |
| hemoglobin, beta adult minor chain | Extracellular Space | other | 4.50E+07 | 4.94E+07 | 5470733.5 | | | | | | |
| hemoglobin, beta adult minor chain | Extracellular Space | other | 2.21E+07 | 1.37E+07 | 838473.312 | | | | | | |
| hemoglobin, beta adult minor chain | Extracellular Space | other | 983132.375 | 1300754.875 | 187932.297 | P02089 | Hbb-b2 | 157146642 | 168123696 | 34318106 | 24 |
| HCK proto-oncogene, Src family tyrosine kinase | Cytoplasm | kinase | 162797.344 | 1180737.5 | 1091383.875 | P08103 | HCK | 162797 | 563687 | 325445 | 3 |
| HECT domain containing E3 ubiquitin protein ligase 3 | Cytoplasm | enzyme | 4022663.75 | 1657754.125 | 1474958 | | | | | | |
| HECT domain containing E3 ubiquitin protein ligase 3 | Cytoplasm | enzyme | 957018.062 | 322876.75 | 491464.406 | Q3U487 | HECTD3 | 2489841 | 1348039 | 550335 | 6 |
| histone cluster 1, H1a | Nucleus | other | 1246035.125 | 4258534 | 4760255.5 | | | | | | |
| histone cluster 1, H1a | Nucleus | other | 514237.625 | 1436217 | 2540163.5 | | | | | | |
| histone cluster 1, H1a | Nucleus | other | 34301.934 | 2506449 | 4429770 | | | | | | |
| histone cluster 1, H1a | Nucleus | other | 0 | 53889.191 | 270500.438 | P43275 | Hist1h1a | 448644 | 1823871 | 526506 | 12 |
| histone cluster 1, H1b | Nucleus | other | 111567.305 | 7882193.5 | 1.24E+07 | | | | | | |
| histone cluster 1, H1b | Nucleus | other | 0 | 284804.469 | 354510.156 | P43276 | Hist1h1b | 55784 | 5336225 | 2178505 | 6 |
| histone cluster 1, H1c | Nucleus | other | 4455556.5 | 6.37E+07 | 8.45E+07 | P15864 | HIST1H1C | 4455557 | 41534564 | 23979992 | 3 |
| histone cluster 1, H1e | Nucleus | other | 2500477.75 | 4.03E+07 | 5.39E+07 | | | | | | |
| histone cluster 1, H1e | Nucleus | other | 941377.062 | 1.09E+07 | 1.95E+07 | | | | | | |
| histone cluster 1, H1e | Nucleus | other | 590612.188 | 2.01E+07 | 2.62E+07 | | | | | | |
| histone cluster 1, H1e | Nucleus | other | 168449.562 | 1.41E+07 | 1.52E+07 | | | | | | |
| histone cluster 1, H1e | Nucleus | other | 4.70E-06 | 78435.844 | 91983.461 | P43274 | Hist1h1e | 840183 | 16387899 | 4231337 | 15 |
| histone cluster 1, H2aj | Nucleus | other | 2.50E+07 | 1.44E+08 | 1.97E+08 | | | | | | |

| | | | | | | | | | | | |
|---|---|---|---|---|---|---|---|---|---|---|---|
| histone cluster 1, H2aj | Nucleus | other | 2.17E+07 | 1.21E+08 | 1.82E+08 | | | | | | |
| histone cluster 1, H2aj | Nucleus | other | 1.07E+07 | 6.14E+07 | 8.01E+07 | | | | | | |
| histone cluster 1, H2aj | Nucleus | other | 5937967 | 2.40E+07 | 4.44E+07 | | | | | | |
| histone cluster 1, H2aj | Nucleus | other | 87322.297 | 405688.312 | 771624.625 | Q8CGP5 | HIST1H2AJ | 12661058 | 68243546 | 17620408 | 15 |
| histone cluster 1, H2ba | Nucleus | other | 2.05E+07 | 1.23E+08 | 1.59E+08 | P70696 | HIST1H2BA | 20470000 | 71928670 | 41528037 | 3 |
| histone cluster 1, H2bb | Nucleus | other | 1.63E+07 | 1.04E+08 | 1.27E+08 | | | | | | |
| histone cluster 1, H2bb | Nucleus | other | 1.34E+07 | 8.69E+07 | 1.30E+08 | | | | | | |
| histone cluster 1, H2bb | Nucleus | other | 1255969.875 | 4659631 | 7469301 | | | | | | |
| histone cluster 1, H2bb | Nucleus | other | 1218408.125 | 1269898.375 | 1909459.75 | | | | | | |
| histone cluster 1, H2bb | Nucleus | other | 525146.125 | 3265834.25 | 3421457.25 | | | | | | |
| histone cluster 1, H2bb | Nucleus | other | 69376.305 | 1506886.625 | 2240874 | Q64475 | HIST1H2BB | 5461483 | 47028448 | 11084711 | 18 |
| histone cluster 2, H2ab | Nucleus | other | 952978.875 | 4760087.5 | 6637428.5 | Q64522 | HIST2H2AB | 952979 | 2896304 | 1672182 | 3 |
| hexokinase 1 | Cytoplasm | kinase | 1994866.625 | 1.70E+07 | 1.08E+07 | | | | | | |
| hexokinase 1 | Cytoplasm | kinase | 1504794.5 | 8273351.5 | 5889428.5 | | | | | | |
| hexokinase 1 | Cytoplasm | kinase | 1450742.875 | 7227266 | 4401408 | | | | | | |
| hexokinase 1 | Cytoplasm | kinase | 1252294.375 | 5972444 | 4122259.25 | | | | | | |
| hexokinase 1 | Cytoplasm | kinase | 920609.125 | 8625723 | 4423819.5 | | | | | | |
| hexokinase 1 | Cytoplasm | kinase | 807702.688 | 7020091.5 | 3758726.25 | | | | | | |
| hexokinase 1 | Cytoplasm | kinase | 744035.125 | 7063577.5 | 5220921.5 | | | | | | |
| hexokinase 1 | Cytoplasm | kinase | 578408.375 | 4505619.5 | 3141926.25 | | | | | | |
| hexokinase 1 | Cytoplasm | kinase | 456849.75 | 4985028.5 | 2914394 | | | | | | |
| hexokinase 1 | Cytoplasm | kinase | 442592.406 | 5189349 | 2316893.25 | | | | | | |
| hexokinase 1 | Cytoplasm | kinase | 175145.812 | 4510892 | 2727881.25 | | | | | | |
| hexokinase 1 | Cytoplasm | kinase | 94833.82 | 5181538 | 2425693.25 | | | | | | |
| hexokinase 1 | Cytoplasm | kinase | 4183.562 | 3623906.75 | 1382249.375 | P17710 | HK1 | 802081 | 3443881 | 551462 | 39 |
| major histocompatibility complex, class I, A | Plasma Membrane | other | 2603452 | 7085832 | 2674377 | | | | | | |
| major histocompatibility complex, class I, A | Plasma Membrane | other | 758700 | 2809982.75 | 1021254.125 | | | | | | |
| major histocompatibility complex, class I, A | Plasma Membrane | other | 184686 | 3221164.25 | 84515.195 | | | | | | |
| major histocompatibility complex, class I, A | Plasma Membrane | other | 143852.812 | 631910.438 | 43149.855 | | | | | | |
| major histocompatibility complex, class I, A | Plasma Membrane | other | 65329.762 | 6270665 | 47633.926 | | | | | | |
| major histocompatibility complex, class I, A | Plasma Membrane | other | 52325.254 | 479812.562 | 4.70E-06 | P01899 | | | | | |
| major histocompatibility complex, class I, A | Plasma Membrane | other | 1486047.5 | 2512539 | 1808907.75 | P01901 | | | | | |
| major histocompatibility complex, class I, A | Plasma Membrane | other | 1494492.75 | 992133.5 | 1198366.625 | | | | | | |
| major histocompatibility complex, class I, A | Plasma Membrane | other | 1152315.625 | 537915.438 | 792039.188 | P01902 | HLA-A | 882356 | 1787034 | 343915 | 27 |
| hydroxymethylbilane synthase | Cytoplasm | enzyme | 1022301.438 | 1234492.75 | 1237455.875 | P22907 | HMBS | 1022301 | 123373 | 71229 | 3 |
| high mobility group box 2 | Nucleus | transcription regulator | 4.70E-06 | 515502.969 | 1532973.875 | P30681 | HMGB2 | 0 | 780064 | 450370 | 3 |
| 3-hydroxymethyl-3-methylglutaryl-CoA lyase | Cytoplasm | enzyme | 712289.875 | 915419 | 1437949 | | | | | | |
| 3-hydroxymethyl-3-methylglutaryl-CoA lyase | Cytoplasm | enzyme | 424128.75 | 269643.156 | 274130.031 | P38060 | HMGCL | 568209 | 453693 | 185219 | 6 |
| Harvey rat sarcoma viral oncogene homolog | Plasma Membrane | enzyme | 2.09E+08 | 1.46E+08 | 1.65E+08 | | | | | | |
| Harvey rat sarcoma viral oncogene homolog | Plasma Membrane | enzyme | 5.01E+07 | 4.53E+07 | 4.64E+07 | | | | | | |
| Harvey rat sarcoma viral oncogene homolog | Plasma Membrane | enzyme | 3.95E+07 | 3.62E+07 | 3.47E+07 | | | | | | |
| Harvey rat sarcoma viral oncogene homolog | Plasma Membrane | enzyme | 3.59E+07 | 2.64E+07 | 3.25E+07 | | | | | | |
| Harvey rat sarcoma viral oncogene homolog | Plasma Membrane | enzyme | 2.13E+07 | 2.20E+07 | 2.09E+07 | | | | | | |
| Harvey rat sarcoma viral oncogene homolog | Plasma Membrane | enzyme | 1.32E+07 | 6288940.5 | 7202512.5 | | | | | | |
| Harvey rat sarcoma viral oncogene homolog | Plasma Membrane | enzyme | 1.26E+07 | 1.57E+07 | 1.74E+07 | | | | | | |
| Harvey rat sarcoma viral oncogene homolog | Plasma Membrane | enzyme | 1.02E+07 | 6354866.5 | 6700284 | | | | | | |
| Harvey rat sarcoma viral oncogene homolog | Plasma Membrane | enzyme | 4809198.5 | 2621091 | 2839681.25 | | | | | | |
| Harvey rat sarcoma viral oncogene homolog | Plasma Membrane | enzyme | 2615958.5 | 2349471 | 1965865 | Q61411 | HRAS | 39941516 | 49950680 | 9119705 | 30 |
| hydroxysteroid (17-beta) dehydrogenase 10 | Cytoplasm | enzyme | 2720509 | 8143952.5 | 5333374 | | | | | | |
| hydroxysteroid (17-beta) dehydrogenase 10 | Cytoplasm | enzyme | 1086403.375 | 2933412 | 1675443.625 | | | | | | |
| hydroxysteroid (17-beta) dehydrogenase 10 | Cytoplasm | enzyme | 1039050.375 | 4155981.5 | 3297196.5 | | | | | | |
| hydroxysteroid (17-beta) dehydrogenase 10 | Cytoplasm | enzyme | 225901.562 | 1448209.875 | 954456.062 | O08756 | HSD17B10 | 1267966 | 2260702 | 652609 | 12 |
| hydroxysteroid (17-beta) dehydrogenase 12 | Cytoplasm | enzyme | 1827658.375 | 3289098.75 | 2373092.75 | | | | | | |
| hydroxysteroid (17-beta) dehydrogenase 12 | Cytoplasm | enzyme | 1400711.125 | 3478531.25 | 1187244.875 | O70503 | HSD17B12 | 1614185 | 962575 | 392969 | 6 |
| hydroxysteroid dehydrogenase like 1 | Cytoplasm | enzyme | 97694.43 | 589540.375 | 359460.906 | Q8BTX9 | HSDL1 | 97694 | 246093 | 142082 | 3 |
| heat shock protein 90kDa alpha (cytosolic), class A member 1 | Cytoplasm | enzyme | 1.17E+07 | 5637457.5 | 6232219 | | | | | | |
| heat shock protein 90kDa alpha (cytosolic), class A member 1 | Cytoplasm | enzyme | 8386179.5 | 4735916.5 | 4914778.5 | | | | | | |
| heat shock protein 90kDa alpha (cytosolic), class A member 1 | Cytoplasm | enzyme | 8236958 | 4297830.5 | 4961761.5 | | | | | | |
| heat shock protein 90kDa alpha (cytosolic), class A member 1 | Cytoplasm | enzyme | 6159140 | 3252448.5 | 3352460.25 | | | | | | |
| heat shock protein 90kDa alpha (cytosolic), class A member 1 | Cytoplasm | enzyme | 5793911 | 2783127 | 3585264 | | | | | | |
| heat shock protein 90kDa alpha (cytosolic), class A member 1 | Cytoplasm | enzyme | 4751321.5 | 2004075.5 | 1930401.25 | P07901 | HSP90AA1 | 7502918 | 2448384 | 577090 | 18 |
| heat shock protein 90kDa alpha (cytosolic), class B member 1 | Cytoplasm | enzyme | 9033925 | 4728932.5 | 4289160 | P11499 | HSP90AB1 | 9033925 | 2621677 | 1513626 | 3 |
| heat shock protein 90kDa beta (Grp94), member 1 | Cytoplasm | other | 4.21E+07 | 4.37E+07 | 3.72E+07 | | | | | | |
| heat shock protein 90kDa beta (Grp94), member 1 | Cytoplasm | other | 4.16E+07 | 4.35E+07 | 3.65E+07 | | | | | | |
| heat shock protein 90kDa beta (Grp94), member 1 | Cytoplasm | other | 3.57E+07 | 3.82E+07 | 3.77E+07 | | | | | | |
| heat shock protein 90kDa beta (Grp94), member 1 | Cytoplasm | other | 3.02E+07 | 2.46E+07 | 2.41E+07 | | | | | | |
| heat shock protein 90kDa beta (Grp94), member 1 | Cytoplasm | other | 2.95E+07 | 3.21E+07 | 3.00E+07 | | | | | | |
| heat shock protein 90kDa beta (Grp94), member 1 | Cytoplasm | other | 2.95E+07 | 3.14E+07 | 2.77E+07 | | | | | | |
| heat shock protein 90kDa beta (Grp94), member 1 | Cytoplasm | other | 2.93E+07 | 3.61E+07 | 3.30E+07 | | | | | | |
| heat shock protein 90kDa beta (Grp94), member 1 | Cytoplasm | other | 2.66E+07 | 2.25E+07 | 2.53E+07 | | | | | | |
| heat shock protein 90kDa beta (Grp94), member 1 | Cytoplasm | other | 2.63E+07 | 2.82E+07 | 2.55E+07 | | | | | | |
| heat shock protein 90kDa beta (Grp94), member 1 | Cytoplasm | other | 2.50E+07 | 2.75E+07 | 2.09E+07 | | | | | | |
| heat shock protein 90kDa beta (Grp94), member 1 | Cytoplasm | other | 2.25E+07 | 2.39E+07 | 2.01E+07 | | | | | | |
| heat shock protein 90kDa beta (Grp94), member 1 | Cytoplasm | other | 2.02E+07 | 2.39E+07 | 1.62E+07 | | | | | | |
| heat shock protein 90kDa beta (Grp94), member 1 | Cytoplasm | other | 1.96E+07 | 2.12E+07 | 1.97E+07 | | | | | | |
| heat shock protein 90kDa beta (Grp94), member 1 | Cytoplasm | other | 1.95E+07 | 3.01E+07 | 2.81E+07 | | | | | | |
| heat shock protein 90kDa beta (Grp94), member 1 | Cytoplasm | other | 1.88E+07 | 1.97E+07 | 1.81E+07 | | | | | | |
| heat shock protein 90kDa beta (Grp94), member 1 | Cytoplasm | other | 1.75E+07 | 2.14E+07 | 1.99E+07 | | | | | | |
| heat shock protein 90kDa beta (Grp94), member 1 | Cytoplasm | other | 1.75E+07 | 2.14E+07 | 1.97E+07 | | | | | | |
| heat shock protein 90kDa beta (Grp94), member 1 | Cytoplasm | other | 1.63E+07 | 1.84E+07 | 2.23E+07 | | | | | | |
| heat shock protein 90kDa beta (Grp94), member 1 | Cytoplasm | other | 1.58E+07 | 1.68E+07 | 1.44E+07 | | | | | | |
| heat shock protein 90kDa beta (Grp94), member 1 | Cytoplasm | other | 1.55E+07 | 1.55E+07 | 1.72E+07 | | | | | | |
| heat shock protein 90kDa beta (Grp94), member 1 | Cytoplasm | other | 1.35E+07 | 1.61E+07 | 1.56E+07 | | | | | | |
| heat shock protein 90kDa beta (Grp94), member 1 | Cytoplasm | other | 9547807 | 1.39E+07 | 1.37E+07 | | | | | | |
| heat shock protein 90kDa beta (Grp94), member 1 | Cytoplasm | other | 8469574 | 9298286 | 8928738 | | | | | | |
| heat shock protein 90kDa beta (Grp94), member 1 | Cytoplasm | other | 6199315 | 9224846 | 8163664.5 | | | | | | |
| heat shock protein 90kDa beta (Grp94), member 1 | Cytoplasm | other | 2801151.5 | 1584290.625 | 1618088.5 | | | | | | |
| heat shock protein 90kDa beta (Grp94), member 1 | Cytoplasm | other | 2155634 | 3340782.5 | 2490278.25 | P08113 | HSP90B1 | 20833595 | 10343991 | 1171226 | 78 |
| heat shock 70kDa protein 4 | Cytoplasm | other | 4.27E+07 | 1.79E+07 | 2.01E+07 | | | | | | |
| heat shock 70kDa protein 4 | Cytoplasm | other | 3.23E+07 | 1.12E+07 | 1.71E+07 | | | | | | |
| heat shock 70kDa protein 4 | Cytoplasm | other | 2.10E+07 | 8292289 | 1.25E+07 | | | | | | |
| heat shock 70kDa protein 4 | Cytoplasm | other | 2.08E+07 | 1.07E+07 | 9527539 | | | | | | |
| heat shock 70kDa protein 4 | Cytoplasm | other | 1.84E+07 | 7522799.5 | 9182183 | | | | | | |
| heat shock 70kDa protein 4 | Cytoplasm | other | 1.83E+07 | 1.03E+07 | 1.12E+07 | | | | | | |
| heat shock 70kDa protein 4 | Cytoplasm | other | 1.76E+07 | 8777780 | 1.20E+07 | | | | | | |

| | | | | | | | | | | | |
|---|---|---|---|---|---|---|---|---|---|---|---|
| heat shock 70kDa protein 4 | Cytoplasm | other | 1.68E+07 | 7607599.5 | 1.08E+07 | | | | | | |
| heat shock 70kDa protein 4 | Cytoplasm | other | 1.65E+07 | 7484095 | 9424219 | | | | | | |
| heat shock 70kDa protein 4 | Cytoplasm | other | 1.65E+07 | 6755663 | 1.02E+07 | | | | | | |
| heat shock 70kDa protein 4 | Cytoplasm | other | 1.30E+07 | 5917325 | 7156666 | | | | | | |
| heat shock 70kDa protein 4 | Cytoplasm | other | 1.28E+07 | 7412529 | 8522039 | | | | | | |
| heat shock 70kDa protein 4 | Cytoplasm | other | 1.23E+07 | 4775279 | 6892899.5 | | | | | | |
| heat shock 70kDa protein 4 | Cytoplasm | other | 1.11E+07 | 4461407.5 | 4666170.5 | | | | | | |
| heat shock 70kDa protein 4 | Cytoplasm | other | 1.05E+07 | 4318487 | 5732691 | | | | | | |
| heat shock 70kDa protein 4 | Cytoplasm | other | 1.02E+07 | 6427205.5 | 6042747 | | | | | | |
| heat shock 70kDa protein 4 | Cytoplasm | other | 7346250 | 2214006.25 | 2532139.5 | | | | | | |
| heat shock 70kDa protein 4 | Cytoplasm | other | 7202887 | 2129332.25 | 3757892.25 | | | | | | |
| heat shock 70kDa protein 4 | Cytoplasm | other | 7094135.5 | 2745300.75 | 4119760 | | | | | | |
| heat shock 70kDa protein 4 | Cytoplasm | other | 6841711 | 3240186.25 | 4414520 | | | | | | |
| heat shock 70kDa protein 4 | Cytoplasm | other | 5702752 | 3292944 | 3832104.75 | | | | | | |
| heat shock 70kDa protein 4 | Cytoplasm | other | 3734399 | 1530062.125 | 1341575.75 | | | | | | |
| heat shock 70kDa protein 4 | Cytoplasm | other | 3017423.25 | 1106287.75 | 1427902.25 | Q61316 | HSPA4 | 14419111 | 7191770 | 865787 | 69 |
| heat shock 70kDa protein 4-like | Cytoplasm | other | 1.16E+07 | 4012661.5 | 6252526.5 | | | | | | |
| heat shock 70kDa protein 4-like | Cytoplasm | other | 5913411 | 2864897.5 | 2991735.5 | | | | | | |
| heat shock 70kDa protein 4-like | Cytoplasm | other | 5056801 | 1829312.5 | 2663811.5 | | | | | | |
| heat shock 70kDa protein 4-like | Cytoplasm | other | 3536953 | 3962956.75 | 1.22E+07 | | | | | | |
| heat shock 70kDa protein 4-like | Cytoplasm | other | 3238201 | 1408478.125 | 1776823.75 | P48722 | HSPA4L | 5873073 | 3283241 | 847729 | 15 |
| heat shock 70kDa protein 5 (glucose-regulated protein, 78kDa) | Cytoplasm | enzyme | 1.99E+08 | 1.55E+08 | 1.56E+08 | | | | | | |
| heat shock 70kDa protein 5 (glucose-regulated protein, 78kDa) | Cytoplasm | enzyme | 7.56E+07 | 7.47E+07 | 6.55E+07 | | | | | | |
| heat shock 70kDa protein 5 (glucose-regulated protein, 78kDa) | Cytoplasm | enzyme | 6.27E+07 | 5.80E+07 | 5.64E+07 | | | | | | |
| heat shock 70kDa protein 5 (glucose-regulated protein, 78kDa) | Cytoplasm | enzyme | 5.34E+07 | 5.60E+07 | 5.30E+07 | | | | | | |
| heat shock 70kDa protein 5 (glucose-regulated protein, 78kDa) | Cytoplasm | enzyme | 5.03E+07 | 6.09E+07 | 6.44E+07 | | | | | | |
| heat shock 70kDa protein 5 (glucose-regulated protein, 78kDa) | Cytoplasm | enzyme | 4.65E+07 | 5.72E+07 | 4.98E+07 | | | | | | |
| heat shock 70kDa protein 5 (glucose-regulated protein, 78kDa) | Cytoplasm | enzyme | 4.53E+07 | 4.41E+07 | 5.11E+07 | | | | | | |
| heat shock 70kDa protein 5 (glucose-regulated protein, 78kDa) | Cytoplasm | enzyme | 4.08E+07 | 4.56E+07 | 4.55E+07 | | | | | | |
| heat shock 70kDa protein 5 (glucose-regulated protein, 78kDa) | Cytoplasm | enzyme | 3.93E+07 | 3.63E+07 | 3.20E+07 | | | | | | |
| heat shock 70kDa protein 5 (glucose-regulated protein, 78kDa) | Cytoplasm | enzyme | 3.38E+07 | 3.21E+07 | 4.23E+07 | | | | | | |
| heat shock 70kDa protein 5 (glucose-regulated protein, 78kDa) | Cytoplasm | enzyme | 3.09E+07 | 3.95E+07 | 3.41E+07 | | | | | | |
| heat shock 70kDa protein 5 (glucose-regulated protein, 78kDa) | Cytoplasm | enzyme | 2.92E+07 | 3.30E+07 | 3.56E+07 | | | | | | |
| heat shock 70kDa protein 5 (glucose-regulated protein, 78kDa) | Cytoplasm | enzyme | 2.60E+07 | 2.21E+07 | 2.32E+07 | | | | | | |
| heat shock 70kDa protein 5 (glucose-regulated protein, 78kDa) | Cytoplasm | enzyme | 2.27E+07 | 1.93E+07 | 2.06E+07 | | | | | | |
| heat shock 70kDa protein 5 (glucose-regulated protein, 78kDa) | Cytoplasm | enzyme | 1.62E+07 | 2.02E+07 | 1.98E+07 | | | | | | |
| heat shock 70kDa protein 5 (glucose-regulated protein, 78kDa) | Cytoplasm | enzyme | 1.50E+07 | 3.09E+07 | 2.07E+07 | | | | | | |
| heat shock 70kDa protein 5 (glucose-regulated protein, 78kDa) | Cytoplasm | enzyme | 1.33E+07 | 1.68E+07 | 1.29E+07 | | | | | | |
| heat shock 70kDa protein 5 (glucose-regulated protein, 78kDa) | Cytoplasm | enzyme | 1.29E+07 | 1.63E+07 | 1.26E+07 | | | | | | |
| heat shock 70kDa protein 5 (glucose-regulated protein, 78kDa) | Cytoplasm | enzyme | 1.29E+07 | 2.78E+07 | 2.61E+07 | | | | | | |
| heat shock 70kDa protein 5 (glucose-regulated protein, 78kDa) | Cytoplasm | enzyme | 7966493 | 7991545.5 | 7783276.5 | | | | | | |
| heat shock 70kDa protein 5 (glucose-regulated protein, 78kDa) | Cytoplasm | enzyme | 6944424.5 | 8800528 | 7425255.5 | | | | | | |
| heat shock 70kDa protein 5 (glucose-regulated protein, 78kDa) | Cytoplasm | enzyme | 5885836 | 7611984.5 | 6929499.5 | | | | | | |
| heat shock 70kDa protein 5 (glucose-regulated protein, 78kDa) | Cytoplasm | enzyme | 5478658.5 | 3765004.5 | 4317528.5 | | | | | | |
| heat shock 70kDa protein 5 (glucose-regulated protein, 78kDa) | Cytoplasm | enzyme | 3059624 | 4620000 | 4774028.5 | | | | | | |
| heat shock 70kDa protein 5 (glucose-regulated protein, 78kDa) | Cytoplasm | enzyme | 2269322.5 | 4000852.25 | 1773035.5 | | | | | | |
| heat shock 70kDa protein 5 (glucose-regulated protein, 78kDa) | Cytoplasm | enzyme | 2138270.75 | 5233971 | 3279475.25 | P20029 | HSPA5 | 33072409 | 34269825 | 3880294 | 78 |
| heat shock 70kDa protein 8 | Cytoplasm | enzyme | 1.77E+08 | 1.12E+08 | 1.10E+08 | | | | | | |
| heat shock 70kDa protein 8 | Cytoplasm | enzyme | 1.69E+08 | 9.52E+07 | 1.08E+08 | | | | | | |
| heat shock 70kDa protein 8 | Cytoplasm | enzyme | 1.58E+08 | 8.82E+07 | 8.18E+07 | | | | | | |
| heat shock 70kDa protein 8 | Cytoplasm | enzyme | 1.35E+08 | 8.99E+07 | 9.32E+07 | | | | | | |
| heat shock 70kDa protein 8 | Cytoplasm | enzyme | 1.33E+08 | 7.96E+07 | 7.95E+07 | | | | | | |
| heat shock 70kDa protein 8 | Cytoplasm | enzyme | 1.25E+08 | 7.92E+07 | 7.17E+07 | | | | | | |
| heat shock 70kDa protein 8 | Cytoplasm | enzyme | 1.22E+08 | 7.58E+07 | 7.36E+07 | | | | | | |
| heat shock 70kDa protein 8 | Cytoplasm | enzyme | 9.73E+07 | 5.72E+07 | 6.09E+07 | | | | | | |
| heat shock 70kDa protein 8 | Cytoplasm | enzyme | 8.60E+07 | 6.05E+07 | 5.53E+07 | | | | | | |
| heat shock 70kDa protein 8 | Cytoplasm | enzyme | 8.30E+07 | 5.44E+07 | 5.45E+07 | | | | | | |
| heat shock 70kDa protein 8 | Cytoplasm | enzyme | 8.18E+07 | 5.84E+07 | 5.63E+07 | | | | | | |
| heat shock 70kDa protein 8 | Cytoplasm | enzyme | 7.70E+07 | 5.27E+07 | 5.43E+07 | | | | | | |
| heat shock 70kDa protein 8 | Cytoplasm | enzyme | 7.66E+07 | 7.24E+07 | 6.86E+07 | | | | | | |
| heat shock 70kDa protein 8 | Cytoplasm | enzyme | 6.84E+07 | 5.03E+07 | 5.02E+07 | | | | | | |
| heat shock 70kDa protein 8 | Cytoplasm | enzyme | 6.43E+07 | 4.51E+07 | 4.42E+07 | | | | | | |
| heat shock 70kDa protein 8 | Cytoplasm | enzyme | 5.63E+07 | 3.68E+07 | 3.43E+07 | | | | | | |
| heat shock 70kDa protein 8 | Cytoplasm | enzyme | 3.64E+07 | 3.10E+07 | 3.18E+07 | | | | | | |
| heat shock 70kDa protein 8 | Cytoplasm | enzyme | 3.61E+07 | 3.55E+07 | 3.20E+07 | | | | | | |
| heat shock 70kDa protein 8 | Cytoplasm | enzyme | 3.44E+07 | 1.98E+07 | 1.62E+07 | | | | | | |
| heat shock 70kDa protein 8 | Cytoplasm | enzyme | 3.39E+07 | 2.16E+07 | 2.23E+07 | | | | | | |
| heat shock 70kDa protein 8 | Cytoplasm | enzyme | 2.11E+07 | 2.24E+07 | 2.40E+07 | | | | | | |
| heat shock 70kDa protein 8 | Cytoplasm | enzyme | 6136084.5 | 3918668.75 | 5797752 | | | | | | |
| heat shock 70kDa protein 8 | Cytoplasm | enzyme | 5586785 | 2994983.25 | 2524495.75 | | | | | | |
| heat shock 70kDa protein 8 | Cytoplasm | enzyme | 5151074 | 2969968.75 | 3173494.75 | | | | | | |
| heat shock 70kDa protein 8 | Cytoplasm | enzyme | 2912935 | 574514.375 | 1642669.875 | | | | | | |
| heat shock 70kDa protein 8 | Cytoplasm | enzyme | 2154573 | 889584.438 | 1755986 | P63017 | HSPA8 | 72835440 | 42234577 | 4782124 | 78 |
| heat shock 70kDa protein 9 (mortalin) | Cytoplasm | other | 4337427 | 5699912.5 | 5931248 | | | | | | |
| heat shock 70kDa protein 9 (mortalin) | Cytoplasm | other | 3713160.75 | 6828648.5 | 5690671 | | | | | | |
| heat shock 70kDa protein 9 (mortalin) | Cytoplasm | other | 3549965 | 7145868.5 | 5896457 | | | | | | |
| heat shock 70kDa protein 9 (mortalin) | Cytoplasm | other | 3382006.75 | 5257284.5 | 5167029 | | | | | | |
| heat shock 70kDa protein 9 (mortalin) | Cytoplasm | other | 3238230.5 | 4776386.5 | 5305319 | | | | | | |
| heat shock 70kDa protein 9 (mortalin) | Cytoplasm | other | 3100445.25 | 4459662 | 4761711.5 | | | | | | |
| heat shock 70kDa protein 9 (mortalin) | Cytoplasm | other | 2564438.5 | 3946036.75 | 3685789.75 | | | | | | |
| heat shock 70kDa protein 9 (mortalin) | Cytoplasm | other | 1697948.375 | 4449104.5 | 3595224.5 | | | | | | |
| heat shock 70kDa protein 9 (mortalin) | Cytoplasm | other | 1595078.25 | 3010841 | 2883207.5 | | | | | | |
| heat shock 70kDa protein 9 (mortalin) | Cytoplasm | other | 1111110.625 | 4281461 | 5824636.5 | | | | | | |
| heat shock 70kDa protein 9 (mortalin) | Cytoplasm | other | 1048501.625 | 2207224.75 | 1253721.75 | | | | | | |
| heat shock 70kDa protein 9 (mortalin) | Cytoplasm | other | 890111.062 | 1772452.5 | 1108003.625 | P38647 | HSPA9 | 2519035 | 1749047 | 291508 | 36 |
| heat shock 60kDa protein 1 (chaperonin) | Cytoplasm | enzyme | 1.21E+07 | 1.68E+07 | 1.63E+07 | | | | | | |
| heat shock 60kDa protein 1 (chaperonin) | Cytoplasm | enzyme | 1.20E+07 | 1.86E+07 | 1.60E+07 | | | | | | |
| heat shock 60kDa protein 1 (chaperonin) | Cytoplasm | enzyme | 1.14E+07 | 1.79E+07 | 1.67E+07 | | | | | | |
| heat shock 60kDa protein 1 (chaperonin) | Cytoplasm | enzyme | 1.10E+07 | 1.30E+07 | 1.56E+07 | | | | | | |
| heat shock 60kDa protein 1 (chaperonin) | Cytoplasm | enzyme | 9564700 | 1.36E+07 | 1.54E+07 | | | | | | |
| heat shock 60kDa protein 1 (chaperonin) | Cytoplasm | enzyme | 9300651 | 1.33E+07 | 1.25E+07 | | | | | | |
| heat shock 60kDa protein 1 (chaperonin) | Cytoplasm | enzyme | 7295339 | 1.46E+07 | 1.37E+07 | | | | | | |
| heat shock 60kDa protein 1 (chaperonin) | Cytoplasm | enzyme | 7191341 | 1.51E+07 | 9362642 | | | | | | |
| heat shock 60kDa protein 1 (chaperonin) | Cytoplasm | enzyme | 6452783 | 9882702 | 1.05E+07 | | | | | | |
| heat shock 60kDa protein 1 (chaperonin) | Cytoplasm | enzyme | 6191913.5 | 8574796 | 8574285 | | | | | | |

| | | | | | | | | | | | |
|---|---|---|---|---|---|---|---|---|---|---|---|
| heat shock 60kDa protein 1 (chaperonin) | Cytoplasm | enzyme | 5911611 | 1.06E+07 | 8967879 | | | | | | |
| heat shock 60kDa protein 1 (chaperonin) | Cytoplasm | enzyme | 5231197 | 7851871 | 7784364.5 | | | | | | |
| heat shock 60kDa protein 1 (chaperonin) | Cytoplasm | enzyme | 5066988.5 | 7791832 | 7297786 | | | | | | |
| heat shock 60kDa protein 1 (chaperonin) | Cytoplasm | enzyme | 4913015 | 7892676 | 1.01E+07 | | | | | | |
| heat shock 60kDa protein 1 (chaperonin) | Cytoplasm | enzyme | 4203513 | 6665120 | 6068853 | | | | | | |
| heat shock 60kDa protein 1 (chaperonin) | Cytoplasm | enzyme | 4072581.75 | 9381749 | 1.34E+07 | | | | | | |
| heat shock 60kDa protein 1 (chaperonin) | Cytoplasm | enzyme | 2485234 | 2771530 | 2343468.75 | | | | | | |
| heat shock 60kDa protein 1 (chaperonin) | Cytoplasm | enzyme | 1070608.75 | 2286894.25 | 2501996.75 | | | | | | |
| heat shock 60kDa protein 1 (chaperonin) | Cytoplasm | enzyme | 1063047.75 | 1889940 | 2289580 | | | | | | |
| heat shock 60kDa protein 1 (chaperonin) | Cytoplasm | enzyme | 841815.625 | 1789061.25 | 2535332.75 | | | | | | |
| heat shock 60kDa protein 1 (chaperonin) | Cytoplasm | enzyme | 547434.812 | 1169715.125 | 964004.938 | | | | | | |
| heat shock 60kDa protein 1 (chaperonin) | Cytoplasm | enzyme | 520541.625 | 795837.188 | 1318951 | | | | | | |
| heat shock 60kDa protein 1 (chaperonin) | Cytoplasm | enzyme | 171063.312 | 1412833.625 | 862089.625 | P63038 | HSPD1 | 5592408 | 5311022 | 639372 | 69 |
| heat shock 10kDa protein 1 | Cytoplasm | enzyme | 1.13E+07 | 1.39E+07 | 1.26E+07 | | | | | | |
| heat shock 10kDa protein 1 | Cytoplasm | enzyme | 8880456 | 1.16E+07 | 9514886 | | | | | | |
| heat shock 10kDa protein 1 | Cytoplasm | enzyme | 5489705.5 | 1.02E+07 | 8320115.5 | | | | | | |
| heat shock 10kDa protein 1 | Cytoplasm | enzyme | 3553638 | 4216599.5 | 3458733.75 | | | | | | |
| heat shock 10kDa protein 1 | Cytoplasm | enzyme | 2712855.75 | 4247193.5 | 2937572.25 | Q64433 | HSPE1 | 6387331 | 3896810 | 1006152 | 15 |
| HIV-1 Tat interactive protein 2, 30kDa | Nucleus | transcription regulator | 727778.688 | 844873.5 | 342860.094 | | | | | | |
| HIV-1 Tat interactive protein 2, 30kDa | Nucleus | transcription regulator | 620662.812 | 302391.781 | 249696.359 | Q9Z2G9 | HTATIP2 | 674221 | 249189 | 101731 | 6 |
| HECT, UBA and WWE domain containing 1, E3 ubiquitin protein ligase | Nucleus | transcription regulator | 3747639.25 | 2547869.75 | 2406845 | Q7TMY8 | HUWE1 | 3747639 | 736779 | 425380 | 3 |
| hyaluronoglucosaminidase 1 | Cytoplasm | enzyme | 3065421.25 | 2793508 | 2139973.5 | Q91ZJ9 | HYAL1 | 3065421 | 475657 | 274621 | 3 |
| hypoxia up-regulated 1 | Cytoplasm | other | 4643533 | 5205090 | 4686312 | | | | | | |
| hypoxia up-regulated 1 | Cytoplasm | other | 4482860 | 3990058.75 | 3973334 | | | | | | |
| hypoxia up-regulated 1 | Cytoplasm | other | 3921883 | 3787954.5 | 4172192 | | | | | | |
| hypoxia up-regulated 1 | Cytoplasm | other | 3492840 | 3107297.5 | 3256489 | | | | | | |
| hypoxia up-regulated 1 | Cytoplasm | other | 3400331.75 | 4302357 | 4484152 | | | | | | |
| hypoxia up-regulated 1 | Cytoplasm | other | 2564755.75 | 2321846 | 1535926.125 | | | | | | |
| hypoxia up-regulated 1 | Cytoplasm | other | 1576167.5 | 2355466.5 | 2273677.25 | Q9JKR6 | HYOU1 | 3440339 | 1054427 | 230095 | 21 |
| isoleucyl-tRNA synthetase 2, mitochondrial | Cytoplasm | enzyme | 1385184.25 | 1447847.75 | 1622318.375 | Q8BIJ6 | IARS2 | 1385184 | 122882 | 70946 | 3 |
| intercellular adhesion molecule 4 (Landsteiner-Wiener blood group) | Plasma Membrane | other | 3228329 | 2868337.5 | 2203116 | | | | | | |
| intercellular adhesion molecule 4 (Landsteiner-Wiener blood group) | Plasma Membrane | other | 3173480.75 | 4407467 | 3618243.5 | Q9ERM2 | ICAM4 | 3200905 | 738010 | 301291 | 6 |
| insulin-degrading enzyme | Extracellular Space | peptidase | 4973240.5 | 2933583 | 2193915.75 | | | | | | |
| insulin-degrading enzyme | Extracellular Space | peptidase | 4552961 | 4188216 | 3291575.25 | Q9JHR7 | IDE | 4763101 | 1059131 | 432389 | 6 |
| isocitrate dehydrogenase 2 (NADP+), mitochondrial | Cytoplasm | enzyme | 3144451.25 | 6268535.5 | 6535187 | | | | | | |
| isocitrate dehydrogenase 2 (NADP+), mitochondrial | Cytoplasm | enzyme | 1051288.875 | 1672152 | 1462094.75 | | | | | | |
| isocitrate dehydrogenase 2 (NADP+), mitochondrial | Cytoplasm | enzyme | 765490.5 | 2546442 | 2310842.25 | | | | | | |
| isocitrate dehydrogenase 2 (NADP+), mitochondrial | Cytoplasm | enzyme | 526056.75 | 1835558 | 2027515.875 | | | | | | |
| isocitrate dehydrogenase 2 (NADP+), mitochondrial | Cytoplasm | enzyme | 454765.312 | 1648229.125 | 1549515.75 | | | | | | |
| isocitrate dehydrogenase 2 (NADP+), mitochondrial | Cytoplasm | enzyme | 144624.578 | 2082088.5 | 947164 | P54071 | IDH2 | 1014446 | 1761251 | 415131 | 18 |
| isocitrate dehydrogenase 3 (NAD+) alpha | Cytoplasm | enzyme | 6049421.5 | 5830681.5 | 8869420 | | | | | | |
| isocitrate dehydrogenase 3 (NAD+) alpha | Cytoplasm | enzyme | 2802443.5 | 3647050 | 3543097.75 | | | | | | |
| isocitrate dehydrogenase 3 (NAD+) alpha | Cytoplasm | enzyme | 2008503.125 | 2453347.5 | 3030324 | | | | | | |
| isocitrate dehydrogenase 3 (NAD+) alpha | Cytoplasm | enzyme | 1938796.625 | 3326628 | 2953305 | | | | | | |
| isocitrate dehydrogenase 3 (NAD+) alpha | Cytoplasm | enzyme | 1805548.875 | 2611541 | 3326688.25 | Q9D6R2 | IDH3A | 2920943 | 1910084 | 493182 | 15 |
| isocitrate dehydrogenase 3 (NAD+) gamma | Cytoplasm | enzyme | 3060005.25 | 4776758 | 4611886 | | | | | | |
| isocitrate dehydrogenase 3 (NAD+) gamma | Cytoplasm | enzyme | 520348.875 | 670527.625 | 897624.062 | P70404 | IDH3G | 1790177 | 1987722 | 811484 | 6 |
| immediate early response 3 interacting protein 1 | Cytoplasm | other | 1.13E+07 | 1.12E+07 | 1.15E+07 | | | | | | |
| immediate early response 3 interacting protein 1 | Cytoplasm | other | 2748630.25 | 4076318.75 | 3509375.5 | Q9CR20 | IER3IP1 | 7004315 | 4337336 | 1770710 | 6 |
| immunoglobulin heavy constant mu | Plasma Membrane | transmembrane receptor | 3856173.75 | 6072276 | 9402836 | | | | | | |
| immunoglobulin heavy constant mu | Plasma Membrane | transmembrane receptor | 927818.562 | 2799667.75 | 2831824.5 | | | | | | |
| immunoglobulin heavy constant mu | Plasma Membrane | transmembrane receptor | 732308.312 | 3158699.75 | 3294943.5 | | | | | | |
| immunoglobulin heavy constant mu | Plasma Membrane | transmembrane receptor | 543437.688 | 2470045.25 | 2849647.25 | P01872 | IGHM | 1514935 | 2456960 | 709263 | 12 |
| integrin-linked kinase | Plasma Membrane | kinase | 2412461.75 | 4129327.5 | 4225369.5 | | | | | | |
| integrin-linked kinase | Plasma Membrane | kinase | 285665.5 | 1498909.625 | 3299992.75 | | | | | | |
| integrin-linked kinase | Plasma Membrane | kinase | 119944.789 | 1775763.125 | 4208115 | | | | | | |
| integrin-linked kinase | Plasma Membrane | kinase | 64539.828 | 1418945.125 | 3224696.25 | | | | | | |
| integrin-linked kinase | Plasma Membrane | kinase | 6199.497 | 785781.562 | 1950048.625 | | | | | | |
| integrin-linked kinase | Plasma Membrane | kinase | 4.70E-06 | 655702.312 | 2047266.625 | O55222 | ILK | 481469 | 1514651 | 357007 | 18 |
| ilvB (bacterial acetolactate synthase)-like | Cytoplasm | enzyme | 876660.375 | 762646.25 | 984080.688 | Q8BU33 | ILVBL | 876660 | 110734 | 63932 | 3 |
| inner membrane protein, mitochondrial | Cytoplasm | other | 2904217.25 | 4568096 | 3147118.25 | | | | | | |
| inner membrane protein, mitochondrial | Cytoplasm | other | 2458898 | 4209607.5 | 4431751.5 | | | | | | |
| inner membrane protein, mitochondrial | Cytoplasm | other | 2328095.75 | 4465179 | 3193026 | | | | | | |
| inner membrane protein, mitochondrial | Cytoplasm | other | 2082687.125 | 3150585.25 | 3612932.25 | | | | | | |
| inner membrane protein, mitochondrial | Cytoplasm | other | 1765967.5 | 2812857.5 | 3140265 | | | | | | |
| inner membrane protein, mitochondrial | Cytoplasm | other | 978959.25 | 1609338.75 | 1365158.5 | | | | | | |
| inner membrane protein, mitochondrial | Cytoplasm | other | 928822.625 | 2058002.75 | 1515777 | | | | | | |
| inner membrane protein, mitochondrial | Cytoplasm | other | 484969.281 | 748418.688 | 499713.625 | | | | | | |
| inner membrane protein, mitochondrial | Cytoplasm | other | 462564.219 | 1300144 | 1317643.625 | Q8CAQ8 | IMMT | 1599465 | 1294962 | 249216 | 27 |
| IMP (inosine 5'-monophosphate) dehydrogenase 2 | Cytoplasm | enzyme | 2736869.75 | 1509013.25 | 1630537.375 | P24547 | IMPDH2 | 2736870 | 676556 | 390610 | 3 |
| inositol polyphosphate-5-phosphatase K | Cytoplasm | phosphatase | 7412012.5 | 5996307 | 5701318.5 | | | | | | |
| inositol polyphosphate-5-phosphatase K | Cytoplasm | phosphatase | 6357432 | 4288642 | 6119295.5 | Q8C5L6 | INPP5K | 6884722 | 1015001 | 414372 | 6 |
| integrator complex subunit 4 | Nucleus | other | 4.46E+07 | 2.57E+07 | 3.09E+07 | Q8CIM8 | INTS4 | 44570000 | 9738220 | 5622364 | 3 |
| importin 5 | Nucleus | transporter | 1.65E+07 | 7317842 | 7516647.5 | | | | | | |
| importin 5 | Nucleus | transporter | 1.32E+07 | 4856586.5 | 3611815.25 | | | | | | |
| importin 5 | Nucleus | transporter | 1.24E+07 | 9793145 | 9871310 | | | | | | |
| importin 5 | Nucleus | transporter | 1.13E+07 | 1.11E+07 | 1.35E+07 | | | | | | |
| importin 5 | Nucleus | transporter | 9439133 | 6045852.5 | 4781252.5 | | | | | | |
| importin 5 | Nucleus | transporter | 9148708 | 6076849.5 | 3854744 | | | | | | |
| importin 5 | Nucleus | transporter | 8755796 | 8428413 | 5098162 | | | | | | |
| importin 5 | Nucleus | transporter | 8513527 | 6537260.5 | 6008526 | | | | | | |
| importin 5 | Nucleus | transporter | 8392086 | 5490968 | 3896957.75 | | | | | | |
| importin 5 | Nucleus | transporter | 8258522 | 4986991.5 | 4990119 | | | | | | |
| importin 5 | Nucleus | transporter | 7092992 | 6086714.5 | 3810982.25 | | | | | | |
| importin 5 | Nucleus | transporter | 6820786 | 4573186 | 3522813.75 | | | | | | |
| importin 5 | Nucleus | transporter | 5737244 | 4533162.5 | 3382016.75 | | | | | | |
| importin 5 | Nucleus | transporter | 5639631 | 6370268 | 3738079.25 | | | | | | |
| importin 5 | Nucleus | transporter | 5513794.5 | 5114498 | 3591645.5 | | | | | | |
| importin 5 | Nucleus | transporter | 5192650.5 | 2945891 | 2560365 | | | | | | |
| importin 5 | Nucleus | transporter | 5149073.5 | 5091360.5 | 2307169.75 | | | | | | |
| importin 5 | Nucleus | transporter | 5055865 | 4252622.5 | 3457311.25 | | | | | | |
| importin 5 | Nucleus | transporter | 3781683.25 | 1781466.875 | 1526512.875 | | | | | | |
| importin 5 | Nucleus | transporter | 3594316.75 | 3268802 | 2688642.75 | Q8BKC5 | IPO5 | 7976790 | 3109988 | 401498 | 60 |
| importin 7 | Nucleus | transporter | 4317617.5 | 4238099.5 | 2744965.5 | | | | | | |
| importin 7 | Nucleus | transporter | 3129323 | 2745932.75 | 2532552.75 | | | | | | |

| | | | | | | | | | | | |
|---|---|---|---|---|---|---|---|---|---|---|---|
| importin 7 | Nucleus | transporter | 2390682.5 | 1955943.25 | 1648153.5 | | | | | | |
| importin 7 | Nucleus | transporter | 2185588 | 2400695.75 | 2241863.75 | | | | | | |
| importin 7 | Nucleus | transporter | 1937654.25 | 1445828.375 | 846123.938 | | | | | | |
| importin 7 | Nucleus | transporter | 1685769.25 | 1379331.375 | 912762.438 | | | | | | |
| importin 7 | Nucleus | transporter | 558498.688 | 712298.75 | 523805.156 | Q9EPL8 | IPO7 | 2315019 | 1067541 | 232956 | 21 |
| immunity-related GTPase family M member 1 | Cytoplasm | other | 1.82E+07 | 1.33E+07 | 1.21E+07 | | | | | | |
| immunity-related GTPase family M member 1 | Cytoplasm | other | 1.60E+07 | 1.73E+07 | 1.17E+07 | | | | | | |
| immunity-related GTPase family M member 1 | Cytoplasm | other | 1.13E+07 | 1.00E+07 | 9955497 | | | | | | |
| immunity-related GTPase family M member 1 | Cytoplasm | other | 1.13E+07 | 6720650.5 | 6686413 | | | | | | |
| immunity-related GTPase family M member 1 | Cytoplasm | other | 9679809 | 6553870 | 5394392 | | | | | | |
| immunity-related GTPase family M member 1 | Cytoplasm | other | 6851140.5 | 5036459 | 6469597.5 | | | | | | |
| immunity-related GTPase family M member 1 | Cytoplasm | other | 5577870.5 | 5280233 | 5900321 | | | | | | |
| immunity-related GTPase family M member 1 | Cytoplasm | other | 4035024 | 2680686.5 | 2583562.25 | | | | | | |
| immunity-related GTPase family M member 1 | Cytoplasm | other | 2150337.5 | 2674550.75 | 3739929.75 | | | | | | |
| immunity-related GTPase family M member 1 | Cytoplasm | other | 1285342.375 | 836303.438 | 1240961 | Q60766 | Irgm1 | 8646952 | 4829752 | 881788 | 30 |
| iron-sulfur cluster assembly 1 | Cytoplasm | other | 1769508.5 | 1356346.875 | 2070368.125 | | | | | | |
| iron-sulfur cluster assembly 1 | Cytoplasm | other | 1107482.375 | 1493457.125 | 1159095.125 | Q9D924 | ISCA1 | 1438495 | 371107 | 151504 | 6 |
| iron-sulfur cluster assembly 2 | Cytoplasm | other | 1534471.75 | 922902.438 | 1851431.375 | Q9DCB8 | ISCA2 | 1534472 | 471990 | 272503 | 3 |
| ISG15 ubiquitin-like modifier | Extracellular Space | other | 5.31E+07 | 3.12E+07 | 4.16E+07 | | | | | | |
| ISG15 ubiquitin-like modifier | Extracellular Space | other | 4.67E+07 | 2.72E+07 | 2.80E+07 | | | | | | |
| ISG15 ubiquitin-like modifier | Extracellular Space | other | 2.57E+07 | 2.52E+07 | 1.88E+07 | | | | | | |
| ISG15 ubiquitin-like modifier | Extracellular Space | other | 2.18E+07 | 2.66E+07 | 1.40E+07 | | | | | | |
| ISG15 ubiquitin-like modifier | Extracellular Space | other | 2.16E+07 | 1.71E+07 | 1.65E+07 | | | | | | |
| ISG15 ubiquitin-like modifier | Extracellular Space | other | 1.40E+07 | 5843803.5 | 6765654.5 | | | | | | |
| ISG15 ubiquitin-like modifier | Extracellular Space | other | 2915585.25 | 6860664.5 | 2915217.75 | | | | | | |
| ISG15 ubiquitin-like modifier | Extracellular Space | other | 2333331 | 2198868.25 | 2311962.75 | Q64339 | ISG15 | 23508615 | 14412867 | 2942014 | 24 |
| interferon stimulated exonuclease gene 20kDa | Nucleus | enzyme | 9033550 | 5705922.5 | 7014156.5 | | | | | | |
| interferon stimulated exonuclease gene 20kDa | Nucleus | enzyme | 6773534.5 | 5270555 | 5815254.5 | | | | | | |
| interferon stimulated exonuclease gene 20kDa | Nucleus | enzyme | 5712277.5 | 5665134 | 4805597 | | | | | | |
| interferon stimulated exonuclease gene 20kDa | Nucleus | enzyme | 4234949.5 | 4657217 | 5136691.5 | Q9JL16 | ISG20 | 6438578 | 1292391 | 373081 | 12 |
| increased sodium tolerance 1 homolog (yeast) | Cytoplasm | other | 4667257 | 2718686 | 2372653 | Q9CX00 | IST1 | 4667257 | 1237058 | 714216 | 3 |
| integrin, alpha 2 (CD49B, alpha 2 subunit of VLA-2 receptor) | Plasma Membrane | transmembrane receptor | 738286.938 | 6800307.5 | 5143662 | | | | | | |
| integrin, alpha 2 (CD49B, alpha 2 subunit of VLA-2 receptor) | Plasma Membrane | transmembrane receptor | 286018.25 | 5341265.5 | 3096993.25 | | | | | | |
| integrin, alpha 2 (CD49B, alpha 2 subunit of VLA-2 receptor) | Plasma Membrane | transmembrane receptor | 279853.25 | 5291953.5 | 3070451.5 | | | | | | |
| integrin, alpha 2 (CD49B, alpha 2 subunit of VLA-2 receptor) | Plasma Membrane | transmembrane receptor | 266110.125 | 3209534.5 | 2589108.75 | | | | | | |
| integrin, alpha 2 (CD49B, alpha 2 subunit of VLA-2 receptor) | Plasma Membrane | transmembrane receptor | 258169.062 | 7952467.5 | 5452576 | | | | | | |
| integrin, alpha 2 (CD49B, alpha 2 subunit of VLA-2 receptor) | Plasma Membrane | transmembrane receptor | 222840.172 | 4534670.5 | 2573695 | | | | | | |
| integrin, alpha 2 (CD49B, alpha 2 subunit of VLA-2 receptor) | Plasma Membrane | transmembrane receptor | 51219.523 | 5067622 | 2961867.75 | | | | | | |
| integrin, alpha 2 (CD49B, alpha 2 subunit of VLA-2 receptor) | Plasma Membrane | transmembrane receptor | 34594.875 | 915637.125 | 1445332.125 | | | | | | |
| integrin, alpha 2 (CD49B, alpha 2 subunit of VLA-2 receptor) | Plasma Membrane | transmembrane receptor | 4.70E-06 | 2781221 | 2082546.875 | | | | | | |
| integrin, alpha 2 (CD49B, alpha 2 subunit of VLA-2 receptor) | Plasma Membrane | transmembrane receptor | 0 | 1194041.5 | 618135.688 | | | | | | |
| integrin, alpha 2 (CD49B, alpha 2 subunit of VLA-2 receptor) | Plasma Membrane | transmembrane receptor | 0 | 4706388 | 2938590.5 | Q62469 | ITGA2 | 194281 | 2271249 | 395374 | 33 |
| integrin, alpha 2b (platelet glycoprotein IIb of IIb/IIIa complex, antigen CD41) | Plasma Membrane | transmembrane receptor | 8218264.5 | 1.79E+08 | 1.34E+08 | | | | | | |
| integrin, alpha 2b (platelet glycoprotein IIb of IIb/IIIa complex, antigen CD41) | Plasma Membrane | transmembrane receptor | 6308755.5 | 6.00E+07 | 3.83E+07 | | | | | | |
| integrin, alpha 2b (platelet glycoprotein IIb of IIb/IIIa complex, antigen CD41) | Plasma Membrane | transmembrane receptor | 3557464.75 | 6.09E+07 | 3.52E+07 | | | | | | |
| integrin, alpha 2b (platelet glycoprotein IIb of IIb/IIIa complex, antigen CD41) | Plasma Membrane | transmembrane receptor | 3138354.5 | 5.19E+07 | 3.32E+07 | | | | | | |
| integrin, alpha 2b (platelet glycoprotein IIb of IIb/IIIa complex, antigen CD41) | Plasma Membrane | transmembrane receptor | 3126101.5 | 7.00E+07 | 5.59E+07 | | | | | | |
| integrin, alpha 2b (platelet glycoprotein IIb of IIb/IIIa complex, antigen CD41) | Plasma Membrane | transmembrane receptor | 3103637.75 | 6.94E+07 | 4.37E+07 | | | | | | |
| integrin, alpha 2b (platelet glycoprotein IIb of IIb/IIIa complex, antigen CD41) | Plasma Membrane | transmembrane receptor | 2205154.75 | 3.83E+07 | 2.77E+07 | | | | | | |
| integrin, alpha 2b (platelet glycoprotein IIb of IIb/IIIa complex, antigen CD41) | Plasma Membrane | transmembrane receptor | 1960731 | 4.13E+07 | 2.86E+07 | | | | | | |
| integrin, alpha 2b (platelet glycoprotein IIb of IIb/IIIa complex, antigen CD41) | Plasma Membrane | transmembrane receptor | 1864715.625 | 1.03E+07 | 9246194 | | | | | | |
| integrin, alpha 2b (platelet glycoprotein IIb of IIb/IIIa complex, antigen CD41) | Plasma Membrane | transmembrane receptor | 1625246.125 | 1.79E+07 | 1.35E+07 | | | | | | |
| integrin, alpha 2b (platelet glycoprotein IIb of IIb/IIIa complex, antigen CD41) | Plasma Membrane | transmembrane receptor | 1571457.75 | 4.17E+07 | 2.22E+07 | | | | | | |
| integrin, alpha 2b (platelet glycoprotein IIb of IIb/IIIa complex, antigen CD41) | Plasma Membrane | transmembrane receptor | 1466583.625 | 2.70E+07 | 2.02E+07 | | | | | | |
| integrin, alpha 2b (platelet glycoprotein IIb of IIb/IIIa complex, antigen CD41) | Plasma Membrane | transmembrane receptor | 1413696.25 | 3.84E+07 | 2.40E+07 | | | | | | |
| integrin, alpha 2b (platelet glycoprotein IIb of IIb/IIIa complex, antigen CD41) | Plasma Membrane | transmembrane receptor | 1366705 | 3.41E+07 | 1.96E+07 | | | | | | |
| integrin, alpha 2b (platelet glycoprotein IIb of IIb/IIIa complex, antigen CD41) | Plasma Membrane | transmembrane receptor | 1284584.625 | 2.94E+07 | 1.99E+07 | | | | | | |
| integrin, alpha 2b (platelet glycoprotein IIb of IIb/IIIa complex, antigen CD41) | Plasma Membrane | transmembrane receptor | 1239702.25 | 3.64E+07 | 2.67E+07 | | | | | | |
| integrin, alpha 2b (platelet glycoprotein IIb of IIb/IIIa complex, antigen CD41) | Plasma Membrane | transmembrane receptor | 1035102.062 | 1.30E+07 | 8369135 | | | | | | |
| integrin, alpha 2b (platelet glycoprotein IIb of IIb/IIIa complex, antigen CD41) | Plasma Membrane | transmembrane receptor | 809862.25 | 2.72E+07 | 1.65E+07 | | | | | | |
| integrin, alpha 2b (platelet glycoprotein IIb of IIb/IIIa complex, antigen CD41) | Plasma Membrane | transmembrane receptor | 653780.312 | 2.34E+07 | 1.22E+07 | | | | | | |
| integrin, alpha 2b (platelet glycoprotein IIb of IIb/IIIa complex, antigen CD41) | Plasma Membrane | transmembrane receptor | 580118.812 | 2.32E+07 | 1.39E+07 | | | | | | |
| integrin, alpha 2b (platelet glycoprotein IIb of IIb/IIIa complex, antigen CD41) | Plasma Membrane | transmembrane receptor | 543821.188 | 1.78E+07 | 1.56E+07 | | | | | | |
| integrin, alpha 2b (platelet glycoprotein IIb of IIb/IIIa complex, antigen CD41) | Plasma Membrane | transmembrane receptor | 490191.094 | 5303606 | 2754162.75 | | | | | | |
| integrin, alpha 2b (platelet glycoprotein IIb of IIb/IIIa complex, antigen CD41) | Plasma Membrane | transmembrane receptor | 258821.312 | 4319330 | 2775155.25 | | | | | | |
| integrin, alpha 2b (platelet glycoprotein IIb of IIb/IIIa complex, antigen CD41) | Plasma Membrane | transmembrane receptor | 133876.906 | 8206484.5 | 5165858.5 | | | | | | |
| integrin, alpha 2b (platelet glycoprotein IIb of IIb/IIIa complex, antigen CD41) | Plasma Membrane | transmembrane receptor | 70608.547 | 3722207.5 | 3227563.5 | | | | | | |
| integrin, alpha 2b (platelet glycoprotein IIb of IIb/IIIa complex, antigen CD41) | Plasma Membrane | transmembrane receptor | 60680.543 | 4530457 | 2358559 | | | | | | |
| integrin, alpha 2b (platelet glycoprotein IIb of IIb/IIIa complex, antigen CD41) | Plasma Membrane | transmembrane receptor | 49820.078 | 2369098.25 | 2131926.75 | Q9QUM0 | ITGA2B | 1782883 | 28564690 | 3173854 | 81 |
| integrin, alpha 5 (fibronectin receptor, alpha polypeptide) | Plasma Membrane | transmembrane receptor | 8364438 | 8049259 | 8410127 | | | | | | |
| integrin, alpha 5 (fibronectin receptor, alpha polypeptide) | Plasma Membrane | transmembrane receptor | 4729253 | 4694239 | 3748131.25 | | | | | | |
| integrin, alpha 5 (fibronectin receptor, alpha polypeptide) | Plasma Membrane | transmembrane receptor | 2526112.75 | 3232993.5 | 3490760 | | | | | | |
| integrin, alpha 5 (fibronectin receptor, alpha polypeptide) | Plasma Membrane | transmembrane receptor | 1223086.25 | 2112071.5 | 1973305.375 | P11688 | ITGA5 | 4210723 | 2569863 | 741856 | 12 |
| integrin, alpha 6 | Plasma Membrane | transmembrane receptor | 314225.625 | 8326964.5 | 5042444 | | | | | | |
| integrin, alpha 6 | Plasma Membrane | transmembrane receptor | 221642.641 | 3104642.25 | 1998900.375 | | | | | | |
| integrin, alpha 6 | Plasma Membrane | transmembrane receptor | 130493.203 | 5735199.5 | 3309107.5 | | | | | | |
| integrin, alpha 6 | Plasma Membrane | transmembrane receptor | 85107.633 | 2299828.25 | 978527.75 | | | | | | |
| integrin, alpha 6 | Plasma Membrane | transmembrane receptor | 4.70E-06 | 1197913 | 1289473.5 | | | | | | |
| integrin, alpha 6 | Plasma Membrane | transmembrane receptor | 4.70E-06 | 2241546.75 | 1502889.5 | | | | | | |
| integrin, alpha 6 | Plasma Membrane | transmembrane receptor | 0 | 5080319.5 | 3119895.75 | Q61739 | ITGA6 | 107353 | 2279413 | 497409 | 21 |
| integrin, beta 1 (fibronectin receptor, beta polypeptide, antigen CD29 includes MDF2, MSK12) | Plasma Membrane | transmembrane receptor | 6371836.5 | 2.70E+07 | 2.07E+07 | | | | | | |
| integrin, beta 1 (fibronectin receptor, beta polypeptide, antigen CD29 includes MDF2, MSK12) | Plasma Membrane | transmembrane receptor | 5569709.5 | 1.73E+07 | 1.27E+07 | | | | | | |
| integrin, beta 1 (fibronectin receptor, beta polypeptide, antigen CD29 includes MDF2, MSK12) | Plasma Membrane | transmembrane receptor | 4705972.5 | 2.22E+07 | 1.52E+07 | | | | | | |
| integrin, beta 1 (fibronectin receptor, beta polypeptide, antigen CD29 includes MDF2, MSK12) | Plasma Membrane | transmembrane receptor | 3452025.25 | 1.47E+07 | 8304354 | | | | | | |
| integrin, beta 1 (fibronectin receptor, beta polypeptide, antigen CD29 includes MDF2, MSK12) | Plasma Membrane | transmembrane receptor | 2663949.75 | 1.02E+07 | 7385932.5 | | | | | | |
| integrin, beta 1 (fibronectin receptor, beta polypeptide, antigen CD29 includes MDF2, MSK12) | Plasma Membrane | transmembrane receptor | 2268335.5 | 8755750 | 6704079 | | | | | | |
| integrin, beta 1 (fibronectin receptor, beta polypeptide, antigen CD29 includes MDF2, MSK12) | Plasma Membrane | transmembrane receptor | 2243173.25 | 9811010 | 7614999 | | | | | | |
| integrin, beta 1 (fibronectin receptor, beta polypeptide, antigen CD29 includes MDF2, MSK12) | Plasma Membrane | transmembrane receptor | 1939862.625 | 5359983 | 4290672.5 | | | | | | |
| integrin, beta 1 (fibronectin receptor, beta polypeptide, antigen CD29 includes MDF2, MSK12) | Plasma Membrane | transmembrane receptor | 1893342.625 | 8759067 | 5716129.5 | | | | | | |
| integrin, beta 1 (fibronectin receptor, beta polypeptide, antigen CD29 includes MDF2, MSK12) | Plasma Membrane | transmembrane receptor | 1579137.375 | 7446702 | 4157389.75 | | | | | | |
| integrin, beta 1 (fibronectin receptor, beta polypeptide, antigen CD29 includes MDF2, MSK12) | Plasma Membrane | transmembrane receptor | 1229182.25 | 5564599 | 3121842.75 | | | | | | |
| integrin, beta 1 (fibronectin receptor, beta polypeptide, antigen CD29 includes MDF2, MSK12) | Plasma Membrane | transmembrane receptor | 755479.375 | 3929082 | 1873193.375 | | | | | | |
| integrin, beta 1 (fibronectin receptor, beta polypeptide, antigen CD29 includes MDF2, MSK12) | Plasma Membrane | transmembrane receptor | 484393.75 | 3361728 | 1578983.875 | P09055 | ITGB1 | 2704338 | 6298776 | 1008611 | 39 |
| integrin, beta 2 (complement component 3 receptor 3 and 4 subunit) | Plasma Membrane | transmembrane receptor | 0 | 1161715.5 | 1158911.125 | P11835 | ITGB2 | | 669909 | 386772 | 3 |
| integrin, beta 3 (platelet glycoprotein IIIa, antigen CD61) | Plasma Membrane | transmembrane receptor | 6613565 | 1.61E+08 | 9.57E+07 | | | | | | |

| | | | | | | | | | | | |
|---|---|---|---|---|---|---|---|---|---|---|---|
| integrin, beta 3 (platelet glycoprotein IIIa, antigen CD61) | Plasma Membrane | transmembrane receptor | 4998616.5 | 1.85E+07 | 1.08E+07 | | | | | | |
| integrin, beta 3 (platelet glycoprotein IIIa, antigen CD61) | Plasma Membrane | transmembrane receptor | 4080349.5 | 4.82E+07 | 2.65E+07 | | | | | | |
| integrin, beta 3 (platelet glycoprotein IIIa, antigen CD61) | Plasma Membrane | transmembrane receptor | 3637579.75 | 6.76E+07 | 3.35E+07 | | | | | | |
| integrin, beta 3 (platelet glycoprotein IIIa, antigen CD61) | Plasma Membrane | transmembrane receptor | 2600505.5 | 5.30E+07 | 3.81E+07 | | | | | | |
| integrin, beta 3 (platelet glycoprotein IIIa, antigen CD61) | Plasma Membrane | transmembrane receptor | 2331472 | 5.17E+07 | 3.07E+07 | | | | | | |
| integrin, beta 3 (platelet glycoprotein IIIa, antigen CD61) | Plasma Membrane | transmembrane receptor | 2182758.5 | 3.42E+07 | 2.04E+07 | | | | | | |
| integrin, beta 3 (platelet glycoprotein IIIa, antigen CD61) | Plasma Membrane | transmembrane receptor | 2092230.5 | 5.99E+07 | 3.29E+07 | | | | | | |
| integrin, beta 3 (platelet glycoprotein IIIa, antigen CD61) | Plasma Membrane | transmembrane receptor | 1514218 | 3.86E+07 | 2.59E+07 | | | | | | |
| integrin, beta 3 (platelet glycoprotein IIIa, antigen CD61) | Plasma Membrane | transmembrane receptor | 1438720.5 | 2.04E+07 | 1.16E+07 | | | | | | |
| integrin, beta 3 (platelet glycoprotein IIIa, antigen CD61) | Plasma Membrane | transmembrane receptor | 1427940 | 2.78E+07 | 1.59E+07 | | | | | | |
| integrin, beta 3 (platelet glycoprotein IIIa, antigen CD61) | Plasma Membrane | transmembrane receptor | 1379546.25 | 3.44E+07 | 1.88E+07 | | | | | | |
| integrin, beta 3 (platelet glycoprotein IIIa, antigen CD61) | Plasma Membrane | transmembrane receptor | 1342123.5 | 2.20E+07 | 1.82E+07 | | | | | | |
| integrin, beta 3 (platelet glycoprotein IIIa, antigen CD61) | Plasma Membrane | transmembrane receptor | 1174390.25 | 3.88E+07 | 2.07E+07 | | | | | | |
| integrin, beta 3 (platelet glycoprotein IIIa, antigen CD61) | Plasma Membrane | transmembrane receptor | 958925.688 | 2.53E+07 | 1.70E+07 | | | | | | |
| integrin, beta 3 (platelet glycoprotein IIIa, antigen CD61) | Plasma Membrane | transmembrane receptor | 858958.625 | 3819475.75 | 3649334.25 | | | | | | |
| integrin, beta 3 (platelet glycoprotein IIIa, antigen CD61) | Plasma Membrane | transmembrane receptor | 807008.938 | 1.77E+07 | 1.24E+07 | | | | | | |
| integrin, beta 3 (platelet glycoprotein IIIa, antigen CD61) | Plasma Membrane | transmembrane receptor | 775514.375 | 1.23E+07 | 9516289 | | | | | | |
| integrin, beta 3 (platelet glycoprotein IIIa, antigen CD61) | Plasma Membrane | transmembrane receptor | 746593.062 | 7782820 | 6782824.5 | | | | | | |
| integrin, beta 3 (platelet glycoprotein IIIa, antigen CD61) | Plasma Membrane | transmembrane receptor | 586541.938 | 2.66E+07 | 1.23E+07 | | | | | | |
| integrin, beta 3 (platelet glycoprotein IIIa, antigen CD61) | Plasma Membrane | transmembrane receptor | 400180.688 | 6540658 | 7164632 | | | | | | |
| integrin, beta 3 (platelet glycoprotein IIIa, antigen CD61) | Plasma Membrane | transmembrane receptor | 378623.719 | 9268383 | 6993771.5 | | | | | | |
| integrin, beta 3 (platelet glycoprotein IIIa, antigen CD61) | Plasma Membrane | transmembrane receptor | 149171.969 | 1.36E+07 | 1.63E+07 | | | | | | |
| integrin, beta 3 (platelet glycoprotein IIIa, antigen CD61) | Plasma Membrane | transmembrane receptor | 4.70E-06 | 3038429.25 | 1101700.625 | O54890 | ITGB3 | 1769814 | 25177179 | 2967159 | 72 |
| integrin, beta 6 | Plasma Membrane | other | 1006529.312 | 1.02E+07 | 5536826.5 | Q9Z0T9 | ITGB6 | 1006529 | 4617005 | 2665629 | 3 |
| jagunal homolog 1 | Cytoplasm | other | 1281434.125 | 1788545.125 | 1471160.625 | Q5XKN4 | JAGN1 | 1281434 | 256220 | 147928 | 3 |
| kalirin, RhoGEF kinase | Cytoplasm | kinase | 47587.082 | 2396046.5 | 1303037.5 | | | | | | |
| kalirin, RhoGEF kinase | Cytoplasm | kinase | 4.70E-06 | 1720799.75 | 518481.094 | A2CG49 | KALRN | 23794 | 969458 | 395780 | 6 |
| potassium channel modulatory factor 1 | Extracellular Space | other | 3927423 | 1634932.5 | 2020418.5 | | | | | | |
| potassium channel modulatory factor 1 | Extracellular Space | other | 1766349.75 | 628729.562 | 1126126.875 | | | | | | |
| potassium channel modulatory factor 1 | Extracellular Space | other | 1400699.75 | 701337.812 | 690535.438 | Q80UY2 | KCMF1 | 2364824 | 1026934 | 342311 | 9 |
| Kell blood group, metallo-endopeptidase | Plasma Membrane | peptidase | 2.02E+08 | 1.89E+08 | 1.44E+08 | | | | | | |
| Kell blood group, metallo-endopeptidase | Plasma Membrane | peptidase | 1.89E+08 | 1.57E+08 | 1.85E+08 | | | | | | |
| Kell blood group, metallo-endopeptidase | Plasma Membrane | peptidase | 1.50E+08 | 1.32E+08 | 1.18E+08 | | | | | | |
| Kell blood group, metallo-endopeptidase | Plasma Membrane | peptidase | 1.47E+08 | 1.33E+08 | 1.09E+08 | | | | | | |
| Kell blood group, metallo-endopeptidase | Plasma Membrane | peptidase | 1.38E+08 | 1.28E+08 | 1.39E+08 | | | | | | |
| Kell blood group, metallo-endopeptidase | Plasma Membrane | peptidase | 1.25E+08 | 8.00E+07 | 7.64E+07 | | | | | | |
| Kell blood group, metallo-endopeptidase | Plasma Membrane | peptidase | 1.07E+08 | 9.64E+07 | 8.39E+07 | | | | | | |
| Kell blood group, metallo-endopeptidase | Plasma Membrane | peptidase | 1.03E+08 | 7.48E+07 | 8.69E+07 | | | | | | |
| Kell blood group, metallo-endopeptidase | Plasma Membrane | peptidase | 8.74E+07 | 8.06E+07 | 8.53E+07 | | | | | | |
| Kell blood group, metallo-endopeptidase | Plasma Membrane | peptidase | 8.33E+07 | 9.29E+07 | 9.14E+07 | | | | | | |
| Kell blood group, metallo-endopeptidase | Plasma Membrane | peptidase | 7.49E+07 | 6.36E+07 | 6.29E+07 | | | | | | |
| Kell blood group, metallo-endopeptidase | Plasma Membrane | peptidase | 7.26E+07 | 6.61E+07 | 6.82E+07 | | | | | | |
| Kell blood group, metallo-endopeptidase | Plasma Membrane | peptidase | 7.05E+07 | 5.54E+07 | 5.86E+07 | | | | | | |
| Kell blood group, metallo-endopeptidase | Plasma Membrane | peptidase | 6.03E+07 | 3.32E+07 | 3.45E+07 | | | | | | |
| Kell blood group, metallo-endopeptidase | Plasma Membrane | peptidase | 5.37E+07 | 6.23E+07 | 4.83E+07 | | | | | | |
| Kell blood group, metallo-endopeptidase | Plasma Membrane | peptidase | 5.28E+07 | 5.19E+07 | 5.37E+07 | | | | | | |
| Kell blood group, metallo-endopeptidase | Plasma Membrane | peptidase | 5.03E+07 | 4.43E+07 | 5.99E+07 | | | | | | |
| Kell blood group, metallo-endopeptidase | Plasma Membrane | peptidase | 4.21E+07 | 3.68E+07 | 3.95E+07 | | | | | | |
| Kell blood group, metallo-endopeptidase | Plasma Membrane | peptidase | 4.03E+07 | 3.45E+07 | 3.26E+07 | | | | | | |
| Kell blood group, metallo-endopeptidase | Plasma Membrane | peptidase | 3.94E+07 | 5.59E+07 | 4.30E+07 | | | | | | |
| Kell blood group, metallo-endopeptidase | Plasma Membrane | peptidase | 3.27E+07 | 3.31E+07 | 2.46E+07 | | | | | | |
| Kell blood group, metallo-endopeptidase | Plasma Membrane | peptidase | 2.58E+07 | 3.15E+07 | 3.60E+07 | | | | | | |
| Kell blood group, metallo-endopeptidase | Plasma Membrane | peptidase | 2.30E+07 | 3.15E+07 | 2.93E+07 | | | | | | |
| Kell blood group, metallo-endopeptidase | Plasma Membrane | peptidase | 2.27E+07 | 1.57E+07 | 1.79E+07 | | | | | | |
| Kell blood group, metallo-endopeptidase | Plasma Membrane | peptidase | 2.14E+07 | 1.76E+07 | 1.34E+07 | | | | | | |
| Kell blood group, metallo-endopeptidase | Plasma Membrane | peptidase | 1.87E+07 | 2.54E+07 | 2.79E+07 | | | | | | |
| Kell blood group, metallo-endopeptidase | Plasma Membrane | peptidase | 1.76E+07 | 1.48E+07 | 1.48E+07 | | | | | | |
| Kell blood group, metallo-endopeptidase | Plasma Membrane | peptidase | 1.70E+07 | 1.09E+07 | 1.34E+07 | | | | | | |
| Kell blood group, metallo-endopeptidase | Plasma Membrane | peptidase | 1.62E+07 | 9456068 | 1.15E+07 | | | | | | |
| Kell blood group, metallo-endopeptidase | Plasma Membrane | peptidase | 1.28E+07 | 2.17E+07 | 1.92E+07 | | | | | | |
| Kell blood group, metallo-endopeptidase | Plasma Membrane | peptidase | 1.12E+07 | 9566789 | 1.08E+07 | | | | | | |
| Kell blood group, metallo-endopeptidase | Plasma Membrane | peptidase | 8826897 | 5061223 | 5529962.5 | | | | | | |
| Kell blood group, metallo-endopeptidase | Plasma Membrane | peptidase | 6202084 | 5623875 | 6498535.5 | | | | | | |
| Kell blood group, metallo-endopeptidase | Plasma Membrane | peptidase | 4532133 | 3032409 | 3893526.25 | | | | | | |
| Kell blood group, metallo-endopeptidase | Plasma Membrane | peptidase | 2943672.75 | 3844896.5 | 3869158.25 | | | | | | |
| Kell blood group, metallo-endopeptidase | Plasma Membrane | peptidase | 2644623.25 | 6127390.5 | 3496035.25 | Q9EQF2 | KEL | 59273039 | 48805655 | 4696326 | 108 |
| KIAA0368 | Cytoplasm | other | 5716296 | 3636969 | 2962953.75 | | | | | | |
| KIAA0368 | Cytoplasm | other | 4892173 | 2276346.5 | 2937322.75 | | | | | | |
| KIAA0368 | Cytoplasm | other | 3461511.25 | 2477341.75 | 3067193.75 | | | | | | |
| KIAA0368 | Cytoplasm | other | 2143001.5 | 1431155.625 | 1421981.5 | | | | | | |
| KIAA0368 | Cytoplasm | other | 2015869.375 | 1105561.625 | 1341799.5 | | | | | | |
| KIAA0368 | Cytoplasm | other | 1549961.25 | 1339832.125 | 1010680.312 | | | | | | |
| KIAA0368 | Cytoplasm | other | 1473742 | 891590.25 | 1197320.375 | | | | | | |
| KIAA0368 | Cytoplasm | other | 1023624.938 | 477741 | 766258.375 | | | | | | |
| KIAA0368 | Cytoplasm | other | 630314.25 | 272323.75 | 312461.844 | Q6PDI5 | KIAA0368 | 2545166 | 1362213 | 262158 | 27 |
| KIAA1715 | Cytoplasm | other | 4280803 | 3227549.75 | 3428491.5 | | | | | | |
| KIAA1715 | Cytoplasm | other | 3293849.25 | 1957446.875 | 2912138.25 | | | | | | |
| KIAA1715 | Cytoplasm | other | 3116863.5 | 2616278 | 2725846.75 | | | | | | |
| KIAA1715 | Cytoplasm | other | 2575665.75 | 2159483.5 | 1795897.625 | | | | | | |
| KIAA1715 | Cytoplasm | other | 2417260.5 | 2162969.5 | 1967434.25 | Q7TQ95 | KIAA1715 | 3136888 | 679663 | 175488 | 15 |
| kinesin family member 1B | Cytoplasm | transporter | 1000077.5 | 2438165 | 3803556.25 | Q60575 | KIF1B | 1000078 | 1401896 | 809385 | 3 |
| kinesin light chain 2 | Cytoplasm | other | 185009.875 | 2122745.25 | 1334273.125 | O88448 | KLC2 | 185010 | 974450 | 562599 | 3 |
| killer cell lectin-like receptor, subfamily A, member 4 | Plasma Membrane | transmembrane receptor | 6822361 | 4867761.5 | 6329052.5 | Q60682 | Klra4 (includes others) | 6822361 | 1016463 | 586855 | 3 |
| karyopherin alpha 3 (importin alpha 4) | Nucleus | transporter | 2753752.5 | 1879679.5 | 1859007.25 | O35344 | KPNA3 | 2753753 | 510718 | 294863 | 3 |
| karyopherin (importin) beta 1 | Nucleus | transporter | 1.89E+07 | 8413057 | 9023283 | | | | | | |
| karyopherin (importin) beta 1 | Nucleus | transporter | 1.71E+07 | 9756209 | 1.19E+07 | | | | | | |
| karyopherin (importin) beta 1 | Nucleus | transporter | 1.43E+07 | 1.79E+07 | 1.53E+07 | | | | | | |
| karyopherin (importin) beta 1 | Nucleus | transporter | 1.34E+07 | 6951557.5 | 6752237.5 | | | | | | |
| karyopherin (importin) beta 1 | Nucleus | transporter | 1.24E+07 | 8331223.5 | 7700783 | | | | | | |
| karyopherin (importin) beta 1 | Nucleus | transporter | 1.13E+07 | 5815096.5 | 5304372.5 | | | | | | |
| karyopherin (importin) beta 1 | Nucleus | transporter | 1.07E+07 | 6388492 | 8256580 | | | | | | |
| karyopherin (importin) beta 1 | Nucleus | transporter | 1.05E+07 | 4674993 | 5340791.5 | | | | | | |
| karyopherin (importin) beta 1 | Nucleus | transporter | 9619471 | 7622809.5 | 7241678 | | | | | | |
| karyopherin (importin) beta 1 | Nucleus | transporter | 9415767 | 7189584.5 | 6933199.5 | | | | | | |
| karyopherin (importin) beta 1 | Nucleus | transporter | 8911918 | 5475177.5 | 5869767.5 | | | | | | |

| | | | | | | | | | | | |
|---|---|---|---|---|---|---|---|---|---|---|---|
| karyopherin (importin) beta 1 | Nucleus | transporter | 6933921 | 4405938.5 | 4708043.5 | | | | | | |
| karyopherin (importin) beta 1 | Nucleus | transporter | 6638495.5 | 4261994 | 3839120.75 | | | | | | |
| karyopherin (importin) beta 1 | Nucleus | transporter | 6636697 | 3153445.5 | 3937754.25 | | | | | | |
| karyopherin (importin) beta 1 | Nucleus | transporter | 6015169 | 4577679.5 | 3706374.75 | | | | | | |
| karyopherin (importin) beta 1 | Nucleus | transporter | 5817691 | 3817425.75 | 3989666.5 | | | | | | |
| karyopherin (importin) beta 1 | Nucleus | transporter | 5472211 | 3062153 | 2800325.5 | | | | | | |
| karyopherin (importin) beta 1 | Nucleus | transporter | 4545192 | 2340147.75 | 2111572.5 | | | | | | |
| karyopherin (importin) beta 1 | Nucleus | transporter | 4412991.5 | 3766724.5 | 3822946.25 | | | | | | |
| karyopherin (importin) beta 1 | Nucleus | transporter | 3881323.25 | 2295580.25 | 1772633.625 | | | | | | |
| karyopherin (importin) beta 1 | Nucleus | transporter | 3208595.5 | 2355440.5 | 2333626 | P70168 | KPNB1 | 9053783 | 4011723 | 505430 | 63 |
| keratin 1, type II | Cytoplasm | other | 1.62E+07 | 3.15E+07 | 2.82E+07 | | | | | | |
| keratin 1, type II | Cytoplasm | other | 4059023.75 | 6507250 | 1425600 | | | | | | |
| keratin 1, type II | Cytoplasm | other | 3983660.25 | 4847393 | 7723854 | | | | | | |
| keratin 1, type II | Cytoplasm | other | 3163564.25 | 7043055.5 | 6700360.5 | P04104 | KRT1 | 6851562 | 9942186 | 2870062 | 12 |
| keratin 10 | Cytoplasm | other | 7679788 | 1.43E+07 | 1.22E+07 | P02535 | Krt10 | 7679788 | 3394538 | 1959837 | 3 |
| keratin 14, type I | Cytoplasm | other | 4194596.5 | 2886050.75 | 1535376.125 | Q61781 | KRT14 | 4194597 | 1329666 | 767683 | 3 |
| keratin 2 | Cytoplasm | other | 4023986.5 | 9870816 | 1.19E+07 | Q3TTY5 | Krt2 | 4023987 | 4085272 | 2358633 | 3 |
| keratin 5, type II | Cytoplasm | other | 2.49E+07 | 2.78E+07 | 2.52E+07 | | | | | | |
| keratin 5, type II | Cytoplasm | other | 8427649 | 7481453.5 | 5920733.5 | | | | | | |
| keratin 5, type II | Cytoplasm | other | 3606912.75 | 1470051.125 | 1707843.375 | | | | | | |
| keratin 5, type II | Cytoplasm | other | 859591.062 | 7706479 | 7641060 | Q922U2 | KRT5 | 9453538 | 9871328 | 2849607 | 12 |
| keratin 73, type II | Extracellular Space | other | 1448962.25 | 1458266.875 | 1915136.75 | Q6NXH9 | KRT73 | 1448962 | 266501 | 153864 | 3 |
| keratin 76, type II | Cytoplasm | other | 2.57E+07 | 2.40E+07 | 1.61E+07 | Q3UV17 | KRT76 | 25740000 | 5126181 | 2959602 | 3 |
| keratin 79, type II | Extracellular Space | other | 1.39E+07 | 7739244.5 | 9014537 | Q8VED5 | KRT79 | 13910000 | 3257559 | 1880753 | 3 |
| keratin 83 | Extracellular Space | other | 1.92E+07 | 1.57E+07 | 1.43E+07 | Q6IMF0 | Krt83 | 19170000 | 2510027 | 1449165 | 3 |
| lysosomal-associated membrane protein 1 | Plasma Membrane | other | 7592876 | 1.25E+07 | 9599633 | | | | | | |
| lysosomal-associated membrane protein 1 | Plasma Membrane | other | 3616334 | 7152215 | 6310950 | | | | | | |
| lysosomal-associated membrane protein 1 | Plasma Membrane | other | 695970.938 | 2973811.5 | 2056775.25 | P11438 | LAMP1 | 3968394 | 3835800 | 1278600 | 9 |
| lysosomal-associated membrane protein 2 | Plasma Membrane | enzyme | 1855237 | 5322791 | 4822066 | | | | | | |
| lysosomal-associated membrane protein 2 | Plasma Membrane | enzyme | 868242.625 | 3536050.25 | 2234236.5 | P17047 | LAMP2 | 1361740 | 1753621 | 715913 | 6 |
| late endosomal/lysosomal adaptor, MAPK and MTOR activator 1 | Plasma Membrane | other | 1.56E+07 | 1.52E+07 | 2.19E+07 | | | | | | |
| late endosomal/lysosomal adaptor, MAPK and MTOR activator 1 | Plasma Membrane | other | 7065188.5 | 5209060.5 | 7154153.5 | | | | | | |
| late endosomal/lysosomal adaptor, MAPK and MTOR activator 1 | Plasma Membrane | other | 5232415.5 | 4440966 | 6747288.5 | | | | | | |
| late endosomal/lysosomal adaptor, MAPK and MTOR activator 1 | Plasma Membrane | other | 2335861 | 2931183.5 | 3169283.25 | Q9CQ22 | LAMTOR1 | 7563366 | 6155612 | 1776972 | 12 |
| late endosomal/lysosomal adaptor, MAPK and MTOR activator 3 | Cytoplasm | other | 9705052 | 8516297 | 1.10E+07 | | | | | | |
| late endosomal/lysosomal adaptor, MAPK and MTOR activator 3 | Cytoplasm | other | 3960483.75 | 3252801 | 5337704 | | | | | | |
| late endosomal/lysosomal adaptor, MAPK and MTOR activator 3 | Cytoplasm | other | 3202432.75 | 2359832.25 | 3617050.5 | O88653 | LAMTOR3 | 5622656 | 3219621 | 1073207 | 9 |
| late endosomal/lysosomal adaptor, MAPK and MTOR activator 5 | Cytoplasm | other | 6685934.5 | 7558508 | 9618549 | | | | | | |
| late endosomal/lysosomal adaptor, MAPK and MTOR activator 5 | Cytoplasm | other | 4384521 | 3024633.25 | 3827502.75 | | | | | | |
| late endosomal/lysosomal adaptor, MAPK and MTOR activator 5 | Cytoplasm | other | 3695318.25 | 2314841 | 3375044 | Q9D1L9 | LAMTOR5 | 4921925 | 2446892 | 815631 | 9 |
| LanC lantibiotic synthetase component C-like 1 (bacterial) | Plasma Membrane | other | 4754973 | 1800464.5 | 2809680 | O89112 | LANCL1 | 4754973 | 1501766 | 867045 | 3 |
| LanC lantibiotic synthetase component C-like 2 (bacterial) | Plasma Membrane | other | 1.50E+07 | 8069866 | 8510782 | | | | | | |
| LanC lantibiotic synthetase component C-like 2 (bacterial) | Plasma Membrane | other | 1.29E+07 | 5876198 | 8058987 | | | | | | |
| LanC lantibiotic synthetase component C-like 2 (bacterial) | Plasma Membrane | other | 1.21E+07 | 7625890 | 7257143.5 | | | | | | |
| LanC lantibiotic synthetase component C-like 2 (bacterial) | Plasma Membrane | other | 1.15E+07 | 3554089.25 | 4456817.5 | | | | | | |
| LanC lantibiotic synthetase component C-like 2 (bacterial) | Plasma Membrane | other | 1.00E+07 | 5080111 | 5984883 | | | | | | |
| LanC lantibiotic synthetase component C-like 2 (bacterial) | Plasma Membrane | other | 8949738 | 3344895.75 | 3451847.25 | | | | | | |
| LanC lantibiotic synthetase component C-like 2 (bacterial) | Plasma Membrane | other | 7223929.5 | 4212736 | 4109655.5 | | | | | | |
| LanC lantibiotic synthetase component C-like 2 (bacterial) | Plasma Membrane | other | 5597943 | 1607883.875 | 2888171.75 | | | | | | |
| LanC lantibiotic synthetase component C-like 2 (bacterial) | Plasma Membrane | other | 5437726.5 | 3861178.75 | 4043252.5 | | | | | | |
| LanC lantibiotic synthetase component C-like 2 (bacterial) | Plasma Membrane | other | 4859961.5 | 4323443.5 | 3048347 | | | | | | |
| LanC lantibiotic synthetase component C-like 2 (bacterial) | Plasma Membrane | other | 4766242 | 5720560 | 2346893 | | | | | | |
| LanC lantibiotic synthetase component C-like 2 (bacterial) | Plasma Membrane | other | 4614056 | 6873122.5 | 3524672 | Q9JJK2 | LANCL2 | 8577466 | 3135713 | 522619 | 36 |
| lactate dehydrogenase A | Cytoplasm | enzyme | 8442347 | 5071964 | 4759581 | | | | | | |
| lactate dehydrogenase A | Cytoplasm | enzyme | 6715481 | 6041222 | 4140436.5 | | | | | | |
| lactate dehydrogenase A | Cytoplasm | enzyme | 5677670 | 3308281.25 | 3514685.5 | | | | | | |
| lactate dehydrogenase A | Cytoplasm | enzyme | 3855549.75 | 2886522 | 3511991 | | | | | | |
| lactate dehydrogenase A | Cytoplasm | enzyme | 3514880.25 | 2608742.25 | 3279173.25 | | | | | | |
| lactate dehydrogenase A | Cytoplasm | enzyme | 2689568.5 | 1872234.5 | 2053028.75 | | | | | | |
| lactate dehydrogenase A | Cytoplasm | enzyme | 2483845 | 1815876.625 | 2229961.5 | | | | | | |
| lactate dehydrogenase A | Cytoplasm | enzyme | 2139432.25 | 1513483.5 | 1641349.25 | | | | | | |
| lactate dehydrogenase A | Cytoplasm | enzyme | 1667594.375 | 2024259.25 | 1211233.5 | P06151 | LDHA | 4131819 | 1777876 | 342152 | 27 |
| leukocyte receptor cluster (LRC) member 8 | Nucleus | other | 161864.859 | 4686392.5 | 7117566 | Q8CBY3 | LENG8 | 161865 | 3529961 | 2038024 | 3 |
| leptin receptor overlapping transcript | Plasma Membrane | other | 4770287 | 3388244.75 | 2430432.5 | | | | | | |
| leptin receptor overlapping transcript | Plasma Membrane | other | 276130.281 | 354086.125 | 303577 | O89013 | LEPROT | 2523209 | 1913512 | 781188 | 6 |
| leucine zipper-EF-hand containing transmembrane protein 1 | Cytoplasm | other | 440723.25 | 2772544 | 1953939 | Q9Z2I0 | LETM1 | 440723 | 1183027 | 683021 | 3 |
| lectin, galactoside-binding, soluble, 9B | Cytoplasm | other | 2.92E+07 | 2.58E+07 | 2.75E+07 | | | | | | |
| lectin, galactoside-binding, soluble, 9B | Cytoplasm | other | 2.45E+07 | 1.58E+07 | 1.66E+07 | | | | | | |
| lectin, galactoside-binding, soluble, 9B | Cytoplasm | other | 2.34E+07 | 1.78E+07 | 2.22E+07 | | | | | | |
| lectin, galactoside-binding, soluble, 9B | Cytoplasm | other | 2.28E+07 | 1.61E+07 | 2.06E+07 | | | | | | |
| lectin, galactoside-binding, soluble, 9B | Cytoplasm | other | 1.25E+07 | 9010934 | 1.12E+07 | | | | | | |
| lectin, galactoside-binding, soluble, 9B | Cytoplasm | other | 1.06E+07 | 1.01E+07 | 7832754.5 | | | | | | |
| lectin, galactoside-binding, soluble, 9B | Cytoplasm | other | 9589569 | 8687899 | 7669614 | | | | | | |
| lectin, galactoside-binding, soluble, 9B | Cytoplasm | other | 5407038.5 | 5685214 | 5973034 | O08573 | LGALS9B | 17240826 | 7587249 | 1548741 | 24 |
| leucine-rich repeat containing G protein-coupled receptor 4 | Plasma Membrane | transmembrane receptor | 453413.812 | 2163059.25 | 1698677.375 | A2ARI4 | LGR4 | 453414 | 884046 | 510404 | 3 |
| LIM and senescent cell antigen-like domains 1 | Plasma Membrane | other | 31175.572 | 1872589.125 | 2339744.5 | Q99JW4 | LIMS1 | 31176 | 1220555 | 704688 | 3 |
| lectin, mannose-binding, 1 | Cytoplasm | other | 1.31E+07 | 9473643 | 1.13E+07 | | | | | | |
| lectin, mannose-binding, 1 | Cytoplasm | other | 1.07E+07 | 9625530 | 1.11E+07 | | | | | | |
| lectin, mannose-binding, 1 | Cytoplasm | other | 9571075 | 9748338 | 1.00E+07 | | | | | | |
| lectin, mannose-binding, 1 | Cytoplasm | other | 8757028 | 8572992 | 9070208 | | | | | | |
| lectin, mannose-binding, 1 | Cytoplasm | other | 8046207 | 8857590 | 8087763 | | | | | | |
| lectin, mannose-binding, 1 | Cytoplasm | other | 7576240 | 5283595 | 4730033.5 | Q9D0F3 | LMAN1 | 9635092 | 2001868 | 471845 | 18 |
| lectin, mannose-binding 2 | Cytoplasm | transporter | 1.26E+07 | 1.22E+07 | 1.15E+07 | | | | | | |
| lectin, mannose-binding 2 | Cytoplasm | transporter | 1.22E+07 | 1.45E+07 | 1.55E+07 | | | | | | |
| lectin, mannose-binding 2 | Cytoplasm | transporter | 4487189.5 | 5510843.5 | 4739517.5 | | | | | | |
| lectin, mannose-binding 2 | Cytoplasm | transporter | 3719739 | 2944733.75 | 3561555.25 | | | | | | |
| lectin, mannose-binding 2 | Cytoplasm | transporter | 3217332 | 3553458.25 | 3645640.5 | Q9DBH5 | LMAN2 | 7234852 | 4765494 | 1230445 | 15 |
| LIM and cysteine-rich domains 1 | Cytoplasm | transcription regulator | 560108.75 | 6680425 | 8661214 | Q8VEE1 | LMCD1 | 560109 | 4223145 | 2438234 | 3 |
| lon peptidase 1, mitochondrial | Cytoplasm | peptidase | 3862734.25 | 3448868.25 | 3598735 | | | | | | |
| lon peptidase 1, mitochondrial | Cytoplasm | peptidase | 1009053 | 3560226.5 | 2434974.25 | Q8CGK3 | LONP1 | 2435894 | 1086282 | 443473 | 6 |
| lysophosphatidylcholine acyltransferase 1 | Cytoplasm | enzyme | 2.31E+07 | 1.58E+07 | 2.11E+07 | | | | | | |
| lysophosphatidylcholine acyltransferase 1 | Cytoplasm | enzyme | 1.30E+07 | 8596600 | 7872716.5 | | | | | | |
| lysophosphatidylcholine acyltransferase 1 | Cytoplasm | enzyme | 1.11E+07 | 9335809 | 8745861 | | | | | | |
| lysophosphatidylcholine acyltransferase 1 | Cytoplasm | enzyme | 1.01E+07 | 6328705.5 | 6614631.5 | | | | | | |
| lysophosphatidylcholine acyltransferase 1 | Cytoplasm | enzyme | 9521133 | 8401479 | 6638948.5 | | | | | | |

| | | | | | | | | | | | |
|---|---|---|---|---|---|---|---|---|---|---|---|
| lysophosphatidylcholine acyltransferase 1 | Cytoplasm | enzyme | 9062505 | 1.23E+07 | 9180725 | | | | | | |
| lysophosphatidylcholine acyltransferase 1 | Cytoplasm | enzyme | 8550265 | 8997225 | 8174852.5 | | | | | | |
| lysophosphatidylcholine acyltransferase 1 | Cytoplasm | enzyme | 5035740.5 | 3357448.25 | 3218054 | Q3TFD2 | LPCAT1 | 11187455 | 4721181 | 963707 | 24 |
| lysophosphatidylcholine acyltransferase 3 | Plasma Membrane | enzyme | 2.53E+07 | 1.80E+07 | 2.06E+07 | | | | | | |
| lysophosphatidylcholine acyltransferase 3 | Plasma Membrane | enzyme | 9989870 | 9402244 | 9123771 | | | | | | |
| lysophosphatidylcholine acyltransferase 3 | Plasma Membrane | enzyme | 9553547 | 7788041.5 | 9457908 | | | | | | |
| lysophosphatidylcholine acyltransferase 3 | Plasma Membrane | enzyme | 2088286.25 | 992205.438 | 2019294.375 | Q91V01 | LPCAT3 | 11742926 | 7533227 | 2174655 | 12 |
| low density lipoprotein receptor-related protein associated protein 1 | Plasma Membrane | transmembrane receptor | 1737223.625 | 1299094.125 | 1301660.625 | P55302 | LRPAP1 | 1737224 | 252217 | 145617 | 3 |
| leucine rich repeat containing 57 | Cytoplasm | other | 9909821 | 6740365 | 9036498 | | | | | | |
| leucine rich repeat containing 57 | Cytoplasm | other | 9553697 | 6297193.5 | 9217553 | | | | | | |
| leucine rich repeat containing 57 | Cytoplasm | other | 8926421 | 6606148 | 9879549 | | | | | | |
| leucine rich repeat containing 57 | Cytoplasm | other | 6695021.5 | 6569384 | 8195660 | | | | | | |
| leucine rich repeat containing 57 | Cytoplasm | other | 6160721 | 5329329 | 6403367 | | | | | | |
| leucine rich repeat containing 57 | Cytoplasm | other | 3004086.5 | 2693110.5 | 3451632.75 | Q9D1G5 | LRRC57 | 7374961 | 2292461 | 540338 | 18 |
| leucine rich repeat containing 59 | Cytoplasm | other | 671108.688 | 2533629.25 | 2594695.75 | Q922Q8 | LRRC59 | 671109 | 1093382 | 631264 | 3 |
| leucine rich repeat containing 7 | Plasma Membrane | other | 887613.188 | 1887676.5 | 1641350.375 | Q80TE7 | LRRC7 | 887613 | 521044 | 300825 | 3 |
| lactotransferrin | Extracellular Space | other | 151456.359 | 2855186.25 | 1812229.625 | | | | | | |
| lactotransferrin | Extracellular Space | other | 77825.352 | 731560.312 | 876370.125 | | | | | | |
| lactotransferrin | Extracellular Space | other | 71204.836 | 1496353.75 | 1566208.125 | | | | | | |
| lactotransferrin | Extracellular Space | other | 66585.469 | 860765.062 | 730541.438 | | | | | | |
| lactotransferrin | Extracellular Space | other | 1332.102 | 1934571.625 | 1444696.125 | P08071 | LTF | 73681 | 854135 | 220537 | 15 |
| LYN proto-oncogene, Src family tyrosine kinase | Cytoplasm | kinase | 1976459.75 | 1.22E+07 | 9887634 | | | | | | |
| LYN proto-oncogene, Src family tyrosine kinase | Cytoplasm | kinase | 1965834.125 | 7835318.5 | 6184925.5 | | | | | | |
| LYN proto-oncogene, Src family tyrosine kinase | Cytoplasm | kinase | 1944894.75 | 8686469 | 6184610 | | | | | | |
| LYN proto-oncogene, Src family tyrosine kinase | Cytoplasm | kinase | 1756887.5 | 5273521 | 4423087.5 | | | | | | |
| LYN proto-oncogene, Src family tyrosine kinase | Cytoplasm | kinase | 519725.719 | 3161360.75 | 1805105.125 | | | | | | |
| LYN proto-oncogene, Src family tyrosine kinase | Cytoplasm | kinase | 322615.062 | 3005571.25 | 2402921.75 | | | | | | |
| LYN proto-oncogene, Src family tyrosine kinase | Cytoplasm | kinase | 191064.844 | 2149598 | 2243291 | | | | | | |
| LYN proto-oncogene, Src family tyrosine kinase | Cytoplasm | kinase | 79078.414 | 1120194.5 | 600926.375 | P25911 | LYN | 1094570 | 3323916 | 678491 | 24 |
| lysozyme | Extracellular Space | enzyme | 1328900.125 | 1.85E+07 | 1.35E+07 | | | | | | |
| lysozyme | Extracellular Space | enzyme | 173001.969 | 1.80E+07 | 5012304 | P08905 | LYZ | 750951 | 8275533 | 3378472 | 6 |
| mannose-6-phosphate receptor (cation dependent) | Cytoplasm | transporter | 2288483 | 2747899.75 | 3924339.25 | | | | | | |
| mannose-6-phosphate receptor (cation dependent) | Cytoplasm | transporter | 1000132.688 | 1156313.625 | 1300396.25 | P24668 | M6PR | 1644308 | 1141717 | 466104 | 6 |
| mesencephalic astrocyte-derived neurotrophic factor | Extracellular Space | other | 1.39E+07 | 1.27E+07 | 1.70E+07 | | | | | | |
| mesencephalic astrocyte-derived neurotrophic factor | Extracellular Space | other | 4092092 | 5051705.5 | 3842605.25 | | | | | | |
| mesencephalic astrocyte-derived neurotrophic factor | Extracellular Space | other | 2533623.25 | 1711180.5 | 1939971.375 | Q9CXI5 | MANF | 6831905 | 5856562 | 1952187 | 9 |
| microtubule-associated protein, RP/EB family, member 1 | Cytoplasm | other | 1583700.75 | 1780166.25 | 1246714.875 | Q61166 | MAPRE1 | 1583701 | 269793 | 155765 | 3 |
| mitochondrial amidoxime reducing component 2 | Cytoplasm | enzyme | 371559.312 | 968976.625 | 515078.031 | Q922Q1 | MARC_2 | 371559 | 311857 | 180051 | 3 |
| membrane-associated ring finger (C3HC4) 8, E3 ubiquitin protein ligase | Cytoplasm | enzyme | 5322463.5 | 3737036 | 4098138.5 | | | | | | |
| membrane-associated ring finger (C3HC4) 8, E3 ubiquitin protein ligase | Cytoplasm | enzyme | 3940972.5 | 2123005.75 | 2920609.5 | Q9DBD2 | MARH8 | 4631718 | 1090334 | 445127 | 6 |
| MAP/microtubule affinity-regulating kinase 2 | Cytoplasm | kinase | 2226624.75 | 2366204.25 | 2008676.75 | | | | | | |
| MAP/microtubule affinity-regulating kinase 2 | Cytoplasm | kinase | 977098.812 | 1023682.875 | 855627.875 | Q05512 | MARK2 | 1601862 | 695358 | 283879 | 6 |
| mitochondrial antiviral signaling protein | Cytoplasm | other | 2337254.75 | 5664388.5 | 5340916.5 | | | | | | |
| mitochondrial antiviral signaling protein | Cytoplasm | other | 918494.188 | 3267819.25 | 2176320 | Q8VCF0 | MAVS | 1627874 | 1877036 | 766297 | 6 |
| metallo-beta-lactamase domain containing 2 | Cytoplasm | other | 1.74E+07 | 1.60E+07 | 1.42E+07 | Q8BL86 | MBLAC2 | 17400000 | 1629305 | 940679 | 3 |
| membrane bound O-acyltransferase domain containing 7 | Plasma Membrane | other | 314667.656 | 4398651 | 3528132.5 | Q8CHK3 | MBOAT7 | 314668 | 2151088 | 1241931 | 3 |
| multiple coagulation factor deficiency 2 | Cytoplasm | other | 1.07E+07 | 7329500 | 9023339 | Q8K5B2 | MCFD2 | 10700000 | 1685257 | 972984 | 3 |
| mitochondrial calcium uniporter | Cytoplasm | ion channel | 409436.875 | 2639371.25 | 1410931.75 | Q3UMR5 | MCU | 409437 | 1116890 | 644837 | 3 |
| malate dehydrogenase 2, NAD (mitochondrial) | Cytoplasm | enzyme | 2.35E+07 | 3.11E+07 | 4.13E+07 | | | | | | |
| malate dehydrogenase 2, NAD (mitochondrial) | Cytoplasm | enzyme | 1.80E+07 | 3.32E+07 | 3.04E+07 | | | | | | |
| malate dehydrogenase 2, NAD (mitochondrial) | Cytoplasm | enzyme | 1.78E+07 | 2.82E+07 | 2.47E+07 | | | | | | |
| malate dehydrogenase 2, NAD (mitochondrial) | Cytoplasm | enzyme | 1.59E+07 | 2.57E+07 | 2.40E+07 | | | | | | |
| malate dehydrogenase 2, NAD (mitochondrial) | Cytoplasm | enzyme | 1.46E+07 | 2.57E+07 | 2.71E+07 | | | | | | |
| malate dehydrogenase 2, NAD (mitochondrial) | Cytoplasm | enzyme | 1.28E+07 | 3.51E+07 | 2.85E+07 | | | | | | |
| malate dehydrogenase 2, NAD (mitochondrial) | Cytoplasm | enzyme | 1.20E+07 | 2.24E+07 | 2.19E+07 | | | | | | |
| malate dehydrogenase 2, NAD (mitochondrial) | Cytoplasm | enzyme | 1.15E+07 | 2.57E+07 | 2.33E+07 | | | | | | |
| malate dehydrogenase 2, NAD (mitochondrial) | Cytoplasm | enzyme | 1.02E+07 | 1.74E+07 | 1.76E+07 | | | | | | |
| malate dehydrogenase 2, NAD (mitochondrial) | Cytoplasm | enzyme | 8108692.5 | 1.40E+07 | 1.09E+07 | | | | | | |
| malate dehydrogenase 2, NAD (mitochondrial) | Cytoplasm | enzyme | 7635212 | 1.95E+07 | 1.39E+07 | | | | | | |
| malate dehydrogenase 2, NAD (mitochondrial) | Cytoplasm | enzyme | 3706547.75 | 8343783 | 6187561 | | | | | | |
| malate dehydrogenase 2, NAD (mitochondrial) | Cytoplasm | enzyme | 1637529.75 | 2331577.75 | 2826691 | P08249 | MDH2 | 12093691 | 9850676 | 1577371 | 39 |
| malic enzyme 2, NAD(+)-dependent, mitochondrial | Cytoplasm | enzyme | 3549060.25 | 4043495.75 | 3662393.75 | | | | | | |
| malic enzyme 2, NAD(+)-dependent, mitochondrial | Cytoplasm | enzyme | 3485035.5 | 3988336 | 4333686.5 | | | | | | |
| malic enzyme 2, NAD(+)-dependent, mitochondrial | Cytoplasm | enzyme | 1008154.5 | 1829072.75 | 1409065.875 | | | | | | |
| malic enzyme 2, NAD(+)-dependent, mitochondrial | Cytoplasm | enzyme | 962973.188 | 2024962.5 | 1783609.125 | Q99KE1 | ME2 | 2251306 | 1278801 | 369158 | 12 |
| mediator of cell motility 1 | Cytoplasm | other | 1.06E+07 | 6340352 | 8466565 | | | | | | |
| mediator of cell motility 1 | Cytoplasm | other | 7651612.5 | 3402479.75 | 3698658.25 | | | | | | |
| mediator of cell motility 1 | Cytoplasm | other | 4954600.5 | 3243581.25 | 4160297.25 | | | | | | |
| mediator of cell motility 1 | Cytoplasm | other | 3326405.5 | 2713671.75 | 2679794.5 | | | | | | |
| mediator of cell motility 1 | Cytoplasm | other | 2450850.5 | 2006531 | 2148788.25 | Q91VH6 | MEMO1 | 5802694 | 2593868 | 669734 | 15 |
| mesoderm development candidate 2 | Extracellular Space | other | 1299740.625 | 1345922.375 | 1305675.25 | Q9ERE7 | MESDC2 | 1299741 | 25126 | 14506 | 3 |
| methionyl aminopeptidase 2 | Cytoplasm | peptidase | 2119599 | 691431.438 | 1961304.25 | O08663 | METAP2 | 2119599 | 782868 | 451989 | 3 |
| mitochondrial fission factor | Cytoplasm | other | 2070086.375 | 3123148.25 | 3100546 | | | | | | |
| mitochondrial fission factor | Cytoplasm | other | 1529773.125 | 4297414 | 3075193.75 | Q6PCP5 | Mff | 1799930 | 962583 | 392973 | 6 |
| major facilitator superfamily domain containing 2B | Plasma Membrane | other | 3.17E+07 | 2.56E+07 | 2.60E+07 | | | | | | |
| major facilitator superfamily domain containing 2B | Plasma Membrane | other | 2088679.25 | 2624222.5 | 2141058 | Q3T9M1 | MFSD2B | 16904340 | 14127339 | 5767462 | 6 |
| microsomal glutathione S-transferase 3 | Cytoplasm | enzyme | 2.55E+07 | 2.32E+07 | 3.15E+07 | | | | | | |
| microsomal glutathione S-transferase 3 | Cytoplasm | enzyme | 7030508 | 3870580.25 | 5845284.5 | | | | | | |
| microsomal glutathione S-transferase 3 | Cytoplasm | enzyme | 2405355 | 3231677.5 | 4254628 | Q9CPU4 | MGST3 | 11655288 | 11427134 | 3809045 | 9 |
| microtubule associated monooxygenase, calponin and LIM domain containing 1 | Cytoplasm | enzyme | 3132395.25 | 2979351.25 | 2883972.75 | Q8VDP3 | MICAL1 | 3132395 | 125322 | 72355 | 3 |
| malectin | Plasma Membrane | other | 1521362.625 | 6279044 | 3904901.5 | | | | | | |
| malectin | Plasma Membrane | other | 1484337.5 | 7849571.5 | 4655335.5 | | | | | | |
| malectin | Plasma Membrane | other | 1401884.375 | 6453207.5 | 4644680.5 | | | | | | |
| malectin | Plasma Membrane | other | 822697.25 | 4653720.5 | 3438300.25 | | | | | | |
| malectin | Plasma Membrane | other | 589419.75 | 4298476.5 | 2605742.75 | Q6ZQI3 | MLEC | 1163940 | 2214710 | 571836 | 15 |
| matrix metallopeptidase 21 | Extracellular Space | peptidase | 1601990.375 | 181950.5 | 881038 | Q8K3F2 | MMP21 | 1601990 | 710048 | 409946 | 3 |
| multimerin 1 | Extracellular Space | other | 325191.219 | 6676892 | 5566795 | | | | | | |
| multimerin 1 | Extracellular Space | other | 290226.531 | 4606605 | 3062927.5 | | | | | | |
| multimerin 1 | Extracellular Space | other | 247999.766 | 4359461 | 3866046.25 | | | | | | |
| multimerin 1 | Extracellular Space | other | 214307.594 | 3875212 | 2371049.5 | | | | | | |
| multimerin 1 | Extracellular Space | other | 174861.469 | 2343371.75 | 2280260.75 | | | | | | |
| multimerin 1 | Extracellular Space | other | 130144.883 | 5415770 | 4253079 | | | | | | |
| multimerin 1 | Extracellular Space | other | 112007.016 | 5915329.5 | 4209734 | | | | | | |
| multimerin 1 | Extracellular Space | other | 83278.789 | 4895722 | 2982581.5 | | | | | | |
| multimerin 1 | Extracellular Space | other | 69119.516 | 2948452.5 | 2542687 | | | | | | |

| | | | | | | | | | | | |
|---|---|---|---|---|---|---|---|---|---|---|---|
| multimerin 1 | Extracellular Space | other | 21561.668 | 3048086.25 | 1878652.375 | | | | | | |
| multimerin 1 | Extracellular Space | other | 0 | 2590642 | 2846383.25 | B2RPV6 | MMRN1 | 151700 | 2036782 | 354558 | 33 |
| membrane protein, palmitoylated 1, 55kDa | Plasma Membrane | kinase | 1.91E+09 | 1.02E+09 | 1.22E+09 | | | | | | |
| membrane protein, palmitoylated 1, 55kDa | Plasma Membrane | kinase | 1.78E+09 | 1.14E+09 | 1.25E+09 | | | | | | |
| membrane protein, palmitoylated 1, 55kDa | Plasma Membrane | kinase | 1.28E+09 | 9.19E+08 | 1.06E+09 | | | | | | |
| membrane protein, palmitoylated 1, 55kDa | Plasma Membrane | kinase | 1.22E+09 | 1.10E+09 | 9.72E+08 | | | | | | |
| membrane protein, palmitoylated 1, 55kDa | Plasma Membrane | kinase | 1.11E+09 | 1.08E+09 | 1.04E+09 | | | | | | |
| membrane protein, palmitoylated 1, 55kDa | Plasma Membrane | kinase | 1.06E+09 | 9.69E+08 | 9.06E+08 | | | | | | |
| membrane protein, palmitoylated 1, 55kDa | Plasma Membrane | kinase | 7.80E+08 | 5.81E+08 | 6.50E+08 | | | | | | |
| membrane protein, palmitoylated 1, 55kDa | Plasma Membrane | kinase | 7.00E+08 | 5.51E+08 | 5.79E+08 | | | | | | |
| membrane protein, palmitoylated 1, 55kDa | Plasma Membrane | kinase | 6.97E+08 | 5.64E+08 | 5.24E+08 | | | | | | |
| membrane protein, palmitoylated 1, 55kDa | Plasma Membrane | kinase | 6.58E+08 | 5.69E+08 | 5.35E+08 | | | | | | |
| membrane protein, palmitoylated 1, 55kDa | Plasma Membrane | kinase | 6.54E+08 | 3.24E+08 | 4.32E+08 | | | | | | |
| membrane protein, palmitoylated 1, 55kDa | Plasma Membrane | kinase | 5.58E+08 | 5.80E+08 | 5.63E+08 | | | | | | |
| membrane protein, palmitoylated 1, 55kDa | Plasma Membrane | kinase | 5.42E+08 | 4.14E+08 | 4.08E+08 | | | | | | |
| membrane protein, palmitoylated 1, 55kDa | Plasma Membrane | kinase | 4.90E+08 | 5.23E+08 | 5.29E+08 | | | | | | |
| membrane protein, palmitoylated 1, 55kDa | Plasma Membrane | kinase | 4.86E+08 | 3.87E+08 | 3.68E+08 | | | | | | |
| membrane protein, palmitoylated 1, 55kDa | Plasma Membrane | kinase | 4.37E+08 | 2.98E+08 | 3.21E+08 | | | | | | |
| membrane protein, palmitoylated 1, 55kDa | Plasma Membrane | kinase | 3.88E+08 | 3.29E+08 | 3.52E+08 | | | | | | |
| membrane protein, palmitoylated 1, 55kDa | Plasma Membrane | kinase | 3.80E+08 | 4.77E+08 | 4.31E+08 | | | | | | |
| membrane protein, palmitoylated 1, 55kDa | Plasma Membrane | kinase | 3.44E+08 | 2.82E+08 | 2.80E+08 | | | | | | |
| membrane protein, palmitoylated 1, 55kDa | Plasma Membrane | kinase | 3.41E+08 | 2.87E+08 | 2.47E+08 | | | | | | |
| membrane protein, palmitoylated 1, 55kDa | Plasma Membrane | kinase | 2.15E+08 | 1.71E+08 | 1.43E+08 | | | | | | |
| membrane protein, palmitoylated 1, 55kDa | Plasma Membrane | kinase | 1.98E+08 | 1.61E+08 | 1.50E+08 | | | | | | |
| membrane protein, palmitoylated 1, 55kDa | Plasma Membrane | kinase | 1.29E+08 | 1.04E+08 | 8.85E+07 | | | | | | |
| membrane protein, palmitoylated 1, 55kDa | Plasma Membrane | kinase | 1.04E+08 | 1.35E+08 | 9.82E+07 | | | | | | |
| membrane protein, palmitoylated 1, 55kDa | Plasma Membrane | kinase | 7.52E+07 | 2.03E+08 | 1.41E+08 | | | | | | |
| membrane protein, palmitoylated 1, 55kDa | Plasma Membrane | kinase | 7.12E+07 | 1.04E+08 | 9.21E+07 | | | | | | |
| membrane protein, palmitoylated 1, 55kDa | Plasma Membrane | kinase | 6.13E+07 | 2.77E+07 | 4.65E+07 | | | | | | |
| membrane protein, palmitoylated 1, 55kDa | Plasma Membrane | kinase | 6.12E+07 | 7.93E+07 | 5.60E+07 | | | | | | |
| membrane protein, palmitoylated 1, 55kDa | Plasma Membrane | kinase | 5.51E+07 | 5.41E+07 | 4.17E+07 | | | | | | |
| membrane protein, palmitoylated 1, 55kDa | Plasma Membrane | kinase | 5.40E+07 | 3.33E+07 | 3.72E+07 | | | | | | |
| membrane protein, palmitoylated 1, 55kDa | Plasma Membrane | kinase | 5.07E+07 | 3.98E+07 | 5.14E+07 | | | | | | |
| membrane protein, palmitoylated 1, 55kDa | Plasma Membrane | kinase | 3.44E+07 | 2.20E+07 | 2.11E+07 | | | | | | |
| membrane protein, palmitoylated 1, 55kDa | Plasma Membrane | kinase | 2.98E+07 | 3.22E+07 | 3.92E+07 | | | | | | |
| membrane protein, palmitoylated 1, 55kDa | Plasma Membrane | kinase | 1.89E+07 | 8.50E+07 | 1.33E+08 | | | | | | |
| membrane protein, palmitoylated 1, 55kDa | Plasma Membrane | kinase | 1.56E+07 | 4.22E+07 | 2.36E+07 | | | | | | |
| membrane protein, palmitoylated 1, 55kDa | Plasma Membrane | kinase | 1.27E+07 | 2.33E+07 | 1.37E+07 | | | | | | |
| membrane protein, palmitoylated 1, 55kDa | Plasma Membrane | kinase | 9253730 | 7137726 | 9229850 | | | | | | |
| membrane protein, palmitoylated 1, 55kDa | Plasma Membrane | kinase | 7653266.5 | 3.01E+07 | 1.56E+07 | | | | | | |
| membrane protein, palmitoylated 1, 55kDa | Plasma Membrane | kinase | 5521123 | 2.81E+07 | 1.46E+07 | | | | | | |
| membrane protein, palmitoylated 1, 55kDa | Plasma Membrane | kinase | 3261180.25 | 2070147.75 | 2157944.5 | | | | | | |
| membrane protein, palmitoylated 1, 55kDa | Plasma Membrane | kinase | 2677349.5 | 2147630 | 2827880 | | | | | | |
| membrane protein, palmitoylated 1, 55kDa | Plasma Membrane | kinase | 276109.656 | 243515.375 | 315340.938 | P70290 | MPP1 | 405636256 | 405153114 | 36093908 | 126 |
| mercaptopyruvate sulfurtransferase | Cytoplasm | enzyme | 8253323.5 | 6819325 | 8891125 | | | | | | |
| mercaptopyruvate sulfurtransferase | Cytoplasm | enzyme | 723238.5 | 627089 | 528980.562 | Q99J99 | MPST | 4488281 | 4087982 | 1668912 | 6 |
| mitochondrial ribosomal protein L12 | Cytoplasm | other | 622013.188 | 1098275.875 | 1239619.875 | Q9DB15 | MRPL12 | 622013 | 323585 | 186822 | 3 |
| mitochondrial ribosomal protein S36 | Cytoplasm | other | 771723.312 | 1518150.125 | 1202710.375 | Q9CQX8 | MRPS36 | 771723 | 374701 | 216334 | 3 |
| murine retrovirus integration site 1 homolog | Cytoplasm | other | 199070.562 | 2849923.5 | 1213693.375 | | | | | | |
| murine retrovirus integration site 1 homolog | Cytoplasm | other | 116345.453 | 1867207.5 | 1082607 | | | | | | |
| murine retrovirus integration site 1 homolog | Cytoplasm | other | 4.70E-06 | 1039973 | 432913.812 | Q9WUX5 | MRVI1 | 105139 | 933267 | 311089 | 9 |
| moesin | Plasma Membrane | other | 9671137 | 7096139 | 8524828 | | | | | | |
| moesin | Plasma Membrane | other | 9267101 | 5711588 | 6271957.5 | | | | | | |
| moesin | Plasma Membrane | other | 8201925.5 | 6872811 | 6201672.5 | | | | | | |
| moesin | Plasma Membrane | other | 7313734 | 4933800.5 | 5796236.5 | | | | | | |
| moesin | Plasma Membrane | other | 5459615 | 3849942 | 3597275.25 | | | | | | |
| moesin | Plasma Membrane | other | 3724817.75 | 1736029.875 | 1765907.625 | | | | | | |
| moesin | Plasma Membrane | other | 3389018 | 4375594 | 4743565 | | | | | | |
| moesin | Plasma Membrane | other | 3323693.75 | 2861913.75 | 2042284.75 | | | | | | |
| moesin | Plasma Membrane | other | 2893368.5 | 2342229.25 | 2328127 | | | | | | |
| moesin | Plasma Membrane | other | 1788869 | 877082.5 | 748966.125 | | | | | | |
| moesin | Plasma Membrane | other | 1607893.5 | 727075.75 | 1119898.375 | | | | | | |
| moesin | Plasma Membrane | other | 1157421 | 1482371.25 | 1718564.875 | P26041 | MSN | 4816550 | 2614896 | 435816 | 36 |
| ATP synthase F0 subunit 8 | Cytoplasm | enzyme | 8771255 | 1.50E+07 | 1.30E+07 | | | | | | |
| ATP synthase F0 subunit 8 | Cytoplasm | enzyme | 2435640.25 | 4838302.5 | 4031384.25 | P03930 | mt-Atp8 | 5603448 | 5126071 | 2092710 | 6 |
| mitochondrial carrier 2 | Cytoplasm | other | 3436243.25 | 5605464 | 5020243 | Q791V5 | MTCH2 | 3436243 | 1122279 | 647948 | 3 |
| cytochrome c oxidase subunit I | Cytoplasm | enzyme | 513720.562 | 1943674.625 | 1392267.125 | P00397 | MT-CO1 | 513721 | 721187 | 416377 | 3 |
| cytochrome c oxidase subunit II | Cytoplasm | enzyme | 1.19E+07 | 2.02E+07 | 2.44E+07 | | | | | | |
| cytochrome c oxidase subunit II | Cytoplasm | enzyme | 9126300 | 1.14E+07 | 1.51E+07 | | | | | | |
| cytochrome c oxidase subunit II | Cytoplasm | enzyme | 1834235 | 3289990.5 | 3245995.75 | | | | | | |
| cytochrome c oxidase subunit II | Cytoplasm | enzyme | 1002725.562 | 1994913.25 | 1923519.375 | | | | | | |
| cytochrome c oxidase subunit II | Cytoplasm | enzyme | 952793.812 | 1230111 | 1205400.875 | | | | | | |
| cytochrome c oxidase subunit II | Cytoplasm | enzyme | 809036.125 | 5967983.5 | 3458296.25 | P00405 | MT-CO2 | 4277515 | 7212496 | 1700002 | 18 |
| cytochrome c oxidase III | Cytoplasm | enzyme | 764169.312 | 1254465.125 | 1337022 | P00416 | MT-CO3 | 764169 | 309668 | 178787 | 3 |
| methylenetetrahydrofolate dehydrogenase (NADP+ dependent) 1, methenyltetrahydrofolate cyclohydrolase, formyltetrahydrofolate synthetase | Cytoplasm | enzyme | 2.40E+07 | 1.57E+07 | 1.63E+07 | | | | | | |
| methylenetetrahydrofolate dehydrogenase (NADP+ dependent) 1, methenyltetrahydrofolate cyclohydrolase, formyltetrahydrofolate synthetase | Cytoplasm | enzyme | 2.34E+07 | 1.09E+07 | 1.35E+07 | | | | | | |
| methylenetetrahydrofolate dehydrogenase (NADP+ dependent) 1, methenyltetrahydrofolate cyclohydrolase, formyltetrahydrofolate synthetase | Cytoplasm | enzyme | 2.22E+07 | 1.23E+07 | 1.54E+07 | | | | | | |
| methylenetetrahydrofolate dehydrogenase (NADP+ dependent) 1, methenyltetrahydrofolate cyclohydrolase, formyltetrahydrofolate synthetase | Cytoplasm | enzyme | 2.04E+07 | 1.04E+07 | 1.30E+07 | | | | | | |
| methylenetetrahydrofolate dehydrogenase (NADP+ dependent) 1, methenyltetrahydrofolate cyclohydrolase, formyltetrahydrofolate synthetase | Cytoplasm | enzyme | 2.02E+07 | 9654595 | 1.06E+07 | | | | | | |
| methylenetetrahydrofolate dehydrogenase (NADP+ dependent) 1, methenyltetrahydrofolate cyclohydrolase, formyltetrahydrofolate synthetase | Cytoplasm | enzyme | 2.01E+07 | 9445357 | 1.17E+07 | | | | | | |
| methylenetetrahydrofolate dehydrogenase (NADP+ dependent) 1, methenyltetrahydrofolate cyclohydrolase, formyltetrahydrofolate synthetase | Cytoplasm | enzyme | 1.84E+07 | 1.05E+07 | 1.59E+07 | | | | | | |
| methylenetetrahydrofolate dehydrogenase (NADP+ dependent) 1, methenyltetrahydrofolate cyclohydrolase, formyltetrahydrofolate synthetase | Cytoplasm | enzyme | 1.83E+07 | 9842771 | 1.25E+07 | | | | | | |
| methylenetetrahydrofolate dehydrogenase (NADP+ dependent) 1, methenyltetrahydrofolate cyclohydrolase, formyltetrahydrofolate synthetase | Cytoplasm | enzyme | 1.66E+07 | 7803742.5 | 4215288 | | | | | | |
| methylenetetrahydrofolate dehydrogenase (NADP+ dependent) 1, methenyltetrahydrofolate cyclohydrolase, formyltetrahydrofolate synthetase | Cytoplasm | enzyme | 1.54E+07 | 8738155 | 8971593 | | | | | | |
| methylenetetrahydrofolate dehydrogenase (NADP+ dependent) 1, methenyltetrahydrofolate cyclohydrolase, formyltetrahydrofolate synthetase | Cytoplasm | enzyme | 1.43E+07 | 9950427 | 7928759 | | | | | | |
| methylenetetrahydrofolate dehydrogenase (NADP+ dependent) 1, methenyltetrahydrofolate cyclohydrolase, formyltetrahydrofolate synthetase | Cytoplasm | enzyme | 1.41E+07 | 5765553.5 | 6047667 | | | | | | |
| methylenetetrahydrofolate dehydrogenase (NADP+ dependent) 1, methenyltetrahydrofolate cyclohydrolase, formyltetrahydrofolate synthetase | Cytoplasm | enzyme | 1.39E+07 | 7108401 | 8382120 | | | | | | |
| methylenetetrahydrofolate dehydrogenase (NADP+ dependent) 1, methenyltetrahydrofolate cyclohydrolase, formyltetrahydrofolate synthetase | Cytoplasm | enzyme | 1.36E+07 | 1.11E+07 | 1.06E+07 | | | | | | |
| methylenetetrahydrofolate dehydrogenase (NADP+ dependent) 1, methenyltetrahydrofolate cyclohydrolase, formyltetrahydrofolate synthetase | Cytoplasm | enzyme | 1.21E+07 | 8763464 | 5818274 | | | | | | |
| methylenetetrahydrofolate dehydrogenase (NADP+ dependent) 1, methenyltetrahydrofolate cyclohydrolase, formyltetrahydrofolate synthetase | Cytoplasm | enzyme | 1.13E+07 | 6205191.5 | 6294091 | | | | | | |
| methylenetetrahydrofolate dehydrogenase (NADP+ dependent) 1, methenyltetrahydrofolate cyclohydrolase, formyltetrahydrofolate synthetase | Cytoplasm | enzyme | 9704487 | 5624972.5 | 7600069 | | | | | | |
| methylenetetrahydrofolate dehydrogenase (NADP+ dependent) 1, methenyltetrahydrofolate cyclohydrolase, formyltetrahydrofolate synthetase | Cytoplasm | enzyme | 8804488 | 4245505.5 | 4707232.5 | | | | | | |
| methylenetetrahydrofolate dehydrogenase (NADP+ dependent) 1, methenyltetrahydrofolate cyclohydrolase, formyltetrahydrofolate synthetase | Cytoplasm | enzyme | 8504414 | 4621014 | 6732983 | | | | | | |
| methylenetetrahydrofolate dehydrogenase (NADP+ dependent) 1, methenyltetrahydrofolate cyclohydrolase, formyltetrahydrofolate synthetase | Cytoplasm | enzyme | 8280399 | 5331470.5 | 6640481.5 | | | | | | |
| methylenetetrahydrofolate dehydrogenase (NADP+ dependent) 1, methenyltetrahydrofolate cyclohydrolase, formyltetrahydrofolate synthetase | Cytoplasm | enzyme | 7432892.5 | 3752940.5 | 4864699 | | | | | | |

| | | | | | | | | | | | |
|---|---|---|---|---|---|---|---|---|---|---|---|
| methylenetetrahydrofolate dehydrogenase (NADP+ dependent) 1, methenyltetrahydrofolate cyclohydrolase, formyltetrahydrofolate synthetase | Cytoplasm | enzyme | 7014220 | 4277197 | 4853748.5 | | | | | | |
| methylenetetrahydrofolate dehydrogenase (NADP+ dependent) 1, methenyltetrahydrofolate cyclohydrolase, formyltetrahydrofolate synthetase | Cytoplasm | enzyme | 6227499.5 | 5450859.5 | 5657879 | | | | | | |
| methylenetetrahydrofolate dehydrogenase (NADP+ dependent) 1, methenyltetrahydrofolate cyclohydrolase, formyltetrahydrofolate synthetase | Cytoplasm | enzyme | 6156822 | 1253304.625 | 966219 | | | | | | |
| methylenetetrahydrofolate dehydrogenase (NADP+ dependent) 1, methenyltetrahydrofolate cyclohydrolase, formyltetrahydrofolate synthetase | Cytoplasm | enzyme | 6148034 | 6509255.5 | 5259866 | | | | | | |
| methylenetetrahydrofolate dehydrogenase (NADP+ dependent) 1, methenyltetrahydrofolate cyclohydrolase, formyltetrahydrofolate synthetase | Cytoplasm | enzyme | 6090534 | 3046370 | 3421762.75 | | | | | | |
| methylenetetrahydrofolate dehydrogenase (NADP+ dependent) 1, methenyltetrahydrofolate cyclohydrolase, formyltetrahydrofolate synthetase | Cytoplasm | enzyme | 4583569.5 | 3057794.5 | 2142037.75 | | | | | | |
| methylenetetrahydrofolate dehydrogenase (NADP+ dependent) 1, methenyltetrahydrofolate cyclohydrolase, formyltetrahydrofolate synthetase | Cytoplasm | enzyme | 4214451.5 | 2383240.5 | 3128903.25 | | | | | | |
| methylenetetrahydrofolate dehydrogenase (NADP+ dependent) 1, methenyltetrahydrofolate cyclohydrolase, formyltetrahydrofolate synthetase | Cytoplasm | enzyme | 3379852.75 | 1237965.25 | 1407511 | Q922D8 | MTHFD1 | 12582126 | 5421849 | 581283 | 87 |
| methylenetetrahydrofolate dehydrogenase (NADP+ dependent) 1-like | Cytoplasm | enzyme | 2122779 | 3856286 | 2092184.5 | | | | | | |
| methylenetetrahydrofolate dehydrogenase (NADP+ dependent) 1-like | Cytoplasm | enzyme | 2006471 | 3722795.5 | 2501537 | | | | | | |
| methylenetetrahydrofolate dehydrogenase (NADP+ dependent) 1-like | Cytoplasm | enzyme | 1615278.5 | 2604859.25 | 1992208.125 | | | | | | |
| methylenetetrahydrofolate dehydrogenase (NADP+ dependent) 1-like | Cytoplasm | enzyme | 1497211 | 1668559.875 | 1809296.125 | | | | | | |
| methylenetetrahydrofolate dehydrogenase (NADP+ dependent) 1-like | Cytoplasm | enzyme | 1276782.375 | 1444249.625 | 1826990.625 | | | | | | |
| methylenetetrahydrofolate dehydrogenase (NADP+ dependent) 1-like | Cytoplasm | enzyme | 1186232.125 | 2370961 | 1758339.625 | | | | | | |
| methylenetetrahydrofolate dehydrogenase (NADP+ dependent) 1-like | Cytoplasm | enzyme | 1142281.875 | 1394505.375 | 1324950.375 | | | | | | |
| methylenetetrahydrofolate dehydrogenase (NADP+ dependent) 1-like | Cytoplasm | enzyme | 447627.812 | 1195112.375 | 1362839.125 | Q3V3R1 | MTHFD1L | 1411833 | 773148 | 157818 | 24 |
| NADH dehydrogenase, subunit 5 (complex I) | Cytoplasm | enzyme | 2348204.75 | 4617148 | 2310097.75 | | | | | | |
| NADH dehydrogenase, subunit 5 (complex I) | Cytoplasm | enzyme | 756502.188 | 1500310.125 | 1247772.625 | | | | | | |
| NADH dehydrogenase, subunit 5 (complex I) | Cytoplasm | enzyme | 739691.438 | 860853.625 | 788514.75 | P03921 | MT-ND5 | 1281466 | 1269447 | 423149 | 9 |
| 5-methyltetrahydrofolate-homocysteine methyltransferase reductase | Cytoplasm | enzyme | 4057716.25 | 3344083.25 | 3264682.5 | Q8C1A3 | MTRR | 4057716 | 436745 | 252155 | 3 |
| murinoglobulin 1 | Extracellular Space | other | 736962.5 | 682248.75 | 1506869.625 | P28665 | Mug1/Mug2 | 736963 | 461113 | 266224 | 3 |
| v-myb avian myeloblastosis viral oncogene homolog-like 2 | Nucleus | transcription regulator | 8026032 | 1.34E+07 | 1.26E+07 | P48972 | MYBL2 | 8026032 | 2903737 | 1676473 | 3 |
| myeloma overexpressed 2 | Nucleus | other | 1.24E+07 | 4767113 | 7223109.5 | Q3U898 | MYEOV2 | 12380000 | 3885475 | 2243280 | 3 |
| myosin, heavy chain 14, non-muscle | Extracellular Space | other | 637943.188 | 5767900.5 | 3805150.5 | | | | | | |
| myosin, heavy chain 14, non-muscle | Extracellular Space | other | 111820.82 | 2914398 | 2018781.5 | Q6URW6 | MYH14 | 374882 | 2095024 | 855290 | 6 |
| myosin, heavy chain 9, non-muscle | Cytoplasm | enzyme | 1.84E+07 | 3.04E+07 | 1.99E+07 | | | | | | |
| myosin, heavy chain 9, non-muscle | Cytoplasm | enzyme | 1.26E+07 | 4.57E+07 | 4.77E+07 | | | | | | |
| myosin, heavy chain 9, non-muscle | Cytoplasm | enzyme | 9957653 | 4.73E+07 | 4.47E+07 | | | | | | |
| myosin, heavy chain 9, non-muscle | Cytoplasm | enzyme | 8707025 | 2.53E+07 | 2.52E+07 | | | | | | |
| myosin, heavy chain 9, non-muscle | Cytoplasm | enzyme | 8602044 | 3.94E+07 | 4.76E+07 | | | | | | |
| myosin, heavy chain 9, non-muscle | Cytoplasm | enzyme | 8210772.5 | 5.53E+07 | 4.70E+07 | | | | | | |
| myosin, heavy chain 9, non-muscle | Cytoplasm | enzyme | 7864087.5 | 3.06E+07 | 3.05E+07 | | | | | | |
| myosin, heavy chain 9, non-muscle | Cytoplasm | enzyme | 7841690 | 4.74E+07 | 3.95E+07 | | | | | | |
| myosin, heavy chain 9, non-muscle | Cytoplasm | enzyme | 7627891 | 4.55E+07 | 4.03E+07 | | | | | | |
| myosin, heavy chain 9, non-muscle | Cytoplasm | enzyme | 7427073 | 3.75E+07 | 2.87E+07 | | | | | | |
| myosin, heavy chain 9, non-muscle | Cytoplasm | enzyme | 7193677.5 | 5.37E+07 | 4.39E+07 | | | | | | |
| myosin, heavy chain 9, non-muscle | Cytoplasm | enzyme | 7124571 | 3.75E+07 | 2.97E+07 | | | | | | |
| myosin, heavy chain 9, non-muscle | Cytoplasm | enzyme | 7024352.5 | 4.29E+07 | 3.29E+07 | | | | | | |
| myosin, heavy chain 9, non-muscle | Cytoplasm | enzyme | 6983599.5 | 4.15E+07 | 3.54E+07 | | | | | | |
| myosin, heavy chain 9, non-muscle | Cytoplasm | enzyme | 6944898.5 | 3.77E+07 | 2.69E+07 | | | | | | |
| myosin, heavy chain 9, non-muscle | Cytoplasm | enzyme | 6197404.5 | 3.39E+07 | 2.72E+07 | | | | | | |
| myosin, heavy chain 9, non-muscle | Cytoplasm | enzyme | 6172533 | 3.39E+07 | 2.60E+07 | | | | | | |
| myosin, heavy chain 9, non-muscle | Cytoplasm | enzyme | 5871991.5 | 3.61E+07 | 2.73E+07 | | | | | | |
| myosin, heavy chain 9, non-muscle | Cytoplasm | enzyme | 5805117 | 3.54E+07 | 2.45E+07 | | | | | | |
| myosin, heavy chain 9, non-muscle | Cytoplasm | enzyme | 5709095 | 2.72E+07 | 2.74E+07 | | | | | | |
| myosin, heavy chain 9, non-muscle | Cytoplasm | enzyme | 5186386 | 4.65E+07 | 3.41E+07 | | | | | | |
| myosin, heavy chain 9, non-muscle | Cytoplasm | enzyme | 5157020 | 2.57E+07 | 2.44E+07 | | | | | | |
| myosin, heavy chain 9, non-muscle | Cytoplasm | enzyme | 5044824 | 3.96E+07 | 2.97E+07 | | | | | | |
| myosin, heavy chain 9, non-muscle | Cytoplasm | enzyme | 5034052 | 1.88E+07 | 2.55E+07 | | | | | | |
| myosin, heavy chain 9, non-muscle | Cytoplasm | enzyme | 5000076.5 | 2.53E+07 | 2.35E+07 | | | | | | |
| myosin, heavy chain 9, non-muscle | Cytoplasm | enzyme | 4910074 | 2.53E+07 | 1.94E+07 | | | | | | |
| myosin, heavy chain 9, non-muscle | Cytoplasm | enzyme | 4815090.5 | 2.45E+07 | 2.23E+07 | | | | | | |
| myosin, heavy chain 9, non-muscle | Cytoplasm | enzyme | 4791312.5 | 3.24E+07 | 2.93E+07 | | | | | | |
| myosin, heavy chain 9, non-muscle | Cytoplasm | enzyme | 4705034 | 2.37E+07 | 2.55E+07 | | | | | | |
| myosin, heavy chain 9, non-muscle | Cytoplasm | enzyme | 4531341.5 | 3.26E+07 | 2.80E+07 | | | | | | |
| myosin, heavy chain 9, non-muscle | Cytoplasm | enzyme | 4517632.5 | 2.94E+07 | 2.45E+07 | | | | | | |
| myosin, heavy chain 9, non-muscle | Cytoplasm | enzyme | 4496785.5 | 2.52E+07 | 2.53E+07 | | | | | | |
| myosin, heavy chain 9, non-muscle | Cytoplasm | enzyme | 4460730.5 | 3.29E+07 | 2.31E+07 | | | | | | |
| myosin, heavy chain 9, non-muscle | Cytoplasm | enzyme | 4275781 | 2.22E+07 | 2.34E+07 | | | | | | |
| myosin, heavy chain 9, non-muscle | Cytoplasm | enzyme | 4111248.25 | 2.69E+07 | 2.05E+07 | | | | | | |
| myosin, heavy chain 9, non-muscle | Cytoplasm | enzyme | 4072403 | 2.70E+07 | 2.33E+07 | | | | | | |
| myosin, heavy chain 9, non-muscle | Cytoplasm | enzyme | 4060996 | 2.06E+07 | 1.97E+07 | | | | | | |
| myosin, heavy chain 9, non-muscle | Cytoplasm | enzyme | 4045315.5 | 2.31E+07 | 2.03E+07 | | | | | | |
| myosin, heavy chain 9, non-muscle | Cytoplasm | enzyme | 3965185.25 | 2.46E+07 | 2.00E+07 | | | | | | |
| myosin, heavy chain 9, non-muscle | Cytoplasm | enzyme | 3916763.75 | 1.77E+07 | 1.52E+07 | | | | | | |
| myosin, heavy chain 9, non-muscle | Cytoplasm | enzyme | 3864115 | 2.10E+07 | 1.82E+07 | | | | | | |
| myosin, heavy chain 9, non-muscle | Cytoplasm | enzyme | 3837734.25 | 2.29E+07 | 2.39E+07 | | | | | | |
| myosin, heavy chain 9, non-muscle | Cytoplasm | enzyme | 3742591.75 | 2.24E+07 | 2.06E+07 | | | | | | |
| myosin, heavy chain 9, non-muscle | Cytoplasm | enzyme | 3555852.25 | 1.64E+07 | 1.54E+07 | | | | | | |
| myosin, heavy chain 9, non-muscle | Cytoplasm | enzyme | 3524653.5 | 2.56E+07 | 2.19E+07 | | | | | | |
| myosin, heavy chain 9, non-muscle | Cytoplasm | enzyme | 3200391.25 | 2.12E+07 | 1.69E+07 | | | | | | |
| myosin, heavy chain 14, non-muscle | Cytoplasm | enzyme | 3141915.75 | 1.92E+07 | 1.88E+07 | | | | | | |
| myosin, heavy chain 9, non-muscle | Cytoplasm | enzyme | 3131051.5 | 2.53E+07 | 1.81E+07 | | | | | | |
| myosin, heavy chain 9, non-muscle | Cytoplasm | enzyme | 2983621 | 1.84E+07 | 1.88E+07 | | | | | | |
| myosin, heavy chain 9, non-muscle | Cytoplasm | enzyme | 2954444.5 | 1.29E+07 | 1.25E+07 | | | | | | |
| myosin, heavy chain 9, non-muscle | Cytoplasm | enzyme | 2851028.5 | 1.65E+07 | 1.80E+07 | | | | | | |
| myosin, heavy chain 9, non-muscle | Cytoplasm | enzyme | 2764340.5 | 1.84E+07 | 1.29E+07 | | | | | | |
| myosin, heavy chain 9, non-muscle | Cytoplasm | enzyme | 2761913.5 | 1.84E+07 | 1.69E+07 | | | | | | |
| myosin, heavy chain 9, non-muscle | Cytoplasm | enzyme | 2745100.25 | 1.52E+07 | 1.46E+07 | | | | | | |
| myosin, heavy chain 9, non-muscle | Cytoplasm | enzyme | 2542464.75 | 1.47E+07 | 1.20E+07 | | | | | | |
| myosin, heavy chain 9, non-muscle | Cytoplasm | enzyme | 2532394.25 | 1.50E+07 | 1.33E+07 | | | | | | |
| myosin, heavy chain 9, non-muscle | Cytoplasm | enzyme | 2388483.75 | 2.06E+07 | 1.38E+07 | | | | | | |
| myosin, heavy chain 9, non-muscle | Cytoplasm | enzyme | 2319219.75 | 1.69E+07 | 1.32E+07 | | | | | | |
| myosin, heavy chain 9, non-muscle | Cytoplasm | enzyme | 2095247.75 | 1.28E+07 | 1.13E+07 | | | | | | |
| myosin, heavy chain 9, non-muscle | Cytoplasm | enzyme | 2021389.125 | 1.32E+07 | 1.56E+07 | | | | | | |
| myosin, heavy chain 9, non-muscle | Cytoplasm | enzyme | 1881755.125 | 1.53E+07 | 1.34E+07 | | | | | | |
| myosin, heavy chain 9, non-muscle | Cytoplasm | enzyme | 1869056 | 4538008 | 6058765 | | | | | | |
| myosin, heavy chain 9, non-muscle | Cytoplasm | enzyme | 1850678.125 | 1.13E+07 | 1.07E+07 | | | | | | |
| myosin, heavy chain 9, non-muscle | Cytoplasm | enzyme | 1775117.75 | 8366149.5 | 9242742 | | | | | | |
| myosin, heavy chain 9, non-muscle | Cytoplasm | enzyme | 1683464.75 | 1.74E+07 | 1.28E+07 | | | | | | |
| myosin, heavy chain 9, non-muscle | Cytoplasm | enzyme | 1357574.5 | 7976045 | 5933860.5 | | | | | | |
| myosin, heavy chain 9, non-muscle | Cytoplasm | enzyme | 1349308.875 | 1.09E+07 | 6963812 | | | | | | |
| myosin, heavy chain 9, non-muscle | Cytoplasm | enzyme | 1303846.875 | 1.35E+07 | 2.19E+07 | | | | | | |
| myosin, heavy chain 9, non-muscle | Cytoplasm | enzyme | 1281842.125 | 8931460 | 7788981.5 | | | | | | |
| myosin, heavy chain 9, non-muscle | Cytoplasm | enzyme | 1238330.5 | 7961607.5 | 6933272.5 | | | | | | |

| | | | | | | | | | | | |
|---|---|---|---|---|---|---|---|---|---|---|---|
| myosin, heavy chain 9, non-muscle | Cytoplasm | enzyme | 1207656.125 | 2.30E+07 | 1.50E+07 | | | | | | |
| myosin, heavy chain 9, non-muscle | Cytoplasm | enzyme | 1176544.875 | 1.63E+07 | 1.62E+07 | | | | | | |
| myosin, heavy chain 9, non-muscle | Cytoplasm | enzyme | 1137683 | 3617815.5 | 1961665.25 | | | | | | |
| myosin, heavy chain 9, non-muscle | Cytoplasm | enzyme | 1114390.875 | 6195583.5 | 6031827.5 | | | | | | |
| myosin, heavy chain 9, non-muscle | Cytoplasm | enzyme | 941624.312 | 7400057 | 5522321.5 | | | | | | |
| myosin, heavy chain 9, non-muscle | Cytoplasm | enzyme | 887042.125 | 4065496.25 | 3876173.5 | | | | | | |
| myosin, heavy chain 9, non-muscle | Cytoplasm | enzyme | 695129.375 | 5837717.5 | 2200005.25 | | | | | | |
| myosin, heavy chain 9, non-muscle | Cytoplasm | enzyme | 646711 | 3447309.75 | 3893004.25 | | | | | | |
| myosin, heavy chain 9, non-muscle | Cytoplasm | enzyme | 637607.188 | 5178305.5 | 3383116 | | | | | | |
| myosin, heavy chain 9, non-muscle | Cytoplasm | enzyme | 581840.938 | 7960722.5 | 4471676 | | | | | | |
| myosin, heavy chain 9, non-muscle | Cytoplasm | enzyme | 571154.188 | 1.10E+07 | 7806166.5 | | | | | | |
| myosin, heavy chain 9, non-muscle | Cytoplasm | enzyme | 556226.688 | 3974216 | 3980050.25 | | | | | | |
| myosin, heavy chain 9, non-muscle | Cytoplasm | enzyme | 405707.406 | 4797896.5 | 3480168.75 | | | | | | |
| myosin, heavy chain 9, non-muscle | Cytoplasm | enzyme | 400142.125 | 2933262 | 3086553.5 | | | | | | |
| myosin, heavy chain 9, non-muscle | Cytoplasm | enzyme | 388734.969 | 1725131.5 | 6055497 | | | | | | |
| myosin, heavy chain 9, non-muscle | Cytoplasm | enzyme | 344933.656 | 6518992 | 3250505.75 | | | | | | |
| myosin, heavy chain 9, non-muscle | Cytoplasm | enzyme | 334966.781 | 1.32E+07 | 9761697 | | | | | | |
| myosin, heavy chain 9, non-muscle | Cytoplasm | enzyme | 297012.594 | 3465440.5 | 3175941.75 | | | | | | |
| myosin, heavy chain 9, non-muscle | Cytoplasm | enzyme | 295611.562 | 1197101.75 | 1622304.75 | | | | | | |
| myosin, heavy chain 9, non-muscle | Cytoplasm | enzyme | 286954.312 | 2813412.5 | 2283885 | | | | | | |
| myosin, heavy chain 9, non-muscle | Cytoplasm | enzyme | 249349.234 | 3852610.75 | 3158736.25 | | | | | | |
| myosin, heavy chain 9, non-muscle | Cytoplasm | enzyme | 205106.547 | 1124753 | 1969238.25 | | | | | | |
| myosin, heavy chain 9, non-muscle | Cytoplasm | enzyme | 125065.867 | 274603.219 | 368151.812 | | | | | | |
| myosin, heavy chain 9, non-muscle | Cytoplasm | enzyme | 118993.273 | 1999102.125 | 1215606.25 | | | | | | |
| myosin, heavy chain 9, non-muscle | Cytoplasm | enzyme | 4.70E-06 | 3215270.5 | 2040119.375 | | | | | | |
| myosin, heavy chain 9, non-muscle | Cytoplasm | enzyme | 0 | 40528.535 | 460327.344 | Q8VDD5 | MYH9 | 3556861 | 12945105 | 762798 | 288 |
| myosin, light chain 1, alkali; skeletal, fast | Cytoplasm | other | 294347.25 | 6456431.5 | 4620410 | P05977 | MYL1 | 294347 | 3163781 | 1826610 | 3 |
| myosin, light chain 12A, regulatory, non-sarcomeric | Cytoplasm | other | 7665141 | 1.61E+07 | 1.34E+07 | | | | | | |
| myosin, light chain 12A, regulatory, non-sarcomeric | Cytoplasm | other | 4245570.5 | 2.30E+07 | 2.54E+07 | | | | | | |
| myosin, light chain 12A, regulatory, non-sarcomeric | Cytoplasm | other | 1577698.5 | 1055128.625 | 1969754.625 | | | | | | |
| myosin, light chain 12A, regulatory, non-sarcomeric | Cytoplasm | other | 773469.812 | 3317610.5 | 3509859.5 | | | | | | |
| myosin, light chain 12A, regulatory, non-sarcomeric | Cytoplasm | other | 290136.031 | 7880182.5 | 5603474.5 | Q3THE2 | MYL12A | 2910403 | 8115308 | 2095363 | 15 |
| myosin, light chain 6, alkali, smooth muscle and non-muscle | Cytoplasm | other | 1.11E+07 | 6.46E+07 | 5.76E+07 | | | | | | |
| myosin, light chain 6, alkali, smooth muscle and non-muscle | Cytoplasm | other | 1.02E+07 | 5.73E+07 | 5.34E+07 | | | | | | |
| myosin, light chain 6, alkali, smooth muscle and non-muscle | Cytoplasm | other | 9180454 | 3.49E+07 | 3.00E+07 | | | | | | |
| myosin, light chain 6, alkali, smooth muscle and non-muscle | Cytoplasm | other | 6972108.5 | 3.86E+07 | 3.00E+07 | | | | | | |
| myosin, light chain 6, alkali, smooth muscle and non-muscle | Cytoplasm | other | 4607807.5 | 2.74E+07 | 2.46E+07 | | | | | | |
| myosin, light chain 6, alkali, smooth muscle and non-muscle | Cytoplasm | other | 4055974 | 2.58E+07 | 2.27E+07 | | | | | | |
| myosin, light chain 6, alkali, smooth muscle and non-muscle | Cytoplasm | other | 1711515.75 | 1.04E+07 | 8716229 | Q60605 | MYL6 | 6826837 | 19603348 | 4277801 | 21 |
| myosin, light chain 9, regulatory | Cytoplasm | other | 3197310 | 2.08E+07 | 1.57E+07 | | | | | | |
| myosin, light chain 9, regulatory | Cytoplasm | other | 2536024 | 2.83E+07 | 2.30E+07 | | | | | | |
| myosin, light chain 9, regulatory | Cytoplasm | other | 2384037 | 6774226 | 7436581 | | | | | | |
| myosin, light chain 9, regulatory | Cytoplasm | other | 1206364.125 | 9586313 | 1.08E+07 | | | | | | |
| myosin, light chain 9, regulatory | Cytoplasm | other | 916139.5 | 1.51E+07 | 1.56E+07 | | | | | | |
| myosin, light chain 9, regulatory | Cytoplasm | other | 868990.375 | 1.17E+07 | 1.18E+07 | | | | | | |
| myosin, light chain 9, regulatory | Cytoplasm | other | 288763.062 | 4696156.5 | 4444023 | | | | | | |
| myosin, light chain 9, regulatory | Cytoplasm | other | 268326.125 | 1.08E+07 | 1.13E+07 | Q9CQ19 | MYL9 | 1458244 | 7721421 | 1576128 | 24 |
| myosin IC | Cytoplasm | other | 332908.625 | 561938.312 | 383464.375 | Q9WTI7 | MYO1C | 332909 | 120321 | 69468 | 3 |
| myosin VI | Cytoplasm | other | 1478718.25 | 964508.688 | 1199879.625 | Q64331 | MYO6 | 1478718 | 257411 | 148616 | 3 |
| nascent polypeptide-associated complex alpha subunit | Cytoplasm | transcription regulator | 1.01E+07 | 5024350 | 5300838 | | | | | | |
| nascent polypeptide-associated complex alpha subunit | Cytoplasm | transcription regulator | 6690468.5 | 3389899.75 | 4071678.5 | | | | | | |
| nascent polypeptide-associated complex alpha subunit | Cytoplasm | transcription regulator | 1289032.75 | 554393.562 | 757586.812 | P70670 | NACA | 6023167 | 3102997 | 1034332 | 9 |
| nucleosome assembly protein 1-like 1 | Nucleus | other | 7.04E+07 | 3.40E+07 | 4.66E+07 | | | | | | |
| nucleosome assembly protein 1-like 1 | Nucleus | other | 4.79E+07 | 1.69E+07 | 2.12E+07 | | | | | | |
| nucleosome assembly protein 1-like 1 | Nucleus | other | 4.39E+07 | 1.90E+07 | 2.77E+07 | | | | | | |
| nucleosome assembly protein 1-like 1 | Nucleus | other | 2.34E+07 | 8975261 | 1.04E+07 | | | | | | |
| nucleosome assembly protein 1-like 1 | Nucleus | other | 1.86E+07 | 1.26E+07 | 1.70E+07 | | | | | | |
| nucleosome assembly protein 1-like 1 | Nucleus | other | 1.47E+07 | 5987662 | 7838537 | | | | | | |
| nucleosome assembly protein 1-like 1 | Nucleus | other | 9754608 | 4581658.5 | 6520506.5 | | | | | | |
| nucleosome assembly protein 1-like 1 | Nucleus | other | 6443283.5 | 2894189.25 | 3502278 | P28656 | NAP1L1 | 29388486 | 17198579 | 3510645 | 24 |
| nucleosome assembly protein 1-like 4 | Cytoplasm | other | 5.05E+07 | 3.04E+07 | 3.12E+07 | | | | | | |
| nucleosome assembly protein 1-like 4 | Cytoplasm | other | 3.30E+07 | 1.73E+07 | 1.98E+07 | | | | | | |
| nucleosome assembly protein 1-like 4 | Cytoplasm | other | 2.63E+07 | 1.03E+07 | 1.24E+07 | | | | | | |
| nucleosome assembly protein 1-like 4 | Cytoplasm | other | 1.85E+07 | 8605380 | 1.23E+07 | | | | | | |
| nucleosome assembly protein 1-like 4 | Cytoplasm | other | 1.80E+07 | 8957187 | 1.11E+07 | | | | | | |
| nucleosome assembly protein 1-like 4 | Cytoplasm | other | 1.55E+07 | 7121108 | 9336494 | | | | | | |
| nucleosome assembly protein 1-like 4 | Cytoplasm | other | 1.41E+07 | 4858081 | 7374328.5 | | | | | | |
| nucleosome assembly protein 1-like 4 | Cytoplasm | other | 1.15E+07 | 4377595 | 7732222.5 | | | | | | |
| nucleosome assembly protein 1-like 4 | Cytoplasm | other | 1.00E+07 | 5296755.5 | 7578452 | | | | | | |
| nucleosome assembly protein 1-like 4 | Cytoplasm | other | 7647452.5 | 2560373.5 | 3686132.75 | | | | | | |
| nucleosome assembly protein 1-like 4 | Cytoplasm | other | 4415725.5 | 1404940.5 | 2326225 | Q78ZA7 | NAP1L4 | 19028471 | 10726705 | 1867280 | 33 |
| N-ethylmaleimide-sensitive factor attachment protein, alpha | Cytoplasm | transporter | 1908906.875 | 2468848.5 | 2671454.75 | Q9DB05 | NAPA | 1908907 | 394982 | 228043 | 3 |
| non-SMC condensin I complex, subunit D2 | Nucleus | other | 872099.125 | 392237.656 | 199387.188 | Q8K2Z4 | NCAPD2 | 872099 | 346410 | 200000 | 3 |
| non-SMC condensin I complex, subunit H | Nucleus | other | 1113911 | 721813.125 | 605673 | Q8C156 | NCAPH | 1113911 | 266313 | 153756 | 3 |
| nicastrin | Plasma Membrane | peptidase | 1.30E+07 | 1.04E+07 | 1.35E+07 | | | | | | |
| nicastrin | Plasma Membrane | peptidase | 1.28E+07 | 9735355 | 1.30E+07 | | | | | | |
| nicastrin | Plasma Membrane | peptidase | 1.13E+07 | 7394520.5 | 7972910.5 | | | | | | |
| nicastrin | Plasma Membrane | peptidase | 8757012 | 3280598 | 5396042 | | | | | | |
| nicastrin | Plasma Membrane | peptidase | 6324018 | 7295732.5 | 1.04E+07 | | | | | | |
| nicastrin | Plasma Membrane | peptidase | 5084752.5 | 5041835 | 4424233.5 | | | | | | |
| nicastrin | Plasma Membrane | peptidase | 4469046 | 3815368 | 5479521 | | | | | | |
| nicastrin | Plasma Membrane | peptidase | 4382926.5 | 3985579 | 5550951 | | | | | | |
| nicastrin | Plasma Membrane | peptidase | 1270834.75 | 311220.969 | 691536.688 | P57716 | NCSTN | 7473177 | 3845591 | 740084 | 27 |
| NADH dehydrogenase (ubiquinone) 1 alpha subcomplex, 10, 42kDa | Cytoplasm | transporter | 2534977 | 4684783 | 4196793 | | | | | | |
| NADH dehydrogenase (ubiquinone) 1 alpha subcomplex, 10, 42kDa | Cytoplasm | transporter | 2130680.25 | 3397775.25 | 3522410.5 | Q99LC3 | NDUFA10 | 2332829 | 965320 | 394090 | 6 |
| NADH dehydrogenase (ubiquinone) 1 alpha subcomplex, 12 | Cytoplasm | enzyme | 359609.625 | 392722.281 | 311634.5 | Q7TMF3 | NDUFA12 | 359610 | 40770 | 23539 | 3 |
| NADH dehydrogenase (ubiquinone) 1 alpha subcomplex, 2, 8kDa | Cytoplasm | enzyme | 1511379.75 | 1882047.5 | 2131184 | Q9CQ75 | NDUFA2 | 1511380 | 311882 | 180065 | 3 |
| NDUFA4, mitochondrial complex associated | Cytoplasm | enzyme | 3163005.75 | 5533499 | 4545474 | | | | | | |
| NDUFA4, mitochondrial complex associated | Cytoplasm | enzyme | 3145937.5 | 3018255.5 | 2916134.75 | Q62425 | NDUFA4 | 3154472 | 1072252 | 437745 | 6 |
| NADH dehydrogenase (ubiquinone) 1 alpha subcomplex, 5 | Cytoplasm | enzyme | 543943.25 | 671431.25 | 876099.25 | Q9CPP6 | NDUFA5 | 543943 | 167566 | 96744 | 3 |
| NADH dehydrogenase (ubiquinone) 1 alpha subcomplex, 8, 19kDa | Cytoplasm | enzyme | 1039292.312 | 1152767.5 | 1532993.625 | Q9DCJ5 | NDUFA8 | 1039292 | 258583 | 149293 | 3 |
| NADH dehydrogenase (ubiquinone) 1 alpha subcomplex, 9, 39kDa | Cytoplasm | enzyme | 5007905.5 | 6616828 | 7426049 | | | | | | |
| NADH dehydrogenase (ubiquinone) 1 alpha subcomplex, 9, 39kDa | Cytoplasm | enzyme | 889203.438 | 2036263.125 | 2275771.75 | | | | | | |
| NADH dehydrogenase (ubiquinone) 1 alpha subcomplex, 9, 39kDa | Cytoplasm | enzyme | 480359.812 | 1191334.75 | 918064.5 | Q9DC69 | NDUFA9 | 2125823 | 2659025 | 886342 | 9 |
| NADH dehydrogenase (ubiquinone) 1, alpha/beta subcomplex, 1, 8kDa | Cytoplasm | enzyme | 2626969.25 | 2721449.5 | 3673292.25 | | | | | | |

| | | | | | | | | | | | |
|---|---|---|---|---|---|---|---|---|---|---|---|
| NADH dehydrogenase (ubiquinone) 1, alpha/beta subcomplex, 1, 8kDa | Cytoplasm | enzyme | 985675.938 | 1695944.125 | 1361631.375 | Q9CR21 | NDUFAB1 | 1806323 | 1005314 | 410418 | 6 |
| NADH dehydrogenase (ubiquinone) Fe-S protein 1, 75kDa (NADH-coenzyme Q reductase) | Cytoplasm | enzyme | 1343870.5 | 1925737.5 | 1705564.125 | | | | | | |
| NADH dehydrogenase (ubiquinone) Fe-S protein 1, 75kDa (NADH-coenzyme Q reductase) | Cytoplasm | enzyme | 1131302.125 | 1474871.5 | 1345564.875 | Q91VD9 | NDUFS1 | 1237586 | 285452 | 116535 | 6 |
| NADH dehydrogenase (ubiquinone) Fe-S protein 2, 49kDa (NADH-coenzyme Q reductase) | Cytoplasm | enzyme | 937118.25 | 1425460.75 | 1516223.125 | Q91WD5 | NDUFS2 | 937118 | 311469 | 179827 | 3 |
| NADH dehydrogenase (ubiquinone) Fe-S protein 3, 30kDa (NADH-coenzyme Q reductase) | Cytoplasm | enzyme | 470151.906 | 1384430 | 1543059.25 | Q9DCT2 | NDUFS3 | 470152 | 579108 | 334348 | 3 |
| NADH dehydrogenase (ubiquinone) Fe-S protein 4, 18kDa (NADH-coenzyme Q reductase) | Cytoplasm | enzyme | 1065595.75 | 2860330.5 | 3147359.75 | | | | | | |
| NADH dehydrogenase (ubiquinone) Fe-S protein 4, 18kDa (NADH-coenzyme Q reductase) | Cytoplasm | enzyme | 849433.562 | 1081861.75 | 917109 | Q9CXZ1 | NDUFS4 | 957515 | 1053494 | 430087 | 6 |
| NADH dehydrogenase (ubiquinone) flavoprotein 1, 51kDa | Cytoplasm | enzyme | 2739606 | 4476642 | 3842466 | | | | | | |
| NADH dehydrogenase (ubiquinone) flavoprotein 1, 51kDa | Cytoplasm | enzyme | 737524.75 | 1385792.125 | 1302117.875 | | | | | | |
| NADH dehydrogenase (ubiquinone) flavoprotein 1, 51kDa | Cytoplasm | enzyme | 371595.562 | 1519713.625 | 844097.938 | Q91YT0 | NDUFV1 | 1282909 | 1444143 | 481381 | 9 |
| NADH dehydrogenase (ubiquinone) flavoprotein 2, 24kDa | Cytoplasm | enzyme | 853393 | 1245930.375 | 1354211.375 | Q9D6J6 | NDUFV2 | 853393 | 263511 | 152138 | 3 |
| NECAP endocytosis associated 2 | Cytoplasm | other | 5691915 | 4123127 | 3798600.75 | Q9D1J1 | NECAP2 | 5691915 | 1012509 | 584573 | 3 |
| sialidase 3 (membrane sialidase) | Plasma Membrane | enzyme | 1.61E+07 | 1.28E+07 | 1.07E+07 | | | | | | |
| sialidase 3 (membrane sialidase) | Plasma Membrane | enzyme | 1.03E+07 | 1.04E+07 | 5481498 | | | | | | |
| sialidase 3 (membrane sialidase) | Plasma Membrane | enzyme | 1202581.5 | 1722249.125 | 959077.812 | Q9JMH7 | NEU3 | 9207527 | 5566996 | 1855665 | 9 |
| nuclear factor of kappa light polypeptide gene enhancer in B-cells inhibitor, zeta | Nucleus | transcription regulator | 3299586.25 | 4476221 | 3701294.5 | Q9EST8 | NFKBIZ | 3299586 | 598101 | 345314 | 3 |
| NLR family member X1 | Cytoplasm | other | 39016.742 | 2142959.5 | 2047958.875 | Q3TL44 | NLRX1 | 39017 | 1188237 | 686029 | 3 |
| NME/NM23 nucleoside diphosphate kinase 3 | Cytoplasm | kinase | 1728207.375 | 1312389 | 1488404.75 | Q9WV85 | NME3 | 1728207 | 208723 | 120506 | 3 |
| neuroplastin | Plasma Membrane | other | 1.22E+08 | 1.07E+08 | 9.36E+07 | | | | | | |
| neuroplastin | Plasma Membrane | other | 4.56E+07 | 3.50E+07 | 3.42E+07 | | | | | | |
| neuroplastin | Plasma Membrane | other | 1.50E+07 | 1.55E+07 | 1.51E+07 | | | | | | |
| neuroplastin | Plasma Membrane | other | 1.48E+07 | 1.07E+07 | 9796420 | | | | | | |
| neuroplastin | Plasma Membrane | other | 1.11E+07 | 6466280.5 | 1.14E+07 | | | | | | |
| neuroplastin | Plasma Membrane | other | 9619563 | 4545159.5 | 5729232 | | | | | | |
| neuroplastin | Plasma Membrane | other | 4147555.25 | 1970788.75 | 2615353.75 | | | | | | |
| neuroplastin | Plasma Membrane | other | 959932.688 | 892007.312 | 563431.125 | | | | | | |
| neuroplastin | Plasma Membrane | other | 240136.891 | 179255.219 | 155883.875 | P97300 | Nptn | 24820799 | 33251908 | 6399333 | 27 |
| neuroblastoma RAS viral (v-ras) oncogene homolog | Plasma Membrane | enzyme | 2.10E+07 | 2.10E+07 | 1.79E+07 | P08556 | NRAS | 20950000 | 1778689 | 1026926 | 3 |
| N-ethylmaleimide-sensitive factor | Cytoplasm | transporter | 1.14E+07 | 7683481.5 | 7749433.5 | | | | | | |
| N-ethylmaleimide-sensitive factor | Cytoplasm | transporter | 9163282 | 6948182 | 6473416.5 | | | | | | |
| N-ethylmaleimide-sensitive factor | Cytoplasm | transporter | 7957305.5 | 6933606.5 | 6303054.5 | | | | | | |
| N-ethylmaleimide-sensitive factor | Cytoplasm | transporter | 7712970 | 4779079 | 7356237.5 | | | | | | |
| N-ethylmaleimide-sensitive factor | Cytoplasm | transporter | 6167164 | 4447875.5 | 5121790 | | | | | | |
| N-ethylmaleimide-sensitive factor | Cytoplasm | transporter | 5702863.5 | 5276254 | 4927730.5 | | | | | | |
| N-ethylmaleimide-sensitive factor | Cytoplasm | transporter | 5314292.5 | 3845008.25 | 4043229.25 | | | | | | |
| N-ethylmaleimide-sensitive factor | Cytoplasm | transporter | 5095448.5 | 2609735 | 3189018.75 | | | | | | |
| N-ethylmaleimide-sensitive factor | Cytoplasm | transporter | 4059112.75 | 3135945.25 | 3404480.75 | | | | | | |
| N-ethylmaleimide-sensitive factor | Cytoplasm | transporter | 3868735.5 | 4559357 | 4639183.5 | | | | | | |
| N-ethylmaleimide-sensitive factor | Cytoplasm | transporter | 3736648.75 | 2677864.75 | 3478978 | | | | | | |
| N-ethylmaleimide-sensitive factor | Cytoplasm | transporter | 3689041.75 | 2869833.25 | 2277427 | | | | | | |
| N-ethylmaleimide-sensitive factor | Cytoplasm | transporter | 1641370.75 | 1073417.625 | 1127250.625 | | | | | | |
| N-ethylmaleimide-sensitive factor | Cytoplasm | transporter | 594309.812 | 210155.469 | 722534.688 | P46460 | NSF | 5435896 | 2456828 | 379097 | 42 |
| NSFL1 (p97) cofactor (p47) | Cytoplasm | other | 6631159 | 2722341 | 3394199.75 | | | | | | |
| NSFL1 (p97) cofactor (p47) | Cytoplasm | other | 4855307.5 | 1771562.875 | 2211649 | | | | | | |
| NSFL1 (p97) cofactor (p47) | Cytoplasm | other | 4573027 | 1145025.25 | 1841923.25 | | | | | | |
| NSFL1 (p97) cofactor (p47) | Cytoplasm | other | 3037179.25 | 422688.719 | 1564338.5 | Q9CZ44 | NSFL1C | 4774168 | 1775415 | 512518 | 12 |
| NACHT and WD repeat domain containing 1 | Cytoplasm | other | 1.49E+07 | 1.51E+07 | 1.37E+07 | A6H603 | NWD1 | 14880000 | 752485 | 434447 | 3 |
| nyctalopin | Extracellular Space | other | 1.07E+08 | 8.45E+07 | 9.33E+07 | P83503 | NYX | 107300000 | 11496157 | 6637310 | 3 |
| ornithine aminotransferase | Cytoplasm | enzyme | 820448.25 | 1112651.375 | 1604581.375 | P29758 | OAT | 820448 | 396283 | 228794 | 3 |
| obscurin, cytoskeletal calmodulin and titin-interacting RhoGEF | Cytoplasm | kinase | 1757516.625 | 1456221.625 | 2353952 | A2AAJ9 | OBSCN | 1757517 | 456880 | 263780 | 3 |
| OCIA domain containing 1 | Cytoplasm | other | 699345.25 | 1161641.625 | 1769092.75 | Q9CRD0 | OCIAD1 | 699345 | 536513 | 309756 | 3 |
| oxoglutarate (alpha-ketoglutarate) dehydrogenase (lipoamide) | Cytoplasm | enzyme | 6007293.5 | 1.13E+07 | 7913947 | | | | | | |
| oxoglutarate (alpha-ketoglutarate) dehydrogenase (lipoamide) | Cytoplasm | enzyme | 4973316.5 | 1.16E+07 | 1.19E+07 | | | | | | |
| oxoglutarate (alpha-ketoglutarate) dehydrogenase (lipoamide) | Cytoplasm | enzyme | 4008244.25 | 5219522 | 5059424.5 | | | | | | |
| oxoglutarate (alpha-ketoglutarate) dehydrogenase (lipoamide) | Cytoplasm | enzyme | 3255471 | 5533839.5 | 6702224.5 | | | | | | |
| oxoglutarate (alpha-ketoglutarate) dehydrogenase (lipoamide) | Cytoplasm | enzyme | 3155304.75 | 6019044 | 5473712.5 | | | | | | |
| oxoglutarate (alpha-ketoglutarate) dehydrogenase (lipoamide) | Cytoplasm | enzyme | 3078706.5 | 5934078.5 | 6354860.5 | | | | | | |
| oxoglutarate (alpha-ketoglutarate) dehydrogenase (lipoamide) | Cytoplasm | enzyme | 2716809 | 5055392.5 | 4975898.5 | | | | | | |
| oxoglutarate (alpha-ketoglutarate) dehydrogenase (lipoamide) | Cytoplasm | enzyme | 2587989.25 | 5515594 | 4328075.5 | | | | | | |
| oxoglutarate (alpha-ketoglutarate) dehydrogenase (lipoamide) | Cytoplasm | enzyme | 2537397.25 | 5060308 | 4065799 | | | | | | |
| oxoglutarate (alpha-ketoglutarate) dehydrogenase (lipoamide) | Cytoplasm | enzyme | 2140798.25 | 3994391 | 4120009.5 | | | | | | |
| oxoglutarate (alpha-ketoglutarate) dehydrogenase (lipoamide) | Cytoplasm | enzyme | 2130368.75 | 4335126 | 3008968.75 | | | | | | |
| oxoglutarate (alpha-ketoglutarate) dehydrogenase (lipoamide) | Cytoplasm | enzyme | 1896731.625 | 3528577 | 3519505 | | | | | | |
| oxoglutarate (alpha-ketoglutarate) dehydrogenase (lipoamide) | Cytoplasm | enzyme | 1404495.625 | 2039682.875 | 2371999.25 | | | | | | |
| oxoglutarate (alpha-ketoglutarate) dehydrogenase (lipoamide) | Cytoplasm | enzyme | 904403.938 | 1716739.625 | 2069652.125 | | | | | | |
| oxoglutarate (alpha-ketoglutarate) dehydrogenase (lipoamide) | Cytoplasm | enzyme | 780671.5 | 1879829.125 | 1389703.25 | | | | | | |
| oxoglutarate (alpha-ketoglutarate) dehydrogenase (lipoamide) | Cytoplasm | enzyme | 691930.062 | 1105274.625 | 835656.875 | | | | | | |
| oxoglutarate (alpha-ketoglutarate) dehydrogenase (lipoamide) | Cytoplasm | enzyme | 659325.688 | 2452960.25 | 1750149.625 | Q60597 | OGDH | 2525250 | 2638675 | 369488 | 51 |
| Obg-like ATPase 1 | Cytoplasm | enzyme | 1833071 | 1075093.125 | 1960115.25 | Q9CZ30 | OLA1 | 1833071 | 478528 | 276278 | 3 |
| osteopetrosis associated transmembrane protein 1 | Cytoplasm | other | 465486.344 | 385042.781 | 491286.938 | Q8BGT0 | OSTM1 | 465486 | 55415 | 31994 | 3 |
| 3-oxoacid CoA transferase 1 | Cytoplasm | enzyme | 634377.062 | 3183776.5 | 1952916.125 | Q9D0K2 | OXCT1 | 634377 | 1274951 | 736093 | 3 |
| prolyl 4-hydroxylase, beta polypeptide | Cytoplasm | enzyme | 2.37E+07 | 2.23E+07 | 2.16E+07 | | | | | | |
| prolyl 4-hydroxylase, beta polypeptide | Cytoplasm | enzyme | 2.26E+07 | 2.10E+07 | 1.89E+07 | | | | | | |
| prolyl 4-hydroxylase, beta polypeptide | Cytoplasm | enzyme | 2.19E+07 | 2.40E+07 | 2.16E+07 | | | | | | |
| prolyl 4-hydroxylase, beta polypeptide | Cytoplasm | enzyme | 2.11E+07 | 2.30E+07 | 1.85E+07 | | | | | | |
| prolyl 4-hydroxylase, beta polypeptide | Cytoplasm | enzyme | 1.96E+07 | 1.88E+07 | 1.98E+07 | | | | | | |
| prolyl 4-hydroxylase, beta polypeptide | Cytoplasm | enzyme | 1.80E+07 | 1.53E+07 | 1.71E+07 | | | | | | |
| prolyl 4-hydroxylase, beta polypeptide | Cytoplasm | enzyme | 1.59E+07 | 1.50E+07 | 1.36E+07 | | | | | | |
| prolyl 4-hydroxylase, beta polypeptide | Cytoplasm | enzyme | 1.43E+07 | 1.70E+07 | 1.48E+07 | | | | | | |
| prolyl 4-hydroxylase, beta polypeptide | Cytoplasm | enzyme | 1.38E+07 | 1.22E+07 | 1.10E+07 | | | | | | |
| prolyl 4-hydroxylase, beta polypeptide | Cytoplasm | enzyme | 1.34E+07 | 1.60E+07 | 1.73E+07 | | | | | | |
| prolyl 4-hydroxylase, beta polypeptide | Cytoplasm | enzyme | 1.33E+07 | 2.20E+07 | 1.59E+07 | | | | | | |
| prolyl 4-hydroxylase, beta polypeptide | Cytoplasm | enzyme | 1.19E+07 | 1.35E+07 | 1.11E+07 | | | | | | |
| prolyl 4-hydroxylase, beta polypeptide | Cytoplasm | enzyme | 1.12E+07 | 1.09E+07 | 1.00E+07 | | | | | | |
| prolyl 4-hydroxylase, beta polypeptide | Cytoplasm | enzyme | 1.10E+07 | 1.21E+07 | 1.12E+07 | | | | | | |
| prolyl 4-hydroxylase, beta polypeptide | Cytoplasm | enzyme | 1.09E+07 | 8803631 | 9871404 | | | | | | |
| prolyl 4-hydroxylase, beta polypeptide | Cytoplasm | enzyme | 8396262 | 7844814 | 9344973 | | | | | | |
| prolyl 4-hydroxylase, beta polypeptide | Cytoplasm | enzyme | 5861632.5 | 4647485 | 5301303.5 | | | | | | |
| prolyl 4-hydroxylase, beta polypeptide | Cytoplasm | enzyme | 5141784.5 | 5858110 | 4954257.5 | | | | | | |
| prolyl 4-hydroxylase, beta polypeptide | Cytoplasm | enzyme | 3273070.25 | 3369644 | 4440091 | P09103 | P4HB | 13960145 | 5901936 | 781730 | 57 |
| proliferation-associated 2G4, 38kDa | Nucleus | transcription regulator | 5558607.5 | 3868665 | 4709881.5 | | | | | | |
| proliferation-associated 2G4, 38kDa | Nucleus | transcription regulator | 4276062.5 | 2598435.5 | 3759354.25 | | | | | | |
| proliferation-associated 2G4, 38kDa | Nucleus | transcription regulator | 4102991.25 | 2557973.75 | 3276865.5 | | | | | | |
| proliferation-associated 2G4, 38kDa | Nucleus | transcription regulator | 3215390.25 | 2457927.5 | 1978584.75 | | | | | | |
| proliferation-associated 2G4, 38kDa | Nucleus | transcription regulator | 2002222.75 | 1336703 | 1764733.375 | P50580 | PA2G4 | 3831055 | 1205530 | 311266 | 15 |

| | | | | | | | | | | | |
|---|---|---|---|---|---|---|---|---|---|---|---|
| poly(A) binding protein, cytoplasmic 1 | Cytoplasm | translation regulator | 2707767.25 | 1219102.125 | 1019493.938 | | | | | | |
| poly(A) binding protein, cytoplasmic 1 | Cytoplasm | translation regulator | 2636947 | 1548713.875 | 1750791.75 | | | | | | |
| poly(A) binding protein, cytoplasmic 1 | Cytoplasm | translation regulator | 2261363.5 | 1155894.625 | 923934.5 | | | | | | |
| poly(A) binding protein, cytoplasmic 1 | Cytoplasm | translation regulator | 2238248.5 | 870084.375 | 680220.938 | | | | | | |
| poly(A) binding protein, cytoplasmic 1 | Cytoplasm | translation regulator | 1841495.375 | 419548.281 | 1147742.875 | P29341 | PABPC1 | 2337164 | 716033 | 184879 | 15 |
| protein kinase C and casein kinase substrate in neurons 2 | Cytoplasm | transporter | 1.21E+07 | 6913788.5 | 7527541.5 | | | | | | |
| protein kinase C and casein kinase substrate in neurons 2 | Cytoplasm | transporter | 1.07E+07 | 4499239 | 6946263.5 | | | | | | |
| protein kinase C and casein kinase substrate in neurons 2 | Cytoplasm | transporter | 9574715 | 4568928.5 | 5276921 | | | | | | |
| protein kinase C and casein kinase substrate in neurons 2 | Cytoplasm | transporter | 9326569 | 4188142 | 5115762.5 | | | | | | |
| protein kinase C and casein kinase substrate in neurons 2 | Cytoplasm | transporter | 6649998.5 | 3018218.25 | 4609406 | | | | | | |
| protein kinase C and casein kinase substrate in neurons 2 | Cytoplasm | transporter | 6242976.5 | 3465209 | 3679071 | | | | | | |
| protein kinase C and casein kinase substrate in neurons 2 | Cytoplasm | transporter | 3687123.25 | 1249676.875 | 1601569.5 | | | | | | |
| protein kinase C and casein kinase substrate in neurons 2 | Cytoplasm | transporter | 1127115.375 | 281503.938 | 709377.75 | Q9WVE8 | PACSIN2 | 7419812 | 3176002 | 648299 | 24 |
| parvin, beta | Cytoplasm | other | 1079500.5 | 2678270.25 | 3305306.5 | | | | | | |
| parvin, beta | Cytoplasm | other | 776674.5 | 4315308.5 | 9378706 | | | | | | |
| parvin, beta | Cytoplasm | other | 126296.883 | 2932230.5 | 4813432 | | | | | | |
| parvin, beta | Cytoplasm | other | 64823.355 | 2287491.75 | 5399513.5 | | | | | | |
| parvin, beta | Cytoplasm | other | 39041.285 | 2049926.625 | 4158536.5 | | | | | | |
| parvin, beta | Cytoplasm | other | 4.70E-06 | 1729070.25 | 2838048.5 | Q9ES46 | PARVB | 347723 | 2392550 | 563930 | 18 |
| pyruvate carboxylase | Cytoplasm | enzyme | 1888640 | 1189606.75 | 1882832.5 | | | | | | |
| pyruvate carboxylase | Cytoplasm | enzyme | 1701950.375 | 2375047 | 1402967.875 | | | | | | |
| pyruvate carboxylase | Cytoplasm | enzyme | 1491031.875 | 1331651.375 | 1918594.125 | | | | | | |
| pyruvate carboxylase | Cytoplasm | enzyme | 713762.812 | 687417.125 | 929360.688 | | | | | | |
| pyruvate carboxylase | Cytoplasm | enzyme | 532138.125 | 693019.25 | 718123.812 | Q05920 | PC | 1265505 | 573163 | 147990 | 15 |
| poly(rC) binding protein 1 | Nucleus | translation regulator | 3200195.5 | 1785305 | 1385207 | | | | | | |
| poly(rC) binding protein 1 | Nucleus | translation regulator | 2490461.75 | 1593494 | 1229018.25 | | | | | | |
| poly(rC) binding protein 1 | Nucleus | translation regulator | 2307923.25 | 934689.312 | 1553723 | | | | | | |
| poly(rC) binding protein 1 | Nucleus | translation regulator | 1979243.125 | 524131.406 | 1229239.25 | | | | | | |
| poly(rC) binding protein 1 | Nucleus | translation regulator | 1425187.125 | 586620.625 | 733570.062 | P60335 | PCBP1 | 2280602 | 741245 | 191389 | 15 |
| phosphoenolpyruvate carboxykinase 2 (mitochondrial) | Cytoplasm | kinase | 9264772 | 8996354 | 9480782 | | | | | | |
| phosphoenolpyruvate carboxykinase 2 (mitochondrial) | Cytoplasm | kinase | 6785180.5 | 8977928 | 9998513 | | | | | | |
| phosphoenolpyruvate carboxykinase 2 (mitochondrial) | Cytoplasm | kinase | 4767254 | 4260690.5 | 4831979 | | | | | | |
| phosphoenolpyruvate carboxykinase 2 (mitochondrial) | Cytoplasm | kinase | 4211675 | 7615678.5 | 6022867.5 | | | | | | |
| phosphoenolpyruvate carboxykinase 2 (mitochondrial) | Cytoplasm | kinase | 3666900.75 | 8452639 | 4142047 | | | | | | |
| phosphoenolpyruvate carboxykinase 2 (mitochondrial) | Cytoplasm | kinase | 2892317.75 | 3720855.75 | 4988324 | | | | | | |
| phosphoenolpyruvate carboxykinase 2 (mitochondrial) | Cytoplasm | kinase | 2148582 | 3555892.75 | 3632505.5 | | | | | | |
| phosphoenolpyruvate carboxykinase 2 (mitochondrial) | Cytoplasm | kinase | 2079954.875 | 2706252.5 | 2665283 | | | | | | |
| phosphoenolpyruvate carboxykinase 2 (mitochondrial) | Cytoplasm | kinase | 2048084.125 | 2261631.25 | 2610024 | | | | | | |
| phosphoenolpyruvate carboxykinase 2 (mitochondrial) | Cytoplasm | kinase | 2042939.25 | 2715368.25 | 3282527.75 | | | | | | |
| phosphoenolpyruvate carboxykinase 2 (mitochondrial) | Cytoplasm | kinase | 1841099.5 | 2897616 | 2619888.75 | | | | | | |
| phosphoenolpyruvate carboxykinase 2 (mitochondrial) | Cytoplasm | kinase | 1797953.75 | 1355406.75 | 1834185.5 | | | | | | |
| phosphoenolpyruvate carboxykinase 2 (mitochondrial) | Cytoplasm | kinase | 1779396 | 2805932.25 | 3099261.75 | | | | | | |
| phosphoenolpyruvate carboxykinase 2 (mitochondrial) | Cytoplasm | kinase | 1705637.5 | 2798647 | 1940109.5 | | | | | | |
| phosphoenolpyruvate carboxykinase 2 (mitochondrial) | Cytoplasm | kinase | 1591140.5 | 1956391.625 | 1572064.875 | | | | | | |
| phosphoenolpyruvate carboxykinase 2 (mitochondrial) | Cytoplasm | kinase | 1475595.5 | 1931705.375 | 2424313.25 | | | | | | |
| phosphoenolpyruvate carboxykinase 2 (mitochondrial) | Cytoplasm | kinase | 762292.812 | 1633976.375 | 1451806 | Q8BH04 | PCK2 | 2991810 | 2471558 | 346087 | 51 |
| prenylcysteine oxidase 1 | Cytoplasm | enzyme | 1.19E+07 | 8394316 | 8504163 | Q9CQF9 | PCYOX1 | 11930000 | 2010368 | 1160687 | 3 |
| phosphodiesterase 6A, cGMP-specific, rod, alpha | Plasma Membrane | enzyme | 1.49E+07 | 1.71E+07 | 1.58E+07 | P27664 | PDE6A | 14870000 | 1133770 | 654582 | 3 |
| pyruvate dehydrogenase (lipoamide) alpha 1 | Cytoplasm | enzyme | 1014873.75 | 980904.125 | 861971.688 | | | | | | |
| pyruvate dehydrogenase (lipoamide) alpha 1 | Cytoplasm | enzyme | 631532.375 | 1091990.5 | 1214376 | | | | | | |
| pyruvate dehydrogenase (lipoamide) alpha 1 | Cytoplasm | enzyme | 274702.844 | 1662304.25 | 1363426 | P35486 | PDHA1 | 640370 | 403977 | 134659 | 9 |
| pyruvate dehydrogenase (lipoamide) beta | Cytoplasm | enzyme | 256243.281 | 2145890.25 | 1507736 | Q9D051 | PDHB | 256243 | 961270 | 554990 | 3 |
| protein disulfide isomerase family A, member 3 | Cytoplasm | peptidase | 1.04E+08 | 7.89E+07 | 1.23E+08 | | | | | | |
| protein disulfide isomerase family A, member 3 | Cytoplasm | peptidase | 4.75E+07 | 4.12E+07 | 5.11E+07 | | | | | | |
| protein disulfide isomerase family A, member 3 | Cytoplasm | peptidase | 4.14E+07 | 3.40E+07 | 3.92E+07 | | | | | | |
| protein disulfide isomerase family A, member 3 | Cytoplasm | peptidase | 2.77E+07 | 2.21E+07 | 2.37E+07 | | | | | | |
| protein disulfide isomerase family A, member 3 | Cytoplasm | peptidase | 2.77E+07 | 1.82E+07 | 2.38E+07 | | | | | | |
| protein disulfide isomerase family A, member 3 | Cytoplasm | peptidase | 2.72E+07 | 2.67E+07 | 3.07E+07 | | | | | | |
| protein disulfide isomerase family A, member 3 | Cytoplasm | peptidase | 2.42E+07 | 1.54E+07 | 2.41E+07 | | | | | | |
| protein disulfide isomerase family A, member 3 | Cytoplasm | peptidase | 2.41E+07 | 1.42E+07 | 2.21E+07 | | | | | | |
| protein disulfide isomerase family A, member 3 | Cytoplasm | peptidase | 2.35E+07 | 1.84E+07 | 3.03E+07 | | | | | | |
| protein disulfide isomerase family A, member 3 | Cytoplasm | peptidase | 2.05E+07 | 1.64E+07 | 2.01E+07 | | | | | | |
| protein disulfide isomerase family A, member 3 | Cytoplasm | peptidase | 2.00E+07 | 1.74E+07 | 1.94E+07 | | | | | | |
| protein disulfide isomerase family A, member 3 | Cytoplasm | peptidase | 1.89E+07 | 1.63E+07 | 2.20E+07 | | | | | | |
| protein disulfide isomerase family A, member 3 | Cytoplasm | peptidase | 1.82E+07 | 1.43E+07 | 2.28E+07 | | | | | | |
| protein disulfide isomerase family A, member 3 | Cytoplasm | peptidase | 1.41E+07 | 1.35E+07 | 1.40E+07 | | | | | | |
| protein disulfide isomerase family A, member 3 | Cytoplasm | peptidase | 1.40E+07 | 1.02E+07 | 1.33E+07 | | | | | | |
| protein disulfide isomerase family A, member 3 | Cytoplasm | peptidase | 1.38E+07 | 1.87E+07 | 1.67E+07 | | | | | | |
| protein disulfide isomerase family A, member 3 | Cytoplasm | peptidase | 1.32E+07 | 1.16E+07 | 1.69E+07 | | | | | | |
| protein disulfide isomerase family A, member 3 | Cytoplasm | peptidase | 1.22E+07 | 6980985 | 1.11E+07 | | | | | | |
| protein disulfide isomerase family A, member 3 | Cytoplasm | peptidase | 1.05E+07 | 6518289 | 1.11E+07 | | | | | | |
| protein disulfide isomerase family A, member 3 | Cytoplasm | peptidase | 5219584 | 4589183 | 8433058 | | | | | | |
| protein disulfide isomerase family A, member 3 | Cytoplasm | peptidase | 5118775.5 | 3464734.5 | 4157229.25 | | | | | | |
| protein disulfide isomerase family A, member 3 | Cytoplasm | peptidase | 3352215.75 | 2801329.75 | 2333585 | | | | | | |
| protein disulfide isomerase family A, member 3 | Cytoplasm | peptidase | 2144656 | 626591.25 | 1640612.625 | P27773 | PDIA3 | 22533706 | 20744834 | 2497385 | 69 |
| protein disulfide isomerase family A, member 4 | Cytoplasm | enzyme | 8996315 | 1.15E+07 | 1.00E+07 | | | | | | |
| protein disulfide isomerase family A, member 4 | Cytoplasm | enzyme | 4309579.5 | 4261178.5 | 4576227 | | | | | | |
| protein disulfide isomerase family A, member 4 | Cytoplasm | enzyme | 4017307.75 | 3060385 | 2996516.25 | | | | | | |
| protein disulfide isomerase family A, member 4 | Cytoplasm | enzyme | 3940981.75 | 4209448.5 | 4045443 | | | | | | |
| protein disulfide isomerase family A, member 4 | Cytoplasm | enzyme | 3393114.25 | 3961788.25 | 3365168 | | | | | | |
| protein disulfide isomerase family A, member 4 | Cytoplasm | enzyme | 3381279 | 3741727.5 | 2870026.25 | | | | | | |
| protein disulfide isomerase family A, member 4 | Cytoplasm | enzyme | 2183269 | 3144347.75 | 2812164.75 | | | | | | |
| protein disulfide isomerase family A, member 4 | Cytoplasm | enzyme | 1723291.5 | 2259982.75 | 1781268.25 | P08003 | PDIA4 | 3993142 | 2463195 | 502798 | 24 |
| protein disulfide isomerase family A, member 6 | Cytoplasm | enzyme | 1.84E+07 | 1.63E+07 | 1.64E+07 | | | | | | |
| protein disulfide isomerase family A, member 6 | Cytoplasm | enzyme | 1.34E+07 | 1.36E+07 | 1.23E+07 | | | | | | |
| protein disulfide isomerase family A, member 6 | Cytoplasm | enzyme | 9328958 | 9599711 | 1.00E+07 | | | | | | |
| protein disulfide isomerase family A, member 6 | Cytoplasm | enzyme | 8734592 | 1.11E+07 | 1.01E+07 | | | | | | |
| protein disulfide isomerase family A, member 6 | Cytoplasm | enzyme | 8380281 | 9862406 | 1.14E+07 | | | | | | |
| protein disulfide isomerase family A, member 6 | Cytoplasm | enzyme | 8230128 | 1.24E+07 | 1.04E+07 | | | | | | |
| protein disulfide isomerase family A, member 6 | Cytoplasm | enzyme | 8158716.5 | 1.08E+07 | 1.09E+07 | | | | | | |
| protein disulfide isomerase family A, member 6 | Cytoplasm | enzyme | 6725224 | 1.06E+07 | 1.09E+07 | | | | | | |
| protein disulfide isomerase family A, member 6 | Cytoplasm | enzyme | 4557269 | 4265135.5 | 7562176 | | | | | | |
| protein disulfide isomerase family A, member 6 | Cytoplasm | enzyme | 3849117 | 1.00E+07 | 5554326 | | | | | | |
| protein disulfide isomerase family A, member 6 | Cytoplasm | enzyme | 3096696 | 4812942.5 | 4611819.5 | Q922R8 | PDIA6 | 8440089 | 3697989 | 643737 | 33 |
| platelet factor 4 | Extracellular Space | cytokine | 3.14E+07 | 1.07E+08 | 7.33E+07 | | | | | | |

| | | | | | | | | | | | |
|---|---|---|---|---|---|---|---|---|---|---|---|
| platelet factor 4 | Extracellular Space | cytokine | 2.49E+07 | 1.50E+08 | 5.87E+07 | | | | | | |
| platelet factor 4 | Extracellular Space | cytokine | 1.31E+07 | 4.34E+07 | 2.82E+07 | Q9Z126 | PF4 | 23133333 | 44655967 | 14885322 | 9 |
| phosphofructokinase, liver | Cytoplasm | kinase | 3098677.25 | 751746.312 | 1821286.625 | P12382 | PFKL | 3098677 | 1174998 | 678386 | 3 |
| phosphofructokinase, muscle | Cytoplasm | kinase | 9579114 | 2520473.5 | 3718070.75 | | | | | | |
| phosphofructokinase, muscle | Cytoplasm | kinase | 5783861 | 3174971.75 | 3202775.25 | | | | | | |
| phosphofructokinase, muscle | Cytoplasm | kinase | 5533937.5 | 2042467.75 | 2469847.25 | | | | | | |
| phosphofructokinase, muscle | Cytoplasm | kinase | 5398756 | 2408501.75 | 2842258.25 | | | | | | |
| phosphofructokinase, muscle | Cytoplasm | kinase | 4031149.75 | 1711294.125 | 1820310.375 | | | | | | |
| phosphofructokinase, muscle | Cytoplasm | kinase | 4016696 | 1682038.625 | 2352571 | P47857 | PFKM | 5723919 | 1985590 | 468008 | 18 |
| phosphogluconate dehydrogenase | Cytoplasm | enzyme | 3947451.25 | 1581356.75 | 2578395.75 | Q9DCD0 | PGD | 3947451 | 1187912 | 685841 | 3 |
| progesterone receptor membrane component 1 | Plasma Membrane | transmembrane receptor | 2351876.25 | 7615170 | 2919543.25 | O55022 | PGRMC1 | 2351876 | 2888870 | 1667890 | 3 |
| progesterone receptor membrane component 2 | Nucleus | other | 2049852.25 | 2135462.5 | 2688610.75 | Q80UU9 | PGRMC2 | 2049852 | 346726 | 200182 | 3 |
| prohibitin | Nucleus | transcription regulator | 6080216 | 5732278 | 5712997 | | | | | | |
| prohibitin | Nucleus | transcription regulator | 3975310.25 | 4898056.5 | 6350039 | | | | | | |
| prohibitin | Nucleus | transcription regulator | 3701790.75 | 4961818.5 | 5081187.5 | | | | | | |
| prohibitin | Nucleus | transcription regulator | 3552802 | 5432167 | 5791219.5 | | | | | | |
| prohibitin | Nucleus | transcription regulator | 2896089.5 | 4271575.5 | 4555479.5 | | | | | | |
| prohibitin | Nucleus | transcription regulator | 2597952.75 | 3173549 | 2567072 | | | | | | |
| prohibitin | Nucleus | transcription regulator | 2113315 | 2010707 | 3493205.25 | | | | | | |
| prohibitin | Nucleus | transcription regulator | 2014381.125 | 2774740.25 | 5096695.5 | | | | | | |
| prohibitin | Nucleus | transcription regulator | 1240822.25 | 1778059.875 | 1881697.5 | | | | | | |
| prohibitin | Nucleus | transcription regulator | 1235570.25 | 1667490 | 2406879.5 | P67778 | PHB | 2940825 | 1593260 | 290888 | 30 |
| prohibitin 2 | Cytoplasm | transcription regulator | 1.15E+07 | 7984477 | 6392178 | | | | | | |
| prohibitin 2 | Cytoplasm | transcription regulator | 7516773 | 1.28E+07 | 8931890 | | | | | | |
| prohibitin 2 | Cytoplasm | transcription regulator | 7041262 | 8256647 | 6939189 | | | | | | |
| prohibitin 2 | Cytoplasm | transcription regulator | 6156346.5 | 1.26E+07 | 1.31E+07 | | | | | | |
| prohibitin 2 | Cytoplasm | transcription regulator | 5639015 | 8161118.5 | 7182043 | | | | | | |
| prohibitin 2 | Cytoplasm | transcription regulator | 5535529.5 | 6900374.5 | 6429711.5 | | | | | | |
| prohibitin 2 | Cytoplasm | transcription regulator | 1354902.75 | 846296.875 | 1132034.25 | | | | | | |
| prohibitin 2 | Cytoplasm | transcription regulator | 1124377.125 | 2614103.25 | 3938030.5 | | | | | | |
| prohibitin 2 | Cytoplasm | transcription regulator | 320740.062 | 533713 | 617662.875 | O35129 | PHB2 | 5128772 | 3928842 | 756106 | 27 |
| phosphoglycerate dehydrogenase | Cytoplasm | enzyme | 6351678 | 3694286.75 | 3264050 | | | | | | |
| phosphoglycerate dehydrogenase | Cytoplasm | enzyme | 5090146 | 3661345 | 2918300.25 | | | | | | |
| phosphoglycerate dehydrogenase | Cytoplasm | enzyme | 4228223.5 | 2453659.25 | 2272070 | | | | | | |
| phosphoglycerate dehydrogenase | Cytoplasm | enzyme | 3823066.5 | 1687760.75 | 1126314.75 | | | | | | |
| phosphoglycerate dehydrogenase | Cytoplasm | enzyme | 2554668.75 | 1882294.375 | 1723005.875 | Q61753 | PHGDH | 4409557 | 1407222 | 363343 | 15 |
| phosphatidylinositol 4-kinase type 2 beta | Cytoplasm | kinase | 2866010.75 | 2266542.75 | 3678382.75 | | | | | | |
| phosphatidylinositol 4-kinase type 2 beta | Cytoplasm | kinase | 2285436 | 1816280.875 | 2774452.75 | | | | | | |
| phosphatidylinositol 4-kinase type 2 beta | Cytoplasm | kinase | 1878765.375 | 1548193.375 | 2521058.5 | | | | | | |
| phosphatidylinositol 4-kinase type 2 beta | Cytoplasm | kinase | 1546207 | 833316.375 | 1033432.562 | Q8CBQ5 | PI4K2B | 2144105 | 812018 | 234410 | 12 |
| phosphatidylinositol binding clathrin assembly protein | Cytoplasm | other | 1.94E+07 | 1.73E+07 | 1.51E+07 | | | | | | |
| phosphatidylinositol binding clathrin assembly protein | Cytoplasm | other | 8143793.5 | 4661635 | 6129475 | | | | | | |
| phosphatidylinositol binding clathrin assembly protein | Cytoplasm | other | 3098461.25 | 2025451.625 | 2592232.75 | | | | | | |
| phosphatidylinositol binding clathrin assembly protein | Cytoplasm | other | 739613.312 | 235778.422 | 809120.562 | Q7M6Y3 | PICALM | 7847967 | 6836178 | 1973435 | 12 |
| piezo-type mechanosensitive ion channel component 1 | Cytoplasm | ion channel | 2.89E+07 | 2.41E+07 | 2.26E+07 | | | | | | |
| piezo-type mechanosensitive ion channel component 1 | Cytoplasm | ion channel | 2.55E+07 | 1.98E+07 | 2.18E+07 | | | | | | |
| piezo-type mechanosensitive ion channel component 1 | Cytoplasm | ion channel | 2.14E+07 | 1.94E+07 | 1.60E+07 | | | | | | |
| piezo-type mechanosensitive ion channel component 1 | Cytoplasm | ion channel | 1.88E+07 | 1.33E+07 | 1.23E+07 | | | | | | |
| piezo-type mechanosensitive ion channel component 1 | Cytoplasm | ion channel | 1.65E+07 | 1.10E+07 | 1.11E+07 | | | | | | |
| piezo-type mechanosensitive ion channel component 1 | Cytoplasm | ion channel | 1.47E+07 | 1.05E+07 | 1.11E+07 | | | | | | |
| piezo-type mechanosensitive ion channel component 1 | Cytoplasm | ion channel | 1.42E+07 | 1.54E+07 | 1.24E+07 | | | | | | |
| piezo-type mechanosensitive ion channel component 1 | Cytoplasm | ion channel | 1.36E+07 | 1.29E+07 | 1.12E+07 | | | | | | |
| piezo-type mechanosensitive ion channel component 1 | Cytoplasm | ion channel | 1.33E+07 | 9426017 | 9558401 | | | | | | |
| piezo-type mechanosensitive ion channel component 1 | Cytoplasm | ion channel | 1.32E+07 | 1.12E+07 | 1.14E+07 | | | | | | |
| piezo-type mechanosensitive ion channel component 1 | Cytoplasm | ion channel | 1.26E+07 | 9298482 | 8270901 | | | | | | |
| piezo-type mechanosensitive ion channel component 1 | Cytoplasm | ion channel | 9108326 | 6189312 | 5187831.5 | | | | | | |
| piezo-type mechanosensitive ion channel component 1 | Cytoplasm | ion channel | 7964057.5 | 6872627.5 | 6907686.5 | | | | | | |
| piezo-type mechanosensitive ion channel component 1 | Cytoplasm | ion channel | 7524113 | 5063208.5 | 5090581 | | | | | | |
| piezo-type mechanosensitive ion channel component 1 | Cytoplasm | ion channel | 7061381 | 6722427 | 6759005.5 | | | | | | |
| piezo-type mechanosensitive ion channel component 1 | Cytoplasm | ion channel | 5583425 | 3989270.5 | 4519599 | | | | | | |
| piezo-type mechanosensitive ion channel component 1 | Cytoplasm | ion channel | 5509056 | 5139332 | 6008572 | | | | | | |
| piezo-type mechanosensitive ion channel component 1 | Cytoplasm | ion channel | 4627987 | 8961199 | 6206044 | | | | | | |
| piezo-type mechanosensitive ion channel component 1 | Cytoplasm | ion channel | 4620231.5 | 3627195.75 | 3940733.5 | | | | | | |
| piezo-type mechanosensitive ion channel component 1 | Cytoplasm | ion channel | 4569672 | 4178141.25 | 3661652.75 | | | | | | |
| piezo-type mechanosensitive ion channel component 1 | Cytoplasm | ion channel | 3756769.75 | 2661980.5 | 3928230.75 | | | | | | |
| piezo-type mechanosensitive ion channel component 1 | Cytoplasm | ion channel | 2707795.5 | 1759572.25 | 2098071.25 | | | | | | |
| piezo-type mechanosensitive ion channel component 1 | Cytoplasm | ion channel | 2702685.25 | 1492049.125 | 1262871 | | | | | | |
| piezo-type mechanosensitive ion channel component 1 | Cytoplasm | ion channel | 2231841.75 | 1693317.875 | 1966264.75 | | | | | | |
| piezo-type mechanosensitive ion channel component 1 | Cytoplasm | ion channel | 2007178.875 | 2420895.25 | 1662770.375 | | | | | | |
| piezo-type mechanosensitive ion channel component 1 | Cytoplasm | ion channel | 1881366.25 | 1519097.75 | 1176562.375 | | | | | | |
| piezo-type mechanosensitive ion channel component 1 | Cytoplasm | ion channel | 725216.5 | 751710.312 | 623496.188 | E2JF22 | PIEZO1 | 9822263 | 6581824 | 731314 | 81 |
| Pim-1 proto-oncogene, serine/threonine kinase | Cytoplasm | kinase | 3.60E+08 | 2.64E+08 | 2.84E+08 | P06803 | PIM1 | 360300000 | 51025876 | 29459803 | 3 |
| phosphatidylinositol-5-phosphate 4-kinase, type II, alpha | Cytoplasm | kinase | 2.07E+07 | 1.39E+07 | 1.37E+07 | | | | | | |
| phosphatidylinositol-5-phosphate 4-kinase, type II, alpha | Cytoplasm | kinase | 6071760.5 | 3496542.75 | 3387701.75 | | | | | | |
| phosphatidylinositol-5-phosphate 4-kinase, type II, alpha | Cytoplasm | kinase | 4980053 | 2665821.25 | 4697611 | | | | | | |
| phosphatidylinositol-5-phosphate 4-kinase, type II, alpha | Cytoplasm | kinase | 4091547.25 | 2492956 | 3967685.5 | O70172 | PIP4K2A | 8970840 | 5822586 | 1680836 | 12 |
| phosphatidylinositol-5-phosphate 4-kinase, type II, beta | Cytoplasm | kinase | 1.11E+07 | 6302471 | 6577173.5 | Q80XI4 | PIP4K2B | 11130000 | 2711357 | 1565402 | 3 |
| phosphatidylinositol-4-phosphate 5-kinase, type I, alpha | Cytoplasm | kinase | 1.51E+07 | 1.48E+07 | 1.38E+07 | | | | | | |
| phosphatidylinositol-4-phosphate 5-kinase, type I, alpha | Cytoplasm | kinase | 9937374 | 1.16E+07 | 8504337 | | | | | | |
| phosphatidylinositol-4-phosphate 5-kinase, type I, alpha | Cytoplasm | kinase | 7779479.5 | 6143407.5 | 7185367 | | | | | | |
| phosphatidylinositol-4-phosphate 5-kinase, type I, alpha | Cytoplasm | kinase | 7160460 | 6903077.5 | 6400206.5 | | | | | | |
| phosphatidylinositol-4-phosphate 5-kinase, type I, alpha | Cytoplasm | kinase | 5908994.5 | 5764159 | 6642665 | | | | | | |
| phosphatidylinositol-4-phosphate 5-kinase, type I, alpha | Cytoplasm | kinase | 5507263.5 | 6786744 | 5798659.5 | | | | | | |
| phosphatidylinositol-4-phosphate 5-kinase, type I, alpha | Cytoplasm | kinase | 3506211.5 | 4687428 | 3608291.75 | | | | | | |
| phosphatidylinositol-4-phosphate 5-kinase, type I, alpha | Cytoplasm | kinase | 3391173.25 | 2897948.25 | 2067801.125 | | | | | | |
| phosphatidylinositol-4-phosphate 5-kinase, type I, alpha | Cytoplasm | kinase | 3357081.25 | 3774173.75 | 3236846 | | | | | | |
| phosphatidylinositol-4-phosphate 5-kinase, type I, alpha | Cytoplasm | kinase | 614231.938 | 133921.125 | 109008.852 | P70182 | PIP5K1A | 6221227 | 3910375 | 713934 | 30 |
| pitrilysin metallopeptidase 1 | Cytoplasm | peptidase | 4410143 | 5878621 | 4995052.5 | | | | | | |
| pitrilysin metallopeptidase 1 | Cytoplasm | peptidase | 2694371 | 3474050.25 | 4929426.5 | | | | | | |
| pitrilysin metallopeptidase 1 | Cytoplasm | peptidase | 2665915.75 | 4450015.5 | 3728908.75 | | | | | | |
| pitrilysin metallopeptidase 1 | Cytoplasm | peptidase | 2241022.75 | 2557190 | 2945960 | | | | | | |
| pitrilysin metallopeptidase 1 | Cytoplasm | peptidase | 1961153.375 | 1956696.25 | 2394135.5 | | | | | | |
| pitrilysin metallopeptidase 1 | Cytoplasm | peptidase | 960526 | 896007 | 846523.25 | | | | | | |
| pitrilysin metallopeptidase 1 | Cytoplasm | peptidase | 558456.5 | 287076.812 | 598237.5 | Q8K411 | PITRM1 | 2213084 | 1642184 | 358354 | 21 |
| polycystic kidney and hepatic disease 1 (autosomal recessive)-like 1 | Extracellular Space | other | 8387783.5 | 7602027.5 | 7502633 | Q80ZA4 | PKHD1L1 | 8387784 | 484903 | 279959 | 3 |

| | | | | | | | | | | | |
|---|---|---|---|---|---|---|---|---|---|---|---|
| pyruvate kinase, liver and RBC | Cytoplasm | kinase | 3.14E+07 | 1.10E+07 | 1.60E+07 | | | | | | |
| pyruvate kinase, liver and RBC | Cytoplasm | kinase | 1.39E+07 | 5652795.5 | 6256975.5 | | | | | | |
| pyruvate kinase, liver and RBC | Cytoplasm | kinase | 1.25E+07 | 5504293 | 9017870 | | | | | | |
| pyruvate kinase, liver and RBC | Cytoplasm | kinase | 1.11E+07 | 4718940 | 6806993.5 | | | | | | |
| pyruvate kinase, liver and RBC | Cytoplasm | kinase | 8633705 | 4837127.5 | 6441510 | | | | | | |
| pyruvate kinase, liver and RBC | Cytoplasm | kinase | 7882479.5 | 3240806.5 | 3884350.75 | | | | | | |
| pyruvate kinase, liver and RBC | Cytoplasm | kinase | 5357516 | 3446748.75 | 3237609.5 | | | | | | |
| pyruvate kinase, liver and RBC | Cytoplasm | kinase | 3929145.5 | 2400023 | 2763914.25 | P53657 | PKLR | 11840356 | 6229870 | 1271667 | 24 |
| pyruvate kinase, muscle | Cytoplasm | kinase | 3346144.5 | 3891566.75 | 5249547 | P52480 | PKM | 3346145 | 980182 | 565908 | 3 |
| phospholipase A2-activating protein | Cytoplasm | other | 1301771.875 | 628511 | 485171.75 | P27612 | PLAA | 1301772 | 436016 | 251734 | 3 |
| pleckstrin | Cytoplasm | other | 157754.688 | 151724.812 | 4388671.5 | Q9JHK5 | PLEK | 157755 | 2444463 | 1411312 | 3 |
| plasminogen | Extracellular Space | peptidase | 639544.812 | 2543604 | 8948364 | | | | | | |
| plasminogen | Extracellular Space | peptidase | 175634.828 | 737136.688 | 4709868 | | | | | | |
| plasminogen | Extracellular Space | peptidase | 82034.938 | 720331.812 | 3157536 | P20918 | PLG | 299072 | 2911528 | 970509 | 9 |
| proteolipid protein 2 (colonic epithelium-enriched) | Cytoplasm | transporter | 1.84E+07 | 1.16E+07 | 1.17E+07 | Q9R1Q7 | PLP2 | 18440000 | 3937846 | 2273517 | 3 |
| phospholipid scramblase 1 | Plasma Membrane | enzyme | 2.90E+07 | 2.04E+07 | 2.24E+07 | | | | | | |
| phospholipid scramblase 1 | Plasma Membrane | enzyme | 1.94E+07 | 1.41E+07 | 1.49E+07 | | | | | | |
| phospholipid scramblase 1 | Plasma Membrane | enzyme | 1.06E+07 | 8692877 | 8294729.5 | Q9JJ00 | PLSCR1 | 19646667 | 6930526 | 2310175 | 9 |
| phospholipid scramblase 3 | Plasma Membrane | enzyme | 1.17E+07 | 8322608 | 8665408 | | | | | | |
| phospholipid scramblase 3 | Plasma Membrane | enzyme | 1.12E+07 | 1.01E+07 | 9824637 | | | | | | |
| phospholipid scramblase 3 | Plasma Membrane | enzyme | 4616824 | 4260383.5 | 3910270 | | | | | | |
| phospholipid scramblase 3 | Plasma Membrane | enzyme | 3915221.75 | 2231684 | 2670993.25 | Q9JIZ9 | PLSCR3 | 7863011 | 3503366 | 1011335 | 12 |
| purine nucleoside phosphorylase | Nucleus | enzyme | 7.05E+07 | 2.33E+07 | 3.21E+07 | | | | | | |
| purine nucleoside phosphorylase | Nucleus | enzyme | 2.45E+07 | 8003570.5 | 8058964 | | | | | | |
| purine nucleoside phosphorylase | Nucleus | enzyme | 2.27E+07 | 7815429.5 | 1.31E+07 | | | | | | |
| purine nucleoside phosphorylase | Nucleus | enzyme | 1.86E+07 | 4735973 | 6473879.5 | | | | | | |
| purine nucleoside phosphorylase | Nucleus | enzyme | 1.28E+07 | 5159264.5 | 7494499 | | | | | | |
| purine nucleoside phosphorylase | Nucleus | enzyme | 3732881 | 1847421.75 | 2079788 | P23492 | PNP | 25480480 | 16368349 | 3858057 | 18 |
| phosphatidic acid phosphatase type 2A | Plasma Membrane | phosphatase | 6.63E+07 | 6.71E+07 | 6.75E+07 | | | | | | |
| phosphatidic acid phosphatase type 2A | Plasma Membrane | phosphatase | 5.66E+07 | 5.11E+07 | 4.38E+07 | | | | | | |
| phosphatidic acid phosphatase type 2A | Plasma Membrane | phosphatase | 5.10E+07 | 2.82E+07 | 4.29E+07 | | | | | | |
| phosphatidic acid phosphatase type 2A | Plasma Membrane | phosphatase | 4.41E+07 | 2.79E+07 | 3.63E+07 | | | | | | |
| phosphatidic acid phosphatase type 2A | Plasma Membrane | phosphatase | 3386180.5 | 4604170.5 | 5245215.5 | Q61469 | Ppap2a | 44263236 | 22104699 | 5707409 | 15 |
| peptidylprolyl isomerase A (cyclophilin A) | Cytoplasm | enzyme | 4620299.5 | 1.83E+07 | 1.82E+07 | | | | | | |
| peptidylprolyl isomerase A (cyclophilin A) | Cytoplasm | enzyme | 2581346.75 | 1.70E+07 | 1.29E+07 | | | | | | |
| peptidylprolyl isomerase A (cyclophilin A) | Cytoplasm | enzyme | 2274293 | 1.32E+07 | 1.44E+07 | | | | | | |
| peptidylprolyl isomerase A (cyclophilin A) | Cytoplasm | enzyme | 1061969.625 | 1.25E+07 | 5342050.5 | | | | | | |
| peptidylprolyl isomerase A (cyclophilin A) | Cytoplasm | enzyme | 771341 | 1.19E+07 | 1.22E+07 | | | | | | |
| peptidylprolyl isomerase A (cyclophilin A) | Cytoplasm | enzyme | 554436.25 | 3773185 | 2582045.25 | | | | | | |
| peptidylprolyl isomerase A (cyclophilin A) | Cytoplasm | enzyme | 260682.547 | 3902883.5 | 2604756 | P17742 | PPIA | 1732053 | 6415683 | 1400017 | 21 |
| peptidylprolyl isomerase B (cyclophilin B) | Cytoplasm | enzyme | 8.39E+07 | 8.85E+07 | 8.35E+07 | | | | | | |
| peptidylprolyl isomerase B (cyclophilin B) | Cytoplasm | enzyme | 4.64E+07 | 5.08E+07 | 4.45E+07 | | | | | | |
| peptidylprolyl isomerase B (cyclophilin B) | Cytoplasm | enzyme | 2.99E+07 | 2.12E+07 | 1.90E+07 | | | | | | |
| peptidylprolyl isomerase B (cyclophilin B) | Cytoplasm | enzyme | 2.51E+07 | 2.42E+07 | 2.48E+07 | | | | | | |
| peptidylprolyl isomerase B (cyclophilin B) | Cytoplasm | enzyme | 2.05E+07 | 2.73E+07 | 2.45E+07 | | | | | | |
| peptidylprolyl isomerase B (cyclophilin B) | Cytoplasm | enzyme | 2.00E+07 | 2.17E+07 | 2.30E+07 | | | | | | |
| peptidylprolyl isomerase B (cyclophilin B) | Cytoplasm | enzyme | 1.76E+07 | 1.63E+07 | 1.71E+07 | | | | | | |
| peptidylprolyl isomerase B (cyclophilin B) | Cytoplasm | enzyme | 1.62E+07 | 2.12E+07 | 1.52E+07 | | | | | | |
| peptidylprolyl isomerase B (cyclophilin B) | Cytoplasm | enzyme | 1.48E+07 | 1.67E+07 | 1.10E+07 | | | | | | |
| peptidylprolyl isomerase B (cyclophilin B) | Cytoplasm | enzyme | 1.42E+07 | 1.40E+07 | 1.15E+07 | | | | | | |
| peptidylprolyl isomerase B (cyclophilin B) | Cytoplasm | enzyme | 1.18E+07 | 8867410 | 1.00E+07 | | | | | | |
| peptidylprolyl isomerase B (cyclophilin B) | Cytoplasm | enzyme | 3543652.75 | 6978621 | 4403660 | P24369 | PPIB | 25333638 | 21114617 | 3519103 | 36 |
| peptidylprolyl isomerase F | Cytoplasm | enzyme | 638831.5 | 1.27E+07 | 5463636 | | | | | | |
| peptidylprolyl isomerase F | Cytoplasm | enzyme | 96668.367 | 5718273.5 | 5339143 | Q99KR7 | PPIF | 367750 | 4546477 | 1856091 | 6 |
| protoporphyrinogen oxidase | Cytoplasm | enzyme | 1.93E+07 | 1.76E+07 | 1.55E+07 | | | | | | |
| protoporphyrinogen oxidase | Cytoplasm | enzyme | 1.68E+07 | 1.55E+07 | 1.72E+07 | | | | | | |
| protoporphyrinogen oxidase | Cytoplasm | enzyme | 6726892.5 | 6210371 | 6942427 | | | | | | |
| protoporphyrinogen oxidase | Cytoplasm | enzyme | 6227639 | 6610401 | 6967832.5 | | | | | | |
| protoporphyrinogen oxidase | Cytoplasm | enzyme | 6087097.5 | 6598516.5 | 7026745.5 | | | | | | |
| protoporphyrinogen oxidase | Cytoplasm | enzyme | 4919424 | 5194487 | 4154863.25 | | | | | | |
| protoporphyrinogen oxidase | Cytoplasm | enzyme | 4074251.5 | 4440514.5 | 4846919 | | | | | | |
| protoporphyrinogen oxidase | Cytoplasm | enzyme | 3949938.75 | 3729944.5 | 4391573 | | | | | | |
| protoporphyrinogen oxidase | Cytoplasm | enzyme | 3858931.25 | 4869517.5 | 5177413.5 | | | | | | |
| protoporphyrinogen oxidase | Cytoplasm | enzyme | 2224063 | 2633613 | 1857557.75 | | | | | | |
| protoporphyrinogen oxidase | Cytoplasm | enzyme | 2221721.25 | 2524505.5 | 2379600.75 | | | | | | |
| protoporphyrinogen oxidase | Cytoplasm | enzyme | 1239045.25 | 1258840.625 | 1832693.25 | | | | | | |
| protoporphyrinogen oxidase | Cytoplasm | enzyme | 1224059.5 | 1870915 | 2207029 | P51175 | PPOX | 6063313 | 5039935 | 807035 | 39 |
| protein phosphatase 1, catalytic subunit, alpha isozyme | Cytoplasm | phosphatase | 3059607.75 | 6040449.5 | 4504338.5 | | | | | | |
| protein phosphatase 1, catalytic subunit, alpha isozyme | Cytoplasm | phosphatase | 2990844.75 | 7684710 | 6480223.5 | | | | | | |
| protein phosphatase 1, catalytic subunit, alpha isozyme | Cytoplasm | phosphatase | 2091906.5 | 2329420.25 | 1817639.125 | | | | | | |
| protein phosphatase 1, catalytic subunit, alpha isozyme | Cytoplasm | phosphatase | 1487836.5 | 3477171.75 | 2710191.75 | | | | | | |
| protein phosphatase 1, catalytic subunit, alpha isozyme | Cytoplasm | phosphatase | 1158544.875 | 1205480.375 | 1449879.125 | | | | | | |
| protein phosphatase 1, catalytic subunit, alpha isozyme | Cytoplasm | phosphatase | 759168.188 | 1742152.875 | 1307738.875 | P62137 | PPP1CA | 1924651 | 2016992 | 475410 | 18 |
| protein phosphatase 1, regulatory subunit 13B | Cytoplasm | phosphatase | 4246512.5 | 1461211.875 | 2100976 | Q62415 | PPP1R13B | 4246513 | 1458911 | 842303 | 3 |
| protein phosphatase 1, regulatory subunit 21 | Cytoplasm | other | 2908225 | 7182241.5 | 8519368 | Q3TDD9 | PPP1R21 | 2908225 | 2930872 | 1692140 | 3 |
| protein phosphatase 2, catalytic subunit, alpha isozyme | Cytoplasm | phosphatase | 9398877 | 4099507.75 | 3714144.5 | | | | | | |
| protein phosphatase 2, catalytic subunit, alpha isozyme | Cytoplasm | phosphatase | 9156457 | 5682358.5 | 5992915 | | | | | | |
| protein phosphatase 2, catalytic subunit, alpha isozyme | Cytoplasm | phosphatase | 6032990.5 | 4453633 | 4938902 | | | | | | |
| protein phosphatase 2, catalytic subunit, alpha isozyme | Cytoplasm | phosphatase | 4838261 | 2236387.5 | 2575200.75 | | | | | | |
| protein phosphatase 2, catalytic subunit, alpha isozyme | Cytoplasm | phosphatase | 4388952 | 2397817.25 | 2415132.25 | P63330 | PPP2CA | 6763108 | 2216863 | 572392 | 15 |
| protein phosphatase 2, regulatory subunit A, alpha | Cytoplasm | phosphatase | 1.32E+07 | 7790672.5 | 1.04E+07 | | | | | | |
| protein phosphatase 2, regulatory subunit A, alpha | Cytoplasm | phosphatase | 9282895 | 2206743.5 | 2669423 | | | | | | |
| protein phosphatase 2, regulatory subunit A, alpha | Cytoplasm | phosphatase | 7494091.5 | 2274587.25 | 3067422.25 | | | | | | |
| protein phosphatase 2, regulatory subunit A, alpha | Cytoplasm | phosphatase | 7370656.5 | 3910781.25 | 5397105 | | | | | | |
| protein phosphatase 2, regulatory subunit A, alpha | Cytoplasm | phosphatase | 5265910.5 | 2304608 | 3860854.5 | | | | | | |
| protein phosphatase 2, regulatory subunit A, alpha | Cytoplasm | phosphatase | 4664187 | 1893206.875 | 2220740 | | | | | | |
| protein phosphatase 2, regulatory subunit A, alpha | Cytoplasm | phosphatase | 2961889 | 1386332.625 | 1828899.25 | Q76MZ3 | PPP2R1A | 7171376 | 3255013 | 710302 | 21 |
| protein phosphatase 2, regulatory subunit B, alpha | Cytoplasm | phosphatase | 7614890 | 1187725.125 | 1040626.938 | Q6P1F6 | PPP2R2A | 7614890 | 3753910 | 2167321 | 3 |
| protein phosphatase 6, catalytic subunit | Nucleus | phosphatase | 1.20E+07 | 5667421.5 | 6656531 | | | | | | |
| protein phosphatase 6, catalytic subunit | Nucleus | phosphatase | 5824406 | 3164461.5 | 2747827.5 | Q9CQR6 | PPP6C | 8917203 | 3325918 | 1357800 | 6 |
| protein phosphatase 6, regulatory subunit 3 | Cytoplasm | other | 2.15E+07 | 1.25E+07 | 1.10E+07 | | | | | | |
| protein phosphatase 6, regulatory subunit 3 | Cytoplasm | other | 1.14E+07 | 7342417 | 9352657 | | | | | | |
| protein phosphatase 6, regulatory subunit 3 | Cytoplasm | other | 9478525 | 3917558.25 | 5121707 | | | | | | |
| protein phosphatase 6, regulatory subunit 3 | Cytoplasm | other | 9391523 | 4245013.5 | 5778201 | | | | | | |
| protein phosphatase 6, regulatory subunit 3 | Cytoplasm | other | 8846557 | 3976904.25 | 3814909 | | | | | | |

| | | | | | | | | | | | |
|---|---|---|---|---|---|---|---|---|---|---|---|
| protein phosphatase 6, regulatory subunit 3 | Cytoplasm | other | 8767766 | 4069572.75 | 3857397.5 | | | | | | |
| protein phosphatase 6, regulatory subunit 3 | Cytoplasm | other | 7743192.5 | 5203848.5 | 5303664 | | | | | | |
| protein phosphatase 6, regulatory subunit 3 | Cytoplasm | other | 6437565.5 | 3619780.75 | 3992295 | | | | | | |
| protein phosphatase 6, regulatory subunit 3 | Cytoplasm | other | 6299008 | 3188652 | 3267970.5 | | | | | | |
| protein phosphatase 6, regulatory subunit 3 | Cytoplasm | other | 5339499.5 | 2287187.25 | 2639049 | | | | | | |
| protein phosphatase 6, regulatory subunit 3 | Cytoplasm | other | 4951365 | 2145855.5 | 2196931.5 | | | | | | |
| protein phosphatase 6, regulatory subunit 3 | Cytoplasm | other | 1493845.125 | 1023324.062 | 1590277.5 | Q922D4 | PPP6R3 | 8468237 | 4013561 | 668927 | 36 |
| prolylcarboxypeptidase (angiotensinase C) | Cytoplasm | peptidase | 1670567.375 | 2729709.25 | 3310859.5 | Q7TMR0 | PRCP | 1670567 | 831673 | 480166 | 3 |
| peroxiredoxin 1 | Cytoplasm | enzyme | 5.29E+07 | 1.31E+07 | 2.56E+07 | | | | | | |
| peroxiredoxin 1 | Cytoplasm | enzyme | 5.23E+07 | 1.85E+07 | 2.41E+07 | | | | | | |
| peroxiredoxin 1 | Cytoplasm | enzyme | 3.75E+07 | 1.27E+07 | 1.75E+07 | | | | | | |
| peroxiredoxin 1 | Cytoplasm | enzyme | 3.59E+07 | 1.57E+07 | 2.44E+07 | | | | | | |
| peroxiredoxin 1 | Cytoplasm | enzyme | 3.46E+07 | 1.44E+07 | 2.32E+07 | | | | | | |
| peroxiredoxin 1 | Cytoplasm | enzyme | 2.67E+07 | 9739484 | 1.54E+07 | | | | | | |
| peroxiredoxin 1 | Cytoplasm | enzyme | 1.80E+07 | 1.24E+07 | 1.10E+07 | | | | | | |
| peroxiredoxin 1 | Cytoplasm | enzyme | 1.47E+07 | 4861626.5 | 7210209.5 | | | | | | |
| peroxiredoxin 1 | Cytoplasm | enzyme | 1.18E+07 | 2335077.25 | 4790418.5 | | | | | | |
| peroxiredoxin 1 | Cytoplasm | enzyme | 9924490 | 3578119 | 2388910.5 | | | | | | |
| peroxiredoxin 1 | Cytoplasm | enzyme | 1872093.75 | 393501.844 | 975510.188 | P35700 | PRDX1 | 26918780 | 13612642 | 2369657 | 33 |
| peroxiredoxin 2 | Cytoplasm | enzyme | 8.85E+08 | 2.07E+08 | 2.95E+08 | | | | | | |
| peroxiredoxin 2 | Cytoplasm | enzyme | 5.02E+08 | 1.63E+08 | 2.21E+08 | | | | | | |
| peroxiredoxin 2 | Cytoplasm | enzyme | 4.56E+08 | 1.49E+08 | 2.20E+08 | | | | | | |
| peroxiredoxin 2 | Cytoplasm | enzyme | 2.89E+08 | 5.48E+07 | 1.03E+08 | | | | | | |
| peroxiredoxin 2 | Cytoplasm | enzyme | 2.27E+08 | 5.84E+07 | 9.16E+07 | | | | | | |
| peroxiredoxin 2 | Cytoplasm | enzyme | 2.00E+08 | 5.93E+07 | 9.47E+07 | | | | | | |
| peroxiredoxin 2 | Cytoplasm | enzyme | 1.96E+08 | 6.35E+07 | 1.06E+08 | | | | | | |
| peroxiredoxin 2 | Cytoplasm | enzyme | 1.41E+08 | 2.66E+07 | 4.65E+07 | | | | | | |
| peroxiredoxin 2 | Cytoplasm | enzyme | 1.03E+08 | 2.66E+07 | 3.61E+07 | | | | | | |
| peroxiredoxin 2 | Cytoplasm | enzyme | 4.08E+07 | 9659971 | 2499952.5 | | | | | | |
| peroxiredoxin 2 | Cytoplasm | enzyme | 4.01E+07 | 9910558 | 1.58E+07 | | | | | | |
| peroxiredoxin 2 | Cytoplasm | enzyme | 3.41E+07 | 3.86E+07 | 3.33E+07 | | | | | | |
| peroxiredoxin 2 | Cytoplasm | enzyme | 2.68E+07 | 1.10E+07 | 8684427 | | | | | | |
| peroxiredoxin 2 | Cytoplasm | enzyme | 8623255 | 1.21E+07 | 8898291 | Q61171 | PRDX2 | 224888804 | 167851530 | 25900053 | 42 |
| peroxiredoxin 3 | Cytoplasm | enzyme | 4.61E+07 | 5.01E+07 | 5.85E+07 | | | | | | |
| peroxiredoxin 3 | Cytoplasm | enzyme | 3.39E+07 | 4.16E+07 | 4.60E+07 | | | | | | |
| peroxiredoxin 3 | Cytoplasm | enzyme | 2.10E+07 | 2.52E+07 | 2.93E+07 | | | | | | |
| peroxiredoxin 3 | Cytoplasm | enzyme | 1.25E+07 | 1.55E+07 | 1.62E+07 | | | | | | |
| peroxiredoxin 3 | Cytoplasm | enzyme | 1.23E+07 | 1.70E+07 | 2.02E+07 | | | | | | |
| peroxiredoxin 3 | Cytoplasm | enzyme | 6988498.5 | 2.61E+07 | 1.69E+07 | | | | | | |
| peroxiredoxin 3 | Cytoplasm | enzyme | 3089876.25 | 2711118.5 | 3583199.5 | P20108 | PRDX3 | 19409768 | 16438247 | 3587120 | 21 |
| peroxiredoxin 4 | Cytoplasm | enzyme | 8968067 | 8264780 | 8129405.5 | | | | | | |
| peroxiredoxin 4 | Cytoplasm | enzyme | 4495310 | 5662518.5 | 5208531 | | | | | | |
| peroxiredoxin 4 | Cytoplasm | enzyme | 1535138.375 | 428795.281 | 962034.125 | O08807 | PRDX4 | 4999505 | 3277573 | 1092524 | 9 |
| peroxiredoxin 5 | Cytoplasm | enzyme | 2326102.5 | 3819527 | 4913880.5 | P99029 | PRDX5 | 2326103 | 1299007 | 749982 | 3 |
| proteoglycan 2, bone marrow (natural killer cell activator, eosinophil granule major basic protein) | Extracellular Space | other | 21925.25 | 3400959.75 | 4272558.5 | Q61878 | PRG2 | 21925 | 2245196 | 1296265 | 3 |
| protein kinase, cAMP-dependent, catalytic, alpha | Cytoplasm | kinase | 3667030.25 | 3353601.25 | 3176094.75 | P05132 | PRKACA | 3667030 | 248584 | 143520 | 3 |
| protein kinase C, alpha | Cytoplasm | kinase | 51849.727 | 2075945.75 | 1307320.25 | P20444 | PRKCA | 51850 | 1021760 | 589913 | 3 |
| protein kinase C substrate 80K-H | Cytoplasm | enzyme | 1.21E+07 | 1.28E+07 | 1.39E+07 | | | | | | |
| protein kinase C substrate 80K-H | Cytoplasm | enzyme | 1.01E+07 | 1.08E+07 | 9787079 | | | | | | |
| protein kinase C substrate 80K-H | Cytoplasm | enzyme | 9224265 | 8732851 | 9042629 | | | | | | |
| protein kinase C substrate 80K-H | Cytoplasm | enzyme | 8231156.5 | 7870223 | 7496519 | | | | | | |
| protein kinase C substrate 80K-H | Cytoplasm | enzyme | 7553271 | 5544860.5 | 6065405.5 | | | | | | |
| protein kinase C substrate 80K-H | Cytoplasm | enzyme | 6852039.5 | 6176407 | 6092170.5 | | | | | | |
| protein kinase C substrate 80K-H | Cytoplasm | enzyme | 6627142.5 | 5943533.5 | 6280497 | | | | | | |
| protein kinase C substrate 80K-H | Cytoplasm | enzyme | 6257360.5 | 3184841.75 | 3547197.5 | | | | | | |
| protein kinase C substrate 80K-H | Cytoplasm | enzyme | 2387720.25 | 4028928.5 | 2596305.25 | | | | | | |
| protein kinase C substrate 80K-H | Cytoplasm | enzyme | 2340806.25 | 3332051.5 | 2675423 | | | | | | |
| protein kinase C substrate 80K-H | Cytoplasm | enzyme | 1350851.25 | 2048984.125 | 1144770.125 | O08795 | PRKCSH | 6645874 | 3404008 | 592562 | 33 |
| protein arginine methyltransferase 5 | Cytoplasm | enzyme | 7362398.5 | 3139734.75 | 4598885.5 | | | | | | |
| protein arginine methyltransferase 5 | Cytoplasm | enzyme | 1373456 | 667128.688 | 824839.062 | Q8CIG8 | PRMT5 | 4367927 | 2623949 | 1071223 | 6 |
| prion protein | Plasma Membrane | other | 1.09E+07 | 1.05E+07 | 6976946.5 | P04925 | PRNP | 10850000 | 2152208 | 1242578 | 3 |
| phosphoribosyl pyrophosphate synthetase 1 | Cytoplasm | kinase | 1.29E+07 | 6549175.5 | 6824323.5 | | | | | | |
| phosphoribosyl pyrophosphate synthetase 1 | Cytoplasm | kinase | 9897321 | 6156220 | 5532343 | | | | | | |
| phosphoribosyl pyrophosphate synthetase 1 | Cytoplasm | kinase | 5907413 | 2150926.75 | 3544283.5 | | | | | | |
| phosphoribosyl pyrophosphate synthetase 1 | Cytoplasm | kinase | 5669409 | 2840207.25 | 3325500.75 | | | | | | |
| phosphoribosyl pyrophosphate synthetase 1 | Cytoplasm | kinase | 3930707.75 | 1906873.875 | 2206896 | | | | | | |
| phosphoribosyl pyrophosphate synthetase 1 | Cytoplasm | kinase | 2821385 | 756363.062 | 1770813.75 | Q9D7G0 | PRPS1 | 6851039 | 3091920 | 728773 | 18 |
| phosphoribosyl pyrophosphate synthetase-associated protein 1 | Extracellular Space | other | 1.13E+07 | 3760221 | 5674166 | | | | | | |
| phosphoribosyl pyrophosphate synthetase-associated protein 1 | Extracellular Space | other | 4218425 | 2787913 | 3159657.5 | | | | | | |
| phosphoribosyl pyrophosphate synthetase-associated protein 1 | Extracellular Space | other | 3932122.5 | 1603227.375 | 2611036.25 | | | | | | |
| phosphoribosyl pyrophosphate synthetase-associated protein 1 | Extracellular Space | other | 3358509.25 | 1578780.625 | 1784086 | | | | | | |
| phosphoribosyl pyrophosphate synthetase-associated protein 1 | Extracellular Space | other | 3115181.75 | 1323137.125 | 1781543.375 | Q9D0M1 | PRPSAP1 | 5186848 | 2478236 | 639878 | 15 |
| phosphoribosyl pyrophosphate synthetase-associated protein 2 | Cytoplasm | other | 4700916.5 | 1404911.625 | 2238337.25 | | | | | | |
| phosphoribosyl pyrophosphate synthetase-associated protein 2 | Cytoplasm | other | 4188525.75 | 2073076.125 | 2913364.75 | Q8R574 | PRPSAP2 | 4444721 | 1285169 | 524668 | 6 |
| prosaposin | Extracellular Space | other | 1502420 | 1844618.75 | 2022984.5 | Q61207 | PSAP | 1502420 | 264544 | 152735 | 3 |
| presenilin 1 | Plasma Membrane | peptidase | 6330520 | 3769265.5 | 4058826.25 | | | | | | |
| presenilin 1 | Plasma Membrane | peptidase | 1545284.5 | 1467642.125 | 1749391.125 | P49769 | PSEN1 | 3937902 | 1933506 | 789351 | 6 |
| presenilin 2 | Cytoplasm | peptidase | 1615870.5 | 1268783.875 | 1174579.5 | Q61144 | PSEN2 | 1615871 | 232408 | 134181 | 3 |
| protein serine kinase H1 | Nucleus | kinase | 3893297.75 | 5360648 | 4042158.5 | Q91YA2 | PSKH1 | 3893298 | 807640 | 466291 | 3 |
| proteasome (prosome, macropain) subunit, alpha type, 1 | Cytoplasm | peptidase | 5.75E+07 | 2.89E+07 | 4.97E+07 | | | | | | |
| proteasome (prosome, macropain) subunit, alpha type, 1 | Cytoplasm | peptidase | 5.63E+07 | 2.42E+07 | 2.56E+07 | | | | | | |
| proteasome (prosome, macropain) subunit, alpha type, 1 | Cytoplasm | peptidase | 4.85E+07 | 2.07E+07 | 2.88E+07 | | | | | | |
| proteasome (prosome, macropain) subunit, alpha type, 1 | Cytoplasm | peptidase | 4.35E+07 | 1.43E+07 | 2.35E+07 | | | | | | |
| proteasome (prosome, macropain) subunit, alpha type, 1 | Cytoplasm | peptidase | 3.93E+07 | 1.54E+07 | 2.16E+07 | | | | | | |
| proteasome (prosome, macropain) subunit, alpha type, 1 | Cytoplasm | peptidase | 3.90E+07 | 2.61E+07 | 3.09E+07 | | | | | | |
| proteasome (prosome, macropain) subunit, alpha type, 1 | Cytoplasm | peptidase | 3.19E+07 | 5723462 | 1.22E+07 | | | | | | |
| proteasome (prosome, macropain) subunit, alpha type, 1 | Cytoplasm | peptidase | 3.00E+07 | 1.26E+07 | 1.75E+07 | | | | | | |
| proteasome (prosome, macropain) subunit, alpha type, 1 | Cytoplasm | peptidase | 2.90E+07 | 1.41E+07 | 1.69E+07 | | | | | | |
| proteasome (prosome, macropain) subunit, alpha type, 1 | Cytoplasm | peptidase | 2.88E+07 | 1.18E+07 | 1.90E+07 | | | | | | |
| proteasome (prosome, macropain) subunit, alpha type, 1 | Cytoplasm | peptidase | 2.81E+07 | 1.14E+07 | 1.77E+07 | | | | | | |
| proteasome (prosome, macropain) subunit, alpha type, 1 | Cytoplasm | peptidase | 1.44E+07 | 7056266 | 7311173.5 | | | | | | |
| proteasome (prosome, macropain) subunit, alpha type, 1 | Cytoplasm | peptidase | 1.24E+07 | 8896428 | 6847732.5 | | | | | | |
| proteasome (prosome, macropain) subunit, alpha type, 1 | Cytoplasm | peptidase | 8351313 | 2382034.5 | 4268395 | | | | | | |
| proteasome (prosome, macropain) subunit, alpha type, 1 | Cytoplasm | peptidase | 6472292.5 | 4331780.5 | 4449911.5 | | | | | | |
| proteasome (prosome, macropain) subunit, alpha type, 1 | Cytoplasm | peptidase | 6201379.5 | 4032125.25 | 9642183 | Q9R1P4 | PSMA1 | 29990937 | 14392914 | 2077438 | 48 |

| | | | | | | | | | | | |
|---|---|---|---|---|---|---|---|---|---|---|---|
| proteasome (prosome, macropain) subunit, alpha type, 2 | Cytoplasm | peptidase | 1.16E+08 | 5.23E+07 | 7.55E+07 | | | | | | |
| proteasome (prosome, macropain) subunit, alpha type, 2 | Cytoplasm | peptidase | 9.17E+07 | 2.61E+07 | 3.28E+07 | | | | | | |
| proteasome (prosome, macropain) subunit, alpha type, 2 | Cytoplasm | peptidase | 4.28E+07 | 1.47E+07 | 2.63E+07 | | | | | | |
| proteasome (prosome, macropain) subunit, alpha type, 2 | Cytoplasm | peptidase | 4.18E+07 | 2.03E+07 | 2.89E+07 | | | | | | |
| proteasome (prosome, macropain) subunit, alpha type, 2 | Cytoplasm | peptidase | 1.94E+07 | 8305368.5 | 1.44E+07 | | | | | | |
| proteasome (prosome, macropain) subunit, alpha type, 2 | Cytoplasm | peptidase | 1.27E+07 | 8485131 | 9360132 | | | | | | |
| proteasome (prosome, macropain) subunit, alpha type, 2 | Cytoplasm | peptidase | 9464894 | 4088194.5 | 4349525 | | | | | | |
| proteasome (prosome, macropain) subunit, alpha type, 2 | Cytoplasm | peptidase | 8099449.5 | 1930695.5 | 3187109.25 | | | | | | |
| proteasome (prosome, macropain) subunit, alpha type, 2 | Cytoplasm | peptidase | 7104912.5 | 2832891.5 | 3684266.75 | | | | | | |
| proteasome (prosome, macropain) subunit, alpha type, 2 | Cytoplasm | peptidase | 3739760.25 | 1314894.875 | 2652098 | P49722 | PSMA2 | 35313902 | 28185169 | 5145884 | 30 |
| proteasome (prosome, macropain) subunit, alpha type, 3 | Cytoplasm | peptidase | 6.39E+07 | 2.27E+07 | 3.10E+07 | | | | | | |
| proteasome (prosome, macropain) subunit, alpha type, 3 | Cytoplasm | peptidase | 4.52E+07 | 2.06E+07 | 2.97E+07 | | | | | | |
| proteasome (prosome, macropain) subunit, alpha type, 3 | Cytoplasm | peptidase | 4.10E+07 | 1.83E+07 | 2.83E+07 | | | | | | |
| proteasome (prosome, macropain) subunit, alpha type, 3 | Cytoplasm | peptidase | 3.32E+07 | 1.16E+07 | 1.87E+07 | | | | | | |
| proteasome (prosome, macropain) subunit, alpha type, 3 | Cytoplasm | peptidase | 3.06E+07 | 1.12E+07 | 1.75E+07 | | | | | | |
| proteasome (prosome, macropain) subunit, alpha type, 3 | Cytoplasm | peptidase | 2.65E+07 | 1.16E+07 | 1.73E+07 | | | | | | |
| proteasome (prosome, macropain) subunit, alpha type, 3 | Cytoplasm | peptidase | 2.25E+07 | 1.01E+07 | 1.33E+07 | | | | | | |
| proteasome (prosome, macropain) subunit, alpha type, 3 | Cytoplasm | peptidase | 1.43E+07 | 1.10E+07 | 1.54E+07 | | | | | | |
| proteasome (prosome, macropain) subunit, alpha type, 3 | Cytoplasm | peptidase | 1.11E+07 | 5993715.5 | 7485630.5 | O70435 | PSMA3 | 32027778 | 13125172 | 2525940 | 27 |
| proteasome (prosome, macropain) subunit, alpha type, 4 | Cytoplasm | peptidase | 9.03E+07 | 6.14E+07 | 3.62E+07 | | | | | | |
| proteasome (prosome, macropain) subunit, alpha type, 4 | Cytoplasm | peptidase | 8.05E+07 | 3.88E+07 | 4.63E+07 | | | | | | |
| proteasome (prosome, macropain) subunit, alpha type, 4 | Cytoplasm | peptidase | 6.36E+07 | 2.92E+07 | 4.33E+07 | | | | | | |
| proteasome (prosome, macropain) subunit, alpha type, 4 | Cytoplasm | peptidase | 4.60E+07 | 2.82E+07 | 3.13E+07 | | | | | | |
| proteasome (prosome, macropain) subunit, alpha type, 4 | Cytoplasm | peptidase | 4.48E+07 | 1.75E+07 | 2.67E+07 | | | | | | |
| proteasome (prosome, macropain) subunit, alpha type, 4 | Cytoplasm | peptidase | 3.63E+07 | 1.40E+07 | 2.12E+07 | | | | | | |
| proteasome (prosome, macropain) subunit, alpha type, 4 | Cytoplasm | peptidase | 3.59E+07 | 1.84E+07 | 1.62E+07 | | | | | | |
| proteasome (prosome, macropain) subunit, alpha type, 4 | Cytoplasm | peptidase | 2.58E+07 | 1.06E+07 | 1.31E+07 | | | | | | |
| proteasome (prosome, macropain) subunit, alpha type, 4 | Cytoplasm | peptidase | 2.13E+07 | 7976471 | 1.00E+07 | | | | | | |
| proteasome (prosome, macropain) subunit, alpha type, 4 | Cytoplasm | peptidase | 1.78E+07 | 6836742 | 1.09E+07 | | | | | | |
| proteasome (prosome, macropain) subunit, alpha type, 4 | Cytoplasm | peptidase | 9182442 | 4186169.75 | 6260164.5 | | | | | | |
| proteasome (prosome, macropain) subunit, alpha type, 4 | Cytoplasm | peptidase | 7127525 | 3978872.75 | 4339141 | | | | | | |
| proteasome (prosome, macropain) subunit, alpha type, 4 | Cytoplasm | peptidase | 1544026.625 | 216929.781 | 875868.188 | Q9R1P0 | PSMA4 | 36941846 | 21855216 | 3499635 | 39 |
| proteasome (prosome, macropain) subunit, alpha type, 5 | Cytoplasm | peptidase | 1.40E+08 | 7.75E+07 | 9.38E+07 | | | | | | |
| proteasome (prosome, macropain) subunit, alpha type, 5 | Cytoplasm | peptidase | 1.11E+08 | 6.04E+07 | 9.21E+07 | | | | | | |
| proteasome (prosome, macropain) subunit, alpha type, 5 | Cytoplasm | peptidase | 1.02E+08 | 5.40E+07 | 7.16E+07 | | | | | | |
| proteasome (prosome, macropain) subunit, alpha type, 5 | Cytoplasm | peptidase | 4.77E+07 | 2.43E+07 | 3.64E+07 | | | | | | |
| proteasome (prosome, macropain) subunit, alpha type, 5 | Cytoplasm | peptidase | 3.87E+07 | 1.79E+07 | 2.50E+07 | | | | | | |
| proteasome (prosome, macropain) subunit, alpha type, 5 | Cytoplasm | peptidase | 3.05E+07 | 1.50E+07 | 2.01E+07 | | | | | | |
| proteasome (prosome, macropain) subunit, alpha type, 5 | Cytoplasm | peptidase | 2.17E+07 | 1.25E+07 | 1.53E+07 | | | | | | |
| proteasome (prosome, macropain) subunit, alpha type, 5 | Cytoplasm | peptidase | 1.93E+07 | 8633281 | 1.04E+07 | | | | | | |
| proteasome (prosome, macropain) subunit, alpha type, 5 | Cytoplasm | peptidase | 1.20E+07 | 1.65E+07 | 1.39E+07 | | | | | | |
| proteasome (prosome, macropain) subunit, alpha type, 5 | Cytoplasm | peptidase | 9836777 | 2619092.25 | 4612519.5 | | | | | | |
| proteasome (prosome, macropain) subunit, alpha type, 5 | Cytoplasm | peptidase | 8287116.5 | 3672109 | 4886350 | | | | | | |
| proteasome (prosome, macropain) subunit, alpha type, 5 | Cytoplasm | peptidase | 7958319 | 4335064 | 5032313 | | | | | | |
| proteasome (prosome, macropain) subunit, alpha type, 5 | Cytoplasm | peptidase | 6330333 | 2177074.75 | 2592408.5 | | | | | | |
| proteasome (prosome, macropain) subunit, alpha type, 5 | Cytoplasm | peptidase | 3804411.25 | 2077820.375 | 1891303.5 | Q9Z2U1 | PSMA5 | 39958354 | 35021880 | 5403993 | 42 |
| proteasome (prosome, macropain) subunit, alpha type, 6 | Cytoplasm | peptidase | 8.48E+07 | 3.70E+07 | 5.16E+07 | | | | | | |
| proteasome (prosome, macropain) subunit, alpha type, 6 | Cytoplasm | peptidase | 7.28E+07 | 3.35E+07 | 5.03E+07 | | | | | | |
| proteasome (prosome, macropain) subunit, alpha type, 6 | Cytoplasm | peptidase | 5.71E+07 | 2.89E+07 | 3.33E+07 | | | | | | |
| proteasome (prosome, macropain) subunit, alpha type, 6 | Cytoplasm | peptidase | 5.15E+07 | 2.40E+07 | 2.80E+07 | | | | | | |
| proteasome (prosome, macropain) subunit, alpha type, 6 | Cytoplasm | peptidase | 5.04E+07 | 2.18E+07 | 2.90E+07 | | | | | | |
| proteasome (prosome, macropain) subunit, alpha type, 6 | Cytoplasm | peptidase | 4.46E+07 | 2.17E+07 | 2.63E+07 | | | | | | |
| proteasome (prosome, macropain) subunit, alpha type, 6 | Cytoplasm | peptidase | 3.69E+07 | 1.46E+07 | 1.66E+07 | | | | | | |
| proteasome (prosome, macropain) subunit, alpha type, 6 | Cytoplasm | peptidase | 3.68E+07 | 1.93E+07 | 2.06E+07 | | | | | | |
| proteasome (prosome, macropain) subunit, alpha type, 6 | Cytoplasm | peptidase | 3.08E+07 | 1.70E+07 | 1.73E+07 | | | | | | |
| proteasome (prosome, macropain) subunit, alpha type, 6 | Cytoplasm | peptidase | 2.34E+07 | 1.14E+07 | 1.51E+07 | | | | | | |
| proteasome (prosome, macropain) subunit, alpha type, 6 | Cytoplasm | peptidase | 3875507.75 | 1724829.125 | 732257.812 | Q9QUM9 | PSMA6 | 44802319 | 19190433 | 3340626 | 33 |
| proteasome (prosome, macropain) subunit, alpha type, 7 | Cytoplasm | peptidase | 1.59E+08 | 5.97E+07 | 8.41E+07 | | | | | | |
| proteasome (prosome, macropain) subunit, alpha type, 7 | Cytoplasm | peptidase | 8.92E+07 | 3.69E+07 | 5.25E+07 | | | | | | |
| proteasome (prosome, macropain) subunit, alpha type, 7 | Cytoplasm | peptidase | 7.87E+07 | 3.84E+07 | 4.26E+07 | | | | | | |
| proteasome (prosome, macropain) subunit, alpha type, 7 | Cytoplasm | peptidase | 6.77E+07 | 2.74E+07 | 3.31E+07 | | | | | | |
| proteasome (prosome, macropain) subunit, alpha type, 7 | Cytoplasm | peptidase | 6.01E+07 | 2.78E+07 | 3.59E+07 | | | | | | |
| proteasome (prosome, macropain) subunit, alpha type, 7 | Cytoplasm | peptidase | 3.75E+07 | 1.61E+07 | 2.13E+07 | | | | | | |
| proteasome (prosome, macropain) subunit, alpha type, 7 | Cytoplasm | peptidase | 2.67E+07 | 9225603 | 1.21E+07 | | | | | | |
| proteasome (prosome, macropain) subunit, alpha type, 7 | Cytoplasm | peptidase | 2.06E+07 | 1.55E+07 | 1.03E+07 | | | | | | |
| proteasome (prosome, macropain) subunit, alpha type, 7 | Cytoplasm | peptidase | 1.52E+07 | 6121722 | 6785595 | | | | | | |
| proteasome (prosome, macropain) subunit, alpha type, 7 | Cytoplasm | peptidase | 9844743 | 6419021 | 5332674 | | | | | | |
| proteasome (prosome, macropain) subunit, alpha type, 7 | Cytoplasm | peptidase | 7928093.5 | 4724099 | 6616354.5 | | | | | | |
| proteasome (prosome, macropain) subunit, alpha type, 7 | Cytoplasm | peptidase | 4105576.5 | 6779761 | 4538920 | Q9Z2U0 | PSMA7 | 48000701 | 32720271 | 5453378 | 36 |
| proteasome (prosome, macropain) subunit, beta type, 1 | Cytoplasm | peptidase | 7.59E+07 | 2.81E+07 | 3.94E+07 | | | | | | |
| proteasome (prosome, macropain) subunit, beta type, 1 | Cytoplasm | peptidase | 6.99E+07 | 2.99E+07 | 3.49E+07 | | | | | | |
| proteasome (prosome, macropain) subunit, beta type, 1 | Cytoplasm | peptidase | 4.87E+07 | 1.96E+07 | 3.31E+07 | | | | | | |
| proteasome (prosome, macropain) subunit, beta type, 1 | Cytoplasm | peptidase | 3.86E+07 | 2.00E+07 | 2.84E+07 | | | | | | |
| proteasome (prosome, macropain) subunit, beta type, 1 | Cytoplasm | peptidase | 2.98E+07 | 1.20E+07 | 2.30E+07 | | | | | | |
| proteasome (prosome, macropain) subunit, beta type, 1 | Cytoplasm | peptidase | 1.67E+07 | 7372633 | 8673179 | | | | | | |
| proteasome (prosome, macropain) subunit, beta type, 1 | Cytoplasm | peptidase | 1.39E+07 | 6125707 | 5552193 | | | | | | |
| proteasome (prosome, macropain) subunit, beta type, 1 | Cytoplasm | peptidase | 1.18E+07 | 3521013.5 | 5606579 | | | | | | |
| proteasome (prosome, macropain) subunit, beta type, 1 | Cytoplasm | peptidase | 1.06E+07 | 2817457.5 | 5764018.5 | | | | | | |
| proteasome (prosome, macropain) subunit, beta type, 1 | Cytoplasm | peptidase | 8632642 | 4288038.5 | 6137856 | | | | | | |
| proteasome (prosome, macropain) subunit, beta type, 1 | Cytoplasm | peptidase | 8488025 | 2266083 | 3838580.5 | | | | | | |
| proteasome (prosome, macropain) subunit, beta type, 1 | Cytoplasm | peptidase | 8045834.5 | 3865764 | 4140928.25 | | | | | | |
| proteasome (prosome, macropain) subunit, beta type, 1 | Cytoplasm | peptidase | 6633206 | 1770831.875 | 3645764 | | | | | | |
| proteasome (prosome, macropain) subunit, beta type, 1 | Cytoplasm | peptidase | 5617945.5 | 2549838.5 | 4506316.5 | | | | | | |
| proteasome (prosome, macropain) subunit, beta type, 1 | Cytoplasm | peptidase | 5210947 | 1267075.875 | 2991457 | | | | | | |
| proteasome (prosome, macropain) subunit, beta type, 1 | Cytoplasm | peptidase | 4181342.5 | 591193.125 | 1770626 | | | | | | |
| proteasome (prosome, macropain) subunit, beta type, 1 | Cytoplasm | peptidase | 3849233 | 4313505 | 2160520 | O09061 | PSMB1 | 21562304 | 16900967 | 2366609 | 51 |
| proteasome (prosome, macropain) subunit, beta type, 2 | Cytoplasm | peptidase | 5.66E+07 | 3.45E+07 | 4.47E+07 | | | | | | |
| proteasome (prosome, macropain) subunit, beta type, 2 | Cytoplasm | peptidase | 5.54E+07 | 2.53E+07 | 3.28E+07 | | | | | | |
| proteasome (prosome, macropain) subunit, beta type, 2 | Cytoplasm | peptidase | 3.52E+07 | 7377967 | 1.15E+07 | | | | | | |
| proteasome (prosome, macropain) subunit, beta type, 2 | Cytoplasm | peptidase | 1.40E+07 | 4972241.5 | 6792260 | | | | | | |
| proteasome (prosome, macropain) subunit, beta type, 2 | Cytoplasm | peptidase | 1.24E+07 | 6077385.5 | 7369556.5 | | | | | | |
| proteasome (prosome, macropain) subunit, beta type, 2 | Cytoplasm | peptidase | 1.15E+07 | 6365529 | 7755657.5 | | | | | | |
| proteasome (prosome, macropain) subunit, beta type, 2 | Cytoplasm | peptidase | 9829915 | 5838400 | 5628709.5 | | | | | | |
| proteasome (prosome, macropain) subunit, beta type, 2 | Cytoplasm | peptidase | 8502954 | 3892220.25 | 3681732 | | | | | | |
| proteasome (prosome, macropain) subunit, beta type, 2 | Cytoplasm | peptidase | 8117803 | 6938159.5 | 5911038.5 | Q9R1P3 | PSMB2 | 23502297 | 15959345 | 3071377 | 27 |

| | | | | | | | | | | | |
|---|---|---|---|---|---|---|---|---|---|---|---|
| proteasome (prosome, macropain) subunit, beta type, 3 | Cytoplasm | peptidase | 5.99E+07 | 2.25E+07 | 3.83E+07 | | | | | | |
| proteasome (prosome, macropain) subunit, beta type, 3 | Cytoplasm | peptidase | 4.01E+07 | 1.99E+07 | 2.51E+07 | | | | | | |
| proteasome (prosome, macropain) subunit, beta type, 3 | Cytoplasm | peptidase | 2.48E+07 | 1.23E+07 | 1.59E+07 | | | | | | |
| proteasome (prosome, macropain) subunit, beta type, 3 | Cytoplasm | peptidase | 2.27E+07 | 1.12E+07 | 1.50E+07 | | | | | | |
| proteasome (prosome, macropain) subunit, beta type, 3 | Cytoplasm | peptidase | 1.49E+07 | 3996940.5 | 8233100.5 | | | | | | |
| proteasome (prosome, macropain) subunit, beta type, 3 | Cytoplasm | peptidase | 1.38E+07 | 5102104.5 | 5196224.5 | | | | | | |
| proteasome (prosome, macropain) subunit, beta type, 3 | Cytoplasm | peptidase | 1.05E+07 | 9644179 | 1.49E+07 | | | | | | |
| proteasome (prosome, macropain) subunit, beta type, 3 | Cytoplasm | peptidase | 1.03E+07 | 2315518.5 | 4697400.5 | | | | | | |
| proteasome (prosome, macropain) subunit, beta type, 3 | Cytoplasm | peptidase | 5085951.5 | 1119373.75 | 3553438.25 | | | | | | |
| proteasome (prosome, macropain) subunit, beta type, 3 | Cytoplasm | peptidase | 165199.781 | 1980816.625 | 1800045.375 | Q9R1P1 | PSMB3 | 20218115 | 13407804 | 2447919 | 30 |
| proteasome (prosome, macropain) subunit, beta type, 4 | Cytoplasm | peptidase | 4.70E+07 | 1.99E+07 | 2.84E+07 | | | | | | |
| proteasome (prosome, macropain) subunit, beta type, 4 | Cytoplasm | peptidase | 4.70E+07 | 2.02E+07 | 2.63E+07 | | | | | | |
| proteasome (prosome, macropain) subunit, beta type, 4 | Cytoplasm | peptidase | 4.59E+07 | 1.86E+07 | 2.29E+07 | | | | | | |
| proteasome (prosome, macropain) subunit, beta type, 4 | Cytoplasm | peptidase | 4.44E+07 | 1.56E+07 | 2.47E+07 | | | | | | |
| proteasome (prosome, macropain) subunit, beta type, 4 | Cytoplasm | peptidase | 3.39E+07 | 1.66E+07 | 2.09E+07 | | | | | | |
| proteasome (prosome, macropain) subunit, beta type, 4 | Cytoplasm | peptidase | 9472627 | 5293842.5 | 6881042.5 | P99026 | PSMB4 | 37922105 | 13497477 | 3181386 | 18 |
| proteasome (prosome, macropain) subunit, beta type, 5 | Cytoplasm | peptidase | 7.95E+07 | 2.55E+07 | 3.59E+07 | | | | | | |
| proteasome (prosome, macropain) subunit, beta type, 5 | Cytoplasm | peptidase | 5.12E+07 | 1.94E+07 | 2.61E+07 | | | | | | |
| proteasome (prosome, macropain) subunit, beta type, 5 | Cytoplasm | peptidase | 4.78E+07 | 2.34E+07 | 2.98E+07 | | | | | | |
| proteasome (prosome, macropain) subunit, beta type, 5 | Cytoplasm | peptidase | 4.63E+07 | 1.81E+07 | 2.58E+07 | | | | | | |
| proteasome (prosome, macropain) subunit, beta type, 5 | Cytoplasm | peptidase | 3.48E+07 | 1.73E+07 | 2.85E+07 | | | | | | |
| proteasome (prosome, macropain) subunit, beta type, 5 | Cytoplasm | peptidase | 2.69E+07 | 9736832 | 1.19E+07 | | | | | | |
| proteasome (prosome, macropain) subunit, beta type, 5 | Cytoplasm | peptidase | 1.67E+07 | 6314964.5 | 9566014 | | | | | | |
| proteasome (prosome, macropain) subunit, beta type, 5 | Cytoplasm | peptidase | 1.58E+07 | 7340942.5 | 1.19E+07 | | | | | | |
| proteasome (prosome, macropain) subunit, beta type, 5 | Cytoplasm | peptidase | 1.42E+07 | 6138245.5 | 7487327 | | | | | | |
| proteasome (prosome, macropain) subunit, beta type, 5 | Cytoplasm | peptidase | 1.09E+07 | 7327175.5 | 4719611.5 | | | | | | |
| proteasome (prosome, macropain) subunit, beta type, 5 | Cytoplasm | peptidase | 6195113.5 | 1580454.5 | 2347181 | O55234 | PSMB5 | 31835919 | 16940178 | 2948906 | 33 |
| proteasome (prosome, macropain) subunit, beta type, 6 | Nucleus | peptidase | 1.74E+08 | 8.69E+07 | 1.12E+08 | | | | | | |
| proteasome (prosome, macropain) subunit, beta type, 6 | Nucleus | peptidase | 5.55E+07 | 2.31E+07 | 3.34E+07 | | | | | | |
| proteasome (prosome, macropain) subunit, beta type, 6 | Nucleus | peptidase | 4.44E+07 | 1.92E+07 | 2.56E+07 | | | | | | |
| proteasome (prosome, macropain) subunit, beta type, 6 | Nucleus | peptidase | 2.69E+07 | 1.05E+07 | 1.35E+07 | | | | | | |
| proteasome (prosome, macropain) subunit, beta type, 6 | Nucleus | peptidase | 9406510 | 3651872.75 | 5797019 | | | | | | |
| proteasome (prosome, macropain) subunit, beta type, 6 | Nucleus | peptidase | 4821672.5 | 2793797.5 | 2031673.5 | Q60692 | PSMB6 | 52539697 | 45750427 | 10783479 | 18 |
| proteasome (prosome, macropain) subunit, beta type, 7 | Cytoplasm | peptidase | 5.51E+07 | 2.14E+07 | 3.93E+07 | | | | | | |
| proteasome (prosome, macropain) subunit, beta type, 7 | Cytoplasm | peptidase | 4.03E+07 | 1.60E+07 | 2.08E+07 | | | | | | |
| proteasome (prosome, macropain) subunit, beta type, 7 | Cytoplasm | peptidase | 3.02E+07 | 1.76E+07 | 2.39E+07 | | | | | | |
| proteasome (prosome, macropain) subunit, beta type, 7 | Cytoplasm | peptidase | 2.98E+07 | 1.26E+07 | 1.58E+07 | | | | | | |
| proteasome (prosome, macropain) subunit, beta type, 7 | Cytoplasm | peptidase | 2.65E+07 | 1.01E+07 | 1.75E+07 | | | | | | |
| proteasome (prosome, macropain) subunit, beta type, 7 | Cytoplasm | peptidase | 1.99E+07 | 9263367 | 1.11E+07 | | | | | | |
| proteasome (prosome, macropain) subunit, beta type, 7 | Cytoplasm | peptidase | 9888502 | 3249629.25 | 4555521 | | | | | | |
| proteasome (prosome, macropain) subunit, beta type, 7 | Cytoplasm | peptidase | 2616689.25 | 350623.281 | 1287907.625 | P70195 | PSMB7 | 26784399 | 13606431 | 2777401 | 24 |
| proteasome (prosome, macropain) subunit, beta type, 9 | Cytoplasm | peptidase | 1278809.125 | 501979.562 | 530969.438 | P28076 | PSMB9 | 1278809 | 440373 | 254249 | 3 |
| proteasome (prosome, macropain) 26S subunit, ATPase, 1 | Nucleus | peptidase | 3.49E+07 | 1.71E+07 | 2.63E+07 | | | | | | |
| proteasome (prosome, macropain) 26S subunit, ATPase, 1 | Nucleus | peptidase | 2.79E+07 | 1.29E+07 | 1.49E+07 | | | | | | |
| proteasome (prosome, macropain) 26S subunit, ATPase, 1 | Nucleus | peptidase | 2.14E+07 | 9199500 | 1.10E+07 | | | | | | |
| proteasome (prosome, macropain) 26S subunit, ATPase, 1 | Nucleus | peptidase | 1.66E+07 | 9488332 | 1.34E+07 | | | | | | |
| proteasome (prosome, macropain) 26S subunit, ATPase, 1 | Nucleus | peptidase | 1.50E+07 | 8221762 | 1.24E+07 | | | | | | |
| proteasome (prosome, macropain) 26S subunit, ATPase, 1 | Nucleus | peptidase | 1.44E+07 | 7859013.5 | 1.04E+07 | | | | | | |
| proteasome (prosome, macropain) 26S subunit, ATPase, 1 | Nucleus | peptidase | 1.43E+07 | 4104926.25 | 7223926 | | | | | | |
| proteasome (prosome, macropain) 26S subunit, ATPase, 1 | Nucleus | peptidase | 1.21E+07 | 6672045.5 | 8087296 | | | | | | |
| proteasome (prosome, macropain) 26S subunit, ATPase, 1 | Nucleus | peptidase | 1.05E+07 | 4985816 | 6246659 | | | | | | |
| proteasome (prosome, macropain) 26S subunit, ATPase, 1 | Nucleus | peptidase | 8829403 | 2544737 | 5128634.5 | | | | | | |
| proteasome (prosome, macropain) 26S subunit, ATPase, 1 | Nucleus | peptidase | 7659943 | 3322735.5 | 4752455 | | | | | | |
| proteasome (prosome, macropain) 26S subunit, ATPase, 1 | Nucleus | peptidase | 4327222.5 | 2305654.25 | 5795210.5 | | | | | | |
| proteasome (prosome, macropain) 26S subunit, ATPase, 1 | Nucleus | peptidase | 2952389.25 | 1562093 | 2811110 | P62192 | PSMC1 | 14685304 | 7402031 | 1185274 | 39 |
| proteasome (prosome, macropain) 26S subunit, ATPase, 2 | Nucleus | peptidase | 3.85E+07 | 1.86E+07 | 2.25E+07 | | | | | | |
| proteasome (prosome, macropain) 26S subunit, ATPase, 2 | Nucleus | peptidase | 2.90E+07 | 1.36E+07 | 1.81E+07 | | | | | | |
| proteasome (prosome, macropain) 26S subunit, ATPase, 2 | Nucleus | peptidase | 2.46E+07 | 9658856 | 1.33E+07 | | | | | | |
| proteasome (prosome, macropain) 26S subunit, ATPase, 2 | Nucleus | peptidase | 1.86E+07 | 9211062 | 1.25E+07 | | | | | | |
| proteasome (prosome, macropain) 26S subunit, ATPase, 2 | Nucleus | peptidase | 1.81E+07 | 7734461.5 | 9446501 | | | | | | |
| proteasome (prosome, macropain) 26S subunit, ATPase, 2 | Nucleus | peptidase | 1.74E+07 | 5027431.5 | 1.11E+07 | | | | | | |
| proteasome (prosome, macropain) 26S subunit, ATPase, 2 | Nucleus | peptidase | 1.74E+07 | 7324124.5 | 9666479 | | | | | | |
| proteasome (prosome, macropain) 26S subunit, ATPase, 2 | Nucleus | peptidase | 1.38E+07 | 8490762 | 1.21E+07 | | | | | | |
| proteasome (prosome, macropain) 26S subunit, ATPase, 2 | Nucleus | peptidase | 1.33E+07 | 6951136 | 9352813 | | | | | | |
| proteasome (prosome, macropain) 26S subunit, ATPase, 2 | Nucleus | peptidase | 1.26E+07 | 7685021.5 | 1.02E+07 | | | | | | |
| proteasome (prosome, macropain) 26S subunit, ATPase, 2 | Nucleus | peptidase | 1.22E+07 | 7479855 | 1.01E+07 | | | | | | |
| proteasome (prosome, macropain) 26S subunit, ATPase, 2 | Nucleus | peptidase | 9434532 | 2714731.75 | 5487445.5 | | | | | | |
| proteasome (prosome, macropain) 26S subunit, ATPase, 2 | Nucleus | peptidase | 8057918.5 | 5485880 | 6030199 | | | | | | |
| proteasome (prosome, macropain) 26S subunit, ATPase, 2 | Nucleus | peptidase | 5474093.5 | 1852698.125 | 2694252.5 | | | | | | |
| proteasome (prosome, macropain) 26S subunit, ATPase, 2 | Nucleus | peptidase | 2812601 | 1665257 | 2500320.25 | P46471 | PSMC2 | 16078610 | 7411189 | 1104795 | 45 |
| proteasome (prosome, macropain) 26S subunit, ATPase, 3 | Nucleus | transcription regulator | 2.85E+07 | 1.48E+07 | 1.68E+07 | | | | | | |
| proteasome (prosome, macropain) 26S subunit, ATPase, 3 | Nucleus | transcription regulator | 2.31E+07 | 8311523.5 | 1.44E+07 | | | | | | |
| proteasome (prosome, macropain) 26S subunit, ATPase, 3 | Nucleus | transcription regulator | 1.90E+07 | 9059324 | 1.25E+07 | | | | | | |
| proteasome (prosome, macropain) 26S subunit, ATPase, 3 | Nucleus | transcription regulator | 1.68E+07 | 6992510.5 | 1.07E+07 | | | | | | |
| proteasome (prosome, macropain) 26S subunit, ATPase, 3 | Nucleus | transcription regulator | 1.38E+07 | 6668819 | 9218666 | | | | | | |
| proteasome (prosome, macropain) 26S subunit, ATPase, 3 | Nucleus | transcription regulator | 1.33E+07 | 5193562.5 | 6601048 | | | | | | |
| proteasome (prosome, macropain) 26S subunit, ATPase, 3 | Nucleus | transcription regulator | 1.33E+07 | 1.01E+07 | 1.17E+07 | | | | | | |
| proteasome (prosome, macropain) 26S subunit, ATPase, 3 | Nucleus | transcription regulator | 1.09E+07 | 6817809 | 9119072 | | | | | | |
| proteasome (prosome, macropain) 26S subunit, ATPase, 3 | Nucleus | transcription regulator | 1.01E+07 | 4936419 | 7682859 | | | | | | |
| proteasome (prosome, macropain) 26S subunit, ATPase, 3 | Nucleus | transcription regulator | 9517274 | 3602881.75 | 5795144 | | | | | | |
| proteasome (prosome, macropain) 26S subunit, ATPase, 3 | Nucleus | transcription regulator | 6065272.5 | 3314764.75 | 3960121 | | | | | | |
| proteasome (prosome, macropain) 26S subunit, ATPase, 3 | Nucleus | transcription regulator | 4973273.5 | 3087760.5 | 3733985.5 | | | | | | |
| proteasome (prosome, macropain) 26S subunit, ATPase, 3 | Nucleus | transcription regulator | 4664830 | 2386640.5 | 2435310 | | | | | | |
| proteasome (prosome, macropain) 26S subunit, ATPase, 3 | Nucleus | transcription regulator | 3116622.5 | 1065822.125 | 1781595 | O88685 | PSMC3 | 12642662 | 5935101 | 915806 | 42 |
| proteasome (prosome, macropain) 26S subunit, ATPase, 4 | Nucleus | peptidase | 2.38E+07 | 1.04E+07 | 1.86E+07 | | | | | | |
| proteasome (prosome, macropain) 26S subunit, ATPase, 4 | Nucleus | peptidase | 2.10E+07 | 1.07E+07 | 1.17E+07 | | | | | | |
| proteasome (prosome, macropain) 26S subunit, ATPase, 4 | Nucleus | peptidase | 1.01E+07 | 4282488.5 | 8605183 | | | | | | |
| proteasome (prosome, macropain) 26S subunit, ATPase, 4 | Nucleus | peptidase | 9062778 | 4893790.5 | 5994819.5 | | | | | | |
| proteasome (prosome, macropain) 26S subunit, ATPase, 4 | Nucleus | peptidase | 8443531 | 4004321.25 | 4851565 | | | | | | |
| proteasome (prosome, macropain) 26S subunit, ATPase, 4 | Nucleus | peptidase | 7995537.5 | 6096595.5 | 7861240.5 | | | | | | |
| proteasome (prosome, macropain) 26S subunit, ATPase, 4 | Nucleus | peptidase | 5891854 | 2327204.75 | 3745731.5 | | | | | | |
| proteasome (prosome, macropain) 26S subunit, ATPase, 4 | Nucleus | peptidase | 5834215 | 3017732.75 | 2508365.75 | | | | | | |
| proteasome (prosome, macropain) 26S subunit, ATPase, 4 | Nucleus | peptidase | 5459271.5 | 2922103 | 4199142.5 | | | | | | |
| proteasome (prosome, macropain) 26S subunit, ATPase, 4 | Nucleus | peptidase | 5295972 | 3691216.75 | 3681791.75 | | | | | | |
| proteasome (prosome, macropain) 26S subunit, ATPase, 4 | Nucleus | peptidase | 4989195 | 1978066.625 | 3603662.25 | | | | | | |

| | | | | | | | | | | | |
|---|---|---|---|---|---|---|---|---|---|---|---|
| proteasome (prosome, macropain) 26S subunit, ATPase, 4 | Nucleus | peptidase | 3553945 | 1871195.25 | 2237922.25 | P54775 | PSMC4 | 9276358 | 5150313 | 858386 | 36 |
| proteasome (prosome, macropain) 26S subunit, ATPase, 5 | Nucleus | transcription regulator | 2.97E+07 | 1.38E+07 | 1.87E+07 | | | | | | |
| proteasome (prosome, macropain) 26S subunit, ATPase, 5 | Nucleus | transcription regulator | 1.95E+07 | 7912580 | 1.20E+07 | | | | | | |
| proteasome (prosome, macropain) 26S subunit, ATPase, 5 | Nucleus | transcription regulator | 1.82E+07 | 7352269.5 | 1.18E+07 | | | | | | |
| proteasome (prosome, macropain) 26S subunit, ATPase, 5 | Nucleus | transcription regulator | 1.73E+07 | 9097310 | 1.05E+07 | | | | | | |
| proteasome (prosome, macropain) 26S subunit, ATPase, 5 | Nucleus | transcription regulator | 1.41E+07 | 6678344.5 | 9186626 | | | | | | |
| proteasome (prosome, macropain) 26S subunit, ATPase, 5 | Nucleus | transcription regulator | 1.28E+07 | 6435487 | 8497651 | | | | | | |
| proteasome (prosome, macropain) 26S subunit, ATPase, 5 | Nucleus | transcription regulator | 1.07E+07 | 4988671.5 | 8173180 | | | | | | |
| proteasome (prosome, macropain) 26S subunit, ATPase, 5 | Nucleus | transcription regulator | 1.05E+07 | 5935650 | 6479024 | | | | | | |
| proteasome (prosome, macropain) 26S subunit, ATPase, 5 | Nucleus | transcription regulator | 9856927 | 5393599 | 6606498 | | | | | | |
| proteasome (prosome, macropain) 26S subunit, ATPase, 5 | Nucleus | transcription regulator | 7535844.5 | 3492980 | 4355633 | | | | | | |
| proteasome (prosome, macropain) 26S subunit, ATPase, 5 | Nucleus | transcription regulator | 7525592 | 4477661 | 5602311.5 | | | | | | |
| proteasome (prosome, macropain) 26S subunit, ATPase, 5 | Nucleus | transcription regulator | 7472180.5 | 2973929.25 | 4571011 | | | | | | |
| proteasome (prosome, macropain) 26S subunit, ATPase, 5 | Nucleus | transcription regulator | 7232715.5 | 5466003.5 | 5163586.5 | | | | | | |
| proteasome (prosome, macropain) 26S subunit, ATPase, 5 | Nucleus | transcription regulator | 6057002 | 3574920.25 | 3448353 | | | | | | |
| proteasome (prosome, macropain) 26S subunit, ATPase, 5 | Nucleus | transcription regulator | 4598040.5 | 1535890.375 | 3894541 | | | | | | |
| proteasome (prosome, macropain) 26S subunit, ATPase, 5 | Nucleus | transcription regulator | 4088481 | 3471940 | 1595124.375 | P62196 | PSMC5 | 11694799 | 5412062 | 781164 | 48 |
| proteasome (prosome, macropain) 26S subunit, ATPase, 6 | Nucleus | peptidase | 2.89E+07 | 1.17E+07 | 1.97E+07 | | | | | | |
| proteasome (prosome, macropain) 26S subunit, ATPase, 6 | Nucleus | peptidase | 1.89E+07 | 8398846 | 1.28E+07 | | | | | | |
| proteasome (prosome, macropain) 26S subunit, ATPase, 6 | Nucleus | peptidase | 1.69E+07 | 5300547.5 | 1.01E+07 | | | | | | |
| proteasome (prosome, macropain) 26S subunit, ATPase, 6 | Nucleus | peptidase | 1.56E+07 | 9043526 | 8274373.5 | | | | | | |
| proteasome (prosome, macropain) 26S subunit, ATPase, 6 | Nucleus | peptidase | 1.51E+07 | 8582071 | 1.21E+07 | | | | | | |
| proteasome (prosome, macropain) 26S subunit, ATPase, 6 | Nucleus | peptidase | 1.50E+07 | 6443197.5 | 8473332 | | | | | | |
| proteasome (prosome, macropain) 26S subunit, ATPase, 6 | Nucleus | peptidase | 1.22E+07 | 4718017 | 6459623.5 | | | | | | |
| proteasome (prosome, macropain) 26S subunit, ATPase, 6 | Nucleus | peptidase | 1.10E+07 | 6036508.5 | 6229341.5 | | | | | | |
| proteasome (prosome, macropain) 26S subunit, ATPase, 6 | Nucleus | peptidase | 1.08E+07 | 5240780 | 6614669 | | | | | | |
| proteasome (prosome, macropain) 26S subunit, ATPase, 6 | Nucleus | peptidase | 1.07E+07 | 4996389.5 | 6757712.5 | | | | | | |
| proteasome (prosome, macropain) 26S subunit, ATPase, 6 | Nucleus | peptidase | 8602080 | 3494956 | 4956118.5 | | | | | | |
| proteasome (prosome, macropain) 26S subunit, ATPase, 6 | Nucleus | peptidase | 7114443 | 4396265 | 5091786 | | | | | | |
| proteasome (prosome, macropain) 26S subunit, ATPase, 6 | Nucleus | peptidase | 3451888.25 | 1280827.875 | 1746375.25 | P62334 | PSMC6 | 13388339 | 5543223 | 887626 | 39 |
| proteasome (prosome, macropain) 26S subunit, non-ATPase, 1 | Cytoplasm | other | 3.15E+07 | 1.98E+07 | 2.11E+07 | | | | | | |
| proteasome (prosome, macropain) 26S subunit, non-ATPase, 1 | Cytoplasm | other | 2.89E+07 | 1.58E+07 | 2.14E+07 | | | | | | |
| proteasome (prosome, macropain) 26S subunit, non-ATPase, 1 | Cytoplasm | other | 2.87E+07 | 1.09E+07 | 1.15E+07 | | | | | | |
| proteasome (prosome, macropain) 26S subunit, non-ATPase, 1 | Cytoplasm | other | 2.60E+07 | 1.50E+07 | 1.77E+07 | | | | | | |
| proteasome (prosome, macropain) 26S subunit, non-ATPase, 1 | Cytoplasm | other | 2.09E+07 | 1.07E+07 | 1.19E+07 | | | | | | |
| proteasome (prosome, macropain) 26S subunit, non-ATPase, 1 | Cytoplasm | other | 1.90E+07 | 9114958 | 1.17E+07 | | | | | | |
| proteasome (prosome, macropain) 26S subunit, non-ATPase, 1 | Cytoplasm | other | 1.73E+07 | 7554723 | 1.02E+07 | | | | | | |
| proteasome (prosome, macropain) 26S subunit, non-ATPase, 1 | Cytoplasm | other | 1.73E+07 | 6893179 | 9718384 | | | | | | |
| proteasome (prosome, macropain) 26S subunit, non-ATPase, 1 | Cytoplasm | other | 1.70E+07 | 1.02E+07 | 1.25E+07 | | | | | | |
| proteasome (prosome, macropain) 26S subunit, non-ATPase, 1 | Cytoplasm | other | 1.38E+07 | 5159001 | 8684004 | | | | | | |
| proteasome (prosome, macropain) 26S subunit, non-ATPase, 1 | Cytoplasm | other | 1.38E+07 | 5861551.5 | 9063175 | | | | | | |
| proteasome (prosome, macropain) 26S subunit, non-ATPase, 1 | Cytoplasm | other | 1.35E+07 | 7192133.5 | 9067084 | | | | | | |
| proteasome (prosome, macropain) 26S subunit, non-ATPase, 1 | Cytoplasm | other | 1.27E+07 | 6524436.5 | 1.01E+07 | | | | | | |
| proteasome (prosome, macropain) 26S subunit, non-ATPase, 1 | Cytoplasm | other | 1.12E+07 | 5279774.5 | 8483342 | | | | | | |
| proteasome (prosome, macropain) 26S subunit, non-ATPase, 1 | Cytoplasm | other | 1.02E+07 | 3681902.5 | 7327972.5 | | | | | | |
| proteasome (prosome, macropain) 26S subunit, non-ATPase, 1 | Cytoplasm | other | 1.01E+07 | 4418362 | 6063614.5 | | | | | | |
| proteasome (prosome, macropain) 26S subunit, non-ATPase, 1 | Cytoplasm | other | 9653481 | 4464974.5 | 5355384.5 | | | | | | |
| proteasome (prosome, macropain) 26S subunit, non-ATPase, 1 | Cytoplasm | other | 9122814 | 4926785.5 | 7060720.5 | | | | | | |
| proteasome (prosome, macropain) 26S subunit, non-ATPase, 1 | Cytoplasm | other | 8286235 | 4200441 | 4686518 | | | | | | |
| proteasome (prosome, macropain) 26S subunit, non-ATPase, 1 | Cytoplasm | other | 7716632 | 4752784 | 5445855.5 | | | | | | |
| proteasome (prosome, macropain) 26S subunit, non-ATPase, 1 | Cytoplasm | other | 5648859.5 | 2704496 | 3457449.75 | | | | | | |
| proteasome (prosome, macropain) 26S subunit, non-ATPase, 1 | Cytoplasm | other | 5467255.5 | 2955313.5 | 3137420.25 | | | | | | |
| proteasome (prosome, macropain) 26S subunit, non-ATPase, 1 | Cytoplasm | other | 5053453.5 | 2299228.75 | 3527686.25 | | | | | | |
| proteasome (prosome, macropain) 26S subunit, non-ATPase, 1 | Cytoplasm | other | 4311117.5 | 1955491.125 | 4396305 | | | | | | |
| proteasome (prosome, macropain) 26S subunit, non-ATPase, 1 | Cytoplasm | other | 2857098 | 2096718.5 | 2144504.75 | | | | | | |
| proteasome (prosome, macropain) 26S subunit, non-ATPase, 1 | Cytoplasm | other | 2656563.75 | 993629.562 | 1428356.75 | | | | | | |
| proteasome (prosome, macropain) 26S subunit, non-ATPase, 1 | Cytoplasm | other | 2620029.25 | 993642.375 | 1753059.75 | | | | | | |
| proteasome (prosome, macropain) 26S subunit, non-ATPase, 1 | Cytoplasm | other | 2137764.5 | 1849336.25 | 1527459.25 | Q3TXS7 | PSMD1 | 12761118 | 6875229 | 750149 | 84 |
| proteasome (prosome, macropain) 26S subunit, non-ATPase, 11 | Cytoplasm | other | 2.39E+07 | 1.01E+07 | 1.53E+07 | | | | | | |
| proteasome (prosome, macropain) 26S subunit, non-ATPase, 11 | Cytoplasm | other | 2.15E+07 | 1.12E+07 | 1.10E+07 | | | | | | |
| proteasome (prosome, macropain) 26S subunit, non-ATPase, 11 | Cytoplasm | other | 2.12E+07 | 1.27E+07 | 1.36E+07 | | | | | | |
| proteasome (prosome, macropain) 26S subunit, non-ATPase, 11 | Cytoplasm | other | 1.86E+07 | 6339478 | 1.61E+07 | | | | | | |
| proteasome (prosome, macropain) 26S subunit, non-ATPase, 11 | Cytoplasm | other | 1.58E+07 | 8103310.5 | 9625975 | | | | | | |
| proteasome (prosome, macropain) 26S subunit, non-ATPase, 11 | Cytoplasm | other | 1.41E+07 | 7824352 | 9543386 | | | | | | |
| proteasome (prosome, macropain) 26S subunit, non-ATPase, 11 | Cytoplasm | other | 1.23E+07 | 6886313 | 9903574 | | | | | | |
| proteasome (prosome, macropain) 26S subunit, non-ATPase, 11 | Cytoplasm | other | 1.21E+07 | 5386925.5 | 6713508 | | | | | | |
| proteasome (prosome, macropain) 26S subunit, non-ATPase, 11 | Cytoplasm | other | 9771932 | 3972405 | 5489027 | | | | | | |
| proteasome (prosome, macropain) 26S subunit, non-ATPase, 11 | Cytoplasm | other | 8197028.5 | 3286001.25 | 6155036.5 | | | | | | |
| proteasome (prosome, macropain) 26S subunit, non-ATPase, 11 | Cytoplasm | other | 7465599.5 | 3375279.25 | 4307128.5 | | | | | | |
| proteasome (prosome, macropain) 26S subunit, non-ATPase, 11 | Cytoplasm | other | 4883830 | 3093166.75 | 3141118 | | | | | | |
| proteasome (prosome, macropain) 26S subunit, non-ATPase, 11 | Cytoplasm | other | 2717633.75 | 1798133.25 | 3352767.5 | | | | | | |
| proteasome (prosome, macropain) 26S subunit, non-ATPase, 11 | Cytoplasm | other | 1058800 | 834167.125 | 2886851.5 | Q8BG32 | PSMD11 | 12401773 | 5829049 | 899442 | 42 |
| proteasome (prosome, macropain) 26S subunit, non-ATPase, 12 | Cytoplasm | other | 2.31E+07 | 9253466 | 1.34E+07 | | | | | | |
| proteasome (prosome, macropain) 26S subunit, non-ATPase, 12 | Cytoplasm | other | 1.84E+07 | 8532574 | 1.17E+07 | | | | | | |
| proteasome (prosome, macropain) 26S subunit, non-ATPase, 12 | Cytoplasm | other | 1.75E+07 | 8397712 | 1.19E+07 | | | | | | |
| proteasome (prosome, macropain) 26S subunit, non-ATPase, 12 | Cytoplasm | other | 1.41E+07 | 6923109 | 8933505 | | | | | | |
| proteasome (prosome, macropain) 26S subunit, non-ATPase, 12 | Cytoplasm | other | 1.37E+07 | 7114263 | 8241422.5 | | | | | | |
| proteasome (prosome, macropain) 26S subunit, non-ATPase, 12 | Cytoplasm | other | 1.25E+07 | 5032243.5 | 7954383.5 | | | | | | |
| proteasome (prosome, macropain) 26S subunit, non-ATPase, 12 | Cytoplasm | other | 1.21E+07 | 5912511 | 7509436.5 | | | | | | |
| proteasome (prosome, macropain) 26S subunit, non-ATPase, 12 | Cytoplasm | other | 1.16E+07 | 7074510.5 | 9044182 | | | | | | |
| proteasome (prosome, macropain) 26S subunit, non-ATPase, 12 | Cytoplasm | other | 1.02E+07 | 6888636 | 7280063 | | | | | | |
| proteasome (prosome, macropain) 26S subunit, non-ATPase, 12 | Cytoplasm | other | 6689080.5 | 5137368 | 4272699 | | | | | | |
| proteasome (prosome, macropain) 26S subunit, non-ATPase, 12 | Cytoplasm | other | 5810997 | 3841774.75 | 4626973.5 | | | | | | |
| proteasome (prosome, macropain) 26S subunit, non-ATPase, 12 | Cytoplasm | other | 5126533 | 2233717.25 | 3417747.75 | | | | | | |
| proteasome (prosome, macropain) 26S subunit, non-ATPase, 12 | Cytoplasm | other | 4208601 | 1577105 | 1795383 | | | | | | |
| proteasome (prosome, macropain) 26S subunit, non-ATPase, 12 | Cytoplasm | other | 3980485 | 2541870.25 | 3455758.75 | | | | | | |
| proteasome (prosome, macropain) 26S subunit, non-ATPase, 12 | Cytoplasm | other | 2581272.5 | 1267921.125 | 991202.625 | Q9D8W5 | PSMD12 | 10768465 | 4837820 | 721180 | 45 |
| proteasome (prosome, macropain) 26S subunit, non-ATPase, 13 | Cytoplasm | peptidase | 3.44E+07 | 1.70E+07 | 2.22E+07 | | | | | | |
| proteasome (prosome, macropain) 26S subunit, non-ATPase, 13 | Cytoplasm | peptidase | 3.13E+07 | 1.32E+07 | 1.79E+07 | | | | | | |
| proteasome (prosome, macropain) 26S subunit, non-ATPase, 13 | Cytoplasm | peptidase | 2.90E+07 | 9358141 | 1.58E+07 | | | | | | |
| proteasome (prosome, macropain) 26S subunit, non-ATPase, 13 | Cytoplasm | peptidase | 1.99E+07 | 9785921 | 1.63E+07 | | | | | | |
| proteasome (prosome, macropain) 26S subunit, non-ATPase, 13 | Cytoplasm | peptidase | 1.85E+07 | 8982958 | 1.22E+07 | | | | | | |
| proteasome (prosome, macropain) 26S subunit, non-ATPase, 13 | Cytoplasm | peptidase | 1.56E+07 | 5086575 | 7960115 | | | | | | |
| proteasome (prosome, macropain) 26S subunit, non-ATPase, 13 | Cytoplasm | peptidase | 1.38E+07 | 8512483 | 8048022.5 | | | | | | |
| proteasome (prosome, macropain) 26S subunit, non-ATPase, 13 | Cytoplasm | peptidase | 1.28E+07 | 7772090.5 | 1.05E+07 | | | | | | |

| | | | | | | | | | | | | |
|---|---|---|---|---|---|---|---|---|---|---|---|---|
| proteasome (prosome, macropain) 26S subunit, non-ATPase, 13 | Cytoplasm | peptidase | 1.28E+07 | 6926264 | 8394060 | | | | | | | |
| proteasome (prosome, macropain) 26S subunit, non-ATPase, 13 | Cytoplasm | peptidase | 1.26E+07 | 7500618.5 | 9801940 | | | | | | | |
| proteasome (prosome, macropain) 26S subunit, non-ATPase, 13 | Cytoplasm | peptidase | 8834298 | 7619352 | 6238314 | | | | | | | |
| proteasome (prosome, macropain) 26S subunit, non-ATPase, 13 | Cytoplasm | peptidase | 7765531 | 4259475 | 5204489.5 | | | | | | | |
| proteasome (prosome, macropain) 26S subunit, non-ATPase, 13 | Cytoplasm | peptidase | 1499190.75 | 2220427.25 | 2575334.5 | Q9WVJ2 | PSMD13 | | 16835309 | 7544281 | 1208052 | 39 |
| proteasome (prosome, macropain) 26S subunit, non-ATPase, 14 | Cytoplasm | peptidase | 3.27E+07 | 1.02E+07 | 1.65E+07 | | | | | | | |
| proteasome (prosome, macropain) 26S subunit, non-ATPase, 14 | Cytoplasm | peptidase | 2.03E+07 | 8406177 | 1.69E+07 | | | | | | | |
| proteasome (prosome, macropain) 26S subunit, non-ATPase, 14 | Cytoplasm | peptidase | 7276973 | 2701946 | 4537577 | | | | | | | |
| proteasome (prosome, macropain) 26S subunit, non-ATPase, 14 | Cytoplasm | peptidase | 3019500.75 | 1253549.75 | 2955087.5 | O35593 | PSMD14 | | 15819118 | 9415440 | 2718003 | 12 |
| proteasome (prosome, macropain) 26S subunit, non-ATPase, 2 | Cytoplasm | other | 4.62E+07 | 1.71E+07 | 2.24E+07 | | | | | | | |
| proteasome (prosome, macropain) 26S subunit, non-ATPase, 2 | Cytoplasm | other | 3.05E+07 | 1.32E+07 | 2.07E+07 | | | | | | | |
| proteasome (prosome, macropain) 26S subunit, non-ATPase, 2 | Cytoplasm | other | 2.60E+07 | 1.00E+07 | 1.18E+07 | | | | | | | |
| proteasome (prosome, macropain) 26S subunit, non-ATPase, 2 | Cytoplasm | other | 2.20E+07 | 1.28E+07 | 1.52E+07 | | | | | | | |
| proteasome (prosome, macropain) 26S subunit, non-ATPase, 2 | Cytoplasm | other | 1.80E+07 | 7901604 | 1.07E+07 | | | | | | | |
| proteasome (prosome, macropain) 26S subunit, non-ATPase, 2 | Cytoplasm | other | 1.60E+07 | 9805438 | 1.03E+07 | | | | | | | |
| proteasome (prosome, macropain) 26S subunit, non-ATPase, 2 | Cytoplasm | other | 1.60E+07 | 1.18E+07 | 1.10E+07 | | | | | | | |
| proteasome (prosome, macropain) 26S subunit, non-ATPase, 2 | Cytoplasm | other | 1.39E+07 | 6099723 | 8717331 | | | | | | | |
| proteasome (prosome, macropain) 26S subunit, non-ATPase, 2 | Cytoplasm | other | 1.30E+07 | 6962280.5 | 8913941 | | | | | | | |
| proteasome (prosome, macropain) 26S subunit, non-ATPase, 2 | Cytoplasm | other | 1.21E+07 | 5678802.5 | 7042253 | | | | | | | |
| proteasome (prosome, macropain) 26S subunit, non-ATPase, 2 | Cytoplasm | other | 1.20E+07 | 6322992.5 | 8991034 | | | | | | | |
| proteasome (prosome, macropain) 26S subunit, non-ATPase, 2 | Cytoplasm | other | 1.12E+07 | 8685798 | 1.07E+07 | | | | | | | |
| proteasome (prosome, macropain) 26S subunit, non-ATPase, 2 | Cytoplasm | other | 1.12E+07 | 6710434 | 6877596.5 | | | | | | | |
| proteasome (prosome, macropain) 26S subunit, non-ATPase, 2 | Cytoplasm | other | 1.08E+07 | 4550854 | 6137314 | | | | | | | |
| proteasome (prosome, macropain) 26S subunit, non-ATPase, 2 | Cytoplasm | other | 1.05E+07 | 4852155 | 6128299.5 | | | | | | | |
| proteasome (prosome, macropain) 26S subunit, non-ATPase, 2 | Cytoplasm | other | 1.05E+07 | 3085273.5 | 5188131 | | | | | | | |
| proteasome (prosome, macropain) 26S subunit, non-ATPase, 2 | Cytoplasm | other | 9526617 | 4904466.5 | 6437019 | | | | | | | |
| proteasome (prosome, macropain) 26S subunit, non-ATPase, 2 | Cytoplasm | other | 9508634 | 1843975.125 | 2442036.5 | | | | | | | |
| proteasome (prosome, macropain) 26S subunit, non-ATPase, 2 | Cytoplasm | other | 8482435 | 3888212.25 | 6076330.5 | | | | | | | |
| proteasome (prosome, macropain) 26S subunit, non-ATPase, 2 | Cytoplasm | other | 7621079 | 3356655 | 5099911.5 | | | | | | | |
| proteasome (prosome, macropain) 26S subunit, non-ATPase, 2 | Cytoplasm | other | 7216854.5 | 3453712.5 | 4417508 | | | | | | | |
| proteasome (prosome, macropain) 26S subunit, non-ATPase, 2 | Cytoplasm | other | 6920141 | 4505488.5 | 5517636.5 | | | | | | | |
| proteasome (prosome, macropain) 26S subunit, non-ATPase, 2 | Cytoplasm | other | 6619682 | 4615496 | 5095594 | | | | | | | |
| proteasome (prosome, macropain) 26S subunit, non-ATPase, 2 | Cytoplasm | other | 5533908 | 2124126.5 | 2809638 | | | | | | | |
| proteasome (prosome, macropain) 26S subunit, non-ATPase, 2 | Cytoplasm | other | 5086271.5 | 2598292.75 | 3182169 | | | | | | | |
| proteasome (prosome, macropain) 26S subunit, non-ATPase, 2 | Cytoplasm | other | 4406073 | 1338470.25 | 2203281.25 | | | | | | | |
| proteasome (prosome, macropain) 26S subunit, non-ATPase, 2 | Cytoplasm | other | 4361806 | 1919758.75 | 3413615 | | | | | | | |
| proteasome (prosome, macropain) 26S subunit, non-ATPase, 2 | Cytoplasm | other | 4198100.5 | 2257581.25 | 3357578.25 | | | | | | | |
| proteasome (prosome, macropain) 26S subunit, non-ATPase, 2 | Cytoplasm | other | 2594818 | 1198572.375 | 2137805 | Q8VDM4 | PSMD2 | | 12477118 | 6968431 | 747094 | 87 |
| proteasome (prosome, macropain) 26S subunit, non-ATPase, 3 | Cytoplasm | other | 3.96E+07 | 1.62E+07 | 2.50E+07 | | | | | | | |
| proteasome (prosome, macropain) 26S subunit, non-ATPase, 3 | Cytoplasm | other | 3.95E+07 | 1.93E+07 | 2.36E+07 | | | | | | | |
| proteasome (prosome, macropain) 26S subunit, non-ATPase, 3 | Cytoplasm | other | 3.01E+07 | 1.44E+07 | 1.74E+07 | | | | | | | |
| proteasome (prosome, macropain) 26S subunit, non-ATPase, 3 | Cytoplasm | other | 1.83E+07 | 9191925 | 1.06E+07 | | | | | | | |
| proteasome (prosome, macropain) 26S subunit, non-ATPase, 3 | Cytoplasm | other | 1.65E+07 | 7952255 | 9361950 | | | | | | | |
| proteasome (prosome, macropain) 26S subunit, non-ATPase, 3 | Cytoplasm | other | 1.53E+07 | 1.19E+07 | 1.27E+07 | | | | | | | |
| proteasome (prosome, macropain) 26S subunit, non-ATPase, 3 | Cytoplasm | other | 1.42E+07 | 8756381 | 9550134 | | | | | | | |
| proteasome (prosome, macropain) 26S subunit, non-ATPase, 3 | Cytoplasm | other | 1.23E+07 | 5304937 | 9595028 | | | | | | | |
| proteasome (prosome, macropain) 26S subunit, non-ATPase, 3 | Cytoplasm | other | 8977552 | 5439927 | 5269466.5 | | | | | | | |
| proteasome (prosome, macropain) 26S subunit, non-ATPase, 3 | Cytoplasm | other | 7500371.5 | 3457786.25 | 4356454 | | | | | | | |
| proteasome (prosome, macropain) 26S subunit, non-ATPase, 3 | Cytoplasm | other | 5843856.5 | 2686647.5 | 4825438 | | | | | | | |
| proteasome (prosome, macropain) 26S subunit, non-ATPase, 3 | Cytoplasm | other | 4786413.5 | 1975544.25 | 1721934.125 | | | | | | | |
| proteasome (prosome, macropain) 26S subunit, non-ATPase, 3 | Cytoplasm | other | 3651733.25 | 1398990.375 | 2505297 | | | | | | | |
| proteasome (prosome, macropain) 26S subunit, non-ATPase, 3 | Cytoplasm | other | 3157252.75 | 1893736 | 1946628.5 | | | | | | | |
| proteasome (prosome, macropain) 26S subunit, non-ATPase, 3 | Cytoplasm | other | 2966590.75 | 5023916 | 4855805.5 | | | | | | | |
| proteasome (prosome, macropain) 26S subunit, non-ATPase, 3 | Cytoplasm | other | 2807921.75 | 1742298.125 | 3219940.75 | P14685 | PSMD3 | | 14089481 | 9170404 | 1323634 | 48 |
| proteasome (prosome, macropain) 26S subunit, non-ATPase, 4 | Cytoplasm | other | 3.22E+07 | 1.95E+07 | 2.05E+07 | | | | | | | |
| proteasome (prosome, macropain) 26S subunit, non-ATPase, 4 | Cytoplasm | other | 2.61E+07 | 1.18E+07 | 1.54E+07 | | | | | | | |
| proteasome (prosome, macropain) 26S subunit, non-ATPase, 4 | Cytoplasm | other | 1.28E+07 | 7710829 | 9741808 | | | | | | | |
| proteasome (prosome, macropain) 26S subunit, non-ATPase, 4 | Cytoplasm | other | 1.09E+07 | 5003792.5 | 6642718.5 | | | | | | | |
| proteasome (prosome, macropain) 26S subunit, non-ATPase, 4 | Cytoplasm | other | 9099155 | 5959519.5 | 8726059 | | | | | | | |
| proteasome (prosome, macropain) 26S subunit, non-ATPase, 4 | Cytoplasm | other | 5928623 | 6205547 | 6115895.5 | | | | | | | |
| proteasome (prosome, macropain) 26S subunit, non-ATPase, 4 | Cytoplasm | other | 4240067.5 | 2273613.25 | 2368465 | | | | | | | |
| proteasome (prosome, macropain) 26S subunit, non-ATPase, 4 | Cytoplasm | other | 2790342.75 | 1816652.625 | 1378623.5 | | | | | | | |
| proteasome (prosome, macropain) 26S subunit, non-ATPase, 4 | Cytoplasm | other | 2532051.25 | 1310988.875 | 1370798 | O35226 | PSMD4 | | 11844471 | 7885443 | 1517554 | 27 |
| proteasome (prosome, macropain) 26S subunit, non-ATPase, 5 | Cytoplasm | other | 6263101 | 3717063.25 | 3365682.25 | | | | | | | |
| proteasome (prosome, macropain) 26S subunit, non-ATPase, 5 | Cytoplasm | other | 4374784 | 3502077.5 | 4292223 | | | | | | | |
| proteasome (prosome, macropain) 26S subunit, non-ATPase, 5 | Cytoplasm | other | 3973959.75 | 2992488.25 | 3357437.25 | | | | | | | |
| proteasome (prosome, macropain) 26S subunit, non-ATPase, 5 | Cytoplasm | other | 3761118.25 | 3412745 | 3879523 | | | | | | | |
| proteasome (prosome, macropain) 26S subunit, non-ATPase, 5 | Cytoplasm | other | 3280917.5 | 2931225.75 | 2369173.25 | | | | | | | |
| proteasome (prosome, macropain) 26S subunit, non-ATPase, 5 | Cytoplasm | other | 3278015.5 | 2346532.5 | 1956382.625 | | | | | | | |
| proteasome (prosome, macropain) 26S subunit, non-ATPase, 5 | Cytoplasm | other | 3072952.75 | 1933596.75 | 2424477.5 | | | | | | | |
| proteasome (prosome, macropain) 26S subunit, non-ATPase, 5 | Cytoplasm | other | 2740406.5 | 1572975.875 | 2276245.5 | | | | | | | |
| proteasome (prosome, macropain) 26S subunit, non-ATPase, 5 | Cytoplasm | other | 2709471.5 | 2127704.25 | 1946224.5 | Q8BJY1 | PSMD5 | | 3717192 | 991053 | 190728 | 27 |
| proteasome (prosome, macropain) 26S subunit, non-ATPase, 6 | Cytoplasm | enzyme | 2.53E+07 | 1.12E+07 | 1.30E+07 | | | | | | | |
| proteasome (prosome, macropain) 26S subunit, non-ATPase, 6 | Cytoplasm | enzyme | 2.15E+07 | 8166882 | 1.27E+07 | | | | | | | |
| proteasome (prosome, macropain) 26S subunit, non-ATPase, 6 | Cytoplasm | enzyme | 1.72E+07 | 9195653 | 9619150 | | | | | | | |
| proteasome (prosome, macropain) 26S subunit, non-ATPase, 6 | Cytoplasm | enzyme | 1.56E+07 | 8231572.5 | 9561379 | | | | | | | |
| proteasome (prosome, macropain) 26S subunit, non-ATPase, 6 | Cytoplasm | enzyme | 1.43E+07 | 6884020.5 | 8379741 | | | | | | | |
| proteasome (prosome, macropain) 26S subunit, non-ATPase, 6 | Cytoplasm | enzyme | 1.07E+07 | 5517147 | 7065037 | | | | | | | |
| proteasome (prosome, macropain) 26S subunit, non-ATPase, 6 | Cytoplasm | enzyme | 9386117 | 5038117.5 | 8199480.5 | | | | | | | |
| proteasome (prosome, macropain) 26S subunit, non-ATPase, 6 | Cytoplasm | enzyme | 5968949 | 3184009 | 3168170.25 | | | | | | | |
| proteasome (prosome, macropain) 26S subunit, non-ATPase, 6 | Cytoplasm | enzyme | 5844068.5 | 4176187.75 | 4269450 | | | | | | | |
| proteasome (prosome, macropain) 26S subunit, non-ATPase, 6 | Cytoplasm | enzyme | 5120140.5 | 2751305.75 | 3137036.75 | | | | | | | |
| proteasome (prosome, macropain) 26S subunit, non-ATPase, 6 | Cytoplasm | enzyme | 5093676.5 | 3243307.25 | 4121663.5 | | | | | | | |
| proteasome (prosome, macropain) 26S subunit, non-ATPase, 6 | Cytoplasm | enzyme | 4871893 | 1682495.5 | 2287365.25 | | | | | | | |
| proteasome (prosome, macropain) 26S subunit, non-ATPase, 6 | Cytoplasm | enzyme | 4572027.5 | 2968678.75 | 6005371 | | | | | | | |
| proteasome (prosome, macropain) 26S subunit, non-ATPase, 6 | Cytoplasm | enzyme | 1538129.625 | 378283.375 | 779191 | Q99JI4 | PSMD6 | | 10500357 | 5422640 | 836732 | 42 |
| proteasome (prosome, macropain) 26S subunit, non-ATPase, 7 | Cytoplasm | other | 3.50E+07 | 1.61E+07 | 2.14E+07 | | | | | | | |
| proteasome (prosome, macropain) 26S subunit, non-ATPase, 7 | Cytoplasm | other | 2.44E+07 | 1.28E+07 | 1.47E+07 | | | | | | | |
| proteasome (prosome, macropain) 26S subunit, non-ATPase, 7 | Cytoplasm | other | 2.38E+07 | 1.50E+07 | 1.50E+07 | | | | | | | |
| proteasome (prosome, macropain) 26S subunit, non-ATPase, 7 | Cytoplasm | other | 2.33E+07 | 1.05E+07 | 1.27E+07 | | | | | | | |
| proteasome (prosome, macropain) 26S subunit, non-ATPase, 7 | Cytoplasm | other | 1.26E+07 | 4954094.5 | 7607966 | | | | | | | |
| proteasome (prosome, macropain) 26S subunit, non-ATPase, 7 | Cytoplasm | other | 8575548 | 3901366 | 4698138 | | | | | | | |
| proteasome (prosome, macropain) 26S subunit, non-ATPase, 7 | Cytoplasm | other | 6407734 | 5334178 | 5429754.5 | | | | | | | |
| proteasome (prosome, macropain) 26S subunit, non-ATPase, 7 | Cytoplasm | other | 4187620.5 | 2689121.75 | 4046717.5 | | | | | | | |
| proteasome (prosome, macropain) 26S subunit, non-ATPase, 7 | Cytoplasm | other | 2734165.25 | 2210621.25 | 2953815.5 | | | | | | | |

| | | | | | | | | | | | |
|---|---|---|---|---|---|---|---|---|---|---|---|
| proteasome (prosome, macropain) 26S subunit, non-ATPase, 7 | Cytoplasm | other | 1899956.375 | 1124022.375 | 1010133.562 | P26516 | PSMD7 | 14290502 | 8560298 | 1562889 | 30 |
| proteasome (prosome, macropain) 26S subunit, non-ATPase, 8 | Cytoplasm | other | 1.96E+07 | 9021820 | 9735671 | | | | | | |
| proteasome (prosome, macropain) 26S subunit, non-ATPase, 8 | Cytoplasm | other | 1.01E+07 | 4948557 | 6593915.5 | | | | | | |
| proteasome (prosome, macropain) 26S subunit, non-ATPase, 8 | Cytoplasm | other | 8891583 | 3444653.5 | 6330849 | Q9CX56 | PSMD8 | 12857194 | 4651411 | 1550470 | 9 |
| proteasome (prosome, macropain) activator subunit 1 (PA28 alpha) | Cytoplasm | other | 3.09E+07 | 1.23E+07 | 1.64E+07 | | | | | | |
| proteasome (prosome, macropain) activator subunit 1 (PA28 alpha) | Cytoplasm | other | 2.46E+07 | 8237567 | 1.37E+07 | | | | | | |
| proteasome (prosome, macropain) activator subunit 1 (PA28 alpha) | Cytoplasm | other | 2.38E+07 | 8036138 | 1.65E+07 | | | | | | |
| proteasome (prosome, macropain) activator subunit 1 (PA28 alpha) | Cytoplasm | other | 2.07E+07 | 7287805 | 1.08E+07 | | | | | | |
| proteasome (prosome, macropain) activator subunit 1 (PA28 alpha) | Cytoplasm | other | 1.76E+07 | 4813925.5 | 9055026 | | | | | | |
| proteasome (prosome, macropain) activator subunit 1 (PA28 alpha) | Cytoplasm | other | 1.39E+07 | 7452509 | 5831518.5 | | | | | | |
| proteasome (prosome, macropain) activator subunit 1 (PA28 alpha) | Cytoplasm | other | 1.26E+07 | 4195629.5 | 5963546.5 | | | | | | |
| proteasome (prosome, macropain) activator subunit 1 (PA28 alpha) | Cytoplasm | other | 1.22E+07 | 4543457.5 | 7227428 | | | | | | |
| proteasome (prosome, macropain) activator subunit 1 (PA28 alpha) | Cytoplasm | other | 9694632 | 4607765.5 | 6722484.5 | | | | | | |
| proteasome (prosome, macropain) activator subunit 1 (PA28 alpha) | Cytoplasm | other | 9397281 | 4385755 | 4864594.5 | | | | | | |
| proteasome (prosome, macropain) activator subunit 1 (PA28 alpha) | Cytoplasm | other | 8629320 | 4605919.5 | 5625208.5 | | | | | | |
| proteasome (prosome, macropain) activator subunit 1 (PA28 alpha) | Cytoplasm | other | 4687396.5 | 3256627.25 | 3161685 | P97371 | PSME1 | 15712386 | 6677402 | 1112900 | 36 |
| proteasome (prosome, macropain) activator subunit 2 (PA28 beta) | Cytoplasm | peptidase | 3.93E+07 | 1.10E+07 | 2.28E+07 | | | | | | |
| proteasome (prosome, macropain) activator subunit 2 (PA28 beta) | Cytoplasm | peptidase | 2.77E+07 | 1.02E+07 | 1.63E+07 | | | | | | |
| proteasome (prosome, macropain) activator subunit 2 (PA28 beta) | Cytoplasm | peptidase | 2.26E+07 | 1.13E+07 | 1.24E+07 | | | | | | |
| proteasome (prosome, macropain) activator subunit 2 (PA28 beta) | Cytoplasm | peptidase | 8444027 | 3625716.25 | 4188993.5 | | | | | | |
| proteasome (prosome, macropain) activator subunit 2 (PA28 beta) | Cytoplasm | peptidase | 5780241.5 | 2058444.75 | 2518038 | | | | | | |
| proteasome (prosome, macropain) activator subunit 2 (PA28 beta) | Cytoplasm | peptidase | 4778445 | 4374867.5 | 2911632.25 | | | | | | |
| proteasome (prosome, macropain) activator subunit 2 (PA28 beta) | Cytoplasm | peptidase | 3482978.75 | 5583116 | 6396348 | P97372 | PSME2 | 16015099 | 9814464 | 2141692 | 21 |
| pentatricopeptide repeat domain 2 | Cytoplasm | other | 2326270.25 | 1904969.5 | 2112359 | Q8R3K3 | PTCD2 | 2326270 | 210659 | 121624 | 3 |
| phosphatase and tensin homolog | Cytoplasm | phosphatase | 2.93E+07 | 2.97E+07 | 3.25E+07 | O08586 | PTEN | 29290000 | 1767739 | 1020604 | 3 |
| protein tyrosine phosphatase type IVA, member 1 | Cytoplasm | phosphatase | 1803146 | 1818368.625 | 1721830.875 | Q63739 | PTP4A1 | 1803146 | 51903 | 29966 | 3 |
| protein tyrosine phosphatase, non-receptor type 1 | Cytoplasm | phosphatase | 629611.562 | 4684242 | 2915153.75 | P35821 | PTPN1 | 629612 | 2032790 | 1173632 | 3 |
| protein tyrosine phosphatase, non-receptor type 21 | Cytoplasm | phosphatase | 2.52E+07 | 2.60E+07 | 2.93E+07 | Q62136 | PTPN21 | 25160000 | 2165741 | 1250391 | 3 |
| protein tyrosine phosphatase, receptor type, A | Plasma Membrane | phosphatase | 9538780 | 1.02E+07 | 7681209.5 | | | | | | |
| protein tyrosine phosphatase, receptor type, A | Plasma Membrane | phosphatase | 5135390.5 | 4511669.5 | 5002310 | | | | | | |
| protein tyrosine phosphatase, receptor type, A | Plasma Membrane | phosphatase | 3773567 | 3551708.75 | 3592530 | | | | | | |
| protein tyrosine phosphatase, receptor type, A | Plasma Membrane | phosphatase | 388253.156 | 863641.125 | 603009.312 | P18052 | PTPRA | 4708998 | 3246512 | 937187 | 12 |
| protein tyrosine phosphatase, receptor type, J | Plasma Membrane | phosphatase | 115111.141 | 8788019 | 6563169 | | | | | | |
| protein tyrosine phosphatase, receptor type, J | Plasma Membrane | phosphatase | 53274.434 | 615742.938 | 775275.312 | | | | | | |
| protein tyrosine phosphatase, receptor type, J | Plasma Membrane | phosphatase | 33700.512 | 984229.438 | 505156.75 | Q64455 | PTPRJ | 67362 | 3255191 | 1085064 | 9 |
| peptidyl-tRNA hydrolase 2 | Cytoplasm | enzyme | 5221813.5 | 6609265.5 | 6189651.5 | | | | | | |
| peptidyl-tRNA hydrolase 2 | Cytoplasm | enzyme | 43739.188 | 266787.531 | 195587.906 | Q8R2Y8 | PTRH2 | 2632776 | 3230032 | 1318655 | 6 |
| purine-rich element binding protein A | Nucleus | transcription regulator | 6927430.5 | 2024609.625 | 3099241 | | | | | | |
| purine-rich element binding protein A | Nucleus | transcription regulator | 4434945.5 | 1109671.125 | 1056446 | P42669 | PURA | 5681188 | 2268252 | 926010 | 6 |
| pyrroline-5-carboxylate reductase family, member 2 | Cytoplasm | enzyme | 1411655.25 | 1825103.125 | 1470433.75 | Q922Q4 | PYCR2 | 1411655 | 223676 | 129139 | 3 |
| pyrroline-5-carboxylate reductase-like | Cytoplasm | enzyme | 8575434 | 3085659.5 | 4166416.25 | | | | | | |
| pyrroline-5-carboxylate reductase-like | Cytoplasm | enzyme | 6890430.5 | 2367439.5 | 3210340.25 | | | | | | |
| pyrroline-5-carboxylate reductase-like | Cytoplasm | enzyme | 4294927 | 2990295 | 2239465 | Q9DCC4 | PYCRL | 6586931 | 2158688 | 719563 | 9 |
| pregnancy zone protein | Extracellular Space | other | 1489379.875 | 1341739.125 | 1981028 | Q61838 | Pzp | 1489380 | 334715 | 193248 | 3 |
| quinoid dihydropteridine reductase | Cytoplasm | enzyme | 1739074.375 | 667210.5 | 989324 | Q8BVI4 | QDPR | 1739074 | 549966 | 317523 | 3 |
| RAB10, member RAS oncogene family | Cytoplasm | enzyme | 2.78E+08 | 2.91E+08 | 2.86E+08 | | | | | | |
| RAB10, member RAS oncogene family | Cytoplasm | enzyme | 1.96E+08 | 1.45E+08 | 1.44E+08 | | | | | | |
| RAB10, member RAS oncogene family | Cytoplasm | enzyme | 1.04E+08 | 1.00E+08 | 1.53E+08 | | | | | | |
| RAB10, member RAS oncogene family | Cytoplasm | enzyme | 8.17E+07 | 4.89E+07 | 3.34E+07 | | | | | | |
| RAB10, member RAS oncogene family | Cytoplasm | enzyme | 4.36E+07 | 3.40E+07 | 3.32E+07 | | | | | | |
| RAB10, member RAS oncogene family | Cytoplasm | enzyme | 4.12E+07 | 2.57E+07 | 3.54E+07 | | | | | | |
| RAB10, member RAS oncogene family | Cytoplasm | enzyme | 3.51E+07 | 3.11E+07 | 3.43E+07 | | | | | | |
| RAB10, member RAS oncogene family | Cytoplasm | enzyme | 2.20E+07 | 2.18E+07 | 2.14E+07 | | | | | | |
| RAB10, member RAS oncogene family | Cytoplasm | enzyme | 1.05E+07 | 2.26E+07 | 1.90E+07 | | | | | | |
| RAB10, member RAS oncogene family | Cytoplasm | enzyme | 8057676 | 1.48E+07 | 1.45E+07 | | | | | | |
| RAB10, member RAS oncogene family | Cytoplasm | enzyme | 4973945 | 4478122.5 | 4671728.5 | P61027 | RAB10 | 75002875 | 83960734 | 14615688 | 33 |
| RAB11B, member RAS oncogene family | Cytoplasm | enzyme | 2.30E+07 | 4.17E+07 | 4.05E+07 | | | | | | |
| RAB11B, member RAS oncogene family | Cytoplasm | enzyme | 1.37E+07 | 2.59E+07 | 2.07E+07 | | | | | | |
| RAB11B, member RAS oncogene family | Cytoplasm | enzyme | 1.36E+07 | 2.62E+07 | 2.17E+07 | | | | | | |
| RAB11B, member RAS oncogene family | Cytoplasm | enzyme | 1.07E+07 | 1.35E+07 | 1.88E+07 | | | | | | |
| RAB11B, member RAS oncogene family | Cytoplasm | enzyme | 5423998.5 | 9400173 | 9347724 | | | | | | |
| RAB11B, member RAS oncogene family | Cytoplasm | enzyme | 4939484 | 9497977 | 7620755.5 | | | | | | |
| RAB11B, member RAS oncogene family | Cytoplasm | enzyme | 4059587.25 | 7242386 | 5873224 | | | | | | |
| RAB11B, member RAS oncogene family | Cytoplasm | enzyme | 2921971.75 | 5393058.5 | 4180011.75 | | | | | | |
| RAB11B, member RAS oncogene family | Cytoplasm | enzyme | 2082270.5 | 9578332 | 6105222.5 | | | | | | |
| RAB11B, member RAS oncogene family | Cytoplasm | enzyme | 659208.812 | 488547.031 | 643739.562 | P46638 | RAB11B | 8105652 | 10834103 | 1978027 | 30 |
| RAB14, member RAS oncogene family | Cytoplasm | enzyme | 2.68E+07 | 1.53E+07 | 2.09E+07 | | | | | | |
| RAB14, member RAS oncogene family | Cytoplasm | enzyme | 1.40E+07 | 1.31E+07 | 1.62E+07 | | | | | | |
| RAB14, member RAS oncogene family | Cytoplasm | enzyme | 9300677 | 1.11E+07 | 1.06E+07 | | | | | | |
| RAB14, member RAS oncogene family | Cytoplasm | enzyme | 8063336 | 8514225 | 7926732 | | | | | | |
| RAB14, member RAS oncogene family | Cytoplasm | enzyme | 7516066.5 | 6717381 | 5116887.5 | | | | | | |
| RAB14, member RAS oncogene family | Cytoplasm | enzyme | 5033540 | 7784565 | 5062327 | | | | | | |
| RAB14, member RAS oncogene family | Cytoplasm | enzyme | 3313755.5 | 6476854.5 | 3530637 | | | | | | |
| RAB14, member RAS oncogene family | Cytoplasm | enzyme | 3034781 | 4111236.75 | 2537781.5 | | | | | | |
| RAB14, member RAS oncogene family | Cytoplasm | enzyme | 1062527.375 | 2936654 | 1584958 | Q91V41 | RAB14 | 8689409 | 6118858 | 1177575 | 27 |
| RAB15, member RAS oncogene family | Cytoplasm | enzyme | 5.12E+08 | 4.53E+08 | 4.08E+08 | Q8K386 | Rab15 | 512200000 | 52256515 | 30170313 | 3 |
| RAB18, member RAS oncogene family | Cytoplasm | enzyme | 2.24E+07 | 2.02E+07 | 1.37E+07 | | | | | | |
| RAB18, member RAS oncogene family | Cytoplasm | enzyme | 2.12E+07 | 1.87E+07 | 1.61E+07 | | | | | | |
| RAB18, member RAS oncogene family | Cytoplasm | enzyme | 2.02E+07 | 1.51E+07 | 1.31E+07 | | | | | | |
| RAB18, member RAS oncogene family | Cytoplasm | enzyme | 1.51E+07 | 1.39E+07 | 1.39E+07 | | | | | | |
| RAB18, member RAS oncogene family | Cytoplasm | enzyme | 1.50E+07 | 1.10E+07 | 1.12E+07 | P35293 | RAB18 | 18766000 | 3614277 | 933202 | 15 |
| RAB1A, member RAS oncogene family | Cytoplasm | enzyme | 8.37E+07 | 6.29E+07 | 7.39E+07 | | | | | | |
| RAB1A, member RAS oncogene family | Cytoplasm | enzyme | 3.67E+07 | 2.38E+07 | 2.95E+07 | | | | | | |
| RAB1A, member RAS oncogene family | Cytoplasm | enzyme | 1.89E+07 | 8497736 | 1.18E+07 | | | | | | |
| RAB1A, member RAS oncogene family | Cytoplasm | enzyme | 4274697.5 | 8188134 | 6878492.5 | | | | | | |
| RAB1A, member RAS oncogene family | Cytoplasm | enzyme | 2963471.5 | 662171.25 | 2114167.75 | | | | | | |
| RAB1A, member RAS oncogene family | Cytoplasm | enzyme | 2678188.5 | 1742111.25 | 2436533.75 | P62821 | RAB1A | 24869393 | 26400127 | 6222570 | 18 |
| RAB1B, member RAS oncogene family | Cytoplasm | other | 4.91E+07 | 6.80E+07 | 5.98E+07 | | | | | | |
| RAB1B, member RAS oncogene family | Cytoplasm | other | 2.71E+07 | 2.31E+07 | 2.59E+07 | | | | | | |
| RAB1B, member RAS oncogene family | Cytoplasm | other | 2.19E+07 | 8269796.5 | 1.16E+07 | | | | | | |
| RAB1B, member RAS oncogene family | Cytoplasm | other | 1.94E+07 | 1.73E+07 | 1.82E+07 | | | | | | |
| RAB1B, member RAS oncogene family | Cytoplasm | other | 6134489.5 | 5475279.5 | 7307990 | Q9D1G1 | RAB1B | 24710898 | 19454964 | 5023250 | 15 |
| RAB21, member RAS oncogene family | Cytoplasm | enzyme | 1.03E+07 | 9154243 | 1.14E+07 | | | | | | |
| RAB21, member RAS oncogene family | Cytoplasm | enzyme | 6661982 | 7346681 | 3757302.25 | | | | | | |
| RAB21, member RAS oncogene family | Cytoplasm | enzyme | 4262913.5 | 3608317.5 | 3166418 | | | | | | |

| | | | | | | | | | | | |
|---|---|---|---|---|---|---|---|---|---|---|---|
| RAB21, member RAS oncogene family | Cytoplasm | enzyme | 4250383 | 2336391 | 3139872.5 | | | | | | |
| RAB21, member RAS oncogene family | Cytoplasm | enzyme | 4040469.25 | 2288452 | 1220812.75 | | | | | | |
| RAB21, member RAS oncogene family | Cytoplasm | enzyme | 3042157.25 | 3191862 | 3824981 | | | | | | |
| RAB21, member RAS oncogene family | Cytoplasm | enzyme | 2449231.75 | 2005967 | 2712678.5 | | | | | | |
| RAB21, member RAS oncogene family | Cytoplasm | enzyme | 1210569.5 | 764784.25 | 687349.25 | P35282 | RAB21 | 4529713 | 2902283 | 592426 | 24 |
| RAB22A, member RAS oncogene family | Cytoplasm | enzyme | 3953155.5 | 3168894.5 | 3513862 | | | | | | |
| RAB22A, member RAS oncogene family | Cytoplasm | enzyme | 2695873.5 | 3051470.75 | 2848749.5 | P35285 | RAB22A | 3324515 | 461764 | 188514 | 6 |
| RAB27A, member RAS oncogene family | Cytoplasm | enzyme | 394758.719 | 2514064.25 | 2149946 | Q9ERI2 | RAB27A | 394759 | 1133190 | 654248 | 3 |
| RAB27B, member RAS oncogene family | Cytoplasm | enzyme | 1243734.5 | 2.34E+07 | 1.54E+07 | | | | | | |
| RAB27B, member RAS oncogene family | Cytoplasm | enzyme | 921927.375 | 1.58E+07 | 1.17E+07 | | | | | | |
| RAB27B, member RAS oncogene family | Cytoplasm | enzyme | 711255 | 2.00E+07 | 1.34E+07 | | | | | | |
| RAB27B, member RAS oncogene family | Cytoplasm | enzyme | 655330.125 | 1.54E+07 | 1.20E+07 | | | | | | |
| RAB27B, member RAS oncogene family | Cytoplasm | enzyme | 439377.719 | 7712113 | 5498877.5 | | | | | | |
| RAB27B, member RAS oncogene family | Cytoplasm | enzyme | 149036.828 | 881193.125 | 1209260 | | | | | | |
| RAB27B, member RAS oncogene family | Cytoplasm | enzyme | 28343.635 | 2514163.75 | 919418.438 | | | | | | |
| RAB27B, member RAS oncogene family | Cytoplasm | enzyme | 3976.828 | 1892160.5 | 1264780.625 | Q99P58 | RAB27B | 519123 | 7402291 | 1510986 | 24 |
| RAB2A, member RAS oncogene family | Cytoplasm | enzyme | 1.24E+07 | 1.20E+07 | 1.20E+07 | | | | | | |
| RAB2A, member RAS oncogene family | Cytoplasm | enzyme | 9575304 | 7332375 | 8351532 | | | | | | |
| RAB2A, member RAS oncogene family | Cytoplasm | enzyme | 4398691 | 4849873 | 3786530 | | | | | | |
| RAB2A, member RAS oncogene family | Cytoplasm | enzyme | 3353529.5 | 3345595.25 | 3045890.25 | | | | | | |
| RAB2A, member RAS oncogene family | Cytoplasm | enzyme | 1888142.75 | 1815031.125 | 2336509.25 | P53994 | RAB2A | 6317133 | 3891901 | 1004885 | 15 |
| RAB35, member RAS oncogene family | Cytoplasm | enzyme | 9.94E+07 | 7.81E+07 | 7.90E+07 | | | | | | |
| RAB35, member RAS oncogene family | Cytoplasm | enzyme | 5.59E+07 | 5.32E+07 | 4.39E+07 | | | | | | |
| RAB35, member RAS oncogene family | Cytoplasm | enzyme | 5.45E+07 | 4.44E+07 | 4.09E+07 | | | | | | |
| RAB35, member RAS oncogene family | Cytoplasm | enzyme | 5.28E+07 | 4.99E+07 | 2.75E+07 | | | | | | |
| RAB35, member RAS oncogene family | Cytoplasm | enzyme | 2.44E+07 | 2.47E+07 | 2.09E+07 | | | | | | |
| RAB35, member RAS oncogene family | Cytoplasm | enzyme | 2.17E+07 | 2.53E+07 | 1.50E+07 | | | | | | |
| RAB35, member RAS oncogene family | Cytoplasm | enzyme | 2743428.75 | 1149010.75 | 2815138.75 | Q6PHN9 | RAB35 | 44481918 | 26327510 | 5745134 | 21 |
| RAB37, member RAS oncogene family | Cytoplasm | enzyme | 1708.659 | 297281.125 | 273965.094 | Q9JKM7 | RAB37 | 1709 | 164332 | 94877 | 3 |
| RAB3 GTPase activating protein subunit 1 (catalytic) | Cytoplasm | other | 7083831 | 5724901.5 | 4489926.5 | | | | | | |
| RAB3 GTPase activating protein subunit 1 (catalytic) | Cytoplasm | other | 4181131 | 6030861 | 2130693.25 | | | | | | |
| RAB3 GTPase activating protein subunit 1 (catalytic) | Cytoplasm | other | 376031.188 | 293674.375 | 219877 | Q80UJ7 | RAB3GAP1 | 3880331 | 2697516 | 899172 | 9 |
| RAB3 GTPase activating protein subunit 2 (non-catalytic) | Cytoplasm | enzyme | 2792122.75 | 2886219 | 1952221.375 | | | | | | |
| RAB3 GTPase activating protein subunit 2 (non-catalytic) | Cytoplasm | enzyme | 1610560.25 | 1105431.625 | 845067.625 | Q8BMG7 | RAB3GAP2 | 2201342 | 847585 | 346025 | 6 |
| RAB3A interacting protein (rabin3)-like 1 | Cytoplasm | other | 1775526 | 1165854.25 | 1014856.438 | Q8VDV3 | RAB3IL1 | 1775526 | 402724 | 232513 | 3 |
| RAB5A, member RAS oncogene family | Cytoplasm | enzyme | 3.04E+07 | 2.55E+07 | 2.66E+07 | | | | | | |
| RAB5A, member RAS oncogene family | Cytoplasm | enzyme | 2.67E+07 | 2.68E+07 | 2.91E+07 | | | | | | |
| RAB5A, member RAS oncogene family | Cytoplasm | enzyme | 1.27E+07 | 7488284.5 | 8931937 | | | | | | |
| RAB5A, member RAS oncogene family | Cytoplasm | enzyme | 9553114 | 6174279.5 | 7544956 | | | | | | |
| RAB5A, member RAS oncogene family | Cytoplasm | enzyme | 7109727 | 8307143.5 | 7527591.5 | Q9CQD1 | RAB5A | 17298568 | 9878662 | 2550660 | 15 |
| RAB5B, member RAS oncogene family | Cytoplasm | enzyme | 2.98E+07 | 2.67E+07 | 3.10E+07 | | | | | | |
| RAB5B, member RAS oncogene family | Cytoplasm | enzyme | 2.00E+07 | 1.66E+07 | 1.71E+07 | | | | | | |
| RAB5B, member RAS oncogene family | Cytoplasm | enzyme | 1.56E+07 | 1.21E+07 | 1.65E+07 | | | | | | |
| RAB5B, member RAS oncogene family | Cytoplasm | enzyme | 1.05E+07 | 8317191 | 7987223.5 | | | | | | |
| RAB5B, member RAS oncogene family | Cytoplasm | enzyme | 9641217 | 7594455.5 | 6624850.5 | | | | | | |
| RAB5B, member RAS oncogene family | Cytoplasm | enzyme | 5128894 | 5148658.5 | 5181511 | P61021 | RAB5B | 15108352 | 8378426 | 1974814 | 18 |
| RAB5C, member RAS oncogene family | Cytoplasm | enzyme | 1.22E+08 | 1.13E+08 | 1.15E+08 | | | | | | |
| RAB5C, member RAS oncogene family | Cytoplasm | enzyme | 1.17E+08 | 8.61E+07 | 9.32E+07 | | | | | | |
| RAB5C, member RAS oncogene family | Cytoplasm | enzyme | 9.56E+07 | 8.29E+07 | 7.80E+07 | | | | | | |
| RAB5C, member RAS oncogene family | Cytoplasm | enzyme | 8.16E+07 | 5.00E+07 | 5.82E+07 | | | | | | |
| RAB5C, member RAS oncogene family | Cytoplasm | enzyme | 6.88E+07 | 5.73E+07 | 7.09E+07 | | | | | | |
| RAB5C, member RAS oncogene family | Cytoplasm | enzyme | 3.19E+07 | 1.93E+07 | 2.63E+07 | | | | | | |
| RAB5C, member RAS oncogene family | Cytoplasm | enzyme | 3.01E+07 | 2.35E+07 | 2.64E+07 | | | | | | |
| RAB5C, member RAS oncogene family | Cytoplasm | enzyme | 2.64E+07 | 1.98E+07 | 2.30E+07 | | | | | | |
| RAB5C, member RAS oncogene family | Cytoplasm | enzyme | 2.17E+07 | 1.37E+07 | 1.17E+07 | | | | | | |
| RAB5C, member RAS oncogene family | Cytoplasm | enzyme | 1.73E+07 | 1.69E+07 | 1.86E+07 | | | | | | |
| RAB5C, member RAS oncogene family | Cytoplasm | enzyme | 1.03E+07 | 4598488 | 6699977 | | | | | | |
| RAB5C, member RAS oncogene family | Cytoplasm | enzyme | 3831715.25 | 2234779.75 | 3685073.75 | | | | | | |
| RAB5C, member RAS oncogene family | Cytoplasm | enzyme | 2754412.5 | 688376.562 | 1347408.5 | P35278 | RAB5C | 48418933 | 38452437 | 6157318 | 39 |
| RAB6A, member RAS oncogene family | Cytoplasm | enzyme | 2727863 | 6243721 | 5673080.5 | | | | | | |
| RAB6A, member RAS oncogene family | Cytoplasm | enzyme | 1376999.25 | 2398189 | 2438682 | P35279 | RAB6A | 2052431 | 1984473 | 810158 | 6 |
| RAB7A, member RAS oncogene family | Cytoplasm | enzyme | 1.88E+07 | 1.75E+07 | 1.98E+07 | | | | | | |
| RAB7A, member RAS oncogene family | Cytoplasm | enzyme | 1.80E+07 | 1.67E+07 | 1.66E+07 | | | | | | |
| RAB7A, member RAS oncogene family | Cytoplasm | enzyme | 1.60E+07 | 1.02E+07 | 1.43E+07 | | | | | | |
| RAB7A, member RAS oncogene family | Cytoplasm | enzyme | 1.49E+07 | 1.25E+07 | 1.49E+07 | | | | | | |
| RAB7A, member RAS oncogene family | Cytoplasm | enzyme | 1.31E+07 | 1.55E+07 | 1.34E+07 | | | | | | |
| RAB7A, member RAS oncogene family | Cytoplasm | enzyme | 1.24E+07 | 1.01E+07 | 1.04E+07 | | | | | | |
| RAB7A, member RAS oncogene family | Cytoplasm | enzyme | 1.19E+07 | 1.32E+07 | 1.27E+07 | | | | | | |
| RAB7A, member RAS oncogene family | Cytoplasm | enzyme | 1.14E+07 | 1.00E+07 | 9673402 | | | | | | |
| RAB7A, member RAS oncogene family | Cytoplasm | enzyme | 9650327 | 9358494 | 9775065 | | | | | | |
| RAB7A, member RAS oncogene family | Cytoplasm | enzyme | 7769770 | 1.14E+07 | 9421563 | | | | | | |
| RAB7A, member RAS oncogene family | Cytoplasm | enzyme | 4814183 | 2932746.75 | 4200802 | P51150 | RAB7A | 12617662 | 4057185 | 706265 | 33 |
| RAB8A, member RAS oncogene family | Plasma Membrane | enzyme | 9.49E+07 | 8.14E+07 | 8.52E+07 | | | | | | |
| RAB8A, member RAS oncogene family | Plasma Membrane | enzyme | 9.37E+07 | 7.76E+07 | 7.20E+07 | | | | | | |
| RAB8A, member RAS oncogene family | Plasma Membrane | enzyme | 8.45E+07 | 6.87E+07 | 6.40E+07 | | | | | | |
| RAB8A, member RAS oncogene family | Plasma Membrane | enzyme | 6.23E+07 | 4.94E+07 | 4.94E+07 | | | | | | |
| RAB8A, member RAS oncogene family | Plasma Membrane | enzyme | 5.12E+07 | 3.78E+07 | 4.55E+07 | | | | | | |
| RAB8A, member RAS oncogene family | Plasma Membrane | enzyme | 1.24E+07 | 7164139.5 | 6894116.5 | | | | | | |
| RAB8A, member RAS oncogene family | Plasma Membrane | enzyme | 6091848 | 2723891.75 | 4479823.5 | | | | | | |
| RAB8A, member RAS oncogene family | Plasma Membrane | enzyme | 340823.812 | 602832.688 | 385023.219 | P55258 | RAB8A | 50676584 | 34463840 | 7034902 | 24 |
| RAB8B, member RAS oncogene family | Cytoplasm | enzyme | 1.32E+08 | 1.11E+08 | 1.02E+08 | | | | | | |
| RAB8B, member RAS oncogene family | Cytoplasm | enzyme | 1.27E+08 | 1.02E+08 | 1.15E+08 | | | | | | |
| RAB8B, member RAS oncogene family | Cytoplasm | enzyme | 6.11E+07 | 5.65E+07 | 5.38E+07 | | | | | | |
| RAB8B, member RAS oncogene family | Cytoplasm | enzyme | 2.77E+07 | 2.94E+07 | 2.54E+07 | | | | | | |
| RAB8B, member RAS oncogene family | Cytoplasm | enzyme | 2814176.75 | 1340557.125 | 3743156.5 | | | | | | |
| RAB8B, member RAS oncogene family | Cytoplasm | enzyme | 706899.812 | 313857.062 | 313100.25 | P61028 | RAB8B | 58626846 | 49543467 | 11677507 | 18 |
| RAB9A, member RAS oncogene family | Cytoplasm | enzyme | 1686128 | 1390088.5 | 2520999.5 | Q9R0M6 | RAB9A | 1686128 | 586460 | 338593 | 3 |
| ras-related C3 botulinum toxin substrate 1 (rho family, small GTP binding protein Rac1) | Plasma Membrane | enzyme | 1.01E+08 | 1.31E+08 | 9.14E+07 | | | | | | |
| ras-related C3 botulinum toxin substrate 1 (rho family, small GTP binding protein Rac1) | Plasma Membrane | enzyme | 9.67E+07 | 1.62E+08 | 8.56E+07 | | | | | | |
| ras-related C3 botulinum toxin substrate 1 (rho family, small GTP binding protein Rac1) | Plasma Membrane | enzyme | 8.32E+07 | 8.80E+07 | 7.44E+07 | | | | | | |
| ras-related C3 botulinum toxin substrate 1 (rho family, small GTP binding protein Rac1) | Plasma Membrane | enzyme | 7.48E+07 | 1.32E+08 | 7.57E+07 | | | | | | |
| ras-related C3 botulinum toxin substrate 1 (rho family, small GTP binding protein Rac1) | Plasma Membrane | enzyme | 6.92E+07 | 9.24E+07 | 6.28E+07 | | | | | | |
| ras-related C3 botulinum toxin substrate 1 (rho family, small GTP binding protein Rac1) | Plasma Membrane | enzyme | 4.95E+07 | 7.81E+07 | 6.63E+07 | | | | | | |
| ras-related C3 botulinum toxin substrate 1 (rho family, small GTP binding protein Rac1) | Plasma Membrane | enzyme | 3.39E+07 | 2.45E+07 | 2.47E+07 | | | | | | |
| ras-related C3 botulinum toxin substrate 1 (rho family, small GTP binding protein Rac1) | Plasma Membrane | enzyme | 2.10E+07 | 3.31E+07 | 2.28E+07 | | | | | | |

| | | | | | | | | | | | |
|---|---|---|---|---|---|---|---|---|---|---|---|
| ras-related C3 botulinum toxin substrate 1 (rho family, small GTP binding protein Rac1) | Plasma Membrane | enzyme | 2.09E+07 | 1.09E+07 | 1.36E+07 | | | | | | |
| ras-related C3 botulinum toxin substrate 1 (rho family, small GTP binding protein Rac1) | Plasma Membrane | enzyme | 1.91E+07 | 1.83E+07 | 1.99E+07 | | | | | | |
| ras-related C3 botulinum toxin substrate 1 (rho family, small GTP binding protein Rac1) | Plasma Membrane | enzyme | 1.03E+07 | 1.56E+07 | 8310663 | | | | | | |
| ras-related C3 botulinum toxin substrate 1 (rho family, small GTP binding protein Rac1) | Plasma Membrane | enzyme | 6091179.5 | 2226751.25 | 3870630 | | | | | | |
| ras-related C3 botulinum toxin substrate 1 (rho family, small GTP binding protein Rac1) | Plasma Membrane | enzyme | 5043485.5 | 4980544.5 | 3496769.5 | | | | | | |
| ras-related C3 botulinum toxin substrate 1 (rho family, small GTP binding protein Rac1) | Plasma Membrane | enzyme | 2671707.5 | 909109.5 | 1326994.125 | P63001 | RAC1 | 42361884 | 42712667 | 6590708 | 42 |
| ras-related C3 botulinum toxin substrate 2 (rho family, small GTP binding protein Rac2) | Cytoplasm | enzyme | 2698724.5 | 5041594.5 | 4104982.75 | Q05144 | RAC2 | 2698725 | 1179254 | 680843 | 3 |
| RAD23 homolog A, nucleotide excision repair protein | Nucleus | other | 1.07E+08 | 3.14E+07 | 5.48E+07 | | | | | | |
| RAD23 homolog A, nucleotide excision repair protein | Nucleus | other | 7.45E+07 | 1.66E+07 | 3.79E+07 | | | | | | |
| RAD23 homolog A, nucleotide excision repair protein | Nucleus | other | 4.76E+07 | 1.28E+07 | 2.23E+07 | | | | | | |
| RAD23 homolog A, nucleotide excision repair protein | Nucleus | other | 1.92E+07 | 6204857 | 7632488.5 | | | | | | |
| RAD23 homolog A, nucleotide excision repair protein | Nucleus | other | 1.28E+07 | 3274545.5 | 7561606 | | | | | | |
| RAD23 homolog A, nucleotide excision repair protein | Nucleus | other | 5249271 | 2458097.25 | 2764306.25 | | | | | | |
| RAD23 homolog A, nucleotide excision repair protein | Nucleus | other | 1841406.875 | 539333.625 | 368062.594 | P54726 | RAD23A | 38271525 | 27904962 | 6089362 | 21 |
| v-ral simian leukemia viral oncogene homolog A (ras related) | Cytoplasm | enzyme | 2280042.75 | 1909859.875 | 2214753.5 | P63321 | RALA | 2280043 | 197593 | 114080 | 3 |
| RAN, member RAS oncogene family | Nucleus | enzyme | 4434035.5 | 3323285.75 | 3081716.25 | P62827 | RAN | 4434036 | 721213 | 416392 | 3 |
| Ran GTPase activating protein 1 | Nucleus | other | 811621.375 | 371071.5 | 383971.281 | P46061 | RANGAP1 | 811621 | 250711 | 144748 | 3 |
| RAP1A, member of RAS oncogene family | Cytoplasm | enzyme | 9.16E+07 | 9.63E+07 | 9.62E+07 | | | | | | |
| RAP1A, member of RAS oncogene family | Cytoplasm | enzyme | 7.25E+07 | 6.22E+07 | 7.36E+07 | | | | | | |
| RAP1A, member of RAS oncogene family | Cytoplasm | enzyme | 4.66E+07 | 8.39E+07 | 3.44E+07 | | | | | | |
| RAP1A, member of RAS oncogene family | Cytoplasm | enzyme | 4.05E+07 | 7.12E+07 | 4.82E+07 | | | | | | |
| RAP1A, member of RAS oncogene family | Cytoplasm | enzyme | 1.25E+07 | 1.66E+07 | 1.17E+07 | | | | | | |
| RAP1A, member of RAS oncogene family | Cytoplasm | enzyme | 5321168.5 | 3831373.5 | 3416601.25 | | | | | | |
| RAP1A, member of RAS oncogene family | Cytoplasm | enzyme | 3122471.5 | 2239837 | 1967123.25 | | | | | | |
| RAP1A, member of RAS oncogene family | Cytoplasm | enzyme | 1666755 | 7290246 | 1775060.875 | P62835 | RAP1A | 34218799 | 35314071 | 7208455 | 24 |
| RAP1B, member of RAS oncogene family | Cytoplasm | enzyme | 1.39E+08 | 1.81E+08 | 1.73E+08 | | | | | | |
| RAP1B, member of RAS oncogene family | Cytoplasm | enzyme | 1.11E+08 | 1.92E+08 | 1.42E+08 | | | | | | |
| RAP1B, member of RAS oncogene family | Cytoplasm | enzyme | 1.00E+08 | 1.76E+08 | 1.53E+08 | | | | | | |
| RAP1B, member of RAS oncogene family | Cytoplasm | enzyme | 1.46E+07 | 7.74E+07 | 7.04E+07 | | | | | | |
| RAP1B, member of RAS oncogene family | Cytoplasm | enzyme | 1.38E+07 | 1.41E+07 | 1.17E+07 | | | | | | |
| RAP1B, member of RAS oncogene family | Cytoplasm | enzyme | 1.09E+07 | 8.06E+07 | 4.30E+07 | | | | | | |
| RAP1B, member of RAS oncogene family | Cytoplasm | enzyme | 6950290.5 | 2.41E+07 | 2.27E+07 | | | | | | |
| RAP1B, member of RAS oncogene family | Cytoplasm | enzyme | 446052.25 | 48142.973 | 470570.781 | Q99JI6 | RAP1B | 49624543 | 68626794 | 14008386 | 24 |
| RAP2B, member of RAS oncogene family | Plasma Membrane | enzyme | 1.21E+07 | 1.15E+07 | 1.06E+07 | | | | | | |
| RAP2B, member of RAS oncogene family | Plasma Membrane | enzyme | 1.10E+07 | 1.03E+07 | 8159837.5 | | | | | | |
| RAP2B, member of RAS oncogene family | Plasma Membrane | enzyme | 1.03E+07 | 1.07E+07 | 9093731 | | | | | | |
| RAP2B, member of RAS oncogene family | Plasma Membrane | enzyme | 9687814 | 7459750.5 | 7758274 | | | | | | |
| RAP2B, member of RAS oncogene family | Plasma Membrane | enzyme | 8290374.5 | 7266763.5 | 5532356 | | | | | | |
| RAP2B, member of RAS oncogene family | Plasma Membrane | enzyme | 2301355.25 | 2739130 | 2369582.75 | P61226 | RAP2B | 8934924 | 3116878 | 734655 | 18 |
| RAP2C, member of RAS oncogene family | Cytoplasm | enzyme | 2227376.5 | 1439105.5 | 1595793.625 | Q8BU31 | RAP2C | 2227377 | 417297 | 240926 | 3 |
| arginyl-tRNA synthetase | Cytoplasm | enzyme | 204597.094 | 83628.617 | 951703.188 | Q9D0I9 | RARS | 204597 | 470169 | 271452 | 3 |
| RAS p21 protein activator 3 | Plasma Membrane | ion channel | 374639.969 | 4886702 | 2872965.25 | Q60790 | RASA3 | 374640 | 2260364 | 1305022 | 3 |
| RAS-like, family 2, locus 9 | Nucleus | transporter | 9372338 | 5291539 | 6272331.5 | | | | | | |
| RAS-like, family 2, locus 9 | Nucleus | transporter | 8491082 | 7557488 | 4031465.25 | Q61820 | Rasl2-9 | 8931710 | 2013025 | 821814 | 6 |
| RNA binding motif (RNP1, RRM) protein 3 | Cytoplasm | other | 7880419 | 3094098.75 | 5356690 | | | | | | |
| RNA binding motif (RNP1, RRM) protein 3 | Cytoplasm | other | 1981448.75 | 714934.25 | 1541324.625 | | | | | | |
| RNA binding motif (RNP1, RRM) protein 3 | Cytoplasm | other | 191773.719 | 74757.539 | 21038.955 | O89086 | RBM3 | 3351214 | 2713203 | 904401 | 9 |
| retinol dehydrogenase 11 (all-trans/9-cis/11-cis) | Cytoplasm | enzyme | 9024213 | 7387683.5 | 7095184.5 | | | | | | |
| retinol dehydrogenase 11 (all-trans/9-cis/11-cis) | Cytoplasm | enzyme | 6114174 | 4710841 | 4837774 | | | | | | |
| retinol dehydrogenase 11 (all-trans/9-cis/11-cis) | Cytoplasm | enzyme | 5954153.5 | 3788195 | 3797986.25 | | | | | | |
| retinol dehydrogenase 11 (all-trans/9-cis/11-cis) | Cytoplasm | enzyme | 5919129.5 | 5205840.5 | 5788750 | | | | | | |
| retinol dehydrogenase 11 (all-trans/9-cis/11-cis) | Cytoplasm | enzyme | 4609866 | 6038755.5 | 5626897 | | | | | | |
| retinol dehydrogenase 11 (all-trans/9-cis/11-cis) | Cytoplasm | enzyme | 3741797.25 | 3940818.5 | 2094647.875 | | | | | | |
| retinol dehydrogenase 11 (all-trans/9-cis/11-cis) | Cytoplasm | enzyme | 3499156.75 | 3416498.25 | 3208996.75 | | | | | | |
| retinol dehydrogenase 11 (all-trans/9-cis/11-cis) | Cytoplasm | enzyme | 3499056.75 | 3812510.25 | 2875035.5 | Q9QYF1 | RDH11 | 5295193 | 1631419 | 333012 | 24 |
| radixin | Cytoplasm | other | 4.62E+07 | 3.21E+07 | 2.93E+07 | | | | | | |
| radixin | Cytoplasm | other | 2.25E+07 | 1.70E+07 | 1.78E+07 | | | | | | |
| radixin | Cytoplasm | other | 2.18E+07 | 1.52E+07 | 1.81E+07 | | | | | | |
| radixin | Cytoplasm | other | 1.44E+07 | 9463084 | 1.10E+07 | | | | | | |
| radixin | Cytoplasm | other | 1.41E+07 | 9460913 | 7819975 | | | | | | |
| radixin | Cytoplasm | other | 1.19E+07 | 5441879 | 7737078 | | | | | | |
| radixin | Cytoplasm | other | 1.16E+07 | 7883348 | 9631895 | | | | | | |
| radixin | Cytoplasm | other | 7638010 | 6120375.5 | 6359883 | | | | | | |
| radixin | Cytoplasm | other | 7166204 | 2906373.75 | 3323968 | | | | | | |
| radixin | Cytoplasm | other | 5893782.5 | 5576923.5 | 4436687.5 | | | | | | |
| radixin | Cytoplasm | other | 5776310.5 | 3574200.25 | 4705236 | | | | | | |
| radixin | Cytoplasm | other | 5028367 | 2006595.25 | 2905075.5 | | | | | | |
| radixin | Cytoplasm | other | 2341612 | 1166537.75 | 1584644.125 | | | | | | |
| radixin | Cytoplasm | other | 1215552.25 | 1582679.125 | 1304346.125 | P26043 | RDX | 12676417 | 9382393 | 1447735 | 42 |
| receptor accessory protein 5 | Extracellular Space | transporter | 3.45E+07 | 4.05E+07 | 2.76E+07 | | | | | | |
| receptor accessory protein 5 | Extracellular Space | transporter | 4037557 | 6141456 | 3384934.75 | | | | | | |
| receptor accessory protein 5 | Extracellular Space | transporter | 574070.25 | 301374.969 | 305688.969 | Q60870 | REEP5 | 13027209 | 16307959 | 5435986 | 9 |
| receptor accessory protein 6 | Plasma Membrane | other | 1.49E+07 | 1.12E+07 | 1.25E+07 | | | | | | |
| receptor accessory protein 6 | Plasma Membrane | other | 6696225.5 | 5937185 | 4667085.5 | | | | | | |
| receptor accessory protein 6 | Plasma Membrane | other | 1883334.875 | 2402333 | 971971.938 | Q9JM62 | REEP6 | 7813187 | 4991467 | 1663822 | 9 |
| RELT-like 1 | Plasma Membrane | other | 6881840 | 2190386 | 2025991.75 | | | | | | |
| RELT-like 1 | Plasma Membrane | other | 4240365 | 3297053.25 | 3464352.5 | | | | | | |
| RELT-like 1 | Plasma Membrane | other | 2722212.25 | 2417603.5 | 1844226.125 | Q8K2J7 | RELL1 | 4614806 | 1572610 | 524203 | 9 |
| Rh-associated glycoprotein | Plasma Membrane | peptidase | 1.20E+09 | 1.09E+09 | 1.14E+09 | | | | | | |
| Rh-associated glycoprotein | Plasma Membrane | peptidase | 7.37E+08 | 3.57E+08 | 4.79E+08 | | | | | | |
| Rh-associated glycoprotein | Plasma Membrane | peptidase | 4.86E+07 | 3.24E+07 | 3.60E+07 | | | | | | |
| Rh-associated glycoprotein | Plasma Membrane | peptidase | 1.03E+07 | 1.56E+07 | 1.62E+07 | Q9QUT0 | RHAG | 499995000 | 487818192 | 140820982 | 12 |
| rhomboid 5 homolog 2 (Drosophila) | Cytoplasm | other | 3755628 | 2885406.5 | 3879717.75 | | | | | | |
| rhomboid 5 homolog 2 (Drosophila) | Cytoplasm | other | 1051167.25 | 651275.062 | 609962.5 | Q80WQ6 | RHBDF2 | 2403398 | 1545016 | 630750 | 6 |
| Rh blood group, D antigen | Plasma Membrane | transporter | 1.73E+09 | 1.14E+09 | 1.37E+09 | | | | | | |
| Rh blood group, D antigen | Plasma Membrane | transporter | 2.16E+08 | 2.00E+08 | 1.63E+08 | | | | | | |
| Rh blood group, D antigen | Plasma Membrane | transporter | 1.19E+08 | 9.04E+07 | 8.82E+07 | | | | | | |
| Rh blood group, D antigen | Plasma Membrane | transporter | 5.45E+07 | 2.30E+07 | 3.40E+07 | | | | | | |
| Rh blood group, D antigen | Plasma Membrane | transporter | 3.47E+07 | 2.01E+07 | 2.32E+07 | | | | | | |
| Rh blood group, D antigen | Plasma Membrane | transporter | 2.24E+07 | 2.49E+07 | 2.27E+07 | | | | | | |
| Rh blood group, D antigen | Plasma Membrane | transporter | 1.97E+07 | 1.33E+08 | 1.43E+08 | | | | | | |
| Rh blood group, D antigen | Plasma Membrane | transporter | 2110711.75 | 1309279.375 | 2944268.75 | Q8CF94 | RHCE/RHD | 274790089 | 467332002 | 95393745 | 24 |
| ras homolog family member A | Cytoplasm | enzyme | 3.16E+07 | 2.92E+07 | 2.94E+07 | | | | | | |
| ras homolog family member A | Cytoplasm | enzyme | 1.88E+07 | 1.45E+07 | 1.44E+07 | | | | | | |
| ras homolog family member A | Cytoplasm | enzyme | 1.74E+07 | 2.63E+07 | 1.79E+07 | | | | | | |

| | | | | | | | | | | | |
|---|---|---|---|---|---|---|---|---|---|---|---|
| ras homolog family member A | Cytoplasm | enzyme | 1.35E+07 | 2.13E+07 | 1.21E+07 | | | | | | |
| ras homolog family member A | Cytoplasm | enzyme | 7941167 | 1.28E+07 | 1.06E+07 | | | | | | |
| ras homolog family member A | Cytoplasm | enzyme | 7263031 | 6048766.5 | 7238732 | | | | | | |
| ras homolog family member A | Cytoplasm | enzyme | 3881673.25 | 2366987.5 | 2827348.25 | | | | | | |
| ras homolog family member A | Cytoplasm | enzyme | 3329385.25 | 5962255 | 3760692.75 | | | | | | |
| ras homolog family member A | Cytoplasm | enzyme | 3136984.25 | 5666699.5 | 9760785 | Q9QUI0 | RHOA | 11872471 | 8843242 | 1701883 | 27 |
| ras homolog family member G | Cytoplasm | enzyme | 5.51E+07 | 4.12E+07 | 5.66E+07 | | | | | | |
| ras homolog family member G | Cytoplasm | enzyme | 3.95E+07 | 2.62E+07 | 2.86E+07 | | | | | | |
| ras homolog family member G | Cytoplasm | enzyme | 3.27E+07 | 2.16E+07 | 3.15E+07 | | | | | | |
| ras homolog family member G | Cytoplasm | enzyme | 2.29E+07 | 2.15E+07 | 2.00E+07 | | | | | | |
| ras homolog family member G | Cytoplasm | enzyme | 2.15E+07 | 1.32E+07 | 2.11E+07 | | | | | | |
| ras homolog family member G | Cytoplasm | enzyme | 1.52E+07 | 8221840.5 | 1.18E+07 | | | | | | |
| ras homolog family member G | Cytoplasm | enzyme | 1.21E+07 | 3497903 | 1.08E+07 | | | | | | |
| ras homolog family member G | Cytoplasm | enzyme | 1.15E+07 | 7235414 | 1.81E+07 | | | | | | |
| ras homolog family member G | Cytoplasm | enzyme | 1.12E+07 | 1.13E+07 | 1.12E+07 | | | | | | |
| ras homolog family member G | Cytoplasm | enzyme | 4581488 | 1445767.625 | 4994640 | | | | | | |
| ras homolog family member G | Cytoplasm | enzyme | 1777031.25 | 564259.562 | 1622161 | P84096 | RHOG | 20732593 | 14509692 | 2525813 | 33 |
| ring finger protein 11 | Cytoplasm | enzyme | 2419634.5 | 1870791.375 | 2416277.25 | Q9QYK7 | RNF11 | 2419635 | 315910 | 182391 | 3 |
| ring finger protein 123 | Cytoplasm | other | 1.45E+07 | 8511001 | 7530303.5 | | | | | | |
| ring finger protein 123 | Cytoplasm | other | 1.29E+07 | 1.01E+07 | 6641255 | | | | | | |
| ring finger protein 123 | Cytoplasm | other | 9886988 | 6133740.5 | 4394331.5 | | | | | | |
| ring finger protein 123 | Cytoplasm | other | 8989893 | 6705630.5 | 6389632 | | | | | | |
| ring finger protein 123 | Cytoplasm | other | 8430178 | 2604909 | 3762297.75 | | | | | | |
| ring finger protein 123 | Cytoplasm | other | 4019359.25 | 2431225.25 | 3022984.75 | | | | | | |
| ring finger protein 123 | Cytoplasm | other | 3601817.5 | 2412847.75 | 2402661 | Q5XPI3 | RNF123 | 8892605 | 3513926 | 766801 | 21 |
| ring finger protein 213 | Cytoplasm | enzyme | 451388.219 | 448432.344 | 410359.031 | E9Q555 | RNF213 | 451388 | 22883 | 13211 | 3 |
| retinitis pigmentosa 2 (X-linked recessive) | Cytoplasm | enzyme | 1.61E+07 | 1.18E+07 | 1.39E+07 | | | | | | |
| retinitis pigmentosa 2 (X-linked recessive) | Cytoplasm | enzyme | 9340735 | 9627995 | 7545845 | | | | | | |
| retinitis pigmentosa 2 (X-linked recessive) | Cytoplasm | enzyme | 8482982 | 8067814.5 | 8536401 | | | | | | |
| retinitis pigmentosa 2 (X-linked recessive) | Cytoplasm | enzyme | 5667764.5 | 5311357 | 4519788.5 | | | | | | |
| retinitis pigmentosa 2 (X-linked recessive) | Cytoplasm | enzyme | 5511251 | 5141801.5 | 5594747 | | | | | | |
| retinitis pigmentosa 2 (X-linked recessive) | Cytoplasm | enzyme | 5280041.5 | 4467834.5 | 5388976.5 | | | | | | |
| retinitis pigmentosa 2 (X-linked recessive) | Cytoplasm | enzyme | 4096161.75 | 5825879 | 2722261.5 | Q9EPK2 | RP2 | 7784134 | 3381337 | 737868 | 21 |
| ribosomal protein L10a | Nucleus | other | 1504374.625 | 2211685.5 | 1747081.125 | | | | | | |
| ribosomal protein L10a | Nucleus | other | 1191237.375 | 784736.375 | 894236.875 | P53026 | RPL10A | 1347806 | 541734 | 221162 | 6 |
| ribosomal protein L10-like | Nucleus | other | 7783871 | 4904106 | 6095894.5 | | | | | | |
| ribosomal protein L10-like | Nucleus | other | 2978705.75 | 2519235.5 | 2623125.25 | P86048 | RPL10L | 5381288 | 2156521 | 880396 | 6 |
| ribosomal protein L11 | Cytoplasm | other | 5053569 | 4235849 | 5430099 | Q9CXW4 | RPL11 | 5053569 | 610556 | 352505 | 3 |
| ribosomal protein L12 | Nucleus | other | 4628550 | 3268376 | 3082400.25 | | | | | | |
| ribosomal protein L12 | Nucleus | other | 4349429.5 | 2198568 | 1991300.875 | | | | | | |
| ribosomal protein L12 | Nucleus | other | 4008711 | 2403696.25 | 3722927 | | | | | | |
| ribosomal protein L12 | Nucleus | other | 3495031 | 2743884.5 | 3277741.5 | | | | | | |
| ribosomal protein L12 | Nucleus | other | 1974580.875 | 1110287.375 | 998563.188 | P35979 | RPL12 | 3691260 | 1099358 | 283853 | 15 |
| ribosomal protein L13 | Nucleus | other | 4329649.5 | 3149442.75 | 3289813 | | | | | | |
| ribosomal protein L13 | Nucleus | other | 1967085.625 | 1750673.5 | 1585483.25 | | | | | | |
| ribosomal protein L13 | Nucleus | other | 1570458.375 | 1515531.75 | 1605324.75 | | | | | | |
| ribosomal protein L13 | Nucleus | other | 1447676.375 | 1789956.25 | 1532232.75 | P47963 | RPL13 | 2328717 | 933917 | 269599 | 12 |
| ribosomal protein L13a | Cytoplasm | other | 2052785.25 | 1691266.875 | 2111869 | P19253 | RPL13A | 2052785 | 227703 | 131465 | 3 |
| ribosomal protein L14 | Cytoplasm | other | 6096668.5 | 4941587 | 5258998 | | | | | | |
| ribosomal protein L14 | Cytoplasm | other | 2725094.5 | 1938036.5 | 2145949.75 | | | | | | |
| ribosomal protein L14 | Cytoplasm | other | 2384943.25 | 2382288 | 2637431.5 | Q9CR57 | RPL14 | 3735569 | 1577838 | 525946 | 9 |
| ribosomal protein L15 | Cytoplasm | other | 3768939.75 | 2717916.75 | 2750624.25 | | | | | | |
| ribosomal protein L15 | Cytoplasm | other | 2687317.25 | 1136243.5 | 2677019.75 | | | | | | |
| ribosomal protein L15 | Cytoplasm | other | 1669693.375 | 1708252.125 | 1696902.75 | Q9CZM2 | RPL15 | 2708650 | 813307 | 271102 | 9 |
| ribosomal protein L17 | Cytoplasm | other | 2490455.5 | 1392214.75 | 2106943 | Q9CPR4 | RPL17 | 2490456 | 557382 | 321805 | 3 |
| ribosomal protein L18 | Cytoplasm | other | 6394480.5 | 4599100 | 6530023 | | | | | | |
| ribosomal protein L18 | Cytoplasm | other | 4206936.5 | 2765605 | 4145647.75 | | | | | | |
| ribosomal protein L18 | Cytoplasm | other | 2974449 | 2829401.75 | 2896650.75 | | | | | | |
| ribosomal protein L18 | Cytoplasm | other | 1955193.25 | 1451747.125 | 1446836.75 | P35980 | RPL18 | 3882765 | 1708455 | 493189 | 12 |
| ribosomal protein L18a | Cytoplasm | other | 2266390 | 2231967.5 | 2130804.25 | P62717 | RPL18A | 2266390 | 70477 | 40690 | 3 |
| ribosomal protein L19 | Cytoplasm | other | 6838673 | 4027540.75 | 5347683.5 | P84099 | RPL19 | 6838673 | 1406431 | 812003 | 3 |
| ribosomal protein L21 | Cytoplasm | other | 4953449 | 4660567 | 5544090.5 | | | | | | |
| ribosomal protein L21 | Cytoplasm | other | 2157780.25 | 2253139.5 | 2468770.75 | O09167 | RPL21 | 3555615 | 1541290 | 629229 | 6 |
| ribosomal protein L22 | Nucleus | other | 1.07E+07 | 7779089.5 | 8201878.5 | | | | | | |
| ribosomal protein L22 | Nucleus | other | 4002699 | 2519956 | 3519214.25 | P67984 | RPL22 | 7326350 | 3219152 | 1314213 | 6 |
| ribosomal protein L23 | Cytoplasm | other | 5221544.5 | 2650713.5 | 3604225 | P62830 | RPL23 | 5221545 | 1299620 | 750336 | 3 |
| ribosomal protein L23A | Nucleus | other | 6631658.5 | 4445106.5 | 5132480.5 | | | | | | |
| ribosomal protein L23A | Nucleus | other | 1111257.75 | 1389985.625 | 1121016 | P62751 | Rpl23a | 3871458 | 2406481 | 982442 | 6 |
| ribosomal protein L24 | Cytoplasm | other | 1891279.125 | 1870091.875 | 1748757 | | | | | | |
| ribosomal protein L24 | Cytoplasm | other | 1861678.75 | 2111950.5 | 1718263.25 | | | | | | |
| ribosomal protein L24 | Cytoplasm | other | 1726355.75 | 1662316.125 | 2216306.5 | Q8BP67 | RPL24 | 1826438 | 187224 | 62408 | 9 |
| ribosomal protein L26 | Cytoplasm | other | 1324738.875 | 889983.812 | 926044.625 | P61255 | RPL26 | 1324739 | 241271 | 139298 | 3 |
| ribosomal protein L27a | Cytoplasm | other | 2497628.5 | 1770078.875 | 1938072.625 | P14115 | RPL27A | 2497629 | 380932 | 219931 | 3 |
| ribosomal protein L28 | Cytoplasm | other | 3325123 | 2184636.25 | 2158204.75 | P41105 | RPL28 | 3325123 | 666222 | 384643 | 3 |
| ribosomal protein L29 | Cytoplasm | other | 373370.156 | 418369.438 | 916547.25 | P47915 | Rpl29 (includes others) | 373370 | 301454 | 174045 | 3 |
| ribosomal protein L3 | Nucleus | other | 8558810 | 5251285.5 | 7372668.5 | | | | | | |
| ribosomal protein L3 | Nucleus | other | 3090547.5 | 5073295 | 5953364 | | | | | | |
| ribosomal protein L3 | Nucleus | other | 2056620 | 2012628.125 | 1838033 | P27659 | RPL3 | 4568659 | 2472491 | 824164 | 9 |
| ribosomal protein L30 | Cytoplasm | other | 4895117 | 4428548.5 | 3908440 | P62889 | RPL30 | 4895117 | 493581 | 284969 | 3 |
| ribosomal protein L31 | Cytoplasm | other | 5337217.5 | 3526393.25 | 3187238.5 | | | | | | |
| ribosomal protein L31 | Cytoplasm | other | 4402473 | 4031438.5 | 4736260 | P62900 | RPL31 | 4869845 | 791152 | 322986 | 6 |
| ribosomal protein L35 | Cytoplasm | other | 4874908.5 | 4736908.5 | 3953429 | | | | | | |
| ribosomal protein L35 | Cytoplasm | other | 1747486.875 | 1272048 | 1649956.125 | Q6ZWV7 | RPL35 | 3311198 | 1661879 | 678459 | 6 |
| ribosomal protein L36 | Nucleus | other | 1805941.375 | 1760513 | 1422836 | | | | | | |
| ribosomal protein L36 | Nucleus | other | 1233600.25 | 1158458.5 | 1067566.75 | P47964 | Rpl36 | 1519771 | 313504 | 127987 | 6 |
| ribosomal protein L4 | Cytoplasm | enzyme | 4108749.5 | 3009495.75 | 3759365 | | | | | | |
| ribosomal protein L4 | Cytoplasm | enzyme | 2889567.5 | 2553644.5 | 2885582.5 | | | | | | |
| ribosomal protein L4 | Cytoplasm | enzyme | 2146040 | 2626146.25 | 2542011 | | | | | | |
| ribosomal protein L4 | Cytoplasm | enzyme | 1249145.375 | 375161.812 | 941872.562 | Q9D8E6 | RPL4 | 2598376 | 1100321 | 317635 | 12 |
| ribosomal protein L5 | Cytoplasm | other | 504075.969 | 1197973.125 | 1329017.25 | P47962 | RPL5 | 504076 | 443320 | 255951 | 3 |
| ribosomal protein L6 | Nucleus | other | 3027717.25 | 2378726.25 | 2786809.75 | | | | | | |
| ribosomal protein L6 | Nucleus | other | 2714354.5 | 2409482.75 | 1849443.625 | | | | | | |
| ribosomal protein L6 | Nucleus | other | 1688661.25 | 573615.5 | 1398971.5 | | | | | | |
| ribosomal protein L6 | Nucleus | other | 854123.188 | 417701.281 | 633223 | P47911 | RPL6 | 2071214 | 944233 | 272577 | 12 |
| ribosomal protein L7 | Nucleus | transcription regulator | 9681199 | 6286726.5 | 6474262.5 | | | | | | |

| | | | | | | | | | | | |
|---|---|---|---|---|---|---|---|---|---|---|---|
| ribosomal protein L7 | Nucleus | transcription regulator | 6029296 | 4129380.75 | 6078136.5 | | | | | | |
| ribosomal protein L7 | Nucleus | transcription regulator | 3833601 | 3205133.25 | 3852075.5 | | | | | | |
| ribosomal protein L7 | Nucleus | transcription regulator | 3580989 | 2655584.75 | 3167186 | | | | | | |
| ribosomal protein L7 | Nucleus | transcription regulator | 499337.781 | 480902.562 | 853508.188 | P14148 | RPL7 | 4724885 | 2534535 | 654414 | 15 |
| ribosomal protein L7a | Cytoplasm | other | 4698297.5 | 3671238.25 | 3736832.75 | | | | | | |
| ribosomal protein L7a | Cytoplasm | other | 3662280 | 3325754.75 | 3880356.25 | | | | | | |
| ribosomal protein L7a | Cytoplasm | other | 1776224.125 | 2301173.75 | 1813598.125 | P12970 | RPL7A | 3378934 | 1012683 | 337561 | 9 |
| ribosomal protein L8 | Cytoplasm | other | 1258992.875 | 883759.812 | 1333925.625 | | | | | | |
| ribosomal protein L8 | Cytoplasm | other | 484713.062 | 425225.312 | 440957.188 | P62918 | RPL8 | 871853 | 417472 | 170432 | 6 |
| ribosomal protein L9 | Nucleus | other | 5504545 | 3239212.25 | 4472036.5 | | | | | | |
| ribosomal protein L9 | Nucleus | other | 2659814.75 | 2323074.5 | 2542638.5 | | | | | | |
| ribosomal protein L9 | Nucleus | other | 2201406.25 | 1889303.75 | 2881766.75 | P51410 | RPL9 | 3455255 | 1178387 | 392796 | 9 |
| ribosomal protein, large, P0 | Cytoplasm | other | 9781133 | 7602796 | 9798406 | | | | | | |
| ribosomal protein, large, P0 | Cytoplasm | other | 4673878 | 2974980 | 4343792 | | | | | | |
| ribosomal protein, large, P0 | Cytoplasm | other | 3905579.75 | 3763273 | 3527607.25 | | | | | | |
| ribosomal protein, large, P0 | Cytoplasm | other | 3257569.25 | 2933479 | 4055842.5 | | | | | | |
| ribosomal protein, large, P0 | Cytoplasm | other | 3012962.75 | 2788714 | 2677703.75 | | | | | | |
| ribosomal protein, large, P0 | Cytoplasm | other | 2106581.75 | 3064545 | 2073702.5 | | | | | | |
| ribosomal protein, large, P0 | Cytoplasm | other | 1833401.625 | 3140671.25 | 3165499.25 | | | | | | |
| ribosomal protein, large, P0 | Cytoplasm | other | 968450.375 | 874031.375 | 1357032.875 | P14869 | RPLP0 | 3692445 | 2332938 | 476209 | 24 |
| ribosomal protein, large, P1 | Nucleus | other | 2.32E+07 | 1.37E+07 | 2.25E+07 | | | | | | |
| ribosomal protein, large, P1 | Nucleus | other | 1.97E+07 | 1.41E+07 | 1.92E+07 | P47955 | Rplp1 (includes others) | 21410000 | 4022078 | 1642006 | 6 |
| ribosomal protein, large, P2 | Cytoplasm | other | 2.29E+07 | 1.39E+07 | 1.62E+07 | | | | | | |
| ribosomal protein, large, P2 | Cytoplasm | other | 2.28E+07 | 1.63E+07 | 2.14E+07 | | | | | | |
| ribosomal protein, large, P2 | Cytoplasm | other | 6150129 | 3725242.75 | 3889275.75 | | | | | | |
| ribosomal protein, large, P2 | Cytoplasm | other | 3001790.25 | 2164612.75 | 2180422 | | | | | | |
| ribosomal protein, large, P2 | Cytoplasm | other | 2861913.5 | 1670527.125 | 2061421.5 | P99027 | RPLP2 | 11540767 | 8426131 | 2175618 | 15 |
| ribophorin I | Cytoplasm | enzyme | 2540542.75 | 5132164.5 | 4160499.5 | | | | | | |
| ribophorin I | Cytoplasm | enzyme | 2392452.75 | 8000925 | 8678252 | | | | | | |
| ribophorin I | Cytoplasm | enzyme | 2389927 | 6707006 | 6618728.5 | | | | | | |
| ribophorin I | Cytoplasm | enzyme | 1908482.875 | 3994751.25 | 3414752.5 | | | | | | |
| ribophorin I | Cytoplasm | enzyme | 1486475.75 | 4601649 | 3429311 | | | | | | |
| ribophorin I | Cytoplasm | enzyme | 1315596.5 | 5345885 | 3971766.25 | | | | | | |
| ribophorin I | Cytoplasm | enzyme | 368684.031 | 2917279.5 | 1945089 | | | | | | |
| ribophorin I | Cytoplasm | enzyme | 234845.844 | 1161832.125 | 1158835.75 | Q91YQ5 | RPN1 | 1579626 | 2323735 | 474330 | 24 |
| ribophorin II | Cytoplasm | enzyme | 4027784.75 | 1.06E+07 | 7765383 | | | | | | |
| ribophorin II | Cytoplasm | enzyme | 3222937.5 | 9018016 | 9081650 | | | | | | |
| ribophorin II | Cytoplasm | enzyme | 1519849.875 | 4901759.5 | 4623755.5 | | | | | | |
| ribophorin II | Cytoplasm | enzyme | 1400848.25 | 4072583.75 | 4107422.5 | | | | | | |
| ribophorin II | Cytoplasm | enzyme | 1301792.375 | 3368237.25 | 3722081.5 | | | | | | |
| ribophorin II | Cytoplasm | enzyme | 1238110.25 | 2725215.5 | 2492988.25 | | | | | | |
| ribophorin II | Cytoplasm | enzyme | 1018680.875 | 3478888.5 | 2892595.5 | | | | | | |
| ribophorin II | Cytoplasm | enzyme | 928619.75 | 2236110.75 | 1909068.25 | | | | | | |
| ribophorin II | Cytoplasm | enzyme | 636167.75 | 2600919 | 2079429.75 | | | | | | |
| ribophorin II | Cytoplasm | enzyme | 8017.086 | 1460686.125 | 1625716.375 | Q9DBG6 | RPN2 | 1530281 | 2635024 | 481087 | 30 |
| ribosomal protein S10 | Cytoplasm | other | 3677242 | 2857758 | 3294171.5 | P63325 | RPS10 | 3677242 | 410031 | 236732 | 3 |
| ribosomal protein S11 | Cytoplasm | other | 2225664.75 | 2486292.5 | 2352748.5 | | | | | | |
| ribosomal protein S11 | Cytoplasm | other | 495268.312 | 129016.344 | 210012.719 | P62281 | RPS11 | 1360467 | 1146966 | 468247 | 6 |
| ribosomal protein S13 | Cytoplasm | other | 1683823.5 | 637288.625 | 979086.625 | | | | | | |
| ribosomal protein S13 | Cytoplasm | other | 957682.188 | 807453 | 995283.062 | P62301 | RPS13 | 1320753 | 357180 | 145818 | 6 |
| ribosomal protein S14 | Cytoplasm | translation regulator | 6586160 | 4572211 | 4376302 | | | | | | |
| ribosomal protein S14 | Cytoplasm | translation regulator | 4620425.5 | 2948824.25 | 3104042 | | | | | | |
| ribosomal protein S14 | Cytoplasm | translation regulator | 2927900 | 1193524.375 | 2307644.5 | P62264 | RPS14 | 4711495 | 1584418 | 528139 | 9 |
| ribosomal protein S15a | Cytoplasm | other | 2271613.25 | 1050682.125 | 1378786.5 | | | | | | |
| ribosomal protein S15a | Cytoplasm | other | 2106915.75 | 1334470.25 | 2233145.5 | P62245 | RPS15A | 2189265 | 534758 | 218314 | 6 |
| ribosomal protein S16 | Cytoplasm | other | 4616897 | 3359011 | 4746060 | | | | | | |
| ribosomal protein S16 | Cytoplasm | other | 1813196.75 | 1939114.25 | 2484486.5 | P14131 | RPS16 | 3215047 | 1299115 | 530362 | 6 |
| ribosomal protein S17 | Cytoplasm | other | 1532752.375 | 1310520.25 | 1551356.375 | P63276 | RPS17 | 1532752 | 134000 | 77365 | 3 |
| ribosomal protein S18 | Cytoplasm | other | 2144836.25 | 2147197.25 | 2373333.75 | | | | | | |
| ribosomal protein S18 | Cytoplasm | other | 2011250.25 | 1478958.5 | 1489474.75 | | | | | | |
| ribosomal protein S18 | Cytoplasm | other | 1346723 | 1457846.625 | 1357167.875 | P62270 | RPS18 | 1834270 | 405238 | 135079 | 9 |
| ribosomal protein S19 | Cytoplasm | other | 4162565 | 2623701.75 | 3281163.25 | | | | | | |
| ribosomal protein S19 | Cytoplasm | other | 3321324.25 | 2391354.75 | 2697213.75 | | | | | | |
| ribosomal protein S19 | Cytoplasm | other | 3273793.75 | 2466842 | 2609272.75 | Q9CZX8 | RPS19 | 3585894 | 575126 | 191709 | 9 |
| ribosomal protein S2 | Cytoplasm | other | 7408788.5 | 3319175.5 | 4787146.5 | | | | | | |
| ribosomal protein S2 | Cytoplasm | other | 3846496.75 | 2336495 | 3492146.75 | | | | | | |
| ribosomal protein S2 | Cytoplasm | other | 3231318.25 | 2375882.5 | 3121240.5 | | | | | | |
| ribosomal protein S2 | Cytoplasm | other | 2468945.25 | 2004435.375 | 3118240 | P25444 | RPS2 | 4238887 | 1456857 | 420559 | 12 |
| ribosomal protein S20 | Cytoplasm | other | 4574746.5 | 3284977.25 | 4139237.25 | | | | | | |
| ribosomal protein S20 | Cytoplasm | other | 2483575.75 | 3009067.75 | 2278358 | P60867 | RPS20 | 3529161 | 908220 | 370779 | 6 |
| ribosomal protein S23 | Cytoplasm | translation regulator | 2956677.75 | 2206375.5 | 3007555.75 | P62267 | RPS23 | 2956678 | 448596 | 258997 | 3 |
| ribosomal protein S24 | Cytoplasm | other | 1257550.5 | 1019934.688 | 1185957.125 | P62849 | RPS24 | 1257551 | 121895 | 70376 | 3 |
| ribosomal protein S25 | Cytoplasm | other | 4723878.5 | 3507892 | 3779716.5 | | | | | | |
| ribosomal protein S25 | Cytoplasm | other | 3665584.75 | 2755243.5 | 3202279.75 | | | | | | |
| ribosomal protein S25 | Cytoplasm | other | 3232740.5 | 1986130.875 | 2706274.5 | P62852 | RPS25 | 3874068 | 776357 | 258786 | 9 |
| ribosomal protein S26 | Cytoplasm | other | 3022712.25 | 1742232.375 | 2175494 | P62855 | RPS26 | 3022712 | 651297 | 376026 | 3 |
| ribosomal protein S27-like | Cytoplasm | translation regulator | 1190044.75 | 777335.812 | 760675.5 | | | | | | |
| ribosomal protein S27-like | Cytoplasm | translation regulator | 497122.844 | 189878.609 | 93298.406 | Q6ZWY3 | RPS27L | 843584 | 409768 | 167287 | 6 |
| ribosomal protein S28 | Cytoplasm | other | 1131259.375 | 340309.406 | 313607.188 | P62858 | RPS28 | 1131259 | 464555 | 268211 | 3 |
| ribosomal protein S3 | Cytoplasm | enzyme | 6223344.5 | 3786604.25 | 3510154.75 | | | | | | |
| ribosomal protein S3 | Cytoplasm | enzyme | 4376300 | 2597402.5 | 2714055.5 | | | | | | |
| ribosomal protein S3 | Cytoplasm | enzyme | 3475406.5 | 2286298.25 | 2531281 | | | | | | |
| ribosomal protein S3 | Cytoplasm | enzyme | 3325245.25 | 2076587.375 | 2817420 | | | | | | |
| ribosomal protein S3 | Cytoplasm | enzyme | 2116357.5 | 1391199.75 | 2215735.25 | | | | | | |
| ribosomal protein S3 | Cytoplasm | enzyme | 1333372.375 | 1900496.5 | 1690682 | | | | | | |
| ribosomal protein S3 | Cytoplasm | enzyme | 1326321.5 | 869044.938 | 687489.188 | | | | | | |
| ribosomal protein S3 | Cytoplasm | enzyme | 1015936.438 | 751907.562 | 504081.281 | P62908 | RPS3 | 2899036 | 1345464 | 274642 | 24 |
| ribosomal protein S3A1 | Cytoplasm | other | 1414804.625 | 1021711.438 | 903056.5 | | | | | | |
| ribosomal protein S3A1 | Cytoplasm | other | 74937.336 | 249220.953 | 241842.625 | P97351 | Rps3a1 | 744871 | 537595 | 219472 | 6 |
| ribosomal protein S4, Y-linked 1 | Cytoplasm | other | 3321758.75 | 3197174.25 | 3633022 | | | | | | |
| ribosomal protein S4, Y-linked 1 | Cytoplasm | other | 2824821.5 | 1917068.625 | 2031586.75 | | | | | | |
| ribosomal protein S4, Y-linked 1 | Cytoplasm | other | 2075388.75 | 1438737.625 | 1153506.25 | | | | | | |
| ribosomal protein S4, Y-linked 1 | Cytoplasm | other | 2042302.25 | 1057678 | 1379773.75 | | | | | | |
| ribosomal protein S4, Y-linked 1 | Cytoplasm | other | 1817565.375 | 753151.062 | 876953.688 | | | | | | |
| ribosomal protein S4, Y-linked 1 | Cytoplasm | other | 1757253.125 | 1844929.75 | 1555767.375 | | | | | | |

| | | | | | | | | | | | |
|---|---|---|---|---|---|---|---|---|---|---|---|
| ribosomal protein S4, Y-linked 1 | Cytoplasm | other | 1020284.062 | 907393.062 | 1256649.75 | | | | | | |
| ribosomal protein S4, Y-linked 1 | Cytoplasm | other | 970473.125 | 816319.438 | 972594.75 | P62702 | RPS4Y1 | 1978731 | 832261 | 169885 | 24 |
| ribosomal protein S6 | Cytoplasm | other | 2900118 | 3154162 | 3172364.75 | | | | | | |
| ribosomal protein S6 | Cytoplasm | other | 497713.812 | 699899.562 | 801094.688 | P62754 | RPS6 | 1698916 | 1326742 | 541640 | 6 |
| ribosomal protein S7 | Cytoplasm | other | 8688807 | 4085739 | 3638340.5 | | | | | | |
| ribosomal protein S7 | Cytoplasm | other | 3731924.5 | 3369677.75 | 3360040.25 | | | | | | |
| ribosomal protein S7 | Cytoplasm | other | 1217111.125 | 1100213.375 | 1621679.375 | P62082 | RPS7 | 4545948 | 2286444 | 762148 | 9 |
| ribosomal protein S8 | Cytoplasm | other | 5395195 | 4294694 | 4896009.5 | | | | | | |
| ribosomal protein S8 | Cytoplasm | other | 4844049.5 | 3146548.75 | 4158062.25 | | | | | | |
| ribosomal protein S8 | Cytoplasm | other | 3679375 | 2573074.75 | 2828346 | | | | | | |
| ribosomal protein S8 | Cytoplasm | other | 1993324.5 | 1679720.375 | 1513851.875 | | | | | | |
| ribosomal protein S8 | Cytoplasm | other | 1168399.375 | 947514.812 | 1126307.75 | P62242 | RPS8 | 3416069 | 1522445 | 393093 | 15 |
| ribosomal protein S9 | Cytoplasm | translation regulator | 4392447 | 3199677.75 | 3602907.5 | | | | | | |
| ribosomal protein S9 | Cytoplasm | translation regulator | 2572944.25 | 1927506.625 | 1952574.75 | | | | | | |
| ribosomal protein S9 | Cytoplasm | translation regulator | 641135.625 | 391436.281 | 859791.438 | Q6ZWN5 | RPS9 | 2535509 | 1393670 | 464557 | 9 |
| ribosomal protein SA | Cytoplasm | translation regulator | 1.10E+07 | 8166999 | 9136963 | | | | | | |
| ribosomal protein SA | Cytoplasm | translation regulator | 5858934.5 | 4592223.5 | 3777361.5 | | | | | | |
| ribosomal protein SA | Cytoplasm | translation regulator | 4251892 | 2957008 | 3448912.5 | | | | | | |
| ribosomal protein SA | Cytoplasm | translation regulator | 2299340.5 | 1216160.125 | 1749342.625 | P14206 | RPSA | 5862542 | 3091826 | 892533 | 12 |
| radical S-adenosyl methionine domain containing 2 | Cytoplasm | enzyme | 7855074 | 8244883 | 1.07E+07 | | | | | | |
| radical S-adenosyl methionine domain containing 2 | Cytoplasm | enzyme | 6765735 | 6038530.5 | 6509373.5 | | | | | | |
| radical S-adenosyl methionine domain containing 2 | Cytoplasm | enzyme | 6060543 | 4975034.5 | 4292286.5 | | | | | | |
| radical S-adenosyl methionine domain containing 2 | Cytoplasm | enzyme | 5055339 | 7569596.5 | 9490193 | | | | | | |
| radical S-adenosyl methionine domain containing 2 | Cytoplasm | enzyme | 3479594 | 3326632.75 | 3428348.75 | | | | | | |
| radical S-adenosyl methionine domain containing 2 | Cytoplasm | enzyme | 2114151.25 | 1678144.375 | 2236948 | | | | | | |
| radical S-adenosyl methionine domain containing 2 | Cytoplasm | enzyme | 2042690.125 | 1479314 | 3073404.75 | | | | | | |
| radical S-adenosyl methionine domain containing 2 | Cytoplasm | enzyme | 2026026.625 | 1468848 | 1753117.5 | Q8CBB9 | RSAD2 | 4424894 | 2751994 | 561749 | 24 |
| Ras suppressor protein 1 | Cytoplasm | other | 835526.125 | 7603209.5 | 1.23E+07 | | | | | | |
| Ras suppressor protein 1 | Cytoplasm | other | 560545.188 | 4440443 | 1.13E+07 | | | | | | |
| Ras suppressor protein 1 | Cytoplasm | other | 4.70E-06 | 2458774.5 | 4267454 | Q01730 | RSU1 | 465357 | 4594424 | 1531475 | 9 |
| reticulon 1 | Cytoplasm | other | 5093824.5 | 8053845 | 9476559 | Q8K0T0 | RTN1 | 5093825 | 2235852 | 1290870 | 3 |
| reticulon 3 | Cytoplasm | other | 6.13E+07 | 8.22E+07 | 6.84E+07 | | | | | | |
| reticulon 3 | Cytoplasm | other | 853444.5 | 245609.484 | 521294.781 | Q9ES97 | RTN3 | 31056722 | 38963594 | 15906821 | 6 |
| reticulon 4 | Cytoplasm | other | 4973468 | 9286726 | 8372285.5 | | | | | | |
| reticulon 4 | Cytoplasm | other | 4580257.5 | 1.40E+07 | 1.03E+07 | | | | | | |
| reticulon 4 | Cytoplasm | other | 2486797.25 | 5925029 | 5066758.5 | | | | | | |
| reticulon 4 | Cytoplasm | other | 1622195.125 | 1543362.75 | 1732280.875 | | | | | | |
| reticulon 4 | Cytoplasm | other | 1345171.375 | 5204246 | 3021340.25 | Q99P72 | RTN4 | 3001578 | 3750368 | 968341 | 15 |
| S100 calcium binding protein A13 | Cytoplasm | other | 2690843.75 | 2440840.5 | 3285264 | P97352 | S100A13 | 2690844 | 433760 | 250432 | 3 |
| S100 calcium binding protein A9 | Cytoplasm | other | 334020.906 | 2711087.25 | 2885823.5 | | | | | | |
| S100 calcium binding protein A9 | Cytoplasm | other | 264457.5 | 1790793 | 1916995.75 | P31725 | S100A9 | 299239 | 1131000 | 461729 | 6 |
| SAC1 suppressor of actin mutations 1-like (yeast) | Cytoplasm | phosphatase | 1.40E+07 | 1.08E+07 | 9915830 | | | | | | |
| SAC1 suppressor of actin mutations 1-like (yeast) | Cytoplasm | phosphatase | 1.11E+07 | 9737984 | 9543877 | | | | | | |
| SAC1 suppressor of actin mutations 1-like (yeast) | Cytoplasm | phosphatase | 1.02E+07 | 1.01E+07 | 1.07E+07 | | | | | | |
| SAC1 suppressor of actin mutations 1-like (yeast) | Cytoplasm | phosphatase | 8503830 | 9537829 | 9911447 | | | | | | |
| SAC1 suppressor of actin mutations 1-like (yeast) | Cytoplasm | phosphatase | 7933863 | 6325717 | 5522490.5 | | | | | | |
| SAC1 suppressor of actin mutations 1-like (yeast) | Cytoplasm | phosphatase | 6766897.5 | 5564190 | 5664650.5 | | | | | | |
| SAC1 suppressor of actin mutations 1-like (yeast) | Cytoplasm | phosphatase | 6529621 | 2896818 | 4104599 | | | | | | |
| SAC1 suppressor of actin mutations 1-like (yeast) | Cytoplasm | phosphatase | 6205615 | 6405087.5 | 5775974 | | | | | | |
| SAC1 suppressor of actin mutations 1-like (yeast) | Cytoplasm | phosphatase | 5596316.5 | 4793045.5 | 4944127 | | | | | | |
| SAC1 suppressor of actin mutations 1-like (yeast) | Cytoplasm | phosphatase | 5338554.5 | 5328546 | 4998538 | | | | | | |
| SAC1 suppressor of actin mutations 1-like (yeast) | Cytoplasm | phosphatase | 5317704 | 4411587 | 4148842 | | | | | | |
| SAC1 suppressor of actin mutations 1-like (yeast) | Cytoplasm | phosphatase | 4176444.75 | 3455973 | 3367898 | | | | | | |
| SAC1 suppressor of actin mutations 1-like (yeast) | Cytoplasm | phosphatase | 3594879.5 | 2950147.5 | 2570152.5 | | | | | | |
| SAC1 suppressor of actin mutations 1-like (yeast) | Cytoplasm | phosphatase | 3044045.75 | 2291871.5 | 2701722.75 | | | | | | |
| SAC1 suppressor of actin mutations 1-like (yeast) | Cytoplasm | phosphatase | 2817713.25 | 2650208.25 | 2136184.5 | | | | | | |
| SAC1 suppressor of actin mutations 1-like (yeast) | Cytoplasm | phosphatase | 2648360.5 | 2545387 | 1871480.5 | | | | | | |
| SAC1 suppressor of actin mutations 1-like (yeast) | Cytoplasm | phosphatase | 2545700.5 | 2870268.75 | 2623776.5 | | | | | | |
| SAC1 suppressor of actin mutations 1-like (yeast) | Cytoplasm | phosphatase | 2252514.5 | 1259431.125 | 1575797.125 | | | | | | |
| SAC1 suppressor of actin mutations 1-like (yeast) | Cytoplasm | phosphatase | 496036.031 | 455862.719 | 1707262.5 | Q9EP69 | SACM1L | 5737268 | 3168423 | 419668 | 57 |
| SAMM50 sorting and assembly machinery component | Cytoplasm | other | 1318309.375 | 1292035.75 | 1335643 | | | | | | |
| SAMM50 sorting and assembly machinery component | Cytoplasm | other | 907609.312 | 2910374.75 | 1856901 | Q8BGH2 | SAMM50 | 1112959 | 708024 | 289050 | 6 |
| secretory carrier membrane protein 1 | Cytoplasm | transporter | 2.51E+07 | 1.78E+07 | 1.52E+07 | Q8K021 | SCAMP1 | 25060000 | 5095550 | 2941917 | 3 |
| secretory carrier membrane protein 2 | Cytoplasm | transporter | 2310915.25 | 5818387.5 | 4151448.5 | | | | | | |
| secretory carrier membrane protein 2 | Cytoplasm | transporter | 1923008.25 | 5315142 | 4820948 | Q9ERN0 | SCAMP2 | 2116962 | 1605034 | 655253 | 6 |
| secretory carrier membrane protein 4 | Cytoplasm | other | 1.07E+07 | 8938398 | 9108230 | | | | | | |
| secretory carrier membrane protein 4 | Cytoplasm | other | 9396277 | 4264584 | 5770647.5 | Q9JKV5 | SCAMP4 | 10038139 | 2456010 | 1002662 | 6 |
| scavenger receptor class B, member 2 | Plasma Membrane | other | 671360.812 | 1510171.625 | 1091739.875 | O35114 | SCARB2 | 671361 | 419406 | 242144 | 3 |
| succinate dehydrogenase complex, subunit A, flavoprotein (Fp) | Cytoplasm | enzyme | 8429917 | 7911210.5 | 1.01E+07 | | | | | | |
| succinate dehydrogenase complex, subunit A, flavoprotein (Fp) | Cytoplasm | enzyme | 4112828 | 4729945.5 | 4872753 | | | | | | |
| succinate dehydrogenase complex, subunit A, flavoprotein (Fp) | Cytoplasm | enzyme | 2404010.5 | 6278583 | 4969988 | | | | | | |
| succinate dehydrogenase complex, subunit A, flavoprotein (Fp) | Cytoplasm | enzyme | 1879509.5 | 3990136.75 | 3260405 | | | | | | |
| succinate dehydrogenase complex, subunit A, flavoprotein (Fp) | Cytoplasm | enzyme | 1856170.375 | 3940053 | 2965282.5 | | | | | | |
| succinate dehydrogenase complex, subunit A, flavoprotein (Fp) | Cytoplasm | enzyme | 1629261.5 | 3552805.75 | 2491720 | | | | | | |
| succinate dehydrogenase complex, subunit A, flavoprotein (Fp) | Cytoplasm | enzyme | 1592518.25 | 2589086.5 | 3129024.25 | | | | | | |
| succinate dehydrogenase complex, subunit A, flavoprotein (Fp) | Cytoplasm | enzyme | 565367.938 | 1528495.5 | 1048145.625 | Q8K2B3 | SDHA | 2808698 | 2413703 | 492695 | 24 |
| succinate dehydrogenase complex, subunit B, iron sulfur (Ip) | Cytoplasm | enzyme | 6449459 | 4797674.5 | 4358725 | | | | | | |
| succinate dehydrogenase complex, subunit B, iron sulfur (Ip) | Cytoplasm | enzyme | 1685644 | 3007637.5 | 2382146 | | | | | | |
| succinate dehydrogenase complex, subunit B, iron sulfur (Ip) | Cytoplasm | enzyme | 934268.625 | 2165674.25 | 1967525.375 | Q9CQA3 | SDHB | 3023124 | 1769637 | 589879 | 9 |
| succinate dehydrogenase complex, subunit C, integral membrane protein, 15kDa | Cytoplasm | enzyme | 1032864.75 | 1603757.875 | 1701245.25 | Q9CZB0 | SDHC | 1032865 | 361053 | 208454 | 3 |
| SEC22 vesicle trafficking protein homolog B (S. cerevisiae) (gene/pseudogene) | Cytoplasm | other | 4840632.5 | 1.06E+07 | 7109689.5 | | | | | | |
| SEC22 vesicle trafficking protein homolog B (S. cerevisiae) (gene/pseudogene) | Cytoplasm | other | 4593113.5 | 8532688 | 6617194 | | | | | | |
| SEC22 vesicle trafficking protein homolog B (S. cerevisiae) (gene/pseudogene) | Cytoplasm | other | 4213450 | 4113114.5 | 4303459.5 | | | | | | |
| SEC22 vesicle trafficking protein homolog B (S. cerevisiae) (gene/pseudogene) | Cytoplasm | other | 3690635.5 | 4491759 | 3402208.5 | | | | | | |
| SEC22 vesicle trafficking protein homolog B (S. cerevisiae) (gene/pseudogene) | Cytoplasm | other | 3023710.75 | 2278862.75 | 2051578.875 | | | | | | |
| SEC22 vesicle trafficking protein homolog B (S. cerevisiae) (gene/pseudogene) | Cytoplasm | other | 1873715 | 1767193.25 | 1752619.25 | | | | | | |
| SEC22 vesicle trafficking protein homolog B (S. cerevisiae) (gene/pseudogene) | Cytoplasm | other | 1865090.25 | 2833661 | 2422054.75 | | | | | | |
| SEC22 vesicle trafficking protein homolog B (S. cerevisiae) (gene/pseudogene) | Cytoplasm | other | 1169714 | 1284662.5 | 1250621.625 | O08547 | SEC22B | 3158758 | 2415016 | 492963 | 24 |
| SEC31 homolog B, COPII coating complex component | Cytoplasm | other | 2170252 | 3.02E+07 | 1.72E+07 | Q3TZ89 | SEC31B | 2170252 | 14008120 | 8087592 | 3 |
| Sec61 alpha 1 subunit (S. cerevisiae) | Cytoplasm | transporter | 1336641.25 | 2253100.75 | 2427252.75 | P61620 | SEC61A1 | 1336641 | 585898 | 338268 | 3 |
| Sec61 beta subunit | Cytoplasm | transporter | 2610136.25 | 5299796.5 | 4403483.5 | | | | | | |
| Sec61 beta subunit | Cytoplasm | transporter | 1216587.625 | 2125141 | 2276430 | Q9CQS8 | SEC61B | 1913362 | 1541398 | 629273 | 6 |
| selenium binding protein 1 | Cytoplasm | other | 799590.25 | 3169003.75 | 909399.25 | P17563 | SELENBP1 | 799590 | 1337410 | 772154 | 3 |
| selectin P (granule membrane protein 140kDa, antigen CD62) | Plasma Membrane | transmembrane receptor | 924869.938 | 4190201 | 2551689.75 | | | | | | |
| selectin P (granule membrane protein 140kDa, antigen CD62) | Plasma Membrane | transmembrane receptor | 445077.062 | 1.73E+07 | 1.09E+07 | | | | | | |

| | | | | | | | | | | | |
|---|---|---|---|---|---|---|---|---|---|---|---|
| selectin P (granule membrane protein 140kDa, antigen CD62) | Plasma Membrane | transmembrane receptor | 290464.156 | 1.06E+07 | 8097630 | | | | | | |
| selectin P (granule membrane protein 140kDa, antigen CD62) | Plasma Membrane | transmembrane receptor | 181035.578 | 1851835.5 | 1047256.75 | | | | | | |
| selectin P (granule membrane protein 140kDa, antigen CD62) | Plasma Membrane | transmembrane receptor | 133712.844 | 2901387.75 | 2941485.75 | | | | | | |
| selectin P (granule membrane protein 140kDa, antigen CD62) | Plasma Membrane | transmembrane receptor | 4.70E-06 | 1009688.188 | 1132056.375 | Q01102 | SELP | 329193 | 4843013 | 1141509 | 18 |
| selenoprotein T | Cytoplasm | other | 2577790.25 | 3808008 | 2457886.25 | P62342 | SELT | 2577790 | 747289 | 431447 | 3 |
| SERPINE1 mRNA binding protein 1 | Cytoplasm | other | 1117366.125 | 503605.781 | 479980.688 | | | | | | |
| SERPINE1 mRNA binding protein 1 | Cytoplasm | other | 675587.562 | 101691.32 | 588871 | | | | | | |
| SERPINE1 mRNA binding protein 1 | Cytoplasm | other | 633420.375 | 94677.414 | 362578.875 | Q9CY58 | SERBP1 | 808791 | 312157 | 104052 | 9 |
| serine incorporator 3 | Cytoplasm | transporter | 3.38E+08 | 3.08E+08 | 3.31E+08 | | | | | | |
| serine incorporator 3 | Cytoplasm | transporter | 1.63E+08 | 1.11E+08 | 1.53E+08 | | | | | | |
| serine incorporator 3 | Cytoplasm | transporter | 1.38E+08 | 1.16E+08 | 1.11E+08 | | | | | | |
| serine incorporator 3 | Cytoplasm | transporter | 3.94E+07 | 2.24E+07 | 3.48E+07 | Q9QZI9 | SERINC3 | 169635000 | 112484989 | 32471619 | 12 |
| serpin peptidase inhibitor, clade A (alpha-1 antiproteinase, antitrypsin), member 1 | Extracellular Space | other | 1.18E+07 | 1.20E+07 | 1.76E+07 | | | | | | |
| serpin peptidase inhibitor, clade A (alpha-1 antiproteinase, antitrypsin), member 1 | Extracellular Space | other | 3885670.75 | 3420469.5 | 5499326 | | | | | | |
| serpin peptidase inhibitor, clade A (alpha-1 antiproteinase, antitrypsin), member 1 | Extracellular Space | other | 1849933.375 | 2725789.75 | 3902896.75 | | | | | | |
| serpin peptidase inhibitor, clade A (alpha-1 antiproteinase, antitrypsin), member 1 | Extracellular Space | other | 1788388.75 | 2255784.5 | 2745287.75 | | | | | | |
| serpin peptidase inhibitor, clade A (alpha-1 antiproteinase, antitrypsin), member 1 | Extracellular Space | other | 1536066.25 | 2626626.75 | 2501110.75 | P07758 | SERPINA1 | 4176012 | 4798081 | 1238859 | 15 |
| serpin peptidase inhibitor, clade A (alpha-1 antiproteinase, antitrypsin), member 3 | Extracellular Space | other | 3.67E+07 | 1.68E+07 | 1.65E+07 | | | | | | |
| serpin peptidase inhibitor, clade A (alpha-1 antiproteinase, antitrypsin), member 3 | Extracellular Space | other | 3.35E+07 | 1.03E+07 | 8893996 | | | | | | |
| serpin peptidase inhibitor, clade A (alpha-1 antiproteinase, antitrypsin), member 3 | Extracellular Space | other | 2.28E+07 | 5303570.5 | 3943760.75 | P07759 | SERPINA3 | 31013333 | 11789834 | 3929945 | 9 |
| sideroflexin 1 | Cytoplasm | transporter | 2362287.5 | 3818222.25 | 3607956.25 | | | | | | |
| sideroflexin 1 | Cytoplasm | transporter | 2177686 | 5418732 | 4264934 | | | | | | |
| sideroflexin 1 | Cytoplasm | transporter | 1702342.75 | 5265043.5 | 2993133.75 | | | | | | |
| sideroflexin 1 | Cytoplasm | transporter | 1617688.875 | 3743342 | 2711996 | | | | | | |
| sideroflexin 1 | Cytoplasm | transporter | 988693.375 | 2940055.25 | 2239088.5 | | | | | | |
| sideroflexin 1 | Cytoplasm | transporter | 358713.031 | 1909490.375 | 520885.562 | Q99JR1 | SFXN1 | 1534569 | 1460794 | 344312 | 18 |
| SH3 domain binding glutamate-rich protein like | Cytoplasm | other | 1.50E+07 | 9882598 | 9546172 | | | | | | |
| SH3 domain binding glutamate-rich protein like | Cytoplasm | other | 8509621 | 6580083 | 6437538 | | | | | | |
| SH3 domain binding glutamate-rich protein like | Cytoplasm | other | 7465450 | 6076608.5 | 8026499 | Q9JJU8 | SH3BGRL | 10308357 | 2729704 | 909901 | 9 |
| SH3 domain binding glutamate-rich protein like 2 | Nucleus | other | 1004238.312 | 2926818.5 | 3329671.5 | Q8BG73 | SH3BGRL2 | 1004238 | 1242728 | 717490 | 3 |
| sialic acid binding Ig-like lectin 9 | Plasma Membrane | other | 4962292 | 4877868.5 | 4808083.5 | Q91Y57 | SIGLEC9 | 4962292 | 77220 | 44583 | 3 |
| src kinase associated phosphoprotein 2 | Cytoplasm | other | 95088.523 | 1630158.875 | 1409242.375 | | | | | | |
| src kinase associated phosphoprotein 2 | Cytoplasm | other | 4.70E-06 | 1734815.375 | 1727130.375 | Q3UND0 | SKAP2 | 47544 | 823751 | 336295 | 6 |
| signaling lymphocytic activation molecule family member 1 | Plasma Membrane | transmembrane receptor | 323918.062 | 5012791.5 | 2922227 | | | | | | |
| signaling lymphocytic activation molecule family member 1 | Plasma Membrane | transmembrane receptor | 257002.922 | 4478447 | 2803544.75 | | | | | | |
| signaling lymphocytic activation molecule family member 1 | Plasma Membrane | transmembrane receptor | 242449.75 | 8082699 | 5019680 | Q9QUM4 | SLAMF1 | 274457 | 2691981 | 897327 | 9 |
| solute carrier family 11 (proton-coupled divalent metal ion transporter), member 2 | Plasma Membrane | transporter | 9149633 | 7580611 | 9609171 | | | | | | |
| solute carrier family 11 (proton-coupled divalent metal ion transporter), member 2 | Plasma Membrane | transporter | 4957790.5 | 2772389 | 3671420.25 | | | | | | |
| solute carrier family 11 (proton-coupled divalent metal ion transporter), member 2 | Plasma Membrane | transporter | 3655913.25 | 2369435.75 | 3210637.75 | | | | | | |
| solute carrier family 11 (proton-coupled divalent metal ion transporter), member 2 | Plasma Membrane | transporter | 1371235.625 | 872912.375 | 1689967 | P49282 | SLC11A2 | 4783643 | 2985632 | 861878 | 12 |
| solute carrier family 14 (urea transporter), member 1 (Kidd blood group) | Plasma Membrane | transporter | 3.26E+08 | 1.97E+08 | 2.58E+08 | | | | | | |
| solute carrier family 14 (urea transporter), member 1 (Kidd blood group) | Plasma Membrane | transporter | 1.97E+08 | 1.52E+08 | 1.24E+08 | | | | | | |
| solute carrier family 14 (urea transporter), member 1 (Kidd blood group) | Plasma Membrane | transporter | 1.76E+08 | 1.88E+08 | 1.39E+08 | | | | | | |
| solute carrier family 14 (urea transporter), member 1 (Kidd blood group) | Plasma Membrane | transporter | 1.52E+08 | 1.15E+08 | 1.37E+08 | | | | | | |
| solute carrier family 14 (urea transporter), member 1 (Kidd blood group) | Plasma Membrane | transporter | 1.49E+08 | 1.13E+08 | 1.37E+08 | | | | | | |
| solute carrier family 14 (urea transporter), member 1 (Kidd blood group) | Plasma Membrane | transporter | 1.29E+08 | 1.08E+08 | 1.08E+08 | | | | | | |
| solute carrier family 14 (urea transporter), member 1 (Kidd blood group) | Plasma Membrane | transporter | 1.24E+08 | 2.79E+08 | 2.00E+08 | | | | | | |
| solute carrier family 14 (urea transporter), member 1 (Kidd blood group) | Plasma Membrane | transporter | 3.65E+07 | 2.46E+07 | 2.59E+07 | | | | | | |
| solute carrier family 14 (urea transporter), member 1 (Kidd blood group) | Plasma Membrane | transporter | 2.52E+07 | 4.66E+07 | 3.04E+07 | | | | | | |
| solute carrier family 14 (urea transporter), member 1 (Kidd blood group) | Plasma Membrane | transporter | 2.22E+07 | 1.38E+07 | 1.31E+07 | | | | | | |
| solute carrier family 14 (urea transporter), member 1 (Kidd blood group) | Plasma Membrane | transporter | 1.97E+07 | 1.04E+07 | 1.99E+07 | | | | | | |
| solute carrier family 14 (urea transporter), member 1 (Kidd blood group) | Plasma Membrane | transporter | 4701699 | 1.02E+07 | 5758068 | Q8VHL0 | SLC14A1 | 113550142 | 86243951 | 14373992 | 36 |
| solute carrier family 16 (monocarboxylate transporter), member 1 | Plasma Membrane | transporter | 7.78E+08 | 5.66E+08 | 4.54E+08 | | | | | | |
| solute carrier family 16 (monocarboxylate transporter), member 1 | Plasma Membrane | transporter | 6.53E+08 | 4.88E+08 | 5.13E+08 | | | | | | |
| solute carrier family 16 (monocarboxylate transporter), member 1 | Plasma Membrane | transporter | 6.08E+08 | 3.78E+08 | 4.65E+08 | | | | | | |
| solute carrier family 16 (monocarboxylate transporter), member 1 | Plasma Membrane | transporter | 3.03E+08 | 2.75E+08 | 2.42E+08 | | | | | | |
| solute carrier family 16 (monocarboxylate transporter), member 1 | Plasma Membrane | transporter | 2.34E+08 | 1.28E+08 | 1.71E+08 | | | | | | |
| solute carrier family 16 (monocarboxylate transporter), member 1 | Plasma Membrane | transporter | 9.48E+07 | 6.22E+07 | 7.35E+07 | | | | | | |
| solute carrier family 16 (monocarboxylate transporter), member 1 | Plasma Membrane | transporter | 3.62E+07 | 2.74E+07 | 2.52E+07 | | | | | | |
| solute carrier family 16 (monocarboxylate transporter), member 1 | Plasma Membrane | transporter | 3.35E+07 | 2.72E+07 | 3.46E+07 | | | | | | |
| solute carrier family 16 (monocarboxylate transporter), member 1 | Plasma Membrane | transporter | 7172651 | 1.03E+07 | 9292629 | P53986 | SLC16A1 | 305165850 | 238287268 | 45858406 | 27 |
| solute carrier family 16 (aromatic amino acid transporter), member 10 | Plasma Membrane | transporter | 2.20E+08 | 1.41E+08 | 1.59E+08 | | | | | | |
| solute carrier family 16 (aromatic amino acid transporter), member 10 | Plasma Membrane | transporter | 1.76E+08 | 1.26E+08 | 1.18E+08 | | | | | | |
| solute carrier family 16 (aromatic amino acid transporter), member 10 | Plasma Membrane | transporter | 1.30E+08 | 6.70E+07 | 9.04E+07 | | | | | | |
| solute carrier family 16 (aromatic amino acid transporter), member 10 | Plasma Membrane | transporter | 7.04E+07 | 4.53E+07 | 5.19E+07 | | | | | | |
| solute carrier family 16 (aromatic amino acid transporter), member 10 | Plasma Membrane | transporter | 5.95E+07 | 5.00E+07 | 4.62E+07 | | | | | | |
| solute carrier family 16 (aromatic amino acid transporter), member 10 | Plasma Membrane | transporter | 4.86E+07 | 2.17E+07 | 3.25E+07 | | | | | | |
| solute carrier family 16 (aromatic amino acid transporter), member 10 | Plasma Membrane | transporter | 2.32E+07 | 1.17E+07 | 1.07E+07 | | | | | | |
| solute carrier family 16 (aromatic amino acid transporter), member 10 | Plasma Membrane | transporter | 1.34E+07 | 7877416 | 1.16E+07 | | | | | | |
| solute carrier family 16 (aromatic amino acid transporter), member 10 | Plasma Membrane | transporter | 1.20E+07 | 4872520.5 | 3809369.5 | | | | | | |
| solute carrier family 16 (aromatic amino acid transporter), member 10 | Plasma Membrane | transporter | 8832718 | 3.21E+07 | 1.38E+07 | Q3U9N9 | SLC16A10 | 76152272 | 58564651 | 10692393 | 30 |
| solute carrier family 18 (vesicular monoamine transporter), member 2 | Plasma Membrane | transporter | 28011.83 | 771169.625 | 431905.562 | Q8BRU6 | SLC18A2 | 28012 | 372047 | 214801 | 3 |
| solute carrier family 19 (folate transporter), member 1 | Plasma Membrane | transporter | 1927803.125 | 1436612 | 1257453.75 | P41438 | SLC19A1 | 1927803 | 347067 | 200379 | 3 |
| solute carrier family 22, member 23 | Plasma Membrane | transporter | 1986577.75 | 1604835.5 | 1422874.125 | Q3UHH2 | SLC22A23 | 1986578 | 287692 | 166099 | 3 |
| solute carrier family 25 (aspartate/glutamate carrier), member 12 | Cytoplasm | transporter | 4350388.5 | 5699455.5 | 5278644 | | | | | | |
| solute carrier family 25 (aspartate/glutamate carrier), member 12 | Cytoplasm | transporter | 1602166.125 | 2287548.75 | 2468042.75 | | | | | | |
| solute carrier family 25 (aspartate/glutamate carrier), member 12 | Cytoplasm | transporter | 1237736.25 | 2014226.625 | 1948921 | | | | | | |
| solute carrier family 25 (aspartate/glutamate carrier), member 12 | Cytoplasm | transporter | 824249.375 | 1181206.125 | 1097041.75 | Q8BH59 | SLC25A12 | 2003635 | 1675551 | 483690 | 12 |
| solute carrier family 25 (mitochondrial oxoadipate carrier), member 21 | Cytoplasm | transporter | 2502447 | 2057766.5 | 3075503.25 | | | | | | |
| solute carrier family 25 (mitochondrial oxoadipate carrier), member 21 | Cytoplasm | transporter | 1607433.875 | 1438651.75 | 1297570.625 | | | | | | |
| solute carrier family 25 (mitochondrial oxoadipate carrier), member 21 | Cytoplasm | transporter | 1380009.125 | 1169856 | 1346040 | | | | | | |
| solute carrier family 25 (mitochondrial oxoadipate carrier), member 21 | Cytoplasm | transporter | 949343.812 | 931296.438 | 970828.312 | | | | | | |
| solute carrier family 25 (mitochondrial oxoadipate carrier), member 21 | Cytoplasm | transporter | 676167.438 | 1118147.625 | 903698.438 | Q8BZ09 | SLC25A21 | 1423080 | 655268 | 169190 | 15 |
| solute carrier family 25 (mitochondrial carrier; phosphate carrier), member 3 | Cytoplasm | transporter | 1.39E+07 | 3.27E+07 | 2.34E+07 | | | | | | |
| solute carrier family 25 (mitochondrial carrier; phosphate carrier), member 3 | Cytoplasm | transporter | 1.08E+07 | 1.96E+07 | 1.56E+07 | | | | | | |
| solute carrier family 25 (mitochondrial carrier; phosphate carrier), member 3 | Cytoplasm | transporter | 7334383 | 2.80E+07 | 1.59E+07 | | | | | | |
| solute carrier family 25 (mitochondrial carrier; phosphate carrier), member 3 | Cytoplasm | transporter | 6074374 | 1.73E+07 | 1.21E+07 | | | | | | |
| solute carrier family 25 (mitochondrial carrier; phosphate carrier), member 3 | Cytoplasm | transporter | 4196712.5 | 1.38E+07 | 1.02E+07 | | | | | | |
| solute carrier family 25 (mitochondrial carrier; phosphate carrier), member 3 | Cytoplasm | transporter | 3308205.5 | 7274713.5 | 7002959 | | | | | | |
| solute carrier family 25 (mitochondrial carrier; phosphate carrier), member 3 | Cytoplasm | transporter | 3145571.25 | 1.09E+07 | 6852866.5 | | | | | | |
| solute carrier family 25 (mitochondrial carrier; phosphate carrier), member 3 | Cytoplasm | transporter | 2152799 | 3423567.75 | 2407864.5 | | | | | | |
| solute carrier family 25 (mitochondrial carrier; phosphate carrier), member 3 | Cytoplasm | transporter | 1418827.25 | 3799550.5 | 2430642.5 | Q8VEM8 | SLC25A3 | 5810097 | 8227171 | 1583320 | 27 |
| solute carrier family 25 (mitochondrial iron transporter), member 37 | Cytoplasm | other | 6166673 | 4059878 | 8440741 | | | | | | |
| solute carrier family 25 (mitochondrial iron transporter), member 37 | Cytoplasm | other | 4122916.5 | 3681753.25 | 3969015 | | | | | | |
| solute carrier family 25 (mitochondrial iron transporter), member 37 | Cytoplasm | other | 4017102.5 | 2795668 | 2752625 | | | | | | |

| | | | | | | | | | | | |
|---|---|---|---|---|---|---|---|---|---|---|---|
| solute carrier family 25 (mitochondrial iron transporter), member 37 | Cytoplasm | other | 3592639.75 | 3997627.25 | 5190302.5 | | | | | | |
| solute carrier family 25 (mitochondrial iron transporter), member 37 | Cytoplasm | other | 3033140.25 | 2507302.75 | 2545992.5 | | | | | | |
| solute carrier family 25 (mitochondrial iron transporter), member 37 | Cytoplasm | other | 2937694.25 | 2514143.5 | 3396398.5 | | | | | | |
| solute carrier family 25 (mitochondrial iron transporter), member 37 | Cytoplasm | other | 2461033 | 1860513.375 | 1629750 | | | | | | |
| solute carrier family 25 (mitochondrial iron transporter), member 37 | Cytoplasm | other | 2164426.25 | 2481307.25 | 2813753.25 | | | | | | |
| solute carrier family 25 (mitochondrial iron transporter), member 37 | Cytoplasm | other | 1613269.5 | 1468265.75 | 1579270.25 | | | | | | |
| solute carrier family 25 (mitochondrial iron transporter), member 37 | Cytoplasm | other | 1191357.5 | 786844.625 | 1083963.75 | | | | | | |
| solute carrier family 25 (mitochondrial iron transporter), member 37 | Cytoplasm | other | 1010028.375 | 475881.344 | 2218605.75 | Q920G8 | SLC25A37 | 2937298 | 1624873 | 282854 | 33 |
| solute carrier family 25 (mitochondrial carrier; adenine nucleotide translocator), member 4 | Cytoplasm | transporter | 910990.75 | 5362308 | 3057652.75 | | | | | | |
| solute carrier family 25 (mitochondrial carrier; adenine nucleotide translocator), member 4 | Cytoplasm | transporter | 675727.375 | 4083961 | 2600228.5 | | | | | | |
| solute carrier family 25 (mitochondrial carrier; adenine nucleotide translocator), member 4 | Cytoplasm | transporter | 50471.242 | 3732701 | 2587454 | P48962 | SLC25A4 | 545730 | 1745281 | 581760 | 9 |
| solute carrier family 25 (mitochondrial carrier; adenine nucleotide translocator), member 5 | Cytoplasm | transporter | 4.04E+07 | 9.23E+07 | 9.06E+07 | | | | | | |
| solute carrier family 25 (mitochondrial carrier; adenine nucleotide translocator), member 5 | Cytoplasm | transporter | 2.94E+07 | 5.90E+07 | 5.13E+07 | | | | | | |
| solute carrier family 25 (mitochondrial carrier; adenine nucleotide translocator), member 5 | Cytoplasm | transporter | 2.42E+07 | 5.48E+07 | 4.43E+07 | | | | | | |
| solute carrier family 25 (mitochondrial carrier; adenine nucleotide translocator), member 5 | Cytoplasm | transporter | 1.33E+07 | 2.90E+07 | 2.40E+07 | | | | | | |
| solute carrier family 25 (mitochondrial carrier; adenine nucleotide translocator), member 5 | Cytoplasm | transporter | 1.16E+07 | 1.66E+07 | 1.26E+07 | | | | | | |
| solute carrier family 25 (mitochondrial carrier; adenine nucleotide translocator), member 5 | Cytoplasm | transporter | 1.15E+07 | 1.56E+07 | 2.04E+07 | | | | | | |
| solute carrier family 25 (mitochondrial carrier; adenine nucleotide translocator), member 5 | Cytoplasm | transporter | 1.03E+07 | 2.80E+07 | 1.72E+07 | | | | | | |
| solute carrier family 25 (mitochondrial carrier; adenine nucleotide translocator), member 5 | Cytoplasm | transporter | 9394171 | 1.96E+07 | 1.72E+07 | | | | | | |
| solute carrier family 25 (mitochondrial carrier; adenine nucleotide translocator), member 5 | Cytoplasm | transporter | 8546387 | 2.35E+07 | 2.01E+07 | | | | | | |
| solute carrier family 25 (mitochondrial carrier; adenine nucleotide translocator), member 5 | Cytoplasm | transporter | 7521726.5 | 1.14E+07 | 1.15E+07 | | | | | | |
| solute carrier family 25 (mitochondrial carrier; adenine nucleotide translocator), member 5 | Cytoplasm | transporter | 4394505.5 | 1.17E+07 | 1.09E+07 | | | | | | |
| solute carrier family 25 (mitochondrial carrier; adenine nucleotide translocator), member 5 | Cytoplasm | transporter | 3298000.75 | 7092020.5 | 5456230 | | | | | | |
| solute carrier family 25 (mitochondrial carrier; adenine nucleotide translocator), member 5 | Cytoplasm | transporter | 1468549 | 1509143.375 | 2966055.75 | | | | | | |
| solute carrier family 25 (mitochondrial carrier; adenine nucleotide translocator), member 5 | Cytoplasm | transporter | 205595.453 | 792001.25 | 277168.188 | | | | | | |
| solute carrier family 25 (mitochondrial carrier; adenine nucleotide translocator), member 5 | Cytoplasm | transporter | 8065.055 | 2517893.75 | 2921990 | P51881 | SLC25A5 | 11700467 | 21477134 | 3201622 | 45 |
| solute carrier family 25, member 51 | Cytoplasm | other | 766489.188 | 1029574.062 | 687377.938 | Q5HZI9 | SLC25A51 | 766489 | 179151 | 103433 | 3 |
| solute carrier family 26 (anion exchanger), member 1 | Plasma Membrane | transporter | 1.20E+07 | 9056866 | 1.00E+07 | | | | | | |
| solute carrier family 26 (anion exchanger), member 1 | Plasma Membrane | transporter | 4954867.5 | 4554023.5 | 3778025 | P58735 | SLC26A1 | 8477434 | 3407982 | 1391303 | 6 |
| solute carrier family 26 (anion exchanger), member 3 | Plasma Membrane | transporter | 2538028.5 | 1974314 | 1624019.375 | Q9WVC8 | SLC26A3 | 2538029 | 461139 | 266239 | 3 |
| solute carrier family 29 (equilibrative nucleoside transporter), member 1 | Plasma Membrane | transporter | 1.62E+08 | 1.33E+08 | 1.50E+08 | | | | | | |
| solute carrier family 29 (equilibrative nucleoside transporter), member 1 | Plasma Membrane | transporter | 8.54E+07 | 6.25E+07 | 6.81E+07 | | | | | | |
| solute carrier family 29 (equilibrative nucleoside transporter), member 1 | Plasma Membrane | transporter | 4.63E+07 | 2.55E+07 | 2.67E+07 | | | | | | |
| solute carrier family 29 (equilibrative nucleoside transporter), member 1 | Plasma Membrane | transporter | 4.36E+07 | 4.00E+07 | 4.49E+07 | | | | | | |
| solute carrier family 29 (equilibrative nucleoside transporter), member 1 | Plasma Membrane | transporter | 3.75E+07 | 4.88E+07 | 3.19E+07 | | | | | | |
| solute carrier family 29 (equilibrative nucleoside transporter), member 1 | Plasma Membrane | transporter | 3.18E+07 | 2.39E+07 | 1.72E+07 | | | | | | |
| solute carrier family 29 (equilibrative nucleoside transporter), member 1 | Plasma Membrane | transporter | 1.44E+07 | 1.60E+07 | 1.05E+07 | | | | | | |
| solute carrier family 29 (equilibrative nucleoside transporter), member 1 | Plasma Membrane | transporter | 4617060 | 3636833.5 | 4352685 | | | | | | |
| solute carrier family 29 (equilibrative nucleoside transporter), member 1 | Plasma Membrane | transporter | 3653377 | 4350789 | 3129514 | | | | | | |
| solute carrier family 29 (equilibrative nucleoside transporter), member 1 | Plasma Membrane | transporter | 3343207.25 | 1605208.625 | 3237337.5 | | | | | | |
| solute carrier family 29 (equilibrative nucleoside transporter), member 1 | Plasma Membrane | transporter | 2212157 | 1527193.625 | 1230393.875 | Q9JIM1 | SLC29A1 | 39516891 | 42562564 | 7409191 | 33 |
| solute carrier family 29 (equilibrative nucleoside transporter), member 2 | Plasma Membrane | transporter | 2.49E+07 | 2.49E+07 | 2.41E+07 | Q61672 | SLC29A2 | 24930000 | 456545 | 263586 | 3 |
| solute carrier family 2 (facilitated glucose transporter), member 3 | Plasma Membrane | transporter | 8400353 | 3.32E+07 | 2.78E+07 | | | | | | |
| solute carrier family 2 (facilitated glucose transporter), member 3 | Plasma Membrane | transporter | 7233695.5 | 3.23E+07 | 1.92E+07 | | | | | | |
| solute carrier family 2 (facilitated glucose transporter), member 3 | Plasma Membrane | transporter | 5277017 | 2.29E+07 | 2.02E+07 | | | | | | |
| solute carrier family 2 (facilitated glucose transporter), member 3 | Plasma Membrane | transporter | 5051474 | 2.31E+07 | 1.51E+07 | | | | | | |
| solute carrier family 2 (facilitated glucose transporter), member 3 | Plasma Membrane | transporter | 1883593.125 | 9129679 | 7588056.5 | | | | | | |
| solute carrier family 2 (facilitated glucose transporter), member 3 | Plasma Membrane | transporter | 997509.125 | 4614210 | 2741831.25 | | | | | | |
| solute carrier family 2 (facilitated glucose transporter), member 3 | Plasma Membrane | transporter | 513119.75 | 1416032.375 | 1618724.875 | P32037 | SLC2A3 | 4193823 | 10851412 | 2367972 | 21 |
| solute carrier family 30 (zinc transporter), member 1 | Plasma Membrane | transporter | 3595088.75 | 3331273.75 | 3304246.25 | | | | | | |
| solute carrier family 30 (zinc transporter), member 1 | Plasma Membrane | transporter | 2592899 | 2701443.25 | 2165986.75 | | | | | | |
| solute carrier family 30 (zinc transporter), member 1 | Plasma Membrane | transporter | 2424198.75 | 2232355.5 | 2186857.25 | | | | | | |
| solute carrier family 30 (zinc transporter), member 1 | Plasma Membrane | transporter | 1770302.375 | 1475499.375 | 1212846.625 | Q60738 | SLC30A1 | 2595622 | 740244 | 213690 | 12 |
| solute carrier family 3 (amino acid transporter heavy chain), member 2 | Plasma Membrane | transporter | 9407040 | 8295683.5 | 8661784 | | | | | | |
| solute carrier family 3 (amino acid transporter heavy chain), member 2 | Plasma Membrane | transporter | 8393475 | 5783866.5 | 5986034 | | | | | | |
| solute carrier family 3 (amino acid transporter heavy chain), member 2 | Plasma Membrane | transporter | 6940880.5 | 5995477 | 6427681 | | | | | | |
| solute carrier family 3 (amino acid transporter heavy chain), member 2 | Plasma Membrane | transporter | 6297527 | 5490742.5 | 5531076 | | | | | | |
| solute carrier family 3 (amino acid transporter heavy chain), member 2 | Plasma Membrane | transporter | 5622987 | 5094240.5 | 5137817 | | | | | | |
| solute carrier family 3 (amino acid transporter heavy chain), member 2 | Plasma Membrane | transporter | 4458240.5 | 3022410.5 | 3560367.5 | | | | | | |
| solute carrier family 3 (amino acid transporter heavy chain), member 2 | Plasma Membrane | transporter | 4086579.25 | 3772916.75 | 4004441.5 | | | | | | |
| solute carrier family 3 (amino acid transporter heavy chain), member 2 | Plasma Membrane | transporter | 3895254.75 | 3454454.5 | 2829718.25 | | | | | | |
| solute carrier family 3 (amino acid transporter heavy chain), member 2 | Plasma Membrane | transporter | 2219523 | 1600469.625 | 1685802.625 | | | | | | |
| solute carrier family 3 (amino acid transporter heavy chain), member 2 | Plasma Membrane | transporter | 1693886.625 | 1026765.125 | 1598479.625 | P10852 | SLC3A2 | 5301539 | 2278486 | 415993 | 30 |
| solute carrier family 40 (iron-regulated transporter), member 1 | Plasma Membrane | transporter | 3.68E+07 | 3.23E+07 | 2.90E+07 | | | | | | |
| solute carrier family 40 (iron-regulated transporter), member 1 | Plasma Membrane | transporter | 3.41E+07 | 1.57E+07 | 1.54E+07 | | | | | | |
| solute carrier family 40 (iron-regulated transporter), member 1 | Plasma Membrane | transporter | 2.81E+07 | 3.10E+07 | 2.73E+07 | | | | | | |
| solute carrier family 40 (iron-regulated transporter), member 1 | Plasma Membrane | transporter | 2.58E+07 | 1.61E+07 | 1.90E+07 | | | | | | |
| solute carrier family 40 (iron-regulated transporter), member 1 | Plasma Membrane | transporter | 1.81E+07 | 3.83E+07 | 2.40E+07 | | | | | | |
| solute carrier family 40 (iron-regulated transporter), member 1 | Plasma Membrane | transporter | 1.71E+07 | 8868785 | 1.10E+07 | | | | | | |
| solute carrier family 40 (iron-regulated transporter), member 1 | Plasma Membrane | transporter | 1.03E+07 | 9638647 | 1.40E+07 | | | | | | |
| solute carrier family 40 (iron-regulated transporter), member 1 | Plasma Membrane | transporter | 1891902.875 | 2910512.5 | 1822286 | Q9JHI9 | SLC40A1 | 21521488 | 11020344 | 2249518 | 24 |
| solute carrier family 43 (amino acid system L transporter), member 1 | Plasma Membrane | transporter | 7.89E+08 | 6.39E+08 | 5.77E+08 | | | | | | |
| solute carrier family 43 (amino acid system L transporter), member 1 | Plasma Membrane | transporter | 4.85E+08 | 4.03E+08 | 3.82E+08 | | | | | | |
| solute carrier family 43 (amino acid system L transporter), member 1 | Plasma Membrane | transporter | 2.37E+08 | 1.63E+08 | 2.24E+08 | | | | | | |
| solute carrier family 43 (amino acid system L transporter), member 1 | Plasma Membrane | transporter | 1.24E+08 | 1.02E+08 | 9.25E+07 | | | | | | |
| solute carrier family 43 (amino acid system L transporter), member 1 | Plasma Membrane | transporter | 9.85E+07 | 5.66E+07 | 5.45E+07 | | | | | | |
| solute carrier family 43 (amino acid system L transporter), member 1 | Plasma Membrane | transporter | 9.60E+07 | 1.08E+08 | 8.34E+07 | | | | | | |
| solute carrier family 43 (amino acid system L transporter), member 1 | Plasma Membrane | transporter | 8.09E+07 | 5.72E+07 | 6.20E+07 | | | | | | |
| solute carrier family 43 (amino acid system L transporter), member 1 | Plasma Membrane | transporter | 7.98E+07 | 1.37E+08 | 8.63E+07 | | | | | | |
| solute carrier family 43 (amino acid system L transporter), member 1 | Plasma Membrane | transporter | 3.77E+07 | 3.34E+07 | 3.01E+07 | | | | | | |
| solute carrier family 43 (amino acid system L transporter), member 1 | Plasma Membrane | transporter | 1.14E+07 | 1.58E+07 | 8099009 | | | | | | |
| solute carrier family 43 (amino acid system L transporter), member 1 | Plasma Membrane | transporter | 9741798 | 5691244 | 6077699 | | | | | | |
| solute carrier family 43 (amino acid system L transporter), member 1 | Plasma Membrane | transporter | 4864703.5 | 3582961.75 | 2660574.5 | | | | | | |
| solute carrier family 43 (amino acid system L transporter), member 1 | Plasma Membrane | transporter | 3287061.5 | 8216759 | 5987862 | | | | | | |
| solute carrier family 43 (amino acid system L transporter), member 1 | Plasma Membrane | transporter | 1880323.875 | 321272.219 | 967332.938 | Q8BSM7 | SLC43A1 | 147037420 | 189519106 | 29243433 | 42 |
| solute carrier family 43, member 3 | Extracellular Space | other | 6.69E+07 | 3.86E+07 | 3.43E+07 | | | | | | |
| solute carrier family 43, member 3 | Extracellular Space | other | 1.72E+07 | 2.68E+07 | 2.51E+07 | | | | | | |
| solute carrier family 43, member 3 | Extracellular Space | other | 1.55E+07 | 1.34E+07 | 1.42E+07 | | | | | | |
| solute carrier family 43, member 3 | Extracellular Space | other | 9029134 | 7045499 | 6376297.5 | A2AVZ9 | SLC43A3 | 27157284 | 17333124 | 5003642 | 12 |
| solute carrier family 44 (choline transporter), member 1 | Plasma Membrane | transporter | 2728212 | 4645957.5 | 3837047.5 | | | | | | |
| solute carrier family 44 (choline transporter), member 1 | Plasma Membrane | transporter | 1959755.5 | 3127938.5 | 2074385 | | | | | | |
| solute carrier family 44 (choline transporter), member 1 | Plasma Membrane | transporter | 1416195.25 | 2592288.75 | 2152986.25 | Q6X893 | SLC44A1 | 2034721 | 1009689 | 336563 | 9 |
| solute carrier family 4 (anion exchanger), member 1 (Diego blood group) | Plasma Membrane | transporter | 3.15E+10 | 2.58E+10 | 2.67E+10 | | | | | | |
| solute carrier family 4 (anion exchanger), member 1 (Diego blood group) | Plasma Membrane | transporter | 1.83E+10 | 1.57E+10 | 1.49E+10 | | | | | | |
| solute carrier family 4 (anion exchanger), member 1 (Diego blood group) | Plasma Membrane | transporter | 1.16E+10 | 9.16E+09 | 8.64E+09 | | | | | | |

| | | | | | | | | | | | |
|---|---|---|---|---|---|---|---|---|---|---|---|
| solute carrier family 4 (anion exchanger), member 1 (Diego blood group) | Plasma Membrane | transporter | 1.11E+10 | 7.88E+09 | 7.91E+09 | | | | | | |
| solute carrier family 4 (anion exchanger), member 1 (Diego blood group) | Plasma Membrane | transporter | 1.08E+10 | 7.22E+09 | 6.90E+09 | | | | | | |
| solute carrier family 4 (anion exchanger), member 1 (Diego blood group) | Plasma Membrane | transporter | 1.01E+10 | 7.79E+09 | 8.10E+09 | | | | | | |
| solute carrier family 4 (anion exchanger), member 1 (Diego blood group) | Plasma Membrane | transporter | 9.78E+09 | 9.28E+09 | 1.02E+10 | | | | | | |
| solute carrier family 4 (anion exchanger), member 1 (Diego blood group) | Plasma Membrane | transporter | 9.45E+09 | 7.41E+09 | 6.49E+09 | | | | | | |
| solute carrier family 4 (anion exchanger), member 1 (Diego blood group) | Plasma Membrane | transporter | 8.09E+09 | 8.00E+09 | 7.18E+09 | | | | | | |
| solute carrier family 4 (anion exchanger), member 1 (Diego blood group) | Plasma Membrane | transporter | 6.54E+09 | 5.02E+09 | 5.05E+09 | | | | | | |
| solute carrier family 4 (anion exchanger), member 1 (Diego blood group) | Plasma Membrane | transporter | 5.52E+09 | 4.22E+09 | 4.71E+09 | | | | | | |
| solute carrier family 4 (anion exchanger), member 1 (Diego blood group) | Plasma Membrane | transporter | 5.50E+09 | 3.71E+09 | 3.50E+09 | | | | | | |
| solute carrier family 4 (anion exchanger), member 1 (Diego blood group) | Plasma Membrane | transporter | 4.71E+09 | 3.57E+09 | 3.89E+09 | | | | | | |
| solute carrier family 4 (anion exchanger), member 1 (Diego blood group) | Plasma Membrane | transporter | 3.97E+09 | 4.62E+09 | 6.79E+09 | | | | | | |
| solute carrier family 4 (anion exchanger), member 1 (Diego blood group) | Plasma Membrane | transporter | 3.72E+09 | 3.64E+09 | 4.45E+09 | | | | | | |
| solute carrier family 4 (anion exchanger), member 1 (Diego blood group) | Plasma Membrane | transporter | 3.32E+09 | 2.17E+09 | 2.61E+09 | | | | | | |
| solute carrier family 4 (anion exchanger), member 1 (Diego blood group) | Plasma Membrane | transporter | 2.67E+09 | 1.06E+09 | 1.47E+09 | | | | | | |
| solute carrier family 4 (anion exchanger), member 1 (Diego blood group) | Plasma Membrane | transporter | 2.12E+09 | 9.54E+08 | 9.18E+08 | | | | | | |
| solute carrier family 4 (anion exchanger), member 1 (Diego blood group) | Plasma Membrane | transporter | 1.94E+09 | 1.30E+09 | 1.54E+09 | | | | | | |
| solute carrier family 4 (anion exchanger), member 1 (Diego blood group) | Plasma Membrane | transporter | 1.05E+09 | 1.15E+09 | 1.67E+09 | | | | | | |
| solute carrier family 4 (anion exchanger), member 1 (Diego blood group) | Plasma Membrane | transporter | 9.07E+08 | 6.51E+08 | 6.77E+08 | | | | | | |
| solute carrier family 4 (anion exchanger), member 1 (Diego blood group) | Plasma Membrane | transporter | 9.05E+08 | 4.27E+08 | 7.08E+08 | | | | | | |
| solute carrier family 4 (anion exchanger), member 1 (Diego blood group) | Plasma Membrane | transporter | 6.80E+08 | 6.20E+08 | 7.24E+08 | | | | | | |
| solute carrier family 4 (anion exchanger), member 1 (Diego blood group) | Plasma Membrane | transporter | 6.57E+08 | 3.21E+08 | 4.52E+08 | | | | | | |
| solute carrier family 4 (anion exchanger), member 1 (Diego blood group) | Plasma Membrane | transporter | 6.32E+08 | 4.28E+08 | 6.49E+08 | | | | | | |
| solute carrier family 4 (anion exchanger), member 1 (Diego blood group) | Plasma Membrane | transporter | 5.61E+08 | 2.33E+08 | 2.97E+08 | | | | | | |
| solute carrier family 4 (anion exchanger), member 1 (Diego blood group) | Plasma Membrane | transporter | 5.02E+08 | 1.58E+08 | 2.61E+08 | | | | | | |
| solute carrier family 4 (anion exchanger), member 1 (Diego blood group) | Plasma Membrane | transporter | 4.61E+08 | 4.93E+08 | 5.99E+08 | | | | | | |
| solute carrier family 4 (anion exchanger), member 1 (Diego blood group) | Plasma Membrane | transporter | 3.90E+08 | 6.61E+08 | 2.49E+08 | | | | | | |
| solute carrier family 4 (anion exchanger), member 1 (Diego blood group) | Plasma Membrane | transporter | 3.72E+08 | 1.27E+08 | 1.93E+08 | | | | | | |
| solute carrier family 4 (anion exchanger), member 1 (Diego blood group) | Plasma Membrane | transporter | 3.50E+08 | 3.29E+08 | 3.03E+08 | | | | | | |
| solute carrier family 4 (anion exchanger), member 1 (Diego blood group) | Plasma Membrane | transporter | 3.21E+08 | 3.65E+08 | 2.62E+08 | | | | | | |
| solute carrier family 4 (anion exchanger), member 1 (Diego blood group) | Plasma Membrane | transporter | 2.44E+08 | 1.84E+08 | 2.94E+08 | | | | | | |
| solute carrier family 4 (anion exchanger), member 1 (Diego blood group) | Plasma Membrane | transporter | 1.73E+08 | 7.39E+07 | 1.36E+08 | | | | | | |
| solute carrier family 4 (anion exchanger), member 1 (Diego blood group) | Plasma Membrane | transporter | 1.57E+08 | 6.09E+07 | 8.21E+07 | | | | | | |
| solute carrier family 4 (anion exchanger), member 1 (Diego blood group) | Plasma Membrane | transporter | 1.47E+08 | 7.92E+07 | 1.07E+08 | | | | | | |
| solute carrier family 4 (anion exchanger), member 1 (Diego blood group) | Plasma Membrane | transporter | 1.19E+08 | 1.65E+08 | 1.23E+08 | | | | | | |
| solute carrier family 4 (anion exchanger), member 1 (Diego blood group) | Plasma Membrane | transporter | 9.33E+07 | 4.29E+07 | 5.95E+07 | | | | | | |
| solute carrier family 4 (anion exchanger), member 1 (Diego blood group) | Plasma Membrane | transporter | 9.19E+07 | 7.60E+07 | 8.42E+07 | | | | | | |
| solute carrier family 4 (anion exchanger), member 1 (Diego blood group) | Plasma Membrane | transporter | 8.74E+07 | 2.77E+07 | 6.02E+07 | | | | | | |
| solute carrier family 4 (anion exchanger), member 1 (Diego blood group) | Plasma Membrane | transporter | 6.71E+07 | 2.20E+08 | 1.11E+08 | | | | | | |
| solute carrier family 4 (anion exchanger), member 1 (Diego blood group) | Plasma Membrane | transporter | 6.28E+07 | 9.32E+07 | 1.45E+08 | | | | | | |
| solute carrier family 4 (anion exchanger), member 1 (Diego blood group) | Plasma Membrane | transporter | 6.22E+07 | 2.96E+07 | 3.83E+07 | | | | | | |
| solute carrier family 4 (anion exchanger), member 1 (Diego blood group) | Plasma Membrane | transporter | 5.78E+07 | 5.71E+07 | 7.65E+07 | | | | | | |
| solute carrier family 4 (anion exchanger), member 1 (Diego blood group) | Plasma Membrane | transporter | 3.76E+07 | 1.37E+07 | 1.88E+07 | | | | | | |
| solute carrier family 4 (anion exchanger), member 1 (Diego blood group) | Plasma Membrane | transporter | 2.80E+07 | 2.87E+07 | 2.67E+07 | | | | | | |
| solute carrier family 4 (anion exchanger), member 1 (Diego blood group) | Plasma Membrane | transporter | 2.49E+07 | 3.97E+07 | 1.20E+07 | | | | | | |
| solute carrier family 4 (anion exchanger), member 1 (Diego blood group) | Plasma Membrane | transporter | 2.35E+07 | 2.85E+07 | 2.94E+07 | | | | | | |
| solute carrier family 4 (anion exchanger), member 1 (Diego blood group) | Plasma Membrane | transporter | 2.27E+07 | 1.36E+07 | 2.19E+07 | | | | | | |
| solute carrier family 4 (anion exchanger), member 1 (Diego blood group) | Plasma Membrane | transporter | 1.68E+07 | 1.96E+07 | 1.92E+07 | | | | | | |
| solute carrier family 4 (anion exchanger), member 1 (Diego blood group) | Plasma Membrane | transporter | 1.60E+07 | 6751294 | 7445888.5 | | | | | | |
| solute carrier family 4 (anion exchanger), member 1 (Diego blood group) | Plasma Membrane | transporter | 1.19E+07 | 6994432 | 7545442 | | | | | | |
| solute carrier family 4 (anion exchanger), member 1 (Diego blood group) | Plasma Membrane | transporter | 7461587 | 6685615.5 | 1.11E+07 | | | | | | |
| solute carrier family 4 (anion exchanger), member 1 (Diego blood group) | Plasma Membrane | transporter | 2912850.25 | 1.16E+07 | 5256331.5 | | | | | | |
| solute carrier family 4 (anion exchanger), member 1 (Diego blood group) | Plasma Membrane | transporter | 2578158 | 1.30E+07 | 2.43E+07 | | | | | | |
| solute carrier family 4 (anion exchanger), member 1 (Diego blood group) | Plasma Membrane | transporter | 1107845.125 | 594502.188 | 896019.938 | | | | | | |
| solute carrier family 4 (anion exchanger), member 1 (Diego blood group) | Plasma Membrane | transporter | 190761.062 | 949166.688 | 2042982.875 | P04919 | SLC4A1 | 2983702477 | 4913738408 | 375762968 | 171 |
| solute carrier family 4 (sodium bicarbonate cotransporter), member 4 | Plasma Membrane | transporter | 1.20E+07 | 1.82E+07 | 1.73E+07 | O88343 | SLC4A4 | 11950000 | 3393705 | 1959356 | 3 |
| solute carrier family 6 (neurotransmitter transporter), member 4 | Plasma Membrane | transporter | 128707.109 | 6085983 | 3374649.75 | Q60857 | SLC6A4 | 128707 | 2982633 | 1722024 | 3 |
| solute carrier family 6 (neurotransmitter transporter, glycine), member 9 | Plasma Membrane | transporter | 4210455 | 4481128 | 4840681 | | | | | | |
| solute carrier family 6 (neurotransmitter transporter, glycine), member 9 | Plasma Membrane | transporter | 1030832 | 1277554.75 | 832345.75 | P28571 | SLC6A9 | 2620644 | 1912941 | 780955 | 6 |
| solute carrier family 7 (cationic amino acid transporter, y+ system), member 1 | Plasma Membrane | transporter | 3526371.75 | 4510210 | 4045111 | Q09143 | SLC7A1 | 3526372 | 492163 | 284150 | 3 |
| solute carrier family 7 (amino acid transporter light chain, L system), member 5 | Plasma Membrane | transporter | 8885757 | 7118544 | 7434527.5 | | | | | | |
| solute carrier family 7 (amino acid transporter light chain, L system), member 5 | Plasma Membrane | transporter | 4281211.5 | 3651837.75 | 4096341.5 | | | | | | |
| solute carrier family 7 (amino acid transporter light chain, L system), member 5 | Plasma Membrane | transporter | 1579649 | 1565904.375 | 1700470.875 | | | | | | |
| solute carrier family 7 (amino acid transporter light chain, L system), member 5 | Plasma Membrane | transporter | 1495952.625 | 1204131.375 | 1157581.5 | Q9Z127 | SLC7A5 | 4060643 | 2757099 | 795906 | 12 |
| solute carrier family 9, subfamily A (NHE1, cation proton antiporter 1), member 1 | Plasma Membrane | ion channel | 1.93E+07 | 1.50E+07 | 1.37E+07 | | | | | | |
| solute carrier family 9, subfamily A (NHE1, cation proton antiporter 1), member 1 | Plasma Membrane | ion channel | 1.23E+07 | 9506748 | 1.04E+07 | | | | | | |
| solute carrier family 9, subfamily A (NHE1, cation proton antiporter 1), member 1 | Plasma Membrane | ion channel | 8952970 | 6316287 | 7639792 | | | | | | |
| solute carrier family 9, subfamily A (NHE1, cation proton antiporter 1), member 1 | Plasma Membrane | ion channel | 7117769.5 | 6326507.5 | 6171452.5 | | | | | | |
| solute carrier family 9, subfamily A (NHE1, cation proton antiporter 1), member 1 | Plasma Membrane | ion channel | 6871188.5 | 5002494 | 5047409.5 | | | | | | |
| solute carrier family 9, subfamily A (NHE1, cation proton antiporter 1), member 1 | Plasma Membrane | ion channel | 6682930.5 | 5081117 | 4983349.5 | | | | | | |
| solute carrier family 9, subfamily A (NHE1, cation proton antiporter 1), member 1 | Plasma Membrane | ion channel | 4493960.5 | 2871887.75 | 3085600.75 | | | | | | |
| solute carrier family 9, subfamily A (NHE1, cation proton antiporter 1), member 1 | Plasma Membrane | ion channel | 4038528.25 | 5250719 | 1659630.5 | Q61165 | SLC9A1 | 8723418 | 4182603 | 853770 | 24 |
| solute carrier family 9, subfamily C (Na+-transporting carboxylic acid decarboxylase), member 1 | Extracellular Space | other | 5.14E+07 | 2.64E+07 | 3.40E+07 | Q6UJY2 | SLC9C1 | 51350000 | 12812323 | 7397198 | 3 |
| structural maintenance of chromosomes 2 | Nucleus | transporter | 1928116.5 | 1691469.25 | 1356761.75 | Q8CG48 | SMC2 | 1928117 | 287076 | 165744 | 3 |
| structural maintenance of chromosomes 4 | Nucleus | transporter | 1.83E+07 | 1.06E+07 | 1.19E+07 | | | | | | |
| structural maintenance of chromosomes 4 | Nucleus | transporter | 934168.062 | 733275.438 | 626047.125 | Q8CG47 | SMC4 | 9602084 | 7487098 | 3056595 | 6 |
| synaptosomal-associated protein, 23kDa | Plasma Membrane | transporter | 3.03E+07 | 3.33E+07 | 2.41E+07 | | | | | | |
| synaptosomal-associated protein, 23kDa | Plasma Membrane | transporter | 2.74E+07 | 3.13E+07 | 2.25E+07 | | | | | | |
| synaptosomal-associated protein, 23kDa | Plasma Membrane | transporter | 1.94E+07 | 2.22E+07 | 1.74E+07 | | | | | | |
| synaptosomal-associated protein, 23kDa | Plasma Membrane | transporter | 1.86E+07 | 2.38E+07 | 2.22E+07 | | | | | | |
| synaptosomal-associated protein, 23kDa | Plasma Membrane | transporter | 7678758.5 | 8472948 | 8530938 | | | | | | |
| synaptosomal-associated protein, 23kDa | Plasma Membrane | transporter | 7372809 | 3759067.25 | 5107424 | | | | | | |
| synaptosomal-associated protein, 23kDa | Plasma Membrane | transporter | 7101334 | 8460234 | 5827274.5 | | | | | | |
| synaptosomal-associated protein, 23kDa | Plasma Membrane | transporter | 3413788 | 2272695.25 | 2159218.5 | | | | | | |
| synaptosomal-associated protein, 23kDa | Plasma Membrane | transporter | 2315954.25 | 1069938.25 | 1318848.875 | | | | | | |
| synaptosomal-associated protein, 23kDa | Plasma Membrane | transporter | 1930070.375 | 860216.625 | 1407577.75 | | | | | | |
| synaptosomal-associated protein, 23kDa | Plasma Membrane | transporter | 1884945.75 | 2679509.25 | 1642124.5 | | | | | | |
| synaptosomal-associated protein, 23kDa | Plasma Membrane | transporter | 479856.75 | 1307489.5 | 896610.875 | O09044 | SNAP23 | 10659793 | 10559516 | 1759919 | 36 |
| synuclein, alpha (non A4 component of amyloid precursor) | Cytoplasm | other | 1.04E+07 | 6083756 | 7778422 | | | | | | |
| synuclein, alpha (non A4 component of amyloid precursor) | Cytoplasm | other | 7775157 | 4981845 | 5208098.5 | | | | | | |
| synuclein, alpha (non A4 component of amyloid precursor) | Cytoplasm | other | 6881085 | 4157534.5 | 5733675.5 | | | | | | |
| synuclein, alpha (non A4 component of amyloid precursor) | Cytoplasm | other | 5371162.5 | 3633062 | 4277619.5 | O55042 | SNCA | 7616851 | 1928166 | 556614 | 12 |
| sorting nexin 12 | Cytoplasm | transporter | 163284.047 | 45852.895 | 291246.781 | O70493 | SNX12 | 163284 | 122735 | 70861 | 3 |
| sine oculis binding protein homolog | Nucleus | other | 582725.5 | 435797.812 | 225828.719 | Q0P5V2 | SOBP | 582726 | 179374 | 103562 | 3 |
| superoxide dismutase 2, mitochondrial | Cytoplasm | enzyme | 1857837 | 3292348.25 | 3502371.5 | | | | | | |
| superoxide dismutase 2, mitochondrial | Cytoplasm | enzyme | 1829399 | 2990004 | 739679.375 | | | | | | |

| | | | | | | | | | | | |
|---|---|---|---|---|---|---|---|---|---|---|---|
| superoxide dismutase 2, mitochondrial | Cytoplasm | enzyme | 694819.812 | 893814.625 | 953357.125 | P09671 | SOD2 | 1460685 | 1139908 | 379969 | 9 |
| signal peptidase complex subunit 2 | Cytoplasm | other | 4317866.5 | 9286873 | 7248815.5 | Q9CYN2 | SPCS2 | 4317867 | 2497838 | 1442127 | 3 |
| signal peptide peptidase like 2A | Plasma Membrane | peptidase | 1.95E+07 | 1.50E+07 | 1.74E+07 | | | | | | |
| signal peptide peptidase like 2A | Plasma Membrane | peptidase | 1.64E+07 | 2.11E+07 | 1.91E+07 | Q9JJF9 | SPPL2A | 17960000 | 2236170 | 912913 | 6 |
| spectrin, alpha, erythrocytic 1 | Cytoplasm | other | 4.26E+09 | 3.53E+09 | 3.54E+09 | | | | | | |
| spectrin, alpha, erythrocytic 1 | Cytoplasm | other | 3.66E+09 | 2.61E+09 | 3.34E+09 | | | | | | |
| spectrin, alpha, erythrocytic 1 | Cytoplasm | other | 3.29E+09 | 2.23E+09 | 3.07E+09 | | | | | | |
| spectrin, alpha, erythrocytic 1 | Cytoplasm | other | 2.95E+09 | 2.09E+09 | 2.56E+09 | | | | | | |
| spectrin, alpha, erythrocytic 1 | Cytoplasm | other | 2.82E+09 | 1.60E+09 | 1.73E+09 | | | | | | |
| spectrin, alpha, erythrocytic 1 | Cytoplasm | other | 2.65E+09 | 2.17E+09 | 2.35E+09 | | | | | | |
| spectrin, alpha, erythrocytic 1 | Cytoplasm | other | 2.62E+09 | 1.75E+09 | 2.41E+09 | | | | | | |
| spectrin, alpha, erythrocytic 1 | Cytoplasm | other | 2.59E+09 | 2.03E+09 | 2.21E+09 | | | | | | |
| spectrin, alpha, erythrocytic 1 | Cytoplasm | other | 2.57E+09 | 1.74E+09 | 1.73E+09 | | | | | | |
| spectrin, alpha, erythrocytic 1 | Cytoplasm | other | 2.49E+09 | 2.43E+09 | 2.16E+09 | | | | | | |
| spectrin, alpha, erythrocytic 1 | Cytoplasm | other | 2.39E+09 | 1.77E+09 | 1.98E+09 | | | | | | |
| spectrin, alpha, erythrocytic 1 | Cytoplasm | other | 2.34E+09 | 1.68E+09 | 1.89E+09 | | | | | | |
| spectrin, alpha, erythrocytic 1 | Cytoplasm | other | 2.30E+09 | 1.71E+09 | 1.74E+09 | | | | | | |
| spectrin, alpha, erythrocytic 1 | Cytoplasm | other | 2.26E+09 | 1.71E+09 | 2.26E+09 | | | | | | |
| spectrin, alpha, erythrocytic 1 | Cytoplasm | other | 2.25E+09 | 1.93E+09 | 2.28E+09 | | | | | | |
| spectrin, alpha, erythrocytic 1 | Cytoplasm | other | 2.24E+09 | 1.68E+09 | 1.86E+09 | | | | | | |
| spectrin, alpha, erythrocytic 1 | Cytoplasm | other | 2.23E+09 | 1.14E+09 | 1.48E+09 | | | | | | |
| spectrin, alpha, erythrocytic 1 | Cytoplasm | other | 2.07E+09 | 1.68E+09 | 1.87E+09 | | | | | | |
| spectrin, alpha, erythrocytic 1 | Cytoplasm | other | 2.01E+09 | 1.49E+09 | 1.86E+09 | | | | | | |
| spectrin, alpha, erythrocytic 1 | Cytoplasm | other | 1.95E+09 | 1.68E+09 | 1.59E+09 | | | | | | |
| spectrin, alpha, erythrocytic 1 | Cytoplasm | other | 1.95E+09 | 1.41E+09 | 1.55E+09 | | | | | | |
| spectrin, alpha, erythrocytic 1 | Cytoplasm | other | 1.92E+09 | 1.37E+09 | 1.45E+09 | | | | | | |
| spectrin, alpha, erythrocytic 1 | Cytoplasm | other | 1.89E+09 | 1.45E+09 | 1.51E+09 | | | | | | |
| spectrin, alpha, erythrocytic 1 | Cytoplasm | other | 1.84E+09 | 1.45E+09 | 1.36E+09 | | | | | | |
| spectrin, alpha, erythrocytic 1 | Cytoplasm | other | 1.80E+09 | 1.23E+09 | 1.18E+09 | | | | | | |
| spectrin, alpha, erythrocytic 1 | Cytoplasm | other | 1.79E+09 | 1.45E+09 | 1.41E+09 | | | | | | |
| spectrin, alpha, erythrocytic 1 | Cytoplasm | other | 1.76E+09 | 1.45E+09 | 1.55E+09 | | | | | | |
| spectrin, alpha, erythrocytic 1 | Cytoplasm | other | 1.76E+09 | 1.77E+09 | 1.73E+09 | | | | | | |
| spectrin, alpha, erythrocytic 1 | Cytoplasm | other | 1.74E+09 | 1.08E+09 | 1.13E+09 | | | | | | |
| spectrin, alpha, erythrocytic 1 | Cytoplasm | other | 1.71E+09 | 1.27E+09 | 1.37E+09 | | | | | | |
| spectrin, alpha, erythrocytic 1 | Cytoplasm | other | 1.71E+09 | 1.20E+09 | 1.29E+09 | | | | | | |
| spectrin, alpha, erythrocytic 1 | Cytoplasm | other | 1.70E+09 | 1.50E+09 | 1.35E+09 | | | | | | |
| spectrin, alpha, erythrocytic 1 | Cytoplasm | other | 1.67E+09 | 1.30E+09 | 1.22E+09 | | | | | | |
| spectrin, alpha, erythrocytic 1 | Cytoplasm | other | 1.67E+09 | 1.19E+09 | 1.15E+09 | | | | | | |
| spectrin, alpha, erythrocytic 1 | Cytoplasm | other | 1.66E+09 | 2.14E+09 | 1.60E+09 | | | | | | |
| spectrin, alpha, erythrocytic 1 | Cytoplasm | other | 1.62E+09 | 1.39E+09 | 1.36E+09 | | | | | | |
| spectrin, alpha, erythrocytic 1 | Cytoplasm | other | 1.55E+09 | 9.34E+08 | 9.96E+08 | | | | | | |
| spectrin, alpha, erythrocytic 1 | Cytoplasm | other | 1.54E+09 | 1.37E+09 | 1.30E+09 | | | | | | |
| spectrin, alpha, erythrocytic 1 | Cytoplasm | other | 1.53E+09 | 1.06E+09 | 1.09E+09 | | | | | | |
| spectrin, alpha, erythrocytic 1 | Cytoplasm | other | 1.52E+09 | 1.26E+09 | 1.25E+09 | | | | | | |
| spectrin, alpha, erythrocytic 1 | Cytoplasm | other | 1.52E+09 | 9.70E+08 | 9.95E+08 | | | | | | |
| spectrin, alpha, erythrocytic 1 | Cytoplasm | other | 1.51E+09 | 1.10E+09 | 1.34E+09 | | | | | | |
| spectrin, alpha, erythrocytic 1 | Cytoplasm | other | 1.50E+09 | 1.25E+09 | 1.34E+09 | | | | | | |
| spectrin, alpha, erythrocytic 1 | Cytoplasm | other | 1.48E+09 | 1.34E+09 | 1.25E+09 | | | | | | |
| spectrin, alpha, erythrocytic 1 | Cytoplasm | other | 1.47E+09 | 1.37E+09 | 1.22E+09 | | | | | | |
| spectrin, alpha, erythrocytic 1 | Cytoplasm | other | 1.45E+09 | 1.16E+09 | 1.15E+09 | | | | | | |
| spectrin, alpha, erythrocytic 1 | Cytoplasm | other | 1.45E+09 | 1.06E+09 | 1.09E+09 | | | | | | |
| spectrin, alpha, erythrocytic 1 | Cytoplasm | other | 1.42E+09 | 1.16E+09 | 1.20E+09 | | | | | | |
| spectrin, alpha, erythrocytic 1 | Cytoplasm | other | 1.42E+09 | 6.46E+08 | 1.12E+09 | | | | | | |
| spectrin, alpha, erythrocytic 1 | Cytoplasm | other | 1.40E+09 | 1.33E+09 | 1.22E+09 | | | | | | |
| spectrin, alpha, erythrocytic 1 | Cytoplasm | other | 1.39E+09 | 1.31E+09 | 1.44E+09 | | | | | | |
| spectrin, alpha, erythrocytic 1 | Cytoplasm | other | 1.37E+09 | 8.69E+08 | 1.13E+09 | | | | | | |
| spectrin, alpha, erythrocytic 1 | Cytoplasm | other | 1.35E+09 | 1.07E+09 | 9.81E+08 | | | | | | |
| spectrin, alpha, erythrocytic 1 | Cytoplasm | other | 1.35E+09 | 8.73E+08 | 9.56E+08 | | | | | | |
| spectrin, alpha, erythrocytic 1 | Cytoplasm | other | 1.33E+09 | 6.49E+08 | 7.74E+08 | | | | | | |
| spectrin, alpha, erythrocytic 1 | Cytoplasm | other | 1.27E+09 | 1.06E+09 | 1.04E+09 | | | | | | |
| spectrin, alpha, erythrocytic 1 | Cytoplasm | other | 1.26E+09 | 1.03E+09 | 9.79E+08 | | | | | | |
| spectrin, alpha, erythrocytic 1 | Cytoplasm | other | 1.25E+09 | 7.62E+08 | 9.34E+08 | | | | | | |
| spectrin, alpha, erythrocytic 1 | Cytoplasm | other | 1.22E+09 | 1.22E+09 | 1.21E+09 | | | | | | |
| spectrin, alpha, erythrocytic 1 | Cytoplasm | other | 1.19E+09 | 8.29E+08 | 9.97E+08 | | | | | | |
| spectrin, alpha, erythrocytic 1 | Cytoplasm | other | 1.18E+09 | 8.03E+08 | 1.13E+09 | | | | | | |
| spectrin, alpha, erythrocytic 1 | Cytoplasm | other | 1.16E+09 | 8.78E+08 | 1.01E+09 | | | | | | |
| spectrin, alpha, erythrocytic 1 | Cytoplasm | other | 1.16E+09 | 9.07E+08 | 9.15E+08 | | | | | | |
| spectrin, alpha, erythrocytic 1 | Cytoplasm | other | 1.15E+09 | 9.49E+08 | 1.02E+09 | | | | | | |
| spectrin, alpha, erythrocytic 1 | Cytoplasm | other | 1.13E+09 | 8.96E+08 | 8.37E+08 | | | | | | |
| spectrin, alpha, erythrocytic 1 | Cytoplasm | other | 1.11E+09 | 1.04E+09 | 1.12E+09 | | | | | | |
| spectrin, alpha, erythrocytic 1 | Cytoplasm | other | 1.11E+09 | 8.37E+08 | 8.28E+08 | | | | | | |
| spectrin, alpha, erythrocytic 1 | Cytoplasm | other | 1.09E+09 | 8.22E+08 | 7.97E+08 | | | | | | |
| spectrin, alpha, erythrocytic 1 | Cytoplasm | other | 1.09E+09 | 7.54E+08 | 9.84E+08 | | | | | | |
| spectrin, alpha, erythrocytic 1 | Cytoplasm | other | 1.07E+09 | 7.65E+08 | 9.29E+08 | | | | | | |
| spectrin, alpha, erythrocytic 1 | Cytoplasm | other | 1.06E+09 | 8.28E+08 | 8.08E+08 | | | | | | |
| spectrin, alpha, erythrocytic 1 | Cytoplasm | other | 1.06E+09 | 1.16E+09 | 1.04E+09 | | | | | | |
| spectrin, alpha, erythrocytic 1 | Cytoplasm | other | 1.06E+09 | 6.61E+08 | 6.52E+08 | | | | | | |
| spectrin, alpha, erythrocytic 1 | Cytoplasm | other | 1.04E+09 | 7.37E+08 | 8.75E+08 | | | | | | |
| spectrin, alpha, erythrocytic 1 | Cytoplasm | other | 1.04E+09 | 7.10E+08 | 7.68E+08 | | | | | | |
| spectrin, alpha, erythrocytic 1 | Cytoplasm | other | 1.03E+09 | 9.09E+08 | 9.74E+08 | | | | | | |
| spectrin, alpha, erythrocytic 1 | Cytoplasm | other | 1.03E+09 | 8.12E+08 | 9.12E+08 | | | | | | |
| spectrin, alpha, erythrocytic 1 | Cytoplasm | other | 1.01E+09 | 8.93E+08 | 7.98E+08 | | | | | | |
| spectrin, alpha, erythrocytic 1 | Cytoplasm | other | 9.88E+08 | 9.92E+08 | 9.15E+08 | | | | | | |
| spectrin, alpha, erythrocytic 1 | Cytoplasm | other | 9.72E+08 | 1.11E+09 | 8.50E+08 | | | | | | |
| spectrin, alpha, erythrocytic 1 | Cytoplasm | other | 9.64E+08 | 6.68E+08 | 7.08E+08 | | | | | | |
| spectrin, alpha, erythrocytic 1 | Cytoplasm | other | 9.58E+08 | 5.43E+08 | 6.23E+08 | | | | | | |
| spectrin, alpha, erythrocytic 1 | Cytoplasm | other | 9.16E+08 | 7.61E+08 | 9.22E+08 | | | | | | |
| spectrin, alpha, erythrocytic 1 | Cytoplasm | other | 8.94E+08 | 6.28E+08 | 6.32E+08 | | | | | | |
| spectrin, alpha, erythrocytic 1 | Cytoplasm | other | 8.86E+08 | 8.59E+08 | 8.41E+08 | | | | | | |
| spectrin, alpha, erythrocytic 1 | Cytoplasm | other | 8.72E+08 | 5.53E+08 | 4.54E+08 | | | | | | |
| spectrin, alpha, erythrocytic 1 | Cytoplasm | other | 8.70E+08 | 6.83E+08 | 6.36E+08 | | | | | | |
| spectrin, alpha, erythrocytic 1 | Cytoplasm | other | 8.50E+08 | 7.29E+08 | 7.63E+08 | | | | | | |
| spectrin, alpha, erythrocytic 1 | Cytoplasm | other | 8.44E+08 | 6.87E+08 | 7.77E+08 | | | | | | |
| spectrin, alpha, erythrocytic 1 | Cytoplasm | other | 8.39E+08 | 6.17E+08 | 6.62E+08 | | | | | | |
| spectrin, alpha, erythrocytic 1 | Cytoplasm | other | 8.37E+08 | 6.44E+08 | 6.44E+08 | | | | | | |

| | | | | | |
|---|---|---|---|---|---|
| spectrin, alpha, erythrocytic 1 | Cytoplasm | other | 8.26E+08 | 7.64E+08 | 8.04E+08 |
| spectrin, alpha, erythrocytic 1 | Cytoplasm | other | 7.76E+08 | 6.32E+08 | 7.58E+08 |
| spectrin, alpha, erythrocytic 1 | Cytoplasm | other | 7.68E+08 | 6.00E+08 | 6.13E+08 |
| spectrin, alpha, erythrocytic 1 | Cytoplasm | other | 7.42E+08 | 4.06E+08 | 4.78E+08 |
| spectrin, alpha, erythrocytic 1 | Cytoplasm | other | 7.17E+08 | 9.31E+08 | 7.70E+08 |
| spectrin, alpha, erythrocytic 1 | Cytoplasm | other | 6.99E+08 | 4.47E+08 | 5.98E+08 |
| spectrin, alpha, erythrocytic 1 | Cytoplasm | other | 6.94E+08 | 5.60E+08 | 6.34E+08 |
| spectrin, alpha, erythrocytic 1 | Cytoplasm | other | 6.65E+08 | 4.52E+08 | 5.45E+08 |
| spectrin, alpha, erythrocytic 1 | Cytoplasm | other | 6.64E+08 | 5.34E+08 | 5.11E+08 |
| spectrin, alpha, erythrocytic 1 | Cytoplasm | other | 6.46E+08 | 3.46E+08 | 4.20E+08 |
| spectrin, alpha, erythrocytic 1 | Cytoplasm | other | 6.34E+08 | 5.30E+08 | 6.39E+08 |
| spectrin, alpha, erythrocytic 1 | Cytoplasm | other | 6.33E+08 | 5.05E+08 | 5.36E+08 |
| spectrin, alpha, erythrocytic 1 | Cytoplasm | other | 6.24E+08 | 1.10E+09 | 8.90E+08 |
| spectrin, alpha, erythrocytic 1 | Cytoplasm | other | 6.18E+08 | 5.13E+08 | 5.31E+08 |
| spectrin, alpha, erythrocytic 1 | Cytoplasm | other | 6.13E+08 | 6.18E+08 | 6.07E+08 |
| spectrin, alpha, erythrocytic 1 | Cytoplasm | other | 6.03E+08 | 4.99E+08 | 3.91E+08 |
| spectrin, alpha, erythrocytic 1 | Cytoplasm | other | 5.74E+08 | 3.86E+08 | 3.25E+08 |
| spectrin, alpha, erythrocytic 1 | Cytoplasm | other | 5.73E+08 | 4.97E+08 | 6.40E+08 |
| spectrin, alpha, erythrocytic 1 | Cytoplasm | other | 5.49E+08 | 4.90E+08 | 3.95E+08 |
| spectrin, alpha, erythrocytic 1 | Cytoplasm | other | 5.40E+08 | 5.23E+08 | 4.38E+08 |
| spectrin, alpha, erythrocytic 1 | Cytoplasm | other | 5.26E+08 | 3.62E+08 | 3.36E+08 |
| spectrin, alpha, erythrocytic 1 | Cytoplasm | other | 5.22E+08 | 5.63E+08 | 3.97E+08 |
| spectrin, alpha, erythrocytic 1 | Cytoplasm | other | 5.21E+08 | 5.99E+08 | 6.02E+08 |
| spectrin, alpha, erythrocytic 1 | Cytoplasm | other | 4.80E+08 | 2.72E+08 | 2.77E+08 |
| spectrin, alpha, erythrocytic 1 | Cytoplasm | other | 4.73E+08 | 3.41E+08 | 3.44E+08 |
| spectrin, alpha, erythrocytic 1 | Cytoplasm | other | 4.64E+08 | 5.33E+08 | 4.91E+08 |
| spectrin, alpha, erythrocytic 1 | Cytoplasm | other | 4.44E+08 | 1.76E+08 | 2.50E+08 |
| spectrin, alpha, erythrocytic 1 | Cytoplasm | other | 4.33E+08 | 1.76E+08 | 2.48E+08 |
| spectrin, alpha, erythrocytic 1 | Cytoplasm | other | 4.33E+08 | 4.48E+08 | 3.88E+08 |
| spectrin, alpha, erythrocytic 1 | Cytoplasm | other | 4.32E+08 | 3.05E+08 | 3.59E+08 |
| spectrin, alpha, erythrocytic 1 | Cytoplasm | other | 4.07E+08 | 3.30E+08 | 4.47E+08 |
| spectrin, alpha, erythrocytic 1 | Cytoplasm | other | 3.96E+08 | 1.85E+08 | 2.57E+08 |
| spectrin, alpha, erythrocytic 1 | Cytoplasm | other | 3.95E+08 | 1.44E+08 | 2.50E+08 |
| spectrin, alpha, erythrocytic 1 | Cytoplasm | other | 3.83E+08 | 2.55E+08 | 2.94E+08 |
| spectrin, alpha, erythrocytic 1 | Cytoplasm | other | 3.67E+08 | 1.83E+08 | 1.72E+08 |
| spectrin, alpha, erythrocytic 1 | Cytoplasm | other | 3.65E+08 | 1.45E+08 | 1.31E+08 |
| spectrin, alpha, erythrocytic 1 | Cytoplasm | other | 3.59E+08 | 1.91E+08 | 2.47E+08 |
| spectrin, alpha, erythrocytic 1 | Cytoplasm | other | 3.57E+08 | 2.17E+08 | 6.93E+08 |
| spectrin, alpha, erythrocytic 1 | Cytoplasm | other | 3.55E+08 | 2.96E+08 | 3.80E+08 |
| spectrin, alpha, erythrocytic 1 | Cytoplasm | other | 3.50E+08 | 6.97E+08 | 4.78E+08 |
| spectrin, alpha, erythrocytic 1 | Cytoplasm | other | 3.36E+08 | 4.74E+08 | 3.59E+08 |
| spectrin, alpha, erythrocytic 1 | Cytoplasm | other | 3.33E+08 | 1.47E+08 | 1.80E+08 |
| spectrin, alpha, erythrocytic 1 | Cytoplasm | other | 3.32E+08 | 2.48E+08 | 3.02E+08 |
| spectrin, alpha, erythrocytic 1 | Cytoplasm | other | 3.29E+08 | 5.28E+08 | 4.15E+08 |
| spectrin, alpha, erythrocytic 1 | Cytoplasm | other | 3.18E+08 | 3.14E+08 | 4.37E+08 |
| spectrin, alpha, erythrocytic 1 | Cytoplasm | other | 3.11E+08 | 1.49E+08 | 2.04E+08 |
| spectrin, alpha, erythrocytic 1 | Cytoplasm | other | 3.09E+08 | 5.04E+08 | 3.96E+08 |
| spectrin, alpha, erythrocytic 1 | Cytoplasm | other | 3.07E+08 | 2.35E+08 | 2.79E+08 |
| spectrin, alpha, erythrocytic 1 | Cytoplasm | other | 3.03E+08 | 2.45E+08 | 1.81E+08 |
| spectrin, alpha, erythrocytic 1 | Cytoplasm | other | 2.97E+08 | 4.11E+08 | 4.22E+08 |
| spectrin, alpha, erythrocytic 1 | Cytoplasm | other | 2.91E+08 | 2.50E+08 | 1.84E+08 |
| spectrin, alpha, erythrocytic 1 | Cytoplasm | other | 2.88E+08 | 1.73E+08 | 2.00E+08 |
| spectrin, alpha, erythrocytic 1 | Cytoplasm | other | 2.82E+08 | 1.98E+08 | 2.22E+08 |
| spectrin, alpha, erythrocytic 1 | Cytoplasm | other | 2.64E+08 | 2.38E+08 | 2.25E+08 |
| spectrin, alpha, erythrocytic 1 | Cytoplasm | other | 2.52E+08 | 1.58E+08 | 1.72E+08 |
| spectrin, alpha, erythrocytic 1 | Cytoplasm | other | 2.48E+08 | 1.44E+08 | 1.42E+08 |
| spectrin, alpha, erythrocytic 1 | Cytoplasm | other | 2.45E+08 | 2.87E+08 | 3.30E+08 |
| spectrin, alpha, erythrocytic 1 | Cytoplasm | other | 2.34E+08 | 2.23E+08 | 1.98E+08 |
| spectrin, alpha, erythrocytic 1 | Cytoplasm | other | 2.34E+08 | 2.51E+08 | 2.95E+08 |
| spectrin, alpha, erythrocytic 1 | Cytoplasm | other | 2.26E+08 | 1.10E+08 | 1.43E+08 |
| spectrin, alpha, erythrocytic 1 | Cytoplasm | other | 2.11E+08 | 1.59E+08 | 1.25E+08 |
| spectrin, alpha, erythrocytic 1 | Cytoplasm | other | 2.08E+08 | 8.75E+07 | 9.86E+07 |
| spectrin, alpha, erythrocytic 1 | Cytoplasm | other | 2.06E+08 | 1.62E+08 | 1.23E+08 |
| spectrin, alpha, erythrocytic 1 | Cytoplasm | other | 2.02E+08 | 2.82E+08 | 1.97E+08 |
| spectrin, alpha, erythrocytic 1 | Cytoplasm | other | 2.01E+08 | 1.76E+08 | 1.63E+08 |
| spectrin, alpha, erythrocytic 1 | Cytoplasm | other | 1.92E+08 | 2.40E+08 | 1.91E+08 |
| spectrin, alpha, erythrocytic 1 | Cytoplasm | other | 1.90E+08 | 9.00E+07 | 1.52E+08 |
| spectrin, alpha, erythrocytic 1 | Cytoplasm | other | 1.88E+08 | 1.24E+08 | 1.21E+08 |
| spectrin, alpha, erythrocytic 1 | Cytoplasm | other | 1.85E+08 | 1.64E+08 | 1.38E+08 |
| spectrin, alpha, erythrocytic 1 | Cytoplasm | other | 1.85E+08 | 1.75E+08 | 2.25E+08 |
| spectrin, alpha, erythrocytic 1 | Cytoplasm | other | 1.78E+08 | 9.69E+07 | 1.07E+08 |
| spectrin, alpha, erythrocytic 1 | Cytoplasm | other | 1.75E+08 | 3.47E+08 | 2.29E+08 |
| spectrin, alpha, erythrocytic 1 | Cytoplasm | other | 1.67E+08 | 1.89E+08 | 2.58E+08 |
| spectrin, alpha, erythrocytic 1 | Cytoplasm | other | 1.65E+08 | 1.10E+08 | 3.39E+07 |
| spectrin, alpha, erythrocytic 1 | Cytoplasm | other | 1.61E+08 | 1.20E+08 | 1.25E+08 |
| spectrin, alpha, erythrocytic 1 | Cytoplasm | other | 1.59E+08 | 2.70E+08 | 2.40E+08 |
| spectrin, alpha, erythrocytic 1 | Cytoplasm | other | 1.56E+08 | 8.21E+07 | 1.11E+08 |
| spectrin, alpha, erythrocytic 1 | Cytoplasm | other | 1.53E+08 | 1.69E+08 | 2.21E+08 |
| spectrin, alpha, erythrocytic 1 | Cytoplasm | other | 1.52E+08 | 2.00E+08 | 1.30E+08 |
| spectrin, alpha, erythrocytic 1 | Cytoplasm | other | 1.51E+08 | 1.37E+08 | 3.44E+08 |
| spectrin, alpha, erythrocytic 1 | Cytoplasm | other | 1.42E+08 | 9.67E+07 | 1.05E+08 |
| spectrin, alpha, erythrocytic 1 | Cytoplasm | other | 1.41E+08 | 1.56E+08 | 3.52E+08 |
| spectrin, alpha, erythrocytic 1 | Cytoplasm | other | 1.39E+08 | 4.47E+08 | 3.05E+08 |
| spectrin, alpha, erythrocytic 1 | Cytoplasm | other | 1.39E+08 | 9.28E+07 | 1.22E+08 |
| spectrin, alpha, erythrocytic 1 | Cytoplasm | other | 1.29E+08 | 9.77E+07 | 9.14E+07 |
| spectrin, alpha, erythrocytic 1 | Cytoplasm | other | 1.28E+08 | 4.87E+07 | 1.54E+08 |
| spectrin, alpha, erythrocytic 1 | Cytoplasm | other | 1.27E+08 | 1.22E+08 | 1.02E+08 |
| spectrin, alpha, erythrocytic 1 | Cytoplasm | other | 1.26E+08 | 1.74E+08 | 1.12E+08 |
| spectrin, alpha, erythrocytic 1 | Cytoplasm | other | 1.25E+08 | 8.89E+07 | 7.57E+07 |
| spectrin, alpha, erythrocytic 1 | Cytoplasm | other | 1.24E+08 | 2.43E+08 | 1.70E+08 |
| spectrin, alpha, erythrocytic 1 | Cytoplasm | other | 1.19E+08 | 7.37E+07 | 8.00E+07 |
| spectrin, alpha, erythrocytic 1 | Cytoplasm | other | 1.18E+08 | 8.78E+07 | 8.47E+07 |
| spectrin, alpha, erythrocytic 1 | Cytoplasm | other | 1.18E+08 | 8.82E+07 | 8.16E+07 |
| spectrin, alpha, erythrocytic 1 | Cytoplasm | other | 1.18E+08 | 6.20E+07 | 1.37E+07 |
| spectrin, alpha, erythrocytic 1 | Cytoplasm | other | 1.14E+08 | 4.81E+07 | 6.50E+07 |

| | | | | | |
|---|---|---|---|---|---|
| spectrin, alpha, erythrocytic 1 | Cytoplasm | other | 1.11E+08 | 6.63E+07 | 7.33E+07 |
| spectrin, alpha, erythrocytic 1 | Cytoplasm | other | 1.09E+08 | 5.75E+07 | 5.59E+07 |
| spectrin, alpha, erythrocytic 1 | Cytoplasm | other | 1.08E+08 | 8.71E+07 | 1.22E+08 |
| spectrin, alpha, erythrocytic 1 | Cytoplasm | other | 1.06E+08 | 5.20E+07 | 7.24E+07 |
| spectrin, alpha, erythrocytic 1 | Cytoplasm | other | 1.05E+08 | 7.84E+07 | 8.20E+07 |
| spectrin, alpha, erythrocytic 1 | Cytoplasm | other | 1.01E+08 | 7.30E+07 | 7.17E+07 |
| spectrin, alpha, erythrocytic 1 | Cytoplasm | other | 9.98E+07 | 5.53E+07 | 7.23E+07 |
| spectrin, alpha, erythrocytic 1 | Cytoplasm | other | 9.78E+07 | 2.58E+08 | 1.15E+08 |
| spectrin, alpha, erythrocytic 1 | Cytoplasm | other | 9.65E+07 | 5.96E+07 | 1.27E+08 |
| spectrin, alpha, erythrocytic 1 | Cytoplasm | other | 8.86E+07 | 3.02E+08 | 2.06E+08 |
| spectrin, alpha, erythrocytic 1 | Cytoplasm | other | 8.73E+07 | 5.00E+07 | 5.40E+07 |
| spectrin, alpha, erythrocytic 1 | Cytoplasm | other | 8.43E+07 | 6.86E+07 | 6.54E+07 |
| spectrin, alpha, erythrocytic 1 | Cytoplasm | other | 8.17E+07 | 2.19E+08 | 2.20E+08 |
| spectrin, alpha, erythrocytic 1 | Cytoplasm | other | 7.91E+07 | 1.27E+08 | 1.41E+08 |
| spectrin, alpha, erythrocytic 1 | Cytoplasm | other | 7.72E+07 | 9.68E+07 | 7.23E+07 |
| spectrin, alpha, erythrocytic 1 | Cytoplasm | other | 7.67E+07 | 5.31E+07 | 5.40E+07 |
| spectrin, alpha, erythrocytic 1 | Cytoplasm | other | 7.45E+07 | 1.11E+08 | 8.14E+07 |
| spectrin, alpha, erythrocytic 1 | Cytoplasm | other | 7.17E+07 | 5.58E+07 | 1.04E+08 |
| spectrin, alpha, erythrocytic 1 | Cytoplasm | other | 6.99E+07 | 3.12E+07 | 3.93E+07 |
| spectrin, alpha, erythrocytic 1 | Cytoplasm | other | 6.68E+07 | 2.94E+07 | 3.82E+07 |
| spectrin, alpha, erythrocytic 1 | Cytoplasm | other | 5.89E+07 | 2.96E+07 | 6.70E+07 |
| spectrin, alpha, erythrocytic 1 | Cytoplasm | other | 5.82E+07 | 4.79E+07 | 3.49E+07 |
| spectrin, alpha, erythrocytic 1 | Cytoplasm | other | 5.50E+07 | 3.24E+07 | 4.32E+07 |
| spectrin, alpha, erythrocytic 1 | Cytoplasm | other | 4.99E+07 | 4.32E+07 | 4.20E+07 |
| spectrin, alpha, erythrocytic 1 | Cytoplasm | other | 4.74E+07 | 1.98E+08 | 1.11E+08 |
| spectrin, alpha, erythrocytic 1 | Cytoplasm | other | 4.74E+07 | 3.08E+07 | 2.67E+07 |
| spectrin, alpha, erythrocytic 1 | Cytoplasm | other | 4.64E+07 | 4.04E+07 | 3.77E+07 |
| spectrin, alpha, erythrocytic 1 | Cytoplasm | other | 4.51E+07 | 2.15E+07 | 3.09E+07 |
| spectrin, alpha, erythrocytic 1 | Cytoplasm | other | 4.50E+07 | 2.34E+07 | 4.83E+07 |
| spectrin, alpha, erythrocytic 1 | Cytoplasm | other | 4.25E+07 | 2.15E+07 | 2.73E+07 |
| spectrin, alpha, erythrocytic 1 | Cytoplasm | other | 4.02E+07 | 1.24E+08 | 9.05E+07 |
| spectrin, alpha, erythrocytic 1 | Cytoplasm | other | 4.00E+07 | 2.07E+07 | 2.31E+07 |
| spectrin, alpha, erythrocytic 1 | Cytoplasm | other | 3.73E+07 | 2.40E+07 | 4.95E+07 |
| spectrin, alpha, erythrocytic 1 | Cytoplasm | other | 3.68E+07 | 8.47E+07 | 4.95E+07 |
| spectrin, alpha, erythrocytic 1 | Cytoplasm | other | 3.56E+07 | 2.94E+07 | 2.65E+07 |
| spectrin, alpha, erythrocytic 1 | Cytoplasm | other | 3.55E+07 | 6.52E+07 | 7.04E+07 |
| spectrin, alpha, erythrocytic 1 | Cytoplasm | other | 3.52E+07 | 5.04E+07 | 4.93E+07 |
| spectrin, alpha, erythrocytic 1 | Cytoplasm | other | 3.52E+07 | 5.96E+07 | 4.25E+07 |
| spectrin, alpha, erythrocytic 1 | Cytoplasm | other | 3.41E+07 | 3.27E+07 | 2.81E+07 |
| spectrin, alpha, erythrocytic 1 | Cytoplasm | other | 3.35E+07 | 2.58E+07 | 3.14E+07 |
| spectrin, alpha, erythrocytic 1 | Cytoplasm | other | 3.26E+07 | 2.05E+07 | 2.57E+07 |
| spectrin, alpha, erythrocytic 1 | Cytoplasm | other | 3.26E+07 | 2.01E+07 | 7141663 |
| spectrin, alpha, erythrocytic 1 | Cytoplasm | other | 3.22E+07 | 4.10E+07 | 4.22E+07 |
| spectrin, alpha, erythrocytic 1 | Cytoplasm | other | 3.06E+07 | 4.21E+07 | 4.67E+07 |
| spectrin, alpha, erythrocytic 1 | Cytoplasm | other | 2.98E+07 | 2.07E+07 | 7225373.5 |
| spectrin, alpha, erythrocytic 1 | Cytoplasm | other | 2.90E+07 | 4.11E+07 | 3.53E+07 |
| spectrin, alpha, erythrocytic 1 | Cytoplasm | other | 2.80E+07 | 8.40E+07 | 3.01E+07 |
| spectrin, alpha, erythrocytic 1 | Cytoplasm | other | 2.77E+07 | 1.85E+07 | 1.97E+07 |
| spectrin, alpha, erythrocytic 1 | Cytoplasm | other | 2.77E+07 | 6.75E+07 | 3.61E+07 |
| spectrin, alpha, erythrocytic 1 | Cytoplasm | other | 2.48E+07 | 2.32E+07 | 1.33E+07 |
| spectrin, alpha, erythrocytic 1 | Cytoplasm | other | 2.48E+07 | 1.41E+07 | 1.91E+07 |
| spectrin, alpha, erythrocytic 1 | Cytoplasm | other | 2.46E+07 | 8730706 | 1.34E+07 |
| spectrin, alpha, erythrocytic 1 | Cytoplasm | other | 2.43E+07 | 7209181.5 | 9182784 |
| spectrin, alpha, erythrocytic 1 | Cytoplasm | other | 2.41E+07 | 4.70E+07 | 2.76E+07 |
| spectrin, alpha, erythrocytic 1 | Cytoplasm | other | 2.35E+07 | 1.44E+08 | 5.63E+07 |
| spectrin, alpha, erythrocytic 1 | Cytoplasm | other | 2.16E+07 | 2.17E+07 | 1.86E+07 |
| spectrin, alpha, erythrocytic 1 | Cytoplasm | other | 2.07E+07 | 1.05E+07 | 1.07E+07 |
| spectrin, alpha, erythrocytic 1 | Cytoplasm | other | 1.89E+07 | 1.04E+07 | 7840668.5 |
| spectrin, alpha, erythrocytic 1 | Cytoplasm | other | 1.88E+07 | 1.42E+07 | 1.53E+07 |
| spectrin, alpha, erythrocytic 1 | Cytoplasm | other | 1.78E+07 | 8919054 | 1.41E+07 |
| spectrin, alpha, erythrocytic 1 | Cytoplasm | other | 1.72E+07 | 1.30E+07 | 1.27E+07 |
| spectrin, alpha, erythrocytic 1 | Cytoplasm | other | 1.68E+07 | 4.04E+07 | 2.33E+07 |
| spectrin, alpha, erythrocytic 1 | Cytoplasm | other | 1.63E+07 | 3.69E+07 | 5.68E+07 |
| spectrin, alpha, erythrocytic 1 | Cytoplasm | other | 1.62E+07 | 8251882.5 | 9543873 |
| spectrin, alpha, erythrocytic 1 | Cytoplasm | other | 1.53E+07 | 2.24E+07 | 1.72E+07 |
| spectrin, alpha, erythrocytic 1 | Cytoplasm | other | 1.48E+07 | 5341309 | 8837666 |
| spectrin, alpha, erythrocytic 1 | Cytoplasm | other | 1.47E+07 | 8714949 | 1.18E+07 |
| spectrin, alpha, erythrocytic 1 | Cytoplasm | other | 1.46E+07 | 8267019.5 | 9142273 |
| spectrin, alpha, erythrocytic 1 | Cytoplasm | other | 1.41E+07 | 1.03E+08 | 5.14E+07 |
| spectrin, alpha, erythrocytic 1 | Cytoplasm | other | 1.35E+07 | 2.95E+07 | 2.95E+07 |
| spectrin, alpha, erythrocytic 1 | Cytoplasm | other | 1.30E+07 | 6981031.5 | 7475257 |
| spectrin, alpha, erythrocytic 1 | Cytoplasm | other | 1.28E+07 | 1.92E+07 | 1.31E+07 |
| spectrin, alpha, erythrocytic 1 | Cytoplasm | other | 1.24E+07 | 7726806 | 8257560.5 |
| spectrin, alpha, erythrocytic 1 | Cytoplasm | other | 1.16E+07 | 7864897.5 | 2738687.75 |
| spectrin, alpha, erythrocytic 1 | Cytoplasm | other | 1.14E+07 | 8732205 | 6907004.5 |
| spectrin, alpha, erythrocytic 1 | Cytoplasm | other | 1.05E+07 | 4622450 | 6354530 |
| spectrin, alpha, erythrocytic 1 | Cytoplasm | other | 1.02E+07 | 1.70E+07 | 1.48E+07 |
| spectrin, alpha, erythrocytic 1 | Cytoplasm | other | 9088375 | 1.50E+07 | 1.28E+07 |
| spectrin, alpha, erythrocytic 1 | Cytoplasm | other | 9023739 | 1.29E+07 | 1.86E+07 |
| spectrin, alpha, erythrocytic 1 | Cytoplasm | other | 8841532 | 2546613.75 | 3329434.75 |
| spectrin, alpha, erythrocytic 1 | Cytoplasm | other | 8461054 | 3.74E+07 | 8896941 |
| spectrin, alpha, erythrocytic 1 | Cytoplasm | other | 8328092.5 | 4868574.5 | 6076855 |
| spectrin, alpha, erythrocytic 1 | Cytoplasm | other | 8303696.5 | 3985183.5 | 1930863.625 |
| spectrin, alpha, erythrocytic 1 | Cytoplasm | other | 7851583 | 4233085 | 1379188.25 |
| spectrin, alpha, erythrocytic 1 | Cytoplasm | other | 7421346 | 6044860.5 | 4769502 |
| spectrin, alpha, erythrocytic 1 | Cytoplasm | other | 6291776 | 1.60E+07 | 1.04E+07 |
| spectrin, alpha, erythrocytic 1 | Cytoplasm | other | 6200211 | 3.52E+07 | 4682939 |
| spectrin, alpha, erythrocytic 1 | Cytoplasm | other | 6124368.5 | 2192564.25 | 2851054 |
| spectrin, alpha, erythrocytic 1 | Cytoplasm | other | 6105301.5 | 4620456 | 3964608 |
| spectrin, alpha, erythrocytic 1 | Cytoplasm | other | 5621353 | 3364675 | 3287020.75 |
| spectrin, alpha, erythrocytic 1 | Cytoplasm | other | 5032289 | 2790141.75 | 4850836 |
| spectrin, alpha, erythrocytic 1 | Cytoplasm | other | 4910489 | 2089769.75 | 4593529.5 |
| spectrin, alpha, erythrocytic 1 | Cytoplasm | other | 4552081 | 2669836 | 2203972.75 |
| spectrin, alpha, erythrocytic 1 | Cytoplasm | other | 3968943 | 2865158.75 | 3931912.25 |
| spectrin, alpha, erythrocytic 1 | Cytoplasm | other | 3920343 | 5830995.5 | 5646173 |

| | | | | | | | | | | | |
|---|---|---|---|---|---|---|---|---|---|---|---|
| spectrin, alpha, erythrocytic 1 | Cytoplasm | other | 2906368.25 | 2.80E+07 | 1.26E+07 | | | | | | |
| spectrin, alpha, erythrocytic 1 | Cytoplasm | other | 2704364.25 | 1951505.375 | 5776820.5 | | | | | | |
| spectrin, alpha, erythrocytic 1 | Cytoplasm | other | 2329428.25 | 1.16E+07 | 6779321 | | | | | | |
| spectrin, alpha, erythrocytic 1 | Cytoplasm | other | 1667181.75 | 1759742.875 | 1210660.375 | | | | | | |
| spectrin, alpha, erythrocytic 1 | Cytoplasm | other | 1655466 | 1877500.875 | 1779020.25 | | | | | | |
| spectrin, alpha, erythrocytic 1 | Cytoplasm | other | 1594036.125 | 3731643.25 | 2297154.75 | | | | | | |
| spectrin, alpha, erythrocytic 1 | Cytoplasm | other | 1579861.125 | 823264.875 | 880268.875 | | | | | | |
| spectrin, alpha, erythrocytic 1 | Cytoplasm | other | 1562770.125 | 4858633 | 6332000 | | | | | | |
| spectrin, alpha, erythrocytic 1 | Cytoplasm | other | 1547656.125 | 3546033 | 2020272.875 | | | | | | |
| spectrin, alpha, erythrocytic 1 | Cytoplasm | other | 1493215.25 | 1307117.875 | 499851.75 | | | | | | |
| spectrin, alpha, erythrocytic 1 | Cytoplasm | other | 1331895.75 | 1206305.875 | 1296583.5 | | | | | | |
| spectrin, alpha, erythrocytic 1 | Cytoplasm | other | 1159140.5 | 8070374 | 2.10E+07 | | | | | | |
| spectrin, alpha, erythrocytic 1 | Cytoplasm | other | 779801.5 | 1.02E+07 | 3757607.25 | | | | | | |
| spectrin, alpha, erythrocytic 1 | Cytoplasm | other | 619085.562 | 1.14E+07 | 4173012.5 | | | | | | |
| spectrin, alpha, erythrocytic 1 | Cytoplasm | other | 524773.938 | 1469237.875 | 1817527.125 | | | | | | |
| spectrin, alpha, erythrocytic 1 | Cytoplasm | other | 399244.906 | 610789.812 | 503928.219 | | | | | | |
| spectrin, alpha, erythrocytic 1 | Cytoplasm | other | 4.70E-06 | 437670.219 | 1024351.75 | P08032 | SPTA1 | 609016916 | 673092989 | 22511597 | 894 |
| spectrin, alpha, non-erythrocytic 1 | Plasma Membrane | other | 1.41E+07 | 4628638.5 | 5985146 | P16546 | SPTAN1 | 14060000 | 5098921 | 2943863 | 3 |
| spectrin, beta, erythrocytic | Plasma Membrane | other | 4.48E+09 | 2.88E+09 | 3.26E+09 | | | | | | |
| spectrin, beta, erythrocytic | Plasma Membrane | other | 3.09E+09 | 2.88E+09 | 2.68E+09 | | | | | | |
| spectrin, beta, erythrocytic | Plasma Membrane | other | 2.90E+09 | 2.29E+09 | 2.40E+09 | | | | | | |
| spectrin, beta, erythrocytic | Plasma Membrane | other | 2.47E+09 | 2.19E+09 | 2.39E+09 | | | | | | |
| spectrin, beta, erythrocytic | Plasma Membrane | other | 2.45E+09 | 1.88E+09 | 1.65E+09 | | | | | | |
| spectrin, beta, erythrocytic | Plasma Membrane | other | 2.39E+09 | 2.15E+09 | 2.02E+09 | | | | | | |
| spectrin, beta, erythrocytic | Plasma Membrane | other | 2.26E+09 | 1.83E+09 | 1.79E+09 | | | | | | |
| spectrin, beta, erythrocytic | Plasma Membrane | other | 2.16E+09 | 1.76E+09 | 1.77E+09 | | | | | | |
| spectrin, beta, erythrocytic | Plasma Membrane | other | 2.05E+09 | 1.48E+09 | 1.52E+09 | | | | | | |
| spectrin, beta, erythrocytic | Plasma Membrane | other | 2.04E+09 | 1.59E+09 | 1.42E+09 | | | | | | |
| spectrin, beta, erythrocytic | Plasma Membrane | other | 2.03E+09 | 9.47E+08 | 1.45E+09 | | | | | | |
| spectrin, beta, erythrocytic | Plasma Membrane | other | 2.03E+09 | 1.64E+09 | 1.70E+09 | | | | | | |
| spectrin, beta, erythrocytic | Plasma Membrane | other | 2.02E+09 | 1.71E+09 | 1.61E+09 | | | | | | |
| spectrin, beta, erythrocytic | Plasma Membrane | other | 2.01E+09 | 2.38E+09 | 2.30E+09 | | | | | | |
| spectrin, beta, erythrocytic | Plasma Membrane | other | 2.00E+09 | 1.57E+09 | 1.45E+09 | | | | | | |
| spectrin, beta, erythrocytic | Plasma Membrane | other | 1.94E+09 | 1.55E+09 | 1.78E+09 | | | | | | |
| spectrin, beta, erythrocytic | Plasma Membrane | other | 1.91E+09 | 1.60E+09 | 1.48E+09 | | | | | | |
| spectrin, beta, erythrocytic | Plasma Membrane | other | 1.80E+09 | 1.18E+09 | 1.17E+09 | | | | | | |
| spectrin, beta, erythrocytic | Plasma Membrane | other | 1.77E+09 | 1.24E+09 | 1.32E+09 | | | | | | |
| spectrin, beta, erythrocytic | Plasma Membrane | other | 1.75E+09 | 1.30E+09 | 1.38E+09 | | | | | | |
| spectrin, beta, erythrocytic | Plasma Membrane | other | 1.74E+09 | 1.35E+09 | 1.32E+09 | | | | | | |
| spectrin, beta, erythrocytic | Plasma Membrane | other | 1.73E+09 | 1.45E+09 | 1.24E+09 | | | | | | |
| spectrin, beta, erythrocytic | Plasma Membrane | other | 1.73E+09 | 1.32E+09 | 1.41E+09 | | | | | | |
| spectrin, beta, erythrocytic | Plasma Membrane | other | 1.72E+09 | 1.37E+09 | 1.43E+09 | | | | | | |
| spectrin, beta, erythrocytic | Plasma Membrane | other | 1.71E+09 | 1.18E+09 | 1.37E+09 | | | | | | |
| spectrin, beta, erythrocytic | Plasma Membrane | other | 1.69E+09 | 1.24E+09 | 1.06E+09 | | | | | | |
| spectrin, beta, erythrocytic | Plasma Membrane | other | 1.68E+09 | 1.49E+09 | 1.45E+09 | | | | | | |
| spectrin, beta, erythrocytic | Plasma Membrane | other | 1.63E+09 | 1.14E+09 | 1.07E+09 | | | | | | |
| spectrin, beta, erythrocytic | Plasma Membrane | other | 1.61E+09 | 1.37E+09 | 1.32E+09 | | | | | | |
| spectrin, beta, erythrocytic | Plasma Membrane | other | 1.59E+09 | 1.17E+09 | 1.21E+09 | | | | | | |
| spectrin, beta, erythrocytic | Plasma Membrane | other | 1.58E+09 | 1.32E+09 | 1.35E+09 | | | | | | |
| spectrin, beta, erythrocytic | Plasma Membrane | other | 1.58E+09 | 1.36E+09 | 1.37E+09 | | | | | | |
| spectrin, beta, erythrocytic | Plasma Membrane | other | 1.54E+09 | 1.13E+09 | 1.14E+09 | | | | | | |
| spectrin, beta, erythrocytic | Plasma Membrane | other | 1.48E+09 | 8.73E+08 | 1.18E+09 | | | | | | |
| spectrin, beta, erythrocytic | Plasma Membrane | other | 1.47E+09 | 1.45E+09 | 1.30E+09 | | | | | | |
| spectrin, beta, erythrocytic | Plasma Membrane | other | 1.46E+09 | 1.02E+09 | 1.10E+09 | | | | | | |
| spectrin, beta, erythrocytic | Plasma Membrane | other | 1.41E+09 | 1.72E+09 | 1.53E+09 | | | | | | |
| spectrin, beta, erythrocytic | Plasma Membrane | other | 1.39E+09 | 9.96E+08 | 9.64E+08 | | | | | | |
| spectrin, beta, erythrocytic | Plasma Membrane | other | 1.37E+09 | 1.16E+09 | 1.07E+09 | | | | | | |
| spectrin, beta, erythrocytic | Plasma Membrane | other | 1.35E+09 | 1.17E+09 | 1.32E+09 | | | | | | |
| spectrin, beta, erythrocytic | Plasma Membrane | other | 1.35E+09 | 1.05E+09 | 1.01E+09 | | | | | | |
| spectrin, beta, erythrocytic | Plasma Membrane | other | 1.34E+09 | 1.16E+09 | 1.18E+09 | | | | | | |
| spectrin, beta, erythrocytic | Plasma Membrane | other | 1.33E+09 | 1.09E+09 | 1.07E+09 | | | | | | |
| spectrin, beta, erythrocytic | Plasma Membrane | other | 1.31E+09 | 7.30E+08 | 8.11E+08 | | | | | | |
| spectrin, beta, erythrocytic | Plasma Membrane | other | 1.30E+09 | 1.16E+09 | 1.13E+09 | | | | | | |
| spectrin, beta, erythrocytic | Plasma Membrane | other | 1.28E+09 | 9.35E+08 | 9.29E+08 | | | | | | |
| spectrin, beta, erythrocytic | Plasma Membrane | other | 1.26E+09 | 1.28E+09 | 1.10E+09 | | | | | | |
| spectrin, beta, erythrocytic | Plasma Membrane | other | 1.23E+09 | 9.44E+08 | 1.15E+09 | | | | | | |
| spectrin, beta, erythrocytic | Plasma Membrane | other | 1.21E+09 | 9.49E+08 | 8.54E+08 | | | | | | |
| spectrin, beta, erythrocytic | Plasma Membrane | other | 1.19E+09 | 9.52E+08 | 9.40E+08 | | | | | | |
| spectrin, beta, erythrocytic | Plasma Membrane | other | 1.17E+09 | 9.95E+08 | 9.27E+08 | | | | | | |
| spectrin, beta, erythrocytic | Plasma Membrane | other | 1.16E+09 | 7.98E+08 | 7.91E+08 | | | | | | |
| spectrin, beta, erythrocytic | Plasma Membrane | other | 1.12E+09 | 1.02E+09 | 1.11E+09 | | | | | | |
| spectrin, beta, erythrocytic | Plasma Membrane | other | 1.11E+09 | 8.87E+08 | 7.76E+08 | | | | | | |
| spectrin, beta, erythrocytic | Plasma Membrane | other | 1.10E+09 | 8.83E+08 | 8.76E+08 | | | | | | |
| spectrin, beta, erythrocytic | Plasma Membrane | other | 1.10E+09 | 6.59E+08 | 7.32E+08 | | | | | | |
| spectrin, beta, erythrocytic | Plasma Membrane | other | 1.07E+09 | 5.67E+08 | 6.89E+08 | | | | | | |
| spectrin, beta, erythrocytic | Plasma Membrane | other | 1.06E+09 | 8.51E+08 | 9.29E+08 | | | | | | |
| spectrin, beta, erythrocytic | Plasma Membrane | other | 1.05E+09 | 1.44E+09 | 1.20E+09 | | | | | | |
| spectrin, beta, erythrocytic | Plasma Membrane | other | 1.04E+09 | 8.21E+08 | 9.26E+08 | | | | | | |
| spectrin, beta, erythrocytic | Plasma Membrane | other | 1.02E+09 | 7.50E+08 | 7.74E+08 | | | | | | |
| spectrin, beta, erythrocytic | Plasma Membrane | other | 1.01E+09 | 4.95E+08 | 6.45E+08 | | | | | | |
| spectrin, beta, erythrocytic | Plasma Membrane | other | 9.97E+08 | 7.79E+08 | 8.84E+08 | | | | | | |
| spectrin, beta, erythrocytic | Plasma Membrane | other | 9.27E+08 | 5.14E+08 | 5.64E+08 | | | | | | |
| spectrin, beta, erythrocytic | Plasma Membrane | other | 9.24E+08 | 8.47E+08 | 8.06E+08 | | | | | | |
| spectrin, beta, erythrocytic | Plasma Membrane | other | 9.09E+08 | 7.06E+08 | 6.36E+08 | | | | | | |
| spectrin, beta, erythrocytic | Plasma Membrane | other | 8.94E+08 | 6.97E+08 | 6.39E+08 | | | | | | |
| spectrin, beta, erythrocytic | Plasma Membrane | other | 8.85E+08 | 6.42E+08 | 6.14E+08 | | | | | | |
| spectrin, beta, erythrocytic | Plasma Membrane | other | 8.56E+08 | 3.60E+08 | 6.82E+08 | | | | | | |
| spectrin, beta, erythrocytic | Plasma Membrane | other | 8.41E+08 | 6.83E+08 | 6.68E+08 | | | | | | |
| spectrin, beta, erythrocytic | Plasma Membrane | other | 8.35E+08 | 1.15E+09 | 1.01E+09 | | | | | | |
| spectrin, beta, erythrocytic | Plasma Membrane | other | 8.28E+08 | 1.04E+09 | 9.84E+08 | | | | | | |
| spectrin, beta, erythrocytic | Plasma Membrane | other | 8.27E+08 | 5.28E+08 | 6.48E+08 | | | | | | |
| spectrin, beta, erythrocytic | Plasma Membrane | other | 8.23E+08 | 8.03E+08 | 8.01E+08 | | | | | | |
| spectrin, beta, erythrocytic | Plasma Membrane | other | 8.21E+08 | 1.09E+09 | 9.05E+08 | | | | | | |
| spectrin, beta, erythrocytic | Plasma Membrane | other | 8.20E+08 | 3.42E+08 | 4.08E+08 | | | | | | |
| spectrin, beta, erythrocytic | Plasma Membrane | other | 7.88E+08 | 5.39E+08 | 6.45E+08 | | | | | | |

| | | | | | |
|---|---|---|---|---|---|
| spectrin, beta, erythrocytic | Plasma Membrane | other | 7.69E+08 | 6.43E+08 | 5.98E+08 |
| spectrin, beta, erythrocytic | Plasma Membrane | other | 7.29E+08 | 4.42E+08 | 4.32E+08 |
| spectrin, beta, erythrocytic | Plasma Membrane | other | 7.10E+08 | 6.98E+08 | 7.55E+08 |
| spectrin, beta, erythrocytic | Plasma Membrane | other | 7.02E+08 | 5.86E+08 | 6.17E+08 |
| spectrin, beta, erythrocytic | Plasma Membrane | other | 6.85E+08 | 5.62E+08 | 6.08E+08 |
| spectrin, beta, erythrocytic | Plasma Membrane | other | 6.80E+08 | 4.43E+08 | 5.35E+08 |
| spectrin, beta, erythrocytic | Plasma Membrane | other | 6.69E+08 | 7.56E+08 | 5.39E+08 |
| spectrin, beta, erythrocytic | Plasma Membrane | other | 6.68E+08 | 6.43E+08 | 5.76E+08 |
| spectrin, beta, erythrocytic | Plasma Membrane | other | 6.41E+08 | 9.44E+08 | 4.97E+08 |
| spectrin, beta, erythrocytic | Plasma Membrane | other | 6.37E+08 | 4.04E+08 | 4.83E+08 |
| spectrin, beta, erythrocytic | Plasma Membrane | other | 6.35E+08 | 2.68E+08 | 3.15E+08 |
| spectrin, beta, erythrocytic | Plasma Membrane | other | 6.32E+08 | 8.27E+08 | 8.61E+08 |
| spectrin, beta, erythrocytic | Plasma Membrane | other | 6.09E+08 | 5.98E+08 | 6.66E+08 |
| spectrin, beta, erythrocytic | Plasma Membrane | other | 5.70E+08 | 4.74E+08 | 5.81E+08 |
| spectrin, beta, erythrocytic | Plasma Membrane | other | 5.53E+08 | 4.18E+08 | 4.15E+08 |
| spectrin, beta, erythrocytic | Plasma Membrane | other | 5.45E+08 | 4.73E+08 | 4.64E+08 |
| spectrin, beta, erythrocytic | Plasma Membrane | other | 5.41E+08 | 1.57E+09 | 1.21E+09 |
| spectrin, beta, erythrocytic | Plasma Membrane | other | 5.38E+08 | 6.06E+08 | 5.21E+08 |
| spectrin, beta, erythrocytic | Plasma Membrane | other | 5.10E+08 | 5.69E+08 | 5.62E+08 |
| spectrin, beta, erythrocytic | Plasma Membrane | other | 5.02E+08 | 3.93E+08 | 3.53E+08 |
| spectrin, beta, erythrocytic | Plasma Membrane | other | 4.82E+08 | 6.18E+08 | 4.83E+08 |
| spectrin, beta, erythrocytic | Plasma Membrane | other | 4.76E+08 | 5.17E+08 | 5.70E+08 |
| spectrin, beta, erythrocytic | Plasma Membrane | other | 4.54E+08 | 2.33E+08 | 2.40E+08 |
| spectrin, beta, erythrocytic | Plasma Membrane | other | 4.52E+08 | 3.70E+08 | 3.75E+08 |
| spectrin, beta, erythrocytic | Plasma Membrane | other | 4.19E+08 | 2.35E+08 | 3.90E+08 |
| spectrin, beta, erythrocytic | Plasma Membrane | other | 4.09E+08 | 4.07E+08 | 4.90E+08 |
| spectrin, beta, erythrocytic | Plasma Membrane | other | 4.06E+08 | 1.85E+08 | 2.11E+08 |
| spectrin, beta, erythrocytic | Plasma Membrane | other | 4.06E+08 | 2.91E+08 | 2.53E+08 |
| spectrin, beta, erythrocytic | Plasma Membrane | other | 4.06E+08 | 3.38E+08 | 2.79E+08 |
| spectrin, beta, erythrocytic | Plasma Membrane | other | 3.88E+08 | 3.15E+08 | 2.91E+08 |
| spectrin, beta, erythrocytic | Plasma Membrane | other | 3.67E+08 | 6.34E+08 | 4.92E+08 |
| spectrin, beta, erythrocytic | Plasma Membrane | other | 3.65E+08 | 2.71E+08 | 2.10E+08 |
| spectrin, beta, erythrocytic | Plasma Membrane | other | 3.63E+08 | 2.35E+08 | 3.02E+08 |
| spectrin, beta, erythrocytic | Plasma Membrane | other | 3.52E+08 | 3.21E+08 | 2.62E+08 |
| spectrin, beta, erythrocytic | Plasma Membrane | other | 3.16E+08 | 4.42E+08 | 3.56E+08 |
| spectrin, beta, erythrocytic | Plasma Membrane | other | 2.86E+08 | 1.96E+08 | 1.44E+08 |
| spectrin, beta, erythrocytic | Plasma Membrane | other | 2.85E+08 | 2.30E+08 | 1.96E+08 |
| spectrin, beta, erythrocytic | Plasma Membrane | other | 2.74E+08 | 1.98E+08 | 2.05E+08 |
| spectrin, beta, erythrocytic | Plasma Membrane | other | 2.72E+08 | 2.48E+08 | 2.34E+08 |
| spectrin, beta, erythrocytic | Plasma Membrane | other | 2.69E+08 | 2.64E+08 | 2.56E+08 |
| spectrin, beta, erythrocytic | Plasma Membrane | other | 2.66E+08 | 2.17E+08 | 2.57E+08 |
| spectrin, beta, erythrocytic | Plasma Membrane | other | 2.66E+08 | 1.29E+08 | 2.25E+08 |
| spectrin, beta, erythrocytic | Plasma Membrane | other | 2.57E+08 | 1.70E+08 | 1.95E+08 |
| spectrin, beta, erythrocytic | Plasma Membrane | other | 2.53E+08 | 4.10E+08 | 4.67E+08 |
| spectrin, beta, erythrocytic | Plasma Membrane | other | 2.43E+08 | 4.37E+08 | 4.02E+08 |
| spectrin, beta, erythrocytic | Plasma Membrane | other | 2.39E+08 | 1.64E+08 | 2.10E+08 |
| spectrin, beta, erythrocytic | Plasma Membrane | other | 2.35E+08 | 2.66E+08 | 2.17E+08 |
| spectrin, beta, erythrocytic | Plasma Membrane | other | 2.32E+08 | 1.54E+08 | 2.38E+08 |
| spectrin, beta, erythrocytic | Plasma Membrane | other | 2.21E+08 | 1.83E+08 | 1.81E+08 |
| spectrin, beta, erythrocytic | Plasma Membrane | other | 2.15E+08 | 1.95E+08 | 2.39E+08 |
| spectrin, beta, erythrocytic | Plasma Membrane | other | 2.00E+08 | 1.90E+08 | 3.21E+08 |
| spectrin, beta, erythrocytic | Plasma Membrane | other | 1.94E+08 | 1.06E+08 | 1.49E+08 |
| spectrin, beta, erythrocytic | Plasma Membrane | other | 1.74E+08 | 1.46E+08 | 2.45E+08 |
| spectrin, beta, erythrocytic | Plasma Membrane | other | 1.70E+08 | 8.67E+07 | 8.75E+07 |
| spectrin, beta, erythrocytic | Plasma Membrane | other | 1.59E+08 | 3.29E+08 | 2.28E+08 |
| spectrin, beta, erythrocytic | Plasma Membrane | other | 1.59E+08 | 9.80E+07 | 1.31E+08 |
| spectrin, beta, erythrocytic | Plasma Membrane | other | 1.58E+08 | 1.13E+08 | 1.13E+08 |
| spectrin, beta, erythrocytic | Plasma Membrane | other | 1.57E+08 | 1.23E+08 | 1.18E+08 |
| spectrin, beta, erythrocytic | Plasma Membrane | other | 1.54E+08 | 8.38E+07 | 6.96E+07 |
| spectrin, beta, erythrocytic | Plasma Membrane | other | 1.53E+08 | 1.03E+08 | 8.10E+07 |
| spectrin, beta, erythrocytic | Plasma Membrane | other | 1.53E+08 | 1.45E+08 | 1.19E+08 |
| spectrin, beta, erythrocytic | Plasma Membrane | other | 1.52E+08 | 2.15E+08 | 1.97E+08 |
| spectrin, beta, erythrocytic | Plasma Membrane | other | 1.46E+08 | 2.73E+08 | 2.19E+08 |
| spectrin, beta, erythrocytic | Plasma Membrane | other | 1.39E+08 | 7.99E+07 | 5.77E+07 |
| spectrin, beta, erythrocytic | Plasma Membrane | other | 1.34E+08 | 2.71E+08 | 1.88E+08 |
| spectrin, beta, erythrocytic | Plasma Membrane | other | 1.29E+08 | 1.75E+08 | 2.07E+08 |
| spectrin, beta, erythrocytic | Plasma Membrane | other | 1.25E+08 | 9.08E+07 | 9.78E+07 |
| spectrin, beta, erythrocytic | Plasma Membrane | other | 1.24E+08 | 6.36E+07 | 8.34E+07 |
| spectrin, beta, erythrocytic | Plasma Membrane | other | 1.22E+08 | 8.28E+07 | 7.60E+07 |
| spectrin, beta, erythrocytic | Plasma Membrane | other | 1.21E+08 | 8.96E+07 | 9.50E+07 |
| spectrin, beta, erythrocytic | Plasma Membrane | other | 1.20E+08 | 7.43E+07 | 8.04E+07 |
| spectrin, beta, erythrocytic | Plasma Membrane | other | 1.19E+08 | 1.00E+08 | 8.88E+07 |
| spectrin, beta, erythrocytic | Plasma Membrane | other | 1.18E+08 | 7.31E+07 | 7.64E+07 |
| spectrin, beta, erythrocytic | Plasma Membrane | other | 1.16E+08 | 1.37E+08 | 8.17E+07 |
| spectrin, beta, erythrocytic | Plasma Membrane | other | 1.15E+08 | 6.57E+07 | 1.17E+08 |
| spectrin, beta, erythrocytic | Plasma Membrane | other | 1.14E+08 | 4.63E+07 | 4.92E+07 |
| spectrin, beta, erythrocytic | Plasma Membrane | other | 1.13E+08 | 1.22E+08 | 1.11E+08 |
| spectrin, beta, erythrocytic | Plasma Membrane | other | 1.06E+08 | 6.96E+07 | 5.85E+07 |
| spectrin, beta, erythrocytic | Plasma Membrane | other | 9.56E+07 | 1.46E+08 | 1.67E+08 |
| spectrin, beta, erythrocytic | Plasma Membrane | other | 9.47E+07 | 6.47E+07 | 6.31E+07 |
| spectrin, beta, erythrocytic | Plasma Membrane | other | 9.37E+07 | 8.39E+07 | 1.16E+08 |
| spectrin, beta, erythrocytic | Plasma Membrane | other | 9.36E+07 | 1.22E+08 | 9.00E+07 |
| spectrin, beta, erythrocytic | Plasma Membrane | other | 9.29E+07 | 1.98E+08 | 1.69E+08 |
| spectrin, beta, erythrocytic | Plasma Membrane | other | 9.15E+07 | 1.64E+08 | 1.58E+08 |
| spectrin, beta, erythrocytic | Plasma Membrane | other | 9.04E+07 | 4.55E+07 | 7.49E+07 |
| spectrin, beta, erythrocytic | Plasma Membrane | other | 9.01E+07 | 2.59E+07 | 5.51E+07 |
| spectrin, beta, erythrocytic | Plasma Membrane | other | 8.65E+07 | 2.68E+08 | 2.57E+08 |
| spectrin, beta, erythrocytic | Plasma Membrane | other | 8.58E+07 | 7.41E+07 | 8.13E+07 |
| spectrin, beta, erythrocytic | Plasma Membrane | other | 8.29E+07 | 1.27E+08 | 8.22E+07 |
| spectrin, beta, erythrocytic | Plasma Membrane | other | 7.83E+07 | 1.88E+08 | 1.46E+08 |
| spectrin, beta, erythrocytic | Plasma Membrane | other | 7.56E+07 | 9.09E+07 | 7.66E+07 |
| spectrin, beta, erythrocytic | Plasma Membrane | other | 7.53E+07 | 2.20E+07 | 6.61E+07 |
| spectrin, beta, erythrocytic | Plasma Membrane | other | 7.44E+07 | 2.40E+08 | 1.24E+08 |
| spectrin, beta, erythrocytic | Plasma Membrane | other | 7.17E+07 | 4.05E+07 | 6.51E+07 |
| spectrin, beta, erythrocytic | Plasma Membrane | other | 6.85E+07 | 2.69E+07 | 2.97E+07 |

| | | | | | | | | | | | |
|---|---|---|---|---|---|---|---|---|---|---|---|
| spectrin, beta, erythrocytic | Plasma Membrane | other | 6.69E+07 | 2.98E+07 | 4.67E+07 | | | | | | |
| spectrin, beta, erythrocytic | Plasma Membrane | other | 6.66E+07 | 4.07E+07 | 4.18E+07 | | | | | | |
| spectrin, beta, erythrocytic | Plasma Membrane | other | 6.64E+07 | 4.39E+07 | 5.21E+07 | | | | | | |
| spectrin, beta, erythrocytic | Plasma Membrane | other | 6.63E+07 | 4.17E+07 | 6.43E+07 | | | | | | |
| spectrin, beta, erythrocytic | Plasma Membrane | other | 6.58E+07 | 1.20E+08 | 6.97E+07 | | | | | | |
| spectrin, beta, erythrocytic | Plasma Membrane | other | 6.52E+07 | 4.69E+07 | 5.19E+07 | | | | | | |
| spectrin, beta, erythrocytic | Plasma Membrane | other | 6.15E+07 | 4.53E+07 | 3.94E+07 | | | | | | |
| spectrin, beta, erythrocytic | Plasma Membrane | other | 5.94E+07 | 4.54E+07 | 5.19E+07 | | | | | | |
| spectrin, beta, erythrocytic | Plasma Membrane | other | 5.88E+07 | 3.62E+07 | 3.54E+07 | | | | | | |
| spectrin, beta, erythrocytic | Plasma Membrane | other | 5.80E+07 | 2.53E+07 | 4.47E+07 | | | | | | |
| spectrin, beta, erythrocytic | Plasma Membrane | other | 5.75E+07 | 2.97E+07 | 4.73E+07 | | | | | | |
| spectrin, beta, erythrocytic | Plasma Membrane | other | 5.71E+07 | 4.04E+07 | 1.06E+08 | | | | | | |
| spectrin, beta, erythrocytic | Plasma Membrane | other | 5.70E+07 | 2.96E+07 | 3.66E+07 | | | | | | |
| spectrin, beta, erythrocytic | Plasma Membrane | other | 5.28E+07 | 3.98E+07 | 4.10E+07 | | | | | | |
| spectrin, beta, erythrocytic | Plasma Membrane | other | 5.02E+07 | 1.85E+07 | 2.73E+07 | | | | | | |
| spectrin, beta, erythrocytic | Plasma Membrane | other | 4.99E+07 | 2.45E+07 | 4.01E+07 | | | | | | |
| spectrin, beta, erythrocytic | Plasma Membrane | other | 4.87E+07 | 4.56E+07 | 4.54E+07 | | | | | | |
| spectrin, beta, erythrocytic | Plasma Membrane | other | 4.84E+07 | 1.31E+07 | 2.49E+07 | | | | | | |
| spectrin, beta, erythrocytic | Plasma Membrane | other | 4.40E+07 | 2.37E+07 | 3.46E+07 | | | | | | |
| spectrin, beta, erythrocytic | Plasma Membrane | other | 4.14E+07 | 2.27E+07 | 3.14E+07 | | | | | | |
| spectrin, beta, erythrocytic | Plasma Membrane | other | 4.07E+07 | 9440227 | 1.70E+07 | | | | | | |
| spectrin, beta, erythrocytic | Plasma Membrane | other | 3.98E+07 | 3.82E+07 | 6.62E+07 | | | | | | |
| spectrin, beta, erythrocytic | Plasma Membrane | other | 3.51E+07 | 4.57E+07 | 3.17E+07 | | | | | | |
| spectrin, beta, erythrocytic | Plasma Membrane | other | 3.28E+07 | 1.42E+07 | 1.64E+07 | | | | | | |
| spectrin, beta, erythrocytic | Plasma Membrane | other | 3.28E+07 | 1.38E+08 | 6.78E+07 | | | | | | |
| spectrin, beta, erythrocytic | Plasma Membrane | other | 3.21E+07 | 2.63E+07 | 2.33E+07 | | | | | | |
| spectrin, beta, erythrocytic | Plasma Membrane | other | 3.08E+07 | 1.49E+07 | 2.05E+07 | | | | | | |
| spectrin, beta, erythrocytic | Plasma Membrane | other | 2.92E+07 | 2.19E+07 | 2.02E+07 | | | | | | |
| spectrin, beta, erythrocytic | Plasma Membrane | other | 2.85E+07 | 1.18E+07 | 1.34E+07 | | | | | | |
| spectrin, beta, erythrocytic | Plasma Membrane | other | 2.73E+07 | 4.96E+07 | 5.71E+07 | | | | | | |
| spectrin, beta, erythrocytic | Plasma Membrane | other | 2.50E+07 | 1.27E+07 | 1.38E+07 | | | | | | |
| spectrin, beta, erythrocytic | Plasma Membrane | other | 2.44E+07 | 1.43E+07 | 1.73E+07 | | | | | | |
| spectrin, beta, erythrocytic | Plasma Membrane | other | 2.39E+07 | 2.83E+07 | 2.63E+07 | | | | | | |
| spectrin, beta, erythrocytic | Plasma Membrane | other | 2.31E+07 | 9202722 | 1.48E+07 | | | | | | |
| spectrin, beta, erythrocytic | Plasma Membrane | other | 2.23E+07 | 1.09E+08 | 5.93E+07 | | | | | | |
| spectrin, beta, erythrocytic | Plasma Membrane | other | 2.12E+07 | 1.06E+07 | 1.84E+07 | | | | | | |
| spectrin, beta, erythrocytic | Plasma Membrane | other | 2.11E+07 | 3.03E+07 | 2.21E+07 | | | | | | |
| spectrin, beta, erythrocytic | Plasma Membrane | other | 2.08E+07 | 9.27E+07 | 4.45E+07 | | | | | | |
| spectrin, beta, erythrocytic | Plasma Membrane | other | 1.96E+07 | 7.21E+07 | 4.12E+07 | | | | | | |
| spectrin, beta, erythrocytic | Plasma Membrane | other | 1.95E+07 | 1.18E+07 | 1.32E+07 | | | | | | |
| spectrin, beta, erythrocytic | Plasma Membrane | other | 1.93E+07 | 1.07E+07 | 9284441 | | | | | | |
| spectrin, beta, erythrocytic | Plasma Membrane | other | 1.91E+07 | 6584300 | 1.26E+07 | | | | | | |
| spectrin, beta, erythrocytic | Plasma Membrane | other | 1.69E+07 | 5.02E+07 | 2.16E+07 | | | | | | |
| spectrin, beta, erythrocytic | Plasma Membrane | other | 1.66E+07 | 1.91E+07 | 2.33E+07 | | | | | | |
| spectrin, beta, erythrocytic | Plasma Membrane | other | 1.62E+07 | 7.30E+07 | 2.03E+07 | | | | | | |
| spectrin, beta, erythrocytic | Plasma Membrane | other | 1.45E+07 | 7583573 | 8726900 | | | | | | |
| spectrin, beta, erythrocytic | Plasma Membrane | other | 1.42E+07 | 2.57E+07 | 5.10E+07 | | | | | | |
| spectrin, beta, erythrocytic | Plasma Membrane | other | 1.36E+07 | 9306575 | 9777560 | | | | | | |
| spectrin, beta, erythrocytic | Plasma Membrane | other | 1.30E+07 | 1.40E+07 | 1.44E+07 | | | | | | |
| spectrin, beta, erythrocytic | Plasma Membrane | other | 1.23E+07 | 9727163 | 7936244 | | | | | | |
| spectrin, beta, erythrocytic | Plasma Membrane | other | 1.13E+07 | 1.33E+07 | 1.16E+07 | | | | | | |
| spectrin, beta, erythrocytic | Plasma Membrane | other | 1.01E+07 | 7092247.5 | 9556386 | | | | | | |
| spectrin, beta, erythrocytic | Plasma Membrane | other | 9851263 | 1.02E+07 | 1.23E+07 | | | | | | |
| spectrin, beta, erythrocytic | Plasma Membrane | other | 9670547 | 2919813.75 | 4702089.5 | | | | | | |
| spectrin, beta, erythrocytic | Plasma Membrane | other | 9181023 | 4.63E+07 | 1.16E+07 | | | | | | |
| spectrin, beta, erythrocytic | Plasma Membrane | other | 8631547 | 9438255 | 8251516.5 | | | | | | |
| spectrin, beta, erythrocytic | Plasma Membrane | other | 8176951 | 4065597.25 | 4575587 | | | | | | |
| spectrin, beta, erythrocytic | Plasma Membrane | other | 7683298.5 | 3.28E+07 | 1.35E+07 | | | | | | |
| spectrin, beta, erythrocytic | Plasma Membrane | other | 7044581.5 | 4.42E+07 | 2.95E+07 | | | | | | |
| spectrin, beta, erythrocytic | Plasma Membrane | other | 6408492.5 | 1.52E+07 | 6346535 | | | | | | |
| spectrin, beta, erythrocytic | Plasma Membrane | other | 5986933.5 | 5350355.5 | 8173324 | | | | | | |
| spectrin, beta, erythrocytic | Plasma Membrane | other | 5961743 | 3.19E+07 | 1.57E+07 | | | | | | |
| spectrin, beta, erythrocytic | Plasma Membrane | other | 5159904.5 | 2354522 | 4793718 | | | | | | |
| spectrin, beta, erythrocytic | Plasma Membrane | other | 5147344.5 | 5565584 | 5562855 | | | | | | |
| spectrin, beta, erythrocytic | Plasma Membrane | other | 5097712.5 | 2.21E+07 | 1.24E+07 | | | | | | |
| spectrin, beta, erythrocytic | Plasma Membrane | other | 4718127.5 | 4205900.5 | 1713789.75 | | | | | | |
| spectrin, beta, erythrocytic | Plasma Membrane | other | 4497307 | 4857263 | 4624020.5 | | | | | | |
| spectrin, beta, erythrocytic | Plasma Membrane | other | 4090346.75 | 3956568 | 5338284.5 | | | | | | |
| spectrin, beta, erythrocytic | Plasma Membrane | other | 3972297 | 2384643.75 | 2304146.5 | | | | | | |
| spectrin, beta, erythrocytic | Plasma Membrane | other | 3467179.25 | 1088291.75 | 3673008 | | | | | | |
| spectrin, beta, erythrocytic | Plasma Membrane | other | 2840383.75 | 2740179 | 2575154.25 | | | | | | |
| spectrin, beta, erythrocytic | Plasma Membrane | other | 2763934.25 | 778263.062 | 1596068.125 | | | | | | |
| spectrin, beta, erythrocytic | Plasma Membrane | other | 2596079.75 | 656939.438 | 1809512.125 | | | | | | |
| spectrin, beta, erythrocytic | Plasma Membrane | other | 2423395.5 | 1.34E+07 | 6475844.5 | | | | | | |
| spectrin, beta, erythrocytic | Plasma Membrane | other | 2379127.25 | 2505003 | 2367243.25 | | | | | | |
| spectrin, beta, erythrocytic | Plasma Membrane | other | 2332556.75 | 1.12E+07 | 2264005.25 | | | | | | |
| spectrin, beta, erythrocytic | Plasma Membrane | other | 2068229.75 | 356894.844 | 1261092.5 | | | | | | |
| spectrin, beta, erythrocytic | Plasma Membrane | other | 2060667.5 | 4.61E+07 | 8.60E+07 | | | | | | |
| spectrin, beta, erythrocytic | Plasma Membrane | other | 1432787.375 | 1.17E+07 | 3686150.5 | | | | | | |
| spectrin, beta, erythrocytic | Plasma Membrane | other | 1387410.375 | 271934.469 | 15690.703 | | | | | | |
| spectrin, beta, erythrocytic | Plasma Membrane | other | 1336797 | 841504.625 | 624090.625 | | | | | | |
| spectrin, beta, erythrocytic | Plasma Membrane | other | 1041942.688 | 5281233.5 | 2401098.25 | | | | | | |
| spectrin, beta, erythrocytic | Plasma Membrane | other | 723983.75 | 4033241.5 | 269771.906 | | | | | | |
| spectrin, beta, erythrocytic | Plasma Membrane | other | 697869.625 | 2051237 | 3077381.25 | | | | | | |
| spectrin, beta, erythrocytic | Plasma Membrane | other | 429840.344 | 5539450.5 | 935962 | | | | | | |
| spectrin, beta, erythrocytic | Plasma Membrane | other | 272864.781 | 916674.875 | 1286247.25 | | | | | | |
| spectrin, beta, erythrocytic | Plasma Membrane | other | 49291.703 | 160635.609 | 164116.812 | | | | | | |
| spectrin, beta, erythrocytic | Plasma Membrane | other | 0 | 61054.828 | 174225.203 | P15508 | SPTB | 559793822 | 631071029 | 22595968 | 780 |
| serglycin | Cytoplasm | other | 277887.25 | 3070085.5 | 3149127.25 | P13609 | Srgn | 277887 | 1635371 | 944182 | 3 |
| signal sequence receptor, alpha | Cytoplasm | other | 1831939.75 | 3583661.25 | 3659655.75 | | | | | | |
| signal sequence receptor, alpha | Cytoplasm | other | 1601641.875 | 1977947.875 | 1796424.375 | Q9CY50 | SSR1 | 1716791 | 947613 | 386861 | 6 |
| signal sequence receptor, gamma (translocon-associated protein gamma) | Cytoplasm | other | 2728234.5 | 6370599.5 | 4786062 | Q9DCF9 | SSR3 | 2728235 | 1826300 | 1054415 | 3 |
| stromal antigen 3 | Nucleus | other | 6166069.5 | 4175968.5 | 4059589.25 | O70576 | STAG3 | 6166070 | 1184012 | 683590 | 3 |
| signal transducing adaptor molecule (SH3 domain and ITAM motif) 1 | Cytoplasm | other | 8653050 | 4511615.5 | 5955188 | | | | | | |
| signal transducing adaptor molecule (SH3 domain and ITAM motif) 1 | Cytoplasm | other | 4791639.5 | 2109555 | 2935694 | | | | | | |

| | | | | | | | | | | | |
|---|---|---|---|---|---|---|---|---|---|---|---|
| signal transducing adaptor molecule (SH3 domain and ITAM motif) 1 | Cytoplasm | other | 3140082.25 | 2153273 | 1820841.375 | P70297 | STAM | 5528257 | 2235135 | 745045 | 9 |
| STEAP family member 3, metalloreductase | Cytoplasm | transporter | 9005202 | 5600681 | 1.04E+07 | | | | | | |
| STEAP family member 3, metalloreductase | Cytoplasm | transporter | 5495389.5 | 3334006.5 | 6070836.5 | | | | | | |
| STEAP family member 3, metalloreductase | Cytoplasm | transporter | 4225756.5 | 3046228.75 | 4593610.5 | | | | | | |
| STEAP family member 3, metalloreductase | Cytoplasm | transporter | 2954660.25 | 1858324.125 | 3330022.75 | | | | | | |
| STEAP family member 3, metalloreductase | Cytoplasm | transporter | 2934896.75 | 2477417.75 | 2677431.75 | Q8CI59 | STEAP3 | 4923181 | 2444818 | 631249 | 15 |
| stromal interaction molecule 1 | Plasma Membrane | ion channel | 329794.844 | 3503194 | 2440671.5 | | | | | | |
| stromal interaction molecule 1 | Plasma Membrane | ion channel | 264631.062 | 3087539.25 | 2872223.25 | | | | | | |
| stromal interaction molecule 1 | Plasma Membrane | ion channel | 254752.391 | 2193743.5 | 1973782.25 | P70302 | STIM1 | 283059 | 1282234 | 427411 | 9 |
| stress-induced phosphoprotein 1 | Cytoplasm | other | 1.12E+07 | 3851257.75 | 4566511.5 | | | | | | |
| stress-induced phosphoprotein 1 | Cytoplasm | other | 9426192 | 5419133 | 5130110.5 | | | | | | |
| stress-induced phosphoprotein 1 | Cytoplasm | other | 8068653.5 | 5500118.5 | 4101820.5 | | | | | | |
| stress-induced phosphoprotein 1 | Cytoplasm | other | 6260908 | 2373041.5 | 2633718.5 | | | | | | |
| stress-induced phosphoprotein 1 | Cytoplasm | other | 5982993.5 | 4004938.5 | 3420280.25 | | | | | | |
| stress-induced phosphoprotein 1 | Cytoplasm | other | 3967539.75 | 2512738.25 | 1855480.625 | | | | | | |
| stress-induced phosphoprotein 1 | Cytoplasm | other | 3867120 | 1623222.75 | 1264958.375 | | | | | | |
| stress-induced phosphoprotein 1 | Cytoplasm | other | 3754855.5 | 1473996.5 | 1594058.5 | | | | | | |
| stress-induced phosphoprotein 1 | Cytoplasm | other | 3294536.25 | 745268.125 | 935796.75 | | | | | | |
| stress-induced phosphoprotein 1 | Cytoplasm | other | 3155226.5 | 2279857.75 | 1759272.125 | Q60864 | STIP1 | 5892803 | 2475608 | 451982 | 30 |
| stomatin | Plasma Membrane | other | 6.92E+08 | 6.04E+08 | 4.58E+08 | | | | | | |
| stomatin | Plasma Membrane | other | 6.75E+08 | 5.66E+08 | 5.49E+08 | | | | | | |
| stomatin | Plasma Membrane | other | 6.18E+08 | 5.47E+08 | 4.62E+08 | | | | | | |
| stomatin | Plasma Membrane | other | 5.75E+08 | 5.23E+08 | 2.92E+08 | | | | | | |
| stomatin | Plasma Membrane | other | 4.75E+08 | 3.47E+08 | 3.59E+08 | | | | | | |
| stomatin | Plasma Membrane | other | 4.74E+08 | 3.68E+08 | 2.39E+08 | | | | | | |
| stomatin | Plasma Membrane | other | 4.69E+08 | 3.78E+08 | 4.59E+08 | | | | | | |
| stomatin | Plasma Membrane | other | 2.80E+08 | 1.79E+08 | 2.22E+08 | | | | | | |
| stomatin | Plasma Membrane | other | 2.19E+08 | 1.85E+08 | 1.77E+08 | | | | | | |
| stomatin | Plasma Membrane | other | 2.07E+08 | 2.34E+08 | 3.59E+08 | | | | | | |
| stomatin | Plasma Membrane | other | 1.99E+08 | 2.35E+08 | 8.94E+08 | | | | | | |
| stomatin | Plasma Membrane | other | 8.86E+07 | 9.03E+07 | 8.27E+07 | | | | | | |
| stomatin | Plasma Membrane | other | 3.07E+07 | 2.73E+07 | 2.21E+07 | | | | | | |
| stomatin | Plasma Membrane | other | 2.90E+07 | 2.20E+07 | 1.50E+07 | | | | | | |
| stomatin | Plasma Membrane | other | 8687961 | 7295523.5 | 5999620.5 | | | | | | |
| stomatin | Plasma Membrane | other | 8482439 | 4940806.5 | 2405167 | | | | | | |
| stomatin | Plasma Membrane | other | 5776484 | 5203781.5 | 7038128.5 | | | | | | |
| stomatin | Plasma Membrane | other | 3628716.25 | 2697695.5 | 1324542.5 | | | | | | |
| stomatin | Plasma Membrane | other | 3559526.5 | 897572.938 | 591724.75 | | | | | | |
| stomatin | Plasma Membrane | other | 2405619.25 | 2068880.375 | 3016710.25 | | | | | | |
| stomatin | Plasma Membrane | other | 1683754.25 | 7069409.5 | 8899089 | | | | | | |
| stomatin | Plasma Membrane | other | 1045633.688 | 391800.312 | 579273.625 | | | | | | |
| stomatin | Plasma Membrane | other | 564140.812 | 530958.188 | 547122.875 | P54116 | STOM | 220258012 | 234826387 | 28269775 | 69 |
| stomatin (EPB72)-like 2 | Plasma Membrane | other | 2921652.75 | 2905684.5 | 3238994.25 | | | | | | |
| stomatin (EPB72)-like 2 | Plasma Membrane | other | 1377458.5 | 2562020.5 | 1762085.875 | | | | | | |
| stomatin (EPB72)-like 2 | Plasma Membrane | other | 1103907.625 | 2211608.75 | 2405006.25 | | | | | | |
| stomatin (EPB72)-like 2 | Plasma Membrane | other | 816028.375 | 1225192.375 | 1477721.5 | | | | | | |
| stomatin (EPB72)-like 2 | Plasma Membrane | other | 572193.25 | 847935.562 | 797284.688 | Q99JB2 | STOML2 | 1358248 | 890978 | 230049 | 15 |
| syntaxin 11 | Plasma Membrane | transporter | 713114.562 | 4595557.5 | 2282529.75 | | | | | | |
| syntaxin 11 | Plasma Membrane | transporter | 708047.25 | 2858058.25 | 1248121.625 | | | | | | |
| syntaxin 11 | Plasma Membrane | transporter | 624061 | 2944448.5 | 1078023.5 | | | | | | |
| syntaxin 11 | Plasma Membrane | transporter | 259283.359 | 2396688.25 | 1539475.125 | | | | | | |
| syntaxin 11 | Plasma Membrane | transporter | 258841.797 | 2203049 | 2048723.25 | | | | | | |
| syntaxin 11 | Plasma Membrane | transporter | 39293.473 | 1266343.25 | 881853.75 | Q9D3G5 | STX11 | 433774 | 1177791 | 277608 | 18 |
| syntaxin 12 | Cytoplasm | other | 1.13E+07 | 1.30E+07 | 1.10E+07 | Q9ER00 | STX12 | 11270000 | 1073732 | 619919 | 3 |
| syntaxin 2 | Cytoplasm | transporter | 4996394.5 | 2974832.5 | 3696763 | | | | | | |
| syntaxin 2 | Cytoplasm | transporter | 4742845.5 | 7080189.5 | 5291847.5 | | | | | | |
| syntaxin 2 | Cytoplasm | transporter | 3494429 | 2482218.75 | 3453191.75 | | | | | | |
| syntaxin 2 | Cytoplasm | transporter | 2111600.75 | 3112292.25 | 1555032.375 | | | | | | |
| syntaxin 2 | Cytoplasm | transporter | 1427190.875 | 1634440.125 | 1308553.125 | | | | | | |
| syntaxin 2 | Cytoplasm | transporter | 1276799 | 1552860.625 | 1110721.75 | Q00262 | STX2 | 3008210 | 1694678 | 399440 | 18 |
| syntaxin 4 | Plasma Membrane | transporter | 1999517.125 | 4056299.25 | 3272125.25 | | | | | | |
| syntaxin 4 | Plasma Membrane | transporter | 992741 | 1647864.625 | 1375674.5 | | | | | | |
| syntaxin 4 | Plasma Membrane | transporter | 940115.938 | 2323006.75 | 1865019 | P70452 | STX4 | 1310791 | 1036589 | 345530 | 9 |
| syntaxin 7 | Plasma Membrane | transporter | 1.34E+07 | 1.38E+07 | 1.23E+07 | | | | | | |
| syntaxin 7 | Plasma Membrane | transporter | 1.31E+07 | 1.31E+07 | 1.10E+07 | | | | | | |
| syntaxin 7 | Plasma Membrane | transporter | 8846038 | 9140734 | 1.03E+07 | | | | | | |
| syntaxin 7 | Plasma Membrane | transporter | 8246318.5 | 8919611 | 7507498.5 | | | | | | |
| syntaxin 7 | Plasma Membrane | transporter | 7810398 | 7230238.5 | 8336821 | | | | | | |
| syntaxin 7 | Plasma Membrane | transporter | 4601965 | 4192291 | 4079233.75 | O70439 | STX7 | 9330787 | 3127184 | 737084 | 18 |
| syntaxin binding protein 1 | Cytoplasm | transporter | 3781331.25 | 3735987.5 | 3164534 | | | | | | |
| syntaxin binding protein 1 | Cytoplasm | transporter | 1084624.25 | 848512.438 | 867082.062 | O08599 | STXBP1 | 2432978 | 1457682 | 595096 | 6 |
| syntaxin binding protein 2 | Plasma Membrane | transporter | 8045458.5 | 2.29E+07 | 1.25E+07 | | | | | | |
| syntaxin binding protein 2 | Plasma Membrane | transporter | 4519911.5 | 8519137 | 5901007.5 | | | | | | |
| syntaxin binding protein 2 | Plasma Membrane | transporter | 4488494 | 1.25E+07 | 6570832 | | | | | | |
| syntaxin binding protein 2 | Plasma Membrane | transporter | 2967497 | 8480016 | 6335145.5 | | | | | | |
| syntaxin binding protein 2 | Plasma Membrane | transporter | 2317372.75 | 6082052.5 | 3985255.75 | | | | | | |
| syntaxin binding protein 2 | Plasma Membrane | transporter | 1954954.5 | 5364350 | 4109926.25 | | | | | | |
| syntaxin binding protein 2 | Plasma Membrane | transporter | 1926344.625 | 5445798.5 | 2795744 | | | | | | |
| syntaxin binding protein 2 | Plasma Membrane | transporter | 1806582.75 | 3401580.25 | 2790902.25 | | | | | | |
| syntaxin binding protein 2 | Plasma Membrane | transporter | 1611784.125 | 2607917.75 | 1916423.25 | | | | | | |
| syntaxin binding protein 2 | Plasma Membrane | transporter | 1376240 | 6047093 | 2444956.5 | | | | | | |
| syntaxin binding protein 2 | Plasma Membrane | transporter | 1166146.25 | 2617302.25 | 2902789.5 | | | | | | |
| syntaxin binding protein 2 | Plasma Membrane | transporter | 928756.562 | 3049059.75 | 2480713.75 | | | | | | |
| syntaxin binding protein 2 | Plasma Membrane | transporter | 567808 | 629901.75 | 458805.781 | | | | | | |
| syntaxin binding protein 2 | Plasma Membrane | transporter | 428044.5 | 1755461.375 | 784894.25 | Q64324 | STXBP2 | 2436100 | 4174148 | 644085 | 42 |
| syntaxin binding protein 3 | Plasma Membrane | transporter | 622879.188 | 1432094.875 | 985406.75 | Q60770 | STXBP3 | 622879 | 405337 | 234021 | 3 |
| syntaxin binding protein 5 (tomosyn) | Plasma Membrane | other | 2.53E+07 | 2.49E+07 | 3.14E+07 | Q8K400 | STXBP5 | 25270000 | 3635991 | 2099241 | 3 |
| succinate-CoA ligase, ADP-forming, beta subunit | Cytoplasm | enzyme | 1686742.375 | 5017786 | 3719952.25 | Q9Z2I9 | SUCLA2 | 1686742 | 1678996 | 969369 | 3 |
| succinate-CoA ligase, alpha subunit | Cytoplasm | enzyme | 1653370.75 | 6240514 | 3449775.25 | | | | | | |
| succinate-CoA ligase, alpha subunit | Cytoplasm | enzyme | 974835.688 | 3416723.25 | 2077090.125 | | | | | | |
| succinate-CoA ligase, alpha subunit | Cytoplasm | enzyme | 649767.5 | 2447093.5 | 845056.75 | Q9WUM5 | SUCLG1 | 1092658 | 1769847 | 589949 | 9 |
| succinate-CoA ligase, GDP-forming, beta subunit | Cytoplasm | enzyme | 483343 | 798823.125 | 679873.375 | Q9Z2I8 | SUCLG2 | 483343 | 159322 | 91985 | 3 |
| surfeit 4 | Cytoplasm | other | 1693064.25 | 4910707 | 3960389.5 | | | | | | |
| surfeit 4 | Cytoplasm | other | 1433180 | 4201467.5 | 3316016.5 | Q64310 | SURF4 | 1563122 | 1406918 | 574372 | 6 |
| sushi domain containing 2 | Extracellular Space | other | 7001626.5 | 5902269 | 3826595 | | | | | | |

| | | | | | | | | | | | |
|---|---|---|---|---|---|---|---|---|---|---|---|
| sushi domain containing 2 | Extracellular Space | other | 4786811 | 2680325.75 | 2485395.5 | | | | | | |
| sushi domain containing 2 | Extracellular Space | other | 4722211 | 3251848 | 2405612.5 | Q9DBX3 | SUSD2 | 5503550 | 1612878 | 537626 | 9 |
| small VCP/p97-interacting protein | Cytoplasm | other | 2.62E+07 | 2.30E+07 | 2.52E+07 | Q3UZP4 | SVIP | 26150000 | 1647463 | 951163 | 3 |
| synaptophysin-like 1 | Plasma Membrane | transporter | 2.02E+07 | 1.76E+07 | 1.89E+07 | | | | | | |
| synaptophysin-like 1 | Plasma Membrane | transporter | 1.77E+07 | 1.83E+07 | 1.70E+07 | | | | | | |
| synaptophysin-like 1 | Plasma Membrane | transporter | 942748.312 | 657624.312 | 574463.438 | O09117 | SYPL1 | 12967583 | 8829481 | 2943160 | 9 |
| T-cell activation RhoGTPase activating protein | Cytoplasm | other | 8773539 | 6754564 | 1.03E+07 | B2RWW0 | TAGAP | 8773539 | 1792697 | 1035014 | 3 |
| transgelin 2 | Cytoplasm | other | 378337.875 | 1527093.625 | 6718576.5 | | | | | | |
| transgelin 2 | Cytoplasm | other | 255157.625 | 13443.513 | 2445726 | | | | | | |
| transgelin 2 | Cytoplasm | other | 252394.109 | 736261.812 | 3575527.75 | | | | | | |
| transgelin 2 | Cytoplasm | other | 128790.703 | 735589.938 | 4042376.5 | Q9WVA4 | TAGLN2 | 253670 | 2085980 | 602171 | 12 |
| TAO kinase 3 | Cytoplasm | kinase | 6473932 | 4398907 | 5889049 | | | | | | |
| TAO kinase 3 | Cytoplasm | kinase | 6039657 | 3601553 | 3985434.5 | | | | | | |
| TAO kinase 3 | Cytoplasm | kinase | 5886484.5 | 4311916 | 6454544 | | | | | | |
| TAO kinase 3 | Cytoplasm | kinase | 4578758.5 | 1838265.625 | 3993358 | | | | | | |
| TAO kinase 3 | Cytoplasm | kinase | 3571789 | 2754892.25 | 3269869.25 | | | | | | |
| TAO kinase 3 | Cytoplasm | kinase | 3057507 | 2087643.125 | 3444998.75 | | | | | | |
| TAO kinase 3 | Cytoplasm | kinase | 2888928.75 | 1328392.25 | 2404810.75 | | | | | | |
| TAO kinase 3 | Cytoplasm | kinase | 1054726.625 | 305971.625 | 993037.5 | Q8BYC6 | TAOK3 | 4193973 | 1779765 | 363293 | 24 |
| TAP binding protein (tapasin) | Cytoplasm | transporter | 790797.188 | 1400068.625 | 1094066.625 | Q9R233 | TAPBP | 790797 | 304637 | 175882 | 3 |
| transmembrane anterior posterior transformation 1 | Plasma Membrane | G-protein coupled receptor | 1151197.625 | 1022226.625 | 1094907.75 | Q4VBD2 | TAPT1 | 1151198 | 64659 | 37331 | 3 |
| threonyl-tRNA synthetase | Nucleus | enzyme | 5530311 | 3640016.5 | 5464728.5 | | | | | | |
| threonyl-tRNA synthetase | Nucleus | enzyme | 1750931.875 | 1125238.75 | 1571252.375 | Q9D0R2 | TARS | 3640621 | 1990372 | 812566 | 6 |
| TBC1 domain family, member 10B | Plasma Membrane | enzyme | 4601084.5 | 5126291 | 3270790.25 | Q8BHL3 | TBC1D10B | 4601085 | 956418 | 552188 | 3 |
| TBC1 domain family, member 24 | Cytoplasm | other | 9618540 | 1.07E+07 | 8759894 | | | | | | |
| TBC1 domain family, member 24 | Cytoplasm | other | 4475418 | 5934628 | 3933194 | Q3UUG6 | TBC1D24 | 7046979 | 2831352 | 1155894 | 6 |
| thromboxane A synthase 1 (platelet) | Plasma Membrane | enzyme | 2279184.25 | 9283476 | 4925575 | | | | | | |
| thromboxane A synthase 1 (platelet) | Plasma Membrane | enzyme | 1253066.75 | 1.86E+07 | 1.12E+07 | | | | | | |
| thromboxane A synthase 1 (platelet) | Plasma Membrane | enzyme | 842403.25 | 4797876 | 3538592.5 | | | | | | |
| thromboxane A synthase 1 (platelet) | Plasma Membrane | enzyme | 779964.688 | 5165792 | 4026864 | | | | | | |
| thromboxane A synthase 1 (platelet) | Plasma Membrane | enzyme | 732055 | 4995577 | 3433495.75 | | | | | | |
| thromboxane A synthase 1 (platelet) | Plasma Membrane | enzyme | 622786.625 | 9134532 | 4804240 | | | | | | |
| thromboxane A synthase 1 (platelet) | Plasma Membrane | enzyme | 600407.562 | 1.24E+07 | 9457502 | | | | | | |
| thromboxane A synthase 1 (platelet) | Plasma Membrane | enzyme | 401824.781 | 4954411.5 | 2435522.75 | | | | | | |
| thromboxane A synthase 1 (platelet) | Plasma Membrane | enzyme | 397605.688 | 7288135 | 5031648 | P36423 | TBXAS1 | 878811 | 4415058 | 849678 | 27 |
| transcription elongation factor A (SII), 1 | Nucleus | transcription regulator | 2.48E+07 | 2.09E+07 | 1.43E+07 | | | | | | |
| transcription elongation factor A (SII), 1 | Nucleus | transcription regulator | 2.41E+07 | 1.60E+07 | 1.63E+07 | | | | | | |
| transcription elongation factor A (SII), 1 | Nucleus | transcription regulator | 1.72E+07 | 1.37E+07 | 1.22E+07 | | | | | | |
| transcription elongation factor A (SII), 1 | Nucleus | transcription regulator | 1.25E+07 | 1.09E+07 | 1.03E+07 | | | | | | |
| transcription elongation factor A (SII), 1 | Nucleus | transcription regulator | 1.12E+07 | 8280727 | 8281868.5 | | | | | | |
| transcription elongation factor A (SII), 1 | Nucleus | transcription regulator | 1.03E+07 | 6510075.5 | 7311208.5 | | | | | | |
| transcription elongation factor A (SII), 1 | Nucleus | transcription regulator | 9088713 | 3407026.25 | 4114594.75 | | | | | | |
| transcription elongation factor A (SII), 1 | Nucleus | transcription regulator | 8999718 | 6215096 | 5115852.5 | | | | | | |
| transcription elongation factor A (SII), 1 | Nucleus | transcription regulator | 8400084 | 6291749 | 8457265 | | | | | | |
| transcription elongation factor A (SII), 1 | Nucleus | transcription regulator | 7802767.5 | 6944799 | 5360267 | | | | | | |
| transcription elongation factor A (SII), 1 | Nucleus | transcription regulator | 3455064.5 | 2071612.375 | 1803222.125 | | | | | | |
| transcription elongation factor A (SII), 1 | Nucleus | transcription regulator | 2022793.25 | 709906 | 1006759.438 | P10711 | TCEA1 | 11659095 | 6086316 | 1014386 | 36 |
| transcription elongation factor A (SII), 2 | Nucleus | transcription regulator | 804365.188 | 9683739 | 4256273 | Q9QVN7 | TCEA2 | 804365 | 4476165 | 2584315 | 3 |
| t-complex 1 | Cytoplasm | other | 4.22E+07 | 1.89E+07 | 2.77E+07 | | | | | | |
| t-complex 1 | Cytoplasm | other | 3.28E+07 | 1.98E+07 | 2.25E+07 | | | | | | |
| t-complex 1 | Cytoplasm | other | 1.74E+07 | 6743469 | 1.11E+07 | | | | | | |
| t-complex 1 | Cytoplasm | other | 1.56E+07 | 7265672 | 1.19E+07 | | | | | | |
| t-complex 1 | Cytoplasm | other | 1.48E+07 | 9606770 | 1.18E+07 | | | | | | |
| t-complex 1 | Cytoplasm | other | 1.32E+07 | 6766113 | 9135586 | | | | | | |
| t-complex 1 | Cytoplasm | other | 1.25E+07 | 5379641.5 | 7820773.5 | | | | | | |
| t-complex 1 | Cytoplasm | other | 1.17E+07 | 6458304.5 | 1.09E+07 | | | | | | |
| t-complex 1 | Cytoplasm | other | 9695881 | 5924924 | 6884365 | | | | | | |
| t-complex 1 | Cytoplasm | other | 9554422 | 6126952 | 6925405 | | | | | | |
| t-complex 1 | Cytoplasm | other | 9130620 | 4146757.75 | 6938454 | | | | | | |
| t-complex 1 | Cytoplasm | other | 8334852.5 | 4507174.5 | 6218144.5 | | | | | | |
| t-complex 1 | Cytoplasm | other | 7932992.5 | 3892354.25 | 5682478 | | | | | | |
| t-complex 1 | Cytoplasm | other | 6188998.5 | 2412226.75 | 3989464.75 | | | | | | |
| t-complex 1 | Cytoplasm | other | 5591363.5 | 2140963 | 1308030.875 | | | | | | |
| t-complex 1 | Cytoplasm | other | 5356245.5 | 2109022.75 | 3507635.75 | | | | | | |
| t-complex 1 | Cytoplasm | other | 5021345 | 4227501 | 3200980.75 | | | | | | |
| t-complex 1 | Cytoplasm | other | 4713501 | 1623427.125 | 1984599 | | | | | | |
| t-complex 1 | Cytoplasm | other | 2975272.75 | 2108414.75 | 2231063 | | | | | | |
| t-complex 1 | Cytoplasm | other | 2109878 | 535778.125 | 762251.25 | | | | | | |
| t-complex 1 | Cytoplasm | other | 115347.695 | 265237.219 | 347060.219 | P11983 | TCP1 | 11277177 | 7798565 | 982527 | 63 |
| transferrin | Extracellular Space | transporter | 2.68E+07 | 3.52E+07 | 3.14E+07 | | | | | | |
| transferrin | Extracellular Space | transporter | 2.39E+07 | 1.34E+07 | 1.63E+07 | | | | | | |
| transferrin | Extracellular Space | transporter | 2.36E+07 | 2.47E+07 | 2.67E+07 | | | | | | |
| transferrin | Extracellular Space | transporter | 2.15E+07 | 1.86E+07 | 1.88E+07 | | | | | | |
| transferrin | Extracellular Space | transporter | 1.92E+07 | 1.29E+07 | 1.67E+07 | | | | | | |
| transferrin | Extracellular Space | transporter | 1.91E+07 | 2.33E+07 | 2.28E+07 | | | | | | |
| transferrin | Extracellular Space | transporter | 1.88E+07 | 1.97E+07 | 2.13E+07 | | | | | | |
| transferrin | Extracellular Space | transporter | 1.84E+07 | 1.89E+07 | 2.17E+07 | | | | | | |
| transferrin | Extracellular Space | transporter | 1.68E+07 | 1.64E+07 | 2.11E+07 | | | | | | |
| transferrin | Extracellular Space | transporter | 1.47E+07 | 1.55E+07 | 1.94E+07 | | | | | | |
| transferrin | Extracellular Space | transporter | 1.46E+07 | 1.13E+07 | 1.76E+07 | | | | | | |
| transferrin | Extracellular Space | transporter | 1.41E+07 | 1.36E+07 | 1.99E+07 | | | | | | |
| transferrin | Extracellular Space | transporter | 1.11E+07 | 1.25E+07 | 1.15E+07 | | | | | | |
| transferrin | Extracellular Space | transporter | 1.08E+07 | 1.42E+07 | 1.46E+07 | | | | | | |
| transferrin | Extracellular Space | transporter | 1.07E+07 | 1.87E+07 | 1.18E+07 | | | | | | |
| transferrin | Extracellular Space | transporter | 9023847 | 8701955 | 1.01E+07 | | | | | | |
| transferrin | Extracellular Space | transporter | 8006892.5 | 1.30E+07 | 9428643 | | | | | | |
| transferrin | Extracellular Space | transporter | 6760157 | 2998782.75 | 4232032.5 | | | | | | |
| transferrin | Extracellular Space | transporter | 6075298.5 | 3654165.5 | 4926863 | | | | | | |
| transferrin | Extracellular Space | transporter | 6001432.5 | 4551972.5 | 6079024 | | | | | | |
| transferrin | Extracellular Space | transporter | 5870899 | 8736892 | 5974881 | | | | | | |
| transferrin | Extracellular Space | transporter | 5404500.5 | 3224413 | 4745795 | | | | | | |
| transferrin | Extracellular Space | transporter | 5238090.5 | 4778458.5 | 5685588.5 | | | | | | |
| transferrin | Extracellular Space | transporter | 4678880.5 | 7629354 | 7261534.5 | | | | | | |
| transferrin | Extracellular Space | transporter | 4433117.5 | 6585625.5 | 6862327.5 | | | | | | |
| transferrin | Extracellular Space | transporter | 4320020 | 6163546.5 | 6329427 | | | | | | |

| | | | | | | | | | | | |
|---|---|---|---|---|---|---|---|---|---|---|---|
| transferrin | Extracellular Space | transporter | 3690952.5 | 3489222.75 | 4252935 | | | | | | |
| transferrin | Extracellular Space | transporter | 3414130 | 2376015.25 | 3284591.5 | | | | | | |
| transferrin | Extracellular Space | transporter | 3309004.75 | 1.92E+07 | 2792458.75 | | | | | | |
| transferrin | Extracellular Space | transporter | 2619226 | 4736924.5 | 3401378.75 | | | | | | |
| transferrin | Extracellular Space | transporter | 2433430.75 | 2271919.5 | 2216344 | | | | | | |
| transferrin | Extracellular Space | transporter | 2182385.25 | 8216547.5 | 1.41E+07 | | | | | | |
| transferrin | Extracellular Space | transporter | 1956895.375 | 2362472.25 | 2431159 | | | | | | |
| transferrin | Extracellular Space | transporter | 965556.375 | 1133066.375 | 1099618 | | | | | | |
| transferrin | Extracellular Space | transporter | 4.70E-06 | 572623.25 | 684514.062 | Q921I1 | TF | 10012420 | 7895304 | 770503 | 105 |
| transferrin receptor | Plasma Membrane | transporter | 4.18E+07 | 3.50E+07 | 3.02E+07 | | | | | | |
| transferrin receptor | Plasma Membrane | transporter | 4.15E+07 | 4.65E+07 | 4.70E+07 | | | | | | |
| transferrin receptor | Plasma Membrane | transporter | 3.94E+07 | 3.30E+07 | 3.17E+07 | | | | | | |
| transferrin receptor | Plasma Membrane | transporter | 3.74E+07 | 3.43E+07 | 5.16E+07 | | | | | | |
| transferrin receptor | Plasma Membrane | transporter | 3.54E+07 | 3.87E+07 | 4.54E+07 | | | | | | |
| transferrin receptor | Plasma Membrane | transporter | 3.50E+07 | 3.22E+07 | 3.79E+07 | | | | | | |
| transferrin receptor | Plasma Membrane | transporter | 3.49E+07 | 3.26E+07 | 2.93E+07 | | | | | | |
| transferrin receptor | Plasma Membrane | transporter | 3.35E+07 | 3.71E+07 | 3.60E+07 | | | | | | |
| transferrin receptor | Plasma Membrane | transporter | 3.32E+07 | 2.99E+07 | 2.86E+07 | | | | | | |
| transferrin receptor | Plasma Membrane | transporter | 3.26E+07 | 3.66E+07 | 3.18E+07 | | | | | | |
| transferrin receptor | Plasma Membrane | transporter | 3.13E+07 | 2.88E+07 | 3.45E+07 | | | | | | |
| transferrin receptor | Plasma Membrane | transporter | 2.66E+07 | 2.21E+07 | 2.16E+07 | | | | | | |
| transferrin receptor | Plasma Membrane | transporter | 2.56E+07 | 3.50E+07 | 3.42E+07 | | | | | | |
| transferrin receptor | Plasma Membrane | transporter | 2.41E+07 | 2.46E+07 | 2.50E+07 | | | | | | |
| transferrin receptor | Plasma Membrane | transporter | 2.28E+07 | 1.88E+07 | 1.73E+07 | | | | | | |
| transferrin receptor | Plasma Membrane | transporter | 2.22E+07 | 2.09E+07 | 2.32E+07 | | | | | | |
| transferrin receptor | Plasma Membrane | transporter | 2.15E+07 | 2.33E+07 | 2.21E+07 | | | | | | |
| transferrin receptor | Plasma Membrane | transporter | 1.91E+07 | 2.01E+07 | 2.50E+07 | | | | | | |
| transferrin receptor | Plasma Membrane | transporter | 1.71E+07 | 1.69E+07 | 2.03E+07 | | | | | | |
| transferrin receptor | Plasma Membrane | transporter | 1.71E+07 | 1.83E+07 | 1.55E+07 | | | | | | |
| transferrin receptor | Plasma Membrane | transporter | 1.64E+07 | 1.70E+07 | 1.47E+07 | | | | | | |
| transferrin receptor | Plasma Membrane | transporter | 1.55E+07 | 1.92E+07 | 2.42E+07 | | | | | | |
| transferrin receptor | Plasma Membrane | transporter | 1.31E+07 | 1.22E+07 | 2.59E+07 | | | | | | |
| transferrin receptor | Plasma Membrane | transporter | 1.04E+07 | 9526707 | 9001165 | | | | | | |
| transferrin receptor | Plasma Membrane | transporter | 1.00E+07 | 8943960 | 1.01E+07 | | | | | | |
| transferrin receptor | Plasma Membrane | transporter | 8889149 | 6843728.5 | 8238176 | | | | | | |
| transferrin receptor | Plasma Membrane | transporter | 8621770 | 7929391 | 7897225.5 | | | | | | |
| transferrin receptor | Plasma Membrane | transporter | 7533637.5 | 1.06E+07 | 8531642 | | | | | | |
| transferrin receptor | Plasma Membrane | transporter | 5560156.5 | 2205688 | 3435905.5 | | | | | | |
| transferrin receptor | Plasma Membrane | transporter | 2893415.5 | 2529319 | 2404025.5 | | | | | | |
| transferrin receptor | Plasma Membrane | transporter | 1837945.25 | 2225625.5 | 1524803.75 | | | | | | |
| transferrin receptor | Plasma Membrane | transporter | 1196992.5 | 978427.125 | 847927.875 | | | | | | |
| transferrin receptor | Plasma Membrane | transporter | 1167238.5 | 444170.094 | 742086 | | | | | | |
| transferrin receptor | Plasma Membrane | transporter | 1050504.875 | 946386.312 | 1707782.125 | | | | | | |
| transferrin receptor | Plasma Membrane | transporter | 918363.688 | 1256595.125 | 1858449.375 | | | | | | |
| transferrin receptor | Plasma Membrane | transporter | 877151.688 | 655965.375 | 1147582.625 | | | | | | |
| transferrin receptor | Plasma Membrane | transporter | 67594.93 | 7431.859 | 41028.578 | Q62351 | TFRC | 18863079 | 13920176 | 1321245 | 111 |
| transforming growth factor, beta receptor II (70/80kDa) | Plasma Membrane | kinase | 2.30E+07 | 1.62E+07 | 1.86E+07 | Q62312 | TGFBR2 | 22990000 | 3435947 | 1983745 | 3 |
| transglutaminase 2 | Cytoplasm | enzyme | 1.19E+07 | 4311795 | 6836487.5 | | | | | | |
| transglutaminase 2 | Cytoplasm | enzyme | 4738098 | 631841.875 | 1540671.25 | P21981 | TGM2 | 8339049 | 4077408 | 1664595 | 6 |
| thrombospondin 1 | Extracellular Space | other | 982663.375 | 2.13E+07 | 2.56E+07 | | | | | | |
| thrombospondin 1 | Extracellular Space | other | 770665.375 | 1.34E+07 | 1.32E+07 | | | | | | |
| thrombospondin 1 | Extracellular Space | other | 527183.875 | 8808306 | 7365265 | | | | | | |
| thrombospondin 1 | Extracellular Space | other | 520716.688 | 1.69E+07 | 1.64E+07 | | | | | | |
| thrombospondin 1 | Extracellular Space | other | 473251.562 | 8751512 | 8176256.5 | | | | | | |
| thrombospondin 1 | Extracellular Space | other | 452276.906 | 5598659 | 5244507.5 | | | | | | |
| thrombospondin 1 | Extracellular Space | other | 398249 | 8447560 | 5863098 | | | | | | |
| thrombospondin 1 | Extracellular Space | other | 362813.938 | 1.31E+07 | 1.03E+07 | | | | | | |
| thrombospondin 1 | Extracellular Space | other | 321261.688 | 8294722 | 8348640.5 | | | | | | |
| thrombospondin 1 | Extracellular Space | other | 277398.094 | 1.13E+07 | 7826356.5 | | | | | | |
| thrombospondin 1 | Extracellular Space | other | 266678.562 | 8359468 | 5394008.5 | | | | | | |
| thrombospondin 1 | Extracellular Space | other | 264726.75 | 2545412.75 | 1613439.5 | | | | | | |
| thrombospondin 1 | Extracellular Space | other | 260687.938 | 2.05E+07 | 8135643.5 | | | | | | |
| thrombospondin 1 | Extracellular Space | other | 239455.359 | 1.23E+07 | 7598608.5 | | | | | | |
| thrombospondin 1 | Extracellular Space | other | 191506.594 | 6329810.5 | 5371580 | | | | | | |
| thrombospondin 1 | Extracellular Space | other | 122902.953 | 7689858.5 | 5335626 | | | | | | |
| thrombospondin 1 | Extracellular Space | other | 113668.367 | 2804286.5 | 1646001.875 | | | | | | |
| thrombospondin 1 | Extracellular Space | other | 90801.422 | 7707769.5 | 5376754.5 | | | | | | |
| thrombospondin 1 | Extracellular Space | other | 85446.32 | 1363446.125 | 1243983 | | | | | | |
| thrombospondin 1 | Extracellular Space | other | 83529.766 | 2645171.25 | 2263023.75 | | | | | | |
| thrombospondin 1 | Extracellular Space | other | 78581.586 | 3228931.5 | 2455091.25 | | | | | | |
| thrombospondin 1 | Extracellular Space | other | 77112.742 | 2887473.25 | 1658897.25 | | | | | | |
| thrombospondin 1 | Extracellular Space | other | 67878.492 | 3550517.25 | 2918719.5 | | | | | | |
| thrombospondin 1 | Extracellular Space | other | 32715.977 | 5209940 | 2652541.25 | | | | | | |
| thrombospondin 1 | Extracellular Space | other | 13352.175 | 5060617 | 2370575 | | | | | | |
| thrombospondin 1 | Extracellular Space | other | 5899.621 | 2289006 | 2768100 | | | | | | |
| thrombospondin 1 | Extracellular Space | other | 4.70E-06 | 2606691.75 | 1732189.5 | | | | | | |
| thrombospondin 1 | Extracellular Space | other | 4.70E-06 | 1579085.375 | 1253850 | | | | | | |
| thrombospondin 1 | Extracellular Space | other | 4.70E-06 | 7322084.5 | 4272748 | | | | | | |
| thrombospondin 1 | Extracellular Space | other | 4.70E-06 | 3294855.5 | 3058728.75 | | | | | | |
| thrombospondin 1 | Extracellular Space | other | 0 | 2957069 | 2065109 | | | | | | |
| thrombospondin 1 | Extracellular Space | other | 0 | 2219998 | 894012.562 | P35441 | THBS1 | 221295 | 5200833 | 530808 | 96 |
| thioesterase superfamily member 6 | Extracellular Space | other | 5072912.5 | 4839591 | 4572826 | | | | | | |
| thioesterase superfamily member 6 | Extracellular Space | other | 5012311 | 4172974.5 | 4196632.5 | | | | | | |
| thioesterase superfamily member 6 | Extracellular Space | other | 3430897.5 | 3600183 | 2960011.75 | | | | | | |
| thioesterase superfamily member 6 | Extracellular Space | other | 2839376.25 | 2857546.5 | 3603211.75 | | | | | | |
| thioesterase superfamily member 6 | Extracellular Space | other | 2101212.75 | 1843425.25 | 1216958.75 | Q80ZW2 | THEM6 | 3691342 | 1186892 | 306454 | 15 |
| tRNA-histidine guanylyltransferase 1-like (S. cerevisiae) | Cytoplasm | enzyme | 1.23E+07 | 1.11E+07 | 7164550 | | | | | | |
| tRNA-histidine guanylyltransferase 1-like (S. cerevisiae) | Cytoplasm | enzyme | 1.17E+07 | 8537956 | 7795431 | | | | | | |
| tRNA-histidine guanylyltransferase 1-like (S. cerevisiae) | Cytoplasm | enzyme | 8963568 | 1.19E+07 | 6924410.5 | | | | | | |
| tRNA-histidine guanylyltransferase 1-like (S. cerevisiae) | Cytoplasm | enzyme | 6698408 | 4411467 | 4188929.5 | | | | | | |
| tRNA-histidine guanylyltransferase 1-like (S. cerevisiae) | Cytoplasm | enzyme | 4808329 | 6278936 | 5407121 | Q9CY52 | THG1L | 8900061 | 2790309 | 720455 | 15 |
| translocase of inner mitochondrial membrane 50 homolog (S. cerevisiae) | Cytoplasm | phosphatase | 1192362.625 | 2039182.25 | 2114462.25 | | | | | | |
| translocase of inner mitochondrial membrane 50 homolog (S. cerevisiae) | Cytoplasm | phosphatase | 1077006.75 | 1653567.875 | 1279119.125 | Q9D880 | TIMM50 | 1134685 | 445675 | 181946 | 6 |
| TIMP metallopeptidase inhibitor 3 | Extracellular Space | other | 557175 | 1.31E+07 | 8897128 | | | | | | |
| TIMP metallopeptidase inhibitor 3 | Extracellular Space | other | 200831.328 | 4223208.5 | 2620236.5 | | | | | | |

| | | | | | | | | | | | |
|---|---|---|---|---|---|---|---|---|---|---|---|
| TIMP metallopeptidase inhibitor 3 | Extracellular Space | other | 112306.836 | 7243450.5 | 4938760.5 | | | | | | |
| TIMP metallopeptidase inhibitor 3 | Extracellular Space | other | 84047.609 | 6438146 | 1289374.75 | P39876 | TIMP3 | 238590 | 4161189 | 1201232 | 12 |
| transketolase | Cytoplasm | enzyme | 2644995.75 | 2917606.5 | 2619685 | | | | | | |
| transketolase | Cytoplasm | enzyme | 22214.49 | 42291.922 | 38373.375 | P40142 | TKT | 1333605 | 1478801 | 603718 | 6 |
| talin 1 | Plasma Membrane | other | 362514 | 1736256.625 | 1692891.25 | | | | | | |
| talin 1 | Plasma Membrane | other | 360567.25 | 3566532.25 | 4195435 | | | | | | |
| talin 1 | Plasma Membrane | other | 170159.953 | 1546744.125 | 2200277.5 | | | | | | |
| talin 1 | Plasma Membrane | other | 117604.102 | 1960786.625 | 2543142.5 | | | | | | |
| talin 1 | Plasma Membrane | other | 108591.578 | 1605592.125 | 3065611.75 | | | | | | |
| talin 1 | Plasma Membrane | other | 28619.428 | 1532546.125 | 3101635.25 | | | | | | |
| talin 1 | Plasma Membrane | other | 725.884 | 182776.766 | 549944.938 | | | | | | |
| talin 1 | Plasma Membrane | other | 4.70E-06 | 954387.062 | 1156498.875 | P26039 | TLN1 | 143598 | 1249897 | 255134 | 24 |
| transmembrane 9 superfamily member 3 | Cytoplasm | transporter | 1039325.125 | 2332531 | 1690056.125 | Q9ET30 | TM9SF3 | 1039325 | 646607 | 373319 | 3 |
| transmembrane channel-like 6 | Cytoplasm | transporter | 2885811.25 | 1746122.125 | 1768989.625 | | | | | | |
| transmembrane channel-like 6 | Cytoplasm | transporter | 1662687.875 | 1367402.125 | 1201397.25 | | | | | | |
| transmembrane channel-like 6 | Cytoplasm | transporter | 536872.688 | 546204.125 | 602721.5 | Q7TN60 | TMC6 | 1695124 | 764353 | 254784 | 9 |
| transmembrane channel-like 8 | Cytoplasm | other | 3371927 | 3600178 | 2899594.75 | | | | | | |
| transmembrane channel-like 8 | Cytoplasm | other | 1582965.75 | 960053.938 | 1636410.625 | | | | | | |
| transmembrane channel-like 8 | Cytoplasm | other | 1198108.875 | 1072234.625 | 1086884 | Q7TN58 | TMC8 | 2051001 | 1057018 | 352339 | 9 |
| transmembrane and coiled-coil domain family 2 | Extracellular Space | other | 1.13E+07 | 7833330 | 1.02E+07 | | | | | | |
| transmembrane and coiled-coil domain family 2 | Extracellular Space | other | 1.11E+07 | 7553757.5 | 1.18E+07 | | | | | | |
| transmembrane and coiled-coil domain family 2 | Extracellular Space | other | 5368131.5 | 3568400 | 4120540.5 | | | | | | |
| transmembrane and coiled-coil domain family 2 | Extracellular Space | other | 4192921.25 | 3088257.25 | 4244649 | | | | | | |
| transmembrane and coiled-coil domain family 2 | Extracellular Space | other | 3305704 | 2314257.75 | 2869678.5 | | | | | | |
| transmembrane and coiled-coil domain family 2 | Extracellular Space | other | 1090553.875 | 767054.688 | 1074834 | Q80W04 | TMCC2 | 6059552 | 3710347 | 874537 | 18 |
| transmembrane and coiled-coil domains 1 | Cytoplasm | other | 2008954 | 6269546.5 | 4318214.5 | Q921L3 | TMCO1 | 2008954 | 2132801 | 1231373 | 3 |
| transmembrane emp24-like trafficking protein 10 (yeast) | Cytoplasm | transporter | 2.38E+07 | 1.70E+07 | 2.07E+07 | | | | | | |
| transmembrane emp24-like trafficking protein 10 (yeast) | Cytoplasm | transporter | 2.02E+07 | 1.68E+07 | 1.91E+07 | | | | | | |
| transmembrane emp24-like trafficking protein 10 (yeast) | Cytoplasm | transporter | 1.67E+07 | 2.04E+07 | 1.26E+07 | | | | | | |
| transmembrane emp24-like trafficking protein 10 (yeast) | Cytoplasm | transporter | 5137650 | 6467024 | 5215800 | | | | | | |
| transmembrane emp24-like trafficking protein 10 (yeast) | Cytoplasm | transporter | 4586068 | 2676392.5 | 3752555.5 | | | | | | |
| transmembrane emp24-like trafficking protein 10 (yeast) | Cytoplasm | transporter | 4359139 | 4999611.5 | 5109313 | Q9D1D4 | TMED10 | 12463810 | 7509414 | 1769986 | 18 |
| transmembrane emp24 domain trafficking protein 2 | Cytoplasm | transporter | 8411628 | 8685371 | 7510536 | | | | | | |
| transmembrane emp24 domain trafficking protein 2 | Cytoplasm | transporter | 6588114.5 | 6376535 | 9914383 | | | | | | |
| transmembrane emp24 domain trafficking protein 2 | Cytoplasm | transporter | 3755796.5 | 3253380.25 | 5254175 | Q9R0Q3 | TMED2 | 6251846 | 2252278 | 750759 | 9 |
| transmembrane emp24 protein transport domain containing 4 | Cytoplasm | transporter | 1.38E+07 | 1.23E+07 | 1.42E+07 | Q8R1V4 | TMED4 | 13780000 | 1011484 | 583981 | 3 |
| transmembrane emp24 protein transport domain containing 5 | Cytoplasm | other | 2070540.875 | 2341339.25 | 3033205.25 | Q9CXE7 | TMED5 | 2070541 | 496443 | 286621 | 3 |
| transmembrane emp24 protein transport domain containing 9 | Cytoplasm | transporter | 2.07E+07 | 1.97E+07 | 2.29E+07 | | | | | | |
| transmembrane emp24 protein transport domain containing 9 | Cytoplasm | transporter | 5298603 | 4128245.75 | 5600459.5 | | | | | | |
| transmembrane emp24 protein transport domain containing 9 | Cytoplasm | transporter | 920568.188 | 2868400.5 | 1794479.75 | Q99KF1 | TMED9 | 8979724 | 8986374 | 2995458 | 9 |
| transmembrane protein 106B | Cytoplasm | other | 267791.938 | 159069.25 | 213300.625 | Q80X71 | TMEM106B | 267792 | 54361 | 31386 | 3 |
| transmembrane protein 14C | Plasma Membrane | other | 8549156 | 9234849 | 8430934 | | | | | | |
| transmembrane protein 14C | Plasma Membrane | other | 3655883.5 | 3027970.25 | 2426021.5 | Q9CQN6 | TMEM14C | 6102520 | 3159021 | 1289665 | 6 |
| transmembrane protein 167A | Extracellular Space | other | 2968664.5 | 3972132 | 3498903 | Q9CR64 | TMEM167A | 2968665 | 502004 | 289832 | 3 |
| transmembrane protein 179B | Extracellular Space | other | 3009648.25 | 2112273.25 | 2720389.25 | Q9CY24 | TMEM179B | 3009648 | 458032 | 264445 | 3 |
| transmembrane protein 206 | Plasma Membrane | other | 1.99E+07 | 1.44E+07 | 1.88E+07 | | | | | | |
| transmembrane protein 206 | Plasma Membrane | other | 7139340 | 3449538 | 5301981.5 | | | | | | |
| transmembrane protein 206 | Plasma Membrane | other | 3746076.25 | 1678647 | 2652218.25 | Q9D771 | TMEM206 | 10258472 | 7160892 | 2386964 | 9 |
| transmembrane protein 256 | Cytoplasm | other | 8816878 | 1.12E+07 | 1.07E+07 | Q5F285 | TMEM256 | 8816878 | 1268618 | 732437 | 3 |
| transmembrane protein 30A | Cytoplasm | other | 1.92E+07 | 1.70E+07 | 1.55E+07 | | | | | | |
| transmembrane protein 30A | Cytoplasm | other | 1.44E+07 | 1.60E+07 | 1.85E+07 | | | | | | |
| transmembrane protein 30A | Cytoplasm | other | 1.10E+07 | 9638619 | 7924422 | | | | | | |
| transmembrane protein 30A | Cytoplasm | other | 8890064 | 8441280 | 1.00E+07 | | | | | | |
| transmembrane protein 30A | Cytoplasm | other | 6502610.5 | 5644565 | 5199830.5 | Q8VEK0 | TMEM30A | 12002535 | 4766710 | 1230759 | 15 |
| transmembrane protein 55A | Cytoplasm | phosphatase | 3205413.25 | 1573197.625 | 2989564.5 | Q9CZX7 | TMEM55A | 3205413 | 886643 | 511904 | 3 |
| transmembrane protein 55B | Cytoplasm | phosphatase | 7863836 | 6032055 | 6356355.5 | | | | | | |
| transmembrane protein 55B | Cytoplasm | phosphatase | 7446244 | 2629729.75 | 4312583 | | | | | | |
| transmembrane protein 55B | Cytoplasm | phosphatase | 5380254.5 | 4232323.5 | 3866177.25 | | | | | | |
| transmembrane protein 55B | Cytoplasm | phosphatase | 4716963.5 | 3517716.25 | 4753146.5 | Q3TWL2 | TMEM55B | 6351825 | 1579319 | 455910 | 12 |
| transmembrane protein 63A | Cytoplasm | other | 373656.094 | 1922842.125 | 1823909.75 | Q91YT8 | TMEM63A | 373656 | 867275 | 500722 | 3 |
| transmembrane protein 9 | Cytoplasm | other | 6952002 | 6126610 | 6679262 | Q9CR23 | TMEM9 | 6952002 | 420532 | 242794 | 3 |
| tropomodulin 1 | Cytoplasm | enzyme | 3.11E+08 | 1.82E+08 | 2.30E+08 | | | | | | |
| tropomodulin 1 | Cytoplasm | enzyme | 2.28E+08 | 1.36E+08 | 1.24E+08 | | | | | | |
| tropomodulin 1 | Cytoplasm | enzyme | 2.16E+08 | 2.13E+08 | 2.16E+08 | | | | | | |
| tropomodulin 1 | Cytoplasm | enzyme | 2.15E+08 | 1.64E+08 | 1.63E+08 | | | | | | |
| tropomodulin 1 | Cytoplasm | enzyme | 2.13E+08 | 7.92E+07 | 1.12E+08 | | | | | | |
| tropomodulin 1 | Cytoplasm | enzyme | 1.90E+08 | 1.61E+08 | 1.55E+08 | | | | | | |
| tropomodulin 1 | Cytoplasm | enzyme | 1.56E+08 | 1.24E+08 | 1.29E+08 | | | | | | |
| tropomodulin 1 | Cytoplasm | enzyme | 1.46E+08 | 1.25E+08 | 1.32E+08 | | | | | | |
| tropomodulin 1 | Cytoplasm | enzyme | 1.42E+08 | 8.74E+07 | 9.20E+07 | | | | | | |
| tropomodulin 1 | Cytoplasm | enzyme | 1.16E+08 | 7.53E+07 | 8.36E+07 | | | | | | |
| tropomodulin 1 | Cytoplasm | enzyme | 6.90E+07 | 5.77E+07 | 5.73E+07 | | | | | | |
| tropomodulin 1 | Cytoplasm | enzyme | 5.05E+07 | 4.45E+07 | 3.77E+07 | | | | | | |
| tropomodulin 1 | Cytoplasm | enzyme | 4.60E+07 | 3.15E+07 | 3.70E+07 | | | | | | |
| tropomodulin 1 | Cytoplasm | enzyme | 2.82E+07 | 5.17E+07 | 1.90E+07 | | | | | | |
| tropomodulin 1 | Cytoplasm | enzyme | 2.35E+07 | 1.54E+07 | 1.95E+07 | | | | | | |
| tropomodulin 1 | Cytoplasm | enzyme | 1.99E+07 | 2.92E+07 | 2.57E+07 | | | | | | |
| tropomodulin 1 | Cytoplasm | enzyme | 1.92E+07 | 7664943.5 | 1.19E+07 | | | | | | |
| tropomodulin 1 | Cytoplasm | enzyme | 1.89E+07 | 1.09E+07 | 1.34E+07 | | | | | | |
| tropomodulin 1 | Cytoplasm | enzyme | 1.12E+07 | 1.11E+07 | 1.24E+07 | | | | | | |
| tropomodulin 1 | Cytoplasm | enzyme | 1.12E+07 | 2.56E+07 | 2.60E+07 | | | | | | |
| tropomodulin 1 | Cytoplasm | enzyme | 7531298.5 | 2179655.75 | 3727798.5 | | | | | | |
| tropomodulin 1 | Cytoplasm | enzyme | 6481760 | 3207729 | 4851148.5 | | | | | | |
| tropomodulin 1 | Cytoplasm | enzyme | 5956478 | 7181858 | 5283721 | | | | | | |
| tropomodulin 1 | Cytoplasm | enzyme | 1719689.625 | 429815.781 | 1075103.75 | P49813 | TMOD1 | 93856634 | 77381676 | 9119518 | 72 |
| thioredoxin-related transmembrane protein 2 | Cytoplasm | enzyme | 448601.688 | 368679.25 | 255888.922 | Q9D710 | TMX2 | 448602 | 96822 | 55900 | 3 |
| thioredoxin-related transmembrane protein 4 | Cytoplasm | enzyme | 3487025.25 | 2141022.25 | 2089425.5 | Q8C0L0 | TMX4 | 3487025 | 792430 | 457510 | 3 |
| target of myb1 membrane trafficking protein | Cytoplasm | transporter | 1962653.875 | 768095.062 | 1264920.25 | O88746 | TOM1 | 1962654 | 600089 | 346461 | 3 |
| target of myb1 like 1 membrane trafficking protein | Cytoplasm | other | 8228039.5 | 2818325.5 | 5254165 | | | | | | |
| target of myb1 like 1 membrane trafficking protein | Cytoplasm | other | 3985153 | 2046856.25 | 1560495.125 | Q923U0 | TOM1L1 | 6106596 | 2474262 | 1010113 | 6 |
| translocase of outer mitochondrial membrane 22 homolog (yeast) | Cytoplasm | transporter | 1310288.5 | 1812920 | 1554349.375 | | | | | | |
| translocase of outer mitochondrial membrane 22 homolog (yeast) | Cytoplasm | transporter | 1278316.5 | 1892479.125 | 1983869.875 | Q9CPQ3 | TOMM22 | 1294303 | 302901 | 123659 | 6 |
| translocase of outer mitochondrial membrane 40 homolog (yeast) | Cytoplasm | ion channel | 2307647 | 4026339.5 | 4070130 | Q9QYA2 | TOMM40 | 2307647 | 1005167 | 580334 | 3 |
| translocase of outer mitochondrial membrane 70 homolog A (S. cerevisiae) | Cytoplasm | transporter | 529870.062 | 1821892.625 | 2054746.25 | Q9CZW5 | TOMM70A | 529870 | 821461 | 474271 | 3 |
| tumor protein D52-like 2 | Cytoplasm | other | 3585962.5 | 2483660.25 | 2985287.75 | Q9CYZ2 | TPD52L2 | 3585963 | 551892 | 318635 | 3 |

| | | | | | | | | | | | |
|---|---|---|---|---|---|---|---|---|---|---|---|
| tropomyosin 1, alpha | Plasma Membrane | other | 4.74E+08 | 3.01E+08 | 3.33E+08 | | | | | | |
| tropomyosin 1, alpha | Plasma Membrane | other | 7.66E+07 | 4.22E+07 | 5.68E+07 | | | | | | |
| tropomyosin 1, alpha | Plasma Membrane | other | 2.38E+07 | 1.05E+07 | 1.79E+07 | | | | | | |
| tropomyosin 1, alpha | Plasma Membrane | other | 9804287 | 8709692 | 1.09E+07 | | | | | | |
| tropomyosin 1, alpha | Plasma Membrane | other | 5570567.5 | 3886687.25 | 3311918.25 | P58771 | Tpm1 | 117906971 | 149137102 | 38507034 | 15 |
| tropomyosin 3 | Cytoplasm | other | 6.54E+08 | 6.87E+08 | 5.41E+08 | | | | | | |
| tropomyosin 3 | Cytoplasm | other | 2.59E+08 | 1.69E+08 | 1.77E+08 | | | | | | |
| tropomyosin 3 | Cytoplasm | other | 2.50E+08 | 2.00E+08 | 1.91E+08 | | | | | | |
| tropomyosin 3 | Cytoplasm | other | 2.45E+08 | 1.73E+08 | 1.95E+08 | | | | | | |
| tropomyosin 3 | Cytoplasm | other | 1.22E+08 | 1.10E+08 | 8.03E+07 | | | | | | |
| tropomyosin 3 | Cytoplasm | other | 1.19E+08 | 5.59E+07 | 8.24E+07 | | | | | | |
| tropomyosin 3 | Cytoplasm | other | 1.17E+08 | 5.21E+07 | 7.83E+07 | | | | | | |
| tropomyosin 3 | Cytoplasm | other | 1.14E+08 | 9.16E+07 | 6.63E+07 | | | | | | |
| tropomyosin 3 | Cytoplasm | other | 8.95E+07 | 5.67E+07 | 5.65E+07 | | | | | | |
| tropomyosin 3 | Cytoplasm | other | 8.08E+07 | 3.30E+07 | 4.80E+07 | | | | | | |
| tropomyosin 3 | Cytoplasm | other | 4.76E+07 | 3.27E+07 | 3.45E+07 | | | | | | |
| tropomyosin 3 | Cytoplasm | other | 1.46E+07 | 5887928 | 9426080 | | | | | | |
| tropomyosin 3 | Cytoplasm | other | 1.34E+07 | 6610543 | 1.19E+07 | | | | | | |
| tropomyosin 3 | Cytoplasm | other | 6977802 | 6069157 | 6137794.5 | P21107 | TPM3 | 152305557 | 158509856 | 24458602 | 42 |
| tripeptidyl peptidase I | Cytoplasm | peptidase | 2284530.5 | 4413206 | 5082897 | O89023 | TPP1 | 2284531 | 1461198 | 843623 | 3 |
| tripeptidyl peptidase II | Cytoplasm | peptidase | 3.78E+07 | 1.55E+07 | 2.03E+07 | | | | | | |
| tripeptidyl peptidase II | Cytoplasm | peptidase | 2.50E+07 | 1.05E+07 | 1.14E+07 | | | | | | |
| tripeptidyl peptidase II | Cytoplasm | peptidase | 2.39E+07 | 8472722 | 1.09E+07 | | | | | | |
| tripeptidyl peptidase II | Cytoplasm | peptidase | 2.22E+07 | 7731330 | 9566329 | | | | | | |
| tripeptidyl peptidase II | Cytoplasm | peptidase | 2.08E+07 | 6748183.5 | 9766811 | | | | | | |
| tripeptidyl peptidase II | Cytoplasm | peptidase | 1.63E+07 | 7413130 | 1.02E+07 | | | | | | |
| tripeptidyl peptidase II | Cytoplasm | peptidase | 1.55E+07 | 7182341 | 7432155 | | | | | | |
| tripeptidyl peptidase II | Cytoplasm | peptidase | 1.45E+07 | 5566670 | 7356630.5 | | | | | | |
| tripeptidyl peptidase II | Cytoplasm | peptidase | 1.36E+07 | 5323060 | 7095733 | | | | | | |
| tripeptidyl peptidase II | Cytoplasm | peptidase | 1.35E+07 | 4583908.5 | 7384914 | | | | | | |
| tripeptidyl peptidase II | Cytoplasm | peptidase | 1.02E+07 | 3724849.75 | 4732636.5 | | | | | | |
| tripeptidyl peptidase II | Cytoplasm | peptidase | 8539036 | 3012813.25 | 3615797.25 | | | | | | |
| tripeptidyl peptidase II | Cytoplasm | peptidase | 8475236 | 1207371 | 1566689.75 | | | | | | |
| tripeptidyl peptidase II | Cytoplasm | peptidase | 8440797 | 3868428.75 | 5183647.5 | | | | | | |
| tripeptidyl peptidase II | Cytoplasm | peptidase | 8415918 | 1820215.625 | 2659214.5 | | | | | | |
| tripeptidyl peptidase II | Cytoplasm | peptidase | 7244141.5 | 2849201 | 3295841.75 | | | | | | |
| tripeptidyl peptidase II | Cytoplasm | peptidase | 6804568 | 2180320.5 | 2954082 | | | | | | |
| tripeptidyl peptidase II | Cytoplasm | peptidase | 6536225.5 | 2631216.25 | 3797775.75 | | | | | | |
| tripeptidyl peptidase II | Cytoplasm | peptidase | 4155005.75 | 1441231.375 | 1863025.5 | | | | | | |
| tripeptidyl peptidase II | Cytoplasm | peptidase | 3029340.5 | 521364.281 | 794951.5 | | | | | | |
| tripeptidyl peptidase II | Cytoplasm | peptidase | 2773610.75 | 1043126.25 | 1268291.5 | | | | | | |
| tripeptidyl peptidase II | Cytoplasm | peptidase | 1994703.375 | 1106708.875 | 1187764.375 | Q64514 | TPP2 | 12707663 | 6999121 | 861532 | 66 |
| TNF receptor-associated protein 1 | Cytoplasm | enzyme | 1.29E+07 | 5490221.5 | 6457602.5 | | | | | | |
| TNF receptor-associated protein 1 | Cytoplasm | enzyme | 597055.562 | 996490.438 | 1110862.5 | Q9CQN1 | TRAP1 | 6733528 | 4771121 | 1947802 | 6 |
| triggering receptor expressed on myeloid cells-like 1 | Plasma Membrane | other | 3999043 | 6.67E+07 | 3.52E+07 | | | | | | |
| triggering receptor expressed on myeloid cells-like 1 | Plasma Membrane | other | 2033835.125 | 4.68E+07 | 2.60E+07 | | | | | | |
| triggering receptor expressed on myeloid cells-like 1 | Plasma Membrane | other | 1666998.625 | 2.80E+07 | 1.73E+07 | | | | | | |
| triggering receptor expressed on myeloid cells-like 1 | Plasma Membrane | other | 1635676.625 | 4.35E+07 | 2.84E+07 | | | | | | |
| triggering receptor expressed on myeloid cells-like 1 | Plasma Membrane | other | 1112229.625 | 3.05E+07 | 1.97E+07 | | | | | | |
| triggering receptor expressed on myeloid cells-like 1 | Plasma Membrane | other | 848164.5 | 1.58E+07 | 1.36E+07 | | | | | | |
| triggering receptor expressed on myeloid cells-like 1 | Plasma Membrane | other | 724682.062 | 2.32E+07 | 1.07E+07 | | | | | | |
| triggering receptor expressed on myeloid cells-like 1 | Plasma Membrane | other | 580427.312 | 4871165.5 | 2564015.25 | Q8K558 | TREML1 | 1575132 | 17764897 | 3626244 | 24 |
| transient receptor potential cation channel, subfamily V, member 2 | Plasma Membrane | ion channel | 2.24E+07 | 2.18E+07 | 1.75E+07 | | | | | | |
| transient receptor potential cation channel, subfamily V, member 2 | Plasma Membrane | ion channel | 1.84E+07 | 1.18E+07 | 1.19E+07 | | | | | | |
| transient receptor potential cation channel, subfamily V, member 2 | Plasma Membrane | ion channel | 9559792 | 7186210.5 | 7890553 | | | | | | |
| transient receptor potential cation channel, subfamily V, member 2 | Plasma Membrane | ion channel | 9113398 | 7627512.5 | 7449693.5 | | | | | | |
| transient receptor potential cation channel, subfamily V, member 2 | Plasma Membrane | ion channel | 8720546 | 1.05E+07 | 7905574 | | | | | | |
| transient receptor potential cation channel, subfamily V, member 2 | Plasma Membrane | ion channel | 8507518 | 7807172 | 8261520.5 | | | | | | |
| transient receptor potential cation channel, subfamily V, member 2 | Plasma Membrane | ion channel | 7980029.5 | 8152402.5 | 5484286.5 | | | | | | |
| transient receptor potential cation channel, subfamily V, member 2 | Plasma Membrane | ion channel | 6924007.5 | 7835852.5 | 6065570.5 | | | | | | |
| transient receptor potential cation channel, subfamily V, member 2 | Plasma Membrane | ion channel | 5936194 | 4236221 | 4680421.5 | | | | | | |
| transient receptor potential cation channel, subfamily V, member 2 | Plasma Membrane | ion channel | 4223797 | 1925650.125 | 1781103.5 | | | | | | |
| transient receptor potential cation channel, subfamily V, member 2 | Plasma Membrane | ion channel | 3858399 | 2983994 | 3474766.5 | | | | | | |
| transient receptor potential cation channel, subfamily V, member 2 | Plasma Membrane | ion channel | 3161141.75 | 2278322.25 | 2319912 | Q9WTR1 | TRPV2 | 9065402 | 5128205 | 854701 | 36 |
| translin | Nucleus | other | 4092619.25 | 2894244.5 | 2952246.5 | | | | | | |
| translin | Nucleus | other | 2016762.625 | 1115747.875 | 866788.75 | Q62348 | TSN | 3054691 | 1226834 | 500853 | 6 |
| tetraspanin 14 | Plasma Membrane | other | 387457.812 | 6565571 | 3825592.5 | Q8QZY6 | TSPAN14 | 387458 | 3095624 | 1787259 | 3 |
| tetraspanin 33 | Plasma Membrane | other | 1.38E+08 | 1.07E+08 | 1.23E+08 | | | | | | |
| tetraspanin 33 | Plasma Membrane | other | 4.43E+07 | 3.38E+07 | 3.81E+07 | | | | | | |
| tetraspanin 33 | Plasma Membrane | other | 2.67E+07 | 3.48E+07 | 2.49E+07 | | | | | | |
| tetraspanin 33 | Plasma Membrane | other | 2.62E+07 | 2.20E+07 | 1.47E+07 | | | | | | |
| tetraspanin 33 | Plasma Membrane | other | 4406335.5 | 1968992.75 | 1977303.375 | | | | | | |
| tetraspanin 33 | Plasma Membrane | other | 2392170.5 | 1676769.625 | 1907299.5 | | | | | | |
| tetraspanin 33 | Plasma Membrane | other | 2079618.125 | 2819225.5 | 1399616.375 | Q8R3S2 | TSPAN33 | 34894018 | 41040173 | 8955700 | 21 |
| tetraspanin 8 | Plasma Membrane | other | 6.18E+07 | 4.62E+07 | 4.64E+07 | | | | | | |
| tetraspanin 8 | Plasma Membrane | other | 3.99E+07 | 3.84E+07 | 2.93E+07 | | | | | | |
| tetraspanin 8 | Plasma Membrane | other | 2.47E+07 | 2.40E+07 | 2.44E+07 | | | | | | |
| tetraspanin 8 | Plasma Membrane | other | 1.90E+07 | 1.07E+07 | 9404191 | | | | | | |
| tetraspanin 8 | Plasma Membrane | other | 7548808 | 9445249 | 6497456.5 | Q8R3G9 | TSPAN8 | 30571762 | 16945893 | 4375411 | 15 |
| translocator protein (18kDa) | Cytoplasm | transmembrane receptor | 4449875.5 | 8206684.5 | 8231730 | P50637 | TSPO | 4449876 | 2176261 | 1256465 | 3 |
| titin | Cytoplasm | kinase | 5224931 | 3082237 | 3326251.5 | A2ASS6 | TTN | 5224931 | 1173006 | 677236 | 3 |
| tubulin, alpha 1a | Cytoplasm | other | 1036830.938 | 2601392.25 | 3122908.25 | P68369 | TUBA1A | 1036831 | 1085630 | 626789 | 3 |
| tubulin, beta 2A class IIa | Cytoplasm | other | 954109.5 | 2213196.25 | 1993829 | | | | | | |
| tubulin, beta 2A class IIa | Cytoplasm | other | 349547.844 | 2471050.25 | 2781472 | Q7TMM9 | TUBB2A | 651829 | 942445 | 384752 | 6 |
| tubulin, beta 4A class IVa | Cytoplasm | other | 1833632.25 | 4007379.75 | 4093326.5 | | | | | | |
| tubulin, beta 4A class IVa | Cytoplasm | other | 868243 | 6103105.5 | 6472481 | Q9D6F9 | TUBB4A | 1350938 | 2235244 | 912535 | 6 |
| Tu translation elongation factor, mitochondrial | Cytoplasm | translation regulator | 2777287.5 | 4320814 | 4375444 | | | | | | |
| Tu translation elongation factor, mitochondrial | Cytoplasm | translation regulator | 2303558.25 | 4824241 | 4074959.5 | | | | | | |
| Tu translation elongation factor, mitochondrial | Cytoplasm | translation regulator | 2155946.25 | 4560447 | 4511831 | | | | | | |
| Tu translation elongation factor, mitochondrial | Cytoplasm | translation regulator | 1971583.375 | 4065335.25 | 2842106.25 | | | | | | |
| Tu translation elongation factor, mitochondrial | Cytoplasm | translation regulator | 1058785.875 | 3709873.25 | 2276280.25 | | | | | | |
| Tu translation elongation factor, mitochondrial | Cytoplasm | translation regulator | 444216.375 | 1812075.5 | 1478699.125 | Q8BFR5 | TUFM | 1785230 | 1357939 | 320069 | 18 |
| thioredoxin domain containing 5 (endoplasmic reticulum) | Cytoplasm | enzyme | 1302285.25 | 722610.688 | 985996.75 | Q91W90 | TXNDC5 | 1302285 | 290239 | 167570 | 3 |
| thioredoxin-like 1 | Cytoplasm | enzyme | 8964218 | 8786208 | 1.21E+07 | | | | | | |
| thioredoxin-like 1 | Cytoplasm | enzyme | 5001140.5 | 3852816.75 | 3537635.25 | | | | | | |

| | | | | | | | | | | | |
|---|---|---|---|---|---|---|---|---|---|---|---|
| thioredoxin-like 1 | Cytoplasm | enzyme | 4574709.5 | 3177182 | 4026771.75 | | | | | | |
| thioredoxin-like 1 | Cytoplasm | enzyme | 4464934 | 1963537.625 | 2379960.75 | | | | | | |
| thioredoxin-like 1 | Cytoplasm | enzyme | 2022193.75 | 448698.344 | 700605.188 | | | | | | |
| thioredoxin-like 1 | Cytoplasm | enzyme | 1429594.875 | 495850.312 | 705001.312 | | | | | | |
| thioredoxin-like 1 | Cytoplasm | enzyme | 1342751 | 86455.617 | 666629.688 | | | | | | |
| thioredoxin-like 1 | Cytoplasm | enzyme | 395769.75 | 177428.547 | 167476.453 | Q8CDN6 | TXNL1 | 3524414 | 3162740 | 645592 | 24 |
| thioredoxin reductase 2 | Cytoplasm | enzyme | 2.26E+07 | 9768311 | 1.33E+07 | | | | | | |
| thioredoxin reductase 2 | Cytoplasm | enzyme | 5965456 | 4761105 | 5646041 | | | | | | |
| thioredoxin reductase 2 | Cytoplasm | enzyme | 1270095.75 | 957268.312 | 815271.375 | Q9JLT4 | TXNRD2 | 9941851 | 7114713 | 2371571 | 9 |
| ubiquitin-like modifier activating enzyme 1 | Cytoplasm | enzyme | 1.28E+07 | 7712013.5 | 6511349 | | | | | | |
| ubiquitin-like modifier activating enzyme 1 | Cytoplasm | enzyme | 1.12E+07 | 5590955.5 | 4585753 | | | | | | |
| ubiquitin-like modifier activating enzyme 1 | Cytoplasm | enzyme | 1.11E+07 | 5875861.5 | 5182661 | | | | | | |
| ubiquitin-like modifier activating enzyme 1 | Cytoplasm | enzyme | 1.03E+07 | 6095688 | 5219776 | | | | | | |
| ubiquitin-like modifier activating enzyme 1 | Cytoplasm | enzyme | 9191075 | 3874681.25 | 4192567.25 | | | | | | |
| ubiquitin-like modifier activating enzyme 1 | Cytoplasm | enzyme | 7784981 | 5414261.5 | 3235215 | | | | | | |
| ubiquitin-like modifier activating enzyme 1 | Cytoplasm | enzyme | 7662501 | 3365023.5 | 2900720.25 | | | | | | |
| ubiquitin-like modifier activating enzyme 1 | Cytoplasm | enzyme | 7362872.5 | 5195431 | 4165768.25 | | | | | | |
| ubiquitin-like modifier activating enzyme 1 | Cytoplasm | enzyme | 5836698.5 | 3601850 | 2036671.25 | | | | | | |
| ubiquitin-like modifier activating enzyme 1 | Cytoplasm | enzyme | 4691941.5 | 1637410.875 | 3054591 | | | | | | |
| ubiquitin-like modifier activating enzyme 1 | Cytoplasm | enzyme | 4333632.5 | 3542673.75 | 2741504.25 | | | | | | |
| ubiquitin-like modifier activating enzyme 1 | Cytoplasm | enzyme | 4044267.5 | 3158356 | 2121965 | | | | | | |
| ubiquitin-like modifier activating enzyme 1 | Cytoplasm | enzyme | 1770721.875 | 1136826.625 | 1158404.75 | | | | | | |
| ubiquitin-like modifier activating enzyme 1 | Cytoplasm | enzyme | 1756163 | 2326392.25 | 799175.75 | Q02053 | UBA1 | 7128204 | 2919458 | 450482 | 42 |
| ubiquitin A-52 residue ribosomal protein fusion product 1 | Cytoplasm | other | 1.56E+09 | 1.05E+09 | 1.06E+09 | | | | | | |
| ubiquitin A-52 residue ribosomal protein fusion product 1 | Cytoplasm | other | 5.78E+08 | 3.71E+08 | 3.88E+08 | | | | | | |
| ubiquitin A-52 residue ribosomal protein fusion product 1 | Cytoplasm | other | 2.77E+08 | 2.10E+08 | 2.28E+08 | | | | | | |
| ubiquitin A-52 residue ribosomal protein fusion product 1 | Cytoplasm | other | 2.64E+08 | 2.54E+08 | 2.18E+08 | | | | | | |
| ubiquitin A-52 residue ribosomal protein fusion product 1 | Cytoplasm | other | 7.71E+07 | 1.15E+08 | 8.44E+07 | | | | | | |
| ubiquitin A-52 residue ribosomal protein fusion product 1 | Cytoplasm | other | 1.26E+07 | 1.67E+07 | 1.05E+07 | | | | | | |
| ubiquitin A-52 residue ribosomal protein fusion product 1 | Cytoplasm | other | 1.22E+07 | 3.20E+07 | 5.00E+07 | | | | | | |
| ubiquitin A-52 residue ribosomal protein fusion product 1 | Cytoplasm | other | 611426.75 | 2113624.75 | 2635262 | P62984 | Uba52 | 347670178 | 400871569 | 81827566 | 24 |
| UBA domain containing 1 | Nucleus | other | 1.50E+07 | 4593902.5 | 5168215.5 | | | | | | |
| UBA domain containing 1 | Nucleus | other | 1.26E+07 | 5023111 | 7499781 | | | | | | |
| UBA domain containing 1 | Nucleus | other | 6924308 | 5066024 | 5106203 | | | | | | |
| UBA domain containing 1 | Nucleus | other | 6152470.5 | 4159693.75 | 4310027.5 | | | | | | |
| UBA domain containing 1 | Nucleus | other | 6022252.5 | 3446579.25 | 3499758 | | | | | | |
| UBA domain containing 1 | Nucleus | other | 5684926.5 | 2988106 | 2720184.25 | | | | | | |
| UBA domain containing 1 | Nucleus | other | 4914896 | 1491105.875 | 1933452.125 | | | | | | |
| UBA domain containing 1 | Nucleus | other | 3556675.25 | 1587708.875 | 1692493 | | | | | | |
| UBA domain containing 1 | Nucleus | other | 2302196.75 | 1839509.375 | 1792051.375 | | | | | | |
| UBA domain containing 1 | Nucleus | other | 609109.438 | 1142454.375 | 2532300 | Q8VDI7 | UBAC1 | 6376683 | 3154723 | 575971 | 30 |
| ubiquitin-conjugating enzyme E2L 3 | Nucleus | enzyme | 1.54E+07 | 6503460.5 | 1.06E+07 | P68037 | UBE2L3 | 15410000 | 4457767 | 2573693 | 3 |
| ubiquitin-conjugating enzyme E2L 6 | Cytoplasm | enzyme | 3.22E+07 | 1.42E+07 | 9334313 | | | | | | |
| ubiquitin-conjugating enzyme E2L 6 | Cytoplasm | enzyme | 7036889.5 | 2435420.5 | 2173713 | | | | | | |
| ubiquitin-conjugating enzyme E2L 6 | Cytoplasm | enzyme | 4235990.5 | 457303.25 | 951598.938 | | | | | | |
| ubiquitin-conjugating enzyme E2L 6 | Cytoplasm | enzyme | 3091884.25 | 1575229.125 | 1206919.125 | Q9QZU9 | UBE2L6 | 11633691 | 9036279 | 2608549 | 12 |
| ubiquitin-conjugating enzyme E2O | Nucleus | enzyme | 5.81E+07 | 1.96E+07 | 3.14E+07 | | | | | | |
| ubiquitin-conjugating enzyme E2O | Nucleus | enzyme | 4.89E+07 | 1.93E+07 | 2.37E+07 | | | | | | |
| ubiquitin-conjugating enzyme E2O | Nucleus | enzyme | 4.78E+07 | 1.88E+07 | 2.83E+07 | | | | | | |
| ubiquitin-conjugating enzyme E2O | Nucleus | enzyme | 4.65E+07 | 1.40E+07 | 2.09E+07 | | | | | | |
| ubiquitin-conjugating enzyme E2O | Nucleus | enzyme | 4.53E+07 | 1.81E+07 | 2.29E+07 | | | | | | |
| ubiquitin-conjugating enzyme E2O | Nucleus | enzyme | 3.59E+07 | 1.13E+07 | 1.52E+07 | | | | | | |
| ubiquitin-conjugating enzyme E2O | Nucleus | enzyme | 2.89E+07 | 1.01E+07 | 1.75E+07 | | | | | | |
| ubiquitin-conjugating enzyme E2O | Nucleus | enzyme | 2.89E+07 | 1.18E+07 | 1.50E+07 | | | | | | |
| ubiquitin-conjugating enzyme E2O | Nucleus | enzyme | 2.88E+07 | 8557099 | 1.71E+07 | | | | | | |
| ubiquitin-conjugating enzyme E2O | Nucleus | enzyme | 2.29E+07 | 8248014 | 1.07E+07 | | | | | | |
| ubiquitin-conjugating enzyme E2O | Nucleus | enzyme | 1.87E+07 | 5474318.5 | 9554135 | | | | | | |
| ubiquitin-conjugating enzyme E2O | Nucleus | enzyme | 1.84E+07 | 8771711 | 1.01E+07 | | | | | | |
| ubiquitin-conjugating enzyme E2O | Nucleus | enzyme | 1.73E+07 | 7262854.5 | 1.07E+07 | | | | | | |
| ubiquitin-conjugating enzyme E2O | Nucleus | enzyme | 1.69E+07 | 8854189 | 1.07E+07 | | | | | | |
| ubiquitin-conjugating enzyme E2O | Nucleus | enzyme | 1.68E+07 | 6420966 | 8435270 | | | | | | |
| ubiquitin-conjugating enzyme E2O | Nucleus | enzyme | 1.65E+07 | 5498177 | 8702412 | | | | | | |
| ubiquitin-conjugating enzyme E2O | Nucleus | enzyme | 1.63E+07 | 6803026.5 | 9294109 | | | | | | |
| ubiquitin-conjugating enzyme E2O | Nucleus | enzyme | 1.52E+07 | 5710539 | 9648744 | | | | | | |
| ubiquitin-conjugating enzyme E2O | Nucleus | enzyme | 1.45E+07 | 3529842 | 7163032.5 | | | | | | |
| ubiquitin-conjugating enzyme E2O | Nucleus | enzyme | 1.39E+07 | 4628634 | 6833986.5 | | | | | | |
| ubiquitin-conjugating enzyme E2O | Nucleus | enzyme | 1.29E+07 | 5372513 | 7437709.5 | | | | | | |
| ubiquitin-conjugating enzyme E2O | Nucleus | enzyme | 1.28E+07 | 3476179.75 | 4194652.5 | | | | | | |
| ubiquitin-conjugating enzyme E2O | Nucleus | enzyme | 1.04E+07 | 3802104.5 | 4723617 | | | | | | |
| ubiquitin-conjugating enzyme E2O | Nucleus | enzyme | 9537897 | 1830091.75 | 3986769.75 | | | | | | |
| ubiquitin-conjugating enzyme E2O | Nucleus | enzyme | 7808289.5 | 2344164.75 | 3264556.25 | | | | | | |
| ubiquitin-conjugating enzyme E2O | Nucleus | enzyme | 7060980 | 2778575 | 3177585.25 | | | | | | |
| ubiquitin-conjugating enzyme E2O | Nucleus | enzyme | 5527240.5 | 4057706.75 | 4478005 | | | | | | |
| ubiquitin-conjugating enzyme E2O | Nucleus | enzyme | 5293163 | 3473093.5 | 3280720 | | | | | | |
| ubiquitin-conjugating enzyme E2O | Nucleus | enzyme | 4591266 | 1495275.125 | 2272969.5 | | | | | | |
| ubiquitin-conjugating enzyme E2O | Nucleus | enzyme | 4513079.5 | 3088917.5 | 1968542.375 | | | | | | |
| ubiquitin-conjugating enzyme E2O | Nucleus | enzyme | 3364655.75 | 1395942.375 | 2103209.5 | | | | | | |
| ubiquitin-conjugating enzyme E2O | Nucleus | enzyme | 3270231.5 | 1942047.625 | 2139381 | | | | | | |
| ubiquitin-conjugating enzyme E2O | Nucleus | enzyme | 2796914.75 | 1378302.625 | 1253528.125 | | | | | | |
| ubiquitin-conjugating enzyme E2O | Nucleus | enzyme | 86223.445 | 167868.234 | 950071.188 | Q6ZPJ3 | UBE2O | 19007057 | 11582120 | 1146801 | 102 |
| ubiquitin-conjugating enzyme E2R 2 | Cytoplasm | enzyme | 4617791.5 | 1422824.375 | 1577062.375 | Q6ZWZ2 | UBE2R2 | 4617792 | 1801742 | 1040236 | 3 |
| ubiquitin protein ligase E3 component n-recognin 4 | Nucleus | enzyme | 2.16E+07 | 8711179 | 1.15E+07 | | | | | | |
| ubiquitin protein ligase E3 component n-recognin 4 | Nucleus | enzyme | 1.18E+07 | 7625348 | 5669658 | | | | | | |
| ubiquitin protein ligase E3 component n-recognin 4 | Nucleus | enzyme | 1.11E+07 | 5531532.5 | 4630280 | | | | | | |
| ubiquitin protein ligase E3 component n-recognin 4 | Nucleus | enzyme | 1.07E+07 | 3513964.5 | 4660863 | | | | | | |
| ubiquitin protein ligase E3 component n-recognin 4 | Nucleus | enzyme | 1.02E+07 | 5360903.5 | 4390181.5 | | | | | | |
| ubiquitin protein ligase E3 component n-recognin 4 | Nucleus | enzyme | 9551360 | 5435864 | 6880333.5 | | | | | | |
| ubiquitin protein ligase E3 component n-recognin 4 | Nucleus | enzyme | 9208158 | 4260170 | 5358656 | | | | | | |
| ubiquitin protein ligase E3 component n-recognin 4 | Nucleus | enzyme | 9100526 | 4012856.75 | 4060190 | | | | | | |
| ubiquitin protein ligase E3 component n-recognin 4 | Nucleus | enzyme | 8937264 | 5475512 | 5295097.5 | | | | | | |
| ubiquitin protein ligase E3 component n-recognin 4 | Nucleus | enzyme | 8481808 | 4005349.25 | 4597312 | | | | | | |
| ubiquitin protein ligase E3 component n-recognin 4 | Nucleus | enzyme | 8418665 | 3576869 | 4475984 | | | | | | |
| ubiquitin protein ligase E3 component n-recognin 4 | Nucleus | enzyme | 8192361 | 5176262.5 | 5867872 | | | | | | |
| ubiquitin protein ligase E3 component n-recognin 4 | Nucleus | enzyme | 8105742 | 4357088 | 6331103 | | | | | | |
| ubiquitin protein ligase E3 component n-recognin 4 | Nucleus | enzyme | 7874598.5 | 2952013 | 3696556.5 | | | | | | |

| | | | | | | | | | | | | |
|---|---|---|---|---|---|---|---|---|---|---|---|---|
| ubiquitin protein ligase E3 component n-recognin 4 | Nucleus | enzyme | 7794014 | 4139865.5 | 4040087.5 | | | | | | | |
| ubiquitin protein ligase E3 component n-recognin 4 | Nucleus | enzyme | 7750631.5 | 4570108 | 5110588 | | | | | | | |
| ubiquitin protein ligase E3 component n-recognin 4 | Nucleus | enzyme | 7646066.5 | 3996911.75 | 4895192 | | | | | | | |
| ubiquitin protein ligase E3 component n-recognin 4 | Nucleus | enzyme | 7592836 | 3383382.25 | 3374820.25 | | | | | | | |
| ubiquitin protein ligase E3 component n-recognin 4 | Nucleus | enzyme | 7571611.5 | 4277587 | 4592425 | | | | | | | |
| ubiquitin protein ligase E3 component n-recognin 4 | Nucleus | enzyme | 7508652 | 5537162 | 5085820.5 | | | | | | | |
| ubiquitin protein ligase E3 component n-recognin 4 | Nucleus | enzyme | 7338001.5 | 2773207 | 4222161 | | | | | | | |
| ubiquitin protein ligase E3 component n-recognin 4 | Nucleus | enzyme | 7145203.5 | 4832288 | 4230609 | | | | | | | |
| ubiquitin protein ligase E3 component n-recognin 4 | Nucleus | enzyme | 6956042.5 | 3907753.5 | 4992970 | | | | | | | |
| ubiquitin protein ligase E3 component n-recognin 4 | Nucleus | enzyme | 6602320 | 4909861 | 4667205.5 | | | | | | | |
| ubiquitin protein ligase E3 component n-recognin 4 | Nucleus | enzyme | 6513120 | 3899247.75 | 4105764.5 | | | | | | | |
| ubiquitin protein ligase E3 component n-recognin 4 | Nucleus | enzyme | 6410951 | 2741152 | 3286976.75 | | | | | | | |
| ubiquitin protein ligase E3 component n-recognin 4 | Nucleus | enzyme | 6357850.5 | 3052107.5 | 3493546.75 | | | | | | | |
| ubiquitin protein ligase E3 component n-recognin 4 | Nucleus | enzyme | 6294871.5 | 2887234.5 | 3190357 | | | | | | | |
| ubiquitin protein ligase E3 component n-recognin 4 | Nucleus | enzyme | 6282822.5 | 2785346.75 | 3888403.5 | | | | | | | |
| ubiquitin protein ligase E3 component n-recognin 4 | Nucleus | enzyme | 6273977.5 | 3944882.75 | 3207115 | | | | | | | |
| ubiquitin protein ligase E3 component n-recognin 4 | Nucleus | enzyme | 6231591.5 | 3961274 | 3769686.25 | | | | | | | |
| ubiquitin protein ligase E3 component n-recognin 4 | Nucleus | enzyme | 6186162 | 2509802.25 | 3922835.75 | | | | | | | |
| ubiquitin protein ligase E3 component n-recognin 4 | Nucleus | enzyme | 6159155 | 3739787 | 4408472.5 | | | | | | | |
| ubiquitin protein ligase E3 component n-recognin 4 | Nucleus | enzyme | 5935706.5 | 1569839.375 | 3232067.75 | | | | | | | |
| ubiquitin protein ligase E3 component n-recognin 4 | Nucleus | enzyme | 5765410.5 | 2472416.75 | 3164105 | | | | | | | |
| ubiquitin protein ligase E3 component n-recognin 4 | Nucleus | enzyme | 5698221 | 2350869 | 3629199.5 | | | | | | | |
| ubiquitin protein ligase E3 component n-recognin 4 | Nucleus | enzyme | 5509447.5 | 3000986.75 | 3138939 | | | | | | | |
| ubiquitin protein ligase E3 component n-recognin 4 | Nucleus | enzyme | 5500777.5 | 3026110.5 | 3648881.5 | | | | | | | |
| ubiquitin protein ligase E3 component n-recognin 4 | Nucleus | enzyme | 5362446 | 2776738.5 | 3768873.25 | | | | | | | |
| ubiquitin protein ligase E3 component n-recognin 4 | Nucleus | enzyme | 5269730.5 | 2244732.25 | 2932576 | | | | | | | |
| ubiquitin protein ligase E3 component n-recognin 4 | Nucleus | enzyme | 5222615.5 | 3301963.5 | 2940108.5 | | | | | | | |
| ubiquitin protein ligase E3 component n-recognin 4 | Nucleus | enzyme | 4638470.5 | 3225958.75 | 3440327.75 | | | | | | | |
| ubiquitin protein ligase E3 component n-recognin 4 | Nucleus | enzyme | 4453097.5 | 1833397.625 | 2472089 | | | | | | | |
| ubiquitin protein ligase E3 component n-recognin 4 | Nucleus | enzyme | 4393593.5 | 1734902.75 | 2923869.5 | | | | | | | |
| ubiquitin protein ligase E3 component n-recognin 4 | Nucleus | enzyme | 4312146.5 | 1906707.125 | 2602279 | | | | | | | |
| ubiquitin protein ligase E3 component n-recognin 4 | Nucleus | enzyme | 4285786 | 2334835.25 | 1884201.375 | | | | | | | |
| ubiquitin protein ligase E3 component n-recognin 4 | Nucleus | enzyme | 4151030.25 | 2366022.5 | 2431314.5 | | | | | | | |
| ubiquitin protein ligase E3 component n-recognin 4 | Nucleus | enzyme | 3519348.75 | 1923742.625 | 2287092.25 | | | | | | | |
| ubiquitin protein ligase E3 component n-recognin 4 | Nucleus | enzyme | 3391616.75 | 1769225.375 | 2484904.5 | | | | | | | |
| ubiquitin protein ligase E3 component n-recognin 4 | Nucleus | enzyme | 3078858 | 2951260.25 | 1490617.5 | | | | | | | |
| ubiquitin protein ligase E3 component n-recognin 4 | Nucleus | enzyme | 2247037 | 1472984.625 | 1633942.75 | | | | | | | |
| ubiquitin protein ligase E3 component n-recognin 4 | Nucleus | enzyme | 1868836.75 | 850700 | 1190504 | | | | | | | |
| ubiquitin protein ligase E3 component n-recognin 4 | Nucleus | enzyme | 1412495.5 | 1112491.75 | 1225416.75 | A2AN08 | UBR4 | | | 6788359 | 2621134 | 207869 | 159 |
| ubiquitin carboxyl-terminal hydrolase L5 | Cytoplasm | peptidase | 1.18E+07 | 4783831 | 5513026.5 | | | | | | | |
| ubiquitin carboxyl-terminal hydrolase L5 | Cytoplasm | peptidase | 7199534 | 2560403 | 4051325.75 | | | | | | | |
| ubiquitin carboxyl-terminal hydrolase L5 | Cytoplasm | peptidase | 3312454.25 | 713652.625 | 1072961 | | | | | | | |
| ubiquitin carboxyl-terminal hydrolase L5 | Cytoplasm | peptidase | 580276.5 | 291276.406 | 725446.75 | Q9WUP7 | UCHL5 | | | 5718066 | 3428650 | 989766 | 12 |
| UDP-glucose glycoprotein glucosyltransferase 1 | Cytoplasm | enzyme | 1.33E+07 | 1.07E+07 | 9824042 | | | | | | | |
| UDP-glucose glycoprotein glucosyltransferase 1 | Cytoplasm | enzyme | 4718512.5 | 4114019.5 | 2603411.25 | | | | | | | |
| UDP-glucose glycoprotein glucosyltransferase 1 | Cytoplasm | enzyme | 3743256.5 | 3319937.75 | 2808768 | | | | | | | |
| UDP-glucose glycoprotein glucosyltransferase 1 | Cytoplasm | enzyme | 3518227 | 4056067 | 4122637.25 | | | | | | | |
| UDP-glucose glycoprotein glucosyltransferase 1 | Cytoplasm | enzyme | 3069949 | 2071321.25 | 2137856.25 | | | | | | | |
| UDP-glucose glycoprotein glucosyltransferase 1 | Cytoplasm | enzyme | 2506517.5 | 2466332.5 | 2387308.75 | | | | | | | |
| UDP-glucose glycoprotein glucosyltransferase 1 | Cytoplasm | enzyme | 2460689 | 2574755.75 | 4072819 | | | | | | | |
| UDP-glucose glycoprotein glucosyltransferase 1 | Cytoplasm | enzyme | 1677157.75 | 1639453 | 2219993.75 | Q6P5E4 | UGGT1 | | | 4371789 | 2975635 | 607399 | 24 |
| ubiquinol-cytochrome c reductase, complex III subunit X | Cytoplasm | enzyme | 905965.75 | 1057003.375 | 1115722.875 | Q8R1I1 | UQCR10 | | | 905966 | 108212 | 62476 | 3 |
| ubiquinol-cytochrome c reductase core protein I | Cytoplasm | enzyme | 8703649 | 1.53E+07 | 1.19E+07 | | | | | | | |
| ubiquinol-cytochrome c reductase core protein I | Cytoplasm | enzyme | 3404963.5 | 4676488 | 3896773.75 | | | | | | | |
| ubiquinol-cytochrome c reductase core protein I | Cytoplasm | enzyme | 3099674.25 | 6000530.5 | 4542620.5 | | | | | | | |
| ubiquinol-cytochrome c reductase core protein I | Cytoplasm | enzyme | 1964462.5 | 4300967 | 3670181.5 | | | | | | | |
| ubiquinol-cytochrome c reductase core protein I | Cytoplasm | enzyme | 1392568.75 | 4713645 | 3849420.25 | Q9CZ13 | UQCRC1 | | | 3713064 | 3770113 | 973439 | 15 |
| ubiquinol-cytochrome c reductase core protein II | Cytoplasm | enzyme | 7035181.5 | 1.11E+07 | 1.12E+07 | | | | | | | |
| ubiquinol-cytochrome c reductase core protein II | Cytoplasm | enzyme | 6716862 | 9262501 | 7581395.5 | | | | | | | |
| ubiquinol-cytochrome c reductase core protein II | Cytoplasm | enzyme | 6512303 | 7244245.5 | 7729958 | | | | | | | |
| ubiquinol-cytochrome c reductase core protein II | Cytoplasm | enzyme | 3601370.5 | 7001957 | 5970754.5 | | | | | | | |
| ubiquinol-cytochrome c reductase core protein II | Cytoplasm | enzyme | 2814043.75 | 5658176 | 3335744.75 | | | | | | | |
| ubiquinol-cytochrome c reductase core protein II | Cytoplasm | enzyme | 2679097.5 | 7359873.5 | 4455027.5 | | | | | | | |
| ubiquinol-cytochrome c reductase core protein II | Cytoplasm | enzyme | 2400491.75 | 2830912.25 | 3273054.75 | | | | | | | |
| ubiquinol-cytochrome c reductase core protein II | Cytoplasm | enzyme | 2163822.75 | 4244035 | 3781046 | | | | | | | |
| ubiquinol-cytochrome c reductase core protein II | Cytoplasm | enzyme | 1462442.5 | 3374651.25 | 3570804 | | | | | | | |
| ubiquinol-cytochrome c reductase core protein II | Cytoplasm | enzyme | 1457774.25 | 2970151.25 | 1937113 | | | | | | | |
| ubiquinol-cytochrome c reductase core protein II | Cytoplasm | enzyme | 799093.938 | 2043540.375 | 1695955.75 | | | | | | | |
| ubiquinol-cytochrome c reductase core protein II | Cytoplasm | enzyme | 212758.328 | 1770641.125 | 1577307.375 | Q9DB77 | UQCRC2 | | | 3154603 | 2862219 | 477037 | 36 |
| ubiquinol-cytochrome c reductase hinge protein like | Cytoplasm | other | 1.60E+07 | 2.19E+07 | 1.84E+07 | | | | | | | |
| ubiquinol-cytochrome c reductase hinge protein like | Cytoplasm | other | 2511540.25 | 5391353 | 6048518.5 | P99028 | UQCRHL | | | 9230770 | 8052003 | 3287216 | 6 |
| uroporphyrinogen decarboxylase | Cytoplasm | enzyme | 4235815 | 1879567.25 | 2155612.75 | P70697 | UROD | | | 4235815 | 1288109 | 743690 | 3 |
| up-regulated during skeletal muscle growth 5 homolog (mouse) | Cytoplasm | other | 3673416.5 | 5522369 | 5348324 | | | | | | | |
| up-regulated during skeletal muscle growth 5 homolog (mouse) | Cytoplasm | other | 2307618 | 5599206.5 | 4252754.5 | Q78IK2 | USMG5 | | | 2990517 | 1304563 | 532586 | 6 |
| ubiquitin specific peptidase 14 (tRNA-guanine transglycosylase) | Cytoplasm | peptidase | 8250871 | 5051807 | 4875069.5 | | | | | | | |
| ubiquitin specific peptidase 14 (tRNA-guanine transglycosylase) | Cytoplasm | peptidase | 5148769 | 2196619 | 2814192.5 | Q9JMA1 | USP14 | | | 6699820 | 2134692 | 871484 | 6 |
| ubiquitin specific peptidase 15 | Cytoplasm | peptidase | 1.37E+07 | 3279010.75 | 3496587.75 | | | | | | | |
| ubiquitin specific peptidase 15 | Cytoplasm | peptidase | 1.34E+07 | 5131413 | 7957404.5 | | | | | | | |
| ubiquitin specific peptidase 15 | Cytoplasm | peptidase | 1.05E+07 | 3919220.5 | 5165521 | | | | | | | |
| ubiquitin specific peptidase 15 | Cytoplasm | peptidase | 9295528 | 2190342 | 4114999.25 | | | | | | | |
| ubiquitin specific peptidase 15 | Cytoplasm | peptidase | 7554391.5 | 2621345.25 | 3713013.5 | | | | | | | |
| ubiquitin specific peptidase 15 | Cytoplasm | peptidase | 6790867 | 3306239 | 4739192 | | | | | | | |
| ubiquitin specific peptidase 15 | Cytoplasm | peptidase | 6081518.5 | 1909574.375 | 3268569.5 | | | | | | | |
| ubiquitin specific peptidase 15 | Cytoplasm | peptidase | 5430106.5 | 1878300.625 | 2575900.25 | | | | | | | |
| ubiquitin specific peptidase 15 | Cytoplasm | peptidase | 4806212 | 1820737.25 | 2767044.75 | | | | | | | |
| ubiquitin specific peptidase 15 | Cytoplasm | peptidase | 4610851.5 | 2726606.75 | 3119870.25 | | | | | | | |
| ubiquitin specific peptidase 15 | Cytoplasm | peptidase | 4068388 | 1259302.875 | 1572492.375 | | | | | | | |
| ubiquitin specific peptidase 15 | Cytoplasm | peptidase | 4044035.75 | 2756717 | 2357109 | | | | | | | |
| ubiquitin specific peptidase 15 | Cytoplasm | peptidase | 3435574 | 2056458 | 2143991.25 | | | | | | | |
| ubiquitin specific peptidase 15 | Cytoplasm | peptidase | 3414667.75 | 1539038.25 | 2000595.5 | | | | | | | |
| ubiquitin specific peptidase 15 | Cytoplasm | peptidase | 2359949 | 566893.188 | 1263124.625 | | | | | | | |
| ubiquitin specific peptidase 15 | Cytoplasm | peptidase | 2295611.75 | 1442983.25 | 1380797 | Q8R5H1 | USP15 | | | 6366731 | 2922306 | 421798 | 48 |
| ubiquitin specific peptidase 19 | Cytoplasm | peptidase | 265051.219 | 90935.836 | 52217.777 | Q3UJD6 | USP19 | | | 265051 | 113368 | 65453 | 3 |
| ubiquitin specific peptidase 22 | Nucleus | peptidase | 1.47E+09 | 9.64E+08 | 1.06E+09 | Q5DU02 | USP22 | | | 1471000000 | 269271090 | 155463736 | 3 |
| ubiquitin specific peptidase 5 (isopeptidase T) | Cytoplasm | peptidase | 2.99E+07 | 6948762.5 | 9696667 | | | | | | | |

| | | | | | | | | | | | |
|---|---|---|---|---|---|---|---|---|---|---|---|
| ubiquitin specific peptidase 5 (isopeptidase T) | Cytoplasm | peptidase | 1.12E+07 | 3508314 | 5973208 | | | | | | |
| ubiquitin specific peptidase 5 (isopeptidase T) | Cytoplasm | peptidase | 1.03E+07 | 3065174.25 | 3707545.75 | | | | | | |
| ubiquitin specific peptidase 5 (isopeptidase T) | Cytoplasm | peptidase | 9419316 | 2825834.5 | 3948297.25 | | | | | | |
| ubiquitin specific peptidase 5 (isopeptidase T) | Cytoplasm | peptidase | 9352179 | 3052377 | 4736260 | | | | | | |
| ubiquitin specific peptidase 5 (isopeptidase T) | Cytoplasm | peptidase | 9289088 | 3516858.75 | 4142428.5 | | | | | | |
| ubiquitin specific peptidase 5 (isopeptidase T) | Cytoplasm | peptidase | 8849405 | 3854094.5 | 5555229.5 | | | | | | |
| ubiquitin specific peptidase 5 (isopeptidase T) | Cytoplasm | peptidase | 7336272.5 | 2536482.75 | 4717107.5 | | | | | | |
| ubiquitin specific peptidase 5 (isopeptidase T) | Cytoplasm | peptidase | 7127297 | 2153128 | 3280464.75 | | | | | | |
| ubiquitin specific peptidase 5 (isopeptidase T) | Cytoplasm | peptidase | 3903479.75 | 1295113.125 | 1424069.25 | | | | | | |
| ubiquitin specific peptidase 5 (isopeptidase T) | Cytoplasm | peptidase | 2416830 | 436848.5 | 1235125.5 | | | | | | |
| ubiquitin specific peptidase 5 (isopeptidase T) | Cytoplasm | peptidase | 2232834.25 | 362266.156 | 1224512.5 | | | | | | |
| ubiquitin specific peptidase 5 (isopeptidase T) | Cytoplasm | peptidase | 1374874.5 | 374738.688 | 1526643 | P56399 | USP5 | 8667044 | 5110957 | 818408 | 39 |
| ubiquitin specific peptidase 7 (herpes virus-associated) | Nucleus | peptidase | 1015498.688 | 815959.25 | 1067404.375 | Q6A4J8 | USP7 | 1015499 | 132750 | 76643 | 3 |
| ubiquitin specific peptidase 9, X-linked | Plasma Membrane | peptidase | 6147368.5 | 3210754.25 | 3678154.75 | | | | | | |
| ubiquitin specific peptidase 9, X-linked | Plasma Membrane | peptidase | 3672221.25 | 2019265.875 | 2301933.25 | | | | | | |
| ubiquitin specific peptidase 9, X-linked | Plasma Membrane | peptidase | 2846538 | 1258053.25 | 1528451.75 | | | | | | |
| ubiquitin specific peptidase 9, X-linked | Plasma Membrane | peptidase | 2138299.5 | 794729.375 | 897114.688 | P70398 | USP9X | 3701107 | 1508616 | 435500 | 12 |
| Vac14 homolog (S. cerevisiae) | Cytoplasm | other | 3370897 | 2189571 | 2203788.5 | Q80WQ2 | VAC14 | 3370897 | 677972 | 391427 | 3 |
| vesicle-associated membrane protein 3 | Plasma Membrane | other | 7.77E+07 | 9.70E+07 | 6.89E+07 | | | | | | |
| vesicle-associated membrane protein 3 | Plasma Membrane | other | 1.39E+07 | 1.34E+07 | 9704473 | | | | | | |
| vesicle-associated membrane protein 3 | Plasma Membrane | other | 6810446 | 5986848 | 4987410 | | | | | | |
| vesicle-associated membrane protein 3 | Plasma Membrane | other | 5906573.5 | 8841654 | 6289297.5 | | | | | | |
| vesicle-associated membrane protein 3 | Plasma Membrane | other | 2540502 | 2633679.75 | 2092830.125 | P63024 | VAMP3 | 21379504 | 31416291 | 8111652 | 15 |
| vesicle-associated membrane protein 5 | Plasma Membrane | transporter | 2.48E+07 | 1.80E+07 | 1.98E+07 | | | | | | |
| vesicle-associated membrane protein 5 | Plasma Membrane | transporter | 8188872 | 7055023.5 | 9439187 | | | | | | |
| vesicle-associated membrane protein 5 | Plasma Membrane | transporter | 6831828.5 | 1.05E+07 | 9389297 | Q9Z2P8 | VAMP5 | 13263567 | 6501527 | 2167176 | 9 |
| vesicle-associated membrane protein 8 | Plasma Membrane | transporter | 9798140 | 1.53E+07 | 1.34E+07 | | | | | | |
| vesicle-associated membrane protein 8 | Plasma Membrane | transporter | 6223299.5 | 1.37E+07 | 9374801 | | | | | | |
| vesicle-associated membrane protein 8 | Plasma Membrane | transporter | 1751968.5 | 7815951 | 2726874.75 | O70404 | VAMP8 | 5924469 | 4790640 | 1596880 | 9 |
| VAMP (vesicle-associated membrane protein)-associated protein A, 33kDa | Plasma Membrane | other | 5.14E+07 | 5.00E+07 | 5.08E+07 | | | | | | |
| VAMP (vesicle-associated membrane protein)-associated protein A, 33kDa | Plasma Membrane | other | 3.16E+07 | 3.28E+07 | 3.31E+07 | | | | | | |
| VAMP (vesicle-associated membrane protein)-associated protein A, 33kDa | Plasma Membrane | other | 2.35E+07 | 2.11E+07 | 1.99E+07 | | | | | | |
| VAMP (vesicle-associated membrane protein)-associated protein A, 33kDa | Plasma Membrane | other | 2.06E+07 | 2.19E+07 | 1.76E+07 | | | | | | |
| VAMP (vesicle-associated membrane protein)-associated protein A, 33kDa | Plasma Membrane | other | 1.03E+07 | 9759346 | 8128284.5 | | | | | | |
| VAMP (vesicle-associated membrane protein)-associated protein A, 33kDa | Plasma Membrane | other | 9661740 | 9572098 | 8765391 | | | | | | |
| VAMP (vesicle-associated membrane protein)-associated protein A, 33kDa | Plasma Membrane | other | 8567506 | 9952677 | 8474108 | | | | | | |
| VAMP (vesicle-associated membrane protein)-associated protein A, 33kDa | Plasma Membrane | other | 6791725.5 | 8485163 | 4833510 | | | | | | |
| VAMP (vesicle-associated membrane protein)-associated protein A, 33kDa | Plasma Membrane | other | 2176985.5 | 2107920.5 | 885508.438 | Q9WV55 | VAPA | 18297551 | 14925523 | 2872418 | 27 |
| VAMP (vesicle-associated membrane protein)-associated protein B and C | Plasma Membrane | other | 4711760.5 | 5608154.5 | 4682829.5 | | | | | | |
| VAMP (vesicle-associated membrane protein)-associated protein B and C | Plasma Membrane | other | 806857.188 | 1586710.5 | 1544517.25 | Q9QY76 | VAPB | 2759309 | 2066043 | 843459 | 6 |
| valyl-tRNA synthetase 2, mitochondrial | Cytoplasm | enzyme | 8609572 | 1.00E+07 | 1.33E+07 | Q3U2A8 | VARS2 | 8609572 | 2384442 | 1376658 | 3 |
| vasodilator-stimulated phosphoprotein | Plasma Membrane | other | 2101634.5 | 5209145 | 4188807.75 | P70460 | VASP | 2101635 | 1583982 | 914513 | 3 |
| valosin containing protein | Cytoplasm | enzyme | 3.59E+08 | 1.21E+08 | 1.51E+08 | | | | | | |
| valosin containing protein | Cytoplasm | enzyme | 3.31E+08 | 2.33E+08 | 2.31E+08 | | | | | | |
| valosin containing protein | Cytoplasm | enzyme | 3.15E+08 | 1.52E+08 | 1.75E+08 | | | | | | |
| valosin containing protein | Cytoplasm | enzyme | 2.86E+08 | 1.19E+08 | 2.35E+08 | | | | | | |
| valosin containing protein | Cytoplasm | enzyme | 2.67E+08 | 1.07E+08 | 1.77E+08 | | | | | | |
| valosin containing protein | Cytoplasm | enzyme | 2.20E+08 | 1.16E+08 | 1.35E+08 | | | | | | |
| valosin containing protein | Cytoplasm | enzyme | 1.96E+08 | 8.87E+07 | 1.48E+08 | | | | | | |
| valosin containing protein | Cytoplasm | enzyme | 1.94E+08 | 8.38E+07 | 1.13E+08 | | | | | | |
| valosin containing protein | Cytoplasm | enzyme | 1.73E+08 | 8.15E+07 | 9.79E+07 | | | | | | |
| valosin containing protein | Cytoplasm | enzyme | 1.69E+08 | 7.97E+07 | 9.89E+07 | | | | | | |
| valosin containing protein | Cytoplasm | enzyme | 1.63E+08 | 8.34E+07 | 7.42E+07 | | | | | | |
| valosin containing protein | Cytoplasm | enzyme | 1.57E+08 | 5.12E+07 | 9.34E+07 | | | | | | |
| valosin containing protein | Cytoplasm | enzyme | 1.54E+08 | 7.75E+07 | 1.02E+08 | | | | | | |
| valosin containing protein | Cytoplasm | enzyme | 1.47E+08 | 5.73E+07 | 6.38E+07 | | | | | | |
| valosin containing protein | Cytoplasm | enzyme | 1.35E+08 | 3.78E+07 | 6.92E+07 | | | | | | |
| valosin containing protein | Cytoplasm | enzyme | 1.29E+08 | 6.31E+07 | 7.16E+07 | | | | | | |
| valosin containing protein | Cytoplasm | enzyme | 1.26E+08 | 4.46E+07 | 6.70E+07 | | | | | | |
| valosin containing protein | Cytoplasm | enzyme | 1.16E+08 | 4.76E+07 | 6.24E+07 | | | | | | |
| valosin containing protein | Cytoplasm | enzyme | 1.15E+08 | 2.39E+07 | 6.17E+07 | | | | | | |
| valosin containing protein | Cytoplasm | enzyme | 1.12E+08 | 5.37E+07 | 6.85E+07 | | | | | | |
| valosin containing protein | Cytoplasm | enzyme | 9.26E+07 | 3.67E+07 | 5.97E+07 | | | | | | |
| valosin containing protein | Cytoplasm | enzyme | 8.29E+07 | 2.89E+07 | 4.97E+07 | | | | | | |
| valosin containing protein | Cytoplasm | enzyme | 7.35E+07 | 3.95E+07 | 4.94E+07 | | | | | | |
| valosin containing protein | Cytoplasm | enzyme | 6.94E+07 | 2.19E+07 | 4.16E+07 | | | | | | |
| valosin containing protein | Cytoplasm | enzyme | 6.51E+07 | 1.83E+07 | 3.86E+07 | | | | | | |
| valosin containing protein | Cytoplasm | enzyme | 6.21E+07 | 3.39E+07 | 4.96E+07 | | | | | | |
| valosin containing protein | Cytoplasm | enzyme | 5.97E+07 | 2.87E+07 | 3.16E+07 | | | | | | |
| valosin containing protein | Cytoplasm | enzyme | 5.73E+07 | 2.86E+07 | 3.43E+07 | | | | | | |
| valosin containing protein | Cytoplasm | enzyme | 5.44E+07 | 2.15E+07 | 3.89E+07 | | | | | | |
| valosin containing protein | Cytoplasm | enzyme | 5.04E+07 | 1.75E+07 | 1.97E+07 | | | | | | |
| valosin containing protein | Cytoplasm | enzyme | 4.87E+07 | 2.16E+07 | 2.57E+07 | | | | | | |
| valosin containing protein | Cytoplasm | enzyme | 4.66E+07 | 3.08E+07 | 4.94E+07 | | | | | | |
| valosin containing protein | Cytoplasm | enzyme | 4.51E+07 | 1.21E+07 | 2.15E+07 | | | | | | |
| valosin containing protein | Cytoplasm | enzyme | 4.40E+07 | 3.83E+07 | 4.49E+07 | | | | | | |
| valosin containing protein | Cytoplasm | enzyme | 4.24E+07 | 2.16E+07 | 2.82E+07 | | | | | | |
| valosin containing protein | Cytoplasm | enzyme | 4.23E+07 | 1.47E+07 | 2.14E+07 | | | | | | |
| valosin containing protein | Cytoplasm | enzyme | 4.06E+07 | 3.51E+07 | 2.87E+07 | | | | | | |
| valosin containing protein | Cytoplasm | enzyme | 4.06E+07 | 1.84E+07 | 2.91E+07 | | | | | | |
| valosin containing protein | Cytoplasm | enzyme | 4.00E+07 | 2.35E+07 | 2.77E+07 | | | | | | |
| valosin containing protein | Cytoplasm | enzyme | 3.97E+07 | 2.87E+07 | 4.66E+07 | | | | | | |
| valosin containing protein | Cytoplasm | enzyme | 3.29E+07 | 1.56E+07 | 1.80E+07 | | | | | | |
| valosin containing protein | Cytoplasm | enzyme | 3.11E+07 | 1.35E+07 | 9824923 | | | | | | |
| valosin containing protein | Cytoplasm | enzyme | 2.41E+07 | 1.16E+07 | 1.24E+07 | | | | | | |
| valosin containing protein | Cytoplasm | enzyme | 1.93E+07 | 3767098.25 | 4859022.5 | | | | | | |
| valosin containing protein | Cytoplasm | enzyme | 1.93E+07 | 1.52E+07 | 7389767 | | | | | | |
| valosin containing protein | Cytoplasm | enzyme | 1.84E+07 | 2.72E+07 | 3.16E+07 | | | | | | |
| valosin containing protein | Cytoplasm | enzyme | 1.75E+07 | 5405044.5 | 7540713.5 | | | | | | |
| valosin containing protein | Cytoplasm | enzyme | 1.71E+07 | 5503121.5 | 7722089.5 | | | | | | |
| valosin containing protein | Cytoplasm | enzyme | 1.71E+07 | 6954456.5 | 1.16E+07 | | | | | | |
| valosin containing protein | Cytoplasm | enzyme | 1.37E+07 | 5147943.5 | 1.06E+07 | | | | | | |
| valosin containing protein | Cytoplasm | enzyme | 1.22E+07 | 6379364.5 | 4679626.5 | | | | | | |
| valosin containing protein | Cytoplasm | enzyme | 1.21E+07 | 9569765 | 1.21E+07 | | | | | | |
| valosin containing protein | Cytoplasm | enzyme | 1.19E+07 | 4575030.5 | 7175142.5 | | | | | | |

| | | | | | | | | | | | |
|---|---|---|---|---|---|---|---|---|---|---|---|
| valosin containing protein | Cytoplasm | enzyme | 1.02E+07 | 4671314 | 8138046 | | | | | | |
| valosin containing protein | Cytoplasm | enzyme | 7667627.5 | 1.28E+07 | 8704454 | | | | | | |
| valosin containing protein | Cytoplasm | enzyme | 6308800.5 | 2455331.75 | 3560066.75 | | | | | | |
| valosin containing protein | Cytoplasm | enzyme | 5127807.5 | 2100115.5 | 2701028 | | | | | | |
| valosin containing protein | Cytoplasm | enzyme | 4659787 | 9343810 | 6795294 | | | | | | |
| valosin containing protein | Cytoplasm | enzyme | 4583773 | 799576.812 | 2019905.25 | | | | | | |
| valosin containing protein | Cytoplasm | enzyme | 3601116.25 | 5687547 | 7860775.5 | | | | | | |
| valosin containing protein | Cytoplasm | enzyme | 3389736.25 | 1673662.875 | 2820368.5 | | | | | | |
| valosin containing protein | Cytoplasm | enzyme | 2960290.25 | 1050003.125 | 1635230.125 | | | | | | |
| valosin containing protein | Cytoplasm | enzyme | 2450707.75 | 625690.812 | 1982508.875 | | | | | | |
| valosin containing protein | Cytoplasm | enzyme | 2275093.75 | 4156755 | 5012108 | | | | | | |
| valosin containing protein | Cytoplasm | enzyme | 1854467.125 | 3290296.5 | 4753573 | Q01853 | VCP | 80983834 | 67387764 | 4825738 | 195 |
| voltage-dependent anion channel 1 | Cytoplasm | ion channel | 1.23E+07 | 2.96E+07 | 1.86E+07 | | | | | | |
| voltage-dependent anion channel 1 | Cytoplasm | ion channel | 1.10E+07 | 9088783 | 8787898 | | | | | | |
| voltage-dependent anion channel 1 | Cytoplasm | ion channel | 6900763 | 1.04E+07 | 1.20E+07 | | | | | | |
| voltage-dependent anion channel 1 | Cytoplasm | ion channel | 6052721 | 1.59E+07 | 1.10E+07 | | | | | | |
| voltage-dependent anion channel 1 | Cytoplasm | ion channel | 6047257.5 | 1.71E+07 | 1.38E+07 | | | | | | |
| voltage-dependent anion channel 1 | Cytoplasm | ion channel | 6030771.5 | 1.08E+07 | 1.03E+07 | | | | | | |
| voltage-dependent anion channel 1 | Cytoplasm | ion channel | 4479381 | 1.05E+07 | 9756926 | | | | | | |
| voltage-dependent anion channel 1 | Cytoplasm | ion channel | 1670113 | 1.13E+07 | 5462020 | Q60932 | VDAC1 | 6812626 | 5657721 | 1154877 | 24 |
| voltage-dependent anion channel 2 | Cytoplasm | ion channel | 1.19E+07 | 1.81E+07 | 1.81E+07 | | | | | | |
| voltage-dependent anion channel 2 | Cytoplasm | ion channel | 1.07E+07 | 1.93E+07 | 1.84E+07 | | | | | | |
| voltage-dependent anion channel 2 | Cytoplasm | ion channel | 7186625 | 1.24E+07 | 1.32E+07 | | | | | | |
| voltage-dependent anion channel 2 | Cytoplasm | ion channel | 6534015 | 1.53E+07 | 1.29E+07 | | | | | | |
| voltage-dependent anion channel 2 | Cytoplasm | ion channel | 6481973.5 | 1.61E+07 | 1.10E+07 | | | | | | |
| voltage-dependent anion channel 2 | Cytoplasm | ion channel | 2354483.75 | 5923451.5 | 4829710.5 | | | | | | |
| voltage-dependent anion channel 2 | Cytoplasm | ion channel | 1601600.5 | 3288569.75 | 4217352.5 | | | | | | |
| voltage-dependent anion channel 2 | Cytoplasm | ion channel | 611147.812 | 1248077.5 | 1085020.125 | | | | | | |
| voltage-dependent anion channel 2 | Cytoplasm | ion channel | 142809.969 | 2312157 | 1221813.375 | Q60930 | VDAC2 | 5274740 | 6432491 | 1237933 | 27 |
| voltage-dependent anion channel 3 | Cytoplasm | ion channel | 1.40E+07 | 2.59E+07 | 2.24E+07 | | | | | | |
| voltage-dependent anion channel 3 | Cytoplasm | ion channel | 8230736.5 | 1.51E+07 | 1.53E+07 | | | | | | |
| voltage-dependent anion channel 3 | Cytoplasm | ion channel | 7379339.5 | 1.64E+07 | 1.69E+07 | | | | | | |
| voltage-dependent anion channel 3 | Cytoplasm | ion channel | 6033180.5 | 1.91E+07 | 1.01E+07 | | | | | | |
| voltage-dependent anion channel 3 | Cytoplasm | ion channel | 3324626.25 | 1.06E+07 | 7283022.5 | | | | | | |
| voltage-dependent anion channel 3 | Cytoplasm | ion channel | 831716.125 | 3762646.25 | 2432972 | Q60931 | VDAC3 | 6631600 | 7164265 | 1688633 | 18 |
| vimentin | Cytoplasm | other | 37215.66 | 621407.188 | 889828.375 | P20152 | VIM | 37216 | 435943 | 251692 | 3 |
| VPS33B interacting protein, apical-basolateral polarity regulator, spe-39 homolog | Plasma Membrane | other | 3.53E+07 | 2.13E+07 | 2.20E+07 | Q8BGQ1 | VIPAS39 | 35280000 | 7865979 | 4541425 | 3 |
| vacuole membrane protein 1 | Plasma Membrane | other | 2.13E+07 | 1.48E+07 | 1.89E+07 | Q99KU0 | VMP1 | 21340000 | 3313734 | 1913185 | 3 |
| vacuolar protein sorting 13 homolog A (S. cerevisiae) | Cytoplasm | transporter | 2.11E+07 | 1.71E+07 | 1.77E+07 | | | | | | |
| vacuolar protein sorting 13 homolog A (S. cerevisiae) | Cytoplasm | transporter | 1.80E+07 | 1.16E+07 | 1.24E+07 | | | | | | |
| vacuolar protein sorting 13 homolog A (S. cerevisiae) | Cytoplasm | transporter | 1.30E+07 | 1.10E+07 | 1.24E+07 | | | | | | |
| vacuolar protein sorting 13 homolog A (S. cerevisiae) | Cytoplasm | transporter | 1.25E+07 | 6450834.5 | 1.05E+07 | | | | | | |
| vacuolar protein sorting 13 homolog A (S. cerevisiae) | Cytoplasm | transporter | 1.22E+07 | 1.12E+07 | 1.09E+07 | | | | | | |
| vacuolar protein sorting 13 homolog A (S. cerevisiae) | Cytoplasm | transporter | 8710360 | 6869428.5 | 8011978 | | | | | | |
| vacuolar protein sorting 13 homolog A (S. cerevisiae) | Cytoplasm | transporter | 7951416.5 | 5520401.5 | 6383352.5 | | | | | | |
| vacuolar protein sorting 13 homolog A (S. cerevisiae) | Cytoplasm | transporter | 7898031.5 | 7401660 | 6808162.5 | | | | | | |
| vacuolar protein sorting 13 homolog A (S. cerevisiae) | Cytoplasm | transporter | 7557549 | 5008041.5 | 6480038 | | | | | | |
| vacuolar protein sorting 13 homolog A (S. cerevisiae) | Cytoplasm | transporter | 7457480.5 | 6274815.5 | 6456256 | | | | | | |
| vacuolar protein sorting 13 homolog A (S. cerevisiae) | Cytoplasm | transporter | 7302933 | 7599061.5 | 7591525 | | | | | | |
| vacuolar protein sorting 13 homolog A (S. cerevisiae) | Cytoplasm | transporter | 7027788 | 4768205 | 3862142.25 | | | | | | |
| vacuolar protein sorting 13 homolog A (S. cerevisiae) | Cytoplasm | transporter | 6838213 | 5992229 | 5863033.5 | | | | | | |
| vacuolar protein sorting 13 homolog A (S. cerevisiae) | Cytoplasm | transporter | 6696580.5 | 5760260.5 | 7420227 | | | | | | |
| vacuolar protein sorting 13 homolog A (S. cerevisiae) | Cytoplasm | transporter | 6399666 | 5612842 | 6151916 | | | | | | |
| vacuolar protein sorting 13 homolog A (S. cerevisiae) | Cytoplasm | transporter | 6079586.5 | 4913517 | 4796081.5 | | | | | | |
| vacuolar protein sorting 13 homolog A (S. cerevisiae) | Cytoplasm | transporter | 5562364 | 5030678 | 6359422 | | | | | | |
| vacuolar protein sorting 13 homolog A (S. cerevisiae) | Cytoplasm | transporter | 4899800 | 5219046 | 4895255 | | | | | | |
| vacuolar protein sorting 13 homolog A (S. cerevisiae) | Cytoplasm | transporter | 4344733 | 3256496.75 | 2747047.25 | | | | | | |
| vacuolar protein sorting 13 homolog A (S. cerevisiae) | Cytoplasm | transporter | 4183945 | 1927707.375 | 2855937.5 | | | | | | |
| vacuolar protein sorting 13 homolog A (S. cerevisiae) | Cytoplasm | transporter | 4063566.5 | 3319197.75 | 3701441.25 | | | | | | |
| vacuolar protein sorting 13 homolog A (S. cerevisiae) | Cytoplasm | transporter | 4054656.25 | 3527182.75 | 3063633.75 | | | | | | |
| vacuolar protein sorting 13 homolog A (S. cerevisiae) | Cytoplasm | transporter | 4023080.25 | 4076702 | 4096482.25 | | | | | | |
| vacuolar protein sorting 13 homolog A (S. cerevisiae) | Cytoplasm | transporter | 3469626 | 3774952.25 | 3901472.25 | | | | | | |
| vacuolar protein sorting 13 homolog A (S. cerevisiae) | Cytoplasm | transporter | 3203424.75 | 3172440.25 | 3248591.75 | | | | | | |
| vacuolar protein sorting 13 homolog A (S. cerevisiae) | Cytoplasm | transporter | 2758936.75 | 2405027.75 | 1814669.75 | | | | | | |
| vacuolar protein sorting 13 homolog A (S. cerevisiae) | Cytoplasm | transporter | 2675969.5 | 4225736.5 | 3983966.5 | | | | | | |
| vacuolar protein sorting 13 homolog A (S. cerevisiae) | Cytoplasm | transporter | 2349008.5 | 2104513.5 | 2326178 | | | | | | |
| vacuolar protein sorting 13 homolog A (S. cerevisiae) | Cytoplasm | transporter | 2091773.625 | 858797.062 | 1001917.062 | | | | | | |
| vacuolar protein sorting 13 homolog A (S. cerevisiae) | Cytoplasm | transporter | 2081943.625 | 1666733 | 1914986.25 | | | | | | |
| vacuolar protein sorting 13 homolog A (S. cerevisiae) | Cytoplasm | transporter | 1747583.25 | 2254966.25 | 1682285.75 | | | | | | |
| vacuolar protein sorting 13 homolog A (S. cerevisiae) | Cytoplasm | transporter | 1279509.75 | 1503196.625 | 1383302.125 | | | | | | |
| vacuolar protein sorting 13 homolog A (S. cerevisiae) | Cytoplasm | transporter | 999617.625 | 988616 | 1151466.875 | | | | | | |
| vacuolar protein sorting 13 homolog A (S. cerevisiae) | Cytoplasm | transporter | 787854.312 | 377793.562 | 2707659.75 | Q5H8C4 | VPS13A | 6211971 | 4002161 | 396273 | 102 |
| vacuolar protein sorting 13 homolog C (S. cerevisiae) | Cytoplasm | other | 1893101.125 | 1444157.375 | 1606485.625 | | | | | | |
| vacuolar protein sorting 13 homolog C (S. cerevisiae) | Cytoplasm | other | 1814432.625 | 1200541.125 | 552803.688 | | | | | | |
| vacuolar protein sorting 13 homolog C (S. cerevisiae) | Cytoplasm | other | 1313201.5 | 720802.688 | 1171527.375 | | | | | | |
| vacuolar protein sorting 13 homolog C (S. cerevisiae) | Cytoplasm | other | 1156654.25 | 1001559.625 | 1018774.312 | | | | | | |
| vacuolar protein sorting 13 homolog C (S. cerevisiae) | Cytoplasm | other | 1082455.75 | 685897.812 | 633206 | Q8BX70 | VPS13C | 1451969 | 412767 | 106576 | 15 |
| vacuolar protein sorting 28 homolog (S. cerevisiae) | Cytoplasm | transporter | 1365972.375 | 1.15E+07 | 6.12E+07 | Q9D1C8 | VPS28 | 1365972 | 32030451 | 18492790 | 3 |
| von Willebrand factor | Extracellular Space | other | 443614.812 | 1.00E+07 | 1.26E+07 | | | | | | |
| von Willebrand factor | Extracellular Space | other | 330451.781 | 2653578.5 | 3670106.25 | | | | | | |
| von Willebrand factor | Extracellular Space | other | 182511.891 | 2950047.75 | 4988640.5 | | | | | | |
| von Willebrand factor | Extracellular Space | other | 168430.188 | 3743776.25 | 6188379 | | | | | | |
| von Willebrand factor | Extracellular Space | other | 154886.109 | 2120266.25 | 4160811.25 | | | | | | |
| von Willebrand factor | Extracellular Space | other | 130058.031 | 1954476.25 | 3546024.5 | | | | | | |
| von Willebrand factor | Extracellular Space | other | 121052.305 | 2959720.75 | 4413031 | | | | | | |
| von Willebrand factor | Extracellular Space | other | 107526.336 | 3360604.75 | 4566921 | | | | | | |
| von Willebrand factor | Extracellular Space | other | 105893.945 | 7192840 | 9314574 | | | | | | |
| von Willebrand factor | Extracellular Space | other | 94770.016 | 3829592.75 | 4930426 | | | | | | |
| von Willebrand factor | Extracellular Space | other | 73244.594 | 3385034.75 | 4807422.5 | | | | | | |
| von Willebrand factor | Extracellular Space | other | 49588.555 | 2022805.125 | 2826350 | | | | | | |
| von Willebrand factor | Extracellular Space | other | 41865.191 | 1638839.875 | 3338603.5 | | | | | | |
| von Willebrand factor | Extracellular Space | other | 6851.076 | 1944394.75 | 3280522.75 | | | | | | |
| von Willebrand factor | Extracellular Space | other | 4.70E-06 | 1302297.25 | 1737251 | Q8CIZ8 | VWF | 134050 | 2861133 | 426513 | 45 |
| WD repeat domain 37 | Nucleus | other | 4146130.5 | 2393424.75 | 2829058.75 | | | | | | |
| WD repeat domain 37 | Nucleus | other | 1706643.625 | 915208.5 | 989501.938 | Q8CBE3 | WDR37 | 2926387 | 1230600 | 502390 | 6 |

| | | | | | | | | | | | |
|---|---|---|---|---|---|---|---|---|---|---|---|
| WD repeat domain 48 | Cytoplasm | other | 974612.5 | 1091039 | 813363.375 | Q8BH57 | WDR48 | 974613 | 139439 | 80505 | 3 |
| WAP four-disulfide core domain 8 | Extracellular Space | other | 1142846.75 | 652230.625 | 649802.625 | Q4KUS1 | WFDC8 | 1142847 | 283961 | 163945 | 3 |
| WNK lysine deficient protein kinase 1 | Cytoplasm | kinase | 2197521.5 | 687628.5 | 1155009.625 | P83741 | WNK1 | 2197522 | 772987 | 446284 | 3 |
| X-linked Kx blood group | Plasma Membrane | transporter | 3.53E+07 | 2.58E+07 | 2.51E+07 | | | | | | |
| X-linked Kx blood group | Plasma Membrane | transporter | 3.20E+07 | 2.09E+07 | 2.43E+07 | | | | | | |
| X-linked Kx blood group | Plasma Membrane | transporter | 2.85E+07 | 3.03E+07 | 2.57E+07 | | | | | | |
| X-linked Kx blood group | Plasma Membrane | transporter | 2.17E+07 | 1.66E+07 | 1.58E+07 | Q9QXY7 | XK | 29377500 | 5889750 | 1700224 | 12 |
| exportin 1 | Nucleus | transporter | 2714303 | 1569724.375 | 1830287.875 | Q6P5F9 | XPO1 | 2714303 | 599922 | 346365 | 3 |
| exportin 7 | Nucleus | transporter | 1961839.5 | 1186313.25 | 919731.938 | | | | | | |
| exportin 7 | Nucleus | transporter | 1823191.125 | 1084843.125 | 829876.25 | Q9EPK7 | XPO7 | 1892515 | 476779 | 194644 | 6 |
| Y box binding protein 1 | Nucleus | transcription regulator | 2662313.75 | 1617272.75 | 1918014.125 | | | | | | |
| Y box binding protein 1 | Nucleus | transcription regulator | 356734.719 | 135046.484 | 135871.484 | P62960 | YBX1 | 1509524 | 1075380 | 439022 | 6 |
| Y box binding protein 3 | Nucleus | transcription regulator | 3526171.25 | 1490498.25 | 1914401.875 | | | | | | |
| Y box binding protein 3 | Nucleus | transcription regulator | 1993480.625 | 1233126.75 | 1836365.75 | Q9JKB3 | YBX3 | 2759826 | 801377 | 327161 | 6 |
| Yip1 interacting factor homolog B (S. cerevisiae) | Cytoplasm | other | 3379546.25 | 2502876.25 | 2713448 | | | | | | |
| Yip1 interacting factor homolog B (S. cerevisiae) | Cytoplasm | other | 2658540.25 | 2095573 | 2402172.5 | Q9CX30 | YIF1B | 3019043 | 429685 | 175418 | 6 |
| tyrosine 3-monooxygenase/tryptophan 5-monooxygenase activation protein, beta | Cytoplasm | transcription regulator | 5171631.5 | 5341714 | 4696224 | | | | | | |
| tyrosine 3-monooxygenase/tryptophan 5-monooxygenase activation protein, beta | Cytoplasm | transcription regulator | 3982280.5 | 5594538.5 | 4809017 | | | | | | |
| tyrosine 3-monooxygenase/tryptophan 5-monooxygenase activation protein, beta | Cytoplasm | transcription regulator | 2481308.5 | 4040403.5 | 3605208.75 | | | | | | |
| tyrosine 3-monooxygenase/tryptophan 5-monooxygenase activation protein, beta | Cytoplasm | transcription regulator | 1563105.75 | 2940354.25 | 2852519 | | | | | | |
| tyrosine 3-monooxygenase/tryptophan 5-monooxygenase activation protein, beta | Cytoplasm | transcription regulator | 446530.562 | 1157733.75 | 679671.125 | | | | | | |
| tyrosine 3-monooxygenase/tryptophan 5-monooxygenase activation protein, beta | Cytoplasm | transcription regulator | 95819.727 | 1137289.25 | 720437.688 | Q9CQV8 | YWHAB | 2290113 | 1876350 | 442260 | 18 |
| tyrosine 3-monooxygenase/tryptophan 5-monooxygenase activation protein, epsilon | Cytoplasm | other | 1.16E+07 | 1.61E+07 | 1.44E+07 | | | | | | |
| tyrosine 3-monooxygenase/tryptophan 5-monooxygenase activation protein, epsilon | Cytoplasm | other | 9167160 | 1.05E+07 | 9978127 | | | | | | |
| tyrosine 3-monooxygenase/tryptophan 5-monooxygenase activation protein, epsilon | Cytoplasm | other | 6874644 | 8904180 | 7550680.5 | | | | | | |
| tyrosine 3-monooxygenase/tryptophan 5-monooxygenase activation protein, epsilon | Cytoplasm | other | 6486524 | 6990309 | 7713308.5 | | | | | | |
| tyrosine 3-monooxygenase/tryptophan 5-monooxygenase activation protein, epsilon | Cytoplasm | other | 4663239 | 6808812 | 6236975 | | | | | | |
| tyrosine 3-monooxygenase/tryptophan 5-monooxygenase activation protein, epsilon | Cytoplasm | other | 4390967.5 | 4799183.5 | 4645330 | | | | | | |
| tyrosine 3-monooxygenase/tryptophan 5-monooxygenase activation protein, epsilon | Cytoplasm | other | 4321038.5 | 6171149 | 7135298.5 | | | | | | |
| tyrosine 3-monooxygenase/tryptophan 5-monooxygenase activation protein, epsilon | Cytoplasm | other | 4013073 | 6788405.5 | 5447768.5 | | | | | | |
| tyrosine 3-monooxygenase/tryptophan 5-monooxygenase activation protein, epsilon | Cytoplasm | other | 775641.375 | 2922554.75 | 3584218.75 | P62259 | YWHAE | 5810254 | 3384502 | 651348 | 27 |
| tyrosine 3-monooxygenase/tryptophan 5-monooxygenase activation protein, gamma | Cytoplasm | other | 2004465.375 | 4853818 | 3436415.5 | | | | | | |
| tyrosine 3-monooxygenase/tryptophan 5-monooxygenase activation protein, gamma | Cytoplasm | other | 885537.375 | 2533060.5 | 1928856.75 | P61982 | YWHAG | 1445001 | 1380431 | 563559 | 6 |
| tyrosine 3-monooxygenase/tryptophan 5-monooxygenase activation protein, eta | Cytoplasm | transcription regulator | 4670984 | 6.34E+07 | 3.55E+07 | | | | | | |
| tyrosine 3-monooxygenase/tryptophan 5-monooxygenase activation protein, eta | Cytoplasm | transcription regulator | 1937206.75 | 1.71E+07 | 1.24E+07 | | | | | | |
| tyrosine 3-monooxygenase/tryptophan 5-monooxygenase activation protein, eta | Cytoplasm | transcription regulator | 1706495.625 | 2.73E+07 | 1.58E+07 | | | | | | |
| tyrosine 3-monooxygenase/tryptophan 5-monooxygenase activation protein, eta | Cytoplasm | transcription regulator | 1319439.75 | 1.46E+07 | 1.01E+07 | | | | | | |
| tyrosine 3-monooxygenase/tryptophan 5-monooxygenase activation protein, eta | Cytoplasm | transcription regulator | 926292.312 | 1.45E+07 | 7694122 | | | | | | |
| tyrosine 3-monooxygenase/tryptophan 5-monooxygenase activation protein, eta | Cytoplasm | transcription regulator | 408307.688 | 3351119.75 | 3070525 | | | | | | |
| tyrosine 3-monooxygenase/tryptophan 5-monooxygenase activation protein, eta | Cytoplasm | transcription regulator | 378693.562 | 8538763 | 6471672 | | | | | | |
| tyrosine 3-monooxygenase/tryptophan 5-monooxygenase activation protein, eta | Cytoplasm | transcription regulator | 316119.875 | 7492050 | 4695957.5 | | | | | | |
| tyrosine 3-monooxygenase/tryptophan 5-monooxygenase activation protein, eta | Cytoplasm | transcription regulator | 0 | 2497416.5 | 1146593.25 | P68510 | YWHAH | 1295949 | 13746149 | 2645448 | 27 |
| tyrosine 3-monooxygenase/tryptophan 5-monooxygenase activation protein, theta | Cytoplasm | other | 1216692.25 | 5801789 | 4230914 | | | | | | |
| tyrosine 3-monooxygenase/tryptophan 5-monooxygenase activation protein, theta | Cytoplasm | other | 673675.625 | 3575237.75 | 2247100.5 | | | | | | |
| tyrosine 3-monooxygenase/tryptophan 5-monooxygenase activation protein, theta | Cytoplasm | other | 371173.75 | 3921849 | 2030625.5 | P68254 | YWHAQ | 753847 | 1823480 | 607827 | 9 |
| tyrosine 3-monooxygenase/tryptophan 5-monooxygenase activation protein, zeta | Cytoplasm | enzyme | 2.26E+07 | 7.04E+07 | 6.31E+07 | | | | | | |
| tyrosine 3-monooxygenase/tryptophan 5-monooxygenase activation protein, zeta | Cytoplasm | enzyme | 1.63E+07 | 3.62E+07 | 3.26E+07 | | | | | | |
| tyrosine 3-monooxygenase/tryptophan 5-monooxygenase activation protein, zeta | Cytoplasm | enzyme | 1.41E+07 | 4.39E+07 | 3.12E+07 | | | | | | |
| tyrosine 3-monooxygenase/tryptophan 5-monooxygenase activation protein, zeta | Cytoplasm | enzyme | 1.32E+07 | 6.36E+07 | 5.09E+07 | | | | | | |
| tyrosine 3-monooxygenase/tryptophan 5-monooxygenase activation protein, zeta | Cytoplasm | enzyme | 1.22E+07 | 6.12E+07 | 4.39E+07 | | | | | | |
| tyrosine 3-monooxygenase/tryptophan 5-monooxygenase activation protein, zeta | Cytoplasm | enzyme | 9525490 | 3.35E+07 | 1.21E+07 | | | | | | |
| tyrosine 3-monooxygenase/tryptophan 5-monooxygenase activation protein, zeta | Cytoplasm | enzyme | 8994783 | 4.12E+07 | 4.21E+07 | | | | | | |
| tyrosine 3-monooxygenase/tryptophan 5-monooxygenase activation protein, zeta | Cytoplasm | enzyme | 7550299 | 2.95E+07 | 2.63E+07 | | | | | | |
| tyrosine 3-monooxygenase/tryptophan 5-monooxygenase activation protein, zeta | Cytoplasm | enzyme | 3949322.25 | 3.37E+07 | 2.15E+07 | | | | | | |
| tyrosine 3-monooxygenase/tryptophan 5-monooxygenase activation protein, zeta | Cytoplasm | enzyme | 2693096.5 | 4449136.5 | 5550694 | | | | | | |
| tyrosine 3-monooxygenase/tryptophan 5-monooxygenase activation protein, zeta | Cytoplasm | enzyme | 2186142.25 | 2.54E+07 | 1.39E+07 | | | | | | |
| tyrosine 3-monooxygenase/tryptophan 5-monooxygenase activation protein, zeta | Cytoplasm | enzyme | 1826668.25 | 7082020 | 6847663.5 | | | | | | |
| tyrosine 3-monooxygenase/tryptophan 5-monooxygenase activation protein, zeta | Cytoplasm | enzyme | 1578915.25 | 8076008.5 | 5760871 | | | | | | |
| tyrosine 3-monooxygenase/tryptophan 5-monooxygenase activation protein, zeta | Cytoplasm | enzyme | 1245976.25 | 7318054.5 | 8062355 | | | | | | |
| tyrosine 3-monooxygenase/tryptophan 5-monooxygenase activation protein, zeta | Cytoplasm | enzyme | 681270 | 6204969.5 | 2946985.25 | | | | | | |
| tyrosine 3-monooxygenase/tryptophan 5-monooxygenase activation protein, zeta | Cytoplasm | enzyme | 295811.062 | 4546467.5 | 1947301.75 | P63101 | YWHAZ | 7429236 | 19514816 | 2816721 | 48 |
| zinc finger protein 524 | Nucleus | other | 4883685.5 | 3928988 | 5100172 | Q9D0B1 | ZNF524 | 4883686 | 623162 | 359783 | 3 |
| zinc finger protein 717 | Nucleus | other | 4.41E+08 | 2.67E+08 | 2.65E+08 | Q02525 | ZNF717 | 440700000 | 100836320 | 58217876 | 3 |
| zinc finger, ZZ-type with EF-hand domain 1 | Cytoplasm | other | 3550270 | 1918355.875 | 1551248.875 | Q5SSH7 | ZZEF1 | 3550270 | 1064111 | 614365 | 3 |

| | # Peptides by Location |
|---|---|
| Plasma Membrane | 1902 |
| Cytoplasm | 4392 |
| Nucleus | 523 |
| Extracellular Space | 347 |
| Other | 24 |
| **Total Peptides** | 7188 |

AQP1 knockout

| ID | Symbol | Entrez Gene Name | Primary Location | Type(s) | Intensity [AQP1KO_1] | Intensity [AQP1KO_2] | Intensity [AQP1KO_3] | ID | Symbol | Mean Intensity | St. Dev | Std. Err | n |
|---|---|---|---|---|---|---|---|---|---|---|---|---|---|
| Q3UHJ0 | AAK1 | AP2 associated kinase 1 | Cytoplasm | kinase | 1315149.25 | 641206.062 | 569730.875 | Q3UHJ0 | AAK1 | 1315149 | 411290 | 237458 | 3 |
| Q9JI39 | ABCB10 | ATP-binding cassette, sub-family B | Cytoplasm | transporter | 1.48E+07 | 1.28E+07 | 2.35E+07 | | | | | | |
| Q9JI39 | ABCB10 | ATP-binding cassette, sub-family B | Cytoplasm | transporter | 1.21E+07 | 1.20E+07 | 1.63E+07 | | | | | | |
| Q9JI39 | ABCB10 | ATP-binding cassette, sub-family B | Cytoplasm | transporter | 5134905 | 6097008.5 | 7486709 | | | | | | |
| Q9JI39 | ABCB10 | ATP-binding cassette, sub-family B | Cytoplasm | transporter | 2767352.5 | 2076733.375 | 1790009.625 | | | | | | |
| Q9JI39 | ABCB10 | ATP-binding cassette, sub-family B | Cytoplasm | transporter | 2965125.25 | 3346896.75 | 4309698.5 | | | | | | |
| Q9JI39 | ABCB10 | ATP-binding cassette, sub-family B | Cytoplasm | transporter | 3482082.5 | 3801826.75 | 5171538 | | | | | | |
| Q9JI39 | ABCB10 | ATP-binding cassette, sub-family B | Cytoplasm | transporter | 3455503 | 2893680.75 | 6031781.5 | | | | | | |
| Q9JI39 | ABCB10 | ATP-binding cassette, sub-family B | Cytoplasm | transporter | 2971433.25 | 3278868 | 4440326 | | | | | | |
| Q9JI39 | ABCB10 | ATP-binding cassette, sub-family B | Cytoplasm | transporter | 1887297.375 | 2486945.75 | 3856554.5 | | | | | | |
| Q9JI39 | ABCB10 | ATP-binding cassette, sub-family B | Cytoplasm | transporter | 2538405.5 | 3991995.5 | 6175970.5 | | | | | | |
| Q9JI39 | ABCB10 | ATP-binding cassette, sub-family B | Cytoplasm | transporter | 1238706.5 | 1098555.25 | 2793082.5 | | | | | | |
| Q9JI39 | ABCB10 | ATP-binding cassette, sub-family B | Cytoplasm | transporter | 945867.812 | 915423.375 | 1316312.125 | Q9JI39 | ABCB10 | 4521390 | 5033113 | 838852 | 36 |
| P21440 | ABCB4 | ATP-binding cassette, sub-family B | Plasma Membrane | transporter | 2.71E+07 | 2.07E+07 | 2.55E+07 | | | | | | |
| P21440 | ABCB4 | ATP-binding cassette, sub-family B | Plasma Membrane | transporter | 1.60E+07 | 1.15E+07 | 1.12E+07 | | | | | | |
| P21440 | ABCB4 | ATP-binding cassette, sub-family B | Plasma Membrane | transporter | 1.49E+07 | 1.49E+07 | 1.24E+07 | | | | | | |
| P21440 | ABCB4 | ATP-binding cassette, sub-family B | Plasma Membrane | transporter | 9780653 | 1.08E+07 | 1.18E+07 | | | | | | |
| P21440 | ABCB4 | ATP-binding cassette, sub-family B | Plasma Membrane | transporter | 1.73E+07 | 1.34E+07 | 1.23E+07 | | | | | | |
| P21440 | ABCB4 | ATP-binding cassette, sub-family B | Plasma Membrane | transporter | 1.15E+07 | 1.11E+07 | 1.48E+07 | | | | | | |
| P21440 | ABCB4 | ATP-binding cassette, sub-family B | Plasma Membrane | transporter | 1.49E+07 | 1.30E+07 | 1.30E+07 | | | | | | |
| P21440 | ABCB4 | ATP-binding cassette, sub-family B | Plasma Membrane | transporter | 1.30E+07 | 1.06E+07 | 1.01E+07 | | | | | | |
| P21440 | ABCB4 | ATP-binding cassette, sub-family B | Plasma Membrane | transporter | 1.28E+07 | 1.01E+07 | 1.04E+07 | | | | | | |
| P21440 | ABCB4 | ATP-binding cassette, sub-family B | Plasma Membrane | transporter | 1.14E+07 | 8578539 | 1.06E+07 | | | | | | |
| P21440 | ABCB4 | ATP-binding cassette, sub-family B | Plasma Membrane | transporter | 9552567 | 9319960 | 9306429 | | | | | | |
| P21440 | ABCB4 | ATP-binding cassette, sub-family B | Plasma Membrane | transporter | 9177367 | 7542390 | 8882015 | | | | | | |
| P21440 | ABCB4 | ATP-binding cassette, sub-family B | Plasma Membrane | transporter | 7792765 | 6725211 | 5455553 | | | | | | |
| P21440 | ABCB4 | ATP-binding cassette, sub-family B | Plasma Membrane | transporter | 7419671.5 | 6667009 | 6861859 | | | | | | |
| P21440 | ABCB4 | ATP-binding cassette, sub-family B | Plasma Membrane | transporter | 8661057 | 7294413.5 | 7978149 | | | | | | |
| P21440 | ABCB4 | ATP-binding cassette, sub-family B | Plasma Membrane | transporter | 8620775 | 6410018 | 8433190 | | | | | | |
| P21440 | ABCB4 | ATP-binding cassette, sub-family B | Plasma Membrane | transporter | 5508740.5 | 4483818.5 | 5447695.5 | | | | | | |
| P21440 | ABCB4 | ATP-binding cassette, sub-family B | Plasma Membrane | transporter | 4113016.25 | 4170613.5 | 5050830.5 | | | | | | |
| P21440 | ABCB4 | ATP-binding cassette, sub-family B | Plasma Membrane | transporter | 4005628.25 | 2923840 | 2982802 | | | | | | |
| P21440 | ABCB4 | ATP-binding cassette, sub-family B | Plasma Membrane | transporter | 3789498.5 | 2222289.75 | 2097144.75 | | | | | | |
| P21440 | ABCB4 | ATP-binding cassette, sub-family B | Plasma Membrane | transporter | 3352497 | 3567431.25 | 3395247.75 | | | | | | |
| P21440 | ABCB4 | ATP-binding cassette, sub-family B | Plasma Membrane | transporter | 3727024.75 | 3882297.75 | 2361420.25 | | | | | | |
| P21440 | ABCB4 | ATP-binding cassette, sub-family B | Plasma Membrane | transporter | 2225233.25 | 1799774.5 | 1563902.75 | | | | | | |
| P21440 | ABCB4 | ATP-binding cassette, sub-family B | Plasma Membrane | transporter | 450019.094 | 1579705.625 | 1967734 | P21440 | ABCB4 | 9453605 | 5342443 | 629613 | 72 |
| Q9DC29 | ABCB6 | ATP-binding cassette, sub-family B | Cytoplasm | transporter | 8.75E+07 | 9.55E+07 | 8.46E+07 | | | | | | |
| Q9DC29 | ABCB6 | ATP-binding cassette, sub-family B | Cytoplasm | transporter | 8.32E+07 | 9.58E+07 | 1.04E+08 | | | | | | |
| Q9DC29 | ABCB6 | ATP-binding cassette, sub-family B | Cytoplasm | transporter | 3.36E+07 | 4.18E+07 | 2.95E+07 | | | | | | |
| Q9DC29 | ABCB6 | ATP-binding cassette, sub-family B | Cytoplasm | transporter | 3.18E+07 | 4.29E+07 | 3.54E+07 | | | | | | |
| Q9DC29 | ABCB6 | ATP-binding cassette, sub-family B | Cytoplasm | transporter | 4.28E+07 | 4.59E+07 | 4.06E+07 | | | | | | |
| Q9DC29 | ABCB6 | ATP-binding cassette, sub-family B | Cytoplasm | transporter | 2.35E+07 | 4.07E+07 | 3.36E+07 | | | | | | |
| Q9DC29 | ABCB6 | ATP-binding cassette, sub-family B | Cytoplasm | transporter | 2.09E+07 | 2.85E+07 | 2.43E+07 | | | | | | |
| Q9DC29 | ABCB6 | ATP-binding cassette, sub-family B | Cytoplasm | transporter | 1.27E+07 | 1.71E+07 | 1.42E+07 | | | | | | |
| Q9DC29 | ABCB6 | ATP-binding cassette, sub-family B | Cytoplasm | transporter | 1.45E+07 | 2.37E+07 | 1.83E+07 | | | | | | |
| Q9DC29 | ABCB6 | ATP-binding cassette, sub-family B | Cytoplasm | transporter | 1.51E+07 | 1.23E+07 | 1.24E+07 | | | | | | |
| Q9DC29 | ABCB6 | ATP-binding cassette, sub-family B | Cytoplasm | transporter | 1.01E+07 | 1.37E+07 | 9591124 | | | | | | |
| Q9DC29 | ABCB6 | ATP-binding cassette, sub-family B | Cytoplasm | transporter | 6505775.5 | 2.08E+07 | 1.57E+07 | | | | | | |
| Q9DC29 | ABCB6 | ATP-binding cassette, sub-family B | Cytoplasm | transporter | 7223106 | 1.25E+07 | 1.09E+07 | | | | | | |
| Q9DC29 | ABCB6 | ATP-binding cassette, sub-family B | Cytoplasm | transporter | 4488753 | 1.20E+07 | 7160934.5 | | | | | | |
| Q9DC29 | ABCB6 | ATP-binding cassette, sub-family B | Cytoplasm | transporter | 3426374.5 | 1.22E+07 | 8845434 | | | | | | |
| Q9DC29 | ABCB6 | ATP-binding cassette, sub-family B | Cytoplasm | transporter | 4943896 | 5511619.5 | 5755294 | | | | | | |
| Q9DC29 | ABCB6 | ATP-binding cassette, sub-family B | Cytoplasm | transporter | 3077742.75 | 3730957.5 | 2783491 | | | | | | |
| Q9DC29 | ABCB6 | ATP-binding cassette, sub-family B | Cytoplasm | transporter | 1602856.75 | 4574552.5 | 3682101.25 | | | | | | |
| Q9DC29 | ABCB6 | ATP-binding cassette, sub-family B | Cytoplasm | transporter | 550334.688 | 1509370.875 | 213271.109 | | | | | | |
| Q9DC29 | ABCB6 | ATP-binding cassette, sub-family B | Cytoplasm | transporter | 257122.781 | 1038832.125 | 2279535.5 | Q9DC29 | ABCB6 | 20386298 | 26347912 | 3401501 | 60 |
| Q61102 | ABCB7 | ATP-binding cassette, sub-family B | Cytoplasm | transporter | 1824504.375 | 1791831.75 | 2696493 | Q61102 | ABCB7 | 1824504 | 513135 | 296258 | 3 |
| Q9JJ59 | ABCB9 | ATP-binding cassette, sub-family B | Cytoplasm | transporter | 8725452 | 7470840 | 9011007 | Q9JJ59 | ABCB9 | 8725452 | 819320 | 473034 | 3 |
| O35379 | ABCC1 | ATP-binding cassette, sub-family C | Plasma Membrane | transporter | 1.53E+07 | 1.36E+07 | 1.50E+07 | | | | | | |
| O35379 | ABCC1 | ATP-binding cassette, sub-family C | Plasma Membrane | transporter | 1.44E+07 | 1.41E+07 | 1.48E+07 | | | | | | |
| O35379 | ABCC1 | ATP-binding cassette, sub-family C | Plasma Membrane | transporter | 1.01E+07 | 1.14E+07 | 1.12E+07 | | | | | | |
| O35379 | ABCC1 | ATP-binding cassette, sub-family C | Plasma Membrane | transporter | 1.21E+07 | 1.22E+07 | 1.38E+07 | | | | | | |
| O35379 | ABCC1 | ATP-binding cassette, sub-family C | Plasma Membrane | transporter | 1.00E+07 | 9390528 | 1.01E+07 | | | | | | |
| O35379 | ABCC1 | ATP-binding cassette, sub-family C | Plasma Membrane | transporter | 7110079.5 | 7067996.5 | 7574361 | | | | | | |
| O35379 | ABCC1 | ATP-binding cassette, sub-family C | Plasma Membrane | transporter | 4972443 | 5429021.5 | 3036780.75 | | | | | | |
| O35379 | ABCC1 | ATP-binding cassette, sub-family C | Plasma Membrane | transporter | 7044457 | 6453937.5 | 7363698 | | | | | | |
| O35379 | ABCC1 | ATP-binding cassette, sub-family C | Plasma Membrane | transporter | 5575399 | 5651611.5 | 5486385.5 | | | | | | |
| O35379 | ABCC1 | ATP-binding cassette, sub-family C | Plasma Membrane | transporter | 4647667 | 4982912 | 4820566 | | | | | | |
| O35379 | ABCC1 | ATP-binding cassette, sub-family C | Plasma Membrane | transporter | 3434105.25 | 4516966.5 | 3701582.5 | | | | | | |
| O35379 | ABCC1 | ATP-binding cassette, sub-family C | Plasma Membrane | transporter | 3295895.25 | 4505546.5 | 4897783 | | | | | | |
| O35379 | ABCC1 | ATP-binding cassette, sub-family C | Plasma Membrane | transporter | 2519416.25 | 2989538.75 | 4368776 | | | | | | |
| O35379 | ABCC1 | ATP-binding cassette, sub-family C | Plasma Membrane | transporter | 3145939.25 | 2502879.75 | 2389394.25 | | | | | | |
| O35379 | ABCC1 | ATP-binding cassette, sub-family C | Plasma Membrane | transporter | 2809037.5 | 3667410 | 3153335.75 | | | | | | |
| O35379 | ABCC1 | ATP-binding cassette, sub-family C | Plasma Membrane | transporter | 3166235.25 | 2999772.25 | 3335512.25 | | | | | | |
| O35379 | ABCC1 | ATP-binding cassette, sub-family C | Plasma Membrane | transporter | 2241797.25 | 2268059.5 | 2476069.5 | | | | | | |
| O35379 | ABCC1 | ATP-binding cassette, sub-family C | Plasma Membrane | transporter | 1411711.375 | 2044942.875 | 2173169.25 | O35379 | ABCC1 | 6289121 | 4175035 | 568150 | 54 |
| Q9R1S7 | ABCC6 | ATP-binding cassette, sub-family C | Plasma Membrane | transporter | 2968568.25 | 7557686 | 8341698 | Q9R1S7 | ABCC6 | 2968568 | 2902447 | 1675729 | 3 |
| P61222 | ABCE1 | ATP-binding cassette, sub-family E | Cytoplasm | transporter | 4314818.5 | 5352311.5 | 2307816.75 | | | | | | |
| P61222 | ABCE1 | ATP-binding cassette, sub-family E | Cytoplasm | transporter | 2670108.5 | 3431863.75 | 1678209.25 | | | | | | |
| P61222 | ABCE1 | ATP-binding cassette, sub-family E | Cytoplasm | transporter | 3716667.5 | 4168675.5 | 2104121 | | | | | | |
| P61222 | ABCE1 | ATP-binding cassette, sub-family E | Cytoplasm | transporter | 1854091.875 | 1925215.125 | 949026.188 | P61222 | ABCE1 | 3138922 | 1312394 | 378855 | 12 |
| Q7TMS5 | ABCG2 | ATP-binding cassette, sub-family G | Plasma Membrane | transporter | 7.97E+07 | 9.91E+07 | 7.29E+07 | | | | | | |
| Q7TMS5 | ABCG2 | ATP-binding cassette, sub-family G | Plasma Membrane | transporter | 6.53E+07 | 7.98E+07 | 6.13E+07 | | | | | | |
| Q7TMS5 | ABCG2 | ATP-binding cassette, sub-family G | Plasma Membrane | transporter | 5.94E+07 | 7.37E+07 | 4.53E+07 | | | | | | |

| | | | | | | | | | | | | | |
|---|---|---|---|---|---|---|---|---|---|---|---|---|---|
| Q7TMS5 | ABCG2 | ATP-binding cassette, sub-family G | Plasma Membrane | transporter | 5.41E+07 | 5.96E+07 | 5.40E+07 | | | | | | |
| Q7TMS5 | ABCG2 | ATP-binding cassette, sub-family G | Plasma Membrane | transporter | 4.04E+07 | 4.56E+07 | 4.12E+07 | | | | | | |
| Q7TMS5 | ABCG2 | ATP-binding cassette, sub-family G | Plasma Membrane | transporter | 2.98E+07 | 2.72E+07 | 2.69E+07 | | | | | | |
| Q7TMS5 | ABCG2 | ATP-binding cassette, sub-family G | Plasma Membrane | transporter | 2.85E+07 | 3.51E+07 | 3.07E+07 | | | | | | |
| Q7TMS5 | ABCG2 | ATP-binding cassette, sub-family G | Plasma Membrane | transporter | 2.17E+07 | 2.67E+07 | 2.00E+07 | | | | | | |
| Q7TMS5 | ABCG2 | ATP-binding cassette, sub-family G | Plasma Membrane | transporter | 2.65E+07 | 3.32E+07 | 2.39E+07 | | | | | | |
| Q7TMS5 | ABCG2 | ATP-binding cassette, sub-family G | Plasma Membrane | transporter | 1.99E+07 | 2.23E+07 | 2.03E+07 | | | | | | |
| Q7TMS5 | ABCG2 | ATP-binding cassette, sub-family G | Plasma Membrane | transporter | 1.34E+07 | 1.98E+07 | 1.29E+07 | | | | | | |
| Q7TMS5 | ABCG2 | ATP-binding cassette, sub-family G | Plasma Membrane | transporter | 9569948 | 1.44E+07 | 1.18E+07 | | | | | | |
| Q7TMS5 | ABCG2 | ATP-binding cassette, sub-family G | Plasma Membrane | transporter | 1.17E+07 | 1.08E+07 | 9239083 | | | | | | |
| Q7TMS5 | ABCG2 | ATP-binding cassette, sub-family G | Plasma Membrane | transporter | 1.36E+07 | 1.58E+07 | 1.58E+07 | | | | | | |
| Q7TMS5 | ABCG2 | ATP-binding cassette, sub-family G | Plasma Membrane | transporter | 5411348.5 | 9329692 | 7469204.5 | | | | | | |
| Q7TMS5 | ABCG2 | ATP-binding cassette, sub-family G | Plasma Membrane | transporter | 6022609 | 8341622.5 | 6983919.5 | | | | | | |
| Q7TMS5 | ABCG2 | ATP-binding cassette, sub-family G | Plasma Membrane | transporter | 3195923.25 | 4193627.75 | 1784041.125 | | | | | | |
| Q7TMS5 | ABCG2 | ATP-binding cassette, sub-family G | Plasma Membrane | transporter | 2581528 | 6011800.5 | 9196885 | | | | | | |
| Q7TMS5 | ABCG2 | ATP-binding cassette, sub-family G | Plasma Membrane | transporter | 4231501.5 | 4409261.5 | 4648851.5 | | | | | | |
| Q7TMS5 | ABCG2 | ATP-binding cassette, sub-family G | Plasma Membrane | transporter | 1021413.562 | 1119728.75 | 593855 | Q7TMS5 | ABCG2 | 24794714 | 23999376 | 3098306 | 60 |
| Q9Z1Q2 | ABHD16A | abhydrolase domain containing 16 | Cytoplasm | enzyme | 5239455.5 | 4237711 | 4334915 | | | | | | |
| Q9Z1Q2 | ABHD16A | abhydrolase domain containing 16 | Cytoplasm | enzyme | 3024969.5 | 5315220.5 | 4630233.5 | | | | | | |
| Q9Z1Q2 | ABHD16A | abhydrolase domain containing 16 | Cytoplasm | enzyme | 710930.438 | 936081.125 | 1080353 | Q9Z1Q2 | ABHD16A | 2991785 | 1898431 | 632810 | 9 |
| Q921H8 | ACAA1 | acetyl-CoA acyltransferase 1 | Cytoplasm | enzyme | 3403160.25 | 3570933.5 | 4651105.5 | Q921H8 | ACAA1 | 3403160 | 677285 | 391030 | 3 |
| Q8BWT1 | ACAA2 | acetyl-CoA acyltransferase 2 | Cytoplasm | enzyme | 8353512.5 | 8423863 | 1.33E+07 | | | | | | |
| Q8BWT1 | ACAA2 | acetyl-CoA acyltransferase 2 | Cytoplasm | enzyme | 6468088.5 | 6025313 | 8508107 | | | | | | |
| Q8BWT1 | ACAA2 | acetyl-CoA acyltransferase 2 | Cytoplasm | enzyme | 4876221 | 3916401 | 7226459.5 | | | | | | |
| Q8BWT1 | ACAA2 | acetyl-CoA acyltransferase 2 | Cytoplasm | enzyme | 5300833.5 | 4615069 | 5117999 | | | | | | |
| Q8BWT1 | ACAA2 | acetyl-CoA acyltransferase 2 | Cytoplasm | enzyme | 3630944 | 3411631.5 | 4481592 | | | | | | |
| Q8BWT1 | ACAA2 | acetyl-CoA acyltransferase 2 | Cytoplasm | enzyme | 3239688.5 | 4774247 | 5557515 | Q8BWT1 | ACAA2 | 5311548 | 2484897 | 585696 | 18 |
| Q8JZN5 | ACAD9 | acyl-CoA dehydrogenase family, m | Cytoplasm | enzyme | 2784238.25 | 2146972 | 2834082 | Q8JZN5 | ACAD9 | 2784238 | 383126 | 221198 | 3 |
| P51174 | ACADL | acyl-CoA dehydrogenase, long cha | Cytoplasm | enzyme | 3.69E+07 | 2.81E+07 | 4.53E+07 | | | | | | |
| P51174 | ACADL | acyl-CoA dehydrogenase, long cha | Cytoplasm | enzyme | 1.21E+07 | 1.19E+07 | 1.37E+07 | | | | | | |
| P51174 | ACADL | acyl-CoA dehydrogenase, long cha | Cytoplasm | enzyme | 1.02E+07 | 8795517 | 1.07E+07 | | | | | | |
| P51174 | ACADL | acyl-CoA dehydrogenase, long cha | Cytoplasm | enzyme | 5382427 | 5062054 | 6047133.5 | | | | | | |
| P51174 | ACADL | acyl-CoA dehydrogenase, long cha | Cytoplasm | enzyme | 8174849 | 3815104 | 4740339 | | | | | | |
| P51174 | ACADL | acyl-CoA dehydrogenase, long cha | Cytoplasm | enzyme | 5440412 | 6683849 | 7718229.5 | | | | | | |
| P51174 | ACADL | acyl-CoA dehydrogenase, long cha | Cytoplasm | enzyme | 1742383 | 6193355.5 | 4535748 | P51174 | ACADL | 11412867 | 11291388 | 2463983 | 21 |
| P45952 | ACADM | acyl-CoA dehydrogenase, C-4 to C- | Cytoplasm | enzyme | 1169462.375 | 1085542.75 | 1455036.375 | P45952 | ACADM | 1169462 | 193701 | 111833 | 3 |
| P50544 | ACADVL | acyl-CoA dehydrogenase, very long | Cytoplasm | enzyme | 3232987.5 | 3446462.5 | 4409895 | | | | | | |
| P50544 | ACADVL | acyl-CoA dehydrogenase, very long | Cytoplasm | enzyme | 2027679.5 | 4117153.75 | 3277306.5 | | | | | | |
| P50544 | ACADVL | acyl-CoA dehydrogenase, very long | Cytoplasm | enzyme | 1547262.25 | 1428928.375 | 1775688.5 | | | | | | |
| P50544 | ACADVL | acyl-CoA dehydrogenase, very long | Cytoplasm | enzyme | 1125155.875 | 1500051.25 | 1850945.125 | | | | | | |
| P50544 | ACADVL | acyl-CoA dehydrogenase, very long | Cytoplasm | enzyme | 772376.312 | 765357.5 | 873456.5 | P50544 | ACADVL | 1741092 | 1228989 | 317324 | 15 |
| Q8QZT1 | ACAT1 | acetyl-CoA acetyltransferase 1 | Cytoplasm | enzyme | 1.53E+07 | 1.76E+07 | 2.02E+07 | | | | | | |
| Q8QZT1 | ACAT1 | acetyl-CoA acetyltransferase 1 | Cytoplasm | enzyme | 1.02E+07 | 1.17E+07 | 1.45E+07 | | | | | | |
| Q8QZT1 | ACAT1 | acetyl-CoA acetyltransferase 1 | Cytoplasm | enzyme | 7180518.5 | 7993669 | 8978147 | | | | | | |
| Q8QZT1 | ACAT1 | acetyl-CoA acetyltransferase 1 | Cytoplasm | enzyme | 1.23E+07 | 1.20E+07 | 1.31E+07 | | | | | | |
| Q8QZT1 | ACAT1 | acetyl-CoA acetyltransferase 1 | Cytoplasm | enzyme | 6599672 | 7100870 | 9515958 | | | | | | |
| Q8QZT1 | ACAT1 | acetyl-CoA acetyltransferase 1 | Cytoplasm | enzyme | 4644215.5 | 5333631 | 7915275.5 | | | | | | |
| Q8QZT1 | ACAT1 | acetyl-CoA acetyltransferase 1 | Cytoplasm | enzyme | 3293615.25 | 5842858.5 | 6112809 | Q8QZT1 | ACAT1 | 8484003 | 4451039 | 971296 | 21 |
| Q9QUI6 | ACKR1 | atypical chemokine receptor 1 (Du | Plasma Membrane | G-protein coupled receptor | 2.45E+07 | 3.06E+07 | 2.88E+07 | | | | | | |
| Q9QUI6 | ACKR1 | atypical chemokine receptor 1 (Du | Plasma Membrane | G-protein coupled receptor | 2.77E+07 | 3.28E+07 | 3.23E+07 | | | | | | |
| Q9QUI6 | ACKR1 | atypical chemokine receptor 1 (Du | Plasma Membrane | G-protein coupled receptor | 6240862 | 8494639 | 8486843 | | | | | | |
| Q9QUI6 | ACKR1 | atypical chemokine receptor 1 (Du | Plasma Membrane | G-protein coupled receptor | 5320406.5 | 480069.844 | 319805.531 | Q9QUI6 | ACKR1 | 15942817 | 13232739 | 3819963 | 12 |
| Q99KI0 | ACO2 | aconitase 2, mitochondrial | Cytoplasm | enzyme | 2.21E+07 | 2.67E+07 | 3.00E+07 | | | | | | |
| Q99KI0 | ACO2 | aconitase 2, mitochondrial | Cytoplasm | enzyme | 1.36E+07 | 1.35E+07 | 1.91E+07 | | | | | | |
| Q99KI0 | ACO2 | aconitase 2, mitochondrial | Cytoplasm | enzyme | 1.07E+07 | 9399966 | 1.51E+07 | | | | | | |
| Q99KI0 | ACO2 | aconitase 2, mitochondrial | Cytoplasm | enzyme | 5342364 | 1.29E+07 | 1.76E+07 | | | | | | |
| Q99KI0 | ACO2 | aconitase 2, mitochondrial | Cytoplasm | enzyme | 1.26E+07 | 8700822 | 9234056 | | | | | | |
| Q99KI0 | ACO2 | aconitase 2, mitochondrial | Cytoplasm | enzyme | 7700695 | 1.10E+07 | 1.12E+07 | | | | | | |
| Q99KI0 | ACO2 | aconitase 2, mitochondrial | Cytoplasm | enzyme | 7570821 | 7957017 | 1.08E+07 | | | | | | |
| Q99KI0 | ACO2 | aconitase 2, mitochondrial | Cytoplasm | enzyme | 5627404.5 | 8332205 | 1.23E+07 | | | | | | |
| Q99KI0 | ACO2 | aconitase 2, mitochondrial | Cytoplasm | enzyme | 8905280 | 8800583 | 1.22E+07 | | | | | | |
| Q99KI0 | ACO2 | aconitase 2, mitochondrial | Cytoplasm | enzyme | 6628208 | 5130331 | 7753342 | | | | | | |
| Q99KI0 | ACO2 | aconitase 2, mitochondrial | Cytoplasm | enzyme | 5922236.5 | 7042861 | 1.09E+07 | | | | | | |
| Q99KI0 | ACO2 | aconitase 2, mitochondrial | Cytoplasm | enzyme | 3038568.75 | 4419363.5 | 4964733.5 | | | | | | |
| Q99KI0 | ACO2 | aconitase 2, mitochondrial | Cytoplasm | enzyme | 4112533.5 | 5305063.5 | 9532339 | | | | | | |
| Q99KI0 | ACO2 | aconitase 2, mitochondrial | Cytoplasm | enzyme | 5671462.5 | 4327980.5 | 5954977.5 | | | | | | |
| Q99KI0 | ACO2 | aconitase 2, mitochondrial | Cytoplasm | enzyme | 3852182.25 | 3903481.5 | 5765372 | | | | | | |
| Q99KI0 | ACO2 | aconitase 2, mitochondrial | Cytoplasm | enzyme | 2626181.5 | 2601220.5 | 3089010.5 | | | | | | |
| Q99KI0 | ACO2 | aconitase 2, mitochondrial | Cytoplasm | enzyme | 1933348.75 | 2388569.75 | 2822915.5 | | | | | | |
| Q99KI0 | ACO2 | aconitase 2, mitochondrial | Cytoplasm | enzyme | 1730987.625 | 6137217.5 | 6248390.5 | | | | | | |
| Q99KI0 | ACO2 | aconitase 2, mitochondrial | Cytoplasm | enzyme | 1113869.875 | 4441517 | 4464518.5 | | | | | | |
| Q99KI0 | ACO2 | aconitase 2, mitochondrial | Cytoplasm | enzyme | 1661490.875 | 1824922.625 | 3742749.25 | | | | | | |
| Q99KI0 | ACO2 | aconitase 2, mitochondrial | Cytoplasm | enzyme | 1146711.125 | 3850089 | 2239378.5 | Q99KI0 | ACO2 | 6360207 | 5878156 | 740578 | 63 |
| P41216 | ACSL1 | acyl-CoA synthetase long-chain far | Cytoplasm | enzyme | 2.12E+07 | 2.15E+07 | 1.85E+07 | | | | | | |
| P41216 | ACSL1 | acyl-CoA synthetase long-chain far | Cytoplasm | enzyme | 1.34E+07 | 1.90E+07 | 1.75E+07 | | | | | | |
| P41216 | ACSL1 | acyl-CoA synthetase long-chain far | Cytoplasm | enzyme | 1.49E+07 | 1.89E+07 | 1.63E+07 | | | | | | |
| P41216 | ACSL1 | acyl-CoA synthetase long-chain far | Cytoplasm | enzyme | 1.17E+07 | 1.35E+07 | 1.13E+07 | | | | | | |
| P41216 | ACSL1 | acyl-CoA synthetase long-chain far | Cytoplasm | enzyme | 9455292 | 2.70E+07 | 2.08E+07 | | | | | | |
| P41216 | ACSL1 | acyl-CoA synthetase long-chain far | Cytoplasm | enzyme | 1.49E+07 | 1.54E+07 | 1.78E+07 | | | | | | |
| P41216 | ACSL1 | acyl-CoA synthetase long-chain far | Cytoplasm | enzyme | 9681499 | 1.15E+07 | 1.06E+07 | | | | | | |
| P41216 | ACSL1 | acyl-CoA synthetase long-chain far | Cytoplasm | enzyme | 8184385 | 8652286 | 9024755 | | | | | | |
| P41216 | ACSL1 | acyl-CoA synthetase long-chain far | Cytoplasm | enzyme | 5942039 | 6993791.5 | 4868333 | | | | | | |
| P41216 | ACSL1 | acyl-CoA synthetase long-chain far | Cytoplasm | enzyme | 5221691.5 | 4976304.5 | 2993884.25 | | | | | | |
| P41216 | ACSL1 | acyl-CoA synthetase long-chain far | Cytoplasm | enzyme | 6870411.5 | 1.01E+07 | 6923664 | | | | | | |
| P41216 | ACSL1 | acyl-CoA synthetase long-chain far | Cytoplasm | enzyme | 5314878 | 6867937.5 | 4768271 | | | | | | |
| P41216 | ACSL1 | acyl-CoA synthetase long-chain far | Cytoplasm | enzyme | 3477832.5 | 3836652 | 3699547.75 | | | | | | |

| | | | | | | | | | | | | | |
|---|---|---|---|---|---|---|---|---|---|---|---|---|---|
| P41216 | ACSL1 | acyl-CoA synthetase long-chain far | Cytoplasm | enzyme | 4377544.5 | 3766194 | 2753496 | | | | | | |
| P41216 | ACSL1 | acyl-CoA synthetase long-chain far | Cytoplasm | enzyme | 3403043 | 4499561 | 3750281.75 | | | | | | |
| P41216 | ACSL1 | acyl-CoA synthetase long-chain far | Cytoplasm | enzyme | 3587876 | 4460751.5 | 4055691.5 | | | | | | |
| P41216 | ACSL1 | acyl-CoA synthetase long-chain far | Cytoplasm | enzyme | 4307706 | 6238236.5 | 4475142.5 | P41216 | ACSL1 | 8580835 | 6206721 | 869115 | 51 |
| Q9QUJ7 | ACSL4 | acyl-CoA synthetase long-chain far | Cytoplasm | enzyme | 1.22E+07 | 1.43E+07 | 1.31E+07 | | | | | | |
| Q9QUJ7 | ACSL4 | acyl-CoA synthetase long-chain far | Cytoplasm | enzyme | 1.44E+07 | 1.45E+07 | 1.44E+07 | | | | | | |
| Q9QUJ7 | ACSL4 | acyl-CoA synthetase long-chain far | Cytoplasm | enzyme | 8199007 | 1.13E+07 | 1.24E+07 | | | | | | |
| Q9QUJ7 | ACSL4 | acyl-CoA synthetase long-chain far | Cytoplasm | enzyme | 4193813.75 | 4992214.5 | 5518231.5 | | | | | | |
| Q9QUJ7 | ACSL4 | acyl-CoA synthetase long-chain far | Cytoplasm | enzyme | 1951520.625 | 1997247 | 2042157 | | | | | | |
| Q9QUJ7 | ACSL4 | acyl-CoA synthetase long-chain far | Cytoplasm | enzyme | 2653015 | 2104527 | 1999470.625 | | | | | | |
| Q9QUJ7 | ACSL4 | acyl-CoA synthetase long-chain far | Cytoplasm | enzyme | 2650751.5 | 2364088.25 | 2822411.5 | | | | | | |
| Q9QUJ7 | ACSL4 | acyl-CoA synthetase long-chain far | Cytoplasm | enzyme | 2992508.25 | 2410333 | 1948890 | | | | | | |
| Q9QUJ7 | ACSL4 | acyl-CoA synthetase long-chain far | Cytoplasm | enzyme | 2316331.75 | 1861156.375 | 1916341.25 | Q9QUJ7 | ACSL4 | 5722994 | 5031521 | 968317 | 27 |
| Q8JZR0 | ACSL5 | acyl-CoA synthetase long-chain far | Cytoplasm | enzyme | 9413270 | 1.08E+07 | 1.20E+07 | | | | | | |
| Q8JZR0 | ACSL5 | acyl-CoA synthetase long-chain far | Cytoplasm | enzyme | 9620572 | 8704911 | 8729794 | | | | | | |
| Q8JZR0 | ACSL5 | acyl-CoA synthetase long-chain far | Cytoplasm | enzyme | 6594268 | 7894585 | 6002260 | | | | | | |
| Q8JZR0 | ACSL5 | acyl-CoA synthetase long-chain far | Cytoplasm | enzyme | 5882927.5 | 6774613.5 | 4382935 | | | | | | |
| Q8JZR0 | ACSL5 | acyl-CoA synthetase long-chain far | Cytoplasm | enzyme | 4759923.5 | 5555086.5 | 5298388 | | | | | | |
| Q8JZR0 | ACSL5 | acyl-CoA synthetase long-chain far | Cytoplasm | enzyme | 3766090 | 3938395 | 4046987.75 | | | | | | |
| Q8JZR0 | ACSL5 | acyl-CoA synthetase long-chain far | Cytoplasm | enzyme | 4904918 | 5280137.5 | 4481269 | | | | | | |
| Q8JZR0 | ACSL5 | acyl-CoA synthetase long-chain far | Cytoplasm | enzyme | 2848003 | 2526100.5 | 2359055.75 | | | | | | |
| Q8JZR0 | ACSL5 | acyl-CoA synthetase long-chain far | Cytoplasm | enzyme | 2919851 | 3429340.5 | 3108521.25 | | | | | | |
| Q8JZR0 | ACSL5 | acyl-CoA synthetase long-chain far | Cytoplasm | enzyme | 2169691.5 | 3249140.5 | 2611426.25 | | | | | | |
| Q8JZR0 | ACSL5 | acyl-CoA synthetase long-chain far | Cytoplasm | enzyme | 3382529.75 | 2580326.25 | 2275947.5 | Q8JZR0 | ACSL5 | 5114731 | 2686380 | 467639 | 33 |
| Q99NB1 | ACSS1 | acyl-CoA synthetase short-chain fa | Cytoplasm | enzyme | 4427494 | 2358082.75 | 3031669.5 | | | | | | |
| Q99NB1 | ACSS1 | acyl-CoA synthetase short-chain fa | Cytoplasm | enzyme | 2136824.5 | 1902518.5 | 2890814.75 | | | | | | |
| Q99NB1 | ACSS1 | acyl-CoA synthetase short-chain fa | Cytoplasm | enzyme | 1137554.625 | 1343116.875 | 2311676.5 | Q99NB1 | ACSS1 | 2567291 | 986617 | 328872 | 9 |
| P60710 | ACTB | actin, beta | Cytoplasm | other | 4.46E+09 | 4.57E+09 | 4.50E+09 | | | | | | |
| P60710 | ACTB | actin, beta | Cytoplasm | other | 3.05E+09 | 2.65E+09 | 2.95E+09 | | | | | | |
| P60710 | ACTB | actin, beta | Cytoplasm | other | 2.71E+09 | 2.47E+09 | 2.56E+09 | | | | | | |
| P60710 | ACTB | actin, beta | Cytoplasm | other | 1.39E+09 | 1.29E+09 | 1.25E+09 | | | | | | |
| P60710 | ACTB | actin, beta | Cytoplasm | other | 1.82E+09 | 1.05E+09 | 1.33E+09 | | | | | | |
| P60710 | ACTB | actin, beta | Cytoplasm | other | 8.30E+08 | 1.61E+09 | 1.19E+09 | | | | | | |
| P60710 | ACTB | actin, beta | Cytoplasm | other | 1.11E+09 | 1.44E+09 | 1.27E+09 | | | | | | |
| P60710 | ACTB | actin, beta | Cytoplasm | other | 9.78E+08 | 1.07E+09 | 1.07E+09 | | | | | | |
| P60710 | ACTB | actin, beta | Cytoplasm | other | 1.28E+09 | 9.53E+08 | 1.08E+09 | | | | | | |
| P60710 | ACTB | actin, beta | Cytoplasm | other | 7.97E+08 | 1.14E+09 | 7.44E+08 | | | | | | |
| P60710 | ACTB | actin, beta | Cytoplasm | other | 1.19E+09 | 9.49E+08 | 1.05E+09 | | | | | | |
| P60710 | ACTB | actin, beta | Cytoplasm | other | 8.75E+08 | 9.29E+08 | 9.95E+08 | | | | | | |
| P60710 | ACTB | actin, beta | Cytoplasm | other | 5.51E+08 | 7.10E+08 | 6.75E+08 | | | | | | |
| P60710 | ACTB | actin, beta | Cytoplasm | other | 5.58E+08 | 5.69E+08 | 5.80E+08 | | | | | | |
| P60710 | ACTB | actin, beta | Cytoplasm | other | 4.85E+08 | 3.12E+08 | 4.41E+08 | | | | | | |
| P60710 | ACTB | actin, beta | Cytoplasm | other | 3.37E+08 | 3.26E+08 | 3.22E+08 | | | | | | |
| P60710 | ACTB | actin, beta | Cytoplasm | other | 2.41E+08 | 3.48E+08 | 3.32E+08 | | | | | | |
| P60710 | ACTB | actin, beta | Cytoplasm | other | 1.78E+08 | 2.50E+08 | 1.55E+08 | | | | | | |
| P60710 | ACTB | actin, beta | Cytoplasm | other | 1.56E+08 | 1.90E+08 | 1.41E+08 | | | | | | |
| P60710 | ACTB | actin, beta | Cytoplasm | other | 2.05E+08 | 3.51E+08 | 2.12E+08 | | | | | | |
| P60710 | ACTB | actin, beta | Cytoplasm | other | 2.13E+08 | 1.26E+08 | 2.12E+08 | | | | | | |
| P60710 | ACTB | actin, beta | Cytoplasm | other | 1.74E+08 | 9.14E+07 | 2.09E+08 | | | | | | |
| P60710 | ACTB | actin, beta | Cytoplasm | other | 3.90E+08 | 1.80E+07 | 1.15E+08 | | | | | | |
| P60710 | ACTB | actin, beta | Cytoplasm | other | 6.52E+07 | 9.89E+07 | 9.57E+07 | | | | | | |
| P60710 | ACTB | actin, beta | Cytoplasm | other | 8.82E+07 | 5.57E+07 | 8.40E+07 | | | | | | |
| P60710 | ACTB | actin, beta | Cytoplasm | other | 2.83E+08 | 1.39E+07 | 8.65E+07 | | | | | | |
| P60710 | ACTB | actin, beta | Cytoplasm | other | 8.14E+07 | 1.33E+08 | 1.03E+08 | | | | | | |
| P60710 | ACTB | actin, beta | Cytoplasm | other | 7.94E+07 | 1.01E+08 | 8.53E+07 | | | | | | |
| P60710 | ACTB | actin, beta | Cytoplasm | other | 7.35E+07 | 6.05E+07 | 6.25E+07 | | | | | | |
| P60710 | ACTB | actin, beta | Cytoplasm | other | 5.34E+07 | 6.92E+07 | 6.86E+07 | | | | | | |
| P60710 | ACTB | actin, beta | Cytoplasm | other | 4.49E+07 | 9.12E+07 | 6.94E+07 | | | | | | |
| P60710 | ACTB | actin, beta | Cytoplasm | other | 1.38E+08 | 1.45E+08 | 1.75E+08 | | | | | | |
| P60710 | ACTB | actin, beta | Cytoplasm | other | 3.40E+07 | 5.23E+07 | 5.09E+07 | | | | | | |
| P60710 | ACTB | actin, beta | Cytoplasm | other | 8.32E+07 | 9261604 | 2.70E+07 | | | | | | |
| P60710 | ACTB | actin, beta | Cytoplasm | other | 2.71E+07 | 1.61E+07 | 2.36E+07 | | | | | | |
| P60710 | ACTB | actin, beta | Cytoplasm | other | 2.73E+07 | 2.98E+07 | 1.74E+07 | | | | | | |
| P60710 | ACTB | actin, beta | Cytoplasm | other | 6.58E+07 | 2794779.25 | 1.10E+07 | | | | | | |
| P60710 | ACTB | actin, beta | Cytoplasm | other | 1.77E+07 | 1.73E+07 | 4.14E+07 | | | | | | |
| P60710 | ACTB | actin, beta | Cytoplasm | other | 5.73E+07 | 2989513 | 2.00E+07 | | | | | | |
| P60710 | ACTB | actin, beta | Cytoplasm | other | 1.22E+07 | 3742182.75 | 2199691.25 | | | | | | |
| P60710 | ACTB | actin, beta | Cytoplasm | other | 1.44E+07 | 8248124.5 | 9482930 | | | | | | |
| P60710 | ACTB | actin, beta | Cytoplasm | other | 1.59E+07 | 3.74E+08 | 3.97E+08 | | | | | | |
| P60710 | ACTB | actin, beta | Cytoplasm | other | 3831635 | 1.01E+07 | 1.55E+07 | | | | | | |
| P60710 | ACTB | actin, beta | Cytoplasm | other | 5105090 | 9403684 | 8724199 | | | | | | |
| P60710 | ACTB | actin, beta | Cytoplasm | other | 3131245.75 | 6066276.5 | 5334903 | | | | | | |
| P60710 | ACTB | actin, beta | Cytoplasm | other | 2586749 | 2461937 | 1720724.5 | | | | | | |
| P60710 | ACTB | actin, beta | Cytoplasm | other | 936325.062 | 6.91E+07 | 7.99E+07 | | | | | | |
| P60710 | ACTB | actin, beta | Cytoplasm | other | 141865.328 | 162757.547 | 341222.625 | P60710 | ACTB | 525967144 | 869684155 | 72473680 | 144 |
| P68033 | ACTC1 | actin, alpha, cardiac muscle 1 | Cytoplasm | enzyme | 2.93E+09 | 3.11E+09 | 2.81E+09 | | | | | | |
| P68033 | ACTC1 | actin, alpha, cardiac muscle 1 | Cytoplasm | enzyme | 4.09E+08 | 5.10E+08 | 3.80E+08 | | | | | | |
| P68033 | ACTC1 | actin, alpha, cardiac muscle 1 | Cytoplasm | enzyme | 1.92E+08 | 2.04E+08 | 1.60E+08 | | | | | | |
| P68033 | ACTC1 | actin, alpha, cardiac muscle 1 | Cytoplasm | enzyme | 2.35E+07 | 2.44E+07 | 8166488.5 | P68033 | ACTC1 | 889175000 | 1250947627 | 361117475 | 12 |
| P61161 | ACTR2 | ARP2 actin-related protein 2 homo | Plasma Membrane | other | 8407380 | 6888169.5 | 5412120.5 | | | | | | |
| P61161 | ACTR2 | ARP2 actin-related protein 2 homo | Plasma Membrane | other | 2739122 | 3390076 | 2354100.25 | | | | | | |
| P61161 | ACTR2 | ARP2 actin-related protein 2 homo | Plasma Membrane | other | 3303913.75 | 3682038 | 2660748.5 | P61161 | ACTR2 | 4816805 | 2118842 | 706281 | 9 |
| Q99JY9 | ACTR3 | ARP3 actin-related protein 3 homo | Plasma Membrane | other | 1.56E+07 | 1.42E+07 | 1.10E+07 | | | | | | |
| Q99JY9 | ACTR3 | ARP3 actin-related protein 3 homo | Plasma Membrane | other | 1941864.375 | 4587637.5 | 2344265.75 | | | | | | |
| Q99JY9 | ACTR3 | ARP3 actin-related protein 3 homo | Plasma Membrane | other | 2476832.25 | 5447100.5 | 4246502.5 | | | | | | |
| Q99JY9 | ACTR3 | ARP3 actin-related protein 3 homo | Plasma Membrane | other | 2326682.25 | 2619361 | 1666422.875 | | | | | | |

| | | | | | | | | | | | | | |
|---|---|---|---|---|---|---|---|---|---|---|---|---|---|
| Q99JY9 | ACTR3 | ARP3 actin-related protein 3 homo | Plasma Membrane | other | 2763004 | 3106105 | 2328722.75 | Q99JY9 | ACTR3 | 5021677 | 4603367 | 1188584 | 15 |
| Q641P0 | ACTR3B | ARP3 actin-related protein 3 homo | Cytoplasm | other | 3382200.5 | 3839868.5 | 4981982.5 | Q641P0 | ACTR3B | 3382201 | 823932 | 475698 | 3 |
| O35598 | ADAM10 | ADAM metallopeptidase domain 1 | Plasma Membrane | peptidase | 6.42E+07 | 3.97E+07 | 4.46E+07 | | | | | | |
| O35598 | ADAM10 | ADAM metallopeptidase domain 1 | Plasma Membrane | peptidase | 2.17E+07 | 3.52E+07 | 2.52E+07 | | | | | | |
| O35598 | ADAM10 | ADAM metallopeptidase domain 1 | Plasma Membrane | peptidase | 3.28E+07 | 3.31E+07 | 2.57E+07 | | | | | | |
| O35598 | ADAM10 | ADAM metallopeptidase domain 1 | Plasma Membrane | peptidase | 4.38E+07 | 4.29E+07 | 4.06E+07 | | | | | | |
| O35598 | ADAM10 | ADAM metallopeptidase domain 1 | Plasma Membrane | peptidase | 3.10E+07 | 3.47E+07 | 2.67E+07 | | | | | | |
| O35598 | ADAM10 | ADAM metallopeptidase domain 1 | Plasma Membrane | peptidase | 2.60E+07 | 3.03E+07 | 2.30E+07 | | | | | | |
| O35598 | ADAM10 | ADAM metallopeptidase domain 1 | Plasma Membrane | peptidase | 3.56E+07 | 3.95E+07 | 3.33E+07 | | | | | | |
| O35598 | ADAM10 | ADAM metallopeptidase domain 1 | Plasma Membrane | peptidase | 2.35E+07 | 2.46E+07 | 2.28E+07 | | | | | | |
| O35598 | ADAM10 | ADAM metallopeptidase domain 1 | Plasma Membrane | peptidase | 1.38E+07 | 1.35E+07 | 1.36E+07 | | | | | | |
| O35598 | ADAM10 | ADAM metallopeptidase domain 1 | Plasma Membrane | peptidase | 2.08E+07 | 1.06E+07 | 9098009 | | | | | | |
| O35598 | ADAM10 | ADAM metallopeptidase domain 1 | Plasma Membrane | peptidase | 1.56E+07 | 1.78E+07 | 1.54E+07 | | | | | | |
| O35598 | ADAM10 | ADAM metallopeptidase domain 1 | Plasma Membrane | peptidase | 1.57E+07 | 5996086.5 | 2442877.25 | | | | | | |
| O35598 | ADAM10 | ADAM metallopeptidase domain 1 | Plasma Membrane | peptidase | 7330035 | 2.45E+07 | 1.35E+07 | | | | | | |
| O35598 | ADAM10 | ADAM metallopeptidase domain 1 | Plasma Membrane | peptidase | 8187766.5 | 9615407 | 8316365.5 | | | | | | |
| O35598 | ADAM10 | ADAM metallopeptidase domain 1 | Plasma Membrane | peptidase | 1.12E+07 | 7132579.5 | 7720465 | | | | | | |
| O35598 | ADAM10 | ADAM metallopeptidase domain 1 | Plasma Membrane | peptidase | 5983181.5 | 5164764 | 5982211.5 | | | | | | |
| O35598 | ADAM10 | ADAM metallopeptidase domain 1 | Plasma Membrane | peptidase | 8614181 | 7270862.5 | 7762127.5 | | | | | | |
| O35598 | ADAM10 | ADAM metallopeptidase domain 1 | Plasma Membrane | peptidase | 3994984 | 2921200.75 | 2569576.25 | | | | | | |
| O35598 | ADAM10 | ADAM metallopeptidase domain 1 | Plasma Membrane | peptidase | 2263817 | 2190632.5 | 1680782.375 | | | | | | |
| O35598 | ADAM10 | ADAM metallopeptidase domain 1 | Plasma Membrane | peptidase | 2887775 | 1541484.875 | 563843.062 | | | | | | |
| O35598 | ADAM10 | ADAM metallopeptidase domain 1 | Plasma Membrane | peptidase | 1162655.625 | 3337437.25 | 2074764.375 | | | | | | |
| O35598 | ADAM10 | ADAM metallopeptidase domain 1 | Plasma Membrane | peptidase | 2441849.75 | 3106256.25 | 2703498.25 | | | | | | |
| O35598 | ADAM10 | ADAM metallopeptidase domain 1 | Plasma Membrane | peptidase | 2853095.25 | 2853672.75 | 2505677.75 | | | | | | |
| O35598 | ADAM10 | ADAM metallopeptidase domain 1 | Plasma Membrane | peptidase | 5012921 | 4914572 | 4585974 | | | | | | |
| O35598 | ADAM10 | ADAM metallopeptidase domain 1 | Plasma Membrane | peptidase | 2445018.5 | 8160040 | 5920143.5 | | | | | | |
| O35598 | ADAM10 | ADAM metallopeptidase domain 1 | Plasma Membrane | peptidase | 1527073.875 | 2140421.5 | 2523011.75 | O35598 | ADAM10 | 15779398 | 14132055 | 1600140 | 78 |
| Q9Z0F8 | ADAM17 | ADAM metallopeptidase domain 1 | Plasma Membrane | peptidase | 5268840.5 | 6245371.5 | 4935689 | | | | | | |
| Q9Z0F8 | ADAM17 | ADAM metallopeptidase domain 1 | Plasma Membrane | peptidase | 170302.438 | 5200505.5 | 6707476.5 | Q9Z0F8 | ADAM17 | 2719571 | 2347253 | 958262 | 6 |
| Q9QYC0 | ADD1 | adducin 1 (alpha) | Cytoplasm | other | 8.19E+08 | 8.26E+08 | 7.92E+08 | | | | | | |
| Q9QYC0 | ADD1 | adducin 1 (alpha) | Cytoplasm | other | 5.01E+08 | 4.46E+08 | 4.78E+08 | | | | | | |
| Q9QYC0 | ADD1 | adducin 1 (alpha) | Cytoplasm | other | 3.25E+08 | 3.36E+08 | 3.60E+08 | | | | | | |
| Q9QYC0 | ADD1 | adducin 1 (alpha) | Cytoplasm | other | 4.45E+08 | 3.04E+08 | 2.26E+08 | | | | | | |
| Q9QYC0 | ADD1 | adducin 1 (alpha) | Cytoplasm | other | 3.64E+08 | 3.61E+08 | 3.30E+08 | | | | | | |
| Q9QYC0 | ADD1 | adducin 1 (alpha) | Cytoplasm | other | 2.41E+08 | 2.17E+08 | 2.21E+08 | | | | | | |
| Q9QYC0 | ADD1 | adducin 1 (alpha) | Cytoplasm | other | 2.31E+08 | 2.04E+08 | 1.80E+08 | | | | | | |
| Q9QYC0 | ADD1 | adducin 1 (alpha) | Cytoplasm | other | 2.45E+08 | 1.87E+08 | 2.34E+08 | | | | | | |
| Q9QYC0 | ADD1 | adducin 1 (alpha) | Cytoplasm | other | 1.62E+08 | 2.77E+08 | 2.79E+08 | | | | | | |
| Q9QYC0 | ADD1 | adducin 1 (alpha) | Cytoplasm | other | 2.20E+08 | 2.16E+08 | 2.02E+08 | | | | | | |
| Q9QYC0 | ADD1 | adducin 1 (alpha) | Cytoplasm | other | 2.21E+08 | 2.12E+08 | 1.85E+08 | | | | | | |
| Q9QYC0 | ADD1 | adducin 1 (alpha) | Cytoplasm | other | 2.49E+08 | 2.09E+08 | 2.26E+08 | | | | | | |
| Q9QYC0 | ADD1 | adducin 1 (alpha) | Cytoplasm | other | 2.01E+08 | 1.77E+08 | 1.62E+08 | | | | | | |
| Q9QYC0 | ADD1 | adducin 1 (alpha) | Cytoplasm | other | 1.90E+08 | 1.97E+08 | 1.78E+08 | | | | | | |
| Q9QYC0 | ADD1 | adducin 1 (alpha) | Cytoplasm | other | 2.14E+08 | 2.26E+08 | 1.97E+08 | | | | | | |
| Q9QYC0 | ADD1 | adducin 1 (alpha) | Cytoplasm | other | 1.18E+08 | 9.31E+07 | 8.54E+07 | | | | | | |
| Q9QYC0 | ADD1 | adducin 1 (alpha) | Cytoplasm | other | 1.41E+08 | 6.91E+07 | 9.58E+07 | | | | | | |
| Q9QYC0 | ADD1 | adducin 1 (alpha) | Cytoplasm | other | 1.31E+08 | 5.20E+07 | 7.88E+07 | | | | | | |
| Q9QYC0 | ADD1 | adducin 1 (alpha) | Cytoplasm | other | 7.33E+07 | 8.25E+07 | 8.79E+07 | | | | | | |
| Q9QYC0 | ADD1 | adducin 1 (alpha) | Cytoplasm | other | 3.27E+07 | 4.87E+07 | 3.31E+07 | | | | | | |
| Q9QYC0 | ADD1 | adducin 1 (alpha) | Cytoplasm | other | 3.30E+07 | 4.25E+07 | 4.08E+07 | | | | | | |
| Q9QYC0 | ADD1 | adducin 1 (alpha) | Cytoplasm | other | 3.25E+07 | 7.39E+07 | 4.65E+07 | | | | | | |
| Q9QYC0 | ADD1 | adducin 1 (alpha) | Cytoplasm | other | 4.29E+07 | 3.49E+07 | 2.92E+07 | | | | | | |
| Q9QYC0 | ADD1 | adducin 1 (alpha) | Cytoplasm | other | 1.81E+07 | 8.24E+07 | 4.57E+07 | | | | | | |
| Q9QYC0 | ADD1 | adducin 1 (alpha) | Cytoplasm | other | 1.34E+07 | 2.71E+07 | 2.04E+07 | | | | | | |
| Q9QYC0 | ADD1 | adducin 1 (alpha) | Cytoplasm | other | 1.67E+07 | 7.42E+07 | 3.16E+07 | | | | | | |
| Q9QYC0 | ADD1 | adducin 1 (alpha) | Cytoplasm | other | 1.24E+07 | 1.95E+07 | 1.61E+07 | | | | | | |
| Q9QYC0 | ADD1 | adducin 1 (alpha) | Cytoplasm | other | 1.55E+07 | 2.74E+07 | 5.19E+07 | | | | | | |
| Q9QYC0 | ADD1 | adducin 1 (alpha) | Cytoplasm | other | 5648123.5 | 1.43E+07 | 1.60E+07 | | | | | | |
| Q9QYC0 | ADD1 | adducin 1 (alpha) | Cytoplasm | other | 1.46E+07 | 1.71E+07 | 1.83E+07 | | | | | | |
| Q9QYC0 | ADD1 | adducin 1 (alpha) | Cytoplasm | other | 8232813.5 | 1.22E+07 | 1.19E+07 | | | | | | |
| Q9QYC0 | ADD1 | adducin 1 (alpha) | Cytoplasm | other | 5190324.5 | 2.64E+07 | 1.13E+07 | | | | | | |
| Q9QYC0 | ADD1 | adducin 1 (alpha) | Cytoplasm | other | 8277516 | 1.10E+07 | 1.06E+07 | | | | | | |
| Q9QYC0 | ADD1 | adducin 1 (alpha) | Cytoplasm | other | 5748543 | 2.29E+07 | 1.37E+07 | | | | | | |
| Q9QYC0 | ADD1 | adducin 1 (alpha) | Cytoplasm | other | 3788363 | 1.24E+07 | 7179107.5 | | | | | | |
| Q9QYC0 | ADD1 | adducin 1 (alpha) | Cytoplasm | other | 1.57E+07 | 7383582.5 | 4231180.5 | | | | | | |
| Q9QYC0 | ADD1 | adducin 1 (alpha) | Cytoplasm | other | 8915132 | 7721025 | 7482874.5 | | | | | | |
| Q9QYC0 | ADD1 | adducin 1 (alpha) | Cytoplasm | other | 4296366 | 3739051.75 | 4632522.5 | | | | | | |
| Q9QYC0 | ADD1 | adducin 1 (alpha) | Cytoplasm | other | 1.36E+07 | 5384533 | 1.37E+07 | | | | | | |
| Q9QYC0 | ADD1 | adducin 1 (alpha) | Cytoplasm | other | 1096185 | 2793745.5 | 2842369.5 | | | | | | |
| Q9QYC0 | ADD1 | adducin 1 (alpha) | Cytoplasm | other | 1083867.75 | 848984.5 | 353210.156 | Q9QYC0 | ADD1 | 131796762 | 163072561 | 14703758 | 123 |
| Q9QYB8 | ADD2 | adducin 2 (beta) | Cytoplasm | other | 3.94E+08 | 4.24E+08 | 3.95E+08 | | | | | | |
| Q9QYB8 | ADD2 | adducin 2 (beta) | Cytoplasm | other | 3.75E+08 | 3.44E+08 | 3.42E+08 | | | | | | |
| Q9QYB8 | ADD2 | adducin 2 (beta) | Cytoplasm | other | 2.99E+08 | 2.43E+08 | 2.48E+08 | | | | | | |
| Q9QYB8 | ADD2 | adducin 2 (beta) | Cytoplasm | other | 4.21E+08 | 4.80E+08 | 3.79E+08 | | | | | | |
| Q9QYB8 | ADD2 | adducin 2 (beta) | Cytoplasm | other | 2.13E+08 | 2.05E+08 | 1.68E+08 | | | | | | |
| Q9QYB8 | ADD2 | adducin 2 (beta) | Cytoplasm | other | 2.25E+08 | 2.12E+08 | 2.40E+08 | | | | | | |
| Q9QYB8 | ADD2 | adducin 2 (beta) | Cytoplasm | other | 1.71E+08 | 1.76E+08 | 1.59E+08 | | | | | | |
| Q9QYB8 | ADD2 | adducin 2 (beta) | Cytoplasm | other | 3.35E+08 | 7.81E+07 | 2.03E+08 | | | | | | |
| Q9QYB8 | ADD2 | adducin 2 (beta) | Cytoplasm | other | 1.93E+08 | 2.13E+08 | 2.07E+08 | | | | | | |
| Q9QYB8 | ADD2 | adducin 2 (beta) | Cytoplasm | other | 2.55E+08 | 2.45E+08 | 2.61E+08 | | | | | | |
| Q9QYB8 | ADD2 | adducin 2 (beta) | Cytoplasm | other | 1.85E+08 | 1.52E+08 | 1.43E+08 | | | | | | |
| Q9QYB8 | ADD2 | adducin 2 (beta) | Cytoplasm | other | 1.49E+08 | 3.15E+08 | 2.07E+08 | | | | | | |
| Q9QYB8 | ADD2 | adducin 2 (beta) | Cytoplasm | other | 1.55E+08 | 1.51E+08 | 1.51E+08 | | | | | | |
| Q9QYB8 | ADD2 | adducin 2 (beta) | Cytoplasm | other | 1.87E+08 | 1.70E+08 | 2.04E+08 | | | | | | |
| Q9QYB8 | ADD2 | adducin 2 (beta) | Cytoplasm | other | 1.64E+08 | 1.56E+08 | 1.56E+08 | | | | | | |

| | | | | | | | | | | | | | |
|---|---|---|---|---|---|---|---|---|---|---|---|---|---|
| Q9QYB8 | ADD2 | adducin 2 (beta) | Cytoplasm | other | 1.59E+08 | 9.13E+07 | 1.45E+08 | | | | | | |
| Q9QYB8 | ADD2 | adducin 2 (beta) | Cytoplasm | other | 1.28E+08 | 1.08E+08 | 1.38E+08 | | | | | | |
| Q9QYB8 | ADD2 | adducin 2 (beta) | Cytoplasm | other | 1.10E+08 | 1.41E+08 | 1.42E+08 | | | | | | |
| Q9QYB8 | ADD2 | adducin 2 (beta) | Cytoplasm | other | 4.26E+07 | 1.20E+08 | 6.90E+07 | | | | | | |
| Q9QYB8 | ADD2 | adducin 2 (beta) | Cytoplasm | other | 8.25E+07 | 8.45E+07 | 6.36E+07 | | | | | | |
| Q9QYB8 | ADD2 | adducin 2 (beta) | Cytoplasm | other | 4.47E+07 | 9.50E+07 | 6.02E+07 | | | | | | |
| Q9QYB8 | ADD2 | adducin 2 (beta) | Cytoplasm | other | 9.07E+07 | 7.52E+07 | 7.36E+07 | | | | | | |
| Q9QYB8 | ADD2 | adducin 2 (beta) | Cytoplasm | other | 4.69E+07 | 8.55E+07 | 6.43E+07 | | | | | | |
| Q9QYB8 | ADD2 | adducin 2 (beta) | Cytoplasm | other | 5.23E+07 | 5.20E+07 | 4.66E+07 | | | | | | |
| Q9QYB8 | ADD2 | adducin 2 (beta) | Cytoplasm | other | 8.95E+07 | 1.64E+08 | 1.47E+08 | | | | | | |
| Q9QYB8 | ADD2 | adducin 2 (beta) | Cytoplasm | other | 3.17E+07 | 3.65E+07 | 1.53E+07 | | | | | | |
| Q9QYB8 | ADD2 | adducin 2 (beta) | Cytoplasm | other | 5.78E+07 | 3.71E+07 | 4.17E+07 | | | | | | |
| Q9QYB8 | ADD2 | adducin 2 (beta) | Cytoplasm | other | 4.40E+07 | 3.42E+07 | 3.84E+07 | | | | | | |
| Q9QYB8 | ADD2 | adducin 2 (beta) | Cytoplasm | other | 4.61E+07 | 6.13E+07 | 3.33E+07 | | | | | | |
| Q9QYB8 | ADD2 | adducin 2 (beta) | Cytoplasm | other | 3.81E+07 | 3.56E+07 | 4.27E+07 | | | | | | |
| Q9QYB8 | ADD2 | adducin 2 (beta) | Cytoplasm | other | 3.48E+07 | 2.22E+07 | 2.19E+07 | | | | | | |
| Q9QYB8 | ADD2 | adducin 2 (beta) | Cytoplasm | other | 2.57E+07 | 2.47E+07 | 1.88E+07 | | | | | | |
| Q9QYB8 | ADD2 | adducin 2 (beta) | Cytoplasm | other | 2.85E+07 | 3.19E+07 | 2.33E+07 | | | | | | |
| Q9QYB8 | ADD2 | adducin 2 (beta) | Cytoplasm | other | 2.17E+07 | 1.93E+07 | 2.01E+07 | | | | | | |
| Q9QYB8 | ADD2 | adducin 2 (beta) | Cytoplasm | other | 1.13E+07 | 3.09E+07 | 1.76E+07 | | | | | | |
| Q9QYB8 | ADD2 | adducin 2 (beta) | Cytoplasm | other | 4.30E+07 | 2.03E+07 | 3.19E+07 | | | | | | |
| Q9QYB8 | ADD2 | adducin 2 (beta) | Cytoplasm | other | 2.25E+07 | 5390743.5 | 1.20E+07 | | | | | | |
| Q9QYB8 | ADD2 | adducin 2 (beta) | Cytoplasm | other | 1.46E+07 | 1.26E+07 | 1.43E+07 | | | | | | |
| Q9QYB8 | ADD2 | adducin 2 (beta) | Cytoplasm | other | 1.40E+07 | 1.43E+07 | 1.71E+07 | | | | | | |
| Q9QYB8 | ADD2 | adducin 2 (beta) | Cytoplasm | other | 2.22E+07 | 7676063.5 | 1.10E+07 | | | | | | |
| Q9QYB8 | ADD2 | adducin 2 (beta) | Cytoplasm | other | 2.42E+07 | 4335709.5 | 3327676 | | | | | | |
| Q9QYB8 | ADD2 | adducin 2 (beta) | Cytoplasm | other | 1.70E+07 | 1.67E+07 | 2.21E+07 | | | | | | |
| Q9QYB8 | ADD2 | adducin 2 (beta) | Cytoplasm | other | 5684348.5 | 1.06E+07 | 1.00E+07 | | | | | | |
| Q9QYB8 | ADD2 | adducin 2 (beta) | Cytoplasm | other | 4944297 | 9854356 | 4767995.5 | | | | | | |
| Q9QYB8 | ADD2 | adducin 2 (beta) | Cytoplasm | other | 6579998 | 5381505 | 9403729 | | | | | | |
| Q9QYB8 | ADD2 | adducin 2 (beta) | Cytoplasm | other | 4990609.5 | 7980524.5 | 5880907 | | | | | | |
| Q9QYB8 | ADD2 | adducin 2 (beta) | Cytoplasm | other | 3844019.25 | 1.41E+07 | 5851492.5 | | | | | | |
| Q9QYB8 | ADD2 | adducin 2 (beta) | Cytoplasm | other | 7004493 | 5571982 | 8272014 | | | | | | |
| Q9QYB8 | ADD2 | adducin 2 (beta) | Cytoplasm | other | 4413006 | 1.16E+07 | 1.18E+07 | | | | | | |
| Q9QYB8 | ADD2 | adducin 2 (beta) | Cytoplasm | other | 3073089.5 | 9531735 | 1.38E+07 | | | | | | |
| Q9QYB8 | ADD2 | adducin 2 (beta) | Cytoplasm | other | 2154647 | 2859970.25 | 2265074.5 | | | | | | |
| Q9QYB8 | ADD2 | adducin 2 (beta) | Cytoplasm | other | 1.79E+07 | 314351.906 | 1551158 | | | | | | |
| Q9QYB8 | ADD2 | adducin 2 (beta) | Cytoplasm | other | 4375765 | 2478405.25 | 6001266.5 | | | | | | |
| Q9QYB8 | ADD2 | adducin 2 (beta) | Cytoplasm | other | 5932405.5 | 4868302.5 | 7816648.5 | | | | | | |
| Q9QYB8 | ADD2 | adducin 2 (beta) | Cytoplasm | other | 3702868.75 | 3866743.75 | 3345370 | | | | | | |
| Q9QYB8 | ADD2 | adducin 2 (beta) | Cytoplasm | other | 1866695.375 | 2896237.25 | 2388733.75 | | | | | | |
| Q9QYB8 | ADD2 | adducin 2 (beta) | Cytoplasm | other | 4494831.5 | 256787.125 | 151124.812 | | | | | | |
| Q9QYB8 | ADD2 | adducin 2 (beta) | Cytoplasm | other | 567824.688 | 2762405.5 | 2270863 | | | | | | |
| Q9QYB8 | ADD2 | adducin 2 (beta) | Cytoplasm | other | 12993.346 | 4562976.5 | 2347534.25 | | | | | | |
| Q9QYB8 | ADD2 | adducin 2 (beta) | Cytoplasm | other | 55218.969 | 2589087.75 | 2001659.625 | | | | | | |
| Q9QYB8 | ADD2 | adducin 2 (beta) | Cytoplasm | other | 2735560 | 1528628.875 | 1249099.875 | | | | | | |
| Q9QYB8 | ADD2 | adducin 2 (beta) | Cytoplasm | other | 1753392.125 | 278.914 | 154519.922 | | | | | | |
| Q9QYB8 | ADD2 | adducin 2 (beta) | Cytoplasm | other | 754129.125 | 803225.938 | 740388.125 | | | | | | |
| Q9QYB8 | ADD2 | adducin 2 (beta) | Cytoplasm | other | 2266929 | 63228.684 | 3170872 | Q9QYB8 | ADD2 | 80505111 | 104963577 | 7575094 | 192 |
| Q9QYB5 | ADD3 | adducin 3 (gamma) | Cytoplasm | other | 2.82E+07 | 3.27E+07 | 2.84E+07 | | | | | | |
| Q9QYB5 | ADD3 | adducin 3 (gamma) | Cytoplasm | other | 1.37E+07 | 1.29E+07 | 1.73E+07 | | | | | | |
| Q9QYB5 | ADD3 | adducin 3 (gamma) | Cytoplasm | other | 7264961.5 | 5453039 | 9046098 | | | | | | |
| Q9QYB5 | ADD3 | adducin 3 (gamma) | Cytoplasm | other | 1.05E+07 | 9288538 | 9470006 | | | | | | |
| Q9QYB5 | ADD3 | adducin 3 (gamma) | Cytoplasm | other | 7758629 | 8027242 | 1.03E+07 | | | | | | |
| Q9QYB5 | ADD3 | adducin 3 (gamma) | Cytoplasm | other | 2611385.25 | 5849840 | 4113377.75 | | | | | | |
| Q9QYB5 | ADD3 | adducin 3 (gamma) | Cytoplasm | other | 7931059 | 3809734.75 | 6433675.5 | | | | | | |
| Q9QYB5 | ADD3 | adducin 3 (gamma) | Cytoplasm | other | 6817955.5 | 5746067 | 7304163 | | | | | | |
| Q9QYB5 | ADD3 | adducin 3 (gamma) | Cytoplasm | other | 4688566.5 | 4865855 | 4636653.5 | | | | | | |
| Q9QYB5 | ADD3 | adducin 3 (gamma) | Cytoplasm | other | 3110467 | 3452544 | 3054580.75 | | | | | | |
| Q9QYB5 | ADD3 | adducin 3 (gamma) | Cytoplasm | other | 4438703.5 | 5104069.5 | 4533634 | | | | | | |
| Q9QYB5 | ADD3 | adducin 3 (gamma) | Cytoplasm | other | 2031433.5 | 5591456.5 | 4236211 | | | | | | |
| Q9QYB5 | ADD3 | adducin 3 (gamma) | Cytoplasm | other | 1153910 | 2667482.5 | 1668164.125 | | | | | | |
| Q9QYB5 | ADD3 | adducin 3 (gamma) | Cytoplasm | other | 2607562 | 4788586 | 2926179 | | | | | | |
| Q9QYB5 | ADD3 | adducin 3 (gamma) | Cytoplasm | other | 1152249.5 | 1919314.125 | 2982873.25 | Q9QYB5 | ADD3 | 6933792 | 6975896 | 1039905 | 45 |
| Q8VDL4 | ADPGK | ADP-dependent glucokinase | Plasma Membrane | kinase | 2780493.75 | 3456847.5 | 2981679 | Q8VDL4 | ADPGK | 2780494 | 347303 | 200515 | 3 |
| P54822 | ADSL | adenylosuccinate lyase | Cytoplasm | enzyme | 9837378 | 9500125 | 1.44E+07 | | | | | | |
| P54822 | ADSL | adenylosuccinate lyase | Cytoplasm | enzyme | 8415421 | 8245532 | 8644312 | | | | | | |
| P54822 | ADSL | adenylosuccinate lyase | Cytoplasm | enzyme | 5426276.5 | 6729849 | 7115159 | | | | | | |
| P54822 | ADSL | adenylosuccinate lyase | Cytoplasm | enzyme | 6871608 | 8531443 | 7591612 | | | | | | |
| P54822 | ADSL | adenylosuccinate lyase | Cytoplasm | enzyme | 9672083 | 5199919.5 | 7458067.5 | | | | | | |
| P54822 | ADSL | adenylosuccinate lyase | Cytoplasm | enzyme | 5648972.5 | 6164599 | 7361727.5 | | | | | | |
| P54822 | ADSL | adenylosuccinate lyase | Cytoplasm | enzyme | 8061605.5 | 3878977 | 4266405.5 | | | | | | |
| P54822 | ADSL | adenylosuccinate lyase | Cytoplasm | enzyme | 7187472 | 6045656.5 | 6863231 | | | | | | |
| P54822 | ADSL | adenylosuccinate lyase | Cytoplasm | enzyme | 3284892.75 | 5735615.5 | 5141061 | | | | | | |
| P54822 | ADSL | adenylosuccinate lyase | Cytoplasm | enzyme | 2763467 | 2658780 | 2589509 | | | | | | |
| P54822 | ADSL | adenylosuccinate lyase | Cytoplasm | enzyme | 1415313 | 1753933.375 | 2524798.75 | | | | | | |
| P54822 | ADSL | adenylosuccinate lyase | Cytoplasm | enzyme | 639473.438 | 715150.875 | 688584.688 | | | | | | |
| P54822 | ADSL | adenylosuccinate lyase | Cytoplasm | enzyme | 651718.312 | 979208.375 | 1691434.125 | P54822 | ADSL | 5375052 | 3235193 | 518046 | 39 |
| Q640N1 | AEBP1 | AE binding protein 1 | Nucleus | peptidase | 8319678.5 | 9407951 | 7985696 | Q640N1 | AEBP1 | 8319679 | 743717 | 429385 | 3 |
| O55112 | Aff2 | AF4/FMR2 family, member 2 | Nucleus | other | 3.73E+07 | 2.08E+07 | 2.17E+07 | O55112 | Aff2 | 37320000 | 9305747 | 5372676 | 3 |
| Q8K2K6 | AGFG1 | ArfGAP with FG repeats 1 | Nucleus | other | 1785666.625 | 3426958.25 | 1802567.25 | Q8K2K6 | AGFG1 | 1785667 | 942759 | 544302 | 3 |
| Q80WC7 | AGFG2 | ArfGAP with FG repeats 2 | Cytoplasm | other | 3788762.75 | 4612348 | 2664434.75 | | | | | | |
| Q80WC7 | AGFG2 | ArfGAP with FG repeats 2 | Cytoplasm | other | 1888354.125 | 4009889.5 | 2706228.25 | | | | | | |
| Q80WC7 | AGFG2 | ArfGAP with FG repeats 2 | Cytoplasm | other | 2266999.5 | 1905713 | 1626327.75 | | | | | | |
| Q80WC7 | AGFG2 | ArfGAP with FG repeats 2 | Cytoplasm | other | 2070786.5 | 4297166 | 2126621.5 | Q80WC7 | AGFG2 | 2503726 | 1055780 | 304777 | 12 |
| Q8CJG0 | AGO2 | argonaute RISC catalytic compone | Cytoplasm | translation regulator | 3.76E+07 | 5.14E+07 | 3.72E+07 | | | | | | |

| | | | | | | | | | | | | | |
|---|---|---|---|---|---|---|---|---|---|---|---|---|---|
| Q8CJG0 | AGO2 | argonaute RISC catalytic compone | Cytoplasm | translation regulator | 3.30E+07 | 3.25E+07 | 2.55E+07 | | | | | | |
| Q8CJG0 | AGO2 | argonaute RISC catalytic compone | Cytoplasm | translation regulator | 3.42E+07 | 3.31E+07 | 2.68E+07 | | | | | | |
| Q8CJG0 | AGO2 | argonaute RISC catalytic compone | Cytoplasm | translation regulator | 2.48E+07 | 2.04E+07 | 1.83E+07 | | | | | | |
| Q8CJG0 | AGO2 | argonaute RISC catalytic compone | Cytoplasm | translation regulator | 2.77E+07 | 2.00E+07 | 2.15E+07 | | | | | | |
| Q8CJG0 | AGO2 | argonaute RISC catalytic compone | Cytoplasm | translation regulator | 2.98E+07 | 3.03E+07 | 2.15E+07 | | | | | | |
| Q8CJG0 | AGO2 | argonaute RISC catalytic compone | Cytoplasm | translation regulator | 2.98E+07 | 2.23E+07 | 2.17E+07 | | | | | | |
| Q8CJG0 | AGO2 | argonaute RISC catalytic compone | Cytoplasm | translation regulator | 2.17E+07 | 1.91E+07 | 1.69E+07 | | | | | | |
| Q8CJG0 | AGO2 | argonaute RISC catalytic compone | Cytoplasm | translation regulator | 2.18E+07 | 2.26E+07 | 2.34E+07 | | | | | | |
| Q8CJG0 | AGO2 | argonaute RISC catalytic compone | Cytoplasm | translation regulator | 2.35E+07 | 2.38E+07 | 3.35E+07 | | | | | | |
| Q8CJG0 | AGO2 | argonaute RISC catalytic compone | Cytoplasm | translation regulator | 2.35E+07 | 2.32E+07 | 1.92E+07 | | | | | | |
| Q8CJG0 | AGO2 | argonaute RISC catalytic compone | Cytoplasm | translation regulator | 1.29E+07 | 1.99E+07 | 1.45E+07 | | | | | | |
| Q8CJG0 | AGO2 | argonaute RISC catalytic compone | Cytoplasm | translation regulator | 2.15E+07 | 1.84E+07 | 1.72E+07 | | | | | | |
| Q8CJG0 | AGO2 | argonaute RISC catalytic compone | Cytoplasm | translation regulator | 2.33E+07 | 2.55E+07 | 2.20E+07 | | | | | | |
| Q8CJG0 | AGO2 | argonaute RISC catalytic compone | Cytoplasm | translation regulator | 2.32E+07 | 1.96E+07 | 1.74E+07 | | | | | | |
| Q8CJG0 | AGO2 | argonaute RISC catalytic compone | Cytoplasm | translation regulator | 2.46E+07 | 2.54E+07 | 2.09E+07 | | | | | | |
| Q8CJG0 | AGO2 | argonaute RISC catalytic compone | Cytoplasm | translation regulator | 1.74E+07 | 1.88E+07 | 1.40E+07 | | | | | | |
| Q8CJG0 | AGO2 | argonaute RISC catalytic compone | Cytoplasm | translation regulator | 1.69E+07 | 2.04E+07 | 1.68E+07 | | | | | | |
| Q8CJG0 | AGO2 | argonaute RISC catalytic compone | Cytoplasm | translation regulator | 8726380 | 2.09E+07 | 1.20E+07 | | | | | | |
| Q8CJG0 | AGO2 | argonaute RISC catalytic compone | Cytoplasm | translation regulator | 1.19E+07 | 1.32E+07 | 9251144 | | | | | | |
| Q8CJG0 | AGO2 | argonaute RISC catalytic compone | Cytoplasm | translation regulator | 2.25E+07 | 2.05E+07 | 1.65E+07 | | | | | | |
| Q8CJG0 | AGO2 | argonaute RISC catalytic compone | Cytoplasm | translation regulator | 1.32E+07 | 1.13E+07 | 1.14E+07 | | | | | | |
| Q8CJG0 | AGO2 | argonaute RISC catalytic compone | Cytoplasm | translation regulator | 1.67E+07 | 9894008 | 1.07E+07 | | | | | | |
| Q8CJG0 | AGO2 | argonaute RISC catalytic compone | Cytoplasm | translation regulator | 1.27E+07 | 1.14E+07 | 1.05E+07 | | | | | | |
| Q8CJG0 | AGO2 | argonaute RISC catalytic compone | Cytoplasm | translation regulator | 8442351 | 1.45E+07 | 9242534 | | | | | | |
| Q8CJG0 | AGO2 | argonaute RISC catalytic compone | Cytoplasm | translation regulator | 8918011 | 8498372 | 7378294 | | | | | | |
| Q8CJG0 | AGO2 | argonaute RISC catalytic compone | Cytoplasm | translation regulator | 1.01E+07 | 7040630.5 | 5658254 | | | | | | |
| Q8CJG0 | AGO2 | argonaute RISC catalytic compone | Cytoplasm | translation regulator | 6012933.5 | 1.07E+07 | 8984684 | | | | | | |
| Q8CJG0 | AGO2 | argonaute RISC catalytic compone | Cytoplasm | translation regulator | 6360207.5 | 7518541 | 7128868 | | | | | | |
| Q8CJG0 | AGO2 | argonaute RISC catalytic compone | Cytoplasm | translation regulator | 9715654 | 6051176.5 | 4696618.5 | | | | | | |
| Q8CJG0 | AGO2 | argonaute RISC catalytic compone | Cytoplasm | translation regulator | 6094734.5 | 4115097.5 | 3599429 | | | | | | |
| Q8CJG0 | AGO2 | argonaute RISC catalytic compone | Cytoplasm | translation regulator | 3419952 | 2641810.5 | 4507916.5 | | | | | | |
| Q8CJG0 | AGO2 | argonaute RISC catalytic compone | Cytoplasm | translation regulator | 5345854.5 | 3368978.5 | 4163584.25 | | | | | | |
| Q8CJG0 | AGO2 | argonaute RISC catalytic compone | Cytoplasm | translation regulator | 2355636.25 | 3728680.75 | 1394007.625 | | | | | | |
| Q8CJG0 | AGO2 | argonaute RISC catalytic compone | Cytoplasm | translation regulator | 3389011.5 | 6948941.5 | 8757759 | | | | | | |
| Q8CJG0 | AGO2 | argonaute RISC catalytic compone | Cytoplasm | translation regulator | 3902068 | 1287329.125 | 471772.125 | | | | | | |
| Q8CJG0 | AGO2 | argonaute RISC catalytic compone | Cytoplasm | translation regulator | 2663749 | 2890068.25 | 2058430 | | | | | | |
| Q8CJG0 | AGO2 | argonaute RISC catalytic compone | Cytoplasm | translation regulator | 1288838.875 | 1.60E+07 | 1.66E+07 | | | | | | |
| Q8CJG0 | AGO2 | argonaute RISC catalytic compone | Cytoplasm | translation regulator | 951237.312 | 1091163.125 | 1000573.688 | | | | | | |
| Q8CJG0 | AGO2 | argonaute RISC catalytic compone | Cytoplasm | translation regulator | 1551735.5 | 269026.531 | 54523.559 | | | | | | |
| Q8CJG0 | AGO2 | argonaute RISC catalytic compone | Cytoplasm | translation regulator | 381667.688 | 259629.578 | 124844.352 | Q8CJG0 | AGO2 | 14962196 | 10210484 | 920648 | 123 |
| O35083 | AGPAT1 | 1-acylglycerol-3-phosphate O-acyl | Cytoplasm | enzyme | 745758.625 | 757730.375 | 1514507.375 | O35083 | AGPAT1 | 745759 | 440422 | 254278 | 3 |
| P50247 | AHCY | adenosylhomocysteinase | Cytoplasm | enzyme | 5051136.5 | 3539937.25 | 4039353 | | | | | | |
| P50247 | AHCY | adenosylhomocysteinase | Cytoplasm | enzyme | 3608322.75 | 2666395.25 | 2957065.75 | | | | | | |
| P50247 | AHCY | adenosylhomocysteinase | Cytoplasm | enzyme | 2246626 | 2771973.25 | 2289973.5 | P50247 | AHCY | 3635362 | 912569 | 304190 | 9 |
| Q8N9S3 | Ahsa2 | AHA1, activator of heat shock prot | Cytoplasm | other | 1.87E+08 | 2.65E+08 | 2.73E+08 | Q8N9S3 | Ahsa2 | 187100000 | 47586167 | 27473887 | 3 |
| Q9Z0X1 | AIFM1 | apoptosis-inducing factor, mitocho | Cytoplasm | enzyme | 2790394.5 | 2563966 | 2820853 | | | | | | |
| Q9Z0X1 | AIFM1 | apoptosis-inducing factor, mitocho | Cytoplasm | enzyme | 1827001.625 | 1830590 | 2069537.5 | Q9Z0X1 | AIFM1 | 2308698 | 464089 | 189464 | 6 |
| P31230 | AIMP1 | aminoacyl tRNA synthetase compl | Extracellular Space | cytokine | 2184634.25 | 1243197.75 | 1784369.75 | | | | | | |
| P31230 | AIMP1 | aminoacyl tRNA synthetase compl | Extracellular Space | cytokine | 1246552.875 | 516241.438 | 260264.969 | P31230 | AIMP1 | 1715594 | 730130 | 298074 | 6 |
| Q9WTP6 | AK2 | adenylate kinase 2 | Cytoplasm | kinase | 1260713.5 | 1137780.875 | 1530561.5 | Q9WTP6 | AK2 | 1260714 | 200917 | 116000 | 3 |
| P10518 | ALAD | aminolevulinate dehydratase | Cytoplasm | enzyme | 1.65E+07 | 5.83E+07 | 5.51E+07 | | | | | | |
| P10518 | ALAD | aminolevulinate dehydratase | Cytoplasm | enzyme | 9407888 | 3.25E+07 | 2.95E+07 | | | | | | |
| P10518 | ALAD | aminolevulinate dehydratase | Cytoplasm | enzyme | 1.07E+07 | 3.00E+07 | 3.24E+07 | | | | | | |
| P10518 | ALAD | aminolevulinate dehydratase | Cytoplasm | enzyme | 5589096.5 | 1.52E+07 | 1.84E+07 | | | | | | |
| P10518 | ALAD | aminolevulinate dehydratase | Cytoplasm | enzyme | 5894901.5 | 1.93E+07 | 1.97E+07 | | | | | | |
| P10518 | ALAD | aminolevulinate dehydratase | Cytoplasm | enzyme | 4483867.5 | 1.74E+07 | 1.21E+07 | | | | | | |
| P10518 | ALAD | aminolevulinate dehydratase | Cytoplasm | enzyme | 3887356.75 | 1.13E+07 | 1.35E+07 | | | | | | |
| P10518 | ALAD | aminolevulinate dehydratase | Cytoplasm | enzyme | 2629900.25 | 1674427.75 | 4801201 | | | | | | |
| P10518 | ALAD | aminolevulinate dehydratase | Cytoplasm | enzyme | 870865.5 | 4005228.75 | 3368272.5 | | | | | | |
| P10518 | ALAD | aminolevulinate dehydratase | Cytoplasm | enzyme | 142108.703 | 5622826 | 2987859.5 | | | | | | |
| P10518 | ALAD | aminolevulinate dehydratase | Cytoplasm | enzyme | 1041868.938 | 9209208 | 1.13E+07 | | | | | | |
| P10518 | ALAD | aminolevulinate dehydratase | Cytoplasm | enzyme | 1155569.75 | 3058779.75 | 3862700.75 | | | | | | |
| P10518 | ALAD | aminolevulinate dehydratase | Cytoplasm | enzyme | 3628496.25 | 4277180.5 | 2901861.25 | | | | | | |
| P10518 | ALAD | aminolevulinate dehydratase | Cytoplasm | enzyme | 1188416.5 | 4024904.5 | 5726811 | | | | | | |
| P10518 | ALAD | aminolevulinate dehydratase | Cytoplasm | enzyme | 741569.5 | 2345646 | 3884250.75 | | | | | | |
| P10518 | ALAD | aminolevulinate dehydratase | Cytoplasm | enzyme | 101642.562 | 376392.781 | 50639.582 | P10518 | ALAD | 4252722 | 13126289 | 1894617 | 48 |
| P08680 | ALAS2 | 5'-aminolevulinate synthase 2 | Cytoplasm | enzyme | 2.60E+07 | 2.98E+07 | 3.67E+07 | | | | | | |
| P08680 | ALAS2 | 5'-aminolevulinate synthase 2 | Cytoplasm | enzyme | 2.17E+07 | 1.89E+07 | 2.62E+07 | | | | | | |
| P08680 | ALAS2 | 5'-aminolevulinate synthase 2 | Cytoplasm | enzyme | 1.49E+07 | 1.36E+07 | 1.45E+07 | | | | | | |
| P08680 | ALAS2 | 5'-aminolevulinate synthase 2 | Cytoplasm | enzyme | 8716587 | 1.14E+07 | 1.08E+07 | | | | | | |
| P08680 | ALAS2 | 5'-aminolevulinate synthase 2 | Cytoplasm | enzyme | 9498997 | 8673981 | 1.32E+07 | | | | | | |
| P08680 | ALAS2 | 5'-aminolevulinate synthase 2 | Cytoplasm | enzyme | 6830284 | 8387055.5 | 1.03E+07 | | | | | | |
| P08680 | ALAS2 | 5'-aminolevulinate synthase 2 | Cytoplasm | enzyme | 7236708.5 | 6918219.5 | 8309505.5 | | | | | | |
| P08680 | ALAS2 | 5'-aminolevulinate synthase 2 | Cytoplasm | enzyme | 5886740 | 6649694 | 8578781 | | | | | | |
| P08680 | ALAS2 | 5'-aminolevulinate synthase 2 | Cytoplasm | enzyme | 3046871.5 | 4771441 | 5555108 | | | | | | |
| P08680 | ALAS2 | 5'-aminolevulinate synthase 2 | Cytoplasm | enzyme | 3808784 | 5052870 | 5321630.5 | | | | | | |
| P08680 | ALAS2 | 5'-aminolevulinate synthase 2 | Cytoplasm | enzyme | 1849602.625 | 2138633.25 | 2080735.75 | | | | | | |
| P08680 | ALAS2 | 5'-aminolevulinate synthase 2 | Cytoplasm | enzyme | 1301882.625 | 1303626.75 | 1282577.5 | P08680 | ALAS2 | 9227205 | 8566270 | 1427712 | 36 |
| P07724 | ALB | albumin | Extracellular Space | transporter | 4.62E+07 | 4.79E+07 | 6.79E+07 | | | | | | |
| P07724 | ALB | albumin | Extracellular Space | transporter | 3.58E+07 | 1.32E+07 | 2.69E+07 | | | | | | |
| P07724 | ALB | albumin | Extracellular Space | transporter | 2.05E+07 | 2.41E+07 | 3.14E+07 | | | | | | |
| P07724 | ALB | albumin | Extracellular Space | transporter | 1.79E+07 | 2.63E+07 | 2.99E+07 | | | | | | |
| P07724 | ALB | albumin | Extracellular Space | transporter | 2.02E+07 | 3.11E+07 | 3.37E+07 | | | | | | |
| P07724 | ALB | albumin | Extracellular Space | transporter | 1.83E+07 | 2.37E+07 | 2.84E+07 | | | | | | |
| P07724 | ALB | albumin | Extracellular Space | transporter | 2.03E+07 | 2.08E+07 | 3.13E+07 | | | | | | |
| P07724 | ALB | albumin | Extracellular Space | transporter | 2.06E+07 | 2.14E+07 | 2.60E+07 | | | | | | |

| | | | | | | | | | | | | | |
|---|---|---|---|---|---|---|---|---|---|---|---|---|---|
| P07724 | ALB | albumin | Extracellular Space | transporter | 2.27E+07 | 2.35E+07 | 2.68E+07 | | | | | | |
| P07724 | ALB | albumin | Extracellular Space | transporter | 2.17E+07 | 2.75E+07 | 3.04E+07 | | | | | | |
| P07724 | ALB | albumin | Extracellular Space | transporter | 1.79E+07 | 1.75E+07 | 2.02E+07 | | | | | | |
| P07724 | ALB | albumin | Extracellular Space | transporter | 2.10E+07 | 8920985 | 1.90E+07 | | | | | | |
| P07724 | ALB | albumin | Extracellular Space | transporter | 1.21E+07 | 1.13E+07 | 1.35E+07 | | | | | | |
| P07724 | ALB | albumin | Extracellular Space | transporter | 1.08E+07 | 1.20E+07 | 1.46E+07 | | | | | | |
| P07724 | ALB | albumin | Extracellular Space | transporter | 1.72E+07 | 1.40E+07 | 1.69E+07 | | | | | | |
| P07724 | ALB | albumin | Extracellular Space | transporter | 1.10E+07 | 1.00E+07 | 1.19E+07 | | | | | | |
| P07724 | ALB | albumin | Extracellular Space | transporter | 9169302 | 1.05E+07 | 1.15E+07 | | | | | | |
| P07724 | ALB | albumin | Extracellular Space | transporter | 3919794.5 | 1.05E+07 | 1.04E+07 | | | | | | |
| P07724 | ALB | albumin | Extracellular Space | transporter | 4467994.5 | 1.03E+07 | 9380657 | | | | | | |
| P07724 | ALB | albumin | Extracellular Space | transporter | 2438365.5 | 3830852.25 | 3871624 | | | | | | |
| P07724 | ALB | albumin | Extracellular Space | transporter | 4061160.75 | 4513344.5 | 8345654 | | | | | | |
| P07724 | ALB | albumin | Extracellular Space | transporter | 2776371.5 | 4265400 | 5462215.5 | | | | | | |
| P07724 | ALB | albumin | Extracellular Space | transporter | 2141511 | 3072863.5 | 3665789.5 | | | | | | |
| P07724 | ALB | albumin | Extracellular Space | transporter | 1269113.25 | 2934954 | 2159790.5 | P07724 | ALB | 15184317 | 12215749 | 1439640 | 72 |
| Q571I9 | ALDH16A1 | aldehyde dehydrogenase 16 famil | Cytoplasm | enzyme | 4786952.5 | 5237560.5 | 2937928.5 | | | | | | |
| Q571I9 | ALDH16A1 | aldehyde dehydrogenase 16 famil | Cytoplasm | enzyme | 5081393.5 | 5874544.5 | 4207153.5 | | | | | | |
| Q571I9 | ALDH16A1 | aldehyde dehydrogenase 16 famil | Cytoplasm | enzyme | 4792474 | 3465606 | 3294319.25 | | | | | | |
| Q571I9 | ALDH16A1 | aldehyde dehydrogenase 16 famil | Cytoplasm | enzyme | 1367870.625 | 1727249.5 | 884324.875 | | | | | | |
| Q571I9 | ALDH16A1 | aldehyde dehydrogenase 16 famil | Cytoplasm | enzyme | 2234805.25 | 2168596.5 | 1827185.75 | | | | | | |
| Q571I9 | ALDH16A1 | aldehyde dehydrogenase 16 famil | Cytoplasm | enzyme | 876341.188 | 581893.625 | 541667 | Q571I9 | ALDH16A1 | 3189973 | 1771296 | 417498 | 18 |
| Q9Z110 | ALDH18A1 | aldehyde dehydrogenase 18 famil | Cytoplasm | kinase | 2966641.5 | 2472409.75 | 3880737.75 | | | | | | |
| Q9Z110 | ALDH18A1 | aldehyde dehydrogenase 18 famil | Cytoplasm | kinase | 1644156.125 | 2453879.5 | 3596096 | | | | | | |
| Q9Z110 | ALDH18A1 | aldehyde dehydrogenase 18 famil | Cytoplasm | kinase | 1240684.625 | 2156729.5 | 2590113.75 | | | | | | |
| Q9Z110 | ALDH18A1 | aldehyde dehydrogenase 18 famil | Cytoplasm | kinase | 1349494.125 | 1236727.25 | 2072940.125 | | | | | | |
| Q9Z110 | ALDH18A1 | aldehyde dehydrogenase 18 famil | Cytoplasm | kinase | 1389067.125 | 1189425.375 | 1953794 | | | | | | |
| Q9Z110 | ALDH18A1 | aldehyde dehydrogenase 18 famil | Cytoplasm | kinase | 677809.062 | 1667504.375 | 2359741.75 | | | | | | |
| Q9Z110 | ALDH18A1 | aldehyde dehydrogenase 18 famil | Cytoplasm | kinase | 819468.625 | 971441.125 | 1471542.5 | | | | | | |
| Q9Z110 | ALDH18A1 | aldehyde dehydrogenase 18 famil | Cytoplasm | kinase | 511060.75 | 946807.875 | 1240574.375 | Q9Z110 | ALDH18A1 | 1324798 | 886819 | 181021 | 24 |
| P24549 | ALDH1A1 | aldehyde dehydrogenase 1 family, | Cytoplasm | enzyme | 3722864.75 | 1370288.5 | 2361204.75 | P24549 | ALDH1A1 | 3722865 | 1181147 | 681935 | 3 |
| P47738 | ALDH2 | aldehyde dehydrogenase 2 family | Cytoplasm | enzyme | 4.96E+07 | 4.55E+07 | 6.32E+07 | | | | | | |
| P47738 | ALDH2 | aldehyde dehydrogenase 2 family | Cytoplasm | enzyme | 3.41E+07 | 3.99E+07 | 5.19E+07 | | | | | | |
| P47738 | ALDH2 | aldehyde dehydrogenase 2 family | Cytoplasm | enzyme | 1.79E+07 | 1.44E+07 | 1.31E+07 | | | | | | |
| P47738 | ALDH2 | aldehyde dehydrogenase 2 family | Cytoplasm | enzyme | 3.12E+07 | 3.58E+07 | 4.49E+07 | | | | | | |
| P47738 | ALDH2 | aldehyde dehydrogenase 2 family | Cytoplasm | enzyme | 2.87E+07 | 3.08E+07 | 4.10E+07 | | | | | | |
| P47738 | ALDH2 | aldehyde dehydrogenase 2 family | Cytoplasm | enzyme | 1.77E+07 | 1.44E+07 | 1.78E+07 | | | | | | |
| P47738 | ALDH2 | aldehyde dehydrogenase 2 family | Cytoplasm | enzyme | 2.73E+07 | 3.13E+07 | 3.57E+07 | | | | | | |
| P47738 | ALDH2 | aldehyde dehydrogenase 2 family | Cytoplasm | enzyme | 1.95E+07 | 2.29E+07 | 2.92E+07 | | | | | | |
| P47738 | ALDH2 | aldehyde dehydrogenase 2 family | Cytoplasm | enzyme | 1.51E+07 | 2.82E+07 | 2.86E+07 | | | | | | |
| P47738 | ALDH2 | aldehyde dehydrogenase 2 family | Cytoplasm | enzyme | 1.70E+07 | 1.90E+07 | 2.32E+07 | | | | | | |
| P47738 | ALDH2 | aldehyde dehydrogenase 2 family | Cytoplasm | enzyme | 1.54E+07 | 2.03E+07 | 1.48E+07 | | | | | | |
| P47738 | ALDH2 | aldehyde dehydrogenase 2 family | Cytoplasm | enzyme | 1.39E+07 | 8524775 | 1.56E+07 | | | | | | |
| P47738 | ALDH2 | aldehyde dehydrogenase 2 family | Cytoplasm | enzyme | 9745019 | 1.64E+07 | 1.49E+07 | | | | | | |
| P47738 | ALDH2 | aldehyde dehydrogenase 2 family | Cytoplasm | enzyme | 1.98E+07 | 2.33E+07 | 2.79E+07 | | | | | | |
| P47738 | ALDH2 | aldehyde dehydrogenase 2 family | Cytoplasm | enzyme | 1.71E+07 | 1.76E+07 | 2.07E+07 | | | | | | |
| P47738 | ALDH2 | aldehyde dehydrogenase 2 family | Cytoplasm | enzyme | 1.87E+07 | 3.40E+07 | 4.29E+07 | | | | | | |
| P47738 | ALDH2 | aldehyde dehydrogenase 2 family | Cytoplasm | enzyme | 1.41E+07 | 1.33E+07 | 1.48E+07 | | | | | | |
| P47738 | ALDH2 | aldehyde dehydrogenase 2 family | Cytoplasm | enzyme | 1.09E+07 | 1.21E+07 | 2.02E+07 | | | | | | |
| P47738 | ALDH2 | aldehyde dehydrogenase 2 family | Cytoplasm | enzyme | 1297922.375 | 2827583 | 5095925 | | | | | | |
| P47738 | ALDH2 | aldehyde dehydrogenase 2 family | Cytoplasm | enzyme | 5140276 | 8646708 | 1.62E+07 | | | | | | |
| P47738 | ALDH2 | aldehyde dehydrogenase 2 family | Cytoplasm | enzyme | 6469986.5 | 5975793 | 1.11E+07 | | | | | | |
| P47738 | ALDH2 | aldehyde dehydrogenase 2 family | Cytoplasm | enzyme | 4724178 | 7136956.5 | 9319231 | | | | | | |
| P47738 | ALDH2 | aldehyde dehydrogenase 2 family | Cytoplasm | enzyme | 7160890 | 4368842.5 | 8459278 | | | | | | |
| P47738 | ALDH2 | aldehyde dehydrogenase 2 family | Cytoplasm | enzyme | 3292871.5 | 6213254.5 | 6389796.5 | P47738 | ALDH2 | 16895464 | 13160268 | 1550952 | 72 |
| P05064 | ALDOA | aldolase A, fructose-bisphosphate | Cytoplasm | enzyme | 5.43E+07 | 7.00E+07 | 5.52E+07 | | | | | | |
| P05064 | ALDOA | aldolase A, fructose-bisphosphate | Cytoplasm | enzyme | 4.50E+07 | 5.05E+07 | 4.37E+07 | | | | | | |
| P05064 | ALDOA | aldolase A, fructose-bisphosphate | Cytoplasm | enzyme | 3.59E+07 | 2.74E+07 | 3.13E+07 | | | | | | |
| P05064 | ALDOA | aldolase A, fructose-bisphosphate | Cytoplasm | enzyme | 2.42E+07 | 3.23E+07 | 3.03E+07 | | | | | | |
| P05064 | ALDOA | aldolase A, fructose-bisphosphate | Cytoplasm | enzyme | 2.65E+07 | 2.75E+07 | 2.72E+07 | | | | | | |
| P05064 | ALDOA | aldolase A, fructose-bisphosphate | Cytoplasm | enzyme | 1.16E+07 | 1.16E+07 | 1.04E+07 | | | | | | |
| P05064 | ALDOA | aldolase A, fructose-bisphosphate | Cytoplasm | enzyme | 3681640.75 | 7224136 | 4332253.5 | | | | | | |
| P05064 | ALDOA | aldolase A, fructose-bisphosphate | Cytoplasm | enzyme | 4621506 | 1.11E+07 | 7163282 | | | | | | |
| P05064 | ALDOA | aldolase A, fructose-bisphosphate | Cytoplasm | enzyme | 4256265 | 5086677 | 3592804.25 | | | | | | |
| P05064 | ALDOA | aldolase A, fructose-bisphosphate | Cytoplasm | enzyme | 4954170 | 3821528.75 | 2993043.75 | | | | | | |
| P05064 | ALDOA | aldolase A, fructose-bisphosphate | Cytoplasm | enzyme | 3466499.5 | 2902949.25 | 2794748.75 | | | | | | |
| P05064 | ALDOA | aldolase A, fructose-bisphosphate | Cytoplasm | enzyme | 553414.125 | 1553180.375 | 848957.25 | P05064 | ALDOA | 18253625 | 19063128 | 3177188 | 36 |
| O08739 | AMPD3 | adenosine monophosphate deami | Cytoplasm | enzyme | 5900140.5 | 3087636.5 | 2789386.25 | | | | | | |
| O08739 | AMPD3 | adenosine monophosphate deami | Cytoplasm | enzyme | 6152447 | 2990607.5 | 2710749.5 | | | | | | |
| O08739 | AMPD3 | adenosine monophosphate deami | Cytoplasm | enzyme | 5389936 | 9248369 | 5723186.5 | | | | | | |
| O08739 | AMPD3 | adenosine monophosphate deami | Cytoplasm | enzyme | 4777191 | 3743310 | 2016271.5 | | | | | | |
| O08739 | AMPD3 | adenosine monophosphate deami | Cytoplasm | enzyme | 3245812.75 | 2565680.25 | 2126933 | | | | | | |
| O08739 | AMPD3 | adenosine monophosphate deami | Cytoplasm | enzyme | 3142058.5 | 2154755.5 | 1443881.125 | | | | | | |
| O08739 | AMPD3 | adenosine monophosphate deami | Cytoplasm | enzyme | 2546690 | 2526683.25 | 1640544.5 | | | | | | |
| O08739 | AMPD3 | adenosine monophosphate deami | Cytoplasm | enzyme | 1154158.875 | 1223209.625 | 463285.156 | O08739 | AMPD3 | 4038554 | 2009447 | 410177 | 24 |
| Q02357 | ANK1 | ankyrin 1, erythrocytic | Plasma Membrane | other | 3.63E+09 | 3.57E+09 | 2.88E+09 | | | | | | |
| Q02357 | ANK1 | ankyrin 1, erythrocytic | Plasma Membrane | other | 3.53E+09 | 3.60E+09 | 3.54E+09 | | | | | | |
| Q02357 | ANK1 | ankyrin 1, erythrocytic | Plasma Membrane | other | 2.04E+09 | 2.48E+09 | 2.46E+09 | | | | | | |
| Q02357 | ANK1 | ankyrin 1, erythrocytic | Plasma Membrane | other | 2.40E+09 | 2.76E+09 | 2.44E+09 | | | | | | |
| Q02357 | ANK1 | ankyrin 1, erythrocytic | Plasma Membrane | other | 1.81E+09 | 2.66E+09 | 2.62E+09 | | | | | | |
| Q02357 | ANK1 | ankyrin 1, erythrocytic | Plasma Membrane | other | 2.54E+09 | 2.32E+09 | 2.55E+09 | | | | | | |
| Q02357 | ANK1 | ankyrin 1, erythrocytic | Plasma Membrane | other | 2.26E+09 | 2.14E+09 | 2.14E+09 | | | | | | |
| Q02357 | ANK1 | ankyrin 1, erythrocytic | Plasma Membrane | other | 2.10E+09 | 1.99E+09 | 2.18E+09 | | | | | | |
| Q02357 | ANK1 | ankyrin 1, erythrocytic | Plasma Membrane | other | 1.56E+09 | 1.86E+09 | 1.84E+09 | | | | | | |
| Q02357 | ANK1 | ankyrin 1, erythrocytic | Plasma Membrane | other | 2.32E+09 | 2.11E+09 | 2.33E+09 | | | | | | |
| Q02357 | ANK1 | ankyrin 1, erythrocytic | Plasma Membrane | other | 1.94E+09 | 2.25E+09 | 2.01E+09 | | | | | | |

| | | | | | | | | |
|---|---|---|---|---|---|---|---|---|
| Q02357 | ANK1 | ankyrin 1, erythrocytic | Plasma Membrane | other | | 1.58E+09 | 1.23E+09 | 1.61E+09 |
| Q02357 | ANK1 | ankyrin 1, erythrocytic | Plasma Membrane | other | | 1.80E+09 | 1.57E+09 | 1.50E+09 |
| Q02357 | ANK1 | ankyrin 1, erythrocytic | Plasma Membrane | other | | 1.59E+09 | 1.30E+09 | 1.59E+09 |
| Q02357 | ANK1 | ankyrin 1, erythrocytic | Plasma Membrane | other | | 1.61E+09 | 1.53E+09 | 1.56E+09 |
| Q02357 | ANK1 | ankyrin 1, erythrocytic | Plasma Membrane | other | | 1.79E+09 | 1.63E+09 | 1.70E+09 |
| Q02357 | ANK1 | ankyrin 1, erythrocytic | Plasma Membrane | other | | 1.56E+09 | 1.60E+09 | 1.47E+09 |
| Q02357 | ANK1 | ankyrin 1, erythrocytic | Plasma Membrane | other | | 1.56E+09 | 1.48E+09 | 1.50E+09 |
| Q02357 | ANK1 | ankyrin 1, erythrocytic | Plasma Membrane | other | | 1.31E+09 | 1.34E+09 | 9.62E+08 |
| Q02357 | ANK1 | ankyrin 1, erythrocytic | Plasma Membrane | other | | 1.34E+09 | 1.29E+09 | 1.45E+09 |
| Q02357 | ANK1 | ankyrin 1, erythrocytic | Plasma Membrane | other | | 1.57E+09 | 1.49E+09 | 1.57E+09 |
| Q02357 | ANK1 | ankyrin 1, erythrocytic | Plasma Membrane | other | | 1.59E+09 | 1.42E+09 | 1.60E+09 |
| Q02357 | ANK1 | ankyrin 1, erythrocytic | Plasma Membrane | other | | 1.56E+09 | 1.25E+09 | 1.24E+09 |
| Q02357 | ANK1 | ankyrin 1, erythrocytic | Plasma Membrane | other | | 1.27E+09 | 1.33E+09 | 1.45E+09 |
| Q02357 | ANK1 | ankyrin 1, erythrocytic | Plasma Membrane | other | | 1.38E+09 | 1.48E+09 | 1.68E+09 |
| Q02357 | ANK1 | ankyrin 1, erythrocytic | Plasma Membrane | other | | 8.84E+08 | 1.02E+09 | 8.88E+08 |
| Q02357 | ANK1 | ankyrin 1, erythrocytic | Plasma Membrane | other | | 1.02E+09 | 1.01E+09 | 9.81E+08 |
| Q02357 | ANK1 | ankyrin 1, erythrocytic | Plasma Membrane | other | | 7.82E+08 | 1.12E+09 | 1.16E+09 |
| Q02357 | ANK1 | ankyrin 1, erythrocytic | Plasma Membrane | other | | 1.18E+09 | 1.26E+09 | 1.04E+09 |
| Q02357 | ANK1 | ankyrin 1, erythrocytic | Plasma Membrane | other | | 7.42E+08 | 9.64E+08 | 9.04E+08 |
| Q02357 | ANK1 | ankyrin 1, erythrocytic | Plasma Membrane | other | | 9.63E+08 | 9.46E+08 | 1.08E+09 |
| Q02357 | ANK1 | ankyrin 1, erythrocytic | Plasma Membrane | other | | 9.38E+08 | 1.28E+09 | 1.38E+09 |
| Q02357 | ANK1 | ankyrin 1, erythrocytic | Plasma Membrane | other | | 1.00E+09 | 1.09E+09 | 1.03E+09 |
| Q02357 | ANK1 | ankyrin 1, erythrocytic | Plasma Membrane | other | | 7.77E+08 | 9.11E+08 | 8.74E+08 |
| Q02357 | ANK1 | ankyrin 1, erythrocytic | Plasma Membrane | other | | 1.04E+09 | 1.25E+09 | 9.09E+08 |
| Q02357 | ANK1 | ankyrin 1, erythrocytic | Plasma Membrane | other | | 7.95E+08 | 8.82E+08 | 7.64E+08 |
| Q02357 | ANK1 | ankyrin 1, erythrocytic | Plasma Membrane | other | | 6.95E+08 | 7.74E+08 | 7.41E+08 |
| Q02357 | ANK1 | ankyrin 1, erythrocytic | Plasma Membrane | other | | 7.32E+08 | 6.51E+08 | 7.62E+08 |
| Q02357 | ANK1 | ankyrin 1, erythrocytic | Plasma Membrane | other | | 1.04E+09 | 6.63E+08 | 7.96E+08 |
| Q02357 | ANK1 | ankyrin 1, erythrocytic | Plasma Membrane | other | | 6.48E+08 | 6.72E+08 | 5.84E+08 |
| Q02357 | ANK1 | ankyrin 1, erythrocytic | Plasma Membrane | other | | 7.59E+08 | 7.09E+08 | 5.14E+08 |
| Q02357 | ANK1 | ankyrin 1, erythrocytic | Plasma Membrane | other | | 7.88E+08 | 6.47E+08 | 5.70E+08 |
| Q02357 | ANK1 | ankyrin 1, erythrocytic | Plasma Membrane | other | | 7.74E+08 | 2.92E+08 | 4.96E+08 |
| Q02357 | ANK1 | ankyrin 1, erythrocytic | Plasma Membrane | other | | 6.59E+08 | 6.87E+08 | 6.65E+08 |
| Q02357 | ANK1 | ankyrin 1, erythrocytic | Plasma Membrane | other | | 5.87E+08 | 6.33E+08 | 5.44E+08 |
| Q02357 | ANK1 | ankyrin 1, erythrocytic | Plasma Membrane | other | | 6.51E+08 | 6.11E+08 | 5.91E+08 |
| Q02357 | ANK1 | ankyrin 1, erythrocytic | Plasma Membrane | other | | 8.10E+08 | 8.69E+08 | 8.80E+08 |
| Q02357 | ANK1 | ankyrin 1, erythrocytic | Plasma Membrane | other | | 7.14E+08 | 4.03E+08 | 2.69E+08 |
| Q02357 | ANK1 | ankyrin 1, erythrocytic | Plasma Membrane | other | | 6.83E+08 | 7.92E+08 | 8.23E+08 |
| Q02357 | ANK1 | ankyrin 1, erythrocytic | Plasma Membrane | other | | 5.38E+08 | 6.03E+08 | 5.06E+08 |
| Q02357 | ANK1 | ankyrin 1, erythrocytic | Plasma Membrane | other | | 5.06E+08 | 5.87E+08 | 5.77E+08 |
| Q02357 | ANK1 | ankyrin 1, erythrocytic | Plasma Membrane | other | | 4.93E+08 | 4.50E+08 | 4.62E+08 |
| Q02357 | ANK1 | ankyrin 1, erythrocytic | Plasma Membrane | other | | 5.84E+08 | 8.92E+08 | 6.67E+08 |
| Q02357 | ANK1 | ankyrin 1, erythrocytic | Plasma Membrane | other | | 4.58E+08 | 5.18E+08 | 4.85E+08 |
| Q02357 | ANK1 | ankyrin 1, erythrocytic | Plasma Membrane | other | | 4.66E+08 | 5.73E+08 | 5.21E+08 |
| Q02357 | ANK1 | ankyrin 1, erythrocytic | Plasma Membrane | other | | 6.14E+08 | 6.25E+08 | 6.41E+08 |
| Q02357 | ANK1 | ankyrin 1, erythrocytic | Plasma Membrane | other | | 4.37E+08 | 5.39E+08 | 5.68E+08 |
| Q02357 | ANK1 | ankyrin 1, erythrocytic | Plasma Membrane | other | | 3.81E+08 | 5.65E+08 | 5.12E+08 |
| Q02357 | ANK1 | ankyrin 1, erythrocytic | Plasma Membrane | other | | 2.52E+08 | 4.68E+08 | 4.59E+08 |
| Q02357 | ANK1 | ankyrin 1, erythrocytic | Plasma Membrane | other | | 5.63E+08 | 4.68E+08 | 3.88E+08 |
| Q02357 | ANK1 | ankyrin 1, erythrocytic | Plasma Membrane | other | | 6.54E+08 | 5.26E+08 | 5.78E+08 |
| Q02357 | ANK1 | ankyrin 1, erythrocytic | Plasma Membrane | other | | 5.40E+08 | 6.36E+08 | 4.25E+08 |
| Q02357 | ANK1 | ankyrin 1, erythrocytic | Plasma Membrane | other | | 3.75E+08 | 3.31E+08 | 2.95E+08 |
| Q02357 | ANK1 | ankyrin 1, erythrocytic | Plasma Membrane | other | | 2.71E+08 | 4.71E+08 | 4.71E+08 |
| Q02357 | ANK1 | ankyrin 1, erythrocytic | Plasma Membrane | other | | 3.15E+08 | 5.71E+08 | 5.42E+08 |
| Q02357 | ANK1 | ankyrin 1, erythrocytic | Plasma Membrane | other | | 3.86E+08 | 3.03E+08 | 4.07E+08 |
| Q02357 | ANK1 | ankyrin 1, erythrocytic | Plasma Membrane | other | | 4.38E+08 | 3.94E+08 | 3.46E+08 |
| Q02357 | ANK1 | ankyrin 1, erythrocytic | Plasma Membrane | other | | 3.76E+08 | 4.21E+08 | 3.27E+08 |
| Q02357 | ANK1 | ankyrin 1, erythrocytic | Plasma Membrane | other | | 2.53E+08 | 3.95E+08 | 3.16E+08 |
| Q02357 | ANK1 | ankyrin 1, erythrocytic | Plasma Membrane | other | | 4.12E+08 | 3.06E+08 | 4.23E+08 |
| Q02357 | ANK1 | ankyrin 1, erythrocytic | Plasma Membrane | other | | 3.66E+08 | 1.81E+08 | 7.57E+07 |
| Q02357 | ANK1 | ankyrin 1, erythrocytic | Plasma Membrane | other | | 5.42E+08 | 3.38E+08 | 4.02E+08 |
| Q02357 | ANK1 | ankyrin 1, erythrocytic | Plasma Membrane | other | | 2.24E+08 | 4.92E+08 | 3.21E+08 |
| Q02357 | ANK1 | ankyrin 1, erythrocytic | Plasma Membrane | other | | 1.16E+08 | 1.95E+08 | 2.31E+08 |
| Q02357 | ANK1 | ankyrin 1, erythrocytic | Plasma Membrane | other | | 5.15E+08 | 6.52E+07 | 2.79E+08 |
| Q02357 | ANK1 | ankyrin 1, erythrocytic | Plasma Membrane | other | | 2.71E+08 | 2.55E+08 | 2.45E+08 |
| Q02357 | ANK1 | ankyrin 1, erythrocytic | Plasma Membrane | other | | 4.46E+08 | 2.01E+08 | 2.29E+08 |
| Q02357 | ANK1 | ankyrin 1, erythrocytic | Plasma Membrane | other | | 2.73E+08 | 2.07E+08 | 1.77E+08 |
| Q02357 | ANK1 | ankyrin 1, erythrocytic | Plasma Membrane | other | | 1.24E+08 | 5.07E+07 | 2.53E+08 |
| Q02357 | ANK1 | ankyrin 1, erythrocytic | Plasma Membrane | other | | 1.46E+08 | 3.81E+08 | 2.76E+08 |
| Q02357 | ANK1 | ankyrin 1, erythrocytic | Plasma Membrane | other | | 2.59E+08 | 1.83E+08 | 1.02E+08 |
| Q02357 | ANK1 | ankyrin 1, erythrocytic | Plasma Membrane | other | | 2.79E+08 | 9.92E+07 | 5.91E+07 |
| Q02357 | ANK1 | ankyrin 1, erythrocytic | Plasma Membrane | other | | 1.57E+08 | 2.38E+08 | 2.61E+08 |
| Q02357 | ANK1 | ankyrin 1, erythrocytic | Plasma Membrane | other | | 2.42E+08 | 1.91E+08 | 2.12E+08 |
| Q02357 | ANK1 | ankyrin 1, erythrocytic | Plasma Membrane | other | | 2.20E+08 | 1.65E+08 | 1.99E+08 |
| Q02357 | ANK1 | ankyrin 1, erythrocytic | Plasma Membrane | other | | 1.91E+08 | 1.88E+08 | 2.26E+08 |
| Q02357 | ANK1 | ankyrin 1, erythrocytic | Plasma Membrane | other | | 2.12E+08 | 9.74E+07 | 5.30E+07 |
| Q02357 | ANK1 | ankyrin 1, erythrocytic | Plasma Membrane | other | | 2.43E+08 | 2.06E+08 | 2.38E+08 |
| Q02357 | ANK1 | ankyrin 1, erythrocytic | Plasma Membrane | other | | 1.40E+08 | 1.83E+08 | 2.62E+08 |
| Q02357 | ANK1 | ankyrin 1, erythrocytic | Plasma Membrane | other | | 1.98E+08 | 2.26E+08 | 2.76E+08 |
| Q02357 | ANK1 | ankyrin 1, erythrocytic | Plasma Membrane | other | | 9.25E+07 | 2.46E+08 | 1.78E+08 |
| Q02357 | ANK1 | ankyrin 1, erythrocytic | Plasma Membrane | other | | 1.02E+08 | 1.01E+08 | 6.87E+07 |
| Q02357 | ANK1 | ankyrin 1, erythrocytic | Plasma Membrane | other | | 1.13E+08 | 1.03E+08 | 1.02E+08 |
| Q02357 | ANK1 | ankyrin 1, erythrocytic | Plasma Membrane | other | | 1.19E+08 | 8.47E+07 | 2.50E+07 |
| Q02357 | ANK1 | ankyrin 1, erythrocytic | Plasma Membrane | other | | 9.19E+07 | 9.56E+07 | 1.12E+08 |
| Q02357 | ANK1 | ankyrin 1, erythrocytic | Plasma Membrane | other | | 1.71E+08 | 1.54E+08 | 2.03E+08 |
| Q02357 | ANK1 | ankyrin 1, erythrocytic | Plasma Membrane | other | | 7.63E+07 | 1.77E+08 | 1.50E+08 |

| | | | | | | | | | | | | | |
|---|---|---|---|---|---|---|---|---|---|---|---|---|---|
| Q02357 | ANK1 | ankyrin 1, erythrocytic | Plasma Membrane | other | 1.36E+08 | 1.23E+08 | 1.20E+08 | | | | | | |
| Q02357 | ANK1 | ankyrin 1, erythrocytic | Plasma Membrane | other | 2.18E+08 | 3.29E+07 | 3.10E+07 | | | | | | |
| Q02357 | ANK1 | ankyrin 1, erythrocytic | Plasma Membrane | other | 7.31E+07 | 9.47E+07 | 6.30E+07 | | | | | | |
| Q02357 | ANK1 | ankyrin 1, erythrocytic | Plasma Membrane | other | 7.20E+07 | 4.33E+07 | 3.02E+07 | | | | | | |
| Q02357 | ANK1 | ankyrin 1, erythrocytic | Plasma Membrane | other | 1.16E+08 | 1.18E+08 | 1.66E+08 | | | | | | |
| Q02357 | ANK1 | ankyrin 1, erythrocytic | Plasma Membrane | other | 1.02E+08 | 1.07E+08 | 8.83E+07 | | | | | | |
| Q02357 | ANK1 | ankyrin 1, erythrocytic | Plasma Membrane | other | 6.53E+07 | 5.28E+07 | 4.70E+07 | | | | | | |
| Q02357 | ANK1 | ankyrin 1, erythrocytic | Plasma Membrane | other | 1.57E+08 | 1.85E+08 | 1.30E+08 | | | | | | |
| Q02357 | ANK1 | ankyrin 1, erythrocytic | Plasma Membrane | other | 5.71E+07 | 4.75E+08 | 5.35E+08 | | | | | | |
| Q02357 | ANK1 | ankyrin 1, erythrocytic | Plasma Membrane | other | 6.02E+07 | 6.29E+07 | 7.25E+07 | | | | | | |
| Q02357 | ANK1 | ankyrin 1, erythrocytic | Plasma Membrane | other | 8.17E+07 | 4.78E+07 | 6.22E+07 | | | | | | |
| Q02357 | ANK1 | ankyrin 1, erythrocytic | Plasma Membrane | other | 8.75E+07 | 6.86E+07 | 6.40E+07 | | | | | | |
| Q02357 | ANK1 | ankyrin 1, erythrocytic | Plasma Membrane | other | 7.16E+07 | 7.32E+07 | 7.13E+07 | | | | | | |
| Q02357 | ANK1 | ankyrin 1, erythrocytic | Plasma Membrane | other | 4.11E+07 | 5.84E+07 | 3.81E+07 | | | | | | |
| Q02357 | ANK1 | ankyrin 1, erythrocytic | Plasma Membrane | other | 2.70E+07 | 1.17E+08 | 6.81E+07 | | | | | | |
| Q02357 | ANK1 | ankyrin 1, erythrocytic | Plasma Membrane | other | 5.59E+07 | 5.95E+07 | 5.59E+07 | | | | | | |
| Q02357 | ANK1 | ankyrin 1, erythrocytic | Plasma Membrane | other | 6.18E+07 | 4.61E+07 | 7.29E+07 | | | | | | |
| Q02357 | ANK1 | ankyrin 1, erythrocytic | Plasma Membrane | other | 2.98E+07 | 1.13E+08 | 6.83E+07 | | | | | | |
| Q02357 | ANK1 | ankyrin 1, erythrocytic | Plasma Membrane | other | 5.50E+07 | 6.08E+07 | 7.23E+07 | | | | | | |
| Q02357 | ANK1 | ankyrin 1, erythrocytic | Plasma Membrane | other | 3.24E+07 | 5.32E+07 | 4.42E+07 | | | | | | |
| Q02357 | ANK1 | ankyrin 1, erythrocytic | Plasma Membrane | other | 9959154 | 8.94E+07 | 9.07E+07 | | | | | | |
| Q02357 | ANK1 | ankyrin 1, erythrocytic | Plasma Membrane | other | 1.46E+08 | 4755778.5 | 3.31E+07 | | | | | | |
| Q02357 | ANK1 | ankyrin 1, erythrocytic | Plasma Membrane | other | 2.97E+07 | 2.04E+07 | 2.67E+07 | | | | | | |
| Q02357 | ANK1 | ankyrin 1, erythrocytic | Plasma Membrane | other | 2.71E+07 | 3.58E+07 | 3.27E+07 | | | | | | |
| Q02357 | ANK1 | ankyrin 1, erythrocytic | Plasma Membrane | other | 3.31E+07 | 3.48E+07 | 3.38E+07 | | | | | | |
| Q02357 | ANK1 | ankyrin 1, erythrocytic | Plasma Membrane | other | 2.96E+07 | 3.33E+07 | 3.06E+07 | | | | | | |
| Q02357 | ANK1 | ankyrin 1, erythrocytic | Plasma Membrane | other | 4.64E+07 | 4.92E+07 | 1.25E+08 | | | | | | |
| Q02357 | ANK1 | ankyrin 1, erythrocytic | Plasma Membrane | other | 3.40E+07 | 1.05E+07 | 2353896 | | | | | | |
| Q02357 | ANK1 | ankyrin 1, erythrocytic | Plasma Membrane | other | 2.81E+07 | 2.54E+07 | 1.44E+07 | | | | | | |
| Q02357 | ANK1 | ankyrin 1, erythrocytic | Plasma Membrane | other | 3.34E+07 | 2.29E+07 | 2.39E+07 | | | | | | |
| Q02357 | ANK1 | ankyrin 1, erythrocytic | Plasma Membrane | other | 1.39E+08 | 5160559.5 | 2.13E+07 | | | | | | |
| Q02357 | ANK1 | ankyrin 1, erythrocytic | Plasma Membrane | other | 2.72E+07 | 2.74E+07 | 2.49E+07 | | | | | | |
| Q02357 | ANK1 | ankyrin 1, erythrocytic | Plasma Membrane | other | 2.52E+07 | 1.87E+07 | 2.15E+07 | | | | | | |
| Q02357 | ANK1 | ankyrin 1, erythrocytic | Plasma Membrane | other | 1.45E+07 | 3.90E+07 | 2.32E+07 | | | | | | |
| Q02357 | ANK1 | ankyrin 1, erythrocytic | Plasma Membrane | other | 5.75E+07 | 1999761.625 | 1.04E+07 | | | | | | |
| Q02357 | ANK1 | ankyrin 1, erythrocytic | Plasma Membrane | other | 2.19E+07 | 7772031 | 2.32E+07 | | | | | | |
| Q02357 | ANK1 | ankyrin 1, erythrocytic | Plasma Membrane | other | 1.58E+07 | 1.48E+07 | 2.05E+07 | | | | | | |
| Q02357 | ANK1 | ankyrin 1, erythrocytic | Plasma Membrane | other | 2.52E+07 | 4.18E+07 | 1.34E+07 | | | | | | |
| Q02357 | ANK1 | ankyrin 1, erythrocytic | Plasma Membrane | other | 1.30E+07 | 1.14E+07 | 9312539 | | | | | | |
| Q02357 | ANK1 | ankyrin 1, erythrocytic | Plasma Membrane | other | 1.57E+07 | 4962595 | 1015270.625 | | | | | | |
| Q02357 | ANK1 | ankyrin 1, erythrocytic | Plasma Membrane | other | 2.24E+07 | 2111955 | 1026147.188 | | | | | | |
| Q02357 | ANK1 | ankyrin 1, erythrocytic | Plasma Membrane | other | 8927128 | 1.83E+07 | 1.30E+07 | | | | | | |
| Q02357 | ANK1 | ankyrin 1, erythrocytic | Plasma Membrane | other | 1.45E+07 | 19158.012 | 28853.453 | | | | | | |
| Q02357 | ANK1 | ankyrin 1, erythrocytic | Plasma Membrane | other | 1.19E+07 | 9787896 | 8817874 | | | | | | |
| Q02357 | ANK1 | ankyrin 1, erythrocytic | Plasma Membrane | other | 1.19E+07 | 7854688.5 | 1937871.125 | | | | | | |
| Q02357 | ANK1 | ankyrin 1, erythrocytic | Plasma Membrane | other | 2.65E+07 | 3388109 | 2621706.75 | | | | | | |
| Q02357 | ANK1 | ankyrin 1, erythrocytic | Plasma Membrane | other | 9147726 | 8767578 | 9221489 | | | | | | |
| Q02357 | ANK1 | ankyrin 1, erythrocytic | Plasma Membrane | other | 7369897.5 | 3675622.5 | 6874296.5 | | | | | | |
| Q02357 | ANK1 | ankyrin 1, erythrocytic | Plasma Membrane | other | 8534919 | 1.40E+07 | 1.31E+07 | | | | | | |
| Q02357 | ANK1 | ankyrin 1, erythrocytic | Plasma Membrane | other | 1.20E+07 | 3634182.25 | 4632564.5 | | | | | | |
| Q02357 | ANK1 | ankyrin 1, erythrocytic | Plasma Membrane | other | 6568046 | 2763343.25 | 4938805.5 | | | | | | |
| Q02357 | ANK1 | ankyrin 1, erythrocytic | Plasma Membrane | other | 6847294.5 | 9905114 | 4795974 | | | | | | |
| Q02357 | ANK1 | ankyrin 1, erythrocytic | Plasma Membrane | other | 1.03E+07 | 7849714 | 8393895 | | | | | | |
| Q02357 | ANK1 | ankyrin 1, erythrocytic | Plasma Membrane | other | 1.01E+07 | 2.84E+07 | 3.40E+07 | | | | | | |
| Q02357 | ANK1 | ankyrin 1, erythrocytic | Plasma Membrane | other | 5652383 | 2111822.75 | 1262824.375 | | | | | | |
| Q02357 | ANK1 | ankyrin 1, erythrocytic | Plasma Membrane | other | 2942223 | 7007828.5 | 4725422.5 | | | | | | |
| Q02357 | ANK1 | ankyrin 1, erythrocytic | Plasma Membrane | other | 2.36E+07 | 1.28E+07 | 1.40E+07 | | | | | | |
| Q02357 | ANK1 | ankyrin 1, erythrocytic | Plasma Membrane | other | 5516465 | 388007.656 | 1634353 | | | | | | |
| Q02357 | ANK1 | ankyrin 1, erythrocytic | Plasma Membrane | other | 1.35E+07 | 1.24E+07 | 1.34E+07 | | | | | | |
| Q02357 | ANK1 | ankyrin 1, erythrocytic | Plasma Membrane | other | 2378345.5 | 2233223.25 | 2536317 | | | | | | |
| Q02357 | ANK1 | ankyrin 1, erythrocytic | Plasma Membrane | other | 6204957.5 | 467507 | 1070772.25 | | | | | | |
| Q02357 | ANK1 | ankyrin 1, erythrocytic | Plasma Membrane | other | 8027058 | 585424.75 | 783726.812 | | | | | | |
| Q02357 | ANK1 | ankyrin 1, erythrocytic | Plasma Membrane | other | 3441592.25 | 2893134 | 2489368 | | | | | | |
| Q02357 | ANK1 | ankyrin 1, erythrocytic | Plasma Membrane | other | 3190180.75 | 2425508.25 | 1462838 | | | | | | |
| Q02357 | ANK1 | ankyrin 1, erythrocytic | Plasma Membrane | other | 2287980.25 | 5724024.5 | 4226783.5 | | | | | | |
| Q02357 | ANK1 | ankyrin 1, erythrocytic | Plasma Membrane | other | 2227868 | 3.34E+07 | 5.75E+07 | | | | | | |
| Q02357 | ANK1 | ankyrin 1, erythrocytic | Plasma Membrane | other | 1.38E+07 | 646366.188 | 1092281.5 | | | | | | |
| Q02357 | ANK1 | ankyrin 1, erythrocytic | Plasma Membrane | other | 1139668.625 | 1105878.375 | 1412249.875 | | | | | | |
| Q02357 | ANK1 | ankyrin 1, erythrocytic | Plasma Membrane | other | 443468.062 | 365219.25 | 559703.375 | | | | | | |
| Q02357 | ANK1 | ankyrin 1, erythrocytic | Plasma Membrane | other | 9524068 | 2429003.5 | 484427 | | | | | | |
| Q02357 | ANK1 | ankyrin 1, erythrocytic | Plasma Membrane | other | 518693.812 | 4748654 | 3424262.25 | | | | | | |
| Q02357 | ANK1 | ankyrin 1, erythrocytic | Plasma Membrane | other | 4.70E-06 | 4706619.5 | 9038282 | Q02357 | ANK1 | 503981474 | 694752565 | 30855045 | 507 |
| Q9JHZ2 | ANKH | ANKH inorganic pyrophosphate tra | Plasma Membrane | transporter | 1860453.375 | 2096608.25 | 1684689.625 | Q9JHZ2 | ANKH | 1860453 | 206696 | 119336 | 3 |
| Q505D1 | ANKRD28 | ankyrin repeat domain 28 | Cytoplasm | other | 1529132.875 | 1104525.5 | 539581.75 | Q505D1 | ANKRD28 | 1529133 | 496431 | 286615 | 3 |
| Q6P9J9 | ANO6 | anoctamin 6 | Plasma Membrane | ion channel | 3703036.75 | 4365237.5 | 3542481 | Q6P9J9 | ANO6 | 3703037 | 436122 | 251795 | 3 |
| O35381 | Anp32a | acidic (leucine-rich) nuclear phosp | Nucleus | other | 1.21E+07 | 3932389 | 9470878 | | | | | | |
| O35381 | Anp32a | acidic (leucine-rich) nuclear phosp | Nucleus | other | 4152429 | 5844542.5 | 3699837.5 | | | | | | |
| O35381 | Anp32a | acidic (leucine-rich) nuclear phosp | Nucleus | other | 3564931.25 | 8239528 | 4918011 | O35381 | Anp32a | 6592453 | 3031007 | 1010336 | 9 |
| O35643 | AP1B1 | adaptor-related protein complex 1 | Cytoplasm | transporter | 2.35E+07 | 1.54E+07 | 1.30E+07 | | | | | | |
| O35643 | AP1B1 | adaptor-related protein complex 1 | Cytoplasm | transporter | 1.13E+07 | 1.31E+07 | 1.23E+07 | | | | | | |
| O35643 | AP1B1 | adaptor-related protein complex 1 | Cytoplasm | transporter | 1.57E+07 | 1.25E+07 | 1.00E+07 | | | | | | |
| O35643 | AP1B1 | adaptor-related protein complex 1 | Cytoplasm | transporter | 9501892 | 1.13E+07 | 1.46E+07 | | | | | | |
| O35643 | AP1B1 | adaptor-related protein complex 1 | Cytoplasm | transporter | 9877692 | 8973001 | 9235629 | | | | | | |
| O35643 | AP1B1 | adaptor-related protein complex 1 | Cytoplasm | transporter | 7521217 | 4990254.5 | 3711286.25 | | | | | | |
| O35643 | AP1B1 | adaptor-related protein complex 1 | Cytoplasm | transporter | 4613006 | 3528009.5 | 2781307.75 | | | | | | |
| O35643 | AP1B1 | adaptor-related protein complex 1 | Cytoplasm | transporter | 8601912 | 6295736.5 | 5016246.5 | | | | | | |

| | | | | | | | | | | | | | |
|---|---|---|---|---|---|---|---|---|---|---|---|---|---|
| O35643 | AP1B1 | adaptor-related protein complex 1 | Cytoplasm | transporter | 4034396.5 | 5697639.5 | 2419467.25 | | | | | | |
| O35643 | AP1B1 | adaptor-related protein complex 1 | Cytoplasm | transporter | 4704773.5 | 4127176.25 | 4002275 | | | | | | |
| O35643 | AP1B1 | adaptor-related protein complex 1 | Cytoplasm | transporter | 3986760.25 | 3270976 | 2890082 | | | | | | |
| O35643 | AP1B1 | adaptor-related protein complex 1 | Cytoplasm | transporter | 1487914.5 | 1914544.5 | 877903.375 | O35643 | AP1B1 | 8728297 | 5078739 | 846457 | 36 |
| P17426 | AP2A1 | adaptor-related protein complex 2 | Cytoplasm | transporter | 5355944 | 6436699.5 | 4211142.5 | | | | | | |
| P17426 | AP2A1 | adaptor-related protein complex 2 | Cytoplasm | transporter | 4832223.5 | 3713233.5 | 3129962.5 | | | | | | |
| P17426 | AP2A1 | adaptor-related protein complex 2 | Cytoplasm | transporter | 1475744.75 | 1588625.875 | 1063900.25 | P17426 | AP2A1 | 3887971 | 1877722 | 625907 | 9 |
| P17427 | AP2A2 | adaptor-related protein complex 2 | Cytoplasm | transporter | 1.32E+07 | 9361003 | 9062224 | | | | | | |
| P17427 | AP2A2 | adaptor-related protein complex 2 | Cytoplasm | transporter | 8668976 | 6438542.5 | 3818344 | | | | | | |
| P17427 | AP2A2 | adaptor-related protein complex 2 | Cytoplasm | transporter | 2.12E+07 | 2.01E+07 | 1.99E+07 | | | | | | |
| P17427 | AP2A2 | adaptor-related protein complex 2 | Cytoplasm | transporter | 2995658 | 3500595 | 2311578.25 | | | | | | |
| P17427 | AP2A2 | adaptor-related protein complex 2 | Cytoplasm | transporter | 3011425.75 | 2659555.75 | 2143583 | | | | | | |
| P17427 | AP2A2 | adaptor-related protein complex 2 | Cytoplasm | transporter | 2650380.75 | 3488027.5 | 1630390.625 | | | | | | |
| P17427 | AP2A2 | adaptor-related protein complex 2 | Cytoplasm | transporter | 2474869.75 | 2195858.5 | 1555648.375 | | | | | | |
| P17427 | AP2A2 | adaptor-related protein complex 2 | Cytoplasm | transporter | 1689719.5 | 1634303.5 | 1387998.75 | P17427 | AP2A2 | 6991379 | 6304664 | 1286934 | 24 |
| Q9DBG3 | Ap2b1 | adaptor-related protein complex 2 | Plasma Membrane | other | 1625470.5 | 1362022.375 | 1177585.5 | | | | | | |
| Q9DBG3 | Ap2b1 | adaptor-related protein complex 2 | Plasma Membrane | other | 85680.539 | 198755.625 | 33597.957 | Q9DBG3 | Ap2b1 | 855576 | 718641 | 293384 | 6 |
| P84091 | AP2M1 | adaptor-related protein complex 2 | Cytoplasm | transporter | 1.13E+07 | 8362507 | 6879939.5 | | | | | | |
| P84091 | AP2M1 | adaptor-related protein complex 2 | Cytoplasm | transporter | 8881405 | 6335698.5 | 5903294 | | | | | | |
| P84091 | AP2M1 | adaptor-related protein complex 2 | Cytoplasm | transporter | 2718282.75 | 2763943.75 | 1998073.125 | | | | | | |
| P84091 | AP2M1 | adaptor-related protein complex 2 | Cytoplasm | transporter | 4228122.5 | 3036514 | 3120022.25 | | | | | | |
| P84091 | AP2M1 | adaptor-related protein complex 2 | Cytoplasm | transporter | 2601850.25 | 2489905 | 2219608.75 | P84091 | AP2M1 | 5949932 | 2917046 | 753178 | 15 |
| Q8R146 | APEH | acylaminoacyl-peptide hydrolase | Cytoplasm | peptidase | 7.22E+07 | 6.14E+07 | 5.55E+07 | | | | | | |
| Q8R146 | APEH | acylaminoacyl-peptide hydrolase | Cytoplasm | peptidase | 3.42E+07 | 2.86E+07 | 3.08E+07 | | | | | | |
| Q8R146 | APEH | acylaminoacyl-peptide hydrolase | Cytoplasm | peptidase | 2.60E+07 | 3.76E+07 | 3.29E+07 | | | | | | |
| Q8R146 | APEH | acylaminoacyl-peptide hydrolase | Cytoplasm | peptidase | 3.23E+07 | 2.67E+07 | 3.26E+07 | | | | | | |
| Q8R146 | APEH | acylaminoacyl-peptide hydrolase | Cytoplasm | peptidase | 3.05E+07 | 2.68E+07 | 2.22E+07 | | | | | | |
| Q8R146 | APEH | acylaminoacyl-peptide hydrolase | Cytoplasm | peptidase | 2.44E+07 | 2.07E+07 | 2.24E+07 | | | | | | |
| Q8R146 | APEH | acylaminoacyl-peptide hydrolase | Cytoplasm | peptidase | 2.64E+07 | 2.37E+07 | 2.70E+07 | | | | | | |
| Q8R146 | APEH | acylaminoacyl-peptide hydrolase | Cytoplasm | peptidase | 2.15E+07 | 1.63E+07 | 1.89E+07 | | | | | | |
| Q8R146 | APEH | acylaminoacyl-peptide hydrolase | Cytoplasm | peptidase | 3.08E+07 | 1.39E+07 | 2.22E+07 | | | | | | |
| Q8R146 | APEH | acylaminoacyl-peptide hydrolase | Cytoplasm | peptidase | 1.90E+07 | 1.68E+07 | 1.82E+07 | | | | | | |
| Q8R146 | APEH | acylaminoacyl-peptide hydrolase | Cytoplasm | peptidase | 1.64E+07 | 1.74E+07 | 1.74E+07 | | | | | | |
| Q8R146 | APEH | acylaminoacyl-peptide hydrolase | Cytoplasm | peptidase | 1.80E+07 | 1.91E+07 | 2.23E+07 | | | | | | |
| Q8R146 | APEH | acylaminoacyl-peptide hydrolase | Cytoplasm | peptidase | 1.91E+07 | 1.62E+07 | 1.34E+07 | | | | | | |
| Q8R146 | APEH | acylaminoacyl-peptide hydrolase | Cytoplasm | peptidase | 1.89E+07 | 1.97E+07 | 2.32E+07 | | | | | | |
| Q8R146 | APEH | acylaminoacyl-peptide hydrolase | Cytoplasm | peptidase | 2.20E+07 | 2.39E+07 | 1.85E+07 | | | | | | |
| Q8R146 | APEH | acylaminoacyl-peptide hydrolase | Cytoplasm | peptidase | 2.00E+07 | 1.66E+07 | 2.04E+07 | | | | | | |
| Q8R146 | APEH | acylaminoacyl-peptide hydrolase | Cytoplasm | peptidase | 1.64E+07 | 1.41E+07 | 1.64E+07 | | | | | | |
| Q8R146 | APEH | acylaminoacyl-peptide hydrolase | Cytoplasm | peptidase | 1.30E+07 | 1.28E+07 | 1.32E+07 | | | | | | |
| Q8R146 | APEH | acylaminoacyl-peptide hydrolase | Cytoplasm | peptidase | 1.07E+07 | 4062645.5 | 7870505 | | | | | | |
| Q8R146 | APEH | acylaminoacyl-peptide hydrolase | Cytoplasm | peptidase | 4769825 | 3825609 | 6655991 | | | | | | |
| Q8R146 | APEH | acylaminoacyl-peptide hydrolase | Cytoplasm | peptidase | 6448770 | 4512175.5 | 3041244 | | | | | | |
| Q8R146 | APEH | acylaminoacyl-peptide hydrolase | Cytoplasm | peptidase | 4343234.5 | 3258639.25 | 3737721 | | | | | | |
| Q8R146 | APEH | acylaminoacyl-peptide hydrolase | Cytoplasm | peptidase | 3319895 | 4494254.5 | 3470328.25 | | | | | | |
| Q8R146 | APEH | acylaminoacyl-peptide hydrolase | Cytoplasm | peptidase | 5254320.5 | 8965642 | 1.04E+07 | | | | | | |
| Q8R146 | APEH | acylaminoacyl-peptide hydrolase | Cytoplasm | peptidase | 1327624.75 | 822147.312 | 849657.562 | Q8R146 | APEH | 19888147 | 13060864 | 1508139 | 75 |
| Q8BVF7 | APH1A | APH1A gamma secretase subunit | Cytoplasm | peptidase | 1747520.25 | 2454240.25 | 1715396.875 | Q8BVF7 | APH1A | 1747520 | 417607 | 241106 | 3 |
| Q00623 | APOA1 | apolipoprotein A-I | Extracellular Space | transporter | 3414599.5 | 1015848.125 | 2494617.25 | | | | | | |
| Q00623 | APOA1 | apolipoprotein A-I | Extracellular Space | transporter | 2188389.75 | 2733733.25 | 2429066 | | | | | | |
| Q00623 | APOA1 | apolipoprotein A-I | Extracellular Space | transporter | 401301.219 | 591548.875 | 623735.562 | Q00623 | APOA1 | 2001430 | 1113136 | 371045 | 9 |
| Q01339 | APOH | apolipoprotein H (beta-2-glycopro | Extracellular Space | transporter | 694399.625 | 630774.75 | 1047230.188 | Q01339 | APOH | 694400 | 224341 | 129523 | 3 |
| Q02013 | AQP1 | aquaporin 1 (Colton blood group) | Plasma Membrane | transporter | 3372884.5 | 573411.125 | 809259.625 | | | | | | |
| Q02013 | AQP1 | aquaporin 1 (Colton blood group) | Plasma Membrane | transporter | 8825466 | 6722767 | 7136432 | | | | | | |
| Q02013 | AQP1 | aquaporin 1 (Colton blood group) | Plasma Membrane | transporter | 1.07E+07 | 1.80E+07 | 2.52E+07 | | | | | | |
| Q02013 | AQP1 | aquaporin 1 (Colton blood group) | Plasma Membrane | transporter | 148100.469 | 323350.906 | 542089.062 | | | | | | |
| Q02013 | AQP1 | aquaporin 1 (Colton blood group) | Plasma Membrane | transporter | 2731997.25 | 5056811 | 4944830 | Q02013 | AQP1 | 5151690 | 7153726 | 1847084 | 15 |
| P84084 | ARF5 | ADP-ribosylation factor 5 | Cytoplasm | enzyme | 2874245.25 | 2110565.5 | 1656176.375 | | | | | | |
| P84084 | ARF5 | ADP-ribosylation factor 5 | Cytoplasm | enzyme | 1539324.25 | 1133838.125 | 1354547.375 | P84084 | ARF5 | 2206785 | 629034 | 256802 | 6 |
| P62331 | ARF6 | ADP-ribosylation factor 6 | Plasma Membrane | transporter | 2.01E+07 | 2.06E+07 | 1.97E+07 | | | | | | |
| P62331 | ARF6 | ADP-ribosylation factor 6 | Plasma Membrane | transporter | 1.15E+07 | 1.53E+07 | 1.71E+07 | | | | | | |
| P62331 | ARF6 | ADP-ribosylation factor 6 | Plasma Membrane | transporter | 5484781 | 7515439 | 8176385.5 | | | | | | |
| P62331 | ARF6 | ADP-ribosylation factor 6 | Plasma Membrane | transporter | 8728586 | 6171332 | 6255396.5 | | | | | | |
| P62331 | ARF6 | ADP-ribosylation factor 6 | Plasma Membrane | transporter | 4956941.5 | 6411957 | 7115261 | | | | | | |
| P62331 | ARF6 | ADP-ribosylation factor 6 | Plasma Membrane | transporter | 5591254 | 5441287 | 6421324.5 | P62331 | ARF6 | 9396927 | 5685930 | 1340187 | 18 |
| Q80YF9 | ARHGAP33 | Rho GTPase activating protein 33 | Plasma Membrane | transporter | 1.12E+07 | 1.52E+07 | 1.58E+07 | Q80YF9 | ARHGAP33 | 11180000 | 2518988 | 1454338 | 3 |
| Q9Z1N7 | ARID3B | AT rich interactive domain 3B (BRI | Nucleus | other | 1.31E+07 | 3938596 | 7680293.5 | Q9Z1N7 | ARID3B | 13110000 | 4611519 | 2662462 | 3 |
| Q9JKW0 | ARL6IP1 | ADP-ribosylation factor-like 6 inte | Cytoplasm | other | 8412707 | 9978464 | 8996203 | Q9JKW0 | ARL6IP1 | 8412707 | 791296 | 456855 | 3 |
| Q8R5J9 | ARL6IP5 | ADP-ribosylation factor-like 6 inte | Cytoplasm | other | 3550395.25 | 4834072.5 | 5057488.5 | Q8R5J9 | ARL6IP5 | 3550395 | 813334 | 469579 | 3 |
| Q8VEH3 | ARL8A | ADP-ribosylation factor-like 8A | Cytoplasm | enzyme | 2590076.5 | 2112034.25 | 1990830.875 | Q8VEH3 | ARL8A | 2590077 | 316836 | 182925 | 3 |
| Q9CQW2 | ARL8B | ADP-ribosylation factor-like 8B | Plasma Membrane | enzyme | 4.72E+07 | 4.48E+07 | 4.72E+07 | | | | | | |
| Q9CQW2 | ARL8B | ADP-ribosylation factor-like 8B | Plasma Membrane | enzyme | 2.52E+07 | 2.84E+07 | 2.87E+07 | | | | | | |
| Q9CQW2 | ARL8B | ADP-ribosylation factor-like 8B | Plasma Membrane | enzyme | 2.90E+07 | 2.70E+07 | 2.67E+07 | | | | | | |
| Q9CQW2 | ARL8B | ADP-ribosylation factor-like 8B | Plasma Membrane | enzyme | 2.40E+07 | 1.70E+07 | 1.39E+07 | | | | | | |
| Q9CQW2 | ARL8B | ADP-ribosylation factor-like 8B | Plasma Membrane | enzyme | 1.33E+07 | 2.69E+07 | 1.77E+07 | | | | | | |
| Q9CQW2 | ARL8B | ADP-ribosylation factor-like 8B | Plasma Membrane | enzyme | 1.66E+07 | 1.32E+07 | 1.03E+07 | | | | | | |
| Q9CQW2 | ARL8B | ADP-ribosylation factor-like 8B | Plasma Membrane | enzyme | 9864236 | 3.03E+07 | 4.00E+07 | | | | | | |
| Q9CQW2 | ARL8B | ADP-ribosylation factor-like 8B | Plasma Membrane | enzyme | 2.04E+07 | 4451655 | 9690786 | | | | | | |
| Q9CQW2 | ARL8B | ADP-ribosylation factor-like 8B | Plasma Membrane | enzyme | 7779071 | 1.25E+07 | 7938409.5 | | | | | | |
| Q9CQW2 | ARL8B | ADP-ribosylation factor-like 8B | Plasma Membrane | enzyme | 1.06E+07 | 2.42E+07 | 3.18E+07 | | | | | | |
| Q9CQW2 | ARL8B | ADP-ribosylation factor-like 8B | Plasma Membrane | enzyme | 6185725 | 2.28E+07 | 1.69E+07 | | | | | | |
| Q9CQW2 | ARL8B | ADP-ribosylation factor-like 8B | Plasma Membrane | enzyme | 1.01E+07 | 9748227 | 9712276 | | | | | | |
| Q9CQW2 | ARL8B | ADP-ribosylation factor-like 8B | Plasma Membrane | enzyme | 1.04E+07 | 9786528 | 9700058 | | | | | | |
| Q9CQW2 | ARL8B | ADP-ribosylation factor-like 8B | Plasma Membrane | enzyme | 1.01E+07 | 1.25E+07 | 9190632 | Q9CQW2 | ARL8B | 17180645 | 11436186 | 1764642 | 42 |
| Q9R0Q6 | ARPC1A | actin related protein 2/3 complex, | Extracellular Space | other | 2869529.25 | 2918672.5 | 2286601 | Q9R0Q6 | ARPC1A | 2869529 | 351600 | 202996 | 3 |
| Q9CVB6 | ARPC2 | actin related protein 2/3 complex, | Cytoplasm | other | 1.47E+07 | 1.60E+07 | 2.03E+07 | | | | | | |

| | | | | | | | | | | | | | |
|---|---|---|---|---|---|---|---|---|---|---|---|---|---|
| Q9CVB6 | ARPC2 | actin related protein 2/3 complex, | Cytoplasm | other | 1.02E+07 | 1.12E+07 | 8793437 | | | | | | |
| Q9CVB6 | ARPC2 | actin related protein 2/3 complex, | Cytoplasm | other | 5324866 | 9055652 | 7449146.5 | Q9CVB6 | ARPC2 | 10098289 | 4719170 | 1573057 | 9 |
| Q9JM76 | ARPC3 | actin related protein 2/3 complex, | Cytoplasm | other | 3852507.75 | 4978972 | 4168160.5 | | | | | | |
| Q9JM76 | ARPC3 | actin related protein 2/3 complex, | Cytoplasm | other | 3079382.75 | 2937548.25 | 2107207.75 | Q9JM76 | ARPC3 | 3465945 | 1018690 | 415878 | 6 |
| P59999 | ARPC4 | actin related protein 2/3 complex, | Cytoplasm | other | 2960291.5 | 4076601 | 2922303.75 | P59999 | ARPC4 | 2960292 | 655743 | 378593 | 3 |
| Q9CPW4 | ARPC5 | actin related protein 2/3 complex, | Cytoplasm | other | 1675400 | 1936957.5 | 1803071 | Q9CPW4 | ARPC5 | 1675400 | 130791 | 75512 | 3 |
| Q9CRA0 | ART4 | ADP-ribosyltransferase 4 (Dombro | Nucleus | enzyme | 2.74E+07 | 3.59E+07 | 3.11E+07 | | | | | | |
| Q9CRA0 | ART4 | ADP-ribosyltransferase 4 (Dombro | Nucleus | enzyme | 2.67E+07 | 2.72E+07 | 2.82E+07 | | | | | | |
| Q9CRA0 | ART4 | ADP-ribosyltransferase 4 (Dombro | Nucleus | enzyme | 9870896 | 8270045 | 6287461.5 | | | | | | |
| Q9CRA0 | ART4 | ADP-ribosyltransferase 4 (Dombro | Nucleus | enzyme | 6105257.5 | 6214657 | 7323918.5 | | | | | | |
| Q9CRA0 | ART4 | ADP-ribosyltransferase 4 (Dombro | Nucleus | enzyme | 4464644.5 | 4018885.75 | 5056130.5 | | | | | | |
| Q9CRA0 | ART4 | ADP-ribosyltransferase 4 (Dombro | Nucleus | enzyme | 2138841.5 | 2521924 | 2803852.5 | | | | | | |
| Q9CRA0 | ART4 | ADP-ribosyltransferase 4 (Dombro | Nucleus | enzyme | 2130033 | 2161355.75 | 3746650.25 | Q9CRA0 | ART4 | 11255668 | 11676985 | 2548127 | 21 |
| Q9QWY8 | ASAP1 | ArfGAP with SH3 domain, ankyrin | Plasma Membrane | other | 3663400.75 | 3450699 | 2709231 | | | | | | |
| Q9QWY8 | ASAP1 | ArfGAP with SH3 domain, ankyrin | Plasma Membrane | other | 1810370.25 | 1381255.125 | 965191.812 | Q9QWY8 | ASAP1 | 2736886 | 1114502 | 454994 | 6 |
| Q7SIG6 | ASAP2 | ArfGAP with SH3 domain, ankyrin | Nucleus | other | 1.54E+07 | 1.24E+07 | 3.07E+07 | Q7SIG6 | ASAP2 | 15390000 | 9824108 | 5671951 | 3 |
| Q9WV71 | ASB4 | ankyrin repeat and SOCS box conta | Nucleus | transcription regulator | 9070696 | 7111267 | 6158828.5 | Q9WV71 | ASB4 | 9070696 | 1484670 | 857175 | 3 |
| Q4QY64 | ATAD5 | ATPase family, AAA domain contai | Extracellular Space | other | 1.82E+07 | 9606087 | 1.64E+07 | Q4QY64 | ATAD5 | 18220000 | 4543657 | 2623282 | 3 |
| P16951 | ATF2 | activating transcription factor 2 | Nucleus | transcription regulator | 7.36E+07 | 5.72E+07 | 4.58E+07 | P16951 | ATF2 | 73630000 | 14015122 | 8091634 | 3 |
| Q811C2 | ATG4C | autophagy related 4C, cysteine pe | Cytoplasm | peptidase | 978144.25 | 565415.938 | 1002484.188 | Q811C2 | ATG4C | 978144 | 245617 | 141807 | 3 |
| Q9CWJ9 | ATIC | 5-aminoimidazole-4-carboxamide | Cytoplasm | enzyme | 8026591 | 9383494 | 9990447 | | | | | | |
| Q9CWJ9 | ATIC | 5-aminoimidazole-4-carboxamide | Cytoplasm | enzyme | 6012429 | 5260898 | 5961675 | | | | | | |
| Q9CWJ9 | ATIC | 5-aminoimidazole-4-carboxamide | Cytoplasm | enzyme | 6979489.5 | 2309548.25 | 3855792.5 | | | | | | |
| Q9CWJ9 | ATIC | 5-aminoimidazole-4-carboxamide | Cytoplasm | enzyme | 4619380.5 | 3628494.5 | 3542098.75 | | | | | | |
| Q9CWJ9 | ATIC | 5-aminoimidazole-4-carboxamide | Cytoplasm | enzyme | 4252438 | 3546395.25 | 4062541 | | | | | | |
| Q9CWJ9 | ATIC | 5-aminoimidazole-4-carboxamide | Cytoplasm | enzyme | 3038481.5 | 4223376 | 5523299.5 | | | | | | |
| Q9CWJ9 | ATIC | 5-aminoimidazole-4-carboxamide | Cytoplasm | enzyme | 3266133.5 | 2073328.25 | 3017099 | | | | | | |
| Q9CWJ9 | ATIC | 5-aminoimidazole-4-carboxamide | Cytoplasm | enzyme | 1676564.875 | 1276415.75 | 1601657.5 | | | | | | |
| Q9CWJ9 | ATIC | 5-aminoimidazole-4-carboxamide | Cytoplasm | enzyme | 443575.406 | 1424389.875 | 1566960.375 | | | | | | |
| Q9CWJ9 | ATIC | 5-aminoimidazole-4-carboxamide | Cytoplasm | enzyme | 648661.625 | 2049686.875 | 2158437.75 | Q9CWJ9 | ATIC | 3896374 | 2444826 | 446362 | 30 |
| Q6PA06 | ATL2 | atlastin GTPase 2 | Cytoplasm | other | 5345405.5 | 5858325 | 6735228.5 | | | | | | |
| Q6PA06 | ATL2 | atlastin GTPase 2 | Cytoplasm | other | 2371042.75 | 2343803.75 | 1873253.25 | | | | | | |
| Q6PA06 | ATL2 | atlastin GTPase 2 | Cytoplasm | other | 1439278.125 | 1777795 | 1659390.625 | Q6PA06 | ATL2 | 3051909 | 2085774 | 695258 | 9 |
| Q91YH5 | ATL3 | atlastin GTPase 3 | Cytoplasm | other | 5.24E+07 | 5.81E+07 | 5.00E+07 | | | | | | |
| Q91YH5 | ATL3 | atlastin GTPase 3 | Cytoplasm | other | 3.35E+07 | 3.42E+07 | 3.05E+07 | | | | | | |
| Q91YH5 | ATL3 | atlastin GTPase 3 | Cytoplasm | other | 1.70E+07 | 1.98E+07 | 1.62E+07 | | | | | | |
| Q91YH5 | ATL3 | atlastin GTPase 3 | Cytoplasm | other | 1.75E+07 | 1.91E+07 | 1.87E+07 | | | | | | |
| Q91YH5 | ATL3 | atlastin GTPase 3 | Cytoplasm | other | 2.27E+07 | 2.78E+07 | 2.08E+07 | | | | | | |
| Q91YH5 | ATL3 | atlastin GTPase 3 | Cytoplasm | other | 1.94E+07 | 2.48E+07 | 2.16E+07 | | | | | | |
| Q91YH5 | ATL3 | atlastin GTPase 3 | Cytoplasm | other | 2.80E+07 | 4.05E+07 | 2.55E+07 | | | | | | |
| Q91YH5 | ATL3 | atlastin GTPase 3 | Cytoplasm | other | 2.15E+07 | 2.04E+07 | 1.82E+07 | | | | | | |
| Q91YH5 | ATL3 | atlastin GTPase 3 | Cytoplasm | other | 1.76E+07 | 2.03E+07 | 1.59E+07 | | | | | | |
| Q91YH5 | ATL3 | atlastin GTPase 3 | Cytoplasm | other | 8817311 | 1.13E+07 | 1.09E+07 | | | | | | |
| Q91YH5 | ATL3 | atlastin GTPase 3 | Cytoplasm | other | 5972772 | 1.25E+07 | 1.03E+07 | | | | | | |
| Q91YH5 | ATL3 | atlastin GTPase 3 | Cytoplasm | other | 3728524.25 | 6313440 | 6334940 | | | | | | |
| Q91YH5 | ATL3 | atlastin GTPase 3 | Cytoplasm | other | 4692125 | 5724483.5 | 5165560 | | | | | | |
| Q91YH5 | ATL3 | atlastin GTPase 3 | Cytoplasm | other | 4415335.5 | 5248775.5 | 4684785 | | | | | | |
| Q91YH5 | ATL3 | atlastin GTPase 3 | Cytoplasm | other | 4641468 | 3381196.5 | 3110471.5 | | | | | | |
| Q91YH5 | ATL3 | atlastin GTPase 3 | Cytoplasm | other | 3353097.25 | 5121145 | 4814366.5 | Q91YH5 | ATL3 | 16575665 | 13393899 | 1933243 | 48 |
| Q9QZW0 | ATP11C | ATPase, class VI, type 11C | Plasma Membrane | transporter | 1.10E+07 | 1.41E+07 | 1.07E+07 | | | | | | |
| Q9QZW0 | ATP11C | ATPase, class VI, type 11C | Plasma Membrane | transporter | 1.02E+07 | 1.23E+07 | 9861329 | | | | | | |
| Q9QZW0 | ATP11C | ATPase, class VI, type 11C | Plasma Membrane | transporter | 5648226 | 5848202 | 6064737 | | | | | | |
| Q9QZW0 | ATP11C | ATPase, class VI, type 11C | Plasma Membrane | transporter | 4291182.5 | 4880578.5 | 5016120 | | | | | | |
| Q9QZW0 | ATP11C | ATPase, class VI, type 11C | Plasma Membrane | transporter | 4388736.5 | 4388060 | 3984852.5 | | | | | | |
| Q9QZW0 | ATP11C | ATPase, class VI, type 11C | Plasma Membrane | transporter | 2362093.25 | 3255539.25 | 3132817 | Q9QZW0 | ATP11C | 6321706 | 3587301 | 845535 | 18 |
| Q9EPE9 | ATP13A1 | ATPase type 13A1 | Extracellular Space | transporter | 1.10E+07 | 1.20E+07 | 1.25E+07 | Q9EPE9 | ATP13A1 | 11040000 | 732143 | 422703 | 3 |
| Q8VDN2 | ATP1A1 | ATPase, Na+/K+ transporting, alph | Plasma Membrane | transporter | 3.32E+07 | 3.71E+07 | 2.83E+07 | | | | | | |
| Q8VDN2 | ATP1A1 | ATPase, Na+/K+ transporting, alph | Plasma Membrane | transporter | 3.09E+07 | 3.31E+07 | 2.37E+07 | | | | | | |
| Q8VDN2 | ATP1A1 | ATPase, Na+/K+ transporting, alph | Plasma Membrane | transporter | 2.96E+07 | 3.21E+07 | 2.63E+07 | | | | | | |
| Q8VDN2 | ATP1A1 | ATPase, Na+/K+ transporting, alph | Plasma Membrane | transporter | 2.75E+07 | 3.02E+07 | 3.25E+07 | | | | | | |
| Q8VDN2 | ATP1A1 | ATPase, Na+/K+ transporting, alph | Plasma Membrane | transporter | 2.69E+07 | 3.10E+07 | 2.69E+07 | | | | | | |
| Q8VDN2 | ATP1A1 | ATPase, Na+/K+ transporting, alph | Plasma Membrane | transporter | 2.58E+07 | 2.63E+07 | 2.31E+07 | | | | | | |
| Q8VDN2 | ATP1A1 | ATPase, Na+/K+ transporting, alph | Plasma Membrane | transporter | 1.98E+07 | 1.80E+07 | 1.83E+07 | | | | | | |
| Q8VDN2 | ATP1A1 | ATPase, Na+/K+ transporting, alph | Plasma Membrane | transporter | 2.18E+07 | 2.01E+07 | 1.27E+07 | | | | | | |
| Q8VDN2 | ATP1A1 | ATPase, Na+/K+ transporting, alph | Plasma Membrane | transporter | 2.06E+07 | 1.86E+07 | 1.62E+07 | | | | | | |
| Q8VDN2 | ATP1A1 | ATPase, Na+/K+ transporting, alph | Plasma Membrane | transporter | 1.86E+07 | 1.66E+07 | 1.59E+07 | | | | | | |
| Q8VDN2 | ATP1A1 | ATPase, Na+/K+ transporting, alph | Plasma Membrane | transporter | 2.00E+07 | 1.69E+07 | 1.49E+07 | | | | | | |
| Q8VDN2 | ATP1A1 | ATPase, Na+/K+ transporting, alph | Plasma Membrane | transporter | 1.26E+07 | 1.18E+07 | 9449345 | | | | | | |
| Q8VDN2 | ATP1A1 | ATPase, Na+/K+ transporting, alph | Plasma Membrane | transporter | 1.08E+07 | 1.20E+07 | 1.06E+07 | | | | | | |
| Q8VDN2 | ATP1A1 | ATPase, Na+/K+ transporting, alph | Plasma Membrane | transporter | 1.30E+07 | 1.00E+07 | 7441013.5 | | | | | | |
| Q8VDN2 | ATP1A1 | ATPase, Na+/K+ transporting, alph | Plasma Membrane | transporter | 9457678 | 1.14E+07 | 1.03E+07 | | | | | | |
| Q8VDN2 | ATP1A1 | ATPase, Na+/K+ transporting, alph | Plasma Membrane | transporter | 9448187 | 1.30E+07 | 1.16E+07 | | | | | | |
| Q8VDN2 | ATP1A1 | ATPase, Na+/K+ transporting, alph | Plasma Membrane | transporter | 9064645 | 1.21E+07 | 9470847 | | | | | | |
| Q8VDN2 | ATP1A1 | ATPase, Na+/K+ transporting, alph | Plasma Membrane | transporter | 4970028.5 | 5515599.5 | 4463771 | | | | | | |
| Q8VDN2 | ATP1A1 | ATPase, Na+/K+ transporting, alph | Plasma Membrane | transporter | 6477543.5 | 6743990 | 5345311 | | | | | | |
| Q8VDN2 | ATP1A1 | ATPase, Na+/K+ transporting, alph | Plasma Membrane | transporter | 3986630.75 | 9119403 | 5549081.5 | | | | | | |
| Q8VDN2 | ATP1A1 | ATPase, Na+/K+ transporting, alph | Plasma Membrane | transporter | 5104016 | 7171029.5 | 6449788.5 | | | | | | |
| Q8VDN2 | ATP1A1 | ATPase, Na+/K+ transporting, alph | Plasma Membrane | transporter | 4173564 | 4532870.5 | 3591052.25 | | | | | | |
| Q8VDN2 | ATP1A1 | ATPase, Na+/K+ transporting, alph | Plasma Membrane | transporter | 4481470 | 5629759.5 | 3956826.5 | | | | | | |
| Q8VDN2 | ATP1A1 | ATPase, Na+/K+ transporting, alph | Plasma Membrane | transporter | 3172186 | 7540850.5 | 7735720 | | | | | | |
| Q8VDN2 | ATP1A1 | ATPase, Na+/K+ transporting, alph | Plasma Membrane | transporter | 4535880 | 6724335.5 | 4420258 | | | | | | |
| Q8VDN2 | ATP1A1 | ATPase, Na+/K+ transporting, alph | Plasma Membrane | transporter | 3298550 | 1972703.25 | 2445918.5 | | | | | | |
| Q8VDN2 | ATP1A1 | ATPase, Na+/K+ transporting, alph | Plasma Membrane | transporter | 2105940 | 2673004.75 | 2216575.5 | | | | | | |
| Q8VDN2 | ATP1A1 | ATPase, Na+/K+ transporting, alph | Plasma Membrane | transporter | 1602253.75 | 1005601 | 750263.312 | | | | | | |
| Q8VDN2 | ATP1A1 | ATPase, Na+/K+ transporting, alph | Plasma Membrane | transporter | 1723359 | 2809041.25 | 3010980.25 | Q8VDN2 | ATP1A1 | 13263170 | 9837141 | 1054652 | 87 |
| P14231 | ATP1B2 | ATPase, Na+/K+ transporting, beta | Plasma Membrane | transporter | 6.71E+07 | 6.29E+07 | 5.30E+07 | | | | | | |

| | | | | | | | | | | | | | |
|---|---|---|---|---|---|---|---|---|---|---|---|---|---|
| P14231 | ATP1B2 | ATPase, Na+/K+ transporting, beta | Plasma Membrane | transporter | 3.59E+07 | 3.40E+07 | 2.74E+07 | | | | | | |
| P14231 | ATP1B2 | ATPase, Na+/K+ transporting, beta | Plasma Membrane | transporter | 8036977.5 | 7820937.5 | 7362742.5 | | | | | | |
| P14231 | ATP1B2 | ATPase, Na+/K+ transporting, beta | Plasma Membrane | transporter | 6057817.5 | 6076965.5 | 4297367 | | | | | | |
| P14231 | ATP1B2 | ATPase, Na+/K+ transporting, beta | Plasma Membrane | transporter | 3094578.5 | 2898047.5 | 2085686 | P14231 | ATP1B2 | 24037875 | 23276435 | 6009950 | 15 |
| P97370 | ATP1B3 | ATPase, Na+/K+ transporting, beta | Plasma Membrane | transporter | 4277534 | 2946381.75 | 4208573 | | | | | | |
| P97370 | ATP1B3 | ATPase, Na+/K+ transporting, beta | Plasma Membrane | transporter | 1719060 | 2129996.75 | 2843763.5 | | | | | | |
| P97370 | ATP1B3 | ATPase, Na+/K+ transporting, beta | Plasma Membrane | transporter | 1707237.25 | 2095547 | 1987249.625 | | | | | | |
| P97370 | ATP1B3 | ATPase, Na+/K+ transporting, beta | Plasma Membrane | transporter | 1504280.5 | 2621272.5 | 2648211.25 | P97370 | ATP1B3 | 2302028 | 913862 | 263809 | 12 |
| O55143 | ATP2A2 | ATPase, Ca++ transporting, cardiac | Cytoplasm | transporter | 1.00E+07 | 1.06E+07 | 1.06E+07 | | | | | | |
| O55143 | ATP2A2 | ATPase, Ca++ transporting, cardiac | Cytoplasm | transporter | 2.05E+07 | 1.99E+07 | 1.81E+07 | | | | | | |
| O55143 | ATP2A2 | ATPase, Ca++ transporting, cardiac | Cytoplasm | transporter | 4182196.75 | 7927473.5 | 6941506.5 | | | | | | |
| O55143 | ATP2A2 | ATPase, Ca++ transporting, cardiac | Cytoplasm | transporter | 3661277 | 4636806.5 | 3266197.5 | | | | | | |
| O55143 | ATP2A2 | ATPase, Ca++ transporting, cardiac | Cytoplasm | transporter | 1999590.625 | 4481465.5 | 2648200 | | | | | | |
| O55143 | ATP2A2 | ATPase, Ca++ transporting, cardiac | Cytoplasm | transporter | 1567434.375 | 3279730.5 | 2212243.25 | O55143 | ATP2A2 | 6988416 | 6211347 | 1464028 | 18 |
| Q64518 | ATP2A3 | ATPase, Ca++ transporting, ubiquit | Cytoplasm | transporter | 3.14E+07 | 3.79E+07 | 3.26E+07 | | | | | | |
| Q64518 | ATP2A3 | ATPase, Ca++ transporting, ubiquit | Cytoplasm | transporter | 1.39E+07 | 1.38E+07 | 1.70E+07 | | | | | | |
| Q64518 | ATP2A3 | ATPase, Ca++ transporting, ubiquit | Cytoplasm | transporter | 8727691 | 9375571 | 1.02E+07 | | | | | | |
| Q64518 | ATP2A3 | ATPase, Ca++ transporting, ubiquit | Cytoplasm | transporter | 8665942 | 9833986 | 8259964 | | | | | | |
| Q64518 | ATP2A3 | ATPase, Ca++ transporting, ubiquit | Cytoplasm | transporter | 2.07E+07 | 2.07E+07 | 2.13E+07 | | | | | | |
| Q64518 | ATP2A3 | ATPase, Ca++ transporting, ubiquit | Cytoplasm | transporter | 5405627.5 | 5935061 | 5752634 | | | | | | |
| Q64518 | ATP2A3 | ATPase, Ca++ transporting, ubiquit | Cytoplasm | transporter | 7742346.5 | 1.02E+07 | 1.03E+07 | | | | | | |
| Q64518 | ATP2A3 | ATPase, Ca++ transporting, ubiquit | Cytoplasm | transporter | 1.36E+07 | 5233159.5 | 1.13E+07 | | | | | | |
| Q64518 | ATP2A3 | ATPase, Ca++ transporting, ubiquit | Cytoplasm | transporter | 6840914.5 | 6783646.5 | 7247861 | | | | | | |
| Q64518 | ATP2A3 | ATPase, Ca++ transporting, ubiquit | Cytoplasm | transporter | 1.19E+07 | 9220781 | 1.28E+07 | | | | | | |
| Q64518 | ATP2A3 | ATPase, Ca++ transporting, ubiquit | Cytoplasm | transporter | 2321967 | 8706.535 | 951268.938 | | | | | | |
| Q64518 | ATP2A3 | ATPase, Ca++ transporting, ubiquit | Cytoplasm | transporter | 1083913.875 | 2263243 | 1655546.375 | | | | | | |
| Q64518 | ATP2A3 | ATPase, Ca++ transporting, ubiquit | Cytoplasm | transporter | 1673188 | 1443952.875 | 1501839.75 | | | | | | |
| Q64518 | ATP2A3 | ATPase, Ca++ transporting, ubiquit | Cytoplasm | transporter | 3487138.25 | 6457057 | 4452122.5 | | | | | | |
| Q64518 | ATP2A3 | ATPase, Ca++ transporting, ubiquit | Cytoplasm | transporter | 1124603.5 | 1212971.875 | 1416023.5 | | | | | | |
| Q64518 | ATP2A3 | ATPase, Ca++ transporting, ubiquit | Cytoplasm | transporter | 5208655.5 | 7543926.5 | 9260906 | Q64518 | ATP2A3 | 8984499 | 8392540 | 1211359 | 48 |
| Q9R0K7 | ATP2B2 | ATPase, Ca++ transporting, plasma | Plasma Membrane | transporter | 1.01E+08 | 1.26E+08 | 1.01E+08 | | | | | | |
| Q9R0K7 | ATP2B2 | ATPase, Ca++ transporting, plasma | Plasma Membrane | transporter | 3.35E+07 | 4.32E+07 | 3.68E+07 | | | | | | |
| Q9R0K7 | ATP2B2 | ATPase, Ca++ transporting, plasma | Plasma Membrane | transporter | 2.49E+07 | 4.01E+07 | 3.23E+07 | | | | | | |
| Q9R0K7 | ATP2B2 | ATPase, Ca++ transporting, plasma | Plasma Membrane | transporter | 3.04E+07 | 2.15E+07 | 1.32E+07 | | | | | | |
| Q9R0K7 | ATP2B2 | ATPase, Ca++ transporting, plasma | Plasma Membrane | transporter | 1.90E+07 | 2.29E+07 | 1.49E+07 | | | | | | |
| Q9R0K7 | ATP2B2 | ATPase, Ca++ transporting, plasma | Plasma Membrane | transporter | 7803906.5 | 8989889 | 7423683 | | | | | | |
| Q9R0K7 | ATP2B2 | ATPase, Ca++ transporting, plasma | Plasma Membrane | transporter | 5906376 | 5220667.5 | 4372962 | | | | | | |
| Q9R0K7 | ATP2B2 | ATPase, Ca++ transporting, plasma | Plasma Membrane | transporter | 9303539 | 2394148.25 | 4832871 | | | | | | |
| Q9R0K7 | ATP2B2 | ATPase, Ca++ transporting, plasma | Plasma Membrane | transporter | 2590890.25 | 2081910.5 | 2697181.5 | | | | | | |
| Q9R0K7 | ATP2B2 | ATPase, Ca++ transporting, plasma | Plasma Membrane | transporter | 1405122.625 | 1661955.625 | 1406877 | Q9R0K7 | ATP2B2 | 23533983 | 31738078 | 5794554 | 30 |
| Q03265 | ATP5A1 | ATP synthase, H+ transporting, mi | Cytoplasm | transporter | 6.75E+07 | 7.31E+07 | 9.71E+07 | | | | | | |
| Q03265 | ATP5A1 | ATP synthase, H+ transporting, mi | Cytoplasm | transporter | 7.23E+07 | 6.24E+07 | 8.85E+07 | | | | | | |
| Q03265 | ATP5A1 | ATP synthase, H+ transporting, mi | Cytoplasm | transporter | 6.31E+07 | 5.62E+07 | 7.36E+07 | | | | | | |
| Q03265 | ATP5A1 | ATP synthase, H+ transporting, mi | Cytoplasm | transporter | 5.46E+07 | 5.35E+07 | 6.55E+07 | | | | | | |
| Q03265 | ATP5A1 | ATP synthase, H+ transporting, mi | Cytoplasm | transporter | 5.11E+07 | 5.43E+07 | 4.94E+07 | | | | | | |
| Q03265 | ATP5A1 | ATP synthase, H+ transporting, mi | Cytoplasm | transporter | 5.70E+07 | 5.37E+07 | 6.43E+07 | | | | | | |
| Q03265 | ATP5A1 | ATP synthase, H+ transporting, mi | Cytoplasm | transporter | 4.97E+07 | 4.82E+07 | 5.79E+07 | | | | | | |
| Q03265 | ATP5A1 | ATP synthase, H+ transporting, mi | Cytoplasm | transporter | 3.72E+07 | 4.20E+07 | 5.64E+07 | | | | | | |
| Q03265 | ATP5A1 | ATP synthase, H+ transporting, mi | Cytoplasm | transporter | 4.09E+07 | 3.70E+07 | 4.86E+07 | | | | | | |
| Q03265 | ATP5A1 | ATP synthase, H+ transporting, mi | Cytoplasm | transporter | 3.57E+07 | 3.46E+07 | 4.24E+07 | | | | | | |
| Q03265 | ATP5A1 | ATP synthase, H+ transporting, mi | Cytoplasm | transporter | 3.43E+07 | 3.85E+07 | 5.53E+07 | | | | | | |
| Q03265 | ATP5A1 | ATP synthase, H+ transporting, mi | Cytoplasm | transporter | 3.64E+07 | 4.42E+07 | 5.16E+07 | | | | | | |
| Q03265 | ATP5A1 | ATP synthase, H+ transporting, mi | Cytoplasm | transporter | 2.51E+07 | 2.61E+07 | 3.04E+07 | | | | | | |
| Q03265 | ATP5A1 | ATP synthase, H+ transporting, mi | Cytoplasm | transporter | 3.37E+07 | 1.72E+07 | 3.37E+07 | | | | | | |
| Q03265 | ATP5A1 | ATP synthase, H+ transporting, mi | Cytoplasm | transporter | 1.94E+07 | 2.03E+07 | 2.50E+07 | | | | | | |
| Q03265 | ATP5A1 | ATP synthase, H+ transporting, mi | Cytoplasm | transporter | 1.37E+07 | 1.34E+07 | 1.62E+07 | | | | | | |
| Q03265 | ATP5A1 | ATP synthase, H+ transporting, mi | Cytoplasm | transporter | 1.68E+07 | 1.66E+07 | 2.18E+07 | | | | | | |
| Q03265 | ATP5A1 | ATP synthase, H+ transporting, mi | Cytoplasm | transporter | 1.22E+07 | 1.19E+07 | 1.57E+07 | | | | | | |
| Q03265 | ATP5A1 | ATP synthase, H+ transporting, mi | Cytoplasm | transporter | 9205461 | 8010911.5 | 8769464 | | | | | | |
| Q03265 | ATP5A1 | ATP synthase, H+ transporting, mi | Cytoplasm | transporter | 7293860 | 8168432.5 | 1.02E+07 | | | | | | |
| Q03265 | ATP5A1 | ATP synthase, H+ transporting, mi | Cytoplasm | transporter | 7329115.5 | 1.07E+07 | 1.40E+07 | | | | | | |
| Q03265 | ATP5A1 | ATP synthase, H+ transporting, mi | Cytoplasm | transporter | 5465716.5 | 1.04E+07 | 1.25E+07 | | | | | | |
| Q03265 | ATP5A1 | ATP synthase, H+ transporting, mi | Cytoplasm | transporter | 661429 | 1.90E+07 | 2.78E+07 | Q03265 | ATP5A1 | 32635460 | 22618731 | 2722975 | 69 |
| P56480 | ATP5B | ATP synthase, H+ transporting, mi | Cytoplasm | transporter | 1.16E+08 | 1.15E+08 | 1.39E+08 | | | | | | |
| P56480 | ATP5B | ATP synthase, H+ transporting, mi | Cytoplasm | transporter | 9.43E+07 | 8.54E+07 | 1.03E+08 | | | | | | |
| P56480 | ATP5B | ATP synthase, H+ transporting, mi | Cytoplasm | transporter | 6.30E+07 | 8.66E+07 | 1.15E+08 | | | | | | |
| P56480 | ATP5B | ATP synthase, H+ transporting, mi | Cytoplasm | transporter | 6.07E+07 | 6.06E+07 | 8.71E+07 | | | | | | |
| P56480 | ATP5B | ATP synthase, H+ transporting, mi | Cytoplasm | transporter | 5.81E+07 | 5.49E+07 | 7.17E+07 | | | | | | |
| P56480 | ATP5B | ATP synthase, H+ transporting, mi | Cytoplasm | transporter | 3.73E+07 | 3.05E+07 | 3.47E+07 | | | | | | |
| P56480 | ATP5B | ATP synthase, H+ transporting, mi | Cytoplasm | transporter | 3.88E+07 | 4.57E+07 | 5.77E+07 | | | | | | |
| P56480 | ATP5B | ATP synthase, H+ transporting, mi | Cytoplasm | transporter | 4.20E+07 | 8.43E+07 | 9.67E+07 | | | | | | |
| P56480 | ATP5B | ATP synthase, H+ transporting, mi | Cytoplasm | transporter | 3.46E+07 | 3.17E+07 | 4.72E+07 | | | | | | |
| P56480 | ATP5B | ATP synthase, H+ transporting, mi | Cytoplasm | transporter | 3.64E+07 | 4.72E+07 | 5.69E+07 | | | | | | |
| P56480 | ATP5B | ATP synthase, H+ transporting, mi | Cytoplasm | transporter | 4.28E+07 | 2.24E+07 | 5.29E+07 | | | | | | |
| P56480 | ATP5B | ATP synthase, H+ transporting, mi | Cytoplasm | transporter | 4.11E+07 | 5.09E+07 | 5.98E+07 | | | | | | |
| P56480 | ATP5B | ATP synthase, H+ transporting, mi | Cytoplasm | transporter | 2.33E+07 | 3.07E+07 | 3.25E+07 | | | | | | |
| P56480 | ATP5B | ATP synthase, H+ transporting, mi | Cytoplasm | transporter | 2.93E+07 | 3.51E+07 | 3.26E+07 | | | | | | |
| P56480 | ATP5B | ATP synthase, H+ transporting, mi | Cytoplasm | transporter | 1.83E+07 | 3.48E+07 | 4.77E+07 | | | | | | |
| P56480 | ATP5B | ATP synthase, H+ transporting, mi | Cytoplasm | transporter | 3.14E+07 | 4.59E+07 | 5.86E+07 | | | | | | |
| P56480 | ATP5B | ATP synthase, H+ transporting, mi | Cytoplasm | transporter | 2.92E+07 | 2.85E+07 | 5.35E+07 | | | | | | |
| P56480 | ATP5B | ATP synthase, H+ transporting, mi | Cytoplasm | transporter | 2.53E+07 | 2.39E+07 | 3.65E+07 | | | | | | |
| P56480 | ATP5B | ATP synthase, H+ transporting, mi | Cytoplasm | transporter | 1.06E+07 | 3.02E+07 | 2.02E+07 | | | | | | |
| P56480 | ATP5B | ATP synthase, H+ transporting, mi | Cytoplasm | transporter | 2.56E+07 | 2.84E+07 | 3.20E+07 | | | | | | |
| P56480 | ATP5B | ATP synthase, H+ transporting, mi | Cytoplasm | transporter | 1.83E+07 | 1.91E+07 | 2.91E+07 | | | | | | |
| P56480 | ATP5B | ATP synthase, H+ transporting, mi | Cytoplasm | transporter | 1.68E+07 | 2.23E+07 | 2.85E+07 | | | | | | |
| P56480 | ATP5B | ATP synthase, H+ transporting, mi | Cytoplasm | transporter | 1.16E+07 | 1.55E+07 | 1.63E+07 | | | | | | |

| | | | | | | | | | | | | | |
|---|---|---|---|---|---|---|---|---|---|---|---|---|---|
| P56480 | ATP5B | ATP synthase, H+ transporting, mi | Cytoplasm | transporter | 8817736 | 1.05E+07 | 1.69E+07 | | | | | | |
| P56480 | ATP5B | ATP synthase, H+ transporting, mi | Cytoplasm | transporter | 8511939 | 8576862 | 9072613 | | | | | | |
| P56480 | ATP5B | ATP synthase, H+ transporting, mi | Cytoplasm | transporter | 6214992 | 1.89E+07 | 1.52E+07 | | | | | | |
| P56480 | ATP5B | ATP synthase, H+ transporting, mi | Cytoplasm | transporter | 3771577.5 | 7197741.5 | 1.08E+07 | | | | | | |
| P56480 | ATP5B | ATP synthase, H+ transporting, mi | Cytoplasm | transporter | 1.42E+07 | 2885088.25 | 3088123.25 | | | | | | |
| P56480 | ATP5B | ATP synthase, H+ transporting, mi | Cytoplasm | transporter | 4895416 | 5284691.5 | 7009677.5 | | | | | | |
| P56480 | ATP5B | ATP synthase, H+ transporting, mi | Cytoplasm | transporter | 5234750 | 3217298.5 | 8038653 | | | | | | |
| P56480 | ATP5B | ATP synthase, H+ transporting, mi | Cytoplasm | transporter | 697452.25 | 1191464.625 | 2793513.5 | | | | | | |
| P56480 | ATP5B | ATP synthase, H+ transporting, mi | Cytoplasm | transporter | 702502.312 | 1549485.875 | 3334920 | P56480 | ATP5B | 29936761 | 30224508 | 3084776 | 96 |
| Q91VR2 | ATP5C1 | ATP synthase, H+ transporting, mi | Cytoplasm | transporter | 1.13E+07 | 1.17E+07 | 1.35E+07 | | | | | | |
| Q91VR2 | ATP5C1 | ATP synthase, H+ transporting, mi | Cytoplasm | transporter | 8338221 | 7302815 | 1.04E+07 | | | | | | |
| Q91VR2 | ATP5C1 | ATP synthase, H+ transporting, mi | Cytoplasm | transporter | 5261766 | 3244014.5 | 2517785.5 | | | | | | |
| Q91VR2 | ATP5C1 | ATP synthase, H+ transporting, mi | Cytoplasm | transporter | 415335.438 | 1728389.875 | 3193501.75 | Q91VR2 | ATP5C1 | 6326331 | 4438068 | 1281160 | 12 |
| Q9D3D9 | ATP5D | ATP synthase, H+ transporting, mi | Cytoplasm | transporter | 1.29E+07 | 1.55E+07 | 1.80E+07 | | | | | | |
| Q9D3D9 | ATP5D | ATP synthase, H+ transporting, mi | Cytoplasm | transporter | 1.56E+07 | 1.64E+07 | 1.92E+07 | Q9D3D9 | ATP5D | 14250000 | 2206790 | 900918 | 6 |
| Q9CQQ7 | ATP5F1 | ATP synthase, H+ transporting, mi | Cytoplasm | transporter | 6702678.5 | 7820262.5 | 9258949 | Q9CQQ7 | ATP5F1 | 6702679 | 1281492 | 739870 | 3 |
| Q9DCX2 | ATP5H | ATP synthase, H+ transporting, mi | Cytoplasm | enzyme | 1.15E+07 | 1.09E+07 | 1.31E+07 | | | | | | |
| Q9DCX2 | ATP5H | ATP synthase, H+ transporting, mi | Cytoplasm | enzyme | 9166241 | 8709675 | 1.05E+07 | | | | | | |
| Q9DCX2 | ATP5H | ATP synthase, H+ transporting, mi | Cytoplasm | enzyme | 7477090 | 7853804 | 8271966.5 | | | | | | |
| Q9DCX2 | ATP5H | ATP synthase, H+ transporting, mi | Cytoplasm | enzyme | 4223950.5 | 5804677 | 5921282 | | | | | | |
| Q9DCX2 | ATP5H | ATP synthase, H+ transporting, mi | Cytoplasm | enzyme | 3428424.25 | 5424014 | 5589976.5 | | | | | | |
| Q9DCX2 | ATP5H | ATP synthase, H+ transporting, mi | Cytoplasm | enzyme | 2614275.5 | 3847692 | 5075958.5 | Q9DCX2 | ATP5H | 6401664 | 3023348 | 712610 | 18 |
| P97450 | ATP5J | ATP synthase, H+ transporting, mi | Cytoplasm | transporter | 5685848.5 | 7318376 | 8154421 | P97450 | ATP5J | 5685849 | 1255519 | 724874 | 3 |
| P56135 | ATP5J2 | ATP synthase, H+ transporting, mi | Cytoplasm | transporter | 6774985.5 | 5457552.5 | 7264535 | P56135 | ATP5J2 | 6774986 | 934565 | 539572 | 3 |
| Q06185 | Atp5k | ATP synthase, H+ transporting, mi | Cytoplasm | enzyme | 1.46E+07 | 1.33E+07 | 1.72E+07 | | | | | | |
| Q06185 | Atp5k | ATP synthase, H+ transporting, mi | Cytoplasm | enzyme | 4346669.5 | 6715727.5 | 6783261.5 | Q06185 | Atp5k | 9483335 | 5204108 | 2124568 | 6 |
| Q9CPQ8 | ATP5L | ATP synthase, H+ transporting, mi | Cytoplasm | enzyme | 6626712.5 | 6988308.5 | 9133412 | | | | | | |
| Q9CPQ8 | ATP5L | ATP synthase, H+ transporting, mi | Cytoplasm | enzyme | 5218832 | 6274444.5 | 6116744.5 | Q9CPQ8 | ATP5L | 5922772 | 1320678 | 539164 | 6 |
| Q9DB20 | ATP5O | ATP synthase, H+ transporting, mi | Cytoplasm | transporter | 1.77E+07 | 1.45E+07 | 1.90E+07 | | | | | | |
| Q9DB20 | ATP5O | ATP synthase, H+ transporting, mi | Cytoplasm | transporter | 1.31E+07 | 1.61E+07 | 1.94E+07 | | | | | | |
| Q9DB20 | ATP5O | ATP synthase, H+ transporting, mi | Cytoplasm | transporter | 1.11E+07 | 9561128 | 1.13E+07 | | | | | | |
| Q9DB20 | ATP5O | ATP synthase, H+ transporting, mi | Cytoplasm | transporter | 1.16E+07 | 1.13E+07 | 1.46E+07 | | | | | | |
| Q9DB20 | ATP5O | ATP synthase, H+ transporting, mi | Cytoplasm | transporter | 1.28E+07 | 1932616.625 | 5853816 | | | | | | |
| Q9DB20 | ATP5O | ATP synthase, H+ transporting, mi | Cytoplasm | transporter | 3661200.25 | 8548997 | 6502058.5 | | | | | | |
| Q9DB20 | ATP5O | ATP synthase, H+ transporting, mi | Cytoplasm | transporter | 6626051.5 | 5425745.5 | 5759692 | | | | | | |
| Q9DB20 | ATP5O | ATP synthase, H+ transporting, mi | Cytoplasm | transporter | 5460766.5 | 5941549 | 7358592.5 | Q9DB20 | ATP5O | 10257252 | 4939444 | 1008260 | 24 |
| Q9R1Q9 | ATP6AP1 | ATPase, H+ transporting, lysosoma | Cytoplasm | transporter | 1547313.25 | 2726216.25 | 1939960.875 | Q9R1Q9 | ATP6AP1 | 1547313 | 600303 | 346585 | 3 |
| Q9CYN9 | ATP6AP2 | ATPase, H+ transporting, lysosoma | Cytoplasm | transporter | 2136095 | 3475192.75 | 3055483.5 | Q9CYN9 | ATP6AP2 | 2136095 | 684910 | 395433 | 3 |
| Q9Z1G4 | ATP6V0A1 | ATPase, H+ transporting, lysosoma | Cytoplasm | transporter | 1.25E+07 | 1.19E+07 | 9793523 | | | | | | |
| Q9Z1G4 | ATP6V0A1 | ATPase, H+ transporting, lysosoma | Cytoplasm | transporter | 5949383.5 | 8953013 | 8196872 | | | | | | |
| Q9Z1G4 | ATP6V0A1 | ATPase, H+ transporting, lysosoma | Cytoplasm | transporter | 6895317 | 9073271 | 7974089.5 | | | | | | |
| Q9Z1G4 | ATP6V0A1 | ATPase, H+ transporting, lysosoma | Cytoplasm | transporter | 6846827.5 | 1.46E+07 | 7515037.5 | | | | | | |
| Q9Z1G4 | ATP6V0A1 | ATPase, H+ transporting, lysosoma | Cytoplasm | transporter | 8837177 | 1.14E+07 | 9176224 | | | | | | |
| Q9Z1G4 | ATP6V0A1 | ATPase, H+ transporting, lysosoma | Cytoplasm | transporter | 4810213 | 4533357 | 4094873.75 | | | | | | |
| Q9Z1G4 | ATP6V0A1 | ATPase, H+ transporting, lysosoma | Cytoplasm | transporter | 5179810.5 | 6126078 | 4884147 | | | | | | |
| Q9Z1G4 | ATP6V0A1 | ATPase, H+ transporting, lysosoma | Cytoplasm | transporter | 3077383 | 3295673.5 | 3531434.25 | | | | | | |
| Q9Z1G4 | ATP6V0A1 | ATPase, H+ transporting, lysosoma | Cytoplasm | transporter | 1982379.5 | 4011784.5 | 2961815.5 | Q9Z1G4 | ATP6V0A1 | 6228721 | 3270013 | 629314 | 27 |
| P63082 | ATP6V0C | ATPase, H+ transporting, lysosoma | Cytoplasm | transporter | 2.21E+07 | 2.40E+07 | 2.59E+07 | | | | | | |
| P63082 | ATP6V0C | ATPase, H+ transporting, lysosoma | Cytoplasm | transporter | 3010375.5 | 3491831.75 | 3051727.5 | P63082 | ATP6V0C | 12575188 | 11472733 | 4683724 | 6 |
| P51863 | ATP6V0D1 | ATPase, H+ transporting, lysosoma | Cytoplasm | transporter | 2.27E+07 | 2.14E+07 | 2.04E+07 | | | | | | |
| P51863 | ATP6V0D1 | ATPase, H+ transporting, lysosoma | Cytoplasm | transporter | 1.11E+07 | 1.35E+07 | 1.10E+07 | | | | | | |
| P51863 | ATP6V0D1 | ATPase, H+ transporting, lysosoma | Cytoplasm | transporter | 8909907 | 1.17E+07 | 9421715 | | | | | | |
| P51863 | ATP6V0D1 | ATPase, H+ transporting, lysosoma | Cytoplasm | transporter | 7129225 | 9797055 | 8435178 | | | | | | |
| P51863 | ATP6V0D1 | ATPase, H+ transporting, lysosoma | Cytoplasm | transporter | 4621148.5 | 4836694 | 5229600 | | | | | | |
| P51863 | ATP6V0D1 | ATPase, H+ transporting, lysosoma | Cytoplasm | transporter | 2690407 | 4039004 | 3289389 | P51863 | ATP6V0D1 | 9535115 | 6143411 | 1448016 | 18 |
| P50516 | ATP6V1A | ATPase, H+ transporting, lysosoma | Plasma Membrane | transporter | 6099126.5 | 4325870.5 | 5829045.5 | | | | | | |
| P50516 | ATP6V1A | ATPase, H+ transporting, lysosoma | Plasma Membrane | transporter | 2289328.75 | 2401698 | 2411103.25 | | | | | | |
| P50516 | ATP6V1A | ATPase, H+ transporting, lysosoma | Plasma Membrane | transporter | 2734840.25 | 2569070.5 | 3334423.25 | | | | | | |
| P50516 | ATP6V1A | ATPase, H+ transporting, lysosoma | Plasma Membrane | transporter | 1914899.875 | 1851381 | 2162322 | | | | | | |
| P50516 | ATP6V1A | ATPase, H+ transporting, lysosoma | Plasma Membrane | transporter | 1067521.75 | 1307942.125 | 726095.312 | P50516 | ATP6V1A | 2821143 | 1577900 | 407412 | 15 |
| P62814 | ATP6V1B2 | ATPase, H+ transporting, lysosoma | Cytoplasm | transporter | 7114942 | 6045933.5 | 8374071 | | | | | | |
| P62814 | ATP6V1B2 | ATPase, H+ transporting, lysosoma | Cytoplasm | transporter | 1962865.75 | 2353262.25 | 2613088.75 | | | | | | |
| P62814 | ATP6V1B2 | ATPase, H+ transporting, lysosoma | Cytoplasm | transporter | 2096123.5 | 1684367.375 | 1650488.25 | P62814 | ATP6V1B2 | 3724644 | 2641525 | 880508 | 9 |
| P50518 | ATP6V1E1 | ATPase, H+ transporting, lysosoma | Cytoplasm | transporter | 1838479.375 | 2102869.25 | 2561311.25 | P50518 | ATP6V1E1 | 1838479 | 365731 | 211155 | 3 |
| Q8BVE3 | ATP6V1H | ATPase, H+ transporting, lysosoma | Cytoplasm | transporter | 1302340.75 | 1078923.875 | 1228138 | | | | | | |
| Q8BVE3 | ATP6V1H | ATPase, H+ transporting, lysosoma | Cytoplasm | transporter | 432603.969 | 234117.359 | 170589.453 | Q8BVE3 | ATP6V1H | 867472 | 518460 | 211661 | 6 |
| Q64446 | ATP7B | ATPase, Cu++ transporting, beta p | Cytoplasm | transporter | 1.47E+07 | 1.85E+07 | 1.36E+07 | | | | | | |
| Q64446 | ATP7B | ATPase, Cu++ transporting, beta p | Cytoplasm | transporter | 2455054 | 3472440.75 | 2567442.5 | | | | | | |
| Q64446 | ATP7B | ATPase, Cu++ transporting, beta p | Cytoplasm | transporter | 2807274.5 | 3214889.75 | 2553506.75 | | | | | | |
| Q64446 | ATP7B | ATPase, Cu++ transporting, beta p | Cytoplasm | transporter | 2769622.75 | 3224437.5 | 3390380 | | | | | | |
| Q64446 | ATP7B | ATPase, Cu++ transporting, beta p | Cytoplasm | transporter | 2304769.5 | 2775208 | 2235583.25 | | | | | | |
| Q64446 | ATP7B | ATPase, Cu++ transporting, beta p | Cytoplasm | transporter | 4755615 | 1186022.25 | 1921958.25 | | | | | | |
| Q64446 | ATP7B | ATPase, Cu++ transporting, beta p | Cytoplasm | transporter | 1820784.75 | 2642831 | 2020396 | | | | | | |
| Q64446 | ATP7B | ATPase, Cu++ transporting, beta p | Cytoplasm | transporter | 1600526.5 | 1695494 | 1354865.125 | Q64446 | ATP7B | 4149206 | 4546370 | 928024 | 24 |
| P70704 | ATP8A1 | ATPase, aminophospholipid transp | Cytoplasm | transporter | 2.20E+07 | 2.02E+07 | 2.42E+07 | | | | | | |
| P70704 | ATP8A1 | ATPase, aminophospholipid transp | Cytoplasm | transporter | 1.45E+07 | 1.71E+07 | 1.64E+07 | | | | | | |
| P70704 | ATP8A1 | ATPase, aminophospholipid transp | Cytoplasm | transporter | 1.67E+07 | 2.07E+07 | 2.16E+07 | | | | | | |
| P70704 | ATP8A1 | ATPase, aminophospholipid transp | Cytoplasm | transporter | 1.23E+07 | 1.20E+07 | 1.17E+07 | | | | | | |
| P70704 | ATP8A1 | ATPase, aminophospholipid transp | Cytoplasm | transporter | 1.18E+07 | 1.27E+07 | 8973433 | | | | | | |
| P70704 | ATP8A1 | ATPase, aminophospholipid transp | Cytoplasm | transporter | 1.12E+07 | 1.15E+07 | 9632686 | | | | | | |
| P70704 | ATP8A1 | ATPase, aminophospholipid transp | Cytoplasm | transporter | 7676151 | 1.11E+07 | 1.04E+07 | | | | | | |
| P70704 | ATP8A1 | ATPase, aminophospholipid transp | Cytoplasm | transporter | 9959953 | 9314191 | 9806149 | | | | | | |
| P70704 | ATP8A1 | ATPase, aminophospholipid transp | Cytoplasm | transporter | 7472743.5 | 7236726 | 8231199.5 | | | | | | |
| P70704 | ATP8A1 | ATPase, aminophospholipid transp | Cytoplasm | transporter | 6859051 | 7881107 | 6561732 | | | | | | |
| P70704 | ATP8A1 | ATPase, aminophospholipid transp | Cytoplasm | transporter | 5540502 | 7395430.5 | 7864826.5 | | | | | | |
| P70704 | ATP8A1 | ATPase, aminophospholipid transp | Cytoplasm | transporter | 5653532 | 4036601.5 | 4587129.5 | | | | | | |

| | | | | | | | | | | | | | |
|---|---|---|---|---|---|---|---|---|---|---|---|---|---|
| P70704 | ATP8A1 | ATPase, aminophospholipid transp | Cytoplasm | transporter | 7563182.5 | 7425358 | 8473061 | | | | | | |
| P70704 | ATP8A1 | ATPase, aminophospholipid transp | Cytoplasm | transporter | 5980679 | 6054506 | 5788164.5 | | | | | | |
| P70704 | ATP8A1 | ATPase, aminophospholipid transp | Cytoplasm | transporter | 5520417 | 7262255 | 9201313 | | | | | | |
| P70704 | ATP8A1 | ATPase, aminophospholipid transp | Cytoplasm | transporter | 4448629 | 3737312.25 | 3478797 | | | | | | |
| P70704 | ATP8A1 | ATPase, aminophospholipid transp | Cytoplasm | transporter | 4795697 | 4529683.5 | 4587035.5 | | | | | | |
| P70704 | ATP8A1 | ATPase, aminophospholipid transp | Cytoplasm | transporter | 4346222 | 4903380.5 | 4957017 | | | | | | |
| P70704 | ATP8A1 | ATPase, aminophospholipid transp | Cytoplasm | transporter | 3010777.5 | 5319368.5 | 4877033 | | | | | | |
| P70704 | ATP8A1 | ATPase, aminophospholipid transp | Cytoplasm | transporter | 3401567 | 4849401 | 5132120.5 | | | | | | |
| P70704 | ATP8A1 | ATPase, aminophospholipid transp | Cytoplasm | transporter | 2459546 | 3292288.5 | 2617806 | | | | | | |
| P70704 | ATP8A1 | ATPase, aminophospholipid transp | Cytoplasm | transporter | 1536271 | 5203407 | 2884602.25 | P70704 | ATP8A1 | 7938405 | 5209400 | 641233 | 66 |
| O35143 | ATPIF1 | ATPase inhibitory factor 1 | Cytoplasm | other | 6282756.5 | 9186363 | 1.43E+07 | O35143 | ATPIF1 | 6282757 | 4075252 | 2352848 | 3 |
| P70295 | AUP1 | ancient ubiquitous protein 1 | Cytoplasm | other | 3786944 | 4277891 | 3536136.5 | P70295 | AUP1 | 3786944 | 377300 | 217834 | 3 |
| P01887 | B2M | beta-2-microglobulin | Plasma Membrane | transmembrane receptor | 9011798 | 8544231 | 1.06E+07 | P01887 | B2M | 9011798 | 1077589 | 622146 | 3 |
| Q09199 | B4GALNT2 | beta-1,4-N-acetyl-galactosaminyl t | Cytoplasm | enzyme | 2.19E+07 | 1.29E+07 | 2.05E+07 | Q09199 | B4GALNT2 | 21870000 | 4809286 | 2776643 | 3 |
| Q9JJ04 | B4GALT4 | UDP-Gal:betaGlcNAc beta 1,4- gal | Cytoplasm | enzyme | 1.07E+08 | 2.91E+08 | 2.03E+08 | Q9JJ04 | B4GALT4 | 107300000 | 91678369 | 52930531 | 3 |
| Q91YN9 | BAG2 | BCL2-associated athanogene 2 | Cytoplasm | other | 4.42E+07 | 3.44E+07 | 3.15E+07 | | | | | | |
| Q91YN9 | BAG2 | BCL2-associated athanogene 2 | Cytoplasm | other | 2.50E+07 | 3.14E+07 | 2.21E+07 | | | | | | |
| Q91YN9 | BAG2 | BCL2-associated athanogene 2 | Cytoplasm | other | 1.62E+07 | 1.48E+07 | 1.61E+07 | | | | | | |
| Q91YN9 | BAG2 | BCL2-associated athanogene 2 | Cytoplasm | other | 1.00E+07 | 1.31E+07 | 1.09E+07 | | | | | | |
| Q91YN9 | BAG2 | BCL2-associated athanogene 2 | Cytoplasm | other | 1.47E+07 | 5380583.5 | 9018829 | | | | | | |
| Q91YN9 | BAG2 | BCL2-associated athanogene 2 | Cytoplasm | other | 8122707 | 1.12E+07 | 1.12E+07 | | | | | | |
| Q91YN9 | BAG2 | BCL2-associated athanogene 2 | Cytoplasm | other | 8957734 | 1.27E+07 | 9609859 | | | | | | |
| Q91YN9 | BAG2 | BCL2-associated athanogene 2 | Cytoplasm | other | 8210220 | 6100741.5 | 6160680 | | | | | | |
| Q91YN9 | BAG2 | BCL2-associated athanogene 2 | Cytoplasm | other | 7755321 | 1783173.875 | 5165686 | | | | | | |
| Q91YN9 | BAG2 | BCL2-associated athanogene 2 | Cytoplasm | other | 1.03E+07 | 2105431.25 | 5606383.5 | | | | | | |
| Q91YN9 | BAG2 | BCL2-associated athanogene 2 | Cytoplasm | other | 1137820.125 | 782320.062 | 770849.625 | Q91YN9 | BAG2 | 14048527 | 10390581 | 1808768 | 33 |
| O08734 | BAK1 | BCL2-antagonist/killer 1 | Cytoplasm | other | 3099165.75 | 4017658.5 | 4868607 | O08734 | BAK1 | 3099166 | 884935 | 510918 | 3 |
| O54962 | BANF1 | barrier to autointegration factor 1 | Nucleus | other | 1.20E+08 | 8.74E+07 | 1.13E+08 | | | | | | |
| O54962 | BANF1 | barrier to autointegration factor 1 | Nucleus | other | 6.17E+07 | 5.26E+07 | 6.26E+07 | | | | | | |
| O54962 | BANF1 | barrier to autointegration factor 1 | Nucleus | other | 2.66E+07 | 1.83E+07 | 2.59E+07 | | | | | | |
| O54962 | BANF1 | barrier to autointegration factor 1 | Nucleus | other | 1.39E+07 | 8149813 | 1.54E+07 | O54962 | BANF1 | 55455000 | 39174649 | 11308747 | 12 |
| Q61335 | BCAP31 | B-cell receptor-associated protein | Cytoplasm | transporter | 4149083.5 | 3978022 | 4115344.75 | Q61335 | BCAP31 | 4149084 | 90607 | 52312 | 3 |
| O35855 | BCAT2 | branched chain amino-acid transa | Cytoplasm | enzyme | 2893576 | 3000787.5 | 2465533 | O35855 | BCAT2 | 2893576 | 283200 | 163505 | 3 |
| P59017 | BCL2L13 | BCL2-like 13 (apoptosis facilitator) | Cytoplasm | other | 4286657.5 | 5505868.5 | 7533824.5 | | | | | | |
| P59017 | BCL2L13 | BCL2-like 13 (apoptosis facilitator) | Cytoplasm | other | 3460822.5 | 3077700 | 3939872.5 | | | | | | |
| P59017 | BCL2L13 | BCL2-like 13 (apoptosis facilitator) | Cytoplasm | other | 3778713.75 | 3207955.25 | 4603293.5 | P59017 | BCL2L13 | 3842065 | 1402788 | 467596 | 9 |
| Q8C6A8 | BHLHE22 | basic helix-loop-helix family, mem | Nucleus | transcription regulator | 1.95E+08 | 1.68E+08 | 1.39E+08 | Q8C6A8 | BHLHE22 | 194700000 | 27660983 | 15970076 | 3 |
| Q8BR07 | BICD1 | bicaudal D homolog 1 (Drosophila) | Cytoplasm | other | 4.16E+07 | 3.13E+07 | 3.37E+07 | Q8BR07 | BICD1 | 41550000 | 5353731 | 3090978 | 3 |
| Q8R016 | BLMH | bleomycin hydrolase | Cytoplasm | peptidase | 7158236.5 | 7067517.5 | 7753486 | | | | | | |
| Q8R016 | BLMH | bleomycin hydrolase | Cytoplasm | peptidase | 9629925 | 4574251 | 5202150.5 | | | | | | |
| Q8R016 | BLMH | bleomycin hydrolase | Cytoplasm | peptidase | 5394507.5 | 5854815.5 | 6117868 | | | | | | |
| Q8R016 | BLMH | bleomycin hydrolase | Cytoplasm | peptidase | 3470053 | 3471090 | 3855496.25 | | | | | | |
| Q8R016 | BLMH | bleomycin hydrolase | Cytoplasm | peptidase | 3096904.75 | 3125498.5 | 2466127.5 | | | | | | |
| Q8R016 | BLMH | bleomycin hydrolase | Cytoplasm | peptidase | 2570468 | 2623985.75 | 3244854 | | | | | | |
| Q8R016 | BLMH | bleomycin hydrolase | Cytoplasm | peptidase | 1927727.875 | 1651694.75 | 1765262.875 | Q8R016 | BLMH | 4749689 | 2208783 | 481996 | 21 |
| Q923D2 | BLVRB | biliverdin reductase B | Cytoplasm | enzyme | 1.42E+07 | 4815083.5 | 7427243 | | | | | | |
| Q923D2 | BLVRB | biliverdin reductase B | Cytoplasm | enzyme | 9806139 | 6123959.5 | 7781940 | | | | | | |
| Q923D2 | BLVRB | biliverdin reductase B | Cytoplasm | enzyme | 2711808.25 | 3474337 | 3965228.75 | | | | | | |
| Q923D2 | BLVRB | biliverdin reductase B | Cytoplasm | enzyme | 1231675.125 | 849611.625 | 1210413.25 | Q923D2 | BLVRB | 6982406 | 3996069 | 1153566 | 12 |
| P15327 | BPGM | 2,3-bisphosphoglycerate mutase | Extracellular Space | phosphatase | 1.48E+07 | 1.24E+07 | 1.84E+07 | | | | | | |
| P15327 | BPGM | 2,3-bisphosphoglycerate mutase | Extracellular Space | phosphatase | 5567254.5 | 4035476.5 | 5701623 | | | | | | |
| P15327 | BPGM | 2,3-bisphosphoglycerate mutase | Extracellular Space | phosphatase | 2416245 | 2263811.5 | 2548512 | | | | | | |
| P15327 | BPGM | 2,3-bisphosphoglycerate mutase | Extracellular Space | phosphatase | 539011.5 | 566956.25 | 846123.5 | P15327 | BPGM | 5833128 | 6036078 | 1742466 | 12 |
| A2AHJ4 | BRWD3 | bromodomain and WD repeat dom | Nucleus | other | 8050664.5 | 1.21E+07 | 1.05E+07 | A2AHJ4 | BRWD3 | 8050665 | 2038773 | 1177086 | 3 |
| P18572 | BSG | basigin (Ok blood group) | Plasma Membrane | transporter | 6.97E+08 | 8.72E+08 | 7.52E+08 | | | | | | |
| P18572 | BSG | basigin (Ok blood group) | Plasma Membrane | transporter | 6.90E+08 | 7.30E+08 | 6.48E+08 | | | | | | |
| P18572 | BSG | basigin (Ok blood group) | Plasma Membrane | transporter | 7.81E+08 | 7.48E+08 | 9.78E+08 | | | | | | |
| P18572 | BSG | basigin (Ok blood group) | Plasma Membrane | transporter | 5.98E+08 | 5.95E+08 | 5.81E+08 | | | | | | |
| P18572 | BSG | basigin (Ok blood group) | Plasma Membrane | transporter | 2.35E+08 | 2.62E+08 | 2.45E+08 | | | | | | |
| P18572 | BSG | basigin (Ok blood group) | Plasma Membrane | transporter | 1.89E+08 | 2.12E+08 | 1.75E+08 | | | | | | |
| P18572 | BSG | basigin (Ok blood group) | Plasma Membrane | transporter | 1.33E+08 | 4.06E+08 | 2.32E+08 | | | | | | |
| P18572 | BSG | basigin (Ok blood group) | Plasma Membrane | transporter | 3.50E+07 | 1.24E+08 | 7.36E+07 | | | | | | |
| P18572 | BSG | basigin (Ok blood group) | Plasma Membrane | transporter | 3.28E+07 | 7.54E+07 | 4.25E+07 | | | | | | |
| P18572 | BSG | basigin (Ok blood group) | Plasma Membrane | transporter | 4.42E+07 | 5.50E+07 | 5.37E+07 | | | | | | |
| P18572 | BSG | basigin (Ok blood group) | Plasma Membrane | transporter | 9339973 | 1.91E+07 | 1.05E+07 | | | | | | |
| P18572 | BSG | basigin (Ok blood group) | Plasma Membrane | transporter | 2113046.25 | 1.44E+07 | 9593504 | | | | | | |
| P18572 | BSG | basigin (Ok blood group) | Plasma Membrane | transporter | 1231934.75 | 1432357 | 1345001.625 | | | | | | |
| P18572 | BSG | basigin (Ok blood group) | Plasma Membrane | transporter | 1791299.5 | 5270113 | 7544049 | P18572 | BSG | 246382590 | 308551160 | 47610478 | 42 |
| Q9D659 | C10orf54 | chromosome 10 open reading frar | Plasma Membrane | other | 9940584 | 9651146 | 5081071 | Q9D659 | C10orf54 | 9940584 | 2725932 | 1573817 | 3 |
| Q91W34 | C16orf58 | chromosome 16 open reading frar | Cytoplasm | other | 764087.375 | 1912164.5 | 960904.938 | Q91W34 | C16orf58 | 764087 | 613964 | 354472 | 3 |
| Q9CX63 | C17orf99 | chromosome 17 open reading frar | Extracellular Space | cytokine | 5.39E+07 | 6.71E+07 | 4.78E+07 | | | | | | |
| Q9CX63 | C17orf99 | chromosome 17 open reading frar | Extracellular Space | cytokine | 3.80E+07 | 5.16E+07 | 3.25E+07 | | | | | | |
| Q9CX63 | C17orf99 | chromosome 17 open reading frar | Extracellular Space | cytokine | 3.94E+07 | 3.58E+07 | 2.64E+07 | | | | | | |
| Q9CX63 | C17orf99 | chromosome 17 open reading frar | Extracellular Space | cytokine | 2.73E+07 | 3.47E+07 | 2.63E+07 | | | | | | |
| Q9CX63 | C17orf99 | chromosome 17 open reading frar | Extracellular Space | cytokine | 1.95E+07 | 2.07E+07 | 1.56E+07 | | | | | | |
| Q9CX63 | C17orf99 | chromosome 17 open reading frar | Extracellular Space | cytokine | 1.32E+07 | 1.24E+07 | 8526329 | | | | | | |
| Q9CX63 | C17orf99 | chromosome 17 open reading frar | Extracellular Space | cytokine | 1.20E+07 | 1.23E+07 | 8408848 | Q9CX63 | C17orf99 | 29054286 | 16590778 | 3620405 | 21 |
| P98086 | C1QA | complement component 1, q subc | Extracellular Space | other | 414262.5 | 1081726.75 | 1409944 | | | | | | |
| P98086 | C1QA | complement component 1, q subc | Extracellular Space | other | 54010.32 | 150453.281 | 683602.438 | P98086 | C1QA | 234136 | 533570 | 217829 | 6 |
| P14106 | C1QB | complement component 1, q subc | Extracellular Space | other | 323430.312 | 773713.5 | 1417271.25 | P14106 | C1QB | 323430 | 549759 | 317403 | 3 |
| O35658 | C1QBP | complement component 1, q subc | Cytoplasm | transcription regulator | 2.46E+07 | 2.29E+07 | 3.76E+07 | | | | | | |
| O35658 | C1QBP | complement component 1, q subc | Cytoplasm | transcription regulator | 905106 | 1.09E+07 | 3.71E+07 | O35658 | C1QBP | 12762553 | 14443517 | 5896541 | 6 |
| Q9D172 | C21orf33/LOC102724023 | chromosome 21 open reading frar | Cytoplasm | other | 7521694 | 7183143 | 9776968 | | | | | | |
| Q9D172 | C21orf33/LOC102724023 | chromosome 21 open reading frar | Cytoplasm | other | 113251.297 | 438667.438 | 620786.75 | Q9D172 | C21orf33/LOC102724023 | 3817473 | 4351111 | 1776334 | 6 |
| Q80X80 | C2CD2L | C2CD2-like | Cytoplasm | other | 2.11E+07 | 1.74E+07 | 1.87E+07 | | | | | | |
| Q80X80 | C2CD2L | C2CD2-like | Cytoplasm | other | 2.27E+07 | 2.76E+07 | 2.41E+07 | | | | | | |

| | | | | | | | | | | | | | |
|---|---|---|---|---|---|---|---|---|---|---|---|---|---|
| Q80X80 | C2CD2L | C2CD2-like | Cytoplasm | other | 1.73E+07 | 1.73E+07 | 1.52E+07 | | | | | | |
| Q80X80 | C2CD2L | C2CD2-like | Cytoplasm | other | 7133998.5 | 6235041 | 6665984.5 | | | | | | |
| Q80X80 | C2CD2L | C2CD2-like | Cytoplasm | other | 6718467.5 | 6562658.5 | 8823049 | | | | | | |
| Q80X80 | C2CD2L | C2CD2-like | Cytoplasm | other | 7872356 | 1.06E+07 | 6493273.5 | | | | | | |
| Q80X80 | C2CD2L | C2CD2-like | Cytoplasm | other | 7989754 | 1.05E+07 | 1.29E+07 | | | | | | |
| Q80X80 | C2CD2L | C2CD2-like | Cytoplasm | other | 8557295 | 1.03E+07 | 7814922 | | | | | | |
| Q80X80 | C2CD2L | C2CD2-like | Cytoplasm | other | 7622689.5 | 6215276.5 | 6904527 | | | | | | |
| Q80X80 | C2CD2L | C2CD2-like | Cytoplasm | other | 7772543 | 7175337.5 | 5760549 | | | | | | |
| Q80X80 | C2CD2L | C2CD2-like | Cytoplasm | other | 3541576.5 | 3672887.25 | 3578040.5 | | | | | | |
| Q80X80 | C2CD2L | C2CD2-like | Cytoplasm | other | 1975102.75 | 1442657.5 | 1777906.625 | | | | | | |
| Q80X80 | C2CD2L | C2CD2-like | Cytoplasm | other | 988847.125 | 447967.75 | 818118.062 | Q80X80 | C2CD2L | 9328664 | 6846689 | 1096348 | 39 |
| P01029 | C4A/C4B | complement component 4B (Chido | Extracellular Space | other | 2068107.25 | 4490996 | 1.58E+07 | | | | | | |
| P01029 | C4A/C4B | complement component 4B (Chido | Extracellular Space | other | 2935743 | 3685761.75 | 1.45E+07 | | | | | | |
| P01029 | C4A/C4B | complement component 4B (Chido | Extracellular Space | other | 2214794.75 | 1638262.125 | 8070655 | | | | | | |
| P01029 | C4A/C4B | complement component 4B (Chido | Extracellular Space | other | 1029642.125 | 871172.625 | 5751907 | | | | | | |
| P01029 | C4A/C4B | complement component 4B (Chido | Extracellular Space | other | 1230660.375 | 701028.438 | 5716454 | P01029 | C4A/C4B | 1895790 | 4745470 | 1225275 | 15 |
| P00920 | CA2 | carbonic anhydrase II | Cytoplasm | enzyme | 3.49E+07 | 3.53E+07 | 3.86E+07 | | | | | | |
| P00920 | CA2 | carbonic anhydrase II | Cytoplasm | enzyme | 1.19E+07 | 1.06E+07 | 1.50E+07 | | | | | | |
| P00920 | CA2 | carbonic anhydrase II | Cytoplasm | enzyme | 8528692 | 6545714.5 | 8902086 | | | | | | |
| P00920 | CA2 | carbonic anhydrase II | Cytoplasm | enzyme | 3566809.75 | 7509178.5 | 1.26E+07 | | | | | | |
| P00920 | CA2 | carbonic anhydrase II | Cytoplasm | enzyme | 7955085.5 | 6826499 | 1.14E+07 | | | | | | |
| P00920 | CA2 | carbonic anhydrase II | Cytoplasm | enzyme | 260157.984 | 193091.328 | 359953.844 | P00920 | CA2 | 11185124 | 11838852 | 2790444 | 18 |
| P62204 | CALM1 | Calmodulin | Cytoplasm | | 2.50E+07 | 3.67E+07 | 3.48E+07 | | | | | | |
| P62204 | CALM1 | Calmodulin | Cytoplasm | | 1.22E+07 | 1.13E+07 | 1.02E+07 | P62204 | CALM1 | 18610000 | 12149192 | 4959887 | 6 |
| Q9D6P8 | CALML3 | calmodulin-like 3 | Cytoplasm | other | 6079201 | 8225202.5 | 6917352 | Q9D6P8 | CALML3 | 6079201 | 1081534 | 624424 | 3 |
| P14211 | CALR | calreticulin | Cytoplasm | transcription regulator | 1.62E+08 | 2.70E+08 | 2.03E+08 | | | | | | |
| P14211 | CALR | calreticulin | Cytoplasm | transcription regulator | 1.08E+08 | 1.78E+08 | 1.24E+08 | | | | | | |
| P14211 | CALR | calreticulin | Cytoplasm | transcription regulator | 1.13E+08 | 2.44E+08 | 2.09E+08 | | | | | | |
| P14211 | CALR | calreticulin | Cytoplasm | transcription regulator | 1.00E+08 | 1.06E+08 | 8.59E+07 | | | | | | |
| P14211 | CALR | calreticulin | Cytoplasm | transcription regulator | 9.15E+07 | 1.32E+08 | 1.13E+08 | | | | | | |
| P14211 | CALR | calreticulin | Cytoplasm | transcription regulator | 6.88E+07 | 1.12E+08 | 6.44E+07 | | | | | | |
| P14211 | CALR | calreticulin | Cytoplasm | transcription regulator | 8.39E+07 | 9.68E+07 | 8.23E+07 | | | | | | |
| P14211 | CALR | calreticulin | Cytoplasm | transcription regulator | 6.46E+07 | 1.35E+08 | 1.25E+08 | | | | | | |
| P14211 | CALR | calreticulin | Cytoplasm | transcription regulator | 4.85E+07 | 5.72E+07 | 4.13E+07 | | | | | | |
| P14211 | CALR | calreticulin | Cytoplasm | transcription regulator | 3.28E+07 | 3.53E+07 | 2.54E+07 | | | | | | |
| P14211 | CALR | calreticulin | Cytoplasm | transcription regulator | 3.92E+07 | 3.68E+07 | 2.68E+07 | | | | | | |
| P14211 | CALR | calreticulin | Cytoplasm | transcription regulator | 1928736.375 | 4391615 | 3192128.75 | | | | | | |
| P14211 | CALR | calreticulin | Cytoplasm | transcription regulator | 5202158 | 9787947 | 6502502 | | | | | | |
| P14211 | CALR | calreticulin | Cytoplasm | transcription regulator | 5792803 | 2.55E+07 | 1.90E+07 | | | | | | |
| P14211 | CALR | calreticulin | Cytoplasm | transcription regulator | 3655941.25 | 6459472.5 | 4261329 | | | | | | |
| P14211 | CALR | calreticulin | Cytoplasm | transcription regulator | 1538676.125 | 1337567.75 | 1458091 | | | | | | |
| P14211 | CALR | calreticulin | Cytoplasm | transcription regulator | 780762.438 | 5319424.5 | 2700517.5 | | | | | | |
| P14211 | CALR | calreticulin | Cytoplasm | transcription regulator | 607189.75 | 374931 | 214599.953 | | | | | | |
| P14211 | CALR | calreticulin | Cytoplasm | transcription regulator | 229343.156 | 8436856 | 5238800 | P14211 | CALR | 49016085 | 68109890 | 9021375 | 57 |
| O35887 | CALU | calumenin | Cytoplasm | other | 8378694 | 1.07E+07 | 9021656 | | | | | | |
| O35887 | CALU | calumenin | Cytoplasm | other | 7071116 | 9528862 | 8196176 | | | | | | |
| O35887 | CALU | calumenin | Cytoplasm | other | 7607828 | 1.11E+07 | 5983144.5 | | | | | | |
| O35887 | CALU | calumenin | Cytoplasm | other | 5466705 | 7364994.5 | 9575208 | | | | | | |
| O35887 | CALU | calumenin | Cytoplasm | other | 3456690.75 | 4238726.5 | 1391900.5 | | | | | | |
| O35887 | CALU | calumenin | Cytoplasm | other | 3063193.75 | 5589861.5 | 2114292.75 | | | | | | |
| O35887 | CALU | calumenin | Cytoplasm | other | 2497752.5 | 4011568 | 3514200.75 | | | | | | |
| O35887 | CALU | calumenin | Cytoplasm | other | 92156.711 | 1315763.75 | 649373.125 | O35887 | CALU | 4704267 | 3327669 | 679258 | 24 |
| P51437 | CAMP | cathelicidin antimicrobial peptide | Cytoplasm | other | 1764320.375 | 883163.188 | 3658241.5 | | | | | | |
| P51437 | CAMP | cathelicidin antimicrobial peptide | Cytoplasm | other | 2135753.5 | 1407295.75 | 5223625 | P51437 | CAMP | 1950037 | 1627320 | 664350 | 6 |
| Q6ZQ38 | CAND1 | cullin-associated and neddylation- | Cytoplasm | transcription regulator | 1.78E+07 | 1.82E+07 | 1.89E+07 | | | | | | |
| Q6ZQ38 | CAND1 | cullin-associated and neddylation- | Cytoplasm | transcription regulator | 1.30E+07 | 9877410 | 1.38E+07 | | | | | | |
| Q6ZQ38 | CAND1 | cullin-associated and neddylation- | Cytoplasm | transcription regulator | 1.53E+07 | 9421603 | 1.13E+07 | | | | | | |
| Q6ZQ38 | CAND1 | cullin-associated and neddylation- | Cytoplasm | transcription regulator | 7980330.5 | 7951508 | 9743936 | | | | | | |
| Q6ZQ38 | CAND1 | cullin-associated and neddylation- | Cytoplasm | transcription regulator | 9841480 | 8989481 | 1.05E+07 | | | | | | |
| Q6ZQ38 | CAND1 | cullin-associated and neddylation- | Cytoplasm | transcription regulator | 1.34E+07 | 1.09E+07 | 1.14E+07 | | | | | | |
| Q6ZQ38 | CAND1 | cullin-associated and neddylation- | Cytoplasm | transcription regulator | 7039834.5 | 7127256 | 6837221.5 | | | | | | |
| Q6ZQ38 | CAND1 | cullin-associated and neddylation- | Cytoplasm | transcription regulator | 8396214 | 8268490.5 | 1.05E+07 | | | | | | |
| Q6ZQ38 | CAND1 | cullin-associated and neddylation- | Cytoplasm | transcription regulator | 1.10E+07 | 7440001.5 | 7747204.5 | | | | | | |
| Q6ZQ38 | CAND1 | cullin-associated and neddylation- | Cytoplasm | transcription regulator | 8190098.5 | 7737118.5 | 7307659 | | | | | | |
| Q6ZQ38 | CAND1 | cullin-associated and neddylation- | Cytoplasm | transcription regulator | 9215377 | 7109340 | 8395257 | | | | | | |
| Q6ZQ38 | CAND1 | cullin-associated and neddylation- | Cytoplasm | transcription regulator | 7138887 | 5473394 | 6944162.5 | | | | | | |
| Q6ZQ38 | CAND1 | cullin-associated and neddylation- | Cytoplasm | transcription regulator | 5027928 | 5319442 | 5578964.5 | | | | | | |
| Q6ZQ38 | CAND1 | cullin-associated and neddylation- | Cytoplasm | transcription regulator | 6221500.5 | 5324889 | 7501191 | | | | | | |
| Q6ZQ38 | CAND1 | cullin-associated and neddylation- | Cytoplasm | transcription regulator | 3910262.5 | 2276032.5 | 3144645 | | | | | | |
| Q6ZQ38 | CAND1 | cullin-associated and neddylation- | Cytoplasm | transcription regulator | 5502756.5 | 4911627.5 | 5521431 | | | | | | |
| Q6ZQ38 | CAND1 | cullin-associated and neddylation- | Cytoplasm | transcription regulator | 4581059.5 | 4340464 | 4516281 | | | | | | |
| Q6ZQ38 | CAND1 | cullin-associated and neddylation- | Cytoplasm | transcription regulator | 4009200.5 | 5132137.5 | 6657029.5 | | | | | | |
| Q6ZQ38 | CAND1 | cullin-associated and neddylation- | Cytoplasm | transcription regulator | 1455949.5 | 2635765 | 2834195.25 | Q6ZQ38 | CAND1 | 8366362 | 3844521 | 509219 | 57 |
| P35564 | CANX | calnexin | Cytoplasm | other | 5.67E+07 | 1.34E+08 | 1.15E+08 | | | | | | |
| P35564 | CANX | calnexin | Cytoplasm | other | 7.90E+07 | 7.74E+07 | 6.13E+07 | | | | | | |
| P35564 | CANX | calnexin | Cytoplasm | other | 6.95E+07 | 8.30E+07 | 6.78E+07 | | | | | | |
| P35564 | CANX | calnexin | Cytoplasm | other | 4.36E+07 | 8.08E+07 | 5.54E+07 | | | | | | |
| P35564 | CANX | calnexin | Cytoplasm | other | 2.27E+07 | 2.05E+07 | 2.23E+07 | | | | | | |
| P35564 | CANX | calnexin | Cytoplasm | other | 4.36E+07 | 5.92E+07 | 5.32E+07 | | | | | | |
| P35564 | CANX | calnexin | Cytoplasm | other | 2.02E+07 | 6.94E+07 | 4.82E+07 | | | | | | |
| P35564 | CANX | calnexin | Cytoplasm | other | 3.44E+07 | 4.70E+07 | 3.60E+07 | | | | | | |
| P35564 | CANX | calnexin | Cytoplasm | other | 3.04E+07 | 4.25E+07 | 3.14E+07 | | | | | | |
| P35564 | CANX | calnexin | Cytoplasm | other | 2.41E+07 | 4.05E+07 | 3.95E+07 | | | | | | |
| P35564 | CANX | calnexin | Cytoplasm | other | 2.86E+07 | 1.58E+07 | 1.34E+07 | | | | | | |
| P35564 | CANX | calnexin | Cytoplasm | other | 2.53E+07 | 1.01E+07 | 7681759.5 | | | | | | |
| P35564 | CANX | calnexin | Cytoplasm | other | 3.60E+07 | 4.22E+07 | 3.32E+07 | | | | | | |

| | | | | | | | | | | | | | |
|---|---|---|---|---|---|---|---|---|---|---|---|---|---|
| P35564 | CANX | calnexin | Cytoplasm | other | 1.34E+07 | 2.93E+07 | 2.51E+07 | | | | | | |
| P35564 | CANX | calnexin | Cytoplasm | other | 8060855.5 | 8864446 | 8394832 | | | | | | |
| P35564 | CANX | calnexin | Cytoplasm | other | 1965594.375 | 5538845 | 4088051.75 | | | | | | |
| P35564 | CANX | calnexin | Cytoplasm | other | 3254419.25 | 1.24E+07 | 1.03E+07 | | | | | | |
| P35564 | CANX | calnexin | Cytoplasm | other | 3152819.5 | 1.36E+07 | 2.44E+07 | | | | | | |
| P35564 | CANX | calnexin | Cytoplasm | other | 5002179 | 931850.25 | 2816453.25 | | | | | | |
| P35564 | CANX | calnexin | Cytoplasm | other | 1360316 | 4859794.5 | 3729060 | | | | | | |
| P35564 | CANX | calnexin | Cytoplasm | other | 776307.625 | 1587217.75 | 1043981.312 | | | | | | |
| P35564 | CANX | calnexin | Cytoplasm | other | 214015.312 | 519527.062 | 514474.5 | P35564 | CANX | 25052114 | 29159622 | 3589301 | 66 |
| P40124 | CAP1 | CAP, adenylate cyclase-associated | Plasma Membrane | other | 3355626.75 | 2598291.75 | 2612254.5 | | | | | | |
| P40124 | CAP1 | CAP, adenylate cyclase-associated | Plasma Membrane | other | 1761439.5 | 2372632.75 | 1676156.125 | | | | | | |
| P40124 | CAP1 | CAP, adenylate cyclase-associated | Plasma Membrane | other | 513324.844 | 1310194.75 | 1275632 | P40124 | CAP1 | 1876797 | 869724 | 289908 | 9 |
| O35350 | CAPN1 | calpain 1, (mu/l) large subunit | Cytoplasm | peptidase | 9806790 | 5970817.5 | 8546147 | | | | | | |
| O35350 | CAPN1 | calpain 1, (mu/l) large subunit | Cytoplasm | peptidase | 6680199 | 4395459.5 | 6016744.5 | | | | | | |
| O35350 | CAPN1 | calpain 1, (mu/l) large subunit | Cytoplasm | peptidase | 4183116.5 | 3609246 | 5860066.5 | | | | | | |
| O35350 | CAPN1 | calpain 1, (mu/l) large subunit | Cytoplasm | peptidase | 5358050 | 4380622.5 | 5851950.5 | | | | | | |
| O35350 | CAPN1 | calpain 1, (mu/l) large subunit | Cytoplasm | peptidase | 3861740.75 | 2071219.625 | 4501328.5 | | | | | | |
| O35350 | CAPN1 | calpain 1, (mu/l) large subunit | Cytoplasm | peptidase | 3000562.75 | 3378435.25 | 4550124.5 | | | | | | |
| O35350 | CAPN1 | calpain 1, (mu/l) large subunit | Cytoplasm | peptidase | 535893 | 456965.25 | 597747.062 | O35350 | CAPN1 | 4775193 | 2409862 | 525875 | 21 |
| O08688 | CAPN5 | calpain 5 | Cytoplasm | peptidase | 3.11E+07 | 4.10E+07 | 3.63E+07 | | | | | | |
| O08688 | CAPN5 | calpain 5 | Cytoplasm | peptidase | 2.80E+07 | 3.02E+07 | 3.37E+07 | | | | | | |
| O08688 | CAPN5 | calpain 5 | Cytoplasm | peptidase | 1.71E+07 | 1.54E+07 | 1.47E+07 | | | | | | |
| O08688 | CAPN5 | calpain 5 | Cytoplasm | peptidase | 1.46E+07 | 1.59E+07 | 1.65E+07 | | | | | | |
| O08688 | CAPN5 | calpain 5 | Cytoplasm | peptidase | 1.80E+07 | 1.44E+07 | 1.55E+07 | | | | | | |
| O08688 | CAPN5 | calpain 5 | Cytoplasm | peptidase | 1.26E+07 | 1.52E+07 | 1.55E+07 | | | | | | |
| O08688 | CAPN5 | calpain 5 | Cytoplasm | peptidase | 1.28E+07 | 1.60E+07 | 1.39E+07 | | | | | | |
| O08688 | CAPN5 | calpain 5 | Cytoplasm | peptidase | 1.05E+07 | 9570984 | 1.08E+07 | | | | | | |
| O08688 | CAPN5 | calpain 5 | Cytoplasm | peptidase | 7775650.5 | 8522807 | 9171315 | | | | | | |
| O08688 | CAPN5 | calpain 5 | Cytoplasm | peptidase | 9314973 | 8123947 | 8573822 | | | | | | |
| O08688 | CAPN5 | calpain 5 | Cytoplasm | peptidase | 6837319.5 | 6540105.5 | 6221750 | | | | | | |
| O08688 | CAPN5 | calpain 5 | Cytoplasm | peptidase | 4889176.5 | 6440570 | 5390064.5 | | | | | | |
| O08688 | CAPN5 | calpain 5 | Cytoplasm | peptidase | 6840275 | 5824675.5 | 6377709 | | | | | | |
| O08688 | CAPN5 | calpain 5 | Cytoplasm | peptidase | 4888987.5 | 4692494.5 | 4660454 | | | | | | |
| O08688 | CAPN5 | calpain 5 | Cytoplasm | peptidase | 3429664.75 | 2184857.5 | 2537567 | | | | | | |
| O08688 | CAPN5 | calpain 5 | Cytoplasm | peptidase | 1652820.75 | 2036098.5 | 1301944.375 | | | | | | |
| O08688 | CAPN5 | calpain 5 | Cytoplasm | peptidase | 2153666 | 4024781.75 | 4610935 | | | | | | |
| O08688 | CAPN5 | calpain 5 | Cytoplasm | peptidase | 416094.156 | 2227791.75 | 1370073.625 | | | | | | |
| O08688 | CAPN5 | calpain 5 | Cytoplasm | peptidase | 519628.5 | 2447027.25 | 3232207 | | | | | | |
| O08688 | CAPN5 | calpain 5 | Cytoplasm | peptidase | 384621.188 | 2854870.25 | 4088805 | O08688 | CAPN5 | 9691644 | 9341172 | 1205940 | 60 |
| O88456 | CAPNS1 | calpain, small subunit 1 | Cytoplasm | peptidase | 1777132.75 | 1143574.25 | 2055161 | O88456 | CAPNS1 | 1777133 | 467206 | 269741 | 3 |
| Q60865 | CAPRIN1 | cell cycle associated protein 1 | Plasma Membrane | other | 1148261.625 | 1343581.5 | 1376109.625 | | | | | | |
| Q60865 | CAPRIN1 | cell cycle associated protein 1 | Plasma Membrane | other | 3038950 | 1383976 | 2456826.75 | Q60865 | CAPRIN1 | 2093606 | 768349 | 313677 | 6 |
| P47753 | CAPZA1 | capping protein (actin filament) m | Cytoplasm | other | 7843657.5 | 9206388 | 7564918 | | | | | | |
| P47753 | CAPZA1 | capping protein (actin filament) m | Cytoplasm | other | 3767375.25 | 4248523.5 | 3166131 | P47753 | CAPZA1 | 5805516 | 2537920 | 1036102 | 6 |
| P47757 | CAPZB | capping protein (actin filament) m | Cytoplasm | other | 4308259 | 4354845.5 | 3399221.75 | | | | | | |
| P47757 | CAPZB | capping protein (actin filament) m | Cytoplasm | other | 2282748.75 | 2584410.75 | 2450504 | P47757 | CAPZB | 3295504 | 935761 | 382023 | 6 |
| Q9CR86 | CARHSP1 | calcium regulated heat stable prot | Cytoplasm | other | 8830022 | 7378209.5 | 7740551.5 | Q9CR86 | CARHSP1 | 8830022 | 755645 | 436272 | 3 |
| Q9WVG6 | CARM1 | coactivator-associated arginine me | Nucleus | transcription regulator | 4418162 | 2008068 | 2135411.75 | | | | | | |
| Q9WVG6 | CARM1 | coactivator-associated arginine me | Nucleus | transcription regulator | 3658456 | 668573.562 | 1345217.625 | | | | | | |
| Q9WVG6 | CARM1 | coactivator-associated arginine me | Nucleus | transcription regulator | 4632235.5 | 2343795.5 | 4503292 | Q9WVG6 | CARM1 | 4236285 | 1479896 | 493299 | 9 |
| Q9ER72 | CARS | cysteinyl-tRNA synthetase | Cytoplasm | enzyme | 1445115.25 | 2434640.5 | 2501196.75 | | | | | | |
| Q9ER72 | CARS | cysteinyl-tRNA synthetase | Cytoplasm | enzyme | 1258937.5 | 1101505.25 | 733296.5 | Q9ER72 | CARS | 1352026 | 727482 | 296993 | 6 |
| P51125 | CAST | calpastatin | Cytoplasm | peptidase | 3.18E+07 | 3.03E+07 | 2.35E+07 | | | | | | |
| P51125 | CAST | calpastatin | Cytoplasm | peptidase | 1.46E+07 | 1.11E+07 | 1.01E+07 | | | | | | |
| P51125 | CAST | calpastatin | Cytoplasm | peptidase | 1.15E+07 | 1.48E+07 | 1.13E+07 | | | | | | |
| P51125 | CAST | calpastatin | Cytoplasm | peptidase | 1.15E+07 | 1.26E+07 | 9215184 | | | | | | |
| P51125 | CAST | calpastatin | Cytoplasm | peptidase | 9172172 | 1.08E+07 | 7554085.5 | | | | | | |
| P51125 | CAST | calpastatin | Cytoplasm | peptidase | 5566117.5 | 1.14E+07 | 8279526 | | | | | | |
| P51125 | CAST | calpastatin | Cytoplasm | peptidase | 1.04E+07 | 3465204.25 | 7001546.5 | | | | | | |
| P51125 | CAST | calpastatin | Cytoplasm | peptidase | 3505624.25 | 7986983.5 | 4899774.5 | | | | | | |
| P51125 | CAST | calpastatin | Cytoplasm | peptidase | 3226337.25 | 8502893 | 5638167.5 | | | | | | |
| P51125 | CAST | calpastatin | Cytoplasm | peptidase | 7244186.5 | 562422 | 776050.438 | | | | | | |
| P51125 | CAST | calpastatin | Cytoplasm | peptidase | 4263579 | 4031084.5 | 3988338 | | | | | | |
| P51125 | CAST | calpastatin | Cytoplasm | peptidase | 1546856.875 | 2959452.5 | 2073698.875 | P51125 | CAST | 9524573 | 7150040 | 1191673 | 36 |
| P24270 | CAT | catalase | Cytoplasm | enzyme | 2.44E+07 | 1.64E+07 | 2.30E+07 | | | | | | |
| P24270 | CAT | catalase | Cytoplasm | enzyme | 2.50E+07 | 1.16E+07 | 2.39E+07 | | | | | | |
| P24270 | CAT | catalase | Cytoplasm | enzyme | 1.37E+07 | 1.10E+07 | 1.67E+07 | | | | | | |
| P24270 | CAT | catalase | Cytoplasm | enzyme | 1.32E+07 | 9520575 | 1.37E+07 | | | | | | |
| P24270 | CAT | catalase | Cytoplasm | enzyme | 1.21E+07 | 6829779 | 1.07E+07 | | | | | | |
| P24270 | CAT | catalase | Cytoplasm | enzyme | 9181951 | 6478267 | 8965458 | | | | | | |
| P24270 | CAT | catalase | Cytoplasm | enzyme | 1.04E+07 | 5792007 | 9088193 | | | | | | |
| P24270 | CAT | catalase | Cytoplasm | enzyme | 7573613 | 6291791 | 7845004.5 | | | | | | |
| P24270 | CAT | catalase | Cytoplasm | enzyme | 6210919.5 | 4548320.5 | 6950891 | | | | | | |
| P24270 | CAT | catalase | Cytoplasm | enzyme | 9634036 | 5157022.5 | 7902037.5 | | | | | | |
| P24270 | CAT | catalase | Cytoplasm | enzyme | 5069222.5 | 3681823.5 | 5488828 | | | | | | |
| P24270 | CAT | catalase | Cytoplasm | enzyme | 4836082.5 | 4825076.5 | 6999913 | | | | | | |
| P24270 | CAT | catalase | Cytoplasm | enzyme | 5518463 | 4496661 | 7089074.5 | | | | | | |
| P24270 | CAT | catalase | Cytoplasm | enzyme | 5350754 | 3070151 | 5776845 | | | | | | |
| P24270 | CAT | catalase | Cytoplasm | enzyme | 3540696 | 2813788.5 | 5670537 | | | | | | |
| P24270 | CAT | catalase | Cytoplasm | enzyme | 4336709 | 2913940.75 | 3968897.75 | | | | | | |
| P24270 | CAT | catalase | Cytoplasm | enzyme | 2562346 | 2013734.125 | 2428483 | | | | | | |
| P24270 | CAT | catalase | Cytoplasm | enzyme | 2803245.25 | 1211161.625 | 2063375.875 | | | | | | |
| P24270 | CAT | catalase | Cytoplasm | enzyme | 840355.562 | 989822.938 | 1076096.5 | | | | | | |
| P24270 | CAT | catalase | Cytoplasm | enzyme | 326856.5 | 1947670.125 | 2189548 | P24270 | CAT | 8333262 | 5907631 | 762672 | 60 |
| Q8C9M2 | CCDC15 | coiled-coil domain containing 15 | Cytoplasm | other | 2375580.75 | 1831373.625 | 2592843 | Q8C9M2 | CCDC15 | 2375581 | 392258 | 226470 | 3 |
| Q3URS9 | CCDC51 | coiled-coil domain containing 51 | Cytoplasm | other | 800367.188 | 735686.438 | 1083628.375 | Q3URS9 | CCDC51 | 800367 | 185060 | 106845 | 3 |

| | | | | | | | | | | | | | |
|---|---|---|---|---|---|---|---|---|---|---|---|---|---|
| Q8BGU5 | CCNY | cyclin Y | Nucleus | other | 3204812 | 3119018.75 | 2655019.5 | | | | | | |
| Q8BGU5 | CCNY | cyclin Y | Nucleus | other | 2648500.75 | 3309736 | 2869259.25 | Q8BGU5 | CCNY | 2926656 | 284765 | 116255 | 6 |
| P80314 | CCT2 | chaperonin containing TCP1, subu | Cytoplasm | kinase | 1.51E+07 | 1.81E+07 | 1.28E+07 | | | | | | |
| P80314 | CCT2 | chaperonin containing TCP1, subu | Cytoplasm | kinase | 1.49E+07 | 1.99E+07 | 1.26E+07 | | | | | | |
| P80314 | CCT2 | chaperonin containing TCP1, subu | Cytoplasm | kinase | 8885532 | 1.28E+07 | 6091377 | | | | | | |
| P80314 | CCT2 | chaperonin containing TCP1, subu | Cytoplasm | kinase | 9811178 | 1.29E+07 | 5447277 | | | | | | |
| P80314 | CCT2 | chaperonin containing TCP1, subu | Cytoplasm | kinase | 8066564 | 9413207 | 9921001 | | | | | | |
| P80314 | CCT2 | chaperonin containing TCP1, subu | Cytoplasm | kinase | 8771966 | 1.25E+07 | 7204162 | | | | | | |
| P80314 | CCT2 | chaperonin containing TCP1, subu | Cytoplasm | kinase | 6817653 | 8768632 | 4459496 | | | | | | |
| P80314 | CCT2 | chaperonin containing TCP1, subu | Cytoplasm | kinase | 7061620 | 9543545 | 5827197 | | | | | | |
| P80314 | CCT2 | chaperonin containing TCP1, subu | Cytoplasm | kinase | 4794070.5 | 5486566 | 3140729.75 | | | | | | |
| P80314 | CCT2 | chaperonin containing TCP1, subu | Cytoplasm | kinase | 5705136 | 7922165.5 | 4476917.5 | | | | | | |
| P80314 | CCT2 | chaperonin containing TCP1, subu | Cytoplasm | kinase | 5435532 | 7442078.5 | 5199498 | | | | | | |
| P80314 | CCT2 | chaperonin containing TCP1, subu | Cytoplasm | kinase | 4510545.5 | 6207741 | 2791102 | | | | | | |
| P80314 | CCT2 | chaperonin containing TCP1, subu | Cytoplasm | kinase | 2737757 | 6827720 | 3738354 | | | | | | |
| P80314 | CCT2 | chaperonin containing TCP1, subu | Cytoplasm | kinase | 2810440.5 | 3959218 | 2145904 | P80314 | CCT2 | 7525571 | 4239645 | 654191 | 42 |
| P80318 | CCT3 | chaperonin containing TCP1, subu | Cytoplasm | other | 2.14E+07 | 2.49E+07 | 1.57E+07 | | | | | | |
| P80318 | CCT3 | chaperonin containing TCP1, subu | Cytoplasm | other | 1.27E+07 | 1.64E+07 | 9134309 | | | | | | |
| P80318 | CCT3 | chaperonin containing TCP1, subu | Cytoplasm | other | 1.30E+07 | 1.79E+07 | 8473324 | | | | | | |
| P80318 | CCT3 | chaperonin containing TCP1, subu | Cytoplasm | other | 1.07E+07 | 1.85E+07 | 1.08E+07 | | | | | | |
| P80318 | CCT3 | chaperonin containing TCP1, subu | Cytoplasm | other | 8618817 | 1.46E+07 | 6519144.5 | | | | | | |
| P80318 | CCT3 | chaperonin containing TCP1, subu | Cytoplasm | other | 7228317.5 | 7391395.5 | 2576161 | | | | | | |
| P80318 | CCT3 | chaperonin containing TCP1, subu | Cytoplasm | other | 8611548 | 9327663 | 4927709.5 | | | | | | |
| P80318 | CCT3 | chaperonin containing TCP1, subu | Cytoplasm | other | 7066240 | 1.16E+07 | 6026949 | | | | | | |
| P80318 | CCT3 | chaperonin containing TCP1, subu | Cytoplasm | other | 5220381 | 7004723.5 | 3517850.75 | | | | | | |
| P80318 | CCT3 | chaperonin containing TCP1, subu | Cytoplasm | other | 4520304 | 1.08E+07 | 6206165.5 | | | | | | |
| P80318 | CCT3 | chaperonin containing TCP1, subu | Cytoplasm | other | 4679785 | 8319766 | 3304118.5 | | | | | | |
| P80318 | CCT3 | chaperonin containing TCP1, subu | Cytoplasm | other | 6028211.5 | 6026851.5 | 3363291 | | | | | | |
| P80318 | CCT3 | chaperonin containing TCP1, subu | Cytoplasm | other | 4991362 | 9128412 | 5008438 | | | | | | |
| P80318 | CCT3 | chaperonin containing TCP1, subu | Cytoplasm | other | 3879752.75 | 5236244 | 1180317 | | | | | | |
| P80318 | CCT3 | chaperonin containing TCP1, subu | Cytoplasm | other | 2629721 | 3472306.5 | 1745894.375 | | | | | | |
| P80318 | CCT3 | chaperonin containing TCP1, subu | Cytoplasm | other | 2641086.75 | 1926089.375 | 1025013.875 | P80318 | CCT3 | 7745970 | 5416834 | 781853 | 48 |
| P80315 | CCT4 | chaperonin containing TCP1, subu | Cytoplasm | other | 2.27E+07 | 3.00E+07 | 1.82E+07 | | | | | | |
| P80315 | CCT4 | chaperonin containing TCP1, subu | Cytoplasm | other | 2.06E+07 | 2.65E+07 | 1.55E+07 | | | | | | |
| P80315 | CCT4 | chaperonin containing TCP1, subu | Cytoplasm | other | 1.36E+07 | 1.73E+07 | 8698934 | | | | | | |
| P80315 | CCT4 | chaperonin containing TCP1, subu | Cytoplasm | other | 1.02E+07 | 1.89E+07 | 9213096 | | | | | | |
| P80315 | CCT4 | chaperonin containing TCP1, subu | Cytoplasm | other | 1.15E+07 | 1.24E+07 | 5733522.5 | | | | | | |
| P80315 | CCT4 | chaperonin containing TCP1, subu | Cytoplasm | other | 7919090 | 9405887 | 5028041 | | | | | | |
| P80315 | CCT4 | chaperonin containing TCP1, subu | Cytoplasm | other | 8319266 | 8788606 | 4943452.5 | | | | | | |
| P80315 | CCT4 | chaperonin containing TCP1, subu | Cytoplasm | other | 7526762 | 8241470 | 4051165.25 | | | | | | |
| P80315 | CCT4 | chaperonin containing TCP1, subu | Cytoplasm | other | 4785399.5 | 8605737 | 3135186.5 | | | | | | |
| P80315 | CCT4 | chaperonin containing TCP1, subu | Cytoplasm | other | 6066467.5 | 6105403.5 | 4320977 | | | | | | |
| P80315 | CCT4 | chaperonin containing TCP1, subu | Cytoplasm | other | 6357050 | 7143200 | 3632739.5 | | | | | | |
| P80315 | CCT4 | chaperonin containing TCP1, subu | Cytoplasm | other | 4407114 | 8798749 | 4634742.5 | | | | | | |
| P80315 | CCT4 | chaperonin containing TCP1, subu | Cytoplasm | other | 3855803.5 | 5147295 | 2467690.25 | | | | | | |
| P80315 | CCT4 | chaperonin containing TCP1, subu | Cytoplasm | other | 2535334.25 | 4787242.5 | 1352017.625 | | | | | | |
| P80315 | CCT4 | chaperonin containing TCP1, subu | Cytoplasm | other | 3502318 | 6911965 | 4016650 | | | | | | |
| P80315 | CCT4 | chaperonin containing TCP1, subu | Cytoplasm | other | 4379228.5 | 660705.312 | 935099.062 | | | | | | |
| P80315 | CCT4 | chaperonin containing TCP1, subu | Cytoplasm | other | 2451755 | 1763038.625 | 525769.938 | | | | | | |
| P80315 | CCT4 | chaperonin containing TCP1, subu | Cytoplasm | other | 1163414.75 | 2447401 | 1558586.375 | P80315 | CCT4 | 7882722 | 6614373 | 900102 | 54 |
| P80316 | CCT5 | chaperonin containing TCP1, subu | Cytoplasm | other | 2.38E+07 | 2.53E+07 | 1.54E+07 | | | | | | |
| P80316 | CCT5 | chaperonin containing TCP1, subu | Cytoplasm | other | 2.23E+07 | 2.76E+07 | 1.45E+07 | | | | | | |
| P80316 | CCT5 | chaperonin containing TCP1, subu | Cytoplasm | other | 9837364 | 1.31E+07 | 8821042 | | | | | | |
| P80316 | CCT5 | chaperonin containing TCP1, subu | Cytoplasm | other | 1.14E+07 | 1.21E+07 | 7744149 | | | | | | |
| P80316 | CCT5 | chaperonin containing TCP1, subu | Cytoplasm | other | 1.45E+07 | 1.57E+07 | 8768239 | | | | | | |
| P80316 | CCT5 | chaperonin containing TCP1, subu | Cytoplasm | other | 7815373.5 | 1.06E+07 | 5152096 | | | | | | |
| P80316 | CCT5 | chaperonin containing TCP1, subu | Cytoplasm | other | 6187142 | 7704115.5 | 4294376 | | | | | | |
| P80316 | CCT5 | chaperonin containing TCP1, subu | Cytoplasm | other | 6297896.5 | 7765671.5 | 4777407.5 | | | | | | |
| P80316 | CCT5 | chaperonin containing TCP1, subu | Cytoplasm | other | 6135961.5 | 6120054.5 | 3545090.5 | | | | | | |
| P80316 | CCT5 | chaperonin containing TCP1, subu | Cytoplasm | other | 4440017 | 5737041.5 | 3020279 | | | | | | |
| P80316 | CCT5 | chaperonin containing TCP1, subu | Cytoplasm | other | 2727907.25 | 2906013 | 1477239.375 | | | | | | |
| P80316 | CCT5 | chaperonin containing TCP1, subu | Cytoplasm | other | 1637171.5 | 2649053.5 | 1423167.875 | | | | | | |
| P80316 | CCT5 | chaperonin containing TCP1, subu | Cytoplasm | other | 1795067.875 | 580933.938 | 257421.781 | P80316 | CCT5 | 9141069 | 6988545 | 1119063 | 39 |
| P80317 | CCT6A | chaperonin containing TCP1, subu | Cytoplasm | other | 2.21E+07 | 2.83E+07 | 1.34E+07 | | | | | | |
| P80317 | CCT6A | chaperonin containing TCP1, subu | Cytoplasm | other | 1.48E+07 | 2.34E+07 | 1.04E+07 | | | | | | |
| P80317 | CCT6A | chaperonin containing TCP1, subu | Cytoplasm | other | 1.45E+07 | 2.25E+07 | 1.21E+07 | | | | | | |
| P80317 | CCT6A | chaperonin containing TCP1, subu | Cytoplasm | other | 1.22E+07 | 1.74E+07 | 1.08E+07 | | | | | | |
| P80317 | CCT6A | chaperonin containing TCP1, subu | Cytoplasm | other | 1.17E+07 | 1.61E+07 | 8576093 | | | | | | |
| P80317 | CCT6A | chaperonin containing TCP1, subu | Cytoplasm | other | 9283715 | 1.14E+07 | 6623921.5 | | | | | | |
| P80317 | CCT6A | chaperonin containing TCP1, subu | Cytoplasm | other | 8660536 | 1.13E+07 | 7969503.5 | | | | | | |
| P80317 | CCT6A | chaperonin containing TCP1, subu | Cytoplasm | other | 8455877 | 9227236 | 5707406.5 | | | | | | |
| P80317 | CCT6A | chaperonin containing TCP1, subu | Cytoplasm | other | 7305027.5 | 7990779 | 3463378.75 | | | | | | |
| P80317 | CCT6A | chaperonin containing TCP1, subu | Cytoplasm | other | 5392518.5 | 7151509 | 3885878.5 | | | | | | |
| P80317 | CCT6A | chaperonin containing TCP1, subu | Cytoplasm | other | 939573.938 | 2343479 | 1075573.875 | | | | | | |
| P80317 | CCT6A | chaperonin containing TCP1, subu | Cytoplasm | other | 904519.062 | 2635015 | 917677.625 | P80317 | CCT6A | 9680981 | 6719175 | 1119862 | 36 |
| P80313 | CCT7 | chaperonin containing TCP1, subu | Cytoplasm | other | 1.52E+07 | 1.89E+07 | 1.20E+07 | | | | | | |
| P80313 | CCT7 | chaperonin containing TCP1, subu | Cytoplasm | other | 1.02E+07 | 2.10E+07 | 9873853 | | | | | | |
| P80313 | CCT7 | chaperonin containing TCP1, subu | Cytoplasm | other | 1.02E+07 | 1.55E+07 | 8249737.5 | | | | | | |
| P80313 | CCT7 | chaperonin containing TCP1, subu | Cytoplasm | other | 1.31E+07 | 1.53E+07 | 7395885.5 | | | | | | |
| P80313 | CCT7 | chaperonin containing TCP1, subu | Cytoplasm | other | 9574792 | 1.73E+07 | 1.35E+07 | | | | | | |
| P80313 | CCT7 | chaperonin containing TCP1, subu | Cytoplasm | other | 8743000 | 1.18E+07 | 5789578.5 | | | | | | |
| P80313 | CCT7 | chaperonin containing TCP1, subu | Cytoplasm | other | 6926171.5 | 9250173 | 4293671 | | | | | | |
| P80313 | CCT7 | chaperonin containing TCP1, subu | Cytoplasm | other | 9307192 | 8538331 | 4386119.5 | | | | | | |
| P80313 | CCT7 | chaperonin containing TCP1, subu | Cytoplasm | other | 6262452.5 | 5587243.5 | 3327766.5 | | | | | | |
| P80313 | CCT7 | chaperonin containing TCP1, subu | Cytoplasm | other | 5338924 | 7422719.5 | 5142704.5 | | | | | | |
| P80313 | CCT7 | chaperonin containing TCP1, subu | Cytoplasm | other | 7618149 | 8359089 | 4846419.5 | | | | | | |

| | | | | | | | | | | | | | |
|---|---|---|---|---|---|---|---|---|---|---|---|---|---|
| P80313 | CCT7 | chaperonin containing TCP1, subu | Cytoplasm | other | 3357656.5 | 3660268.75 | 1838290.25 | | | | | | |
| P80313 | CCT7 | chaperonin containing TCP1, subu | Cytoplasm | other | 3409276 | 4756777 | 2124478.5 | | | | | | |
| P80313 | CCT7 | chaperonin containing TCP1, subu | Cytoplasm | other | 2525517.25 | 8294559.5 | 2571084.25 | | | | | | |
| P80313 | CCT7 | chaperonin containing TCP1, subu | Cytoplasm | other | 2465708 | 2893854.75 | 1666814.625 | P80313 | CCT7 | 7614589 | 4846604 | 722489 | 45 |
| P42932 | CCT8 | chaperonin containing TCP1, subu | Cytoplasm | enzyme | 1.60E+07 | 1.77E+07 | 8989833 | | | | | | |
| P42932 | CCT8 | chaperonin containing TCP1, subu | Cytoplasm | enzyme | 1.04E+07 | 1.35E+07 | 8028463.5 | | | | | | |
| P42932 | CCT8 | chaperonin containing TCP1, subu | Cytoplasm | enzyme | 9135368 | 1.32E+07 | 6295169 | | | | | | |
| P42932 | CCT8 | chaperonin containing TCP1, subu | Cytoplasm | enzyme | 1.05E+07 | 1.42E+07 | 7436482 | | | | | | |
| P42932 | CCT8 | chaperonin containing TCP1, subu | Cytoplasm | enzyme | 8593275 | 1.30E+07 | 6952781 | | | | | | |
| P42932 | CCT8 | chaperonin containing TCP1, subu | Cytoplasm | enzyme | 1.08E+07 | 1.11E+07 | 4386192.5 | | | | | | |
| P42932 | CCT8 | chaperonin containing TCP1, subu | Cytoplasm | enzyme | 8897531 | 9717174 | 6107981.5 | | | | | | |
| P42932 | CCT8 | chaperonin containing TCP1, subu | Cytoplasm | enzyme | 8509329 | 1.18E+07 | 5653667 | | | | | | |
| P42932 | CCT8 | chaperonin containing TCP1, subu | Cytoplasm | enzyme | 5447456 | 1.02E+07 | 3778638 | | | | | | |
| P42932 | CCT8 | chaperonin containing TCP1, subu | Cytoplasm | enzyme | 8037414.5 | 7474922 | 4848602.5 | | | | | | |
| P42932 | CCT8 | chaperonin containing TCP1, subu | Cytoplasm | enzyme | 7125250.5 | 7832908 | 3572496.5 | | | | | | |
| P42932 | CCT8 | chaperonin containing TCP1, subu | Cytoplasm | enzyme | 5965689.5 | 1.14E+07 | 4907689.5 | | | | | | |
| P42932 | CCT8 | chaperonin containing TCP1, subu | Cytoplasm | enzyme | 6296334.5 | 5995992 | 4419958.5 | | | | | | |
| P42932 | CCT8 | chaperonin containing TCP1, subu | Cytoplasm | enzyme | 4559983.5 | 5372231 | 3508536.5 | | | | | | |
| P42932 | CCT8 | chaperonin containing TCP1, subu | Cytoplasm | enzyme | 3523859.75 | 4550622 | 2400622.5 | | | | | | |
| P42932 | CCT8 | chaperonin containing TCP1, subu | Cytoplasm | enzyme | 8480243 | 1851895.125 | 2846421.25 | | | | | | |
| P42932 | CCT8 | chaperonin containing TCP1, subu | Cytoplasm | enzyme | 6125847 | 9313769 | 6164316.5 | | | | | | |
| P42932 | CCT8 | chaperonin containing TCP1, subu | Cytoplasm | enzyme | 4417093 | 3600432.25 | 1956879.5 | | | | | | |
| P42932 | CCT8 | chaperonin containing TCP1, subu | Cytoplasm | enzyme | 2355372 | 2874046 | 1421185.375 | | | | | | |
| P42932 | CCT8 | chaperonin containing TCP1, subu | Cytoplasm | enzyme | 2621290 | 3006136.25 | 1865195.375 | | | | | | |
| P42932 | CCT8 | chaperonin containing TCP1, subu | Cytoplasm | enzyme | 2149898.75 | 2231524 | 1267011.75 | | | | | | |
| P42932 | CCT8 | chaperonin containing TCP1, subu | Cytoplasm | enzyme | 3664601.75 | 4549152 | 2845835.5 | | | | | | |
| P42932 | CCT8 | chaperonin containing TCP1, subu | Cytoplasm | enzyme | 2764777.75 | 2664480.5 | 1051176.875 | | | | | | |
| P42932 | CCT8 | chaperonin containing TCP1, subu | Cytoplasm | enzyme | 525827.438 | 3776420.5 | 1388313.25 | P42932 | CCT8 | 6536102 | 3879297 | 457179 | 72 |
| Q8K4F0 | CD226 | CD226 molecule | Plasma Membrane | other | 3256935 | 3159416 | 2905226.5 | Q8K4F0 | CD226 | 3256935 | 181577 | 104833 | 3 |
| Q08857 | CD36 | CD36 molecule (thrombospondin r | Plasma Membrane | transmembrane receptor | 6.16E+08 | 5.90E+08 | 5.53E+08 | | | | | | |
| Q08857 | CD36 | CD36 molecule (thrombospondin r | Plasma Membrane | transmembrane receptor | 2.45E+08 | 2.23E+08 | 2.61E+08 | | | | | | |
| Q08857 | CD36 | CD36 molecule (thrombospondin r | Plasma Membrane | transmembrane receptor | 8.42E+07 | 1.76E+08 | 1.52E+08 | | | | | | |
| Q08857 | CD36 | CD36 molecule (thrombospondin r | Plasma Membrane | transmembrane receptor | 1.11E+08 | 1.42E+08 | 1.37E+08 | | | | | | |
| Q08857 | CD36 | CD36 molecule (thrombospondin r | Plasma Membrane | transmembrane receptor | 6.65E+07 | 2.86E+07 | 1.83E+07 | | | | | | |
| Q08857 | CD36 | CD36 molecule (thrombospondin r | Plasma Membrane | transmembrane receptor | 4.62E+07 | 1.49E+08 | 1.22E+08 | | | | | | |
| Q08857 | CD36 | CD36 molecule (thrombospondin r | Plasma Membrane | transmembrane receptor | 8.49E+07 | 6.49E+07 | 4.97E+07 | | | | | | |
| Q08857 | CD36 | CD36 molecule (thrombospondin r | Plasma Membrane | transmembrane receptor | 7.17E+07 | 1.04E+08 | 1.09E+08 | | | | | | |
| Q08857 | CD36 | CD36 molecule (thrombospondin r | Plasma Membrane | transmembrane receptor | 7.67E+07 | 7.48E+07 | 6.86E+07 | | | | | | |
| Q08857 | CD36 | CD36 molecule (thrombospondin r | Plasma Membrane | transmembrane receptor | 1.02E+08 | 1.71E+08 | 1.60E+08 | | | | | | |
| Q08857 | CD36 | CD36 molecule (thrombospondin r | Plasma Membrane | transmembrane receptor | 1.01E+07 | 4.85E+07 | 3.56E+07 | | | | | | |
| Q08857 | CD36 | CD36 molecule (thrombospondin r | Plasma Membrane | transmembrane receptor | 1.17E+07 | 2.44E+07 | 2.26E+07 | | | | | | |
| Q08857 | CD36 | CD36 molecule (thrombospondin r | Plasma Membrane | transmembrane receptor | 2.49E+07 | 2.10E+07 | 2.23E+07 | | | | | | |
| Q08857 | CD36 | CD36 molecule (thrombospondin r | Plasma Membrane | transmembrane receptor | 1.11E+07 | 1.14E+07 | 1.54E+07 | | | | | | |
| Q08857 | CD36 | CD36 molecule (thrombospondin r | Plasma Membrane | transmembrane receptor | 3561800 | 2280475 | 2257873 | | | | | | |
| Q08857 | CD36 | CD36 molecule (thrombospondin r | Plasma Membrane | transmembrane receptor | 4.70E-06 | 3294649.5 | 5944497.5 | Q08857 | CD36 | 97836363 | 142149624 | 20517531 | 48 |
| P15379 | CD44 | CD44 molecule (Indian blood grou | Plasma Membrane | enzyme | 1.65E+07 | 1.49E+07 | 1.74E+07 | | | | | | |
| P15379 | CD44 | CD44 molecule (Indian blood grou | Plasma Membrane | enzyme | 5524198 | 8106175 | 6657902.5 | | | | | | |
| P15379 | CD44 | CD44 molecule (Indian blood grou | Plasma Membrane | enzyme | 2355496 | 4207019 | 3651010.25 | | | | | | |
| P15379 | CD44 | CD44 molecule (Indian blood grou | Plasma Membrane | enzyme | 966277 | 1432368 | 1011714.688 | P15379 | CD44 | 6343993 | 6096442 | 1759891 | 12 |
| Q61735 | CD47 | CD47 molecule | Plasma Membrane | transmembrane receptor | 5.18E+08 | 5.38E+08 | 5.76E+08 | | | | | | |
| Q61735 | CD47 | CD47 molecule | Plasma Membrane | transmembrane receptor | 3.40E+08 | 2.67E+08 | 3.28E+08 | | | | | | |
| Q61735 | CD47 | CD47 molecule | Plasma Membrane | transmembrane receptor | 1.37E+08 | 2.03E+08 | 1.96E+08 | | | | | | |
| Q61735 | CD47 | CD47 molecule | Plasma Membrane | transmembrane receptor | 1.29E+08 | 1.18E+08 | 1.25E+08 | | | | | | |
| Q61735 | CD47 | CD47 molecule | Plasma Membrane | transmembrane receptor | 7.10E+07 | 1.06E+08 | 9.50E+07 | | | | | | |
| Q61735 | CD47 | CD47 molecule | Plasma Membrane | transmembrane receptor | 7037595 | 8485070 | 9719181 | Q61735 | CD47 | 200282933 | 182139107 | 42930599 | 18 |
| P18181 | CD48 | CD48 molecule | Plasma Membrane | other | 1945206.625 | 2560806 | 1965509.875 | P18181 | CD48 | 1945207 | 349703 | 201901 | 3 |
| Q61475 | CD55 | CD55 molecule, decay accelerating | Plasma Membrane | other | 1.13E+07 | 1.30E+07 | 1.22E+07 | | | | | | |
| Q61475 | CD55 | CD55 molecule, decay accelerating | Plasma Membrane | other | 1.15E+07 | 1.19E+07 | 1.07E+07 | | | | | | |
| Q61475 | CD55 | CD55 molecule, decay accelerating | Plasma Membrane | other | 4254778.5 | 7672391.5 | 6772439 | | | | | | |
| Q61475 | CD55 | CD55 molecule, decay accelerating | Plasma Membrane | other | 4249687.5 | 6309682.5 | 6458394 | | | | | | |
| Q61475 | CD55 | CD55 molecule, decay accelerating | Plasma Membrane | other | 5695951.5 | 5088053 | 5900782.5 | | | | | | |
| Q61475 | CD55 | CD55 molecule, decay accelerating | Plasma Membrane | other | 1978302.75 | 3995142 | 3553894.25 | Q61475 | CD55 | 6483120 | 3469958 | 817877 | 18 |
| O55186 | Cd59a | CD59a antigen | Plasma Membrane | other | 9.90E+07 | 8.88E+07 | 1.12E+08 | | | | | | |
| O55186 | Cd59a | CD59a antigen | Plasma Membrane | other | 7721246 | 7681169.5 | 4977050.5 | | | | | | |
| O55186 | Cd59a | CD59a antigen | Plasma Membrane | other | 2516504.25 | 1904868.5 | 1474524.875 | O55186 | Cd59a | 36395917 | 48112234 | 16037411 | 9 |
| P40237 | CD82 | CD82 molecule | Plasma Membrane | other | 2694257.75 | 9644478 | 7733070.5 | | | | | | |
| P40237 | CD82 | CD82 molecule | Plasma Membrane | other | 3517701.25 | 8371678.5 | 3054009.25 | P40237 | CD82 | 3105980 | 3082777 | 1258538 | 6 |
| P40240 | CD9 | CD9 molecule | Plasma Membrane | other | 3.37E+07 | 5.12E+07 | 3.81E+07 | | | | | | |
| P40240 | CD9 | CD9 molecule | Plasma Membrane | other | 1993519.625 | 6459962.5 | 7167189 | | | | | | |
| P40240 | CD9 | CD9 molecule | Plasma Membrane | other | 2341904.25 | 1090500.25 | 2541798.75 | P40240 | CD9 | 12685141 | 19353296 | 6451099 | 9 |
| Q8CII2 | CDC123 | cell division cycle 123 | Cytoplasm | other | 5244124.5 | 9347171 | 7731145.5 | Q8CII2 | CDC123 | 5244125 | 2066874 | 1193310 | 3 |
| P60766 | Cdc42 | cell division cycle 42 | Plasma Membrane | enzyme | 1.21E+07 | 7848287.5 | 6730854 | | | | | | |
| P60766 | Cdc42 | cell division cycle 42 | Plasma Membrane | enzyme | 1.16E+07 | 4999144 | 4974296.5 | | | | | | |
| P60766 | Cdc42 | cell division cycle 42 | Plasma Membrane | enzyme | 8035463.5 | 7772131 | 7072205.5 | | | | | | |
| P60766 | Cdc42 | cell division cycle 42 | Plasma Membrane | enzyme | 3996927 | 4382540 | 4607143.5 | | | | | | |
| P60766 | Cdc42 | cell division cycle 42 | Plasma Membrane | enzyme | 2491397.25 | 2871424 | 2676925.25 | P60766 | Cdc42 | 7634758 | 2974321 | 767966 | 15 |
| Q922M5 | CDCA7L | cell division cycle associated 7-like | Nucleus | other | 3.06E+07 | 1.57E+07 | 2.34E+07 | Q922M5 | CDCA7L | 30630000 | 7466233 | 4310632 | 3 |
| Q8VDP6 | CDIPT | CDP-diacylglycerol--inositol 3-phos | Cytoplasm | enzyme | 2469102 | 2857943.5 | 2412590.75 | Q8VDP6 | CDIPT | 2469102 | 242463 | 139986 | 3 |
| Q99L43 | CDS2 | CDP-diacylglycerol synthase (phos | Cytoplasm | enzyme | 5690125.5 | 3384527.75 | 4837291.5 | Q99L43 | CDS2 | 5690126 | 1165735 | 673037 | 3 |
| Q4VAA2 | CDV3 | CDV3 homolog (mouse) | Cytoplasm | other | 3448284 | 2526248 | 1510580.5 | Q4VAA2 | CDV3 | 3448284 | 969229 | 559584 | 3 |
| Q6AW46 | CES5A | carboxylesterase 5A | Extracellular Space | enzyme | 1.42E+07 | 1.68E+07 | 1.57E+07 | Q6AW46 | CES5A | 14230000 | 1268161 | 732173 | 3 |
| P18760 | CFL1 | cofilin 1 (non-muscle) | Nucleus | other | 1.19E+07 | 1.23E+07 | 1.36E+07 | | | | | | |
| P18760 | CFL1 | cofilin 1 (non-muscle) | Nucleus | other | 2630517 | 3873394.5 | 3686277 | | | | | | |
| P18760 | CFL1 | cofilin 1 (non-muscle) | Nucleus | other | 3524848.75 | 4648995.5 | 5176880 | | | | | | |
| P18760 | CFL1 | cofilin 1 (non-muscle) | Nucleus | other | 4201902.5 | 5810939 | 5396878.5 | | | | | | |
| P18760 | CFL1 | cofilin 1 (non-muscle) | Nucleus | other | 1366941.375 | 2073288 | 2289586.75 | P18760 | CFL1 | 4728842 | 3888628 | 1004040 | 15 |

| | | | | | | | | | | | | | |
|---|---|---|---|---|---|---|---|---|---|---|---|---|---|
| Q9CRB9 | CHCHD3 | coiled-coil-helix-coiled-coil-helix d | Cytoplasm | other | 4032050 | 4656372.5 | 5861668.5 | Q9CRB9 | CHCHD3 | 4032050 | 930056 | 536968 | 3 |
| O35744 | Chil3/Chil4 | chitinase-like 3 | Cytoplasm | enzyme | 1246010.125 | 562210.875 | 1641805.625 | | | | | | |
| O35744 | Chil3/Chil4 | chitinase-like 3 | Cytoplasm | enzyme | 3477832.5 | 1049261.125 | 4476415.5 | | | | | | |
| O35744 | Chil3/Chil4 | chitinase-like 3 | Cytoplasm | enzyme | 1557064.25 | 1322376.125 | 2495042.25 | | | | | | |
| O35744 | Chil3/Chil4 | chitinase-like 3 | Cytoplasm | enzyme | 1446459.75 | 610616.438 | 2297749.25 | | | | | | |
| O35744 | Chil3/Chil4 | chitinase-like 3 | Cytoplasm | enzyme | 876255.25 | 356071.781 | 927074.875 | | | | | | |
| O35744 | Chil3/Chil4 | chitinase-like 3 | Cytoplasm | enzyme | 891114.812 | 307173.219 | 1058234.5 | O35744 | Chil3/Chil4 | 1582456 | 1095417 | 258192 | 18 |
| Q9DB34 | CHMP2A | charged multivesicular body prote | Cytoplasm | other | 2097610.5 | 1924367.125 | 1332703.375 | Q9DB34 | CHMP2A | 2097611 | 401074 | 231560 | 3 |
| P0C0A3 | CHMP6 | charged multivesicular body prote | Cytoplasm | other | 4365233.5 | 4401979.5 | 4012180.75 | P0C0A3 | CHMP6 | 4365234 | 215228 | 124262 | 3 |
| P61022 | CHP1 | calcineurin-like EF-hand protein 1 | Cytoplasm | transporter | 7555505.5 | 5243058.5 | 6308054 | | | | | | |
| P61022 | CHP1 | calcineurin-like EF-hand protein 1 | Cytoplasm | transporter | 4613776.5 | 4378443 | 5001064.5 | | | | | | |
| P61022 | CHP1 | calcineurin-like EF-hand protein 1 | Cytoplasm | transporter | 3099908 | 2285772 | 1886909.375 | | | | | | |
| P61022 | CHP1 | calcineurin-like EF-hand protein 1 | Cytoplasm | transporter | 2890442 | 4721830 | 3263617 | | | | | | |
| P61022 | CHP1 | calcineurin-like EF-hand protein 1 | Cytoplasm | transporter | 3988872.75 | 4240395.5 | 3982605.5 | | | | | | |
| P61022 | CHP1 | calcineurin-like EF-hand protein 1 | Cytoplasm | transporter | 3522623.75 | 3103679 | 3329919.25 | P61022 | CHP1 | 4278521 | 1392656 | 328252 | 18 |
| Q9Z0F4 | CIB1 | calcium and integrin binding 1 (cal | Nucleus | other | 4185508.75 | 1.36E+07 | 8345757.5 | | | | | | |
| Q9Z0F4 | CIB1 | calcium and integrin binding 1 (cal | Nucleus | other | 4949985.5 | 7844570.5 | 6958545 | | | | | | |
| Q9Z0F4 | CIB1 | calcium and integrin binding 1 (cal | Nucleus | other | 2609994.25 | 4874616 | 3627083.5 | | | | | | |
| Q9Z0F4 | CIB1 | calcium and integrin binding 1 (cal | Nucleus | other | 219152.719 | 1121864.375 | 454037.062 | Q9Z0F4 | CIB1 | 2991160 | 3867089 | 1116333 | 12 |
| Q91WS0 | CISD1 | CDGSH iron sulfur domain 1 | Cytoplasm | other | 2192525.75 | 2238026 | 3228037.75 | | | | | | |
| Q91WS0 | CISD1 | CDGSH iron sulfur domain 1 | Cytoplasm | other | 3859662.25 | 3876293.75 | 3550380 | Q91WS0 | CISD1 | 3026094 | 767546 | 313349 | 6 |
| Q9CQB5 | CISD2 | CDGSH iron sulfur domain 2 | Cytoplasm | other | 5594330.5 | 8973193 | 6974573.5 | | | | | | |
| Q9CQB5 | CISD2 | CDGSH iron sulfur domain 2 | Cytoplasm | other | 5713730 | 8247278.5 | 4780801 | | | | | | |
| Q9CQB5 | CISD2 | CDGSH iron sulfur domain 2 | Cytoplasm | other | 5526238.5 | 5789076 | 5411041 | | | | | | |
| Q9CQB5 | CISD2 | CDGSH iron sulfur domain 2 | Cytoplasm | other | 3415505.5 | 4014273.75 | 3159858 | | | | | | |
| Q9CQB5 | CISD2 | CDGSH iron sulfur domain 2 | Cytoplasm | other | 1897754.75 | 4849794.5 | 3521539.75 | Q9CQB5 | CISD2 | 4429512 | 1896637 | 489709 | 15 |
| Q9Z0S4 | Cldn13 | claudin 13 | Plasma Membrane | other | 1.83E+08 | 2.28E+08 | 1.91E+08 | | | | | | |
| Q9Z0S4 | Cldn13 | claudin 13 | Plasma Membrane | other | 5.41E+07 | 5.29E+07 | 4.79E+07 | | | | | | |
| Q9Z0S4 | Cldn13 | claudin 13 | Plasma Membrane | other | 4.07E+07 | 7.26E+07 | 5.67E+07 | | | | | | |
| Q9Z0S4 | Cldn13 | claudin 13 | Plasma Membrane | other | 3.88E+07 | 3.32E+07 | 2.81E+07 | Q9Z0S4 | Cldn13 | 79097500 | 71122971 | 20531433 | 12 |
| Q9JL99 | CLEC1B | C-type lectin domain family 1, mer | Plasma Membrane | transmembrane receptor | 8850786 | 1.01E+07 | 1.02E+07 | Q9JL99 | CLEC1B | 8850786 | 757771 | 437499 | 3 |
| Q9QUK3 | CLN8 | ceroid-lipofuscinosis, neuronal 8 | Cytoplasm | other | 5365502 | 6937252.5 | 5581264.5 | Q9QUK3 | CLN8 | 5365502 | 852023 | 491916 | 3 |
| Q61189 | CLNS1A | chloride channel, nucleotide-sensi | Plasma Membrane | ion channel | 3243941.75 | 2277121.5 | 2890515.25 | Q61189 | CLNS1A | 3243942 | 489201 | 282440 | 3 |
| Q8VBZ3 | CLPTM1 | cleft lip and palate associated tran | Plasma Membrane | other | 1478961.25 | 936176.062 | 1118500.5 | Q8VBZ3 | CLPTM1 | 1478961 | 276222 | 159477 | 3 |
| Q68FD5 | CLTC | clathrin, heavy chain (Hc) | Plasma Membrane | other | 1.45E+07 | 1.41E+07 | 1.24E+07 | | | | | | |
| Q68FD5 | CLTC | clathrin, heavy chain (Hc) | Plasma Membrane | other | 1.85E+07 | 1.92E+07 | 1.40E+07 | | | | | | |
| Q68FD5 | CLTC | clathrin, heavy chain (Hc) | Plasma Membrane | other | 1.89E+07 | 2.14E+07 | 1.30E+07 | | | | | | |
| Q68FD5 | CLTC | clathrin, heavy chain (Hc) | Plasma Membrane | other | 1.77E+07 | 1.86E+07 | 1.22E+07 | | | | | | |
| Q68FD5 | CLTC | clathrin, heavy chain (Hc) | Plasma Membrane | other | 1.20E+07 | 1.22E+07 | 9048821 | | | | | | |
| Q68FD5 | CLTC | clathrin, heavy chain (Hc) | Plasma Membrane | other | 1.35E+07 | 1.64E+07 | 1.02E+07 | | | | | | |
| Q68FD5 | CLTC | clathrin, heavy chain (Hc) | Plasma Membrane | other | 1.62E+07 | 1.70E+07 | 1.21E+07 | | | | | | |
| Q68FD5 | CLTC | clathrin, heavy chain (Hc) | Plasma Membrane | other | 8842186 | 1.20E+07 | 7533483 | | | | | | |
| Q68FD5 | CLTC | clathrin, heavy chain (Hc) | Plasma Membrane | other | 9491274 | 9861780 | 6955853 | | | | | | |
| Q68FD5 | CLTC | clathrin, heavy chain (Hc) | Plasma Membrane | other | 1.24E+07 | 1.08E+07 | 9144868 | | | | | | |
| Q68FD5 | CLTC | clathrin, heavy chain (Hc) | Plasma Membrane | other | 1.39E+07 | 1.56E+07 | 1.20E+07 | | | | | | |
| Q68FD5 | CLTC | clathrin, heavy chain (Hc) | Plasma Membrane | other | 8426825 | 1.27E+07 | 5489888.5 | | | | | | |
| Q68FD5 | CLTC | clathrin, heavy chain (Hc) | Plasma Membrane | other | 6882171.5 | 8410103 | 4713951 | | | | | | |
| Q68FD5 | CLTC | clathrin, heavy chain (Hc) | Plasma Membrane | other | 7054054.5 | 6213219 | 3763197.25 | | | | | | |
| Q68FD5 | CLTC | clathrin, heavy chain (Hc) | Plasma Membrane | other | 8086061.5 | 9415210 | 7105471.5 | | | | | | |
| Q68FD5 | CLTC | clathrin, heavy chain (Hc) | Plasma Membrane | other | 8384867.5 | 9603956 | 4674295 | | | | | | |
| Q68FD5 | CLTC | clathrin, heavy chain (Hc) | Plasma Membrane | other | 5921677 | 9530503 | 7381385 | | | | | | |
| Q68FD5 | CLTC | clathrin, heavy chain (Hc) | Plasma Membrane | other | 6251545.5 | 8972885 | 5356281.5 | | | | | | |
| Q68FD5 | CLTC | clathrin, heavy chain (Hc) | Plasma Membrane | other | 6549985 | 8214639 | 5483572 | | | | | | |
| Q68FD5 | CLTC | clathrin, heavy chain (Hc) | Plasma Membrane | other | 7518781 | 8954679 | 3364538.5 | | | | | | |
| Q68FD5 | CLTC | clathrin, heavy chain (Hc) | Plasma Membrane | other | 5711531 | 9204579 | 4864520 | | | | | | |
| Q68FD5 | CLTC | clathrin, heavy chain (Hc) | Plasma Membrane | other | 3583864.75 | 4596680.5 | 1872656.625 | | | | | | |
| Q68FD5 | CLTC | clathrin, heavy chain (Hc) | Plasma Membrane | other | 8678592 | 8562370 | 5300732 | | | | | | |
| Q68FD5 | CLTC | clathrin, heavy chain (Hc) | Plasma Membrane | other | 6733542 | 5755319 | 1.08E+07 | | | | | | |
| Q68FD5 | CLTC | clathrin, heavy chain (Hc) | Plasma Membrane | other | 4467513 | 6142853.5 | 3699595.75 | | | | | | |
| Q68FD5 | CLTC | clathrin, heavy chain (Hc) | Plasma Membrane | other | 7343243.5 | 6479466 | 4769131 | | | | | | |
| Q68FD5 | CLTC | clathrin, heavy chain (Hc) | Plasma Membrane | other | 6697040.5 | 7447416 | 5872201.5 | | | | | | |
| Q68FD5 | CLTC | clathrin, heavy chain (Hc) | Plasma Membrane | other | 4892590.5 | 7472765 | 5035565 | | | | | | |
| Q68FD5 | CLTC | clathrin, heavy chain (Hc) | Plasma Membrane | other | 6417912.5 | 2476182 | 4157802.5 | | | | | | |
| Q68FD5 | CLTC | clathrin, heavy chain (Hc) | Plasma Membrane | other | 5318280 | 5261728.5 | 4096239.75 | | | | | | |
| Q68FD5 | CLTC | clathrin, heavy chain (Hc) | Plasma Membrane | other | 5087893 | 4787772.5 | 3340763.5 | | | | | | |
| Q68FD5 | CLTC | clathrin, heavy chain (Hc) | Plasma Membrane | other | 2264078.75 | 7115600.5 | 3660709.25 | | | | | | |
| Q68FD5 | CLTC | clathrin, heavy chain (Hc) | Plasma Membrane | other | 2765869.25 | 3261195.5 | 1912517 | | | | | | |
| Q68FD5 | CLTC | clathrin, heavy chain (Hc) | Plasma Membrane | other | 3897835.75 | 7207926 | 3838789.25 | | | | | | |
| Q68FD5 | CLTC | clathrin, heavy chain (Hc) | Plasma Membrane | other | 2880762.75 | 3394228.75 | 1942343.5 | | | | | | |
| Q68FD5 | CLTC | clathrin, heavy chain (Hc) | Plasma Membrane | other | 2558413.25 | 2501072.25 | 1645639.75 | | | | | | |
| Q68FD5 | CLTC | clathrin, heavy chain (Hc) | Plasma Membrane | other | 2313668.5 | 3846175.5 | 1246857.375 | | | | | | |
| Q68FD5 | CLTC | clathrin, heavy chain (Hc) | Plasma Membrane | other | 2852382.75 | 3296167.25 | 1568998.875 | | | | | | |
| Q68FD5 | CLTC | clathrin, heavy chain (Hc) | Plasma Membrane | other | 2460011.25 | 4632207 | 2514822 | | | | | | |
| Q68FD5 | CLTC | clathrin, heavy chain (Hc) | Plasma Membrane | other | 2286780.75 | 3616384.25 | 1695637.875 | | | | | | |
| Q68FD5 | CLTC | clathrin, heavy chain (Hc) | Plasma Membrane | other | 1204940.5 | 1859683.125 | 1315732.75 | | | | | | |
| Q68FD5 | CLTC | clathrin, heavy chain (Hc) | Plasma Membrane | other | 2324553.5 | 577869.312 | 1077479.625 | | | | | | |
| Q68FD5 | CLTC | clathrin, heavy chain (Hc) | Plasma Membrane | other | 3039955.25 | 721147.062 | 664433.188 | | | | | | |
| Q68FD5 | CLTC | clathrin, heavy chain (Hc) | Plasma Membrane | other | 1593463.5 | 1663354 | 1564925 | | | | | | |
| Q68FD5 | CLTC | clathrin, heavy chain (Hc) | Plasma Membrane | other | 4.70E-06 | 1302683.875 | 997017.312 | | | | | | |
| Q68FD5 | CLTC | clathrin, heavy chain (Hc) | Plasma Membrane | other | 71882.078 | 360222.906 | 281677.156 | Q68FD5 | CLTC | 6921218 | 4789236 | 407687 | 138 |
| Q06890 | CLU | clusterin | Cytoplasm | other | 1.33E+07 | 1.18E+07 | 1.45E+07 | | | | | | |
| Q06890 | CLU | clusterin | Cytoplasm | other | 2484128.25 | 3756156.75 | 4995753 | | | | | | |
| Q06890 | CLU | clusterin | Cytoplasm | other | 3232436.75 | 220260.891 | 1433220.875 | Q06890 | CLU | 6342188 | 5466225 | 1822075 | 9 |
| Q8R4N0 | CLYBL | citrate lyase beta like | Cytoplasm | enzyme | 3081254.5 | 2923675.5 | 5454803.5 | Q8R4N0 | CLYBL | 3081255 | 1418049 | 818711 | 3 |
| O35372 | CNIH1 | cornichon family AMPA receptor a | Plasma Membrane | other | 3495308 | 4335982.5 | 4462046 | O35372 | CNIH1 | 3495308 | 525549 | 303426 | 3 |
| Q08093 | CNN2 | calponin 2 | Cytoplasm | other | 2568861 | 4499479.5 | 4376801 | Q08093 | CNN2 | 2568861 | 1080971 | 624099 | 3 |

| | | | | | | | | | | | | | |
|---|---|---|---|---|---|---|---|---|---|---|---|---|---|
| Q3TWN3 | CNNM2 | cyclin and CBS domain divalent me | Plasma Membrane | other | 4014787.5 | 4675386 | 3756273.5 | | | | | | |
| Q3TWN3 | CNNM2 | cyclin and CBS domain divalent me | Plasma Membrane | other | 3661906 | 3831089.75 | 3706825.5 | Q3TWN3 | CNNM2 | 3838347 | 380448 | 155317 | 6 |
| P16330 | CNP | 2',3'-cyclic nucleotide 3' phosphod | Cytoplasm | enzyme | 2.97E+07 | 2.26E+07 | 3.73E+07 | | | | | | |
| P16330 | CNP | 2',3'-cyclic nucleotide 3' phosphod | Cytoplasm | enzyme | 1.41E+07 | 1.13E+07 | 1.57E+07 | | | | | | |
| P16330 | CNP | 2',3'-cyclic nucleotide 3' phosphod | Cytoplasm | enzyme | 8867757 | 7315468 | 1.05E+07 | | | | | | |
| P16330 | CNP | 2',3'-cyclic nucleotide 3' phosphod | Cytoplasm | enzyme | 7727071 | 5862656 | 8057757 | | | | | | |
| P16330 | CNP | 2',3'-cyclic nucleotide 3' phosphod | Cytoplasm | enzyme | 3966282.75 | 4066734.25 | 4902753.5 | | | | | | |
| P16330 | CNP | 2',3'-cyclic nucleotide 3' phosphod | Cytoplasm | enzyme | 4190769 | 2737421.25 | 3872331.5 | | | | | | |
| P16330 | CNP | 2',3'-cyclic nucleotide 3' phosphod | Cytoplasm | enzyme | 5665685.5 | 1774229.625 | 2671656.5 | | | | | | |
| P16330 | CNP | 2',3'-cyclic nucleotide 3' phosphod | Cytoplasm | enzyme | 930044.312 | 3985902.25 | 5627157.5 | P16330 | CNP | 9390951 | 9008552 | 1838863 | 24 |
| Q9QXT0 | CNPY2 | canopy FGF signaling regulator 2 | Plasma Membrane | other | 5901154 | 6794639 | 5625834 | | | | | | |
| Q9QXT0 | CNPY2 | canopy FGF signaling regulator 2 | Plasma Membrane | other | 6527204.5 | 5670263.5 | 5351300 | | | | | | |
| Q9QXT0 | CNPY2 | canopy FGF signaling regulator 2 | Plasma Membrane | other | 2524755 | 2873009 | 3158998.25 | Q9QXT0 | CNPY2 | 4984371 | 1632990 | 544330 | 9 |
| Q8BQ47 | CNPY4 | canopy FGF signaling regulator 4 | Extracellular Space | other | 1848511.375 | 2354804.5 | 1938583 | Q8BQ47 | CNPY4 | 1848511 | 270088 | 155936 | 3 |
| O88587 | COMT | catechol-O-methyltransferase | Cytoplasm | enzyme | 4626540.5 | 8139940.5 | 4855803 | | | | | | |
| O88587 | COMT | catechol-O-methyltransferase | Cytoplasm | enzyme | 7679575.5 | 8511996 | 4749793.5 | | | | | | |
| O88587 | COMT | catechol-O-methyltransferase | Cytoplasm | enzyme | 4670467 | 4961631.5 | 3457996 | | | | | | |
| O88587 | COMT | catechol-O-methyltransferase | Cytoplasm | enzyme | 4385249 | 5484482.5 | 3443729.75 | O88587 | COMT | 5340458 | 1733696 | 500475 | 12 |
| Q8BIG7 | COMTD1 | catechol-O-methyltransferase don | Plasma Membrane | enzyme | 3808231.5 | 4761323 | 5664664 | Q8BIG7 | COMTD1 | 3808232 | 928327 | 535970 | 3 |
| P61202 | Cops2 | COP9 (constitutive photomorphog | Nucleus | transcription regulator | 6952522.5 | 7028388.5 | 5773690 | | | | | | |
| P61202 | Cops2 | COP9 (constitutive photomorphog | Nucleus | transcription regulator | 4587460 | 3174559.75 | 3282222.25 | | | | | | |
| P61202 | Cops2 | COP9 (constitutive photomorphog | Nucleus | transcription regulator | 1920382.375 | 2281121.75 | 2431571.25 | | | | | | |
| P61202 | Cops2 | COP9 (constitutive photomorphog | Nucleus | transcription regulator | 1588685.375 | 1178956.625 | 1057676.875 | | | | | | |
| P61202 | Cops2 | COP9 (constitutive photomorphog | Nucleus | transcription regulator | 308192.312 | 170755.719 | 25411.814 | P61202 | Cops2 | 3071449 | 2340016 | 604189 | 15 |
| O88543 | COPS3 | COP9 signalosome subunit 3 | Cytoplasm | other | 4436958.5 | 2910861 | 2876908 | | | | | | |
| O88543 | COPS3 | COP9 signalosome subunit 3 | Cytoplasm | other | 3758886 | 3228887 | 3357941.75 | | | | | | |
| O88543 | COPS3 | COP9 signalosome subunit 3 | Cytoplasm | other | 3563053.25 | 3254514.25 | 2274703.75 | | | | | | |
| O88543 | COPS3 | COP9 signalosome subunit 3 | Cytoplasm | other | 1957344 | 1393189.125 | 1250548.125 | | | | | | |
| O88543 | COPS3 | COP9 signalosome subunit 3 | Cytoplasm | other | 1467290.625 | 1105153.625 | 1051533.375 | | | | | | |
| O88543 | COPS3 | COP9 signalosome subunit 3 | Cytoplasm | other | 1460229.5 | 2152395.25 | 3522970.25 | O88543 | COPS3 | 2773960 | 1060081 | 249863 | 18 |
| O88544 | COPS4 | COP9 signalosome subunit 4 | Cytoplasm | other | 3568870.25 | 3902295.25 | 3522533.25 | | | | | | |
| O88544 | COPS4 | COP9 signalosome subunit 4 | Cytoplasm | other | 2731126 | 2612636 | 3008047 | | | | | | |
| O88544 | COPS4 | COP9 signalosome subunit 4 | Cytoplasm | other | 1664429.875 | 1801137.625 | 1753703.375 | | | | | | |
| O88544 | COPS4 | COP9 signalosome subunit 4 | Cytoplasm | other | 2972311.5 | 376594 | 521136.75 | | | | | | |
| O88544 | COPS4 | COP9 signalosome subunit 4 | Cytoplasm | other | 1350527.125 | 2068845.75 | 2444567.75 | | | | | | |
| O88544 | COPS4 | COP9 signalosome subunit 4 | Cytoplasm | other | 1339976.25 | 1741681.5 | 1643933 | O88544 | COPS4 | 2271207 | 1001472 | 236049 | 18 |
| O35864 | COPS5 | COP9 signalosome subunit 5 | Nucleus | transcription regulator | 7712161 | 7598358 | 6502452.5 | | | | | | |
| O35864 | COPS5 | COP9 signalosome subunit 5 | Nucleus | transcription regulator | 2297771 | 1499379.875 | 2016605.625 | O35864 | COPS5 | 5004966 | 2962530 | 1209448 | 6 |
| O88545 | COPS6 | COP9 signalosome subunit 6 | Nucleus | other | 3842455 | 6273184.5 | 9197324 | | | | | | |
| O88545 | COPS6 | COP9 signalosome subunit 6 | Nucleus | other | 5195441 | 4245291 | 6171823.5 | | | | | | |
| O88545 | COPS6 | COP9 signalosome subunit 6 | Nucleus | other | 4510388.5 | 4637864 | 4513805.5 | | | | | | |
| O88545 | COPS6 | COP9 signalosome subunit 6 | Nucleus | other | 1306674.875 | 1267146.125 | 1070793.125 | | | | | | |
| O88545 | COPS6 | COP9 signalosome subunit 6 | Nucleus | other | 1497018.875 | 1351258 | 1038680.125 | O88545 | COPS6 | 3270396 | 2444975 | 631290 | 15 |
| Q9CZ04 | COPS7A | COP9 signalosome subunit 7A | Cytoplasm | other | 743125.188 | 613611.125 | 657370.438 | Q9CZ04 | COPS7A | 743125 | 65882 | 38037 | 3 |
| Q8BV13 | COPS7B | COP9 signalosome subunit 7B | Cytoplasm | other | 1877084 | 1452714.25 | 1890317.75 | Q8BV13 | COPS7B | 1877084 | 248918 | 143713 | 3 |
| Q8VBV7 | COPS8 | COP9 signalosome subunit 8 | Nucleus | other | 1864468.875 | 2383142.75 | 2072595 | Q8VBV7 | COPS8 | 1864469 | 261017 | 150698 | 3 |
| O89053 | CORO1A | coronin, actin binding protein, 1A | Cytoplasm | other | 4.09E+07 | 5.36E+07 | 6.14E+07 | | | | | | |
| O89053 | CORO1A | coronin, actin binding protein, 1A | Cytoplasm | other | 6388574.5 | 1.01E+07 | 7945649.5 | | | | | | |
| O89053 | CORO1A | coronin, actin binding protein, 1A | Cytoplasm | other | 2.07E+07 | 3.16E+07 | 3.03E+07 | | | | | | |
| O89053 | CORO1A | coronin, actin binding protein, 1A | Cytoplasm | other | 2.33E+07 | 1.31E+07 | 9422515 | | | | | | |
| O89053 | CORO1A | coronin, actin binding protein, 1A | Cytoplasm | other | 6217753 | 5374874.5 | 6537513.5 | | | | | | |
| O89053 | CORO1A | coronin, actin binding protein, 1A | Cytoplasm | other | 1.25E+07 | 1.55E+07 | 1.46E+07 | | | | | | |
| O89053 | CORO1A | coronin, actin binding protein, 1A | Cytoplasm | other | 8530389 | 948742 | 1198191.875 | | | | | | |
| O89053 | CORO1A | coronin, actin binding protein, 1A | Cytoplasm | other | 5127017.5 | 5124542.5 | 6130738.5 | | | | | | |
| O89053 | CORO1A | coronin, actin binding protein, 1A | Cytoplasm | other | 5495924.5 | 1.02E+07 | 7668930 | | | | | | |
| O89053 | CORO1A | coronin, actin binding protein, 1A | Cytoplasm | other | 6031617.5 | 7517163.5 | 7236073.5 | | | | | | |
| O89053 | CORO1A | coronin, actin binding protein, 1A | Cytoplasm | other | 1.42E+07 | 1.58E+07 | 1.58E+07 | | | | | | |
| O89053 | CORO1A | coronin, actin binding protein, 1A | Cytoplasm | other | 4620410 | 9934545 | 2.06E+07 | | | | | | |
| O89053 | CORO1A | coronin, actin binding protein, 1A | Cytoplasm | other | 6208398.5 | 5811629.5 | 5784330 | O89053 | CORO1A | 12318468 | 13351955 | 2138024 | 39 |
| Q9WUM4 | CORO1C | coronin, actin binding protein, 1C | Cytoplasm | other | 9615806 | 7263166.5 | 8725381 | | | | | | |
| Q9WUM4 | CORO1C | coronin, actin binding protein, 1C | Cytoplasm | other | 1668155 | 3906909 | 3901321.5 | Q9WUM4 | CORO1C | 5641981 | 3146645 | 1284612 | 6 |
| P19783 | COX4I1 | cytochrome c oxidase subunit IV is | Cytoplasm | enzyme | 2.31E+07 | 2.37E+07 | 2.93E+07 | | | | | | |
| P19783 | COX4I1 | cytochrome c oxidase subunit IV is | Cytoplasm | enzyme | 1.15E+07 | 1.23E+07 | 1.50E+07 | | | | | | |
| P19783 | COX4I1 | cytochrome c oxidase subunit IV is | Cytoplasm | enzyme | 5059801 | 5975705 | 6686390.5 | P19783 | COX4I1 | 13213267 | 8760659 | 2920220 | 9 |
| P12787 | COX5A | cytochrome c oxidase subunit Va | Cytoplasm | enzyme | 1.33E+07 | 1.34E+07 | 1.72E+07 | | | | | | |
| P12787 | COX5A | cytochrome c oxidase subunit Va | Cytoplasm | enzyme | 7102689.5 | 1.49E+07 | 2.09E+07 | | | | | | |
| P12787 | COX5A | cytochrome c oxidase subunit Va | Cytoplasm | enzyme | 6282337 | 5764117.5 | 7085656.5 | P12787 | COX5A | 8905009 | 5453673 | 1817891 | 9 |
| P19536 | Cox5b | cytochrome c oxidase subunit Vb | Cytoplasm | other | 1.37E+07 | 1.21E+07 | 1.64E+07 | P19536 | Cox5b | 13670000 | 2203845 | 1272391 | 3 |
| P56391 | COX6B1 | cytochrome c oxidase subunit VIb | Cytoplasm | enzyme | 6822950 | 6807406 | 5293774 | P56391 | COX6B1 | 6822950 | 878417 | 507155 | 3 |
| Q9CPQ1 | Cox6c | cytochrome c oxidase subunit VIc | Cytoplasm | enzyme | 1898905 | 2419565.25 | 3397303.25 | Q9CPQ1 | Cox6c | 1898905 | 760729 | 439207 | 3 |
| P48771 | COX7A2 | cytochrome c oxidase subunit VIIa | Cytoplasm | enzyme | 6604115.5 | 7081054 | 8655708 | P48771 | COX7A2 | 6604116 | 1073626 | 619858 | 3 |
| P59108 | CPNE2 | copine II | Cytoplasm | other | 3506303.75 | 4371944 | 2994200 | | | | | | |
| P59108 | CPNE2 | copine II | Cytoplasm | other | 2751151 | 4706449 | 2335119.75 | P59108 | CPNE2 | 3128727 | 934956 | 381694 | 6 |
| P36552 | CPOX | coproporphyrinogen oxidase | Cytoplasm | enzyme | 1.46E+07 | 9145915 | 1.37E+07 | | | | | | |
| P36552 | CPOX | coproporphyrinogen oxidase | Cytoplasm | enzyme | 1.15E+07 | 1.17E+07 | 1.58E+07 | | | | | | |
| P36552 | CPOX | coproporphyrinogen oxidase | Cytoplasm | enzyme | 1.24E+07 | 1.19E+07 | 1.69E+07 | | | | | | |
| P36552 | CPOX | coproporphyrinogen oxidase | Cytoplasm | enzyme | 1.30E+07 | 1.05E+07 | 1.24E+07 | | | | | | |
| P36552 | CPOX | coproporphyrinogen oxidase | Cytoplasm | enzyme | 8017276.5 | 1.05E+07 | 1.31E+07 | | | | | | |
| P36552 | CPOX | coproporphyrinogen oxidase | Cytoplasm | enzyme | 8844763 | 6860959 | 1.06E+07 | | | | | | |
| P36552 | CPOX | coproporphyrinogen oxidase | Cytoplasm | enzyme | 6785015.5 | 6061150.5 | 8949435 | | | | | | |
| P36552 | CPOX | coproporphyrinogen oxidase | Cytoplasm | enzyme | 5581438 | 7683496.5 | 1.03E+07 | | | | | | |
| P36552 | CPOX | coproporphyrinogen oxidase | Cytoplasm | enzyme | 7458334.5 | 5909135.5 | 1.22E+07 | | | | | | |
| P36552 | CPOX | coproporphyrinogen oxidase | Cytoplasm | enzyme | 5800488.5 | 4039626.75 | 8378078.5 | | | | | | |
| P36552 | CPOX | coproporphyrinogen oxidase | Cytoplasm | enzyme | 3298880.75 | 2515323.5 | 2594550.5 | | | | | | |
| P36552 | CPOX | coproporphyrinogen oxidase | Cytoplasm | enzyme | 2144650 | 1769435.875 | 2204012.75 | P36552 | CPOX | 8288404 | 4133894 | 688982 | 36 |
| P52825 | CPT2 | carnitine palmitoyltransferase 2 | Cytoplasm | enzyme | 5334923.5 | 2584711 | 5230418.5 | | | | | | |

| | | | | | | | | | | | | | |
|---|---|---|---|---|---|---|---|---|---|---|---|---|---|
| P52825 | CPT2 | carnitine palmitoyltransferase 2 | Cytoplasm | enzyme | 1894105 | 3179392.25 | 3789095.25 | | | | | | |
| P52825 | CPT2 | carnitine palmitoyltransferase 2 | Cytoplasm | enzyme | 850690.25 | 1046899.312 | 1247968.875 | P52825 | CPT2 | 2693240 | 1717694 | 572565 | 9 |
| Q64735 | CR1L | complement component (3b/4b) r | Plasma Membrane | other | 5.01E+07 | 6.29E+07 | 4.58E+07 | | | | | | |
| Q64735 | CR1L | complement component (3b/4b) r | Plasma Membrane | other | 3.37E+07 | 3.26E+07 | 2.55E+07 | | | | | | |
| Q64735 | CR1L | complement component (3b/4b) r | Plasma Membrane | other | 2.84E+07 | 2.58E+07 | 2.20E+07 | | | | | | |
| Q64735 | CR1L | complement component (3b/4b) r | Plasma Membrane | other | 2.14E+07 | 2.35E+07 | 2.08E+07 | | | | | | |
| Q64735 | CR1L | complement component (3b/4b) r | Plasma Membrane | other | 7115841.5 | 1.00E+07 | 6700578 | | | | | | |
| Q64735 | CR1L | complement component (3b/4b) r | Plasma Membrane | other | 5762688 | 7086900.5 | 5488267.5 | | | | | | |
| Q64735 | CR1L | complement component (3b/4b) r | Plasma Membrane | other | 9659487 | 5663534 | 5469700 | | | | | | |
| Q64735 | CR1L | complement component (3b/4b) r | Plasma Membrane | other | 5201632 | 3762518.75 | 2484131 | | | | | | |
| Q64735 | CR1L | complement component (3b/4b) r | Plasma Membrane | other | 1188715.875 | 1304358 | 1303521 | Q64735 | CR1L | 18060929 | 16550413 | 3185129 | 27 |
| Q9Z125 | CREB3L1 | cAMP responsive element binding | Nucleus | transcription regulator | 3431786.75 | 3776714.25 | 4974322.5 | Q9Z125 | CREB3L1 | 3431787 | 809594 | 467419 | 3 |
| O88668 | CREG1 | cellular repressor of E1A-stimulate | Nucleus | transcription regulator | 5101440 | 5696968 | 6499286 | | | | | | |
| O88668 | CREG1 | cellular repressor of E1A-stimulate | Nucleus | transcription regulator | 4030366.75 | 4599445 | 5301513.5 | O88668 | CREG1 | 4565903 | 858337 | 350414 | 6 |
| Q9DCT8 | Crip2 | cysteine rich protein 2 | Plasma Membrane | other | 3.33E+07 | 3.85E+07 | 3.25E+07 | | | | | | |
| Q9DCT8 | Crip2 | cysteine rich protein 2 | Plasma Membrane | other | 2.47E+07 | 5.26E+07 | 3.33E+07 | | | | | | |
| Q9DCT8 | Crip2 | cysteine rich protein 2 | Plasma Membrane | other | 2.51E+07 | 3.40E+07 | 2.10E+07 | | | | | | |
| Q9DCT8 | Crip2 | cysteine rich protein 2 | Plasma Membrane | other | 2.45E+07 | 4.50E+07 | 3.28E+07 | | | | | | |
| Q9DCT8 | Crip2 | cysteine rich protein 2 | Plasma Membrane | other | 1.75E+07 | 2.03E+07 | 1.06E+07 | | | | | | |
| Q9DCT8 | Crip2 | cysteine rich protein 2 | Plasma Membrane | other | 1.26E+07 | 2.17E+07 | 8120762 | | | | | | |
| Q9DCT8 | Crip2 | cysteine rich protein 2 | Plasma Membrane | other | 1.01E+07 | 1.26E+07 | 6794574 | | | | | | |
| Q9DCT8 | Crip2 | cysteine rich protein 2 | Plasma Membrane | other | 9909173 | 1.49E+07 | 6521038.5 | | | | | | |
| Q9DCT8 | Crip2 | cysteine rich protein 2 | Plasma Membrane | other | 1.19E+07 | 822657.062 | 0 | Q9DCT8 | Crip2 | 18833241 | 13469988 | 2592300 | 27 |
| Q9Z2L7 | CRLF3 | cytokine receptor-like factor 3 | Cytoplasm | transcription regulator | 6335565.5 | 3310453.5 | 3569567 | Q9Z2L7 | CRLF3 | 6335566 | 1676762 | 968079 | 3 |
| Q9CZU6 | CS | citrate synthase | Cytoplasm | enzyme | 3.28E+07 | 3.10E+07 | 4.33E+07 | | | | | | |
| Q9CZU6 | CS | citrate synthase | Cytoplasm | enzyme | 1.26E+07 | 1.61E+07 | 1.15E+07 | | | | | | |
| Q9CZU6 | CS | citrate synthase | Cytoplasm | enzyme | 1.71E+07 | 2.90E+07 | 2.90E+07 | | | | | | |
| Q9CZU6 | CS | citrate synthase | Cytoplasm | enzyme | 8256615.5 | 9764743 | 1.37E+07 | | | | | | |
| Q9CZU6 | CS | citrate synthase | Cytoplasm | enzyme | 7860374.5 | 1.05E+07 | 1.45E+07 | | | | | | |
| Q9CZU6 | CS | citrate synthase | Cytoplasm | enzyme | 1.09E+07 | 8862336 | 1.32E+07 | | | | | | |
| Q9CZU6 | CS | citrate synthase | Cytoplasm | enzyme | 1.29E+07 | 2871938.25 | 1.06E+07 | | | | | | |
| Q9CZU6 | CS | citrate synthase | Cytoplasm | enzyme | 6430969 | 3565052 | 4169557 | | | | | | |
| Q9CZU6 | CS | citrate synthase | Cytoplasm | enzyme | 4957674.5 | 4624188 | 4946220 | | | | | | |
| Q9CZU6 | CS | citrate synthase | Cytoplasm | enzyme | 3753019.25 | 4149556.75 | 6418525 | Q9CZU6 | CS | 11750865 | 10154675 | 1853981 | 30 |
| Q8BK63 | CSNK1A1 | casein kinase 1, alpha 1 | Cytoplasm | kinase | 9546768 | 7250752 | 9366224 | | | | | | |
| Q8BK63 | CSNK1A1 | casein kinase 1, alpha 1 | Cytoplasm | kinase | 8895995 | 1371285 | 543719.188 | | | | | | |
| Q8BK63 | CSNK1A1 | casein kinase 1, alpha 1 | Cytoplasm | kinase | 6221738 | 6011301.5 | 9973706 | | | | | | |
| Q8BK63 | CSNK1A1 | casein kinase 1, alpha 1 | Cytoplasm | kinase | 4813845 | 4139588.75 | 5679398.5 | | | | | | |
| Q8BK63 | CSNK1A1 | casein kinase 1, alpha 1 | Cytoplasm | kinase | 4450981.5 | 5361100.5 | 5691733 | | | | | | |
| Q8BK63 | CSNK1A1 | casein kinase 1, alpha 1 | Cytoplasm | kinase | 3170399.75 | 2990389 | 3417284.5 | | | | | | |
| Q8BK63 | CSNK1A1 | casein kinase 1, alpha 1 | Cytoplasm | kinase | 3257123 | 4057838.25 | 3827402 | | | | | | |
| Q8BK63 | CSNK1A1 | casein kinase 1, alpha 1 | Cytoplasm | kinase | 1875700.75 | 2198996 | 1243093.125 | Q8BK63 | CSNK1A1 | 5279069 | 2706045 | 552369 | 24 |
| Q60737 | CSNK2A1 | casein kinase 2, alpha 1 polypeptic | Nucleus | kinase | 2369045.25 | 2216705.5 | 1994346.125 | Q60737 | CSNK2A1 | 2369045 | 188437 | 108794 | 3 |
| P67871 | CSNK2B | casein kinase 2, beta polypeptide | Cytoplasm | kinase | 2876129.5 | 3480966 | 2370355.5 | P67871 | CSNK2B | 2876130 | 556041 | 321030 | 3 |
| P97315 | CSRP1 | cysteine and glycine-rich protein 1 | Nucleus | other | 2.65E+07 | 3.15E+07 | 2.50E+07 | | | | | | |
| P97315 | CSRP1 | cysteine and glycine-rich protein 1 | Nucleus | other | 2.38E+07 | 3.10E+07 | 3.02E+07 | | | | | | |
| P97315 | CSRP1 | cysteine and glycine-rich protein 1 | Nucleus | other | 2.22E+07 | 2.66E+07 | 2.22E+07 | | | | | | |
| P97315 | CSRP1 | cysteine and glycine-rich protein 1 | Nucleus | other | 1.78E+07 | 2.14E+07 | 1.65E+07 | | | | | | |
| P97315 | CSRP1 | cysteine and glycine-rich protein 1 | Nucleus | other | 4791477 | 3545879 | 3708586.25 | P97315 | CSRP1 | 19044295 | 9547731 | 2465213 | 15 |
| P10605 | CTSB | cathepsin B | Cytoplasm | peptidase | 1.12E+07 | 1.55E+07 | 1.30E+07 | | | | | | |
| P10605 | CTSB | cathepsin B | Cytoplasm | peptidase | 9615856 | 1.60E+07 | 1.25E+07 | | | | | | |
| P10605 | CTSB | cathepsin B | Cytoplasm | peptidase | 9895461 | 1.15E+07 | 1.35E+07 | | | | | | |
| P10605 | CTSB | cathepsin B | Cytoplasm | peptidase | 5366646.5 | 1.06E+07 | 8487001 | | | | | | |
| P10605 | CTSB | cathepsin B | Cytoplasm | peptidase | 4549577.5 | 4814929.5 | 4355186 | P10605 | CTSB | 8129508 | 3881783 | 1002272 | 15 |
| P18242 | CTSD | cathepsin D | Cytoplasm | peptidase | 2.83E+07 | 1.91E+07 | 2.42E+07 | | | | | | |
| P18242 | CTSD | cathepsin D | Cytoplasm | peptidase | 1.41E+07 | 1.12E+07 | 1.35E+07 | | | | | | |
| P18242 | CTSD | cathepsin D | Cytoplasm | peptidase | 1.56E+07 | 1.57E+07 | 1.60E+07 | | | | | | |
| P18242 | CTSD | cathepsin D | Cytoplasm | peptidase | 1.18E+07 | 1.22E+07 | 1.70E+07 | | | | | | |
| P18242 | CTSD | cathepsin D | Cytoplasm | peptidase | 1.02E+07 | 9933251 | 1.16E+07 | | | | | | |
| P18242 | CTSD | cathepsin D | Cytoplasm | peptidase | 5068468 | 3891263.75 | 6010799.5 | | | | | | |
| P18242 | CTSD | cathepsin D | Cytoplasm | peptidase | 3067513.25 | 3596340 | 5880688 | P18242 | CTSD | 12573712 | 6638405 | 1448619 | 21 |
| P70269 | CTSE | cathepsin E | Cytoplasm | peptidase | 8.35E+07 | 1.25E+08 | 1.18E+08 | | | | | | |
| P70269 | CTSE | cathepsin E | Cytoplasm | peptidase | 8.49E+07 | 1.21E+08 | 9.24E+07 | | | | | | |
| P70269 | CTSE | cathepsin E | Cytoplasm | peptidase | 2.86E+07 | 1021903.188 | 8966125 | P70269 | CTSE | 65663333 | 48618132 | 16206044 | 9 |
| Q9WTX6 | CUL1 | cullin 1 | Nucleus | enzyme | 4146118.75 | 2438598 | 2742725.5 | | | | | | |
| Q9WTX6 | CUL1 | cullin 1 | Nucleus | enzyme | 3432104.75 | 3229452.75 | 3573004.75 | | | | | | |
| Q9WTX6 | CUL1 | cullin 1 | Nucleus | enzyme | 2629931.75 | 2619131.25 | 2451114 | | | | | | |
| Q9WTX6 | CUL1 | cullin 1 | Nucleus | enzyme | 2005105.625 | 2549619.5 | 1996323.5 | | | | | | |
| Q9WTX6 | CUL1 | cullin 1 | Nucleus | enzyme | 2102078.75 | 1506404.5 | 1916968 | | | | | | |
| Q9WTX6 | CUL1 | cullin 1 | Nucleus | enzyme | 1337877.25 | 1469142.375 | 1530862.125 | | | | | | |
| Q9WTX6 | CUL1 | cullin 1 | Nucleus | enzyme | 1272488.75 | 29141.814 | 497727.562 | | | | | | |
| Q9WTX6 | CUL1 | cullin 1 | Nucleus | enzyme | 1296734.875 | 355387.5 | 203469.125 | | | | | | |
| Q9WTX6 | CUL1 | cullin 1 | Nucleus | enzyme | 1202067.625 | 1176646.75 | 784029 | Q9WTX6 | CUL1 | 2158279 | 1061728 | 204330 | 27 |
| Q9JLV5 | CUL3 | cullin 3 | Nucleus | enzyme | 3572892 | 2791793.5 | 3070787.25 | | | | | | |
| Q9JLV5 | CUL3 | cullin 3 | Nucleus | enzyme | 2669824.5 | 2847742.25 | 2341640.25 | Q9JLV5 | CUL3 | 3121358 | 414557 | 169242 | 6 |
| P70658 | CXCR4 | chemokine (C-X-C motif) receptor | Plasma Membrane | G-protein coupled receptor | 3760118.5 | 3910230.75 | 4131016.5 | P70658 | CXCR4 | 3760119 | 186568 | 107715 | 3 |
| P56395 | CYB5A | cytochrome b5 type A (microsoma | Cytoplasm | enzyme | 1.20E+07 | 2.64E+07 | 1.06E+07 | | | | | | |
| P56395 | CYB5A | cytochrome b5 type A (microsoma | Cytoplasm | enzyme | 1.14E+07 | 1.74E+07 | 1.08E+07 | | | | | | |
| P56395 | CYB5A | cytochrome b5 type A (microsoma | Cytoplasm | enzyme | 2240951.75 | 5468194.5 | 3632904.25 | | | | | | |
| P56395 | CYB5A | cytochrome b5 type A (microsoma | Cytoplasm | enzyme | 2492779 | 6140474.5 | 2306018 | | | | | | |
| P56395 | CYB5A | cytochrome b5 type A (microsoma | Cytoplasm | enzyme | 5397509.5 | 3227022.75 | 3366382.75 | P56395 | CYB5A | 6714248 | 6790760 | 1753367 | 15 |
| Q9CQX2 | CYB5B | cytochrome b5 type B (outer mito | Cytoplasm | enzyme | 2.33E+07 | 2.09E+07 | 2.20E+07 | | | | | | |
| Q9CQX2 | CYB5B | cytochrome b5 type B (outer mito | Cytoplasm | enzyme | 2062294.875 | 995262.375 | 71574.094 | | | | | | |
| Q9CQX2 | CYB5B | cytochrome b5 type B (outer mito | Cytoplasm | enzyme | 930944.938 | 1221699 | 612793.938 | Q9CQX2 | CYB5B | 8767747 | 10573495 | 3524498 | 9 |
| Q9DCN2 | Cyb5r3 | cytochrome b5 reductase 3 | Cytoplasm | enzyme | 3.54E+07 | 4.56E+07 | 3.18E+07 | | | | | | |
| Q9DCN2 | Cyb5r3 | cytochrome b5 reductase 3 | Cytoplasm | enzyme | 8376768 | 3.21E+07 | 1.37E+07 | | | | | | |

| | | | | | | | | | | | | | |
|---|---|---|---|---|---|---|---|---|---|---|---|---|---|
| Q9DCN2 | Cyb5r3 | cytochrome b5 reductase 3 | Cytoplasm | enzyme | 2.29E+07 | 2.76E+07 | 2.20E+07 | | | | | | |
| Q9DCN2 | Cyb5r3 | cytochrome b5 reductase 3 | Cytoplasm | enzyme | 1.24E+07 | 1.40E+07 | 1.81E+07 | | | | | | |
| Q9DCN2 | Cyb5r3 | cytochrome b5 reductase 3 | Cytoplasm | enzyme | 1.27E+07 | 1.73E+07 | 1.28E+07 | | | | | | |
| Q9DCN2 | Cyb5r3 | cytochrome b5 reductase 3 | Cytoplasm | enzyme | 1.40E+07 | 1.37E+07 | 1.06E+07 | | | | | | |
| Q9DCN2 | Cyb5r3 | cytochrome b5 reductase 3 | Cytoplasm | enzyme | 7204791 | 1.05E+07 | 7371050.5 | | | | | | |
| Q9DCN2 | Cyb5r3 | cytochrome b5 reductase 3 | Cytoplasm | enzyme | 6218165.5 | 1.07E+07 | 7932531 | | | | | | |
| Q9DCN2 | Cyb5r3 | cytochrome b5 reductase 3 | Cytoplasm | enzyme | 7398934.5 | 8136567 | 7367756.5 | | | | | | |
| Q9DCN2 | Cyb5r3 | cytochrome b5 reductase 3 | Cytoplasm | enzyme | 5164653 | 5734619.5 | 5121971 | | | | | | |
| Q9DCN2 | Cyb5r3 | cytochrome b5 reductase 3 | Cytoplasm | enzyme | 3460829.25 | 5817843.5 | 6142900.5 | | | | | | |
| Q9DCN2 | Cyb5r3 | cytochrome b5 reductase 3 | Cytoplasm | enzyme | 1453004.375 | 8774100 | 4082188 | Q9DCN2 | Cyb5r3 | 11393095 | 10110483 | 1685080 | 36 |
| Q9D0M3 | CYC1 | cytochrome c-1 | Cytoplasm | enzyme | 1.83E+07 | 2.12E+07 | 1.97E+07 | Q9D0M3 | CYC1 | 18280000 | 1455644 | 840417 | 3 |
| P62897 | CYCS | cytochrome c, somatic | Cytoplasm | transporter | 1.04E+07 | 1.11E+07 | 1.15E+07 | | | | | | |
| P62897 | CYCS | cytochrome c, somatic | Cytoplasm | transporter | 4669116.5 | 4727594.5 | 4393803 | | | | | | |
| P62897 | CYCS | cytochrome c, somatic | Cytoplasm | transporter | 7495929 | 2112938.25 | 4262661 | | | | | | |
| P62897 | CYCS | cytochrome c, somatic | Cytoplasm | transporter | 3357050.75 | 4524184 | 4394696.5 | | | | | | |
| P62897 | CYCS | cytochrome c, somatic | Cytoplasm | transporter | 1061836.375 | 1529867 | 1631860.625 | P62897 | CYCS | 5400787 | 3441065 | 888479 | 15 |
| Q8BPM0 | DAAM1 | dishevelled associated activator of | Cytoplasm | other | 8639254 | 5044710.5 | 7681557.5 | | | | | | |
| Q8BPM0 | DAAM1 | dishevelled associated activator of | Cytoplasm | other | 5598811 | 6133310 | 5098763 | | | | | | |
| Q8BPM0 | DAAM1 | dishevelled associated activator of | Cytoplasm | other | 7524181 | 6240811.5 | 6476802 | | | | | | |
| Q8BPM0 | DAAM1 | dishevelled associated activator of | Cytoplasm | other | 7348454.5 | 4793036.5 | 5532783.5 | | | | | | |
| Q8BPM0 | DAAM1 | dishevelled associated activator of | Cytoplasm | other | 4084929.75 | 5651835.5 | 6301975 | | | | | | |
| Q8BPM0 | DAAM1 | dishevelled associated activator of | Cytoplasm | other | 2965355 | 2328104 | 2205108.75 | | | | | | |
| Q8BPM0 | DAAM1 | dishevelled associated activator of | Cytoplasm | other | 4220285.5 | 6553310.5 | 6756549 | | | | | | |
| Q8BPM0 | DAAM1 | dishevelled associated activator of | Cytoplasm | other | 2752719.25 | 3068545.5 | 3848318 | | | | | | |
| Q8BPM0 | DAAM1 | dishevelled associated activator of | Cytoplasm | other | 3517056.5 | 3362162.75 | 3820996.25 | | | | | | |
| Q8BPM0 | DAAM1 | dishevelled associated activator of | Cytoplasm | other | 2490289 | 2254185.5 | 2883079.25 | | | | | | |
| Q8BPM0 | DAAM1 | dishevelled associated activator of | Cytoplasm | other | 1323871.125 | 2085804.625 | 2006083.5 | | | | | | |
| Q8BPM0 | DAAM1 | dishevelled associated activator of | Cytoplasm | other | 2563478.75 | 148317.203 | 430878.625 | Q8BPM0 | DAAM1 | 4419057 | 2158284 | 359714 | 36 |
| P61804 | DAD1 | defender against cell death 1 | Cytoplasm | other | 2929263 | 3246854 | 3217554.75 | P61804 | DAD1 | 2929263 | 175516 | 101334 | 3 |
| Q91WC9 | DAGLB | diacylglycerol lipase, beta | Plasma Membrane | enzyme | 6792364 | 9976432 | 6558921.5 | | | | | | |
| Q91WC9 | DAGLB | diacylglycerol lipase, beta | Plasma Membrane | enzyme | 3824705.5 | 7462523 | 4986748.5 | | | | | | |
| Q91WC9 | DAGLB | diacylglycerol lipase, beta | Plasma Membrane | enzyme | 2514495.25 | 3922088.75 | 2654056 | | | | | | |
| Q91WC9 | DAGLB | diacylglycerol lipase, beta | Plasma Membrane | enzyme | 2118313 | 8211638 | 3941407 | | | | | | |
| Q91WC9 | DAGLB | diacylglycerol lipase, beta | Plasma Membrane | enzyme | 3544058.5 | 3237836.25 | 2911531 | | | | | | |
| Q91WC9 | DAGLB | diacylglycerol lipase, beta | Plasma Membrane | enzyme | 2036505 | 1327921.875 | 1952991.625 | Q91WC9 | DAGLB | 3471740 | 2467900 | 581690 | 18 |
| Q922B2 | DARS | aspartyl-tRNA synthetase | Cytoplasm | enzyme | 4724412.5 | 6683472.5 | 6621849 | | | | | | |
| Q922B2 | DARS | aspartyl-tRNA synthetase | Cytoplasm | enzyme | 5658694 | 5734165.5 | 3461623.75 | | | | | | |
| Q922B2 | DARS | aspartyl-tRNA synthetase | Cytoplasm | enzyme | 4376437.5 | 3512346.5 | 2680590.25 | | | | | | |
| Q922B2 | DARS | aspartyl-tRNA synthetase | Cytoplasm | enzyme | 3810120.25 | 3725306.75 | 2641949.25 | | | | | | |
| Q922B2 | DARS | aspartyl-tRNA synthetase | Cytoplasm | enzyme | 2378001.5 | 2316683 | 1531146 | | | | | | |
| Q922B2 | DARS | aspartyl-tRNA synthetase | Cytoplasm | enzyme | 2100875.5 | 1965309.875 | 1833242.25 | | | | | | |
| Q922B2 | DARS | aspartyl-tRNA synthetase | Cytoplasm | enzyme | 2408204 | 3299109.25 | 1716973.75 | | | | | | |
| Q922B2 | DARS | aspartyl-tRNA synthetase | Cytoplasm | enzyme | 2417758 | 2554718.75 | 1557631.25 | Q922B2 | DARS | 3484313 | 1566836 | 319829 | 24 |
| Q9JII5 | Dazap1 | DAZ associated protein 1 | Cytoplasm | other | 2252456.75 | 1779254 | 945128.25 | Q9JII5 | Dazap1 | 2252457 | 661916 | 382157 | 3 |
| Q9QZ73 | DCUN1D1 | DCN1, defective in cullin neddylati | Nucleus | other | 2672045.75 | 3220432.75 | 2175506.75 | Q9QZ73 | DCUN1D1 | 2672046 | 522677 | 301768 | 3 |
| A2ADY9 | DDI2 | DNA-damage inducible 1 homolog | Plasma Membrane | transporter | 6006832 | 5052378 | 4965124.5 | | | | | | |
| A2ADY9 | DDI2 | DNA-damage inducible 1 homolog | Plasma Membrane | transporter | 3148434.25 | 2936792.75 | 5020707.5 | A2ADY9 | DDI2 | 4577633 | 1210839 | 494323 | 6 |
| O54734 | DDOST | dolichyl-diphosphooligosaccharide | Cytoplasm | enzyme | 3791752.75 | 6199886 | 5334717.5 | | | | | | |
| O54734 | DDOST | dolichyl-diphosphooligosaccharide | Cytoplasm | enzyme | 2860424.75 | 4382703 | 4412328 | | | | | | |
| O54734 | DDOST | dolichyl-diphosphooligosaccharide | Cytoplasm | enzyme | 2440902.25 | 4857059 | 4427378 | O54734 | DDOST | 3031027 | 1162601 | 387534 | 9 |
| Q8C4S8 | DENND2A | DENN/MADD domain containing 2 | Cytoplasm | other | 2441461.5 | 2718762.5 | 3080972 | Q8C4S8 | DENND2A | 2441462 | 320693 | 185152 | 3 |
| Q91YP3 | DERA | deoxyribose-phosphate aldolase (p | Cytoplasm | enzyme | 1951351.625 | 2515591.5 | 2365102 | | | | | | |
| Q91YP3 | DERA | deoxyribose-phosphate aldolase (p | Cytoplasm | enzyme | 1399423.75 | 1202934 | 1213010.75 | | | | | | |
| Q91YP3 | DERA | deoxyribose-phosphate aldolase (p | Cytoplasm | enzyme | 1395540.75 | 1483250.875 | 880472.375 | Q91YP3 | DERA | 1582105 | 555421 | 185140 | 9 |
| Q99J56 | DERL1 | derlin 1 | Cytoplasm | other | 1.08E+07 | 1.59E+07 | 1.40E+07 | | | | | | |
| Q99J56 | DERL1 | derlin 1 | Cytoplasm | other | 4445162 | 5755934 | 5858501 | Q99J56 | DERL1 | 7602581 | 4796516 | 1958169 | 6 |
| Q8BNI4 | DERL2 | derlin 2 | Cytoplasm | other | 4145268.25 | 4710614 | 2785606.25 | Q8BNI4 | DERL2 | 4145268 | 989440 | 571254 | 3 |
| Q3U0B3 | DHRS11 | dehydrogenase/reductase (SDR fa | Cytoplasm | other | 5877451.5 | 2714917.25 | 5295416 | Q3U0B3 | DHRS11 | 5877452 | 1683219 | 971807 | 3 |
| A2ATU0 | DHTKD1 | dehydrogenase E1 and transketola | Cytoplasm | enzyme | 512878.812 | 260300.266 | 204787.453 | A2ATU0 | DHTKD1 | 512879 | 164214 | 94809 | 3 |
| Q80VY9 | DHX33 | DEAH (Asp-Glu-Ala-His) box polype | Nucleus | enzyme | 3971806 | 5848249.5 | 5750296 | Q80VY9 | DHX33 | 3971806 | 1056225 | 609812 | 3 |
| Q8BMF4 | DLAT | dihydrolipoamide S-acetyltransfer | Cytoplasm | enzyme | 2972198 | 4079477.25 | 4051451.5 | | | | | | |
| Q8BMF4 | DLAT | dihydrolipoamide S-acetyltransfer | Cytoplasm | enzyme | 2552736.25 | 2721324.5 | 3879354.5 | Q8BMF4 | DLAT | 2762467 | 703406 | 287164 | 6 |
| O08749 | DLD | dihydrolipoamide dehydrogenase | Cytoplasm | enzyme | 1.59E+07 | 2.03E+07 | 2.56E+07 | | | | | | |
| O08749 | DLD | dihydrolipoamide dehydrogenase | Cytoplasm | enzyme | 1.12E+07 | 1.21E+07 | 1.50E+07 | | | | | | |
| O08749 | DLD | dihydrolipoamide dehydrogenase | Cytoplasm | enzyme | 7559651.5 | 7210495 | 7590468 | O08749 | DLD | 11556551 | 6305246 | 2101749 | 9 |
| Q811D0 | DLG1 | discs, large homolog 1 (Drosophila | Plasma Membrane | kinase | 1817706.875 | 2487432.5 | 1219697.875 | | | | | | |
| Q811D0 | DLG1 | discs, large homolog 1 (Drosophila | Plasma Membrane | kinase | 1602549.625 | 967116.562 | 438444 | | | | | | |
| Q811D0 | DLG1 | discs, large homolog 1 (Drosophila | Plasma Membrane | kinase | 584208.812 | 787116.938 | 292578.406 | Q811D0 | DLG1 | 1334822 | 721833 | 240611 | 9 |
| Q9D2G2 | DLST | dihydrolipoamide S-succinyltransfe | Cytoplasm | enzyme | 1.43E+07 | 1.28E+07 | 1.76E+07 | | | | | | |
| Q9D2G2 | DLST | dihydrolipoamide S-succinyltransfe | Cytoplasm | enzyme | 7747031 | 8623312 | 1.10E+07 | | | | | | |
| Q9D2G2 | DLST | dihydrolipoamide S-succinyltransfe | Cytoplasm | enzyme | 8332336.5 | 7517615.5 | 1.35E+07 | | | | | | |
| Q9D2G2 | DLST | dihydrolipoamide S-succinyltransfe | Cytoplasm | enzyme | 1.07E+07 | 3274680 | 9413495 | | | | | | |
| Q9D2G2 | DLST | dihydrolipoamide S-succinyltransfe | Cytoplasm | enzyme | 4701282 | 1.07E+07 | 9947100 | | | | | | |
| Q9D2G2 | DLST | dihydrolipoamide S-succinyltransfe | Cytoplasm | enzyme | 2933641.5 | 8629439 | 9215421 | | | | | | |
| Q9D2G2 | DLST | dihydrolipoamide S-succinyltransfe | Cytoplasm | enzyme | 3709165 | 4723281 | 6041340 | Q9D2G2 | DLST | 7489065 | 3831894 | 836188 | 21 |
| Q80WT2 | DMRT3 | doublesex and mab-3 related tran | Nucleus | transcription regulator | 4519629 | 5749171.5 | 4702136.5 | Q80WT2 | DMRT3 | 4519629 | 663497 | 383070 | 3 |
| Q9WV69 | DMTN | dematin actin binding protein | Plasma Membrane | other | 3.56E+08 | 6.36E+08 | 4.86E+08 | | | | | | |
| Q9WV69 | DMTN | dematin actin binding protein | Plasma Membrane | other | 4.49E+08 | 6.73E+08 | 6.01E+08 | | | | | | |
| Q9WV69 | DMTN | dematin actin binding protein | Plasma Membrane | other | 3.77E+08 | 3.56E+08 | 2.94E+08 | | | | | | |
| Q9WV69 | DMTN | dematin actin binding protein | Plasma Membrane | other | 3.42E+08 | 2.91E+08 | 2.70E+08 | | | | | | |
| Q9WV69 | DMTN | dematin actin binding protein | Plasma Membrane | other | 4.10E+08 | 3.66E+08 | 3.47E+08 | | | | | | |
| Q9WV69 | DMTN | dematin actin binding protein | Plasma Membrane | other | 2.99E+08 | 3.10E+08 | 3.19E+08 | | | | | | |
| Q9WV69 | DMTN | dematin actin binding protein | Plasma Membrane | other | 3.95E+08 | 3.23E+08 | 3.25E+08 | | | | | | |
| Q9WV69 | DMTN | dematin actin binding protein | Plasma Membrane | other | 3.25E+08 | 3.16E+08 | 3.12E+08 | | | | | | |
| Q9WV69 | DMTN | dematin actin binding protein | Plasma Membrane | other | 3.06E+08 | 1.66E+08 | 1.55E+08 | | | | | | |
| Q9WV69 | DMTN | dematin actin binding protein | Plasma Membrane | other | 2.06E+08 | 2.04E+08 | 1.96E+08 | | | | | | |

| | | | | | | | | | | | | | |
|---|---|---|---|---|---|---|---|---|---|---|---|---|---|
| Q9WV69 | DMTN | dematin actin binding protein | Plasma Membrane | other | 2.07E+08 | 1.65E+08 | 1.73E+08 | | | | | | |
| Q9WV69 | DMTN | dematin actin binding protein | Plasma Membrane | other | 8.64E+07 | 1.80E+08 | 1.14E+08 | | | | | | |
| Q9WV69 | DMTN | dematin actin binding protein | Plasma Membrane | other | 1.18E+08 | 6.05E+07 | 7.23E+07 | | | | | | |
| Q9WV69 | DMTN | dematin actin binding protein | Plasma Membrane | other | 6.22E+07 | 1.14E+08 | 2.16E+08 | | | | | | |
| Q9WV69 | DMTN | dematin actin binding protein | Plasma Membrane | other | 5.65E+07 | 1.04E+08 | 1.96E+08 | | | | | | |
| Q9WV69 | DMTN | dematin actin binding protein | Plasma Membrane | other | 3.47E+07 | 6.36E+07 | 6.93E+07 | | | | | | |
| Q9WV69 | DMTN | dematin actin binding protein | Plasma Membrane | other | 3981590.75 | 4.18E+07 | 2.62E+07 | | | | | | |
| Q9WV69 | DMTN | dematin actin binding protein | Plasma Membrane | other | 1.82E+07 | 1.84E+07 | 2.13E+07 | | | | | | |
| Q9WV69 | DMTN | dematin actin binding protein | Plasma Membrane | other | 1.51E+07 | 1.88E+07 | 1.38E+07 | | | | | | |
| Q9WV69 | DMTN | dematin actin binding protein | Plasma Membrane | other | 1.27E+07 | 1.19E+07 | 1.22E+07 | | | | | | |
| Q9WV69 | DMTN | dematin actin binding protein | Plasma Membrane | other | 1.04E+07 | 2639490 | 2871756 | | | | | | |
| Q9WV69 | DMTN | dematin actin binding protein | Plasma Membrane | other | 8029880.5 | 8481590 | 8204292.5 | | | | | | |
| Q9WV69 | DMTN | dematin actin binding protein | Plasma Membrane | other | 5078337.5 | 5247606 | 5472428 | | | | | | |
| Q9WV69 | DMTN | dematin actin binding protein | Plasma Membrane | other | 1.05E+07 | 184766.594 | 3166357.75 | | | | | | |
| Q9WV69 | DMTN | dematin actin binding protein | Plasma Membrane | other | 1724589.75 | 2327390.25 | 723629.875 | Q9WV69 | DMTN | 164605776 | 171780252 | 19835475 | 75 |
| Q9JMC3 | DNAJA4 | DnaJ (Hsp40) homolog, subfamily | Nucleus | other | 3190072.5 | 1620414.375 | 1979343.625 | Q9JMC3 | DNAJA4 | 3190073 | 822448 | 474840 | 3 |
| Q9QYJ3 | DNAJB1 | DnaJ (Hsp40) homolog, subfamily | Nucleus | other | 1.50E+07 | 1.85E+07 | 1.65E+07 | | | | | | |
| Q9QYJ3 | DNAJB1 | DnaJ (Hsp40) homolog, subfamily | Nucleus | other | 1.06E+07 | 9776371 | 1.08E+07 | | | | | | |
| Q9QYJ3 | DNAJB1 | DnaJ (Hsp40) homolog, subfamily | Nucleus | other | 9731864 | 7139609 | 5267453 | | | | | | |
| Q9QYJ3 | DNAJB1 | DnaJ (Hsp40) homolog, subfamily | Nucleus | other | 8777566 | 1.48E+07 | 1.17E+07 | | | | | | |
| Q9QYJ3 | DNAJB1 | DnaJ (Hsp40) homolog, subfamily | Nucleus | other | 4295023.5 | 3940321.75 | 3775667.25 | | | | | | |
| Q9QYJ3 | DNAJB1 | DnaJ (Hsp40) homolog, subfamily | Nucleus | other | 4129323.5 | 6628533.5 | 4592604.5 | | | | | | |
| Q9QYJ3 | DNAJB1 | DnaJ (Hsp40) homolog, subfamily | Nucleus | other | 3266367.5 | 3957051.25 | 3279129.5 | | | | | | |
| Q9QYJ3 | DNAJB1 | DnaJ (Hsp40) homolog, subfamily | Nucleus | other | 2778645.25 | 2032567 | 2843271.75 | | | | | | |
| Q9QYJ3 | DNAJB1 | DnaJ (Hsp40) homolog, subfamily | Nucleus | other | 4.70E-06 | 2324982 | 2900253.5 | Q9QYJ3 | DNAJB1 | 6505421 | 4971292 | 956726 | 27 |
| Q99KV1 | DNAJB11 | DnaJ (Hsp40) homolog, subfamily | Cytoplasm | other | 4991597.5 | 5720503 | 4761393.5 | | | | | | |
| Q99KV1 | DNAJB11 | DnaJ (Hsp40) homolog, subfamily | Cytoplasm | other | 3293813.25 | 4059748.25 | 2654896.5 | | | | | | |
| Q99KV1 | DNAJB11 | DnaJ (Hsp40) homolog, subfamily | Cytoplasm | other | 2054983.375 | 3073833 | 1830203.625 | Q99KV1 | DNAJB11 | 3446798 | 1359052 | 453017 | 9 |
| Q9D832 | DNAJB4 | DnaJ (Hsp40) homolog, subfamily | Nucleus | other | 2.19E+07 | 2.11E+07 | 2.33E+07 | | | | | | |
| Q9D832 | DNAJB4 | DnaJ (Hsp40) homolog, subfamily | Nucleus | other | 1.60E+07 | 1.67E+07 | 1.43E+07 | | | | | | |
| Q9D832 | DNAJB4 | DnaJ (Hsp40) homolog, subfamily | Nucleus | other | 1.23E+07 | 1.38E+07 | 1.38E+07 | | | | | | |
| Q9D832 | DNAJB4 | DnaJ (Hsp40) homolog, subfamily | Nucleus | other | 5738530.5 | 4330931 | 3810376.75 | | | | | | |
| Q9D832 | DNAJB4 | DnaJ (Hsp40) homolog, subfamily | Nucleus | other | 8326420 | 7299986.5 | 8692086 | | | | | | |
| Q9D832 | DNAJB4 | DnaJ (Hsp40) homolog, subfamily | Nucleus | other | 7787154 | 4362018.5 | 6823230 | | | | | | |
| Q9D832 | DNAJB4 | DnaJ (Hsp40) homolog, subfamily | Nucleus | other | 6496087 | 5825170.5 | 5573912 | | | | | | |
| Q9D832 | DNAJB4 | DnaJ (Hsp40) homolog, subfamily | Nucleus | other | 5045401.5 | 4535337 | 6102351.5 | | | | | | |
| Q9D832 | DNAJB4 | DnaJ (Hsp40) homolog, subfamily | Nucleus | other | 2694049.25 | 5449718 | 2972911.5 | | | | | | |
| Q9D832 | DNAJB4 | DnaJ (Hsp40) homolog, subfamily | Nucleus | other | 920827.625 | 3839994.75 | 4785630 | Q9D832 | DNAJB4 | 8717847 | 6066801 | 1107641 | 30 |
| Q9DC23 | DNAJC10 | DnaJ (Hsp40) homolog, subfamily | Cytoplasm | enzyme | 526934.625 | 506716.125 | 220092.984 | Q9DC23 | DNAJC10 | 526935 | 171617 | 99083 | 3 |
| Q5U458 | DNAJC11 | DnaJ (Hsp40) homolog, subfamily | Cytoplasm | other | 2674096.25 | 2491481 | 3357158.25 | | | | | | |
| Q5U458 | DNAJC11 | DnaJ (Hsp40) homolog, subfamily | Cytoplasm | other | 1437341 | 1875623.125 | 2743525.75 | Q5U458 | DNAJC11 | 2055719 | 680292 | 277728 | 6 |
| P60904 | DNAJC5 | DnaJ (Hsp40) homolog, subfamily | Plasma Membrane | other | 3.28E+07 | 4.68E+07 | 4.29E+07 | | | | | | |
| P60904 | DNAJC5 | DnaJ (Hsp40) homolog, subfamily | Plasma Membrane | other | 2.70E+07 | 3.74E+07 | 4.20E+07 | | | | | | |
| P60904 | DNAJC5 | DnaJ (Hsp40) homolog, subfamily | Plasma Membrane | other | 1.26E+07 | 1.66E+07 | 1.51E+07 | | | | | | |
| P60904 | DNAJC5 | DnaJ (Hsp40) homolog, subfamily | Plasma Membrane | other | 7716076 | 2071698.125 | 321718.344 | | | | | | |
| P60904 | DNAJC5 | DnaJ (Hsp40) homolog, subfamily | Plasma Membrane | other | 712703.125 | 61086.445 | 0 | P60904 | DNAJC5 | 16151756 | 17606429 | 4545960 | 15 |
| Q9D7J6 | DNASE1L1 | deoxyribonuclease I-like 1 | Cytoplasm | enzyme | 374793.844 | 517202.25 | 495204.625 | Q9D7J6 | DNASE1L1 | 374794 | 76662 | 44261 | 3 |
| P39054 | DNM2 | dynamin 2 | Plasma Membrane | enzyme | 4.09E+07 | 4.38E+07 | 2.56E+07 | | | | | | |
| P39054 | DNM2 | dynamin 2 | Plasma Membrane | enzyme | 2.83E+07 | 3.30E+07 | 1.62E+07 | | | | | | |
| P39054 | DNM2 | dynamin 2 | Plasma Membrane | enzyme | 2.98E+07 | 3.59E+07 | 2.33E+07 | | | | | | |
| P39054 | DNM2 | dynamin 2 | Plasma Membrane | enzyme | 2.42E+07 | 2.43E+07 | 1.25E+07 | | | | | | |
| P39054 | DNM2 | dynamin 2 | Plasma Membrane | enzyme | 1.81E+07 | 2.02E+07 | 1.16E+07 | | | | | | |
| P39054 | DNM2 | dynamin 2 | Plasma Membrane | enzyme | 2.03E+07 | 2.49E+07 | 9658635 | | | | | | |
| P39054 | DNM2 | dynamin 2 | Plasma Membrane | enzyme | 1.21E+07 | 1.38E+07 | 1.14E+07 | | | | | | |
| P39054 | DNM2 | dynamin 2 | Plasma Membrane | enzyme | 9435401 | 1.33E+07 | 7814624 | | | | | | |
| P39054 | DNM2 | dynamin 2 | Plasma Membrane | enzyme | 1.24E+07 | 1.14E+07 | 8045128.5 | | | | | | |
| P39054 | DNM2 | dynamin 2 | Plasma Membrane | enzyme | 8820659 | 1.10E+07 | 6934880 | | | | | | |
| P39054 | DNM2 | dynamin 2 | Plasma Membrane | enzyme | 8984620 | 9622472 | 6064899 | | | | | | |
| P39054 | DNM2 | dynamin 2 | Plasma Membrane | enzyme | 5769959 | 1.32E+07 | 6478622 | | | | | | |
| P39054 | DNM2 | dynamin 2 | Plasma Membrane | enzyme | 6617645.5 | 7585194 | 4427460.5 | | | | | | |
| P39054 | DNM2 | dynamin 2 | Plasma Membrane | enzyme | 5676773.5 | 9141628 | 4175367 | | | | | | |
| P39054 | DNM2 | dynamin 2 | Plasma Membrane | enzyme | 3916608.5 | 463738.5 | 362813.5 | | | | | | |
| P39054 | DNM2 | dynamin 2 | Plasma Membrane | enzyme | 7628546 | 7921924 | 6264179.5 | | | | | | |
| P39054 | DNM2 | dynamin 2 | Plasma Membrane | enzyme | 5652669 | 5365993.5 | 3211065 | | | | | | |
| P39054 | DNM2 | dynamin 2 | Plasma Membrane | enzyme | 3917698.25 | 3979140.25 | 2227934 | | | | | | |
| P39054 | DNM2 | dynamin 2 | Plasma Membrane | enzyme | 2733633 | 3226055.75 | 2205696.25 | | | | | | |
| P39054 | DNM2 | dynamin 2 | Plasma Membrane | enzyme | 1918344.875 | 2891496.5 | 1593026.625 | | | | | | |
| P39054 | DNM2 | dynamin 2 | Plasma Membrane | enzyme | 1929816.25 | 2293235.75 | 413549.188 | | | | | | |
| P39054 | DNM2 | dynamin 2 | Plasma Membrane | enzyme | 1506151.25 | 1958180.875 | 1433710.875 | | | | | | |
| P39054 | DNM2 | dynamin 2 | Plasma Membrane | enzyme | 1288032 | 3726629.75 | 941632.438 | P39054 | DNM2 | 11381589 | 10156489 | 1222698 | 69 |
| Q8BZ98 | DNM3 | dynamin 3 | Cytoplasm | enzyme | 8650986 | 9896302 | 5422412.5 | | | | | | |
| Q8BZ98 | DNM3 | dynamin 3 | Cytoplasm | enzyme | 9881335 | 1.06E+07 | 7516149 | | | | | | |
| Q8BZ98 | DNM3 | dynamin 3 | Cytoplasm | enzyme | 2492611.25 | 4418332.5 | 2140138.25 | Q8BZ98 | DNM3 | 7008311 | 3266718 | 1088906 | 9 |
| Q9Z2W0 | DNPEP | aspartyl aminopeptidase | Cytoplasm | peptidase | 2.86E+07 | 2.28E+07 | 2.58E+07 | | | | | | |
| Q9Z2W0 | DNPEP | aspartyl aminopeptidase | Cytoplasm | peptidase | 7473613.5 | 5685976 | 6902659.5 | | | | | | |
| Q9Z2W0 | DNPEP | aspartyl aminopeptidase | Cytoplasm | peptidase | 5885886.5 | 5756475 | 5842465.5 | | | | | | |
| Q9Z2W0 | DNPEP | aspartyl aminopeptidase | Cytoplasm | peptidase | 6391217.5 | 5169061 | 4321096.5 | | | | | | |
| Q9Z2W0 | DNPEP | aspartyl aminopeptidase | Cytoplasm | peptidase | 5119491 | 4668388.5 | 4186583.5 | | | | | | |
| Q9Z2W0 | DNPEP | aspartyl aminopeptidase | Cytoplasm | peptidase | 5651927.5 | 3699193.25 | 4904871.5 | | | | | | |
| Q9Z2W0 | DNPEP | aspartyl aminopeptidase | Cytoplasm | peptidase | 4468770.5 | 4024203.5 | 4093625.25 | | | | | | |
| Q9Z2W0 | DNPEP | aspartyl aminopeptidase | Cytoplasm | peptidase | 3218975.75 | 2817408 | 4377394.5 | | | | | | |
| Q9Z2W0 | DNPEP | aspartyl aminopeptidase | Cytoplasm | peptidase | 2044725.125 | 1531169.375 | 1823863.5 | Q9Z2W0 | DNPEP | 7646067 | 6966385 | 1340681 | 27 |
| Q9DD18 | DTD1 | D-tyrosyl-tRNA deacylase 1 | Cytoplasm | enzyme | 125637.555 | 39224.18 | 185965.922 | Q9DD18 | DTD1 | 125638 | 73756 | 42583 | 3 |
| Q9JHU4 | DYNC1H1 | dynein, cytoplasmic 1, heavy chain | Cytoplasm | peptidase | 1911889.625 | 1374517.75 | 1940199 | Q9JHU4 | DYNC1H1 | 1911890 | 318738 | 184024 | 3 |
| P42125 | ECI1 | enoyl-CoA delta isomerase 1 | Cytoplasm | enzyme | 5671852 | 9153889 | 7620152 | | | | | | |
| P42125 | ECI1 | enoyl-CoA delta isomerase 1 | Cytoplasm | enzyme | 2739973.25 | 3810391 | 3837285.5 | P42125 | ECI1 | 4205913 | 2494314 | 1018299 | 6 |

| | | | | | | | | | | | | | |
|---|---|---|---|---|---|---|---|---|---|---|---|---|---|
| Q8BL66 | EEA1 | early endosome antigen 1 | Cytoplasm | other | 1536229.5 | 1417311.25 | 974902.375 | Q8BL66 | EEA1 | 1536230 | 295791 | 170775 | 3 |
| P10126 | EEF1A1 | eukaryotic translation elongation f | Cytoplasm | translation regulator | 3.07E+08 | 3.43E+08 | 2.52E+08 | | | | | | |
| P10126 | EEF1A1 | eukaryotic translation elongation f | Cytoplasm | translation regulator | 1.20E+08 | 1.11E+08 | 8.47E+07 | | | | | | |
| P10126 | EEF1A1 | eukaryotic translation elongation f | Cytoplasm | translation regulator | 1.33E+08 | 4.03E+07 | 2.72E+07 | | | | | | |
| P10126 | EEF1A1 | eukaryotic translation elongation f | Cytoplasm | translation regulator | 6.41E+07 | 1.14E+08 | 7.01E+07 | | | | | | |
| P10126 | EEF1A1 | eukaryotic translation elongation f | Cytoplasm | translation regulator | 5.52E+07 | 4.61E+07 | 3.35E+07 | | | | | | |
| P10126 | EEF1A1 | eukaryotic translation elongation f | Cytoplasm | translation regulator | 7.35E+07 | 6.07E+07 | 4.52E+07 | | | | | | |
| P10126 | EEF1A1 | eukaryotic translation elongation f | Cytoplasm | translation regulator | 6.89E+07 | 1.04E+08 | 6.95E+07 | | | | | | |
| P10126 | EEF1A1 | eukaryotic translation elongation f | Cytoplasm | translation regulator | 1.19E+08 | 3.06E+07 | 2.00E+07 | | | | | | |
| P10126 | EEF1A1 | eukaryotic translation elongation f | Cytoplasm | translation regulator | 7.43E+07 | 4.82E+07 | 5.02E+07 | | | | | | |
| P10126 | EEF1A1 | eukaryotic translation elongation f | Cytoplasm | translation regulator | 4.33E+07 | 6.60E+07 | 5.73E+07 | | | | | | |
| P10126 | EEF1A1 | eukaryotic translation elongation f | Cytoplasm | translation regulator | 6.26E+07 | 6.36E+07 | 5.10E+07 | | | | | | |
| P10126 | EEF1A1 | eukaryotic translation elongation f | Cytoplasm | translation regulator | 6.16E+07 | 5.81E+07 | 4.60E+07 | | | | | | |
| P10126 | EEF1A1 | eukaryotic translation elongation f | Cytoplasm | translation regulator | 3.56E+07 | 5.69E+07 | 3.42E+07 | | | | | | |
| P10126 | EEF1A1 | eukaryotic translation elongation f | Cytoplasm | translation regulator | 4.24E+07 | 3.39E+07 | 3.33E+07 | | | | | | |
| P10126 | EEF1A1 | eukaryotic translation elongation f | Cytoplasm | translation regulator | 2.70E+07 | 5.88E+07 | 3.74E+07 | | | | | | |
| P10126 | EEF1A1 | eukaryotic translation elongation f | Cytoplasm | translation regulator | 7.29E+07 | 1.56E+07 | 2.66E+07 | | | | | | |
| P10126 | EEF1A1 | eukaryotic translation elongation f | Cytoplasm | translation regulator | 1.40E+07 | 1.23E+07 | 8267749.5 | | | | | | |
| P10126 | EEF1A1 | eukaryotic translation elongation f | Cytoplasm | translation regulator | 3121294.25 | 4890039.5 | 3750722 | | | | | | |
| P10126 | EEF1A1 | eukaryotic translation elongation f | Cytoplasm | translation regulator | 6094816 | 4852769 | 3552826 | | | | | | |
| P10126 | EEF1A1 | eukaryotic translation elongation f | Cytoplasm | translation regulator | 2979980.25 | 2391281 | 3176060.25 | | | | | | |
| P10126 | EEF1A1 | eukaryotic translation elongation f | Cytoplasm | translation regulator | 5031019 | 2604874.5 | 2291877.25 | | | | | | |
| P10126 | EEF1A1 | eukaryotic translation elongation f | Cytoplasm | translation regulator | 9214503 | 1376816.125 | 2116453.75 | | | | | | |
| P10126 | EEF1A1 | eukaryotic translation elongation f | Cytoplasm | translation regulator | 851872.625 | 844732 | 648768.875 | P10126 | EEF1A1 | 60906238 | 63434330 | 7636596 | 69 |
| O70251 | EEF1B2 | eukaryotic translation elongation f | Cytoplasm | translation regulator | 1837833.25 | 1463542.125 | 1714686.125 | O70251 | EEF1B2 | 1837833 | 190758 | 110134 | 3 |
| P57776 | EEF1D | eukaryotic translation elongation f | Cytoplasm | translation regulator | 3161082.75 | 3331283.25 | 2413484.75 | P57776 | EEF1D | 3161083 | 488232 | 281881 | 3 |
| Q9D8N0 | EEF1G | eukaryotic translation elongation f | Cytoplasm | translation regulator | 4972986.5 | 9478326 | 6794120.5 | | | | | | |
| Q9D8N0 | EEF1G | eukaryotic translation elongation f | Cytoplasm | translation regulator | 2213297.75 | 5219380 | 3947377.75 | | | | | | |
| Q9D8N0 | EEF1G | eukaryotic translation elongation f | Cytoplasm | translation regulator | 4146653 | 3636651.25 | 3616915.75 | Q9D8N0 | EEF1G | 3777646 | 2138669 | 712890 | 9 |
| P58252 | EEF2 | eukaryotic translation elongation f | Cytoplasm | translation regulator | 1298874.25 | 1211264.75 | 1056520.875 | | | | | | |
| P58252 | EEF2 | eukaryotic translation elongation f | Cytoplasm | translation regulator | 2283442.25 | 2273775.25 | 1859268.25 | | | | | | |
| P58252 | EEF2 | eukaryotic translation elongation f | Cytoplasm | translation regulator | 2749070 | 2852876 | 2368795.5 | | | | | | |
| P58252 | EEF2 | eukaryotic translation elongation f | Cytoplasm | translation regulator | 1381917.875 | 1383774.875 | 1393734.375 | | | | | | |
| P58252 | EEF2 | eukaryotic translation elongation f | Cytoplasm | translation regulator | 502334.688 | 1131375.625 | 748689.688 | P58252 | EEF2 | 1643128 | 720924 | 186142 | 15 |
| Q8BG67 | EFR3A | EFR3 homolog A | Plasma Membrane | other | 3979258.25 | 3018869.75 | 3344109 | | | | | | |
| Q8BG67 | EFR3A | EFR3 homolog A | Plasma Membrane | other | 3450658 | 4402275.5 | 4146722.25 | | | | | | |
| Q8BG67 | EFR3A | EFR3 homolog A | Plasma Membrane | other | 4429582 | 4913976.5 | 3799287.5 | | | | | | |
| Q8BG67 | EFR3A | EFR3 homolog A | Plasma Membrane | other | 3466285.75 | 5639488.5 | 5521744.5 | | | | | | |
| Q8BG67 | EFR3A | EFR3 homolog A | Plasma Membrane | other | 3523856.25 | 4322472 | 3096391.5 | | | | | | |
| Q8BG67 | EFR3A | EFR3 homolog A | Plasma Membrane | other | 3410548.5 | 3779902.25 | 3043874.25 | | | | | | |
| Q8BG67 | EFR3A | EFR3 homolog A | Plasma Membrane | other | 2513532.5 | 3206634.25 | 2295740 | Q8BG67 | EFR3A | 3539103 | 876877 | 191350 | 21 |
| Q9WVK4 | EHD1 | EH-domain containing 1 | Cytoplasm | other | 4319125.5 | 6316362.5 | 6374703.5 | | | | | | |
| Q9WVK4 | EHD1 | EH-domain containing 1 | Cytoplasm | other | 7362156.5 | 9617278 | 6392467.5 | | | | | | |
| Q9WVK4 | EHD1 | EH-domain containing 1 | Cytoplasm | other | 2816135.75 | 2585187 | 1660331.125 | | | | | | |
| Q9WVK4 | EHD1 | EH-domain containing 1 | Cytoplasm | other | 2479802.25 | 1156220.75 | 794512.125 | Q9WVK4 | EHD1 | 4244305 | 2828236 | 816441 | 12 |
| Q9EQP2 | EHD4 | EH-domain containing 4 | Plasma Membrane | enzyme | 1631020.875 | 2304138 | 1153805.25 | | | | | | |
| Q9EQP2 | EHD4 | EH-domain containing 4 | Plasma Membrane | enzyme | 1555556.25 | 1831492.625 | 847173 | Q9EQP2 | EHD4 | 1593289 | 510961 | 208599 | 6 |
| Q6ZWX6 | EIF2S1 | eukaryotic translation initiation fa | Cytoplasm | translation regulator | 6747063 | 5801578.5 | 3415225 | | | | | | |
| Q6ZWX6 | EIF2S1 | eukaryotic translation initiation fa | Cytoplasm | translation regulator | 3252838.5 | 4491664.5 | 3199756 | | | | | | |
| Q6ZWX6 | EIF2S1 | eukaryotic translation initiation fa | Cytoplasm | translation regulator | 3854646.25 | 4387677 | 2488608.5 | | | | | | |
| Q6ZWX6 | EIF2S1 | eukaryotic translation initiation fa | Cytoplasm | translation regulator | 2939088.5 | 3529008 | 1495446.625 | | | | | | |
| Q6ZWX6 | EIF2S1 | eukaryotic translation initiation fa | Cytoplasm | translation regulator | 3437772 | 2837590.75 | 1579949.125 | | | | | | |
| Q6ZWX6 | EIF2S1 | eukaryotic translation initiation fa | Cytoplasm | translation regulator | 2876967.25 | 3443376.5 | 2476043.75 | Q6ZWX6 | EIF2S1 | 3851396 | 1301381 | 306738 | 18 |
| Q99L45 | EIF2S2 | eukaryotic translation initiation fa | Cytoplasm | translation regulator | 5173058 | 5449325.5 | 2984821.75 | | | | | | |
| Q99L45 | EIF2S2 | eukaryotic translation initiation fa | Cytoplasm | translation regulator | 3467619.5 | 6629178.5 | 3096137.25 | | | | | | |
| Q99L45 | EIF2S2 | eukaryotic translation initiation fa | Cytoplasm | translation regulator | 4045265.5 | 5257819 | 1849577.375 | Q99L45 | EIF2S2 | 4228648 | 1512652 | 504217 | 9 |
| Q9Z0N1 | EIF2S3 | eukaryotic translation initiation fa | Cytoplasm | translation regulator | 8696938 | 9460128 | 5880031.5 | | | | | | |
| Q9Z0N1 | EIF2S3 | eukaryotic translation initiation fa | Cytoplasm | translation regulator | 1.24E+07 | 1.28E+07 | 8509856 | | | | | | |
| Q9Z0N1 | EIF2S3 | eukaryotic translation initiation fa | Cytoplasm | translation regulator | 7704891 | 9203832 | 6826967.5 | | | | | | |
| Q9Z0N1 | EIF2S3 | eukaryotic translation initiation fa | Cytoplasm | translation regulator | 5626763 | 5208494 | 4355901.5 | | | | | | |
| Q9Z0N1 | EIF2S3 | eukaryotic translation initiation fa | Cytoplasm | translation regulator | 6551041 | 7787878.5 | 3595660.5 | | | | | | |
| Q9Z0N1 | EIF2S3 | eukaryotic translation initiation fa | Cytoplasm | translation regulator | 5990962.5 | 6195376.5 | 3156989 | | | | | | |
| Q9Z0N1 | EIF2S3 | eukaryotic translation initiation fa | Cytoplasm | translation regulator | 5043444.5 | 4419188.5 | 2800773.25 | | | | | | |
| Q9Z0N1 | EIF2S3 | eukaryotic translation initiation fa | Cytoplasm | translation regulator | 3417731.25 | 4070858.25 | 2334295.75 | | | | | | |
| Q9Z0N1 | EIF2S3 | eukaryotic translation initiation fa | Cytoplasm | translation regulator | 1981319 | 1943326.25 | 1840312.625 | | | | | | |
| Q9Z0N1 | EIF2S3 | eukaryotic translation initiation fa | Cytoplasm | translation regulator | 1750625.375 | 2146327.25 | 1359593.25 | | | | | | |
| Q9Z0N1 | EIF2S3 | eukaryotic translation initiation fa | Cytoplasm | translation regulator | 1681510 | 1535263.125 | 986005.562 | Q9Z0N1 | EIF2S3 | 5526839 | 3181142 | 553766 | 33 |
| Q3UGC7 | EIF3J | eukaryotic translation initiation fa | Cytoplasm | translation regulator | 8585740 | 1.10E+07 | 6723998.5 | | | | | | |
| Q3UGC7 | EIF3J | eukaryotic translation initiation fa | Cytoplasm | translation regulator | 6927638.5 | 1.15E+07 | 5760730.5 | | | | | | |
| Q3UGC7 | EIF3J | eukaryotic translation initiation fa | Cytoplasm | translation regulator | 4356683 | 6768899 | 4301193 | | | | | | |
| Q3UGC7 | EIF3J | eukaryotic translation initiation fa | Cytoplasm | translation regulator | 31891.195 | 90518.828 | 30533.439 | | | | | | |
| Q3UGC7 | EIF3J | eukaryotic translation initiation fa | Cytoplasm | translation regulator | 3242039.75 | 2655795.25 | 3070602.75 | Q3UGC7 | EIF3J | 4628798 | 3664271 | 946111 | 15 |
| Q8BGD9 | EIF4B | eukaryotic translation initiation fa | Cytoplasm | translation regulator | 7727001 | 1.02E+07 | 7089852.5 | | | | | | |
| Q8BGD9 | EIF4B | eukaryotic translation initiation fa | Cytoplasm | translation regulator | 5709912.5 | 6164796 | 4371587 | | | | | | |
| Q8BGD9 | EIF4B | eukaryotic translation initiation fa | Cytoplasm | translation regulator | 5989930.5 | 6137209 | 4276846 | | | | | | |
| Q8BGD9 | EIF4B | eukaryotic translation initiation fa | Cytoplasm | translation regulator | 2279199.5 | 6871018 | 2868365.25 | | | | | | |
| Q8BGD9 | EIF4B | eukaryotic translation initiation fa | Cytoplasm | translation regulator | 3467000.5 | 3780246.75 | 2901306 | | | | | | |
| Q8BGD9 | EIF4B | eukaryotic translation initiation fa | Cytoplasm | translation regulator | 2039905.375 | 2067166.25 | 1732756.75 | Q8BGD9 | EIF4B | 4535492 | 2353355 | 554691 | 18 |
| Q6NZJ6 | EIF4G1 | eukaryotic translation initiation fa | Cytoplasm | translation regulator | 4458645 | 4461330.5 | 2065506.125 | | | | | | |
| Q6NZJ6 | EIF4G1 | eukaryotic translation initiation fa | Cytoplasm | translation regulator | 1464945.75 | 1866065.5 | 1212597.875 | | | | | | |
| Q6NZJ6 | EIF4G1 | eukaryotic translation initiation fa | Cytoplasm | translation regulator | 854097.438 | 676413.312 | 469666.969 | Q6NZJ6 | EIF4G1 | 2259229 | 1517147 | 505716 | 9 |
| P59325 | EIF5 | eukaryotic translation initiation fa | Cytoplasm | translation regulator | 2501011.5 | 1768433.375 | 2126373.25 | P59325 | EIF5 | 2501012 | 366321 | 211495 | 3 |
| P63242 | EIF5A | eukaryotic translation initiation fa | Cytoplasm | translation regulator | 1919710.5 | 3468346.25 | 2329364.75 | P63242 | EIF5A | 1919711 | 802430 | 463283 | 3 |
| Q99K41 | EMILIN1 | elastin microfibril interfacer 1 | Extracellular Space | other | 6275711 | 6234883.5 | 6558458.5 | | | | | | |
| Q99K41 | EMILIN1 | elastin microfibril interfacer 1 | Extracellular Space | other | 1352048.625 | 880207.312 | 976718.375 | Q99K41 | EMILIN1 | 3813880 | 2902070 | 1184765 | 6 |
| Q8C522 | ENDOD1 | endonuclease domain containing 1 | Extracellular Space | enzyme | 2.24E+07 | 2.67E+07 | 2.34E+07 | | | | | | |

| | | | | | | | | | | | | | |
|---|---|---|---|---|---|---|---|---|---|---|---|---|---|
| Q8C522 | ENDOD1 | endonuclease domain containing 1 | Extracellular Space | enzyme | 1.64E+07 | 9010818 | 1.21E+07 | | | | | | |
| Q8C522 | ENDOD1 | endonuclease domain containing 1 | Extracellular Space | enzyme | 1.29E+07 | 8180743 | 8786929 | | | | | | |
| Q8C522 | ENDOD1 | endonuclease domain containing 1 | Extracellular Space | enzyme | 6447936 | 7424288.5 | 7583102.5 | | | | | | |
| Q8C522 | ENDOD1 | endonuclease domain containing 1 | Extracellular Space | enzyme | 8234954 | 1.00E+07 | 1.06E+07 | | | | | | |
| Q8C522 | ENDOD1 | endonuclease domain containing 1 | Extracellular Space | enzyme | 6695785 | 9986402 | 5053827 | | | | | | |
| Q8C522 | ENDOD1 | endonuclease domain containing 1 | Extracellular Space | enzyme | 8895452 | 5374318 | 4247524 | | | | | | |
| Q8C522 | ENDOD1 | endonuclease domain containing 1 | Extracellular Space | enzyme | 6164167 | 9745568 | 4721280.5 | Q8C522 | ENDOD1 | 11016037 | 6008343 | 1226448 | 24 |
| Q8BTJ4 | ENPP4 | ectonucleotide pyrophosphatase/p | Cytoplasm | enzyme | 996231.188 | 551155.812 | 1125308.625 | Q8BTJ4 | ENPP4 | 996231 | 301221 | 173910 | 3 |
| P10404 | ENV1 | MLV-related proviral Env polyprot | other | | 2.27E+07 | 3.34E+07 | 1.82E+07 | | | | | | |
| P10404 | ENV1 | MLV-related proviral Env polyprot | other | | 1.55E+07 | 3.11E+07 | 1.07E+07 | | | | | | |
| P10404 | ENV1 | MLV-related proviral Env polyprot | other | | 2.47E+07 | 3.46E+07 | 1.23E+07 | | | | | | |
| P10404 | ENV1 | MLV-related proviral Env polyprot | other | | 1.33E+07 | 1.76E+07 | 6123258.5 | | | | | | |
| P10404 | ENV1 | MLV-related proviral Env polyprot | other | | 9461509 | 2.30E+07 | 9317924 | | | | | | |
| P10404 | ENV1 | MLV-related proviral Env polyprot | other | | 1.10E+07 | 2.20E+07 | 1.05E+07 | | | | | | |
| P10404 | ENV1 | MLV-related proviral Env polyprot | other | | 1.35E+07 | 2.05E+07 | 8136012 | | | | | | |
| P10404 | ENV1 | MLV-related proviral Env polyprot | other | | 1.05E+07 | 1.69E+07 | 6047378.5 | | | | | | |
| P10404 | ENV1 | MLV-related proviral Env polyprot | other | | 5166782.5 | 1.24E+07 | 4901673 | | | | | | |
| P10404 | ENV1 | MLV-related proviral Env polyprot | other | | 9796593 | 6516193.5 | 5222518.5 | | | | | | |
| P10404 | ENV1 | MLV-related proviral Env polyprot | other | | 6734599 | 1.16E+07 | 4207081 | | | | | | |
| P10404 | ENV1 | MLV-related proviral Env polyprot | other | | 6387114.5 | 9367882 | 3617615 | | | | | | |
| P10404 | ENV1 | MLV-related proviral Env polyprot | other | | 3964426.5 | 4836250.5 | 2406155.5 | | | | | | |
| P10404 | ENV1 | MLV-related proviral Env polyprot | other | | 3133159.5 | 4988595.5 | 3344187 | | | | | | |
| P10404 | ENV1 | MLV-related proviral Env polyprot | other | | 3198925 | 2758184.75 | 954879.438 | | | | | | |
| P10404 | ENV1 | MLV-related proviral Env polyprot | other | | 2397833 | 3599545.75 | 1548478.875 | | | | | | |
| P10404 | ENV1 | MLV-related proviral Env polyprot | other | | 2140997 | 2792752.25 | 1434990.375 | | | | | | |
| P10404 | ENV1 | MLV-related proviral Env polyprot | other | | 1404115.5 | 3580326 | 1068360.5 | | | | | | |
| P10404 | ENV1 | MLV-related proviral Env polyprot | other | | 491024.594 | 2178493.25 | 1045986.312 | | | | | | |
| P10404 | ENV1 | MLV-related proviral Env polyprot | other | | 1042105 | 4463025.5 | 1348839.625 | | | | | | |
| P10404 | ENV1 | MLV-related proviral Env polyprot | other | | 381148.469 | 3455058 | 379422.062 | | | | | | |
| P10404 | ENV1 | MLV-related proviral Env polyprot | other | | 712557.375 | 1145594.75 | 491375.719 | | | | | | |
| P10404 | ENV1 | MLV-related proviral Env polyprot | other | | 833675.812 | 1008150.812 | 0 | | | | | | |
| P10404 | ENV1 | MLV-related proviral Env polyprot | other | | 276292.531 | 760459.938 | 645139.625 | P10404 | ENV1 | 7024702 | 8297940 | 977922 | 72 |
| P48193 | EPB41 | erythrocyte membrane protein ba | Plasma Membrane | other | 1.28E+09 | 2.83E+09 | 2.28E+09 | | | | | | |
| P48193 | EPB41 | erythrocyte membrane protein ba | Plasma Membrane | other | 1.73E+09 | 1.50E+09 | 1.49E+09 | | | | | | |
| P48193 | EPB41 | erythrocyte membrane protein ba | Plasma Membrane | other | 1.83E+09 | 1.55E+09 | 1.50E+09 | | | | | | |
| P48193 | EPB41 | erythrocyte membrane protein ba | Plasma Membrane | other | 1.31E+09 | 1.73E+09 | 1.70E+09 | | | | | | |
| P48193 | EPB41 | erythrocyte membrane protein ba | Plasma Membrane | other | 1.24E+09 | 1.20E+09 | 1.03E+09 | | | | | | |
| P48193 | EPB41 | erythrocyte membrane protein ba | Plasma Membrane | other | 1.09E+09 | 1.36E+09 | 1.42E+09 | | | | | | |
| P48193 | EPB41 | erythrocyte membrane protein ba | Plasma Membrane | other | 1.21E+09 | 1.40E+09 | 1.16E+09 | | | | | | |
| P48193 | EPB41 | erythrocyte membrane protein ba | Plasma Membrane | other | 9.47E+08 | 1.45E+09 | 1.28E+09 | | | | | | |
| P48193 | EPB41 | erythrocyte membrane protein ba | Plasma Membrane | other | 1.11E+09 | 1.05E+09 | 9.62E+08 | | | | | | |
| P48193 | EPB41 | erythrocyte membrane protein ba | Plasma Membrane | other | 9.71E+08 | 8.23E+08 | 9.28E+08 | | | | | | |
| P48193 | EPB41 | erythrocyte membrane protein ba | Plasma Membrane | other | 1.05E+09 | 9.61E+08 | 9.78E+08 | | | | | | |
| P48193 | EPB41 | erythrocyte membrane protein ba | Plasma Membrane | other | 7.85E+08 | 8.51E+08 | 8.20E+08 | | | | | | |
| P48193 | EPB41 | erythrocyte membrane protein ba | Plasma Membrane | other | 7.70E+08 | 8.48E+08 | 9.25E+08 | | | | | | |
| P48193 | EPB41 | erythrocyte membrane protein ba | Plasma Membrane | other | 7.54E+08 | 8.34E+08 | 7.61E+08 | | | | | | |
| P48193 | EPB41 | erythrocyte membrane protein ba | Plasma Membrane | other | 6.51E+08 | 7.96E+08 | 7.31E+08 | | | | | | |
| P48193 | EPB41 | erythrocyte membrane protein ba | Plasma Membrane | other | 5.35E+08 | 5.68E+08 | 5.24E+08 | | | | | | |
| P48193 | EPB41 | erythrocyte membrane protein ba | Plasma Membrane | other | 3.81E+08 | 5.25E+08 | 5.32E+08 | | | | | | |
| P48193 | EPB41 | erythrocyte membrane protein ba | Plasma Membrane | other | 3.29E+08 | 4.21E+08 | 4.03E+08 | | | | | | |
| P48193 | EPB41 | erythrocyte membrane protein ba | Plasma Membrane | other | 3.79E+08 | 6.56E+08 | 4.49E+08 | | | | | | |
| P48193 | EPB41 | erythrocyte membrane protein ba | Plasma Membrane | other | 3.89E+08 | 3.14E+08 | 2.96E+08 | | | | | | |
| P48193 | EPB41 | erythrocyte membrane protein ba | Plasma Membrane | other | 2.14E+08 | 5.92E+08 | 3.90E+08 | | | | | | |
| P48193 | EPB41 | erythrocyte membrane protein ba | Plasma Membrane | other | 2.03E+08 | 2.59E+08 | 2.12E+08 | | | | | | |
| P48193 | EPB41 | erythrocyte membrane protein ba | Plasma Membrane | other | 3.13E+08 | 2.07E+08 | 2.09E+08 | | | | | | |
| P48193 | EPB41 | erythrocyte membrane protein ba | Plasma Membrane | other | 2.05E+08 | 1.86E+08 | 1.79E+08 | | | | | | |
| P48193 | EPB41 | erythrocyte membrane protein ba | Plasma Membrane | other | 2.74E+08 | 2.21E+08 | 2.34E+08 | | | | | | |
| P48193 | EPB41 | erythrocyte membrane protein ba | Plasma Membrane | other | 1.32E+08 | 1.11E+08 | 1.20E+08 | | | | | | |
| P48193 | EPB41 | erythrocyte membrane protein ba | Plasma Membrane | other | 5.18E+07 | 1.22E+08 | 1.28E+08 | | | | | | |
| P48193 | EPB41 | erythrocyte membrane protein ba | Plasma Membrane | other | 7.70E+07 | 9.86E+07 | 9.96E+07 | | | | | | |
| P48193 | EPB41 | erythrocyte membrane protein ba | Plasma Membrane | other | 1.72E+08 | 2.08E+08 | 1.26E+08 | | | | | | |
| P48193 | EPB41 | erythrocyte membrane protein ba | Plasma Membrane | other | 4.04E+07 | 1.67E+08 | 7.52E+07 | | | | | | |
| P48193 | EPB41 | erythrocyte membrane protein ba | Plasma Membrane | other | 5.79E+07 | 5.63E+07 | 5.44E+07 | | | | | | |
| P48193 | EPB41 | erythrocyte membrane protein ba | Plasma Membrane | other | 4.33E+07 | 8.11E+07 | 7.42E+07 | | | | | | |
| P48193 | EPB41 | erythrocyte membrane protein ba | Plasma Membrane | other | 5.50E+07 | 6.89E+07 | 6.10E+07 | | | | | | |
| P48193 | EPB41 | erythrocyte membrane protein ba | Plasma Membrane | other | 5.62E+07 | 6.07E+07 | 5.31E+07 | | | | | | |
| P48193 | EPB41 | erythrocyte membrane protein ba | Plasma Membrane | other | 4.83E+07 | 3.31E+07 | 2.70E+07 | | | | | | |
| P48193 | EPB41 | erythrocyte membrane protein ba | Plasma Membrane | other | 1.95E+07 | 4.92E+07 | 2.05E+07 | | | | | | |
| P48193 | EPB41 | erythrocyte membrane protein ba | Plasma Membrane | other | 3.56E+07 | 3.38E+07 | 3.08E+07 | | | | | | |
| P48193 | EPB41 | erythrocyte membrane protein ba | Plasma Membrane | other | 3.11E+07 | 1.94E+07 | 2.06E+07 | | | | | | |
| P48193 | EPB41 | erythrocyte membrane protein ba | Plasma Membrane | other | 3.76E+07 | 2.96E+07 | 3.19E+07 | | | | | | |
| P48193 | EPB41 | erythrocyte membrane protein ba | Plasma Membrane | other | 3.49E+07 | 3.33E+07 | 3.26E+07 | | | | | | |
| P48193 | EPB41 | erythrocyte membrane protein ba | Plasma Membrane | other | 1.39E+07 | 6.90E+07 | 3.53E+07 | | | | | | |
| P48193 | EPB41 | erythrocyte membrane protein ba | Plasma Membrane | other | 1.35E+07 | 3.73E+07 | 2.61E+07 | | | | | | |
| P48193 | EPB41 | erythrocyte membrane protein ba | Plasma Membrane | other | 2.31E+07 | 2.04E+07 | 1.86E+07 | | | | | | |
| P48193 | EPB41 | erythrocyte membrane protein ba | Plasma Membrane | other | 2.05E+07 | 1.79E+07 | 1.22E+07 | | | | | | |
| P48193 | EPB41 | erythrocyte membrane protein ba | Plasma Membrane | other | 1.68E+07 | 1.78E+07 | 6123518.5 | | | | | | |
| P48193 | EPB41 | erythrocyte membrane protein ba | Plasma Membrane | other | 1.16E+07 | 1.84E+07 | 1.34E+07 | | | | | | |
| P48193 | EPB41 | erythrocyte membrane protein ba | Plasma Membrane | other | 6953195.5 | 8060517 | 7084270.5 | | | | | | |
| P48193 | EPB41 | erythrocyte membrane protein ba | Plasma Membrane | other | 8984573 | 6.63E+07 | 6.43E+07 | | | | | | |
| P48193 | EPB41 | erythrocyte membrane protein ba | Plasma Membrane | other | 9924210 | 1.10E+07 | 1.03E+07 | | | | | | |
| P48193 | EPB41 | erythrocyte membrane protein ba | Plasma Membrane | other | 5.03E+07 | 7.26E+07 | 6.52E+07 | | | | | | |
| P48193 | EPB41 | erythrocyte membrane protein ba | Plasma Membrane | other | 2.52E+07 | 898600 | 986245.625 | | | | | | |
| P48193 | EPB41 | erythrocyte membrane protein ba | Plasma Membrane | other | 9360744 | 6556566 | 9118898 | | | | | | |
| P48193 | EPB41 | erythrocyte membrane protein ba | Plasma Membrane | other | 5317280 | 5457115 | 5610539 | | | | | | |
| P48193 | EPB41 | erythrocyte membrane protein ba | Plasma Membrane | other | 6251769 | 4880038 | 3895424 | | | | | | |

| | | | | | | | | | | | | | |
|---|---|---|---|---|---|---|---|---|---|---|---|---|---|
| P48193 | EPB41 | erythrocyte membrane protein ba | Plasma Membrane | other | 2495758.5 | 3573380 | 2561711.25 | | | | | | |
| P48193 | EPB41 | erythrocyte membrane protein ba | Plasma Membrane | other | 2993665.5 | 3428167.5 | 2825705 | | | | | | |
| P48193 | EPB41 | erythrocyte membrane protein ba | Plasma Membrane | other | 2886766.25 | 4303108.5 | 3117911.25 | | | | | | |
| P48193 | EPB41 | erythrocyte membrane protein ba | Plasma Membrane | other | 5623236.5 | 277727.562 | 314186.281 | | | | | | |
| P48193 | EPB41 | erythrocyte membrane protein ba | Plasma Membrane | other | 925819.812 | 1453049.5 | 1259618.25 | | | | | | |
| P48193 | EPB41 | erythrocyte membrane protein ba | Plasma Membrane | other | 83717.43 | 262414.406 | 108034.273 | | | | | | |
| P48193 | EPB41 | erythrocyte membrane protein ba | Plasma Membrane | other | 564741.562 | 449439.281 | 9064386 | | | | | | |
| P48193 | EPB41 | erythrocyte membrane protein ba | Plasma Membrane | other | 4157.404 | 482019.812 | 130002.398 | P48193 | EPB41 | 340004349 | 531328238 | 38958877 | 186 |
| P49222 | EPB42 | erythrocyte membrane protein ba | Plasma Membrane | transporter | 1.77E+09 | 1.98E+09 | 1.77E+09 | | | | | | |
| P49222 | EPB42 | erythrocyte membrane protein ba | Plasma Membrane | transporter | 1.81E+09 | 1.69E+09 | 1.56E+09 | | | | | | |
| P49222 | EPB42 | erythrocyte membrane protein ba | Plasma Membrane | transporter | 1.90E+09 | 1.36E+09 | 1.40E+09 | | | | | | |
| P49222 | EPB42 | erythrocyte membrane protein ba | Plasma Membrane | transporter | 1.71E+09 | 1.78E+09 | 1.57E+09 | | | | | | |
| P49222 | EPB42 | erythrocyte membrane protein ba | Plasma Membrane | transporter | 1.29E+09 | 1.39E+09 | 1.28E+09 | | | | | | |
| P49222 | EPB42 | erythrocyte membrane protein ba | Plasma Membrane | transporter | 1.31E+09 | 1.24E+09 | 1.25E+09 | | | | | | |
| P49222 | EPB42 | erythrocyte membrane protein ba | Plasma Membrane | transporter | 1.28E+09 | 1.08E+09 | 1.16E+09 | | | | | | |
| P49222 | EPB42 | erythrocyte membrane protein ba | Plasma Membrane | transporter | 1.00E+09 | 9.35E+08 | 9.60E+08 | | | | | | |
| P49222 | EPB42 | erythrocyte membrane protein ba | Plasma Membrane | transporter | 8.59E+08 | 9.29E+08 | 1.03E+09 | | | | | | |
| P49222 | EPB42 | erythrocyte membrane protein ba | Plasma Membrane | transporter | 8.38E+08 | 1.06E+09 | 6.21E+08 | | | | | | |
| P49222 | EPB42 | erythrocyte membrane protein ba | Plasma Membrane | transporter | 3.38E+08 | 4.83E+08 | 5.24E+08 | | | | | | |
| P49222 | EPB42 | erythrocyte membrane protein ba | Plasma Membrane | transporter | 8.30E+08 | 9.80E+08 | 7.95E+08 | | | | | | |
| P49222 | EPB42 | erythrocyte membrane protein ba | Plasma Membrane | transporter | 7.05E+08 | 5.91E+08 | 6.85E+08 | | | | | | |
| P49222 | EPB42 | erythrocyte membrane protein ba | Plasma Membrane | transporter | 7.46E+08 | 6.62E+08 | 6.05E+08 | | | | | | |
| P49222 | EPB42 | erythrocyte membrane protein ba | Plasma Membrane | transporter | 5.70E+08 | 5.51E+08 | 5.16E+08 | | | | | | |
| P49222 | EPB42 | erythrocyte membrane protein ba | Plasma Membrane | transporter | 5.05E+08 | 4.51E+08 | 3.97E+08 | | | | | | |
| P49222 | EPB42 | erythrocyte membrane protein ba | Plasma Membrane | transporter | 5.76E+08 | 6.11E+08 | 5.81E+08 | | | | | | |
| P49222 | EPB42 | erythrocyte membrane protein ba | Plasma Membrane | transporter | 4.88E+08 | 5.31E+08 | 4.96E+08 | | | | | | |
| P49222 | EPB42 | erythrocyte membrane protein ba | Plasma Membrane | transporter | 3.93E+08 | 4.68E+08 | 4.19E+08 | | | | | | |
| P49222 | EPB42 | erythrocyte membrane protein ba | Plasma Membrane | transporter | 3.79E+08 | 3.93E+08 | 3.33E+08 | | | | | | |
| P49222 | EPB42 | erythrocyte membrane protein ba | Plasma Membrane | transporter | 2.00E+08 | 2.88E+08 | 2.15E+08 | | | | | | |
| P49222 | EPB42 | erythrocyte membrane protein ba | Plasma Membrane | transporter | 3.39E+08 | 2.42E+08 | 2.44E+08 | | | | | | |
| P49222 | EPB42 | erythrocyte membrane protein ba | Plasma Membrane | transporter | 2.55E+08 | 2.77E+08 | 2.21E+08 | | | | | | |
| P49222 | EPB42 | erythrocyte membrane protein ba | Plasma Membrane | transporter | 2.70E+08 | 2.65E+08 | 3.38E+08 | | | | | | |
| P49222 | EPB42 | erythrocyte membrane protein ba | Plasma Membrane | transporter | 2.83E+08 | 2.73E+08 | 2.39E+08 | | | | | | |
| P49222 | EPB42 | erythrocyte membrane protein ba | Plasma Membrane | transporter | 1.04E+08 | 2.70E+08 | 1.49E+08 | | | | | | |
| P49222 | EPB42 | erythrocyte membrane protein ba | Plasma Membrane | transporter | 1.64E+08 | 1.49E+08 | 1.33E+08 | | | | | | |
| P49222 | EPB42 | erythrocyte membrane protein ba | Plasma Membrane | transporter | 1.28E+08 | 2.41E+08 | 1.80E+08 | | | | | | |
| P49222 | EPB42 | erythrocyte membrane protein ba | Plasma Membrane | transporter | 1.22E+08 | 1.35E+08 | 1.74E+08 | | | | | | |
| P49222 | EPB42 | erythrocyte membrane protein ba | Plasma Membrane | transporter | 6.56E+07 | 2.77E+08 | 1.54E+08 | | | | | | |
| P49222 | EPB42 | erythrocyte membrane protein ba | Plasma Membrane | transporter | 1.26E+08 | 1.24E+08 | 1.29E+08 | | | | | | |
| P49222 | EPB42 | erythrocyte membrane protein ba | Plasma Membrane | transporter | 7.53E+07 | 8.91E+07 | 8.80E+07 | | | | | | |
| P49222 | EPB42 | erythrocyte membrane protein ba | Plasma Membrane | transporter | 1.08E+08 | 1.49E+08 | 1.33E+08 | | | | | | |
| P49222 | EPB42 | erythrocyte membrane protein ba | Plasma Membrane | transporter | 1.32E+08 | 3.96E+07 | 6.55E+07 | | | | | | |
| P49222 | EPB42 | erythrocyte membrane protein ba | Plasma Membrane | transporter | 6.25E+07 | 4.12E+07 | 3.94E+07 | | | | | | |
| P49222 | EPB42 | erythrocyte membrane protein ba | Plasma Membrane | transporter | 5.32E+07 | 5.53E+07 | 5.61E+07 | | | | | | |
| P49222 | EPB42 | erythrocyte membrane protein ba | Plasma Membrane | transporter | 4.40E+07 | 9.72E+07 | 8.98E+07 | | | | | | |
| P49222 | EPB42 | erythrocyte membrane protein ba | Plasma Membrane | transporter | 4.16E+07 | 4.05E+07 | 3.20E+07 | | | | | | |
| P49222 | EPB42 | erythrocyte membrane protein ba | Plasma Membrane | transporter | 2.92E+07 | 3.19E+07 | 3.00E+07 | | | | | | |
| P49222 | EPB42 | erythrocyte membrane protein ba | Plasma Membrane | transporter | 2.23E+07 | 5870426 | 1913708 | | | | | | |
| P49222 | EPB42 | erythrocyte membrane protein ba | Plasma Membrane | transporter | 2.94E+07 | 3.05E+07 | 3.65E+07 | | | | | | |
| P49222 | EPB42 | erythrocyte membrane protein ba | Plasma Membrane | transporter | 2.59E+07 | 3.42E+07 | 3.49E+07 | | | | | | |
| P49222 | EPB42 | erythrocyte membrane protein ba | Plasma Membrane | transporter | 2.80E+07 | 5.72E+07 | 4.09E+07 | | | | | | |
| P49222 | EPB42 | erythrocyte membrane protein ba | Plasma Membrane | transporter | 1.41E+07 | 1.38E+07 | 1.60E+07 | | | | | | |
| P49222 | EPB42 | erythrocyte membrane protein ba | Plasma Membrane | transporter | 1.59E+07 | 1.52E+07 | 1.58E+07 | | | | | | |
| P49222 | EPB42 | erythrocyte membrane protein ba | Plasma Membrane | transporter | 1.38E+07 | 2091658.75 | 1864383.125 | | | | | | |
| P49222 | EPB42 | erythrocyte membrane protein ba | Plasma Membrane | transporter | 5.38E+08 | 4.77E+08 | 9.69E+08 | | | | | | |
| P49222 | EPB42 | erythrocyte membrane protein ba | Plasma Membrane | transporter | 8198479 | 1.28E+07 | 1.40E+07 | | | | | | |
| P49222 | EPB42 | erythrocyte membrane protein ba | Plasma Membrane | transporter | 4351443 | 5312249.5 | 4914757.5 | | | | | | |
| P49222 | EPB42 | erythrocyte membrane protein ba | Plasma Membrane | transporter | 4249105 | 2919739.5 | 3596001.75 | | | | | | |
| P49222 | EPB42 | erythrocyte membrane protein ba | Plasma Membrane | transporter | 7499838.5 | 1389735.875 | 1744639.25 | | | | | | |
| P49222 | EPB42 | erythrocyte membrane protein ba | Plasma Membrane | transporter | 2832658.75 | 6733664 | 6620415 | | | | | | |
| P49222 | EPB42 | erythrocyte membrane protein ba | Plasma Membrane | transporter | 2465894.5 | 5999824 | 9161748 | | | | | | |
| P49222 | EPB42 | erythrocyte membrane protein ba | Plasma Membrane | transporter | 7101147.5 | 2086883.625 | 2038423.75 | | | | | | |
| P49222 | EPB42 | erythrocyte membrane protein ba | Plasma Membrane | transporter | 8499913 | 6315492 | 6671196 | | | | | | |
| P49222 | EPB42 | erythrocyte membrane protein ba | Plasma Membrane | transporter | 1582831.875 | 7907494 | 5170186 | | | | | | |
| P49222 | EPB42 | erythrocyte membrane protein ba | Plasma Membrane | transporter | 1552872.875 | 404205.375 | 437736.656 | | | | | | |
| P49222 | EPB42 | erythrocyte membrane protein ba | Plasma Membrane | transporter | 1597340.625 | 1.46E+07 | 1.12E+07 | | | | | | |
| P49222 | EPB42 | erythrocyte membrane protein ba | Plasma Membrane | transporter | 778387.5 | 1086715 | 942262.875 | | | | | | |
| P49222 | EPB42 | erythrocyte membrane protein ba | Plasma Membrane | transporter | 2223086.25 | 1979542.125 | 1319575.125 | | | | | | |
| P49222 | EPB42 | erythrocyte membrane protein ba | Plasma Membrane | transporter | 1095109.25 | 799605.875 | 421981.906 | | | | | | |
| P49222 | EPB42 | erythrocyte membrane protein ba | Plasma Membrane | transporter | 2.35E+07 | 2.22E+07 | 4.81E+07 | | | | | | |
| P49222 | EPB42 | erythrocyte membrane protein ba | Plasma Membrane | transporter | 329315.281 | 32425.238 | 4.70E-06 | | | | | | |
| P49222 | EPB42 | erythrocyte membrane protein ba | Plasma Membrane | transporter | 213854.562 | 762206.125 | 2017906.5 | P49222 | EPB42 | 354220489 | 490159731 | 35374232 | 192 |
| Q80VP1 | EPN1 | epsin 1 | Plasma Membrane | other | 3074678.25 | 3209511.5 | 2910248.5 | | | | | | |
| Q80VP1 | EPN1 | epsin 1 | Plasma Membrane | other | 3198878 | 3456181 | 3044758 | | | | | | |
| Q80VP1 | EPN1 | epsin 1 | Plasma Membrane | other | 1271567.125 | 1391191.75 | 1576015.375 | | | | | | |
| Q80VP1 | EPN1 | epsin 1 | Plasma Membrane | other | 1557001.75 | 1050364.125 | 1455096.125 | Q80VP1 | EPN1 | 2275531 | 940056 | 271371 | 12 |
| Q8CGC7 | EPRS | glutamyl-prolyl-tRNA synthetase | Cytoplasm | enzyme | 1590843.75 | 1903661 | 1323373.125 | | | | | | |
| Q8CGC7 | EPRS | glutamyl-prolyl-tRNA synthetase | Cytoplasm | enzyme | 1854836 | 2776266.75 | 1400779.875 | Q8CGC7 | EPRS | 1722840 | 528532 | 215772 | 6 |
| P42567 | EPS15 | epidermal growth factor receptor | Cytoplasm | other | 4428392.5 | 4408967 | 3089497.25 | | | | | | |
| P42567 | EPS15 | epidermal growth factor receptor | Cytoplasm | other | 2252748.25 | 5723847 | 3849743.75 | | | | | | |
| P42567 | EPS15 | epidermal growth factor receptor | Cytoplasm | other | 3347740.25 | 3282547.75 | 1680185.125 | | | | | | |
| P42567 | EPS15 | epidermal growth factor receptor | Cytoplasm | other | 3442351 | 3718526.75 | 2471321.75 | | | | | | |
| P42567 | EPS15 | epidermal growth factor receptor | Cytoplasm | other | 2863769.75 | 2950439.25 | 1706799.75 | P42567 | EPS15 | 3267000 | 1074886 | 277534 | 15 |
| Q60902 | EPS15L1 | epidermal growth factor receptor | Plasma Membrane | other | 424061.719 | 374051.375 | 239655.609 | Q60902 | EPS15L1 | 424062 | 95367 | 55060 | 3 |
| Q9EQH2 | ERAP1 | endoplasmic reticulum aminopept | Extracellular Space | peptidase | 3569555 | 2661818 | 2799567.5 | Q9EQH2 | ERAP1 | 3569555 | 489190 | 282434 | 3 |
| Q80TH2 | ERBB2IP | erbb2 interacting protein | Plasma Membrane | other | 6496586 | 6384522 | 7789238 | | | | | | |

| | | | | | | | | | | | | | |
|---|---|---|---|---|---|---|---|---|---|---|---|---|---|
| Q80TH2 | ERBB2IP | erbb2 interacting protein | Plasma Membrane | other | 4531058.5 | 7446027 | 7644192 | | | | | | |
| Q80TH2 | ERBB2IP | erbb2 interacting protein | Plasma Membrane | other | 4694660 | 5494571.5 | 4823953 | | | | | | |
| Q80TH2 | ERBB2IP | erbb2 interacting protein | Plasma Membrane | other | 5244299.5 | 4679256 | 4719665.5 | | | | | | |
| Q80TH2 | ERBB2IP | erbb2 interacting protein | Plasma Membrane | other | 5171471 | 4380430.5 | 4786479.5 | | | | | | |
| Q80TH2 | ERBB2IP | erbb2 interacting protein | Plasma Membrane | other | 2908854 | 3862225.25 | 2663043.25 | | | | | | |
| Q80TH2 | ERBB2IP | erbb2 interacting protein | Plasma Membrane | other | 4692924.5 | 4216996 | 4029228.5 | | | | | | |
| Q80TH2 | ERBB2IP | erbb2 interacting protein | Plasma Membrane | other | 2514742.5 | 2741256.75 | 2595410.25 | Q80TH2 | ERBB2IP | 4531825 | 1543342 | 315033 | 24 |
| Q8BFZ9 | ERLIN2 | ER lipid raft associated 2 | Plasma Membrane | other | 2346011.5 | 4102537 | 3668749.25 | Q8BFZ9 | ERLIN2 | 2346012 | 914985 | 528267 | 3 |
| Q9JLN5 | ERMAP | erythroblast membrane-associate | Cytoplasm | other | 7.88E+07 | 9.82E+07 | 1.09E+08 | | | | | | |
| Q9JLN5 | ERMAP | erythroblast membrane-associate | Cytoplasm | other | 5.32E+07 | 6.42E+07 | 6.63E+07 | | | | | | |
| Q9JLN5 | ERMAP | erythroblast membrane-associate | Cytoplasm | other | 4.81E+07 | 4.90E+07 | 6.06E+07 | | | | | | |
| Q9JLN5 | ERMAP | erythroblast membrane-associate | Cytoplasm | other | 4.69E+07 | 5.29E+07 | 6.07E+07 | | | | | | |
| Q9JLN5 | ERMAP | erythroblast membrane-associate | Cytoplasm | other | 3.77E+07 | 4.10E+07 | 4.20E+07 | | | | | | |
| Q9JLN5 | ERMAP | erythroblast membrane-associate | Cytoplasm | other | 3.95E+07 | 4.79E+07 | 5.29E+07 | | | | | | |
| Q9JLN5 | ERMAP | erythroblast membrane-associate | Cytoplasm | other | 2.82E+07 | 2.52E+07 | 3.23E+07 | | | | | | |
| Q9JLN5 | ERMAP | erythroblast membrane-associate | Cytoplasm | other | 2.11E+07 | 2.11E+07 | 2.03E+07 | | | | | | |
| Q9JLN5 | ERMAP | erythroblast membrane-associate | Cytoplasm | other | 1.74E+07 | 1.43E+07 | 1.92E+07 | | | | | | |
| Q9JLN5 | ERMAP | erythroblast membrane-associate | Cytoplasm | other | 1.00E+07 | 1.85E+07 | 1.41E+07 | | | | | | |
| Q9JLN5 | ERMAP | erythroblast membrane-associate | Cytoplasm | other | 9461333 | 1.17E+07 | 1.39E+07 | | | | | | |
| Q9JLN5 | ERMAP | erythroblast membrane-associate | Cytoplasm | other | 6152004 | 6341429 | 6648828 | | | | | | |
| Q9JLN5 | ERMAP | erythroblast membrane-associate | Cytoplasm | other | 5248537.5 | 7372679 | 9277955 | | | | | | |
| Q9JLN5 | ERMAP | erythroblast membrane-associate | Cytoplasm | other | 4045528 | 8875865 | 8417030 | | | | | | |
| Q9JLN5 | ERMAP | erythroblast membrane-associate | Cytoplasm | other | 5722872.5 | 6326670.5 | 7858361 | | | | | | |
| Q9JLN5 | ERMAP | erythroblast membrane-associate | Cytoplasm | other | 4772693.5 | 4565914.5 | 5653812.5 | | | | | | |
| Q9JLN5 | ERMAP | erythroblast membrane-associate | Cytoplasm | other | 3293371.25 | 3502216 | 3991641.25 | | | | | | |
| Q9JLN5 | ERMAP | erythroblast membrane-associate | Cytoplasm | other | 3257238.25 | 1857475.25 | 4656555 | | | | | | |
| Q9JLN5 | ERMAP | erythroblast membrane-associate | Cytoplasm | other | 869391.812 | 5925718.5 | 7463648 | | | | | | |
| Q9JLN5 | ERMAP | erythroblast membrane-associate | Cytoplasm | other | 3175102.25 | 3201650.25 | 4322072 | Q9JLN5 | ERMAP | 21342904 | 25479732 | 3289419 | 60 |
| Q8R180 | ERO1A | endoplasmic reticulum oxidoreduc | Cytoplasm | enzyme | 1349002.625 | 1899309.875 | 1497061.5 | Q8R180 | ERO1A | 1349003 | 284770 | 164412 | 3 |
| P57759 | ERP29 | endoplasmic reticulum protein 29 | Cytoplasm | transporter | 9443199 | 1.35E+07 | 8289512.5 | | | | | | |
| P57759 | ERP29 | endoplasmic reticulum protein 29 | Cytoplasm | transporter | 5519861.5 | 8204504 | 5050724.5 | | | | | | |
| P57759 | ERP29 | endoplasmic reticulum protein 29 | Cytoplasm | transporter | 4567806 | 6284336.5 | 4397755 | | | | | | |
| P57759 | ERP29 | endoplasmic reticulum protein 29 | Cytoplasm | transporter | 3123415 | 3830536 | 2864182.25 | P57759 | ERP29 | 5663570 | 3109437 | 897617 | 12 |
| Q9D1Q6 | ERP44 | endoplasmic reticulum protein 44 | Cytoplasm | enzyme | 1.75E+07 | 1.84E+07 | 1.58E+07 | | | | | | |
| Q9D1Q6 | ERP44 | endoplasmic reticulum protein 44 | Cytoplasm | enzyme | 1.53E+07 | 1.72E+07 | 1.50E+07 | | | | | | |
| Q9D1Q6 | ERP44 | endoplasmic reticulum protein 44 | Cytoplasm | enzyme | 1.01E+07 | 1.28E+07 | 1.05E+07 | | | | | | |
| Q9D1Q6 | ERP44 | endoplasmic reticulum protein 44 | Cytoplasm | enzyme | 8993530 | 1.01E+07 | 9135856 | | | | | | |
| Q9D1Q6 | ERP44 | endoplasmic reticulum protein 44 | Cytoplasm | enzyme | 2825769.5 | 7997460 | 7340018 | | | | | | |
| Q9D1Q6 | ERP44 | endoplasmic reticulum protein 44 | Cytoplasm | enzyme | 4519463 | 3183056.25 | 1832453.75 | Q9D1Q6 | ERP44 | 9853127 | 5298345 | 1248832 | 18 |
| Q3U7R1 | ESYT1 | extended synaptotagmin-like prot | Cytoplasm | other | 2.28E+07 | 3.10E+07 | 2.02E+07 | | | | | | |
| Q3U7R1 | ESYT1 | extended synaptotagmin-like prot | Cytoplasm | other | 3.19E+07 | 2.76E+07 | 2.41E+07 | | | | | | |
| Q3U7R1 | ESYT1 | extended synaptotagmin-like prot | Cytoplasm | other | 2.82E+07 | 2.57E+07 | 2.07E+07 | | | | | | |
| Q3U7R1 | ESYT1 | extended synaptotagmin-like prot | Cytoplasm | other | 1.62E+07 | 1.58E+07 | 1.58E+07 | | | | | | |
| Q3U7R1 | ESYT1 | extended synaptotagmin-like prot | Cytoplasm | other | 1.62E+07 | 2.22E+07 | 1.60E+07 | | | | | | |
| Q3U7R1 | ESYT1 | extended synaptotagmin-like prot | Cytoplasm | other | 1.33E+07 | 1.37E+07 | 1.30E+07 | | | | | | |
| Q3U7R1 | ESYT1 | extended synaptotagmin-like prot | Cytoplasm | other | 1.12E+07 | 1.72E+07 | 1.34E+07 | | | | | | |
| Q3U7R1 | ESYT1 | extended synaptotagmin-like prot | Cytoplasm | other | 1.49E+07 | 1.31E+07 | 1.41E+07 | | | | | | |
| Q3U7R1 | ESYT1 | extended synaptotagmin-like prot | Cytoplasm | other | 1.38E+07 | 1.53E+07 | 1.29E+07 | | | | | | |
| Q3U7R1 | ESYT1 | extended synaptotagmin-like prot | Cytoplasm | other | 1.37E+07 | 1.31E+07 | 1.17E+07 | | | | | | |
| Q3U7R1 | ESYT1 | extended synaptotagmin-like prot | Cytoplasm | other | 1.09E+07 | 8727358 | 8377239.5 | | | | | | |
| Q3U7R1 | ESYT1 | extended synaptotagmin-like prot | Cytoplasm | other | 1.15E+07 | 1.25E+07 | 1.09E+07 | | | | | | |
| Q3U7R1 | ESYT1 | extended synaptotagmin-like prot | Cytoplasm | other | 1.01E+07 | 1.04E+07 | 9776888 | | | | | | |
| Q3U7R1 | ESYT1 | extended synaptotagmin-like prot | Cytoplasm | other | 9312672 | 9227535 | 6046565 | | | | | | |
| Q3U7R1 | ESYT1 | extended synaptotagmin-like prot | Cytoplasm | other | 6321894.5 | 9717902 | 6809082 | | | | | | |
| Q3U7R1 | ESYT1 | extended synaptotagmin-like prot | Cytoplasm | other | 7439710.5 | 7976855.5 | 6877747.5 | | | | | | |
| Q3U7R1 | ESYT1 | extended synaptotagmin-like prot | Cytoplasm | other | 8127792.5 | 1.07E+07 | 8258439 | | | | | | |
| Q3U7R1 | ESYT1 | extended synaptotagmin-like prot | Cytoplasm | other | 9410608 | 6967364 | 5557965.5 | | | | | | |
| Q3U7R1 | ESYT1 | extended synaptotagmin-like prot | Cytoplasm | other | 6200666.5 | 6438008 | 5866702 | | | | | | |
| Q3U7R1 | ESYT1 | extended synaptotagmin-like prot | Cytoplasm | other | 1.19E+07 | 3614246.5 | 3360112.75 | | | | | | |
| Q3U7R1 | ESYT1 | extended synaptotagmin-like prot | Cytoplasm | other | 6533781 | 5699515.5 | 4535616 | | | | | | |
| Q3U7R1 | ESYT1 | extended synaptotagmin-like prot | Cytoplasm | other | 6393230 | 6324009 | 5576888.5 | | | | | | |
| Q3U7R1 | ESYT1 | extended synaptotagmin-like prot | Cytoplasm | other | 4566718.5 | 6802488 | 6150052.5 | | | | | | |
| Q3U7R1 | ESYT1 | extended synaptotagmin-like prot | Cytoplasm | other | 6776340 | 6974367.5 | 6637390.5 | | | | | | |
| Q3U7R1 | ESYT1 | extended synaptotagmin-like prot | Cytoplasm | other | 4533210 | 1.04E+07 | 7696427.5 | | | | | | |
| Q3U7R1 | ESYT1 | extended synaptotagmin-like prot | Cytoplasm | other | 4279095.5 | 5034080.5 | 3616720.75 | | | | | | |
| Q3U7R1 | ESYT1 | extended synaptotagmin-like prot | Cytoplasm | other | 2453169.75 | 6230503.5 | 2844765.5 | | | | | | |
| Q3U7R1 | ESYT1 | extended synaptotagmin-like prot | Cytoplasm | other | 1692820.5 | 3762371.25 | 3713660.5 | | | | | | |
| Q3U7R1 | ESYT1 | extended synaptotagmin-like prot | Cytoplasm | other | 1388939.75 | 1459014.75 | 1520388.375 | Q3U7R1 | ESYT1 | 10750022 | 6792515 | 728234 | 87 |
| Q3TZZ7 | ESYT2 | extended synaptotagmin-like prot | Plasma Membrane | other | 9460137 | 1.16E+07 | 1.02E+07 | | | | | | |
| Q3TZZ7 | ESYT2 | extended synaptotagmin-like prot | Plasma Membrane | other | 9679195 | 9910724 | 9742347 | | | | | | |
| Q3TZZ7 | ESYT2 | extended synaptotagmin-like prot | Plasma Membrane | other | 9721866 | 1.01E+07 | 1.00E+07 | | | | | | |
| Q3TZZ7 | ESYT2 | extended synaptotagmin-like prot | Plasma Membrane | other | 1.02E+07 | 1.02E+07 | 7702671.5 | | | | | | |
| Q3TZZ7 | ESYT2 | extended synaptotagmin-like prot | Plasma Membrane | other | 8414859 | 5933656.5 | 6161388.5 | | | | | | |
| Q3TZZ7 | ESYT2 | extended synaptotagmin-like prot | Plasma Membrane | other | 9006091 | 9966853 | 8356485.5 | | | | | | |
| Q3TZZ7 | ESYT2 | extended synaptotagmin-like prot | Plasma Membrane | other | 7533350 | 7836918 | 7493562.5 | | | | | | |
| Q3TZZ7 | ESYT2 | extended synaptotagmin-like prot | Plasma Membrane | other | 9059510 | 6189388.5 | 7511765 | | | | | | |
| Q3TZZ7 | ESYT2 | extended synaptotagmin-like prot | Plasma Membrane | other | 8603426 | 9480121 | 6531764.5 | | | | | | |
| Q3TZZ7 | ESYT2 | extended synaptotagmin-like prot | Plasma Membrane | other | 4384611 | 8956900 | 6245991.5 | | | | | | |
| Q3TZZ7 | ESYT2 | extended synaptotagmin-like prot | Plasma Membrane | other | 5813859 | 9925905 | 8839993 | | | | | | |
| Q3TZZ7 | ESYT2 | extended synaptotagmin-like prot | Plasma Membrane | other | 5342682.5 | 5532335 | 4079453.75 | | | | | | |
| Q3TZZ7 | ESYT2 | extended synaptotagmin-like prot | Plasma Membrane | other | 5290820 | 5305524.5 | 4828360.5 | | | | | | |
| Q3TZZ7 | ESYT2 | extended synaptotagmin-like prot | Plasma Membrane | other | 4302819 | 5220747 | 3711226 | | | | | | |
| Q3TZZ7 | ESYT2 | extended synaptotagmin-like prot | Plasma Membrane | other | 8015319 | 8541610 | 7108658.5 | | | | | | |
| Q3TZZ7 | ESYT2 | extended synaptotagmin-like prot | Plasma Membrane | other | 5736751.5 | 6111255 | 4533517.5 | | | | | | |
| Q3TZZ7 | ESYT2 | extended synaptotagmin-like prot | Plasma Membrane | other | 4381302 | 3705055.25 | 3017814.75 | | | | | | |
| Q3TZZ7 | ESYT2 | extended synaptotagmin-like prot | Plasma Membrane | other | 1476097.375 | 5738221 | 3247939.25 | | | | | | |

| | | | | | | | | | | | | | |
|---|---|---|---|---|---|---|---|---|---|---|---|---|---|
| Q3TZZ7 | ESYT2 | extended synaptotagmin-like prot | Plasma Membrane | other | 1675556.875 | 1026561.812 | 1042735.188 | | | | | | |
| Q3TZZ7 | ESYT2 | extended synaptotagmin-like prot | Plasma Membrane | other | 68873.234 | 301548.969 | 28201.098 | Q3TZZ7 | ESYT2 | 6410356 | 2995885 | 386767 | 60 |
| Q99LC5 | ETFA | electron-transfer-flavoprotein, alp | Cytoplasm | transporter | 8035747 | 9307556 | 1.28E+07 | | | | | | |
| Q99LC5 | ETFA | electron-transfer-flavoprotein, alp | Cytoplasm | transporter | 6193012.5 | 8369626.5 | 1.04E+07 | | | | | | |
| Q99LC5 | ETFA | electron-transfer-flavoprotein, alp | Cytoplasm | transporter | 6733326.5 | 8834294 | 1.04E+07 | | | | | | |
| Q99LC5 | ETFA | electron-transfer-flavoprotein, alp | Cytoplasm | transporter | 5764593.5 | 4747419 | 6275438.5 | | | | | | |
| Q99LC5 | ETFA | electron-transfer-flavoprotein, alp | Cytoplasm | transporter | 2236386.25 | 2526070.5 | 3033434 | | | | | | |
| Q99LC5 | ETFA | electron-transfer-flavoprotein, alp | Cytoplasm | transporter | 3368055.5 | 2738068.25 | 4860300 | | | | | | |
| Q99LC5 | ETFA | electron-transfer-flavoprotein, alp | Cytoplasm | transporter | 2068649.25 | 2406191.25 | 2789394.25 | | | | | | |
| Q99LC5 | ETFA | electron-transfer-flavoprotein, alp | Cytoplasm | transporter | 503673.438 | 2122089.5 | 3409352.25 | | | | | | |
| Q99LC5 | ETFA | electron-transfer-flavoprotein, alp | Cytoplasm | transporter | 431254.219 | 682685.188 | 1259843.625 | Q99LC5 | ETFA | 3926078 | 3444953 | 662981 | 27 |
| Q9DCW4 | ETFB | electron-transfer-flavoprotein, bet | Cytoplasm | transporter | 1.19E+07 | 1.34E+07 | 1.74E+07 | | | | | | |
| Q9DCW4 | ETFB | electron-transfer-flavoprotein, bet | Cytoplasm | transporter | 3723932.75 | 3894503.75 | 5329181.5 | | | | | | |
| Q9DCW4 | ETFB | electron-transfer-flavoprotein, bet | Cytoplasm | transporter | 2845449.75 | 3122613 | 3707050.25 | | | | | | |
| Q9DCW4 | ETFB | electron-transfer-flavoprotein, bet | Cytoplasm | transporter | 1945531.125 | 2156190.25 | 2784102.5 | Q9DCW4 | ETFB | 5111228 | 5175346 | 1493994 | 12 |
| Q6DIA2 | EXOC3L4 | exocyst complex component 3-like | Cytoplasm | other | 1355715.625 | 1828232.875 | 2056587.875 | Q6DIA2 | EXOC3L4 | 1355716 | 357454 | 206376 | 3 |
| P26040 | EZR | ezrin | Plasma Membrane | other | 2.18E+07 | 1.61E+07 | 2.18E+07 | | | | | | |
| P26040 | EZR | ezrin | Plasma Membrane | other | 1.93E+07 | 9604126 | 1.56E+07 | | | | | | |
| P26040 | EZR | ezrin | Plasma Membrane | other | 1.20E+07 | 8608775 | 1.06E+07 | | | | | | |
| P26040 | EZR | ezrin | Plasma Membrane | other | 7226976.5 | 5838726 | 6354096.5 | | | | | | |
| P26040 | EZR | ezrin | Plasma Membrane | other | 8908526 | 4980604 | 6055170.5 | | | | | | |
| P26040 | EZR | ezrin | Plasma Membrane | other | 3526147.5 | 1156896.5 | 1231024.625 | | | | | | |
| P26040 | EZR | ezrin | Plasma Membrane | other | 3320829.75 | 4990207 | 2666932.5 | | | | | | |
| P26040 | EZR | ezrin | Plasma Membrane | other | 3743520.75 | 1279057.875 | 1825898.875 | | | | | | |
| P26040 | EZR | ezrin | Plasma Membrane | other | 2936668 | 1900130.625 | 1758859.625 | | | | | | |
| P26040 | EZR | ezrin | Plasma Membrane | other | 713208.562 | 1053.391 | 4.70E-06 | P26040 | EZR | 8348588 | 6421040 | 1172316 | 30 |
| O88792 | F11R | F11 receptor | Plasma Membrane | other | 1.22E+07 | 1.29E+07 | 1.22E+07 | | | | | | |
| O88792 | F11R | F11 receptor | Plasma Membrane | other | 3035525.5 | 3734481 | 4124935.5 | | | | | | |
| O88792 | F11R | F11 receptor | Plasma Membrane | other | 1824203.75 | 3445952.75 | 3772173.5 | | | | | | |
| O88792 | F11R | F11 receptor | Plasma Membrane | other | 2726828.25 | 4246342.5 | 4131972.75 | | | | | | |
| O88792 | F11R | F11 receptor | Plasma Membrane | other | 3776267.25 | 3934604 | 3678293.25 | O88792 | F11R | 4702565 | 3730288 | 963156 | 15 |
| P19221 | F2 | coagulation factor II (thrombin) | Extracellular Space | peptidase | 5544167 | 5075234 | 4580836.5 | P19221 | F2 | 5544167 | 481721 | 278122 | 3 |
| O88783 | F5 | coagulation factor V (proaccelerin, | Plasma Membrane | enzyme | 5005538 | 6241425 | 6863110 | | | | | | |
| O88783 | F5 | coagulation factor V (proaccelerin, | Plasma Membrane | enzyme | 1234816.75 | 2006699 | 1562193 | | | | | | |
| O88783 | F5 | coagulation factor V (proaccelerin, | Plasma Membrane | enzyme | 788572.75 | 804893.625 | 1697153.25 | O88783 | F5 | 2342976 | 2422768 | 807589 | 9 |
| Q920L1 | FADS1 | fatty acid desaturase 1 | Plasma Membrane | enzyme | 2711816 | 3305691.25 | 3853430.75 | Q920L1 | FADS1 | 2711816 | 570963 | 329645 | 3 |
| P54731 | FAF1 | Fas (TNFRSF6) associated factor 1 | Nucleus | other | 3536700 | 3005020.25 | 2761874.25 | | | | | | |
| P54731 | FAF1 | Fas (TNFRSF6) associated factor 1 | Nucleus | other | 2886942.25 | 2573590.5 | 2367038.25 | | | | | | |
| P54731 | FAF1 | Fas (TNFRSF6) associated factor 1 | Nucleus | other | 2546588.25 | 2235147 | 2614592.5 | P54731 | FAF1 | 2990077 | 387281 | 129094 | 9 |
| Q8R0F8 | FAHD1 | fumarylacetoacetate hydrolase do | Cytoplasm | enzyme | 2360784.25 | 2825126.25 | 2667014 | | | | | | |
| Q8R0F8 | FAHD1 | fumarylacetoacetate hydrolase do | Cytoplasm | enzyme | 1834203.375 | 2753119 | 3368764.5 | Q8R0F8 | FAHD1 | 2097494 | 511078 | 208647 | 6 |
| Q6NS82 | FAM134A | family with sequence similarity 13 | Extracellular Space | other | 2645562.75 | 2973780.25 | 2615209 | Q6NS82 | FAM134A | 2645563 | 198839 | 114800 | 3 |
| Q9D8B6 | FAM210B | family with sequence similarity 21 | Extracellular Space | other | 3357324.75 | 2626754.5 | 3134622.5 | | | | | | |
| Q9D8B6 | FAM210B | family with sequence similarity 21 | Extracellular Space | other | 145395.984 | 501119.719 | 327647.531 | Q9D8B6 | FAM210B | 1751360 | 1509920 | 616422 | 6 |
| Q9CYH2 | FAM213A | family with sequence similarity 21 | Extracellular Space | other | 1.38E+08 | 2.25E+08 | 1.50E+08 | | | | | | |
| Q9CYH2 | FAM213A | family with sequence similarity 21 | Extracellular Space | other | 9.02E+07 | 1.85E+08 | 1.34E+08 | | | | | | |
| Q9CYH2 | FAM213A | family with sequence similarity 21 | Extracellular Space | other | 5.69E+07 | 1.78E+08 | 1.10E+08 | | | | | | |
| Q9CYH2 | FAM213A | family with sequence similarity 21 | Extracellular Space | other | 6.52E+07 | 1.46E+08 | 1.13E+08 | | | | | | |
| Q9CYH2 | FAM213A | family with sequence similarity 21 | Extracellular Space | other | 8.36E+07 | 3.23E+07 | 6.13E+07 | | | | | | |
| Q9CYH2 | FAM213A | family with sequence similarity 21 | Extracellular Space | other | 3.93E+07 | 6.63E+07 | 4.89E+07 | | | | | | |
| Q9CYH2 | FAM213A | family with sequence similarity 21 | Extracellular Space | other | 2.36E+07 | 1.69E+07 | 1.15E+07 | | | | | | |
| Q9CYH2 | FAM213A | family with sequence similarity 21 | Extracellular Space | other | 1.08E+07 | 3.74E+07 | 2.50E+07 | | | | | | |
| Q9CYH2 | FAM213A | family with sequence similarity 21 | Extracellular Space | other | 9455955 | 1.86E+07 | 1.29E+07 | | | | | | |
| Q9CYH2 | FAM213A | family with sequence similarity 21 | Extracellular Space | other | 1.87E+07 | 7042330 | 1.36E+07 | | | | | | |
| Q9CYH2 | FAM213A | family with sequence similarity 21 | Extracellular Space | other | 9847650 | 1.24E+07 | 9066777 | | | | | | |
| Q9CYH2 | FAM213A | family with sequence similarity 21 | Extracellular Space | other | 792973.938 | 2885447.5 | 1060182.25 | Q9CYH2 | FAM213A | 45508882 | 61509338 | 10251556 | 36 |
| Q9WUA2 | FARSB | phenylalanyl-tRNA synthetase, bet | Cytoplasm | enzyme | 868776.75 | 1262312.125 | 950108.938 | Q9WUA2 | FARSB | 868777 | 207748 | 119943 | 3 |
| P19096 | FASN | fatty acid synthase | Cytoplasm | enzyme | 7655979.5 | 7653737.5 | 7637121 | | | | | | |
| P19096 | FASN | fatty acid synthase | Cytoplasm | enzyme | 1.03E+07 | 6094851 | 7565622 | | | | | | |
| P19096 | FASN | fatty acid synthase | Cytoplasm | enzyme | 6793245.5 | 7354767 | 8217931.5 | | | | | | |
| P19096 | FASN | fatty acid synthase | Cytoplasm | enzyme | 4216690.5 | 3528668 | 4224384 | | | | | | |
| P19096 | FASN | fatty acid synthase | Cytoplasm | enzyme | 4205126 | 4449465.5 | 6250015.5 | | | | | | |
| P19096 | FASN | fatty acid synthase | Cytoplasm | enzyme | 5233571.5 | 5275224.5 | 3627180.5 | | | | | | |
| P19096 | FASN | fatty acid synthase | Cytoplasm | enzyme | 5010391 | 5138775 | 5490646 | | | | | | |
| P19096 | FASN | fatty acid synthase | Cytoplasm | enzyme | 5231616.5 | 3742458 | 4746687.5 | | | | | | |
| P19096 | FASN | fatty acid synthase | Cytoplasm | enzyme | 5222780.5 | 2849228.5 | 4129589.75 | | | | | | |
| P19096 | FASN | fatty acid synthase | Cytoplasm | enzyme | 2812386.5 | 2273708 | 1984131.75 | | | | | | |
| P19096 | FASN | fatty acid synthase | Cytoplasm | enzyme | 3191225.25 | 2771212.5 | 2815074 | | | | | | |
| P19096 | FASN | fatty acid synthase | Cytoplasm | enzyme | 3011871.5 | 1743911 | 2478897.5 | | | | | | |
| P19096 | FASN | fatty acid synthase | Cytoplasm | enzyme | 3422886.75 | 3286853 | 2689876.75 | | | | | | |
| P19096 | FASN | fatty acid synthase | Cytoplasm | enzyme | 3623405 | 2382800 | 3271339.75 | | | | | | |
| P19096 | FASN | fatty acid synthase | Cytoplasm | enzyme | 2811795.75 | 2870720 | 1733331.375 | | | | | | |
| P19096 | FASN | fatty acid synthase | Cytoplasm | enzyme | 1981257.5 | 2122621.5 | 1854110.125 | | | | | | |
| P19096 | FASN | fatty acid synthase | Cytoplasm | enzyme | 2536140.25 | 2164701.25 | 2396225 | | | | | | |
| P19096 | FASN | fatty acid synthase | Cytoplasm | enzyme | 1625369.125 | 1347710 | 1259265.625 | | | | | | |
| P19096 | FASN | fatty acid synthase | Cytoplasm | enzyme | 996300.938 | 893416 | 314591.156 | | | | | | |
| P19096 | FASN | fatty acid synthase | Cytoplasm | enzyme | 2450187 | 1538968.375 | 1417630.25 | | | | | | |
| P19096 | FASN | fatty acid synthase | Cytoplasm | enzyme | 1078480.25 | 1510853.75 | 1821528.5 | | | | | | |
| P19096 | FASN | fatty acid synthase | Cytoplasm | enzyme | 244464.094 | 347999.219 | 357231.531 | | | | | | |
| P19096 | FASN | fatty acid synthase | Cytoplasm | enzyme | 372400.344 | 748165.375 | 691649.375 | P19096 | FASN | 3654242 | 2243964 | 270141 | 69 |
| P62862 | FAU | Finkel-Biskis-Reilly murine sarcoma | Cytoplasm | other | 5375533.5 | 6835382 | 6186480.5 | P62862 | FAU | 5375534 | 731422 | 422286 | 3 |
| Q3U7U3 | FBXO7 | F-box protein 7 | Cytoplasm | enzyme | 1991970.5 | 1462085.375 | 2447470 | Q3U7U3 | FBXO7 | 1991971 | 493160 | 284726 | 3 |
| P22315 | FECH | ferrochelatase | Cytoplasm | enzyme | 5.06E+07 | 5.57E+07 | 7.63E+07 | | | | | | |
| P22315 | FECH | ferrochelatase | Cytoplasm | enzyme | 3.43E+07 | 3.91E+07 | 4.49E+07 | | | | | | |
| P22315 | FECH | ferrochelatase | Cytoplasm | enzyme | 2.54E+07 | 2.02E+07 | 3.69E+07 | | | | | | |
| P22315 | FECH | ferrochelatase | Cytoplasm | enzyme | 2.12E+07 | 1.95E+07 | 3.20E+07 | | | | | | |

| | | | | | | | | | | | | | |
|---|---|---|---|---|---|---|---|---|---|---|---|---|---|
| P22315 | FECH | ferrochelatase | Cytoplasm | enzyme | 1.16E+07 | 1.13E+07 | 1.76E+07 | | | | | | |
| P22315 | FECH | ferrochelatase | Cytoplasm | enzyme | 7718520.5 | 9087027 | 1.43E+07 | | | | | | |
| P22315 | FECH | ferrochelatase | Cytoplasm | enzyme | 2740537 | 1.24E+07 | 1.79E+07 | | | | | | |
| P22315 | FECH | ferrochelatase | Cytoplasm | enzyme | 3932284 | 2.39E+07 | 4.10E+07 | | | | | | |
| P22315 | FECH | ferrochelatase | Cytoplasm | enzyme | 7996716 | 7469610.5 | 9070013 | | | | | | |
| P22315 | FECH | ferrochelatase | Cytoplasm | enzyme | 4983112 | 1738377.125 | 1905578.625 | | | | | | |
| P22315 | FECH | ferrochelatase | Cytoplasm | enzyme | 2171674.5 | 3441871.25 | 3789252.5 | | | | | | |
| P22315 | FECH | ferrochelatase | Cytoplasm | enzyme | 3684330 | 2819758.75 | 5427936 | | | | | | |
| P22315 | FECH | ferrochelatase | Cytoplasm | enzyme | 3478173 | 2656599.25 | 5228465 | | | | | | |
| P22315 | FECH | ferrochelatase | Cytoplasm | enzyme | 1947074.875 | 7485091.5 | 6734811 | | | | | | |
| P22315 | FECH | ferrochelatase | Cytoplasm | enzyme | 3048668.25 | 3002429.25 | 3299676 | | | | | | |
| P22315 | FECH | ferrochelatase | Cytoplasm | enzyme | 2409667.75 | 1650530.875 | 2115246 | | | | | | |
| P22315 | FECH | ferrochelatase | Cytoplasm | enzyme | 2792644 | 2209900.5 | 3482900.25 | | | | | | |
| P22315 | FECH | ferrochelatase | Cytoplasm | enzyme | 2448349 | 481592.219 | 904301.688 | | | | | | |
| P22315 | FECH | ferrochelatase | Cytoplasm | enzyme | 471144.406 | 2310885.5 | 3282914.75 | P22315 | FECH | | 10153837 | 16343027 | 2164687 | 57 |
| Q8K1B8 | FERMT3 | fermitin family member 3 | Cytoplasm | enzyme | 2.39E+07 | 2.34E+07 | 1.97E+07 | | | | | | |
| Q8K1B8 | FERMT3 | fermitin family member 3 | Cytoplasm | enzyme | 2.89E+07 | 2.73E+07 | 2.54E+07 | | | | | | |
| Q8K1B8 | FERMT3 | fermitin family member 3 | Cytoplasm | enzyme | 1.71E+07 | 1.46E+07 | 1.49E+07 | | | | | | |
| Q8K1B8 | FERMT3 | fermitin family member 3 | Cytoplasm | enzyme | 1.78E+07 | 1.54E+07 | 1.58E+07 | | | | | | |
| Q8K1B8 | FERMT3 | fermitin family member 3 | Cytoplasm | enzyme | 1.66E+07 | 9334277 | 1.49E+07 | | | | | | |
| Q8K1B8 | FERMT3 | fermitin family member 3 | Cytoplasm | enzyme | 1.67E+07 | 1.48E+07 | 1.33E+07 | | | | | | |
| Q8K1B8 | FERMT3 | fermitin family member 3 | Cytoplasm | enzyme | 1.36E+07 | 1.21E+07 | 1.21E+07 | | | | | | |
| Q8K1B8 | FERMT3 | fermitin family member 3 | Cytoplasm | enzyme | 1.48E+07 | 1.30E+07 | 1.37E+07 | | | | | | |
| Q8K1B8 | FERMT3 | fermitin family member 3 | Cytoplasm | enzyme | 1.07E+07 | 8497557 | 1.07E+07 | | | | | | |
| Q8K1B8 | FERMT3 | fermitin family member 3 | Cytoplasm | enzyme | 1.29E+07 | 9560043 | 9893366 | | | | | | |
| Q8K1B8 | FERMT3 | fermitin family member 3 | Cytoplasm | enzyme | 8837906 | 8530405 | 1.06E+07 | | | | | | |
| Q8K1B8 | FERMT3 | fermitin family member 3 | Cytoplasm | enzyme | 4685458 | 5380604 | 5179721.5 | | | | | | |
| Q8K1B8 | FERMT3 | fermitin family member 3 | Cytoplasm | enzyme | 5396698.5 | 4330893.5 | 5701378.5 | | | | | | |
| Q8K1B8 | FERMT3 | fermitin family member 3 | Cytoplasm | enzyme | 4990288 | 4271559.5 | 4507247.5 | | | | | | |
| Q8K1B8 | FERMT3 | fermitin family member 3 | Cytoplasm | enzyme | 5457932.5 | 4276801 | 5129545 | | | | | | |
| Q8K1B8 | FERMT3 | fermitin family member 3 | Cytoplasm | enzyme | 7297761.5 | 4114824.5 | 3203496.75 | Q8K1B8 | FERMT3 | | 13104128 | 6565013 | 947578 | 48 |
| Q8K0E8 | FGB | fibrinogen beta chain | Extracellular Space | other | 2.18E+07 | 2.03E+07 | 3.92E+07 | | | | | | |
| Q8K0E8 | FGB | fibrinogen beta chain | Extracellular Space | other | 3.48E+07 | 6.20E+07 | 6.69E+07 | | | | | | |
| Q8K0E8 | FGB | fibrinogen beta chain | Extracellular Space | other | 3.63E+07 | 3.43E+07 | 3.90E+07 | | | | | | |
| Q8K0E8 | FGB | fibrinogen beta chain | Extracellular Space | other | 2.02E+07 | 2.20E+07 | 2.24E+07 | | | | | | |
| Q8K0E8 | FGB | fibrinogen beta chain | Extracellular Space | other | 2.47E+07 | 2.72E+07 | 3.08E+07 | | | | | | |
| Q8K0E8 | FGB | fibrinogen beta chain | Extracellular Space | other | 3.34E+07 | 9764030 | 1.08E+07 | | | | | | |
| Q8K0E8 | FGB | fibrinogen beta chain | Extracellular Space | other | 2.31E+07 | 2.39E+07 | 2.58E+07 | | | | | | |
| Q8K0E8 | FGB | fibrinogen beta chain | Extracellular Space | other | 2.22E+07 | 2.03E+07 | 2.23E+07 | | | | | | |
| Q8K0E8 | FGB | fibrinogen beta chain | Extracellular Space | other | 2.47E+07 | 2.36E+07 | 3.04E+07 | | | | | | |
| Q8K0E8 | FGB | fibrinogen beta chain | Extracellular Space | other | 1.99E+07 | 2.27E+07 | 2.03E+07 | | | | | | |
| Q8K0E8 | FGB | fibrinogen beta chain | Extracellular Space | other | 1.35E+07 | 1.17E+07 | 1.08E+07 | | | | | | |
| Q8K0E8 | FGB | fibrinogen beta chain | Extracellular Space | other | 8077256.5 | 1.62E+07 | 1.88E+07 | | | | | | |
| Q8K0E8 | FGB | fibrinogen beta chain | Extracellular Space | other | 9277229 | 6226813 | 4977670 | | | | | | |
| Q8K0E8 | FGB | fibrinogen beta chain | Extracellular Space | other | 1.37E+07 | 9520470 | 1.22E+07 | | | | | | |
| Q8K0E8 | FGB | fibrinogen beta chain | Extracellular Space | other | 7255641 | 7110649.5 | 7072506 | | | | | | |
| Q8K0E8 | FGB | fibrinogen beta chain | Extracellular Space | other | 8375577 | 7159199.5 | 1.08E+07 | | | | | | |
| Q8K0E8 | FGB | fibrinogen beta chain | Extracellular Space | other | 3043518.5 | 4384970.5 | 8211383.5 | | | | | | |
| Q8K0E8 | FGB | fibrinogen beta chain | Extracellular Space | other | 3096045.75 | 1932189.875 | 3365568 | | | | | | |
| Q8K0E8 | FGB | fibrinogen beta chain | Extracellular Space | other | 1686733.125 | 3082329.75 | 6022827 | | | | | | |
| Q8K0E8 | FGB | fibrinogen beta chain | Extracellular Space | other | 2227769 | 695039.812 | 676055.062 | | | | | | |
| Q8K0E8 | FGB | fibrinogen beta chain | Extracellular Space | other | 1188139.25 | 1712182.25 | 1757164.5 | | | | | | |
| Q8K0E8 | FGB | fibrinogen beta chain | Extracellular Space | other | 1577884 | 3393650.25 | 2457710 | | | | | | |
| Q8K0E8 | FGB | fibrinogen beta chain | Extracellular Space | other | 953951.188 | 2347827.75 | 2911405.75 | | | | | | |
| Q8K0E8 | FGB | fibrinogen beta chain | Extracellular Space | other | 1198106 | 497776.875 | 244445.203 | Q8K0E8 | FGB | | 14012827 | 13937828 | 1642589 | 72 |
| Q8VCM7 | FGG | fibrinogen gamma chain | Extracellular Space | other | 5.75E+07 | 3.40E+07 | 3.12E+07 | | | | | | |
| Q8VCM7 | FGG | fibrinogen gamma chain | Extracellular Space | other | 7.07E+07 | 6.28E+07 | 5.74E+07 | | | | | | |
| Q8VCM7 | FGG | fibrinogen gamma chain | Extracellular Space | other | 3.48E+07 | 2.96E+07 | 2.71E+07 | | | | | | |
| Q8VCM7 | FGG | fibrinogen gamma chain | Extracellular Space | other | 1.31E+07 | 2.05E+07 | 1.92E+07 | | | | | | |
| Q8VCM7 | FGG | fibrinogen gamma chain | Extracellular Space | other | 2.03E+07 | 2.28E+07 | 1.71E+07 | | | | | | |
| Q8VCM7 | FGG | fibrinogen gamma chain | Extracellular Space | other | 1.94E+07 | 1.64E+07 | 1.73E+07 | | | | | | |
| Q8VCM7 | FGG | fibrinogen gamma chain | Extracellular Space | other | 1.50E+07 | 1.53E+07 | 1.77E+07 | | | | | | |
| Q8VCM7 | FGG | fibrinogen gamma chain | Extracellular Space | other | 1.45E+07 | 1.35E+07 | 1.85E+07 | | | | | | |
| Q8VCM7 | FGG | fibrinogen gamma chain | Extracellular Space | other | 7149292 | 1.54E+07 | 1.61E+07 | | | | | | |
| Q8VCM7 | FGG | fibrinogen gamma chain | Extracellular Space | other | 1.46E+07 | 1.06E+07 | 1.39E+07 | | | | | | |
| Q8VCM7 | FGG | fibrinogen gamma chain | Extracellular Space | other | 6860746.5 | 6766016 | 7426084.5 | | | | | | |
| Q8VCM7 | FGG | fibrinogen gamma chain | Extracellular Space | other | 9908530 | 8838382 | 1.20E+07 | | | | | | |
| Q8VCM7 | FGG | fibrinogen gamma chain | Extracellular Space | other | 5628048.5 | 1.29E+07 | 1.31E+07 | | | | | | |
| Q8VCM7 | FGG | fibrinogen gamma chain | Extracellular Space | other | 1.11E+07 | 1.12E+07 | 1.23E+07 | | | | | | |
| Q8VCM7 | FGG | fibrinogen gamma chain | Extracellular Space | other | 1.25E+07 | 1.25E+07 | 1.69E+07 | | | | | | |
| Q8VCM7 | FGG | fibrinogen gamma chain | Extracellular Space | other | 7752828 | 7844088 | 8866916 | | | | | | |
| Q8VCM7 | FGG | fibrinogen gamma chain | Extracellular Space | other | 1.46E+07 | 1.55E+07 | 1.66E+07 | | | | | | |
| Q8VCM7 | FGG | fibrinogen gamma chain | Extracellular Space | other | 1689483.875 | 2721804 | 3143319.75 | | | | | | |
| Q8VCM7 | FGG | fibrinogen gamma chain | Extracellular Space | other | 1327187.125 | 1.04E+07 | 1.52E+07 | | | | | | |
| Q8VCM7 | FGG | fibrinogen gamma chain | Extracellular Space | other | 5150983 | 4709998.5 | 3651385 | | | | | | |
| Q8VCM7 | FGG | fibrinogen gamma chain | Extracellular Space | other | 3545973.75 | 2417287.25 | 3146277 | | | | | | |
| Q8VCM7 | FGG | fibrinogen gamma chain | Extracellular Space | other | 475589.281 | 482461.375 | 870113.938 | | | | | | |
| Q8VCM7 | FGG | fibrinogen gamma chain | Extracellular Space | other | 468746.062 | 918233.062 | 1166126.25 | Q8VCM7 | FGG | | 15129018 | 14277186 | 1718771 | 69 |
| P97807 | FH | fumarate hydratase | Cytoplasm | enzyme | 1.31E+07 | 1.16E+07 | 1.45E+07 | | | | | | |
| P97807 | FH | fumarate hydratase | Cytoplasm | enzyme | 6813218.5 | 6517636 | 7973911 | | | | | | |
| P97807 | FH | fumarate hydratase | Cytoplasm | enzyme | 3650026 | 3785881.5 | 4067724.25 | | | | | | |
| P97807 | FH | fumarate hydratase | Cytoplasm | enzyme | 4223345.5 | 3643714.75 | 4023854 | | | | | | |
| P97807 | FH | fumarate hydratase | Cytoplasm | enzyme | 2273235.75 | 2432419 | 3035863.5 | | | | | | |
| P97807 | FH | fumarate hydratase | Cytoplasm | enzyme | 1159329.5 | 2766900.5 | 2707856.25 | | | | | | |
| P97807 | FH | fumarate hydratase | Cytoplasm | enzyme | 4072024.25 | 2711002 | 4120326.25 | P97807 | FH | | 5047311 | 3685021 | 804138 | 21 |
| Q9CQ92 | FIS1 | fission, mitochondrial 1 | Cytoplasm | other | 6066951 | 4961726 | 5813609.5 | | | | | | |

| | | | | | | | | | | | | | | |
|---|---|---|---|---|---|---|---|---|---|---|---|---|---|---|
| Q9CQ92 | FIS1 | fission, mitochondrial 1 | Cytoplasm | other | | 5555874 | 5763053.5 | 6729528 | Q9CQ92 | FIS1 | 5811413 | 582712 | 237891 | 6 |
| Q6P9Q6 | FKBP15 | FK506 binding protein 15, 133kDa | Plasma Membrane | enzyme | | 1264216.75 | 1299533.875 | 614303.312 | | | | | | |
| Q6P9Q6 | FKBP15 | FK506 binding protein 15, 133kDa | Plasma Membrane | enzyme | | 503245.875 | 545696.188 | 209179.297 | Q6P9Q6 | FKBP15 | 883731 | 442551 | 180671 | 6 |
| P45878 | FKBP2 | FK506 binding protein 2, 13kDa | Cytoplasm | enzyme | | 1.65E+07 | 2.24E+07 | 1.76E+07 | | | | | | |
| P45878 | FKBP2 | FK506 binding protein 2, 13kDa | Cytoplasm | enzyme | | 1.37E+07 | 1.92E+07 | 1.54E+07 | P45878 | FKBP2 | 15080000 | 3057775 | 1248331 | 6 |
| O35450 | FKBPL | FK506 binding protein like | Nucleus | other | | 1.92E+07 | 2.47E+07 | 1.15E+07 | O35450 | FKBPL | 19240000 | 6663740 | 3847312 | 3 |
| Q2VIS4 | Flg2 | filaggrin family member 2 | Extracellular Space | other | | 4.66E+07 | 2.72E+07 | 3.91E+07 | Q2VIS4 | Flg2 | 46640000 | 9818637 | 5668793 | 3 |
| Q8BTM8 | FLNA | filamin A, alpha | Cytoplasm | other | | 1.89E+08 | 1.79E+08 | 1.63E+08 | | | | | | |
| Q8BTM8 | FLNA | filamin A, alpha | Cytoplasm | other | | 1.33E+08 | 1.39E+08 | 1.22E+08 | | | | | | |
| Q8BTM8 | FLNA | filamin A, alpha | Cytoplasm | other | | 1.64E+08 | 8.81E+07 | 1.18E+08 | | | | | | |
| Q8BTM8 | FLNA | filamin A, alpha | Cytoplasm | other | | 1.16E+08 | 1.41E+08 | 1.55E+08 | | | | | | |
| Q8BTM8 | FLNA | filamin A, alpha | Cytoplasm | other | | 1.04E+08 | 1.18E+08 | 1.16E+08 | | | | | | |
| Q8BTM8 | FLNA | filamin A, alpha | Cytoplasm | other | | 2.28E+07 | 3.66E+07 | 2.70E+07 | | | | | | |
| Q8BTM8 | FLNA | filamin A, alpha | Cytoplasm | other | | 8.44E+07 | 1.05E+08 | 7.60E+07 | | | | | | |
| Q8BTM8 | FLNA | filamin A, alpha | Cytoplasm | other | | 7.71E+07 | 9.61E+07 | 9.83E+07 | | | | | | |
| Q8BTM8 | FLNA | filamin A, alpha | Cytoplasm | other | | 5.97E+07 | 4.69E+07 | 6.05E+07 | | | | | | |
| Q8BTM8 | FLNA | filamin A, alpha | Cytoplasm | other | | 6.47E+07 | 7.04E+07 | 6.71E+07 | | | | | | |
| Q8BTM8 | FLNA | filamin A, alpha | Cytoplasm | other | | 4.45E+07 | 5.04E+07 | 5.76E+07 | | | | | | |
| Q8BTM8 | FLNA | filamin A, alpha | Cytoplasm | other | | 6.25E+07 | 5.98E+07 | 6.48E+07 | | | | | | |
| Q8BTM8 | FLNA | filamin A, alpha | Cytoplasm | other | | 4.94E+07 | 4.20E+07 | 4.35E+07 | | | | | | |
| Q8BTM8 | FLNA | filamin A, alpha | Cytoplasm | other | | 4.67E+07 | 6.30E+07 | 6.00E+07 | | | | | | |
| Q8BTM8 | FLNA | filamin A, alpha | Cytoplasm | other | | 7.55E+07 | 8.79E+07 | 8.45E+07 | | | | | | |
| Q8BTM8 | FLNA | filamin A, alpha | Cytoplasm | other | | 6.77E+07 | 7.60E+07 | 6.89E+07 | | | | | | |
| Q8BTM8 | FLNA | filamin A, alpha | Cytoplasm | other | | 4.55E+07 | 5.34E+07 | 6.04E+07 | | | | | | |
| Q8BTM8 | FLNA | filamin A, alpha | Cytoplasm | other | | 7.97E+07 | 8.18E+07 | 8.68E+07 | | | | | | |
| Q8BTM8 | FLNA | filamin A, alpha | Cytoplasm | other | | 6.67E+07 | 6.11E+07 | 5.78E+07 | | | | | | |
| Q8BTM8 | FLNA | filamin A, alpha | Cytoplasm | other | | 4.55E+07 | 6.01E+07 | 5.98E+07 | | | | | | |
| Q8BTM8 | FLNA | filamin A, alpha | Cytoplasm | other | | 5.38E+07 | 6.82E+07 | 6.40E+07 | | | | | | |
| Q8BTM8 | FLNA | filamin A, alpha | Cytoplasm | other | | 5.46E+07 | 6.22E+07 | 5.98E+07 | | | | | | |
| Q8BTM8 | FLNA | filamin A, alpha | Cytoplasm | other | | 5.96E+07 | 6.32E+07 | 5.95E+07 | | | | | | |
| Q8BTM8 | FLNA | filamin A, alpha | Cytoplasm | other | | 6.83E+07 | 7.79E+07 | 1.12E+08 | | | | | | |
| Q8BTM8 | FLNA | filamin A, alpha | Cytoplasm | other | | 6.47E+07 | 6.43E+07 | 6.24E+07 | | | | | | |
| Q8BTM8 | FLNA | filamin A, alpha | Cytoplasm | other | | 2.51E+07 | 4.12E+07 | 3.56E+07 | | | | | | |
| Q8BTM8 | FLNA | filamin A, alpha | Cytoplasm | other | | 3.99E+07 | 8385359.5 | 1.84E+07 | | | | | | |
| Q8BTM8 | FLNA | filamin A, alpha | Cytoplasm | other | | 5.75E+07 | 6.38E+07 | 7.22E+07 | | | | | | |
| Q8BTM8 | FLNA | filamin A, alpha | Cytoplasm | other | | 5.26E+07 | 6.28E+07 | 5.62E+07 | | | | | | |
| Q8BTM8 | FLNA | filamin A, alpha | Cytoplasm | other | | 5.65E+07 | 6.02E+07 | 5.45E+07 | | | | | | |
| Q8BTM8 | FLNA | filamin A, alpha | Cytoplasm | other | | 5.12E+07 | 5.63E+07 | 4.86E+07 | | | | | | |
| Q8BTM8 | FLNA | filamin A, alpha | Cytoplasm | other | | 4.86E+07 | 5.30E+07 | 5.80E+07 | | | | | | |
| Q8BTM8 | FLNA | filamin A, alpha | Cytoplasm | other | | 2.16E+07 | 2.61E+07 | 2.30E+07 | | | | | | |
| Q8BTM8 | FLNA | filamin A, alpha | Cytoplasm | other | | 3.95E+07 | 7.50E+07 | 6.24E+07 | | | | | | |
| Q8BTM8 | FLNA | filamin A, alpha | Cytoplasm | other | | 3.74E+07 | 3.19E+07 | 4.21E+07 | | | | | | |
| Q8BTM8 | FLNA | filamin A, alpha | Cytoplasm | other | | 4.75E+07 | 3.71E+07 | 3.15E+07 | | | | | | |
| Q8BTM8 | FLNA | filamin A, alpha | Cytoplasm | other | | 3.23E+07 | 5.85E+07 | 5.60E+07 | | | | | | |
| Q8BTM8 | FLNA | filamin A, alpha | Cytoplasm | other | | 4.93E+07 | 5.72E+07 | 6.06E+07 | | | | | | |
| Q8BTM8 | FLNA | filamin A, alpha | Cytoplasm | other | | 3.48E+07 | 4.71E+07 | 5.02E+07 | | | | | | |
| Q8BTM8 | FLNA | filamin A, alpha | Cytoplasm | other | | 3.73E+07 | 4.22E+07 | 4.97E+07 | | | | | | |
| Q8BTM8 | FLNA | filamin A, alpha | Cytoplasm | other | | 4.60E+07 | 4.59E+07 | 5.99E+07 | | | | | | |
| Q8BTM8 | FLNA | filamin A, alpha | Cytoplasm | other | | 1.53E+07 | 1.50E+07 | 1.41E+07 | | | | | | |
| Q8BTM8 | FLNA | filamin A, alpha | Cytoplasm | other | | 4.13E+07 | 5.00E+07 | 5.90E+07 | | | | | | |
| Q8BTM8 | FLNA | filamin A, alpha | Cytoplasm | other | | 4.04E+07 | 3.74E+07 | 3.72E+07 | | | | | | |
| Q8BTM8 | FLNA | filamin A, alpha | Cytoplasm | other | | 2.38E+07 | 3.06E+07 | 3.29E+07 | | | | | | |
| Q8BTM8 | FLNA | filamin A, alpha | Cytoplasm | other | | 4.68E+07 | 5.47E+07 | 6.19E+07 | | | | | | |
| Q8BTM8 | FLNA | filamin A, alpha | Cytoplasm | other | | 3.73E+07 | 6.60E+07 | 6.46E+07 | | | | | | |
| Q8BTM8 | FLNA | filamin A, alpha | Cytoplasm | other | | 4.24E+07 | 5.72E+07 | 5.35E+07 | | | | | | |
| Q8BTM8 | FLNA | filamin A, alpha | Cytoplasm | other | | 9169626 | 1.15E+07 | 1.10E+07 | | | | | | |
| Q8BTM8 | FLNA | filamin A, alpha | Cytoplasm | other | | 4.37E+07 | 4.02E+07 | 4.42E+07 | | | | | | |
| Q8BTM8 | FLNA | filamin A, alpha | Cytoplasm | other | | 2.87E+07 | 3.02E+07 | 3.20E+07 | | | | | | |
| Q8BTM8 | FLNA | filamin A, alpha | Cytoplasm | other | | 3.58E+07 | 4.53E+07 | 3.70E+07 | | | | | | |
| Q8BTM8 | FLNA | filamin A, alpha | Cytoplasm | other | | 3.68E+07 | 4.36E+07 | 5.09E+07 | | | | | | |
| Q8BTM8 | FLNA | filamin A, alpha | Cytoplasm | other | | 2.94E+07 | 3.05E+07 | 3.09E+07 | | | | | | |
| Q8BTM8 | FLNA | filamin A, alpha | Cytoplasm | other | | 3.89E+07 | 3.59E+07 | 3.79E+07 | | | | | | |
| Q8BTM8 | FLNA | filamin A, alpha | Cytoplasm | other | | 3.67E+07 | 4.97E+07 | 5.11E+07 | | | | | | |
| Q8BTM8 | FLNA | filamin A, alpha | Cytoplasm | other | | 2.98E+07 | 3.36E+07 | 4.20E+07 | | | | | | |
| Q8BTM8 | FLNA | filamin A, alpha | Cytoplasm | other | | 3.02E+07 | 4.22E+07 | 3.86E+07 | | | | | | |
| Q8BTM8 | FLNA | filamin A, alpha | Cytoplasm | other | | 1.04E+07 | 1.23E+07 | 9579644 | | | | | | |
| Q8BTM8 | FLNA | filamin A, alpha | Cytoplasm | other | | 6458924 | 1309278.25 | 6818596.5 | | | | | | |
| Q8BTM8 | FLNA | filamin A, alpha | Cytoplasm | other | | 1.73E+07 | 2.67E+07 | 2.64E+07 | | | | | | |
| Q8BTM8 | FLNA | filamin A, alpha | Cytoplasm | other | | 2.43E+07 | 3.05E+07 | 3.08E+07 | | | | | | |
| Q8BTM8 | FLNA | filamin A, alpha | Cytoplasm | other | | 1.96E+07 | 1.94E+07 | 1.81E+07 | | | | | | |
| Q8BTM8 | FLNA | filamin A, alpha | Cytoplasm | other | | 1.12E+07 | 1.32E+07 | 1.41E+07 | | | | | | |
| Q8BTM8 | FLNA | filamin A, alpha | Cytoplasm | other | | 2.98E+07 | 2.50E+07 | 2.69E+07 | | | | | | |
| Q8BTM8 | FLNA | filamin A, alpha | Cytoplasm | other | | 2.98E+07 | 2.49E+07 | 2.72E+07 | | | | | | |
| Q8BTM8 | FLNA | filamin A, alpha | Cytoplasm | other | | 2.68E+07 | 3.18E+07 | 3.23E+07 | | | | | | |
| Q8BTM8 | FLNA | filamin A, alpha | Cytoplasm | other | | 1.21E+07 | 4.48E+07 | 2.43E+07 | | | | | | |
| Q8BTM8 | FLNA | filamin A, alpha | Cytoplasm | other | | 1.82E+07 | 1.33E+07 | 1.85E+07 | | | | | | |
| Q8BTM8 | FLNA | filamin A, alpha | Cytoplasm | other | | 2.96E+07 | 2.01E+07 | 2.48E+07 | | | | | | |
| Q8BTM8 | FLNA | filamin A, alpha | Cytoplasm | other | | 2.15E+07 | 2.75E+07 | 2.28E+07 | | | | | | |
| Q8BTM8 | FLNA | filamin A, alpha | Cytoplasm | other | | 2.19E+07 | 2.25E+07 | 2.56E+07 | | | | | | |
| Q8BTM8 | FLNA | filamin A, alpha | Cytoplasm | other | | 2.58E+07 | 2.75E+07 | 2.82E+07 | | | | | | |
| Q8BTM8 | FLNA | filamin A, alpha | Cytoplasm | other | | 2.18E+07 | 2.10E+07 | 2.55E+07 | | | | | | |
| Q8BTM8 | FLNA | filamin A, alpha | Cytoplasm | other | | 5582868 | 8500602 | 1.18E+07 | | | | | | |
| Q8BTM8 | FLNA | filamin A, alpha | Cytoplasm | other | | 1.88E+07 | 1.63E+07 | 1.53E+07 | | | | | | |
| Q8BTM8 | FLNA | filamin A, alpha | Cytoplasm | other | | 2.51E+07 | 1.54E+07 | 1.45E+07 | | | | | | |
| Q8BTM8 | FLNA | filamin A, alpha | Cytoplasm | other | | 8444284 | 8203471 | 1.07E+07 | | | | | | |
| Q8BTM8 | FLNA | filamin A, alpha | Cytoplasm | other | | 9945540 | 1.34E+07 | 1.36E+07 | | | | | | |

| | | | | | | | | | | | | | |
|---|---|---|---|---|---|---|---|---|---|---|---|---|---|
| Q8BTM8 | FLNA | filamin A, alpha | Cytoplasm | other | 2.15E+07 | 2.01E+07 | 2.33E+07 | | | | | | |
| Q8BTM8 | FLNA | filamin A, alpha | Cytoplasm | other | 1.73E+07 | 1.53E+07 | 1.66E+07 | | | | | | |
| Q8BTM8 | FLNA | filamin A, alpha | Cytoplasm | other | 1.30E+07 | 1.19E+07 | 1.65E+07 | | | | | | |
| Q8BTM8 | FLNA | filamin A, alpha | Cytoplasm | other | 2.76E+07 | 2.11E+07 | 1.76E+07 | | | | | | |
| Q8BTM8 | FLNA | filamin A, alpha | Cytoplasm | other | 1.75E+07 | 1.64E+07 | 1.66E+07 | | | | | | |
| Q8BTM8 | FLNA | filamin A, alpha | Cytoplasm | other | 2.62E+07 | 4396680.5 | 4509579.5 | | | | | | |
| Q8BTM8 | FLNA | filamin A, alpha | Cytoplasm | other | 1.37E+07 | 1.40E+07 | 1.75E+07 | | | | | | |
| Q8BTM8 | FLNA | filamin A, alpha | Cytoplasm | other | 8687348 | 8571790 | 5613251.5 | | | | | | |
| Q8BTM8 | FLNA | filamin A, alpha | Cytoplasm | other | 2366147.75 | 5494977.5 | 4113187 | | | | | | |
| Q8BTM8 | FLNA | filamin A, alpha | Cytoplasm | other | 1.04E+07 | 9863384 | 1.00E+07 | | | | | | |
| Q8BTM8 | FLNA | filamin A, alpha | Cytoplasm | other | 1.37E+07 | 5.22E+07 | 4.71E+07 | | | | | | |
| Q8BTM8 | FLNA | filamin A, alpha | Cytoplasm | other | 1.58E+07 | 2145966.5 | 8220601.5 | | | | | | |
| Q8BTM8 | FLNA | filamin A, alpha | Cytoplasm | other | 9838715 | 1.36E+07 | 1.43E+07 | | | | | | |
| Q8BTM8 | FLNA | filamin A, alpha | Cytoplasm | other | 1.36E+07 | 2.12E+07 | 1.70E+07 | | | | | | |
| Q8BTM8 | FLNA | filamin A, alpha | Cytoplasm | other | 1.19E+07 | 1.04E+07 | 9852565 | | | | | | |
| Q8BTM8 | FLNA | filamin A, alpha | Cytoplasm | other | 9364723 | 1.17E+07 | 9512667 | | | | | | |
| Q8BTM8 | FLNA | filamin A, alpha | Cytoplasm | other | 1.21E+07 | 1.38E+07 | 1.39E+07 | | | | | | |
| Q8BTM8 | FLNA | filamin A, alpha | Cytoplasm | other | 1.07E+07 | 1.46E+07 | 1.45E+07 | | | | | | |
| Q8BTM8 | FLNA | filamin A, alpha | Cytoplasm | other | 1.14E+07 | 1.02E+07 | 1.07E+07 | | | | | | |
| Q8BTM8 | FLNA | filamin A, alpha | Cytoplasm | other | 6855021 | 6570192.5 | 6347064.5 | | | | | | |
| Q8BTM8 | FLNA | filamin A, alpha | Cytoplasm | other | 1.11E+07 | 8991531 | 8966249 | | | | | | |
| Q8BTM8 | FLNA | filamin A, alpha | Cytoplasm | other | 2.57E+07 | 5279112 | 1.46E+07 | | | | | | |
| Q8BTM8 | FLNA | filamin A, alpha | Cytoplasm | other | 5926830 | 1.01E+07 | 9103691 | | | | | | |
| Q8BTM8 | FLNA | filamin A, alpha | Cytoplasm | other | 6595419.5 | 6833299 | 7337291 | | | | | | |
| Q8BTM8 | FLNA | filamin A, alpha | Cytoplasm | other | 3495952 | 1922689 | 2779197.25 | | | | | | |
| Q8BTM8 | FLNA | filamin A, alpha | Cytoplasm | other | 3199244.25 | 9809297 | 1.11E+07 | | | | | | |
| Q8BTM8 | FLNA | filamin A, alpha | Cytoplasm | other | 4576484.5 | 4632403.5 | 3542966.25 | | | | | | |
| Q8BTM8 | FLNA | filamin A, alpha | Cytoplasm | other | 1.01E+07 | 1.85E+07 | 2.11E+07 | | | | | | |
| Q8BTM8 | FLNA | filamin A, alpha | Cytoplasm | other | 2715728.75 | 5182097 | 4058103.75 | | | | | | |
| Q8BTM8 | FLNA | filamin A, alpha | Cytoplasm | other | 6037983.5 | 5122051.5 | 5201656.5 | | | | | | |
| Q8BTM8 | FLNA | filamin A, alpha | Cytoplasm | other | 5680504.5 | 983862.5 | 661331.125 | | | | | | |
| Q8BTM8 | FLNA | filamin A, alpha | Cytoplasm | other | 2670514 | 1378881.875 | 1563249.5 | | | | | | |
| Q8BTM8 | FLNA | filamin A, alpha | Cytoplasm | other | 8142496 | 363274.375 | 3333562.5 | | | | | | |
| Q8BTM8 | FLNA | filamin A, alpha | Cytoplasm | other | 2122227.75 | 5631922 | 4290337 | | | | | | |
| Q8BTM8 | FLNA | filamin A, alpha | Cytoplasm | other | 3796576.25 | 2024127.75 | 3402911.5 | | | | | | |
| Q8BTM8 | FLNA | filamin A, alpha | Cytoplasm | other | 4554810 | 2202861.75 | 2276152 | | | | | | |
| Q8BTM8 | FLNA | filamin A, alpha | Cytoplasm | other | 2431862.75 | 3108589.75 | 4698046 | | | | | | |
| Q8BTM8 | FLNA | filamin A, alpha | Cytoplasm | other | 733752.062 | 1533521.875 | 2850267.5 | | | | | | |
| Q8BTM8 | FLNA | filamin A, alpha | Cytoplasm | other | 2556491.5 | 883214.438 | 1808486.125 | Q8BTM8 | FLNA | 33247204 | 31918329 | 1696440 | 354 |
| O08917 | FLOT1 | flotillin 1 | Plasma Membrane | other | 1.41E+07 | 1.41E+07 | 1.71E+07 | | | | | | |
| O08917 | FLOT1 | flotillin 1 | Plasma Membrane | other | 1.86E+07 | 2.08E+07 | 1.91E+07 | | | | | | |
| O08917 | FLOT1 | flotillin 1 | Plasma Membrane | other | 1.89E+07 | 1.23E+07 | 1.01E+07 | | | | | | |
| O08917 | FLOT1 | flotillin 1 | Plasma Membrane | other | 1.40E+07 | 1.13E+07 | 1.27E+07 | | | | | | |
| O08917 | FLOT1 | flotillin 1 | Plasma Membrane | other | 9996720 | 1.04E+07 | 1.05E+07 | | | | | | |
| O08917 | FLOT1 | flotillin 1 | Plasma Membrane | other | 1.02E+07 | 9919506 | 8500205 | | | | | | |
| O08917 | FLOT1 | flotillin 1 | Plasma Membrane | other | 1.36E+07 | 1.02E+07 | 7668915 | | | | | | |
| O08917 | FLOT1 | flotillin 1 | Plasma Membrane | other | 1.05E+07 | 5515052 | 7670119 | | | | | | |
| O08917 | FLOT1 | flotillin 1 | Plasma Membrane | other | 1.03E+07 | 1.10E+07 | 1.05E+07 | | | | | | |
| O08917 | FLOT1 | flotillin 1 | Plasma Membrane | other | 5575284.5 | 6061116 | 5628352 | | | | | | |
| O08917 | FLOT1 | flotillin 1 | Plasma Membrane | other | 5678699.5 | 6669003.5 | 5734712.5 | | | | | | |
| O08917 | FLOT1 | flotillin 1 | Plasma Membrane | other | 3338006 | 7541102 | 6427733 | | | | | | |
| O08917 | FLOT1 | flotillin 1 | Plasma Membrane | other | 3194886 | 3684352.75 | 3321693.75 | | | | | | |
| O08917 | FLOT1 | flotillin 1 | Plasma Membrane | other | 4740157.5 | 5513286.5 | 4575770.5 | | | | | | |
| O08917 | FLOT1 | flotillin 1 | Plasma Membrane | other | 2182096.5 | 3484667.75 | 4551225.5 | | | | | | |
| O08917 | FLOT1 | flotillin 1 | Plasma Membrane | other | 2907334 | 59832.258 | 244698.562 | O08917 | FLOT1 | 9237074 | 5034241 | 726630 | 48 |
| Q60634 | FLOT2 | flotillin 2 | Plasma Membrane | other | 3.28E+07 | 2.11E+07 | 2.86E+07 | | | | | | |
| Q60634 | FLOT2 | flotillin 2 | Plasma Membrane | other | 2.22E+07 | 2.15E+07 | 2.70E+07 | | | | | | |
| Q60634 | FLOT2 | flotillin 2 | Plasma Membrane | other | 1.66E+07 | 1.57E+07 | 1.60E+07 | | | | | | |
| Q60634 | FLOT2 | flotillin 2 | Plasma Membrane | other | 1.75E+07 | 1.75E+07 | 2.01E+07 | | | | | | |
| Q60634 | FLOT2 | flotillin 2 | Plasma Membrane | other | 1.64E+07 | 1.49E+07 | 1.38E+07 | | | | | | |
| Q60634 | FLOT2 | flotillin 2 | Plasma Membrane | other | 1.49E+07 | 1.20E+07 | 1.18E+07 | | | | | | |
| Q60634 | FLOT2 | flotillin 2 | Plasma Membrane | other | 1.07E+07 | 1.28E+07 | 1.19E+07 | | | | | | |
| Q60634 | FLOT2 | flotillin 2 | Plasma Membrane | other | 9704218 | 9876025 | 1.06E+07 | | | | | | |
| Q60634 | FLOT2 | flotillin 2 | Plasma Membrane | other | 7490574.5 | 7721533 | 8548283 | | | | | | |
| Q60634 | FLOT2 | flotillin 2 | Plasma Membrane | other | 8270696 | 1.03E+07 | 9373429 | | | | | | |
| Q60634 | FLOT2 | flotillin 2 | Plasma Membrane | other | 7700761 | 6412771 | 6189686.5 | | | | | | |
| Q60634 | FLOT2 | flotillin 2 | Plasma Membrane | other | 3943632.75 | 5676397 | 5840548 | | | | | | |
| Q60634 | FLOT2 | flotillin 2 | Plasma Membrane | other | 3172066.5 | 3042449.25 | 3020421.75 | | | | | | |
| Q60634 | FLOT2 | flotillin 2 | Plasma Membrane | other | 3554432 | 3091235 | 3748990 | | | | | | |
| Q60634 | FLOT2 | flotillin 2 | Plasma Membrane | other | 2520578.25 | 3868166.75 | 3867352.75 | | | | | | |
| Q60634 | FLOT2 | flotillin 2 | Plasma Membrane | other | 763160.5 | 4360351.5 | 2749959.75 | | | | | | |
| Q60634 | FLOT2 | flotillin 2 | Plasma Membrane | other | 1861339.125 | 442749.5 | 171868.578 | Q60634 | FLOT2 | 10597733 | 7681896 | 1075681 | 51 |
| P11276 | FN1 | fibronectin 1 | Extracellular Space | enzyme | 299415.875 | 315601.594 | 416168.344 | | | | | | |
| P11276 | FN1 | fibronectin 1 | Extracellular Space | enzyme | 303568.125 | 349461.594 | 358762.219 | | | | | | |
| P11276 | FN1 | fibronectin 1 | Extracellular Space | enzyme | 735051.75 | 0 | 300489.688 | P11276 | FN1 | 446012 | 188083 | 62694 | 9 |
| Q9ER35 | FN3K | fructosamine 3 kinase | Cytoplasm | kinase | 1.23E+07 | 1.23E+07 | 1.21E+07 | | | | | | |
| Q9ER35 | FN3K | fructosamine 3 kinase | Cytoplasm | kinase | 1.16E+07 | 1.11E+07 | 1.06E+07 | | | | | | |
| Q9ER35 | FN3K | fructosamine 3 kinase | Cytoplasm | kinase | 1.12E+07 | 1.41E+07 | 1.07E+07 | | | | | | |
| Q9ER35 | FN3K | fructosamine 3 kinase | Cytoplasm | kinase | 7618483.5 | 6419934 | 4014542.25 | | | | | | |
| Q9ER35 | FN3K | fructosamine 3 kinase | Cytoplasm | kinase | 1.31E+07 | 9754650 | 8184408.5 | | | | | | |
| Q9ER35 | FN3K | fructosamine 3 kinase | Cytoplasm | kinase | 1918846.125 | 3469046.5 | 1482148.875 | | | | | | |
| Q9ER35 | FN3K | fructosamine 3 kinase | Cytoplasm | kinase | 1952655.25 | 2540693.5 | 1470780.875 | Q9ER35 | FN3K | 8531426 | 4439638 | 968808 | 21 |
| Q8K274 | FN3KRP | fructosamine 3 kinase related prot | Cytoplasm | kinase | 2.28E+07 | 2.94E+07 | 2.09E+07 | | | | | | |
| Q8K274 | FN3KRP | fructosamine 3 kinase related prot | Cytoplasm | kinase | 1.62E+07 | 1.55E+07 | 1.18E+07 | | | | | | |
| Q8K274 | FN3KRP | fructosamine 3 kinase related prot | Cytoplasm | kinase | 9975366 | 1.07E+07 | 8110553.5 | | | | | | |
| Q8K274 | FN3KRP | fructosamine 3 kinase related prot | Cytoplasm | kinase | 8222028.5 | 9102641 | 7603507 | | | | | | |

| | | | | | | | | | | | | | |
|---|---|---|---|---|---|---|---|---|---|---|---|---|---|
| Q8K274 | FN3KRP | fructosamine 3 kinase related prot | Cytoplasm | kinase | 6147588.5 | 5181165 | 7274182.5 | | | | | | |
| Q8K274 | FN3KRP | fructosamine 3 kinase related prot | Cytoplasm | kinase | 4544268 | 5116258 | 6804519.5 | Q8K274 | FN3KRP | 11318209 | 6943473 | 1636592 | 18 |
| Q3UFK8 | FRMD8 | FERM domain containing 8 | Cytoplasm | other | 4527805.5 | 7208731 | 6142874 | Q3UFK8 | FRMD8 | 4527806 | 1349806 | 779311 | 3 |
| P09528 | FTH1 | ferritin, heavy polypeptide 1 | Cytoplasm | enzyme | 1836814.875 | 1873934 | 3172387.25 | P09528 | FTH1 | 1836815 | 760604 | 439135 | 3 |
| P29391 | FTL | ferritin, light polypeptide | Cytoplasm | enzyme | 5908476 | 4501758 | 1.06E+07 | | | | | | |
| P29391 | FTL | ferritin, light polypeptide | Cytoplasm | enzyme | 2480834.5 | 2902324 | 4733930.5 | | | | | | |
| P29391 | FTL | ferritin, light polypeptide | Cytoplasm | enzyme | 1203320 | 855761.438 | 1446902.5 | P29391 | FTL | 3197544 | 3075285 | 1025095 | 9 |
| Q9D6K8 | FUNDC2 | FUN14 domain containing 2 | Cytoplasm | other | 6934546 | 8313268 | 9271200 | | | | | | |
| Q9D6K8 | FUNDC2 | FUN14 domain containing 2 | Cytoplasm | other | 3071093.5 | 6055881 | 7709968.5 | Q9D6K8 | FUNDC2 | 5002820 | 2174852 | 887879 | 6 |
| O35943 | FXN | frataxin | Cytoplasm | kinase | 3066425.25 | 2725103.75 | 3270231.5 | | | | | | |
| O35943 | FXN | frataxin | Cytoplasm | kinase | 3271099.25 | 2708996.5 | 3035137.25 | O35943 | FXN | 3168762 | 249572 | 101887 | 6 |
| O35601 | FYB | FYN binding protein | Nucleus | other | 3531200 | 2704041.5 | 3127478.75 | | | | | | |
| O35601 | FYB | FYN binding protein | Nucleus | other | 1348019.75 | 1305440 | 1385614.625 | | | | | | |
| O35601 | FYB | FYN binding protein | Nucleus | other | 1761404.625 | 1481920.375 | 1707286.25 | O35601 | FYB | 2213541 | 851002 | 283667 | 9 |
| P97855 | G3BP1 | GTPase activating protein (SH3 do | Nucleus | enzyme | 8508107 | 6950029 | 4481356.5 | | | | | | |
| P97855 | G3BP1 | GTPase activating protein (SH3 do | Nucleus | enzyme | 6649377 | 5728587 | 4688580 | P97855 | G3BP1 | 7578742 | 1519600 | 620374 | 6 |
| Q6NSQ9 | G6PC3 | glucose 6 phosphatase, catalytic, 3 | Cytoplasm | phosphatase | 2439961 | 2642534.75 | 3353619 | Q6NSQ9 | G6PC3 | 2439961 | 479835 | 277033 | 3 |
| Q00612 | G6PD | glucose-6-phosphate dehydrogena | Cytoplasm | enzyme | 2343171.25 | 1039685.562 | 1899593.5 | | | | | | |
| Q00612 | G6PD | glucose-6-phosphate dehydrogena | Cytoplasm | enzyme | 1464955.125 | 619535.938 | 1177824.25 | | | | | | |
| Q00612 | G6PD | glucose-6-phosphate dehydrogena | Cytoplasm | enzyme | 1058909.875 | 690812.688 | 766609.562 | | | | | | |
| Q00612 | G6PD | glucose-6-phosphate dehydrogena | Cytoplasm | enzyme | 1327496 | 517922.312 | 696405.75 | Q00612 | G6PD | 1548633 | 554220 | 159990 | 12 |
| P60521 | GABARAPL2 | GABA(A) receptor-associated prot | Cytoplasm | other | 1.71E+07 | 1.67E+07 | 1.22E+07 | | | | | | |
| P60521 | GABARAPL2 | GABA(A) receptor-associated prot | Cytoplasm | other | 1.33E+07 | 1.36E+07 | 9063504 | | | | | | |
| P60521 | GABARAPL2 | GABA(A) receptor-associated prot | Cytoplasm | other | 1.14E+07 | 1.22E+07 | 7356434 | | | | | | |
| P60521 | GABARAPL2 | GABA(A) receptor-associated prot | Cytoplasm | other | 6398918 | 9020012 | 7058171 | | | | | | |
| P60521 | GABARAPL2 | GABA(A) receptor-associated prot | Cytoplasm | other | 1505132.875 | 1803430.625 | 699368.688 | P60521 | GABARAPL2 | 9940810 | 5202782 | 1343352 | 15 |
| Q8BHN3 | GANAB | glucosidase, alpha; neutral AB | Cytoplasm | enzyme | 1.66E+07 | 1.49E+07 | 1.42E+07 | | | | | | |
| Q8BHN3 | GANAB | glucosidase, alpha; neutral AB | Cytoplasm | enzyme | 1.60E+07 | 1.65E+07 | 1.62E+07 | | | | | | |
| Q8BHN3 | GANAB | glucosidase, alpha; neutral AB | Cytoplasm | enzyme | 1.53E+07 | 1.52E+07 | 1.59E+07 | | | | | | |
| Q8BHN3 | GANAB | glucosidase, alpha; neutral AB | Cytoplasm | enzyme | 9885804 | 8800784 | 5593895.5 | | | | | | |
| Q8BHN3 | GANAB | glucosidase, alpha; neutral AB | Cytoplasm | enzyme | 1.43E+07 | 1.38E+07 | 1.21E+07 | | | | | | |
| Q8BHN3 | GANAB | glucosidase, alpha; neutral AB | Cytoplasm | enzyme | 9363919 | 1.10E+07 | 1.13E+07 | | | | | | |
| Q8BHN3 | GANAB | glucosidase, alpha; neutral AB | Cytoplasm | enzyme | 1.02E+07 | 1.31E+07 | 1.14E+07 | | | | | | |
| Q8BHN3 | GANAB | glucosidase, alpha; neutral AB | Cytoplasm | enzyme | 7959747 | 8324732.5 | 9082837 | | | | | | |
| Q8BHN3 | GANAB | glucosidase, alpha; neutral AB | Cytoplasm | enzyme | 8027487 | 9611591 | 7993428.5 | | | | | | |
| Q8BHN3 | GANAB | glucosidase, alpha; neutral AB | Cytoplasm | enzyme | 5917396 | 9404079 | 8023180 | | | | | | |
| Q8BHN3 | GANAB | glucosidase, alpha; neutral AB | Cytoplasm | enzyme | 5680743 | 7659928.5 | 7181964 | | | | | | |
| Q8BHN3 | GANAB | glucosidase, alpha; neutral AB | Cytoplasm | enzyme | 1.24E+07 | 1985620.625 | 6141558.5 | | | | | | |
| Q8BHN3 | GANAB | glucosidase, alpha; neutral AB | Cytoplasm | enzyme | 3968895.75 | 7455456.5 | 4866101 | | | | | | |
| Q8BHN3 | GANAB | glucosidase, alpha; neutral AB | Cytoplasm | enzyme | 3120135.25 | 6406685.5 | 5749909 | | | | | | |
| Q8BHN3 | GANAB | glucosidase, alpha; neutral AB | Cytoplasm | enzyme | 3908400.5 | 4090864 | 4829710.5 | | | | | | |
| Q8BHN3 | GANAB | glucosidase, alpha; neutral AB | Cytoplasm | enzyme | 3491683.25 | 4169950.75 | 3131320.5 | | | | | | |
| Q8BHN3 | GANAB | glucosidase, alpha; neutral AB | Cytoplasm | enzyme | 2636197.75 | 3832730.25 | 2914938 | | | | | | |
| Q8BHN3 | GANAB | glucosidase, alpha; neutral AB | Cytoplasm | enzyme | 2712380.75 | 2760556 | 2794348.5 | | | | | | |
| Q8BHN3 | GANAB | glucosidase, alpha; neutral AB | Cytoplasm | enzyme | 1261655.75 | 1517669.5 | 1523661.75 | | | | | | |
| Q8BHN3 | GANAB | glucosidase, alpha; neutral AB | Cytoplasm | enzyme | 868007 | 1386274.375 | 1065581.125 | Q8BHN3 | GANAB | 7674623 | 4788103 | 618141 | 60 |
| P16858 | GAPDH | glyceraldehyde-3-phosphate dehy | Cytoplasm | enzyme | 2.57E+08 | 2.49E+08 | 2.26E+08 | | | | | | |
| P16858 | GAPDH | glyceraldehyde-3-phosphate dehy | Cytoplasm | enzyme | 1.82E+08 | 1.13E+08 | 2.06E+08 | | | | | | |
| P16858 | GAPDH | glyceraldehyde-3-phosphate dehy | Cytoplasm | enzyme | 1.88E+08 | 1.81E+08 | 1.98E+08 | | | | | | |
| P16858 | GAPDH | glyceraldehyde-3-phosphate dehy | Cytoplasm | enzyme | 2.06E+08 | 6.42E+07 | 1.40E+08 | | | | | | |
| P16858 | GAPDH | glyceraldehyde-3-phosphate dehy | Cytoplasm | enzyme | 1.06E+08 | 2.52E+08 | 1.78E+08 | | | | | | |
| P16858 | GAPDH | glyceraldehyde-3-phosphate dehy | Cytoplasm | enzyme | 8.40E+07 | 1.15E+08 | 9.43E+07 | | | | | | |
| P16858 | GAPDH | glyceraldehyde-3-phosphate dehy | Cytoplasm | enzyme | 1.41E+08 | 1.36E+08 | 1.52E+08 | | | | | | |
| P16858 | GAPDH | glyceraldehyde-3-phosphate dehy | Cytoplasm | enzyme | 1.15E+08 | 1.04E+08 | 1.02E+08 | | | | | | |
| P16858 | GAPDH | glyceraldehyde-3-phosphate dehy | Cytoplasm | enzyme | 7.24E+07 | 8.79E+07 | 1.12E+08 | | | | | | |
| P16858 | GAPDH | glyceraldehyde-3-phosphate dehy | Cytoplasm | enzyme | 6.41E+07 | 1.45E+08 | 1.69E+08 | | | | | | |
| P16858 | GAPDH | glyceraldehyde-3-phosphate dehy | Cytoplasm | enzyme | 8.09E+07 | 6.34E+07 | 6.11E+07 | | | | | | |
| P16858 | GAPDH | glyceraldehyde-3-phosphate dehy | Cytoplasm | enzyme | 4.45E+07 | 3.34E+07 | 2.68E+07 | | | | | | |
| P16858 | GAPDH | glyceraldehyde-3-phosphate dehy | Cytoplasm | enzyme | 5.71E+07 | 5.82E+07 | 6.89E+07 | | | | | | |
| P16858 | GAPDH | glyceraldehyde-3-phosphate dehy | Cytoplasm | enzyme | 3.61E+07 | 3.94E+07 | 7.35E+07 | | | | | | |
| P16858 | GAPDH | glyceraldehyde-3-phosphate dehy | Cytoplasm | enzyme | 5.16E+07 | 7956148 | 8122013.5 | | | | | | |
| P16858 | GAPDH | glyceraldehyde-3-phosphate dehy | Cytoplasm | enzyme | 2.64E+07 | 2.18E+07 | 2.68E+07 | | | | | | |
| P16858 | GAPDH | glyceraldehyde-3-phosphate dehy | Cytoplasm | enzyme | 1.23E+07 | 1.66E+07 | 1.76E+07 | | | | | | |
| P16858 | GAPDH | glyceraldehyde-3-phosphate dehy | Cytoplasm | enzyme | 7805172.5 | 1.64E+07 | 1.23E+07 | | | | | | |
| P16858 | GAPDH | glyceraldehyde-3-phosphate dehy | Cytoplasm | enzyme | 1.33E+07 | 2800979.25 | 6738873.5 | | | | | | |
| P16858 | GAPDH | glyceraldehyde-3-phosphate dehy | Cytoplasm | enzyme | 8883443 | 3.27E+07 | 7.19E+07 | | | | | | |
| P16858 | GAPDH | glyceraldehyde-3-phosphate dehy | Cytoplasm | enzyme | 1.04E+07 | 1.00E+07 | 1.15E+07 | | | | | | |
| P16858 | GAPDH | glyceraldehyde-3-phosphate dehy | Cytoplasm | enzyme | 1.04E+07 | 9158458 | 9531339 | | | | | | |
| P16858 | GAPDH | glyceraldehyde-3-phosphate dehy | Cytoplasm | enzyme | 5286719.5 | 7883254 | 5703879 | | | | | | |
| P16858 | GAPDH | glyceraldehyde-3-phosphate dehy | Cytoplasm | enzyme | 2612309 | 1.68E+07 | 8746592 | | | | | | |
| P16858 | GAPDH | glyceraldehyde-3-phosphate dehy | Cytoplasm | enzyme | 6808510 | 7613924 | 7587817.5 | | | | | | |
| P16858 | GAPDH | glyceraldehyde-3-phosphate dehy | Cytoplasm | enzyme | 3622883 | 7621188 | 6900769.5 | | | | | | |
| P16858 | GAPDH | glyceraldehyde-3-phosphate dehy | Cytoplasm | enzyme | 2923327.75 | 4220734.5 | 4229450.5 | | | | | | |
| P16858 | GAPDH | glyceraldehyde-3-phosphate dehy | Cytoplasm | enzyme | 2259685 | 3072199 | 3275403.5 | | | | | | |
| P16858 | GAPDH | glyceraldehyde-3-phosphate dehy | Cytoplasm | enzyme | 1206913.375 | 221169.391 | 241503.859 | P16858 | GAPDH | 62075137 | 71598783 | 7676195 | 87 |
| Q6PAR5 | GAPVD1 | GTPase activating protein and VPS | Cytoplasm | other | 9933166 | 6693638.5 | 6673529 | | | | | | |
| Q6PAR5 | GAPVD1 | GTPase activating protein and VPS | Cytoplasm | other | 8048963.5 | 8977145 | 8081130 | | | | | | |
| Q6PAR5 | GAPVD1 | GTPase activating protein and VPS | Cytoplasm | other | 8561378 | 6385101 | 5707227 | | | | | | |
| Q6PAR5 | GAPVD1 | GTPase activating protein and VPS | Cytoplasm | other | 5437890 | 6669802 | 5170612.5 | | | | | | |
| Q6PAR5 | GAPVD1 | GTPase activating protein and VPS | Cytoplasm | other | 6364065.5 | 5979848.5 | 6796652 | | | | | | |
| Q6PAR5 | GAPVD1 | GTPase activating protein and VPS | Cytoplasm | other | 5973353.5 | 5882767.5 | 6086726 | | | | | | |
| Q6PAR5 | GAPVD1 | GTPase activating protein and VPS | Cytoplasm | other | 5904052 | 4976544.5 | 5539490 | | | | | | |
| Q6PAR5 | GAPVD1 | GTPase activating protein and VPS | Cytoplasm | other | 4200186.5 | 4645988.5 | 3282320.75 | | | | | | |
| Q6PAR5 | GAPVD1 | GTPase activating protein and VPS | Cytoplasm | other | 4703328 | 4219357 | 3789916.25 | | | | | | |
| Q6PAR5 | GAPVD1 | GTPase activating protein and VPS | Cytoplasm | other | 2696421 | 3245509.25 | 3718133 | | | | | | |
| Q6PAR5 | GAPVD1 | GTPase activating protein and VPS | Cytoplasm | other | 2869520.75 | 2537731.75 | 3754760.5 | | | | | | |

| | | | | | | | | | | | | | |
|---|---|---|---|---|---|---|---|---|---|---|---|---|---|
| Q6PAR5 | GAPVD1 | GTPase activating protein and VPS | Cytoplasm | other | 3404094 | 2893604.75 | 2870504.5 | | | | | | |
| Q6PAR5 | GAPVD1 | GTPase activating protein and VPS | Cytoplasm | other | 2646034 | 3040281.75 | 2677486.25 | | | | | | |
| Q6PAR5 | GAPVD1 | GTPase activating protein and VPS | Cytoplasm | other | 1994134.75 | 2140704.25 | 728771.75 | | | | | | |
| Q6PAR5 | GAPVD1 | GTPase activating protein and VPS | Cytoplasm | other | 2532744.75 | 1788647.625 | 1836072.75 | | | | | | |
| Q6PAR5 | GAPVD1 | GTPase activating protein and VPS | Cytoplasm | other | 1339533.875 | 1796719.25 | 1540314.625 | | | | | | |
| Q6PAR5 | GAPVD1 | GTPase activating protein and VPS | Cytoplasm | other | 912000.375 | 1228980.125 | 964737.125 | Q6PAR5 | GAPVD1 | 4560051 | 2313495 | 323954 | 51 |
| Q3V0G7 | GARNL3 | GTPase activating Rap/RanGAP do | Cytoplasm | other | 3704114.75 | 1418055.125 | 3732596 | Q3V0G7 | GARNL3 | 3704115 | 1328155 | 766811 | 3 |
| O09172 | GCLM | glutamate-cysteine ligase, modifie | Cytoplasm | enzyme | 7119186 | 4892882.5 | 6728850.5 | | | | | | |
| O09172 | GCLM | glutamate-cysteine ligase, modifie | Cytoplasm | enzyme | 6781637 | 3501103 | 5081570.5 | | | | | | |
| O09172 | GCLM | glutamate-cysteine ligase, modifie | Cytoplasm | enzyme | 4477492 | 2865684 | 3531658.25 | O09172 | GCLM | 6126105 | 1577309 | 525770 | 9 |
| Q9R111 | GDA | guanine deaminase | Cytoplasm | enzyme | 2.95E+07 | 2.00E+07 | 2.80E+07 | | | | | | |
| Q9R111 | GDA | guanine deaminase | Cytoplasm | enzyme | 1.68E+07 | 1.29E+07 | 2.18E+07 | | | | | | |
| Q9R111 | GDA | guanine deaminase | Cytoplasm | enzyme | 2.32E+07 | 1.49E+07 | 2.01E+07 | | | | | | |
| Q9R111 | GDA | guanine deaminase | Cytoplasm | enzyme | 1.45E+07 | 8052166 | 9109360 | | | | | | |
| Q9R111 | GDA | guanine deaminase | Cytoplasm | enzyme | 1.60E+07 | 1.18E+07 | 1.34E+07 | | | | | | |
| Q9R111 | GDA | guanine deaminase | Cytoplasm | enzyme | 8690465 | 5011854.5 | 7531341.5 | | | | | | |
| Q9R111 | GDA | guanine deaminase | Cytoplasm | enzyme | 1.17E+07 | 7114979 | 1.20E+07 | | | | | | |
| Q9R111 | GDA | guanine deaminase | Cytoplasm | enzyme | 7548862 | 5357142.5 | 6274301 | | | | | | |
| Q9R111 | GDA | guanine deaminase | Cytoplasm | enzyme | 9637286 | 5317186.5 | 9982816 | | | | | | |
| Q9R111 | GDA | guanine deaminase | Cytoplasm | enzyme | 3359286.75 | 2599137.5 | 2630936.25 | | | | | | |
| Q9R111 | GDA | guanine deaminase | Cytoplasm | enzyme | 3051915.25 | 1485276.875 | 2453104.25 | Q9R111 | GDA | 13078892 | 7412422 | 1290337 | 33 |
| Q9JL56 | GDE1 | glycerophosphodiester phosphodi | Plasma Membrane | enzyme | 5943148 | 7971161 | 6112167.5 | | | | | | |
| Q9JL56 | GDE1 | glycerophosphodiester phosphodi | Plasma Membrane | enzyme | 2882257.75 | 2393472.75 | 2239904.5 | | | | | | |
| Q9JL56 | GDE1 | glycerophosphodiester phosphodi | Plasma Membrane | enzyme | 671754.875 | 745082.375 | 365445.75 | Q9JL56 | GDE1 | 3165720 | 2758296 | 919432 | 9 |
| Q9CRY7 | GDPD1 | glycerophosphodiester phosphodi | Plasma Membrane | enzyme | 2441463 | 2690917 | 2721309.25 | Q9CRY7 | GDPD1 | 2441463 | 153550 | 88652 | 3 |
| Q91VC9 | GHITM | growth hormone inducible transm | Cytoplasm | other | 2151141.25 | 1799230.5 | 3226479 | | | | | | |
| Q91VC9 | GHITM | growth hormone inducible transm | Cytoplasm | other | 1554343 | 1597478 | 1909693.75 | Q91VC9 | GHITM | 1852742 | 620761 | 253425 | 6 |
| Q9CYL5 | GLIPR2 | GLI pathogenesis-related 2 | Cytoplasm | other | 3620591 | 3195244.25 | 2838973.75 | Q9CYL5 | GLIPR2 | 3620591 | 391317 | 225927 | 3 |
| Q80Y14 | GLRX5 | glutaredoxin 5 | Cytoplasm | other | 2.37E+07 | 2.33E+07 | 3.59E+07 | | | | | | |
| Q80Y14 | GLRX5 | glutaredoxin 5 | Cytoplasm | other | 1.58E+07 | 1.24E+07 | 2.03E+07 | Q80Y14 | GLRX5 | 19740000 | 8139831 | 3323072 | 6 |
| P26443 | GLUD1 | glutamate dehydrogenase 1 | Cytoplasm | enzyme | 2886873.5 | 3007232.75 | 2987508.5 | | | | | | |
| P26443 | GLUD1 | glutamate dehydrogenase 1 | Cytoplasm | enzyme | 2790341.5 | 2621868.5 | 2820065.5 | P26443 | GLUD1 | 2838608 | 142448 | 58154 | 6 |
| P63158 | Gm21596/Hmgb1 | high mobility group box 1 | Nucleus | transcription regulator | 325784.344 | 119385.023 | 742926.5 | | | | | | |
| P63158 | Gm21596/Hmgb1 | high mobility group box 1 | Nucleus | transcription regulator | 222034.625 | 384033.844 | 682741.438 | P63158 | Gm21596/Hmgb1 | 273909 | 250094 | 102100 | 6 |
| P21278 | GNA11 | guanine nucleotide binding protei | Plasma Membrane | enzyme | 3321035.75 | 3796911.25 | 3071435 | P21278 | GNA11 | 3321036 | 368572 | 212795 | 3 |
| P27600 | GNA12 | guanine nucleotide binding protei | Plasma Membrane | enzyme | 5.60E+07 | 6.88E+07 | 3.68E+07 | P27600 | GNA12 | 55970000 | 16144733 | 9321166 | 3 |
| P08752 | GNAI2 | guanine nucleotide binding protei | Plasma Membrane | other | 9.16E+07 | 1.14E+08 | 6.51E+07 | | | | | | |
| P08752 | GNAI2 | guanine nucleotide binding protei | Plasma Membrane | other | 8.90E+07 | 9.51E+07 | 6.20E+07 | | | | | | |
| P08752 | GNAI2 | guanine nucleotide binding protei | Plasma Membrane | other | 5.32E+07 | 8.00E+07 | 3.94E+07 | | | | | | |
| P08752 | GNAI2 | guanine nucleotide binding protei | Plasma Membrane | other | 7.96E+07 | 5.42E+07 | 4.17E+07 | | | | | | |
| P08752 | GNAI2 | guanine nucleotide binding protei | Plasma Membrane | other | 4.57E+07 | 8.63E+07 | 5.13E+07 | | | | | | |
| P08752 | GNAI2 | guanine nucleotide binding protei | Plasma Membrane | other | 5.79E+07 | 7.37E+07 | 4.13E+07 | | | | | | |
| P08752 | GNAI2 | guanine nucleotide binding protei | Plasma Membrane | other | 5.31E+07 | 5.86E+07 | 3.70E+07 | | | | | | |
| P08752 | GNAI2 | guanine nucleotide binding protei | Plasma Membrane | other | 3.46E+07 | 4.92E+07 | 2.78E+07 | | | | | | |
| P08752 | GNAI2 | guanine nucleotide binding protei | Plasma Membrane | other | 3.68E+07 | 5.28E+07 | 3.08E+07 | | | | | | |
| P08752 | GNAI2 | guanine nucleotide binding protei | Plasma Membrane | other | 3.61E+07 | 4.37E+07 | 3.13E+07 | | | | | | |
| P08752 | GNAI2 | guanine nucleotide binding protei | Plasma Membrane | other | 2.15E+07 | 2.77E+07 | 1.62E+07 | | | | | | |
| P08752 | GNAI2 | guanine nucleotide binding protei | Plasma Membrane | other | 1.71E+07 | 2.52E+07 | 1.46E+07 | | | | | | |
| P08752 | GNAI2 | guanine nucleotide binding protei | Plasma Membrane | other | 1.43E+07 | 1.40E+07 | 6971051 | | | | | | |
| P08752 | GNAI2 | guanine nucleotide binding protei | Plasma Membrane | other | 1.22E+07 | 2.64E+07 | 1.91E+07 | | | | | | |
| P08752 | GNAI2 | guanine nucleotide binding protei | Plasma Membrane | other | 1.42E+07 | 1.68E+07 | 9277668 | | | | | | |
| P08752 | GNAI2 | guanine nucleotide binding protei | Plasma Membrane | other | 1.55E+07 | 2.29E+07 | 1.16E+07 | | | | | | |
| P08752 | GNAI2 | guanine nucleotide binding protei | Plasma Membrane | other | 9239506 | 1.70E+07 | 1.17E+07 | | | | | | |
| P08752 | GNAI2 | guanine nucleotide binding protei | Plasma Membrane | other | 1.37E+07 | 1.52E+07 | 7943777.5 | | | | | | |
| P08752 | GNAI2 | guanine nucleotide binding protei | Plasma Membrane | other | 1.29E+07 | 1.54E+07 | 1.10E+07 | | | | | | |
| P08752 | GNAI2 | guanine nucleotide binding protei | Plasma Membrane | other | 1.95E+07 | 5115230.5 | 7859546 | | | | | | |
| P08752 | GNAI2 | guanine nucleotide binding protei | Plasma Membrane | other | 1.17E+07 | 2.76E+07 | 1.18E+07 | | | | | | |
| P08752 | GNAI2 | guanine nucleotide binding protei | Plasma Membrane | other | 6314747.5 | 1.16E+07 | 5268458.5 | | | | | | |
| P08752 | GNAI2 | guanine nucleotide binding protei | Plasma Membrane | other | 7378185 | 7191231 | 6947420 | | | | | | |
| P08752 | GNAI2 | guanine nucleotide binding protei | Plasma Membrane | other | 1.11E+07 | 819462.75 | 1649673.125 | | | | | | |
| P08752 | GNAI2 | guanine nucleotide binding protei | Plasma Membrane | other | 5515783.5 | 4308286 | 3229150.5 | | | | | | |
| P08752 | GNAI2 | guanine nucleotide binding protei | Plasma Membrane | other | 4200403 | 4944485.5 | 2885950.5 | P08752 | GNAI2 | 29762255 | 26420145 | 2991492 | 78 |
| Q9DC51 | GNAI3 | guanine nucleotide binding protei | Cytoplasm | enzyme | 3362029.75 | 2236774.25 | 3164621.75 | Q9DC51 | GNAI3 | 3362030 | 600843 | 346897 | 3 |
| P21279 | GNAQ | guanine nucleotide binding protei | Plasma Membrane | enzyme | 8623020 | 4944568.5 | 1.13E+07 | P21279 | GNAQ | 8623020 | 3181411 | 1836788 | 3 |
| Q6R0H7 | GNAS | GNAS complex locus | Plasma Membrane | enzyme | 3417284.5 | 3605587.5 | 3166889.5 | | | | | | |
| Q6R0H7 | GNAS | GNAS complex locus | Plasma Membrane | enzyme | 4042151 | 4568885 | 3815825 | Q6R0H7 | GNAS | 3729718 | 495896 | 202449 | 6 |
| O70443 | GNAZ | guanine nucleotide binding protei | Plasma Membrane | enzyme | 2534133.75 | 3149255 | 3644871.75 | | | | | | |
| O70443 | GNAZ | guanine nucleotide binding protei | Plasma Membrane | enzyme | 2504030.5 | 3143996.5 | 4397740 | O70443 | GNAZ | 2519082 | 715669 | 292170 | 6 |
| P62874 | GNB1 | guanine nucleotide binding protei | Plasma Membrane | enzyme | 8.22E+07 | 1.03E+08 | 4.98E+07 | | | | | | |
| P62874 | GNB1 | guanine nucleotide binding protei | Plasma Membrane | enzyme | 6.59E+07 | 7.23E+07 | 4.53E+07 | | | | | | |
| P62874 | GNB1 | guanine nucleotide binding protei | Plasma Membrane | enzyme | 2.50E+07 | 2.71E+07 | 1.82E+07 | | | | | | |
| P62874 | GNB1 | guanine nucleotide binding protei | Plasma Membrane | enzyme | 2.71E+07 | 3.37E+07 | 2.60E+07 | | | | | | |
| P62874 | GNB1 | guanine nucleotide binding protei | Plasma Membrane | enzyme | 2.18E+07 | 2.52E+07 | 1.50E+07 | | | | | | |
| P62874 | GNB1 | guanine nucleotide binding protei | Plasma Membrane | enzyme | 1.89E+07 | 2.82E+07 | 2.12E+07 | | | | | | |
| P62874 | GNB1 | guanine nucleotide binding protei | Plasma Membrane | enzyme | 1.54E+07 | 1.39E+07 | 8702611 | | | | | | |
| P62874 | GNB1 | guanine nucleotide binding protei | Plasma Membrane | enzyme | 1.75E+07 | 1.65E+07 | 1.62E+07 | | | | | | |
| P62874 | GNB1 | guanine nucleotide binding protei | Plasma Membrane | enzyme | 1.39E+07 | 1.47E+07 | 1.02E+07 | | | | | | |
| P62874 | GNB1 | guanine nucleotide binding protei | Plasma Membrane | enzyme | 6213462 | 1.16E+07 | 7902354 | | | | | | |
| P62874 | GNB1 | guanine nucleotide binding protei | Plasma Membrane | enzyme | 6774646 | 5499090 | 3511399.5 | | | | | | |
| P62874 | GNB1 | guanine nucleotide binding protei | Plasma Membrane | enzyme | 5181313.5 | 3239413.25 | 2036503.375 | | | | | | |
| P62874 | GNB1 | guanine nucleotide binding protei | Plasma Membrane | enzyme | 2669202.25 | 2281528.25 | 2304168.5 | P62874 | GNB1 | 23733740 | 23240146 | 3721402 | 39 |
| P62880 | GNB2 | guanine nucleotide binding protei | Plasma Membrane | enzyme | 5.59E+07 | 7.98E+07 | 4.75E+07 | | | | | | |
| P62880 | GNB2 | guanine nucleotide binding protei | Plasma Membrane | enzyme | 5.16E+07 | 5.64E+07 | 3.44E+07 | | | | | | |
| P62880 | GNB2 | guanine nucleotide binding protei | Plasma Membrane | enzyme | 3.53E+07 | 4.99E+07 | 2.85E+07 | | | | | | |
| P62880 | GNB2 | guanine nucleotide binding protei | Plasma Membrane | enzyme | 4.21E+07 | 6.20E+07 | 2.92E+07 | | | | | | |
| P62880 | GNB2 | guanine nucleotide binding protei | Plasma Membrane | enzyme | 3.24E+07 | 3.51E+07 | 1.96E+07 | | | | | | |

| | | | | | | | | | | | | | |
|---|---|---|---|---|---|---|---|---|---|---|---|---|---|
| P62880 | GNB2 | guanine nucleotide binding protei | Plasma Membrane | enzyme | 3.45E+07 | 2.93E+07 | 2.10E+07 | | | | | | |
| P62880 | GNB2 | guanine nucleotide binding protei | Plasma Membrane | enzyme | 1.20E+07 | 2.03E+07 | 1.36E+07 | | | | | | |
| P62880 | GNB2 | guanine nucleotide binding protei | Plasma Membrane | enzyme | 1.30E+07 | 1.93E+07 | 7708001 | | | | | | |
| P62880 | GNB2 | guanine nucleotide binding protei | Plasma Membrane | enzyme | 1.03E+07 | 1.23E+07 | 5524735.5 | | | | | | |
| P62880 | GNB2 | guanine nucleotide binding protei | Plasma Membrane | enzyme | 5381762 | 4978672.5 | 3780860.25 | | | | | | |
| P62880 | GNB2 | guanine nucleotide binding protei | Plasma Membrane | enzyme | 1439956.25 | 4653072.5 | 3999947.75 | | | | | | |
| P62880 | GNB2 | guanine nucleotide binding protei | Plasma Membrane | enzyme | 3419518 | 4138662.5 | 4489631 | P62880 | GNB2 | 24775103 | 20347452 | 3391242 | 36 |
| P68040 | GNB2L1 | guanine nucleotide binding protei | Cytoplasm | enzyme | 7609490.5 | 8018110.5 | 6679332 | | | | | | |
| P68040 | GNB2L1 | guanine nucleotide binding protei | Cytoplasm | enzyme | 6393703 | 6549832 | 5671137.5 | | | | | | |
| P68040 | GNB2L1 | guanine nucleotide binding protei | Cytoplasm | enzyme | 6499320 | 4770813 | 5322501.5 | | | | | | |
| P68040 | GNB2L1 | guanine nucleotide binding protei | Cytoplasm | enzyme | 5243116 | 3464727.75 | 4361090 | | | | | | |
| P68040 | GNB2L1 | guanine nucleotide binding protei | Cytoplasm | enzyme | 2081679.875 | 2064774.375 | 1934669.25 | P68040 | GNB2L1 | 5565462 | 1983972 | 512259 | 15 |
| Q9DAS9 | GNG12 | guanine nucleotide binding protei | Plasma Membrane | enzyme | 7285862 | 8133148.5 | 6213474 | Q9DAS9 | GNG12 | 7285862 | 962034 | 555431 | 3 |
| Q91W53 | GOLGA7 | golgin A7 | Cytoplasm | other | 4505359.5 | 4567899.5 | 3975703 | | | | | | |
| Q91W53 | GOLGA7 | golgin A7 | Cytoplasm | other | 3341158.75 | 3539065.5 | 3456162.75 | Q91W53 | GOLGA7 | 3923259 | 539966 | 220440 | 6 |
| Q9CR60 | GOLT1B | golgi transport 1B | Cytoplasm | other | 3193742.5 | 3560726 | 3878508.75 | Q9CR60 | GOLT1B | 3193743 | 342678 | 197845 | 3 |
| P05202 | GOT2 | glutamic-oxaloacetic transaminase | Cytoplasm | enzyme | 2.13E+07 | 1.99E+07 | 3.19E+07 | | | | | | |
| P05202 | GOT2 | glutamic-oxaloacetic transaminase | Cytoplasm | enzyme | 1.62E+07 | 1.24E+07 | 1.63E+07 | | | | | | |
| P05202 | GOT2 | glutamic-oxaloacetic transaminase | Cytoplasm | enzyme | 1.12E+07 | 1.07E+07 | 1.40E+07 | | | | | | |
| P05202 | GOT2 | glutamic-oxaloacetic transaminase | Cytoplasm | enzyme | 8013560.5 | 9050876 | 9443911 | | | | | | |
| P05202 | GOT2 | glutamic-oxaloacetic transaminase | Cytoplasm | enzyme | 5556846.5 | 6113305 | 6917866.5 | | | | | | |
| P05202 | GOT2 | glutamic-oxaloacetic transaminase | Cytoplasm | enzyme | 7617517 | 9865681 | 1.14E+07 | | | | | | |
| P05202 | GOT2 | glutamic-oxaloacetic transaminase | Cytoplasm | enzyme | 6731126.5 | 6566316.5 | 8454151 | | | | | | |
| P05202 | GOT2 | glutamic-oxaloacetic transaminase | Cytoplasm | enzyme | 2297087.5 | 4250569.5 | 4382780.5 | | | | | | |
| P05202 | GOT2 | glutamic-oxaloacetic transaminase | Cytoplasm | enzyme | 2288431.5 | 2614696.25 | 1675602.875 | | | | | | |
| P05202 | GOT2 | glutamic-oxaloacetic transaminase | Cytoplasm | enzyme | 2102528 | 2107348.75 | 3779265.75 | | | | | | |
| P05202 | GOT2 | glutamic-oxaloacetic transaminase | Cytoplasm | enzyme | 1747425.875 | 1132391.5 | 1389587.75 | P05202 | GOT2 | 7734048 | 6852876 | 1192933 | 33 |
| O35930 | GP1BA | glycoprotein Ib (platelet), alpha pc | Plasma Membrane | other | 5.71E+07 | 6.78E+07 | 6.64E+07 | | | | | | |
| O35930 | GP1BA | glycoprotein Ib (platelet), alpha pc | Plasma Membrane | other | 7.49E+07 | 4.83E+07 | 8.38E+07 | | | | | | |
| O35930 | GP1BA | glycoprotein Ib (platelet), alpha pc | Plasma Membrane | other | 3.55E+07 | 4.36E+07 | 4.71E+07 | | | | | | |
| O35930 | GP1BA | glycoprotein Ib (platelet), alpha pc | Plasma Membrane | other | 3.02E+07 | 3.59E+07 | 3.42E+07 | | | | | | |
| O35930 | GP1BA | glycoprotein Ib (platelet), alpha pc | Plasma Membrane | other | 3.37E+07 | 3.01E+07 | 3.79E+07 | | | | | | |
| O35930 | GP1BA | glycoprotein Ib (platelet), alpha pc | Plasma Membrane | other | 2.77E+07 | 3.34E+07 | 3.30E+07 | | | | | | |
| O35930 | GP1BA | glycoprotein Ib (platelet), alpha pc | Plasma Membrane | other | 1.78E+07 | 1.90E+07 | 2.22E+07 | | | | | | |
| O35930 | GP1BA | glycoprotein Ib (platelet), alpha pc | Plasma Membrane | other | 8290655.5 | 1.74E+07 | 1.56E+07 | | | | | | |
| O35930 | GP1BA | glycoprotein Ib (platelet), alpha pc | Plasma Membrane | other | 5985780 | 6794046 | 9224617 | | | | | | |
| O35930 | GP1BA | glycoprotein Ib (platelet), alpha pc | Plasma Membrane | other | 6493933.5 | 7352469.5 | 7144055 | | | | | | |
| O35930 | GP1BA | glycoprotein Ib (platelet), alpha pc | Plasma Membrane | other | 5986625 | 1.15E+07 | 8851630 | | | | | | |
| O35930 | GP1BA | glycoprotein Ib (platelet), alpha pc | Plasma Membrane | other | 3345855.25 | 1704347.75 | 1.43E+07 | | | | | | |
| O35930 | GP1BA | glycoprotein Ib (platelet), alpha pc | Plasma Membrane | other | 211117.641 | 1430655.75 | 2438158 | O35930 | GP1BA | 23617997 | 22153667 | 3547426 | 39 |
| P56400 | GP1BB | glycoprotein Ib (platelet), beta pol | Plasma Membrane | other | 9.74E+07 | 1.19E+08 | 1.21E+08 | | | | | | |
| P56400 | GP1BB | glycoprotein Ib (platelet), beta pol | Plasma Membrane | other | 6.30E+07 | 6.16E+07 | 6.66E+07 | | | | | | |
| P56400 | GP1BB | glycoprotein Ib (platelet), beta pol | Plasma Membrane | other | 2.83E+07 | 2.59E+07 | 3.01E+07 | | | | | | |
| P56400 | GP1BB | glycoprotein Ib (platelet), beta pol | Plasma Membrane | other | 2.92E+07 | 3.88E+07 | 3.38E+07 | | | | | | |
| P56400 | GP1BB | glycoprotein Ib (platelet), beta pol | Plasma Membrane | other | 1.44E+07 | 1.30E+07 | 1.36E+07 | | | | | | |
| P56400 | GP1BB | glycoprotein Ib (platelet), beta pol | Plasma Membrane | other | 1.14E+07 | 1.61E+07 | 1.58E+07 | P56400 | GP1BB | 40611667 | 36084051 | 8505092 | 18 |
| O08742 | GP5 | glycoprotein V (platelet) | Plasma Membrane | other | 5.35E+07 | 5.55E+07 | 6.66E+07 | | | | | | |
| O08742 | GP5 | glycoprotein V (platelet) | Plasma Membrane | other | 4.10E+07 | 1.80E+07 | 4.48E+07 | | | | | | |
| O08742 | GP5 | glycoprotein V (platelet) | Plasma Membrane | other | 2.78E+07 | 2.57E+07 | 3.68E+07 | | | | | | |
| O08742 | GP5 | glycoprotein V (platelet) | Plasma Membrane | other | 2.84E+07 | 4.84E+07 | 5.01E+07 | | | | | | |
| O08742 | GP5 | glycoprotein V (platelet) | Plasma Membrane | other | 2.37E+07 | 3.01E+07 | 3.48E+07 | | | | | | |
| O08742 | GP5 | glycoprotein V (platelet) | Plasma Membrane | other | 3.79E+07 | 3.53E+07 | 3.87E+07 | | | | | | |
| O08742 | GP5 | glycoprotein V (platelet) | Plasma Membrane | other | 1.16E+07 | 1.33E+07 | 1.70E+07 | | | | | | |
| O08742 | GP5 | glycoprotein V (platelet) | Plasma Membrane | other | 9501131 | 9821189 | 1.27E+07 | | | | | | |
| O08742 | GP5 | glycoprotein V (platelet) | Plasma Membrane | other | 9454052 | 1.01E+07 | 1.26E+07 | | | | | | |
| O08742 | GP5 | glycoprotein V (platelet) | Plasma Membrane | other | 3635714.25 | 7463242.5 | 7741091 | O08742 | GP5 | 24641090 | 17224694 | 3144785 | 30 |
| O88186 | GP9 | glycoprotein IX (platelet) | Plasma Membrane | other | 7.71E+07 | 8.86E+07 | 9.42E+07 | | | | | | |
| O88186 | GP9 | glycoprotein IX (platelet) | Plasma Membrane | other | 2.26E+07 | 3.32E+07 | 2.57E+07 | | | | | | |
| O88186 | GP9 | glycoprotein IX (platelet) | Plasma Membrane | other | 4746031 | 7282877 | 6941900.5 | | | | | | |
| O88186 | GP9 | glycoprotein IX (platelet) | Plasma Membrane | other | 3625994.75 | 5567480.5 | 5350274 | O88186 | GP9 | 27008006 | 34924421 | 10081812 | 12 |
| P51655 | GPC4 | glypican 4 | Plasma Membrane | transmembrane receptor | 5485597.5 | 5389203 | 6668570.5 | | | | | | |
| P51655 | GPC4 | glypican 4 | Plasma Membrane | transmembrane receptor | 3625294 | 5925255 | 5817713 | | | | | | |
| P51655 | GPC4 | glypican 4 | Plasma Membrane | transmembrane receptor | 3447102.25 | 2396072.75 | 2919381.5 | P51655 | GPC4 | 4185998 | 1536143 | 512048 | 9 |
| Q64521 | GPD2 | glycerol-3-phosphate dehydrogena | Cytoplasm | enzyme | 6197499.5 | 9039041 | 1.14E+07 | | | | | | |
| Q64521 | GPD2 | glycerol-3-phosphate dehydrogena | Cytoplasm | enzyme | 8907043 | 8381348.5 | 1.23E+07 | | | | | | |
| Q64521 | GPD2 | glycerol-3-phosphate dehydrogena | Cytoplasm | enzyme | 3750067 | 4479439 | 4362391 | | | | | | |
| Q64521 | GPD2 | glycerol-3-phosphate dehydrogena | Cytoplasm | enzyme | 8248486.5 | 6501782.5 | 8900188 | | | | | | |
| Q64521 | GPD2 | glycerol-3-phosphate dehydrogena | Cytoplasm | enzyme | 1491634 | 1154847.25 | 1893962.5 | | | | | | |
| Q64521 | GPD2 | glycerol-3-phosphate dehydrogena | Cytoplasm | enzyme | 5486754.5 | 5811163.5 | 6414481.5 | | | | | | |
| Q64521 | GPD2 | glycerol-3-phosphate dehydrogena | Cytoplasm | enzyme | 4200096.5 | 3985550.5 | 4550865 | | | | | | |
| Q64521 | GPD2 | glycerol-3-phosphate dehydrogena | Cytoplasm | enzyme | 2760187 | 3198154.25 | 4492790 | | | | | | |
| Q64521 | GPD2 | glycerol-3-phosphate dehydrogena | Cytoplasm | enzyme | 3631242 | 5616177 | 5837102.5 | | | | | | |
| Q64521 | GPD2 | glycerol-3-phosphate dehydrogena | Cytoplasm | enzyme | 3961493.25 | 4042251.25 | 6356513 | Q64521 | GPD2 | 4863450 | 2722122 | 496989 | 30 |
| Q99LD4 | GPS1 | G protein pathway suppressor 1 | Cytoplasm | other | 4831571 | 4623800 | 3561360 | | | | | | |
| Q99LD4 | GPS1 | G protein pathway suppressor 1 | Cytoplasm | other | 4917654 | 4280235 | 5199696.5 | | | | | | |
| Q99LD4 | GPS1 | G protein pathway suppressor 1 | Cytoplasm | other | 3401891.75 | 1418794.625 | 1946622 | | | | | | |
| Q99LD4 | GPS1 | G protein pathway suppressor 1 | Cytoplasm | other | 1003797 | 1465738.625 | 1261980.25 | | | | | | |
| Q99LD4 | GPS1 | G protein pathway suppressor 1 | Cytoplasm | other | 510459.219 | 1040040.312 | 548201.25 | Q99LD4 | GPS1 | 2933075 | 1772492 | 457655 | 15 |
| P11352 | GPX1 | glutathione peroxidase 1 | Cytoplasm | enzyme | 4.27E+07 | 3.61E+07 | 5.39E+07 | | | | | | |
| P11352 | GPX1 | glutathione peroxidase 1 | Cytoplasm | enzyme | 3.73E+07 | 1.57E+07 | 3.36E+07 | | | | | | |
| P11352 | GPX1 | glutathione peroxidase 1 | Cytoplasm | enzyme | 2.96E+07 | 1.92E+07 | 2.72E+07 | | | | | | |
| P11352 | GPX1 | glutathione peroxidase 1 | Cytoplasm | enzyme | 1.67E+07 | 1.19E+07 | 2.07E+07 | | | | | | |
| P11352 | GPX1 | glutathione peroxidase 1 | Cytoplasm | enzyme | 1.16E+07 | 7577516 | 9305190 | | | | | | |
| P11352 | GPX1 | glutathione peroxidase 1 | Cytoplasm | enzyme | 1.33E+07 | 9204702 | 1.08E+07 | | | | | | |
| P11352 | GPX1 | glutathione peroxidase 1 | Cytoplasm | enzyme | 6862717 | 7564745 | 1.11E+07 | | | | | | |
| P11352 | GPX1 | glutathione peroxidase 1 | Cytoplasm | enzyme | 3779254.75 | 8720488 | 9732667 | | | | | | |

| | | | | | | | | | | | | | |
|---|---|---|---|---|---|---|---|---|---|---|---|---|---|
| P11352 | GPX1 | glutathione peroxidase 1 | Cytoplasm | enzyme | 3941651.5 | 4202031 | 3369708.5 | | | | | | |
| P11352 | GPX1 | glutathione peroxidase 1 | Cytoplasm | enzyme | 4229140 | 4591336 | 5278251 | P11352 | GPX1 | 17010276 | 13316025 | 2431162 | 30 |
| O70325 | GPX4 | glutathione peroxidase 4 | Cytoplasm | enzyme | 3.21E+07 | 3.87E+07 | 4.56E+07 | | | | | | |
| O70325 | GPX4 | glutathione peroxidase 4 | Cytoplasm | enzyme | 2.89E+07 | 2.99E+07 | 2.74E+07 | | | | | | |
| O70325 | GPX4 | glutathione peroxidase 4 | Cytoplasm | enzyme | 1.77E+07 | 1.64E+07 | 1.27E+07 | | | | | | |
| O70325 | GPX4 | glutathione peroxidase 4 | Cytoplasm | enzyme | 1.38E+07 | 1.22E+07 | 1.35E+07 | | | | | | |
| O70325 | GPX4 | glutathione peroxidase 4 | Cytoplasm | enzyme | 1.28E+07 | 1.19E+07 | 1.04E+07 | | | | | | |
| O70325 | GPX4 | glutathione peroxidase 4 | Cytoplasm | enzyme | 6114915 | 4715174.5 | 3306055.75 | | | | | | |
| O70325 | GPX4 | glutathione peroxidase 4 | Cytoplasm | enzyme | 2927530 | 2344242.25 | 2566688.75 | O70325 | GPX4 | 16346064 | 12556118 | 2739970 | 21 |
| Q8R050 | GSPT1 | G1 to S phase transition 1 | Cytoplasm | translation regulator | 2863654.5 | 2442336 | 2263419.25 | Q8R050 | GSPT1 | 2863655 | 308167 | 177921 | 3 |
| P19157 | GSTP1 | glutathione S-transferase pi 1 | Cytoplasm | enzyme | 2020213 | 2224867 | 1880344.875 | P19157 | GSTP1 | 2020213 | 173273 | 100039 | 3 |
| Q8BZI6 | GUCD1 | guanylyl cyclase domain containin | Nucleus | other | 1500639.625 | 1777200.125 | 1746785.25 | Q8BZI6 | GUCD1 | 1500640 | 151657 | 87559 | 3 |
| P14220 | Gypa | glycophorin A | Plasma Membrane | other | 5.11E+09 | 3.65E+09 | 2.56E+09 | | | | | | |
| P14220 | Gypa | glycophorin A | Plasma Membrane | other | 5.31E+07 | 1.51E+07 | 1.84E+07 | P14220 | Gypa | 2582540000 | 2205128573 | 900239970 | 6 |
| Q78HU7 | GYPC | glycophorin C (Gerbich blood grou | Plasma Membrane | other | 1.18E+09 | 6.73E+08 | 5.91E+08 | | | | | | |
| Q78HU7 | GYPC | glycophorin C (Gerbich blood grou | Plasma Membrane | other | 6.08E+08 | 1.46E+09 | 1.18E+09 | | | | | | |
| Q78HU7 | GYPC | glycophorin C (Gerbich blood grou | Plasma Membrane | other | 3.78E+07 | 2.67E+07 | 2.38E+07 | Q78HU7 | GYPC | 607916667 | 545208667 | 181736222 | 9 |
| Q3THW5 | H2AFV | H2A histone family, member V | Nucleus | other | 4248813.5 | 2734254 | 2406870.25 | Q3THW5 | H2AFV | 4248814 | 982669 | 567344 | 3 |
| P27661 | H2AFX | H2A histone family, member X | Nucleus | transcription regulator | 1.09E+07 | 7966861.5 | 7241246.5 | | | | | | |
| P27661 | H2AFX | H2A histone family, member X | Nucleus | transcription regulator | 1526033 | 219336.281 | 453939.781 | P27661 | H2AFX | 6188017 | 4541623 | 1854110 | 6 |
| Q9QZQ8 | H2AFY | H2A histone family, member Y | Nucleus | other | 4860416 | 2058076.125 | 2033998.25 | Q9QZQ8 | H2AFY | 4860416 | 1624927 | 938152 | 3 |
| P68433 | H31 | Histone H3.1 | Nucleus | | 1.21E+08 | 7.56E+07 | 6.31E+07 | | | | | | |
| P68433 | H31 | Histone H3.1 | Nucleus | | 1.10E+08 | 6.62E+07 | 5.75E+07 | | | | | | |
| P68433 | H31 | Histone H3.1 | Nucleus | | 3.83E+07 | 2.09E+07 | 1.55E+07 | | | | | | |
| P68433 | H31 | Histone H3.1 | Nucleus | | 5.87E+07 | 2.80E+07 | 2.86E+07 | | | | | | |
| P68433 | H31 | Histone H3.1 | Nucleus | | 9235880 | 7940239 | 1.30E+07 | | | | | | |
| P68433 | H31 | Histone H3.1 | Nucleus | | 1.90E+07 | 6917381.5 | 2.04E+07 | P68433 | H31 | 59395980 | 34636698 | 8163948 | 18 |
| P62806 | H4 | Histone H4 | Nucleus | | 2.78E+08 | 1.70E+08 | 1.70E+08 | | | | | | |
| P62806 | H4 | Histone H4 | Nucleus | | 2.95E+08 | 1.94E+08 | 1.88E+08 | | | | | | |
| P62806 | H4 | Histone H4 | Nucleus | | 7.30E+07 | 4.07E+07 | 2.91E+07 | | | | | | |
| P62806 | H4 | Histone H4 | Nucleus | | 8.57E+07 | 4.24E+07 | 4.39E+07 | | | | | | |
| P62806 | H4 | Histone H4 | Nucleus | | 3.96E+07 | 1.76E+07 | 1.75E+07 | | | | | | |
| P62806 | H4 | Histone H4 | Nucleus | | 2040859.875 | 5956247.5 | 1.37E+07 | | | | | | |
| P62806 | H4 | Histone H4 | Nucleus | | 413639.812 | 256751.734 | 175141.547 | | | | | | |
| P62806 | H4 | Histone H4 | Nucleus | | 334848.75 | 3387573.75 | 3466689.5 | P62806 | H4 | 96726169 | 91826281 | 18743961 | 24 |
| Q3U0D9 | HACE1 | HECT domain and ankyrin repeat c | Cytoplasm | enzyme | 853676.938 | 920746.125 | 657783.438 | Q3U0D9 | HACE1 | 853677 | 136639 | 78889 | 3 |
| Q61425 | HADH | hydroxyacyl-CoA dehydrogenase | Cytoplasm | enzyme | 2666216 | 3790676 | 5050756 | | | | | | |
| Q61425 | HADH | hydroxyacyl-CoA dehydrogenase | Cytoplasm | enzyme | 684393.375 | 1081496.5 | 1257443.5 | Q61425 | HADH | 1675305 | 1732957 | 707477 | 6 |
| Q8BMS1 | HADHA | hydroxyacyl-CoA dehydrogenase/3 | Cytoplasm | enzyme | 6046268 | 8746041 | 1.24E+07 | | | | | | |
| Q8BMS1 | HADHA | hydroxyacyl-CoA dehydrogenase/3 | Cytoplasm | enzyme | 5728928.5 | 6633778.5 | 7144069 | | | | | | |
| Q8BMS1 | HADHA | hydroxyacyl-CoA dehydrogenase/3 | Cytoplasm | enzyme | 5509668.5 | 6620067.5 | 8658814 | | | | | | |
| Q8BMS1 | HADHA | hydroxyacyl-CoA dehydrogenase/3 | Cytoplasm | enzyme | 5885535.5 | 5595542.5 | 6978161.5 | | | | | | |
| Q8BMS1 | HADHA | hydroxyacyl-CoA dehydrogenase/3 | Cytoplasm | enzyme | 5730664.5 | 4754251.5 | 8787671 | | | | | | |
| Q8BMS1 | HADHA | hydroxyacyl-CoA dehydrogenase/3 | Cytoplasm | enzyme | 3752976.5 | 4358540 | 5752217 | | | | | | |
| Q8BMS1 | HADHA | hydroxyacyl-CoA dehydrogenase/3 | Cytoplasm | enzyme | 4012971.5 | 4077334.5 | 5906049.5 | | | | | | |
| Q8BMS1 | HADHA | hydroxyacyl-CoA dehydrogenase/3 | Cytoplasm | enzyme | 4535425.5 | 3704268.5 | 5581169 | | | | | | |
| Q8BMS1 | HADHA | hydroxyacyl-CoA dehydrogenase/3 | Cytoplasm | enzyme | 4127751.25 | 3200820.5 | 5258473.5 | | | | | | |
| Q8BMS1 | HADHA | hydroxyacyl-CoA dehydrogenase/3 | Cytoplasm | enzyme | 3656155.5 | 3711895.5 | 4600044 | | | | | | |
| Q8BMS1 | HADHA | hydroxyacyl-CoA dehydrogenase/3 | Cytoplasm | enzyme | 1787460.875 | 1487625.125 | 2484804.75 | | | | | | |
| Q8BMS1 | HADHA | hydroxyacyl-CoA dehydrogenase/3 | Cytoplasm | enzyme | 1803965.25 | 1881047 | 2474068.75 | | | | | | |
| Q8BMS1 | HADHA | hydroxyacyl-CoA dehydrogenase/3 | Cytoplasm | enzyme | 1263631.875 | 1423343.375 | 1898828.75 | | | | | | |
| Q8BMS1 | HADHA | hydroxyacyl-CoA dehydrogenase/3 | Cytoplasm | enzyme | 539701.875 | 760536.5 | 804233.188 | Q8BMS1 | HADHA | 3884365 | 2547633 | 393108 | 42 |
| Q99JY0 | HADHB | hydroxyacyl-CoA dehydrogenase/3 | Cytoplasm | enzyme | 8021201.5 | 1.32E+07 | 1.36E+07 | | | | | | |
| Q99JY0 | HADHB | hydroxyacyl-CoA dehydrogenase/3 | Cytoplasm | enzyme | 7595412 | 7565171 | 1.15E+07 | | | | | | |
| Q99JY0 | HADHB | hydroxyacyl-CoA dehydrogenase/3 | Cytoplasm | enzyme | 7524156 | 7743107.5 | 1.17E+07 | | | | | | |
| Q99JY0 | HADHB | hydroxyacyl-CoA dehydrogenase/3 | Cytoplasm | enzyme | 3983322 | 4220754.5 | 5378745.5 | | | | | | |
| Q99JY0 | HADHB | hydroxyacyl-CoA dehydrogenase/3 | Cytoplasm | enzyme | 1130646.625 | 1306982.875 | 2249539.25 | | | | | | |
| Q99JY0 | HADHB | hydroxyacyl-CoA dehydrogenase/3 | Cytoplasm | enzyme | 2499026.5 | 2292022.5 | 3279700.75 | | | | | | |
| Q99JY0 | HADHB | hydroxyacyl-CoA dehydrogenase/3 | Cytoplasm | enzyme | 1370128.75 | 2730414 | 4547088 | Q99JY0 | HADHB | 4589128 | 4012995 | 875707 | 21 |
| Q99KB8 | HAGH | hydroxyacylglutathione hydrolase | Cytoplasm | enzyme | 9370782 | 7462772.5 | 7919645.5 | | | | | | |
| Q99KB8 | HAGH | hydroxyacylglutathione hydrolase | Cytoplasm | enzyme | 2605013.5 | 3907401 | 5886243 | | | | | | |
| Q99KB8 | HAGH | hydroxyacylglutathione hydrolase | Cytoplasm | enzyme | 622089.062 | 702734.875 | 829910.75 | Q99KB8 | HAGH | 4199295 | 3406410 | 1135470 | 9 |
| P01942 | HBA1/HBA2 | hemoglobin, alpha 1 | Extracellular Space | transporter | 2.59E+09 | 2.18E+09 | 3.08E+09 | | | | | | |
| P01942 | HBA1/HBA2 | hemoglobin, alpha 1 | Extracellular Space | transporter | 3.06E+09 | 1.78E+09 | 2.03E+09 | | | | | | |
| P01942 | HBA1/HBA2 | hemoglobin, alpha 1 | Extracellular Space | transporter | 2.22E+09 | 1.32E+09 | 1.71E+09 | | | | | | |
| P01942 | HBA1/HBA2 | hemoglobin, alpha 1 | Extracellular Space | transporter | 1.96E+09 | 8.70E+08 | 1.29E+09 | | | | | | |
| P01942 | HBA1/HBA2 | hemoglobin, alpha 1 | Extracellular Space | transporter | 2.79E+09 | 1.26E+09 | 2.09E+09 | | | | | | |
| P01942 | HBA1/HBA2 | hemoglobin, alpha 1 | Extracellular Space | transporter | 1.92E+09 | 1.01E+09 | 1.09E+09 | | | | | | |
| P01942 | HBA1/HBA2 | hemoglobin, alpha 1 | Extracellular Space | transporter | 5.58E+08 | 2.62E+08 | 4.14E+08 | | | | | | |
| P01942 | HBA1/HBA2 | hemoglobin, alpha 1 | Extracellular Space | transporter | 1.38E+08 | 2.44E+08 | 2.07E+08 | | | | | | |
| P01942 | HBA1/HBA2 | hemoglobin, alpha 1 | Extracellular Space | transporter | 2.53E+08 | 6.84E+08 | 1.04E+09 | | | | | | |
| P01942 | HBA1/HBA2 | hemoglobin, alpha 1 | Extracellular Space | transporter | 3.00E+08 | 1.52E+08 | 2.43E+08 | | | | | | |
| P01942 | HBA1/HBA2 | hemoglobin, alpha 1 | Extracellular Space | transporter | 1.78E+08 | 1.20E+08 | 1.22E+08 | P01942 | HBA1/HBA2 | 1450963636 | 953873008 | 166047977 | 33 |
| P02088 | Hbb-b1 | hemoglobin, beta adult major chai | Extracellular Space | transporter | 5.28E+09 | 3.24E+09 | 4.61E+09 | | | | | | |
| P02088 | Hbb-b1 | hemoglobin, beta adult major chai | Extracellular Space | transporter | 4.15E+09 | 2.31E+09 | 3.57E+09 | | | | | | |
| P02088 | Hbb-b1 | hemoglobin, beta adult major chai | Extracellular Space | transporter | 4.39E+09 | 2.22E+09 | 2.99E+09 | | | | | | |
| P02088 | Hbb-b1 | hemoglobin, beta adult major chai | Extracellular Space | transporter | 3.06E+09 | 1.77E+09 | 2.63E+09 | | | | | | |
| P02088 | Hbb-b1 | hemoglobin, beta adult major chai | Extracellular Space | transporter | 4.06E+09 | 1.47E+09 | 1.94E+09 | | | | | | |
| P02088 | Hbb-b1 | hemoglobin, beta adult major chai | Extracellular Space | transporter | 2.24E+09 | 1.41E+09 | 1.80E+09 | | | | | | |
| P02088 | Hbb-b1 | hemoglobin, beta adult major chai | Extracellular Space | transporter | 3.76E+09 | 1.08E+09 | 1.46E+09 | | | | | | |
| P02088 | Hbb-b1 | hemoglobin, beta adult major chai | Extracellular Space | transporter | 2.06E+09 | 1.02E+09 | 1.54E+09 | | | | | | |
| P02088 | Hbb-b1 | hemoglobin, beta adult major chai | Extracellular Space | transporter | 1.03E+09 | 4.03E+08 | 5.20E+08 | | | | | | |
| P02088 | Hbb-b1 | hemoglobin, beta adult major chai | Extracellular Space | transporter | 9.56E+08 | 5.96E+08 | 8.26E+08 | | | | | | |
| P02088 | Hbb-b1 | hemoglobin, beta adult major chai | Extracellular Space | transporter | 3.86E+08 | 2.14E+08 | 3.06E+08 | | | | | | |
| P02088 | Hbb-b1 | hemoglobin, beta adult major chai | Extracellular Space | transporter | 4.82E+08 | 2.72E+08 | 5.28E+08 | | | | | | |
| P02088 | Hbb-b1 | hemoglobin, beta adult major chai | Extracellular Space | transporter | 2.16E+08 | 9.64E+07 | 1.63E+08 | | | | | | |

| | | | | | | | | | | | | | |
|---|---|---|---|---|---|---|---|---|---|---|---|---|---|
| P02088 | Hbb-b1 | hemoglobin, beta adult major chai | Extracellular Space | transporter | 5.56E+07 | 1.11E+08 | 2.15E+08 | | | | | | |
| P02088 | Hbb-b1 | hemoglobin, beta adult major chai | Extracellular Space | transporter | 1.29E+08 | 7551379.5 | 7462437.5 | | | | | | |
| P02088 | Hbb-b1 | hemoglobin, beta adult major chai | Extracellular Space | transporter | 2.45E+07 | 9363816 | 1.52E+07 | | | | | | |
| P02088 | Hbb-b1 | hemoglobin, beta adult major chai | Extracellular Space | transporter | 2.65E+07 | 1.13E+07 | 2.36E+07 | | | | | | |
| P02088 | Hbb-b1 | hemoglobin, beta adult major chai | Extracellular Space | transporter | 9400307 | 1.27E+07 | 2.09E+07 | | | | | | |
| P02088 | Hbb-b1 | hemoglobin, beta adult major chai | Extracellular Space | transporter | 1.04E+07 | 110109.953 | 211313.188 | P02088 | Hbb-b1 | 1701731595 | 1460357643 | 193429095 | 57 |
| P02089 | Hbb-b2 | hemoglobin, beta adult minor cha | Extracellular Space | other | 1.10E+09 | 3.36E+08 | 4.46E+08 | | | | | | |
| P02089 | Hbb-b2 | hemoglobin, beta adult minor cha | Extracellular Space | other | 7.77E+08 | 3.03E+08 | 4.33E+08 | | | | | | |
| P02089 | Hbb-b2 | hemoglobin, beta adult minor cha | Extracellular Space | other | 7.30E+08 | 2.09E+08 | 3.14E+08 | | | | | | |
| P02089 | Hbb-b2 | hemoglobin, beta adult minor cha | Extracellular Space | other | 2.35E+08 | 8.51E+07 | 1.14E+08 | | | | | | |
| P02089 | Hbb-b2 | hemoglobin, beta adult minor cha | Extracellular Space | other | 2.40E+08 | 6.69E+07 | 1.30E+08 | | | | | | |
| P02089 | Hbb-b2 | hemoglobin, beta adult minor cha | Extracellular Space | other | 1.06E+08 | 3.06E+07 | 4.63E+07 | | | | | | |
| P02089 | Hbb-b2 | hemoglobin, beta adult minor cha | Extracellular Space | other | 6.63E+07 | 1.72E+07 | 2.93E+07 | | | | | | |
| P02089 | Hbb-b2 | hemoglobin, beta adult minor cha | Extracellular Space | other | 717281.312 | 4927152.5 | 8840465 | P02089 | Hbb-b2 | 406682160 | 282609004 | 57687321 | 24 |
| P08103 | HCK | HCK proto-oncogene, Src family ty | Cytoplasm | kinase | 1691686.25 | 1712436.25 | 1666567.875 | P08103 | HCK | 1691686 | 22969 | 13261 | 3 |
| Q3U487 | HECTD3 | HECT domain containing E3 ubiqui | Cytoplasm | enzyme | 2373932.5 | 4318969 | 1175052.375 | | | | | | |
| Q3U487 | HECTD3 | HECT domain containing E3 ubiqui | Cytoplasm | enzyme | 464103.969 | 703648.938 | 430373.219 | Q3U487 | HECTD3 | 1419018 | 1525194 | 622658 | 6 |
| P43275 | Hist1h1a | histone cluster 1, H1a | Nucleus | other | 9668471 | 4786397.5 | 5432454.5 | | | | | | |
| P43275 | Hist1h1a | histone cluster 1, H1a | Nucleus | other | 5619315.5 | 3545001 | 4413115.5 | | | | | | |
| P43275 | Hist1h1a | histone cluster 1, H1a | Nucleus | other | 9343993 | 4569286 | 5268280 | | | | | | |
| P43275 | Hist1h1a | histone cluster 1, H1a | Nucleus | other | 922036.312 | 306757.5 | 130477.211 | P43275 | Hist1h1a | 6388454 | 3065559 | 884951 | 12 |
| P43276 | Hist1h1b | histone cluster 1, H1b | Nucleus | other | 2.21E+07 | 1.35E+07 | 1.23E+07 | | | | | | |
| P43276 | Hist1h1b | histone cluster 1, H1b | Nucleus | other | 1344401.25 | 669765.5 | 399616.281 | P43276 | Hist1h1b | 11732201 | 8976633 | 3664695 | 6 |
| P15864 | HIST1H1C | histone cluster 1, H1c | Nucleus | other | 1.74E+08 | 1.06E+08 | 8.96E+07 | P15864 | HIST1H1C | 174000000 | 44847826 | 25892904 | 3 |
| P43274 | Hist1h1e | histone cluster 1, H1e | Nucleus | other | 1.09E+08 | 5.89E+07 | 5.03E+07 | | | | | | |
| P43274 | Hist1h1e | histone cluster 1, H1e | Nucleus | other | 3.25E+07 | 2.86E+07 | 2.23E+07 | | | | | | |
| P43274 | Hist1h1e | histone cluster 1, H1e | Nucleus | other | 4.97E+07 | 2.69E+07 | 2.62E+07 | | | | | | |
| P43274 | Hist1h1e | histone cluster 1, H1e | Nucleus | other | 2.96E+07 | 6265240.5 | 1.04E+07 | | | | | | |
| P43274 | Hist1h1e | histone cluster 1, H1e | Nucleus | other | 260509.375 | 229714.938 | 147716.094 | P43274 | Hist1h1e | 44150102 | 28714246 | 7413986 | 15 |
| Q8CGP5 | HIST1H2AJ | histone cluster 1, H2aj | Nucleus | other | 3.37E+08 | 1.85E+08 | 2.20E+08 | | | | | | |
| Q8CGP5 | HIST1H2AJ | histone cluster 1, H2aj | Nucleus | other | 3.08E+08 | 1.71E+08 | 1.85E+08 | | | | | | |
| Q8CGP5 | HIST1H2AJ | histone cluster 1, H2aj | Nucleus | other | 1.22E+08 | 6.54E+07 | 6.60E+07 | | | | | | |
| Q8CGP5 | HIST1H2AJ | histone cluster 1, H2aj | Nucleus | other | 5.77E+07 | 7.08E+07 | 5.38E+07 | | | | | | |
| Q8CGP5 | HIST1H2AJ | histone cluster 1, H2aj | Nucleus | other | 523523.562 | 431897.688 | 321290.656 | Q8CGP5 | HIST1H2AJ | 165168705 | 107875923 | 27853444 | 15 |
| P70696 | HIST1H2BA | histone cluster 1, H2ba | Nucleus | other | 2.61E+08 | 1.25E+08 | 1.54E+08 | P70696 | HIST1H2BA | 260800000 | 71476080 | 41266734 | 3 |
| Q64475 | HIST1H2BB | histone cluster 1, H2bb | Nucleus | other | 2.23E+08 | 1.16E+08 | 1.16E+08 | | | | | | |
| Q64475 | HIST1H2BB | histone cluster 1, H2bb | Nucleus | other | 2.12E+08 | 1.15E+08 | 1.09E+08 | | | | | | |
| Q64475 | HIST1H2BB | histone cluster 1, H2bb | Nucleus | other | 1.31E+07 | 1.12E+07 | 1.61E+07 | | | | | | |
| Q64475 | HIST1H2BB | histone cluster 1, H2bb | Nucleus | other | 4349996 | 2955838.25 | 2504720.75 | | | | | | |
| Q64475 | HIST1H2BB | histone cluster 1, H2bb | Nucleus | other | 5596274 | 3501828.25 | 3889179 | | | | | | |
| Q64475 | HIST1H2BB | histone cluster 1, H2bb | Nucleus | other | 4999416 | 2496077.75 | 3589037.5 | Q64475 | HIST1H2BB | 77087614 | 74898636 | 17653778 | 18 |
| Q64522 | HIST2H2AB | histone cluster 2, H2ab | Nucleus | other | 1.31E+07 | 6330031 | 6827609.5 | Q64522 | HIST2H2AB | 13050000 | 3744412 | 2161837 | 3 |
| P17710 | HK1 | hexokinase 1 | Cytoplasm | kinase | 1.69E+07 | 1.88E+07 | 2.17E+07 | | | | | | |
| P17710 | HK1 | hexokinase 1 | Cytoplasm | kinase | 7291285.5 | 8959883 | 8077118 | | | | | | |
| P17710 | HK1 | hexokinase 1 | Cytoplasm | kinase | 6816020.5 | 6746155.5 | 9181286 | | | | | | |
| P17710 | HK1 | hexokinase 1 | Cytoplasm | kinase | 5619546 | 4956286 | 7983186 | | | | | | |
| P17710 | HK1 | hexokinase 1 | Cytoplasm | kinase | 5562520.5 | 6399198.5 | 8149716 | | | | | | |
| P17710 | HK1 | hexokinase 1 | Cytoplasm | kinase | 6724941.5 | 7403920 | 7084530.5 | | | | | | |
| P17710 | HK1 | hexokinase 1 | Cytoplasm | kinase | 8109741 | 7965668.5 | 8608295 | | | | | | |
| P17710 | HK1 | hexokinase 1 | Cytoplasm | kinase | 5546270.5 | 5469277 | 5106946 | | | | | | |
| P17710 | HK1 | hexokinase 1 | Cytoplasm | kinase | 4530453 | 5719171 | 6711298.5 | | | | | | |
| P17710 | HK1 | hexokinase 1 | Cytoplasm | kinase | 3832786.75 | 3080220.5 | 3816974 | | | | | | |
| P17710 | HK1 | hexokinase 1 | Cytoplasm | kinase | 4149265.75 | 5487801.5 | 5564607 | | | | | | |
| P17710 | HK1 | hexokinase 1 | Cytoplasm | kinase | 2762487 | 3057371.25 | 3130894.5 | | | | | | |
| P17710 | HK1 | hexokinase 1 | Cytoplasm | kinase | 2040684.875 | 2256359 | 3214842.5 | P17710 | HK1 | 6141231 | 4137304 | 662499 | 39 |
| P01899 | HLA-A | major histocompatibility complex, | Plasma Membrane | other | 6228991 | 5377593.5 | 6212030.5 | | | | | | |
| P01899 | HLA-A | major histocompatibility complex, | Plasma Membrane | other | 3937605.5 | 2129539.75 | 2096933.375 | | | | | | |
| P01899 | HLA-A | major histocompatibility complex, | Plasma Membrane | other | 258502.578 | 108649.969 | 133933 | | | | | | |
| P01899 | HLA-A | major histocompatibility complex, | Plasma Membrane | other | 303108.812 | 1019184.188 | 732622.562 | | | | | | |
| P01899 | HLA-A | major histocompatibility complex, | Plasma Membrane | other | 3213.335 | 50565.055 | 1343.74 | | | | | | |
| P01899 | HLA-A | major histocompatibility complex, | Plasma Membrane | other | 4.70E-06 | 25188.891 | 46297.035 | P01899 | | | | | |
| P01901 | HLA-A | major histocompatibility complex, | Plasma Membrane | other | 5770374.5 | 3295016 | 5528991.5 | P01901 | | | | | |
| P01902 | HLA-A | major histocompatibility complex, | Plasma Membrane | other | 3650154.75 | 4281487.5 | 5158373.5 | | | | | | |
| P01902 | HLA-A | major histocompatibility complex, | Plasma Membrane | other | 3411813.75 | 2691620 | 3008605.5 | P01902 | HLA-A | 2618196 | 2276353 | 438084 | 27 |
| P22907 | HMBS | hydroxymethylbilane synthase | Cytoplasm | enzyme | 1336936.375 | 1083789 | 1140986 | P22907 | HMBS | 1336936 | 132760 | 76649 | 3 |
| P30681 | HMGB2 | high mobility group box 2 | Nucleus | transcription regulator | 2611339 | 1572713.25 | 1613524.625 | P30681 | HMGB2 | 2611339 | 588224 | 339611 | 3 |
| P38060 | HMGCL | 3-hydroxymethyl-3-methylglutaryl | Cytoplasm | enzyme | 1890439.625 | 1859930 | 2355849.25 | | | | | | |
| P38060 | HMGCL | 3-hydroxymethyl-3-methylglutaryl | Cytoplasm | enzyme | 1036851.625 | 373083.938 | 646072.875 | P38060 | HMGCL | 1463646 | 788813 | 322032 | 6 |
| Q61411 | HRAS | Harvey rat sarcoma viral oncogene | Plasma Membrane | enzyme | 1.77E+08 | 1.33E+08 | 1.47E+08 | | | | | | |
| Q61411 | HRAS | Harvey rat sarcoma viral oncogene | Plasma Membrane | enzyme | 4.94E+07 | 4.49E+07 | 4.43E+07 | | | | | | |
| Q61411 | HRAS | Harvey rat sarcoma viral oncogene | Plasma Membrane | enzyme | 4.00E+07 | 1.03E+07 | 1.85E+07 | | | | | | |
| Q61411 | HRAS | Harvey rat sarcoma viral oncogene | Plasma Membrane | enzyme | 2.75E+07 | 4.95E+07 | 3.52E+07 | | | | | | |
| Q61411 | HRAS | Harvey rat sarcoma viral oncogene | Plasma Membrane | enzyme | 1.61E+07 | 2.24E+07 | 2.07E+07 | | | | | | |
| Q61411 | HRAS | Harvey rat sarcoma viral oncogene | Plasma Membrane | enzyme | 7737845.5 | 1.68E+07 | 8720072 | | | | | | |
| Q61411 | HRAS | Harvey rat sarcoma viral oncogene | Plasma Membrane | enzyme | 1.52E+07 | 8530157 | 1.07E+07 | | | | | | |
| Q61411 | HRAS | Harvey rat sarcoma viral oncogene | Plasma Membrane | enzyme | 7718226 | 1.29E+07 | 8717605 | | | | | | |
| Q61411 | HRAS | Harvey rat sarcoma viral oncogene | Plasma Membrane | enzyme | 2607767.25 | 5389660.5 | 3375972 | | | | | | |
| Q61411 | HRAS | Harvey rat sarcoma viral oncogene | Plasma Membrane | enzyme | 2487787.75 | 1245755.125 | 2665392.75 | Q61411 | HRAS | 34577163 | 43811008 | 7998759 | 30 |
| O08756 | HSD17B10 | hydroxysteroid (17-beta) dehydro | Cytoplasm | enzyme | 8610517 | 7178700 | 1.24E+07 | | | | | | |
| O08756 | HSD17B10 | hydroxysteroid (17-beta) dehydro | Cytoplasm | enzyme | 3896580.5 | 2770960 | 3109116.5 | | | | | | |
| O08756 | HSD17B10 | hydroxysteroid (17-beta) dehydro | Cytoplasm | enzyme | 4299401.5 | 5344760.5 | 7793177.5 | | | | | | |
| O08756 | HSD17B10 | hydroxysteroid (17-beta) dehydro | Cytoplasm | enzyme | 1131364.25 | 947309.938 | 1937223 | O08756 | HSD17B10 | 4484466 | 3449169 | 995689 | 12 |
| O70503 | HSD17B12 | hydroxysteroid (17-beta) dehydro | Cytoplasm | enzyme | 3928757.75 | 4154274.25 | 3207075.25 | | | | | | |
| O70503 | HSD17B12 | hydroxysteroid (17-beta) dehydro | Cytoplasm | enzyme | 1190245.125 | 1623335.125 | 903432.938 | O70503 | HSD17B12 | 2559501 | 1436032 | 586258 | 6 |
| Q8BTX9 | HSDL1 | hydroxysteroid dehydrogenase like | Cytoplasm | enzyme | 498573.094 | 718247.688 | 612421.312 | Q8BTX9 | HSDL1 | 498573 | 109862 | 63429 | 3 |
| P07901 | HSP90AA1 | heat shock protein 90kDa alpha (c | Cytoplasm | enzyme | 1.23E+07 | 1.23E+07 | 7567834.5 | | | | | | |

| | | | | | | | | | | | | | |
|---|---|---|---|---|---|---|---|---|---|---|---|---|---|
| P07901 | HSP90AA1 | heat shock protein 90kDa alpha (c | Cytoplasm | enzyme | 1.06E+07 | 7482357 | 4080978.5 | | | | | | |
| P07901 | HSP90AA1 | heat shock protein 90kDa alpha (c | Cytoplasm | enzyme | 9789329 | 7915819.5 | 5515475.5 | | | | | | |
| P07901 | HSP90AA1 | heat shock protein 90kDa alpha (c | Cytoplasm | enzyme | 7675979 | 6216860 | 3354204.75 | | | | | | |
| P07901 | HSP90AA1 | heat shock protein 90kDa alpha (c | Cytoplasm | enzyme | 6238814 | 5667586 | 3591215 | | | | | | |
| P07901 | HSP90AA1 | heat shock protein 90kDa alpha (c | Cytoplasm | enzyme | 4853482 | 4329296 | 1765622.5 | P07901 | HSP90AA1 | 8564601 | 3009741 | 709403 | 18 |
| P11499 | HSP90AB1 | heat shock protein 90kDa alpha (c | Cytoplasm | enzyme | 8950438 | 8229099.5 | 5486545 | P11499 | HSP90AB1 | 8950438 | 1827589 | 1055159 | 3 |
| P08113 | HSP90B1 | heat shock protein 90kDa beta (Gr | Cytoplasm | other | 4.11E+07 | 5.08E+07 | 4.25E+07 | | | | | | |
| P08113 | HSP90B1 | heat shock protein 90kDa beta (Gr | Cytoplasm | other | 4.55E+07 | 4.84E+07 | 3.80E+07 | | | | | | |
| P08113 | HSP90B1 | heat shock protein 90kDa beta (Gr | Cytoplasm | other | 3.90E+07 | 3.95E+07 | 3.58E+07 | | | | | | |
| P08113 | HSP90B1 | heat shock protein 90kDa beta (Gr | Cytoplasm | other | 2.20E+07 | 2.28E+07 | 1.99E+07 | | | | | | |
| P08113 | HSP90B1 | heat shock protein 90kDa beta (Gr | Cytoplasm | other | 2.60E+07 | 3.50E+07 | 2.77E+07 | | | | | | |
| P08113 | HSP90B1 | heat shock protein 90kDa beta (Gr | Cytoplasm | other | 3.48E+07 | 4.30E+07 | 3.04E+07 | | | | | | |
| P08113 | HSP90B1 | heat shock protein 90kDa beta (Gr | Cytoplasm | other | 3.27E+07 | 4.24E+07 | 3.05E+07 | | | | | | |
| P08113 | HSP90B1 | heat shock protein 90kDa beta (Gr | Cytoplasm | other | 2.84E+07 | 2.39E+07 | 2.38E+07 | | | | | | |
| P08113 | HSP90B1 | heat shock protein 90kDa beta (Gr | Cytoplasm | other | 2.64E+07 | 2.82E+07 | 2.35E+07 | | | | | | |
| P08113 | HSP90B1 | heat shock protein 90kDa beta (Gr | Cytoplasm | other | 3.05E+07 | 3.25E+07 | 2.30E+07 | | | | | | |
| P08113 | HSP90B1 | heat shock protein 90kDa beta (Gr | Cytoplasm | other | 2.28E+07 | 2.49E+07 | 2.00E+07 | | | | | | |
| P08113 | HSP90B1 | heat shock protein 90kDa beta (Gr | Cytoplasm | other | 2.14E+07 | 2.19E+07 | 1.35E+07 | | | | | | |
| P08113 | HSP90B1 | heat shock protein 90kDa beta (Gr | Cytoplasm | other | 2.31E+07 | 3.73E+07 | 2.76E+07 | | | | | | |
| P08113 | HSP90B1 | heat shock protein 90kDa beta (Gr | Cytoplasm | other | 2.67E+07 | 2.43E+07 | 2.41E+07 | | | | | | |
| P08113 | HSP90B1 | heat shock protein 90kDa beta (Gr | Cytoplasm | other | 1.84E+07 | 2.66E+07 | 2.05E+07 | | | | | | |
| P08113 | HSP90B1 | heat shock protein 90kDa beta (Gr | Cytoplasm | other | 1.84E+07 | 2.16E+07 | 1.73E+07 | | | | | | |
| P08113 | HSP90B1 | heat shock protein 90kDa beta (Gr | Cytoplasm | other | 1.73E+07 | 2.09E+07 | 2.03E+07 | | | | | | |
| P08113 | HSP90B1 | heat shock protein 90kDa beta (Gr | Cytoplasm | other | 1.78E+07 | 2.03E+07 | 1.70E+07 | | | | | | |
| P08113 | HSP90B1 | heat shock protein 90kDa beta (Gr | Cytoplasm | other | 1.65E+07 | 1.89E+07 | 1.47E+07 | | | | | | |
| P08113 | HSP90B1 | heat shock protein 90kDa beta (Gr | Cytoplasm | other | 1.89E+07 | 2.34E+07 | 2.09E+07 | | | | | | |
| P08113 | HSP90B1 | heat shock protein 90kDa beta (Gr | Cytoplasm | other | 1.38E+07 | 1.57E+07 | 1.71E+07 | | | | | | |
| P08113 | HSP90B1 | heat shock protein 90kDa beta (Gr | Cytoplasm | other | 1.34E+07 | 1.27E+07 | 1.28E+07 | | | | | | |
| P08113 | HSP90B1 | heat shock protein 90kDa beta (Gr | Cytoplasm | other | 9318226 | 1.09E+07 | 9136209 | | | | | | |
| P08113 | HSP90B1 | heat shock protein 90kDa beta (Gr | Cytoplasm | other | 7209460 | 9177327 | 7950298 | | | | | | |
| P08113 | HSP90B1 | heat shock protein 90kDa beta (Gr | Cytoplasm | other | 1626277.25 | 2213544 | 2041220.625 | | | | | | |
| P08113 | HSP90B1 | heat shock protein 90kDa beta (Gr | Cytoplasm | other | 2701221.25 | 2983049.75 | 2312862.25 | P08113 | HSP90B1 | 22139430 | 11402184 | 1291043 | 78 |
| Q61316 | HSPA4 | heat shock 70kDa protein 4 | Cytoplasm | other | 2.71E+07 | 2.28E+07 | 1.91E+07 | | | | | | |
| Q61316 | HSPA4 | heat shock 70kDa protein 4 | Cytoplasm | other | 1.39E+07 | 1.33E+07 | 1.32E+07 | | | | | | |
| Q61316 | HSPA4 | heat shock 70kDa protein 4 | Cytoplasm | other | 1.17E+07 | 1.27E+07 | 9555393 | | | | | | |
| Q61316 | HSPA4 | heat shock 70kDa protein 4 | Cytoplasm | other | 9278628 | 6609921.5 | 5354919.5 | | | | | | |
| Q61316 | HSPA4 | heat shock 70kDa protein 4 | Cytoplasm | other | 1.07E+07 | 9508163 | 7762594 | | | | | | |
| Q61316 | HSPA4 | heat shock 70kDa protein 4 | Cytoplasm | other | 1.22E+07 | 4875313.5 | 5176053 | | | | | | |
| Q61316 | HSPA4 | heat shock 70kDa protein 4 | Cytoplasm | other | 1.12E+07 | 9104994 | 9551228 | | | | | | |
| Q61316 | HSPA4 | heat shock 70kDa protein 4 | Cytoplasm | other | 8294282.5 | 9714708 | 1.13E+07 | | | | | | |
| Q61316 | HSPA4 | heat shock 70kDa protein 4 | Cytoplasm | other | 8374762 | 8368608.5 | 8504033 | | | | | | |
| Q61316 | HSPA4 | heat shock 70kDa protein 4 | Cytoplasm | other | 8951958 | 8740219 | 1.00E+07 | | | | | | |
| Q61316 | HSPA4 | heat shock 70kDa protein 4 | Cytoplasm | other | 5737764 | 5248410 | 5206816.5 | | | | | | |
| Q61316 | HSPA4 | heat shock 70kDa protein 4 | Cytoplasm | other | 1.02E+07 | 6122288.5 | 7494359.5 | | | | | | |
| Q61316 | HSPA4 | heat shock 70kDa protein 4 | Cytoplasm | other | 6244713 | 5139327 | 7607726.5 | | | | | | |
| Q61316 | HSPA4 | heat shock 70kDa protein 4 | Cytoplasm | other | 3681923.75 | 3224864.25 | 2781860 | | | | | | |
| Q61316 | HSPA4 | heat shock 70kDa protein 4 | Cytoplasm | other | 5408922 | 4892681.5 | 5133421 | | | | | | |
| Q61316 | HSPA4 | heat shock 70kDa protein 4 | Cytoplasm | other | 6825649 | 4589540 | 4354276 | | | | | | |
| Q61316 | HSPA4 | heat shock 70kDa protein 4 | Cytoplasm | other | 3951620.5 | 3077073 | 3437211 | | | | | | |
| Q61316 | HSPA4 | heat shock 70kDa protein 4 | Cytoplasm | other | 3100384.5 | 4245216 | 4286082.5 | | | | | | |
| Q61316 | HSPA4 | heat shock 70kDa protein 4 | Cytoplasm | other | 3512633 | 2753894.25 | 3350822.25 | | | | | | |
| Q61316 | HSPA4 | heat shock 70kDa protein 4 | Cytoplasm | other | 5048245.5 | 4009393.75 | 4659693.5 | | | | | | |
| Q61316 | HSPA4 | heat shock 70kDa protein 4 | Cytoplasm | other | 2723538 | 3991919.5 | 4066202.75 | | | | | | |
| Q61316 | HSPA4 | heat shock 70kDa protein 4 | Cytoplasm | other | 1859309.75 | 1353331.375 | 802361.188 | | | | | | |
| Q61316 | HSPA4 | heat shock 70kDa protein 4 | Cytoplasm | other | 3237130.5 | 3591568.25 | 1182662.25 | Q61316 | HSPA4 | 7970933 | 4755289 | 572469 | 69 |
| P48722 | HSPA4L | heat shock 70kDa protein 4-like | Cytoplasm | other | 5690185 | 6192820.5 | 6212961.5 | | | | | | |
| P48722 | HSPA4L | heat shock 70kDa protein 4-like | Cytoplasm | other | 3541567.75 | 2417497 | 3448864.75 | | | | | | |
| P48722 | HSPA4L | heat shock 70kDa protein 4-like | Cytoplasm | other | 2904995.25 | 2721300 | 2826161 | | | | | | |
| P48722 | HSPA4L | heat shock 70kDa protein 4-like | Cytoplasm | other | 4372774 | 3970682.25 | 3842238.5 | | | | | | |
| P48722 | HSPA4L | heat shock 70kDa protein 4-like | Cytoplasm | other | 1846061.625 | 1604610.125 | 1559781.375 | P48722 | HSPA4L | 3671117 | 1539097 | 397393 | 15 |
| P20029 | HSPA5 | heat shock 70kDa protein 5 (gluco | Cytoplasm | enzyme | 2.00E+08 | 1.95E+08 | 1.81E+08 | | | | | | |
| P20029 | HSPA5 | heat shock 70kDa protein 5 (gluco | Cytoplasm | enzyme | 8.21E+07 | 1.11E+08 | 7.20E+07 | | | | | | |
| P20029 | HSPA5 | heat shock 70kDa protein 5 (gluco | Cytoplasm | enzyme | 5.67E+07 | 8.44E+07 | 6.65E+07 | | | | | | |
| P20029 | HSPA5 | heat shock 70kDa protein 5 (gluco | Cytoplasm | enzyme | 5.87E+07 | 7.44E+07 | 6.36E+07 | | | | | | |
| P20029 | HSPA5 | heat shock 70kDa protein 5 (gluco | Cytoplasm | enzyme | 6.38E+07 | 8.57E+07 | 7.01E+07 | | | | | | |
| P20029 | HSPA5 | heat shock 70kDa protein 5 (gluco | Cytoplasm | enzyme | 5.12E+07 | 6.35E+07 | 4.98E+07 | | | | | | |
| P20029 | HSPA5 | heat shock 70kDa protein 5 (gluco | Cytoplasm | enzyme | 5.28E+07 | 8.20E+07 | 7.48E+07 | | | | | | |
| P20029 | HSPA5 | heat shock 70kDa protein 5 (gluco | Cytoplasm | enzyme | 4.53E+07 | 6.59E+07 | 4.46E+07 | | | | | | |
| P20029 | HSPA5 | heat shock 70kDa protein 5 (gluco | Cytoplasm | enzyme | 3.84E+07 | 4.61E+07 | 3.60E+07 | | | | | | |
| P20029 | HSPA5 | heat shock 70kDa protein 5 (gluco | Cytoplasm | enzyme | 3.69E+07 | 5.23E+07 | 4.72E+07 | | | | | | |
| P20029 | HSPA5 | heat shock 70kDa protein 5 (gluco | Cytoplasm | enzyme | 4.67E+07 | 5.44E+07 | 3.74E+07 | | | | | | |
| P20029 | HSPA5 | heat shock 70kDa protein 5 (gluco | Cytoplasm | enzyme | 3.24E+07 | 3.85E+07 | 3.50E+07 | | | | | | |
| P20029 | HSPA5 | heat shock 70kDa protein 5 (gluco | Cytoplasm | enzyme | 2.71E+07 | 2.87E+07 | 2.58E+07 | | | | | | |
| P20029 | HSPA5 | heat shock 70kDa protein 5 (gluco | Cytoplasm | enzyme | 1.98E+07 | 2.62E+07 | 2.65E+07 | | | | | | |
| P20029 | HSPA5 | heat shock 70kDa protein 5 (gluco | Cytoplasm | enzyme | 2.23E+07 | 1.86E+07 | 2.08E+07 | | | | | | |
| P20029 | HSPA5 | heat shock 70kDa protein 5 (gluco | Cytoplasm | enzyme | 2.66E+07 | 9038222 | 7150298 | | | | | | |
| P20029 | HSPA5 | heat shock 70kDa protein 5 (gluco | Cytoplasm | enzyme | 1.40E+07 | 1.84E+07 | 1.55E+07 | | | | | | |
| P20029 | HSPA5 | heat shock 70kDa protein 5 (gluco | Cytoplasm | enzyme | 1.59E+07 | 1.19E+07 | 9061766 | | | | | | |
| P20029 | HSPA5 | heat shock 70kDa protein 5 (gluco | Cytoplasm | enzyme | 1.38E+07 | 3.83E+07 | 2.91E+07 | | | | | | |
| P20029 | HSPA5 | heat shock 70kDa protein 5 (gluco | Cytoplasm | enzyme | 7766572 | 1.52E+07 | 1.17E+07 | | | | | | |
| P20029 | HSPA5 | heat shock 70kDa protein 5 (gluco | Cytoplasm | enzyme | 7441908 | 8017693.5 | 7389009 | | | | | | |
| P20029 | HSPA5 | heat shock 70kDa protein 5 (gluco | Cytoplasm | enzyme | 7535258.5 | 7668139.5 | 3698291.25 | | | | | | |
| P20029 | HSPA5 | heat shock 70kDa protein 5 (gluco | Cytoplasm | enzyme | 3995851.25 | 5200450.5 | 5083481.5 | | | | | | |
| P20029 | HSPA5 | heat shock 70kDa protein 5 (gluco | Cytoplasm | enzyme | 5807824 | 8847291 | 5290083 | | | | | | |
| P20029 | HSPA5 | heat shock 70kDa protein 5 (gluco | Cytoplasm | enzyme | 2465874 | 718329.562 | 399813.625 | | | | | | |
| P20029 | HSPA5 | heat shock 70kDa protein 5 (gluco | Cytoplasm | enzyme | 5267480.5 | 2075306.5 | 3038906.25 | P20029 | HSPA5 | 36350030 | 40352257 | 4568993 | 78 |

| | | | | | | | | | | | | | |
|---|---|---|---|---|---|---|---|---|---|---|---|---|---|
| P63017 | HSPA8 | heat shock 70kDa protein 8 | Cytoplasm | enzyme | 1.42E+08 | 1.37E+08 | 1.25E+08 | | | | | | |
| P63017 | HSPA8 | heat shock 70kDa protein 8 | Cytoplasm | enzyme | 1.23E+08 | 1.28E+08 | 1.26E+08 | | | | | | |
| P63017 | HSPA8 | heat shock 70kDa protein 8 | Cytoplasm | enzyme | 1.16E+08 | 1.06E+08 | 9.87E+07 | | | | | | |
| P63017 | HSPA8 | heat shock 70kDa protein 8 | Cytoplasm | enzyme | 1.07E+08 | 1.00E+08 | 1.03E+08 | | | | | | |
| P63017 | HSPA8 | heat shock 70kDa protein 8 | Cytoplasm | enzyme | 9.84E+07 | 9.30E+07 | 8.35E+07 | | | | | | |
| P63017 | HSPA8 | heat shock 70kDa protein 8 | Cytoplasm | enzyme | 9.61E+07 | 8.11E+07 | 9.18E+07 | | | | | | |
| P63017 | HSPA8 | heat shock 70kDa protein 8 | Cytoplasm | enzyme | 9.35E+07 | 9.57E+07 | 9.34E+07 | | | | | | |
| P63017 | HSPA8 | heat shock 70kDa protein 8 | Cytoplasm | enzyme | 7.69E+07 | 7.04E+07 | 6.23E+07 | | | | | | |
| P63017 | HSPA8 | heat shock 70kDa protein 8 | Cytoplasm | enzyme | 8.75E+07 | 6.11E+07 | 6.62E+07 | | | | | | |
| P63017 | HSPA8 | heat shock 70kDa protein 8 | Cytoplasm | enzyme | 6.22E+07 | 5.75E+07 | 4.93E+07 | | | | | | |
| P63017 | HSPA8 | heat shock 70kDa protein 8 | Cytoplasm | enzyme | 5.76E+07 | 6.97E+07 | 4.67E+07 | | | | | | |
| P63017 | HSPA8 | heat shock 70kDa protein 8 | Cytoplasm | enzyme | 6.97E+07 | 6.16E+07 | 4.88E+07 | | | | | | |
| P63017 | HSPA8 | heat shock 70kDa protein 8 | Cytoplasm | enzyme | 7.76E+07 | 7.50E+07 | 7.59E+07 | | | | | | |
| P63017 | HSPA8 | heat shock 70kDa protein 8 | Cytoplasm | enzyme | 5.98E+07 | 5.85E+07 | 5.24E+07 | | | | | | |
| P63017 | HSPA8 | heat shock 70kDa protein 8 | Cytoplasm | enzyme | 4.84E+07 | 5.05E+07 | 4.63E+07 | | | | | | |
| P63017 | HSPA8 | heat shock 70kDa protein 8 | Cytoplasm | enzyme | 3.58E+07 | 3.54E+07 | 3.48E+07 | | | | | | |
| P63017 | HSPA8 | heat shock 70kDa protein 8 | Cytoplasm | enzyme | 3.06E+07 | 2.96E+07 | 2.46E+07 | | | | | | |
| P63017 | HSPA8 | heat shock 70kDa protein 8 | Cytoplasm | enzyme | 4.95E+07 | 3.11E+07 | 3.71E+07 | | | | | | |
| P63017 | HSPA8 | heat shock 70kDa protein 8 | Cytoplasm | enzyme | 2.42E+07 | 1.97E+07 | 1.83E+07 | | | | | | |
| P63017 | HSPA8 | heat shock 70kDa protein 8 | Cytoplasm | enzyme | 2.93E+07 | 2.13E+07 | 2.32E+07 | | | | | | |
| P63017 | HSPA8 | heat shock 70kDa protein 8 | Cytoplasm | enzyme | 2.03E+07 | 2.68E+07 | 2.70E+07 | | | | | | |
| P63017 | HSPA8 | heat shock 70kDa protein 8 | Cytoplasm | enzyme | 3764585.75 | 4712091 | 5774065 | | | | | | |
| P63017 | HSPA8 | heat shock 70kDa protein 8 | Cytoplasm | enzyme | 3059788.75 | 2634359.75 | 2149803.25 | | | | | | |
| P63017 | HSPA8 | heat shock 70kDa protein 8 | Cytoplasm | enzyme | 3460789.75 | 3784996.5 | 3742749.25 | | | | | | |
| P63017 | HSPA8 | heat shock 70kDa protein 8 | Cytoplasm | enzyme | 1905285.25 | 2034988.375 | 2197112 | | | | | | |
| P63017 | HSPA8 | heat shock 70kDa protein 8 | Cytoplasm | enzyme | 1211799.875 | 1585052.25 | 1613932.5 | P63017 | HSPA8 | 58405471 | 39506319 | 4473210 | 78 |
| P38647 | HSPA9 | heat shock 70kDa protein 9 (morta | Cytoplasm | other | 1.21E+07 | 1.11E+07 | 1.44E+07 | | | | | | |
| P38647 | HSPA9 | heat shock 70kDa protein 9 (morta | Cytoplasm | other | 9927600 | 8240593 | 1.17E+07 | | | | | | |
| P38647 | HSPA9 | heat shock 70kDa protein 9 (morta | Cytoplasm | other | 1.10E+07 | 8951685 | 1.36E+07 | | | | | | |
| P38647 | HSPA9 | heat shock 70kDa protein 9 (morta | Cytoplasm | other | 8444272 | 8656869 | 9790030 | | | | | | |
| P38647 | HSPA9 | heat shock 70kDa protein 9 (morta | Cytoplasm | other | 7595241.5 | 6960055.5 | 1.07E+07 | | | | | | |
| P38647 | HSPA9 | heat shock 70kDa protein 9 (morta | Cytoplasm | other | 7399764 | 6847049.5 | 7524992.5 | | | | | | |
| P38647 | HSPA9 | heat shock 70kDa protein 9 (morta | Cytoplasm | other | 6532328.5 | 5273267 | 7148504 | | | | | | |
| P38647 | HSPA9 | heat shock 70kDa protein 9 (morta | Cytoplasm | other | 6226795 | 4595310 | 6443071.5 | | | | | | |
| P38647 | HSPA9 | heat shock 70kDa protein 9 (morta | Cytoplasm | other | 4674821.5 | 4326754 | 5849608.5 | | | | | | |
| P38647 | HSPA9 | heat shock 70kDa protein 9 (morta | Cytoplasm | other | 1791531.875 | 9532176 | 1.22E+07 | | | | | | |
| P38647 | HSPA9 | heat shock 70kDa protein 9 (morta | Cytoplasm | other | 2569209.25 | 2020281.25 | 2178628.25 | | | | | | |
| P38647 | HSPA9 | heat shock 70kDa protein 9 (morta | Cytoplasm | other | 2493573 | 1599179 | 2088739.375 | P38647 | HSPA9 | 6727928 | 3558412 | 593069 | 36 |
| P63038 | HSPD1 | heat shock 60kDa protein 1 (chape | Cytoplasm | enzyme | 2.40E+07 | 3.00E+07 | 4.52E+07 | | | | | | |
| P63038 | HSPD1 | heat shock 60kDa protein 1 (chape | Cytoplasm | enzyme | 2.82E+07 | 2.01E+07 | 2.70E+07 | | | | | | |
| P63038 | HSPD1 | heat shock 60kDa protein 1 (chape | Cytoplasm | enzyme | 2.33E+07 | 3.07E+07 | 4.31E+07 | | | | | | |
| P63038 | HSPD1 | heat shock 60kDa protein 1 (chape | Cytoplasm | enzyme | 2.59E+07 | 2.76E+07 | 3.10E+07 | | | | | | |
| P63038 | HSPD1 | heat shock 60kDa protein 1 (chape | Cytoplasm | enzyme | 2.02E+07 | 2.07E+07 | 3.02E+07 | | | | | | |
| P63038 | HSPD1 | heat shock 60kDa protein 1 (chape | Cytoplasm | enzyme | 1.83E+07 | 2.31E+07 | 2.49E+07 | | | | | | |
| P63038 | HSPD1 | heat shock 60kDa protein 1 (chape | Cytoplasm | enzyme | 1.66E+07 | 2.05E+07 | 2.76E+07 | | | | | | |
| P63038 | HSPD1 | heat shock 60kDa protein 1 (chape | Cytoplasm | enzyme | 2.35E+07 | 2.15E+07 | 2.33E+07 | | | | | | |
| P63038 | HSPD1 | heat shock 60kDa protein 1 (chape | Cytoplasm | enzyme | 1.52E+07 | 1.82E+07 | 2.37E+07 | | | | | | |
| P63038 | HSPD1 | heat shock 60kDa protein 1 (chape | Cytoplasm | enzyme | 1.62E+07 | 1.46E+07 | 2.00E+07 | | | | | | |
| P63038 | HSPD1 | heat shock 60kDa protein 1 (chape | Cytoplasm | enzyme | 1.44E+07 | 1.33E+07 | 1.84E+07 | | | | | | |
| P63038 | HSPD1 | heat shock 60kDa protein 1 (chape | Cytoplasm | enzyme | 1.36E+07 | 1.12E+07 | 1.63E+07 | | | | | | |
| P63038 | HSPD1 | heat shock 60kDa protein 1 (chape | Cytoplasm | enzyme | 1.29E+07 | 1.16E+07 | 1.53E+07 | | | | | | |
| P63038 | HSPD1 | heat shock 60kDa protein 1 (chape | Cytoplasm | enzyme | 1.37E+07 | 1.78E+07 | 2.15E+07 | | | | | | |
| P63038 | HSPD1 | heat shock 60kDa protein 1 (chape | Cytoplasm | enzyme | 8615636 | 9725966 | 1.26E+07 | | | | | | |
| P63038 | HSPD1 | heat shock 60kDa protein 1 (chape | Cytoplasm | enzyme | 1.44E+07 | 1.59E+07 | 3.24E+07 | | | | | | |
| P63038 | HSPD1 | heat shock 60kDa protein 1 (chape | Cytoplasm | enzyme | 4242698.5 | 4110846 | 6757377.5 | | | | | | |
| P63038 | HSPD1 | heat shock 60kDa protein 1 (chape | Cytoplasm | enzyme | 2153025 | 3817941.5 | 5778496 | | | | | | |
| P63038 | HSPD1 | heat shock 60kDa protein 1 (chape | Cytoplasm | enzyme | 4232464.5 | 2431139.25 | 3223532.75 | | | | | | |
| P63038 | HSPD1 | heat shock 60kDa protein 1 (chape | Cytoplasm | enzyme | 2197084.75 | 4928020 | 6115504.5 | | | | | | |
| P63038 | HSPD1 | heat shock 60kDa protein 1 (chape | Cytoplasm | enzyme | 1190399.875 | 1729058.75 | 1818260.875 | | | | | | |
| P63038 | HSPD1 | heat shock 60kDa protein 1 (chape | Cytoplasm | enzyme | 2487381.25 | 2214867.75 | 3335937.25 | | | | | | |
| P63038 | HSPD1 | heat shock 60kDa protein 1 (chape | Cytoplasm | enzyme | 1922628.5 | 1868139.75 | 2080550.25 | P63038 | HSPD1 | 13364405 | 10530160 | 1267682 | 69 |
| Q64433 | HSPE1 | heat shock 10kDa protein 1 | Cytoplasm | enzyme | 2.47E+07 | 2.43E+07 | 3.24E+07 | | | | | | |
| Q64433 | HSPE1 | heat shock 10kDa protein 1 | Cytoplasm | enzyme | 1.63E+07 | 1.58E+07 | 2.05E+07 | | | | | | |
| Q64433 | HSPE1 | heat shock 10kDa protein 1 | Cytoplasm | enzyme | 1.32E+07 | 1.51E+07 | 2.02E+07 | | | | | | |
| Q64433 | HSPE1 | heat shock 10kDa protein 1 | Cytoplasm | enzyme | 5962272 | 6614189.5 | 9686783 | | | | | | |
| Q64433 | HSPE1 | heat shock 10kDa protein 1 | Cytoplasm | enzyme | 5819537 | 4856438 | 6353781.5 | Q64433 | HSPE1 | 13184362 | 8391497 | 2166675 | 15 |
| Q9Z2G9 | HTATIP2 | HIV-1 Tat interactive protein 2, 30 | Nucleus | transcription regulator | 2424699 | 2897010.5 | 1728998.75 | | | | | | |
| Q9Z2G9 | HTATIP2 | HIV-1 Tat interactive protein 2, 30 | Nucleus | transcription regulator | 1683613.5 | 917770.688 | 924633.625 | Q9Z2G9 | HTATIP2 | 2054156 | 793543 | 323963 | 6 |
| Q7TMY8 | HUWE1 | HECT, UBA and WWE domain cont | Nucleus | transcription regulator | 2301852.75 | 2505018.25 | 2406527.75 | Q7TMY8 | HUWE1 | 2301853 | 101598 | 58658 | 3 |
| Q91ZJ9 | HYAL1 | hyaluronoglucosaminidase 1 | Cytoplasm | enzyme | 5463096.5 | 4963992 | 2748070 | Q91ZJ9 | HYAL1 | 5463097 | 1445152 | 834359 | 3 |
| Q9JKR6 | HYOU1 | hypoxia up-regulated 1 | Cytoplasm | other | 4066113.25 | 5206332 | 4308557.5 | | | | | | |
| Q9JKR6 | HYOU1 | hypoxia up-regulated 1 | Cytoplasm | other | 4549985.5 | 5103611.5 | 2998100 | | | | | | |
| Q9JKR6 | HYOU1 | hypoxia up-regulated 1 | Cytoplasm | other | 3079378.5 | 3199830.75 | 4624097.5 | | | | | | |
| Q9JKR6 | HYOU1 | hypoxia up-regulated 1 | Cytoplasm | other | 2906988.5 | 3970726 | 2613316 | | | | | | |
| Q9JKR6 | HYOU1 | hypoxia up-regulated 1 | Cytoplasm | other | 4066707.25 | 3476692.25 | 2136279.5 | | | | | | |
| Q9JKR6 | HYOU1 | hypoxia up-regulated 1 | Cytoplasm | other | 2842710.25 | 2882108.25 | 1986871.25 | | | | | | |
| Q9JKR6 | HYOU1 | hypoxia up-regulated 1 | Cytoplasm | other | 1454590.75 | 3588794 | 3114076.75 | Q9JKR6 | HYOU1 | 3280925 | 1006073 | 219543 | 21 |
| Q8BIJ6 | IARS2 | isoleucyl-tRNA synthetase 2, mitoc | Cytoplasm | enzyme | 2239343.25 | 2352547.75 | 3519249.75 | Q8BIJ6 | IARS2 | 2239343 | 708540 | 409075 | 3 |
| Q9ERM2 | ICAM4 | intercellular adhesion molecule 4 ( | Plasma Membrane | other | 1960857.375 | 2722378.25 | 1568382 | | | | | | |
| Q9ERM2 | ICAM4 | intercellular adhesion molecule 4 ( | Plasma Membrane | other | 2981533.75 | 5425608.5 | 3496267.25 | Q9ERM2 | ICAM4 | 2471196 | 1366207 | 557752 | 6 |
| Q9JHR7 | IDE | insulin-degrading enzyme | Extracellular Space | peptidase | 4046053.25 | 3980428 | 4199595.5 | | | | | | |
| Q9JHR7 | IDE | insulin-degrading enzyme | Extracellular Space | peptidase | 3668903.25 | 5142967 | 3346414.25 | Q9JHR7 | IDE | 3857478 | 610064 | 249058 | 6 |
| P54071 | IDH2 | isocitrate dehydrogenase 2 (NADP | Cytoplasm | enzyme | 7270838 | 1.11E+07 | 1.17E+07 | | | | | | |
| P54071 | IDH2 | isocitrate dehydrogenase 2 (NADP | Cytoplasm | enzyme | 1693440 | 2063846.625 | 2410051 | | | | | | |
| P54071 | IDH2 | isocitrate dehydrogenase 2 (NADP | Cytoplasm | enzyme | 2816638.5 | 3329069.25 | 3935486.5 | | | | | | |
| P54071 | IDH2 | isocitrate dehydrogenase 2 (NADP | Cytoplasm | enzyme | 2126544 | 2641640.25 | 3363997.25 | | | | | | |

| | | | | | | | | | | | | | |
|---|---|---|---|---|---|---|---|---|---|---|---|---|---|
| P54071 | IDH2 | isocitrate dehydrogenase 2 (NADP | Cytoplasm | enzyme | 1873739.75 | 2116863.75 | 3569601 | | | | | | |
| P54071 | IDH2 | isocitrate dehydrogenase 2 (NADP | Cytoplasm | enzyme | 1061076 | 1252094.5 | 889870.562 | P54071 | IDH2 | 2807046 | 3169896 | 747152 | 18 |
| Q9D6R2 | IDH3A | isocitrate dehydrogenase 3 (NAD+ | Cytoplasm | enzyme | 1.24E+07 | 1.89E+07 | 2.19E+07 | | | | | | |
| Q9D6R2 | IDH3A | isocitrate dehydrogenase 3 (NAD+ | Cytoplasm | enzyme | 6528680.5 | 6101443 | 7768954.5 | | | | | | |
| Q9D6R2 | IDH3A | isocitrate dehydrogenase 3 (NAD+ | Cytoplasm | enzyme | 4530409.5 | 6450803.5 | 8826493 | | | | | | |
| Q9D6R2 | IDH3A | isocitrate dehydrogenase 3 (NAD+ | Cytoplasm | enzyme | 4986705.5 | 5660057.5 | 6257235.5 | | | | | | |
| Q9D6R2 | IDH3A | isocitrate dehydrogenase 3 (NAD+ | Cytoplasm | enzyme | 4481949 | 5074982.5 | 6174954 | Q9D6R2 | IDH3A | 6587549 | 5298221 | 1367995 | 15 |
| P70404 | IDH3G | isocitrate dehydrogenase 3 (NAD+ | Cytoplasm | enzyme | 6315953 | 5762569.5 | 7779190 | | | | | | |
| P70404 | IDH3G | isocitrate dehydrogenase 3 (NAD+ | Cytoplasm | enzyme | 1109845.125 | 1523808.125 | 2537206 | P70404 | IDH3G | 3712899 | 2800019 | 1143103 | 6 |
| Q9CR20 | IER3IP1 | immediate early response 3 intera | Cytoplasm | other | 1.13E+07 | 1.14E+07 | 1.26E+07 | | | | | | |
| Q9CR20 | IER3IP1 | immediate early response 3 intera | Cytoplasm | other | 3217843.5 | 3729690 | 3583334.5 | Q9CR20 | IER3IP1 | 7278922 | 4537366 | 1852372 | 6 |
| P01872 | IGHM | immunoglobulin heavy constant m | Plasma Membrane | transmembrane receptor | 4953099.5 | 8468592 | 6180731 | | | | | | |
| P01872 | IGHM | immunoglobulin heavy constant m | Plasma Membrane | transmembrane receptor | 1385706.875 | 2197517 | 1614153 | | | | | | |
| P01872 | IGHM | immunoglobulin heavy constant m | Plasma Membrane | transmembrane receptor | 1185242.875 | 3373852.25 | 2476574.25 | | | | | | |
| P01872 | IGHM | immunoglobulin heavy constant m | Plasma Membrane | transmembrane receptor | 1285393.5 | 2864807.25 | 2084157.75 | P01872 | IGHM | 2202361 | 2259417 | 652238 | 12 |
| O55222 | ILK | integrin-linked kinase | Plasma Membrane | kinase | 5392863 | 5770958.5 | 7589131.5 | | | | | | |
| O55222 | ILK | integrin-linked kinase | Plasma Membrane | kinase | 2927401.5 | 2086775.25 | 2288501.5 | | | | | | |
| O55222 | ILK | integrin-linked kinase | Plasma Membrane | kinase | 3699708.5 | 2627609.75 | 3318926.75 | | | | | | |
| O55222 | ILK | integrin-linked kinase | Plasma Membrane | kinase | 2494715.75 | 2054374 | 1969326.625 | | | | | | |
| O55222 | ILK | integrin-linked kinase | Plasma Membrane | kinase | 1101525.25 | 1262510 | 1073257.125 | | | | | | |
| O55222 | ILK | integrin-linked kinase | Plasma Membrane | kinase | 1258869.5 | 878933.312 | 1079750.75 | O55222 | ILK | 2812514 | 1860511 | 438527 | 18 |
| Q8BU33 | ILVBL | ilvB (bacterial acetolactate syntha | Cytoplasm | enzyme | 612862.625 | 1266760.375 | 1059112.375 | Q8BU33 | ILVBL | 612863 | 334125 | 192907 | 3 |
| Q8CAQ8 | IMMT | inner membrane protein, mitocho | Cytoplasm | other | 4582109 | 5198645 | 6846504 | | | | | | |
| Q8CAQ8 | IMMT | inner membrane protein, mitocho | Cytoplasm | other | 5864228 | 6042441.5 | 7128072.5 | | | | | | |
| Q8CAQ8 | IMMT | inner membrane protein, mitocho | Cytoplasm | other | 5202128.5 | 5053414.5 | 6574798.5 | | | | | | |
| Q8CAQ8 | IMMT | inner membrane protein, mitocho | Cytoplasm | other | 3272253.25 | 5363452 | 5871330.5 | | | | | | |
| Q8CAQ8 | IMMT | inner membrane protein, mitocho | Cytoplasm | other | 5060498.5 | 4167289.25 | 5278709 | | | | | | |
| Q8CAQ8 | IMMT | inner membrane protein, mitocho | Cytoplasm | other | 2317940 | 2545216.75 | 2059716.125 | | | | | | |
| Q8CAQ8 | IMMT | inner membrane protein, mitocho | Cytoplasm | other | 2763007 | 2908934.75 | 3080095.75 | | | | | | |
| Q8CAQ8 | IMMT | inner membrane protein, mitocho | Cytoplasm | other | 862609.688 | 877190.812 | 1316698 | | | | | | |
| Q8CAQ8 | IMMT | inner membrane protein, mitocho | Cytoplasm | other | 1900481.875 | 2132150.75 | 2624421.75 | Q8CAQ8 | IMMT | 3536140 | 1901303 | 365906 | 27 |
| P24547 | IMPDH2 | IMP (inosine 5'-monophosphate) c | Cytoplasm | enzyme | 1459235 | 353662.938 | 296958.5 | P24547 | IMPDH2 | 1459235 | 655285 | 378329 | 3 |
| Q8C5L6 | INPP5K | inositol polyphosphate-5-phospha | Cytoplasm | phosphatase | 5000372.5 | 7438032 | 5068357 | | | | | | |
| Q8C5L6 | INPP5K | inositol polyphosphate-5-phospha | Cytoplasm | phosphatase | 7620359 | 7591970.5 | 7761065 | Q8C5L6 | INPP5K | 6310366 | 1330499 | 543174 | 6 |
| Q8CIM8 | INTS4 | integrator complex subunit 4 | Nucleus | other | 2.91E+07 | 3.92E+07 | 3.30E+07 | Q8CIM8 | INTS4 | 29070000 | 5103375 | 2946435 | 3 |
| Q8BKC5 | IPO5 | importin 5 | Nucleus | transporter | 1.24E+07 | 1.27E+07 | 8415714 | | | | | | |
| Q8BKC5 | IPO5 | importin 5 | Nucleus | transporter | 5943029 | 5580365 | 2191288.75 | | | | | | |
| Q8BKC5 | IPO5 | importin 5 | Nucleus | transporter | 1.17E+07 | 1.39E+07 | 1.09E+07 | | | | | | |
| Q8BKC5 | IPO5 | importin 5 | Nucleus | transporter | 1.16E+07 | 1.93E+07 | 1.24E+07 | | | | | | |
| Q8BKC5 | IPO5 | importin 5 | Nucleus | transporter | 7434531 | 6901704.5 | 4990154.5 | | | | | | |
| Q8BKC5 | IPO5 | importin 5 | Nucleus | transporter | 7089757.5 | 7319842.5 | 5434992.5 | | | | | | |
| Q8BKC5 | IPO5 | importin 5 | Nucleus | transporter | 5875096.5 | 5875887 | 4856220 | | | | | | |
| Q8BKC5 | IPO5 | importin 5 | Nucleus | transporter | 7213849.5 | 7114096.5 | 6137955.5 | | | | | | |
| Q8BKC5 | IPO5 | importin 5 | Nucleus | transporter | 5982713.5 | 5466681.5 | 2690187.25 | | | | | | |
| Q8BKC5 | IPO5 | importin 5 | Nucleus | transporter | 5451148.5 | 6080674.5 | 5067682.5 | | | | | | |
| Q8BKC5 | IPO5 | importin 5 | Nucleus | transporter | 4876079 | 5171893.5 | 4013199.5 | | | | | | |
| Q8BKC5 | IPO5 | importin 5 | Nucleus | transporter | 5608331.5 | 5074408.5 | 4396825.5 | | | | | | |
| Q8BKC5 | IPO5 | importin 5 | Nucleus | transporter | 5225567 | 4795775 | 2965519 | | | | | | |
| Q8BKC5 | IPO5 | importin 5 | Nucleus | transporter | 3859793.25 | 4773149 | 2261876.5 | | | | | | |
| Q8BKC5 | IPO5 | importin 5 | Nucleus | transporter | 4692118.5 | 4547294.5 | 2659219.5 | | | | | | |
| Q8BKC5 | IPO5 | importin 5 | Nucleus | transporter | 2924790.75 | 2923171.75 | 2806827.25 | | | | | | |
| Q8BKC5 | IPO5 | importin 5 | Nucleus | transporter | 3056729 | 2420054 | 1833617 | | | | | | |
| Q8BKC5 | IPO5 | importin 5 | Nucleus | transporter | 5086203.5 | 3909354 | 3015298 | | | | | | |
| Q8BKC5 | IPO5 | importin 5 | Nucleus | transporter | 2876356.25 | 2462810.5 | 2564689.75 | | | | | | |
| Q8BKC5 | IPO5 | importin 5 | Nucleus | transporter | 2907302 | 3384453.25 | 2171796.75 | Q8BKC5 | IPO5 | 6091670 | 3433668 | 443285 | 60 |
| Q9EPL8 | IPO7 | importin 7 | Nucleus | transporter | 3376429.75 | 4231865 | 2815822.75 | | | | | | |
| Q9EPL8 | IPO7 | importin 7 | Nucleus | transporter | 3209510 | 2868366.5 | 1194013.5 | | | | | | |
| Q9EPL8 | IPO7 | importin 7 | Nucleus | transporter | 2353387.75 | 2258729.25 | 1383345.25 | | | | | | |
| Q9EPL8 | IPO7 | importin 7 | Nucleus | transporter | 2298732.25 | 2416714.75 | 1578855.25 | | | | | | |
| Q9EPL8 | IPO7 | importin 7 | Nucleus | transporter | 1534051 | 1602273.625 | 917798.375 | | | | | | |
| Q9EPL8 | IPO7 | importin 7 | Nucleus | transporter | 1613034 | 1800612.5 | 1130925.125 | | | | | | |
| Q9EPL8 | IPO7 | importin 7 | Nucleus | transporter | 591010.062 | 818914.562 | 329086.844 | Q9EPL8 | IPO7 | 2139451 | 998745 | 217944 | 21 |
| Q60766 | Irgm1 | immunity-related GTPase family M | Cytoplasm | other | 1.83E+07 | 1.75E+07 | 1.45E+07 | | | | | | |
| Q60766 | Irgm1 | immunity-related GTPase family M | Cytoplasm | other | 1.40E+07 | 1.31E+07 | 8031236.5 | | | | | | |
| Q60766 | Irgm1 | immunity-related GTPase family M | Cytoplasm | other | 1.31E+07 | 1.36E+07 | 1.57E+07 | | | | | | |
| Q60766 | Irgm1 | immunity-related GTPase family M | Cytoplasm | other | 1.28E+07 | 1.04E+07 | 1.04E+07 | | | | | | |
| Q60766 | Irgm1 | immunity-related GTPase family M | Cytoplasm | other | 9705876 | 7714240 | 9003922 | | | | | | |
| Q60766 | Irgm1 | immunity-related GTPase family M | Cytoplasm | other | 7765086.5 | 7970590.5 | 9560002 | | | | | | |
| Q60766 | Irgm1 | immunity-related GTPase family M | Cytoplasm | other | 6581995 | 7648288 | 7760017 | | | | | | |
| Q60766 | Irgm1 | immunity-related GTPase family M | Cytoplasm | other | 5493310.5 | 4002281 | 3792199.5 | | | | | | |
| Q60766 | Irgm1 | immunity-related GTPase family M | Cytoplasm | other | 3869169.25 | 3878416.25 | 1.17E+07 | | | | | | |
| Q60766 | Irgm1 | immunity-related GTPase family M | Cytoplasm | other | 962032.5 | 1976564.5 | 3040298 | Q60766 | Irgm1 | 9260747 | 4622363 | 843924 | 30 |
| Q9D924 | ISCA1 | iron-sulfur cluster assembly 1 | Cytoplasm | other | 4259231 | 4379684.5 | 4889806.5 | | | | | | |
| Q9D924 | ISCA1 | iron-sulfur cluster assembly 1 | Cytoplasm | other | 2528644.25 | 2077860 | 1875859.25 | Q9D924 | ISCA1 | 3393938 | 1320810 | 539218 | 6 |
| Q9DCB8 | ISCA2 | iron-sulfur cluster assembly 2 | Cytoplasm | other | 2142904.25 | 1911728.875 | 3364377.75 | Q9DCB8 | ISCA2 | 2142904 | 780558 | 450656 | 3 |
| Q64339 | ISG15 | ISG15 ubiquitin-like modifier | Extracellular Space | other | 4.88E+07 | 7.40E+07 | 7.64E+07 | | | | | | |
| Q64339 | ISG15 | ISG15 ubiquitin-like modifier | Extracellular Space | other | 4.97E+07 | 5.34E+07 | 5.91E+07 | | | | | | |
| Q64339 | ISG15 | ISG15 ubiquitin-like modifier | Extracellular Space | other | 3.42E+07 | 3.56E+07 | 2.96E+07 | | | | | | |
| Q64339 | ISG15 | ISG15 ubiquitin-like modifier | Extracellular Space | other | 2.62E+07 | 1.34E+07 | 1.25E+07 | | | | | | |
| Q64339 | ISG15 | ISG15 ubiquitin-like modifier | Extracellular Space | other | 2.42E+07 | 4.14E+07 | 3.19E+07 | | | | | | |
| Q64339 | ISG15 | ISG15 ubiquitin-like modifier | Extracellular Space | other | 8493878 | 1.92E+07 | 1.33E+07 | | | | | | |
| Q64339 | ISG15 | ISG15 ubiquitin-like modifier | Extracellular Space | other | 4489121.5 | 2322165.5 | 2060433.625 | | | | | | |
| Q64339 | ISG15 | ISG15 ubiquitin-like modifier | Extracellular Space | other | 2621000 | 3761046.5 | 4927869.5 | Q64339 | ISG15 | 24839250 | 22893718 | 4673161 | 24 |
| Q9JL16 | ISG20 | interferon stimulated exonuclease | Nucleus | enzyme | 7666691 | 9927449 | 5329461.5 | | | | | | |
| Q9JL16 | ISG20 | interferon stimulated exonuclease | Nucleus | enzyme | 6671558.5 | 7443374 | 3551483.25 | | | | | | |
| Q9JL16 | ISG20 | interferon stimulated exonuclease | Nucleus | enzyme | 4479526.5 | 7060090.5 | 3259494.75 | | | | | | |

| | | | | | | | | | | | | | |
|---|---|---|---|---|---|---|---|---|---|---|---|---|---|
| Q9JL16 | ISG20 | interferon stimulated exonuclease | Nucleus | enzyme | 5180075 | 5468152.5 | 4824883 | Q9JL16 | ISG20 | 5999463 | 1915393 | 552926 | 12 |
| Q9CX00 | IST1 | increased sodium tolerance 1 hom | Cytoplasm | other | 1728472.375 | 1910528.625 | 1581908.75 | Q9CX00 | IST1 | 1728472 | 164629 | 95049 | 3 |
| Q62469 | ITGA2 | integrin, alpha 2 (CD49B, alpha 2 s | Plasma Membrane | transmembrane receptor | 6568184 | 6820219.5 | 9007617 | | | | | | |
| Q62469 | ITGA2 | integrin, alpha 2 (CD49B, alpha 2 s | Plasma Membrane | transmembrane receptor | 5437062.5 | 5759584.5 | 5553343 | | | | | | |
| Q62469 | ITGA2 | integrin, alpha 2 (CD49B, alpha 2 s | Plasma Membrane | transmembrane receptor | 5941747 | 4554486 | 6022922 | | | | | | |
| Q62469 | ITGA2 | integrin, alpha 2 (CD49B, alpha 2 s | Plasma Membrane | transmembrane receptor | 3807219.5 | 3469540 | 4397553.5 | | | | | | |
| Q62469 | ITGA2 | integrin, alpha 2 (CD49B, alpha 2 s | Plasma Membrane | transmembrane receptor | 1.02E+07 | 7590424 | 7699141 | | | | | | |
| Q62469 | ITGA2 | integrin, alpha 2 (CD49B, alpha 2 s | Plasma Membrane | transmembrane receptor | 3160577.5 | 4598521 | 4577889 | | | | | | |
| Q62469 | ITGA2 | integrin, alpha 2 (CD49B, alpha 2 s | Plasma Membrane | transmembrane receptor | 4978131 | 4156552.75 | 4839665 | | | | | | |
| Q62469 | ITGA2 | integrin, alpha 2 (CD49B, alpha 2 s | Plasma Membrane | transmembrane receptor | 1193846.875 | 2907796.25 | 3139526.5 | | | | | | |
| Q62469 | ITGA2 | integrin, alpha 2 (CD49B, alpha 2 s | Plasma Membrane | transmembrane receptor | 2147050.75 | 2449122.75 | 2715241 | | | | | | |
| Q62469 | ITGA2 | integrin, alpha 2 (CD49B, alpha 2 s | Plasma Membrane | transmembrane receptor | 1369697.75 | 1053646.625 | 1209547.25 | | | | | | |
| Q62469 | ITGA2 | integrin, alpha 2 (CD49B, alpha 2 s | Plasma Membrane | transmembrane receptor | 4680939.5 | 4556127.5 | 5068545.5 | Q62469 | ITGA2 | 4500405 | 2201477 | 383228 | 33 |
| Q9QUM0 | ITGA2B | integrin, alpha 2b (platelet glycopr | Plasma Membrane | transmembrane receptor | 1.91E+08 | 1.89E+08 | 2.18E+08 | | | | | | |
| Q9QUM0 | ITGA2B | integrin, alpha 2b (platelet glycopr | Plasma Membrane | transmembrane receptor | 5.63E+07 | 5.30E+07 | 5.40E+07 | | | | | | |
| Q9QUM0 | ITGA2B | integrin, alpha 2b (platelet glycopr | Plasma Membrane | transmembrane receptor | 5.48E+07 | 6.63E+07 | 6.25E+07 | | | | | | |
| Q9QUM0 | ITGA2B | integrin, alpha 2b (platelet glycopr | Plasma Membrane | transmembrane receptor | 4.54E+07 | 4.96E+07 | 5.55E+07 | | | | | | |
| Q9QUM0 | ITGA2B | integrin, alpha 2b (platelet glycopr | Plasma Membrane | transmembrane receptor | 8.20E+07 | 8.71E+07 | 8.75E+07 | | | | | | |
| Q9QUM0 | ITGA2B | integrin, alpha 2b (platelet glycopr | Plasma Membrane | transmembrane receptor | 6.81E+07 | 7.96E+07 | 7.79E+07 | | | | | | |
| Q9QUM0 | ITGA2B | integrin, alpha 2b (platelet glycopr | Plasma Membrane | transmembrane receptor | 3.30E+07 | 3.94E+07 | 3.51E+07 | | | | | | |
| Q9QUM0 | ITGA2B | integrin, alpha 2b (platelet glycopr | Plasma Membrane | transmembrane receptor | 4.02E+07 | 3.89E+07 | 4.99E+07 | | | | | | |
| Q9QUM0 | ITGA2B | integrin, alpha 2b (platelet glycopr | Plasma Membrane | transmembrane receptor | 8894281 | 2.48E+07 | 2.00E+07 | | | | | | |
| Q9QUM0 | ITGA2B | integrin, alpha 2b (platelet glycopr | Plasma Membrane | transmembrane receptor | 1.38E+07 | 1.96E+07 | 2.29E+07 | | | | | | |
| Q9QUM0 | ITGA2B | integrin, alpha 2b (platelet glycopr | Plasma Membrane | transmembrane receptor | 4.25E+07 | 3.80E+07 | 4.96E+07 | | | | | | |
| Q9QUM0 | ITGA2B | integrin, alpha 2b (platelet glycopr | Plasma Membrane | transmembrane receptor | 2.36E+07 | 5.57E+07 | 3.85E+07 | | | | | | |
| Q9QUM0 | ITGA2B | integrin, alpha 2b (platelet glycopr | Plasma Membrane | transmembrane receptor | 2.76E+07 | 3.42E+07 | 2.56E+07 | | | | | | |
| Q9QUM0 | ITGA2B | integrin, alpha 2b (platelet glycopr | Plasma Membrane | transmembrane receptor | 2.35E+07 | 3.21E+07 | 3.89E+07 | | | | | | |
| Q9QUM0 | ITGA2B | integrin, alpha 2b (platelet glycopr | Plasma Membrane | transmembrane receptor | 2.54E+07 | 2.55E+07 | 3.64E+07 | | | | | | |
| Q9QUM0 | ITGA2B | integrin, alpha 2b (platelet glycopr | Plasma Membrane | transmembrane receptor | 3.17E+07 | 2.96E+07 | 3.77E+07 | | | | | | |
| Q9QUM0 | ITGA2B | integrin, alpha 2b (platelet glycopr | Plasma Membrane | transmembrane receptor | 1.22E+07 | 1.66E+07 | 1.53E+07 | | | | | | |
| Q9QUM0 | ITGA2B | integrin, alpha 2b (platelet glycopr | Plasma Membrane | transmembrane receptor | 2.43E+07 | 2.94E+07 | 3.23E+07 | | | | | | |
| Q9QUM0 | ITGA2B | integrin, alpha 2b (platelet glycopr | Plasma Membrane | transmembrane receptor | 1.85E+07 | 1.84E+07 | 1.32E+07 | | | | | | |
| Q9QUM0 | ITGA2B | integrin, alpha 2b (platelet glycopr | Plasma Membrane | transmembrane receptor | 2.39E+07 | 3.13E+07 | 2.69E+07 | | | | | | |
| Q9QUM0 | ITGA2B | integrin, alpha 2b (platelet glycopr | Plasma Membrane | transmembrane receptor | 1.49E+07 | 2.28E+07 | 2.02E+07 | | | | | | |
| Q9QUM0 | ITGA2B | integrin, alpha 2b (platelet glycopr | Plasma Membrane | transmembrane receptor | 4905725.5 | 4535847.5 | 4224033 | | | | | | |
| Q9QUM0 | ITGA2B | integrin, alpha 2b (platelet glycopr | Plasma Membrane | transmembrane receptor | 6012110.5 | 3223479.25 | 2943137.5 | | | | | | |
| Q9QUM0 | ITGA2B | integrin, alpha 2b (platelet glycopr | Plasma Membrane | transmembrane receptor | 8100265 | 1.17E+07 | 8538616 | | | | | | |
| Q9QUM0 | ITGA2B | integrin, alpha 2b (platelet glycopr | Plasma Membrane | transmembrane receptor | 3462114.5 | 1.21E+07 | 8267619 | | | | | | |
| Q9QUM0 | ITGA2B | integrin, alpha 2b (platelet glycopr | Plasma Membrane | transmembrane receptor | 3343784.75 | 4186066 | 4332989.5 | | | | | | |
| Q9QUM0 | ITGA2B | integrin, alpha 2b (platelet glycopr | Plasma Membrane | transmembrane receptor | 2456399.75 | 8816369 | 4634625.5 | Q9QUM0 | ITGA2B | 32939433 | 38792670 | 4310297 | 81 |
| P11688 | ITGA5 | integrin, alpha 5 (fibronectin recep | Plasma Membrane | transmembrane receptor | 1.02E+07 | 1.00E+07 | 9750413 | | | | | | |
| P11688 | ITGA5 | integrin, alpha 5 (fibronectin recep | Plasma Membrane | transmembrane receptor | 4973710 | 5045558 | 4670783.5 | | | | | | |
| P11688 | ITGA5 | integrin, alpha 5 (fibronectin recep | Plasma Membrane | transmembrane receptor | 4048336 | 3733946.5 | 2966272 | | | | | | |
| P11688 | ITGA5 | integrin, alpha 5 (fibronectin recep | Plasma Membrane | transmembrane receptor | 2111668.25 | 2726765.5 | 2815375 | P11688 | ITGA5 | 5343429 | 3009853 | 868870 | 12 |
| Q61739 | ITGA6 | integrin, alpha 6 | Plasma Membrane | transmembrane receptor | 8763647 | 7270436 | 8823719 | | | | | | |
| Q61739 | ITGA6 | integrin, alpha 6 | Plasma Membrane | transmembrane receptor | 2491602 | 2701744.75 | 3445758.5 | | | | | | |
| Q61739 | ITGA6 | integrin, alpha 6 | Plasma Membrane | transmembrane receptor | 4448816 | 4558323.5 | 5484841 | | | | | | |
| Q61739 | ITGA6 | integrin, alpha 6 | Plasma Membrane | transmembrane receptor | 2002234.25 | 1981887.25 | 2479726.5 | | | | | | |
| Q61739 | ITGA6 | integrin, alpha 6 | Plasma Membrane | transmembrane receptor | 132117.094 | 1853277 | 74544.875 | | | | | | |
| Q61739 | ITGA6 | integrin, alpha 6 | Plasma Membrane | transmembrane receptor | 2295372.75 | 2164582.25 | 2149717 | | | | | | |
| Q61739 | ITGA6 | integrin, alpha 6 | Plasma Membrane | transmembrane receptor | 5109291 | 5141789.5 | 5552617 | Q61739 | ITGA6 | 3606154 | 2473988 | 539868 | 21 |
| P09055 | ITGB1 | integrin, beta 1 (fibronectin recept | Plasma Membrane | transmembrane receptor | 2.65E+07 | 2.37E+07 | 2.99E+07 | | | | | | |
| P09055 | ITGB1 | integrin, beta 1 (fibronectin recept | Plasma Membrane | transmembrane receptor | 1.79E+07 | 1.78E+07 | 2.31E+07 | | | | | | |
| P09055 | ITGB1 | integrin, beta 1 (fibronectin recept | Plasma Membrane | transmembrane receptor | 1.80E+07 | 2.22E+07 | 2.39E+07 | | | | | | |
| P09055 | ITGB1 | integrin, beta 1 (fibronectin recept | Plasma Membrane | transmembrane receptor | 1.72E+07 | 1.94E+07 | 2.08E+07 | | | | | | |
| P09055 | ITGB1 | integrin, beta 1 (fibronectin recept | Plasma Membrane | transmembrane receptor | 8589947 | 1.12E+07 | 1.39E+07 | | | | | | |
| P09055 | ITGB1 | integrin, beta 1 (fibronectin recept | Plasma Membrane | transmembrane receptor | 1.03E+07 | 9688770 | 9580617 | | | | | | |
| P09055 | ITGB1 | integrin, beta 1 (fibronectin recept | Plasma Membrane | transmembrane receptor | 1.05E+07 | 1.03E+07 | 1.06E+07 | | | | | | |
| P09055 | ITGB1 | integrin, beta 1 (fibronectin recept | Plasma Membrane | transmembrane receptor | 6382390.5 | 9504313 | 8712920 | | | | | | |
| P09055 | ITGB1 | integrin, beta 1 (fibronectin recept | Plasma Membrane | transmembrane receptor | 8876776 | 8674333 | 9237730 | | | | | | |
| P09055 | ITGB1 | integrin, beta 1 (fibronectin recept | Plasma Membrane | transmembrane receptor | 6723581.5 | 6314548.5 | 6587435.5 | | | | | | |
| P09055 | ITGB1 | integrin, beta 1 (fibronectin recept | Plasma Membrane | transmembrane receptor | 4507120.5 | 5434173.5 | 4163526.5 | | | | | | |
| P09055 | ITGB1 | integrin, beta 1 (fibronectin recept | Plasma Membrane | transmembrane receptor | 2914721 | 2962562 | 3145368.75 | | | | | | |
| P09055 | ITGB1 | integrin, beta 1 (fibronectin recept | Plasma Membrane | transmembrane receptor | 2577891 | 1727600.25 | 2002875.875 | P09055 | ITGB1 | 10841725 | 7603079 | 1217467 | 39 |
| P11835 | ITGB2 | integrin, beta 2 (complement com | Plasma Membrane | transmembrane receptor | 857676.125 | 1515175.875 | 2308708.5 | P11835 | ITGB2 | 857676 | 726578 | 419490 | 3 |
| O54890 | ITGB3 | integrin, beta 3 (platelet glycoprot | Plasma Membrane | transmembrane receptor | 1.52E+08 | 1.41E+08 | 1.55E+08 | | | | | | |
| O54890 | ITGB3 | integrin, beta 3 (platelet glycoprot | Plasma Membrane | transmembrane receptor | 1.69E+07 | 1.85E+07 | 2.00E+07 | | | | | | |
| O54890 | ITGB3 | integrin, beta 3 (platelet glycoprot | Plasma Membrane | transmembrane receptor | 5.83E+07 | 4.75E+07 | 4.95E+07 | | | | | | |
| O54890 | ITGB3 | integrin, beta 3 (platelet glycoprot | Plasma Membrane | transmembrane receptor | 5.02E+07 | 6.22E+07 | 5.09E+07 | | | | | | |
| O54890 | ITGB3 | integrin, beta 3 (platelet glycoprot | Plasma Membrane | transmembrane receptor | 5.07E+07 | 5.41E+07 | 6.50E+07 | | | | | | |
| O54890 | ITGB3 | integrin, beta 3 (platelet glycoprot | Plasma Membrane | transmembrane receptor | 5.60E+07 | 5.23E+07 | 4.79E+07 | | | | | | |
| O54890 | ITGB3 | integrin, beta 3 (platelet glycoprot | Plasma Membrane | transmembrane receptor | 3.11E+07 | 2.73E+07 | 2.93E+07 | | | | | | |
| O54890 | ITGB3 | integrin, beta 3 (platelet glycoprot | Plasma Membrane | transmembrane receptor | 6.21E+07 | 5.72E+07 | 5.44E+07 | | | | | | |
| O54890 | ITGB3 | integrin, beta 3 (platelet glycoprot | Plasma Membrane | transmembrane receptor | 2.94E+07 | 3.25E+07 | 3.15E+07 | | | | | | |
| O54890 | ITGB3 | integrin, beta 3 (platelet glycoprot | Plasma Membrane | transmembrane receptor | 2.06E+07 | 1.59E+07 | 1.99E+07 | | | | | | |
| O54890 | ITGB3 | integrin, beta 3 (platelet glycoprot | Plasma Membrane | transmembrane receptor | 2.28E+07 | 2.49E+07 | 2.27E+07 | | | | | | |
| O54890 | ITGB3 | integrin, beta 3 (platelet glycoprot | Plasma Membrane | transmembrane receptor | 2.70E+07 | 2.94E+07 | 2.57E+07 | | | | | | |
| O54890 | ITGB3 | integrin, beta 3 (platelet glycoprot | Plasma Membrane | transmembrane receptor | 2.39E+07 | 3.45E+07 | 4.17E+07 | | | | | | |
| O54890 | ITGB3 | integrin, beta 3 (platelet glycoprot | Plasma Membrane | transmembrane receptor | 3.54E+07 | 3.18E+07 | 3.40E+07 | | | | | | |
| O54890 | ITGB3 | integrin, beta 3 (platelet glycoprot | Plasma Membrane | transmembrane receptor | 1.93E+07 | 3.70E+07 | 3.89E+07 | | | | | | |
| O54890 | ITGB3 | integrin, beta 3 (platelet glycoprot | Plasma Membrane | transmembrane receptor | 7127314 | 2605901.5 | 4643639 | | | | | | |
| O54890 | ITGB3 | integrin, beta 3 (platelet glycoprot | Plasma Membrane | transmembrane receptor | 1.48E+07 | 2.68E+07 | 3.61E+07 | | | | | | |
| O54890 | ITGB3 | integrin, beta 3 (platelet glycoprot | Plasma Membrane | transmembrane receptor | 1.57E+07 | 2.41E+07 | 2.33E+07 | | | | | | |
| O54890 | ITGB3 | integrin, beta 3 (platelet glycoprot | Plasma Membrane | transmembrane receptor | 8714321 | 3.28E+07 | 1.93E+07 | | | | | | |
| O54890 | ITGB3 | integrin, beta 3 (platelet glycoprot | Plasma Membrane | transmembrane receptor | 2.20E+07 | 2.16E+07 | 2.07E+07 | | | | | | |
| O54890 | ITGB3 | integrin, beta 3 (platelet glycoprot | Plasma Membrane | transmembrane receptor | 3328484.75 | 1.88E+07 | 2.15E+07 | | | | | | |

| | | | | | | | | | | | | | |
|---|---|---|---|---|---|---|---|---|---|---|---|---|---|
| O54890 | ITGB3 | integrin, beta 3 (platelet glycoprot | Plasma Membrane | transmembrane receptor | 8966690 | 1.44E+07 | 1.52E+07 | | | | | | |
| O54890 | ITGB3 | integrin, beta 3 (platelet glycoprot | Plasma Membrane | transmembrane receptor | 8730548 | 3.22E+07 | 3.60E+07 | | | | | | |
| O54890 | ITGB3 | integrin, beta 3 (platelet glycoprot | Plasma Membrane | transmembrane receptor | 1549454.375 | 396002.812 | 455963.188 | O54890 | ITGB3 | 31092784 | 29243451 | 3446374 | 72 |
| Q9Z0T9 | ITGB6 | integrin, beta 6 | Plasma Membrane | other | 7580600 | 8385315.5 | 1.14E+07 | Q9Z0T9 | ITGB6 | 7580600 | 2030417 | 1172262 | 3 |
| Q5XKN4 | JAGN1 | jagunal homolog 1 | Cytoplasm | other | 2418748.5 | 2185706.75 | 2051816 | Q5XKN4 | JAGN1 | 2418749 | 185686 | 107206 | 3 |
| A2CG49 | KALRN | kalirin, RhoGEF kinase | Cytoplasm | kinase | 2827465.75 | 2042635.5 | 2288858.5 | | | | | | |
| A2CG49 | KALRN | kalirin, RhoGEF kinase | Cytoplasm | kinase | 1040330.188 | 985055.125 | 851109.312 | A2CG49 | KALRN | 1933898 | 824348 | 336539 | 6 |
| Q80UY2 | KCMF1 | potassium channel modulatory fac | Extracellular Space | other | 1679760.125 | 2855132 | 2573378.25 | | | | | | |
| Q80UY2 | KCMF1 | potassium channel modulatory fac | Extracellular Space | other | 574234 | 662730.75 | 1203447.75 | | | | | | |
| Q80UY2 | KCMF1 | potassium channel modulatory fac | Extracellular Space | other | 565790.125 | 1079955.25 | 760195.25 | Q80UY2 | KCMF1 | 939928 | 865880 | 288627 | 9 |
| Q9EQF2 | KEL | Kell blood group, metallo-endopep | Plasma Membrane | peptidase | 1.39E+08 | 1.59E+08 | 1.23E+08 | | | | | | |
| Q9EQF2 | KEL | Kell blood group, metallo-endopep | Plasma Membrane | peptidase | 1.77E+08 | 1.73E+08 | 1.62E+08 | | | | | | |
| Q9EQF2 | KEL | Kell blood group, metallo-endopep | Plasma Membrane | peptidase | 1.23E+08 | 1.51E+08 | 1.22E+08 | | | | | | |
| Q9EQF2 | KEL | Kell blood group, metallo-endopep | Plasma Membrane | peptidase | 9.08E+07 | 1.59E+08 | 1.21E+08 | | | | | | |
| Q9EQF2 | KEL | Kell blood group, metallo-endopep | Plasma Membrane | peptidase | 1.23E+08 | 1.14E+08 | 1.23E+08 | | | | | | |
| Q9EQF2 | KEL | Kell blood group, metallo-endopep | Plasma Membrane | peptidase | 7.80E+07 | 1.05E+08 | 8.63E+07 | | | | | | |
| Q9EQF2 | KEL | Kell blood group, metallo-endopep | Plasma Membrane | peptidase | 7.30E+07 | 8.00E+07 | 8.45E+07 | | | | | | |
| Q9EQF2 | KEL | Kell blood group, metallo-endopep | Plasma Membrane | peptidase | 8.25E+07 | 1.17E+08 | 9.72E+07 | | | | | | |
| Q9EQF2 | KEL | Kell blood group, metallo-endopep | Plasma Membrane | peptidase | 7.79E+07 | 6.32E+07 | 7.52E+07 | | | | | | |
| Q9EQF2 | KEL | Kell blood group, metallo-endopep | Plasma Membrane | peptidase | 7.17E+07 | 8.34E+07 | 7.42E+07 | | | | | | |
| Q9EQF2 | KEL | Kell blood group, metallo-endopep | Plasma Membrane | peptidase | 4.30E+07 | 4.89E+07 | 5.27E+07 | | | | | | |
| Q9EQF2 | KEL | Kell blood group, metallo-endopep | Plasma Membrane | peptidase | 5.99E+07 | 5.54E+07 | 5.83E+07 | | | | | | |
| Q9EQF2 | KEL | Kell blood group, metallo-endopep | Plasma Membrane | peptidase | 4.80E+07 | 6.48E+07 | 7.38E+07 | | | | | | |
| Q9EQF2 | KEL | Kell blood group, metallo-endopep | Plasma Membrane | peptidase | 2.67E+07 | 4.71E+07 | 3.44E+07 | | | | | | |
| Q9EQF2 | KEL | Kell blood group, metallo-endopep | Plasma Membrane | peptidase | 4.10E+07 | 3.91E+07 | 3.84E+07 | | | | | | |
| Q9EQF2 | KEL | Kell blood group, metallo-endopep | Plasma Membrane | peptidase | 6.33E+07 | 3.02E+07 | 5.24E+07 | | | | | | |
| Q9EQF2 | KEL | Kell blood group, metallo-endopep | Plasma Membrane | peptidase | 5.98E+07 | 2.93E+07 | 5.23E+07 | | | | | | |
| Q9EQF2 | KEL | Kell blood group, metallo-endopep | Plasma Membrane | peptidase | 2.86E+07 | 3.15E+07 | 2.64E+07 | | | | | | |
| Q9EQF2 | KEL | Kell blood group, metallo-endopep | Plasma Membrane | peptidase | 3.35E+07 | 3.88E+07 | 3.67E+07 | | | | | | |
| Q9EQF2 | KEL | Kell blood group, metallo-endopep | Plasma Membrane | peptidase | 3.47E+07 | 3.52E+07 | 3.30E+07 | | | | | | |
| Q9EQF2 | KEL | Kell blood group, metallo-endopep | Plasma Membrane | peptidase | 2.19E+07 | 2.08E+07 | 1.71E+07 | | | | | | |
| Q9EQF2 | KEL | Kell blood group, metallo-endopep | Plasma Membrane | peptidase | 4.11E+07 | 1.21E+07 | 3.21E+07 | | | | | | |
| Q9EQF2 | KEL | Kell blood group, metallo-endopep | Plasma Membrane | peptidase | 1.17E+07 | 2.12E+07 | 2.29E+07 | | | | | | |
| Q9EQF2 | KEL | Kell blood group, metallo-endopep | Plasma Membrane | peptidase | 1.85E+07 | 2.74E+07 | 2.20E+07 | | | | | | |
| Q9EQF2 | KEL | Kell blood group, metallo-endopep | Plasma Membrane | peptidase | 1.32E+07 | 1.59E+07 | 1.14E+07 | | | | | | |
| Q9EQF2 | KEL | Kell blood group, metallo-endopep | Plasma Membrane | peptidase | 2.02E+07 | 2.78E+07 | 2.52E+07 | | | | | | |
| Q9EQF2 | KEL | Kell blood group, metallo-endopep | Plasma Membrane | peptidase | 1.23E+07 | 1.16E+07 | 1.25E+07 | | | | | | |
| Q9EQF2 | KEL | Kell blood group, metallo-endopep | Plasma Membrane | peptidase | 1.37E+07 | 2.08E+07 | 1.76E+07 | | | | | | |
| Q9EQF2 | KEL | Kell blood group, metallo-endopep | Plasma Membrane | peptidase | 1.46E+07 | 1.58E+07 | 1.52E+07 | | | | | | |
| Q9EQF2 | KEL | Kell blood group, metallo-endopep | Plasma Membrane | peptidase | 1.78E+07 | 1.78E+07 | 1.45E+07 | | | | | | |
| Q9EQF2 | KEL | Kell blood group, metallo-endopep | Plasma Membrane | peptidase | 8031516 | 1.51E+07 | 1.97E+07 | | | | | | |
| Q9EQF2 | KEL | Kell blood group, metallo-endopep | Plasma Membrane | peptidase | 4099016.5 | 8637664 | 5200842.5 | | | | | | |
| Q9EQF2 | KEL | Kell blood group, metallo-endopep | Plasma Membrane | peptidase | 5422231.5 | 8744719 | 7998727 | | | | | | |
| Q9EQF2 | KEL | Kell blood group, metallo-endopep | Plasma Membrane | peptidase | 1779120.25 | 5526707.5 | 3907222 | | | | | | |
| Q9EQF2 | KEL | Kell blood group, metallo-endopep | Plasma Membrane | peptidase | 4142412.75 | 945452.062 | 2329634 | | | | | | |
| Q9EQF2 | KEL | Kell blood group, metallo-endopep | Plasma Membrane | peptidase | 3920539.25 | 647018.062 | 494754.281 | Q9EQF2 | KEL | 46857357 | 44711953 | 4302410 | 108 |
| Q6PDI5 | KIAA0368 | KIAA0368 | Cytoplasm | other | 5369574 | 5643590.5 | 3392756.5 | | | | | | |
| Q6PDI5 | KIAA0368 | KIAA0368 | Cytoplasm | other | 2508246.25 | 5831961 | 2919094.5 | | | | | | |
| Q6PDI5 | KIAA0368 | KIAA0368 | Cytoplasm | other | 4200515 | 4284279 | 2212069 | | | | | | |
| Q6PDI5 | KIAA0368 | KIAA0368 | Cytoplasm | other | 2262887.5 | 2646883 | 2089405.5 | | | | | | |
| Q6PDI5 | KIAA0368 | KIAA0368 | Cytoplasm | other | 1451643.5 | 2349508 | 1024597.938 | | | | | | |
| Q6PDI5 | KIAA0368 | KIAA0368 | Cytoplasm | other | 1073350 | 1534537.5 | 1185727.5 | | | | | | |
| Q6PDI5 | KIAA0368 | KIAA0368 | Cytoplasm | other | 1220312 | 2054648.125 | 1266729.375 | | | | | | |
| Q6PDI5 | KIAA0368 | KIAA0368 | Cytoplasm | other | 521391.562 | 905211.812 | 443148.125 | | | | | | |
| Q6PDI5 | KIAA0368 | KIAA0368 | Cytoplasm | other | 629481.438 | 594804 | 327753.844 | Q6PDI5 | KIAA0368 | 2137489 | 1621583 | 312074 | 27 |
| Q7TQ95 | KIAA1715 | KIAA1715 | Cytoplasm | other | 2736201.25 | 5242061 | 3381471.25 | | | | | | |
| Q7TQ95 | KIAA1715 | KIAA1715 | Cytoplasm | other | 2538626 | 5001952 | 4148206.5 | | | | | | |
| Q7TQ95 | KIAA1715 | KIAA1715 | Cytoplasm | other | 2401283.75 | 3967856 | 2821906.25 | | | | | | |
| Q7TQ95 | KIAA1715 | KIAA1715 | Cytoplasm | other | 2497670.5 | 2618175 | 2340797.5 | | | | | | |
| Q7TQ95 | KIAA1715 | KIAA1715 | Cytoplasm | other | 1856539.125 | 2314950.5 | 1809993.25 | Q7TQ95 | KIAA1715 | 2406064 | 1072477 | 276912 | 15 |
| Q60575 | KIF1B | kinesin family member 1B | Cytoplasm | transporter | 912794.062 | 1898198.625 | 6491471 | Q60575 | KIF1B | 912794 | 2977438 | 1719024 | 3 |
| O88448 | KLC2 | kinesin light chain 2 | Cytoplasm | other | 2564679.75 | 2702397.5 | 2337230 | O88448 | KLC2 | 2564680 | 184412 | 106470 | 3 |
| Q60682 | Klra4 (includes others) | killer cell lectin-like receptor, subfa | Plasma Membrane | transmembrane receptor | 5048062.5 | 6782827.5 | 7176049.5 | Q60682 | Klra4 (includes others) | 5048063 | 1132281 | 653723 | 3 |
| O35344 | KPNA3 | karyopherin alpha 3 (importin alph | Nucleus | transporter | 2544795.5 | 2505181.25 | 1810937.125 | O35344 | KPNA3 | 2544796 | 412733 | 238292 | 3 |
| P70168 | KPNB1 | karyopherin (importin) beta 1 | Nucleus | transporter | 8925267 | 9423450 | 8364246.5 | | | | | | |
| P70168 | KPNB1 | karyopherin (importin) beta 1 | Nucleus | transporter | 1.18E+07 | 1.18E+07 | 9594327 | | | | | | |
| P70168 | KPNB1 | karyopherin (importin) beta 1 | Nucleus | transporter | 1.64E+07 | 1.35E+07 | 1.34E+07 | | | | | | |
| P70168 | KPNB1 | karyopherin (importin) beta 1 | Nucleus | transporter | 1.01E+07 | 6300111 | 5349181.5 | | | | | | |
| P70168 | KPNB1 | karyopherin (importin) beta 1 | Nucleus | transporter | 1.02E+07 | 8169200.5 | 6949777 | | | | | | |
| P70168 | KPNB1 | karyopherin (importin) beta 1 | Nucleus | transporter | 7786477.5 | 5847097 | 4324993 | | | | | | |
| P70168 | KPNB1 | karyopherin (importin) beta 1 | Nucleus | transporter | 5572021.5 | 7957882.5 | 6730157 | | | | | | |
| P70168 | KPNB1 | karyopherin (importin) beta 1 | Nucleus | transporter | 5547967 | 5609998.5 | 3729328.5 | | | | | | |
| P70168 | KPNB1 | karyopherin (importin) beta 1 | Nucleus | transporter | 8047589 | 6811369 | 5336385 | | | | | | |
| P70168 | KPNB1 | karyopherin (importin) beta 1 | Nucleus | transporter | 7950343.5 | 5968818 | 5971267.5 | | | | | | |
| P70168 | KPNB1 | karyopherin (importin) beta 1 | Nucleus | transporter | 5636481 | 5866998.5 | 4739678 | | | | | | |
| P70168 | KPNB1 | karyopherin (importin) beta 1 | Nucleus | transporter | 4774885 | 4671104.5 | 4421146 | | | | | | |
| P70168 | KPNB1 | karyopherin (importin) beta 1 | Nucleus | transporter | 3399079 | 5641477 | 4994280 | | | | | | |
| P70168 | KPNB1 | karyopherin (importin) beta 1 | Nucleus | transporter | 3420628 | 5429231 | 3617231.75 | | | | | | |
| P70168 | KPNB1 | karyopherin (importin) beta 1 | Nucleus | transporter | 5532979 | 1063247.625 | 2215361 | | | | | | |
| P70168 | KPNB1 | karyopherin (importin) beta 1 | Nucleus | transporter | 4001383.25 | 4321298.5 | 3085646.25 | | | | | | |
| P70168 | KPNB1 | karyopherin (importin) beta 1 | Nucleus | transporter | 4002466.25 | 3573825.25 | 3277477.25 | | | | | | |
| P70168 | KPNB1 | karyopherin (importin) beta 1 | Nucleus | transporter | 2809956 | 2913912.75 | 1818800.375 | | | | | | |
| P70168 | KPNB1 | karyopherin (importin) beta 1 | Nucleus | transporter | 4132700.5 | 4207687.5 | 4075080 | | | | | | |
| P70168 | KPNB1 | karyopherin (importin) beta 1 | Nucleus | transporter | 2673352.25 | 2448450.75 | 1459042.375 | | | | | | |
| P70168 | KPNB1 | karyopherin (importin) beta 1 | Nucleus | transporter | 2923008.25 | 2658459 | 2398387 | P70168 | KPNB1 | 6456028 | 3145278 | 396268 | 63 |
| P04104 | KRT1 | keratin 1, type II | Cytoplasm | other | 1.86E+07 | 2.93E+07 | 5.43E+07 | | | | | | |

| | | | | | | | | | | | | | |
|---|---|---|---|---|---|---|---|---|---|---|---|---|---|
| P04104 | KRT1 | keratin 1, type II | Cytoplasm | other | 728554.375 | 5793029.5 | 8734984 | | | | | | |
| P04104 | KRT1 | keratin 1, type II | Cytoplasm | other | 5445471 | 7190517.5 | 1.62E+07 | | | | | | |
| P04104 | KRT1 | keratin 1, type II | Cytoplasm | other | 3074894 | 4260847 | 9554317 | P04104 | KRT1 | 6962230 | 15097299 | 4358215 | 12 |
| P02535 | Krt10 | keratin 10 | Cytoplasm | other | 8510603 | 7786043 | 1.00E+07 | P02535 | Krt10 | 8510603 | 1134252 | 654861 | 3 |
| Q61781 | KRT14 | keratin 14, type I | Cytoplasm | other | 1181707.125 | 2675373 | 6412235.5 | Q61781 | KRT14 | 1181707 | 2694241 | 1555521 | 3 |
| Q3TTY5 | Krt2 | keratin 2 | Cytoplasm | other | 8006854.5 | 2966207 | 9919837 | Q3TTY5 | Krt2 | 8006855 | 3592135 | 2073920 | 3 |
| Q922U2 | KRT5 | keratin 5, type II | Cytoplasm | other | 1.84E+07 | 3.36E+07 | 6.28E+07 | | | | | | |
| Q922U2 | KRT5 | keratin 5, type II | Cytoplasm | other | 4629462.5 | 6903183 | 6356000.5 | | | | | | |
| Q922U2 | KRT5 | keratin 5, type II | Cytoplasm | other | 1036743 | 1603281.375 | 2548817.5 | | | | | | |
| Q922U2 | KRT5 | keratin 5, type II | Cytoplasm | other | 2449629 | 6826824.5 | 1.78E+07 | Q922U2 | KRT5 | 6626459 | 18165304 | 5243872 | 12 |
| Q6NXH9 | KRT73 | keratin 73, type II | Extracellular Space | other | 2295151.5 | 1423709.25 | 2009131.5 | Q6NXH9 | KRT73 | 2295152 | 444211 | 256465 | 3 |
| Q3UV17 | KRT76 | keratin 76, type II | Cytoplasm | other | 1.50E+07 | 2.32E+07 | 4.19E+07 | Q3UV17 | KRT76 | 15040000 | 13790737 | 7962086 | 3 |
| Q8VED5 | KRT79 | keratin 79, type II | Extracellular Space | other | 4602531.5 | 9970124 | 1.49E+07 | Q8VED5 | KRT79 | 4602532 | 5165075 | 2982058 | 3 |
| Q6IMF0 | Krt83 | keratin 83 | Extracellular Space | other | 9697870 | 1.56E+07 | 1.47E+07 | Q6IMF0 | Krt83 | 9697870 | 3174076 | 1832554 | 3 |
| P11438 | LAMP1 | lysosomal-associated membrane p | Plasma Membrane | other | 1.29E+07 | 1.28E+07 | 1.46E+07 | | | | | | |
| P11438 | LAMP1 | lysosomal-associated membrane p | Plasma Membrane | other | 9377829 | 8905004 | 8455812 | | | | | | |
| P11438 | LAMP1 | lysosomal-associated membrane p | Plasma Membrane | other | 2925377 | 2866953.75 | 2874710.25 | P11438 | LAMP1 | 8401069 | 4622800 | 1540933 | 9 |
| P17047 | LAMP2 | lysosomal-associated membrane p | Plasma Membrane | enzyme | 6342400 | 7841323 | 1.02E+07 | | | | | | |
| P17047 | LAMP2 | lysosomal-associated membrane p | Plasma Membrane | enzyme | 3671101.25 | 4220821 | 4949607.5 | P17047 | LAMP2 | 5006751 | 2470232 | 1008468 | 6 |
| Q9CQ22 | LAMTOR1 | late endosomal/lysosomal adapto | Plasma Membrane | other | 2.37E+07 | 2.40E+07 | 2.13E+07 | | | | | | |
| Q9CQ22 | LAMTOR1 | late endosomal/lysosomal adapto | Plasma Membrane | other | 8386723.5 | 7741674.5 | 6036022 | | | | | | |
| Q9CQ22 | LAMTOR1 | late endosomal/lysosomal adapto | Plasma Membrane | other | 6535530 | 7946768 | 6608594 | | | | | | |
| Q9CQ22 | LAMTOR1 | late endosomal/lysosomal adapto | Plasma Membrane | other | 3679884 | 3668323.75 | 3057155.25 | Q9CQ22 | LAMTOR1 | 10575534 | 7919101 | 2286048 | 12 |
| O88653 | LAMTOR3 | late endosomal/lysosomal adapto | Cytoplasm | other | 1.29E+07 | 1.72E+07 | 1.18E+07 | | | | | | |
| O88653 | LAMTOR3 | late endosomal/lysosomal adapto | Cytoplasm | other | 4882059.5 | 6845018 | 5180250.5 | | | | | | |
| O88653 | LAMTOR3 | late endosomal/lysosomal adapto | Cytoplasm | other | 3770611.25 | 5880766.5 | 3371452.5 | O88653 | LAMTOR3 | 7197557 | 4818844 | 1606281 | 9 |
| Q9D1L9 | LAMTOR5 | late endosomal/lysosomal adapto | Cytoplasm | other | 9692616 | 1.23E+07 | 8945184 | | | | | | |
| Q9D1L9 | LAMTOR5 | late endosomal/lysosomal adapto | Cytoplasm | other | 4134335.75 | 5885013 | 5200453.5 | | | | | | |
| Q9D1L9 | LAMTOR5 | late endosomal/lysosomal adapto | Cytoplasm | other | 3849297 | 3866871 | 3569827.5 | Q9D1L9 | LAMTOR5 | 5892083 | 3157894 | 1052631 | 9 |
| O89112 | LANCL1 | LanC lantibiotic synthetase compo | Plasma Membrane | other | 1643422.25 | 3431020.5 | 2500488.75 | O89112 | LANCL1 | 1643422 | 894051 | 516180 | 3 |
| Q9JJK2 | LANCL2 | LanC lantibiotic synthetase compo | Plasma Membrane | other | 7485180.5 | 8916319 | 8902757 | | | | | | |
| Q9JJK2 | LANCL2 | LanC lantibiotic synthetase compo | Plasma Membrane | other | 7639340 | 9645720 | 8666649 | | | | | | |
| Q9JJK2 | LANCL2 | LanC lantibiotic synthetase compo | Plasma Membrane | other | 7622882.5 | 7859366 | 8830077 | | | | | | |
| Q9JJK2 | LANCL2 | LanC lantibiotic synthetase compo | Plasma Membrane | other | 4828839.5 | 5687031.5 | 5677968 | | | | | | |
| Q9JJK2 | LANCL2 | LanC lantibiotic synthetase compo | Plasma Membrane | other | 6420868 | 6221776.5 | 6163858 | | | | | | |
| Q9JJK2 | LANCL2 | LanC lantibiotic synthetase compo | Plasma Membrane | other | 3277388.25 | 6109625.5 | 4094788 | | | | | | |
| Q9JJK2 | LANCL2 | LanC lantibiotic synthetase compo | Plasma Membrane | other | 3559154.25 | 4468664 | 3595945 | | | | | | |
| Q9JJK2 | LANCL2 | LanC lantibiotic synthetase compo | Plasma Membrane | other | 2659601.5 | 3662475.75 | 3134860.25 | | | | | | |
| Q9JJK2 | LANCL2 | LanC lantibiotic synthetase compo | Plasma Membrane | other | 3755158.5 | 3454804 | 2915042.5 | | | | | | |
| Q9JJK2 | LANCL2 | LanC lantibiotic synthetase compo | Plasma Membrane | other | 2059397.75 | 2914313.5 | 2306916.5 | | | | | | |
| Q9JJK2 | LANCL2 | LanC lantibiotic synthetase compo | Plasma Membrane | other | 1707243.75 | 2294268.25 | 1765982.75 | | | | | | |
| Q9JJK2 | LANCL2 | LanC lantibiotic synthetase compo | Plasma Membrane | other | 4679329.5 | 494387.375 | 338759.406 | Q9JJK2 | LANCL2 | 4641199 | 2577974 | 429662 | 36 |
| P06151 | LDHA | lactate dehydrogenase A | Cytoplasm | enzyme | 9009249 | 7232073.5 | 6691969 | | | | | | |
| P06151 | LDHA | lactate dehydrogenase A | Cytoplasm | enzyme | 7389710.5 | 6292556.5 | 6399678 | | | | | | |
| P06151 | LDHA | lactate dehydrogenase A | Cytoplasm | enzyme | 5726461 | 5803865.5 | 5608251 | | | | | | |
| P06151 | LDHA | lactate dehydrogenase A | Cytoplasm | enzyme | 4290281.5 | 4388661 | 4455493 | | | | | | |
| P06151 | LDHA | lactate dehydrogenase A | Cytoplasm | enzyme | 5572537.5 | 5069426 | 5800704 | | | | | | |
| P06151 | LDHA | lactate dehydrogenase A | Cytoplasm | enzyme | 3088030.25 | 4003516.75 | 2737011.25 | | | | | | |
| P06151 | LDHA | lactate dehydrogenase A | Cytoplasm | enzyme | 3155836.5 | 1983064.75 | 2184980.5 | | | | | | |
| P06151 | LDHA | lactate dehydrogenase A | Cytoplasm | enzyme | 2803245.25 | 1507493.75 | 1676670.25 | | | | | | |
| P06151 | LDHA | lactate dehydrogenase A | Cytoplasm | enzyme | 1739234.75 | 1282715.625 | 2003486 | P06151 | LDHA | 4752732 | 2128659 | 409661 | 27 |
| Q8CBY3 | LENG8 | leukocyte receptor cluster (LRC) m | Nucleus | other | 5963671.5 | 7931490.5 | 5358648 | Q8CBY3 | LENG8 | 5963672 | 1345231 | 776670 | 3 |
| O89013 | LEPROT | leptin receptor overlapping transc | Plasma Membrane | other | 3163981 | 3431416.5 | 2896319 | | | | | | |
| O89013 | LEPROT | leptin receptor overlapping transc | Plasma Membrane | other | 224755.25 | 254874.5 | 183906.016 | O89013 | LEPROT | 1694368 | 1620812 | 661694 | 6 |
| Q9Z2I0 | LETM1 | leucine zipper-EF-hand containing | Cytoplasm | other | 2091175.625 | 1876937.875 | 3103020 | Q9Z2I0 | LETM1 | 2091176 | 654854 | 378080 | 3 |
| O08573 | LGALS9B | lectin, galactoside-binding, soluble | Cytoplasm | other | 2.72E+07 | 2.31E+07 | 2.88E+07 | | | | | | |
| O08573 | LGALS9B | lectin, galactoside-binding, soluble | Cytoplasm | other | 2.12E+07 | 1.44E+07 | 2.33E+07 | | | | | | |
| O08573 | LGALS9B | lectin, galactoside-binding, soluble | Cytoplasm | other | 2.08E+07 | 1.67E+07 | 2.30E+07 | | | | | | |
| O08573 | LGALS9B | lectin, galactoside-binding, soluble | Cytoplasm | other | 2.11E+07 | 1.74E+07 | 2.62E+07 | | | | | | |
| O08573 | LGALS9B | lectin, galactoside-binding, soluble | Cytoplasm | other | 1.05E+07 | 9375106 | 1.74E+07 | | | | | | |
| O08573 | LGALS9B | lectin, galactoside-binding, soluble | Cytoplasm | other | 6925709 | 6061538 | 9658257 | | | | | | |
| O08573 | LGALS9B | lectin, galactoside-binding, soluble | Cytoplasm | other | 7134386 | 5916255.5 | 1.05E+07 | | | | | | |
| O08573 | LGALS9B | lectin, galactoside-binding, soluble | Cytoplasm | other | 4243555 | 3703261 | 5673581.5 | O08573 | LGALS9B | 14882956 | 8101619 | 1653736 | 24 |
| A2ARI4 | LGR4 | leucine-rich repeat containing G p | Plasma Membrane | transmembrane receptor | 2202717.25 | 5385931 | 5063711 | A2ARI4 | LGR4 | 2202717 | 1752235 | 1011653 | 3 |
| Q99JW4 | LIMS1 | LIM and senescent cell antigen-like | Plasma Membrane | other | 1642042.125 | 1779912.5 | 1281245.5 | Q99JW4 | LIMS1 | 1642042 | 257504 | 148670 | 3 |
| Q9D0F3 | LMAN1 | lectin, mannose-binding, 1 | Cytoplasm | other | 1.62E+07 | 1.81E+07 | 1.42E+07 | | | | | | |
| Q9D0F3 | LMAN1 | lectin, mannose-binding, 1 | Cytoplasm | other | 9381315 | 1.03E+07 | 9884229 | | | | | | |
| Q9D0F3 | LMAN1 | lectin, mannose-binding, 1 | Cytoplasm | other | 1.15E+07 | 1.44E+07 | 1.26E+07 | | | | | | |
| Q9D0F3 | LMAN1 | lectin, mannose-binding, 1 | Cytoplasm | other | 1.32E+07 | 1.49E+07 | 9378116 | | | | | | |
| Q9D0F3 | LMAN1 | lectin, mannose-binding, 1 | Cytoplasm | other | 1.03E+07 | 1.14E+07 | 8732609 | | | | | | |
| Q9D0F3 | LMAN1 | lectin, mannose-binding, 1 | Cytoplasm | other | 6079566.5 | 6006574.5 | 2684540 | Q9D0F3 | LMAN1 | 11091814 | 3848110 | 907008 | 18 |
| Q9DBH5 | LMAN2 | lectin, mannose-binding 2 | Cytoplasm | transporter | 1.85E+07 | 1.59E+07 | 1.32E+07 | | | | | | |
| Q9DBH5 | LMAN2 | lectin, mannose-binding 2 | Cytoplasm | transporter | 2.16E+07 | 2.32E+07 | 1.83E+07 | | | | | | |
| Q9DBH5 | LMAN2 | lectin, mannose-binding 2 | Cytoplasm | transporter | 7534399.5 | 5432143.5 | 5695087.5 | | | | | | |
| Q9DBH5 | LMAN2 | lectin, mannose-binding 2 | Cytoplasm | transporter | 4062370.25 | 5472289.5 | 4552289.5 | | | | | | |
| Q9DBH5 | LMAN2 | lectin, mannose-binding 2 | Cytoplasm | transporter | 4429013.5 | 5077106.5 | 4645017.5 | Q9DBH5 | LMAN2 | 11239157 | 7099754 | 1833149 | 15 |
| Q8VEE1 | LMCD1 | LIM and cysteine-rich domains 1 | Cytoplasm | transcription regulator | 1061291.125 | 1.40E+07 | 1.51E+07 | Q8VEE1 | LMCD1 | 1061291 | 7818626 | 4514086 | 3 |
| Q8CGK3 | LONP1 | lon peptidase 1, mitochondrial | Cytoplasm | peptidase | 5868709 | 4848781 | 7360516.5 | | | | | | |
| Q8CGK3 | LONP1 | lon peptidase 1, mitochondrial | Cytoplasm | peptidase | 4497764.5 | 1187963.125 | 2223371.75 | Q8CGK3 | LONP1 | 5183237 | 2287322 | 933795 | 6 |
| Q3TFD2 | LPCAT1 | lysophosphatidylcholine acyltransf | Cytoplasm | enzyme | 1.93E+07 | 2.29E+07 | 2.31E+07 | | | | | | |
| Q3TFD2 | LPCAT1 | lysophosphatidylcholine acyltransf | Cytoplasm | enzyme | 8554295 | 9886663 | 8471776 | | | | | | |
| Q3TFD2 | LPCAT1 | lysophosphatidylcholine acyltransf | Cytoplasm | enzyme | 5959752.5 | 9401975 | 8633717 | | | | | | |
| Q3TFD2 | LPCAT1 | lysophosphatidylcholine acyltransf | Cytoplasm | enzyme | 5790161.5 | 7535722.5 | 6213438.5 | | | | | | |
| Q3TFD2 | LPCAT1 | lysophosphatidylcholine acyltransf | Cytoplasm | enzyme | 7342189.5 | 9302874 | 6477676.5 | | | | | | |
| Q3TFD2 | LPCAT1 | lysophosphatidylcholine acyltransf | Cytoplasm | enzyme | 6390339.5 | 2736266.5 | 3305530.5 | | | | | | |
| Q3TFD2 | LPCAT1 | lysophosphatidylcholine acyltransf | Cytoplasm | enzyme | 6851192.5 | 8073701 | 8183174.5 | | | | | | |

| | | | | | | | | | | | | | |
|---|---|---|---|---|---|---|---|---|---|---|---|---|---|
| Q3TFD2 | LPCAT1 | lysophosphatidylcholine acyltransf | Cytoplasm | enzyme | 2817473.75 | 4659177.5 | 3553338 | Q3TFD2 | LPCAT1 | 7874426 | 5533398 | 1129500 | 24 |
| Q91V01 | LPCAT3 | lysophosphatidylcholine acyltransf | Plasma Membrane | enzyme | 2.44E+07 | 2.69E+07 | 2.16E+07 | | | | | | |
| Q91V01 | LPCAT3 | lysophosphatidylcholine acyltransf | Plasma Membrane | enzyme | 1.02E+07 | 1.21E+07 | 1.01E+07 | | | | | | |
| Q91V01 | LPCAT3 | lysophosphatidylcholine acyltransf | Plasma Membrane | enzyme | 7096677.5 | 9915513 | 8900116 | | | | | | |
| Q91V01 | LPCAT3 | lysophosphatidylcholine acyltransf | Plasma Membrane | enzyme | 1710440.5 | 2938819 | 2067479.875 | Q91V01 | LPCAT3 | 10846780 | 8515698 | 2458270 | 12 |
| P55302 | LRPAP1 | low density lipoprotein receptor-r | Plasma Membrane | transmembrane receptor | 1043917.75 | 2521343.75 | 1363890.125 | P55302 | LRPAP1 | 1043918 | 777268 | 448756 | 3 |
| Q9D1G5 | LRRC57 | leucine rich repeat containing 57 | Cytoplasm | other | 8193463 | 6768324 | 5276380 | | | | | | |
| Q9D1G5 | LRRC57 | leucine rich repeat containing 57 | Cytoplasm | other | 7924278.5 | 6477896 | 5859708.5 | | | | | | |
| Q9D1G5 | LRRC57 | leucine rich repeat containing 57 | Cytoplasm | other | 7297156 | 6791524.5 | 5766736 | | | | | | |
| Q9D1G5 | LRRC57 | leucine rich repeat containing 57 | Cytoplasm | other | 7226304 | 5326139 | 4093734.75 | | | | | | |
| Q9D1G5 | LRRC57 | leucine rich repeat containing 57 | Cytoplasm | other | 5431073 | 5291693 | 3391331.75 | | | | | | |
| Q9D1G5 | LRRC57 | leucine rich repeat containing 57 | Cytoplasm | other | 3289186.5 | 2521898.75 | 1570399.125 | Q9D1G5 | LRRC57 | 6560244 | 1871021 | 441004 | 18 |
| Q922Q8 | LRRC59 | leucine rich repeat containing 59 | Cytoplasm | other | 3773615.75 | 5418593 | 4838914.5 | Q922Q8 | LRRC59 | 3773616 | 834350 | 481712 | 3 |
| Q80TE7 | LRRC7 | leucine rich repeat containing 7 | Plasma Membrane | other | 3003918.75 | 2445968.75 | 2602715 | Q80TE7 | LRRC7 | 3003919 | 287762 | 166140 | 3 |
| P08071 | LTF | lactotransferrin | Extracellular Space | other | 941995.625 | 478881.031 | 1965274.25 | | | | | | |
| P08071 | LTF | lactotransferrin | Extracellular Space | other | 616683.5 | 238704.547 | 1241443.5 | | | | | | |
| P08071 | LTF | lactotransferrin | Extracellular Space | other | 1005929.125 | 456228.156 | 1831395.5 | | | | | | |
| P08071 | LTF | lactotransferrin | Extracellular Space | other | 610742.562 | 160087.125 | 801214.812 | | | | | | |
| P08071 | LTF | lactotransferrin | Extracellular Space | other | 1371890.5 | 406033.688 | 2238528.75 | P08071 | LTF | 909448 | 647693 | 167234 | 15 |
| P25911 | LYN | LYN proto-oncogene, Src family ty | Cytoplasm | kinase | 1.04E+07 | 1.68E+07 | 1.43E+07 | | | | | | |
| P25911 | LYN | LYN proto-oncogene, Src family ty | Cytoplasm | kinase | 8230163 | 8876759 | 8432579 | | | | | | |
| P25911 | LYN | LYN proto-oncogene, Src family ty | Cytoplasm | kinase | 8933160 | 1.14E+07 | 1.22E+07 | | | | | | |
| P25911 | LYN | LYN proto-oncogene, Src family ty | Cytoplasm | kinase | 8510766 | 7415217 | 6282882.5 | | | | | | |
| P25911 | LYN | LYN proto-oncogene, Src family ty | Cytoplasm | kinase | 3380017.25 | 3447566.5 | 3684968.5 | | | | | | |
| P25911 | LYN | LYN proto-oncogene, Src family ty | Cytoplasm | kinase | 2058268.125 | 3401465 | 2913506.75 | | | | | | |
| P25911 | LYN | LYN proto-oncogene, Src family ty | Cytoplasm | kinase | 2377713 | 3158020 | 3419746.5 | | | | | | |
| P25911 | LYN | LYN proto-oncogene, Src family ty | Cytoplasm | kinase | 1216934.875 | 1145463.375 | 740916.75 | P25911 | LYN | 5632128 | 4457022 | 909786 | 24 |
| P08905 | LYZ | lysozyme | Extracellular Space | enzyme | 1.46E+07 | 7035654.5 | 1.80E+07 | | | | | | |
| P08905 | LYZ | lysozyme | Extracellular Space | enzyme | 3073346.25 | 5086126 | 3646810 | P08905 | LYZ | 8831673 | 6218601 | 2538733 | 6 |
| P24668 | M6PR | mannose-6-phosphate receptor (c | Cytoplasm | transporter | 4545170.5 | 5544477 | 5441084.5 | | | | | | |
| P24668 | M6PR | mannose-6-phosphate receptor (c | Cytoplasm | transporter | 976703.938 | 1209125.75 | 1253245.875 | P24668 | M6PR | 2760937 | 2236791 | 913166 | 6 |
| Q9CXI5 | MANF | mesencephalic astrocyte-derived r | Extracellular Space | other | 1.68E+07 | 2.15E+07 | 2.20E+07 | | | | | | |
| Q9CXI5 | MANF | mesencephalic astrocyte-derived r | Extracellular Space | other | 4354286.5 | 5705748.5 | 4865383.5 | | | | | | |
| Q9CXI5 | MANF | mesencephalic astrocyte-derived r | Extracellular Space | other | 1607872.75 | 3688181 | 2406843 | Q9CXI5 | MANF | 7597386 | 8394587 | 2798196 | 9 |
| Q61166 | MAPRE1 | microtubule-associated protein, R | Cytoplasm | other | 735372.5 | 630731.625 | 1050573.75 | Q61166 | MAPRE1 | 735373 | 218544 | 126176 | 3 |
| Q922Q1 | MARC_2 | mitochondrial amidoxime reducing | Cytoplasm | enzyme | 994026.312 | 1424500.125 | 1335464.25 | Q922Q1 | MARC_2 | 994026 | 227235 | 131194 | 3 |
| Q9DBD2 | MARH8 | membrane-associated ring finger ( | Cytoplasm | enzyme | 3715649.5 | 5521990 | 3971199.75 | | | | | | |
| Q9DBD2 | MARH8 | membrane-associated ring finger ( | Cytoplasm | enzyme | 2823939.75 | 4597027 | 3492806.5 | Q9DBD2 | MARH8 | 3269795 | 937277 | 382642 | 6 |
| Q05512 | MARK2 | MAP/microtubule affinity-regulati | Cytoplasm | kinase | 2615019.25 | 3461860 | 2023495.75 | | | | | | |
| Q05512 | MARK2 | MAP/microtubule affinity-regulati | Cytoplasm | kinase | 1271435.75 | 1217969.625 | 457986.781 | Q05512 | MARK2 | 1943228 | 1084886 | 442903 | 6 |
| Q8VCF0 | MAVS | mitochondrial antiviral signaling pi | Cytoplasm | other | 5891561.5 | 7108618 | 8467756 | | | | | | |
| Q8VCF0 | MAVS | mitochondrial antiviral signaling pi | Cytoplasm | other | 3907423.25 | 3303639.25 | 4105966.25 | Q8VCF0 | MAVS | 4899492 | 2041787 | 833556 | 6 |
| Q8BL86 | MBLAC2 | metallo-beta-lactamase domain cc | Cytoplasm | other | 1.38E+07 | 1.25E+07 | 1.21E+07 | Q8BL86 | MBLAC2 | 13760000 | 877097 | 506392 | 3 |
| Q8CHK3 | MBOAT7 | membrane bound O-acyltransfera | Plasma Membrane | other | 4253361.5 | 6698066 | 7020343 | Q8CHK3 | MBOAT7 | 4253362 | 1513089 | 873582 | 3 |
| Q8K5B2 | MCFD2 | multiple coagulation factor deficie | Cytoplasm | other | 9107340 | 1.18E+07 | 1.08E+07 | Q8K5B2 | MCFD2 | 9107340 | 1354902 | 782253 | 3 |
| Q3UMR5 | MCU | mitochondrial calcium uniporter | Cytoplasm | ion channel | 1706888.25 | 1915372.75 | 2477308.75 | Q3UMR5 | MCU | 1706888 | 398494 | 230071 | 3 |
| P08249 | MDH2 | malate dehydrogenase 2, NAD (mi | Cytoplasm | enzyme | 5.27E+07 | 5.05E+07 | 7.34E+07 | | | | | | |
| P08249 | MDH2 | malate dehydrogenase 2, NAD (mi | Cytoplasm | enzyme | 4.16E+07 | 4.46E+07 | 5.01E+07 | | | | | | |
| P08249 | MDH2 | malate dehydrogenase 2, NAD (mi | Cytoplasm | enzyme | 4.10E+07 | 4.44E+07 | 5.51E+07 | | | | | | |
| P08249 | MDH2 | malate dehydrogenase 2, NAD (mi | Cytoplasm | enzyme | 4.18E+07 | 3.82E+07 | 4.70E+07 | | | | | | |
| P08249 | MDH2 | malate dehydrogenase 2, NAD (mi | Cytoplasm | enzyme | 3.94E+07 | 3.82E+07 | 4.68E+07 | | | | | | |
| P08249 | MDH2 | malate dehydrogenase 2, NAD (mi | Cytoplasm | enzyme | 3.61E+07 | 3.88E+07 | 3.57E+07 | | | | | | |
| P08249 | MDH2 | malate dehydrogenase 2, NAD (mi | Cytoplasm | enzyme | 3.35E+07 | 3.01E+07 | 4.27E+07 | | | | | | |
| P08249 | MDH2 | malate dehydrogenase 2, NAD (mi | Cytoplasm | enzyme | 3.62E+07 | 3.00E+07 | 4.24E+07 | | | | | | |
| P08249 | MDH2 | malate dehydrogenase 2, NAD (mi | Cytoplasm | enzyme | 2.42E+07 | 2.88E+07 | 3.31E+07 | | | | | | |
| P08249 | MDH2 | malate dehydrogenase 2, NAD (mi | Cytoplasm | enzyme | 1.61E+07 | 1.33E+07 | 1.70E+07 | | | | | | |
| P08249 | MDH2 | malate dehydrogenase 2, NAD (mi | Cytoplasm | enzyme | 2.01E+07 | 1.79E+07 | 1.93E+07 | | | | | | |
| P08249 | MDH2 | malate dehydrogenase 2, NAD (mi | Cytoplasm | enzyme | 1.01E+07 | 1.10E+07 | 1.37E+07 | | | | | | |
| P08249 | MDH2 | malate dehydrogenase 2, NAD (mi | Cytoplasm | enzyme | 3378704.25 | 3642226.75 | 6197413.5 | P08249 | MDH2 | 30467593 | 16023768 | 2565856 | 39 |
| Q99KE1 | ME2 | malic enzyme 2, NAD(+)-depender | Cytoplasm | enzyme | 4279390 | 5969635.5 | 6039562 | | | | | | |
| Q99KE1 | ME2 | malic enzyme 2, NAD(+)-depender | Cytoplasm | enzyme | 7200939 | 8675442 | 1.08E+07 | | | | | | |
| Q99KE1 | ME2 | malic enzyme 2, NAD(+)-depender | Cytoplasm | enzyme | 1995517.125 | 2064055.625 | 2538166.75 | | | | | | |
| Q99KE1 | ME2 | malic enzyme 2, NAD(+)-depender | Cytoplasm | enzyme | 3116802.75 | 2453854 | 3525685 | Q99KE1 | ME2 | 4148162 | 2870683 | 828695 | 12 |
| Q91VH6 | MEMO1 | mediator of cell motility 1 | Cytoplasm | other | 1.08E+07 | 1.51E+07 | 8776444 | | | | | | |
| Q91VH6 | MEMO1 | mediator of cell motility 1 | Cytoplasm | other | 3884284.25 | 6217201.5 | 3459440.5 | | | | | | |
| Q91VH6 | MEMO1 | mediator of cell motility 1 | Cytoplasm | other | 3102986 | 7160750 | 5443639.5 | | | | | | |
| Q91VH6 | MEMO1 | mediator of cell motility 1 | Cytoplasm | other | 3062804.75 | 4837252 | 3696053.75 | | | | | | |
| Q91VH6 | MEMO1 | mediator of cell motility 1 | Cytoplasm | other | 1402457.25 | 4553869 | 3251691.25 | Q91VH6 | MEMO1 | 4452506 | 3571139 | 922064 | 15 |
| Q9ERE7 | MESDC2 | mesoderm development candidat | Extracellular Space | other | 1702291.125 | 2128421.75 | 1136357 | Q9ERE7 | MESDC2 | 1702291 | 497671 | 287331 | 3 |
| O08663 | METAP2 | methionyl aminopeptidase 2 | Cytoplasm | peptidase | 3353268.5 | 3178404.5 | 2585673.25 | O08663 | METAP2 | 3353269 | 402308 | 232273 | 3 |
| Q6PCP5 | Mff | mitochondrial fission factor | Cytoplasm | other | 6330668 | 8002119.5 | 1.17E+07 | | | | | | |
| Q6PCP5 | Mff | mitochondrial fission factor | Cytoplasm | other | 6076220.5 | 5054427.5 | 7227414.5 | Q6PCP5 | Mff | 6203444 | 2339487 | 955092 | 6 |
| Q3T9M1 | MFSD2B | major facilitator superfamily doma | Plasma Membrane | other | 2.94E+07 | 3.32E+07 | 3.48E+07 | | | | | | |
| Q3T9M1 | MFSD2B | major facilitator superfamily doma | Plasma Membrane | other | 2072056.25 | 2681949.75 | 2233270.25 | Q3T9M1 | MFSD2B | 15746028 | 16598770 | 6776419 | 6 |
| Q9CPU4 | MGST3 | microsomal glutathione S-transfer | Cytoplasm | enzyme | 2.69E+07 | 4.81E+07 | 6.66E+07 | | | | | | |
| Q9CPU4 | MGST3 | microsomal glutathione S-transfer | Cytoplasm | enzyme | 7435247.5 | 1.37E+07 | 1.36E+07 | | | | | | |
| Q9CPU4 | MGST3 | microsomal glutathione S-transfer | Cytoplasm | enzyme | 4541155.5 | 7161126 | 9240939 | Q9CPU4 | MGST3 | 12965468 | 21598335 | 7199445 | 9 |
| Q8VDP3 | MICAL1 | microtubule associated monooxyg | Cytoplasm | enzyme | 3003252 | 1650966.125 | 2198075.25 | Q8VDP3 | MICAL1 | 3003252 | 680235 | 392734 | 3 |
| Q6ZQI3 | MLEC | malectin | Plasma Membrane | other | 5964870 | 7342410 | 5869219 | | | | | | |
| Q6ZQI3 | MLEC | malectin | Plasma Membrane | other | 5983715.5 | 8293618 | 8981732 | | | | | | |
| Q6ZQI3 | MLEC | malectin | Plasma Membrane | other | 6938891 | 6187059 | 5516781 | | | | | | |
| Q6ZQI3 | MLEC | malectin | Plasma Membrane | other | 3362631.25 | 4674127.5 | 4790802.5 | | | | | | |
| Q6ZQI3 | MLEC | malectin | Plasma Membrane | other | 3181173 | 4052975.25 | 3852597.75 | Q6ZQI3 | MLEC | 5086256 | 1731521 | 447077 | 15 |
| Q8K3F2 | MMP21 | matrix metallopeptidase 21 | Extracellular Space | peptidase | 3936884 | 1811750.125 | 3471080 | Q8K3F2 | MMP21 | 3936884 | 1117031 | 644918 | 3 |
| B2RPV6 | MMRN1 | multimerin 1 | Extracellular Space | other | 5478235 | 4335868.5 | 5238590 | | | | | | |
| B2RPV6 | MMRN1 | multimerin 1 | Extracellular Space | other | 4035276.25 | 3786604.25 | 4114473 | | | | | | |

| | | | | | | | | | | | | | |
|---|---|---|---|---|---|---|---|---|---|---|---|---|---|
| B2RPV6 | MMRN1 | multimerin 1 | Extracellular Space | other | 4861288 | 5974365 | 6025244 | | | | | | |
| B2RPV6 | MMRN1 | multimerin 1 | Extracellular Space | other | 3459970.5 | 4135547.25 | 4565969 | | | | | | |
| B2RPV6 | MMRN1 | multimerin 1 | Extracellular Space | other | 2408550 | 2568145 | 4048670.25 | | | | | | |
| B2RPV6 | MMRN1 | multimerin 1 | Extracellular Space | other | 4151779 | 5909674 | 5952543 | | | | | | |
| B2RPV6 | MMRN1 | multimerin 1 | Extracellular Space | other | 5319389 | 5475974.5 | 5687669 | | | | | | |
| B2RPV6 | MMRN1 | multimerin 1 | Extracellular Space | other | 4197007 | 4120644.75 | 4059570 | | | | | | |
| B2RPV6 | MMRN1 | multimerin 1 | Extracellular Space | other | 3395523.25 | 2856234.5 | 3693592 | | | | | | |
| B2RPV6 | MMRN1 | multimerin 1 | Extracellular Space | other | 2572519.25 | 2836371.75 | 2830070.5 | | | | | | |
| B2RPV6 | MMRN1 | multimerin 1 | Extracellular Space | other | 2651540 | 2174927.25 | 4423630 | B2RPV6 | MMRN1 | 3866462 | 1161177 | 202135 | 33 |
| P70290 | MPP1 | membrane protein, palmitoylated | Plasma Membrane | kinase | 1.33E+09 | 1.49E+09 | 1.55E+09 | | | | | | |
| P70290 | MPP1 | membrane protein, palmitoylated | Plasma Membrane | kinase | 1.43E+09 | 1.46E+09 | 1.42E+09 | | | | | | |
| P70290 | MPP1 | membrane protein, palmitoylated | Plasma Membrane | kinase | 1.07E+09 | 1.00E+09 | 9.49E+08 | | | | | | |
| P70290 | MPP1 | membrane protein, palmitoylated | Plasma Membrane | kinase | 8.72E+08 | 9.27E+08 | 1.00E+09 | | | | | | |
| P70290 | MPP1 | membrane protein, palmitoylated | Plasma Membrane | kinase | 1.12E+09 | 1.03E+09 | 1.15E+09 | | | | | | |
| P70290 | MPP1 | membrane protein, palmitoylated | Plasma Membrane | kinase | 9.33E+08 | 1.02E+09 | 1.02E+09 | | | | | | |
| P70290 | MPP1 | membrane protein, palmitoylated | Plasma Membrane | kinase | 7.22E+08 | 6.06E+08 | 6.78E+08 | | | | | | |
| P70290 | MPP1 | membrane protein, palmitoylated | Plasma Membrane | kinase | 5.75E+08 | 6.18E+08 | 5.36E+08 | | | | | | |
| P70290 | MPP1 | membrane protein, palmitoylated | Plasma Membrane | kinase | 6.53E+08 | 5.50E+08 | 3.71E+08 | | | | | | |
| P70290 | MPP1 | membrane protein, palmitoylated | Plasma Membrane | kinase | 5.22E+08 | 6.97E+08 | 4.93E+08 | | | | | | |
| P70290 | MPP1 | membrane protein, palmitoylated | Plasma Membrane | kinase | 3.35E+08 | 5.66E+08 | 6.15E+08 | | | | | | |
| P70290 | MPP1 | membrane protein, palmitoylated | Plasma Membrane | kinase | 6.51E+08 | 4.92E+08 | 4.80E+08 | | | | | | |
| P70290 | MPP1 | membrane protein, palmitoylated | Plasma Membrane | kinase | 5.22E+08 | 4.83E+08 | 4.32E+08 | | | | | | |
| P70290 | MPP1 | membrane protein, palmitoylated | Plasma Membrane | kinase | 5.40E+08 | 4.58E+08 | 5.56E+08 | | | | | | |
| P70290 | MPP1 | membrane protein, palmitoylated | Plasma Membrane | kinase | 3.92E+08 | 3.73E+08 | 3.54E+08 | | | | | | |
| P70290 | MPP1 | membrane protein, palmitoylated | Plasma Membrane | kinase | 3.24E+08 | 4.43E+08 | 3.66E+08 | | | | | | |
| P70290 | MPP1 | membrane protein, palmitoylated | Plasma Membrane | kinase | 3.45E+08 | 2.82E+08 | 2.69E+08 | | | | | | |
| P70290 | MPP1 | membrane protein, palmitoylated | Plasma Membrane | kinase | 4.50E+08 | 1.60E+08 | 1.37E+08 | | | | | | |
| P70290 | MPP1 | membrane protein, palmitoylated | Plasma Membrane | kinase | 2.98E+08 | 2.62E+08 | 2.86E+08 | | | | | | |
| P70290 | MPP1 | membrane protein, palmitoylated | Plasma Membrane | kinase | 2.51E+08 | 2.58E+08 | 2.47E+08 | | | | | | |
| P70290 | MPP1 | membrane protein, palmitoylated | Plasma Membrane | kinase | 1.54E+08 | 1.73E+08 | 1.41E+08 | | | | | | |
| P70290 | MPP1 | membrane protein, palmitoylated | Plasma Membrane | kinase | 1.55E+08 | 1.53E+08 | 1.41E+08 | | | | | | |
| P70290 | MPP1 | membrane protein, palmitoylated | Plasma Membrane | kinase | 1.02E+08 | 1.13E+08 | 9.11E+07 | | | | | | |
| P70290 | MPP1 | membrane protein, palmitoylated | Plasma Membrane | kinase | 1.23E+08 | 4.17E+07 | 3.28E+07 | | | | | | |
| P70290 | MPP1 | membrane protein, palmitoylated | Plasma Membrane | kinase | 1.76E+08 | 8072326 | 5.97E+07 | | | | | | |
| P70290 | MPP1 | membrane protein, palmitoylated | Plasma Membrane | kinase | 7.83E+07 | 8.55E+07 | 9.24E+07 | | | | | | |
| P70290 | MPP1 | membrane protein, palmitoylated | Plasma Membrane | kinase | 5.25E+07 | 5.24E+07 | 5.07E+07 | | | | | | |
| P70290 | MPP1 | membrane protein, palmitoylated | Plasma Membrane | kinase | 8.17E+07 | 2.43E+07 | 1.79E+07 | | | | | | |
| P70290 | MPP1 | membrane protein, palmitoylated | Plasma Membrane | kinase | 3.39E+07 | 4.22E+07 | 3.84E+07 | | | | | | |
| P70290 | MPP1 | membrane protein, palmitoylated | Plasma Membrane | kinase | 3.92E+07 | 4.81E+07 | 3.26E+07 | | | | | | |
| P70290 | MPP1 | membrane protein, palmitoylated | Plasma Membrane | kinase | 6.03E+07 | 4.89E+07 | 5.32E+07 | | | | | | |
| P70290 | MPP1 | membrane protein, palmitoylated | Plasma Membrane | kinase | 3.00E+07 | 3.13E+07 | 4.26E+07 | | | | | | |
| P70290 | MPP1 | membrane protein, palmitoylated | Plasma Membrane | kinase | 2.52E+07 | 3.37E+07 | 3.64E+07 | | | | | | |
| P70290 | MPP1 | membrane protein, palmitoylated | Plasma Membrane | kinase | 1.07E+07 | 1.39E+08 | 1.02E+08 | | | | | | |
| P70290 | MPP1 | membrane protein, palmitoylated | Plasma Membrane | kinase | 3.45E+07 | 1312491.75 | 9812919 | | | | | | |
| P70290 | MPP1 | membrane protein, palmitoylated | Plasma Membrane | kinase | 2.02E+07 | 2082198.875 | 2805036 | | | | | | |
| P70290 | MPP1 | membrane protein, palmitoylated | Plasma Membrane | kinase | 1.13E+07 | 9621256 | 1.11E+07 | | | | | | |
| P70290 | MPP1 | membrane protein, palmitoylated | Plasma Membrane | kinase | 2.71E+07 | 3421160.25 | 4014337.25 | | | | | | |
| P70290 | MPP1 | membrane protein, palmitoylated | Plasma Membrane | kinase | 2.16E+07 | 3167574.25 | 3498014.75 | | | | | | |
| P70290 | MPP1 | membrane protein, palmitoylated | Plasma Membrane | kinase | 2624611.25 | 3302285.25 | 3075280.25 | | | | | | |
| P70290 | MPP1 | membrane protein, palmitoylated | Plasma Membrane | kinase | 2790944.75 | 3572012.25 | 2747178 | | | | | | |
| P70290 | MPP1 | membrane protein, palmitoylated | Plasma Membrane | kinase | 278077.25 | 590828.812 | 562646.062 | P70290 | MPP1 | 347045563 | 399603092 | 35599473 | 126 |
| Q99J99 | MPST | mercaptopyruvate sulfurtransfera | Cytoplasm | enzyme | 1.24E+07 | 1.27E+07 | 1.77E+07 | | | | | | |
| Q99J99 | MPST | mercaptopyruvate sulfurtransfera | Cytoplasm | enzyme | 963457.25 | 986521.188 | 1439812.625 | Q99J99 | MPST | 6696729 | 7439289 | 3037077 | 6 |
| Q9DB15 | MRPL12 | mitochondrial ribosomal protein L | Cytoplasm | other | 2017793.5 | 2099498.5 | 2682623.25 | Q9DB15 | MRPL12 | 2017794 | 362562 | 209325 | 3 |
| Q9CQX8 | MRPS36 | mitochondrial ribosomal protein S | Cytoplasm | other | 1660903.5 | 2067050.75 | 2295475 | Q9CQX8 | MRPS36 | 1660904 | 321407 | 185564 | 3 |
| Q9WUX5 | MRVI1 | murine retrovirus integration site | Cytoplasm | other | 3211010.25 | 3018594.5 | 3483419.25 | | | | | | |
| Q9WUX5 | MRVI1 | murine retrovirus integration site | Cytoplasm | other | 1742890.375 | 2343308 | 2175135 | | | | | | |
| Q9WUX5 | MRVI1 | murine retrovirus integration site | Cytoplasm | other | 955513.688 | 985262.938 | 1019843.938 | Q9WUX5 | MRVI1 | 1969805 | 993948 | 331316 | 9 |
| P26041 | MSN | moesin | Plasma Membrane | other | 9034239 | 8613240 | 9261789 | | | | | | |
| P26041 | MSN | moesin | Plasma Membrane | other | 8835756 | 7821986 | 7659442.5 | | | | | | |
| P26041 | MSN | moesin | Plasma Membrane | other | 8408042 | 6888678.5 | 5362988.5 | | | | | | |
| P26041 | MSN | moesin | Plasma Membrane | other | 7918757.5 | 6945912 | 6435210.5 | | | | | | |
| P26041 | MSN | moesin | Plasma Membrane | other | 5792463 | 3857903.75 | 4075190.75 | | | | | | |
| P26041 | MSN | moesin | Plasma Membrane | other | 3420419 | 1500252 | 2077998 | | | | | | |
| P26041 | MSN | moesin | Plasma Membrane | other | 4113120.25 | 4993846.5 | 3625899.5 | | | | | | |
| P26041 | MSN | moesin | Plasma Membrane | other | 3990649.25 | 2316621.5 | 2348151 | | | | | | |
| P26041 | MSN | moesin | Plasma Membrane | other | 2208722 | 2589449.75 | 2581692.75 | | | | | | |
| P26041 | MSN | moesin | Plasma Membrane | other | 1284993 | 2381340.75 | 1077799.75 | | | | | | |
| P26041 | MSN | moesin | Plasma Membrane | other | 1130772.125 | 1233655.625 | 621598.875 | | | | | | |
| P26041 | MSN | moesin | Plasma Membrane | other | 2900947.5 | 4547229 | 2304030.25 | P26041 | MSN | 4919907 | 2702816 | 450469 | 36 |
| P03930 | mt-Atp8 | ATP synthase F0 subunit 8 | Cytoplasm | enzyme | 1.73E+07 | 1.51E+07 | 1.96E+07 | | | | | | |
| P03930 | mt-Atp8 | ATP synthase F0 subunit 8 | Cytoplasm | enzyme | 4580932.5 | 5889836 | 6781058 | P03930 | mt-Atp8 | 10950466 | 6536081 | 2668344 | 6 |
| Q791V5 | MTCH2 | mitochondrial carrier 2 | Cytoplasm | other | 7871702.5 | 7254315.5 | 6966696 | Q791V5 | MTCH2 | 7871703 | 462408 | 266972 | 3 |
| P00397 | MT-CO1 | cytochrome c oxidase subunit I | Cytoplasm | enzyme | 3193346.5 | 2753959 | 2672826.5 | P00397 | MT-CO1 | 3193347 | 280055 | 161690 | 3 |
| P00405 | MT-CO2 | cytochrome c oxidase subunit II | Cytoplasm | enzyme | 2.99E+07 | 3.17E+07 | 4.24E+07 | | | | | | |
| P00405 | MT-CO2 | cytochrome c oxidase subunit II | Cytoplasm | enzyme | 1.90E+07 | 2.19E+07 | 2.67E+07 | | | | | | |
| P00405 | MT-CO2 | cytochrome c oxidase subunit II | Cytoplasm | enzyme | 4732546.5 | 6097724.5 | 7824393.5 | | | | | | |
| P00405 | MT-CO2 | cytochrome c oxidase subunit II | Cytoplasm | enzyme | 3173276.5 | 1953703.75 | 3650614.75 | | | | | | |
| P00405 | MT-CO2 | cytochrome c oxidase subunit II | Cytoplasm | enzyme | 1631807.25 | 3070067.75 | 4197325.5 | | | | | | |
| P00405 | MT-CO2 | cytochrome c oxidase subunit II | Cytoplasm | enzyme | 6715385 | 203069.422 | 2435487.75 | P00405 | MT-CO2 | 10852169 | 12950325 | 3052421 | 18 |
| P00416 | MT-CO3 | cytochrome c oxidase III | Cytoplasm | enzyme | 1409032.875 | 2470053.5 | 2671319.75 | P00416 | MT-CO3 | 1409033 | 678189 | 391553 | 3 |
| Q922D8 | MTHFD1 | methylenetetrahydrofolate dehyd | Cytoplasm | enzyme | 2.22E+07 | 3.07E+07 | 1.64E+07 | | | | | | |
| Q922D8 | MTHFD1 | methylenetetrahydrofolate dehyd | Cytoplasm | enzyme | 1.79E+07 | 2.28E+07 | 1.14E+07 | | | | | | |
| Q922D8 | MTHFD1 | methylenetetrahydrofolate dehyd | Cytoplasm | enzyme | 1.93E+07 | 3.49E+07 | 1.53E+07 | | | | | | |
| Q922D8 | MTHFD1 | methylenetetrahydrofolate dehyd | Cytoplasm | enzyme | 1.95E+07 | 2.25E+07 | 1.16E+07 | | | | | | |
| Q922D8 | MTHFD1 | methylenetetrahydrofolate dehyd | Cytoplasm | enzyme | 1.77E+07 | 2.38E+07 | 1.04E+07 | | | | | | |

| | | | | | | | | | | | | | |
|---|---|---|---|---|---|---|---|---|---|---|---|---|---|
| Q922D8 | MTHFD1 | methylenetetrahydrofolate dehyd | Cytoplasm | enzyme | 1.48E+07 | 2.26E+07 | 1.08E+07 | | | | | | |
| Q922D8 | MTHFD1 | methylenetetrahydrofolate dehyd | Cytoplasm | enzyme | 1.42E+07 | 2.27E+07 | 1.23E+07 | | | | | | |
| Q922D8 | MTHFD1 | methylenetetrahydrofolate dehyd | Cytoplasm | enzyme | 1.63E+07 | 1.96E+07 | 9429781 | | | | | | |
| Q922D8 | MTHFD1 | methylenetetrahydrofolate dehyd | Cytoplasm | enzyme | 1.48E+07 | 8964333 | 2505419 | | | | | | |
| Q922D8 | MTHFD1 | methylenetetrahydrofolate dehyd | Cytoplasm | enzyme | 1.29E+07 | 1.84E+07 | 8382116 | | | | | | |
| Q922D8 | MTHFD1 | methylenetetrahydrofolate dehyd | Cytoplasm | enzyme | 1.15E+07 | 1.58E+07 | 7024738 | | | | | | |
| Q922D8 | MTHFD1 | methylenetetrahydrofolate dehyd | Cytoplasm | enzyme | 7172408.5 | 1.50E+07 | 7674767 | | | | | | |
| Q922D8 | MTHFD1 | methylenetetrahydrofolate dehyd | Cytoplasm | enzyme | 1.16E+07 | 1.79E+07 | 7809429.5 | | | | | | |
| Q922D8 | MTHFD1 | methylenetetrahydrofolate dehyd | Cytoplasm | enzyme | 1.38E+07 | 1.62E+07 | 6595199 | | | | | | |
| Q922D8 | MTHFD1 | methylenetetrahydrofolate dehyd | Cytoplasm | enzyme | 8146497.5 | 1.18E+07 | 3903360.5 | | | | | | |
| Q922D8 | MTHFD1 | methylenetetrahydrofolate dehyd | Cytoplasm | enzyme | 1.01E+07 | 1.22E+07 | 6679667 | | | | | | |
| Q922D8 | MTHFD1 | methylenetetrahydrofolate dehyd | Cytoplasm | enzyme | 7391656.5 | 1.08E+07 | 6184518 | | | | | | |
| Q922D8 | MTHFD1 | methylenetetrahydrofolate dehyd | Cytoplasm | enzyme | 7967336.5 | 7997288.5 | 5379120 | | | | | | |
| Q922D8 | MTHFD1 | methylenetetrahydrofolate dehyd | Cytoplasm | enzyme | 5412025 | 1.29E+07 | 7588559.5 | | | | | | |
| Q922D8 | MTHFD1 | methylenetetrahydrofolate dehyd | Cytoplasm | enzyme | 9781110 | 1.01E+07 | 7274553.5 | | | | | | |
| Q922D8 | MTHFD1 | methylenetetrahydrofolate dehyd | Cytoplasm | enzyme | 6936766 | 8990095 | 4557623.5 | | | | | | |
| Q922D8 | MTHFD1 | methylenetetrahydrofolate dehyd | Cytoplasm | enzyme | 6302026 | 8566370 | 3969073.75 | | | | | | |
| Q922D8 | MTHFD1 | methylenetetrahydrofolate dehyd | Cytoplasm | enzyme | 7751278.5 | 3900493 | 2017172.375 | | | | | | |
| Q922D8 | MTHFD1 | methylenetetrahydrofolate dehyd | Cytoplasm | enzyme | 2426291.25 | 7766750 | 3512289.5 | | | | | | |
| Q922D8 | MTHFD1 | methylenetetrahydrofolate dehyd | Cytoplasm | enzyme | 7751249 | 8808928 | 4211606.5 | | | | | | |
| Q922D8 | MTHFD1 | methylenetetrahydrofolate dehyd | Cytoplasm | enzyme | 4852231.5 | 5614215.5 | 3082778.5 | | | | | | |
| Q922D8 | MTHFD1 | methylenetetrahydrofolate dehyd | Cytoplasm | enzyme | 3184372.25 | 4448204.5 | 2001668.875 | | | | | | |
| Q922D8 | MTHFD1 | methylenetetrahydrofolate dehyd | Cytoplasm | enzyme | 2692846 | 6217186.5 | 2549387.25 | | | | | | |
| Q922D8 | MTHFD1 | methylenetetrahydrofolate dehyd | Cytoplasm | enzyme | 1934642.75 | 3797870.25 | 798617.812 | Q922D8 | MTHFD1 | 10559405 | 6794261 | 728421 | 87 |
| Q3V3R1 | MTHFD1L | methylenetetrahydrofolate dehyd | Cytoplasm | enzyme | 6429950.5 | 6120408 | 7171272.5 | | | | | | |
| Q3V3R1 | MTHFD1L | methylenetetrahydrofolate dehyd | Cytoplasm | enzyme | 3827541 | 5064627 | 6088384.5 | | | | | | |
| Q3V3R1 | MTHFD1L | methylenetetrahydrofolate dehyd | Cytoplasm | enzyme | 3391571.25 | 3307016.75 | 4332793 | | | | | | |
| Q3V3R1 | MTHFD1L | methylenetetrahydrofolate dehyd | Cytoplasm | enzyme | 3265965.5 | 4212629.5 | 4141404.75 | | | | | | |
| Q3V3R1 | MTHFD1L | methylenetetrahydrofolate dehyd | Cytoplasm | enzyme | 2315732.25 | 2879898 | 3811322.5 | | | | | | |
| Q3V3R1 | MTHFD1L | methylenetetrahydrofolate dehyd | Cytoplasm | enzyme | 3092840.5 | 2691277.75 | 3579201.75 | | | | | | |
| Q3V3R1 | MTHFD1L | methylenetetrahydrofolate dehyd | Cytoplasm | enzyme | 2892232.25 | 1877484.375 | 3230732.25 | | | | | | |
| Q3V3R1 | MTHFD1L | methylenetetrahydrofolate dehyd | Cytoplasm | enzyme | 1693986.125 | 2191757.25 | 2355066 | Q3V3R1 | MTHFD1L | 3363727 | 1481582 | 302427 | 24 |
| P03921 | MT-ND5 | NADH dehydrogenase, subunit 5 ( | Cytoplasm | enzyme | 4049607.5 | 3794866.25 | 6671727.5 | | | | | | |
| P03921 | MT-ND5 | NADH dehydrogenase, subunit 5 ( | Cytoplasm | enzyme | 1890953.125 | 1961267.625 | 2909122 | | | | | | |
| P03921 | MT-ND5 | NADH dehydrogenase, subunit 5 ( | Cytoplasm | enzyme | 1294767 | 1194789.5 | 1929345.375 | P03921 | MT-ND5 | 2411776 | 1755576 | 585192 | 9 |
| Q8C1A3 | MTRR | 5-methyltetrahydrofolate-homocy | Cytoplasm | enzyme | 6141403 | 5255807 | 4686732.5 | Q8C1A3 | MTRR | 6141403 | 733052 | 423228 | 3 |
| P28665 | Mug1/Mug2 | murinoglobulin 1 | Extracellular Space | other | 66672.594 | 4.70E-06 | 128878.688 | P28665 | Mug1/Mug2 | 66673 | 64452 | 37212 | 3 |
| P48972 | MYBL2 | v-myb avian myeloblastosis viral o | Nucleus | transcription regulator | 9290160 | 8010369 | 1.18E+07 | P48972 | MYBL2 | 9290160 | 1900593 | 1097308 | 3 |
| Q3U898 | MYEOV2 | myeloma overexpressed 2 | Nucleus | other | 4020938.25 | 6556566 | 6458684 | Q3U898 | MYEOV2 | 4020938 | 1436523 | 829377 | 3 |
| Q6URW6 | MYH14 | myosin, heavy chain 14, non-musc | Extracellular Space | other | 7970271.5 | 4181157 | 5835465 | | | | | | |
| Q6URW6 | MYH14 | myosin, heavy chain 14, non-musc | Extracellular Space | other | 2413293.5 | 1119490.375 | 2987699.75 | Q6URW6 | MYH14 | 5191783 | 2488438 | 1015901 | 6 |
| Q8VDD5 | MYH9 | myosin, heavy chain 9, non-muscl | Cytoplasm | enzyme | 2.76E+07 | 3.81E+07 | 2.97E+07 | | | | | | |
| Q8VDD5 | MYH9 | myosin, heavy chain 9, non-muscl | Cytoplasm | enzyme | 6.40E+07 | 7.28E+07 | 6.51E+07 | | | | | | |
| Q8VDD5 | MYH9 | myosin, heavy chain 9, non-muscl | Cytoplasm | enzyme | 5.95E+07 | 5.57E+07 | 4.93E+07 | | | | | | |
| Q8VDD5 | MYH9 | myosin, heavy chain 9, non-muscl | Cytoplasm | enzyme | 3.41E+07 | 4.57E+07 | 3.72E+07 | | | | | | |
| Q8VDD5 | MYH9 | myosin, heavy chain 9, non-muscl | Cytoplasm | enzyme | 6.15E+07 | 7.20E+07 | 5.21E+07 | | | | | | |
| Q8VDD5 | MYH9 | myosin, heavy chain 9, non-muscl | Cytoplasm | enzyme | 7.05E+07 | 7.36E+07 | 5.02E+07 | | | | | | |
| Q8VDD5 | MYH9 | myosin, heavy chain 9, non-muscl | Cytoplasm | enzyme | 3.80E+07 | 3.83E+07 | 3.10E+07 | | | | | | |
| Q8VDD5 | MYH9 | myosin, heavy chain 9, non-muscl | Cytoplasm | enzyme | 5.56E+07 | 5.03E+07 | 3.82E+07 | | | | | | |
| Q8VDD5 | MYH9 | myosin, heavy chain 9, non-muscl | Cytoplasm | enzyme | 5.37E+07 | 5.10E+07 | 4.94E+07 | | | | | | |
| Q8VDD5 | MYH9 | myosin, heavy chain 9, non-muscl | Cytoplasm | enzyme | 3.46E+07 | 4.16E+07 | 4.03E+07 | | | | | | |
| Q8VDD5 | MYH9 | myosin, heavy chain 9, non-muscl | Cytoplasm | enzyme | 5.13E+07 | 3.18E+07 | 3.95E+07 | | | | | | |
| Q8VDD5 | MYH9 | myosin, heavy chain 9, non-muscl | Cytoplasm | enzyme | 4.51E+07 | 4.57E+07 | 3.77E+07 | | | | | | |
| Q8VDD5 | MYH9 | myosin, heavy chain 9, non-muscl | Cytoplasm | enzyme | 5.09E+07 | 4.93E+07 | 3.88E+07 | | | | | | |
| Q8VDD5 | MYH9 | myosin, heavy chain 9, non-muscl | Cytoplasm | enzyme | 5.41E+07 | 4.94E+07 | 4.21E+07 | | | | | | |
| Q8VDD5 | MYH9 | myosin, heavy chain 9, non-muscl | Cytoplasm | enzyme | 4.96E+07 | 4.35E+07 | 4.21E+07 | | | | | | |
| Q8VDD5 | MYH9 | myosin, heavy chain 9, non-muscl | Cytoplasm | enzyme | 4.90E+07 | 3.87E+07 | 3.63E+07 | | | | | | |
| Q8VDD5 | MYH9 | myosin, heavy chain 9, non-muscl | Cytoplasm | enzyme | 3.99E+07 | 4.15E+07 | 3.64E+07 | | | | | | |
| Q8VDD5 | MYH9 | myosin, heavy chain 9, non-muscl | Cytoplasm | enzyme | 4.23E+07 | 3.97E+07 | 3.76E+07 | | | | | | |
| Q8VDD5 | MYH9 | myosin, heavy chain 9, non-muscl | Cytoplasm | enzyme | 3.04E+07 | 2.48E+07 | 1.73E+07 | | | | | | |
| Q8VDD5 | MYH9 | myosin, heavy chain 9, non-muscl | Cytoplasm | enzyme | 3.60E+07 | 4.43E+07 | 3.84E+07 | | | | | | |
| Q8VDD5 | MYH9 | myosin, heavy chain 9, non-muscl | Cytoplasm | enzyme | 4.76E+07 | 4.29E+07 | 4.65E+07 | | | | | | |
| Q8VDD5 | MYH9 | myosin, heavy chain 9, non-muscl | Cytoplasm | enzyme | 3.06E+07 | 3.46E+07 | 2.48E+07 | | | | | | |
| Q8VDD5 | MYH9 | myosin, heavy chain 9, non-muscl | Cytoplasm | enzyme | 3.34E+07 | 3.98E+07 | 3.25E+07 | | | | | | |
| Q8VDD5 | MYH9 | myosin, heavy chain 9, non-muscl | Cytoplasm | enzyme | 2.85E+07 | 5.74E+07 | 3.55E+07 | | | | | | |
| Q8VDD5 | MYH9 | myosin, heavy chain 9, non-muscl | Cytoplasm | enzyme | 3.19E+07 | 2.59E+07 | 2.57E+07 | | | | | | |
| Q8VDD5 | MYH9 | myosin, heavy chain 9, non-muscl | Cytoplasm | enzyme | 2.95E+07 | 2.78E+07 | 2.09E+07 | | | | | | |
| Q8VDD5 | MYH9 | myosin, heavy chain 9, non-muscl | Cytoplasm | enzyme | 3.21E+07 | 3.28E+07 | 3.36E+07 | | | | | | |
| Q8VDD5 | MYH9 | myosin, heavy chain 9, non-muscl | Cytoplasm | enzyme | 3.79E+07 | 2.95E+07 | 3.82E+07 | | | | | | |
| Q8VDD5 | MYH9 | myosin, heavy chain 9, non-muscl | Cytoplasm | enzyme | 2.89E+07 | 4.97E+07 | 3.39E+07 | | | | | | |
| Q8VDD5 | MYH9 | myosin, heavy chain 9, non-muscl | Cytoplasm | enzyme | 3.65E+07 | 3.22E+07 | 2.87E+07 | | | | | | |
| Q8VDD5 | MYH9 | myosin, heavy chain 9, non-muscl | Cytoplasm | enzyme | 3.72E+07 | 3.02E+07 | 2.54E+07 | | | | | | |
| Q8VDD5 | MYH9 | myosin, heavy chain 9, non-muscl | Cytoplasm | enzyme | 3.06E+07 | 5.81E+07 | 3.79E+07 | | | | | | |
| Q8VDD5 | MYH9 | myosin, heavy chain 9, non-muscl | Cytoplasm | enzyme | 3.94E+07 | 3.18E+07 | 2.31E+07 | | | | | | |
| Q8VDD5 | MYH9 | myosin, heavy chain 9, non-muscl | Cytoplasm | enzyme | 2.52E+07 | 4.11E+07 | 3.18E+07 | | | | | | |
| Q8VDD5 | MYH9 | myosin, heavy chain 9, non-muscl | Cytoplasm | enzyme | 3.01E+07 | 2.76E+07 | 2.07E+07 | | | | | | |
| Q8VDD5 | MYH9 | myosin, heavy chain 9, non-muscl | Cytoplasm | enzyme | 3.12E+07 | 2.42E+07 | 2.54E+07 | | | | | | |
| Q8VDD5 | MYH9 | myosin, heavy chain 9, non-muscl | Cytoplasm | enzyme | 3.00E+07 | 2.85E+07 | 3.21E+07 | | | | | | |
| Q8VDD5 | MYH9 | myosin, heavy chain 9, non-muscl | Cytoplasm | enzyme | 2.99E+07 | 2.74E+07 | 2.16E+07 | | | | | | |
| Q8VDD5 | MYH9 | myosin, heavy chain 9, non-muscl | Cytoplasm | enzyme | 3.68E+07 | 2.73E+07 | 2.69E+07 | | | | | | |
| Q8VDD5 | MYH9 | myosin, heavy chain 9, non-muscl | Cytoplasm | enzyme | 2.25E+07 | 2.71E+07 | 2.16E+07 | | | | | | |
| Q8VDD5 | MYH9 | myosin, heavy chain 9, non-muscl | Cytoplasm | enzyme | 2.07E+07 | 2.93E+07 | 2.24E+07 | | | | | | |
| Q8VDD5 | MYH9 | myosin, heavy chain 9, non-muscl | Cytoplasm | enzyme | 3.10E+07 | 3.57E+07 | 3.13E+07 | | | | | | |
| Q8VDD5 | MYH9 | myosin, heavy chain 9, non-muscl | Cytoplasm | enzyme | 2.71E+07 | 2.84E+07 | 2.00E+07 | | | | | | |
| Q8VDD5 | MYH9 | myosin, heavy chain 9, non-muscl | Cytoplasm | enzyme | 2.18E+07 | 3.11E+07 | 2.47E+07 | | | | | | |
| Q8VDD5 | MYH9 | myosin, heavy chain 9, non-muscl | Cytoplasm | enzyme | 2.93E+07 | 2.53E+07 | 2.08E+07 | | | | | | |

| | | | | | | | | | | | | | | | |
|---|---|---|---|---|---|---|---|---|---|---|---|---|---|---|---|
| Q8VDD5 | MYH9 | myosin, heavy chain 9, non-muscl | Cytoplasm | enzyme | | 2.76E+07 | 2.07E+07 | 1.86E+07 | | | | | | | |
| Q8VDD5 | MYH9 | myosin, heavy chain 9, non-muscl | Cytoplasm | enzyme | | 1.93E+07 | 2.55E+07 | 1.86E+07 | | | | | | | |
| Q8VDD5 | MYH9 | myosin, heavy chain 9, non-muscl | Cytoplasm | enzyme | | 1.46E+07 | 1.54E+07 | 2.45E+07 | | | | | | | |
| Q8VDD5 | MYH9 | myosin, heavy chain 9, non-muscl | Cytoplasm | enzyme | | 2.20E+07 | 2.50E+07 | 1.96E+07 | | | | | | | |
| Q8VDD5 | MYH9 | myosin, heavy chain 9, non-muscl | Cytoplasm | enzyme | | 1.25E+07 | 1.31E+07 | 9765139 | | | | | | | |
| Q8VDD5 | MYH9 | myosin, heavy chain 9, non-muscl | Cytoplasm | enzyme | | 1.86E+07 | 2.96E+07 | 2.02E+07 | | | | | | | |
| Q8VDD5 | MYH9 | myosin, heavy chain 9, non-muscl | Cytoplasm | enzyme | | 2.02E+07 | 1.65E+07 | 1.48E+07 | | | | | | | |
| Q8VDD5 | MYH9 | myosin, heavy chain 9, non-muscl | Cytoplasm | enzyme | | 1.88E+07 | 1.60E+07 | 1.56E+07 | | | | | | | |
| Q8VDD5 | MYH9 | myosin, heavy chain 9, non-muscl | Cytoplasm | enzyme | | 1.88E+07 | 1.88E+07 | 1.73E+07 | | | | | | | |
| Q8VDD5 | MYH9 | myosin, heavy chain 9, non-muscl | Cytoplasm | enzyme | | 1.78E+07 | 1.71E+07 | 1.49E+07 | | | | | | | |
| Q8VDD5 | MYH9 | myosin, heavy chain 9, non-muscl | Cytoplasm | enzyme | | 2.12E+07 | 1.60E+07 | 1.56E+07 | | | | | | | |
| Q8VDD5 | MYH9 | myosin, heavy chain 9, non-muscl | Cytoplasm | enzyme | | 1.97E+07 | 1.53E+07 | 1.18E+07 | | | | | | | |
| Q8VDD5 | MYH9 | myosin, heavy chain 9, non-muscl | Cytoplasm | enzyme | | 1.73E+07 | 2.17E+07 | 2.13E+07 | | | | | | | |
| Q8VDD5 | MYH9 | myosin, heavy chain 9, non-muscl | Cytoplasm | enzyme | | 1.40E+07 | 1.76E+07 | 1.48E+07 | | | | | | | |
| Q8VDD5 | MYH9 | myosin, heavy chain 9, non-muscl | Cytoplasm | enzyme | | 1.59E+07 | 1.85E+07 | 1.56E+07 | | | | | | | |
| Q8VDD5 | MYH9 | myosin, heavy chain 9, non-muscl | Cytoplasm | enzyme | | 1.38E+07 | 1.56E+07 | 1.36E+07 | | | | | | | |
| Q8VDD5 | MYH9 | myosin, heavy chain 9, non-muscl | Cytoplasm | enzyme | | 7327521.5 | 1.73E+07 | 1.28E+07 | | | | | | | |
| Q8VDD5 | MYH9 | myosin, heavy chain 9, non-muscl | Cytoplasm | enzyme | | 1.26E+07 | 1.30E+07 | 6278636.5 | | | | | | | |
| Q8VDD5 | MYH9 | myosin, heavy chain 9, non-muscl | Cytoplasm | enzyme | | 8231788.5 | 1.52E+07 | 1.18E+07 | | | | | | | |
| Q8VDD5 | MYH9 | myosin, heavy chain 9, non-muscl | Cytoplasm | enzyme | | 1.86E+07 | 6304705.5 | 1.05E+07 | | | | | | | |
| Q8VDD5 | MYH9 | myosin, heavy chain 9, non-muscl | Cytoplasm | enzyme | | 8183245 | 7378963 | 7398652 | | | | | | | |
| Q8VDD5 | MYH9 | myosin, heavy chain 9, non-muscl | Cytoplasm | enzyme | | 1.11E+07 | 1.03E+07 | 7478595.5 | | | | | | | |
| Q8VDD5 | MYH9 | myosin, heavy chain 9, non-muscl | Cytoplasm | enzyme | | 2208347.5 | 2.36E+07 | 2.94E+07 | | | | | | | |
| Q8VDD5 | MYH9 | myosin, heavy chain 9, non-muscl | Cytoplasm | enzyme | | 9207714 | 1.16E+07 | 1.05E+07 | | | | | | | |
| Q8VDD5 | MYH9 | myosin, heavy chain 9, non-muscl | Cytoplasm | enzyme | | 9812334 | 1.05E+07 | 7191659.5 | | | | | | | |
| Q8VDD5 | MYH9 | myosin, heavy chain 9, non-muscl | Cytoplasm | enzyme | | 2.72E+07 | 2676994.5 | 9648104 | | | | | | | |
| Q8VDD5 | MYH9 | myosin, heavy chain 9, non-muscl | Cytoplasm | enzyme | | 1.84E+07 | 1.40E+07 | 1.29E+07 | | | | | | | |
| Q8VDD5 | MYH9 | myosin, heavy chain 9, non-muscl | Cytoplasm | enzyme | | 3112438 | 4019248.25 | 2731075.25 | | | | | | | |
| Q8VDD5 | MYH9 | myosin, heavy chain 9, non-muscl | Cytoplasm | enzyme | | 7049331.5 | 8732372 | 8515675 | | | | | | | |
| Q8VDD5 | MYH9 | myosin, heavy chain 9, non-muscl | Cytoplasm | enzyme | | 7129626.5 | 9656912 | 7595959 | | | | | | | |
| Q8VDD5 | MYH9 | myosin, heavy chain 9, non-muscl | Cytoplasm | enzyme | | 6326854.5 | 8094853 | 5547324 | | | | | | | |
| Q8VDD5 | MYH9 | myosin, heavy chain 9, non-muscl | Cytoplasm | enzyme | | 4000556.5 | 94775.758 | 175534.359 | | | | | | | |
| Q8VDD5 | MYH9 | myosin, heavy chain 9, non-muscl | Cytoplasm | enzyme | | 4446857 | 7763260.5 | 5930843.5 | | | | | | | |
| Q8VDD5 | MYH9 | myosin, heavy chain 9, non-muscl | Cytoplasm | enzyme | | 6626718.5 | 9527171 | 8898410 | | | | | | | |
| Q8VDD5 | MYH9 | myosin, heavy chain 9, non-muscl | Cytoplasm | enzyme | | 5689227 | 4328026 | 3738958.75 | | | | | | | |
| Q8VDD5 | MYH9 | myosin, heavy chain 9, non-muscl | Cytoplasm | enzyme | | 1.32E+07 | 1.06E+07 | 9254688 | | | | | | | |
| Q8VDD5 | MYH9 | myosin, heavy chain 9, non-muscl | Cytoplasm | enzyme | | 5622080 | 6595265.5 | 7521353.5 | | | | | | | |
| Q8VDD5 | MYH9 | myosin, heavy chain 9, non-muscl | Cytoplasm | enzyme | | 5599703.5 | 4575323 | 3269028.5 | | | | | | | |
| Q8VDD5 | MYH9 | myosin, heavy chain 9, non-muscl | Cytoplasm | enzyme | | 3352660.25 | 4014557.5 | 4348597 | | | | | | | |
| Q8VDD5 | MYH9 | myosin, heavy chain 9, non-muscl | Cytoplasm | enzyme | | 667487.062 | 9685231 | 1.09E+07 | | | | | | | |
| Q8VDD5 | MYH9 | myosin, heavy chain 9, non-muscl | Cytoplasm | enzyme | | 6476589 | 1167807.875 | 1972899.5 | | | | | | | |
| Q8VDD5 | MYH9 | myosin, heavy chain 9, non-muscl | Cytoplasm | enzyme | | 1.73E+07 | 1296078.5 | 6820054 | | | | | | | |
| Q8VDD5 | MYH9 | myosin, heavy chain 9, non-muscl | Cytoplasm | enzyme | | 1963494.5 | 5602017.5 | 4042810.5 | | | | | | | |
| Q8VDD5 | MYH9 | myosin, heavy chain 9, non-muscl | Cytoplasm | enzyme | | 1692185 | 2318117.25 | 1604262.125 | | | | | | | |
| Q8VDD5 | MYH9 | myosin, heavy chain 9, non-muscl | Cytoplasm | enzyme | | 3884868.25 | 2695579.25 | 2616264.5 | | | | | | | |
| Q8VDD5 | MYH9 | myosin, heavy chain 9, non-muscl | Cytoplasm | enzyme | | 4036734.25 | 4433310 | 3963346.5 | | | | | | | |
| Q8VDD5 | MYH9 | myosin, heavy chain 9, non-muscl | Cytoplasm | enzyme | | 1555861 | 6165852 | 3664622.75 | | | | | | | |
| Q8VDD5 | MYH9 | myosin, heavy chain 9, non-muscl | Cytoplasm | enzyme | | 436683.406 | 862731.875 | 303895.594 | | | | | | | |
| Q8VDD5 | MYH9 | myosin, heavy chain 9, non-muscl | Cytoplasm | enzyme | | 808535.938 | 1823546 | 897498.938 | | | | | | | |
| Q8VDD5 | MYH9 | myosin, heavy chain 9, non-muscl | Cytoplasm | enzyme | | 212778.719 | 2545760 | 2256717.5 | | | | | | | |
| Q8VDD5 | MYH9 | myosin, heavy chain 9, non-muscl | Cytoplasm | enzyme | | 23609.678 | 2153095.25 | 1571074.125 | Q8VDD5 | MYH9 | | 23575373 | 16610194 | 978765 | 288 |
| P05977 | MYL1 | myosin, light chain 1, alkali; skelet | Cytoplasm | other | | 7633071 | 5634120.5 | 6287818 | P05977 | MYL1 | | 7633071 | 1019218 | 588446 | 3 |
| Q3THE2 | MYL12A | myosin, light chain 12A, regulatory | Cytoplasm | other | | 1.94E+07 | 2.01E+07 | 1.95E+07 | | | | | | | |
| Q3THE2 | MYL12A | myosin, light chain 12A, regulatory | Cytoplasm | other | | 3.18E+07 | 2.82E+07 | 3.06E+07 | | | | | | | |
| Q3THE2 | MYL12A | myosin, light chain 12A, regulatory | Cytoplasm | other | | 1867484.625 | 9126514 | 3470854.75 | | | | | | | |
| Q3THE2 | MYL12A | myosin, light chain 12A, regulatory | Cytoplasm | other | | 4265107 | 1.02E+07 | 5445707.5 | | | | | | | |
| Q3THE2 | MYL12A | myosin, light chain 12A, regulatory | Cytoplasm | other | | 8483835 | 3278776 | 5635585.5 | Q3THE2 | MYL12A | | 13175285 | 10572994 | 2729935 | 15 |
| Q60605 | MYL6 | myosin, light chain 6, alkali, smoot | Cytoplasm | other | | 6.78E+07 | 6.57E+07 | 5.86E+07 | | | | | | | |
| Q60605 | MYL6 | myosin, light chain 6, alkali, smoot | Cytoplasm | other | | 5.93E+07 | 6.00E+07 | 6.47E+07 | | | | | | | |
| Q60605 | MYL6 | myosin, light chain 6, alkali, smoot | Cytoplasm | other | | 4.49E+07 | 5.04E+07 | 5.06E+07 | | | | | | | |
| Q60605 | MYL6 | myosin, light chain 6, alkali, smoot | Cytoplasm | other | | 4.03E+07 | 3.60E+07 | 3.40E+07 | | | | | | | |
| Q60605 | MYL6 | myosin, light chain 6, alkali, smoot | Cytoplasm | other | | 2.94E+07 | 4.05E+07 | 4.09E+07 | | | | | | | |
| Q60605 | MYL6 | myosin, light chain 6, alkali, smoot | Cytoplasm | other | | 2.98E+07 | 2.82E+07 | 2.79E+07 | | | | | | | |
| Q60605 | MYL6 | myosin, light chain 6, alkali, smoot | Cytoplasm | other | | 1.24E+07 | 1.01E+07 | 1.01E+07 | Q60605 | MYL6 | | 40557143 | 18014409 | 3931066 | 21 |
| Q9CQ19 | MYL9 | myosin, light chain 9, regulatory | Cytoplasm | other | | 2.18E+07 | 1.95E+07 | 1.97E+07 | | | | | | | |
| Q9CQ19 | MYL9 | myosin, light chain 9, regulatory | Cytoplasm | other | | 3.31E+07 | 2.69E+07 | 2.88E+07 | | | | | | | |
| Q9CQ19 | MYL9 | myosin, light chain 9, regulatory | Cytoplasm | other | | 8293784 | 1.82E+07 | 9010122 | | | | | | | |
| Q9CQ19 | MYL9 | myosin, light chain 9, regulatory | Cytoplasm | other | | 1.34E+07 | 1.18E+07 | 1.06E+07 | | | | | | | |
| Q9CQ19 | MYL9 | myosin, light chain 9, regulatory | Cytoplasm | other | | 1.47E+07 | 1.61E+07 | 1.24E+07 | | | | | | | |
| Q9CQ19 | MYL9 | myosin, light chain 9, regulatory | Cytoplasm | other | | 1.33E+07 | 1.90E+07 | 1.35E+07 | | | | | | | |
| Q9CQ19 | MYL9 | myosin, light chain 9, regulatory | Cytoplasm | other | | 5571623 | 4915742.5 | 5643744 | | | | | | | |
| Q9CQ19 | MYL9 | myosin, light chain 9, regulatory | Cytoplasm | other | | 1.37E+07 | 1.59E+07 | 1.29E+07 | Q9CQ19 | MYL9 | | 15469426 | 7166971 | 1462952 | 24 |
| Q9WTI7 | MYO1C | myosin IC | Cytoplasm | other | | 231818.344 | 180854.453 | 256722 | Q9WTI7 | MYO1C | | 231818 | 38673 | 22328 | 3 |
| Q64331 | MYO6 | myosin VI | Cytoplasm | other | | 2771127.75 | 2401289.5 | 2830669 | Q64331 | MYO6 | | 2771128 | 232627 | 134307 | 3 |
| P70670 | NACA | nascent polypeptide-associated cc | Cytoplasm | transcription regulator | | 6607207 | 6594661 | 7195493 | | | | | | | |
| P70670 | NACA | nascent polypeptide-associated cc | Cytoplasm | transcription regulator | | 4505549 | 5718268 | 5125562 | | | | | | | |
| P70670 | NACA | nascent polypeptide-associated cc | Cytoplasm | transcription regulator | | 654833.375 | 891786.5 | 1173918.125 | P70670 | NACA | | 3922530 | 2654651 | 884884 | 9 |
| P28656 | NAP1L1 | nucleosome assembly protein 1-lik | Nucleus | other | | 5.28E+07 | 4.81E+07 | 2.76E+07 | | | | | | | |
| P28656 | NAP1L1 | nucleosome assembly protein 1-lik | Nucleus | other | | 2.83E+07 | 3.26E+07 | 1.68E+07 | | | | | | | |
| P28656 | NAP1L1 | nucleosome assembly protein 1-lik | Nucleus | other | | 2.76E+07 | 2.50E+07 | 1.63E+07 | | | | | | | |
| P28656 | NAP1L1 | nucleosome assembly protein 1-lik | Nucleus | other | | 1.73E+07 | 1.21E+07 | 3464157.5 | | | | | | | |
| P28656 | NAP1L1 | nucleosome assembly protein 1-lik | Nucleus | other | | 1.65E+07 | 1.39E+07 | 1.12E+07 | | | | | | | |
| P28656 | NAP1L1 | nucleosome assembly protein 1-lik | Nucleus | other | | 8558797 | 8735805 | 4935105 | | | | | | | |
| P28656 | NAP1L1 | nucleosome assembly protein 1-lik | Nucleus | other | | 7189848.5 | 7835710.5 | 5284671.5 | | | | | | | |
| P28656 | NAP1L1 | nucleosome assembly protein 1-lik | Nucleus | other | | 5149415 | 4023149.5 | 2053130.75 | P28656 | NAP1L1 | | 20431008 | 13661182 | 2788577 | 24 |
| Q78ZA7 | NAP1L4 | nucleosome assembly protein 1-lik | Cytoplasm | other | | 3.52E+07 | 2.76E+07 | 1.75E+07 | | | | | | | |

| | | | | | | | | | | | | | |
|---|---|---|---|---|---|---|---|---|---|---|---|---|---|
| Q78ZA7 | NAP1L4 | nucleosome assembly protein 1-lik | Cytoplasm | other | 2.17E+07 | 2.17E+07 | 1.20E+07 | | | | | | |
| Q78ZA7 | NAP1L4 | nucleosome assembly protein 1-lik | Cytoplasm | other | 1.74E+07 | 1.18E+07 | 5282932 | | | | | | |
| Q78ZA7 | NAP1L4 | nucleosome assembly protein 1-lik | Cytoplasm | other | 1.19E+07 | 1.18E+07 | 9638284 | | | | | | |
| Q78ZA7 | NAP1L4 | nucleosome assembly protein 1-lik | Cytoplasm | other | 1.23E+07 | 1.26E+07 | 7786689.5 | | | | | | |
| Q78ZA7 | NAP1L4 | nucleosome assembly protein 1-lik | Cytoplasm | other | 1.06E+07 | 9334393 | 6391528.5 | | | | | | |
| Q78ZA7 | NAP1L4 | nucleosome assembly protein 1-lik | Cytoplasm | other | 5431306.5 | 1.45E+07 | 5518726.5 | | | | | | |
| Q78ZA7 | NAP1L4 | nucleosome assembly protein 1-lik | Cytoplasm | other | 7163405 | 6439473 | 6264165 | | | | | | |
| Q78ZA7 | NAP1L4 | nucleosome assembly protein 1-lik | Cytoplasm | other | 5530714.5 | 5703361.5 | 4900302.5 | | | | | | |
| Q78ZA7 | NAP1L4 | nucleosome assembly protein 1-lik | Cytoplasm | other | 2522517.75 | 4540566.5 | 3206672.75 | | | | | | |
| Q78ZA7 | NAP1L4 | nucleosome assembly protein 1-lik | Cytoplasm | other | 2826007.25 | 4498678.5 | 2502606.25 | Q78ZA7 | NAP1L4 | 12045814 | 7537781 | 1312159 | 33 |
| Q9DB05 | NAPA | N-ethylmaleimide-sensitive factor | Cytoplasm | transporter | 4234248.5 | 3868877.5 | 2619433.75 | Q9DB05 | NAPA | 4234249 | 846781 | 488889 | 3 |
| Q8K2Z4 | NCAPD2 | non-SMC condensin I complex, sub | Nucleus | other | 1907278.25 | 4.70E-06 | 338506.5 | Q8K2Z4 | NCAPD2 | 1907278 | 1017623 | 587525 | 3 |
| Q8C156 | NCAPH | non-SMC condensin I complex, sub | Nucleus | other | 1577030.5 | 3281309 | 909884.562 | Q8C156 | NCAPH | 1577031 | 1222927 | 706057 | 3 |
| P57716 | NCSTN | nicastrin | Plasma Membrane | peptidase | 8869394 | 1.35E+07 | 1.30E+07 | | | | | | |
| P57716 | NCSTN | nicastrin | Plasma Membrane | peptidase | 1.17E+07 | 1.18E+07 | 1.12E+07 | | | | | | |
| P57716 | NCSTN | nicastrin | Plasma Membrane | peptidase | 7912353.5 | 8314529 | 6922346.5 | | | | | | |
| P57716 | NCSTN | nicastrin | Plasma Membrane | peptidase | 4408447.5 | 6109503 | 4336030 | | | | | | |
| P57716 | NCSTN | nicastrin | Plasma Membrane | peptidase | 6568086.5 | 7693388 | 9606087 | | | | | | |
| P57716 | NCSTN | nicastrin | Plasma Membrane | peptidase | 4032975.75 | 2853065 | 3657815.25 | | | | | | |
| P57716 | NCSTN | nicastrin | Plasma Membrane | peptidase | 4299912.5 | 4635045.5 | 5491277.5 | | | | | | |
| P57716 | NCSTN | nicastrin | Plasma Membrane | peptidase | 4553617 | 5037648.5 | 6396000 | | | | | | |
| P57716 | NCSTN | nicastrin | Plasma Membrane | peptidase | 101937.305 | 1691129.125 | 1558047.75 | P57716 | NCSTN | 5830747 | 3570012 | 687049 | 27 |
| Q99LC3 | NDUFA10 | NADH dehydrogenase (ubiquinone | Cytoplasm | transporter | 6618094 | 5294729 | 9554007 | | | | | | |
| Q99LC3 | NDUFA10 | NADH dehydrogenase (ubiquinone | Cytoplasm | transporter | 5317399.5 | 4482370.5 | 6649592.5 | Q99LC3 | NDUFA10 | 5967747 | 1793854 | 732338 | 6 |
| Q7TMF3 | NDUFA12 | NADH dehydrogenase (ubiquinone | Cytoplasm | enzyme | 724440.312 | 860207.938 | 767423 | Q7TMF3 | NDUFA12 | 724440 | 69389 | 40062 | 3 |
| Q9CQ75 | NDUFA2 | NADH dehydrogenase (ubiquinone | Cytoplasm | enzyme | 3644316.75 | 2807309.5 | 4074119.5 | Q9CQ75 | NDUFA2 | 3644317 | 644220 | 371941 | 3 |
| Q62425 | NDUFA4 | NDUFA4, mitochondrial complex a | Cytoplasm | enzyme | 7611323.5 | 7063449.5 | 9537552 | | | | | | |
| Q62425 | NDUFA4 | NDUFA4, mitochondrial complex a | Cytoplasm | enzyme | 3572912.5 | 6518457 | 7518831.5 | Q62425 | NDUFA4 | 5592118 | 1952315 | 797029 | 6 |
| Q9CPP6 | NDUFA5 | NADH dehydrogenase (ubiquinone | Cytoplasm | enzyme | 1575730.875 | 1690581 | 2257635.75 | Q9CPP6 | NDUFA5 | 1575731 | 365088 | 210784 | 3 |
| Q9DCJ5 | NDUFA8 | NADH dehydrogenase (ubiquinone | Cytoplasm | enzyme | 1418298.375 | 2544134.5 | 3909092.5 | Q9DCJ5 | NDUFA8 | 1418298 | 1247309 | 720134 | 3 |
| Q9DC69 | NDUFA9 | NADH dehydrogenase (ubiquinone | Cytoplasm | enzyme | 9699494 | 8295715 | 1.40E+07 | | | | | | |
| Q9DC69 | NDUFA9 | NADH dehydrogenase (ubiquinone | Cytoplasm | enzyme | 2598429.5 | 3386074 | 4846933 | | | | | | |
| Q9DC69 | NDUFA9 | NADH dehydrogenase (ubiquinone | Cytoplasm | enzyme | 1710041.875 | 1369869.5 | 2796172.25 | Q9DC69 | NDUFA9 | 4669322 | 4315556 | 1438519 | 9 |
| Q9CR21 | NDUFAB1 | NADH dehydrogenase (ubiquinone | Cytoplasm | enzyme | 4458726 | 8246444.5 | 1.17E+07 | | | | | | |
| Q9CR21 | NDUFAB1 | NADH dehydrogenase (ubiquinone | Cytoplasm | enzyme | 1776120.25 | 2013719.375 | 2304217 | Q9CR21 | NDUFAB1 | 3117423 | 4065885 | 1659891 | 6 |
| Q91VD9 | NDUFS1 | NADH dehydrogenase (ubiquinone | Cytoplasm | enzyme | 2922718.25 | 3185013.75 | 3577122 | | | | | | |
| Q91VD9 | NDUFS1 | NADH dehydrogenase (ubiquinone | Cytoplasm | enzyme | 2023782.75 | 3686227.75 | 3976222.75 | Q91VD9 | NDUFS1 | 2473251 | 698123 | 285008 | 6 |
| Q91WD5 | NDUFS2 | NADH dehydrogenase (ubiquinone | Cytoplasm | enzyme | 2209990.5 | 2029543.875 | 3003149 | Q91WD5 | NDUFS2 | 2209991 | 517940 | 299033 | 3 |
| Q9DCT2 | NDUFS3 | NADH dehydrogenase (ubiquinone | Cytoplasm | enzyme | 1307556.5 | 2837315.5 | 3707749.25 | Q9DCT2 | NDUFS3 | 1307557 | 1215096 | 701536 | 3 |
| Q9CXZ1 | NDUFS4 | NADH dehydrogenase (ubiquinone | Cytoplasm | enzyme | 3725406.25 | 4333426 | 6208508 | | | | | | |
| Q9CXZ1 | NDUFS4 | NADH dehydrogenase (ubiquinone | Cytoplasm | enzyme | 2038823.25 | 1723850.125 | 1190102.875 | Q9CXZ1 | NDUFS4 | 2882115 | 1906767 | 778434 | 6 |
| Q91YT0 | NDUFV1 | NADH dehydrogenase (ubiquinone | Cytoplasm | enzyme | 5182977.5 | 4247929.5 | 8180825 | | | | | | |
| Q91YT0 | NDUFV1 | NADH dehydrogenase (ubiquinone | Cytoplasm | enzyme | 2301337.5 | 2550539 | 2691871.75 | | | | | | |
| Q91YT0 | NDUFV1 | NADH dehydrogenase (ubiquinone | Cytoplasm | enzyme | 1096766.375 | 1916080 | 2216742.75 | Q91YT0 | NDUFV1 | 2860360 | 2181921 | 727307 | 9 |
| Q9D6J6 | NDUFV2 | NADH dehydrogenase (ubiquinone | Cytoplasm | enzyme | 1887800 | 2185429.25 | 3748803.75 | Q9D6J6 | NDUFV2 | 1887800 | 999671 | 577161 | 3 |
| Q9D1J1 | NECAP2 | NECAP endocytosis associated 2 | Cytoplasm | other | 8065948 | 9033231 | 5265769.5 | Q9D1J1 | NECAP2 | 8065948 | 1956630 | 1129661 | 3 |
| Q9JMH7 | NEU3 | sialidase 3 (membrane sialidase) | Plasma Membrane | enzyme | 1.25E+07 | 1.48E+07 | 1.54E+07 | | | | | | |
| Q9JMH7 | NEU3 | sialidase 3 (membrane sialidase) | Plasma Membrane | enzyme | 7401035 | 7230245.5 | 5936213.5 | | | | | | |
| Q9JMH7 | NEU3 | sialidase 3 (membrane sialidase) | Plasma Membrane | enzyme | 1111462.375 | 1823856.875 | 1135567.875 | Q9JMH7 | NEU3 | 7004166 | 5659921 | 1886640 | 9 |
| Q9EST8 | NFKBIZ | nuclear factor of kappa light polyp | Nucleus | transcription regulator | 5605728.5 | 2115466.25 | 3213595 | Q9EST8 | NFKBIZ | 5605729 | 1784662 | 1030375 | 3 |
| Q3TL44 | NLRX1 | NLR family member X1 | Cytoplasm | other | 309516.938 | 3466426.5 | 4681243.5 | Q3TL44 | NLRX1 | 309517 | 2256614 | 1302857 | 3 |
| Q9WV85 | NME3 | NME/NM23 nucleoside diphospha | Cytoplasm | kinase | 1292491.625 | 1772156.625 | 1683546 | Q9WV85 | NME3 | 1292492 | 255230 | 147357 | 3 |
| P97300 | Nptn | neuroplastin | Plasma Membrane | other | 8.97E+07 | 9.91E+07 | 7.99E+07 | | | | | | |
| P97300 | Nptn | neuroplastin | Plasma Membrane | other | 3.88E+07 | 3.27E+07 | 2.62E+07 | | | | | | |
| P97300 | Nptn | neuroplastin | Plasma Membrane | other | 9947931 | 1.85E+07 | 1.63E+07 | | | | | | |
| P97300 | Nptn | neuroplastin | Plasma Membrane | other | 9384497 | 8790438 | 7121624.5 | | | | | | |
| P97300 | Nptn | neuroplastin | Plasma Membrane | other | 7332658 | 9486830 | 1.20E+07 | | | | | | |
| P97300 | Nptn | neuroplastin | Plasma Membrane | other | 5203046.5 | 1.04E+07 | 9452302 | | | | | | |
| P97300 | Nptn | neuroplastin | Plasma Membrane | other | 2705595 | 4484841 | 3739953 | | | | | | |
| P97300 | Nptn | neuroplastin | Plasma Membrane | other | 615575.625 | 363307.25 | 285082.25 | | | | | | |
| P97300 | Nptn | neuroplastin | Plasma Membrane | other | 72731.602 | 109746.852 | 182047.516 | P97300 | Nptn | 18200226 | 27464062 | 5285461 | 27 |
| P08556 | NRAS | neuroblastoma RAS viral (v-ras) or | Plasma Membrane | enzyme | 1.96E+07 | 1.97E+07 | 1.87E+07 | P08556 | NRAS | 19560000 | 526530 | 303992 | 3 |
| P46460 | NSF | N-ethylmaleimide-sensitive factor | Cytoplasm | transporter | 1.02E+07 | 9598291 | 7719846 | | | | | | |
| P46460 | NSF | N-ethylmaleimide-sensitive factor | Cytoplasm | transporter | 6818524.5 | 7206427 | 5614673.5 | | | | | | |
| P46460 | NSF | N-ethylmaleimide-sensitive factor | Cytoplasm | transporter | 6798627 | 9331174 | 6782811.5 | | | | | | |
| P46460 | NSF | N-ethylmaleimide-sensitive factor | Cytoplasm | transporter | 8122792.5 | 6185486 | 5701169 | | | | | | |
| P46460 | NSF | N-ethylmaleimide-sensitive factor | Cytoplasm | transporter | 6511526 | 4969689 | 4246091 | | | | | | |
| P46460 | NSF | N-ethylmaleimide-sensitive factor | Cytoplasm | transporter | 5839718 | 6773851 | 6175036.5 | | | | | | |
| P46460 | NSF | N-ethylmaleimide-sensitive factor | Cytoplasm | transporter | 3944462.75 | 3872250.25 | 4039194.75 | | | | | | |
| P46460 | NSF | N-ethylmaleimide-sensitive factor | Cytoplasm | transporter | 3795603.25 | 5494589.5 | 4490610 | | | | | | |
| P46460 | NSF | N-ethylmaleimide-sensitive factor | Cytoplasm | transporter | 3011226 | 4197700 | 2912783.5 | | | | | | |
| P46460 | NSF | N-ethylmaleimide-sensitive factor | Cytoplasm | transporter | 5485082 | 1541277.25 | 3210490.75 | | | | | | |
| P46460 | NSF | N-ethylmaleimide-sensitive factor | Cytoplasm | transporter | 3273679.5 | 3640587.75 | 3638736.25 | | | | | | |
| P46460 | NSF | N-ethylmaleimide-sensitive factor | Cytoplasm | transporter | 2185531.5 | 2423217.25 | 2032099.125 | | | | | | |
| P46460 | NSF | N-ethylmaleimide-sensitive factor | Cytoplasm | transporter | 1342496.25 | 1205277 | 1348802.25 | | | | | | |
| P46460 | NSF | N-ethylmaleimide-sensitive factor | Cytoplasm | transporter | 161564.328 | 499636.188 | 161667.359 | P46460 | NSF | 4822202 | 2565397 | 395849 | 42 |
| Q9CZ44 | NSFL1C | NSFL1 (p97) cofactor (p47) | Cytoplasm | other | 3752260 | 2727509 | 3258620.75 | | | | | | |
| Q9CZ44 | NSFL1C | NSFL1 (p97) cofactor (p47) | Cytoplasm | other | 1515657.5 | 1616920.5 | 2614172 | | | | | | |
| Q9CZ44 | NSFL1C | NSFL1 (p97) cofactor (p47) | Cytoplasm | other | 2660986 | 2135025.25 | 1658199.25 | | | | | | |
| Q9CZ44 | NSFL1C | NSFL1 (p97) cofactor (p47) | Cytoplasm | other | 1253205.25 | 764508.562 | 1773512.5 | Q9CZ44 | NSFL1C | 2295527 | 873872 | 252265 | 12 |
| A6H603 | NWD1 | NACHT and WD repeat domain co | Cytoplasm | other | 1.22E+07 | 1.14E+07 | 1.10E+07 | A6H603 | NWD1 | 12200000 | 612454 | 353601 | 3 |
| P83503 | NYX | nyctalopin | Extracellular Space | other | 5.98E+07 | 9.32E+07 | 7.98E+07 | P83503 | NYX | 59800000 | 16818922 | 9710409 | 3 |
| P29758 | OAT | ornithine aminotransferase | Cytoplasm | enzyme | 2315021.5 | 2210867.5 | 3578515.25 | P29758 | OAT | 2315022 | 761328 | 439553 | 3 |
| A2AAJ9 | OBSCN | obscurin, cytoskeletal calmodulin | Cytoplasm | kinase | 2261898 | 2660884 | 1905338 | A2AAJ9 | OBSCN | 2261898 | 377971 | 218222 | 3 |
| Q9CRD0 | OCIAD1 | OCIA domain containing 1 | Cytoplasm | other | 2649512.75 | 2416597.25 | 3236625.25 | Q9CRD0 | OCIAD1 | 2649513 | 422571 | 243971 | 3 |
| Q60597 | OGDH | oxoglutarate (alpha-ketoglutarate | Cytoplasm | enzyme | 1.27E+07 | 1.20E+07 | 1.43E+07 | | | | | | |

| | | | | | | | | | | | | | |
|---|---|---|---|---|---|---|---|---|---|---|---|---|---|
| Q60597 | OGDH | oxoglutarate (alpha-ketoglutarate | Cytoplasm | enzyme | 1.84E+07 | 1.45E+07 | 2.55E+07 | | | | | | |
| Q60597 | OGDH | oxoglutarate (alpha-ketoglutarate | Cytoplasm | enzyme | 6108346 | 7103309.5 | 1.05E+07 | | | | | | |
| Q60597 | OGDH | oxoglutarate (alpha-ketoglutarate | Cytoplasm | enzyme | 7277865 | 8569492 | 1.17E+07 | | | | | | |
| Q60597 | OGDH | oxoglutarate (alpha-ketoglutarate | Cytoplasm | enzyme | 6267952.5 | 1.10E+07 | 1.93E+07 | | | | | | |
| Q60597 | OGDH | oxoglutarate (alpha-ketoglutarate | Cytoplasm | enzyme | 8123986 | 7853800.5 | 9567944 | | | | | | |
| Q60597 | OGDH | oxoglutarate (alpha-ketoglutarate | Cytoplasm | enzyme | 5610665 | 8725165 | 9427060 | | | | | | |
| Q60597 | OGDH | oxoglutarate (alpha-ketoglutarate | Cytoplasm | enzyme | 5069426 | 5395537.5 | 8343178.5 | | | | | | |
| Q60597 | OGDH | oxoglutarate (alpha-ketoglutarate | Cytoplasm | enzyme | 6797699.5 | 5257530 | 7254810.5 | | | | | | |
| Q60597 | OGDH | oxoglutarate (alpha-ketoglutarate | Cytoplasm | enzyme | 6124582 | 6048250 | 9139369 | | | | | | |
| Q60597 | OGDH | oxoglutarate (alpha-ketoglutarate | Cytoplasm | enzyme | 5500774.5 | 5053783.5 | 6407835 | | | | | | |
| Q60597 | OGDH | oxoglutarate (alpha-ketoglutarate | Cytoplasm | enzyme | 3747807.5 | 4177547.25 | 7760254 | | | | | | |
| Q60597 | OGDH | oxoglutarate (alpha-ketoglutarate | Cytoplasm | enzyme | 3141116.5 | 2930742.75 | 4697429.5 | | | | | | |
| Q60597 | OGDH | oxoglutarate (alpha-ketoglutarate | Cytoplasm | enzyme | 2275073.75 | 2884717.5 | 5177097.5 | | | | | | |
| Q60597 | OGDH | oxoglutarate (alpha-ketoglutarate | Cytoplasm | enzyme | 1798930.5 | 2104545.75 | 2644113.25 | | | | | | |
| Q60597 | OGDH | oxoglutarate (alpha-ketoglutarate | Cytoplasm | enzyme | 1554447 | 1939626.375 | 2257260.75 | | | | | | |
| Q60597 | OGDH | oxoglutarate (alpha-ketoglutarate | Cytoplasm | enzyme | 3549533 | 3162391.5 | 3956687 | Q60597 | OGDH | 6119306 | 4825645 | 675725 | 51 |
| Q9CZ30 | OLA1 | Obg-like ATPase 1 | Cytoplasm | enzyme | 4407166.5 | 1808724.125 | 3187786.5 | Q9CZ30 | OLA1 | 4407167 | 1300039 | 750578 | 3 |
| Q8BGT0 | OSTM1 | osteopetrosis associated transmer | Cytoplasm | other | 707289.75 | 992712 | 710934.562 | Q8BGT0 | OSTM1 | 707290 | 163747 | 94539 | 3 |
| Q9D0K2 | OXCT1 | 3-oxoacid CoA transferase 1 | Cytoplasm | enzyme | 3549663.25 | 3186742.5 | 3526755.25 | Q9D0K2 | OXCT1 | 3549663 | 203242 | 117342 | 3 |
| P09103 | P4HB | prolyl 4-hydroxylase, beta polypep | Cytoplasm | enzyme | 2.34E+07 | 2.48E+07 | 2.09E+07 | | | | | | |
| P09103 | P4HB | prolyl 4-hydroxylase, beta polypep | Cytoplasm | enzyme | 1.88E+07 | 2.13E+07 | 1.55E+07 | | | | | | |
| P09103 | P4HB | prolyl 4-hydroxylase, beta polypep | Cytoplasm | enzyme | 2.15E+07 | 2.54E+07 | 2.02E+07 | | | | | | |
| P09103 | P4HB | prolyl 4-hydroxylase, beta polypep | Cytoplasm | enzyme | 1.94E+07 | 2.45E+07 | 1.88E+07 | | | | | | |
| P09103 | P4HB | prolyl 4-hydroxylase, beta polypep | Cytoplasm | enzyme | 2.17E+07 | 2.29E+07 | 2.28E+07 | | | | | | |
| P09103 | P4HB | prolyl 4-hydroxylase, beta polypep | Cytoplasm | enzyme | 1.45E+07 | 2.41E+07 | 1.89E+07 | | | | | | |
| P09103 | P4HB | prolyl 4-hydroxylase, beta polypep | Cytoplasm | enzyme | 1.43E+07 | 1.79E+07 | 1.16E+07 | | | | | | |
| P09103 | P4HB | prolyl 4-hydroxylase, beta polypep | Cytoplasm | enzyme | 1.39E+07 | 1.80E+07 | 1.25E+07 | | | | | | |
| P09103 | P4HB | prolyl 4-hydroxylase, beta polypep | Cytoplasm | enzyme | 9316413 | 1.63E+07 | 1.02E+07 | | | | | | |
| P09103 | P4HB | prolyl 4-hydroxylase, beta polypep | Cytoplasm | enzyme | 1.35E+07 | 2.19E+07 | 1.61E+07 | | | | | | |
| P09103 | P4HB | prolyl 4-hydroxylase, beta polypep | Cytoplasm | enzyme | 1.74E+07 | 1.34E+07 | 8555103 | | | | | | |
| P09103 | P4HB | prolyl 4-hydroxylase, beta polypep | Cytoplasm | enzyme | 8529660 | 1.28E+07 | 1.01E+07 | | | | | | |
| P09103 | P4HB | prolyl 4-hydroxylase, beta polypep | Cytoplasm | enzyme | 8986690 | 1.34E+07 | 9937876 | | | | | | |
| P09103 | P4HB | prolyl 4-hydroxylase, beta polypep | Cytoplasm | enzyme | 1.16E+07 | 1.65E+07 | 1.19E+07 | | | | | | |
| P09103 | P4HB | prolyl 4-hydroxylase, beta polypep | Cytoplasm | enzyme | 1.06E+07 | 1.13E+07 | 1.02E+07 | | | | | | |
| P09103 | P4HB | prolyl 4-hydroxylase, beta polypep | Cytoplasm | enzyme | 7129735.5 | 1.31E+07 | 7845285.5 | | | | | | |
| P09103 | P4HB | prolyl 4-hydroxylase, beta polypep | Cytoplasm | enzyme | 4784201 | 1.04E+07 | 5151263 | | | | | | |
| P09103 | P4HB | prolyl 4-hydroxylase, beta polypep | Cytoplasm | enzyme | 4871191 | 5855402 | 4720604.5 | | | | | | |
| P09103 | P4HB | prolyl 4-hydroxylase, beta polypep | Cytoplasm | enzyme | 2253117 | 6054254.5 | 5722184.5 | P09103 | P4HB | 12970579 | 6204932 | 821863 | 57 |
| P50580 | PA2G4 | proliferation-associated 2G4, 38kD | Nucleus | transcription regulator | 8434244 | 8475883 | 5629211.5 | | | | | | |
| P50580 | PA2G4 | proliferation-associated 2G4, 38kD | Nucleus | transcription regulator | 6241401 | 5794168.5 | 5502712 | | | | | | |
| P50580 | PA2G4 | proliferation-associated 2G4, 38kD | Nucleus | transcription regulator | 5650781.5 | 5902587.5 | 3965588.25 | | | | | | |
| P50580 | PA2G4 | proliferation-associated 2G4, 38kD | Nucleus | transcription regulator | 3913604.25 | 3666225.25 | 3010295 | | | | | | |
| P50580 | PA2G4 | proliferation-associated 2G4, 38kD | Nucleus | transcription regulator | 3802986.5 | 3978136 | 3235582.5 | P50580 | PA2G4 | 5608603 | 1715248 | 442875 | 15 |
| P29341 | PABPC1 | poly(A) binding protein, cytoplasm | Cytoplasm | translation regulator | 3564099.25 | 2766117.5 | 2349239.75 | | | | | | |
| P29341 | PABPC1 | poly(A) binding protein, cytoplasm | Cytoplasm | translation regulator | 5427255 | 5200609.5 | 3351771 | | | | | | |
| P29341 | PABPC1 | poly(A) binding protein, cytoplasm | Cytoplasm | translation regulator | 3467339.75 | 3796467.5 | 1947742.625 | | | | | | |
| P29341 | PABPC1 | poly(A) binding protein, cytoplasm | Cytoplasm | translation regulator | 3575261.5 | 2383962.25 | 2356724.5 | | | | | | |
| P29341 | PABPC1 | poly(A) binding protein, cytoplasm | Cytoplasm | translation regulator | 1900421.125 | 1853871 | 1150170.5 | P29341 | PABPC1 | 3586875 | 1215455 | 313829 | 15 |
| Q9WVE8 | PACSIN2 | protein kinase C and casein kinase | Cytoplasm | transporter | 9939706 | 6943738.5 | 7673156.5 | | | | | | |
| Q9WVE8 | PACSIN2 | protein kinase C and casein kinase | Cytoplasm | transporter | 9081191 | 7162240 | 5635096.5 | | | | | | |
| Q9WVE8 | PACSIN2 | protein kinase C and casein kinase | Cytoplasm | transporter | 4404471 | 4206870 | 3521500.75 | | | | | | |
| Q9WVE8 | PACSIN2 | protein kinase C and casein kinase | Cytoplasm | transporter | 5748225 | 3531833.5 | 3860778.75 | | | | | | |
| Q9WVE8 | PACSIN2 | protein kinase C and casein kinase | Cytoplasm | transporter | 5145009 | 3703165.5 | 3963783.25 | | | | | | |
| Q9WVE8 | PACSIN2 | protein kinase C and casein kinase | Cytoplasm | transporter | 5510499 | 4132718 | 3613206.5 | | | | | | |
| Q9WVE8 | PACSIN2 | protein kinase C and casein kinase | Cytoplasm | transporter | 1481168.625 | 1206507.75 | 1666930.875 | | | | | | |
| Q9WVE8 | PACSIN2 | protein kinase C and casein kinase | Cytoplasm | transporter | 657401.312 | 2698919.25 | 1991565.375 | Q9WVE8 | PACSIN2 | 5245959 | 2418309 | 493635 | 24 |
| Q9ES46 | PARVB | parvin, beta | Cytoplasm | other | 3371652 | 2967253.75 | 2330634.25 | | | | | | |
| Q9ES46 | PARVB | parvin, beta | Cytoplasm | other | 6546995 | 5780288.5 | 6913953.5 | | | | | | |
| Q9ES46 | PARVB | parvin, beta | Cytoplasm | other | 4343021 | 4576128.5 | 4549859.5 | | | | | | |
| Q9ES46 | PARVB | parvin, beta | Cytoplasm | other | 2862643.25 | 3766607.5 | 2658162 | | | | | | |
| Q9ES46 | PARVB | parvin, beta | Cytoplasm | other | 2270484.5 | 2840362 | 2588627.75 | | | | | | |
| Q9ES46 | PARVB | parvin, beta | Cytoplasm | other | 2264292.75 | 1980331.375 | 2751661.75 | Q9ES46 | PARVB | 3609848 | 1511276 | 356211 | 18 |
| Q05920 | PC | pyruvate carboxylase | Cytoplasm | enzyme | 2334675 | 2342261 | 3545963.75 | | | | | | |
| Q05920 | PC | pyruvate carboxylase | Cytoplasm | enzyme | 2750092.5 | 3516524.75 | 3368219.5 | | | | | | |
| Q05920 | PC | pyruvate carboxylase | Cytoplasm | enzyme | 2699439.75 | 2088577.75 | 3877121 | | | | | | |
| Q05920 | PC | pyruvate carboxylase | Cytoplasm | enzyme | 1414082.75 | 920321.125 | 1244074.625 | | | | | | |
| Q05920 | PC | pyruvate carboxylase | Cytoplasm | enzyme | 1275707.5 | 1173607.75 | 1718078.125 | Q05920 | PC | 2094800 | 982959 | 253799 | 15 |
| P60335 | PCBP1 | poly(rC) binding protein 1 | Nucleus | translation regulator | 3335072.75 | 2938706.75 | 1725767 | | | | | | |
| P60335 | PCBP1 | poly(rC) binding protein 1 | Nucleus | translation regulator | 3217412.25 | 3633929.25 | 3012669.5 | | | | | | |
| P60335 | PCBP1 | poly(rC) binding protein 1 | Nucleus | translation regulator | 5383025 | 3369922.25 | 2290150.75 | | | | | | |
| P60335 | PCBP1 | poly(rC) binding protein 1 | Nucleus | translation regulator | 2555061.5 | 1962175.5 | 1296088.5 | | | | | | |
| P60335 | PCBP1 | poly(rC) binding protein 1 | Nucleus | translation regulator | 2349609.75 | 1545995 | 1244755.125 | P60335 | PCBP1 | 3368036 | 1087062 | 280678 | 15 |
| Q8BH04 | PCK2 | phosphoenolpyruvate carboxykina | Cytoplasm | kinase | 1.07E+07 | 1.42E+07 | 1.50E+07 | | | | | | |
| Q8BH04 | PCK2 | phosphoenolpyruvate carboxykina | Cytoplasm | kinase | 1.59E+07 | 1.33E+07 | 2.15E+07 | | | | | | |
| Q8BH04 | PCK2 | phosphoenolpyruvate carboxykina | Cytoplasm | kinase | 6447016.5 | 7928150.5 | 8186107 | | | | | | |
| Q8BH04 | PCK2 | phosphoenolpyruvate carboxykina | Cytoplasm | kinase | 1.07E+07 | 1.18E+07 | 1.38E+07 | | | | | | |
| Q8BH04 | PCK2 | phosphoenolpyruvate carboxykina | Cytoplasm | kinase | 7228090 | 6293895.5 | 7856483 | | | | | | |
| Q8BH04 | PCK2 | phosphoenolpyruvate carboxykina | Cytoplasm | kinase | 7096843.5 | 7764427 | 1.19E+07 | | | | | | |
| Q8BH04 | PCK2 | phosphoenolpyruvate carboxykina | Cytoplasm | kinase | 4434262 | 5605025 | 8857336 | | | | | | |
| Q8BH04 | PCK2 | phosphoenolpyruvate carboxykina | Cytoplasm | kinase | 4435601.5 | 3885758 | 5952864 | | | | | | |
| Q8BH04 | PCK2 | phosphoenolpyruvate carboxykina | Cytoplasm | kinase | 4545853.5 | 3292684.5 | 5283199 | | | | | | |
| Q8BH04 | PCK2 | phosphoenolpyruvate carboxykina | Cytoplasm | kinase | 4693719.5 | 4703952.5 | 6539428.5 | | | | | | |
| Q8BH04 | PCK2 | phosphoenolpyruvate carboxykina | Cytoplasm | kinase | 5922499.5 | 4926809 | 7342014 | | | | | | |
| Q8BH04 | PCK2 | phosphoenolpyruvate carboxykina | Cytoplasm | kinase | 1939304.125 | 3456682.5 | 3629722.25 | | | | | | |
| Q8BH04 | PCK2 | phosphoenolpyruvate carboxykina | Cytoplasm | kinase | 5099466.5 | 4813372 | 7877393.5 | | | | | | |
| Q8BH04 | PCK2 | phosphoenolpyruvate carboxykina | Cytoplasm | kinase | 4306391 | 5073244 | 8469663 | | | | | | |

| | | | | | | | | | | | | | |
|---|---|---|---|---|---|---|---|---|---|---|---|---|---|
| Q8BH04 | PCK2 | phosphoenolpyruvate carboxykina | Cytoplasm | kinase | 4101356.5 | 3470137.5 | 5433162 | | | | | | |
| Q8BH04 | PCK2 | phosphoenolpyruvate carboxykina | Cytoplasm | kinase | 3105178.75 | 2358556.75 | 3707043.25 | | | | | | |
| Q8BH04 | PCK2 | phosphoenolpyruvate carboxykina | Cytoplasm | kinase | 2314298.25 | 1804108.25 | 2225950.75 | Q8BH04 | PCK2 | 6060581 | 4126474 | 577822 | 51 |
| Q9CQF9 | PCYOX1 | prenylcysteine oxidase 1 | Cytoplasm | enzyme | 6911731 | 8784289 | 8224500 | Q9CQF9 | PCYOX1 | 6911731 | 961180 | 554937 | 3 |
| P27664 | PDE6A | phosphodiesterase 6A, cGMP-spec | Plasma Membrane | enzyme | 1.52E+07 | 8734271 | 1.12E+07 | P27664 | PDE6A | 15240000 | 3283663 | 1895824 | 3 |
| P35486 | PDHA1 | pyruvate dehydrogenase (lipoamid | Cytoplasm | enzyme | 1148707.5 | 2020520.5 | 1760111.125 | | | | | | |
| P35486 | PDHA1 | pyruvate dehydrogenase (lipoamid | Cytoplasm | enzyme | 1859658.75 | 1804157.375 | 1982430.25 | | | | | | |
| P35486 | PDHA1 | pyruvate dehydrogenase (lipoamid | Cytoplasm | enzyme | 1617933.875 | 2221313.75 | 2109603 | P35486 | PDHA1 | 1542100 | 317425 | 105808 | 9 |
| Q9D051 | PDHB | pyruvate dehydrogenase (lipoamid | Cytoplasm | enzyme | 3465574.5 | 3017368.25 | 3583440.5 | Q9D051 | PDHB | 3465575 | 298669 | 172437 | 3 |
| P27773 | PDIA3 | protein disulfide isomerase family | Cytoplasm | peptidase | 1.27E+08 | 1.34E+08 | 1.17E+08 | | | | | | |
| P27773 | PDIA3 | protein disulfide isomerase family | Cytoplasm | peptidase | 5.51E+07 | 6.99E+07 | 5.43E+07 | | | | | | |
| P27773 | PDIA3 | protein disulfide isomerase family | Cytoplasm | peptidase | 4.31E+07 | 5.84E+07 | 4.21E+07 | | | | | | |
| P27773 | PDIA3 | protein disulfide isomerase family | Cytoplasm | peptidase | 2.77E+07 | 3.53E+07 | 3.00E+07 | | | | | | |
| P27773 | PDIA3 | protein disulfide isomerase family | Cytoplasm | peptidase | 2.22E+07 | 3.09E+07 | 2.93E+07 | | | | | | |
| P27773 | PDIA3 | protein disulfide isomerase family | Cytoplasm | peptidase | 3.71E+07 | 2.98E+07 | 3.07E+07 | | | | | | |
| P27773 | PDIA3 | protein disulfide isomerase family | Cytoplasm | peptidase | 2.35E+07 | 2.58E+07 | 2.45E+07 | | | | | | |
| P27773 | PDIA3 | protein disulfide isomerase family | Cytoplasm | peptidase | 2.26E+07 | 3.52E+07 | 2.84E+07 | | | | | | |
| P27773 | PDIA3 | protein disulfide isomerase family | Cytoplasm | peptidase | 2.47E+07 | 3.72E+07 | 3.60E+07 | | | | | | |
| P27773 | PDIA3 | protein disulfide isomerase family | Cytoplasm | peptidase | 1.78E+07 | 3.02E+07 | 2.30E+07 | | | | | | |
| P27773 | PDIA3 | protein disulfide isomerase family | Cytoplasm | peptidase | 1.87E+07 | 2.00E+07 | 1.96E+07 | | | | | | |
| P27773 | PDIA3 | protein disulfide isomerase family | Cytoplasm | peptidase | 2.03E+07 | 2.25E+07 | 2.02E+07 | | | | | | |
| P27773 | PDIA3 | protein disulfide isomerase family | Cytoplasm | peptidase | 1.72E+07 | 2.37E+07 | 2.27E+07 | | | | | | |
| P27773 | PDIA3 | protein disulfide isomerase family | Cytoplasm | peptidase | 1.54E+07 | 1.75E+07 | 1.30E+07 | | | | | | |
| P27773 | PDIA3 | protein disulfide isomerase family | Cytoplasm | peptidase | 1.49E+07 | 1.91E+07 | 1.61E+07 | | | | | | |
| P27773 | PDIA3 | protein disulfide isomerase family | Cytoplasm | peptidase | 1.52E+07 | 1.75E+07 | 1.27E+07 | | | | | | |
| P27773 | PDIA3 | protein disulfide isomerase family | Cytoplasm | peptidase | 1.96E+07 | 1.79E+07 | 1.89E+07 | | | | | | |
| P27773 | PDIA3 | protein disulfide isomerase family | Cytoplasm | peptidase | 1.02E+07 | 1.26E+07 | 1.11E+07 | | | | | | |
| P27773 | PDIA3 | protein disulfide isomerase family | Cytoplasm | peptidase | 9481311 | 1.15E+07 | 1.13E+07 | | | | | | |
| P27773 | PDIA3 | protein disulfide isomerase family | Cytoplasm | peptidase | 8451800 | 1.75E+07 | 1.29E+07 | | | | | | |
| P27773 | PDIA3 | protein disulfide isomerase family | Cytoplasm | peptidase | 3750391 | 7305459.5 | 5211367.5 | | | | | | |
| P27773 | PDIA3 | protein disulfide isomerase family | Cytoplasm | peptidase | 2964163.25 | 2358107.75 | 1551000.625 | | | | | | |
| P27773 | PDIA3 | protein disulfide isomerase family | Cytoplasm | peptidase | 1957151.5 | 4445136.5 | 2513350 | P27773 | PDIA3 | 24306296 | 25406267 | 3058555 | 69 |
| P08003 | PDIA4 | protein disulfide isomerase family | Cytoplasm | enzyme | 1.17E+07 | 8489337 | 8949751 | | | | | | |
| P08003 | PDIA4 | protein disulfide isomerase family | Cytoplasm | enzyme | 6094086 | 4622253.5 | 6612993.5 | | | | | | |
| P08003 | PDIA4 | protein disulfide isomerase family | Cytoplasm | enzyme | 2962935.5 | 4732286.5 | 3474610.75 | | | | | | |
| P08003 | PDIA4 | protein disulfide isomerase family | Cytoplasm | enzyme | 4216674 | 4333744.5 | 3172290.5 | | | | | | |
| P08003 | PDIA4 | protein disulfide isomerase family | Cytoplasm | enzyme | 3546026.25 | 4619792.5 | 4222181 | | | | | | |
| P08003 | PDIA4 | protein disulfide isomerase family | Cytoplasm | enzyme | 3076404.75 | 3135879.5 | 2908890 | | | | | | |
| P08003 | PDIA4 | protein disulfide isomerase family | Cytoplasm | enzyme | 2407984 | 3437261.75 | 3007703.5 | | | | | | |
| P08003 | PDIA4 | protein disulfide isomerase family | Cytoplasm | enzyme | 1650012.875 | 2190207.75 | 1934451.125 | P08003 | PDIA4 | 4454265 | 2418723 | 493720 | 24 |
| Q922R8 | PDIA6 | protein disulfide isomerase family | Cytoplasm | enzyme | 2.00E+07 | 2.53E+07 | 2.82E+07 | | | | | | |
| Q922R8 | PDIA6 | protein disulfide isomerase family | Cytoplasm | enzyme | 1.48E+07 | 1.86E+07 | 1.62E+07 | | | | | | |
| Q922R8 | PDIA6 | protein disulfide isomerase family | Cytoplasm | enzyme | 1.28E+07 | 1.60E+07 | 1.37E+07 | | | | | | |
| Q922R8 | PDIA6 | protein disulfide isomerase family | Cytoplasm | enzyme | 1.37E+07 | 1.32E+07 | 1.13E+07 | | | | | | |
| Q922R8 | PDIA6 | protein disulfide isomerase family | Cytoplasm | enzyme | 1.19E+07 | 1.36E+07 | 1.22E+07 | | | | | | |
| Q922R8 | PDIA6 | protein disulfide isomerase family | Cytoplasm | enzyme | 1.57E+07 | 7074409 | 1.25E+07 | | | | | | |
| Q922R8 | PDIA6 | protein disulfide isomerase family | Cytoplasm | enzyme | 9470693 | 1.52E+07 | 1.36E+07 | | | | | | |
| Q922R8 | PDIA6 | protein disulfide isomerase family | Cytoplasm | enzyme | 1.18E+07 | 1.25E+07 | 1.42E+07 | | | | | | |
| Q922R8 | PDIA6 | protein disulfide isomerase family | Cytoplasm | enzyme | 6058089 | 1.07E+07 | 1.04E+07 | | | | | | |
| Q922R8 | PDIA6 | protein disulfide isomerase family | Cytoplasm | enzyme | 5114922.5 | 7032101 | 5317511 | | | | | | |
| Q922R8 | PDIA6 | protein disulfide isomerase family | Cytoplasm | enzyme | 4897488.5 | 5796385.5 | 4915449 | Q922R8 | PDIA6 | 11470108 | 5433369 | 945828 | 33 |
| Q9Z126 | PF4 | platelet factor 4 | Extracellular Space | cytokine | 1.14E+08 | 1.28E+08 | 1.26E+08 | | | | | | |
| Q9Z126 | PF4 | platelet factor 4 | Extracellular Space | cytokine | 9.49E+07 | 7.89E+07 | 9.69E+07 | | | | | | |
| Q9Z126 | PF4 | platelet factor 4 | Extracellular Space | cytokine | 4.50E+07 | 4.95E+07 | 5.65E+07 | Q9Z126 | PF4 | 84490000 | 32069310 | 10689770 | 9 |
| P12382 | PFKL | phosphofructokinase, liver | Cytoplasm | kinase | 2471148.25 | 2809863.5 | 2352370 | P12382 | PFKL | 2471148 | 237394 | 137060 | 3 |
| P47857 | PFKM | phosphofructokinase, muscle | Cytoplasm | kinase | 3423432.25 | 5242638.5 | 3466259.25 | | | | | | |
| P47857 | PFKM | phosphofructokinase, muscle | Cytoplasm | kinase | 2667754 | 2726985.75 | 2483338.75 | | | | | | |
| P47857 | PFKM | phosphofructokinase, muscle | Cytoplasm | kinase | 2355797 | 2886184.5 | 2176624.5 | | | | | | |
| P47857 | PFKM | phosphofructokinase, muscle | Cytoplasm | kinase | 2608076.25 | 2853994.25 | 2846134.5 | | | | | | |
| P47857 | PFKM | phosphofructokinase, muscle | Cytoplasm | kinase | 1975478.25 | 2205549 | 1396223.75 | | | | | | |
| P47857 | PFKM | phosphofructokinase, muscle | Cytoplasm | kinase | 2177655.75 | 2622653.25 | 2613736.25 | P47857 | PFKM | 2534699 | 799243 | 188383 | 18 |
| Q9DCD0 | PGD | phosphogluconate dehydrogenase | Cytoplasm | enzyme | 4228926 | 3084118 | 3917102 | Q9DCD0 | PGD | 4228926 | 591845 | 341702 | 3 |
| O55022 | PGRMC1 | progesterone receptor membrane | Plasma Membrane | transmembrane receptor | 6001659 | 4494147.5 | 4851743.5 | O55022 | PGRMC1 | 6001659 | 787694 | 454775 | 3 |
| Q80UU9 | PGRMC2 | progesterone receptor membrane | Nucleus | other | 2441065.25 | 2704707.75 | 2624746.25 | Q80UU9 | PGRMC2 | 2441065 | 135179 | 78046 | 3 |
| P67778 | PHB | prohibitin | Nucleus | transcription regulator | 9280761 | 9813682 | 1.06E+07 | | | | | | |
| P67778 | PHB | prohibitin | Nucleus | transcription regulator | 7285524.5 | 1.09E+07 | 1.59E+07 | | | | | | |
| P67778 | PHB | prohibitin | Nucleus | transcription regulator | 9011183 | 7824277.5 | 1.01E+07 | | | | | | |
| P67778 | PHB | prohibitin | Nucleus | transcription regulator | 5434619 | 1.19E+07 | 1.10E+07 | | | | | | |
| P67778 | PHB | prohibitin | Nucleus | transcription regulator | 5678759 | 8668670 | 9752101 | | | | | | |
| P67778 | PHB | prohibitin | Nucleus | transcription regulator | 5375849 | 4676041.5 | 6073429 | | | | | | |
| P67778 | PHB | prohibitin | Nucleus | transcription regulator | 3484837.5 | 5551292.5 | 8236329.5 | | | | | | |
| P67778 | PHB | prohibitin | Nucleus | transcription regulator | 6341096 | 6900615 | 1.29E+07 | | | | | | |
| P67778 | PHB | prohibitin | Nucleus | transcription regulator | 1246986.75 | 3545848.5 | 4194608.5 | | | | | | |
| P67778 | PHB | prohibitin | Nucleus | transcription regulator | 2505568.5 | 2435627.5 | 3193919.25 | P67778 | PHB | 5564518 | 3494791 | 638059 | 30 |
| O35129 | PHB2 | prohibitin 2 | Cytoplasm | transcription regulator | 1.22E+07 | 9200707 | 1.13E+07 | | | | | | |
| O35129 | PHB2 | prohibitin 2 | Cytoplasm | transcription regulator | 1.23E+07 | 1.35E+07 | 1.49E+07 | | | | | | |
| O35129 | PHB2 | prohibitin 2 | Cytoplasm | transcription regulator | 8147310 | 9290834 | 1.14E+07 | | | | | | |
| O35129 | PHB2 | prohibitin 2 | Cytoplasm | transcription regulator | 1.29E+07 | 2.29E+07 | 2.14E+07 | | | | | | |
| O35129 | PHB2 | prohibitin 2 | Cytoplasm | transcription regulator | 1.04E+07 | 1.34E+07 | 1.36E+07 | | | | | | |
| O35129 | PHB2 | prohibitin 2 | Cytoplasm | transcription regulator | 1.18E+07 | 1.15E+07 | 1.40E+07 | | | | | | |
| O35129 | PHB2 | prohibitin 2 | Cytoplasm | transcription regulator | 1331151.375 | 1856165 | 1849117.625 | | | | | | |
| O35129 | PHB2 | prohibitin 2 | Cytoplasm | transcription regulator | 5000329.5 | 3981167 | 6032294 | | | | | | |
| O35129 | PHB2 | prohibitin 2 | Cytoplasm | transcription regulator | 1143903.5 | 1343053.25 | 1295531.75 | O35129 | PHB2 | 8355855 | 5920680 | 1139435 | 27 |
| Q61753 | PHGDH | phosphoglycerate dehydrogenase | Cytoplasm | enzyme | 4523063.5 | 2827418.5 | 2451302.25 | | | | | | |
| Q61753 | PHGDH | phosphoglycerate dehydrogenase | Cytoplasm | enzyme | 3980397.5 | 3056180.5 | 2572733 | | | | | | |
| Q61753 | PHGDH | phosphoglycerate dehydrogenase | Cytoplasm | enzyme | 2734376.75 | 1990769 | 1760303.625 | | | | | | |

| | | | | | | | | | | | | | |
|---|---|---|---|---|---|---|---|---|---|---|---|---|---|
| Q61753 | PHGDH | phosphoglycerate dehydrogenase | Cytoplasm | enzyme | 3215448.5 | 352609 | 126976.492 | | | | | | |
| Q61753 | PHGDH | phosphoglycerate dehydrogenase | Cytoplasm | enzyme | 2061868.5 | 1832423.625 | 1500842.875 | Q61753 | PHGDH | 3303031 | 1181379 | 305031 | 15 |
| Q8CBQ5 | PI4K2B | phosphatidylinositol 4-kinase type | Cytoplasm | kinase | 2004866.125 | 2196781.5 | 3465399.25 | | | | | | |
| Q8CBQ5 | PI4K2B | phosphatidylinositol 4-kinase type | Cytoplasm | kinase | 1914376.75 | 2395704.5 | 1928998.625 | | | | | | |
| Q8CBQ5 | PI4K2B | phosphatidylinositol 4-kinase type | Cytoplasm | kinase | 2083987.5 | 3009632.25 | 2119269.25 | | | | | | |
| Q8CBQ5 | PI4K2B | phosphatidylinositol 4-kinase type | Cytoplasm | kinase | 1120159.25 | 1096358.5 | 730650.375 | Q8CBQ5 | PI4K2B | 1780847 | 774438 | 223561 | 12 |
| Q7M6Y3 | PICALM | phosphatidylinositol binding clath | Cytoplasm | other | 2.56E+07 | 2.71E+07 | 2.29E+07 | | | | | | |
| Q7M6Y3 | PICALM | phosphatidylinositol binding clath | Cytoplasm | other | 7749740.5 | 9977498 | 6987532 | | | | | | |
| Q7M6Y3 | PICALM | phosphatidylinositol binding clath | Cytoplasm | other | 3450606.75 | 3426981.25 | 2744270.75 | | | | | | |
| Q7M6Y3 | PICALM | phosphatidylinositol binding clath | Cytoplasm | other | 791693 | 3148580 | 2187214.5 | Q7M6Y3 | PICALM | 9393010 | 9752223 | 2815224 | 12 |
| E2JF22 | PIEZO1 | piezo-type mechanosensitive ion c | Cytoplasm | ion channel | 2.62E+07 | 2.36E+07 | 2.05E+07 | | | | | | |
| E2JF22 | PIEZO1 | piezo-type mechanosensitive ion c | Cytoplasm | ion channel | 2.27E+07 | 2.05E+07 | 2.49E+07 | | | | | | |
| E2JF22 | PIEZO1 | piezo-type mechanosensitive ion c | Cytoplasm | ion channel | 1.54E+07 | 1.56E+07 | 1.55E+07 | | | | | | |
| E2JF22 | PIEZO1 | piezo-type mechanosensitive ion c | Cytoplasm | ion channel | 1.33E+07 | 1.23E+07 | 1.21E+07 | | | | | | |
| E2JF22 | PIEZO1 | piezo-type mechanosensitive ion c | Cytoplasm | ion channel | 8866911 | 1.25E+07 | 1.21E+07 | | | | | | |
| E2JF22 | PIEZO1 | piezo-type mechanosensitive ion c | Cytoplasm | ion channel | 1.11E+07 | 1.21E+07 | 9744405 | | | | | | |
| E2JF22 | PIEZO1 | piezo-type mechanosensitive ion c | Cytoplasm | ion channel | 1.30E+07 | 1.28E+07 | 1.08E+07 | | | | | | |
| E2JF22 | PIEZO1 | piezo-type mechanosensitive ion c | Cytoplasm | ion channel | 1.04E+07 | 1.00E+07 | 9042146 | | | | | | |
| E2JF22 | PIEZO1 | piezo-type mechanosensitive ion c | Cytoplasm | ion channel | 8596448 | 8543526 | 1.05E+07 | | | | | | |
| E2JF22 | PIEZO1 | piezo-type mechanosensitive ion c | Cytoplasm | ion channel | 1.12E+07 | 1.08E+07 | 1.18E+07 | | | | | | |
| E2JF22 | PIEZO1 | piezo-type mechanosensitive ion c | Cytoplasm | ion channel | 9930309 | 8301443 | 8617391 | | | | | | |
| E2JF22 | PIEZO1 | piezo-type mechanosensitive ion c | Cytoplasm | ion channel | 7137831.5 | 5248262 | 5890698.5 | | | | | | |
| E2JF22 | PIEZO1 | piezo-type mechanosensitive ion c | Cytoplasm | ion channel | 6350875 | 5487851.5 | 6739731.5 | | | | | | |
| E2JF22 | PIEZO1 | piezo-type mechanosensitive ion c | Cytoplasm | ion channel | 5378635 | 4419802 | 4491229.5 | | | | | | |
| E2JF22 | PIEZO1 | piezo-type mechanosensitive ion c | Cytoplasm | ion channel | 5647851.5 | 5240435 | 7164980.5 | | | | | | |
| E2JF22 | PIEZO1 | piezo-type mechanosensitive ion c | Cytoplasm | ion channel | 4485472.5 | 4853095.5 | 4045790.5 | | | | | | |
| E2JF22 | PIEZO1 | piezo-type mechanosensitive ion c | Cytoplasm | ion channel | 5122264 | 4659580 | 6350193.5 | | | | | | |
| E2JF22 | PIEZO1 | piezo-type mechanosensitive ion c | Cytoplasm | ion channel | 6747504 | 4280178 | 6146729 | | | | | | |
| E2JF22 | PIEZO1 | piezo-type mechanosensitive ion c | Cytoplasm | ion channel | 3293074.25 | 4409842 | 3994507.25 | | | | | | |
| E2JF22 | PIEZO1 | piezo-type mechanosensitive ion c | Cytoplasm | ion channel | 3245744.75 | 3379670.5 | 3868482.5 | | | | | | |
| E2JF22 | PIEZO1 | piezo-type mechanosensitive ion c | Cytoplasm | ion channel | 3438290.5 | 4504237.5 | 4383385 | | | | | | |
| E2JF22 | PIEZO1 | piezo-type mechanosensitive ion c | Cytoplasm | ion channel | 1833769.5 | 2304366.75 | 2845305.75 | | | | | | |
| E2JF22 | PIEZO1 | piezo-type mechanosensitive ion c | Cytoplasm | ion channel | 1063881.25 | 1514075.5 | 1154121.375 | | | | | | |
| E2JF22 | PIEZO1 | piezo-type mechanosensitive ion c | Cytoplasm | ion channel | 1223858.875 | 1068981.375 | 1727776.5 | | | | | | |
| E2JF22 | PIEZO1 | piezo-type mechanosensitive ion c | Cytoplasm | ion channel | 1728590.5 | 1334579.875 | 1206536.625 | | | | | | |
| E2JF22 | PIEZO1 | piezo-type mechanosensitive ion c | Cytoplasm | ion channel | 1132251.875 | 1227758.125 | 1108563.75 | | | | | | |
| E2JF22 | PIEZO1 | piezo-type mechanosensitive ion c | Cytoplasm | ion channel | 195482.469 | 385062.219 | 932857.25 | E2JF22 | PIEZO1 | 7727742 | 6019935 | 668882 | 81 |
| P06803 | PIM1 | Pim-1 proto-oncogene, serine/thre | Cytoplasm | kinase | 2.49E+08 | 2.05E+08 | 2.06E+08 | P06803 | PIM1 | 248700000 | 25117922 | 14501839 | 3 |
| O70172 | PIP4K2A | phosphatidylinositol-5-phosphate | Cytoplasm | kinase | 1.45E+07 | 1.31E+07 | 9740130 | | | | | | |
| O70172 | PIP4K2A | phosphatidylinositol-5-phosphate | Cytoplasm | kinase | 3839355.5 | 2346031.75 | 2881875.75 | | | | | | |
| O70172 | PIP4K2A | phosphatidylinositol-5-phosphate | Cytoplasm | kinase | 1821807.875 | 3303331.75 | 4293751 | | | | | | |
| O70172 | PIP4K2A | phosphatidylinositol-5-phosphate | Cytoplasm | kinase | 3181651 | 2193873.5 | 2964028.75 | O70172 | PIP4K2A | 5845704 | 4468151 | 1289844 | 12 |
| Q80XI4 | PIP4K2B | phosphatidylinositol-5-phosphate | Cytoplasm | kinase | 6737272.5 | 5057698.5 | 5229794.5 | Q80XI4 | PIP4K2B | 6737273 | 924038 | 533494 | 3 |
| P70182 | PIP5K1A | phosphatidylinositol-4-phosphate | Cytoplasm | kinase | 1.32E+07 | 1.13E+07 | 1.18E+07 | | | | | | |
| P70182 | PIP5K1A | phosphatidylinositol-4-phosphate | Cytoplasm | kinase | 8457769 | 9233637 | 8270664.5 | | | | | | |
| P70182 | PIP5K1A | phosphatidylinositol-4-phosphate | Cytoplasm | kinase | 7443015.5 | 6526936.5 | 5687902.5 | | | | | | |
| P70182 | PIP5K1A | phosphatidylinositol-4-phosphate | Cytoplasm | kinase | 6504264 | 5952364.5 | 5023274 | | | | | | |
| P70182 | PIP5K1A | phosphatidylinositol-4-phosphate | Cytoplasm | kinase | 7423594.5 | 5891983 | 6873322.5 | | | | | | |
| P70182 | PIP5K1A | phosphatidylinositol-4-phosphate | Cytoplasm | kinase | 5712018.5 | 5794008 | 5914819 | | | | | | |
| P70182 | PIP5K1A | phosphatidylinositol-4-phosphate | Cytoplasm | kinase | 2831062.25 | 2780289.25 | 2341545 | | | | | | |
| P70182 | PIP5K1A | phosphatidylinositol-4-phosphate | Cytoplasm | kinase | 2566708.25 | 2073569.125 | 1838599.75 | | | | | | |
| P70182 | PIP5K1A | phosphatidylinositol-4-phosphate | Cytoplasm | kinase | 4463911.5 | 3963893 | 2786201.75 | | | | | | |
| P70182 | PIP5K1A | phosphatidylinositol-4-phosphate | Cytoplasm | kinase | 306210.938 | 624740.562 | 378424.625 | P70182 | PIP5K1A | 5890855 | 3331071 | 608168 | 30 |
| Q8K411 | PITRM1 | pitrilysin metallopeptidase 1 | Cytoplasm | peptidase | 6999743 | 8239642 | 1.04E+07 | | | | | | |
| Q8K411 | PITRM1 | pitrilysin metallopeptidase 1 | Cytoplasm | peptidase | 6607601 | 6268117 | 9094512 | | | | | | |
| Q8K411 | PITRM1 | pitrilysin metallopeptidase 1 | Cytoplasm | peptidase | 4063446.5 | 3629270.5 | 6890864.5 | | | | | | |
| Q8K411 | PITRM1 | pitrilysin metallopeptidase 1 | Cytoplasm | peptidase | 4131720.5 | 4870163.5 | 5627705 | | | | | | |
| Q8K411 | PITRM1 | pitrilysin metallopeptidase 1 | Cytoplasm | peptidase | 3296287 | 3311463 | 4628294 | | | | | | |
| Q8K411 | PITRM1 | pitrilysin metallopeptidase 1 | Cytoplasm | peptidase | 1439562.75 | 925567.812 | 940385.188 | | | | | | |
| Q8K411 | PITRM1 | pitrilysin metallopeptidase 1 | Cytoplasm | peptidase | 421447.094 | 658046.312 | 765673.438 | Q8K411 | PITRM1 | 3851401 | 2956048 | 645063 | 21 |
| Q80ZA4 | PKHD1L1 | polycystic kidney and hepatic dise | Extracellular Space | other | 7814419 | 9039636 | 7164840.5 | Q80ZA4 | PKHD1L1 | 7814419 | 952013 | 549645 | 3 |
| P53657 | PKLR | pyruvate kinase, liver and RBC | Cytoplasm | kinase | 1.73E+07 | 1.50E+07 | 1.39E+07 | | | | | | |
| P53657 | PKLR | pyruvate kinase, liver and RBC | Cytoplasm | kinase | 9490740 | 7553145 | 7704387.5 | | | | | | |
| P53657 | PKLR | pyruvate kinase, liver and RBC | Cytoplasm | kinase | 8996504 | 8070062.5 | 7467384 | | | | | | |
| P53657 | PKLR | pyruvate kinase, liver and RBC | Cytoplasm | kinase | 7594893.5 | 5175166.5 | 6700172 | | | | | | |
| P53657 | PKLR | pyruvate kinase, liver and RBC | Cytoplasm | kinase | 6461063 | 5344625 | 3897381.75 | | | | | | |
| P53657 | PKLR | pyruvate kinase, liver and RBC | Cytoplasm | kinase | 4474320.5 | 3108576 | 3858522.25 | | | | | | |
| P53657 | PKLR | pyruvate kinase, liver and RBC | Cytoplasm | kinase | 2898611 | 1960781.875 | 1509906.5 | | | | | | |
| P53657 | PKLR | pyruvate kinase, liver and RBC | Cytoplasm | kinase | 2921702.5 | 3281083.25 | 2987222 | P53657 | PKLR | 7518479 | 4126445 | 842307 | 24 |
| P52480 | PKM | pyruvate kinase, muscle | Cytoplasm | kinase | 4411644 | 5653402.5 | 4715132 | P52480 | PKM | 4411644 | 647356 | 373751 | 3 |
| P27612 | PLAA | phospholipase A2-activating prote | Cytoplasm | other | 1260387.375 | 753212.875 | 579190.938 | P27612 | PLAA | 1260387 | 353916 | 204333 | 3 |
| Q9JHK5 | PLEK | pleckstrin | Cytoplasm | other | 234741.172 | 870978.375 | 357072.562 | Q9JHK5 | PLEK | 234741 | 337605 | 194916 | 3 |
| P20918 | PLG | plasminogen | Extracellular Space | peptidase | 2776720.25 | 1332215.375 | 1100833.375 | | | | | | |
| P20918 | PLG | plasminogen | Extracellular Space | peptidase | 773900.75 | 107960.484 | 169213.312 | | | | | | |
| P20918 | PLG | plasminogen | Extracellular Space | peptidase | 490774.406 | 509263.719 | 311519.562 | P20918 | PLG | 1347132 | 833818 | 277939 | 9 |
| Q9R1Q7 | PLP2 | proteolipid protein 2 (colonic epith | Cytoplasm | transporter | 1.33E+07 | 1.65E+07 | 1.03E+07 | Q9R1Q7 | PLP2 | 13290000 | 3100650 | 1790161 | 3 |
| Q9JJ00 | PLSCR1 | phospholipid scramblase 1 | Plasma Membrane | enzyme | 1.94E+07 | 2.07E+07 | 2.04E+07 | | | | | | |
| Q9JJ00 | PLSCR1 | phospholipid scramblase 1 | Plasma Membrane | enzyme | 1.39E+07 | 1.37E+07 | 1.35E+07 | | | | | | |
| Q9JJ00 | PLSCR1 | phospholipid scramblase 1 | Plasma Membrane | enzyme | 8251292 | 6991796.5 | 8996160 | Q9JJ00 | PLSCR1 | 13857097 | 5271987 | 1757329 | 9 |
| Q9JIZ9 | PLSCR3 | phospholipid scramblase 3 | Plasma Membrane | enzyme | 8030237 | 8531251 | 9503302 | | | | | | |
| Q9JIZ9 | PLSCR3 | phospholipid scramblase 3 | Plasma Membrane | enzyme | 1.03E+07 | 9951884 | 1.12E+07 | | | | | | |
| Q9JIZ9 | PLSCR3 | phospholipid scramblase 3 | Plasma Membrane | enzyme | 4447657 | 3895693.5 | 3343736.75 | | | | | | |
| Q9JIZ9 | PLSCR3 | phospholipid scramblase 3 | Plasma Membrane | enzyme | 2442090 | 2771318.25 | 2436711.5 | Q9JIZ9 | PLSCR3 | 6302496 | 3457355 | 998052 | 12 |
| P23492 | PNP | purine nucleoside phosphorylase | Nucleus | enzyme | 4.46E+07 | 3.62E+07 | 3.76E+07 | | | | | | |
| P23492 | PNP | purine nucleoside phosphorylase | Nucleus | enzyme | 1.25E+07 | 8560725 | 8867753 | | | | | | |
| P23492 | PNP | purine nucleoside phosphorylase | Nucleus | enzyme | 2.07E+07 | 8477213 | 1.33E+07 | | | | | | |

| | | | | | | | | | | | | | |
|---|---|---|---|---|---|---|---|---|---|---|---|---|---|
| P23492 | PNP | purine nucleoside phosphorylase | Nucleus | enzyme | 8186915 | 6743597.5 | 8104752 | | | | | | |
| P23492 | PNP | purine nucleoside phosphorylase | Nucleus | enzyme | 7504856 | 6445620 | 8718624 | | | | | | |
| P23492 | PNP | purine nucleoside phosphorylase | Nucleus | enzyme | 3002137 | 2052046.25 | 2510962.25 | P23492 | PNP | 16070651 | 12739418 | 3002710 | 18 |
| Q61469 | Ppap2a | phosphatidic acid phosphatase typ | Plasma Membrane | phosphatase | 3.52E+07 | 5.38E+07 | 5.74E+07 | | | | | | |
| Q61469 | Ppap2a | phosphatidic acid phosphatase typ | Plasma Membrane | phosphatase | 2.72E+07 | 3.36E+07 | 3.93E+07 | | | | | | |
| Q61469 | Ppap2a | phosphatidic acid phosphatase typ | Plasma Membrane | phosphatase | 2.79E+07 | 7.30E+07 | 5.88E+07 | | | | | | |
| Q61469 | Ppap2a | phosphatidic acid phosphatase typ | Plasma Membrane | phosphatase | 2.09E+07 | 3.76E+07 | 3.36E+07 | | | | | | |
| Q61469 | Ppap2a | phosphatidic acid phosphatase typ | Plasma Membrane | phosphatase | 1912553.875 | 8889195 | 4068403.75 | Q61469 | Ppap2a | 22640511 | 20570246 | 5311215 | 15 |
| P17742 | PPIA | peptidylprolyl isomerase A (cyclop | Cytoplasm | enzyme | 1.94E+07 | 2.56E+07 | 3.02E+07 | | | | | | |
| P17742 | PPIA | peptidylprolyl isomerase A (cyclop | Cytoplasm | enzyme | 1.60E+07 | 1.54E+07 | 1.70E+07 | | | | | | |
| P17742 | PPIA | peptidylprolyl isomerase A (cyclop | Cytoplasm | enzyme | 2.05E+07 | 2.02E+07 | 2.32E+07 | | | | | | |
| P17742 | PPIA | peptidylprolyl isomerase A (cyclop | Cytoplasm | enzyme | 1.46E+07 | 5308132.5 | 6771483 | | | | | | |
| P17742 | PPIA | peptidylprolyl isomerase A (cyclop | Cytoplasm | enzyme | 1.65E+07 | 1.37E+07 | 1.52E+07 | | | | | | |
| P17742 | PPIA | peptidylprolyl isomerase A (cyclop | Cytoplasm | enzyme | 3905361.5 | 1403989.625 | 778208.25 | | | | | | |
| P17742 | PPIA | peptidylprolyl isomerase A (cyclop | Cytoplasm | enzyme | 1040245.125 | 4978204.5 | 4469293 | P17742 | PPIA | 13126515 | 8662344 | 1890278 | 21 |
| P24369 | PPIB | peptidylprolyl isomerase B (cyclop | Cytoplasm | enzyme | 8.61E+07 | 9.10E+07 | 6.81E+07 | | | | | | |
| P24369 | PPIB | peptidylprolyl isomerase B (cyclop | Cytoplasm | enzyme | 5.03E+07 | 3.75E+07 | 4.76E+07 | | | | | | |
| P24369 | PPIB | peptidylprolyl isomerase B (cyclop | Cytoplasm | enzyme | 1.63E+07 | 3.54E+07 | 2.50E+07 | | | | | | |
| P24369 | PPIB | peptidylprolyl isomerase B (cyclop | Cytoplasm | enzyme | 2.28E+07 | 3.01E+07 | 2.85E+07 | | | | | | |
| P24369 | PPIB | peptidylprolyl isomerase B (cyclop | Cytoplasm | enzyme | 2.67E+07 | 2.93E+07 | 2.56E+07 | | | | | | |
| P24369 | PPIB | peptidylprolyl isomerase B (cyclop | Cytoplasm | enzyme | 2.08E+07 | 2.20E+07 | 2.82E+07 | | | | | | |
| P24369 | PPIB | peptidylprolyl isomerase B (cyclop | Cytoplasm | enzyme | 1.52E+07 | 1.91E+07 | 2.02E+07 | | | | | | |
| P24369 | PPIB | peptidylprolyl isomerase B (cyclop | Cytoplasm | enzyme | 1.31E+07 | 1.52E+07 | 1.15E+07 | | | | | | |
| P24369 | PPIB | peptidylprolyl isomerase B (cyclop | Cytoplasm | enzyme | 1.33E+07 | 9921896 | 1.11E+07 | | | | | | |
| P24369 | PPIB | peptidylprolyl isomerase B (cyclop | Cytoplasm | enzyme | 1.13E+07 | 1.45E+07 | 1.22E+07 | | | | | | |
| P24369 | PPIB | peptidylprolyl isomerase B (cyclop | Cytoplasm | enzyme | 1.28E+07 | 1.19E+07 | 9592740 | | | | | | |
| P24369 | PPIB | peptidylprolyl isomerase B (cyclop | Cytoplasm | enzyme | 5620618.5 | 981702.312 | 1276272.875 | P24369 | PPIB | 24525885 | 20837127 | 3472854 | 36 |
| Q99KR7 | PPIF | peptidylprolyl isomerase F | Cytoplasm | enzyme | 9056425 | 7205942.5 | 9766434 | | | | | | |
| Q99KR7 | PPIF | peptidylprolyl isomerase F | Cytoplasm | enzyme | 7431932 | 7692969.5 | 8135632 | Q99KR7 | PPIF | 8244179 | 1002512 | 409274 | 6 |
| P51175 | PPOX | protoporphyrinogen oxidase | Cytoplasm | enzyme | 2.61E+07 | 2.47E+07 | 3.27E+07 | | | | | | |
| P51175 | PPOX | protoporphyrinogen oxidase | Cytoplasm | enzyme | 2.42E+07 | 2.94E+07 | 4.50E+07 | | | | | | |
| P51175 | PPOX | protoporphyrinogen oxidase | Cytoplasm | enzyme | 8341769.5 | 9379310 | 1.37E+07 | | | | | | |
| P51175 | PPOX | protoporphyrinogen oxidase | Cytoplasm | enzyme | 1.01E+07 | 9570450 | 9806935 | | | | | | |
| P51175 | PPOX | protoporphyrinogen oxidase | Cytoplasm | enzyme | 9192157 | 6088050.5 | 1.03E+07 | | | | | | |
| P51175 | PPOX | protoporphyrinogen oxidase | Cytoplasm | enzyme | 7337245.5 | 5903179 | 6520435 | | | | | | |
| P51175 | PPOX | protoporphyrinogen oxidase | Cytoplasm | enzyme | 6246510 | 7518881.5 | 1.03E+07 | | | | | | |
| P51175 | PPOX | protoporphyrinogen oxidase | Cytoplasm | enzyme | 7028596.5 | 6511529 | 8685110 | | | | | | |
| P51175 | PPOX | protoporphyrinogen oxidase | Cytoplasm | enzyme | 7824759 | 5942665.5 | 1.05E+07 | | | | | | |
| P51175 | PPOX | protoporphyrinogen oxidase | Cytoplasm | enzyme | 4272554.5 | 3096060.75 | 4042417.25 | | | | | | |
| P51175 | PPOX | protoporphyrinogen oxidase | Cytoplasm | enzyme | 3392175 | 3868535.75 | 4257085.5 | | | | | | |
| P51175 | PPOX | protoporphyrinogen oxidase | Cytoplasm | enzyme | 2231129.75 | 3676619 | 5770283.5 | | | | | | |
| P51175 | PPOX | protoporphyrinogen oxidase | Cytoplasm | enzyme | 4150202.5 | 2597489.25 | 3960918.75 | P51175 | PPOX | 9260546 | 9458815 | 1514623 | 39 |
| P62137 | PPP1CA | protein phosphatase 1, catalytic su | Cytoplasm | phosphatase | 6674726 | 4859755 | 6049465 | | | | | | |
| P62137 | PPP1CA | protein phosphatase 1, catalytic su | Cytoplasm | phosphatase | 6797826 | 8170354 | 8429217 | | | | | | |
| P62137 | PPP1CA | protein phosphatase 1, catalytic su | Cytoplasm | phosphatase | 2457921.25 | 2759237 | 2330823 | | | | | | |
| P62137 | PPP1CA | protein phosphatase 1, catalytic su | Cytoplasm | phosphatase | 3145301 | 4025778.5 | 3998978 | | | | | | |
| P62137 | PPP1CA | protein phosphatase 1, catalytic su | Cytoplasm | phosphatase | 2543595.5 | 4064297.75 | 2725487.75 | | | | | | |
| P62137 | PPP1CA | protein phosphatase 1, catalytic su | Cytoplasm | phosphatase | 1655064.375 | 1477246.875 | 1510883 | P62137 | PPP1CA | 3879072 | 2249424 | 530194 | 18 |
| Q62415 | PPP1R13B | protein phosphatase 1, regulatory | Cytoplasm | phosphatase | 2195394 | 4882094.5 | 2240164.25 | Q62415 | PPP1R13B | 2195394 | 1538406 | 888199 | 3 |
| Q3TDD9 | PPP1R21 | protein phosphatase 1, regulatory | Cytoplasm | other | 5202436 | 1.64E+07 | 1.15E+07 | Q3TDD9 | PPP1R21 | 5202436 | 5605035 | 3236068 | 3 |
| P63330 | PPP2CA | protein phosphatase 2, catalytic su | Cytoplasm | phosphatase | 5474830.5 | 4301646.5 | 3964303.75 | | | | | | |
| P63330 | PPP2CA | protein phosphatase 2, catalytic su | Cytoplasm | phosphatase | 6652956 | 2482689.25 | 4404763.5 | | | | | | |
| P63330 | PPP2CA | protein phosphatase 2, catalytic su | Cytoplasm | phosphatase | 4303116.5 | 3188531.75 | 4216847 | | | | | | |
| P63330 | PPP2CA | protein phosphatase 2, catalytic su | Cytoplasm | phosphatase | 2821342 | 1737945.25 | 2272787.5 | | | | | | |
| P63330 | PPP2CA | protein phosphatase 2, catalytic su | Cytoplasm | phosphatase | 2277861.25 | 2453966.5 | 2567628.75 | P63330 | PPP2CA | 4306021 | 1370911 | 353968 | 15 |
| Q76MZ3 | PPP2R1A | protein phosphatase 2, regulatory | Cytoplasm | phosphatase | 7323656.5 | 6997105.5 | 7799388.5 | | | | | | |
| Q76MZ3 | PPP2R1A | protein phosphatase 2, regulatory | Cytoplasm | phosphatase | 2495863.25 | 2464713.75 | 2723815 | | | | | | |
| Q76MZ3 | PPP2R1A | protein phosphatase 2, regulatory | Cytoplasm | phosphatase | 2411287.25 | 1876159.5 | 2695625.25 | | | | | | |
| Q76MZ3 | PPP2R1A | protein phosphatase 2, regulatory | Cytoplasm | phosphatase | 4045784.75 | 3845146 | 5671078 | | | | | | |
| Q76MZ3 | PPP2R1A | protein phosphatase 2, regulatory | Cytoplasm | phosphatase | 3724732.75 | 3086577.25 | 3222579.25 | | | | | | |
| Q76MZ3 | PPP2R1A | protein phosphatase 2, regulatory | Cytoplasm | phosphatase | 2915050.75 | 1952239 | 2408255.5 | | | | | | |
| Q76MZ3 | PPP2R1A | protein phosphatase 2, regulatory | Cytoplasm | phosphatase | 1411055.625 | 1346413.25 | 1870990.125 | Q76MZ3 | PPP2R1A | 3475347 | 1916122 | 418132 | 21 |
| Q6P1F6 | PPP2R2A | protein phosphatase 2, regulatory | Cytoplasm | phosphatase | 3170106.5 | 1870585.375 | 838424.75 | Q6P1F6 | PPP2R2A | 3170107 | 1168393 | 674572 | 3 |
| Q9CQR6 | PPP6C | protein phosphatase 6, catalytic su | Nucleus | phosphatase | 4737376.5 | 7165172 | 5258285.5 | | | | | | |
| Q9CQR6 | PPP6C | protein phosphatase 6, catalytic su | Nucleus | phosphatase | 3489206.25 | 2914968.75 | 2342288.75 | Q9CQR6 | PPP6C | 4113291 | 1773428 | 723999 | 6 |
| Q922D4 | PPP6R3 | protein phosphatase 6, regulatory | Cytoplasm | other | 1.01E+07 | 1.24E+07 | 6888912 | | | | | | |
| Q922D4 | PPP6R3 | protein phosphatase 6, regulatory | Cytoplasm | other | 6363194 | 7196522.5 | 8106051.5 | | | | | | |
| Q922D4 | PPP6R3 | protein phosphatase 6, regulatory | Cytoplasm | other | 6056505.5 | 5449819.5 | 3835950.75 | | | | | | |
| Q922D4 | PPP6R3 | protein phosphatase 6, regulatory | Cytoplasm | other | 5307691.5 | 5928167.5 | 5329052 | | | | | | |
| Q922D4 | PPP6R3 | protein phosphatase 6, regulatory | Cytoplasm | other | 4359103.5 | 4097286 | 3296022.75 | | | | | | |
| Q922D4 | PPP6R3 | protein phosphatase 6, regulatory | Cytoplasm | other | 5234555.5 | 4501141.5 | 3049530 | | | | | | |
| Q922D4 | PPP6R3 | protein phosphatase 6, regulatory | Cytoplasm | other | 4778921.5 | 4743576.5 | 4451323.5 | | | | | | |
| Q922D4 | PPP6R3 | protein phosphatase 6, regulatory | Cytoplasm | other | 3455595.5 | 5137525 | 2964696.75 | | | | | | |
| Q922D4 | PPP6R3 | protein phosphatase 6, regulatory | Cytoplasm | other | 3793700 | 3879693.5 | 2620262.75 | | | | | | |
| Q922D4 | PPP6R3 | protein phosphatase 6, regulatory | Cytoplasm | other | 2414281.5 | 3195282.75 | 2178085.75 | | | | | | |
| Q922D4 | PPP6R3 | protein phosphatase 6, regulatory | Cytoplasm | other | 2827226.75 | 1953777.125 | 1881191.625 | | | | | | |
| Q922D4 | PPP6R3 | protein phosphatase 6, regulatory | Cytoplasm | other | 747797.375 | 1034199.562 | 1288733.375 | Q922D4 | PPP6R3 | 4618214 | 2429699 | 404950 | 36 |
| Q7TMR0 | PRCP | prolylcarboxypeptidase (angiotens | Cytoplasm | peptidase | 4788476.5 | 5068364 | 8311376.5 | Q7TMR0 | PRCP | 4788477 | 1958158 | 1130543 | 3 |
| P35700 | PRDX1 | peroxiredoxin 1 | Cytoplasm | enzyme | 2.41E+07 | 2.41E+07 | 2.65E+07 | | | | | | |
| P35700 | PRDX1 | peroxiredoxin 1 | Cytoplasm | enzyme | 3.14E+07 | 1.92E+07 | 2.23E+07 | | | | | | |
| P35700 | PRDX1 | peroxiredoxin 1 | Cytoplasm | enzyme | 1.59E+07 | 1.50E+07 | 1.89E+07 | | | | | | |
| P35700 | PRDX1 | peroxiredoxin 1 | Cytoplasm | enzyme | 2.22E+07 | 1.96E+07 | 2.42E+07 | | | | | | |
| P35700 | PRDX1 | peroxiredoxin 1 | Cytoplasm | enzyme | 1.62E+07 | 1.70E+07 | 2.43E+07 | | | | | | |
| P35700 | PRDX1 | peroxiredoxin 1 | Cytoplasm | enzyme | 1.57E+07 | 1.48E+07 | 1.64E+07 | | | | | | |
| P35700 | PRDX1 | peroxiredoxin 1 | Cytoplasm | enzyme | 8181855.5 | 1.02E+07 | 1.18E+07 | | | | | | |
| P35700 | PRDX1 | peroxiredoxin 1 | Cytoplasm | enzyme | 7867783.5 | 6063163.5 | 8201006 | | | | | | |

| | | | | | | | | | | | | | |
|---|---|---|---|---|---|---|---|---|---|---|---|---|---|
| P35700 | PRDX1 | peroxiredoxin 1 | Cytoplasm | enzyme | 4250009.5 | 9449534 | 1.02E+07 | | | | | | |
| P35700 | PRDX1 | peroxiredoxin 1 | Cytoplasm | enzyme | 4708434.5 | 2318089.5 | 1833555.75 | | | | | | |
| P35700 | PRDX1 | peroxiredoxin 1 | Cytoplasm | enzyme | 377857.125 | 1819660.625 | 1489583.625 | P35700 | PRDX1 | 13723267 | 8472956 | 1474952 | 33 |
| Q61171 | PRDX2 | peroxiredoxin 2 | Cytoplasm | enzyme | 3.67E+08 | 3.78E+08 | 4.51E+08 | | | | | | |
| Q61171 | PRDX2 | peroxiredoxin 2 | Cytoplasm | enzyme | 2.39E+08 | 1.82E+08 | 2.51E+08 | | | | | | |
| Q61171 | PRDX2 | peroxiredoxin 2 | Cytoplasm | enzyme | 1.92E+08 | 1.80E+08 | 1.99E+08 | | | | | | |
| Q61171 | PRDX2 | peroxiredoxin 2 | Cytoplasm | enzyme | 9.52E+07 | 1.71E+08 | 2.68E+08 | | | | | | |
| Q61171 | PRDX2 | peroxiredoxin 2 | Cytoplasm | enzyme | 1.07E+08 | 9.04E+07 | 1.02E+08 | | | | | | |
| Q61171 | PRDX2 | peroxiredoxin 2 | Cytoplasm | enzyme | 9.28E+07 | 8.36E+07 | 1.07E+08 | | | | | | |
| Q61171 | PRDX2 | peroxiredoxin 2 | Cytoplasm | enzyme | 9.01E+07 | 1.29E+08 | 1.65E+08 | | | | | | |
| Q61171 | PRDX2 | peroxiredoxin 2 | Cytoplasm | enzyme | 5.17E+07 | 6.98E+07 | 8.74E+07 | | | | | | |
| Q61171 | PRDX2 | peroxiredoxin 2 | Cytoplasm | enzyme | 4.46E+07 | 4.06E+07 | 4.81E+07 | | | | | | |
| Q61171 | PRDX2 | peroxiredoxin 2 | Cytoplasm | enzyme | 1.04E+07 | 1.18E+07 | 1.52E+07 | | | | | | |
| Q61171 | PRDX2 | peroxiredoxin 2 | Cytoplasm | enzyme | 1.48E+07 | 2.57E+07 | 2.75E+07 | | | | | | |
| Q61171 | PRDX2 | peroxiredoxin 2 | Cytoplasm | enzyme | 4.79E+07 | 6360842 | 1.58E+07 | | | | | | |
| Q61171 | PRDX2 | peroxiredoxin 2 | Cytoplasm | enzyme | 6503333.5 | 2982649.75 | 1035687.875 | | | | | | |
| Q61171 | PRDX2 | peroxiredoxin 2 | Cytoplasm | enzyme | 1.35E+07 | 1781295.75 | 5218921.5 | Q61171 | PRDX2 | 97934524 | 111381773 | 17186581 | 42 |
| P20108 | PRDX3 | peroxiredoxin 3 | Cytoplasm | enzyme | 7.69E+07 | 1.14E+08 | 1.21E+08 | | | | | | |
| P20108 | PRDX3 | peroxiredoxin 3 | Cytoplasm | enzyme | 6.74E+07 | 5.51E+07 | 8.11E+07 | | | | | | |
| P20108 | PRDX3 | peroxiredoxin 3 | Cytoplasm | enzyme | 4.14E+07 | 3.37E+07 | 5.36E+07 | | | | | | |
| P20108 | PRDX3 | peroxiredoxin 3 | Cytoplasm | enzyme | 2.75E+07 | 1.96E+07 | 2.87E+07 | | | | | | |
| P20108 | PRDX3 | peroxiredoxin 3 | Cytoplasm | enzyme | 2.77E+07 | 2.95E+07 | 4.58E+07 | | | | | | |
| P20108 | PRDX3 | peroxiredoxin 3 | Cytoplasm | enzyme | 2.67E+07 | 2151590.5 | 1948040.625 | | | | | | |
| P20108 | PRDX3 | peroxiredoxin 3 | Cytoplasm | enzyme | 5604672 | 3309819.75 | 5463803 | P20108 | PRDX3 | 39024953 | 34561916 | 7542028 | 21 |
| O08807 | PRDX4 | peroxiredoxin 4 | Cytoplasm | enzyme | 1.04E+07 | 1.10E+07 | 7877912 | | | | | | |
| O08807 | PRDX4 | peroxiredoxin 4 | Cytoplasm | enzyme | 4994104 | 4797290 | 4329228 | | | | | | |
| O08807 | PRDX4 | peroxiredoxin 4 | Cytoplasm | enzyme | 805148.75 | 2901754.75 | 2017127.625 | O08807 | PRDX4 | 5413084 | 3595998 | 1198666 | 9 |
| P99029 | PRDX5 | peroxiredoxin 5 | Cytoplasm | enzyme | 6296442.5 | 6275124 | 5905649 | P99029 | PRDX5 | 6296443 | 219729 | 126861 | 3 |
| Q61878 | PRG2 | proteoglycan 2, bone marrow (nat | Extracellular Space | other | 1047241.188 | 577979.875 | 2662797.75 | Q61878 | PRG2 | 1047241 | 1093671 | 631431 | 3 |
| P05132 | PRKACA | protein kinase, cAMP-dependent, | Cytoplasm | kinase | 4749551 | 6027553 | 5898198.5 | P05132 | PRKACA | 4749551 | 703493 | 406162 | 3 |
| P20444 | PRKCA | protein kinase C, alpha | Cytoplasm | kinase | 2548198.25 | 1677726.75 | 1686389 | P20444 | PRKCA | 2548198 | 500085 | 288724 | 3 |
| O08795 | PRKCSH | protein kinase C substrate 80K-H | Cytoplasm | enzyme | 1.18E+07 | 1.40E+07 | 1.24E+07 | | | | | | |
| O08795 | PRKCSH | protein kinase C substrate 80K-H | Cytoplasm | enzyme | 8444385 | 1.11E+07 | 1.16E+07 | | | | | | |
| O08795 | PRKCSH | protein kinase C substrate 80K-H | Cytoplasm | enzyme | 8571569 | 1.17E+07 | 9712350 | | | | | | |
| O08795 | PRKCSH | protein kinase C substrate 80K-H | Cytoplasm | enzyme | 6412826 | 8002196 | 7190521 | | | | | | |
| O08795 | PRKCSH | protein kinase C substrate 80K-H | Cytoplasm | enzyme | 4982049 | 7101073 | 3727389 | | | | | | |
| O08795 | PRKCSH | protein kinase C substrate 80K-H | Cytoplasm | enzyme | 6139311.5 | 6443815.5 | 5354659 | | | | | | |
| O08795 | PRKCSH | protein kinase C substrate 80K-H | Cytoplasm | enzyme | 5364025 | 6075089.5 | 5405031.5 | | | | | | |
| O08795 | PRKCSH | protein kinase C substrate 80K-H | Cytoplasm | enzyme | 4137916 | 7172877 | 4739515.5 | | | | | | |
| O08795 | PRKCSH | protein kinase C substrate 80K-H | Cytoplasm | enzyme | 3111438.25 | 4893980 | 2821006.75 | | | | | | |
| O08795 | PRKCSH | protein kinase C substrate 80K-H | Cytoplasm | enzyme | 3949804.75 | 4134760 | 2372085.75 | | | | | | |
| O08795 | PRKCSH | protein kinase C substrate 80K-H | Cytoplasm | enzyme | 1515423.875 | 970914.25 | 407888.781 | O08795 | PRKCSH | 5854432 | 3474134 | 604769 | 33 |
| Q8CIG8 | PRMT5 | protein arginine methyltransferas | Cytoplasm | enzyme | 3903779.75 | 5475551.5 | 2921040.5 | | | | | | |
| Q8CIG8 | PRMT5 | protein arginine methyltransferas | Cytoplasm | enzyme | 1090564.25 | 1167929.625 | 498324.844 | Q8CIG8 | PRMT5 | 2497172 | 1937465 | 790967 | 6 |
| P04925 | PRNP | prion protein | Plasma Membrane | other | 1.47E+07 | 1.44E+07 | 1.82E+07 | P04925 | PRNP | 14680000 | 2104978 | 1215310 | 3 |
| Q9D7G0 | PRPS1 | phosphoribosyl pyrophosphate sy | Cytoplasm | kinase | 7318023 | 8479850 | 4753001.5 | | | | | | |
| Q9D7G0 | PRPS1 | phosphoribosyl pyrophosphate sy | Cytoplasm | kinase | 7628619 | 6882611.5 | 4921057.5 | | | | | | |
| Q9D7G0 | PRPS1 | phosphoribosyl pyrophosphate sy | Cytoplasm | kinase | 3792092.75 | 4723626 | 3014143.75 | | | | | | |
| Q9D7G0 | PRPS1 | phosphoribosyl pyrophosphate sy | Cytoplasm | kinase | 5017204 | 4776063 | 3525430.75 | | | | | | |
| Q9D7G0 | PRPS1 | phosphoribosyl pyrophosphate sy | Cytoplasm | kinase | 2668094 | 2938203.25 | 2028765.375 | | | | | | |
| Q9D7G0 | PRPS1 | phosphoribosyl pyrophosphate sy | Cytoplasm | kinase | 2260736.25 | 3664456.75 | 2316220.75 | Q9D7G0 | PRPS1 | 4780795 | 1969144 | 464132 | 18 |
| Q9D0M1 | PRPSAP1 | phosphoribosyl pyrophosphate sy | Extracellular Space | other | 5894989 | 5053140 | 5514191.5 | | | | | | |
| Q9D0M1 | PRPSAP1 | phosphoribosyl pyrophosphate sy | Extracellular Space | other | 3557149 | 2787629.5 | 2164900 | | | | | | |
| Q9D0M1 | PRPSAP1 | phosphoribosyl pyrophosphate sy | Extracellular Space | other | 2417834.5 | 2585287.25 | 1680484.125 | | | | | | |
| Q9D0M1 | PRPSAP1 | phosphoribosyl pyrophosphate sy | Extracellular Space | other | 2656913.5 | 2647564 | 1959121.875 | | | | | | |
| Q9D0M1 | PRPSAP1 | phosphoribosyl pyrophosphate sy | Extracellular Space | other | 1604872.75 | 1751611.625 | 1154599.375 | Q9D0M1 | PRPSAP1 | 3226352 | 1471002 | 379811 | 15 |
| Q8R574 | PRPSAP2 | phosphoribosyl pyrophosphate sy | Cytoplasm | other | 2461619.75 | 1774981.625 | 1920905.625 | | | | | | |
| Q8R574 | PRPSAP2 | phosphoribosyl pyrophosphate sy | Cytoplasm | other | 3477231.75 | 3583333 | 2832749.5 | Q8R574 | PRPSAP2 | 2969426 | 763905 | 311863 | 6 |
| Q61207 | PSAP | prosaposin | Extracellular Space | other | 4561437 | 4560932 | 5852375 | Q61207 | PSAP | 4561437 | 745469 | 430397 | 3 |
| P49769 | PSEN1 | presenilin 1 | Plasma Membrane | peptidase | 3316067.25 | 3683090.75 | 5149223.5 | | | | | | |
| P49769 | PSEN1 | presenilin 1 | Plasma Membrane | peptidase | 1307447.75 | 2602542.25 | 1443844 | P49769 | PSEN1 | 2311758 | 1455246 | 594102 | 6 |
| Q61144 | PSEN2 | presenilin 2 | Cytoplasm | peptidase | 1596527.25 | 1680302.25 | 1010088.625 | Q61144 | PSEN2 | 1596527 | 365175 | 210834 | 3 |
| Q91YA2 | PSKH1 | protein serine kinase H1 | Nucleus | kinase | 5211059 | 2266722.25 | 3199305.25 | Q91YA2 | PSKH1 | 5211059 | 1504769 | 868779 | 3 |
| Q9R1P4 | PSMA1 | proteasome (prosome, macropain | Cytoplasm | peptidase | 3.64E+07 | 3.59E+07 | 4.41E+07 | | | | | | |
| Q9R1P4 | PSMA1 | proteasome (prosome, macropain | Cytoplasm | peptidase | 2.54E+07 | 2.07E+07 | 7374712 | | | | | | |
| Q9R1P4 | PSMA1 | proteasome (prosome, macropain | Cytoplasm | peptidase | 2.08E+07 | 2.16E+07 | 1.93E+07 | | | | | | |
| Q9R1P4 | PSMA1 | proteasome (prosome, macropain | Cytoplasm | peptidase | 1.69E+07 | 2.96E+07 | 2.07E+07 | | | | | | |
| Q9R1P4 | PSMA1 | proteasome (prosome, macropain | Cytoplasm | peptidase | 2.05E+07 | 1.81E+07 | 1.77E+07 | | | | | | |
| Q9R1P4 | PSMA1 | proteasome (prosome, macropain | Cytoplasm | peptidase | 3.24E+07 | 8304655 | 1.57E+07 | | | | | | |
| Q9R1P4 | PSMA1 | proteasome (prosome, macropain | Cytoplasm | peptidase | 8809058 | 2.57E+07 | 1.48E+07 | | | | | | |
| Q9R1P4 | PSMA1 | proteasome (prosome, macropain | Cytoplasm | peptidase | 1.40E+07 | 1.39E+07 | 1.49E+07 | | | | | | |
| Q9R1P4 | PSMA1 | proteasome (prosome, macropain | Cytoplasm | peptidase | 1.30E+07 | 1.44E+07 | 1.24E+07 | | | | | | |
| Q9R1P4 | PSMA1 | proteasome (prosome, macropain | Cytoplasm | peptidase | 1.41E+07 | 1.40E+07 | 1.46E+07 | | | | | | |
| Q9R1P4 | PSMA1 | proteasome (prosome, macropain | Cytoplasm | peptidase | 1.21E+07 | 1.07E+07 | 9621609 | | | | | | |
| Q9R1P4 | PSMA1 | proteasome (prosome, macropain | Cytoplasm | peptidase | 5520356.5 | 5825765 | 4859018 | | | | | | |
| Q9R1P4 | PSMA1 | proteasome (prosome, macropain | Cytoplasm | peptidase | 4157086.25 | 3950909.25 | 2754880.5 | | | | | | |
| Q9R1P4 | PSMA1 | proteasome (prosome, macropain | Cytoplasm | peptidase | 3767391.25 | 2867870.75 | 2430373.25 | | | | | | |
| Q9R1P4 | PSMA1 | proteasome (prosome, macropain | Cytoplasm | peptidase | 4519437 | 805471.812 | 1974512.375 | | | | | | |
| Q9R1P4 | PSMA1 | proteasome (prosome, macropain | Cytoplasm | peptidase | 2866042.5 | 9095783 | 1.21E+07 | Q9R1P4 | PSMA1 | 14701836 | 9940799 | 1434831 | 48 |
| P49722 | PSMA2 | proteasome (prosome, macropain | Cytoplasm | peptidase | 6.56E+07 | 6.79E+07 | 5.15E+07 | | | | | | |
| P49722 | PSMA2 | proteasome (prosome, macropain | Cytoplasm | peptidase | 4.28E+07 | 2.17E+07 | 1.86E+07 | | | | | | |
| P49722 | PSMA2 | proteasome (prosome, macropain | Cytoplasm | peptidase | 2.05E+07 | 1.29E+07 | 1.29E+07 | | | | | | |
| P49722 | PSMA2 | proteasome (prosome, macropain | Cytoplasm | peptidase | 2.14E+07 | 2.13E+07 | 2.19E+07 | | | | | | |
| P49722 | PSMA2 | proteasome (prosome, macropain | Cytoplasm | peptidase | 9584571 | 1.17E+07 | 1.35E+07 | | | | | | |
| P49722 | PSMA2 | proteasome (prosome, macropain | Cytoplasm | peptidase | 1.09E+07 | 1135729.625 | 4663014 | | | | | | |
| P49722 | PSMA2 | proteasome (prosome, macropain | Cytoplasm | peptidase | 4526637 | 4252484.5 | 1961705.5 | | | | | | |

| | | | | | | | | | | | | | |
|---|---|---|---|---|---|---|---|---|---|---|---|---|---|
| P49722 | PSMA2 | proteasome (prosome, macropain | Cytoplasm | peptidase | 2892154.75 | 6372809.5 | 4981022.5 | | | | | | |
| P49722 | PSMA2 | proteasome (prosome, macropain | Cytoplasm | peptidase | 3300374 | 3780916.25 | 5322273 | | | | | | |
| P49722 | PSMA2 | proteasome (prosome, macropain | Cytoplasm | peptidase | 2281631.75 | 2164740.5 | 2325357.25 | P49722 | PSMA2 | 18370537 | 18174560 | 3318206 | 30 |
| O70435 | PSMA3 | proteasome (prosome, macropain | Cytoplasm | peptidase | 3.02E+07 | 2.65E+07 | 2.25E+07 | | | | | | |
| O70435 | PSMA3 | proteasome (prosome, macropain | Cytoplasm | peptidase | 2.08E+07 | 2.45E+07 | 2.23E+07 | | | | | | |
| O70435 | PSMA3 | proteasome (prosome, macropain | Cytoplasm | peptidase | 2.32E+07 | 2.15E+07 | 1.98E+07 | | | | | | |
| O70435 | PSMA3 | proteasome (prosome, macropain | Cytoplasm | peptidase | 1.39E+07 | 1.37E+07 | 1.32E+07 | | | | | | |
| O70435 | PSMA3 | proteasome (prosome, macropain | Cytoplasm | peptidase | 1.50E+07 | 1.47E+07 | 1.21E+07 | | | | | | |
| O70435 | PSMA3 | proteasome (prosome, macropain | Cytoplasm | peptidase | 1.38E+07 | 1.53E+07 | 1.49E+07 | | | | | | |
| O70435 | PSMA3 | proteasome (prosome, macropain | Cytoplasm | peptidase | 1.18E+07 | 1.15E+07 | 6103385 | | | | | | |
| O70435 | PSMA3 | proteasome (prosome, macropain | Cytoplasm | peptidase | 1.50E+07 | 2746369 | 5939080.5 | | | | | | |
| O70435 | PSMA3 | proteasome (prosome, macropain | Cytoplasm | peptidase | 9219431 | 1754711 | 4576323 | O70435 | PSMA3 | 16972159 | 7323852 | 1409476 | 27 |
| Q9R1P0 | PSMA4 | proteasome (prosome, macropain | Cytoplasm | peptidase | 2.61E+07 | 2.97E+07 | 2.53E+07 | | | | | | |
| Q9R1P0 | PSMA4 | proteasome (prosome, macropain | Cytoplasm | peptidase | 3.51E+07 | 3.32E+07 | 3.09E+07 | | | | | | |
| Q9R1P0 | PSMA4 | proteasome (prosome, macropain | Cytoplasm | peptidase | 3.59E+07 | 3.60E+07 | 3.32E+07 | | | | | | |
| Q9R1P0 | PSMA4 | proteasome (prosome, macropain | Cytoplasm | peptidase | 2.14E+07 | 2.44E+07 | 2.49E+07 | | | | | | |
| Q9R1P0 | PSMA4 | proteasome (prosome, macropain | Cytoplasm | peptidase | 2.14E+07 | 1.93E+07 | 2.09E+07 | | | | | | |
| Q9R1P0 | PSMA4 | proteasome (prosome, macropain | Cytoplasm | peptidase | 2.07E+07 | 1.58E+07 | 1.62E+07 | | | | | | |
| Q9R1P0 | PSMA4 | proteasome (prosome, macropain | Cytoplasm | peptidase | 1.46E+07 | 1.54E+07 | 1.31E+07 | | | | | | |
| Q9R1P0 | PSMA4 | proteasome (prosome, macropain | Cytoplasm | peptidase | 1.15E+07 | 1.14E+07 | 9476473 | | | | | | |
| Q9R1P0 | PSMA4 | proteasome (prosome, macropain | Cytoplasm | peptidase | 8722942 | 7491961 | 7062812.5 | | | | | | |
| Q9R1P0 | PSMA4 | proteasome (prosome, macropain | Cytoplasm | peptidase | 9177402 | 9644552 | 7744987.5 | | | | | | |
| Q9R1P0 | PSMA4 | proteasome (prosome, macropain | Cytoplasm | peptidase | 3904495 | 3778859.75 | 4796731.5 | | | | | | |
| Q9R1P0 | PSMA4 | proteasome (prosome, macropain | Cytoplasm | peptidase | 3124220.25 | 2621375.5 | 1100480.75 | | | | | | |
| Q9R1P0 | PSMA4 | proteasome (prosome, macropain | Cytoplasm | peptidase | 763023.25 | 617588.438 | 696977 | Q9R1P0 | PSMA4 | 16334006 | 11190926 | 1791982 | 39 |
| Q9Z2U1 | PSMA5 | proteasome (prosome, macropain | Cytoplasm | peptidase | 9.22E+07 | 8.24E+07 | 7.21E+07 | | | | | | |
| Q9Z2U1 | PSMA5 | proteasome (prosome, macropain | Cytoplasm | peptidase | 6.13E+07 | 6.60E+07 | 5.55E+07 | | | | | | |
| Q9Z2U1 | PSMA5 | proteasome (prosome, macropain | Cytoplasm | peptidase | 6.13E+07 | 6.31E+07 | 5.42E+07 | | | | | | |
| Q9Z2U1 | PSMA5 | proteasome (prosome, macropain | Cytoplasm | peptidase | 3.37E+07 | 2.56E+07 | 2.68E+07 | | | | | | |
| Q9Z2U1 | PSMA5 | proteasome (prosome, macropain | Cytoplasm | peptidase | 1.75E+07 | 1.86E+07 | 1.70E+07 | | | | | | |
| Q9Z2U1 | PSMA5 | proteasome (prosome, macropain | Cytoplasm | peptidase | 1.83E+07 | 1.76E+07 | 1.53E+07 | | | | | | |
| Q9Z2U1 | PSMA5 | proteasome (prosome, macropain | Cytoplasm | peptidase | 1.04E+07 | 1.16E+07 | 1.11E+07 | | | | | | |
| Q9Z2U1 | PSMA5 | proteasome (prosome, macropain | Cytoplasm | peptidase | 9667567 | 9039809 | 7436641.5 | | | | | | |
| Q9Z2U1 | PSMA5 | proteasome (prosome, macropain | Cytoplasm | peptidase | 1.56E+07 | 821227.375 | 4963982.5 | | | | | | |
| Q9Z2U1 | PSMA5 | proteasome (prosome, macropain | Cytoplasm | peptidase | 3373451.25 | 7520858 | 2949328 | | | | | | |
| Q9Z2U1 | PSMA5 | proteasome (prosome, macropain | Cytoplasm | peptidase | 3653047.75 | 4203044 | 3914809 | | | | | | |
| Q9Z2U1 | PSMA5 | proteasome (prosome, macropain | Cytoplasm | peptidase | 5760725 | 4726052 | 3114502.25 | | | | | | |
| Q9Z2U1 | PSMA5 | proteasome (prosome, macropain | Cytoplasm | peptidase | 2775364.25 | 2936337 | 4011738.25 | | | | | | |
| Q9Z2U1 | PSMA5 | proteasome (prosome, macropain | Cytoplasm | peptidase | 2509397 | 2989961 | 1867394.5 | Q9Z2U1 | PSMA5 | 24147111 | 25642249 | 3956685 | 42 |
| Q9QUM9 | PSMA6 | proteasome (prosome, macropain | Cytoplasm | peptidase | 4.26E+07 | 4.08E+07 | 4.25E+07 | | | | | | |
| Q9QUM9 | PSMA6 | proteasome (prosome, macropain | Cytoplasm | peptidase | 3.63E+07 | 3.67E+07 | 3.99E+07 | | | | | | |
| Q9QUM9 | PSMA6 | proteasome (prosome, macropain | Cytoplasm | peptidase | 2.72E+07 | 2.78E+07 | 2.74E+07 | | | | | | |
| Q9QUM9 | PSMA6 | proteasome (prosome, macropain | Cytoplasm | peptidase | 2.79E+07 | 2.26E+07 | 1.88E+07 | | | | | | |
| Q9QUM9 | PSMA6 | proteasome (prosome, macropain | Cytoplasm | peptidase | 2.50E+07 | 2.21E+07 | 1.68E+07 | | | | | | |
| Q9QUM9 | PSMA6 | proteasome (prosome, macropain | Cytoplasm | peptidase | 1.77E+07 | 2.25E+07 | 1.80E+07 | | | | | | |
| Q9QUM9 | PSMA6 | proteasome (prosome, macropain | Cytoplasm | peptidase | 1.52E+07 | 1.45E+07 | 1.18E+07 | | | | | | |
| Q9QUM9 | PSMA6 | proteasome (prosome, macropain | Cytoplasm | peptidase | 1.55E+07 | 1.56E+07 | 1.27E+07 | | | | | | |
| Q9QUM9 | PSMA6 | proteasome (prosome, macropain | Cytoplasm | peptidase | 1.48E+07 | 1.55E+07 | 1.28E+07 | | | | | | |
| Q9QUM9 | PSMA6 | proteasome (prosome, macropain | Cytoplasm | peptidase | 1.22E+07 | 1.72E+07 | 1.49E+07 | | | | | | |
| Q9QUM9 | PSMA6 | proteasome (prosome, macropain | Cytoplasm | peptidase | 1959672.125 | 338115.906 | 134968.844 | Q9QUM9 | PSMA6 | 21475425 | 11419894 | 1987948 | 33 |
| Q9Z2U0 | PSMA7 | proteasome (prosome, macropain | Cytoplasm | peptidase | 8.89E+07 | 7.83E+07 | 7.68E+07 | | | | | | |
| Q9Z2U0 | PSMA7 | proteasome (prosome, macropain | Cytoplasm | peptidase | 4.54E+07 | 4.17E+07 | 4.33E+07 | | | | | | |
| Q9Z2U0 | PSMA7 | proteasome (prosome, macropain | Cytoplasm | peptidase | 3.87E+07 | 4.42E+07 | 3.72E+07 | | | | | | |
| Q9Z2U0 | PSMA7 | proteasome (prosome, macropain | Cytoplasm | peptidase | 2.94E+07 | 2.94E+07 | 2.36E+07 | | | | | | |
| Q9Z2U0 | PSMA7 | proteasome (prosome, macropain | Cytoplasm | peptidase | 3.21E+07 | 3.27E+07 | 3.07E+07 | | | | | | |
| Q9Z2U0 | PSMA7 | proteasome (prosome, macropain | Cytoplasm | peptidase | 1.87E+07 | 1.77E+07 | 1.20E+07 | | | | | | |
| Q9Z2U0 | PSMA7 | proteasome (prosome, macropain | Cytoplasm | peptidase | 9814885 | 8680476 | 6073528 | | | | | | |
| Q9Z2U0 | PSMA7 | proteasome (prosome, macropain | Cytoplasm | peptidase | 7682826 | 6489981.5 | 6370742.5 | | | | | | |
| Q9Z2U0 | PSMA7 | proteasome (prosome, macropain | Cytoplasm | peptidase | 6180433 | 5845164.5 | 4811553.5 | | | | | | |
| Q9Z2U0 | PSMA7 | proteasome (prosome, macropain | Cytoplasm | peptidase | 4374176.5 | 3353273.25 | 3036896.5 | | | | | | |
| Q9Z2U0 | PSMA7 | proteasome (prosome, macropain | Cytoplasm | peptidase | 4663125.5 | 6046772.5 | 5658799.5 | | | | | | |
| Q9Z2U0 | PSMA7 | proteasome (prosome, macropain | Cytoplasm | peptidase | 5946670.5 | 623584.812 | 478667.312 | Q9Z2U0 | PSMA7 | 24321843 | 23085100 | 3847517 | 36 |
| O09061 | PSMB1 | proteasome (prosome, macropain | Cytoplasm | peptidase | 4.31E+07 | 3.65E+07 | 3.32E+07 | | | | | | |
| O09061 | PSMB1 | proteasome (prosome, macropain | Cytoplasm | peptidase | 2.87E+07 | 2.94E+07 | 2.83E+07 | | | | | | |
| O09061 | PSMB1 | proteasome (prosome, macropain | Cytoplasm | peptidase | 2.83E+07 | 2.51E+07 | 2.31E+07 | | | | | | |
| O09061 | PSMB1 | proteasome (prosome, macropain | Cytoplasm | peptidase | 2.29E+07 | 1.40E+07 | 1.97E+07 | | | | | | |
| O09061 | PSMB1 | proteasome (prosome, macropain | Cytoplasm | peptidase | 1.64E+07 | 1.46E+07 | 1.28E+07 | | | | | | |
| O09061 | PSMB1 | proteasome (prosome, macropain | Cytoplasm | peptidase | 7839670 | 6219355 | 5437788.5 | | | | | | |
| O09061 | PSMB1 | proteasome (prosome, macropain | Cytoplasm | peptidase | 6191459 | 4959544 | 5311958 | | | | | | |
| O09061 | PSMB1 | proteasome (prosome, macropain | Cytoplasm | peptidase | 3934491.25 | 8065251.5 | 4824590.5 | | | | | | |
| O09061 | PSMB1 | proteasome (prosome, macropain | Cytoplasm | peptidase | 3660148.25 | 8350831 | 6090473 | | | | | | |
| O09061 | PSMB1 | proteasome (prosome, macropain | Cytoplasm | peptidase | 5691708.5 | 2573065 | 4072799.75 | | | | | | |
| O09061 | PSMB1 | proteasome (prosome, macropain | Cytoplasm | peptidase | 4386774.5 | 3873621.75 | 3996130 | | | | | | |
| O09061 | PSMB1 | proteasome (prosome, macropain | Cytoplasm | peptidase | 3499711.25 | 2604507.5 | 3579762 | | | | | | |
| O09061 | PSMB1 | proteasome (prosome, macropain | Cytoplasm | peptidase | 2717378.25 | 4381530 | 5055990.5 | | | | | | |
| O09061 | PSMB1 | proteasome (prosome, macropain | Cytoplasm | peptidase | 3286445 | 3228591 | 3893790.25 | | | | | | |
| O09061 | PSMB1 | proteasome (prosome, macropain | Cytoplasm | peptidase | 2163568.5 | 3728876.5 | 3752911.75 | | | | | | |
| O09061 | PSMB1 | proteasome (prosome, macropain | Cytoplasm | peptidase | 1763111.875 | 2199093.75 | 2414234 | | | | | | |
| O09061 | PSMB1 | proteasome (prosome, macropain | Cytoplasm | peptidase | 1020499.938 | 911351.562 | 606418.062 | O09061 | PSMB1 | 10914410 | 10829740 | 1516467 | 51 |
| Q9R1P3 | PSMB2 | proteasome (prosome, macropain | Cytoplasm | peptidase | 2.94E+07 | 3.86E+07 | 3.91E+07 | | | | | | |
| Q9R1P3 | PSMB2 | proteasome (prosome, macropain | Cytoplasm | peptidase | 2.44E+07 | 2.49E+07 | 2.34E+07 | | | | | | |
| Q9R1P3 | PSMB2 | proteasome (prosome, macropain | Cytoplasm | peptidase | 1.25E+07 | 1.17E+07 | 6220684.5 | | | | | | |
| Q9R1P3 | PSMB2 | proteasome (prosome, macropain | Cytoplasm | peptidase | 5146796.5 | 7483313.5 | 5749031.5 | | | | | | |
| Q9R1P3 | PSMB2 | proteasome (prosome, macropain | Cytoplasm | peptidase | 5597371.5 | 7632168 | 4660578.5 | | | | | | |
| Q9R1P3 | PSMB2 | proteasome (prosome, macropain | Cytoplasm | peptidase | 7574465.5 | 5880441 | 7660169.5 | | | | | | |
| Q9R1P3 | PSMB2 | proteasome (prosome, macropain | Cytoplasm | peptidase | 4368924 | 4538930.5 | 4096007.25 | | | | | | |

| | | | | | | | | | | | | | |
|---|---|---|---|---|---|---|---|---|---|---|---|---|---|
| Q9R1P3 | PSMB2 | proteasome (prosome, macropain | Cytoplasm | peptidase | 2835689.25 | 2338877.75 | 1721526.625 | | | | | | |
| Q9R1P3 | PSMB2 | proteasome (prosome, macropain | Cytoplasm | peptidase | 5824336.5 | 1692092 | 3237970.75 | Q9R1P3 | PSMB2 | 10841954 | 11055365 | 2127606 | 27 |
| Q9R1P1 | PSMB3 | proteasome (prosome, macropain | Cytoplasm | peptidase | 3.19E+07 | 3.10E+07 | 3.13E+07 | | | | | | |
| Q9R1P1 | PSMB3 | proteasome (prosome, macropain | Cytoplasm | peptidase | 2.29E+07 | 1.54E+07 | 1.95E+07 | | | | | | |
| Q9R1P1 | PSMB3 | proteasome (prosome, macropain | Cytoplasm | peptidase | 1.21E+07 | 1.21E+07 | 1.29E+07 | | | | | | |
| Q9R1P1 | PSMB3 | proteasome (prosome, macropain | Cytoplasm | peptidase | 1.45E+07 | 1.22E+07 | 1.21E+07 | | | | | | |
| Q9R1P1 | PSMB3 | proteasome (prosome, macropain | Cytoplasm | peptidase | 6215988.5 | 5891463 | 5317113 | | | | | | |
| Q9R1P1 | PSMB3 | proteasome (prosome, macropain | Cytoplasm | peptidase | 5389511.5 | 5031746.5 | 2771633.5 | | | | | | |
| Q9R1P1 | PSMB3 | proteasome (prosome, macropain | Cytoplasm | peptidase | 8036602 | 6336811 | 9824206 | | | | | | |
| Q9R1P1 | PSMB3 | proteasome (prosome, macropain | Cytoplasm | peptidase | 2992518.25 | 5675143.5 | 3318784.25 | | | | | | |
| Q9R1P1 | PSMB3 | proteasome (prosome, macropain | Cytoplasm | peptidase | 2455117.25 | 4057319.5 | 5107750.5 | | | | | | |
| Q9R1P1 | PSMB3 | proteasome (prosome, macropain | Cytoplasm | peptidase | 4.70E-06 | 1308418 | 1200647.875 | Q9R1P1 | PSMB3 | 10644974 | 9008415 | 1644704 | 30 |
| P99026 | PSMB4 | proteasome (prosome, macropain | Cytoplasm | peptidase | 2.13E+07 | 2.15E+07 | 1.95E+07 | | | | | | |
| P99026 | PSMB4 | proteasome (prosome, macropain | Cytoplasm | peptidase | 2.47E+07 | 2.07E+07 | 1.90E+07 | | | | | | |
| P99026 | PSMB4 | proteasome (prosome, macropain | Cytoplasm | peptidase | 2.36E+07 | 1.99E+07 | 1.79E+07 | | | | | | |
| P99026 | PSMB4 | proteasome (prosome, macropain | Cytoplasm | peptidase | 1.90E+07 | 2.15E+07 | 1.85E+07 | | | | | | |
| P99026 | PSMB4 | proteasome (prosome, macropain | Cytoplasm | peptidase | 1.89E+07 | 1.98E+07 | 1.68E+07 | | | | | | |
| P99026 | PSMB4 | proteasome (prosome, macropain | Cytoplasm | peptidase | 5667664.5 | 5728177 | 5364631.5 | P99026 | PSMB4 | 18846277 | 5904475 | 1391698 | 18 |
| O55234 | PSMB5 | proteasome (prosome, macropain | Cytoplasm | peptidase | 4.00E+07 | 2.91E+07 | 3.06E+07 | | | | | | |
| O55234 | PSMB5 | proteasome (prosome, macropain | Cytoplasm | peptidase | 2.44E+07 | 2.48E+07 | 2.07E+07 | | | | | | |
| O55234 | PSMB5 | proteasome (prosome, macropain | Cytoplasm | peptidase | 2.10E+07 | 2.53E+07 | 2.18E+07 | | | | | | |
| O55234 | PSMB5 | proteasome (prosome, macropain | Cytoplasm | peptidase | 2.28E+07 | 2.11E+07 | 2.01E+07 | | | | | | |
| O55234 | PSMB5 | proteasome (prosome, macropain | Cytoplasm | peptidase | 1.82E+07 | 2.00E+07 | 1.98E+07 | | | | | | |
| O55234 | PSMB5 | proteasome (prosome, macropain | Cytoplasm | peptidase | 1.07E+07 | 1.18E+07 | 1.03E+07 | | | | | | |
| O55234 | PSMB5 | proteasome (prosome, macropain | Cytoplasm | peptidase | 7844548 | 8055854.5 | 6751793 | | | | | | |
| O55234 | PSMB5 | proteasome (prosome, macropain | Cytoplasm | peptidase | 7745006 | 8096853.5 | 7675744.5 | | | | | | |
| O55234 | PSMB5 | proteasome (prosome, macropain | Cytoplasm | peptidase | 6331302.5 | 7167517 | 5713553 | | | | | | |
| O55234 | PSMB5 | proteasome (prosome, macropain | Cytoplasm | peptidase | 3437841 | 4117061.75 | 2906564 | | | | | | |
| O55234 | PSMB5 | proteasome (prosome, macropain | Cytoplasm | peptidase | 1631595.625 | 2025647 | 1666574.25 | O55234 | PSMB5 | 14915481 | 9936927 | 1729797 | 33 |
| Q60692 | PSMB6 | proteasome (prosome, macropain | Nucleus | peptidase | 9.67E+07 | 9.27E+07 | 9.51E+07 | | | | | | |
| Q60692 | PSMB6 | proteasome (prosome, macropain | Nucleus | peptidase | 2.99E+07 | 3.04E+07 | 2.38E+07 | | | | | | |
| Q60692 | PSMB6 | proteasome (prosome, macropain | Nucleus | peptidase | 2.40E+07 | 2.44E+07 | 2.11E+07 | | | | | | |
| Q60692 | PSMB6 | proteasome (prosome, macropain | Nucleus | peptidase | 1.50E+07 | 1.19E+07 | 1.18E+07 | | | | | | |
| Q60692 | PSMB6 | proteasome (prosome, macropain | Nucleus | peptidase | 3136122 | 5090112 | 4418446.5 | | | | | | |
| Q60692 | PSMB6 | proteasome (prosome, macropain | Nucleus | peptidase | 2625882 | 1241745.5 | 534758.625 | Q60692 | PSMB6 | 28563667 | 32550379 | 7672198 | 18 |
| P70195 | PSMB7 | proteasome (prosome, macropain | Cytoplasm | peptidase | 2.65E+07 | 3.76E+07 | 2.65E+07 | | | | | | |
| P70195 | PSMB7 | proteasome (prosome, macropain | Cytoplasm | peptidase | 2.14E+07 | 2.02E+07 | 1.72E+07 | | | | | | |
| P70195 | PSMB7 | proteasome (prosome, macropain | Cytoplasm | peptidase | 1.83E+07 | 2.11E+07 | 1.99E+07 | | | | | | |
| P70195 | PSMB7 | proteasome (prosome, macropain | Cytoplasm | peptidase | 1.35E+07 | 1.65E+07 | 1.44E+07 | | | | | | |
| P70195 | PSMB7 | proteasome (prosome, macropain | Cytoplasm | peptidase | 1.39E+07 | 1.43E+07 | 1.65E+07 | | | | | | |
| P70195 | PSMB7 | proteasome (prosome, macropain | Cytoplasm | peptidase | 9986811 | 9828636 | 8020658 | | | | | | |
| P70195 | PSMB7 | proteasome (prosome, macropain | Cytoplasm | peptidase | 4495258 | 4732943.5 | 4706415 | | | | | | |
| P70195 | PSMB7 | proteasome (prosome, macropain | Cytoplasm | peptidase | 796211.062 | 1517773.625 | 1306831 | P70195 | PSMB7 | 13601035 | 9113879 | 1860363 | 24 |
| P28076 | PSMB9 | proteasome (prosome, macropain | Cytoplasm | peptidase | 905486.875 | 540917.25 | 1147362.125 | P28076 | PSMB9 | 905487 | 305284 | 176256 | 3 |
| P62192 | PSMC1 | proteasome (prosome, macropain | Nucleus | peptidase | 2.96E+07 | 2.99E+07 | 2.47E+07 | | | | | | |
| P62192 | PSMC1 | proteasome (prosome, macropain | Nucleus | peptidase | 1.47E+07 | 1.55E+07 | 1.23E+07 | | | | | | |
| P62192 | PSMC1 | proteasome (prosome, macropain | Nucleus | peptidase | 1.22E+07 | 1.53E+07 | 9959482 | | | | | | |
| P62192 | PSMC1 | proteasome (prosome, macropain | Nucleus | peptidase | 1.41E+07 | 1.28E+07 | 8665470 | | | | | | |
| P62192 | PSMC1 | proteasome (prosome, macropain | Nucleus | peptidase | 8439267 | 1.11E+07 | 6972917 | | | | | | |
| P62192 | PSMC1 | proteasome (prosome, macropain | Nucleus | peptidase | 7686671 | 1.10E+07 | 8941136 | | | | | | |
| P62192 | PSMC1 | proteasome (prosome, macropain | Nucleus | peptidase | 6404131 | 1.27E+07 | 8385115.5 | | | | | | |
| P62192 | PSMC1 | proteasome (prosome, macropain | Nucleus | peptidase | 7442788.5 | 9453191 | 6754419 | | | | | | |
| P62192 | PSMC1 | proteasome (prosome, macropain | Nucleus | peptidase | 6059722 | 6431276 | 4957547 | | | | | | |
| P62192 | PSMC1 | proteasome (prosome, macropain | Nucleus | peptidase | 3154440.5 | 9001895 | 3762703.5 | | | | | | |
| P62192 | PSMC1 | proteasome (prosome, macropain | Nucleus | peptidase | 4420745 | 6135511 | 4085703.75 | | | | | | |
| P62192 | PSMC1 | proteasome (prosome, macropain | Nucleus | peptidase | 2657421 | 6645007 | 5582212.5 | | | | | | |
| P62192 | PSMC1 | proteasome (prosome, macropain | Nucleus | peptidase | 1339745.625 | 2945494.5 | 2852880 | P62192 | PSMC1 | 9091918 | 6582926 | 1054112 | 39 |
| P46471 | PSMC2 | proteasome (prosome, macropain | Nucleus | peptidase | 3.38E+07 | 3.17E+07 | 1.63E+07 | | | | | | |
| P46471 | PSMC2 | proteasome (prosome, macropain | Nucleus | peptidase | 1.85E+07 | 1.97E+07 | 1.61E+07 | | | | | | |
| P46471 | PSMC2 | proteasome (prosome, macropain | Nucleus | peptidase | 1.71E+07 | 1.60E+07 | 1.17E+07 | | | | | | |
| P46471 | PSMC2 | proteasome (prosome, macropain | Nucleus | peptidase | 1.22E+07 | 1.39E+07 | 8869352 | | | | | | |
| P46471 | PSMC2 | proteasome (prosome, macropain | Nucleus | peptidase | 1.22E+07 | 1.03E+07 | 8332527 | | | | | | |
| P46471 | PSMC2 | proteasome (prosome, macropain | Nucleus | peptidase | 1.06E+07 | 1.11E+07 | 7825580.5 | | | | | | |
| P46471 | PSMC2 | proteasome (prosome, macropain | Nucleus | peptidase | 9896000 | 1.09E+07 | 7223688 | | | | | | |
| P46471 | PSMC2 | proteasome (prosome, macropain | Nucleus | peptidase | 1.03E+07 | 1.29E+07 | 6894435.5 | | | | | | |
| P46471 | PSMC2 | proteasome (prosome, macropain | Nucleus | peptidase | 8584188 | 1.27E+07 | 8907714 | | | | | | |
| P46471 | PSMC2 | proteasome (prosome, macropain | Nucleus | peptidase | 7843156.5 | 9144955 | 7256562 | | | | | | |
| P46471 | PSMC2 | proteasome (prosome, macropain | Nucleus | peptidase | 1.15E+07 | 1.17E+07 | 8942036 | | | | | | |
| P46471 | PSMC2 | proteasome (prosome, macropain | Nucleus | peptidase | 5058978 | 5723317.5 | 5229892 | | | | | | |
| P46471 | PSMC2 | proteasome (prosome, macropain | Nucleus | peptidase | 3827420.25 | 5536113.5 | 4342032.5 | | | | | | |
| P46471 | PSMC2 | proteasome (prosome, macropain | Nucleus | peptidase | 3169380 | 3290610.75 | 2143498 | | | | | | |
| P46471 | PSMC2 | proteasome (prosome, macropain | Nucleus | peptidase | 1655268 | 6324498 | 2853450.5 | P46471 | PSMC2 | 11074293 | 6558628 | 977703 | 45 |
| O88685 | PSMC3 | proteasome (prosome, macropain | Nucleus | transcription regulator | 1.90E+07 | 2.60E+07 | 1.52E+07 | | | | | | |
| O88685 | PSMC3 | proteasome (prosome, macropain | Nucleus | transcription regulator | 1.52E+07 | 1.62E+07 | 1.36E+07 | | | | | | |
| O88685 | PSMC3 | proteasome (prosome, macropain | Nucleus | transcription regulator | 1.09E+07 | 1.73E+07 | 1.17E+07 | | | | | | |
| O88685 | PSMC3 | proteasome (prosome, macropain | Nucleus | transcription regulator | 1.09E+07 | 1.51E+07 | 9713411 | | | | | | |
| O88685 | PSMC3 | proteasome (prosome, macropain | Nucleus | transcription regulator | 1.05E+07 | 1.17E+07 | 7961133 | | | | | | |
| O88685 | PSMC3 | proteasome (prosome, macropain | Nucleus | transcription regulator | 9851996 | 9110677 | 5914562 | | | | | | |
| O88685 | PSMC3 | proteasome (prosome, macropain | Nucleus | transcription regulator | 1.46E+07 | 7633810 | 9363749 | | | | | | |
| O88685 | PSMC3 | proteasome (prosome, macropain | Nucleus | transcription regulator | 8655890 | 1.06E+07 | 7012089 | | | | | | |
| O88685 | PSMC3 | proteasome (prosome, macropain | Nucleus | transcription regulator | 5319262 | 7543764.5 | 4878764.5 | | | | | | |
| O88685 | PSMC3 | proteasome (prosome, macropain | Nucleus | transcription regulator | 3536934.5 | 6002291.5 | 5037355.5 | | | | | | |
| O88685 | PSMC3 | proteasome (prosome, macropain | Nucleus | transcription regulator | 3263827.25 | 5157950 | 3533931.25 | | | | | | |
| O88685 | PSMC3 | proteasome (prosome, macropain | Nucleus | transcription regulator | 3370245.5 | 4355963.5 | 3620546 | | | | | | |
| O88685 | PSMC3 | proteasome (prosome, macropain | Nucleus | transcription regulator | 2952500.25 | 3120184.5 | 2458335.5 | | | | | | |
| O88685 | PSMC3 | proteasome (prosome, macropain | Nucleus | transcription regulator | 1472245.125 | 4228294 | 1893202.375 | O88685 | PSMC3 | 8538064 | 5428833 | 837687 | 42 |

| | | | | | | | | | | | | | |
|---|---|---|---|---|---|---|---|---|---|---|---|---|---|
| P54775 | PSMC4 | proteasome (prosome, macropain | Nucleus | peptidase | 1.28E+07 | 1.97E+07 | 1.30E+07 | | | | | | |
| P54775 | PSMC4 | proteasome (prosome, macropain | Nucleus | peptidase | 1.11E+07 | 1.60E+07 | 1.33E+07 | | | | | | |
| P54775 | PSMC4 | proteasome (prosome, macropain | Nucleus | peptidase | 5829641 | 6124979.5 | 5531197.5 | | | | | | |
| P54775 | PSMC4 | proteasome (prosome, macropain | Nucleus | peptidase | 7659420.5 | 7058191.5 | 3328722.75 | | | | | | |
| P54775 | PSMC4 | proteasome (prosome, macropain | Nucleus | peptidase | 4955922 | 6077719 | 4695547.5 | | | | | | |
| P54775 | PSMC4 | proteasome (prosome, macropain | Nucleus | peptidase | 9893621 | 6404326.5 | 6018943.5 | | | | | | |
| P54775 | PSMC4 | proteasome (prosome, macropain | Nucleus | peptidase | 3794176.25 | 6115323.5 | 4339731 | | | | | | |
| P54775 | PSMC4 | proteasome (prosome, macropain | Nucleus | peptidase | 3392136.25 | 2757156.5 | 1914040.375 | | | | | | |
| P54775 | PSMC4 | proteasome (prosome, macropain | Nucleus | peptidase | 3232398 | 3888896.75 | 1676196.5 | | | | | | |
| P54775 | PSMC4 | proteasome (prosome, macropain | Nucleus | peptidase | 2795288.75 | 3549578.5 | 4303090 | | | | | | |
| P54775 | PSMC4 | proteasome (prosome, macropain | Nucleus | peptidase | 2754676.75 | 4880156.5 | 3427110.5 | | | | | | |
| P54775 | PSMC4 | proteasome (prosome, macropain | Nucleus | peptidase | 2774332.25 | 2690015.25 | 1267647.875 | P54775 | PSMC4 | 5910134 | 4287490 | 714582 | 36 |
| P62196 | PSMC5 | proteasome (prosome, macropain | Nucleus | transcription regulator | 2.32E+07 | 1.78E+07 | 1.69E+07 | | | | | | |
| P62196 | PSMC5 | proteasome (prosome, macropain | Nucleus | transcription regulator | 1.23E+07 | 1.23E+07 | 9792425 | | | | | | |
| P62196 | PSMC5 | proteasome (prosome, macropain | Nucleus | transcription regulator | 8322648 | 1.36E+07 | 1.08E+07 | | | | | | |
| P62196 | PSMC5 | proteasome (prosome, macropain | Nucleus | transcription regulator | 1.22E+07 | 1.47E+07 | 1.02E+07 | | | | | | |
| P62196 | PSMC5 | proteasome (prosome, macropain | Nucleus | transcription regulator | 9996644 | 9599120 | 7035396 | | | | | | |
| P62196 | PSMC5 | proteasome (prosome, macropain | Nucleus | transcription regulator | 8482294 | 9437354 | 5707227 | | | | | | |
| P62196 | PSMC5 | proteasome (prosome, macropain | Nucleus | transcription regulator | 6872204.5 | 7820896.5 | 4560862.5 | | | | | | |
| P62196 | PSMC5 | proteasome (prosome, macropain | Nucleus | transcription regulator | 6991029.5 | 7194641.5 | 5222962 | | | | | | |
| P62196 | PSMC5 | proteasome (prosome, macropain | Nucleus | transcription regulator | 6473382.5 | 7540181 | 5856285 | | | | | | |
| P62196 | PSMC5 | proteasome (prosome, macropain | Nucleus | transcription regulator | 4839108 | 6254638.5 | 2591058.5 | | | | | | |
| P62196 | PSMC5 | proteasome (prosome, macropain | Nucleus | transcription regulator | 4945653.5 | 5426905.5 | 3003381 | | | | | | |
| P62196 | PSMC5 | proteasome (prosome, macropain | Nucleus | transcription regulator | 3745985.25 | 6556003 | 4821875 | | | | | | |
| P62196 | PSMC5 | proteasome (prosome, macropain | Nucleus | transcription regulator | 5910167.5 | 3068592.25 | 3198032 | | | | | | |
| P62196 | PSMC5 | proteasome (prosome, macropain | Nucleus | transcription regulator | 4008140.75 | 4837935.5 | 3594920.75 | | | | | | |
| P62196 | PSMC5 | proteasome (prosome, macropain | Nucleus | transcription regulator | 1948265.5 | 6046253.5 | 2947401.5 | | | | | | |
| P62196 | PSMC5 | proteasome (prosome, macropain | Nucleus | transcription regulator | 2873586.5 | 522448.594 | 67084.234 | P62196 | PSMC5 | 7690569 | 4602219 | 664273 | 48 |
| P62334 | PSMC6 | proteasome (prosome, macropain | Nucleus | peptidase | 1.64E+07 | 2.18E+07 | 1.82E+07 | | | | | | |
| P62334 | PSMC6 | proteasome (prosome, macropain | Nucleus | peptidase | 9819114 | 1.20E+07 | 7527020.5 | | | | | | |
| P62334 | PSMC6 | proteasome (prosome, macropain | Nucleus | peptidase | 8108283 | 1.61E+07 | 1.03E+07 | | | | | | |
| P62334 | PSMC6 | proteasome (prosome, macropain | Nucleus | peptidase | 6197759.5 | 6464698 | 5165878 | | | | | | |
| P62334 | PSMC6 | proteasome (prosome, macropain | Nucleus | peptidase | 1.30E+07 | 5794967 | 6987808.5 | | | | | | |
| P62334 | PSMC6 | proteasome (prosome, macropain | Nucleus | peptidase | 8132139.5 | 9475329 | 7134124 | | | | | | |
| P62334 | PSMC6 | proteasome (prosome, macropain | Nucleus | peptidase | 7024068 | 8050399.5 | 5705375.5 | | | | | | |
| P62334 | PSMC6 | proteasome (prosome, macropain | Nucleus | peptidase | 4799453.5 | 6984829 | 3348096.5 | | | | | | |
| P62334 | PSMC6 | proteasome (prosome, macropain | Nucleus | peptidase | 5598146 | 7341811 | 6492737.5 | | | | | | |
| P62334 | PSMC6 | proteasome (prosome, macropain | Nucleus | peptidase | 6942592.5 | 6423542.5 | 5086458.5 | | | | | | |
| P62334 | PSMC6 | proteasome (prosome, macropain | Nucleus | peptidase | 5538795 | 5081342.5 | 3346131.5 | | | | | | |
| P62334 | PSMC6 | proteasome (prosome, macropain | Nucleus | peptidase | 5065770 | 5619256.5 | 4395258.5 | | | | | | |
| P62334 | PSMC6 | proteasome (prosome, macropain | Nucleus | peptidase | 1712970.5 | 1558806.125 | 1132962.875 | P62334 | PSMC6 | 7564546 | 4435891 | 710311 | 39 |
| Q3TXS7 | PSMD1 | proteasome (prosome, macropain | Cytoplasm | other | 1.75E+07 | 2.53E+07 | 1.38E+07 | | | | | | |
| Q3TXS7 | PSMD1 | proteasome (prosome, macropain | Cytoplasm | other | 2.18E+07 | 2.94E+07 | 1.41E+07 | | | | | | |
| Q3TXS7 | PSMD1 | proteasome (prosome, macropain | Cytoplasm | other | 1.40E+07 | 1.20E+07 | 8313644.5 | | | | | | |
| Q3TXS7 | PSMD1 | proteasome (prosome, macropain | Cytoplasm | other | 1.80E+07 | 2.02E+07 | 1.39E+07 | | | | | | |
| Q3TXS7 | PSMD1 | proteasome (prosome, macropain | Cytoplasm | other | 1.13E+07 | 1.60E+07 | 9091176 | | | | | | |
| Q3TXS7 | PSMD1 | proteasome (prosome, macropain | Cytoplasm | other | 1.35E+07 | 1.31E+07 | 7943000.5 | | | | | | |
| Q3TXS7 | PSMD1 | proteasome (prosome, macropain | Cytoplasm | other | 1.09E+07 | 1.15E+07 | 7269756 | | | | | | |
| Q3TXS7 | PSMD1 | proteasome (prosome, macropain | Cytoplasm | other | 9615246 | 1.36E+07 | 7624642.5 | | | | | | |
| Q3TXS7 | PSMD1 | proteasome (prosome, macropain | Cytoplasm | other | 1.15E+07 | 1.88E+07 | 1.11E+07 | | | | | | |
| Q3TXS7 | PSMD1 | proteasome (prosome, macropain | Cytoplasm | other | 7773384.5 | 1.06E+07 | 9570897 | | | | | | |
| Q3TXS7 | PSMD1 | proteasome (prosome, macropain | Cytoplasm | other | 8100964.5 | 8568234 | 6902386 | | | | | | |
| Q3TXS7 | PSMD1 | proteasome (prosome, macropain | Cytoplasm | other | 9120112 | 1.19E+07 | 7427572.5 | | | | | | |
| Q3TXS7 | PSMD1 | proteasome (prosome, macropain | Cytoplasm | other | 8176147.5 | 9504313 | 7523599.5 | | | | | | |
| Q3TXS7 | PSMD1 | proteasome (prosome, macropain | Cytoplasm | other | 6457525.5 | 8633100 | 6304714.5 | | | | | | |
| Q3TXS7 | PSMD1 | proteasome (prosome, macropain | Cytoplasm | other | 5661967.5 | 1.04E+07 | 4317535 | | | | | | |
| Q3TXS7 | PSMD1 | proteasome (prosome, macropain | Cytoplasm | other | 5715079.5 | 6607787 | 4250827 | | | | | | |
| Q3TXS7 | PSMD1 | proteasome (prosome, macropain | Cytoplasm | other | 6185084.5 | 6131165.5 | 4452139.5 | | | | | | |
| Q3TXS7 | PSMD1 | proteasome (prosome, macropain | Cytoplasm | other | 4981158 | 6816358.5 | 4995307.5 | | | | | | |
| Q3TXS7 | PSMD1 | proteasome (prosome, macropain | Cytoplasm | other | 4371553.5 | 4188287.75 | 4002038.25 | | | | | | |
| Q3TXS7 | PSMD1 | proteasome (prosome, macropain | Cytoplasm | other | 5397527.5 | 2885737 | 1977163 | | | | | | |
| Q3TXS7 | PSMD1 | proteasome (prosome, macropain | Cytoplasm | other | 3593785 | 3344049.75 | 2166027.75 | | | | | | |
| Q3TXS7 | PSMD1 | proteasome (prosome, macropain | Cytoplasm | other | 3185587 | 4311424.5 | 2634785.25 | | | | | | |
| Q3TXS7 | PSMD1 | proteasome (prosome, macropain | Cytoplasm | other | 3246373.75 | 5795338 | 3442921.25 | | | | | | |
| Q3TXS7 | PSMD1 | proteasome (prosome, macropain | Cytoplasm | other | 2974335.25 | 4005339.75 | 3019915.75 | | | | | | |
| Q3TXS7 | PSMD1 | proteasome (prosome, macropain | Cytoplasm | other | 2992889.75 | 3617684.25 | 889455.812 | | | | | | |
| Q3TXS7 | PSMD1 | proteasome (prosome, macropain | Cytoplasm | other | 1547749.25 | 1469667.375 | 1228359.875 | | | | | | |
| Q3TXS7 | PSMD1 | proteasome (prosome, macropain | Cytoplasm | other | 1123651.625 | 3216905.5 | 2665900.5 | | | | | | |
| Q3TXS7 | PSMD1 | proteasome (prosome, macropain | Cytoplasm | other | 1266132.5 | 1824791.125 | 1711017.125 | Q3TXS7 | PSMD1 | 7856652 | 5706499 | 622630 | 84 |
| Q8BG32 | PSMD11 | proteasome (prosome, macropain | Cytoplasm | other | 1.44E+07 | 1.70E+07 | 1.30E+07 | | | | | | |
| Q8BG32 | PSMD11 | proteasome (prosome, macropain | Cytoplasm | other | 1.04E+07 | 1.27E+07 | 9609556 | | | | | | |
| Q8BG32 | PSMD11 | proteasome (prosome, macropain | Cytoplasm | other | 1.35E+07 | 1.74E+07 | 1.18E+07 | | | | | | |
| Q8BG32 | PSMD11 | proteasome (prosome, macropain | Cytoplasm | other | 8589427 | 1.27E+07 | 1.41E+07 | | | | | | |
| Q8BG32 | PSMD11 | proteasome (prosome, macropain | Cytoplasm | other | 1.07E+07 | 1.21E+07 | 1.08E+07 | | | | | | |
| Q8BG32 | PSMD11 | proteasome (prosome, macropain | Cytoplasm | other | 1.07E+07 | 9095783 | 6277708 | | | | | | |
| Q8BG32 | PSMD11 | proteasome (prosome, macropain | Cytoplasm | other | 6533279.5 | 5796872.5 | 3661007.75 | | | | | | |
| Q8BG32 | PSMD11 | proteasome (prosome, macropain | Cytoplasm | other | 5917362 | 1.31E+07 | 6501286 | | | | | | |
| Q8BG32 | PSMD11 | proteasome (prosome, macropain | Cytoplasm | other | 6038830 | 5777983.5 | 4505858.5 | | | | | | |
| Q8BG32 | PSMD11 | proteasome (prosome, macropain | Cytoplasm | other | 4223362 | 7830662.5 | 5782986.5 | | | | | | |
| Q8BG32 | PSMD11 | proteasome (prosome, macropain | Cytoplasm | other | 4113477.5 | 3909467.5 | 2595521.75 | | | | | | |
| Q8BG32 | PSMD11 | proteasome (prosome, macropain | Cytoplasm | other | 3298850.75 | 4097418.75 | 2491279.5 | | | | | | |
| Q8BG32 | PSMD11 | proteasome (prosome, macropain | Cytoplasm | other | 2540071 | 3554883.25 | 2231677.5 | | | | | | |
| Q8BG32 | PSMD11 | proteasome (prosome, macropain | Cytoplasm | other | 589279.625 | 2560735.25 | 2367664.5 | Q8BG32 | PSMD11 | 7253853 | 4513843 | 696501 | 42 |
| Q9D8W5 | PSMD12 | proteasome (prosome, macropain | Cytoplasm | other | 1.43E+07 | 2.02E+07 | 1.26E+07 | | | | | | |
| Q9D8W5 | PSMD12 | proteasome (prosome, macropain | Cytoplasm | other | 1.07E+07 | 1.14E+07 | 1.01E+07 | | | | | | |
| Q9D8W5 | PSMD12 | proteasome (prosome, macropain | Cytoplasm | other | 1.31E+07 | 1.32E+07 | 1.10E+07 | | | | | | |

| | | | | | | | | | | | | | |
|---|---|---|---|---|---|---|---|---|---|---|---|---|---|
| Q9D8W5 | PSMD12 | proteasome (prosome, macropain | Cytoplasm | other | 1.01E+07 | 8886690 | 7059185 | | | | | | |
| Q9D8W5 | PSMD12 | proteasome (prosome, macropain | Cytoplasm | other | 6833892 | 9782062 | 5369330.5 | | | | | | |
| Q9D8W5 | PSMD12 | proteasome (prosome, macropain | Cytoplasm | other | 4839993 | 8847658 | 6101766.5 | | | | | | |
| Q9D8W5 | PSMD12 | proteasome (prosome, macropain | Cytoplasm | other | 6904490.5 | 1.03E+07 | 4977354 | | | | | | |
| Q9D8W5 | PSMD12 | proteasome (prosome, macropain | Cytoplasm | other | 5123075.5 | 5717657 | 4262045 | | | | | | |
| Q9D8W5 | PSMD12 | proteasome (prosome, macropain | Cytoplasm | other | 7018240 | 7689337.5 | 3862586.5 | | | | | | |
| Q9D8W5 | PSMD12 | proteasome (prosome, macropain | Cytoplasm | other | 3364566 | 4586085.5 | 2239402 | | | | | | |
| Q9D8W5 | PSMD12 | proteasome (prosome, macropain | Cytoplasm | other | 4461445.5 | 3659390.25 | 2968284.75 | | | | | | |
| Q9D8W5 | PSMD12 | proteasome (prosome, macropain | Cytoplasm | other | 3379189.75 | 4787165 | 3215025 | | | | | | |
| Q9D8W5 | PSMD12 | proteasome (prosome, macropain | Cytoplasm | other | 1774317.875 | 3536251.25 | 1488533.875 | | | | | | |
| Q9D8W5 | PSMD12 | proteasome (prosome, macropain | Cytoplasm | other | 2478870.5 | 3232004.5 | 2085544.625 | | | | | | |
| Q9D8W5 | PSMD12 | proteasome (prosome, macropain | Cytoplasm | other | 1151189.375 | 1187361.25 | 749970.938 | Q9D8W5 | PSMD12 | 6365951 | 4250026 | 633556 | 45 |
| Q9WVJ2 | PSMD13 | proteasome (prosome, macropain | Cytoplasm | peptidase | 1.65E+07 | 2.40E+07 | 1.73E+07 | | | | | | |
| Q9WVJ2 | PSMD13 | proteasome (prosome, macropain | Cytoplasm | peptidase | 1.69E+07 | 3.13E+07 | 1.57E+07 | | | | | | |
| Q9WVJ2 | PSMD13 | proteasome (prosome, macropain | Cytoplasm | peptidase | 1.31E+07 | 1.38E+07 | 8328867.5 | | | | | | |
| Q9WVJ2 | PSMD13 | proteasome (prosome, macropain | Cytoplasm | peptidase | 1.50E+07 | 1.83E+07 | 1.38E+07 | | | | | | |
| Q9WVJ2 | PSMD13 | proteasome (prosome, macropain | Cytoplasm | peptidase | 1.16E+07 | 1.48E+07 | 8947741 | | | | | | |
| Q9WVJ2 | PSMD13 | proteasome (prosome, macropain | Cytoplasm | peptidase | 8078813.5 | 1.02E+07 | 7276667.5 | | | | | | |
| Q9WVJ2 | PSMD13 | proteasome (prosome, macropain | Cytoplasm | peptidase | 7917870 | 1.08E+07 | 6523123.5 | | | | | | |
| Q9WVJ2 | PSMD13 | proteasome (prosome, macropain | Cytoplasm | peptidase | 8458144 | 1.18E+07 | 9029255 | | | | | | |
| Q9WVJ2 | PSMD13 | proteasome (prosome, macropain | Cytoplasm | peptidase | 7957582.5 | 8664400 | 5035152 | | | | | | |
| Q9WVJ2 | PSMD13 | proteasome (prosome, macropain | Cytoplasm | peptidase | 8152231.5 | 1.12E+07 | 6611555 | | | | | | |
| Q9WVJ2 | PSMD13 | proteasome (prosome, macropain | Cytoplasm | peptidase | 6506123 | 6919599.5 | 4099045.75 | | | | | | |
| Q9WVJ2 | PSMD13 | proteasome (prosome, macropain | Cytoplasm | peptidase | 3382518.5 | 5557234 | 3299742 | | | | | | |
| Q9WVJ2 | PSMD13 | proteasome (prosome, macropain | Cytoplasm | peptidase | 1771322.125 | 1515922.125 | 1217996.5 | Q9WVJ2 | PSMD13 | 9639585 | 6165777 | 987315 | 39 |
| O35593 | PSMD14 | proteasome (prosome, macropain | Cytoplasm | peptidase | 1.27E+07 | 1.80E+07 | 1.24E+07 | | | | | | |
| O35593 | PSMD14 | proteasome (prosome, macropain | Cytoplasm | peptidase | 9669818 | 1.99E+07 | 1.12E+07 | | | | | | |
| O35593 | PSMD14 | proteasome (prosome, macropain | Cytoplasm | peptidase | 3582166.75 | 5643038.5 | 3503995 | | | | | | |
| O35593 | PSMD14 | proteasome (prosome, macropain | Cytoplasm | peptidase | 1357548 | 4055968.25 | 2056987.375 | O35593 | PSMD14 | 6824883 | 6265101 | 1808579 | 12 |
| Q8VDM4 | PSMD2 | proteasome (prosome, macropain | Cytoplasm | other | 2.14E+07 | 2.13E+07 | 8946558 | | | | | | |
| Q8VDM4 | PSMD2 | proteasome (prosome, macropain | Cytoplasm | other | 1.72E+07 | 1.94E+07 | 1.30E+07 | | | | | | |
| Q8VDM4 | PSMD2 | proteasome (prosome, macropain | Cytoplasm | other | 1.53E+07 | 1.36E+07 | 1.02E+07 | | | | | | |
| Q8VDM4 | PSMD2 | proteasome (prosome, macropain | Cytoplasm | other | 1.37E+07 | 8731764 | 5611990 | | | | | | |
| Q8VDM4 | PSMD2 | proteasome (prosome, macropain | Cytoplasm | other | 1.02E+07 | 1.04E+07 | 8561987 | | | | | | |
| Q8VDM4 | PSMD2 | proteasome (prosome, macropain | Cytoplasm | other | 9687980 | 6807029.5 | 8463294 | | | | | | |
| Q8VDM4 | PSMD2 | proteasome (prosome, macropain | Cytoplasm | other | 9237113 | 1.37E+07 | 1.16E+07 | | | | | | |
| Q8VDM4 | PSMD2 | proteasome (prosome, macropain | Cytoplasm | other | 8738510 | 1.02E+07 | 6881685.5 | | | | | | |
| Q8VDM4 | PSMD2 | proteasome (prosome, macropain | Cytoplasm | other | 6888780.5 | 9717744 | 7037608 | | | | | | |
| Q8VDM4 | PSMD2 | proteasome (prosome, macropain | Cytoplasm | other | 4773308.5 | 1.14E+07 | 5088169.5 | | | | | | |
| Q8VDM4 | PSMD2 | proteasome (prosome, macropain | Cytoplasm | other | 6164649 | 9317257 | 7080186 | | | | | | |
| Q8VDM4 | PSMD2 | proteasome (prosome, macropain | Cytoplasm | other | 7169141.5 | 1.12E+07 | 8556740 | | | | | | |
| Q8VDM4 | PSMD2 | proteasome (prosome, macropain | Cytoplasm | other | 7335691.5 | 5972270.5 | 4407818.5 | | | | | | |
| Q8VDM4 | PSMD2 | proteasome (prosome, macropain | Cytoplasm | other | 6461146 | 7846775 | 6071094.5 | | | | | | |
| Q8VDM4 | PSMD2 | proteasome (prosome, macropain | Cytoplasm | other | 6314792.5 | 8041449 | 5124652.5 | | | | | | |
| Q8VDM4 | PSMD2 | proteasome (prosome, macropain | Cytoplasm | other | 5340042 | 1.16E+07 | 7879363 | | | | | | |
| Q8VDM4 | PSMD2 | proteasome (prosome, macropain | Cytoplasm | other | 5437062 | 5928685 | 4391350.5 | | | | | | |
| Q8VDM4 | PSMD2 | proteasome (prosome, macropain | Cytoplasm | other | 3283284.5 | 4250198.5 | 2241704.5 | | | | | | |
| Q8VDM4 | PSMD2 | proteasome (prosome, macropain | Cytoplasm | other | 6019549.5 | 6708792 | 4711306.5 | | | | | | |
| Q8VDM4 | PSMD2 | proteasome (prosome, macropain | Cytoplasm | other | 3891731.75 | 5003931 | 4580100 | | | | | | |
| Q8VDM4 | PSMD2 | proteasome (prosome, macropain | Cytoplasm | other | 5409722 | 4313356.5 | 3274075 | | | | | | |
| Q8VDM4 | PSMD2 | proteasome (prosome, macropain | Cytoplasm | other | 4197742 | 4533247 | 4256527 | | | | | | |
| Q8VDM4 | PSMD2 | proteasome (prosome, macropain | Cytoplasm | other | 4483228 | 4664531.5 | 2979226.25 | | | | | | |
| Q8VDM4 | PSMD2 | proteasome (prosome, macropain | Cytoplasm | other | 2357948.75 | 2169082.25 | 1471924.375 | | | | | | |
| Q8VDM4 | PSMD2 | proteasome (prosome, macropain | Cytoplasm | other | 3617226.5 | 2240001.75 | 1898168.625 | | | | | | |
| Q8VDM4 | PSMD2 | proteasome (prosome, macropain | Cytoplasm | other | 1592364.125 | 3613308.25 | 2170092.5 | | | | | | |
| Q8VDM4 | PSMD2 | proteasome (prosome, macropain | Cytoplasm | other | 2172506.25 | 3080301.5 | 2221188.75 | | | | | | |
| Q8VDM4 | PSMD2 | proteasome (prosome, macropain | Cytoplasm | other | 3132459.75 | 2507620 | 2435127.75 | | | | | | |
| Q8VDM4 | PSMD2 | proteasome (prosome, macropain | Cytoplasm | other | 1937417.375 | 2682313.5 | 2309687.25 | Q8VDM4 | PSMD2 | 7012876 | 4359536 | 467391 | 87 |
| P14685 | PSMD3 | proteasome (prosome, macropain | Cytoplasm | other | 2.14E+07 | 2.67E+07 | 1.80E+07 | | | | | | |
| P14685 | PSMD3 | proteasome (prosome, macropain | Cytoplasm | other | 2.36E+07 | 2.91E+07 | 1.54E+07 | | | | | | |
| P14685 | PSMD3 | proteasome (prosome, macropain | Cytoplasm | other | 1.89E+07 | 2.21E+07 | 1.52E+07 | | | | | | |
| P14685 | PSMD3 | proteasome (prosome, macropain | Cytoplasm | other | 1.15E+07 | 1.32E+07 | 1.00E+07 | | | | | | |
| P14685 | PSMD3 | proteasome (prosome, macropain | Cytoplasm | other | 7852358 | 1.11E+07 | 6823474 | | | | | | |
| P14685 | PSMD3 | proteasome (prosome, macropain | Cytoplasm | other | 1.29E+07 | 8871565 | 6762353.5 | | | | | | |
| P14685 | PSMD3 | proteasome (prosome, macropain | Cytoplasm | other | 7841603.5 | 1.05E+07 | 6860412.5 | | | | | | |
| P14685 | PSMD3 | proteasome (prosome, macropain | Cytoplasm | other | 5376849.5 | 9598355 | 7194415 | | | | | | |
| P14685 | PSMD3 | proteasome (prosome, macropain | Cytoplasm | other | 5386488.5 | 3943926.5 | 2258972.25 | | | | | | |
| P14685 | PSMD3 | proteasome (prosome, macropain | Cytoplasm | other | 3552912.25 | 5167626 | 3739330 | | | | | | |
| P14685 | PSMD3 | proteasome (prosome, macropain | Cytoplasm | other | 2812146 | 6530845.5 | 4651372 | | | | | | |
| P14685 | PSMD3 | proteasome (prosome, macropain | Cytoplasm | other | 2991675.75 | 2024195.75 | 1178438.125 | | | | | | |
| P14685 | PSMD3 | proteasome (prosome, macropain | Cytoplasm | other | 2505555.25 | 3732651 | 1380753.5 | | | | | | |
| P14685 | PSMD3 | proteasome (prosome, macropain | Cytoplasm | other | 1645909.375 | 2538852.5 | 1201006.75 | | | | | | |
| P14685 | PSMD3 | proteasome (prosome, macropain | Cytoplasm | other | 4384031 | 299901 | 1082890.625 | | | | | | |
| P14685 | PSMD3 | proteasome (prosome, macropain | Cytoplasm | other | 3201023.5 | 3326216.75 | 2467891.75 | P14685 | PSMD3 | 8495660 | 7280628 | 1050868 | 48 |
| O35226 | PSMD4 | proteasome (prosome, macropain | Cytoplasm | other | 2.18E+07 | 2.92E+07 | 1.81E+07 | | | | | | |
| O35226 | PSMD4 | proteasome (prosome, macropain | Cytoplasm | other | 1.96E+07 | 2.15E+07 | 1.61E+07 | | | | | | |
| O35226 | PSMD4 | proteasome (prosome, macropain | Cytoplasm | other | 9003510 | 1.63E+07 | 8348688.5 | | | | | | |
| O35226 | PSMD4 | proteasome (prosome, macropain | Cytoplasm | other | 8696984 | 1.03E+07 | 5565695.5 | | | | | | |
| O35226 | PSMD4 | proteasome (prosome, macropain | Cytoplasm | other | 1.11E+07 | 1.36E+07 | 7333319.5 | | | | | | |
| O35226 | PSMD4 | proteasome (prosome, macropain | Cytoplasm | other | 9833025 | 1086284.375 | 3782587.5 | | | | | | |
| O35226 | PSMD4 | proteasome (prosome, macropain | Cytoplasm | other | 3858093.25 | 3251934.75 | 2226975.5 | | | | | | |
| O35226 | PSMD4 | proteasome (prosome, macropain | Cytoplasm | other | 2254339.75 | 3424249.5 | 1822953.75 | | | | | | |
| O35226 | PSMD4 | proteasome (prosome, macropain | Cytoplasm | other | 1604897.25 | 2108978.75 | 1342676.375 | O35226 | PSMD4 | 9743428 | 7721955 | 1486091 | 27 |
| Q8BJY1 | PSMD5 | proteasome (prosome, macropain | Cytoplasm | other | 3528060 | 2307534.75 | 3646261.75 | | | | | | |
| Q8BJY1 | PSMD5 | proteasome (prosome, macropain | Cytoplasm | other | 6805496.5 | 2264184.75 | 2238975.5 | | | | | | |
| Q8BJY1 | PSMD5 | proteasome (prosome, macropain | Cytoplasm | other | 2962420.75 | 2528675.75 | 2461569.25 | | | | | | |

| | | | | | | | | | | | | | |
|---|---|---|---|---|---|---|---|---|---|---|---|---|---|
| Q8BJY1 | PSMD5 | proteasome (prosome, macropain | Cytoplasm | other | 2997051.25 | 3757680.75 | 3284961.5 | | | | | | |
| Q8BJY1 | PSMD5 | proteasome (prosome, macropain | Cytoplasm | other | 2617561.25 | 2257018.5 | 1997355.75 | | | | | | |
| Q8BJY1 | PSMD5 | proteasome (prosome, macropain | Cytoplasm | other | 2519059 | 872242.188 | 1486788.75 | | | | | | |
| Q8BJY1 | PSMD5 | proteasome (prosome, macropain | Cytoplasm | other | 2250362.25 | 1418606.125 | 2026101.75 | | | | | | |
| Q8BJY1 | PSMD5 | proteasome (prosome, macropain | Cytoplasm | other | 2183927.25 | 1723832.875 | 1892318.625 | | | | | | |
| Q8BJY1 | PSMD5 | proteasome (prosome, macropain | Cytoplasm | other | 975730.688 | 1005900.938 | 1742058 | Q8BJY1 | PSMD5 | 2982185 | 1164821 | 224170 | 27 |
| Q99JI4 | PSMD6 | proteasome (prosome, macropain | Cytoplasm | enzyme | 1.60E+07 | 1.57E+07 | 1.14E+07 | | | | | | |
| Q99JI4 | PSMD6 | proteasome (prosome, macropain | Cytoplasm | enzyme | 1.37E+07 | 1.74E+07 | 1.16E+07 | | | | | | |
| Q99JI4 | PSMD6 | proteasome (prosome, macropain | Cytoplasm | enzyme | 1.14E+07 | 1.25E+07 | 7680803 | | | | | | |
| Q99JI4 | PSMD6 | proteasome (prosome, macropain | Cytoplasm | enzyme | 9085094 | 1.12E+07 | 6600676 | | | | | | |
| Q99JI4 | PSMD6 | proteasome (prosome, macropain | Cytoplasm | enzyme | 8823942 | 1.12E+07 | 7139895 | | | | | | |
| Q99JI4 | PSMD6 | proteasome (prosome, macropain | Cytoplasm | enzyme | 6985085.5 | 8568789 | 5461181 | | | | | | |
| Q99JI4 | PSMD6 | proteasome (prosome, macropain | Cytoplasm | enzyme | 6398768.5 | 7837681 | 6126128 | | | | | | |
| Q99JI4 | PSMD6 | proteasome (prosome, macropain | Cytoplasm | enzyme | 3036535.75 | 4395806 | 2596642.5 | | | | | | |
| Q99JI4 | PSMD6 | proteasome (prosome, macropain | Cytoplasm | enzyme | 3111727.75 | 4113962.5 | 2624925.5 | | | | | | |
| Q99JI4 | PSMD6 | proteasome (prosome, macropain | Cytoplasm | enzyme | 2401810 | 3992346.25 | 1977937.125 | | | | | | |
| Q99JI4 | PSMD6 | proteasome (prosome, macropain | Cytoplasm | enzyme | 3692779.75 | 5555603.5 | 3128675 | | | | | | |
| Q99JI4 | PSMD6 | proteasome (prosome, macropain | Cytoplasm | enzyme | 2982415 | 2268539.5 | 1992779.375 | | | | | | |
| Q99JI4 | PSMD6 | proteasome (prosome, macropain | Cytoplasm | enzyme | 2492141 | 6007118.5 | 4706774.5 | | | | | | |
| Q99JI4 | PSMD6 | proteasome (prosome, macropain | Cytoplasm | enzyme | 1046484.688 | 2599730.75 | 768958.125 | Q99JI4 | PSMD6 | 6505485 | 4403145 | 679420 | 42 |
| P26516 | PSMD7 | proteasome (prosome, macropain | Cytoplasm | other | 2.22E+07 | 3.30E+07 | 1.97E+07 | | | | | | |
| P26516 | PSMD7 | proteasome (prosome, macropain | Cytoplasm | other | 1.71E+07 | 1.89E+07 | 1.24E+07 | | | | | | |
| P26516 | PSMD7 | proteasome (prosome, macropain | Cytoplasm | other | 1.96E+07 | 1.85E+07 | 1.15E+07 | | | | | | |
| P26516 | PSMD7 | proteasome (prosome, macropain | Cytoplasm | other | 1.37E+07 | 1.41E+07 | 4438701.5 | | | | | | |
| P26516 | PSMD7 | proteasome (prosome, macropain | Cytoplasm | other | 7725704 | 7557726 | 5159350.5 | | | | | | |
| P26516 | PSMD7 | proteasome (prosome, macropain | Cytoplasm | other | 3870242 | 5277047 | 3315158.75 | | | | | | |
| P26516 | PSMD7 | proteasome (prosome, macropain | Cytoplasm | other | 6615256 | 6565643 | 5452407 | | | | | | |
| P26516 | PSMD7 | proteasome (prosome, macropain | Cytoplasm | other | 3860729.25 | 3398660.75 | 2834788.25 | | | | | | |
| P26516 | PSMD7 | proteasome (prosome, macropain | Cytoplasm | other | 2475482 | 2246336.25 | 1932845.125 | | | | | | |
| P26516 | PSMD7 | proteasome (prosome, macropain | Cytoplasm | other | 1066068.25 | 1062820 | 93168.68 | P26516 | PSMD7 | 9818348 | 8030048 | 1466080 | 30 |
| Q9CX56 | PSMD8 | proteasome (prosome, macropain | Cytoplasm | other | 1.23E+07 | 1.33E+07 | 1.14E+07 | | | | | | |
| Q9CX56 | PSMD8 | proteasome (prosome, macropain | Cytoplasm | other | 5552376 | 6823386 | 5726756 | | | | | | |
| Q9CX56 | PSMD8 | proteasome (prosome, macropain | Cytoplasm | other | 5346017.5 | 6556606.5 | 4211170.5 | Q9CX56 | PSMD8 | 7716131 | 3425575 | 1141858 | 9 |
| P97371 | PSME1 | proteasome (prosome, macropain | Cytoplasm | other | 1.50E+07 | 1.58E+07 | 1.22E+07 | | | | | | |
| P97371 | PSME1 | proteasome (prosome, macropain | Cytoplasm | other | 1.11E+07 | 1.31E+07 | 1.26E+07 | | | | | | |
| P97371 | PSME1 | proteasome (prosome, macropain | Cytoplasm | other | 1.36E+07 | 1.09E+07 | 1.15E+07 | | | | | | |
| P97371 | PSME1 | proteasome (prosome, macropain | Cytoplasm | other | 1.07E+07 | 9733568 | 9550435 | | | | | | |
| P97371 | PSME1 | proteasome (prosome, macropain | Cytoplasm | other | 1.04E+07 | 8260318.5 | 1.06E+07 | | | | | | |
| P97371 | PSME1 | proteasome (prosome, macropain | Cytoplasm | other | 5532570 | 5373297.5 | 4542729 | | | | | | |
| P97371 | PSME1 | proteasome (prosome, macropain | Cytoplasm | other | 6503584.5 | 5750707.5 | 2441130.75 | | | | | | |
| P97371 | PSME1 | proteasome (prosome, macropain | Cytoplasm | other | 5987359.5 | 5652944 | 7134355.5 | | | | | | |
| P97371 | PSME1 | proteasome (prosome, macropain | Cytoplasm | other | 4710953.5 | 5317074.5 | 6657299 | | | | | | |
| P97371 | PSME1 | proteasome (prosome, macropain | Cytoplasm | other | 4261841.5 | 4111838.75 | 4747063.5 | | | | | | |
| P97371 | PSME1 | proteasome (prosome, macropain | Cytoplasm | other | 5326998 | 6378296.5 | 5836247.5 | | | | | | |
| P97371 | PSME1 | proteasome (prosome, macropain | Cytoplasm | other | 3287038 | 729185.438 | 1670090.625 | P97371 | PSME1 | 8031695 | 3860734 | 643456 | 36 |
| P97372 | PSME2 | proteasome (prosome, macropain | Cytoplasm | peptidase | 1.87E+07 | 2.00E+07 | 2.62E+07 | | | | | | |
| P97372 | PSME2 | proteasome (prosome, macropain | Cytoplasm | peptidase | 1.41E+07 | 1.43E+07 | 1.16E+07 | | | | | | |
| P97372 | PSME2 | proteasome (prosome, macropain | Cytoplasm | peptidase | 1.32E+07 | 9935919 | 1.07E+07 | | | | | | |
| P97372 | PSME2 | proteasome (prosome, macropain | Cytoplasm | peptidase | 2978634.75 | 4597733.5 | 5004497 | | | | | | |
| P97372 | PSME2 | proteasome (prosome, macropain | Cytoplasm | peptidase | 2484012.25 | 2028853.5 | 2475815.25 | | | | | | |
| P97372 | PSME2 | proteasome (prosome, macropain | Cytoplasm | peptidase | 3140470.5 | 1821275.375 | 2341671.25 | | | | | | |
| P97372 | PSME2 | proteasome (prosome, macropain | Cytoplasm | peptidase | 4862394.5 | 936667.375 | 4240080 | P97372 | PSME2 | 8479359 | 7154242 | 1561184 | 21 |
| Q8R3K3 | PTCD2 | pentatricopeptide repeat domain | Cytoplasm | other | 2480456.75 | 2964617.25 | 2654817.75 | Q8R3K3 | PTCD2 | 2480457 | 245217 | 141576 | 3 |
| O08586 | PTEN | phosphatase and tensin homolog | Cytoplasm | phosphatase | 2.40E+07 | 2.67E+07 | 3.35E+07 | O08586 | PTEN | 24010000 | 4885622 | 2820715 | 3 |
| Q63739 | PTP4A1 | protein tyrosine phosphatase type | Cytoplasm | phosphatase | 1692810 | 2166138.5 | 1806317.5 | Q63739 | PTP4A1 | 1692810 | 247115 | 142672 | 3 |
| P35821 | PTPN1 | protein tyrosine phosphatase, non | Cytoplasm | phosphatase | 4248675.5 | 4892303.5 | 5359384.5 | P35821 | PTPN1 | 4248676 | 557688 | 321981 | 3 |
| Q62136 | PTPN21 | protein tyrosine phosphatase, non | Cytoplasm | phosphatase | 2.01E+07 | 2.28E+07 | 2.54E+07 | Q62136 | PTPN21 | 20060000 | 2675126 | 1544485 | 3 |
| P18052 | PTPRA | protein tyrosine phosphatase, rec | Plasma Membrane | phosphatase | 9371909 | 6921999.5 | 7843246 | | | | | | |
| P18052 | PTPRA | protein tyrosine phosphatase, rec | Plasma Membrane | phosphatase | 3899082.25 | 3315872.5 | 4502851.5 | | | | | | |
| P18052 | PTPRA | protein tyrosine phosphatase, rec | Plasma Membrane | phosphatase | 4158890 | 2933444.25 | 3075189.25 | | | | | | |
| P18052 | PTPRA | protein tyrosine phosphatase, rec | Plasma Membrane | phosphatase | 270460.188 | 625240.875 | 242013.578 | P18052 | PTPRA | 4425085 | 2927570 | 845117 | 12 |
| Q64455 | PTPRJ | protein tyrosine phosphatase, rec | Plasma Membrane | phosphatase | 8178031.5 | 1.10E+07 | 8593435 | | | | | | |
| Q64455 | PTPRJ | protein tyrosine phosphatase, rec | Plasma Membrane | phosphatase | 77416.773 | 581853.438 | 831952.25 | | | | | | |
| Q64455 | PTPRJ | protein tyrosine phosphatase, rec | Plasma Membrane | phosphatase | 1227887.25 | 927593.562 | 917555.125 | Q64455 | PTPRJ | 3161112 | 4316507 | 1438836 | 9 |
| Q8R2Y8 | PTRH2 | peptidyl-tRNA hydrolase 2 | Cytoplasm | enzyme | 8283445 | 8649934 | 1.10E+07 | | | | | | |
| Q8R2Y8 | PTRH2 | peptidyl-tRNA hydrolase 2 | Cytoplasm | enzyme | 258894.875 | 220090.859 | 324614.812 | Q8R2Y8 | PTRH2 | 4271170 | 5032894 | 2054670 | 6 |
| P42669 | PURA | purine-rich element binding prote | Nucleus | transcription regulator | 4518533.5 | 3830503.25 | 4081720.5 | | | | | | |
| P42669 | PURA | purine-rich element binding prote | Nucleus | transcription regulator | 1182787.25 | 293266.469 | 654.293 | P42669 | PURA | 2850660 | 2049345 | 836641 | 6 |
| Q922Q4 | PYCR2 | pyrroline-5-carboxylate reductase | Cytoplasm | enzyme | 3728353.5 | 2760037 | 3669306 | Q922Q4 | PYCR2 | 3728354 | 542816 | 313395 | 3 |
| Q9DCC4 | PYCRL | pyrroline-5-carboxylate reductase | Cytoplasm | enzyme | 3600107.5 | 3996604.5 | 3812377.25 | | | | | | |
| Q9DCC4 | PYCRL | pyrroline-5-carboxylate reductase | Cytoplasm | enzyme | 2901583 | 2444525.75 | 2271283.5 | | | | | | |
| Q9DCC4 | PYCRL | pyrroline-5-carboxylate reductase | Cytoplasm | enzyme | 2176803.25 | 1942002.125 | 1935251.5 | Q9DCC4 | PYCRL | 2892831 | 820358 | 273453 | 9 |
| Q61838 | Pzp | pregnancy zone protein | Extracellular Space | other | 1349723 | 2257681.5 | 2129047 | Q61838 | Pzp | 1349723 | 491305 | 283655 | 3 |
| Q8BVI4 | QDPR | quinoid dihydropteridine reductas | Cytoplasm | enzyme | 687325.875 | 637392.75 | 1015503.562 | Q8BVI4 | QDPR | 687326 | 205411 | 118594 | 3 |
| P61027 | RAB10 | RAB10, member RAS oncogene far | Cytoplasm | enzyme | 2.26E+08 | 2.64E+08 | 2.95E+08 | | | | | | |
| P61027 | RAB10 | RAB10, member RAS oncogene far | Cytoplasm | enzyme | 1.51E+08 | 1.44E+08 | 1.43E+08 | | | | | | |
| P61027 | RAB10 | RAB10, member RAS oncogene far | Cytoplasm | enzyme | 1.71E+08 | 1.13E+08 | 1.63E+08 | | | | | | |
| P61027 | RAB10 | RAB10, member RAS oncogene far | Cytoplasm | enzyme | 2.90E+07 | 4.51E+07 | 3.20E+07 | | | | | | |
| P61027 | RAB10 | RAB10, member RAS oncogene far | Cytoplasm | enzyme | 3.37E+07 | 4.13E+07 | 4.23E+07 | | | | | | |
| P61027 | RAB10 | RAB10, member RAS oncogene far | Cytoplasm | enzyme | 2.97E+07 | 4.95E+07 | 4.61E+07 | | | | | | |
| P61027 | RAB10 | RAB10, member RAS oncogene far | Cytoplasm | enzyme | 3.15E+07 | 2.89E+07 | 3.00E+07 | | | | | | |
| P61027 | RAB10 | RAB10, member RAS oncogene far | Cytoplasm | enzyme | 1.93E+07 | 2.14E+07 | 1.87E+07 | | | | | | |
| P61027 | RAB10 | RAB10, member RAS oncogene far | Cytoplasm | enzyme | 8905072 | 2.84E+07 | 3.08E+07 | | | | | | |
| P61027 | RAB10 | RAB10, member RAS oncogene far | Cytoplasm | enzyme | 9810654 | 1.99E+07 | 1.97E+07 | | | | | | |
| P61027 | RAB10 | RAB10, member RAS oncogene far | Cytoplasm | enzyme | 5865434 | 4072381.75 | 4722341.5 | P61027 | RAB10 | 65081015 | 79213903 | 13789371 | 33 |
| P46638 | RAB11B | RAB11B, member RAS oncogene fa | Cytoplasm | enzyme | 3.72E+07 | 4.14E+07 | 4.44E+07 | | | | | | |

| | | | | | | | | | | | | | |
|---|---|---|---|---|---|---|---|---|---|---|---|---|---|
| P46638 | RAB11B | RAB11B, member RAS oncogene fa | Cytoplasm | enzyme | 2.25E+07 | 2.78E+07 | 2.72E+07 | | | | | | |
| P46638 | RAB11B | RAB11B, member RAS oncogene fa | Cytoplasm | enzyme | 2.34E+07 | 2.73E+07 | 3.19E+07 | | | | | | |
| P46638 | RAB11B | RAB11B, member RAS oncogene fa | Cytoplasm | enzyme | 1.98E+07 | 1.89E+07 | 2.29E+07 | | | | | | |
| P46638 | RAB11B | RAB11B, member RAS oncogene fa | Cytoplasm | enzyme | 6931873.5 | 8633495 | 8387027.5 | | | | | | |
| P46638 | RAB11B | RAB11B, member RAS oncogene fa | Cytoplasm | enzyme | 9574623 | 9120264 | 1.05E+07 | | | | | | |
| P46638 | RAB11B | RAB11B, member RAS oncogene fa | Cytoplasm | enzyme | 6730780 | 6948378 | 6505096 | | | | | | |
| P46638 | RAB11B | RAB11B, member RAS oncogene fa | Cytoplasm | enzyme | 5601255.5 | 3213188.5 | 3463758.75 | | | | | | |
| P46638 | RAB11B | RAB11B, member RAS oncogene fa | Cytoplasm | enzyme | 9270842 | 1120997.25 | 3872984 | | | | | | |
| P46638 | RAB11B | RAB11B, member RAS oncogene fa | Cytoplasm | enzyme | 515430.5 | 2231549.75 | 1393456.125 | P46638 | RAB11B | 14152480 | 12781810 | 2333629 | 30 |
| Q91V41 | RAB14 | RAB14, member RAS oncogene far | Cytoplasm | enzyme | 2.20E+07 | 3.65E+07 | 3.89E+07 | | | | | | |
| Q91V41 | RAB14 | RAB14, member RAS oncogene far | Cytoplasm | enzyme | 1.56E+07 | 2.39E+07 | 2.06E+07 | | | | | | |
| Q91V41 | RAB14 | RAB14, member RAS oncogene far | Cytoplasm | enzyme | 1.31E+07 | 5557788 | 7884491 | | | | | | |
| Q91V41 | RAB14 | RAB14, member RAS oncogene far | Cytoplasm | enzyme | 9699586 | 1.14E+07 | 1.16E+07 | | | | | | |
| Q91V41 | RAB14 | RAB14, member RAS oncogene far | Cytoplasm | enzyme | 6823282 | 6468514 | 8912381 | | | | | | |
| Q91V41 | RAB14 | RAB14, member RAS oncogene far | Cytoplasm | enzyme | 4254616 | 5884592 | 5149926 | | | | | | |
| Q91V41 | RAB14 | RAB14, member RAS oncogene far | Cytoplasm | enzyme | 5082838.5 | 624718.25 | 321976.188 | | | | | | |
| Q91V41 | RAB14 | RAB14, member RAS oncogene far | Cytoplasm | enzyme | 3903772.25 | 660764.812 | 2847636.25 | | | | | | |
| Q91V41 | RAB14 | RAB14, member RAS oncogene far | Cytoplasm | enzyme | 1763299.25 | 139271.406 | 0 | Q91V41 | RAB14 | 9125266 | 10377521 | 1997155 | 27 |
| Q8K386 | Rab15 | RAB15, member RAS oncogene far | Cytoplasm | enzyme | 4.25E+08 | 4.26E+08 | 4.21E+08 | Q8K386 | Rab15 | 424700000 | 2497999 | 1442221 | 3 |
| P35293 | RAB18 | RAB18, member RAS oncogene far | Cytoplasm | enzyme | 1.49E+07 | 1.74E+07 | 1.22E+07 | | | | | | |
| P35293 | RAB18 | RAB18, member RAS oncogene far | Cytoplasm | enzyme | 2.15E+07 | 2.51E+07 | 1.84E+07 | | | | | | |
| P35293 | RAB18 | RAB18, member RAS oncogene far | Cytoplasm | enzyme | 1.93E+07 | 2.08E+07 | 1.47E+07 | | | | | | |
| P35293 | RAB18 | RAB18, member RAS oncogene far | Cytoplasm | enzyme | 1.51E+07 | 1.60E+07 | 1.56E+07 | | | | | | |
| P35293 | RAB18 | RAB18, member RAS oncogene far | Cytoplasm | enzyme | 1.10E+07 | 1.23E+07 | 1.15E+07 | P35293 | RAB18 | 16350000 | 4036599 | 1042246 | 15 |
| P62821 | RAB1A | RAB1A, member RAS oncogene fa | Cytoplasm | enzyme | 6.34E+07 | 7.90E+07 | 8.47E+07 | | | | | | |
| P62821 | RAB1A | RAB1A, member RAS oncogene fa | Cytoplasm | enzyme | 2.93E+07 | 3.38E+07 | 3.64E+07 | | | | | | |
| P62821 | RAB1A | RAB1A, member RAS oncogene fa | Cytoplasm | enzyme | 1.40E+07 | 1.34E+07 | 1.47E+07 | | | | | | |
| P62821 | RAB1A | RAB1A, member RAS oncogene fa | Cytoplasm | enzyme | 5705776 | 3187351.25 | 4978603.5 | | | | | | |
| P62821 | RAB1A | RAB1A, member RAS oncogene fa | Cytoplasm | enzyme | 2690138.75 | 2607049.25 | 3047866.75 | | | | | | |
| P62821 | RAB1A | RAB1A, member RAS oncogene fa | Cytoplasm | enzyme | 2693837.25 | 2683092.25 | 2184895.25 | P62821 | RAB1A | 19629959 | 27267194 | 6426939 | 18 |
| Q9D1G1 | RAB1B | RAB1B, member RAS oncogene far | Cytoplasm | other | 5.62E+07 | 3.01E+07 | 5.21E+07 | | | | | | |
| Q9D1G1 | RAB1B | RAB1B, member RAS oncogene far | Cytoplasm | other | 2.76E+07 | 2.53E+07 | 3.17E+07 | | | | | | |
| Q9D1G1 | RAB1B | RAB1B, member RAS oncogene far | Cytoplasm | other | 8142927 | 2.52E+07 | 1.83E+07 | | | | | | |
| Q9D1G1 | RAB1B | RAB1B, member RAS oncogene far | Cytoplasm | other | 1.31E+07 | 1.31E+07 | 1.49E+07 | | | | | | |
| Q9D1G1 | RAB1B | RAB1B, member RAS oncogene far | Cytoplasm | other | 4407160 | 7539440.5 | 8083755.5 | Q9D1G1 | RAB1B | 21888017 | 15602073 | 4028438 | 15 |
| P35282 | RAB21 | RAB21, member RAS oncogene far | Cytoplasm | enzyme | 1.08E+07 | 1.31E+07 | 1.37E+07 | | | | | | |
| P35282 | RAB21 | RAB21, member RAS oncogene far | Cytoplasm | enzyme | 9862067 | 3382809.25 | 727536.688 | | | | | | |
| P35282 | RAB21 | RAB21, member RAS oncogene far | Cytoplasm | enzyme | 4497174 | 3895541 | 3710477 | | | | | | |
| P35282 | RAB21 | RAB21, member RAS oncogene far | Cytoplasm | enzyme | 2073404.5 | 5778887.5 | 4668281.5 | | | | | | |
| P35282 | RAB21 | RAB21, member RAS oncogene far | Cytoplasm | enzyme | 3989272.25 | 1299042 | 1147947.625 | | | | | | |
| P35282 | RAB21 | RAB21, member RAS oncogene far | Cytoplasm | enzyme | 4012309 | 3469854.5 | 2974260 | | | | | | |
| P35282 | RAB21 | RAB21, member RAS oncogene far | Cytoplasm | enzyme | 3174987 | 2287976 | 2926955.75 | | | | | | |
| P35282 | RAB21 | RAB21, member RAS oncogene far | Cytoplasm | enzyme | 948608.812 | 794333.688 | 569843.625 | P35282 | RAB21 | 4919728 | 3772526 | 770064 | 24 |
| P35285 | RAB22A | RAB22A, member RAS oncogene f | Cytoplasm | enzyme | 3382791.5 | 3282616.75 | 3550781.5 | | | | | | |
| P35285 | RAB22A | RAB22A, member RAS oncogene f | Cytoplasm | enzyme | 3543209.5 | 3309653.75 | 2879749.5 | P35285 | RAB22A | 3463001 | 245850 | 100368 | 6 |
| Q9ERI2 | RAB27A | RAB27A, member RAS oncogene f | Cytoplasm | enzyme | 2046176.75 | 3213713 | 2846172.5 | Q9ERI2 | RAB27A | 2046177 | 596967 | 344659 | 3 |
| Q99P58 | RAB27B | RAB27B, member RAS oncogene f | Cytoplasm | enzyme | 2.20E+07 | 2.62E+07 | 2.61E+07 | | | | | | |
| Q99P58 | RAB27B | RAB27B, member RAS oncogene f | Cytoplasm | enzyme | 1.79E+07 | 1.74E+07 | 1.94E+07 | | | | | | |
| Q99P58 | RAB27B | RAB27B, member RAS oncogene f | Cytoplasm | enzyme | 1.64E+07 | 1.66E+07 | 1.97E+07 | | | | | | |
| Q99P58 | RAB27B | RAB27B, member RAS oncogene f | Cytoplasm | enzyme | 1.47E+07 | 1.72E+07 | 2.11E+07 | | | | | | |
| Q99P58 | RAB27B | RAB27B, member RAS oncogene f | Cytoplasm | enzyme | 1.03E+07 | 8770853 | 1.23E+07 | | | | | | |
| Q99P58 | RAB27B | RAB27B, member RAS oncogene f | Cytoplasm | enzyme | 2178335 | 2520457.75 | 2821056.25 | | | | | | |
| Q99P58 | RAB27B | RAB27B, member RAS oncogene f | Cytoplasm | enzyme | 1925625.875 | 1565709.125 | 1329514 | | | | | | |
| Q99P58 | RAB27B | RAB27B, member RAS oncogene f | Cytoplasm | enzyme | 782989.25 | 2219995 | 2071190 | Q99P58 | RAB27B | 10772119 | 8752582 | 1786613 | 24 |
| P53994 | RAB2A | RAB2A, member RAS oncogene fa | Cytoplasm | enzyme | 1.40E+07 | 1.27E+07 | 1.37E+07 | | | | | | |
| P53994 | RAB2A | RAB2A, member RAS oncogene fa | Cytoplasm | enzyme | 7305316.5 | 8542894 | 8077214 | | | | | | |
| P53994 | RAB2A | RAB2A, member RAS oncogene fa | Cytoplasm | enzyme | 3208690.5 | 4964984.5 | 4856841 | | | | | | |
| P53994 | RAB2A | RAB2A, member RAS oncogene fa | Cytoplasm | enzyme | 4132327.75 | 4608579.5 | 4326271 | | | | | | |
| P53994 | RAB2A | RAB2A, member RAS oncogene fa | Cytoplasm | enzyme | 2012277.375 | 3279600.5 | 2883924.5 | P53994 | RAB2A | 6121722 | 4016868 | 1037151 | 15 |
| Q6PHN9 | RAB35 | RAB35, member RAS oncogene far | Cytoplasm | enzyme | 8.52E+07 | 7.87E+07 | 6.05E+07 | | | | | | |
| Q6PHN9 | RAB35 | RAB35, member RAS oncogene far | Cytoplasm | enzyme | 4.91E+07 | 4.59E+07 | 4.64E+07 | | | | | | |
| Q6PHN9 | RAB35 | RAB35, member RAS oncogene far | Cytoplasm | enzyme | 4.25E+07 | 4.28E+07 | 4.26E+07 | | | | | | |
| Q6PHN9 | RAB35 | RAB35, member RAS oncogene far | Cytoplasm | enzyme | 2.99E+07 | 2.34E+07 | 1.30E+07 | | | | | | |
| Q6PHN9 | RAB35 | RAB35, member RAS oncogene far | Cytoplasm | enzyme | 2.04E+07 | 2.60E+07 | 2.37E+07 | | | | | | |
| Q6PHN9 | RAB35 | RAB35, member RAS oncogene far | Cytoplasm | enzyme | 1.60E+07 | 2129463.5 | 1856147.25 | | | | | | |
| Q6PHN9 | RAB35 | RAB35, member RAS oncogene far | Cytoplasm | enzyme | 3417433 | 5845661 | 5550736.5 | Q6PHN9 | RAB35 | 35231062 | 24277556 | 5297797 | 21 |
| Q9JKM7 | RAB37 | RAB37, member RAS oncogene far | Cytoplasm | enzyme | 523765.969 | 788694.062 | 735762.062 | Q9JKM7 | RAB37 | 523766 | 140197 | 80943 | 3 |
| Q80UJ7 | RAB3GAP1 | RAB3 GTPase activating protein su | Cytoplasm | other | 5325445.5 | 5063981.5 | 3886234.5 | | | | | | |
| Q80UJ7 | RAB3GAP1 | RAB3 GTPase activating protein su | Cytoplasm | other | 5769989 | 1752462.375 | 1139442 | | | | | | |
| Q80UJ7 | RAB3GAP1 | RAB3 GTPase activating protein su | Cytoplasm | other | 145736.328 | 82181.453 | 33317.652 | Q80UJ7 | RAB3GAP1 | 3747057 | 2423846 | 807949 | 9 |
| Q8BMG7 | RAB3GAP2 | RAB3 GTPase activating protein su | Cytoplasm | enzyme | 1257669.625 | 1556937 | 1171103.375 | | | | | | |
| Q8BMG7 | RAB3GAP2 | RAB3 GTPase activating protein su | Cytoplasm | enzyme | 1582070.5 | 781675.625 | 553398.438 | Q8BMG7 | RAB3GAP2 | 1419870 | 413743 | 168910 | 6 |
| Q8VDV3 | RAB3IL1 | RAB3A interacting protein (rabin3) | Cytoplasm | other | 1644742.5 | 2424754.5 | 2365647.75 | Q8VDV3 | RAB3IL1 | 1644743 | 434284 | 250734 | 3 |
| Q9CQD1 | RAB5A | RAB5A, member RAS oncogene fa | Cytoplasm | enzyme | 2.99E+07 | 3.14E+07 | 2.67E+07 | | | | | | |
| Q9CQD1 | RAB5A | RAB5A, member RAS oncogene fa | Cytoplasm | enzyme | 3.14E+07 | 2.79E+07 | 3.08E+07 | | | | | | |
| Q9CQD1 | RAB5A | RAB5A, member RAS oncogene fa | Cytoplasm | enzyme | 9164165 | 1.15E+07 | 1.10E+07 | | | | | | |
| Q9CQD1 | RAB5A | RAB5A, member RAS oncogene fa | Cytoplasm | enzyme | 5609444.5 | 1.64E+07 | 1.07E+07 | | | | | | |
| Q9CQD1 | RAB5A | RAB5A, member RAS oncogene fa | Cytoplasm | enzyme | 6802958 | 8133629.5 | 7765708.5 | Q9CQD1 | RAB5A | 16579314 | 10498118 | 2710602 | 15 |
| P61021 | RAB5B | RAB5B, member RAS oncogene fa | Cytoplasm | enzyme | 2.55E+07 | 2.61E+07 | 2.30E+07 | | | | | | |
| P61021 | RAB5B | RAB5B, member RAS oncogene fa | Cytoplasm | enzyme | 2.02E+07 | 1.79E+07 | 1.80E+07 | | | | | | |
| P61021 | RAB5B | RAB5B, member RAS oncogene fa | Cytoplasm | enzyme | 1.41E+07 | 1.41E+07 | 1.56E+07 | | | | | | |
| P61021 | RAB5B | RAB5B, member RAS oncogene fa | Cytoplasm | enzyme | 8402729 | 8653974 | 9329964 | | | | | | |
| P61021 | RAB5B | RAB5B, member RAS oncogene fa | Cytoplasm | enzyme | 6209930 | 7719154 | 8897213 | | | | | | |
| P61021 | RAB5B | RAB5B, member RAS oncogene fa | Cytoplasm | enzyme | 4813046 | 5255223 | 6352726.5 | P61021 | RAB5B | 13197618 | 7079287 | 1668604 | 18 |
| P35278 | RAB5C | RAB5C, member RAS oncogene fa | Cytoplasm | enzyme | 1.20E+08 | 1.10E+08 | 1.13E+08 | | | | | | |
| P35278 | RAB5C | RAB5C, member RAS oncogene fa | Cytoplasm | enzyme | 8.35E+07 | 1.15E+08 | 1.12E+08 | | | | | | |

| | | | | | | | | | | | | | |
|---|---|---|---|---|---|---|---|---|---|---|---|---|---|
| P35278 | RAB5C | RAB5C, member RAS oncogene far | Cytoplasm | enzyme | 6.26E+07 | 7.62E+07 | 5.70E+07 | | | | | | |
| P35278 | RAB5C | RAB5C, member RAS oncogene far | Cytoplasm | enzyme | 5.55E+07 | 8.82E+07 | 6.61E+07 | | | | | | |
| P35278 | RAB5C | RAB5C, member RAS oncogene far | Cytoplasm | enzyme | 6.17E+07 | 6.94E+07 | 6.59E+07 | | | | | | |
| P35278 | RAB5C | RAB5C, member RAS oncogene far | Cytoplasm | enzyme | 2.44E+07 | 2.44E+07 | 2.38E+07 | | | | | | |
| P35278 | RAB5C | RAB5C, member RAS oncogene far | Cytoplasm | enzyme | 2.74E+07 | 3.13E+07 | 3.05E+07 | | | | | | |
| P35278 | RAB5C | RAB5C, member RAS oncogene far | Cytoplasm | enzyme | 2.84E+07 | 2.44E+07 | 2.55E+07 | | | | | | |
| P35278 | RAB5C | RAB5C, member RAS oncogene far | Cytoplasm | enzyme | 1.93E+07 | 1.44E+07 | 1.02E+07 | | | | | | |
| P35278 | RAB5C | RAB5C, member RAS oncogene far | Cytoplasm | enzyme | 1.48E+07 | 1.69E+07 | 1.48E+07 | | | | | | |
| P35278 | RAB5C | RAB5C, member RAS oncogene far | Cytoplasm | enzyme | 7455104.5 | 7983654.5 | 9640403 | | | | | | |
| P35278 | RAB5C | RAB5C, member RAS oncogene far | Cytoplasm | enzyme | 2629173.25 | 2715691 | 2201089.75 | | | | | | |
| P35278 | RAB5C | RAB5C, member RAS oncogene far | Cytoplasm | enzyme | 1771571.375 | 2276051 | 2250279.75 | P35278 | RAB5C | 39195065 | 37526112 | 6008987 | 39 |
| P35279 | RAB6A | RAB6A, member RAS oncogene far | Cytoplasm | enzyme | 4946973 | 5670561.5 | 5473754 | | | | | | |
| P35279 | RAB6A | RAB6A, member RAS oncogene far | Cytoplasm | enzyme | 2254453.5 | 2530349.25 | 3054475.75 | P35279 | RAB6A | 3600713 | 1546577 | 631387 | 6 |
| P51150 | RAB7A | RAB7A, member RAS oncogene far | Cytoplasm | enzyme | 1.99E+07 | 2.26E+07 | 2.35E+07 | | | | | | |
| P51150 | RAB7A | RAB7A, member RAS oncogene far | Cytoplasm | enzyme | 1.97E+07 | 2.03E+07 | 1.96E+07 | | | | | | |
| P51150 | RAB7A | RAB7A, member RAS oncogene far | Cytoplasm | enzyme | 1.51E+07 | 2.72E+07 | 1.99E+07 | | | | | | |
| P51150 | RAB7A | RAB7A, member RAS oncogene far | Cytoplasm | enzyme | 1.57E+07 | 1.53E+07 | 1.38E+07 | | | | | | |
| P51150 | RAB7A | RAB7A, member RAS oncogene far | Cytoplasm | enzyme | 1.48E+07 | 1.39E+07 | 1.08E+07 | | | | | | |
| P51150 | RAB7A | RAB7A, member RAS oncogene far | Cytoplasm | enzyme | 1.00E+07 | 1.25E+07 | 1.25E+07 | | | | | | |
| P51150 | RAB7A | RAB7A, member RAS oncogene far | Cytoplasm | enzyme | 1.45E+07 | 1.29E+07 | 1.41E+07 | | | | | | |
| P51150 | RAB7A | RAB7A, member RAS oncogene far | Cytoplasm | enzyme | 8104984 | 1.73E+07 | 1.41E+07 | | | | | | |
| P51150 | RAB7A | RAB7A, member RAS oncogene far | Cytoplasm | enzyme | 1.04E+07 | 8309398.5 | 1.15E+07 | | | | | | |
| P51150 | RAB7A | RAB7A, member RAS oncogene far | Cytoplasm | enzyme | 9729116 | 3418157.25 | 6504658.5 | | | | | | |
| P51150 | RAB7A | RAB7A, member RAS oncogene far | Cytoplasm | enzyme | 3950975.25 | 3285809.75 | 3435329.75 | P51150 | RAB7A | 12883189 | 6006469 | 1045592 | 33 |
| P55258 | RAB8A | RAB8A, member RAS oncogene far | Plasma Membrane | enzyme | 7.94E+07 | 8.50E+07 | 8.51E+07 | | | | | | |
| P55258 | RAB8A | RAB8A, member RAS oncogene far | Plasma Membrane | enzyme | 6.87E+07 | 8.06E+07 | 6.69E+07 | | | | | | |
| P55258 | RAB8A | RAB8A, member RAS oncogene far | Plasma Membrane | enzyme | 6.59E+07 | 6.82E+07 | 6.34E+07 | | | | | | |
| P55258 | RAB8A | RAB8A, member RAS oncogene far | Plasma Membrane | enzyme | 5.40E+07 | 5.41E+07 | 5.85E+07 | | | | | | |
| P55258 | RAB8A | RAB8A, member RAS oncogene far | Plasma Membrane | enzyme | 4.28E+07 | 4.54E+07 | 4.64E+07 | | | | | | |
| P55258 | RAB8A | RAB8A, member RAS oncogene far | Plasma Membrane | enzyme | 3204590 | 6028841.5 | 7092870 | | | | | | |
| P55258 | RAB8A | RAB8A, member RAS oncogene far | Plasma Membrane | enzyme | 4436694 | 6055260 | 7811381.5 | | | | | | |
| P55258 | RAB8A | RAB8A, member RAS oncogene far | Plasma Membrane | enzyme | 467166 | 312805.594 | 586724.688 | P55258 | RAB8A | 39846056 | 31828808 | 6497028 | 24 |
| P61028 | RAB8B | RAB8B, member RAS oncogene far | Cytoplasm | enzyme | 1.05E+08 | 1.05E+08 | 1.04E+08 | | | | | | |
| P61028 | RAB8B | RAB8B, member RAS oncogene far | Cytoplasm | enzyme | 1.13E+08 | 1.24E+08 | 1.20E+08 | | | | | | |
| P61028 | RAB8B | RAB8B, member RAS oncogene far | Cytoplasm | enzyme | 4.64E+07 | 5.68E+07 | 5.70E+07 | | | | | | |
| P61028 | RAB8B | RAB8B, member RAS oncogene far | Cytoplasm | enzyme | 1.82E+07 | 2.30E+07 | 2.21E+07 | | | | | | |
| P61028 | RAB8B | RAB8B, member RAS oncogene far | Cytoplasm | enzyme | 3057532 | 4450663 | 4926999.5 | | | | | | |
| P61028 | RAB8B | RAB8B, member RAS oncogene far | Cytoplasm | enzyme | 235203.5 | 448248.312 | 570165.625 | P61028 | RAB8B | 47705456 | 48255780 | 11373996 | 18 |
| Q9R0M6 | RAB9A | RAB9A, member RAS oncogene far | Cytoplasm | enzyme | 2628907.25 | 1708490.375 | 2320591.25 | Q9R0M6 | RAB9A | 2628907 | 468489 | 270482 | 3 |
| P63001 | RAC1 | ras-related C3 botulinum toxin sub | Plasma Membrane | enzyme | 1.03E+08 | 9.85E+07 | 1.10E+08 | | | | | | |
| P63001 | RAC1 | ras-related C3 botulinum toxin sub | Plasma Membrane | enzyme | 1.08E+08 | 4.19E+07 | 3.66E+07 | | | | | | |
| P63001 | RAC1 | ras-related C3 botulinum toxin sub | Plasma Membrane | enzyme | 8.14E+07 | 8.48E+07 | 9.26E+07 | | | | | | |
| P63001 | RAC1 | ras-related C3 botulinum toxin sub | Plasma Membrane | enzyme | 7.61E+07 | 6.14E+07 | 5.80E+07 | | | | | | |
| P63001 | RAC1 | ras-related C3 botulinum toxin sub | Plasma Membrane | enzyme | 6.07E+07 | 6.55E+07 | 7.78E+07 | | | | | | |
| P63001 | RAC1 | ras-related C3 botulinum toxin sub | Plasma Membrane | enzyme | 6.50E+07 | 4.62E+07 | 6.01E+07 | | | | | | |
| P63001 | RAC1 | ras-related C3 botulinum toxin sub | Plasma Membrane | enzyme | 3.17E+07 | 4.91E+07 | 4.86E+07 | | | | | | |
| P63001 | RAC1 | ras-related C3 botulinum toxin sub | Plasma Membrane | enzyme | 2.03E+07 | 1.84E+07 | 2.20E+07 | | | | | | |
| P63001 | RAC1 | ras-related C3 botulinum toxin sub | Plasma Membrane | enzyme | 1.32E+07 | 4.37E+07 | 4.20E+07 | | | | | | |
| P63001 | RAC1 | ras-related C3 botulinum toxin sub | Plasma Membrane | enzyme | 1.48E+07 | 2.19E+07 | 2.85E+07 | | | | | | |
| P63001 | RAC1 | ras-related C3 botulinum toxin sub | Plasma Membrane | enzyme | 1.08E+07 | 6221981.5 | 6965965 | | | | | | |
| P63001 | RAC1 | ras-related C3 botulinum toxin sub | Plasma Membrane | enzyme | 1895392.875 | 1.07E+07 | 9848237 | | | | | | |
| P63001 | RAC1 | ras-related C3 botulinum toxin sub | Plasma Membrane | enzyme | 6566200.5 | 3621793.75 | 5556184 | | | | | | |
| P63001 | RAC1 | ras-related C3 botulinum toxin sub | Plasma Membrane | enzyme | 2098989.5 | 3052058 | 3851180.5 | P63001 | RAC1 | 42540756 | 33560848 | 5178551 | 42 |
| Q05144 | RAC2 | ras-related C3 botulinum toxin sub | Cytoplasm | enzyme | 7338551 | 5916233 | 6391553 | Q05144 | RAC2 | 7338551 | 724077 | 418046 | 3 |
| P54726 | RAD23A | RAD23 homolog A, nucleotide exci | Nucleus | other | 5.16E+07 | 3.66E+07 | 6.74E+07 | | | | | | |
| P54726 | RAD23A | RAD23 homolog A, nucleotide exci | Nucleus | other | 3.56E+07 | 3.30E+07 | 4.09E+07 | | | | | | |
| P54726 | RAD23A | RAD23 homolog A, nucleotide exci | Nucleus | other | 2.15E+07 | 2.61E+07 | 2.93E+07 | | | | | | |
| P54726 | RAD23A | RAD23 homolog A, nucleotide exci | Nucleus | other | 9312454 | 8617551 | 9306544 | | | | | | |
| P54726 | RAD23A | RAD23 homolog A, nucleotide exci | Nucleus | other | 5674948.5 | 5761019.5 | 9259638 | | | | | | |
| P54726 | RAD23A | RAD23 homolog A, nucleotide exci | Nucleus | other | 3497709.25 | 2770925.5 | 3093515.25 | | | | | | |
| P54726 | RAD23A | RAD23 homolog A, nucleotide exci | Nucleus | other | 882459 | 451813.438 | 595300 | P54726 | RAD23A | 18293939 | 19159292 | 4180900 | 21 |
| P63321 | RALA | v-ral simian leukemia viral oncoge | Cytoplasm | enzyme | 2928393.5 | 3065971.5 | 3312765.5 | P63321 | RALA | 2928394 | 194755 | 112442 | 3 |
| P62827 | RAN | RAN, member RAS oncogene famil | Nucleus | enzyme | 2771569.75 | 2111100.5 | 1880555.625 | P62827 | RAN | 2771570 | 462471 | 267008 | 3 |
| P46061 | RANGAP1 | Ran GTPase activating protein 1 | Nucleus | other | 471362.188 | 321755.188 | 180698.922 | P46061 | RANGAP1 | 471362 | 145353 | 83919 | 3 |
| P62835 | RAP1A | RAP1A, member of RAS oncogene | Cytoplasm | enzyme | 1.03E+08 | 1.23E+08 | 1.13E+08 | | | | | | |
| P62835 | RAP1A | RAP1A, member of RAS oncogene | Cytoplasm | enzyme | 5.79E+07 | 8.17E+07 | 7.48E+07 | | | | | | |
| P62835 | RAP1A | RAP1A, member of RAS oncogene | Cytoplasm | enzyme | 3.26E+07 | 3.70E+07 | 2.30E+07 | | | | | | |
| P62835 | RAP1A | RAP1A, member of RAS oncogene | Cytoplasm | enzyme | 6.68E+07 | 5.68E+07 | 5.71E+07 | | | | | | |
| P62835 | RAP1A | RAP1A, member of RAS oncogene | Cytoplasm | enzyme | 1.92E+07 | 1.51E+07 | 2.08E+07 | | | | | | |
| P62835 | RAP1A | RAP1A, member of RAS oncogene | Cytoplasm | enzyme | 5978245.5 | 5963871 | 5383739.5 | | | | | | |
| P62835 | RAP1A | RAP1A, member of RAS oncogene | Cytoplasm | enzyme | 3321806.25 | 2548234.75 | 1895832.875 | | | | | | |
| P62835 | RAP1A | RAP1A, member of RAS oncogene | Cytoplasm | enzyme | 3845476.25 | 234768.578 | 180787 | P62835 | RAP1A | 36643191 | 38818058 | 7923703 | 24 |
| Q99JI6 | RAP1B | RAP1B, member of RAS oncogene | Cytoplasm | enzyme | 2.03E+08 | 1.96E+08 | 2.29E+08 | | | | | | |
| Q99JI6 | RAP1B | RAP1B, member of RAS oncogene | Cytoplasm | enzyme | 2.02E+08 | 1.46E+08 | 1.53E+08 | | | | | | |
| Q99JI6 | RAP1B | RAP1B, member of RAS oncogene | Cytoplasm | enzyme | 1.54E+08 | 1.65E+08 | 1.73E+08 | | | | | | |
| Q99JI6 | RAP1B | RAP1B, member of RAS oncogene | Cytoplasm | enzyme | 6.06E+07 | 8.45E+07 | 7.36E+07 | | | | | | |
| Q99JI6 | RAP1B | RAP1B, member of RAS oncogene | Cytoplasm | enzyme | 1.43E+07 | 2.61E+07 | 1.62E+07 | | | | | | |
| Q99JI6 | RAP1B | RAP1B, member of RAS oncogene | Cytoplasm | enzyme | 4.62E+07 | 5.76E+07 | 4.81E+07 | | | | | | |
| Q99JI6 | RAP1B | RAP1B, member of RAS oncogene | Cytoplasm | enzyme | 2.52E+07 | 3.39E+07 | 3.82E+07 | | | | | | |
| Q99JI6 | RAP1B | RAP1B, member of RAS oncogene | Cytoplasm | enzyme | 93522.141 | 397810.125 | 504480.594 | Q99JI6 | RAP1B | 88101690 | 76543011 | 15624277 | 24 |
| P61226 | RAP2B | RAP2B, member of RAS oncogene | Plasma Membrane | enzyme | 1.15E+07 | 1.26E+07 | 1.13E+07 | | | | | | |
| P61226 | RAP2B | RAP2B, member of RAS oncogene | Plasma Membrane | enzyme | 9505863 | 1.03E+07 | 7443370.5 | | | | | | |
| P61226 | RAP2B | RAP2B, member of RAS oncogene | Plasma Membrane | enzyme | 8815456 | 1.10E+07 | 8324136.5 | | | | | | |
| P61226 | RAP2B | RAP2B, member of RAS oncogene | Plasma Membrane | enzyme | 1.18E+07 | 1.30E+07 | 7605770.5 | | | | | | |
| P61226 | RAP2B | RAP2B, member of RAS oncogene | Plasma Membrane | enzyme | 7398987.5 | 6896869.5 | 7732938 | | | | | | |
| P61226 | RAP2B | RAP2B, member of RAS oncogene | Plasma Membrane | enzyme | 2439259.75 | 2759759 | 3276836 | P61226 | RAP2B | 8576594 | 3235988 | 762730 | 18 |

| | | | | | | | | | | | | | |
|---|---|---|---|---|---|---|---|---|---|---|---|---|---|
| Q8BU31 | RAP2C | RAP2C, member of RAS oncogene | Cytoplasm | enzyme | 2198911 | 2583110.75 | 1268771.125 | Q8BU31 | RAP2C | 2198911 | 675803 | 390175 | 3 |
| Q9D0I9 | RARS | arginyl-tRNA synthetase | Cytoplasm | enzyme | 435604.719 | 460139.312 | 474362.312 | Q9D0I9 | RARS | 435605 | 19606 | 11320 | 3 |
| Q60790 | RASA3 | RAS p21 protein activator 3 | Plasma Membrane | ion channel | 3643680 | 5455813.5 | 4408693.5 | Q60790 | RASA3 | 3643680 | 909719 | 525227 | 3 |
| Q61820 | Rasl2-9 | RAS-like, family 2, locus 9 | Nucleus | transporter | 5933563.5 | 4727594.5 | 5024211 | | | | | | |
| Q61820 | Rasl2-9 | RAS-like, family 2, locus 9 | Nucleus | transporter | 3825651 | 3362004 | 3530959 | Q61820 | Rasl2-9 | 4879607 | 1001131 | 408710 | 6 |
| O89086 | RBM3 | RNA binding motif (RNP1, RRM) pr | Cytoplasm | other | 9606875 | 8924684 | 5836832.5 | | | | | | |
| O89086 | RBM3 | RNA binding motif (RNP1, RRM) pr | Cytoplasm | other | 3316022.75 | 2729763.75 | 2059345.375 | | | | | | |
| O89086 | RBM3 | RNA binding motif (RNP1, RRM) pr | Cytoplasm | other | 269717.969 | 114773.5 | 228189.484 | O89086 | RBM3 | 4397539 | 3660672 | 1220224 | 9 |
| Q9QYF1 | RDH11 | retinol dehydrogenase 11 (all-tran | Cytoplasm | enzyme | 6568788.5 | 8920241 | 8741616 | | | | | | |
| Q9QYF1 | RDH11 | retinol dehydrogenase 11 (all-tran | Cytoplasm | enzyme | 4212782.5 | 6745727.5 | 4504098 | | | | | | |
| Q9QYF1 | RDH11 | retinol dehydrogenase 11 (all-tran | Cytoplasm | enzyme | 5011424 | 6990979.5 | 4487491.5 | | | | | | |
| Q9QYF1 | RDH11 | retinol dehydrogenase 11 (all-tran | Cytoplasm | enzyme | 4740341 | 5384805.5 | 4892672.5 | | | | | | |
| Q9QYF1 | RDH11 | retinol dehydrogenase 11 (all-tran | Cytoplasm | enzyme | 3871550 | 5685984 | 5330817 | | | | | | |
| Q9QYF1 | RDH11 | retinol dehydrogenase 11 (all-tran | Cytoplasm | enzyme | 2307045.5 | 3535385.5 | 2042179.375 | | | | | | |
| Q9QYF1 | RDH11 | retinol dehydrogenase 11 (all-tran | Cytoplasm | enzyme | 4482026 | 4396385 | 4713429 | | | | | | |
| Q9QYF1 | RDH11 | retinol dehydrogenase 11 (all-tran | Cytoplasm | enzyme | 2043755.75 | 2914303.5 | 2015809.375 | Q9QYF1 | RDH11 | 4154714 | 1884290 | 384629 | 24 |
| P26043 | RDX | radixin | Cytoplasm | other | 4.21E+07 | 2.54E+07 | 3.18E+07 | | | | | | |
| P26043 | RDX | radixin | Cytoplasm | other | 1.80E+07 | 1.76E+07 | 1.84E+07 | | | | | | |
| P26043 | RDX | radixin | Cytoplasm | other | 1.64E+07 | 2.45E+07 | 3.11E+07 | | | | | | |
| P26043 | RDX | radixin | Cytoplasm | other | 1.06E+07 | 7935237 | 1.15E+07 | | | | | | |
| P26043 | RDX | radixin | Cytoplasm | other | 1.25E+07 | 9449823 | 9721764 | | | | | | |
| P26043 | RDX | radixin | Cytoplasm | other | 9404788 | 8328883 | 9833076 | | | | | | |
| P26043 | RDX | radixin | Cytoplasm | other | 9670150 | 7496769.5 | 8506512 | | | | | | |
| P26043 | RDX | radixin | Cytoplasm | other | 8179467 | 2744728 | 5105304 | | | | | | |
| P26043 | RDX | radixin | Cytoplasm | other | 5562600 | 2729112.25 | 2570676.5 | | | | | | |
| P26043 | RDX | radixin | Cytoplasm | other | 7646945.5 | 2160417.75 | 1845548.5 | | | | | | |
| P26043 | RDX | radixin | Cytoplasm | other | 3196211.5 | 3201175 | 3510864.75 | | | | | | |
| P26043 | RDX | radixin | Cytoplasm | other | 2465206.75 | 4808652.5 | 3035840 | | | | | | |
| P26043 | RDX | radixin | Cytoplasm | other | 2800144.75 | 1042395.125 | 1137627 | | | | | | |
| P26043 | RDX | radixin | Cytoplasm | other | 1622573 | 4.70E-06 | 28292.666 | P26043 | RDX | 10722006 | 9545193 | 1472855 | 42 |
| Q60870 | REEP5 | receptor accessory protein 5 | Extracellular Space | transporter | 1.49E+07 | 2.32E+07 | 1.42E+07 | | | | | | |
| Q60870 | REEP5 | receptor accessory protein 5 | Extracellular Space | transporter | 1251270.5 | 1329452.25 | 991907.625 | | | | | | |
| Q60870 | REEP5 | receptor accessory protein 5 | Extracellular Space | transporter | 220608.031 | 413996.281 | 155396.859 | Q60870 | REEP5 | 5453960 | 8733837 | 2911279 | 9 |
| Q9JM62 | REEP6 | receptor accessory protein 6 | Plasma Membrane | other | 1.01E+07 | 1.23E+07 | 1.02E+07 | | | | | | |
| Q9JM62 | REEP6 | receptor accessory protein 6 | Plasma Membrane | other | 4652799 | 5401922.5 | 4734857 | | | | | | |
| Q9JM62 | REEP6 | receptor accessory protein 6 | Plasma Membrane | other | 1661099.125 | 228226.328 | 128409.5 | Q9JM62 | REEP6 | 5454633 | 4483382 | 1494461 | 9 |
| Q8K2J7 | RELL1 | RELT-like 1 | Plasma Membrane | other | 4827678.5 | 1409878 | 921657.875 | | | | | | |
| Q8K2J7 | RELL1 | RELT-like 1 | Plasma Membrane | other | 3748191.75 | 3555427.75 | 4148240.25 | | | | | | |
| Q8K2J7 | RELL1 | RELT-like 1 | Plasma Membrane | other | 2429654.25 | 2284267.75 | 1886511.5 | Q8K2J7 | RELL1 | 3668508 | 1327458 | 442486 | 9 |
| Q9QUT0 | RHAG | Rh-associated glycoprotein | Plasma Membrane | peptidase | 9.70E+08 | 1.16E+09 | 1.25E+09 | | | | | | |
| Q9QUT0 | RHAG | Rh-associated glycoprotein | Plasma Membrane | peptidase | 5.17E+08 | 6.55E+08 | 6.42E+08 | | | | | | |
| Q9QUT0 | RHAG | Rh-associated glycoprotein | Plasma Membrane | peptidase | 3.22E+07 | 4.72E+07 | 7.51E+07 | | | | | | |
| Q9QUT0 | RHAG | Rh-associated glycoprotein | Plasma Membrane | peptidase | 1.56E+07 | 6195332.5 | 1.52E+07 | Q9QUT0 | RHAG | 383490000 | 480977189 | 138846155 | 12 |
| Q80WQ6 | RHBDF2 | rhomboid 5 homolog 2 (Drosophila | Cytoplasm | other | 1972032.875 | 3238994.25 | 3127019.25 | | | | | | |
| Q80WQ6 | RHBDF2 | rhomboid 5 homolog 2 (Drosophila | Cytoplasm | other | 564829 | 800225.062 | 600243.25 | Q80WQ6 | RHBDF2 | 1268431 | 1247781 | 509404 | 6 |
| Q8CF94 | RHCE/RHD | Rh blood group, D antigen | Plasma Membrane | transporter | 1.04E+09 | 1.30E+09 | 1.43E+09 | | | | | | |
| Q8CF94 | RHCE/RHD | Rh blood group, D antigen | Plasma Membrane | transporter | 1.97E+08 | 1.46E+08 | 1.44E+08 | | | | | | |
| Q8CF94 | RHCE/RHD | Rh blood group, D antigen | Plasma Membrane | transporter | 7.92E+07 | 1.39E+08 | 8.03E+07 | | | | | | |
| Q8CF94 | RHCE/RHD | Rh blood group, D antigen | Plasma Membrane | transporter | 3.94E+07 | 8.59E+07 | 5.68E+07 | | | | | | |
| Q8CF94 | RHCE/RHD | Rh blood group, D antigen | Plasma Membrane | transporter | 3.72E+07 | 4.01E+07 | 3.71E+07 | | | | | | |
| Q8CF94 | RHCE/RHD | Rh blood group, D antigen | Plasma Membrane | transporter | 2.72E+07 | 1.85E+07 | 3.27E+07 | | | | | | |
| Q8CF94 | RHCE/RHD | Rh blood group, D antigen | Plasma Membrane | transporter | 2.80E+07 | 1.03E+08 | 1.31E+08 | | | | | | |
| Q8CF94 | RHCE/RHD | Rh blood group, D antigen | Plasma Membrane | transporter | 2259952.75 | 3720215 | 5791219.5 | Q8CF94 | RHCE/RHD | 180638744 | 408003139 | 83283292 | 24 |
| Q9QUI0 | RHOA | ras homolog family member A | Cytoplasm | enzyme | 3.30E+07 | 3.80E+07 | 3.00E+07 | | | | | | |
| Q9QUI0 | RHOA | ras homolog family member A | Cytoplasm | enzyme | 2.03E+07 | 3.52E+07 | 2.00E+07 | | | | | | |
| Q9QUI0 | RHOA | ras homolog family member A | Cytoplasm | enzyme | 2.77E+07 | 1.15E+07 | 9601048 | | | | | | |
| Q9QUI0 | RHOA | ras homolog family member A | Cytoplasm | enzyme | 1.65E+07 | 8367905.5 | 8027165 | | | | | | |
| Q9QUI0 | RHOA | ras homolog family member A | Cytoplasm | enzyme | 1.24E+07 | 9618544 | 9601423 | | | | | | |
| Q9QUI0 | RHOA | ras homolog family member A | Cytoplasm | enzyme | 7765560.5 | 1.29E+07 | 1.49E+07 | | | | | | |
| Q9QUI0 | RHOA | ras homolog family member A | Cytoplasm | enzyme | 4467063 | 5422648 | 2561719.5 | | | | | | |
| Q9QUI0 | RHOA | ras homolog family member A | Cytoplasm | enzyme | 6322241.5 | 1778957 | 3728727 | | | | | | |
| Q9QUI0 | RHOA | ras homolog family member A | Cytoplasm | enzyme | 2429861.75 | 1.71E+07 | 1.41E+07 | Q9QUI0 | RHOA | 14538303 | 10449115 | 2010933 | 27 |
| P84096 | RHOG | ras homolog family member G | Cytoplasm | enzyme | 6.72E+07 | 5.19E+07 | 5.48E+07 | | | | | | |
| P84096 | RHOG | ras homolog family member G | Cytoplasm | enzyme | 3.69E+07 | 3.84E+07 | 2.67E+07 | | | | | | |
| P84096 | RHOG | ras homolog family member G | Cytoplasm | enzyme | 4.03E+07 | 3.79E+07 | 3.26E+07 | | | | | | |
| P84096 | RHOG | ras homolog family member G | Cytoplasm | enzyme | 2.23E+07 | 2.44E+07 | 1.72E+07 | | | | | | |
| P84096 | RHOG | ras homolog family member G | Cytoplasm | enzyme | 2.28E+07 | 3.14E+07 | 2.41E+07 | | | | | | |
| P84096 | RHOG | ras homolog family member G | Cytoplasm | enzyme | 1.43E+07 | 1.50E+07 | 1.29E+07 | | | | | | |
| P84096 | RHOG | ras homolog family member G | Cytoplasm | enzyme | 1.01E+07 | 1.18E+07 | 9979688 | | | | | | |
| P84096 | RHOG | ras homolog family member G | Cytoplasm | enzyme | 4171397.75 | 2.13E+07 | 1.38E+07 | | | | | | |
| P84096 | RHOG | ras homolog family member G | Cytoplasm | enzyme | 9456835 | 8384211.5 | 8208771 | | | | | | |
| P84096 | RHOG | ras homolog family member G | Cytoplasm | enzyme | 3270417.25 | 2007634.25 | 652955.75 | | | | | | |
| P84096 | RHOG | ras homolog family member G | Cytoplasm | enzyme | 2076044.25 | 5823636.5 | 2737564 | P84096 | RHOG | 21172245 | 16802804 | 2924993 | 33 |
| Q9QYK7 | RNF11 | ring finger protein 11 | Cytoplasm | enzyme | 2872603.25 | 2731386.5 | 2934878.25 | Q9QYK7 | RNF11 | 2872603 | 104267 | 60198 | 3 |
| Q5XPI3 | RNF123 | ring finger protein 123 | Cytoplasm | other | 6289276.5 | 6454876.5 | 6852866.5 | | | | | | |
| Q5XPI3 | RNF123 | ring finger protein 123 | Cytoplasm | other | 6909838.5 | 7276893 | 5679932.5 | | | | | | |
| Q5XPI3 | RNF123 | ring finger protein 123 | Cytoplasm | other | 6250442 | 5717788 | 4404662.5 | | | | | | |
| Q5XPI3 | RNF123 | ring finger protein 123 | Cytoplasm | other | 9644423 | 7543448 | 6072551.5 | | | | | | |
| Q5XPI3 | RNF123 | ring finger protein 123 | Cytoplasm | other | 3989478 | 5783140.5 | 4060984.25 | | | | | | |
| Q5XPI3 | RNF123 | ring finger protein 123 | Cytoplasm | other | 2129960.75 | 2241247.25 | 2498242.75 | | | | | | |
| Q5XPI3 | RNF123 | ring finger protein 123 | Cytoplasm | other | 2798399.75 | 2080207.125 | 2073986.75 | Q5XPI3 | RNF123 | 5430260 | 2174164 | 474442 | 21 |
| E9Q555 | RNF213 | ring finger protein 213 | Cytoplasm | enzyme | 1551312.625 | 1813187.625 | 1508798.75 | E9Q555 | RNF213 | 1551313 | 164843 | 95172 | 3 |
| Q9EPK2 | RP2 | retinitis pigmentosa 2 (X-linked rec | Cytoplasm | enzyme | 1.45E+07 | 1.83E+07 | 1.75E+07 | | | | | | |
| Q9EPK2 | RP2 | retinitis pigmentosa 2 (X-linked rec | Cytoplasm | enzyme | 1.03E+07 | 6059106.5 | 8086702 | | | | | | |
| Q9EPK2 | RP2 | retinitis pigmentosa 2 (X-linked rec | Cytoplasm | enzyme | 6672431 | 8361972.5 | 6262868 | | | | | | |
| Q9EPK2 | RP2 | retinitis pigmentosa 2 (X-linked rec | Cytoplasm | enzyme | 6547773 | 5794359 | 4502851.5 | | | | | | |

| | | | | | | | | | | | | | |
|---|---|---|---|---|---|---|---|---|---|---|---|---|---|
| Q9EPK2 | RP2 | retinitis pigmentosa 2 (X-linked re | Cytoplasm | enzyme | 5273828 | 6763986 | 5910311 | | | | | | |
| Q9EPK2 | RP2 | retinitis pigmentosa 2 (X-linked re | Cytoplasm | enzyme | 5773819.5 | 6874250.5 | 5903077.5 | | | | | | |
| Q9EPK2 | RP2 | retinitis pigmentosa 2 (X-linked re | Cytoplasm | enzyme | 2513884.25 | 3087241.75 | 1922608.875 | Q9EPK2 | RP2 | 7364534 | 4371667 | 953976 | 21 |
| P53026 | RPL10A | ribosomal protein L10a | Nucleus | other | 4037986.5 | 4232933.5 | 3177364 | | | | | | |
| P53026 | RPL10A | ribosomal protein L10a | Nucleus | other | 1934049.375 | 1603336.5 | 1584522.25 | P53026 | RPL10A | 2986018 | 1214806 | 495943 | 6 |
| P86048 | RPL10L | ribosomal protein L10-like | Nucleus | other | 1.02E+07 | 1.39E+07 | 1.41E+07 | | | | | | |
| P86048 | RPL10L | ribosomal protein L10-like | Nucleus | other | 5093302 | 4731934.5 | 4437674 | P86048 | RPL10L | 7641651 | 4578213 | 1869048 | 6 |
| Q9CXW4 | RPL11 | ribosomal protein L11 | Cytoplasm | other | 9962034 | 9587035 | 9954381 | Q9CXW4 | RPL11 | 9962034 | 214331 | 123744 | 3 |
| P35979 | RPL12 | ribosomal protein L12 | Nucleus | other | 6666608 | 6672243 | 5277948.5 | | | | | | |
| P35979 | RPL12 | ribosomal protein L12 | Nucleus | other | 4297744.5 | 8138663.5 | 6724778 | | | | | | |
| P35979 | RPL12 | ribosomal protein L12 | Nucleus | other | 7394704 | 6188299 | 8642036 | | | | | | |
| P35979 | RPL12 | ribosomal protein L12 | Nucleus | other | 6169818.5 | 7508802.5 | 6357107.5 | | | | | | |
| P35979 | RPL12 | ribosomal protein L12 | Nucleus | other | 3293506.5 | 2749631.5 | 3148593.5 | P35979 | RPL12 | 5564476 | 1828141 | 472024 | 15 |
| P47963 | RPL13 | ribosomal protein L13 | Nucleus | other | 6947410 | 5923952 | 6049121.5 | | | | | | |
| P47963 | RPL13 | ribosomal protein L13 | Nucleus | other | 2973456.5 | 3110925.75 | 3682623 | | | | | | |
| P47963 | RPL13 | ribosomal protein L13 | Nucleus | other | 2771666.25 | 2701321.75 | 4201547.5 | | | | | | |
| P47963 | RPL13 | ribosomal protein L13 | Nucleus | other | 3085537.25 | 2455222.5 | 3625989.5 | P47963 | RPL13 | 3944518 | 1510949 | 436173 | 12 |
| P19253 | RPL13A | ribosomal protein L13a | Cytoplasm | other | 3244016.25 | 4477838 | 4787745 | P19253 | RPL13A | 3244016 | 816645 | 471490 | 3 |
| Q9CR57 | RPL14 | ribosomal protein L14 | Cytoplasm | other | 8977736 | 1.04E+07 | 9884404 | | | | | | |
| Q9CR57 | RPL14 | ribosomal protein L14 | Cytoplasm | other | 2341542.75 | 3799193.25 | 4051405 | | | | | | |
| Q9CR57 | RPL14 | ribosomal protein L14 | Cytoplasm | other | 4730713 | 5707232.5 | 5496333.5 | Q9CR57 | RPL14 | 5349997 | 2890127 | 963376 | 9 |
| Q9CZM2 | RPL15 | ribosomal protein L15 | Cytoplasm | other | 4851127.5 | 4758815.5 | 5627168 | | | | | | |
| Q9CZM2 | RPL15 | ribosomal protein L15 | Cytoplasm | other | 3638656.25 | 3821806 | 3270768.25 | | | | | | |
| Q9CZM2 | RPL15 | ribosomal protein L15 | Cytoplasm | other | 4343207.5 | 3740672.25 | 3170793.25 | Q9CZM2 | RPL15 | 4277664 | 817651 | 272550 | 9 |
| Q9CPR4 | RPL17 | ribosomal protein L17 | Cytoplasm | other | 3481732 | 5622112.5 | 4544749.5 | Q9CPR4 | RPL17 | 3481732 | 1070198 | 617879 | 3 |
| P35980 | RPL18 | ribosomal protein L18 | Cytoplasm | other | 1.19E+07 | 1.25E+07 | 1.43E+07 | | | | | | |
| P35980 | RPL18 | ribosomal protein L18 | Cytoplasm | other | 7173961.5 | 7682869.5 | 7809064 | | | | | | |
| P35980 | RPL18 | ribosomal protein L18 | Cytoplasm | other | 5744422.5 | 5217562 | 7023109.5 | | | | | | |
| P35980 | RPL18 | ribosomal protein L18 | Cytoplasm | other | 3764454.75 | 3370162 | 3252237.5 | P35980 | RPL18 | 7143210 | 3677354 | 1061561 | 12 |
| P62717 | RPL18A | ribosomal protein L18a | Cytoplasm | other | 3271303.75 | 3981119.5 | 4304248 | P62717 | RPL18A | 3271304 | 528398 | 305071 | 3 |
| P84099 | RPL19 | ribosomal protein L19 | Cytoplasm | other | 9529166 | 8777190 | 1.11E+07 | P84099 | RPL19 | 9529166 | 1201655 | 693776 | 3 |
| O09167 | RPL21 | ribosomal protein L21 | Cytoplasm | other | 9674164 | 8385851.5 | 8027935 | | | | | | |
| O09167 | RPL21 | ribosomal protein L21 | Cytoplasm | other | 2942602 | 2739106.25 | 3902084.75 | O09167 | RPL21 | 6308383 | 3087675 | 1260538 | 6 |
| P67984 | RPL22 | ribosomal protein L22 | Nucleus | other | 1.65E+07 | 1.63E+07 | 1.36E+07 | | | | | | |
| P67984 | RPL22 | ribosomal protein L22 | Nucleus | other | 6436460 | 6348055 | 6204182 | P67984 | RPL22 | 11458230 | 5109814 | 2086073 | 6 |
| P62830 | RPL23 | ribosomal protein L23 | Cytoplasm | other | 7868211.5 | 7522885.5 | 8087331 | P62830 | RPL23 | 7868212 | 284565 | 164293 | 3 |
| P62751 | Rpl23a | ribosomal protein L23A | Nucleus | other | 1.24E+07 | 1.05E+07 | 1.19E+07 | | | | | | |
| P62751 | Rpl23a | ribosomal protein L23A | Nucleus | other | 2121482.5 | 2031962.25 | 2273332 | P62751 | Rpl23a | 7275741 | 5215938 | 2129398 | 6 |
| Q8BP67 | RPL24 | ribosomal protein L24 | Cytoplasm | other | 2887315.75 | 3981305.5 | 4265508 | | | | | | |
| Q8BP67 | RPL24 | ribosomal protein L24 | Cytoplasm | other | 2144789 | 3100201 | 3573431 | | | | | | |
| Q8BP67 | RPL24 | ribosomal protein L24 | Cytoplasm | other | 2743956.5 | 3091288 | 4036291 | Q8BP67 | RPL24 | 2592020 | 699021 | 233007 | 9 |
| P61255 | RPL26 | ribosomal protein L26 | Cytoplasm | other | 2439459.75 | 2217154.5 | 2101608 | P61255 | RPL26 | 2439460 | 171714 | 99139 | 3 |
| P14115 | RPL27A | ribosomal protein L27a | Cytoplasm | other | 3992592 | 4223087.5 | 4182064.75 | P14115 | RPL27A | 3992592 | 122957 | 70989 | 3 |
| P41105 | RPL28 | ribosomal protein L28 | Cytoplasm | other | 5595847 | 4714549 | 4607308.5 | P41105 | RPL28 | 5595847 | 542432 | 313173 | 3 |
| P47915 | Rpl29 (includes others) | ribosomal protein L29 | Cytoplasm | other | 122581.828 | 1565912.625 | 1127949.375 | P47915 | Rpl29 (includes others) | 122582 | 740020 | 427251 | 3 |
| P27659 | RPL3 | ribosomal protein L3 | Nucleus | other | 1.52E+07 | 1.19E+07 | 1.28E+07 | | | | | | |
| P27659 | RPL3 | ribosomal protein L3 | Nucleus | other | 3781690.5 | 5866240 | 7448791 | | | | | | |
| P27659 | RPL3 | ribosomal protein L3 | Nucleus | other | 3879710 | 4014304.5 | 3697020.5 | P27659 | RPL3 | 7613800 | 4508690 | 1502897 | 9 |
| P62889 | RPL30 | ribosomal protein L30 | Cytoplasm | other | 6810384.5 | 8346940 | 7056670 | P62889 | RPL30 | 6810385 | 825273 | 476472 | 3 |
| P62900 | RPL31 | ribosomal protein L31 | Cytoplasm | other | 6968451 | 6262333 | 7862365.5 | | | | | | |
| P62900 | RPL31 | ribosomal protein L31 | Cytoplasm | other | 9530439 | 8021527 | 8280687.5 | P62900 | RPL31 | 8249445 | 1125318 | 459409 | 6 |
| Q6ZWV7 | RPL35 | ribosomal protein L35 | Cytoplasm | other | 7717195 | 6338226 | 8515773 | | | | | | |
| Q6ZWV7 | RPL35 | ribosomal protein L35 | Cytoplasm | other | 2699497.5 | 3335916.5 | 3748821.5 | Q6ZWV7 | RPL35 | 5208346 | 2459143 | 1003941 | 6 |
| P47964 | Rpl36 | ribosomal protein L36 | Nucleus | other | 2519982.5 | 3049821 | 3291994.75 | | | | | | |
| P47964 | Rpl36 | ribosomal protein L36 | Nucleus | other | 2019662.875 | 2201015 | 2148927.75 | P47964 | Rpl36 | 2269823 | 522379 | 213260 | 6 |
| Q9D8E6 | RPL4 | ribosomal protein L4 | Cytoplasm | enzyme | 5622847.5 | 5176724.5 | 6946121 | | | | | | |
| Q9D8E6 | RPL4 | ribosomal protein L4 | Cytoplasm | enzyme | 5304775 | 4502956.5 | 4896941.5 | | | | | | |
| Q9D8E6 | RPL4 | ribosomal protein L4 | Cytoplasm | enzyme | 4996640 | 2575687.5 | 5509965.5 | | | | | | |
| Q9D8E6 | RPL4 | ribosomal protein L4 | Cytoplasm | enzyme | 1351749 | 3880086 | 2659081 | Q9D8E6 | RPL4 | 4319003 | 1570588 | 453390 | 12 |
| P47962 | RPL5 | ribosomal protein L5 | Cytoplasm | other | 1945173.125 | 1615141.125 | 1882075 | P47962 | RPL5 | 1945173 | 175193 | 101148 | 3 |
| P47911 | RPL6 | ribosomal protein L6 | Nucleus | other | 4609309.5 | 5790277.5 | 6601413 | | | | | | |
| P47911 | RPL6 | ribosomal protein L6 | Nucleus | other | 4133231 | 4820317.5 | 3881199 | | | | | | |
| P47911 | RPL6 | ribosomal protein L6 | Nucleus | other | 2185864.75 | 4299124.5 | 4033884.25 | | | | | | |
| P47911 | RPL6 | ribosomal protein L6 | Nucleus | other | 1247558.625 | 1027211.625 | 522894.625 | P47911 | RPL6 | 3043991 | 1930038 | 557154 | 12 |
| P14148 | RPL7 | ribosomal protein L7 | Nucleus | transcription regulator | 2.02E+07 | 1.37E+07 | 1.50E+07 | | | | | | |
| P14148 | RPL7 | ribosomal protein L7 | Nucleus | transcription regulator | 6331036.5 | 9845236 | 1.08E+07 | | | | | | |
| P14148 | RPL7 | ribosomal protein L7 | Nucleus | transcription regulator | 7162950.5 | 6988442 | 6592689 | | | | | | |
| P14148 | RPL7 | ribosomal protein L7 | Nucleus | transcription regulator | 6174097 | 5320011 | 6524188 | | | | | | |
| P14148 | RPL7 | ribosomal protein L7 | Nucleus | transcription regulator | 944332.062 | 1185415.75 | 2188046 | P14148 | RPL7 | 8158483 | 5276772 | 1362457 | 15 |
| P12970 | RPL7A | ribosomal protein L7a | Cytoplasm | other | 9265541 | 8159989.5 | 8501632 | | | | | | |
| P12970 | RPL7A | ribosomal protein L7a | Cytoplasm | other | 6684104 | 7520126.5 | 7062149 | | | | | | |
| P12970 | RPL7A | ribosomal protein L7a | Cytoplasm | other | 3449209 | 3980494.5 | 3781865.5 | P12970 | RPL7A | 6466285 | 2203502 | 734501 | 9 |
| P62918 | RPL8 | ribosomal protein L8 | Cytoplasm | other | 2044224.25 | 1712569.625 | 2454808.75 | | | | | | |
| P62918 | RPL8 | ribosomal protein L8 | Cytoplasm | other | 1423086.75 | 1137511.75 | 1411297 | P62918 | RPL8 | 1733656 | 482652 | 197042 | 6 |
| P51410 | RPL9 | ribosomal protein L9 | Nucleus | other | 8497193 | 8631625 | 7836338.5 | | | | | | |
| P51410 | RPL9 | ribosomal protein L9 | Nucleus | other | 3568899 | 4496571 | 4824986.5 | | | | | | |
| P51410 | RPL9 | ribosomal protein L9 | Nucleus | other | 4755202 | 5778725 | 6672507 | P51410 | RPL9 | 5607098 | 1871169 | 623723 | 9 |
| P14869 | RPLP0 | ribosomal protein, large, P0 | Cytoplasm | other | 1.50E+07 | 2.07E+07 | 1.78E+07 | | | | | | |
| P14869 | RPLP0 | ribosomal protein, large, P0 | Cytoplasm | other | 9380375 | 7862669.5 | 3251975.25 | | | | | | |
| P14869 | RPLP0 | ribosomal protein, large, P0 | Cytoplasm | other | 7294963.5 | 8962901 | 7887820.5 | | | | | | |
| P14869 | RPLP0 | ribosomal protein, large, P0 | Cytoplasm | other | 8416910 | 5396672.5 | 5390527.5 | | | | | | |
| P14869 | RPLP0 | ribosomal protein, large, P0 | Cytoplasm | other | 8065198 | 6896583.5 | 6922155 | | | | | | |
| P14869 | RPLP0 | ribosomal protein, large, P0 | Cytoplasm | other | 4184328.5 | 4267102 | 3776845.75 | | | | | | |
| P14869 | RPLP0 | ribosomal protein, large, P0 | Cytoplasm | other | 4128464.5 | 7199266.5 | 6965533 | | | | | | |
| P14869 | RPLP0 | ribosomal protein, large, P0 | Cytoplasm | other | 2704504 | 1663201.375 | 2209777.25 | P14869 | RPLP0 | 7399343 | 4685054 | 956333 | 24 |
| P47955 | Rplp1 (includes others) | ribosomal protein, large, P1 | Nucleus | other | 3.71E+07 | 3.73E+07 | 3.28E+07 | | | | | | |

| | | | | | | | | | | | | | |
|---|---|---|---|---|---|---|---|---|---|---|---|---|---|
| P47955 | Rplp1 (includes others) | ribosomal protein, large, P1 | Nucleus | other | 3.05E+07 | 3.34E+07 | 3.57E+07 | P47955 | Rplp1 (includes others) | 33770000 | 2702071 | 1103116 | 6 |
| P99027 | RPLP2 | ribosomal protein, large, P2 | Cytoplasm | other | 3.79E+07 | 4.72E+07 | 3.83E+07 | | | | | | |
| P99027 | RPLP2 | ribosomal protein, large, P2 | Cytoplasm | other | 3.64E+07 | 4.67E+07 | 4.15E+07 | | | | | | |
| P99027 | RPLP2 | ribosomal protein, large, P2 | Cytoplasm | other | 9638546 | 9100444 | 8643459 | | | | | | |
| P99027 | RPLP2 | ribosomal protein, large, P2 | Cytoplasm | other | 5007948.5 | 5306537 | 4590309 | | | | | | |
| P99027 | RPLP2 | ribosomal protein, large, P2 | Cytoplasm | other | 3625977.5 | 3994749.25 | 3960737 | P99027 | RPLP2 | 18524494 | 18232113 | 4707511 | 15 |
| Q91YQ5 | RPN1 | ribophorin I | Cytoplasm | enzyme | 5327427.5 | 5555895 | 6728359.5 | | | | | | |
| Q91YQ5 | RPN1 | ribophorin I | Cytoplasm | enzyme | 7760461.5 | 8579112 | 1.04E+07 | | | | | | |
| Q91YQ5 | RPN1 | ribophorin I | Cytoplasm | enzyme | 9141091 | 9982273 | 9927894 | | | | | | |
| Q91YQ5 | RPN1 | ribophorin I | Cytoplasm | enzyme | 4779382 | 4578249.5 | 4925840.5 | | | | | | |
| Q91YQ5 | RPN1 | ribophorin I | Cytoplasm | enzyme | 4009283 | 4419368 | 4496637.5 | | | | | | |
| Q91YQ5 | RPN1 | ribophorin I | Cytoplasm | enzyme | 4442613 | 4488506.5 | 5024849 | | | | | | |
| Q91YQ5 | RPN1 | ribophorin I | Cytoplasm | enzyme | 1620677.625 | 2852212.75 | 2889825.5 | | | | | | |
| Q91YQ5 | RPN1 | ribophorin I | Cytoplasm | enzyme | 1680535.875 | 2187104.75 | 2555043.25 | Q91YQ5 | RPN1 | 4845184 | 2670395 | 545092 | 24 |
| Q9DBG6 | RPN2 | ribophorin II | Cytoplasm | enzyme | 1.05E+07 | 1.26E+07 | 1.38E+07 | | | | | | |
| Q9DBG6 | RPN2 | ribophorin II | Cytoplasm | enzyme | 1.08E+07 | 1.11E+07 | 1.58E+07 | | | | | | |
| Q9DBG6 | RPN2 | ribophorin II | Cytoplasm | enzyme | 5235477 | 4859458.5 | 7349990 | | | | | | |
| Q9DBG6 | RPN2 | ribophorin II | Cytoplasm | enzyme | 5650107.5 | 5271679.5 | 7315904.5 | | | | | | |
| Q9DBG6 | RPN2 | ribophorin II | Cytoplasm | enzyme | 3456247 | 5585500.5 | 6567573 | | | | | | |
| Q9DBG6 | RPN2 | ribophorin II | Cytoplasm | enzyme | 3502468.5 | 3794415.25 | 4014410 | | | | | | |
| Q9DBG6 | RPN2 | ribophorin II | Cytoplasm | enzyme | 3842183.75 | 4579918.5 | 4676468 | | | | | | |
| Q9DBG6 | RPN2 | ribophorin II | Cytoplasm | enzyme | 2115894.75 | 2509261.5 | 3348803 | | | | | | |
| Q9DBG6 | RPN2 | ribophorin II | Cytoplasm | enzyme | 3370842.5 | 4352502 | 4218491.5 | | | | | | |
| Q9DBG6 | RPN2 | ribophorin II | Cytoplasm | enzyme | 671168.188 | 1387096.25 | 1629335.875 | Q9DBG6 | RPN2 | 4913439 | 3810119 | 695629 | 30 |
| P63325 | RPS10 | ribosomal protein S10 | Cytoplasm | other | 6315160 | 5685892 | 8140162 | P63325 | RPS10 | 6315160 | 1274758 | 735982 | 3 |
| P62281 | RPS11 | ribosomal protein S11 | Cytoplasm | other | 3626410 | 5774792 | 6442573.5 | | | | | | |
| P62281 | RPS11 | ribosomal protein S11 | Cytoplasm | other | 425802.812 | 1387723.5 | 928184.875 | P62281 | RPS11 | 2026106 | 2584739 | 1055215 | 6 |
| P62301 | RPS13 | ribosomal protein S13 | Cytoplasm | other | 2005376.625 | 2649089 | 2651782.75 | | | | | | |
| P62301 | RPS13 | ribosomal protein S13 | Cytoplasm | other | 2199053 | 2681025.75 | 2335263.75 | P62301 | RPS13 | 2102215 | 283637 | 115794 | 6 |
| P62264 | RPS14 | ribosomal protein S14 | Cytoplasm | translation regulator | 1.17E+07 | 1.34E+07 | 1.08E+07 | | | | | | |
| P62264 | RPS14 | ribosomal protein S14 | Cytoplasm | translation regulator | 7994271.5 | 6388385.5 | 7045939 | | | | | | |
| P62264 | RPS14 | ribosomal protein S14 | Cytoplasm | translation regulator | 3570158 | 7782734.5 | 5990862.5 | P62264 | RPS14 | 7768143 | 3108230 | 1036077 | 9 |
| P62245 | RPS15A | ribosomal protein S15a | Cytoplasm | other | 2520437.5 | 2945630.75 | 2968847.25 | | | | | | |
| P62245 | RPS15A | ribosomal protein S15a | Cytoplasm | other | 3159274.25 | 3629679 | 4840611.5 | P62245 | RPS15A | 2839856 | 816489 | 333330 | 6 |
| P14131 | RPS16 | ribosomal protein S16 | Cytoplasm | other | 8643001 | 8774732 | 9062743 | | | | | | |
| P14131 | RPS16 | ribosomal protein S16 | Cytoplasm | other | 4387335.5 | 4765234.5 | 4921689.5 | P14131 | RPS16 | 6515168 | 2275762 | 929076 | 6 |
| P63276 | RPS17 | ribosomal protein S17 | Cytoplasm | other | 2920988.75 | 2687229 | 2737328.75 | P63276 | RPS17 | 2920989 | 123075 | 71057 | 3 |
| P62270 | RPS18 | ribosomal protein S18 | Cytoplasm | other | 5895118 | 5695343 | 5734094 | | | | | | |
| P62270 | RPS18 | ribosomal protein S18 | Cytoplasm | other | 3737815.25 | 3965968.5 | 3979930.25 | | | | | | |
| P62270 | RPS18 | ribosomal protein S18 | Cytoplasm | other | 3141127 | 4232212.5 | 5198525.5 | P62270 | RPS18 | 4258020 | 1019272 | 339757 | 9 |
| Q9CZX8 | RPS19 | ribosomal protein S19 | Cytoplasm | other | 5882127.5 | 6082319.5 | 6839443 | | | | | | |
| Q9CZX8 | RPS19 | ribosomal protein S19 | Cytoplasm | other | 5845608 | 4643716.5 | 5383456.5 | | | | | | |
| Q9CZX8 | RPS19 | ribosomal protein S19 | Cytoplasm | other | 5288220.5 | 6269017 | 6703983 | Q9CZX8 | RPS19 | 5671985 | 699826 | 233275 | 9 |
| P25444 | RPS2 | ribosomal protein S2 | Cytoplasm | other | 9333378 | 9840537 | 8857184 | | | | | | |
| P25444 | RPS2 | ribosomal protein S2 | Cytoplasm | other | 6582711 | 7755939 | 8691887 | | | | | | |
| P25444 | RPS2 | ribosomal protein S2 | Cytoplasm | other | 5398539.5 | 6761953.5 | 6082639 | | | | | | |
| P25444 | RPS2 | ribosomal protein S2 | Cytoplasm | other | 4207850 | 5775742.5 | 4609663.5 | P25444 | RPS2 | 6380620 | 1883930 | 543844 | 12 |
| P60867 | RPS20 | ribosomal protein S20 | Cytoplasm | other | 7591449 | 6911249.5 | 7656406.5 | | | | | | |
| P60867 | RPS20 | ribosomal protein S20 | Cytoplasm | other | 6213752.5 | 4817062 | 5764709 | P60867 | RPS20 | 6902601 | 1109251 | 452850 | 6 |
| P62267 | RPS23 | ribosomal protein S23 | Cytoplasm | translation regulator | 4146498.5 | 4723091.5 | 5171010 | P62267 | RPS23 | 4146499 | 513601 | 296528 | 3 |
| P62849 | RPS24 | ribosomal protein S24 | Cytoplasm | other | 2979128 | 2766393.25 | 2675179 | P62849 | RPS24 | 2979128 | 155971 | 90050 | 3 |
| P62852 | RPS25 | ribosomal protein S25 | Cytoplasm | other | 8506127 | 8971297 | 9264352 | | | | | | |
| P62852 | RPS25 | ribosomal protein S25 | Cytoplasm | other | 7056208.5 | 7442955 | 7136745 | | | | | | |
| P62852 | RPS25 | ribosomal protein S25 | Cytoplasm | other | 5231329.5 | 5831485 | 5048101 | P62852 | RPS25 | 6931222 | 1563534 | 521178 | 9 |
| P62855 | RPS26 | ribosomal protein S26 | Cytoplasm | other | 4220052 | 3895005.5 | 4288775 | P62855 | RPS26 | 4220052 | 210330 | 121434 | 3 |
| Q6ZWY3 | RPS27L | ribosomal protein S27-like | Cytoplasm | translation regulator | 1371564.25 | 2003789.25 | 1962893.5 | | | | | | |
| Q6ZWY3 | RPS27L | ribosomal protein S27-like | Cytoplasm | translation regulator | 537130.5 | 363973.219 | 480864.031 | Q6ZWY3 | RPS27L | 954347 | 758241 | 309551 | 6 |
| P62858 | RPS28 | ribosomal protein S28 | Cytoplasm | other | 564345.5 | 2441046.75 | 1279476.75 | P62858 | RPS28 | 564346 | 947159 | 546843 | 3 |
| P62908 | RPS3 | ribosomal protein S3 | Cytoplasm | enzyme | 9344693 | 7573859 | 8763033 | | | | | | |
| P62908 | RPS3 | ribosomal protein S3 | Cytoplasm | enzyme | 7495171 | 7657681.5 | 6106504.5 | | | | | | |
| P62908 | RPS3 | ribosomal protein S3 | Cytoplasm | enzyme | 3470319.75 | 4043636.5 | 3570831.25 | | | | | | |
| P62908 | RPS3 | ribosomal protein S3 | Cytoplasm | enzyme | 6956696 | 5358883.5 | 5475851.5 | | | | | | |
| P62908 | RPS3 | ribosomal protein S3 | Cytoplasm | enzyme | 4029132.25 | 4107679.25 | 3874682.75 | | | | | | |
| P62908 | RPS3 | ribosomal protein S3 | Cytoplasm | enzyme | 4125329.25 | 1158232.5 | 2996071.25 | | | | | | |
| P62908 | RPS3 | ribosomal protein S3 | Cytoplasm | enzyme | 1503306.875 | 1880615.75 | 2659522.75 | | | | | | |
| P62908 | RPS3 | ribosomal protein S3 | Cytoplasm | enzyme | 1055300.125 | 1607006.125 | 2015977.75 | P62908 | RPS3 | 4747494 | 2488118 | 507885 | 24 |
| P97351 | Rps3a1 | ribosomal protein S3A1 | Cytoplasm | other | 2309796.5 | 2459178.5 | 2092304.125 | | | | | | |
| P97351 | Rps3a1 | ribosomal protein S3A1 | Cytoplasm | other | 85298.562 | 643210.812 | 1024591.375 | P97351 | Rps3a1 | 1197548 | 986237 | 402629 | 6 |
| P62702 | RPS4Y1 | ribosomal protein S4, Y-linked 1 | Cytoplasm | other | 7990398 | 5863548.5 | 7000177 | | | | | | |
| P62702 | RPS4Y1 | ribosomal protein S4, Y-linked 1 | Cytoplasm | other | 4714335.5 | 5294567 | 3258679.5 | | | | | | |
| P62702 | RPS4Y1 | ribosomal protein S4, Y-linked 1 | Cytoplasm | other | 4305263 | 1472187.375 | 1362561 | | | | | | |
| P62702 | RPS4Y1 | ribosomal protein S4, Y-linked 1 | Cytoplasm | other | 2112140.5 | 3756454.5 | 2454356.5 | | | | | | |
| P62702 | RPS4Y1 | ribosomal protein S4, Y-linked 1 | Cytoplasm | other | 2158161.5 | 3171071.75 | 2545457.5 | | | | | | |
| P62702 | RPS4Y1 | ribosomal protein S4, Y-linked 1 | Cytoplasm | other | 1968425.75 | 2213589.5 | 1756202.5 | | | | | | |
| P62702 | RPS4Y1 | ribosomal protein S4, Y-linked 1 | Cytoplasm | other | 2300697.75 | 2627158.25 | 2509077.25 | | | | | | |
| P62702 | RPS4Y1 | ribosomal protein S4, Y-linked 1 | Cytoplasm | other | 2002193.75 | 2494810 | 1707738.125 | P62702 | RPS4Y1 | 3443952 | 1777389 | 362808 | 24 |
| P62754 | RPS6 | ribosomal protein S6 | Cytoplasm | other | 5623660 | 5597652 | 4087581.75 | | | | | | |
| P62754 | RPS6 | ribosomal protein S6 | Cytoplasm | other | 2926484.75 | 825408.938 | 1088745.125 | P62754 | RPS6 | 4275072 | 2118079 | 864702 | 6 |
| P62082 | RPS7 | ribosomal protein S7 | Cytoplasm | other | 1.28E+07 | 1.29E+07 | 1.32E+07 | | | | | | |
| P62082 | RPS7 | ribosomal protein S7 | Cytoplasm | other | 7640149 | 6674506.5 | 8352110 | | | | | | |
| P62082 | RPS7 | ribosomal protein S7 | Cytoplasm | other | 2497235 | 2449321.5 | 3236646.5 | P62082 | RPS7 | 7629128 | 4459894 | 1486631 | 9 |
| P62242 | RPS8 | ribosomal protein S8 | Cytoplasm | other | 7685853.5 | 1.06E+07 | 8849224 | | | | | | |
| P62242 | RPS8 | ribosomal protein S8 | Cytoplasm | other | 7751396.5 | 9163492 | 8135713 | | | | | | |
| P62242 | RPS8 | ribosomal protein S8 | Cytoplasm | other | 5736776 | 6129378.5 | 7162226 | | | | | | |
| P62242 | RPS8 | ribosomal protein S8 | Cytoplasm | other | 3000655.5 | 3205656.25 | 3569948.5 | | | | | | |
| P62242 | RPS8 | ribosomal protein S8 | Cytoplasm | other | 2445617.25 | 2001427 | 2513155.75 | P62242 | RPS8 | 5324060 | 2858611 | 738090 | 15 |

| | | | | | | | | | | | | | |
|---|---|---|---|---|---|---|---|---|---|---|---|---|---|
| Q6ZWN5 | RPS9 | ribosomal protein S9 | Cytoplasm | translation regulator | 7638436.5 | 6759360.5 | 7949960.5 | | | | | | |
| Q6ZWN5 | RPS9 | ribosomal protein S9 | Cytoplasm | translation regulator | 4667221 | 4816892 | 4314141 | | | | | | |
| Q6ZWN5 | RPS9 | ribosomal protein S9 | Cytoplasm | translation regulator | 2731007.25 | 2651957.5 | 2262186.5 | Q6ZWN5 | RPS9 | 5012222 | 2161265 | 720422 | 9 |
| P14206 | RPSA | ribosomal protein SA | Cytoplasm | translation regulator | 1.89E+07 | 1.92E+07 | 1.95E+07 | | | | | | |
| P14206 | RPSA | ribosomal protein SA | Cytoplasm | translation regulator | 9827595 | 9676845 | 8843464 | | | | | | |
| P14206 | RPSA | ribosomal protein SA | Cytoplasm | translation regulator | 6016896.5 | 7672548.5 | 7152986.5 | | | | | | |
| P14206 | RPSA | ribosomal protein SA | Cytoplasm | translation regulator | 4836437.5 | 3671169.75 | 4217455 | P14206 | RPSA | 9902732 | 5910780 | 1706295 | 12 |
| Q8CBB9 | RSAD2 | radical S-adenosyl methionine dor | Cytoplasm | enzyme | 1.12E+07 | 5734912 | 1.48E+07 | | | | | | |
| Q8CBB9 | RSAD2 | radical S-adenosyl methionine dor | Cytoplasm | enzyme | 7628633.5 | 1.06E+07 | 2.06E+07 | | | | | | |
| Q8CBB9 | RSAD2 | radical S-adenosyl methionine dor | Cytoplasm | enzyme | 5977738 | 5125498.5 | 7956007.5 | | | | | | |
| Q8CBB9 | RSAD2 | radical S-adenosyl methionine dor | Cytoplasm | enzyme | 6570424.5 | 7321833 | 1.91E+07 | | | | | | |
| Q8CBB9 | RSAD2 | radical S-adenosyl methionine dor | Cytoplasm | enzyme | 3127774.25 | 4546387 | 9145352 | | | | | | |
| Q8CBB9 | RSAD2 | radical S-adenosyl methionine dor | Cytoplasm | enzyme | 2321619.75 | 2212410.25 | 4696587 | | | | | | |
| Q8CBB9 | RSAD2 | radical S-adenosyl methionine dor | Cytoplasm | enzyme | 4478998.5 | 2064752.625 | 6602436.5 | | | | | | |
| Q8CBB9 | RSAD2 | radical S-adenosyl methionine dor | Cytoplasm | enzyme | 2950840.75 | 2038789.25 | 4634167.5 | Q8CBB9 | RSAD2 | 5533254 | 5033657 | 1027491 | 24 |
| Q01730 | RSU1 | Ras suppressor protein 1 | Cytoplasm | other | 1.02E+07 | 9665898 | 1.07E+07 | | | | | | |
| Q01730 | RSU1 | Ras suppressor protein 1 | Cytoplasm | other | 9802048 | 6589966 | 7673270 | | | | | | |
| Q01730 | RSU1 | Ras suppressor protein 1 | Cytoplasm | other | 3963509 | 700149 | 1719471.875 | Q01730 | RSU1 | 7971852 | 3805257 | 1268419 | 9 |
| Q8K0T0 | RTN1 | reticulon 1 | Cytoplasm | other | 6433126.5 | 1.65E+07 | 1.56E+07 | Q8K0T0 | RTN1 | 6433127 | 5573794 | 3218031 | 3 |
| Q9ES97 | RTN3 | reticulon 3 | Cytoplasm | other | 7.41E+07 | 9.03E+07 | 8.36E+07 | | | | | | |
| Q9ES97 | RTN3 | reticulon 3 | Cytoplasm | other | 424607.156 | 1153544.875 | 1279254.75 | Q9ES97 | RTN3 | 37267304 | 45060083 | 18395702 | 6 |
| Q99P72 | RTN4 | reticulon 4 | Cytoplasm | other | 1.20E+07 | 2.39E+07 | 2.61E+07 | | | | | | |
| Q99P72 | RTN4 | reticulon 4 | Cytoplasm | other | 1.42E+07 | 1.75E+07 | 1.29E+07 | | | | | | |
| Q99P72 | RTN4 | reticulon 4 | Cytoplasm | other | 7913878.5 | 1.73E+07 | 9714843 | | | | | | |
| Q99P72 | RTN4 | reticulon 4 | Cytoplasm | other | 4276852.5 | 4758359.5 | 3768407.5 | | | | | | |
| Q99P72 | RTN4 | reticulon 4 | Cytoplasm | other | 4398865 | 4745254 | 4475939 | Q99P72 | RTN4 | 8569919 | 7383083 | 1906304 | 15 |
| P97352 | S100A13 | S100 calcium binding protein A13 | Cytoplasm | other | 4837035 | 5819831 | 7875154 | P97352 | S100A13 | 4837035 | 1550291 | 895061 | 3 |
| P31725 | S100A9 | S100 calcium binding protein A9 | Cytoplasm | other | 1005405.25 | 732410 | 665457 | | | | | | |
| P31725 | S100A9 | S100 calcium binding protein A9 | Cytoplasm | other | 985756.75 | 582468.312 | 283005.875 | P31725 | S100A9 | 995581 | 269954 | 110208 | 6 |
| Q9EP69 | SACM1L | SAC1 suppressor of actin mutation | Cytoplasm | phosphatase | 1.12E+07 | 1.05E+07 | 1.12E+07 | | | | | | |
| Q9EP69 | SACM1L | SAC1 suppressor of actin mutation | Cytoplasm | phosphatase | 8953491 | 1.10E+07 | 8908258 | | | | | | |
| Q9EP69 | SACM1L | SAC1 suppressor of actin mutation | Cytoplasm | phosphatase | 1.14E+07 | 1.09E+07 | 8934025 | | | | | | |
| Q9EP69 | SACM1L | SAC1 suppressor of actin mutation | Cytoplasm | phosphatase | 8383459.5 | 9391290 | 1.00E+07 | | | | | | |
| Q9EP69 | SACM1L | SAC1 suppressor of actin mutation | Cytoplasm | phosphatase | 5269326 | 5445250 | 4041369.75 | | | | | | |
| Q9EP69 | SACM1L | SAC1 suppressor of actin mutation | Cytoplasm | phosphatase | 4852021 | 7091412 | 3649893.5 | | | | | | |
| Q9EP69 | SACM1L | SAC1 suppressor of actin mutation | Cytoplasm | phosphatase | 3766490.75 | 7253443.5 | 5497602.5 | | | | | | |
| Q9EP69 | SACM1L | SAC1 suppressor of actin mutation | Cytoplasm | phosphatase | 4201170.5 | 4726115 | 4352101 | | | | | | |
| Q9EP69 | SACM1L | SAC1 suppressor of actin mutation | Cytoplasm | phosphatase | 3146090.75 | 5670750.5 | 3641175.25 | | | | | | |
| Q9EP69 | SACM1L | SAC1 suppressor of actin mutation | Cytoplasm | phosphatase | 4933654.5 | 4552554.5 | 3844841.25 | | | | | | |
| Q9EP69 | SACM1L | SAC1 suppressor of actin mutation | Cytoplasm | phosphatase | 5501420.5 | 4673795.5 | 3637277.75 | | | | | | |
| Q9EP69 | SACM1L | SAC1 suppressor of actin mutation | Cytoplasm | phosphatase | 3633939.5 | 3539156.75 | 3751108.75 | | | | | | |
| Q9EP69 | SACM1L | SAC1 suppressor of actin mutation | Cytoplasm | phosphatase | 2375966.5 | 2362638 | 2730204.25 | | | | | | |
| Q9EP69 | SACM1L | SAC1 suppressor of actin mutation | Cytoplasm | phosphatase | 1723596.125 | 2608028.5 | 2444937.75 | | | | | | |
| Q9EP69 | SACM1L | SAC1 suppressor of actin mutation | Cytoplasm | phosphatase | 1961883.125 | 1957387 | 2043749.875 | | | | | | |
| Q9EP69 | SACM1L | SAC1 suppressor of actin mutation | Cytoplasm | phosphatase | 1853391.125 | 2274905.5 | 2015381 | | | | | | |
| Q9EP69 | SACM1L | SAC1 suppressor of actin mutation | Cytoplasm | phosphatase | 2221976.75 | 2978439.75 | 2313458.75 | | | | | | |
| Q9EP69 | SACM1L | SAC1 suppressor of actin mutation | Cytoplasm | phosphatase | 1436883.125 | 2409421.75 | 1660849.25 | | | | | | |
| Q9EP69 | SACM1L | SAC1 suppressor of actin mutation | Cytoplasm | phosphatase | 356223.938 | 985312.938 | 2135591 | Q9EP69 | SACM1L | 4590052 | 3108043 | 411670 | 57 |
| Q8BGH2 | SAMM50 | SAMM50 sorting and assembly ma | Cytoplasm | other | 1386888.125 | 2127904.25 | 3229070.25 | | | | | | |
| Q8BGH2 | SAMM50 | SAMM50 sorting and assembly ma | Cytoplasm | other | 3099982 | 1117195.75 | 1890879 | Q8BGH2 | SAMM50 | 2243435 | 869842 | 355112 | 6 |
| Q8K021 | SCAMP1 | secretory carrier membrane prote | Cytoplasm | transporter | 1.39E+07 | 1.32E+07 | 1.07E+07 | Q8K021 | SCAMP1 | 13890000 | 1673091 | 965960 | 3 |
| Q9ERN0 | SCAMP2 | secretory carrier membrane prote | Cytoplasm | transporter | 8217590 | 7121869.5 | 7510368 | | | | | | |
| Q9ERN0 | SCAMP2 | secretory carrier membrane prote | Cytoplasm | transporter | 5449707.5 | 7378670.5 | 6581866 | Q9ERN0 | SCAMP2 | 6833649 | 945327 | 385928 | 6 |
| Q9JKV5 | SCAMP4 | secretory carrier membrane prote | Cytoplasm | other | 8875946 | 8480040 | 9138619 | | | | | | |
| Q9JKV5 | SCAMP4 | secretory carrier membrane prote | Cytoplasm | other | 4635136.5 | 1.08E+07 | 8246047 | Q9JKV5 | SCAMP4 | 6755541 | 2028917 | 828302 | 6 |
| O35114 | SCARB2 | scavenger receptor class B, memb | Plasma Membrane | other | 1270428.5 | 2600042.25 | 4445639 | O35114 | SCARB2 | 1270429 | 1594577 | 920630 | 3 |
| Q8K2B3 | SDHA | succinate dehydrogenase complex | Cytoplasm | enzyme | 1.10E+07 | 1.42E+07 | 1.66E+07 | | | | | | |
| Q8K2B3 | SDHA | succinate dehydrogenase complex | Cytoplasm | enzyme | 5605102.5 | 1.22E+07 | 1.30E+07 | | | | | | |
| Q8K2B3 | SDHA | succinate dehydrogenase complex | Cytoplasm | enzyme | 6120840 | 6878308.5 | 9338156 | | | | | | |
| Q8K2B3 | SDHA | succinate dehydrogenase complex | Cytoplasm | enzyme | 4857469 | 5463276 | 6034409.5 | | | | | | |
| Q8K2B3 | SDHA | succinate dehydrogenase complex | Cytoplasm | enzyme | 3707837.75 | 4481517 | 4931750.5 | | | | | | |
| Q8K2B3 | SDHA | succinate dehydrogenase complex | Cytoplasm | enzyme | 4573035.5 | 4218983 | 4946409 | | | | | | |
| Q8K2B3 | SDHA | succinate dehydrogenase complex | Cytoplasm | enzyme | 3962331 | 4817512.5 | 7159254 | | | | | | |
| Q8K2B3 | SDHA | succinate dehydrogenase complex | Cytoplasm | enzyme | 2125842.25 | 1677645.75 | 2385862.25 | Q8K2B3 | SDHA | 5241557 | 3968699 | 810107 | 24 |
| Q9CQA3 | SDHB | succinate dehydrogenase complex | Cytoplasm | enzyme | 4052483.75 | 4582918 | 5821650 | | | | | | |
| Q9CQA3 | SDHB | succinate dehydrogenase complex | Cytoplasm | enzyme | 3438047.5 | 3085569.5 | 5236711 | | | | | | |
| Q9CQA3 | SDHB | succinate dehydrogenase complex | Cytoplasm | enzyme | 2729262 | 2642012.25 | 3596961.25 | Q9CQA3 | SDHB | 3406598 | 1111870 | 370623 | 9 |
| Q9CZB0 | SDHC | succinate dehydrogenase complex | Cytoplasm | enzyme | 2051191.875 | 2907629.5 | 4299752.5 | Q9CZB0 | SDHC | 2051192 | 1134865 | 655215 | 3 |
| O08547 | SEC22B | SEC22 vesicle trafficking protein h | Cytoplasm | other | 1.07E+07 | 3639120.25 | 6417511.5 | | | | | | |
| O08547 | SEC22B | SEC22 vesicle trafficking protein h | Cytoplasm | other | 9339982 | 1604281 | 4740748 | | | | | | |
| O08547 | SEC22B | SEC22 vesicle trafficking protein h | Cytoplasm | other | 4590503.5 | 6723267 | 4739253 | | | | | | |
| O08547 | SEC22B | SEC22 vesicle trafficking protein h | Cytoplasm | other | 5051006.5 | 5439680.5 | 5390638 | | | | | | |
| O08547 | SEC22B | SEC22 vesicle trafficking protein h | Cytoplasm | other | 2633198 | 2603387.25 | 2184493.25 | | | | | | |
| O08547 | SEC22B | SEC22 vesicle trafficking protein h | Cytoplasm | other | 2827070.5 | 3707162 | 3620492.5 | | | | | | |
| O08547 | SEC22B | SEC22 vesicle trafficking protein h | Cytoplasm | other | 2508006.5 | 2016801.125 | 2625901.75 | | | | | | |
| O08547 | SEC22B | SEC22 vesicle trafficking protein h | Cytoplasm | other | 839339 | 2648187.5 | 1776998.125 | O08547 | SEC22B | 4809888 | 2397932 | 489476 | 24 |
| Q3TZ89 | SEC31B | SEC31 homolog B, COPII coating co | Cytoplasm | other | 2.85E+07 | 2.78E+07 | 2.92E+07 | Q3TZ89 | SEC31B | 28520000 | 715285 | 412970 | 3 |
| P61620 | SEC61A1 | Sec61 alpha 1 subunit (S. cerevisia | Cytoplasm | transporter | 3385534 | 3710562 | 3458428.25 | P61620 | SEC61A1 | 3385534 | 170552 | 98468 | 3 |
| Q9CQS8 | SEC61B | Sec61 beta subunit | Cytoplasm | transporter | 6388044 | 8535371 | 6621065.5 | | | | | | |
| Q9CQS8 | SEC61B | Sec61 beta subunit | Cytoplasm | transporter | 2895015.5 | 4414535 | 3347356.75 | Q9CQS8 | SEC61B | 4641530 | 2179479 | 889769 | 6 |
| P17563 | SELENBP1 | selenium binding protein 1 | Cytoplasm | other | 1845548 | 489758.688 | 967821.438 | P17563 | SELENBP1 | 1845548 | 687642 | 397011 | 3 |
| Q01102 | SELP | selectin P (granule membrane pro | Plasma Membrane | transmembrane receptor | 4290287.5 | 3831943.75 | 5108532.5 | | | | | | |
| Q01102 | SELP | selectin P (granule membrane pro | Plasma Membrane | transmembrane receptor | 1.41E+07 | 1.90E+07 | 2.09E+07 | | | | | | |
| Q01102 | SELP | selectin P (granule membrane pro | Plasma Membrane | transmembrane receptor | 1.12E+07 | 1.21E+07 | 1.26E+07 | | | | | | |
| Q01102 | SELP | selectin P (granule membrane pro | Plasma Membrane | transmembrane receptor | 1914582.5 | 1834187.25 | 2160133 | | | | | | |
| Q01102 | SELP | selectin P (granule membrane pro | Plasma Membrane | transmembrane receptor | 3253821.5 | 4231435 | 5299419.5 | | | | | | |

| | | | | | | | | | | | | | |
|---|---|---|---|---|---|---|---|---|---|---|---|---|---|
| Q01102 | SELP | selectin P (granule membrane pro | Plasma Membrane | transmembrane receptor | 580034 | 1483228.875 | 1253307.625 | Q01102 | SELP | 5883121 | 6358799 | 1498783 | 18 |
| P62342 | SELT | selenoprotein T | Cytoplasm | other | 2510359.5 | 4114979.75 | 3303024.5 | P62342 | SELT | 2510360 | 802329 | 463225 | 3 |
| Q9CY58 | SERBP1 | SERPINE1 mRNA binding protein 1 | Cytoplasm | other | 1308044.125 | 1241915.875 | 1472231 | | | | | | |
| Q9CY58 | SERBP1 | SERPINE1 mRNA binding protein 1 | Cytoplasm | other | 1038839.562 | 3125274.75 | 2119011 | | | | | | |
| Q9CY58 | SERBP1 | SERPINE1 mRNA binding protein 1 | Cytoplasm | other | 702951.625 | 2254752 | 1531411.25 | Q9CY58 | SERBP1 | 1016612 | 738354 | 246118 | 9 |
| Q9QZI9 | SERINC3 | serine incorporator 3 | Cytoplasm | transporter | 2.78E+08 | 2.49E+08 | 2.67E+08 | | | | | | |
| Q9QZI9 | SERINC3 | serine incorporator 3 | Cytoplasm | transporter | 1.21E+08 | 1.48E+08 | 1.19E+08 | | | | | | |
| Q9QZI9 | SERINC3 | serine incorporator 3 | Cytoplasm | transporter | 1.09E+08 | 1.67E+08 | 1.58E+08 | | | | | | |
| Q9QZI9 | SERINC3 | serine incorporator 3 | Cytoplasm | transporter | 1.98E+07 | 4.54E+07 | 4.60E+07 | Q9QZI9 | SERINC3 | 131992500 | 86263830 | 24902223 | 12 |
| P07758 | SERPINA1 | serpin peptidase inhibitor, clade A | Extracellular Space | other | 6082473.5 | 6539182 | 8455595 | | | | | | |
| P07758 | SERPINA1 | serpin peptidase inhibitor, clade A | Extracellular Space | other | 1898297.375 | 1776959.5 | 3749687.5 | | | | | | |
| P07758 | SERPINA1 | serpin peptidase inhibitor, clade A | Extracellular Space | other | 2205641.5 | 2305887 | 2562615.25 | | | | | | |
| P07758 | SERPINA1 | serpin peptidase inhibitor, clade A | Extracellular Space | other | 1666126.875 | 1215576 | 1932670.75 | | | | | | |
| P07758 | SERPINA1 | serpin peptidase inhibitor, clade A | Extracellular Space | other | 1752973.5 | 1176314.875 | 2447959.75 | P07758 | SERPINA1 | 2721103 | 2197654 | 567432 | 15 |
| P07759 | SERPINA3 | serpin peptidase inhibitor, clade A | Extracellular Space | other | 2.03E+07 | 2.05E+07 | 1.95E+07 | | | | | | |
| P07759 | SERPINA3 | serpin peptidase inhibitor, clade A | Extracellular Space | other | 1.63E+07 | 1.50E+07 | 1.29E+07 | | | | | | |
| P07759 | SERPINA3 | serpin peptidase inhibitor, clade A | Extracellular Space | other | 1.21E+07 | 6817383 | 6653267 | P07759 | SERPINA3 | 16240000 | 5330625 | 1776875 | 9 |
| Q99JR1 | SFXN1 | sideroflexin 1 | Cytoplasm | transporter | 5784730 | 5258456 | 7787154 | | | | | | |
| Q99JR1 | SFXN1 | sideroflexin 1 | Cytoplasm | transporter | 6075591 | 5457623 | 8173566 | | | | | | |
| Q99JR1 | SFXN1 | sideroflexin 1 | Cytoplasm | transporter | 4897163.5 | 7696445.5 | 8178090 | | | | | | |
| Q99JR1 | SFXN1 | sideroflexin 1 | Cytoplasm | transporter | 4420846.5 | 3822552 | 5203014.5 | | | | | | |
| Q99JR1 | SFXN1 | sideroflexin 1 | Cytoplasm | transporter | 3557490 | 3063372 | 4260060.5 | | | | | | |
| Q99JR1 | SFXN1 | sideroflexin 1 | Cytoplasm | transporter | 1254969.5 | 1345987.5 | 1708456.125 | Q99JR1 | SFXN1 | 4331798 | 2214777 | 522028 | 18 |
| Q9JJU8 | SH3BGRL | SH3 domain binding glutamate-ric | Cytoplasm | other | 1.58E+07 | 1.74E+07 | 1.57E+07 | | | | | | |
| Q9JJU8 | SH3BGRL | SH3 domain binding glutamate-ric | Cytoplasm | other | 1.23E+07 | 7911682 | 8656154 | | | | | | |
| Q9JJU8 | SH3BGRL | SH3 domain binding glutamate-ric | Cytoplasm | other | 1.15E+07 | 1.05E+07 | 1.07E+07 | Q9JJU8 | SH3BGRL | 13200000 | 3326441 | 1108814 | 9 |
| Q8BG73 | SH3BGRL2 | SH3 domain binding glutamate-ric | Nucleus | other | 2345647.25 | 2768708 | 4296632.5 | Q8BG73 | SH3BGRL2 | 2345647 | 1026310 | 592541 | 3 |
| Q91Y57 | SIGLEC9 | sialic acid binding Ig-like lectin 9 | Plasma Membrane | other | 3427575 | 5269841 | 5441905 | Q91Y57 | SIGLEC9 | 3427575 | 1116623 | 644682 | 3 |
| Q3UND0 | SKAP2 | src kinase associated phosphoprot | Cytoplasm | other | 1971408.25 | 1861083.125 | 1631692.25 | | | | | | |
| Q3UND0 | SKAP2 | src kinase associated phosphoprot | Cytoplasm | other | 1201027.375 | 1700947.75 | 2149601 | Q3UND0 | SKAP2 | 1586218 | 328260 | 134012 | 6 |
| Q9QUM4 | SLAMF1 | signaling lymphocytic activation m | Plasma Membrane | transmembrane receptor | 4283933.5 | 5782313 | 4727357.5 | | | | | | |
| Q9QUM4 | SLAMF1 | signaling lymphocytic activation m | Plasma Membrane | transmembrane receptor | 4343980.5 | 3523134.75 | 4575755 | | | | | | |
| Q9QUM4 | SLAMF1 | signaling lymphocytic activation m | Plasma Membrane | transmembrane receptor | 8222409 | 8071798.5 | 8847645 | Q9QUM4 | SLAMF1 | 5616774 | 2017609 | 672536 | 9 |
| P49282 | SLC11A2 | solute carrier family 11 (proton-co | Plasma Membrane | transporter | 7576084.5 | 9894036 | 8974246 | | | | | | |
| P49282 | SLC11A2 | solute carrier family 11 (proton-co | Plasma Membrane | transporter | 3820476.75 | 6031779 | 4329674.5 | | | | | | |
| P49282 | SLC11A2 | solute carrier family 11 (proton-co | Plasma Membrane | transporter | 7914909.5 | 7176875 | 7291759 | | | | | | |
| P49282 | SLC11A2 | solute carrier family 11 (proton-co | Plasma Membrane | transporter | 1879278.25 | 4204985.5 | 3039513 | P49282 | SLC11A2 | 5297687 | 2517592 | 726766 | 12 |
| Q8VHL0 | SLC14A1 | solute carrier family 14 (urea trans | Plasma Membrane | transporter | 2.01E+08 | 3.07E+08 | 2.93E+08 | | | | | | |
| Q8VHL0 | SLC14A1 | solute carrier family 14 (urea trans | Plasma Membrane | transporter | 1.29E+08 | 1.70E+08 | 1.57E+08 | | | | | | |
| Q8VHL0 | SLC14A1 | solute carrier family 14 (urea trans | Plasma Membrane | transporter | 1.38E+08 | 1.40E+08 | 1.09E+08 | | | | | | |
| Q8VHL0 | SLC14A1 | solute carrier family 14 (urea trans | Plasma Membrane | transporter | 9.14E+07 | 1.49E+08 | 1.73E+08 | | | | | | |
| Q8VHL0 | SLC14A1 | solute carrier family 14 (urea trans | Plasma Membrane | transporter | 1.19E+08 | 1.22E+08 | 1.14E+08 | | | | | | |
| Q8VHL0 | SLC14A1 | solute carrier family 14 (urea trans | Plasma Membrane | transporter | 8.93E+07 | 1.01E+08 | 1.15E+08 | | | | | | |
| Q8VHL0 | SLC14A1 | solute carrier family 14 (urea trans | Plasma Membrane | transporter | 2.22E+08 | 2.34E+07 | 8.91E+07 | | | | | | |
| Q8VHL0 | SLC14A1 | solute carrier family 14 (urea trans | Plasma Membrane | transporter | 2.70E+07 | 3.37E+07 | 3.42E+07 | | | | | | |
| Q8VHL0 | SLC14A1 | solute carrier family 14 (urea trans | Plasma Membrane | transporter | 3.81E+07 | 2679587.5 | 6066928 | | | | | | |
| Q8VHL0 | SLC14A1 | solute carrier family 14 (urea trans | Plasma Membrane | transporter | 1.14E+07 | 2.58E+07 | 1.37E+07 | | | | | | |
| Q8VHL0 | SLC14A1 | solute carrier family 14 (urea trans | Plasma Membrane | transporter | 1.09E+07 | 3.20E+07 | 3.07E+07 | | | | | | |
| Q8VHL0 | SLC14A1 | solute carrier family 14 (urea trans | Plasma Membrane | transporter | 6540539 | 369657.219 | 409503.969 | Q8VHL0 | SLC14A1 | 90300878 | 81683555 | 13613926 | 36 |
| P53986 | SLC16A1 | solute carrier family 16 (monocarb | Plasma Membrane | transporter | 6.41E+08 | 5.59E+08 | 4.65E+08 | | | | | | |
| P53986 | SLC16A1 | solute carrier family 16 (monocarb | Plasma Membrane | transporter | 5.28E+08 | 5.48E+08 | 4.09E+08 | | | | | | |
| P53986 | SLC16A1 | solute carrier family 16 (monocarb | Plasma Membrane | transporter | 3.85E+08 | 4.76E+08 | 3.66E+08 | | | | | | |
| P53986 | SLC16A1 | solute carrier family 16 (monocarb | Plasma Membrane | transporter | 1.68E+08 | 1.97E+08 | 2.15E+08 | | | | | | |
| P53986 | SLC16A1 | solute carrier family 16 (monocarb | Plasma Membrane | transporter | 1.46E+08 | 1.43E+08 | 1.24E+08 | | | | | | |
| P53986 | SLC16A1 | solute carrier family 16 (monocarb | Plasma Membrane | transporter | 5.88E+07 | 9.71E+07 | 8.58E+07 | | | | | | |
| P53986 | SLC16A1 | solute carrier family 16 (monocarb | Plasma Membrane | transporter | 2.18E+07 | 3.22E+07 | 3.14E+07 | | | | | | |
| P53986 | SLC16A1 | solute carrier family 16 (monocarb | Plasma Membrane | transporter | 3.85E+07 | 5.13E+07 | 6.12E+07 | | | | | | |
| P53986 | SLC16A1 | solute carrier family 16 (monocarb | Plasma Membrane | transporter | 2741919 | 8019418.5 | 7998795.5 | P53986 | SLC16A1 | 221161324 | 207646509 | 39961589 | 27 |
| Q3U9N9 | SLC16A10 | solute carrier family 16 (aromatic a | Plasma Membrane | transporter | 1.71E+08 | 1.81E+08 | 1.82E+08 | | | | | | |
| Q3U9N9 | SLC16A10 | solute carrier family 16 (aromatic a | Plasma Membrane | transporter | 1.35E+08 | 1.39E+08 | 1.35E+08 | | | | | | |
| Q3U9N9 | SLC16A10 | solute carrier family 16 (aromatic a | Plasma Membrane | transporter | 9.03E+07 | 1.06E+08 | 1.23E+08 | | | | | | |
| Q3U9N9 | SLC16A10 | solute carrier family 16 (aromatic a | Plasma Membrane | transporter | 4.29E+07 | 8.16E+07 | 7.11E+07 | | | | | | |
| Q3U9N9 | SLC16A10 | solute carrier family 16 (aromatic a | Plasma Membrane | transporter | 4.91E+07 | 4.93E+07 | 4.58E+07 | | | | | | |
| Q3U9N9 | SLC16A10 | solute carrier family 16 (aromatic a | Plasma Membrane | transporter | 2.18E+07 | 9.82E+07 | 6.01E+07 | | | | | | |
| Q3U9N9 | SLC16A10 | solute carrier family 16 (aromatic a | Plasma Membrane | transporter | 1.80E+07 | 1.86E+07 | 1.58E+07 | | | | | | |
| Q3U9N9 | SLC16A10 | solute carrier family 16 (aromatic a | Plasma Membrane | transporter | 6988398.5 | 8558846 | 1.06E+07 | | | | | | |
| Q3U9N9 | SLC16A10 | solute carrier family 16 (aromatic a | Plasma Membrane | transporter | 6209284 | 8456841 | 8514644 | | | | | | |
| Q3U9N9 | SLC16A10 | solute carrier family 16 (aromatic a | Plasma Membrane | transporter | 1.72E+07 | 255857.188 | 251714.719 | Q3U9N9 | SLC16A10 | 55802768 | 58868383 | 10747847 | 30 |
| Q8BRU6 | SLC18A2 | solute carrier family 18 (vesicular r | Plasma Membrane | transporter | 901223.125 | 933646 | 1251318.875 | Q8BRU6 | SLC18A2 | 901223 | 193449 | 111688 | 3 |
| P41438 | SLC19A1 | solute carrier family 19 (folate trar | Plasma Membrane | transporter | 1105503 | 1767621.75 | 1433079.375 | P41438 | SLC19A1 | 1105503 | 331065 | 191141 | 3 |
| Q3UHH2 | SLC22A23 | solute carrier family 22, member 2 | Plasma Membrane | transporter | 1606564 | 1592139 | 1463613.75 | Q3UHH2 | SLC22A23 | 1606564 | 78699 | 45437 | 3 |
| Q8BH59 | SLC25A12 | solute carrier family 25 (aspartate, | Cytoplasm | transporter | 1.08E+07 | 7582708.5 | 1.09E+07 | | | | | | |
| Q8BH59 | SLC25A12 | solute carrier family 25 (aspartate, | Cytoplasm | transporter | 3318938 | 3331779.5 | 5416284.5 | | | | | | |
| Q8BH59 | SLC25A12 | solute carrier family 25 (aspartate, | Cytoplasm | transporter | 3740997.25 | 2540084.75 | 4990278.5 | | | | | | |
| Q8BH59 | SLC25A12 | solute carrier family 25 (aspartate, | Cytoplasm | transporter | 1876163.125 | 1765578.75 | 2267573.25 | Q8BH59 | SLC25A12 | 4939025 | 3256383 | 940037 | 12 |
| Q8BZ09 | SLC25A21 | solute carrier family 25 (mitochon | Cytoplasm | transporter | 3554116.5 | 4369966.5 | 5848645.5 | | | | | | |
| Q8BZ09 | SLC25A21 | solute carrier family 25 (mitochon | Cytoplasm | transporter | 2295811.25 | 2872590.75 | 3565298.5 | | | | | | |
| Q8BZ09 | SLC25A21 | solute carrier family 25 (mitochon | Cytoplasm | transporter | 1399925.25 | 1484239.625 | 2480951 | | | | | | |
| Q8BZ09 | SLC25A21 | solute carrier family 25 (mitochon | Cytoplasm | transporter | 1434708.875 | 1418551.875 | 2081177.875 | | | | | | |
| Q8BZ09 | SLC25A21 | solute carrier family 25 (mitochon | Cytoplasm | transporter | 1257612.5 | 1648811.125 | 1794802.125 | Q8BZ09 | SLC25A21 | 1988435 | 1326224 | 342430 | 15 |
| Q8VEM8 | SLC25A3 | solute carrier family 25 (mitochon | Cytoplasm | transporter | 3.71E+07 | 3.33E+07 | 4.32E+07 | | | | | | |
| Q8VEM8 | SLC25A3 | solute carrier family 25 (mitochon | Cytoplasm | transporter | 2.58E+07 | 2.31E+07 | 3.04E+07 | | | | | | |
| Q8VEM8 | SLC25A3 | solute carrier family 25 (mitochon | Cytoplasm | transporter | 1.35E+07 | 1.40E+07 | 1.91E+07 | | | | | | |
| Q8VEM8 | SLC25A3 | solute carrier family 25 (mitochon | Cytoplasm | transporter | 1.57E+07 | 1.84E+07 | 2.53E+07 | | | | | | |
| Q8VEM8 | SLC25A3 | solute carrier family 25 (mitochon | Cytoplasm | transporter | 1.09E+07 | 1.57E+07 | 1.98E+07 | | | | | | |
| Q8VEM8 | SLC25A3 | solute carrier family 25 (mitochon | Cytoplasm | transporter | 8050276.5 | 7846633 | 1.03E+07 | | | | | | |

| | | | | | | | | | | | | | |
|---|---|---|---|---|---|---|---|---|---|---|---|---|---|
| Q8VEM8 | SLC25A3 | solute carrier family 25 (mitochon | Cytoplasm | transporter | 7313866.5 | 7770289 | 9619852 | | | | | | |
| Q8VEM8 | SLC25A3 | solute carrier family 25 (mitochon | Cytoplasm | transporter | 4972410 | 2817976.75 | 4718715 | | | | | | |
| Q8VEM8 | SLC25A3 | solute carrier family 25 (mitochon | Cytoplasm | transporter | 4511602.5 | 2642138.5 | 4476043.5 | Q8VEM8 | SLC25A3 | 14197573 | 11084164 | 2133148 | 27 |
| Q920G8 | SLC25A37 | solute carrier family 25 (mitochon | Cytoplasm | other | 7590297.5 | 1.10E+07 | 1.25E+07 | | | | | | |
| Q920G8 | SLC25A37 | solute carrier family 25 (mitochon | Cytoplasm | other | 4511958 | 5113266 | 8627785 | | | | | | |
| Q920G8 | SLC25A37 | solute carrier family 25 (mitochon | Cytoplasm | other | 6799010 | 4258219 | 4476620.5 | | | | | | |
| Q920G8 | SLC25A37 | solute carrier family 25 (mitochon | Cytoplasm | other | 4418798.5 | 4944693.5 | 7530249.5 | | | | | | |
| Q920G8 | SLC25A37 | solute carrier family 25 (mitochon | Cytoplasm | other | 4739002 | 3966544.25 | 5520098.5 | | | | | | |
| Q920G8 | SLC25A37 | solute carrier family 25 (mitochon | Cytoplasm | other | 4101117.75 | 4277150 | 4693363.5 | | | | | | |
| Q920G8 | SLC25A37 | solute carrier family 25 (mitochon | Cytoplasm | other | 3758401.75 | 2997278.5 | 4325059 | | | | | | |
| Q920G8 | SLC25A37 | solute carrier family 25 (mitochon | Cytoplasm | other | 3589174.25 | 3563833.75 | 4686994 | | | | | | |
| Q920G8 | SLC25A37 | solute carrier family 25 (mitochon | Cytoplasm | other | 2095302.75 | 1903008 | 2441629.5 | | | | | | |
| Q920G8 | SLC25A37 | solute carrier family 25 (mitochon | Cytoplasm | other | 1326926.875 | 1164306.125 | 1502040 | | | | | | |
| Q920G8 | SLC25A37 | solute carrier family 25 (mitochon | Cytoplasm | other | 1539980.25 | 2692465.5 | 7338670 | Q920G8 | SLC25A37 | 4042725 | 2622369 | 456496 | 33 |
| P48962 | SLC25A4 | solute carrier family 25 (mitochon | Cytoplasm | transporter | 5007334.5 | 6322313.5 | 7463320.5 | | | | | | |
| P48962 | SLC25A4 | solute carrier family 25 (mitochon | Cytoplasm | transporter | 3746705.75 | 4640886 | 5288104 | | | | | | |
| P48962 | SLC25A4 | solute carrier family 25 (mitochon | Cytoplasm | transporter | 3300989.75 | 2931397.75 | 3872782.5 | P48962 | SLC25A4 | 4018343 | 1474007 | 491336 | 9 |
| P51881 | SLC25A5 | solute carrier family 25 (mitochon | Cytoplasm | transporter | 1.23E+08 | 1.11E+08 | 1.63E+08 | | | | | | |
| P51881 | SLC25A5 | solute carrier family 25 (mitochon | Cytoplasm | transporter | 7.98E+07 | 7.42E+07 | 9.29E+07 | | | | | | |
| P51881 | SLC25A5 | solute carrier family 25 (mitochon | Cytoplasm | transporter | 6.90E+07 | 7.31E+07 | 7.75E+07 | | | | | | |
| P51881 | SLC25A5 | solute carrier family 25 (mitochon | Cytoplasm | transporter | 3.09E+07 | 3.83E+07 | 4.19E+07 | | | | | | |
| P51881 | SLC25A5 | solute carrier family 25 (mitochon | Cytoplasm | transporter | 1.32E+07 | 3.09E+07 | 2.55E+07 | | | | | | |
| P51881 | SLC25A5 | solute carrier family 25 (mitochon | Cytoplasm | transporter | 2.11E+07 | 4.64E+07 | 5.31E+07 | | | | | | |
| P51881 | SLC25A5 | solute carrier family 25 (mitochon | Cytoplasm | transporter | 2.48E+07 | 1.20E+07 | 1.05E+07 | | | | | | |
| P51881 | SLC25A5 | solute carrier family 25 (mitochon | Cytoplasm | transporter | 1.95E+07 | 2.38E+07 | 3.19E+07 | | | | | | |
| P51881 | SLC25A5 | solute carrier family 25 (mitochon | Cytoplasm | transporter | 1.88E+07 | 8867508 | 2.11E+07 | | | | | | |
| P51881 | SLC25A5 | solute carrier family 25 (mitochon | Cytoplasm | transporter | 9271080 | 2.01E+07 | 3.03E+07 | | | | | | |
| P51881 | SLC25A5 | solute carrier family 25 (mitochon | Cytoplasm | transporter | 2.22E+07 | 2.40E+07 | 2.40E+07 | | | | | | |
| P51881 | SLC25A5 | solute carrier family 25 (mitochon | Cytoplasm | transporter | 7948204.5 | 7689223.5 | 9699780 | | | | | | |
| P51881 | SLC25A5 | solute carrier family 25 (mitochon | Cytoplasm | transporter | 1791973 | 4050776.75 | 7219557.5 | | | | | | |
| P51881 | SLC25A5 | solute carrier family 25 (mitochon | Cytoplasm | transporter | 1197386.75 | 2970051 | 2839810.25 | | | | | | |
| P51881 | SLC25A5 | solute carrier family 25 (mitochon | Cytoplasm | transporter | 251195.734 | 7570680 | 1.06E+07 | P51881 | SLC25A5 | 29519323 | 36089087 | 5379843 | 45 |
| Q5HZI9 | SLC25A51 | solute carrier family 25, member 5 | Cytoplasm | other | 868693.562 | 1148930.5 | 2351979.25 | Q5HZI9 | SLC25A51 | 868694 | 788035 | 454972 | 3 |
| P58735 | SLC26A1 | solute carrier family 26 (anion excl | Plasma Membrane | transporter | 9132061 | 8247003 | 8092143 | | | | | | |
| P58735 | SLC26A1 | solute carrier family 26 (anion excl | Plasma Membrane | transporter | 3622243.25 | 3173335.5 | 3205756 | P58735 | SLC26A1 | 6377152 | 2851003 | 1163917 | 6 |
| Q9WVC8 | SLC26A3 | solute carrier family 26 (anion excl | Plasma Membrane | transporter | 1252418.875 | 2425488.5 | 1521566.5 | Q9WVC8 | SLC26A3 | 1252419 | 614493 | 354778 | 3 |
| Q9JIM1 | SLC29A1 | solute carrier family 29 (equilibrat | Plasma Membrane | transporter | 1.30E+08 | 1.27E+08 | 1.27E+08 | | | | | | |
| Q9JIM1 | SLC29A1 | solute carrier family 29 (equilibrat | Plasma Membrane | transporter | 6.17E+07 | 6.67E+07 | 5.53E+07 | | | | | | |
| Q9JIM1 | SLC29A1 | solute carrier family 29 (equilibrat | Plasma Membrane | transporter | 2.97E+07 | 5.00E+07 | 3.01E+07 | | | | | | |
| Q9JIM1 | SLC29A1 | solute carrier family 29 (equilibrat | Plasma Membrane | transporter | 2.50E+07 | 4.32E+07 | 3.82E+07 | | | | | | |
| Q9JIM1 | SLC29A1 | solute carrier family 29 (equilibrat | Plasma Membrane | transporter | 1.38E+07 | 2.01E+07 | 2.89E+07 | | | | | | |
| Q9JIM1 | SLC29A1 | solute carrier family 29 (equilibrat | Plasma Membrane | transporter | 2.24E+07 | 1.65E+07 | 6421879 | | | | | | |
| Q9JIM1 | SLC29A1 | solute carrier family 29 (equilibrat | Plasma Membrane | transporter | 1.06E+07 | 1.46E+07 | 8246192.5 | | | | | | |
| Q9JIM1 | SLC29A1 | solute carrier family 29 (equilibrat | Plasma Membrane | transporter | 3204379.25 | 4482071 | 5950181 | | | | | | |
| Q9JIM1 | SLC29A1 | solute carrier family 29 (equilibrat | Plasma Membrane | transporter | 672873.438 | 1393674 | 2669691.75 | | | | | | |
| Q9JIM1 | SLC29A1 | solute carrier family 29 (equilibrat | Plasma Membrane | transporter | 3034197.5 | 3779554.25 | 3186029.5 | | | | | | |
| Q9JIM1 | SLC29A1 | solute carrier family 29 (equilibrat | Plasma Membrane | transporter | 931077.5 | 1680854.25 | 1433172.25 | Q9JIM1 | SLC29A1 | 27392048 | 36945891 | 6431454 | 33 |
| Q61672 | SLC29A2 | solute carrier family 29 (equilibrat | Plasma Membrane | transporter | 1.71E+07 | 2.80E+07 | 2.68E+07 | Q61672 | SLC29A2 | 17090000 | 5984455 | 3455127 | 3 |
| P32037 | SLC2A3 | solute carrier family 2 (facilitated g | Plasma Membrane | transporter | 3.55E+07 | 4.40E+07 | 4.04E+07 | | | | | | |
| P32037 | SLC2A3 | solute carrier family 2 (facilitated g | Plasma Membrane | transporter | 3.03E+07 | 3.12E+07 | 2.95E+07 | | | | | | |
| P32037 | SLC2A3 | solute carrier family 2 (facilitated g | Plasma Membrane | transporter | 2.60E+07 | 2.65E+07 | 2.97E+07 | | | | | | |
| P32037 | SLC2A3 | solute carrier family 2 (facilitated g | Plasma Membrane | transporter | 2.12E+07 | 2.59E+07 | 1.92E+07 | | | | | | |
| P32037 | SLC2A3 | solute carrier family 2 (facilitated g | Plasma Membrane | transporter | 1.12E+07 | 1.17E+07 | 1.19E+07 | | | | | | |
| P32037 | SLC2A3 | solute carrier family 2 (facilitated g | Plasma Membrane | transporter | 4704138.5 | 5163751.5 | 3639842.5 | | | | | | |
| P32037 | SLC2A3 | solute carrier family 2 (facilitated g | Plasma Membrane | transporter | 1429119.625 | 1880948.125 | 2479465.75 | P32037 | SLC2A3 | 18620465 | 13551048 | 2957081 | 21 |
| Q60738 | SLC30A1 | solute carrier family 30 (zinc transp | Plasma Membrane | transporter | 3925352.25 | 4462327 | 4331885.5 | | | | | | |
| Q60738 | SLC30A1 | solute carrier family 30 (zinc transp | Plasma Membrane | transporter | 3065347 | 3123949 | 1534371.75 | | | | | | |
| Q60738 | SLC30A1 | solute carrier family 30 (zinc transp | Plasma Membrane | transporter | 2895839.25 | 2888526.25 | 2405123 | | | | | | |
| Q60738 | SLC30A1 | solute carrier family 30 (zinc transp | Plasma Membrane | transporter | 2185743.75 | 1897604.25 | 1946351.875 | Q60738 | SLC30A1 | 3018071 | 960720 | 277336 | 12 |
| P10852 | SLC3A2 | solute carrier family 3 (amino acid | Plasma Membrane | transporter | 1.50E+07 | 1.45E+07 | 1.30E+07 | | | | | | |
| P10852 | SLC3A2 | solute carrier family 3 (amino acid | Plasma Membrane | transporter | 1.06E+07 | 1.11E+07 | 1.36E+07 | | | | | | |
| P10852 | SLC3A2 | solute carrier family 3 (amino acid | Plasma Membrane | transporter | 1.10E+07 | 9283746 | 1.02E+07 | | | | | | |
| P10852 | SLC3A2 | solute carrier family 3 (amino acid | Plasma Membrane | transporter | 7669475 | 8157576.5 | 9093709 | | | | | | |
| P10852 | SLC3A2 | solute carrier family 3 (amino acid | Plasma Membrane | transporter | 8953282 | 7265772.5 | 9469830 | | | | | | |
| P10852 | SLC3A2 | solute carrier family 3 (amino acid | Plasma Membrane | transporter | 6436438.5 | 4606851 | 6056930 | | | | | | |
| P10852 | SLC3A2 | solute carrier family 3 (amino acid | Plasma Membrane | transporter | 4784415.5 | 5356258 | 5553594.5 | | | | | | |
| P10852 | SLC3A2 | solute carrier family 3 (amino acid | Plasma Membrane | transporter | 6884762.5 | 7392312.5 | 8399743 | | | | | | |
| P10852 | SLC3A2 | solute carrier family 3 (amino acid | Plasma Membrane | transporter | 3825413.5 | 3246889.75 | 3787610.75 | | | | | | |
| P10852 | SLC3A2 | solute carrier family 3 (amino acid | Plasma Membrane | transporter | 3355481 | 1522746.125 | 1975618.5 | P10852 | SLC3A2 | 7855927 | 3635136 | 663682 | 30 |
| Q9JHI9 | SLC40A1 | solute carrier family 40 (iron-regul | Plasma Membrane | transporter | 2.58E+07 | 3.05E+07 | 2.65E+07 | | | | | | |
| Q9JHI9 | SLC40A1 | solute carrier family 40 (iron-regul | Plasma Membrane | transporter | 1.84E+07 | 2.80E+07 | 2.32E+07 | | | | | | |
| Q9JHI9 | SLC40A1 | solute carrier family 40 (iron-regul | Plasma Membrane | transporter | 2.72E+07 | 1.29E+07 | 1.99E+07 | | | | | | |
| Q9JHI9 | SLC40A1 | solute carrier family 40 (iron-regul | Plasma Membrane | transporter | 1.58E+07 | 1.74E+07 | 1.46E+07 | | | | | | |
| Q9JHI9 | SLC40A1 | solute carrier family 40 (iron-regul | Plasma Membrane | transporter | 2.52E+07 | 4936577 | 4404528 | | | | | | |
| Q9JHI9 | SLC40A1 | solute carrier family 40 (iron-regul | Plasma Membrane | transporter | 1.01E+07 | 1.93E+07 | 1.19E+07 | | | | | | |
| Q9JHI9 | SLC40A1 | solute carrier family 40 (iron-regul | Plasma Membrane | transporter | 6125262.5 | 1.70E+07 | 1.37E+07 | | | | | | |
| Q9JHI9 | SLC40A1 | solute carrier family 40 (iron-regul | Plasma Membrane | transporter | 2319755 | 230935.078 | 63381.707 | Q9JHI9 | SLC40A1 | 16365627 | 9243930 | 1886909 | 24 |
| Q8BSM7 | SLC43A1 | solute carrier family 43 (amino aci | Plasma Membrane | transporter | 5.88E+08 | 7.02E+08 | 6.33E+08 | | | | | | |
| Q8BSM7 | SLC43A1 | solute carrier family 43 (amino aci | Plasma Membrane | transporter | 3.27E+08 | 4.25E+08 | 4.07E+08 | | | | | | |
| Q8BSM7 | SLC43A1 | solute carrier family 43 (amino aci | Plasma Membrane | transporter | 1.57E+08 | 2.19E+08 | 2.34E+08 | | | | | | |
| Q8BSM7 | SLC43A1 | solute carrier family 43 (amino aci | Plasma Membrane | transporter | 9.58E+07 | 1.27E+08 | 1.09E+08 | | | | | | |
| Q8BSM7 | SLC43A1 | solute carrier family 43 (amino aci | Plasma Membrane | transporter | 6.43E+07 | 6.98E+07 | 5.20E+07 | | | | | | |
| Q8BSM7 | SLC43A1 | solute carrier family 43 (amino aci | Plasma Membrane | transporter | 6.59E+07 | 5.43E+07 | 8.78E+07 | | | | | | |
| Q8BSM7 | SLC43A1 | solute carrier family 43 (amino aci | Plasma Membrane | transporter | 5.38E+07 | 7.52E+07 | 6.66E+07 | | | | | | |
| Q8BSM7 | SLC43A1 | solute carrier family 43 (amino aci | Plasma Membrane | transporter | 9.39E+07 | 2.29E+07 | 4.84E+07 | | | | | | |
| Q8BSM7 | SLC43A1 | solute carrier family 43 (amino aci | Plasma Membrane | transporter | 2.42E+07 | 2.28E+07 | 2.97E+07 | | | | | | |

| | | | | | | | | | | | | | |
|---|---|---|---|---|---|---|---|---|---|---|---|---|---|
| Q8BSM7 | SLC43A1 | solute carrier family 43 (amino aci | Plasma Membrane | transporter | 1.21E+07 | 1535684.625 | 2973955 | | | | | | |
| Q8BSM7 | SLC43A1 | solute carrier family 43 (amino aci | Plasma Membrane | transporter | 4576279 | 1.29E+07 | 8032823 | | | | | | |
| Q8BSM7 | SLC43A1 | solute carrier family 43 (amino aci | Plasma Membrane | transporter | 4103322.25 | 4299851 | 2580633.25 | | | | | | |
| Q8BSM7 | SLC43A1 | solute carrier family 43 (amino aci | Plasma Membrane | transporter | 1.05E+07 | 725504.438 | 1894236.25 | | | | | | |
| Q8BSM7 | SLC43A1 | solute carrier family 43 (amino aci | Plasma Membrane | transporter | 805099.812 | 3759549.75 | 2066641.75 | Q8BSM7 | SLC43A1 | 107263193 | 180275150 | 27817059 | 42 |
| A2AVZ9 | SLC43A3 | solute carrier family 43, member 3 | Extracellular Space | other | 4.50E+07 | 9.51E+07 | 4.73E+07 | | | | | | |
| A2AVZ9 | SLC43A3 | solute carrier family 43, member 3 | Extracellular Space | other | 1.71E+07 | 1.70E+07 | 1.34E+07 | | | | | | |
| A2AVZ9 | SLC43A3 | solute carrier family 43, member 3 | Extracellular Space | other | 1.01E+07 | 1.91E+07 | 1.85E+07 | | | | | | |
| A2AVZ9 | SLC43A3 | solute carrier family 43, member 3 | Extracellular Space | other | 6707812.5 | 7616579.5 | 7202925 | A2AVZ9 | SLC43A3 | 19734453 | 25806694 | 7449751 | 12 |
| Q6X893 | SLC44A1 | solute carrier family 44 (choline tr | Plasma Membrane | transporter | 3985816.75 | 3993045.5 | 3502486.75 | | | | | | |
| Q6X893 | SLC44A1 | solute carrier family 44 (choline tr | Plasma Membrane | transporter | 1330585.875 | 2300267.25 | 2055873.75 | | | | | | |
| Q6X893 | SLC44A1 | solute carrier family 44 (choline tr | Plasma Membrane | transporter | 1866481.75 | 2541546.5 | 1663946.875 | Q6X893 | SLC44A1 | 2394295 | 1005466 | 335155 | 9 |
| P04919 | SLC4A1 | solute carrier family 4 (anion exch | Plasma Membrane | transporter | 2.45E+10 | 2.98E+10 | 3.00E+10 | | | | | | |
| P04919 | SLC4A1 | solute carrier family 4 (anion exch | Plasma Membrane | transporter | 1.46E+10 | 1.42E+10 | 1.58E+10 | | | | | | |
| P04919 | SLC4A1 | solute carrier family 4 (anion exch | Plasma Membrane | transporter | 9.26E+09 | 8.47E+09 | 8.33E+09 | | | | | | |
| P04919 | SLC4A1 | solute carrier family 4 (anion exch | Plasma Membrane | transporter | 7.78E+09 | 8.35E+09 | 7.08E+09 | | | | | | |
| P04919 | SLC4A1 | solute carrier family 4 (anion exch | Plasma Membrane | transporter | 6.46E+09 | 7.95E+09 | 5.66E+09 | | | | | | |
| P04919 | SLC4A1 | solute carrier family 4 (anion exch | Plasma Membrane | transporter | 7.57E+09 | 8.14E+09 | 8.42E+09 | | | | | | |
| P04919 | SLC4A1 | solute carrier family 4 (anion exch | Plasma Membrane | transporter | 8.80E+09 | 9.19E+09 | 9.99E+09 | | | | | | |
| P04919 | SLC4A1 | solute carrier family 4 (anion exch | Plasma Membrane | transporter | 6.31E+09 | 6.25E+09 | 5.73E+09 | | | | | | |
| P04919 | SLC4A1 | solute carrier family 4 (anion exch | Plasma Membrane | transporter | 5.70E+09 | 6.85E+09 | 6.53E+09 | | | | | | |
| P04919 | SLC4A1 | solute carrier family 4 (anion exch | Plasma Membrane | transporter | 4.12E+09 | 6.17E+09 | 5.74E+09 | | | | | | |
| P04919 | SLC4A1 | solute carrier family 4 (anion exch | Plasma Membrane | transporter | 4.32E+09 | 4.26E+09 | 4.44E+09 | | | | | | |
| P04919 | SLC4A1 | solute carrier family 4 (anion exch | Plasma Membrane | transporter | 4.42E+09 | 3.35E+09 | 3.31E+09 | | | | | | |
| P04919 | SLC4A1 | solute carrier family 4 (anion exch | Plasma Membrane | transporter | 2.65E+09 | 3.98E+09 | 3.47E+09 | | | | | | |
| P04919 | SLC4A1 | solute carrier family 4 (anion exch | Plasma Membrane | transporter | 2.71E+09 | 7.18E+09 | 7.33E+09 | | | | | | |
| P04919 | SLC4A1 | solute carrier family 4 (anion exch | Plasma Membrane | transporter | 2.88E+09 | 4.23E+09 | 4.55E+09 | | | | | | |
| P04919 | SLC4A1 | solute carrier family 4 (anion exch | Plasma Membrane | transporter | 2.07E+09 | 3.09E+09 | 2.95E+09 | | | | | | |
| P04919 | SLC4A1 | solute carrier family 4 (anion exch | Plasma Membrane | transporter | 1.16E+09 | 3.54E+09 | 1.90E+09 | | | | | | |
| P04919 | SLC4A1 | solute carrier family 4 (anion exch | Plasma Membrane | transporter | 7.84E+08 | 1.90E+09 | 1.21E+09 | | | | | | |
| P04919 | SLC4A1 | solute carrier family 4 (anion exch | Plasma Membrane | transporter | 1.52E+09 | 1.87E+09 | 1.82E+09 | | | | | | |
| P04919 | SLC4A1 | solute carrier family 4 (anion exch | Plasma Membrane | transporter | 4.72E+08 | 1.82E+09 | 1.76E+09 | | | | | | |
| P04919 | SLC4A1 | solute carrier family 4 (anion exch | Plasma Membrane | transporter | 6.58E+08 | 7.76E+08 | 6.11E+08 | | | | | | |
| P04919 | SLC4A1 | solute carrier family 4 (anion exch | Plasma Membrane | transporter | 9.55E+08 | 8.04E+08 | 8.52E+08 | | | | | | |
| P04919 | SLC4A1 | solute carrier family 4 (anion exch | Plasma Membrane | transporter | 5.80E+08 | 1.07E+09 | 8.49E+08 | | | | | | |
| P04919 | SLC4A1 | solute carrier family 4 (anion exch | Plasma Membrane | transporter | 5.88E+08 | 5.94E+08 | 5.91E+08 | | | | | | |
| P04919 | SLC4A1 | solute carrier family 4 (anion exch | Plasma Membrane | transporter | 3.73E+08 | 7.42E+08 | 6.16E+08 | | | | | | |
| P04919 | SLC4A1 | solute carrier family 4 (anion exch | Plasma Membrane | transporter | 3.33E+08 | 6.40E+08 | 6.27E+08 | | | | | | |
| P04919 | SLC4A1 | solute carrier family 4 (anion exch | Plasma Membrane | transporter | 1.68E+08 | 5.96E+08 | 3.85E+08 | | | | | | |
| P04919 | SLC4A1 | solute carrier family 4 (anion exch | Plasma Membrane | transporter | 3.03E+08 | 6.08E+08 | 6.63E+08 | | | | | | |
| P04919 | SLC4A1 | solute carrier family 4 (anion exch | Plasma Membrane | transporter | 2.96E+08 | 4.07E+07 | 1.31E+07 | | | | | | |
| P04919 | SLC4A1 | solute carrier family 4 (anion exch | Plasma Membrane | transporter | 1.46E+08 | 5.11E+08 | 1.86E+08 | | | | | | |
| P04919 | SLC4A1 | solute carrier family 4 (anion exch | Plasma Membrane | transporter | 2.14E+08 | 3.11E+08 | 3.01E+08 | | | | | | |
| P04919 | SLC4A1 | solute carrier family 4 (anion exch | Plasma Membrane | transporter | 6.65E+08 | 6.72E+08 | 3.56E+08 | | | | | | |
| P04919 | SLC4A1 | solute carrier family 4 (anion exch | Plasma Membrane | transporter | 3.24E+08 | 5.90E+08 | 2.53E+08 | | | | | | |
| P04919 | SLC4A1 | solute carrier family 4 (anion exch | Plasma Membrane | transporter | 9.93E+07 | 2.63E+08 | 2.43E+08 | | | | | | |
| P04919 | SLC4A1 | solute carrier family 4 (anion exch | Plasma Membrane | transporter | 7.47E+07 | 1.31E+08 | 7.72E+07 | | | | | | |
| P04919 | SLC4A1 | solute carrier family 4 (anion exch | Plasma Membrane | transporter | 7.39E+07 | 1.86E+08 | 1.60E+08 | | | | | | |
| P04919 | SLC4A1 | solute carrier family 4 (anion exch | Plasma Membrane | transporter | 1.92E+08 | 3.34E+07 | 7.67E+07 | | | | | | |
| P04919 | SLC4A1 | solute carrier family 4 (anion exch | Plasma Membrane | transporter | 1.02E+08 | 3.78E+08 | 1.79E+08 | | | | | | |
| P04919 | SLC4A1 | solute carrier family 4 (anion exch | Plasma Membrane | transporter | 6.81E+07 | 9.54E+07 | 9.76E+07 | | | | | | |
| P04919 | SLC4A1 | solute carrier family 4 (anion exch | Plasma Membrane | transporter | 5.11E+07 | 6.40E+07 | 7.06E+07 | | | | | | |
| P04919 | SLC4A1 | solute carrier family 4 (anion exch | Plasma Membrane | transporter | 1.01E+08 | 1.95E+07 | 1.40E+07 | | | | | | |
| P04919 | SLC4A1 | solute carrier family 4 (anion exch | Plasma Membrane | transporter | 4.55E+07 | 3.26E+08 | 3.32E+08 | | | | | | |
| P04919 | SLC4A1 | solute carrier family 4 (anion exch | Plasma Membrane | transporter | 4.08E+07 | 4.77E+07 | 3.37E+07 | | | | | | |
| P04919 | SLC4A1 | solute carrier family 4 (anion exch | Plasma Membrane | transporter | 4.78E+07 | 7.87E+07 | 8.07E+07 | | | | | | |
| P04919 | SLC4A1 | solute carrier family 4 (anion exch | Plasma Membrane | transporter | 1.33E+07 | 3.62E+07 | 2.48E+07 | | | | | | |
| P04919 | SLC4A1 | solute carrier family 4 (anion exch | Plasma Membrane | transporter | 1.64E+07 | 2.78E+07 | 2.70E+07 | | | | | | |
| P04919 | SLC4A1 | solute carrier family 4 (anion exch | Plasma Membrane | transporter | 1.71E+07 | 1686859.125 | 128122.227 | | | | | | |
| P04919 | SLC4A1 | solute carrier family 4 (anion exch | Plasma Membrane | transporter | 3.24E+07 | 1.14E+07 | 9733201 | | | | | | |
| P04919 | SLC4A1 | solute carrier family 4 (anion exch | Plasma Membrane | transporter | 1.04E+07 | 1.58E+07 | 1786807.375 | | | | | | |
| P04919 | SLC4A1 | solute carrier family 4 (anion exch | Plasma Membrane | transporter | 9337038 | 5.28E+07 | 2.55E+07 | | | | | | |
| P04919 | SLC4A1 | solute carrier family 4 (anion exch | Plasma Membrane | transporter | 8347099.5 | 4357202.5 | 3166129.25 | | | | | | |
| P04919 | SLC4A1 | solute carrier family 4 (anion exch | Plasma Membrane | transporter | 1.64E+07 | 1.82E+07 | 8165351 | | | | | | |
| P04919 | SLC4A1 | solute carrier family 4 (anion exch | Plasma Membrane | transporter | 7945289.5 | 2.43E+07 | 1.69E+07 | | | | | | |
| P04919 | SLC4A1 | solute carrier family 4 (anion exch | Plasma Membrane | transporter | 6354545.5 | 1369266.5 | 957939.25 | | | | | | |
| P04919 | SLC4A1 | solute carrier family 4 (anion exch | Plasma Membrane | transporter | 1917388.25 | 9.72E+07 | 1.19E+08 | | | | | | |
| P04919 | SLC4A1 | solute carrier family 4 (anion exch | Plasma Membrane | transporter | 579565.312 | 4147171.25 | 2850403.75 | | | | | | |
| P04919 | SLC4A1 | solute carrier family 4 (anion exch | Plasma Membrane | transporter | 3762708.75 | 1.41E+07 | 6504326 | P04919 | SLC4A1 | 2188020415 | 4673329602 | 357378447 | 171 |
| O88343 | SLC4A4 | solute carrier family 4 (sodium bic | Plasma Membrane | transporter | 1.66E+07 | 7966550 | 1.19E+07 | O88343 | SLC4A4 | 16580000 | 4311552 | 2489276 | 3 |
| Q60857 | SLC6A4 | solute carrier family 6 (neurotrans | Plasma Membrane | transporter | 4986422.5 | 5089500.5 | 5997992 | Q60857 | SLC6A4 | 4986423 | 556665 | 321391 | 3 |
| P28571 | SLC6A9 | solute carrier family 6 (neurotrans | Plasma Membrane | transporter | 6397144.5 | 6817679 | 9777826 | | | | | | |
| P28571 | SLC6A9 | solute carrier family 6 (neurotrans | Plasma Membrane | transporter | 1358096.375 | 1973295.875 | 2113745.5 | P28571 | SLC6A9 | 3877620 | 3418529 | 1395609 | 6 |
| Q09143 | SLC7A1 | solute carrier family 7 (cationic am | Plasma Membrane | transporter | 6701965.5 | 5731017.5 | 6593752 | Q09143 | SLC7A1 | 6701966 | 532097 | 307206 | 3 |
| Q9Z127 | SLC7A5 | solute carrier family 7 (amino acid | Plasma Membrane | transporter | 1.39E+07 | 1.17E+07 | 1.34E+07 | | | | | | |
| Q9Z127 | SLC7A5 | solute carrier family 7 (amino acid | Plasma Membrane | transporter | 8806105 | 6168444 | 6269301 | | | | | | |
| Q9Z127 | SLC7A5 | solute carrier family 7 (amino acid | Plasma Membrane | transporter | 2663014 | 2605462.5 | 2423130.5 | | | | | | |
| Q9Z127 | SLC7A5 | solute carrier family 7 (amino acid | Plasma Membrane | transporter | 2115276.25 | 1765948.125 | 2203359 | Q9Z127 | SLC7A5 | 6866099 | 4664119 | 1346415 | 12 |
| Q61165 | SLC9A1 | solute carrier family 9, subfamily A | Plasma Membrane | ion channel | 1.52E+07 | 1.68E+07 | 1.04E+07 | | | | | | |
| Q61165 | SLC9A1 | solute carrier family 9, subfamily A | Plasma Membrane | ion channel | 9006895 | 1.07E+07 | 8392630 | | | | | | |
| Q61165 | SLC9A1 | solute carrier family 9, subfamily A | Plasma Membrane | ion channel | 5994141.5 | 7454301 | 7716260 | | | | | | |
| Q61165 | SLC9A1 | solute carrier family 9, subfamily A | Plasma Membrane | ion channel | 6732363 | 6722061.5 | 7062897 | | | | | | |
| Q61165 | SLC9A1 | solute carrier family 9, subfamily A | Plasma Membrane | ion channel | 6004603 | 5323611 | 4679959 | | | | | | |
| Q61165 | SLC9A1 | solute carrier family 9, subfamily A | Plasma Membrane | ion channel | 4032627.25 | 5070826.5 | 4913505 | | | | | | |
| Q61165 | SLC9A1 | solute carrier family 9, subfamily A | Plasma Membrane | ion channel | 2246253 | 2886074.5 | 2796933.5 | | | | | | |
| Q61165 | SLC9A1 | solute carrier family 9, subfamily A | Plasma Membrane | ion channel | 2240945.75 | 89692.188 | 65633.062 | Q61165 | SLC9A1 | 6425979 | 4097935 | 836487 | 24 |

| | | | | | | | | | | | | | |
|---|---|---|---|---|---|---|---|---|---|---|---|---|---|
| Q6UJY2 | SLC9C1 | solute carrier family 9, subfamily C | Extracellular Space | other | 2.72E+07 | 6.57E+07 | 4.76E+07 | Q6UJY2 | SLC9C1 | 27150000 | 19277978 | 11130146 | 3 |
| Q8CG48 | SMC2 | structural maintenance of chromo | Nucleus | transporter | 1974976.375 | 2353706.25 | 1400498.75 | Q8CG48 | SMC2 | 1974976 | 479942 | 277095 | 3 |
| Q8CG47 | SMC4 | structural maintenance of chromo | Nucleus | transporter | 1.15E+07 | 1.47E+07 | 1.07E+07 | | | | | | |
| Q8CG47 | SMC4 | structural maintenance of chromo | Nucleus | transporter | 1015288.875 | 1176904.375 | 423362.969 | Q8CG47 | SMC4 | 6257644 | 6395723 | 2611043 | 6 |
| O09044 | SNAP23 | synaptosomal-associated protein, | Plasma Membrane | transporter | 4.12E+07 | 3.13E+07 | 2.99E+07 | | | | | | |
| O09044 | SNAP23 | synaptosomal-associated protein, | Plasma Membrane | transporter | 2.95E+07 | 2.94E+07 | 2.67E+07 | | | | | | |
| O09044 | SNAP23 | synaptosomal-associated protein, | Plasma Membrane | transporter | 2.14E+07 | 2.56E+07 | 1.86E+07 | | | | | | |
| O09044 | SNAP23 | synaptosomal-associated protein, | Plasma Membrane | transporter | 2.50E+07 | 2.65E+07 | 2.77E+07 | | | | | | |
| O09044 | SNAP23 | synaptosomal-associated protein, | Plasma Membrane | transporter | 1.28E+07 | 1.03E+07 | 5681951.5 | | | | | | |
| O09044 | SNAP23 | synaptosomal-associated protein, | Plasma Membrane | transporter | 6956168.5 | 2.17E+07 | 1.18E+07 | | | | | | |
| O09044 | SNAP23 | synaptosomal-associated protein, | Plasma Membrane | transporter | 1.05E+07 | 1.09E+07 | 1.10E+07 | | | | | | |
| O09044 | SNAP23 | synaptosomal-associated protein, | Plasma Membrane | transporter | 2386154.25 | 7511138 | 4600786 | | | | | | |
| O09044 | SNAP23 | synaptosomal-associated protein, | Plasma Membrane | transporter | 1739485.625 | 3446391.75 | 1791182.75 | | | | | | |
| O09044 | SNAP23 | synaptosomal-associated protein, | Plasma Membrane | transporter | 1814134.125 | 4311424.5 | 1842950.75 | | | | | | |
| O09044 | SNAP23 | synaptosomal-associated protein, | Plasma Membrane | transporter | 3106796 | 880120.625 | 1207051.25 | | | | | | |
| O09044 | SNAP23 | synaptosomal-associated protein, | Plasma Membrane | transporter | 413335.656 | 1202225.125 | 950026.438 | O09044 | SNAP23 | 13060506 | 11764904 | 1960817 | 36 |
| O55042 | SNCA | synuclein, alpha (non A4 compone | Cytoplasm | other | 1.41E+07 | 1.39E+07 | 2.09E+07 | | | | | | |
| O55042 | SNCA | synuclein, alpha (non A4 compone | Cytoplasm | other | 1.04E+07 | 9817531 | 1.45E+07 | | | | | | |
| O55042 | SNCA | synuclein, alpha (non A4 compone | Cytoplasm | other | 8755391 | 8483924 | 1.27E+07 | | | | | | |
| O55042 | SNCA | synuclein, alpha (non A4 compone | Cytoplasm | other | 7443899.5 | 6124959 | 8793831 | O55042 | SNCA | 10167323 | 4091654 | 1181159 | 12 |
| O70493 | SNX12 | sorting nexin 12 | Cytoplasm | transporter | 603701.062 | 777064.562 | 547303.562 | O70493 | SNX12 | 603701 | 119740 | 69132 | 3 |
| Q0P5V2 | SOBP | sine oculis binding protein homolo | Nucleus | other | 76708.055 | 795946.125 | 979048.188 | Q0P5V2 | SOBP | 76708 | 476978 | 275383 | 3 |
| P09671 | SOD2 | superoxide dismutase 2, mitochon | Cytoplasm | enzyme | 6850911.5 | 6313931 | 8655015 | | | | | | |
| P09671 | SOD2 | superoxide dismutase 2, mitochon | Cytoplasm | enzyme | 5942966.5 | 1085916.125 | 904752.688 | | | | | | |
| P09671 | SOD2 | superoxide dismutase 2, mitochon | Cytoplasm | enzyme | 1389597.75 | 2077333.25 | 2275822.5 | P09671 | SOD2 | 4727825 | 2967620 | 989207 | 9 |
| Q9CYN2 | SPCS2 | signal peptidase complex subunit | Cytoplasm | other | 7844769 | 1.13E+07 | 7891151 | Q9CYN2 | SPCS2 | 7844769 | 1998944 | 1154091 | 3 |
| Q9JJF9 | SPPL2A | signal peptide peptidase like 2A | Plasma Membrane | peptidase | 1.18E+07 | 1.49E+07 | 1.35E+07 | | | | | | |
| Q9JJF9 | SPPL2A | signal peptide peptidase like 2A | Plasma Membrane | peptidase | 1.20E+07 | 1.55E+07 | 1.46E+07 | Q9JJF9 | SPPL2A | 11870000 | 1566390 | 639476 | 6 |
| P08032 | SPTA1 | spectrin, alpha, erythrocytic 1 | Cytoplasm | other | 3.62E+09 | 3.86E+09 | 3.61E+09 | | | | | | |
| P08032 | SPTA1 | spectrin, alpha, erythrocytic 1 | Cytoplasm | other | 2.40E+09 | 2.65E+09 | 2.76E+09 | | | | | | |
| P08032 | SPTA1 | spectrin, alpha, erythrocytic 1 | Cytoplasm | other | 2.14E+09 | 2.90E+09 | 2.76E+09 | | | | | | |
| P08032 | SPTA1 | spectrin, alpha, erythrocytic 1 | Cytoplasm | other | 2.05E+09 | 2.55E+09 | 2.56E+09 | | | | | | |
| P08032 | SPTA1 | spectrin, alpha, erythrocytic 1 | Cytoplasm | other | 1.54E+09 | 3.03E+09 | 2.13E+09 | | | | | | |
| P08032 | SPTA1 | spectrin, alpha, erythrocytic 1 | Cytoplasm | other | 2.39E+09 | 2.31E+09 | 2.20E+09 | | | | | | |
| P08032 | SPTA1 | spectrin, alpha, erythrocytic 1 | Cytoplasm | other | 2.36E+09 | 2.14E+09 | 2.48E+09 | | | | | | |
| P08032 | SPTA1 | spectrin, alpha, erythrocytic 1 | Cytoplasm | other | 2.13E+09 | 2.22E+09 | 1.83E+09 | | | | | | |
| P08032 | SPTA1 | spectrin, alpha, erythrocytic 1 | Cytoplasm | other | 1.74E+09 | 1.58E+09 | 1.41E+09 | | | | | | |
| P08032 | SPTA1 | spectrin, alpha, erythrocytic 1 | Cytoplasm | other | 2.20E+09 | 1.85E+09 | 2.09E+09 | | | | | | |
| P08032 | SPTA1 | spectrin, alpha, erythrocytic 1 | Cytoplasm | other | 1.60E+09 | 1.87E+09 | 1.71E+09 | | | | | | |
| P08032 | SPTA1 | spectrin, alpha, erythrocytic 1 | Cytoplasm | other | 2.05E+09 | 2.16E+09 | 1.86E+09 | | | | | | |
| P08032 | SPTA1 | spectrin, alpha, erythrocytic 1 | Cytoplasm | other | 1.80E+09 | 1.72E+09 | 1.83E+09 | | | | | | |
| P08032 | SPTA1 | spectrin, alpha, erythrocytic 1 | Cytoplasm | other | 1.58E+09 | 2.27E+09 | 2.42E+09 | | | | | | |
| P08032 | SPTA1 | spectrin, alpha, erythrocytic 1 | Cytoplasm | other | 2.13E+09 | 1.25E+09 | 2.02E+09 | | | | | | |
| P08032 | SPTA1 | spectrin, alpha, erythrocytic 1 | Cytoplasm | other | 1.73E+09 | 1.77E+09 | 1.52E+09 | | | | | | |
| P08032 | SPTA1 | spectrin, alpha, erythrocytic 1 | Cytoplasm | other | 1.23E+09 | 1.13E+09 | 1.29E+09 | | | | | | |
| P08032 | SPTA1 | spectrin, alpha, erythrocytic 1 | Cytoplasm | other | 1.87E+09 | 1.96E+09 | 1.83E+09 | | | | | | |
| P08032 | SPTA1 | spectrin, alpha, erythrocytic 1 | Cytoplasm | other | 1.38E+09 | 1.81E+09 | 1.64E+09 | | | | | | |
| P08032 | SPTA1 | spectrin, alpha, erythrocytic 1 | Cytoplasm | other | 1.53E+09 | 1.46E+09 | 1.32E+09 | | | | | | |
| P08032 | SPTA1 | spectrin, alpha, erythrocytic 1 | Cytoplasm | other | 1.78E+09 | 1.49E+09 | 1.52E+09 | | | | | | |
| P08032 | SPTA1 | spectrin, alpha, erythrocytic 1 | Cytoplasm | other | 1.49E+09 | 1.46E+09 | 1.43E+09 | | | | | | |
| P08032 | SPTA1 | spectrin, alpha, erythrocytic 1 | Cytoplasm | other | 1.45E+09 | 1.46E+09 | 1.33E+09 | | | | | | |
| P08032 | SPTA1 | spectrin, alpha, erythrocytic 1 | Cytoplasm | other | 1.54E+09 | 1.36E+09 | 1.24E+09 | | | | | | |
| P08032 | SPTA1 | spectrin, alpha, erythrocytic 1 | Cytoplasm | other | 1.25E+09 | 1.31E+09 | 1.38E+09 | | | | | | |
| P08032 | SPTA1 | spectrin, alpha, erythrocytic 1 | Cytoplasm | other | 1.18E+09 | 1.29E+09 | 1.32E+09 | | | | | | |
| P08032 | SPTA1 | spectrin, alpha, erythrocytic 1 | Cytoplasm | other | 1.54E+09 | 1.56E+09 | 1.55E+09 | | | | | | |
| P08032 | SPTA1 | spectrin, alpha, erythrocytic 1 | Cytoplasm | other | 1.66E+09 | 1.55E+09 | 1.63E+09 | | | | | | |
| P08032 | SPTA1 | spectrin, alpha, erythrocytic 1 | Cytoplasm | other | 1.12E+09 | 1.46E+09 | 1.21E+09 | | | | | | |
| P08032 | SPTA1 | spectrin, alpha, erythrocytic 1 | Cytoplasm | other | 1.36E+09 | 1.34E+09 | 1.26E+09 | | | | | | |
| P08032 | SPTA1 | spectrin, alpha, erythrocytic 1 | Cytoplasm | other | 1.21E+09 | 1.09E+09 | 1.14E+09 | | | | | | |
| P08032 | SPTA1 | spectrin, alpha, erythrocytic 1 | Cytoplasm | other | 1.41E+09 | 1.27E+09 | 1.26E+09 | | | | | | |
| P08032 | SPTA1 | spectrin, alpha, erythrocytic 1 | Cytoplasm | other | 1.25E+09 | 1.22E+09 | 1.08E+09 | | | | | | |
| P08032 | SPTA1 | spectrin, alpha, erythrocytic 1 | Cytoplasm | other | 1.14E+09 | 1.16E+09 | 9.67E+08 | | | | | | |
| P08032 | SPTA1 | spectrin, alpha, erythrocytic 1 | Cytoplasm | other | 1.85E+09 | 4.00E+08 | 1.12E+09 | | | | | | |
| P08032 | SPTA1 | spectrin, alpha, erythrocytic 1 | Cytoplasm | other | 1.52E+09 | 1.42E+09 | 1.40E+09 | | | | | | |
| P08032 | SPTA1 | spectrin, alpha, erythrocytic 1 | Cytoplasm | other | 9.70E+08 | 1.53E+09 | 1.13E+09 | | | | | | |
| P08032 | SPTA1 | spectrin, alpha, erythrocytic 1 | Cytoplasm | other | 1.24E+09 | 1.28E+09 | 1.00E+09 | | | | | | |
| P08032 | SPTA1 | spectrin, alpha, erythrocytic 1 | Cytoplasm | other | 1.15E+09 | 1.11E+09 | 1.13E+09 | | | | | | |
| P08032 | SPTA1 | spectrin, alpha, erythrocytic 1 | Cytoplasm | other | 1.13E+09 | 1.33E+09 | 1.27E+09 | | | | | | |
| P08032 | SPTA1 | spectrin, alpha, erythrocytic 1 | Cytoplasm | other | 1.01E+09 | 1.18E+09 | 1.09E+09 | | | | | | |
| P08032 | SPTA1 | spectrin, alpha, erythrocytic 1 | Cytoplasm | other | 1.27E+09 | 1.16E+09 | 1.14E+09 | | | | | | |
| P08032 | SPTA1 | spectrin, alpha, erythrocytic 1 | Cytoplasm | other | 1.03E+09 | 1.23E+09 | 1.23E+09 | | | | | | |
| P08032 | SPTA1 | spectrin, alpha, erythrocytic 1 | Cytoplasm | other | 1.30E+09 | 1.11E+09 | 1.20E+09 | | | | | | |
| P08032 | SPTA1 | spectrin, alpha, erythrocytic 1 | Cytoplasm | other | 1.22E+09 | 1.20E+09 | 1.22E+09 | | | | | | |
| P08032 | SPTA1 | spectrin, alpha, erythrocytic 1 | Cytoplasm | other | 1.10E+09 | 1.21E+09 | 1.01E+09 | | | | | | |
| P08032 | SPTA1 | spectrin, alpha, erythrocytic 1 | Cytoplasm | other | 1.06E+09 | 1.10E+09 | 9.67E+08 | | | | | | |
| P08032 | SPTA1 | spectrin, alpha, erythrocytic 1 | Cytoplasm | other | 1.22E+09 | 1.29E+09 | 1.06E+09 | | | | | | |
| P08032 | SPTA1 | spectrin, alpha, erythrocytic 1 | Cytoplasm | other | 6.09E+08 | 1.80E+09 | 1.38E+09 | | | | | | |
| P08032 | SPTA1 | spectrin, alpha, erythrocytic 1 | Cytoplasm | other | 1.20E+09 | 1.24E+09 | 1.28E+09 | | | | | | |
| P08032 | SPTA1 | spectrin, alpha, erythrocytic 1 | Cytoplasm | other | 1.44E+09 | 9.08E+08 | 9.23E+08 | | | | | | |
| P08032 | SPTA1 | spectrin, alpha, erythrocytic 1 | Cytoplasm | other | 1.08E+09 | 9.40E+08 | 9.46E+08 | | | | | | |
| P08032 | SPTA1 | spectrin, alpha, erythrocytic 1 | Cytoplasm | other | 1.09E+09 | 1.02E+09 | 9.75E+08 | | | | | | |
| P08032 | SPTA1 | spectrin, alpha, erythrocytic 1 | Cytoplasm | other | 8.43E+08 | 1.39E+09 | 1.14E+09 | | | | | | |
| P08032 | SPTA1 | spectrin, alpha, erythrocytic 1 | Cytoplasm | other | 7.19E+08 | 1.01E+09 | 7.91E+08 | | | | | | |
| P08032 | SPTA1 | spectrin, alpha, erythrocytic 1 | Cytoplasm | other | 9.07E+08 | 1.08E+09 | 1.07E+09 | | | | | | |
| P08032 | SPTA1 | spectrin, alpha, erythrocytic 1 | Cytoplasm | other | 1.10E+09 | 8.62E+08 | 8.75E+08 | | | | | | |
| P08032 | SPTA1 | spectrin, alpha, erythrocytic 1 | Cytoplasm | other | 7.64E+08 | 1.38E+09 | 1.21E+09 | | | | | | |

| | | | | | | | |
|---|---|---|---|---|---|---|---|
| P08032 | SPTA1 | spectrin, alpha, erythrocytic 1 | Cytoplasm | other | 8.71E+08 | 1.14E+09 | 7.79E+08 |
| P08032 | SPTA1 | spectrin, alpha, erythrocytic 1 | Cytoplasm | other | 1.03E+09 | 1.47E+09 | 1.22E+09 |
| P08032 | SPTA1 | spectrin, alpha, erythrocytic 1 | Cytoplasm | other | 7.57E+08 | 9.98E+08 | 1.12E+09 |
| P08032 | SPTA1 | spectrin, alpha, erythrocytic 1 | Cytoplasm | other | 1.10E+09 | 1.04E+09 | 1.04E+09 |
| P08032 | SPTA1 | spectrin, alpha, erythrocytic 1 | Cytoplasm | other | 9.63E+08 | 9.57E+08 | 8.46E+08 |
| P08032 | SPTA1 | spectrin, alpha, erythrocytic 1 | Cytoplasm | other | 1.16E+09 | 8.60E+08 | 1.11E+09 |
| P08032 | SPTA1 | spectrin, alpha, erythrocytic 1 | Cytoplasm | other | 8.46E+08 | 8.33E+08 | 8.13E+08 |
| P08032 | SPTA1 | spectrin, alpha, erythrocytic 1 | Cytoplasm | other | 1.01E+09 | 8.17E+08 | 9.23E+08 |
| P08032 | SPTA1 | spectrin, alpha, erythrocytic 1 | Cytoplasm | other | 7.65E+08 | 8.26E+08 | 7.88E+08 |
| P08032 | SPTA1 | spectrin, alpha, erythrocytic 1 | Cytoplasm | other | 8.01E+08 | 7.99E+08 | 7.83E+08 |
| P08032 | SPTA1 | spectrin, alpha, erythrocytic 1 | Cytoplasm | other | 6.89E+08 | 1.13E+09 | 8.79E+08 |
| P08032 | SPTA1 | spectrin, alpha, erythrocytic 1 | Cytoplasm | other | 7.08E+08 | 1.00E+09 | 9.22E+08 |
| P08032 | SPTA1 | spectrin, alpha, erythrocytic 1 | Cytoplasm | other | 7.70E+08 | 7.45E+08 | 5.33E+08 |
| P08032 | SPTA1 | spectrin, alpha, erythrocytic 1 | Cytoplasm | other | 1.03E+09 | 5.19E+08 | 6.40E+08 |
| P08032 | SPTA1 | spectrin, alpha, erythrocytic 1 | Cytoplasm | other | 6.63E+08 | 9.48E+08 | 7.79E+08 |
| P08032 | SPTA1 | spectrin, alpha, erythrocytic 1 | Cytoplasm | other | 9.22E+08 | 8.08E+08 | 8.95E+08 |
| P08032 | SPTA1 | spectrin, alpha, erythrocytic 1 | Cytoplasm | other | 8.03E+08 | 7.28E+08 | 6.70E+08 |
| P08032 | SPTA1 | spectrin, alpha, erythrocytic 1 | Cytoplasm | other | 1.01E+09 | 1.04E+09 | 1.04E+09 |
| P08032 | SPTA1 | spectrin, alpha, erythrocytic 1 | Cytoplasm | other | 7.74E+08 | 8.23E+08 | 7.82E+08 |
| P08032 | SPTA1 | spectrin, alpha, erythrocytic 1 | Cytoplasm | other | 7.40E+08 | 7.08E+08 | 9.03E+08 |
| P08032 | SPTA1 | spectrin, alpha, erythrocytic 1 | Cytoplasm | other | 7.96E+08 | 8.77E+08 | 9.41E+08 |
| P08032 | SPTA1 | spectrin, alpha, erythrocytic 1 | Cytoplasm | other | 9.46E+08 | 6.81E+08 | 6.21E+08 |
| P08032 | SPTA1 | spectrin, alpha, erythrocytic 1 | Cytoplasm | other | 6.89E+08 | 7.30E+08 | 6.31E+08 |
| P08032 | SPTA1 | spectrin, alpha, erythrocytic 1 | Cytoplasm | other | 5.49E+08 | 7.43E+08 | 5.73E+08 |
| P08032 | SPTA1 | spectrin, alpha, erythrocytic 1 | Cytoplasm | other | 8.00E+08 | 8.30E+08 | 8.85E+08 |
| P08032 | SPTA1 | spectrin, alpha, erythrocytic 1 | Cytoplasm | other | 6.31E+08 | 6.03E+08 | 5.94E+08 |
| P08032 | SPTA1 | spectrin, alpha, erythrocytic 1 | Cytoplasm | other | 8.80E+08 | 7.92E+08 | 8.48E+08 |
| P08032 | SPTA1 | spectrin, alpha, erythrocytic 1 | Cytoplasm | other | 5.42E+08 | 6.71E+08 | 5.15E+08 |
| P08032 | SPTA1 | spectrin, alpha, erythrocytic 1 | Cytoplasm | other | 6.02E+08 | 6.29E+08 | 5.77E+08 |
| P08032 | SPTA1 | spectrin, alpha, erythrocytic 1 | Cytoplasm | other | 5.69E+08 | 7.03E+08 | 7.39E+08 |
| P08032 | SPTA1 | spectrin, alpha, erythrocytic 1 | Cytoplasm | other | 7.82E+08 | 7.78E+08 | 7.30E+08 |
| P08032 | SPTA1 | spectrin, alpha, erythrocytic 1 | Cytoplasm | other | 6.65E+08 | 5.64E+08 | 5.72E+08 |
| P08032 | SPTA1 | spectrin, alpha, erythrocytic 1 | Cytoplasm | other | 6.44E+08 | 6.68E+08 | 6.53E+08 |
| P08032 | SPTA1 | spectrin, alpha, erythrocytic 1 | Cytoplasm | other | 8.27E+08 | 6.75E+08 | 7.06E+08 |
| P08032 | SPTA1 | spectrin, alpha, erythrocytic 1 | Cytoplasm | other | 4.94E+08 | 9.40E+08 | 6.12E+08 |
| P08032 | SPTA1 | spectrin, alpha, erythrocytic 1 | Cytoplasm | other | 5.60E+08 | 6.47E+08 | 5.75E+08 |
| P08032 | SPTA1 | spectrin, alpha, erythrocytic 1 | Cytoplasm | other | 4.38E+08 | 6.91E+08 | 6.22E+08 |
| P08032 | SPTA1 | spectrin, alpha, erythrocytic 1 | Cytoplasm | other | 9.15E+08 | 2.73E+08 | 6.01E+08 |
| P08032 | SPTA1 | spectrin, alpha, erythrocytic 1 | Cytoplasm | other | 4.83E+08 | 8.96E+08 | 7.55E+08 |
| P08032 | SPTA1 | spectrin, alpha, erythrocytic 1 | Cytoplasm | other | 7.09E+08 | 5.17E+08 | 7.45E+08 |
| P08032 | SPTA1 | spectrin, alpha, erythrocytic 1 | Cytoplasm | other | 4.99E+08 | 7.03E+08 | 4.26E+08 |
| P08032 | SPTA1 | spectrin, alpha, erythrocytic 1 | Cytoplasm | other | 5.25E+08 | 3.75E+08 | 4.63E+08 |
| P08032 | SPTA1 | spectrin, alpha, erythrocytic 1 | Cytoplasm | other | 3.77E+08 | 7.06E+08 | 5.14E+08 |
| P08032 | SPTA1 | spectrin, alpha, erythrocytic 1 | Cytoplasm | other | 4.59E+08 | 5.02E+08 | 6.37E+08 |
| P08032 | SPTA1 | spectrin, alpha, erythrocytic 1 | Cytoplasm | other | 4.98E+08 | 4.93E+08 | 4.89E+08 |
| P08032 | SPTA1 | spectrin, alpha, erythrocytic 1 | Cytoplasm | other | 9.44E+08 | 2.46E+08 | 5.20E+08 |
| P08032 | SPTA1 | spectrin, alpha, erythrocytic 1 | Cytoplasm | other | 5.39E+08 | 5.63E+08 | 5.70E+08 |
| P08032 | SPTA1 | spectrin, alpha, erythrocytic 1 | Cytoplasm | other | 5.65E+08 | 6.62E+08 | 5.58E+08 |
| P08032 | SPTA1 | spectrin, alpha, erythrocytic 1 | Cytoplasm | other | 4.78E+08 | 4.43E+08 | 3.66E+08 |
| P08032 | SPTA1 | spectrin, alpha, erythrocytic 1 | Cytoplasm | other | 3.27E+08 | 3.85E+08 | 3.08E+08 |
| P08032 | SPTA1 | spectrin, alpha, erythrocytic 1 | Cytoplasm | other | 6.18E+08 | 4.41E+08 | 5.64E+08 |
| P08032 | SPTA1 | spectrin, alpha, erythrocytic 1 | Cytoplasm | other | 4.45E+08 | 4.09E+08 | 3.79E+08 |
| P08032 | SPTA1 | spectrin, alpha, erythrocytic 1 | Cytoplasm | other | 3.84E+08 | 4.61E+08 | 3.73E+08 |
| P08032 | SPTA1 | spectrin, alpha, erythrocytic 1 | Cytoplasm | other | 3.21E+08 | 3.40E+08 | 2.96E+08 |
| P08032 | SPTA1 | spectrin, alpha, erythrocytic 1 | Cytoplasm | other | 3.13E+08 | 3.11E+08 | 2.12E+08 |
| P08032 | SPTA1 | spectrin, alpha, erythrocytic 1 | Cytoplasm | other | 4.94E+08 | 5.40E+08 | 6.38E+08 |
| P08032 | SPTA1 | spectrin, alpha, erythrocytic 1 | Cytoplasm | other | 2.51E+08 | 4.96E+08 | 3.32E+08 |
| P08032 | SPTA1 | spectrin, alpha, erythrocytic 1 | Cytoplasm | other | 4.02E+08 | 3.13E+08 | 3.11E+08 |
| P08032 | SPTA1 | spectrin, alpha, erythrocytic 1 | Cytoplasm | other | 5.77E+08 | 4.86E+08 | 5.11E+08 |
| P08032 | SPTA1 | spectrin, alpha, erythrocytic 1 | Cytoplasm | other | 1.67E+08 | 6.56E+08 | 3.77E+08 |
| P08032 | SPTA1 | spectrin, alpha, erythrocytic 1 | Cytoplasm | other | 2.28E+08 | 4.90E+08 | 2.94E+08 |
| P08032 | SPTA1 | spectrin, alpha, erythrocytic 1 | Cytoplasm | other | 3.57E+08 | 4.20E+08 | 3.66E+08 |
| P08032 | SPTA1 | spectrin, alpha, erythrocytic 1 | Cytoplasm | other | 2.92E+08 | 3.44E+08 | 3.69E+08 |
| P08032 | SPTA1 | spectrin, alpha, erythrocytic 1 | Cytoplasm | other | 3.87E+08 | 3.92E+08 | 5.02E+08 |
| P08032 | SPTA1 | spectrin, alpha, erythrocytic 1 | Cytoplasm | other | 2.15E+08 | 4.14E+08 | 2.93E+08 |
| P08032 | SPTA1 | spectrin, alpha, erythrocytic 1 | Cytoplasm | other | 1.83E+08 | 4.45E+08 | 2.46E+08 |
| P08032 | SPTA1 | spectrin, alpha, erythrocytic 1 | Cytoplasm | other | 2.59E+08 | 3.44E+08 | 2.94E+08 |
| P08032 | SPTA1 | spectrin, alpha, erythrocytic 1 | Cytoplasm | other | 1.96E+08 | 4.45E+08 | 2.53E+08 |
| P08032 | SPTA1 | spectrin, alpha, erythrocytic 1 | Cytoplasm | other | 1.74E+08 | 2.13E+08 | 1.05E+08 |
| P08032 | SPTA1 | spectrin, alpha, erythrocytic 1 | Cytoplasm | other | 2.07E+08 | 5.11E+08 | 3.19E+08 |
| P08032 | SPTA1 | spectrin, alpha, erythrocytic 1 | Cytoplasm | other | 7.48E+07 | 3.12E+08 | 6.90E+08 |
| P08032 | SPTA1 | spectrin, alpha, erythrocytic 1 | Cytoplasm | other | 4.18E+08 | 4.52E+08 | 4.73E+08 |
| P08032 | SPTA1 | spectrin, alpha, erythrocytic 1 | Cytoplasm | other | 6.01E+08 | 5.59E+07 | 2.27E+08 |
| P08032 | SPTA1 | spectrin, alpha, erythrocytic 1 | Cytoplasm | other | 4.55E+08 | 1.26E+08 | 2.71E+08 |
| P08032 | SPTA1 | spectrin, alpha, erythrocytic 1 | Cytoplasm | other | 1.86E+08 | 4.69E+08 | 2.40E+08 |
| P08032 | SPTA1 | spectrin, alpha, erythrocytic 1 | Cytoplasm | other | 3.12E+08 | 2.81E+08 | 3.08E+08 |
| P08032 | SPTA1 | spectrin, alpha, erythrocytic 1 | Cytoplasm | other | 3.87E+08 | 8.57E+07 | 2.42E+08 |
| P08032 | SPTA1 | spectrin, alpha, erythrocytic 1 | Cytoplasm | other | 1.79E+08 | 6.15E+08 | 4.10E+08 |
| P08032 | SPTA1 | spectrin, alpha, erythrocytic 1 | Cytoplasm | other | 1.97E+08 | 4.86E+08 | 3.88E+08 |
| P08032 | SPTA1 | spectrin, alpha, erythrocytic 1 | Cytoplasm | other | 4.67E+08 | 5.71E+07 | 1.40E+08 |
| P08032 | SPTA1 | spectrin, alpha, erythrocytic 1 | Cytoplasm | other | 2.57E+08 | 2.06E+08 | 1.96E+08 |
| P08032 | SPTA1 | spectrin, alpha, erythrocytic 1 | Cytoplasm | other | 2.27E+08 | 2.26E+08 | 1.78E+08 |
| P08032 | SPTA1 | spectrin, alpha, erythrocytic 1 | Cytoplasm | other | 4.73E+08 | 3.72E+07 | 1.91E+08 |
| P08032 | SPTA1 | spectrin, alpha, erythrocytic 1 | Cytoplasm | other | 2.55E+08 | 1.45E+08 | 6.20E+07 |
| P08032 | SPTA1 | spectrin, alpha, erythrocytic 1 | Cytoplasm | other | 2.20E+08 | 4.25E+08 | 2.80E+08 |
| P08032 | SPTA1 | spectrin, alpha, erythrocytic 1 | Cytoplasm | other | 1.83E+08 | 2.21E+08 | 2.13E+08 |

| | | | | | | |
|---|---|---|---|---|---|---|
| P08032 | SPTA1 | spectrin, alpha, erythrocytic 1 | Cytoplasm | other | 2.38E+08 | 2.10E+08 | 1.97E+08 |
| P08032 | SPTA1 | spectrin, alpha, erythrocytic 1 | Cytoplasm | other | 1.40E+08 | 2.33E+08 | 1.75E+08 |
| P08032 | SPTA1 | spectrin, alpha, erythrocytic 1 | Cytoplasm | other | 1.43E+08 | 2.46E+08 | 1.58E+08 |
| P08032 | SPTA1 | spectrin, alpha, erythrocytic 1 | Cytoplasm | other | 3.79E+08 | 3.76E+08 | 3.83E+08 |
| P08032 | SPTA1 | spectrin, alpha, erythrocytic 1 | Cytoplasm | other | 2.42E+08 | 2.56E+08 | 2.41E+08 |
| P08032 | SPTA1 | spectrin, alpha, erythrocytic 1 | Cytoplasm | other | 3.43E+08 | 2.13E+08 | 3.13E+08 |
| P08032 | SPTA1 | spectrin, alpha, erythrocytic 1 | Cytoplasm | other | 7.99E+07 | 3.53E+08 | 1.64E+08 |
| P08032 | SPTA1 | spectrin, alpha, erythrocytic 1 | Cytoplasm | other | 1.57E+08 | 1.29E+08 | 4.90E+07 |
| P08032 | SPTA1 | spectrin, alpha, erythrocytic 1 | Cytoplasm | other | 1.62E+08 | 8.80E+07 | 8.81E+07 |
| P08032 | SPTA1 | spectrin, alpha, erythrocytic 1 | Cytoplasm | other | 1.65E+08 | 1.26E+08 | 1.06E+08 |
| P08032 | SPTA1 | spectrin, alpha, erythrocytic 1 | Cytoplasm | other | 2.89E+08 | 3.16E+07 | 1.20E+08 |
| P08032 | SPTA1 | spectrin, alpha, erythrocytic 1 | Cytoplasm | other | 1.82E+08 | 1.81E+08 | 1.77E+08 |
| P08032 | SPTA1 | spectrin, alpha, erythrocytic 1 | Cytoplasm | other | 2.20E+08 | 4.27E+07 | 1.29E+08 |
| P08032 | SPTA1 | spectrin, alpha, erythrocytic 1 | Cytoplasm | other | 1.03E+08 | 2.73E+08 | 2.93E+08 |
| P08032 | SPTA1 | spectrin, alpha, erythrocytic 1 | Cytoplasm | other | 1.21E+08 | 1.86E+08 | 1.77E+08 |
| P08032 | SPTA1 | spectrin, alpha, erythrocytic 1 | Cytoplasm | other | 1.57E+08 | 1.36E+08 | 1.27E+08 |
| P08032 | SPTA1 | spectrin, alpha, erythrocytic 1 | Cytoplasm | other | 1.22E+08 | 3.51E+08 | 2.73E+08 |
| P08032 | SPTA1 | spectrin, alpha, erythrocytic 1 | Cytoplasm | other | 6.89E+07 | 8.99E+07 | 1.08E+08 |
| P08032 | SPTA1 | spectrin, alpha, erythrocytic 1 | Cytoplasm | other | 3.17E+08 | 2.54E+07 | 9.79E+07 |
| P08032 | SPTA1 | spectrin, alpha, erythrocytic 1 | Cytoplasm | other | 1.03E+08 | 3.11E+08 | 2.27E+08 |
| P08032 | SPTA1 | spectrin, alpha, erythrocytic 1 | Cytoplasm | other | 5.93E+07 | 2.73E+07 | 1.46E+07 |
| P08032 | SPTA1 | spectrin, alpha, erythrocytic 1 | Cytoplasm | other | 1.42E+08 | 1.02E+08 | 9.55E+07 |
| P08032 | SPTA1 | spectrin, alpha, erythrocytic 1 | Cytoplasm | other | 2.70E+08 | 3.90E+07 | 1.08E+08 |
| P08032 | SPTA1 | spectrin, alpha, erythrocytic 1 | Cytoplasm | other | 1.03E+08 | 1.10E+08 | 1.04E+08 |
| P08032 | SPTA1 | spectrin, alpha, erythrocytic 1 | Cytoplasm | other | 1.23E+08 | 1.91E+08 | 1.09E+08 |
| P08032 | SPTA1 | spectrin, alpha, erythrocytic 1 | Cytoplasm | other | 1.64E+08 | 3.29E+07 | 7.91E+07 |
| P08032 | SPTA1 | spectrin, alpha, erythrocytic 1 | Cytoplasm | other | 1.07E+08 | 1.44E+08 | 3.07E+08 |
| P08032 | SPTA1 | spectrin, alpha, erythrocytic 1 | Cytoplasm | other | 7.49E+07 | 1.37E+08 | 9.43E+07 |
| P08032 | SPTA1 | spectrin, alpha, erythrocytic 1 | Cytoplasm | other | 1.17E+08 | 3.29E+08 | 7.09E+08 |
| P08032 | SPTA1 | spectrin, alpha, erythrocytic 1 | Cytoplasm | other | 3.52E+08 | 3.25E+07 | 1.14E+08 |
| P08032 | SPTA1 | spectrin, alpha, erythrocytic 1 | Cytoplasm | other | 1.20E+08 | 1.45E+08 | 1.22E+08 |
| P08032 | SPTA1 | spectrin, alpha, erythrocytic 1 | Cytoplasm | other | 9.10E+07 | 9.59E+07 | 1.09E+08 |
| P08032 | SPTA1 | spectrin, alpha, erythrocytic 1 | Cytoplasm | other | 5.55E+07 | 3.46E+08 | 5.05E+08 |
| P08032 | SPTA1 | spectrin, alpha, erythrocytic 1 | Cytoplasm | other | 1.28E+08 | 4.16E+07 | 2.02E+07 |
| P08032 | SPTA1 | spectrin, alpha, erythrocytic 1 | Cytoplasm | other | 1.45E+08 | 3.19E+07 | 1.50E+07 |
| P08032 | SPTA1 | spectrin, alpha, erythrocytic 1 | Cytoplasm | other | 9.02E+07 | 6.74E+07 | 9.86E+07 |
| P08032 | SPTA1 | spectrin, alpha, erythrocytic 1 | Cytoplasm | other | 1.81E+08 | 1.24E+08 | 1.11E+08 |
| P08032 | SPTA1 | spectrin, alpha, erythrocytic 1 | Cytoplasm | other | 8.06E+07 | 1.55E+08 | 1.05E+08 |
| P08032 | SPTA1 | spectrin, alpha, erythrocytic 1 | Cytoplasm | other | 8.86E+07 | 9.37E+07 | 1.16E+08 |
| P08032 | SPTA1 | spectrin, alpha, erythrocytic 1 | Cytoplasm | other | 3.51E+07 | 7.17E+07 | 6.37E+07 |
| P08032 | SPTA1 | spectrin, alpha, erythrocytic 1 | Cytoplasm | other | 6.15E+07 | 1664601.25 | 1572178 |
| P08032 | SPTA1 | spectrin, alpha, erythrocytic 1 | Cytoplasm | other | 5.25E+07 | 1.30E+08 | 8.76E+07 |
| P08032 | SPTA1 | spectrin, alpha, erythrocytic 1 | Cytoplasm | other | 6.20E+07 | 1.16E+08 | 1.07E+08 |
| P08032 | SPTA1 | spectrin, alpha, erythrocytic 1 | Cytoplasm | other | 6.96E+07 | 7.29E+07 | 8.70E+07 |
| P08032 | SPTA1 | spectrin, alpha, erythrocytic 1 | Cytoplasm | other | 4.96E+07 | 7.56E+07 | 7.84E+07 |
| P08032 | SPTA1 | spectrin, alpha, erythrocytic 1 | Cytoplasm | other | 6.17E+07 | 1.59E+08 | 9.06E+07 |
| P08032 | SPTA1 | spectrin, alpha, erythrocytic 1 | Cytoplasm | other | 6.38E+07 | 6.00E+07 | 8.73E+07 |
| P08032 | SPTA1 | spectrin, alpha, erythrocytic 1 | Cytoplasm | other | 7.64E+07 | 6.67E+07 | 5.99E+07 |
| P08032 | SPTA1 | spectrin, alpha, erythrocytic 1 | Cytoplasm | other | 9.21E+07 | 9.33E+07 | 8.11E+07 |
| P08032 | SPTA1 | spectrin, alpha, erythrocytic 1 | Cytoplasm | other | 2.80E+08 | 4109204.75 | 3.27E+07 |
| P08032 | SPTA1 | spectrin, alpha, erythrocytic 1 | Cytoplasm | other | 5.69E+07 | 1.53E+08 | 2.93E+08 |
| P08032 | SPTA1 | spectrin, alpha, erythrocytic 1 | Cytoplasm | other | 2.60E+08 | 1.34E+07 | 6.32E+07 |
| P08032 | SPTA1 | spectrin, alpha, erythrocytic 1 | Cytoplasm | other | 6.27E+07 | 6.43E+07 | 7.47E+07 |
| P08032 | SPTA1 | spectrin, alpha, erythrocytic 1 | Cytoplasm | other | 6.85E+07 | 1.02E+08 | 9.81E+07 |
| P08032 | SPTA1 | spectrin, alpha, erythrocytic 1 | Cytoplasm | other | 1.88E+08 | 1.74E+07 | 1.04E+08 |
| P08032 | SPTA1 | spectrin, alpha, erythrocytic 1 | Cytoplasm | other | 7.63E+07 | 1.64E+08 | 1.50E+08 |
| P08032 | SPTA1 | spectrin, alpha, erythrocytic 1 | Cytoplasm | other | 9.46E+07 | 2.29E+07 | 4.48E+07 |
| P08032 | SPTA1 | spectrin, alpha, erythrocytic 1 | Cytoplasm | other | 6.21E+07 | 5.78E+07 | 6.14E+07 |
| P08032 | SPTA1 | spectrin, alpha, erythrocytic 1 | Cytoplasm | other | 1.01E+08 | 3.21E+07 | 4.67E+07 |
| P08032 | SPTA1 | spectrin, alpha, erythrocytic 1 | Cytoplasm | other | 4.43E+07 | 5.04E+07 | 4.56E+07 |
| P08032 | SPTA1 | spectrin, alpha, erythrocytic 1 | Cytoplasm | other | 2.32E+07 | 1.16E+08 | 5.20E+07 |
| P08032 | SPTA1 | spectrin, alpha, erythrocytic 1 | Cytoplasm | other | 3.04E+07 | 5.51E+07 | 3.88E+07 |
| P08032 | SPTA1 | spectrin, alpha, erythrocytic 1 | Cytoplasm | other | 2.47E+07 | 1.63E+08 | 1.76E+08 |
| P08032 | SPTA1 | spectrin, alpha, erythrocytic 1 | Cytoplasm | other | 4.85E+07 | 2.34E+07 | 2.52E+07 |
| P08032 | SPTA1 | spectrin, alpha, erythrocytic 1 | Cytoplasm | other | 5.25E+07 | 5.09E+07 | 4.23E+07 |
| P08032 | SPTA1 | spectrin, alpha, erythrocytic 1 | Cytoplasm | other | 5.48E+07 | 4.60E+07 | 5.27E+07 |
| P08032 | SPTA1 | spectrin, alpha, erythrocytic 1 | Cytoplasm | other | 1.60E+08 | 8648845 | 4.19E+07 |
| P08032 | SPTA1 | spectrin, alpha, erythrocytic 1 | Cytoplasm | other | 3.65E+07 | 3.85E+07 | 3.89E+07 |
| P08032 | SPTA1 | spectrin, alpha, erythrocytic 1 | Cytoplasm | other | 3.40E+07 | 3.99E+07 | 4.06E+07 |
| P08032 | SPTA1 | spectrin, alpha, erythrocytic 1 | Cytoplasm | other | 2.85E+07 | 6.25E+07 | 6.31E+07 |
| P08032 | SPTA1 | spectrin, alpha, erythrocytic 1 | Cytoplasm | other | 1.53E+07 | 1.47E+08 | 4.78E+07 |
| P08032 | SPTA1 | spectrin, alpha, erythrocytic 1 | Cytoplasm | other | 2.43E+07 | 5.47E+07 | 3.63E+07 |
| P08032 | SPTA1 | spectrin, alpha, erythrocytic 1 | Cytoplasm | other | 1.00E+08 | 9914274 | 3.12E+07 |
| P08032 | SPTA1 | spectrin, alpha, erythrocytic 1 | Cytoplasm | other | 2.18E+07 | 4.16E+07 | 2.94E+07 |
| P08032 | SPTA1 | spectrin, alpha, erythrocytic 1 | Cytoplasm | other | 1.41E+07 | 1.27E+08 | 1.24E+08 |
| P08032 | SPTA1 | spectrin, alpha, erythrocytic 1 | Cytoplasm | other | 5.38E+07 | 6199820.5 | 2755513.25 |
| P08032 | SPTA1 | spectrin, alpha, erythrocytic 1 | Cytoplasm | other | 1.58E+07 | 2.68E+07 | 2.77E+07 |
| P08032 | SPTA1 | spectrin, alpha, erythrocytic 1 | Cytoplasm | other | 7.06E+07 | 1.10E+08 | 8.58E+07 |
| P08032 | SPTA1 | spectrin, alpha, erythrocytic 1 | Cytoplasm | other | 6.01E+07 | 3812748.75 | 2.17E+07 |
| P08032 | SPTA1 | spectrin, alpha, erythrocytic 1 | Cytoplasm | other | 5.24E+07 | 8762556 | 1.80E+07 |
| P08032 | SPTA1 | spectrin, alpha, erythrocytic 1 | Cytoplasm | other | 3.08E+07 | 3.55E+07 | 3.15E+07 |
| P08032 | SPTA1 | spectrin, alpha, erythrocytic 1 | Cytoplasm | other | 2.01E+07 | 2.43E+07 | 2.38E+07 |
| P08032 | SPTA1 | spectrin, alpha, erythrocytic 1 | Cytoplasm | other | 2.42E+07 | 4.21E+07 | 3.97E+07 |
| P08032 | SPTA1 | spectrin, alpha, erythrocytic 1 | Cytoplasm | other | 9324562 | 3181473.5 | 2823992.25 |
| P08032 | SPTA1 | spectrin, alpha, erythrocytic 1 | Cytoplasm | other | 6.44E+07 | 6.77E+07 | 5.03E+07 |
| P08032 | SPTA1 | spectrin, alpha, erythrocytic 1 | Cytoplasm | other | 2.97E+07 | 6.91E+07 | 4.30E+07 |

| | | | | | | | | | | | | | |
|---|---|---|---|---|---|---|---|---|---|---|---|---|---|
| P08032 | SPTA1 | spectrin, alpha, erythrocytic 1 | Cytoplasm | other | 8855858 | 1833242.25 | 1487010.625 | | | | | | |
| P08032 | SPTA1 | spectrin, alpha, erythrocytic 1 | Cytoplasm | other | 2.02E+07 | 7315335.5 | 2.31E+07 | | | | | | |
| P08032 | SPTA1 | spectrin, alpha, erythrocytic 1 | Cytoplasm | other | 4.82E+07 | 1725184.25 | 1283034.5 | | | | | | |
| P08032 | SPTA1 | spectrin, alpha, erythrocytic 1 | Cytoplasm | other | 2.83E+07 | 2.02E+07 | 2.26E+07 | | | | | | |
| P08032 | SPTA1 | spectrin, alpha, erythrocytic 1 | Cytoplasm | other | 6.07E+07 | 4501483 | 2.13E+07 | | | | | | |
| P08032 | SPTA1 | spectrin, alpha, erythrocytic 1 | Cytoplasm | other | 2.99E+07 | 8974486 | 8981814 | | | | | | |
| P08032 | SPTA1 | spectrin, alpha, erythrocytic 1 | Cytoplasm | other | 2.04E+07 | 2.65E+07 | 2.33E+07 | | | | | | |
| P08032 | SPTA1 | spectrin, alpha, erythrocytic 1 | Cytoplasm | other | 1.08E+07 | 2.61E+07 | 2.47E+07 | | | | | | |
| P08032 | SPTA1 | spectrin, alpha, erythrocytic 1 | Cytoplasm | other | 9356664 | 2.23E+07 | 9967633 | | | | | | |
| P08032 | SPTA1 | spectrin, alpha, erythrocytic 1 | Cytoplasm | other | 4.71E+07 | 2949384.75 | 9295249 | | | | | | |
| P08032 | SPTA1 | spectrin, alpha, erythrocytic 1 | Cytoplasm | other | 9.69E+07 | 152036.078 | 9596044 | | | | | | |
| P08032 | SPTA1 | spectrin, alpha, erythrocytic 1 | Cytoplasm | other | 1.56E+07 | 1.45E+07 | 1.82E+07 | | | | | | |
| P08032 | SPTA1 | spectrin, alpha, erythrocytic 1 | Cytoplasm | other | 1.76E+07 | 1.54E+07 | 1.44E+07 | | | | | | |
| P08032 | SPTA1 | spectrin, alpha, erythrocytic 1 | Cytoplasm | other | 9702409 | 1.41E+07 | 8747964 | | | | | | |
| P08032 | SPTA1 | spectrin, alpha, erythrocytic 1 | Cytoplasm | other | 1.55E+07 | 1.66E+07 | 1.72E+07 | | | | | | |
| P08032 | SPTA1 | spectrin, alpha, erythrocytic 1 | Cytoplasm | other | 8540535 | 1.83E+07 | 1.38E+07 | | | | | | |
| P08032 | SPTA1 | spectrin, alpha, erythrocytic 1 | Cytoplasm | other | 1.37E+07 | 1.95E+07 | 1.53E+07 | | | | | | |
| P08032 | SPTA1 | spectrin, alpha, erythrocytic 1 | Cytoplasm | other | 4.94E+07 | 3618033 | 9308307 | | | | | | |
| P08032 | SPTA1 | spectrin, alpha, erythrocytic 1 | Cytoplasm | other | 2.92E+07 | 8640045 | 5.89E+07 | | | | | | |
| P08032 | SPTA1 | spectrin, alpha, erythrocytic 1 | Cytoplasm | other | 1.04E+07 | 1.22E+07 | 1.48E+07 | | | | | | |
| P08032 | SPTA1 | spectrin, alpha, erythrocytic 1 | Cytoplasm | other | 2.08E+07 | 3015230.5 | 1.04E+07 | | | | | | |
| P08032 | SPTA1 | spectrin, alpha, erythrocytic 1 | Cytoplasm | other | 7378058.5 | 3.23E+07 | 1.48E+07 | | | | | | |
| P08032 | SPTA1 | spectrin, alpha, erythrocytic 1 | Cytoplasm | other | 1.05E+07 | 7264594 | 9301085 | | | | | | |
| P08032 | SPTA1 | spectrin, alpha, erythrocytic 1 | Cytoplasm | other | 9649011 | 1.16E+07 | 1.09E+07 | | | | | | |
| P08032 | SPTA1 | spectrin, alpha, erythrocytic 1 | Cytoplasm | other | 6.20E+07 | 385444.312 | 7828851 | | | | | | |
| P08032 | SPTA1 | spectrin, alpha, erythrocytic 1 | Cytoplasm | other | 1.89E+07 | 2.00E+07 | 2.26E+07 | | | | | | |
| P08032 | SPTA1 | spectrin, alpha, erythrocytic 1 | Cytoplasm | other | 8264263.5 | 2.39E+07 | 1.34E+07 | | | | | | |
| P08032 | SPTA1 | spectrin, alpha, erythrocytic 1 | Cytoplasm | other | 1.70E+07 | 1101402.5 | 4847233.5 | | | | | | |
| P08032 | SPTA1 | spectrin, alpha, erythrocytic 1 | Cytoplasm | other | 1.02E+07 | 1.08E+07 | 8019721 | | | | | | |
| P08032 | SPTA1 | spectrin, alpha, erythrocytic 1 | Cytoplasm | other | 4098023 | 1368507.75 | 851750.312 | | | | | | |
| P08032 | SPTA1 | spectrin, alpha, erythrocytic 1 | Cytoplasm | other | 7862527 | 7996983.5 | 6017103.5 | | | | | | |
| P08032 | SPTA1 | spectrin, alpha, erythrocytic 1 | Cytoplasm | other | 7253935 | 6824258.5 | 5806325 | | | | | | |
| P08032 | SPTA1 | spectrin, alpha, erythrocytic 1 | Cytoplasm | other | 2.08E+07 | 1.48E+07 | 1.25E+07 | | | | | | |
| P08032 | SPTA1 | spectrin, alpha, erythrocytic 1 | Cytoplasm | other | 1.55E+07 | 1386108.25 | 8485248 | | | | | | |
| P08032 | SPTA1 | spectrin, alpha, erythrocytic 1 | Cytoplasm | other | 4154315.75 | 3.05E+07 | 2.33E+07 | | | | | | |
| P08032 | SPTA1 | spectrin, alpha, erythrocytic 1 | Cytoplasm | other | 2920841 | 6798834.5 | 4247753 | | | | | | |
| P08032 | SPTA1 | spectrin, alpha, erythrocytic 1 | Cytoplasm | other | 1.43E+07 | 16813.092 | 6074.841 | | | | | | |
| P08032 | SPTA1 | spectrin, alpha, erythrocytic 1 | Cytoplasm | other | 4786706 | 4435007 | 5158230.5 | | | | | | |
| P08032 | SPTA1 | spectrin, alpha, erythrocytic 1 | Cytoplasm | other | 7751711 | 1953186.875 | 2070397.875 | | | | | | |
| P08032 | SPTA1 | spectrin, alpha, erythrocytic 1 | Cytoplasm | other | 1895319 | 537406.625 | 278149.219 | | | | | | |
| P08032 | SPTA1 | spectrin, alpha, erythrocytic 1 | Cytoplasm | other | 5793126.5 | 6935912.5 | 5942402 | | | | | | |
| P08032 | SPTA1 | spectrin, alpha, erythrocytic 1 | Cytoplasm | other | 1.40E+07 | 1813033.125 | 5947439.5 | | | | | | |
| P08032 | SPTA1 | spectrin, alpha, erythrocytic 1 | Cytoplasm | other | 1.55E+07 | 4.70E-06 | 4.70E-06 | | | | | | |
| P08032 | SPTA1 | spectrin, alpha, erythrocytic 1 | Cytoplasm | other | 2617689.75 | 9998465 | 4522854 | | | | | | |
| P08032 | SPTA1 | spectrin, alpha, erythrocytic 1 | Cytoplasm | other | 3448459.75 | 4504923 | 3140520 | | | | | | |
| P08032 | SPTA1 | spectrin, alpha, erythrocytic 1 | Cytoplasm | other | 4096232 | 3438249.25 | 4719960 | | | | | | |
| P08032 | SPTA1 | spectrin, alpha, erythrocytic 1 | Cytoplasm | other | 6656435.5 | 6747163 | 7341867 | | | | | | |
| P08032 | SPTA1 | spectrin, alpha, erythrocytic 1 | Cytoplasm | other | 4611412.5 | 8270384 | 7263779.5 | | | | | | |
| P08032 | SPTA1 | spectrin, alpha, erythrocytic 1 | Cytoplasm | other | 4420521.5 | 4063644.25 | 3208018.25 | | | | | | |
| P08032 | SPTA1 | spectrin, alpha, erythrocytic 1 | Cytoplasm | other | 2120206.5 | 4922203.5 | 7060458 | | | | | | |
| P08032 | SPTA1 | spectrin, alpha, erythrocytic 1 | Cytoplasm | other | 3799378.5 | 9196574 | 5313580 | | | | | | |
| P08032 | SPTA1 | spectrin, alpha, erythrocytic 1 | Cytoplasm | other | 1.38E+07 | 348206.594 | 2987176.5 | | | | | | |
| P08032 | SPTA1 | spectrin, alpha, erythrocytic 1 | Cytoplasm | other | 703025.438 | 1.27E+07 | 1.32E+07 | | | | | | |
| P08032 | SPTA1 | spectrin, alpha, erythrocytic 1 | Cytoplasm | other | 8437689 | 21760.238 | 398956 | | | | | | |
| P08032 | SPTA1 | spectrin, alpha, erythrocytic 1 | Cytoplasm | other | 2554799.25 | 765292.25 | 1790400.875 | | | | | | |
| P08032 | SPTA1 | spectrin, alpha, erythrocytic 1 | Cytoplasm | other | 1486555 | 2050586 | 1954723 | | | | | | |
| P08032 | SPTA1 | spectrin, alpha, erythrocytic 1 | Cytoplasm | other | 2937276.5 | 275685.062 | 1134278.5 | | | | | | |
| P08032 | SPTA1 | spectrin, alpha, erythrocytic 1 | Cytoplasm | other | 844488.688 | 1040174.938 | 804740.688 | | | | | | |
| P08032 | SPTA1 | spectrin, alpha, erythrocytic 1 | Cytoplasm | other | 3183611 | 6960248 | 6592626 | | | | | | |
| P08032 | SPTA1 | spectrin, alpha, erythrocytic 1 | Cytoplasm | other | 3519757 | 466104.25 | 931391.125 | | | | | | |
| P08032 | SPTA1 | spectrin, alpha, erythrocytic 1 | Cytoplasm | other | 1266498.875 | 1536788.875 | 1241274.375 | | | | | | |
| P08032 | SPTA1 | spectrin, alpha, erythrocytic 1 | Cytoplasm | other | 400462.906 | 1676655.875 | 2069122.75 | | | | | | |
| P08032 | SPTA1 | spectrin, alpha, erythrocytic 1 | Cytoplasm | other | 974162.062 | 4.08E+07 | 5.88E+07 | | | | | | |
| P08032 | SPTA1 | spectrin, alpha, erythrocytic 1 | Cytoplasm | other | 7216989 | 4.70E-06 | 396491.469 | | | | | | |
| P08032 | SPTA1 | spectrin, alpha, erythrocytic 1 | Cytoplasm | other | 3272147.25 | 0 | 0 | | | | | | |
| P08032 | SPTA1 | spectrin, alpha, erythrocytic 1 | Cytoplasm | other | 2861398.5 | 1686447 | 1929042.75 | | | | | | |
| P08032 | SPTA1 | spectrin, alpha, erythrocytic 1 | Cytoplasm | other | 100901.539 | 340460.25 | 797790.5 | | | | | | |
| P08032 | SPTA1 | spectrin, alpha, erythrocytic 1 | Cytoplasm | other | 109567.188 | 1417408.75 | 1800574.25 | P08032 | SPTA1 | 480941468 | 616440184 | 20616844 | 894 |
| P16546 | SPTAN1 | spectrin, alpha, non-erythrocytic 1 | Plasma Membrane | other | 5150154.5 | 1.09E+07 | 7431198 | P16546 | SPTAN1 | 5150155 | 2911225 | 1680797 | 3 |
| P15508 | SPTB | spectrin, beta, erythrocytic | Plasma Membrane | other | 2.77E+09 | 3.49E+09 | 2.83E+09 | | | | | | |
| P15508 | SPTB | spectrin, beta, erythrocytic | Plasma Membrane | other | 2.58E+09 | 2.30E+09 | 2.07E+09 | | | | | | |
| P15508 | SPTB | spectrin, beta, erythrocytic | Plasma Membrane | other | 2.57E+09 | 2.17E+09 | 2.14E+09 | | | | | | |
| P15508 | SPTB | spectrin, beta, erythrocytic | Plasma Membrane | other | 2.42E+09 | 2.18E+09 | 2.32E+09 | | | | | | |
| P15508 | SPTB | spectrin, beta, erythrocytic | Plasma Membrane | other | 1.87E+09 | 1.92E+09 | 1.55E+09 | | | | | | |
| P15508 | SPTB | spectrin, beta, erythrocytic | Plasma Membrane | other | 1.92E+09 | 2.19E+09 | 2.19E+09 | | | | | | |
| P15508 | SPTB | spectrin, beta, erythrocytic | Plasma Membrane | other | 1.67E+09 | 1.75E+09 | 1.82E+09 | | | | | | |
| P15508 | SPTB | spectrin, beta, erythrocytic | Plasma Membrane | other | 1.74E+09 | 1.51E+09 | 1.32E+09 | | | | | | |
| P15508 | SPTB | spectrin, beta, erythrocytic | Plasma Membrane | other | 1.59E+09 | 1.51E+09 | 1.62E+09 | | | | | | |
| P15508 | SPTB | spectrin, beta, erythrocytic | Plasma Membrane | other | 1.53E+09 | 1.65E+09 | 1.66E+09 | | | | | | |
| P15508 | SPTB | spectrin, beta, erythrocytic | Plasma Membrane | other | 9.94E+08 | 2.58E+09 | 1.70E+09 | | | | | | |
| P15508 | SPTB | spectrin, beta, erythrocytic | Plasma Membrane | other | 1.90E+09 | 1.53E+09 | 1.50E+09 | | | | | | |
| P15508 | SPTB | spectrin, beta, erythrocytic | Plasma Membrane | other | 1.60E+09 | 1.39E+09 | 1.32E+09 | | | | | | |
| P15508 | SPTB | spectrin, beta, erythrocytic | Plasma Membrane | other | 2.09E+09 | 2.07E+09 | 2.42E+09 | | | | | | |
| P15508 | SPTB | spectrin, beta, erythrocytic | Plasma Membrane | other | 1.39E+09 | 1.56E+09 | 1.52E+09 | | | | | | |
| P15508 | SPTB | spectrin, beta, erythrocytic | Plasma Membrane | other | 1.59E+09 | 1.70E+09 | 1.64E+09 | | | | | | |
| P15508 | SPTB | spectrin, beta, erythrocytic | Plasma Membrane | other | 1.45E+09 | 1.46E+09 | 1.36E+09 | | | | | | |

| | | | | | | | |
|---|---|---|---|---|---|---|---|
| P15508 | SPTB | spectrin, beta, erythrocytic | Plasma Membrane | other | 1.18E+09 | 1.57E+09 | 1.11E+09 |
| P15508 | SPTB | spectrin, beta, erythrocytic | Plasma Membrane | other | 1.30E+09 | 1.31E+09 | 1.16E+09 |
| P15508 | SPTB | spectrin, beta, erythrocytic | Plasma Membrane | other | 1.19E+09 | 1.20E+09 | 1.40E+09 |
| P15508 | SPTB | spectrin, beta, erythrocytic | Plasma Membrane | other | 1.40E+09 | 1.64E+09 | 1.29E+09 |
| P15508 | SPTB | spectrin, beta, erythrocytic | Plasma Membrane | other | 1.51E+09 | 1.30E+09 | 1.06E+09 |
| P15508 | SPTB | spectrin, beta, erythrocytic | Plasma Membrane | other | 1.32E+09 | 1.30E+09 | 1.20E+09 |
| P15508 | SPTB | spectrin, beta, erythrocytic | Plasma Membrane | other | 1.31E+09 | 1.49E+09 | 1.42E+09 |
| P15508 | SPTB | spectrin, beta, erythrocytic | Plasma Membrane | other | 1.05E+09 | 1.07E+09 | 8.05E+08 |
| P15508 | SPTB | spectrin, beta, erythrocytic | Plasma Membrane | other | 1.10E+09 | 1.14E+09 | 1.06E+09 |
| P15508 | SPTB | spectrin, beta, erythrocytic | Plasma Membrane | other | 1.39E+09 | 1.35E+09 | 1.22E+09 |
| P15508 | SPTB | spectrin, beta, erythrocytic | Plasma Membrane | other | 1.31E+09 | 1.15E+09 | 1.09E+09 |
| P15508 | SPTB | spectrin, beta, erythrocytic | Plasma Membrane | other | 1.32E+09 | 1.33E+09 | 1.25E+09 |
| P15508 | SPTB | spectrin, beta, erythrocytic | Plasma Membrane | other | 1.05E+09 | 1.02E+09 | 8.90E+08 |
| P15508 | SPTB | spectrin, beta, erythrocytic | Plasma Membrane | other | 1.48E+09 | 1.52E+09 | 1.40E+09 |
| P15508 | SPTB | spectrin, beta, erythrocytic | Plasma Membrane | other | 1.23E+09 | 1.41E+09 | 9.66E+08 |
| P15508 | SPTB | spectrin, beta, erythrocytic | Plasma Membrane | other | 9.94E+08 | 1.62E+09 | 1.23E+09 |
| P15508 | SPTB | spectrin, beta, erythrocytic | Plasma Membrane | other | 1.06E+09 | 1.62E+09 | 1.25E+09 |
| P15508 | SPTB | spectrin, beta, erythrocytic | Plasma Membrane | other | 1.28E+09 | 3.43E+08 | 5.35E+08 |
| P15508 | SPTB | spectrin, beta, erythrocytic | Plasma Membrane | other | 1.20E+09 | 9.90E+08 | 1.10E+09 |
| P15508 | SPTB | spectrin, beta, erythrocytic | Plasma Membrane | other | 1.41E+09 | 1.36E+09 | 1.43E+09 |
| P15508 | SPTB | spectrin, beta, erythrocytic | Plasma Membrane | other | 1.10E+09 | 1.06E+09 | 1.06E+09 |
| P15508 | SPTB | spectrin, beta, erythrocytic | Plasma Membrane | other | 9.25E+08 | 1.09E+09 | 1.04E+09 |
| P15508 | SPTB | spectrin, beta, erythrocytic | Plasma Membrane | other | 1.08E+09 | 1.20E+09 | 1.06E+09 |
| P15508 | SPTB | spectrin, beta, erythrocytic | Plasma Membrane | other | 1.07E+09 | 9.40E+08 | 8.80E+08 |
| P15508 | SPTB | spectrin, beta, erythrocytic | Plasma Membrane | other | 1.22E+09 | 1.36E+09 | 1.29E+09 |
| P15508 | SPTB | spectrin, beta, erythrocytic | Plasma Membrane | other | 9.18E+08 | 1.27E+09 | 8.09E+08 |
| P15508 | SPTB | spectrin, beta, erythrocytic | Plasma Membrane | other | 7.82E+08 | 1.09E+09 | 1.02E+09 |
| P15508 | SPTB | spectrin, beta, erythrocytic | Plasma Membrane | other | 1.06E+09 | 1.16E+09 | 1.12E+09 |
| P15508 | SPTB | spectrin, beta, erythrocytic | Plasma Membrane | other | 9.20E+08 | 9.47E+08 | 8.08E+08 |
| P15508 | SPTB | spectrin, beta, erythrocytic | Plasma Membrane | other | 1.23E+09 | 1.08E+09 | 1.10E+09 |
| P15508 | SPTB | spectrin, beta, erythrocytic | Plasma Membrane | other | 9.48E+08 | 1.14E+09 | 1.24E+09 |
| P15508 | SPTB | spectrin, beta, erythrocytic | Plasma Membrane | other | 9.47E+08 | 9.98E+08 | 8.10E+08 |
| P15508 | SPTB | spectrin, beta, erythrocytic | Plasma Membrane | other | 8.59E+08 | 9.31E+08 | 9.36E+08 |
| P15508 | SPTB | spectrin, beta, erythrocytic | Plasma Membrane | other | 8.66E+08 | 9.36E+08 | 8.24E+08 |
| P15508 | SPTB | spectrin, beta, erythrocytic | Plasma Membrane | other | 7.79E+08 | 1.11E+09 | 9.91E+08 |
| P15508 | SPTB | spectrin, beta, erythrocytic | Plasma Membrane | other | 9.52E+08 | 9.42E+08 | 1.05E+09 |
| P15508 | SPTB | spectrin, beta, erythrocytic | Plasma Membrane | other | 1.00E+09 | 7.55E+08 | 7.31E+08 |
| P15508 | SPTB | spectrin, beta, erythrocytic | Plasma Membrane | other | 9.07E+08 | 8.34E+08 | 8.11E+08 |
| P15508 | SPTB | spectrin, beta, erythrocytic | Plasma Membrane | other | 6.45E+08 | 6.89E+08 | 4.50E+08 |
| P15508 | SPTB | spectrin, beta, erythrocytic | Plasma Membrane | other | 5.86E+08 | 8.17E+08 | 6.27E+08 |
| P15508 | SPTB | spectrin, beta, erythrocytic | Plasma Membrane | other | 8.58E+08 | 9.26E+08 | 9.15E+08 |
| P15508 | SPTB | spectrin, beta, erythrocytic | Plasma Membrane | other | 1.22E+09 | 4.00E+08 | 7.83E+08 |
| P15508 | SPTB | spectrin, beta, erythrocytic | Plasma Membrane | other | 8.17E+08 | 8.33E+08 | 8.48E+08 |
| P15508 | SPTB | spectrin, beta, erythrocytic | Plasma Membrane | other | 6.99E+08 | 8.21E+08 | 7.74E+08 |
| P15508 | SPTB | spectrin, beta, erythrocytic | Plasma Membrane | other | 5.58E+08 | 9.57E+08 | 7.52E+08 |
| P15508 | SPTB | spectrin, beta, erythrocytic | Plasma Membrane | other | 7.20E+08 | 9.37E+08 | 8.87E+08 |
| P15508 | SPTB | spectrin, beta, erythrocytic | Plasma Membrane | other | 5.77E+08 | 6.87E+08 | 3.61E+08 |
| P15508 | SPTB | spectrin, beta, erythrocytic | Plasma Membrane | other | 7.05E+08 | 8.23E+08 | 7.45E+08 |
| P15508 | SPTB | spectrin, beta, erythrocytic | Plasma Membrane | other | 7.15E+08 | 5.85E+08 | 6.16E+08 |
| P15508 | SPTB | spectrin, beta, erythrocytic | Plasma Membrane | other | 5.54E+08 | 6.32E+08 | 5.96E+08 |
| P15508 | SPTB | spectrin, beta, erythrocytic | Plasma Membrane | other | 6.16E+08 | 5.85E+08 | 5.74E+08 |
| P15508 | SPTB | spectrin, beta, erythrocytic | Plasma Membrane | other | 5.03E+08 | 9.06E+08 | 1.46E+09 |
| P15508 | SPTB | spectrin, beta, erythrocytic | Plasma Membrane | other | 6.78E+08 | 7.20E+08 | 7.10E+08 |
| P15508 | SPTB | spectrin, beta, erythrocytic | Plasma Membrane | other | 1.19E+09 | 1.43E+08 | 5.16E+08 |
| P15508 | SPTB | spectrin, beta, erythrocytic | Plasma Membrane | other | 1.04E+09 | 2.95E+08 | 6.38E+08 |
| P15508 | SPTB | spectrin, beta, erythrocytic | Plasma Membrane | other | 5.86E+08 | 6.51E+08 | 5.90E+08 |
| P15508 | SPTB | spectrin, beta, erythrocytic | Plasma Membrane | other | 6.40E+08 | 7.18E+08 | 6.75E+08 |
| P15508 | SPTB | spectrin, beta, erythrocytic | Plasma Membrane | other | 7.83E+08 | 8.47E+08 | 6.97E+08 |
| P15508 | SPTB | spectrin, beta, erythrocytic | Plasma Membrane | other | 3.26E+08 | 1.26E+09 | 5.26E+08 |
| P15508 | SPTB | spectrin, beta, erythrocytic | Plasma Membrane | other | 5.01E+08 | 6.13E+08 | 7.39E+08 |
| P15508 | SPTB | spectrin, beta, erythrocytic | Plasma Membrane | other | 5.44E+08 | 9.52E+08 | 6.17E+08 |
| P15508 | SPTB | spectrin, beta, erythrocytic | Plasma Membrane | other | 4.41E+08 | 6.06E+08 | 5.22E+08 |
| P15508 | SPTB | spectrin, beta, erythrocytic | Plasma Membrane | other | 5.84E+08 | 7.56E+08 | 7.48E+08 |
| P15508 | SPTB | spectrin, beta, erythrocytic | Plasma Membrane | other | 5.87E+08 | 5.67E+08 | 6.25E+08 |
| P15508 | SPTB | spectrin, beta, erythrocytic | Plasma Membrane | other | 5.12E+08 | 5.48E+08 | 5.08E+08 |
| P15508 | SPTB | spectrin, beta, erythrocytic | Plasma Membrane | other | 4.92E+08 | 1.08E+09 | 7.45E+08 |
| P15508 | SPTB | spectrin, beta, erythrocytic | Plasma Membrane | other | 4.52E+08 | 4.56E+08 | 3.89E+08 |
| P15508 | SPTB | spectrin, beta, erythrocytic | Plasma Membrane | other | 5.84E+08 | 6.60E+08 | 6.34E+08 |
| P15508 | SPTB | spectrin, beta, erythrocytic | Plasma Membrane | other | 4.01E+08 | 4.88E+08 | 4.21E+08 |
| P15508 | SPTB | spectrin, beta, erythrocytic | Plasma Membrane | other | 3.22E+08 | 5.05E+08 | 4.57E+08 |
| P15508 | SPTB | spectrin, beta, erythrocytic | Plasma Membrane | other | 3.02E+08 | 1.08E+09 | 4.72E+08 |
| P15508 | SPTB | spectrin, beta, erythrocytic | Plasma Membrane | other | 5.14E+08 | 8.80E+08 | 8.73E+08 |
| P15508 | SPTB | spectrin, beta, erythrocytic | Plasma Membrane | other | 6.48E+08 | 6.22E+08 | 7.49E+08 |
| P15508 | SPTB | spectrin, beta, erythrocytic | Plasma Membrane | other | 4.33E+08 | 5.14E+08 | 5.08E+08 |
| P15508 | SPTB | spectrin, beta, erythrocytic | Plasma Membrane | other | 4.17E+08 | 6.05E+08 | 4.72E+08 |
| P15508 | SPTB | spectrin, beta, erythrocytic | Plasma Membrane | other | 4.31E+08 | 4.34E+08 | 3.66E+08 |
| P15508 | SPTB | spectrin, beta, erythrocytic | Plasma Membrane | other | 5.90E+08 | 5.63E+08 | 8.09E+08 |
| P15508 | SPTB | spectrin, beta, erythrocytic | Plasma Membrane | other | 5.57E+08 | 3.58E+08 | 3.95E+08 |
| P15508 | SPTB | spectrin, beta, erythrocytic | Plasma Membrane | other | 4.82E+08 | 4.31E+08 | 4.01E+08 |
| P15508 | SPTB | spectrin, beta, erythrocytic | Plasma Membrane | other | 4.05E+08 | 3.42E+08 | 2.71E+08 |
| P15508 | SPTB | spectrin, beta, erythrocytic | Plasma Membrane | other | 4.56E+08 | 2.58E+08 | 4.01E+08 |
| P15508 | SPTB | spectrin, beta, erythrocytic | Plasma Membrane | other | 4.61E+08 | 5.45E+08 | 5.29E+08 |
| P15508 | SPTB | spectrin, beta, erythrocytic | Plasma Membrane | other | 2.40E+08 | 4.35E+08 | 2.29E+08 |
| P15508 | SPTB | spectrin, beta, erythrocytic | Plasma Membrane | other | 5.40E+08 | 5.24E+08 | 4.96E+08 |
| P15508 | SPTB | spectrin, beta, erythrocytic | Plasma Membrane | other | 4.29E+08 | 3.97E+08 | 5.04E+08 |
| P15508 | SPTB | spectrin, beta, erythrocytic | Plasma Membrane | other | 3.56E+08 | 4.52E+08 | 5.25E+08 |

| | | | | | | | |
|---|---|---|---|---|---|---|---|
| P15508 | SPTB | spectrin, beta, erythrocytic | Plasma Membrane | other | 1.90E+08 | 4.02E+08 | 2.92E+08 |
| P15508 | SPTB | spectrin, beta, erythrocytic | Plasma Membrane | other | 3.22E+08 | 2.51E+08 | 2.46E+08 |
| P15508 | SPTB | spectrin, beta, erythrocytic | Plasma Membrane | other | 2.58E+08 | 2.74E+08 | 2.68E+08 |
| P15508 | SPTB | spectrin, beta, erythrocytic | Plasma Membrane | other | 2.52E+08 | 2.64E+08 | 2.47E+08 |
| P15508 | SPTB | spectrin, beta, erythrocytic | Plasma Membrane | other | 6.15E+08 | 4.78E+07 | 2.28E+08 |
| P15508 | SPTB | spectrin, beta, erythrocytic | Plasma Membrane | other | 2.41E+08 | 2.08E+08 | 1.89E+08 |
| P15508 | SPTB | spectrin, beta, erythrocytic | Plasma Membrane | other | 2.41E+08 | 3.30E+08 | 2.96E+08 |
| P15508 | SPTB | spectrin, beta, erythrocytic | Plasma Membrane | other | 2.60E+08 | 2.33E+08 | 1.15E+08 |
| P15508 | SPTB | spectrin, beta, erythrocytic | Plasma Membrane | other | 3.61E+08 | 6.95E+07 | 1.90E+08 |
| P15508 | SPTB | spectrin, beta, erythrocytic | Plasma Membrane | other | 1.64E+08 | 1.37E+08 | 1.00E+08 |
| P15508 | SPTB | spectrin, beta, erythrocytic | Plasma Membrane | other | 1.60E+08 | 1.92E+08 | 1.81E+08 |
| P15508 | SPTB | spectrin, beta, erythrocytic | Plasma Membrane | other | 1.70E+08 | 1.74E+08 | 1.34E+08 |
| P15508 | SPTB | spectrin, beta, erythrocytic | Plasma Membrane | other | 1.70E+08 | 2.84E+08 | 1.20E+08 |
| P15508 | SPTB | spectrin, beta, erythrocytic | Plasma Membrane | other | 2.23E+08 | 2.54E+08 | 2.64E+08 |
| P15508 | SPTB | spectrin, beta, erythrocytic | Plasma Membrane | other | 2.00E+08 | 3.93E+08 | 1.84E+08 |
| P15508 | SPTB | spectrin, beta, erythrocytic | Plasma Membrane | other | 1.92E+08 | 3.55E+08 | 4.73E+08 |
| P15508 | SPTB | spectrin, beta, erythrocytic | Plasma Membrane | other | 1.75E+08 | 1.89E+08 | 1.16E+08 |
| P15508 | SPTB | spectrin, beta, erythrocytic | Plasma Membrane | other | 2.59E+08 | 5.53E+08 | 4.55E+08 |
| P15508 | SPTB | spectrin, beta, erythrocytic | Plasma Membrane | other | 4.60E+08 | 3.72E+08 | 3.85E+08 |
| P15508 | SPTB | spectrin, beta, erythrocytic | Plasma Membrane | other | 2.42E+08 | 1.55E+08 | 1.57E+08 |
| P15508 | SPTB | spectrin, beta, erythrocytic | Plasma Membrane | other | 2.06E+08 | 2.15E+08 | 1.89E+08 |
| P15508 | SPTB | spectrin, beta, erythrocytic | Plasma Membrane | other | 2.30E+08 | 2.91E+08 | 2.74E+08 |
| P15508 | SPTB | spectrin, beta, erythrocytic | Plasma Membrane | other | 2.20E+08 | 1.80E+08 | 1.75E+08 |
| P15508 | SPTB | spectrin, beta, erythrocytic | Plasma Membrane | other | 2.22E+08 | 1.59E+08 | 1.81E+08 |
| P15508 | SPTB | spectrin, beta, erythrocytic | Plasma Membrane | other | 1.31E+08 | 3.81E+08 | 8.57E+08 |
| P15508 | SPTB | spectrin, beta, erythrocytic | Plasma Membrane | other | 8.44E+07 | 2.26E+08 | 2.19E+08 |
| P15508 | SPTB | spectrin, beta, erythrocytic | Plasma Membrane | other | 1.63E+08 | 2.58E+08 | 5.31E+08 |
| P15508 | SPTB | spectrin, beta, erythrocytic | Plasma Membrane | other | 1.02E+08 | 2.36E+08 | 1.58E+08 |
| P15508 | SPTB | spectrin, beta, erythrocytic | Plasma Membrane | other | 3.50E+08 | 6224695.5 | 7.22E+07 |
| P15508 | SPTB | spectrin, beta, erythrocytic | Plasma Membrane | other | 8.70E+07 | 7.20E+07 | 1.06E+08 |
| P15508 | SPTB | spectrin, beta, erythrocytic | Plasma Membrane | other | 1.05E+08 | 1.91E+08 | 1.87E+08 |
| P15508 | SPTB | spectrin, beta, erythrocytic | Plasma Membrane | other | 1.30E+08 | 1.60E+08 | 1.39E+08 |
| P15508 | SPTB | spectrin, beta, erythrocytic | Plasma Membrane | other | 8.50E+07 | 1.15E+08 | 9.75E+07 |
| P15508 | SPTB | spectrin, beta, erythrocytic | Plasma Membrane | other | 1.06E+08 | 1.38E+08 | 8.79E+07 |
| P15508 | SPTB | spectrin, beta, erythrocytic | Plasma Membrane | other | 1.37E+08 | 8.98E+07 | 9.37E+07 |
| P15508 | SPTB | spectrin, beta, erythrocytic | Plasma Membrane | other | 1.62E+08 | 7.08E+07 | 2.96E+08 |
| P15508 | SPTB | spectrin, beta, erythrocytic | Plasma Membrane | other | 1.79E+08 | 2.13E+08 | 2.03E+08 |
| P15508 | SPTB | spectrin, beta, erythrocytic | Plasma Membrane | other | 7.66E+07 | 4.68E+07 | 4.04E+07 |
| P15508 | SPTB | spectrin, beta, erythrocytic | Plasma Membrane | other | 2.03E+08 | 4.55E+07 | 5.01E+07 |
| P15508 | SPTB | spectrin, beta, erythrocytic | Plasma Membrane | other | 1.69E+08 | 5.93E+07 | 2.76E+08 |
| P15508 | SPTB | spectrin, beta, erythrocytic | Plasma Membrane | other | 1.13E+08 | 1.15E+08 | 9.74E+07 |
| P15508 | SPTB | spectrin, beta, erythrocytic | Plasma Membrane | other | 7.31E+07 | 1.28E+08 | 6.47E+07 |
| P15508 | SPTB | spectrin, beta, erythrocytic | Plasma Membrane | other | 9.17E+07 | 1.13E+08 | 1.01E+08 |
| P15508 | SPTB | spectrin, beta, erythrocytic | Plasma Membrane | other | 9.29E+07 | 1.01E+08 | 9.42E+07 |
| P15508 | SPTB | spectrin, beta, erythrocytic | Plasma Membrane | other | 1.12E+08 | 1.05E+08 | 1.01E+08 |
| P15508 | SPTB | spectrin, beta, erythrocytic | Plasma Membrane | other | 8.24E+07 | 1.06E+08 | 9.91E+07 |
| P15508 | SPTB | spectrin, beta, erythrocytic | Plasma Membrane | other | 7.38E+07 | 8.83E+07 | 7.73E+07 |
| P15508 | SPTB | spectrin, beta, erythrocytic | Plasma Membrane | other | 6.57E+07 | 1.02E+08 | 7.61E+07 |
| P15508 | SPTB | spectrin, beta, erythrocytic | Plasma Membrane | other | 7.12E+07 | 8.91E+07 | 9.83E+07 |
| P15508 | SPTB | spectrin, beta, erythrocytic | Plasma Membrane | other | 6.27E+07 | 8.80E+07 | 7.22E+07 |
| P15508 | SPTB | spectrin, beta, erythrocytic | Plasma Membrane | other | 1.08E+08 | 7.64E+07 | 9.02E+07 |
| P15508 | SPTB | spectrin, beta, erythrocytic | Plasma Membrane | other | 8.15E+07 | 8.45E+07 | 6.79E+07 |
| P15508 | SPTB | spectrin, beta, erythrocytic | Plasma Membrane | other | 1.29E+08 | 1.26E+08 | 1.20E+08 |
| P15508 | SPTB | spectrin, beta, erythrocytic | Plasma Membrane | other | 7.50E+07 | 8.79E+07 | 9.07E+07 |
| P15508 | SPTB | spectrin, beta, erythrocytic | Plasma Membrane | other | 6.23E+07 | 1.64E+08 | 3.70E+08 |
| P15508 | SPTB | spectrin, beta, erythrocytic | Plasma Membrane | other | 1.11E+08 | 2.99E+07 | 3.46E+07 |
| P15508 | SPTB | spectrin, beta, erythrocytic | Plasma Membrane | other | 1.93E+08 | 1.66E+08 | 1.70E+08 |
| P15508 | SPTB | spectrin, beta, erythrocytic | Plasma Membrane | other | 8.41E+07 | 1.92E+08 | 2.21E+08 |
| P15508 | SPTB | spectrin, beta, erythrocytic | Plasma Membrane | other | 9.34E+07 | 9.12E+07 | 9.54E+07 |
| P15508 | SPTB | spectrin, beta, erythrocytic | Plasma Membrane | other | 9.60E+07 | 9.09E+07 | 5.25E+07 |
| P15508 | SPTB | spectrin, beta, erythrocytic | Plasma Membrane | other | 8.72E+07 | 3.43E+08 | 3.13E+08 |
| P15508 | SPTB | spectrin, beta, erythrocytic | Plasma Membrane | other | 1.12E+08 | 4.41E+07 | 6.69E+07 |
| P15508 | SPTB | spectrin, beta, erythrocytic | Plasma Membrane | other | 1.33E+08 | 4.14E+07 | 7.55E+07 |
| P15508 | SPTB | spectrin, beta, erythrocytic | Plasma Membrane | other | 1.96E+08 | 2074150.5 | 5.05E+07 |
| P15508 | SPTB | spectrin, beta, erythrocytic | Plasma Membrane | other | 1.07E+08 | 1.82E+07 | 8.87E+07 |
| P15508 | SPTB | spectrin, beta, erythrocytic | Plasma Membrane | other | 1.08E+08 | 2.90E+07 | 5.14E+07 |
| P15508 | SPTB | spectrin, beta, erythrocytic | Plasma Membrane | other | 1.28E+08 | 7177984 | 4173398.75 |
| P15508 | SPTB | spectrin, beta, erythrocytic | Plasma Membrane | other | 6.39E+07 | 7.71E+07 | 7.80E+07 |
| P15508 | SPTB | spectrin, beta, erythrocytic | Plasma Membrane | other | 3.51E+07 | 4.84E+07 | 2.60E+07 |
| P15508 | SPTB | spectrin, beta, erythrocytic | Plasma Membrane | other | 3.87E+07 | 8.27E+07 | 6.43E+07 |
| P15508 | SPTB | spectrin, beta, erythrocytic | Plasma Membrane | other | 6.68E+07 | 5.13E+07 | 4.87E+07 |
| P15508 | SPTB | spectrin, beta, erythrocytic | Plasma Membrane | other | 2.43E+07 | 4.46E+07 | 5.45E+07 |
| P15508 | SPTB | spectrin, beta, erythrocytic | Plasma Membrane | other | 4.89E+07 | 7.36E+07 | 8.89E+07 |
| P15508 | SPTB | spectrin, beta, erythrocytic | Plasma Membrane | other | 1.06E+08 | 6388117.5 | 3.38E+07 |
| P15508 | SPTB | spectrin, beta, erythrocytic | Plasma Membrane | other | 3.00E+07 | 6.62E+07 | 6.37E+07 |
| P15508 | SPTB | spectrin, beta, erythrocytic | Plasma Membrane | other | 4.39E+07 | 4.49E+07 | 4.19E+07 |
| P15508 | SPTB | spectrin, beta, erythrocytic | Plasma Membrane | other | 5.32E+07 | 6.23E+07 | 6.05E+07 |
| P15508 | SPTB | spectrin, beta, erythrocytic | Plasma Membrane | other | 3.76E+07 | 3.21E+07 | 2.15E+07 |
| P15508 | SPTB | spectrin, beta, erythrocytic | Plasma Membrane | other | 2.04E+07 | 1.62E+08 | 6.79E+07 |
| P15508 | SPTB | spectrin, beta, erythrocytic | Plasma Membrane | other | 5.86E+07 | 5.64E+07 | 7.08E+07 |
| P15508 | SPTB | spectrin, beta, erythrocytic | Plasma Membrane | other | 2.65E+07 | 1.69E+08 | 1.44E+08 |
| P15508 | SPTB | spectrin, beta, erythrocytic | Plasma Membrane | other | 3.23E+07 | 8.04E+07 | 5.38E+07 |
| P15508 | SPTB | spectrin, beta, erythrocytic | Plasma Membrane | other | 3.32E+07 | 6.27E+07 | 7.08E+07 |
| P15508 | SPTB | spectrin, beta, erythrocytic | Plasma Membrane | other | 3.26E+07 | 3.42E+07 | 3.39E+07 |
| P15508 | SPTB | spectrin, beta, erythrocytic | Plasma Membrane | other | 4.07E+07 | 6.14E+07 | 8.81E+07 |
| P15508 | SPTB | spectrin, beta, erythrocytic | Plasma Membrane | other | 5.19E+07 | 4.62E+07 | 4.89E+07 |

| | | | | | | | | | | | | | |
|---|---|---|---|---|---|---|---|---|---|---|---|---|---|
| P15508 | SPTB | spectrin, beta, erythrocytic | Plasma Membrane | other | 1.89E+07 | 1.36E+08 | 7.87E+07 | | | | | | |
| P15508 | SPTB | spectrin, beta, erythrocytic | Plasma Membrane | other | 2.82E+07 | 4.16E+07 | 5.90E+07 | | | | | | |
| P15508 | SPTB | spectrin, beta, erythrocytic | Plasma Membrane | other | 2.71E+07 | 3.94E+07 | 3.82E+07 | | | | | | |
| P15508 | SPTB | spectrin, beta, erythrocytic | Plasma Membrane | other | 2.18E+07 | 3.60E+07 | 2.34E+07 | | | | | | |
| P15508 | SPTB | spectrin, beta, erythrocytic | Plasma Membrane | other | 3.05E+07 | 7.22E+07 | 9.50E+07 | | | | | | |
| P15508 | SPTB | spectrin, beta, erythrocytic | Plasma Membrane | other | 3.46E+07 | 1.67E+07 | 2.45E+07 | | | | | | |
| P15508 | SPTB | spectrin, beta, erythrocytic | Plasma Membrane | other | 1.49E+07 | 3.80E+07 | 3.06E+07 | | | | | | |
| P15508 | SPTB | spectrin, beta, erythrocytic | Plasma Membrane | other | 1.07E+08 | 1171325.75 | 1.73E+07 | | | | | | |
| P15508 | SPTB | spectrin, beta, erythrocytic | Plasma Membrane | other | 2.22E+07 | 1.78E+07 | 1.83E+07 | | | | | | |
| P15508 | SPTB | spectrin, beta, erythrocytic | Plasma Membrane | other | 1.72E+07 | 3.61E+07 | 3.21E+07 | | | | | | |
| P15508 | SPTB | spectrin, beta, erythrocytic | Plasma Membrane | other | 2.72E+07 | 1.57E+07 | 1.29E+07 | | | | | | |
| P15508 | SPTB | spectrin, beta, erythrocytic | Plasma Membrane | other | 1.17E+07 | 4.07E+07 | 2.17E+07 | | | | | | |
| P15508 | SPTB | spectrin, beta, erythrocytic | Plasma Membrane | other | 7.19E+07 | 6.48E+07 | 6.66E+07 | | | | | | |
| P15508 | SPTB | spectrin, beta, erythrocytic | Plasma Membrane | other | 1.68E+07 | 2.52E+07 | 3.21E+07 | | | | | | |
| P15508 | SPTB | spectrin, beta, erythrocytic | Plasma Membrane | other | 1.84E+07 | 2.11E+07 | 1.92E+07 | | | | | | |
| P15508 | SPTB | spectrin, beta, erythrocytic | Plasma Membrane | other | 3.20E+07 | 9638141 | 1.91E+07 | | | | | | |
| P15508 | SPTB | spectrin, beta, erythrocytic | Plasma Membrane | other | 1.65E+07 | 2.78E+07 | 2.42E+07 | | | | | | |
| P15508 | SPTB | spectrin, beta, erythrocytic | Plasma Membrane | other | 7.93E+07 | 2073742.125 | 1.39E+07 | | | | | | |
| P15508 | SPTB | spectrin, beta, erythrocytic | Plasma Membrane | other | 1.68E+07 | 3.71E+07 | 2.28E+07 | | | | | | |
| P15508 | SPTB | spectrin, beta, erythrocytic | Plasma Membrane | other | 1.04E+07 | 3.03E+07 | 2.50E+07 | | | | | | |
| P15508 | SPTB | spectrin, beta, erythrocytic | Plasma Membrane | other | 6.78E+07 | 1163641.75 | 1.16E+07 | | | | | | |
| P15508 | SPTB | spectrin, beta, erythrocytic | Plasma Membrane | other | 5.39E+07 | 703003.25 | 1.44E+07 | | | | | | |
| P15508 | SPTB | spectrin, beta, erythrocytic | Plasma Membrane | other | 1.74E+07 | 1.72E+07 | 1.24E+07 | | | | | | |
| P15508 | SPTB | spectrin, beta, erythrocytic | Plasma Membrane | other | 1.28E+07 | 7786871.5 | 7060444.5 | | | | | | |
| P15508 | SPTB | spectrin, beta, erythrocytic | Plasma Membrane | other | 1.38E+07 | 2.07E+07 | 3.26E+07 | | | | | | |
| P15508 | SPTB | spectrin, beta, erythrocytic | Plasma Membrane | other | 4.33E+07 | 206597.328 | 5050628 | | | | | | |
| P15508 | SPTB | spectrin, beta, erythrocytic | Plasma Membrane | other | 1.63E+07 | 1.14E+07 | 1.24E+08 | | | | | | |
| P15508 | SPTB | spectrin, beta, erythrocytic | Plasma Membrane | other | 3.58E+07 | 1546864.625 | 579689 | | | | | | |
| P15508 | SPTB | spectrin, beta, erythrocytic | Plasma Membrane | other | 1.04E+07 | 8983163 | 6171193.5 | | | | | | |
| P15508 | SPTB | spectrin, beta, erythrocytic | Plasma Membrane | other | 2.04E+07 | 5.41E+07 | 6.80E+07 | | | | | | |
| P15508 | SPTB | spectrin, beta, erythrocytic | Plasma Membrane | other | 8473344 | 6861417.5 | 3708524.5 | | | | | | |
| P15508 | SPTB | spectrin, beta, erythrocytic | Plasma Membrane | other | 2.08E+07 | 7616525 | 1.12E+07 | | | | | | |
| P15508 | SPTB | spectrin, beta, erythrocytic | Plasma Membrane | other | 1.14E+07 | 1.07E+07 | 1.07E+07 | | | | | | |
| P15508 | SPTB | spectrin, beta, erythrocytic | Plasma Membrane | other | 1.21E+07 | 1.67E+07 | 2.18E+07 | | | | | | |
| P15508 | SPTB | spectrin, beta, erythrocytic | Plasma Membrane | other | 9062929 | 1.02E+07 | 1.21E+07 | | | | | | |
| P15508 | SPTB | spectrin, beta, erythrocytic | Plasma Membrane | other | 4573587.5 | 1.65E+07 | 1.26E+07 | | | | | | |
| P15508 | SPTB | spectrin, beta, erythrocytic | Plasma Membrane | other | 4244391.5 | 7861874.5 | 6148043 | | | | | | |
| P15508 | SPTB | spectrin, beta, erythrocytic | Plasma Membrane | other | 2.11E+07 | 2511547.25 | 1108441.875 | | | | | | |
| P15508 | SPTB | spectrin, beta, erythrocytic | Plasma Membrane | other | 7494699 | 8781767 | 1.37E+07 | | | | | | |
| P15508 | SPTB | spectrin, beta, erythrocytic | Plasma Membrane | other | 3070151 | 5550212 | 3971070.5 | | | | | | |
| P15508 | SPTB | spectrin, beta, erythrocytic | Plasma Membrane | other | 2.33E+07 | 67561.172 | 2578132.25 | | | | | | |
| P15508 | SPTB | spectrin, beta, erythrocytic | Plasma Membrane | other | 1.20E+07 | 1.16E+07 | 1.54E+07 | | | | | | |
| P15508 | SPTB | spectrin, beta, erythrocytic | Plasma Membrane | other | 1.82E+07 | 3253161.5 | 4466223.5 | | | | | | |
| P15508 | SPTB | spectrin, beta, erythrocytic | Plasma Membrane | other | 4918893.5 | 1.16E+07 | 1.19E+07 | | | | | | |
| P15508 | SPTB | spectrin, beta, erythrocytic | Plasma Membrane | other | 2.72E+07 | 156048.875 | 3496414.25 | | | | | | |
| P15508 | SPTB | spectrin, beta, erythrocytic | Plasma Membrane | other | 2776509.25 | 8926485 | 8369299 | | | | | | |
| P15508 | SPTB | spectrin, beta, erythrocytic | Plasma Membrane | other | 5255576.5 | 9177174 | 5199612 | | | | | | |
| P15508 | SPTB | spectrin, beta, erythrocytic | Plasma Membrane | other | 1.29E+07 | 3209476.25 | 2043057.375 | | | | | | |
| P15508 | SPTB | spectrin, beta, erythrocytic | Plasma Membrane | other | 3930139.5 | 607433.688 | 170547.953 | | | | | | |
| P15508 | SPTB | spectrin, beta, erythrocytic | Plasma Membrane | other | 4079163.75 | 2346150.25 | 2891699.25 | | | | | | |
| P15508 | SPTB | spectrin, beta, erythrocytic | Plasma Membrane | other | 3694193.25 | 2.07E+07 | 9280328 | | | | | | |
| P15508 | SPTB | spectrin, beta, erythrocytic | Plasma Membrane | other | 4477795.5 | 4040870.25 | 3249351.5 | | | | | | |
| P15508 | SPTB | spectrin, beta, erythrocytic | Plasma Membrane | other | 539273.688 | 2580806.75 | 4396857 | | | | | | |
| P15508 | SPTB | spectrin, beta, erythrocytic | Plasma Membrane | other | 3934057.75 | 1101482.125 | 1786445.875 | | | | | | |
| P15508 | SPTB | spectrin, beta, erythrocytic | Plasma Membrane | other | 1554854 | 5509450 | 3066435.5 | | | | | | |
| P15508 | SPTB | spectrin, beta, erythrocytic | Plasma Membrane | other | 1643542.375 | 3480715.25 | 5477487.5 | | | | | | |
| P15508 | SPTB | spectrin, beta, erythrocytic | Plasma Membrane | other | 8087033.5 | 1145815.5 | 2081918.5 | | | | | | |
| P15508 | SPTB | spectrin, beta, erythrocytic | Plasma Membrane | other | 2974868 | 981905.188 | 1948057.375 | | | | | | |
| P15508 | SPTB | spectrin, beta, erythrocytic | Plasma Membrane | other | 3947247.75 | 905548.688 | 793417.5 | | | | | | |
| P15508 | SPTB | spectrin, beta, erythrocytic | Plasma Membrane | other | 658727.062 | 4686887 | 3416663.25 | | | | | | |
| P15508 | SPTB | spectrin, beta, erythrocytic | Plasma Membrane | other | 2016128 | 1.21E+08 | 1.61E+08 | | | | | | |
| P15508 | SPTB | spectrin, beta, erythrocytic | Plasma Membrane | other | 7736277 | 455258.062 | 810280.625 | | | | | | |
| P15508 | SPTB | spectrin, beta, erythrocytic | Plasma Membrane | other | 718565.875 | 60598.316 | 1252.724 | | | | | | |
| P15508 | SPTB | spectrin, beta, erythrocytic | Plasma Membrane | other | 481577.906 | 3003774 | 564354 | | | | | | |
| P15508 | SPTB | spectrin, beta, erythrocytic | Plasma Membrane | other | 3883213 | 63133.312 | 706674.375 | | | | | | |
| P15508 | SPTB | spectrin, beta, erythrocytic | Plasma Membrane | other | 2433752.75 | 3262.843 | 0 | | | | | | |
| P15508 | SPTB | spectrin, beta, erythrocytic | Plasma Membrane | other | 464343.438 | 6095874.5 | 6246995.5 | | | | | | |
| P15508 | SPTB | spectrin, beta, erythrocytic | Plasma Membrane | other | 1933747.25 | 7738.142 | 4.70E-06 | | | | | | |
| P15508 | SPTB | spectrin, beta, erythrocytic | Plasma Membrane | other | 210561.672 | 3370274.5 | 3717025.75 | | | | | | |
| P15508 | SPTB | spectrin, beta, erythrocytic | Plasma Membrane | other | 4.70E-06 | 223502.312 | 294774.25 | | | | | | |
| P15508 | SPTB | spectrin, beta, erythrocytic | Plasma Membrane | other | 64331.84 | 2449911.25 | 1793720.875 | P15508 | SPTB | 442589860 | 566334452 | 20278026 | 780 |
| P13609 | Srgn | serglycin | Cytoplasm | other | 2638056.25 | 3942436 | 3503545.25 | P13609 | Srgn | 2638056 | 663715 | 383196 | 3 |
| Q9CY50 | SSR1 | signal sequence receptor, alpha | Cytoplasm | other | 5251889 | 4475174.5 | 4184801.75 | | | | | | |
| Q9CY50 | SSR1 | signal sequence receptor, alpha | Cytoplasm | other | 1878389.75 | 3927091.25 | 3948263.25 | Q9CY50 | SSR1 | 3565139 | 1124078 | 458903 | 6 |
| Q9DCF9 | SSR3 | signal sequence receptor, gamma | Cytoplasm | other | 6620209.5 | 8742775 | 6989332 | Q9DCF9 | SSR3 | 6620210 | 1134027 | 654731 | 3 |
| O70576 | STAG3 | stromal antigen 3 | Nucleus | other | 2339020.75 | 4653640.5 | 4425912 | O70576 | STAG3 | 2339021 | 1275698 | 736525 | 3 |
| P70297 | STAM | signal transducing adaptor molecu | Cytoplasm | other | 6301157.5 | 4321552 | 5145755.5 | | | | | | |
| P70297 | STAM | signal transducing adaptor molecu | Cytoplasm | other | 4174247.25 | 2542584 | 2645513.75 | | | | | | |
| P70297 | STAM | signal transducing adaptor molecu | Cytoplasm | other | 1235384.75 | 1362336.375 | 1861254.75 | P70297 | STAM | 3903597 | 1778675 | 592892 | 9 |
| Q8CI59 | STEAP3 | STEAP family member 3, metallore | Cytoplasm | transporter | 1.20E+07 | 1.32E+07 | 1.34E+07 | | | | | | |
| Q8CI59 | STEAP3 | STEAP family member 3, metallore | Cytoplasm | transporter | 7185953.5 | 7113408 | 7074237 | | | | | | |
| Q8CI59 | STEAP3 | STEAP family member 3, metallore | Cytoplasm | transporter | 6065608 | 7212242.5 | 5252383 | | | | | | |
| Q8CI59 | STEAP3 | STEAP family member 3, metallore | Cytoplasm | transporter | 4378133.5 | 4980369 | 4468235.5 | | | | | | |
| Q8CI59 | STEAP3 | STEAP family member 3, metallore | Cytoplasm | transporter | 3077419.75 | 3410447.75 | 3030214.25 | Q8CI59 | STEAP3 | 6543423 | 3479681 | 898450 | 15 |
| P70302 | STIM1 | stromal interaction molecule 1 | Plasma Membrane | ion channel | 2269632.5 | 3343057 | 4376517 | | | | | | |
| P70302 | STIM1 | stromal interaction molecule 1 | Plasma Membrane | ion channel | 3221075 | 3863563.75 | 3927079.75 | | | | | | |

| | | | | | | | | | | | | | |
|---|---|---|---|---|---|---|---|---|---|---|---|---|---|
| P70302 | STIM1 | stromal interaction molecule 1 | Plasma Membrane | ion channel | 2523459.5 | 1524777.5 | 1787256.375 | P70302 | STIM1 | 2671389 | 1004438 | 334813 | 9 |
| Q60864 | STIP1 | stress-induced phosphoprotein 1 | Cytoplasm | other | 7322049.5 | 7831888 | 4542731.5 | | | | | | |
| Q60864 | STIP1 | stress-induced phosphoprotein 1 | Cytoplasm | other | 7534087 | 9122723 | 6659853 | | | | | | |
| Q60864 | STIP1 | stress-induced phosphoprotein 1 | Cytoplasm | other | 8289160.5 | 6270052 | 5553618.5 | | | | | | |
| Q60864 | STIP1 | stress-induced phosphoprotein 1 | Cytoplasm | other | 6378171.5 | 5014464 | 3346187.5 | | | | | | |
| Q60864 | STIP1 | stress-induced phosphoprotein 1 | Cytoplasm | other | 6298563.5 | 5787477 | 3413950.75 | | | | | | |
| Q60864 | STIP1 | stress-induced phosphoprotein 1 | Cytoplasm | other | 2661009.75 | 2216941.75 | 2561126.5 | | | | | | |
| Q60864 | STIP1 | stress-induced phosphoprotein 1 | Cytoplasm | other | 3463547.25 | 2449852.5 | 1963763.625 | | | | | | |
| Q60864 | STIP1 | stress-induced phosphoprotein 1 | Cytoplasm | other | 3100406.75 | 2229162 | 2052250.125 | | | | | | |
| Q60864 | STIP1 | stress-induced phosphoprotein 1 | Cytoplasm | other | 1888546.75 | 2470326 | 2187428.75 | | | | | | |
| Q60864 | STIP1 | stress-induced phosphoprotein 1 | Cytoplasm | other | 3649445.75 | 3246321 | 2304601 | Q60864 | STIP1 | 5058499 | 2230951 | 407314 | 30 |
| P54116 | STOM | stomatin | Plasma Membrane | other | 4.82E+08 | 4.59E+08 | 4.29E+08 | | | | | | |
| P54116 | STOM | stomatin | Plasma Membrane | other | 4.61E+08 | 5.38E+08 | 5.77E+08 | | | | | | |
| P54116 | STOM | stomatin | Plasma Membrane | other | 4.91E+08 | 4.85E+08 | 4.91E+08 | | | | | | |
| P54116 | STOM | stomatin | Plasma Membrane | other | 2.68E+08 | 2.10E+08 | 1.76E+08 | | | | | | |
| P54116 | STOM | stomatin | Plasma Membrane | other | 3.44E+08 | 3.42E+08 | 2.99E+08 | | | | | | |
| P54116 | STOM | stomatin | Plasma Membrane | other | 1.92E+08 | 1.81E+08 | 1.95E+08 | | | | | | |
| P54116 | STOM | stomatin | Plasma Membrane | other | 3.57E+08 | 4.67E+08 | 4.34E+08 | | | | | | |
| P54116 | STOM | stomatin | Plasma Membrane | other | 1.94E+08 | 1.68E+08 | 1.43E+08 | | | | | | |
| P54116 | STOM | stomatin | Plasma Membrane | other | 1.28E+08 | 2.02E+08 | 1.84E+08 | | | | | | |
| P54116 | STOM | stomatin | Plasma Membrane | other | 1.56E+08 | 3.84E+08 | 4.10E+08 | | | | | | |
| P54116 | STOM | stomatin | Plasma Membrane | other | 6.24E+07 | 8.69E+08 | 1.46E+09 | | | | | | |
| P54116 | STOM | stomatin | Plasma Membrane | other | 7.41E+07 | 7.18E+07 | 8.85E+07 | | | | | | |
| P54116 | STOM | stomatin | Plasma Membrane | other | 2.18E+07 | 1.40E+07 | 1.70E+07 | | | | | | |
| P54116 | STOM | stomatin | Plasma Membrane | other | 1.79E+07 | 1.92E+07 | 1.84E+07 | | | | | | |
| P54116 | STOM | stomatin | Plasma Membrane | other | 4607933 | 4892060.5 | 5835793.5 | | | | | | |
| P54116 | STOM | stomatin | Plasma Membrane | other | 2634719.75 | 2632534.5 | 3262944.25 | | | | | | |
| P54116 | STOM | stomatin | Plasma Membrane | other | 2160075.5 | 8146902 | 2902496 | | | | | | |
| P54116 | STOM | stomatin | Plasma Membrane | other | 1510476.625 | 1466495 | 1246071.375 | | | | | | |
| P54116 | STOM | stomatin | Plasma Membrane | other | 2014107 | 386410.656 | 255800.25 | | | | | | |
| P54116 | STOM | stomatin | Plasma Membrane | other | 1041701.875 | 5538477.5 | 2851836.75 | | | | | | |
| P54116 | STOM | stomatin | Plasma Membrane | other | 5226624 | 6047932.5 | 8147753 | | | | | | |
| P54116 | STOM | stomatin | Plasma Membrane | other | 869537.062 | 662413.625 | 451113.125 | | | | | | |
| P54116 | STOM | stomatin | Plasma Membrane | other | 393265.656 | 591828.25 | 485887.812 | P54116 | STOM | 142131237 | 254160027 | 30597272 | 69 |
| Q99JB2 | STOML2 | stomatin (EPB72)-like 2 | Plasma Membrane | other | 4707983 | 5396330 | 7963119 | | | | | | |
| Q99JB2 | STOML2 | stomatin (EPB72)-like 2 | Plasma Membrane | other | 2598708.5 | 3063733.5 | 3328303.5 | | | | | | |
| Q99JB2 | STOML2 | stomatin (EPB72)-like 2 | Plasma Membrane | other | 3738200.75 | 3565475.75 | 5219674 | | | | | | |
| Q99JB2 | STOML2 | stomatin (EPB72)-like 2 | Plasma Membrane | other | 2554128.5 | 2481947 | 2862617.25 | | | | | | |
| Q99JB2 | STOML2 | stomatin (EPB72)-like 2 | Plasma Membrane | other | 1372094.5 | 1465447.75 | 2015867.375 | Q99JB2 | STOML2 | 2994223 | 1736801 | 448440 | 15 |
| Q9D3G5 | STX11 | syntaxin 11 | Plasma Membrane | transporter | 3345210.5 | 5357953 | 3927548.5 | | | | | | |
| Q9D3G5 | STX11 | syntaxin 11 | Plasma Membrane | transporter | 3177216.75 | 2733238.75 | 2808494.25 | | | | | | |
| Q9D3G5 | STX11 | syntaxin 11 | Plasma Membrane | transporter | 3417075.75 | 520770.406 | 189972.156 | | | | | | |
| Q9D3G5 | STX11 | syntaxin 11 | Plasma Membrane | transporter | 2335420.5 | 2281138.25 | 2592292.25 | | | | | | |
| Q9D3G5 | STX11 | syntaxin 11 | Plasma Membrane | transporter | 2044862.125 | 5041633 | 5554649.5 | | | | | | |
| Q9D3G5 | STX11 | syntaxin 11 | Plasma Membrane | transporter | 1507306.75 | 1081581.25 | 1622788.5 | Q9D3G5 | STX11 | 2637849 | 1540179 | 363024 | 18 |
| Q9ER00 | STX12 | syntaxin 12 | Cytoplasm | other | 1.58E+07 | 1.91E+07 | 1.45E+07 | Q9ER00 | STX12 | 15780000 | 2360388 | 1362771 | 3 |
| Q00262 | STX2 | syntaxin 2 | Cytoplasm | transporter | 3399988 | 5647003 | 5849232 | | | | | | |
| Q00262 | STX2 | syntaxin 2 | Cytoplasm | transporter | 7646019 | 973720.125 | 1043941.375 | | | | | | |
| Q00262 | STX2 | syntaxin 2 | Cytoplasm | transporter | 5029466.5 | 5717098 | 4955475.5 | | | | | | |
| Q00262 | STX2 | syntaxin 2 | Cytoplasm | transporter | 2343395 | 3121857.5 | 2113471 | | | | | | |
| Q00262 | STX2 | syntaxin 2 | Cytoplasm | transporter | 1487893.375 | 1483345.875 | 1519367.375 | | | | | | |
| Q00262 | STX2 | syntaxin 2 | Cytoplasm | transporter | 1823481.125 | 1579293.625 | 1690960.625 | Q00262 | STX2 | 3621707 | 2066580 | 487098 | 18 |
| P70452 | STX4 | syntaxin 4 | Plasma Membrane | transporter | 3357062 | 4700225.5 | 4125949.5 | | | | | | |
| P70452 | STX4 | syntaxin 4 | Plasma Membrane | transporter | 1714834 | 2185773.75 | 2110429.25 | | | | | | |
| P70452 | STX4 | syntaxin 4 | Plasma Membrane | transporter | 2787101 | 2903016.75 | 2735259.25 | P70452 | STX4 | 2619666 | 968292 | 322764 | 9 |
| O70439 | STX7 | syntaxin 7 | Plasma Membrane | transporter | 1.52E+07 | 1.53E+07 | 1.31E+07 | | | | | | |
| O70439 | STX7 | syntaxin 7 | Plasma Membrane | transporter | 1.27E+07 | 1.60E+07 | 1.56E+07 | | | | | | |
| O70439 | STX7 | syntaxin 7 | Plasma Membrane | transporter | 1.12E+07 | 1.14E+07 | 1.12E+07 | | | | | | |
| O70439 | STX7 | syntaxin 7 | Plasma Membrane | transporter | 9713467 | 7616467 | 7798339.5 | | | | | | |
| O70439 | STX7 | syntaxin 7 | Plasma Membrane | transporter | 7459483.5 | 9464682 | 1.29E+07 | | | | | | |
| O70439 | STX7 | syntaxin 7 | Plasma Membrane | transporter | 4535709 | 4769024 | 5782900.5 | O70439 | STX7 | 10133110 | 3732056 | 879654 | 18 |
| O08599 | STXBP1 | syntaxin binding protein 1 | Cytoplasm | transporter | 3839240 | 3084054.75 | 3127753.5 | | | | | | |
| O08599 | STXBP1 | syntaxin binding protein 1 | Cytoplasm | transporter | 1256601.125 | 830418.562 | 1024245 | O08599 | STXBP1 | 2547921 | 1302099 | 531580 | 6 |
| Q64324 | STXBP2 | syntaxin binding protein 2 | Plasma Membrane | transporter | 2.15E+07 | 2.38E+07 | 1.52E+07 | | | | | | |
| Q64324 | STXBP2 | syntaxin binding protein 2 | Plasma Membrane | transporter | 8893088 | 1.25E+07 | 9823381 | | | | | | |
| Q64324 | STXBP2 | syntaxin binding protein 2 | Plasma Membrane | transporter | 8955734 | 1.04E+07 | 7847647.5 | | | | | | |
| Q64324 | STXBP2 | syntaxin binding protein 2 | Plasma Membrane | transporter | 1.03E+07 | 8914728 | 6911167 | | | | | | |
| Q64324 | STXBP2 | syntaxin binding protein 2 | Plasma Membrane | transporter | 6121243 | 8172112 | 5259226.5 | | | | | | |
| Q64324 | STXBP2 | syntaxin binding protein 2 | Plasma Membrane | transporter | 6923149 | 6799165.5 | 7536039 | | | | | | |
| Q64324 | STXBP2 | syntaxin binding protein 2 | Plasma Membrane | transporter | 4628667 | 5364585.5 | 3670710.75 | | | | | | |
| Q64324 | STXBP2 | syntaxin binding protein 2 | Plasma Membrane | transporter | 3988105.75 | 4380719 | 4058274.25 | | | | | | |
| Q64324 | STXBP2 | syntaxin binding protein 2 | Plasma Membrane | transporter | 3050218.5 | 3300408.25 | 2873601.5 | | | | | | |
| Q64324 | STXBP2 | syntaxin binding protein 2 | Plasma Membrane | transporter | 2442647.5 | 3966125.75 | 2008140.625 | | | | | | |
| Q64324 | STXBP2 | syntaxin binding protein 2 | Plasma Membrane | transporter | 3870157 | 4086123 | 3711562.5 | | | | | | |
| Q64324 | STXBP2 | syntaxin binding protein 2 | Plasma Membrane | transporter | 2918659.75 | 3527485.75 | 3053083.75 | | | | | | |
| Q64324 | STXBP2 | syntaxin binding protein 2 | Plasma Membrane | transporter | 842024.75 | 2540059 | 1503487.125 | | | | | | |
| Q64324 | STXBP2 | syntaxin binding protein 2 | Plasma Membrane | transporter | 1210706 | 2991815.75 | 1499237.75 | Q64324 | STXBP2 | 6121743 | 4941888 | 762550 | 42 |
| Q60770 | STXBP3 | syntaxin binding protein 3 | Plasma Membrane | transporter | 1245225.375 | 1371598.25 | 1459691.875 | Q60770 | STXBP3 | 1245225 | 107801 | 62239 | 3 |
| Q8K400 | STXBP5 | syntaxin binding protein 5 (tomosy | Plasma Membrane | other | 2.51E+07 | 2.88E+07 | 3.92E+07 | Q8K400 | STXBP5 | 25080000 | 7337911 | 4236545 | 3 |
| Q9Z2I9 | SUCLA2 | succinate-CoA ligase, ADP-forming | Cytoplasm | enzyme | 5888555 | 6778923.5 | 8771870 | Q9Z2I9 | SUCLA2 | 5888555 | 1476375 | 852385 | 3 |
| Q9WUM5 | SUCLG1 | succinate-CoA ligase, alpha subuni | Cytoplasm | enzyme | 5197895.5 | 5414359.5 | 6580990 | | | | | | |
| Q9WUM5 | SUCLG1 | succinate-CoA ligase, alpha subuni | Cytoplasm | enzyme | 3025436 | 2417131.25 | 4845136 | | | | | | |
| Q9WUM5 | SUCLG1 | succinate-CoA ligase, alpha subuni | Cytoplasm | enzyme | 3346141.25 | 2576473.5 | 2946106 | Q9WUM5 | SUCLG1 | 3856491 | 1492374 | 497458 | 9 |
| Q9Z2I8 | SUCLG2 | succinate-CoA ligase, GDP-forming | Cytoplasm | enzyme | 1436711.5 | 1235663.625 | 1197302.625 | Q9Z2I8 | SUCLG2 | 1436712 | 128587 | 74240 | 3 |
| Q64310 | SURF4 | surfeit 4 | Cytoplasm | other | 2846088.5 | 7320163.5 | 4949112 | | | | | | |
| Q64310 | SURF4 | surfeit 4 | Cytoplasm | other | 3248858.5 | 3512271 | 3628885.75 | Q64310 | SURF4 | 3047474 | 1662404 | 678674 | 6 |

| | | | | | | | | | | | | | |
|---|---|---|---|---|---|---|---|---|---|---|---|---|---|
| Q9DBX3 | SUSD2 | sushi domain containing 2 | Extracellular Space | other | 7692988 | 6369648.5 | 3635624 | | | | | | |
| Q9DBX3 | SUSD2 | sushi domain containing 2 | Extracellular Space | other | 3962684.5 | 5432530 | 2523558.5 | | | | | | |
| Q9DBX3 | SUSD2 | sushi domain containing 2 | Extracellular Space | other | 5533602 | 6635105 | 3038035 | Q9DBX3 | SUSD2 | 5729758 | 1774460 | 591487 | 9 |
| Q3UZP4 | SVIP | small VCP/p97-interacting protein | Cytoplasm | other | 2.05E+07 | 2.10E+07 | 2.46E+07 | Q3UZP4 | SVIP | 20520000 | 2222169 | 1282970 | 3 |
| O09117 | SYPL1 | synaptophysin-like 1 | Plasma Membrane | transporter | 2.41E+07 | 3.34E+07 | 2.35E+07 | | | | | | |
| O09117 | SYPL1 | synaptophysin-like 1 | Plasma Membrane | transporter | 1.70E+07 | 2.05E+07 | 1.72E+07 | | | | | | |
| O09117 | SYPL1 | synaptophysin-like 1 | Plasma Membrane | transporter | 1334209.25 | 1160685.625 | 1291236.125 | O09117 | SYPL1 | 14131403 | 11709022 | 3903007 | 9 |
| B2RWW0 | TAGAP | T-cell activation RhoGTPase activa | Cytoplasm | other | 6829202.5 | 7554773.5 | 7915695 | B2RWW0 | TAGAP | 6829203 | 553351 | 319477 | 3 |
| Q9WVA4 | TAGLN2 | transgelin 2 | Cytoplasm | other | 2020051.875 | 3495940.5 | 3437847.5 | | | | | | |
| Q9WVA4 | TAGLN2 | transgelin 2 | Cytoplasm | other | 956412.125 | 1261429.5 | 1801047.75 | | | | | | |
| Q9WVA4 | TAGLN2 | transgelin 2 | Cytoplasm | other | 1582623.625 | 1283122.125 | 1804027.625 | | | | | | |
| Q9WVA4 | TAGLN2 | transgelin 2 | Cytoplasm | other | 1825802 | 1730680.875 | 1435014.625 | Q9WVA4 | TAGLN2 | 1596222 | 796178 | 229837 | 12 |
| Q8BYC6 | TAOK3 | TAO kinase 3 | Cytoplasm | kinase | 6019064 | 5155639.5 | 4600441.5 | | | | | | |
| Q8BYC6 | TAOK3 | TAO kinase 3 | Cytoplasm | kinase | 4078410.75 | 4154444.75 | 3186557.25 | | | | | | |
| Q8BYC6 | TAOK3 | TAO kinase 3 | Cytoplasm | kinase | 6071856 | 2777045 | 2685178.5 | | | | | | |
| Q8BYC6 | TAOK3 | TAO kinase 3 | Cytoplasm | kinase | 2970790.75 | 3358514 | 3078102.75 | | | | | | |
| Q8BYC6 | TAOK3 | TAO kinase 3 | Cytoplasm | kinase | 3158838.5 | 2443147.75 | 1752623.125 | | | | | | |
| Q8BYC6 | TAOK3 | TAO kinase 3 | Cytoplasm | kinase | 2708117.25 | 2017041 | 1856421.25 | | | | | | |
| Q8BYC6 | TAOK3 | TAO kinase 3 | Cytoplasm | kinase | 1912047.625 | 2737043.5 | 1766246.75 | | | | | | |
| Q8BYC6 | TAOK3 | TAO kinase 3 | Cytoplasm | kinase | 562879.25 | 624467.25 | 365458.75 | Q8BYC6 | TAOK3 | 3435251 | 1531736 | 312664 | 24 |
| Q9R233 | TAPBP | TAP binding protein (tapasin) | Cytoplasm | transporter | 2042470.625 | 1915811.875 | 2038044.125 | Q9R233 | TAPBP | 2042471 | 71883 | 41502 | 3 |
| Q4VBD2 | TAPT1 | transmembrane anterior posterior | Plasma Membrane | G-protein coupled receptor | 1042711.75 | 1605604 | 1138860.25 | Q4VBD2 | TAPT1 | 1042712 | 301093 | 173836 | 3 |
| Q9D0R2 | TARS | threonyl-tRNA synthetase | Nucleus | enzyme | 8483326 | 8490240 | 4786340.5 | | | | | | |
| Q9D0R2 | TARS | threonyl-tRNA synthetase | Nucleus | enzyme | 2532939.5 | 2485352.5 | 1088200.25 | Q9D0R2 | TARS | 5508133 | 3203570 | 1307852 | 6 |
| Q8BHL3 | TBC1D10B | TBC1 domain family, member 10B | Plasma Membrane | enzyme | 6340603 | 1399097.375 | 836967 | Q8BHL3 | TBC1D10B | 6340603 | 3028324 | 1748404 | 3 |
| Q3UUG6 | TBC1D24 | TBC1 domain family, member 24 | Cytoplasm | other | 5553555 | 9674671 | 4155029.75 | | | | | | |
| Q3UUG6 | TBC1D24 | TBC1 domain family, member 24 | Cytoplasm | other | 2235175.5 | 3711922.25 | 1859380.75 | Q3UUG6 | TBC1D24 | 3894365 | 2853895 | 1165098 | 6 |
| P36423 | TBXAS1 | thromboxane A synthase 1 (platel | Plasma Membrane | enzyme | 6380965.5 | 9029866 | 1.05E+07 | | | | | | |
| P36423 | TBXAS1 | thromboxane A synthase 1 (platel | Plasma Membrane | enzyme | 1.16E+07 | 1.42E+07 | 1.49E+07 | | | | | | |
| P36423 | TBXAS1 | thromboxane A synthase 1 (platel | Plasma Membrane | enzyme | 5130051.5 | 5343164 | 6744122 | | | | | | |
| P36423 | TBXAS1 | thromboxane A synthase 1 (platel | Plasma Membrane | enzyme | 4697979 | 5451015.5 | 5581169 | | | | | | |
| P36423 | TBXAS1 | thromboxane A synthase 1 (platel | Plasma Membrane | enzyme | 3595734.25 | 7652222.5 | 5110117.5 | | | | | | |
| P36423 | TBXAS1 | thromboxane A synthase 1 (platel | Plasma Membrane | enzyme | 8447020 | 8591492 | 7887798 | | | | | | |
| P36423 | TBXAS1 | thromboxane A synthase 1 (platel | Plasma Membrane | enzyme | 1.01E+07 | 1.35E+07 | 1.87E+07 | | | | | | |
| P36423 | TBXAS1 | thromboxane A synthase 1 (platel | Plasma Membrane | enzyme | 2808686 | 3964292 | 4205367 | | | | | | |
| P36423 | TBXAS1 | thromboxane A synthase 1 (platel | Plasma Membrane | enzyme | 6485843 | 6903828.5 | 6663250.5 | P36423 | TBXAS1 | 6575142 | 3866449 | 744098 | 27 |
| P10711 | TCEA1 | transcription elongation factor A (S | Nucleus | transcription regulator | 1.70E+07 | 2.75E+07 | 1.65E+07 | | | | | | |
| P10711 | TCEA1 | transcription elongation factor A (S | Nucleus | transcription regulator | 1.48E+07 | 1.73E+07 | 1.69E+07 | | | | | | |
| P10711 | TCEA1 | transcription elongation factor A (S | Nucleus | transcription regulator | 1.42E+07 | 1.35E+07 | 1.22E+07 | | | | | | |
| P10711 | TCEA1 | transcription elongation factor A (S | Nucleus | transcription regulator | 9546823 | 7736085 | 1.13E+07 | | | | | | |
| P10711 | TCEA1 | transcription elongation factor A (S | Nucleus | transcription regulator | 8169434 | 9683508 | 9436454 | | | | | | |
| P10711 | TCEA1 | transcription elongation factor A (S | Nucleus | transcription regulator | 6377243.5 | 8612505 | 7265120.5 | | | | | | |
| P10711 | TCEA1 | transcription elongation factor A (S | Nucleus | transcription regulator | 5146929 | 7853624 | 6968976 | | | | | | |
| P10711 | TCEA1 | transcription elongation factor A (S | Nucleus | transcription regulator | 7053920 | 5109766 | 5321282.5 | | | | | | |
| P10711 | TCEA1 | transcription elongation factor A (S | Nucleus | transcription regulator | 7889962 | 7182975 | 9164117 | | | | | | |
| P10711 | TCEA1 | transcription elongation factor A (S | Nucleus | transcription regulator | 6452709 | 6866810.5 | 3890135 | | | | | | |
| P10711 | TCEA1 | transcription elongation factor A (S | Nucleus | transcription regulator | 2333288.75 | 2170969.75 | 1537092.625 | | | | | | |
| P10711 | TCEA1 | transcription elongation factor A (S | Nucleus | transcription regulator | 631625.875 | 1573346 | 1835988.5 | P10711 | TCEA1 | 8303495 | 5655619 | 942603 | 36 |
| Q9QVN7 | TCEA2 | transcription elongation factor A (S | Nucleus | transcription regulator | 4890035 | 4851558 | 3792431 | Q9QVN7 | TCEA2 | 4890035 | 622892 | 359627 | 3 |
| P11983 | TCP1 | t-complex 1 | Cytoplasm | other | 2.75E+07 | 2.64E+07 | 1.65E+07 | | | | | | |
| P11983 | TCP1 | t-complex 1 | Cytoplasm | other | 2.19E+07 | 1.81E+07 | 1.12E+07 | | | | | | |
| P11983 | TCP1 | t-complex 1 | Cytoplasm | other | 9911277 | 1.24E+07 | 6874119.5 | | | | | | |
| P11983 | TCP1 | t-complex 1 | Cytoplasm | other | 9479981 | 1.20E+07 | 5436634 | | | | | | |
| P11983 | TCP1 | t-complex 1 | Cytoplasm | other | 1.29E+07 | 1.75E+07 | 9573504 | | | | | | |
| P11983 | TCP1 | t-complex 1 | Cytoplasm | other | 6889303 | 9362789 | 7641938.5 | | | | | | |
| P11983 | TCP1 | t-complex 1 | Cytoplasm | other | 7645191.5 | 8276580.5 | 5123804 | | | | | | |
| P11983 | TCP1 | t-complex 1 | Cytoplasm | other | 7928645.5 | 1.09E+07 | 7337473 | | | | | | |
| P11983 | TCP1 | t-complex 1 | Cytoplasm | other | 8739539 | 9527480 | 5254483.5 | | | | | | |
| P11983 | TCP1 | t-complex 1 | Cytoplasm | other | 5390069.5 | 8027490.5 | 4014783.5 | | | | | | |
| P11983 | TCP1 | t-complex 1 | Cytoplasm | other | 6927913 | 7336622.5 | 3764731.25 | | | | | | |
| P11983 | TCP1 | t-complex 1 | Cytoplasm | other | 6703363 | 6428506 | 3955490 | | | | | | |
| P11983 | TCP1 | t-complex 1 | Cytoplasm | other | 6467453 | 7699600 | 4483765 | | | | | | |
| P11983 | TCP1 | t-complex 1 | Cytoplasm | other | 5316803.5 | 5876520.5 | 2270484.5 | | | | | | |
| P11983 | TCP1 | t-complex 1 | Cytoplasm | other | 3056630 | 1905077.75 | 151753.406 | | | | | | |
| P11983 | TCP1 | t-complex 1 | Cytoplasm | other | 3201425.5 | 3330750.75 | 2209461.5 | | | | | | |
| P11983 | TCP1 | t-complex 1 | Cytoplasm | other | 2827264.5 | 3831247 | 1192614.625 | | | | | | |
| P11983 | TCP1 | t-complex 1 | Cytoplasm | other | 2849575.25 | 3087877 | 1343804.375 | | | | | | |
| P11983 | TCP1 | t-complex 1 | Cytoplasm | other | 2990667.75 | 2157102.75 | 1424139.875 | | | | | | |
| P11983 | TCP1 | t-complex 1 | Cytoplasm | other | 774263.062 | 2493456 | 595838.75 | | | | | | |
| P11983 | TCP1 | t-complex 1 | Cytoplasm | other | 24404.488 | 498894.125 | 21253.744 | P11983 | TCP1 | 7593037 | 5894497 | 742637 | 63 |
| Q921I1 | TF | transferrin | Extracellular Space | transporter | 5.59E+07 | 5.27E+07 | 6.71E+07 | | | | | | |
| Q921I1 | TF | transferrin | Extracellular Space | transporter | 2.34E+07 | 4.31E+07 | 3.96E+07 | | | | | | |
| Q921I1 | TF | transferrin | Extracellular Space | transporter | 4.54E+07 | 6.53E+07 | 5.68E+07 | | | | | | |
| Q921I1 | TF | transferrin | Extracellular Space | transporter | 3.53E+07 | 3.89E+07 | 5.03E+07 | | | | | | |
| Q921I1 | TF | transferrin | Extracellular Space | transporter | 2.99E+07 | 4.78E+07 | 4.90E+07 | | | | | | |
| Q921I1 | TF | transferrin | Extracellular Space | transporter | 3.53E+07 | 4.57E+07 | 5.55E+07 | | | | | | |
| Q921I1 | TF | transferrin | Extracellular Space | transporter | 3.24E+07 | 4.54E+07 | 5.17E+07 | | | | | | |
| Q921I1 | TF | transferrin | Extracellular Space | transporter | 3.98E+07 | 3.98E+07 | 5.29E+07 | | | | | | |
| Q921I1 | TF | transferrin | Extracellular Space | transporter | 3.52E+07 | 3.61E+07 | 4.49E+07 | | | | | | |
| Q921I1 | TF | transferrin | Extracellular Space | transporter | 2.64E+07 | 3.36E+07 | 4.73E+07 | | | | | | |
| Q921I1 | TF | transferrin | Extracellular Space | transporter | 1.70E+07 | 2.65E+07 | 3.32E+07 | | | | | | |
| Q921I1 | TF | transferrin | Extracellular Space | transporter | 1.96E+07 | 3.34E+07 | 5.45E+07 | | | | | | |
| Q921I1 | TF | transferrin | Extracellular Space | transporter | 2.14E+07 | 2.12E+07 | 2.58E+07 | | | | | | |
| Q921I1 | TF | transferrin | Extracellular Space | transporter | 2.36E+07 | 2.08E+07 | 3.39E+07 | | | | | | |
| Q921I1 | TF | transferrin | Extracellular Space | transporter | 1.51E+07 | 2.19E+07 | 2.07E+07 | | | | | | |
| Q921I1 | TF | transferrin | Extracellular Space | transporter | 1.46E+07 | 1.99E+07 | 2.46E+07 | | | | | | |

| | | | | | | | | | | | | | |
|---|---|---|---|---|---|---|---|---|---|---|---|---|---|
| Q921I1 | TF | transferrin | Extracellular Space | transporter | 1.27E+07 | 1.46E+07 | 1.99E+07 | | | | | | |
| Q921I1 | TF | transferrin | Extracellular Space | transporter | 5975294.5 | 9883626 | 7864012.5 | | | | | | |
| Q921I1 | TF | transferrin | Extracellular Space | transporter | 6327685 | 1.84E+07 | 1.66E+07 | | | | | | |
| Q921I1 | TF | transferrin | Extracellular Space | transporter | 1.09E+07 | 1.13E+07 | 1.82E+07 | | | | | | |
| Q921I1 | TF | transferrin | Extracellular Space | transporter | 8741724 | 1.20E+07 | 1.19E+07 | | | | | | |
| Q921I1 | TF | transferrin | Extracellular Space | transporter | 7048077 | 1.53E+07 | 1.63E+07 | | | | | | |
| Q921I1 | TF | transferrin | Extracellular Space | transporter | 9538234 | 1.14E+07 | 1.36E+07 | | | | | | |
| Q921I1 | TF | transferrin | Extracellular Space | transporter | 6438149 | 8126059.5 | 7991373.5 | | | | | | |
| Q921I1 | TF | transferrin | Extracellular Space | transporter | 9191526 | 1.25E+07 | 1.59E+07 | | | | | | |
| Q921I1 | TF | transferrin | Extracellular Space | transporter | 1.25E+07 | 1.10E+07 | 1.43E+07 | | | | | | |
| Q921I1 | TF | transferrin | Extracellular Space | transporter | 7966014 | 7796650.5 | 1.09E+07 | | | | | | |
| Q921I1 | TF | transferrin | Extracellular Space | transporter | 4156856 | 1.18E+07 | 1.52E+07 | | | | | | |
| Q921I1 | TF | transferrin | Extracellular Space | transporter | 2943450.75 | 3268984.75 | 3188475.75 | | | | | | |
| Q921I1 | TF | transferrin | Extracellular Space | transporter | 4981388.5 | 6768414.5 | 8076547.5 | | | | | | |
| Q921I1 | TF | transferrin | Extracellular Space | transporter | 5257796 | 6919890 | 3561416.25 | | | | | | |
| Q921I1 | TF | transferrin | Extracellular Space | transporter | 3531444.25 | 2.39E+07 | 3.37E+07 | | | | | | |
| Q921I1 | TF | transferrin | Extracellular Space | transporter | 4897439.5 | 4204799 | 4281794 | | | | | | |
| Q921I1 | TF | transferrin | Extracellular Space | transporter | 2049729.25 | 2675921.25 | 2286854.25 | | | | | | |
| Q921I1 | TF | transferrin | Extracellular Space | transporter | 44208.539 | 3289968.5 | 3817770 | Q921I1 | TF | 17008829 | 17095267 | 1668327 | 105 |
| Q62351 | TFRC | transferrin receptor | Plasma Membrane | transporter | 7.25E+07 | 7.44E+07 | 9.12E+07 | | | | | | |
| Q62351 | TFRC | transferrin receptor | Plasma Membrane | transporter | 8.46E+07 | 7.64E+07 | 1.01E+08 | | | | | | |
| Q62351 | TFRC | transferrin receptor | Plasma Membrane | transporter | 6.62E+07 | 5.28E+07 | 6.93E+07 | | | | | | |
| Q62351 | TFRC | transferrin receptor | Plasma Membrane | transporter | 6.45E+07 | 8.78E+07 | 1.10E+08 | | | | | | |
| Q62351 | TFRC | transferrin receptor | Plasma Membrane | transporter | 8.98E+07 | 5.92E+07 | 1.01E+08 | | | | | | |
| Q62351 | TFRC | transferrin receptor | Plasma Membrane | transporter | 7.67E+07 | 6.05E+07 | 9.01E+07 | | | | | | |
| Q62351 | TFRC | transferrin receptor | Plasma Membrane | transporter | 7.13E+07 | 5.50E+07 | 7.47E+07 | | | | | | |
| Q62351 | TFRC | transferrin receptor | Plasma Membrane | transporter | 6.28E+07 | 6.60E+07 | 9.71E+07 | | | | | | |
| Q62351 | TFRC | transferrin receptor | Plasma Membrane | transporter | 7.85E+07 | 6.58E+07 | 6.84E+07 | | | | | | |
| Q62351 | TFRC | transferrin receptor | Plasma Membrane | transporter | 5.96E+07 | 6.66E+07 | 5.16E+07 | | | | | | |
| Q62351 | TFRC | transferrin receptor | Plasma Membrane | transporter | 6.66E+07 | 5.60E+07 | 7.56E+07 | | | | | | |
| Q62351 | TFRC | transferrin receptor | Plasma Membrane | transporter | 6.13E+07 | 4.80E+07 | 6.01E+07 | | | | | | |
| Q62351 | TFRC | transferrin receptor | Plasma Membrane | transporter | 7.23E+07 | 4.08E+07 | 5.19E+07 | | | | | | |
| Q62351 | TFRC | transferrin receptor | Plasma Membrane | transporter | 5.09E+07 | 3.49E+07 | 5.72E+07 | | | | | | |
| Q62351 | TFRC | transferrin receptor | Plasma Membrane | transporter | 3.00E+07 | 2.53E+07 | 3.89E+07 | | | | | | |
| Q62351 | TFRC | transferrin receptor | Plasma Membrane | transporter | 5.06E+07 | 4.23E+07 | 4.91E+07 | | | | | | |
| Q62351 | TFRC | transferrin receptor | Plasma Membrane | transporter | 5.14E+07 | 4.32E+07 | 5.83E+07 | | | | | | |
| Q62351 | TFRC | transferrin receptor | Plasma Membrane | transporter | 5.24E+07 | 4.15E+07 | 6.25E+07 | | | | | | |
| Q62351 | TFRC | transferrin receptor | Plasma Membrane | transporter | 3.61E+07 | 3.80E+07 | 4.48E+07 | | | | | | |
| Q62351 | TFRC | transferrin receptor | Plasma Membrane | transporter | 2.99E+07 | 2.81E+07 | 3.34E+07 | | | | | | |
| Q62351 | TFRC | transferrin receptor | Plasma Membrane | transporter | 3.45E+07 | 2.61E+07 | 3.40E+07 | | | | | | |
| Q62351 | TFRC | transferrin receptor | Plasma Membrane | transporter | 4.82E+07 | 3.89E+07 | 6.15E+07 | | | | | | |
| Q62351 | TFRC | transferrin receptor | Plasma Membrane | transporter | 1.22E+07 | 4.13E+07 | 5.27E+07 | | | | | | |
| Q62351 | TFRC | transferrin receptor | Plasma Membrane | transporter | 1.69E+07 | 1.44E+07 | 2.13E+07 | | | | | | |
| Q62351 | TFRC | transferrin receptor | Plasma Membrane | transporter | 9757810 | 1.48E+07 | 1.33E+07 | | | | | | |
| Q62351 | TFRC | transferrin receptor | Plasma Membrane | transporter | 1.39E+07 | 1.15E+07 | 1.85E+07 | | | | | | |
| Q62351 | TFRC | transferrin receptor | Plasma Membrane | transporter | 1.65E+07 | 1.48E+07 | 1.64E+07 | | | | | | |
| Q62351 | TFRC | transferrin receptor | Plasma Membrane | transporter | 1.47E+07 | 1.58E+07 | 1.65E+07 | | | | | | |
| Q62351 | TFRC | transferrin receptor | Plasma Membrane | transporter | 7753453 | 1.20E+07 | 2.01E+07 | | | | | | |
| Q62351 | TFRC | transferrin receptor | Plasma Membrane | transporter | 5094707 | 4789362.5 | 6106396.5 | | | | | | |
| Q62351 | TFRC | transferrin receptor | Plasma Membrane | transporter | 4280272 | 3158053 | 3647535.5 | | | | | | |
| Q62351 | TFRC | transferrin receptor | Plasma Membrane | transporter | 3420251 | 2595682 | 3839703.75 | | | | | | |
| Q62351 | TFRC | transferrin receptor | Plasma Membrane | transporter | 2530696 | 1055523.75 | 2127760.75 | | | | | | |
| Q62351 | TFRC | transferrin receptor | Plasma Membrane | transporter | 3028731.75 | 2884374.75 | 3521845.5 | | | | | | |
| Q62351 | TFRC | transferrin receptor | Plasma Membrane | transporter | 2347165.25 | 2099093.5 | 3543224.75 | | | | | | |
| Q62351 | TFRC | transferrin receptor | Plasma Membrane | transporter | 1195174.625 | 2518594.75 | 5535255 | | | | | | |
| Q62351 | TFRC | transferrin receptor | Plasma Membrane | transporter | 111171.867 | 133937.984 | 258336.609 | Q62351 | TFRC | 38490525 | 29576509 | 2807278 | 111 |
| Q62312 | TGFBR2 | transforming growth factor, beta r | Plasma Membrane | kinase | 1.63E+07 | 1.85E+07 | 2.04E+07 | Q62312 | TGFBR2 | 16310000 | 2050984 | 1184136 | 3 |
| P21981 | TGM2 | transglutaminase 2 | Cytoplasm | enzyme | 6425449 | 5569592.5 | 6379221.5 | | | | | | |
| P21981 | TGM2 | transglutaminase 2 | Cytoplasm | enzyme | 1367709.875 | 1411902.75 | 2356146 | P21981 | TGM2 | 3896579 | 2461567 | 1004931 | 6 |
| P35441 | THBS1 | thrombospondin 1 | Extracellular Space | other | 3.27E+07 | 3.42E+07 | 2.78E+07 | | | | | | |
| P35441 | THBS1 | thrombospondin 1 | Extracellular Space | other | 1.69E+07 | 1.73E+07 | 1.68E+07 | | | | | | |
| P35441 | THBS1 | thrombospondin 1 | Extracellular Space | other | 1.13E+07 | 1.22E+07 | 1.22E+07 | | | | | | |
| P35441 | THBS1 | thrombospondin 1 | Extracellular Space | other | 2.44E+07 | 2.34E+07 | 2.11E+07 | | | | | | |
| P35441 | THBS1 | thrombospondin 1 | Extracellular Space | other | 1.10E+07 | 1.14E+07 | 1.07E+07 | | | | | | |
| P35441 | THBS1 | thrombospondin 1 | Extracellular Space | other | 7902116.5 | 6391507 | 6088648.5 | | | | | | |
| P35441 | THBS1 | thrombospondin 1 | Extracellular Space | other | 8923198 | 1.09E+07 | 1.06E+07 | | | | | | |
| P35441 | THBS1 | thrombospondin 1 | Extracellular Space | other | 1.69E+07 | 1.98E+07 | 1.72E+07 | | | | | | |
| P35441 | THBS1 | thrombospondin 1 | Extracellular Space | other | 1.36E+07 | 1.69E+07 | 1.41E+07 | | | | | | |
| P35441 | THBS1 | thrombospondin 1 | Extracellular Space | other | 1.33E+07 | 1.69E+07 | 1.47E+07 | | | | | | |
| P35441 | THBS1 | thrombospondin 1 | Extracellular Space | other | 1.05E+07 | 1.09E+07 | 1.05E+07 | | | | | | |
| P35441 | THBS1 | thrombospondin 1 | Extracellular Space | other | 2908940.25 | 3384894.25 | 2958061.5 | | | | | | |
| P35441 | THBS1 | thrombospondin 1 | Extracellular Space | other | 3.24E+07 | 4.98E+07 | 3.82E+07 | | | | | | |
| P35441 | THBS1 | thrombospondin 1 | Extracellular Space | other | 1.56E+07 | 1.65E+07 | 1.48E+07 | | | | | | |
| P35441 | THBS1 | thrombospondin 1 | Extracellular Space | other | 7791850.5 | 8708553 | 9086602 | | | | | | |
| P35441 | THBS1 | thrombospondin 1 | Extracellular Space | other | 1.04E+07 | 1.05E+07 | 9094815 | | | | | | |
| P35441 | THBS1 | thrombospondin 1 | Extracellular Space | other | 3848896.75 | 3106719.25 | 3577613.75 | | | | | | |
| P35441 | THBS1 | thrombospondin 1 | Extracellular Space | other | 8413181 | 9993812 | 1.01E+07 | | | | | | |
| P35441 | THBS1 | thrombospondin 1 | Extracellular Space | other | 2044198.25 | 5590814 | 4822922 | | | | | | |
| P35441 | THBS1 | thrombospondin 1 | Extracellular Space | other | 2292098 | 4813452.5 | 3352481 | | | | | | |
| P35441 | THBS1 | thrombospondin 1 | Extracellular Space | other | 3243496 | 3942697.5 | 3466449.5 | | | | | | |
| P35441 | THBS1 | thrombospondin 1 | Extracellular Space | other | 3360802.25 | 4118604.25 | 3310012.5 | | | | | | |
| P35441 | THBS1 | thrombospondin 1 | Extracellular Space | other | 6781805.5 | 1.25E+07 | 1.05E+07 | | | | | | |
| P35441 | THBS1 | thrombospondin 1 | Extracellular Space | other | 5456633.5 | 6272586 | 5794798.5 | | | | | | |
| P35441 | THBS1 | thrombospondin 1 | Extracellular Space | other | 6323149 | 72534.305 | 1809721.875 | | | | | | |
| P35441 | THBS1 | thrombospondin 1 | Extracellular Space | other | 2759771.75 | 2903842.75 | 3696718.75 | | | | | | |
| P35441 | THBS1 | thrombospondin 1 | Extracellular Space | other | 3304210 | 1781958.375 | 1345062.75 | | | | | | |

| | | | | | | | | | | | | | |
|---|---|---|---|---|---|---|---|---|---|---|---|---|---|
| P35441 | THBS1 | thrombospondin 1 | Extracellular Space | other | 2387528.5 | 6449271 | 3599571.5 | | | | | | |
| P35441 | THBS1 | thrombospondin 1 | Extracellular Space | other | 8825019 | 1.20E+07 | 1.02E+07 | | | | | | |
| P35441 | THBS1 | thrombospondin 1 | Extracellular Space | other | 4287860.5 | 3366278 | 4128998.25 | | | | | | |
| P35441 | THBS1 | thrombospondin 1 | Extracellular Space | other | 3512661.75 | 7632510 | 3945310 | | | | | | |
| P35441 | THBS1 | thrombospondin 1 | Extracellular Space | other | 2337302.75 | 2240969.25 | 2296277.5 | P35441 | THBS1 | 9556397 | 8765874 | 894663 | 96 |
| Q80ZW2 | THEM6 | thioesterase superfamily member | Extracellular Space | other | 4341800.5 | 5671297 | 4594895.5 | | | | | | |
| Q80ZW2 | THEM6 | thioesterase superfamily member | Extracellular Space | other | 3393477 | 3267947 | 3170962.75 | | | | | | |
| Q80ZW2 | THEM6 | thioesterase superfamily member | Extracellular Space | other | 1893075.875 | 1653417 | 1666739.875 | | | | | | |
| Q80ZW2 | THEM6 | thioesterase superfamily member | Extracellular Space | other | 1869081.875 | 2148915.5 | 2241359.5 | | | | | | |
| Q80ZW2 | THEM6 | thioesterase superfamily member | Extracellular Space | other | 900302.188 | 1835840.375 | 1356763.25 | Q80ZW2 | THEM6 | 2479547 | 1360994 | 351407 | 15 |
| Q9CY52 | THG1L | tRNA-histidine guanylyltransferase | Cytoplasm | enzyme | 7055491.5 | 5317270 | 5286825.5 | | | | | | |
| Q9CY52 | THG1L | tRNA-histidine guanylyltransferase | Cytoplasm | enzyme | 8271804.5 | 6833628 | 6394401.5 | | | | | | |
| Q9CY52 | THG1L | tRNA-histidine guanylyltransferase | Cytoplasm | enzyme | 4819998 | 4908952 | 2970989.25 | | | | | | |
| Q9CY52 | THG1L | tRNA-histidine guanylyltransferase | Cytoplasm | enzyme | 4528544.5 | 3388354.75 | 2580014.75 | | | | | | |
| Q9CY52 | THG1L | tRNA-histidine guanylyltransferase | Cytoplasm | enzyme | 4071167.25 | 3783880 | 2465346.75 | Q9CY52 | THG1L | 5749401 | 1725007 | 445395 | 15 |
| Q9D880 | TIMM50 | translocase of inner mitochondrial | Cytoplasm | phosphatase | 3207750.25 | 3314587.5 | 3998376.75 | | | | | | |
| Q9D880 | TIMM50 | translocase of inner mitochondrial | Cytoplasm | phosphatase | 2499502.5 | 2071856.375 | 2974771.25 | Q9D880 | TIMM50 | 2853626 | 670928 | 273905 | 6 |
| P39876 | TIMP3 | TIMP metallopeptidase inhibitor 3 | Extracellular Space | other | 1.04E+07 | 1.34E+07 | 1.49E+07 | | | | | | |
| P39876 | TIMP3 | TIMP metallopeptidase inhibitor 3 | Extracellular Space | other | 2350507.5 | 3927389 | 3124995.75 | | | | | | |
| P39876 | TIMP3 | TIMP metallopeptidase inhibitor 3 | Extracellular Space | other | 3899309.25 | 5793220 | 6264442.5 | | | | | | |
| P39876 | TIMP3 | TIMP metallopeptidase inhibitor 3 | Extracellular Space | other | 1973564.875 | 1365426.5 | 1141776.75 | P39876 | TIMP3 | 4665845 | 4713701 | 1360728 | 12 |
| P40142 | TKT | transketolase | Cytoplasm | enzyme | 2875734 | 1820130.875 | 2568667 | | | | | | |
| P40142 | TKT | transketolase | Cytoplasm | enzyme | 14892.106 | 1512709.125 | 852383.562 | P40142 | TKT | 1445313 | 1067057 | 435624 | 6 |
| P26039 | TLN1 | talin 1 | Plasma Membrane | other | 1224424.75 | 1569906.625 | 2522188 | | | | | | |
| P26039 | TLN1 | talin 1 | Plasma Membrane | other | 2195235.75 | 2064338.5 | 4920182 | | | | | | |
| P26039 | TLN1 | talin 1 | Plasma Membrane | other | 1576123.125 | 1729677.5 | 1975995 | | | | | | |
| P26039 | TLN1 | talin 1 | Plasma Membrane | other | 979475.625 | 1065965.875 | 1768497.125 | | | | | | |
| P26039 | TLN1 | talin 1 | Plasma Membrane | other | 1969462.375 | 1947835.125 | 2526858.25 | | | | | | |
| P26039 | TLN1 | talin 1 | Plasma Membrane | other | 1685262.375 | 1526845.125 | 2356522.75 | | | | | | |
| P26039 | TLN1 | talin 1 | Plasma Membrane | other | 1.05 | 223916.812 | 726402.312 | | | | | | |
| P26039 | TLN1 | talin 1 | Plasma Membrane | other | 885502.625 | 582788.938 | 745643.625 | P26039 | TLN1 | 1314436 | 991947 | 202480 | 24 |
| Q9ET30 | TM9SF3 | transmembrane 9 superfamily me | Cytoplasm | transporter | 2147728.5 | 2258495.25 | 2285141.25 | Q9ET30 | TM9SF3 | 2147729 | 72872 | 42072 | 3 |
| Q7TN60 | TMC6 | transmembrane channel-like 6 | Cytoplasm | transporter | 1789201.75 | 3453556 | 2073191.875 | | | | | | |
| Q7TN60 | TMC6 | transmembrane channel-like 6 | Cytoplasm | transporter | 1298353.625 | 1544165.625 | 1251048 | | | | | | |
| Q7TN60 | TMC6 | transmembrane channel-like 6 | Cytoplasm | transporter | 460177.656 | 753664.062 | 797456.438 | Q7TN60 | TMC6 | 1182578 | 898907 | 299636 | 9 |
| Q7TN58 | TMC8 | transmembrane channel-like 8 | Cytoplasm | other | 2986038.75 | 3603801.75 | 2310555.25 | | | | | | |
| Q7TN58 | TMC8 | transmembrane channel-like 8 | Cytoplasm | other | 1522181 | 1777466.375 | 1354635.375 | | | | | | |
| Q7TN58 | TMC8 | transmembrane channel-like 8 | Cytoplasm | other | 1834327.875 | 1642879 | 933971.25 | Q7TN58 | TMC8 | 2114183 | 838637 | 279546 | 9 |
| Q80W04 | TMCC2 | transmembrane and coiled-coil do | Extracellular Space | other | 1.05E+07 | 1.16E+07 | 9892158 | | | | | | |
| Q80W04 | TMCC2 | transmembrane and coiled-coil do | Extracellular Space | other | 1.16E+07 | 1.67E+07 | 1.63E+07 | | | | | | |
| Q80W04 | TMCC2 | transmembrane and coiled-coil do | Extracellular Space | other | 4342312 | 4351164.5 | 5692319.5 | | | | | | |
| Q80W04 | TMCC2 | transmembrane and coiled-coil do | Extracellular Space | other | 3327868.25 | 4147139.75 | 3302157.25 | | | | | | |
| Q80W04 | TMCC2 | transmembrane and coiled-coil do | Extracellular Space | other | 2732438 | 4018997 | 3273104.75 | | | | | | |
| Q80W04 | TMCC2 | transmembrane and coiled-coil do | Extracellular Space | other | 873368.25 | 1837862.125 | 1385056.5 | Q80W04 | TMCC2 | 5565998 | 5004750 | 1179631 | 18 |
| Q921L3 | TMCO1 | transmembrane and coiled-coil do | Cytoplasm | other | 5853522.5 | 7905966.5 | 6842830.5 | Q921L3 | TMCO1 | 5853523 | 1026443 | 592617 | 3 |
| Q9D1D4 | TMED10 | transmembrane emp24-like traffic | Cytoplasm | transporter | 2.60E+07 | 3.05E+07 | 2.31E+07 | | | | | | |
| Q9D1D4 | TMED10 | transmembrane emp24-like traffic | Cytoplasm | transporter | 2.46E+07 | 2.12E+07 | 2.15E+07 | | | | | | |
| Q9D1D4 | TMED10 | transmembrane emp24-like traffic | Cytoplasm | transporter | 2.08E+07 | 1.91E+07 | 1.71E+07 | | | | | | |
| Q9D1D4 | TMED10 | transmembrane emp24-like traffic | Cytoplasm | transporter | 6652400.5 | 5333226 | 3170524 | | | | | | |
| Q9D1D4 | TMED10 | transmembrane emp24-like traffic | Cytoplasm | transporter | 5058364.5 | 7699361.5 | 7804561.5 | | | | | | |
| Q9D1D4 | TMED10 | transmembrane emp24-like traffic | Cytoplasm | transporter | 7001119 | 6641590.5 | 5099999 | Q9D1D4 | TMED10 | 15025314 | 9027928 | 2127903 | 18 |
| Q9R0Q3 | TMED2 | transmembrane emp24 domain tr | Cytoplasm | transporter | 8806088 | 1.05E+07 | 8824388 | | | | | | |
| Q9R0Q3 | TMED2 | transmembrane emp24 domain tr | Cytoplasm | transporter | 6437974 | 1.67E+07 | 1.28E+07 | | | | | | |
| Q9R0Q3 | TMED2 | transmembrane emp24 domain tr | Cytoplasm | transporter | 3566144.25 | 5769692 | 5215924.5 | Q9R0Q3 | TMED2 | 6270069 | 4120576 | 1373525 | 9 |
| Q8R1V4 | TMED4 | transmembrane emp24 protein tra | Cytoplasm | transporter | 1.83E+07 | 1.92E+07 | 1.51E+07 | Q8R1V4 | TMED4 | 18310000 | 2150752 | 1241737 | 3 |
| Q9CXE7 | TMED5 | transmembrane emp24 protein tra | Cytoplasm | other | 2211473.25 | 4284784 | 2829041.25 | Q9CXE7 | TMED5 | 2211473 | 1064518 | 614600 | 3 |
| Q99KF1 | TMED9 | transmembrane emp24 protein tra | Cytoplasm | transporter | 2.50E+07 | 2.72E+07 | 2.29E+07 | | | | | | |
| Q99KF1 | TMED9 | transmembrane emp24 protein tra | Cytoplasm | transporter | 6861872.5 | 1.56E+07 | 9642330 | | | | | | |
| Q99KF1 | TMED9 | transmembrane emp24 protein tra | Cytoplasm | transporter | 2881021.25 | 74258.406 | 952470.438 | Q99KF1 | TMED9 | 11564298 | 10659244 | 3553081 | 9 |
| Q80X71 | TMEM106B | transmembrane protein 106B | Cytoplasm | other | 308053.094 | 371788.781 | 339937.844 | Q80X71 | TMEM106B | 308053 | 31868 | 18399 | 3 |
| Q9CQN6 | TMEM14C | transmembrane protein 14C | Plasma Membrane | other | 8918875 | 1.47E+07 | 1.62E+07 | | | | | | |
| Q9CQN6 | TMEM14C | transmembrane protein 14C | Plasma Membrane | other | 2628118.25 | 5335203.5 | 5542215.5 | Q9CQN6 | TMEM14C | 5773497 | 5480194 | 2237280 | 6 |
| Q9CR64 | TMEM167A | transmembrane protein 167A | Extracellular Space | other | 2668662 | 3751461.25 | 2899603 | Q9CR64 | TMEM167A | 2668662 | 570300 | 329263 | 3 |
| Q9CY24 | TMEM179B | transmembrane protein 179B | Extracellular Space | other | 2441768.25 | 3675917.25 | 2305936.25 | Q9CY24 | TMEM179B | 2441768 | 754809 | 435789 | 3 |
| Q9D771 | TMEM206 | transmembrane protein 206 | Plasma Membrane | other | 1.01E+07 | 1.21E+07 | 1.20E+07 | | | | | | |
| Q9D771 | TMEM206 | transmembrane protein 206 | Plasma Membrane | other | 2151361.25 | 4844190.5 | 3696798 | | | | | | |
| Q9D771 | TMEM206 | transmembrane protein 206 | Plasma Membrane | other | 1207869.25 | 2490610 | 2528297.75 | Q9D771 | TMEM206 | 4479744 | 4445375 | 1481792 | 9 |
| Q5F285 | TMEM256 | transmembrane protein 256 | Cytoplasm | other | 2.39E+07 | 1.94E+07 | 2.67E+07 | Q5F285 | TMEM256 | 23920000 | 3650128 | 2107402 | 3 |
| Q8VEK0 | TMEM30A | transmembrane protein 30A | Cytoplasm | other | 2.18E+07 | 1.86E+07 | 1.54E+07 | | | | | | |
| Q8VEK0 | TMEM30A | transmembrane protein 30A | Cytoplasm | other | 1.63E+07 | 1.81E+07 | 1.81E+07 | | | | | | |
| Q8VEK0 | TMEM30A | transmembrane protein 30A | Cytoplasm | other | 9533499 | 1.18E+07 | 8429297 | | | | | | |
| Q8VEK0 | TMEM30A | transmembrane protein 30A | Cytoplasm | other | 8628896 | 9876458 | 9548818 | | | | | | |
| Q8VEK0 | TMEM30A | transmembrane protein 30A | Cytoplasm | other | 3704125.25 | 8145087 | 5771313.5 | Q8VEK0 | TMEM30A | 11995304 | 5390362 | 1391786 | 15 |
| Q9CZX7 | TMEM55A | transmembrane protein 55A | Cytoplasm | phosphatase | 2673326.75 | 4136471 | 3523589 | Q9CZX7 | TMEM55A | 2673327 | 734774 | 424222 | 3 |
| Q3TWL2 | TMEM55B | transmembrane protein 55B | Cytoplasm | phosphatase | 5243036 | 6635516 | 4999652 | | | | | | |
| Q3TWL2 | TMEM55B | transmembrane protein 55B | Cytoplasm | phosphatase | 3747133.25 | 4398091 | 2977742 | | | | | | |
| Q3TWL2 | TMEM55B | transmembrane protein 55B | Cytoplasm | phosphatase | 2982086 | 3852005.75 | 2759990.5 | | | | | | |
| Q3TWL2 | TMEM55B | transmembrane protein 55B | Cytoplasm | phosphatase | 4206689.5 | 3495349.5 | 3325583.25 | Q3TWL2 | TMEM55B | 4044736 | 1131105 | 326522 | 12 |
| Q91YT8 | TMEM63A | transmembrane protein 63A | Cytoplasm | other | 1621454 | 1511327.375 | 2485857 | Q91YT8 | TMEM63A | 1621454 | 533702 | 308133 | 3 |
| Q9CR23 | TMEM9 | transmembrane protein 9 | Cytoplasm | other | 5639510 | 6859784 | 4161216.5 | Q9CR23 | TMEM9 | 5639510 | 1351338 | 780195 | 3 |
| P49813 | TMOD1 | tropomodulin 1 | Cytoplasm | enzyme | 2.31E+08 | 3.24E+08 | 2.58E+08 | | | | | | |
| P49813 | TMOD1 | tropomodulin 1 | Cytoplasm | enzyme | 1.36E+08 | 2.38E+08 | 2.17E+08 | | | | | | |
| P49813 | TMOD1 | tropomodulin 1 | Cytoplasm | enzyme | 2.12E+08 | 1.69E+08 | 2.30E+08 | | | | | | |
| P49813 | TMOD1 | tropomodulin 1 | Cytoplasm | enzyme | 1.57E+08 | 1.90E+08 | 1.98E+08 | | | | | | |
| P49813 | TMOD1 | tropomodulin 1 | Cytoplasm | enzyme | 9.59E+07 | 2.46E+08 | 1.73E+08 | | | | | | |
| P49813 | TMOD1 | tropomodulin 1 | Cytoplasm | enzyme | 1.93E+08 | 1.75E+08 | 1.62E+08 | | | | | | |

| | | | | | | | | | | | | | |
|---|---|---|---|---|---|---|---|---|---|---|---|---|---|
| P49813 | TMOD1 | tropomodulin 1 | Cytoplasm | enzyme | 1.28E+08 | 1.15E+08 | 1.26E+08 | | | | | | |
| P49813 | TMOD1 | tropomodulin 1 | Cytoplasm | enzyme | 1.31E+08 | 1.27E+08 | 1.13E+08 | | | | | | |
| P49813 | TMOD1 | tropomodulin 1 | Cytoplasm | enzyme | 9.65E+07 | 1.08E+08 | 1.02E+08 | | | | | | |
| P49813 | TMOD1 | tropomodulin 1 | Cytoplasm | enzyme | 8.04E+07 | 1.02E+08 | 8.97E+07 | | | | | | |
| P49813 | TMOD1 | tropomodulin 1 | Cytoplasm | enzyme | 5.71E+07 | 6.02E+07 | 5.00E+07 | | | | | | |
| P49813 | TMOD1 | tropomodulin 1 | Cytoplasm | enzyme | 3.60E+07 | 3.84E+07 | 4.01E+07 | | | | | | |
| P49813 | TMOD1 | tropomodulin 1 | Cytoplasm | enzyme | 3.24E+07 | 4.78E+07 | 5.05E+07 | | | | | | |
| P49813 | TMOD1 | tropomodulin 1 | Cytoplasm | enzyme | 4.40E+07 | 1495476.375 | 916586.125 | | | | | | |
| P49813 | TMOD1 | tropomodulin 1 | Cytoplasm | enzyme | 1.60E+07 | 1.94E+07 | 1.83E+07 | | | | | | |
| P49813 | TMOD1 | tropomodulin 1 | Cytoplasm | enzyme | 2.87E+07 | 7087924.5 | 1.56E+07 | | | | | | |
| P49813 | TMOD1 | tropomodulin 1 | Cytoplasm | enzyme | 1.03E+07 | 2.50E+07 | 2.44E+07 | | | | | | |
| P49813 | TMOD1 | tropomodulin 1 | Cytoplasm | enzyme | 1.12E+07 | 8368895.5 | 1.03E+07 | | | | | | |
| P49813 | TMOD1 | tropomodulin 1 | Cytoplasm | enzyme | 1.24E+07 | 1.13E+07 | 1.22E+07 | | | | | | |
| P49813 | TMOD1 | tropomodulin 1 | Cytoplasm | enzyme | 7464075.5 | 4.29E+07 | 3.49E+07 | | | | | | |
| P49813 | TMOD1 | tropomodulin 1 | Cytoplasm | enzyme | 3432073.5 | 1.39E+07 | 1.16E+07 | | | | | | |
| P49813 | TMOD1 | tropomodulin 1 | Cytoplasm | enzyme | 6367682.5 | 6250728 | 5787424.5 | | | | | | |
| P49813 | TMOD1 | tropomodulin 1 | Cytoplasm | enzyme | 7377604.5 | 1121515.75 | 2558096 | | | | | | |
| P49813 | TMOD1 | tropomodulin 1 | Cytoplasm | enzyme | 744878.188 | 1370763.375 | 1021570.25 | P49813 | TMOD1 | 72207346 | 82459447 | 9717939 | 72 |
| Q9D710 | TMX2 | thioredoxin-related transmembrar | Cytoplasm | enzyme | 309907.344 | 515765.906 | 665560.938 | Q9D710 | TMX2 | 309907 | 178562 | 103093 | 3 |
| Q8C0L0 | TMX4 | thioredoxin-related transmembrar | Cytoplasm | enzyme | 3567266 | 4436865.5 | 3388417.75 | Q8C0L0 | TMX4 | 3567266 | 560867 | 323817 | 3 |
| O88746 | TOM1 | target of myb1 membrane traffick | Cytoplasm | transporter | 898543.562 | 1524180.625 | 1215987.5 | O88746 | TOM1 | 898544 | 312830 | 180612 | 3 |
| Q923U0 | TOM1L1 | target of myb1 like 1 membrane tr | Cytoplasm | other | 3588080 | 3668070 | 4132984.5 | | | | | | |
| Q923U0 | TOM1L1 | target of myb1 like 1 membrane tr | Cytoplasm | other | 1359818.75 | 1394647.125 | 1401444.75 | Q923U0 | TOM1L1 | 2473949 | 1333722 | 544490 | 6 |
| Q9CPQ3 | TOMM22 | translocase of outer mitochondria | Cytoplasm | transporter | 2564950.5 | 2545382.5 | 2947512.75 | | | | | | |
| Q9CPQ3 | TOMM22 | translocase of outer mitochondria | Cytoplasm | transporter | 2846582.75 | 2632378.5 | 3600007.75 | Q9CPQ3 | TOMM22 | 2705767 | 398299 | 162605 | 6 |
| Q9QYA2 | TOMM40 | translocase of outer mitochondria | Cytoplasm | ion channel | 6306005 | 5950976 | 6383581 | Q9QYA2 | TOMM40 | 6306005 | 230655 | 133169 | 3 |
| Q9CZW5 | TOMM70A | translocase of outer mitochondria | Cytoplasm | transporter | 2027571.375 | 2956885.5 | 2901484.75 | Q9CZW5 | TOMM70A | 2027571 | 521283 | 300963 | 3 |
| Q9CYZ2 | TPD52L2 | tumor protein D52-like 2 | Cytoplasm | other | 3124535 | 4401557 | 3046043 | Q9CYZ2 | TPD52L2 | 3124535 | 760960 | 439341 | 3 |
| P58771 | Tpm1 | tropomyosin 1, alpha | Plasma Membrane | other | 2.73E+08 | 2.70E+08 | 2.61E+08 | | | | | | |
| P58771 | Tpm1 | tropomyosin 1, alpha | Plasma Membrane | other | 5.39E+07 | 5.92E+07 | 6.84E+07 | | | | | | |
| P58771 | Tpm1 | tropomyosin 1, alpha | Plasma Membrane | other | 8624579 | 2.22E+07 | 1.72E+07 | | | | | | |
| P58771 | Tpm1 | tropomyosin 1, alpha | Plasma Membrane | other | 9670722 | 1.27E+07 | 1.44E+07 | | | | | | |
| P58771 | Tpm1 | tropomyosin 1, alpha | Plasma Membrane | other | 4361061 | 2616755 | 3048012 | P58771 | Tpm1 | 69801272 | 103564375 | 26740207 | 15 |
| P21107 | TPM3 | tropomyosin 3 | Cytoplasm | other | 5.75E+08 | 4.34E+08 | 4.80E+08 | | | | | | |
| P21107 | TPM3 | tropomyosin 3 | Cytoplasm | other | 1.77E+08 | 1.75E+08 | 1.87E+08 | | | | | | |
| P21107 | TPM3 | tropomyosin 3 | Cytoplasm | other | 2.02E+08 | 1.45E+08 | 1.74E+08 | | | | | | |
| P21107 | TPM3 | tropomyosin 3 | Cytoplasm | other | 1.46E+08 | 2.17E+08 | 1.87E+08 | | | | | | |
| P21107 | TPM3 | tropomyosin 3 | Cytoplasm | other | 9.75E+07 | 6.67E+07 | 3.88E+07 | | | | | | |
| P21107 | TPM3 | tropomyosin 3 | Cytoplasm | other | 5.64E+07 | 8.38E+07 | 9.84E+07 | | | | | | |
| P21107 | TPM3 | tropomyosin 3 | Cytoplasm | other | 4.78E+07 | 1.11E+08 | 8.71E+07 | | | | | | |
| P21107 | TPM3 | tropomyosin 3 | Cytoplasm | other | 8.80E+07 | 4.06E+07 | 3.60E+07 | | | | | | |
| P21107 | TPM3 | tropomyosin 3 | Cytoplasm | other | 5.23E+07 | 8.29E+07 | 6.58E+07 | | | | | | |
| P21107 | TPM3 | tropomyosin 3 | Cytoplasm | other | 3.48E+07 | 7.66E+07 | 4.10E+07 | | | | | | |
| P21107 | TPM3 | tropomyosin 3 | Cytoplasm | other | 2.62E+07 | 4.28E+07 | 2.40E+07 | | | | | | |
| P21107 | TPM3 | tropomyosin 3 | Cytoplasm | other | 5401969 | 1.68E+07 | 1.13E+07 | | | | | | |
| P21107 | TPM3 | tropomyosin 3 | Cytoplasm | other | 5784222.5 | 1.05E+07 | 9682547 | | | | | | |
| P21107 | TPM3 | tropomyosin 3 | Cytoplasm | other | 5540415 | 6368348 | 8666392 | P21107 | TPM3 | 108529758 | 126812523 | 19567597 | 42 |
| O89023 | TPP1 | tripeptidyl peptidase I | Cytoplasm | peptidase | 6968973 | 8278886.5 | 8313573 | O89023 | TPP1 | 6968973 | 766488 | 442532 | 3 |
| Q64514 | TPP2 | tripeptidyl peptidase II | Cytoplasm | peptidase | 1.75E+07 | 1.34E+07 | 1.59E+07 | | | | | | |
| Q64514 | TPP2 | tripeptidyl peptidase II | Cytoplasm | peptidase | 1.00E+07 | 1.09E+07 | 1.14E+07 | | | | | | |
| Q64514 | TPP2 | tripeptidyl peptidase II | Cytoplasm | peptidase | 8763183 | 9512522 | 7952456 | | | | | | |
| Q64514 | TPP2 | tripeptidyl peptidase II | Cytoplasm | peptidase | 7477989 | 7586221.5 | 7853377 | | | | | | |
| Q64514 | TPP2 | tripeptidyl peptidase II | Cytoplasm | peptidase | 9982140 | 6920028.5 | 6498966.5 | | | | | | |
| Q64514 | TPP2 | tripeptidyl peptidase II | Cytoplasm | peptidase | 8372996.5 | 7757663 | 8526459 | | | | | | |
| Q64514 | TPP2 | tripeptidyl peptidase II | Cytoplasm | peptidase | 6058250.5 | 5158604.5 | 4761998 | | | | | | |
| Q64514 | TPP2 | tripeptidyl peptidase II | Cytoplasm | peptidase | 5538208 | 5066111 | 5859885 | | | | | | |
| Q64514 | TPP2 | tripeptidyl peptidase II | Cytoplasm | peptidase | 6459001 | 5574494.5 | 7027962.5 | | | | | | |
| Q64514 | TPP2 | tripeptidyl peptidase II | Cytoplasm | peptidase | 4894960.5 | 4883166 | 4994461.5 | | | | | | |
| Q64514 | TPP2 | tripeptidyl peptidase II | Cytoplasm | peptidase | 3876532.5 | 2942818.5 | 3571833.25 | | | | | | |
| Q64514 | TPP2 | tripeptidyl peptidase II | Cytoplasm | peptidase | 3121216.75 | 2618204.75 | 2281068.5 | | | | | | |
| Q64514 | TPP2 | tripeptidyl peptidase II | Cytoplasm | peptidase | 2484147.5 | 3852919.5 | 1278844.5 | | | | | | |
| Q64514 | TPP2 | tripeptidyl peptidase II | Cytoplasm | peptidase | 4088428.25 | 3645819.5 | 2993149.5 | | | | | | |
| Q64514 | TPP2 | tripeptidyl peptidase II | Cytoplasm | peptidase | 3885424.5 | 3114133.5 | 2644989.75 | | | | | | |
| Q64514 | TPP2 | tripeptidyl peptidase II | Cytoplasm | peptidase | 3438037.75 | 1029471.625 | 2298011.25 | | | | | | |
| Q64514 | TPP2 | tripeptidyl peptidase II | Cytoplasm | peptidase | 2465787 | 3072791.5 | 2715759.75 | | | | | | |
| Q64514 | TPP2 | tripeptidyl peptidase II | Cytoplasm | peptidase | 2891368 | 2412340.75 | 2002588.625 | | | | | | |
| Q64514 | TPP2 | tripeptidyl peptidase II | Cytoplasm | peptidase | 1352144.375 | 343474.438 | 453417.75 | | | | | | |
| Q64514 | TPP2 | tripeptidyl peptidase II | Cytoplasm | peptidase | 706293.812 | 919195 | 880716.812 | | | | | | |
| Q64514 | TPP2 | tripeptidyl peptidase II | Cytoplasm | peptidase | 1218426.875 | 1054154.625 | 1133551.375 | | | | | | |
| Q64514 | TPP2 | tripeptidyl peptidase II | Cytoplasm | peptidase | 688382.562 | 574397.875 | 817390.75 | Q64514 | TPP2 | 5240133 | 3744281 | 460889 | 66 |
| Q9CQN1 | TRAP1 | TNF receptor-associated protein 1 | Cytoplasm | enzyme | 1.57E+07 | 1.56E+07 | 1.18E+07 | | | | | | |
| Q9CQN1 | TRAP1 | TNF receptor-associated protein 1 | Cytoplasm | enzyme | 1525429 | 1698989.875 | 2424710.25 | Q9CQN1 | TRAP1 | 8632715 | 7001230 | 2858240 | 6 |
| Q8K558 | TREML1 | triggering receptor expressed on n | Plasma Membrane | other | 4.14E+07 | 6.78E+07 | 5.50E+07 | | | | | | |
| Q8K558 | TREML1 | triggering receptor expressed on n | Plasma Membrane | other | 3.48E+07 | 4.19E+07 | 4.31E+07 | | | | | | |
| Q8K558 | TREML1 | triggering receptor expressed on n | Plasma Membrane | other | 2.45E+07 | 3.01E+07 | 2.64E+07 | | | | | | |
| Q8K558 | TREML1 | triggering receptor expressed on n | Plasma Membrane | other | 3.07E+07 | 4.91E+07 | 4.67E+07 | | | | | | |
| Q8K558 | TREML1 | triggering receptor expressed on n | Plasma Membrane | other | 2.12E+07 | 2.90E+07 | 2.85E+07 | | | | | | |
| Q8K558 | TREML1 | triggering receptor expressed on n | Plasma Membrane | other | 1.65E+07 | 2.42E+07 | 2.24E+07 | | | | | | |
| Q8K558 | TREML1 | triggering receptor expressed on n | Plasma Membrane | other | 1.99E+07 | 2.38E+07 | 1.95E+07 | | | | | | |
| Q8K558 | TREML1 | triggering receptor expressed on n | Plasma Membrane | other | 3719061.25 | 6463622 | 6984899 | Q8K558 | TREML1 | 24079883 | 15591272 | 3182555 | 24 |
| Q9WTR1 | TRPV2 | transient receptor potential catior | Plasma Membrane | ion channel | 1.53E+07 | 1.94E+07 | 1.49E+07 | | | | | | |
| Q9WTR1 | TRPV2 | transient receptor potential catior | Plasma Membrane | ion channel | 1.20E+07 | 1.63E+07 | 1.28E+07 | | | | | | |
| Q9WTR1 | TRPV2 | transient receptor potential catior | Plasma Membrane | ion channel | 5106310.5 | 7787763 | 8513121 | | | | | | |
| Q9WTR1 | TRPV2 | transient receptor potential catior | Plasma Membrane | ion channel | 5937216 | 9373804 | 5345558.5 | | | | | | |
| Q9WTR1 | TRPV2 | transient receptor potential catior | Plasma Membrane | ion channel | 4731650 | 7506929.5 | 5976420.5 | | | | | | |
| Q9WTR1 | TRPV2 | transient receptor potential catior | Plasma Membrane | ion channel | 8122277 | 8744940 | 6895902.5 | | | | | | |

| | | | | | | | | | | | | | |
|---|---|---|---|---|---|---|---|---|---|---|---|---|---|
| Q9WTR1 | TRPV2 | transient receptor potential cation | Plasma Membrane | ion channel | 5186458 | 6462663 | 4728948 | | | | | | |
| Q9WTR1 | TRPV2 | transient receptor potential cation | Plasma Membrane | ion channel | 3511451.25 | 4249381 | 4067574.75 | | | | | | |
| Q9WTR1 | TRPV2 | transient receptor potential cation | Plasma Membrane | ion channel | 4454513 | 6985465.5 | 4562669.5 | | | | | | |
| Q9WTR1 | TRPV2 | transient receptor potential cation | Plasma Membrane | ion channel | 2951370.5 | 5554933 | 2561984 | | | | | | |
| Q9WTR1 | TRPV2 | transient receptor potential cation | Plasma Membrane | ion channel | 2686818.25 | 3418292.5 | 2943596.75 | | | | | | |
| Q9WTR1 | TRPV2 | transient receptor potential cation | Plasma Membrane | ion channel | 1515240.625 | 3328973.75 | 2617431 | Q9WTR1 | TRPV2 | 5956109 | 4320365 | 720061 | 36 |
| Q62348 | TSN | translin | Nucleus | other | 2763525.25 | 3371436 | 2593688.5 | | | | | | |
| Q62348 | TSN | translin | Nucleus | other | 1431522.875 | 1422447.75 | 634138.188 | Q62348 | TSN | 2097524 | 1032546 | 421535 | 6 |
| Q8QZY6 | TSPAN14 | tetraspanin 14 | Plasma Membrane | other | 5838634 | 5513484 | 6005494 | Q8QZY6 | TSPAN14 | 5838634 | 250213 | 144460 | 3 |
| Q8R3S2 | TSPAN33 | tetraspanin 33 | Plasma Membrane | other | 8.77E+07 | 1.19E+08 | 1.03E+08 | | | | | | |
| Q8R3S2 | TSPAN33 | tetraspanin 33 | Plasma Membrane | other | 3.68E+07 | 3.89E+07 | 3.44E+07 | | | | | | |
| Q8R3S2 | TSPAN33 | tetraspanin 33 | Plasma Membrane | other | 1.71E+07 | 1.61E+07 | 1.12E+07 | | | | | | |
| Q8R3S2 | TSPAN33 | tetraspanin 33 | Plasma Membrane | other | 1.13E+07 | 1.39E+07 | 9562086 | | | | | | |
| Q8R3S2 | TSPAN33 | tetraspanin 33 | Plasma Membrane | other | 2211281 | 6493797 | 4010207 | | | | | | |
| Q8R3S2 | TSPAN33 | tetraspanin 33 | Plasma Membrane | other | 1631449.5 | 4079787 | 3175681.25 | | | | | | |
| Q8R3S2 | TSPAN33 | tetraspanin 33 | Plasma Membrane | other | 1127262.5 | 1284261.625 | 1283956 | Q8R3S2 | TSPAN33 | 22544285 | 35068408 | 7652554 | 21 |
| Q8R3G9 | TSPAN8 | tetraspanin 8 | Plasma Membrane | other | 4.40E+07 | 4.35E+07 | 4.49E+07 | | | | | | |
| Q8R3G9 | TSPAN8 | tetraspanin 8 | Plasma Membrane | other | 3.22E+07 | 2.52E+07 | 2.89E+07 | | | | | | |
| Q8R3G9 | TSPAN8 | tetraspanin 8 | Plasma Membrane | other | 2.42E+07 | 1.75E+07 | 2.14E+07 | | | | | | |
| Q8R3G9 | TSPAN8 | tetraspanin 8 | Plasma Membrane | other | 1.10E+07 | 1.11E+07 | 1.19E+07 | | | | | | |
| Q8R3G9 | TSPAN8 | tetraspanin 8 | Plasma Membrane | other | 7079662.5 | 5395444.5 | 4405635.5 | Q8R3G9 | TSPAN8 | 23691933 | 14162193 | 3656663 | 15 |
| P50637 | TSPO | translocator protein (18kDa) | Cytoplasm | transmembrane receptor | 9668490 | 1.36E+07 | 1.50E+07 | P50637 | TSPO | 9668490 | 2768170 | 1598204 | 3 |
| A2ASS6 | TTN | titin | Cytoplasm | kinase | 3600968.25 | 4059754 | 3375587 | A2ASS6 | TTN | 3600968 | 348656 | 201297 | 3 |
| P68369 | TUBA1A | tubulin, alpha 1a | Cytoplasm | other | 5078839 | 4446952.5 | 3168852.25 | P68369 | TUBA1A | 5078839 | 973042 | 561786 | 3 |
| Q7TMM9 | TUBB2A | tubulin, beta 2A class IIa | Cytoplasm | other | 2930894 | 3481713.75 | 1878840.75 | | | | | | |
| Q7TMM9 | TUBB2A | tubulin, beta 2A class IIa | Cytoplasm | other | 3609109.75 | 4019234.5 | 2895792.25 | Q7TMM9 | TUBB2A | 3270002 | 748822 | 305705 | 6 |
| Q9D6F9 | TUBB4A | tubulin, beta 4A class IVa | Cytoplasm | other | 7532430.5 | 6708404.5 | 3598812.5 | | | | | | |
| Q9D6F9 | TUBB4A | tubulin, beta 4A class IVa | Cytoplasm | other | 8237755.5 | 7867284 | 5081743 | Q9D6F9 | TUBB4A | 7885093 | 1812558 | 739974 | 6 |
| Q8BFR5 | TUFM | Tu translation elongation factor, m | Cytoplasm | translation regulator | 4791779 | 8735634 | 8114659.5 | | | | | | |
| Q8BFR5 | TUFM | Tu translation elongation factor, m | Cytoplasm | translation regulator | 7311397 | 7818129 | 9582153 | | | | | | |
| Q8BFR5 | TUFM | Tu translation elongation factor, m | Cytoplasm | translation regulator | 6465019 | 6304636.5 | 9599587 | | | | | | |
| Q8BFR5 | TUFM | Tu translation elongation factor, m | Cytoplasm | translation regulator | 5664234 | 4684774 | 5301339 | | | | | | |
| Q8BFR5 | TUFM | Tu translation elongation factor, m | Cytoplasm | translation regulator | 4111856.75 | 3850881 | 4726110.5 | | | | | | |
| Q8BFR5 | TUFM | Tu translation elongation factor, m | Cytoplasm | translation regulator | 1773899.625 | 2107498.5 | 3732455.25 | Q8BFR5 | TUFM | 5019698 | 2354161 | 554881 | 18 |
| Q91W90 | TXNDC5 | thioredoxin domain containing 5 ( | Cytoplasm | enzyme | 1365890.625 | 1495870.875 | 680904.562 | Q91W90 | TXNDC5 | 1365891 | 437849 | 252792 | 3 |
| Q8CDN6 | TXNL1 | thioredoxin-like 1 | Cytoplasm | enzyme | 1.04E+07 | 6416060.5 | 2316546 | | | | | | |
| Q8CDN6 | TXNL1 | thioredoxin-like 1 | Cytoplasm | enzyme | 7159855.5 | 8504772 | 4274708 | | | | | | |
| Q8CDN6 | TXNL1 | thioredoxin-like 1 | Cytoplasm | enzyme | 6552691.5 | 7126732.5 | 3520938 | | | | | | |
| Q8CDN6 | TXNL1 | thioredoxin-like 1 | Cytoplasm | enzyme | 4878371 | 5362044.5 | 2416599.25 | | | | | | |
| Q8CDN6 | TXNL1 | thioredoxin-like 1 | Cytoplasm | enzyme | 1278237 | 4687909 | 2537953.75 | | | | | | |
| Q8CDN6 | TXNL1 | thioredoxin-like 1 | Cytoplasm | enzyme | 2062093.375 | 1589547.5 | 602057.625 | | | | | | |
| Q8CDN6 | TXNL1 | thioredoxin-like 1 | Cytoplasm | enzyme | 2540192.25 | 1498665 | 672535.75 | | | | | | |
| Q8CDN6 | TXNL1 | thioredoxin-like 1 | Cytoplasm | enzyme | 248965.734 | 444096.719 | 256320.953 | Q8CDN6 | TXNL1 | 4395051 | 2877075 | 587281 | 24 |
| Q9JLT4 | TXNRD2 | thioredoxin reductase 2 | Cytoplasm | enzyme | 1.81E+07 | 1.27E+07 | 1.90E+07 | | | | | | |
| Q9JLT4 | TXNRD2 | thioredoxin reductase 2 | Cytoplasm | enzyme | 6490396.5 | 4988459.5 | 6707688 | | | | | | |
| Q9JLT4 | TXNRD2 | thioredoxin reductase 2 | Cytoplasm | enzyme | 1064957.375 | 846287.938 | 1052451 | Q9JLT4 | TXNRD2 | 8538451 | 7114876 | 2371625 | 9 |
| Q02053 | UBA1 | ubiquitin-like modifier activating e | Cytoplasm | enzyme | 8415605 | 5426955 | 8033087.5 | | | | | | |
| Q02053 | UBA1 | ubiquitin-like modifier activating e | Cytoplasm | enzyme | 6330985 | 4385933.5 | 3900640 | | | | | | |
| Q02053 | UBA1 | ubiquitin-like modifier activating e | Cytoplasm | enzyme | 7118985.5 | 4434607 | 6248623 | | | | | | |
| Q02053 | UBA1 | ubiquitin-like modifier activating e | Cytoplasm | enzyme | 6451425.5 | 5580713.5 | 4975758.5 | | | | | | |
| Q02053 | UBA1 | ubiquitin-like modifier activating e | Cytoplasm | enzyme | 5298727 | 3332143.75 | 4621764 | | | | | | |
| Q02053 | UBA1 | ubiquitin-like modifier activating e | Cytoplasm | enzyme | 5986988 | 3595360 | 4201507.5 | | | | | | |
| Q02053 | UBA1 | ubiquitin-like modifier activating e | Cytoplasm | enzyme | 3730567.5 | 3218585.5 | 3057492.25 | | | | | | |
| Q02053 | UBA1 | ubiquitin-like modifier activating e | Cytoplasm | enzyme | 4925626.5 | 3737865.25 | 4315874.5 | | | | | | |
| Q02053 | UBA1 | ubiquitin-like modifier activating e | Cytoplasm | enzyme | 4017558.75 | 1676633.75 | 2594216 | | | | | | |
| Q02053 | UBA1 | ubiquitin-like modifier activating e | Cytoplasm | enzyme | 2660761 | 2972830.75 | 3967928 | | | | | | |
| Q02053 | UBA1 | ubiquitin-like modifier activating e | Cytoplasm | enzyme | 2836840.5 | 2317399.25 | 2541434.75 | | | | | | |
| Q02053 | UBA1 | ubiquitin-like modifier activating e | Cytoplasm | enzyme | 1769465.875 | 2461784 | 3081157.25 | | | | | | |
| Q02053 | UBA1 | ubiquitin-like modifier activating e | Cytoplasm | enzyme | 1277355 | 531756.438 | 1257833.625 | | | | | | |
| Q02053 | UBA1 | ubiquitin-like modifier activating e | Cytoplasm | enzyme | 1333265.875 | 261194.797 | 166.49 | Q02053 | UBA1 | 4439583 | 1999312 | 308500 | 42 |
| P62984 | Uba52 | ubiquitin A-52 residue ribosomal p | Cytoplasm | other | 1.06E+09 | 1.05E+09 | 1.07E+09 | | | | | | |
| P62984 | Uba52 | ubiquitin A-52 residue ribosomal p | Cytoplasm | other | 3.73E+08 | 3.70E+08 | 4.31E+08 | | | | | | |
| P62984 | Uba52 | ubiquitin A-52 residue ribosomal p | Cytoplasm | other | 2.03E+08 | 1.58E+08 | 2.05E+08 | | | | | | |
| P62984 | Uba52 | ubiquitin A-52 residue ribosomal p | Cytoplasm | other | 2.16E+08 | 1.99E+08 | 2.32E+08 | | | | | | |
| P62984 | Uba52 | ubiquitin A-52 residue ribosomal p | Cytoplasm | other | 1.25E+08 | 1.15E+07 | 3.19E+07 | | | | | | |
| P62984 | Uba52 | ubiquitin A-52 residue ribosomal p | Cytoplasm | other | 1.63E+07 | 1513898.625 | 3364339.5 | | | | | | |
| P62984 | Uba52 | ubiquitin A-52 residue ribosomal p | Cytoplasm | other | 1.71E+07 | 3.84E+07 | 1.80E+07 | | | | | | |
| P62984 | Uba52 | ubiquitin A-52 residue ribosomal p | Cytoplasm | other | 974435.875 | 354904.844 | 1583835.375 | P62984 | Uba52 | 251598054 | 342471523 | 69906707 | 24 |
| Q8VDI7 | UBAC1 | UBA domain containing 1 | Nucleus | other | 4887490 | 1.13E+07 | 7915472 | | | | | | |
| Q8VDI7 | UBAC1 | UBA domain containing 1 | Nucleus | other | 6244901 | 6059702 | 6871680 | | | | | | |
| Q8VDI7 | UBAC1 | UBA domain containing 1 | Nucleus | other | 4000026 | 5258870 | 5614312 | | | | | | |
| Q8VDI7 | UBAC1 | UBA domain containing 1 | Nucleus | other | 4074737.75 | 4097602.5 | 3355734 | | | | | | |
| Q8VDI7 | UBAC1 | UBA domain containing 1 | Nucleus | other | 4492901.5 | 3120850.25 | 3459385.75 | | | | | | |
| Q8VDI7 | UBAC1 | UBA domain containing 1 | Nucleus | other | 2964674 | 2738610.75 | 2886979.5 | | | | | | |
| Q8VDI7 | UBAC1 | UBA domain containing 1 | Nucleus | other | 2842217.5 | 2485774.25 | 2605521.25 | | | | | | |
| Q8VDI7 | UBAC1 | UBA domain containing 1 | Nucleus | other | 1964183.875 | 974297.688 | 1325395 | | | | | | |
| Q8VDI7 | UBAC1 | UBA domain containing 1 | Nucleus | other | 2079818.75 | 1421406.125 | 1619045 | | | | | | |
| Q8VDI7 | UBAC1 | UBA domain containing 1 | Nucleus | other | 263599.469 | 2308478.75 | 2891275.5 | Q8VDI7 | UBAC1 | 3381455 | 2305799 | 420979 | 30 |
| P68037 | UBE2L3 | ubiquitin-conjugating enzyme E2L | Nucleus | enzyme | 8471877 | 7663174 | 1.02E+07 | P68037 | UBE2L3 | 8471877 | 1306860 | 754516 | 3 |
| Q9QZU9 | UBE2L6 | ubiquitin-conjugating enzyme E2L | Cytoplasm | enzyme | 1.63E+07 | 2.30E+07 | 2.20E+07 | | | | | | |
| Q9QZU9 | UBE2L6 | ubiquitin-conjugating enzyme E2L | Cytoplasm | enzyme | 4074189.5 | 6504546.5 | 4929913 | | | | | | |
| Q9QZU9 | UBE2L6 | ubiquitin-conjugating enzyme E2L | Cytoplasm | enzyme | 1203137 | 6619382 | 4412823 | | | | | | |
| Q9QZU9 | UBE2L6 | ubiquitin-conjugating enzyme E2L | Cytoplasm | enzyme | 1939635.125 | 1517554.625 | 1674431.375 | Q9QZU9 | UBE2L6 | 5866740 | 7945301 | 2293611 | 12 |
| Q6ZPJ3 | UBE2O | ubiquitin-conjugating enzyme E2O | Nucleus | enzyme | 2.99E+07 | 4.00E+07 | 2.44E+07 | | | | | | |
| Q6ZPJ3 | UBE2O | ubiquitin-conjugating enzyme E2O | Nucleus | enzyme | 2.71E+07 | 3.89E+07 | 1.60E+07 | | | | | | |
| Q6ZPJ3 | UBE2O | ubiquitin-conjugating enzyme E2O | Nucleus | enzyme | 2.05E+07 | 2.72E+07 | 1.60E+07 | | | | | | |

| | | | | | | | | | | | | | |
|---|---|---|---|---|---|---|---|---|---|---|---|---|---|
| Q6ZPJ3 | UBE2O | ubiquitin-conjugating enzyme E2O | Nucleus | enzyme | 2.58E+07 | 2.71E+07 | 2.11E+07 | | | | | | |
| Q6ZPJ3 | UBE2O | ubiquitin-conjugating enzyme E2O | Nucleus | enzyme | 1.84E+07 | 2.28E+07 | 1.64E+07 | | | | | | |
| Q6ZPJ3 | UBE2O | ubiquitin-conjugating enzyme E2O | Nucleus | enzyme | 1.86E+07 | 2.25E+07 | 1.18E+07 | | | | | | |
| Q6ZPJ3 | UBE2O | ubiquitin-conjugating enzyme E2O | Nucleus | enzyme | 1.73E+07 | 2.10E+07 | 1.40E+07 | | | | | | |
| Q6ZPJ3 | UBE2O | ubiquitin-conjugating enzyme E2O | Nucleus | enzyme | 1.26E+07 | 1.81E+07 | 1.21E+07 | | | | | | |
| Q6ZPJ3 | UBE2O | ubiquitin-conjugating enzyme E2O | Nucleus | enzyme | 1.35E+07 | 1.75E+07 | 1.25E+07 | | | | | | |
| Q6ZPJ3 | UBE2O | ubiquitin-conjugating enzyme E2O | Nucleus | enzyme | 1.20E+07 | 1.31E+07 | 8318095.5 | | | | | | |
| Q6ZPJ3 | UBE2O | ubiquitin-conjugating enzyme E2O | Nucleus | enzyme | 1.02E+07 | 8465741 | 6156149.5 | | | | | | |
| Q6ZPJ3 | UBE2O | ubiquitin-conjugating enzyme E2O | Nucleus | enzyme | 1.00E+07 | 1.06E+07 | 7336724 | | | | | | |
| Q6ZPJ3 | UBE2O | ubiquitin-conjugating enzyme E2O | Nucleus | enzyme | 8390949 | 1.10E+07 | 7902501 | | | | | | |
| Q6ZPJ3 | UBE2O | ubiquitin-conjugating enzyme E2O | Nucleus | enzyme | 1.23E+07 | 1.32E+07 | 6840598.5 | | | | | | |
| Q6ZPJ3 | UBE2O | ubiquitin-conjugating enzyme E2O | Nucleus | enzyme | 9221885 | 9881557 | 5915727.5 | | | | | | |
| Q6ZPJ3 | UBE2O | ubiquitin-conjugating enzyme E2O | Nucleus | enzyme | 9830473 | 1.06E+07 | 6360235 | | | | | | |
| Q6ZPJ3 | UBE2O | ubiquitin-conjugating enzyme E2O | Nucleus | enzyme | 8365762.5 | 1.05E+07 | 6763944 | | | | | | |
| Q6ZPJ3 | UBE2O | ubiquitin-conjugating enzyme E2O | Nucleus | enzyme | 9284401 | 1.43E+07 | 7310591.5 | | | | | | |
| Q6ZPJ3 | UBE2O | ubiquitin-conjugating enzyme E2O | Nucleus | enzyme | 8077665 | 1.20E+07 | 5631051.5 | | | | | | |
| Q6ZPJ3 | UBE2O | ubiquitin-conjugating enzyme E2O | Nucleus | enzyme | 6367424 | 8226599 | 6651442 | | | | | | |
| Q6ZPJ3 | UBE2O | ubiquitin-conjugating enzyme E2O | Nucleus | enzyme | 6325403 | 9591664 | 5835373 | | | | | | |
| Q6ZPJ3 | UBE2O | ubiquitin-conjugating enzyme E2O | Nucleus | enzyme | 4591417 | 4045302.25 | 1059831.5 | | | | | | |
| Q6ZPJ3 | UBE2O | ubiquitin-conjugating enzyme E2O | Nucleus | enzyme | 5529289.5 | 6182235 | 2975799.5 | | | | | | |
| Q6ZPJ3 | UBE2O | ubiquitin-conjugating enzyme E2O | Nucleus | enzyme | 3028120.25 | 7205086.5 | 4711048 | | | | | | |
| Q6ZPJ3 | UBE2O | ubiquitin-conjugating enzyme E2O | Nucleus | enzyme | 3165881.5 | 3916397.25 | 2655852 | | | | | | |
| Q6ZPJ3 | UBE2O | ubiquitin-conjugating enzyme E2O | Nucleus | enzyme | 3755905.75 | 4394314.5 | 2659136.75 | | | | | | |
| Q6ZPJ3 | UBE2O | ubiquitin-conjugating enzyme E2O | Nucleus | enzyme | 4759133.5 | 1001470 | 2032972.75 | | | | | | |
| Q6ZPJ3 | UBE2O | ubiquitin-conjugating enzyme E2O | Nucleus | enzyme | 3214790 | 4657779.5 | 2246965 | | | | | | |
| Q6ZPJ3 | UBE2O | ubiquitin-conjugating enzyme E2O | Nucleus | enzyme | 1700971.375 | 1786594.875 | 1311232.125 | | | | | | |
| Q6ZPJ3 | UBE2O | ubiquitin-conjugating enzyme E2O | Nucleus | enzyme | 3154505.25 | 264032.656 | 92282.484 | | | | | | |
| Q6ZPJ3 | UBE2O | ubiquitin-conjugating enzyme E2O | Nucleus | enzyme | 1514393.125 | 2185008.75 | 1912898 | | | | | | |
| Q6ZPJ3 | UBE2O | ubiquitin-conjugating enzyme E2O | Nucleus | enzyme | 1849378.25 | 2930751.25 | 1980219.875 | | | | | | |
| Q6ZPJ3 | UBE2O | ubiquitin-conjugating enzyme E2O | Nucleus | enzyme | 1327686.25 | 1112686 | 732652.625 | | | | | | |
| Q6ZPJ3 | UBE2O | ubiquitin-conjugating enzyme E2O | Nucleus | enzyme | 4.70E-06 | 1111038.875 | 885795.875 | Q6ZPJ3 | UBE2O | 9755748 | 8356433 | 827410 | 102 |
| Q6ZWZ2 | UBE2R2 | ubiquitin-conjugating enzyme E2R | Cytoplasm | enzyme | 1886794.125 | 1832225.875 | 1612992 | Q6ZWZ2 | UBE2R2 | 1886794 | 144919 | 83669 | 3 |
| A2AN08 | UBR4 | ubiquitin protein ligase E3 compor | Nucleus | enzyme | 1.16E+07 | 1.30E+07 | 9298406 | | | | | | |
| A2AN08 | UBR4 | ubiquitin protein ligase E3 compor | Nucleus | enzyme | 5732885 | 7693098 | 5063501 | | | | | | |
| A2AN08 | UBR4 | ubiquitin protein ligase E3 compor | Nucleus | enzyme | 8237704.5 | 3139803.75 | 1574075.125 | | | | | | |
| A2AN08 | UBR4 | ubiquitin protein ligase E3 compor | Nucleus | enzyme | 4056938.25 | 9586477 | 5096092.5 | | | | | | |
| A2AN08 | UBR4 | ubiquitin protein ligase E3 compor | Nucleus | enzyme | 4591274.5 | 5060016 | 4463622 | | | | | | |
| A2AN08 | UBR4 | ubiquitin protein ligase E3 compor | Nucleus | enzyme | 5593244.5 | 6183432.5 | 4278260.5 | | | | | | |
| A2AN08 | UBR4 | ubiquitin protein ligase E3 compor | Nucleus | enzyme | 5897619 | 4888019.5 | 4120845.25 | | | | | | |
| A2AN08 | UBR4 | ubiquitin protein ligase E3 compor | Nucleus | enzyme | 3850831.5 | 4387884.5 | 4096210.75 | | | | | | |
| A2AN08 | UBR4 | ubiquitin protein ligase E3 compor | Nucleus | enzyme | 5098770.5 | 4065940.75 | 3653108.5 | | | | | | |
| A2AN08 | UBR4 | ubiquitin protein ligase E3 compor | Nucleus | enzyme | 5503799 | 4133359 | 3276499.5 | | | | | | |
| A2AN08 | UBR4 | ubiquitin protein ligase E3 compor | Nucleus | enzyme | 4264592 | 3605282.75 | 2873058.5 | | | | | | |
| A2AN08 | UBR4 | ubiquitin protein ligase E3 compor | Nucleus | enzyme | 5997259.5 | 5469141.5 | 4414897.5 | | | | | | |
| A2AN08 | UBR4 | ubiquitin protein ligase E3 compor | Nucleus | enzyme | 4572933 | 6583376.5 | 4899311.5 | | | | | | |
| A2AN08 | UBR4 | ubiquitin protein ligase E3 compor | Nucleus | enzyme | 3932188.25 | 3464084.75 | 2981538.25 | | | | | | |
| A2AN08 | UBR4 | ubiquitin protein ligase E3 compor | Nucleus | enzyme | 3896385.25 | 5353966.5 | 2871442.25 | | | | | | |
| A2AN08 | UBR4 | ubiquitin protein ligase E3 compor | Nucleus | enzyme | 4626975.5 | 5337388 | 3944236.75 | | | | | | |
| A2AN08 | UBR4 | ubiquitin protein ligase E3 compor | Nucleus | enzyme | 5155696.5 | 5553844 | 4264256.5 | | | | | | |
| A2AN08 | UBR4 | ubiquitin protein ligase E3 compor | Nucleus | enzyme | 5316839 | 4028399.75 | 3462299.5 | | | | | | |
| A2AN08 | UBR4 | ubiquitin protein ligase E3 compor | Nucleus | enzyme | 4814837.5 | 6118565.5 | 3340401.5 | | | | | | |
| A2AN08 | UBR4 | ubiquitin protein ligase E3 compor | Nucleus | enzyme | 3967852.5 | 4262262.5 | 3054900.25 | | | | | | |
| A2AN08 | UBR4 | ubiquitin protein ligase E3 compor | Nucleus | enzyme | 3851287.25 | 3756065.5 | 3751182.25 | | | | | | |
| A2AN08 | UBR4 | ubiquitin protein ligase E3 compor | Nucleus | enzyme | 5118585 | 5544178 | 2947616.75 | | | | | | |
| A2AN08 | UBR4 | ubiquitin protein ligase E3 compor | Nucleus | enzyme | 4780203.5 | 4659873.5 | 4082813.5 | | | | | | |
| A2AN08 | UBR4 | ubiquitin protein ligase E3 compor | Nucleus | enzyme | 4172524.5 | 4892014 | 3302431.75 | | | | | | |
| A2AN08 | UBR4 | ubiquitin protein ligase E3 compor | Nucleus | enzyme | 3570832.75 | 4342345 | 2965790.75 | | | | | | |
| A2AN08 | UBR4 | ubiquitin protein ligase E3 compor | Nucleus | enzyme | 3519874.25 | 3733160.5 | 2898170.75 | | | | | | |
| A2AN08 | UBR4 | ubiquitin protein ligase E3 compor | Nucleus | enzyme | 4297570 | 3544114.25 | 2831239.25 | | | | | | |
| A2AN08 | UBR4 | ubiquitin protein ligase E3 compor | Nucleus | enzyme | 3050840.25 | 3542360.75 | 3239572.25 | | | | | | |
| A2AN08 | UBR4 | ubiquitin protein ligase E3 compor | Nucleus | enzyme | 4087193.75 | 4655728.5 | 4114581 | | | | | | |
| A2AN08 | UBR4 | ubiquitin protein ligase E3 compor | Nucleus | enzyme | 2947194.75 | 3759914.25 | 2049455.125 | | | | | | |
| A2AN08 | UBR4 | ubiquitin protein ligase E3 compor | Nucleus | enzyme | 4485796 | 4397329 | 2775836 | | | | | | |
| A2AN08 | UBR4 | ubiquitin protein ligase E3 compor | Nucleus | enzyme | 2852264.5 | 4192617.25 | 2560846.75 | | | | | | |
| A2AN08 | UBR4 | ubiquitin protein ligase E3 compor | Nucleus | enzyme | 5468557 | 5048609 | 4233054.5 | | | | | | |
| A2AN08 | UBR4 | ubiquitin protein ligase E3 compor | Nucleus | enzyme | 2645579.25 | 4298527.5 | 4041427.5 | | | | | | |
| A2AN08 | UBR4 | ubiquitin protein ligase E3 compor | Nucleus | enzyme | 3619775.5 | 2781963.5 | 1673258.75 | | | | | | |
| A2AN08 | UBR4 | ubiquitin protein ligase E3 compor | Nucleus | enzyme | 3606751 | 3548374.5 | 3567550.5 | | | | | | |
| A2AN08 | UBR4 | ubiquitin protein ligase E3 compor | Nucleus | enzyme | 1962124.375 | 2673963.5 | 2114536.25 | | | | | | |
| A2AN08 | UBR4 | ubiquitin protein ligase E3 compor | Nucleus | enzyme | 3192709 | 4257648 | 2825061.5 | | | | | | |
| A2AN08 | UBR4 | ubiquitin protein ligase E3 compor | Nucleus | enzyme | 3297851.25 | 3916184.25 | 2572245.5 | | | | | | |
| A2AN08 | UBR4 | ubiquitin protein ligase E3 compor | Nucleus | enzyme | 3020388.5 | 3130458.25 | 1889184.625 | | | | | | |
| A2AN08 | UBR4 | ubiquitin protein ligase E3 compor | Nucleus | enzyme | 2562851 | 3113436.25 | 2547656.75 | | | | | | |
| A2AN08 | UBR4 | ubiquitin protein ligase E3 compor | Nucleus | enzyme | 3329905.25 | 1462327.125 | 1066946.125 | | | | | | |
| A2AN08 | UBR4 | ubiquitin protein ligase E3 compor | Nucleus | enzyme | 2345331.75 | 3903016 | 2486307 | | | | | | |
| A2AN08 | UBR4 | ubiquitin protein ligase E3 compor | Nucleus | enzyme | 2997577.75 | 2861194 | 2405963 | | | | | | |
| A2AN08 | UBR4 | ubiquitin protein ligase E3 compor | Nucleus | enzyme | 2106501.25 | 2452099.25 | 1417882.375 | | | | | | |
| A2AN08 | UBR4 | ubiquitin protein ligase E3 compor | Nucleus | enzyme | 1685346.125 | 2243435.75 | 1280065.875 | | | | | | |
| A2AN08 | UBR4 | ubiquitin protein ligase E3 compor | Nucleus | enzyme | 2660031.75 | 2790201.5 | 2189602.25 | | | | | | |
| A2AN08 | UBR4 | ubiquitin protein ligase E3 compor | Nucleus | enzyme | 1602723.5 | 2222117.25 | 1664159.125 | | | | | | |
| A2AN08 | UBR4 | ubiquitin protein ligase E3 compor | Nucleus | enzyme | 1564703.625 | 2682007.5 | 1589209.625 | | | | | | |
| A2AN08 | UBR4 | ubiquitin protein ligase E3 compor | Nucleus | enzyme | 2351091 | 557784.5 | 823154.562 | | | | | | |
| A2AN08 | UBR4 | ubiquitin protein ligase E3 compor | Nucleus | enzyme | 1579845.25 | 3476709 | 1683852.375 | | | | | | |
| A2AN08 | UBR4 | ubiquitin protein ligase E3 compor | Nucleus | enzyme | 1072772.25 | 1491574.125 | 955417.438 | | | | | | |
| A2AN08 | UBR4 | ubiquitin protein ligase E3 compor | Nucleus | enzyme | 768976.562 | 1400285.375 | 856501.812 | A2AN08 | UBR4 | 3903853 | 1814503 | 143899 | 159 |
| Q9WUP7 | UCHL5 | ubiquitin carboxyl-terminal hydrol | Cytoplasm | peptidase | 8560937 | 1.03E+07 | 4762598 | | | | | | |

| | | | | | | | | | | | | | |
|---|---|---|---|---|---|---|---|---|---|---|---|---|---|
| Q9WUP7 | UCHL5 | ubiquitin carboxyl-terminal hydrol | Cytoplasm | peptidase | 5077591 | 1.17E+07 | 4270332.5 | | | | | | |
| Q9WUP7 | UCHL5 | ubiquitin carboxyl-terminal hydrol | Cytoplasm | peptidase | 2033094.375 | 4029749.5 | 1170888.375 | | | | | | |
| Q9WUP7 | UCHL5 | ubiquitin carboxyl-terminal hydrol | Cytoplasm | peptidase | 372082.625 | 1067419.125 | 1875277.875 | Q9WUP7 | UCHL5 | 4010926 | 3754582 | 1083854 | 12 |
| Q6P5E4 | UGGT1 | UDP-glucose glycoprotein glucosyl | Cytoplasm | enzyme | 9471913 | 1.21E+07 | 1.06E+07 | | | | | | |
| Q6P5E4 | UGGT1 | UDP-glucose glycoprotein glucosyl | Cytoplasm | enzyme | 4664683 | 3938641.25 | 3869210 | | | | | | |
| Q6P5E4 | UGGT1 | UDP-glucose glycoprotein glucosyl | Cytoplasm | enzyme | 2964766 | 2732263.25 | 2256360 | | | | | | |
| Q6P5E4 | UGGT1 | UDP-glucose glycoprotein glucosyl | Cytoplasm | enzyme | 4944901 | 5336881.5 | 5276418 | | | | | | |
| Q6P5E4 | UGGT1 | UDP-glucose glycoprotein glucosyl | Cytoplasm | enzyme | 2378827 | 2335144 | 2511770.25 | | | | | | |
| Q6P5E4 | UGGT1 | UDP-glucose glycoprotein glucosyl | Cytoplasm | enzyme | 2342580.75 | 2140275.25 | 2369859 | | | | | | |
| Q6P5E4 | UGGT1 | UDP-glucose glycoprotein glucosyl | Cytoplasm | enzyme | 2844579.25 | 4079853 | 3254595 | | | | | | |
| Q6P5E4 | UGGT1 | UDP-glucose glycoprotein glucosyl | Cytoplasm | enzyme | 2017373.375 | 2518546.5 | 1482351.75 | Q6P5E4 | UGGT1 | 3953703 | 2790033 | 569513 | 24 |
| Q8R1I1 | UQCR10 | ubiquinol-cytochrome c reductase | Cytoplasm | enzyme | 1299579.25 | 2313134 | 2990624.75 | Q8R1I1 | UQCR10 | 1299579 | 851070 | 491366 | 3 |
| Q9CZ13 | UQCRC1 | ubiquinol-cytochrome c reductase | Cytoplasm | enzyme | 2.02E+07 | 1.54E+07 | 1.64E+07 | | | | | | |
| Q9CZ13 | UQCRC1 | ubiquinol-cytochrome c reductase | Cytoplasm | enzyme | 6504072 | 6233756.5 | 9970381 | | | | | | |
| Q9CZ13 | UQCRC1 | ubiquinol-cytochrome c reductase | Cytoplasm | enzyme | 7964578 | 6514568 | 1.00E+07 | | | | | | |
| Q9CZ13 | UQCRC1 | ubiquinol-cytochrome c reductase | Cytoplasm | enzyme | 5200696.5 | 4248639 | 6622863 | | | | | | |
| Q9CZ13 | UQCRC1 | ubiquinol-cytochrome c reductase | Cytoplasm | enzyme | 6340757.5 | 6727637 | 1.12E+07 | Q9CZ13 | UQCRC1 | 9244021 | 4649461 | 1200486 | 15 |
| Q9DB77 | UQCRC2 | ubiquinol-cytochrome c reductase | Cytoplasm | enzyme | 1.52E+07 | 1.96E+07 | 2.39E+07 | | | | | | |
| Q9DB77 | UQCRC2 | ubiquinol-cytochrome c reductase | Cytoplasm | enzyme | 1.32E+07 | 1.34E+07 | 1.50E+07 | | | | | | |
| Q9DB77 | UQCRC2 | ubiquinol-cytochrome c reductase | Cytoplasm | enzyme | 1.23E+07 | 1.19E+07 | 1.56E+07 | | | | | | |
| Q9DB77 | UQCRC2 | ubiquinol-cytochrome c reductase | Cytoplasm | enzyme | 9619604 | 9678687 | 9852034 | | | | | | |
| Q9DB77 | UQCRC2 | ubiquinol-cytochrome c reductase | Cytoplasm | enzyme | 4872444 | 5888434 | 6852127.5 | | | | | | |
| Q9DB77 | UQCRC2 | ubiquinol-cytochrome c reductase | Cytoplasm | enzyme | 5795465 | 7197807 | 7183057 | | | | | | |
| Q9DB77 | UQCRC2 | ubiquinol-cytochrome c reductase | Cytoplasm | enzyme | 3470435.75 | 3145544 | 5054410.5 | | | | | | |
| Q9DB77 | UQCRC2 | ubiquinol-cytochrome c reductase | Cytoplasm | enzyme | 4604908.5 | 5240970.5 | 6421781 | | | | | | |
| Q9DB77 | UQCRC2 | ubiquinol-cytochrome c reductase | Cytoplasm | enzyme | 5434881 | 4032881.5 | 6257050.5 | | | | | | |
| Q9DB77 | UQCRC2 | ubiquinol-cytochrome c reductase | Cytoplasm | enzyme | 3525964.25 | 3451904.75 | 4926160.5 | | | | | | |
| Q9DB77 | UQCRC2 | ubiquinol-cytochrome c reductase | Cytoplasm | enzyme | 2346315 | 2448241.5 | 3519132 | | | | | | |
| Q9DB77 | UQCRC2 | ubiquinol-cytochrome c reductase | Cytoplasm | enzyme | 558999.562 | 2636983.25 | 4469630 | Q9DB77 | UQCRC2 | 6744918 | 5327248 | 887875 | 36 |
| P99028 | UQCRHL | ubiquinol-cytochrome c reductase | Cytoplasm | other | 3.40E+07 | 2.86E+07 | 3.80E+07 | | | | | | |
| P99028 | UQCRHL | ubiquinol-cytochrome c reductase | Cytoplasm | other | 5225769 | 1.33E+07 | 1.75E+07 | P99028 | UQCRHL | 19622885 | 12782641 | 5218491 | 6 |
| P70697 | UROD | uroporphyrinogen decarboxylase | Cytoplasm | enzyme | 3921286 | 2523676.75 | 3432986 | P70697 | UROD | 3921286 | 709294 | 409511 | 3 |
| Q78IK2 | USMG5 | up-regulated during skeletal musc | Cytoplasm | other | 8308740.5 | 9941793 | 1.27E+07 | | | | | | |
| Q78IK2 | USMG5 | up-regulated during skeletal musc | Cytoplasm | other | 4772359 | 5994779.5 | 7522720.5 | Q78IK2 | USMG5 | 6540550 | 2847662 | 1162553 | 6 |
| Q9JMA1 | USP14 | ubiquitin specific peptidase 14 (tR | Cytoplasm | peptidase | 6184188 | 6962699 | 5345964 | | | | | | |
| Q9JMA1 | USP14 | ubiquitin specific peptidase 14 (tR | Cytoplasm | peptidase | 3825236.5 | 3209012 | 3010815 | Q9JMA1 | USP14 | 5004712 | 1646965 | 672370 | 6 |
| Q8R5H1 | USP15 | ubiquitin specific peptidase 15 | Cytoplasm | peptidase | 2748986 | 2662557.75 | 2554234.75 | | | | | | |
| Q8R5H1 | USP15 | ubiquitin specific peptidase 15 | Cytoplasm | peptidase | 8344615 | 6684939.5 | 6673277.5 | | | | | | |
| Q8R5H1 | USP15 | ubiquitin specific peptidase 15 | Cytoplasm | peptidase | 4768412 | 4066439.5 | 3923178.25 | | | | | | |
| Q8R5H1 | USP15 | ubiquitin specific peptidase 15 | Cytoplasm | peptidase | 3504476.75 | 4034610 | 5005435.5 | | | | | | |
| Q8R5H1 | USP15 | ubiquitin specific peptidase 15 | Cytoplasm | peptidase | 4032030.75 | 2735117 | 1953347 | | | | | | |
| Q8R5H1 | USP15 | ubiquitin specific peptidase 15 | Cytoplasm | peptidase | 4019175.25 | 4015918 | 3494696 | | | | | | |
| Q8R5H1 | USP15 | ubiquitin specific peptidase 15 | Cytoplasm | peptidase | 3307223.5 | 1690709.5 | 2282222.25 | | | | | | |
| Q8R5H1 | USP15 | ubiquitin specific peptidase 15 | Cytoplasm | peptidase | 2815380.5 | 2670952.5 | 2771621.25 | | | | | | |
| Q8R5H1 | USP15 | ubiquitin specific peptidase 15 | Cytoplasm | peptidase | 2750333 | 2273954.5 | 2438278.25 | | | | | | |
| Q8R5H1 | USP15 | ubiquitin specific peptidase 15 | Cytoplasm | peptidase | 3142841.25 | 1452648.5 | 1962418.25 | | | | | | |
| Q8R5H1 | USP15 | ubiquitin specific peptidase 15 | Cytoplasm | peptidase | 2273386.25 | 2255640.25 | 1649582.5 | | | | | | |
| Q8R5H1 | USP15 | ubiquitin specific peptidase 15 | Cytoplasm | peptidase | 2265482.25 | 2329440.25 | 2645314 | | | | | | |
| Q8R5H1 | USP15 | ubiquitin specific peptidase 15 | Cytoplasm | peptidase | 2565356.75 | 2058710.125 | 1988124.875 | | | | | | |
| Q8R5H1 | USP15 | ubiquitin specific peptidase 15 | Cytoplasm | peptidase | 1649066.375 | 1622117.25 | 2151340.75 | | | | | | |
| Q8R5H1 | USP15 | ubiquitin specific peptidase 15 | Cytoplasm | peptidase | 1018519 | 982577.375 | 948416.812 | | | | | | |
| Q8R5H1 | USP15 | ubiquitin specific peptidase 15 | Cytoplasm | peptidase | 1327133.875 | 1230845.75 | 1329111.875 | Q8R5H1 | USP15 | 3158276 | 1518853 | 219228 | 48 |
| Q3UJD6 | USP19 | ubiquitin specific peptidase 19 | Cytoplasm | peptidase | 196195.484 | 109112.617 | 135099.297 | Q3UJD6 | USP19 | 196195 | 44705 | 25811 | 3 |
| Q5DU02 | USP22 | ubiquitin specific peptidase 22 | Nucleus | peptidase | 9.62E+08 | 1.16E+09 | 1.10E+09 | Q5DU02 | USP22 | 961600000 | 99932577 | 57696100 | 3 |
| P56399 | USP5 | ubiquitin specific peptidase 5 (isop | Cytoplasm | peptidase | 7800813.5 | 7030165.5 | 6121202 | | | | | | |
| P56399 | USP5 | ubiquitin specific peptidase 5 (isop | Cytoplasm | peptidase | 5157618 | 4535255 | 4496005 | | | | | | |
| P56399 | USP5 | ubiquitin specific peptidase 5 (isop | Cytoplasm | peptidase | 4622743 | 3680990.75 | 3896556.25 | | | | | | |
| P56399 | USP5 | ubiquitin specific peptidase 5 (isop | Cytoplasm | peptidase | 4039491.75 | 3716052.25 | 2977952.5 | | | | | | |
| P56399 | USP5 | ubiquitin specific peptidase 5 (isop | Cytoplasm | peptidase | 3895191.5 | 2684816.75 | 3767163 | | | | | | |
| P56399 | USP5 | ubiquitin specific peptidase 5 (isop | Cytoplasm | peptidase | 4637076.5 | 4198784 | 3881471.5 | | | | | | |
| P56399 | USP5 | ubiquitin specific peptidase 5 (isop | Cytoplasm | peptidase | 4670977.5 | 3178853.5 | 3555799.5 | | | | | | |
| P56399 | USP5 | ubiquitin specific peptidase 5 (isop | Cytoplasm | peptidase | 4073610 | 2925652.25 | 4255720.5 | | | | | | |
| P56399 | USP5 | ubiquitin specific peptidase 5 (isop | Cytoplasm | peptidase | 2205477 | 1984081.25 | 1396079.75 | | | | | | |
| P56399 | USP5 | ubiquitin specific peptidase 5 (isop | Cytoplasm | peptidase | 1418381.125 | 796029.938 | 153999.906 | | | | | | |
| P56399 | USP5 | ubiquitin specific peptidase 5 (isop | Cytoplasm | peptidase | 1201139.375 | 858960.25 | 1557024.125 | | | | | | |
| P56399 | USP5 | ubiquitin specific peptidase 5 (isop | Cytoplasm | peptidase | 664102.875 | 1235707.25 | 1386466.5 | | | | | | |
| P56399 | USP5 | ubiquitin specific peptidase 5 (isop | Cytoplasm | peptidase | 228383.266 | 1071494.875 | 2756844.75 | P56399 | USP5 | 3431923 | 1831168 | 293221 | 39 |
| Q6A4J8 | USP7 | ubiquitin specific peptidase 7 (her | Nucleus | peptidase | 1127051.25 | 793845.25 | 888376.312 | Q6A4J8 | USP7 | 1127051 | 171721 | 99143 | 3 |
| P70398 | USP9X | ubiquitin specific peptidase 9, X-lir | Plasma Membrane | peptidase | 5924161 | 2941099.75 | 3507540.5 | | | | | | |
| P70398 | USP9X | ubiquitin specific peptidase 9, X-lir | Plasma Membrane | peptidase | 2220426.5 | 2876279.25 | 1936353.875 | | | | | | |
| P70398 | USP9X | ubiquitin specific peptidase 9, X-lir | Plasma Membrane | peptidase | 2241759.75 | 1991076.125 | 851730.75 | | | | | | |
| P70398 | USP9X | ubiquitin specific peptidase 9, X-lir | Plasma Membrane | peptidase | 956649.125 | 1488572.25 | 765768.5 | P70398 | USP9X | 2835749 | 1429771 | 412739 | 12 |
| Q80WQ2 | VAC14 | Vac14 homolog (S. cerevisiae) | Cytoplasm | other | 2131856.5 | 2642373.25 | 2596140.25 | Q80WQ2 | VAC14 | 2131857 | 282349 | 163014 | 3 |
| P63024 | VAMP3 | vesicle-associated membrane prot | Plasma Membrane | other | 5.72E+07 | 8.03E+07 | 8.06E+07 | | | | | | |
| P63024 | VAMP3 | vesicle-associated membrane prot | Plasma Membrane | other | 7398147.5 | 1.31E+07 | 1.07E+07 | | | | | | |
| P63024 | VAMP3 | vesicle-associated membrane prot | Plasma Membrane | other | 4435115 | 7373867.5 | 6722218.5 | | | | | | |
| P63024 | VAMP3 | vesicle-associated membrane prot | Plasma Membrane | other | 4383845 | 6309872 | 4334073 | | | | | | |
| P63024 | VAMP3 | vesicle-associated membrane prot | Plasma Membrane | other | 1519038.25 | 6221055.5 | 3037563.75 | P63024 | VAMP3 | 14985229 | 28099891 | 7255361 | 15 |
| Q9Z2P8 | VAMP5 | vesicle-associated membrane prot | Plasma Membrane | transporter | 2.76E+07 | 3.13E+07 | 1.97E+07 | | | | | | |
| Q9Z2P8 | VAMP5 | vesicle-associated membrane prot | Plasma Membrane | transporter | 6170534 | 1.31E+07 | 1.58E+07 | | | | | | |
| Q9Z2P8 | VAMP5 | vesicle-associated membrane prot | Plasma Membrane | transporter | 7807775.5 | 1.26E+07 | 1.48E+07 | Q9Z2P8 | VAMP5 | 13862770 | 8408758 | 2802919 | 9 |
| O70404 | VAMP8 | vesicle-associated membrane prot | Plasma Membrane | transporter | 2.06E+07 | 2.78E+07 | 2.97E+07 | | | | | | |
| O70404 | VAMP8 | vesicle-associated membrane prot | Plasma Membrane | transporter | 1.41E+07 | 1.76E+07 | 1.78E+07 | | | | | | |
| O70404 | VAMP8 | vesicle-associated membrane prot | Plasma Membrane | transporter | 5740452.5 | 486836.406 | 155596.562 | O70404 | VAMP8 | 13463484 | 10851875 | 3617292 | 9 |
| Q9WV55 | VAPA | VAMP (vesicle-associated membra | Plasma Membrane | other | 4.85E+07 | 5.03E+07 | 5.25E+07 | | | | | | |
| Q9WV55 | VAPA | VAMP (vesicle-associated membra | Plasma Membrane | other | 3.56E+07 | 2.91E+07 | 2.62E+07 | | | | | | |

| | | | | | | | | | | | | | |
|---|---|---|---|---|---|---|---|---|---|---|---|---|---|
| Q9WV55 | VAPA | VAMP (vesicle-associated membra | Plasma Membrane | other | 2.37E+07 | 2.59E+07 | 2.01E+07 | | | | | | |
| Q9WV55 | VAPA | VAMP (vesicle-associated membra | Plasma Membrane | other | 1.93E+07 | 1.23E+07 | 8408812 | | | | | | |
| Q9WV55 | VAPA | VAMP (vesicle-associated membra | Plasma Membrane | other | 9219136 | 7773788.5 | 6271269.5 | | | | | | |
| Q9WV55 | VAPA | VAMP (vesicle-associated membra | Plasma Membrane | other | 1.04E+07 | 1.13E+07 | 9890620 | | | | | | |
| Q9WV55 | VAPA | VAMP (vesicle-associated membra | Plasma Membrane | other | 8912581 | 1.00E+07 | 7875785.5 | | | | | | |
| Q9WV55 | VAPA | VAMP (vesicle-associated membra | Plasma Membrane | other | 6146902 | 4369893.5 | 1618806 | | | | | | |
| Q9WV55 | VAPA | VAMP (vesicle-associated membra | Plasma Membrane | other | 1816115.625 | 541541.938 | 426441.75 | Q9WV55 | VAPA | 18172748 | 15243706 | 2933653 | 27 |
| Q9QY76 | VAPB | VAMP (vesicle-associated membra | Plasma Membrane | other | 5904795.5 | 7369567 | 6052565.5 | | | | | | |
| Q9QY76 | VAPB | VAMP (vesicle-associated membra | Plasma Membrane | other | 1029484.562 | 2094076.75 | 1387463.375 | Q9QY76 | VAPB | 3467140 | 2773904 | 1132442 | 6 |
| Q3U2A8 | VARS2 | valyl-tRNA synthetase 2, mitochon | Cytoplasm | enzyme | 7537085 | 1.55E+07 | 1.27E+07 | Q3U2A8 | VARS2 | 7537085 | 4016308 | 2318816 | 3 |
| P70460 | VASP | vasodilator-stimulated phosphopr | Plasma Membrane | other | 5146602.5 | 7074794 | 3232512.25 | P70460 | VASP | 5146603 | 1921145 | 1109174 | 3 |
| Q01853 | VCP | valosin containing protein | Cytoplasm | enzyme | 1.62E+08 | 2.34E+08 | 1.46E+08 | | | | | | |
| Q01853 | VCP | valosin containing protein | Cytoplasm | enzyme | 2.20E+08 | 8.61E+07 | 1.84E+08 | | | | | | |
| Q01853 | VCP | valosin containing protein | Cytoplasm | enzyme | 1.82E+08 | 1.02E+08 | 9.13E+07 | | | | | | |
| Q01853 | VCP | valosin containing protein | Cytoplasm | enzyme | 1.24E+08 | 3.31E+08 | 2.65E+08 | | | | | | |
| Q01853 | VCP | valosin containing protein | Cytoplasm | enzyme | 1.47E+08 | 1.52E+08 | 1.78E+08 | | | | | | |
| Q01853 | VCP | valosin containing protein | Cytoplasm | enzyme | 1.25E+08 | 1.05E+08 | 6.53E+07 | | | | | | |
| Q01853 | VCP | valosin containing protein | Cytoplasm | enzyme | 1.10E+08 | 1.20E+08 | 1.21E+08 | | | | | | |
| Q01853 | VCP | valosin containing protein | Cytoplasm | enzyme | 1.05E+08 | 1.04E+08 | 9.81E+07 | | | | | | |
| Q01853 | VCP | valosin containing protein | Cytoplasm | enzyme | 8.82E+07 | 9.79E+07 | 9.47E+07 | | | | | | |
| Q01853 | VCP | valosin containing protein | Cytoplasm | enzyme | 9.27E+07 | 9.71E+07 | 7.30E+07 | | | | | | |
| Q01853 | VCP | valosin containing protein | Cytoplasm | enzyme | 7.75E+07 | 8.29E+07 | 7.71E+07 | | | | | | |
| Q01853 | VCP | valosin containing protein | Cytoplasm | enzyme | 7.19E+07 | 9.69E+07 | 1.02E+08 | | | | | | |
| Q01853 | VCP | valosin containing protein | Cytoplasm | enzyme | 7.77E+07 | 8.30E+07 | 7.98E+07 | | | | | | |
| Q01853 | VCP | valosin containing protein | Cytoplasm | enzyme | 6.18E+07 | 6.32E+07 | 6.17E+07 | | | | | | |
| Q01853 | VCP | valosin containing protein | Cytoplasm | enzyme | 5.30E+07 | 9.90E+07 | 7.87E+07 | | | | | | |
| Q01853 | VCP | valosin containing protein | Cytoplasm | enzyme | 7.54E+07 | 5.13E+07 | 5.64E+07 | | | | | | |
| Q01853 | VCP | valosin containing protein | Cytoplasm | enzyme | 6.48E+07 | 8.71E+07 | 6.61E+07 | | | | | | |
| Q01853 | VCP | valosin containing protein | Cytoplasm | enzyme | 4.46E+07 | 5.05E+07 | 5.73E+07 | | | | | | |
| Q01853 | VCP | valosin containing protein | Cytoplasm | enzyme | 5.70E+07 | 9.84E+07 | 9.92E+07 | | | | | | |
| Q01853 | VCP | valosin containing protein | Cytoplasm | enzyme | 6.35E+07 | 5.31E+07 | 4.77E+07 | | | | | | |
| Q01853 | VCP | valosin containing protein | Cytoplasm | enzyme | 3.56E+07 | 6.32E+07 | 7.22E+07 | | | | | | |
| Q01853 | VCP | valosin containing protein | Cytoplasm | enzyme | 3.23E+07 | 4.66E+07 | 4.72E+07 | | | | | | |
| Q01853 | VCP | valosin containing protein | Cytoplasm | enzyme | 4.31E+07 | 2.55E+07 | 3.85E+07 | | | | | | |
| Q01853 | VCP | valosin containing protein | Cytoplasm | enzyme | 3.39E+07 | 5.15E+07 | 4.60E+07 | | | | | | |
| Q01853 | VCP | valosin containing protein | Cytoplasm | enzyme | 2.68E+07 | 6.83E+07 | 7.95E+07 | | | | | | |
| Q01853 | VCP | valosin containing protein | Cytoplasm | enzyme | 4.44E+07 | 4.00E+07 | 4.27E+07 | | | | | | |
| Q01853 | VCP | valosin containing protein | Cytoplasm | enzyme | 3.39E+07 | 2.78E+07 | 1.17E+07 | | | | | | |
| Q01853 | VCP | valosin containing protein | Cytoplasm | enzyme | 2.91E+07 | 2.63E+07 | 2.58E+07 | | | | | | |
| Q01853 | VCP | valosin containing protein | Cytoplasm | enzyme | 2.64E+07 | 3.60E+07 | 3.47E+07 | | | | | | |
| Q01853 | VCP | valosin containing protein | Cytoplasm | enzyme | 1.83E+07 | 2.58E+07 | 1.41E+07 | | | | | | |
| Q01853 | VCP | valosin containing protein | Cytoplasm | enzyme | 2.36E+07 | 2.47E+07 | 2.19E+07 | | | | | | |
| Q01853 | VCP | valosin containing protein | Cytoplasm | enzyme | 3.59E+07 | 3.34E+07 | 3.81E+07 | | | | | | |
| Q01853 | VCP | valosin containing protein | Cytoplasm | enzyme | 1.63E+07 | 4.15E+07 | 2.71E+07 | | | | | | |
| Q01853 | VCP | valosin containing protein | Cytoplasm | enzyme | 2.91E+07 | 2.42E+07 | 2.94E+07 | | | | | | |
| Q01853 | VCP | valosin containing protein | Cytoplasm | enzyme | 2.78E+07 | 1.89E+07 | 2.24E+07 | | | | | | |
| Q01853 | VCP | valosin containing protein | Cytoplasm | enzyme | 2.14E+07 | 3.19E+07 | 2.18E+07 | | | | | | |
| Q01853 | VCP | valosin containing protein | Cytoplasm | enzyme | 2.33E+07 | 9311499 | 5587899 | | | | | | |
| Q01853 | VCP | valosin containing protein | Cytoplasm | enzyme | 2.73E+07 | 2.27E+07 | 2.86E+07 | | | | | | |
| Q01853 | VCP | valosin containing protein | Cytoplasm | enzyme | 2.59E+07 | 2.25E+07 | 2.34E+07 | | | | | | |
| Q01853 | VCP | valosin containing protein | Cytoplasm | enzyme | 3.62E+07 | 3.53E+07 | 4.20E+07 | | | | | | |
| Q01853 | VCP | valosin containing protein | Cytoplasm | enzyme | 1.97E+07 | 1.44E+07 | 1.48E+07 | | | | | | |
| Q01853 | VCP | valosin containing protein | Cytoplasm | enzyme | 8610500 | 4501745 | 1995439 | | | | | | |
| Q01853 | VCP | valosin containing protein | Cytoplasm | enzyme | 9088874 | 1.86E+07 | 1.30E+07 | | | | | | |
| Q01853 | VCP | valosin containing protein | Cytoplasm | enzyme | 5396296.5 | 1.26E+07 | 7118452 | | | | | | |
| Q01853 | VCP | valosin containing protein | Cytoplasm | enzyme | 7362785 | 295686.219 | 336850.188 | | | | | | |
| Q01853 | VCP | valosin containing protein | Cytoplasm | enzyme | 2.77E+07 | 1313963.125 | 1.03E+07 | | | | | | |
| Q01853 | VCP | valosin containing protein | Cytoplasm | enzyme | 6574344 | 7351961 | 7655417 | | | | | | |
| Q01853 | VCP | valosin containing protein | Cytoplasm | enzyme | 6006955.5 | 1.43E+07 | 1.10E+07 | | | | | | |
| Q01853 | VCP | valosin containing protein | Cytoplasm | enzyme | 9465608 | 1.45E+07 | 1.01E+07 | | | | | | |
| Q01853 | VCP | valosin containing protein | Cytoplasm | enzyme | 3776384.75 | 2.77E+07 | 1.62E+07 | | | | | | |
| Q01853 | VCP | valosin containing protein | Cytoplasm | enzyme | 6411835 | 5679518 | 4839540 | | | | | | |
| Q01853 | VCP | valosin containing protein | Cytoplasm | enzyme | 1.05E+07 | 5707913 | 8988795 | | | | | | |
| Q01853 | VCP | valosin containing protein | Cytoplasm | enzyme | 6235445.5 | 7653459.5 | 7286758.5 | | | | | | |
| Q01853 | VCP | valosin containing protein | Cytoplasm | enzyme | 4995929.5 | 1.25E+07 | 9056257 | | | | | | |
| Q01853 | VCP | valosin containing protein | Cytoplasm | enzyme | 9481953 | 1636527.875 | 1870421.5 | | | | | | |
| Q01853 | VCP | valosin containing protein | Cytoplasm | enzyme | 3668917.25 | 5125811.5 | 4030345.75 | | | | | | |
| Q01853 | VCP | valosin containing protein | Cytoplasm | enzyme | 1985749 | 3876262.5 | 2280329.25 | | | | | | |
| Q01853 | VCP | valosin containing protein | Cytoplasm | enzyme | 9827885 | 369601.938 | 1127649.625 | | | | | | |
| Q01853 | VCP | valosin containing protein | Cytoplasm | enzyme | 1415847.375 | 3257207.25 | 3953631.25 | | | | | | |
| Q01853 | VCP | valosin containing protein | Cytoplasm | enzyme | 2056871.75 | 3883250 | 5031962.5 | | | | | | |
| Q01853 | VCP | valosin containing protein | Cytoplasm | enzyme | 2457288 | 2089192.25 | 3971899 | | | | | | |
| Q01853 | VCP | valosin containing protein | Cytoplasm | enzyme | 1364690 | 1272494.875 | 1182293.625 | | | | | | |
| Q01853 | VCP | valosin containing protein | Cytoplasm | enzyme | 1267669 | 1434735.625 | 1865216.75 | | | | | | |
| Q01853 | VCP | valosin containing protein | Cytoplasm | enzyme | 5462218 | 96626.641 | 1097085.375 | | | | | | |
| Q01853 | VCP | valosin containing protein | Cytoplasm | enzyme | 2297836.75 | 1284696.375 | 2828764.5 | Q01853 | VCP | 42576337 | 51950283 | 3720237 | 195 |
| Q60932 | VDAC1 | voltage-dependent anion channel | Cytoplasm | ion channel | 3.28E+07 | 3.13E+07 | 3.54E+07 | | | | | | |
| Q60932 | VDAC1 | voltage-dependent anion channel | Cytoplasm | ion channel | 1.35E+07 | 2.06E+07 | 1.60E+07 | | | | | | |
| Q60932 | VDAC1 | voltage-dependent anion channel | Cytoplasm | ion channel | 1.59E+07 | 2.95E+07 | 2.99E+07 | | | | | | |
| Q60932 | VDAC1 | voltage-dependent anion channel | Cytoplasm | ion channel | 1.76E+07 | 1.17E+07 | 1.88E+07 | | | | | | |
| Q60932 | VDAC1 | voltage-dependent anion channel | Cytoplasm | ion channel | 2.01E+07 | 2.34E+07 | 3.13E+07 | | | | | | |
| Q60932 | VDAC1 | voltage-dependent anion channel | Cytoplasm | ion channel | 1.30E+07 | 1.99E+07 | 1.65E+07 | | | | | | |
| Q60932 | VDAC1 | voltage-dependent anion channel | Cytoplasm | ion channel | 1.53E+07 | 2.04E+07 | 2.38E+07 | | | | | | |
| Q60932 | VDAC1 | voltage-dependent anion channel | Cytoplasm | ion channel | 1.35E+07 | 1791976.875 | 2112458 | Q60932 | VDAC1 | 17721250 | 8843355 | 1805142 | 24 |
| Q60930 | VDAC2 | voltage-dependent anion channel | Cytoplasm | ion channel | 2.63E+07 | 3.36E+07 | 3.93E+07 | | | | | | |
| Q60930 | VDAC2 | voltage-dependent anion channel | Cytoplasm | ion channel | 3.05E+07 | 2.55E+07 | 3.88E+07 | | | | | | |

| | | | | | | | | | | | | | |
|---|---|---|---|---|---|---|---|---|---|---|---|---|---|
| Q60930 | VDAC2 | voltage-dependent anion channel | Cytoplasm | ion channel | 2.03E+07 | 2.81E+07 | 3.03E+07 | | | | | | |
| Q60930 | VDAC2 | voltage-dependent anion channel | Cytoplasm | ion channel | 1.91E+07 | 2.40E+07 | 1.81E+07 | | | | | | |
| Q60930 | VDAC2 | voltage-dependent anion channel | Cytoplasm | ion channel | 1.62E+07 | 2.01E+07 | 1.40E+07 | | | | | | |
| Q60930 | VDAC2 | voltage-dependent anion channel | Cytoplasm | ion channel | 5569595 | 8719531 | 6572396.5 | | | | | | |
| Q60930 | VDAC2 | voltage-dependent anion channel | Cytoplasm | ion channel | 3090408.75 | 6736401.5 | 9576551 | | | | | | |
| Q60930 | VDAC2 | voltage-dependent anion channel | Cytoplasm | ion channel | 1827876.625 | 1728839.375 | 1826136.5 | | | | | | |
| Q60930 | VDAC2 | voltage-dependent anion channel | Cytoplasm | ion channel | 3071977.75 | 2433012.5 | 3938987 | Q60930 | VDAC2 | 13991095 | 12223509 | 2352415 | 27 |
| Q60931 | VDAC3 | voltage-dependent anion channel | Cytoplasm | ion channel | 3.59E+07 | 3.78E+07 | 4.67E+07 | | | | | | |
| Q60931 | VDAC3 | voltage-dependent anion channel | Cytoplasm | ion channel | 2.38E+07 | 3.29E+07 | 4.25E+07 | | | | | | |
| Q60931 | VDAC3 | voltage-dependent anion channel | Cytoplasm | ion channel | 2.06E+07 | 2.38E+07 | 2.88E+07 | | | | | | |
| Q60931 | VDAC3 | voltage-dependent anion channel | Cytoplasm | ion channel | 1.55E+07 | 1.95E+07 | 1.88E+07 | | | | | | |
| Q60931 | VDAC3 | voltage-dependent anion channel | Cytoplasm | ion channel | 1.37E+07 | 6484018 | 8523336 | | | | | | |
| Q60931 | VDAC3 | voltage-dependent anion channel | Cytoplasm | ion channel | 4045302.25 | 3795755.75 | 5327977 | Q60931 | VDAC3 | 18905884 | 13560352 | 3196206 | 18 |
| P20152 | VIM | vimentin | Cytoplasm | other | 1859551.25 | 762751 | 710586 | P20152 | VIM | 1859551 | 648821 | 374597 | 3 |
| Q8BGQ1 | VIPAS39 | VPS33B interacting protein, apical | Plasma Membrane | other | 2.81E+07 | 2.03E+07 | 2.25E+07 | Q8BGQ1 | VIPAS39 | 28060000 | 4017474 | 2319490 | 3 |
| Q99KU0 | VMP1 | vacuole membrane protein 1 | Plasma Membrane | other | 1.94E+07 | 2.82E+07 | 1.92E+07 | Q99KU0 | VMP1 | 19390000 | 5125059 | 2958954 | 3 |
| Q5H8C4 | VPS13A | vacuolar protein sorting 13 homol | Cytoplasm | transporter | 1.83E+07 | 1.65E+07 | 1.68E+07 | | | | | | |
| Q5H8C4 | VPS13A | vacuolar protein sorting 13 homol | Cytoplasm | transporter | 1.26E+07 | 1.77E+07 | 1.29E+07 | | | | | | |
| Q5H8C4 | VPS13A | vacuolar protein sorting 13 homol | Cytoplasm | transporter | 1.45E+07 | 1.18E+07 | 1.34E+07 | | | | | | |
| Q5H8C4 | VPS13A | vacuolar protein sorting 13 homol | Cytoplasm | transporter | 8766821 | 1.37E+07 | 1.45E+07 | | | | | | |
| Q5H8C4 | VPS13A | vacuolar protein sorting 13 homol | Cytoplasm | transporter | 1.09E+07 | 9176119 | 1.26E+07 | | | | | | |
| Q5H8C4 | VPS13A | vacuolar protein sorting 13 homol | Cytoplasm | transporter | 9617925 | 9585526 | 9897921 | | | | | | |
| Q5H8C4 | VPS13A | vacuolar protein sorting 13 homol | Cytoplasm | transporter | 5596375.5 | 7052661.5 | 7170201.5 | | | | | | |
| Q5H8C4 | VPS13A | vacuolar protein sorting 13 homol | Cytoplasm | transporter | 6932293.5 | 8079549.5 | 5755338 | | | | | | |
| Q5H8C4 | VPS13A | vacuolar protein sorting 13 homol | Cytoplasm | transporter | 6411636 | 9010831 | 7264892 | | | | | | |
| Q5H8C4 | VPS13A | vacuolar protein sorting 13 homol | Cytoplasm | transporter | 7643641.5 | 8795245 | 5896184 | | | | | | |
| Q5H8C4 | VPS13A | vacuolar protein sorting 13 homol | Cytoplasm | transporter | 8393431 | 9156771 | 8128851 | | | | | | |
| Q5H8C4 | VPS13A | vacuolar protein sorting 13 homol | Cytoplasm | transporter | 5267244.5 | 6460884 | 4709281.5 | | | | | | |
| Q5H8C4 | VPS13A | vacuolar protein sorting 13 homol | Cytoplasm | transporter | 6969731 | 6390089.5 | 7255862.5 | | | | | | |
| Q5H8C4 | VPS13A | vacuolar protein sorting 13 homol | Cytoplasm | transporter | 7219857 | 7918259 | 6618615 | | | | | | |
| Q5H8C4 | VPS13A | vacuolar protein sorting 13 homol | Cytoplasm | transporter | 6414255 | 6240960.5 | 6175033.5 | | | | | | |
| Q5H8C4 | VPS13A | vacuolar protein sorting 13 homol | Cytoplasm | transporter | 5751130 | 5448043.5 | 5735705.5 | | | | | | |
| Q5H8C4 | VPS13A | vacuolar protein sorting 13 homol | Cytoplasm | transporter | 7296073.5 | 5626958.5 | 6341465.5 | | | | | | |
| Q5H8C4 | VPS13A | vacuolar protein sorting 13 homol | Cytoplasm | transporter | 5588232.5 | 6523051.5 | 5898657.5 | | | | | | |
| Q5H8C4 | VPS13A | vacuolar protein sorting 13 homol | Cytoplasm | transporter | 4331642 | 3948054.25 | 3087461.5 | | | | | | |
| Q5H8C4 | VPS13A | vacuolar protein sorting 13 homol | Cytoplasm | transporter | 2633042 | 5178498.5 | 3886577.75 | | | | | | |
| Q5H8C4 | VPS13A | vacuolar protein sorting 13 homol | Cytoplasm | transporter | 4437164 | 5558687 | 5106101 | | | | | | |
| Q5H8C4 | VPS13A | vacuolar protein sorting 13 homol | Cytoplasm | transporter | 3825234.5 | 2413540.25 | 3870223.5 | | | | | | |
| Q5H8C4 | VPS13A | vacuolar protein sorting 13 homol | Cytoplasm | transporter | 4537280.5 | 5155175 | 5364406.5 | | | | | | |
| Q5H8C4 | VPS13A | vacuolar protein sorting 13 homol | Cytoplasm | transporter | 4346838 | 4305723 | 5300886.5 | | | | | | |
| Q5H8C4 | VPS13A | vacuolar protein sorting 13 homol | Cytoplasm | transporter | 3088190.75 | 3285209.25 | 2914310.5 | | | | | | |
| Q5H8C4 | VPS13A | vacuolar protein sorting 13 homol | Cytoplasm | transporter | 2979815 | 3047347.25 | 2250325.75 | | | | | | |
| Q5H8C4 | VPS13A | vacuolar protein sorting 13 homol | Cytoplasm | transporter | 5619101 | 3829455.75 | 4621955.5 | | | | | | |
| Q5H8C4 | VPS13A | vacuolar protein sorting 13 homol | Cytoplasm | transporter | 2523652.25 | 2607127.25 | 2367848.75 | | | | | | |
| Q5H8C4 | VPS13A | vacuolar protein sorting 13 homol | Cytoplasm | transporter | 1424660.75 | 2413760.25 | 1680869.125 | | | | | | |
| Q5H8C4 | VPS13A | vacuolar protein sorting 13 homol | Cytoplasm | transporter | 1915172.75 | 2111961 | 2333621.75 | | | | | | |
| Q5H8C4 | VPS13A | vacuolar protein sorting 13 homol | Cytoplasm | transporter | 1420702.375 | 2148810 | 1699467.5 | | | | | | |
| Q5H8C4 | VPS13A | vacuolar protein sorting 13 homol | Cytoplasm | transporter | 1654327.625 | 1980393.25 | 1579004.25 | | | | | | |
| Q5H8C4 | VPS13A | vacuolar protein sorting 13 homol | Cytoplasm | transporter | 1315632.75 | 1180566.375 | 1239527.375 | | | | | | |
| Q5H8C4 | VPS13A | vacuolar protein sorting 13 homol | Cytoplasm | transporter | 2926314.25 | 4673815.5 | 4188301.75 | Q5H8C4 | VPS13A | 5976395 | 3907874 | 386937 | 102 |
| Q8BX70 | VPS13C | vacuolar protein sorting 13 homol | Cytoplasm | other | 2673860 | 2245489.25 | 1747553.625 | | | | | | |
| Q8BX70 | VPS13C | vacuolar protein sorting 13 homol | Cytoplasm | other | 2705862.5 | 2076721.5 | 1600244.375 | | | | | | |
| Q8BX70 | VPS13C | vacuolar protein sorting 13 homol | Cytoplasm | other | 2003213.625 | 3012537.25 | 2811630.75 | | | | | | |
| Q8BX70 | VPS13C | vacuolar protein sorting 13 homol | Cytoplasm | other | 1756132 | 1712218.625 | 1306574.375 | | | | | | |
| Q8BX70 | VPS13C | vacuolar protein sorting 13 homol | Cytoplasm | other | 1350604 | 1278752.25 | 831528.625 | Q8BX70 | VPS13C | 2097934 | 643867 | 166246 | 15 |
| Q9D1C8 | VPS28 | vacuolar protein sorting 28 homol | Cytoplasm | transporter | 7095614.5 | 8643999 | 9131434 | Q9D1C8 | VPS28 | 7095615 | 1062987 | 613716 | 3 |
| Q8CIZ8 | VWF | von Willebrand factor | Extracellular Space | other | 9427182 | 6690513 | 6418020 | | | | | | |
| Q8CIZ8 | VWF | von Willebrand factor | Extracellular Space | other | 2371522.5 | 2259730.25 | 2609104.5 | | | | | | |
| Q8CIZ8 | VWF | von Willebrand factor | Extracellular Space | other | 3745615.25 | 2424603 | 3376919.75 | | | | | | |
| Q8CIZ8 | VWF | von Willebrand factor | Extracellular Space | other | 4274430.5 | 3764153 | 3471479.5 | | | | | | |
| Q8CIZ8 | VWF | von Willebrand factor | Extracellular Space | other | 2160840.75 | 1028414.062 | 1886386.75 | | | | | | |
| Q8CIZ8 | VWF | von Willebrand factor | Extracellular Space | other | 2613705 | 1122567.75 | 1685314.625 | | | | | | |
| Q8CIZ8 | VWF | von Willebrand factor | Extracellular Space | other | 2725630.75 | 3053832.75 | 3393580.75 | | | | | | |
| Q8CIZ8 | VWF | von Willebrand factor | Extracellular Space | other | 3979563.75 | 2740490 | 3420587 | | | | | | |
| Q8CIZ8 | VWF | von Willebrand factor | Extracellular Space | other | 7395765.5 | 5016735 | 5854056 | | | | | | |
| Q8CIZ8 | VWF | von Willebrand factor | Extracellular Space | other | 3449235.25 | 3448321.5 | 2886721.75 | | | | | | |
| Q8CIZ8 | VWF | von Willebrand factor | Extracellular Space | other | 4180237.25 | 3629294.5 | 3673055.25 | | | | | | |
| Q8CIZ8 | VWF | von Willebrand factor | Extracellular Space | other | 2056399.375 | 1643389.25 | 1494115 | | | | | | |
| Q8CIZ8 | VWF | von Willebrand factor | Extracellular Space | other | 2063656.5 | 2013802.125 | 2040501.25 | | | | | | |
| Q8CIZ8 | VWF | von Willebrand factor | Extracellular Space | other | 2120355.25 | 1835231.875 | 2187728.5 | | | | | | |
| Q8CIZ8 | VWF | von Willebrand factor | Extracellular Space | other | 1045746.188 | 938092.75 | 907883.375 | Q8CIZ8 | VWF | 3573992 | 1786048 | 266248 | 45 |
| Q8CBE3 | WDR37 | WD repeat domain 37 | Nucleus | other | 3675106.75 | 2076966.125 | 2259966.5 | | | | | | |
| Q8CBE3 | WDR37 | WD repeat domain 37 | Nucleus | other | 1142177 | 891901.875 | 845442.25 | Q8CBE3 | WDR37 | 2408642 | 1092837 | 446149 | 6 |
| Q8BH57 | WDR48 | WD repeat domain 48 | Cytoplasm | other | 2029056.375 | 1736782.625 | 556300.812 | Q8BH57 | WDR48 | 2029056 | 779740 | 450183 | 3 |
| Q4KUS1 | WFDC8 | WAP four-disulfide core domain 8 | Extracellular Space | other | 923766.062 | 880746.312 | 434907.344 | Q4KUS1 | WFDC8 | 923766 | 270680 | 156277 | 3 |
| P83741 | WNK1 | WNK lysine deficient protein kinas | Cytoplasm | kinase | 1285073.375 | 1874160.75 | 1117084.75 | P83741 | WNK1 | 1285073 | 397578 | 229542 | 3 |
| Q9QXY7 | XK | X-linked Kx blood group | Plasma Membrane | transporter | 2.91E+07 | 2.61E+07 | 2.33E+07 | | | | | | |
| Q9QXY7 | XK | X-linked Kx blood group | Plasma Membrane | transporter | 1.72E+07 | 2.43E+07 | 2.68E+07 | | | | | | |
| Q9QXY7 | XK | X-linked Kx blood group | Plasma Membrane | transporter | 1.87E+07 | 2.67E+07 | 2.96E+07 | | | | | | |
| Q9QXY7 | XK | X-linked Kx blood group | Plasma Membrane | transporter | 1.81E+07 | 1.68E+07 | 1.55E+07 | Q9QXY7 | XK | 20780000 | 5115138 | 1476613 | 12 |
| Q6P5F9 | XPO1 | exportin 1 | Nucleus | transporter | 2664938.5 | 1175208.625 | 1769888.125 | Q6P5F9 | XPO1 | 2664939 | 749895 | 432952 | 3 |
| Q9EPK7 | XPO7 | exportin 7 | Nucleus | transporter | 2766321 | 2531358.5 | 1469543.75 | | | | | | |
| Q9EPK7 | XPO7 | exportin 7 | Nucleus | transporter | 2591624 | 1764742.25 | 863732.875 | Q9EPK7 | XPO7 | 2678973 | 754641 | 308081 | 6 |
| P62960 | YBX1 | Y box binding protein 1 | Nucleus | transcription regulator | 5283270 | 4504480.5 | 2708131.75 | | | | | | |
| P62960 | YBX1 | Y box binding protein 1 | Nucleus | transcription regulator | 597584.062 | 329062.625 | 262983.25 | P62960 | YBX1 | 2940427 | 2229631 | 910243 | 6 |
| Q9JKB3 | YBX3 | Y box binding protein 3 | Nucleus | transcription regulator | 2192187.75 | 3081788.5 | 1994655.125 | | | | | | |

| | | | | | | | | | | | | | |
|---|---|---|---|---|---|---|---|---|---|---|---|---|---|
| Q9JKB3 | YBX3 | Y box binding protein 3 | Nucleus | transcription regulator | 2431539.75 | 3028406.5 | 1883388 | Q9JKB3 | YBX3 | 2311864 | 515289 | 210366 | 6 |
| Q9CX30 | YIF1B | Yip1 interacting factor homolog B | Cytoplasm | other | 2851247.5 | 3619108.75 | 3540371.5 | | | | | | |
| Q9CX30 | YIF1B | Yip1 interacting factor homolog B | Cytoplasm | other | 2616878.75 | 3872693.75 | 2917321 | Q9CX30 | YIF1B | 2734063 | 505509 | 206373 | 6 |
| Q9CQV8 | YWHAB | tyrosine 3-monooxygenase/trypto | Cytoplasm | transcription regulator | 6221014 | 8336678 | 7970062.5 | | | | | | |
| Q9CQV8 | YWHAB | tyrosine 3-monooxygenase/trypto | Cytoplasm | transcription regulator | 6648454 | 7570416.5 | 7080696 | | | | | | |
| Q9CQV8 | YWHAB | tyrosine 3-monooxygenase/trypto | Cytoplasm | transcription regulator | 3492179.75 | 4875441.5 | 5386640 | | | | | | |
| Q9CQV8 | YWHAB | tyrosine 3-monooxygenase/trypto | Cytoplasm | transcription regulator | 3539425.25 | 3562383.25 | 3620160.75 | | | | | | |
| Q9CQV8 | YWHAB | tyrosine 3-monooxygenase/trypto | Cytoplasm | transcription regulator | 623969.812 | 1232228.625 | 736570.375 | | | | | | |
| Q9CQV8 | YWHAB | tyrosine 3-monooxygenase/trypto | Cytoplasm | transcription regulator | 4.70E-06 | 52864.539 | 1191035.75 | Q9CQV8 | YWHAB | 3420840 | 2884299 | 679836 | 18 |
| P62259 | YWHAE | tyrosine 3-monooxygenase/trypto | Cytoplasm | other | 1.89E+07 | 1.60E+07 | 1.77E+07 | | | | | | |
| P62259 | YWHAE | tyrosine 3-monooxygenase/trypto | Cytoplasm | other | 1.05E+07 | 1.25E+07 | 1.27E+07 | | | | | | |
| P62259 | YWHAE | tyrosine 3-monooxygenase/trypto | Cytoplasm | other | 8962127 | 1.10E+07 | 9405761 | | | | | | |
| P62259 | YWHAE | tyrosine 3-monooxygenase/trypto | Cytoplasm | other | 8875671 | 9102966 | 9534713 | | | | | | |
| P62259 | YWHAE | tyrosine 3-monooxygenase/trypto | Cytoplasm | other | 8297393 | 9691992 | 8606862 | | | | | | |
| P62259 | YWHAE | tyrosine 3-monooxygenase/trypto | Cytoplasm | other | 5002355.5 | 5323812 | 5976546 | | | | | | |
| P62259 | YWHAE | tyrosine 3-monooxygenase/trypto | Cytoplasm | other | 5759994 | 7746587.5 | 8001478.5 | | | | | | |
| P62259 | YWHAE | tyrosine 3-monooxygenase/trypto | Cytoplasm | other | 8160180 | 6295526.5 | 5882770.5 | | | | | | |
| P62259 | YWHAE | tyrosine 3-monooxygenase/trypto | Cytoplasm | other | 1362944.625 | 4363717.5 | 3864790 | P62259 | YWHAE | 8422296 | 4078796 | 784965 | 27 |
| P61982 | YWHAG | tyrosine 3-monooxygenase/trypto | Cytoplasm | other | 4243353 | 6259881 | 5317536.5 | | | | | | |
| P61982 | YWHAG | tyrosine 3-monooxygenase/trypto | Cytoplasm | other | 2704403 | 3184908.75 | 2992044 | P61982 | YWHAG | 3473878 | 1426811 | 582493 | 6 |
| P68510 | YWHAH | tyrosine 3-monooxygenase/trypto | Cytoplasm | transcription regulator | 6.23E+07 | 4.49E+07 | 5.54E+07 | | | | | | |
| P68510 | YWHAH | tyrosine 3-monooxygenase/trypto | Cytoplasm | transcription regulator | 2.11E+07 | 1.92E+07 | 2.06E+07 | | | | | | |
| P68510 | YWHAH | tyrosine 3-monooxygenase/trypto | Cytoplasm | transcription regulator | 1.97E+07 | 2.44E+07 | 2.69E+07 | | | | | | |
| P68510 | YWHAH | tyrosine 3-monooxygenase/trypto | Cytoplasm | transcription regulator | 1.49E+07 | 1.69E+07 | 1.61E+07 | | | | | | |
| P68510 | YWHAH | tyrosine 3-monooxygenase/trypto | Cytoplasm | transcription regulator | 1.32E+07 | 5151929 | 9129705 | | | | | | |
| P68510 | YWHAH | tyrosine 3-monooxygenase/trypto | Cytoplasm | transcription regulator | 3262553.5 | 1.06E+07 | 6557144.5 | | | | | | |
| P68510 | YWHAH | tyrosine 3-monooxygenase/trypto | Cytoplasm | transcription regulator | 9043160 | 7344018 | 1.02E+07 | | | | | | |
| P68510 | YWHAH | tyrosine 3-monooxygenase/trypto | Cytoplasm | transcription regulator | 6352308.5 | 6566896 | 6462912.5 | | | | | | |
| P68510 | YWHAH | tyrosine 3-monooxygenase/trypto | Cytoplasm | transcription regulator | 2638784.5 | 1140812 | 1425427.375 | P68510 | YWHAH | 16957423 | 15553472 | 2993267 | 27 |
| P68254 | YWHAQ | tyrosine 3-monooxygenase/trypto | Cytoplasm | other | 4486239 | 5746264.5 | 6099251.5 | | | | | | |
| P68254 | YWHAQ | tyrosine 3-monooxygenase/trypto | Cytoplasm | other | 5583782 | 5826557.5 | 4413490.5 | | | | | | |
| P68254 | YWHAQ | tyrosine 3-monooxygenase/trypto | Cytoplasm | other | 3152432.5 | 3706236.5 | 3284773.25 | P68254 | YWHAQ | 4407485 | 1151952 | 383984 | 9 |
| P63101 | YWHAZ | tyrosine 3-monooxygenase/trypto | Cytoplasm | enzyme | 6.92E+07 | 7.83E+07 | 8.95E+07 | | | | | | |
| P63101 | YWHAZ | tyrosine 3-monooxygenase/trypto | Cytoplasm | enzyme | 4.39E+07 | 6.70E+07 | 6.43E+07 | | | | | | |
| P63101 | YWHAZ | tyrosine 3-monooxygenase/trypto | Cytoplasm | enzyme | 5.19E+07 | 5.73E+07 | 5.11E+07 | | | | | | |
| P63101 | YWHAZ | tyrosine 3-monooxygenase/trypto | Cytoplasm | enzyme | 6.42E+07 | 4.02E+07 | 5.83E+07 | | | | | | |
| P63101 | YWHAZ | tyrosine 3-monooxygenase/trypto | Cytoplasm | enzyme | 4.99E+07 | 6.16E+07 | 6.43E+07 | | | | | | |
| P63101 | YWHAZ | tyrosine 3-monooxygenase/trypto | Cytoplasm | enzyme | 3.96E+07 | 3.82E+07 | 4.56E+07 | | | | | | |
| P63101 | YWHAZ | tyrosine 3-monooxygenase/trypto | Cytoplasm | enzyme | 5.30E+07 | 6.17E+07 | 5.85E+07 | | | | | | |
| P63101 | YWHAZ | tyrosine 3-monooxygenase/trypto | Cytoplasm | enzyme | 3.36E+07 | 3.67E+07 | 4.33E+07 | | | | | | |
| P63101 | YWHAZ | tyrosine 3-monooxygenase/trypto | Cytoplasm | enzyme | 2.57E+07 | 1.22E+07 | 2.23E+07 | | | | | | |
| P63101 | YWHAZ | tyrosine 3-monooxygenase/trypto | Cytoplasm | enzyme | 4603827.5 | 7333357.5 | 1.00E+07 | | | | | | |
| P63101 | YWHAZ | tyrosine 3-monooxygenase/trypto | Cytoplasm | enzyme | 2.03E+07 | 2980735.25 | 4326192.5 | | | | | | |
| P63101 | YWHAZ | tyrosine 3-monooxygenase/trypto | Cytoplasm | enzyme | 9139888 | 1.37E+07 | 1.41E+07 | | | | | | |
| P63101 | YWHAZ | tyrosine 3-monooxygenase/trypto | Cytoplasm | enzyme | 6861663 | 4826110.5 | 6810950 | | | | | | |
| P63101 | YWHAZ | tyrosine 3-monooxygenase/trypto | Cytoplasm | enzyme | 6305529.5 | 1.88E+07 | 1.35E+07 | | | | | | |
| P63101 | YWHAZ | tyrosine 3-monooxygenase/trypto | Cytoplasm | enzyme | 5740513 | 2414505 | 2950649.25 | | | | | | |
| P63101 | YWHAZ | tyrosine 3-monooxygenase/trypto | Cytoplasm | enzyme | 3503581.75 | 648070.125 | 471976.875 | P63101 | YWHAZ | 30460313 | 25604834 | 3695739 | 48 |
| Q9D0B1 | ZNF524 | zinc finger protein 524 | Nucleus | other | 2337472.25 | 9636721 | 6622503 | Q9D0B1 | ZNF524 | 2337472 | 3668016 | 2117730 | 3 |
| Q02525 | ZNF717 | zinc finger protein 717 | Nucleus | other | 3.01E+08 | 2.73E+08 | 2.57E+08 | Q02525 | ZNF717 | 300500000 | 22199625 | 12816959 | 3 |
| Q5SSH7 | ZZEF1 | zinc finger, ZZ-type with EF-hand d | Cytoplasm | other | 1348374.75 | 2750872.5 | 1241030.375 | Q5SSH7 | ZZEF1 | 1348375 | 842432 | 486378 | 3 |

| # Peptides by Location | |
|---|---|
| Plasma Membrane | 1902 |
| Cytoplasm | 4392 |
| Nucleus | 523 |
| Extracellular Space | 347 |
| Other | 24 |
| **Total Peptides** | **7188** |

### RhAG knockout

| ID | Symbol | Entrez Gene Name | Primary Location | Type(s) | Intensity [RhAGKO_1] | Intensity [RhAGKO_2] | Intensity [RhAGKO_3] | ID | Symbol | Mean Intensity | St. Dev | Std. Err | n |
|---|---|---|---|---|---|---|---|---|---|---|---|---|---|
| Q3UHJ0 | AAK1 | AP2 associated kinase 1 | Cytoplasm | kinase | 1101591.125 | 872663.438 | 758731.375 | Q3UHJ0 | AAK1 | 1101591 | 174614 | 100814 | 3 |
| Q9JI39 | ABCB10 | ATP-binding cassette, sub-family B (MDR/TAP), member 10 | Cytoplasm | transporter | 1.07E+07 | 1.68E+07 | 1.81E+07 | | | | | | |
| Q9JI39 | ABCB10 | ATP-binding cassette, sub-family B (MDR/TAP), member 10 | Cytoplasm | transporter | 9621738 | 1.44E+07 | 1.55E+07 | | | | | | |
| Q9JI39 | ABCB10 | ATP-binding cassette, sub-family B (MDR/TAP), member 10 | Cytoplasm | transporter | 3691963.75 | 7599960.5 | 7532876 | | | | | | |
| Q9JI39 | ABCB10 | ATP-binding cassette, sub-family B (MDR/TAP), member 10 | Cytoplasm | transporter | 2215662.5 | 2250750.5 | 2641814.5 | | | | | | |
| Q9JI39 | ABCB10 | ATP-binding cassette, sub-family B (MDR/TAP), member 10 | Cytoplasm | transporter | 2719897.25 | 5039134.5 | 5116280 | | | | | | |
| Q9JI39 | ABCB10 | ATP-binding cassette, sub-family B (MDR/TAP), member 10 | Cytoplasm | transporter | 2394133.25 | 5452794.5 | 5765157 | | | | | | |
| Q9JI39 | ABCB10 | ATP-binding cassette, sub-family B (MDR/TAP), member 10 | Cytoplasm | transporter | 2608369.75 | 4341638.5 | 5416581.5 | | | | | | |
| Q9JI39 | ABCB10 | ATP-binding cassette, sub-family B (MDR/TAP), member 10 | Cytoplasm | transporter | 1847704.125 | 4225914 | 4726169 | | | | | | |
| Q9JI39 | ABCB10 | ATP-binding cassette, sub-family B (MDR/TAP), member 10 | Cytoplasm | transporter | 1909124.125 | 2744847.5 | 3237393 | | | | | | |
| Q9JI39 | ABCB10 | ATP-binding cassette, sub-family B (MDR/TAP), member 10 | Cytoplasm | transporter | 2761271.5 | 5047171 | 5684115.5 | | | | | | |
| Q9JI39 | ABCB10 | ATP-binding cassette, sub-family B (MDR/TAP), member 10 | Cytoplasm | transporter | 1345937.25 | 1636913.375 | 2260400.5 | | | | | | |
| Q9JI39 | ABCB10 | ATP-binding cassette, sub-family B (MDR/TAP), member 10 | Cytoplasm | transporter | 725046.5 | 1078015.25 | 1292958.125 | Q9JI39 | ABCB10 | 3540904 | 4564730 | 760788 | 36 |
| P21440 | ABCB4 | ATP-binding cassette, sub-family B (MDR/TAP), member 4 | Plasma Membrane | transporter | 2.90E+07 | 2.57E+07 | 2.42E+07 | | | | | | |
| P21440 | ABCB4 | ATP-binding cassette, sub-family B (MDR/TAP), member 4 | Plasma Membrane | transporter | 1.57E+07 | 1.24E+07 | 1.07E+07 | | | | | | |
| P21440 | ABCB4 | ATP-binding cassette, sub-family B (MDR/TAP), member 4 | Plasma Membrane | transporter | 1.55E+07 | 1.80E+07 | 1.45E+07 | | | | | | |
| P21440 | ABCB4 | ATP-binding cassette, sub-family B (MDR/TAP), member 4 | Plasma Membrane | transporter | 1.28E+07 | 1.40E+07 | 1.20E+07 | | | | | | |
| P21440 | ABCB4 | ATP-binding cassette, sub-family B (MDR/TAP), member 4 | Plasma Membrane | transporter | 1.47E+07 | 1.47E+07 | 1.16E+07 | | | | | | |
| P21440 | ABCB4 | ATP-binding cassette, sub-family B (MDR/TAP), member 4 | Plasma Membrane | transporter | 1.19E+07 | 1.41E+07 | 1.25E+07 | | | | | | |
| P21440 | ABCB4 | ATP-binding cassette, sub-family B (MDR/TAP), member 4 | Plasma Membrane | transporter | 1.87E+07 | 1.52E+07 | 1.54E+07 | | | | | | |
| P21440 | ABCB4 | ATP-binding cassette, sub-family B (MDR/TAP), member 4 | Plasma Membrane | transporter | 1.26E+07 | 1.53E+07 | 6689358 | | | | | | |
| P21440 | ABCB4 | ATP-binding cassette, sub-family B (MDR/TAP), member 4 | Plasma Membrane | transporter | 1.17E+07 | 1.09E+07 | 1.03E+07 | | | | | | |
| P21440 | ABCB4 | ATP-binding cassette, sub-family B (MDR/TAP), member 4 | Plasma Membrane | transporter | 1.14E+07 | 8664280 | 7927802.5 | | | | | | |
| P21440 | ABCB4 | ATP-binding cassette, sub-family B (MDR/TAP), member 4 | Plasma Membrane | transporter | 9868161 | 8502768 | 8821640 | | | | | | |
| P21440 | ABCB4 | ATP-binding cassette, sub-family B (MDR/TAP), member 4 | Plasma Membrane | transporter | 1.05E+07 | 7459775.5 | 7902644.5 | | | | | | |
| P21440 | ABCB4 | ATP-binding cassette, sub-family B (MDR/TAP), member 4 | Plasma Membrane | transporter | 6200758.5 | 6147119 | 5345316.5 | | | | | | |
| P21440 | ABCB4 | ATP-binding cassette, sub-family B (MDR/TAP), member 4 | Plasma Membrane | transporter | 6999255.5 | 6248495 | 5500906 | | | | | | |
| P21440 | ABCB4 | ATP-binding cassette, sub-family B (MDR/TAP), member 4 | Plasma Membrane | transporter | 6202433 | 9398491 | 7333127 | | | | | | |
| P21440 | ABCB4 | ATP-binding cassette, sub-family B (MDR/TAP), member 4 | Plasma Membrane | transporter | 7034590.5 | 6801267.5 | 6082633 | | | | | | |
| P21440 | ABCB4 | ATP-binding cassette, sub-family B (MDR/TAP), member 4 | Plasma Membrane | transporter | 4690289.5 | 5250238 | 4308785.5 | | | | | | |
| P21440 | ABCB4 | ATP-binding cassette, sub-family B (MDR/TAP), member 4 | Plasma Membrane | transporter | 5875553 | 4916612.5 | 4507142 | | | | | | |
| P21440 | ABCB4 | ATP-binding cassette, sub-family B (MDR/TAP), member 4 | Plasma Membrane | transporter | 4030943.75 | 3111476.75 | 3095747.25 | | | | | | |
| P21440 | ABCB4 | ATP-binding cassette, sub-family B (MDR/TAP), member 4 | Plasma Membrane | transporter | 3514818 | 2554964 | 3034778 | | | | | | |
| P21440 | ABCB4 | ATP-binding cassette, sub-family B (MDR/TAP), member 4 | Plasma Membrane | transporter | 3869112 | 3859194.5 | 2722052.25 | | | | | | |
| P21440 | ABCB4 | ATP-binding cassette, sub-family B (MDR/TAP), member 4 | Plasma Membrane | transporter | 4413075.5 | 4294058.5 | 3252051.5 | | | | | | |
| P21440 | ABCB4 | ATP-binding cassette, sub-family B (MDR/TAP), member 4 | Plasma Membrane | transporter | 3649639 | 2710339.25 | 2183691.25 | | | | | | |
| P21440 | ABCB4 | ATP-binding cassette, sub-family B (MDR/TAP), member 4 | Plasma Membrane | transporter | 1371637 | 1954752.5 | 2024310.5 | P21440 | ABCB4 | 9671261 | 5780924 | 681288 | 72 |
| Q9DC29 | ABCB6 | ATP-binding cassette, sub-family B (MDR/TAP), member 6 (Langereis blood group) | Cytoplasm | transporter | 8.39E+07 | 8.19E+07 | 1.04E+08 | | | | | | |
| Q9DC29 | ABCB6 | ATP-binding cassette, sub-family B (MDR/TAP), member 6 (Langereis blood group) | Cytoplasm | transporter | 8.33E+07 | 1.02E+08 | 1.03E+08 | | | | | | |
| Q9DC29 | ABCB6 | ATP-binding cassette, sub-family B (MDR/TAP), member 6 (Langereis blood group) | Cytoplasm | transporter | 4.06E+07 | 4.08E+07 | 3.83E+07 | | | | | | |
| Q9DC29 | ABCB6 | ATP-binding cassette, sub-family B (MDR/TAP), member 6 (Langereis blood group) | Cytoplasm | transporter | 3.26E+07 | 3.75E+07 | 4.21E+07 | | | | | | |
| Q9DC29 | ABCB6 | ATP-binding cassette, sub-family B (MDR/TAP), member 6 (Langereis blood group) | Cytoplasm | transporter | 4.07E+07 | 4.86E+07 | 4.99E+07 | | | | | | |
| Q9DC29 | ABCB6 | ATP-binding cassette, sub-family B (MDR/TAP), member 6 (Langereis blood group) | Cytoplasm | transporter | 3.41E+07 | 3.17E+07 | 3.97E+07 | | | | | | |
| Q9DC29 | ABCB6 | ATP-binding cassette, sub-family B (MDR/TAP), member 6 (Langereis blood group) | Cytoplasm | transporter | 2.60E+07 | 2.76E+07 | 3.10E+07 | | | | | | |
| Q9DC29 | ABCB6 | ATP-binding cassette, sub-family B (MDR/TAP), member 6 (Langereis blood group) | Cytoplasm | transporter | 1.76E+07 | 1.75E+07 | 2.09E+07 | | | | | | |
| Q9DC29 | ABCB6 | ATP-binding cassette, sub-family B (MDR/TAP), member 6 (Langereis blood group) | Cytoplasm | transporter | 1.98E+07 | 2.01E+07 | 2.18E+07 | | | | | | |
| Q9DC29 | ABCB6 | ATP-binding cassette, sub-family B (MDR/TAP), member 6 (Langereis blood group) | Cytoplasm | transporter | 1.32E+07 | 1.79E+07 | 1.20E+07 | | | | | | |
| Q9DC29 | ABCB6 | ATP-binding cassette, sub-family B (MDR/TAP), member 6 (Langereis blood group) | Cytoplasm | transporter | 1.33E+07 | 1.44E+07 | 1.63E+07 | | | | | | |
| Q9DC29 | ABCB6 | ATP-binding cassette, sub-family B (MDR/TAP), member 6 (Langereis blood group) | Cytoplasm | transporter | 1.52E+07 | 1.82E+07 | 1.45E+07 | | | | | | |
| Q9DC29 | ABCB6 | ATP-binding cassette, sub-family B (MDR/TAP), member 6 (Langereis blood group) | Cytoplasm | transporter | 1.08E+07 | 1.00E+07 | 1.15E+07 | | | | | | |
| Q9DC29 | ABCB6 | ATP-binding cassette, sub-family B (MDR/TAP), member 6 (Langereis blood group) | Cytoplasm | transporter | 8234651.5 | 1.06E+07 | 7678055.5 | | | | | | |
| Q9DC29 | ABCB6 | ATP-binding cassette, sub-family B (MDR/TAP), member 6 (Langereis blood group) | Cytoplasm | transporter | 8100268.5 | 9513779 | 9555593 | | | | | | |
| Q9DC29 | ABCB6 | ATP-binding cassette, sub-family B (MDR/TAP), member 6 (Langereis blood group) | Cytoplasm | transporter | 3895307 | 5667289 | 9318729 | | | | | | |
| Q9DC29 | ABCB6 | ATP-binding cassette, sub-family B (MDR/TAP), member 6 (Langereis blood group) | Cytoplasm | transporter | 4004374.5 | 3068810.5 | 2928204.75 | | | | | | |
| Q9DC29 | ABCB6 | ATP-binding cassette, sub-family B (MDR/TAP), member 6 (Langereis blood group) | Cytoplasm | transporter | 3048911.5 | 3893749.5 | 3980988.5 | | | | | | |
| Q9DC29 | ABCB6 | ATP-binding cassette, sub-family B (MDR/TAP), member 6 (Langereis blood group) | Cytoplasm | transporter | 2429824.5 | 932137.125 | 363525.531 | | | | | | |
| Q9DC29 | ABCB6 | ATP-binding cassette, sub-family B (MDR/TAP), member 6 (Langereis blood group) | Cytoplasm | transporter | 3185954.75 | 3251284.5 | 3514383.75 | Q9DC29 | ABCB6 | 23190465 | 26565422 | 3429581 | 60 |
| Q61102 | ABCB7 | ATP-binding cassette, sub-family B (MDR/TAP), member 7 | Cytoplasm | transporter | 1285864.125 | 2088254.625 | 3198073.25 | Q61102 | ABCB7 | 1285864 | 960215 | 554380 | 3 |
| Q9JJ59 | ABCB9 | ATP-binding cassette, sub-family B (MDR/TAP), member 9 | Cytoplasm | transporter | 3971241.25 | 3865634.75 | 2.17E+07 | Q9JJ59 | ABCB9 | 3971241 | 10237458 | 5910599 | 3 |
| O35379 | ABCC1 | ATP-binding cassette, sub-family C (CFTR/MRP), member 1 | Plasma Membrane | transporter | 1.48E+07 | 1.66E+07 | 1.50E+07 | | | | | | |
| O35379 | ABCC1 | ATP-binding cassette, sub-family C (CFTR/MRP), member 1 | Plasma Membrane | transporter | 1.26E+07 | 1.58E+07 | 1.48E+07 | | | | | | |
| O35379 | ABCC1 | ATP-binding cassette, sub-family C (CFTR/MRP), member 1 | Plasma Membrane | transporter | 1.14E+07 | 1.26E+07 | 9956485 | | | | | | |
| O35379 | ABCC1 | ATP-binding cassette, sub-family C (CFTR/MRP), member 1 | Plasma Membrane | transporter | 1.16E+07 | 1.24E+07 | 8819103 | | | | | | |
| O35379 | ABCC1 | ATP-binding cassette, sub-family C (CFTR/MRP), member 1 | Plasma Membrane | transporter | 8441256 | 8980383 | 8961841 | | | | | | |
| O35379 | ABCC1 | ATP-binding cassette, sub-family C (CFTR/MRP), member 1 | Plasma Membrane | transporter | 7546892.5 | 7178152 | 5602325 | | | | | | |
| O35379 | ABCC1 | ATP-binding cassette, sub-family C (CFTR/MRP), member 1 | Plasma Membrane | transporter | 6975342.5 | 5205438 | 3544792.75 | | | | | | |
| O35379 | ABCC1 | ATP-binding cassette, sub-family C (CFTR/MRP), member 1 | Plasma Membrane | transporter | 5973929 | 6399028 | 6390397.5 | | | | | | |
| O35379 | ABCC1 | ATP-binding cassette, sub-family C (CFTR/MRP), member 1 | Plasma Membrane | transporter | 5837191.5 | 5161817.5 | 4614613 | | | | | | |
| O35379 | ABCC1 | ATP-binding cassette, sub-family C (CFTR/MRP), member 1 | Plasma Membrane | transporter | 4716741 | 4705216 | 3855684 | | | | | | |
| O35379 | ABCC1 | ATP-binding cassette, sub-family C (CFTR/MRP), member 1 | Plasma Membrane | transporter | 4366592 | 4634296 | 3392243.25 | | | | | | |
| O35379 | ABCC1 | ATP-binding cassette, sub-family C (CFTR/MRP), member 1 | Plasma Membrane | transporter | 3525126.25 | 5996264 | 5167687.5 | | | | | | |
| O35379 | ABCC1 | ATP-binding cassette, sub-family C (CFTR/MRP), member 1 | Plasma Membrane | transporter | 3975111.25 | 4470059 | 4651216.5 | | | | | | |
| O35379 | ABCC1 | ATP-binding cassette, sub-family C (CFTR/MRP), member 1 | Plasma Membrane | transporter | 3066203 | 3415052.5 | 2591467 | | | | | | |
| O35379 | ABCC1 | ATP-binding cassette, sub-family C (CFTR/MRP), member 1 | Plasma Membrane | transporter | 2451720.25 | 3997100.75 | 2134533 | | | | | | |
| O35379 | ABCC1 | ATP-binding cassette, sub-family C (CFTR/MRP), member 1 | Plasma Membrane | transporter | 2459440.5 | 3615507.75 | 2617426.25 | | | | | | |
| O35379 | ABCC1 | ATP-binding cassette, sub-family C (CFTR/MRP), member 1 | Plasma Membrane | transporter | 2510386.75 | 2063901.875 | 2604212.75 | | | | | | |
| O35379 | ABCC1 | ATP-binding cassette, sub-family C (CFTR/MRP), member 1 | Plasma Membrane | transporter | 1740471.375 | 2335973.75 | 2263636.5 | O35379 | ABCC1 | 6328689 | 4101030 | 558079 | 54 |
| Q9R1S7 | ABCC6 | ATP-binding cassette, sub-family C (CFTR/MRP), member 6 | Plasma Membrane | transporter | 4761171 | 8132295 | 7780860 | Q9R1S7 | ABCC6 | 4761171 | 1853218 | 1069956 | 3 |
| P61222 | ABCE1 | ATP-binding cassette, sub-family E (OABP), member 1 | Cytoplasm | transporter | 3966254.5 | 4507437 | 4060850.75 | | | | | | |
| P61222 | ABCE1 | ATP-binding cassette, sub-family E (OABP), member 1 | Cytoplasm | transporter | 2473629.5 | 2357840.75 | 2039743.25 | | | | | | |
| P61222 | ABCE1 | ATP-binding cassette, sub-family E (OABP), member 1 | Cytoplasm | transporter | 2412858.25 | 2946005 | 2434740.5 | | | | | | |
| P61222 | ABCE1 | ATP-binding cassette, sub-family E (OABP), member 1 | Cytoplasm | transporter | 1212791 | 1405815 | 1659792.25 | P61222 | ABCE1 | 2516383 | 1063339 | 306960 | 12 |
| Q7TMS5 | ABCG2 | ATP-binding cassette, sub-family G (WHITE), member 2 (Junior blood group) | Plasma Membrane | transporter | 1.26E+08 | 1.29E+08 | 1.12E+08 | | | | | | |
| Q7TMS5 | ABCG2 | ATP-binding cassette, sub-family G (WHITE), member 2 (Junior blood group) | Plasma Membrane | transporter | 9.93E+07 | 9.40E+07 | 8.62E+07 | | | | | | |
| Q7TMS5 | ABCG2 | ATP-binding cassette, sub-family G (WHITE), member 2 (Junior blood group) | Plasma Membrane | transporter | 9.48E+07 | 9.80E+07 | 8.48E+07 | | | | | | |
| Q7TMS5 | ABCG2 | ATP-binding cassette, sub-family G (WHITE), member 2 (Junior blood group) | Plasma Membrane | transporter | 7.34E+07 | 7.91E+07 | 7.15E+07 | | | | | | |
| Q7TMS5 | ABCG2 | ATP-binding cassette, sub-family G (WHITE), member 2 (Junior blood group) | Plasma Membrane | transporter | 5.67E+07 | 5.68E+07 | 5.65E+07 | | | | | | |
| Q7TMS5 | ABCG2 | ATP-binding cassette, sub-family G (WHITE), member 2 (Junior blood group) | Plasma Membrane | transporter | 3.55E+07 | 3.94E+07 | 3.93E+07 | | | | | | |
| Q7TMS5 | ABCG2 | ATP-binding cassette, sub-family G (WHITE), member 2 (Junior blood group) | Plasma Membrane | transporter | 4.36E+07 | 4.79E+07 | 3.85E+07 | | | | | | |
| Q7TMS5 | ABCG2 | ATP-binding cassette, sub-family G (WHITE), member 2 (Junior blood group) | Plasma Membrane | transporter | 2.95E+07 | 3.08E+07 | 2.88E+07 | | | | | | |
| Q7TMS5 | ABCG2 | ATP-binding cassette, sub-family G (WHITE), member 2 (Junior blood group) | Plasma Membrane | transporter | 4.27E+07 | 4.81E+07 | 3.08E+07 | | | | | | |
| Q7TMS5 | ABCG2 | ATP-binding cassette, sub-family G (WHITE), member 2 (Junior blood group) | Plasma Membrane | transporter | 2.48E+07 | 2.86E+07 | 2.74E+07 | | | | | | |
| Q7TMS5 | ABCG2 | ATP-binding cassette, sub-family G (WHITE), member 2 (Junior blood group) | Plasma Membrane | transporter | 2.46E+07 | 2.35E+07 | 2.12E+07 | | | | | | |
| Q7TMS5 | ABCG2 | ATP-binding cassette, sub-family G (WHITE), member 2 (Junior blood group) | Plasma Membrane | transporter | 1.77E+07 | 1.78E+07 | 1.45E+07 | | | | | | |
| Q7TMS5 | ABCG2 | ATP-binding cassette, sub-family G (WHITE), member 2 (Junior blood group) | Plasma Membrane | transporter | 1.27E+07 | 1.49E+07 | 1.27E+07 | | | | | | |
| Q7TMS5 | ABCG2 | ATP-binding cassette, sub-family G (WHITE), member 2 (Junior blood group) | Plasma Membrane | transporter | 2.19E+07 | 2.04E+07 | 1.96E+07 | | | | | | |
| Q7TMS5 | ABCG2 | ATP-binding cassette, sub-family G (WHITE), member 2 (Junior blood group) | Plasma Membrane | transporter | 1.30E+07 | 1.31E+07 | 9509156 | | | | | | |
| Q7TMS5 | ABCG2 | ATP-binding cassette, sub-family G (WHITE), member 2 (Junior blood group) | Plasma Membrane | transporter | 8927720 | 1.06E+07 | 9750185 | | | | | | |
| Q7TMS5 | ABCG2 | ATP-binding cassette, sub-family G (WHITE), member 2 (Junior blood group) | Plasma Membrane | transporter | 3828686.5 | 4798760 | 4750544 | | | | | | |
| Q7TMS5 | ABCG2 | ATP-binding cassette, sub-family G (WHITE), member 2 (Junior blood group) | Plasma Membrane | transporter | 6076298.5 | 1.02E+07 | 9874654 | | | | | | |
| Q7TMS5 | ABCG2 | ATP-binding cassette, sub-family G (WHITE), member 2 (Junior blood group) | Plasma Membrane | transporter | 6078470 | 6761047 | 6204644 | | | | | | |
| Q7TMS5 | ABCG2 | ATP-binding cassette, sub-family G (WHITE), member 2 (Junior blood group) | Plasma Membrane | transporter | 1314429.125 | 1085062.25 | 814962 | Q7TMS5 | ABCG2 | 37143280 | 33929875 | 4380328 | 60 |
| Q9Z1Q2 | ABHD16A | abhydrolase domain containing 16A | Cytoplasm | enzyme | 4284823 | 5624232 | 5688654.5 | | | | | | |
| Q9Z1Q2 | ABHD16A | abhydrolase domain containing 16A | Cytoplasm | enzyme | 4219069 | 5597855 | 4651507.5 | | | | | | |
| Q9Z1Q2 | ABHD16A | abhydrolase domain containing 16A | Cytoplasm | enzyme | 788688.312 | 919572.375 | 1158311.75 | Q9Z1Q2 | ABHD16A | 3097527 | 2104394 | 701465 | 9 |
| Q921H8 | ACAA1 | acetyl-CoA acyltransferase 1 | Cytoplasm | enzyme | 2893885.5 | 2988005 | 5172271 | Q921H8 | ACAA1 | 2893886 | 1289116 | 744271 | 3 |
| Q8BWT1 | ACAA2 | acetyl-CoA acyltransferase 2 | Cytoplasm | enzyme | 6196118.5 | 1.06E+07 | 1.43E+07 | | | | | | |
| Q8BWT1 | ACAA2 | acetyl-CoA acyltransferase 2 | Cytoplasm | enzyme | 4197395.5 | 7099633.5 | 8645298 | | | | | | |

| | | | | | | | | | | | | | |
|---|---|---|---|---|---|---|---|---|---|---|---|---|---|
| Q8BWT1 | ACAA2 | acetyl-CoA acyltransferase 2 | Cytoplasm | enzyme | 3057260.25 | 5146860.5 | 5945297.5 | | | | | | |
| Q8BWT1 | ACAA2 | acetyl-CoA acyltransferase 2 | Cytoplasm | enzyme | 2653808 | 5661116.5 | 6009948 | | | | | | |
| Q8BWT1 | ACAA2 | acetyl-CoA acyltransferase 2 | Cytoplasm | enzyme | 1477558.125 | 3469064.75 | 5590259 | | | | | | |
| Q8BWT1 | ACAA2 | acetyl-CoA acyltransferase 2 | Cytoplasm | enzyme | 1867031.875 | 4257934.5 | 5056492.5 | Q8BWT1 | ACAA2 | 3241529 | 3133707 | 738622 | 18 |
| Q8JZN5 | ACAD9 | acyl-CoA dehydrogenase family, member 9 | Cytoplasm | enzyme | 1789085.5 | 2163761 | 3428151 | Q8JZN5 | ACAD9 | 1789086 | 858836 | 495849 | 3 |
| P51174 | ACADL | acyl-CoA dehydrogenase, long chain | Cytoplasm | enzyme | 1.97E+07 | 3.04E+07 | 4.29E+07 | | | | | | |
| P51174 | ACADL | acyl-CoA dehydrogenase, long chain | Cytoplasm | enzyme | 8102927.5 | 1.26E+07 | 1.41E+07 | | | | | | |
| P51174 | ACADL | acyl-CoA dehydrogenase, long chain | Cytoplasm | enzyme | 6332003 | 1.09E+07 | 1.10E+07 | | | | | | |
| P51174 | ACADL | acyl-CoA dehydrogenase, long chain | Cytoplasm | enzyme | 3765376.5 | 5706222.5 | 8118376.5 | | | | | | |
| P51174 | ACADL | acyl-CoA dehydrogenase, long chain | Cytoplasm | enzyme | 3698395.25 | 4028103.75 | 3583493.75 | | | | | | |
| P51174 | ACADL | acyl-CoA dehydrogenase, long chain | Cytoplasm | enzyme | 4175099.75 | 7330080.5 | 1.06E+07 | | | | | | |
| P51174 | ACADL | acyl-CoA dehydrogenase, long chain | Cytoplasm | enzyme | 2428315.75 | 5392485 | 4444865 | P51174 | ACADL | 6887445 | 9893301 | 2158895 | 21 |
| P45952 | ACADM | acyl-CoA dehydrogenase, C-4 to C-12 straight chain | Cytoplasm | enzyme | 821226.938 | 1070649.75 | 1775297.5 | P45952 | ACADM | 821227 | 494805 | 285676 | 3 |
| P50544 | ACADVL | acyl-CoA dehydrogenase, very long chain | Cytoplasm | enzyme | 2065991.75 | 4657697.5 | 5153030.5 | | | | | | |
| P50544 | ACADVL | acyl-CoA dehydrogenase, very long chain | Cytoplasm | enzyme | 1908065.5 | 3583086.5 | 3351230 | | | | | | |
| P50544 | ACADVL | acyl-CoA dehydrogenase, very long chain | Cytoplasm | enzyme | 974240.375 | 1603273.75 | 2559655.75 | | | | | | |
| P50544 | ACADVL | acyl-CoA dehydrogenase, very long chain | Cytoplasm | enzyme | 1050075.875 | 2251417.75 | 1643020.625 | | | | | | |
| P50544 | ACADVL | acyl-CoA dehydrogenase, very long chain | Cytoplasm | enzyme | 467639.812 | 772689.938 | 1026903.625 | P50544 | ACADVL | 1293203 | 1417729 | 366056 | 15 |
| Q8QZT1 | ACAT1 | acetyl-CoA acetyltransferase 1 | Cytoplasm | enzyme | 9861399 | 2.03E+07 | 2.11E+07 | | | | | | |
| Q8QZT1 | ACAT1 | acetyl-CoA acetyltransferase 1 | Cytoplasm | enzyme | 6764851 | 1.09E+07 | 1.50E+07 | | | | | | |
| Q8QZT1 | ACAT1 | acetyl-CoA acetyltransferase 1 | Cytoplasm | enzyme | 4687725.5 | 8124648.5 | 1.09E+07 | | | | | | |
| Q8QZT1 | ACAT1 | acetyl-CoA acetyltransferase 1 | Cytoplasm | enzyme | 8215136.5 | 9998823 | 1.37E+07 | | | | | | |
| Q8QZT1 | ACAT1 | acetyl-CoA acetyltransferase 1 | Cytoplasm | enzyme | 4736350.5 | 6600695 | 8634755 | | | | | | |
| Q8QZT1 | ACAT1 | acetyl-CoA acetyltransferase 1 | Cytoplasm | enzyme | 3403127.75 | 4650966 | 7610692 | | | | | | |
| Q8QZT1 | ACAT1 | acetyl-CoA acetyltransferase 1 | Cytoplasm | enzyme | 2872545.75 | 4300661.5 | 7384713.5 | Q8QZT1 | ACAT1 | 5791591 | 5013455 | 1094025 | 21 |
| Q9QUI6 | ACKR1 | atypical chemokine receptor 1 (Duffy blood group) | Plasma Membrane | G-protein coupled receptor | 3.73E+07 | 3.74E+07 | 2.34E+07 | | | | | | |
| Q9QUI6 | ACKR1 | atypical chemokine receptor 1 (Duffy blood group) | Plasma Membrane | G-protein coupled receptor | 3.43E+07 | 3.64E+07 | 3.03E+07 | | | | | | |
| Q9QUI6 | ACKR1 | atypical chemokine receptor 1 (Duffy blood group) | Plasma Membrane | G-protein coupled receptor | 9735634 | 1.11E+07 | 8596145 | | | | | | |
| Q9QUI6 | ACKR1 | atypical chemokine receptor 1 (Duffy blood group) | Plasma Membrane | G-protein coupled receptor | 1154238.875 | 925885.125 | 3087784 | Q9QUI6 | ACKR1 | 20617468 | 15104210 | 4360210 | 12 |
| Q99KI0 | ACO2 | aconitase 2, mitochondrial | Cytoplasm | enzyme | 1.67E+07 | 2.58E+07 | 2.48E+07 | | | | | | |
| Q99KI0 | ACO2 | aconitase 2, mitochondrial | Cytoplasm | enzyme | 7736579.5 | 1.38E+07 | 1.60E+07 | | | | | | |
| Q99KI0 | ACO2 | aconitase 2, mitochondrial | Cytoplasm | enzyme | 6844038.5 | 1.08E+07 | 1.25E+07 | | | | | | |
| Q99KI0 | ACO2 | aconitase 2, mitochondrial | Cytoplasm | enzyme | 7990550.5 | 1.43E+07 | 8283662.5 | | | | | | |
| Q99KI0 | ACO2 | aconitase 2, mitochondrial | Cytoplasm | enzyme | 7174218.5 | 9343600 | 1.35E+07 | | | | | | |
| Q99KI0 | ACO2 | aconitase 2, mitochondrial | Cytoplasm | enzyme | 5222633.5 | 9600828 | 1.28E+07 | | | | | | |
| Q99KI0 | ACO2 | aconitase 2, mitochondrial | Cytoplasm | enzyme | 4006125.5 | 7933632 | 1.04E+07 | | | | | | |
| Q99KI0 | ACO2 | aconitase 2, mitochondrial | Cytoplasm | enzyme | 5363646.5 | 1.11E+07 | 1.10E+07 | | | | | | |
| Q99KI0 | ACO2 | aconitase 2, mitochondrial | Cytoplasm | enzyme | 6234146 | 9677584 | 1.01E+07 | | | | | | |
| Q99KI0 | ACO2 | aconitase 2, mitochondrial | Cytoplasm | enzyme | 4306839 | 6049857.5 | 1.06E+07 | | | | | | |
| Q99KI0 | ACO2 | aconitase 2, mitochondrial | Cytoplasm | enzyme | 4386912.5 | 7470377 | 1.05E+07 | | | | | | |
| Q99KI0 | ACO2 | aconitase 2, mitochondrial | Cytoplasm | enzyme | 2233532.75 | 5021766.5 | 6068960 | | | | | | |
| Q99KI0 | ACO2 | aconitase 2, mitochondrial | Cytoplasm | enzyme | 3971745.25 | 5845429.5 | 7735856.5 | | | | | | |
| Q99KI0 | ACO2 | aconitase 2, mitochondrial | Cytoplasm | enzyme | 3498108.25 | 4816708 | 4859374.5 | | | | | | |
| Q99KI0 | ACO2 | aconitase 2, mitochondrial | Cytoplasm | enzyme | 2364834.25 | 4414387.5 | 5174033 | | | | | | |
| Q99KI0 | ACO2 | aconitase 2, mitochondrial | Cytoplasm | enzyme | 1547509.25 | 2651457.5 | 2239256.5 | | | | | | |
| Q99KI0 | ACO2 | aconitase 2, mitochondrial | Cytoplasm | enzyme | 1564816.625 | 2597304.5 | 2984468.5 | | | | | | |
| Q99KI0 | ACO2 | aconitase 2, mitochondrial | Cytoplasm | enzyme | 2905327.75 | 6054566.5 | 3729898.25 | | | | | | |
| Q99KI0 | ACO2 | aconitase 2, mitochondrial | Cytoplasm | enzyme | 1682276.5 | 4242059 | 2598629 | | | | | | |
| Q99KI0 | ACO2 | aconitase 2, mitochondrial | Cytoplasm | enzyme | 1585944.25 | 2219149.25 | 3713439 | | | | | | |
| Q99KI0 | ACO2 | aconitase 2, mitochondrial | Cytoplasm | enzyme | 2107452.5 | 2905477.75 | 1667184.625 | Q99KI0 | ACO2 | 4734154 | 5141602 | 647781 | 63 |
| P41216 | ACSL1 | acyl-CoA synthetase long-chain family member 1 | Cytoplasm | enzyme | 1.86E+07 | 2.03E+07 | 1.65E+07 | | | | | | |
| P41216 | ACSL1 | acyl-CoA synthetase long-chain family member 1 | Cytoplasm | enzyme | 1.79E+07 | 1.73E+07 | 1.61E+07 | | | | | | |
| P41216 | ACSL1 | acyl-CoA synthetase long-chain family member 1 | Cytoplasm | enzyme | 1.84E+07 | 1.73E+07 | 1.84E+07 | | | | | | |
| P41216 | ACSL1 | acyl-CoA synthetase long-chain family member 1 | Cytoplasm | enzyme | 1.31E+07 | 1.27E+07 | 1.24E+07 | | | | | | |
| P41216 | ACSL1 | acyl-CoA synthetase long-chain family member 1 | Cytoplasm | enzyme | 1.87E+07 | 2.25E+07 | 1.53E+07 | | | | | | |
| P41216 | ACSL1 | acyl-CoA synthetase long-chain family member 1 | Cytoplasm | enzyme | 1.39E+07 | 1.53E+07 | 1.68E+07 | | | | | | |
| P41216 | ACSL1 | acyl-CoA synthetase long-chain family member 1 | Cytoplasm | enzyme | 9135525 | 1.10E+07 | 1.32E+07 | | | | | | |
| P41216 | ACSL1 | acyl-CoA synthetase long-chain family member 1 | Cytoplasm | enzyme | 8547671 | 8931860 | 9913636 | | | | | | |
| P41216 | ACSL1 | acyl-CoA synthetase long-chain family member 1 | Cytoplasm | enzyme | 6135434.5 | 6771773.5 | 6687060.5 | | | | | | |
| P41216 | ACSL1 | acyl-CoA synthetase long-chain family member 1 | Cytoplasm | enzyme | 5459532 | 5947277.5 | 5265624 | | | | | | |
| P41216 | ACSL1 | acyl-CoA synthetase long-chain family member 1 | Cytoplasm | enzyme | 6897682.5 | 8767256 | 9665460 | | | | | | |
| P41216 | ACSL1 | acyl-CoA synthetase long-chain family member 1 | Cytoplasm | enzyme | 5695799.5 | 6895201.5 | 6810443 | | | | | | |
| P41216 | ACSL1 | acyl-CoA synthetase long-chain family member 1 | Cytoplasm | enzyme | 3447805 | 4096292.5 | 3531936.5 | | | | | | |
| P41216 | ACSL1 | acyl-CoA synthetase long-chain family member 1 | Cytoplasm | enzyme | 3052027.25 | 3809931 | 4870096 | | | | | | |
| P41216 | ACSL1 | acyl-CoA synthetase long-chain family member 1 | Cytoplasm | enzyme | 3651635.75 | 4815878.5 | 4319239 | | | | | | |
| P41216 | ACSL1 | acyl-CoA synthetase long-chain family member 1 | Cytoplasm | enzyme | 3381137 | 4534499.5 | 4629027 | | | | | | |
| P41216 | ACSL1 | acyl-CoA synthetase long-chain family member 1 | Cytoplasm | enzyme | 4750165.5 | 5790971 | 6591943.5 | P41216 | ACSL1 | 9457907 | 5641118 | 789914 | 51 |
| Q9QUJ7 | ACSL4 | acyl-CoA synthetase long-chain family member 4 | Cytoplasm | enzyme | 1.26E+07 | 1.21E+07 | 1.30E+07 | | | | | | |
| Q9QUJ7 | ACSL4 | acyl-CoA synthetase long-chain family member 4 | Cytoplasm | enzyme | 1.27E+07 | 1.55E+07 | 1.36E+07 | | | | | | |
| Q9QUJ7 | ACSL4 | acyl-CoA synthetase long-chain family member 4 | Cytoplasm | enzyme | 1.11E+07 | 1.35E+07 | 1.21E+07 | | | | | | |
| Q9QUJ7 | ACSL4 | acyl-CoA synthetase long-chain family member 4 | Cytoplasm | enzyme | 5505640 | 4844066 | 5626921 | | | | | | |
| Q9QUJ7 | ACSL4 | acyl-CoA synthetase long-chain family member 4 | Cytoplasm | enzyme | 2457823 | 2353661.25 | 2489198 | | | | | | |
| Q9QUJ7 | ACSL4 | acyl-CoA synthetase long-chain family member 4 | Cytoplasm | enzyme | 1707130.75 | 2240006.25 | 2216840.25 | | | | | | |
| Q9QUJ7 | ACSL4 | acyl-CoA synthetase long-chain family member 4 | Cytoplasm | enzyme | 2705980 | 2834920.75 | 3955518.25 | | | | | | |
| Q9QUJ7 | ACSL4 | acyl-CoA synthetase long-chain family member 4 | Cytoplasm | enzyme | 2874234 | 2216901.5 | 2263299.25 | | | | | | |
| Q9QUJ7 | ACSL4 | acyl-CoA synthetase long-chain family member 4 | Cytoplasm | enzyme | 2418201.5 | 1643336.625 | 1632346 | Q9QUJ7 | ACSL4 | 6002112 | 4956849 | 953946 | 27 |
| Q8JZR0 | ACSL5 | acyl-CoA synthetase long-chain family member 5 | Cytoplasm | enzyme | 1.29E+07 | 1.33E+07 | 1.32E+07 | | | | | | |
| Q8JZR0 | ACSL5 | acyl-CoA synthetase long-chain family member 5 | Cytoplasm | enzyme | 9699091 | 7374624 | 8153309 | | | | | | |
| Q8JZR0 | ACSL5 | acyl-CoA synthetase long-chain family member 5 | Cytoplasm | enzyme | 6226064.5 | 7916854 | 7051911.5 | | | | | | |
| Q8JZR0 | ACSL5 | acyl-CoA synthetase long-chain family member 5 | Cytoplasm | enzyme | 6064719.5 | 5091953.5 | 4969020.5 | | | | | | |
| Q8JZR0 | ACSL5 | acyl-CoA synthetase long-chain family member 5 | Cytoplasm | enzyme | 5739404 | 5572367 | 4692637.5 | | | | | | |
| Q8JZR0 | ACSL5 | acyl-CoA synthetase long-chain family member 5 | Cytoplasm | enzyme | 4315034.5 | 4758280 | 4958370 | | | | | | |
| Q8JZR0 | ACSL5 | acyl-CoA synthetase long-chain family member 5 | Cytoplasm | enzyme | 5149123 | 5269190 | 5916086 | | | | | | |
| Q8JZR0 | ACSL5 | acyl-CoA synthetase long-chain family member 5 | Cytoplasm | enzyme | 3896600.75 | 2637216.25 | 3159321 | | | | | | |
| Q8JZR0 | ACSL5 | acyl-CoA synthetase long-chain family member 5 | Cytoplasm | enzyme | 3547422.5 | 3158977.25 | 3437151.75 | | | | | | |
| Q8JZR0 | ACSL5 | acyl-CoA synthetase long-chain family member 5 | Cytoplasm | enzyme | 2617016.25 | 2636054.5 | 2741781.5 | | | | | | |
| Q8JZR0 | ACSL5 | acyl-CoA synthetase long-chain family member 5 | Cytoplasm | enzyme | 3223479 | 3084609.5 | 2549193.5 | Q8JZR0 | ACSL5 | 5758905 | 3003944 | 522919 | 33 |
| Q99NB1 | ACSS1 | acyl-CoA synthetase short-chain family member 1 | Cytoplasm | enzyme | 1729559.375 | 3518974 | 5673662.5 | | | | | | |
| Q99NB1 | ACSS1 | acyl-CoA synthetase short-chain family member 1 | Cytoplasm | enzyme | 1348604.875 | 2305774.75 | 3448860 | | | | | | |
| Q99NB1 | ACSS1 | acyl-CoA synthetase short-chain family member 1 | Cytoplasm | enzyme | 1043189.625 | 2179105 | 1900535.5 | Q99NB1 | ACSS1 | 1373785 | 1436390 | 478797 | 9 |
| P60710 | ACTB | actin, beta | Cytoplasm | other | 4.93E+09 | 4.68E+09 | 4.33E+09 | | | | | | |
| P60710 | ACTB | actin, beta | Cytoplasm | other | 2.37E+09 | 2.78E+09 | 2.82E+09 | | | | | | |
| P60710 | ACTB | actin, beta | Cytoplasm | other | 2.82E+09 | 2.71E+09 | 2.53E+09 | | | | | | |
| P60710 | ACTB | actin, beta | Cytoplasm | other | 1.37E+09 | 1.36E+09 | 1.30E+09 | | | | | | |
| P60710 | ACTB | actin, beta | Cytoplasm | other | 1.31E+09 | 1.10E+09 | 1.03E+09 | | | | | | |
| P60710 | ACTB | actin, beta | Cytoplasm | other | 1.06E+09 | 1.43E+09 | 1.04E+09 | | | | | | |
| P60710 | ACTB | actin, beta | Cytoplasm | other | 1.38E+09 | 1.43E+09 | 1.25E+09 | | | | | | |
| P60710 | ACTB | actin, beta | Cytoplasm | other | 1.11E+09 | 1.06E+09 | 9.79E+08 | | | | | | |
| P60710 | ACTB | actin, beta | Cytoplasm | other | 1.24E+09 | 1.06E+09 | 9.73E+08 | | | | | | |
| P60710 | ACTB | actin, beta | Cytoplasm | other | 9.81E+08 | 1.01E+09 | 7.11E+08 | | | | | | |
| P60710 | ACTB | actin, beta | Cytoplasm | other | 1.13E+09 | 9.99E+08 | 1.03E+09 | | | | | | |
| P60710 | ACTB | actin, beta | Cytoplasm | other | 9.65E+08 | 9.64E+08 | 9.45E+08 | | | | | | |
| P60710 | ACTB | actin, beta | Cytoplasm | other | 7.79E+08 | 7.21E+08 | 6.26E+08 | | | | | | |
| P60710 | ACTB | actin, beta | Cytoplasm | other | 6.49E+08 | 5.75E+08 | 5.13E+08 | | | | | | |
| P60710 | ACTB | actin, beta | Cytoplasm | other | 5.37E+08 | 3.76E+08 | 3.76E+08 | | | | | | |
| P60710 | ACTB | actin, beta | Cytoplasm | other | 3.70E+08 | 4.59E+08 | 3.11E+08 | | | | | | |
| P60710 | ACTB | actin, beta | Cytoplasm | other | 2.81E+08 | 3.35E+08 | 3.64E+08 | | | | | | |
| P60710 | ACTB | actin, beta | Cytoplasm | other | 2.05E+08 | 2.30E+08 | 1.72E+08 | | | | | | |
| P60710 | ACTB | actin, beta | Cytoplasm | other | 1.84E+08 | 1.99E+08 | 1.45E+08 | | | | | | |

| | | | | | | | | | | | | | |
|---|---|---|---|---|---|---|---|---|---|---|---|---|---|
| P60710 | ACTB | actin, beta | Cytoplasm | other | 3.54E+08 | 4.18E+08 | 2.92E+08 | | | | | | |
| P60710 | ACTB | actin, beta | Cytoplasm | other | 1.82E+08 | 1.56E+08 | 1.84E+08 | | | | | | |
| P60710 | ACTB | actin, beta | Cytoplasm | other | 1.34E+08 | 1.59E+08 | 1.79E+08 | | | | | | |
| P60710 | ACTB | actin, beta | Cytoplasm | other | 1.17E+08 | 3.56E+07 | 1.58E+08 | | | | | | |
| P60710 | ACTB | actin, beta | Cytoplasm | other | 1.02E+08 | 1.05E+08 | 7.82E+07 | | | | | | |
| P60710 | ACTB | actin, beta | Cytoplasm | other | 1.07E+08 | 6.70E+07 | 7.27E+07 | | | | | | |
| P60710 | ACTB | actin, beta | Cytoplasm | other | 9.17E+07 | 3.32E+07 | 1.29E+08 | | | | | | |
| P60710 | ACTB | actin, beta | Cytoplasm | other | 9.46E+07 | 1.30E+08 | 7.99E+07 | | | | | | |
| P60710 | ACTB | actin, beta | Cytoplasm | other | 8.65E+07 | 8.98E+07 | 7.50E+07 | | | | | | |
| P60710 | ACTB | actin, beta | Cytoplasm | other | 8.69E+07 | 6.29E+07 | 5.80E+07 | | | | | | |
| P60710 | ACTB | actin, beta | Cytoplasm | other | 7.49E+07 | 6.77E+07 | 6.05E+07 | | | | | | |
| P60710 | ACTB | actin, beta | Cytoplasm | other | 7.19E+07 | 9.84E+07 | 4.63E+07 | | | | | | |
| P60710 | ACTB | actin, beta | Cytoplasm | other | 1.57E+08 | 2.17E+08 | 1.51E+08 | | | | | | |
| P60710 | ACTB | actin, beta | Cytoplasm | other | 5.10E+07 | 5.47E+07 | 3.97E+07 | | | | | | |
| P60710 | ACTB | actin, beta | Cytoplasm | other | 3.62E+07 | 1.48E+07 | 3.20E+07 | | | | | | |
| P60710 | ACTB | actin, beta | Cytoplasm | other | 1.92E+07 | 2.17E+07 | 1.79E+07 | | | | | | |
| P60710 | ACTB | actin, beta | Cytoplasm | other | 2.59E+07 | 2.79E+07 | 2.33E+07 | | | | | | |
| P60710 | ACTB | actin, beta | Cytoplasm | other | 1.99E+07 | 5861383.5 | 2.47E+07 | | | | | | |
| P60710 | ACTB | actin, beta | Cytoplasm | other | 1.62E+07 | 3.70E+07 | 2.54E+07 | | | | | | |
| P60710 | ACTB | actin, beta | Cytoplasm | other | 2.19E+07 | 5400546.5 | 2.73E+07 | | | | | | |
| P60710 | ACTB | actin, beta | Cytoplasm | other | 1.03E+07 | 5287849.5 | 1613013.625 | | | | | | |
| P60710 | ACTB | actin, beta | Cytoplasm | other | 9563600 | 6729707.5 | 1763952.625 | | | | | | |
| P60710 | ACTB | actin, beta | Cytoplasm | other | 1.83E+08 | 3.99E+08 | 5.57E+08 | | | | | | |
| P60710 | ACTB | actin, beta | Cytoplasm | other | 6777736.5 | 1.16E+07 | 1.46E+07 | | | | | | |
| P60710 | ACTB | actin, beta | Cytoplasm | other | 6686072 | 1.16E+07 | 5055884.5 | | | | | | |
| P60710 | ACTB | actin, beta | Cytoplasm | other | 4994997.5 | 5650425.5 | 6303084.5 | | | | | | |
| P60710 | ACTB | actin, beta | Cytoplasm | other | 3038101.75 | 2898810.25 | 2492030.5 | | | | | | |
| P60710 | ACTB | actin, beta | Cytoplasm | other | 3.99E+07 | 6.86E+07 | 8.24E+07 | | | | | | |
| P60710 | ACTB | actin, beta | Cytoplasm | other | 153940.797 | 165873.969 | 356620.5 | P60710 | ACTB | 537215718 | 870960448 | 72580037 | 144 |
| P68033 | ACTC1 | actin, alpha, cardiac muscle 1 | Cytoplasm | enzyme | 3.16E+09 | 3.06E+09 | 2.55E+09 | | | | | | |
| P68033 | ACTC1 | actin, alpha, cardiac muscle 1 | Cytoplasm | enzyme | 5.19E+08 | 4.88E+08 | 4.46E+08 | | | | | | |
| P68033 | ACTC1 | actin, alpha, cardiac muscle 1 | Cytoplasm | enzyme | 2.46E+08 | 2.09E+08 | 1.60E+08 | | | | | | |
| P68033 | ACTC1 | actin, alpha, cardiac muscle 1 | Cytoplasm | enzyme | 2.88E+07 | 2.19E+07 | 1.76E+07 | P68033 | ACTC1 | 987790000 | 1233924993 | 356203464 | 12 |
| P61161 | ACTR2 | ARP2 actin-related protein 2 homolog (yeast) | Plasma Membrane | other | 6048916.5 | 4993587 | 7032741.5 | | | | | | |
| P61161 | ACTR2 | ARP2 actin-related protein 2 homolog (yeast) | Plasma Membrane | other | 2362344.75 | 3327558.5 | 3404515 | | | | | | |
| P61161 | ACTR2 | ARP2 actin-related protein 2 homolog (yeast) | Plasma Membrane | other | 2734565 | 3266924.25 | 4108600.75 | P61161 | ACTR2 | 3715275 | 1574803 | 524934 | 9 |
| Q99JY9 | ACTR3 | ARP3 actin-related protein 3 homolog (yeast) | Plasma Membrane | other | 1.08E+07 | 1.02E+07 | 1.80E+07 | | | | | | |
| Q99JY9 | ACTR3 | ARP3 actin-related protein 3 homolog (yeast) | Plasma Membrane | other | 2508569.5 | 3615419.75 | 2887375.25 | | | | | | |
| Q99JY9 | ACTR3 | ARP3 actin-related protein 3 homolog (yeast) | Plasma Membrane | other | 3854013.75 | 3893645.25 | 4155091.25 | | | | | | |
| Q99JY9 | ACTR3 | ARP3 actin-related protein 3 homolog (yeast) | Plasma Membrane | other | 2375804.25 | 2372478.5 | 2933500 | | | | | | |
| Q99JY9 | ACTR3 | ARP3 actin-related protein 3 homolog (yeast) | Plasma Membrane | other | 2379104.25 | 2251113.5 | 3555259.75 | Q99JY9 | ACTR3 | 4377498 | 4475188 | 1155489 | 15 |
| Q641P0 | ACTR3B | ARP3 actin-related protein 3 homolog B (yeast) | Cytoplasm | other | 4018302.5 | 3800045.5 | 6817445 | Q641P0 | ACTR3B | 4018303 | 1682634 | 971469 | 3 |
| O35598 | ADAM10 | ADAM metallopeptidase domain 10 | Plasma Membrane | peptidase | 9.04E+07 | 7.22E+07 | 8.34E+07 | | | | | | |
| O35598 | ADAM10 | ADAM metallopeptidase domain 10 | Plasma Membrane | peptidase | 5.11E+07 | 5.31E+07 | 5.32E+07 | | | | | | |
| O35598 | ADAM10 | ADAM metallopeptidase domain 10 | Plasma Membrane | peptidase | 5.69E+07 | 5.81E+07 | 5.33E+07 | | | | | | |
| O35598 | ADAM10 | ADAM metallopeptidase domain 10 | Plasma Membrane | peptidase | 6.59E+07 | 6.35E+07 | 6.03E+07 | | | | | | |
| O35598 | ADAM10 | ADAM metallopeptidase domain 10 | Plasma Membrane | peptidase | 5.69E+07 | 5.36E+07 | 5.05E+07 | | | | | | |
| O35598 | ADAM10 | ADAM metallopeptidase domain 10 | Plasma Membrane | peptidase | 4.84E+07 | 4.71E+07 | 4.54E+07 | | | | | | |
| O35598 | ADAM10 | ADAM metallopeptidase domain 10 | Plasma Membrane | peptidase | 7.63E+07 | 6.37E+07 | 6.89E+07 | | | | | | |
| O35598 | ADAM10 | ADAM metallopeptidase domain 10 | Plasma Membrane | peptidase | 3.71E+07 | 4.37E+07 | 3.99E+07 | | | | | | |
| O35598 | ADAM10 | ADAM metallopeptidase domain 10 | Plasma Membrane | peptidase | 2.56E+07 | 2.27E+07 | 2.58E+07 | | | | | | |
| O35598 | ADAM10 | ADAM metallopeptidase domain 10 | Plasma Membrane | peptidase | 2.30E+07 | 2.01E+07 | 2.08E+07 | | | | | | |
| O35598 | ADAM10 | ADAM metallopeptidase domain 10 | Plasma Membrane | peptidase | 2.71E+07 | 2.84E+07 | 2.17E+07 | | | | | | |
| O35598 | ADAM10 | ADAM metallopeptidase domain 10 | Plasma Membrane | peptidase | 1.42E+07 | 9163557 | 1.50E+07 | | | | | | |
| O35598 | ADAM10 | ADAM metallopeptidase domain 10 | Plasma Membrane | peptidase | 2.24E+07 | 3.20E+07 | 1.63E+07 | | | | | | |
| O35598 | ADAM10 | ADAM metallopeptidase domain 10 | Plasma Membrane | peptidase | 1.94E+07 | 1.74E+07 | 1.55E+07 | | | | | | |
| O35598 | ADAM10 | ADAM metallopeptidase domain 10 | Plasma Membrane | peptidase | 1.53E+07 | 1.14E+07 | 1.33E+07 | | | | | | |
| O35598 | ADAM10 | ADAM metallopeptidase domain 10 | Plasma Membrane | peptidase | 1.22E+07 | 7733473 | 6740789.5 | | | | | | |
| O35598 | ADAM10 | ADAM metallopeptidase domain 10 | Plasma Membrane | peptidase | 1.39E+07 | 1.14E+07 | 1.12E+07 | | | | | | |
| O35598 | ADAM10 | ADAM metallopeptidase domain 10 | Plasma Membrane | peptidase | 6708961.5 | 4421507 | 5591659.5 | | | | | | |
| O35598 | ADAM10 | ADAM metallopeptidase domain 10 | Plasma Membrane | peptidase | 4874260 | 4348194.5 | 4289162 | | | | | | |
| O35598 | ADAM10 | ADAM metallopeptidase domain 10 | Plasma Membrane | peptidase | 1905413 | 893566.875 | 512217.406 | | | | | | |
| O35598 | ADAM10 | ADAM metallopeptidase domain 10 | Plasma Membrane | peptidase | 3952855.5 | 5150894.5 | 2919041.5 | | | | | | |
| O35598 | ADAM10 | ADAM metallopeptidase domain 10 | Plasma Membrane | peptidase | 4567139 | 5599671 | 3638257 | | | | | | |
| O35598 | ADAM10 | ADAM metallopeptidase domain 10 | Plasma Membrane | peptidase | 4054962 | 3264238.75 | 3498419 | | | | | | |
| O35598 | ADAM10 | ADAM metallopeptidase domain 10 | Plasma Membrane | peptidase | 1.07E+07 | 7987498 | 9006156 | | | | | | |
| O35598 | ADAM10 | ADAM metallopeptidase domain 10 | Plasma Membrane | peptidase | 7987620 | 1.02E+07 | 8051267.5 | | | | | | |
| O35598 | ADAM10 | ADAM metallopeptidase domain 10 | Plasma Membrane | peptidase | 3465024 | 2320323.25 | 2437507.75 | O35598 | ADAM10 | 27082932 | 23851369 | 2700636 | 78 |
| Q9Z0F8 | ADAM17 | ADAM metallopeptidase domain 17 | Plasma Membrane | peptidase | 5406827 | 7139299 | 6287563.5 | | | | | | |
| Q9Z0F8 | ADAM17 | ADAM metallopeptidase domain 17 | Plasma Membrane | peptidase | 824233.625 | 4942306 | 6155157 | Q9Z0F8 | ADAM17 | 3115530 | 2239624 | 914322 | 6 |
| Q9QYC0 | ADD1 | adducin 1 (alpha) | Cytoplasm | other | 8.25E+08 | 1.05E+09 | 7.01E+08 | | | | | | |
| Q9QYC0 | ADD1 | adducin 1 (alpha) | Cytoplasm | other | 6.24E+08 | 4.81E+08 | 4.63E+08 | | | | | | |
| Q9QYC0 | ADD1 | adducin 1 (alpha) | Cytoplasm | other | 3.78E+08 | 3.53E+08 | 3.26E+08 | | | | | | |
| Q9QYC0 | ADD1 | adducin 1 (alpha) | Cytoplasm | other | 4.03E+08 | 3.45E+08 | 4.42E+08 | | | | | | |
| Q9QYC0 | ADD1 | adducin 1 (alpha) | Cytoplasm | other | 4.11E+08 | 3.64E+08 | 3.45E+08 | | | | | | |
| Q9QYC0 | ADD1 | adducin 1 (alpha) | Cytoplasm | other | 2.93E+08 | 2.42E+08 | 2.30E+08 | | | | | | |
| Q9QYC0 | ADD1 | adducin 1 (alpha) | Cytoplasm | other | 2.57E+08 | 2.24E+08 | 1.93E+08 | | | | | | |
| Q9QYC0 | ADD1 | adducin 1 (alpha) | Cytoplasm | other | 3.41E+08 | 2.75E+08 | 2.32E+08 | | | | | | |
| Q9QYC0 | ADD1 | adducin 1 (alpha) | Cytoplasm | other | 3.12E+08 | 3.00E+08 | 1.77E+08 | | | | | | |
| Q9QYC0 | ADD1 | adducin 1 (alpha) | Cytoplasm | other | 2.79E+08 | 2.31E+08 | 2.08E+08 | | | | | | |
| Q9QYC0 | ADD1 | adducin 1 (alpha) | Cytoplasm | other | 2.51E+08 | 2.26E+08 | 1.80E+08 | | | | | | |
| Q9QYC0 | ADD1 | adducin 1 (alpha) | Cytoplasm | other | 2.74E+08 | 2.51E+08 | 2.36E+08 | | | | | | |
| Q9QYC0 | ADD1 | adducin 1 (alpha) | Cytoplasm | other | 2.22E+08 | 2.09E+08 | 1.92E+08 | | | | | | |
| Q9QYC0 | ADD1 | adducin 1 (alpha) | Cytoplasm | other | 2.42E+08 | 2.32E+08 | 2.08E+08 | | | | | | |
| Q9QYC0 | ADD1 | adducin 1 (alpha) | Cytoplasm | other | 2.70E+08 | 2.88E+08 | 1.88E+08 | | | | | | |
| Q9QYC0 | ADD1 | adducin 1 (alpha) | Cytoplasm | other | 1.68E+08 | 1.05E+08 | 8.99E+07 | | | | | | |
| Q9QYC0 | ADD1 | adducin 1 (alpha) | Cytoplasm | other | 1.18E+08 | 9.24E+07 | 1.23E+08 | | | | | | |
| Q9QYC0 | ADD1 | adducin 1 (alpha) | Cytoplasm | other | 1.16E+08 | 7.16E+07 | 8.64E+07 | | | | | | |
| Q9QYC0 | ADD1 | adducin 1 (alpha) | Cytoplasm | other | 1.16E+08 | 9.82E+07 | 9.37E+07 | | | | | | |
| Q9QYC0 | ADD1 | adducin 1 (alpha) | Cytoplasm | other | 4.49E+07 | 5.58E+07 | 3.94E+07 | | | | | | |
| Q9QYC0 | ADD1 | adducin 1 (alpha) | Cytoplasm | other | 5.28E+07 | 4.68E+07 | 4.26E+07 | | | | | | |
| Q9QYC0 | ADD1 | adducin 1 (alpha) | Cytoplasm | other | 6.17E+07 | 6.87E+07 | 3.69E+07 | | | | | | |
| Q9QYC0 | ADD1 | adducin 1 (alpha) | Cytoplasm | other | 4.56E+07 | 4.09E+07 | 3.68E+07 | | | | | | |
| Q9QYC0 | ADD1 | adducin 1 (alpha) | Cytoplasm | other | 4.85E+07 | 7.06E+07 | 3.73E+07 | | | | | | |
| Q9QYC0 | ADD1 | adducin 1 (alpha) | Cytoplasm | other | 3.26E+07 | 3.67E+07 | 2.64E+07 | | | | | | |
| Q9QYC0 | ADD1 | adducin 1 (alpha) | Cytoplasm | other | 3.52E+07 | 5.27E+07 | 3.07E+07 | | | | | | |
| Q9QYC0 | ADD1 | adducin 1 (alpha) | Cytoplasm | other | 1.94E+07 | 1.87E+07 | 1.76E+07 | | | | | | |
| Q9QYC0 | ADD1 | adducin 1 (alpha) | Cytoplasm | other | 3.15E+07 | 4.14E+07 | 3.65E+07 | | | | | | |
| Q9QYC0 | ADD1 | adducin 1 (alpha) | Cytoplasm | other | 2.50E+07 | 1.76E+07 | 1.74E+07 | | | | | | |
| Q9QYC0 | ADD1 | adducin 1 (alpha) | Cytoplasm | other | 1.98E+07 | 1.96E+07 | 1.43E+07 | | | | | | |
| Q9QYC0 | ADD1 | adducin 1 (alpha) | Cytoplasm | other | 1.08E+07 | 1.48E+07 | 9795654 | | | | | | |
| Q9QYC0 | ADD1 | adducin 1 (alpha) | Cytoplasm | other | 1.36E+07 | 2.34E+07 | 9939777 | | | | | | |
| Q9QYC0 | ADD1 | adducin 1 (alpha) | Cytoplasm | other | 1.53E+07 | 1.32E+07 | 1.17E+07 | | | | | | |
| Q9QYC0 | ADD1 | adducin 1 (alpha) | Cytoplasm | other | 1.53E+07 | 1.78E+07 | 9556815 | | | | | | |
| Q9QYC0 | ADD1 | adducin 1 (alpha) | Cytoplasm | other | 9222928 | 1.15E+07 | 5533454 | | | | | | |
| Q9QYC0 | ADD1 | adducin 1 (alpha) | Cytoplasm | other | 1.49E+07 | 8195324 | 1.06E+07 | | | | | | |
| Q9QYC0 | ADD1 | adducin 1 (alpha) | Cytoplasm | other | 9345606 | 6097171.5 | 6060844 | | | | | | |
| Q9QYC0 | ADD1 | adducin 1 (alpha) | Cytoplasm | other | 7292552 | 5618235 | 3946569.5 | | | | | | |
| Q9QYC0 | ADD1 | adducin 1 (alpha) | Cytoplasm | other | 1.80E+07 | 1.33E+07 | 1.46E+07 | | | | | | |

| | | | | | | | | | | | | | |
|---|---|---|---|---|---|---|---|---|---|---|---|---|---|
| Q9QYC0 | ADD1 | adducin 1 (alpha) | Cytoplasm | other | 1573066.125 | 1935572.5 | 2057542.75 | | | | | | |
| Q9QYC0 | ADD1 | adducin 1 (alpha) | Cytoplasm | other | 520216.094 | 377822.781 | 444964.031 | Q9QYC0 | ADD1 | 156847911 | 178172720 | 16065293 | 123 |
| Q9QYB8 | ADD2 | adducin 2 (beta) | Cytoplasm | other | 4.35E+08 | 4.37E+08 | 4.41E+08 | | | | | | |
| Q9QYB8 | ADD2 | adducin 2 (beta) | Cytoplasm | other | 3.66E+08 | 4.42E+08 | 3.47E+08 | | | | | | |
| Q9QYB8 | ADD2 | adducin 2 (beta) | Cytoplasm | other | 2.86E+08 | 2.39E+08 | 2.14E+08 | | | | | | |
| Q9QYB8 | ADD2 | adducin 2 (beta) | Cytoplasm | other | 4.79E+08 | 4.91E+08 | 4.38E+08 | | | | | | |
| Q9QYB8 | ADD2 | adducin 2 (beta) | Cytoplasm | other | 2.30E+08 | 2.36E+08 | 1.69E+08 | | | | | | |
| Q9QYB8 | ADD2 | adducin 2 (beta) | Cytoplasm | other | 2.48E+08 | 2.51E+08 | 2.15E+08 | | | | | | |
| Q9QYB8 | ADD2 | adducin 2 (beta) | Cytoplasm | other | 2.31E+08 | 2.01E+08 | 1.79E+08 | | | | | | |
| Q9QYB8 | ADD2 | adducin 2 (beta) | Cytoplasm | other | 2.38E+08 | 1.33E+08 | 2.00E+08 | | | | | | |
| Q9QYB8 | ADD2 | adducin 2 (beta) | Cytoplasm | other | 2.28E+08 | 2.09E+08 | 1.93E+08 | | | | | | |
| Q9QYB8 | ADD2 | adducin 2 (beta) | Cytoplasm | other | 2.40E+08 | 2.70E+08 | 2.40E+08 | | | | | | |
| Q9QYB8 | ADD2 | adducin 2 (beta) | Cytoplasm | other | 2.08E+08 | 1.67E+08 | 1.32E+08 | | | | | | |
| Q9QYB8 | ADD2 | adducin 2 (beta) | Cytoplasm | other | 2.64E+08 | 2.75E+08 | 1.93E+08 | | | | | | |
| Q9QYB8 | ADD2 | adducin 2 (beta) | Cytoplasm | other | 1.84E+08 | 1.74E+08 | 1.62E+08 | | | | | | |
| Q9QYB8 | ADD2 | adducin 2 (beta) | Cytoplasm | other | 2.11E+08 | 1.67E+08 | 1.76E+08 | | | | | | |
| Q9QYB8 | ADD2 | adducin 2 (beta) | Cytoplasm | other | 1.85E+08 | 1.72E+08 | 1.30E+08 | | | | | | |
| Q9QYB8 | ADD2 | adducin 2 (beta) | Cytoplasm | other | 1.69E+08 | 1.06E+08 | 1.29E+08 | | | | | | |
| Q9QYB8 | ADD2 | adducin 2 (beta) | Cytoplasm | other | 1.63E+08 | 1.47E+08 | 1.37E+08 | | | | | | |
| Q9QYB8 | ADD2 | adducin 2 (beta) | Cytoplasm | other | 1.53E+08 | 1.47E+08 | 1.19E+08 | | | | | | |
| Q9QYB8 | ADD2 | adducin 2 (beta) | Cytoplasm | other | 8.36E+07 | 1.19E+08 | 5.97E+07 | | | | | | |
| Q9QYB8 | ADD2 | adducin 2 (beta) | Cytoplasm | other | 9.14E+07 | 9.37E+07 | 7.77E+07 | | | | | | |
| Q9QYB8 | ADD2 | adducin 2 (beta) | Cytoplasm | other | 8.22E+07 | 8.69E+07 | 4.56E+07 | | | | | | |
| Q9QYB8 | ADD2 | adducin 2 (beta) | Cytoplasm | other | 9.49E+07 | 7.91E+07 | 8.52E+07 | | | | | | |
| Q9QYB8 | ADD2 | adducin 2 (beta) | Cytoplasm | other | 6.73E+07 | 8.71E+07 | 4.72E+07 | | | | | | |
| Q9QYB8 | ADD2 | adducin 2 (beta) | Cytoplasm | other | 4.70E+07 | 5.06E+07 | 4.81E+07 | | | | | | |
| Q9QYB8 | ADD2 | adducin 2 (beta) | Cytoplasm | other | 1.60E+08 | 1.69E+08 | 1.20E+08 | | | | | | |
| Q9QYB8 | ADD2 | adducin 2 (beta) | Cytoplasm | other | 3.80E+07 | 3.00E+07 | 1.45E+07 | | | | | | |
| Q9QYB8 | ADD2 | adducin 2 (beta) | Cytoplasm | other | 6.56E+07 | 4.83E+07 | 4.16E+07 | | | | | | |
| Q9QYB8 | ADD2 | adducin 2 (beta) | Cytoplasm | other | 5.00E+07 | 4.12E+07 | 1.86E+07 | | | | | | |
| Q9QYB8 | ADD2 | adducin 2 (beta) | Cytoplasm | other | 4.67E+07 | 5.24E+07 | 4.19E+07 | | | | | | |
| Q9QYB8 | ADD2 | adducin 2 (beta) | Cytoplasm | other | 3.98E+07 | 4.15E+07 | 3.17E+07 | | | | | | |
| Q9QYB8 | ADD2 | adducin 2 (beta) | Cytoplasm | other | 3.70E+07 | 2.68E+07 | 2.21E+07 | | | | | | |
| Q9QYB8 | ADD2 | adducin 2 (beta) | Cytoplasm | other | 3.08E+07 | 2.17E+07 | 2.19E+07 | | | | | | |
| Q9QYB8 | ADD2 | adducin 2 (beta) | Cytoplasm | other | 3.22E+07 | 3.26E+07 | 2.70E+07 | | | | | | |
| Q9QYB8 | ADD2 | adducin 2 (beta) | Cytoplasm | other | 2.43E+07 | 2.22E+07 | 1.58E+07 | | | | | | |
| Q9QYB8 | ADD2 | adducin 2 (beta) | Cytoplasm | other | 2.03E+07 | 2.64E+07 | 1.53E+07 | | | | | | |
| Q9QYB8 | ADD2 | adducin 2 (beta) | Cytoplasm | other | 4.22E+07 | 2.42E+07 | 2.98E+07 | | | | | | |
| Q9QYB8 | ADD2 | adducin 2 (beta) | Cytoplasm | other | 1.88E+07 | 1.05E+07 | 9174531 | | | | | | |
| Q9QYB8 | ADD2 | adducin 2 (beta) | Cytoplasm | other | 1.57E+07 | 1.64E+07 | 9977641 | | | | | | |
| Q9QYB8 | ADD2 | adducin 2 (beta) | Cytoplasm | other | 2.19E+07 | 1.94E+07 | 1.31E+07 | | | | | | |
| Q9QYB8 | ADD2 | adducin 2 (beta) | Cytoplasm | other | 1.46E+07 | 1.10E+07 | 8711923 | | | | | | |
| Q9QYB8 | ADD2 | adducin 2 (beta) | Cytoplasm | other | 5358952.5 | 4400370.5 | 1.39E+07 | | | | | | |
| Q9QYB8 | ADD2 | adducin 2 (beta) | Cytoplasm | other | 2.17E+07 | 2.03E+07 | 2.28E+07 | | | | | | |
| Q9QYB8 | ADD2 | adducin 2 (beta) | Cytoplasm | other | 1.40E+07 | 1.49E+07 | 7754359 | | | | | | |
| Q9QYB8 | ADD2 | adducin 2 (beta) | Cytoplasm | other | 1.02E+07 | 8900753 | 3968502 | | | | | | |
| Q9QYB8 | ADD2 | adducin 2 (beta) | Cytoplasm | other | 6162067.5 | 7702256 | 5728694 | | | | | | |
| Q9QYB8 | ADD2 | adducin 2 (beta) | Cytoplasm | other | 7611897.5 | 8031451 | 4542042 | | | | | | |
| Q9QYB8 | ADD2 | adducin 2 (beta) | Cytoplasm | other | 7036648 | 1.11E+07 | 4713469.5 | | | | | | |
| Q9QYB8 | ADD2 | adducin 2 (beta) | Cytoplasm | other | 7527897 | 7915581.5 | 4876323.5 | | | | | | |
| Q9QYB8 | ADD2 | adducin 2 (beta) | Cytoplasm | other | 1.05E+07 | 1.22E+07 | 6342675.5 | | | | | | |
| Q9QYB8 | ADD2 | adducin 2 (beta) | Cytoplasm | other | 4668709 | 1.40E+07 | 1.07E+07 | | | | | | |
| Q9QYB8 | ADD2 | adducin 2 (beta) | Cytoplasm | other | 3250324 | 3356080 | 2850175 | | | | | | |
| Q9QYB8 | ADD2 | adducin 2 (beta) | Cytoplasm | other | 3787549.5 | 1468526.125 | 1231861.125 | | | | | | |
| Q9QYB8 | ADD2 | adducin 2 (beta) | Cytoplasm | other | 5068417.5 | 3540657 | 3209401 | | | | | | |
| Q9QYB8 | ADD2 | adducin 2 (beta) | Cytoplasm | other | 7231915 | 6717073.5 | 5298638.5 | | | | | | |
| Q9QYB8 | ADD2 | adducin 2 (beta) | Cytoplasm | other | 4931602.5 | 3439034 | 2435480 | | | | | | |
| Q9QYB8 | ADD2 | adducin 2 (beta) | Cytoplasm | other | 2664395 | 2935447.25 | 1588189.125 | | | | | | |
| Q9QYB8 | ADD2 | adducin 2 (beta) | Cytoplasm | other | 309796.906 | 377295.188 | 2271398.75 | | | | | | |
| Q9QYB8 | ADD2 | adducin 2 (beta) | Cytoplasm | other | 1272465.625 | 2801361.75 | 1099650.125 | | | | | | |
| Q9QYB8 | ADD2 | adducin 2 (beta) | Cytoplasm | other | 6438238.5 | 3211894.5 | 2127628 | | | | | | |
| Q9QYB8 | ADD2 | adducin 2 (beta) | Cytoplasm | other | 3486037 | 2249087.5 | 1317561 | | | | | | |
| Q9QYB8 | ADD2 | adducin 2 (beta) | Cytoplasm | other | 1090409.5 | 1272313 | 1063095.625 | | | | | | |
| Q9QYB8 | ADD2 | adducin 2 (beta) | Cytoplasm | other | 256429.562 | 4014.461 | 1201.406 | | | | | | |
| Q9QYB8 | ADD2 | adducin 2 (beta) | Cytoplasm | other | 843295.938 | 679675.688 | 410986.5 | | | | | | |
| Q9QYB8 | ADD2 | adducin 2 (beta) | Cytoplasm | other | 3529473 | 3160702.5 | 3132990.5 | Q9QYB8 | ADD2 | 90340883 | 110979207 | 8009234 | 192 |
| Q9QYB5 | ADD3 | adducin 3 (gamma) | Cytoplasm | other | 2.67E+07 | 2.84E+07 | 3.12E+07 | | | | | | |
| Q9QYB5 | ADD3 | adducin 3 (gamma) | Cytoplasm | other | 1.74E+07 | 1.63E+07 | 1.75E+07 | | | | | | |
| Q9QYB5 | ADD3 | adducin 3 (gamma) | Cytoplasm | other | 1.22E+07 | 1.14E+07 | 8273959 | | | | | | |
| Q9QYB5 | ADD3 | adducin 3 (gamma) | Cytoplasm | other | 1.00E+07 | 9660925 | 1.09E+07 | | | | | | |
| Q9QYB5 | ADD3 | adducin 3 (gamma) | Cytoplasm | other | 7378421 | 8050196 | 9892545 | | | | | | |
| Q9QYB5 | ADD3 | adducin 3 (gamma) | Cytoplasm | other | 3975171.75 | 5345742.5 | 3771775.5 | | | | | | |
| Q9QYB5 | ADD3 | adducin 3 (gamma) | Cytoplasm | other | 6318984.5 | 5806884.5 | 6750247.5 | | | | | | |
| Q9QYB5 | ADD3 | adducin 3 (gamma) | Cytoplasm | other | 8136190.5 | 6148632.5 | 6374758.5 | | | | | | |
| Q9QYB5 | ADD3 | adducin 3 (gamma) | Cytoplasm | other | 5719460.5 | 5479565.5 | 4723977.5 | | | | | | |
| Q9QYB5 | ADD3 | adducin 3 (gamma) | Cytoplasm | other | 4301917.5 | 3324393 | 3175100.5 | | | | | | |
| Q9QYB5 | ADD3 | adducin 3 (gamma) | Cytoplasm | other | 7327965.5 | 6862088.5 | 6432230 | | | | | | |
| Q9QYB5 | ADD3 | adducin 3 (gamma) | Cytoplasm | other | 4525520.5 | 5372446.5 | 3467123 | | | | | | |
| Q9QYB5 | ADD3 | adducin 3 (gamma) | Cytoplasm | other | 2341210.75 | 2553067.75 | 1541815.25 | | | | | | |
| Q9QYB5 | ADD3 | adducin 3 (gamma) | Cytoplasm | other | 3640553.25 | 3961605 | 4184568.25 | | | | | | |
| Q9QYB5 | ADD3 | adducin 3 (gamma) | Cytoplasm | other | 3151766 | 2628863.25 | 2555534.75 | Q9QYB5 | ADD3 | 8207811 | 6774175 | 1009834 | 45 |
| Q8VDL4 | ADPGK | ADP-dependent glucokinase | Plasma Membrane | kinase | 3317001.25 | 3329964 | 3969130.75 | Q8VDL4 | ADPGK | 3317001 | 372821 | 215249 | 3 |
| P54822 | ADSL | adenylosuccinate lyase | Cytoplasm | enzyme | 1.27E+07 | 1.18E+07 | 9150192 | | | | | | |
| P54822 | ADSL | adenylosuccinate lyase | Cytoplasm | enzyme | 9878107 | 7472444 | 7390828 | | | | | | |
| P54822 | ADSL | adenylosuccinate lyase | Cytoplasm | enzyme | 1.11E+07 | 8015655.5 | 6705346 | | | | | | |
| P54822 | ADSL | adenylosuccinate lyase | Cytoplasm | enzyme | 1.14E+07 | 8516285 | 7982698.5 | | | | | | |
| P54822 | ADSL | adenylosuccinate lyase | Cytoplasm | enzyme | 1.04E+07 | 7469276 | 6615426.5 | | | | | | |
| P54822 | ADSL | adenylosuccinate lyase | Cytoplasm | enzyme | 8208743.5 | 5438328.5 | 5820597.5 | | | | | | |
| P54822 | ADSL | adenylosuccinate lyase | Cytoplasm | enzyme | 6527842.5 | 4773190 | 4073011.25 | | | | | | |
| P54822 | ADSL | adenylosuccinate lyase | Cytoplasm | enzyme | 8023524.5 | 6402558 | 5056224.5 | | | | | | |
| P54822 | ADSL | adenylosuccinate lyase | Cytoplasm | enzyme | 5534463 | 5492836.5 | 3828677.5 | | | | | | |
| P54822 | ADSL | adenylosuccinate lyase | Cytoplasm | enzyme | 2108966.5 | 2813392 | 3014369.75 | | | | | | |
| P54822 | ADSL | adenylosuccinate lyase | Cytoplasm | enzyme | 2200541.5 | 2592604.25 | 1887907.75 | | | | | | |
| P54822 | ADSL | adenylosuccinate lyase | Cytoplasm | enzyme | 891923.5 | 790334.375 | 763010.125 | | | | | | |
| P54822 | ADSL | adenylosuccinate lyase | Cytoplasm | enzyme | 2237012 | 1408587.125 | 1162310 | P54822 | ADSL | 7015471 | 3385721 | 542149 | 39 |
| Q640N1 | AEBP1 | AE binding protein 1 | Nucleus | peptidase | 9436323 | 1.02E+07 | 7899814 | Q640N1 | AEBP1 | 9436323 | 1175896 | 678904 | 3 |
| O55112 | Aff2 | AF4/FMR2 family, member 2 | Nucleus | other | 2.90E+07 | 1.90E+07 | 6816319.5 | O55112 | Aff2 | 28960000 | 11090433 | 6403065 | 3 |
| Q8K2K6 | AGFG1 | ArfGAP with FG repeats 1 | Nucleus | other | 2337962.25 | 2528213.25 | 2265261.5 | Q8K2K6 | AGFG1 | 2337962 | 135784 | 78395 | 3 |
| Q80WC7 | AGFG2 | ArfGAP with FG repeats 2 | Cytoplasm | other | 3478510 | 3907921.5 | 3224673.25 | | | | | | |
| Q80WC7 | AGFG2 | ArfGAP with FG repeats 2 | Cytoplasm | other | 2798751.25 | 3298583.25 | 2239008.75 | | | | | | |
| Q80WC7 | AGFG2 | ArfGAP with FG repeats 2 | Cytoplasm | other | 2161893.5 | 1808166.5 | 2367622.75 | | | | | | |
| Q80WC7 | AGFG2 | ArfGAP with FG repeats 2 | Cytoplasm | other | 2364910.25 | 4165229.5 | 2806955.75 | Q80WC7 | AGFG2 | 2701016 | 737039 | 212765 | 12 |
| Q8CJG0 | AGO2 | argonaute RISC catalytic component 2 | Cytoplasm | translation regulator | 4.63E+07 | 4.27E+07 | 3.92E+07 | | | | | | |
| Q8CJG0 | AGO2 | argonaute RISC catalytic component 2 | Cytoplasm | translation regulator | 3.28E+07 | 3.17E+07 | 3.36E+07 | | | | | | |
| Q8CJG0 | AGO2 | argonaute RISC catalytic component 2 | Cytoplasm | translation regulator | 3.34E+07 | 3.09E+07 | 2.41E+07 | | | | | | |
| Q8CJG0 | AGO2 | argonaute RISC catalytic component 2 | Cytoplasm | translation regulator | 2.22E+07 | 1.96E+07 | 1.86E+07 | | | | | | |
| Q8CJG0 | AGO2 | argonaute RISC catalytic component 2 | Cytoplasm | translation regulator | 2.69E+07 | 2.12E+07 | 1.76E+07 | | | | | | |
| Q8CJG0 | AGO2 | argonaute RISC catalytic component 2 | Cytoplasm | translation regulator | 2.92E+07 | 2.75E+07 | 2.04E+07 | | | | | | |
| Q8CJG0 | AGO2 | argonaute RISC catalytic component 2 | Cytoplasm | translation regulator | 3.11E+07 | 2.12E+07 | 1.79E+07 | | | | | | |

| | | | | | | | | | | | | | |
|---|---|---|---|---|---|---|---|---|---|---|---|---|---|
| Q8CJG0 | AGO2 | argonaute RISC catalytic component 2 | Cytoplasm | translation regulator | 2.32E+07 | 2.14E+07 | 1.88E+07 | | | | | | |
| Q8CJG0 | AGO2 | argonaute RISC catalytic component 2 | Cytoplasm | translation regulator | 2.44E+07 | 2.21E+07 | 2.48E+07 | | | | | | |
| Q8CJG0 | AGO2 | argonaute RISC catalytic component 2 | Cytoplasm | translation regulator | 2.78E+07 | 2.83E+07 | 2.09E+07 | | | | | | |
| Q8CJG0 | AGO2 | argonaute RISC catalytic component 2 | Cytoplasm | translation regulator | 2.26E+07 | 2.06E+07 | 1.71E+07 | | | | | | |
| Q8CJG0 | AGO2 | argonaute RISC catalytic component 2 | Cytoplasm | translation regulator | 1.58E+07 | 1.80E+07 | 1.46E+07 | | | | | | |
| Q8CJG0 | AGO2 | argonaute RISC catalytic component 2 | Cytoplasm | translation regulator | 2.39E+07 | 2.10E+07 | 1.46E+07 | | | | | | |
| Q8CJG0 | AGO2 | argonaute RISC catalytic component 2 | Cytoplasm | translation regulator | 2.40E+07 | 2.19E+07 | 2.59E+07 | | | | | | |
| Q8CJG0 | AGO2 | argonaute RISC catalytic component 2 | Cytoplasm | translation regulator | 2.37E+07 | 2.00E+07 | 1.73E+07 | | | | | | |
| Q8CJG0 | AGO2 | argonaute RISC catalytic component 2 | Cytoplasm | translation regulator | 2.65E+07 | 2.50E+07 | 2.06E+07 | | | | | | |
| Q8CJG0 | AGO2 | argonaute RISC catalytic component 2 | Cytoplasm | translation regulator | 2.21E+07 | 1.52E+07 | 1.50E+07 | | | | | | |
| Q8CJG0 | AGO2 | argonaute RISC catalytic component 2 | Cytoplasm | translation regulator | 1.94E+07 | 1.74E+07 | 1.66E+07 | | | | | | |
| Q8CJG0 | AGO2 | argonaute RISC catalytic component 2 | Cytoplasm | translation regulator | 1.34E+07 | 1.62E+07 | 1.25E+07 | | | | | | |
| Q8CJG0 | AGO2 | argonaute RISC catalytic component 2 | Cytoplasm | translation regulator | 1.48E+07 | 1.20E+07 | 1.23E+07 | | | | | | |
| Q8CJG0 | AGO2 | argonaute RISC catalytic component 2 | Cytoplasm | translation regulator | 2.10E+07 | 1.72E+07 | 1.56E+07 | | | | | | |
| Q8CJG0 | AGO2 | argonaute RISC catalytic component 2 | Cytoplasm | translation regulator | 1.39E+07 | 1.10E+07 | 1.10E+07 | | | | | | |
| Q8CJG0 | AGO2 | argonaute RISC catalytic component 2 | Cytoplasm | translation regulator | 1.48E+07 | 1.28E+07 | 1.25E+07 | | | | | | |
| Q8CJG0 | AGO2 | argonaute RISC catalytic component 2 | Cytoplasm | translation regulator | 1.27E+07 | 1.11E+07 | 8915387 | | | | | | |
| Q8CJG0 | AGO2 | argonaute RISC catalytic component 2 | Cytoplasm | translation regulator | 1.37E+07 | 1.52E+07 | 1.14E+07 | | | | | | |
| Q8CJG0 | AGO2 | argonaute RISC catalytic component 2 | Cytoplasm | translation regulator | 1.05E+07 | 8655733 | 6258172.5 | | | | | | |
| Q8CJG0 | AGO2 | argonaute RISC catalytic component 2 | Cytoplasm | translation regulator | 8465535 | 6458545 | 6039389 | | | | | | |
| Q8CJG0 | AGO2 | argonaute RISC catalytic component 2 | Cytoplasm | translation regulator | 9008025 | 9494820 | 8252988.5 | | | | | | |
| Q8CJG0 | AGO2 | argonaute RISC catalytic component 2 | Cytoplasm | translation regulator | 7526827 | 7087502 | 6436638 | | | | | | |
| Q8CJG0 | AGO2 | argonaute RISC catalytic component 2 | Cytoplasm | translation regulator | 8825394 | 6387569 | 4825546 | | | | | | |
| Q8CJG0 | AGO2 | argonaute RISC catalytic component 2 | Cytoplasm | translation regulator | 5740494 | 4360112.5 | 3056859.25 | | | | | | |
| Q8CJG0 | AGO2 | argonaute RISC catalytic component 2 | Cytoplasm | translation regulator | 6067006 | 5406409 | 4087414 | | | | | | |
| Q8CJG0 | AGO2 | argonaute RISC catalytic component 2 | Cytoplasm | translation regulator | 5129719 | 3284165.25 | 5160808 | | | | | | |
| Q8CJG0 | AGO2 | argonaute RISC catalytic component 2 | Cytoplasm | translation regulator | 7355525 | 2713577.25 | 1165767.25 | | | | | | |
| Q8CJG0 | AGO2 | argonaute RISC catalytic component 2 | Cytoplasm | translation regulator | 6848764.5 | 7844084 | 3620629 | | | | | | |
| Q8CJG0 | AGO2 | argonaute RISC catalytic component 2 | Cytoplasm | translation regulator | 3347451 | 888215.688 | 5014.692 | | | | | | |
| Q8CJG0 | AGO2 | argonaute RISC catalytic component 2 | Cytoplasm | translation regulator | 2162385 | 2122484.75 | 1520947.125 | | | | | | |
| Q8CJG0 | AGO2 | argonaute RISC catalytic component 2 | Cytoplasm | translation regulator | 5008307 | 1.56E+07 | 1.54E+07 | | | | | | |
| Q8CJG0 | AGO2 | argonaute RISC catalytic component 2 | Cytoplasm | translation regulator | 789816.875 | 1221051.375 | 456869.719 | | | | | | |
| Q8CJG0 | AGO2 | argonaute RISC catalytic component 2 | Cytoplasm | translation regulator | 842303.688 | 322887 | 361247.875 | | | | | | |
| Q8CJG0 | AGO2 | argonaute RISC catalytic component 2 | Cytoplasm | translation regulator | 344270.969 | 129178.469 | 47274.703 | Q8CJG0 | AGO2 | 16273703 | 10019259 | 903406 | 123 |
| O35083 | AGPAT1 | 1-acylglycerol-3-phosphate O-acyltransferase 1 | Cytoplasm | enzyme | 1215250.625 | 1292447.75 | 1019387.938 | O35083 | AGPAT1 | 1215251 | 140762 | 81269 | 3 |
| P50247 | AHCY | adenosylhomocysteinase | Cytoplasm | enzyme | 3969678 | 3199980.25 | 1942590.375 | | | | | | |
| P50247 | AHCY | adenosylhomocysteinase | Cytoplasm | enzyme | 3190363.25 | 2237612.5 | 1376298.5 | | | | | | |
| P50247 | AHCY | adenosylhomocysteinase | Cytoplasm | enzyme | 2577634.75 | 1687509.25 | 1381129 | P50247 | AHCY | 3245892 | 907554 | 302518 | 9 |
| Q8N9S3 | Ahsa2 | AHA1, activator of heat shock protein ATPase 2 | Cytoplasm | other | 2.68E+08 | 3.32E+08 | 3.36E+08 | Q8N9S3 | Ahsa2 | 268000000 | 37963974 | 21918511 | 3 |
| Q9Z0X1 | AIFM1 | apoptosis-inducing factor, mitochondrion-associated, 1 | Cytoplasm | enzyme | 1563899.75 | 2194170 | 3009359.5 | | | | | | |
| Q9Z0X1 | AIFM1 | apoptosis-inducing factor, mitochondrion-associated, 1 | Cytoplasm | enzyme | 1338441.375 | 1617862.75 | 2663592.5 | Q9Z0X1 | AIFM1 | 1451171 | 670184 | 273601 | 6 |
| P31230 | AIMP1 | aminoacyl tRNA synthetase complex-interacting multifunctional protein 1 | Extracellular Space | cytokine | 974152.25 | 918895.938 | 1140852.25 | | | | | | |
| P31230 | AIMP1 | aminoacyl tRNA synthetase complex-interacting multifunctional protein 1 | Extracellular Space | cytokine | 119396.977 | 135728.922 | 276183.031 | P31230 | AIMP1 | 546775 | 465916 | 190209 | 6 |
| Q9WTP6 | AK2 | adenylate kinase 2 | Cytoplasm | kinase | 503571.688 | 1161536.125 | 1361762.875 | Q9WTP6 | AK2 | 503572 | 448980 | 259219 | 3 |
| P10518 | ALAD | aminolevulinate dehydratase | Cytoplasm | enzyme | 4.90E+07 | 4.52E+07 | 5.54E+07 | | | | | | |
| P10518 | ALAD | aminolevulinate dehydratase | Cytoplasm | enzyme | 2.76E+07 | 2.21E+07 | 2.94E+07 | | | | | | |
| P10518 | ALAD | aminolevulinate dehydratase | Cytoplasm | enzyme | 3.06E+07 | 2.73E+07 | 4.07E+07 | | | | | | |
| P10518 | ALAD | aminolevulinate dehydratase | Cytoplasm | enzyme | 1.69E+07 | 1.24E+07 | 1.51E+07 | | | | | | |
| P10518 | ALAD | aminolevulinate dehydratase | Cytoplasm | enzyme | 1.77E+07 | 1.51E+07 | 2.04E+07 | | | | | | |
| P10518 | ALAD | aminolevulinate dehydratase | Cytoplasm | enzyme | 1.40E+07 | 1.30E+07 | 1.31E+07 | | | | | | |
| P10518 | ALAD | aminolevulinate dehydratase | Cytoplasm | enzyme | 9583515 | 9884927 | 1.28E+07 | | | | | | |
| P10518 | ALAD | aminolevulinate dehydratase | Cytoplasm | enzyme | 4332776.5 | 2152690 | 6373767 | | | | | | |
| P10518 | ALAD | aminolevulinate dehydratase | Cytoplasm | enzyme | 2809695.75 | 2118090.5 | 3145117.75 | | | | | | |
| P10518 | ALAD | aminolevulinate dehydratase | Cytoplasm | enzyme | 2655672.25 | 3448560.5 | 2481018.5 | | | | | | |
| P10518 | ALAD | aminolevulinate dehydratase | Cytoplasm | enzyme | 2582229.25 | 8172322.5 | 1.07E+07 | | | | | | |
| P10518 | ALAD | aminolevulinate dehydratase | Cytoplasm | enzyme | 3168481.5 | 2319668.5 | 4009929.5 | | | | | | |
| P10518 | ALAD | aminolevulinate dehydratase | Cytoplasm | enzyme | 2827081 | 2849801 | 2436483.5 | | | | | | |
| P10518 | ALAD | aminolevulinate dehydratase | Cytoplasm | enzyme | 2908472.25 | 3853250.5 | 3572344.75 | | | | | | |
| P10518 | ALAD | aminolevulinate dehydratase | Cytoplasm | enzyme | 1940079.5 | 1903299.75 | 2396405.75 | | | | | | |
| P10518 | ALAD | aminolevulinate dehydratase | Cytoplasm | enzyme | 462799.219 | 158929.766 | 57674.664 | P10518 | ALAD | 11818175 | 13682782 | 1974939 | 48 |
| P08680 | ALAS2 | 5'-aminolevulinate synthase 2 | Cytoplasm | enzyme | 1.67E+07 | 3.32E+07 | 3.04E+07 | | | | | | |
| P08680 | ALAS2 | 5'-aminolevulinate synthase 2 | Cytoplasm | enzyme | 1.13E+07 | 2.17E+07 | 2.63E+07 | | | | | | |
| P08680 | ALAS2 | 5'-aminolevulinate synthase 2 | Cytoplasm | enzyme | 9723976 | 1.50E+07 | 1.46E+07 | | | | | | |
| P08680 | ALAS2 | 5'-aminolevulinate synthase 2 | Cytoplasm | enzyme | 6010375.5 | 1.04E+07 | 1.16E+07 | | | | | | |
| P08680 | ALAS2 | 5'-aminolevulinate synthase 2 | Cytoplasm | enzyme | 5978599 | 9997349 | 1.01E+07 | | | | | | |
| P08680 | ALAS2 | 5'-aminolevulinate synthase 2 | Cytoplasm | enzyme | 4527643.5 | 1.05E+07 | 9600008 | | | | | | |
| P08680 | ALAS2 | 5'-aminolevulinate synthase 2 | Cytoplasm | enzyme | 4241632 | 7813166.5 | 7824568.5 | | | | | | |
| P08680 | ALAS2 | 5'-aminolevulinate synthase 2 | Cytoplasm | enzyme | 3785110 | 7271660 | 8597958 | | | | | | |
| P08680 | ALAS2 | 5'-aminolevulinate synthase 2 | Cytoplasm | enzyme | 2724358.5 | 5812752.5 | 6056852 | | | | | | |
| P08680 | ALAS2 | 5'-aminolevulinate synthase 2 | Cytoplasm | enzyme | 2635171.25 | 4423271 | 6297337.5 | | | | | | |
| P08680 | ALAS2 | 5'-aminolevulinate synthase 2 | Cytoplasm | enzyme | 1000031.562 | 3732945 | 2477891.75 | | | | | | |
| P08680 | ALAS2 | 5'-aminolevulinate synthase 2 | Cytoplasm | enzyme | 883215.625 | 2163229 | 1695177.375 | P08680 | ALAS2 | 5787509 | 7867759 | 1311293 | 36 |
| P07724 | ALB | albumin | Extracellular Space | transporter | 6.81E+07 | 5.75E+07 | 7.20E+07 | | | | | | |
| P07724 | ALB | albumin | Extracellular Space | transporter | 3.35E+07 | 2.11E+07 | 4.45E+07 | | | | | | |
| P07724 | ALB | albumin | Extracellular Space | transporter | 3.57E+07 | 3.42E+07 | 4.64E+07 | | | | | | |
| P07724 | ALB | albumin | Extracellular Space | transporter | 3.64E+07 | 3.70E+07 | 3.77E+07 | | | | | | |
| P07724 | ALB | albumin | Extracellular Space | transporter | 3.08E+07 | 4.20E+07 | 5.09E+07 | | | | | | |
| P07724 | ALB | albumin | Extracellular Space | transporter | 3.53E+07 | 3.74E+07 | 3.91E+07 | | | | | | |
| P07724 | ALB | albumin | Extracellular Space | transporter | 3.75E+07 | 3.98E+07 | 4.24E+07 | | | | | | |
| P07724 | ALB | albumin | Extracellular Space | transporter | 3.24E+07 | 2.76E+07 | 3.22E+07 | | | | | | |
| P07724 | ALB | albumin | Extracellular Space | transporter | 3.27E+07 | 3.21E+07 | 3.87E+07 | | | | | | |
| P07724 | ALB | albumin | Extracellular Space | transporter | 3.80E+07 | 3.33E+07 | 4.00E+07 | | | | | | |
| P07724 | ALB | albumin | Extracellular Space | transporter | 2.19E+07 | 2.26E+07 | 3.20E+07 | | | | | | |
| P07724 | ALB | albumin | Extracellular Space | transporter | 1.83E+07 | 1.24E+07 | 3.18E+07 | | | | | | |
| P07724 | ALB | albumin | Extracellular Space | transporter | 1.72E+07 | 1.92E+07 | 2.05E+07 | | | | | | |
| P07724 | ALB | albumin | Extracellular Space | transporter | 1.44E+07 | 1.77E+07 | 1.84E+07 | | | | | | |
| P07724 | ALB | albumin | Extracellular Space | transporter | 1.87E+07 | 1.97E+07 | 2.69E+07 | | | | | | |
| P07724 | ALB | albumin | Extracellular Space | transporter | 1.66E+07 | 1.58E+07 | 1.48E+07 | | | | | | |
| P07724 | ALB | albumin | Extracellular Space | transporter | 1.27E+07 | 1.41E+07 | 1.78E+07 | | | | | | |
| P07724 | ALB | albumin | Extracellular Space | transporter | 9533958 | 1.39E+07 | 1.26E+07 | | | | | | |
| P07724 | ALB | albumin | Extracellular Space | transporter | 9974515 | 1.41E+07 | 1.13E+07 | | | | | | |
| P07724 | ALB | albumin | Extracellular Space | transporter | 4974498 | 5074709 | 5992043 | | | | | | |
| P07724 | ALB | albumin | Extracellular Space | transporter | 7330773 | 7472782.5 | 1.19E+07 | | | | | | |
| P07724 | ALB | albumin | Extracellular Space | transporter | 6731448 | 6068010.5 | 8589628 | | | | | | |
| P07724 | ALB | albumin | Extracellular Space | transporter | 3890983.5 | 5165713 | 6798491 | | | | | | |
| P07724 | ALB | albumin | Extracellular Space | transporter | 2714709.75 | 3353191.5 | 3404989.5 | P07724 | ALB | 22710037 | 15704282 | 1850767 | 72 |
| Q571I9 | ALDH16A1 | aldehyde dehydrogenase 16 family, member A1 | Cytoplasm | enzyme | 7308150.5 | 4837861.5 | 3802387.5 | | | | | | |
| Q571I9 | ALDH16A1 | aldehyde dehydrogenase 16 family, member A1 | Cytoplasm | enzyme | 6337745.5 | 4991131 | 4097360.25 | | | | | | |
| Q571I9 | ALDH16A1 | aldehyde dehydrogenase 16 family, member A1 | Cytoplasm | enzyme | 4492792 | 4170088 | 3538685.5 | | | | | | |
| Q571I9 | ALDH16A1 | aldehyde dehydrogenase 16 family, member A1 | Cytoplasm | enzyme | 2012636.125 | 1053683.5 | 733780.375 | | | | | | |
| Q571I9 | ALDH16A1 | aldehyde dehydrogenase 16 family, member A1 | Cytoplasm | enzyme | 2716825.5 | 2285325.75 | 1815091.875 | | | | | | |
| Q571I9 | ALDH16A1 | aldehyde dehydrogenase 16 family, member A1 | Cytoplasm | enzyme | 962456.688 | 591055.125 | 453687.969 | Q571I9 | ALDH16A1 | 3971768 | 2041363 | 481154 | 18 |
| Q9Z110 | ALDH18A1 | aldehyde dehydrogenase 18 family, member A1 | Cytoplasm | kinase | 2326732.25 | 2520650.5 | 3389013 | | | | | | |
| Q9Z110 | ALDH18A1 | aldehyde dehydrogenase 18 family, member A1 | Cytoplasm | kinase | 1733196.75 | 3327248.75 | 3174167 | | | | | | |
| Q9Z110 | ALDH18A1 | aldehyde dehydrogenase 18 family, member A1 | Cytoplasm | kinase | 1597110.875 | 2349414.5 | 2936747.75 | | | | | | |
| Q9Z110 | ALDH18A1 | aldehyde dehydrogenase 18 family, member A1 | Cytoplasm | kinase | 1248273.875 | 1752536 | 2246161.5 | | | | | | |
| Q9Z110 | ALDH18A1 | aldehyde dehydrogenase 18 family, member A1 | Cytoplasm | kinase | 1280769.125 | 1710052.5 | 2170269.75 | | | | | | |
| Q9Z110 | ALDH18A1 | aldehyde dehydrogenase 18 family, member A1 | Cytoplasm | kinase | 1216824.875 | 1360755.75 | 1889356.125 | | | | | | |
| Q9Z110 | ALDH18A1 | aldehyde dehydrogenase 18 family, member A1 | Cytoplasm | kinase | 492262.281 | 1182164.25 | 1464935.5 | | | | | | |

| | | | | | | | | | | | | | |
|---|---|---|---|---|---|---|---|---|---|---|---|---|---|
| Q9Z110 | ALDH18A1 | aldehyde dehydrogenase 18 family, member A1 | Cytoplasm | kinase | 803042.5 | 1348801 | 932221 | Q9Z110 | ALDH18A1 | 1337277 | 800356 | 163372 | 24 |
| P24549 | ALDH1A1 | aldehyde dehydrogenase 1 family, member A1 | Cytoplasm | enzyme | 853954.375 | 589726.938 | 2165928.5 | P24549 | ALDH1A1 | 853954 | 844147 | 487368 | 3 |
| P47738 | ALDH2 | aldehyde dehydrogenase 2 family (mitochondrial) | Cytoplasm | enzyme | 2.52E+07 | 3.54E+07 | 7.10E+07 | | | | | | |
| P47738 | ALDH2 | aldehyde dehydrogenase 2 family (mitochondrial) | Cytoplasm | enzyme | 2.31E+07 | 3.01E+07 | 5.77E+07 | | | | | | |
| P47738 | ALDH2 | aldehyde dehydrogenase 2 family (mitochondrial) | Cytoplasm | enzyme | 1.46E+07 | 1.35E+07 | 2.18E+07 | | | | | | |
| P47738 | ALDH2 | aldehyde dehydrogenase 2 family (mitochondrial) | Cytoplasm | enzyme | 2.10E+07 | 2.77E+07 | 4.82E+07 | | | | | | |
| P47738 | ALDH2 | aldehyde dehydrogenase 2 family (mitochondrial) | Cytoplasm | enzyme | 1.65E+07 | 2.35E+07 | 3.94E+07 | | | | | | |
| P47738 | ALDH2 | aldehyde dehydrogenase 2 family (mitochondrial) | Cytoplasm | enzyme | 1.06E+07 | 1.05E+07 | 1.76E+07 | | | | | | |
| P47738 | ALDH2 | aldehyde dehydrogenase 2 family (mitochondrial) | Cytoplasm | enzyme | 2.02E+07 | 2.53E+07 | 3.72E+07 | | | | | | |
| P47738 | ALDH2 | aldehyde dehydrogenase 2 family (mitochondrial) | Cytoplasm | enzyme | 1.72E+07 | 1.92E+07 | 2.96E+07 | | | | | | |
| P47738 | ALDH2 | aldehyde dehydrogenase 2 family (mitochondrial) | Cytoplasm | enzyme | 1.50E+07 | 1.90E+07 | 2.76E+07 | | | | | | |
| P47738 | ALDH2 | aldehyde dehydrogenase 2 family (mitochondrial) | Cytoplasm | enzyme | 1.08E+07 | 1.49E+07 | 2.12E+07 | | | | | | |
| P47738 | ALDH2 | aldehyde dehydrogenase 2 family (mitochondrial) | Cytoplasm | enzyme | 1.16E+07 | 1.28E+07 | 1.93E+07 | | | | | | |
| P47738 | ALDH2 | aldehyde dehydrogenase 2 family (mitochondrial) | Cytoplasm | enzyme | 8030095 | 9046568 | 1.94E+07 | | | | | | |
| P47738 | ALDH2 | aldehyde dehydrogenase 2 family (mitochondrial) | Cytoplasm | enzyme | 7462875.5 | 1.32E+07 | 1.42E+07 | | | | | | |
| P47738 | ALDH2 | aldehyde dehydrogenase 2 family (mitochondrial) | Cytoplasm | enzyme | 1.30E+07 | 1.98E+07 | 1.86E+07 | | | | | | |
| P47738 | ALDH2 | aldehyde dehydrogenase 2 family (mitochondrial) | Cytoplasm | enzyme | 1.10E+07 | 1.51E+07 | 2.36E+07 | | | | | | |
| P47738 | ALDH2 | aldehyde dehydrogenase 2 family (mitochondrial) | Cytoplasm | enzyme | 1.69E+07 | 2.69E+07 | 3.78E+07 | | | | | | |
| P47738 | ALDH2 | aldehyde dehydrogenase 2 family (mitochondrial) | Cytoplasm | enzyme | 7828597 | 8977192 | 1.35E+07 | | | | | | |
| P47738 | ALDH2 | aldehyde dehydrogenase 2 family (mitochondrial) | Cytoplasm | enzyme | 8914847 | 1.08E+07 | 1.82E+07 | | | | | | |
| P47738 | ALDH2 | aldehyde dehydrogenase 2 family (mitochondrial) | Cytoplasm | enzyme | 1917598 | 2219490.25 | 4135401 | | | | | | |
| P47738 | ALDH2 | aldehyde dehydrogenase 2 family (mitochondrial) | Cytoplasm | enzyme | 4812958.5 | 9527630 | 2.00E+07 | | | | | | |
| P47738 | ALDH2 | aldehyde dehydrogenase 2 family (mitochondrial) | Cytoplasm | enzyme | 5534497 | 6884230.5 | 1.10E+07 | | | | | | |
| P47738 | ALDH2 | aldehyde dehydrogenase 2 family (mitochondrial) | Cytoplasm | enzyme | 4289243.5 | 4966607.5 | 7699861 | | | | | | |
| P47738 | ALDH2 | aldehyde dehydrogenase 2 family (mitochondrial) | Cytoplasm | enzyme | 3803812.25 | 3814058.75 | 7922899 | | | | | | |
| P47738 | ALDH2 | aldehyde dehydrogenase 2 family (mitochondrial) | Cytoplasm | enzyme | 2453605.25 | 3271488 | 5323776.5 | P47738 | ALDH2 | 11736172 | 12703566 | 1497130 | 72 |
| P05064 | ALDOA | aldolase A, fructose-bisphosphate | Cytoplasm | enzyme | 6.96E+07 | 6.27E+07 | 5.27E+07 | | | | | | |
| P05064 | ALDOA | aldolase A, fructose-bisphosphate | Cytoplasm | enzyme | 5.26E+07 | 4.68E+07 | 3.82E+07 | | | | | | |
| P05064 | ALDOA | aldolase A, fructose-bisphosphate | Cytoplasm | enzyme | 3.92E+07 | 3.35E+07 | 2.97E+07 | | | | | | |
| P05064 | ALDOA | aldolase A, fructose-bisphosphate | Cytoplasm | enzyme | 3.08E+07 | 3.52E+07 | 3.20E+07 | | | | | | |
| P05064 | ALDOA | aldolase A, fructose-bisphosphate | Cytoplasm | enzyme | 2.71E+07 | 2.52E+07 | 2.01E+07 | | | | | | |
| P05064 | ALDOA | aldolase A, fructose-bisphosphate | Cytoplasm | enzyme | 1.40E+07 | 1.16E+07 | 8869712 | | | | | | |
| P05064 | ALDOA | aldolase A, fructose-bisphosphate | Cytoplasm | enzyme | 8093073.5 | 4984229.5 | 2938669 | | | | | | |
| P05064 | ALDOA | aldolase A, fructose-bisphosphate | Cytoplasm | enzyme | 7172449.5 | 8882291 | 7136445.5 | | | | | | |
| P05064 | ALDOA | aldolase A, fructose-bisphosphate | Cytoplasm | enzyme | 4178663.75 | 3667406.5 | 3602799.25 | | | | | | |
| P05064 | ALDOA | aldolase A, fructose-bisphosphate | Cytoplasm | enzyme | 3597781.75 | 3389136 | 3536019.75 | | | | | | |
| P05064 | ALDOA | aldolase A, fructose-bisphosphate | Cytoplasm | enzyme | 3280612 | 2678552.75 | 2668392.25 | | | | | | |
| P05064 | ALDOA | aldolase A, fructose-bisphosphate | Cytoplasm | enzyme | 1234626.125 | 1601297.125 | 582986.5 | P05064 | ALDOA | 21748934 | 19683544 | 3280591 | 36 |
| O08739 | AMPD3 | adenosine monophosphate deaminase 3 | Cytoplasm | enzyme | 4032084.75 | 3529306.25 | 2388519.25 | | | | | | |
| O08739 | AMPD3 | adenosine monophosphate deaminase 3 | Cytoplasm | enzyme | 3978711.25 | 4145038.5 | 2278370 | | | | | | |
| O08739 | AMPD3 | adenosine monophosphate deaminase 3 | Cytoplasm | enzyme | 4573622.5 | 8136738 | 4071876.5 | | | | | | |
| O08739 | AMPD3 | adenosine monophosphate deaminase 3 | Cytoplasm | enzyme | 3544156.5 | 3420698 | 1952354.625 | | | | | | |
| O08739 | AMPD3 | adenosine monophosphate deaminase 3 | Cytoplasm | enzyme | 2598970 | 2908724.75 | 2059163.375 | | | | | | |
| O08739 | AMPD3 | adenosine monophosphate deaminase 3 | Cytoplasm | enzyme | 1708992.875 | 2035930.625 | 1537814 | | | | | | |
| O08739 | AMPD3 | adenosine monophosphate deaminase 3 | Cytoplasm | enzyme | 1882860.875 | 2833483.75 | 2063969.875 | | | | | | |
| O08739 | AMPD3 | adenosine monophosphate deaminase 3 | Cytoplasm | enzyme | 833122.562 | 1040898.062 | 421571.625 | O08739 | AMPD3 | 2894065 | 1595355 | 325651 | 24 |
| Q02357 | ANK1 | ankyrin 1, erythrocytic | Plasma Membrane | other | 3.12E+09 | 3.19E+09 | 3.05E+09 | | | | | | |
| Q02357 | ANK1 | ankyrin 1, erythrocytic | Plasma Membrane | other | 3.29E+09 | 3.32E+09 | 3.20E+09 | | | | | | |
| Q02357 | ANK1 | ankyrin 1, erythrocytic | Plasma Membrane | other | 2.28E+09 | 2.27E+09 | 1.89E+09 | | | | | | |
| Q02357 | ANK1 | ankyrin 1, erythrocytic | Plasma Membrane | other | 2.61E+09 | 2.51E+09 | 2.39E+09 | | | | | | |
| Q02357 | ANK1 | ankyrin 1, erythrocytic | Plasma Membrane | other | 2.36E+09 | 2.79E+09 | 1.75E+09 | | | | | | |
| Q02357 | ANK1 | ankyrin 1, erythrocytic | Plasma Membrane | other | 2.39E+09 | 2.41E+09 | 2.41E+09 | | | | | | |
| Q02357 | ANK1 | ankyrin 1, erythrocytic | Plasma Membrane | other | 2.04E+09 | 2.26E+09 | 1.77E+09 | | | | | | |
| Q02357 | ANK1 | ankyrin 1, erythrocytic | Plasma Membrane | other | 1.98E+09 | 2.09E+09 | 2.15E+09 | | | | | | |
| Q02357 | ANK1 | ankyrin 1, erythrocytic | Plasma Membrane | other | 2.03E+09 | 2.11E+09 | 1.65E+09 | | | | | | |
| Q02357 | ANK1 | ankyrin 1, erythrocytic | Plasma Membrane | other | 1.92E+09 | 2.14E+09 | 2.05E+09 | | | | | | |
| Q02357 | ANK1 | ankyrin 1, erythrocytic | Plasma Membrane | other | 2.00E+09 | 1.86E+09 | 1.79E+09 | | | | | | |
| Q02357 | ANK1 | ankyrin 1, erythrocytic | Plasma Membrane | other | 1.66E+09 | 1.24E+09 | 1.25E+09 | | | | | | |
| Q02357 | ANK1 | ankyrin 1, erythrocytic | Plasma Membrane | other | 1.65E+09 | 1.53E+09 | 1.56E+09 | | | | | | |
| Q02357 | ANK1 | ankyrin 1, erythrocytic | Plasma Membrane | other | 1.50E+09 | 1.30E+09 | 1.39E+09 | | | | | | |
| Q02357 | ANK1 | ankyrin 1, erythrocytic | Plasma Membrane | other | 1.52E+09 | 1.64E+09 | 1.45E+09 | | | | | | |
| Q02357 | ANK1 | ankyrin 1, erythrocytic | Plasma Membrane | other | 1.39E+09 | 1.50E+09 | 1.54E+09 | | | | | | |
| Q02357 | ANK1 | ankyrin 1, erythrocytic | Plasma Membrane | other | 1.69E+09 | 1.58E+09 | 1.32E+09 | | | | | | |
| Q02357 | ANK1 | ankyrin 1, erythrocytic | Plasma Membrane | other | 1.69E+09 | 1.49E+09 | 1.25E+09 | | | | | | |
| Q02357 | ANK1 | ankyrin 1, erythrocytic | Plasma Membrane | other | 1.22E+09 | 1.11E+09 | 9.99E+08 | | | | | | |
| Q02357 | ANK1 | ankyrin 1, erythrocytic | Plasma Membrane | other | 1.49E+09 | 1.46E+09 | 1.39E+09 | | | | | | |
| Q02357 | ANK1 | ankyrin 1, erythrocytic | Plasma Membrane | other | 1.55E+09 | 1.53E+09 | 1.34E+09 | | | | | | |
| Q02357 | ANK1 | ankyrin 1, erythrocytic | Plasma Membrane | other | 1.25E+09 | 1.51E+09 | 1.39E+09 | | | | | | |
| Q02357 | ANK1 | ankyrin 1, erythrocytic | Plasma Membrane | other | 1.38E+09 | 1.47E+09 | 1.16E+09 | | | | | | |
| Q02357 | ANK1 | ankyrin 1, erythrocytic | Plasma Membrane | other | 1.27E+09 | 1.68E+09 | 1.28E+09 | | | | | | |
| Q02357 | ANK1 | ankyrin 1, erythrocytic | Plasma Membrane | other | 1.42E+09 | 1.49E+09 | 1.16E+09 | | | | | | |
| Q02357 | ANK1 | ankyrin 1, erythrocytic | Plasma Membrane | other | 9.91E+08 | 1.04E+09 | 1.05E+09 | | | | | | |
| Q02357 | ANK1 | ankyrin 1, erythrocytic | Plasma Membrane | other | 1.04E+09 | 1.12E+09 | 8.89E+08 | | | | | | |
| Q02357 | ANK1 | ankyrin 1, erythrocytic | Plasma Membrane | other | 1.16E+09 | 1.19E+09 | 8.24E+08 | | | | | | |
| Q02357 | ANK1 | ankyrin 1, erythrocytic | Plasma Membrane | other | 1.28E+09 | 9.85E+08 | 8.98E+08 | | | | | | |
| Q02357 | ANK1 | ankyrin 1, erythrocytic | Plasma Membrane | other | 9.50E+08 | 9.23E+08 | 8.68E+08 | | | | | | |
| Q02357 | ANK1 | ankyrin 1, erythrocytic | Plasma Membrane | other | 1.22E+09 | 1.00E+09 | 9.11E+08 | | | | | | |
| Q02357 | ANK1 | ankyrin 1, erythrocytic | Plasma Membrane | other | 1.14E+09 | 1.34E+09 | 1.07E+09 | | | | | | |
| Q02357 | ANK1 | ankyrin 1, erythrocytic | Plasma Membrane | other | 1.04E+09 | 1.09E+09 | 1.08E+09 | | | | | | |
| Q02357 | ANK1 | ankyrin 1, erythrocytic | Plasma Membrane | other | 8.70E+08 | 8.52E+08 | 7.19E+08 | | | | | | |
| Q02357 | ANK1 | ankyrin 1, erythrocytic | Plasma Membrane | other | 1.17E+09 | 1.20E+09 | 1.11E+09 | | | | | | |
| Q02357 | ANK1 | ankyrin 1, erythrocytic | Plasma Membrane | other | 8.32E+08 | 8.39E+08 | 6.79E+08 | | | | | | |
| Q02357 | ANK1 | ankyrin 1, erythrocytic | Plasma Membrane | other | 7.90E+08 | 8.06E+08 | 7.01E+08 | | | | | | |
| Q02357 | ANK1 | ankyrin 1, erythrocytic | Plasma Membrane | other | 7.16E+08 | 7.89E+08 | 9.50E+08 | | | | | | |
| Q02357 | ANK1 | ankyrin 1, erythrocytic | Plasma Membrane | other | 9.17E+08 | 7.63E+08 | 7.66E+08 | | | | | | |
| Q02357 | ANK1 | ankyrin 1, erythrocytic | Plasma Membrane | other | 6.37E+08 | 6.23E+08 | 6.30E+08 | | | | | | |
| Q02357 | ANK1 | ankyrin 1, erythrocytic | Plasma Membrane | other | 7.92E+08 | 7.24E+08 | 6.84E+08 | | | | | | |
| Q02357 | ANK1 | ankyrin 1, erythrocytic | Plasma Membrane | other | 8.00E+08 | 6.50E+08 | 7.37E+08 | | | | | | |
| Q02357 | ANK1 | ankyrin 1, erythrocytic | Plasma Membrane | other | 3.55E+08 | 3.84E+08 | 5.70E+08 | | | | | | |
| Q02357 | ANK1 | ankyrin 1, erythrocytic | Plasma Membrane | other | 6.30E+08 | 6.69E+08 | 6.43E+08 | | | | | | |
| Q02357 | ANK1 | ankyrin 1, erythrocytic | Plasma Membrane | other | 6.38E+08 | 6.15E+08 | 5.74E+08 | | | | | | |
| Q02357 | ANK1 | ankyrin 1, erythrocytic | Plasma Membrane | other | 6.91E+08 | 5.94E+08 | 5.46E+08 | | | | | | |
| Q02357 | ANK1 | ankyrin 1, erythrocytic | Plasma Membrane | other | 9.62E+08 | 8.89E+08 | 7.50E+08 | | | | | | |
| Q02357 | ANK1 | ankyrin 1, erythrocytic | Plasma Membrane | other | 4.10E+08 | 4.08E+08 | 5.37E+08 | | | | | | |
| Q02357 | ANK1 | ankyrin 1, erythrocytic | Plasma Membrane | other | 7.20E+08 | 7.22E+08 | 4.30E+08 | | | | | | |
| Q02357 | ANK1 | ankyrin 1, erythrocytic | Plasma Membrane | other | 5.18E+08 | 6.76E+08 | 5.49E+08 | | | | | | |
| Q02357 | ANK1 | ankyrin 1, erythrocytic | Plasma Membrane | other | 5.68E+08 | 5.36E+08 | 4.71E+08 | | | | | | |
| Q02357 | ANK1 | ankyrin 1, erythrocytic | Plasma Membrane | other | 4.89E+08 | 4.43E+08 | 4.50E+08 | | | | | | |
| Q02357 | ANK1 | ankyrin 1, erythrocytic | Plasma Membrane | other | 5.72E+08 | 7.73E+08 | 4.76E+08 | | | | | | |
| Q02357 | ANK1 | ankyrin 1, erythrocytic | Plasma Membrane | other | 5.02E+08 | 4.41E+08 | 3.76E+08 | | | | | | |
| Q02357 | ANK1 | ankyrin 1, erythrocytic | Plasma Membrane | other | 5.00E+08 | 5.18E+08 | 4.51E+08 | | | | | | |
| Q02357 | ANK1 | ankyrin 1, erythrocytic | Plasma Membrane | other | 6.29E+08 | 6.51E+08 | 5.71E+08 | | | | | | |
| Q02357 | ANK1 | ankyrin 1, erythrocytic | Plasma Membrane | other | 5.11E+08 | 5.54E+08 | 4.43E+08 | | | | | | |
| Q02357 | ANK1 | ankyrin 1, erythrocytic | Plasma Membrane | other | 4.40E+08 | 5.29E+08 | 3.82E+08 | | | | | | |
| Q02357 | ANK1 | ankyrin 1, erythrocytic | Plasma Membrane | other | 3.56E+08 | 4.69E+08 | 3.65E+08 | | | | | | |
| Q02357 | ANK1 | ankyrin 1, erythrocytic | Plasma Membrane | other | 4.83E+08 | 4.63E+08 | 4.35E+08 | | | | | | |
| Q02357 | ANK1 | ankyrin 1, erythrocytic | Plasma Membrane | other | 6.00E+08 | 5.05E+08 | 4.75E+08 | | | | | | |
| Q02357 | ANK1 | ankyrin 1, erythrocytic | Plasma Membrane | other | 5.62E+08 | 5.98E+08 | 5.47E+08 | | | | | | |
| Q02357 | ANK1 | ankyrin 1, erythrocytic | Plasma Membrane | other | 4.07E+08 | 3.84E+08 | 3.73E+08 | | | | | | |

| | | | | | | | | | | | | | |
|---|---|---|---|---|---|---|---|---|---|---|---|---|---|
| Q02357 | ANK1 | ankyrin 1, erythrocytic | Plasma Membrane | other | 4.48E+08 | 4.44E+08 | 2.85E+08 | | | | | | |
| Q02357 | ANK1 | ankyrin 1, erythrocytic | Plasma Membrane | other | 5.78E+08 | 5.20E+08 | 3.59E+08 | | | | | | |
| Q02357 | ANK1 | ankyrin 1, erythrocytic | Plasma Membrane | other | 3.48E+08 | 3.49E+08 | 3.24E+08 | | | | | | |
| Q02357 | ANK1 | ankyrin 1, erythrocytic | Plasma Membrane | other | 3.82E+08 | 3.92E+08 | 3.46E+08 | | | | | | |
| Q02357 | ANK1 | ankyrin 1, erythrocytic | Plasma Membrane | other | 3.24E+08 | 3.04E+08 | 3.60E+08 | | | | | | |
| Q02357 | ANK1 | ankyrin 1, erythrocytic | Plasma Membrane | other | 2.81E+08 | 3.69E+08 | 3.03E+08 | | | | | | |
| Q02357 | ANK1 | ankyrin 1, erythrocytic | Plasma Membrane | other | 2.98E+08 | 3.30E+08 | 2.81E+08 | | | | | | |
| Q02357 | ANK1 | ankyrin 1, erythrocytic | Plasma Membrane | other | 2.60E+08 | 1.10E+08 | 7.21E+07 | | | | | | |
| Q02357 | ANK1 | ankyrin 1, erythrocytic | Plasma Membrane | other | 4.11E+08 | 3.58E+08 | 3.81E+08 | | | | | | |
| Q02357 | ANK1 | ankyrin 1, erythrocytic | Plasma Membrane | other | 3.65E+08 | 4.64E+08 | 2.66E+08 | | | | | | |
| Q02357 | ANK1 | ankyrin 1, erythrocytic | Plasma Membrane | other | 2.36E+08 | 2.43E+08 | 2.37E+08 | | | | | | |
| Q02357 | ANK1 | ankyrin 1, erythrocytic | Plasma Membrane | other | 3.72E+08 | 1.33E+08 | 3.42E+08 | | | | | | |
| Q02357 | ANK1 | ankyrin 1, erythrocytic | Plasma Membrane | other | 2.49E+08 | 2.66E+08 | 2.26E+08 | | | | | | |
| Q02357 | ANK1 | ankyrin 1, erythrocytic | Plasma Membrane | other | 3.36E+08 | 2.11E+08 | 2.16E+08 | | | | | | |
| Q02357 | ANK1 | ankyrin 1, erythrocytic | Plasma Membrane | other | 2.31E+08 | 1.79E+08 | 1.63E+08 | | | | | | |
| Q02357 | ANK1 | ankyrin 1, erythrocytic | Plasma Membrane | other | 2.15E+08 | 1.18E+08 | 3.31E+08 | | | | | | |
| Q02357 | ANK1 | ankyrin 1, erythrocytic | Plasma Membrane | other | 2.47E+08 | 3.31E+08 | 1.94E+08 | | | | | | |
| Q02357 | ANK1 | ankyrin 1, erythrocytic | Plasma Membrane | other | 2.21E+08 | 1.57E+08 | 1.44E+08 | | | | | | |
| Q02357 | ANK1 | ankyrin 1, erythrocytic | Plasma Membrane | other | 1.03E+08 | 8.58E+07 | 2.13E+08 | | | | | | |
| Q02357 | ANK1 | ankyrin 1, erythrocytic | Plasma Membrane | other | 2.10E+08 | 2.17E+08 | 1.60E+08 | | | | | | |
| Q02357 | ANK1 | ankyrin 1, erythrocytic | Plasma Membrane | other | 2.38E+08 | 2.12E+08 | 2.17E+08 | | | | | | |
| Q02357 | ANK1 | ankyrin 1, erythrocytic | Plasma Membrane | other | 1.99E+08 | 1.50E+08 | 1.84E+08 | | | | | | |
| Q02357 | ANK1 | ankyrin 1, erythrocytic | Plasma Membrane | other | 2.01E+08 | 1.76E+08 | 1.93E+08 | | | | | | |
| Q02357 | ANK1 | ankyrin 1, erythrocytic | Plasma Membrane | other | 1.57E+08 | 6.35E+07 | 2.41E+07 | | | | | | |
| Q02357 | ANK1 | ankyrin 1, erythrocytic | Plasma Membrane | other | 2.37E+08 | 2.55E+08 | 3.12E+08 | | | | | | |
| Q02357 | ANK1 | ankyrin 1, erythrocytic | Plasma Membrane | other | 1.60E+08 | 2.62E+08 | 1.61E+08 | | | | | | |
| Q02357 | ANK1 | ankyrin 1, erythrocytic | Plasma Membrane | other | 2.15E+08 | 2.49E+08 | 2.07E+08 | | | | | | |
| Q02357 | ANK1 | ankyrin 1, erythrocytic | Plasma Membrane | other | 1.92E+08 | 2.08E+08 | 1.12E+08 | | | | | | |
| Q02357 | ANK1 | ankyrin 1, erythrocytic | Plasma Membrane | other | 1.32E+08 | 7.33E+07 | 5.98E+07 | | | | | | |
| Q02357 | ANK1 | ankyrin 1, erythrocytic | Plasma Membrane | other | 1.17E+08 | 1.09E+08 | 9.43E+07 | | | | | | |
| Q02357 | ANK1 | ankyrin 1, erythrocytic | Plasma Membrane | other | 1.10E+08 | 9.55E+07 | 1.06E+08 | | | | | | |
| Q02357 | ANK1 | ankyrin 1, erythrocytic | Plasma Membrane | other | 1.05E+08 | 1.14E+08 | 1.16E+08 | | | | | | |
| Q02357 | ANK1 | ankyrin 1, erythrocytic | Plasma Membrane | other | 1.52E+08 | 1.82E+08 | 1.44E+08 | | | | | | |
| Q02357 | ANK1 | ankyrin 1, erythrocytic | Plasma Membrane | other | 1.44E+08 | 1.38E+08 | 1.08E+08 | | | | | | |
| Q02357 | ANK1 | ankyrin 1, erythrocytic | Plasma Membrane | other | 1.15E+08 | 1.12E+08 | 8.77E+07 | | | | | | |
| Q02357 | ANK1 | ankyrin 1, erythrocytic | Plasma Membrane | other | 4.14E+07 | 4.54E+07 | 1.33E+08 | | | | | | |
| Q02357 | ANK1 | ankyrin 1, erythrocytic | Plasma Membrane | other | 6.78E+07 | 7.11E+07 | 6.54E+07 | | | | | | |
| Q02357 | ANK1 | ankyrin 1, erythrocytic | Plasma Membrane | other | 5.25E+07 | 4.30E+07 | 6.07E+07 | | | | | | |
| Q02357 | ANK1 | ankyrin 1, erythrocytic | Plasma Membrane | other | 1.39E+08 | 1.57E+08 | 1.11E+08 | | | | | | |
| Q02357 | ANK1 | ankyrin 1, erythrocytic | Plasma Membrane | other | 1.02E+08 | 9.84E+07 | 8.14E+07 | | | | | | |
| Q02357 | ANK1 | ankyrin 1, erythrocytic | Plasma Membrane | other | 6.66E+07 | 4.62E+07 | 3.95E+07 | | | | | | |
| Q02357 | ANK1 | ankyrin 1, erythrocytic | Plasma Membrane | other | 9.51E+07 | 1.79E+08 | 1.33E+08 | | | | | | |
| Q02357 | ANK1 | ankyrin 1, erythrocytic | Plasma Membrane | other | 2.17E+08 | 4.97E+08 | 4.92E+08 | | | | | | |
| Q02357 | ANK1 | ankyrin 1, erythrocytic | Plasma Membrane | other | 6.79E+07 | 6.91E+07 | 5.65E+07 | | | | | | |
| Q02357 | ANK1 | ankyrin 1, erythrocytic | Plasma Membrane | other | 7.36E+07 | 5.60E+07 | 6.19E+07 | | | | | | |
| Q02357 | ANK1 | ankyrin 1, erythrocytic | Plasma Membrane | other | 7.28E+07 | 7.32E+07 | 4.33E+07 | | | | | | |
| Q02357 | ANK1 | ankyrin 1, erythrocytic | Plasma Membrane | other | 5.91E+07 | 6.22E+07 | 5.28E+07 | | | | | | |
| Q02357 | ANK1 | ankyrin 1, erythrocytic | Plasma Membrane | other | 7.58E+07 | 6.42E+07 | 5.75E+07 | | | | | | |
| Q02357 | ANK1 | ankyrin 1, erythrocytic | Plasma Membrane | other | 4.59E+07 | 9.10E+07 | 4.63E+07 | | | | | | |
| Q02357 | ANK1 | ankyrin 1, erythrocytic | Plasma Membrane | other | 5.87E+07 | 6.03E+07 | 4.74E+07 | | | | | | |
| Q02357 | ANK1 | ankyrin 1, erythrocytic | Plasma Membrane | other | 7.41E+07 | 6.35E+07 | 7.54E+07 | | | | | | |
| Q02357 | ANK1 | ankyrin 1, erythrocytic | Plasma Membrane | other | 5.39E+07 | 7.41E+07 | 4.88E+07 | | | | | | |
| Q02357 | ANK1 | ankyrin 1, erythrocytic | Plasma Membrane | other | 5.43E+07 | 6.44E+07 | 5.03E+07 | | | | | | |
| Q02357 | ANK1 | ankyrin 1, erythrocytic | Plasma Membrane | other | 4.28E+07 | 4.87E+07 | 3.69E+07 | | | | | | |
| Q02357 | ANK1 | ankyrin 1, erythrocytic | Plasma Membrane | other | 5.90E+07 | 8.63E+07 | 8.54E+07 | | | | | | |
| Q02357 | ANK1 | ankyrin 1, erythrocytic | Plasma Membrane | other | 2.97E+07 | 1.03E+07 | 3.28E+07 | | | | | | |
| Q02357 | ANK1 | ankyrin 1, erythrocytic | Plasma Membrane | other | 1.95E+07 | 2.06E+07 | 1.97E+07 | | | | | | |
| Q02357 | ANK1 | ankyrin 1, erythrocytic | Plasma Membrane | other | 3.04E+07 | 3.38E+07 | 2.43E+07 | | | | | | |
| Q02357 | ANK1 | ankyrin 1, erythrocytic | Plasma Membrane | other | 2.99E+07 | 3.34E+07 | 2.39E+07 | | | | | | |
| Q02357 | ANK1 | ankyrin 1, erythrocytic | Plasma Membrane | other | 2.84E+07 | 2.56E+07 | 2.68E+07 | | | | | | |
| Q02357 | ANK1 | ankyrin 1, erythrocytic | Plasma Membrane | other | 4.67E+07 | 8.11E+07 | 1.02E+08 | | | | | | |
| Q02357 | ANK1 | ankyrin 1, erythrocytic | Plasma Membrane | other | 2.17E+07 | 8150715 | 7376175.5 | | | | | | |
| Q02357 | ANK1 | ankyrin 1, erythrocytic | Plasma Membrane | other | 2.85E+07 | 2.56E+07 | 2.38E+07 | | | | | | |
| Q02357 | ANK1 | ankyrin 1, erythrocytic | Plasma Membrane | other | 2.67E+07 | 2.04E+07 | 2.62E+07 | | | | | | |
| Q02357 | ANK1 | ankyrin 1, erythrocytic | Plasma Membrane | other | 2.08E+07 | 1.84E+07 | 3.57E+07 | | | | | | |
| Q02357 | ANK1 | ankyrin 1, erythrocytic | Plasma Membrane | other | 2.72E+07 | 2.59E+07 | 2.18E+07 | | | | | | |
| Q02357 | ANK1 | ankyrin 1, erythrocytic | Plasma Membrane | other | 1.93E+07 | 2.13E+07 | 2.07E+07 | | | | | | |
| Q02357 | ANK1 | ankyrin 1, erythrocytic | Plasma Membrane | other | 2.72E+07 | 3.09E+07 | 1.61E+07 | | | | | | |
| Q02357 | ANK1 | ankyrin 1, erythrocytic | Plasma Membrane | other | 9727868 | 5264501 | 9161411 | | | | | | |
| Q02357 | ANK1 | ankyrin 1, erythrocytic | Plasma Membrane | other | 1.66E+07 | 1.35E+07 | 2.17E+07 | | | | | | |
| Q02357 | ANK1 | ankyrin 1, erythrocytic | Plasma Membrane | other | 2.00E+07 | 1.47E+07 | 1.57E+07 | | | | | | |
| Q02357 | ANK1 | ankyrin 1, erythrocytic | Plasma Membrane | other | 3.67E+07 | 2.96E+07 | 3.09E+07 | | | | | | |
| Q02357 | ANK1 | ankyrin 1, erythrocytic | Plasma Membrane | other | 1.13E+07 | 1.09E+07 | 1.03E+07 | | | | | | |
| Q02357 | ANK1 | ankyrin 1, erythrocytic | Plasma Membrane | other | 6988498.5 | 4483055 | 1.21E+07 | | | | | | |
| Q02357 | ANK1 | ankyrin 1, erythrocytic | Plasma Membrane | other | 3142156 | 2925762.75 | 1.55E+07 | | | | | | |
| Q02357 | ANK1 | ankyrin 1, erythrocytic | Plasma Membrane | other | 1.50E+07 | 1.72E+07 | 7613190 | | | | | | |
| Q02357 | ANK1 | ankyrin 1, erythrocytic | Plasma Membrane | other | 508670.844 | 53533.207 | 828587.5 | | | | | | |
| Q02357 | ANK1 | ankyrin 1, erythrocytic | Plasma Membrane | other | 1.05E+07 | 8474533 | 6420525 | | | | | | |
| Q02357 | ANK1 | ankyrin 1, erythrocytic | Plasma Membrane | other | 1.14E+07 | 5221041 | 5858291.5 | | | | | | |
| Q02357 | ANK1 | ankyrin 1, erythrocytic | Plasma Membrane | other | 3928399.5 | 4379774 | 1.76E+07 | | | | | | |
| Q02357 | ANK1 | ankyrin 1, erythrocytic | Plasma Membrane | other | 1.20E+07 | 1.19E+07 | 7576677.5 | | | | | | |
| Q02357 | ANK1 | ankyrin 1, erythrocytic | Plasma Membrane | other | 6237064 | 4528384.5 | 5056012 | | | | | | |
| Q02357 | ANK1 | ankyrin 1, erythrocytic | Plasma Membrane | other | 8185630.5 | 1.42E+07 | 8597178 | | | | | | |
| Q02357 | ANK1 | ankyrin 1, erythrocytic | Plasma Membrane | other | 7189457.5 | 4927063 | 7345370.5 | | | | | | |
| Q02357 | ANK1 | ankyrin 1, erythrocytic | Plasma Membrane | other | 4502415.5 | 3729396.5 | 1835297.125 | | | | | | |
| Q02357 | ANK1 | ankyrin 1, erythrocytic | Plasma Membrane | other | 9755018 | 7392432.5 | 6841437 | | | | | | |
| Q02357 | ANK1 | ankyrin 1, erythrocytic | Plasma Membrane | other | 9112606 | 7527293.5 | 5259969 | | | | | | |
| Q02357 | ANK1 | ankyrin 1, erythrocytic | Plasma Membrane | other | 3.04E+07 | 2.74E+07 | 2.02E+07 | | | | | | |
| Q02357 | ANK1 | ankyrin 1, erythrocytic | Plasma Membrane | other | 4685055.5 | 2050820 | 1980965.5 | | | | | | |
| Q02357 | ANK1 | ankyrin 1, erythrocytic | Plasma Membrane | other | 3556750 | 6184456.5 | 4767340.5 | | | | | | |
| Q02357 | ANK1 | ankyrin 1, erythrocytic | Plasma Membrane | other | 1.80E+07 | 1.33E+07 | 1.46E+07 | | | | | | |
| Q02357 | ANK1 | ankyrin 1, erythrocytic | Plasma Membrane | other | 1369518.375 | 8048687 | 1.39E+07 | | | | | | |
| Q02357 | ANK1 | ankyrin 1, erythrocytic | Plasma Membrane | other | 1.07E+07 | 1.27E+07 | 1.49E+07 | | | | | | |
| Q02357 | ANK1 | ankyrin 1, erythrocytic | Plasma Membrane | other | 3217381.25 | 2508184.75 | 2225880.5 | | | | | | |
| Q02357 | ANK1 | ankyrin 1, erythrocytic | Plasma Membrane | other | 1440430.375 | 624256.625 | 698742.875 | | | | | | |
| Q02357 | ANK1 | ankyrin 1, erythrocytic | Plasma Membrane | other | 363522.906 | 630457.125 | 2207992.25 | | | | | | |
| Q02357 | ANK1 | ankyrin 1, erythrocytic | Plasma Membrane | other | 3652746.25 | 2728307.5 | 2845827.5 | | | | | | |
| Q02357 | ANK1 | ankyrin 1, erythrocytic | Plasma Membrane | other | 2935944.75 | 2185790.75 | 2509174.5 | | | | | | |
| Q02357 | ANK1 | ankyrin 1, erythrocytic | Plasma Membrane | other | 3029392.75 | 5580618 | 3701231 | | | | | | |
| Q02357 | ANK1 | ankyrin 1, erythrocytic | Plasma Membrane | other | 6286534.5 | 3.67E+07 | 5.69E+07 | | | | | | |
| Q02357 | ANK1 | ankyrin 1, erythrocytic | Plasma Membrane | other | 1992983 | 854041.375 | 2171284.75 | | | | | | |
| Q02357 | ANK1 | ankyrin 1, erythrocytic | Plasma Membrane | other | 1203368.875 | 965540.5 | 1583166.125 | | | | | | |
| Q02357 | ANK1 | ankyrin 1, erythrocytic | Plasma Membrane | other | 912670.438 | 360549.719 | 894354.938 | | | | | | |
| Q02357 | ANK1 | ankyrin 1, erythrocytic | Plasma Membrane | other | 846983.25 | 502971.969 | 1900778.25 | | | | | | |
| Q02357 | ANK1 | ankyrin 1, erythrocytic | Plasma Membrane | other | 3590137 | 4164957.25 | 1998111 | | | | | | |
| Q02357 | ANK1 | ankyrin 1, erythrocytic | Plasma Membrane | other | 314502.719 | 8378166.5 | 8458669 | Q02357 | ANK1 | 504901809 | 663806943 | 29480701 | 507 |
| Q9JHZ2 | ANKH | ANKH inorganic pyrophosphate transport regulator | Plasma Membrane | transporter | 1489273.75 | 2027331.25 | 1951905.625 | Q9JHZ2 | ANKH | 1489274 | 291325 | 168197 | 3 |
| Q505D1 | ANKRD28 | ankyrin repeat domain 28 | Cytoplasm | other | 1334152.5 | 1194770.125 | 788385.625 | Q505D1 | ANKRD28 | 1334153 | 283560 | 163713 | 3 |
| Q6P9J9 | ANO6 | anoctamin 6 | Plasma Membrane | ion channel | 2163863.25 | 2366027.25 | 4111870.5 | Q6P9J9 | ANO6 | 2163863 | 1071103 | 618402 | 3 |

| | | | | | | | | | | | | | |
|---|---|---|---|---|---|---|---|---|---|---|---|---|---|
| O35381 | Anp32a | acidic (leucine-rich) nuclear phosphoprotein 32 family, member A | Nucleus | other | 9542653 | 7712367 | 1.26E+07 | | | | | | |
| O35381 | Anp32a | acidic (leucine-rich) nuclear phosphoprotein 32 family, member A | Nucleus | other | 5959920 | 4972232 | 5033151 | | | | | | |
| O35381 | Anp32a | acidic (leucine-rich) nuclear phosphoprotein 32 family, member A | Nucleus | other | 5793789.5 | 5328800.5 | 6091002 | O35381 | Anp32a | 7098788 | 2547333 | 849111 | 9 |
| O35643 | AP1B1 | adaptor-related protein complex 1, beta 1 subunit | Cytoplasm | transporter | 1.25E+07 | 1.55E+07 | 1.24E+07 | | | | | | |
| O35643 | AP1B1 | adaptor-related protein complex 1, beta 1 subunit | Cytoplasm | transporter | 1.34E+07 | 1.72E+07 | 1.44E+07 | | | | | | |
| O35643 | AP1B1 | adaptor-related protein complex 1, beta 1 subunit | Cytoplasm | transporter | 1.07E+07 | 1.14E+07 | 1.16E+07 | | | | | | |
| O35643 | AP1B1 | adaptor-related protein complex 1, beta 1 subunit | Cytoplasm | transporter | 1.61E+07 | 1.62E+07 | 1.51E+07 | | | | | | |
| O35643 | AP1B1 | adaptor-related protein complex 1, beta 1 subunit | Cytoplasm | transporter | 6998507.5 | 9389445 | 8509398 | | | | | | |
| O35643 | AP1B1 | adaptor-related protein complex 1, beta 1 subunit | Cytoplasm | transporter | 6699181 | 5398189.5 | 3619269.25 | | | | | | |
| O35643 | AP1B1 | adaptor-related protein complex 1, beta 1 subunit | Cytoplasm | transporter | 4712406 | 3863635.75 | 2201216.25 | | | | | | |
| O35643 | AP1B1 | adaptor-related protein complex 1, beta 1 subunit | Cytoplasm | transporter | 6675423.5 | 7091790.5 | 6236427.5 | | | | | | |
| O35643 | AP1B1 | adaptor-related protein complex 1, beta 1 subunit | Cytoplasm | transporter | 4051565.5 | 4066592.75 | 2927333 | | | | | | |
| O35643 | AP1B1 | adaptor-related protein complex 1, beta 1 subunit | Cytoplasm | transporter | 4217082.5 | 4690806.5 | 4160462 | | | | | | |
| O35643 | AP1B1 | adaptor-related protein complex 1, beta 1 subunit | Cytoplasm | transporter | 4456832.5 | 3493743.5 | 3661250.75 | | | | | | |
| O35643 | AP1B1 | adaptor-related protein complex 1, beta 1 subunit | Cytoplasm | transporter | 1676568.5 | 2144944 | 1593125.5 | O35643 | AP1B1 | 7678131 | 4851200 | 808533 | 36 |
| P17426 | AP2A1 | adaptor-related protein complex 2, alpha 1 subunit | Cytoplasm | transporter | 4055637.5 | 5836657 | 4509519.5 | | | | | | |
| P17426 | AP2A1 | adaptor-related protein complex 2, alpha 1 subunit | Cytoplasm | transporter | 2978363.25 | 4169831.5 | 4956515.5 | | | | | | |
| P17426 | AP2A1 | adaptor-related protein complex 2, alpha 1 subunit | Cytoplasm | transporter | 859352.062 | 1500464 | 1156158.875 | P17426 | AP2A1 | 2631118 | 1797646 | 599215 | 9 |
| P17427 | AP2A2 | adaptor-related protein complex 2, alpha 2 subunit | Cytoplasm | transporter | 1.18E+07 | 1.43E+07 | 9721907 | | | | | | |
| P17427 | AP2A2 | adaptor-related protein complex 2, alpha 2 subunit | Cytoplasm | transporter | 6098405 | 5801119.5 | 4199892 | | | | | | |
| P17427 | AP2A2 | adaptor-related protein complex 2, alpha 2 subunit | Cytoplasm | transporter | 1.23E+07 | 1.46E+07 | 2.15E+07 | | | | | | |
| P17427 | AP2A2 | adaptor-related protein complex 2, alpha 2 subunit | Cytoplasm | transporter | 3447041.5 | 3472815 | 3934530.75 | | | | | | |
| P17427 | AP2A2 | adaptor-related protein complex 2, alpha 2 subunit | Cytoplasm | transporter | 2708916.5 | 3197884 | 2547784.5 | | | | | | |
| P17427 | AP2A2 | adaptor-related protein complex 2, alpha 2 subunit | Cytoplasm | transporter | 2925226.5 | 3448354.75 | 3166611.5 | | | | | | |
| P17427 | AP2A2 | adaptor-related protein complex 2, alpha 2 subunit | Cytoplasm | transporter | 1551385.625 | 2097114.625 | 1185219.125 | | | | | | |
| P17427 | AP2A2 | adaptor-related protein complex 2, alpha 2 subunit | Cytoplasm | transporter | 1504674.375 | 1881565.375 | 1455073.125 | P17427 | AP2A2 | 5295706 | 5357753 | 1093647 | 24 |
| Q9DBG3 | Ap2b1 | adaptor-related protein complex 2, beta 1 subunit | Plasma Membrane | other | 1355484.875 | 1801935.75 | 2540124.75 | | | | | | |
| Q9DBG3 | Ap2b1 | adaptor-related protein complex 2, beta 1 subunit | Plasma Membrane | other | 121308.477 | 192752.328 | 163799.438 | Q9DBG3 | Ap2b1 | 738397 | 1025603 | 418701 | 6 |
| P84091 | AP2M1 | adaptor-related protein complex 2, mu 1 subunit | Cytoplasm | transporter | 1.04E+07 | 7970613.5 | 8264252 | | | | | | |
| P84091 | AP2M1 | adaptor-related protein complex 2, mu 1 subunit | Cytoplasm | transporter | 6896521 | 6866361.5 | 6121009.5 | | | | | | |
| P84091 | AP2M1 | adaptor-related protein complex 2, mu 1 subunit | Cytoplasm | transporter | 2611409 | 2974180.75 | 3292706.75 | | | | | | |
| P84091 | AP2M1 | adaptor-related protein complex 2, mu 1 subunit | Cytoplasm | transporter | 4079177.5 | 4132173.75 | 4110801 | | | | | | |
| P84091 | AP2M1 | adaptor-related protein complex 2, mu 1 subunit | Cytoplasm | transporter | 2501777 | 2408068 | 2463020.75 | P84091 | AP2M1 | 5291777 | 2551815 | 658876 | 15 |
| Q8R146 | APEH | acylaminoacyl-peptide hydrolase | Cytoplasm | peptidase | 8.35E+07 | 5.04E+07 | 4.15E+07 | | | | | | |
| Q8R146 | APEH | acylaminoacyl-peptide hydrolase | Cytoplasm | peptidase | 4.61E+07 | 3.23E+07 | 2.57E+07 | | | | | | |
| Q8R146 | APEH | acylaminoacyl-peptide hydrolase | Cytoplasm | peptidase | 4.04E+07 | 3.11E+07 | 2.34E+07 | | | | | | |
| Q8R146 | APEH | acylaminoacyl-peptide hydrolase | Cytoplasm | peptidase | 4.11E+07 | 2.74E+07 | 2.32E+07 | | | | | | |
| Q8R146 | APEH | acylaminoacyl-peptide hydrolase | Cytoplasm | peptidase | 4.06E+07 | 2.32E+07 | 1.74E+07 | | | | | | |
| Q8R146 | APEH | acylaminoacyl-peptide hydrolase | Cytoplasm | peptidase | 2.96E+07 | 2.00E+07 | 1.64E+07 | | | | | | |
| Q8R146 | APEH | acylaminoacyl-peptide hydrolase | Cytoplasm | peptidase | 3.36E+07 | 2.23E+07 | 1.88E+07 | | | | | | |
| Q8R146 | APEH | acylaminoacyl-peptide hydrolase | Cytoplasm | peptidase | 2.77E+07 | 1.62E+07 | 1.32E+07 | | | | | | |
| Q8R146 | APEH | acylaminoacyl-peptide hydrolase | Cytoplasm | peptidase | 2.70E+07 | 1.42E+07 | 1.78E+07 | | | | | | |
| Q8R146 | APEH | acylaminoacyl-peptide hydrolase | Cytoplasm | peptidase | 2.19E+07 | 1.74E+07 | 1.57E+07 | | | | | | |
| Q8R146 | APEH | acylaminoacyl-peptide hydrolase | Cytoplasm | peptidase | 2.27E+07 | 1.70E+07 | 1.33E+07 | | | | | | |
| Q8R146 | APEH | acylaminoacyl-peptide hydrolase | Cytoplasm | peptidase | 2.58E+07 | 1.81E+07 | 1.49E+07 | | | | | | |
| Q8R146 | APEH | acylaminoacyl-peptide hydrolase | Cytoplasm | peptidase | 2.22E+07 | 1.54E+07 | 1.46E+07 | | | | | | |
| Q8R146 | APEH | acylaminoacyl-peptide hydrolase | Cytoplasm | peptidase | 2.95E+07 | 2.78E+07 | 1.74E+07 | | | | | | |
| Q8R146 | APEH | acylaminoacyl-peptide hydrolase | Cytoplasm | peptidase | 3.64E+07 | 1.97E+07 | 1.32E+07 | | | | | | |
| Q8R146 | APEH | acylaminoacyl-peptide hydrolase | Cytoplasm | peptidase | 2.31E+07 | 1.71E+07 | 1.47E+07 | | | | | | |
| Q8R146 | APEH | acylaminoacyl-peptide hydrolase | Cytoplasm | peptidase | 2.08E+07 | 1.56E+07 | 1.26E+07 | | | | | | |
| Q8R146 | APEH | acylaminoacyl-peptide hydrolase | Cytoplasm | peptidase | 1.53E+07 | 1.22E+07 | 1.06E+07 | | | | | | |
| Q8R146 | APEH | acylaminoacyl-peptide hydrolase | Cytoplasm | peptidase | 9918006 | 5369604.5 | 5255205.5 | | | | | | |
| Q8R146 | APEH | acylaminoacyl-peptide hydrolase | Cytoplasm | peptidase | 4670616.5 | 4453102 | 4274695.5 | | | | | | |
| Q8R146 | APEH | acylaminoacyl-peptide hydrolase | Cytoplasm | peptidase | 6098733.5 | 3826224.25 | 3115003 | | | | | | |
| Q8R146 | APEH | acylaminoacyl-peptide hydrolase | Cytoplasm | peptidase | 4738751 | 2967949.25 | 2353600.5 | | | | | | |
| Q8R146 | APEH | acylaminoacyl-peptide hydrolase | Cytoplasm | peptidase | 4780427 | 4220930 | 2789016.5 | | | | | | |
| Q8R146 | APEH | acylaminoacyl-peptide hydrolase | Cytoplasm | peptidase | 1.05E+07 | 8570874 | 6591305 | | | | | | |
| Q8R146 | APEH | acylaminoacyl-peptide hydrolase | Cytoplasm | peptidase | 1061100.5 | 1097693.625 | 917620.812 | Q8R146 | APEH | 25155505 | 13865953 | 1601102 | 75 |
| Q8BVF7 | APH1A | APH1A gamma secretase subunit | Cytoplasm | peptidase | 1532686.75 | 1665300 | 794187.312 | Q8BVF7 | APH1A | 1532687 | 469362 | 270986 | 3 |
| Q00623 | APOA1 | apolipoprotein A-I | Extracellular Space | transporter | 3311861.5 | 2726521 | 5729565.5 | | | | | | |
| Q00623 | APOA1 | apolipoprotein A-I | Extracellular Space | transporter | 3247598 | 3136833 | 6276432.5 | | | | | | |
| Q00623 | APOA1 | apolipoprotein A-I | Extracellular Space | transporter | 1069230.375 | 732036.25 | 1393660 | Q00623 | APOA1 | 2542897 | 1931523 | 643841 | 9 |
| Q01339 | APOH | apolipoprotein H (beta-2-glycoprotein I) | Extracellular Space | transporter | 620430.562 | 590856.562 | 724717.25 | Q01339 | APOH | 620431 | 70320 | 40599 | 3 |
| Q02013 | AQP1 | aquaporin 1 (Colton blood group) | Plasma Membrane | transporter | 2.66E+09 | 2.67E+09 | 2.30E+09 | | | | | | |
| Q02013 | AQP1 | aquaporin 1 (Colton blood group) | Plasma Membrane | transporter | 2.38E+09 | 2.54E+09 | 2.03E+09 | | | | | | |
| Q02013 | AQP1 | aquaporin 1 (Colton blood group) | Plasma Membrane | transporter | 1.02E+09 | 6.77E+08 | 9.30E+08 | | | | | | |
| Q02013 | AQP1 | aquaporin 1 (Colton blood group) | Plasma Membrane | transporter | 4.62E+08 | 4.02E+08 | 3.41E+08 | | | | | | |
| Q02013 | AQP1 | aquaporin 1 (Colton blood group) | Plasma Membrane | transporter | 9.57E+07 | 4.81E+07 | 5.58E+07 | Q02013 | AQP1 | 1321930000 | 1050278029 | 271180621 | 15 |
| P84084 | ARF5 | ADP-ribosylation factor 5 | Cytoplasm | enzyme | 1975531.125 | 998292.125 | 1754387.5 | | | | | | |
| P84084 | ARF5 | ADP-ribosylation factor 5 | Cytoplasm | enzyme | 943219.5 | 917268.062 | 1325547.75 | P84084 | ARF5 | 1459375 | 453010 | 184941 | 6 |
| P62331 | ARF6 | ADP-ribosylation factor 6 | Plasma Membrane | transporter | 1.23E+07 | 1.48E+07 | 1.71E+07 | | | | | | |
| P62331 | ARF6 | ADP-ribosylation factor 6 | Plasma Membrane | transporter | 1.21E+07 | 1.39E+07 | 1.24E+07 | | | | | | |
| P62331 | ARF6 | ADP-ribosylation factor 6 | Plasma Membrane | transporter | 4298733 | 7630992 | 6626134 | | | | | | |
| P62331 | ARF6 | ADP-ribosylation factor 6 | Plasma Membrane | transporter | 4478267.5 | 5024511 | 6012985 | | | | | | |
| P62331 | ARF6 | ADP-ribosylation factor 6 | Plasma Membrane | transporter | 3857399.25 | 4631539 | 5844701.5 | | | | | | |
| P62331 | ARF6 | ADP-ribosylation factor 6 | Plasma Membrane | transporter | 3896236.25 | 4627415 | 4578924.5 | P62331 | ARF6 | 6815106 | 4409792 | 1039398 | 18 |
| Q80YF9 | ARHGAP33 | Rho GTPase activating protein 33 | Plasma Membrane | transporter | 1.60E+07 | 1.45E+07 | 1.21E+07 | Q80YF9 | ARHGAP33 | 15970000 | 1983742 | 1145314 | 3 |
| Q9Z1N7 | ARID3B | AT rich interactive domain 3B (BRIGHT-like) | Nucleus | other | 8983707 | 4422222 | 8522608 | Q9Z1N7 | ARID3B | 8983707 | 2511073 | 1449769 | 3 |
| Q9JKW0 | ARL6IP1 | ADP-ribosylation factor-like 6 interacting protein 1 | Cytoplasm | other | 9615544 | 1.01E+07 | 9494553 | Q9JKW0 | ARL6IP1 | 9615544 | 337416 | 194807 | 3 |
| Q8R5J9 | ARL6IP5 | ADP-ribosylation factor-like 6 interacting protein 5 | Cytoplasm | other | 3651065.75 | 5294352.5 | 3515531.25 | Q8R5J9 | ARL6IP5 | 3651066 | 990199 | 571692 | 3 |
| Q8VEH3 | ARL8A | ADP-ribosylation factor-like 8A | Cytoplasm | enzyme | 2531382.5 | 1795716.25 | 2042820.875 | Q8VEH3 | ARL8A | 2531383 | 374379 | 216148 | 3 |
| Q9CQW2 | ARL8B | ADP-ribosylation factor-like 8B | Plasma Membrane | enzyme | 4.68E+07 | 4.86E+07 | 4.16E+07 | | | | | | |
| Q9CQW2 | ARL8B | ADP-ribosylation factor-like 8B | Plasma Membrane | enzyme | 2.34E+07 | 2.75E+07 | 2.60E+07 | | | | | | |
| Q9CQW2 | ARL8B | ADP-ribosylation factor-like 8B | Plasma Membrane | enzyme | 2.50E+07 | 2.67E+07 | 2.06E+07 | | | | | | |
| Q9CQW2 | ARL8B | ADP-ribosylation factor-like 8B | Plasma Membrane | enzyme | 1.56E+07 | 1.88E+07 | 1.82E+07 | | | | | | |
| Q9CQW2 | ARL8B | ADP-ribosylation factor-like 8B | Plasma Membrane | enzyme | 1.64E+07 | 1.90E+07 | 1.72E+07 | | | | | | |
| Q9CQW2 | ARL8B | ADP-ribosylation factor-like 8B | Plasma Membrane | enzyme | 1.34E+07 | 1.35E+07 | 1.69E+07 | | | | | | |
| Q9CQW2 | ARL8B | ADP-ribosylation factor-like 8B | Plasma Membrane | enzyme | 2.97E+07 | 2.80E+07 | 1.24E+07 | | | | | | |
| Q9CQW2 | ARL8B | ADP-ribosylation factor-like 8B | Plasma Membrane | enzyme | 9884795 | 3675636.75 | 1.17E+07 | | | | | | |
| Q9CQW2 | ARL8B | ADP-ribosylation factor-like 8B | Plasma Membrane | enzyme | 9563253 | 9763066 | 6292268.5 | | | | | | |
| Q9CQW2 | ARL8B | ADP-ribosylation factor-like 8B | Plasma Membrane | enzyme | 2.28E+07 | 1.97E+07 | 1.08E+07 | | | | | | |
| Q9CQW2 | ARL8B | ADP-ribosylation factor-like 8B | Plasma Membrane | enzyme | 1.28E+07 | 1.95E+07 | 9576875 | | | | | | |
| Q9CQW2 | ARL8B | ADP-ribosylation factor-like 8B | Plasma Membrane | enzyme | 8477824 | 9505324 | 1.08E+07 | | | | | | |
| Q9CQW2 | ARL8B | ADP-ribosylation factor-like 8B | Plasma Membrane | enzyme | 8984633 | 1.07E+07 | 8116839 | | | | | | |
| Q9CQW2 | ARL8B | ADP-ribosylation factor-like 8B | Plasma Membrane | enzyme | 9479298 | 1.03E+07 | 1.18E+07 | Q9CQW2 | ARL8B | 18014986 | 10282581 | 1586637 | 42 |
| Q9R0Q6 | ARPC1A | actin related protein 2/3 complex, subunit 1A, 41kDa | Extracellular Space | other | 2170913.75 | 3347403.25 | 4821753 | Q9R0Q6 | ARPC1A | 2170914 | 1328206 | 766840 | 3 |
| Q9CVB6 | ARPC2 | actin related protein 2/3 complex, subunit 2, 34kDa | Cytoplasm | other | 1.73E+07 | 1.86E+07 | 2.34E+07 | | | | | | |
| Q9CVB6 | ARPC2 | actin related protein 2/3 complex, subunit 2, 34kDa | Cytoplasm | other | 9350217 | 9779645 | 1.14E+07 | | | | | | |
| Q9CVB6 | ARPC2 | actin related protein 2/3 complex, subunit 2, 34kDa | Cytoplasm | other | 6343547 | 6323433 | 8193369.5 | Q9CVB6 | ARPC2 | 11004588 | 6035998 | 2011999 | 9 |
| Q9JM76 | ARPC3 | actin related protein 2/3 complex, subunit 3, 21kDa | Cytoplasm | other | 2724609.75 | 3902422 | 4001578 | | | | | | |
| Q9JM76 | ARPC3 | actin related protein 2/3 complex, subunit 3, 21kDa | Cytoplasm | other | 1889520.375 | 2316590.25 | 3122318 | Q9JM76 | ARPC3 | 2307065 | 849498 | 346806 | 6 |
| P59999 | ARPC4 | actin related protein 2/3 complex, subunit 4, 20kDa | Cytoplasm | other | 2508965.75 | 3466707.75 | 4538621 | P59999 | ARPC4 | 2508966 | 1015363 | 586220 | 3 |
| Q9CPW4 | ARPC5 | actin related protein 2/3 complex, subunit 5, 16kDa | Cytoplasm | other | 1440001.25 | 1533064 | 2363661.5 | Q9CPW4 | ARPC5 | 1440001 | 508544 | 293608 | 3 |
| Q9CRA0 | ART4 | ADP-ribosyltransferase 4 (Dombrock blood group) | Nucleus | enzyme | 3.32E+07 | 3.85E+07 | 3.05E+07 | | | | | | |
| Q9CRA0 | ART4 | ADP-ribosyltransferase 4 (Dombrock blood group) | Nucleus | enzyme | 3.38E+07 | 3.14E+07 | 2.86E+07 | | | | | | |
| Q9CRA0 | ART4 | ADP-ribosyltransferase 4 (Dombrock blood group) | Nucleus | enzyme | 9367681 | 8855587 | 7016137.5 | | | | | | |
| Q9CRA0 | ART4 | ADP-ribosyltransferase 4 (Dombrock blood group) | Nucleus | enzyme | 7334498.5 | 6724710.5 | 7895462.5 | | | | | | |
| Q9CRA0 | ART4 | ADP-ribosyltransferase 4 (Dombrock blood group) | Nucleus | enzyme | 6035187 | 4347425 | 4367436 | | | | | | |
| Q9CRA0 | ART4 | ADP-ribosyltransferase 4 (Dombrock blood group) | Nucleus | enzyme | 3857009 | 3708361.5 | 3192380 | | | | | | |

| | | | | | | | | | | | | | |
|---|---|---|---|---|---|---|---|---|---|---|---|---|---|
| Q9CRA0 | ART4 | ADP-ribosyltransferase 4 (Dombrock blood group) | Nucleus | enzyme | 2719010.5 | 4468148 | 3333255.5 | Q9CRA0 | ART4 | 13761912 | 12793894 | 2791857 | 21 |
| Q9QWY8 | ASAP1 | ArfGAP with SH3 domain, ankyrin repeat and PH domain 1 | Plasma Membrane | other | 3560855.25 | 2598876 | 3682820 | | | | | | |
| Q9QWY8 | ASAP1 | ArfGAP with SH3 domain, ankyrin repeat and PH domain 1 | Plasma Membrane | other | 1787224 | 987655.812 | 1360411.75 | Q9QWY8 | ASAP1 | 2674040 | 1136135 | 463825 | 6 |
| Q7SIG6 | ASAP2 | ArfGAP with SH3 domain, ankyrin repeat and PH domain 2 | Nucleus | other | 1.07E+07 | 5155600.5 | 1.19E+07 | Q7SIG6 | ASAP2 | 10680000 | 3579114 | 2066403 | 3 |
| Q9WV71 | ASB4 | ankyrin repeat and SOCS box containing 4 | Nucleus | transcription regulator | 6185663 | 8235535.5 | 7079676 | Q9WV71 | ASB4 | 6185663 | 1027720 | 593354 | 3 |
| Q4QY64 | ATAD5 | ATPase family, AAA domain containing 5 | Extracellular Space | other | 1.57E+07 | 1.06E+07 | 1.29E+07 | Q4QY64 | ATAD5 | 15680000 | 2534798 | 1463466 | 3 |
| P16951 | ATF2 | activating transcription factor 2 | Nucleus | transcription regulator | 6.08E+07 | 6.67E+07 | 7.06E+07 | P16951 | ATF2 | 60820000 | 4896798 | 2827168 | 3 |
| Q811C2 | ATG4C | autophagy related 4C, cysteine peptidase | Cytoplasm | peptidase | 912470 | 895156 | 850212.062 | Q811C2 | ATG4C | 912470 | 32135 | 18553 | 3 |
| Q9CWJ9 | ATIC | 5-aminoimidazole-4-carboxamide ribonucleotide formyltransferase/IMP cyclohydrolase | Cytoplasm | enzyme | 1.18E+07 | 7623802.5 | 5119889 | | | | | | |
| Q9CWJ9 | ATIC | 5-aminoimidazole-4-carboxamide ribonucleotide formyltransferase/IMP cyclohydrolase | Cytoplasm | enzyme | 7308569 | 4986261 | 6418966 | | | | | | |
| Q9CWJ9 | ATIC | 5-aminoimidazole-4-carboxamide ribonucleotide formyltransferase/IMP cyclohydrolase | Cytoplasm | enzyme | 5263305.5 | 2306283 | 2988615.25 | | | | | | |
| Q9CWJ9 | ATIC | 5-aminoimidazole-4-carboxamide ribonucleotide formyltransferase/IMP cyclohydrolase | Cytoplasm | enzyme | 4603485 | 3842946.25 | 2384423 | | | | | | |
| Q9CWJ9 | ATIC | 5-aminoimidazole-4-carboxamide ribonucleotide formyltransferase/IMP cyclohydrolase | Cytoplasm | enzyme | 3926471 | 3621062.75 | 2375582.25 | | | | | | |
| Q9CWJ9 | ATIC | 5-aminoimidazole-4-carboxamide ribonucleotide formyltransferase/IMP cyclohydrolase | Cytoplasm | enzyme | 4985675.5 | 4257770 | 2717767.5 | | | | | | |
| Q9CWJ9 | ATIC | 5-aminoimidazole-4-carboxamide ribonucleotide formyltransferase/IMP cyclohydrolase | Cytoplasm | enzyme | 3486692.25 | 2817058.25 | 1225427.25 | | | | | | |
| Q9CWJ9 | ATIC | 5-aminoimidazole-4-carboxamide ribonucleotide formyltransferase/IMP cyclohydrolase | Cytoplasm | enzyme | 1991232.125 | 1273849.625 | 959946 | | | | | | |
| Q9CWJ9 | ATIC | 5-aminoimidazole-4-carboxamide ribonucleotide formyltransferase/IMP cyclohydrolase | Cytoplasm | enzyme | 1395428.75 | 886129.562 | 584746.25 | | | | | | |
| Q9CWJ9 | ATIC | 5-aminoimidazole-4-carboxamide ribonucleotide formyltransferase/IMP cyclohydrolase | Cytoplasm | enzyme | 1838588 | 2018852.75 | 1982221 | Q9CWJ9 | ATIC | 4662945 | 2440245 | 445526 | 30 |
| Q6PA06 | ATL2 | atlastin GTPase 2 | Cytoplasm | other | 4578341 | 5335934.5 | 5530440 | | | | | | |
| Q6PA06 | ATL2 | atlastin GTPase 2 | Cytoplasm | other | 1710224.375 | 1974426.5 | 2670339.5 | | | | | | |
| Q6PA06 | ATL2 | atlastin GTPase 2 | Cytoplasm | other | 1101638 | 1493251.625 | 2198995 | Q6PA06 | ATL2 | 2463401 | 1720182 | 573394 | 9 |
| Q91YH5 | ATL3 | atlastin GTPase 3 | Cytoplasm | other | 6.52E+07 | 5.82E+07 | 5.52E+07 | | | | | | |
| Q91YH5 | ATL3 | atlastin GTPase 3 | Cytoplasm | other | 3.12E+07 | 3.24E+07 | 3.62E+07 | | | | | | |
| Q91YH5 | ATL3 | atlastin GTPase 3 | Cytoplasm | other | 2.48E+07 | 2.18E+07 | 2.41E+07 | | | | | | |
| Q91YH5 | ATL3 | atlastin GTPase 3 | Cytoplasm | other | 2.16E+07 | 1.80E+07 | 1.95E+07 | | | | | | |
| Q91YH5 | ATL3 | atlastin GTPase 3 | Cytoplasm | other | 2.71E+07 | 2.63E+07 | 2.84E+07 | | | | | | |
| Q91YH5 | ATL3 | atlastin GTPase 3 | Cytoplasm | other | 2.47E+07 | 2.26E+07 | 2.39E+07 | | | | | | |
| Q91YH5 | ATL3 | atlastin GTPase 3 | Cytoplasm | other | 2.85E+07 | 3.42E+07 | 3.39E+07 | | | | | | |
| Q91YH5 | ATL3 | atlastin GTPase 3 | Cytoplasm | other | 2.12E+07 | 1.92E+07 | 2.01E+07 | | | | | | |
| Q91YH5 | ATL3 | atlastin GTPase 3 | Cytoplasm | other | 2.08E+07 | 1.66E+07 | 1.93E+07 | | | | | | |
| Q91YH5 | ATL3 | atlastin GTPase 3 | Cytoplasm | other | 1.01E+07 | 9950593 | 1.05E+07 | | | | | | |
| Q91YH5 | ATL3 | atlastin GTPase 3 | Cytoplasm | other | 1.15E+07 | 1.16E+07 | 7859714.5 | | | | | | |
| Q91YH5 | ATL3 | atlastin GTPase 3 | Cytoplasm | other | 6235776 | 6455172 | 5559989 | | | | | | |
| Q91YH5 | ATL3 | atlastin GTPase 3 | Cytoplasm | other | 6020405.5 | 5661997 | 7371430.5 | | | | | | |
| Q91YH5 | ATL3 | atlastin GTPase 3 | Cytoplasm | other | 5578581 | 5020796 | 4359095 | | | | | | |
| Q91YH5 | ATL3 | atlastin GTPase 3 | Cytoplasm | other | 5322697 | 3019124.5 | 3940410.25 | | | | | | |
| Q91YH5 | ATL3 | atlastin GTPase 3 | Cytoplasm | other | 5390285.5 | 5137613 | 4659071 | Q91YH5 | ATL3 | 19692984 | 14443625 | 2084758 | 48 |
| Q9QZW0 | ATP11C | ATPase, class VI, type 11C | Plasma Membrane | transporter | 1.11E+07 | 1.06E+07 | 9624194 | | | | | | |
| Q9QZW0 | ATP11C | ATPase, class VI, type 11C | Plasma Membrane | transporter | 9520511 | 9908568 | 9989875 | | | | | | |
| Q9QZW0 | ATP11C | ATPase, class VI, type 11C | Plasma Membrane | transporter | 6791158.5 | 5632330.5 | 5268114 | | | | | | |
| Q9QZW0 | ATP11C | ATPase, class VI, type 11C | Plasma Membrane | transporter | 3538648.5 | 4722889 | 4103860.75 | | | | | | |
| Q9QZW0 | ATP11C | ATPase, class VI, type 11C | Plasma Membrane | transporter | 4339954.5 | 4799327.5 | 4128391.5 | | | | | | |
| Q9QZW0 | ATP11C | ATPase, class VI, type 11C | Plasma Membrane | transporter | 2873017.25 | 2630099.75 | 3015096.25 | Q9QZW0 | ATP11C | 6360548 | 2997114 | 706427 | 18 |
| Q9EPE9 | ATP13A1 | ATPase type 13A1 | Extracellular Space | transporter | 1.21E+07 | 1.19E+07 | 9807084 | Q9EPE9 | ATP13A1 | 12070000 | 1268014 | 732088 | 3 |
| Q8VDN2 | ATP1A1 | ATPase, Na+/K+ transporting, alpha 1 polypeptide | Plasma Membrane | transporter | 3.04E+07 | 3.21E+07 | 3.37E+07 | | | | | | |
| Q8VDN2 | ATP1A1 | ATPase, Na+/K+ transporting, alpha 1 polypeptide | Plasma Membrane | transporter | 3.26E+07 | 4.06E+07 | 3.24E+07 | | | | | | |
| Q8VDN2 | ATP1A1 | ATPase, Na+/K+ transporting, alpha 1 polypeptide | Plasma Membrane | transporter | 2.83E+07 | 3.08E+07 | 2.88E+07 | | | | | | |
| Q8VDN2 | ATP1A1 | ATPase, Na+/K+ transporting, alpha 1 polypeptide | Plasma Membrane | transporter | 2.96E+07 | 3.41E+07 | 3.91E+07 | | | | | | |
| Q8VDN2 | ATP1A1 | ATPase, Na+/K+ transporting, alpha 1 polypeptide | Plasma Membrane | transporter | 2.30E+07 | 2.86E+07 | 2.97E+07 | | | | | | |
| Q8VDN2 | ATP1A1 | ATPase, Na+/K+ transporting, alpha 1 polypeptide | Plasma Membrane | transporter | 2.19E+07 | 2.92E+07 | 2.95E+07 | | | | | | |
| Q8VDN2 | ATP1A1 | ATPase, Na+/K+ transporting, alpha 1 polypeptide | Plasma Membrane | transporter | 2.09E+07 | 1.78E+07 | 2.02E+07 | | | | | | |
| Q8VDN2 | ATP1A1 | ATPase, Na+/K+ transporting, alpha 1 polypeptide | Plasma Membrane | transporter | 1.98E+07 | 2.18E+07 | 1.88E+07 | | | | | | |
| Q8VDN2 | ATP1A1 | ATPase, Na+/K+ transporting, alpha 1 polypeptide | Plasma Membrane | transporter | 1.77E+07 | 1.82E+07 | 1.87E+07 | | | | | | |
| Q8VDN2 | ATP1A1 | ATPase, Na+/K+ transporting, alpha 1 polypeptide | Plasma Membrane | transporter | 1.80E+07 | 1.76E+07 | 2.00E+07 | | | | | | |
| Q8VDN2 | ATP1A1 | ATPase, Na+/K+ transporting, alpha 1 polypeptide | Plasma Membrane | transporter | 1.76E+07 | 1.65E+07 | 1.76E+07 | | | | | | |
| Q8VDN2 | ATP1A1 | ATPase, Na+/K+ transporting, alpha 1 polypeptide | Plasma Membrane | transporter | 1.10E+07 | 1.13E+07 | 1.08E+07 | | | | | | |
| Q8VDN2 | ATP1A1 | ATPase, Na+/K+ transporting, alpha 1 polypeptide | Plasma Membrane | transporter | 1.21E+07 | 1.33E+07 | 1.18E+07 | | | | | | |
| Q8VDN2 | ATP1A1 | ATPase, Na+/K+ transporting, alpha 1 polypeptide | Plasma Membrane | transporter | 1.05E+07 | 9599230 | 9196227 | | | | | | |
| Q8VDN2 | ATP1A1 | ATPase, Na+/K+ transporting, alpha 1 polypeptide | Plasma Membrane | transporter | 1.06E+07 | 1.22E+07 | 1.18E+07 | | | | | | |
| Q8VDN2 | ATP1A1 | ATPase, Na+/K+ transporting, alpha 1 polypeptide | Plasma Membrane | transporter | 1.29E+07 | 1.44E+07 | 1.28E+07 | | | | | | |
| Q8VDN2 | ATP1A1 | ATPase, Na+/K+ transporting, alpha 1 polypeptide | Plasma Membrane | transporter | 9397707 | 1.36E+07 | 1.25E+07 | | | | | | |
| Q8VDN2 | ATP1A1 | ATPase, Na+/K+ transporting, alpha 1 polypeptide | Plasma Membrane | transporter | 5833054.5 | 6249016.5 | 6205455 | | | | | | |
| Q8VDN2 | ATP1A1 | ATPase, Na+/K+ transporting, alpha 1 polypeptide | Plasma Membrane | transporter | 6659386 | 6761176 | 6578482 | | | | | | |
| Q8VDN2 | ATP1A1 | ATPase, Na+/K+ transporting, alpha 1 polypeptide | Plasma Membrane | transporter | 5959317.5 | 8644353 | 6016756 | | | | | | |
| Q8VDN2 | ATP1A1 | ATPase, Na+/K+ transporting, alpha 1 polypeptide | Plasma Membrane | transporter | 5808957 | 7019770 | 5326004.5 | | | | | | |
| Q8VDN2 | ATP1A1 | ATPase, Na+/K+ transporting, alpha 1 polypeptide | Plasma Membrane | transporter | 2714836.75 | 3667984 | 3683509 | | | | | | |
| Q8VDN2 | ATP1A1 | ATPase, Na+/K+ transporting, alpha 1 polypeptide | Plasma Membrane | transporter | 5694625.5 | 5690236.5 | 5234962.5 | | | | | | |
| Q8VDN2 | ATP1A1 | ATPase, Na+/K+ transporting, alpha 1 polypeptide | Plasma Membrane | transporter | 6490176.5 | 7789256.5 | 5255165 | | | | | | |
| Q8VDN2 | ATP1A1 | ATPase, Na+/K+ transporting, alpha 1 polypeptide | Plasma Membrane | transporter | 5662178 | 6928967.5 | 7750435.5 | | | | | | |
| Q8VDN2 | ATP1A1 | ATPase, Na+/K+ transporting, alpha 1 polypeptide | Plasma Membrane | transporter | 2790942 | 2266281.75 | 2950854.5 | | | | | | |
| Q8VDN2 | ATP1A1 | ATPase, Na+/K+ transporting, alpha 1 polypeptide | Plasma Membrane | transporter | 2953167 | 3445490.5 | 2857204.25 | | | | | | |
| Q8VDN2 | ATP1A1 | ATPase, Na+/K+ transporting, alpha 1 polypeptide | Plasma Membrane | transporter | 1195311.75 | 971405.312 | 1207517 | | | | | | |
| Q8VDN2 | ATP1A1 | ATPase, Na+/K+ transporting, alpha 1 polypeptide | Plasma Membrane | transporter | 2626659.25 | 3410976.75 | 3104105.75 | Q8VDN2 | ATP1A1 | 13124356 | 10299913 | 1104267 | 87 |
| P14231 | ATP1B2 | ATPase, Na+/K+ transporting, beta 2 polypeptide | Plasma Membrane | transporter | 6.29E+07 | 5.18E+07 | 4.92E+07 | | | | | | |
| P14231 | ATP1B2 | ATPase, Na+/K+ transporting, beta 2 polypeptide | Plasma Membrane | transporter | 3.42E+07 | 3.18E+07 | 2.76E+07 | | | | | | |
| P14231 | ATP1B2 | ATPase, Na+/K+ transporting, beta 2 polypeptide | Plasma Membrane | transporter | 7695976 | 7117454 | 6610249.5 | | | | | | |
| P14231 | ATP1B2 | ATPase, Na+/K+ transporting, beta 2 polypeptide | Plasma Membrane | transporter | 5938627 | 5319653 | 5339910 | | | | | | |
| P14231 | ATP1B2 | ATPase, Na+/K+ transporting, beta 2 polypeptide | Plasma Membrane | transporter | 2528254.25 | 2707380.5 | 2731706 | P14231 | ATP1B2 | 22644571 | 20918420 | 5401113 | 15 |
| P97370 | ATP1B3 | ATPase, Na+/K+ transporting, beta 3 polypeptide | Plasma Membrane | transporter | 3384697.25 | 5133068 | 7502350.5 | | | | | | |
| P97370 | ATP1B3 | ATPase, Na+/K+ transporting, beta 3 polypeptide | Plasma Membrane | transporter | 2833127 | 3050116.75 | 5138442 | | | | | | |
| P97370 | ATP1B3 | ATPase, Na+/K+ transporting, beta 3 polypeptide | Plasma Membrane | transporter | 2301252.75 | 2776378 | 3193913.25 | | | | | | |
| P97370 | ATP1B3 | ATPase, Na+/K+ transporting, beta 3 polypeptide | Plasma Membrane | transporter | 2464791.25 | 4383864 | 4485995 | P97370 | ATP1B3 | 2745967 | 1510860 | 436148 | 12 |
| O55143 | ATP2A2 | ATPase, Ca++ transporting, cardiac muscle, slow twitch 2 | Cytoplasm | transporter | 5750271.5 | 9571304 | 1.11E+07 | | | | | | |
| O55143 | ATP2A2 | ATPase, Ca++ transporting, cardiac muscle, slow twitch 2 | Cytoplasm | transporter | 1.16E+07 | 1.40E+07 | 2.28E+07 | | | | | | |
| O55143 | ATP2A2 | ATPase, Ca++ transporting, cardiac muscle, slow twitch 2 | Cytoplasm | transporter | 3230310 | 4516647.5 | 6243152.5 | | | | | | |
| O55143 | ATP2A2 | ATPase, Ca++ transporting, cardiac muscle, slow twitch 2 | Cytoplasm | transporter | 2564219.25 | 3286296 | 4018114.75 | | | | | | |
| O55143 | ATP2A2 | ATPase, Ca++ transporting, cardiac muscle, slow twitch 2 | Cytoplasm | transporter | 1377423.75 | 2521755.5 | 2550339 | | | | | | |
| O55143 | ATP2A2 | ATPase, Ca++ transporting, cardiac muscle, slow twitch 2 | Cytoplasm | transporter | 1079762.625 | 2190519.5 | 2212523.25 | O55143 | ATP2A2 | 4258665 | 5668882 | 1336168 | 18 |
| Q64518 | ATP2A3 | ATPase, Ca++ transporting, ubiquitous | Cytoplasm | transporter | 1.84E+07 | 2.54E+07 | 4.02E+07 | | | | | | |
| Q64518 | ATP2A3 | ATPase, Ca++ transporting, ubiquitous | Cytoplasm | transporter | 9722097 | 1.02E+07 | 1.36E+07 | | | | | | |
| Q64518 | ATP2A3 | ATPase, Ca++ transporting, ubiquitous | Cytoplasm | transporter | 5609867.5 | 7471670.5 | 9426381 | | | | | | |
| Q64518 | ATP2A3 | ATPase, Ca++ transporting, ubiquitous | Cytoplasm | transporter | 6240040.5 | 6450668 | 1.23E+07 | | | | | | |
| Q64518 | ATP2A3 | ATPase, Ca++ transporting, ubiquitous | Cytoplasm | transporter | 1.22E+07 | 1.72E+07 | 2.72E+07 | | | | | | |
| Q64518 | ATP2A3 | ATPase, Ca++ transporting, ubiquitous | Cytoplasm | transporter | 3295121.5 | 5990451 | 6756423.5 | | | | | | |
| Q64518 | ATP2A3 | ATPase, Ca++ transporting, ubiquitous | Cytoplasm | transporter | 5126516 | 6742275.5 | 1.24E+07 | | | | | | |
| Q64518 | ATP2A3 | ATPase, Ca++ transporting, ubiquitous | Cytoplasm | transporter | 5565116.5 | 4907088 | 1.14E+07 | | | | | | |
| Q64518 | ATP2A3 | ATPase, Ca++ transporting, ubiquitous | Cytoplasm | transporter | 4357770 | 5145127 | 8298976.5 | | | | | | |
| Q64518 | ATP2A3 | ATPase, Ca++ transporting, ubiquitous | Cytoplasm | transporter | 5992374.5 | 7969165 | 1.29E+07 | | | | | | |
| Q64518 | ATP2A3 | ATPase, Ca++ transporting, ubiquitous | Cytoplasm | transporter | 258071.062 | 73976.656 | 2303870.75 | | | | | | |
| Q64518 | ATP2A3 | ATPase, Ca++ transporting, ubiquitous | Cytoplasm | transporter | 984976.438 | 1790053.625 | 1466182 | | | | | | |
| Q64518 | ATP2A3 | ATPase, Ca++ transporting, ubiquitous | Cytoplasm | transporter | 638709.062 | 934806.312 | 1707628.375 | | | | | | |
| Q64518 | ATP2A3 | ATPase, Ca++ transporting, ubiquitous | Cytoplasm | transporter | 3108770.5 | 4128281.25 | 3831503 | | | | | | |
| Q64518 | ATP2A3 | ATPase, Ca++ transporting, ubiquitous | Cytoplasm | transporter | 452231.938 | 724507.25 | 1612270.875 | | | | | | |
| Q64518 | ATP2A3 | ATPase, Ca++ transporting, ubiquitous | Cytoplasm | transporter | 3679664.75 | 6219331.5 | 8416866 | Q64518 | ATP2A3 | 5356333 | 7682412 | 1108861 | 48 |
| Q9R0K7 | ATP2B2 | ATPase, Ca++ transporting, plasma membrane 2 | Plasma Membrane | transporter | 1.15E+08 | 1.22E+08 | 1.07E+08 | | | | | | |
| Q9R0K7 | ATP2B2 | ATPase, Ca++ transporting, plasma membrane 2 | Plasma Membrane | transporter | 3.98E+07 | 4.74E+07 | 3.74E+07 | | | | | | |
| Q9R0K7 | ATP2B2 | ATPase, Ca++ transporting, plasma membrane 2 | Plasma Membrane | transporter | 3.31E+07 | 4.06E+07 | 3.27E+07 | | | | | | |
| Q9R0K7 | ATP2B2 | ATPase, Ca++ transporting, plasma membrane 2 | Plasma Membrane | transporter | 2.23E+07 | 1.74E+07 | 1.47E+07 | | | | | | |
| Q9R0K7 | ATP2B2 | ATPase, Ca++ transporting, plasma membrane 2 | Plasma Membrane | transporter | 2.41E+07 | 2.09E+07 | 1.86E+07 | | | | | | |

| | | | | | | | | | | | | | |
|---|---|---|---|---|---|---|---|---|---|---|---|---|---|
| Q9R0K7 | ATP2B2 | ATPase, Ca++ transporting, plasma membrane 2 | Plasma Membrane | transporter | 9004919 | 9207908 | 8076875 | | | | | | |
| Q9R0K7 | ATP2B2 | ATPase, Ca++ transporting, plasma membrane 2 | Plasma Membrane | transporter | 8178421.5 | 6159310.5 | 4935114.5 | | | | | | |
| Q9R0K7 | ATP2B2 | ATPase, Ca++ transporting, plasma membrane 2 | Plasma Membrane | transporter | 6161362 | 3447566.25 | 5865756 | | | | | | |
| Q9R0K7 | ATP2B2 | ATPase, Ca++ transporting, plasma membrane 2 | Plasma Membrane | transporter | 3775788 | 3075426.75 | 2350872.25 | | | | | | |
| Q9R0K7 | ATP2B2 | ATPase, Ca++ transporting, plasma membrane 2 | Plasma Membrane | transporter | 1869784.625 | 1584558.625 | 1352532.25 | Q9R0K7 | ATP2B2 | 26281028 | 33006078 | 6026058 | 30 |
| Q03265 | ATP5A1 | ATP synthase, H+ transporting, mitochondrial F1 complex, alpha subunit 1, cardiac muscle | Cytoplasm | transporter | 4.85E+07 | 7.82E+07 | 9.52E+07 | | | | | | |
| Q03265 | ATP5A1 | ATP synthase, H+ transporting, mitochondrial F1 complex, alpha subunit 1, cardiac muscle | Cytoplasm | transporter | 4.20E+07 | 6.91E+07 | 1.05E+08 | | | | | | |
| Q03265 | ATP5A1 | ATP synthase, H+ transporting, mitochondrial F1 complex, alpha subunit 1, cardiac muscle | Cytoplasm | transporter | 3.35E+07 | 5.40E+07 | 8.10E+07 | | | | | | |
| Q03265 | ATP5A1 | ATP synthase, H+ transporting, mitochondrial F1 complex, alpha subunit 1, cardiac muscle | Cytoplasm | transporter | 3.71E+07 | 5.17E+07 | 6.02E+07 | | | | | | |
| Q03265 | ATP5A1 | ATP synthase, H+ transporting, mitochondrial F1 complex, alpha subunit 1, cardiac muscle | Cytoplasm | transporter | 3.33E+07 | 4.88E+07 | 5.77E+07 | | | | | | |
| Q03265 | ATP5A1 | ATP synthase, H+ transporting, mitochondrial F1 complex, alpha subunit 1, cardiac muscle | Cytoplasm | transporter | 3.67E+07 | 4.93E+07 | 7.99E+07 | | | | | | |
| Q03265 | ATP5A1 | ATP synthase, H+ transporting, mitochondrial F1 complex, alpha subunit 1, cardiac muscle | Cytoplasm | transporter | 3.01E+07 | 5.06E+07 | 6.34E+07 | | | | | | |
| Q03265 | ATP5A1 | ATP synthase, H+ transporting, mitochondrial F1 complex, alpha subunit 1, cardiac muscle | Cytoplasm | transporter | 2.85E+07 | 4.61E+07 | 6.29E+07 | | | | | | |
| Q03265 | ATP5A1 | ATP synthase, H+ transporting, mitochondrial F1 complex, alpha subunit 1, cardiac muscle | Cytoplasm | transporter | 2.28E+07 | 3.62E+07 | 5.02E+07 | | | | | | |
| Q03265 | ATP5A1 | ATP synthase, H+ transporting, mitochondrial F1 complex, alpha subunit 1, cardiac muscle | Cytoplasm | transporter | 2.38E+07 | 3.83E+07 | 4.27E+07 | | | | | | |
| Q03265 | ATP5A1 | ATP synthase, H+ transporting, mitochondrial F1 complex, alpha subunit 1, cardiac muscle | Cytoplasm | transporter | 2.35E+07 | 3.88E+07 | 4.74E+07 | | | | | | |
| Q03265 | ATP5A1 | ATP synthase, H+ transporting, mitochondrial F1 complex, alpha subunit 1, cardiac muscle | Cytoplasm | transporter | 2.49E+07 | 3.81E+07 | 5.31E+07 | | | | | | |
| Q03265 | ATP5A1 | ATP synthase, H+ transporting, mitochondrial F1 complex, alpha subunit 1, cardiac muscle | Cytoplasm | transporter | 1.67E+07 | 2.43E+07 | 3.75E+07 | | | | | | |
| Q03265 | ATP5A1 | ATP synthase, H+ transporting, mitochondrial F1 complex, alpha subunit 1, cardiac muscle | Cytoplasm | transporter | 1.59E+07 | 2.05E+07 | 3.67E+07 | | | | | | |
| Q03265 | ATP5A1 | ATP synthase, H+ transporting, mitochondrial F1 complex, alpha subunit 1, cardiac muscle | Cytoplasm | transporter | 1.28E+07 | 2.11E+07 | 2.82E+07 | | | | | | |
| Q03265 | ATP5A1 | ATP synthase, H+ transporting, mitochondrial F1 complex, alpha subunit 1, cardiac muscle | Cytoplasm | transporter | 7526870 | 1.35E+07 | 2.24E+07 | | | | | | |
| Q03265 | ATP5A1 | ATP synthase, H+ transporting, mitochondrial F1 complex, alpha subunit 1, cardiac muscle | Cytoplasm | transporter | 1.03E+07 | 1.79E+07 | 2.55E+07 | | | | | | |
| Q03265 | ATP5A1 | ATP synthase, H+ transporting, mitochondrial F1 complex, alpha subunit 1, cardiac muscle | Cytoplasm | transporter | 8194565.5 | 1.21E+07 | 1.54E+07 | | | | | | |
| Q03265 | ATP5A1 | ATP synthase, H+ transporting, mitochondrial F1 complex, alpha subunit 1, cardiac muscle | Cytoplasm | transporter | 4025152.25 | 8636753 | 9291033 | | | | | | |
| Q03265 | ATP5A1 | ATP synthase, H+ transporting, mitochondrial F1 complex, alpha subunit 1, cardiac muscle | Cytoplasm | transporter | 3499070 | 5696044 | 9084145 | | | | | | |
| Q03265 | ATP5A1 | ATP synthase, H+ transporting, mitochondrial F1 complex, alpha subunit 1, cardiac muscle | Cytoplasm | transporter | 6253794.5 | 1.10E+07 | 1.47E+07 | | | | | | |
| Q03265 | ATP5A1 | ATP synthase, H+ transporting, mitochondrial F1 complex, alpha subunit 1, cardiac muscle | Cytoplasm | transporter | 4673699.5 | 1.07E+07 | 1.24E+07 | | | | | | |
| Q03265 | ATP5A1 | ATP synthase, H+ transporting, mitochondrial F1 complex, alpha subunit 1, cardiac muscle | Cytoplasm | transporter | 3806138.25 | 1.69E+07 | 2.63E+07 | Q03265 | ATP5A1 | 20797795 | 23333188 | 2808986 | 69 |
| P56480 | ATP5B | ATP synthase, H+ transporting, mitochondrial F1 complex, beta polypeptide | Cytoplasm | transporter | 8.21E+07 | 1.38E+08 | 1.53E+08 | | | | | | |
| P56480 | ATP5B | ATP synthase, H+ transporting, mitochondrial F1 complex, beta polypeptide | Cytoplasm | transporter | 6.97E+07 | 9.16E+07 | 8.59E+07 | | | | | | |
| P56480 | ATP5B | ATP synthase, H+ transporting, mitochondrial F1 complex, beta polypeptide | Cytoplasm | transporter | 5.02E+07 | 7.49E+07 | 1.18E+08 | | | | | | |
| P56480 | ATP5B | ATP synthase, H+ transporting, mitochondrial F1 complex, beta polypeptide | Cytoplasm | transporter | 3.71E+07 | 6.35E+07 | 8.45E+07 | | | | | | |
| P56480 | ATP5B | ATP synthase, H+ transporting, mitochondrial F1 complex, beta polypeptide | Cytoplasm | transporter | 3.44E+07 | 5.70E+07 | 8.44E+07 | | | | | | |
| P56480 | ATP5B | ATP synthase, H+ transporting, mitochondrial F1 complex, beta polypeptide | Cytoplasm | transporter | 2.24E+07 | 3.26E+07 | 4.07E+07 | | | | | | |
| P56480 | ATP5B | ATP synthase, H+ transporting, mitochondrial F1 complex, beta polypeptide | Cytoplasm | transporter | 3.04E+07 | 4.39E+07 | 5.42E+07 | | | | | | |
| P56480 | ATP5B | ATP synthase, H+ transporting, mitochondrial F1 complex, beta polypeptide | Cytoplasm | transporter | 3.55E+07 | 8.24E+07 | 8.93E+07 | | | | | | |
| P56480 | ATP5B | ATP synthase, H+ transporting, mitochondrial F1 complex, beta polypeptide | Cytoplasm | transporter | 2.38E+07 | 4.09E+07 | 4.88E+07 | | | | | | |
| P56480 | ATP5B | ATP synthase, H+ transporting, mitochondrial F1 complex, beta polypeptide | Cytoplasm | transporter | 3.44E+07 | 4.95E+07 | 6.03E+07 | | | | | | |
| P56480 | ATP5B | ATP synthase, H+ transporting, mitochondrial F1 complex, beta polypeptide | Cytoplasm | transporter | 3.35E+07 | 4.16E+07 | 5.91E+07 | | | | | | |
| P56480 | ATP5B | ATP synthase, H+ transporting, mitochondrial F1 complex, beta polypeptide | Cytoplasm | transporter | 3.16E+07 | 5.39E+07 | 7.21E+07 | | | | | | |
| P56480 | ATP5B | ATP synthase, H+ transporting, mitochondrial F1 complex, beta polypeptide | Cytoplasm | transporter | 1.85E+07 | 3.16E+07 | 2.94E+07 | | | | | | |
| P56480 | ATP5B | ATP synthase, H+ transporting, mitochondrial F1 complex, beta polypeptide | Cytoplasm | transporter | 2.45E+07 | 3.88E+07 | 5.33E+07 | | | | | | |
| P56480 | ATP5B | ATP synthase, H+ transporting, mitochondrial F1 complex, beta polypeptide | Cytoplasm | transporter | 2.24E+07 | 3.80E+07 | 5.11E+07 | | | | | | |
| P56480 | ATP5B | ATP synthase, H+ transporting, mitochondrial F1 complex, beta polypeptide | Cytoplasm | transporter | 2.89E+07 | 3.97E+07 | 5.28E+07 | | | | | | |
| P56480 | ATP5B | ATP synthase, H+ transporting, mitochondrial F1 complex, beta polypeptide | Cytoplasm | transporter | 2.51E+07 | 4.13E+07 | 4.26E+07 | | | | | | |
| P56480 | ATP5B | ATP synthase, H+ transporting, mitochondrial F1 complex, beta polypeptide | Cytoplasm | transporter | 1.60E+07 | 2.33E+07 | 2.86E+07 | | | | | | |
| P56480 | ATP5B | ATP synthase, H+ transporting, mitochondrial F1 complex, beta polypeptide | Cytoplasm | transporter | 1.19E+07 | 2.67E+07 | 1.95E+07 | | | | | | |
| P56480 | ATP5B | ATP synthase, H+ transporting, mitochondrial F1 complex, beta polypeptide | Cytoplasm | transporter | 1.64E+07 | 2.86E+07 | 3.13E+07 | | | | | | |
| P56480 | ATP5B | ATP synthase, H+ transporting, mitochondrial F1 complex, beta polypeptide | Cytoplasm | transporter | 1.30E+07 | 2.09E+07 | 2.82E+07 | | | | | | |
| P56480 | ATP5B | ATP synthase, H+ transporting, mitochondrial F1 complex, beta polypeptide | Cytoplasm | transporter | 1.46E+07 | 2.13E+07 | 3.37E+07 | | | | | | |
| P56480 | ATP5B | ATP synthase, H+ transporting, mitochondrial F1 complex, beta polypeptide | Cytoplasm | transporter | 8961665 | 1.60E+07 | 1.58E+07 | | | | | | |
| P56480 | ATP5B | ATP synthase, H+ transporting, mitochondrial F1 complex, beta polypeptide | Cytoplasm | transporter | 7024567 | 1.02E+07 | 1.65E+07 | | | | | | |
| P56480 | ATP5B | ATP synthase, H+ transporting, mitochondrial F1 complex, beta polypeptide | Cytoplasm | transporter | 6096296 | 8108464.5 | 1.03E+07 | | | | | | |
| P56480 | ATP5B | ATP synthase, H+ transporting, mitochondrial F1 complex, beta polypeptide | Cytoplasm | transporter | 5957131.5 | 1.45E+07 | 1.37E+07 | | | | | | |
| P56480 | ATP5B | ATP synthase, H+ transporting, mitochondrial F1 complex, beta polypeptide | Cytoplasm | transporter | 3096393.25 | 7511991 | 7683251 | | | | | | |
| P56480 | ATP5B | ATP synthase, H+ transporting, mitochondrial F1 complex, beta polypeptide | Cytoplasm | transporter | 2305192.25 | 4151266.25 | 1.30E+07 | | | | | | |
| P56480 | ATP5B | ATP synthase, H+ transporting, mitochondrial F1 complex, beta polypeptide | Cytoplasm | transporter | 3890734.5 | 6137487 | 7673698 | | | | | | |
| P56480 | ATP5B | ATP synthase, H+ transporting, mitochondrial F1 complex, beta polypeptide | Cytoplasm | transporter | 3797004 | 4690359 | 8175438 | | | | | | |
| P56480 | ATP5B | ATP synthase, H+ transporting, mitochondrial F1 complex, beta polypeptide | Cytoplasm | transporter | 630987.062 | 1911819.625 | 1391715 | | | | | | |
| P56480 | ATP5B | ATP synthase, H+ transporting, mitochondrial F1 complex, beta polypeptide | Cytoplasm | transporter | 282489.781 | 2138093.25 | 1648805.25 | P56480 | ATP5B | 22447577 | 30254000 | 3087786 | 96 |
| Q91VR2 | ATP5C1 | ATP synthase, H+ transporting, mitochondrial F1 complex, gamma polypeptide 1 | Cytoplasm | transporter | 7159643.5 | 1.23E+07 | 1.59E+07 | | | | | | |
| Q91VR2 | ATP5C1 | ATP synthase, H+ transporting, mitochondrial F1 complex, gamma polypeptide 1 | Cytoplasm | transporter | 4956149 | 7134107 | 1.03E+07 | | | | | | |
| Q91VR2 | ATP5C1 | ATP synthase, H+ transporting, mitochondrial F1 complex, gamma polypeptide 1 | Cytoplasm | transporter | 2838847.75 | 2943328.25 | 2878718.5 | | | | | | |
| Q91VR2 | ATP5C1 | ATP synthase, H+ transporting, mitochondrial F1 complex, gamma polypeptide 1 | Cytoplasm | transporter | 352249.062 | 1531177.5 | 3582867.75 | Q91VR2 | ATP5C1 | 3826722 | 4738194 | 1367799 | 12 |
| Q9D3D9 | ATP5D | ATP synthase, H+ transporting, mitochondrial F1 complex, delta subunit | Cytoplasm | transporter | 8887071 | 1.49E+07 | 1.75E+07 | | | | | | |
| Q9D3D9 | ATP5D | ATP synthase, H+ transporting, mitochondrial F1 complex, delta subunit | Cytoplasm | transporter | 1.16E+07 | 1.83E+07 | 1.87E+07 | Q9D3D9 | ATP5D | 10253536 | 3984359 | 1626608 | 6 |
| Q9CQQ7 | ATP5F1 | ATP synthase, H+ transporting, mitochondrial Fo complex, subunit B1 | Cytoplasm | transporter | 3797618 | 6783287 | 8913346 | Q9CQQ7 | ATP5F1 | 3797618 | 2569761 | 1483652 | 3 |
| Q9DCX2 | ATP5H | ATP synthase, H+ transporting, mitochondrial Fo complex, subunit d | Cytoplasm | enzyme | 7958790 | 1.23E+07 | 1.46E+07 | | | | | | |
| Q9DCX2 | ATP5H | ATP synthase, H+ transporting, mitochondrial Fo complex, subunit d | Cytoplasm | enzyme | 4899435.5 | 9728889 | 1.19E+07 | | | | | | |
| Q9DCX2 | ATP5H | ATP synthase, H+ transporting, mitochondrial Fo complex, subunit d | Cytoplasm | enzyme | 5098576 | 7388198.5 | 9380151 | | | | | | |
| Q9DCX2 | ATP5H | ATP synthase, H+ transporting, mitochondrial Fo complex, subunit d | Cytoplasm | enzyme | 3215875.25 | 4288057 | 5858411.5 | | | | | | |
| Q9DCX2 | ATP5H | ATP synthase, H+ transporting, mitochondrial Fo complex, subunit d | Cytoplasm | enzyme | 3163378.75 | 4471238.5 | 6868897.5 | | | | | | |
| Q9DCX2 | ATP5H | ATP synthase, H+ transporting, mitochondrial Fo complex, subunit d | Cytoplasm | enzyme | 2280767 | 3604175 | 4143521.75 | Q9DCX2 | ATP5H | 4436137 | 3564026 | 840049 | 18 |
| P97450 | ATP5J | ATP synthase, H+ transporting, mitochondrial Fo complex, subunit F6 | Cytoplasm | transporter | 4015837.25 | 6741564.5 | 8641863 | P97450 | ATP5J | 4015837 | 2325254 | 1342486 | 3 |
| P56135 | ATP5J2 | ATP synthase, H+ transporting, mitochondrial Fo complex, subunit F2 | Cytoplasm | transporter | 3814308.25 | 5741099 | 7566499 | P56135 | ATP5J2 | 3814308 | 1876324 | 1083296 | 3 |
| Q06185 | Atp5k | ATP synthase, H+ transporting, mitochondrial F1F0 complex, subunit E | Cytoplasm | enzyme | 8852504 | 1.54E+07 | 1.92E+07 | | | | | | |
| Q06185 | Atp5k | ATP synthase, H+ transporting, mitochondrial F1F0 complex, subunit E | Cytoplasm | enzyme | 4331670.5 | 7314885.5 | 9020116 | Q06185 | Atp5k | 6592087 | 5525996 | 2255978 | 6 |
| Q9CPQ8 | ATP5L | ATP synthase, H+ transporting, mitochondrial Fo complex, subunit G | Cytoplasm | enzyme | 5204802 | 9896165 | 1.02E+07 | | | | | | |
| Q9CPQ8 | ATP5L | ATP synthase, H+ transporting, mitochondrial Fo complex, subunit G | Cytoplasm | enzyme | 2706220.25 | 5109295.5 | 4981017.5 | Q9CPQ8 | ATP5L | 3955511 | 3015856 | 1231218 | 6 |
| Q9DB20 | ATP5O | ATP synthase, H+ transporting, mitochondrial F1 complex, O subunit | Cytoplasm | transporter | 1.12E+07 | 1.82E+07 | 2.17E+07 | | | | | | |
| Q9DB20 | ATP5O | ATP synthase, H+ transporting, mitochondrial F1 complex, O subunit | Cytoplasm | transporter | 1.18E+07 | 1.52E+07 | 2.32E+07 | | | | | | |
| Q9DB20 | ATP5O | ATP synthase, H+ transporting, mitochondrial F1 complex, O subunit | Cytoplasm | transporter | 6561519 | 9521329 | 1.19E+07 | | | | | | |
| Q9DB20 | ATP5O | ATP synthase, H+ transporting, mitochondrial F1 complex, O subunit | Cytoplasm | transporter | 6702691.5 | 1.18E+07 | 2.02E+07 | | | | | | |
| Q9DB20 | ATP5O | ATP synthase, H+ transporting, mitochondrial F1 complex, O subunit | Cytoplasm | transporter | 3717004.75 | 3536840.25 | 9253112 | | | | | | |
| Q9DB20 | ATP5O | ATP synthase, H+ transporting, mitochondrial F1 complex, O subunit | Cytoplasm | transporter | 2873246.5 | 6772365 | 8134529.5 | | | | | | |
| Q9DB20 | ATP5O | ATP synthase, H+ transporting, mitochondrial F1 complex, O subunit | Cytoplasm | transporter | 3266815 | 5210522 | 5505198.5 | | | | | | |
| Q9DB20 | ATP5O | ATP synthase, H+ transporting, mitochondrial F1 complex, O subunit | Cytoplasm | transporter | 3666445.75 | 5681905.5 | 8854857 | Q9DB20 | ATP5O | 6219715 | 6037297 | 1232358 | 24 |
| Q9R1Q9 | ATP6AP1 | ATPase, H+ transporting, lysosomal accessory protein 1 | Cytoplasm | transporter | 1849522.25 | 2606769 | 1779266.375 | Q9R1Q9 | ATP6AP1 | 1849522 | 458824 | 264902 | 3 |
| Q9CYN9 | ATP6AP2 | ATPase, H+ transporting, lysosomal accessory protein 2 | Cytoplasm | transporter | 2390660 | 2815771.75 | 2992928.25 | Q9CYN9 | ATP6AP2 | 2390660 | 309524 | 178704 | 3 |
| Q9Z1G4 | ATP6V0A1 | ATPase, H+ transporting, lysosomal V0 subunit a1 | Cytoplasm | transporter | 9313085 | 9090244 | 1.08E+07 | | | | | | |
| Q9Z1G4 | ATP6V0A1 | ATPase, H+ transporting, lysosomal V0 subunit a1 | Cytoplasm | transporter | 5782677.5 | 7718800.5 | 8036214.5 | | | | | | |
| Q9Z1G4 | ATP6V0A1 | ATPase, H+ transporting, lysosomal V0 subunit a1 | Cytoplasm | transporter | 8063640 | 8721598 | 8599820 | | | | | | |
| Q9Z1G4 | ATP6V0A1 | ATPase, H+ transporting, lysosomal V0 subunit a1 | Cytoplasm | transporter | 8933594 | 1.14E+07 | 8080902.5 | | | | | | |
| Q9Z1G4 | ATP6V0A1 | ATPase, H+ transporting, lysosomal V0 subunit a1 | Cytoplasm | transporter | 8347955.5 | 9562255 | 1.08E+07 | | | | | | |
| Q9Z1G4 | ATP6V0A1 | ATPase, H+ transporting, lysosomal V0 subunit a1 | Cytoplasm | transporter | 3926232.75 | 4181889.25 | 3460845.5 | | | | | | |
| Q9Z1G4 | ATP6V0A1 | ATPase, H+ transporting, lysosomal V0 subunit a1 | Cytoplasm | transporter | 4673108.5 | 5103436 | 4836354.5 | | | | | | |
| Q9Z1G4 | ATP6V0A1 | ATPase, H+ transporting, lysosomal V0 subunit a1 | Cytoplasm | transporter | 3080351.75 | 3995453 | 3816276.5 | | | | | | |
| Q9Z1G4 | ATP6V0A1 | ATPase, H+ transporting, lysosomal V0 subunit a1 | Cytoplasm | transporter | 2376795 | 3530077.75 | 2889411.75 | Q9Z1G4 | ATP6V0A1 | 6055271 | 2811047 | 540986 | 27 |
| P63082 | ATP6V0C | ATPase, H+ transporting, lysosomal 16kDa, V0 subunit c | Cytoplasm | transporter | 2.07E+07 | 2.43E+07 | 2.30E+07 | | | | | | |
| P63082 | ATP6V0C | ATPase, H+ transporting, lysosomal 16kDa, V0 subunit c | Cytoplasm | transporter | 2422707.5 | 2944693.25 | 2837473 | P63082 | ATP6V0C | 11581354 | 10982642 | 4483645 | 6 |
| P51863 | ATP6V0D1 | ATPase, H+ transporting, lysosomal 38kDa, V0 subunit d1 | Cytoplasm | transporter | 1.40E+07 | 1.95E+07 | 2.44E+07 | | | | | | |
| P51863 | ATP6V0D1 | ATPase, H+ transporting, lysosomal 38kDa, V0 subunit d1 | Cytoplasm | transporter | 8697125 | 1.15E+07 | 1.06E+07 | | | | | | |
| P51863 | ATP6V0D1 | ATPase, H+ transporting, lysosomal 38kDa, V0 subunit d1 | Cytoplasm | transporter | 7122739 | 8843354 | 7509737 | | | | | | |
| P51863 | ATP6V0D1 | ATPase, H+ transporting, lysosomal 38kDa, V0 subunit d1 | Cytoplasm | transporter | 5632513.5 | 9099011 | 6876717.5 | | | | | | |
| P51863 | ATP6V0D1 | ATPase, H+ transporting, lysosomal 38kDa, V0 subunit d1 | Cytoplasm | transporter | 3915401.25 | 5727300 | 6702502.5 | | | | | | |
| P51863 | ATP6V0D1 | ATPase, H+ transporting, lysosomal 38kDa, V0 subunit d1 | Cytoplasm | transporter | 2487589.25 | 3808157 | 3237252.25 | P51863 | ATP6V0D1 | 6974228 | 5673634 | 1337288 | 18 |
| P50516 | ATP6V1A | ATPase, H+ transporting, lysosomal 70kDa, V1 subunit A | Plasma Membrane | transporter | 4972815.5 | 4751806.5 | 4562477.5 | | | | | | |
| P50516 | ATP6V1A | ATPase, H+ transporting, lysosomal 70kDa, V1 subunit A | Plasma Membrane | transporter | 1532059 | 1997988.25 | 2185196.75 | | | | | | |
| P50516 | ATP6V1A | ATPase, H+ transporting, lysosomal 70kDa, V1 subunit A | Plasma Membrane | transporter | 2982291 | 2677891.5 | 1947464.5 | | | | | | |

| | | | | | | | | | | | | | |
|---|---|---|---|---|---|---|---|---|---|---|---|---|---|
| P50516 | ATP6V1A | ATPase, H+ transporting, lysosomal 70kDa, V1 subunit A | Plasma Membrane | transporter | 1689096.5 | 1385471.125 | 1522540.25 | | | | | | |
| P50516 | ATP6V1A | ATPase, H+ transporting, lysosomal 70kDa, V1 subunit A | Plasma Membrane | transporter | 775010 | 619480.125 | 838558.812 | P50516 | ATP6V1A | 2390254 | 1434278 | 370329 | 15 |
| P62814 | ATP6V1B2 | ATPase, H+ transporting, lysosomal 56/58kDa, V1 subunit B2 | Cytoplasm | transporter | 8427954 | 7508566 | 5486508.5 | | | | | | |
| P62814 | ATP6V1B2 | ATPase, H+ transporting, lysosomal 56/58kDa, V1 subunit B2 | Cytoplasm | transporter | 2021776.625 | 2413760.5 | 2762019.25 | | | | | | |
| P62814 | ATP6V1B2 | ATPase, H+ transporting, lysosomal 56/58kDa, V1 subunit B2 | Cytoplasm | transporter | 1616064.375 | 1813197.625 | 1741096.75 | P62814 | ATP6V1B2 | 4021932 | 2671945 | 890648 | 9 |
| P50518 | ATP6V1E1 | ATPase, H+ transporting, lysosomal 31kDa, V1 subunit E1 | Cytoplasm | transporter | 2049214.375 | 1792960.125 | 2242533 | P50518 | ATP6V1E1 | 2049214 | 225519 | 130204 | 3 |
| Q8BVE3 | ATP6V1H | ATPase, H+ transporting, lysosomal 50/57kDa, V1 subunit H | Cytoplasm | transporter | 1138858.625 | 898819.75 | 669603.812 | | | | | | |
| Q8BVE3 | ATP6V1H | ATPase, H+ transporting, lysosomal 50/57kDa, V1 subunit H | Cytoplasm | transporter | 452165.219 | 249448.828 | 121440.844 | Q8BVE3 | ATP6V1H | 795512 | 389219 | 158898 | 6 |
| Q64446 | ATP7B | ATPase, Cu++ transporting, beta polypeptide | Cytoplasm | transporter | 2.02E+07 | 2.05E+07 | 2.12E+07 | | | | | | |
| Q64446 | ATP7B | ATPase, Cu++ transporting, beta polypeptide | Cytoplasm | transporter | 3957478.25 | 3452193.25 | 3042909.75 | | | | | | |
| Q64446 | ATP7B | ATPase, Cu++ transporting, beta polypeptide | Cytoplasm | transporter | 3282812.5 | 3794893.25 | 3709492.5 | | | | | | |
| Q64446 | ATP7B | ATPase, Cu++ transporting, beta polypeptide | Cytoplasm | transporter | 4384387 | 4385950 | 4507641 | | | | | | |
| Q64446 | ATP7B | ATPase, Cu++ transporting, beta polypeptide | Cytoplasm | transporter | 2949543.5 | 2829939.5 | 3083183 | | | | | | |
| Q64446 | ATP7B | ATPase, Cu++ transporting, beta polypeptide | Cytoplasm | transporter | 2071186 | 2030927.5 | 3531306.25 | | | | | | |
| Q64446 | ATP7B | ATPase, Cu++ transporting, beta polypeptide | Cytoplasm | transporter | 2708648.75 | 2744577.5 | 2423926.75 | | | | | | |
| Q64446 | ATP7B | ATPase, Cu++ transporting, beta polypeptide | Cytoplasm | transporter | 1545683 | 1795250 | 1952342.875 | Q64446 | ATP7B | 5134967 | 5986980 | 1222087 | 24 |
| P70704 | ATP8A1 | ATPase, aminophospholipid transporter (APLT), class I, type 8A, member 1 | Cytoplasm | transporter | 2.33E+07 | 2.34E+07 | 2.47E+07 | | | | | | |
| P70704 | ATP8A1 | ATPase, aminophospholipid transporter (APLT), class I, type 8A, member 1 | Cytoplasm | transporter | 1.40E+07 | 1.48E+07 | 1.37E+07 | | | | | | |
| P70704 | ATP8A1 | ATPase, aminophospholipid transporter (APLT), class I, type 8A, member 1 | Cytoplasm | transporter | 1.82E+07 | 1.85E+07 | 1.93E+07 | | | | | | |
| P70704 | ATP8A1 | ATPase, aminophospholipid transporter (APLT), class I, type 8A, member 1 | Cytoplasm | transporter | 1.19E+07 | 1.16E+07 | 1.16E+07 | | | | | | |
| P70704 | ATP8A1 | ATPase, aminophospholipid transporter (APLT), class I, type 8A, member 1 | Cytoplasm | transporter | 1.19E+07 | 9586052 | 1.06E+07 | | | | | | |
| P70704 | ATP8A1 | ATPase, aminophospholipid transporter (APLT), class I, type 8A, member 1 | Cytoplasm | transporter | 1.02E+07 | 9837101 | 1.12E+07 | | | | | | |
| P70704 | ATP8A1 | ATPase, aminophospholipid transporter (APLT), class I, type 8A, member 1 | Cytoplasm | transporter | 1.07E+07 | 1.04E+07 | 8899365 | | | | | | |
| P70704 | ATP8A1 | ATPase, aminophospholipid transporter (APLT), class I, type 8A, member 1 | Cytoplasm | transporter | 8228699 | 9360275 | 8146148 | | | | | | |
| P70704 | ATP8A1 | ATPase, aminophospholipid transporter (APLT), class I, type 8A, member 1 | Cytoplasm | transporter | 6932965 | 7412744.5 | 8080116 | | | | | | |
| P70704 | ATP8A1 | ATPase, aminophospholipid transporter (APLT), class I, type 8A, member 1 | Cytoplasm | transporter | 6756326.5 | 7135884 | 5927090 | | | | | | |
| P70704 | ATP8A1 | ATPase, aminophospholipid transporter (APLT), class I, type 8A, member 1 | Cytoplasm | transporter | 6397773 | 6558186 | 7762334.5 | | | | | | |
| P70704 | ATP8A1 | ATPase, aminophospholipid transporter (APLT), class I, type 8A, member 1 | Cytoplasm | transporter | 4845753.5 | 4699671.5 | 3664206.75 | | | | | | |
| P70704 | ATP8A1 | ATPase, aminophospholipid transporter (APLT), class I, type 8A, member 1 | Cytoplasm | transporter | 7009978.5 | 7500819 | 7042471 | | | | | | |
| P70704 | ATP8A1 | ATPase, aminophospholipid transporter (APLT), class I, type 8A, member 1 | Cytoplasm | transporter | 6953521 | 6345875 | 4779240.5 | | | | | | |
| P70704 | ATP8A1 | ATPase, aminophospholipid transporter (APLT), class I, type 8A, member 1 | Cytoplasm | transporter | 7029022 | 6894820 | 7809749.5 | | | | | | |
| P70704 | ATP8A1 | ATPase, aminophospholipid transporter (APLT), class I, type 8A, member 1 | Cytoplasm | transporter | 4698555 | 3252363.25 | 3023663 | | | | | | |
| P70704 | ATP8A1 | ATPase, aminophospholipid transporter (APLT), class I, type 8A, member 1 | Cytoplasm | transporter | 5661767.5 | 5394031.5 | 5119329.5 | | | | | | |
| P70704 | ATP8A1 | ATPase, aminophospholipid transporter (APLT), class I, type 8A, member 1 | Cytoplasm | transporter | 5323268.5 | 3621050.5 | 4540733 | | | | | | |
| P70704 | ATP8A1 | ATPase, aminophospholipid transporter (APLT), class I, type 8A, member 1 | Cytoplasm | transporter | 4569436.5 | 4727107 | 4183865.25 | | | | | | |
| P70704 | ATP8A1 | ATPase, aminophospholipid transporter (APLT), class I, type 8A, member 1 | Cytoplasm | transporter | 3951897.5 | 4720620 | 4760898 | | | | | | |
| P70704 | ATP8A1 | ATPase, aminophospholipid transporter (APLT), class I, type 8A, member 1 | Cytoplasm | transporter | 2691592.75 | 2441743.75 | 2396808.5 | | | | | | |
| P70704 | ATP8A1 | ATPase, aminophospholipid transporter (APLT), class I, type 8A, member 1 | Cytoplasm | transporter | 2515644.5 | 3392528.25 | 1606980.375 | P70704 | ATP8A1 | 8353464 | 5186122 | 638367 | 66 |
| O35143 | ATPIF1 | ATPase inhibitory factor 1 | Cytoplasm | other | 6829398 | 1.07E+07 | 1.54E+07 | O35143 | ATPIF1 | 6829398 | 4271080 | 2465909 | 3 |
| P70295 | AUP1 | ancient ubiquitous protein 1 | Cytoplasm | other | 2412859.5 | 3568655.75 | 4563396.5 | P70295 | AUP1 | 2412860 | 1076273 | 621387 | 3 |
| P01887 | B2M | beta-2-microglobulin | Plasma Membrane | transmembrane receptor | 7290124 | 1.09E+07 | 1.21E+07 | P01887 | B2M | 7290124 | 2522753 | 1456512 | 3 |
| Q09199 | B4GALNT2 | beta-1,4-N-acetyl-galactosaminyl transferase 2 | Cytoplasm | enzyme | 2.02E+07 | 1.50E+07 | 2.03E+07 | Q09199 | B4GALNT2 | 20240000 | 3045527 | 1758336 | 3 |
| Q9JJ04 | B4GALT4 | UDP-Gal:betaGlcNAc beta 1,4- galactosyltransferase, polypeptide 4 | Cytoplasm | enzyme | 2.10E+08 | 2.80E+08 | 1.86E+08 | Q9JJ04 | B4GALT4 | 210400000 | 48864541 | 28211956 | 3 |
| Q91YN9 | BAG2 | BCL2-associated athanogene 2 | Cytoplasm | other | 3.76E+07 | 3.00E+07 | 2.70E+07 | | | | | | |
| Q91YN9 | BAG2 | BCL2-associated athanogene 2 | Cytoplasm | other | 3.06E+07 | 2.72E+07 | 1.99E+07 | | | | | | |
| Q91YN9 | BAG2 | BCL2-associated athanogene 2 | Cytoplasm | other | 2.00E+07 | 1.49E+07 | 1.46E+07 | | | | | | |
| Q91YN9 | BAG2 | BCL2-associated athanogene 2 | Cytoplasm | other | 1.28E+07 | 1.25E+07 | 7915713.5 | | | | | | |
| Q91YN9 | BAG2 | BCL2-associated athanogene 2 | Cytoplasm | other | 9458084 | 7514697 | 8461292 | | | | | | |
| Q91YN9 | BAG2 | BCL2-associated athanogene 2 | Cytoplasm | other | 9409279 | 9991209 | 1.16E+07 | | | | | | |
| Q91YN9 | BAG2 | BCL2-associated athanogene 2 | Cytoplasm | other | 1.00E+07 | 9815522 | 1.03E+07 | | | | | | |
| Q91YN9 | BAG2 | BCL2-associated athanogene 2 | Cytoplasm | other | 6628562 | 6276432.5 | 7979850.5 | | | | | | |
| Q91YN9 | BAG2 | BCL2-associated athanogene 2 | Cytoplasm | other | 5323931.5 | 3759532 | 4673599.5 | | | | | | |
| Q91YN9 | BAG2 | BCL2-associated athanogene 2 | Cytoplasm | other | 6238582 | 3097472 | 5494825.5 | | | | | | |
| Q91YN9 | BAG2 | BCL2-associated athanogene 2 | Cytoplasm | other | 848270.438 | 928148.25 | 1359816.75 | Q91YN9 | BAG2 | 13535155 | 9288438 | 1616910 | 33 |
| O08734 | BAK1 | BCL2-antagonist/killer 1 | Cytoplasm | other | 2477329 | 3953823.5 | 5093494 | O08734 | BAK1 | 2477329 | 1311691 | 757305 | 3 |
| O54962 | BANF1 | barrier to autointegration factor 1 | Nucleus | other | 5.53E+07 | 4.27E+07 | 2.02E+08 | | | | | | |
| O54962 | BANF1 | barrier to autointegration factor 1 | Nucleus | other | 3.34E+07 | 2.36E+07 | 8.47E+07 | | | | | | |
| O54962 | BANF1 | barrier to autointegration factor 1 | Nucleus | other | 1.24E+07 | 9015960 | 4.52E+07 | | | | | | |
| O54962 | BANF1 | barrier to autointegration factor 1 | Nucleus | other | 5234358 | 5787018.5 | 2.08E+07 | O54962 | BANF1 | 26606090 | 54843454 | 15831942 | 12 |
| Q61335 | BCAP31 | B-cell receptor-associated protein 31 | Cytoplasm | transporter | 2818634.75 | 2994327 | 3981309.5 | Q61335 | BCAP31 | 2818635 | 626740 | 361848 | 3 |
| O35855 | BCAT2 | branched chain amino-acid transaminase 2, mitochondrial | Cytoplasm | enzyme | 1586062.5 | 2829608.75 | 5381199.5 | O35855 | BCAT2 | 1586063 | 1934773 | 1117042 | 3 |
| P59017 | BCL2L13 | BCL2-like 13 (apoptosis facilitator) | Cytoplasm | other | 3072372.25 | 7728332 | 8456970 | | | | | | |
| P59017 | BCL2L13 | BCL2-like 13 (apoptosis facilitator) | Cytoplasm | other | 2232400.25 | 3258852.25 | 4037514.5 | | | | | | |
| P59017 | BCL2L13 | BCL2-like 13 (apoptosis facilitator) | Cytoplasm | other | 2336357.5 | 3455019.75 | 4252953.5 | P59017 | BCL2L13 | 2547043 | 2251083 | 750361 | 9 |
| Q8C6A8 | BHLHE22 | basic helix-loop-helix family, member e22 | Nucleus | transcription regulator | 1.67E+08 | 1.71E+08 | 1.57E+08 | Q8C6A8 | BHLHE22 | 167100000 | 7502666 | 4331666 | 3 |
| Q8BR07 | BICD1 | bicaudal D homolog 1 (Drosophila) | Cytoplasm | other | 4.67E+07 | 3.04E+07 | 2.43E+07 | Q8BR07 | BICD1 | 46660000 | 11563604 | 6676250 | 3 |
| Q8R016 | BLMH | bleomycin hydrolase | Cytoplasm | peptidase | 8658946 | 5464910.5 | 4767843 | | | | | | |
| Q8R016 | BLMH | bleomycin hydrolase | Cytoplasm | peptidase | 7172209.5 | 4856315 | 4456683.5 | | | | | | |
| Q8R016 | BLMH | bleomycin hydrolase | Cytoplasm | peptidase | 7398532 | 5661186.5 | 3754363 | | | | | | |
| Q8R016 | BLMH | bleomycin hydrolase | Cytoplasm | peptidase | 4346821 | 3376128.25 | 2026729.75 | | | | | | |
| Q8R016 | BLMH | bleomycin hydrolase | Cytoplasm | peptidase | 4366612.5 | 3343566.25 | 1772472 | | | | | | |
| Q8R016 | BLMH | bleomycin hydrolase | Cytoplasm | peptidase | 3435451 | 2697141.5 | 1744925.125 | | | | | | |
| Q8R016 | BLMH | bleomycin hydrolase | Cytoplasm | peptidase | 1647373.875 | 1417687.625 | 1451466.625 | Q8R016 | BLMH | 5289421 | 2068146 | 451307 | 21 |
| Q923D2 | BLVRB | biliverdin reductase B | Cytoplasm | enzyme | 6045310 | 4394749 | 4881183.5 | | | | | | |
| Q923D2 | BLVRB | biliverdin reductase B | Cytoplasm | enzyme | 5485095 | 5085007.5 | 4815839.5 | | | | | | |
| Q923D2 | BLVRB | biliverdin reductase B | Cytoplasm | enzyme | 2022191.5 | 2371571.25 | 1691425.25 | | | | | | |
| Q923D2 | BLVRB | biliverdin reductase B | Cytoplasm | enzyme | 852409.688 | 741277.312 | 480903.719 | Q923D2 | BLVRB | 3601252 | 2067422 | 596813 | 12 |
| P15327 | BPGM | 2,3-bisphosphoglycerate mutase | Extracellular Space | phosphatase | 1.01E+07 | 8495713 | 5678006 | | | | | | |
| P15327 | BPGM | 2,3-bisphosphoglycerate mutase | Extracellular Space | phosphatase | 3151322.25 | 2428996.25 | 1687606.75 | | | | | | |
| P15327 | BPGM | 2,3-bisphosphoglycerate mutase | Extracellular Space | phosphatase | 1233087.875 | 1493156.75 | 838527.562 | | | | | | |
| P15327 | BPGM | 2,3-bisphosphoglycerate mutase | Extracellular Space | phosphatase | 425956.25 | 372369.531 | 49515.602 | P15327 | BPGM | 3720092 | 3326619 | 960312 | 12 |
| A2AHJ4 | BRWD3 | bromodomain and WD repeat domain containing 3 | Nucleus | other | 1.26E+07 | 1.20E+07 | 9975071 | A2AHJ4 | BRWD3 | 12620000 | 1383256 | 798623 | 3 |
| P18572 | BSG | basigin (Ok blood group) | Plasma Membrane | transporter | 8.55E+08 | 8.29E+08 | 8.13E+08 | | | | | | |
| P18572 | BSG | basigin (Ok blood group) | Plasma Membrane | transporter | 6.61E+08 | 6.85E+08 | 6.29E+08 | | | | | | |
| P18572 | BSG | basigin (Ok blood group) | Plasma Membrane | transporter | 8.94E+08 | 7.86E+08 | 1.04E+09 | | | | | | |
| P18572 | BSG | basigin (Ok blood group) | Plasma Membrane | transporter | 5.89E+08 | 5.64E+08 | 4.78E+08 | | | | | | |
| P18572 | BSG | basigin (Ok blood group) | Plasma Membrane | transporter | 2.29E+08 | 2.72E+08 | 2.99E+08 | | | | | | |
| P18572 | BSG | basigin (Ok blood group) | Plasma Membrane | transporter | 2.20E+08 | 1.95E+08 | 1.92E+08 | | | | | | |
| P18572 | BSG | basigin (Ok blood group) | Plasma Membrane | transporter | 2.27E+08 | 2.95E+08 | 2.10E+08 | | | | | | |
| P18572 | BSG | basigin (Ok blood group) | Plasma Membrane | transporter | 6.49E+07 | 1.01E+08 | 6.04E+07 | | | | | | |
| P18572 | BSG | basigin (Ok blood group) | Plasma Membrane | transporter | 5.76E+07 | 7.35E+07 | 4.62E+07 | | | | | | |
| P18572 | BSG | basigin (Ok blood group) | Plasma Membrane | transporter | 4.33E+07 | 5.48E+07 | 5.80E+07 | | | | | | |
| P18572 | BSG | basigin (Ok blood group) | Plasma Membrane | transporter | 1.50E+07 | 1.63E+07 | 9400585 | | | | | | |
| P18572 | BSG | basigin (Ok blood group) | Plasma Membrane | transporter | 8081916 | 9491021 | 7821352 | | | | | | |
| P18572 | BSG | basigin (Ok blood group) | Plasma Membrane | transporter | 2109684.5 | 1654580.875 | 1810117 | | | | | | |
| P18572 | BSG | basigin (Ok blood group) | Plasma Membrane | transporter | 5729828 | 4983304.5 | 1684611.5 | P18572 | BSG | 276570816 | 317346238 | 48967588 | 42 |
| Q9D659 | C10orf54 | chromosome 10 open reading frame 54 | Plasma Membrane | other | 9342308 | 1.07E+07 | 8308237 | Q9D659 | C10orf54 | 9342308 | 1220404 | 704601 | 3 |
| Q91W34 | C16orf58 | chromosome 16 open reading frame 58 | Cytoplasm | other | 899071.812 | 1411220.375 | 1176406.5 | Q91W34 | C16orf58 | 899072 | 256368 | 148014 | 3 |
| Q9CX63 | C17orf99 | chromosome 17 open reading frame 99 | Extracellular Space | cytokine | 393037.594 | 2175781 | 1229781.75 | | | | | | |
| Q9CX63 | C17orf99 | chromosome 17 open reading frame 99 | Extracellular Space | cytokine | 76100.117 | 0 | 373510.719 | | | | | | |
| Q9CX63 | C17orf99 | chromosome 17 open reading frame 99 | Extracellular Space | cytokine | 83021.305 | 14481.472 | 0 | | | | | | |
| Q9CX63 | C17orf99 | chromosome 17 open reading frame 99 | Extracellular Space | cytokine | 51642.344 | 4.70E-06 | 49485.91 | | | | | | |
| Q9CX63 | C17orf99 | chromosome 17 open reading frame 99 | Extracellular Space | cytokine | 605911.438 | 354088.031 | 733894.375 | | | | | | |
| Q9CX63 | C17orf99 | chromosome 17 open reading frame 99 | Extracellular Space | cytokine | 624940.75 | 485583.875 | 120158.977 | | | | | | |
| Q9CX63 | C17orf99 | chromosome 17 open reading frame 99 | Extracellular Space | cytokine | 46673.957 | 0 | 0 | Q9CX63 | C17orf99 | 268761 | 529720 | 115594 | 21 |
| P98086 | C1QA | complement component 1, q subcomponent, A chain | Extracellular Space | other | 59353.238 | 1944655.125 | 355959.938 | | | | | | |
| P98086 | C1QA | complement component 1, q subcomponent, A chain | Extracellular Space | other | 4021.414 | 672844.125 | 177005.609 | P98086 | C1QA | 31687 | 731371 | 298581 | 6 |
| P14106 | C1QB | complement component 1, q subcomponent, B chain | Extracellular Space | other | 43892.996 | 2110693.25 | 394637.5 | P14106 | C1QB | 43893 | 1106009 | 638554 | 3 |
| O35658 | C1QBP | complement component 1, q subcomponent binding protein | Cytoplasm | transcription regulator | 1.39E+07 | 2.73E+07 | 2.91E+07 | | | | | | |

| | | | | | | | | | | | | | |
|---|---|---|---|---|---|---|---|---|---|---|---|---|---|
| O35658 | C1QBP | complement component 1, q subcomponent binding protein | Cytoplasm | transcription regulator | 2935978.5 | 1.77E+07 | 1.97E+07 | O35658 | C1QBP | 8397989 | 9555997 | 3901219 | 6 |
| Q9D172 | C21orf33/LOC102724023 | chromosome 21 open reading frame 33 | Cytoplasm | other | 6011795 | 7672625.5 | 1.08E+07 | | | | | | |
| Q9D172 | C21orf33/LOC102724023 | chromosome 21 open reading frame 33 | Cytoplasm | other | 255358.703 | 332039.094 | 304419.969 | Q9D172 | C21orf33/LOC102724023 | 3133577 | 4567952 | 1864859 | 6 |
| Q80X80 | C2CD2L | C2CD2-like | Cytoplasm | other | 1.96E+07 | 2.01E+07 | 1.80E+07 | | | | | | |
| Q80X80 | C2CD2L | C2CD2-like | Cytoplasm | other | 2.95E+07 | 2.62E+07 | 2.44E+07 | | | | | | |
| Q80X80 | C2CD2L | C2CD2-like | Cytoplasm | other | 1.77E+07 | 1.80E+07 | 1.54E+07 | | | | | | |
| Q80X80 | C2CD2L | C2CD2-like | Cytoplasm | other | 9009714 | 7870677.5 | 5558116.5 | | | | | | |
| Q80X80 | C2CD2L | C2CD2-like | Cytoplasm | other | 8296577.5 | 6826323 | 6787168.5 | | | | | | |
| Q80X80 | C2CD2L | C2CD2-like | Cytoplasm | other | 8802103 | 1.06E+07 | 6617784.5 | | | | | | |
| Q80X80 | C2CD2L | C2CD2-like | Cytoplasm | other | 1.02E+07 | 1.28E+07 | 1.21E+07 | | | | | | |
| Q80X80 | C2CD2L | C2CD2-like | Cytoplasm | other | 9853345 | 9513316 | 8379361.5 | | | | | | |
| Q80X80 | C2CD2L | C2CD2-like | Cytoplasm | other | 8923897 | 7502010.5 | 5553242.5 | | | | | | |
| Q80X80 | C2CD2L | C2CD2-like | Cytoplasm | other | 8061847.5 | 6808393 | 5088490 | | | | | | |
| Q80X80 | C2CD2L | C2CD2-like | Cytoplasm | other | 4678483.5 | 3649036.75 | 3498078.25 | | | | | | |
| Q80X80 | C2CD2L | C2CD2-like | Cytoplasm | other | 2846535.5 | 2187681.75 | 1614892.375 | | | | | | |
| Q80X80 | C2CD2L | C2CD2-like | Cytoplasm | other | 918340.812 | 367404.562 | 956070.125 | Q80X80 | C2CD2L | 10650065 | 7178859 | 1149537 | 39 |
| P01029 | C4A/C4B | complement component 4B (Chido blood group) | Extracellular Space | other | 3944685 | 5456631 | 4487911 | | | | | | |
| P01029 | C4A/C4B | complement component 4B (Chido blood group) | Extracellular Space | other | 3697200.5 | 4516296 | 6224707 | | | | | | |
| P01029 | C4A/C4B | complement component 4B (Chido blood group) | Extracellular Space | other | 2162343.5 | 2586442.5 | 3127834 | | | | | | |
| P01029 | C4A/C4B | complement component 4B (Chido blood group) | Extracellular Space | other | 1338617.25 | 1513588.375 | 1250334.75 | | | | | | |
| P01029 | C4A/C4B | complement component 4B (Chido blood group) | Extracellular Space | other | 1842305.125 | 2112292.75 | 1319799.125 | P01029 | C4A/C4B | 2597030 | 1607442 | 415040 | 15 |
| P00920 | CA2 | carbonic anhydrase II | Cytoplasm | enzyme | 2.18E+07 | 2.47E+07 | 2.26E+07 | | | | | | |
| P00920 | CA2 | carbonic anhydrase II | Cytoplasm | enzyme | 7959352 | 7917062 | 8420870 | | | | | | |
| P00920 | CA2 | carbonic anhydrase II | Cytoplasm | enzyme | 5024997.5 | 5510415 | 5570634 | | | | | | |
| P00920 | CA2 | carbonic anhydrase II | Cytoplasm | enzyme | 2357153 | 2379432.5 | 2290430.75 | | | | | | |
| P00920 | CA2 | carbonic anhydrase II | Cytoplasm | enzyme | 4039945 | 6173655.5 | 7820878 | | | | | | |
| P00920 | CA2 | carbonic anhydrase II | Cytoplasm | enzyme | 222670.188 | 195759.969 | 480888.969 | P00920 | CA2 | 6895686 | 7641857 | 1801203 | 18 |
| P62204 | CALM1 | Calmodulin | Cytoplasm | | 3.79E+07 | 3.31E+07 | 2.88E+07 | | | | | | |
| P62204 | CALM1 | Calmodulin | Cytoplasm | | 1.01E+07 | 1.16E+07 | 1.07E+07 | P62204 | CALM1 | 23980000 | 12655297 | 5166503 | 6 |
| Q9D6P8 | CALML3 | calmodulin-like 3 | Cytoplasm | other | 7524256.5 | 5956864.5 | 6634228 | Q9D6P8 | CALML3 | 7524257 | 786097 | 453853 | 3 |
| P14211 | CALR | calreticulin | Cytoplasm | transcription regulator | 2.30E+08 | 2.19E+08 | 1.70E+08 | | | | | | |
| P14211 | CALR | calreticulin | Cytoplasm | transcription regulator | 1.38E+08 | 1.46E+08 | 1.17E+08 | | | | | | |
| P14211 | CALR | calreticulin | Cytoplasm | transcription regulator | 1.68E+08 | 1.90E+08 | 1.49E+08 | | | | | | |
| P14211 | CALR | calreticulin | Cytoplasm | transcription regulator | 1.21E+08 | 9.14E+07 | 7.55E+07 | | | | | | |
| P14211 | CALR | calreticulin | Cytoplasm | transcription regulator | 1.11E+08 | 1.05E+08 | 9.95E+07 | | | | | | |
| P14211 | CALR | calreticulin | Cytoplasm | transcription regulator | 7.44E+07 | 8.10E+07 | 7.80E+07 | | | | | | |
| P14211 | CALR | calreticulin | Cytoplasm | transcription regulator | 8.51E+07 | 8.07E+07 | 7.52E+07 | | | | | | |
| P14211 | CALR | calreticulin | Cytoplasm | transcription regulator | 9.72E+07 | 1.37E+08 | 8.28E+07 | | | | | | |
| P14211 | CALR | calreticulin | Cytoplasm | transcription regulator | 4.75E+07 | 4.60E+07 | 4.71E+07 | | | | | | |
| P14211 | CALR | calreticulin | Cytoplasm | transcription regulator | 3.03E+07 | 2.95E+07 | 2.59E+07 | | | | | | |
| P14211 | CALR | calreticulin | Cytoplasm | transcription regulator | 3.32E+07 | 2.71E+07 | 3.65E+07 | | | | | | |
| P14211 | CALR | calreticulin | Cytoplasm | transcription regulator | 4320183 | 1915211.25 | 2058160 | | | | | | |
| P14211 | CALR | calreticulin | Cytoplasm | transcription regulator | 8342685 | 6785640.5 | 5075832.5 | | | | | | |
| P14211 | CALR | calreticulin | Cytoplasm | transcription regulator | 1.13E+07 | 2.00E+07 | 2.23E+07 | | | | | | |
| P14211 | CALR | calreticulin | Cytoplasm | transcription regulator | 7011945 | 4587399 | 4133262.5 | | | | | | |
| P14211 | CALR | calreticulin | Cytoplasm | transcription regulator | 1329874.25 | 1405293.75 | 1019678.25 | | | | | | |
| P14211 | CALR | calreticulin | Cytoplasm | transcription regulator | 2636360.5 | 2841428 | 1371098.75 | | | | | | |
| P14211 | CALR | calreticulin | Cytoplasm | transcription regulator | 454257.312 | 263213.594 | 235623.984 | | | | | | |
| P14211 | CALR | calreticulin | Cytoplasm | transcription regulator | 2058878.25 | 3566813.25 | 1371292.875 | P14211 | CALR | 61768641 | 62748132 | 8311193 | 57 |
| O35887 | CALU | calumenin | Cytoplasm | other | 9552239 | 8518393 | 6945332 | | | | | | |
| O35887 | CALU | calumenin | Cytoplasm | other | 8130464 | 8425129 | 7354721.5 | | | | | | |
| O35887 | CALU | calumenin | Cytoplasm | other | 8540123 | 8058725.5 | 5303793 | | | | | | |
| O35887 | CALU | calumenin | Cytoplasm | other | 7712521.5 | 7451055 | 6894274.5 | | | | | | |
| O35887 | CALU | calumenin | Cytoplasm | other | 3869270.75 | 5015402 | 3348128.75 | | | | | | |
| O35887 | CALU | calumenin | Cytoplasm | other | 3419731.75 | 4931209.5 | 2910091.5 | | | | | | |
| O35887 | CALU | calumenin | Cytoplasm | other | 3331882.75 | 3426363 | 2921935.5 | | | | | | |
| O35887 | CALU | calumenin | Cytoplasm | other | 1285984.75 | 712219 | 148170.297 | O35887 | CALU | 5730277 | 2768328 | 565083 | 24 |
| P51437 | CAMP | cathelicidin antimicrobial peptide | Cytoplasm | other | 1157775.25 | 635396.688 | 1.10E+07 | | | | | | |
| P51437 | CAMP | cathelicidin antimicrobial peptide | Cytoplasm | other | 1376070.75 | 1061631.375 | 1.31E+07 | P51437 | CAMP | 1266923 | 5720919 | 2335555 | 6 |
| Q6ZQ38 | CAND1 | cullin-associated and neddylation-dissociated 1 | Cytoplasm | transcription regulator | 2.08E+07 | 1.48E+07 | 1.30E+07 | | | | | | |
| Q6ZQ38 | CAND1 | cullin-associated and neddylation-dissociated 1 | Cytoplasm | transcription regulator | 1.64E+07 | 1.15E+07 | 1.29E+07 | | | | | | |
| Q6ZQ38 | CAND1 | cullin-associated and neddylation-dissociated 1 | Cytoplasm | transcription regulator | 1.49E+07 | 1.16E+07 | 1.40E+07 | | | | | | |
| Q6ZQ38 | CAND1 | cullin-associated and neddylation-dissociated 1 | Cytoplasm | transcription regulator | 1.01E+07 | 6601029 | 7837210 | | | | | | |
| Q6ZQ38 | CAND1 | cullin-associated and neddylation-dissociated 1 | Cytoplasm | transcription regulator | 1.10E+07 | 9673047 | 5808012 | | | | | | |
| Q6ZQ38 | CAND1 | cullin-associated and neddylation-dissociated 1 | Cytoplasm | transcription regulator | 1.59E+07 | 9392383 | 9008326 | | | | | | |
| Q6ZQ38 | CAND1 | cullin-associated and neddylation-dissociated 1 | Cytoplasm | transcription regulator | 8458519 | 7283929.5 | 6127820 | | | | | | |
| Q6ZQ38 | CAND1 | cullin-associated and neddylation-dissociated 1 | Cytoplasm | transcription regulator | 1.07E+07 | 9287471 | 1.03E+07 | | | | | | |
| Q6ZQ38 | CAND1 | cullin-associated and neddylation-dissociated 1 | Cytoplasm | transcription regulator | 9675917 | 7769658.5 | 6568259 | | | | | | |
| Q6ZQ38 | CAND1 | cullin-associated and neddylation-dissociated 1 | Cytoplasm | transcription regulator | 1.01E+07 | 6563745 | 7499659.5 | | | | | | |
| Q6ZQ38 | CAND1 | cullin-associated and neddylation-dissociated 1 | Cytoplasm | transcription regulator | 1.23E+07 | 7952255 | 8238435.5 | | | | | | |
| Q6ZQ38 | CAND1 | cullin-associated and neddylation-dissociated 1 | Cytoplasm | transcription regulator | 8611145 | 5270371.5 | 4987141 | | | | | | |
| Q6ZQ38 | CAND1 | cullin-associated and neddylation-dissociated 1 | Cytoplasm | transcription regulator | 6202081 | 3760506.25 | 4034975.75 | | | | | | |
| Q6ZQ38 | CAND1 | cullin-associated and neddylation-dissociated 1 | Cytoplasm | transcription regulator | 7699181.5 | 5512624 | 6454516 | | | | | | |
| Q6ZQ38 | CAND1 | cullin-associated and neddylation-dissociated 1 | Cytoplasm | transcription regulator | 3525109.5 | 2701772 | 3426508.75 | | | | | | |
| Q6ZQ38 | CAND1 | cullin-associated and neddylation-dissociated 1 | Cytoplasm | transcription regulator | 6058580 | 4360863.5 | 5090875 | | | | | | |
| Q6ZQ38 | CAND1 | cullin-associated and neddylation-dissociated 1 | Cytoplasm | transcription regulator | 4587490.5 | 3330245.25 | 4079864.5 | | | | | | |
| Q6ZQ38 | CAND1 | cullin-associated and neddylation-dissociated 1 | Cytoplasm | transcription regulator | 5000296 | 5965498.5 | 4672306 | | | | | | |
| Q6ZQ38 | CAND1 | cullin-associated and neddylation-dissociated 1 | Cytoplasm | transcription regulator | 2822550.25 | 2399451.25 | 1841759.25 | Q6ZQ38 | CAND1 | 9731625 | 4012532 | 531473 | 57 |
| P35564 | CANX | calnexin | Cytoplasm | other | 9.50E+07 | 1.01E+08 | 9.19E+07 | | | | | | |
| P35564 | CANX | calnexin | Cytoplasm | other | 6.83E+07 | 6.79E+07 | 6.33E+07 | | | | | | |
| P35564 | CANX | calnexin | Cytoplasm | other | 7.28E+07 | 7.52E+07 | 6.49E+07 | | | | | | |
| P35564 | CANX | calnexin | Cytoplasm | other | 5.33E+07 | 6.38E+07 | 5.63E+07 | | | | | | |
| P35564 | CANX | calnexin | Cytoplasm | other | 3.25E+07 | 2.10E+07 | 1.58E+07 | | | | | | |
| P35564 | CANX | calnexin | Cytoplasm | other | 5.36E+07 | 4.62E+07 | 4.94E+07 | | | | | | |
| P35564 | CANX | calnexin | Cytoplasm | other | 4.36E+07 | 4.98E+07 | 2.61E+07 | | | | | | |
| P35564 | CANX | calnexin | Cytoplasm | other | 3.81E+07 | 3.99E+07 | 3.05E+07 | | | | | | |
| P35564 | CANX | calnexin | Cytoplasm | other | 3.06E+07 | 3.63E+07 | 3.29E+07 | | | | | | |
| P35564 | CANX | calnexin | Cytoplasm | other | 3.46E+07 | 3.48E+07 | 3.34E+07 | | | | | | |
| P35564 | CANX | calnexin | Cytoplasm | other | 2.01E+07 | 1.75E+07 | 2.19E+07 | | | | | | |
| P35564 | CANX | calnexin | Cytoplasm | other | 8188907 | 1.10E+07 | 1.45E+07 | | | | | | |
| P35564 | CANX | calnexin | Cytoplasm | other | 3.64E+07 | 3.44E+07 | 3.27E+07 | | | | | | |
| P35564 | CANX | calnexin | Cytoplasm | other | 1.97E+07 | 2.50E+07 | 2.59E+07 | | | | | | |
| P35564 | CANX | calnexin | Cytoplasm | other | 7284659 | 7256309 | 7594484.5 | | | | | | |
| P35564 | CANX | calnexin | Cytoplasm | other | 4255543.5 | 3984350.5 | 2855104.5 | | | | | | |
| P35564 | CANX | calnexin | Cytoplasm | other | 4701986.5 | 6220399.5 | 3599585.25 | | | | | | |
| P35564 | CANX | calnexin | Cytoplasm | other | 4802237 | 1.71E+07 | 1.55E+07 | | | | | | |
| P35564 | CANX | calnexin | Cytoplasm | other | 2632254.25 | 1026052.438 | 3308472.5 | | | | | | |
| P35564 | CANX | calnexin | Cytoplasm | other | 3510851.5 | 4025292.5 | 1389280.75 | | | | | | |
| P35564 | CANX | calnexin | Cytoplasm | other | 1538985.75 | 1362376.5 | 1038749.812 | | | | | | |
| P35564 | CANX | calnexin | Cytoplasm | other | 621225.688 | 388607.938 | 165686.344 | P35564 | CANX | 28906211 | 26195811 | 3224481 | 66 |
| P40124 | CAP1 | CAP, adenylate cyclase-associated protein 1 (yeast) | Plasma Membrane | other | 3387313.75 | 2779769.25 | 2785727.25 | | | | | | |
| P40124 | CAP1 | CAP, adenylate cyclase-associated protein 1 (yeast) | Plasma Membrane | other | 2529460.75 | 1846150.125 | 2168658 | | | | | | |
| P40124 | CAP1 | CAP, adenylate cyclase-associated protein 1 (yeast) | Plasma Membrane | other | 1887265.5 | 374168.5 | 1329108 | P40124 | CAP1 | 2601347 | 898414 | 299471 | 9 |
| O35350 | CAPN1 | calpain 1, (mu/I) large subunit | Cytoplasm | peptidase | 6443640.5 | 5313856.5 | 7105190.5 | | | | | | |
| O35350 | CAPN1 | calpain 1, (mu/I) large subunit | Cytoplasm | peptidase | 5715349.5 | 3734746.5 | 4526993 | | | | | | |
| O35350 | CAPN1 | calpain 1, (mu/I) large subunit | Cytoplasm | peptidase | 5300636 | 3327353.5 | 3770213.25 | | | | | | |
| O35350 | CAPN1 | calpain 1, (mu/I) large subunit | Cytoplasm | peptidase | 4340938.5 | 3222125.25 | 3598498 | | | | | | |
| O35350 | CAPN1 | calpain 1, (mu/I) large subunit | Cytoplasm | peptidase | 3888889.5 | 2833612.5 | 4383523 | | | | | | |
| O35350 | CAPN1 | calpain 1, (mu/I) large subunit | Cytoplasm | peptidase | 3707666.25 | 2613682.75 | 2380945.5 | | | | | | |

| | | | | | | | | | | | | | |
|---|---|---|---|---|---|---|---|---|---|---|---|---|---|
| O35350 | CAPN1 | calpain 1, (mu/I) large subunit | Cytoplasm | peptidase | 751971.812 | 529772 | 650544.75 | O35350 | CAPN1 | 4307013 | 1761167 | 384318 | 21 |
| O08688 | CAPN5 | calpain 5 | Cytoplasm | peptidase | 2.06E+07 | 2.55E+07 | 2.24E+07 | | | | | | |
| O08688 | CAPN5 | calpain 5 | Cytoplasm | peptidase | 2.12E+07 | 2.23E+07 | 2.06E+07 | | | | | | |
| O08688 | CAPN5 | calpain 5 | Cytoplasm | peptidase | 1.12E+07 | 1.17E+07 | 9772221 | | | | | | |
| O08688 | CAPN5 | calpain 5 | Cytoplasm | peptidase | 1.24E+07 | 1.26E+07 | 9756209 | | | | | | |
| O08688 | CAPN5 | calpain 5 | Cytoplasm | peptidase | 1.37E+07 | 1.26E+07 | 1.23E+07 | | | | | | |
| O08688 | CAPN5 | calpain 5 | Cytoplasm | peptidase | 1.02E+07 | 9280027 | 9003510 | | | | | | |
| O08688 | CAPN5 | calpain 5 | Cytoplasm | peptidase | 1.27E+07 | 1.24E+07 | 9252486 | | | | | | |
| O08688 | CAPN5 | calpain 5 | Cytoplasm | peptidase | 6298089 | 6160712.5 | 5386103 | | | | | | |
| O08688 | CAPN5 | calpain 5 | Cytoplasm | peptidase | 5834073 | 6240350 | 4440044.5 | | | | | | |
| O08688 | CAPN5 | calpain 5 | Cytoplasm | peptidase | 6371046 | 5780832 | 5545879 | | | | | | |
| O08688 | CAPN5 | calpain 5 | Cytoplasm | peptidase | 4993131.5 | 4149671.5 | 3493972 | | | | | | |
| O08688 | CAPN5 | calpain 5 | Cytoplasm | peptidase | 4806711.5 | 3898462.5 | 4370433.5 | | | | | | |
| O08688 | CAPN5 | calpain 5 | Cytoplasm | peptidase | 5590686 | 5474901 | 4039609.25 | | | | | | |
| O08688 | CAPN5 | calpain 5 | Cytoplasm | peptidase | 2623609 | 2648799.5 | 1938918.25 | | | | | | |
| O08688 | CAPN5 | calpain 5 | Cytoplasm | peptidase | 1695165.625 | 1960942.625 | 2008644.125 | | | | | | |
| O08688 | CAPN5 | calpain 5 | Cytoplasm | peptidase | 1565357.5 | 1076637.5 | 818370.25 | | | | | | |
| O08688 | CAPN5 | calpain 5 | Cytoplasm | peptidase | 2264181.5 | 2950756.25 | 2056240.25 | | | | | | |
| O08688 | CAPN5 | calpain 5 | Cytoplasm | peptidase | 1206991.75 | 1756383.25 | 1319927.75 | | | | | | |
| O08688 | CAPN5 | calpain 5 | Cytoplasm | peptidase | 1062245.375 | 1827221.875 | 1826099 | | | | | | |
| O08688 | CAPN5 | calpain 5 | Cytoplasm | peptidase | 1488966 | 1973819 | 2195001 | O08688 | CAPN5 | 7386013 | 6285998 | 811519 | 60 |
| O88456 | CAPNS1 | calpain, small subunit 1 | Cytoplasm | peptidase | 1944353.875 | 1397406.75 | 1575136.875 | O88456 | CAPNS1 | 1944354 | 279004 | 161083 | 3 |
| Q60865 | CAPRIN1 | cell cycle associated protein 1 | Plasma Membrane | other | 368206.531 | 1034291.062 | 1148631.125 | | | | | | |
| Q60865 | CAPRIN1 | cell cycle associated protein 1 | Plasma Membrane | other | 1064370.5 | 1437308.75 | 1290917.75 | Q60865 | CAPRIN1 | 716289 | 369528 | 150859 | 6 |
| P47753 | CAPZA1 | capping protein (actin filament) muscle Z-line, alpha 1 | Cytoplasm | other | 6941258 | 7774393 | 7298513.5 | | | | | | |
| P47753 | CAPZA1 | capping protein (actin filament) muscle Z-line, alpha 1 | Cytoplasm | other | 2735183.75 | 2777417.5 | 3874026.75 | P47753 | CAPZA1 | 4838221 | 2356224 | 961924 | 6 |
| P47757 | CAPZB | capping protein (actin filament) muscle Z-line, beta | Cytoplasm | other | 3758986.5 | 4158294.5 | 4617573.5 | | | | | | |
| P47757 | CAPZB | capping protein (actin filament) muscle Z-line, beta | Cytoplasm | other | 1854456.5 | 1899523.625 | 2061515 | P47757 | CAPZB | 2806722 | 1258401 | 513740 | 6 |
| Q9CR86 | CARHSP1 | calcium regulated heat stable protein 1, 24kDa | Cytoplasm | other | 9158615 | 8004643 | 7756109 | Q9CR86 | CARHSP1 | 9158615 | 748381 | 432078 | 3 |
| Q9WVG6 | CARM1 | coactivator-associated arginine methyltransferase 1 | Nucleus | transcription regulator | 3170639.25 | 2158243.25 | 1767328.25 | | | | | | |
| Q9WVG6 | CARM1 | coactivator-associated arginine methyltransferase 1 | Nucleus | transcription regulator | 2183221 | 902185.812 | 1069528 | | | | | | |
| Q9WVG6 | CARM1 | coactivator-associated arginine methyltransferase 1 | Nucleus | transcription regulator | 4382960 | 2864803 | 3074427.25 | Q9WVG6 | CARM1 | 3245607 | 1102052 | 367351 | 9 |
| Q9ER72 | CARS | cysteinyl-tRNA synthetase | Cytoplasm | enzyme | 3628228 | 3244333.75 | 2520993.75 | | | | | | |
| Q9ER72 | CARS | cysteinyl-tRNA synthetase | Cytoplasm | enzyme | 1383500 | 1152811.5 | 619323.938 | Q9ER72 | CARS | 2505864 | 1218583 | 497485 | 6 |
| P51125 | CAST | calpastatin | Cytoplasm | peptidase | 2.63E+07 | 2.09E+07 | 2.92E+07 | | | | | | |
| P51125 | CAST | calpastatin | Cytoplasm | peptidase | 1.40E+07 | 1.04E+07 | 1.51E+07 | | | | | | |
| P51125 | CAST | calpastatin | Cytoplasm | peptidase | 1.30E+07 | 9623923 | 1.26E+07 | | | | | | |
| P51125 | CAST | calpastatin | Cytoplasm | peptidase | 8573111 | 6882661 | 1.02E+07 | | | | | | |
| P51125 | CAST | calpastatin | Cytoplasm | peptidase | 6558658.5 | 5615678.5 | 8624744 | | | | | | |
| P51125 | CAST | calpastatin | Cytoplasm | peptidase | 8993286 | 7034772 | 6670858.5 | | | | | | |
| P51125 | CAST | calpastatin | Cytoplasm | peptidase | 8090900 | 3536887.5 | 7285080 | | | | | | |
| P51125 | CAST | calpastatin | Cytoplasm | peptidase | 5619519 | 5030354.5 | 5494165 | | | | | | |
| P51125 | CAST | calpastatin | Cytoplasm | peptidase | 6079059 | 5057300.5 | 3484341.75 | | | | | | |
| P51125 | CAST | calpastatin | Cytoplasm | peptidase | 1605883.625 | 718169.5 | 5376718.5 | | | | | | |
| P51125 | CAST | calpastatin | Cytoplasm | peptidase | 4310570.5 | 3036692.5 | 4346813 | | | | | | |
| P51125 | CAST | calpastatin | Cytoplasm | peptidase | 2281787.75 | 1676080.875 | 1903095.5 | P51125 | CAST | 8788565 | 6421306 | 1070218 | 36 |
| P24270 | CAT | catalase | Cytoplasm | enzyme | 1.20E+07 | 9437683 | 7701561.5 | | | | | | |
| P24270 | CAT | catalase | Cytoplasm | enzyme | 1.37E+07 | 9458743 | 6266553 | | | | | | |
| P24270 | CAT | catalase | Cytoplasm | enzyme | 6374646 | 7125359.5 | 5136736 | | | | | | |
| P24270 | CAT | catalase | Cytoplasm | enzyme | 6804432 | 5630533 | 6132140 | | | | | | |
| P24270 | CAT | catalase | Cytoplasm | enzyme | 5285218.5 | 3938154.5 | 3772241.75 | | | | | | |
| P24270 | CAT | catalase | Cytoplasm | enzyme | 4328061 | 3899316.5 | 3288717 | | | | | | |
| P24270 | CAT | catalase | Cytoplasm | enzyme | 4549662 | 4279786 | 3748751.75 | | | | | | |
| P24270 | CAT | catalase | Cytoplasm | enzyme | 3762802.25 | 3203937 | 2255013.5 | | | | | | |
| P24270 | CAT | catalase | Cytoplasm | enzyme | 4299939 | 3226799.5 | 3678293.5 | | | | | | |
| P24270 | CAT | catalase | Cytoplasm | enzyme | 3736982.25 | 3520393.5 | 2822946.25 | | | | | | |
| P24270 | CAT | catalase | Cytoplasm | enzyme | 2653330.25 | 2419552.5 | 1926305.625 | | | | | | |
| P24270 | CAT | catalase | Cytoplasm | enzyme | 2917689 | 2629008 | 2369355.5 | | | | | | |
| P24270 | CAT | catalase | Cytoplasm | enzyme | 2981612 | 2790128 | 2096081.375 | | | | | | |
| P24270 | CAT | catalase | Cytoplasm | enzyme | 2689013.75 | 2404006 | 2325503.75 | | | | | | |
| P24270 | CAT | catalase | Cytoplasm | enzyme | 1845302 | 1780820.25 | 1045351.312 | | | | | | |
| P24270 | CAT | catalase | Cytoplasm | enzyme | 1584400.75 | 1330040.125 | 1556107.5 | | | | | | |
| P24270 | CAT | catalase | Cytoplasm | enzyme | 1786232.625 | 1031625.25 | 849588.062 | | | | | | |
| P24270 | CAT | catalase | Cytoplasm | enzyme | 940335.25 | 740705.125 | 620546.375 | | | | | | |
| P24270 | CAT | catalase | Cytoplasm | enzyme | 448981.594 | 532630.25 | 135150.531 | | | | | | |
| P24270 | CAT | catalase | Cytoplasm | enzyme | 535798.188 | 1056293.375 | 698187.625 | P24270 | CAT | 4160722 | 2776804 | 358484 | 60 |
| Q8C9M2 | CCDC15 | coiled-coil domain containing 15 | Cytoplasm | other | 2074735.625 | 2963469.75 | 2977470.25 | Q8C9M2 | CCDC15 | 2074736 | 517200 | 298605 | 3 |
| Q3URS9 | CCDC51 | coiled-coil domain containing 51 | Cytoplasm | other | 690002.5 | 1014224.5 | 1046860.125 | Q3URS9 | CCDC51 | 690003 | 197287 | 113904 | 3 |
| Q8BGU5 | CCNY | cyclin Y | Nucleus | other | 3417044.75 | 3832366.25 | 3126367 | | | | | | |
| Q8BGU5 | CCNY | cyclin Y | Nucleus | other | 3583570.5 | 3950157.25 | 2998914.25 | Q8BGU5 | CCNY | 3500308 | 378488 | 154517 | 6 |
| P80314 | CCT2 | chaperonin containing TCP1, subunit 2 (beta) | Cytoplasm | kinase | 1.79E+07 | 1.68E+07 | 1.33E+07 | | | | | | |
| P80314 | CCT2 | chaperonin containing TCP1, subunit 2 (beta) | Cytoplasm | kinase | 1.72E+07 | 1.38E+07 | 9929409 | | | | | | |
| P80314 | CCT2 | chaperonin containing TCP1, subunit 2 (beta) | Cytoplasm | kinase | 1.09E+07 | 9543873 | 8752572 | | | | | | |
| P80314 | CCT2 | chaperonin containing TCP1, subunit 2 (beta) | Cytoplasm | kinase | 1.02E+07 | 8086393.5 | 5526470.5 | | | | | | |
| P80314 | CCT2 | chaperonin containing TCP1, subunit 2 (beta) | Cytoplasm | kinase | 9665843 | 9238788 | 8084052.5 | | | | | | |
| P80314 | CCT2 | chaperonin containing TCP1, subunit 2 (beta) | Cytoplasm | kinase | 9110351 | 1.08E+07 | 7561902 | | | | | | |
| P80314 | CCT2 | chaperonin containing TCP1, subunit 2 (beta) | Cytoplasm | kinase | 8337891 | 6006050 | 4514548.5 | | | | | | |
| P80314 | CCT2 | chaperonin containing TCP1, subunit 2 (beta) | Cytoplasm | kinase | 8892155 | 7833151 | 5621364 | | | | | | |
| P80314 | CCT2 | chaperonin containing TCP1, subunit 2 (beta) | Cytoplasm | kinase | 5579502 | 4773053.5 | 3140515.5 | | | | | | |
| P80314 | CCT2 | chaperonin containing TCP1, subunit 2 (beta) | Cytoplasm | kinase | 5237493 | 4989578.5 | 3546677.75 | | | | | | |
| P80314 | CCT2 | chaperonin containing TCP1, subunit 2 (beta) | Cytoplasm | kinase | 5486380 | 6533463.5 | 4661728.5 | | | | | | |
| P80314 | CCT2 | chaperonin containing TCP1, subunit 2 (beta) | Cytoplasm | kinase | 4918980 | 4181651.75 | 2756817 | | | | | | |
| P80314 | CCT2 | chaperonin containing TCP1, subunit 2 (beta) | Cytoplasm | kinase | 4007429.25 | 5300600.5 | 3625920.25 | | | | | | |
| P80314 | CCT2 | chaperonin containing TCP1, subunit 2 (beta) | Cytoplasm | kinase | 3591232.25 | 2845625.25 | 2642061.75 | P80314 | CCT2 | 8641233 | 3946094 | 608896 | 42 |
| P80318 | CCT3 | chaperonin containing TCP1, subunit 3 (gamma) | Cytoplasm | other | 1.99E+07 | 2.33E+07 | 2.27E+07 | | | | | | |
| P80318 | CCT3 | chaperonin containing TCP1, subunit 3 (gamma) | Cytoplasm | other | 1.48E+07 | 1.13E+07 | 1.04E+07 | | | | | | |
| P80318 | CCT3 | chaperonin containing TCP1, subunit 3 (gamma) | Cytoplasm | other | 1.50E+07 | 1.10E+07 | 8455768 | | | | | | |
| P80318 | CCT3 | chaperonin containing TCP1, subunit 3 (gamma) | Cytoplasm | other | 1.42E+07 | 1.34E+07 | 8901428 | | | | | | |
| P80318 | CCT3 | chaperonin containing TCP1, subunit 3 (gamma) | Cytoplasm | other | 1.05E+07 | 1.04E+07 | 7643674 | | | | | | |
| P80318 | CCT3 | chaperonin containing TCP1, subunit 3 (gamma) | Cytoplasm | other | 7937963 | 5383572.5 | 2118037.75 | | | | | | |
| P80318 | CCT3 | chaperonin containing TCP1, subunit 3 (gamma) | Cytoplasm | other | 8496848 | 7548754 | 5226554.5 | | | | | | |
| P80318 | CCT3 | chaperonin containing TCP1, subunit 3 (gamma) | Cytoplasm | other | 8627669 | 8191290.5 | 5065656.5 | | | | | | |
| P80318 | CCT3 | chaperonin containing TCP1, subunit 3 (gamma) | Cytoplasm | other | 6000900 | 5001272 | 3827411.25 | | | | | | |
| P80318 | CCT3 | chaperonin containing TCP1, subunit 3 (gamma) | Cytoplasm | other | 7122776.5 | 6716695.5 | 4104479.5 | | | | | | |
| P80318 | CCT3 | chaperonin containing TCP1, subunit 3 (gamma) | Cytoplasm | other | 5170237.5 | 4314505 | 3371780.5 | | | | | | |
| P80318 | CCT3 | chaperonin containing TCP1, subunit 3 (gamma) | Cytoplasm | other | 5429832 | 5653799 | 3563167 | | | | | | |
| P80318 | CCT3 | chaperonin containing TCP1, subunit 3 (gamma) | Cytoplasm | other | 5975094.5 | 6519483.5 | 5113802.5 | | | | | | |
| P80318 | CCT3 | chaperonin containing TCP1, subunit 3 (gamma) | Cytoplasm | other | 3928309.5 | 2767265 | 2216149 | | | | | | |
| P80318 | CCT3 | chaperonin containing TCP1, subunit 3 (gamma) | Cytoplasm | other | 3078050 | 3063003.75 | 2367287.25 | | | | | | |
| P80318 | CCT3 | chaperonin containing TCP1, subunit 3 (gamma) | Cytoplasm | other | 2397692 | 1426624.25 | 997479.5 | P80318 | CCT3 | 8660336 | 5174187 | 746830 | 48 |
| P80315 | CCT4 | chaperonin containing TCP1, subunit 4 (delta) | Cytoplasm | other | 2.25E+07 | 2.09E+07 | 1.50E+07 | | | | | | |
| P80315 | CCT4 | chaperonin containing TCP1, subunit 4 (delta) | Cytoplasm | other | 1.99E+07 | 1.78E+07 | 1.37E+07 | | | | | | |
| P80315 | CCT4 | chaperonin containing TCP1, subunit 4 (delta) | Cytoplasm | other | 1.57E+07 | 1.47E+07 | 9044290 | | | | | | |
| P80315 | CCT4 | chaperonin containing TCP1, subunit 4 (delta) | Cytoplasm | other | 1.29E+07 | 1.14E+07 | 1.25E+07 | | | | | | |
| P80315 | CCT4 | chaperonin containing TCP1, subunit 4 (delta) | Cytoplasm | other | 8451292 | 9330137 | 7300282.5 | | | | | | |
| P80315 | CCT4 | chaperonin containing TCP1, subunit 4 (delta) | Cytoplasm | other | 8068611 | 7260512.5 | 5175749 | | | | | | |
| P80315 | CCT4 | chaperonin containing TCP1, subunit 4 (delta) | Cytoplasm | other | 7753094 | 7019425 | 7563471 | | | | | | |
| P80315 | CCT4 | chaperonin containing TCP1, subunit 4 (delta) | Cytoplasm | other | 8716025 | 6022597.5 | 3885659.75 | | | | | | |
| P80315 | CCT4 | chaperonin containing TCP1, subunit 4 (delta) | Cytoplasm | other | 6013791.5 | 5787761.5 | 4022550.25 | | | | | | |

| | | | | | | | | | | | | | |
|---|---|---|---|---|---|---|---|---|---|---|---|---|---|
| P80315 | CCT4 | chaperonin containing TCP1, subunit 4 (delta) | Cytoplasm | other | 5727625 | 4880403.5 | 5952489.5 | | | | | | |
| P80315 | CCT4 | chaperonin containing TCP1, subunit 4 (delta) | Cytoplasm | other | 6357401.5 | 5620967 | 4650073.5 | | | | | | |
| P80315 | CCT4 | chaperonin containing TCP1, subunit 4 (delta) | Cytoplasm | other | 5292115 | 6288688.5 | 3587205.25 | | | | | | |
| P80315 | CCT4 | chaperonin containing TCP1, subunit 4 (delta) | Cytoplasm | other | 4503872 | 4158760.75 | 3062413 | | | | | | |
| P80315 | CCT4 | chaperonin containing TCP1, subunit 4 (delta) | Cytoplasm | other | 3199230.5 | 2453373.75 | 2006251.5 | | | | | | |
| P80315 | CCT4 | chaperonin containing TCP1, subunit 4 (delta) | Cytoplasm | other | 3305456.25 | 4708063.5 | 2238651.25 | | | | | | |
| P80315 | CCT4 | chaperonin containing TCP1, subunit 4 (delta) | Cytoplasm | other | 1305756.5 | 1256576.5 | 2212206.25 | | | | | | |
| P80315 | CCT4 | chaperonin containing TCP1, subunit 4 (delta) | Cytoplasm | other | 2249266.5 | 748595.062 | 463874.469 | | | | | | |
| P80315 | CCT4 | chaperonin containing TCP1, subunit 4 (delta) | Cytoplasm | other | 2456754 | 2011197.5 | 1664888.75 | P80315 | CCT4 | 8018349 | 5342641 | 727041 | 54 |
| P80316 | CCT5 | chaperonin containing TCP1, subunit 5 (epsilon) | Cytoplasm | other | 2.38E+07 | 2.00E+07 | 1.40E+07 | | | | | | |
| P80316 | CCT5 | chaperonin containing TCP1, subunit 5 (epsilon) | Cytoplasm | other | 1.91E+07 | 1.59E+07 | 1.39E+07 | | | | | | |
| P80316 | CCT5 | chaperonin containing TCP1, subunit 5 (epsilon) | Cytoplasm | other | 1.19E+07 | 1.28E+07 | 7415033 | | | | | | |
| P80316 | CCT5 | chaperonin containing TCP1, subunit 5 (epsilon) | Cytoplasm | other | 1.14E+07 | 9575633 | 8644201 | | | | | | |
| P80316 | CCT5 | chaperonin containing TCP1, subunit 5 (epsilon) | Cytoplasm | other | 1.02E+07 | 1.13E+07 | 1.01E+07 | | | | | | |
| P80316 | CCT5 | chaperonin containing TCP1, subunit 5 (epsilon) | Cytoplasm | other | 9221507 | 8027330 | 5781317 | | | | | | |
| P80316 | CCT5 | chaperonin containing TCP1, subunit 5 (epsilon) | Cytoplasm | other | 7602672.5 | 5638498 | 3964369.75 | | | | | | |
| P80316 | CCT5 | chaperonin containing TCP1, subunit 5 (epsilon) | Cytoplasm | other | 6974913 | 4745505.5 | 3753965 | | | | | | |
| P80316 | CCT5 | chaperonin containing TCP1, subunit 5 (epsilon) | Cytoplasm | other | 6374448.5 | 4228857 | 2990990 | | | | | | |
| P80316 | CCT5 | chaperonin containing TCP1, subunit 5 (epsilon) | Cytoplasm | other | 4823182 | 4010172.75 | 3077419.5 | | | | | | |
| P80316 | CCT5 | chaperonin containing TCP1, subunit 5 (epsilon) | Cytoplasm | other | 2382005 | 2100812.25 | 1517137.375 | | | | | | |
| P80316 | CCT5 | chaperonin containing TCP1, subunit 5 (epsilon) | Cytoplasm | other | 2567572.25 | 2527188.75 | 1606440.375 | | | | | | |
| P80316 | CCT5 | chaperonin containing TCP1, subunit 5 (epsilon) | Cytoplasm | other | 277305.281 | 360121.656 | 695343.875 | P80316 | CCT5 | 8975662 | 5729106 | 917391 | 39 |
| P80317 | CCT6A | chaperonin containing TCP1, subunit 6A (zeta 1) | Cytoplasm | other | 2.07E+07 | 1.88E+07 | 1.31E+07 | | | | | | |
| P80317 | CCT6A | chaperonin containing TCP1, subunit 6A (zeta 1) | Cytoplasm | other | 1.67E+07 | 2.01E+07 | 1.36E+07 | | | | | | |
| P80317 | CCT6A | chaperonin containing TCP1, subunit 6A (zeta 1) | Cytoplasm | other | 1.77E+07 | 1.66E+07 | 1.17E+07 | | | | | | |
| P80317 | CCT6A | chaperonin containing TCP1, subunit 6A (zeta 1) | Cytoplasm | other | 1.79E+07 | 1.40E+07 | 9219259 | | | | | | |
| P80317 | CCT6A | chaperonin containing TCP1, subunit 6A (zeta 1) | Cytoplasm | other | 1.21E+07 | 1.18E+07 | 8046460.5 | | | | | | |
| P80317 | CCT6A | chaperonin containing TCP1, subunit 6A (zeta 1) | Cytoplasm | other | 1.08E+07 | 8434385 | 5735333.5 | | | | | | |
| P80317 | CCT6A | chaperonin containing TCP1, subunit 6A (zeta 1) | Cytoplasm | other | 1.10E+07 | 8517804 | 7091171.5 | | | | | | |
| P80317 | CCT6A | chaperonin containing TCP1, subunit 6A (zeta 1) | Cytoplasm | other | 8094513.5 | 7101700 | 5508278 | | | | | | |
| P80317 | CCT6A | chaperonin containing TCP1, subunit 6A (zeta 1) | Cytoplasm | other | 6782397.5 | 6370341 | 3919901.25 | | | | | | |
| P80317 | CCT6A | chaperonin containing TCP1, subunit 6A (zeta 1) | Cytoplasm | other | 6275654 | 5243821.5 | 3873579 | | | | | | |
| P80317 | CCT6A | chaperonin containing TCP1, subunit 6A (zeta 1) | Cytoplasm | other | 1939863 | 1279219.25 | 888367.312 | | | | | | |
| P80317 | CCT6A | chaperonin containing TCP1, subunit 6A (zeta 1) | Cytoplasm | other | 2076234.25 | 1384066.125 | 699022.125 | P80317 | CCT6A | 10999055 | 5820384 | 970064 | 36 |
| P80313 | CCT7 | chaperonin containing TCP1, subunit 7 (eta) | Cytoplasm | other | 1.43E+07 | 1.45E+07 | 1.10E+07 | | | | | | |
| P80313 | CCT7 | chaperonin containing TCP1, subunit 7 (eta) | Cytoplasm | other | 1.27E+07 | 1.68E+07 | 9472934 | | | | | | |
| P80313 | CCT7 | chaperonin containing TCP1, subunit 7 (eta) | Cytoplasm | other | 1.09E+07 | 1.27E+07 | 9921162 | | | | | | |
| P80313 | CCT7 | chaperonin containing TCP1, subunit 7 (eta) | Cytoplasm | other | 1.23E+07 | 1.00E+07 | 8111246 | | | | | | |
| P80313 | CCT7 | chaperonin containing TCP1, subunit 7 (eta) | Cytoplasm | other | 1.46E+07 | 1.49E+07 | 1.34E+07 | | | | | | |
| P80313 | CCT7 | chaperonin containing TCP1, subunit 7 (eta) | Cytoplasm | other | 1.04E+07 | 1.03E+07 | 5610325 | | | | | | |
| P80313 | CCT7 | chaperonin containing TCP1, subunit 7 (eta) | Cytoplasm | other | 8330734.5 | 7298663.5 | 4936492 | | | | | | |
| P80313 | CCT7 | chaperonin containing TCP1, subunit 7 (eta) | Cytoplasm | other | 9785982 | 7243996.5 | 4496296.5 | | | | | | |
| P80313 | CCT7 | chaperonin containing TCP1, subunit 7 (eta) | Cytoplasm | other | 6809260.5 | 4153953 | 3092419.75 | | | | | | |
| P80313 | CCT7 | chaperonin containing TCP1, subunit 7 (eta) | Cytoplasm | other | 5746369 | 6122350 | 5779789.5 | | | | | | |
| P80313 | CCT7 | chaperonin containing TCP1, subunit 7 (eta) | Cytoplasm | other | 6463507.5 | 5837225 | 4798821.5 | | | | | | |
| P80313 | CCT7 | chaperonin containing TCP1, subunit 7 (eta) | Cytoplasm | other | 3485509.5 | 2916725 | 2649827.5 | | | | | | |
| P80313 | CCT7 | chaperonin containing TCP1, subunit 7 (eta) | Cytoplasm | other | 4112576.75 | 2876082.75 | 2351508.75 | | | | | | |
| P80313 | CCT7 | chaperonin containing TCP1, subunit 7 (eta) | Cytoplasm | other | 3659830.5 | 4432641.5 | 2219581.5 | | | | | | |
| P80313 | CCT7 | chaperonin containing TCP1, subunit 7 (eta) | Cytoplasm | other | 3086611 | 2115782.5 | 1071210 | P80313 | CCT7 | 8447359 | 4248018 | 633257 | 45 |
| P42932 | CCT8 | chaperonin containing TCP1, subunit 8 (theta) | Cytoplasm | enzyme | 1.78E+07 | 1.47E+07 | 8748357 | | | | | | |
| P42932 | CCT8 | chaperonin containing TCP1, subunit 8 (theta) | Cytoplasm | enzyme | 1.26E+07 | 9402065 | 6869281 | | | | | | |
| P42932 | CCT8 | chaperonin containing TCP1, subunit 8 (theta) | Cytoplasm | enzyme | 1.17E+07 | 8493116 | 8426125 | | | | | | |
| P42932 | CCT8 | chaperonin containing TCP1, subunit 8 (theta) | Cytoplasm | enzyme | 1.25E+07 | 1.12E+07 | 7651112.5 | | | | | | |
| P42932 | CCT8 | chaperonin containing TCP1, subunit 8 (theta) | Cytoplasm | enzyme | 1.24E+07 | 1.03E+07 | 6732723 | | | | | | |
| P42932 | CCT8 | chaperonin containing TCP1, subunit 8 (theta) | Cytoplasm | enzyme | 9817845 | 8003704 | 5386311 | | | | | | |
| P42932 | CCT8 | chaperonin containing TCP1, subunit 8 (theta) | Cytoplasm | enzyme | 9156885 | 8041537.5 | 6751632 | | | | | | |
| P42932 | CCT8 | chaperonin containing TCP1, subunit 8 (theta) | Cytoplasm | enzyme | 8272593.5 | 7251288 | 5006135 | | | | | | |
| P42932 | CCT8 | chaperonin containing TCP1, subunit 8 (theta) | Cytoplasm | enzyme | 7554363 | 7801185.5 | 4090594.5 | | | | | | |
| P42932 | CCT8 | chaperonin containing TCP1, subunit 8 (theta) | Cytoplasm | enzyme | 7007460.5 | 5429568 | 5214777.5 | | | | | | |
| P42932 | CCT8 | chaperonin containing TCP1, subunit 8 (theta) | Cytoplasm | enzyme | 7246194.5 | 6162323 | 4300546.5 | | | | | | |
| P42932 | CCT8 | chaperonin containing TCP1, subunit 8 (theta) | Cytoplasm | enzyme | 8044944.5 | 8099971 | 5206791.5 | | | | | | |
| P42932 | CCT8 | chaperonin containing TCP1, subunit 8 (theta) | Cytoplasm | enzyme | 7500557.5 | 6076893 | 3442896.5 | | | | | | |
| P42932 | CCT8 | chaperonin containing TCP1, subunit 8 (theta) | Cytoplasm | enzyme | 5734999.5 | 4425644 | 3566437 | | | | | | |
| P42932 | CCT8 | chaperonin containing TCP1, subunit 8 (theta) | Cytoplasm | enzyme | 3483800 | 2848327 | 2849980.75 | | | | | | |
| P42932 | CCT8 | chaperonin containing TCP1, subunit 8 (theta) | Cytoplasm | enzyme | 4285491.5 | 1882658.875 | 3306628.75 | | | | | | |
| P42932 | CCT8 | chaperonin containing TCP1, subunit 8 (theta) | Cytoplasm | enzyme | 6294382 | 7164984 | 7277174 | | | | | | |
| P42932 | CCT8 | chaperonin containing TCP1, subunit 8 (theta) | Cytoplasm | enzyme | 4844146.5 | 4239422 | 2982063.25 | | | | | | |
| P42932 | CCT8 | chaperonin containing TCP1, subunit 8 (theta) | Cytoplasm | enzyme | 3314152.5 | 2706372.5 | 2639518.75 | | | | | | |
| P42932 | CCT8 | chaperonin containing TCP1, subunit 8 (theta) | Cytoplasm | enzyme | 3580924 | 2759441.25 | 1789817.75 | | | | | | |
| P42932 | CCT8 | chaperonin containing TCP1, subunit 8 (theta) | Cytoplasm | enzyme | 2541666.25 | 2033573.75 | 1780858.125 | | | | | | |
| P42932 | CCT8 | chaperonin containing TCP1, subunit 8 (theta) | Cytoplasm | enzyme | 3575089 | 3798598.75 | 3248356 | | | | | | |
| P42932 | CCT8 | chaperonin containing TCP1, subunit 8 (theta) | Cytoplasm | enzyme | 3105558.25 | 2260801.25 | 1928972.25 | | | | | | |
| P42932 | CCT8 | chaperonin containing TCP1, subunit 8 (theta) | Cytoplasm | enzyme | 1188145.5 | 2678412 | 1394157.625 | P42932 | CCT8 | 7229133 | 3418641 | 402891 | 72 |
| Q8K4F0 | CD226 | CD226 molecule | Plasma Membrane | other | 2570814 | 2649828.75 | 3307212.5 | Q8K4F0 | CD226 | 2570814 | 404285 | 233414 | 3 |
| Q08857 | CD36 | CD36 molecule (thrombospondin receptor) | Plasma Membrane | transmembrane receptor | 5.32E+08 | 5.40E+08 | 4.51E+08 | | | | | | |
| Q08857 | CD36 | CD36 molecule (thrombospondin receptor) | Plasma Membrane | transmembrane receptor | 2.09E+08 | 2.39E+08 | 1.93E+08 | | | | | | |
| Q08857 | CD36 | CD36 molecule (thrombospondin receptor) | Plasma Membrane | transmembrane receptor | 1.30E+08 | 1.42E+08 | 1.02E+08 | | | | | | |
| Q08857 | CD36 | CD36 molecule (thrombospondin receptor) | Plasma Membrane | transmembrane receptor | 1.07E+08 | 1.02E+08 | 1.06E+08 | | | | | | |
| Q08857 | CD36 | CD36 molecule (thrombospondin receptor) | Plasma Membrane | transmembrane receptor | 2.49E+07 | 2.50E+07 | 2.21E+07 | | | | | | |
| Q08857 | CD36 | CD36 molecule (thrombospondin receptor) | Plasma Membrane | transmembrane receptor | 9.64E+07 | 1.05E+08 | 4.57E+07 | | | | | | |
| Q08857 | CD36 | CD36 molecule (thrombospondin receptor) | Plasma Membrane | transmembrane receptor | 5.87E+07 | 5.93E+07 | 6.74E+07 | | | | | | |
| Q08857 | CD36 | CD36 molecule (thrombospondin receptor) | Plasma Membrane | transmembrane receptor | 8.16E+07 | 9.20E+07 | 8.78E+07 | | | | | | |
| Q08857 | CD36 | CD36 molecule (thrombospondin receptor) | Plasma Membrane | transmembrane receptor | 7.41E+07 | 7.42E+07 | 5.82E+07 | | | | | | |
| Q08857 | CD36 | CD36 molecule (thrombospondin receptor) | Plasma Membrane | transmembrane receptor | 1.33E+08 | 1.21E+08 | 9.38E+07 | | | | | | |
| Q08857 | CD36 | CD36 molecule (thrombospondin receptor) | Plasma Membrane | transmembrane receptor | 2.57E+07 | 3.16E+07 | 1.44E+07 | | | | | | |
| Q08857 | CD36 | CD36 molecule (thrombospondin receptor) | Plasma Membrane | transmembrane receptor | 2.19E+07 | 2.24E+07 | 1.52E+07 | | | | | | |
| Q08857 | CD36 | CD36 molecule (thrombospondin receptor) | Plasma Membrane | transmembrane receptor | 2.18E+07 | 2.61E+07 | 1.58E+07 | | | | | | |
| Q08857 | CD36 | CD36 molecule (thrombospondin receptor) | Plasma Membrane | transmembrane receptor | 1.14E+07 | 1.35E+07 | 6387666 | | | | | | |
| Q08857 | CD36 | CD36 molecule (thrombospondin receptor) | Plasma Membrane | transmembrane receptor | 1725860.25 | 2034529.625 | 2250784.75 | | | | | | |
| Q08857 | CD36 | CD36 molecule (thrombospondin receptor) | Plasma Membrane | transmembrane receptor | 2706700.75 | 2500758.25 | 4814061 | Q08857 | CD36 | 95685160 | 122839782 | 17730395 | 48 |
| P15379 | CD44 | CD44 molecule (Indian blood group) | Plasma Membrane | enzyme | 1.34E+07 | 1.78E+07 | 2.16E+07 | | | | | | |
| P15379 | CD44 | CD44 molecule (Indian blood group) | Plasma Membrane | enzyme | 3818282 | 6688802.5 | 9313152 | | | | | | |
| P15379 | CD44 | CD44 molecule (Indian blood group) | Plasma Membrane | enzyme | 3300277.75 | 4206760 | 4257892 | | | | | | |
| P15379 | CD44 | CD44 molecule (Indian blood group) | Plasma Membrane | enzyme | 1331573.875 | 1722060.375 | 1369544.75 | P15379 | CD44 | 5467533 | 6789733 | 1960027 | 12 |
| Q61735 | CD47 | CD47 molecule | Plasma Membrane | transmembrane receptor | 5.70E+08 | 5.59E+08 | 4.72E+08 | | | | | | |
| Q61735 | CD47 | CD47 molecule | Plasma Membrane | transmembrane receptor | 3.25E+08 | 3.48E+08 | 3.03E+08 | | | | | | |
| Q61735 | CD47 | CD47 molecule | Plasma Membrane | transmembrane receptor | 1.63E+08 | 2.01E+08 | 1.92E+08 | | | | | | |
| Q61735 | CD47 | CD47 molecule | Plasma Membrane | transmembrane receptor | 1.17E+08 | 1.37E+08 | 1.33E+08 | | | | | | |
| Q61735 | CD47 | CD47 molecule | Plasma Membrane | transmembrane receptor | 1.05E+08 | 1.12E+08 | 7.79E+07 | | | | | | |
| Q61735 | CD47 | CD47 molecule | Plasma Membrane | transmembrane receptor | 7560798 | 8181114 | 7826991.5 | Q61735 | CD47 | 214593466 | 178919708 | 42171780 | 18 |
| P18181 | CD48 | CD48 molecule | Plasma Membrane | other | 1258305.75 | 1607800.25 | 2344229.75 | P18181 | CD48 | 1258306 | 554332 | 320044 | 3 |
| Q61475 | CD55 | CD55 molecule, decay accelerating factor for complement (Cromer blood group) | Plasma Membrane | other | 1.26E+07 | 1.34E+07 | 1.40E+07 | | | | | | |
| Q61475 | CD55 | CD55 molecule, decay accelerating factor for complement (Cromer blood group) | Plasma Membrane | other | 8159055.5 | 1.21E+07 | 1.09E+07 | | | | | | |
| Q61475 | CD55 | CD55 molecule, decay accelerating factor for complement (Cromer blood group) | Plasma Membrane | other | 6360854 | 6458798 | 6267109 | | | | | | |
| Q61475 | CD55 | CD55 molecule, decay accelerating factor for complement (Cromer blood group) | Plasma Membrane | other | 4599210 | 6284195.5 | 5411397.5 | | | | | | |
| Q61475 | CD55 | CD55 molecule, decay accelerating factor for complement (Cromer blood group) | Plasma Membrane | other | 5671013 | 4801980 | 4940483.5 | | | | | | |
| Q61475 | CD55 | CD55 molecule, decay accelerating factor for complement (Cromer blood group) | Plasma Membrane | other | 3525959.25 | 4179417.5 | 3905672.75 | Q61475 | CD55 | 6812682 | 3515158 | 828531 | 18 |
| O55186 | Cd59a | CD59a antigen | Plasma Membrane | other | 1.08E+08 | 9.89E+07 | 9.78E+07 | | | | | | |
| O55186 | Cd59a | CD59a antigen | Plasma Membrane | other | 8022166 | 6560070 | 7654858.5 | | | | | | |

| | | | | | | | | | | | | | |
|---|---|---|---|---|---|---|---|---|---|---|---|---|---|
| O55186 | Cd59a | CD59a antigen | Plasma Membrane | other | 2487252 | 2194727.5 | 3275597.25 | O55186 | Cd59a | 39536473 | 48420598 | 16140199 | 9 |
| P40237 | CD82 | CD82 molecule | Plasma Membrane | other | 8383307.5 | 1.17E+07 | 9943614 | | | | | | |
| P40237 | CD82 | CD82 molecule | Plasma Membrane | other | 6684652.5 | 1.05E+07 | 6930825 | P40237 | CD82 | 7533980 | 2010060 | 820604 | 6 |
| P40240 | CD9 | CD9 molecule | Plasma Membrane | other | 2.75E+07 | 3.77E+07 | 4.79E+07 | | | | | | |
| P40240 | CD9 | CD9 molecule | Plasma Membrane | other | 4596347 | 4946335.5 | 4013369.75 | | | | | | |
| P40240 | CD9 | CD9 molecule | Plasma Membrane | other | 1116697 | 1229952.25 | 3747528.5 | P40240 | CD9 | 11084348 | 18008602 | 6002867 | 9 |
| Q8CII2 | CDC123 | cell division cycle 123 | Cytoplasm | other | 6530004.5 | 7033496.5 | 6583125.5 | Q8CII2 | CDC123 | 6530005 | 276635 | 159715 | 3 |
| P60766 | Cdc42 | cell division cycle 42 | Plasma Membrane | enzyme | 9624855 | 8411592 | 9450332 | | | | | | |
| P60766 | Cdc42 | cell division cycle 42 | Plasma Membrane | enzyme | 5970282 | 4448608 | 6192605 | | | | | | |
| P60766 | Cdc42 | cell division cycle 42 | Plasma Membrane | enzyme | 9402487 | 8023876.5 | 1.02E+07 | | | | | | |
| P60766 | Cdc42 | cell division cycle 42 | Plasma Membrane | enzyme | 3832307.75 | 5019483.5 | 4257839 | | | | | | |
| P60766 | Cdc42 | cell division cycle 42 | Plasma Membrane | enzyme | 3224693 | 4279700 | 2375484.5 | P60766 | Cdc42 | 6410925 | 2639711 | 681571 | 15 |
| Q922M5 | CDCA7L | cell division cycle associated 7-like | Nucleus | other | 2.98E+07 | 1.85E+07 | 2.19E+07 | Q922M5 | CDCA7L | 29750000 | 5762369 | 3326905 | 3 |
| Q8VDP6 | CDIPT | CDP-diacylglycerol--inositol 3-phosphatidyltransferase | Cytoplasm | enzyme | 2267210.5 | 2383142.5 | 2321506.5 | Q8VDP6 | CDIPT | 2267211 | 58005 | 33489 | 3 |
| Q99L43 | CDS2 | CDP-diacylglycerol synthase (phosphatidate cytidylyltransferase) 2 | Cytoplasm | enzyme | 3278182.75 | 2133408 | 4425885 | Q99L43 | CDS2 | 3278183 | 1146239 | 661781 | 3 |
| Q4VAA2 | CDV3 | CDV3 homolog (mouse) | Cytoplasm | other | 1739132.875 | 1469491 | 2836168.75 | Q4VAA2 | CDV3 | 1739133 | 723879 | 417932 | 3 |
| Q6AW46 | CES5A | carboxylesterase 5A | Extracellular Space | enzyme | 1.64E+07 | 1.65E+07 | 1.76E+07 | Q6AW46 | CES5A | 16440000 | 661992 | 382201 | 3 |
| P18760 | CFL1 | cofilin 1 (non-muscle) | Nucleus | other | 7376042 | 1.25E+07 | 1.76E+07 | | | | | | |
| P18760 | CFL1 | cofilin 1 (non-muscle) | Nucleus | other | 2288383.5 | 3455600.5 | 4346993.5 | | | | | | |
| P18760 | CFL1 | cofilin 1 (non-muscle) | Nucleus | other | 2894903.5 | 4016078.75 | 5908648 | | | | | | |
| P18760 | CFL1 | cofilin 1 (non-muscle) | Nucleus | other | 3422559.75 | 4362922 | 6144152 | | | | | | |
| P18760 | CFL1 | cofilin 1 (non-muscle) | Nucleus | other | 1216603.25 | 2056935.625 | 3439989.5 | P18760 | CFL1 | 3439698 | 4345908 | 1122109 | 15 |
| Q9CRB9 | CHCHD3 | coiled-coil-helix-coiled-coil-helix domain containing 3 | Cytoplasm | other | 2647526.25 | 5538504 | 6241862.5 | Q9CRB9 | CHCHD3 | 2647526 | 1904894 | 1099791 | 3 |
| O35744 | Chil3/Chil4 | chitinase-like 3 | Cytoplasm | enzyme | 773427 | 546694.312 | 4597617 | | | | | | |
| O35744 | Chil3/Chil4 | chitinase-like 3 | Cytoplasm | enzyme | 1600697.5 | 1456287 | 1.10E+07 | | | | | | |
| O35744 | Chil3/Chil4 | chitinase-like 3 | Cytoplasm | enzyme | 1308379.25 | 813542.062 | 7574928 | | | | | | |
| O35744 | Chil3/Chil4 | chitinase-like 3 | Cytoplasm | enzyme | 953110.188 | 626012.188 | 8516992 | | | | | | |
| O35744 | Chil3/Chil4 | chitinase-like 3 | Cytoplasm | enzyme | 439674.938 | 252569.719 | 2767748.75 | | | | | | |
| O35744 | Chil3/Chil4 | chitinase-like 3 | Cytoplasm | enzyme | 430732.562 | 458096.406 | 4410050.5 | O35744 | Chil3/Chil4 | 917670 | 3236580 | 762869 | 18 |
| Q9DB34 | CHMP2A | charged multivesicular body protein 2A | Cytoplasm | other | 2124946.25 | 1769442.25 | 1979991.875 | Q9DB34 | CHMP2A | 2124946 | 178758 | 103206 | 3 |
| P0C0A3 | CHMP6 | charged multivesicular body protein 6 | Cytoplasm | other | 3250180 | 4015186 | 3648005.75 | P0C0A3 | CHMP6 | 3250180 | 382605 | 220897 | 3 |
| P61022 | CHP1 | calcineurin-like EF-hand protein 1 | Cytoplasm | transporter | 6075162 | 5986913.5 | 6072734 | | | | | | |
| P61022 | CHP1 | calcineurin-like EF-hand protein 1 | Cytoplasm | transporter | 4518585.5 | 4827805.5 | 5108706 | | | | | | |
| P61022 | CHP1 | calcineurin-like EF-hand protein 1 | Cytoplasm | transporter | 2643472 | 2080192.25 | 2211380.25 | | | | | | |
| P61022 | CHP1 | calcineurin-like EF-hand protein 1 | Cytoplasm | transporter | 3526795.5 | 3594227.5 | 3259210 | | | | | | |
| P61022 | CHP1 | calcineurin-like EF-hand protein 1 | Cytoplasm | transporter | 2507532.5 | 2515526.75 | 4332919.5 | | | | | | |
| P61022 | CHP1 | calcineurin-like EF-hand protein 1 | Cytoplasm | transporter | 3791904.5 | 2188258.25 | 3257933.25 | P61022 | CHP1 | 3843909 | 1374850 | 324055 | 18 |
| Q9Z0F4 | CIB1 | calcium and integrin binding 1 (calmyrin) | Nucleus | other | 7161737.5 | 1.09E+07 | 7114246 | | | | | | |
| Q9Z0F4 | CIB1 | calcium and integrin binding 1 (calmyrin) | Nucleus | other | 6882004.5 | 6900025.5 | 7119800.5 | | | | | | |
| Q9Z0F4 | CIB1 | calcium and integrin binding 1 (calmyrin) | Nucleus | other | 3686141.75 | 3149663.75 | 3152736.75 | | | | | | |
| Q9Z0F4 | CIB1 | calcium and integrin binding 1 (calmyrin) | Nucleus | other | 844583.25 | 875755.938 | 961244.688 | Q9Z0F4 | CIB1 | 4643617 | 3221236 | 929891 | 12 |
| Q91WS0 | CISD1 | CDGSH iron sulfur domain 1 | Cytoplasm | other | 1376088.75 | 2892096.75 | 2892283.25 | | | | | | |
| Q91WS0 | CISD1 | CDGSH iron sulfur domain 1 | Cytoplasm | other | 2492242 | 5033084 | 5003694.5 | Q91WS0 | CISD1 | 1934165 | 1455379 | 594156 | 6 |
| Q9CQB5 | CISD2 | CDGSH iron sulfur domain 2 | Cytoplasm | other | 7272087 | 7157642.5 | 5880424.5 | | | | | | |
| Q9CQB5 | CISD2 | CDGSH iron sulfur domain 2 | Cytoplasm | other | 6024405 | 6447207 | 6885879.5 | | | | | | |
| Q9CQB5 | CISD2 | CDGSH iron sulfur domain 2 | Cytoplasm | other | 7198562 | 6303833.5 | 6213130 | | | | | | |
| Q9CQB5 | CISD2 | CDGSH iron sulfur domain 2 | Cytoplasm | other | 3920219.25 | 4091145 | 3894855 | | | | | | |
| Q9CQB5 | CISD2 | CDGSH iron sulfur domain 2 | Cytoplasm | other | 3558541.5 | 3224624 | 3573448.25 | Q9CQB5 | CISD2 | 5594763 | 1531389 | 395403 | 15 |
| Q9Z0S4 | Cldn13 | claudin 13 | Plasma Membrane | other | 2.09E+08 | 2.54E+08 | 1.96E+08 | | | | | | |
| Q9Z0S4 | Cldn13 | claudin 13 | Plasma Membrane | other | 6.27E+07 | 6.06E+07 | 5.43E+07 | | | | | | |
| Q9Z0S4 | Cldn13 | claudin 13 | Plasma Membrane | other | 6.26E+07 | 6.36E+07 | 6.36E+07 | | | | | | |
| Q9Z0S4 | Cldn13 | claudin 13 | Plasma Membrane | other | 3.79E+07 | 3.60E+07 | 4.04E+07 | Q9Z0S4 | Cldn13 | 93040000 | 76934616 | 22209111 | 12 |
| Q9JL99 | CLEC1B | C-type lectin domain family 1, member B | Plasma Membrane | transmembrane receptor | 4611527 | 6689993 | 8782647 | Q9JL99 | CLEC1B | 4611527 | 2085564 | 1204101 | 3 |
| Q9QUK3 | CLN8 | ceroid-lipofuscinosis, neuronal 8 | Cytoplasm | other | 6942023 | 7619708.5 | 5993787 | Q9QUK3 | CLN8 | 6942023 | 816704 | 471524 | 3 |
| Q61189 | CLNS1A | chloride channel, nucleotide-sensitive, 1A | Plasma Membrane | ion channel | 5317866.5 | 3388209.25 | 2313204.5 | Q61189 | CLNS1A | 5317867 | 1522454 | 878990 | 3 |
| Q8VBZ3 | CLPTM1 | cleft lip and palate associated transmembrane protein 1 | Plasma Membrane | other | 1139244.25 | 896571.938 | 1198186.25 | Q8VBZ3 | CLPTM1 | 1139244 | 159862 | 92296 | 3 |
| Q68FD5 | CLTC | clathrin, heavy chain (Hc) | Plasma Membrane | other | 1.27E+07 | 1.11E+07 | 1.07E+07 | | | | | | |
| Q68FD5 | CLTC | clathrin, heavy chain (Hc) | Plasma Membrane | other | 1.44E+07 | 1.19E+07 | 9339176 | | | | | | |
| Q68FD5 | CLTC | clathrin, heavy chain (Hc) | Plasma Membrane | other | 1.15E+07 | 1.42E+07 | 1.13E+07 | | | | | | |
| Q68FD5 | CLTC | clathrin, heavy chain (Hc) | Plasma Membrane | other | 1.04E+07 | 1.27E+07 | 1.28E+07 | | | | | | |
| Q68FD5 | CLTC | clathrin, heavy chain (Hc) | Plasma Membrane | other | 9840025 | 8067653 | 8235013 | | | | | | |
| Q68FD5 | CLTC | clathrin, heavy chain (Hc) | Plasma Membrane | other | 8924041 | 1.10E+07 | 9856730 | | | | | | |
| Q68FD5 | CLTC | clathrin, heavy chain (Hc) | Plasma Membrane | other | 1.17E+07 | 1.26E+07 | 1.02E+07 | | | | | | |
| Q68FD5 | CLTC | clathrin, heavy chain (Hc) | Plasma Membrane | other | 7932519.5 | 8407324 | 6916418 | | | | | | |
| Q68FD5 | CLTC | clathrin, heavy chain (Hc) | Plasma Membrane | other | 7923862.5 | 7199160 | 6602972 | | | | | | |
| Q68FD5 | CLTC | clathrin, heavy chain (Hc) | Plasma Membrane | other | 8090456.5 | 7351820.5 | 7010329.5 | | | | | | |
| Q68FD5 | CLTC | clathrin, heavy chain (Hc) | Plasma Membrane | other | 1.05E+07 | 9526108 | 7119692 | | | | | | |
| Q68FD5 | CLTC | clathrin, heavy chain (Hc) | Plasma Membrane | other | 7933889.5 | 8436900 | 6759586 | | | | | | |
| Q68FD5 | CLTC | clathrin, heavy chain (Hc) | Plasma Membrane | other | 5044903 | 3826759.25 | 4225482.5 | | | | | | |
| Q68FD5 | CLTC | clathrin, heavy chain (Hc) | Plasma Membrane | other | 5001718 | 4395124 | 3992729 | | | | | | |
| Q68FD5 | CLTC | clathrin, heavy chain (Hc) | Plasma Membrane | other | 5970883 | 6097247 | 6826495.5 | | | | | | |
| Q68FD5 | CLTC | clathrin, heavy chain (Hc) | Plasma Membrane | other | 7118062 | 4777193.5 | 4799298 | | | | | | |
| Q68FD5 | CLTC | clathrin, heavy chain (Hc) | Plasma Membrane | other | 6018828.5 | 5836517.5 | 4063044.75 | | | | | | |
| Q68FD5 | CLTC | clathrin, heavy chain (Hc) | Plasma Membrane | other | 4594702.5 | 5942796.5 | 3136379.5 | | | | | | |
| Q68FD5 | CLTC | clathrin, heavy chain (Hc) | Plasma Membrane | other | 5224520 | 5000942.5 | 4312741 | | | | | | |
| Q68FD5 | CLTC | clathrin, heavy chain (Hc) | Plasma Membrane | other | 4554571 | 4987867 | 3727241.5 | | | | | | |
| Q68FD5 | CLTC | clathrin, heavy chain (Hc) | Plasma Membrane | other | 4384598 | 5027355 | 4236271.5 | | | | | | |
| Q68FD5 | CLTC | clathrin, heavy chain (Hc) | Plasma Membrane | other | 2905251.75 | 3003883 | 2509974.75 | | | | | | |
| Q68FD5 | CLTC | clathrin, heavy chain (Hc) | Plasma Membrane | other | 4966385 | 5063646 | 4506279.5 | | | | | | |
| Q68FD5 | CLTC | clathrin, heavy chain (Hc) | Plasma Membrane | other | 6447324 | 6374369.5 | 1.07E+07 | | | | | | |
| Q68FD5 | CLTC | clathrin, heavy chain (Hc) | Plasma Membrane | other | 3715958.25 | 4399808 | 4024729.75 | | | | | | |
| Q68FD5 | CLTC | clathrin, heavy chain (Hc) | Plasma Membrane | other | 4847696 | 4147254.5 | 3647298.75 | | | | | | |
| Q68FD5 | CLTC | clathrin, heavy chain (Hc) | Plasma Membrane | other | 5327178.5 | 4400274 | 5061479 | | | | | | |
| Q68FD5 | CLTC | clathrin, heavy chain (Hc) | Plasma Membrane | other | 4051859.25 | 4475430.5 | 3567490.75 | | | | | | |
| Q68FD5 | CLTC | clathrin, heavy chain (Hc) | Plasma Membrane | other | 2507565 | 2309317 | 2900971 | | | | | | |
| Q68FD5 | CLTC | clathrin, heavy chain (Hc) | Plasma Membrane | other | 3782706.5 | 3750364 | 3130268.5 | | | | | | |
| Q68FD5 | CLTC | clathrin, heavy chain (Hc) | Plasma Membrane | other | 2627229.75 | 2764546.5 | 2812711.25 | | | | | | |
| Q68FD5 | CLTC | clathrin, heavy chain (Hc) | Plasma Membrane | other | 3071244.75 | 4082373.25 | 3228015 | | | | | | |
| Q68FD5 | CLTC | clathrin, heavy chain (Hc) | Plasma Membrane | other | 1503931.875 | 2382641.75 | 2352453.25 | | | | | | |
| Q68FD5 | CLTC | clathrin, heavy chain (Hc) | Plasma Membrane | other | 3597189.5 | 4393434 | 3128391 | | | | | | |
| Q68FD5 | CLTC | clathrin, heavy chain (Hc) | Plasma Membrane | other | 2157719.5 | 2270286 | 2164988.5 | | | | | | |
| Q68FD5 | CLTC | clathrin, heavy chain (Hc) | Plasma Membrane | other | 2277485.75 | 2197765.75 | 2256434.25 | | | | | | |
| Q68FD5 | CLTC | clathrin, heavy chain (Hc) | Plasma Membrane | other | 2309996 | 2004845.125 | 1445452.625 | | | | | | |
| Q68FD5 | CLTC | clathrin, heavy chain (Hc) | Plasma Membrane | other | 1939808 | 2309828.25 | 1479215 | | | | | | |
| Q68FD5 | CLTC | clathrin, heavy chain (Hc) | Plasma Membrane | other | 2527482 | 1991059.375 | 1587642.5 | | | | | | |
| Q68FD5 | CLTC | clathrin, heavy chain (Hc) | Plasma Membrane | other | 2007093.625 | 2292213 | 1633286.375 | | | | | | |
| Q68FD5 | CLTC | clathrin, heavy chain (Hc) | Plasma Membrane | other | 1404374.625 | 1201926.25 | 1390210.625 | | | | | | |
| Q68FD5 | CLTC | clathrin, heavy chain (Hc) | Plasma Membrane | other | 465039.656 | 471123.344 | 401597.562 | | | | | | |
| Q68FD5 | CLTC | clathrin, heavy chain (Hc) | Plasma Membrane | other | 792853.625 | 712955.188 | 603458.062 | | | | | | |
| Q68FD5 | CLTC | clathrin, heavy chain (Hc) | Plasma Membrane | other | 1349200 | 492019.656 | 1207920 | | | | | | |
| Q68FD5 | CLTC | clathrin, heavy chain (Hc) | Plasma Membrane | other | 925199.188 | 4.70E-06 | 800787.625 | | | | | | |
| Q68FD5 | CLTC | clathrin, heavy chain (Hc) | Plasma Membrane | other | 109093.727 | 238612.281 | 251061.812 | Q68FD5 | CLTC | 5158358 | 3457793 | 294347 | 138 |
| Q06890 | CLU | clusterin | Cytoplasm | other | 7702682.5 | 1.21E+07 | 1.27E+07 | | | | | | |
| Q06890 | CLU | clusterin | Cytoplasm | other | 3035259 | 3523557 | 3537069.5 | | | | | | |
| Q06890 | CLU | clusterin | Cytoplasm | other | 1060697.125 | 684650.75 | 1692648.625 | Q06890 | CLU | 3932880 | 4598162 | 1532721 | 9 |
| Q8R4N0 | CLYBL | citrate lyase beta like | Cytoplasm | enzyme | 1560418.375 | 2475548.5 | 2444430.25 | Q8R4N0 | CLYBL | 1560418 | 519601 | 299992 | 3 |
| O35372 | CNIH1 | cornichon family AMPA receptor auxiliary protein 1 | Plasma Membrane | other | 5212837 | 5960719 | 4069739.75 | O35372 | CNIH1 | 5212837 | 952348 | 549838 | 3 |
| Q08093 | CNN2 | calponin 2 | Cytoplasm | other | 3151003.75 | 3990424.5 | 6186720 | Q08093 | CNN2 | 3151004 | 1567584 | 905045 | 3 |
| Q3TWN3 | CNNM2 | cyclin and CBS domain divalent metal cation transport mediator 2 | Plasma Membrane | other | 4645139.5 | 4357032 | 3700692.25 | | | | | | |

| | | | | | | | | | | | | | |
|---|---|---|---|---|---|---|---|---|---|---|---|---|---|
| Q3TWN3 | CNNM2 | cyclin and CBS domain divalent metal cation transport mediator 2 | Plasma Membrane | other | 3834084 | 4493083.5 | 4106654.25 | Q3TWN3 | CNNM2 | 4239612 | 374154 | 152748 | 6 |
| P16330 | CNP | 2',3'-cyclic nucleotide 3' phosphodiesterase | Cytoplasm | enzyme | 3.02E+07 | 3.27E+07 | 3.50E+07 | | | | | | |
| P16330 | CNP | 2',3'-cyclic nucleotide 3' phosphodiesterase | Cytoplasm | enzyme | 1.51E+07 | 1.32E+07 | 1.47E+07 | | | | | | |
| P16330 | CNP | 2',3'-cyclic nucleotide 3' phosphodiesterase | Cytoplasm | enzyme | 8502601 | 9097093 | 8681109 | | | | | | |
| P16330 | CNP | 2',3'-cyclic nucleotide 3' phosphodiesterase | Cytoplasm | enzyme | 7076650.5 | 6876589 | 8000306.5 | | | | | | |
| P16330 | CNP | 2',3'-cyclic nucleotide 3' phosphodiesterase | Cytoplasm | enzyme | 6359792 | 4660818.5 | 4467167.5 | | | | | | |
| P16330 | CNP | 2',3'-cyclic nucleotide 3' phosphodiesterase | Cytoplasm | enzyme | 3746421.25 | 3485652.5 | 3563400 | | | | | | |
| P16330 | CNP | 2',3'-cyclic nucleotide 3' phosphodiesterase | Cytoplasm | enzyme | 3780853.25 | 4441214 | 3674810.5 | | | | | | |
| P16330 | CNP | 2',3'-cyclic nucleotide 3' phosphodiesterase | Cytoplasm | enzyme | 2126450.5 | 5060272 | 5044624 | P16330 | CNP | 9611596 | 9431592 | 1925216 | 24 |
| Q9QXT0 | CNPY2 | canopy FGF signaling regulator 2 | Plasma Membrane | other | 6216400.5 | 5827013.5 | 5121958.5 | | | | | | |
| Q9QXT0 | CNPY2 | canopy FGF signaling regulator 2 | Plasma Membrane | other | 5717381.5 | 5623046 | 4901736 | | | | | | |
| Q9QXT0 | CNPY2 | canopy FGF signaling regulator 2 | Plasma Membrane | other | 2323956.5 | 3040678 | 2705429.5 | Q9QXT0 | CNPY2 | 4752580 | 1499064 | 499688 | 9 |
| Q8BQ47 | CNPY4 | canopy FGF signaling regulator 4 | Extracellular Space | other | 2514924 | 2550849.5 | 2193612 | Q8BQ47 | CNPY4 | 2514924 | 196702 | 113566 | 3 |
| O88587 | COMT | catechol-O-methyltransferase | Cytoplasm | enzyme | 4250421.5 | 4935512 | 3817138 | | | | | | |
| O88587 | COMT | catechol-O-methyltransferase | Cytoplasm | enzyme | 4490338 | 5558079.5 | 3587708.25 | | | | | | |
| O88587 | COMT | catechol-O-methyltransferase | Cytoplasm | enzyme | 3330332.75 | 4901140 | 3383763.75 | | | | | | |
| O88587 | COMT | catechol-O-methyltransferase | Cytoplasm | enzyme | 3541571.25 | 3884182.25 | 3167154 | O88587 | COMT | 3903166 | 757207 | 218587 | 12 |
| Q8BIG7 | COMTD1 | catechol-O-methyltransferase domain containing 1 | Plasma Membrane | enzyme | 2280336 | 4031544.25 | 5000613.5 | Q8BIG7 | COMTD1 | 2280336 | 1378752 | 796023 | 3 |
| P61202 | Cops2 | COP9 (constitutive photomorphogenic) homolog, subunit 2 (Arabidopsis thaliana) | Nucleus | transcription regulator | 8587202 | 5401198.5 | 3451076 | | | | | | |
| P61202 | Cops2 | COP9 (constitutive photomorphogenic) homolog, subunit 2 (Arabidopsis thaliana) | Nucleus | transcription regulator | 4818195.5 | 3230134.25 | 2971406.5 | | | | | | |
| P61202 | Cops2 | COP9 (constitutive photomorphogenic) homolog, subunit 2 (Arabidopsis thaliana) | Nucleus | transcription regulator | 2684948.75 | 1992824.75 | 1010120 | | | | | | |
| P61202 | Cops2 | COP9 (constitutive photomorphogenic) homolog, subunit 2 (Arabidopsis thaliana) | Nucleus | transcription regulator | 2517149.75 | 1221877.125 | 1323681.875 | | | | | | |
| P61202 | Cops2 | COP9 (constitutive photomorphogenic) homolog, subunit 2 (Arabidopsis thaliana) | Nucleus | transcription regulator | 465622.469 | 192221.469 | 202357.188 | P61202 | Cops2 | 3814624 | 2273088 | 586909 | 15 |
| O88543 | COPS3 | COP9 signalosome subunit 3 | Cytoplasm | other | 4898248 | 3080686.75 | 2314407.75 | | | | | | |
| O88543 | COPS3 | COP9 signalosome subunit 3 | Cytoplasm | other | 3148707.75 | 3042314.25 | 3096656 | | | | | | |
| O88543 | COPS3 | COP9 signalosome subunit 3 | Cytoplasm | other | 3412372.25 | 2947203.25 | 2257184.5 | | | | | | |
| O88543 | COPS3 | COP9 signalosome subunit 3 | Cytoplasm | other | 2483885.25 | 1180492.5 | 670314.375 | | | | | | |
| O88543 | COPS3 | COP9 signalosome subunit 3 | Cytoplasm | other | 1793536.625 | 870644.562 | 867372.125 | | | | | | |
| O88543 | COPS3 | COP9 signalosome subunit 3 | Cytoplasm | other | 2272506.5 | 2841272 | 2339213 | O88543 | COPS3 | 3001543 | 1064560 | 250919 | 18 |
| O88544 | COPS4 | COP9 signalosome subunit 4 | Cytoplasm | other | 3922556.75 | 3498120 | 3211939 | | | | | | |
| O88544 | COPS4 | COP9 signalosome subunit 4 | Cytoplasm | other | 2868814.25 | 2384142 | 2552054.25 | | | | | | |
| O88544 | COPS4 | COP9 signalosome subunit 4 | Cytoplasm | other | 1860610.25 | 1613553.125 | 1187731.375 | | | | | | |
| O88544 | COPS4 | COP9 signalosome subunit 4 | Cytoplasm | other | 1482271.5 | 391100.562 | 423240.5 | | | | | | |
| O88544 | COPS4 | COP9 signalosome subunit 4 | Cytoplasm | other | 2695025.75 | 2073336.125 | 1495230.125 | | | | | | |
| O88544 | COPS4 | COP9 signalosome subunit 4 | Cytoplasm | other | 2536541 | 1866983.375 | 1705012.25 | O88544 | COPS4 | 2560970 | 962533 | 226871 | 18 |
| O35864 | COPS5 | COP9 signalosome subunit 5 | Nucleus | transcription regulator | 1.09E+07 | 6936107.5 | 5214635.5 | | | | | | |
| O35864 | COPS5 | COP9 signalosome subunit 5 | Nucleus | transcription regulator | 2405201 | 1557280 | 1341490.5 | O35864 | COPS5 | 6667601 | 3754853 | 1532912 | 6 |
| O88545 | COPS6 | COP9 signalosome subunit 6 | Nucleus | other | 5748351.5 | 5642061 | 5883817 | | | | | | |
| O88545 | COPS6 | COP9 signalosome subunit 6 | Nucleus | other | 5304456 | 5805361 | 4211371.5 | | | | | | |
| O88545 | COPS6 | COP9 signalosome subunit 6 | Nucleus | other | 3456374 | 3579671.75 | 3245669 | | | | | | |
| O88545 | COPS6 | COP9 signalosome subunit 6 | Nucleus | other | 1529241.375 | 1000159.75 | 706556.375 | | | | | | |
| O88545 | COPS6 | COP9 signalosome subunit 6 | Nucleus | other | 1726478.625 | 1291278.875 | 734738 | O88545 | COPS6 | 3552980 | 2028724 | 523814 | 15 |
| Q9CZ04 | COPS7A | COP9 signalosome subunit 7A | Cytoplasm | other | 1019223.312 | 696694.625 | 856178.812 | Q9CZ04 | COPS7A | 1019223 | 161268 | 93108 | 3 |
| Q8BV13 | COPS7B | COP9 signalosome subunit 7B | Cytoplasm | other | 2252633.25 | 1645424.75 | 1917492.125 | Q8BV13 | COPS7B | 2252633 | 304150 | 175601 | 3 |
| Q8VBV7 | COPS8 | COP9 signalosome subunit 8 | Nucleus | other | 2045619.125 | 1784984.75 | 1658651.625 | Q8VBV7 | COPS8 | 2045619 | 197330 | 113928 | 3 |
| O89053 | CORO1A | coronin, actin binding protein, 1A | Cytoplasm | other | 3.10E+07 | 4.36E+07 | 7.79E+07 | | | | | | |
| O89053 | CORO1A | coronin, actin binding protein, 1A | Cytoplasm | other | 4933144 | 6662481 | 1.05E+07 | | | | | | |
| O89053 | CORO1A | coronin, actin binding protein, 1A | Cytoplasm | other | 1.67E+07 | 2.41E+07 | 3.74E+07 | | | | | | |
| O89053 | CORO1A | coronin, actin binding protein, 1A | Cytoplasm | other | 1.09E+07 | 1.01E+07 | 1.99E+07 | | | | | | |
| O89053 | CORO1A | coronin, actin binding protein, 1A | Cytoplasm | other | 3779162.75 | 4512093.5 | 8910039 | | | | | | |
| O89053 | CORO1A | coronin, actin binding protein, 1A | Cytoplasm | other | 8637540 | 1.22E+07 | 2.25E+07 | | | | | | |
| O89053 | CORO1A | coronin, actin binding protein, 1A | Cytoplasm | other | 1174257.75 | 2372037 | 5192675.5 | | | | | | |
| O89053 | CORO1A | coronin, actin binding protein, 1A | Cytoplasm | other | 3165898 | 4369576.5 | 6918081 | | | | | | |
| O89053 | CORO1A | coronin, actin binding protein, 1A | Cytoplasm | other | 5037947 | 6748312.5 | 8678567 | | | | | | |
| O89053 | CORO1A | coronin, actin binding protein, 1A | Cytoplasm | other | 4270752.5 | 6150108 | 1.04E+07 | | | | | | |
| O89053 | CORO1A | coronin, actin binding protein, 1A | Cytoplasm | other | 1.04E+07 | 1.43E+07 | 2.36E+07 | | | | | | |
| O89053 | CORO1A | coronin, actin binding protein, 1A | Cytoplasm | other | 3956866.25 | 1.25E+07 | 2.52E+07 | | | | | | |
| O89053 | CORO1A | coronin, actin binding protein, 1A | Cytoplasm | other | 3592121.75 | 4447888 | 7775164.5 | O89053 | CORO1A | 8273668 | 14478330 | 2318388 | 39 |
| Q9WUM4 | CORO1C | coronin, actin binding protein, 1C | Cytoplasm | other | 5405937 | 7090698 | 1.03E+07 | | | | | | |
| Q9WUM4 | CORO1C | coronin, actin binding protein, 1C | Cytoplasm | other | 1656582.375 | 3435557.75 | 4748910 | Q9WUM4 | CORO1C | 3531260 | 2995474 | 1222897 | 6 |
| P19783 | COX4I1 | cytochrome c oxidase subunit IV isoform 1 | Cytoplasm | enzyme | 1.68E+07 | 2.30E+07 | 2.94E+07 | | | | | | |
| P19783 | COX4I1 | cytochrome c oxidase subunit IV isoform 1 | Cytoplasm | enzyme | 7106248 | 1.36E+07 | 1.52E+07 | | | | | | |
| P19783 | COX4I1 | cytochrome c oxidase subunit IV isoform 1 | Cytoplasm | enzyme | 3535954.25 | 5279117.5 | 6292286.5 | P19783 | COX4I1 | 9140734 | 8773711 | 2924570 | 9 |
| P12787 | COX5A | cytochrome c oxidase subunit Va | Cytoplasm | enzyme | 8005819.5 | 1.20E+07 | 1.76E+07 | | | | | | |
| P12787 | COX5A | cytochrome c oxidase subunit Va | Cytoplasm | enzyme | 7312506 | 1.44E+07 | 9851263 | | | | | | |
| P12787 | COX5A | cytochrome c oxidase subunit Va | Cytoplasm | enzyme | 4145616.25 | 6186138 | 6723899 | P12787 | COX5A | 6487981 | 4339234 | 1446411 | 9 |
| P19536 | Cox5b | cytochrome c oxidase subunit Vb | Cytoplasm | other | 8487300 | 1.33E+07 | 1.60E+07 | P19536 | Cox5b | 8487300 | 3784176 | 2184795 | 3 |
| P56391 | COX6B1 | cytochrome c oxidase subunit VIb polypeptide 1 (ubiquitous) | Cytoplasm | enzyme | 3922030.5 | 7664658 | 1.07E+07 | P56391 | COX6B1 | 3922031 | 3385501 | 1954620 | 3 |
| Q9CPQ1 | Cox6c | cytochrome c oxidase subunit VIc | Cytoplasm | enzyme | 1276566.5 | 2517721.75 | 2563170.5 | Q9CPQ1 | Cox6c | 1276567 | 730055 | 421497 | 3 |
| P48771 | COX7A2 | cytochrome c oxidase subunit VIIa polypeptide 2 (liver) | Cytoplasm | enzyme | 4594713.5 | 8052200.5 | 9855310 | P48771 | COX7A2 | 4594714 | 2673303 | 1543432 | 3 |
| P59108 | CPNE2 | copine II | Cytoplasm | other | 2452470.75 | 4948569 | 2874497.75 | | | | | | |
| P59108 | CPNE2 | copine II | Cytoplasm | other | 2549870.5 | 4419943.5 | 2944752.25 | P59108 | CPNE2 | 2501171 | 1052135 | 429532 | 6 |
| P36552 | CPOX | coproporphyrinogen oxidase | Cytoplasm | enzyme | 8679797 | 1.75E+07 | 1.50E+07 | | | | | | |
| P36552 | CPOX | coproporphyrinogen oxidase | Cytoplasm | enzyme | 7587749 | 1.69E+07 | 1.67E+07 | | | | | | |
| P36552 | CPOX | coproporphyrinogen oxidase | Cytoplasm | enzyme | 6091417.5 | 1.42E+07 | 1.86E+07 | | | | | | |
| P36552 | CPOX | coproporphyrinogen oxidase | Cytoplasm | enzyme | 8478665 | 1.53E+07 | 1.60E+07 | | | | | | |
| P36552 | CPOX | coproporphyrinogen oxidase | Cytoplasm | enzyme | 5065091 | 1.49E+07 | 1.12E+07 | | | | | | |
| P36552 | CPOX | coproporphyrinogen oxidase | Cytoplasm | enzyme | 3972636.25 | 1.10E+07 | 1.19E+07 | | | | | | |
| P36552 | CPOX | coproporphyrinogen oxidase | Cytoplasm | enzyme | 4624512.5 | 9437476 | 1.07E+07 | | | | | | |
| P36552 | CPOX | coproporphyrinogen oxidase | Cytoplasm | enzyme | 4324075 | 1.01E+07 | 1.36E+07 | | | | | | |
| P36552 | CPOX | coproporphyrinogen oxidase | Cytoplasm | enzyme | 3710829.25 | 9402244 | 1.11E+07 | | | | | | |
| P36552 | CPOX | coproporphyrinogen oxidase | Cytoplasm | enzyme | 2817986.25 | 6222204 | 8721241 | | | | | | |
| P36552 | CPOX | coproporphyrinogen oxidase | Cytoplasm | enzyme | 1627211.375 | 3750090.25 | 3200108.5 | | | | | | |
| P36552 | CPOX | coproporphyrinogen oxidase | Cytoplasm | enzyme | 1217242.625 | 2376924.5 | 2792215 | P36552 | CPOX | 4849768 | 5194062 | 865677 | 36 |
| P52825 | CPT2 | carnitine palmitoyltransferase 2 | Cytoplasm | enzyme | 2218758.25 | 3243773.25 | 5008531.5 | | | | | | |
| P52825 | CPT2 | carnitine palmitoyltransferase 2 | Cytoplasm | enzyme | 1110369.625 | 3035808 | 3555921.75 | | | | | | |
| P52825 | CPT2 | carnitine palmitoyltransferase 2 | Cytoplasm | enzyme | 413933.219 | 1019424.188 | 891876.562 | P52825 | CPT2 | 1247687 | 1538784 | 512928 | 9 |
| Q64735 | CR1L | complement component (3b/4b) receptor 1-like | Plasma Membrane | other | 5.75E+07 | 6.14E+07 | 5.50E+07 | | | | | | |
| Q64735 | CR1L | complement component (3b/4b) receptor 1-like | Plasma Membrane | other | 3.27E+07 | 3.08E+07 | 3.32E+07 | | | | | | |
| Q64735 | CR1L | complement component (3b/4b) receptor 1-like | Plasma Membrane | other | 3.25E+07 | 2.60E+07 | 2.13E+07 | | | | | | |
| Q64735 | CR1L | complement component (3b/4b) receptor 1-like | Plasma Membrane | other | 2.85E+07 | 2.35E+07 | 2.20E+07 | | | | | | |
| Q64735 | CR1L | complement component (3b/4b) receptor 1-like | Plasma Membrane | other | 8502313 | 8885566 | 7109462.5 | | | | | | |
| Q64735 | CR1L | complement component (3b/4b) receptor 1-like | Plasma Membrane | other | 7676883.5 | 8230406.5 | 8650178 | | | | | | |
| Q64735 | CR1L | complement component (3b/4b) receptor 1-like | Plasma Membrane | other | 6950473 | 7307317 | 6212214.5 | | | | | | |
| Q64735 | CR1L | complement component (3b/4b) receptor 1-like | Plasma Membrane | other | 5453528.5 | 3204204.75 | 2921790.75 | | | | | | |
| Q64735 | CR1L | complement component (3b/4b) receptor 1-like | Plasma Membrane | other | 1009608.5 | 1258331.625 | 693796.812 | Q64735 | CR1L | 20080312 | 17845107 | 3434292 | 27 |
| Q9Z125 | CREB3L1 | cAMP responsive element binding protein 3-like 1 | Nucleus | transcription regulator | 2930324.5 | 3947345.5 | 4552552.5 | Q9Z125 | CREB3L1 | 2930325 | 819780 | 473300 | 3 |
| O88668 | CREG1 | cellular repressor of E1A-stimulated genes 1 | Nucleus | transcription regulator | 3508161.5 | 4277632 | 5580902.5 | | | | | | |
| O88668 | CREG1 | cellular repressor of E1A-stimulated genes 1 | Nucleus | transcription regulator | 3134799 | 3199022.75 | 3540700.75 | O88668 | CREG1 | 3321480 | 929879 | 379621 | 6 |
| Q9DCT8 | Crip2 | cysteine rich protein 2 | Plasma Membrane | other | 4.77E+07 | 4.11E+07 | 2.84E+07 | | | | | | |
| Q9DCT8 | Crip2 | cysteine rich protein 2 | Plasma Membrane | other | 3.75E+07 | 4.58E+07 | 3.59E+07 | | | | | | |
| Q9DCT8 | Crip2 | cysteine rich protein 2 | Plasma Membrane | other | 3.61E+07 | 2.95E+07 | 2.78E+07 | | | | | | |
| Q9DCT8 | Crip2 | cysteine rich protein 2 | Plasma Membrane | other | 4.50E+07 | 4.02E+07 | 2.55E+07 | | | | | | |
| Q9DCT8 | Crip2 | cysteine rich protein 2 | Plasma Membrane | other | 2.21E+07 | 1.81E+07 | 1.63E+07 | | | | | | |
| Q9DCT8 | Crip2 | cysteine rich protein 2 | Plasma Membrane | other | 2.18E+07 | 2.03E+07 | 1.60E+07 | | | | | | |
| Q9DCT8 | Crip2 | cysteine rich protein 2 | Plasma Membrane | other | 1.46E+07 | 1.35E+07 | 1.16E+07 | | | | | | |
| Q9DCT8 | Crip2 | cysteine rich protein 2 | Plasma Membrane | other | 1.42E+07 | 1.18E+07 | 7460996 | | | | | | |
| Q9DCT8 | Crip2 | cysteine rich protein 2 | Plasma Membrane | other | 3359280.5 | 1862207.375 | 5607004.5 | Q9DCT8 | Crip2 | 26921031 | 13695015 | 2635607 | 27 |
| Q9Z2L7 | CRLF3 | cytokine receptor-like factor 3 | Cytoplasm | transcription regulator | 3686224.25 | 3255907.75 | 3869339.25 | Q9Z2L7 | CRLF3 | 3686224 | 314908 | 181812 | 3 |

| | | | | | | | | | | | | | |
|---|---|---|---|---|---|---|---|---|---|---|---|---|---|
| Q9CZU6 | CS | citrate synthase | Cytoplasm | enzyme | 2.16E+07 | 3.88E+07 | 4.69E+07 | | | | | | |
| Q9CZU6 | CS | citrate synthase | Cytoplasm | enzyme | 8430266 | 1.80E+07 | 1.58E+07 | | | | | | |
| Q9CZU6 | CS | citrate synthase | Cytoplasm | enzyme | 1.49E+07 | 2.68E+07 | 2.98E+07 | | | | | | |
| Q9CZU6 | CS | citrate synthase | Cytoplasm | enzyme | 6772978.5 | 1.13E+07 | 1.51E+07 | | | | | | |
| Q9CZU6 | CS | citrate synthase | Cytoplasm | enzyme | 6344367.5 | 1.05E+07 | 1.47E+07 | | | | | | |
| Q9CZU6 | CS | citrate synthase | Cytoplasm | enzyme | 6038735 | 1.14E+07 | 1.52E+07 | | | | | | |
| Q9CZU6 | CS | citrate synthase | Cytoplasm | enzyme | 4678458.5 | 3765700 | 1.50E+07 | | | | | | |
| Q9CZU6 | CS | citrate synthase | Cytoplasm | enzyme | 3068762.5 | 3710778.25 | 4632231 | | | | | | |
| Q9CZU6 | CS | citrate synthase | Cytoplasm | enzyme | 2838731.25 | 4454827.5 | 5312335.5 | | | | | | |
| Q9CZU6 | CS | citrate synthase | Cytoplasm | enzyme | 2986936.75 | 5531553.5 | 8032719.5 | Q9CZU6 | CS | 7768924 | 10829860 | 1977253 | 30 |
| Q8BK63 | CSNK1A1 | casein kinase 1, alpha 1 | Cytoplasm | kinase | 1.28E+07 | 1.03E+07 | 8330814 | | | | | | |
| Q8BK63 | CSNK1A1 | casein kinase 1, alpha 1 | Cytoplasm | kinase | 9971751 | 725946.5 | 72585.742 | | | | | | |
| Q8BK63 | CSNK1A1 | casein kinase 1, alpha 1 | Cytoplasm | kinase | 1.14E+07 | 1.09E+07 | 6515777 | | | | | | |
| Q8BK63 | CSNK1A1 | casein kinase 1, alpha 1 | Cytoplasm | kinase | 6218471 | 8157117.5 | 6228236 | | | | | | |
| Q8BK63 | CSNK1A1 | casein kinase 1, alpha 1 | Cytoplasm | kinase | 7890447.5 | 6441279.5 | 6090839.5 | | | | | | |
| Q8BK63 | CSNK1A1 | casein kinase 1, alpha 1 | Cytoplasm | kinase | 5096686 | 3326008.75 | 3047805.5 | | | | | | |
| Q8BK63 | CSNK1A1 | casein kinase 1, alpha 1 | Cytoplasm | kinase | 7042108.5 | 4879030 | 3164154.75 | | | | | | |
| Q8BK63 | CSNK1A1 | casein kinase 1, alpha 1 | Cytoplasm | kinase | 4266407.5 | 1881663.375 | 925112.875 | Q8BK63 | CSNK1A1 | 8086984 | 3509248 | 716322 | 24 |
| Q60737 | CSNK2A1 | casein kinase 2, alpha 1 polypeptide | Nucleus | kinase | 3221596.25 | 2070492.5 | 1987580.875 | Q60737 | CSNK2A1 | 3221596 | 689771 | 398240 | 3 |
| P67871 | CSNK2B | casein kinase 2, beta polypeptide | Cytoplasm | kinase | 2795629 | 2614764.5 | 2422632 | P67871 | CSNK2B | 2795629 | 186527 | 107691 | 3 |
| P97315 | CSRP1 | cysteine and glycine-rich protein 1 | Nucleus | other | 1.77E+07 | 2.24E+07 | 2.69E+07 | | | | | | |
| P97315 | CSRP1 | cysteine and glycine-rich protein 1 | Nucleus | other | 1.77E+07 | 2.42E+07 | 2.98E+07 | | | | | | |
| P97315 | CSRP1 | cysteine and glycine-rich protein 1 | Nucleus | other | 1.34E+07 | 2.14E+07 | 2.47E+07 | | | | | | |
| P97315 | CSRP1 | cysteine and glycine-rich protein 1 | Nucleus | other | 1.06E+07 | 1.56E+07 | 2.30E+07 | | | | | | |
| P97315 | CSRP1 | cysteine and glycine-rich protein 1 | Nucleus | other | 1209763 | 2137665.25 | 6027783 | P97315 | CSRP1 | 12107953 | 8882019 | 2293327 | 15 |
| P10605 | CTSB | cathepsin B | Cytoplasm | peptidase | 1.03E+07 | 1.01E+07 | 9450030 | | | | | | |
| P10605 | CTSB | cathepsin B | Cytoplasm | peptidase | 7905668.5 | 1.05E+07 | 9885771 | | | | | | |
| P10605 | CTSB | cathepsin B | Cytoplasm | peptidase | 6304137 | 9043716 | 1.15E+07 | | | | | | |
| P10605 | CTSB | cathepsin B | Cytoplasm | peptidase | 5133825 | 7134209.5 | 3598625 | | | | | | |
| P10605 | CTSB | cathepsin B | Cytoplasm | peptidase | 2866161.25 | 3555412.5 | 2922009.5 | P10605 | CTSB | 6497958 | 3057788 | 789517 | 15 |
| P18242 | CTSD | cathepsin D | Cytoplasm | peptidase | 1.09E+07 | 1.94E+07 | 2.22E+07 | | | | | | |
| P18242 | CTSD | cathepsin D | Cytoplasm | peptidase | 7847771 | 1.34E+07 | 1.41E+07 | | | | | | |
| P18242 | CTSD | cathepsin D | Cytoplasm | peptidase | 7777731.5 | 1.28E+07 | 1.62E+07 | | | | | | |
| P18242 | CTSD | cathepsin D | Cytoplasm | peptidase | 6114518.5 | 1.30E+07 | 1.35E+07 | | | | | | |
| P18242 | CTSD | cathepsin D | Cytoplasm | peptidase | 6911292.5 | 1.12E+07 | 1.35E+07 | | | | | | |
| P18242 | CTSD | cathepsin D | Cytoplasm | peptidase | 2387163.75 | 4100321.5 | 6319982.5 | | | | | | |
| P18242 | CTSD | cathepsin D | Cytoplasm | peptidase | 1556228.75 | 3097198.75 | 3477675 | P18242 | CTSD | 6206387 | 5690621 | 1241795 | 21 |
| P70269 | CTSE | cathepsin E | Cytoplasm | peptidase | 1.29E+08 | 1.27E+08 | 1.01E+08 | | | | | | |
| P70269 | CTSE | cathepsin E | Cytoplasm | peptidase | 1.21E+08 | 1.27E+08 | 1.07E+08 | | | | | | |
| P70269 | CTSE | cathepsin E | Cytoplasm | peptidase | 9858220 | 3738866.25 | 1.70E+07 | P70269 | CTSE | 86586073 | 55115526 | 18371842 | 9 |
| Q9WTX6 | CUL1 | cullin 1 | Nucleus | enzyme | 3645819.5 | 2203142.25 | 1872192.125 | | | | | | |
| Q9WTX6 | CUL1 | cullin 1 | Nucleus | enzyme | 3906709.5 | 3137576 | 2572474.25 | | | | | | |
| Q9WTX6 | CUL1 | cullin 1 | Nucleus | enzyme | 2795708 | 2311640.25 | 3151555.75 | | | | | | |
| Q9WTX6 | CUL1 | cullin 1 | Nucleus | enzyme | 2283827 | 2788473 | 2444852.5 | | | | | | |
| Q9WTX6 | CUL1 | cullin 1 | Nucleus | enzyme | 2705420.5 | 2708276.25 | 2044567.875 | | | | | | |
| Q9WTX6 | CUL1 | cullin 1 | Nucleus | enzyme | 2448127.75 | 1497770.125 | 1576958.5 | | | | | | |
| Q9WTX6 | CUL1 | cullin 1 | Nucleus | enzyme | 819435.312 | 133374.422 | 795140.75 | | | | | | |
| Q9WTX6 | CUL1 | cullin 1 | Nucleus | enzyme | 1320136.75 | 114614.531 | 122363.25 | | | | | | |
| Q9WTX6 | CUL1 | cullin 1 | Nucleus | enzyme | 995783.062 | 1107071 | 565632.375 | Q9WTX6 | CUL1 | 2324552 | 1077936 | 207449 | 27 |
| Q9JLV5 | CUL3 | cullin 3 | Nucleus | enzyme | 4145877.25 | 2765071.75 | 2566559.75 | | | | | | |
| Q9JLV5 | CUL3 | cullin 3 | Nucleus | enzyme | 2392106.5 | 2222200 | 3020079 | Q9JLV5 | CUL3 | 3268992 | 692756 | 282816 | 6 |
| P70658 | CXCR4 | chemokine (C-X-C motif) receptor 4 | Plasma Membrane | G-protein coupled receptor | 4768582.5 | 4311566 | 2846991.75 | P70658 | CXCR4 | 4768583 | 1003855 | 579576 | 3 |
| P56395 | CYB5A | cytochrome b5 type A (microsomal) | Cytoplasm | enzyme | 1.16E+07 | 1.43E+07 | 1.03E+07 | | | | | | |
| P56395 | CYB5A | cytochrome b5 type A (microsomal) | Cytoplasm | enzyme | 8136648.5 | 1.10E+07 | 1.21E+07 | | | | | | |
| P56395 | CYB5A | cytochrome b5 type A (microsomal) | Cytoplasm | enzyme | 2836025 | 3510633.5 | 3041026.25 | | | | | | |
| P56395 | CYB5A | cytochrome b5 type A (microsomal) | Cytoplasm | enzyme | 2796169.75 | 3061299.5 | 3072060 | | | | | | |
| P56395 | CYB5A | cytochrome b5 type A (microsomal) | Cytoplasm | enzyme | 2579994 | 2129098.75 | 4914147.5 | P56395 | CYB5A | 5581767 | 4339908 | 1120559 | 15 |
| Q9CQX2 | CYB5B | cytochrome b5 type B (outer mitochondrial membrane) | Cytoplasm | enzyme | 1.29E+07 | 1.96E+07 | 1.79E+07 | | | | | | |
| Q9CQX2 | CYB5B | cytochrome b5 type B (outer mitochondrial membrane) | Cytoplasm | enzyme | 1985788.5 | 508315.875 | 18230.643 | | | | | | |
| Q9CQX2 | CYB5B | cytochrome b5 type B (outer mitochondrial membrane) | Cytoplasm | enzyme | 1392307.875 | 752591.562 | 85096.273 | Q9CQX2 | CYB5B | 5429365 | 8216946 | 2738982 | 9 |
| Q9DCN2 | Cyb5r3 | cytochrome b5 reductase 3 | Cytoplasm | enzyme | 2.81E+07 | 3.90E+07 | 3.62E+07 | | | | | | |
| Q9DCN2 | Cyb5r3 | cytochrome b5 reductase 3 | Cytoplasm | enzyme | 1.95E+07 | 2.83E+07 | 1.35E+07 | | | | | | |
| Q9DCN2 | Cyb5r3 | cytochrome b5 reductase 3 | Cytoplasm | enzyme | 2.22E+07 | 2.95E+07 | 2.70E+07 | | | | | | |
| Q9DCN2 | Cyb5r3 | cytochrome b5 reductase 3 | Cytoplasm | enzyme | 1.32E+07 | 1.41E+07 | 2.46E+07 | | | | | | |
| Q9DCN2 | Cyb5r3 | cytochrome b5 reductase 3 | Cytoplasm | enzyme | 1.28E+07 | 1.53E+07 | 1.76E+07 | | | | | | |
| Q9DCN2 | Cyb5r3 | cytochrome b5 reductase 3 | Cytoplasm | enzyme | 8896831 | 1.10E+07 | 1.36E+07 | | | | | | |
| Q9DCN2 | Cyb5r3 | cytochrome b5 reductase 3 | Cytoplasm | enzyme | 7283679.5 | 7687338.5 | 8851191 | | | | | | |
| Q9DCN2 | Cyb5r3 | cytochrome b5 reductase 3 | Cytoplasm | enzyme | 7632634 | 8708063 | 1.01E+07 | | | | | | |
| Q9DCN2 | Cyb5r3 | cytochrome b5 reductase 3 | Cytoplasm | enzyme | 8327008 | 7877983.5 | 8639209 | | | | | | |
| Q9DCN2 | Cyb5r3 | cytochrome b5 reductase 3 | Cytoplasm | enzyme | 5355555.5 | 5617382.5 | 5861719 | | | | | | |
| Q9DCN2 | Cyb5r3 | cytochrome b5 reductase 3 | Cytoplasm | enzyme | 5971387.5 | 5624787.5 | 9253929 | | | | | | |
| Q9DCN2 | Cyb5r3 | cytochrome b5 reductase 3 | Cytoplasm | enzyme | 3769411.25 | 5269280.5 | 3239696.25 | Q9DCN2 | Cyb5r3 | 11918876 | 9489061 | 1581510 | 36 |
| Q9D0M3 | CYC1 | cytochrome c-1 | Cytoplasm | enzyme | 1.21E+07 | 1.62E+07 | 1.87E+07 | Q9D0M3 | CYC1 | 12090000 | 3349632 | 1933911 | 3 |
| P62897 | CYCS | cytochrome c, somatic | Cytoplasm | transporter | 7096501.5 | 1.20E+07 | 1.21E+07 | | | | | | |
| P62897 | CYCS | cytochrome c, somatic | Cytoplasm | transporter | 3881615.75 | 6486994.5 | 7057602 | | | | | | |
| P62897 | CYCS | cytochrome c, somatic | Cytoplasm | transporter | 3527107 | 3298299.75 | 5902089 | | | | | | |
| P62897 | CYCS | cytochrome c, somatic | Cytoplasm | transporter | 3191921.25 | 5313658.5 | 4326663 | | | | | | |
| P62897 | CYCS | cytochrome c, somatic | Cytoplasm | transporter | 1165917.75 | 2044009.5 | 1892296.625 | P62897 | CYCS | 3772613 | 3299391 | 851899 | 15 |
| Q8BPM0 | DAAM1 | dishevelled associated activator of morphogenesis 1 | Cytoplasm | other | 7405075 | 7217350.5 | 7082777 | | | | | | |
| Q8BPM0 | DAAM1 | dishevelled associated activator of morphogenesis 1 | Cytoplasm | other | 5117415 | 4198972 | 4029093.75 | | | | | | |
| Q8BPM0 | DAAM1 | dishevelled associated activator of morphogenesis 1 | Cytoplasm | other | 4457428 | 4858781.5 | 5715878.5 | | | | | | |
| Q8BPM0 | DAAM1 | dishevelled associated activator of morphogenesis 1 | Cytoplasm | other | 4592734 | 4548288 | 4034436.75 | | | | | | |
| Q8BPM0 | DAAM1 | dishevelled associated activator of morphogenesis 1 | Cytoplasm | other | 4689141.5 | 4761900 | 3369071.5 | | | | | | |
| Q8BPM0 | DAAM1 | dishevelled associated activator of morphogenesis 1 | Cytoplasm | other | 1351304.375 | 1673301.5 | 2208912.25 | | | | | | |
| Q8BPM0 | DAAM1 | dishevelled associated activator of morphogenesis 1 | Cytoplasm | other | 4989978.5 | 5467795 | 5823208.5 | | | | | | |
| Q8BPM0 | DAAM1 | dishevelled associated activator of morphogenesis 1 | Cytoplasm | other | 2114359.25 | 3590632.25 | 2641463.75 | | | | | | |
| Q8BPM0 | DAAM1 | dishevelled associated activator of morphogenesis 1 | Cytoplasm | other | 3623380.75 | 2982080.25 | 2856315 | | | | | | |
| Q8BPM0 | DAAM1 | dishevelled associated activator of morphogenesis 1 | Cytoplasm | other | 3089293.5 | 2812676.25 | 1851666.625 | | | | | | |
| Q8BPM0 | DAAM1 | dishevelled associated activator of morphogenesis 1 | Cytoplasm | other | 1861591.625 | 1767349 | 1167037.625 | | | | | | |
| Q8BPM0 | DAAM1 | dishevelled associated activator of morphogenesis 1 | Cytoplasm | other | 240330.812 | 371180.906 | 760635.438 | Q8BPM0 | DAAM1 | 3627669 | 1890860 | 315143 | 36 |
| P61804 | DAD1 | defender against cell death 1 | Cytoplasm | other | 2360395.25 | 2366586.25 | 3887394 | P61804 | DAD1 | 2360395 | 879831 | 507971 | 3 |
| Q91WC9 | DAGLB | diacylglycerol lipase, beta | Plasma Membrane | enzyme | 7782616 | 7487445 | 7669903 | | | | | | |
| Q91WC9 | DAGLB | diacylglycerol lipase, beta | Plasma Membrane | enzyme | 4016473.75 | 6154035.5 | 4670122 | | | | | | |
| Q91WC9 | DAGLB | diacylglycerol lipase, beta | Plasma Membrane | enzyme | 3349424.75 | 3937466.5 | 2958894.25 | | | | | | |
| Q91WC9 | DAGLB | diacylglycerol lipase, beta | Plasma Membrane | enzyme | 5233444 | 7023391 | 3510082.5 | | | | | | |
| Q91WC9 | DAGLB | diacylglycerol lipase, beta | Plasma Membrane | enzyme | 3232263.75 | 3326429.5 | 2409424 | | | | | | |
| Q91WC9 | DAGLB | diacylglycerol lipase, beta | Plasma Membrane | enzyme | 2536817 | 2071382.5 | 2109509.25 | Q91WC9 | DAGLB | 4358507 | 1990286 | 469115 | 18 |
| Q922B2 | DARS | aspartyl-tRNA synthetase | Cytoplasm | enzyme | 4562965.5 | 6790258 | 6862772.5 | | | | | | |
| Q922B2 | DARS | aspartyl-tRNA synthetase | Cytoplasm | enzyme | 2687626.5 | 2819061.25 | 4382274 | | | | | | |
| Q922B2 | DARS | aspartyl-tRNA synthetase | Cytoplasm | enzyme | 1653962.875 | 1658584.625 | 3157622 | | | | | | |
| Q922B2 | DARS | aspartyl-tRNA synthetase | Cytoplasm | enzyme | 2301639.5 | 1928666.875 | 3241244.25 | | | | | | |
| Q922B2 | DARS | aspartyl-tRNA synthetase | Cytoplasm | enzyme | 1280898.875 | 983791.25 | 1623353 | | | | | | |
| Q922B2 | DARS | aspartyl-tRNA synthetase | Cytoplasm | enzyme | 1198542 | 1008074.438 | 1742233.25 | | | | | | |
| Q922B2 | DARS | aspartyl-tRNA synthetase | Cytoplasm | enzyme | 1312966.75 | 1546461.5 | 2902080.5 | | | | | | |
| Q922B2 | DARS | aspartyl-tRNA synthetase | Cytoplasm | enzyme | 2005687.875 | 1203000.125 | 1906862.875 | Q922B2 | DARS | 2125536 | 1640904 | 334948 | 24 |
| Q9JII5 | Dazap1 | DAZ associated protein 1 | Cytoplasm | other | 799815.312 | 1245831.375 | 945069.438 | Q9JII5 | Dazap1 | 799815 | 227481 | 131336 | 3 |
| Q9QZ73 | DCUN1D1 | DCN1, defective in cullin neddylation 1, domain containing 1 | Nucleus | other | 1991872.625 | 1893198.25 | 2113595.5 | Q9QZ73 | DCUN1D1 | 1991873 | 110399 | 63739 | 3 |
| A2ADY9 | DDI2 | DNA-damage inducible 1 homolog 2 | Plasma Membrane | transporter | 7656695 | 5624057.5 | 5211775 | | | | | | |
| A2ADY9 | DDI2 | DNA-damage inducible 1 homolog 2 | Plasma Membrane | transporter | 5643014 | 3016366.25 | 3143632 | A2ADY9 | DDI2 | 6649855 | 1747692 | 713492 | 6 |

| | | | | | | | | | | | | | |
|---|---|---|---|---|---|---|---|---|---|---|---|---|---|
| O54734 | DDOST | dolichyl-diphosphooligosaccharide--protein glycosyltransferase subunit (non-catalytic) | Cytoplasm | enzyme | 2904372.75 | 4623760 | 6834143 | | | | | | |
| O54734 | DDOST | dolichyl-diphosphooligosaccharide--protein glycosyltransferase subunit (non-catalytic) | Cytoplasm | enzyme | 1794374.375 | 3305992.5 | 4346761 | | | | | | |
| O54734 | DDOST | dolichyl-diphosphooligosaccharide--protein glycosyltransferase subunit (non-catalytic) | Cytoplasm | enzyme | 2350908 | 2817658 | 4868871.5 | O54734 | DDOST | 2349885 | 1559077 | 519692 | 9 |
| Q8C4S8 | DENND2A | DENN/MADD domain containing 2A | Cytoplasm | other | 2342002.25 | 3078540.5 | 2801746.75 | Q8C4S8 | DENND2A | 2342002 | 372037 | 214796 | 3 |
| Q91YP3 | DERA | deoxyribose-phosphate aldolase (putative) | Cytoplasm | enzyme | 4023468 | 2288961.25 | 2846425 | | | | | | |
| Q91YP3 | DERA | deoxyribose-phosphate aldolase (putative) | Cytoplasm | enzyme | 1754867.375 | 1178331.125 | 1574823.125 | | | | | | |
| Q91YP3 | DERA | deoxyribose-phosphate aldolase (putative) | Cytoplasm | enzyme | 2068565.25 | 1470503.5 | 1365110.875 | Q91YP3 | DERA | 2615634 | 898769 | 299590 | 9 |
| Q99J56 | DERL1 | derlin 1 | Cytoplasm | other | 1.02E+07 | 1.11E+07 | 1.40E+07 | | | | | | |
| Q99J56 | DERL1 | derlin 1 | Cytoplasm | other | 4103424 | 4689685.5 | 8798291 | Q99J56 | DERL1 | 7146712 | 3813359 | 1556797 | 6 |
| Q8BNI4 | DERL2 | derlin 2 | Cytoplasm | other | 2801483.5 | 2856196.75 | 3952840.75 | Q8BNI4 | DERL2 | 2801484 | 649518 | 375000 | 3 |
| Q3U0B3 | DHRS11 | dehydrogenase/reductase (SDR family) member 11 | Cytoplasm | other | 2441957.25 | 2188129 | 2366125.5 | Q3U0B3 | DHRS11 | 2441957 | 130296 | 75226 | 3 |
| A2ATU0 | DHTKD1 | dehydrogenase E1 and transketolase domain containing 1 | Cytoplasm | enzyme | 1.21E+07 | 8458144 | 1.12E+07 | A2ATU0 | DHTKD1 | 12120000 | 1904966 | 1099832 | 3 |
| Q80VY9 | DHX33 | DEAH (Asp-Glu-Ala-His) box polypeptide 33 | Nucleus | enzyme | 7030876.5 | 5875758 | 5735566 | Q80VY9 | DHX33 | 7030877 | 710842 | 410405 | 3 |
| Q8BMF4 | DLAT | dihydrolipoamide S-acetyltransferase | Cytoplasm | enzyme | 1669196.875 | 3498691.25 | 4138814.75 | | | | | | |
| Q8BMF4 | DLAT | dihydrolipoamide S-acetyltransferase | Cytoplasm | enzyme | 1589597.25 | 4150578.75 | 3960457.5 | Q8BMF4 | DLAT | 1629397 | 1215159 | 496086 | 6 |
| O08749 | DLD | dihydrolipoamide dehydrogenase | Cytoplasm | enzyme | 1.20E+07 | 2.28E+07 | 2.27E+07 | | | | | | |
| O08749 | DLD | dihydrolipoamide dehydrogenase | Cytoplasm | enzyme | 8735992 | 1.34E+07 | 1.54E+07 | | | | | | |
| O08749 | DLD | dihydrolipoamide dehydrogenase | Cytoplasm | enzyme | 4865966.5 | 7987284.5 | 1.09E+07 | O08749 | DLD | 8547320 | 6225232 | 2075077 | 9 |
| Q811D0 | DLG1 | discs, large homolog 1 (Drosophila) | Plasma Membrane | kinase | 2169088.5 | 2491181 | 2545542.5 | | | | | | |
| Q811D0 | DLG1 | discs, large homolog 1 (Drosophila) | Plasma Membrane | kinase | 1739806.75 | 932699.938 | 747906.938 | | | | | | |
| Q811D0 | DLG1 | discs, large homolog 1 (Drosophila) | Plasma Membrane | kinase | 736352.625 | 729185.438 | 722970.125 | Q811D0 | DLG1 | 1548416 | 806006 | 268669 | 9 |
| Q9D2G2 | DLST | dihydrolipoamide S-succinyltransferase (E2 component of 2-oxo-glutarate complex) | Cytoplasm | enzyme | 8770594 | 1.50E+07 | 1.67E+07 | | | | | | |
| Q9D2G2 | DLST | dihydrolipoamide S-succinyltransferase (E2 component of 2-oxo-glutarate complex) | Cytoplasm | enzyme | 5907633 | 8609806 | 9428310 | | | | | | |
| Q9D2G2 | DLST | dihydrolipoamide S-succinyltransferase (E2 component of 2-oxo-glutarate complex) | Cytoplasm | enzyme | 5158075.5 | 1.10E+07 | 1.16E+07 | | | | | | |
| Q9D2G2 | DLST | dihydrolipoamide S-succinyltransferase (E2 component of 2-oxo-glutarate complex) | Cytoplasm | enzyme | 6336605.5 | 4971717 | 1.06E+07 | | | | | | |
| Q9D2G2 | DLST | dihydrolipoamide S-succinyltransferase (E2 component of 2-oxo-glutarate complex) | Cytoplasm | enzyme | 4887933 | 1.00E+07 | 9124032 | | | | | | |
| Q9D2G2 | DLST | dihydrolipoamide S-succinyltransferase (E2 component of 2-oxo-glutarate complex) | Cytoplasm | enzyme | 5356938 | 8226630.5 | 6304841 | | | | | | |
| Q9D2G2 | DLST | dihydrolipoamide S-succinyltransferase (E2 component of 2-oxo-glutarate complex) | Cytoplasm | enzyme | 3324245.5 | 5129902.5 | 7693509 | Q9D2G2 | DLST | 5677432 | 3415425 | 745307 | 21 |
| Q80WT2 | DMRT3 | doublesex and mab-3 related transcription factor 3 | Nucleus | transcription regulator | 5161620.5 | 5171816.5 | 4990812 | Q80WT2 | DMRT3 | 5161621 | 101688 | 58709 | 3 |
| Q9WV69 | DMTN | dematin actin binding protein | Plasma Membrane | other | 5.11E+08 | 6.65E+08 | 4.30E+08 | | | | | | |
| Q9WV69 | DMTN | dematin actin binding protein | Plasma Membrane | other | 7.02E+08 | 7.21E+08 | 5.97E+08 | | | | | | |
| Q9WV69 | DMTN | dematin actin binding protein | Plasma Membrane | other | 3.93E+08 | 3.76E+08 | 3.07E+08 | | | | | | |
| Q9WV69 | DMTN | dematin actin binding protein | Plasma Membrane | other | 3.52E+08 | 3.44E+08 | 2.85E+08 | | | | | | |
| Q9WV69 | DMTN | dematin actin binding protein | Plasma Membrane | other | 4.18E+08 | 4.10E+08 | 3.80E+08 | | | | | | |
| Q9WV69 | DMTN | dematin actin binding protein | Plasma Membrane | other | 3.18E+08 | 3.16E+08 | 2.85E+08 | | | | | | |
| Q9WV69 | DMTN | dematin actin binding protein | Plasma Membrane | other | 3.80E+08 | 3.87E+08 | 3.09E+08 | | | | | | |
| Q9WV69 | DMTN | dematin actin binding protein | Plasma Membrane | other | 3.45E+08 | 3.44E+08 | 3.22E+08 | | | | | | |
| Q9WV69 | DMTN | dematin actin binding protein | Plasma Membrane | other | 2.86E+08 | 2.62E+08 | 1.44E+08 | | | | | | |
| Q9WV69 | DMTN | dematin actin binding protein | Plasma Membrane | other | 2.23E+08 | 2.08E+08 | 1.92E+08 | | | | | | |
| Q9WV69 | DMTN | dematin actin binding protein | Plasma Membrane | other | 2.06E+08 | 1.99E+08 | 1.73E+08 | | | | | | |
| Q9WV69 | DMTN | dematin actin binding protein | Plasma Membrane | other | 1.45E+08 | 1.76E+08 | 1.00E+08 | | | | | | |
| Q9WV69 | DMTN | dematin actin binding protein | Plasma Membrane | other | 9.07E+07 | 7.07E+07 | 7.11E+07 | | | | | | |
| Q9WV69 | DMTN | dematin actin binding protein | Plasma Membrane | other | 9.02E+07 | 1.88E+08 | 1.81E+08 | | | | | | |
| Q9WV69 | DMTN | dematin actin binding protein | Plasma Membrane | other | 8.21E+07 | 1.63E+08 | 1.59E+08 | | | | | | |
| Q9WV69 | DMTN | dematin actin binding protein | Plasma Membrane | other | 4.55E+07 | 7.53E+07 | 7.37E+07 | | | | | | |
| Q9WV69 | DMTN | dematin actin binding protein | Plasma Membrane | other | 5.18E+07 | 3.39E+07 | 1.54E+07 | | | | | | |
| Q9WV69 | DMTN | dematin actin binding protein | Plasma Membrane | other | 2.10E+07 | 2.29E+07 | 1.90E+07 | | | | | | |
| Q9WV69 | DMTN | dematin actin binding protein | Plasma Membrane | other | 1.96E+07 | 1.74E+07 | 1.63E+07 | | | | | | |
| Q9WV69 | DMTN | dematin actin binding protein | Plasma Membrane | other | 1.84E+07 | 1.47E+07 | 1.30E+07 | | | | | | |
| Q9WV69 | DMTN | dematin actin binding protein | Plasma Membrane | other | 6623839.5 | 2533080 | 3179430.25 | | | | | | |
| Q9WV69 | DMTN | dematin actin binding protein | Plasma Membrane | other | 8958887 | 9628712 | 7206437.5 | | | | | | |
| Q9WV69 | DMTN | dematin actin binding protein | Plasma Membrane | other | 4366713 | 4656064 | 4249144 | | | | | | |
| Q9WV69 | DMTN | dematin actin binding protein | Plasma Membrane | other | 3656688 | 1057676.5 | 4675981.5 | | | | | | |
| Q9WV69 | DMTN | dematin actin binding protein | Plasma Membrane | other | 725960.312 | 1710968.875 | 1065295.25 | Q9WV69 | DMTN | 188882884 | 186336882 | 21516330 | 75 |
| Q9JMC3 | DNAJA4 | DnaJ (Hsp40) homolog, subfamily A, member 4 | Nucleus | other | 1794445 | 1488136.375 | 1058272.625 | Q9JMC3 | DNAJA4 | 1794445 | 369810 | 213510 | 3 |
| Q9QYJ3 | DNAJB1 | DnaJ (Hsp40) homolog, subfamily B, member 1 | Nucleus | other | 1.68E+07 | 1.72E+07 | 1.89E+07 | | | | | | |
| Q9QYJ3 | DNAJB1 | DnaJ (Hsp40) homolog, subfamily B, member 1 | Nucleus | other | 9497692 | 8266913.5 | 1.17E+07 | | | | | | |
| Q9QYJ3 | DNAJB1 | DnaJ (Hsp40) homolog, subfamily B, member 1 | Nucleus | other | 9355169 | 6168553 | 6104532.5 | | | | | | |
| Q9QYJ3 | DNAJB1 | DnaJ (Hsp40) homolog, subfamily B, member 1 | Nucleus | other | 1.24E+07 | 1.15E+07 | 1.19E+07 | | | | | | |
| Q9QYJ3 | DNAJB1 | DnaJ (Hsp40) homolog, subfamily B, member 1 | Nucleus | other | 4277048 | 4565542 | 3956377 | | | | | | |
| Q9QYJ3 | DNAJB1 | DnaJ (Hsp40) homolog, subfamily B, member 1 | Nucleus | other | 4361458.5 | 6032343 | 4901542 | | | | | | |
| Q9QYJ3 | DNAJB1 | DnaJ (Hsp40) homolog, subfamily B, member 1 | Nucleus | other | 4136474.75 | 3667112.5 | 3168384.75 | | | | | | |
| Q9QYJ3 | DNAJB1 | DnaJ (Hsp40) homolog, subfamily B, member 1 | Nucleus | other | 2977783 | 2366491.25 | 2578438.5 | | | | | | |
| Q9QYJ3 | DNAJB1 | DnaJ (Hsp40) homolog, subfamily B, member 1 | Nucleus | other | 336420.031 | 2331450.75 | 3499407.25 | Q9QYJ3 | DNAJB1 | 7119116 | 5016152 | 965359 | 27 |
| Q99KV1 | DNAJB11 | DnaJ (Hsp40) homolog, subfamily B, member 11 | Cytoplasm | other | 5909561 | 6354627 | 5084221.5 | | | | | | |
| Q99KV1 | DNAJB11 | DnaJ (Hsp40) homolog, subfamily B, member 11 | Cytoplasm | other | 3316803.25 | 3401654.75 | 3261441.75 | | | | | | |
| Q99KV1 | DNAJB11 | DnaJ (Hsp40) homolog, subfamily B, member 11 | Cytoplasm | other | 2231176.75 | 2425344.75 | 1964894.125 | Q99KV1 | DNAJB11 | 3819180 | 1620931 | 540310 | 9 |
| Q9D832 | DNAJB4 | DnaJ (Hsp40) homolog, subfamily B, member 4 | Nucleus | other | 1.71E+07 | 2.08E+07 | 1.83E+07 | | | | | | |
| Q9D832 | DNAJB4 | DnaJ (Hsp40) homolog, subfamily B, member 4 | Nucleus | other | 1.79E+07 | 1.66E+07 | 1.93E+07 | | | | | | |
| Q9D832 | DNAJB4 | DnaJ (Hsp40) homolog, subfamily B, member 4 | Nucleus | other | 1.26E+07 | 1.21E+07 | 1.03E+07 | | | | | | |
| Q9D832 | DNAJB4 | DnaJ (Hsp40) homolog, subfamily B, member 4 | Nucleus | other | 5293855 | 4238356.5 | 5415264 | | | | | | |
| Q9D832 | DNAJB4 | DnaJ (Hsp40) homolog, subfamily B, member 4 | Nucleus | other | 5411895.5 | 7564070 | 7915374 | | | | | | |
| Q9D832 | DNAJB4 | DnaJ (Hsp40) homolog, subfamily B, member 4 | Nucleus | other | 3814029.75 | 5418859 | 6671087.5 | | | | | | |
| Q9D832 | DNAJB4 | DnaJ (Hsp40) homolog, subfamily B, member 4 | Nucleus | other | 5923709 | 5701830 | 4244499 | | | | | | |
| Q9D832 | DNAJB4 | DnaJ (Hsp40) homolog, subfamily B, member 4 | Nucleus | other | 5298737 | 4550485 | 5238962.5 | | | | | | |
| Q9D832 | DNAJB4 | DnaJ (Hsp40) homolog, subfamily B, member 4 | Nucleus | other | 3686408.75 | 3731933.25 | 1999797.875 | | | | | | |
| Q9D832 | DNAJB4 | DnaJ (Hsp40) homolog, subfamily B, member 4 | Nucleus | other | 3185856 | 3063261 | 2100041 | Q9D832 | DNAJB4 | 8021449 | 5751133 | 1050008 | 30 |
| Q9DC23 | DNAJC10 | DnaJ (Hsp40) homolog, subfamily C, member 10 | Cytoplasm | enzyme | 471543.906 | 355508.719 | 304404.906 | Q9DC23 | DNAJC10 | 471544 | 85646 | 49448 | 3 |
| Q5U458 | DNAJC11 | DnaJ (Hsp40) homolog, subfamily C, member 11 | Cytoplasm | other | 1265604.875 | 2579882 | 3014407.25 | | | | | | |
| Q5U458 | DNAJC11 | DnaJ (Hsp40) homolog, subfamily C, member 11 | Cytoplasm | other | 1013225.312 | 2532323.25 | 2062891.125 | Q5U458 | DNAJC11 | 1139415 | 791089 | 322961 | 6 |
| P60904 | DNAJC5 | DnaJ (Hsp40) homolog, subfamily C, member 5 | Plasma Membrane | other | 4.38E+07 | 3.83E+07 | 3.55E+07 | | | | | | |
| P60904 | DNAJC5 | DnaJ (Hsp40) homolog, subfamily C, member 5 | Plasma Membrane | other | 2.78E+07 | 3.92E+07 | 3.09E+07 | | | | | | |
| P60904 | DNAJC5 | DnaJ (Hsp40) homolog, subfamily C, member 5 | Plasma Membrane | other | 1.46E+07 | 1.33E+07 | 1.26E+07 | | | | | | |
| P60904 | DNAJC5 | DnaJ (Hsp40) homolog, subfamily C, member 5 | Plasma Membrane | other | 1645440.125 | 2472415.75 | 4749805 | | | | | | |
| P60904 | DNAJC5 | DnaJ (Hsp40) homolog, subfamily C, member 5 | Plasma Membrane | other | 44530.609 | 0 | 75036.859 | P60904 | DNAJC5 | 17583994 | 16496122 | 4259280 | 15 |
| Q9D7J6 | DNASE1L1 | deoxyribonuclease I-like 1 | Cytoplasm | enzyme | 499889.156 | 648978.75 | 794006.375 | Q9D7J6 | DNASE1L1 | 499889 | 147063 | 84907 | 3 |
| P39054 | DNM2 | dynamin 2 | Plasma Membrane | enzyme | 4.89E+07 | 5.02E+07 | 5.41E+07 | | | | | | |
| P39054 | DNM2 | dynamin 2 | Plasma Membrane | enzyme | 2.93E+07 | 2.83E+07 | 3.24E+07 | | | | | | |
| P39054 | DNM2 | dynamin 2 | Plasma Membrane | enzyme | 4.16E+07 | 3.35E+07 | 4.20E+07 | | | | | | |
| P39054 | DNM2 | dynamin 2 | Plasma Membrane | enzyme | 2.51E+07 | 2.60E+07 | 2.76E+07 | | | | | | |
| P39054 | DNM2 | dynamin 2 | Plasma Membrane | enzyme | 1.70E+07 | 2.10E+07 | 2.14E+07 | | | | | | |
| P39054 | DNM2 | dynamin 2 | Plasma Membrane | enzyme | 1.77E+07 | 1.77E+07 | 1.93E+07 | | | | | | |
| P39054 | DNM2 | dynamin 2 | Plasma Membrane | enzyme | 1.41E+07 | 1.51E+07 | 1.81E+07 | | | | | | |
| P39054 | DNM2 | dynamin 2 | Plasma Membrane | enzyme | 1.32E+07 | 1.26E+07 | 1.43E+07 | | | | | | |
| P39054 | DNM2 | dynamin 2 | Plasma Membrane | enzyme | 1.11E+07 | 1.26E+07 | 1.41E+07 | | | | | | |
| P39054 | DNM2 | dynamin 2 | Plasma Membrane | enzyme | 1.02E+07 | 1.01E+07 | 1.16E+07 | | | | | | |
| P39054 | DNM2 | dynamin 2 | Plasma Membrane | enzyme | 9666184 | 8945991 | 1.09E+07 | | | | | | |
| P39054 | DNM2 | dynamin 2 | Plasma Membrane | enzyme | 1.13E+07 | 1.39E+07 | 1.23E+07 | | | | | | |
| P39054 | DNM2 | dynamin 2 | Plasma Membrane | enzyme | 7719143 | 7322231 | 7772494.5 | | | | | | |
| P39054 | DNM2 | dynamin 2 | Plasma Membrane | enzyme | 7661185.5 | 7747559.5 | 9270643 | | | | | | |
| P39054 | DNM2 | dynamin 2 | Plasma Membrane | enzyme | 2180592.75 | 303901.844 | 48774.723 | | | | | | |
| P39054 | DNM2 | dynamin 2 | Plasma Membrane | enzyme | 5692414.5 | 6607787 | 8095432.5 | | | | | | |
| P39054 | DNM2 | dynamin 2 | Plasma Membrane | enzyme | 4126636.5 | 6082374.5 | 5988973.5 | | | | | | |
| P39054 | DNM2 | dynamin 2 | Plasma Membrane | enzyme | 4699528 | 3644779.25 | 3645647.25 | | | | | | |
| P39054 | DNM2 | dynamin 2 | Plasma Membrane | enzyme | 3697887 | 2720272.75 | 3771003.5 | | | | | | |
| P39054 | DNM2 | dynamin 2 | Plasma Membrane | enzyme | 2793899 | 3241307.75 | 3795196 | | | | | | |
| P39054 | DNM2 | dynamin 2 | Plasma Membrane | enzyme | 2267243.25 | 2281827 | 2448247.25 | | | | | | |
| P39054 | DNM2 | dynamin 2 | Plasma Membrane | enzyme | 1898958.5 | 2724110 | 4183124.5 | | | | | | |
| P39054 | DNM2 | dynamin 2 | Plasma Membrane | enzyme | 2117906.25 | 2915711.75 | 1486878.5 | P39054 | DNM2 | 12783112 | 12643575 | 1522108 | 69 |

| | | | | | | | | | | | | | |
|---|---|---|---|---|---|---|---|---|---|---|---|---|---|
| Q8BZ98 | DNM3 | dynamin 3 | Cytoplasm | enzyme | 8269772.5 | 8347955.5 | 9267804 | | | | | | |
| Q8BZ98 | DNM3 | dynamin 3 | Cytoplasm | enzyme | 1.17E+07 | 1.28E+07 | 1.15E+07 | | | | | | |
| Q8BZ98 | DNM3 | dynamin 3 | Cytoplasm | enzyme | 4332644.5 | 3742586.75 | 5068862.5 | Q8BZ98 | DNM3 | 8087472 | 3347285 | 1115762 | 9 |
| Q9Z2W0 | DNPEP | aspartyl aminopeptidase | Cytoplasm | peptidase | 2.70E+07 | 2.38E+07 | 1.96E+07 | | | | | | |
| Q9Z2W0 | DNPEP | aspartyl aminopeptidase | Cytoplasm | peptidase | 6207708.5 | 5145024 | 4642274.5 | | | | | | |
| Q9Z2W0 | DNPEP | aspartyl aminopeptidase | Cytoplasm | peptidase | 6332335.5 | 5729462 | 5086262 | | | | | | |
| Q9Z2W0 | DNPEP | aspartyl aminopeptidase | Cytoplasm | peptidase | 6041219 | 5017740.5 | 4078550.75 | | | | | | |
| Q9Z2W0 | DNPEP | aspartyl aminopeptidase | Cytoplasm | peptidase | 5174685 | 3982118.75 | 3449887 | | | | | | |
| Q9Z2W0 | DNPEP | aspartyl aminopeptidase | Cytoplasm | peptidase | 3844810.25 | 3062664.75 | 4407553.5 | | | | | | |
| Q9Z2W0 | DNPEP | aspartyl aminopeptidase | Cytoplasm | peptidase | 5296103.5 | 3724117.5 | 2754519 | | | | | | |
| Q9Z2W0 | DNPEP | aspartyl aminopeptidase | Cytoplasm | peptidase | 3478742.25 | 2330438.25 | 3659605.25 | | | | | | |
| Q9Z2W0 | DNPEP | aspartyl aminopeptidase | Cytoplasm | peptidase | 1845236.5 | 1089883.375 | 1262632 | Q9Z2W0 | DNPEP | 7244538 | 6449917 | 1241287 | 27 |
| Q9DD18 | DTD1 | D-tyrosyl-tRNA deacylase 1 | Cytoplasm | enzyme | 174792.438 | 46949.656 | 182425.875 | Q9DD18 | DTD1 | 174792 | 76109 | 43942 | 3 |
| Q9JHU4 | DYNC1H1 | dynein, cytoplasmic 1, heavy chain 1 | Cytoplasm | peptidase | 1862330.75 | 1815545.375 | 1617580.625 | Q9JHU4 | DYNC1H1 | 1862331 | 129924 | 75012 | 3 |
| P42125 | ECI1 | enoyl-CoA delta isomerase 1 | Cytoplasm | enzyme | 3494202 | 8673692 | 1.01E+07 | | | | | | |
| P42125 | ECI1 | enoyl-CoA delta isomerase 1 | Cytoplasm | enzyme | 2240498.25 | 3873643.75 | 4357672.5 | P42125 | ECI1 | 2867350 | 3168323 | 1293462 | 6 |
| Q8BL66 | EEA1 | early endosome antigen 1 | Cytoplasm | other | 1752384.5 | 1246630.375 | 718254.625 | Q8BL66 | EEA1 | 1752385 | 517106 | 298551 | 3 |
| P10126 | EEF1A1 | eukaryotic translation elongation factor 1 alpha 1 | Cytoplasm | translation regulator | 1.89E+08 | 2.34E+08 | 2.81E+08 | | | | | | |
| P10126 | EEF1A1 | eukaryotic translation elongation factor 1 alpha 1 | Cytoplasm | translation regulator | 7.71E+07 | 9.81E+07 | 8.65E+07 | | | | | | |
| P10126 | EEF1A1 | eukaryotic translation elongation factor 1 alpha 1 | Cytoplasm | translation regulator | 3.51E+07 | 3.51E+07 | 8.26E+07 | | | | | | |
| P10126 | EEF1A1 | eukaryotic translation elongation factor 1 alpha 1 | Cytoplasm | translation regulator | 5.24E+07 | 7.40E+07 | 6.22E+07 | | | | | | |
| P10126 | EEF1A1 | eukaryotic translation elongation factor 1 alpha 1 | Cytoplasm | translation regulator | 2.91E+07 | 3.57E+07 | 3.91E+07 | | | | | | |
| P10126 | EEF1A1 | eukaryotic translation elongation factor 1 alpha 1 | Cytoplasm | translation regulator | 4.69E+07 | 4.48E+07 | 4.45E+07 | | | | | | |
| P10126 | EEF1A1 | eukaryotic translation elongation factor 1 alpha 1 | Cytoplasm | translation regulator | 6.01E+07 | 7.77E+07 | 8.21E+07 | | | | | | |
| P10126 | EEF1A1 | eukaryotic translation elongation factor 1 alpha 1 | Cytoplasm | translation regulator | 2.73E+07 | 2.50E+07 | 6.99E+07 | | | | | | |
| P10126 | EEF1A1 | eukaryotic translation elongation factor 1 alpha 1 | Cytoplasm | translation regulator | 3.76E+07 | 4.54E+07 | 6.33E+07 | | | | | | |
| P10126 | EEF1A1 | eukaryotic translation elongation factor 1 alpha 1 | Cytoplasm | translation regulator | 4.48E+07 | 4.85E+07 | 4.14E+07 | | | | | | |
| P10126 | EEF1A1 | eukaryotic translation elongation factor 1 alpha 1 | Cytoplasm | translation regulator | 3.76E+07 | 5.00E+07 | 5.56E+07 | | | | | | |
| P10126 | EEF1A1 | eukaryotic translation elongation factor 1 alpha 1 | Cytoplasm | translation regulator | 3.24E+07 | 3.91E+07 | 4.33E+07 | | | | | | |
| P10126 | EEF1A1 | eukaryotic translation elongation factor 1 alpha 1 | Cytoplasm | translation regulator | 3.80E+07 | 5.52E+07 | 2.73E+07 | | | | | | |
| P10126 | EEF1A1 | eukaryotic translation elongation factor 1 alpha 1 | Cytoplasm | translation regulator | 2.92E+07 | 2.91E+07 | 2.98E+07 | | | | | | |
| P10126 | EEF1A1 | eukaryotic translation elongation factor 1 alpha 1 | Cytoplasm | translation regulator | 2.39E+07 | 4.07E+07 | 3.56E+07 | | | | | | |
| P10126 | EEF1A1 | eukaryotic translation elongation factor 1 alpha 1 | Cytoplasm | translation regulator | 2.74E+07 | 2.01E+07 | 3.28E+07 | | | | | | |
| P10126 | EEF1A1 | eukaryotic translation elongation factor 1 alpha 1 | Cytoplasm | translation regulator | 9737050 | 8142014.5 | 8236337 | | | | | | |
| P10126 | EEF1A1 | eukaryotic translation elongation factor 1 alpha 1 | Cytoplasm | translation regulator | 3391775.25 | 3919102.5 | 2967260.5 | | | | | | |
| P10126 | EEF1A1 | eukaryotic translation elongation factor 1 alpha 1 | Cytoplasm | translation regulator | 2631799.5 | 4149444 | 2232408 | | | | | | |
| P10126 | EEF1A1 | eukaryotic translation elongation factor 1 alpha 1 | Cytoplasm | translation regulator | 3137225.5 | 3548542 | 4305246.5 | | | | | | |
| P10126 | EEF1A1 | eukaryotic translation elongation factor 1 alpha 1 | Cytoplasm | translation regulator | 1871878.625 | 2365690.75 | 3082444.5 | | | | | | |
| P10126 | EEF1A1 | eukaryotic translation elongation factor 1 alpha 1 | Cytoplasm | translation regulator | 3319665.25 | 902175.312 | 2787155.75 | | | | | | |
| P10126 | EEF1A1 | eukaryotic translation elongation factor 1 alpha 1 | Cytoplasm | translation regulator | 462342 | 468958 | 608989.188 | P10126 | EEF1A1 | 35337467 | 49184761 | 5921149 | 69 |
| O70251 | EEF1B2 | eukaryotic translation elongation factor 1 beta 2 | Cytoplasm | translation regulator | 432794.938 | 935456.812 | 1543679.875 | O70251 | EEF1B2 | 432795 | 556278 | 321167 | 3 |
| P57776 | EEF1D | eukaryotic translation elongation factor 1 delta (guanine nucleotide exchange protein) | Cytoplasm | translation regulator | 826614.375 | 1677891.75 | 1578533.125 | P57776 | EEF1D | 826614 | 465462 | 268734 | 3 |
| Q9D8N0 | EEF1G | eukaryotic translation elongation factor 1 gamma | Cytoplasm | translation regulator | 1649914.75 | 4317963.5 | 4824894.5 | | | | | | |
| Q9D8N0 | EEF1G | eukaryotic translation elongation factor 1 gamma | Cytoplasm | translation regulator | 663487.188 | 1799003.625 | 3349639 | | | | | | |
| Q9D8N0 | EEF1G | eukaryotic translation elongation factor 1 gamma | Cytoplasm | translation regulator | 1168226.125 | 1988089.75 | 2420666.5 | Q9D8N0 | EEF1G | 1160543 | 1416402 | 472134 | 9 |
| P58252 | EEF2 | eukaryotic translation elongation factor 2 | Cytoplasm | translation regulator | 763794.875 | 1171591.25 | 1527353.375 | | | | | | |
| P58252 | EEF2 | eukaryotic translation elongation factor 2 | Cytoplasm | translation regulator | 1154812.5 | 1846016.125 | 2940583.25 | | | | | | |
| P58252 | EEF2 | eukaryotic translation elongation factor 2 | Cytoplasm | translation regulator | 1613558.125 | 2523840.5 | 3744046.25 | | | | | | |
| P58252 | EEF2 | eukaryotic translation elongation factor 2 | Cytoplasm | translation regulator | 718879.75 | 1226632.875 | 1268449 | | | | | | |
| P58252 | EEF2 | eukaryotic translation elongation factor 2 | Cytoplasm | translation regulator | 349626.719 | 754692.875 | 838368.625 | P58252 | EEF2 | 920134 | 930304 | 240203 | 15 |
| Q8BG67 | EFR3A | EFR3 homolog A | Plasma Membrane | other | 5048575.5 | 4383079.5 | 4058406 | | | | | | |
| Q8BG67 | EFR3A | EFR3 homolog A | Plasma Membrane | other | 4938973 | 4539316.5 | 4763563.5 | | | | | | |
| Q8BG67 | EFR3A | EFR3 homolog A | Plasma Membrane | other | 4467611 | 4506419.5 | 3922249.5 | | | | | | |
| Q8BG67 | EFR3A | EFR3 homolog A | Plasma Membrane | other | 3284314 | 4237994 | 4630827.5 | | | | | | |
| Q8BG67 | EFR3A | EFR3 homolog A | Plasma Membrane | other | 3538415.25 | 4221548.5 | 3144550.25 | | | | | | |
| Q8BG67 | EFR3A | EFR3 homolog A | Plasma Membrane | other | 4260751.5 | 4439618.5 | 3312093.75 | | | | | | |
| Q8BG67 | EFR3A | EFR3 homolog A | Plasma Membrane | other | 2429215.5 | 4051687.25 | 2359854.5 | Q8BG67 | EFR3A | 3995408 | 752591 | 164229 | 21 |
| Q9WVK4 | EHD1 | EH-domain containing 1 | Cytoplasm | other | 8075977.5 | 7952494 | 6319723.5 | | | | | | |
| Q9WVK4 | EHD1 | EH-domain containing 1 | Cytoplasm | other | 5253949.5 | 9054594 | 1.10E+07 | | | | | | |
| Q9WVK4 | EHD1 | EH-domain containing 1 | Cytoplasm | other | 2199486.5 | 2021225 | 2923322.25 | | | | | | |
| Q9WVK4 | EHD1 | EH-domain containing 1 | Cytoplasm | other | 2045907.375 | 2285550.75 | 3486835.75 | Q9WVK4 | EHD1 | 4393830 | 3175477 | 916681 | 12 |
| Q9EQP2 | EHD4 | EH-domain containing 4 | Plasma Membrane | enzyme | 1442031.75 | 1525026.375 | 1999818.25 | | | | | | |
| Q9EQP2 | EHD4 | EH-domain containing 4 | Plasma Membrane | enzyme | 1214781.875 | 916226.75 | 1702821 | Q9EQP2 | EHD4 | 1328407 | 376952 | 153890 | 6 |
| Q6ZWX6 | EIF2S1 | eukaryotic translation initiation factor 2, subunit 1 alpha, 35kDa | Cytoplasm | translation regulator | 3817633.5 | 4312041.5 | 4640731 | | | | | | |
| Q6ZWX6 | EIF2S1 | eukaryotic translation initiation factor 2, subunit 1 alpha, 35kDa | Cytoplasm | translation regulator | 2442662.75 | 3428372 | 4527417 | | | | | | |
| Q6ZWX6 | EIF2S1 | eukaryotic translation initiation factor 2, subunit 1 alpha, 35kDa | Cytoplasm | translation regulator | 2452323 | 3039514.25 | 2693752.25 | | | | | | |
| Q6ZWX6 | EIF2S1 | eukaryotic translation initiation factor 2, subunit 1 alpha, 35kDa | Cytoplasm | translation regulator | 2207646.5 | 2298474.25 | 3043771 | | | | | | |
| Q6ZWX6 | EIF2S1 | eukaryotic translation initiation factor 2, subunit 1 alpha, 35kDa | Cytoplasm | translation regulator | 666991.938 | 1567699.875 | 2873874.75 | | | | | | |
| Q6ZWX6 | EIF2S1 | eukaryotic translation initiation factor 2, subunit 1 alpha, 35kDa | Cytoplasm | translation regulator | 1711317.875 | 2639582 | 2890616.25 | Q6ZWX6 | EIF2S1 | 2216429 | 1038109 | 244685 | 18 |
| Q99L45 | EIF2S2 | eukaryotic translation initiation factor 2, subunit 2 beta, 38kDa | Cytoplasm | translation regulator | 2684192.5 | 2946945.5 | 3631574.75 | | | | | | |
| Q99L45 | EIF2S2 | eukaryotic translation initiation factor 2, subunit 2 beta, 38kDa | Cytoplasm | translation regulator | 2861298.75 | 3676289.25 | 3716988.5 | | | | | | |
| Q99L45 | EIF2S2 | eukaryotic translation initiation factor 2, subunit 2 beta, 38kDa | Cytoplasm | translation regulator | 1972048.875 | 2229596.5 | 3824928 | Q99L45 | EIF2S2 | 2505847 | 688291 | 229430 | 9 |
| Q9Z0N1 | EIF2S3 | eukaryotic translation initiation factor 2, subunit 3 gamma, 52kDa | Cytoplasm | translation regulator | 6544587.5 | 6445322 | 9543085 | | | | | | |
| Q9Z0N1 | EIF2S3 | eukaryotic translation initiation factor 2, subunit 3 gamma, 52kDa | Cytoplasm | translation regulator | 6504255 | 8102270.5 | 1.01E+07 | | | | | | |
| Q9Z0N1 | EIF2S3 | eukaryotic translation initiation factor 2, subunit 3 gamma, 52kDa | Cytoplasm | translation regulator | 4938798.5 | 6246429.5 | 6443434 | | | | | | |
| Q9Z0N1 | EIF2S3 | eukaryotic translation initiation factor 2, subunit 3 gamma, 52kDa | Cytoplasm | translation regulator | 4887446 | 4774420 | 4779190.5 | | | | | | |
| Q9Z0N1 | EIF2S3 | eukaryotic translation initiation factor 2, subunit 3 gamma, 52kDa | Cytoplasm | translation regulator | 5219479.5 | 5654322.5 | 4225543 | | | | | | |
| Q9Z0N1 | EIF2S3 | eukaryotic translation initiation factor 2, subunit 3 gamma, 52kDa | Cytoplasm | translation regulator | 4139371.5 | 4951854 | 3895171 | | | | | | |
| Q9Z0N1 | EIF2S3 | eukaryotic translation initiation factor 2, subunit 3 gamma, 52kDa | Cytoplasm | translation regulator | 2629438.25 | 3450129.25 | 3371550.25 | | | | | | |
| Q9Z0N1 | EIF2S3 | eukaryotic translation initiation factor 2, subunit 3 gamma, 52kDa | Cytoplasm | translation regulator | 2226550.25 | 2689600.5 | 3157528.5 | | | | | | |
| Q9Z0N1 | EIF2S3 | eukaryotic translation initiation factor 2, subunit 3 gamma, 52kDa | Cytoplasm | translation regulator | 1334550.625 | 1523601.375 | 1781910.75 | | | | | | |
| Q9Z0N1 | EIF2S3 | eukaryotic translation initiation factor 2, subunit 3 gamma, 52kDa | Cytoplasm | translation regulator | 1159040.25 | 1041870.938 | 1431074 | | | | | | |
| Q9Z0N1 | EIF2S3 | eukaryotic translation initiation factor 2, subunit 3 gamma, 52kDa | Cytoplasm | translation regulator | 1247120.25 | 1218547.375 | 918756.188 | Q9Z0N1 | EIF2S3 | 3711876 | 2474992 | 430841 | 33 |
| Q3UGC7 | EIF3J | eukaryotic translation initiation factor 3, subunit J | Cytoplasm | translation regulator | 9105090 | 6797261.5 | 7170266.5 | | | | | | |
| Q3UGC7 | EIF3J | eukaryotic translation initiation factor 3, subunit J | Cytoplasm | translation regulator | 1.04E+07 | 8714242 | 7141431.5 | | | | | | |
| Q3UGC7 | EIF3J | eukaryotic translation initiation factor 3, subunit J | Cytoplasm | translation regulator | 6728532.5 | 4610044 | 3685031.5 | | | | | | |
| Q3UGC7 | EIF3J | eukaryotic translation initiation factor 3, subunit J | Cytoplasm | translation regulator | 107774 | 40404.645 | 38283.883 | | | | | | |
| Q3UGC7 | EIF3J | eukaryotic translation initiation factor 3, subunit J | Cytoplasm | translation regulator | 6647087 | 4469442.5 | 2937782.75 | Q3UGC7 | EIF3J | 6603697 | 3349302 | 864786 | 15 |
| Q8BGD9 | EIF4B | eukaryotic translation initiation factor 4B | Cytoplasm | translation regulator | 6860517.5 | 6673812.5 | 7465343 | | | | | | |
| Q8BGD9 | EIF4B | eukaryotic translation initiation factor 4B | Cytoplasm | translation regulator | 6277894 | 4140106.5 | 5316179.5 | | | | | | |
| Q8BGD9 | EIF4B | eukaryotic translation initiation factor 4B | Cytoplasm | translation regulator | 7316302.5 | 4894444.5 | 5080249.5 | | | | | | |
| Q8BGD9 | EIF4B | eukaryotic translation initiation factor 4B | Cytoplasm | translation regulator | 4614875 | 3984761.25 | 3307530 | | | | | | |
| Q8BGD9 | EIF4B | eukaryotic translation initiation factor 4B | Cytoplasm | translation regulator | 3308155 | 2873000.75 | 3379662.25 | | | | | | |
| Q8BGD9 | EIF4B | eukaryotic translation initiation factor 4B | Cytoplasm | translation regulator | 1789854.875 | 1509214.125 | 1677665.375 | Q8BGD9 | EIF4B | 5027933 | 1921002 | 452785 | 18 |
| Q6NZJ6 | EIF4G1 | eukaryotic translation initiation factor 4 gamma, 1 | Cytoplasm | translation regulator | 2691425.5 | 3229662.25 | 2399735.25 | | | | | | |
| Q6NZJ6 | EIF4G1 | eukaryotic translation initiation factor 4 gamma, 1 | Cytoplasm | translation regulator | 1068552.625 | 1746404 | 1642932.375 | | | | | | |
| Q6NZJ6 | EIF4G1 | eukaryotic translation initiation factor 4 gamma, 1 | Cytoplasm | translation regulator | 560755.562 | 518135.5 | 442589.219 | Q6NZJ6 | EIF4G1 | 1440245 | 1023551 | 341184 | 9 |
| P59325 | EIF5 | eukaryotic translation initiation factor 5 | Cytoplasm | translation regulator | 1331449.75 | 1662838 | 1172798.875 | P59325 | EIF5 | 1331450 | 250042 | 144362 | 3 |
| P63242 | EIF5A | eukaryotic translation initiation factor 5A | Cytoplasm | translation regulator | 1798071 | 2694153.75 | 2895170.5 | P63242 | EIF5A | 1798071 | 584095 | 337227 | 3 |
| Q99K41 | EMILIN1 | elastin microfibril interfacer 1 | Extracellular Space | other | 3814661.5 | 4373970 | 8251363 | | | | | | |
| Q99K41 | EMILIN1 | elastin microfibril interfacer 1 | Extracellular Space | other | 727432.375 | 586590 | 1070627.875 | Q99K41 | EMILIN1 | 2271047 | 2990870 | 1221018 | 6 |
| Q8C522 | ENDOD1 | endonuclease domain containing 1 | Extracellular Space | enzyme | 1.65E+07 | 1.74E+07 | 2.03E+07 | | | | | | |
| Q8C522 | ENDOD1 | endonuclease domain containing 1 | Extracellular Space | enzyme | 1.28E+07 | 8201581 | 1.12E+07 | | | | | | |
| Q8C522 | ENDOD1 | endonuclease domain containing 1 | Extracellular Space | enzyme | 7204756.5 | 5637040.5 | 1.12E+07 | | | | | | |
| Q8C522 | ENDOD1 | endonuclease domain containing 1 | Extracellular Space | enzyme | 5450230 | 6737150.5 | 5846068 | | | | | | |
| Q8C522 | ENDOD1 | endonuclease domain containing 1 | Extracellular Space | enzyme | 5628669 | 8695370 | 8896933 | | | | | | |
| Q8C522 | ENDOD1 | endonuclease domain containing 1 | Extracellular Space | enzyme | 5008120.5 | 6176837 | 5532261 | | | | | | |
| Q8C522 | ENDOD1 | endonuclease domain containing 1 | Extracellular Space | enzyme | 3936164.75 | 4064709 | 6766000 | | | | | | |
| Q8C522 | ENDOD1 | endonuclease domain containing 1 | Extracellular Space | enzyme | 5065893.5 | 6108561.5 | 6235098 | Q8C522 | ENDOD1 | 7689229 | 4394723 | 897069 | 24 |

| | | | | | | | | | | | | | |
|---|---|---|---|---|---|---|---|---|---|---|---|---|---|
| Q8BTJ4 | ENPP4 | ectonucleotide pyrophosphatase/phosphodiesterase 4 (putative) | Cytoplasm | enzyme | 1220418.25 | 2012575.25 | 2372657.5 | Q8BTJ4 | ENPP4 | 1220418 | 589467 | 340329 | 3 |
| P10404 | ENV1 | MLV-related proviral Env polyprotein | other | | 9.38E+07 | 1.04E+08 | 8.80E+07 | | | | | | |
| P10404 | ENV1 | MLV-related proviral Env polyprotein | other | | 6.32E+07 | 7.21E+07 | 6.26E+07 | | | | | | |
| P10404 | ENV1 | MLV-related proviral Env polyprotein | other | | 7.74E+07 | 6.57E+07 | 7.38E+07 | | | | | | |
| P10404 | ENV1 | MLV-related proviral Env polyprotein | other | | 3.95E+07 | 4.57E+07 | 4.40E+07 | | | | | | |
| P10404 | ENV1 | MLV-related proviral Env polyprotein | other | | 4.00E+07 | 5.33E+07 | 4.13E+07 | | | | | | |
| P10404 | ENV1 | MLV-related proviral Env polyprotein | other | | 4.19E+07 | 5.26E+07 | 6.34E+07 | | | | | | |
| P10404 | ENV1 | MLV-related proviral Env polyprotein | other | | 5.38E+07 | 5.62E+07 | 4.74E+07 | | | | | | |
| P10404 | ENV1 | MLV-related proviral Env polyprotein | other | | 3.75E+07 | 3.80E+07 | 3.96E+07 | | | | | | |
| P10404 | ENV1 | MLV-related proviral Env polyprotein | other | | 2.89E+07 | 3.61E+07 | 3.45E+07 | | | | | | |
| P10404 | ENV1 | MLV-related proviral Env polyprotein | other | | 3.60E+07 | 2.28E+07 | 3.69E+07 | | | | | | |
| P10404 | ENV1 | MLV-related proviral Env polyprotein | other | | 2.27E+07 | 2.57E+07 | 2.72E+07 | | | | | | |
| P10404 | ENV1 | MLV-related proviral Env polyprotein | other | | 2.20E+07 | 2.44E+07 | 2.52E+07 | | | | | | |
| P10404 | ENV1 | MLV-related proviral Env polyprotein | other | | 1.40E+07 | 1.43E+07 | 1.29E+07 | | | | | | |
| P10404 | ENV1 | MLV-related proviral Env polyprotein | other | | 1.47E+07 | 2.03E+07 | 1.87E+07 | | | | | | |
| P10404 | ENV1 | MLV-related proviral Env polyprotein | other | | 8564380 | 1.03E+07 | 1.29E+07 | | | | | | |
| P10404 | ENV1 | MLV-related proviral Env polyprotein | other | | 8677979 | 9578625 | 1.06E+07 | | | | | | |
| P10404 | ENV1 | MLV-related proviral Env polyprotein | other | | 8125784 | 8288646.5 | 8849026 | | | | | | |
| P10404 | ENV1 | MLV-related proviral Env polyprotein | other | | 6571926 | 7582242 | 7690717 | | | | | | |
| P10404 | ENV1 | MLV-related proviral Env polyprotein | other | | 5277430 | 5439148.5 | 6035480.5 | | | | | | |
| P10404 | ENV1 | MLV-related proviral Env polyprotein | other | | 7437177 | 1.20E+07 | 7547277.5 | | | | | | |
| P10404 | ENV1 | MLV-related proviral Env polyprotein | other | | 4703912 | 8145013.5 | 5723334 | | | | | | |
| P10404 | ENV1 | MLV-related proviral Env polyprotein | other | | 4534019 | 3715293.25 | 3407966.5 | | | | | | |
| P10404 | ENV1 | MLV-related proviral Env polyprotein | other | | 3054719.5 | 2312189 | 2705956.75 | | | | | | |
| P10404 | ENV1 | MLV-related proviral Env polyprotein | other | | 3167379.25 | 2846122 | 1851163.5 | P10404 | ENV1 | 26897696 | 25250767 | 2975831 | 72 |
| P48193 | EPB41 | erythrocyte membrane protein band 4.1 | Plasma Membrane | other | 2.60E+09 | 2.89E+09 | 1.83E+09 | | | | | | |
| P48193 | EPB41 | erythrocyte membrane protein band 4.1 | Plasma Membrane | other | 1.78E+09 | 1.76E+09 | 1.66E+09 | | | | | | |
| P48193 | EPB41 | erythrocyte membrane protein band 4.1 | Plasma Membrane | other | 2.05E+09 | 1.73E+09 | 1.44E+09 | | | | | | |
| P48193 | EPB41 | erythrocyte membrane protein band 4.1 | Plasma Membrane | other | 1.77E+09 | 1.84E+09 | 1.30E+09 | | | | | | |
| P48193 | EPB41 | erythrocyte membrane protein band 4.1 | Plasma Membrane | other | 1.36E+09 | 1.33E+09 | 1.58E+09 | | | | | | |
| P48193 | EPB41 | erythrocyte membrane protein band 4.1 | Plasma Membrane | other | 1.64E+09 | 1.45E+09 | 1.34E+09 | | | | | | |
| P48193 | EPB41 | erythrocyte membrane protein band 4.1 | Plasma Membrane | other | 1.47E+09 | 1.46E+09 | 1.27E+09 | | | | | | |
| P48193 | EPB41 | erythrocyte membrane protein band 4.1 | Plasma Membrane | other | 1.41E+09 | 1.62E+09 | 1.47E+09 | | | | | | |
| P48193 | EPB41 | erythrocyte membrane protein band 4.1 | Plasma Membrane | other | 1.30E+09 | 1.27E+09 | 9.40E+08 | | | | | | |
| P48193 | EPB41 | erythrocyte membrane protein band 4.1 | Plasma Membrane | other | 9.40E+08 | 9.58E+08 | 1.10E+09 | | | | | | |
| P48193 | EPB41 | erythrocyte membrane protein band 4.1 | Plasma Membrane | other | 1.20E+09 | 1.14E+09 | 1.01E+09 | | | | | | |
| P48193 | EPB41 | erythrocyte membrane protein band 4.1 | Plasma Membrane | other | 9.69E+08 | 9.11E+08 | 8.14E+08 | | | | | | |
| P48193 | EPB41 | erythrocyte membrane protein band 4.1 | Plasma Membrane | other | 1.14E+09 | 1.04E+09 | 9.09E+08 | | | | | | |
| P48193 | EPB41 | erythrocyte membrane protein band 4.1 | Plasma Membrane | other | 8.93E+08 | 9.13E+08 | 7.72E+08 | | | | | | |
| P48193 | EPB41 | erythrocyte membrane protein band 4.1 | Plasma Membrane | other | 9.65E+08 | 9.57E+08 | 8.67E+08 | | | | | | |
| P48193 | EPB41 | erythrocyte membrane protein band 4.1 | Plasma Membrane | other | 5.74E+08 | 5.91E+08 | 5.69E+08 | | | | | | |
| P48193 | EPB41 | erythrocyte membrane protein band 4.1 | Plasma Membrane | other | 6.02E+08 | 5.46E+08 | 4.60E+08 | | | | | | |
| P48193 | EPB41 | erythrocyte membrane protein band 4.1 | Plasma Membrane | other | 5.03E+08 | 4.69E+08 | 4.81E+08 | | | | | | |
| P48193 | EPB41 | erythrocyte membrane protein band 4.1 | Plasma Membrane | other | 6.86E+08 | 7.03E+08 | 4.28E+08 | | | | | | |
| P48193 | EPB41 | erythrocyte membrane protein band 4.1 | Plasma Membrane | other | 4.20E+08 | 3.22E+08 | 3.06E+08 | | | | | | |
| P48193 | EPB41 | erythrocyte membrane protein band 4.1 | Plasma Membrane | other | 4.49E+08 | 5.84E+08 | 2.70E+08 | | | | | | |
| P48193 | EPB41 | erythrocyte membrane protein band 4.1 | Plasma Membrane | other | 2.87E+08 | 2.71E+08 | 1.61E+08 | | | | | | |
| P48193 | EPB41 | erythrocyte membrane protein band 4.1 | Plasma Membrane | other | 2.83E+08 | 2.08E+08 | 2.54E+08 | | | | | | |
| P48193 | EPB41 | erythrocyte membrane protein band 4.1 | Plasma Membrane | other | 2.19E+08 | 2.23E+08 | 2.47E+08 | | | | | | |
| P48193 | EPB41 | erythrocyte membrane protein band 4.1 | Plasma Membrane | other | 2.84E+08 | 2.58E+08 | 2.70E+08 | | | | | | |
| P48193 | EPB41 | erythrocyte membrane protein band 4.1 | Plasma Membrane | other | 1.25E+08 | 1.23E+08 | 1.12E+08 | | | | | | |
| P48193 | EPB41 | erythrocyte membrane protein band 4.1 | Plasma Membrane | other | 1.73E+08 | 1.41E+08 | 1.34E+08 | | | | | | |
| P48193 | EPB41 | erythrocyte membrane protein band 4.1 | Plasma Membrane | other | 1.01E+08 | 1.17E+08 | 1.18E+08 | | | | | | |
| P48193 | EPB41 | erythrocyte membrane protein band 4.1 | Plasma Membrane | other | 1.31E+08 | 1.79E+08 | 1.78E+08 | | | | | | |
| P48193 | EPB41 | erythrocyte membrane protein band 4.1 | Plasma Membrane | other | 9.94E+07 | 1.33E+08 | 6.29E+07 | | | | | | |
| P48193 | EPB41 | erythrocyte membrane protein band 4.1 | Plasma Membrane | other | 7.91E+07 | 5.97E+07 | 4.87E+07 | | | | | | |
| P48193 | EPB41 | erythrocyte membrane protein band 4.1 | Plasma Membrane | other | 8.44E+07 | 9.16E+07 | 5.34E+07 | | | | | | |
| P48193 | EPB41 | erythrocyte membrane protein band 4.1 | Plasma Membrane | other | 5.97E+07 | 9.67E+07 | 8.33E+07 | | | | | | |
| P48193 | EPB41 | erythrocyte membrane protein band 4.1 | Plasma Membrane | other | 5.76E+07 | 6.49E+07 | 5.15E+07 | | | | | | |
| P48193 | EPB41 | erythrocyte membrane protein band 4.1 | Plasma Membrane | other | 5.29E+07 | 3.66E+07 | 3.98E+07 | | | | | | |
| P48193 | EPB41 | erythrocyte membrane protein band 4.1 | Plasma Membrane | other | 6.11E+07 | 4.77E+07 | 1.69E+07 | | | | | | |
| P48193 | EPB41 | erythrocyte membrane protein band 4.1 | Plasma Membrane | other | 5.01E+07 | 5.22E+07 | 4.14E+07 | | | | | | |
| P48193 | EPB41 | erythrocyte membrane protein band 4.1 | Plasma Membrane | other | 3.23E+07 | 2.41E+07 | 2.37E+07 | | | | | | |
| P48193 | EPB41 | erythrocyte membrane protein band 4.1 | Plasma Membrane | other | 5.47E+07 | 3.63E+07 | 3.98E+07 | | | | | | |
| P48193 | EPB41 | erythrocyte membrane protein band 4.1 | Plasma Membrane | other | 3.65E+07 | 3.79E+07 | 2.78E+07 | | | | | | |
| P48193 | EPB41 | erythrocyte membrane protein band 4.1 | Plasma Membrane | other | 4.05E+07 | 5.97E+07 | 2.52E+07 | | | | | | |
| P48193 | EPB41 | erythrocyte membrane protein band 4.1 | Plasma Membrane | other | 3.14E+07 | 3.48E+07 | 1.83E+07 | | | | | | |
| P48193 | EPB41 | erythrocyte membrane protein band 4.1 | Plasma Membrane | other | 2.59E+07 | 2.32E+07 | 2.49E+07 | | | | | | |
| P48193 | EPB41 | erythrocyte membrane protein band 4.1 | Plasma Membrane | other | 2.39E+07 | 2.12E+07 | 1.87E+07 | | | | | | |
| P48193 | EPB41 | erythrocyte membrane protein band 4.1 | Plasma Membrane | other | 1.45E+07 | 1.62E+07 | 1.82E+07 | | | | | | |
| P48193 | EPB41 | erythrocyte membrane protein band 4.1 | Plasma Membrane | other | 1.97E+07 | 2.12E+07 | 1.38E+07 | | | | | | |
| P48193 | EPB41 | erythrocyte membrane protein band 4.1 | Plasma Membrane | other | 1.04E+07 | 8295735 | 5410083.5 | | | | | | |
| P48193 | EPB41 | erythrocyte membrane protein band 4.1 | Plasma Membrane | other | 3.29E+07 | 7.70E+07 | 6.56E+07 | | | | | | |
| P48193 | EPB41 | erythrocyte membrane protein band 4.1 | Plasma Membrane | other | 1.33E+07 | 9765265 | 8047228 | | | | | | |
| P48193 | EPB41 | erythrocyte membrane protein band 4.1 | Plasma Membrane | other | 6.97E+07 | 8.57E+07 | 8.76E+07 | | | | | | |
| P48193 | EPB41 | erythrocyte membrane protein band 4.1 | Plasma Membrane | other | 2055439.75 | 1776146.5 | 1.41E+07 | | | | | | |
| P48193 | EPB41 | erythrocyte membrane protein band 4.1 | Plasma Membrane | other | 9087729 | 8664830 | 1.12E+07 | | | | | | |
| P48193 | EPB41 | erythrocyte membrane protein band 4.1 | Plasma Membrane | other | 4267767.5 | 5695468 | 3125826.75 | | | | | | |
| P48193 | EPB41 | erythrocyte membrane protein band 4.1 | Plasma Membrane | other | 5574013.5 | 5788949 | 5674285.5 | | | | | | |
| P48193 | EPB41 | erythrocyte membrane protein band 4.1 | Plasma Membrane | other | 4938247 | 3733253.5 | 3115462.75 | | | | | | |
| P48193 | EPB41 | erythrocyte membrane protein band 4.1 | Plasma Membrane | other | 2856644 | 4268831 | 5714109 | | | | | | |
| P48193 | EPB41 | erythrocyte membrane protein band 4.1 | Plasma Membrane | other | 4373928.5 | 4741010.5 | 5039716.5 | | | | | | |
| P48193 | EPB41 | erythrocyte membrane protein band 4.1 | Plasma Membrane | other | 1194701.625 | 340311.812 | 1892888.125 | | | | | | |
| P48193 | EPB41 | erythrocyte membrane protein band 4.1 | Plasma Membrane | other | 1943912.25 | 1740095.25 | 1719547.875 | | | | | | |
| P48193 | EPB41 | erythrocyte membrane protein band 4.1 | Plasma Membrane | other | 427306.312 | 375895.969 | 205579.156 | | | | | | |
| P48193 | EPB41 | erythrocyte membrane protein band 4.1 | Plasma Membrane | other | 651020.5 | 1719151.625 | 1.85E+07 | | | | | | |
| P48193 | EPB41 | erythrocyte membrane protein band 4.1 | Plasma Membrane | other | 420015.531 | 669377.688 | 608365.438 | P48193 | EPB41 | 440612270 | 593966695 | 43551751 | 186 |
| P49222 | EPB42 | erythrocyte membrane protein band 4.2 | Plasma Membrane | transporter | 1.80E+09 | 1.73E+09 | 1.78E+09 | | | | | | |
| P49222 | EPB42 | erythrocyte membrane protein band 4.2 | Plasma Membrane | transporter | 1.73E+09 | 1.65E+09 | 1.44E+09 | | | | | | |
| P49222 | EPB42 | erythrocyte membrane protein band 4.2 | Plasma Membrane | transporter | 1.49E+09 | 1.43E+09 | 1.40E+09 | | | | | | |
| P49222 | EPB42 | erythrocyte membrane protein band 4.2 | Plasma Membrane | transporter | 1.63E+09 | 1.60E+09 | 1.66E+09 | | | | | | |
| P49222 | EPB42 | erythrocyte membrane protein band 4.2 | Plasma Membrane | transporter | 1.46E+09 | 1.59E+09 | 1.04E+09 | | | | | | |
| P49222 | EPB42 | erythrocyte membrane protein band 4.2 | Plasma Membrane | transporter | 1.30E+09 | 1.32E+09 | 1.26E+09 | | | | | | |
| P49222 | EPB42 | erythrocyte membrane protein band 4.2 | Plasma Membrane | transporter | 1.32E+09 | 1.23E+09 | 1.26E+09 | | | | | | |
| P49222 | EPB42 | erythrocyte membrane protein band 4.2 | Plasma Membrane | transporter | 8.93E+08 | 9.04E+08 | 8.29E+08 | | | | | | |
| P49222 | EPB42 | erythrocyte membrane protein band 4.2 | Plasma Membrane | transporter | 8.32E+08 | 1.18E+09 | 9.52E+08 | | | | | | |
| P49222 | EPB42 | erythrocyte membrane protein band 4.2 | Plasma Membrane | transporter | 8.40E+08 | 9.66E+08 | 9.07E+08 | | | | | | |
| P49222 | EPB42 | erythrocyte membrane protein band 4.2 | Plasma Membrane | transporter | 4.42E+08 | 6.39E+08 | 3.92E+08 | | | | | | |
| P49222 | EPB42 | erythrocyte membrane protein band 4.2 | Plasma Membrane | transporter | 8.87E+08 | 1.03E+09 | 8.85E+08 | | | | | | |
| P49222 | EPB42 | erythrocyte membrane protein band 4.2 | Plasma Membrane | transporter | 7.20E+08 | 6.80E+08 | 5.84E+08 | | | | | | |
| P49222 | EPB42 | erythrocyte membrane protein band 4.2 | Plasma Membrane | transporter | 6.39E+08 | 6.87E+08 | 6.24E+08 | | | | | | |
| P49222 | EPB42 | erythrocyte membrane protein band 4.2 | Plasma Membrane | transporter | 5.31E+08 | 5.64E+08 | 4.92E+08 | | | | | | |
| P49222 | EPB42 | erythrocyte membrane protein band 4.2 | Plasma Membrane | transporter | 4.70E+08 | 5.20E+08 | 4.91E+08 | | | | | | |
| P49222 | EPB42 | erythrocyte membrane protein band 4.2 | Plasma Membrane | transporter | 4.83E+08 | 5.49E+08 | 4.58E+08 | | | | | | |
| P49222 | EPB42 | erythrocyte membrane protein band 4.2 | Plasma Membrane | transporter | 4.49E+08 | 4.97E+08 | 5.31E+08 | | | | | | |
| P49222 | EPB42 | erythrocyte membrane protein band 4.2 | Plasma Membrane | transporter | 4.51E+08 | 4.52E+08 | 3.85E+08 | | | | | | |
| P49222 | EPB42 | erythrocyte membrane protein band 4.2 | Plasma Membrane | transporter | 4.04E+08 | 3.78E+08 | 3.57E+08 | | | | | | |
| P49222 | EPB42 | erythrocyte membrane protein band 4.2 | Plasma Membrane | transporter | 2.58E+08 | 2.74E+08 | 2.33E+08 | | | | | | |
| P49222 | EPB42 | erythrocyte membrane protein band 4.2 | Plasma Membrane | transporter | 2.79E+08 | 2.32E+08 | 2.63E+08 | | | | | | |

| | | | | | | | | | | | | | |
|---|---|---|---|---|---|---|---|---|---|---|---|---|---|
| P49222 | EPB42 | erythrocyte membrane protein band 4.2 | Plasma Membrane | transporter | 2.78E+08 | 2.65E+08 | 2.59E+08 | | | | | | |
| P49222 | EPB42 | erythrocyte membrane protein band 4.2 | Plasma Membrane | transporter | 2.65E+08 | 3.28E+08 | 3.04E+08 | | | | | | |
| P49222 | EPB42 | erythrocyte membrane protein band 4.2 | Plasma Membrane | transporter | 2.61E+08 | 2.44E+08 | 1.98E+08 | | | | | | |
| P49222 | EPB42 | erythrocyte membrane protein band 4.2 | Plasma Membrane | transporter | 1.83E+08 | 2.05E+08 | 9.62E+07 | | | | | | |
| P49222 | EPB42 | erythrocyte membrane protein band 4.2 | Plasma Membrane | transporter | 1.70E+08 | 1.62E+08 | 1.26E+08 | | | | | | |
| P49222 | EPB42 | erythrocyte membrane protein band 4.2 | Plasma Membrane | transporter | 1.85E+08 | 2.33E+08 | 1.81E+08 | | | | | | |
| P49222 | EPB42 | erythrocyte membrane protein band 4.2 | Plasma Membrane | transporter | 1.62E+08 | 1.49E+08 | 1.24E+08 | | | | | | |
| P49222 | EPB42 | erythrocyte membrane protein band 4.2 | Plasma Membrane | transporter | 1.47E+08 | 2.14E+08 | 1.01E+08 | | | | | | |
| P49222 | EPB42 | erythrocyte membrane protein band 4.2 | Plasma Membrane | transporter | 1.43E+08 | 1.23E+08 | 1.04E+08 | | | | | | |
| P49222 | EPB42 | erythrocyte membrane protein band 4.2 | Plasma Membrane | transporter | 7.80E+07 | 9.86E+07 | 9.04E+07 | | | | | | |
| P49222 | EPB42 | erythrocyte membrane protein band 4.2 | Plasma Membrane | transporter | 1.33E+08 | 1.41E+08 | 1.16E+08 | | | | | | |
| P49222 | EPB42 | erythrocyte membrane protein band 4.2 | Plasma Membrane | transporter | 8.15E+07 | 4.25E+07 | 7.23E+07 | | | | | | |
| P49222 | EPB42 | erythrocyte membrane protein band 4.2 | Plasma Membrane | transporter | 4.95E+07 | 4.06E+07 | 4.20E+07 | | | | | | |
| P49222 | EPB42 | erythrocyte membrane protein band 4.2 | Plasma Membrane | transporter | 4.70E+07 | 5.32E+07 | 5.26E+07 | | | | | | |
| P49222 | EPB42 | erythrocyte membrane protein band 4.2 | Plasma Membrane | transporter | 7.18E+07 | 8.76E+07 | 7.94E+07 | | | | | | |
| P49222 | EPB42 | erythrocyte membrane protein band 4.2 | Plasma Membrane | transporter | 3.55E+07 | 3.42E+07 | 2.42E+07 | | | | | | |
| P49222 | EPB42 | erythrocyte membrane protein band 4.2 | Plasma Membrane | transporter | 2.85E+07 | 3.07E+07 | 3.38E+07 | | | | | | |
| P49222 | EPB42 | erythrocyte membrane protein band 4.2 | Plasma Membrane | transporter | 1.19E+07 | 6755447 | 1974267.25 | | | | | | |
| P49222 | EPB42 | erythrocyte membrane protein band 4.2 | Plasma Membrane | transporter | 2.80E+07 | 3.47E+07 | 2.84E+07 | | | | | | |
| P49222 | EPB42 | erythrocyte membrane protein band 4.2 | Plasma Membrane | transporter | 2.42E+07 | 2.95E+07 | 2.67E+07 | | | | | | |
| P49222 | EPB42 | erythrocyte membrane protein band 4.2 | Plasma Membrane | transporter | 4.48E+07 | 7.16E+07 | 4.22E+07 | | | | | | |
| P49222 | EPB42 | erythrocyte membrane protein band 4.2 | Plasma Membrane | transporter | 1.72E+07 | 1.86E+07 | 9541988 | | | | | | |
| P49222 | EPB42 | erythrocyte membrane protein band 4.2 | Plasma Membrane | transporter | 1.07E+07 | 1.40E+07 | 1.37E+07 | | | | | | |
| P49222 | EPB42 | erythrocyte membrane protein band 4.2 | Plasma Membrane | transporter | 4060449.5 | 2975318 | 7258552.5 | | | | | | |
| P49222 | EPB42 | erythrocyte membrane protein band 4.2 | Plasma Membrane | transporter | 1.32E+09 | 1.21E+09 | 1.03E+09 | | | | | | |
| P49222 | EPB42 | erythrocyte membrane protein band 4.2 | Plasma Membrane | transporter | 1.46E+07 | 1.58E+07 | 9555397 | | | | | | |
| P49222 | EPB42 | erythrocyte membrane protein band 4.2 | Plasma Membrane | transporter | 6076739 | 5099111 | 4229430.5 | | | | | | |
| P49222 | EPB42 | erythrocyte membrane protein band 4.2 | Plasma Membrane | transporter | 6016695.5 | 4694270.5 | 3803527.25 | | | | | | |
| P49222 | EPB42 | erythrocyte membrane protein band 4.2 | Plasma Membrane | transporter | 1772228.625 | 2040819 | 5108664.5 | | | | | | |
| P49222 | EPB42 | erythrocyte membrane protein band 4.2 | Plasma Membrane | transporter | 7925272.5 | 7086474.5 | 2836033.25 | | | | | | |
| P49222 | EPB42 | erythrocyte membrane protein band 4.2 | Plasma Membrane | transporter | 5011641.5 | 8000532 | 5091356 | | | | | | |
| P49222 | EPB42 | erythrocyte membrane protein band 4.2 | Plasma Membrane | transporter | 2384879.5 | 2620421.75 | 4180480.5 | | | | | | |
| P49222 | EPB42 | erythrocyte membrane protein band 4.2 | Plasma Membrane | transporter | 3512438.75 | 5599302.5 | 7812000 | | | | | | |
| P49222 | EPB42 | erythrocyte membrane protein band 4.2 | Plasma Membrane | transporter | 6128472 | 6602603.5 | 3410312.5 | | | | | | |
| P49222 | EPB42 | erythrocyte membrane protein band 4.2 | Plasma Membrane | transporter | 747364.938 | 624013.75 | 1294581.375 | | | | | | |
| P49222 | EPB42 | erythrocyte membrane protein band 4.2 | Plasma Membrane | transporter | 3412476.25 | 1.29E+07 | 1.40E+07 | | | | | | |
| P49222 | EPB42 | erythrocyte membrane protein band 4.2 | Plasma Membrane | transporter | 1196974.75 | 988609.688 | 1231336 | | | | | | |
| P49222 | EPB42 | erythrocyte membrane protein band 4.2 | Plasma Membrane | transporter | 708563.125 | 966515.812 | 919138.5 | | | | | | |
| P49222 | EPB42 | erythrocyte membrane protein band 4.2 | Plasma Membrane | transporter | 112654.086 | 423632.312 | 1300569.75 | | | | | | |
| P49222 | EPB42 | erythrocyte membrane protein band 4.2 | Plasma Membrane | transporter | 7.10E+07 | 5.38E+07 | 4.87E+07 | | | | | | |
| P49222 | EPB42 | erythrocyte membrane protein band 4.2 | Plasma Membrane | transporter | 87985.336 | 263409.25 | 116579.578 | | | | | | |
| P49222 | EPB42 | erythrocyte membrane protein band 4.2 | Plasma Membrane | transporter | 457167.594 | 1804197.375 | 721293.062 | P49222 | EPB42 | 362571281 | 490807979 | 35421015 | 192 |
| Q80VP1 | EPN1 | epsin 1 | Plasma Membrane | other | 4028695.75 | 2913067 | 2386856 | | | | | | |
| Q80VP1 | EPN1 | epsin 1 | Plasma Membrane | other | 4160350.75 | 2957316 | 2851064 | | | | | | |
| Q80VP1 | EPN1 | epsin 1 | Plasma Membrane | other | 2070377.75 | 1474414.25 | 1426051.75 | | | | | | |
| Q80VP1 | EPN1 | epsin 1 | Plasma Membrane | other | 1746810 | 1109578.75 | 1090358.25 | Q80VP1 | EPN1 | 3001559 | 1054056 | 304280 | 12 |
| Q8CGC7 | EPRS | glutamyl-prolyl-tRNA synthetase | Cytoplasm | enzyme | 580045.938 | 853797.812 | 904485.188 | | | | | | |
| Q8CGC7 | EPRS | glutamyl-prolyl-tRNA synthetase | Cytoplasm | enzyme | 447463.375 | 614942.125 | 562425.188 | Q8CGC7 | EPRS | 513755 | 179134 | 73131 | 6 |
| P42567 | EPS15 | epidermal growth factor receptor pathway substrate 15 | Cytoplasm | other | 4301683.5 | 3586755 | 3083073 | | | | | | |
| P42567 | EPS15 | epidermal growth factor receptor pathway substrate 15 | Cytoplasm | other | 3529004.75 | 4939946.5 | 2763447.5 | | | | | | |
| P42567 | EPS15 | epidermal growth factor receptor pathway substrate 15 | Cytoplasm | other | 3022593.75 | 2496215.75 | 1976153.5 | | | | | | |
| P42567 | EPS15 | epidermal growth factor receptor pathway substrate 15 | Cytoplasm | other | 2738414.5 | 3339975.75 | 2457638.5 | | | | | | |
| P42567 | EPS15 | epidermal growth factor receptor pathway substrate 15 | Cytoplasm | other | 2422218.25 | 2197910.75 | 1727883.375 | P42567 | EPS15 | 3202783 | 866411 | 223706 | 15 |
| Q60902 | EPS15L1 | epidermal growth factor receptor pathway substrate 15-like 1 | Plasma Membrane | other | 292821.906 | 272481.938 | 229884.766 | Q60902 | EPS15L1 | 292822 | 32118 | 18543 | 3 |
| Q9EQH2 | ERAP1 | endoplasmic reticulum aminopeptidase 1 | Extracellular Space | peptidase | 2855649.75 | 3149293.75 | 2295018.75 | Q9EQH2 | ERAP1 | 2855650 | 434035 | 250590 | 3 |
| Q80TH2 | ERBB2IP | erbb2 interacting protein | Plasma Membrane | other | 5724877 | 5588555 | 9258428 | | | | | | |
| Q80TH2 | ERBB2IP | erbb2 interacting protein | Plasma Membrane | other | 4654929 | 6984066 | 8101810.5 | | | | | | |
| Q80TH2 | ERBB2IP | erbb2 interacting protein | Plasma Membrane | other | 2960161.75 | 5049645 | 4515044 | | | | | | |
| Q80TH2 | ERBB2IP | erbb2 interacting protein | Plasma Membrane | other | 3472207 | 3779732.75 | 5432809.5 | | | | | | |
| Q80TH2 | ERBB2IP | erbb2 interacting protein | Plasma Membrane | other | 3034279.75 | 3658045.75 | 4073915.25 | | | | | | |
| Q80TH2 | ERBB2IP | erbb2 interacting protein | Plasma Membrane | other | 1759593.25 | 2732183.5 | 2743938 | | | | | | |
| Q80TH2 | ERBB2IP | erbb2 interacting protein | Plasma Membrane | other | 2473421.75 | 3379080.25 | 4502982.5 | | | | | | |
| Q80TH2 | ERBB2IP | erbb2 interacting protein | Plasma Membrane | other | 1552079.5 | 2467896.25 | 2913075.5 | Q80TH2 | ERBB2IP | 3203944 | 1921514 | 392227 | 24 |
| Q8BFZ9 | ERLIN2 | ER lipid raft associated 2 | Plasma Membrane | other | 2741868.25 | 2688059 | 3153670 | Q8BFZ9 | ERLIN2 | 2741868 | 254712 | 147058 | 3 |
| Q9JLN5 | ERMAP | erythroblast membrane-associated protein (Scianna blood group) | Cytoplasm | other | 1.70E+08 | 1.73E+08 | 1.67E+08 | | | | | | |
| Q9JLN5 | ERMAP | erythroblast membrane-associated protein (Scianna blood group) | Cytoplasm | other | 1.04E+08 | 1.10E+08 | 1.11E+08 | | | | | | |
| Q9JLN5 | ERMAP | erythroblast membrane-associated protein (Scianna blood group) | Cytoplasm | other | 8.40E+07 | 8.23E+07 | 7.20E+07 | | | | | | |
| Q9JLN5 | ERMAP | erythroblast membrane-associated protein (Scianna blood group) | Cytoplasm | other | 1.08E+08 | 1.11E+08 | 7.90E+07 | | | | | | |
| Q9JLN5 | ERMAP | erythroblast membrane-associated protein (Scianna blood group) | Cytoplasm | other | 7.80E+07 | 7.09E+07 | 6.40E+07 | | | | | | |
| Q9JLN5 | ERMAP | erythroblast membrane-associated protein (Scianna blood group) | Cytoplasm | other | 1.02E+08 | 9.10E+07 | 7.47E+07 | | | | | | |
| Q9JLN5 | ERMAP | erythroblast membrane-associated protein (Scianna blood group) | Cytoplasm | other | 6.00E+07 | 6.49E+07 | 5.34E+07 | | | | | | |
| Q9JLN5 | ERMAP | erythroblast membrane-associated protein (Scianna blood group) | Cytoplasm | other | 4.94E+07 | 4.15E+07 | 2.86E+07 | | | | | | |
| Q9JLN5 | ERMAP | erythroblast membrane-associated protein (Scianna blood group) | Cytoplasm | other | 3.51E+07 | 2.85E+07 | 2.49E+07 | | | | | | |
| Q9JLN5 | ERMAP | erythroblast membrane-associated protein (Scianna blood group) | Cytoplasm | other | 2.95E+07 | 3.32E+07 | 2.34E+07 | | | | | | |
| Q9JLN5 | ERMAP | erythroblast membrane-associated protein (Scianna blood group) | Cytoplasm | other | 2.64E+07 | 2.29E+07 | 1.99E+07 | | | | | | |
| Q9JLN5 | ERMAP | erythroblast membrane-associated protein (Scianna blood group) | Cytoplasm | other | 1.75E+07 | 1.45E+07 | 1.01E+07 | | | | | | |
| Q9JLN5 | ERMAP | erythroblast membrane-associated protein (Scianna blood group) | Cytoplasm | other | 1.35E+07 | 1.54E+07 | 1.39E+07 | | | | | | |
| Q9JLN5 | ERMAP | erythroblast membrane-associated protein (Scianna blood group) | Cytoplasm | other | 1.16E+07 | 1.47E+07 | 9876797 | | | | | | |
| Q9JLN5 | ERMAP | erythroblast membrane-associated protein (Scianna blood group) | Cytoplasm | other | 1.12E+07 | 9538066 | 9488826 | | | | | | |
| Q9JLN5 | ERMAP | erythroblast membrane-associated protein (Scianna blood group) | Cytoplasm | other | 9281846 | 9000547 | 4179876 | | | | | | |
| Q9JLN5 | ERMAP | erythroblast membrane-associated protein (Scianna blood group) | Cytoplasm | other | 7206918.5 | 6170681 | 4501264 | | | | | | |
| Q9JLN5 | ERMAP | erythroblast membrane-associated protein (Scianna blood group) | Cytoplasm | other | 5147238.5 | 5186596.5 | 6006909.5 | | | | | | |
| Q9JLN5 | ERMAP | erythroblast membrane-associated protein (Scianna blood group) | Cytoplasm | other | 3245845.25 | 1.20E+07 | 9652675 | | | | | | |
| Q9JLN5 | ERMAP | erythroblast membrane-associated protein (Scianna blood group) | Cytoplasm | other | 8561586 | 7501441.5 | 5646973.5 | Q9JLN5 | ERMAP | 46658172 | 44860872 | 5791514 | 60 |
| Q8R180 | ERO1A | endoplasmic reticulum oxidoreductase alpha | Cytoplasm | enzyme | 1476509.375 | 1778271 | 1699305.125 | Q8R180 | ERO1A | 1476509 | 156489 | 90349 | 3 |
| P57759 | ERP29 | endoplasmic reticulum protein 29 | Cytoplasm | transporter | 1.04E+07 | 1.19E+07 | 9188409 | | | | | | |
| P57759 | ERP29 | endoplasmic reticulum protein 29 | Cytoplasm | transporter | 5967798.5 | 7328283.5 | 5835671 | | | | | | |
| P57759 | ERP29 | endoplasmic reticulum protein 29 | Cytoplasm | transporter | 3866552 | 4627795 | 3943600.75 | | | | | | |
| P57759 | ERP29 | endoplasmic reticulum protein 29 | Cytoplasm | transporter | 2411618.75 | 4395569 | 4597645.5 | P57759 | ERP29 | 5651492 | 2911661 | 840524 | 12 |
| Q9D1Q6 | ERP44 | endoplasmic reticulum protein 44 | Cytoplasm | enzyme | 1.73E+07 | 1.56E+07 | 1.41E+07 | | | | | | |
| Q9D1Q6 | ERP44 | endoplasmic reticulum protein 44 | Cytoplasm | enzyme | 1.44E+07 | 1.39E+07 | 1.05E+07 | | | | | | |
| Q9D1Q6 | ERP44 | endoplasmic reticulum protein 44 | Cytoplasm | enzyme | 9913130 | 1.02E+07 | 1.03E+07 | | | | | | |
| Q9D1Q6 | ERP44 | endoplasmic reticulum protein 44 | Cytoplasm | enzyme | 9377481 | 9382930 | 7157188 | | | | | | |
| Q9D1Q6 | ERP44 | endoplasmic reticulum protein 44 | Cytoplasm | enzyme | 5991122.5 | 5842462.5 | 2726032.75 | | | | | | |
| Q9D1Q6 | ERP44 | endoplasmic reticulum protein 44 | Cytoplasm | enzyme | 2876630.75 | 2640084.75 | 3553580.5 | Q9D1Q6 | ERP44 | 9968061 | 4645364 | 1094923 | 18 |
| Q3U7R1 | ESYT1 | extended synaptotagmin-like protein 1 | Cytoplasm | other | 1.98E+07 | 2.36E+07 | 2.03E+07 | | | | | | |
| Q3U7R1 | ESYT1 | extended synaptotagmin-like protein 1 | Cytoplasm | other | 2.45E+07 | 2.83E+07 | 2.98E+07 | | | | | | |
| Q3U7R1 | ESYT1 | extended synaptotagmin-like protein 1 | Cytoplasm | other | 2.56E+07 | 2.25E+07 | 2.63E+07 | | | | | | |
| Q3U7R1 | ESYT1 | extended synaptotagmin-like protein 1 | Cytoplasm | other | 1.92E+07 | 1.63E+07 | 1.71E+07 | | | | | | |
| Q3U7R1 | ESYT1 | extended synaptotagmin-like protein 1 | Cytoplasm | other | 1.74E+07 | 2.07E+07 | 1.92E+07 | | | | | | |
| Q3U7R1 | ESYT1 | extended synaptotagmin-like protein 1 | Cytoplasm | other | 1.39E+07 | 1.49E+07 | 1.70E+07 | | | | | | |
| Q3U7R1 | ESYT1 | extended synaptotagmin-like protein 1 | Cytoplasm | other | 1.32E+07 | 1.47E+07 | 1.57E+07 | | | | | | |
| Q3U7R1 | ESYT1 | extended synaptotagmin-like protein 1 | Cytoplasm | other | 1.29E+07 | 1.85E+07 | 1.58E+07 | | | | | | |
| Q3U7R1 | ESYT1 | extended synaptotagmin-like protein 1 | Cytoplasm | other | 1.18E+07 | 1.33E+07 | 1.39E+07 | | | | | | |
| Q3U7R1 | ESYT1 | extended synaptotagmin-like protein 1 | Cytoplasm | other | 1.33E+07 | 1.14E+07 | 1.20E+07 | | | | | | |
| Q3U7R1 | ESYT1 | extended synaptotagmin-like protein 1 | Cytoplasm | other | 9880600 | 8498186 | 8215940 | | | | | | |
| Q3U7R1 | ESYT1 | extended synaptotagmin-like protein 1 | Cytoplasm | other | 1.19E+07 | 1.34E+07 | 1.40E+07 | | | | | | |
| Q3U7R1 | ESYT1 | extended synaptotagmin-like protein 1 | Cytoplasm | other | 8605651 | 9720484 | 8262285 | | | | | | |
| Q3U7R1 | ESYT1 | extended synaptotagmin-like protein 1 | Cytoplasm | other | 7510188.5 | 7994172.5 | 9956936 | | | | | | |

| | | | | | | | | | | | | | |
|---|---|---|---|---|---|---|---|---|---|---|---|---|---|
| Q3U7R1 | ESYT1 | extended synaptotagmin-like protein 1 | Cytoplasm | other | 7273450.5 | 8195492 | 8129638 | | | | | | |
| Q3U7R1 | ESYT1 | extended synaptotagmin-like protein 1 | Cytoplasm | other | 7604276 | 7635605.5 | 8981008 | | | | | | |
| Q3U7R1 | ESYT1 | extended synaptotagmin-like protein 1 | Cytoplasm | other | 1.04E+07 | 8891939 | 9946768 | | | | | | |
| Q3U7R1 | ESYT1 | extended synaptotagmin-like protein 1 | Cytoplasm | other | 7888080 | 6663441 | 7005034 | | | | | | |
| Q3U7R1 | ESYT1 | extended synaptotagmin-like protein 1 | Cytoplasm | other | 6430892.5 | 6480607 | 6905765.5 | | | | | | |
| Q3U7R1 | ESYT1 | extended synaptotagmin-like protein 1 | Cytoplasm | other | 6043364 | 4555192.5 | 7704935 | | | | | | |
| Q3U7R1 | ESYT1 | extended synaptotagmin-like protein 1 | Cytoplasm | other | 5411333 | 5574912 | 6579229 | | | | | | |
| Q3U7R1 | ESYT1 | extended synaptotagmin-like protein 1 | Cytoplasm | other | 7737325 | 6131385 | 7137712 | | | | | | |
| Q3U7R1 | ESYT1 | extended synaptotagmin-like protein 1 | Cytoplasm | other | 6217296.5 | 5912550.5 | 7289717.5 | | | | | | |
| Q3U7R1 | ESYT1 | extended synaptotagmin-like protein 1 | Cytoplasm | other | 6510724.5 | 7624355 | 7047196.5 | | | | | | |
| Q3U7R1 | ESYT1 | extended synaptotagmin-like protein 1 | Cytoplasm | other | 6329052.5 | 1.09E+07 | 6096275.5 | | | | | | |
| Q3U7R1 | ESYT1 | extended synaptotagmin-like protein 1 | Cytoplasm | other | 4295202 | 3639668.75 | 4251494.5 | | | | | | |
| Q3U7R1 | ESYT1 | extended synaptotagmin-like protein 1 | Cytoplasm | other | 3261648.75 | 4935044 | 2891615.25 | | | | | | |
| Q3U7R1 | ESYT1 | extended synaptotagmin-like protein 1 | Cytoplasm | other | 3768017 | 3731306.5 | 3782139.75 | | | | | | |
| Q3U7R1 | ESYT1 | extended synaptotagmin-like protein 1 | Cytoplasm | other | 1015543.375 | 1391945.25 | 808485.188 | Q3U7R1 | ESYT1 | 10333539 | 6487684 | 695553 | 87 |
| Q3TZZ7 | ESYT2 | extended synaptotagmin-like protein 2 | Plasma Membrane | other | 1.03E+07 | 1.15E+07 | 9747404 | | | | | | |
| Q3TZZ7 | ESYT2 | extended synaptotagmin-like protein 2 | Plasma Membrane | other | 1.01E+07 | 1.14E+07 | 9183769 | | | | | | |
| Q3TZZ7 | ESYT2 | extended synaptotagmin-like protein 2 | Plasma Membrane | other | 1.03E+07 | 1.13E+07 | 1.20E+07 | | | | | | |
| Q3TZZ7 | ESYT2 | extended synaptotagmin-like protein 2 | Plasma Membrane | other | 1.01E+07 | 9530075 | 9517415 | | | | | | |
| Q3TZZ7 | ESYT2 | extended synaptotagmin-like protein 2 | Plasma Membrane | other | 8120774 | 8209738.5 | 5935367 | | | | | | |
| Q3TZZ7 | ESYT2 | extended synaptotagmin-like protein 2 | Plasma Membrane | other | 8833599 | 1.12E+07 | 6513365.5 | | | | | | |
| Q3TZZ7 | ESYT2 | extended synaptotagmin-like protein 2 | Plasma Membrane | other | 7060855.5 | 8362259.5 | 7486787.5 | | | | | | |
| Q3TZZ7 | ESYT2 | extended synaptotagmin-like protein 2 | Plasma Membrane | other | 8194389.5 | 8564057 | 7175734.5 | | | | | | |
| Q3TZZ7 | ESYT2 | extended synaptotagmin-like protein 2 | Plasma Membrane | other | 7389492 | 8278831 | 7005034 | | | | | | |
| Q3TZZ7 | ESYT2 | extended synaptotagmin-like protein 2 | Plasma Membrane | other | 5459235 | 8371670.5 | 6032463.5 | | | | | | |
| Q3TZZ7 | ESYT2 | extended synaptotagmin-like protein 2 | Plasma Membrane | other | 8086173.5 | 1.15E+07 | 7507248 | | | | | | |
| Q3TZZ7 | ESYT2 | extended synaptotagmin-like protein 2 | Plasma Membrane | other | 5797389.5 | 5234672.5 | 4828358 | | | | | | |
| Q3TZZ7 | ESYT2 | extended synaptotagmin-like protein 2 | Plasma Membrane | other | 5864443.5 | 5712193 | 5023135 | | | | | | |
| Q3TZZ7 | ESYT2 | extended synaptotagmin-like protein 2 | Plasma Membrane | other | 4493066.5 | 6075950.5 | 3967933.75 | | | | | | |
| Q3TZZ7 | ESYT2 | extended synaptotagmin-like protein 2 | Plasma Membrane | other | 8170241 | 6998243.5 | 6918444 | | | | | | |
| Q3TZZ7 | ESYT2 | extended synaptotagmin-like protein 2 | Plasma Membrane | other | 5296063 | 5098099 | 5926476.5 | | | | | | |
| Q3TZZ7 | ESYT2 | extended synaptotagmin-like protein 2 | Plasma Membrane | other | 4064769 | 4159854.5 | 1928746.125 | | | | | | |
| Q3TZZ7 | ESYT2 | extended synaptotagmin-like protein 2 | Plasma Membrane | other | 4818981.5 | 4664066.5 | 1501564.875 | | | | | | |
| Q3TZZ7 | ESYT2 | extended synaptotagmin-like protein 2 | Plasma Membrane | other | 1352101.75 | 1200695 | 936052 | | | | | | |
| Q3TZZ7 | ESYT2 | extended synaptotagmin-like protein 2 | Plasma Membrane | other | 140422.922 | 271645.281 | 264047.844 | Q3TZZ7 | ESYT2 | 6693100 | 3096717 | 399784 | 60 |
| Q99LC5 | ETFA | electron-transfer-flavoprotein, alpha polypeptide | Cytoplasm | transporter | 5550040 | 1.02E+07 | 1.25E+07 | | | | | | |
| Q99LC5 | ETFA | electron-transfer-flavoprotein, alpha polypeptide | Cytoplasm | transporter | 4249357 | 7343489 | 9993740 | | | | | | |
| Q99LC5 | ETFA | electron-transfer-flavoprotein, alpha polypeptide | Cytoplasm | transporter | 4733361.5 | 8995821 | 7962785 | | | | | | |
| Q99LC5 | ETFA | electron-transfer-flavoprotein, alpha polypeptide | Cytoplasm | transporter | 4240424 | 4657730.5 | 6688091 | | | | | | |
| Q99LC5 | ETFA | electron-transfer-flavoprotein, alpha polypeptide | Cytoplasm | transporter | 1830117.125 | 2170353.5 | 3000960.75 | | | | | | |
| Q99LC5 | ETFA | electron-transfer-flavoprotein, alpha polypeptide | Cytoplasm | transporter | 2097573.5 | 3258376.25 | 3964095.5 | | | | | | |
| Q99LC5 | ETFA | electron-transfer-flavoprotein, alpha polypeptide | Cytoplasm | transporter | 1636135.5 | 2800080.75 | 3538550.25 | | | | | | |
| Q99LC5 | ETFA | electron-transfer-flavoprotein, alpha polypeptide | Cytoplasm | transporter | 1474707.375 | 2439519.25 | 2989286 | | | | | | |
| Q99LC5 | ETFA | electron-transfer-flavoprotein, alpha polypeptide | Cytoplasm | transporter | 268561.938 | 568142.438 | 1016027.75 | Q99LC5 | ETFA | 2897809 | 3213044 | 618351 | 27 |
| Q9DCW4 | ETFB | electron-transfer-flavoprotein, beta polypeptide | Cytoplasm | transporter | 8481501 | 1.36E+07 | 1.42E+07 | | | | | | |
| Q9DCW4 | ETFB | electron-transfer-flavoprotein, beta polypeptide | Cytoplasm | transporter | 2466027.25 | 4173518.25 | 4067792 | | | | | | |
| Q9DCW4 | ETFB | electron-transfer-flavoprotein, beta polypeptide | Cytoplasm | transporter | 1414197 | 3485712.25 | 4249330.5 | | | | | | |
| Q9DCW4 | ETFB | electron-transfer-flavoprotein, beta polypeptide | Cytoplasm | transporter | 1026661.125 | 1969038.25 | 3658961 | Q9DCW4 | ETFB | 3347097 | 4479742 | 1293190 | 12 |
| Q6DIA2 | EXOC3L4 | exocyst complex component 3-like 4 | Cytoplasm | other | 878378.625 | 1459046.5 | 2241659.5 | Q6DIA2 | EXOC3L4 | 878379 | 684129 | 394982 | 3 |
| P26040 | EZR | ezrin | Plasma Membrane | other | 2.76E+07 | 2.08E+07 | 2.38E+07 | | | | | | |
| P26040 | EZR | ezrin | Plasma Membrane | other | 1.77E+07 | 1.20E+07 | 1.56E+07 | | | | | | |
| P26040 | EZR | ezrin | Plasma Membrane | other | 1.26E+07 | 7981891 | 1.06E+07 | | | | | | |
| P26040 | EZR | ezrin | Plasma Membrane | other | 7468004 | 5088179.5 | 6469600.5 | | | | | | |
| P26040 | EZR | ezrin | Plasma Membrane | other | 5893689.5 | 4778205.5 | 7580914.5 | | | | | | |
| P26040 | EZR | ezrin | Plasma Membrane | other | 2168498.25 | 882800.375 | 2265269.25 | | | | | | |
| P26040 | EZR | ezrin | Plasma Membrane | other | 4507114 | 3944302.75 | 3819408.5 | | | | | | |
| P26040 | EZR | ezrin | Plasma Membrane | other | 3043582.25 | 1709547.625 | 2924927.5 | | | | | | |
| P26040 | EZR | ezrin | Plasma Membrane | other | 2967388 | 1630160.375 | 1959287.5 | | | | | | |
| P26040 | EZR | ezrin | Plasma Membrane | other | 727147.625 | 3528.697 | 4.70E-06 | P26040 | EZR | 8464542 | 7283222 | 1329728 | 30 |
| O88792 | F11R | F11 receptor | Plasma Membrane | other | 6565956.5 | 1.02E+07 | 1.69E+07 | | | | | | |
| O88792 | F11R | F11 receptor | Plasma Membrane | other | 2876309.25 | 3654706 | 3318494.25 | | | | | | |
| O88792 | F11R | F11 receptor | Plasma Membrane | other | 1826303.125 | 2046165.375 | 4712939 | | | | | | |
| O88792 | F11R | F11 receptor | Plasma Membrane | other | 2006861 | 2414134 | 3785524 | | | | | | |
| O88792 | F11R | F11 receptor | Plasma Membrane | other | 2134455.25 | 3192884.5 | 4433388.5 | O88792 | F11R | 3081977 | 4017264 | 1037253 | 15 |
| P19221 | F2 | coagulation factor II (thrombin) | Extracellular Space | peptidase | 5027635.5 | 5294832.5 | 5935409.5 | P19221 | F2 | 5027636 | 466510 | 269339 | 3 |
| O88783 | F5 | coagulation factor V (proaccelerin, labile factor) | Plasma Membrane | enzyme | 3537995 | 3720309 | 5422293.5 | | | | | | |
| O88783 | F5 | coagulation factor V (proaccelerin, labile factor) | Plasma Membrane | enzyme | 924933.938 | 1117454.375 | 1880631.375 | | | | | | |
| O88783 | F5 | coagulation factor V (proaccelerin, labile factor) | Plasma Membrane | enzyme | 553688.625 | 913192.875 | 1394331.5 | O88783 | F5 | 1672206 | 1672845 | 557615 | 9 |
| Q920L1 | FADS1 | fatty acid desaturase 1 | Plasma Membrane | enzyme | 2971988 | 2758369 | 2574420 | Q920L1 | FADS1 | 2971988 | 198968 | 114874 | 3 |
| P54731 | FAF1 | Fas (TNFRSF6) associated factor 1 | Nucleus | other | 3241586.25 | 1928099.875 | 1640114.625 | | | | | | |
| P54731 | FAF1 | Fas (TNFRSF6) associated factor 1 | Nucleus | other | 3114123 | 1971746.25 | 1967640 | | | | | | |
| P54731 | FAF1 | Fas (TNFRSF6) associated factor 1 | Nucleus | other | 3353566.25 | 2369058 | 2423303.25 | P54731 | FAF1 | 3236425 | 640398 | 213466 | 9 |
| Q8R0F8 | FAHD1 | fumarylacetoacetate hydrolase domain containing 1 | Cytoplasm | enzyme | 1541488.875 | 2859317.75 | 3181116.5 | | | | | | |
| Q8R0F8 | FAHD1 | fumarylacetoacetate hydrolase domain containing 1 | Cytoplasm | enzyme | 1384807 | 3449344 | 4368459 | Q8R0F8 | FAHD1 | 1463148 | 1150397 | 469647 | 6 |
| Q6NS82 | FAM134A | family with sequence similarity 134, member A | Extracellular Space | other | 2662777.5 | 2579938.25 | 2164750.75 | Q6NS82 | FAM134A | 2662778 | 266856 | 154070 | 3 |
| Q9D8B6 | FAM210B | family with sequence similarity 210, member B | Extracellular Space | other | 2408904 | 3407964.75 | 3299209.75 | | | | | | |
| Q9D8B6 | FAM210B | family with sequence similarity 210, member B | Extracellular Space | other | 205797.125 | 528804.75 | 323369.438 | Q9D8B6 | FAM210B | 1307351 | 1515025 | 618506 | 6 |
| Q9CYH2 | FAM213A | family with sequence similarity 213, member A | Extracellular Space | other | 1.57E+08 | 1.96E+08 | 1.36E+08 | | | | | | |
| Q9CYH2 | FAM213A | family with sequence similarity 213, member A | Extracellular Space | other | 1.27E+08 | 1.31E+08 | 1.03E+08 | | | | | | |
| Q9CYH2 | FAM213A | family with sequence similarity 213, member A | Extracellular Space | other | 9.19E+07 | 1.19E+08 | 7.39E+07 | | | | | | |
| Q9CYH2 | FAM213A | family with sequence similarity 213, member A | Extracellular Space | other | 1.12E+08 | 1.18E+08 | 7.06E+07 | | | | | | |
| Q9CYH2 | FAM213A | family with sequence similarity 213, member A | Extracellular Space | other | 6.90E+07 | 4.26E+07 | 5.09E+07 | | | | | | |
| Q9CYH2 | FAM213A | family with sequence similarity 213, member A | Extracellular Space | other | 4.70E+07 | 5.18E+07 | 4.02E+07 | | | | | | |
| Q9CYH2 | FAM213A | family with sequence similarity 213, member A | Extracellular Space | other | 1.63E+07 | 1.62E+07 | 1.88E+07 | | | | | | |
| Q9CYH2 | FAM213A | family with sequence similarity 213, member A | Extracellular Space | other | 2.43E+07 | 2.69E+07 | 1.36E+07 | | | | | | |
| Q9CYH2 | FAM213A | family with sequence similarity 213, member A | Extracellular Space | other | 1.48E+07 | 1.55E+07 | 1.02E+07 | | | | | | |
| Q9CYH2 | FAM213A | family with sequence similarity 213, member A | Extracellular Space | other | 1.47E+07 | 8519458 | 1.44E+07 | | | | | | |
| Q9CYH2 | FAM213A | family with sequence similarity 213, member A | Extracellular Space | other | 1.09E+07 | 9668822 | 7768984 | | | | | | |
| Q9CYH2 | FAM213A | family with sequence similarity 213, member A | Extracellular Space | other | 1597062.125 | 1713113.75 | 1090729.625 | Q9CYH2 | FAM213A | 57148922 | 52614874 | 8769146 | 36 |
| Q9WUA2 | FARSB | phenylalanyl-tRNA synthetase, beta subunit | Cytoplasm | enzyme | 442159.688 | 770174.375 | 620420.125 | Q9WUA2 | FARSB | 442160 | 164214 | 94809 | 3 |
| P19096 | FASN | fatty acid synthase | Cytoplasm | enzyme | 8857227 | 7116551 | 6613886.5 | | | | | | |
| P19096 | FASN | fatty acid synthase | Cytoplasm | enzyme | 9197122 | 4690101.5 | 7458480.5 | | | | | | |
| P19096 | FASN | fatty acid synthase | Cytoplasm | enzyme | 8896551 | 6719253 | 6781857 | | | | | | |
| P19096 | FASN | fatty acid synthase | Cytoplasm | enzyme | 4989352.5 | 3883277.75 | 3155416.5 | | | | | | |
| P19096 | FASN | fatty acid synthase | Cytoplasm | enzyme | 6500380.5 | 5020645.5 | 4491926 | | | | | | |
| P19096 | FASN | fatty acid synthase | Cytoplasm | enzyme | 6238394.5 | 3775429.5 | 4257229.5 | | | | | | |
| P19096 | FASN | fatty acid synthase | Cytoplasm | enzyme | 7168272.5 | 4823787 | 2988352.75 | | | | | | |
| P19096 | FASN | fatty acid synthase | Cytoplasm | enzyme | 5046054 | 4067941.75 | 4236213 | | | | | | |
| P19096 | FASN | fatty acid synthase | Cytoplasm | enzyme | 3996871.75 | 3759740.25 | 2341941 | | | | | | |
| P19096 | FASN | fatty acid synthase | Cytoplasm | enzyme | 2583530.25 | 1401632.25 | 1981094.25 | | | | | | |
| P19096 | FASN | fatty acid synthase | Cytoplasm | enzyme | 3897541.75 | 2362324.5 | 2349639 | | | | | | |
| P19096 | FASN | fatty acid synthase | Cytoplasm | enzyme | 2655156.25 | 1802514.75 | 1599935.875 | | | | | | |
| P19096 | FASN | fatty acid synthase | Cytoplasm | enzyme | 3678829 | 2716361.5 | 2128977.75 | | | | | | |
| P19096 | FASN | fatty acid synthase | Cytoplasm | enzyme | 3702393.25 | 2966449 | 2100772.25 | | | | | | |
| P19096 | FASN | fatty acid synthase | Cytoplasm | enzyme | 2498139.25 | 2298366 | 2023606.25 | | | | | | |
| P19096 | FASN | fatty acid synthase | Cytoplasm | enzyme | 2166402.25 | 1844624 | 1326576.375 | | | | | | |
| P19096 | FASN | fatty acid synthase | Cytoplasm | enzyme | 3091586 | 2084775.875 | 1773009.25 | | | | | | |
| P19096 | FASN | fatty acid synthase | Cytoplasm | enzyme | 1283106.25 | 1189951.5 | 291197.938 | | | | | | |
| P19096 | FASN | fatty acid synthase | Cytoplasm | enzyme | 965870.562 | 567442.562 | 160882.859 | | | | | | |

| | | | | | | | | | | | | | |
|---|---|---|---|---|---|---|---|---|---|---|---|---|---|
| P19096 | FASN | fatty acid synthase | Cytoplasm | enzyme | 1916905.75 | 1177060.875 | 999884.25 | | | | | | |
| P19096 | FASN | fatty acid synthase | Cytoplasm | enzyme | 999126.938 | 1410060.875 | 1125571.875 | | | | | | |
| P19096 | FASN | fatty acid synthase | Cytoplasm | enzyme | 490509.062 | 321549.688 | 304296.281 | | | | | | |
| P19096 | FASN | fatty acid synthase | Cytoplasm | enzyme | 588888.812 | 478640.688 | 405284.719 | P19096 | FASN | 3974270 | 2307769 | 277823 | 69 |
| P62862 | FAU | Finkel-Biskis-Reilly murine sarcoma virus (FBR-MuSV) ubiquitously expressed | Cytoplasm | other | 3194959.25 | 5389002.5 | 6301449 | P62862 | FAU | 3194959 | 1596698 | 921854 | 3 |
| Q3U7U3 | FBXO7 | F-box protein 7 | Cytoplasm | enzyme | 2102671.5 | 1541185.875 | 1866399.75 | Q3U7U3 | FBXO7 | 2102672 | 281914 | 162763 | 3 |
| P22315 | FECH | ferrochelatase | Cytoplasm | enzyme | 3.87E+07 | 7.00E+07 | 9.96E+07 | | | | | | |
| P22315 | FECH | ferrochelatase | Cytoplasm | enzyme | 2.50E+07 | 5.33E+07 | 5.02E+07 | | | | | | |
| P22315 | FECH | ferrochelatase | Cytoplasm | enzyme | 1.65E+07 | 3.32E+07 | 3.57E+07 | | | | | | |
| P22315 | FECH | ferrochelatase | Cytoplasm | enzyme | 1.37E+07 | 2.85E+07 | 3.80E+07 | | | | | | |
| P22315 | FECH | ferrochelatase | Cytoplasm | enzyme | 7295433 | 1.52E+07 | 2.78E+07 | | | | | | |
| P22315 | FECH | ferrochelatase | Cytoplasm | enzyme | 6063525 | 1.22E+07 | 1.46E+07 | | | | | | |
| P22315 | FECH | ferrochelatase | Cytoplasm | enzyme | 7880847.5 | 2.01E+07 | 2.40E+07 | | | | | | |
| P22315 | FECH | ferrochelatase | Cytoplasm | enzyme | 8227494 | 3.37E+07 | 5.14E+07 | | | | | | |
| P22315 | FECH | ferrochelatase | Cytoplasm | enzyme | 5635771 | 7810655 | 1.14E+07 | | | | | | |
| P22315 | FECH | ferrochelatase | Cytoplasm | enzyme | 1189052.375 | 2396997.5 | 4348450 | | | | | | |
| P22315 | FECH | ferrochelatase | Cytoplasm | enzyme | 2397378.5 | 4889298 | 3813158 | | | | | | |
| P22315 | FECH | ferrochelatase | Cytoplasm | enzyme | 2137009.25 | 3677426.5 | 6126832.5 | | | | | | |
| P22315 | FECH | ferrochelatase | Cytoplasm | enzyme | 2501384.25 | 5100683 | 3375601.75 | | | | | | |
| P22315 | FECH | ferrochelatase | Cytoplasm | enzyme | 3933227.75 | 6324166 | 4481350 | | | | | | |
| P22315 | FECH | ferrochelatase | Cytoplasm | enzyme | 1701572.75 | 3756343.25 | 5085218.5 | | | | | | |
| P22315 | FECH | ferrochelatase | Cytoplasm | enzyme | 1747086.125 | 2738175.25 | 2981313 | | | | | | |
| P22315 | FECH | ferrochelatase | Cytoplasm | enzyme | 1903953.5 | 3131387.75 | 4048297.25 | | | | | | |
| P22315 | FECH | ferrochelatase | Cytoplasm | enzyme | 721887.125 | 412438.812 | 4321175 | | | | | | |
| P22315 | FECH | ferrochelatase | Cytoplasm | enzyme | 927030.812 | 3046290.25 | 4535876 | P22315 | FECH | 7797508 | 19625488 | 2599459 | 57 |
| Q8K1B8 | FERMT3 | fermitin family member 3 | Cytoplasm | enzyme | 1.70E+07 | 1.25E+07 | 4.32E+07 | | | | | | |
| Q8K1B8 | FERMT3 | fermitin family member 3 | Cytoplasm | enzyme | 1.84E+07 | 1.45E+07 | 4.23E+07 | | | | | | |
| Q8K1B8 | FERMT3 | fermitin family member 3 | Cytoplasm | enzyme | 1.06E+07 | 9044290 | 2.50E+07 | | | | | | |
| Q8K1B8 | FERMT3 | fermitin family member 3 | Cytoplasm | enzyme | 1.08E+07 | 8676386 | 3.15E+07 | | | | | | |
| Q8K1B8 | FERMT3 | fermitin family member 3 | Cytoplasm | enzyme | 1.18E+07 | 7435956.5 | 2.80E+07 | | | | | | |
| Q8K1B8 | FERMT3 | fermitin family member 3 | Cytoplasm | enzyme | 1.23E+07 | 9772007 | 2.72E+07 | | | | | | |
| Q8K1B8 | FERMT3 | fermitin family member 3 | Cytoplasm | enzyme | 1.12E+07 | 7441560 | 2.02E+07 | | | | | | |
| Q8K1B8 | FERMT3 | fermitin family member 3 | Cytoplasm | enzyme | 1.05E+07 | 9625580 | 2.64E+07 | | | | | | |
| Q8K1B8 | FERMT3 | fermitin family member 3 | Cytoplasm | enzyme | 7811936.5 | 5678594 | 1.91E+07 | | | | | | |
| Q8K1B8 | FERMT3 | fermitin family member 3 | Cytoplasm | enzyme | 7473945 | 5955210.5 | 1.72E+07 | | | | | | |
| Q8K1B8 | FERMT3 | fermitin family member 3 | Cytoplasm | enzyme | 6964021 | 5222718 | 1.88E+07 | | | | | | |
| Q8K1B8 | FERMT3 | fermitin family member 3 | Cytoplasm | enzyme | 4186183.75 | 2732029.5 | 5994968 | | | | | | |
| Q8K1B8 | FERMT3 | fermitin family member 3 | Cytoplasm | enzyme | 3885785.75 | 3042703.5 | 1.04E+07 | | | | | | |
| Q8K1B8 | FERMT3 | fermitin family member 3 | Cytoplasm | enzyme | 3597467.75 | 2877594.75 | 8534218 | | | | | | |
| Q8K1B8 | FERMT3 | fermitin family member 3 | Cytoplasm | enzyme | 4467587.5 | 3165884.5 | 7188857.5 | | | | | | |
| Q8K1B8 | FERMT3 | fermitin family member 3 | Cytoplasm | enzyme | 3787255 | 1726257.25 | 9342816 | Q8K1B8 | FERMT3 | 9038386 | 9777479 | 1411258 | 48 |
| Q8K0E8 | FGB | fibrinogen beta chain | Extracellular Space | other | 1.31E+07 | 1.50E+07 | 5.06E+07 | | | | | | |
| Q8K0E8 | FGB | fibrinogen beta chain | Extracellular Space | other | 3.03E+07 | 3.08E+07 | 8.47E+07 | | | | | | |
| Q8K0E8 | FGB | fibrinogen beta chain | Extracellular Space | other | 2.01E+07 | 1.84E+07 | 1.04E+08 | | | | | | |
| Q8K0E8 | FGB | fibrinogen beta chain | Extracellular Space | other | 1.17E+07 | 1.24E+07 | 8.08E+07 | | | | | | |
| Q8K0E8 | FGB | fibrinogen beta chain | Extracellular Space | other | 1.45E+07 | 1.40E+07 | 8.01E+07 | | | | | | |
| Q8K0E8 | FGB | fibrinogen beta chain | Extracellular Space | other | 9596955 | 5741241.5 | 5.45E+07 | | | | | | |
| Q8K0E8 | FGB | fibrinogen beta chain | Extracellular Space | other | 1.18E+07 | 1.22E+07 | 6.44E+07 | | | | | | |
| Q8K0E8 | FGB | fibrinogen beta chain | Extracellular Space | other | 1.10E+07 | 1.13E+07 | 4.98E+07 | | | | | | |
| Q8K0E8 | FGB | fibrinogen beta chain | Extracellular Space | other | 1.21E+07 | 1.28E+07 | 7.37E+07 | | | | | | |
| Q8K0E8 | FGB | fibrinogen beta chain | Extracellular Space | other | 1.06E+07 | 1.05E+07 | 5.05E+07 | | | | | | |
| Q8K0E8 | FGB | fibrinogen beta chain | Extracellular Space | other | 6603334.5 | 6004560 | 3.47E+07 | | | | | | |
| Q8K0E8 | FGB | fibrinogen beta chain | Extracellular Space | other | 6822670 | 7891125 | 2.64E+07 | | | | | | |
| Q8K0E8 | FGB | fibrinogen beta chain | Extracellular Space | other | 4076801.5 | 2952806.25 | 2.27E+07 | | | | | | |
| Q8K0E8 | FGB | fibrinogen beta chain | Extracellular Space | other | 5978801.5 | 5319660.5 | 2.86E+07 | | | | | | |
| Q8K0E8 | FGB | fibrinogen beta chain | Extracellular Space | other | 4074069 | 3352006 | 1.78E+07 | | | | | | |
| Q8K0E8 | FGB | fibrinogen beta chain | Extracellular Space | other | 3381078.75 | 4931014 | 1.91E+07 | | | | | | |
| Q8K0E8 | FGB | fibrinogen beta chain | Extracellular Space | other | 1950645.625 | 3098477.75 | 1.77E+07 | | | | | | |
| Q8K0E8 | FGB | fibrinogen beta chain | Extracellular Space | other | 2079699.625 | 1180275.25 | 8209299.5 | | | | | | |
| Q8K0E8 | FGB | fibrinogen beta chain | Extracellular Space | other | 1758080.125 | 1208792.375 | 7379691.5 | | | | | | |
| Q8K0E8 | FGB | fibrinogen beta chain | Extracellular Space | other | 326554.062 | 581600.812 | 3888787.5 | | | | | | |
| Q8K0E8 | FGB | fibrinogen beta chain | Extracellular Space | other | 885985.688 | 716944.438 | 2482782.75 | | | | | | |
| Q8K0E8 | FGB | fibrinogen beta chain | Extracellular Space | other | 1248857 | 2030147.5 | 9780214 | | | | | | |
| Q8K0E8 | FGB | fibrinogen beta chain | Extracellular Space | other | 1199022.875 | 1405113.875 | 8677528 | | | | | | |
| Q8K0E8 | FGB | fibrinogen beta chain | Extracellular Space | other | 217146.516 | 190966.609 | 2365067 | Q8K0E8 | FGB | 7727904 | 23231673 | 2737879 | 72 |
| Q8VCM7 | FGG | fibrinogen gamma chain | Extracellular Space | other | 2.10E+07 | 2.10E+07 | 1.21E+08 | | | | | | |
| Q8VCM7 | FGG | fibrinogen gamma chain | Extracellular Space | other | 3.58E+07 | 3.82E+07 | 1.46E+08 | | | | | | |
| Q8VCM7 | FGG | fibrinogen gamma chain | Extracellular Space | other | 1.44E+07 | 1.23E+07 | 7.80E+07 | | | | | | |
| Q8VCM7 | FGG | fibrinogen gamma chain | Extracellular Space | other | 8840271 | 9395694 | 3.85E+07 | | | | | | |
| Q8VCM7 | FGG | fibrinogen gamma chain | Extracellular Space | other | 1.04E+07 | 1.18E+07 | 6.11E+07 | | | | | | |
| Q8VCM7 | FGG | fibrinogen gamma chain | Extracellular Space | other | 8705588 | 8996688 | 5.58E+07 | | | | | | |
| Q8VCM7 | FGG | fibrinogen gamma chain | Extracellular Space | other | 1.03E+07 | 8873723 | 4.33E+07 | | | | | | |
| Q8VCM7 | FGG | fibrinogen gamma chain | Extracellular Space | other | 8832600 | 8903241 | 6.23E+07 | | | | | | |
| Q8VCM7 | FGG | fibrinogen gamma chain | Extracellular Space | other | 7014668.5 | 6808952 | 2.18E+07 | | | | | | |
| Q8VCM7 | FGG | fibrinogen gamma chain | Extracellular Space | other | 7849174.5 | 5887847 | 3.72E+07 | | | | | | |
| Q8VCM7 | FGG | fibrinogen gamma chain | Extracellular Space | other | 4095503 | 3580818 | 1.91E+07 | | | | | | |
| Q8VCM7 | FGG | fibrinogen gamma chain | Extracellular Space | other | 5018782 | 5835326 | 3.29E+07 | | | | | | |
| Q8VCM7 | FGG | fibrinogen gamma chain | Extracellular Space | other | 7176440 | 7134897 | 2.48E+07 | | | | | | |
| Q8VCM7 | FGG | fibrinogen gamma chain | Extracellular Space | other | 6786747.5 | 6551391 | 1.59E+07 | | | | | | |
| Q8VCM7 | FGG | fibrinogen gamma chain | Extracellular Space | other | 8244863.5 | 6076565.5 | 3.77E+07 | | | | | | |
| Q8VCM7 | FGG | fibrinogen gamma chain | Extracellular Space | other | 4845605.5 | 4406910 | 2.13E+07 | | | | | | |
| Q8VCM7 | FGG | fibrinogen gamma chain | Extracellular Space | other | 8710693 | 7159377 | 3.25E+07 | | | | | | |
| Q8VCM7 | FGG | fibrinogen gamma chain | Extracellular Space | other | 1360232.875 | 1140821.75 | 3193968 | | | | | | |
| Q8VCM7 | FGG | fibrinogen gamma chain | Extracellular Space | other | 3030082.5 | 6871847.5 | 4.27E+07 | | | | | | |
| Q8VCM7 | FGG | fibrinogen gamma chain | Extracellular Space | other | 2515057.25 | 2408048.75 | 1.35E+07 | | | | | | |
| Q8VCM7 | FGG | fibrinogen gamma chain | Extracellular Space | other | 1626792 | 1277910.75 | 9072536 | | | | | | |
| Q8VCM7 | FGG | fibrinogen gamma chain | Extracellular Space | other | 244075.453 | 100212.656 | 1696869.875 | | | | | | |
| Q8VCM7 | FGG | fibrinogen gamma chain | Extracellular Space | other | 268512.688 | 487602.594 | 1680834.125 | Q8VCM7 | FGG | 8129378 | 26218978 | 3156394 | 69 |
| P97807 | FH | fumarate hydratase | Cytoplasm | enzyme | 8138205 | 1.26E+07 | 1.36E+07 | | | | | | |
| P97807 | FH | fumarate hydratase | Cytoplasm | enzyme | 4236023 | 6114130.5 | 8247396.5 | | | | | | |
| P97807 | FH | fumarate hydratase | Cytoplasm | enzyme | 1907064.5 | 3119575.75 | 5362968.5 | | | | | | |
| P97807 | FH | fumarate hydratase | Cytoplasm | enzyme | 2455282.25 | 4029470.5 | 4561663.5 | | | | | | |
| P97807 | FH | fumarate hydratase | Cytoplasm | enzyme | 1939246.125 | 3015288 | 3842429.5 | | | | | | |
| P97807 | FH | fumarate hydratase | Cytoplasm | enzyme | 987962.938 | 2794052 | 2635052.75 | | | | | | |
| P97807 | FH | fumarate hydratase | Cytoplasm | enzyme | 2183082.75 | 3323362 | 4966283 | P97807 | FH | 3120981 | 3351925 | 731450 | 21 |
| Q9CQ92 | FIS1 | fission, mitochondrial 1 | Cytoplasm | other | 4688497 | 6801274 | 6198454.5 | | | | | | |
| Q9CQ92 | FIS1 | fission, mitochondrial 1 | Cytoplasm | other | 4565733.5 | 6290827.5 | 6656118 | Q9CQ92 | FIS1 | 4627115 | 986634 | 402792 | 6 |
| Q6P9Q6 | FKBP15 | FK506 binding protein 15, 133kDa | Plasma Membrane | enzyme | 330949.844 | 332883.469 | 92176.938 | | | | | | |
| Q6P9Q6 | FKBP15 | FK506 binding protein 15, 133kDa | Plasma Membrane | enzyme | 130802.492 | 222478.188 | 300123.156 | Q6P9Q6 | FKBP15 | 230876 | 104346 | 42599 | 6 |
| P45878 | FKBP2 | FK506 binding protein 2, 13kDa | Cytoplasm | enzyme | 1.82E+07 | 2.14E+07 | 1.56E+07 | | | | | | |
| P45878 | FKBP2 | FK506 binding protein 2, 13kDa | Cytoplasm | enzyme | 1.43E+07 | 1.86E+07 | 1.81E+07 | P45878 | FKBP2 | 16250000 | 2469540 | 1008185 | 6 |
| O35450 | FKBPL | FK506 binding protein like | Nucleus | other | 2.49E+07 | 2.65E+07 | 2.28E+07 | O35450 | FKBPL | 24890000 | 1859543 | 1073608 | 3 |
| Q2VIS4 | Flg2 | filaggrin family member 2 | Extracellular Space | other | 4.29E+07 | 3.26E+07 | 3.83E+07 | Q2VIS4 | Flg2 | 42850000 | 5160875 | 2979633 | 3 |
| Q8BTM8 | FLNA | filamin A, alpha | Cytoplasm | other | 9.07E+07 | 1.06E+08 | 1.95E+08 | | | | | | |
| Q8BTM8 | FLNA | filamin A, alpha | Cytoplasm | other | 7.64E+07 | 1.01E+08 | 1.33E+08 | | | | | | |
| Q8BTM8 | FLNA | filamin A, alpha | Cytoplasm | other | 7.42E+07 | 8.97E+07 | 1.63E+08 | | | | | | |
| Q8BTM8 | FLNA | filamin A, alpha | Cytoplasm | other | 8.43E+07 | 1.03E+08 | 1.76E+08 | | | | | | |
| Q8BTM8 | FLNA | filamin A, alpha | Cytoplasm | other | 6.54E+07 | 9.09E+07 | 1.66E+08 | | | | | | |
| Q8BTM8 | FLNA | filamin A, alpha | Cytoplasm | other | 1.87E+07 | 2.75E+07 | 4.27E+07 | | | | | | |

| | | | | | | | |
|---|---|---|---|---|---|---|---|
| Q8BTM8 | FLNA | filamin A, alpha | Cytoplasm | other | 4.84E+07 | 6.29E+07 | 1.04E+08 |
| Q8BTM8 | FLNA | filamin A, alpha | Cytoplasm | other | 4.75E+07 | 6.89E+07 | 1.23E+08 |
| Q8BTM8 | FLNA | filamin A, alpha | Cytoplasm | other | 3.33E+07 | 4.66E+07 | 6.53E+07 |
| Q8BTM8 | FLNA | filamin A, alpha | Cytoplasm | other | 3.56E+07 | 5.24E+07 | 8.66E+07 |
| Q8BTM8 | FLNA | filamin A, alpha | Cytoplasm | other | 2.38E+07 | 4.11E+07 | 4.67E+07 |
| Q8BTM8 | FLNA | filamin A, alpha | Cytoplasm | other | 3.28E+07 | 4.33E+07 | 7.96E+07 |
| Q8BTM8 | FLNA | filamin A, alpha | Cytoplasm | other | 3.79E+07 | 2.63E+07 | 3.96E+07 |
| Q8BTM8 | FLNA | filamin A, alpha | Cytoplasm | other | 3.00E+07 | 4.64E+07 | 6.39E+07 |
| Q8BTM8 | FLNA | filamin A, alpha | Cytoplasm | other | 4.12E+07 | 5.60E+07 | 1.05E+08 |
| Q8BTM8 | FLNA | filamin A, alpha | Cytoplasm | other | 4.51E+07 | 4.97E+07 | 7.93E+07 |
| Q8BTM8 | FLNA | filamin A, alpha | Cytoplasm | other | 3.40E+07 | 4.03E+07 | 6.28E+07 |
| Q8BTM8 | FLNA | filamin A, alpha | Cytoplasm | other | 3.78E+07 | 5.22E+07 | 8.86E+07 |
| Q8BTM8 | FLNA | filamin A, alpha | Cytoplasm | other | 3.72E+07 | 4.50E+07 | 6.16E+07 |
| Q8BTM8 | FLNA | filamin A, alpha | Cytoplasm | other | 3.06E+07 | 4.23E+07 | 7.34E+07 |
| Q8BTM8 | FLNA | filamin A, alpha | Cytoplasm | other | 2.96E+07 | 4.05E+07 | 6.45E+07 |
| Q8BTM8 | FLNA | filamin A, alpha | Cytoplasm | other | 3.55E+07 | 4.95E+07 | 7.21E+07 |
| Q8BTM8 | FLNA | filamin A, alpha | Cytoplasm | other | 3.77E+07 | 4.48E+07 | 7.39E+07 |
| Q8BTM8 | FLNA | filamin A, alpha | Cytoplasm | other | 5.04E+07 | 6.49E+07 | 7.20E+07 |
| Q8BTM8 | FLNA | filamin A, alpha | Cytoplasm | other | 3.68E+07 | 4.10E+07 | 5.53E+07 |
| Q8BTM8 | FLNA | filamin A, alpha | Cytoplasm | other | 1.97E+07 | 3.41E+07 | 4.23E+07 |
| Q8BTM8 | FLNA | filamin A, alpha | Cytoplasm | other | 1.13E+07 | 6485011 | 1.63E+07 |
| Q8BTM8 | FLNA | filamin A, alpha | Cytoplasm | other | 3.34E+07 | 4.24E+07 | 7.59E+07 |
| Q8BTM8 | FLNA | filamin A, alpha | Cytoplasm | other | 3.22E+07 | 3.67E+07 | 6.19E+07 |
| Q8BTM8 | FLNA | filamin A, alpha | Cytoplasm | other | 3.42E+07 | 4.21E+07 | 5.88E+07 |
| Q8BTM8 | FLNA | filamin A, alpha | Cytoplasm | other | 3.10E+07 | 3.87E+07 | 5.79E+07 |
| Q8BTM8 | FLNA | filamin A, alpha | Cytoplasm | other | 2.97E+07 | 3.78E+07 | 5.95E+07 |
| Q8BTM8 | FLNA | filamin A, alpha | Cytoplasm | other | 1.57E+07 | 1.78E+07 | 2.50E+07 |
| Q8BTM8 | FLNA | filamin A, alpha | Cytoplasm | other | 3.24E+07 | 4.70E+07 | 6.17E+07 |
| Q8BTM8 | FLNA | filamin A, alpha | Cytoplasm | other | 2.68E+07 | 3.01E+07 | 5.08E+07 |
| Q8BTM8 | FLNA | filamin A, alpha | Cytoplasm | other | 2.05E+07 | 2.54E+07 | 4.89E+07 |
| Q8BTM8 | FLNA | filamin A, alpha | Cytoplasm | other | 2.94E+07 | 4.06E+07 | 5.91E+07 |
| Q8BTM8 | FLNA | filamin A, alpha | Cytoplasm | other | 3.05E+07 | 4.02E+07 | 6.95E+07 |
| Q8BTM8 | FLNA | filamin A, alpha | Cytoplasm | other | 2.47E+07 | 3.15E+07 | 4.73E+07 |
| Q8BTM8 | FLNA | filamin A, alpha | Cytoplasm | other | 2.60E+07 | 3.07E+07 | 5.40E+07 |
| Q8BTM8 | FLNA | filamin A, alpha | Cytoplasm | other | 2.91E+07 | 3.45E+07 | 5.55E+07 |
| Q8BTM8 | FLNA | filamin A, alpha | Cytoplasm | other | 9887941 | 1.40E+07 | 1.73E+07 |
| Q8BTM8 | FLNA | filamin A, alpha | Cytoplasm | other | 2.92E+07 | 3.33E+07 | 5.30E+07 |
| Q8BTM8 | FLNA | filamin A, alpha | Cytoplasm | other | 2.00E+07 | 2.82E+07 | 4.25E+07 |
| Q8BTM8 | FLNA | filamin A, alpha | Cytoplasm | other | 1.55E+07 | 2.20E+07 | 3.32E+07 |
| Q8BTM8 | FLNA | filamin A, alpha | Cytoplasm | other | 3.27E+07 | 3.93E+07 | 7.55E+07 |
| Q8BTM8 | FLNA | filamin A, alpha | Cytoplasm | other | 3.06E+07 | 4.23E+07 | 6.82E+07 |
| Q8BTM8 | FLNA | filamin A, alpha | Cytoplasm | other | 2.99E+07 | 3.63E+07 | 5.65E+07 |
| Q8BTM8 | FLNA | filamin A, alpha | Cytoplasm | other | 8268388 | 7831432 | 1.34E+07 |
| Q8BTM8 | FLNA | filamin A, alpha | Cytoplasm | other | 2.42E+07 | 3.06E+07 | 4.46E+07 |
| Q8BTM8 | FLNA | filamin A, alpha | Cytoplasm | other | 1.59E+07 | 2.20E+07 | 4.06E+07 |
| Q8BTM8 | FLNA | filamin A, alpha | Cytoplasm | other | 2.56E+07 | 2.75E+07 | 3.45E+07 |
| Q8BTM8 | FLNA | filamin A, alpha | Cytoplasm | other | 2.76E+07 | 3.51E+07 | 5.03E+07 |
| Q8BTM8 | FLNA | filamin A, alpha | Cytoplasm | other | 2.11E+07 | 2.47E+07 | 4.35E+07 |
| Q8BTM8 | FLNA | filamin A, alpha | Cytoplasm | other | 2.16E+07 | 3.09E+07 | 4.84E+07 |
| Q8BTM8 | FLNA | filamin A, alpha | Cytoplasm | other | 2.80E+07 | 3.68E+07 | 5.71E+07 |
| Q8BTM8 | FLNA | filamin A, alpha | Cytoplasm | other | 2.27E+07 | 2.41E+07 | 3.86E+07 |
| Q8BTM8 | FLNA | filamin A, alpha | Cytoplasm | other | 1.40E+07 | 2.27E+07 | 4.84E+07 |
| Q8BTM8 | FLNA | filamin A, alpha | Cytoplasm | other | 5819764.5 | 7428872.5 | 1.32E+07 |
| Q8BTM8 | FLNA | filamin A, alpha | Cytoplasm | other | 3454357 | 1089382.625 | 8282275.5 |
| Q8BTM8 | FLNA | filamin A, alpha | Cytoplasm | other | 1.56E+07 | 1.82E+07 | 2.35E+07 |
| Q8BTM8 | FLNA | filamin A, alpha | Cytoplasm | other | 1.89E+07 | 2.32E+07 | 3.35E+07 |
| Q8BTM8 | FLNA | filamin A, alpha | Cytoplasm | other | 1.18E+07 | 1.37E+07 | 2.12E+07 |
| Q8BTM8 | FLNA | filamin A, alpha | Cytoplasm | other | 8779911 | 9807295 | 1.56E+07 |
| Q8BTM8 | FLNA | filamin A, alpha | Cytoplasm | other | 1.39E+07 | 2.03E+07 | 2.74E+07 |
| Q8BTM8 | FLNA | filamin A, alpha | Cytoplasm | other | 1.91E+07 | 1.77E+07 | 2.99E+07 |
| Q8BTM8 | FLNA | filamin A, alpha | Cytoplasm | other | 1.55E+07 | 2.28E+07 | 3.37E+07 |
| Q8BTM8 | FLNA | filamin A, alpha | Cytoplasm | other | 1.31E+07 | 2.71E+07 | 3.13E+07 |
| Q8BTM8 | FLNA | filamin A, alpha | Cytoplasm | other | 8991030 | 1.11E+07 | 2.11E+07 |
| Q8BTM8 | FLNA | filamin A, alpha | Cytoplasm | other | 1.44E+07 | 1.79E+07 | 2.91E+07 |
| Q8BTM8 | FLNA | filamin A, alpha | Cytoplasm | other | 1.38E+07 | 1.85E+07 | 2.62E+07 |
| Q8BTM8 | FLNA | filamin A, alpha | Cytoplasm | other | 1.39E+07 | 1.72E+07 | 2.80E+07 |
| Q8BTM8 | FLNA | filamin A, alpha | Cytoplasm | other | 1.46E+07 | 1.85E+07 | 3.47E+07 |
| Q8BTM8 | FLNA | filamin A, alpha | Cytoplasm | other | 1.36E+07 | 1.53E+07 | 2.58E+07 |
| Q8BTM8 | FLNA | filamin A, alpha | Cytoplasm | other | 5105681.5 | 6105187.5 | 1.28E+07 |
| Q8BTM8 | FLNA | filamin A, alpha | Cytoplasm | other | 1.02E+07 | 1.04E+07 | 1.18E+07 |
| Q8BTM8 | FLNA | filamin A, alpha | Cytoplasm | other | 1.00E+07 | 1.21E+07 | 2.69E+07 |
| Q8BTM8 | FLNA | filamin A, alpha | Cytoplasm | other | 5241113 | 6971147.5 | 1.23E+07 |
| Q8BTM8 | FLNA | filamin A, alpha | Cytoplasm | other | 6724483 | 1.04E+07 | 1.38E+07 |
| Q8BTM8 | FLNA | filamin A, alpha | Cytoplasm | other | 1.34E+07 | 1.31E+07 | 2.36E+07 |
| Q8BTM8 | FLNA | filamin A, alpha | Cytoplasm | other | 1.13E+07 | 1.35E+07 | 2.57E+07 |
| Q8BTM8 | FLNA | filamin A, alpha | Cytoplasm | other | 7080858 | 9762810 | 1.96E+07 |
| Q8BTM8 | FLNA | filamin A, alpha | Cytoplasm | other | 1.11E+07 | 1.40E+07 | 2.26E+07 |
| Q8BTM8 | FLNA | filamin A, alpha | Cytoplasm | other | 9475257 | 1.18E+07 | 2.01E+07 |
| Q8BTM8 | FLNA | filamin A, alpha | Cytoplasm | other | 4773600 | 5447453.5 | 2.10E+07 |
| Q8BTM8 | FLNA | filamin A, alpha | Cytoplasm | other | 8417560 | 1.14E+07 | 1.97E+07 |
| Q8BTM8 | FLNA | filamin A, alpha | Cytoplasm | other | 3807670 | 7082790.5 | 1.21E+07 |
| Q8BTM8 | FLNA | filamin A, alpha | Cytoplasm | other | 2446745.5 | 3101628 | 3687513.75 |
| Q8BTM8 | FLNA | filamin A, alpha | Cytoplasm | other | 6781572.5 | 7206076.5 | 8770916 |
| Q8BTM8 | FLNA | filamin A, alpha | Cytoplasm | other | 1.45E+07 | 3.67E+07 | 5.65E+07 |
| Q8BTM8 | FLNA | filamin A, alpha | Cytoplasm | other | 5034928.5 | 2074704.25 | 1.39E+07 |
| Q8BTM8 | FLNA | filamin A, alpha | Cytoplasm | other | 6524545.5 | 9346997 | 1.52E+07 |
| Q8BTM8 | FLNA | filamin A, alpha | Cytoplasm | other | 1.11E+07 | 1.49E+07 | 1.73E+07 |
| Q8BTM8 | FLNA | filamin A, alpha | Cytoplasm | other | 6417995.5 | 6884066.5 | 1.03E+07 |
| Q8BTM8 | FLNA | filamin A, alpha | Cytoplasm | other | 7035503.5 | 7231353 | 1.27E+07 |
| Q8BTM8 | FLNA | filamin A, alpha | Cytoplasm | other | 7005274.5 | 1.03E+07 | 1.64E+07 |
| Q8BTM8 | FLNA | filamin A, alpha | Cytoplasm | other | 8286227 | 1.15E+07 | 1.50E+07 |
| Q8BTM8 | FLNA | filamin A, alpha | Cytoplasm | other | 5494081 | 7641672.5 | 1.45E+07 |
| Q8BTM8 | FLNA | filamin A, alpha | Cytoplasm | other | 2905946.5 | 3797036.25 | 6071879.5 |
| Q8BTM8 | FLNA | filamin A, alpha | Cytoplasm | other | 5155728.5 | 5771355 | 1.20E+07 |
| Q8BTM8 | FLNA | filamin A, alpha | Cytoplasm | other | 8556511 | 5303193.5 | 2.16E+07 |
| Q8BTM8 | FLNA | filamin A, alpha | Cytoplasm | other | 4768259.5 | 6667727.5 | 1.02E+07 |
| Q8BTM8 | FLNA | filamin A, alpha | Cytoplasm | other | 4587493 | 4818678 | 8016053 |
| Q8BTM8 | FLNA | filamin A, alpha | Cytoplasm | other | 1943058.375 | 1552906.25 | 3215172.25 |
| Q8BTM8 | FLNA | filamin A, alpha | Cytoplasm | other | 5247616 | 7557531 | 1.38E+07 |
| Q8BTM8 | FLNA | filamin A, alpha | Cytoplasm | other | 3193960.75 | 3241321.75 | 6282628 |
| Q8BTM8 | FLNA | filamin A, alpha | Cytoplasm | other | 8612854 | 1.41E+07 | 1.61E+07 |
| Q8BTM8 | FLNA | filamin A, alpha | Cytoplasm | other | 2794419.25 | 3247530 | 4738050.5 |
| Q8BTM8 | FLNA | filamin A, alpha | Cytoplasm | other | 4516761.5 | 4068048.25 | 9323144 |
| Q8BTM8 | FLNA | filamin A, alpha | Cytoplasm | other | 814450.312 | 641484.812 | 4610092.5 |
| Q8BTM8 | FLNA | filamin A, alpha | Cytoplasm | other | 970286.75 | 1114468.25 | 2763265.5 |
| Q8BTM8 | FLNA | filamin A, alpha | Cytoplasm | other | 1317808 | 583982 | 3930263.25 |
| Q8BTM8 | FLNA | filamin A, alpha | Cytoplasm | other | 1417262.25 | 2747709.5 | 3664993.75 |
| Q8BTM8 | FLNA | filamin A, alpha | Cytoplasm | other | 5797085.5 | 2321405.75 | 4157917.5 |
| Q8BTM8 | FLNA | filamin A, alpha | Cytoplasm | other | 1438144.875 | 1289297.125 | 3084718.75 |

| | | | | | | | | | | | | | |
|---|---|---|---|---|---|---|---|---|---|---|---|---|---|
| Q8BTM8 | FLNA | filamin A, alpha | Cytoplasm | other | 1570613.375 | 1806655.25 | 5926239 | | | | | | |
| Q8BTM8 | FLNA | filamin A, alpha | Cytoplasm | other | 1028783.5 | 1174781.625 | 1475107 | | | | | | |
| Q8BTM8 | FLNA | filamin A, alpha | Cytoplasm | other | 1524430 | 767407.188 | 2025930.5 | Q8BTM8 | FLNA | 20160983 | 28570622 | 1518511 | 354 |
| O08917 | FLOT1 | flotillin 1 | Plasma Membrane | other | 1.49E+07 | 1.79E+07 | 1.85E+07 | | | | | | |
| O08917 | FLOT1 | flotillin 1 | Plasma Membrane | other | 2.14E+07 | 2.18E+07 | 1.88E+07 | | | | | | |
| O08917 | FLOT1 | flotillin 1 | Plasma Membrane | other | 1.99E+07 | 1.42E+07 | 9581453 | | | | | | |
| O08917 | FLOT1 | flotillin 1 | Plasma Membrane | other | 1.22E+07 | 1.25E+07 | 1.14E+07 | | | | | | |
| O08917 | FLOT1 | flotillin 1 | Plasma Membrane | other | 1.17E+07 | 1.26E+07 | 9786500 | | | | | | |
| O08917 | FLOT1 | flotillin 1 | Plasma Membrane | other | 9779911 | 1.10E+07 | 1.04E+07 | | | | | | |
| O08917 | FLOT1 | flotillin 1 | Plasma Membrane | other | 1.19E+07 | 1.04E+07 | 1.01E+07 | | | | | | |
| O08917 | FLOT1 | flotillin 1 | Plasma Membrane | other | 8212766 | 7072836.5 | 8842371 | | | | | | |
| O08917 | FLOT1 | flotillin 1 | Plasma Membrane | other | 1.12E+07 | 1.01E+07 | 1.06E+07 | | | | | | |
| O08917 | FLOT1 | flotillin 1 | Plasma Membrane | other | 5887187 | 6321225 | 7463890.5 | | | | | | |
| O08917 | FLOT1 | flotillin 1 | Plasma Membrane | other | 7518849.5 | 6376173 | 6192043.5 | | | | | | |
| O08917 | FLOT1 | flotillin 1 | Plasma Membrane | other | 5619216 | 8654317 | 5711432.5 | | | | | | |
| O08917 | FLOT1 | flotillin 1 | Plasma Membrane | other | 3233084.5 | 4246259.5 | 3206669.5 | | | | | | |
| O08917 | FLOT1 | flotillin 1 | Plasma Membrane | other | 4698833 | 6649361 | 5768497.5 | | | | | | |
| O08917 | FLOT1 | flotillin 1 | Plasma Membrane | other | 3314239.5 | 4084791.25 | 4094076.5 | | | | | | |
| O08917 | FLOT1 | flotillin 1 | Plasma Membrane | other | 870251 | 442733.344 | 1473239.5 | O08917 | FLOT1 | 9517146 | 5248621 | 757573 | 48 |
| Q60634 | FLOT2 | flotillin 2 | Plasma Membrane | other | 2.68E+07 | 2.66E+07 | 3.09E+07 | | | | | | |
| Q60634 | FLOT2 | flotillin 2 | Plasma Membrane | other | 2.53E+07 | 2.26E+07 | 2.37E+07 | | | | | | |
| Q60634 | FLOT2 | flotillin 2 | Plasma Membrane | other | 1.83E+07 | 1.73E+07 | 1.50E+07 | | | | | | |
| Q60634 | FLOT2 | flotillin 2 | Plasma Membrane | other | 2.16E+07 | 2.08E+07 | 1.87E+07 | | | | | | |
| Q60634 | FLOT2 | flotillin 2 | Plasma Membrane | other | 1.59E+07 | 1.62E+07 | 1.32E+07 | | | | | | |
| Q60634 | FLOT2 | flotillin 2 | Plasma Membrane | other | 1.40E+07 | 1.29E+07 | 1.20E+07 | | | | | | |
| Q60634 | FLOT2 | flotillin 2 | Plasma Membrane | other | 1.14E+07 | 1.25E+07 | 1.35E+07 | | | | | | |
| Q60634 | FLOT2 | flotillin 2 | Plasma Membrane | other | 1.02E+07 | 1.04E+07 | 1.17E+07 | | | | | | |
| Q60634 | FLOT2 | flotillin 2 | Plasma Membrane | other | 8572039 | 8874600 | 8131042 | | | | | | |
| Q60634 | FLOT2 | flotillin 2 | Plasma Membrane | other | 9594936 | 1.05E+07 | 8277334.5 | | | | | | |
| Q60634 | FLOT2 | flotillin 2 | Plasma Membrane | other | 8005911 | 6294739.5 | 6559162.5 | | | | | | |
| Q60634 | FLOT2 | flotillin 2 | Plasma Membrane | other | 5130514 | 5550220 | 4900155.5 | | | | | | |
| Q60634 | FLOT2 | flotillin 2 | Plasma Membrane | other | 4767929.5 | 3956333.75 | 3667684.75 | | | | | | |
| Q60634 | FLOT2 | flotillin 2 | Plasma Membrane | other | 3007643.5 | 3748045.25 | 4195657.5 | | | | | | |
| Q60634 | FLOT2 | flotillin 2 | Plasma Membrane | other | 4797976.5 | 4466986 | 4258347 | | | | | | |
| Q60634 | FLOT2 | flotillin 2 | Plasma Membrane | other | 2407176 | 3486321.25 | 2412047.25 | | | | | | |
| Q60634 | FLOT2 | flotillin 2 | Plasma Membrane | other | 730872.938 | 487504.375 | 529378.562 | Q60634 | FLOT2 | 11213235 | 7726213 | 1081886 | 51 |
| P11276 | FN1 | fibronectin 1 | Extracellular Space | enzyme | 362149.969 | 159601.266 | 330077.125 | | | | | | |
| P11276 | FN1 | fibronectin 1 | Extracellular Space | enzyme | 336343.469 | 286694.719 | 456202.969 | | | | | | |
| P11276 | FN1 | fibronectin 1 | Extracellular Space | enzyme | 516128.312 | 376694.875 | 557366.75 | P11276 | FN1 | 404874 | 121264 | 40421 | 9 |
| Q9ER35 | FN3K | fructosamine 3 kinase | Cytoplasm | kinase | 1.61E+07 | 1.27E+07 | 1.28E+07 | | | | | | |
| Q9ER35 | FN3K | fructosamine 3 kinase | Cytoplasm | kinase | 1.56E+07 | 1.20E+07 | 1.13E+07 | | | | | | |
| Q9ER35 | FN3K | fructosamine 3 kinase | Cytoplasm | kinase | 1.48E+07 | 1.02E+07 | 1.06E+07 | | | | | | |
| Q9ER35 | FN3K | fructosamine 3 kinase | Cytoplasm | kinase | 9832396 | 6114364 | 7709049 | | | | | | |
| Q9ER35 | FN3K | fructosamine 3 kinase | Cytoplasm | kinase | 1.71E+07 | 1.05E+07 | 1.05E+07 | | | | | | |
| Q9ER35 | FN3K | fructosamine 3 kinase | Cytoplasm | kinase | 3182974.25 | 3005643 | 1890632 | | | | | | |
| Q9ER35 | FN3K | fructosamine 3 kinase | Cytoplasm | kinase | 2404157.75 | 1848352.375 | 2586439 | Q9ER35 | FN3K | 11282790 | 5046871 | 1101318 | 21 |
| Q8K274 | FN3KRP | fructosamine 3 kinase related protein | Cytoplasm | kinase | 2.02E+07 | 2.17E+07 | 1.89E+07 | | | | | | |
| Q8K274 | FN3KRP | fructosamine 3 kinase related protein | Cytoplasm | kinase | 1.60E+07 | 1.33E+07 | 8626550 | | | | | | |
| Q8K274 | FN3KRP | fructosamine 3 kinase related protein | Cytoplasm | kinase | 1.13E+07 | 8175675.5 | 7828720.5 | | | | | | |
| Q8K274 | FN3KRP | fructosamine 3 kinase related protein | Cytoplasm | kinase | 7445224.5 | 7183674 | 6552672.5 | | | | | | |
| Q8K274 | FN3KRP | fructosamine 3 kinase related protein | Cytoplasm | kinase | 5566954.5 | 5900321 | 4428155.5 | | | | | | |
| Q8K274 | FN3KRP | fructosamine 3 kinase related protein | Cytoplasm | kinase | 4297266.5 | 4250594 | 3796280.75 | Q8K274 | FN3KRP | 10799908 | 5804871 | 1368221 | 18 |
| Q3UFK8 | FRMD8 | FERM domain containing 8 | Cytoplasm | other | 4236518.5 | 5779147 | 6007973 | Q3UFK8 | FRMD8 | 4236519 | 963511 | 556283 | 3 |
| P09528 | FTH1 | ferritin, heavy polypeptide 1 | Cytoplasm | enzyme | 659294.062 | 1321630.5 | 1685636.75 | P09528 | FTH1 | 659294 | 520348 | 300423 | 3 |
| P29391 | FTL | ferritin, light polypeptide | Cytoplasm | enzyme | 2121547.5 | 3901738.25 | 5121587 | | | | | | |
| P29391 | FTL | ferritin, light polypeptide | Cytoplasm | enzyme | 933229.062 | 2133887.5 | 2498690 | | | | | | |
| P29391 | FTL | ferritin, light polypeptide | Cytoplasm | enzyme | 345334.281 | 1131369.125 | 446090.625 | P29391 | FTL | 1133370 | 1606346 | 535449 | 9 |
| Q9D6K8 | FUNDC2 | FUN14 domain containing 2 | Cytoplasm | other | 4593676.5 | 6768398.5 | 8095069.5 | | | | | | |
| Q9D6K8 | FUNDC2 | FUN14 domain containing 2 | Cytoplasm | other | 3186227 | 5537370.5 | 4503094 | Q9D6K8 | FUNDC2 | 3889952 | 1760497 | 718720 | 6 |
| O35943 | FXN | frataxin | Cytoplasm | kinase | 1873778.25 | 4232872.5 | 3229788.5 | | | | | | |
| O35943 | FXN | frataxin | Cytoplasm | kinase | 2494311 | 3061863.5 | 3349789.25 | O35943 | FXN | 2184045 | 801656 | 327275 | 6 |
| O35601 | FYB | FYN binding protein | Nucleus | other | 2098798.75 | 1553570.875 | 4875346 | | | | | | |
| O35601 | FYB | FYN binding protein | Nucleus | other | 996189.438 | 655609.688 | 1990548.875 | | | | | | |
| O35601 | FYB | FYN binding protein | Nucleus | other | 1055873.625 | 805445.938 | 2538462.25 | O35601 | FYB | 1383621 | 1307326 | 435775 | 9 |
| P97855 | G3BP1 | GTPase activating protein (SH3 domain) binding protein 1 | Nucleus | enzyme | 3361002.75 | 4927855.5 | 5428757 | | | | | | |
| P97855 | G3BP1 | GTPase activating protein (SH3 domain) binding protein 1 | Nucleus | enzyme | 3222140.75 | 4561539.5 | 4721485.5 | P97855 | G3BP1 | 3291572 | 886347 | 361850 | 6 |
| Q6NSQ9 | G6PC3 | glucose 6 phosphatase, catalytic, 3 | Cytoplasm | phosphatase | 1583802.125 | 2424460.5 | 2986413.75 | Q6NSQ9 | G6PC3 | 1583802 | 705906 | 407555 | 3 |
| Q00612 | G6PD | glucose-6-phosphate dehydrogenase | Cytoplasm | enzyme | 1167803.375 | 805018.5 | 826599.375 | | | | | | |
| Q00612 | G6PD | glucose-6-phosphate dehydrogenase | Cytoplasm | enzyme | 820053.688 | 984958.562 | 656376.812 | | | | | | |
| Q00612 | G6PD | glucose-6-phosphate dehydrogenase | Cytoplasm | enzyme | 635896.688 | 957910.125 | 691533.75 | | | | | | |
| Q00612 | G6PD | glucose-6-phosphate dehydrogenase | Cytoplasm | enzyme | 694437.438 | 432404.969 | 130379.289 | Q00612 | G6PD | 829548 | 269116 | 77687 | 12 |
| P60521 | GABARAPL2 | GABA(A) receptor-associated protein like 2 | Cytoplasm | other | 1.26E+07 | 1.75E+07 | 1.68E+07 | | | | | | |
| P60521 | GABARAPL2 | GABA(A) receptor-associated protein like 2 | Cytoplasm | other | 9684649 | 1.10E+07 | 1.19E+07 | | | | | | |
| P60521 | GABARAPL2 | GABA(A) receptor-associated protein like 2 | Cytoplasm | other | 9258075 | 9424201 | 9492963 | | | | | | |
| P60521 | GABARAPL2 | GABA(A) receptor-associated protein like 2 | Cytoplasm | other | 5614288 | 7661668 | 1.02E+07 | | | | | | |
| P60521 | GABARAPL2 | GABA(A) receptor-associated protein like 2 | Cytoplasm | other | 1152258.25 | 1340589.25 | 1255938 | P60521 | GABARAPL2 | 7663854 | 5019255 | 1295966 | 15 |
| Q8BHN3 | GANAB | glucosidase, alpha; neutral AB | Cytoplasm | enzyme | 1.62E+07 | 2.02E+07 | 1.57E+07 | | | | | | |
| Q8BHN3 | GANAB | glucosidase, alpha; neutral AB | Cytoplasm | enzyme | 1.88E+07 | 1.52E+07 | 1.42E+07 | | | | | | |
| Q8BHN3 | GANAB | glucosidase, alpha; neutral AB | Cytoplasm | enzyme | 1.66E+07 | 1.61E+07 | 1.31E+07 | | | | | | |
| Q8BHN3 | GANAB | glucosidase, alpha; neutral AB | Cytoplasm | enzyme | 1.06E+07 | 7249884 | 4939131 | | | | | | |
| Q8BHN3 | GANAB | glucosidase, alpha; neutral AB | Cytoplasm | enzyme | 1.57E+07 | 1.51E+07 | 1.12E+07 | | | | | | |
| Q8BHN3 | GANAB | glucosidase, alpha; neutral AB | Cytoplasm | enzyme | 1.02E+07 | 9700127 | 8543318 | | | | | | |
| Q8BHN3 | GANAB | glucosidase, alpha; neutral AB | Cytoplasm | enzyme | 1.23E+07 | 1.21E+07 | 1.37E+07 | | | | | | |
| Q8BHN3 | GANAB | glucosidase, alpha; neutral AB | Cytoplasm | enzyme | 7720321.5 | 7384164 | 7823516 | | | | | | |
| Q8BHN3 | GANAB | glucosidase, alpha; neutral AB | Cytoplasm | enzyme | 1.27E+07 | 1.07E+07 | 8069862 | | | | | | |
| Q8BHN3 | GANAB | glucosidase, alpha; neutral AB | Cytoplasm | enzyme | 1.09E+07 | 1.03E+07 | 6734682.5 | | | | | | |
| Q8BHN3 | GANAB | glucosidase, alpha; neutral AB | Cytoplasm | enzyme | 7348170.5 | 8034486 | 6380898.5 | | | | | | |
| Q8BHN3 | GANAB | glucosidase, alpha; neutral AB | Cytoplasm | enzyme | 6933080.5 | 2877909.25 | 7289818 | | | | | | |
| Q8BHN3 | GANAB | glucosidase, alpha; neutral AB | Cytoplasm | enzyme | 6279412.5 | 6059407 | 3934812.25 | | | | | | |
| Q8BHN3 | GANAB | glucosidase, alpha; neutral AB | Cytoplasm | enzyme | 5060136.5 | 6438038.5 | 4409011 | | | | | | |
| Q8BHN3 | GANAB | glucosidase, alpha; neutral AB | Cytoplasm | enzyme | 4562917.5 | 5702036.5 | 5848124 | | | | | | |
| Q8BHN3 | GANAB | glucosidase, alpha; neutral AB | Cytoplasm | enzyme | 4058711.75 | 3964553.25 | 3361038 | | | | | | |
| Q8BHN3 | GANAB | glucosidase, alpha; neutral AB | Cytoplasm | enzyme | 3594522.75 | 3184504.25 | 2230492.75 | | | | | | |
| Q8BHN3 | GANAB | glucosidase, alpha; neutral AB | Cytoplasm | enzyme | 3235680 | 2763681 | 2284314.75 | | | | | | |
| Q8BHN3 | GANAB | glucosidase, alpha; neutral AB | Cytoplasm | enzyme | 1773294.75 | 1814253.625 | 1117105.5 | | | | | | |
| Q8BHN3 | GANAB | glucosidase, alpha; neutral AB | Cytoplasm | enzyme | 971551 | 949040.25 | 738823.062 | Q8BHN3 | GANAB | 8773390 | 5030028 | 649374 | 60 |
| P16858 | GAPDH | glyceraldehyde-3-phosphate dehydrogenase | Cytoplasm | enzyme | 3.64E+08 | 2.39E+08 | 2.17E+08 | | | | | | |
| P16858 | GAPDH | glyceraldehyde-3-phosphate dehydrogenase | Cytoplasm | enzyme | 3.48E+08 | 2.28E+08 | 1.62E+08 | | | | | | |
| P16858 | GAPDH | glyceraldehyde-3-phosphate dehydrogenase | Cytoplasm | enzyme | 2.73E+08 | 1.68E+08 | 1.65E+08 | | | | | | |
| P16858 | GAPDH | glyceraldehyde-3-phosphate dehydrogenase | Cytoplasm | enzyme | 2.15E+08 | 9.64E+07 | 1.44E+08 | | | | | | |
| P16858 | GAPDH | glyceraldehyde-3-phosphate dehydrogenase | Cytoplasm | enzyme | 2.37E+08 | 1.84E+08 | 1.49E+08 | | | | | | |
| P16858 | GAPDH | glyceraldehyde-3-phosphate dehydrogenase | Cytoplasm | enzyme | 1.57E+08 | 1.02E+08 | 1.18E+08 | | | | | | |
| P16858 | GAPDH | glyceraldehyde-3-phosphate dehydrogenase | Cytoplasm | enzyme | 2.44E+08 | 1.53E+08 | 1.47E+08 | | | | | | |
| P16858 | GAPDH | glyceraldehyde-3-phosphate dehydrogenase | Cytoplasm | enzyme | 1.89E+08 | 9.68E+07 | 1.05E+08 | | | | | | |
| P16858 | GAPDH | glyceraldehyde-3-phosphate dehydrogenase | Cytoplasm | enzyme | 1.54E+08 | 9.32E+07 | 6.97E+07 | | | | | | |
| P16858 | GAPDH | glyceraldehyde-3-phosphate dehydrogenase | Cytoplasm | enzyme | 2.31E+08 | 1.40E+08 | 7.51E+07 | | | | | | |
| P16858 | GAPDH | glyceraldehyde-3-phosphate dehydrogenase | Cytoplasm | enzyme | 1.02E+08 | 5.82E+07 | 5.55E+07 | | | | | | |
| P16858 | GAPDH | glyceraldehyde-3-phosphate dehydrogenase | Cytoplasm | enzyme | 6.11E+07 | 2.65E+07 | 2.65E+07 | | | | | | |
| P16858 | GAPDH | glyceraldehyde-3-phosphate dehydrogenase | Cytoplasm | enzyme | 6.71E+07 | 5.31E+07 | 6.18E+07 | | | | | | |

| | | | | | | | | | | | | | |
|---|---|---|---|---|---|---|---|---|---|---|---|---|---|
| P16858 | GAPDH | glyceraldehyde-3-phosphate dehydrogenase | Cytoplasm | enzyme | 5.34E+07 | 5.37E+07 | 4.09E+07 | | | | | | |
| P16858 | GAPDH | glyceraldehyde-3-phosphate dehydrogenase | Cytoplasm | enzyme | 2.31E+07 | 1.05E+07 | 3.06E+07 | | | | | | |
| P16858 | GAPDH | glyceraldehyde-3-phosphate dehydrogenase | Cytoplasm | enzyme | 3.55E+07 | 2.15E+07 | 2.08E+07 | | | | | | |
| P16858 | GAPDH | glyceraldehyde-3-phosphate dehydrogenase | Cytoplasm | enzyme | 1.69E+07 | 1.60E+07 | 2.09E+07 | | | | | | |
| P16858 | GAPDH | glyceraldehyde-3-phosphate dehydrogenase | Cytoplasm | enzyme | 1.62E+07 | 1.39E+07 | 9740864 | | | | | | |
| P16858 | GAPDH | glyceraldehyde-3-phosphate dehydrogenase | Cytoplasm | enzyme | 1.35E+07 | 4248296.5 | 7452082.5 | | | | | | |
| P16858 | GAPDH | glyceraldehyde-3-phosphate dehydrogenase | Cytoplasm | enzyme | 2.31E+07 | 4.08E+07 | 3.67E+07 | | | | | | |
| P16858 | GAPDH | glyceraldehyde-3-phosphate dehydrogenase | Cytoplasm | enzyme | 1.59E+07 | 1.07E+07 | 9768656 | | | | | | |
| P16858 | GAPDH | glyceraldehyde-3-phosphate dehydrogenase | Cytoplasm | enzyme | 1.63E+07 | 8172607 | 6979313.5 | | | | | | |
| P16858 | GAPDH | glyceraldehyde-3-phosphate dehydrogenase | Cytoplasm | enzyme | 1.75E+07 | 7360492 | 4274190 | | | | | | |
| P16858 | GAPDH | glyceraldehyde-3-phosphate dehydrogenase | Cytoplasm | enzyme | 1.12E+07 | 1.20E+07 | 3563614.25 | | | | | | |
| P16858 | GAPDH | glyceraldehyde-3-phosphate dehydrogenase | Cytoplasm | enzyme | 1.07E+07 | 7370825.5 | 4406336 | | | | | | |
| P16858 | GAPDH | glyceraldehyde-3-phosphate dehydrogenase | Cytoplasm | enzyme | 8325634 | 7370917 | 5762072 | | | | | | |
| P16858 | GAPDH | glyceraldehyde-3-phosphate dehydrogenase | Cytoplasm | enzyme | 6004374 | 4015973.5 | 5706203 | | | | | | |
| P16858 | GAPDH | glyceraldehyde-3-phosphate dehydrogenase | Cytoplasm | enzyme | 5486403.5 | 2569676.5 | 1586897.375 | | | | | | |
| P16858 | GAPDH | glyceraldehyde-3-phosphate dehydrogenase | Cytoplasm | enzyme | 655009.938 | 545805.312 | 1187984.75 | P16858 | GAPDH | 100477980 | 86817670 | 9307831 | 87 |
| Q6PAR5 | GAPVD1 | GTPase activating protein and VPS9 domains 1 | Cytoplasm | other | 1.03E+07 | 7587010.5 | 6061821.5 | | | | | | |
| Q6PAR5 | GAPVD1 | GTPase activating protein and VPS9 domains 1 | Cytoplasm | other | 9134157 | 6477043 | 8707718 | | | | | | |
| Q6PAR5 | GAPVD1 | GTPase activating protein and VPS9 domains 1 | Cytoplasm | other | 8255528.5 | 5553547 | 5685135 | | | | | | |
| Q6PAR5 | GAPVD1 | GTPase activating protein and VPS9 domains 1 | Cytoplasm | other | 6714065 | 5905832.5 | 5556658.5 | | | | | | |
| Q6PAR5 | GAPVD1 | GTPase activating protein and VPS9 domains 1 | Cytoplasm | other | 9728996 | 5277294 | 6071960.5 | | | | | | |
| Q6PAR5 | GAPVD1 | GTPase activating protein and VPS9 domains 1 | Cytoplasm | other | 6850326.5 | 5452038 | 6219310.5 | | | | | | |
| Q6PAR5 | GAPVD1 | GTPase activating protein and VPS9 domains 1 | Cytoplasm | other | 5973661 | 5660020 | 6295700.5 | | | | | | |
| Q6PAR5 | GAPVD1 | GTPase activating protein and VPS9 domains 1 | Cytoplasm | other | 5512442.5 | 3628975.75 | 2983156.5 | | | | | | |
| Q6PAR5 | GAPVD1 | GTPase activating protein and VPS9 domains 1 | Cytoplasm | other | 5920878 | 4457398.5 | 3417400.25 | | | | | | |
| Q6PAR5 | GAPVD1 | GTPase activating protein and VPS9 domains 1 | Cytoplasm | other | 3894085.75 | 2887673 | 3913654.75 | | | | | | |
| Q6PAR5 | GAPVD1 | GTPase activating protein and VPS9 domains 1 | Cytoplasm | other | 4873025 | 2992529.5 | 3085675.75 | | | | | | |
| Q6PAR5 | GAPVD1 | GTPase activating protein and VPS9 domains 1 | Cytoplasm | other | 4271777.5 | 2813216 | 3203087 | | | | | | |
| Q6PAR5 | GAPVD1 | GTPase activating protein and VPS9 domains 1 | Cytoplasm | other | 4190341 | 3294525.25 | 2060389.125 | | | | | | |
| Q6PAR5 | GAPVD1 | GTPase activating protein and VPS9 domains 1 | Cytoplasm | other | 3425910 | 1637856 | 762250.5 | | | | | | |
| Q6PAR5 | GAPVD1 | GTPase activating protein and VPS9 domains 1 | Cytoplasm | other | 2634696.75 | 1873877.625 | 2036683 | | | | | | |
| Q6PAR5 | GAPVD1 | GTPase activating protein and VPS9 domains 1 | Cytoplasm | other | 2179706.5 | 1222024.25 | 1158959.375 | | | | | | |
| Q6PAR5 | GAPVD1 | GTPase activating protein and VPS9 domains 1 | Cytoplasm | other | 1454806.25 | 1105437.875 | 1055203.75 | Q6PAR5 | GAPVD1 | 5605553 | 2393113 | 335103 | 51 |
| Q3V0G7 | GARNL3 | GTPase activating Rap/RanGAP domain-like 3 | Cytoplasm | other | 2243666 | 1690126.375 | 3183252.25 | Q3V0G7 | GARNL3 | 2243666 | 754835 | 435804 | 3 |
| O09172 | GCLM | glutamate-cysteine ligase, modifier subunit | Cytoplasm | enzyme | 4560906.5 | 4103056 | 3349535.25 | | | | | | |
| O09172 | GCLM | glutamate-cysteine ligase, modifier subunit | Cytoplasm | enzyme | 3431698.25 | 2340555.75 | 2539110.75 | | | | | | |
| O09172 | GCLM | glutamate-cysteine ligase, modifier subunit | Cytoplasm | enzyme | 2953781.75 | 2035458.75 | 2902688.5 | O09172 | GCLM | 3648796 | 820550 | 273517 | 9 |
| Q9R111 | GDA | guanine deaminase | Cytoplasm | enzyme | 2.15E+07 | 1.44E+07 | 9383405 | | | | | | |
| Q9R111 | GDA | guanine deaminase | Cytoplasm | enzyme | 1.90E+07 | 1.32E+07 | 8625365 | | | | | | |
| Q9R111 | GDA | guanine deaminase | Cytoplasm | enzyme | 1.79E+07 | 1.36E+07 | 7576193 | | | | | | |
| Q9R111 | GDA | guanine deaminase | Cytoplasm | enzyme | 9353964 | 7307090.5 | 3337316.5 | | | | | | |
| Q9R111 | GDA | guanine deaminase | Cytoplasm | enzyme | 1.30E+07 | 8910332 | 4259568.5 | | | | | | |
| Q9R111 | GDA | guanine deaminase | Cytoplasm | enzyme | 5873171 | 4654182.5 | 2290171.75 | | | | | | |
| Q9R111 | GDA | guanine deaminase | Cytoplasm | enzyme | 9365701 | 6631573 | 4000629 | | | | | | |
| Q9R111 | GDA | guanine deaminase | Cytoplasm | enzyme | 6007253.5 | 4038876.75 | 2359447.75 | | | | | | |
| Q9R111 | GDA | guanine deaminase | Cytoplasm | enzyme | 7825673.5 | 5648450 | 4188573.75 | | | | | | |
| Q9R111 | GDA | guanine deaminase | Cytoplasm | enzyme | 2485784.5 | 1855766.125 | 849706.375 | | | | | | |
| Q9R111 | GDA | guanine deaminase | Cytoplasm | enzyme | 2569705 | 2050725 | 1261230.625 | Q9R111 | GDA | 10447387 | 5379619 | 936471 | 33 |
| Q9JL56 | GDE1 | glycerophosphodiester phosphodiesterase 1 | Plasma Membrane | enzyme | 6517148 | 6027377.5 | 6780307.5 | | | | | | |
| Q9JL56 | GDE1 | glycerophosphodiester phosphodiesterase 1 | Plasma Membrane | enzyme | 3060069.25 | 3250721.25 | 2953860.5 | | | | | | |
| Q9JL56 | GDE1 | glycerophosphodiester phosphodiesterase 1 | Plasma Membrane | enzyme | 974157.125 | 622221.438 | 926717.688 | Q9JL56 | GDE1 | 3517125 | 2451321 | 817107 | 9 |
| Q9CRY7 | GDPD1 | glycerophosphodiester phosphodiesterase domain containing 1 | Plasma Membrane | enzyme | 3532595.75 | 2684773.25 | 2127639.25 | Q9CRY7 | GDPD1 | 3532596 | 707473 | 408459 | 3 |
| Q91VC9 | GHITM | growth hormone inducible transmembrane protein | Cytoplasm | other | 1473610.5 | 2393240.75 | 2443287.75 | | | | | | |
| Q91VC9 | GHITM | growth hormone inducible transmembrane protein | Cytoplasm | other | 1516920.625 | 1751984.375 | 1248074.5 | Q91VC9 | GHITM | 1495266 | 501841 | 204876 | 6 |
| Q9CYL5 | GLIPR2 | GLI pathogenesis-related 2 | Cytoplasm | other | 2634693 | 3462504.25 | 3240116.75 | Q9CYL5 | GLIPR2 | 2634693 | 428421 | 247349 | 3 |
| Q80Y14 | GLRX5 | glutaredoxin 5 | Cytoplasm | other | 1.42E+07 | 2.75E+07 | 2.75E+07 | | | | | | |
| Q80Y14 | GLRX5 | glutaredoxin 5 | Cytoplasm | other | 8741724 | 1.49E+07 | 1.54E+07 | Q80Y14 | GLRX5 | 11460862 | 7704321 | 3145276 | 6 |
| P26443 | GLUD1 | glutamate dehydrogenase 1 | Cytoplasm | enzyme | 2249323.5 | 2696496.5 | 3682746 | | | | | | |
| P26443 | GLUD1 | glutamate dehydrogenase 1 | Cytoplasm | enzyme | 1221264.5 | 1836312.5 | 3616601.75 | P26443 | GLUD1 | 1735294 | 980562 | 400313 | 6 |
| P63158 | Gm21596/Hmgb1 | high mobility group box 1 | Nucleus | transcription regulator | 213980.375 | 51483.703 | 4454073 | | | | | | |
| P63158 | Gm21596/Hmgb1 | high mobility group box 1 | Nucleus | transcription regulator | 183234.5 | 20054.871 | 2604563.25 | P63158 | Gm21596/Hmgb1 | 198607 | 1858028 | 758537 | 6 |
| P21278 | GNA11 | guanine nucleotide binding protein (G protein), alpha 11 (Gq class) | Plasma Membrane | enzyme | 2162282.75 | 2764972.75 | 3760488.25 | P21278 | GNA11 | 2162283 | 807109 | 465984 | 3 |
| P27600 | GNA12 | guanine nucleotide binding protein (G protein) alpha 12 | Plasma Membrane | enzyme | 4.28E+07 | 5.24E+07 | 5.87E+07 | P27600 | GNA12 | 42760000 | 8016117 | 4628107 | 3 |
| P08752 | GNAI2 | guanine nucleotide binding protein (G protein), alpha inhibiting activity polypeptide 2 | Plasma Membrane | other | 7.43E+07 | 9.31E+07 | 1.07E+08 | | | | | | |
| P08752 | GNAI2 | guanine nucleotide binding protein (G protein), alpha inhibiting activity polypeptide 2 | Plasma Membrane | other | 6.22E+07 | 7.63E+07 | 9.27E+07 | | | | | | |
| P08752 | GNAI2 | guanine nucleotide binding protein (G protein), alpha inhibiting activity polypeptide 2 | Plasma Membrane | other | 5.42E+07 | 6.12E+07 | 5.04E+07 | | | | | | |
| P08752 | GNAI2 | guanine nucleotide binding protein (G protein), alpha inhibiting activity polypeptide 2 | Plasma Membrane | other | 5.33E+07 | 4.79E+07 | 4.92E+07 | | | | | | |
| P08752 | GNAI2 | guanine nucleotide binding protein (G protein), alpha inhibiting activity polypeptide 2 | Plasma Membrane | other | 5.37E+07 | 6.15E+07 | 8.71E+07 | | | | | | |
| P08752 | GNAI2 | guanine nucleotide binding protein (G protein), alpha inhibiting activity polypeptide 2 | Plasma Membrane | other | 4.78E+07 | 5.78E+07 | 5.47E+07 | | | | | | |
| P08752 | GNAI2 | guanine nucleotide binding protein (G protein), alpha inhibiting activity polypeptide 2 | Plasma Membrane | other | 4.21E+07 | 5.05E+07 | 5.31E+07 | | | | | | |
| P08752 | GNAI2 | guanine nucleotide binding protein (G protein), alpha inhibiting activity polypeptide 2 | Plasma Membrane | other | 3.03E+07 | 4.75E+07 | 4.16E+07 | | | | | | |
| P08752 | GNAI2 | guanine nucleotide binding protein (G protein), alpha inhibiting activity polypeptide 2 | Plasma Membrane | other | 2.94E+07 | 3.78E+07 | 4.48E+07 | | | | | | |
| P08752 | GNAI2 | guanine nucleotide binding protein (G protein), alpha inhibiting activity polypeptide 2 | Plasma Membrane | other | 2.96E+07 | 3.90E+07 | 4.19E+07 | | | | | | |
| P08752 | GNAI2 | guanine nucleotide binding protein (G protein), alpha inhibiting activity polypeptide 2 | Plasma Membrane | other | 1.62E+07 | 2.25E+07 | 2.28E+07 | | | | | | |
| P08752 | GNAI2 | guanine nucleotide binding protein (G protein), alpha inhibiting activity polypeptide 2 | Plasma Membrane | other | 1.62E+07 | 1.88E+07 | 1.62E+07 | | | | | | |
| P08752 | GNAI2 | guanine nucleotide binding protein (G protein), alpha inhibiting activity polypeptide 2 | Plasma Membrane | other | 1.10E+07 | 1.08E+07 | 1.04E+07 | | | | | | |
| P08752 | GNAI2 | guanine nucleotide binding protein (G protein), alpha inhibiting activity polypeptide 2 | Plasma Membrane | other | 1.89E+07 | 1.99E+07 | 2.66E+07 | | | | | | |
| P08752 | GNAI2 | guanine nucleotide binding protein (G protein), alpha inhibiting activity polypeptide 2 | Plasma Membrane | other | 1.12E+07 | 1.49E+07 | 1.55E+07 | | | | | | |
| P08752 | GNAI2 | guanine nucleotide binding protein (G protein), alpha inhibiting activity polypeptide 2 | Plasma Membrane | other | 1.38E+07 | 1.62E+07 | 1.81E+07 | | | | | | |
| P08752 | GNAI2 | guanine nucleotide binding protein (G protein), alpha inhibiting activity polypeptide 2 | Plasma Membrane | other | 1.14E+07 | 1.87E+07 | 1.57E+07 | | | | | | |
| P08752 | GNAI2 | guanine nucleotide binding protein (G protein), alpha inhibiting activity polypeptide 2 | Plasma Membrane | other | 1.09E+07 | 1.08E+07 | 1.20E+07 | | | | | | |
| P08752 | GNAI2 | guanine nucleotide binding protein (G protein), alpha inhibiting activity polypeptide 2 | Plasma Membrane | other | 1.09E+07 | 1.34E+07 | 1.15E+07 | | | | | | |
| P08752 | GNAI2 | guanine nucleotide binding protein (G protein), alpha inhibiting activity polypeptide 2 | Plasma Membrane | other | 8230273 | 3677982.75 | 1.14E+07 | | | | | | |
| P08752 | GNAI2 | guanine nucleotide binding protein (G protein), alpha inhibiting activity polypeptide 2 | Plasma Membrane | other | 1.26E+07 | 1.63E+07 | 1.53E+07 | | | | | | |
| P08752 | GNAI2 | guanine nucleotide binding protein (G protein), alpha inhibiting activity polypeptide 2 | Plasma Membrane | other | 8135232 | 8728665 | 7905649.5 | | | | | | |
| P08752 | GNAI2 | guanine nucleotide binding protein (G protein), alpha inhibiting activity polypeptide 2 | Plasma Membrane | other | 6579257 | 8501271 | 7029186.5 | | | | | | |
| P08752 | GNAI2 | guanine nucleotide binding protein (G protein), alpha inhibiting activity polypeptide 2 | Plasma Membrane | other | 3790162.25 | 1318588 | 3440933.5 | | | | | | |
| P08752 | GNAI2 | guanine nucleotide binding protein (G protein), alpha inhibiting activity polypeptide 2 | Plasma Membrane | other | 3595356.5 | 3341825.5 | 3731037.5 | | | | | | |
| P08752 | GNAI2 | guanine nucleotide binding protein (G protein), alpha inhibiting activity polypeptide 2 | Plasma Membrane | other | 3106566.5 | 3735081.75 | 4192663.25 | P08752 | GNAI2 | 24745263 | 25069064 | 2838513 | 78 |
| Q9DC51 | GNAI3 | guanine nucleotide binding protein (G protein), alpha inhibiting activity polypeptide 3 | Cytoplasm | enzyme | 1674573.75 | 2507897.75 | 2853512 | Q9DC51 | GNAI3 | 1674574 | 606049 | 349903 | 3 |
| P21279 | GNAQ | guanine nucleotide binding protein (G protein), q polypeptide | Plasma Membrane | enzyme | 5513429 | 5794693.5 | 1.17E+07 | P21279 | GNAQ | 5513429 | 3510763 | 2026940 | 3 |
| Q6R0H7 | GNAS | GNAS complex locus | Plasma Membrane | enzyme | 3135129.75 | 5258215 | 3646195.5 | | | | | | |
| Q6R0H7 | GNAS | GNAS complex locus | Plasma Membrane | enzyme | 4401681 | 5225173 | 5421931.5 | Q6R0H7 | GNAS | 3768405 | 954038 | 389484 | 6 |
| O70443 | GNAZ | guanine nucleotide binding protein (G protein), alpha z polypeptide | Plasma Membrane | enzyme | 1719548.625 | 2544087.25 | 3524559.5 | | | | | | |
| O70443 | GNAZ | guanine nucleotide binding protein (G protein), alpha z polypeptide | Plasma Membrane | enzyme | 1589121.125 | 2453285.5 | 3864371.25 | O70443 | GNAZ | 1654335 | 924518 | 377433 | 6 |
| P62874 | GNB1 | guanine nucleotide binding protein (G protein), beta polypeptide 1 | Plasma Membrane | enzyme | 5.94E+07 | 7.81E+07 | 8.76E+07 | | | | | | |
| P62874 | GNB1 | guanine nucleotide binding protein (G protein), beta polypeptide 1 | Plasma Membrane | enzyme | 4.92E+07 | 5.27E+07 | 5.94E+07 | | | | | | |
| P62874 | GNB1 | guanine nucleotide binding protein (G protein), beta polypeptide 1 | Plasma Membrane | enzyme | 2.44E+07 | 2.38E+07 | 2.45E+07 | | | | | | |
| P62874 | GNB1 | guanine nucleotide binding protein (G protein), beta polypeptide 1 | Plasma Membrane | enzyme | 2.56E+07 | 2.96E+07 | 3.05E+07 | | | | | | |
| P62874 | GNB1 | guanine nucleotide binding protein (G protein), beta polypeptide 1 | Plasma Membrane | enzyme | 1.45E+07 | 1.96E+07 | 2.09E+07 | | | | | | |
| P62874 | GNB1 | guanine nucleotide binding protein (G protein), beta polypeptide 1 | Plasma Membrane | enzyme | 1.50E+07 | 2.22E+07 | 2.54E+07 | | | | | | |
| P62874 | GNB1 | guanine nucleotide binding protein (G protein), beta polypeptide 1 | Plasma Membrane | enzyme | 1.15E+07 | 1.32E+07 | 1.38E+07 | | | | | | |
| P62874 | GNB1 | guanine nucleotide binding protein (G protein), beta polypeptide 1 | Plasma Membrane | enzyme | 1.29E+07 | 1.42E+07 | 1.65E+07 | | | | | | |
| P62874 | GNB1 | guanine nucleotide binding protein (G protein), beta polypeptide 1 | Plasma Membrane | enzyme | 1.16E+07 | 1.28E+07 | 1.68E+07 | | | | | | |
| P62874 | GNB1 | guanine nucleotide binding protein (G protein), beta polypeptide 1 | Plasma Membrane | enzyme | 9240996 | 1.15E+07 | 9286288 | | | | | | |
| P62874 | GNB1 | guanine nucleotide binding protein (G protein), beta polypeptide 1 | Plasma Membrane | enzyme | 5323507 | 4278003.5 | 7190860.5 | | | | | | |
| P62874 | GNB1 | guanine nucleotide binding protein (G protein), beta polypeptide 1 | Plasma Membrane | enzyme | 3040913.25 | 2673014 | 3518616.5 | | | | | | |
| P62874 | GNB1 | guanine nucleotide binding protein (G protein), beta polypeptide 1 | Plasma Membrane | enzyme | 2749547.5 | 2688827.5 | 3626626.5 | P62874 | GNB1 | 18794997 | 20687471 | 3312647 | 39 |
| P62880 | GNB2 | guanine nucleotide binding protein (G protein), beta polypeptide 2 | Plasma Membrane | enzyme | 4.71E+07 | 7.08E+07 | 6.31E+07 | | | | | | |

| | | | | | | | | | | | | | |
|---|---|---|---|---|---|---|---|---|---|---|---|---|---|
| P62880 | GNB2 | guanine nucleotide binding protein (G protein), beta polypeptide 2 | Plasma Membrane | enzyme | 3.32E+07 | 4.54E+07 | 4.98E+07 | | | | | | |
| P62880 | GNB2 | guanine nucleotide binding protein (G protein), beta polypeptide 2 | Plasma Membrane | enzyme | 2.89E+07 | 3.99E+07 | 4.50E+07 | | | | | | |
| P62880 | GNB2 | guanine nucleotide binding protein (G protein), beta polypeptide 2 | Plasma Membrane | enzyme | 2.76E+07 | 4.27E+07 | 4.86E+07 | | | | | | |
| P62880 | GNB2 | guanine nucleotide binding protein (G protein), beta polypeptide 2 | Plasma Membrane | enzyme | 2.25E+07 | 3.02E+07 | 3.53E+07 | | | | | | |
| P62880 | GNB2 | guanine nucleotide binding protein (G protein), beta polypeptide 2 | Plasma Membrane | enzyme | 2.11E+07 | 2.72E+07 | 2.85E+07 | | | | | | |
| P62880 | GNB2 | guanine nucleotide binding protein (G protein), beta polypeptide 2 | Plasma Membrane | enzyme | 1.35E+07 | 1.80E+07 | 2.01E+07 | | | | | | |
| P62880 | GNB2 | guanine nucleotide binding protein (G protein), beta polypeptide 2 | Plasma Membrane | enzyme | 1.03E+07 | 1.82E+07 | 1.58E+07 | | | | | | |
| P62880 | GNB2 | guanine nucleotide binding protein (G protein), beta polypeptide 2 | Plasma Membrane | enzyme | 6958173 | 8460751 | 1.10E+07 | | | | | | |
| P62880 | GNB2 | guanine nucleotide binding protein (G protein), beta polypeptide 2 | Plasma Membrane | enzyme | 3866254.75 | 5422663.5 | 5639873 | | | | | | |
| P62880 | GNB2 | guanine nucleotide binding protein (G protein), beta polypeptide 2 | Plasma Membrane | enzyme | 3461866.75 | 5021987 | 4617428 | | | | | | |
| P62880 | GNB2 | guanine nucleotide binding protein (G protein), beta polypeptide 2 | Plasma Membrane | enzyme | 2294525 | 4920665.5 | 3992557.75 | P62880 | GNB2 | 18404235 | 18513068 | 3085511 | 36 |
| P68040 | GNB2L1 | guanine nucleotide binding protein (G protein), beta polypeptide 2-like 1 | Cytoplasm | enzyme | 4922001.5 | 7560297 | 8575843 | | | | | | |
| P68040 | GNB2L1 | guanine nucleotide binding protein (G protein), beta polypeptide 2-like 1 | Cytoplasm | enzyme | 2866577.5 | 5386278 | 5895399.5 | | | | | | |
| P68040 | GNB2L1 | guanine nucleotide binding protein (G protein), beta polypeptide 2-like 1 | Cytoplasm | enzyme | 2981672 | 4738464 | 4918661 | | | | | | |
| P68040 | GNB2L1 | guanine nucleotide binding protein (G protein), beta polypeptide 2-like 1 | Cytoplasm | enzyme | 2801885.75 | 3609070.25 | 4108916.25 | | | | | | |
| P68040 | GNB2L1 | guanine nucleotide binding protein (G protein), beta polypeptide 2-like 1 | Cytoplasm | enzyme | 1332276.625 | 2107436 | 2555706.75 | P68040 | GNB2L1 | 2980883 | 2014350 | 520103 | 15 |
| Q9DAS9 | GNG12 | guanine nucleotide binding protein (G protein), gamma 12 | Plasma Membrane | enzyme | 6392937.5 | 8006254.5 | 7210371.5 | Q9DAS9 | GNG12 | 6392938 | 806682 | 465738 | 3 |
| Q91W53 | GOLGA7 | golgin A7 | Cytoplasm | other | 4883522.5 | 4910374 | 4040303.5 | | | | | | |
| Q91W53 | GOLGA7 | golgin A7 | Cytoplasm | other | 4222574 | 4699420.5 | 3742977.75 | Q91W53 | GOLGA7 | 4553048 | 484733 | 197891 | 6 |
| Q9CR60 | GOLT1B | golgi transport 1B | Cytoplasm | other | 4010362 | 2633215.5 | 3336932.5 | Q9CR60 | GOLT1B | 4010362 | 688629 | 397580 | 3 |
| P05202 | GOT2 | glutamic-oxaloacetic transaminase 2, mitochondrial | Cytoplasm | enzyme | 1.81E+07 | 2.11E+07 | 2.89E+07 | | | | | | |
| P05202 | GOT2 | glutamic-oxaloacetic transaminase 2, mitochondrial | Cytoplasm | enzyme | 8824316 | 1.28E+07 | 1.81E+07 | | | | | | |
| P05202 | GOT2 | glutamic-oxaloacetic transaminase 2, mitochondrial | Cytoplasm | enzyme | 7631727.5 | 1.20E+07 | 1.71E+07 | | | | | | |
| P05202 | GOT2 | glutamic-oxaloacetic transaminase 2, mitochondrial | Cytoplasm | enzyme | 7076367 | 1.11E+07 | 1.28E+07 | | | | | | |
| P05202 | GOT2 | glutamic-oxaloacetic transaminase 2, mitochondrial | Cytoplasm | enzyme | 5194846 | 6202403.5 | 7996830.5 | | | | | | |
| P05202 | GOT2 | glutamic-oxaloacetic transaminase 2, mitochondrial | Cytoplasm | enzyme | 4689629.5 | 8127711 | 1.16E+07 | | | | | | |
| P05202 | GOT2 | glutamic-oxaloacetic transaminase 2, mitochondrial | Cytoplasm | enzyme | 5238438 | 5372326 | 8213275.5 | | | | | | |
| P05202 | GOT2 | glutamic-oxaloacetic transaminase 2, mitochondrial | Cytoplasm | enzyme | 1767622.625 | 5073968 | 3649377.75 | | | | | | |
| P05202 | GOT2 | glutamic-oxaloacetic transaminase 2, mitochondrial | Cytoplasm | enzyme | 2348408.75 | 2122360.5 | 1793982.25 | | | | | | |
| P05202 | GOT2 | glutamic-oxaloacetic transaminase 2, mitochondrial | Cytoplasm | enzyme | 1283794.5 | 3005153.75 | 3166415.25 | | | | | | |
| P05202 | GOT2 | glutamic-oxaloacetic transaminase 2, mitochondrial | Cytoplasm | enzyme | 912894.938 | 1250826.75 | 1516295.25 | P05202 | GOT2 | 5730731 | 6656285 | 1158711 | 33 |
| O35930 | GP1BA | glycoprotein Ib (platelet), alpha polypeptide | Plasma Membrane | other | 3.09E+07 | 5.12E+07 | 6.71E+07 | | | | | | |
| O35930 | GP1BA | glycoprotein Ib (platelet), alpha polypeptide | Plasma Membrane | other | 3.49E+07 | 4.16E+07 | 8.43E+07 | | | | | | |
| O35930 | GP1BA | glycoprotein Ib (platelet), alpha polypeptide | Plasma Membrane | other | 2.07E+07 | 3.17E+07 | 4.72E+07 | | | | | | |
| O35930 | GP1BA | glycoprotein Ib (platelet), alpha polypeptide | Plasma Membrane | other | 1.82E+07 | 2.34E+07 | 3.46E+07 | | | | | | |
| O35930 | GP1BA | glycoprotein Ib (platelet), alpha polypeptide | Plasma Membrane | other | 1.79E+07 | 2.17E+07 | 4.70E+07 | | | | | | |
| O35930 | GP1BA | glycoprotein Ib (platelet), alpha polypeptide | Plasma Membrane | other | 1.73E+07 | 2.13E+07 | 3.00E+07 | | | | | | |
| O35930 | GP1BA | glycoprotein Ib (platelet), alpha polypeptide | Plasma Membrane | other | 1.01E+07 | 1.46E+07 | 1.97E+07 | | | | | | |
| O35930 | GP1BA | glycoprotein Ib (platelet), alpha polypeptide | Plasma Membrane | other | 6501221 | 1.23E+07 | 1.40E+07 | | | | | | |
| O35930 | GP1BA | glycoprotein Ib (platelet), alpha polypeptide | Plasma Membrane | other | 3964757.75 | 6911055 | 9903923 | | | | | | |
| O35930 | GP1BA | glycoprotein Ib (platelet), alpha polypeptide | Plasma Membrane | other | 4098412 | 4951859 | 8309014 | | | | | | |
| O35930 | GP1BA | glycoprotein Ib (platelet), alpha polypeptide | Plasma Membrane | other | 4673059.5 | 7495990 | 1.00E+07 | | | | | | |
| O35930 | GP1BA | glycoprotein Ib (platelet), alpha polypeptide | Plasma Membrane | other | 5905547.5 | 7906709.5 | 1.54E+07 | | | | | | |
| O35930 | GP1BA | glycoprotein Ib (platelet), alpha polypeptide | Plasma Membrane | other | 78213.164 | 573268.75 | 3001369 | O35930 | GP1BA | 13479324 | 18872850 | 3022075 | 39 |
| P56400 | GP1BB | glycoprotein Ib (platelet), beta polypeptide | Plasma Membrane | other | 5.58E+07 | 8.21E+07 | 1.29E+08 | | | | | | |
| P56400 | GP1BB | glycoprotein Ib (platelet), beta polypeptide | Plasma Membrane | other | 3.35E+07 | 3.93E+07 | 7.64E+07 | | | | | | |
| P56400 | GP1BB | glycoprotein Ib (platelet), beta polypeptide | Plasma Membrane | other | 1.50E+07 | 1.98E+07 | 3.44E+07 | | | | | | |
| P56400 | GP1BB | glycoprotein Ib (platelet), beta polypeptide | Plasma Membrane | other | 2.14E+07 | 2.53E+07 | 3.67E+07 | | | | | | |
| P56400 | GP1BB | glycoprotein Ib (platelet), beta polypeptide | Plasma Membrane | other | 7739300 | 9100270 | 1.48E+07 | | | | | | |
| P56400 | GP1BB | glycoprotein Ib (platelet), beta polypeptide | Plasma Membrane | other | 8863718 | 1.25E+07 | 1.56E+07 | P56400 | GP1BB | 23718836 | 32093975 | 7564622 | 18 |
| O08742 | GP5 | glycoprotein V (platelet) | Plasma Membrane | other | 3.09E+07 | 4.37E+07 | 7.17E+07 | | | | | | |
| O08742 | GP5 | glycoprotein V (platelet) | Plasma Membrane | other | 2.24E+07 | 1.73E+07 | 4.62E+07 | | | | | | |
| O08742 | GP5 | glycoprotein V (platelet) | Plasma Membrane | other | 1.79E+07 | 3.12E+07 | 3.47E+07 | | | | | | |
| O08742 | GP5 | glycoprotein V (platelet) | Plasma Membrane | other | 2.11E+07 | 3.26E+07 | 4.54E+07 | | | | | | |
| O08742 | GP5 | glycoprotein V (platelet) | Plasma Membrane | other | 1.83E+07 | 1.97E+07 | 3.34E+07 | | | | | | |
| O08742 | GP5 | glycoprotein V (platelet) | Plasma Membrane | other | 2.01E+07 | 2.86E+07 | 4.70E+07 | | | | | | |
| O08742 | GP5 | glycoprotein V (platelet) | Plasma Membrane | other | 8216191 | 9119834 | 1.56E+07 | | | | | | |
| O08742 | GP5 | glycoprotein V (platelet) | Plasma Membrane | other | 6151619.5 | 7330392 | 1.12E+07 | | | | | | |
| O08742 | GP5 | glycoprotein V (platelet) | Plasma Membrane | other | 4686455 | 7454340 | 1.20E+07 | | | | | | |
| O08742 | GP5 | glycoprotein V (platelet) | Plasma Membrane | other | 2987792.5 | 4657739.5 | 7614196.5 | O08742 | GP5 | 15271206 | 16424223 | 2998639 | 30 |
| O88186 | GP9 | glycoprotein IX (platelet) | Plasma Membrane | other | 5.57E+07 | 7.35E+07 | 9.46E+07 | | | | | | |
| O88186 | GP9 | glycoprotein IX (platelet) | Plasma Membrane | other | 1.89E+07 | 2.15E+07 | 3.95E+07 | | | | | | |
| O88186 | GP9 | glycoprotein IX (platelet) | Plasma Membrane | other | 3929432.75 | 4279849.5 | 7362792 | | | | | | |
| O88186 | GP9 | glycoprotein IX (platelet) | Plasma Membrane | other | 3072049.75 | 3876403 | 7802883 | O88186 | GP9 | 20382871 | 31193958 | 9004920 | 12 |
| P51655 | GPC4 | glypican 4 | Plasma Membrane | transmembrane receptor | 8178230.5 | 7274276 | 5545704.5 | | | | | | |
| P51655 | GPC4 | glypican 4 | Plasma Membrane | transmembrane receptor | 7399294.5 | 7067949.5 | 4815501.5 | | | | | | |
| P51655 | GPC4 | glypican 4 | Plasma Membrane | transmembrane receptor | 3327375.75 | 4265060 | 2529343.25 | P51655 | GPC4 | 6301634 | 1994900 | 664967 | 9 |
| Q64521 | GPD2 | glycerol-3-phosphate dehydrogenase 2 (mitochondrial) | Cytoplasm | enzyme | 5593556.5 | 7805794 | 1.26E+07 | | | | | | |
| Q64521 | GPD2 | glycerol-3-phosphate dehydrogenase 2 (mitochondrial) | Cytoplasm | enzyme | 5055428.5 | 7793308 | 1.28E+07 | | | | | | |
| Q64521 | GPD2 | glycerol-3-phosphate dehydrogenase 2 (mitochondrial) | Cytoplasm | enzyme | 2555216.5 | 3184221.75 | 4572433.5 | | | | | | |
| Q64521 | GPD2 | glycerol-3-phosphate dehydrogenase 2 (mitochondrial) | Cytoplasm | enzyme | 4743719 | 5257109 | 6336167 | | | | | | |
| Q64521 | GPD2 | glycerol-3-phosphate dehydrogenase 2 (mitochondrial) | Cytoplasm | enzyme | 903061.562 | 1402757.375 | 1890004.5 | | | | | | |
| Q64521 | GPD2 | glycerol-3-phosphate dehydrogenase 2 (mitochondrial) | Cytoplasm | enzyme | 2876663.75 | 4811976 | 7666029.5 | | | | | | |
| Q64521 | GPD2 | glycerol-3-phosphate dehydrogenase 2 (mitochondrial) | Cytoplasm | enzyme | 2697444.25 | 3465012.25 | 5073815.5 | | | | | | |
| Q64521 | GPD2 | glycerol-3-phosphate dehydrogenase 2 (mitochondrial) | Cytoplasm | enzyme | 1931842.375 | 2258850.25 | 4268155 | | | | | | |
| Q64521 | GPD2 | glycerol-3-phosphate dehydrogenase 2 (mitochondrial) | Cytoplasm | enzyme | 3208012.25 | 3396162 | 6345402.5 | | | | | | |
| Q64521 | GPD2 | glycerol-3-phosphate dehydrogenase 2 (mitochondrial) | Cytoplasm | enzyme | 1940079.5 | 3626524.25 | 4549434 | Q64521 | GPD2 | 3150502 | 2863635 | 522826 | 30 |
| Q99LD4 | GPS1 | G protein pathway suppressor 1 | Cytoplasm | other | 5629491 | 4580952.5 | 3635792.25 | | | | | | |
| Q99LD4 | GPS1 | G protein pathway suppressor 1 | Cytoplasm | other | 6080468.5 | 4343273.5 | 3658225.75 | | | | | | |
| Q99LD4 | GPS1 | G protein pathway suppressor 1 | Cytoplasm | other | 2992231 | 1431067.375 | 1728614.375 | | | | | | |
| Q99LD4 | GPS1 | G protein pathway suppressor 1 | Cytoplasm | other | 1618182.75 | 1399353.5 | 964564.5 | | | | | | |
| Q99LD4 | GPS1 | G protein pathway suppressor 1 | Cytoplasm | other | 1046976.875 | 818964.375 | 409907.375 | Q99LD4 | GPS1 | 3473470 | 1852955 | 478431 | 15 |
| P11352 | GPX1 | glutathione peroxidase 1 | Cytoplasm | enzyme | 3.59E+07 | 3.49E+07 | 2.75E+07 | | | | | | |
| P11352 | GPX1 | glutathione peroxidase 1 | Cytoplasm | enzyme | 1.85E+07 | 1.69E+07 | 2.60E+07 | | | | | | |
| P11352 | GPX1 | glutathione peroxidase 1 | Cytoplasm | enzyme | 1.85E+07 | 1.92E+07 | 1.66E+07 | | | | | | |
| P11352 | GPX1 | glutathione peroxidase 1 | Cytoplasm | enzyme | 1.10E+07 | 1.24E+07 | 1.18E+07 | | | | | | |
| P11352 | GPX1 | glutathione peroxidase 1 | Cytoplasm | enzyme | 8040344.5 | 8120041.5 | 7726990 | | | | | | |
| P11352 | GPX1 | glutathione peroxidase 1 | Cytoplasm | enzyme | 8653013 | 9002947 | 7221135 | | | | | | |
| P11352 | GPX1 | glutathione peroxidase 1 | Cytoplasm | enzyme | 8336359.5 | 7408188 | 6659745 | | | | | | |
| P11352 | GPX1 | glutathione peroxidase 1 | Cytoplasm | enzyme | 4683039.5 | 9122431 | 3786712.5 | | | | | | |
| P11352 | GPX1 | glutathione peroxidase 1 | Cytoplasm | enzyme | 3723428.25 | 2652597.25 | 1381060.875 | | | | | | |
| P11352 | GPX1 | glutathione peroxidase 1 | Cytoplasm | enzyme | 3915162.25 | 3487115.25 | 3420430.5 | P11352 | GPX1 | 12122135 | 9237445 | 1686519 | 30 |
| O70325 | GPX4 | glutathione peroxidase 4 | Cytoplasm | enzyme | 4.20E+07 | 4.60E+07 | 3.70E+07 | | | | | | |
| O70325 | GPX4 | glutathione peroxidase 4 | Cytoplasm | enzyme | 3.03E+07 | 2.98E+07 | 3.19E+07 | | | | | | |
| O70325 | GPX4 | glutathione peroxidase 4 | Cytoplasm | enzyme | 1.81E+07 | 1.54E+07 | 1.38E+07 | | | | | | |
| O70325 | GPX4 | glutathione peroxidase 4 | Cytoplasm | enzyme | 1.72E+07 | 1.34E+07 | 1.41E+07 | | | | | | |
| O70325 | GPX4 | glutathione peroxidase 4 | Cytoplasm | enzyme | 1.47E+07 | 1.20E+07 | 1.12E+07 | | | | | | |
| O70325 | GPX4 | glutathione peroxidase 4 | Cytoplasm | enzyme | 4143626.5 | 4180638.25 | 3593653 | | | | | | |
| O70325 | GPX4 | glutathione peroxidase 4 | Cytoplasm | enzyme | 3298970.75 | 2559667.25 | 2940751.5 | O70325 | GPX4 | 18527514 | 13455723 | 2936279 | 21 |
| Q8R050 | GSPT1 | G1 to S phase transition 1 | Cytoplasm | translation regulator | 2018492.25 | 1788336.75 | 1410670.75 | Q8R050 | GSPT1 | 2018492 | 306879 | 177177 | 3 |
| P19157 | GSTP1 | glutathione S-transferase pi 1 | Cytoplasm | enzyme | 1536555.125 | 1734331.875 | 1807948.375 | P19157 | GSTP1 | 1536555 | 140350 | 81031 | 3 |
| Q8BZI6 | GUCD1 | guanylyl cyclase domain containing 1 | Nucleus | other | 1726199.5 | 1859574.75 | 1867556.25 | Q8BZI6 | GUCD1 | 1726200 | 79409 | 45847 | 3 |
| P14220 | Gypa | glycophorin A | Plasma Membrane | other | 5.14E+09 | 4.06E+09 | 1.86E+09 | | | | | | |
| P14220 | Gypa | glycophorin A | Plasma Membrane | other | 2.14E+07 | 2.05E+07 | 2.42E+07 | P14220 | Gypa | 2581215000 | 2268103360 | 925949319 | 6 |
| Q78HU7 | GYPC | glycophorin C (Gerbich blood group) | Plasma Membrane | other | 8.39E+08 | 8.83E+08 | 1.34E+09 | | | | | | |
| Q78HU7 | GYPC | glycophorin C (Gerbich blood group) | Plasma Membrane | other | 1.09E+09 | 1.31E+09 | 9.47E+08 | | | | | | |
| Q78HU7 | GYPC | glycophorin C (Gerbich blood group) | Plasma Membrane | other | 3.81E+07 | 3.69E+07 | 5.72E+07 | Q78HU7 | GYPC | 653976667 | 539902431 | 179967477 | 9 |
| Q3THW5 | H2AFV | H2A histone family, member V | Nucleus | other | 1551983.875 | 509067.344 | 1.16E+07 | Q3THW5 | H2AFV | 1551984 | 6147539 | 3549283 | 3 |
| P27661 | H2AFX | H2A histone family, member X | Nucleus | transcription regulator | 5430402 | 1908238.125 | 3.05E+07 | | | | | | |

| | | | | | | | | | | | | | |
|---|---|---|---|---|---|---|---|---|---|---|---|---|---|
| P27661 | H2AFX | H2A histone family, member X | Nucleus | transcription regulator | 434227.406 | 71921.461 | 2289583.5 | P27661 | H2AFX | 2932315 | 11786212 | 4811701 | 6 |
| Q9QZQ8 | H2AFY | H2A histone family, member Y | Nucleus | other | 1577580.125 | 234881.469 | 1.12E+07 | Q9QZQ8 | H2AFY | 1577580 | 5969435 | 3446455 | 3 |
| P68433 | H31 | Histone H3.1 | Nucleus | | 5.44E+07 | 1.75E+07 | 3.21E+08 | | | | | | |
| P68433 | H31 | Histone H3.1 | Nucleus | | 5.08E+07 | 1.67E+07 | 2.66E+08 | | | | | | |
| P68433 | H31 | Histone H3.1 | Nucleus | | 2.05E+07 | 7040311.5 | 8.08E+07 | | | | | | |
| P68433 | H31 | Histone H3.1 | Nucleus | | 2.24E+07 | 7256929 | 1.37E+08 | | | | | | |
| P68433 | H31 | Histone H3.1 | Nucleus | | 5483930.5 | 3600011.5 | 4.28E+07 | | | | | | |
| P68433 | H31 | Histone H3.1 | Nucleus | | 1.37E+07 | 2061714.875 | 7.91E+07 | P68433 | H31 | 27873988 | 91146365 | 21483404 | 18 |
| P62806 | H4 | Histone H4 | Nucleus | | 1.24E+08 | 4.54E+07 | 6.85E+08 | | | | | | |
| P62806 | H4 | Histone H4 | Nucleus | | 1.47E+08 | 4.63E+07 | 7.21E+08 | | | | | | |
| P62806 | H4 | Histone H4 | Nucleus | | 2.14E+07 | 8898138 | 1.94E+08 | | | | | | |
| P62806 | H4 | Histone H4 | Nucleus | | 3.90E+07 | 1.14E+07 | 1.64E+08 | | | | | | |
| P62806 | H4 | Histone H4 | Nucleus | | 1.30E+07 | 4017267.25 | 6.06E+07 | | | | | | |
| P62806 | H4 | Histone H4 | Nucleus | | 7006498 | 1718341.125 | 1.08E+08 | | | | | | |
| P62806 | H4 | Histone H4 | Nucleus | | 216929.094 | 22684.6 | 753448.625 | | | | | | |
| P62806 | H4 | Histone H4 | Nucleus | | 372962.5 | 808218.25 | 1.72E+07 | P62806 | H4 | 43977049 | 194301484 | 39661624 | 24 |
| Q3U0D9 | HACE1 | HECT domain and ankyrin repeat containing E3 ubiquitin protein ligase 1 | Cytoplasm | enzyme | 594981.938 | 396785.031 | 482441.25 | Q3U0D9 | HACE1 | 594982 | 99402 | 57390 | 3 |
| Q61425 | HADH | hydroxyacyl-CoA dehydrogenase | Cytoplasm | enzyme | 2682919.5 | 3613560 | 5257006 | | | | | | |
| Q61425 | HADH | hydroxyacyl-CoA dehydrogenase | Cytoplasm | enzyme | 517999.094 | 874682.812 | 1502177 | Q61425 | HADH | 1600459 | 1810496 | 739132 | 6 |
| Q8BMS1 | HADHA | hydroxyacyl-CoA dehydrogenase/3-ketoacyl-CoA thiolase/enoyl-CoA hydratase (trifunctional protein), alpha subunit | Cytoplasm | enzyme | 6150167 | 9696972 | 1.31E+07 | | | | | | |
| Q8BMS1 | HADHA | hydroxyacyl-CoA dehydrogenase/3-ketoacyl-CoA thiolase/enoyl-CoA hydratase (trifunctional protein), alpha subunit | Cytoplasm | enzyme | 3941649.5 | 6729319 | 8055478 | | | | | | |
| Q8BMS1 | HADHA | hydroxyacyl-CoA dehydrogenase/3-ketoacyl-CoA thiolase/enoyl-CoA hydratase (trifunctional protein), alpha subunit | Cytoplasm | enzyme | 4044194.25 | 7533602 | 8467215 | | | | | | |
| Q8BMS1 | HADHA | hydroxyacyl-CoA dehydrogenase/3-ketoacyl-CoA thiolase/enoyl-CoA hydratase (trifunctional protein), alpha subunit | Cytoplasm | enzyme | 3864559.5 | 5927319 | 7361654 | | | | | | |
| Q8BMS1 | HADHA | hydroxyacyl-CoA dehydrogenase/3-ketoacyl-CoA thiolase/enoyl-CoA hydratase (trifunctional protein), alpha subunit | Cytoplasm | enzyme | 4198305 | 8193408.5 | 1.01E+07 | | | | | | |
| Q8BMS1 | HADHA | hydroxyacyl-CoA dehydrogenase/3-ketoacyl-CoA thiolase/enoyl-CoA hydratase (trifunctional protein), alpha subunit | Cytoplasm | enzyme | 2591551.25 | 4363228 | 6419980 | | | | | | |
| Q8BMS1 | HADHA | hydroxyacyl-CoA dehydrogenase/3-ketoacyl-CoA thiolase/enoyl-CoA hydratase (trifunctional protein), alpha subunit | Cytoplasm | enzyme | 3149764.5 | 3989070.5 | 5689476.5 | | | | | | |
| Q8BMS1 | HADHA | hydroxyacyl-CoA dehydrogenase/3-ketoacyl-CoA thiolase/enoyl-CoA hydratase (trifunctional protein), alpha subunit | Cytoplasm | enzyme | 2231937.5 | 4560464.5 | 5451229 | | | | | | |
| Q8BMS1 | HADHA | hydroxyacyl-CoA dehydrogenase/3-ketoacyl-CoA thiolase/enoyl-CoA hydratase (trifunctional protein), alpha subunit | Cytoplasm | enzyme | 2310564 | 4441987.5 | 7440640.5 | | | | | | |
| Q8BMS1 | HADHA | hydroxyacyl-CoA dehydrogenase/3-ketoacyl-CoA thiolase/enoyl-CoA hydratase (trifunctional protein), alpha subunit | Cytoplasm | enzyme | 2326741.25 | 5322506.5 | 5046398 | | | | | | |
| Q8BMS1 | HADHA | hydroxyacyl-CoA dehydrogenase/3-ketoacyl-CoA thiolase/enoyl-CoA hydratase (trifunctional protein), alpha subunit | Cytoplasm | enzyme | 1359637.375 | 2045075.125 | 2803261.25 | | | | | | |
| Q8BMS1 | HADHA | hydroxyacyl-CoA dehydrogenase/3-ketoacyl-CoA thiolase/enoyl-CoA hydratase (trifunctional protein), alpha subunit | Cytoplasm | enzyme | 993245.875 | 2347293 | 2075492.75 | | | | | | |
| Q8BMS1 | HADHA | hydroxyacyl-CoA dehydrogenase/3-ketoacyl-CoA thiolase/enoyl-CoA hydratase (trifunctional protein), alpha subunit | Cytoplasm | enzyme | 906696.938 | 1314003.375 | 1791774.875 | | | | | | |
| Q8BMS1 | HADHA | hydroxyacyl-CoA dehydrogenase/3-ketoacyl-CoA thiolase/enoyl-CoA hydratase (trifunctional protein), alpha subunit | Cytoplasm | enzyme | 596031.5 | 1087182.375 | 1071663.625 | Q8BMS1 | HADHA | 2761789 | 2915166 | 449820 | 42 |
| Q99JY0 | HADHB | hydroxyacyl-CoA dehydrogenase/3-ketoacyl-CoA thiolase/enoyl-CoA hydratase (trifunctional protein), beta subunit | Cytoplasm | enzyme | 7378706 | 1.42E+07 | 1.30E+07 | | | | | | |
| Q99JY0 | HADHB | hydroxyacyl-CoA dehydrogenase/3-ketoacyl-CoA thiolase/enoyl-CoA hydratase (trifunctional protein), beta subunit | Cytoplasm | enzyme | 6143847 | 8465640 | 1.24E+07 | | | | | | |
| Q99JY0 | HADHB | hydroxyacyl-CoA dehydrogenase/3-ketoacyl-CoA thiolase/enoyl-CoA hydratase (trifunctional protein), beta subunit | Cytoplasm | enzyme | 5299111 | 9231449 | 9231990 | | | | | | |
| Q99JY0 | HADHB | hydroxyacyl-CoA dehydrogenase/3-ketoacyl-CoA thiolase/enoyl-CoA hydratase (trifunctional protein), beta subunit | Cytoplasm | enzyme | 2169872.75 | 4178619.75 | 6915807.5 | | | | | | |
| Q99JY0 | HADHB | hydroxyacyl-CoA dehydrogenase/3-ketoacyl-CoA thiolase/enoyl-CoA hydratase (trifunctional protein), beta subunit | Cytoplasm | enzyme | 1163996.125 | 1821164.875 | 1813727.75 | | | | | | |
| Q99JY0 | HADHB | hydroxyacyl-CoA dehydrogenase/3-ketoacyl-CoA thiolase/enoyl-CoA hydratase (trifunctional protein), beta subunit | Cytoplasm | enzyme | 1504761.375 | 2818642.75 | 3806376.25 | | | | | | |
| Q99JY0 | HADHB | hydroxyacyl-CoA dehydrogenase/3-ketoacyl-CoA thiolase/enoyl-CoA hydratase (trifunctional protein), beta subunit | Cytoplasm | enzyme | 1610760.25 | 3882934.75 | 6217785.5 | Q99JY0 | HADHB | 3610151 | 4010345 | 875129 | 21 |
| Q99KB8 | HAGH | hydroxyacylglutathione hydrolase | Cytoplasm | enzyme | 7006257.5 | 7489381 | 5813104.5 | | | | | | |
| Q99KB8 | HAGH | hydroxyacylglutathione hydrolase | Cytoplasm | enzyme | 3508859.75 | 3868312.5 | 3667252.5 | | | | | | |
| Q99KB8 | HAGH | hydroxyacylglutathione hydrolase | Cytoplasm | enzyme | 545167.938 | 416429.75 | 395998.375 | Q99KB8 | HAGH | 3686762 | 2771158 | 923719 | 9 |
| P01942 | HBA1/HBA2 | hemoglobin, alpha 1 | Extracellular Space | transporter | 2.07E+09 | 2.10E+09 | 1.94E+09 | | | | | | |
| P01942 | HBA1/HBA2 | hemoglobin, alpha 1 | Extracellular Space | transporter | 1.60E+09 | 1.46E+09 | 1.19E+09 | | | | | | |
| P01942 | HBA1/HBA2 | hemoglobin, alpha 1 | Extracellular Space | transporter | 1.22E+09 | 1.09E+09 | 9.59E+08 | | | | | | |
| P01942 | HBA1/HBA2 | hemoglobin, alpha 1 | Extracellular Space | transporter | 1.31E+09 | 8.72E+08 | 9.34E+08 | | | | | | |
| P01942 | HBA1/HBA2 | hemoglobin, alpha 1 | Extracellular Space | transporter | 1.29E+09 | 1.15E+09 | 1.15E+09 | | | | | | |
| P01942 | HBA1/HBA2 | hemoglobin, alpha 1 | Extracellular Space | transporter | 9.54E+08 | 9.70E+08 | 1.03E+09 | | | | | | |
| P01942 | HBA1/HBA2 | hemoglobin, alpha 1 | Extracellular Space | transporter | 3.91E+08 | 2.67E+08 | 4.51E+08 | | | | | | |
| P01942 | HBA1/HBA2 | hemoglobin, alpha 1 | Extracellular Space | transporter | 2.07E+08 | 1.68E+08 | 1.42E+08 | | | | | | |
| P01942 | HBA1/HBA2 | hemoglobin, alpha 1 | Extracellular Space | transporter | 4.06E+08 | 6.42E+08 | 5.75E+08 | | | | | | |
| P01942 | HBA1/HBA2 | hemoglobin, alpha 1 | Extracellular Space | transporter | 1.35E+08 | 1.47E+08 | 1.28E+08 | | | | | | |
| P01942 | HBA1/HBA2 | hemoglobin, alpha 1 | Extracellular Space | transporter | 7.84E+07 | 8.69E+07 | 7.31E+07 | P01942 | HBA1/HBA2 | 878189091 | 608175744 | 105869808 | 33 |
| P02088 | Hbb-b1 | hemoglobin, beta adult major chain | Extracellular Space | transporter | 3.16E+09 | 2.88E+09 | 2.45E+09 | | | | | | |
| P02088 | Hbb-b1 | hemoglobin, beta adult major chain | Extracellular Space | transporter | 2.28E+09 | 2.09E+09 | 1.94E+09 | | | | | | |
| P02088 | Hbb-b1 | hemoglobin, beta adult major chain | Extracellular Space | transporter | 2.19E+09 | 2.05E+09 | 2.41E+09 | | | | | | |
| P02088 | Hbb-b1 | hemoglobin, beta adult major chain | Extracellular Space | transporter | 1.58E+09 | 1.32E+09 | 1.46E+09 | | | | | | |
| P02088 | Hbb-b1 | hemoglobin, beta adult major chain | Extracellular Space | transporter | 2.20E+09 | 2.09E+09 | 1.97E+09 | | | | | | |
| P02088 | Hbb-b1 | hemoglobin, beta adult major chain | Extracellular Space | transporter | 1.32E+09 | 1.45E+09 | 1.33E+09 | | | | | | |
| P02088 | Hbb-b1 | hemoglobin, beta adult major chain | Extracellular Space | transporter | 1.80E+09 | 1.64E+09 | 1.49E+09 | | | | | | |
| P02088 | Hbb-b1 | hemoglobin, beta adult major chain | Extracellular Space | transporter | 1.16E+09 | 1.10E+09 | 1.02E+09 | | | | | | |
| P02088 | Hbb-b1 | hemoglobin, beta adult major chain | Extracellular Space | transporter | 6.72E+08 | 6.07E+08 | 6.55E+08 | | | | | | |
| P02088 | Hbb-b1 | hemoglobin, beta adult major chain | Extracellular Space | transporter | 5.57E+08 | 4.79E+08 | 4.57E+08 | | | | | | |
| P02088 | Hbb-b1 | hemoglobin, beta adult major chain | Extracellular Space | transporter | 1.85E+08 | 1.55E+08 | 1.58E+08 | | | | | | |
| P02088 | Hbb-b1 | hemoglobin, beta adult major chain | Extracellular Space | transporter | 2.42E+08 | 2.58E+08 | 2.91E+08 | | | | | | |
| P02088 | Hbb-b1 | hemoglobin, beta adult major chain | Extracellular Space | transporter | 1.02E+08 | 8.24E+07 | 7.75E+07 | | | | | | |
| P02088 | Hbb-b1 | hemoglobin, beta adult major chain | Extracellular Space | transporter | 1.28E+08 | 1.37E+08 | 1.68E+08 | | | | | | |
| P02088 | Hbb-b1 | hemoglobin, beta adult major chain | Extracellular Space | transporter | 1.01E+07 | 8667116 | 2.00E+07 | | | | | | |
| P02088 | Hbb-b1 | hemoglobin, beta adult major chain | Extracellular Space | transporter | 9705908 | 1.00E+07 | 1.24E+07 | | | | | | |
| P02088 | Hbb-b1 | hemoglobin, beta adult major chain | Extracellular Space | transporter | 1.25E+07 | 1.38E+07 | 1.06E+07 | | | | | | |
| P02088 | Hbb-b1 | hemoglobin, beta adult major chain | Extracellular Space | transporter | 9213896 | 1.09E+07 | 1.51E+07 | | | | | | |
| P02088 | Hbb-b1 | hemoglobin, beta adult major chain | Extracellular Space | transporter | 923531.688 | 216121.078 | 1179325.25 | P02088 | Hbb-b1 | 926008070 | 924202808 | 122413652 | 57 |
| P02089 | Hbb-b2 | hemoglobin, beta adult minor chain | Extracellular Space | other | 5.31E+08 | 4.97E+08 | 4.19E+08 | | | | | | |
| P02089 | Hbb-b2 | hemoglobin, beta adult minor chain | Extracellular Space | other | 4.27E+08 | 4.26E+08 | 3.77E+08 | | | | | | |
| P02089 | Hbb-b2 | hemoglobin, beta adult minor chain | Extracellular Space | other | 3.25E+08 | 3.13E+08 | 3.14E+08 | | | | | | |
| P02089 | Hbb-b2 | hemoglobin, beta adult minor chain | Extracellular Space | other | 1.48E+08 | 1.39E+08 | 1.37E+08 | | | | | | |
| P02089 | Hbb-b2 | hemoglobin, beta adult minor chain | Extracellular Space | other | 1.32E+08 | 1.45E+08 | 1.30E+08 | | | | | | |
| P02089 | Hbb-b2 | hemoglobin, beta adult minor chain | Extracellular Space | other | 5.26E+07 | 4.67E+07 | 4.56E+07 | | | | | | |
| P02089 | Hbb-b2 | hemoglobin, beta adult minor chain | Extracellular Space | other | 2.97E+07 | 2.50E+07 | 1.95E+07 | | | | | | |
| P02089 | Hbb-b2 | hemoglobin, beta adult minor chain | Extracellular Space | other | 2879511.75 | 7730035.5 | 9794098 | P02089 | Hbb-b2 | 205867439 | 176437991 | 36015254 | 24 |
| P08103 | HCK | HCK proto-oncogene, Src family tyrosine kinase | Cytoplasm | kinase | 977344.438 | 1365189.125 | 1542227.125 | P08103 | HCK | 977344 | 288923 | 166810 | 3 |
| Q3U487 | HECTD3 | HECT domain containing E3 ubiquitin protein ligase 3 | Cytoplasm | enzyme | 2884096.75 | 2558523.5 | 2382354.25 | | | | | | |
| Q3U487 | HECTD3 | HECT domain containing E3 ubiquitin protein ligase 3 | Cytoplasm | enzyme | 802001.438 | 840738.125 | 874349.688 | Q3U487 | HECTD3 | 1843049 | 982631 | 401158 | 6 |
| P43275 | Hist1h1a | histone cluster 1, H1a | Nucleus | other | 4300288 | 1780005.125 | 2.45E+07 | | | | | | |
| P43275 | Hist1h1a | histone cluster 1, H1a | Nucleus | other | 2293242.25 | 1391045.375 | 2.01E+07 | | | | | | |
| P43275 | Hist1h1a | histone cluster 1, H1a | Nucleus | other | 4081278.25 | 1197500.125 | 3.05E+07 | | | | | | |
| P43275 | Hist1h1a | histone cluster 1, H1a | Nucleus | other | 256813.391 | 0 | 2406590.25 | P43275 | Hist1h1a | 2732905 | 10756272 | 3105068 | 12 |
| P43276 | Hist1h1b | histone cluster 1, H1b | Nucleus | other | 8096135.5 | 2481366.5 | 7.34E+07 | | | | | | |
| P43276 | Hist1h1b | histone cluster 1, H1b | Nucleus | other | 317886.438 | 89424.867 | 3176227 | P43276 | Hist1h1b | 4207011 | 28945768 | 11817060 | 6 |
| P15864 | HIST1H1C | histone cluster 1, H1c | Nucleus | other | 7.77E+07 | 2.36E+07 | 3.69E+08 | P15864 | HIST1H1C | 77650000 | 185575785 | 107142230 | 3 |
| P43274 | Hist1h1e | histone cluster 1, H1e | Nucleus | other | 4.62E+07 | 1.27E+07 | 2.45E+08 | | | | | | |
| P43274 | Hist1h1e | histone cluster 1, H1e | Nucleus | other | 1.72E+07 | 6290509.5 | 1.05E+08 | | | | | | |
| P43274 | Hist1h1e | histone cluster 1, H1e | Nucleus | other | 2.44E+07 | 7305717.5 | 1.31E+08 | | | | | | |
| P43274 | Hist1h1e | histone cluster 1, H1e | Nucleus | other | 8265186 | 1832451.625 | 5.87E+07 | | | | | | |
| P43274 | Hist1h1e | histone cluster 1, H1e | Nucleus | other | 53772.727 | 4.70E-06 | 1654301.875 | P43274 | Hist1h1e | 19215792 | 68290663 | 17632573 | 15 |
| Q8CGP5 | HIST1H2AJ | histone cluster 1, H2aj | Nucleus | other | 1.28E+08 | 5.38E+07 | 9.86E+08 | | | | | | |
| Q8CGP5 | HIST1H2AJ | histone cluster 1, H2aj | Nucleus | other | 1.25E+08 | 4.41E+07 | 7.11E+08 | | | | | | |
| Q8CGP5 | HIST1H2AJ | histone cluster 1, H2aj | Nucleus | other | 5.86E+07 | 1.70E+07 | 2.96E+08 | | | | | | |
| Q8CGP5 | HIST1H2AJ | histone cluster 1, H2aj | Nucleus | other | 4.23E+07 | 1.82E+07 | 2.42E+08 | | | | | | |
| Q8CGP5 | HIST1H2AJ | histone cluster 1, H2aj | Nucleus | other | 712171.5 | 273193.094 | 7911191.5 | Q8CGP5 | HIST1H2AJ | 70910434 | 288941575 | 74604394 | 15 |
| P70696 | HIST1H2BA | histone cluster 1, H2ba | Nucleus | other | 1.05E+08 | 3.76E+07 | 6.75E+08 | P70696 | HIST1H2BA | 105000000 | 349874350 | 202000050 | 3 |
| Q64475 | HIST1H2BB | histone cluster 1, H2bb | Nucleus | other | 7.98E+07 | 3.02E+07 | 5.99E+08 | | | | | | |
| Q64475 | HIST1H2BB | histone cluster 1, H2bb | Nucleus | other | 8.74E+07 | 2.95E+07 | 4.77E+08 | | | | | | |
| Q64475 | HIST1H2BB | histone cluster 1, H2bb | Nucleus | other | 6828603 | 3298320.5 | 5.33E+07 | | | | | | |
| Q64475 | HIST1H2BB | histone cluster 1, H2bb | Nucleus | other | 1943098.25 | 2035996.75 | 8303756 | | | | | | |
| Q64475 | HIST1H2BB | histone cluster 1, H2bb | Nucleus | other | 2847940.5 | 869703.5 | 2.15E+07 | | | | | | |
| Q64475 | HIST1H2BB | histone cluster 1, H2bb | Nucleus | other | 1407131 | 594132.938 | 1.19E+07 | Q64475 | HIST1H2BB | 30027795 | 170335325 | 40148421 | 18 |
| Q64522 | HIST2H2AB | histone cluster 2, H2ab | Nucleus | other | 4306913 | 1629472.5 | 3.28E+07 | Q64522 | HIST2H2AB | 4306913 | 17281107 | 9977252 | 3 |

| | | | | | | | | | | | | | |
|---|---|---|---|---|---|---|---|---|---|---|---|---|---|
| P17710 | HK1 | hexokinase 1 | Cytoplasm | kinase | 9695534 | 1.24E+07 | 1.90E+07 | | | | | | |
| P17710 | HK1 | hexokinase 1 | Cytoplasm | kinase | 5736559.5 | 5986825 | 7313929.5 | | | | | | |
| P17710 | HK1 | hexokinase 1 | Cytoplasm | kinase | 4180221.25 | 5692053 | 7644961.5 | | | | | | |
| P17710 | HK1 | hexokinase 1 | Cytoplasm | kinase | 3015696.5 | 4103382.75 | 7970313 | | | | | | |
| P17710 | HK1 | hexokinase 1 | Cytoplasm | kinase | 4809416.5 | 5691567 | 8802334 | | | | | | |
| P17710 | HK1 | hexokinase 1 | Cytoplasm | kinase | 4068164.75 | 5296406.5 | 6386109 | | | | | | |
| P17710 | HK1 | hexokinase 1 | Cytoplasm | kinase | 4574458.5 | 5449029 | 6699373 | | | | | | |
| P17710 | HK1 | hexokinase 1 | Cytoplasm | kinase | 3033240 | 3829384.5 | 4828959.5 | | | | | | |
| P17710 | HK1 | hexokinase 1 | Cytoplasm | kinase | 3593792 | 4795701.5 | 5698071.5 | | | | | | |
| P17710 | HK1 | hexokinase 1 | Cytoplasm | kinase | 2093316.5 | 2196550.75 | 5581198 | | | | | | |
| P17710 | HK1 | hexokinase 1 | Cytoplasm | kinase | 3103366.5 | 4071039 | 5492985.5 | | | | | | |
| P17710 | HK1 | hexokinase 1 | Cytoplasm | kinase | 1293562.75 | 2145191.5 | 4522351.5 | | | | | | |
| P17710 | HK1 | hexokinase 1 | Cytoplasm | kinase | 1198169.625 | 2051955.5 | 3037218.5 | P17710 | HK1 | 3876577 | 3233408 | 517760 | 39 |
| P01899 | HLA-A | major histocompatibility complex, class I, A | Plasma Membrane | other | 1.10E+07 | 1.43E+07 | 1.45E+07 | | | | | | |
| P01899 | HLA-A | major histocompatibility complex, class I, A | Plasma Membrane | other | 2949752 | 3664262.5 | 5453778 | | | | | | |
| P01899 | HLA-A | major histocompatibility complex, class I, A | Plasma Membrane | other | 5337126 | 6249666.5 | 7127283.5 | | | | | | |
| P01899 | HLA-A | major histocompatibility complex, class I, A | Plasma Membrane | other | 1643867 | 3152774.25 | 3049269.25 | | | | | | |
| P01899 | HLA-A | major histocompatibility complex, class I, A | Plasma Membrane | other | 1.13E+07 | 1.61E+07 | 1.69E+07 | | | | | | |
| P01899 | HLA-A | major histocompatibility complex, class I, A | Plasma Membrane | other | 1073087.625 | 1479022.125 | 1066965 | P01899 | | | | | |
| P01901 | HLA-A | major histocompatibility complex, class I, A | Plasma Membrane | other | 921725.5 | 960069.562 | 1650317.875 | P01901 | | | | | |
| P01902 | HLA-A | major histocompatibility complex, class I, A | Plasma Membrane | other | 19120.637 | 26086.707 | 222194.484 | | | | | | |
| P01902 | HLA-A | major histocompatibility complex, class I, A | Plasma Membrane | other | 0 | 0 | 0 | P01902 | HLA-A | 3810520 | 5484751 | 1055541 | 27 |
| P22907 | HMBS | hydroxymethylbilane synthase | Cytoplasm | enzyme | 1402017 | 1016018.562 | 1289933.375 | P22907 | HMBS | 1402017 | 198573 | 114646 | 3 |
| P30681 | HMGB2 | high mobility group box 2 | Nucleus | transcription regulator | 1232145.625 | 128847.688 | 4320523 | P30681 | HMGB2 | 1232146 | 2172766 | 1254447 | 3 |
| P38060 | HMGCL | 3-hydroxymethyl-3-methylglutaryl-CoA lyase | Cytoplasm | enzyme | 1338612.125 | 1713712.375 | 3077842.75 | | | | | | |
| P38060 | HMGCL | 3-hydroxymethyl-3-methylglutaryl-CoA lyase | Cytoplasm | enzyme | 355301.5 | 745162.5 | 446963.281 | P38060 | HMGCL | 846957 | 1025558 | 418682 | 6 |
| Q61411 | HRAS | Harvey rat sarcoma viral oncogene homolog | Plasma Membrane | enzyme | 1.37E+08 | 1.21E+08 | 1.08E+08 | | | | | | |
| Q61411 | HRAS | Harvey rat sarcoma viral oncogene homolog | Plasma Membrane | enzyme | 3.23E+07 | 3.26E+07 | 3.46E+07 | | | | | | |
| Q61411 | HRAS | Harvey rat sarcoma viral oncogene homolog | Plasma Membrane | enzyme | 2.05E+07 | 1.46E+07 | 1.37E+07 | | | | | | |
| Q61411 | HRAS | Harvey rat sarcoma viral oncogene homolog | Plasma Membrane | enzyme | 2.56E+07 | 3.31E+07 | 2.11E+07 | | | | | | |
| Q61411 | HRAS | Harvey rat sarcoma viral oncogene homolog | Plasma Membrane | enzyme | 2.34E+07 | 2.11E+07 | 1.59E+07 | | | | | | |
| Q61411 | HRAS | Harvey rat sarcoma viral oncogene homolog | Plasma Membrane | enzyme | 8614366 | 1.04E+07 | 5953597 | | | | | | |
| Q61411 | HRAS | Harvey rat sarcoma viral oncogene homolog | Plasma Membrane | enzyme | 1.43E+07 | 8733355 | 7551052 | | | | | | |
| Q61411 | HRAS | Harvey rat sarcoma viral oncogene homolog | Plasma Membrane | enzyme | 8657092 | 1.07E+07 | 5691149 | | | | | | |
| Q61411 | HRAS | Harvey rat sarcoma viral oncogene homolog | Plasma Membrane | enzyme | 3741031 | 5183848 | 3059219.25 | | | | | | |
| Q61411 | HRAS | Harvey rat sarcoma viral oncogene homolog | Plasma Membrane | enzyme | 1519362 | 1703043.5 | 1713723 | Q61411 | HRAS | 27591185 | 34570255 | 6311636 | 30 |
| O08756 | HSD17B10 | hydroxysteroid (17-beta) dehydrogenase 10 | Cytoplasm | enzyme | 6146025.5 | 7965212.5 | 1.37E+07 | | | | | | |
| O08756 | HSD17B10 | hydroxysteroid (17-beta) dehydrogenase 10 | Cytoplasm | enzyme | 2047681.5 | 2662561.75 | 3352879.75 | | | | | | |
| O08756 | HSD17B10 | hydroxysteroid (17-beta) dehydrogenase 10 | Cytoplasm | enzyme | 3007497 | 5570408.5 | 7054024.5 | | | | | | |
| O08756 | HSD17B10 | hydroxysteroid (17-beta) dehydrogenase 10 | Cytoplasm | enzyme | 886305.938 | 1257101.625 | 2709700.25 | O08756 | HSD17B10 | 3021877 | 3637645 | 1050098 | 12 |
| O70503 | HSD17B12 | hydroxysteroid (17-beta) dehydrogenase 12 | Cytoplasm | enzyme | 2795058.25 | 2432745.5 | 2929264.75 | | | | | | |
| O70503 | HSD17B12 | hydroxysteroid (17-beta) dehydrogenase 12 | Cytoplasm | enzyme | 2009126.75 | 1670598.875 | 2029428.25 | O70503 | HSD17B12 | 2402093 | 492323 | 200990 | 6 |
| Q8BTX9 | HSDL1 | hydroxysteroid dehydrogenase like 1 | Cytoplasm | enzyme | 423698.656 | 504185.531 | 715889.5 | Q8BTX9 | HSDL1 | 423699 | 150926 | 87137 | 3 |
| P07901 | HSP90AA1 | heat shock protein 90kDa alpha (cytosolic), class A member 1 | Cytoplasm | enzyme | 9971199 | 9549574 | 5538945.5 | | | | | | |
| P07901 | HSP90AA1 | heat shock protein 90kDa alpha (cytosolic), class A member 1 | Cytoplasm | enzyme | 7338467 | 4610194 | 2779825 | | | | | | |
| P07901 | HSP90AA1 | heat shock protein 90kDa alpha (cytosolic), class A member 1 | Cytoplasm | enzyme | 7476270 | 5530097 | 3700240 | | | | | | |
| P07901 | HSP90AA1 | heat shock protein 90kDa alpha (cytosolic), class A member 1 | Cytoplasm | enzyme | 5413210 | 3967538 | 1794754.125 | | | | | | |
| P07901 | HSP90AA1 | heat shock protein 90kDa alpha (cytosolic), class A member 1 | Cytoplasm | enzyme | 4777143 | 4237642.5 | 2089391.5 | | | | | | |
| P07901 | HSP90AA1 | heat shock protein 90kDa alpha (cytosolic), class A member 1 | Cytoplasm | enzyme | 3952619.75 | 2624192.5 | 1834210.125 | P07901 | HSP90AA1 | 6488151 | 2440952 | 575338 | 18 |
| P11499 | HSP90AB1 | heat shock protein 90kDa alpha (cytosolic), class B member 1 | Cytoplasm | enzyme | 6855057 | 5464243.5 | 4039497.5 | P11499 | HSP90AB1 | 6855057 | 1407814 | 812802 | 3 |
| P08113 | HSP90B1 | heat shock protein 90kDa beta (Grp94), member 1 | Cytoplasm | other | 4.00E+07 | 4.35E+07 | 4.18E+07 | | | | | | |
| P08113 | HSP90B1 | heat shock protein 90kDa beta (Grp94), member 1 | Cytoplasm | other | 4.55E+07 | 4.38E+07 | 3.61E+07 | | | | | | |
| P08113 | HSP90B1 | heat shock protein 90kDa beta (Grp94), member 1 | Cytoplasm | other | 3.86E+07 | 3.74E+07 | 3.42E+07 | | | | | | |
| P08113 | HSP90B1 | heat shock protein 90kDa beta (Grp94), member 1 | Cytoplasm | other | 2.50E+07 | 2.38E+07 | 2.33E+07 | | | | | | |
| P08113 | HSP90B1 | heat shock protein 90kDa beta (Grp94), member 1 | Cytoplasm | other | 2.59E+07 | 2.74E+07 | 2.82E+07 | | | | | | |
| P08113 | HSP90B1 | heat shock protein 90kDa beta (Grp94), member 1 | Cytoplasm | other | 3.31E+07 | 3.16E+07 | 3.55E+07 | | | | | | |
| P08113 | HSP90B1 | heat shock protein 90kDa beta (Grp94), member 1 | Cytoplasm | other | 3.39E+07 | 3.39E+07 | 3.25E+07 | | | | | | |
| P08113 | HSP90B1 | heat shock protein 90kDa beta (Grp94), member 1 | Cytoplasm | other | 2.30E+07 | 2.55E+07 | 2.67E+07 | | | | | | |
| P08113 | HSP90B1 | heat shock protein 90kDa beta (Grp94), member 1 | Cytoplasm | other | 2.62E+07 | 2.83E+07 | 2.26E+07 | | | | | | |
| P08113 | HSP90B1 | heat shock protein 90kDa beta (Grp94), member 1 | Cytoplasm | other | 2.42E+07 | 2.51E+07 | 2.44E+07 | | | | | | |
| P08113 | HSP90B1 | heat shock protein 90kDa beta (Grp94), member 1 | Cytoplasm | other | 2.23E+07 | 2.24E+07 | 1.86E+07 | | | | | | |
| P08113 | HSP90B1 | heat shock protein 90kDa beta (Grp94), member 1 | Cytoplasm | other | 1.95E+07 | 1.73E+07 | 1.40E+07 | | | | | | |
| P08113 | HSP90B1 | heat shock protein 90kDa beta (Grp94), member 1 | Cytoplasm | other | 2.50E+07 | 3.09E+07 | 2.96E+07 | | | | | | |
| P08113 | HSP90B1 | heat shock protein 90kDa beta (Grp94), member 1 | Cytoplasm | other | 2.99E+07 | 2.53E+07 | 2.17E+07 | | | | | | |
| P08113 | HSP90B1 | heat shock protein 90kDa beta (Grp94), member 1 | Cytoplasm | other | 2.03E+07 | 2.22E+07 | 1.96E+07 | | | | | | |
| P08113 | HSP90B1 | heat shock protein 90kDa beta (Grp94), member 1 | Cytoplasm | other | 1.99E+07 | 2.06E+07 | 1.96E+07 | | | | | | |
| P08113 | HSP90B1 | heat shock protein 90kDa beta (Grp94), member 1 | Cytoplasm | other | 1.98E+07 | 2.04E+07 | 2.05E+07 | | | | | | |
| P08113 | HSP90B1 | heat shock protein 90kDa beta (Grp94), member 1 | Cytoplasm | other | 1.80E+07 | 2.24E+07 | 1.55E+07 | | | | | | |
| P08113 | HSP90B1 | heat shock protein 90kDa beta (Grp94), member 1 | Cytoplasm | other | 1.77E+07 | 1.74E+07 | 1.69E+07 | | | | | | |
| P08113 | HSP90B1 | heat shock protein 90kDa beta (Grp94), member 1 | Cytoplasm | other | 1.68E+07 | 2.06E+07 | 2.19E+07 | | | | | | |
| P08113 | HSP90B1 | heat shock protein 90kDa beta (Grp94), member 1 | Cytoplasm | other | 1.47E+07 | 1.67E+07 | 1.56E+07 | | | | | | |
| P08113 | HSP90B1 | heat shock protein 90kDa beta (Grp94), member 1 | Cytoplasm | other | 1.26E+07 | 1.29E+07 | 1.27E+07 | | | | | | |
| P08113 | HSP90B1 | heat shock protein 90kDa beta (Grp94), member 1 | Cytoplasm | other | 9513516 | 9440582 | 9686797 | | | | | | |
| P08113 | HSP90B1 | heat shock protein 90kDa beta (Grp94), member 1 | Cytoplasm | other | 8414248 | 8106473 | 7751726 | | | | | | |
| P08113 | HSP90B1 | heat shock protein 90kDa beta (Grp94), member 1 | Cytoplasm | other | 2042544.875 | 1825999.125 | 1938065.125 | | | | | | |
| P08113 | HSP90B1 | heat shock protein 90kDa beta (Grp94), member 1 | Cytoplasm | other | 2711073 | 2252896.5 | 1822552 | P08113 | HSP90B1 | 22094669 | 10433464 | 1181357 | 78 |
| Q61316 | HSPA4 | heat shock 70kDa protein 4 | Cytoplasm | other | 2.52E+07 | 1.82E+07 | 1.48E+07 | | | | | | |
| Q61316 | HSPA4 | heat shock 70kDa protein 4 | Cytoplasm | other | 1.55E+07 | 1.22E+07 | 8530458 | | | | | | |
| Q61316 | HSPA4 | heat shock 70kDa protein 4 | Cytoplasm | other | 1.39E+07 | 9723229 | 7200805 | | | | | | |
| Q61316 | HSPA4 | heat shock 70kDa protein 4 | Cytoplasm | other | 9148642 | 5497070.5 | 5491468.5 | | | | | | |
| Q61316 | HSPA4 | heat shock 70kDa protein 4 | Cytoplasm | other | 1.04E+07 | 7959253 | 7150615.5 | | | | | | |
| Q61316 | HSPA4 | heat shock 70kDa protein 4 | Cytoplasm | other | 8527606 | 6421539 | 6285458 | | | | | | |
| Q61316 | HSPA4 | heat shock 70kDa protein 4 | Cytoplasm | other | 1.19E+07 | 8446758 | 7149520.5 | | | | | | |
| Q61316 | HSPA4 | heat shock 70kDa protein 4 | Cytoplasm | other | 1.22E+07 | 8895481 | 6860468 | | | | | | |
| Q61316 | HSPA4 | heat shock 70kDa protein 4 | Cytoplasm | other | 1.08E+07 | 8173659.5 | 5246579.5 | | | | | | |
| Q61316 | HSPA4 | heat shock 70kDa protein 4 | Cytoplasm | other | 1.21E+07 | 9542780 | 8831707 | | | | | | |
| Q61316 | HSPA4 | heat shock 70kDa protein 4 | Cytoplasm | other | 7662567 | 5631941 | 4387875.5 | | | | | | |
| Q61316 | HSPA4 | heat shock 70kDa protein 4 | Cytoplasm | other | 7609523 | 5928467.5 | 5218272 | | | | | | |
| Q61316 | HSPA4 | heat shock 70kDa protein 4 | Cytoplasm | other | 7519240 | 5202605 | 3755106.5 | | | | | | |
| Q61316 | HSPA4 | heat shock 70kDa protein 4 | Cytoplasm | other | 5047275 | 3612511.75 | 3307528.5 | | | | | | |
| Q61316 | HSPA4 | heat shock 70kDa protein 4 | Cytoplasm | other | 6832493.5 | 3824504.75 | 3892538 | | | | | | |
| Q61316 | HSPA4 | heat shock 70kDa protein 4 | Cytoplasm | other | 7433052 | 3412276 | 3460231.25 | | | | | | |
| Q61316 | HSPA4 | heat shock 70kDa protein 4 | Cytoplasm | other | 5229974.5 | 2778361.75 | 2341909.75 | | | | | | |
| Q61316 | HSPA4 | heat shock 70kDa protein 4 | Cytoplasm | other | 5688065.5 | 3969534 | 2231247 | | | | | | |
| Q61316 | HSPA4 | heat shock 70kDa protein 4 | Cytoplasm | other | 5083780 | 3467993.5 | 2478519 | | | | | | |
| Q61316 | HSPA4 | heat shock 70kDa protein 4 | Cytoplasm | other | 4819236.5 | 3269451 | 2709030.25 | | | | | | |
| Q61316 | HSPA4 | heat shock 70kDa protein 4 | Cytoplasm | other | 3438101.75 | 3476687.5 | 3681340.5 | | | | | | |
| Q61316 | HSPA4 | heat shock 70kDa protein 4 | Cytoplasm | other | 1654286.125 | 970715.5 | 296075.594 | | | | | | |
| Q61316 | HSPA4 | heat shock 70kDa protein 4 | Cytoplasm | other | 3554369.25 | 2639331.25 | 2090079.5 | Q61316 | HSPA4 | 8746879 | 4296950 | 517292 | 69 |
| P48722 | HSPA4L | heat shock 70kDa protein 4-like | Cytoplasm | other | 6520155.5 | 5171227 | 3186653 | | | | | | |
| P48722 | HSPA4L | heat shock 70kDa protein 4-like | Cytoplasm | other | 3676210.25 | 2133230.25 | 1270611.375 | | | | | | |
| P48722 | HSPA4L | heat shock 70kDa protein 4-like | Cytoplasm | other | 2665464 | 2321236.5 | 1657292.375 | | | | | | |
| P48722 | HSPA4L | heat shock 70kDa protein 4-like | Cytoplasm | other | 3424636.5 | 2920647 | 5816719.5 | | | | | | |
| P48722 | HSPA4L | heat shock 70kDa protein 4-like | Cytoplasm | other | 1908585.875 | 1730235.25 | 1034048.562 | P48722 | HSPA4L | 3639010 | 1658132 | 428128 | 15 |
| P20029 | HSPA5 | heat shock 70kDa protein 5 (glucose-regulated protein, 78kDa) | Cytoplasm | enzyme | 1.74E+08 | 1.61E+08 | 1.61E+08 | | | | | | |
| P20029 | HSPA5 | heat shock 70kDa protein 5 (glucose-regulated protein, 78kDa) | Cytoplasm | enzyme | 7.17E+07 | 7.46E+07 | 8.04E+07 | | | | | | |
| P20029 | HSPA5 | heat shock 70kDa protein 5 (glucose-regulated protein, 78kDa) | Cytoplasm | enzyme | 6.85E+07 | 6.92E+07 | 7.27E+07 | | | | | | |
| P20029 | HSPA5 | heat shock 70kDa protein 5 (glucose-regulated protein, 78kDa) | Cytoplasm | enzyme | 4.94E+07 | 6.25E+07 | 7.22E+07 | | | | | | |
| P20029 | HSPA5 | heat shock 70kDa protein 5 (glucose-regulated protein, 78kDa) | Cytoplasm | enzyme | 6.73E+07 | 7.72E+07 | 6.45E+07 | | | | | | |

| | | | | | | | | | | | | | |
|---|---|---|---|---|---|---|---|---|---|---|---|---|---|
| P20029 | HSPA5 | heat shock 70kDa protein 5 (glucose-regulated protein, 78kDa) | Cytoplasm | enzyme | 5.24E+07 | 6.20E+07 | 5.19E+07 | | | | | | |
| P20029 | HSPA5 | heat shock 70kDa protein 5 (glucose-regulated protein, 78kDa) | Cytoplasm | enzyme | 7.33E+07 | 7.87E+07 | 5.31E+07 | | | | | | |
| P20029 | HSPA5 | heat shock 70kDa protein 5 (glucose-regulated protein, 78kDa) | Cytoplasm | enzyme | 4.73E+07 | 5.40E+07 | 4.75E+07 | | | | | | |
| P20029 | HSPA5 | heat shock 70kDa protein 5 (glucose-regulated protein, 78kDa) | Cytoplasm | enzyme | 3.48E+07 | 4.16E+07 | 3.80E+07 | | | | | | |
| P20029 | HSPA5 | heat shock 70kDa protein 5 (glucose-regulated protein, 78kDa) | Cytoplasm | enzyme | 3.98E+07 | 4.10E+07 | 4.14E+07 | | | | | | |
| P20029 | HSPA5 | heat shock 70kDa protein 5 (glucose-regulated protein, 78kDa) | Cytoplasm | enzyme | 3.93E+07 | 4.21E+07 | 3.89E+07 | | | | | | |
| P20029 | HSPA5 | heat shock 70kDa protein 5 (glucose-regulated protein, 78kDa) | Cytoplasm | enzyme | 3.25E+07 | 3.52E+07 | 3.18E+07 | | | | | | |
| P20029 | HSPA5 | heat shock 70kDa protein 5 (glucose-regulated protein, 78kDa) | Cytoplasm | enzyme | 2.47E+07 | 2.53E+07 | 2.33E+07 | | | | | | |
| P20029 | HSPA5 | heat shock 70kDa protein 5 (glucose-regulated protein, 78kDa) | Cytoplasm | enzyme | 2.22E+07 | 2.65E+07 | 2.45E+07 | | | | | | |
| P20029 | HSPA5 | heat shock 70kDa protein 5 (glucose-regulated protein, 78kDa) | Cytoplasm | enzyme | 2.35E+07 | 1.75E+07 | 1.74E+07 | | | | | | |
| P20029 | HSPA5 | heat shock 70kDa protein 5 (glucose-regulated protein, 78kDa) | Cytoplasm | enzyme | 1.07E+07 | 1.04E+07 | 1.64E+07 | | | | | | |
| P20029 | HSPA5 | heat shock 70kDa protein 5 (glucose-regulated protein, 78kDa) | Cytoplasm | enzyme | 1.11E+07 | 1.57E+07 | 1.38E+07 | | | | | | |
| P20029 | HSPA5 | heat shock 70kDa protein 5 (glucose-regulated protein, 78kDa) | Cytoplasm | enzyme | 9555862 | 9425428 | 1.20E+07 | | | | | | |
| P20029 | HSPA5 | heat shock 70kDa protein 5 (glucose-regulated protein, 78kDa) | Cytoplasm | enzyme | 1.89E+07 | 3.42E+07 | 3.18E+07 | | | | | | |
| P20029 | HSPA5 | heat shock 70kDa protein 5 (glucose-regulated protein, 78kDa) | Cytoplasm | enzyme | 1.06E+07 | 1.16E+07 | 9892668 | | | | | | |
| P20029 | HSPA5 | heat shock 70kDa protein 5 (glucose-regulated protein, 78kDa) | Cytoplasm | enzyme | 7553480 | 8819515 | 6811330 | | | | | | |
| P20029 | HSPA5 | heat shock 70kDa protein 5 (glucose-regulated protein, 78kDa) | Cytoplasm | enzyme | 7074723 | 6673153.5 | 6994726 | | | | | | |
| P20029 | HSPA5 | heat shock 70kDa protein 5 (glucose-regulated protein, 78kDa) | Cytoplasm | enzyme | 2921681.75 | 4149820 | 4531406.5 | | | | | | |
| P20029 | HSPA5 | heat shock 70kDa protein 5 (glucose-regulated protein, 78kDa) | Cytoplasm | enzyme | 7779713 | 6706085 | 4635899.5 | | | | | | |
| P20029 | HSPA5 | heat shock 70kDa protein 5 (glucose-regulated protein, 78kDa) | Cytoplasm | enzyme | 837712 | 1050258.125 | 1947471 | | | | | | |
| P20029 | HSPA5 | heat shock 70kDa protein 5 (glucose-regulated protein, 78kDa) | Cytoplasm | enzyme | 2409021.25 | 1640544.5 | 2917875.25 | P20029 | HSPA5 | 34998931 | 35359460 | 4003670 | 78 |
| P63017 | HSPA8 | heat shock 70kDa protein 8 | Cytoplasm | enzyme | 1.33E+08 | 1.25E+08 | 9.61E+07 | | | | | | |
| P63017 | HSPA8 | heat shock 70kDa protein 8 | Cytoplasm | enzyme | 1.30E+08 | 1.14E+08 | 1.13E+08 | | | | | | |
| P63017 | HSPA8 | heat shock 70kDa protein 8 | Cytoplasm | enzyme | 1.06E+08 | 9.37E+07 | 8.10E+07 | | | | | | |
| P63017 | HSPA8 | heat shock 70kDa protein 8 | Cytoplasm | enzyme | 1.08E+08 | 8.97E+07 | 7.47E+07 | | | | | | |
| P63017 | HSPA8 | heat shock 70kDa protein 8 | Cytoplasm | enzyme | 8.62E+07 | 7.87E+07 | 8.81E+07 | | | | | | |
| P63017 | HSPA8 | heat shock 70kDa protein 8 | Cytoplasm | enzyme | 1.01E+08 | 8.34E+07 | 7.99E+07 | | | | | | |
| P63017 | HSPA8 | heat shock 70kDa protein 8 | Cytoplasm | enzyme | 9.06E+07 | 8.63E+07 | 7.42E+07 | | | | | | |
| P63017 | HSPA8 | heat shock 70kDa protein 8 | Cytoplasm | enzyme | 7.27E+07 | 5.92E+07 | 5.02E+07 | | | | | | |
| P63017 | HSPA8 | heat shock 70kDa protein 8 | Cytoplasm | enzyme | 6.53E+07 | 4.93E+07 | 5.15E+07 | | | | | | |
| P63017 | HSPA8 | heat shock 70kDa protein 8 | Cytoplasm | enzyme | 5.83E+07 | 5.30E+07 | 4.55E+07 | | | | | | |
| P63017 | HSPA8 | heat shock 70kDa protein 8 | Cytoplasm | enzyme | 6.17E+07 | 6.18E+07 | 5.65E+07 | | | | | | |
| P63017 | HSPA8 | heat shock 70kDa protein 8 | Cytoplasm | enzyme | 6.49E+07 | 5.97E+07 | 4.37E+07 | | | | | | |
| P63017 | HSPA8 | heat shock 70kDa protein 8 | Cytoplasm | enzyme | 7.44E+07 | 6.21E+07 | 6.46E+07 | | | | | | |
| P63017 | HSPA8 | heat shock 70kDa protein 8 | Cytoplasm | enzyme | 5.30E+07 | 4.90E+07 | 4.23E+07 | | | | | | |
| P63017 | HSPA8 | heat shock 70kDa protein 8 | Cytoplasm | enzyme | 5.04E+07 | 4.44E+07 | 4.39E+07 | | | | | | |
| P63017 | HSPA8 | heat shock 70kDa protein 8 | Cytoplasm | enzyme | 3.75E+07 | 3.29E+07 | 2.77E+07 | | | | | | |
| P63017 | HSPA8 | heat shock 70kDa protein 8 | Cytoplasm | enzyme | 3.13E+07 | 3.00E+07 | 2.75E+07 | | | | | | |
| P63017 | HSPA8 | heat shock 70kDa protein 8 | Cytoplasm | enzyme | 3.64E+07 | 2.95E+07 | 2.84E+07 | | | | | | |
| P63017 | HSPA8 | heat shock 70kDa protein 8 | Cytoplasm | enzyme | 2.31E+07 | 1.75E+07 | 1.57E+07 | | | | | | |
| P63017 | HSPA8 | heat shock 70kDa protein 8 | Cytoplasm | enzyme | 2.71E+07 | 2.13E+07 | 2.44E+07 | | | | | | |
| P63017 | HSPA8 | heat shock 70kDa protein 8 | Cytoplasm | enzyme | 2.01E+07 | 2.32E+07 | 2.23E+07 | | | | | | |
| P63017 | HSPA8 | heat shock 70kDa protein 8 | Cytoplasm | enzyme | 5021091 | 4418474 | 6527715 | | | | | | |
| P63017 | HSPA8 | heat shock 70kDa protein 8 | Cytoplasm | enzyme | 3649646 | 2695325.5 | 2244739.75 | | | | | | |
| P63017 | HSPA8 | heat shock 70kDa protein 8 | Cytoplasm | enzyme | 4153200 | 3701245 | 2674030 | | | | | | |
| P63017 | HSPA8 | heat shock 70kDa protein 8 | Cytoplasm | enzyme | 2490687.25 | 2083116.875 | 814181.375 | | | | | | |
| P63017 | HSPA8 | heat shock 70kDa protein 8 | Cytoplasm | enzyme | 1827395.25 | 1850382 | 917080.188 | P63017 | HSPA8 | 55678155 | 35816283 | 4055396 | 78 |
| P38647 | HSPA9 | heat shock 70kDa protein 9 (mortalin) | Cytoplasm | other | 6483343.5 | 1.16E+07 | 1.20E+07 | | | | | | |
| P38647 | HSPA9 | heat shock 70kDa protein 9 (mortalin) | Cytoplasm | other | 5189284.5 | 9074223 | 1.10E+07 | | | | | | |
| P38647 | HSPA9 | heat shock 70kDa protein 9 (mortalin) | Cytoplasm | other | 5849145.5 | 9496686 | 1.29E+07 | | | | | | |
| P38647 | HSPA9 | heat shock 70kDa protein 9 (mortalin) | Cytoplasm | other | 5370724 | 8869411 | 1.08E+07 | | | | | | |
| P38647 | HSPA9 | heat shock 70kDa protein 9 (mortalin) | Cytoplasm | other | 3817065 | 8646927 | 1.04E+07 | | | | | | |
| P38647 | HSPA9 | heat shock 70kDa protein 9 (mortalin) | Cytoplasm | other | 4689996.5 | 8368919.5 | 8761023 | | | | | | |
| P38647 | HSPA9 | heat shock 70kDa protein 9 (mortalin) | Cytoplasm | other | 3696540.5 | 7134747 | 7500690 | | | | | | |
| P38647 | HSPA9 | heat shock 70kDa protein 9 (mortalin) | Cytoplasm | other | 3230195.75 | 5258920 | 8217915.5 | | | | | | |
| P38647 | HSPA9 | heat shock 70kDa protein 9 (mortalin) | Cytoplasm | other | 2820095.25 | 4482285 | 6790099.5 | | | | | | |
| P38647 | HSPA9 | heat shock 70kDa protein 9 (mortalin) | Cytoplasm | other | 2946677 | 9242834 | 1.16E+07 | | | | | | |
| P38647 | HSPA9 | heat shock 70kDa protein 9 (mortalin) | Cytoplasm | other | 1727562.75 | 1599968.75 | 1726275 | | | | | | |
| P38647 | HSPA9 | heat shock 70kDa protein 9 (mortalin) | Cytoplasm | other | 1389503.875 | 1549453.25 | 2372368.75 | P38647 | HSPA9 | 3934178 | 3487685 | 581281 | 36 |
| P63038 | HSPD1 | heat shock 60kDa protein 1 (chaperonin) | Cytoplasm | enzyme | 1.79E+07 | 3.53E+07 | 3.71E+07 | | | | | | |
| P63038 | HSPD1 | heat shock 60kDa protein 1 (chaperonin) | Cytoplasm | enzyme | 1.57E+07 | 2.58E+07 | 2.57E+07 | | | | | | |
| P63038 | HSPD1 | heat shock 60kDa protein 1 (chaperonin) | Cytoplasm | enzyme | 1.59E+07 | 2.70E+07 | 2.83E+07 | | | | | | |
| P63038 | HSPD1 | heat shock 60kDa protein 1 (chaperonin) | Cytoplasm | enzyme | 1.43E+07 | 2.78E+07 | 3.49E+07 | | | | | | |
| P63038 | HSPD1 | heat shock 60kDa protein 1 (chaperonin) | Cytoplasm | enzyme | 1.34E+07 | 2.59E+07 | 2.32E+07 | | | | | | |
| P63038 | HSPD1 | heat shock 60kDa protein 1 (chaperonin) | Cytoplasm | enzyme | 1.26E+07 | 2.43E+07 | 2.52E+07 | | | | | | |
| P63038 | HSPD1 | heat shock 60kDa protein 1 (chaperonin) | Cytoplasm | enzyme | 1.12E+07 | 2.18E+07 | 2.62E+07 | | | | | | |
| P63038 | HSPD1 | heat shock 60kDa protein 1 (chaperonin) | Cytoplasm | enzyme | 1.08E+07 | 2.01E+07 | 2.54E+07 | | | | | | |
| P63038 | HSPD1 | heat shock 60kDa protein 1 (chaperonin) | Cytoplasm | enzyme | 1.04E+07 | 1.90E+07 | 2.21E+07 | | | | | | |
| P63038 | HSPD1 | heat shock 60kDa protein 1 (chaperonin) | Cytoplasm | enzyme | 9017203 | 1.71E+07 | 1.72E+07 | | | | | | |
| P63038 | HSPD1 | heat shock 60kDa protein 1 (chaperonin) | Cytoplasm | enzyme | 1.05E+07 | 1.59E+07 | 1.51E+07 | | | | | | |
| P63038 | HSPD1 | heat shock 60kDa protein 1 (chaperonin) | Cytoplasm | enzyme | 7527642 | 1.51E+07 | 1.50E+07 | | | | | | |
| P63038 | HSPD1 | heat shock 60kDa protein 1 (chaperonin) | Cytoplasm | enzyme | 7124101.5 | 1.39E+07 | 1.48E+07 | | | | | | |
| P63038 | HSPD1 | heat shock 60kDa protein 1 (chaperonin) | Cytoplasm | enzyme | 1.00E+07 | 1.92E+07 | 1.96E+07 | | | | | | |
| P63038 | HSPD1 | heat shock 60kDa protein 1 (chaperonin) | Cytoplasm | enzyme | 6820636 | 1.01E+07 | 1.47E+07 | | | | | | |
| P63038 | HSPD1 | heat shock 60kDa protein 1 (chaperonin) | Cytoplasm | enzyme | 1.29E+07 | 2.09E+07 | 2.55E+07 | | | | | | |
| P63038 | HSPD1 | heat shock 60kDa protein 1 (chaperonin) | Cytoplasm | enzyme | 3047443.25 | 4971610.5 | 5010020.5 | | | | | | |
| P63038 | HSPD1 | heat shock 60kDa protein 1 (chaperonin) | Cytoplasm | enzyme | 2048179 | 4158102 | 5709710.5 | | | | | | |
| P63038 | HSPD1 | heat shock 60kDa protein 1 (chaperonin) | Cytoplasm | enzyme | 2285461 | 2344145.75 | 3465253.75 | | | | | | |
| P63038 | HSPD1 | heat shock 60kDa protein 1 (chaperonin) | Cytoplasm | enzyme | 1729018.25 | 4980704 | 7131762.5 | | | | | | |
| P63038 | HSPD1 | heat shock 60kDa protein 1 (chaperonin) | Cytoplasm | enzyme | 789650.188 | 2037172.25 | 2147175.75 | | | | | | |
| P63038 | HSPD1 | heat shock 60kDa protein 1 (chaperonin) | Cytoplasm | enzyme | 1851754.125 | 3187042.25 | 2244398 | | | | | | |
| P63038 | HSPD1 | heat shock 60kDa protein 1 (chaperonin) | Cytoplasm | enzyme | 596412.75 | 1868688.625 | 2297803 | P63038 | HSPD1 | 8627717 | 9620743 | 1158201 | 69 |
| Q64433 | HSPE1 | heat shock 10kDa protein 1 | Cytoplasm | enzyme | 1.45E+07 | 2.41E+07 | 2.70E+07 | | | | | | |
| Q64433 | HSPE1 | heat shock 10kDa protein 1 | Cytoplasm | enzyme | 9717058 | 1.75E+07 | 1.97E+07 | | | | | | |
| Q64433 | HSPE1 | heat shock 10kDa protein 1 | Cytoplasm | enzyme | 9239453 | 1.57E+07 | 1.61E+07 | | | | | | |
| Q64433 | HSPE1 | heat shock 10kDa protein 1 | Cytoplasm | enzyme | 3795380.5 | 7085274.5 | 8950917 | | | | | | |
| Q64433 | HSPE1 | heat shock 10kDa protein 1 | Cytoplasm | enzyme | 3328127 | 5493106 | 6229264 | Q64433 | HSPE1 | 8120004 | 7357997 | 1899827 | 15 |
| Q9Z2G9 | HTATIP2 | HIV-1 Tat interactive protein 2, 30kDa | Nucleus | transcription regulator | 861454.5 | 2359234 | 464610.406 | | | | | | |
| Q9Z2G9 | HTATIP2 | HIV-1 Tat interactive protein 2, 30kDa | Nucleus | transcription regulator | 601318.812 | 1072760.25 | 465039.875 | Q9Z2G9 | HTATIP2 | 731387 | 720921 | 294315 | 6 |
| Q7TMY8 | HUWE1 | HECT, UBA and WWE domain containing 1, E3 ubiquitin protein ligase | Nucleus | transcription regulator | 2515339.25 | 2036116 | 2372839.75 | Q7TMY8 | HUWE1 | 2515339 | 246084 | 142077 | 3 |
| Q91ZJ9 | HYAL1 | hyaluronoglucosaminidase 1 | Cytoplasm | enzyme | 4353636 | 4373475.5 | 4395573 | Q91ZJ9 | HYAL1 | 4353636 | 20979 | 12112 | 3 |
| Q9JKR6 | HYOU1 | hypoxia up-regulated 1 | Cytoplasm | other | 5092016.5 | 4941721.5 | 4549477.5 | | | | | | |
| Q9JKR6 | HYOU1 | hypoxia up-regulated 1 | Cytoplasm | other | 3882054.75 | 4096191 | 3554116.5 | | | | | | |
| Q9JKR6 | HYOU1 | hypoxia up-regulated 1 | Cytoplasm | other | 4310322 | 3504324.5 | 4224077.5 | | | | | | |
| Q9JKR6 | HYOU1 | hypoxia up-regulated 1 | Cytoplasm | other | 3778647 | 3336709.75 | 3294568 | | | | | | |
| Q9JKR6 | HYOU1 | hypoxia up-regulated 1 | Cytoplasm | other | 4062300.5 | 2643294.25 | 2536274.5 | | | | | | |
| Q9JKR6 | HYOU1 | hypoxia up-regulated 1 | Cytoplasm | other | 2918322.75 | 2318986.5 | 1834444.375 | | | | | | |
| Q9JKR6 | HYOU1 | hypoxia up-regulated 1 | Cytoplasm | other | 2144380.5 | 2280043.75 | 2755744.75 | Q9JKR6 | HYOU1 | 3741149 | 943750 | 205943 | 21 |
| Q8BIJ6 | IARS2 | isoleucyl-tRNA synthetase 2, mitochondrial | Cytoplasm | enzyme | 1157895.875 | 2680691.75 | 3650097.25 | Q8BIJ6 | IARS2 | 1157896 | 1256299 | 725325 | 3 |
| Q9ERM2 | ICAM4 | intercellular adhesion molecule 4 (Landsteiner-Wiener blood group) | Plasma Membrane | other | 34660.59 | 22197.473 | 1880.717 | | | | | | |
| Q9ERM2 | ICAM4 | intercellular adhesion molecule 4 (Landsteiner-Wiener blood group) | Plasma Membrane | other | 113493.664 | 147105.484 | 187040.5 | Q9ERM2 | ICAM4 | 74077 | 75453 | 30804 | 6 |
| Q9JHR7 | IDE | insulin-degrading enzyme | Extracellular Space | peptidase | 2564229.25 | 1508484.5 | 1707192 | | | | | | |
| Q9JHR7 | IDE | insulin-degrading enzyme | Extracellular Space | peptidase | 2937950.75 | 1996102.375 | 1220719.875 | Q9JHR7 | IDE | 2751090 | 652995 | 266584 | 6 |
| P54071 | IDH2 | isocitrate dehydrogenase 2 (NADP+), mitochondrial | Cytoplasm | enzyme | 3901503.75 | 9061801 | 1.04E+07 | | | | | | |
| P54071 | IDH2 | isocitrate dehydrogenase 2 (NADP+), mitochondrial | Cytoplasm | enzyme | 2049340.375 | 1806593.625 | 1761601.625 | | | | | | |
| P54071 | IDH2 | isocitrate dehydrogenase 2 (NADP+), mitochondrial | Cytoplasm | enzyme | 1502038.75 | 3260822 | 3730419.75 | | | | | | |
| P54071 | IDH2 | isocitrate dehydrogenase 2 (NADP+), mitochondrial | Cytoplasm | enzyme | 1341358.5 | 2123936.75 | 3282206.25 | | | | | | |
| P54071 | IDH2 | isocitrate dehydrogenase 2 (NADP+), mitochondrial | Cytoplasm | enzyme | 1124359.375 | 2275571.5 | 3332940.5 | | | | | | |
| P54071 | IDH2 | isocitrate dehydrogenase 2 (NADP+), mitochondrial | Cytoplasm | enzyme | 936319.875 | 612591 | 1834256.5 | P54071 | IDH2 | 1809153 | 2630658 | 620052 | 18 |

| | | | | | | | | | | | | | |
|---|---|---|---|---|---|---|---|---|---|---|---|---|---|
| Q9D6R2 | IDH3A | isocitrate dehydrogenase 3 (NAD+) alpha | Cytoplasm | enzyme | 1.00E+07 | 2.18E+07 | 2.03E+07 | | | | | | |
| Q9D6R2 | IDH3A | isocitrate dehydrogenase 3 (NAD+) alpha | Cytoplasm | enzyme | 3831614.5 | 6346280.5 | 7122358.5 | | | | | | |
| Q9D6R2 | IDH3A | isocitrate dehydrogenase 3 (NAD+) alpha | Cytoplasm | enzyme | 2977808.75 | 6579637 | 6600607 | | | | | | |
| Q9D6R2 | IDH3A | isocitrate dehydrogenase 3 (NAD+) alpha | Cytoplasm | enzyme | 3048099.25 | 4965901.5 | 7094504.5 | | | | | | |
| Q9D6R2 | IDH3A | isocitrate dehydrogenase 3 (NAD+) alpha | Cytoplasm | enzyme | 4046145.75 | 6099889 | 6276342.5 | Q9D6R2 | IDH3A | 4788734 | 5673212 | 1464817 | 15 |
| P70404 | IDH3G | isocitrate dehydrogenase 3 (NAD+) gamma | Cytoplasm | enzyme | 4212406.5 | 7694460 | 8180325.5 | | | | | | |
| P70404 | IDH3G | isocitrate dehydrogenase 3 (NAD+) gamma | Cytoplasm | enzyme | 1145929.5 | 1912141.625 | 2130943.5 | P70404 | IDH3G | 2679168 | 3062550 | 1250281 | 6 |
| Q9CR20 | IER3IP1 | immediate early response 3 interacting protein 1 | Cytoplasm | other | 1.24E+07 | 1.03E+07 | 1.13E+07 | | | | | | |
| Q9CR20 | IER3IP1 | immediate early response 3 interacting protein 1 | Cytoplasm | other | 3239512 | 3603702 | 2891366.75 | Q9CR20 | IER3IP1 | 7829756 | 4485536 | 1831213 | 6 |
| P01872 | IGHM | immunoglobulin heavy constant mu | Plasma Membrane | transmembrane receptor | 6124520.5 | 4460866.5 | 2936522.25 | | | | | | |
| P01872 | IGHM | immunoglobulin heavy constant mu | Plasma Membrane | transmembrane receptor | 1926157.5 | 687003.625 | 1597856.625 | | | | | | |
| P01872 | IGHM | immunoglobulin heavy constant mu | Plasma Membrane | transmembrane receptor | 2581957.75 | 1284220.5 | 1933100.875 | | | | | | |
| P01872 | IGHM | immunoglobulin heavy constant mu | Plasma Membrane | transmembrane receptor | 2290530.25 | 1338306.375 | 2316778.25 | P01872 | IGHM | 3230792 | 1501916 | 433566 | 12 |
| O55222 | ILK | integrin-linked kinase | Plasma Membrane | kinase | 4327573.5 | 7009787.5 | 9323869 | | | | | | |
| O55222 | ILK | integrin-linked kinase | Plasma Membrane | kinase | 1579860.75 | 1215503.875 | 6765277 | | | | | | |
| O55222 | ILK | integrin-linked kinase | Plasma Membrane | kinase | 2153092 | 1049947.75 | 6417971 | | | | | | |
| O55222 | ILK | integrin-linked kinase | Plasma Membrane | kinase | 1303600.125 | 861037.875 | 5213324.5 | | | | | | |
| O55222 | ILK | integrin-linked kinase | Plasma Membrane | kinase | 1239628.75 | 239648.031 | 4379400 | | | | | | |
| O55222 | ILK | integrin-linked kinase | Plasma Membrane | kinase | 895776.062 | 574790.375 | 3790185.75 | O55222 | ILK | 1916589 | 2746906 | 647452 | 18 |
| Q8BU33 | ILVBL | ilvB (bacterial acetolactate synthase)-like | Cytoplasm | enzyme | 825252.125 | 1612855.625 | 1031487.938 | Q8BU33 | ILVBL | 825252 | 408420 | 235801 | 3 |
| Q8CAQ8 | IMMT | inner membrane protein, mitochondrial | Cytoplasm | other | 3649041.75 | 5802334.5 | 7487509 | | | | | | |
| Q8CAQ8 | IMMT | inner membrane protein, mitochondrial | Cytoplasm | other | 3992517.75 | 5733894.5 | 8164572 | | | | | | |
| Q8CAQ8 | IMMT | inner membrane protein, mitochondrial | Cytoplasm | other | 3106274.25 | 5689493 | 6286669.5 | | | | | | |
| Q8CAQ8 | IMMT | inner membrane protein, mitochondrial | Cytoplasm | other | 3708418 | 6318918.5 | 7217643 | | | | | | |
| Q8CAQ8 | IMMT | inner membrane protein, mitochondrial | Cytoplasm | other | 2735014 | 5268858 | 6279993.5 | | | | | | |
| Q8CAQ8 | IMMT | inner membrane protein, mitochondrial | Cytoplasm | other | 1780623.375 | 2095921.25 | 2542375 | | | | | | |
| Q8CAQ8 | IMMT | inner membrane protein, mitochondrial | Cytoplasm | other | 1616222.25 | 3112739.5 | 4351472 | | | | | | |
| Q8CAQ8 | IMMT | inner membrane protein, mitochondrial | Cytoplasm | other | 757739.062 | 1130740.125 | 1175396.875 | | | | | | |
| Q8CAQ8 | IMMT | inner membrane protein, mitochondrial | Cytoplasm | other | 1195922.25 | 1945868.125 | 2383804.75 | Q8CAQ8 | IMMT | 2504641 | 2214040 | 426092 | 27 |
| P24547 | IMPDH2 | IMP (inosine 5'-monophosphate) dehydrogenase 2 | Cytoplasm | enzyme | 2341658 | 236163.672 | 31239.066 | P24547 | IMPDH2 | 2341658 | 1278876 | 738359 | 3 |
| Q8C5L6 | INPP5K | inositol polyphosphate-5-phosphatase K | Cytoplasm | phosphatase | 5344923.5 | 6161206 | 6555521 | | | | | | |
| Q8C5L6 | INPP5K | inositol polyphosphate-5-phosphatase K | Cytoplasm | phosphatase | 6710520.5 | 6684407 | 7682356.5 | Q8C5L6 | INPP5K | 6027722 | 764871 | 312257 | 6 |
| Q8CIM8 | INTS4 | integrator complex subunit 4 | Nucleus | other | 3.39E+07 | 3.51E+07 | 3.56E+07 | Q8CIM8 | INTS4 | 33930000 | 855628 | 493997 | 3 |
| Q8BKC5 | IPO5 | importin 5 | Nucleus | transporter | 9079783 | 9453700 | 7747500.5 | | | | | | |
| Q8BKC5 | IPO5 | importin 5 | Nucleus | transporter | 4735467 | 3812868.75 | 1819942.625 | | | | | | |
| Q8BKC5 | IPO5 | importin 5 | Nucleus | transporter | 1.26E+07 | 1.41E+07 | 9886582 | | | | | | |
| Q8BKC5 | IPO5 | importin 5 | Nucleus | transporter | 1.64E+07 | 1.95E+07 | 1.05E+07 | | | | | | |
| Q8BKC5 | IPO5 | importin 5 | Nucleus | transporter | 6498200.5 | 6071966 | 4428265.5 | | | | | | |
| Q8BKC5 | IPO5 | importin 5 | Nucleus | transporter | 7507173 | 6434667 | 4964674.5 | | | | | | |
| Q8BKC5 | IPO5 | importin 5 | Nucleus | transporter | 6790384 | 6595347 | 5831329.5 | | | | | | |
| Q8BKC5 | IPO5 | importin 5 | Nucleus | transporter | 7118836 | 6441270.5 | 5909070.5 | | | | | | |
| Q8BKC5 | IPO5 | importin 5 | Nucleus | transporter | 5974080 | 4796521 | 3304800 | | | | | | |
| Q8BKC5 | IPO5 | importin 5 | Nucleus | transporter | 5923587.5 | 5142695 | 4842640 | | | | | | |
| Q8BKC5 | IPO5 | importin 5 | Nucleus | transporter | 5561520 | 4736545 | 4338114 | | | | | | |
| Q8BKC5 | IPO5 | importin 5 | Nucleus | transporter | 5161081.5 | 5102542.5 | 5367460.5 | | | | | | |
| Q8BKC5 | IPO5 | importin 5 | Nucleus | transporter | 5810160 | 4426850 | 3371383.25 | | | | | | |
| Q8BKC5 | IPO5 | importin 5 | Nucleus | transporter | 4963265 | 4229892.5 | 3254417.75 | | | | | | |
| Q8BKC5 | IPO5 | importin 5 | Nucleus | transporter | 3257157.5 | 3891204.5 | 2895043.25 | | | | | | |
| Q8BKC5 | IPO5 | importin 5 | Nucleus | transporter | 2804063 | 3672804.75 | 3203573.25 | | | | | | |
| Q8BKC5 | IPO5 | importin 5 | Nucleus | transporter | 3589229 | 2449790.5 | 1761578.875 | | | | | | |
| Q8BKC5 | IPO5 | importin 5 | Nucleus | transporter | 4562787 | 3925472.5 | 3190395.5 | | | | | | |
| Q8BKC5 | IPO5 | importin 5 | Nucleus | transporter | 2625923.5 | 2524777 | 2160356 | | | | | | |
| Q8BKC5 | IPO5 | importin 5 | Nucleus | transporter | 4115582.5 | 3054831.5 | 2541893.25 | Q8BKC5 | IPO5 | 6254914 | 3396424 | 438476 | 60 |
| Q9EPL8 | IPO7 | importin 7 | Nucleus | transporter | 3885832.25 | 3385403.25 | 2770563.25 | | | | | | |
| Q9EPL8 | IPO7 | importin 7 | Nucleus | transporter | 2824935 | 2403393.25 | 2251412.75 | | | | | | |
| Q9EPL8 | IPO7 | importin 7 | Nucleus | transporter | 1864391.125 | 2254239.5 | 1555955.375 | | | | | | |
| Q9EPL8 | IPO7 | importin 7 | Nucleus | transporter | 2789116 | 2449803.25 | 2178958.25 | | | | | | |
| Q9EPL8 | IPO7 | importin 7 | Nucleus | transporter | 1718543.125 | 1199616.875 | 1295468.5 | | | | | | |
| Q9EPL8 | IPO7 | importin 7 | Nucleus | transporter | 1411155.375 | 1488267.875 | 1360862.875 | | | | | | |
| Q9EPL8 | IPO7 | importin 7 | Nucleus | transporter | 817903.875 | 671674.75 | 897909.188 | Q9EPL8 | IPO7 | 2187411 | 857607 | 187145 | 21 |
| Q60766 | Irgm1 | immunity-related GTPase family M member 1 | Cytoplasm | other | 1.71E+07 | 2.03E+07 | 1.37E+07 | | | | | | |
| Q60766 | Irgm1 | immunity-related GTPase family M member 1 | Cytoplasm | other | 1.22E+07 | 1.55E+07 | 1.23E+07 | | | | | | |
| Q60766 | Irgm1 | immunity-related GTPase family M member 1 | Cytoplasm | other | 1.07E+07 | 1.65E+07 | 1.30E+07 | | | | | | |
| Q60766 | Irgm1 | immunity-related GTPase family M member 1 | Cytoplasm | other | 1.32E+07 | 1.41E+07 | 8121761.5 | | | | | | |
| Q60766 | Irgm1 | immunity-related GTPase family M member 1 | Cytoplasm | other | 9372718 | 9533886 | 8902991 | | | | | | |
| Q60766 | Irgm1 | immunity-related GTPase family M member 1 | Cytoplasm | other | 8793516 | 9572778 | 6864180.5 | | | | | | |
| Q60766 | Irgm1 | immunity-related GTPase family M member 1 | Cytoplasm | other | 7399368.5 | 1.07E+07 | 6470715 | | | | | | |
| Q60766 | Irgm1 | immunity-related GTPase family M member 1 | Cytoplasm | other | 4011108.75 | 4475433 | 3293443.75 | | | | | | |
| Q60766 | Irgm1 | immunity-related GTPase family M member 1 | Cytoplasm | other | 3513070.75 | 9065847 | 8266191.5 | | | | | | |
| Q60766 | Irgm1 | immunity-related GTPase family M member 1 | Cytoplasm | other | 1459981.875 | 3832688.25 | 2506644.5 | Q60766 | Irgm1 | 8765976 | 4700102 | 858117 | 30 |
| Q9D924 | ISCA1 | iron-sulfur cluster assembly 1 | Cytoplasm | other | 2063059.625 | 3945987.75 | 6495777 | | | | | | |
| Q9D924 | ISCA1 | iron-sulfur cluster assembly 1 | Cytoplasm | other | 980395.125 | 2020973.125 | 2897574.5 | Q9D924 | ISCA1 | 1521727 | 1950068 | 796112 | 6 |
| Q9DCB8 | ISCA2 | iron-sulfur cluster assembly 2 | Cytoplasm | other | 1764914.25 | 2729117.5 | 3871330 | Q9DCB8 | ISCA2 | 1764914 | 1054461 | 608793 | 3 |
| Q64339 | ISG15 | ISG15 ubiquitin-like modifier | Extracellular Space | other | 5.85E+07 | 5.97E+07 | 3.45E+07 | | | | | | |
| Q64339 | ISG15 | ISG15 ubiquitin-like modifier | Extracellular Space | other | 4.83E+07 | 4.75E+07 | 2.97E+07 | | | | | | |
| Q64339 | ISG15 | ISG15 ubiquitin-like modifier | Extracellular Space | other | 3.68E+07 | 2.58E+07 | 1.65E+07 | | | | | | |
| Q64339 | ISG15 | ISG15 ubiquitin-like modifier | Extracellular Space | other | 1.56E+07 | 1.07E+07 | 1.34E+07 | | | | | | |
| Q64339 | ISG15 | ISG15 ubiquitin-like modifier | Extracellular Space | other | 2.68E+07 | 2.74E+07 | 2.35E+07 | | | | | | |
| Q64339 | ISG15 | ISG15 ubiquitin-like modifier | Extracellular Space | other | 1.53E+07 | 1.42E+07 | 6378454.5 | | | | | | |
| Q64339 | ISG15 | ISG15 ubiquitin-like modifier | Extracellular Space | other | 3586660.75 | 2360906.25 | 2868704.75 | | | | | | |
| Q64339 | ISG15 | ISG15 ubiquitin-like modifier | Extracellular Space | other | 3820883.5 | 4553743 | 3380199.75 | Q64339 | ISG15 | 26083443 | 17806216 | 3634679 | 24 |
| Q9JL16 | ISG20 | interferon stimulated exonuclease gene 20kDa | Nucleus | enzyme | 5503358 | 6121339.5 | 5917974.5 | | | | | | |
| Q9JL16 | ISG20 | interferon stimulated exonuclease gene 20kDa | Nucleus | enzyme | 4333680 | 4547830 | 2723742 | | | | | | |
| Q9JL16 | ISG20 | interferon stimulated exonuclease gene 20kDa | Nucleus | enzyme | 3845810.25 | 3973127.25 | 3243901.5 | | | | | | |
| Q9JL16 | ISG20 | interferon stimulated exonuclease gene 20kDa | Nucleus | enzyme | 3655293.75 | 4571009 | 4174552 | Q9JL16 | ISG20 | 4334536 | 1034775 | 298714 | 12 |
| Q9CX00 | IST1 | increased sodium tolerance 1 homolog (yeast) | Cytoplasm | other | 2000069.125 | 1734286.75 | 2099090.5 | Q9CX00 | IST1 | 2000069 | 188648 | 108916 | 3 |
| Q62469 | ITGA2 | integrin, alpha 2 (CD49B, alpha 2 subunit of VLA-2 receptor) | Plasma Membrane | transmembrane receptor | 4044244.5 | 6340297.5 | 8240762 | | | | | | |
| Q62469 | ITGA2 | integrin, alpha 2 (CD49B, alpha 2 subunit of VLA-2 receptor) | Plasma Membrane | transmembrane receptor | 3223726.75 | 3650616.5 | 6720243.5 | | | | | | |
| Q62469 | ITGA2 | integrin, alpha 2 (CD49B, alpha 2 subunit of VLA-2 receptor) | Plasma Membrane | transmembrane receptor | 3123252.5 | 4971750.5 | 6627275 | | | | | | |
| Q62469 | ITGA2 | integrin, alpha 2 (CD49B, alpha 2 subunit of VLA-2 receptor) | Plasma Membrane | transmembrane receptor | 1938685.75 | 2713822.25 | 5342170 | | | | | | |
| Q62469 | ITGA2 | integrin, alpha 2 (CD49B, alpha 2 subunit of VLA-2 receptor) | Plasma Membrane | transmembrane receptor | 4379960 | 6128799.5 | 7772001.5 | | | | | | |
| Q62469 | ITGA2 | integrin, alpha 2 (CD49B, alpha 2 subunit of VLA-2 receptor) | Plasma Membrane | transmembrane receptor | 2211952.75 | 3045288.25 | 4858514.5 | | | | | | |
| Q62469 | ITGA2 | integrin, alpha 2 (CD49B, alpha 2 subunit of VLA-2 receptor) | Plasma Membrane | transmembrane receptor | 2381423 | 2444757 | 5069111.5 | | | | | | |
| Q62469 | ITGA2 | integrin, alpha 2 (CD49B, alpha 2 subunit of VLA-2 receptor) | Plasma Membrane | transmembrane receptor | 1397577.5 | 2566306.5 | 2841085 | | | | | | |
| Q62469 | ITGA2 | integrin, alpha 2 (CD49B, alpha 2 subunit of VLA-2 receptor) | Plasma Membrane | transmembrane receptor | 1438021.375 | 1806868.875 | 3906899.5 | | | | | | |
| Q62469 | ITGA2 | integrin, alpha 2 (CD49B, alpha 2 subunit of VLA-2 receptor) | Plasma Membrane | transmembrane receptor | 1009687.375 | 835672.062 | 1501067.75 | | | | | | |
| Q62469 | ITGA2 | integrin, alpha 2 (CD49B, alpha 2 subunit of VLA-2 receptor) | Plasma Membrane | transmembrane receptor | 2800163.75 | 3276623.25 | 5273762.5 | Q62469 | ITGA2 | 2540790 | 1985734 | 345672 | 33 |
| Q9QUM0 | ITGA2B | integrin, alpha 2b (platelet glycoprotein IIb of IIb/IIIa complex, antigen CD41) | Plasma Membrane | transmembrane receptor | 9.38E+07 | 1.67E+08 | 2.57E+08 | | | | | | |
| Q9QUM0 | ITGA2B | integrin, alpha 2b (platelet glycoprotein IIb of IIb/IIIa complex, antigen CD41) | Plasma Membrane | transmembrane receptor | 3.10E+07 | 3.82E+07 | 5.43E+07 | | | | | | |
| Q9QUM0 | ITGA2B | integrin, alpha 2b (platelet glycoprotein IIb of IIb/IIIa complex, antigen CD41) | Plasma Membrane | transmembrane receptor | 3.67E+07 | 4.51E+07 | 7.29E+07 | | | | | | |
| Q9QUM0 | ITGA2B | integrin, alpha 2b (platelet glycoprotein IIb of IIb/IIIa complex, antigen CD41) | Plasma Membrane | transmembrane receptor | 2.80E+07 | 4.11E+07 | 5.73E+07 | | | | | | |
| Q9QUM0 | ITGA2B | integrin, alpha 2b (platelet glycoprotein IIb of IIb/IIIa complex, antigen CD41) | Plasma Membrane | transmembrane receptor | 4.67E+07 | 6.26E+07 | 1.00E+08 | | | | | | |
| Q9QUM0 | ITGA2B | integrin, alpha 2b (platelet glycoprotein IIb of IIb/IIIa complex, antigen CD41) | Plasma Membrane | transmembrane receptor | 4.12E+07 | 5.39E+07 | 9.02E+07 | | | | | | |
| Q9QUM0 | ITGA2B | integrin, alpha 2b (platelet glycoprotein IIb of IIb/IIIa complex, antigen CD41) | Plasma Membrane | transmembrane receptor | 2.33E+07 | 3.46E+07 | 4.44E+07 | | | | | | |
| Q9QUM0 | ITGA2B | integrin, alpha 2b (platelet glycoprotein IIb of IIb/IIIa complex, antigen CD41) | Plasma Membrane | transmembrane receptor | 2.50E+07 | 2.94E+07 | 5.48E+07 | | | | | | |
| Q9QUM0 | ITGA2B | integrin, alpha 2b (platelet glycoprotein IIb of IIb/IIIa complex, antigen CD41) | Plasma Membrane | transmembrane receptor | 7681711.5 | 1.92E+07 | 2.00E+07 | | | | | | |
| Q9QUM0 | ITGA2B | integrin, alpha 2b (platelet glycoprotein IIb of IIb/IIIa complex, antigen CD41) | Plasma Membrane | transmembrane receptor | 1.05E+07 | 1.38E+07 | 2.06E+07 | | | | | | |
| Q9QUM0 | ITGA2B | integrin, alpha 2b (platelet glycoprotein IIb of IIb/IIIa complex, antigen CD41) | Plasma Membrane | transmembrane receptor | 2.30E+07 | 3.12E+07 | 5.37E+07 | | | | | | |
| Q9QUM0 | ITGA2B | integrin, alpha 2b (platelet glycoprotein IIb of IIb/IIIa complex, antigen CD41) | Plasma Membrane | transmembrane receptor | 2.04E+07 | 3.19E+07 | 3.87E+07 | | | | | | |

| | | | | | | | | | | | | | |
|---|---|---|---|---|---|---|---|---|---|---|---|---|---|
| Q9QUM0 | ITGA2B | integrin, alpha 2b (platelet glycoprotein IIb of IIb/IIIa complex, antigen CD41) | Plasma Membrane | transmembrane receptor | 1.95E+07 | 2.64E+07 | 4.57E+07 | | | | | | |
| Q9QUM0 | ITGA2B | integrin, alpha 2b (platelet glycoprotein IIb of IIb/IIIa complex, antigen CD41) | Plasma Membrane | transmembrane receptor | 1.75E+07 | 2.47E+07 | 3.93E+07 | | | | | | |
| Q9QUM0 | ITGA2B | integrin, alpha 2b (platelet glycoprotein IIb of IIb/IIIa complex, antigen CD41) | Plasma Membrane | transmembrane receptor | 1.66E+07 | 2.47E+07 | 3.19E+07 | | | | | | |
| Q9QUM0 | ITGA2B | integrin, alpha 2b (platelet glycoprotein IIb of IIb/IIIa complex, antigen CD41) | Plasma Membrane | transmembrane receptor | 1.76E+07 | 2.22E+07 | 3.79E+07 | | | | | | |
| Q9QUM0 | ITGA2B | integrin, alpha 2b (platelet glycoprotein IIb of IIb/IIIa complex, antigen CD41) | Plasma Membrane | transmembrane receptor | 9550367 | 1.01E+07 | 1.49E+07 | | | | | | |
| Q9QUM0 | ITGA2B | integrin, alpha 2b (platelet glycoprotein IIb of IIb/IIIa complex, antigen CD41) | Plasma Membrane | transmembrane receptor | 1.54E+07 | 2.16E+07 | 4.05E+07 | | | | | | |
| Q9QUM0 | ITGA2B | integrin, alpha 2b (platelet glycoprotein IIb of IIb/IIIa complex, antigen CD41) | Plasma Membrane | transmembrane receptor | 1.11E+07 | 1.40E+07 | 2.07E+07 | | | | | | |
| Q9QUM0 | ITGA2B | integrin, alpha 2b (platelet glycoprotein IIb of IIb/IIIa complex, antigen CD41) | Plasma Membrane | transmembrane receptor | 1.50E+07 | 1.93E+07 | 2.61E+07 | | | | | | |
| Q9QUM0 | ITGA2B | integrin, alpha 2b (platelet glycoprotein IIb of IIb/IIIa complex, antigen CD41) | Plasma Membrane | transmembrane receptor | 8965201 | 1.49E+07 | 2.29E+07 | | | | | | |
| Q9QUM0 | ITGA2B | integrin, alpha 2b (platelet glycoprotein IIb of IIb/IIIa complex, antigen CD41) | Plasma Membrane | transmembrane receptor | 3053610 | 2709021.5 | 5014112.5 | | | | | | |
| Q9QUM0 | ITGA2B | integrin, alpha 2b (platelet glycoprotein IIb of IIb/IIIa complex, antigen CD41) | Plasma Membrane | transmembrane receptor | 3676057.75 | 2432141.5 | 4000235.75 | | | | | | |
| Q9QUM0 | ITGA2B | integrin, alpha 2b (platelet glycoprotein IIb of IIb/IIIa complex, antigen CD41) | Plasma Membrane | transmembrane receptor | 4534889 | 7266857.5 | 8532480 | | | | | | |
| Q9QUM0 | ITGA2B | integrin, alpha 2b (platelet glycoprotein IIb of IIb/IIIa complex, antigen CD41) | Plasma Membrane | transmembrane receptor | 4047719.75 | 8547003 | 7532840 | | | | | | |
| Q9QUM0 | ITGA2B | integrin, alpha 2b (platelet glycoprotein IIb of IIb/IIIa complex, antigen CD41) | Plasma Membrane | transmembrane receptor | 2395463 | 3291996.25 | 4193701.5 | | | | | | |
| Q9QUM0 | ITGA2B | integrin, alpha 2b (platelet glycoprotein IIb of IIb/IIIa complex, antigen CD41) | Plasma Membrane | transmembrane receptor | 1928995.75 | 4765312 | 4603170.5 | Q9QUM0 | ITGA2B | 19924223 | 36657471 | 4073052 | 81 |
| P11688 | ITGA5 | integrin, alpha 5 (fibronectin receptor, alpha polypeptide) | Plasma Membrane | transmembrane receptor | 7062533 | 8854962 | 8210098.5 | | | | | | |
| P11688 | ITGA5 | integrin, alpha 5 (fibronectin receptor, alpha polypeptide) | Plasma Membrane | transmembrane receptor | 4607770.5 | 4525196.5 | 4358604.5 | | | | | | |
| P11688 | ITGA5 | integrin, alpha 5 (fibronectin receptor, alpha polypeptide) | Plasma Membrane | transmembrane receptor | 3144527.75 | 3011368.25 | 3020387.25 | | | | | | |
| P11688 | ITGA5 | integrin, alpha 5 (fibronectin receptor, alpha polypeptide) | Plasma Membrane | transmembrane receptor | 1813302.375 | 1881624.75 | 2016385.125 | P11688 | ITGA5 | 4157033 | 2442590 | 705115 | 12 |
| Q61739 | ITGA6 | integrin, alpha 6 | Plasma Membrane | transmembrane receptor | 3888214.25 | 5645540 | 1.03E+07 | | | | | | |
| Q61739 | ITGA6 | integrin, alpha 6 | Plasma Membrane | transmembrane receptor | 1209990.875 | 1706407.25 | 2995389.25 | | | | | | |
| Q61739 | ITGA6 | integrin, alpha 6 | Plasma Membrane | transmembrane receptor | 2541169 | 3431431.5 | 6289582.5 | | | | | | |
| Q61739 | ITGA6 | integrin, alpha 6 | Plasma Membrane | transmembrane receptor | 1232231.625 | 1535648 | 2205867 | | | | | | |
| Q61739 | ITGA6 | integrin, alpha 6 | Plasma Membrane | transmembrane receptor | 1913608.875 | 2202353.5 | 3718026.25 | | | | | | |
| Q61739 | ITGA6 | integrin, alpha 6 | Plasma Membrane | transmembrane receptor | 989193.875 | 1771261.25 | 2970008.25 | | | | | | |
| Q61739 | ITGA6 | integrin, alpha 6 | Plasma Membrane | transmembrane receptor | 2556011 | 3711931.25 | 6423448 | Q61739 | ITGA6 | 2047203 | 2246910 | 490316 | 21 |
| P09055 | ITGB1 | integrin, beta 1 (fibronectin receptor, beta polypeptide, antigen CD29 includes MDF2, MSK12) | Plasma Membrane | transmembrane receptor | 2.06E+07 | 2.06E+07 | 3.23E+07 | | | | | | |
| P09055 | ITGB1 | integrin, beta 1 (fibronectin receptor, beta polypeptide, antigen CD29 includes MDF2, MSK12) | Plasma Membrane | transmembrane receptor | 1.15E+07 | 1.54E+07 | 2.61E+07 | | | | | | |
| P09055 | ITGB1 | integrin, beta 1 (fibronectin receptor, beta polypeptide, antigen CD29 includes MDF2, MSK12) | Plasma Membrane | transmembrane receptor | 1.21E+07 | 1.92E+07 | 2.34E+07 | | | | | | |
| P09055 | ITGB1 | integrin, beta 1 (fibronectin receptor, beta polypeptide, antigen CD29 includes MDF2, MSK12) | Plasma Membrane | transmembrane receptor | 9338975 | 1.62E+07 | 2.20E+07 | | | | | | |
| P09055 | ITGB1 | integrin, beta 1 (fibronectin receptor, beta polypeptide, antigen CD29 includes MDF2, MSK12) | Plasma Membrane | transmembrane receptor | 7800713 | 9550048 | 1.34E+07 | | | | | | |
| P09055 | ITGB1 | integrin, beta 1 (fibronectin receptor, beta polypeptide, antigen CD29 includes MDF2, MSK12) | Plasma Membrane | transmembrane receptor | 5844004.5 | 6728028.5 | 9672175 | | | | | | |
| P09055 | ITGB1 | integrin, beta 1 (fibronectin receptor, beta polypeptide, antigen CD29 includes MDF2, MSK12) | Plasma Membrane | transmembrane receptor | 6145512 | 8042439 | 1.10E+07 | | | | | | |
| P09055 | ITGB1 | integrin, beta 1 (fibronectin receptor, beta polypeptide, antigen CD29 includes MDF2, MSK12) | Plasma Membrane | transmembrane receptor | 6187682 | 7145885.5 | 7901559 | | | | | | |
| P09055 | ITGB1 | integrin, beta 1 (fibronectin receptor, beta polypeptide, antigen CD29 includes MDF2, MSK12) | Plasma Membrane | transmembrane receptor | 5658149 | 6660326.5 | 8603176 | | | | | | |
| P09055 | ITGB1 | integrin, beta 1 (fibronectin receptor, beta polypeptide, antigen CD29 includes MDF2, MSK12) | Plasma Membrane | transmembrane receptor | 4459145 | 4973646 | 7021287 | | | | | | |
| P09055 | ITGB1 | integrin, beta 1 (fibronectin receptor, beta polypeptide, antigen CD29 includes MDF2, MSK12) | Plasma Membrane | transmembrane receptor | 2028332.625 | 3408677.25 | 6477065 | | | | | | |
| P09055 | ITGB1 | integrin, beta 1 (fibronectin receptor, beta polypeptide, antigen CD29 includes MDF2, MSK12) | Plasma Membrane | transmembrane receptor | 1563716.375 | 2440173 | 3103857 | | | | | | |
| P09055 | ITGB1 | integrin, beta 1 (fibronectin receptor, beta polypeptide, antigen CD29 includes MDF2, MSK12) | Plasma Membrane | transmembrane receptor | 1446485.375 | 1447447.125 | 2646641.75 | P09055 | ITGB1 | 7283286 | 7517998 | 1203843 | 39 |
| P11835 | ITGB2 | integrin, beta 2 (complement component 3 receptor 3 and 4 subunit) | Plasma Membrane | transmembrane receptor | 370733.875 | 1214443.375 | 2607503.5 | P11835 | ITGB2 | 370734 | 1129572 | 652159 | 3 |
| O54890 | ITGB3 | integrin, beta 3 (platelet glycoprotein IIIa, antigen CD61) | Plasma Membrane | transmembrane receptor | 7.10E+07 | 9.38E+07 | 1.65E+08 | | | | | | |
| O54890 | ITGB3 | integrin, beta 3 (platelet glycoprotein IIIa, antigen CD61) | Plasma Membrane | transmembrane receptor | 1.05E+07 | 1.46E+07 | 2.03E+07 | | | | | | |
| O54890 | ITGB3 | integrin, beta 3 (platelet glycoprotein IIIa, antigen CD61) | Plasma Membrane | transmembrane receptor | 2.74E+07 | 3.28E+07 | 5.70E+07 | | | | | | |
| O54890 | ITGB3 | integrin, beta 3 (platelet glycoprotein IIIa, antigen CD61) | Plasma Membrane | transmembrane receptor | 3.03E+07 | 4.14E+07 | 6.04E+07 | | | | | | |
| O54890 | ITGB3 | integrin, beta 3 (platelet glycoprotein IIIa, antigen CD61) | Plasma Membrane | transmembrane receptor | 2.87E+07 | 3.91E+07 | 6.18E+07 | | | | | | |
| O54890 | ITGB3 | integrin, beta 3 (platelet glycoprotein IIIa, antigen CD61) | Plasma Membrane | transmembrane receptor | 2.98E+07 | 3.59E+07 | 5.51E+07 | | | | | | |
| O54890 | ITGB3 | integrin, beta 3 (platelet glycoprotein IIIa, antigen CD61) | Plasma Membrane | transmembrane receptor | 1.74E+07 | 1.94E+07 | 2.90E+07 | | | | | | |
| O54890 | ITGB3 | integrin, beta 3 (platelet glycoprotein IIIa, antigen CD61) | Plasma Membrane | transmembrane receptor | 2.45E+07 | 3.65E+07 | 7.31E+07 | | | | | | |
| O54890 | ITGB3 | integrin, beta 3 (platelet glycoprotein IIIa, antigen CD61) | Plasma Membrane | transmembrane receptor | 1.69E+07 | 2.31E+07 | 3.55E+07 | | | | | | |
| O54890 | ITGB3 | integrin, beta 3 (platelet glycoprotein IIIa, antigen CD61) | Plasma Membrane | transmembrane receptor | 9870204 | 1.30E+07 | 2.48E+07 | | | | | | |
| O54890 | ITGB3 | integrin, beta 3 (platelet glycoprotein IIIa, antigen CD61) | Plasma Membrane | transmembrane receptor | 1.45E+07 | 1.90E+07 | 2.54E+07 | | | | | | |
| O54890 | ITGB3 | integrin, beta 3 (platelet glycoprotein IIIa, antigen CD61) | Plasma Membrane | transmembrane receptor | 1.47E+07 | 2.12E+07 | 3.67E+07 | | | | | | |
| O54890 | ITGB3 | integrin, beta 3 (platelet glycoprotein IIIa, antigen CD61) | Plasma Membrane | transmembrane receptor | 1.82E+07 | 2.53E+07 | 3.82E+07 | | | | | | |
| O54890 | ITGB3 | integrin, beta 3 (platelet glycoprotein IIIa, antigen CD61) | Plasma Membrane | transmembrane receptor | 1.61E+07 | 2.12E+07 | 3.40E+07 | | | | | | |
| O54890 | ITGB3 | integrin, beta 3 (platelet glycoprotein IIIa, antigen CD61) | Plasma Membrane | transmembrane receptor | 1.56E+07 | 2.63E+07 | 4.19E+07 | | | | | | |
| O54890 | ITGB3 | integrin, beta 3 (platelet glycoprotein IIIa, antigen CD61) | Plasma Membrane | transmembrane receptor | 1827829.125 | 3136216.25 | 4197824 | | | | | | |
| O54890 | ITGB3 | integrin, beta 3 (platelet glycoprotein IIIa, antigen CD61) | Plasma Membrane | transmembrane receptor | 1.73E+07 | 2.16E+07 | 2.80E+07 | | | | | | |
| O54890 | ITGB3 | integrin, beta 3 (platelet glycoprotein IIIa, antigen CD61) | Plasma Membrane | transmembrane receptor | 1.16E+07 | 1.67E+07 | 1.86E+07 | | | | | | |
| O54890 | ITGB3 | integrin, beta 3 (platelet glycoprotein IIIa, antigen CD61) | Plasma Membrane | transmembrane receptor | 7790877 | 1.64E+07 | 1.55E+07 | | | | | | |
| O54890 | ITGB3 | integrin, beta 3 (platelet glycoprotein IIIa, antigen CD61) | Plasma Membrane | transmembrane receptor | 9830183 | 1.35E+07 | 2.22E+07 | | | | | | |
| O54890 | ITGB3 | integrin, beta 3 (platelet glycoprotein IIIa, antigen CD61) | Plasma Membrane | transmembrane receptor | 4607983.5 | 1.36E+07 | 2.24E+07 | | | | | | |
| O54890 | ITGB3 | integrin, beta 3 (platelet glycoprotein IIIa, antigen CD61) | Plasma Membrane | transmembrane receptor | 6365359 | 1.00E+07 | 1.50E+07 | | | | | | |
| O54890 | ITGB3 | integrin, beta 3 (platelet glycoprotein IIIa, antigen CD61) | Plasma Membrane | transmembrane receptor | 1.18E+07 | 2.35E+07 | 4.33E+07 | | | | | | |
| O54890 | ITGB3 | integrin, beta 3 (platelet glycoprotein IIIa, antigen CD61) | Plasma Membrane | transmembrane receptor | 568662.938 | 285953.25 | 1444237.375 | O54890 | ITGB3 | 17389212 | 24405880 | 2876261 | 72 |
| Q9Z0T9 | ITGB6 | integrin, beta 6 | Plasma Membrane | other | 5548433 | 6721471.5 | 1.01E+07 | Q9Z0T9 | ITGB6 | 5548433 | 2385541 | 1377293 | 3 |
| Q5XKN4 | JAGN1 | jagunal homolog 1 | Cytoplasm | other | 2014990.25 | 2387643.5 | 2353975.5 | Q5XKN4 | JAGN1 | 2014990 | 206121 | 119004 | 3 |
| A2CG49 | KALRN | kalirin, RhoGEF kinase | Cytoplasm | kinase | 1230005.75 | 1824368 | 2145729.5 | | | | | | |
| A2CG49 | KALRN | kalirin, RhoGEF kinase | Cytoplasm | kinase | 517539.531 | 490062.5 | 980580.375 | A2CG49 | KALRN | 873773 | 678679 | 277070 | 6 |
| Q80UY2 | KCMF1 | potassium channel modulatory factor 1 | Extracellular Space | other | 3079216.75 | 2693609.75 | 2973567.5 | | | | | | |
| Q80UY2 | KCMF1 | potassium channel modulatory factor 1 | Extracellular Space | other | 1672099.25 | 1413959.75 | 1366678.25 | | | | | | |
| Q80UY2 | KCMF1 | potassium channel modulatory factor 1 | Extracellular Space | other | 822833.875 | 1002378.188 | 536065.75 | Q80UY2 | KCMF1 | 1858050 | 955803 | 318601 | 9 |
| Q9EQF2 | KEL | Kell blood group, metallo-endopeptidase | Plasma Membrane | peptidase | 1.73E+08 | 1.58E+08 | 1.21E+08 | | | | | | |
| Q9EQF2 | KEL | Kell blood group, metallo-endopeptidase | Plasma Membrane | peptidase | 1.75E+08 | 1.81E+08 | 1.61E+08 | | | | | | |
| Q9EQF2 | KEL | Kell blood group, metallo-endopeptidase | Plasma Membrane | peptidase | 1.29E+08 | 1.41E+08 | 1.31E+08 | | | | | | |
| Q9EQF2 | KEL | Kell blood group, metallo-endopeptidase | Plasma Membrane | peptidase | 1.14E+08 | 1.31E+08 | 9.77E+07 | | | | | | |
| Q9EQF2 | KEL | Kell blood group, metallo-endopeptidase | Plasma Membrane | peptidase | 1.45E+08 | 1.43E+08 | 1.22E+08 | | | | | | |
| Q9EQF2 | KEL | Kell blood group, metallo-endopeptidase | Plasma Membrane | peptidase | 9.40E+07 | 1.05E+08 | 1.09E+08 | | | | | | |
| Q9EQF2 | KEL | Kell blood group, metallo-endopeptidase | Plasma Membrane | peptidase | 9.27E+07 | 9.59E+07 | 7.31E+07 | | | | | | |
| Q9EQF2 | KEL | Kell blood group, metallo-endopeptidase | Plasma Membrane | peptidase | 1.13E+08 | 1.08E+08 | 7.76E+07 | | | | | | |
| Q9EQF2 | KEL | Kell blood group, metallo-endopeptidase | Plasma Membrane | peptidase | 9.37E+07 | 7.54E+07 | 7.33E+07 | | | | | | |
| Q9EQF2 | KEL | Kell blood group, metallo-endopeptidase | Plasma Membrane | peptidase | 1.01E+08 | 8.32E+07 | 8.12E+07 | | | | | | |
| Q9EQF2 | KEL | Kell blood group, metallo-endopeptidase | Plasma Membrane | peptidase | 6.07E+07 | 6.05E+07 | 6.10E+07 | | | | | | |
| Q9EQF2 | KEL | Kell blood group, metallo-endopeptidase | Plasma Membrane | peptidase | 7.54E+07 | 6.50E+07 | 4.76E+07 | | | | | | |
| Q9EQF2 | KEL | Kell blood group, metallo-endopeptidase | Plasma Membrane | peptidase | 8.02E+07 | 7.57E+07 | 6.58E+07 | | | | | | |
| Q9EQF2 | KEL | Kell blood group, metallo-endopeptidase | Plasma Membrane | peptidase | 4.09E+07 | 5.39E+07 | 3.40E+07 | | | | | | |
| Q9EQF2 | KEL | Kell blood group, metallo-endopeptidase | Plasma Membrane | peptidase | 4.85E+07 | 5.06E+07 | 4.65E+07 | | | | | | |
| Q9EQF2 | KEL | Kell blood group, metallo-endopeptidase | Plasma Membrane | peptidase | 4.92E+07 | 3.67E+07 | 4.50E+07 | | | | | | |
| Q9EQF2 | KEL | Kell blood group, metallo-endopeptidase | Plasma Membrane | peptidase | 6.32E+07 | 4.83E+07 | 5.52E+07 | | | | | | |
| Q9EQF2 | KEL | Kell blood group, metallo-endopeptidase | Plasma Membrane | peptidase | 3.49E+07 | 3.58E+07 | 2.66E+07 | | | | | | |
| Q9EQF2 | KEL | Kell blood group, metallo-endopeptidase | Plasma Membrane | peptidase | 3.29E+07 | 3.88E+07 | 3.97E+07 | | | | | | |
| Q9EQF2 | KEL | Kell blood group, metallo-endopeptidase | Plasma Membrane | peptidase | 5.02E+07 | 4.96E+07 | 4.18E+07 | | | | | | |
| Q9EQF2 | KEL | Kell blood group, metallo-endopeptidase | Plasma Membrane | peptidase | 2.69E+07 | 2.25E+07 | 1.94E+07 | | | | | | |
| Q9EQF2 | KEL | Kell blood group, metallo-endopeptidase | Plasma Membrane | peptidase | 3.73E+07 | 2.62E+07 | 3.13E+07 | | | | | | |
| Q9EQF2 | KEL | Kell blood group, metallo-endopeptidase | Plasma Membrane | peptidase | 2.75E+07 | 2.67E+07 | 2.38E+07 | | | | | | |
| Q9EQF2 | KEL | Kell blood group, metallo-endopeptidase | Plasma Membrane | peptidase | 2.55E+07 | 3.27E+07 | 1.96E+07 | | | | | | |
| Q9EQF2 | KEL | Kell blood group, metallo-endopeptidase | Plasma Membrane | peptidase | 1.74E+07 | 1.58E+07 | 1.29E+07 | | | | | | |
| Q9EQF2 | KEL | Kell blood group, metallo-endopeptidase | Plasma Membrane | peptidase | 2.78E+07 | 2.75E+07 | 2.66E+07 | | | | | | |
| Q9EQF2 | KEL | Kell blood group, metallo-endopeptidase | Plasma Membrane | peptidase | 1.61E+07 | 1.33E+07 | 1.54E+07 | | | | | | |
| Q9EQF2 | KEL | Kell blood group, metallo-endopeptidase | Plasma Membrane | peptidase | 1.79E+07 | 2.27E+07 | 1.56E+07 | | | | | | |
| Q9EQF2 | KEL | Kell blood group, metallo-endopeptidase | Plasma Membrane | peptidase | 1.62E+07 | 1.47E+07 | 1.49E+07 | | | | | | |
| Q9EQF2 | KEL | Kell blood group, metallo-endopeptidase | Plasma Membrane | peptidase | 2.18E+07 | 2.02E+07 | 1.78E+07 | | | | | | |
| Q9EQF2 | KEL | Kell blood group, metallo-endopeptidase | Plasma Membrane | peptidase | 1.53E+07 | 1.95E+07 | 1.70E+07 | | | | | | |
| Q9EQF2 | KEL | Kell blood group, metallo-endopeptidase | Plasma Membrane | peptidase | 7225925 | 9004180 | 4787187.5 | | | | | | |
| Q9EQF2 | KEL | Kell blood group, metallo-endopeptidase | Plasma Membrane | peptidase | 9538011 | 7803419 | 6281270.5 | | | | | | |
| Q9EQF2 | KEL | Kell blood group, metallo-endopeptidase | Plasma Membrane | peptidase | 5228544.5 | 5099665.5 | 3364795.5 | | | | | | |
| Q9EQF2 | KEL | Kell blood group, metallo-endopeptidase | Plasma Membrane | peptidase | 3378069.25 | 1645854.75 | 2479366.5 | | | | | | |
| Q9EQF2 | KEL | Kell blood group, metallo-endopeptidase | Plasma Membrane | peptidase | 865258.25 | 765490.188 | 2096175.5 | Q9EQF2 | KEL | 56709328 | 45909705 | 4417663 | 108 |
| Q6PDI5 | KIAA0368 | KIAA0368 | Cytoplasm | other | 4924604.5 | 5534146 | 4537166 | | | | | | |
| Q6PDI5 | KIAA0368 | KIAA0368 | Cytoplasm | other | 5006211.5 | 3267117.5 | 3415836.5 | | | | | | |

| | | | | | | | | | | | | | |
|---|---|---|---|---|---|---|---|---|---|---|---|---|---|
| Q6PDI5 | KIAA0368 | KIAA0368 | Cytoplasm | other | 3424301.5 | 3843342.5 | 3308925.75 | | | | | | |
| Q6PDI5 | KIAA0368 | KIAA0368 | Cytoplasm | other | 2187800.5 | 2097964 | 1573242.375 | | | | | | |
| Q6PDI5 | KIAA0368 | KIAA0368 | Cytoplasm | other | 1907449.875 | 1564025 | 1717603.75 | | | | | | |
| Q6PDI5 | KIAA0368 | KIAA0368 | Cytoplasm | other | 1550240.5 | 1689417.25 | 1730725 | | | | | | |
| Q6PDI5 | KIAA0368 | KIAA0368 | Cytoplasm | other | 2035328.875 | 1377796.25 | 2030224.375 | | | | | | |
| Q6PDI5 | KIAA0368 | KIAA0368 | Cytoplasm | other | 873347.875 | 828925.062 | 820235 | | | | | | |
| Q6PDI5 | KIAA0368 | KIAA0368 | Cytoplasm | other | 600527.562 | 479711 | 364426.719 | Q6PDI5 | KIAA0368 | 2501090 | 1476927 | 284235 | 27 |
| Q7TQ95 | KIAA1715 | KIAA1715 | Cytoplasm | other | 4587359.5 | 4164862 | 4972901 | | | | | | |
| Q7TQ95 | KIAA1715 | KIAA1715 | Cytoplasm | other | 5439859.5 | 5465197.5 | 3818774.25 | | | | | | |
| Q7TQ95 | KIAA1715 | KIAA1715 | Cytoplasm | other | 4414530.5 | 4111111 | 3238437.5 | | | | | | |
| Q7TQ95 | KIAA1715 | KIAA1715 | Cytoplasm | other | 2107291.5 | 3170076.25 | 2955023 | | | | | | |
| Q7TQ95 | KIAA1715 | KIAA1715 | Cytoplasm | other | 3035064.75 | 2189954.75 | 1996704.875 | Q7TQ95 | KIAA1715 | 3916821 | 1154735 | 298151 | 15 |
| Q60575 | KIF1B | kinesin family member 1B | Cytoplasm | transporter | 1564541 | 3865684.5 | 7614599.5 | Q60575 | KIF1B | 1564541 | 3053764 | 1763091 | 3 |
| O88448 | KLC2 | kinesin light chain 2 | Cytoplasm | other | 1099895.625 | 1866875.5 | 3316736.75 | O88448 | KLC2 | 1099896 | 1125814 | 649989 | 3 |
| Q60682 | Klra4 (includes others) | killer cell lectin-like receptor, subfamily A, member 4 | Plasma Membrane | transmembrane receptor | 7317925.5 | 5845014 | 4624837 | Q60682 | Klra4 (includes others) | 7317926 | 1348519 | 778568 | 3 |
| O35344 | KPNA3 | karyopherin alpha 3 (importin alpha 4) | Nucleus | transporter | 2129632.5 | 1612393 | 2038688 | O35344 | KPNA3 | 2129633 | 276145 | 159432 | 3 |
| P70168 | KPNB1 | karyopherin (importin) beta 1 | Nucleus | transporter | 1.02E+07 | 8262092 | 8348322 | | | | | | |
| P70168 | KPNB1 | karyopherin (importin) beta 1 | Nucleus | transporter | 1.15E+07 | 1.02E+07 | 1.21E+07 | | | | | | |
| P70168 | KPNB1 | karyopherin (importin) beta 1 | Nucleus | transporter | 1.39E+07 | 1.22E+07 | 1.34E+07 | | | | | | |
| P70168 | KPNB1 | karyopherin (importin) beta 1 | Nucleus | transporter | 9562524 | 6208514 | 5195277.5 | | | | | | |
| P70168 | KPNB1 | karyopherin (importin) beta 1 | Nucleus | transporter | 9352969 | 7005996.5 | 8406794 | | | | | | |
| P70168 | KPNB1 | karyopherin (importin) beta 1 | Nucleus | transporter | 6884697 | 4906509.5 | 3912614.25 | | | | | | |
| P70168 | KPNB1 | karyopherin (importin) beta 1 | Nucleus | transporter | 9699415 | 7256863 | 5575369.5 | | | | | | |
| P70168 | KPNB1 | karyopherin (importin) beta 1 | Nucleus | transporter | 6766500 | 4505899 | 4218880 | | | | | | |
| P70168 | KPNB1 | karyopherin (importin) beta 1 | Nucleus | transporter | 8431662 | 7229866 | 6361761.5 | | | | | | |
| P70168 | KPNB1 | karyopherin (importin) beta 1 | Nucleus | transporter | 7189622 | 5421072.5 | 6787715.5 | | | | | | |
| P70168 | KPNB1 | karyopherin (importin) beta 1 | Nucleus | transporter | 5465980 | 4291751.5 | 4500686 | | | | | | |
| P70168 | KPNB1 | karyopherin (importin) beta 1 | Nucleus | transporter | 5242756 | 4447455.5 | 5220779.5 | | | | | | |
| P70168 | KPNB1 | karyopherin (importin) beta 1 | Nucleus | transporter | 5016086.5 | 4745227 | 3779339.5 | | | | | | |
| P70168 | KPNB1 | karyopherin (importin) beta 1 | Nucleus | transporter | 5345842 | 4074920.75 | 3707577.75 | | | | | | |
| P70168 | KPNB1 | karyopherin (importin) beta 1 | Nucleus | transporter | 3791432.25 | 1536137.5 | 2665067 | | | | | | |
| P70168 | KPNB1 | karyopherin (importin) beta 1 | Nucleus | transporter | 5400788.5 | 3848233.5 | 3334340.5 | | | | | | |
| P70168 | KPNB1 | karyopherin (importin) beta 1 | Nucleus | transporter | 4233895 | 3041309.75 | 3002545.5 | | | | | | |
| P70168 | KPNB1 | karyopherin (importin) beta 1 | Nucleus | transporter | 2898159.75 | 1806399.375 | 1882866.25 | | | | | | |
| P70168 | KPNB1 | karyopherin (importin) beta 1 | Nucleus | transporter | 4148701.5 | 4028177 | 5289166.5 | | | | | | |
| P70168 | KPNB1 | karyopherin (importin) beta 1 | Nucleus | transporter | 2794099 | 1255879.875 | 1747754.375 | | | | | | |
| P70168 | KPNB1 | karyopherin (importin) beta 1 | Nucleus | transporter | 3285025.75 | 2103934.75 | 1634836 | P70168 | KPNB1 | 6721436 | 3040152 | 383023 | 63 |
| P04104 | KRT1 | keratin 1, type II | Cytoplasm | other | 1.78E+07 | 9698517 | 1.76E+07 | | | | | | |
| P04104 | KRT1 | keratin 1, type II | Cytoplasm | other | 152116.859 | 777976.188 | 2840527.25 | | | | | | |
| P04104 | KRT1 | keratin 1, type II | Cytoplasm | other | 4984650.5 | 2470215.25 | 3901701 | | | | | | |
| P04104 | KRT1 | keratin 1, type II | Cytoplasm | other | 1333066.5 | 1435697.875 | 2178515 | P04104 | KRT1 | 6054958 | 6246188 | 1803119 | 12 |
| P02535 | Krt10 | keratin 10 | Cytoplasm | other | 6953222.5 | 2701601.5 | 6595765.5 | P02535 | Krt10 | 6953223 | 2358268 | 1361547 | 3 |
| Q61781 | KRT14 | keratin 14, type I | Cytoplasm | other | 1013572.188 | 312469.531 | 2252061.75 | Q61781 | KRT14 | 1013572 | 982125 | 567030 | 3 |
| Q3TTY5 | Krt2 | keratin 2 | Cytoplasm | other | 2731431 | 662889.5 | 3082014.75 | Q3TTY5 | Krt2 | 2731431 | 1307283 | 754760 | 3 |
| Q922U2 | KRT5 | keratin 5, type II | Cytoplasm | other | 1.13E+07 | 1.38E+07 | 2.11E+07 | | | | | | |
| Q922U2 | KRT5 | keratin 5, type II | Cytoplasm | other | 6400310 | 7911655.5 | 7569929 | | | | | | |
| Q922U2 | KRT5 | keratin 5, type II | Cytoplasm | other | 904725.125 | 412650.906 | 1394854 | | | | | | |
| Q922U2 | KRT5 | keratin 5, type II | Cytoplasm | other | 3978481.5 | 2455832 | 4592442.5 | Q922U2 | KRT5 | 5638379 | 6123794 | 1767787 | 12 |
| Q6NXH9 | KRT73 | keratin 73, type II | Extracellular Space | other | 945537.75 | 579997 | 556948.5 | Q6NXH9 | KRT73 | 945538 | 218003 | 125864 | 3 |
| Q3UV17 | KRT76 | keratin 76, type II | Cytoplasm | other | 1.14E+07 | 7976265.5 | 1.53E+07 | Q3UV17 | KRT76 | 11360000 | 3660266 | 2113255 | 3 |
| Q8VED5 | KRT79 | keratin 79, type II | Extracellular Space | other | 3529611 | 2731505.25 | 8904074 | Q8VED5 | KRT79 | 3529611 | 3357142 | 1938247 | 3 |
| Q6IMF0 | Krt83 | keratin 83 | Extracellular Space | other | 1.56E+07 | 1.75E+07 | 1.39E+07 | Q6IMF0 | Krt83 | 15630000 | 1775284 | 1024961 | 3 |
| P11438 | LAMP1 | lysosomal-associated membrane protein 1 | Plasma Membrane | other | 9595046 | 1.31E+07 | 1.28E+07 | | | | | | |
| P11438 | LAMP1 | lysosomal-associated membrane protein 1 | Plasma Membrane | other | 6100686.5 | 7805440.5 | 8418845 | | | | | | |
| P11438 | LAMP1 | lysosomal-associated membrane protein 1 | Plasma Membrane | other | 2147185 | 2276442 | 2916466.25 | P11438 | LAMP1 | 5947639 | 4230798 | 1410266 | 9 |
| P17047 | LAMP2 | lysosomal-associated membrane protein 2 | Plasma Membrane | enzyme | 4094000.25 | 7381600 | 9977546 | | | | | | |
| P17047 | LAMP2 | lysosomal-associated membrane protein 2 | Plasma Membrane | enzyme | 2365894.75 | 4039304.75 | 5436860 | P17047 | LAMP2 | 3229948 | 2739028 | 1118204 | 6 |
| Q9CQ22 | LAMTOR1 | late endosomal/lysosomal adaptor, MAPK and MTOR activator 1 | Plasma Membrane | other | 2.05E+07 | 2.64E+07 | 2.61E+07 | | | | | | |
| Q9CQ22 | LAMTOR1 | late endosomal/lysosomal adaptor, MAPK and MTOR activator 1 | Plasma Membrane | other | 5941463.5 | 7345598 | 7825509.5 | | | | | | |
| Q9CQ22 | LAMTOR1 | late endosomal/lysosomal adaptor, MAPK and MTOR activator 1 | Plasma Membrane | other | 5898182 | 7391970.5 | 7960229 | | | | | | |
| Q9CQ22 | LAMTOR1 | late endosomal/lysosomal adaptor, MAPK and MTOR activator 1 | Plasma Membrane | other | 2428308.75 | 3304542.75 | 3884708.5 | Q9CQ22 | LAMTOR1 | 8696989 | 8700398 | 2511589 | 12 |
| O88653 | LAMTOR3 | late endosomal/lysosomal adaptor, MAPK and MTOR activator 3 | Cytoplasm | other | 1.02E+07 | 1.27E+07 | 1.29E+07 | | | | | | |
| O88653 | LAMTOR3 | late endosomal/lysosomal adaptor, MAPK and MTOR activator 3 | Cytoplasm | other | 3879712 | 6271616.5 | 4966617 | | | | | | |
| O88653 | LAMTOR3 | late endosomal/lysosomal adaptor, MAPK and MTOR activator 3 | Cytoplasm | other | 2954408 | 3644104.5 | 5081801 | O88653 | LAMTOR3 | 5684707 | 3936197 | 1312066 | 9 |
| Q9D1L9 | LAMTOR5 | late endosomal/lysosomal adaptor, MAPK and MTOR activator 5 | Cytoplasm | other | 8495041 | 1.10E+07 | 9549870 | | | | | | |
| Q9D1L9 | LAMTOR5 | late endosomal/lysosomal adaptor, MAPK and MTOR activator 5 | Cytoplasm | other | 3994353 | 5068703 | 5899327.5 | | | | | | |
| Q9D1L9 | LAMTOR5 | late endosomal/lysosomal adaptor, MAPK and MTOR activator 5 | Cytoplasm | other | 3108059.75 | 3918855.75 | 3897359.5 | Q9D1L9 | LAMTOR5 | 5199151 | 2860355 | 953452 | 9 |
| O89112 | LANCL1 | LanC lantibiotic synthetase component C-like 1 (bacterial) | Plasma Membrane | other | 2612747 | 2672468 | 2188691 | O89112 | LANCL1 | 2612747 | 263765 | 152285 | 3 |
| Q9JJK2 | LANCL2 | LanC lantibiotic synthetase component C-like 2 (bacterial) | Plasma Membrane | other | 8862518 | 7533921.5 | 8002314.5 | | | | | | |
| Q9JJK2 | LANCL2 | LanC lantibiotic synthetase component C-like 2 (bacterial) | Plasma Membrane | other | 8974751 | 8335031.5 | 7903104.5 | | | | | | |
| Q9JJK2 | LANCL2 | LanC lantibiotic synthetase component C-like 2 (bacterial) | Plasma Membrane | other | 8871772 | 7076677.5 | 8652175 | | | | | | |
| Q9JJK2 | LANCL2 | LanC lantibiotic synthetase component C-like 2 (bacterial) | Plasma Membrane | other | 7108682.5 | 5042792.5 | 3918203 | | | | | | |
| Q9JJK2 | LANCL2 | LanC lantibiotic synthetase component C-like 2 (bacterial) | Plasma Membrane | other | 6278445 | 5821730.5 | 5868446 | | | | | | |
| Q9JJK2 | LANCL2 | LanC lantibiotic synthetase component C-like 2 (bacterial) | Plasma Membrane | other | 4866503 | 4737211.5 | 3787365.25 | | | | | | |
| Q9JJK2 | LANCL2 | LanC lantibiotic synthetase component C-like 2 (bacterial) | Plasma Membrane | other | 4830118.5 | 4910685.5 | 4002307.5 | | | | | | |
| Q9JJK2 | LANCL2 | LanC lantibiotic synthetase component C-like 2 (bacterial) | Plasma Membrane | other | 4037830.5 | 3763671.5 | 2117524 | | | | | | |
| Q9JJK2 | LANCL2 | LanC lantibiotic synthetase component C-like 2 (bacterial) | Plasma Membrane | other | 3647877 | 3331704.75 | 3657520.5 | | | | | | |
| Q9JJK2 | LANCL2 | LanC lantibiotic synthetase component C-like 2 (bacterial) | Plasma Membrane | other | 3077431.5 | 2737737.75 | 2516908.25 | | | | | | |
| Q9JJK2 | LANCL2 | LanC lantibiotic synthetase component C-like 2 (bacterial) | Plasma Membrane | other | 2452342.75 | 2603480.5 | 2406588.75 | | | | | | |
| Q9JJK2 | LANCL2 | LanC lantibiotic synthetase component C-like 2 (bacterial) | Plasma Membrane | other | 1107402.875 | 1156001.875 | 2284376.5 | Q9JJK2 | LANCL2 | 5342973 | 2376385 | 396064 | 36 |
| P06151 | LDHA | lactate dehydrogenase A | Cytoplasm | enzyme | 6445608 | 6020394 | 5181147.5 | | | | | | |
| P06151 | LDHA | lactate dehydrogenase A | Cytoplasm | enzyme | 5886982 | 6293553.5 | 6599379 | | | | | | |
| P06151 | LDHA | lactate dehydrogenase A | Cytoplasm | enzyme | 5154515.5 | 5235149.5 | 3774604.25 | | | | | | |
| P06151 | LDHA | lactate dehydrogenase A | Cytoplasm | enzyme | 4149105.25 | 3486076.75 | 4055345 | | | | | | |
| P06151 | LDHA | lactate dehydrogenase A | Cytoplasm | enzyme | 4652886 | 5053887 | 4993370 | | | | | | |
| P06151 | LDHA | lactate dehydrogenase A | Cytoplasm | enzyme | 2715680.75 | 2981783 | 2089782 | | | | | | |
| P06151 | LDHA | lactate dehydrogenase A | Cytoplasm | enzyme | 2474549.5 | 1512510.75 | 1905204.25 | | | | | | |
| P06151 | LDHA | lactate dehydrogenase A | Cytoplasm | enzyme | 1818938 | 1956808.75 | 1930067.875 | | | | | | |
| P06151 | LDHA | lactate dehydrogenase A | Cytoplasm | enzyme | 1442826.75 | 1362414.5 | 1498521.125 | P06151 | LDHA | 3860121 | 1786452 | 343803 | 27 |
| Q8CBY3 | LENG8 | leukocyte receptor cluster (LRC) member 8 | Nucleus | other | 4663103 | 1790803.75 | 2.28E+07 | Q8CBY3 | LENG8 | 4663103 | 11414306 | 6590053 | 3 |
| O89013 | LEPROT | leptin receptor overlapping transcript | Plasma Membrane | other | 2994608.75 | 3106142.25 | 2985363 | | | | | | |
| O89013 | LEPROT | leptin receptor overlapping transcript | Plasma Membrane | other | 248785.234 | 168006.609 | 208409.641 | O89013 | LEPROT | 1621697 | 1545540 | 630964 | 6 |
| Q9Z2I0 | LETM1 | leucine zipper-EF-hand containing transmembrane protein 1 | Cytoplasm | other | 1218856.75 | 2183263 | 3306562.5 | Q9Z2I0 | LETM1 | 1218857 | 1044860 | 603250 | 3 |
| O08573 | LGALS9B | lectin, galactoside-binding, soluble, 9B | Cytoplasm | other | 1.86E+07 | 2.29E+07 | 2.08E+07 | | | | | | |
| O08573 | LGALS9B | lectin, galactoside-binding, soluble, 9B | Cytoplasm | other | 1.12E+07 | 1.71E+07 | 1.86E+07 | | | | | | |
| O08573 | LGALS9B | lectin, galactoside-binding, soluble, 9B | Cytoplasm | other | 1.59E+07 | 1.69E+07 | 1.51E+07 | | | | | | |
| O08573 | LGALS9B | lectin, galactoside-binding, soluble, 9B | Cytoplasm | other | 1.48E+07 | 1.93E+07 | 1.98E+07 | | | | | | |
| O08573 | LGALS9B | lectin, galactoside-binding, soluble, 9B | Cytoplasm | other | 8637696 | 1.18E+07 | 9862547 | | | | | | |
| O08573 | LGALS9B | lectin, galactoside-binding, soluble, 9B | Cytoplasm | other | 6574842.5 | 7519007 | 7030088.5 | | | | | | |
| O08573 | LGALS9B | lectin, galactoside-binding, soluble, 9B | Cytoplasm | other | 7701620.5 | 7919196 | 6081083.5 | | | | | | |
| O08573 | LGALS9B | lectin, galactoside-binding, soluble, 9B | Cytoplasm | other | 3679362.5 | 5319592 | 4332745.5 | O08573 | LGALS9B | 10876690 | 5936855 | 1211855 | 24 |
| A2ARI4 | LGR4 | leucine-rich repeat containing G protein-coupled receptor 4 | Plasma Membrane | transmembrane receptor | 2087767.75 | 3725326.25 | 4489564.5 | A2ARI4 | LGR4 | 2087768 | 1227075 | 708452 | 3 |
| Q99JW4 | LIMS1 | LIM and senescent cell antigen-like domains 1 | Plasma Membrane | other | 1167797.875 | 856635.5 | 4879383.5 | Q99JW4 | LIMS1 | 1167798 | 2238124 | 1292181 | 3 |
| Q9D0F3 | LMAN1 | lectin, mannose-binding, 1 | Cytoplasm | other | 1.85E+07 | 1.54E+07 | 1.04E+07 | | | | | | |
| Q9D0F3 | LMAN1 | lectin, mannose-binding, 1 | Cytoplasm | other | 1.04E+07 | 9384649 | 6402884.5 | | | | | | |
| Q9D0F3 | LMAN1 | lectin, mannose-binding, 1 | Cytoplasm | other | 1.15E+07 | 1.28E+07 | 8595456 | | | | | | |
| Q9D0F3 | LMAN1 | lectin, mannose-binding, 1 | Cytoplasm | other | 1.21E+07 | 1.15E+07 | 8597506 | | | | | | |
| Q9D0F3 | LMAN1 | lectin, mannose-binding, 1 | Cytoplasm | other | 1.19E+07 | 8757789 | 5473075.5 | | | | | | |
| Q9D0F3 | LMAN1 | lectin, mannose-binding, 1 | Cytoplasm | other | 5071076 | 5910328 | 3856332 | Q9D0F3 | LMAN1 | 11558513 | 3742129 | 882028 | 18 |

| | | | | | | | | | | | | | |
|---|---|---|---|---|---|---|---|---|---|---|---|---|---|
| Q9DBH5 | LMAN2 | lectin, mannose-binding 2 | Cytoplasm | transporter | 1.26E+07 | 1.27E+07 | 1.39E+07 | | | | | | |
| Q9DBH5 | LMAN2 | lectin, mannose-binding 2 | Cytoplasm | transporter | 1.79E+07 | 1.93E+07 | 1.69E+07 | | | | | | |
| Q9DBH5 | LMAN2 | lectin, mannose-binding 2 | Cytoplasm | transporter | 5174559 | 4999358.5 | 4273559.5 | | | | | | |
| Q9DBH5 | LMAN2 | lectin, mannose-binding 2 | Cytoplasm | transporter | 5192400.5 | 5195716 | 3677177 | | | | | | |
| Q9DBH5 | LMAN2 | lectin, mannose-binding 2 | Cytoplasm | transporter | 4066949.75 | 3852430.5 | 3437023.75 | Q9DBH5 | LMAN2 | 8986782 | 5929899 | 1531093 | 15 |
| Q8VEE1 | LMCD1 | LIM and cysteine-rich domains 1 | Cytoplasm | transcription regulator | 3230655 | 1.34E+07 | 1.46E+07 | Q8VEE1 | LMCD1 | 3230655 | 6252314 | 3609775 | 3 |
| Q8CGK3 | LONP1 | lon peptidase 1, mitochondrial | Cytoplasm | peptidase | 4289565 | 6245943.5 | 8953586 | | | | | | |
| Q8CGK3 | LONP1 | lon peptidase 1, mitochondrial | Cytoplasm | peptidase | 1287190 | 2984344.5 | 5250418.5 | Q8CGK3 | LONP1 | 2788378 | 2662159 | 1086822 | 6 |
| Q3TFD2 | LPCAT1 | lysophosphatidylcholine acyltransferase 1 | Cytoplasm | enzyme | 2.23E+07 | 2.55E+07 | 2.80E+07 | | | | | | |
| Q3TFD2 | LPCAT1 | lysophosphatidylcholine acyltransferase 1 | Cytoplasm | enzyme | 1.23E+07 | 1.02E+07 | 1.12E+07 | | | | | | |
| Q3TFD2 | LPCAT1 | lysophosphatidylcholine acyltransferase 1 | Cytoplasm | enzyme | 1.15E+07 | 9747883 | 1.04E+07 | | | | | | |
| Q3TFD2 | LPCAT1 | lysophosphatidylcholine acyltransferase 1 | Cytoplasm | enzyme | 8037104 | 1.01E+07 | 7442412 | | | | | | |
| Q3TFD2 | LPCAT1 | lysophosphatidylcholine acyltransferase 1 | Cytoplasm | enzyme | 7234969 | 8846236 | 8944519 | | | | | | |
| Q3TFD2 | LPCAT1 | lysophosphatidylcholine acyltransferase 1 | Cytoplasm | enzyme | 3150627.75 | 3353836.75 | 7943216.5 | | | | | | |
| Q3TFD2 | LPCAT1 | lysophosphatidylcholine acyltransferase 1 | Cytoplasm | enzyme | 8668658 | 1.03E+07 | 7696258.5 | | | | | | |
| Q3TFD2 | LPCAT1 | lysophosphatidylcholine acyltransferase 1 | Cytoplasm | enzyme | 4430930.5 | 3933595.75 | 3359277 | Q3TFD2 | LPCAT1 | 9709036 | 6458278 | 1318290 | 24 |
| Q91V01 | LPCAT3 | lysophosphatidylcholine acyltransferase 3 | Plasma Membrane | enzyme | 2.48E+07 | 2.33E+07 | 1.78E+07 | | | | | | |
| Q91V01 | LPCAT3 | lysophosphatidylcholine acyltransferase 3 | Plasma Membrane | enzyme | 1.04E+07 | 1.09E+07 | 9295315 | | | | | | |
| Q91V01 | LPCAT3 | lysophosphatidylcholine acyltransferase 3 | Plasma Membrane | enzyme | 9041270 | 8319520 | 8289180.5 | | | | | | |
| Q91V01 | LPCAT3 | lysophosphatidylcholine acyltransferase 3 | Plasma Membrane | enzyme | 2065881 | 2284708.25 | 1545023.875 | Q91V01 | LPCAT3 | 11571788 | 7694277 | 2221146 | 12 |
| P55302 | LRPAP1 | low density lipoprotein receptor-related protein associated protein 1 | Plasma Membrane | transmembrane receptor | 1934062.25 | 2214844.5 | 1477991.25 | P55302 | LRPAP1 | 1934062 | 371885 | 214708 | 3 |
| Q9D1G5 | LRRC57 | leucine rich repeat containing 57 | Cytoplasm | other | 6989492 | 7078088.5 | 4466916 | | | | | | |
| Q9D1G5 | LRRC57 | leucine rich repeat containing 57 | Cytoplasm | other | 6019868.5 | 5251112.5 | 4540269.5 | | | | | | |
| Q9D1G5 | LRRC57 | leucine rich repeat containing 57 | Cytoplasm | other | 6483340.5 | 5842660.5 | 4924442.5 | | | | | | |
| Q9D1G5 | LRRC57 | leucine rich repeat containing 57 | Cytoplasm | other | 3993432.25 | 4558380 | 2906329.25 | | | | | | |
| Q9D1G5 | LRRC57 | leucine rich repeat containing 57 | Cytoplasm | other | 3712443 | 3784017.25 | 3248242.75 | | | | | | |
| Q9D1G5 | LRRC57 | leucine rich repeat containing 57 | Cytoplasm | other | 2165470.5 | 1979526.75 | 1638205.75 | Q9D1G5 | LRRC57 | 4894008 | 1663061 | 391987 | 18 |
| Q922Q8 | LRRC59 | leucine rich repeat containing 59 | Cytoplasm | other | 2651198.25 | 3773797.5 | 3852344.25 | Q922Q8 | LRRC59 | 2651198 | 671956 | 387954 | 3 |
| Q80TE7 | LRRC7 | leucine rich repeat containing 7 | Plasma Membrane | other | 1679054.5 | 2219717 | 2690160.25 | Q80TE7 | LRRC7 | 1679055 | 505959 | 292116 | 3 |
| P08071 | LTF | lactotransferrin | Extracellular Space | other | 729337 | 106522.203 | 8370732 | | | | | | |
| P08071 | LTF | lactotransferrin | Extracellular Space | other | 512833.781 | 470072.031 | 3785402.75 | | | | | | |
| P08071 | LTF | lactotransferrin | Extracellular Space | other | 808559.125 | 368222.469 | 3474877.5 | | | | | | |
| P08071 | LTF | lactotransferrin | Extracellular Space | other | 129202.398 | 215484.469 | 2904648.5 | | | | | | |
| P08071 | LTF | lactotransferrin | Extracellular Space | other | 711110.875 | 689814.375 | 5902249.5 | P08071 | LTF | 578209 | 2476270 | 639370 | 15 |
| P25911 | LYN | LYN proto-oncogene, Src family tyrosine kinase | Cytoplasm | kinase | 6315190.5 | 1.26E+07 | 1.74E+07 | | | | | | |
| P25911 | LYN | LYN proto-oncogene, Src family tyrosine kinase | Cytoplasm | kinase | 4342329 | 6812659 | 1.01E+07 | | | | | | |
| P25911 | LYN | LYN proto-oncogene, Src family tyrosine kinase | Cytoplasm | kinase | 5387519 | 9004227 | 1.22E+07 | | | | | | |
| P25911 | LYN | LYN proto-oncogene, Src family tyrosine kinase | Cytoplasm | kinase | 4189391.5 | 5451408.5 | 8790581 | | | | | | |
| P25911 | LYN | LYN proto-oncogene, Src family tyrosine kinase | Cytoplasm | kinase | 1106129.125 | 2227753.25 | 3375132.75 | | | | | | |
| P25911 | LYN | LYN proto-oncogene, Src family tyrosine kinase | Cytoplasm | kinase | 2014827 | 2421297.75 | 3230390 | | | | | | |
| P25911 | LYN | LYN proto-oncogene, Src family tyrosine kinase | Cytoplasm | kinase | 1481487.875 | 2122151.25 | 3171273 | | | | | | |
| P25911 | LYN | LYN proto-oncogene, Src family tyrosine kinase | Cytoplasm | kinase | 763689.062 | 832077.25 | 926753.375 | P25911 | LYN | 3200070 | 4387557 | 895606 | 24 |
| P08905 | LYZ | lysozyme | Extracellular Space | enzyme | 1.06E+07 | 7285166.5 | 1.69E+07 | | | | | | |
| P08905 | LYZ | lysozyme | Extracellular Space | enzyme | 3601360.5 | 4485543.5 | 7162192 | P08905 | LYZ | 7120680 | 4862177 | 1984976 | 6 |
| P24668 | M6PR | mannose-6-phosphate receptor (cation dependent) | Cytoplasm | transporter | 3779345 | 4571931.5 | 6928590.5 | | | | | | |
| P24668 | M6PR | mannose-6-phosphate receptor (cation dependent) | Cytoplasm | transporter | 665955.562 | 1532303.5 | 2421783.5 | P24668 | M6PR | 2222650 | 2273622 | 928202 | 6 |
| Q9CXI5 | MANF | mesencephalic astrocyte-derived neurotrophic factor | Extracellular Space | other | 1.77E+07 | 2.44E+07 | 2.13E+07 | | | | | | |
| Q9CXI5 | MANF | mesencephalic astrocyte-derived neurotrophic factor | Extracellular Space | other | 5213981 | 5303466.5 | 5138853.5 | | | | | | |
| Q9CXI5 | MANF | mesencephalic astrocyte-derived neurotrophic factor | Extracellular Space | other | 1868764.5 | 2562632.25 | 3252118.25 | Q9CXI5 | MANF | 8260915 | 8862574 | 2954191 | 9 |
| Q61166 | MAPRE1 | microtubule-associated protein, RP/EB family, member 1 | Cytoplasm | other | 708648 | 626281.625 | 565409.75 | Q61166 | MAPRE1 | 708648 | 71887 | 41504 | 3 |
| Q922Q1 | MARC_2 | mitochondrial amidoxime reducing component 2 | Cytoplasm | enzyme | 744978.75 | 1173694.25 | 1750946.125 | Q922Q1 | MARC_2 | 744979 | 504808 | 291451 | 3 |
| Q9DBD2 | MARH8 | membrane-associated ring finger (C3HC4) 8, E3 ubiquitin protein ligase | Cytoplasm | enzyme | 4662395.5 | 6014425.5 | 5157399 | | | | | | |
| Q9DBD2 | MARH8 | membrane-associated ring finger (C3HC4) 8, E3 ubiquitin protein ligase | Cytoplasm | enzyme | 4426163.5 | 4645210.5 | 3766823.25 | Q9DBD2 | MARH8 | 4544280 | 754977 | 308218 | 6 |
| Q05512 | MARK2 | MAP/microtubule affinity-regulating kinase 2 | Cytoplasm | kinase | 2341315.75 | 2562654 | 2300770.25 | | | | | | |
| Q05512 | MARK2 | MAP/microtubule affinity-regulating kinase 2 | Cytoplasm | kinase | 1167903.375 | 1520030.75 | 886318.125 | Q05512 | MARK2 | 1754610 | 698299 | 285079 | 6 |
| Q8VCF0 | MAVS | mitochondrial antiviral signaling protein | Cytoplasm | other | 5077433.5 | 9077973 | 1.17E+07 | | | | | | |
| Q8VCF0 | MAVS | mitochondrial antiviral signaling protein | Cytoplasm | other | 2295766.25 | 3143055.75 | 3599267.5 | Q8VCF0 | MAVS | 3686600 | 3748546 | 1530337 | 6 |
| Q8BL86 | MBLAC2 | metallo-beta-lactamase domain containing 2 | Cytoplasm | other | 1.33E+07 | 1.31E+07 | 1.51E+07 | Q8BL86 | MBLAC2 | 13310000 | 1121844 | 647697 | 3 |
| Q8CHK3 | MBOAT7 | membrane bound O-acyltransferase domain containing 7 | Plasma Membrane | enzyme | 3490547 | 4961503.5 | 9374158 | Q8CHK3 | MBOAT7 | 3490547 | 3061920 | 1767800 | 3 |
| Q8K5B2 | MCFD2 | multiple coagulation factor deficiency 2 | Cytoplasm | other | 1.17E+07 | 1.16E+07 | 9367337 | Q8K5B2 | MCFD2 | 11670000 | 1318048 | 760975 | 3 |
| Q3UMR5 | MCU | mitochondrial calcium uniporter | Cytoplasm | ion channel | 1168810 | 1657710.875 | 2274434 | Q3UMR5 | MCU | 1168810 | 554042 | 319876 | 3 |
| P08249 | MDH2 | malate dehydrogenase 2, NAD (mitochondrial) | Cytoplasm | enzyme | 3.41E+07 | 6.57E+07 | 7.75E+07 | | | | | | |
| P08249 | MDH2 | malate dehydrogenase 2, NAD (mitochondrial) | Cytoplasm | enzyme | 2.78E+07 | 4.06E+07 | 5.17E+07 | | | | | | |
| P08249 | MDH2 | malate dehydrogenase 2, NAD (mitochondrial) | Cytoplasm | enzyme | 2.50E+07 | 4.22E+07 | 6.30E+07 | | | | | | |
| P08249 | MDH2 | malate dehydrogenase 2, NAD (mitochondrial) | Cytoplasm | enzyme | 2.69E+07 | 3.98E+07 | 4.64E+07 | | | | | | |
| P08249 | MDH2 | malate dehydrogenase 2, NAD (mitochondrial) | Cytoplasm | enzyme | 2.44E+07 | 4.39E+07 | 5.67E+07 | | | | | | |
| P08249 | MDH2 | malate dehydrogenase 2, NAD (mitochondrial) | Cytoplasm | enzyme | 2.55E+07 | 3.72E+07 | 4.68E+07 | | | | | | |
| P08249 | MDH2 | malate dehydrogenase 2, NAD (mitochondrial) | Cytoplasm | enzyme | 1.84E+07 | 3.37E+07 | 5.12E+07 | | | | | | |
| P08249 | MDH2 | malate dehydrogenase 2, NAD (mitochondrial) | Cytoplasm | enzyme | 2.25E+07 | 3.53E+07 | 3.90E+07 | | | | | | |
| P08249 | MDH2 | malate dehydrogenase 2, NAD (mitochondrial) | Cytoplasm | enzyme | 1.54E+07 | 2.93E+07 | 3.66E+07 | | | | | | |
| P08249 | MDH2 | malate dehydrogenase 2, NAD (mitochondrial) | Cytoplasm | enzyme | 9339550 | 1.37E+07 | 1.71E+07 | | | | | | |
| P08249 | MDH2 | malate dehydrogenase 2, NAD (mitochondrial) | Cytoplasm | enzyme | 1.15E+07 | 1.77E+07 | 3.05E+07 | | | | | | |
| P08249 | MDH2 | malate dehydrogenase 2, NAD (mitochondrial) | Cytoplasm | enzyme | 6165690.5 | 1.07E+07 | 1.25E+07 | | | | | | |
| P08249 | MDH2 | malate dehydrogenase 2, NAD (mitochondrial) | Cytoplasm | enzyme | 1916325.375 | 4457966 | 7083017 | P08249 | MDH2 | 19138582 | 18233179 | 2919645 | 39 |
| Q99KE1 | ME2 | malic enzyme 2, NAD(+)-dependent, mitochondrial | Cytoplasm | enzyme | 4536516.5 | 6151696 | 5411322.5 | | | | | | |
| Q99KE1 | ME2 | malic enzyme 2, NAD(+)-dependent, mitochondrial | Cytoplasm | enzyme | 4647254.5 | 9194504 | 1.10E+07 | | | | | | |
| Q99KE1 | ME2 | malic enzyme 2, NAD(+)-dependent, mitochondrial | Cytoplasm | enzyme | 1315572.5 | 2528319.5 | 2946824.75 | | | | | | |
| Q99KE1 | ME2 | malic enzyme 2, NAD(+)-dependent, mitochondrial | Cytoplasm | enzyme | 1867745.375 | 3069597.5 | 3669195.75 | Q99KE1 | ME2 | 3091772 | 2923660 | 843988 | 12 |
| Q91VH6 | MEMO1 | mediator of cell motility 1 | Cytoplasm | other | 7912286 | 1.19E+07 | 8471097 | | | | | | |
| Q91VH6 | MEMO1 | mediator of cell motility 1 | Cytoplasm | other | 3149707.5 | 5571458 | 5464407.5 | | | | | | |
| Q91VH6 | MEMO1 | mediator of cell motility 1 | Cytoplasm | other | 4518643.5 | 7426137.5 | 4572813 | | | | | | |
| Q91VH6 | MEMO1 | mediator of cell motility 1 | Cytoplasm | other | 3254004.5 | 6160830 | 4964544 | | | | | | |
| Q91VH6 | MEMO1 | mediator of cell motility 1 | Cytoplasm | other | 3287659.25 | 5403874 | 2914050.25 | Q91VH6 | MEMO1 | 4424460 | 2445781 | 631498 | 15 |
| Q9ERE7 | MESDC2 | mesoderm development candidate 2 | Extracellular Space | other | 1796553.125 | 1574378.125 | 1788859.125 | Q9ERE7 | MESDC2 | 1796553 | 126110 | 72810 | 3 |
| O08663 | METAP2 | methionyl aminopeptidase 2 | Cytoplasm | peptidase | 1177878.25 | 2348141.75 | 2326204.5 | O08663 | METAP2 | 1177878 | 669409 | 386484 | 3 |
| Q6PCP5 | Mff | mitochondrial fission factor | Cytoplasm | other | 5973752.5 | 9150484 | 5814250.5 | | | | | | |
| Q6PCP5 | Mff | mitochondrial fission factor | Cytoplasm | other | 3594500.5 | 6648695 | 7715145 | Q6PCP5 | Mff | 4784127 | 1882078 | 768355 | 6 |
| Q3T9M1 | MFSD2B | major facilitator superfamily domain containing 2B | Plasma Membrane | other | 2.64E+07 | 3.14E+07 | 3.42E+07 | | | | | | |
| Q3T9M1 | MFSD2B | major facilitator superfamily domain containing 2B | Plasma Membrane | other | 2647994 | 1989022.375 | 2013383.75 | Q3T9M1 | MFSD2B | 14513997 | 15764065 | 6435653 | 6 |
| Q9CPU4 | MGST3 | microsomal glutathione S-transferase 3 | Cytoplasm | enzyme | 3.11E+07 | 6.63E+07 | 6.20E+07 | | | | | | |
| Q9CPU4 | MGST3 | microsomal glutathione S-transferase 3 | Cytoplasm | enzyme | 9136545 | 2.03E+07 | 1.38E+07 | | | | | | |
| Q9CPU4 | MGST3 | microsomal glutathione S-transferase 3 | Cytoplasm | enzyme | 5094344.5 | 8966942 | 9598548 | Q9CPU4 | MGST3 | 15103630 | 23455564 | 7818521 | 9 |
| Q8VDP3 | MICAL1 | microtubule associated monooxygenase, calponin and LIM domain containing 1 | Cytoplasm | enzyme | 2825949 | 1758767.25 | 1847946 | Q8VDP3 | MICAL1 | 2825949 | 592075 | 341835 | 3 |
| Q6ZQI3 | MLEC | malectin | Plasma Membrane | other | 3748710.75 | 6174344.5 | 7072843.5 | | | | | | |
| Q6ZQI3 | MLEC | malectin | Plasma Membrane | other | 4987231.5 | 5719880.5 | 8953931 | | | | | | |
| Q6ZQI3 | MLEC | malectin | Plasma Membrane | other | 4449612 | 5300383 | 6061240 | | | | | | |
| Q6ZQI3 | MLEC | malectin | Plasma Membrane | other | 3426197.75 | 3656658.25 | 4064202.5 | | | | | | |
| Q6ZQI3 | MLEC | malectin | Plasma Membrane | other | 2008302 | 3115807.75 | 5231958.5 | Q6ZQI3 | MLEC | 3724011 | 1743430 | 450152 | 15 |
| Q8K3F2 | MMP21 | matrix metallopeptidase 21 | Extracellular Space | peptidase | 1978087.375 | 1996304.5 | 1651151.5 | Q8K3F2 | MMP21 | 1978087 | 194229 | 112138 | 3 |
| B2RPV6 | MMRN1 | multimerin 1 | Extracellular Space | other | 3725726.25 | 3222256 | 5505803 | | | | | | |
| B2RPV6 | MMRN1 | multimerin 1 | Extracellular Space | other | 1989100.25 | 3071889.75 | 4443121.5 | | | | | | |
| B2RPV6 | MMRN1 | multimerin 1 | Extracellular Space | other | 3992237.5 | 4811815 | 5619650.5 | | | | | | |
| B2RPV6 | MMRN1 | multimerin 1 | Extracellular Space | other | 2998377.75 | 3442822.75 | 5132769.5 | | | | | | |
| B2RPV6 | MMRN1 | multimerin 1 | Extracellular Space | other | 1726212.625 | 2570748 | 2987547.25 | | | | | | |
| B2RPV6 | MMRN1 | multimerin 1 | Extracellular Space | other | 3246902 | 4127686.25 | 4917135.5 | | | | | | |
| B2RPV6 | MMRN1 | multimerin 1 | Extracellular Space | other | 3382728.5 | 3699226.5 | 5551721.5 | | | | | | |
| B2RPV6 | MMRN1 | multimerin 1 | Extracellular Space | other | 2913452.5 | 3475350.25 | 4096373 | | | | | | |
| B2RPV6 | MMRN1 | multimerin 1 | Extracellular Space | other | 2116627.25 | 2569955 | 3714091 | | | | | | |
| B2RPV6 | MMRN1 | multimerin 1 | Extracellular Space | other | 1843919.25 | 1601988.875 | 2771207.25 | | | | | | |

| | | | | | | | | | | | | | |
|---|---|---|---|---|---|---|---|---|---|---|---|---|---|
| B2RPV6 | MMRN1 | multimerin 1 | Extracellular Space | other | 1438518.75 | 1846577 | 3834969.75 | B2RPV6 | MMRN1 | 2670346 | 1187026 | 206635 | 33 |
| P70290 | MPP1 | membrane protein, palmitoylated 1, 55kDa | Plasma Membrane | kinase | 1.50E+09 | 1.65E+09 | 1.57E+09 | | | | | | |
| P70290 | MPP1 | membrane protein, palmitoylated 1, 55kDa | Plasma Membrane | kinase | 1.35E+09 | 1.49E+09 | 1.55E+09 | | | | | | |
| P70290 | MPP1 | membrane protein, palmitoylated 1, 55kDa | Plasma Membrane | kinase | 1.21E+09 | 1.17E+09 | 1.09E+09 | | | | | | |
| P70290 | MPP1 | membrane protein, palmitoylated 1, 55kDa | Plasma Membrane | kinase | 1.26E+09 | 1.12E+09 | 9.71E+08 | | | | | | |
| P70290 | MPP1 | membrane protein, palmitoylated 1, 55kDa | Plasma Membrane | kinase | 1.41E+09 | 1.32E+09 | 1.04E+09 | | | | | | |
| P70290 | MPP1 | membrane protein, palmitoylated 1, 55kDa | Plasma Membrane | kinase | 1.11E+09 | 1.09E+09 | 1.12E+09 | | | | | | |
| P70290 | MPP1 | membrane protein, palmitoylated 1, 55kDa | Plasma Membrane | kinase | 7.80E+08 | 8.14E+08 | 6.22E+08 | | | | | | |
| P70290 | MPP1 | membrane protein, palmitoylated 1, 55kDa | Plasma Membrane | kinase | 6.80E+08 | 6.47E+08 | 5.85E+08 | | | | | | |
| P70290 | MPP1 | membrane protein, palmitoylated 1, 55kDa | Plasma Membrane | kinase | 6.20E+08 | 6.05E+08 | 4.81E+08 | | | | | | |
| P70290 | MPP1 | membrane protein, palmitoylated 1, 55kDa | Plasma Membrane | kinase | 5.58E+08 | 6.06E+08 | 5.61E+08 | | | | | | |
| P70290 | MPP1 | membrane protein, palmitoylated 1, 55kDa | Plasma Membrane | kinase | 6.00E+08 | 6.86E+08 | 4.02E+08 | | | | | | |
| P70290 | MPP1 | membrane protein, palmitoylated 1, 55kDa | Plasma Membrane | kinase | 8.36E+08 | 5.35E+08 | 4.53E+08 | | | | | | |
| P70290 | MPP1 | membrane protein, palmitoylated 1, 55kDa | Plasma Membrane | kinase | 4.66E+08 | 4.01E+08 | 4.66E+08 | | | | | | |
| P70290 | MPP1 | membrane protein, palmitoylated 1, 55kDa | Plasma Membrane | kinase | 6.56E+08 | 6.12E+08 | 5.33E+08 | | | | | | |
| P70290 | MPP1 | membrane protein, palmitoylated 1, 55kDa | Plasma Membrane | kinase | 4.22E+08 | 4.70E+08 | 3.82E+08 | | | | | | |
| P70290 | MPP1 | membrane protein, palmitoylated 1, 55kDa | Plasma Membrane | kinase | 4.62E+08 | 4.71E+08 | 3.61E+08 | | | | | | |
| P70290 | MPP1 | membrane protein, palmitoylated 1, 55kDa | Plasma Membrane | kinase | 3.52E+08 | 3.38E+08 | 3.46E+08 | | | | | | |
| P70290 | MPP1 | membrane protein, palmitoylated 1, 55kDa | Plasma Membrane | kinase | 1.87E+08 | 2.24E+08 | 3.77E+08 | | | | | | |
| P70290 | MPP1 | membrane protein, palmitoylated 1, 55kDa | Plasma Membrane | kinase | 3.22E+08 | 3.20E+08 | 2.88E+08 | | | | | | |
| P70290 | MPP1 | membrane protein, palmitoylated 1, 55kDa | Plasma Membrane | kinase | 3.09E+08 | 2.86E+08 | 2.74E+08 | | | | | | |
| P70290 | MPP1 | membrane protein, palmitoylated 1, 55kDa | Plasma Membrane | kinase | 2.01E+08 | 1.71E+08 | 1.53E+08 | | | | | | |
| P70290 | MPP1 | membrane protein, palmitoylated 1, 55kDa | Plasma Membrane | kinase | 1.92E+08 | 1.71E+08 | 1.43E+08 | | | | | | |
| P70290 | MPP1 | membrane protein, palmitoylated 1, 55kDa | Plasma Membrane | kinase | 1.21E+08 | 1.04E+08 | 1.02E+08 | | | | | | |
| P70290 | MPP1 | membrane protein, palmitoylated 1, 55kDa | Plasma Membrane | kinase | 5.44E+07 | 5.32E+07 | 1.08E+08 | | | | | | |
| P70290 | MPP1 | membrane protein, palmitoylated 1, 55kDa | Plasma Membrane | kinase | 6.71E+07 | 2.01E+07 | 8.34E+07 | | | | | | |
| P70290 | MPP1 | membrane protein, palmitoylated 1, 55kDa | Plasma Membrane | kinase | 1.06E+08 | 1.02E+08 | 8.88E+07 | | | | | | |
| P70290 | MPP1 | membrane protein, palmitoylated 1, 55kDa | Plasma Membrane | kinase | 5.86E+07 | 6.47E+07 | 4.24E+07 | | | | | | |
| P70290 | MPP1 | membrane protein, palmitoylated 1, 55kDa | Plasma Membrane | kinase | 3.51E+07 | 3.23E+07 | 6.13E+07 | | | | | | |
| P70290 | MPP1 | membrane protein, palmitoylated 1, 55kDa | Plasma Membrane | kinase | 4.41E+07 | 4.63E+07 | 3.91E+07 | | | | | | |
| P70290 | MPP1 | membrane protein, palmitoylated 1, 55kDa | Plasma Membrane | kinase | 4.34E+07 | 4.45E+07 | 3.79E+07 | | | | | | |
| P70290 | MPP1 | membrane protein, palmitoylated 1, 55kDa | Plasma Membrane | kinase | 7.46E+07 | 7.04E+07 | 5.09E+07 | | | | | | |
| P70290 | MPP1 | membrane protein, palmitoylated 1, 55kDa | Plasma Membrane | kinase | 2.83E+07 | 3.69E+07 | 3.65E+07 | | | | | | |
| P70290 | MPP1 | membrane protein, palmitoylated 1, 55kDa | Plasma Membrane | kinase | 3.73E+07 | 4.22E+07 | 4.10E+07 | | | | | | |
| P70290 | MPP1 | membrane protein, palmitoylated 1, 55kDa | Plasma Membrane | kinase | 5.53E+07 | 1.52E+08 | 1.24E+08 | | | | | | |
| P70290 | MPP1 | membrane protein, palmitoylated 1, 55kDa | Plasma Membrane | kinase | 1.31E+07 | 3159508.25 | 1.45E+07 | | | | | | |
| P70290 | MPP1 | membrane protein, palmitoylated 1, 55kDa | Plasma Membrane | kinase | 5100305.5 | 4521080.5 | 1.03E+07 | | | | | | |
| P70290 | MPP1 | membrane protein, palmitoylated 1, 55kDa | Plasma Membrane | kinase | 1.29E+07 | 1.49E+07 | 8254721 | | | | | | |
| P70290 | MPP1 | membrane protein, palmitoylated 1, 55kDa | Plasma Membrane | kinase | 5044260.5 | 4097184.25 | 5659396.5 | | | | | | |
| P70290 | MPP1 | membrane protein, palmitoylated 1, 55kDa | Plasma Membrane | kinase | 5286699 | 4407940 | 5163889.5 | | | | | | |
| P70290 | MPP1 | membrane protein, palmitoylated 1, 55kDa | Plasma Membrane | kinase | 3478340.5 | 3225604.5 | 2579958.25 | | | | | | |
| P70290 | MPP1 | membrane protein, palmitoylated 1, 55kDa | Plasma Membrane | kinase | 2484416.5 | 5112392.5 | 2283285 | | | | | | |
| P70290 | MPP1 | membrane protein, palmitoylated 1, 55kDa | Plasma Membrane | kinase | 288988.781 | 569744.5 | 268524.25 | P70290 | MPP1 | 387043643 | 438506504 | 39065264 | 126 |
| Q99J99 | MPST | mercaptopyruvate sulfurtransferase | Cytoplasm | enzyme | 6996895 | 1.47E+07 | 2.03E+07 | | | | | | |
| Q99J99 | MPST | mercaptopyruvate sulfurtransferase | Cytoplasm | enzyme | 664256.812 | 1283741.75 | 1584083.25 | Q99J99 | MPST | 3830576 | 8176488 | 3338037 | 6 |
| Q9DB15 | MRPL12 | mitochondrial ribosomal protein L12 | Cytoplasm | other | 1448071.5 | 2007567.125 | 2603950.5 | Q9DB15 | MRPL12 | 1448072 | 578038 | 333730 | 3 |
| Q9CQX8 | MRPS36 | mitochondrial ribosomal protein S36 | Cytoplasm | other | 1366380.875 | 2729921.5 | 2661925 | Q9CQX8 | MRPS36 | 1366381 | 768364 | 443615 | 3 |
| Q9WUX5 | MRVI1 | murine retrovirus integration site 1 homolog | Cytoplasm | other | 1500630.25 | 1786088.375 | 2390271.75 | | | | | | |
| Q9WUX5 | MRVI1 | murine retrovirus integration site 1 homolog | Cytoplasm | other | 758763.375 | 1417906.5 | 2553711.75 | | | | | | |
| Q9WUX5 | MRVI1 | murine retrovirus integration site 1 homolog | Cytoplasm | other | 540275.062 | 795506.562 | 974992.812 | Q9WUX5 | MRVI1 | 933223 | 721630 | 240543 | 9 |
| P26041 | MSN | moesin | Plasma Membrane | other | 1.12E+07 | 9465531 | 1.01E+07 | | | | | | |
| P26041 | MSN | moesin | Plasma Membrane | other | 9091666 | 6850464 | 1.02E+07 | | | | | | |
| P26041 | MSN | moesin | Plasma Membrane | other | 8046368 | 5649342 | 7725339 | | | | | | |
| P26041 | MSN | moesin | Plasma Membrane | other | 6320549.5 | 6156687 | 8839091 | | | | | | |
| P26041 | MSN | moesin | Plasma Membrane | other | 5136255.5 | 3982844.75 | 6143375 | | | | | | |
| P26041 | MSN | moesin | Plasma Membrane | other | 2063989.625 | 2077887.875 | 3106455 | | | | | | |
| P26041 | MSN | moesin | Plasma Membrane | other | 4372811.5 | 3863396 | 6457448.5 | | | | | | |
| P26041 | MSN | moesin | Plasma Membrane | other | 2545809 | 1910061.75 | 3857526.5 | | | | | | |
| P26041 | MSN | moesin | Plasma Membrane | other | 3190990.75 | 3094196 | 3998546.75 | | | | | | |
| P26041 | MSN | moesin | Plasma Membrane | other | 1856356.5 | 2105912 | 2388779.25 | | | | | | |
| P26041 | MSN | moesin | Plasma Membrane | other | 913375.375 | 733953.312 | 1726289.375 | | | | | | |
| P26041 | MSN | moesin | Plasma Membrane | other | 2471464.75 | 2505803.25 | 4724261.5 | P26041 | MSN | 4768303 | 2931136 | 488523 | 36 |
| P03930 | mt-Atp8 | ATP synthase F0 subunit 8 | Cytoplasm | enzyme | 1.18E+07 | 1.68E+07 | 2.29E+07 | | | | | | |
| P03930 | mt-Atp8 | ATP synthase F0 subunit 8 | Cytoplasm | enzyme | 3459712.75 | 6267052.5 | 8603049 | P03930 | mt-Atp8 | 7649856 | 7199416 | 2939149 | 6 |
| Q791V5 | MTCH2 | mitochondrial carrier 2 | Cytoplasm | other | 5045582 | 9776874 | 9683818 | Q791V5 | MTCH2 | 5045582 | 2705150 | 1561819 | 3 |
| P00397 | MT-CO1 | cytochrome c oxidase subunit I | Cytoplasm | enzyme | 1321108 | 2764693.25 | 3928530.75 | P00397 | MT-CO1 | 1321108 | 1306210 | 754141 | 3 |
| P00405 | MT-CO2 | cytochrome c oxidase subunit II | Cytoplasm | enzyme | 2.13E+07 | 3.67E+07 | 3.30E+07 | | | | | | |
| P00405 | MT-CO2 | cytochrome c oxidase subunit II | Cytoplasm | enzyme | 1.29E+07 | 2.51E+07 | 2.51E+07 | | | | | | |
| P00405 | MT-CO2 | cytochrome c oxidase subunit II | Cytoplasm | enzyme | 3593162.75 | 6050310.5 | 6967311 | | | | | | |
| P00405 | MT-CO2 | cytochrome c oxidase subunit II | Cytoplasm | enzyme | 1475808.875 | 2532006.5 | 1345711.375 | | | | | | |
| P00405 | MT-CO2 | cytochrome c oxidase subunit II | Cytoplasm | enzyme | 2084139.125 | 3278715 | 3736151.5 | | | | | | |
| P00405 | MT-CO2 | cytochrome c oxidase subunit II | Cytoplasm | enzyme | 1568572.375 | 494532.812 | 3396080.75 | P00405 | MT-CO2 | 7146947 | 12002385 | 2828989 | 18 |
| P00416 | MT-CO3 | cytochrome c oxidase III | Cytoplasm | enzyme | 1297281.625 | 3013216 | 2459819.5 | P00416 | MT-CO3 | 1297282 | 875802 | 505644 | 3 |
| Q922D8 | MTHFD1 | methylenetetrahydrofolate dehydrogenase (NADP+ dependent) 1, methenyltetrahydrofolate cyclohydrolase, formyltetrahydrofolate synthetase | Cytoplasm | enzyme | 1.88E+07 | 2.20E+07 | 2.32E+07 | | | | | | |
| Q922D8 | MTHFD1 | methylenetetrahydrofolate dehydrogenase (NADP+ dependent) 1, methenyltetrahydrofolate cyclohydrolase, formyltetrahydrofolate synthetase | Cytoplasm | enzyme | 1.43E+07 | 1.61E+07 | 1.70E+07 | | | | | | |
| Q922D8 | MTHFD1 | methylenetetrahydrofolate dehydrogenase (NADP+ dependent) 1, methenyltetrahydrofolate cyclohydrolase, formyltetrahydrofolate synthetase | Cytoplasm | enzyme | 1.52E+07 | 2.08E+07 | 2.84E+07 | | | | | | |
| Q922D8 | MTHFD1 | methylenetetrahydrofolate dehydrogenase (NADP+ dependent) 1, methenyltetrahydrofolate cyclohydrolase, formyltetrahydrofolate synthetase | Cytoplasm | enzyme | 1.49E+07 | 1.71E+07 | 1.65E+07 | | | | | | |
| Q922D8 | MTHFD1 | methylenetetrahydrofolate dehydrogenase (NADP+ dependent) 1, methenyltetrahydrofolate cyclohydrolase, formyltetrahydrofolate synthetase | Cytoplasm | enzyme | 1.20E+07 | 1.50E+07 | 1.61E+07 | | | | | | |
| Q922D8 | MTHFD1 | methylenetetrahydrofolate dehydrogenase (NADP+ dependent) 1, methenyltetrahydrofolate cyclohydrolase, formyltetrahydrofolate synthetase | Cytoplasm | enzyme | 1.23E+07 | 1.39E+07 | 1.78E+07 | | | | | | |
| Q922D8 | MTHFD1 | methylenetetrahydrofolate dehydrogenase (NADP+ dependent) 1, methenyltetrahydrofolate cyclohydrolase, formyltetrahydrofolate synthetase | Cytoplasm | enzyme | 1.49E+07 | 2.06E+07 | 1.63E+07 | | | | | | |
| Q922D8 | MTHFD1 | methylenetetrahydrofolate dehydrogenase (NADP+ dependent) 1, methenyltetrahydrofolate cyclohydrolase, formyltetrahydrofolate synthetase | Cytoplasm | enzyme | 1.21E+07 | 1.37E+07 | 1.35E+07 | | | | | | |
| Q922D8 | MTHFD1 | methylenetetrahydrofolate dehydrogenase (NADP+ dependent) 1, methenyltetrahydrofolate cyclohydrolase, formyltetrahydrofolate synthetase | Cytoplasm | enzyme | 7886206.5 | 4337129 | 2245557.75 | | | | | | |
| Q922D8 | MTHFD1 | methylenetetrahydrofolate dehydrogenase (NADP+ dependent) 1, methenyltetrahydrofolate cyclohydrolase, formyltetrahydrofolate synthetase | Cytoplasm | enzyme | 1.15E+07 | 1.27E+07 | 1.27E+07 | | | | | | |
| Q922D8 | MTHFD1 | methylenetetrahydrofolate dehydrogenase (NADP+ dependent) 1, methenyltetrahydrofolate cyclohydrolase, formyltetrahydrofolate synthetase | Cytoplasm | enzyme | 1.15E+07 | 1.07E+07 | 1.19E+07 | | | | | | |
| Q922D8 | MTHFD1 | methylenetetrahydrofolate dehydrogenase (NADP+ dependent) 1, methenyltetrahydrofolate cyclohydrolase, formyltetrahydrofolate synthetase | Cytoplasm | enzyme | 7973675 | 1.01E+07 | 1.09E+07 | | | | | | |
| Q922D8 | MTHFD1 | methylenetetrahydrofolate dehydrogenase (NADP+ dependent) 1, methenyltetrahydrofolate cyclohydrolase, formyltetrahydrofolate synthetase | Cytoplasm | enzyme | 8004895 | 9084847 | 1.22E+07 | | | | | | |
| Q922D8 | MTHFD1 | methylenetetrahydrofolate dehydrogenase (NADP+ dependent) 1, methenyltetrahydrofolate cyclohydrolase, formyltetrahydrofolate synthetase | Cytoplasm | enzyme | 1.09E+07 | 1.06E+07 | 1.21E+07 | | | | | | |
| Q922D8 | MTHFD1 | methylenetetrahydrofolate dehydrogenase (NADP+ dependent) 1, methenyltetrahydrofolate cyclohydrolase, formyltetrahydrofolate synthetase | Cytoplasm | enzyme | 7263232 | 7592068.5 | 1.05E+07 | | | | | | |
| Q922D8 | MTHFD1 | methylenetetrahydrofolate dehydrogenase (NADP+ dependent) 1, methenyltetrahydrofolate cyclohydrolase, formyltetrahydrofolate synthetase | Cytoplasm | enzyme | 6153744.5 | 7641133 | 1.46E+07 | | | | | | |
| Q922D8 | MTHFD1 | methylenetetrahydrofolate dehydrogenase (NADP+ dependent) 1, methenyltetrahydrofolate cyclohydrolase, formyltetrahydrofolate synthetase | Cytoplasm | enzyme | 5648670.5 | 8110898 | 9766075 | | | | | | |
| Q922D8 | MTHFD1 | methylenetetrahydrofolate dehydrogenase (NADP+ dependent) 1, methenyltetrahydrofolate cyclohydrolase, formyltetrahydrofolate synthetase | Cytoplasm | enzyme | 5595732 | 6163455 | 6783853.5 | | | | | | |
| Q922D8 | MTHFD1 | methylenetetrahydrofolate dehydrogenase (NADP+ dependent) 1, methenyltetrahydrofolate cyclohydrolase, formyltetrahydrofolate synthetase | Cytoplasm | enzyme | 7826091.5 | 8406999 | 9944026 | | | | | | |
| Q922D8 | MTHFD1 | methylenetetrahydrofolate dehydrogenase (NADP+ dependent) 1, methenyltetrahydrofolate cyclohydrolase, formyltetrahydrofolate synthetase | Cytoplasm | enzyme | 7659062.5 | 7417513 | 9554900 | | | | | | |
| Q922D8 | MTHFD1 | methylenetetrahydrofolate dehydrogenase (NADP+ dependent) 1, methenyltetrahydrofolate cyclohydrolase, formyltetrahydrofolate synthetase | Cytoplasm | enzyme | 5181086 | 6381014.5 | 7620115.5 | | | | | | |
| Q922D8 | MTHFD1 | methylenetetrahydrofolate dehydrogenase (NADP+ dependent) 1, methenyltetrahydrofolate cyclohydrolase, formyltetrahydrofolate synthetase | Cytoplasm | enzyme | 5267619 | 5486450.5 | 6976224 | | | | | | |
| Q922D8 | MTHFD1 | methylenetetrahydrofolate dehydrogenase (NADP+ dependent) 1, methenyltetrahydrofolate cyclohydrolase, formyltetrahydrofolate synthetase | Cytoplasm | enzyme | 3054550.25 | 3174493 | 8768419 | | | | | | |
| Q922D8 | MTHFD1 | methylenetetrahydrofolate dehydrogenase (NADP+ dependent) 1, methenyltetrahydrofolate cyclohydrolase, formyltetrahydrofolate synthetase | Cytoplasm | enzyme | 5802445.5 | 4615166 | 3269757 | | | | | | |
| Q922D8 | MTHFD1 | methylenetetrahydrofolate dehydrogenase (NADP+ dependent) 1, methenyltetrahydrofolate cyclohydrolase, formyltetrahydrofolate synthetase | Cytoplasm | enzyme | 4770278 | 7115681.5 | 6841982 | | | | | | |
| Q922D8 | MTHFD1 | methylenetetrahydrofolate dehydrogenase (NADP+ dependent) 1, methenyltetrahydrofolate cyclohydrolase, formyltetrahydrofolate synthetase | Cytoplasm | enzyme | 3524386 | 4213255 | 4967292.5 | | | | | | |
| Q922D8 | MTHFD1 | methylenetetrahydrofolate dehydrogenase (NADP+ dependent) 1, methenyltetrahydrofolate cyclohydrolase, formyltetrahydrofolate synthetase | Cytoplasm | enzyme | 3208091.75 | 3384474.25 | 4109749.75 | | | | | | |
| Q922D8 | MTHFD1 | methylenetetrahydrofolate dehydrogenase (NADP+ dependent) 1, methenyltetrahydrofolate cyclohydrolase, formyltetrahydrofolate synthetase | Cytoplasm | enzyme | 3035863.5 | 3892372.75 | 5614746 | | | | | | |
| Q922D8 | MTHFD1 | methylenetetrahydrofolate dehydrogenase (NADP+ dependent) 1, methenyltetrahydrofolate cyclohydrolase, formyltetrahydrofolate synthetase | Cytoplasm | enzyme | 1574903.25 | 2632287 | 2761393 | Q922D8 | MTHFD1 | 8542777 | 5461338 | 585517 | 87 |
| Q3V3R1 | MTHFD1L | methylenetetrahydrofolate dehydrogenase (NADP+ dependent) 1-like | Cytoplasm | enzyme | 2601290.5 | 5633706.5 | 7905234.5 | | | | | | |
| Q3V3R1 | MTHFD1L | methylenetetrahydrofolate dehydrogenase (NADP+ dependent) 1-like | Cytoplasm | enzyme | 2873027 | 6016971.5 | 7478799 | | | | | | |
| Q3V3R1 | MTHFD1L | methylenetetrahydrofolate dehydrogenase (NADP+ dependent) 1-like | Cytoplasm | enzyme | 2320450.5 | 3679894.5 | 5234290.5 | | | | | | |
| Q3V3R1 | MTHFD1L | methylenetetrahydrofolate dehydrogenase (NADP+ dependent) 1-like | Cytoplasm | enzyme | 1954528.875 | 3948003.25 | 4448657 | | | | | | |
| Q3V3R1 | MTHFD1L | methylenetetrahydrofolate dehydrogenase (NADP+ dependent) 1-like | Cytoplasm | enzyme | 1234058.5 | 2909826.25 | 5081519.5 | | | | | | |
| Q3V3R1 | MTHFD1L | methylenetetrahydrofolate dehydrogenase (NADP+ dependent) 1-like | Cytoplasm | enzyme | 2434905.25 | 3060826 | 3880824.75 | | | | | | |
| Q3V3R1 | MTHFD1L | methylenetetrahydrofolate dehydrogenase (NADP+ dependent) 1-like | Cytoplasm | enzyme | 1564209.875 | 2911950.75 | 3361084.5 | | | | | | |

| | | | | | | | | | | | | | |
|---|---|---|---|---|---|---|---|---|---|---|---|---|---|
| Q3V3R1 | MTHFD1L | methylenetetrahydrofolate dehydrogenase (NADP+ dependent) 1-like | Cytoplasm | enzyme | 1619897.625 | 2705947.5 | 2912862.75 | Q3V3R1 | MTHFD1L | 2075296 | 1778273 | 362988 | 24 |
| P03921 | MT-ND5 | NADH dehydrogenase, subunit 5 (complex I) | Cytoplasm | enzyme | 3705999.75 | 4365215 | 9824689 | | | | | | |
| P03921 | MT-ND5 | NADH dehydrogenase, subunit 5 (complex I) | Cytoplasm | enzyme | 2028564.875 | 2426000.25 | 2609938 | | | | | | |
| P03921 | MT-ND5 | NADH dehydrogenase, subunit 5 (complex I) | Cytoplasm | enzyme | 912968.625 | 1601103.25 | 2216804.25 | P03921 | MT-ND5 | 2215844 | 2657188 | 885729 | 9 |
| Q8C1A3 | MTRR | 5-methyltetrahydrofolate-homocysteine methyltransferase reductase | Cytoplasm | enzyme | 4744178.5 | 4499586.5 | 4057565.25 | Q8C1A3 | MTRR | 4744179 | 348005 | 200921 | 3 |
| P28665 | Mug1/Mug2 | murinoglobulin 1 | Extracellular Space | other | 280181.75 | 547336.625 | 1799022.5 | P28665 | Mug1/Mug2 | 280182 | 810860 | 468150 | 3 |
| P48972 | MYBL2 | v-myb avian myeloblastosis viral oncogene homolog-like 2 | Nucleus | transcription regulator | 7967390 | 7924849.5 | 1.01E+07 | P48972 | MYBL2 | 7967390 | 1249498 | 721398 | 3 |
| Q3U898 | MYEOV2 | myeloma overexpressed 2 | Nucleus | other | 6039029 | 6664566.5 | 4429836 | Q3U898 | MYEOV2 | 6039029 | 1152882 | 665617 | 3 |
| Q6URW6 | MYH14 | myosin, heavy chain 14, non-muscle | Extracellular Space | other | 3063958.75 | 2850361.75 | 8057830 | | | | | | |
| Q6URW6 | MYH14 | myosin, heavy chain 14, non-muscle | Extracellular Space | other | 1358345.75 | 1460769.25 | 3604116.5 | Q6URW6 | MYH14 | 2211152 | 2452796 | 1001350 | 6 |
| Q8VDD5 | MYH9 | myosin, heavy chain 9, non-muscle | Cytoplasm | enzyme | 1.92E+07 | 2.13E+07 | 3.41E+07 | | | | | | |
| Q8VDD5 | MYH9 | myosin, heavy chain 9, non-muscle | Cytoplasm | enzyme | 3.84E+07 | 5.10E+07 | 8.26E+07 | | | | | | |
| Q8VDD5 | MYH9 | myosin, heavy chain 9, non-muscle | Cytoplasm | enzyme | 2.82E+07 | 4.21E+07 | 7.73E+07 | | | | | | |
| Q8VDD5 | MYH9 | myosin, heavy chain 9, non-muscle | Cytoplasm | enzyme | 2.33E+07 | 2.60E+07 | 4.72E+07 | | | | | | |
| Q8VDD5 | MYH9 | myosin, heavy chain 9, non-muscle | Cytoplasm | enzyme | 3.42E+07 | 4.53E+07 | 7.30E+07 | | | | | | |
| Q8VDD5 | MYH9 | myosin, heavy chain 9, non-muscle | Cytoplasm | enzyme | 3.29E+07 | 4.10E+07 | 6.81E+07 | | | | | | |
| Q8VDD5 | MYH9 | myosin, heavy chain 9, non-muscle | Cytoplasm | enzyme | 1.96E+07 | 2.50E+07 | 4.70E+07 | | | | | | |
| Q8VDD5 | MYH9 | myosin, heavy chain 9, non-muscle | Cytoplasm | enzyme | 2.76E+07 | 3.12E+07 | 5.82E+07 | | | | | | |
| Q8VDD5 | MYH9 | myosin, heavy chain 9, non-muscle | Cytoplasm | enzyme | 2.70E+07 | 3.22E+07 | 7.05E+07 | | | | | | |
| Q8VDD5 | MYH9 | myosin, heavy chain 9, non-muscle | Cytoplasm | enzyme | 2.32E+07 | 2.79E+07 | 6.32E+07 | | | | | | |
| Q8VDD5 | MYH9 | myosin, heavy chain 9, non-muscle | Cytoplasm | enzyme | 2.65E+07 | 2.38E+07 | 4.63E+07 | | | | | | |
| Q8VDD5 | MYH9 | myosin, heavy chain 9, non-muscle | Cytoplasm | enzyme | 2.74E+07 | 2.79E+07 | 5.05E+07 | | | | | | |
| Q8VDD5 | MYH9 | myosin, heavy chain 9, non-muscle | Cytoplasm | enzyme | 2.75E+07 | 3.03E+07 | 5.57E+07 | | | | | | |
| Q8VDD5 | MYH9 | myosin, heavy chain 9, non-muscle | Cytoplasm | enzyme | 2.37E+07 | 3.33E+07 | 6.31E+07 | | | | | | |
| Q8VDD5 | MYH9 | myosin, heavy chain 9, non-muscle | Cytoplasm | enzyme | 2.65E+07 | 3.03E+07 | 5.25E+07 | | | | | | |
| Q8VDD5 | MYH9 | myosin, heavy chain 9, non-muscle | Cytoplasm | enzyme | 2.26E+07 | 2.85E+07 | 4.17E+07 | | | | | | |
| Q8VDD5 | MYH9 | myosin, heavy chain 9, non-muscle | Cytoplasm | enzyme | 2.02E+07 | 2.62E+07 | 5.29E+07 | | | | | | |
| Q8VDD5 | MYH9 | myosin, heavy chain 9, non-muscle | Cytoplasm | enzyme | 1.97E+07 | 2.50E+07 | 4.66E+07 | | | | | | |
| Q8VDD5 | MYH9 | myosin, heavy chain 9, non-muscle | Cytoplasm | enzyme | 1.57E+07 | 2.27E+07 | 3.42E+07 | | | | | | |
| Q8VDD5 | MYH9 | myosin, heavy chain 9, non-muscle | Cytoplasm | enzyme | 2.06E+07 | 2.70E+07 | 4.65E+07 | | | | | | |
| Q8VDD5 | MYH9 | myosin, heavy chain 9, non-muscle | Cytoplasm | enzyme | 2.80E+07 | 3.31E+07 | 5.93E+07 | | | | | | |
| Q8VDD5 | MYH9 | myosin, heavy chain 9, non-muscle | Cytoplasm | enzyme | 1.52E+07 | 2.36E+07 | 4.04E+07 | | | | | | |
| Q8VDD5 | MYH9 | myosin, heavy chain 9, non-muscle | Cytoplasm | enzyme | 2.37E+07 | 2.80E+07 | 4.79E+07 | | | | | | |
| Q8VDD5 | MYH9 | myosin, heavy chain 9, non-muscle | Cytoplasm | enzyme | 2.21E+07 | 3.16E+07 | 5.00E+07 | | | | | | |
| Q8VDD5 | MYH9 | myosin, heavy chain 9, non-muscle | Cytoplasm | enzyme | 7924944 | 1.75E+07 | 2.98E+07 | | | | | | |
| Q8VDD5 | MYH9 | myosin, heavy chain 9, non-muscle | Cytoplasm | enzyme | 1.63E+07 | 1.75E+07 | 3.00E+07 | | | | | | |
| Q8VDD5 | MYH9 | myosin, heavy chain 9, non-muscle | Cytoplasm | enzyme | 1.50E+07 | 2.18E+07 | 4.86E+07 | | | | | | |
| Q8VDD5 | MYH9 | myosin, heavy chain 9, non-muscle | Cytoplasm | enzyme | 2.66E+07 | 2.52E+07 | 4.34E+07 | | | | | | |
| Q8VDD5 | MYH9 | myosin, heavy chain 9, non-muscle | Cytoplasm | enzyme | 2.11E+07 | 2.86E+07 | 3.33E+07 | | | | | | |
| Q8VDD5 | MYH9 | myosin, heavy chain 9, non-muscle | Cytoplasm | enzyme | 2.07E+07 | 2.02E+07 | 4.45E+07 | | | | | | |
| Q8VDD5 | MYH9 | myosin, heavy chain 9, non-muscle | Cytoplasm | enzyme | 1.72E+07 | 2.13E+07 | 3.31E+07 | | | | | | |
| Q8VDD5 | MYH9 | myosin, heavy chain 9, non-muscle | Cytoplasm | enzyme | 2.45E+07 | 3.41E+07 | 4.56E+07 | | | | | | |
| Q8VDD5 | MYH9 | myosin, heavy chain 9, non-muscle | Cytoplasm | enzyme | 1.73E+07 | 1.80E+07 | 3.33E+07 | | | | | | |
| Q8VDD5 | MYH9 | myosin, heavy chain 9, non-muscle | Cytoplasm | enzyme | 1.61E+07 | 2.45E+07 | 4.52E+07 | | | | | | |
| Q8VDD5 | MYH9 | myosin, heavy chain 9, non-muscle | Cytoplasm | enzyme | 1.52E+07 | 1.95E+07 | 3.32E+07 | | | | | | |
| Q8VDD5 | MYH9 | myosin, heavy chain 9, non-muscle | Cytoplasm | enzyme | 1.82E+07 | 1.61E+07 | 3.08E+07 | | | | | | |
| Q8VDD5 | MYH9 | myosin, heavy chain 9, non-muscle | Cytoplasm | enzyme | 1.52E+07 | 1.79E+07 | 3.58E+07 | | | | | | |
| Q8VDD5 | MYH9 | myosin, heavy chain 9, non-muscle | Cytoplasm | enzyme | 1.50E+07 | 1.78E+07 | 3.16E+07 | | | | | | |
| Q8VDD5 | MYH9 | myosin, heavy chain 9, non-muscle | Cytoplasm | enzyme | 1.93E+07 | 1.81E+07 | 2.89E+07 | | | | | | |
| Q8VDD5 | MYH9 | myosin, heavy chain 9, non-muscle | Cytoplasm | enzyme | 1.47E+07 | 1.48E+07 | 2.84E+07 | | | | | | |
| Q8VDD5 | MYH9 | myosin, heavy chain 9, non-muscle | Cytoplasm | enzyme | 1.64E+07 | 1.86E+07 | 2.78E+07 | | | | | | |
| Q8VDD5 | MYH9 | myosin, heavy chain 9, non-muscle | Cytoplasm | enzyme | 2.00E+07 | 2.41E+07 | 3.49E+07 | | | | | | |
| Q8VDD5 | MYH9 | myosin, heavy chain 9, non-muscle | Cytoplasm | enzyme | 1.25E+07 | 1.64E+07 | 2.93E+07 | | | | | | |
| Q8VDD5 | MYH9 | myosin, heavy chain 9, non-muscle | Cytoplasm | enzyme | 1.63E+07 | 1.74E+07 | 2.73E+07 | | | | | | |
| Q8VDD5 | MYH9 | myosin, heavy chain 9, non-muscle | Cytoplasm | enzyme | 1.35E+07 | 1.67E+07 | 3.14E+07 | | | | | | |
| Q8VDD5 | MYH9 | myosin, heavy chain 9, non-muscle | Cytoplasm | enzyme | 1.18E+07 | 1.43E+07 | 2.86E+07 | | | | | | |
| Q8VDD5 | MYH9 | myosin, heavy chain 9, non-muscle | Cytoplasm | enzyme | 1.39E+07 | 1.60E+07 | 2.79E+07 | | | | | | |
| Q8VDD5 | MYH9 | myosin, heavy chain 9, non-muscle | Cytoplasm | enzyme | 1.44E+07 | 1.90E+07 | 3.76E+07 | | | | | | |
| Q8VDD5 | MYH9 | myosin, heavy chain 9, non-muscle | Cytoplasm | enzyme | 1.20E+07 | 1.44E+07 | 2.27E+07 | | | | | | |
| Q8VDD5 | MYH9 | myosin, heavy chain 9, non-muscle | Cytoplasm | enzyme | 7607650 | 8714001 | 1.39E+07 | | | | | | |
| Q8VDD5 | MYH9 | myosin, heavy chain 9, non-muscle | Cytoplasm | enzyme | 1.26E+07 | 2.19E+07 | 2.40E+07 | | | | | | |
| Q8VDD5 | MYH9 | myosin, heavy chain 9, non-muscle | Cytoplasm | enzyme | 9610372 | 1.08E+07 | 1.98E+07 | | | | | | |
| Q8VDD5 | MYH9 | myosin, heavy chain 9, non-muscle | Cytoplasm | enzyme | 1.12E+07 | 1.18E+07 | 1.98E+07 | | | | | | |
| Q8VDD5 | MYH9 | myosin, heavy chain 9, non-muscle | Cytoplasm | enzyme | 8662743 | 1.19E+07 | 1.59E+07 | | | | | | |
| Q8VDD5 | MYH9 | myosin, heavy chain 9, non-muscle | Cytoplasm | enzyme | 9708811 | 1.11E+07 | 1.82E+07 | | | | | | |
| Q8VDD5 | MYH9 | myosin, heavy chain 9, non-muscle | Cytoplasm | enzyme | 1.09E+07 | 1.30E+07 | 1.81E+07 | | | | | | |
| Q8VDD5 | MYH9 | myosin, heavy chain 9, non-muscle | Cytoplasm | enzyme | 8104427.5 | 1.17E+07 | 2.48E+07 | | | | | | |
| Q8VDD5 | MYH9 | myosin, heavy chain 9, non-muscle | Cytoplasm | enzyme | 1.05E+07 | 1.53E+07 | 2.73E+07 | | | | | | |
| Q8VDD5 | MYH9 | myosin, heavy chain 9, non-muscle | Cytoplasm | enzyme | 8585822 | 1.22E+07 | 1.81E+07 | | | | | | |
| Q8VDD5 | MYH9 | myosin, heavy chain 9, non-muscle | Cytoplasm | enzyme | 1.18E+07 | 1.37E+07 | 1.98E+07 | | | | | | |
| Q8VDD5 | MYH9 | myosin, heavy chain 9, non-muscle | Cytoplasm | enzyme | 1.08E+07 | 9857990 | 1.85E+07 | | | | | | |
| Q8VDD5 | MYH9 | myosin, heavy chain 9, non-muscle | Cytoplasm | enzyme | 8234483 | 9831627 | 1.08E+07 | | | | | | |
| Q8VDD5 | MYH9 | myosin, heavy chain 9, non-muscle | Cytoplasm | enzyme | 8683462 | 9735773 | 1.35E+07 | | | | | | |
| Q8VDD5 | MYH9 | myosin, heavy chain 9, non-muscle | Cytoplasm | enzyme | 6540966.5 | 8167992.5 | 1.69E+07 | | | | | | |
| Q8VDD5 | MYH9 | myosin, heavy chain 9, non-muscle | Cytoplasm | enzyme | 6011990 | 4342341 | 1.57E+07 | | | | | | |
| Q8VDD5 | MYH9 | myosin, heavy chain 9, non-muscle | Cytoplasm | enzyme | 5516631 | 5037093.5 | 1.11E+07 | | | | | | |
| Q8VDD5 | MYH9 | myosin, heavy chain 9, non-muscle | Cytoplasm | enzyme | 5878974 | 5917842 | 1.05E+07 | | | | | | |
| Q8VDD5 | MYH9 | myosin, heavy chain 9, non-muscle | Cytoplasm | enzyme | 1.12E+07 | 2.11E+07 | 4.10E+07 | | | | | | |
| Q8VDD5 | MYH9 | myosin, heavy chain 9, non-muscle | Cytoplasm | enzyme | 6102779.5 | 6964911.5 | 1.20E+07 | | | | | | |
| Q8VDD5 | MYH9 | myosin, heavy chain 9, non-muscle | Cytoplasm | enzyme | 5424242 | 6575175 | 1.15E+07 | | | | | | |
| Q8VDD5 | MYH9 | myosin, heavy chain 9, non-muscle | Cytoplasm | enzyme | 4758198 | 2670405.75 | 1.64E+07 | | | | | | |
| Q8VDD5 | MYH9 | myosin, heavy chain 9, non-muscle | Cytoplasm | enzyme | 9436035 | 8776620 | 1.93E+07 | | | | | | |
| Q8VDD5 | MYH9 | myosin, heavy chain 9, non-muscle | Cytoplasm | enzyme | 1888012.25 | 2385295.5 | 3098341.75 | | | | | | |
| Q8VDD5 | MYH9 | myosin, heavy chain 9, non-muscle | Cytoplasm | enzyme | 4741399.5 | 5687677 | 8811088 | | | | | | |
| Q8VDD5 | MYH9 | myosin, heavy chain 9, non-muscle | Cytoplasm | enzyme | 4034954.5 | 5964522.5 | 8616733 | | | | | | |
| Q8VDD5 | MYH9 | myosin, heavy chain 9, non-muscle | Cytoplasm | enzyme | 4090465.75 | 4372947.5 | 5953415 | | | | | | |
| Q8VDD5 | MYH9 | myosin, heavy chain 9, non-muscle | Cytoplasm | enzyme | 213366.484 | 278320.875 | 1984723.75 | | | | | | |
| Q8VDD5 | MYH9 | myosin, heavy chain 9, non-muscle | Cytoplasm | enzyme | 3796166.75 | 4291395.5 | 5753776 | | | | | | |
| Q8VDD5 | MYH9 | myosin, heavy chain 9, non-muscle | Cytoplasm | enzyme | 3437227.25 | 5100595.5 | 1.06E+07 | | | | | | |
| Q8VDD5 | MYH9 | myosin, heavy chain 9, non-muscle | Cytoplasm | enzyme | 3035640 | 3506827.25 | 7128120 | | | | | | |
| Q8VDD5 | MYH9 | myosin, heavy chain 9, non-muscle | Cytoplasm | enzyme | 7152048 | 5264059 | 1.01E+07 | | | | | | |
| Q8VDD5 | MYH9 | myosin, heavy chain 9, non-muscle | Cytoplasm | enzyme | 3348066.25 | 4054194 | 8843059 | | | | | | |
| Q8VDD5 | MYH9 | myosin, heavy chain 9, non-muscle | Cytoplasm | enzyme | 2246217.5 | 2861407 | 4609301 | | | | | | |
| Q8VDD5 | MYH9 | myosin, heavy chain 9, non-muscle | Cytoplasm | enzyme | 1828691.875 | 3167414 | 6733204.5 | | | | | | |
| Q8VDD5 | MYH9 | myosin, heavy chain 9, non-muscle | Cytoplasm | enzyme | 5458346.5 | 6944683 | 1.11E+07 | | | | | | |
| Q8VDD5 | MYH9 | myosin, heavy chain 9, non-muscle | Cytoplasm | enzyme | 2038162.25 | 834856.938 | 3621453.5 | | | | | | |
| Q8VDD5 | MYH9 | myosin, heavy chain 9, non-muscle | Cytoplasm | enzyme | 4765564.5 | 1222276.625 | 1.00E+07 | | | | | | |
| Q8VDD5 | MYH9 | myosin, heavy chain 9, non-muscle | Cytoplasm | enzyme | 2134569.25 | 3117708.75 | 2970429.25 | | | | | | |
| Q8VDD5 | MYH9 | myosin, heavy chain 9, non-muscle | Cytoplasm | enzyme | 837604.562 | 1444776.5 | 1872965.75 | | | | | | |
| Q8VDD5 | MYH9 | myosin, heavy chain 9, non-muscle | Cytoplasm | enzyme | 1602249.5 | 2030806.25 | 3723115.5 | | | | | | |
| Q8VDD5 | MYH9 | myosin, heavy chain 9, non-muscle | Cytoplasm | enzyme | 2170616.25 | 2810294.25 | 5568173.5 | | | | | | |
| Q8VDD5 | MYH9 | myosin, heavy chain 9, non-muscle | Cytoplasm | enzyme | 2279055.5 | 3477110.5 | 3583266 | | | | | | |
| Q8VDD5 | MYH9 | myosin, heavy chain 9, non-muscle | Cytoplasm | enzyme | 262102.344 | 412152.562 | 317949.156 | | | | | | |
| Q8VDD5 | MYH9 | myosin, heavy chain 9, non-muscle | Cytoplasm | enzyme | 535860.312 | 1183942 | 1997828 | | | | | | |
| Q8VDD5 | MYH9 | myosin, heavy chain 9, non-muscle | Cytoplasm | enzyme | 400817.469 | 1701635.375 | 3329418.75 | | | | | | |
| Q8VDD5 | MYH9 | myosin, heavy chain 9, non-muscle | Cytoplasm | enzyme | 198168.469 | 718117.938 | 1733155 | Q8VDD5 | MYH9 | 13166228 | 15709944 | 925717 | 288 |
| P05977 | MYL1 | myosin, light chain 1, alkali; skeletal, fast | Cytoplasm | other | 3529228.75 | 3908296 | 7750683 | P05977 | MYL1 | 3529229 | 2335534 | 1348421 | 3 |
| Q3THE2 | MYL12A | myosin, light chain 12A, regulatory, non-sarcomeric | Cytoplasm | other | 1.29E+07 | 1.63E+07 | 2.50E+07 | | | | | | |
| Q3THE2 | MYL12A | myosin, light chain 12A, regulatory, non-sarcomeric | Cytoplasm | other | 1.69E+07 | 2.07E+07 | 3.95E+07 | | | | | | |

| | | | | | | | | | | | | | |
|---|---|---|---|---|---|---|---|---|---|---|---|---|---|
| Q3THE2 | MYL12A | myosin, light chain 12A, regulatory, non-sarcomeric | Cytoplasm | other | 2944683 | 3972543.25 | 3564815.25 | | | | | | |
| Q3THE2 | MYL12A | myosin, light chain 12A, regulatory, non-sarcomeric | Cytoplasm | other | 3233237.5 | 5649606 | 5307180 | | | | | | |
| Q3THE2 | MYL12A | myosin, light chain 12A, regulatory, non-sarcomeric | Cytoplasm | other | 3468300 | 2313068 | 5966244.5 | Q3THE2 | MYL12A | 7891244 | 10752860 | 2776377 | 15 |
| Q60605 | MYL6 | myosin, light chain 6, alkali, smooth muscle and non-muscle | Cytoplasm | other | 4.49E+07 | 4.78E+07 | 7.50E+07 | | | | | | |
| Q60605 | MYL6 | myosin, light chain 6, alkali, smooth muscle and non-muscle | Cytoplasm | other | 3.41E+07 | 4.53E+07 | 8.01E+07 | | | | | | |
| Q60605 | MYL6 | myosin, light chain 6, alkali, smooth muscle and non-muscle | Cytoplasm | other | 2.85E+07 | 3.75E+07 | 5.86E+07 | | | | | | |
| Q60605 | MYL6 | myosin, light chain 6, alkali, smooth muscle and non-muscle | Cytoplasm | other | 2.19E+07 | 2.69E+07 | 4.26E+07 | | | | | | |
| Q60605 | MYL6 | myosin, light chain 6, alkali, smooth muscle and non-muscle | Cytoplasm | other | 2.00E+07 | 2.50E+07 | 4.57E+07 | | | | | | |
| Q60605 | MYL6 | myosin, light chain 6, alkali, smooth muscle and non-muscle | Cytoplasm | other | 1.60E+07 | 1.65E+07 | 3.03E+07 | | | | | | |
| Q60605 | MYL6 | myosin, light chain 6, alkali, smooth muscle and non-muscle | Cytoplasm | other | 5814874.5 | 7377840 | 1.34E+07 | Q60605 | MYL6 | 24456411 | 20182420 | 4404165 | 21 |
| Q9CQ19 | MYL9 | myosin, light chain 9, regulatory | Cytoplasm | other | 1.25E+07 | 1.23E+07 | 2.15E+07 | | | | | | |
| Q9CQ19 | MYL9 | myosin, light chain 9, regulatory | Cytoplasm | other | 1.70E+07 | 1.90E+07 | 3.29E+07 | | | | | | |
| Q9CQ19 | MYL9 | myosin, light chain 9, regulatory | Cytoplasm | other | 7459565.5 | 1.08E+07 | 9514355 | | | | | | |
| Q9CQ19 | MYL9 | myosin, light chain 9, regulatory | Cytoplasm | other | 6153518.5 | 6050354 | 1.11E+07 | | | | | | |
| Q9CQ19 | MYL9 | myosin, light chain 9, regulatory | Cytoplasm | other | 8604087 | 1.01E+07 | 2.02E+07 | | | | | | |
| Q9CQ19 | MYL9 | myosin, light chain 9, regulatory | Cytoplasm | other | 8405026 | 1.15E+07 | 2.08E+07 | | | | | | |
| Q9CQ19 | MYL9 | myosin, light chain 9, regulatory | Cytoplasm | other | 3154756.75 | 3737897.25 | 5902379 | | | | | | |
| Q9CQ19 | MYL9 | myosin, light chain 9, regulatory | Cytoplasm | other | 8128374 | 1.14E+07 | 1.68E+07 | Q9CQ19 | MYL9 | 8923166 | 6888372 | 1406083 | 24 |
| Q9WTI7 | MYO1C | myosin IC | Cytoplasm | other | 491182.719 | 500801.938 | 607389.438 | Q9WTI7 | MYO1C | 491183 | 64495 | 37236 | 3 |
| Q64331 | MYO6 | myosin VI | Cytoplasm | other | 1079455 | 2174298.5 | 2301601 | Q64331 | MYO6 | 1079455 | 671879 | 387910 | 3 |
| P70670 | NACA | nascent polypeptide-associated complex alpha subunit | Cytoplasm | transcription regulator | 5230840 | 5091110.5 | 3703861.75 | | | | | | |
| P70670 | NACA | nascent polypeptide-associated complex alpha subunit | Cytoplasm | transcription regulator | 3352284.5 | 3852039 | 3744198 | | | | | | |
| P70670 | NACA | nascent polypeptide-associated complex alpha subunit | Cytoplasm | transcription regulator | 752830.312 | 941160.25 | 871582.5 | P70670 | NACA | 3111985 | 1769007 | 589669 | 9 |
| P28656 | NAP1L1 | nucleosome assembly protein 1-like 1 | Nucleus | other | 4.71E+07 | 4.17E+07 | 2.64E+07 | | | | | | |
| P28656 | NAP1L1 | nucleosome assembly protein 1-like 1 | Nucleus | other | 3.28E+07 | 2.71E+07 | 1.82E+07 | | | | | | |
| P28656 | NAP1L1 | nucleosome assembly protein 1-like 1 | Nucleus | other | 2.88E+07 | 2.11E+07 | 1.68E+07 | | | | | | |
| P28656 | NAP1L1 | nucleosome assembly protein 1-like 1 | Nucleus | other | 1.27E+07 | 8553908 | 4080706 | | | | | | |
| P28656 | NAP1L1 | nucleosome assembly protein 1-like 1 | Nucleus | other | 1.59E+07 | 1.47E+07 | 1.19E+07 | | | | | | |
| P28656 | NAP1L1 | nucleosome assembly protein 1-like 1 | Nucleus | other | 8533535 | 7420319.5 | 4489989 | | | | | | |
| P28656 | NAP1L1 | nucleosome assembly protein 1-like 1 | Nucleus | other | 6313497.5 | 5314706 | 3460366.75 | | | | | | |
| P28656 | NAP1L1 | nucleosome assembly protein 1-like 1 | Nucleus | other | 4023374 | 3135830.25 | 1608236.25 | P28656 | NAP1L1 | 19516301 | 12685742 | 2589466 | 24 |
| Q78ZA7 | NAP1L4 | nucleosome assembly protein 1-like 4 | Cytoplasm | other | 3.65E+07 | 2.22E+07 | 1.84E+07 | | | | | | |
| Q78ZA7 | NAP1L4 | nucleosome assembly protein 1-like 4 | Cytoplasm | other | 2.62E+07 | 1.84E+07 | 1.22E+07 | | | | | | |
| Q78ZA7 | NAP1L4 | nucleosome assembly protein 1-like 4 | Cytoplasm | other | 1.48E+07 | 1.03E+07 | 5444847.5 | | | | | | |
| Q78ZA7 | NAP1L4 | nucleosome assembly protein 1-like 4 | Cytoplasm | other | 1.28E+07 | 1.08E+07 | 7173024 | | | | | | |
| Q78ZA7 | NAP1L4 | nucleosome assembly protein 1-like 4 | Cytoplasm | other | 1.44E+07 | 1.00E+07 | 5828398 | | | | | | |
| Q78ZA7 | NAP1L4 | nucleosome assembly protein 1-like 4 | Cytoplasm | other | 1.11E+07 | 7901604 | 5136532.5 | | | | | | |
| Q78ZA7 | NAP1L4 | nucleosome assembly protein 1-like 4 | Cytoplasm | other | 1.04E+07 | 1.03E+07 | 6074646.5 | | | | | | |
| Q78ZA7 | NAP1L4 | nucleosome assembly protein 1-like 4 | Cytoplasm | other | 6308659 | 6059141 | 3687283.5 | | | | | | |
| Q78ZA7 | NAP1L4 | nucleosome assembly protein 1-like 4 | Cytoplasm | other | 7773718 | 5506533 | 4201880.5 | | | | | | |
| Q78ZA7 | NAP1L4 | nucleosome assembly protein 1-like 4 | Cytoplasm | other | 4538452 | 4069860.25 | 973457 | | | | | | |
| Q78ZA7 | NAP1L4 | nucleosome assembly protein 1-like 4 | Cytoplasm | other | 4052078 | 3794160 | 1019383.75 | Q78ZA7 | NAP1L4 | 13533901 | 7566371 | 1317136 | 33 |
| Q9DB05 | NAPA | N-ethylmaleimide-sensitive factor attachment protein, alpha | Cytoplasm | transporter | 3880934 | 3592358.25 | 3455539.25 | Q9DB05 | NAPA | 3880934 | 217162 | 125379 | 3 |
| Q8K2Z4 | NCAPD2 | non-SMC condensin I complex, subunit D2 | Nucleus | other | 4.70E-06 | 383381.594 | 1307004.75 | Q8K2Z4 | NCAPD2 | 0 | 671853 | 387895 | 3 |
| Q8C156 | NCAPH | non-SMC condensin I complex, subunit H | Nucleus | other | 1533286.375 | 1082914.625 | 897890.25 | Q8C156 | NCAPH | 1533286 | 326802 | 188679 | 3 |
| P57716 | NCSTN | nicastrin | Plasma Membrane | peptidase | 1.25E+07 | 1.16E+07 | 1.27E+07 | | | | | | |
| P57716 | NCSTN | nicastrin | Plasma Membrane | peptidase | 1.22E+07 | 1.16E+07 | 1.00E+07 | | | | | | |
| P57716 | NCSTN | nicastrin | Plasma Membrane | peptidase | 6629134.5 | 6714696 | 6454479 | | | | | | |
| P57716 | NCSTN | nicastrin | Plasma Membrane | peptidase | 4828487.5 | 4582336.5 | 3698958.25 | | | | | | |
| P57716 | NCSTN | nicastrin | Plasma Membrane | peptidase | 7787454.5 | 8244017.5 | 7560701 | | | | | | |
| P57716 | NCSTN | nicastrin | Plasma Membrane | peptidase | 3428021.5 | 3506517.75 | 3958201.5 | | | | | | |
| P57716 | NCSTN | nicastrin | Plasma Membrane | peptidase | 4447105.5 | 4485121.5 | 4840891 | | | | | | |
| P57716 | NCSTN | nicastrin | Plasma Membrane | peptidase | 5038160.5 | 5307403 | 6467804.5 | | | | | | |
| P57716 | NCSTN | nicastrin | Plasma Membrane | peptidase | 490731.469 | 1681071.875 | 1535380.875 | P57716 | NCSTN | 6371011 | 3476696 | 669090 | 27 |
| Q99LC3 | NDUFA10 | NADH dehydrogenase (ubiquinone) 1 alpha subcomplex, 10, 42kDa | Cytoplasm | transporter | 4938157.5 | 7725729.5 | 8632828 | | | | | | |
| Q99LC3 | NDUFA10 | NADH dehydrogenase (ubiquinone) 1 alpha subcomplex, 10, 42kDa | Cytoplasm | transporter | 2664587.25 | 4261437 | 6061966 | Q99LC3 | NDUFA10 | 3801372 | 2222543 | 907349 | 6 |
| Q7TMF3 | NDUFA12 | NADH dehydrogenase (ubiquinone) 1 alpha subcomplex, 12 | Cytoplasm | enzyme | 494517.875 | 660185.062 | 863663.438 | Q7TMF3 | NDUFA12 | 494518 | 184895 | 106749 | 3 |
| Q9CQ75 | NDUFA2 | NADH dehydrogenase (ubiquinone) 1 alpha subcomplex, 2, 8kDa | Cytoplasm | enzyme | 2209930.25 | 3173004 | 3827740.25 | Q9CQ75 | NDUFA2 | 2209930 | 813787 | 469840 | 3 |
| Q62425 | NDUFA4 | NDUFA4, mitochondrial complex associated | Cytoplasm | enzyme | 4453592.5 | 7810230 | 9126318 | | | | | | |
| Q62425 | NDUFA4 | NDUFA4, mitochondrial complex associated | Cytoplasm | enzyme | 3087104.75 | 8540380 | 6260334.5 | Q62425 | NDUFA4 | 3770349 | 2394059 | 977371 | 6 |
| Q9CPP6 | NDUFA5 | NADH dehydrogenase (ubiquinone) 1 alpha subcomplex, 5 | Cytoplasm | enzyme | 1003694.688 | 1778212.875 | 1870020.375 | Q9CPP6 | NDUFA5 | 1003695 | 475890 | 274755 | 3 |
| Q9DCJ5 | NDUFA8 | NADH dehydrogenase (ubiquinone) 1 alpha subcomplex, 8, 19kDa | Cytoplasm | enzyme | 1707030.125 | 3010572 | 2852039.75 | Q9DCJ5 | NDUFA8 | 1707030 | 711267 | 410650 | 3 |
| Q9DC69 | NDUFA9 | NADH dehydrogenase (ubiquinone) 1 alpha subcomplex, 9, 39kDa | Cytoplasm | enzyme | 6245371.5 | 1.02E+07 | 1.11E+07 | | | | | | |
| Q9DC69 | NDUFA9 | NADH dehydrogenase (ubiquinone) 1 alpha subcomplex, 9, 39kDa | Cytoplasm | enzyme | 1920468 | 3254023.25 | 4721336.5 | | | | | | |
| Q9DC69 | NDUFA9 | NADH dehydrogenase (ubiquinone) 1 alpha subcomplex, 9, 39kDa | Cytoplasm | enzyme | 1275682 | 2256877 | 3372382.5 | Q9DC69 | NDUFA9 | 3147174 | 3581528 | 1193843 | 9 |
| Q9CR21 | NDUFAB1 | NADH dehydrogenase (ubiquinone) 1, alpha/beta subcomplex, 1, 8kDa | Cytoplasm | enzyme | 5227222.5 | 8785940 | 7683368 | | | | | | |
| Q9CR21 | NDUFAB1 | NADH dehydrogenase (ubiquinone) 1, alpha/beta subcomplex, 1, 8kDa | Cytoplasm | enzyme | 1339050.5 | 1956000.25 | 2370748.25 | Q9CR21 | NDUFAB1 | 3283137 | 3162510 | 1291089 | 6 |
| Q91VD9 | NDUFS1 | NADH dehydrogenase (ubiquinone) Fe-S protein 1, 75kDa (NADH-coenzyme Q reductase) | Cytoplasm | enzyme | 2058877.625 | 3384274 | 4050052 | | | | | | |
| Q91VD9 | NDUFS1 | NADH dehydrogenase (ubiquinone) Fe-S protein 1, 75kDa (NADH-coenzyme Q reductase) | Cytoplasm | enzyme | 2110466.5 | 3832166.75 | 3515964 | Q91VD9 | NDUFS1 | 2084672 | 864290 | 352845 | 6 |
| Q91WD5 | NDUFS2 | NADH dehydrogenase (ubiquinone) Fe-S protein 2, 49kDa (NADH-coenzyme Q reductase) | Cytoplasm | enzyme | 1474739.5 | 2515205 | 2949715.5 | Q91WD5 | NDUFS2 | 1474740 | 757949 | 437602 | 3 |
| Q9DCT2 | NDUFS3 | NADH dehydrogenase (ubiquinone) Fe-S protein 3, 30kDa (NADH-coenzyme Q reductase) | Cytoplasm | enzyme | 1454796.5 | 3841019.25 | 3640586 | Q9DCT2 | NDUFS3 | 1454797 | 1323626 | 764196 | 3 |
| Q9CXZ1 | NDUFS4 | NADH dehydrogenase (ubiquinone) Fe-S protein 4, 18kDa (NADH-coenzyme Q reductase) | Cytoplasm | enzyme | 2201290.5 | 5598683 | 5883118 | | | | | | |
| Q9CXZ1 | NDUFS4 | NADH dehydrogenase (ubiquinone) Fe-S protein 4, 18kDa (NADH-coenzyme Q reductase) | Cytoplasm | enzyme | 1064314.5 | 1471295.375 | 1658791.125 | Q9CXZ1 | NDUFS4 | 1632803 | 2171802 | 886634 | 6 |
| Q91YT0 | NDUFV1 | NADH dehydrogenase (ubiquinone) flavoprotein 1, 51kDa | Cytoplasm | enzyme | 2807524 | 5586108 | 7170916.5 | | | | | | |
| Q91YT0 | NDUFV1 | NADH dehydrogenase (ubiquinone) flavoprotein 1, 51kDa | Cytoplasm | enzyme | 1368440.875 | 2597475.75 | 3603683.25 | | | | | | |
| Q91YT0 | NDUFV1 | NADH dehydrogenase (ubiquinone) flavoprotein 1, 51kDa | Cytoplasm | enzyme | 1055803 | 1595876 | 2317176.5 | Q91YT0 | NDUFV1 | 1743923 | 2043053 | 681018 | 9 |
| Q9D6J6 | NDUFV2 | NADH dehydrogenase (ubiquinone) flavoprotein 2, 24kDa | Cytoplasm | enzyme | 1284396.75 | 2278060 | 2840619.5 | Q9D6J6 | NDUFV2 | 1284397 | 788001 | 454952 | 3 |
| Q9D1J1 | NECAP2 | NECAP endocytosis associated 2 | Cytoplasm | other | 5519324.5 | 7411220.5 | 6179728.5 | Q9D1J1 | NECAP2 | 5519325 | 960206 | 554375 | 3 |
| Q9JMH7 | NEU3 | sialidase 3 (membrane sialidase) | Plasma Membrane | enzyme | 1.88E+07 | 2.13E+07 | 1.37E+07 | | | | | | |
| Q9JMH7 | NEU3 | sialidase 3 (membrane sialidase) | Plasma Membrane | enzyme | 1.02E+07 | 8341893 | 6729829.5 | | | | | | |
| Q9JMH7 | NEU3 | sialidase 3 (membrane sialidase) | Plasma Membrane | enzyme | 1335108 | 1933367.125 | 1360245.125 | Q9JMH7 | NEU3 | 10105036 | 7440781 | 2480260 | 9 |
| Q9EST8 | NFKBIZ | nuclear factor of kappa light polypeptide gene enhancer in B-cells inhibitor, zeta | Nucleus | transcription regulator | 3276521.5 | 2595559 | 2842403.25 | Q9EST8 | NFKBIZ | 3276522 | 344746 | 199039 | 3 |
| Q3TL44 | NLRX1 | NLR family member X1 | Cytoplasm | other | 859026 | 2662050.5 | 4630761.5 | Q3TL44 | NLRX1 | 859026 | 1886474 | 1089156 | 3 |
| Q9WV85 | NME3 | NME/NM23 nucleoside diphosphate kinase 3 | Cytoplasm | kinase | 1165099.125 | 1852561 | 1778786.25 | Q9WV85 | NME3 | 1165099 | 377416 | 217901 | 3 |
| P97300 | Nptn | neuroplastin | Plasma Membrane | other | 1.03E+08 | 9.44E+07 | 8.33E+07 | | | | | | |
| P97300 | Nptn | neuroplastin | Plasma Membrane | other | 3.77E+07 | 3.23E+07 | 2.56E+07 | | | | | | |
| P97300 | Nptn | neuroplastin | Plasma Membrane | other | 1.52E+07 | 1.60E+07 | 1.70E+07 | | | | | | |
| P97300 | Nptn | neuroplastin | Plasma Membrane | other | 9158881 | 6778939.5 | 6704757.5 | | | | | | |
| P97300 | Nptn | neuroplastin | Plasma Membrane | other | 7783514 | 9512876 | 1.18E+07 | | | | | | |
| P97300 | Nptn | neuroplastin | Plasma Membrane | other | 9931048 | 1.12E+07 | 5173073 | | | | | | |
| P97300 | Nptn | neuroplastin | Plasma Membrane | other | 3958124 | 4142780.25 | 2776535.75 | | | | | | |
| P97300 | Nptn | neuroplastin | Plasma Membrane | other | 506013.281 | 279689.5 | 379318.562 | | | | | | |
| P97300 | Nptn | neuroplastin | Plasma Membrane | other | 151164.672 | 346912.25 | 40915.688 | P97300 | Nptn | 20766527 | 28556976 | 5495793 | 27 |
| P08556 | NRAS | neuroblastoma RAS viral (v-ras) oncogene homolog | Plasma Membrane | enzyme | 1.17E+07 | 1.27E+07 | 1.11E+07 | P08556 | NRAS | 11690000 | 768722 | 443822 | 3 |
| P46460 | NSF | N-ethylmaleimide-sensitive factor | Cytoplasm | transporter | 9351586 | 7387856.5 | 6459676 | | | | | | |
| P46460 | NSF | N-ethylmaleimide-sensitive factor | Cytoplasm | transporter | 7242096 | 7112556 | 5918423.5 | | | | | | |
| P46460 | NSF | N-ethylmaleimide-sensitive factor | Cytoplasm | transporter | 7730843 | 7736801 | 7047156 | | | | | | |
| P46460 | NSF | N-ethylmaleimide-sensitive factor | Cytoplasm | transporter | 7905163 | 5139677.5 | 4961536.5 | | | | | | |
| P46460 | NSF | N-ethylmaleimide-sensitive factor | Cytoplasm | transporter | 4866270.5 | 5022190.5 | 4608812.5 | | | | | | |
| P46460 | NSF | N-ethylmaleimide-sensitive factor | Cytoplasm | transporter | 7946692 | 6454812 | 5706080.5 | | | | | | |
| P46460 | NSF | N-ethylmaleimide-sensitive factor | Cytoplasm | transporter | 4830655.5 | 4120459.75 | 3217814.25 | | | | | | |
| P46460 | NSF | N-ethylmaleimide-sensitive factor | Cytoplasm | transporter | 5210092 | 5254290 | 3485534.25 | | | | | | |
| P46460 | NSF | N-ethylmaleimide-sensitive factor | Cytoplasm | transporter | 3750210 | 3332910.25 | 2904789.75 | | | | | | |
| P46460 | NSF | N-ethylmaleimide-sensitive factor | Cytoplasm | transporter | 2851736.25 | 2233955.25 | 3781963 | | | | | | |
| P46460 | NSF | N-ethylmaleimide-sensitive factor | Cytoplasm | transporter | 3369864.25 | 3758419.5 | 3289514.5 | | | | | | |
| P46460 | NSF | N-ethylmaleimide-sensitive factor | Cytoplasm | transporter | 2806328.5 | 2084528 | 1813113.625 | | | | | | |
| P46460 | NSF | N-ethylmaleimide-sensitive factor | Cytoplasm | transporter | 1154037.5 | 1504278.25 | 991454.938 | | | | | | |
| P46460 | NSF | N-ethylmaleimide-sensitive factor | Cytoplasm | transporter | 489277.656 | 441172.781 | 170363.109 | P46460 | NSF | 4964632 | 2362329 | 364515 | 42 |
| Q9CZ44 | NSFL1C | NSFL1 (p97) cofactor (p47) | Cytoplasm | other | 3691190.5 | 2547029.5 | 1819012.625 | | | | | | |

| | | | | | | | | | | | | | |
|---|---|---|---|---|---|---|---|---|---|---|---|---|---|
| Q9CZ44 | NSFL1C | NSFL1 (p97) cofactor (p47) | Cytoplasm | other | 3647490.5 | 1195204.875 | 884990.375 | | | | | | |
| Q9CZ44 | NSFL1C | NSFL1 (p97) cofactor (p47) | Cytoplasm | other | 2276093.25 | 1552143 | 1634428 | | | | | | |
| Q9CZ44 | NSFL1C | NSFL1 (p97) cofactor (p47) | Cytoplasm | other | 1475685 | 987916.188 | 979277.75 | Q9CZ44 | NSFL1C | 2772615 | 972622 | 280772 | 12 |
| A6H603 | NWD1 | NACHT and WD repeat domain containing 1 | Cytoplasm | other | 1.37E+07 | 1.15E+07 | 1.16E+07 | A6H603 | NWD1 | 13740000 | 1285392 | 742122 | 3 |
| P83503 | NYX | nyctalopin | Extracellular Space | other | 7.87E+07 | 1.07E+08 | 8.27E+07 | P83503 | NYX | 78740000 | 15191344 | 8770727 | 3 |
| P29758 | OAT | ornithine aminotransferase | Cytoplasm | enzyme | 1316869.625 | 2559017 | 3106231.25 | P29758 | OAT | 1316870 | 916896 | 529370 | 3 |
| A2AAJ9 | OBSCN | obscurin, cytoskeletal calmodulin and titin-interacting RhoGEF | Cytoplasm | kinase | 1725108.5 | 2349601.75 | 3320617.25 | A2AAJ9 | OBSCN | 1725109 | 804002 | 464191 | 3 |
| Q9CRD0 | OCIAD1 | OCIA domain containing 1 | Cytoplasm | other | 1002143.438 | 2622467 | 3115721.5 | Q9CRD0 | OCIAD1 | 1002143 | 1105740 | 638399 | 3 |
| Q60597 | OGDH | oxoglutarate (alpha-ketoglutarate) dehydrogenase (lipoamide) | Cytoplasm | enzyme | 1.05E+07 | 1.37E+07 | 1.81E+07 | | | | | | |
| Q60597 | OGDH | oxoglutarate (alpha-ketoglutarate) dehydrogenase (lipoamide) | Cytoplasm | enzyme | 1.34E+07 | 2.15E+07 | 1.93E+07 | | | | | | |
| Q60597 | OGDH | oxoglutarate (alpha-ketoglutarate) dehydrogenase (lipoamide) | Cytoplasm | enzyme | 3878529.25 | 8378178.5 | 9552066 | | | | | | |
| Q60597 | OGDH | oxoglutarate (alpha-ketoglutarate) dehydrogenase (lipoamide) | Cytoplasm | enzyme | 5100546.5 | 9168480 | 1.04E+07 | | | | | | |
| Q60597 | OGDH | oxoglutarate (alpha-ketoglutarate) dehydrogenase (lipoamide) | Cytoplasm | enzyme | 8540832 | 1.60E+07 | 1.26E+07 | | | | | | |
| Q60597 | OGDH | oxoglutarate (alpha-ketoglutarate) dehydrogenase (lipoamide) | Cytoplasm | enzyme | 6581090.5 | 1.10E+07 | 1.11E+07 | | | | | | |
| Q60597 | OGDH | oxoglutarate (alpha-ketoglutarate) dehydrogenase (lipoamide) | Cytoplasm | enzyme | 4804627.5 | 9708006 | 1.09E+07 | | | | | | |
| Q60597 | OGDH | oxoglutarate (alpha-ketoglutarate) dehydrogenase (lipoamide) | Cytoplasm | enzyme | 4734897.5 | 6623672 | 9670228 | | | | | | |
| Q60597 | OGDH | oxoglutarate (alpha-ketoglutarate) dehydrogenase (lipoamide) | Cytoplasm | enzyme | 4221550.5 | 6515307.5 | 8864298 | | | | | | |
| Q60597 | OGDH | oxoglutarate (alpha-ketoglutarate) dehydrogenase (lipoamide) | Cytoplasm | enzyme | 3936166.5 | 7994797.5 | 7472508 | | | | | | |
| Q60597 | OGDH | oxoglutarate (alpha-ketoglutarate) dehydrogenase (lipoamide) | Cytoplasm | enzyme | 3697242.75 | 5673080.5 | 6216557.5 | | | | | | |
| Q60597 | OGDH | oxoglutarate (alpha-ketoglutarate) dehydrogenase (lipoamide) | Cytoplasm | enzyme | 3243558 | 6447155 | 8490017 | | | | | | |
| Q60597 | OGDH | oxoglutarate (alpha-ketoglutarate) dehydrogenase (lipoamide) | Cytoplasm | enzyme | 2308574.75 | 3829631.25 | 4707158.5 | | | | | | |
| Q60597 | OGDH | oxoglutarate (alpha-ketoglutarate) dehydrogenase (lipoamide) | Cytoplasm | enzyme | 1982415.125 | 3373944 | 4138739.5 | | | | | | |
| Q60597 | OGDH | oxoglutarate (alpha-ketoglutarate) dehydrogenase (lipoamide) | Cytoplasm | enzyme | 1686410 | 2290159.5 | 2314363.5 | | | | | | |
| Q60597 | OGDH | oxoglutarate (alpha-ketoglutarate) dehydrogenase (lipoamide) | Cytoplasm | enzyme | 1477158.125 | 1922268.125 | 1850531.375 | | | | | | |
| Q60597 | OGDH | oxoglutarate (alpha-ketoglutarate) dehydrogenase (lipoamide) | Cytoplasm | enzyme | 2175807 | 3833445.5 | 5769970 | Q60597 | OGDH | 4837024 | 4752180 | 665438 | 51 |
| Q9CZ30 | OLA1 | Obg-like ATPase 1 | Cytoplasm | enzyme | 1688510.375 | 1766274.125 | 2482882.5 | Q9CZ30 | OLA1 | 1688510 | 437912 | 252829 | 3 |
| Q8BGT0 | OSTM1 | osteopetrosis associated transmembrane protein 1 | Cytoplasm | other | 528001.312 | 820266.438 | 707347 | Q8BGT0 | OSTM1 | 528001 | 147385 | 85093 | 3 |
| Q9D0K2 | OXCT1 | 3-oxoacid CoA transferase 1 | Cytoplasm | enzyme | 2398076.75 | 2593522.5 | 3096056 | Q9D0K2 | OXCT1 | 2398077 | 360073 | 207888 | 3 |
| P09103 | P4HB | prolyl 4-hydroxylase, beta polypeptide | Cytoplasm | enzyme | 2.46E+07 | 2.76E+07 | 2.14E+07 | | | | | | |
| P09103 | P4HB | prolyl 4-hydroxylase, beta polypeptide | Cytoplasm | enzyme | 2.18E+07 | 1.88E+07 | 1.81E+07 | | | | | | |
| P09103 | P4HB | prolyl 4-hydroxylase, beta polypeptide | Cytoplasm | enzyme | 2.26E+07 | 2.70E+07 | 2.50E+07 | | | | | | |
| P09103 | P4HB | prolyl 4-hydroxylase, beta polypeptide | Cytoplasm | enzyme | 1.74E+07 | 2.03E+07 | 2.05E+07 | | | | | | |
| P09103 | P4HB | prolyl 4-hydroxylase, beta polypeptide | Cytoplasm | enzyme | 2.17E+07 | 2.22E+07 | 2.27E+07 | | | | | | |
| P09103 | P4HB | prolyl 4-hydroxylase, beta polypeptide | Cytoplasm | enzyme | 1.77E+07 | 1.98E+07 | 2.45E+07 | | | | | | |
| P09103 | P4HB | prolyl 4-hydroxylase, beta polypeptide | Cytoplasm | enzyme | 1.36E+07 | 1.34E+07 | 1.26E+07 | | | | | | |
| P09103 | P4HB | prolyl 4-hydroxylase, beta polypeptide | Cytoplasm | enzyme | 1.49E+07 | 1.58E+07 | 1.38E+07 | | | | | | |
| P09103 | P4HB | prolyl 4-hydroxylase, beta polypeptide | Cytoplasm | enzyme | 1.06E+07 | 1.49E+07 | 1.48E+07 | | | | | | |
| P09103 | P4HB | prolyl 4-hydroxylase, beta polypeptide | Cytoplasm | enzyme | 1.73E+07 | 1.90E+07 | 1.84E+07 | | | | | | |
| P09103 | P4HB | prolyl 4-hydroxylase, beta polypeptide | Cytoplasm | enzyme | 1.19E+07 | 1.17E+07 | 1.49E+07 | | | | | | |
| P09103 | P4HB | prolyl 4-hydroxylase, beta polypeptide | Cytoplasm | enzyme | 9391303 | 1.11E+07 | 1.25E+07 | | | | | | |
| P09103 | P4HB | prolyl 4-hydroxylase, beta polypeptide | Cytoplasm | enzyme | 1.10E+07 | 1.13E+07 | 1.04E+07 | | | | | | |
| P09103 | P4HB | prolyl 4-hydroxylase, beta polypeptide | Cytoplasm | enzyme | 1.21E+07 | 1.13E+07 | 1.26E+07 | | | | | | |
| P09103 | P4HB | prolyl 4-hydroxylase, beta polypeptide | Cytoplasm | enzyme | 7681198.5 | 8809701 | 1.32E+07 | | | | | | |
| P09103 | P4HB | prolyl 4-hydroxylase, beta polypeptide | Cytoplasm | enzyme | 8452026 | 1.14E+07 | 1.04E+07 | | | | | | |
| P09103 | P4HB | prolyl 4-hydroxylase, beta polypeptide | Cytoplasm | enzyme | 6912595 | 7549578.5 | 5771798 | | | | | | |
| P09103 | P4HB | prolyl 4-hydroxylase, beta polypeptide | Cytoplasm | enzyme | 5217978 | 4813105.5 | 3726869.75 | | | | | | |
| P09103 | P4HB | prolyl 4-hydroxylase, beta polypeptide | Cytoplasm | enzyme | 3496584.25 | 3999565.75 | 4082466.5 | P09103 | P4HB | 13597457 | 6373505 | 844191 | 57 |
| P50580 | PA2G4 | proliferation-associated 2G4, 38kDa | Nucleus | transcription regulator | 4635698 | 5847753 | 5769290 | | | | | | |
| P50580 | PA2G4 | proliferation-associated 2G4, 38kDa | Nucleus | transcription regulator | 3400950 | 4407681.5 | 5295246.5 | | | | | | |
| P50580 | PA2G4 | proliferation-associated 2G4, 38kDa | Nucleus | transcription regulator | 2761120 | 4053290.25 | 3513548.5 | | | | | | |
| P50580 | PA2G4 | proliferation-associated 2G4, 38kDa | Nucleus | transcription regulator | 2468313.75 | 2841193.5 | 3768010 | | | | | | |
| P50580 | PA2G4 | proliferation-associated 2G4, 38kDa | Nucleus | transcription regulator | 1834508.375 | 2847711 | 3421702.5 | P50580 | PA2G4 | 3020118 | 1206092 | 311411 | 15 |
| P29341 | PABPC1 | poly(A) binding protein, cytoplasmic 1 | Cytoplasm | translation regulator | 1782897 | 2750708.5 | 2105647.75 | | | | | | |
| P29341 | PABPC1 | poly(A) binding protein, cytoplasmic 1 | Cytoplasm | translation regulator | 3410345.25 | 3331919.5 | 3401679.25 | | | | | | |
| P29341 | PABPC1 | poly(A) binding protein, cytoplasmic 1 | Cytoplasm | translation regulator | 1871385.75 | 3140133.25 | 3222616 | | | | | | |
| P29341 | PABPC1 | poly(A) binding protein, cytoplasmic 1 | Cytoplasm | translation regulator | 2399615.25 | 2154642.25 | 3352335.75 | | | | | | |
| P29341 | PABPC1 | poly(A) binding protein, cytoplasmic 1 | Cytoplasm | translation regulator | 1110165.25 | 1743478.75 | 1804143.625 | P29341 | PABPC1 | 2114882 | 767050 | 198051 | 15 |
| Q9WVE8 | PACSIN2 | protein kinase C and casein kinase substrate in neurons 2 | Cytoplasm | transporter | 5908163 | 5071342 | 6873857 | | | | | | |
| Q9WVE8 | PACSIN2 | protein kinase C and casein kinase substrate in neurons 2 | Cytoplasm | transporter | 6762266.5 | 5413414.5 | 6940092.5 | | | | | | |
| Q9WVE8 | PACSIN2 | protein kinase C and casein kinase substrate in neurons 2 | Cytoplasm | transporter | 5209147.5 | 4074450 | 4667838 | | | | | | |
| Q9WVE8 | PACSIN2 | protein kinase C and casein kinase substrate in neurons 2 | Cytoplasm | transporter | 4303603 | 3564240.25 | 4725321 | | | | | | |
| Q9WVE8 | PACSIN2 | protein kinase C and casein kinase substrate in neurons 2 | Cytoplasm | transporter | 3523784 | 3641081.25 | 4175311 | | | | | | |
| Q9WVE8 | PACSIN2 | protein kinase C and casein kinase substrate in neurons 2 | Cytoplasm | transporter | 4729370 | 3731913.75 | 3785285.25 | | | | | | |
| Q9WVE8 | PACSIN2 | protein kinase C and casein kinase substrate in neurons 2 | Cytoplasm | transporter | 1306407.625 | 1215062.375 | 1643113.25 | | | | | | |
| Q9WVE8 | PACSIN2 | protein kinase C and casein kinase substrate in neurons 2 | Cytoplasm | transporter | 1650600.625 | 1487406.875 | 407628.844 | Q9WVE8 | PACSIN2 | 4174168 | 1868773 | 381462 | 24 |
| Q9ES46 | PARVB | parvin, beta | Cytoplasm | other | 2926227.75 | 1582477.625 | 7554673 | | | | | | |
| Q9ES46 | PARVB | parvin, beta | Cytoplasm | other | 4865430 | 2647927.25 | 1.69E+07 | | | | | | |
| Q9ES46 | PARVB | parvin, beta | Cytoplasm | other | 3220870.75 | 1810657.75 | 1.08E+07 | | | | | | |
| Q9ES46 | PARVB | parvin, beta | Cytoplasm | other | 3296091.75 | 1189815 | 9591339 | | | | | | |
| Q9ES46 | PARVB | parvin, beta | Cytoplasm | other | 3166851.75 | 1503285.25 | 7512833 | | | | | | |
| Q9ES46 | PARVB | parvin, beta | Cytoplasm | other | 1742855.875 | 1052498.25 | 6043981 | Q9ES46 | PARVB | 3203055 | 4233501 | 997846 | 18 |
| Q05920 | PC | pyruvate carboxylase | Cytoplasm | enzyme | 1984392.375 | 3528853.25 | 3494824.25 | | | | | | |
| Q05920 | PC | pyruvate carboxylase | Cytoplasm | enzyme | 1786731.875 | 3558690.75 | 5161108.5 | | | | | | |
| Q05920 | PC | pyruvate carboxylase | Cytoplasm | enzyme | 1837748 | 2667768 | 3235632 | | | | | | |
| Q05920 | PC | pyruvate carboxylase | Cytoplasm | enzyme | 863768.812 | 1069563.75 | 1085088.625 | | | | | | |
| Q05920 | PC | pyruvate carboxylase | Cytoplasm | enzyme | 728740.562 | 1544527.625 | 1828466 | Q05920 | PC | 1440276 | 1270798 | 328119 | 15 |
| P60335 | PCBP1 | poly(rC) binding protein 1 | Nucleus | translation regulator | 1805633.375 | 2622912.5 | 2501640.75 | | | | | | |
| P60335 | PCBP1 | poly(rC) binding protein 1 | Nucleus | translation regulator | 2269483.25 | 3446311.25 | 3153793.25 | | | | | | |
| P60335 | PCBP1 | poly(rC) binding protein 1 | Nucleus | translation regulator | 2586466 | 2521479.75 | 3708290.75 | | | | | | |
| P60335 | PCBP1 | poly(rC) binding protein 1 | Nucleus | translation regulator | 1201213.5 | 1842430.5 | 1632452.5 | | | | | | |
| P60335 | PCBP1 | poly(rC) binding protein 1 | Nucleus | translation regulator | 1243943.875 | 1948573.5 | 1292693.75 | P60335 | PCBP1 | 1821348 | 785065 | 202703 | 15 |
| Q8BH04 | PCK2 | phosphoenolpyruvate carboxykinase 2 (mitochondrial) | Cytoplasm | kinase | 7138972 | 1.94E+07 | 1.82E+07 | | | | | | |
| Q8BH04 | PCK2 | phosphoenolpyruvate carboxykinase 2 (mitochondrial) | Cytoplasm | kinase | 1.00E+07 | 1.88E+07 | 2.35E+07 | | | | | | |
| Q8BH04 | PCK2 | phosphoenolpyruvate carboxykinase 2 (mitochondrial) | Cytoplasm | kinase | 4637393 | 9632309 | 1.13E+07 | | | | | | |
| Q8BH04 | PCK2 | phosphoenolpyruvate carboxykinase 2 (mitochondrial) | Cytoplasm | kinase | 7362145.5 | 1.14E+07 | 1.42E+07 | | | | | | |
| Q8BH04 | PCK2 | phosphoenolpyruvate carboxykinase 2 (mitochondrial) | Cytoplasm | kinase | 5403051.5 | 9376572 | 8862497 | | | | | | |
| Q8BH04 | PCK2 | phosphoenolpyruvate carboxykinase 2 (mitochondrial) | Cytoplasm | kinase | 5135103.5 | 9334762 | 1.04E+07 | | | | | | |
| Q8BH04 | PCK2 | phosphoenolpyruvate carboxykinase 2 (mitochondrial) | Cytoplasm | kinase | 4097113.75 | 6332133 | 8889695 | | | | | | |
| Q8BH04 | PCK2 | phosphoenolpyruvate carboxykinase 2 (mitochondrial) | Cytoplasm | kinase | 2745155.25 | 4979694 | 6643485.5 | | | | | | |
| Q8BH04 | PCK2 | phosphoenolpyruvate carboxykinase 2 (mitochondrial) | Cytoplasm | kinase | 3386358.25 | 4777606 | 5240910 | | | | | | |
| Q8BH04 | PCK2 | phosphoenolpyruvate carboxykinase 2 (mitochondrial) | Cytoplasm | kinase | 3561965.25 | 5768594 | 5466726 | | | | | | |
| Q8BH04 | PCK2 | phosphoenolpyruvate carboxykinase 2 (mitochondrial) | Cytoplasm | kinase | 3034130.5 | 5866674 | 7427005.5 | | | | | | |
| Q8BH04 | PCK2 | phosphoenolpyruvate carboxykinase 2 (mitochondrial) | Cytoplasm | kinase | 2330362.5 | 4989357 | 2641224.25 | | | | | | |
| Q8BH04 | PCK2 | phosphoenolpyruvate carboxykinase 2 (mitochondrial) | Cytoplasm | kinase | 4568582 | 7016807 | 7153986.5 | | | | | | |
| Q8BH04 | PCK2 | phosphoenolpyruvate carboxykinase 2 (mitochondrial) | Cytoplasm | kinase | 3788227.5 | 5585524.5 | 7729427 | | | | | | |
| Q8BH04 | PCK2 | phosphoenolpyruvate carboxykinase 2 (mitochondrial) | Cytoplasm | kinase | 2916971.5 | 4263723 | 4794750 | | | | | | |
| Q8BH04 | PCK2 | phosphoenolpyruvate carboxykinase 2 (mitochondrial) | Cytoplasm | kinase | 2498353.5 | 3645602 | 3903291.5 | | | | | | |
| Q8BH04 | PCK2 | phosphoenolpyruvate carboxykinase 2 (mitochondrial) | Cytoplasm | kinase | 1721255.25 | 2472459.25 | 2475203 | Q8BH04 | PCK2 | 4374420 | 4744206 | 664322 | 51 |
| Q9CQF9 | PCYOX1 | prenylcysteine oxidase 1 | Cytoplasm | enzyme | 8786581 | 1.01E+07 | 9336762 | Q9CQF9 | PCYOX1 | 8786581 | 670095 | 386880 | 3 |
| P27664 | PDE6A | phosphodiesterase 6A, cGMP-specific, rod, alpha | Plasma Membrane | enzyme | 1.43E+07 | 9887469 | 1.28E+07 | P27664 | PDE6A | 14270000 | 2232667 | 1289031 | 3 |
| P35486 | PDHA1 | pyruvate dehydrogenase (lipoamide) alpha 1 | Cytoplasm | enzyme | 911735.125 | 1986885.125 | 1270318.625 | | | | | | |
| P35486 | PDHA1 | pyruvate dehydrogenase (lipoamide) alpha 1 | Cytoplasm | enzyme | 1323088.625 | 1534406.5 | 2209444.5 | | | | | | |
| P35486 | PDHA1 | pyruvate dehydrogenase (lipoamide) alpha 1 | Cytoplasm | enzyme | 757588.125 | 2181687 | 3374625.25 | P35486 | PDHA1 | 997471 | 808927 | 269642 | 9 |
| Q9D051 | PDHB | pyruvate dehydrogenase (lipoamide) beta | Cytoplasm | enzyme | 1619890.75 | 3092157 | 4825827 | Q9D051 | PDHB | 1619891 | 1604743 | 926499 | 3 |
| P27773 | PDIA3 | protein disulfide isomerase family A, member 3 | Cytoplasm | peptidase | 1.11E+08 | 1.25E+08 | 1.13E+08 | | | | | | |
| P27773 | PDIA3 | protein disulfide isomerase family A, member 3 | Cytoplasm | peptidase | 5.44E+07 | 5.61E+07 | 5.47E+07 | | | | | | |
| P27773 | PDIA3 | protein disulfide isomerase family A, member 3 | Cytoplasm | peptidase | 3.95E+07 | 4.80E+07 | 4.18E+07 | | | | | | |
| P27773 | PDIA3 | protein disulfide isomerase family A, member 3 | Cytoplasm | peptidase | 2.64E+07 | 2.85E+07 | 2.81E+07 | | | | | | |
| P27773 | PDIA3 | protein disulfide isomerase family A, member 3 | Cytoplasm | peptidase | 2.73E+07 | 2.79E+07 | 2.48E+07 | | | | | | |

| | | | | | | | | | | | | | |
|---|---|---|---|---|---|---|---|---|---|---|---|---|---|
| P27773 | PDIA3 | protein disulfide isomerase family A, member 3 | Cytoplasm | peptidase | 3.15E+07 | 2.80E+07 | 2.72E+07 | | | | | | |
| P27773 | PDIA3 | protein disulfide isomerase family A, member 3 | Cytoplasm | peptidase | 2.54E+07 | 2.63E+07 | 2.53E+07 | | | | | | |
| P27773 | PDIA3 | protein disulfide isomerase family A, member 3 | Cytoplasm | peptidase | 2.75E+07 | 2.98E+07 | 2.28E+07 | | | | | | |
| P27773 | PDIA3 | protein disulfide isomerase family A, member 3 | Cytoplasm | peptidase | 2.93E+07 | 3.12E+07 | 3.21E+07 | | | | | | |
| P27773 | PDIA3 | protein disulfide isomerase family A, member 3 | Cytoplasm | peptidase | 2.50E+07 | 2.26E+07 | 2.12E+07 | | | | | | |
| P27773 | PDIA3 | protein disulfide isomerase family A, member 3 | Cytoplasm | peptidase | 1.82E+07 | 2.28E+07 | 1.66E+07 | | | | | | |
| P27773 | PDIA3 | protein disulfide isomerase family A, member 3 | Cytoplasm | peptidase | 1.96E+07 | 2.05E+07 | 2.05E+07 | | | | | | |
| P27773 | PDIA3 | protein disulfide isomerase family A, member 3 | Cytoplasm | peptidase | 1.93E+07 | 2.17E+07 | 2.22E+07 | | | | | | |
| P27773 | PDIA3 | protein disulfide isomerase family A, member 3 | Cytoplasm | peptidase | 1.58E+07 | 1.51E+07 | 1.32E+07 | | | | | | |
| P27773 | PDIA3 | protein disulfide isomerase family A, member 3 | Cytoplasm | peptidase | 1.44E+07 | 1.55E+07 | 1.57E+07 | | | | | | |
| P27773 | PDIA3 | protein disulfide isomerase family A, member 3 | Cytoplasm | peptidase | 1.52E+07 | 1.72E+07 | 1.75E+07 | | | | | | |
| P27773 | PDIA3 | protein disulfide isomerase family A, member 3 | Cytoplasm | peptidase | 1.49E+07 | 1.63E+07 | 1.80E+07 | | | | | | |
| P27773 | PDIA3 | protein disulfide isomerase family A, member 3 | Cytoplasm | peptidase | 1.10E+07 | 1.18E+07 | 1.14E+07 | | | | | | |
| P27773 | PDIA3 | protein disulfide isomerase family A, member 3 | Cytoplasm | peptidase | 1.10E+07 | 9836008 | 1.25E+07 | | | | | | |
| P27773 | PDIA3 | protein disulfide isomerase family A, member 3 | Cytoplasm | peptidase | 1.08E+07 | 1.54E+07 | 1.34E+07 | | | | | | |
| P27773 | PDIA3 | protein disulfide isomerase family A, member 3 | Cytoplasm | peptidase | 5092454 | 5312138 | 4127462 | | | | | | |
| P27773 | PDIA3 | protein disulfide isomerase family A, member 3 | Cytoplasm | peptidase | 2712821 | 1645921.625 | 1330230.25 | | | | | | |
| P27773 | PDIA3 | protein disulfide isomerase family A, member 3 | Cytoplasm | peptidase | 2304899.25 | 3128591 | 1710379.25 | P27773 | PDIA3 | 24233921 | 23105394 | 2781563 | 69 |
| P08003 | PDIA4 | protein disulfide isomerase family A, member 4 | Cytoplasm | enzyme | 8385551.5 | 6695031 | 8926298 | | | | | | |
| P08003 | PDIA4 | protein disulfide isomerase family A, member 4 | Cytoplasm | enzyme | 6271170.5 | 6019432 | 4472352.5 | | | | | | |
| P08003 | PDIA4 | protein disulfide isomerase family A, member 4 | Cytoplasm | enzyme | 3434570.75 | 4185113.5 | 3472788.75 | | | | | | |
| P08003 | PDIA4 | protein disulfide isomerase family A, member 4 | Cytoplasm | enzyme | 4627351 | 3717034.5 | 3513528.25 | | | | | | |
| P08003 | PDIA4 | protein disulfide isomerase family A, member 4 | Cytoplasm | enzyme | 3541941.5 | 3460991 | 3515487.5 | | | | | | |
| P08003 | PDIA4 | protein disulfide isomerase family A, member 4 | Cytoplasm | enzyme | 2602419.25 | 3282776.5 | 2511203.25 | | | | | | |
| P08003 | PDIA4 | protein disulfide isomerase family A, member 4 | Cytoplasm | enzyme | 2406137.25 | 2813647.25 | 2384039.25 | | | | | | |
| P08003 | PDIA4 | protein disulfide isomerase family A, member 4 | Cytoplasm | enzyme | 1721300.5 | 1788949.25 | 1763383.875 | P08003 | PDIA4 | 4123805 | 1962901 | 400675 | 24 |
| Q922R8 | PDIA6 | protein disulfide isomerase family A, member 6 | Cytoplasm | enzyme | 2.58E+07 | 2.75E+07 | 2.23E+07 | | | | | | |
| Q922R8 | PDIA6 | protein disulfide isomerase family A, member 6 | Cytoplasm | enzyme | 1.66E+07 | 1.73E+07 | 1.53E+07 | | | | | | |
| Q922R8 | PDIA6 | protein disulfide isomerase family A, member 6 | Cytoplasm | enzyme | 1.34E+07 | 1.30E+07 | 1.33E+07 | | | | | | |
| Q922R8 | PDIA6 | protein disulfide isomerase family A, member 6 | Cytoplasm | enzyme | 1.19E+07 | 1.14E+07 | 9118533 | | | | | | |
| Q922R8 | PDIA6 | protein disulfide isomerase family A, member 6 | Cytoplasm | enzyme | 1.08E+07 | 1.19E+07 | 1.10E+07 | | | | | | |
| Q922R8 | PDIA6 | protein disulfide isomerase family A, member 6 | Cytoplasm | enzyme | 4881400.5 | 5812503 | 1.43E+07 | | | | | | |
| Q922R8 | PDIA6 | protein disulfide isomerase family A, member 6 | Cytoplasm | enzyme | 1.20E+07 | 1.19E+07 | 1.05E+07 | | | | | | |
| Q922R8 | PDIA6 | protein disulfide isomerase family A, member 6 | Cytoplasm | enzyme | 1.08E+07 | 1.28E+07 | 1.07E+07 | | | | | | |
| Q922R8 | PDIA6 | protein disulfide isomerase family A, member 6 | Cytoplasm | enzyme | 6295664.5 | 1.03E+07 | 9859184 | | | | | | |
| Q922R8 | PDIA6 | protein disulfide isomerase family A, member 6 | Cytoplasm | enzyme | 5745650.5 | 6639275 | 6356167.5 | | | | | | |
| Q922R8 | PDIA6 | protein disulfide isomerase family A, member 6 | Cytoplasm | enzyme | 4509112.5 | 4691144.5 | 4929405 | Q922R8 | PDIA6 | 11171075 | 5606664 | 975995 | 33 |
| Q9Z126 | PF4 | platelet factor 4 | Extracellular Space | cytokine | 1.07E+08 | 9.13E+07 | 1.61E+08 | | | | | | |
| Q9Z126 | PF4 | platelet factor 4 | Extracellular Space | cytokine | 7.70E+07 | 7.05E+07 | 1.20E+08 | | | | | | |
| Q9Z126 | PF4 | platelet factor 4 | Extracellular Space | cytokine | 4.43E+07 | 4.23E+07 | 6.31E+07 | Q9Z126 | PF4 | 75963333 | 38244671 | 12748224 | 9 |
| P12382 | PFKL | phosphofructokinase, liver | Cytoplasm | kinase | 2278612.5 | 2614568.5 | 1953714.625 | P12382 | PFKL | 2278613 | 330442 | 190781 | 3 |
| P47857 | PFKM | phosphofructokinase, muscle | Cytoplasm | kinase | 6437381.5 | 6033813.5 | 3563112.75 | | | | | | |
| P47857 | PFKM | phosphofructokinase, muscle | Cytoplasm | kinase | 3278172 | 2631848.5 | 3603084.75 | | | | | | |
| P47857 | PFKM | phosphofructokinase, muscle | Cytoplasm | kinase | 4085391.75 | 2641326.25 | 2837512 | | | | | | |
| P47857 | PFKM | phosphofructokinase, muscle | Cytoplasm | kinase | 3226895 | 3655084.5 | 3072300.5 | | | | | | |
| P47857 | PFKM | phosphofructokinase, muscle | Cytoplasm | kinase | 1802519.375 | 1617652.875 | 2008540.875 | | | | | | |
| P47857 | PFKM | phosphofructokinase, muscle | Cytoplasm | kinase | 4409354 | 3088378 | 2886987.75 | P47857 | PFKM | 3873286 | 1270429 | 299443 | 18 |
| Q9DCD0 | PGD | phosphogluconate dehydrogenase | Cytoplasm | enzyme | 3069281 | 2741838 | 2310292 | Q9DCD0 | PGD | 3069281 | 380683 | 219787 | 3 |
| O55022 | PGRMC1 | progesterone receptor membrane component 1 | Plasma Membrane | transmembrane receptor | 3118293.75 | 3661429.25 | 7178121 | O55022 | PGRMC1 | 3118294 | 2203948 | 1272450 | 3 |
| Q80UU9 | PGRMC2 | progesterone receptor membrane component 2 | Nucleus | other | 2345972 | 2309242.75 | 3679984.25 | Q80UU9 | PGRMC2 | 2345972 | 781011 | 450917 | 3 |
| P67778 | PHB | prohibitin | Nucleus | transcription regulator | 7894355.5 | 1.04E+07 | 1.17E+07 | | | | | | |
| P67778 | PHB | prohibitin | Nucleus | transcription regulator | 7074676 | 1.40E+07 | 1.43E+07 | | | | | | |
| P67778 | PHB | prohibitin | Nucleus | transcription regulator | 5898061 | 8706809 | 1.18E+07 | | | | | | |
| P67778 | PHB | prohibitin | Nucleus | transcription regulator | 5945135.5 | 1.26E+07 | 1.13E+07 | | | | | | |
| P67778 | PHB | prohibitin | Nucleus | transcription regulator | 4954872.5 | 9559974 | 9773322 | | | | | | |
| P67778 | PHB | prohibitin | Nucleus | transcription regulator | 4264506.5 | 5579505 | 6069910 | | | | | | |
| P67778 | PHB | prohibitin | Nucleus | transcription regulator | 3272972 | 7612653 | 8716803 | | | | | | |
| P67778 | PHB | prohibitin | Nucleus | transcription regulator | 5346788 | 8203522 | 1.17E+07 | | | | | | |
| P67778 | PHB | prohibitin | Nucleus | transcription regulator | 1489570.125 | 3913757.25 | 4688620 | | | | | | |
| P67778 | PHB | prohibitin | Nucleus | transcription regulator | 3493428.25 | 4307015.5 | 3763725.25 | P67778 | PHB | 4963437 | 3478944 | 635165 | 30 |
| O35129 | PHB2 | prohibitin 2 | Cytoplasm | transcription regulator | 8595505 | 9600558 | 8340349 | | | | | | |
| O35129 | PHB2 | prohibitin 2 | Cytoplasm | transcription regulator | 1.02E+07 | 1.38E+07 | 1.83E+07 | | | | | | |
| O35129 | PHB2 | prohibitin 2 | Cytoplasm | transcription regulator | 7089676.5 | 1.13E+07 | 1.11E+07 | | | | | | |
| O35129 | PHB2 | prohibitin 2 | Cytoplasm | transcription regulator | 1.45E+07 | 2.30E+07 | 2.72E+07 | | | | | | |
| O35129 | PHB2 | prohibitin 2 | Cytoplasm | transcription regulator | 7852553 | 1.46E+07 | 1.83E+07 | | | | | | |
| O35129 | PHB2 | prohibitin 2 | Cytoplasm | transcription regulator | 7993043.5 | 1.18E+07 | 1.38E+07 | | | | | | |
| O35129 | PHB2 | prohibitin 2 | Cytoplasm | transcription regulator | 1428428 | 2134906.75 | 2379611 | | | | | | |
| O35129 | PHB2 | prohibitin 2 | Cytoplasm | transcription regulator | 3343919 | 5119833 | 7132936.5 | | | | | | |
| O35129 | PHB2 | prohibitin 2 | Cytoplasm | transcription regulator | 885256.062 | 1461194.375 | 1441168 | O35129 | PHB2 | 6880931 | 6784164 | 1305613 | 27 |
| Q61753 | PHGDH | phosphoglycerate dehydrogenase | Cytoplasm | enzyme | 3268781.75 | 2045599.25 | 1590345.5 | | | | | | |
| Q61753 | PHGDH | phosphoglycerate dehydrogenase | Cytoplasm | enzyme | 3020312.25 | 1688470.375 | 1104044.875 | | | | | | |
| Q61753 | PHGDH | phosphoglycerate dehydrogenase | Cytoplasm | enzyme | 1955282 | 1370573.5 | 1315890.125 | | | | | | |
| Q61753 | PHGDH | phosphoglycerate dehydrogenase | Cytoplasm | enzyme | 1753066 | 440311.562 | 69856.93 | | | | | | |
| Q61753 | PHGDH | phosphoglycerate dehydrogenase | Cytoplasm | enzyme | 1416636.75 | 1084511.75 | 1125911.5 | Q61753 | PHGDH | 2282816 | 832825 | 215034 | 15 |
| Q8CBQ5 | PI4K2B | phosphatidylinositol 4-kinase type 2 beta | Cytoplasm | kinase | 2618754.75 | 2029774.25 | 2301200 | | | | | | |
| Q8CBQ5 | PI4K2B | phosphatidylinositol 4-kinase type 2 beta | Cytoplasm | kinase | 1859311.5 | 1902024.75 | 1921090.625 | | | | | | |
| Q8CBQ5 | PI4K2B | phosphatidylinositol 4-kinase type 2 beta | Cytoplasm | kinase | 2008829.25 | 2351499.75 | 2410299.5 | | | | | | |
| Q8CBQ5 | PI4K2B | phosphatidylinositol 4-kinase type 2 beta | Cytoplasm | kinase | 849162.062 | 1044697.938 | 1091383.875 | Q8CBQ5 | PI4K2B | 1834014 | 575889 | 166245 | 12 |
| Q7M6Y3 | PICALM | phosphatidylinositol binding clathrin assembly protein | Cytoplasm | other | 2.07E+07 | 2.28E+07 | 2.29E+07 | | | | | | |
| Q7M6Y3 | PICALM | phosphatidylinositol binding clathrin assembly protein | Cytoplasm | other | 7594999 | 8845632 | 8225127.5 | | | | | | |
| Q7M6Y3 | PICALM | phosphatidylinositol binding clathrin assembly protein | Cytoplasm | other | 2876717 | 3121887.25 | 4147410.75 | | | | | | |
| Q7M6Y3 | PICALM | phosphatidylinositol binding clathrin assembly protein | Cytoplasm | other | 1474738.375 | 2506196.5 | 1675251.125 | Q7M6Y3 | PICALM | 8151614 | 8364280 | 2414560 | 12 |
| E2JF22 | PIEZO1 | piezo-type mechanosensitive ion channel component 1 | Cytoplasm | ion channel | 2.79E+07 | 2.84E+07 | 2.09E+07 | | | | | | |
| E2JF22 | PIEZO1 | piezo-type mechanosensitive ion channel component 1 | Cytoplasm | ion channel | 1.90E+07 | 2.14E+07 | 1.99E+07 | | | | | | |
| E2JF22 | PIEZO1 | piezo-type mechanosensitive ion channel component 1 | Cytoplasm | ion channel | 1.81E+07 | 1.79E+07 | 1.11E+07 | | | | | | |
| E2JF22 | PIEZO1 | piezo-type mechanosensitive ion channel component 1 | Cytoplasm | ion channel | 1.21E+07 | 1.26E+07 | 1.21E+07 | | | | | | |
| E2JF22 | PIEZO1 | piezo-type mechanosensitive ion channel component 1 | Cytoplasm | ion channel | 1.29E+07 | 1.27E+07 | 1.09E+07 | | | | | | |
| E2JF22 | PIEZO1 | piezo-type mechanosensitive ion channel component 1 | Cytoplasm | ion channel | 1.38E+07 | 1.38E+07 | 9592410 | | | | | | |
| E2JF22 | PIEZO1 | piezo-type mechanosensitive ion channel component 1 | Cytoplasm | ion channel | 1.28E+07 | 1.18E+07 | 1.23E+07 | | | | | | |
| E2JF22 | PIEZO1 | piezo-type mechanosensitive ion channel component 1 | Cytoplasm | ion channel | 1.13E+07 | 1.08E+07 | 9304152 | | | | | | |
| E2JF22 | PIEZO1 | piezo-type mechanosensitive ion channel component 1 | Cytoplasm | ion channel | 9417335 | 9954381 | 9744754 | | | | | | |
| E2JF22 | PIEZO1 | piezo-type mechanosensitive ion channel component 1 | Cytoplasm | ion channel | 1.20E+07 | 1.19E+07 | 1.08E+07 | | | | | | |
| E2JF22 | PIEZO1 | piezo-type mechanosensitive ion channel component 1 | Cytoplasm | ion channel | 9307987 | 8212621 | 8622683 | | | | | | |
| E2JF22 | PIEZO1 | piezo-type mechanosensitive ion channel component 1 | Cytoplasm | ion channel | 8699266 | 6800005.5 | 5989199.5 | | | | | | |
| E2JF22 | PIEZO1 | piezo-type mechanosensitive ion channel component 1 | Cytoplasm | ion channel | 8178687 | 6775703 | 6538009 | | | | | | |
| E2JF22 | PIEZO1 | piezo-type mechanosensitive ion channel component 1 | Cytoplasm | ion channel | 6059508 | 5872863 | 4779167.5 | | | | | | |
| E2JF22 | PIEZO1 | piezo-type mechanosensitive ion channel component 1 | Cytoplasm | ion channel | 7527663.5 | 7045334 | 6938447.5 | | | | | | |
| E2JF22 | PIEZO1 | piezo-type mechanosensitive ion channel component 1 | Cytoplasm | ion channel | 5204929 | 4890728 | 3760522.5 | | | | | | |
| E2JF22 | PIEZO1 | piezo-type mechanosensitive ion channel component 1 | Cytoplasm | ion channel | 7245424 | 7867881 | 7042192.5 | | | | | | |
| E2JF22 | PIEZO1 | piezo-type mechanosensitive ion channel component 1 | Cytoplasm | ion channel | 3857392 | 5024237.5 | 6428432.5 | | | | | | |
| E2JF22 | PIEZO1 | piezo-type mechanosensitive ion channel component 1 | Cytoplasm | ion channel | 3869246.75 | 4484366 | 3440431 | | | | | | |
| E2JF22 | PIEZO1 | piezo-type mechanosensitive ion channel component 1 | Cytoplasm | ion channel | 5071126.5 | 5156848 | 3649588.75 | | | | | | |
| E2JF22 | PIEZO1 | piezo-type mechanosensitive ion channel component 1 | Cytoplasm | ion channel | 5186116.5 | 5446798.5 | 4270481 | | | | | | |
| E2JF22 | PIEZO1 | piezo-type mechanosensitive ion channel component 1 | Cytoplasm | ion channel | 4493887.5 | 2799629 | 2274474.25 | | | | | | |
| E2JF22 | PIEZO1 | piezo-type mechanosensitive ion channel component 1 | Cytoplasm | ion channel | 2045214.375 | 2012334 | 698444.125 | | | | | | |
| E2JF22 | PIEZO1 | piezo-type mechanosensitive ion channel component 1 | Cytoplasm | ion channel | 1254619.75 | 1372842.375 | 1203488 | | | | | | |
| E2JF22 | PIEZO1 | piezo-type mechanosensitive ion channel component 1 | Cytoplasm | ion channel | 1976139.375 | 1399694.625 | 1623235.125 | | | | | | |
| E2JF22 | PIEZO1 | piezo-type mechanosensitive ion channel component 1 | Cytoplasm | ion channel | 1415601.125 | 1299809.625 | 1232295.25 | | | | | | |
| E2JF22 | PIEZO1 | piezo-type mechanosensitive ion channel component 1 | Cytoplasm | ion channel | 397314.188 | 764370.562 | 939407.812 | E2JF22 | PIEZO1 | 8558795 | 6013334 | 668148 | 81 |

| | | | | | | | | | | | | | |
|---|---|---|---|---|---|---|---|---|---|---|---|---|---|
| P06803 | PIM1 | Pim-1 proto-oncogene, serine/threonine kinase | Cytoplasm | kinase | 2.30E+08 | 2.02E+08 | 2.07E+08 | P06803 | PIM1 | 230200000 | 14990108 | 8654543 | 3 |
| O70172 | PIP4K2A | phosphatidylinositol-5-phosphate 4-kinase, type II, alpha | Cytoplasm | kinase | 1.30E+07 | 1.15E+07 | 1.39E+07 | | | | | | |
| O70172 | PIP4K2A | phosphatidylinositol-5-phosphate 4-kinase, type II, alpha | Cytoplasm | kinase | 3371035.75 | 2441108.75 | 2441496.75 | | | | | | |
| O70172 | PIP4K2A | phosphatidylinositol-5-phosphate 4-kinase, type II, alpha | Cytoplasm | kinase | 2787331.25 | 3717572.25 | 3895299.25 | | | | | | |
| O70172 | PIP4K2A | phosphatidylinositol-5-phosphate 4-kinase, type II, alpha | Cytoplasm | kinase | 3376797 | 2611089.5 | 2547824.75 | O70172 | PIP4K2A | 5633791 | 4483124 | 1294166 | 12 |
| Q80XI4 | PIP4K2B | phosphatidylinositol-5-phosphate 4-kinase, type II, beta | Cytoplasm | kinase | 6637074 | 4789091 | 4823027.5 | Q80XI4 | PIP4K2B | 6637074 | 1057273 | 610417 | 3 |
| P70182 | PIP5K1A | phosphatidylinositol-4-phosphate 5-kinase, type I, alpha | Cytoplasm | kinase | 1.55E+07 | 1.21E+07 | 1.03E+07 | | | | | | |
| P70182 | PIP5K1A | phosphatidylinositol-4-phosphate 5-kinase, type I, alpha | Cytoplasm | kinase | 9450540 | 6489350 | 5743491 | | | | | | |
| P70182 | PIP5K1A | phosphatidylinositol-4-phosphate 5-kinase, type I, alpha | Cytoplasm | kinase | 6520040 | 5730785 | 5480339.5 | | | | | | |
| P70182 | PIP5K1A | phosphatidylinositol-4-phosphate 5-kinase, type I, alpha | Cytoplasm | kinase | 7082412 | 4726435 | 4867030 | | | | | | |
| P70182 | PIP5K1A | phosphatidylinositol-4-phosphate 5-kinase, type I, alpha | Cytoplasm | kinase | 6399492 | 5199552.5 | 6137168 | | | | | | |
| P70182 | PIP5K1A | phosphatidylinositol-4-phosphate 5-kinase, type I, alpha | Cytoplasm | kinase | 5527931.5 | 4233301 | 3977786.75 | | | | | | |
| P70182 | PIP5K1A | phosphatidylinositol-4-phosphate 5-kinase, type I, alpha | Cytoplasm | kinase | 2656881.75 | 2379420 | 3067381.25 | | | | | | |
| P70182 | PIP5K1A | phosphatidylinositol-4-phosphate 5-kinase, type I, alpha | Cytoplasm | kinase | 2765127.5 | 1684063.625 | 1849732.875 | | | | | | |
| P70182 | PIP5K1A | phosphatidylinositol-4-phosphate 5-kinase, type I, alpha | Cytoplasm | kinase | 3930828 | 3418335 | 3029728 | | | | | | |
| P70182 | PIP5K1A | phosphatidylinositol-4-phosphate 5-kinase, type I, alpha | Cytoplasm | kinase | 467103.875 | 365839.281 | 137965.234 | P70182 | PIP5K1A | 6033036 | 3421328 | 624646 | 30 |
| Q8K411 | PITRM1 | pitrilysin metallopeptidase 1 | Cytoplasm | peptidase | 4685833.5 | 8940842 | 1.22E+07 | | | | | | |
| Q8K411 | PITRM1 | pitrilysin metallopeptidase 1 | Cytoplasm | peptidase | 3697440.5 | 8324244 | 9200325 | | | | | | |
| Q8K411 | PITRM1 | pitrilysin metallopeptidase 1 | Cytoplasm | peptidase | 3437271.75 | 6540779.5 | 8658261 | | | | | | |
| Q8K411 | PITRM1 | pitrilysin metallopeptidase 1 | Cytoplasm | peptidase | 2632705.5 | 5690685 | 7246288 | | | | | | |
| Q8K411 | PITRM1 | pitrilysin metallopeptidase 1 | Cytoplasm | peptidase | 2107579 | 3788090 | 4849266.5 | | | | | | |
| Q8K411 | PITRM1 | pitrilysin metallopeptidase 1 | Cytoplasm | peptidase | 880455.875 | 1341856.5 | 1425808.875 | | | | | | |
| Q8K411 | PITRM1 | pitrilysin metallopeptidase 1 | Cytoplasm | peptidase | 259516.094 | 1043800.438 | 1153913.875 | Q8K411 | PITRM1 | 2528686 | 3399438 | 741818 | 21 |
| Q80ZA4 | PKHD1L1 | polycystic kidney and hepatic disease 1 (autosomal recessive)-like 1 | Extracellular Space | other | 6258432.5 | 7528277.5 | 7549027.5 | Q80ZA4 | PKHD1L1 | 6258433 | 739208 | 426782 | 3 |
| P53657 | PKLR | pyruvate kinase, liver and RBC | Cytoplasm | kinase | 1.44E+07 | 1.20E+07 | 9940569 | | | | | | |
| P53657 | PKLR | pyruvate kinase, liver and RBC | Cytoplasm | kinase | 1.05E+07 | 7164081.5 | 5417734.5 | | | | | | |
| P53657 | PKLR | pyruvate kinase, liver and RBC | Cytoplasm | kinase | 9498471 | 9055462 | 4874164.5 | | | | | | |
| P53657 | PKLR | pyruvate kinase, liver and RBC | Cytoplasm | kinase | 8191439 | 5105896 | 4588898.5 | | | | | | |
| P53657 | PKLR | pyruvate kinase, liver and RBC | Cytoplasm | kinase | 5766505 | 4156033 | 3662703 | | | | | | |
| P53657 | PKLR | pyruvate kinase, liver and RBC | Cytoplasm | kinase | 5117285.5 | 3254044.75 | 2277874 | | | | | | |
| P53657 | PKLR | pyruvate kinase, liver and RBC | Cytoplasm | kinase | 2590644.5 | 1787549.625 | 2120679.75 | | | | | | |
| P53657 | PKLR | pyruvate kinase, liver and RBC | Cytoplasm | kinase | 3330560 | 2758726 | 2075648.375 | P53657 | PKLR | 7419363 | 3501832 | 714808 | 24 |
| P52480 | PKM | pyruvate kinase, muscle | Cytoplasm | kinase | 4802922 | 6121418.5 | 6522028 | P52480 | PKM | 4802922 | 899467 | 519308 | 3 |
| P27612 | PLAA | phospholipase A2-activating protein | Cytoplasm | other | 998427.25 | 567924.625 | 395231.938 | P27612 | PLAA | 998427 | 310644 | 179351 | 3 |
| Q9JHK5 | PLEK | pleckstrin | Cytoplasm | other | 109695.281 | 305657.25 | 604348.625 | Q9JHK5 | PLEK | 109695 | 249098 | 143817 | 3 |
| P20918 | PLG | plasminogen | Extracellular Space | peptidase | 960215.875 | 529341.688 | 4465952.5 | | | | | | |
| P20918 | PLG | plasminogen | Extracellular Space | peptidase | 2776.57 | 0 | 1932590.75 | | | | | | |
| P20918 | PLG | plasminogen | Extracellular Space | peptidase | 124887.844 | 67296.227 | 1666022.5 | P20918 | PLG | 362627 | 1462750 | 487583 | 9 |
| Q9R1Q7 | PLP2 | proteolipid protein 2 (colonic epithelium-enriched) | Cytoplasm | transporter | 1.40E+07 | 1.34E+07 | 1.08E+07 | Q9R1Q7 | PLP2 | 13960000 | 1661214 | 959103 | 3 |
| Q9JJ00 | PLSCR1 | phospholipid scramblase 1 | Plasma Membrane | enzyme | 2.04E+07 | 2.00E+07 | 1.98E+07 | | | | | | |
| Q9JJ00 | PLSCR1 | phospholipid scramblase 1 | Plasma Membrane | enzyme | 1.47E+07 | 1.41E+07 | 1.19E+07 | | | | | | |
| Q9JJ00 | PLSCR1 | phospholipid scramblase 1 | Plasma Membrane | enzyme | 7155703 | 7185572.5 | 7224432.5 | Q9JJ00 | PLSCR1 | 14061901 | 5619708 | 1873236 | 9 |
| Q9JIZ9 | PLSCR3 | phospholipid scramblase 3 | Plasma Membrane | enzyme | 8240562 | 8193940 | 7598097 | | | | | | |
| Q9JIZ9 | PLSCR3 | phospholipid scramblase 3 | Plasma Membrane | enzyme | 9829020 | 9637939 | 9946318 | | | | | | |
| Q9JIZ9 | PLSCR3 | phospholipid scramblase 3 | Plasma Membrane | enzyme | 3687089.75 | 3627997.5 | 2928849.25 | | | | | | |
| Q9JIZ9 | PLSCR3 | phospholipid scramblase 3 | Plasma Membrane | enzyme | 2805137.75 | 2056142.75 | 2302920.25 | Q9JIZ9 | PLSCR3 | 6140452 | 3241560 | 935758 | 12 |
| P23492 | PNP | purine nucleoside phosphorylase | Nucleus | enzyme | 3.72E+07 | 3.18E+07 | 2.19E+07 | | | | | | |
| P23492 | PNP | purine nucleoside phosphorylase | Nucleus | enzyme | 1.07E+07 | 8582395 | 5732546 | | | | | | |
| P23492 | PNP | purine nucleoside phosphorylase | Nucleus | enzyme | 1.41E+07 | 9726244 | 8316980.5 | | | | | | |
| P23492 | PNP | purine nucleoside phosphorylase | Nucleus | enzyme | 9409777 | 6553173 | 5071588.5 | | | | | | |
| P23492 | PNP | purine nucleoside phosphorylase | Nucleus | enzyme | 9019290 | 6473534 | 4283574 | | | | | | |
| P23492 | PNP | purine nucleoside phosphorylase | Nucleus | enzyme | 2486727 | 2008669.125 | 1249298.25 | P23492 | PNP | 13824299 | 9897708 | 2332912 | 18 |
| Q61469 | Ppap2a | phosphatidic acid phosphatase type 2A | Plasma Membrane | phosphatase | 6.36E+07 | 5.47E+07 | 6.01E+07 | | | | | | |
| Q61469 | Ppap2a | phosphatidic acid phosphatase type 2A | Plasma Membrane | phosphatase | 4.32E+07 | 3.20E+07 | 3.63E+07 | | | | | | |
| Q61469 | Ppap2a | phosphatidic acid phosphatase type 2A | Plasma Membrane | phosphatase | 6.20E+07 | 6.56E+07 | 3.90E+07 | | | | | | |
| Q61469 | Ppap2a | phosphatidic acid phosphatase type 2A | Plasma Membrane | phosphatase | 3.82E+07 | 3.95E+07 | 2.68E+07 | | | | | | |
| Q61469 | Ppap2a | phosphatidic acid phosphatase type 2A | Plasma Membrane | phosphatase | 4293835 | 7768799 | 5012318.5 | Q61469 | Ppap2a | 42234767 | 20849007 | 5383191 | 15 |
| P17742 | PPIA | peptidylprolyl isomerase A (cyclophilin A) | Cytoplasm | enzyme | 1.62E+07 | 2.30E+07 | 3.28E+07 | | | | | | |
| P17742 | PPIA | peptidylprolyl isomerase A (cyclophilin A) | Cytoplasm | enzyme | 6260266 | 1.08E+07 | 2.64E+07 | | | | | | |
| P17742 | PPIA | peptidylprolyl isomerase A (cyclophilin A) | Cytoplasm | enzyme | 1.08E+07 | 1.69E+07 | 2.52E+07 | | | | | | |
| P17742 | PPIA | peptidylprolyl isomerase A (cyclophilin A) | Cytoplasm | enzyme | 4971643.5 | 4351273 | 9547333 | | | | | | |
| P17742 | PPIA | peptidylprolyl isomerase A (cyclophilin A) | Cytoplasm | enzyme | 7972199.5 | 1.38E+07 | 2.12E+07 | | | | | | |
| P17742 | PPIA | peptidylprolyl isomerase A (cyclophilin A) | Cytoplasm | enzyme | 674374.688 | 1070588.875 | 4715930.5 | | | | | | |
| P17742 | PPIA | peptidylprolyl isomerase A (cyclophilin A) | Cytoplasm | enzyme | 1665230.75 | 3055482 | 6303734 | P17742 | PPIA | 6933388 | 9348562 | 2040023 | 21 |
| P24369 | PPIB | peptidylprolyl isomerase B (cyclophilin B) | Cytoplasm | enzyme | 8.56E+07 | 8.58E+07 | 8.34E+07 | | | | | | |
| P24369 | PPIB | peptidylprolyl isomerase B (cyclophilin B) | Cytoplasm | enzyme | 4.87E+07 | 3.56E+07 | 4.53E+07 | | | | | | |
| P24369 | PPIB | peptidylprolyl isomerase B (cyclophilin B) | Cytoplasm | enzyme | 3.14E+07 | 3.21E+07 | 2.20E+07 | | | | | | |
| P24369 | PPIB | peptidylprolyl isomerase B (cyclophilin B) | Cytoplasm | enzyme | 3.10E+07 | 2.81E+07 | 2.47E+07 | | | | | | |
| P24369 | PPIB | peptidylprolyl isomerase B (cyclophilin B) | Cytoplasm | enzyme | 2.78E+07 | 2.52E+07 | 2.68E+07 | | | | | | |
| P24369 | PPIB | peptidylprolyl isomerase B (cyclophilin B) | Cytoplasm | enzyme | 3.03E+07 | 2.43E+07 | 2.36E+07 | | | | | | |
| P24369 | PPIB | peptidylprolyl isomerase B (cyclophilin B) | Cytoplasm | enzyme | 1.85E+07 | 1.66E+07 | 2.19E+07 | | | | | | |
| P24369 | PPIB | peptidylprolyl isomerase B (cyclophilin B) | Cytoplasm | enzyme | 1.75E+07 | 1.37E+07 | 1.16E+07 | | | | | | |
| P24369 | PPIB | peptidylprolyl isomerase B (cyclophilin B) | Cytoplasm | enzyme | 1.30E+07 | 8496500 | 1.16E+07 | | | | | | |
| P24369 | PPIB | peptidylprolyl isomerase B (cyclophilin B) | Cytoplasm | enzyme | 1.53E+07 | 1.34E+07 | 1.46E+07 | | | | | | |
| P24369 | PPIB | peptidylprolyl isomerase B (cyclophilin B) | Cytoplasm | enzyme | 9470616 | 1.12E+07 | 1.48E+07 | | | | | | |
| P24369 | PPIB | peptidylprolyl isomerase B (cyclophilin B) | Cytoplasm | enzyme | 3586190 | 1325989.875 | 3585996.75 | P24369 | PPIB | 27680567 | 21018715 | 3503119 | 36 |
| Q99KR7 | PPIF | peptidylprolyl isomerase F | Cytoplasm | enzyme | 4540588 | 5376775 | 1.23E+07 | | | | | | |
| Q99KR7 | PPIF | peptidylprolyl isomerase F | Cytoplasm | enzyme | 4181111 | 6364539 | 8617605 | Q99KR7 | PPIF | 4360850 | 3101929 | 1266357 | 6 |
| P51175 | PPOX | protoporphyrinogen oxidase | Cytoplasm | enzyme | 1.71E+07 | 3.37E+07 | 3.69E+07 | | | | | | |
| P51175 | PPOX | protoporphyrinogen oxidase | Cytoplasm | enzyme | 1.77E+07 | 3.63E+07 | 4.90E+07 | | | | | | |
| P51175 | PPOX | protoporphyrinogen oxidase | Cytoplasm | enzyme | 6765599.5 | 1.38E+07 | 1.47E+07 | | | | | | |
| P51175 | PPOX | protoporphyrinogen oxidase | Cytoplasm | enzyme | 6894547 | 1.30E+07 | 1.66E+07 | | | | | | |
| P51175 | PPOX | protoporphyrinogen oxidase | Cytoplasm | enzyme | 5288238 | 1.12E+07 | 1.37E+07 | | | | | | |
| P51175 | PPOX | protoporphyrinogen oxidase | Cytoplasm | enzyme | 4373223 | 7910153.5 | 9035584 | | | | | | |
| P51175 | PPOX | protoporphyrinogen oxidase | Cytoplasm | enzyme | 4446700 | 1.00E+07 | 9518427 | | | | | | |
| P51175 | PPOX | protoporphyrinogen oxidase | Cytoplasm | enzyme | 3674116.25 | 8018818 | 1.01E+07 | | | | | | |
| P51175 | PPOX | protoporphyrinogen oxidase | Cytoplasm | enzyme | 5715450.5 | 7562490 | 1.09E+07 | | | | | | |
| P51175 | PPOX | protoporphyrinogen oxidase | Cytoplasm | enzyme | 2747329.25 | 4194434.5 | 6316718 | | | | | | |
| P51175 | PPOX | protoporphyrinogen oxidase | Cytoplasm | enzyme | 2240380.5 | 3818453.5 | 6539597 | | | | | | |
| P51175 | PPOX | protoporphyrinogen oxidase | Cytoplasm | enzyme | 2083885 | 4717810 | 5366298 | | | | | | |
| P51175 | PPOX | protoporphyrinogen oxidase | Cytoplasm | enzyme | 1933961.875 | 2976745.75 | 3518791.25 | P51175 | PPOX | 6227956 | 10672621 | 1708987 | 39 |
| P62137 | PPP1CA | protein phosphatase 1, catalytic subunit, alpha isozyme | Cytoplasm | phosphatase | 3996930.75 | 4495477 | 7213226.5 | | | | | | |
| P62137 | PPP1CA | protein phosphatase 1, catalytic subunit, alpha isozyme | Cytoplasm | phosphatase | 5583987 | 6073461 | 1.08E+07 | | | | | | |
| P62137 | PPP1CA | protein phosphatase 1, catalytic subunit, alpha isozyme | Cytoplasm | phosphatase | 2209498.5 | 2928681.5 | 3133917.75 | | | | | | |
| P62137 | PPP1CA | protein phosphatase 1, catalytic subunit, alpha isozyme | Cytoplasm | phosphatase | 2676177.75 | 3388747.75 | 3938643 | | | | | | |
| P62137 | PPP1CA | protein phosphatase 1, catalytic subunit, alpha isozyme | Cytoplasm | phosphatase | 1634635 | 3377411.5 | 3250574 | | | | | | |
| P62137 | PPP1CA | protein phosphatase 1, catalytic subunit, alpha isozyme | Cytoplasm | phosphatase | 1222137 | 1274629.875 | 1256108 | P62137 | PPP1CA | 2887228 | 2421124 | 570664 | 18 |
| Q62415 | PPP1R13B | protein phosphatase 1, regulatory subunit 13B | Cytoplasm | phosphatase | 3473405 | 2932948.5 | 2720658.25 | Q62415 | PPP1R13B | 3473405 | 388113 | 224077 | 3 |
| Q3TDD9 | PPP1R21 | protein phosphatase 1, regulatory subunit 21 | Cytoplasm | other | 1.22E+07 | 1.62E+07 | 1.04E+07 | Q3TDD9 | PPP1R21 | 12200000 | 2974228 | 1717171 | 3 |
| P63330 | PPP2CA | protein phosphatase 2, catalytic subunit, alpha isozyme | Cytoplasm | phosphatase | 4468159 | 3250988.25 | 2633904.75 | | | | | | |
| P63330 | PPP2CA | protein phosphatase 2, catalytic subunit, alpha isozyme | Cytoplasm | phosphatase | 4070813.75 | 1909186.125 | 3037808.75 | | | | | | |
| P63330 | PPP2CA | protein phosphatase 2, catalytic subunit, alpha isozyme | Cytoplasm | phosphatase | 3301492.25 | 2768392.25 | 1985130.5 | | | | | | |
| P63330 | PPP2CA | protein phosphatase 2, catalytic subunit, alpha isozyme | Cytoplasm | phosphatase | 1903314.625 | 1167744.75 | 674396.312 | | | | | | |
| P63330 | PPP2CA | protein phosphatase 2, catalytic subunit, alpha isozyme | Cytoplasm | phosphatase | 2291794 | 1760533.625 | 1550835.375 | P63330 | PPP2CA | 3207115 | 1050870 | 271333 | 15 |
| Q76MZ3 | PPP2R1A | protein phosphatase 2, regulatory subunit A, alpha | Cytoplasm | phosphatase | 8321207 | 4792437.5 | 5722864 | | | | | | |
| Q76MZ3 | PPP2R1A | protein phosphatase 2, regulatory subunit A, alpha | Cytoplasm | phosphatase | 2690185 | 2218629 | 2105396.5 | | | | | | |
| Q76MZ3 | PPP2R1A | protein phosphatase 2, regulatory subunit A, alpha | Cytoplasm | phosphatase | 3037057.5 | 2086234.875 | 1828991.5 | | | | | | |
| Q76MZ3 | PPP2R1A | protein phosphatase 2, regulatory subunit A, alpha | Cytoplasm | phosphatase | 6021548.5 | 4174352.75 | 3675741.75 | | | | | | |
| Q76MZ3 | PPP2R1A | protein phosphatase 2, regulatory subunit A, alpha | Cytoplasm | phosphatase | 3328867.5 | 1888188.125 | 1726150.875 | | | | | | |

| | | | | | | | | | | | | | |
|---|---|---|---|---|---|---|---|---|---|---|---|---|---|
| Q76MZ3 | PPP2R1A | protein phosphatase 2, regulatory subunit A, alpha | Cytoplasm | phosphatase | 2754989.75 | 1595577.875 | 1450122.5 | | | | | | |
| Q76MZ3 | PPP2R1A | protein phosphatase 2, regulatory subunit A, alpha | Cytoplasm | phosphatase | 1656815.875 | 1231772.75 | 1253682.125 | Q76MZ3 | PPP2R1A | 3972953 | 1857269 | 405289 | 21 |
| Q6P1F6 | PPP2R2A | protein phosphatase 2, regulatory subunit B, alpha | Cytoplasm | phosphatase | 2695864.75 | 1585220 | 1144624.625 | Q6P1F6 | PPP2R2A | 2695865 | 799375 | 461519 | 3 |
| Q9CQR6 | PPP6C | protein phosphatase 6, catalytic subunit | Nucleus | phosphatase | 6910877 | 5368362.5 | 5647824.5 | | | | | | |
| Q9CQR6 | PPP6C | protein phosphatase 6, catalytic subunit | Nucleus | phosphatase | 3881979 | 2690143.75 | 3198270.25 | Q9CQR6 | PPP6C | 5396428 | 1622026 | 662189 | 6 |
| Q922D4 | PPP6R3 | protein phosphatase 6, regulatory subunit 3 | Cytoplasm | other | 1.38E+07 | 8449132 | 8641220 | | | | | | |
| Q922D4 | PPP6R3 | protein phosphatase 6, regulatory subunit 3 | Cytoplasm | other | 1.19E+07 | 7825490.5 | 8209703 | | | | | | |
| Q922D4 | PPP6R3 | protein phosphatase 6, regulatory subunit 3 | Cytoplasm | other | 5543794.5 | 3944054.5 | 4464261 | | | | | | |
| Q922D4 | PPP6R3 | protein phosphatase 6, regulatory subunit 3 | Cytoplasm | other | 6684011.5 | 5649838 | 4022895.75 | | | | | | |
| Q922D4 | PPP6R3 | protein phosphatase 6, regulatory subunit 3 | Cytoplasm | other | 4506882 | 3463610.25 | 3890712.5 | | | | | | |
| Q922D4 | PPP6R3 | protein phosphatase 6, regulatory subunit 3 | Cytoplasm | other | 5536459.5 | 3815144 | 3568781.75 | | | | | | |
| Q922D4 | PPP6R3 | protein phosphatase 6, regulatory subunit 3 | Cytoplasm | other | 5582074 | 4048677.75 | 4049659.5 | | | | | | |
| Q922D4 | PPP6R3 | protein phosphatase 6, regulatory subunit 3 | Cytoplasm | other | 5067293 | 3506280.25 | 3362915.5 | | | | | | |
| Q922D4 | PPP6R3 | protein phosphatase 6, regulatory subunit 3 | Cytoplasm | other | 4505351 | 2821926.5 | 2401825 | | | | | | |
| Q922D4 | PPP6R3 | protein phosphatase 6, regulatory subunit 3 | Cytoplasm | other | 4482244 | 2991281.5 | 2067339 | | | | | | |
| Q922D4 | PPP6R3 | protein phosphatase 6, regulatory subunit 3 | Cytoplasm | other | 3304667.5 | 1999199.125 | 1798201.125 | | | | | | |
| Q922D4 | PPP6R3 | protein phosphatase 6, regulatory subunit 3 | Cytoplasm | other | 1478910.125 | 875096.5 | 1142961.5 | Q922D4 | PPP6R3 | 6028474 | 2809494 | 468249 | 36 |
| Q7TMR0 | PRCP | prolylcarboxypeptidase (angiotensinase C) | Cytoplasm | peptidase | 3048165 | 5172221.5 | 5897917.5 | Q7TMR0 | PRCP | 3048165 | 1480954 | 855029 | 3 |
| P35700 | PRDX1 | peroxiredoxin 1 | Cytoplasm | enzyme | 2.47E+07 | 2.09E+07 | 1.91E+07 | | | | | | |
| P35700 | PRDX1 | peroxiredoxin 1 | Cytoplasm | enzyme | 2.66E+07 | 1.91E+07 | 1.97E+07 | | | | | | |
| P35700 | PRDX1 | peroxiredoxin 1 | Cytoplasm | enzyme | 1.74E+07 | 1.17E+07 | 1.07E+07 | | | | | | |
| P35700 | PRDX1 | peroxiredoxin 1 | Cytoplasm | enzyme | 2.28E+07 | 1.52E+07 | 1.44E+07 | | | | | | |
| P35700 | PRDX1 | peroxiredoxin 1 | Cytoplasm | enzyme | 2.09E+07 | 1.52E+07 | 1.35E+07 | | | | | | |
| P35700 | PRDX1 | peroxiredoxin 1 | Cytoplasm | enzyme | 1.27E+07 | 1.36E+07 | 1.28E+07 | | | | | | |
| P35700 | PRDX1 | peroxiredoxin 1 | Cytoplasm | enzyme | 1.36E+07 | 9337684 | 9672166 | | | | | | |
| P35700 | PRDX1 | peroxiredoxin 1 | Cytoplasm | enzyme | 6710837.5 | 5054061 | 3744466 | | | | | | |
| P35700 | PRDX1 | peroxiredoxin 1 | Cytoplasm | enzyme | 9028467 | 8142608.5 | 4156641.75 | | | | | | |
| P35700 | PRDX1 | peroxiredoxin 1 | Cytoplasm | enzyme | 3273521.75 | 1613397.375 | 665124.25 | | | | | | |
| P35700 | PRDX1 | peroxiredoxin 1 | Cytoplasm | enzyme | 918108.438 | 1377260.5 | 653717.938 | P35700 | PRDX1 | 14424630 | 7483760 | 1302755 | 33 |
| Q61171 | PRDX2 | peroxiredoxin 2 | Cytoplasm | enzyme | 3.84E+08 | 2.82E+08 | 2.95E+08 | | | | | | |
| Q61171 | PRDX2 | peroxiredoxin 2 | Cytoplasm | enzyme | 2.21E+08 | 1.51E+08 | 1.34E+08 | | | | | | |
| Q61171 | PRDX2 | peroxiredoxin 2 | Cytoplasm | enzyme | 2.15E+08 | 1.53E+08 | 1.42E+08 | | | | | | |
| Q61171 | PRDX2 | peroxiredoxin 2 | Cytoplasm | enzyme | 1.78E+08 | 1.46E+08 | 9.06E+07 | | | | | | |
| Q61171 | PRDX2 | peroxiredoxin 2 | Cytoplasm | enzyme | 9.68E+07 | 7.01E+07 | 8.34E+07 | | | | | | |
| Q61171 | PRDX2 | peroxiredoxin 2 | Cytoplasm | enzyme | 1.05E+08 | 7.24E+07 | 6.52E+07 | | | | | | |
| Q61171 | PRDX2 | peroxiredoxin 2 | Cytoplasm | enzyme | 1.33E+08 | 1.12E+08 | 1.24E+08 | | | | | | |
| Q61171 | PRDX2 | peroxiredoxin 2 | Cytoplasm | enzyme | 8.20E+07 | 5.67E+07 | 2.77E+07 | | | | | | |
| Q61171 | PRDX2 | peroxiredoxin 2 | Cytoplasm | enzyme | 3.76E+07 | 3.21E+07 | 2.85E+07 | | | | | | |
| Q61171 | PRDX2 | peroxiredoxin 2 | Cytoplasm | enzyme | 1.39E+07 | 3245152.75 | 7937357 | | | | | | |
| Q61171 | PRDX2 | peroxiredoxin 2 | Cytoplasm | enzyme | 2.61E+07 | 1.90E+07 | 1.17E+07 | | | | | | |
| Q61171 | PRDX2 | peroxiredoxin 2 | Cytoplasm | enzyme | 2.05E+07 | 6324274.5 | 1.63E+07 | | | | | | |
| Q61171 | PRDX2 | peroxiredoxin 2 | Cytoplasm | enzyme | 2510543.75 | 2333408.75 | 3765452 | | | | | | |
| Q61171 | PRDX2 | peroxiredoxin 2 | Cytoplasm | enzyme | 5218237 | 2066630.25 | 5077836 | Q61171 | PRDX2 | 108634199 | 90880978 | 14023239 | 42 |
| P20108 | PRDX3 | peroxiredoxin 3 | Cytoplasm | enzyme | 6.98E+07 | 1.15E+08 | 1.17E+08 | | | | | | |
| P20108 | PRDX3 | peroxiredoxin 3 | Cytoplasm | enzyme | 3.75E+07 | 6.62E+07 | 7.25E+07 | | | | | | |
| P20108 | PRDX3 | peroxiredoxin 3 | Cytoplasm | enzyme | 2.39E+07 | 4.86E+07 | 4.92E+07 | | | | | | |
| P20108 | PRDX3 | peroxiredoxin 3 | Cytoplasm | enzyme | 1.34E+07 | 2.18E+07 | 2.73E+07 | | | | | | |
| P20108 | PRDX3 | peroxiredoxin 3 | Cytoplasm | enzyme | 1.84E+07 | 3.47E+07 | 3.34E+07 | | | | | | |
| P20108 | PRDX3 | peroxiredoxin 3 | Cytoplasm | enzyme | 2180135.5 | 1880794.875 | 2.23E+07 | | | | | | |
| P20108 | PRDX3 | peroxiredoxin 3 | Cytoplasm | enzyme | 2412488.5 | 3902198.5 | 5857914.5 | P20108 | PRDX3 | 23928946 | 34188318 | 7460503 | 21 |
| O08807 | PRDX4 | peroxiredoxin 4 | Cytoplasm | enzyme | 8182878 | 9846908 | 9849280 | | | | | | |
| O08807 | PRDX4 | peroxiredoxin 4 | Cytoplasm | enzyme | 3990447.25 | 4167052.75 | 6212961.5 | | | | | | |
| O08807 | PRDX4 | peroxiredoxin 4 | Cytoplasm | enzyme | 1918724.875 | 2275792 | 892722.125 | O08807 | PRDX4 | 4697350 | 3418728 | 1139576 | 9 |
| P99029 | PRDX5 | peroxiredoxin 5 | Cytoplasm | enzyme | 3032419.25 | 4571868 | 5613385.5 | P99029 | PRDX5 | 3032419 | 1298464 | 749668 | 3 |
| Q61878 | PRG2 | proteoglycan 2, bone marrow (natural killer cell activator, eosinophil granule major basic protein) | Extracellular Space | other | 610044.062 | 239482.25 | 1.46E+07 | Q61878 | PRG2 | 610044 | 8168858 | 4716292 | 3 |
| P05132 | PRKACA | protein kinase, cAMP-dependent, catalytic, alpha | Cytoplasm | kinase | 5526523 | 5078092.5 | 4651809 | P05132 | PRKACA | 5526523 | 437404 | 252535 | 3 |
| P20444 | PRKCA | protein kinase C, alpha | Cytoplasm | kinase | 1026311.938 | 1187758.625 | 1880619.375 | P20444 | PRKCA | 1026312 | 453865 | 262039 | 3 |
| O08795 | PRKCSH | protein kinase C substrate 80K-H | Cytoplasm | enzyme | 1.50E+07 | 1.34E+07 | 9282466 | | | | | | |
| O08795 | PRKCSH | protein kinase C substrate 80K-H | Cytoplasm | enzyme | 1.18E+07 | 1.05E+07 | 9110225 | | | | | | |
| O08795 | PRKCSH | protein kinase C substrate 80K-H | Cytoplasm | enzyme | 9388652 | 8274871 | 8453816 | | | | | | |
| O08795 | PRKCSH | protein kinase C substrate 80K-H | Cytoplasm | enzyme | 7899135.5 | 7100243.5 | 5604177.5 | | | | | | |
| O08795 | PRKCSH | protein kinase C substrate 80K-H | Cytoplasm | enzyme | 5582218 | 7912810.5 | 5070161 | | | | | | |
| O08795 | PRKCSH | protein kinase C substrate 80K-H | Cytoplasm | enzyme | 6115373 | 5588776.5 | 5608893.5 | | | | | | |
| O08795 | PRKCSH | protein kinase C substrate 80K-H | Cytoplasm | enzyme | 7222399.5 | 6396439.5 | 6015059.5 | | | | | | |
| O08795 | PRKCSH | protein kinase C substrate 80K-H | Cytoplasm | enzyme | 5868745.5 | 5492802 | 3558485.25 | | | | | | |
| O08795 | PRKCSH | protein kinase C substrate 80K-H | Cytoplasm | enzyme | 2915180.25 | 3828551.25 | 3436074.25 | | | | | | |
| O08795 | PRKCSH | protein kinase C substrate 80K-H | Cytoplasm | enzyme | 4532877 | 4434522.5 | 3455219.5 | | | | | | |
| O08795 | PRKCSH | protein kinase C substrate 80K-H | Cytoplasm | enzyme | 796526.625 | 827428.25 | 1229530.25 | O08795 | PRKCSH | 7005555 | 3321218 | 578150 | 33 |
| Q8CIG8 | PRMT5 | protein arginine methyltransferase 5 | Cytoplasm | enzyme | 6184441.5 | 6226979.5 | 4983385.5 | | | | | | |
| Q8CIG8 | PRMT5 | protein arginine methyltransferase 5 | Cytoplasm | enzyme | 1430815.75 | 1034146 | 761846.625 | Q8CIG8 | PRMT5 | 3807629 | 2633578 | 1075154 | 6 |
| P04925 | PRNP | prion protein | Plasma Membrane | other | 7945009 | 8698013 | 5863268.5 | P04925 | PRNP | 7945009 | 1468357 | 847756 | 3 |
| Q9D7G0 | PRPS1 | phosphoribosyl pyrophosphate synthetase 1 | Cytoplasm | kinase | 8438571 | 7340673 | 7134315 | | | | | | |
| Q9D7G0 | PRPS1 | phosphoribosyl pyrophosphate synthetase 1 | Cytoplasm | kinase | 7555300 | 6324863 | 5026486.5 | | | | | | |
| Q9D7G0 | PRPS1 | phosphoribosyl pyrophosphate synthetase 1 | Cytoplasm | kinase | 3490985 | 3323741 | 3608391.5 | | | | | | |
| Q9D7G0 | PRPS1 | phosphoribosyl pyrophosphate synthetase 1 | Cytoplasm | kinase | 4333529 | 3968465.75 | 3928414.5 | | | | | | |
| Q9D7G0 | PRPS1 | phosphoribosyl pyrophosphate synthetase 1 | Cytoplasm | kinase | 2887129.75 | 2662214.5 | 2137642.75 | | | | | | |
| Q9D7G0 | PRPS1 | phosphoribosyl pyrophosphate synthetase 1 | Cytoplasm | kinase | 2588491.75 | 2513673 | 2232079.25 | Q9D7G0 | PRPS1 | 4882334 | 2050447 | 483295 | 18 |
| Q9D0M1 | PRPSAP1 | phosphoribosyl pyrophosphate synthetase-associated protein 1 | Extracellular Space | other | 7781550.5 | 5496315 | 4382142.5 | | | | | | |
| Q9D0M1 | PRPSAP1 | phosphoribosyl pyrophosphate synthetase-associated protein 1 | Extracellular Space | other | 3828730.5 | 2918176.5 | 2679785.75 | | | | | | |
| Q9D0M1 | PRPSAP1 | phosphoribosyl pyrophosphate synthetase-associated protein 1 | Extracellular Space | other | 2351894 | 1952898.75 | 1714061.25 | | | | | | |
| Q9D0M1 | PRPSAP1 | phosphoribosyl pyrophosphate synthetase-associated protein 1 | Extracellular Space | other | 2636127.5 | 2031297.625 | 1699923.5 | | | | | | |
| Q9D0M1 | PRPSAP1 | phosphoribosyl pyrophosphate synthetase-associated protein 1 | Extracellular Space | other | 1286150.625 | 1355704.5 | 1826577.875 | Q9D0M1 | PRPSAP1 | 3576891 | 1786366 | 461238 | 15 |
| Q8R574 | PRPSAP2 | phosphoribosyl pyrophosphate synthetase-associated protein 2 | Cytoplasm | other | 2774308.75 | 1718446.625 | 1651574 | | | | | | |
| Q8R574 | PRPSAP2 | phosphoribosyl pyrophosphate synthetase-associated protein 2 | Cytoplasm | other | 3473080.5 | 2619860.25 | 2709123.5 | Q8R574 | PRPSAP2 | 3123695 | 694320 | 283455 | 6 |
| Q61207 | PSAP | prosaposin | Extracellular Space | other | 2270406.25 | 3520329.5 | 4989666.5 | Q61207 | PSAP | 2270406 | 1361105 | 785834 | 3 |
| P49769 | PSEN1 | presenilin 1 | Plasma Membrane | peptidase | 5246239.5 | 4913111.5 | 5668935.5 | | | | | | |
| P49769 | PSEN1 | presenilin 1 | Plasma Membrane | peptidase | 1951237 | 2477260.25 | 1750041.375 | P49769 | PSEN1 | 3598738 | 1793800 | 732316 | 6 |
| Q61144 | PSEN2 | presenilin 2 | Cytoplasm | peptidase | 1128021.125 | 1262355 | 1256911.75 | Q61144 | PSEN2 | 1128021 | 76035 | 43899 | 3 |
| Q91YA2 | PSKH1 | protein serine kinase H1 | Nucleus | kinase | 2723934.5 | 3211400.75 | 2912402.5 | Q91YA2 | PSKH1 | 2723935 | 245813 | 141920 | 3 |
| Q9R1P4 | PSMA1 | proteasome (prosome, macropain) subunit, alpha type, 1 | Cytoplasm | peptidase | 4.17E+07 | 3.49E+07 | 3.73E+07 | | | | | | |
| Q9R1P4 | PSMA1 | proteasome (prosome, macropain) subunit, alpha type, 1 | Cytoplasm | peptidase | 2.66E+07 | 1.30E+07 | 9091276 | | | | | | |
| Q9R1P4 | PSMA1 | proteasome (prosome, macropain) subunit, alpha type, 1 | Cytoplasm | peptidase | 2.94E+07 | 2.12E+07 | 1.86E+07 | | | | | | |
| Q9R1P4 | PSMA1 | proteasome (prosome, macropain) subunit, alpha type, 1 | Cytoplasm | peptidase | 2.88E+07 | 2.60E+07 | 1.71E+07 | | | | | | |
| Q9R1P4 | PSMA1 | proteasome (prosome, macropain) subunit, alpha type, 1 | Cytoplasm | peptidase | 2.24E+07 | 1.81E+07 | 1.73E+07 | | | | | | |
| Q9R1P4 | PSMA1 | proteasome (prosome, macropain) subunit, alpha type, 1 | Cytoplasm | peptidase | 2.36E+07 | 1.10E+07 | 1.64E+07 | | | | | | |
| Q9R1P4 | PSMA1 | proteasome (prosome, macropain) subunit, alpha type, 1 | Cytoplasm | peptidase | 2.01E+07 | 2.06E+07 | 9609047 | | | | | | |
| Q9R1P4 | PSMA1 | proteasome (prosome, macropain) subunit, alpha type, 1 | Cytoplasm | peptidase | 1.85E+07 | 1.50E+07 | 1.28E+07 | | | | | | |
| Q9R1P4 | PSMA1 | proteasome (prosome, macropain) subunit, alpha type, 1 | Cytoplasm | peptidase | 1.95E+07 | 1.43E+07 | 1.25E+07 | | | | | | |
| Q9R1P4 | PSMA1 | proteasome (prosome, macropain) subunit, alpha type, 1 | Cytoplasm | peptidase | 1.58E+07 | 1.34E+07 | 1.38E+07 | | | | | | |
| Q9R1P4 | PSMA1 | proteasome (prosome, macropain) subunit, alpha type, 1 | Cytoplasm | peptidase | 1.74E+07 | 1.07E+07 | 9823297 | | | | | | |
| Q9R1P4 | PSMA1 | proteasome (prosome, macropain) subunit, alpha type, 1 | Cytoplasm | peptidase | 7941102.5 | 5220837 | 4294903 | | | | | | |
| Q9R1P4 | PSMA1 | proteasome (prosome, macropain) subunit, alpha type, 1 | Cytoplasm | peptidase | 5084680 | 4712160.5 | 3961052.75 | | | | | | |
| Q9R1P4 | PSMA1 | proteasome (prosome, macropain) subunit, alpha type, 1 | Cytoplasm | peptidase | 4046416 | 3048306 | 2203799 | | | | | | |
| Q9R1P4 | PSMA1 | proteasome (prosome, macropain) subunit, alpha type, 1 | Cytoplasm | peptidase | 2870810.25 | 1118994.875 | 1447575.875 | | | | | | |
| Q9R1P4 | PSMA1 | proteasome (prosome, macropain) subunit, alpha type, 1 | Cytoplasm | peptidase | 6473348.5 | 1.04E+07 | 1.07E+07 | Q9R1P4 | PSMA1 | 18138522 | 9613501 | 1387589 | 48 |
| P49722 | PSMA2 | proteasome (prosome, macropain) subunit, alpha type, 2 | Cytoplasm | peptidase | 6.56E+07 | 6.66E+07 | 6.30E+07 | | | | | | |
| P49722 | PSMA2 | proteasome (prosome, macropain) subunit, alpha type, 2 | Cytoplasm | peptidase | 5.13E+07 | 1.98E+07 | 1.23E+07 | | | | | | |
| P49722 | PSMA2 | proteasome (prosome, macropain) subunit, alpha type, 2 | Cytoplasm | peptidase | 2.68E+07 | 1.47E+07 | 1.01E+07 | | | | | | |
| P49722 | PSMA2 | proteasome (prosome, macropain) subunit, alpha type, 2 | Cytoplasm | peptidase | 3.14E+07 | 2.03E+07 | 2.09E+07 | | | | | | |

| | | | | | | | | | | | | | |
|---|---|---|---|---|---|---|---|---|---|---|---|---|---|
| P49722 | PSMA2 | proteasome (prosome, macropain) subunit, alpha type, 2 | Cytoplasm | peptidase | 1.17E+07 | 1.25E+07 | 1.22E+07 | | | | | | |
| P49722 | PSMA2 | proteasome (prosome, macropain) subunit, alpha type, 2 | Cytoplasm | peptidase | 5412613.5 | 2109723.25 | 6117839 | | | | | | |
| P49722 | PSMA2 | proteasome (prosome, macropain) subunit, alpha type, 2 | Cytoplasm | peptidase | 4391485 | 2933985 | 3165931.5 | | | | | | |
| P49722 | PSMA2 | proteasome (prosome, macropain) subunit, alpha type, 2 | Cytoplasm | peptidase | 4486226 | 6135610.5 | 2603408.5 | | | | | | |
| P49722 | PSMA2 | proteasome (prosome, macropain) subunit, alpha type, 2 | Cytoplasm | peptidase | 4802287.5 | 4405104 | 3445556.25 | | | | | | |
| P49722 | PSMA2 | proteasome (prosome, macropain) subunit, alpha type, 2 | Cytoplasm | peptidase | 3356251.25 | 2537720.75 | 1673273.875 | P49722 | PSMA2 | 20922886 | 19668094 | 3590886 | 30 |
| O70435 | PSMA3 | proteasome (prosome, macropain) subunit, alpha type, 3 | Cytoplasm | peptidase | 3.65E+07 | 2.80E+07 | 2.43E+07 | | | | | | |
| O70435 | PSMA3 | proteasome (prosome, macropain) subunit, alpha type, 3 | Cytoplasm | peptidase | 2.67E+07 | 2.17E+07 | 2.21E+07 | | | | | | |
| O70435 | PSMA3 | proteasome (prosome, macropain) subunit, alpha type, 3 | Cytoplasm | peptidase | 2.91E+07 | 2.11E+07 | 1.87E+07 | | | | | | |
| O70435 | PSMA3 | proteasome (prosome, macropain) subunit, alpha type, 3 | Cytoplasm | peptidase | 2.14E+07 | 1.35E+07 | 1.26E+07 | | | | | | |
| O70435 | PSMA3 | proteasome (prosome, macropain) subunit, alpha type, 3 | Cytoplasm | peptidase | 1.77E+07 | 1.54E+07 | 1.12E+07 | | | | | | |
| O70435 | PSMA3 | proteasome (prosome, macropain) subunit, alpha type, 3 | Cytoplasm | peptidase | 1.69E+07 | 1.41E+07 | 1.27E+07 | | | | | | |
| O70435 | PSMA3 | proteasome (prosome, macropain) subunit, alpha type, 3 | Cytoplasm | peptidase | 1.09E+07 | 9170951 | 8144224.5 | | | | | | |
| O70435 | PSMA3 | proteasome (prosome, macropain) subunit, alpha type, 3 | Cytoplasm | peptidase | 1.04E+07 | 3573434.5 | 8761336 | | | | | | |
| O70435 | PSMA3 | proteasome (prosome, macropain) subunit, alpha type, 3 | Cytoplasm | peptidase | 6163681.5 | 2024142.625 | 6375360.5 | O70435 | PSMA3 | 19531520 | 8476202 | 1631246 | 27 |
| Q9R1P0 | PSMA4 | proteasome (prosome, macropain) subunit, alpha type, 4 | Cytoplasm | peptidase | 3.25E+07 | 2.77E+07 | 3.14E+07 | | | | | | |
| Q9R1P0 | PSMA4 | proteasome (prosome, macropain) subunit, alpha type, 4 | Cytoplasm | peptidase | 4.07E+07 | 3.93E+07 | 3.19E+07 | | | | | | |
| Q9R1P0 | PSMA4 | proteasome (prosome, macropain) subunit, alpha type, 4 | Cytoplasm | peptidase | 4.86E+07 | 3.27E+07 | 3.04E+07 | | | | | | |
| Q9R1P0 | PSMA4 | proteasome (prosome, macropain) subunit, alpha type, 4 | Cytoplasm | peptidase | 3.55E+07 | 2.36E+07 | 2.57E+07 | | | | | | |
| Q9R1P0 | PSMA4 | proteasome (prosome, macropain) subunit, alpha type, 4 | Cytoplasm | peptidase | 2.50E+07 | 2.15E+07 | 2.01E+07 | | | | | | |
| Q9R1P0 | PSMA4 | proteasome (prosome, macropain) subunit, alpha type, 4 | Cytoplasm | peptidase | 2.18E+07 | 1.80E+07 | 1.56E+07 | | | | | | |
| Q9R1P0 | PSMA4 | proteasome (prosome, macropain) subunit, alpha type, 4 | Cytoplasm | peptidase | 1.86E+07 | 1.40E+07 | 1.53E+07 | | | | | | |
| Q9R1P0 | PSMA4 | proteasome (prosome, macropain) subunit, alpha type, 4 | Cytoplasm | peptidase | 1.31E+07 | 9501027 | 9119146 | | | | | | |
| Q9R1P0 | PSMA4 | proteasome (prosome, macropain) subunit, alpha type, 4 | Cytoplasm | peptidase | 1.25E+07 | 8011366.5 | 7726525.5 | | | | | | |
| Q9R1P0 | PSMA4 | proteasome (prosome, macropain) subunit, alpha type, 4 | Cytoplasm | peptidase | 1.09E+07 | 8085521.5 | 8155471.5 | | | | | | |
| Q9R1P0 | PSMA4 | proteasome (prosome, macropain) subunit, alpha type, 4 | Cytoplasm | peptidase | 6105091.5 | 5191082.5 | 5181783 | | | | | | |
| Q9R1P0 | PSMA4 | proteasome (prosome, macropain) subunit, alpha type, 4 | Cytoplasm | peptidase | 4172244 | 2286947 | 2043583.25 | | | | | | |
| Q9R1P0 | PSMA4 | proteasome (prosome, macropain) subunit, alpha type, 4 | Cytoplasm | peptidase | 706790.125 | 604045.438 | 149761.547 | Q9R1P0 | PSMA4 | 20777240 | 12761420 | 2043463 | 39 |
| Q9Z2U1 | PSMA5 | proteasome (prosome, macropain) subunit, alpha type, 5 | Cytoplasm | peptidase | 8.96E+07 | 7.41E+07 | 6.70E+07 | | | | | | |
| Q9Z2U1 | PSMA5 | proteasome (prosome, macropain) subunit, alpha type, 5 | Cytoplasm | peptidase | 7.88E+07 | 5.89E+07 | 4.68E+07 | | | | | | |
| Q9Z2U1 | PSMA5 | proteasome (prosome, macropain) subunit, alpha type, 5 | Cytoplasm | peptidase | 7.83E+07 | 4.81E+07 | 4.16E+07 | | | | | | |
| Q9Z2U1 | PSMA5 | proteasome (prosome, macropain) subunit, alpha type, 5 | Cytoplasm | peptidase | 3.44E+07 | 2.70E+07 | 2.02E+07 | | | | | | |
| Q9Z2U1 | PSMA5 | proteasome (prosome, macropain) subunit, alpha type, 5 | Cytoplasm | peptidase | 2.25E+07 | 1.92E+07 | 1.61E+07 | | | | | | |
| Q9Z2U1 | PSMA5 | proteasome (prosome, macropain) subunit, alpha type, 5 | Cytoplasm | peptidase | 2.38E+07 | 1.33E+07 | 1.13E+07 | | | | | | |
| Q9Z2U1 | PSMA5 | proteasome (prosome, macropain) subunit, alpha type, 5 | Cytoplasm | peptidase | 1.32E+07 | 1.06E+07 | 9888870 | | | | | | |
| Q9Z2U1 | PSMA5 | proteasome (prosome, macropain) subunit, alpha type, 5 | Cytoplasm | peptidase | 1.23E+07 | 6769887.5 | 6178576 | | | | | | |
| Q9Z2U1 | PSMA5 | proteasome (prosome, macropain) subunit, alpha type, 5 | Cytoplasm | peptidase | 7088682 | 2736250.5 | 6293142 | | | | | | |
| Q9Z2U1 | PSMA5 | proteasome (prosome, macropain) subunit, alpha type, 5 | Cytoplasm | peptidase | 5247355.5 | 4632719.5 | 2455635.25 | | | | | | |
| Q9Z2U1 | PSMA5 | proteasome (prosome, macropain) subunit, alpha type, 5 | Cytoplasm | peptidase | 4487900.5 | 3724467.75 | 3399486.5 | | | | | | |
| Q9Z2U1 | PSMA5 | proteasome (prosome, macropain) subunit, alpha type, 5 | Cytoplasm | peptidase | 6182076 | 2914829.75 | 5287337.5 | | | | | | |
| Q9Z2U1 | PSMA5 | proteasome (prosome, macropain) subunit, alpha type, 5 | Cytoplasm | peptidase | 3218273.5 | 2183915.5 | 2566352.75 | | | | | | |
| Q9Z2U1 | PSMA5 | proteasome (prosome, macropain) subunit, alpha type, 5 | Cytoplasm | peptidase | 3181374.75 | 2087351 | 1511964.375 | Q9Z2U1 | PSMA5 | 27303976 | 25207229 | 3889560 | 42 |
| Q9QUM9 | PSMA6 | proteasome (prosome, macropain) subunit, alpha type, 6 | Cytoplasm | peptidase | 4.68E+07 | 3.70E+07 | 3.34E+07 | | | | | | |
| Q9QUM9 | PSMA6 | proteasome (prosome, macropain) subunit, alpha type, 6 | Cytoplasm | peptidase | 5.54E+07 | 3.33E+07 | 3.13E+07 | | | | | | |
| Q9QUM9 | PSMA6 | proteasome (prosome, macropain) subunit, alpha type, 6 | Cytoplasm | peptidase | 3.53E+07 | 2.59E+07 | 2.21E+07 | | | | | | |
| Q9QUM9 | PSMA6 | proteasome (prosome, macropain) subunit, alpha type, 6 | Cytoplasm | peptidase | 3.24E+07 | 2.14E+07 | 1.83E+07 | | | | | | |
| Q9QUM9 | PSMA6 | proteasome (prosome, macropain) subunit, alpha type, 6 | Cytoplasm | peptidase | 3.33E+07 | 2.27E+07 | 2.21E+07 | | | | | | |
| Q9QUM9 | PSMA6 | proteasome (prosome, macropain) subunit, alpha type, 6 | Cytoplasm | peptidase | 2.85E+07 | 1.91E+07 | 1.68E+07 | | | | | | |
| Q9QUM9 | PSMA6 | proteasome (prosome, macropain) subunit, alpha type, 6 | Cytoplasm | peptidase | 2.08E+07 | 1.33E+07 | 1.09E+07 | | | | | | |
| Q9QUM9 | PSMA6 | proteasome (prosome, macropain) subunit, alpha type, 6 | Cytoplasm | peptidase | 1.95E+07 | 1.54E+07 | 1.28E+07 | | | | | | |
| Q9QUM9 | PSMA6 | proteasome (prosome, macropain) subunit, alpha type, 6 | Cytoplasm | peptidase | 1.86E+07 | 1.25E+07 | 1.34E+07 | | | | | | |
| Q9QUM9 | PSMA6 | proteasome (prosome, macropain) subunit, alpha type, 6 | Cytoplasm | peptidase | 1.75E+07 | 1.45E+07 | 1.19E+07 | | | | | | |
| Q9QUM9 | PSMA6 | proteasome (prosome, macropain) subunit, alpha type, 6 | Cytoplasm | peptidase | 1532144.25 | 373069.75 | 70049.32 | Q9QUM9 | PSMA6 | 28130195 | 12340815 | 2148260 | 33 |
| Q9Z2U0 | PSMA7 | proteasome (prosome, macropain) subunit, alpha type, 7 | Cytoplasm | peptidase | 9.43E+07 | 7.86E+07 | 7.74E+07 | | | | | | |
| Q9Z2U0 | PSMA7 | proteasome (prosome, macropain) subunit, alpha type, 7 | Cytoplasm | peptidase | 4.56E+07 | 3.78E+07 | 3.57E+07 | | | | | | |
| Q9Z2U0 | PSMA7 | proteasome (prosome, macropain) subunit, alpha type, 7 | Cytoplasm | peptidase | 4.59E+07 | 3.94E+07 | 4.22E+07 | | | | | | |
| Q9Z2U0 | PSMA7 | proteasome (prosome, macropain) subunit, alpha type, 7 | Cytoplasm | peptidase | 3.43E+07 | 2.81E+07 | 2.68E+07 | | | | | | |
| Q9Z2U0 | PSMA7 | proteasome (prosome, macropain) subunit, alpha type, 7 | Cytoplasm | peptidase | 4.18E+07 | 3.34E+07 | 3.12E+07 | | | | | | |
| Q9Z2U0 | PSMA7 | proteasome (prosome, macropain) subunit, alpha type, 7 | Cytoplasm | peptidase | 1.88E+07 | 2.06E+07 | 1.39E+07 | | | | | | |
| Q9Z2U0 | PSMA7 | proteasome (prosome, macropain) subunit, alpha type, 7 | Cytoplasm | peptidase | 1.43E+07 | 1.21E+07 | 6985009 | | | | | | |
| Q9Z2U0 | PSMA7 | proteasome (prosome, macropain) subunit, alpha type, 7 | Cytoplasm | peptidase | 1.04E+07 | 7195280 | 7446364.5 | | | | | | |
| Q9Z2U0 | PSMA7 | proteasome (prosome, macropain) subunit, alpha type, 7 | Cytoplasm | peptidase | 8332869 | 6132321 | 4842460 | | | | | | |
| Q9Z2U0 | PSMA7 | proteasome (prosome, macropain) subunit, alpha type, 7 | Cytoplasm | peptidase | 6180999.5 | 3191485.75 | 4344177.5 | | | | | | |
| Q9Z2U0 | PSMA7 | proteasome (prosome, macropain) subunit, alpha type, 7 | Cytoplasm | peptidase | 6254662.5 | 5288374 | 5445102 | | | | | | |
| Q9Z2U0 | PSMA7 | proteasome (prosome, macropain) subunit, alpha type, 7 | Cytoplasm | peptidase | 1283192.625 | 675003 | 3018565.5 | Q9Z2U0 | PSMA7 | 27281810 | 23414294 | 3902382 | 36 |
| O09061 | PSMB1 | proteasome (prosome, macropain) subunit, beta type, 1 | Cytoplasm | peptidase | 4.49E+07 | 3.36E+07 | 3.12E+07 | | | | | | |
| O09061 | PSMB1 | proteasome (prosome, macropain) subunit, beta type, 1 | Cytoplasm | peptidase | 3.98E+07 | 2.91E+07 | 2.61E+07 | | | | | | |
| O09061 | PSMB1 | proteasome (prosome, macropain) subunit, beta type, 1 | Cytoplasm | peptidase | 3.33E+07 | 2.49E+07 | 2.13E+07 | | | | | | |
| O09061 | PSMB1 | proteasome (prosome, macropain) subunit, beta type, 1 | Cytoplasm | peptidase | 2.29E+07 | 1.64E+07 | 1.86E+07 | | | | | | |
| O09061 | PSMB1 | proteasome (prosome, macropain) subunit, beta type, 1 | Cytoplasm | peptidase | 1.97E+07 | 1.55E+07 | 1.05E+07 | | | | | | |
| O09061 | PSMB1 | proteasome (prosome, macropain) subunit, beta type, 1 | Cytoplasm | peptidase | 9578972 | 5723527.5 | 6297869.5 | | | | | | |
| O09061 | PSMB1 | proteasome (prosome, macropain) subunit, beta type, 1 | Cytoplasm | peptidase | 7890436 | 3961952.75 | 6451773 | | | | | | |
| O09061 | PSMB1 | proteasome (prosome, macropain) subunit, beta type, 1 | Cytoplasm | peptidase | 7996571.5 | 6540679.5 | 3404594.5 | | | | | | |
| O09061 | PSMB1 | proteasome (prosome, macropain) subunit, beta type, 1 | Cytoplasm | peptidase | 7121176 | 7663828 | 5054946 | | | | | | |
| O09061 | PSMB1 | proteasome (prosome, macropain) subunit, beta type, 1 | Cytoplasm | peptidase | 5679661.5 | 3333115.5 | 4806922.5 | | | | | | |
| O09061 | PSMB1 | proteasome (prosome, macropain) subunit, beta type, 1 | Cytoplasm | peptidase | 4624005 | 4179471.5 | 3768666.5 | | | | | | |
| O09061 | PSMB1 | proteasome (prosome, macropain) subunit, beta type, 1 | Cytoplasm | peptidase | 5144736.5 | 2511969.25 | 4307472 | | | | | | |
| O09061 | PSMB1 | proteasome (prosome, macropain) subunit, beta type, 1 | Cytoplasm | peptidase | 5686554 | 4211393.5 | 4427076 | | | | | | |
| O09061 | PSMB1 | proteasome (prosome, macropain) subunit, beta type, 1 | Cytoplasm | peptidase | 4374364.5 | 5093499 | 3811822.5 | | | | | | |
| O09061 | PSMB1 | proteasome (prosome, macropain) subunit, beta type, 1 | Cytoplasm | peptidase | 4279802 | 3693586.75 | 3238715.75 | | | | | | |
| O09061 | PSMB1 | proteasome (prosome, macropain) subunit, beta type, 1 | Cytoplasm | peptidase | 2774456.75 | 1812131.75 | 1363581.125 | | | | | | |
| O09061 | PSMB1 | proteasome (prosome, macropain) subunit, beta type, 1 | Cytoplasm | peptidase | 1684453 | 1235338.625 | 1426341.125 | O09061 | PSMB1 | 13371482 | 11122951 | 1557525 | 51 |
| Q9R1P3 | PSMB2 | proteasome (prosome, macropain) subunit, beta type, 2 | Cytoplasm | peptidase | 4.94E+07 | 3.82E+07 | 3.76E+07 | | | | | | |
| Q9R1P3 | PSMB2 | proteasome (prosome, macropain) subunit, beta type, 2 | Cytoplasm | peptidase | 3.28E+07 | 2.42E+07 | 2.29E+07 | | | | | | |
| Q9R1P3 | PSMB2 | proteasome (prosome, macropain) subunit, beta type, 2 | Cytoplasm | peptidase | 1.47E+07 | 1.29E+07 | 4334433 | | | | | | |
| Q9R1P3 | PSMB2 | proteasome (prosome, macropain) subunit, beta type, 2 | Cytoplasm | peptidase | 8628810 | 6995550 | 6875284 | | | | | | |
| Q9R1P3 | PSMB2 | proteasome (prosome, macropain) subunit, beta type, 2 | Cytoplasm | peptidase | 5797461.5 | 7546129.5 | 5320203.5 | | | | | | |
| Q9R1P3 | PSMB2 | proteasome (prosome, macropain) subunit, beta type, 2 | Cytoplasm | peptidase | 9109551 | 6510265 | 7972575.5 | | | | | | |
| Q9R1P3 | PSMB2 | proteasome (prosome, macropain) subunit, beta type, 2 | Cytoplasm | peptidase | 5078792.5 | 4853794.5 | 3943634.5 | | | | | | |
| Q9R1P3 | PSMB2 | proteasome (prosome, macropain) subunit, beta type, 2 | Cytoplasm | peptidase | 3740079.5 | 2429562.25 | 1655770.125 | | | | | | |
| Q9R1P3 | PSMB2 | proteasome (prosome, macropain) subunit, beta type, 2 | Cytoplasm | peptidase | 5504489.5 | 2385477.25 | 3657731.5 | Q9R1P3 | PSMB2 | 14974354 | 12940733 | 2490445 | 27 |
| Q9R1P1 | PSMB3 | proteasome (prosome, macropain) subunit, beta type, 3 | Cytoplasm | peptidase | 4.12E+07 | 3.65E+07 | 2.81E+07 | | | | | | |
| Q9R1P1 | PSMB3 | proteasome (prosome, macropain) subunit, beta type, 3 | Cytoplasm | peptidase | 2.46E+07 | 2.21E+07 | 1.62E+07 | | | | | | |
| Q9R1P1 | PSMB3 | proteasome (prosome, macropain) subunit, beta type, 3 | Cytoplasm | peptidase | 1.50E+07 | 1.20E+07 | 1.30E+07 | | | | | | |
| Q9R1P1 | PSMB3 | proteasome (prosome, macropain) subunit, beta type, 3 | Cytoplasm | peptidase | 1.54E+07 | 1.45E+07 | 1.29E+07 | | | | | | |
| Q9R1P1 | PSMB3 | proteasome (prosome, macropain) subunit, beta type, 3 | Cytoplasm | peptidase | 8902383 | 5300524.5 | 4874888 | | | | | | |
| Q9R1P1 | PSMB3 | proteasome (prosome, macropain) subunit, beta type, 3 | Cytoplasm | peptidase | 3774910.75 | 4093221 | 3391176.5 | | | | | | |
| Q9R1P1 | PSMB3 | proteasome (prosome, macropain) subunit, beta type, 3 | Cytoplasm | peptidase | 9930669 | 7406184.5 | 9928164 | | | | | | |
| Q9R1P1 | PSMB3 | proteasome (prosome, macropain) subunit, beta type, 3 | Cytoplasm | peptidase | 5915589.5 | 4948141.5 | 4031195.75 | | | | | | |
| Q9R1P1 | PSMB3 | proteasome (prosome, macropain) subunit, beta type, 3 | Cytoplasm | peptidase | 3232600.25 | 4201183 | 3424043.5 | | | | | | |
| Q9R1P1 | PSMB3 | proteasome (prosome, macropain) subunit, beta type, 3 | Cytoplasm | peptidase | 549440 | 1553841 | 1331798.5 | Q9R1P1 | PSMB3 | 12854559 | 10272258 | 1875449 | 30 |
| P99026 | PSMB4 | proteasome (prosome, macropain) subunit, beta type, 4 | Cytoplasm | peptidase | 2.73E+07 | 2.25E+07 | 1.59E+07 | | | | | | |
| P99026 | PSMB4 | proteasome (prosome, macropain) subunit, beta type, 4 | Cytoplasm | peptidase | 2.67E+07 | 1.81E+07 | 1.97E+07 | | | | | | |
| P99026 | PSMB4 | proteasome (prosome, macropain) subunit, beta type, 4 | Cytoplasm | peptidase | 2.75E+07 | 1.98E+07 | 1.59E+07 | | | | | | |
| P99026 | PSMB4 | proteasome (prosome, macropain) subunit, beta type, 4 | Cytoplasm | peptidase | 2.38E+07 | 2.00E+07 | 2.05E+07 | | | | | | |
| P99026 | PSMB4 | proteasome (prosome, macropain) subunit, beta type, 4 | Cytoplasm | peptidase | 2.30E+07 | 1.77E+07 | 1.61E+07 | | | | | | |
| P99026 | PSMB4 | proteasome (prosome, macropain) subunit, beta type, 4 | Cytoplasm | peptidase | 5533657.5 | 4037295 | 5223914 | P99026 | PSMB4 | 22283943 | 7153657 | 1686133 | 18 |
| O55234 | PSMB5 | proteasome (prosome, macropain) subunit, beta type, 5 | Cytoplasm | peptidase | 4.60E+07 | 2.69E+07 | 2.33E+07 | | | | | | |
| O55234 | PSMB5 | proteasome (prosome, macropain) subunit, beta type, 5 | Cytoplasm | peptidase | 3.25E+07 | 2.19E+07 | 1.93E+07 | | | | | | |

| | | | | | | | | | | | | | |
|---|---|---|---|---|---|---|---|---|---|---|---|---|---|
| O55234 | PSMB5 | proteasome (prosome, macropain) subunit, beta type, 5 | Cytoplasm | peptidase | 2.93E+07 | 2.16E+07 | 2.09E+07 | | | | | | |
| O55234 | PSMB5 | proteasome (prosome, macropain) subunit, beta type, 5 | Cytoplasm | peptidase | 2.72E+07 | 1.91E+07 | 1.57E+07 | | | | | | |
| O55234 | PSMB5 | proteasome (prosome, macropain) subunit, beta type, 5 | Cytoplasm | peptidase | 2.56E+07 | 1.76E+07 | 2.00E+07 | | | | | | |
| O55234 | PSMB5 | proteasome (prosome, macropain) subunit, beta type, 5 | Cytoplasm | peptidase | 1.50E+07 | 1.08E+07 | 9180138 | | | | | | |
| O55234 | PSMB5 | proteasome (prosome, macropain) subunit, beta type, 5 | Cytoplasm | peptidase | 9477191 | 7223402 | 6788787.5 | | | | | | |
| O55234 | PSMB5 | proteasome (prosome, macropain) subunit, beta type, 5 | Cytoplasm | peptidase | 1.14E+07 | 6678134 | 5585607 | | | | | | |
| O55234 | PSMB5 | proteasome (prosome, macropain) subunit, beta type, 5 | Cytoplasm | peptidase | 8108306 | 5772894.5 | 5647663 | | | | | | |
| O55234 | PSMB5 | proteasome (prosome, macropain) subunit, beta type, 5 | Cytoplasm | peptidase | 5347668.5 | 3892872.5 | 3132884.5 | | | | | | |
| O55234 | PSMB5 | proteasome (prosome, macropain) subunit, beta type, 5 | Cytoplasm | peptidase | 2013904 | 2202674.25 | 1524178 | O55234 | PSMB5 | 19254279 | 10611663 | 1847253 | 33 |
| Q60692 | PSMB6 | proteasome (prosome, macropain) subunit, beta type, 6 | Nucleus | peptidase | 9.38E+07 | 9.30E+07 | 8.64E+07 | | | | | | |
| Q60692 | PSMB6 | proteasome (prosome, macropain) subunit, beta type, 6 | Nucleus | peptidase | 2.99E+07 | 2.52E+07 | 1.96E+07 | | | | | | |
| Q60692 | PSMB6 | proteasome (prosome, macropain) subunit, beta type, 6 | Nucleus | peptidase | 2.66E+07 | 2.18E+07 | 1.67E+07 | | | | | | |
| Q60692 | PSMB6 | proteasome (prosome, macropain) subunit, beta type, 6 | Nucleus | peptidase | 1.35E+07 | 1.05E+07 | 9817414 | | | | | | |
| Q60692 | PSMB6 | proteasome (prosome, macropain) subunit, beta type, 6 | Nucleus | peptidase | 4822033.5 | 3820801.25 | 4077830.5 | | | | | | |
| Q60692 | PSMB6 | proteasome (prosome, macropain) subunit, beta type, 6 | Nucleus | peptidase | 2591365.5 | 798008.688 | 199307.031 | Q60692 | PSMB6 | 28540567 | 31447225 | 7412182 | 18 |
| P70195 | PSMB7 | proteasome (prosome, macropain) subunit, beta type, 7 | Cytoplasm | peptidase | 4.14E+07 | 3.28E+07 | 2.70E+07 | | | | | | |
| P70195 | PSMB7 | proteasome (prosome, macropain) subunit, beta type, 7 | Cytoplasm | peptidase | 2.52E+07 | 1.86E+07 | 1.46E+07 | | | | | | |
| P70195 | PSMB7 | proteasome (prosome, macropain) subunit, beta type, 7 | Cytoplasm | peptidase | 2.43E+07 | 2.42E+07 | 1.80E+07 | | | | | | |
| P70195 | PSMB7 | proteasome (prosome, macropain) subunit, beta type, 7 | Cytoplasm | peptidase | 1.61E+07 | 1.61E+07 | 1.55E+07 | | | | | | |
| P70195 | PSMB7 | proteasome (prosome, macropain) subunit, beta type, 7 | Cytoplasm | peptidase | 1.84E+07 | 1.43E+07 | 1.37E+07 | | | | | | |
| P70195 | PSMB7 | proteasome (prosome, macropain) subunit, beta type, 7 | Cytoplasm | peptidase | 1.20E+07 | 8069954.5 | 7399644 | | | | | | |
| P70195 | PSMB7 | proteasome (prosome, macropain) subunit, beta type, 7 | Cytoplasm | peptidase | 6403947.5 | 4035661.25 | 3334838.75 | | | | | | |
| P70195 | PSMB7 | proteasome (prosome, macropain) subunit, beta type, 7 | Cytoplasm | peptidase | 969266.125 | 1158734.375 | 1201362.125 | P70195 | PSMB7 | 18087902 | 10408734 | 2124674 | 24 |
| P28076 | PSMB9 | proteasome (prosome, macropain) subunit, beta type, 9 | Cytoplasm | peptidase | 845317.25 | 556225.75 | 587133.5 | P28076 | PSMB9 | 845317 | 158739 | 91648 | 3 |
| P62192 | PSMC1 | proteasome (prosome, macropain) 26S subunit, ATPase, 1 | Nucleus | peptidase | 2.91E+07 | 2.91E+07 | 2.85E+07 | | | | | | |
| P62192 | PSMC1 | proteasome (prosome, macropain) 26S subunit, ATPase, 1 | Nucleus | peptidase | 1.74E+07 | 1.39E+07 | 1.53E+07 | | | | | | |
| P62192 | PSMC1 | proteasome (prosome, macropain) 26S subunit, ATPase, 1 | Nucleus | peptidase | 1.45E+07 | 1.18E+07 | 1.06E+07 | | | | | | |
| P62192 | PSMC1 | proteasome (prosome, macropain) 26S subunit, ATPase, 1 | Nucleus | peptidase | 1.45E+07 | 1.21E+07 | 9450972 | | | | | | |
| P62192 | PSMC1 | proteasome (prosome, macropain) 26S subunit, ATPase, 1 | Nucleus | peptidase | 1.01E+07 | 8036789.5 | 7687844.5 | | | | | | |
| P62192 | PSMC1 | proteasome (prosome, macropain) 26S subunit, ATPase, 1 | Nucleus | peptidase | 1.10E+07 | 9679869 | 8319782 | | | | | | |
| P62192 | PSMC1 | proteasome (prosome, macropain) 26S subunit, ATPase, 1 | Nucleus | peptidase | 9769434 | 1.12E+07 | 7184993.5 | | | | | | |
| P62192 | PSMC1 | proteasome (prosome, macropain) 26S subunit, ATPase, 1 | Nucleus | peptidase | 9953104 | 8474504 | 6944630 | | | | | | |
| P62192 | PSMC1 | proteasome (prosome, macropain) 26S subunit, ATPase, 1 | Nucleus | peptidase | 7961972 | 5944639.5 | 5297531 | | | | | | |
| P62192 | PSMC1 | proteasome (prosome, macropain) 26S subunit, ATPase, 1 | Nucleus | peptidase | 5455220 | 7796740 | 4282092 | | | | | | |
| P62192 | PSMC1 | proteasome (prosome, macropain) 26S subunit, ATPase, 1 | Nucleus | peptidase | 6314814 | 6179392.5 | 3491658.25 | | | | | | |
| P62192 | PSMC1 | proteasome (prosome, macropain) 26S subunit, ATPase, 1 | Nucleus | peptidase | 6270300 | 6160162.5 | 4717328 | | | | | | |
| P62192 | PSMC1 | proteasome (prosome, macropain) 26S subunit, ATPase, 1 | Nucleus | peptidase | 2681823.25 | 2670133 | 3096285.25 | P62192 | PSMC1 | 11141282 | 6553953 | 1049472 | 39 |
| P46471 | PSMC2 | proteasome (prosome, macropain) 26S subunit, ATPase, 2 | Nucleus | peptidase | 2.40E+07 | 1.90E+07 | 1.96E+07 | | | | | | |
| P46471 | PSMC2 | proteasome (prosome, macropain) 26S subunit, ATPase, 2 | Nucleus | peptidase | 1.77E+07 | 1.82E+07 | 1.65E+07 | | | | | | |
| P46471 | PSMC2 | proteasome (prosome, macropain) 26S subunit, ATPase, 2 | Nucleus | peptidase | 1.70E+07 | 1.34E+07 | 1.07E+07 | | | | | | |
| P46471 | PSMC2 | proteasome (prosome, macropain) 26S subunit, ATPase, 2 | Nucleus | peptidase | 1.35E+07 | 1.02E+07 | 1.07E+07 | | | | | | |
| P46471 | PSMC2 | proteasome (prosome, macropain) 26S subunit, ATPase, 2 | Nucleus | peptidase | 1.43E+07 | 8539203 | 7268303 | | | | | | |
| P46471 | PSMC2 | proteasome (prosome, macropain) 26S subunit, ATPase, 2 | Nucleus | peptidase | 1.20E+07 | 1.03E+07 | 1.05E+07 | | | | | | |
| P46471 | PSMC2 | proteasome (prosome, macropain) 26S subunit, ATPase, 2 | Nucleus | peptidase | 1.01E+07 | 9149890 | 7839285 | | | | | | |
| P46471 | PSMC2 | proteasome (prosome, macropain) 26S subunit, ATPase, 2 | Nucleus | peptidase | 1.30E+07 | 1.04E+07 | 1.01E+07 | | | | | | |
| P46471 | PSMC2 | proteasome (prosome, macropain) 26S subunit, ATPase, 2 | Nucleus | peptidase | 1.11E+07 | 8446037 | 7957336 | | | | | | |
| P46471 | PSMC2 | proteasome (prosome, macropain) 26S subunit, ATPase, 2 | Nucleus | peptidase | 1.13E+07 | 8712862 | 8623337 | | | | | | |
| P46471 | PSMC2 | proteasome (prosome, macropain) 26S subunit, ATPase, 2 | Nucleus | peptidase | 1.06E+07 | 9589720 | 8352456.5 | | | | | | |
| P46471 | PSMC2 | proteasome (prosome, macropain) 26S subunit, ATPase, 2 | Nucleus | peptidase | 5218102.5 | 5682621.5 | 5502680.5 | | | | | | |
| P46471 | PSMC2 | proteasome (prosome, macropain) 26S subunit, ATPase, 2 | Nucleus | peptidase | 6099170 | 4626265 | 4561911.5 | | | | | | |
| P46471 | PSMC2 | proteasome (prosome, macropain) 26S subunit, ATPase, 2 | Nucleus | peptidase | 4741399.5 | 2497882.5 | 2302206.75 | | | | | | |
| P46471 | PSMC2 | proteasome (prosome, macropain) 26S subunit, ATPase, 2 | Nucleus | peptidase | 2319963 | 3617604.5 | 2295893.5 | P46471 | PSMC2 | 11527909 | 5028182 | 749557 | 45 |
| O88685 | PSMC3 | proteasome (prosome, macropain) 26S subunit, ATPase, 3 | Nucleus | transcription regulator | 2.03E+07 | 1.66E+07 | 1.46E+07 | | | | | | |
| O88685 | PSMC3 | proteasome (prosome, macropain) 26S subunit, ATPase, 3 | Nucleus | transcription regulator | 1.36E+07 | 1.49E+07 | 1.29E+07 | | | | | | |
| O88685 | PSMC3 | proteasome (prosome, macropain) 26S subunit, ATPase, 3 | Nucleus | transcription regulator | 1.43E+07 | 1.28E+07 | 1.29E+07 | | | | | | |
| O88685 | PSMC3 | proteasome (prosome, macropain) 26S subunit, ATPase, 3 | Nucleus | transcription regulator | 1.14E+07 | 9949862 | 1.01E+07 | | | | | | |
| O88685 | PSMC3 | proteasome (prosome, macropain) 26S subunit, ATPase, 3 | Nucleus | transcription regulator | 9267123 | 7659544.5 | 9841701 | | | | | | |
| O88685 | PSMC3 | proteasome (prosome, macropain) 26S subunit, ATPase, 3 | Nucleus | transcription regulator | 7252800 | 6471026.5 | 6532534.5 | | | | | | |
| O88685 | PSMC3 | proteasome (prosome, macropain) 26S subunit, ATPase, 3 | Nucleus | transcription regulator | 1.09E+07 | 7530899.5 | 1.02E+07 | | | | | | |
| O88685 | PSMC3 | proteasome (prosome, macropain) 26S subunit, ATPase, 3 | Nucleus | transcription regulator | 9967899 | 8280221.5 | 7689300.5 | | | | | | |
| O88685 | PSMC3 | proteasome (prosome, macropain) 26S subunit, ATPase, 3 | Nucleus | transcription regulator | 6849121 | 5442777.5 | 5021381 | | | | | | |
| O88685 | PSMC3 | proteasome (prosome, macropain) 26S subunit, ATPase, 3 | Nucleus | transcription regulator | 6448726.5 | 6803987 | 4787299.5 | | | | | | |
| O88685 | PSMC3 | proteasome (prosome, macropain) 26S subunit, ATPase, 3 | Nucleus | transcription regulator | 4771480 | 3978684.5 | 3503677.25 | | | | | | |
| O88685 | PSMC3 | proteasome (prosome, macropain) 26S subunit, ATPase, 3 | Nucleus | transcription regulator | 4547865 | 3858154 | 3346040.5 | | | | | | |
| O88685 | PSMC3 | proteasome (prosome, macropain) 26S subunit, ATPase, 3 | Nucleus | transcription regulator | 2561154.5 | 2592157.25 | 2823625.5 | | | | | | |
| O88685 | PSMC3 | proteasome (prosome, macropain) 26S subunit, ATPase, 3 | Nucleus | transcription regulator | 3407818.25 | 2989149 | 1714491.5 | O88685 | PSMC3 | 8967428 | 4429476 | 683483 | 42 |
| P54775 | PSMC4 | proteasome (prosome, macropain) 26S subunit, ATPase, 4 | Nucleus | peptidase | 1.54E+07 | 1.43E+07 | 1.14E+07 | | | | | | |
| P54775 | PSMC4 | proteasome (prosome, macropain) 26S subunit, ATPase, 4 | Nucleus | peptidase | 1.52E+07 | 1.38E+07 | 1.50E+07 | | | | | | |
| P54775 | PSMC4 | proteasome (prosome, macropain) 26S subunit, ATPase, 4 | Nucleus | peptidase | 6227909.5 | 5134844 | 5947450.5 | | | | | | |
| P54775 | PSMC4 | proteasome (prosome, macropain) 26S subunit, ATPase, 4 | Nucleus | peptidase | 5371516 | 6069131 | 4724187 | | | | | | |
| P54775 | PSMC4 | proteasome (prosome, macropain) 26S subunit, ATPase, 4 | Nucleus | peptidase | 5646229.5 | 5363019.5 | 5761739.5 | | | | | | |
| P54775 | PSMC4 | proteasome (prosome, macropain) 26S subunit, ATPase, 4 | Nucleus | peptidase | 8153787 | 6867819.5 | 7199881 | | | | | | |
| P54775 | PSMC4 | proteasome (prosome, macropain) 26S subunit, ATPase, 4 | Nucleus | peptidase | 4929998 | 4659200 | 4220942 | | | | | | |
| P54775 | PSMC4 | proteasome (prosome, macropain) 26S subunit, ATPase, 4 | Nucleus | peptidase | 3900116.75 | 2564878 | 1591164.875 | | | | | | |
| P54775 | PSMC4 | proteasome (prosome, macropain) 26S subunit, ATPase, 4 | Nucleus | peptidase | 3581575.25 | 3947031 | 3551259.5 | | | | | | |
| P54775 | PSMC4 | proteasome (prosome, macropain) 26S subunit, ATPase, 4 | Nucleus | peptidase | 5200934.5 | 4595224 | 4315306 | | | | | | |
| P54775 | PSMC4 | proteasome (prosome, macropain) 26S subunit, ATPase, 4 | Nucleus | peptidase | 4613935 | 3195666.75 | 2592475.25 | | | | | | |
| P54775 | PSMC4 | proteasome (prosome, macropain) 26S subunit, ATPase, 4 | Nucleus | peptidase | 3166812.5 | 2201433.5 | 2395043.25 | P54775 | PSMC4 | 6777735 | 3929202 | 654867 | 36 |
| P62196 | PSMC5 | proteasome (prosome, macropain) 26S subunit, ATPase, 5 | Nucleus | transcription regulator | 2.14E+07 | 2.03E+07 | 1.98E+07 | | | | | | |
| P62196 | PSMC5 | proteasome (prosome, macropain) 26S subunit, ATPase, 5 | Nucleus | transcription regulator | 1.42E+07 | 1.35E+07 | 9346106 | | | | | | |
| P62196 | PSMC5 | proteasome (prosome, macropain) 26S subunit, ATPase, 5 | Nucleus | transcription regulator | 1.36E+07 | 1.52E+07 | 1.19E+07 | | | | | | |
| P62196 | PSMC5 | proteasome (prosome, macropain) 26S subunit, ATPase, 5 | Nucleus | transcription regulator | 1.38E+07 | 1.20E+07 | 1.21E+07 | | | | | | |
| P62196 | PSMC5 | proteasome (prosome, macropain) 26S subunit, ATPase, 5 | Nucleus | transcription regulator | 1.08E+07 | 8187266.5 | 8178609 | | | | | | |
| P62196 | PSMC5 | proteasome (prosome, macropain) 26S subunit, ATPase, 5 | Nucleus | transcription regulator | 1.07E+07 | 8057299.5 | 7021126.5 | | | | | | |
| P62196 | PSMC5 | proteasome (prosome, macropain) 26S subunit, ATPase, 5 | Nucleus | transcription regulator | 7768669 | 6957871 | 5821428 | | | | | | |
| P62196 | PSMC5 | proteasome (prosome, macropain) 26S subunit, ATPase, 5 | Nucleus | transcription regulator | 9933829 | 5809328.5 | 5159116.5 | | | | | | |
| P62196 | PSMC5 | proteasome (prosome, macropain) 26S subunit, ATPase, 5 | Nucleus | transcription regulator | 8283370 | 6684824.5 | 7018414 | | | | | | |
| P62196 | PSMC5 | proteasome (prosome, macropain) 26S subunit, ATPase, 5 | Nucleus | transcription regulator | 4810403.5 | 3954340.5 | 4381965 | | | | | | |
| P62196 | PSMC5 | proteasome (prosome, macropain) 26S subunit, ATPase, 5 | Nucleus | transcription regulator | 5975665 | 4780080 | 4590744.5 | | | | | | |
| P62196 | PSMC5 | proteasome (prosome, macropain) 26S subunit, ATPase, 5 | Nucleus | transcription regulator | 5280460 | 6302173.5 | 5806341.5 | | | | | | |
| P62196 | PSMC5 | proteasome (prosome, macropain) 26S subunit, ATPase, 5 | Nucleus | transcription regulator | 6349345 | 3474209.25 | 3962711 | | | | | | |
| P62196 | PSMC5 | proteasome (prosome, macropain) 26S subunit, ATPase, 5 | Nucleus | transcription regulator | 4637293 | 4509898 | 3494467.25 | | | | | | |
| P62196 | PSMC5 | proteasome (prosome, macropain) 26S subunit, ATPase, 5 | Nucleus | transcription regulator | 4334364.5 | 4687054.5 | 2954589.75 | | | | | | |
| P62196 | PSMC5 | proteasome (prosome, macropain) 26S subunit, ATPase, 5 | Nucleus | transcription regulator | 1838154.125 | 698864.812 | 20012.359 | P62196 | PSMC5 | 8984472 | 4877231 | 703968 | 48 |
| P62334 | PSMC6 | proteasome (prosome, macropain) 26S subunit, ATPase, 6 | Nucleus | peptidase | 2.12E+07 | 1.66E+07 | 1.79E+07 | | | | | | |
| P62334 | PSMC6 | proteasome (prosome, macropain) 26S subunit, ATPase, 6 | Nucleus | peptidase | 1.29E+07 | 1.14E+07 | 6138646.5 | | | | | | |
| P62334 | PSMC6 | proteasome (prosome, macropain) 26S subunit, ATPase, 6 | Nucleus | peptidase | 1.43E+07 | 1.40E+07 | 9995409 | | | | | | |
| P62334 | PSMC6 | proteasome (prosome, macropain) 26S subunit, ATPase, 6 | Nucleus | peptidase | 1.01E+07 | 7575210 | 7804614 | | | | | | |
| P62334 | PSMC6 | proteasome (prosome, macropain) 26S subunit, ATPase, 6 | Nucleus | peptidase | 1.14E+07 | 6709256 | 8887571 | | | | | | |
| P62334 | PSMC6 | proteasome (prosome, macropain) 26S subunit, ATPase, 6 | Nucleus | peptidase | 1.12E+07 | 7621529.5 | 7065933.5 | | | | | | |
| P62334 | PSMC6 | proteasome (prosome, macropain) 26S subunit, ATPase, 6 | Nucleus | peptidase | 9540300 | 6993094 | 5965706.5 | | | | | | |
| P62334 | PSMC6 | proteasome (prosome, macropain) 26S subunit, ATPase, 6 | Nucleus | peptidase | 7955837 | 5254082 | 6220316.5 | | | | | | |
| P62334 | PSMC6 | proteasome (prosome, macropain) 26S subunit, ATPase, 6 | Nucleus | peptidase | 7895248 | 7214152.5 | 6865362.5 | | | | | | |
| P62334 | PSMC6 | proteasome (prosome, macropain) 26S subunit, ATPase, 6 | Nucleus | peptidase | 8149160 | 6155826.5 | 5505220 | | | | | | |
| P62334 | PSMC6 | proteasome (prosome, macropain) 26S subunit, ATPase, 6 | Nucleus | peptidase | 6112864.5 | 5407918 | 3894050.25 | | | | | | |
| P62334 | PSMC6 | proteasome (prosome, macropain) 26S subunit, ATPase, 6 | Nucleus | peptidase | 6581385.5 | 5803057 | 5308269 | | | | | | |
| P62334 | PSMC6 | proteasome (prosome, macropain) 26S subunit, ATPase, 6 | Nucleus | peptidase | 2343894 | 1367608.75 | 1347840.125 | P62334 | PSMC6 | 9971438 | 4236918 | 678450 | 39 |
| Q3TXS7 | PSMD1 | proteasome (prosome, macropain) 26S subunit, non-ATPase, 1 | Cytoplasm | other | 2.37E+07 | 1.94E+07 | 2.38E+07 | | | | | | |
| Q3TXS7 | PSMD1 | proteasome (prosome, macropain) 26S subunit, non-ATPase, 1 | Cytoplasm | other | 2.82E+07 | 2.16E+07 | 1.85E+07 | | | | | | |

| | | | | | | | | | | | | | |
|---|---|---|---|---|---|---|---|---|---|---|---|---|---|
| Q3TXS7 | PSMD1 | proteasome (prosome, macropain) 26S subunit, non-ATPase, 1 | Cytoplasm | other | 1.39E+07 | 1.09E+07 | 1.25E+07 | | | | | | |
| Q3TXS7 | PSMD1 | proteasome (prosome, macropain) 26S subunit, non-ATPase, 1 | Cytoplasm | other | 2.20E+07 | 1.91E+07 | 1.53E+07 | | | | | | |
| Q3TXS7 | PSMD1 | proteasome (prosome, macropain) 26S subunit, non-ATPase, 1 | Cytoplasm | other | 1.52E+07 | 1.20E+07 | 1.16E+07 | | | | | | |
| Q3TXS7 | PSMD1 | proteasome (prosome, macropain) 26S subunit, non-ATPase, 1 | Cytoplasm | other | 1.30E+07 | 1.02E+07 | 8241930 | | | | | | |
| Q3TXS7 | PSMD1 | proteasome (prosome, macropain) 26S subunit, non-ATPase, 1 | Cytoplasm | other | 1.22E+07 | 9738128 | 8618225 | | | | | | |
| Q3TXS7 | PSMD1 | proteasome (prosome, macropain) 26S subunit, non-ATPase, 1 | Cytoplasm | other | 9395632 | 1.14E+07 | 1.01E+07 | | | | | | |
| Q3TXS7 | PSMD1 | proteasome (prosome, macropain) 26S subunit, non-ATPase, 1 | Cytoplasm | other | 1.63E+07 | 1.44E+07 | 1.46E+07 | | | | | | |
| Q3TXS7 | PSMD1 | proteasome (prosome, macropain) 26S subunit, non-ATPase, 1 | Cytoplasm | other | 1.06E+07 | 1.18E+07 | 6987968.5 | | | | | | |
| Q3TXS7 | PSMD1 | proteasome (prosome, macropain) 26S subunit, non-ATPase, 1 | Cytoplasm | other | 7848980 | 7106705.5 | 7590859 | | | | | | |
| Q3TXS7 | PSMD1 | proteasome (prosome, macropain) 26S subunit, non-ATPase, 1 | Cytoplasm | other | 1.11E+07 | 1.09E+07 | 6505468.5 | | | | | | |
| Q3TXS7 | PSMD1 | proteasome (prosome, macropain) 26S subunit, non-ATPase, 1 | Cytoplasm | other | 1.06E+07 | 7969655.5 | 8597367 | | | | | | |
| Q3TXS7 | PSMD1 | proteasome (prosome, macropain) 26S subunit, non-ATPase, 1 | Cytoplasm | other | 9073851 | 7799905.5 | 6252476 | | | | | | |
| Q3TXS7 | PSMD1 | proteasome (prosome, macropain) 26S subunit, non-ATPase, 1 | Cytoplasm | other | 6090819 | 6288187 | 5808583 | | | | | | |
| Q3TXS7 | PSMD1 | proteasome (prosome, macropain) 26S subunit, non-ATPase, 1 | Cytoplasm | other | 6586496 | 5462001.5 | 4782651 | | | | | | |
| Q3TXS7 | PSMD1 | proteasome (prosome, macropain) 26S subunit, non-ATPase, 1 | Cytoplasm | other | 6263621 | 4988400 | 4493924 | | | | | | |
| Q3TXS7 | PSMD1 | proteasome (prosome, macropain) 26S subunit, non-ATPase, 1 | Cytoplasm | other | 7650152.5 | 6446232 | 6554836.5 | | | | | | |
| Q3TXS7 | PSMD1 | proteasome (prosome, macropain) 26S subunit, non-ATPase, 1 | Cytoplasm | other | 5215384.5 | 4856164.5 | 4540802.5 | | | | | | |
| Q3TXS7 | PSMD1 | proteasome (prosome, macropain) 26S subunit, non-ATPase, 1 | Cytoplasm | other | 4088922.25 | 3425697.75 | 3473848 | | | | | | |
| Q3TXS7 | PSMD1 | proteasome (prosome, macropain) 26S subunit, non-ATPase, 1 | Cytoplasm | other | 4601198.5 | 2454703.25 | 2425849.75 | | | | | | |
| Q3TXS7 | PSMD1 | proteasome (prosome, macropain) 26S subunit, non-ATPase, 1 | Cytoplasm | other | 3477343 | 4040209 | 3605013 | | | | | | |
| Q3TXS7 | PSMD1 | proteasome (prosome, macropain) 26S subunit, non-ATPase, 1 | Cytoplasm | other | 5451585 | 4281585.5 | 4257382 | | | | | | |
| Q3TXS7 | PSMD1 | proteasome (prosome, macropain) 26S subunit, non-ATPase, 1 | Cytoplasm | other | 4271614.5 | 4515246.5 | 2753342.25 | | | | | | |
| Q3TXS7 | PSMD1 | proteasome (prosome, macropain) 26S subunit, non-ATPase, 1 | Cytoplasm | other | 3998102.25 | 2489982.5 | 1805109.375 | | | | | | |
| Q3TXS7 | PSMD1 | proteasome (prosome, macropain) 26S subunit, non-ATPase, 1 | Cytoplasm | other | 1849735.5 | 1775738 | 1314738.875 | | | | | | |
| Q3TXS7 | PSMD1 | proteasome (prosome, macropain) 26S subunit, non-ATPase, 1 | Cytoplasm | other | 2359128 | 2676160.25 | 1380503.875 | | | | | | |
| Q3TXS7 | PSMD1 | proteasome (prosome, macropain) 26S subunit, non-ATPase, 1 | Cytoplasm | other | 2974839.5 | 1982221.375 | 1681598.75 | Q3TXS7 | PSMD1 | 9568122 | 5892532 | 642928 | 84 |
| Q8BG32 | PSMD11 | proteasome (prosome, macropain) 26S subunit, non-ATPase, 11 | Cytoplasm | other | 1.83E+07 | 1.59E+07 | 1.42E+07 | | | | | | |
| Q8BG32 | PSMD11 | proteasome (prosome, macropain) 26S subunit, non-ATPase, 11 | Cytoplasm | other | 1.60E+07 | 1.19E+07 | 1.16E+07 | | | | | | |
| Q8BG32 | PSMD11 | proteasome (prosome, macropain) 26S subunit, non-ATPase, 11 | Cytoplasm | other | 1.44E+07 | 1.55E+07 | 1.24E+07 | | | | | | |
| Q8BG32 | PSMD11 | proteasome (prosome, macropain) 26S subunit, non-ATPase, 11 | Cytoplasm | other | 1.30E+07 | 1.31E+07 | 1.91E+07 | | | | | | |
| Q8BG32 | PSMD11 | proteasome (prosome, macropain) 26S subunit, non-ATPase, 11 | Cytoplasm | other | 1.52E+07 | 1.05E+07 | 1.33E+07 | | | | | | |
| Q8BG32 | PSMD11 | proteasome (prosome, macropain) 26S subunit, non-ATPase, 11 | Cytoplasm | other | 1.18E+07 | 9687786 | 9544783 | | | | | | |
| Q8BG32 | PSMD11 | proteasome (prosome, macropain) 26S subunit, non-ATPase, 11 | Cytoplasm | other | 1.18E+07 | 5430366 | 4776115.5 | | | | | | |
| Q8BG32 | PSMD11 | proteasome (prosome, macropain) 26S subunit, non-ATPase, 11 | Cytoplasm | other | 1.29E+07 | 1.16E+07 | 8125908 | | | | | | |
| Q8BG32 | PSMD11 | proteasome (prosome, macropain) 26S subunit, non-ATPase, 11 | Cytoplasm | other | 6219726 | 5760739 | 5124109.5 | | | | | | |
| Q8BG32 | PSMD11 | proteasome (prosome, macropain) 26S subunit, non-ATPase, 11 | Cytoplasm | other | 3273490.75 | 6072024 | 5543062 | | | | | | |
| Q8BG32 | PSMD11 | proteasome (prosome, macropain) 26S subunit, non-ATPase, 11 | Cytoplasm | other | 6377058 | 3540875 | 3328673.5 | | | | | | |
| Q8BG32 | PSMD11 | proteasome (prosome, macropain) 26S subunit, non-ATPase, 11 | Cytoplasm | other | 4722587 | 3401383.5 | 2517062 | | | | | | |
| Q8BG32 | PSMD11 | proteasome (prosome, macropain) 26S subunit, non-ATPase, 11 | Cytoplasm | other | 3164314.75 | 2824549 | 2514496.25 | | | | | | |
| Q8BG32 | PSMD11 | proteasome (prosome, macropain) 26S subunit, non-ATPase, 11 | Cytoplasm | other | 1465853.5 | 2722047 | 3057714.25 | Q8BG32 | PSMD11 | 9889502 | 5086464 | 784858 | 42 |
| Q9D8W5 | PSMD12 | proteasome (prosome, macropain) 26S subunit, non-ATPase, 12 | Cytoplasm | other | 1.91E+07 | 1.49E+07 | 1.48E+07 | | | | | | |
| Q9D8W5 | PSMD12 | proteasome (prosome, macropain) 26S subunit, non-ATPase, 12 | Cytoplasm | other | 1.36E+07 | 1.14E+07 | 1.25E+07 | | | | | | |
| Q9D8W5 | PSMD12 | proteasome (prosome, macropain) 26S subunit, non-ATPase, 12 | Cytoplasm | other | 1.45E+07 | 1.47E+07 | 9133333 | | | | | | |
| Q9D8W5 | PSMD12 | proteasome (prosome, macropain) 26S subunit, non-ATPase, 12 | Cytoplasm | other | 1.08E+07 | 8201886 | 6719140.5 | | | | | | |
| Q9D8W5 | PSMD12 | proteasome (prosome, macropain) 26S subunit, non-ATPase, 12 | Cytoplasm | other | 9232250 | 8004341.5 | 8306850 | | | | | | |
| Q9D8W5 | PSMD12 | proteasome (prosome, macropain) 26S subunit, non-ATPase, 12 | Cytoplasm | other | 7246986.5 | 7940455 | 6894988 | | | | | | |
| Q9D8W5 | PSMD12 | proteasome (prosome, macropain) 26S subunit, non-ATPase, 12 | Cytoplasm | other | 8408759 | 7821661.5 | 6562943.5 | | | | | | |
| Q9D8W5 | PSMD12 | proteasome (prosome, macropain) 26S subunit, non-ATPase, 12 | Cytoplasm | other | 8191188.5 | 8455631 | 7926758.5 | | | | | | |
| Q9D8W5 | PSMD12 | proteasome (prosome, macropain) 26S subunit, non-ATPase, 12 | Cytoplasm | other | 7623435 | 6581718.5 | 6585996.5 | | | | | | |
| Q9D8W5 | PSMD12 | proteasome (prosome, macropain) 26S subunit, non-ATPase, 12 | Cytoplasm | other | 4357111 | 4150378.75 | 3486714.25 | | | | | | |
| Q9D8W5 | PSMD12 | proteasome (prosome, macropain) 26S subunit, non-ATPase, 12 | Cytoplasm | other | 6134949 | 3098170 | 3187783.25 | | | | | | |
| Q9D8W5 | PSMD12 | proteasome (prosome, macropain) 26S subunit, non-ATPase, 12 | Cytoplasm | other | 3868800 | 4134679 | 4084904.5 | | | | | | |
| Q9D8W5 | PSMD12 | proteasome (prosome, macropain) 26S subunit, non-ATPase, 12 | Cytoplasm | other | 1805256.875 | 2098831.25 | 1822791.75 | | | | | | |
| Q9D8W5 | PSMD12 | proteasome (prosome, macropain) 26S subunit, non-ATPase, 12 | Cytoplasm | other | 3672527.75 | 3248444.5 | 2619954 | | | | | | |
| Q9D8W5 | PSMD12 | proteasome (prosome, macropain) 26S subunit, non-ATPase, 12 | Cytoplasm | other | 1094317.25 | 1283616 | 1160526.875 | Q9D8W5 | PSMD12 | 7973039 | 4323642 | 644530 | 45 |
| Q9WVJ2 | PSMD13 | proteasome (prosome, macropain) 26S subunit, non-ATPase, 13 | Cytoplasm | peptidase | 2.70E+07 | 2.08E+07 | 2.35E+07 | | | | | | |
| Q9WVJ2 | PSMD13 | proteasome (prosome, macropain) 26S subunit, non-ATPase, 13 | Cytoplasm | peptidase | 2.58E+07 | 2.22E+07 | 1.93E+07 | | | | | | |
| Q9WVJ2 | PSMD13 | proteasome (prosome, macropain) 26S subunit, non-ATPase, 13 | Cytoplasm | peptidase | 1.88E+07 | 1.34E+07 | 9156134 | | | | | | |
| Q9WVJ2 | PSMD13 | proteasome (prosome, macropain) 26S subunit, non-ATPase, 13 | Cytoplasm | peptidase | 1.64E+07 | 1.58E+07 | 1.61E+07 | | | | | | |
| Q9WVJ2 | PSMD13 | proteasome (prosome, macropain) 26S subunit, non-ATPase, 13 | Cytoplasm | peptidase | 1.36E+07 | 1.17E+07 | 1.37E+07 | | | | | | |
| Q9WVJ2 | PSMD13 | proteasome (prosome, macropain) 26S subunit, non-ATPase, 13 | Cytoplasm | peptidase | 1.02E+07 | 8217688 | 9312454 | | | | | | |
| Q9WVJ2 | PSMD13 | proteasome (prosome, macropain) 26S subunit, non-ATPase, 13 | Cytoplasm | peptidase | 1.21E+07 | 8570817 | 8576518 | | | | | | |
| Q9WVJ2 | PSMD13 | proteasome (prosome, macropain) 26S subunit, non-ATPase, 13 | Cytoplasm | peptidase | 1.30E+07 | 9556992 | 9830291 | | | | | | |
| Q9WVJ2 | PSMD13 | proteasome (prosome, macropain) 26S subunit, non-ATPase, 13 | Cytoplasm | peptidase | 9541078 | 7979020.5 | 7828380.5 | | | | | | |
| Q9WVJ2 | PSMD13 | proteasome (prosome, macropain) 26S subunit, non-ATPase, 13 | Cytoplasm | peptidase | 1.18E+07 | 1.03E+07 | 9187778 | | | | | | |
| Q9WVJ2 | PSMD13 | proteasome (prosome, macropain) 26S subunit, non-ATPase, 13 | Cytoplasm | peptidase | 8537423 | 5678320.5 | 7004125 | | | | | | |
| Q9WVJ2 | PSMD13 | proteasome (prosome, macropain) 26S subunit, non-ATPase, 13 | Cytoplasm | peptidase | 5217888.5 | 4749340.5 | 4235879.5 | | | | | | |
| Q9WVJ2 | PSMD13 | proteasome (prosome, macropain) 26S subunit, non-ATPase, 13 | Cytoplasm | peptidase | 1616408 | 1401996.25 | 1807897.125 | Q9WVJ2 | PSMD13 | 13355600 | 6408420 | 1026169 | 39 |
| O35593 | PSMD14 | proteasome (prosome, macropain) 26S subunit, non-ATPase, 14 | Cytoplasm | peptidase | 2.12E+07 | 2.08E+07 | 1.68E+07 | | | | | | |
| O35593 | PSMD14 | proteasome (prosome, macropain) 26S subunit, non-ATPase, 14 | Cytoplasm | peptidase | 1.47E+07 | 1.40E+07 | 1.34E+07 | | | | | | |
| O35593 | PSMD14 | proteasome (prosome, macropain) 26S subunit, non-ATPase, 14 | Cytoplasm | peptidase | 5108735 | 3841567.5 | 3683029 | | | | | | |
| O35593 | PSMD14 | proteasome (prosome, macropain) 26S subunit, non-ATPase, 14 | Cytoplasm | peptidase | 3760761 | 3583917.75 | 2304154.25 | O35593 | PSMD14 | 11189874 | 7244049 | 2091177 | 12 |
| Q8VDM4 | PSMD2 | proteasome (prosome, macropain) 26S subunit, non-ATPase, 2 | Cytoplasm | other | 2.60E+07 | 1.47E+07 | 8731980 | | | | | | |
| Q8VDM4 | PSMD2 | proteasome (prosome, macropain) 26S subunit, non-ATPase, 2 | Cytoplasm | other | 2.39E+07 | 1.68E+07 | 1.42E+07 | | | | | | |
| Q8VDM4 | PSMD2 | proteasome (prosome, macropain) 26S subunit, non-ATPase, 2 | Cytoplasm | other | 1.94E+07 | 1.13E+07 | 1.06E+07 | | | | | | |
| Q8VDM4 | PSMD2 | proteasome (prosome, macropain) 26S subunit, non-ATPase, 2 | Cytoplasm | other | 9150205 | 7292684 | 1.06E+07 | | | | | | |
| Q8VDM4 | PSMD2 | proteasome (prosome, macropain) 26S subunit, non-ATPase, 2 | Cytoplasm | other | 1.11E+07 | 8128463 | 8901602 | | | | | | |
| Q8VDM4 | PSMD2 | proteasome (prosome, macropain) 26S subunit, non-ATPase, 2 | Cytoplasm | other | 1.26E+07 | 7164632 | 1.10E+07 | | | | | | |
| Q8VDM4 | PSMD2 | proteasome (prosome, macropain) 26S subunit, non-ATPase, 2 | Cytoplasm | other | 1.30E+07 | 1.21E+07 | 1.15E+07 | | | | | | |
| Q8VDM4 | PSMD2 | proteasome (prosome, macropain) 26S subunit, non-ATPase, 2 | Cytoplasm | other | 9785585 | 8561966 | 7963719.5 | | | | | | |
| Q8VDM4 | PSMD2 | proteasome (prosome, macropain) 26S subunit, non-ATPase, 2 | Cytoplasm | other | 1.00E+07 | 9045213 | 7807220.5 | | | | | | |
| Q8VDM4 | PSMD2 | proteasome (prosome, macropain) 26S subunit, non-ATPase, 2 | Cytoplasm | other | 8577803 | 8421605 | 5924073.5 | | | | | | |
| Q8VDM4 | PSMD2 | proteasome (prosome, macropain) 26S subunit, non-ATPase, 2 | Cytoplasm | other | 7135240.5 | 6815549 | 7643328 | | | | | | |
| Q8VDM4 | PSMD2 | proteasome (prosome, macropain) 26S subunit, non-ATPase, 2 | Cytoplasm | other | 1.13E+07 | 8672935 | 8174938.5 | | | | | | |
| Q8VDM4 | PSMD2 | proteasome (prosome, macropain) 26S subunit, non-ATPase, 2 | Cytoplasm | other | 7064147 | 4911855 | 4469502 | | | | | | |
| Q8VDM4 | PSMD2 | proteasome (prosome, macropain) 26S subunit, non-ATPase, 2 | Cytoplasm | other | 7216951 | 8256347.5 | 5656143.5 | | | | | | |
| Q8VDM4 | PSMD2 | proteasome (prosome, macropain) 26S subunit, non-ATPase, 2 | Cytoplasm | other | 7068394.5 | 6960845.5 | 5771602.5 | | | | | | |
| Q8VDM4 | PSMD2 | proteasome (prosome, macropain) 26S subunit, non-ATPase, 2 | Cytoplasm | other | 1.06E+07 | 8869919 | 4420215.5 | | | | | | |
| Q8VDM4 | PSMD2 | proteasome (prosome, macropain) 26S subunit, non-ATPase, 2 | Cytoplasm | other | 7343054.5 | 5015440 | 5702975 | | | | | | |
| Q8VDM4 | PSMD2 | proteasome (prosome, macropain) 26S subunit, non-ATPase, 2 | Cytoplasm | other | 4473940.5 | 5002577.5 | 3456011.25 | | | | | | |
| Q8VDM4 | PSMD2 | proteasome (prosome, macropain) 26S subunit, non-ATPase, 2 | Cytoplasm | other | 6244349.5 | 4821946.5 | 6372830.5 | | | | | | |
| Q8VDM4 | PSMD2 | proteasome (prosome, macropain) 26S subunit, non-ATPase, 2 | Cytoplasm | other | 3541623.75 | 4620546.5 | 4831036 | | | | | | |
| Q8VDM4 | PSMD2 | proteasome (prosome, macropain) 26S subunit, non-ATPase, 2 | Cytoplasm | other | 6022063 | 3820950.75 | 4100550.25 | | | | | | |
| Q8VDM4 | PSMD2 | proteasome (prosome, macropain) 26S subunit, non-ATPase, 2 | Cytoplasm | other | 4464131 | 3747557 | 5543493 | | | | | | |
| Q8VDM4 | PSMD2 | proteasome (prosome, macropain) 26S subunit, non-ATPase, 2 | Cytoplasm | other | 4529065.5 | 3611482.5 | 3108828.25 | | | | | | |
| Q8VDM4 | PSMD2 | proteasome (prosome, macropain) 26S subunit, non-ATPase, 2 | Cytoplasm | other | 2303883.75 | 2064598.875 | 2826147.25 | | | | | | |
| Q8VDM4 | PSMD2 | proteasome (prosome, macropain) 26S subunit, non-ATPase, 2 | Cytoplasm | other | 3137045.75 | 2015589.875 | 2138513.75 | | | | | | |
| Q8VDM4 | PSMD2 | proteasome (prosome, macropain) 26S subunit, non-ATPase, 2 | Cytoplasm | other | 2335905.75 | 2548930.5 | 1751827.125 | | | | | | |
| Q8VDM4 | PSMD2 | proteasome (prosome, macropain) 26S subunit, non-ATPase, 2 | Cytoplasm | other | 3182854 | 2230761.25 | 2947740.5 | | | | | | |
| Q8VDM4 | PSMD2 | proteasome (prosome, macropain) 26S subunit, non-ATPase, 2 | Cytoplasm | other | 2377347.75 | 2403252.25 | 3157228.75 | | | | | | |
| Q8VDM4 | PSMD2 | proteasome (prosome, macropain) 26S subunit, non-ATPase, 2 | Cytoplasm | other | 2837019 | 3040518.25 | 1630709.5 | Q8VDM4 | PSMD2 | 8505883 | 4573031 | 490280 | 87 |
| P14685 | PSMD3 | proteasome (prosome, macropain) 26S subunit, non-ATPase, 3 | Cytoplasm | other | 2.37E+07 | 2.04E+07 | 1.78E+07 | | | | | | |
| P14685 | PSMD3 | proteasome (prosome, macropain) 26S subunit, non-ATPase, 3 | Cytoplasm | other | 2.62E+07 | 2.09E+07 | 2.09E+07 | | | | | | |
| P14685 | PSMD3 | proteasome (prosome, macropain) 26S subunit, non-ATPase, 3 | Cytoplasm | other | 1.92E+07 | 1.65E+07 | 1.98E+07 | | | | | | |
| P14685 | PSMD3 | proteasome (prosome, macropain) 26S subunit, non-ATPase, 3 | Cytoplasm | other | 1.26E+07 | 9693606 | 1.21E+07 | | | | | | |
| P14685 | PSMD3 | proteasome (prosome, macropain) 26S subunit, non-ATPase, 3 | Cytoplasm | other | 1.22E+07 | 8913818 | 7873047 | | | | | | |
| P14685 | PSMD3 | proteasome (prosome, macropain) 26S subunit, non-ATPase, 3 | Cytoplasm | other | 1.05E+07 | 7656377.5 | 9429808 | | | | | | |
| P14685 | PSMD3 | proteasome (prosome, macropain) 26S subunit, non-ATPase, 3 | Cytoplasm | other | 1.05E+07 | 8731847 | 8046978.5 | | | | | | |
| P14685 | PSMD3 | proteasome (prosome, macropain) 26S subunit, non-ATPase, 3 | Cytoplasm | other | 9607154 | 7404432 | 9108504 | | | | | | |

| | | | | | | | | | | | | | |
|---|---|---|---|---|---|---|---|---|---|---|---|---|---|
| P14685 | PSMD3 | proteasome (prosome, macropain) 26S subunit, non-ATPase, 3 | Cytoplasm | other | 3184846.5 | 3442980.5 | 4638962 | | | | | | |
| P14685 | PSMD3 | proteasome (prosome, macropain) 26S subunit, non-ATPase, 3 | Cytoplasm | other | 5349932 | 4521488 | 4802610.5 | | | | | | |
| P14685 | PSMD3 | proteasome (prosome, macropain) 26S subunit, non-ATPase, 3 | Cytoplasm | other | 5563393.5 | 5323329 | 5395292.5 | | | | | | |
| P14685 | PSMD3 | proteasome (prosome, macropain) 26S subunit, non-ATPase, 3 | Cytoplasm | other | 2459812.75 | 1462298.5 | 1055291.25 | | | | | | |
| P14685 | PSMD3 | proteasome (prosome, macropain) 26S subunit, non-ATPase, 3 | Cytoplasm | other | 2839503.75 | 2800186.25 | 2342212.5 | | | | | | |
| P14685 | PSMD3 | proteasome (prosome, macropain) 26S subunit, non-ATPase, 3 | Cytoplasm | other | 2385118.5 | 1885519.375 | 1934572.25 | | | | | | |
| P14685 | PSMD3 | proteasome (prosome, macropain) 26S subunit, non-ATPase, 3 | Cytoplasm | other | 2959928.5 | 861514.375 | 2737396.5 | | | | | | |
| P14685 | PSMD3 | proteasome (prosome, macropain) 26S subunit, non-ATPase, 3 | Cytoplasm | other | 3416495 | 3300098 | 2960228 | P14685 | PSMD3 | 9545387 | 6769419 | 977081 | 48 |
| O35226 | PSMD4 | proteasome (prosome, macropain) 26S subunit, non-ATPase, 4 | Cytoplasm | other | 2.85E+07 | 2.54E+07 | 2.22E+07 | | | | | | |
| O35226 | PSMD4 | proteasome (prosome, macropain) 26S subunit, non-ATPase, 4 | Cytoplasm | other | 1.90E+07 | 1.79E+07 | 2.23E+07 | | | | | | |
| O35226 | PSMD4 | proteasome (prosome, macropain) 26S subunit, non-ATPase, 4 | Cytoplasm | other | 1.00E+07 | 1.08E+07 | 1.33E+07 | | | | | | |
| O35226 | PSMD4 | proteasome (prosome, macropain) 26S subunit, non-ATPase, 4 | Cytoplasm | other | 1.12E+07 | 7427937 | 7751822 | | | | | | |
| O35226 | PSMD4 | proteasome (prosome, macropain) 26S subunit, non-ATPase, 4 | Cytoplasm | other | 1.28E+07 | 1.11E+07 | 7675408 | | | | | | |
| O35226 | PSMD4 | proteasome (prosome, macropain) 26S subunit, non-ATPase, 4 | Cytoplasm | other | 5099646.5 | 2362481.5 | 4850961 | | | | | | |
| O35226 | PSMD4 | proteasome (prosome, macropain) 26S subunit, non-ATPase, 4 | Cytoplasm | other | 3812601 | 2725604.5 | 2553482.25 | | | | | | |
| O35226 | PSMD4 | proteasome (prosome, macropain) 26S subunit, non-ATPase, 4 | Cytoplasm | other | 3311291 | 2006488.25 | 1755327.25 | | | | | | |
| O35226 | PSMD4 | proteasome (prosome, macropain) 26S subunit, non-ATPase, 4 | Cytoplasm | other | 2160906 | 1444249.625 | 1739113.75 | O35226 | PSMD4 | 10644938 | 8075945 | 1554216 | 27 |
| Q8BJY1 | PSMD5 | proteasome (prosome, macropain) 26S subunit, non-ATPase, 5 | Cytoplasm | other | 3196492.25 | 1805807.125 | 2207654.25 | | | | | | |
| Q8BJY1 | PSMD5 | proteasome (prosome, macropain) 26S subunit, non-ATPase, 5 | Cytoplasm | other | 2901529.25 | 2140291.75 | 2950031 | | | | | | |
| Q8BJY1 | PSMD5 | proteasome (prosome, macropain) 26S subunit, non-ATPase, 5 | Cytoplasm | other | 1957108.75 | 1231100.5 | 1733061.375 | | | | | | |
| Q8BJY1 | PSMD5 | proteasome (prosome, macropain) 26S subunit, non-ATPase, 5 | Cytoplasm | other | 3009849.25 | 2122030.75 | 2434554.25 | | | | | | |
| Q8BJY1 | PSMD5 | proteasome (prosome, macropain) 26S subunit, non-ATPase, 5 | Cytoplasm | other | 1735811.375 | 1184212.625 | 1819086.5 | | | | | | |
| Q8BJY1 | PSMD5 | proteasome (prosome, macropain) 26S subunit, non-ATPase, 5 | Cytoplasm | other | 1080263.75 | 745328.938 | 1055457.75 | | | | | | |
| Q8BJY1 | PSMD5 | proteasome (prosome, macropain) 26S subunit, non-ATPase, 5 | Cytoplasm | other | 1208270 | 1076995.375 | 1079534.5 | | | | | | |
| Q8BJY1 | PSMD5 | proteasome (prosome, macropain) 26S subunit, non-ATPase, 5 | Cytoplasm | other | 1619140.625 | 1260639.25 | 1787680.25 | | | | | | |
| Q8BJY1 | PSMD5 | proteasome (prosome, macropain) 26S subunit, non-ATPase, 5 | Cytoplasm | other | 990683.312 | 482469.938 | 634100.625 | Q8BJY1 | PSMD5 | 1966572 | 753394 | 144991 | 27 |
| Q99JI4 | PSMD6 | proteasome (prosome, macropain) 26S subunit, non-ATPase, 6 | Cytoplasm | enzyme | 2.06E+07 | 1.51E+07 | 1.18E+07 | | | | | | |
| Q99JI4 | PSMD6 | proteasome (prosome, macropain) 26S subunit, non-ATPase, 6 | Cytoplasm | enzyme | 1.71E+07 | 1.47E+07 | 1.29E+07 | | | | | | |
| Q99JI4 | PSMD6 | proteasome (prosome, macropain) 26S subunit, non-ATPase, 6 | Cytoplasm | enzyme | 1.56E+07 | 1.02E+07 | 8546860 | | | | | | |
| Q99JI4 | PSMD6 | proteasome (prosome, macropain) 26S subunit, non-ATPase, 6 | Cytoplasm | enzyme | 1.23E+07 | 9605024 | 7620126.5 | | | | | | |
| Q99JI4 | PSMD6 | proteasome (prosome, macropain) 26S subunit, non-ATPase, 6 | Cytoplasm | enzyme | 1.06E+07 | 8751412 | 8575205 | | | | | | |
| Q99JI4 | PSMD6 | proteasome (prosome, macropain) 26S subunit, non-ATPase, 6 | Cytoplasm | enzyme | 8960793 | 6010940.5 | 7022406 | | | | | | |
| Q99JI4 | PSMD6 | proteasome (prosome, macropain) 26S subunit, non-ATPase, 6 | Cytoplasm | enzyme | 7171789 | 7929061.5 | 7444052.5 | | | | | | |
| Q99JI4 | PSMD6 | proteasome (prosome, macropain) 26S subunit, non-ATPase, 6 | Cytoplasm | enzyme | 4789742.5 | 3558811.5 | 3195595.25 | | | | | | |
| Q99JI4 | PSMD6 | proteasome (prosome, macropain) 26S subunit, non-ATPase, 6 | Cytoplasm | enzyme | 4696649.5 | 4168514.25 | 3431921 | | | | | | |
| Q99JI4 | PSMD6 | proteasome (prosome, macropain) 26S subunit, non-ATPase, 6 | Cytoplasm | enzyme | 4292223 | 2787425.5 | 4011650.25 | | | | | | |
| Q99JI4 | PSMD6 | proteasome (prosome, macropain) 26S subunit, non-ATPase, 6 | Cytoplasm | enzyme | 4761182.5 | 4275875 | 4825152.5 | | | | | | |
| Q99JI4 | PSMD6 | proteasome (prosome, macropain) 26S subunit, non-ATPase, 6 | Cytoplasm | enzyme | 2480046.25 | 1848070 | 2416337.75 | | | | | | |
| Q99JI4 | PSMD6 | proteasome (prosome, macropain) 26S subunit, non-ATPase, 6 | Cytoplasm | enzyme | 5078582 | 5292175.5 | 5777016 | | | | | | |
| Q99JI4 | PSMD6 | proteasome (prosome, macropain) 26S subunit, non-ATPase, 6 | Cytoplasm | enzyme | 1833850 | 1836526.25 | 1518164.125 | Q99JI4 | PSMD6 | 8588204 | 4632722 | 714845 | 42 |
| P26516 | PSMD7 | proteasome (prosome, macropain) 26S subunit, non-ATPase, 7 | Cytoplasm | other | 2.81E+07 | 2.48E+07 | 2.42E+07 | | | | | | |
| P26516 | PSMD7 | proteasome (prosome, macropain) 26S subunit, non-ATPase, 7 | Cytoplasm | other | 1.94E+07 | 1.46E+07 | 1.38E+07 | | | | | | |
| P26516 | PSMD7 | proteasome (prosome, macropain) 26S subunit, non-ATPase, 7 | Cytoplasm | other | 1.99E+07 | 1.47E+07 | 1.72E+07 | | | | | | |
| P26516 | PSMD7 | proteasome (prosome, macropain) 26S subunit, non-ATPase, 7 | Cytoplasm | other | 1.37E+07 | 1.01E+07 | 7546316.5 | | | | | | |
| P26516 | PSMD7 | proteasome (prosome, macropain) 26S subunit, non-ATPase, 7 | Cytoplasm | other | 9728156 | 7014414 | 6453158.5 | | | | | | |
| P26516 | PSMD7 | proteasome (prosome, macropain) 26S subunit, non-ATPase, 7 | Cytoplasm | other | 5737452 | 3449969.5 | 3925506 | | | | | | |
| P26516 | PSMD7 | proteasome (prosome, macropain) 26S subunit, non-ATPase, 7 | Cytoplasm | other | 6803779.5 | 7791048 | 6654797 | | | | | | |
| P26516 | PSMD7 | proteasome (prosome, macropain) 26S subunit, non-ATPase, 7 | Cytoplasm | other | 4373417 | 3742084.75 | 4152199.25 | | | | | | |
| P26516 | PSMD7 | proteasome (prosome, macropain) 26S subunit, non-ATPase, 7 | Cytoplasm | other | 3612494.25 | 2339119 | 2704909.25 | | | | | | |
| P26516 | PSMD7 | proteasome (prosome, macropain) 26S subunit, non-ATPase, 7 | Cytoplasm | other | 1540486.125 | 537810.5 | 24553.502 | P26516 | PSMD7 | 11288578 | 7703849 | 1406524 | 30 |
| Q9CX56 | PSMD8 | proteasome (prosome, macropain) 26S subunit, non-ATPase, 8 | Cytoplasm | other | 1.38E+07 | 1.16E+07 | 1.38E+07 | | | | | | |
| Q9CX56 | PSMD8 | proteasome (prosome, macropain) 26S subunit, non-ATPase, 8 | Cytoplasm | other | 7476230.5 | 5836211 | 6329502.5 | | | | | | |
| Q9CX56 | PSMD8 | proteasome (prosome, macropain) 26S subunit, non-ATPase, 8 | Cytoplasm | other | 7191097 | 5524024 | 6184155.5 | Q9CX56 | PSMD8 | 9499109 | 3444115 | 1148038 | 9 |
| P97371 | PSME1 | proteasome (prosome, macropain) activator subunit 1 (PA28 alpha) | Cytoplasm | other | 1.79E+07 | 1.51E+07 | 1.03E+07 | | | | | | |
| P97371 | PSME1 | proteasome (prosome, macropain) activator subunit 1 (PA28 alpha) | Cytoplasm | other | 1.37E+07 | 1.08E+07 | 9569131 | | | | | | |
| P97371 | PSME1 | proteasome (prosome, macropain) activator subunit 1 (PA28 alpha) | Cytoplasm | other | 1.67E+07 | 1.17E+07 | 1.12E+07 | | | | | | |
| P97371 | PSME1 | proteasome (prosome, macropain) activator subunit 1 (PA28 alpha) | Cytoplasm | other | 1.23E+07 | 8979813 | 7500235.5 | | | | | | |
| P97371 | PSME1 | proteasome (prosome, macropain) activator subunit 1 (PA28 alpha) | Cytoplasm | other | 1.32E+07 | 9536314 | 7432403.5 | | | | | | |
| P97371 | PSME1 | proteasome (prosome, macropain) activator subunit 1 (PA28 alpha) | Cytoplasm | other | 6946932.5 | 5036468.5 | 4552146 | | | | | | |
| P97371 | PSME1 | proteasome (prosome, macropain) activator subunit 1 (PA28 alpha) | Cytoplasm | other | 6882710 | 4967364 | 4109677.25 | | | | | | |
| P97371 | PSME1 | proteasome (prosome, macropain) activator subunit 1 (PA28 alpha) | Cytoplasm | other | 7441411 | 5401634 | 5169714.5 | | | | | | |
| P97371 | PSME1 | proteasome (prosome, macropain) activator subunit 1 (PA28 alpha) | Cytoplasm | other | 8049408.5 | 5433844 | 4379826 | | | | | | |
| P97371 | PSME1 | proteasome (prosome, macropain) activator subunit 1 (PA28 alpha) | Cytoplasm | other | 5842245.5 | 4157455.25 | 3225870.75 | | | | | | |
| P97371 | PSME1 | proteasome (prosome, macropain) activator subunit 1 (PA28 alpha) | Cytoplasm | other | 6221563 | 5427053 | 5009148 | | | | | | |
| P97371 | PSME1 | proteasome (prosome, macropain) activator subunit 1 (PA28 alpha) | Cytoplasm | other | 2772350.5 | 919109.188 | 1426273.875 | P97371 | PSME1 | 9828885 | 4161142 | 693524 | 36 |
| P97372 | PSME2 | proteasome (prosome, macropain) activator subunit 2 (PA28 beta) | Cytoplasm | peptidase | 2.62E+07 | 2.50E+07 | 1.97E+07 | | | | | | |
| P97372 | PSME2 | proteasome (prosome, macropain) activator subunit 2 (PA28 beta) | Cytoplasm | peptidase | 1.74E+07 | 1.19E+07 | 8677698 | | | | | | |
| P97372 | PSME2 | proteasome (prosome, macropain) activator subunit 2 (PA28 beta) | Cytoplasm | peptidase | 1.42E+07 | 9430663 | 8105889 | | | | | | |
| P97372 | PSME2 | proteasome (prosome, macropain) activator subunit 2 (PA28 beta) | Cytoplasm | peptidase | 5785290.5 | 4096345.5 | 4060207.5 | | | | | | |
| P97372 | PSME2 | proteasome (prosome, macropain) activator subunit 2 (PA28 beta) | Cytoplasm | peptidase | 2267869.75 | 1382488.75 | 1559257.5 | | | | | | |
| P97372 | PSME2 | proteasome (prosome, macropain) activator subunit 2 (PA28 beta) | Cytoplasm | peptidase | 3467326.5 | 2648583 | 1650906.25 | | | | | | |
| P97372 | PSME2 | proteasome (prosome, macropain) activator subunit 2 (PA28 beta) | Cytoplasm | peptidase | 4546964 | 1509023.75 | 2961534.25 | P97372 | PSME2 | 10546779 | 7811415 | 1704590 | 21 |
| Q8R3K3 | PTCD2 | pentatricopeptide repeat domain 2 | Cytoplasm | other | 2706366.25 | 2771458.5 | 2998600.75 | Q8R3K3 | PTCD2 | 2706366 | 153423 | 88579 | 3 |
| O08586 | PTEN | phosphatase and tensin homolog | Cytoplasm | phosphatase | 3.28E+07 | 3.01E+07 | 2.56E+07 | O08586 | PTEN | 32760000 | 3602254 | 2079762 | 3 |
| Q63739 | PTP4A1 | protein tyrosine phosphatase type IVA, member 1 | Cytoplasm | phosphatase | 1912708 | 2272255.5 | 1974969.5 | Q63739 | PTP4A1 | 1912708 | 192150 | 110938 | 3 |
| P35821 | PTPN1 | protein tyrosine phosphatase, non-receptor type 1 | Cytoplasm | phosphatase | 3248160.75 | 3347452.75 | 4242251.5 | P35821 | PTPN1 | 3248161 | 547531 | 316117 | 3 |
| Q62136 | PTPN21 | protein tyrosine phosphatase, non-receptor type 21 | Cytoplasm | phosphatase | 2.18E+07 | 2.14E+07 | 2.47E+07 | Q62136 | PTPN21 | 21760000 | 1783003 | 1029417 | 3 |
| P18052 | PTPRA | protein tyrosine phosphatase, receptor type, A | Plasma Membrane | phosphatase | 7485452 | 7071959.5 | 9678669 | | | | | | |
| P18052 | PTPRA | protein tyrosine phosphatase, receptor type, A | Plasma Membrane | phosphatase | 3812381.25 | 5290587.5 | 3822461 | | | | | | |
| P18052 | PTPRA | protein tyrosine phosphatase, receptor type, A | Plasma Membrane | phosphatase | 2973252.5 | 3111866 | 2919579.5 | | | | | | |
| P18052 | PTPRA | protein tyrosine phosphatase, receptor type, A | Plasma Membrane | phosphatase | 275440.438 | 650665.312 | 684913.188 | P18052 | PTPRA | 3636632 | 2933370 | 846791 | 12 |
| Q64455 | PTPRJ | protein tyrosine phosphatase, receptor type, J | Plasma Membrane | phosphatase | 5385900 | 6960553.5 | 7261877.5 | | | | | | |
| Q64455 | PTPRJ | protein tyrosine phosphatase, receptor type, J | Plasma Membrane | phosphatase | 538985.938 | 532723.25 | 1209420.625 | | | | | | |
| Q64455 | PTPRJ | protein tyrosine phosphatase, receptor type, J | Plasma Membrane | phosphatase | 851782.938 | 838991.188 | 1184631 | Q64455 | PTPRJ | 2258890 | 2892179 | 964060 | 9 |
| Q8R2Y8 | PTRH2 | peptidyl-tRNA hydrolase 2 | Cytoplasm | enzyme | 5258049.5 | 1.06E+07 | 8957165 | | | | | | |
| Q8R2Y8 | PTRH2 | peptidyl-tRNA hydrolase 2 | Cytoplasm | enzyme | 173171.781 | 271731.594 | 252732.547 | Q8R2Y8 | PTRH2 | 2715611 | 4725839 | 1929316 | 6 |
| P42669 | PURA | purine-rich element binding protein A | Nucleus | transcription regulator | 4774978.5 | 4452424 | 2422487.75 | | | | | | |
| P42669 | PURA | purine-rich element binding protein A | Nucleus | transcription regulator | 321694.812 | 538548.375 | 493613.781 | P42669 | PURA | 2548337 | 2046813 | 835608 | 6 |
| Q922Q4 | PYCR2 | pyrroline-5-carboxylate reductase family, member 2 | Cytoplasm | enzyme | 1861998.375 | 4757415 | 4489693.5 | Q922Q4 | PYCR2 | 1861998 | 1599994 | 923757 | 3 |
| Q9DCC4 | PYCRL | pyrroline-5-carboxylate reductase-like | Cytoplasm | enzyme | 5266076 | 3366959.25 | 2056721.125 | | | | | | |
| Q9DCC4 | PYCRL | pyrroline-5-carboxylate reductase-like | Cytoplasm | enzyme | 2854371.5 | 2298212 | 1961869.125 | | | | | | |
| Q9DCC4 | PYCRL | pyrroline-5-carboxylate reductase-like | Cytoplasm | enzyme | 2580171 | 1965594.375 | 1571030.25 | Q9DCC4 | PYCRL | 3566873 | 1117241 | 372414 | 9 |
| Q61838 | Pzp | pregnancy zone protein | Extracellular Space | other | 2784258 | 2161144.5 | 5717119.5 | Q61838 | Pzp | 2784258 | 1898899 | 1096330 | 3 |
| Q8BVI4 | QDPR | quinoid dihydropteridine reductase | Cytoplasm | enzyme | 1443077 | 887214.438 | 898254.812 | Q8BVI4 | QDPR | 1443077 | 317788 | 183475 | 3 |
| P61027 | RAB10 | RAB10, member RAS oncogene family | Cytoplasm | enzyme | 2.78E+08 | 2.85E+08 | 2.71E+08 | | | | | | |
| P61027 | RAB10 | RAB10, member RAS oncogene family | Cytoplasm | enzyme | 1.50E+08 | 1.47E+08 | 1.19E+08 | | | | | | |
| P61027 | RAB10 | RAB10, member RAS oncogene family | Cytoplasm | enzyme | 1.39E+08 | 1.56E+08 | 1.34E+08 | | | | | | |
| P61027 | RAB10 | RAB10, member RAS oncogene family | Cytoplasm | enzyme | 3.43E+07 | 4.87E+07 | 2.66E+07 | | | | | | |
| P61027 | RAB10 | RAB10, member RAS oncogene family | Cytoplasm | enzyme | 4.59E+07 | 4.34E+07 | 3.46E+07 | | | | | | |
| P61027 | RAB10 | RAB10, member RAS oncogene family | Cytoplasm | enzyme | 4.55E+07 | 5.13E+07 | 3.60E+07 | | | | | | |
| P61027 | RAB10 | RAB10, member RAS oncogene family | Cytoplasm | enzyme | 3.50E+07 | 3.38E+07 | 2.87E+07 | | | | | | |
| P61027 | RAB10 | RAB10, member RAS oncogene family | Cytoplasm | enzyme | 2.29E+07 | 2.32E+07 | 1.79E+07 | | | | | | |
| P61027 | RAB10 | RAB10, member RAS oncogene family | Cytoplasm | enzyme | 1.76E+07 | 2.97E+07 | 3.34E+07 | | | | | | |
| P61027 | RAB10 | RAB10, member RAS oncogene family | Cytoplasm | enzyme | 1.52E+07 | 1.77E+07 | 1.58E+07 | | | | | | |
| P61027 | RAB10 | RAB10, member RAS oncogene family | Cytoplasm | enzyme | 4401996 | 4603608 | 4197872.5 | P61027 | RAB10 | 71611091 | 80505462 | 14014202 | 33 |
| P46638 | RAB11B | RAB11B, member RAS oncogene family | Cytoplasm | enzyme | 3.15E+07 | 3.87E+07 | 4.42E+07 | | | | | | |
| P46638 | RAB11B | RAB11B, member RAS oncogene family | Cytoplasm | enzyme | 2.22E+07 | 2.61E+07 | 2.58E+07 | | | | | | |
| P46638 | RAB11B | RAB11B, member RAS oncogene family | Cytoplasm | enzyme | 2.03E+07 | 2.56E+07 | 2.99E+07 | | | | | | |
| P46638 | RAB11B | RAB11B, member RAS oncogene family | Cytoplasm | enzyme | 1.41E+07 | 2.02E+07 | 2.41E+07 | | | | | | |

| | | | | | | | | | | | | | |
|---|---|---|---|---|---|---|---|---|---|---|---|---|---|
| P46638 | RAB11B | RAB11B, member RAS oncogene family | Cytoplasm | enzyme | 7324627.5 | 9983049 | 1.12E+07 | | | | | | |
| P46638 | RAB11B | RAB11B, member RAS oncogene family | Cytoplasm | enzyme | 7053062 | 9562323 | 1.06E+07 | | | | | | |
| P46638 | RAB11B | RAB11B, member RAS oncogene family | Cytoplasm | enzyme | 4641087.5 | 6192903.5 | 7445512 | | | | | | |
| P46638 | RAB11B | RAB11B, member RAS oncogene family | Cytoplasm | enzyme | 3248382.25 | 4401712.5 | 3109779.75 | | | | | | |
| P46638 | RAB11B | RAB11B, member RAS oncogene family | Cytoplasm | enzyme | 3762949.5 | 2221010.5 | 5084267.5 | | | | | | |
| P46638 | RAB11B | RAB11B, member RAS oncogene family | Cytoplasm | enzyme | 690904.375 | 1185962.75 | 1299534.5 | P46638 | RAB11B | 11475101 | 12088572 | 2207061 | 30 |
| Q91V41 | RAB14 | RAB14, member RAS oncogene family | Cytoplasm | enzyme | 3.52E+07 | 3.21E+07 | 3.14E+07 | | | | | | |
| Q91V41 | RAB14 | RAB14, member RAS oncogene family | Cytoplasm | enzyme | 2.05E+07 | 2.27E+07 | 2.15E+07 | | | | | | |
| Q91V41 | RAB14 | RAB14, member RAS oncogene family | Cytoplasm | enzyme | 7736982 | 6939225 | 8417978 | | | | | | |
| Q91V41 | RAB14 | RAB14, member RAS oncogene family | Cytoplasm | enzyme | 1.15E+07 | 1.14E+07 | 9684935 | | | | | | |
| Q91V41 | RAB14 | RAB14, member RAS oncogene family | Cytoplasm | enzyme | 8827954 | 8442762 | 7190394 | | | | | | |
| Q91V41 | RAB14 | RAB14, member RAS oncogene family | Cytoplasm | enzyme | 4917053.5 | 5390288 | 5915979 | | | | | | |
| Q91V41 | RAB14 | RAB14, member RAS oncogene family | Cytoplasm | enzyme | 898749.188 | 1122974.25 | 3362801.5 | | | | | | |
| Q91V41 | RAB14 | RAB14, member RAS oncogene family | Cytoplasm | enzyme | 2226338 | 1992132.25 | 3402729.75 | | | | | | |
| Q91V41 | RAB14 | RAB14, member RAS oncogene family | Cytoplasm | enzyme | 287472.5 | 296036.312 | 1078657.75 | Q91V41 | RAB14 | 10232728 | 10231522 | 1969057 | 27 |
| Q8K386 | Rab15 | RAB15, member RAS oncogene family | Cytoplasm | enzyme | 4.42E+08 | 4.33E+08 | 3.84E+08 | Q8K386 | Rab15 | 442100000 | 31557144 | 18219526 | 3 |
| P35293 | RAB18 | RAB18, member RAS oncogene family | Cytoplasm | enzyme | 1.51E+07 | 1.63E+07 | 1.13E+07 | | | | | | |
| P35293 | RAB18 | RAB18, member RAS oncogene family | Cytoplasm | enzyme | 2.05E+07 | 1.86E+07 | 1.53E+07 | | | | | | |
| P35293 | RAB18 | RAB18, member RAS oncogene family | Cytoplasm | enzyme | 1.34E+07 | 1.31E+07 | 1.16E+07 | | | | | | |
| P35293 | RAB18 | RAB18, member RAS oncogene family | Cytoplasm | enzyme | 1.47E+07 | 1.38E+07 | 1.07E+07 | | | | | | |
| P35293 | RAB18 | RAB18, member RAS oncogene family | Cytoplasm | enzyme | 1.17E+07 | 1.13E+07 | 7502332.5 | P35293 | RAB18 | 15078000 | 3278661 | 846547 | 15 |
| P62821 | RAB1A | RAB1A, member RAS oncogene family | Cytoplasm | enzyme | 7.12E+07 | 7.73E+07 | 7.30E+07 | | | | | | |
| P62821 | RAB1A | RAB1A, member RAS oncogene family | Cytoplasm | enzyme | 2.77E+07 | 3.28E+07 | 3.15E+07 | | | | | | |
| P62821 | RAB1A | RAB1A, member RAS oncogene family | Cytoplasm | enzyme | 1.12E+07 | 1.35E+07 | 1.09E+07 | | | | | | |
| P62821 | RAB1A | RAB1A, member RAS oncogene family | Cytoplasm | enzyme | 5648557.5 | 2831041 | 5592292 | | | | | | |
| P62821 | RAB1A | RAB1A, member RAS oncogene family | Cytoplasm | enzyme | 2892336 | 3054661 | 2027784.5 | | | | | | |
| P62821 | RAB1A | RAB1A, member RAS oncogene family | Cytoplasm | enzyme | 1909386.75 | 2426831.25 | 1974903.75 | P62821 | RAB1A | 20086713 | 26371053 | 6215717 | 18 |
| Q9D1G1 | RAB1B | RAB1B, member RAS oncogene family | Cytoplasm | other | 5.45E+07 | 3.96E+07 | 5.57E+07 | | | | | | |
| Q9D1G1 | RAB1B | RAB1B, member RAS oncogene family | Cytoplasm | other | 2.63E+07 | 2.73E+07 | 3.22E+07 | | | | | | |
| Q9D1G1 | RAB1B | RAB1B, member RAS oncogene family | Cytoplasm | other | 1.70E+07 | 2.03E+07 | 1.44E+07 | | | | | | |
| Q9D1G1 | RAB1B | RAB1B, member RAS oncogene family | Cytoplasm | other | 1.32E+07 | 1.52E+07 | 1.71E+07 | | | | | | |
| Q9D1G1 | RAB1B | RAB1B, member RAS oncogene family | Cytoplasm | other | 7457672.5 | 6716951.5 | 8731959 | Q9D1G1 | RAB1B | 23687535 | 15741630 | 4064471 | 15 |
| P35282 | RAB21 | RAB21, member RAS oncogene family | Cytoplasm | enzyme | 8973467 | 1.16E+07 | 8805529 | | | | | | |
| P35282 | RAB21 | RAB21, member RAS oncogene family | Cytoplasm | enzyme | 1.07E+07 | 1993078.875 | 742059.312 | | | | | | |
| P35282 | RAB21 | RAB21, member RAS oncogene family | Cytoplasm | enzyme | 3755588.5 | 2828705 | 3534707 | | | | | | |
| P35282 | RAB21 | RAB21, member RAS oncogene family | Cytoplasm | enzyme | 597785.5 | 4661608 | 2527745 | | | | | | |
| P35282 | RAB21 | RAB21, member RAS oncogene family | Cytoplasm | enzyme | 4914886 | 1297782.125 | 301817.875 | | | | | | |
| P35282 | RAB21 | RAB21, member RAS oncogene family | Cytoplasm | enzyme | 3715202.5 | 3396033.75 | 3161137.25 | | | | | | |
| P35282 | RAB21 | RAB21, member RAS oncogene family | Cytoplasm | enzyme | 1695936.375 | 2083274.375 | 2762571.75 | | | | | | |
| P35282 | RAB21 | RAB21, member RAS oncogene family | Cytoplasm | enzyme | 837594.125 | 868314.5 | 346179.594 | P35282 | RAB21 | 4392558 | 3247973 | 662990 | 24 |
| P35285 | RAB22A | RAB22A, member RAS oncogene family | Cytoplasm | enzyme | 3451059.75 | 3977230.5 | 3708972 | | | | | | |
| P35285 | RAB22A | RAB22A, member RAS oncogene family | Cytoplasm | enzyme | 2746898.75 | 3509742.25 | 3587985.75 | P35285 | RAB22A | 3098979 | 412009 | 168202 | 6 |
| Q9ERI2 | RAB27A | RAB27A, member RAS oncogene family | Cytoplasm | enzyme | 2047174.625 | 2529750.25 | 2370241 | Q9ERI2 | RAB27A | 2047175 | 245864 | 141950 | 3 |
| Q99P58 | RAB27B | RAB27B, member RAS oncogene family | Cytoplasm | enzyme | 1.34E+07 | 1.78E+07 | 2.66E+07 | | | | | | |
| Q99P58 | RAB27B | RAB27B, member RAS oncogene family | Cytoplasm | enzyme | 9937602 | 1.35E+07 | 2.00E+07 | | | | | | |
| Q99P58 | RAB27B | RAB27B, member RAS oncogene family | Cytoplasm | enzyme | 1.02E+07 | 1.46E+07 | 2.36E+07 | | | | | | |
| Q99P58 | RAB27B | RAB27B, member RAS oncogene family | Cytoplasm | enzyme | 9119172 | 1.26E+07 | 2.01E+07 | | | | | | |
| Q99P58 | RAB27B | RAB27B, member RAS oncogene family | Cytoplasm | enzyme | 6889733.5 | 6773738 | 8216034 | | | | | | |
| Q99P58 | RAB27B | RAB27B, member RAS oncogene family | Cytoplasm | enzyme | 950459.438 | 1906640.75 | 1981585 | | | | | | |
| Q99P58 | RAB27B | RAB27B, member RAS oncogene family | Cytoplasm | enzyme | 688124.062 | 1183365.875 | 1201152.625 | | | | | | |
| Q99P58 | RAB27B | RAB27B, member RAS oncogene family | Cytoplasm | enzyme | 783876.25 | 1726663.75 | 2413825 | Q99P58 | RAB27B | 6491121 | 7947234 | 1622222 | 24 |
| P53994 | RAB2A | RAB2A, member RAS oncogene family | Cytoplasm | enzyme | 1.31E+07 | 1.19E+07 | 1.09E+07 | | | | | | |
| P53994 | RAB2A | RAB2A, member RAS oncogene family | Cytoplasm | enzyme | 8514473 | 8994955 | 7457352.5 | | | | | | |
| P53994 | RAB2A | RAB2A, member RAS oncogene family | Cytoplasm | enzyme | 4697237 | 4593803.5 | 5177819 | | | | | | |
| P53994 | RAB2A | RAB2A, member RAS oncogene family | Cytoplasm | enzyme | 3087094.5 | 2926239 | 3594752.75 | | | | | | |
| P53994 | RAB2A | RAB2A, member RAS oncogene family | Cytoplasm | enzyme | 2505227.5 | 2435963.5 | 2290776.25 | P53994 | RAB2A | 6382806 | 3714398 | 959053 | 15 |
| Q6PHN9 | RAB35 | RAB35, member RAS oncogene family | Cytoplasm | enzyme | 8.62E+07 | 9.06E+07 | 8.02E+07 | | | | | | |
| Q6PHN9 | RAB35 | RAB35, member RAS oncogene family | Cytoplasm | enzyme | 5.11E+07 | 4.61E+07 | 5.15E+07 | | | | | | |
| Q6PHN9 | RAB35 | RAB35, member RAS oncogene family | Cytoplasm | enzyme | 4.96E+07 | 4.86E+07 | 4.26E+07 | | | | | | |
| Q6PHN9 | RAB35 | RAB35, member RAS oncogene family | Cytoplasm | enzyme | 2.01E+07 | 2.35E+07 | 2.10E+07 | | | | | | |
| Q6PHN9 | RAB35 | RAB35, member RAS oncogene family | Cytoplasm | enzyme | 2.48E+07 | 2.64E+07 | 2.24E+07 | | | | | | |
| Q6PHN9 | RAB35 | RAB35, member RAS oncogene family | Cytoplasm | enzyme | 1.95E+07 | 2941909.75 | 579931.625 | | | | | | |
| Q6PHN9 | RAB35 | RAB35, member RAS oncogene family | Cytoplasm | enzyme | 2983977 | 3627931.75 | 4343817 | Q6PHN9 | RAB35 | 36316282 | 27518864 | 6005108 | 21 |
| Q9JKM7 | RAB37 | RAB37, member RAS oncogene family | Cytoplasm | enzyme | 405528.5 | 532021.5 | 734267.562 | Q9JKM7 | RAB37 | 405529 | 165818 | 95735 | 3 |
| Q80UJ7 | RAB3GAP1 | RAB3 GTPase activating protein subunit 1 (catalytic) | Cytoplasm | other | 7426754 | 5248515 | 3840119.5 | | | | | | |
| Q80UJ7 | RAB3GAP1 | RAB3 GTPase activating protein subunit 1 (catalytic) | Cytoplasm | other | 3016639.5 | 1813886.25 | 720741.75 | | | | | | |
| Q80UJ7 | RAB3GAP1 | RAB3 GTPase activating protein subunit 1 (catalytic) | Cytoplasm | other | 269110.375 | 191839.156 | 335449.781 | Q80UJ7 | RAB3GAP1 | 3570835 | 2561779 | 853926 | 9 |
| Q8BMG7 | RAB3GAP2 | RAB3 GTPase activating protein subunit 2 (non-catalytic) | Cytoplasm | enzyme | 1810280.375 | 2008127.125 | 942939.312 | | | | | | |
| Q8BMG7 | RAB3GAP2 | RAB3 GTPase activating protein subunit 2 (non-catalytic) | Cytoplasm | enzyme | 1532721.25 | 925283.312 | 710231.438 | Q8BMG7 | RAB3GAP2 | 1671501 | 534588 | 218245 | 6 |
| Q8VDV3 | RAB3IL1 | RAB3A interacting protein (rabin3)-like 1 | Cytoplasm | other | 1492717.125 | 1585136 | 2236927 | Q8VDV3 | RAB3IL1 | 1492717 | 405631 | 234191 | 3 |
| Q9CQD1 | RAB5A | RAB5A, member RAS oncogene family | Cytoplasm | enzyme | 3.29E+07 | 2.77E+07 | 2.45E+07 | | | | | | |
| Q9CQD1 | RAB5A | RAB5A, member RAS oncogene family | Cytoplasm | enzyme | 2.90E+07 | 3.15E+07 | 2.58E+07 | | | | | | |
| Q9CQD1 | RAB5A | RAB5A, member RAS oncogene family | Cytoplasm | enzyme | 9886186 | 1.11E+07 | 1.06E+07 | | | | | | |
| Q9CQD1 | RAB5A | RAB5A, member RAS oncogene family | Cytoplasm | enzyme | 1.09E+07 | 1.49E+07 | 8425920 | | | | | | |
| Q9CQD1 | RAB5A | RAB5A, member RAS oncogene family | Cytoplasm | enzyme | 7303372 | 1.02E+07 | 8018072 | Q9CQD1 | RAB5A | 17993912 | 9682983 | 2500135 | 15 |
| P61021 | RAB5B | RAB5B, member RAS oncogene family | Cytoplasm | enzyme | 3.13E+07 | 2.66E+07 | 2.14E+07 | | | | | | |
| P61021 | RAB5B | RAB5B, member RAS oncogene family | Cytoplasm | enzyme | 2.06E+07 | 1.92E+07 | 1.83E+07 | | | | | | |
| P61021 | RAB5B | RAB5B, member RAS oncogene family | Cytoplasm | enzyme | 1.40E+07 | 1.46E+07 | 1.12E+07 | | | | | | |
| P61021 | RAB5B | RAB5B, member RAS oncogene family | Cytoplasm | enzyme | 9787494 | 9683674 | 7923836 | | | | | | |
| P61021 | RAB5B | RAB5B, member RAS oncogene family | Cytoplasm | enzyme | 8707403 | 8809361 | 6482183.5 | | | | | | |
| P61021 | RAB5B | RAB5B, member RAS oncogene family | Cytoplasm | enzyme | 6171879.5 | 5763463.5 | 5586515.5 | P61021 | RAB5B | 15091129 | 7643836 | 1801669 | 18 |
| P35278 | RAB5C | RAB5C, member RAS oncogene family | Cytoplasm | enzyme | 1.31E+08 | 1.27E+08 | 1.06E+08 | | | | | | |
| P35278 | RAB5C | RAB5C, member RAS oncogene family | Cytoplasm | enzyme | 1.26E+08 | 1.25E+08 | 9.53E+07 | | | | | | |
| P35278 | RAB5C | RAB5C, member RAS oncogene family | Cytoplasm | enzyme | 1.05E+08 | 8.43E+07 | 9.00E+07 | | | | | | |
| P35278 | RAB5C | RAB5C, member RAS oncogene family | Cytoplasm | enzyme | 7.75E+07 | 8.89E+07 | 5.71E+07 | | | | | | |
| P35278 | RAB5C | RAB5C, member RAS oncogene family | Cytoplasm | enzyme | 6.28E+07 | 6.90E+07 | 5.49E+07 | | | | | | |
| P35278 | RAB5C | RAB5C, member RAS oncogene family | Cytoplasm | enzyme | 2.59E+07 | 2.77E+07 | 2.96E+07 | | | | | | |
| P35278 | RAB5C | RAB5C, member RAS oncogene family | Cytoplasm | enzyme | 3.12E+07 | 3.38E+07 | 3.49E+07 | | | | | | |
| P35278 | RAB5C | RAB5C, member RAS oncogene family | Cytoplasm | enzyme | 2.55E+07 | 2.68E+07 | 2.81E+07 | | | | | | |
| P35278 | RAB5C | RAB5C, member RAS oncogene family | Cytoplasm | enzyme | 1.66E+07 | 1.40E+07 | 1.03E+07 | | | | | | |
| P35278 | RAB5C | RAB5C, member RAS oncogene family | Cytoplasm | enzyme | 1.85E+07 | 1.62E+07 | 1.31E+07 | | | | | | |
| P35278 | RAB5C | RAB5C, member RAS oncogene family | Cytoplasm | enzyme | 8910940 | 9758443 | 7334897 | | | | | | |
| P35278 | RAB5C | RAB5C, member RAS oncogene family | Cytoplasm | enzyme | 2625391.75 | 2471022 | 626838.438 | | | | | | |
| P35278 | RAB5C | RAB5C, member RAS oncogene family | Cytoplasm | enzyme | 2416220 | 2037683.375 | 1727733.875 | P35278 | RAB5C | 48810966 | 42197045 | 6756935 | 39 |
| P35279 | RAB6A | RAB6A, member RAS oncogene family | Cytoplasm | enzyme | 4569033 | 4933807.5 | 7168461 | | | | | | |
| P35279 | RAB6A | RAB6A, member RAS oncogene family | Cytoplasm | enzyme | 2671022.5 | 2023497.625 | 2818326.5 | P35279 | RAB6A | 3620028 | 1913028 | 780990 | 6 |
| P51150 | RAB7A | RAB7A, member RAS oncogene family | Cytoplasm | enzyme | 1.70E+07 | 2.46E+07 | 1.97E+07 | | | | | | |
| P51150 | RAB7A | RAB7A, member RAS oncogene family | Cytoplasm | enzyme | 1.51E+07 | 1.97E+07 | 2.17E+07 | | | | | | |
| P51150 | RAB7A | RAB7A, member RAS oncogene family | Cytoplasm | enzyme | 1.70E+07 | 2.18E+07 | 1.72E+07 | | | | | | |
| P51150 | RAB7A | RAB7A, member RAS oncogene family | Cytoplasm | enzyme | 1.48E+07 | 1.74E+07 | 1.60E+07 | | | | | | |
| P51150 | RAB7A | RAB7A, member RAS oncogene family | Cytoplasm | enzyme | 1.25E+07 | 1.33E+07 | 1.59E+07 | | | | | | |
| P51150 | RAB7A | RAB7A, member RAS oncogene family | Cytoplasm | enzyme | 1.11E+07 | 1.20E+07 | 1.12E+07 | | | | | | |
| P51150 | RAB7A | RAB7A, member RAS oncogene family | Cytoplasm | enzyme | 9757084 | 1.33E+07 | 1.47E+07 | | | | | | |
| P51150 | RAB7A | RAB7A, member RAS oncogene family | Cytoplasm | enzyme | 1.19E+07 | 1.54E+07 | 1.15E+07 | | | | | | |
| P51150 | RAB7A | RAB7A, member RAS oncogene family | Cytoplasm | enzyme | 8453893 | 1.00E+07 | 9792028 | | | | | | |
| P51150 | RAB7A | RAB7A, member RAS oncogene family | Cytoplasm | enzyme | 7109696.5 | 3530544.5 | 7282456.5 | | | | | | |
| P51150 | RAB7A | RAB7A, member RAS oncogene family | Cytoplasm | enzyme | 2476529.25 | 3431801.5 | 3506199.75 | P51150 | RAB7A | 11549746 | 5562105 | 968238 | 33 |
| P55258 | RAB8A | RAB8A, member RAS oncogene family | Plasma Membrane | enzyme | 9.31E+07 | 9.96E+07 | 1.06E+08 | | | | | | |
| P55258 | RAB8A | RAB8A, member RAS oncogene family | Plasma Membrane | enzyme | 9.30E+07 | 9.43E+07 | 8.10E+07 | | | | | | |

| | | | | | | | | | | | | | |
|---|---|---|---|---|---|---|---|---|---|---|---|---|---|
| P55258 | RAB8A | RAB8A, member RAS oncogene family | Plasma Membrane | enzyme | 7.87E+07 | 8.17E+07 | 6.42E+07 | | | | | | |
| P55258 | RAB8A | RAB8A, member RAS oncogene family | Plasma Membrane | enzyme | 5.75E+07 | 6.19E+07 | 5.02E+07 | | | | | | |
| P55258 | RAB8A | RAB8A, member RAS oncogene family | Plasma Membrane | enzyme | 5.05E+07 | 5.37E+07 | 4.10E+07 | | | | | | |
| P55258 | RAB8A | RAB8A, member RAS oncogene family | Plasma Membrane | enzyme | 8243573.5 | 7431453 | 6306784 | | | | | | |
| P55258 | RAB8A | RAB8A, member RAS oncogene family | Plasma Membrane | enzyme | 5192673 | 7973135 | 6312955.5 | | | | | | |
| P55258 | RAB8A | RAB8A, member RAS oncogene family | Plasma Membrane | enzyme | 555832.312 | 624385.312 | 567725.562 | P55258 | RAB8A | 48359010 | 37803568 | 7716621 | 24 |
| P61028 | RAB8B | RAB8B, member RAS oncogene family | Cytoplasm | enzyme | 1.22E+08 | 1.26E+08 | 1.03E+08 | | | | | | |
| P61028 | RAB8B | RAB8B, member RAS oncogene family | Cytoplasm | enzyme | 1.27E+08 | 1.50E+08 | 1.55E+08 | | | | | | |
| P61028 | RAB8B | RAB8B, member RAS oncogene family | Cytoplasm | enzyme | 6.32E+07 | 7.05E+07 | 5.64E+07 | | | | | | |
| P61028 | RAB8B | RAB8B, member RAS oncogene family | Cytoplasm | enzyme | 2.92E+07 | 2.97E+07 | 2.84E+07 | | | | | | |
| P61028 | RAB8B | RAB8B, member RAS oncogene family | Cytoplasm | enzyme | 3283494.5 | 5154516 | 4406746 | | | | | | |
| P61028 | RAB8B | RAB8B, member RAS oncogene family | Cytoplasm | enzyme | 572444.062 | 523952.938 | 557230.688 | P61028 | RAB8B | 57477656 | 56589248 | 13338214 | 18 |
| Q9R0M6 | RAB9A | RAB9A, member RAS oncogene family | Cytoplasm | enzyme | 2355987.25 | 2561135 | 2449271.25 | Q9R0M6 | RAB9A | 2355987 | 102714 | 59302 | 3 |
| P63001 | RAC1 | ras-related C3 botulinum toxin substrate 1 (rho family, small GTP binding protein Rac1) | Plasma Membrane | enzyme | 9.02E+07 | 9.99E+07 | 1.04E+08 | | | | | | |
| P63001 | RAC1 | ras-related C3 botulinum toxin substrate 1 (rho family, small GTP binding protein Rac1) | Plasma Membrane | enzyme | 6.81E+07 | 4.75E+07 | 2.33E+07 | | | | | | |
| P63001 | RAC1 | ras-related C3 botulinum toxin substrate 1 (rho family, small GTP binding protein Rac1) | Plasma Membrane | enzyme | 8.77E+07 | 8.42E+07 | 7.63E+07 | | | | | | |
| P63001 | RAC1 | ras-related C3 botulinum toxin substrate 1 (rho family, small GTP binding protein Rac1) | Plasma Membrane | enzyme | 6.45E+07 | 6.26E+07 | 7.86E+07 | | | | | | |
| P63001 | RAC1 | ras-related C3 botulinum toxin substrate 1 (rho family, small GTP binding protein Rac1) | Plasma Membrane | enzyme | 7.55E+07 | 7.54E+07 | 7.60E+07 | | | | | | |
| P63001 | RAC1 | ras-related C3 botulinum toxin substrate 1 (rho family, small GTP binding protein Rac1) | Plasma Membrane | enzyme | 6.39E+07 | 6.01E+07 | 6.94E+07 | | | | | | |
| P63001 | RAC1 | ras-related C3 botulinum toxin substrate 1 (rho family, small GTP binding protein Rac1) | Plasma Membrane | enzyme | 4.36E+07 | 4.79E+07 | 3.67E+07 | | | | | | |
| P63001 | RAC1 | ras-related C3 botulinum toxin substrate 1 (rho family, small GTP binding protein Rac1) | Plasma Membrane | enzyme | 2.27E+07 | 2.25E+07 | 2.57E+07 | | | | | | |
| P63001 | RAC1 | ras-related C3 botulinum toxin substrate 1 (rho family, small GTP binding protein Rac1) | Plasma Membrane | enzyme | 3.01E+07 | 3.79E+07 | 2.08E+07 | | | | | | |
| P63001 | RAC1 | ras-related C3 botulinum toxin substrate 1 (rho family, small GTP binding protein Rac1) | Plasma Membrane | enzyme | 2.00E+07 | 3.14E+07 | 2.20E+07 | | | | | | |
| P63001 | RAC1 | ras-related C3 botulinum toxin substrate 1 (rho family, small GTP binding protein Rac1) | Plasma Membrane | enzyme | 7125033 | 8540580 | 8588571 | | | | | | |
| P63001 | RAC1 | ras-related C3 botulinum toxin substrate 1 (rho family, small GTP binding protein Rac1) | Plasma Membrane | enzyme | 8564425 | 1.06E+07 | 3958577.25 | | | | | | |
| P63001 | RAC1 | ras-related C3 botulinum toxin substrate 1 (rho family, small GTP binding protein Rac1) | Plasma Membrane | enzyme | 5017920 | 4997190.5 | 4610242 | | | | | | |
| P63001 | RAC1 | ras-related C3 botulinum toxin substrate 1 (rho family, small GTP binding protein Rac1) | Plasma Membrane | enzyme | 3085870.25 | 3096931.25 | 1779133.375 | P63001 | RAC1 | 42155946 | 31753992 | 4899747 | 42 |
| Q05144 | RAC2 | ras-related C3 botulinum toxin substrate 2 (rho family, small GTP binding protein Rac2) | Cytoplasm | enzyme | 3213351 | 5085575 | 7559240 | Q05144 | RAC2 | 3213351 | 2179870 | 1258548 | 3 |
| P54726 | RAD23A | RAD23 homolog A, nucleotide excision repair protein | Nucleus | other | 4.40E+07 | 4.33E+07 | 3.37E+07 | | | | | | |
| P54726 | RAD23A | RAD23 homolog A, nucleotide excision repair protein | Nucleus | other | 3.47E+07 | 3.00E+07 | 1.35E+07 | | | | | | |
| P54726 | RAD23A | RAD23 homolog A, nucleotide excision repair protein | Nucleus | other | 2.19E+07 | 1.69E+07 | 1.06E+07 | | | | | | |
| P54726 | RAD23A | RAD23 homolog A, nucleotide excision repair protein | Nucleus | other | 9537770 | 7513378 | 4278656.5 | | | | | | |
| P54726 | RAD23A | RAD23 homolog A, nucleotide excision repair protein | Nucleus | other | 6424391.5 | 5968855 | 5770226 | | | | | | |
| P54726 | RAD23A | RAD23 homolog A, nucleotide excision repair protein | Nucleus | other | 2672734.25 | 2661699.75 | 1518881.75 | | | | | | |
| P54726 | RAD23A | RAD23 homolog A, nucleotide excision repair protein | Nucleus | other | 796313.562 | 241030.953 | 197155 | P54726 | RAD23A | 17154458 | 14539649 | 3172812 | 21 |
| P63321 | RALA | v-ral simian leukemia viral oncogene homolog A (ras related) | Cytoplasm | enzyme | 4077575.75 | 2574178 | 1639120.25 | P63321 | RALA | 4077576 | 1230217 | 710266 | 3 |
| P62827 | RAN | RAN, member RAS oncogene family | Nucleus | enzyme | 1810447.25 | 1729210.75 | 2198945.5 | P62827 | RAN | 1810447 | 251058 | 144948 | 3 |
| P46061 | RANGAP1 | Ran GTPase activating protein 1 | Nucleus | other | 410984.281 | 93570.68 | 121729.562 | P46061 | RANGAP1 | 410984 | 175695 | 101438 | 3 |
| P62835 | RAP1A | RAP1A, member of RAS oncogene family | Cytoplasm | enzyme | 1.17E+08 | 1.34E+08 | 1.19E+08 | | | | | | |
| P62835 | RAP1A | RAP1A, member of RAS oncogene family | Cytoplasm | enzyme | 6.62E+07 | 7.57E+07 | 7.92E+07 | | | | | | |
| P62835 | RAP1A | RAP1A, member of RAS oncogene family | Cytoplasm | enzyme | 3.84E+07 | 3.94E+07 | 4.77E+07 | | | | | | |
| P62835 | RAP1A | RAP1A, member of RAS oncogene family | Cytoplasm | enzyme | 5.85E+07 | 5.76E+07 | 5.90E+07 | | | | | | |
| P62835 | RAP1A | RAP1A, member of RAS oncogene family | Cytoplasm | enzyme | 1.29E+07 | 1.53E+07 | 2.11E+07 | | | | | | |
| P62835 | RAP1A | RAP1A, member of RAS oncogene family | Cytoplasm | enzyme | 5445276 | 5793007.5 | 4436738.5 | | | | | | |
| P62835 | RAP1A | RAP1A, member of RAS oncogene family | Cytoplasm | enzyme | 3725047 | 2450127.25 | 1286223.875 | | | | | | |
| P62835 | RAP1A | RAP1A, member of RAS oncogene family | Cytoplasm | enzyme | 147037.859 | 169438.141 | 1048212.312 | P62835 | RAP1A | 37732170 | 41547112 | 8480769 | 24 |
| Q99JI6 | RAP1B | RAP1B, member of RAS oncogene family | Cytoplasm | enzyme | 1.62E+08 | 1.75E+08 | 1.99E+08 | | | | | | |
| Q99JI6 | RAP1B | RAP1B, member of RAS oncogene family | Cytoplasm | enzyme | 1.51E+08 | 1.47E+08 | 1.69E+08 | | | | | | |
| Q99JI6 | RAP1B | RAP1B, member of RAS oncogene family | Cytoplasm | enzyme | 1.41E+08 | 1.94E+08 | 1.88E+08 | | | | | | |
| Q99JI6 | RAP1B | RAP1B, member of RAS oncogene family | Cytoplasm | enzyme | 5.38E+07 | 7.31E+07 | 9.08E+07 | | | | | | |
| Q99JI6 | RAP1B | RAP1B, member of RAS oncogene family | Cytoplasm | enzyme | 1.35E+07 | 2.06E+07 | 1.57E+07 | | | | | | |
| Q99JI6 | RAP1B | RAP1B, member of RAS oncogene family | Cytoplasm | enzyme | 3.47E+07 | 4.10E+07 | 7.60E+07 | | | | | | |
| Q99JI6 | RAP1B | RAP1B, member of RAS oncogene family | Cytoplasm | enzyme | 1.81E+07 | 2.85E+07 | 3.10E+07 | | | | | | |
| Q99JI6 | RAP1B | RAP1B, member of RAS oncogene family | Cytoplasm | enzyme | 504980.188 | 340213.094 | 101220.133 | Q99JI6 | RAP1B | 71928123 | 72172851 | 14732222 | 24 |
| P61226 | RAP2B | RAP2B, member of RAS oncogene family | Plasma Membrane | enzyme | 1.07E+07 | 1.12E+07 | 1.01E+07 | | | | | | |
| P61226 | RAP2B | RAP2B, member of RAS oncogene family | Plasma Membrane | enzyme | 9637557 | 8371690.5 | 7153594 | | | | | | |
| P61226 | RAP2B | RAP2B, member of RAS oncogene family | Plasma Membrane | enzyme | 1.01E+07 | 8750873 | 9371672 | | | | | | |
| P61226 | RAP2B | RAP2B, member of RAS oncogene family | Plasma Membrane | enzyme | 9384018 | 1.16E+07 | 7989948 | | | | | | |
| P61226 | RAP2B | RAP2B, member of RAS oncogene family | Plasma Membrane | enzyme | 7113750.5 | 5644743 | 6854298.5 | | | | | | |
| P61226 | RAP2B | RAP2B, member of RAS oncogene family | Plasma Membrane | enzyme | 2103754.25 | 2775251.75 | 2691314 | P61226 | RAP2B | 8169847 | 2917709 | 687711 | 18 |
| Q8BU31 | RAP2C | RAP2C, member of RAS oncogene family | Cytoplasm | enzyme | 2702585.75 | 2669434.25 | 1939152.125 | Q8BU31 | RAP2C | 2702586 | 431517 | 249136 | 3 |
| Q9D0I9 | RARS | arginyl-tRNA synthetase | Cytoplasm | enzyme | 123954.734 | 304779.625 | 191098.25 | Q9D0I9 | RARS | 123955 | 91405 | 52773 | 3 |
| Q60790 | RASA3 | RAS p21 protein activator 3 | Plasma Membrane | ion channel | 3033212.5 | 3497729.25 | 4907170 | Q60790 | RASA3 | 3033213 | 975877 | 563423 | 3 |
| Q61820 | Rasl2-9 | RAS-like, family 2, locus 9 | Nucleus | transporter | 4573982.5 | 3565058.5 | 4169234.5 | | | | | | |
| Q61820 | Rasl2-9 | RAS-like, family 2, locus 9 | Nucleus | transporter | 3270818.5 | 2777283.5 | 3310859.5 | Q61820 | Rasl2-9 | 3922401 | 654517 | 267205 | 6 |
| O89086 | RBM3 | RNA binding motif (RNP1, RRM) protein 3 | Cytoplasm | other | 4282852.5 | 7856491 | 1.50E+07 | | | | | | |
| O89086 | RBM3 | RNA binding motif (RNP1, RRM) protein 3 | Cytoplasm | other | 1520560.625 | 2793405.25 | 4830916 | | | | | | |
| O89086 | RBM3 | RNA binding motif (RNP1, RRM) protein 3 | Cytoplasm | other | 94026.773 | 52761.672 | 553367.438 | O89086 | RBM3 | 1965813 | 4835717 | 1611906 | 9 |
| Q9QYF1 | RDH11 | retinol dehydrogenase 11 (all-trans/9-cis/11-cis) | Cytoplasm | enzyme | 7211792 | 7520836.5 | 7524479 | | | | | | |
| Q9QYF1 | RDH11 | retinol dehydrogenase 11 (all-trans/9-cis/11-cis) | Cytoplasm | enzyme | 4718589 | 5441027.5 | 4362474.5 | | | | | | |
| Q9QYF1 | RDH11 | retinol dehydrogenase 11 (all-trans/9-cis/11-cis) | Cytoplasm | enzyme | 4119587 | 6837120.5 | 6217180.5 | | | | | | |
| Q9QYF1 | RDH11 | retinol dehydrogenase 11 (all-trans/9-cis/11-cis) | Cytoplasm | enzyme | 4385577.5 | 5308864.5 | 5364701 | | | | | | |
| Q9QYF1 | RDH11 | retinol dehydrogenase 11 (all-trans/9-cis/11-cis) | Cytoplasm | enzyme | 5011125 | 4689806 | 5184797.5 | | | | | | |
| Q9QYF1 | RDH11 | retinol dehydrogenase 11 (all-trans/9-cis/11-cis) | Cytoplasm | enzyme | 3108570.5 | 2792413.5 | 2308463.75 | | | | | | |
| Q9QYF1 | RDH11 | retinol dehydrogenase 11 (all-trans/9-cis/11-cis) | Cytoplasm | enzyme | 3507913.75 | 5076215 | 5158538 | | | | | | |
| Q9QYF1 | RDH11 | retinol dehydrogenase 11 (all-trans/9-cis/11-cis) | Cytoplasm | enzyme | 3434249.25 | 2946108.75 | 3007812.75 | Q9QYF1 | RDH11 | 4437176 | 1508159 | 307852 | 24 |
| P26043 | RDX | radixin | Cytoplasm | other | 4.14E+07 | 2.91E+07 | 3.96E+07 | | | | | | |
| P26043 | RDX | radixin | Cytoplasm | other | 2.25E+07 | 1.80E+07 | 2.11E+07 | | | | | | |
| P26043 | RDX | radixin | Cytoplasm | other | 3.04E+07 | 2.63E+07 | 2.40E+07 | | | | | | |
| P26043 | RDX | radixin | Cytoplasm | other | 1.57E+07 | 1.03E+07 | 1.04E+07 | | | | | | |
| P26043 | RDX | radixin | Cytoplasm | other | 1.33E+07 | 1.01E+07 | 9335920 | | | | | | |
| P26043 | RDX | radixin | Cytoplasm | other | 1.47E+07 | 1.09E+07 | 7406541 | | | | | | |
| P26043 | RDX | radixin | Cytoplasm | other | 1.24E+07 | 7858627 | 7115196.5 | | | | | | |
| P26043 | RDX | radixin | Cytoplasm | other | 6964944.5 | 3650497.75 | 4069648.25 | | | | | | |
| P26043 | RDX | radixin | Cytoplasm | other | 5844347.5 | 2212189.5 | 2020955.75 | | | | | | |
| P26043 | RDX | radixin | Cytoplasm | other | 3692538 | 2608760.75 | 6002108.5 | | | | | | |
| P26043 | RDX | radixin | Cytoplasm | other | 5994113 | 3577077.25 | 3782184.75 | | | | | | |
| P26043 | RDX | radixin | Cytoplasm | other | 5374625.5 | 5080118.5 | 2952572.25 | | | | | | |
| P26043 | RDX | radixin | Cytoplasm | other | 1966377.375 | 1857375.625 | 1760159.875 | | | | | | |
| P26043 | RDX | radixin | Cytoplasm | other | 482636.25 | 167392.391 | 577747.125 | P26043 | RDX | 12919970 | 10497364 | 1619778 | 42 |
| Q60870 | REEP5 | receptor accessory protein 5 | Extracellular Space | transporter | 2.11E+07 | 2.73E+07 | 2.04E+07 | | | | | | |
| Q60870 | REEP5 | receptor accessory protein 5 | Extracellular Space | transporter | 2152416.75 | 1442358.375 | 1641095.125 | | | | | | |
| Q60870 | REEP5 | receptor accessory protein 5 | Extracellular Space | transporter | 273769.75 | 317370.625 | 109970.375 | Q60870 | REEP5 | 7842062 | 11163441 | 3721147 | 9 |
| Q9JM62 | REEP6 | receptor accessory protein 6 | Plasma Membrane | other | 1.27E+07 | 1.20E+07 | 9692963 | | | | | | |
| Q9JM62 | REEP6 | receptor accessory protein 6 | Plasma Membrane | other | 4879954 | 4478932 | 4726658.5 | | | | | | |
| Q9JM62 | REEP6 | receptor accessory protein 6 | Plasma Membrane | other | 1420031.25 | 251692.719 | 116356.789 | Q9JM62 | REEP6 | 6339995 | 4842579 | 1614193 | 9 |
| Q8K2J7 | RELL1 | RELT-like 1 | Plasma Membrane | other | 3411253.75 | 1353141.125 | 626014.25 | | | | | | |
| Q8K2J7 | RELL1 | RELT-like 1 | Plasma Membrane | other | 3799503.25 | 4315668.5 | 3480388 | | | | | | |
| Q8K2J7 | RELL1 | RELT-like 1 | Plasma Membrane | other | 2121066.5 | 2896982.75 | 2480800.5 | Q8K2J7 | RELL1 | 3110608 | 1197280 | 399093 | 9 |
| Q9QUT0 | RHAG | Rh-associated glycoprotein | Plasma Membrane | peptidase | 1533138.75 | 771479.375 | 323789.594 | | | | | | |
| Q9QUT0 | RHAG | Rh-associated glycoprotein | Plasma Membrane | peptidase | 866557.688 | 1021592.375 | 1605249.25 | | | | | | |
| Q9QUT0 | RHAG | Rh-associated glycoprotein | Plasma Membrane | peptidase | 7982226.5 | 9717568 | 1.40E+07 | | | | | | |
| Q9QUT0 | RHAG | Rh-associated glycoprotein | Plasma Membrane | peptidase | 2013103.375 | 1778619.25 | 2376991.5 | Q9QUT0 | RHAG | 3098757 | 4406468 | 1272038 | 12 |
| Q80WQ6 | RHBDF2 | rhomboid 5 homolog 2 (Drosophila) | Cytoplasm | other | 3316793.75 | 3896928.25 | 3947826.25 | | | | | | |
| Q80WQ6 | RHBDF2 | rhomboid 5 homolog 2 (Drosophila) | Cytoplasm | other | 822816.125 | 609477.062 | 892431.25 | Q80WQ6 | RHBDF2 | 2069805 | 1631206 | 665937 | 6 |
| Q8CF94 | RHCE/RHD | Rh blood group, D antigen | Plasma Membrane | transporter | 3607203.75 | 3367064 | 3504522 | | | | | | |
| Q8CF94 | RHCE/RHD | Rh blood group, D antigen | Plasma Membrane | transporter | 1.13E+07 | 9545812 | 1.06E+07 | | | | | | |
| Q8CF94 | RHCE/RHD | Rh blood group, D antigen | Plasma Membrane | transporter | 137700.234 | 197322.734 | 489011.219 | | | | | | |
| Q8CF94 | RHCE/RHD | Rh blood group, D antigen | Plasma Membrane | transporter | 7831.782 | 69026.508 | 4310.843 | | | | | | |

| | | | | | | | | | | | | | |
|---|---|---|---|---|---|---|---|---|---|---|---|---|---|
| Q8CF94 | RHCE/RHD | Rh blood group, D antigen | Plasma Membrane | transporter | 843191.438 | 1391747.5 | 647970.875 | | | | | | |
| Q8CF94 | RHCE/RHD | Rh blood group, D antigen | Plasma Membrane | transporter | 1413533.875 | 803431.625 | 1386314.75 | | | | | | |
| Q8CF94 | RHCE/RHD | Rh blood group, D antigen | Plasma Membrane | transporter | 163514.859 | 111333.141 | 137358.766 | | | | | | |
| Q8CF94 | RHCE/RHD | Rh blood group, D antigen | Plasma Membrane | transporter | 0 | 0 | 0 | Q8CF94 | RHCE/RHD | 2182872 | 3444695 | 703145 | 24 |
| Q9QUI0 | RHOA | ras homolog family member A | Cytoplasm | enzyme | 3.68E+07 | 3.80E+07 | 3.34E+07 | | | | | | |
| Q9QUI0 | RHOA | ras homolog family member A | Cytoplasm | enzyme | 2.59E+07 | 2.95E+07 | 2.09E+07 | | | | | | |
| Q9QUI0 | RHOA | ras homolog family member A | Cytoplasm | enzyme | 1.40E+07 | 1.30E+07 | 2.08E+07 | | | | | | |
| Q9QUI0 | RHOA | ras homolog family member A | Cytoplasm | enzyme | 1.24E+07 | 1.03E+07 | 1.40E+07 | | | | | | |
| Q9QUI0 | RHOA | ras homolog family member A | Cytoplasm | enzyme | 1.25E+07 | 1.15E+07 | 1.56E+07 | | | | | | |
| Q9QUI0 | RHOA | ras homolog family member A | Cytoplasm | enzyme | 1.35E+07 | 1.34E+07 | 7615406 | | | | | | |
| Q9QUI0 | RHOA | ras homolog family member A | Cytoplasm | enzyme | 6600619.5 | 5709806 | 4078200.25 | | | | | | |
| Q9QUI0 | RHOA | ras homolog family member A | Cytoplasm | enzyme | 5363490 | 3231893.5 | 3869577.25 | | | | | | |
| Q9QUI0 | RHOA | ras homolog family member A | Cytoplasm | enzyme | 8211254 | 1.88E+07 | 1.85E+07 | Q9QUI0 | RHOA | 15010596 | 9899801 | 1905218 | 27 |
| P84096 | RHOG | ras homolog family member G | Cytoplasm | enzyme | 4.59E+07 | 4.56E+07 | 4.46E+07 | | | | | | |
| P84096 | RHOG | ras homolog family member G | Cytoplasm | enzyme | 2.76E+07 | 3.00E+07 | 2.79E+07 | | | | | | |
| P84096 | RHOG | ras homolog family member G | Cytoplasm | enzyme | 2.76E+07 | 3.18E+07 | 2.85E+07 | | | | | | |
| P84096 | RHOG | ras homolog family member G | Cytoplasm | enzyme | 1.52E+07 | 1.75E+07 | 1.93E+07 | | | | | | |
| P84096 | RHOG | ras homolog family member G | Cytoplasm | enzyme | 1.61E+07 | 2.16E+07 | 1.99E+07 | | | | | | |
| P84096 | RHOG | ras homolog family member G | Cytoplasm | enzyme | 9127955 | 1.03E+07 | 9278132 | | | | | | |
| P84096 | RHOG | ras homolog family member G | Cytoplasm | enzyme | 9228706 | 1.21E+07 | 8274772.5 | | | | | | |
| P84096 | RHOG | ras homolog family member G | Cytoplasm | enzyme | 1.16E+07 | 1.85E+07 | 1.86E+07 | | | | | | |
| P84096 | RHOG | ras homolog family member G | Cytoplasm | enzyme | 8188692.5 | 8254745 | 9709455 | | | | | | |
| P84096 | RHOG | ras homolog family member G | Cytoplasm | enzyme | 2324894.25 | 1454740.125 | 413368.438 | | | | | | |
| P84096 | RHOG | ras homolog family member G | Cytoplasm | enzyme | 2269572 | 3063096 | 2031593.875 | P84096 | RHOG | 15919984 | 12813955 | 2230623 | 33 |
| Q9QYK7 | RNF11 | ring finger protein 11 | Cytoplasm | enzyme | 2140665.5 | 3043894.5 | 3263247.75 | Q9QYK7 | RNF11 | 2140666 | 594997 | 343522 | 3 |
| Q5XPI3 | RNF123 | ring finger protein 123 | Cytoplasm | other | 1.03E+07 | 6471493 | 7438270 | | | | | | |
| Q5XPI3 | RNF123 | ring finger protein 123 | Cytoplasm | other | 8301974 | 8250706 | 4938591 | | | | | | |
| Q5XPI3 | RNF123 | ring finger protein 123 | Cytoplasm | other | 7385855 | 4833051 | 4288978 | | | | | | |
| Q5XPI3 | RNF123 | ring finger protein 123 | Cytoplasm | other | 8172275.5 | 6442865.5 | 5548393 | | | | | | |
| Q5XPI3 | RNF123 | ring finger protein 123 | Cytoplasm | other | 6259388 | 5668235 | 2719892 | | | | | | |
| Q5XPI3 | RNF123 | ring finger protein 123 | Cytoplasm | other | 3167799.75 | 2524345.25 | 2138762 | | | | | | |
| Q5XPI3 | RNF123 | ring finger protein 123 | Cytoplasm | other | 3047412.75 | 2624195 | 2127856.5 | Q5XPI3 | RNF123 | 6662101 | 2410075 | 525921 | 21 |
| E9Q555 | RNF213 | ring finger protein 213 | Cytoplasm | enzyme | 817903.875 | 1098732.375 | 1295809.625 | E9Q555 | RNF213 | 817904 | 240173 | 138664 | 3 |
| Q9EPK2 | RP2 | retinitis pigmentosa 2 (X-linked recessive) | Cytoplasm | enzyme | 1.49E+07 | 1.56E+07 | 1.66E+07 | | | | | | |
| Q9EPK2 | RP2 | retinitis pigmentosa 2 (X-linked recessive) | Cytoplasm | enzyme | 9416131 | 7379191.5 | 7639230.5 | | | | | | |
| Q9EPK2 | RP2 | retinitis pigmentosa 2 (X-linked recessive) | Cytoplasm | enzyme | 8129037 | 8404473 | 8057357 | | | | | | |
| Q9EPK2 | RP2 | retinitis pigmentosa 2 (X-linked recessive) | Cytoplasm | enzyme | 6460656 | 6825590 | 5561631.5 | | | | | | |
| Q9EPK2 | RP2 | retinitis pigmentosa 2 (X-linked recessive) | Cytoplasm | enzyme | 6385707 | 6243629 | 5978068.5 | | | | | | |
| Q9EPK2 | RP2 | retinitis pigmentosa 2 (X-linked recessive) | Cytoplasm | enzyme | 5308340 | 5877653 | 5682513 | | | | | | |
| Q9EPK2 | RP2 | retinitis pigmentosa 2 (X-linked recessive) | Cytoplasm | enzyme | 3458184 | 3182041.5 | 2469465.25 | Q9EPK2 | RP2 | 7724008 | 3807949 | 830962 | 21 |
| P53026 | RPL10A | ribosomal protein L10a | Nucleus | other | 1659320.125 | 3124869 | 3541968.5 | | | | | | |
| P53026 | RPL10A | ribosomal protein L10a | Nucleus | other | 1125866.75 | 1436548.125 | 1874855.25 | P53026 | RPL10A | 1392593 | 975620 | 398295 | 6 |
| P86048 | RPL10L | ribosomal protein L10-like | Nucleus | other | 6484175.5 | 1.07E+07 | 1.07E+07 | | | | | | |
| P86048 | RPL10L | ribosomal protein L10-like | Nucleus | other | 2835395.5 | 3976547.25 | 3945936.75 | P86048 | RPL10L | 4659786 | 3507083 | 1431761 | 6 |
| Q9CXW4 | RPL11 | ribosomal protein L11 | Cytoplasm | other | 5643787 | 9740590 | 9646319 | Q9CXW4 | RPL11 | 5643787 | 2338552 | 1350163 | 3 |
| P35979 | RPL12 | ribosomal protein L12 | Nucleus | other | 3224254.5 | 5536253.5 | 4656006 | | | | | | |
| P35979 | RPL12 | ribosomal protein L12 | Nucleus | other | 2885507 | 7216992.5 | 6667336.5 | | | | | | |
| P35979 | RPL12 | ribosomal protein L12 | Nucleus | other | 3529358.5 | 6773075.5 | 7020483.5 | | | | | | |
| P35979 | RPL12 | ribosomal protein L12 | Nucleus | other | 2915220.5 | 5309859.5 | 5868947 | | | | | | |
| P35979 | RPL12 | ribosomal protein L12 | Nucleus | other | 1706580 | 1909670.125 | 2874428.75 | P35979 | RPL12 | 2852184 | 1928117 | 497838 | 15 |
| P47963 | RPL13 | ribosomal protein L13 | Nucleus | other | 3439577.25 | 5570535.5 | 5738656.5 | | | | | | |
| P47963 | RPL13 | ribosomal protein L13 | Nucleus | other | 1478918.875 | 3662503.75 | 3327614 | | | | | | |
| P47963 | RPL13 | ribosomal protein L13 | Nucleus | other | 1457157.875 | 3030073.75 | 2921649.5 | | | | | | |
| P47963 | RPL13 | ribosomal protein L13 | Nucleus | other | 1610958.125 | 2160802.5 | 3137255.25 | P47963 | RPL13 | 1996653 | 1414357 | 408290 | 12 |
| P19253 | RPL13A | ribosomal protein L13a | Cytoplasm | other | 1718581.75 | 3945441.75 | 4029295.75 | P19253 | RPL13A | 1718582 | 1310556 | 756650 | 3 |
| Q9CR57 | RPL14 | ribosomal protein L14 | Cytoplasm | other | 6368861 | 8760057 | 1.16E+07 | | | | | | |
| Q9CR57 | RPL14 | ribosomal protein L14 | Cytoplasm | other | 1727585.875 | 3128429.75 | 4189995 | | | | | | |
| Q9CR57 | RPL14 | ribosomal protein L14 | Cytoplasm | other | 3121194.5 | 4345216 | 5734165.5 | Q9CR57 | RPL14 | 3739214 | 3103584 | 1034528 | 9 |
| Q9CZM2 | RPL15 | ribosomal protein L15 | Cytoplasm | other | 3071175.75 | 4590585 | 6033169 | | | | | | |
| Q9CZM2 | RPL15 | ribosomal protein L15 | Cytoplasm | other | 2084153.75 | 2973336 | 3437350.25 | | | | | | |
| Q9CZM2 | RPL15 | ribosomal protein L15 | Cytoplasm | other | 2189806.25 | 2769780.25 | 3133419.75 | Q9CZM2 | RPL15 | 2448379 | 1239744 | 413248 | 9 |
| Q9CPR4 | RPL17 | ribosomal protein L17 | Cytoplasm | other | 2663541.5 | 4111372 | 4163439 | Q9CPR4 | RPL17 | 2663542 | 851334 | 491518 | 3 |
| P35980 | RPL18 | ribosomal protein L18 | Cytoplasm | other | 7108404 | 9955554 | 1.03E+07 | | | | | | |
| P35980 | RPL18 | ribosomal protein L18 | Cytoplasm | other | 3632836.5 | 6393218 | 6842537 | | | | | | |
| P35980 | RPL18 | ribosomal protein L18 | Cytoplasm | other | 3017218.5 | 4536289 | 5114068.5 | | | | | | |
| P35980 | RPL18 | ribosomal protein L18 | Cytoplasm | other | 1799957.625 | 2843972.25 | 3791942.5 | P35980 | RPL18 | 3889604 | 2735887 | 789783 | 12 |
| P62717 | RPL18A | ribosomal protein L18a | Cytoplasm | other | 1708978.625 | 4096419.75 | 4901271 | P62717 | RPL18A | 1708979 | 1660241 | 958540 | 3 |
| P84099 | RPL19 | ribosomal protein L19 | Cytoplasm | other | 3246301 | 7979222.5 | 1.02E+07 | P84099 | RPL19 | 3246301 | 3534487 | 2040637 | 3 |
| O09167 | RPL21 | ribosomal protein L21 | Cytoplasm | other | 4136038.5 | 8181020.5 | 9031615 | | | | | | |
| O09167 | RPL21 | ribosomal protein L21 | Cytoplasm | other | 1762466 | 2786803 | 2989458.5 | O09167 | RPL21 | 2949252 | 3044177 | 1242780 | 6 |
| P67984 | RPL22 | ribosomal protein L22 | Nucleus | other | 7592626 | 1.18E+07 | 1.41E+07 | | | | | | |
| P67984 | RPL22 | ribosomal protein L22 | Nucleus | other | 3671351.75 | 5492105 | 6341486.5 | P67984 | RPL22 | 5631989 | 3991621 | 1629572 | 6 |
| P62830 | RPL23 | ribosomal protein L23 | Cytoplasm | other | 2901537.5 | 5315773.5 | 8828771 | P62830 | RPL23 | 2901538 | 2980542 | 1720817 | 3 |
| P62751 | Rpl23a | ribosomal protein L23A | Nucleus | other | 6121310 | 9952392 | 1.09E+07 | | | | | | |
| P62751 | Rpl23a | ribosomal protein L23A | Nucleus | other | 1382155.75 | 1709502 | 2113661.75 | P62751 | Rpl23a | 3751733 | 4282939 | 1748502 | 6 |
| Q8BP67 | RPL24 | ribosomal protein L24 | Cytoplasm | other | 1893788.5 | 3183647.25 | 4044777 | | | | | | |
| Q8BP67 | RPL24 | ribosomal protein L24 | Cytoplasm | other | 1341480.875 | 2500060 | 2678533.5 | | | | | | |
| Q8BP67 | RPL24 | ribosomal protein L24 | Cytoplasm | other | 2259352.75 | 2860706.25 | 3706146.25 | Q8BP67 | RPL24 | 1831541 | 852020 | 284007 | 9 |
| P61255 | RPL26 | ribosomal protein L26 | Cytoplasm | other | 1374454.125 | 2101280 | 1859184.5 | P61255 | RPL26 | 1374454 | 370101 | 213678 | 3 |
| P14115 | RPL27A | ribosomal protein L27a | Cytoplasm | other | 1727243.375 | 3791821.25 | 4478588 | P14115 | RPL27A | 1727243 | 1432017 | 826775 | 3 |
| P41105 | RPL28 | ribosomal protein L28 | Cytoplasm | other | 3018294.5 | 4029916.75 | 4001030 | P41105 | RPL28 | 3018295 | 575903 | 332498 | 3 |
| P47915 | Rpl29 (includes others) | ribosomal protein L29 | Cytoplasm | other | 843466.312 | 972340.25 | 755440.125 | P47915 | Rpl29 (includes others) | 843466 | 109089 | 62983 | 3 |
| P27659 | RPL3 | ribosomal protein L3 | Nucleus | other | 6624493.5 | 1.17E+07 | 1.13E+07 | | | | | | |
| P27659 | RPL3 | ribosomal protein L3 | Nucleus | other | 3808569.75 | 3982546.75 | 9221089 | | | | | | |
| P27659 | RPL3 | ribosomal protein L3 | Nucleus | other | 2042860.875 | 3147643.75 | 4177880 | P27659 | RPL3 | 4158641 | 3649629 | 1216543 | 9 |
| P62889 | RPL30 | ribosomal protein L30 | Cytoplasm | other | 3551497 | 6501069 | 7815951.5 | P62889 | RPL30 | 3551497 | 2183822 | 1260830 | 3 |
| P62900 | RPL31 | ribosomal protein L31 | Cytoplasm | other | 3210524.25 | 5572311.5 | 6661562.5 | | | | | | |
| P62900 | RPL31 | ribosomal protein L31 | Cytoplasm | other | 3989362 | 8017682 | 1.06E+07 | P62900 | RPL31 | 3599943 | 2716210 | 1108888 | 6 |
| Q6ZWV7 | RPL35 | ribosomal protein L35 | Cytoplasm | other | 4004976.5 | 6381191 | 6463825 | | | | | | |
| Q6ZWV7 | RPL35 | ribosomal protein L35 | Cytoplasm | other | 1531139.375 | 2960483.75 | 3453158.75 | Q6ZWV7 | RPL35 | 2768058 | 1955001 | 798126 | 6 |
| P47964 | Rpl36 | ribosomal protein L36 | Nucleus | other | 1809137.625 | 2909503.75 | 3295739.5 | | | | | | |
| P47964 | Rpl36 | ribosomal protein L36 | Nucleus | other | 1564069.375 | 2034331.5 | 1593752.375 | P47964 | Rpl36 | 1686604 | 728862 | 297557 | 6 |
| Q9D8E6 | RPL4 | ribosomal protein L4 | Cytoplasm | enzyme | 3715516.5 | 4428635 | 7323097.5 | | | | | | |
| Q9D8E6 | RPL4 | ribosomal protein L4 | Cytoplasm | enzyme | 2251863 | 4462544 | 4860372 | | | | | | |
| Q9D8E6 | RPL4 | ribosomal protein L4 | Cytoplasm | enzyme | 2170865.75 | 3038963 | 4394470 | | | | | | |
| Q9D8E6 | RPL4 | ribosomal protein L4 | Cytoplasm | enzyme | 1338192 | 3191079 | 2481136.75 | Q9D8E6 | RPL4 | 2369109 | 1601626 | 462350 | 12 |
| P47962 | RPL5 | ribosomal protein L5 | Cytoplasm | other | 858456.438 | 1176604.5 | 3981923.25 | P47962 | RPL5 | 858456 | 1718870 | 992390 | 3 |
| P47911 | RPL6 | ribosomal protein L6 | Nucleus | other | 2777193.5 | 5579656.5 | 6151745.5 | | | | | | |
| P47911 | RPL6 | ribosomal protein L6 | Nucleus | other | 487017.75 | 4011489.5 | 3694450.5 | | | | | | |
| P47911 | RPL6 | ribosomal protein L6 | Nucleus | other | 1102774.375 | 3656227.25 | 2906784.5 | | | | | | |
| P47911 | RPL6 | ribosomal protein L6 | Nucleus | other | 2716304.25 | 316673.719 | 19613.844 | P47911 | RPL6 | 1770822 | 2002401 | 578043 | 12 |
| P14148 | RPL7 | ribosomal protein L7 | Nucleus | transcription regulator | 7726153.5 | 1.28E+07 | 1.24E+07 | | | | | | |
| P14148 | RPL7 | ribosomal protein L7 | Nucleus | transcription regulator | 6376312.5 | 9366439 | 8156105.5 | | | | | | |
| P14148 | RPL7 | ribosomal protein L7 | Nucleus | transcription regulator | 3249277 | 7713478 | 8012081 | | | | | | |
| P14148 | RPL7 | ribosomal protein L7 | Nucleus | transcription regulator | 2746319.5 | 5144825 | 7799504 | | | | | | |
| P14148 | RPL7 | ribosomal protein L7 | Nucleus | transcription regulator | 757285.688 | 938091 | 1169875 | P14148 | RPL7 | 4171070 | 3878441 | 1001409 | 15 |
| P12970 | RPL7A | ribosomal protein L7a | Cytoplasm | other | 4548924 | 7259858 | 8032003 | | | | | | |
| P12970 | RPL7A | ribosomal protein L7a | Cytoplasm | other | 2897563.5 | 6154505 | 6274423.5 | | | | | | |
| P12970 | RPL7A | ribosomal protein L7a | Cytoplasm | other | 1311591.875 | 3550166.5 | 4476266 | P12970 | RPL7A | 2919360 | 2174690 | 724897 | 9 |

| | | | | | | | | | | | | | |
|---|---|---|---|---|---|---|---|---|---|---|---|---|---|
| P62918 | RPL8 | ribosomal protein L8 | Cytoplasm | other | 1197353 | 1750346.5 | 2219811.25 | | | | | | |
| P62918 | RPL8 | ribosomal protein L8 | Cytoplasm | other | 694143.5 | 924698.5 | 784321.438 | P62918 | RPL8 | 945748 | 604065 | 246609 | 6 |
| P51410 | RPL9 | ribosomal protein L9 | Nucleus | other | 3925789 | 6057149.5 | 7301853.5 | | | | | | |
| P51410 | RPL9 | ribosomal protein L9 | Nucleus | other | 2280661.25 | 4354955.5 | 4346387.5 | | | | | | |
| P51410 | RPL9 | ribosomal protein L9 | Nucleus | other | 3020165.25 | 6009036.5 | 5843700.5 | P51410 | RPL9 | 3075539 | 1623011 | 541004 | 9 |
| P14869 | RPLP0 | ribosomal protein, large, P0 | Cytoplasm | other | 9931114 | 1.43E+07 | 1.63E+07 | | | | | | |
| P14869 | RPLP0 | ribosomal protein, large, P0 | Cytoplasm | other | 3343930 | 5963495.5 | 5276161 | | | | | | |
| P14869 | RPLP0 | ribosomal protein, large, P0 | Cytoplasm | other | 4080770.25 | 5731630 | 6158441 | | | | | | |
| P14869 | RPLP0 | ribosomal protein, large, P0 | Cytoplasm | other | 3463947.25 | 4596621 | 5978710.5 | | | | | | |
| P14869 | RPLP0 | ribosomal protein, large, P0 | Cytoplasm | other | 2281030.5 | 5287670.5 | 5036132 | | | | | | |
| P14869 | RPLP0 | ribosomal protein, large, P0 | Cytoplasm | other | 2323533.75 | 4171584.75 | 4600588.5 | | | | | | |
| P14869 | RPLP0 | ribosomal protein, large, P0 | Cytoplasm | other | 3453152 | 5424350.5 | 6111295.5 | | | | | | |
| P14869 | RPLP0 | ribosomal protein, large, P0 | Cytoplasm | other | 1028261.688 | 1061584.625 | 1890048.75 | P14869 | RPLP0 | 3738217 | 3648028 | 744651 | 24 |
| P47955 | Rplp1 (includes others) | ribosomal protein, large, P1 | Nucleus | other | 1.76E+07 | 3.01E+07 | 3.20E+07 | | | | | | |
| P47955 | Rplp1 (includes others) | ribosomal protein, large, P1 | Nucleus | other | 1.94E+07 | 2.40E+07 | 3.86E+07 | P47955 | Rplp1 (includes others) | 18485000 | 8057153 | 3289319 | 6 |
| P99027 | RPLP2 | ribosomal protein, large, P2 | Cytoplasm | other | 1.67E+07 | 2.65E+07 | 2.53E+07 | | | | | | |
| P99027 | RPLP2 | ribosomal protein, large, P2 | Cytoplasm | other | 2.23E+07 | 3.34E+07 | 3.13E+07 | | | | | | |
| P99027 | RPLP2 | ribosomal protein, large, P2 | Cytoplasm | other | 3979015.25 | 6613139 | 9508925 | | | | | | |
| P99027 | RPLP2 | ribosomal protein, large, P2 | Cytoplasm | other | 2715412.5 | 4192549.25 | 4455312.5 | | | | | | |
| P99027 | RPLP2 | ribosomal protein, large, P2 | Cytoplasm | other | 2050366.25 | 4008538.75 | 3708860.5 | P99027 | RPLP2 | 9546959 | 11525878 | 2975969 | 15 |
| Q91YQ5 | RPN1 | ribophorin I | Cytoplasm | enzyme | 4089318 | 3769860.75 | 6058493.5 | | | | | | |
| Q91YQ5 | RPN1 | ribophorin I | Cytoplasm | enzyme | 5384088.5 | 6510352 | 1.17E+07 | | | | | | |
| Q91YQ5 | RPN1 | ribophorin I | Cytoplasm | enzyme | 5337861.5 | 9744480 | 1.12E+07 | | | | | | |
| Q91YQ5 | RPN1 | ribophorin I | Cytoplasm | enzyme | 2967293 | 4009124 | 5239420 | | | | | | |
| Q91YQ5 | RPN1 | ribophorin I | Cytoplasm | enzyme | 2850968.5 | 3793276.5 | 5029960.5 | | | | | | |
| Q91YQ5 | RPN1 | ribophorin I | Cytoplasm | enzyme | 3040367.5 | 4754489.5 | 5488655 | | | | | | |
| Q91YQ5 | RPN1 | ribophorin I | Cytoplasm | enzyme | 1513894.25 | 2046234.75 | 3053922 | | | | | | |
| Q91YQ5 | RPN1 | ribophorin I | Cytoplasm | enzyme | 1565928.25 | 1709186.75 | 1995997.5 | Q91YQ5 | RPN1 | 3343715 | 2805473 | 572665 | 24 |
| Q9DBG6 | RPN2 | ribophorin II | Cytoplasm | enzyme | 7486205.5 | 1.04E+07 | 1.44E+07 | | | | | | |
| Q9DBG6 | RPN2 | ribophorin II | Cytoplasm | enzyme | 8774603 | 1.07E+07 | 1.53E+07 | | | | | | |
| Q9DBG6 | RPN2 | ribophorin II | Cytoplasm | enzyme | 4101732.5 | 4741375 | 7420188.5 | | | | | | |
| Q9DBG6 | RPN2 | ribophorin II | Cytoplasm | enzyme | 3763299.75 | 5442208.5 | 5711547 | | | | | | |
| Q9DBG6 | RPN2 | ribophorin II | Cytoplasm | enzyme | 3248178 | 4206756 | 5758153 | | | | | | |
| Q9DBG6 | RPN2 | ribophorin II | Cytoplasm | enzyme | 2627729 | 2931464.75 | 4154639 | | | | | | |
| Q9DBG6 | RPN2 | ribophorin II | Cytoplasm | enzyme | 3143680.25 | 3516524.75 | 5493630.5 | | | | | | |
| Q9DBG6 | RPN2 | ribophorin II | Cytoplasm | enzyme | 1867489.875 | 2852366.5 | 3447910.25 | | | | | | |
| Q9DBG6 | RPN2 | ribophorin II | Cytoplasm | enzyme | 1994716.125 | 2576894.25 | 4988621.5 | | | | | | |
| Q9DBG6 | RPN2 | ribophorin II | Cytoplasm | enzyme | 1032170.438 | 1589285.125 | 2642609 | Q9DBG6 | RPN2 | 3803980 | 3562402 | 650403 | 30 |
| P63325 | RPS10 | ribosomal protein S10 | Cytoplasm | other | 3623062.75 | 5521776.5 | 7375697 | P63325 | RPS10 | 3623063 | 1876362 | 1083318 | 3 |
| P62281 | RPS11 | ribosomal protein S11 | Cytoplasm | other | 2315234.5 | 5238335.5 | 6286789.5 | | | | | | |
| P62281 | RPS11 | ribosomal protein S11 | Cytoplasm | other | 290773.406 | 1201564.375 | 970425.75 | P62281 | RPS11 | 1303004 | 2469639 | 1008226 | 6 |
| P62301 | RPS13 | ribosomal protein S13 | Cytoplasm | other | 1073442.875 | 2303256.75 | 2138453.5 | | | | | | |
| P62301 | RPS13 | ribosomal protein S13 | Cytoplasm | other | 990028.125 | 2173923 | 2596769 | P62301 | RPS13 | 1031736 | 676561 | 276205 | 6 |
| P62264 | RPS14 | ribosomal protein S14 | Cytoplasm | translation regulator | 5042586 | 9677639 | 1.05E+07 | | | | | | |
| P62264 | RPS14 | ribosomal protein S14 | Cytoplasm | translation regulator | 3763738 | 6300487 | 6321071 | | | | | | |
| P62264 | RPS14 | ribosomal protein S14 | Cytoplasm | translation regulator | 3099316.25 | 6768169 | 4634694 | P62264 | RPS14 | 3968547 | 2504376 | 834792 | 9 |
| P62245 | RPS15A | ribosomal protein S15a | Cytoplasm | other | 1581128.25 | 2429526.5 | 2377600.75 | | | | | | |
| P62245 | RPS15A | ribosomal protein S15a | Cytoplasm | other | 2956512.5 | 3436328.25 | 3805735 | P62245 | RPS15A | 2268820 | 804321 | 328363 | 6 |
| P14131 | RPS16 | ribosomal protein S16 | Cytoplasm | other | 4692116 | 9058793 | 8718957 | | | | | | |
| P14131 | RPS16 | ribosomal protein S16 | Cytoplasm | other | 1675151.875 | 4768851 | 5557045.5 | P14131 | RPS16 | 3183634 | 2774385 | 1132638 | 6 |
| P63276 | RPS17 | ribosomal protein S17 | Cytoplasm | other | 1387655 | 2653349.5 | 3279802.75 | P63276 | RPS17 | 1387655 | 963903 | 556509 | 3 |
| P62270 | RPS18 | ribosomal protein S18 | Cytoplasm | other | 2772845.5 | 5326812.5 | 5200629.5 | | | | | | |
| P62270 | RPS18 | ribosomal protein S18 | Cytoplasm | other | 1584777.375 | 2825684.5 | 2850557.5 | | | | | | |
| P62270 | RPS18 | ribosomal protein S18 | Cytoplasm | other | 1644815.25 | 2180901.75 | 3244643.25 | P62270 | RPS18 | 2000813 | 1364137 | 454712 | 9 |
| Q9CZX8 | RPS19 | ribosomal protein S19 | Cytoplasm | other | 3174702 | 6308057 | 8037330.5 | | | | | | |
| Q9CZX8 | RPS19 | ribosomal protein S19 | Cytoplasm | other | 2739788.75 | 5605701.5 | 4920569.5 | | | | | | |
| Q9CZX8 | RPS19 | ribosomal protein S19 | Cytoplasm | other | 3170035.25 | 4815947 | 5378296.5 | Q9CZX8 | RPS19 | 3028175 | 1702770 | 567590 | 9 |
| P25444 | RPS2 | ribosomal protein S2 | Cytoplasm | other | 6066427 | 1.01E+07 | 9056823 | | | | | | |
| P25444 | RPS2 | ribosomal protein S2 | Cytoplasm | other | 3567961 | 9204592 | 7606932 | | | | | | |
| P25444 | RPS2 | ribosomal protein S2 | Cytoplasm | other | 2849184.75 | 5849622.5 | 5565600 | | | | | | |
| P25444 | RPS2 | ribosomal protein S2 | Cytoplasm | other | 2534005.25 | 5017419.5 | 5508927 | P25444 | RPS2 | 3754395 | 2487267 | 718012 | 12 |
| P60867 | RPS20 | ribosomal protein S20 | Cytoplasm | other | 4454651 | 7553746.5 | 7431559.5 | | | | | | |
| P60867 | RPS20 | ribosomal protein S20 | Cytoplasm | other | 2791961.5 | 5391677 | 6112185 | P60867 | RPS20 | 3623306 | 1825996 | 745460 | 6 |
| P62267 | RPS23 | ribosomal protein S23 | Cytoplasm | translation regulator | 2395434.5 | 4226668 | 4967581.5 | P62267 | RPS23 | 2395435 | 1324029 | 764428 | 3 |
| P62849 | RPS24 | ribosomal protein S24 | Cytoplasm | other | 1111784.5 | 1830181.75 | 2048230.75 | P62849 | RPS24 | 1111785 | 489995 | 282899 | 3 |
| P62852 | RPS25 | ribosomal protein S25 | Cytoplasm | other | 3923324.25 | 8001318 | 8121754 | | | | | | |
| P62852 | RPS25 | ribosomal protein S25 | Cytoplasm | other | 3588962 | 5752754.5 | 7782712.5 | | | | | | |
| P62852 | RPS25 | ribosomal protein S25 | Cytoplasm | other | 2763421 | 5541924.5 | 5351706 | P62852 | RPS25 | 3425236 | 1995605 | 665202 | 9 |
| P62855 | RPS26 | ribosomal protein S26 | Cytoplasm | other | 2131146 | 2873987 | 3685881 | P62855 | RPS26 | 2131146 | 777623 | 448961 | 3 |
| Q6ZWY3 | RPS27L | ribosomal protein S27-like | Cytoplasm | translation regulator | 1168203.25 | 1961442.875 | 1784173.75 | | | | | | |
| Q6ZWY3 | RPS27L | ribosomal protein S27-like | Cytoplasm | translation regulator | 440794.156 | 603191.438 | 444727.562 | Q6ZWY3 | RPS27L | 804499 | 681041 | 278034 | 6 |
| P62858 | RPS28 | ribosomal protein S28 | Cytoplasm | other | 697472.688 | 1850285.625 | 1465111.875 | P62858 | RPS28 | 697473 | 586885 | 338838 | 3 |
| P62908 | RPS3 | ribosomal protein S3 | Cytoplasm | enzyme | 4667001 | 7263991 | 7954987 | | | | | | |
| P62908 | RPS3 | ribosomal protein S3 | Cytoplasm | enzyme | 3785020 | 7077406.5 | 6474503.5 | | | | | | |
| P62908 | RPS3 | ribosomal protein S3 | Cytoplasm | enzyme | 2299506.25 | 3253214 | 3498263.5 | | | | | | |
| P62908 | RPS3 | ribosomal protein S3 | Cytoplasm | enzyme | 2725736 | 6198114.5 | 5632147.5 | | | | | | |
| P62908 | RPS3 | ribosomal protein S3 | Cytoplasm | enzyme | 2146494 | 3942010.75 | 3332030.5 | | | | | | |
| P62908 | RPS3 | ribosomal protein S3 | Cytoplasm | enzyme | 1305686.625 | 1192319.875 | 2565158.5 | | | | | | |
| P62908 | RPS3 | ribosomal protein S3 | Cytoplasm | enzyme | 1136573.875 | 2049875.75 | 2091053.5 | | | | | | |
| P62908 | RPS3 | ribosomal protein S3 | Cytoplasm | enzyme | 793652.188 | 1585622.625 | 1315865 | P62908 | RPS3 | 2357459 | 2182245 | 445449 | 24 |
| P97351 | Rps3a1 | ribosomal protein S3A1 | Cytoplasm | other | 1200540.625 | 2041206.875 | 2723222 | | | | | | |
| P97351 | Rps3a1 | ribosomal protein S3A1 | Cytoplasm | other | 292042.094 | 805601.75 | 1092540.5 | P97351 | Rps3a1 | 746291 | 879462 | 359039 | 6 |
| P62702 | RPS4Y1 | ribosomal protein S4, Y-linked 1 | Cytoplasm | other | 3749904 | 6546923.5 | 7633395 | | | | | | |
| P62702 | RPS4Y1 | ribosomal protein S4, Y-linked 1 | Cytoplasm | other | 2567382.25 | 5028636 | 5940965 | | | | | | |
| P62702 | RPS4Y1 | ribosomal protein S4, Y-linked 1 | Cytoplasm | other | 955269.125 | 1075186 | 147472.359 | | | | | | |
| P62702 | RPS4Y1 | ribosomal protein S4, Y-linked 1 | Cytoplasm | other | 1255148.875 | 3005452 | 2494126.5 | | | | | | |
| P62702 | RPS4Y1 | ribosomal protein S4, Y-linked 1 | Cytoplasm | other | 1411701.25 | 2617407.25 | 2596209.5 | | | | | | |
| P62702 | RPS4Y1 | ribosomal protein S4, Y-linked 1 | Cytoplasm | other | 1370639.625 | 1903263.375 | 2088619.375 | | | | | | |
| P62702 | RPS4Y1 | ribosomal protein S4, Y-linked 1 | Cytoplasm | other | 1214319.75 | 2783844.5 | 2745172.25 | | | | | | |
| P62702 | RPS4Y1 | ribosomal protein S4, Y-linked 1 | Cytoplasm | other | 1173602.75 | 1915749.75 | 1833189.625 | P62702 | RPS4Y1 | 1712246 | 1876036 | 382944 | 24 |
| P62754 | RPS6 | ribosomal protein S6 | Cytoplasm | other | 3181610 | 4756275.5 | 5989793.5 | | | | | | |
| P62754 | RPS6 | ribosomal protein S6 | Cytoplasm | other | 710959.375 | 860212.875 | 1671898.75 | P62754 | RPS6 | 1946285 | 2169055 | 885513 | 6 |
| P62082 | RPS7 | ribosomal protein S7 | Cytoplasm | other | 6581166 | 1.19E+07 | 1.27E+07 | | | | | | |
| P62082 | RPS7 | ribosomal protein S7 | Cytoplasm | other | 3265638 | 7308248 | 5429290.5 | | | | | | |
| P62082 | RPS7 | ribosomal protein S7 | Cytoplasm | other | 1579277 | 2958699.5 | 2938706.75 | P62082 | RPS7 | 3808694 | 3998690 | 1332897 | 9 |
| P62242 | RPS8 | ribosomal protein S8 | Cytoplasm | other | 4813436.5 | 8555079 | 9953057 | | | | | | |
| P62242 | RPS8 | ribosomal protein S8 | Cytoplasm | other | 4416758 | 7796814.5 | 6993361 | | | | | | |
| P62242 | RPS8 | ribosomal protein S8 | Cytoplasm | other | 3317968.75 | 6742182 | 6380375 | | | | | | |
| P62242 | RPS8 | ribosomal protein S8 | Cytoplasm | other | 2073389.625 | 3068935 | 3115754 | | | | | | |
| P62242 | RPS8 | ribosomal protein S8 | Cytoplasm | other | 1616867.5 | 2347440 | 2309284.75 | P62242 | RPS8 | 3247684 | 2652870 | 684968 | 15 |
| Q6ZWN5 | RPS9 | ribosomal protein S9 | Cytoplasm | translation regulator | 4490659.5 | 6344873 | 6669273.5 | | | | | | |
| Q6ZWN5 | RPS9 | ribosomal protein S9 | Cytoplasm | translation regulator | 2180479 | 4098003.25 | 3935133.5 | | | | | | |
| Q6ZWN5 | RPS9 | ribosomal protein S9 | Cytoplasm | translation regulator | 1480021.75 | 3120122 | 2809239.75 | Q6ZWN5 | RPS9 | 2717053 | 1755225 | 585075 | 9 |
| P14206 | RPSA | ribosomal protein SA | Cytoplasm | translation regulator | 9278539 | 1.66E+07 | 1.88E+07 | | | | | | |
| P14206 | RPSA | ribosomal protein SA | Cytoplasm | translation regulator | 4554788.5 | 8788665 | 9482125 | | | | | | |
| P14206 | RPSA | ribosomal protein SA | Cytoplasm | translation regulator | 4113265.5 | 8077788.5 | 7780685.5 | | | | | | |
| P14206 | RPSA | ribosomal protein SA | Cytoplasm | translation regulator | 1833838.625 | 3695898.5 | 3389595.5 | P14206 | RPSA | 4945108 | 5209258 | 1503783 | 12 |
| Q8CBB9 | RSAD2 | radical S-adenosyl methionine domain containing 2 | Cytoplasm | enzyme | 7429294.5 | 8752827 | 2.08E+07 | | | | | | |
| Q8CBB9 | RSAD2 | radical S-adenosyl methionine domain containing 2 | Cytoplasm | enzyme | 5899065.5 | 1.27E+07 | 2.28E+07 | | | | | | |

| | | | | | | | | | | | | | |
|---|---|---|---|---|---|---|---|---|---|---|---|---|---|
| Q8CBB9 | RSAD2 | radical S-adenosyl methionine domain containing 2 | Cytoplasm | enzyme | 3978718.75 | 6068045 | 1.10E+07 | | | | | | |
| Q8CBB9 | RSAD2 | radical S-adenosyl methionine domain containing 2 | Cytoplasm | enzyme | 4246875.5 | 1.08E+07 | 2.85E+07 | | | | | | |
| Q8CBB9 | RSAD2 | radical S-adenosyl methionine domain containing 2 | Cytoplasm | enzyme | 2529234.5 | 5955818.5 | 9958679 | | | | | | |
| Q8CBB9 | RSAD2 | radical S-adenosyl methionine domain containing 2 | Cytoplasm | enzyme | 1901504.25 | 3610476 | 5893847 | | | | | | |
| Q8CBB9 | RSAD2 | radical S-adenosyl methionine domain containing 2 | Cytoplasm | enzyme | 1938794.75 | 3345896.75 | 6503079 | | | | | | |
| Q8CBB9 | RSAD2 | radical S-adenosyl methionine domain containing 2 | Cytoplasm | enzyme | 1556315.75 | 2537627.75 | 5531543 | Q8CBB9 | RSAD2 | 3684975 | 6963913 | 1421503 | 24 |
| Q01730 | RSU1 | Ras suppressor protein 1 | Cytoplasm | other | 9689436 | 4675914.5 | 2.86E+07 | | | | | | |
| Q01730 | RSU1 | Ras suppressor protein 1 | Cytoplasm | other | 5927087 | 3920951 | 2.10E+07 | | | | | | |
| Q01730 | RSU1 | Ras suppressor protein 1 | Cytoplasm | other | 1427508.125 | 328425.875 | 5348645.5 | Q01730 | RSU1 | 5681344 | 9538580 | 3179527 | 9 |
| Q8K0T0 | RTN1 | reticulon 1 | Cytoplasm | other | 8623448 | 1.27E+07 | 1.03E+07 | Q8K0T0 | RTN1 | 8623448 | 2025780 | 1169585 | 3 |
| Q9ES97 | RTN3 | reticulon 3 | Cytoplasm | other | 6.38E+07 | 6.95E+07 | 7.22E+07 | | | | | | |
| Q9ES97 | RTN3 | reticulon 3 | Cytoplasm | other | 781623.625 | 1018793.312 | 579428.625 | Q9ES97 | RTN3 | 32285812 | 37178931 | 15178235 | 6 |
| Q99P72 | RTN4 | reticulon 4 | Cytoplasm | other | 1.57E+07 | 1.79E+07 | 1.28E+07 | | | | | | |
| Q99P72 | RTN4 | reticulon 4 | Cytoplasm | other | 9268905 | 1.07E+07 | 1.46E+07 | | | | | | |
| Q99P72 | RTN4 | reticulon 4 | Cytoplasm | other | 7607004 | 1.02E+07 | 8505534 | | | | | | |
| Q99P72 | RTN4 | reticulon 4 | Cytoplasm | other | 2854565 | 3686421.25 | 2738447.5 | | | | | | |
| Q99P72 | RTN4 | reticulon 4 | Cytoplasm | other | 3818679.5 | 3376771.25 | 3974762 | Q99P72 | RTN4 | 7851831 | 5091979 | 1314743 | 15 |
| P97352 | S100A13 | S100 calcium binding protein A13 | Cytoplasm | other | 2403123.5 | 5844609.5 | 2971692.75 | P97352 | S100A13 | 2403124 | 1844846 | 1065123 | 3 |
| P31725 | S100A9 | S100 calcium binding protein A9 | Cytoplasm | other | 730405.5 | 392219.562 | 2277821.75 | | | | | | |
| P31725 | S100A9 | S100 calcium binding protein A9 | Cytoplasm | other | 464014.188 | 312309.594 | 2356633.25 | P31725 | S100A9 | 597210 | 962096 | 392774 | 6 |
| Q9EP69 | SACM1L | SAC1 suppressor of actin mutations 1-like (yeast) | Cytoplasm | phosphatase | 1.20E+07 | 1.22E+07 | 9498625 | | | | | | |
| Q9EP69 | SACM1L | SAC1 suppressor of actin mutations 1-like (yeast) | Cytoplasm | phosphatase | 1.24E+07 | 9157684 | 9752524 | | | | | | |
| Q9EP69 | SACM1L | SAC1 suppressor of actin mutations 1-like (yeast) | Cytoplasm | phosphatase | 1.18E+07 | 1.06E+07 | 9612655 | | | | | | |
| Q9EP69 | SACM1L | SAC1 suppressor of actin mutations 1-like (yeast) | Cytoplasm | phosphatase | 9979902 | 9435018 | 6775577 | | | | | | |
| Q9EP69 | SACM1L | SAC1 suppressor of actin mutations 1-like (yeast) | Cytoplasm | phosphatase | 4302948.5 | 5732510.5 | 6317568 | | | | | | |
| Q9EP69 | SACM1L | SAC1 suppressor of actin mutations 1-like (yeast) | Cytoplasm | phosphatase | 7637540 | 5882568.5 | 6320088 | | | | | | |
| Q9EP69 | SACM1L | SAC1 suppressor of actin mutations 1-like (yeast) | Cytoplasm | phosphatase | 7695080 | 7919672 | 4961378 | | | | | | |
| Q9EP69 | SACM1L | SAC1 suppressor of actin mutations 1-like (yeast) | Cytoplasm | phosphatase | 6885528 | 6547048 | 6031433.5 | | | | | | |
| Q9EP69 | SACM1L | SAC1 suppressor of actin mutations 1-like (yeast) | Cytoplasm | phosphatase | 5285070 | 4085869.5 | 4013653.25 | | | | | | |
| Q9EP69 | SACM1L | SAC1 suppressor of actin mutations 1-like (yeast) | Cytoplasm | phosphatase | 5748038.5 | 5335565 | 4404862 | | | | | | |
| Q9EP69 | SACM1L | SAC1 suppressor of actin mutations 1-like (yeast) | Cytoplasm | phosphatase | 5576638.5 | 4322981.5 | 3668742 | | | | | | |
| Q9EP69 | SACM1L | SAC1 suppressor of actin mutations 1-like (yeast) | Cytoplasm | phosphatase | 3685214.5 | 3550590 | 3155594 | | | | | | |
| Q9EP69 | SACM1L | SAC1 suppressor of actin mutations 1-like (yeast) | Cytoplasm | phosphatase | 3055534.5 | 2924236.5 | 3028882.25 | | | | | | |
| Q9EP69 | SACM1L | SAC1 suppressor of actin mutations 1-like (yeast) | Cytoplasm | phosphatase | 2735024.25 | 2916062.5 | 2476519.75 | | | | | | |
| Q9EP69 | SACM1L | SAC1 suppressor of actin mutations 1-like (yeast) | Cytoplasm | phosphatase | 2985424.5 | 2193359.25 | 1631730.5 | | | | | | |
| Q9EP69 | SACM1L | SAC1 suppressor of actin mutations 1-like (yeast) | Cytoplasm | phosphatase | 1931206.375 | 2194187.75 | 1728848.125 | | | | | | |
| Q9EP69 | SACM1L | SAC1 suppressor of actin mutations 1-like (yeast) | Cytoplasm | phosphatase | 2370159.75 | 2442642.75 | 2120026.25 | | | | | | |
| Q9EP69 | SACM1L | SAC1 suppressor of actin mutations 1-like (yeast) | Cytoplasm | phosphatase | 2401606 | 2619103.5 | 1757256.375 | | | | | | |
| Q9EP69 | SACM1L | SAC1 suppressor of actin mutations 1-like (yeast) | Cytoplasm | phosphatase | 2066586.5 | 1193707 | 1666294 | Q9EP69 | SACM1L | 5816395 | 3171003 | 420010 | 57 |
| Q8BGH2 | SAMM50 | SAMM50 sorting and assembly machinery component | Cytoplasm | other | 1694474.25 | 2207687.5 | 3023696.25 | | | | | | |
| Q8BGH2 | SAMM50 | SAMM50 sorting and assembly machinery component | Cytoplasm | other | 1349682.5 | 1188022.75 | 3880052.5 | Q8BGH2 | SAMM50 | 1522078 | 1048844 | 428189 | 6 |
| Q8K021 | SCAMP1 | secretory carrier membrane protein 1 | Cytoplasm | transporter | 1.73E+07 | 1.24E+07 | 8287543.5 | Q8K021 | SCAMP1 | 17260000 | 4492584 | 2593795 | 3 |
| Q9ERN0 | SCAMP2 | secretory carrier membrane protein 2 | Cytoplasm | transporter | 5122369.5 | 6606403.5 | 7976300 | | | | | | |
| Q9ERN0 | SCAMP2 | secretory carrier membrane protein 2 | Cytoplasm | transporter | 4187476.5 | 5653399.5 | 6727024 | Q9ERN0 | SCAMP2 | 4654923 | 1339047 | 546664 | 6 |
| Q9JKV5 | SCAMP4 | secretory carrier membrane protein 4 | Cytoplasm | other | 1.06E+07 | 1.04E+07 | 8269603 | | | | | | |
| Q9JKV5 | SCAMP4 | secretory carrier membrane protein 4 | Cytoplasm | other | 9258022 | 1.10E+07 | 5923805 | Q9JKV5 | SCAMP4 | 9924011 | 1902672 | 776762 | 6 |
| O35114 | SCARB2 | scavenger receptor class B, member 2 | Plasma Membrane | other | 1136375 | 3583750 | 2191144 | O35114 | SCARB2 | 1136375 | 1227568 | 708736 | 3 |
| Q8K2B3 | SDHA | succinate dehydrogenase complex, subunit A, flavoprotein (Fp) | Cytoplasm | enzyme | 8989923 | 1.62E+07 | 1.46E+07 | | | | | | |
| Q8K2B3 | SDHA | succinate dehydrogenase complex, subunit A, flavoprotein (Fp) | Cytoplasm | enzyme | 5887150.5 | 1.32E+07 | 1.11E+07 | | | | | | |
| Q8K2B3 | SDHA | succinate dehydrogenase complex, subunit A, flavoprotein (Fp) | Cytoplasm | enzyme | 4544732.5 | 8205447.5 | 8478058 | | | | | | |
| Q8K2B3 | SDHA | succinate dehydrogenase complex, subunit A, flavoprotein (Fp) | Cytoplasm | enzyme | 3606714.75 | 5349046 | 6074072.5 | | | | | | |
| Q8K2B3 | SDHA | succinate dehydrogenase complex, subunit A, flavoprotein (Fp) | Cytoplasm | enzyme | 3253784 | 4855759 | 5258882.5 | | | | | | |
| Q8K2B3 | SDHA | succinate dehydrogenase complex, subunit A, flavoprotein (Fp) | Cytoplasm | enzyme | 2270964 | 4726717 | 7199953.5 | | | | | | |
| Q8K2B3 | SDHA | succinate dehydrogenase complex, subunit A, flavoprotein (Fp) | Cytoplasm | enzyme | 2444314.75 | 5118741 | 8247211.5 | | | | | | |
| Q8K2B3 | SDHA | succinate dehydrogenase complex, subunit A, flavoprotein (Fp) | Cytoplasm | enzyme | 812178.312 | 2008181.875 | 2174269.5 | Q8K2B3 | SDHA | 3976220 | 4073454 | 831490 | 24 |
| Q9CQA3 | SDHB | succinate dehydrogenase complex, subunit B, iron sulfur (Ip) | Cytoplasm | enzyme | 3320652.25 | 3713389.25 | 3531457.75 | | | | | | |
| Q9CQA3 | SDHB | succinate dehydrogenase complex, subunit B, iron sulfur (Ip) | Cytoplasm | enzyme | 2715545.75 | 4469248.5 | 5924675.5 | | | | | | |
| Q9CQA3 | SDHB | succinate dehydrogenase complex, subunit B, iron sulfur (Ip) | Cytoplasm | enzyme | 1378679 | 3038317.5 | 3600030.25 | Q9CQA3 | SDHB | 2471626 | 1236508 | 412169 | 9 |
| Q9CZB0 | SDHC | succinate dehydrogenase complex, subunit C, integral membrane protein, 15kDa | Cytoplasm | enzyme | 1491520.25 | 3366406.5 | 3906549 | Q9CZB0 | SDHC | 1491520 | 1267499 | 731791 | 3 |
| O08547 | SEC22B | SEC22 vesicle trafficking protein homolog B (S. cerevisiae) (gene/pseudogene) | Cytoplasm | other | 6769412.5 | 4693813.5 | 7177987.5 | | | | | | |
| O08547 | SEC22B | SEC22 vesicle trafficking protein homolog B (S. cerevisiae) (gene/pseudogene) | Cytoplasm | other | 4115623.75 | 2126965.75 | 6085460 | | | | | | |
| O08547 | SEC22B | SEC22 vesicle trafficking protein homolog B (S. cerevisiae) (gene/pseudogene) | Cytoplasm | other | 5227232.5 | 6050521 | 4373991 | | | | | | |
| O08547 | SEC22B | SEC22 vesicle trafficking protein homolog B (S. cerevisiae) (gene/pseudogene) | Cytoplasm | other | 4620086 | 5177194 | 4398477 | | | | | | |
| O08547 | SEC22B | SEC22 vesicle trafficking protein homolog B (S. cerevisiae) (gene/pseudogene) | Cytoplasm | other | 2074072.875 | 2899461.75 | 2497834.75 | | | | | | |
| O08547 | SEC22B | SEC22 vesicle trafficking protein homolog B (S. cerevisiae) (gene/pseudogene) | Cytoplasm | other | 2641069.25 | 3139493.5 | 2327773.75 | | | | | | |
| O08547 | SEC22B | SEC22 vesicle trafficking protein homolog B (S. cerevisiae) (gene/pseudogene) | Cytoplasm | other | 2027924.5 | 1870474.25 | 2262826 | | | | | | |
| O08547 | SEC22B | SEC22 vesicle trafficking protein homolog B (S. cerevisiae) (gene/pseudogene) | Cytoplasm | other | 1504695 | 1719166.75 | 1192078.25 | O08547 | SEC22B | 3622515 | 1791545 | 365698 | 24 |
| Q3TZ89 | SEC31B | SEC31 homolog B, COPII coating complex component | Cytoplasm | other | 1.55E+07 | 2.00E+07 | 2.87E+07 | Q3TZ89 | SEC31B | 15490000 | 6711738 | 3875024 | 3 |
| P61620 | SEC61A1 | Sec61 alpha 1 subunit (S. cerevisiae) | Cytoplasm | transporter | 1398913.5 | 2845068 | 3699629 | P61620 | SEC61A1 | 1398914 | 1162965 | 671438 | 3 |
| Q9CQS8 | SEC61B | Sec61 beta subunit | Cytoplasm | transporter | 4226295 | 5851328 | 6482125 | | | | | | |
| Q9CQS8 | SEC61B | Sec61 beta subunit | Cytoplasm | transporter | 2238666.25 | 2420477 | 3036961.75 | Q9CQS8 | SEC61B | 3232481 | 1797403 | 733787 | 6 |
| P17563 | SELENBP1 | selenium binding protein 1 | Cytoplasm | other | 170060.016 | 221247.328 | 373456.125 | P17563 | SELENBP1 | 170060 | 105797 | 61082 | 3 |
| Q01102 | SELP | selectin P (granule membrane protein 140kDa, antigen CD62) | Plasma Membrane | transmembrane receptor | 3311956.25 | 3121441.75 | 4054834.25 | | | | | | |
| Q01102 | SELP | selectin P (granule membrane protein 140kDa, antigen CD62) | Plasma Membrane | transmembrane receptor | 1.26E+07 | 1.19E+07 | 1.46E+07 | | | | | | |
| Q01102 | SELP | selectin P (granule membrane protein 140kDa, antigen CD62) | Plasma Membrane | transmembrane receptor | 6997192 | 9831946 | 8910430 | | | | | | |
| Q01102 | SELP | selectin P (granule membrane protein 140kDa, antigen CD62) | Plasma Membrane | transmembrane receptor | 1216340.375 | 1361860.5 | 1646485.125 | | | | | | |
| Q01102 | SELP | selectin P (granule membrane protein 140kDa, antigen CD62) | Plasma Membrane | transmembrane receptor | 2900269 | 3862231 | 4077898.75 | | | | | | |
| Q01102 | SELP | selectin P (granule membrane protein 140kDa, antigen CD62) | Plasma Membrane | transmembrane receptor | 681425 | 654780.562 | 939368.688 | Q01102 | SELP | 4609530 | 4505182 | 1061882 | 18 |
| P62342 | SELT | selenoprotein T | Cytoplasm | other | 3101478.75 | 3359492 | 2834916.75 | P62342 | SELT | 3101479 | 262299 | 151439 | 3 |
| Q9CY58 | SERBP1 | SERPINE1 mRNA binding protein 1 | Cytoplasm | other | 636843.812 | 1068359.875 | 1151568.75 | | | | | | |
| Q9CY58 | SERBP1 | SERPINE1 mRNA binding protein 1 | Cytoplasm | other | 1161560.5 | 2734101.5 | 1707738.125 | | | | | | |
| Q9CY58 | SERBP1 | SERPINE1 mRNA binding protein 1 | Cytoplasm | other | 790015.875 | 1989956.625 | 1052898.125 | Q9CY58 | SERBP1 | 862807 | 662913 | 220971 | 9 |
| Q9QZI9 | SERINC3 | serine incorporator 3 | Cytoplasm | transporter | 2.52E+08 | 2.16E+08 | 2.25E+08 | | | | | | |
| Q9QZI9 | SERINC3 | serine incorporator 3 | Cytoplasm | transporter | 1.32E+08 | 1.39E+08 | 1.09E+08 | | | | | | |
| Q9QZI9 | SERINC3 | serine incorporator 3 | Cytoplasm | transporter | 1.22E+08 | 1.39E+08 | 8.49E+07 | | | | | | |
| Q9QZI9 | SERINC3 | serine incorporator 3 | Cytoplasm | transporter | 3.81E+07 | 3.75E+07 | 2.81E+07 | Q9QZI9 | SERINC3 | 135805000 | 74655255 | 21551116 | 12 |
| P07758 | SERPINA1 | serpin peptidase inhibitor, clade A (alpha-1 antiproteinase, antitrypsin), member 1 | Extracellular Space | other | 9331446 | 1.01E+07 | 2.02E+07 | | | | | | |
| P07758 | SERPINA1 | serpin peptidase inhibitor, clade A (alpha-1 antiproteinase, antitrypsin), member 1 | Extracellular Space | other | 4749422 | 3549751.25 | 7202540 | | | | | | |
| P07758 | SERPINA1 | serpin peptidase inhibitor, clade A (alpha-1 antiproteinase, antitrypsin), member 1 | Extracellular Space | other | 2877501.5 | 2926279.5 | 5557472.5 | | | | | | |
| P07758 | SERPINA1 | serpin peptidase inhibitor, clade A (alpha-1 antiproteinase, antitrypsin), member 1 | Extracellular Space | other | 2903226 | 2363235 | 4045918 | | | | | | |
| P07758 | SERPINA1 | serpin peptidase inhibitor, clade A (alpha-1 antiproteinase, antitrypsin), member 1 | Extracellular Space | other | 2245516.25 | 1819203 | 4589743.5 | P07758 | SERPINA1 | 4421422 | 4748729 | 1226117 | 15 |
| P07759 | SERPINA3 | serpin peptidase inhibitor, clade A (alpha-1 antiproteinase, antitrypsin), member 3 | Extracellular Space | other | 2.69E+07 | 2.31E+07 | 2.55E+07 | | | | | | |
| P07759 | SERPINA3 | serpin peptidase inhibitor, clade A (alpha-1 antiproteinase, antitrypsin), member 3 | Extracellular Space | other | 1.72E+07 | 1.94E+07 | 1.51E+07 | | | | | | |
| P07759 | SERPINA3 | serpin peptidase inhibitor, clade A (alpha-1 antiproteinase, antitrypsin), member 3 | Extracellular Space | other | 1.02E+07 | 7322867 | 6987122 | P07759 | SERPINA3 | 18103333 | 7549624 | 2516541 | 9 |
| Q99JR1 | SFXN1 | sideroflexin 1 | Cytoplasm | transporter | 4074154.5 | 5712111 | 7730208.5 | | | | | | |
| Q99JR1 | SFXN1 | sideroflexin 1 | Cytoplasm | transporter | 3467197.5 | 5161288 | 6942731.5 | | | | | | |
| Q99JR1 | SFXN1 | sideroflexin 1 | Cytoplasm | transporter | 3788711.75 | 5411302 | 7083314 | | | | | | |
| Q99JR1 | SFXN1 | sideroflexin 1 | Cytoplasm | transporter | 3093519.75 | 4364429.5 | 4312130 | | | | | | |
| Q99JR1 | SFXN1 | sideroflexin 1 | Cytoplasm | transporter | 2355424.75 | 3867559.25 | 4484922.5 | | | | | | |
| Q99JR1 | SFXN1 | sideroflexin 1 | Cytoplasm | transporter | 1158472.375 | 1761461.875 | 2627835.75 | Q99JR1 | SFXN1 | 2989580 | 1815716 | 427968 | 18 |
| Q9JJU8 | SH3BGRL | SH3 domain binding glutamate-rich protein like | Cytoplasm | other | 1.45E+07 | 1.23E+07 | 1.05E+07 | | | | | | |
| Q9JJU8 | SH3BGRL | SH3 domain binding glutamate-rich protein like | Cytoplasm | other | 7078737 | 8337242.5 | 8040206.5 | | | | | | |
| Q9JJU8 | SH3BGRL | SH3 domain binding glutamate-rich protein like | Cytoplasm | other | 6794538.5 | 1.10E+07 | 1.07E+07 | Q9JJU8 | SH3BGRL | 9441092 | 2558950 | 852983 | 9 |
| Q8BG73 | SH3BGRL2 | SH3 domain binding glutamate-rich protein like 2 | Nucleus | other | 3941489.75 | 3718727.25 | 3714084 | Q8BG73 | SH3BGRL2 | 3941490 | 129973 | 75040 | 3 |
| Q91Y57 | SIGLEC9 | sialic acid binding Ig-like lectin 9 | Plasma Membrane | other | 5146995.5 | 5294559.5 | 4245420.5 | Q91Y57 | SIGLEC9 | 5146996 | 567936 | 327898 | 3 |
| Q3UND0 | SKAP2 | src kinase associated phosphoprotein 2 | Cytoplasm | other | 1317143.625 | 1075448.875 | 3125053.75 | | | | | | |
| Q3UND0 | SKAP2 | src kinase associated phosphoprotein 2 | Cytoplasm | other | 1441862.375 | 863661.75 | 3497400.5 | Q3UND0 | SKAP2 | 1379503 | 1127376 | 460249 | 6 |
| Q9QUM4 | SLAMF1 | signaling lymphocytic activation molecule family member 1 | Plasma Membrane | transmembrane receptor | 2569642.25 | 3832148.5 | 4563246 | | | | | | |
| Q9QUM4 | SLAMF1 | signaling lymphocytic activation molecule family member 1 | Plasma Membrane | transmembrane receptor | 2140733 | 2570048.25 | 4045109 | | | | | | |

| | | | | | | | | | | | | | |
|---|---|---|---|---|---|---|---|---|---|---|---|---|---|
| Q9QUM4 | SLAMF1 | signaling lymphocytic activation molecule family member 1 | Plasma Membrane | transmembrane receptor | 5258345.5 | 6028361 | 6199445 | Q9QUM4 | SLAMF1 | 3322907 | 1511140 | 503713 | 9 |
| P49282 | SLC11A2 | solute carrier family 11 (proton-coupled divalent metal ion transporter), member 2 | Plasma Membrane | transporter | 9331650 | 1.14E+07 | 1.11E+07 | | | | | | |
| P49282 | SLC11A2 | solute carrier family 11 (proton-coupled divalent metal ion transporter), member 2 | Plasma Membrane | transporter | 3914093.5 | 5970712 | 7041618 | | | | | | |
| P49282 | SLC11A2 | solute carrier family 11 (proton-coupled divalent metal ion transporter), member 2 | Plasma Membrane | transporter | 3750292.5 | 6847216.5 | 6635320 | | | | | | |
| P49282 | SLC11A2 | solute carrier family 11 (proton-coupled divalent metal ion transporter), member 2 | Plasma Membrane | transporter | 1315429.125 | 3853752.75 | 3878823.5 | P49282 | SLC11A2 | 4577866 | 3132977 | 904413 | 12 |
| Q8VHL0 | SLC14A1 | solute carrier family 14 (urea transporter), member 1 (Kidd blood group) | Plasma Membrane | transporter | 4.04E+08 | 4.18E+08 | 3.39E+08 | | | | | | |
| Q8VHL0 | SLC14A1 | solute carrier family 14 (urea transporter), member 1 (Kidd blood group) | Plasma Membrane | transporter | 1.80E+08 | 2.36E+08 | 2.83E+08 | | | | | | |
| Q8VHL0 | SLC14A1 | solute carrier family 14 (urea transporter), member 1 (Kidd blood group) | Plasma Membrane | transporter | 2.28E+08 | 2.18E+08 | 1.83E+08 | | | | | | |
| Q8VHL0 | SLC14A1 | solute carrier family 14 (urea transporter), member 1 (Kidd blood group) | Plasma Membrane | transporter | 2.07E+08 | 2.17E+08 | 1.74E+08 | | | | | | |
| Q8VHL0 | SLC14A1 | solute carrier family 14 (urea transporter), member 1 (Kidd blood group) | Plasma Membrane | transporter | 1.55E+08 | 1.56E+08 | 1.45E+08 | | | | | | |
| Q8VHL0 | SLC14A1 | solute carrier family 14 (urea transporter), member 1 (Kidd blood group) | Plasma Membrane | transporter | 1.67E+08 | 1.52E+08 | 1.26E+08 | | | | | | |
| Q8VHL0 | SLC14A1 | solute carrier family 14 (urea transporter), member 1 (Kidd blood group) | Plasma Membrane | transporter | 1.47E+08 | 5.68E+07 | 1.56E+08 | | | | | | |
| Q8VHL0 | SLC14A1 | solute carrier family 14 (urea transporter), member 1 (Kidd blood group) | Plasma Membrane | transporter | 4.17E+07 | 4.37E+07 | 3.89E+07 | | | | | | |
| Q8VHL0 | SLC14A1 | solute carrier family 14 (urea transporter), member 1 (Kidd blood group) | Plasma Membrane | transporter | 2.20E+07 | 7099545.5 | 2.31E+07 | | | | | | |
| Q8VHL0 | SLC14A1 | solute carrier family 14 (urea transporter), member 1 (Kidd blood group) | Plasma Membrane | transporter | 2.32E+07 | 3.10E+07 | 2.01E+07 | | | | | | |
| Q8VHL0 | SLC14A1 | solute carrier family 14 (urea transporter), member 1 (Kidd blood group) | Plasma Membrane | transporter | 3.41E+07 | 4.53E+07 | 2.59E+07 | | | | | | |
| Q8VHL0 | SLC14A1 | solute carrier family 14 (urea transporter), member 1 (Kidd blood group) | Plasma Membrane | transporter | 1713555.625 | 1321509.875 | 6535732.5 | Q8VHL0 | SLC14A1 | 134199463 | 113971577 | 18995263 | 36 |
| P53986 | SLC16A1 | solute carrier family 16 (monocarboxylate transporter), member 1 | Plasma Membrane | transporter | 5.90E+08 | 4.90E+08 | 4.69E+08 | | | | | | |
| P53986 | SLC16A1 | solute carrier family 16 (monocarboxylate transporter), member 1 | Plasma Membrane | transporter | 4.84E+08 | 5.01E+08 | 4.14E+08 | | | | | | |
| P53986 | SLC16A1 | solute carrier family 16 (monocarboxylate transporter), member 1 | Plasma Membrane | transporter | 5.01E+08 | 5.59E+08 | 3.89E+08 | | | | | | |
| P53986 | SLC16A1 | solute carrier family 16 (monocarboxylate transporter), member 1 | Plasma Membrane | transporter | 2.32E+08 | 2.30E+08 | 2.32E+08 | | | | | | |
| P53986 | SLC16A1 | solute carrier family 16 (monocarboxylate transporter), member 1 | Plasma Membrane | transporter | 1.41E+08 | 1.31E+08 | 1.50E+08 | | | | | | |
| P53986 | SLC16A1 | solute carrier family 16 (monocarboxylate transporter), member 1 | Plasma Membrane | transporter | 8.33E+07 | 9.04E+07 | 8.29E+07 | | | | | | |
| P53986 | SLC16A1 | solute carrier family 16 (monocarboxylate transporter), member 1 | Plasma Membrane | transporter | 3.54E+07 | 2.87E+07 | 2.15E+07 | | | | | | |
| P53986 | SLC16A1 | solute carrier family 16 (monocarboxylate transporter), member 1 | Plasma Membrane | transporter | 5.20E+07 | 4.66E+07 | 3.80E+07 | | | | | | |
| P53986 | SLC16A1 | solute carrier family 16 (monocarboxylate transporter), member 1 | Plasma Membrane | transporter | 9808871 | 9902232 | 8696731 | P53986 | SLC16A1 | 236499875 | 204586948 | 39372776 | 27 |
| Q3U9N9 | SLC16A10 | solute carrier family 16 (aromatic amino acid transporter), member 10 | Plasma Membrane | transporter | 1.99E+08 | 1.76E+08 | 1.40E+08 | | | | | | |
| Q3U9N9 | SLC16A10 | solute carrier family 16 (aromatic amino acid transporter), member 10 | Plasma Membrane | transporter | 1.30E+08 | 1.35E+08 | 1.09E+08 | | | | | | |
| Q3U9N9 | SLC16A10 | solute carrier family 16 (aromatic amino acid transporter), member 10 | Plasma Membrane | transporter | 8.76E+07 | 9.02E+07 | 9.37E+07 | | | | | | |
| Q3U9N9 | SLC16A10 | solute carrier family 16 (aromatic amino acid transporter), member 10 | Plasma Membrane | transporter | 7.53E+07 | 6.44E+07 | 5.22E+07 | | | | | | |
| Q3U9N9 | SLC16A10 | solute carrier family 16 (aromatic amino acid transporter), member 10 | Plasma Membrane | transporter | 4.93E+07 | 4.48E+07 | 4.04E+07 | | | | | | |
| Q3U9N9 | SLC16A10 | solute carrier family 16 (aromatic amino acid transporter), member 10 | Plasma Membrane | transporter | 4.41E+07 | 7.03E+07 | 3.78E+07 | | | | | | |
| Q3U9N9 | SLC16A10 | solute carrier family 16 (aromatic amino acid transporter), member 10 | Plasma Membrane | transporter | 1.80E+07 | 1.55E+07 | 1.37E+07 | | | | | | |
| Q3U9N9 | SLC16A10 | solute carrier family 16 (aromatic amino acid transporter), member 10 | Plasma Membrane | transporter | 1.02E+07 | 8158125 | 7565048 | | | | | | |
| Q3U9N9 | SLC16A10 | solute carrier family 16 (aromatic amino acid transporter), member 10 | Plasma Membrane | transporter | 9722979 | 8528367 | 5557536 | | | | | | |
| Q3U9N9 | SLC16A10 | solute carrier family 16 (aromatic amino acid transporter), member 10 | Plasma Membrane | transporter | 1575778.25 | 1495925.5 | 1.07E+07 | Q3U9N9 | SLC16A10 | 62476876 | 55059131 | 10052376 | 30 |
| Q8BRU6 | SLC18A2 | solute carrier family 18 (vesicular monoamine transporter), member 2 | Plasma Membrane | transporter | 634636.562 | 686259.938 | 1265266.125 | Q8BRU6 | SLC18A2 | 634637 | 350144 | 202156 | 3 |
| P41438 | SLC19A1 | solute carrier family 19 (folate transporter), member 1 | Plasma Membrane | transporter | 1594346.5 | 1866612.375 | 1514453.125 | P41438 | SLC19A1 | 1594347 | 184629 | 106596 | 3 |
| Q3UHH2 | SLC22A23 | solute carrier family 22, member 23 | Plasma Membrane | transporter | 2254844.5 | 2185773.75 | 1860231.75 | Q3UHH2 | SLC22A23 | 2254845 | 210740 | 121671 | 3 |
| Q8BH59 | SLC25A12 | solute carrier family 25 (aspartate/glutamate carrier), member 12 | Cytoplasm | transporter | 5338740.5 | 7400385.5 | 9696908 | | | | | | |
| Q8BH59 | SLC25A12 | solute carrier family 25 (aspartate/glutamate carrier), member 12 | Cytoplasm | transporter | 2559288.25 | 4981317 | 6659523 | | | | | | |
| Q8BH59 | SLC25A12 | solute carrier family 25 (aspartate/glutamate carrier), member 12 | Cytoplasm | transporter | 1604396.625 | 3189109 | 4260186.5 | | | | | | |
| Q8BH59 | SLC25A12 | solute carrier family 25 (aspartate/glutamate carrier), member 12 | Cytoplasm | transporter | 802322.812 | 2032628.125 | 2759387.25 | Q8BH59 | SLC25A12 | 2576187 | 2647837 | 764365 | 12 |
| Q8BZ09 | SLC25A21 | solute carrier family 25 (mitochondrial oxoadipate carrier), member 21 | Cytoplasm | transporter | 2791264.5 | 5618562 | 5722108 | | | | | | |
| Q8BZ09 | SLC25A21 | solute carrier family 25 (mitochondrial oxoadipate carrier), member 21 | Cytoplasm | transporter | 2230128.75 | 3722292.75 | 3924747.25 | | | | | | |
| Q8BZ09 | SLC25A21 | solute carrier family 25 (mitochondrial oxoadipate carrier), member 21 | Cytoplasm | transporter | 1467843.25 | 2364562.5 | 3289172.5 | | | | | | |
| Q8BZ09 | SLC25A21 | solute carrier family 25 (mitochondrial oxoadipate carrier), member 21 | Cytoplasm | transporter | 1235938.875 | 1826260 | 1837950.5 | | | | | | |
| Q8BZ09 | SLC25A21 | solute carrier family 25 (mitochondrial oxoadipate carrier), member 21 | Cytoplasm | transporter | 833768.375 | 2030535.375 | 1542284.625 | Q8BZ09 | SLC25A21 | 1711789 | 1495810 | 386216 | 15 |
| Q8VEM8 | SLC25A3 | solute carrier family 25 (mitochondrial carrier; phosphate carrier), member 3 | Cytoplasm | transporter | 2.39E+07 | 3.58E+07 | 4.03E+07 | | | | | | |
| Q8VEM8 | SLC25A3 | solute carrier family 25 (mitochondrial carrier; phosphate carrier), member 3 | Cytoplasm | transporter | 1.80E+07 | 2.58E+07 | 2.91E+07 | | | | | | |
| Q8VEM8 | SLC25A3 | solute carrier family 25 (mitochondrial carrier; phosphate carrier), member 3 | Cytoplasm | transporter | 1.14E+07 | 2.17E+07 | 2.43E+07 | | | | | | |
| Q8VEM8 | SLC25A3 | solute carrier family 25 (mitochondrial carrier; phosphate carrier), member 3 | Cytoplasm | transporter | 1.27E+07 | 2.08E+07 | 3.08E+07 | | | | | | |
| Q8VEM8 | SLC25A3 | solute carrier family 25 (mitochondrial carrier; phosphate carrier), member 3 | Cytoplasm | transporter | 1.11E+07 | 1.58E+07 | 1.91E+07 | | | | | | |
| Q8VEM8 | SLC25A3 | solute carrier family 25 (mitochondrial carrier; phosphate carrier), member 3 | Cytoplasm | transporter | 5647197 | 7842572.5 | 9307268 | | | | | | |
| Q8VEM8 | SLC25A3 | solute carrier family 25 (mitochondrial carrier; phosphate carrier), member 3 | Cytoplasm | transporter | 6293935 | 8601215 | 1.29E+07 | | | | | | |
| Q8VEM8 | SLC25A3 | solute carrier family 25 (mitochondrial carrier; phosphate carrier), member 3 | Cytoplasm | transporter | 2244798.75 | 3257368.75 | 3802560 | | | | | | |
| Q8VEM8 | SLC25A3 | solute carrier family 25 (mitochondrial carrier; phosphate carrier), member 3 | Cytoplasm | transporter | 2320994.5 | 3692895.75 | 5635290 | Q8VEM8 | SLC25A3 | 10398547 | 10812851 | 2080934 | 27 |
| Q920G8 | SLC25A37 | solute carrier family 25 (mitochondrial iron transporter), member 37 | Cytoplasm | other | 6081707.5 | 1.71E+07 | 1.50E+07 | | | | | | |
| Q920G8 | SLC25A37 | solute carrier family 25 (mitochondrial iron transporter), member 37 | Cytoplasm | other | 4089640 | 7719868.5 | 9244796 | | | | | | |
| Q920G8 | SLC25A37 | solute carrier family 25 (mitochondrial iron transporter), member 37 | Cytoplasm | other | 4142770.5 | 5452404.5 | 4633774 | | | | | | |
| Q920G8 | SLC25A37 | solute carrier family 25 (mitochondrial iron transporter), member 37 | Cytoplasm | other | 3372239.25 | 7392697 | 1.09E+07 | | | | | | |
| Q920G8 | SLC25A37 | solute carrier family 25 (mitochondrial iron transporter), member 37 | Cytoplasm | other | 3027048.25 | 4929915.5 | 6837949 | | | | | | |
| Q920G8 | SLC25A37 | solute carrier family 25 (mitochondrial iron transporter), member 37 | Cytoplasm | other | 2467929.25 | 4809939.5 | 5076697 | | | | | | |
| Q920G8 | SLC25A37 | solute carrier family 25 (mitochondrial iron transporter), member 37 | Cytoplasm | other | 2314835.5 | 4847733 | 4261943 | | | | | | |
| Q920G8 | SLC25A37 | solute carrier family 25 (mitochondrial iron transporter), member 37 | Cytoplasm | other | 2452718.75 | 4222565.5 | 5109795.5 | | | | | | |
| Q920G8 | SLC25A37 | solute carrier family 25 (mitochondrial iron transporter), member 37 | Cytoplasm | other | 1943237.125 | 3240328.5 | 3612625.25 | | | | | | |
| Q920G8 | SLC25A37 | solute carrier family 25 (mitochondrial iron transporter), member 37 | Cytoplasm | other | 930125.125 | 2034553.25 | 1948832.625 | | | | | | |
| Q920G8 | SLC25A37 | solute carrier family 25 (mitochondrial iron transporter), member 37 | Cytoplasm | other | 1715387.375 | 5104907 | 5688860.5 | Q920G8 | SLC25A37 | 2957967 | 3556874 | 619172 | 33 |
| P48962 | SLC25A4 | solute carrier family 25 (mitochondrial carrier; adenine nucleotide translocator), member 4 | Cytoplasm | transporter | 2717540.25 | 4650016 | 6194283 | | | | | | |
| P48962 | SLC25A4 | solute carrier family 25 (mitochondrial carrier; adenine nucleotide translocator), member 4 | Cytoplasm | transporter | 2476181.75 | 3607511.75 | 4668764.5 | | | | | | |
| P48962 | SLC25A4 | solute carrier family 25 (mitochondrial carrier; adenine nucleotide translocator), member 4 | Cytoplasm | transporter | 1717506.625 | 2728102.75 | 3554737.25 | P48962 | SLC25A4 | 2303743 | 1383929 | 461310 | 9 |
| P51881 | SLC25A5 | solute carrier family 25 (mitochondrial carrier; adenine nucleotide translocator), member 5 | Cytoplasm | transporter | 7.92E+07 | 1.21E+08 | 1.64E+08 | | | | | | |
| P51881 | SLC25A5 | solute carrier family 25 (mitochondrial carrier; adenine nucleotide translocator), member 5 | Cytoplasm | transporter | 5.25E+07 | 7.84E+07 | 1.11E+08 | | | | | | |
| P51881 | SLC25A5 | solute carrier family 25 (mitochondrial carrier; adenine nucleotide translocator), member 5 | Cytoplasm | transporter | 4.96E+07 | 6.34E+07 | 8.14E+07 | | | | | | |
| P51881 | SLC25A5 | solute carrier family 25 (mitochondrial carrier; adenine nucleotide translocator), member 5 | Cytoplasm | transporter | 2.24E+07 | 3.57E+07 | 5.12E+07 | | | | | | |
| P51881 | SLC25A5 | solute carrier family 25 (mitochondrial carrier; adenine nucleotide translocator), member 5 | Cytoplasm | transporter | 1.51E+07 | 3.02E+07 | 3.40E+07 | | | | | | |
| P51881 | SLC25A5 | solute carrier family 25 (mitochondrial carrier; adenine nucleotide translocator), member 5 | Cytoplasm | transporter | 3.06E+07 | 4.23E+07 | 4.07E+07 | | | | | | |
| P51881 | SLC25A5 | solute carrier family 25 (mitochondrial carrier; adenine nucleotide translocator), member 5 | Cytoplasm | transporter | 7926021.5 | 1.51E+07 | 3.08E+07 | | | | | | |
| P51881 | SLC25A5 | solute carrier family 25 (mitochondrial carrier; adenine nucleotide translocator), member 5 | Cytoplasm | transporter | 1.47E+07 | 2.26E+07 | 3.11E+07 | | | | | | |
| P51881 | SLC25A5 | solute carrier family 25 (mitochondrial carrier; adenine nucleotide translocator), member 5 | Cytoplasm | transporter | 1.56E+07 | 1.36E+07 | 2.68E+07 | | | | | | |
| P51881 | SLC25A5 | solute carrier family 25 (mitochondrial carrier; adenine nucleotide translocator), member 5 | Cytoplasm | transporter | 1.06E+07 | 2.81E+07 | 3.52E+07 | | | | | | |
| P51881 | SLC25A5 | solute carrier family 25 (mitochondrial carrier; adenine nucleotide translocator), member 5 | Cytoplasm | transporter | 1.22E+07 | 2.50E+07 | 3.52E+07 | | | | | | |
| P51881 | SLC25A5 | solute carrier family 25 (mitochondrial carrier; adenine nucleotide translocator), member 5 | Cytoplasm | transporter | 5567209.5 | 7769922 | 9128325 | | | | | | |
| P51881 | SLC25A5 | solute carrier family 25 (mitochondrial carrier; adenine nucleotide translocator), member 5 | Cytoplasm | transporter | 2585125.5 | 4633314.5 | 6491660 | | | | | | |
| P51881 | SLC25A5 | solute carrier family 25 (mitochondrial carrier; adenine nucleotide translocator), member 5 | Cytoplasm | transporter | 537974 | 2947773 | 1336909.375 | | | | | | |
| P51881 | SLC25A5 | solute carrier family 25 (mitochondrial carrier; adenine nucleotide translocator), member 5 | Cytoplasm | transporter | 777049.688 | 6920963 | 1.09E+07 | P51881 | SLC25A5 | 21324892 | 34656536 | 5166291 | 45 |
| Q5HZI9 | SLC25A51 | solute carrier family 25, member 51 | Cytoplasm | other | 1307824.625 | 909399.812 | 2280701.5 | Q5HZI9 | SLC25A51 | 1307825 | 705420 | 407274 | 3 |
| P58735 | SLC26A1 | solute carrier family 26 (anion exchanger), member 1 | Plasma Membrane | transporter | 9937488 | 1.02E+07 | 1.20E+07 | | | | | | |
| P58735 | SLC26A1 | solute carrier family 26 (anion exchanger), member 1 | Plasma Membrane | transporter | 3797966.25 | 3336216 | 4678809.5 | P58735 | SLC26A1 | 6867727 | 3816050 | 1557896 | 6 |
| Q9WVC8 | SLC26A3 | solute carrier family 26 (anion exchanger), member 3 | Plasma Membrane | transporter | 1987176.5 | 1599809 | 1621706.5 | Q9WVC8 | SLC26A3 | 1987177 | 217601 | 125632 | 3 |
| Q9JIM1 | SLC29A1 | solute carrier family 29 (equilibrative nucleoside transporter), member 1 | Plasma Membrane | transporter | 1.60E+08 | 1.53E+08 | 1.28E+08 | | | | | | |
| Q9JIM1 | SLC29A1 | solute carrier family 29 (equilibrative nucleoside transporter), member 1 | Plasma Membrane | transporter | 7.89E+07 | 7.96E+07 | 7.85E+07 | | | | | | |
| Q9JIM1 | SLC29A1 | solute carrier family 29 (equilibrative nucleoside transporter), member 1 | Plasma Membrane | transporter | 4.60E+07 | 5.02E+07 | 3.95E+07 | | | | | | |
| Q9JIM1 | SLC29A1 | solute carrier family 29 (equilibrative nucleoside transporter), member 1 | Plasma Membrane | transporter | 4.84E+07 | 4.39E+07 | 4.75E+07 | | | | | | |
| Q9JIM1 | SLC29A1 | solute carrier family 29 (equilibrative nucleoside transporter), member 1 | Plasma Membrane | transporter | 3.70E+07 | 3.68E+07 | 5.09E+07 | | | | | | |
| Q9JIM1 | SLC29A1 | solute carrier family 29 (equilibrative nucleoside transporter), member 1 | Plasma Membrane | transporter | 2.29E+07 | 1.64E+07 | 1.70E+07 | | | | | | |
| Q9JIM1 | SLC29A1 | solute carrier family 29 (equilibrative nucleoside transporter), member 1 | Plasma Membrane | transporter | 1.24E+07 | 1.62E+07 | 1.26E+07 | | | | | | |
| Q9JIM1 | SLC29A1 | solute carrier family 29 (equilibrative nucleoside transporter), member 1 | Plasma Membrane | transporter | 4768198 | 7046702.5 | 5653467.5 | | | | | | |
| Q9JIM1 | SLC29A1 | solute carrier family 29 (equilibrative nucleoside transporter), member 1 | Plasma Membrane | transporter | 3067928.75 | 3714206.25 | 4299284.5 | | | | | | |
| Q9JIM1 | SLC29A1 | solute carrier family 29 (equilibrative nucleoside transporter), member 1 | Plasma Membrane | transporter | 3611534.25 | 4637331 | 2983978.5 | | | | | | |
| Q9JIM1 | SLC29A1 | solute carrier family 29 (equilibrative nucleoside transporter), member 1 | Plasma Membrane | transporter | 1800541 | 2010730.125 | 1379314.125 | Q9JIM1 | SLC29A1 | 38024382 | 42819607 | 7453937 | 33 |
| Q61672 | SLC29A2 | solute carrier family 29 (equilibrative nucleoside transporter), member 2 | Plasma Membrane | transporter | 2.88E+07 | 2.69E+07 | 2.88E+07 | Q61672 | SLC29A2 | 28830000 | 1096966 | 633333 | 3 |
| P32037 | SLC2A3 | solute carrier family 2 (facilitated glucose transporter), member 3 | Plasma Membrane | transporter | 2.26E+07 | 2.82E+07 | 4.28E+07 | | | | | | |
| P32037 | SLC2A3 | solute carrier family 2 (facilitated glucose transporter), member 3 | Plasma Membrane | transporter | 1.98E+07 | 2.38E+07 | 2.66E+07 | | | | | | |
| P32037 | SLC2A3 | solute carrier family 2 (facilitated glucose transporter), member 3 | Plasma Membrane | transporter | 1.66E+07 | 2.06E+07 | 2.68E+07 | | | | | | |
| P32037 | SLC2A3 | solute carrier family 2 (facilitated glucose transporter), member 3 | Plasma Membrane | transporter | 1.33E+07 | 1.75E+07 | 1.77E+07 | | | | | | |
| P32037 | SLC2A3 | solute carrier family 2 (facilitated glucose transporter), member 3 | Plasma Membrane | transporter | 6628195 | 9060832 | 1.14E+07 | | | | | | |
| P32037 | SLC2A3 | solute carrier family 2 (facilitated glucose transporter), member 3 | Plasma Membrane | transporter | 2878830 | 3046982.25 | 3326474 | | | | | | |
| P32037 | SLC2A3 | solute carrier family 2 (facilitated glucose transporter), member 3 | Plasma Membrane | transporter | 1139371.75 | 1603800.75 | 1919837.625 | P32037 | SLC2A3 | 11835200 | 11217463 | 2447851 | 21 |

| | | | | | | | | | | | | | |
|---|---|---|---|---|---|---|---|---|---|---|---|---|---|
| Q60738 | SLC30A1 | solute carrier family 30 (zinc transporter), member 1 | Plasma Membrane | transporter | 6454987.5 | 5999228.5 | 5120399.5 | | | | | | |
| Q60738 | SLC30A1 | solute carrier family 30 (zinc transporter), member 1 | Plasma Membrane | transporter | 4176427 | 3841927 | 2861296.25 | | | | | | |
| Q60738 | SLC30A1 | solute carrier family 30 (zinc transporter), member 1 | Plasma Membrane | transporter | 4330513 | 4124075.5 | 3331496.25 | | | | | | |
| Q60738 | SLC30A1 | solute carrier family 30 (zinc transporter), member 1 | Plasma Membrane | transporter | 2672057 | 2920270.75 | 3129487.25 | Q60738 | SLC30A1 | 4408496 | 1236320 | 356895 | 12 |
| P10852 | SLC3A2 | solute carrier family 3 (amino acid transporter heavy chain), member 2 | Plasma Membrane | transporter | 9085246 | 1.25E+07 | 1.07E+07 | | | | | | |
| P10852 | SLC3A2 | solute carrier family 3 (amino acid transporter heavy chain), member 2 | Plasma Membrane | transporter | 7138557 | 1.08E+07 | 9944226 | | | | | | |
| P10852 | SLC3A2 | solute carrier family 3 (amino acid transporter heavy chain), member 2 | Plasma Membrane | transporter | 7570629.5 | 8099044 | 7523994.5 | | | | | | |
| P10852 | SLC3A2 | solute carrier family 3 (amino acid transporter heavy chain), member 2 | Plasma Membrane | transporter | 5860821.5 | 7401275 | 6330103.5 | | | | | | |
| P10852 | SLC3A2 | solute carrier family 3 (amino acid transporter heavy chain), member 2 | Plasma Membrane | transporter | 7187366 | 7319158 | 7138410.5 | | | | | | |
| P10852 | SLC3A2 | solute carrier family 3 (amino acid transporter heavy chain), member 2 | Plasma Membrane | transporter | 3837748.75 | 4126776.5 | 4858111 | | | | | | |
| P10852 | SLC3A2 | solute carrier family 3 (amino acid transporter heavy chain), member 2 | Plasma Membrane | transporter | 3422225 | 4926999.5 | 3979269.5 | | | | | | |
| P10852 | SLC3A2 | solute carrier family 3 (amino acid transporter heavy chain), member 2 | Plasma Membrane | transporter | 4700044 | 8415364 | 5726193.5 | | | | | | |
| P10852 | SLC3A2 | solute carrier family 3 (amino acid transporter heavy chain), member 2 | Plasma Membrane | transporter | 2513976.5 | 3019996.5 | 2457386.25 | | | | | | |
| P10852 | SLC3A2 | solute carrier family 3 (amino acid transporter heavy chain), member 2 | Plasma Membrane | transporter | 1651368.5 | 965014.062 | 1180450.75 | P10852 | SLC3A2 | 5296798 | 2987185 | 545383 | 30 |
| Q9JHI9 | SLC40A1 | solute carrier family 40 (iron-regulated transporter), member 1 | Plasma Membrane | transporter | 3.08E+07 | 2.72E+07 | 2.55E+07 | | | | | | |
| Q9JHI9 | SLC40A1 | solute carrier family 40 (iron-regulated transporter), member 1 | Plasma Membrane | transporter | 3.00E+07 | 2.42E+07 | 1.56E+07 | | | | | | |
| Q9JHI9 | SLC40A1 | solute carrier family 40 (iron-regulated transporter), member 1 | Plasma Membrane | transporter | 2.18E+07 | 1.48E+07 | 2.04E+07 | | | | | | |
| Q9JHI9 | SLC40A1 | solute carrier family 40 (iron-regulated transporter), member 1 | Plasma Membrane | transporter | 1.49E+07 | 1.52E+07 | 1.51E+07 | | | | | | |
| Q9JHI9 | SLC40A1 | solute carrier family 40 (iron-regulated transporter), member 1 | Plasma Membrane | transporter | 6934526 | 5469979 | 1.67E+07 | | | | | | |
| Q9JHI9 | SLC40A1 | solute carrier family 40 (iron-regulated transporter), member 1 | Plasma Membrane | transporter | 1.03E+07 | 1.58E+07 | 1.05E+07 | | | | | | |
| Q9JHI9 | SLC40A1 | solute carrier family 40 (iron-regulated transporter), member 1 | Plasma Membrane | transporter | 1.50E+07 | 1.55E+07 | 1.00E+07 | | | | | | |
| Q9JHI9 | SLC40A1 | solute carrier family 40 (iron-regulated transporter), member 1 | Plasma Membrane | transporter | 445663.031 | 289938.656 | 1189807 | Q9JHI9 | SLC40A1 | 16270024 | 8700608 | 1776004 | 24 |
| Q8BSM7 | SLC43A1 | solute carrier family 43 (amino acid system L transporter), member 1 | Plasma Membrane | transporter | 6.80E+08 | 6.80E+08 | 5.75E+08 | | | | | | |
| Q8BSM7 | SLC43A1 | solute carrier family 43 (amino acid system L transporter), member 1 | Plasma Membrane | transporter | 4.62E+08 | 4.20E+08 | 3.67E+08 | | | | | | |
| Q8BSM7 | SLC43A1 | solute carrier family 43 (amino acid system L transporter), member 1 | Plasma Membrane | transporter | 2.84E+08 | 2.70E+08 | 1.82E+08 | | | | | | |
| Q8BSM7 | SLC43A1 | solute carrier family 43 (amino acid system L transporter), member 1 | Plasma Membrane | transporter | 1.17E+08 | 1.21E+08 | 1.13E+08 | | | | | | |
| Q8BSM7 | SLC43A1 | solute carrier family 43 (amino acid system L transporter), member 1 | Plasma Membrane | transporter | 8.84E+07 | 7.01E+07 | 4.37E+07 | | | | | | |
| Q8BSM7 | SLC43A1 | solute carrier family 43 (amino acid system L transporter), member 1 | Plasma Membrane | transporter | 8.24E+07 | 7.03E+07 | 7.60E+07 | | | | | | |
| Q8BSM7 | SLC43A1 | solute carrier family 43 (amino acid system L transporter), member 1 | Plasma Membrane | transporter | 6.60E+07 | 9.57E+07 | 6.61E+07 | | | | | | |
| Q8BSM7 | SLC43A1 | solute carrier family 43 (amino acid system L transporter), member 1 | Plasma Membrane | transporter | 7.29E+07 | 3.39E+07 | 5.27E+07 | | | | | | |
| Q8BSM7 | SLC43A1 | solute carrier family 43 (amino acid system L transporter), member 1 | Plasma Membrane | transporter | 2.98E+07 | 2.75E+07 | 3.48E+07 | | | | | | |
| Q8BSM7 | SLC43A1 | solute carrier family 43 (amino acid system L transporter), member 1 | Plasma Membrane | transporter | 6821147 | 1641285.625 | 3332924.75 | | | | | | |
| Q8BSM7 | SLC43A1 | solute carrier family 43 (amino acid system L transporter), member 1 | Plasma Membrane | transporter | 1.07E+07 | 1.07E+07 | 6576332.5 | | | | | | |
| Q8BSM7 | SLC43A1 | solute carrier family 43 (amino acid system L transporter), member 1 | Plasma Membrane | transporter | 5441269 | 4435091.5 | 3252660 | | | | | | |
| Q8BSM7 | SLC43A1 | solute carrier family 43 (amino acid system L transporter), member 1 | Plasma Membrane | transporter | 2308070 | 1895370.25 | 2490669.5 | | | | | | |
| Q8BSM7 | SLC43A1 | solute carrier family 43 (amino acid system L transporter), member 1 | Plasma Membrane | transporter | 2657441.25 | 3767425.5 | 969400.375 | Q8BSM7 | SLC43A1 | 136488423 | 185754915 | 28662606 | 42 |
| A2AVZ9 | SLC43A3 | solute carrier family 43, member 3 | Extracellular Space | other | 8.15E+07 | 8.45E+07 | 3.85E+07 | | | | | | |
| A2AVZ9 | SLC43A3 | solute carrier family 43, member 3 | Extracellular Space | other | 2.16E+07 | 2.25E+07 | 1.92E+07 | | | | | | |
| A2AVZ9 | SLC43A3 | solute carrier family 43, member 3 | Extracellular Space | other | 1.81E+07 | 1.98E+07 | 1.65E+07 | | | | | | |
| A2AVZ9 | SLC43A3 | solute carrier family 43, member 3 | Extracellular Space | other | 8926259 | 9760622 | 6704143 | A2AVZ9 | SLC43A3 | 32506565 | 26524058 | 7656836 | 12 |
| Q6X893 | SLC44A1 | solute carrier family 44 (choline transporter), member 1 | Plasma Membrane | transporter | 4962436 | 5307279 | 5528374.5 | | | | | | |
| Q6X893 | SLC44A1 | solute carrier family 44 (choline transporter), member 1 | Plasma Membrane | transporter | 2269383.25 | 2982948.75 | 3187815 | | | | | | |
| Q6X893 | SLC44A1 | solute carrier family 44 (choline transporter), member 1 | Plasma Membrane | transporter | 2162502 | 2162205.5 | 2638261.75 | Q6X893 | SLC44A1 | 3131440 | 1401328 | 467109 | 9 |
| P04919 | SLC4A1 | solute carrier family 4 (anion exchanger), member 1 (Diego blood group) | Plasma Membrane | transporter | 2.97E+10 | 2.82E+10 | 2.54E+10 | | | | | | |
| P04919 | SLC4A1 | solute carrier family 4 (anion exchanger), member 1 (Diego blood group) | Plasma Membrane | transporter | 1.72E+10 | 1.53E+10 | 1.33E+10 | | | | | | |
| P04919 | SLC4A1 | solute carrier family 4 (anion exchanger), member 1 (Diego blood group) | Plasma Membrane | transporter | 1.03E+10 | 8.67E+09 | 7.50E+09 | | | | | | |
| P04919 | SLC4A1 | solute carrier family 4 (anion exchanger), member 1 (Diego blood group) | Plasma Membrane | transporter | 8.19E+09 | 8.35E+09 | 6.50E+09 | | | | | | |
| P04919 | SLC4A1 | solute carrier family 4 (anion exchanger), member 1 (Diego blood group) | Plasma Membrane | transporter | 7.55E+09 | 9.09E+09 | 6.85E+09 | | | | | | |
| P04919 | SLC4A1 | solute carrier family 4 (anion exchanger), member 1 (Diego blood group) | Plasma Membrane | transporter | 8.18E+09 | 8.17E+09 | 7.07E+09 | | | | | | |
| P04919 | SLC4A1 | solute carrier family 4 (anion exchanger), member 1 (Diego blood group) | Plasma Membrane | transporter | 1.03E+10 | 9.99E+09 | 8.43E+09 | | | | | | |
| P04919 | SLC4A1 | solute carrier family 4 (anion exchanger), member 1 (Diego blood group) | Plasma Membrane | transporter | 7.13E+09 | 6.20E+09 | 5.07E+09 | | | | | | |
| P04919 | SLC4A1 | solute carrier family 4 (anion exchanger), member 1 (Diego blood group) | Plasma Membrane | transporter | 7.97E+09 | 7.27E+09 | 7.09E+09 | | | | | | |
| P04919 | SLC4A1 | solute carrier family 4 (anion exchanger), member 1 (Diego blood group) | Plasma Membrane | transporter | 6.39E+09 | 6.22E+09 | 5.13E+09 | | | | | | |
| P04919 | SLC4A1 | solute carrier family 4 (anion exchanger), member 1 (Diego blood group) | Plasma Membrane | transporter | 4.85E+09 | 4.81E+09 | 4.28E+09 | | | | | | |
| P04919 | SLC4A1 | solute carrier family 4 (anion exchanger), member 1 (Diego blood group) | Plasma Membrane | transporter | 3.96E+09 | 3.39E+09 | 3.19E+09 | | | | | | |
| P04919 | SLC4A1 | solute carrier family 4 (anion exchanger), member 1 (Diego blood group) | Plasma Membrane | transporter | 4.43E+09 | 3.86E+09 | 3.57E+09 | | | | | | |
| P04919 | SLC4A1 | solute carrier family 4 (anion exchanger), member 1 (Diego blood group) | Plasma Membrane | transporter | 6.24E+09 | 7.41E+09 | 7.20E+09 | | | | | | |
| P04919 | SLC4A1 | solute carrier family 4 (anion exchanger), member 1 (Diego blood group) | Plasma Membrane | transporter | 4.09E+09 | 4.48E+09 | 4.57E+09 | | | | | | |
| P04919 | SLC4A1 | solute carrier family 4 (anion exchanger), member 1 (Diego blood group) | Plasma Membrane | transporter | 3.01E+09 | 2.96E+09 | 2.34E+09 | | | | | | |
| P04919 | SLC4A1 | solute carrier family 4 (anion exchanger), member 1 (Diego blood group) | Plasma Membrane | transporter | 2.09E+09 | 2.87E+09 | 1.59E+09 | | | | | | |
| P04919 | SLC4A1 | solute carrier family 4 (anion exchanger), member 1 (Diego blood group) | Plasma Membrane | transporter | 2.03E+08 | 1.73E+09 | 9.18E+08 | | | | | | |
| P04919 | SLC4A1 | solute carrier family 4 (anion exchanger), member 1 (Diego blood group) | Plasma Membrane | transporter | 1.86E+09 | 1.89E+09 | 1.78E+09 | | | | | | |
| P04919 | SLC4A1 | solute carrier family 4 (anion exchanger), member 1 (Diego blood group) | Plasma Membrane | transporter | 1.78E+09 | 1.81E+09 | 1.69E+09 | | | | | | |
| P04919 | SLC4A1 | solute carrier family 4 (anion exchanger), member 1 (Diego blood group) | Plasma Membrane | transporter | 7.86E+08 | 7.42E+08 | 5.91E+08 | | | | | | |
| P04919 | SLC4A1 | solute carrier family 4 (anion exchanger), member 1 (Diego blood group) | Plasma Membrane | transporter | 8.70E+08 | 8.25E+08 | 6.59E+08 | | | | | | |
| P04919 | SLC4A1 | solute carrier family 4 (anion exchanger), member 1 (Diego blood group) | Plasma Membrane | transporter | 9.76E+08 | 9.50E+08 | 6.42E+08 | | | | | | |
| P04919 | SLC4A1 | solute carrier family 4 (anion exchanger), member 1 (Diego blood group) | Plasma Membrane | transporter | 6.36E+08 | 5.71E+08 | 4.71E+08 | | | | | | |
| P04919 | SLC4A1 | solute carrier family 4 (anion exchanger), member 1 (Diego blood group) | Plasma Membrane | transporter | 5.90E+08 | 6.97E+08 | 6.91E+08 | | | | | | |
| P04919 | SLC4A1 | solute carrier family 4 (anion exchanger), member 1 (Diego blood group) | Plasma Membrane | transporter | 6.76E+08 | 5.72E+08 | 3.49E+08 | | | | | | |
| P04919 | SLC4A1 | solute carrier family 4 (anion exchanger), member 1 (Diego blood group) | Plasma Membrane | transporter | 3.37E+08 | 3.77E+08 | 2.18E+08 | | | | | | |
| P04919 | SLC4A1 | solute carrier family 4 (anion exchanger), member 1 (Diego blood group) | Plasma Membrane | transporter | 6.03E+08 | 6.80E+08 | 6.95E+08 | | | | | | |
| P04919 | SLC4A1 | solute carrier family 4 (anion exchanger), member 1 (Diego blood group) | Plasma Membrane | transporter | 1.45E+07 | 4.23E+07 | 1.87E+08 | | | | | | |
| P04919 | SLC4A1 | solute carrier family 4 (anion exchanger), member 1 (Diego blood group) | Plasma Membrane | transporter | 2.85E+08 | 3.88E+08 | 1.14E+08 | | | | | | |
| P04919 | SLC4A1 | solute carrier family 4 (anion exchanger), member 1 (Diego blood group) | Plasma Membrane | transporter | 3.88E+08 | 3.60E+08 | 3.89E+08 | | | | | | |
| P04919 | SLC4A1 | solute carrier family 4 (anion exchanger), member 1 (Diego blood group) | Plasma Membrane | transporter | 3.07E+08 | 5.01E+08 | 5.87E+08 | | | | | | |
| P04919 | SLC4A1 | solute carrier family 4 (anion exchanger), member 1 (Diego blood group) | Plasma Membrane | transporter | 3.29E+08 | 5.11E+08 | 4.44E+08 | | | | | | |
| P04919 | SLC4A1 | solute carrier family 4 (anion exchanger), member 1 (Diego blood group) | Plasma Membrane | transporter | 2.28E+08 | 2.21E+08 | 1.45E+08 | | | | | | |
| P04919 | SLC4A1 | solute carrier family 4 (anion exchanger), member 1 (Diego blood group) | Plasma Membrane | transporter | 1.24E+08 | 1.56E+08 | 3.76E+07 | | | | | | |
| P04919 | SLC4A1 | solute carrier family 4 (anion exchanger), member 1 (Diego blood group) | Plasma Membrane | transporter | 1.92E+08 | 2.81E+08 | 1.17E+08 | | | | | | |
| P04919 | SLC4A1 | solute carrier family 4 (anion exchanger), member 1 (Diego blood group) | Plasma Membrane | transporter | 7.83E+07 | 5.99E+07 | 8.93E+07 | | | | | | |
| P04919 | SLC4A1 | solute carrier family 4 (anion exchanger), member 1 (Diego blood group) | Plasma Membrane | transporter | 1.53E+08 | 1.99E+08 | 1.13E+08 | | | | | | |
| P04919 | SLC4A1 | solute carrier family 4 (anion exchanger), member 1 (Diego blood group) | Plasma Membrane | transporter | 8.45E+07 | 1.13E+08 | 9.76E+07 | | | | | | |
| P04919 | SLC4A1 | solute carrier family 4 (anion exchanger), member 1 (Diego blood group) | Plasma Membrane | transporter | 6.68E+07 | 6.47E+07 | 5.28E+07 | | | | | | |
| P04919 | SLC4A1 | solute carrier family 4 (anion exchanger), member 1 (Diego blood group) | Plasma Membrane | transporter | 2.56E+07 | 2.47E+07 | 6.63E+07 | | | | | | |
| P04919 | SLC4A1 | solute carrier family 4 (anion exchanger), member 1 (Diego blood group) | Plasma Membrane | transporter | 2.09E+08 | 3.22E+08 | 1.41E+08 | | | | | | |
| P04919 | SLC4A1 | solute carrier family 4 (anion exchanger), member 1 (Diego blood group) | Plasma Membrane | transporter | 5.51E+07 | 7.42E+07 | 1.98E+07 | | | | | | |
| P04919 | SLC4A1 | solute carrier family 4 (anion exchanger), member 1 (Diego blood group) | Plasma Membrane | transporter | 7.18E+07 | 7.66E+07 | 9.73E+07 | | | | | | |
| P04919 | SLC4A1 | solute carrier family 4 (anion exchanger), member 1 (Diego blood group) | Plasma Membrane | transporter | 2.48E+07 | 3.70E+07 | 1.47E+07 | | | | | | |
| P04919 | SLC4A1 | solute carrier family 4 (anion exchanger), member 1 (Diego blood group) | Plasma Membrane | transporter | 2.69E+07 | 3.30E+07 | 2.36E+07 | | | | | | |
| P04919 | SLC4A1 | solute carrier family 4 (anion exchanger), member 1 (Diego blood group) | Plasma Membrane | transporter | 539540.375 | 1898416.5 | 8007966 | | | | | | |
| P04919 | SLC4A1 | solute carrier family 4 (anion exchanger), member 1 (Diego blood group) | Plasma Membrane | transporter | 1.15E+07 | 1.49E+07 | 2.95E+07 | | | | | | |
| P04919 | SLC4A1 | solute carrier family 4 (anion exchanger), member 1 (Diego blood group) | Plasma Membrane | transporter | 1.52E+07 | 2.49E+07 | 1.76E+07 | | | | | | |
| P04919 | SLC4A1 | solute carrier family 4 (anion exchanger), member 1 (Diego blood group) | Plasma Membrane | transporter | 3.15E+07 | 3.23E+07 | 3.08E+07 | | | | | | |
| P04919 | SLC4A1 | solute carrier family 4 (anion exchanger), member 1 (Diego blood group) | Plasma Membrane | transporter | 5515562.5 | 4266428 | 6490350 | | | | | | |
| P04919 | SLC4A1 | solute carrier family 4 (anion exchanger), member 1 (Diego blood group) | Plasma Membrane | transporter | 1.42E+07 | 1.26E+07 | 1.32E+07 | | | | | | |
| P04919 | SLC4A1 | solute carrier family 4 (anion exchanger), member 1 (Diego blood group) | Plasma Membrane | transporter | 1.96E+07 | 3.10E+07 | 1.45E+07 | | | | | | |
| P04919 | SLC4A1 | solute carrier family 4 (anion exchanger), member 1 (Diego blood group) | Plasma Membrane | transporter | 863289.375 | 864395.625 | 2030366.875 | | | | | | |
| P04919 | SLC4A1 | solute carrier family 4 (anion exchanger), member 1 (Diego blood group) | Plasma Membrane | transporter | 1.57E+07 | 9.54E+07 | 5.36E+07 | | | | | | |
| P04919 | SLC4A1 | solute carrier family 4 (anion exchanger), member 1 (Diego blood group) | Plasma Membrane | transporter | 3335341.75 | 2876349 | 1193218 | | | | | | |
| P04919 | SLC4A1 | solute carrier family 4 (anion exchanger), member 1 (Diego blood group) | Plasma Membrane | transporter | 6006611.5 | 7090167 | 3563619.5 | P04919 | SLC4A1 | 2697304743 | 4722322098 | 361124997 | 171 |
| O88343 | SLC4A4 | solute carrier family 4 (sodium bicarbonate cotransporter), member 4 | Plasma Membrane | transporter | 1.44E+07 | 8058994.5 | 1.29E+07 | O88343 | SLC4A4 | 14400000 | 3312277 | 1912344 | 3 |
| Q60857 | SLC6A4 | solute carrier family 6 (neurotransmitter transporter), member 4 | Plasma Membrane | transporter | 3199162 | 4112086.25 | 5815848.5 | Q60857 | SLC6A4 | 3199162 | 1328112 | 766786 | 3 |
| P28571 | SLC6A9 | solute carrier family 6 (neurotransmitter transporter, glycine), member 9 | Plasma Membrane | transporter | 8456809 | 1.05E+07 | 1.16E+07 | | | | | | |
| P28571 | SLC6A9 | solute carrier family 6 (neurotransmitter transporter, glycine), member 9 | Plasma Membrane | transporter | 2412600 | 3425795.5 | 2545543.75 | P28571 | SLC6A9 | 5434705 | 4192258 | 1711482 | 6 |
| Q09143 | SLC7A1 | solute carrier family 7 (cationic amino acid transporter, y+ system), member 1 | Plasma Membrane | transporter | 4207713.5 | 6520880 | 9241503 | Q09143 | SLC7A1 | 4207714 | 2519642 | 1454716 | 3 |
| Q9Z127 | SLC7A5 | solute carrier family 7 (amino acid transporter light chain, L system), member 5 | Plasma Membrane | transporter | 1.02E+07 | 1.19E+07 | 8853210 | | | | | | |
| Q9Z127 | SLC7A5 | solute carrier family 7 (amino acid transporter light chain, L system), member 5 | Plasma Membrane | transporter | 4970054.5 | 5785492 | 4898893 | | | | | | |
| Q9Z127 | SLC7A5 | solute carrier family 7 (amino acid transporter light chain, L system), member 5 | Plasma Membrane | transporter | 2460477.25 | 2372015.25 | 3071668.25 | | | | | | |
| Q9Z127 | SLC7A5 | solute carrier family 7 (amino acid transporter light chain, L system), member 5 | Plasma Membrane | transporter | 1875221.5 | 2188735.75 | 1589873.75 | Q9Z127 | SLC7A5 | 4881438 | 3532363 | 1019705 | 12 |

| | | | | | | | | | | | | | |
|---|---|---|---|---|---|---|---|---|---|---|---|---|---|
| Q61165 | SLC9A1 | solute carrier family 9, subfamily A (NHE1, cation proton antiporter 1), member 1 | Plasma Membrane | ion channel | 1.39E+07 | 1.32E+07 | 1.21E+07 | | | | | | |
| Q61165 | SLC9A1 | solute carrier family 9, subfamily A (NHE1, cation proton antiporter 1), member 1 | Plasma Membrane | ion channel | 1.15E+07 | 7759721 | 7852583 | | | | | | |
| Q61165 | SLC9A1 | solute carrier family 9, subfamily A (NHE1, cation proton antiporter 1), member 1 | Plasma Membrane | ion channel | 6910109 | 8068388 | 6310898.5 | | | | | | |
| Q61165 | SLC9A1 | solute carrier family 9, subfamily A (NHE1, cation proton antiporter 1), member 1 | Plasma Membrane | ion channel | 6192368.5 | 7229473 | 6614303 | | | | | | |
| Q61165 | SLC9A1 | solute carrier family 9, subfamily A (NHE1, cation proton antiporter 1), member 1 | Plasma Membrane | ion channel | 4675524 | 4953274.5 | 3868057.75 | | | | | | |
| Q61165 | SLC9A1 | solute carrier family 9, subfamily A (NHE1, cation proton antiporter 1), member 1 | Plasma Membrane | ion channel | 5293718.5 | 4535166 | 5358569 | | | | | | |
| Q61165 | SLC9A1 | solute carrier family 9, subfamily A (NHE1, cation proton antiporter 1), member 1 | Plasma Membrane | ion channel | 3315035.5 | 2748600.25 | 2345483 | | | | | | |
| Q61165 | SLC9A1 | solute carrier family 9, subfamily A (NHE1, cation proton antiporter 1), member 1 | Plasma Membrane | ion channel | 116999.219 | 286684.969 | 663509.75 | Q61165 | SLC9A1 | 6492969 | 3809465 | 777604 | 24 |
| Q6UJY2 | SLC9C1 | solute carrier family 9, subfamily C (Na+-transporting carboxylic acid decarboxylase), member 1 | Extracellular Space | other | 2.92E+07 | 5.91E+07 | 4.23E+07 | Q6UJY2 | SLC9C1 | 29200000 | 15008056 | 8664905 | 3 |
| Q8CG48 | SMC2 | structural maintenance of chromosomes 2 | Nucleus | transporter | 1105809.625 | 1282572 | 1754694.25 | Q8CG48 | SMC2 | 1105810 | 335459 | 193677 | 3 |
| Q8CG47 | SMC4 | structural maintenance of chromosomes 4 | Nucleus | transporter | 1.17E+07 | 1.31E+07 | 1.01E+07 | | | | | | |
| Q8CG47 | SMC4 | structural maintenance of chromosomes 4 | Nucleus | transporter | 815558.625 | 600136.812 | 516061.688 | Q8CG47 | SMC4 | 6272779 | 6110263 | 2494505 | 6 |
| O09044 | SNAP23 | synaptosomal-associated protein, 23kDa | Plasma Membrane | transporter | 3.92E+07 | 3.80E+07 | 3.54E+07 | | | | | | |
| O09044 | SNAP23 | synaptosomal-associated protein, 23kDa | Plasma Membrane | transporter | 4.16E+07 | 3.67E+07 | 2.71E+07 | | | | | | |
| O09044 | SNAP23 | synaptosomal-associated protein, 23kDa | Plasma Membrane | transporter | 2.46E+07 | 2.66E+07 | 2.67E+07 | | | | | | |
| O09044 | SNAP23 | synaptosomal-associated protein, 23kDa | Plasma Membrane | transporter | 3.17E+07 | 3.42E+07 | 3.17E+07 | | | | | | |
| O09044 | SNAP23 | synaptosomal-associated protein, 23kDa | Plasma Membrane | transporter | 1.32E+07 | 1.24E+07 | 8465155 | | | | | | |
| O09044 | SNAP23 | synaptosomal-associated protein, 23kDa | Plasma Membrane | transporter | 1.27E+07 | 2.12E+07 | 1.01E+07 | | | | | | |
| O09044 | SNAP23 | synaptosomal-associated protein, 23kDa | Plasma Membrane | transporter | 1.23E+07 | 1.15E+07 | 9991548 | | | | | | |
| O09044 | SNAP23 | synaptosomal-associated protein, 23kDa | Plasma Membrane | transporter | 5346732 | 6679322.5 | 3739260.5 | | | | | | |
| O09044 | SNAP23 | synaptosomal-associated protein, 23kDa | Plasma Membrane | transporter | 3177065.25 | 3943433.25 | 2003813.25 | | | | | | |
| O09044 | SNAP23 | synaptosomal-associated protein, 23kDa | Plasma Membrane | transporter | 2420694.25 | 3815249.5 | 1722619.875 | | | | | | |
| O09044 | SNAP23 | synaptosomal-associated protein, 23kDa | Plasma Membrane | transporter | 2300562.75 | 1643258.5 | 1172855.625 | | | | | | |
| O09044 | SNAP23 | synaptosomal-associated protein, 23kDa | Plasma Membrane | transporter | 789621.875 | 1465905.75 | 1380522.375 | O09044 | SNAP23 | 15760390 | 13716352 | 2286059 | 36 |
| O55042 | SNCA | synuclein, alpha (non A4 component of amyloid precursor) | Cytoplasm | other | 1.13E+07 | 1.42E+07 | 1.02E+07 | | | | | | |
| O55042 | SNCA | synuclein, alpha (non A4 component of amyloid precursor) | Cytoplasm | other | 9645854 | 1.02E+07 | 5781394.5 | | | | | | |
| O55042 | SNCA | synuclein, alpha (non A4 component of amyloid precursor) | Cytoplasm | other | 7345882 | 9580846 | 5627997.5 | | | | | | |
| O55042 | SNCA | synuclein, alpha (non A4 component of amyloid precursor) | Cytoplasm | other | 5380237 | 6981328 | 4397354 | O55042 | SNCA | 8415493 | 2927673 | 845146 | 12 |
| O70493 | SNX12 | sorting nexin 12 | Cytoplasm | transporter | 373096.125 | 1064836.75 | 687827.188 | O70493 | SNX12 | 373096 | 346337 | 199958 | 3 |
| Q0P5V2 | SOBP | sine oculis binding protein homolog | Nucleus | other | 474929.344 | 697743.312 | 1005310.562 | Q0P5V2 | SOBP | 474929 | 266317 | 153758 | 3 |
| P09671 | SOD2 | superoxide dismutase 2, mitochondrial | Cytoplasm | enzyme | 3915434.75 | 6898561 | 6532188.5 | | | | | | |
| P09671 | SOD2 | superoxide dismutase 2, mitochondrial | Cytoplasm | enzyme | 2346719.5 | 1210794.625 | 605663.688 | | | | | | |
| P09671 | SOD2 | superoxide dismutase 2, mitochondrial | Cytoplasm | enzyme | 1323632.625 | 2279333 | 2117248 | P09671 | SOD2 | 2528596 | 2290660 | 763553 | 9 |
| Q9CYN2 | SPCS2 | signal peptidase complex subunit 2 | Cytoplasm | other | 7185998 | 8760688 | 9378787 | Q9CYN2 | SPCS2 | 7185998 | 1130635 | 652773 | 3 |
| Q9JJF9 | SPPL2A | signal peptide peptidase like 2A | Plasma Membrane | peptidase | 1.87E+07 | 1.39E+07 | 1.56E+07 | | | | | | |
| Q9JJF9 | SPPL2A | signal peptide peptidase like 2A | Plasma Membrane | peptidase | 1.72E+07 | 1.57E+07 | 1.59E+07 | Q9JJF9 | SPPL2A | 17940000 | 1629650 | 665302 | 6 |
| P08032 | SPTA1 | spectrin, alpha, erythrocytic 1 | Cytoplasm | other | 3.45E+09 | 3.93E+09 | 3.34E+09 | | | | | | |
| P08032 | SPTA1 | spectrin, alpha, erythrocytic 1 | Cytoplasm | other | 2.92E+09 | 2.63E+09 | 2.64E+09 | | | | | | |
| P08032 | SPTA1 | spectrin, alpha, erythrocytic 1 | Cytoplasm | other | 3.12E+09 | 3.03E+09 | 2.29E+09 | | | | | | |
| P08032 | SPTA1 | spectrin, alpha, erythrocytic 1 | Cytoplasm | other | 2.41E+09 | 2.83E+09 | 2.31E+09 | | | | | | |
| P08032 | SPTA1 | spectrin, alpha, erythrocytic 1 | Cytoplasm | other | 2.27E+09 | 2.77E+09 | 2.18E+09 | | | | | | |
| P08032 | SPTA1 | spectrin, alpha, erythrocytic 1 | Cytoplasm | other | 2.39E+09 | 2.68E+09 | 2.18E+09 | | | | | | |
| P08032 | SPTA1 | spectrin, alpha, erythrocytic 1 | Cytoplasm | other | 2.05E+09 | 2.48E+09 | 1.94E+09 | | | | | | |
| P08032 | SPTA1 | spectrin, alpha, erythrocytic 1 | Cytoplasm | other | 2.28E+09 | 2.02E+09 | 1.78E+09 | | | | | | |
| P08032 | SPTA1 | spectrin, alpha, erythrocytic 1 | Cytoplasm | other | 1.72E+09 | 1.79E+09 | 1.68E+09 | | | | | | |
| P08032 | SPTA1 | spectrin, alpha, erythrocytic 1 | Cytoplasm | other | 2.44E+09 | 2.10E+09 | 1.86E+09 | | | | | | |
| P08032 | SPTA1 | spectrin, alpha, erythrocytic 1 | Cytoplasm | other | 1.82E+09 | 1.81E+09 | 1.62E+09 | | | | | | |
| P08032 | SPTA1 | spectrin, alpha, erythrocytic 1 | Cytoplasm | other | 1.94E+09 | 1.93E+09 | 2.01E+09 | | | | | | |
| P08032 | SPTA1 | spectrin, alpha, erythrocytic 1 | Cytoplasm | other | 1.74E+09 | 1.97E+09 | 1.70E+09 | | | | | | |
| P08032 | SPTA1 | spectrin, alpha, erythrocytic 1 | Cytoplasm | other | 2.44E+09 | 2.30E+09 | 1.78E+09 | | | | | | |
| P08032 | SPTA1 | spectrin, alpha, erythrocytic 1 | Cytoplasm | other | 1.96E+09 | 1.87E+09 | 2.00E+09 | | | | | | |
| P08032 | SPTA1 | spectrin, alpha, erythrocytic 1 | Cytoplasm | other | 2.09E+09 | 1.81E+09 | 1.84E+09 | | | | | | |
| P08032 | SPTA1 | spectrin, alpha, erythrocytic 1 | Cytoplasm | other | 1.07E+09 | 1.40E+09 | 1.07E+09 | | | | | | |
| P08032 | SPTA1 | spectrin, alpha, erythrocytic 1 | Cytoplasm | other | 2.06E+09 | 2.29E+09 | 1.91E+09 | | | | | | |
| P08032 | SPTA1 | spectrin, alpha, erythrocytic 1 | Cytoplasm | other | 1.71E+09 | 1.77E+09 | 1.70E+09 | | | | | | |
| P08032 | SPTA1 | spectrin, alpha, erythrocytic 1 | Cytoplasm | other | 1.71E+09 | 1.38E+09 | 1.18E+09 | | | | | | |
| P08032 | SPTA1 | spectrin, alpha, erythrocytic 1 | Cytoplasm | other | 1.72E+09 | 1.50E+09 | 1.44E+09 | | | | | | |
| P08032 | SPTA1 | spectrin, alpha, erythrocytic 1 | Cytoplasm | other | 1.63E+09 | 1.70E+09 | 1.23E+09 | | | | | | |
| P08032 | SPTA1 | spectrin, alpha, erythrocytic 1 | Cytoplasm | other | 1.45E+09 | 1.43E+09 | 1.25E+09 | | | | | | |
| P08032 | SPTA1 | spectrin, alpha, erythrocytic 1 | Cytoplasm | other | 1.44E+09 | 1.50E+09 | 1.29E+09 | | | | | | |
| P08032 | SPTA1 | spectrin, alpha, erythrocytic 1 | Cytoplasm | other | 1.27E+09 | 1.26E+09 | 1.24E+09 | | | | | | |
| P08032 | SPTA1 | spectrin, alpha, erythrocytic 1 | Cytoplasm | other | 1.48E+09 | 1.35E+09 | 1.24E+09 | | | | | | |
| P08032 | SPTA1 | spectrin, alpha, erythrocytic 1 | Cytoplasm | other | 1.80E+09 | 1.62E+09 | 1.49E+09 | | | | | | |
| P08032 | SPTA1 | spectrin, alpha, erythrocytic 1 | Cytoplasm | other | 1.95E+09 | 1.57E+09 | 1.41E+09 | | | | | | |
| P08032 | SPTA1 | spectrin, alpha, erythrocytic 1 | Cytoplasm | other | 1.41E+09 | 1.40E+09 | 1.23E+09 | | | | | | |
| P08032 | SPTA1 | spectrin, alpha, erythrocytic 1 | Cytoplasm | other | 1.48E+09 | 1.45E+09 | 1.28E+09 | | | | | | |
| P08032 | SPTA1 | spectrin, alpha, erythrocytic 1 | Cytoplasm | other | 1.29E+09 | 1.11E+09 | 9.68E+08 | | | | | | |
| P08032 | SPTA1 | spectrin, alpha, erythrocytic 1 | Cytoplasm | other | 1.45E+09 | 1.30E+09 | 1.12E+09 | | | | | | |
| P08032 | SPTA1 | spectrin, alpha, erythrocytic 1 | Cytoplasm | other | 1.44E+09 | 1.23E+09 | 1.00E+09 | | | | | | |
| P08032 | SPTA1 | spectrin, alpha, erythrocytic 1 | Cytoplasm | other | 1.17E+09 | 1.23E+09 | 1.01E+09 | | | | | | |
| P08032 | SPTA1 | spectrin, alpha, erythrocytic 1 | Cytoplasm | other | 1.40E+09 | 6.79E+08 | 1.43E+09 | | | | | | |
| P08032 | SPTA1 | spectrin, alpha, erythrocytic 1 | Cytoplasm | other | 1.54E+09 | 1.55E+09 | 1.35E+09 | | | | | | |
| P08032 | SPTA1 | spectrin, alpha, erythrocytic 1 | Cytoplasm | other | 1.28E+09 | 1.52E+09 | 1.07E+09 | | | | | | |
| P08032 | SPTA1 | spectrin, alpha, erythrocytic 1 | Cytoplasm | other | 1.38E+09 | 1.30E+09 | 1.11E+09 | | | | | | |
| P08032 | SPTA1 | spectrin, alpha, erythrocytic 1 | Cytoplasm | other | 1.26E+09 | 1.16E+09 | 9.71E+08 | | | | | | |
| P08032 | SPTA1 | spectrin, alpha, erythrocytic 1 | Cytoplasm | other | 1.25E+09 | 1.28E+09 | 1.20E+09 | | | | | | |
| P08032 | SPTA1 | spectrin, alpha, erythrocytic 1 | Cytoplasm | other | 1.35E+09 | 1.27E+09 | 9.24E+08 | | | | | | |
| P08032 | SPTA1 | spectrin, alpha, erythrocytic 1 | Cytoplasm | other | 1.24E+09 | 1.16E+09 | 1.10E+09 | | | | | | |
| P08032 | SPTA1 | spectrin, alpha, erythrocytic 1 | Cytoplasm | other | 1.40E+09 | 1.42E+09 | 1.18E+09 | | | | | | |
| P08032 | SPTA1 | spectrin, alpha, erythrocytic 1 | Cytoplasm | other | 1.38E+09 | 1.19E+09 | 9.32E+08 | | | | | | |
| P08032 | SPTA1 | spectrin, alpha, erythrocytic 1 | Cytoplasm | other | 1.45E+09 | 1.25E+09 | 1.09E+09 | | | | | | |
| P08032 | SPTA1 | spectrin, alpha, erythrocytic 1 | Cytoplasm | other | 1.14E+09 | 1.06E+09 | 1.08E+09 | | | | | | |
| P08032 | SPTA1 | spectrin, alpha, erythrocytic 1 | Cytoplasm | other | 1.13E+09 | 1.12E+09 | 9.58E+08 | | | | | | |
| P08032 | SPTA1 | spectrin, alpha, erythrocytic 1 | Cytoplasm | other | 1.48E+09 | 1.45E+09 | 1.15E+09 | | | | | | |
| P08032 | SPTA1 | spectrin, alpha, erythrocytic 1 | Cytoplasm | other | 1.23E+09 | 1.85E+09 | 1.18E+09 | | | | | | |
| P08032 | SPTA1 | spectrin, alpha, erythrocytic 1 | Cytoplasm | other | 1.43E+09 | 1.52E+09 | 1.20E+09 | | | | | | |
| P08032 | SPTA1 | spectrin, alpha, erythrocytic 1 | Cytoplasm | other | 1.11E+09 | 9.90E+08 | 8.88E+08 | | | | | | |
| P08032 | SPTA1 | spectrin, alpha, erythrocytic 1 | Cytoplasm | other | 1.15E+09 | 9.69E+08 | 8.50E+08 | | | | | | |
| P08032 | SPTA1 | spectrin, alpha, erythrocytic 1 | Cytoplasm | other | 1.06E+09 | 9.93E+08 | 9.12E+08 | | | | | | |
| P08032 | SPTA1 | spectrin, alpha, erythrocytic 1 | Cytoplasm | other | 1.31E+09 | 1.56E+09 | 1.02E+09 | | | | | | |
| P08032 | SPTA1 | spectrin, alpha, erythrocytic 1 | Cytoplasm | other | 8.74E+08 | 1.11E+09 | 6.94E+08 | | | | | | |
| P08032 | SPTA1 | spectrin, alpha, erythrocytic 1 | Cytoplasm | other | 1.09E+09 | 1.12E+09 | 9.88E+08 | | | | | | |
| P08032 | SPTA1 | spectrin, alpha, erythrocytic 1 | Cytoplasm | other | 1.18E+09 | 9.31E+08 | 7.88E+08 | | | | | | |
| P08032 | SPTA1 | spectrin, alpha, erythrocytic 1 | Cytoplasm | other | 1.27E+09 | 1.29E+09 | 1.05E+09 | | | | | | |
| P08032 | SPTA1 | spectrin, alpha, erythrocytic 1 | Cytoplasm | other | 1.17E+09 | 1.18E+09 | 1.03E+09 | | | | | | |
| P08032 | SPTA1 | spectrin, alpha, erythrocytic 1 | Cytoplasm | other | 1.29E+09 | 1.45E+09 | 9.56E+08 | | | | | | |
| P08032 | SPTA1 | spectrin, alpha, erythrocytic 1 | Cytoplasm | other | 1.23E+09 | 1.05E+09 | 7.94E+08 | | | | | | |
| P08032 | SPTA1 | spectrin, alpha, erythrocytic 1 | Cytoplasm | other | 9.34E+08 | 1.08E+09 | 1.03E+09 | | | | | | |
| P08032 | SPTA1 | spectrin, alpha, erythrocytic 1 | Cytoplasm | other | 9.19E+08 | 9.60E+08 | 7.93E+08 | | | | | | |
| P08032 | SPTA1 | spectrin, alpha, erythrocytic 1 | Cytoplasm | other | 9.35E+08 | 1.13E+09 | 1.11E+09 | | | | | | |
| P08032 | SPTA1 | spectrin, alpha, erythrocytic 1 | Cytoplasm | other | 9.49E+08 | 8.98E+08 | 7.59E+08 | | | | | | |
| P08032 | SPTA1 | spectrin, alpha, erythrocytic 1 | Cytoplasm | other | 9.78E+08 | 9.53E+08 | 1.00E+09 | | | | | | |
| P08032 | SPTA1 | spectrin, alpha, erythrocytic 1 | Cytoplasm | other | 8.69E+08 | 8.24E+08 | 7.53E+08 | | | | | | |
| P08032 | SPTA1 | spectrin, alpha, erythrocytic 1 | Cytoplasm | other | 8.33E+08 | 8.06E+08 | 7.52E+08 | | | | | | |
| P08032 | SPTA1 | spectrin, alpha, erythrocytic 1 | Cytoplasm | other | 9.99E+08 | 1.21E+09 | 7.69E+08 | | | | | | |
| P08032 | SPTA1 | spectrin, alpha, erythrocytic 1 | Cytoplasm | other | 9.16E+08 | 9.40E+08 | 8.13E+08 | | | | | | |
| P08032 | SPTA1 | spectrin, alpha, erythrocytic 1 | Cytoplasm | other | 8.06E+08 | 6.87E+08 | 5.67E+08 | | | | | | |
| P08032 | SPTA1 | spectrin, alpha, erythrocytic 1 | Cytoplasm | other | 9.61E+08 | 7.08E+08 | 8.51E+08 | | | | | | |
| P08032 | SPTA1 | spectrin, alpha, erythrocytic 1 | Cytoplasm | other | 8.66E+08 | 9.03E+08 | 6.73E+08 | | | | | | |

| | | | | | | | |
|---|---|---|---|---|---|---|---|
| P08032 | SPTA1 | spectrin, alpha, erythrocytic 1 | Cytoplasm | other | 7.34E+08 | 8.15E+08 | 7.98E+08 |
| P08032 | SPTA1 | spectrin, alpha, erythrocytic 1 | Cytoplasm | other | 7.70E+08 | 7.71E+08 | 7.16E+08 |
| P08032 | SPTA1 | spectrin, alpha, erythrocytic 1 | Cytoplasm | other | 1.17E+09 | 1.08E+09 | 9.06E+08 |
| P08032 | SPTA1 | spectrin, alpha, erythrocytic 1 | Cytoplasm | other | 9.04E+08 | 8.83E+08 | 8.37E+08 |
| P08032 | SPTA1 | spectrin, alpha, erythrocytic 1 | Cytoplasm | other | 8.86E+08 | 8.28E+08 | 7.65E+08 |
| P08032 | SPTA1 | spectrin, alpha, erythrocytic 1 | Cytoplasm | other | 8.89E+08 | 8.17E+08 | 8.21E+08 |
| P08032 | SPTA1 | spectrin, alpha, erythrocytic 1 | Cytoplasm | other | 8.24E+08 | 8.53E+08 | 8.22E+08 |
| P08032 | SPTA1 | spectrin, alpha, erythrocytic 1 | Cytoplasm | other | 6.76E+08 | 6.95E+08 | 7.32E+08 |
| P08032 | SPTA1 | spectrin, alpha, erythrocytic 1 | Cytoplasm | other | 7.05E+08 | 7.21E+08 | 5.19E+08 |
| P08032 | SPTA1 | spectrin, alpha, erythrocytic 1 | Cytoplasm | other | 9.09E+08 | 9.03E+08 | 7.86E+08 |
| P08032 | SPTA1 | spectrin, alpha, erythrocytic 1 | Cytoplasm | other | 6.62E+08 | 6.08E+08 | 5.70E+08 |
| P08032 | SPTA1 | spectrin, alpha, erythrocytic 1 | Cytoplasm | other | 8.33E+08 | 9.02E+08 | 7.39E+08 |
| P08032 | SPTA1 | spectrin, alpha, erythrocytic 1 | Cytoplasm | other | 5.98E+08 | 5.87E+08 | 5.10E+08 |
| P08032 | SPTA1 | spectrin, alpha, erythrocytic 1 | Cytoplasm | other | 6.33E+08 | 7.12E+08 | 6.10E+08 |
| P08032 | SPTA1 | spectrin, alpha, erythrocytic 1 | Cytoplasm | other | 8.30E+08 | 6.77E+08 | 7.22E+08 |
| P08032 | SPTA1 | spectrin, alpha, erythrocytic 1 | Cytoplasm | other | 7.70E+08 | 8.64E+08 | 6.19E+08 |
| P08032 | SPTA1 | spectrin, alpha, erythrocytic 1 | Cytoplasm | other | 6.52E+08 | 6.30E+08 | 5.77E+08 |
| P08032 | SPTA1 | spectrin, alpha, erythrocytic 1 | Cytoplasm | other | 7.03E+08 | 7.07E+08 | 6.68E+08 |
| P08032 | SPTA1 | spectrin, alpha, erythrocytic 1 | Cytoplasm | other | 8.25E+08 | 7.50E+08 | 5.80E+08 |
| P08032 | SPTA1 | spectrin, alpha, erythrocytic 1 | Cytoplasm | other | 8.48E+08 | 9.06E+08 | 6.72E+08 |
| P08032 | SPTA1 | spectrin, alpha, erythrocytic 1 | Cytoplasm | other | 7.26E+08 | 6.54E+08 | 5.11E+08 |
| P08032 | SPTA1 | spectrin, alpha, erythrocytic 1 | Cytoplasm | other | 6.98E+08 | 7.29E+08 | 4.70E+08 |
| P08032 | SPTA1 | spectrin, alpha, erythrocytic 1 | Cytoplasm | other | 7.08E+08 | 5.01E+08 | 6.59E+08 |
| P08032 | SPTA1 | spectrin, alpha, erythrocytic 1 | Cytoplasm | other | 6.38E+08 | 7.73E+08 | 5.80E+08 |
| P08032 | SPTA1 | spectrin, alpha, erythrocytic 1 | Cytoplasm | other | 9.02E+08 | 5.45E+08 | 4.74E+08 |
| P08032 | SPTA1 | spectrin, alpha, erythrocytic 1 | Cytoplasm | other | 5.53E+08 | 8.08E+08 | 6.49E+08 |
| P08032 | SPTA1 | spectrin, alpha, erythrocytic 1 | Cytoplasm | other | 5.67E+08 | 4.75E+08 | 5.27E+08 |
| P08032 | SPTA1 | spectrin, alpha, erythrocytic 1 | Cytoplasm | other | 5.17E+08 | 6.02E+08 | 4.31E+08 |
| P08032 | SPTA1 | spectrin, alpha, erythrocytic 1 | Cytoplasm | other | 6.33E+08 | 5.86E+08 | 4.93E+08 |
| P08032 | SPTA1 | spectrin, alpha, erythrocytic 1 | Cytoplasm | other | 4.68E+08 | 4.93E+08 | 5.73E+08 |
| P08032 | SPTA1 | spectrin, alpha, erythrocytic 1 | Cytoplasm | other | 6.22E+08 | 3.68E+08 | 5.49E+08 |
| P08032 | SPTA1 | spectrin, alpha, erythrocytic 1 | Cytoplasm | other | 6.07E+08 | 5.34E+08 | 5.79E+08 |
| P08032 | SPTA1 | spectrin, alpha, erythrocytic 1 | Cytoplasm | other | 5.44E+08 | 6.26E+08 | 5.62E+08 |
| P08032 | SPTA1 | spectrin, alpha, erythrocytic 1 | Cytoplasm | other | 4.53E+08 | 3.82E+08 | 3.33E+08 |
| P08032 | SPTA1 | spectrin, alpha, erythrocytic 1 | Cytoplasm | other | 4.25E+08 | 3.47E+08 | 3.17E+08 |
| P08032 | SPTA1 | spectrin, alpha, erythrocytic 1 | Cytoplasm | other | 5.21E+08 | 6.10E+08 | 5.24E+08 |
| P08032 | SPTA1 | spectrin, alpha, erythrocytic 1 | Cytoplasm | other | 4.81E+08 | 3.96E+08 | 3.73E+08 |
| P08032 | SPTA1 | spectrin, alpha, erythrocytic 1 | Cytoplasm | other | 4.94E+08 | 4.73E+08 | 3.58E+08 |
| P08032 | SPTA1 | spectrin, alpha, erythrocytic 1 | Cytoplasm | other | 3.76E+08 | 3.04E+08 | 3.04E+08 |
| P08032 | SPTA1 | spectrin, alpha, erythrocytic 1 | Cytoplasm | other | 3.54E+08 | 2.81E+08 | 2.43E+08 |
| P08032 | SPTA1 | spectrin, alpha, erythrocytic 1 | Cytoplasm | other | 5.73E+08 | 5.78E+08 | 6.34E+08 |
| P08032 | SPTA1 | spectrin, alpha, erythrocytic 1 | Cytoplasm | other | 3.68E+08 | 4.15E+08 | 3.30E+08 |
| P08032 | SPTA1 | spectrin, alpha, erythrocytic 1 | Cytoplasm | other | 3.39E+08 | 3.19E+08 | 2.74E+08 |
| P08032 | SPTA1 | spectrin, alpha, erythrocytic 1 | Cytoplasm | other | 4.99E+08 | 5.33E+08 | 6.06E+08 |
| P08032 | SPTA1 | spectrin, alpha, erythrocytic 1 | Cytoplasm | other | 3.98E+08 | 5.54E+08 | 3.02E+08 |
| P08032 | SPTA1 | spectrin, alpha, erythrocytic 1 | Cytoplasm | other | 3.04E+08 | 4.48E+08 | 2.43E+08 |
| P08032 | SPTA1 | spectrin, alpha, erythrocytic 1 | Cytoplasm | other | 3.76E+08 | 3.85E+08 | 3.61E+08 |
| P08032 | SPTA1 | spectrin, alpha, erythrocytic 1 | Cytoplasm | other | 3.61E+08 | 3.75E+08 | 4.19E+08 |
| P08032 | SPTA1 | spectrin, alpha, erythrocytic 1 | Cytoplasm | other | 3.81E+08 | 4.13E+08 | 4.21E+08 |
| P08032 | SPTA1 | spectrin, alpha, erythrocytic 1 | Cytoplasm | other | 2.71E+08 | 3.70E+08 | 2.52E+08 |
| P08032 | SPTA1 | spectrin, alpha, erythrocytic 1 | Cytoplasm | other | 3.27E+08 | 4.57E+08 | 2.12E+08 |
| P08032 | SPTA1 | spectrin, alpha, erythrocytic 1 | Cytoplasm | other | 3.38E+08 | 3.37E+08 | 2.64E+08 |
| P08032 | SPTA1 | spectrin, alpha, erythrocytic 1 | Cytoplasm | other | 3.50E+08 | 3.78E+08 | 2.48E+08 |
| P08032 | SPTA1 | spectrin, alpha, erythrocytic 1 | Cytoplasm | other | 2.39E+08 | 1.61E+08 | 1.05E+08 |
| P08032 | SPTA1 | spectrin, alpha, erythrocytic 1 | Cytoplasm | other | 3.07E+08 | 4.74E+08 | 2.64E+08 |
| P08032 | SPTA1 | spectrin, alpha, erythrocytic 1 | Cytoplasm | other | 5.37E+08 | 5.02E+08 | 7.27E+08 |
| P08032 | SPTA1 | spectrin, alpha, erythrocytic 1 | Cytoplasm | other | 4.72E+08 | 4.14E+08 | 3.51E+08 |
| P08032 | SPTA1 | spectrin, alpha, erythrocytic 1 | Cytoplasm | other | 2.84E+08 | 9.68E+07 | 3.16E+08 |
| P08032 | SPTA1 | spectrin, alpha, erythrocytic 1 | Cytoplasm | other | 3.32E+08 | 2.11E+08 | 3.16E+08 |
| P08032 | SPTA1 | spectrin, alpha, erythrocytic 1 | Cytoplasm | other | 2.77E+08 | 3.65E+08 | 2.13E+08 |
| P08032 | SPTA1 | spectrin, alpha, erythrocytic 1 | Cytoplasm | other | 3.26E+08 | 3.05E+08 | 2.38E+08 |
| P08032 | SPTA1 | spectrin, alpha, erythrocytic 1 | Cytoplasm | other | 2.73E+08 | 1.73E+08 | 3.27E+08 |
| P08032 | SPTA1 | spectrin, alpha, erythrocytic 1 | Cytoplasm | other | 3.84E+08 | 4.37E+08 | 2.20E+08 |
| P08032 | SPTA1 | spectrin, alpha, erythrocytic 1 | Cytoplasm | other | 3.96E+08 | 3.96E+08 | 2.39E+08 |
| P08032 | SPTA1 | spectrin, alpha, erythrocytic 1 | Cytoplasm | other | 2.17E+08 | 9.59E+07 | 1.37E+08 |
| P08032 | SPTA1 | spectrin, alpha, erythrocytic 1 | Cytoplasm | other | 2.79E+08 | 2.43E+08 | 2.30E+08 |
| P08032 | SPTA1 | spectrin, alpha, erythrocytic 1 | Cytoplasm | other | 2.44E+08 | 2.07E+08 | 1.81E+08 |
| P08032 | SPTA1 | spectrin, alpha, erythrocytic 1 | Cytoplasm | other | 2.13E+08 | 9.40E+07 | 1.93E+08 |
| P08032 | SPTA1 | spectrin, alpha, erythrocytic 1 | Cytoplasm | other | 2.08E+08 | 1.35E+08 | 1.26E+08 |
| P08032 | SPTA1 | spectrin, alpha, erythrocytic 1 | Cytoplasm | other | 2.76E+08 | 3.48E+08 | 2.08E+08 |
| P08032 | SPTA1 | spectrin, alpha, erythrocytic 1 | Cytoplasm | other | 1.96E+08 | 2.51E+08 | 1.79E+08 |
| P08032 | SPTA1 | spectrin, alpha, erythrocytic 1 | Cytoplasm | other | 2.43E+08 | 2.01E+08 | 1.73E+08 |
| P08032 | SPTA1 | spectrin, alpha, erythrocytic 1 | Cytoplasm | other | 1.98E+08 | 2.24E+08 | 1.73E+08 |
| P08032 | SPTA1 | spectrin, alpha, erythrocytic 1 | Cytoplasm | other | 2.06E+08 | 2.28E+08 | 1.45E+08 |
| P08032 | SPTA1 | spectrin, alpha, erythrocytic 1 | Cytoplasm | other | 4.02E+08 | 3.26E+08 | 2.68E+08 |
| P08032 | SPTA1 | spectrin, alpha, erythrocytic 1 | Cytoplasm | other | 2.92E+08 | 2.54E+08 | 2.03E+08 |
| P08032 | SPTA1 | spectrin, alpha, erythrocytic 1 | Cytoplasm | other | 2.91E+08 | 3.08E+08 | 3.01E+08 |
| P08032 | SPTA1 | spectrin, alpha, erythrocytic 1 | Cytoplasm | other | 1.82E+08 | 2.46E+08 | 1.29E+08 |
| P08032 | SPTA1 | spectrin, alpha, erythrocytic 1 | Cytoplasm | other | 1.45E+08 | 1.19E+08 | 1.08E+08 |
| P08032 | SPTA1 | spectrin, alpha, erythrocytic 1 | Cytoplasm | other | 1.56E+08 | 8.93E+07 | 5.21E+07 |
| P08032 | SPTA1 | spectrin, alpha, erythrocytic 1 | Cytoplasm | other | 1.41E+08 | 1.10E+08 | 1.01E+08 |
| P08032 | SPTA1 | spectrin, alpha, erythrocytic 1 | Cytoplasm | other | 1.22E+08 | 5.68E+07 | 1.50E+08 |
| P08032 | SPTA1 | spectrin, alpha, erythrocytic 1 | Cytoplasm | other | 2.00E+08 | 1.78E+08 | 1.58E+08 |
| P08032 | SPTA1 | spectrin, alpha, erythrocytic 1 | Cytoplasm | other | 1.51E+08 | 6.31E+07 | 1.23E+08 |
| P08032 | SPTA1 | spectrin, alpha, erythrocytic 1 | Cytoplasm | other | 2.12E+08 | 2.67E+08 | 1.34E+08 |
| P08032 | SPTA1 | spectrin, alpha, erythrocytic 1 | Cytoplasm | other | 2.05E+08 | 1.95E+08 | 1.42E+08 |
| P08032 | SPTA1 | spectrin, alpha, erythrocytic 1 | Cytoplasm | other | 1.58E+08 | 1.28E+08 | 1.09E+08 |
| P08032 | SPTA1 | spectrin, alpha, erythrocytic 1 | Cytoplasm | other | 2.77E+08 | 2.65E+08 | 1.97E+08 |
| P08032 | SPTA1 | spectrin, alpha, erythrocytic 1 | Cytoplasm | other | 9.08E+07 | 1.03E+08 | 6.61E+07 |
| P08032 | SPTA1 | spectrin, alpha, erythrocytic 1 | Cytoplasm | other | 1.40E+08 | 4.46E+07 | 9.19E+07 |
| P08032 | SPTA1 | spectrin, alpha, erythrocytic 1 | Cytoplasm | other | 2.31E+08 | 2.50E+08 | 1.17E+08 |
| P08032 | SPTA1 | spectrin, alpha, erythrocytic 1 | Cytoplasm | other | 2.07E+07 | 2.93E+07 | 3.97E+07 |
| P08032 | SPTA1 | spectrin, alpha, erythrocytic 1 | Cytoplasm | other | 1.23E+08 | 9.12E+07 | 9.09E+07 |
| P08032 | SPTA1 | spectrin, alpha, erythrocytic 1 | Cytoplasm | other | 1.22E+08 | 5.87E+07 | 1.08E+08 |
| P08032 | SPTA1 | spectrin, alpha, erythrocytic 1 | Cytoplasm | other | 9.41E+07 | 1.06E+08 | 9.42E+07 |
| P08032 | SPTA1 | spectrin, alpha, erythrocytic 1 | Cytoplasm | other | 1.97E+08 | 2.03E+08 | 1.39E+08 |
| P08032 | SPTA1 | spectrin, alpha, erythrocytic 1 | Cytoplasm | other | 9.29E+07 | 4.50E+07 | 8.25E+07 |
| P08032 | SPTA1 | spectrin, alpha, erythrocytic 1 | Cytoplasm | other | 1.93E+08 | 2.20E+08 | 1.83E+08 |
| P08032 | SPTA1 | spectrin, alpha, erythrocytic 1 | Cytoplasm | other | 1.18E+08 | 1.21E+08 | 8.89E+07 |
| P08032 | SPTA1 | spectrin, alpha, erythrocytic 1 | Cytoplasm | other | 1.96E+08 | 5.01E+08 | 4.61E+08 |
| P08032 | SPTA1 | spectrin, alpha, erythrocytic 1 | Cytoplasm | other | 9.93E+07 | 3.70E+07 | 1.23E+08 |
| P08032 | SPTA1 | spectrin, alpha, erythrocytic 1 | Cytoplasm | other | 1.26E+08 | 1.54E+08 | 8.85E+07 |
| P08032 | SPTA1 | spectrin, alpha, erythrocytic 1 | Cytoplasm | other | 1.25E+08 | 1.06E+08 | 8.77E+07 |
| P08032 | SPTA1 | spectrin, alpha, erythrocytic 1 | Cytoplasm | other | 1.72E+08 | 5.45E+08 | 2.61E+08 |
| P08032 | SPTA1 | spectrin, alpha, erythrocytic 1 | Cytoplasm | other | 9.92E+07 | 5.15E+07 | 6.20E+07 |
| P08032 | SPTA1 | spectrin, alpha, erythrocytic 1 | Cytoplasm | other | 4.03E+07 | 3.93E+07 | 9.41E+07 |
| P08032 | SPTA1 | spectrin, alpha, erythrocytic 1 | Cytoplasm | other | 1.12E+08 | 6.42E+07 | 5.71E+07 |
| P08032 | SPTA1 | spectrin, alpha, erythrocytic 1 | Cytoplasm | other | 1.57E+08 | 1.39E+08 | 1.36E+08 |
| P08032 | SPTA1 | spectrin, alpha, erythrocytic 1 | Cytoplasm | other | 1.10E+08 | 1.27E+08 | 9.34E+07 |

| | | | | | | | |
|---|---|---|---|---|---|---|---|
| P08032 | SPTA1 | spectrin, alpha, erythrocytic 1 | Cytoplasm | other | 1.09E+08 | 1.07E+08 | 1.00E+08 |
| P08032 | SPTA1 | spectrin, alpha, erythrocytic 1 | Cytoplasm | other | 9.18E+07 | 6.07E+07 | 5.45E+07 |
| P08032 | SPTA1 | spectrin, alpha, erythrocytic 1 | Cytoplasm | other | 9563600 | 5729218.5 | 3.45E+07 |
| P08032 | SPTA1 | spectrin, alpha, erythrocytic 1 | Cytoplasm | other | 9.78E+07 | 1.24E+08 | 7.04E+07 |
| P08032 | SPTA1 | spectrin, alpha, erythrocytic 1 | Cytoplasm | other | 5.96E+07 | 1.02E+08 | 6.44E+07 |
| P08032 | SPTA1 | spectrin, alpha, erythrocytic 1 | Cytoplasm | other | 7.96E+07 | 7.76E+07 | 6.66E+07 |
| P08032 | SPTA1 | spectrin, alpha, erythrocytic 1 | Cytoplasm | other | 1.02E+08 | 7.85E+07 | 7.17E+07 |
| P08032 | SPTA1 | spectrin, alpha, erythrocytic 1 | Cytoplasm | other | 9.46E+07 | 1.26E+08 | 7.06E+07 |
| P08032 | SPTA1 | spectrin, alpha, erythrocytic 1 | Cytoplasm | other | 8.86E+07 | 7.47E+07 | 8.23E+07 |
| P08032 | SPTA1 | spectrin, alpha, erythrocytic 1 | Cytoplasm | other | 7.63E+07 | 6.89E+07 | 6.64E+07 |
| P08032 | SPTA1 | spectrin, alpha, erythrocytic 1 | Cytoplasm | other | 9.64E+07 | 9.53E+07 | 6.85E+07 |
| P08032 | SPTA1 | spectrin, alpha, erythrocytic 1 | Cytoplasm | other | 5.56E+07 | 9453335 | 4.66E+07 |
| P08032 | SPTA1 | spectrin, alpha, erythrocytic 1 | Cytoplasm | other | 8.96E+07 | 2.20E+08 | 1.56E+08 |
| P08032 | SPTA1 | spectrin, alpha, erythrocytic 1 | Cytoplasm | other | 7.64E+07 | 2.52E+07 | 9.93E+07 |
| P08032 | SPTA1 | spectrin, alpha, erythrocytic 1 | Cytoplasm | other | 6.74E+07 | 6.14E+07 | 5.93E+07 |
| P08032 | SPTA1 | spectrin, alpha, erythrocytic 1 | Cytoplasm | other | 9.58E+07 | 9.31E+07 | 8.04E+07 |
| P08032 | SPTA1 | spectrin, alpha, erythrocytic 1 | Cytoplasm | other | 1.06E+08 | 3.68E+07 | 1.25E+08 |
| P08032 | SPTA1 | spectrin, alpha, erythrocytic 1 | Cytoplasm | other | 1.18E+08 | 1.61E+08 | 1.50E+08 |
| P08032 | SPTA1 | spectrin, alpha, erythrocytic 1 | Cytoplasm | other | 6.58E+07 | 3.47E+07 | 4.95E+07 |
| P08032 | SPTA1 | spectrin, alpha, erythrocytic 1 | Cytoplasm | other | 6.21E+07 | 5.73E+07 | 4.57E+07 |
| P08032 | SPTA1 | spectrin, alpha, erythrocytic 1 | Cytoplasm | other | 5.91E+07 | 3.61E+07 | 4.92E+07 |
| P08032 | SPTA1 | spectrin, alpha, erythrocytic 1 | Cytoplasm | other | 2.89E+07 | 8.10E+07 | 1.50E+08 |
| P08032 | SPTA1 | spectrin, alpha, erythrocytic 1 | Cytoplasm | other | 5.62E+07 | 7.53E+07 | 4.25E+07 |
| P08032 | SPTA1 | spectrin, alpha, erythrocytic 1 | Cytoplasm | other | 3.88E+07 | 5.01E+07 | 4.00E+07 |
| P08032 | SPTA1 | spectrin, alpha, erythrocytic 1 | Cytoplasm | other | 7.44E+07 | 1.85E+08 | 1.26E+08 |
| P08032 | SPTA1 | spectrin, alpha, erythrocytic 1 | Cytoplasm | other | 3.56E+07 | 2.12E+07 | 2.42E+07 |
| P08032 | SPTA1 | spectrin, alpha, erythrocytic 1 | Cytoplasm | other | 6.01E+07 | 5.07E+07 | 4.30E+07 |
| P08032 | SPTA1 | spectrin, alpha, erythrocytic 1 | Cytoplasm | other | 6.09E+07 | 5.66E+07 | 4.11E+07 |
| P08032 | SPTA1 | spectrin, alpha, erythrocytic 1 | Cytoplasm | other | 4.03E+07 | 1.27E+07 | 4.26E+07 |
| P08032 | SPTA1 | spectrin, alpha, erythrocytic 1 | Cytoplasm | other | 4.27E+07 | 4.17E+07 | 3.57E+07 |
| P08032 | SPTA1 | spectrin, alpha, erythrocytic 1 | Cytoplasm | other | 3.86E+07 | 3.68E+07 | 3.80E+07 |
| P08032 | SPTA1 | spectrin, alpha, erythrocytic 1 | Cytoplasm | other | 4.38E+07 | 5.53E+07 | 3.17E+07 |
| P08032 | SPTA1 | spectrin, alpha, erythrocytic 1 | Cytoplasm | other | 4.21E+07 | 7.46E+07 | 2.52E+07 |
| P08032 | SPTA1 | spectrin, alpha, erythrocytic 1 | Cytoplasm | other | 3.47E+07 | 5.00E+07 | 3.16E+07 |
| P08032 | SPTA1 | spectrin, alpha, erythrocytic 1 | Cytoplasm | other | 2.97E+07 | 1.01E+07 | 2.16E+07 |
| P08032 | SPTA1 | spectrin, alpha, erythrocytic 1 | Cytoplasm | other | 2.90E+07 | 3.46E+07 | 2.59E+07 |
| P08032 | SPTA1 | spectrin, alpha, erythrocytic 1 | Cytoplasm | other | 5.61E+07 | 1.44E+08 | 8.01E+07 |
| P08032 | SPTA1 | spectrin, alpha, erythrocytic 1 | Cytoplasm | other | 1.20E+07 | 9671188 | 3.76E+07 |
| P08032 | SPTA1 | spectrin, alpha, erythrocytic 1 | Cytoplasm | other | 3.18E+07 | 2.97E+07 | 2.34E+07 |
| P08032 | SPTA1 | spectrin, alpha, erythrocytic 1 | Cytoplasm | other | 9.96E+07 | 1.03E+08 | 6.97E+07 |
| P08032 | SPTA1 | spectrin, alpha, erythrocytic 1 | Cytoplasm | other | 2.35E+07 | 9895405 | 2.28E+07 |
| P08032 | SPTA1 | spectrin, alpha, erythrocytic 1 | Cytoplasm | other | 2.80E+07 | 1.23E+07 | 2.34E+07 |
| P08032 | SPTA1 | spectrin, alpha, erythrocytic 1 | Cytoplasm | other | 3.91E+07 | 3.54E+07 | 2.81E+07 |
| P08032 | SPTA1 | spectrin, alpha, erythrocytic 1 | Cytoplasm | other | 2.40E+07 | 2.43E+07 | 1.97E+07 |
| P08032 | SPTA1 | spectrin, alpha, erythrocytic 1 | Cytoplasm | other | 4.06E+07 | 4.11E+07 | 2.62E+07 |
| P08032 | SPTA1 | spectrin, alpha, erythrocytic 1 | Cytoplasm | other | 3029119.25 | 4625589.5 | 4055859.75 |
| P08032 | SPTA1 | spectrin, alpha, erythrocytic 1 | Cytoplasm | other | 4.46E+07 | 6.06E+07 | 7.42E+07 |
| P08032 | SPTA1 | spectrin, alpha, erythrocytic 1 | Cytoplasm | other | 4.20E+07 | 3.93E+07 | 2.33E+07 |
| P08032 | SPTA1 | spectrin, alpha, erythrocytic 1 | Cytoplasm | other | 2316695.25 | 2313812.25 | 4889930 |
| P08032 | SPTA1 | spectrin, alpha, erythrocytic 1 | Cytoplasm | other | 2.71E+07 | 1.15E+07 | 2.95E+07 |
| P08032 | SPTA1 | spectrin, alpha, erythrocytic 1 | Cytoplasm | other | 9809957 | 2662025 | 5507773.5 |
| P08032 | SPTA1 | spectrin, alpha, erythrocytic 1 | Cytoplasm | other | 2.74E+07 | 2.20E+07 | 1.75E+07 |
| P08032 | SPTA1 | spectrin, alpha, erythrocytic 1 | Cytoplasm | other | 2.11E+07 | 8520031 | 2.69E+07 |
| P08032 | SPTA1 | spectrin, alpha, erythrocytic 1 | Cytoplasm | other | 1.64E+07 | 7155508.5 | 5417489 |
| P08032 | SPTA1 | spectrin, alpha, erythrocytic 1 | Cytoplasm | other | 2.50E+07 | 2.54E+07 | 1.90E+07 |
| P08032 | SPTA1 | spectrin, alpha, erythrocytic 1 | Cytoplasm | other | 2.69E+07 | 2.27E+07 | 1.47E+07 |
| P08032 | SPTA1 | spectrin, alpha, erythrocytic 1 | Cytoplasm | other | 1.10E+07 | 1.74E+07 | 9350525 |
| P08032 | SPTA1 | spectrin, alpha, erythrocytic 1 | Cytoplasm | other | 1.69E+07 | 4774440.5 | 8770652 |
| P08032 | SPTA1 | spectrin, alpha, erythrocytic 1 | Cytoplasm | other | 1.29E+07 | 2184150.5 | 1.99E+07 |
| P08032 | SPTA1 | spectrin, alpha, erythrocytic 1 | Cytoplasm | other | 1.92E+07 | 1.53E+07 | 1.38E+07 |
| P08032 | SPTA1 | spectrin, alpha, erythrocytic 1 | Cytoplasm | other | 2.00E+07 | 1.37E+07 | 1.08E+07 |
| P08032 | SPTA1 | spectrin, alpha, erythrocytic 1 | Cytoplasm | other | 1.01E+07 | 1.10E+07 | 7350691.5 |
| P08032 | SPTA1 | spectrin, alpha, erythrocytic 1 | Cytoplasm | other | 1.74E+07 | 1.61E+07 | 1.39E+07 |
| P08032 | SPTA1 | spectrin, alpha, erythrocytic 1 | Cytoplasm | other | 1.40E+07 | 1.63E+07 | 7586844 |
| P08032 | SPTA1 | spectrin, alpha, erythrocytic 1 | Cytoplasm | other | 1.76E+07 | 1.69E+07 | 1.50E+07 |
| P08032 | SPTA1 | spectrin, alpha, erythrocytic 1 | Cytoplasm | other | 1.10E+07 | 8373883.5 | 1.15E+07 |
| P08032 | SPTA1 | spectrin, alpha, erythrocytic 1 | Cytoplasm | other | 1.79E+07 | 1.70E+07 | 4.98E+07 |
| P08032 | SPTA1 | spectrin, alpha, erythrocytic 1 | Cytoplasm | other | 1.38E+07 | 1.51E+07 | 1.24E+07 |
| P08032 | SPTA1 | spectrin, alpha, erythrocytic 1 | Cytoplasm | other | 1.32E+07 | 5577812 | 1.16E+07 |
| P08032 | SPTA1 | spectrin, alpha, erythrocytic 1 | Cytoplasm | other | 1.35E+07 | 2.61E+07 | 1.28E+07 |
| P08032 | SPTA1 | spectrin, alpha, erythrocytic 1 | Cytoplasm | other | 7455037 | 9099328 | 8606041 |
| P08032 | SPTA1 | spectrin, alpha, erythrocytic 1 | Cytoplasm | other | 7528152 | 1.46E+07 | 1.27E+07 |
| P08032 | SPTA1 | spectrin, alpha, erythrocytic 1 | Cytoplasm | other | 6342675.5 | 4969063.5 | 1.48E+07 |
| P08032 | SPTA1 | spectrin, alpha, erythrocytic 1 | Cytoplasm | other | 2.87E+07 | 2.50E+07 | 2.24E+07 |
| P08032 | SPTA1 | spectrin, alpha, erythrocytic 1 | Cytoplasm | other | 1.50E+07 | 1.87E+07 | 6873939 |
| P08032 | SPTA1 | spectrin, alpha, erythrocytic 1 | Cytoplasm | other | 8204660.5 | 2837632.5 | 9314009 |
| P08032 | SPTA1 | spectrin, alpha, erythrocytic 1 | Cytoplasm | other | 1.14E+07 | 9431185 | 6764896 |
| P08032 | SPTA1 | spectrin, alpha, erythrocytic 1 | Cytoplasm | other | 1772833.125 | 2258027.25 | 2848305 |
| P08032 | SPTA1 | spectrin, alpha, erythrocytic 1 | Cytoplasm | other | 7750587 | 7068232.5 | 6719169.5 |
| P08032 | SPTA1 | spectrin, alpha, erythrocytic 1 | Cytoplasm | other | 7762942 | 7914199.5 | 5710590.5 |
| P08032 | SPTA1 | spectrin, alpha, erythrocytic 1 | Cytoplasm | other | 1.69E+07 | 1.22E+07 | 1.18E+07 |
| P08032 | SPTA1 | spectrin, alpha, erythrocytic 1 | Cytoplasm | other | 8545710 | 2620246.5 | 7220911 |
| P08032 | SPTA1 | spectrin, alpha, erythrocytic 1 | Cytoplasm | other | 2.31E+07 | 2.89E+07 | 2.01E+07 |
| P08032 | SPTA1 | spectrin, alpha, erythrocytic 1 | Cytoplasm | other | 4781526.5 | 5060856 | 2981817 |
| P08032 | SPTA1 | spectrin, alpha, erythrocytic 1 | Cytoplasm | other | 258589.547 | 181728.312 | 6448628 |
| P08032 | SPTA1 | spectrin, alpha, erythrocytic 1 | Cytoplasm | other | 5281042 | 4461179.5 | 4712963.5 |
| P08032 | SPTA1 | spectrin, alpha, erythrocytic 1 | Cytoplasm | other | 3327700 | 2558774 | 3024886 |
| P08032 | SPTA1 | spectrin, alpha, erythrocytic 1 | Cytoplasm | other | 375618.969 | 818243.375 | 673387.125 |
| P08032 | SPTA1 | spectrin, alpha, erythrocytic 1 | Cytoplasm | other | 8085583.5 | 6217524.5 | 4312823.5 |
| P08032 | SPTA1 | spectrin, alpha, erythrocytic 1 | Cytoplasm | other | 8022594.5 | 3774982.5 | 6553560.5 |
| P08032 | SPTA1 | spectrin, alpha, erythrocytic 1 | Cytoplasm | other | 446162.812 | 58811.836 | 6098716.5 |
| P08032 | SPTA1 | spectrin, alpha, erythrocytic 1 | Cytoplasm | other | 4036162 | 7069912 | 3385592.25 |
| P08032 | SPTA1 | spectrin, alpha, erythrocytic 1 | Cytoplasm | other | 4281857 | 3003564.5 | 4766828.5 |
| P08032 | SPTA1 | spectrin, alpha, erythrocytic 1 | Cytoplasm | other | 5848687.5 | 4501173.5 | 3555402.25 |
| P08032 | SPTA1 | spectrin, alpha, erythrocytic 1 | Cytoplasm | other | 7239885.5 | 6441940.5 | 5374264.5 |
| P08032 | SPTA1 | spectrin, alpha, erythrocytic 1 | Cytoplasm | other | 5302806 | 9060435 | 5672377 |
| P08032 | SPTA1 | spectrin, alpha, erythrocytic 1 | Cytoplasm | other | 3808755 | 2809465.25 | 2227291.75 |
| P08032 | SPTA1 | spectrin, alpha, erythrocytic 1 | Cytoplasm | other | 4681516 | 5522511.5 | 5637839 |
| P08032 | SPTA1 | spectrin, alpha, erythrocytic 1 | Cytoplasm | other | 6609997.5 | 4254035.5 | 4056555 |
| P08032 | SPTA1 | spectrin, alpha, erythrocytic 1 | Cytoplasm | other | 4598426.5 | 1363132.75 | 4933085 |
| P08032 | SPTA1 | spectrin, alpha, erythrocytic 1 | Cytoplasm | other | 4750061 | 1.40E+07 | 1.43E+07 |
| P08032 | SPTA1 | spectrin, alpha, erythrocytic 1 | Cytoplasm | other | 2428219.75 | 451366.125 | 2834670.5 |
| P08032 | SPTA1 | spectrin, alpha, erythrocytic 1 | Cytoplasm | other | 1610116.25 | 1312031.5 | 2043558.5 |
| P08032 | SPTA1 | spectrin, alpha, erythrocytic 1 | Cytoplasm | other | 1934358.25 | 1848545.75 | 1884421.25 |
| P08032 | SPTA1 | spectrin, alpha, erythrocytic 1 | Cytoplasm | other | 1765209.75 | 282158.188 | 1168677.5 |
| P08032 | SPTA1 | spectrin, alpha, erythrocytic 1 | Cytoplasm | other | 993257.312 | 1130803.5 | 750991.125 |
| P08032 | SPTA1 | spectrin, alpha, erythrocytic 1 | Cytoplasm | other | 7222389 | 5155728.5 | 5333081 |
| P08032 | SPTA1 | spectrin, alpha, erythrocytic 1 | Cytoplasm | other | 508095.781 | 609396.938 | 1337163.75 |
| P08032 | SPTA1 | spectrin, alpha, erythrocytic 1 | Cytoplasm | other | 1257865 | 1468801.25 | 1115126.5 |

| | | | | | | | | | | | | | |
|---|---|---|---|---|---|---|---|---|---|---|---|---|---|
| P08032 | SPTA1 | spectrin, alpha, erythrocytic 1 | Cytoplasm | other | 967304 | 351815.156 | 1555477.125 | | | | | | |
| P08032 | SPTA1 | spectrin, alpha, erythrocytic 1 | Cytoplasm | other | 4211232.5 | 4.24E+07 | 7.09E+07 | | | | | | |
| P08032 | SPTA1 | spectrin, alpha, erythrocytic 1 | Cytoplasm | other | 382895.719 | 125588.984 | 893483.188 | | | | | | |
| P08032 | SPTA1 | spectrin, alpha, erythrocytic 1 | Cytoplasm | other | 120010.539 | 0 | 1780274.25 | | | | | | |
| P08032 | SPTA1 | spectrin, alpha, erythrocytic 1 | Cytoplasm | other | 2095704.125 | 2573059.75 | 3528336 | | | | | | |
| P08032 | SPTA1 | spectrin, alpha, erythrocytic 1 | Cytoplasm | other | 287272.688 | 514750.719 | 686904.625 | | | | | | |
| P08032 | SPTA1 | spectrin, alpha, erythrocytic 1 | Cytoplasm | other | 752513.75 | 1994267.125 | 1901949.375 | P08032 | SPTA1 | 525877770 | 634585593 | 21223717 | 894 |
| P16546 | SPTAN1 | spectrin, alpha, non-erythrocytic 1 | Plasma Membrane | other | 9566639 | 1.04E+07 | 6598359 | P16546 | SPTAN1 | 9566639 | 1986679 | 1147009 | 3 |
| P15508 | SPTB | spectrin, beta, erythrocytic | Plasma Membrane | other | 3.02E+09 | 3.08E+09 | 3.09E+09 | | | | | | |
| P15508 | SPTB | spectrin, beta, erythrocytic | Plasma Membrane | other | 2.06E+09 | 2.57E+09 | 2.35E+09 | | | | | | |
| P15508 | SPTB | spectrin, beta, erythrocytic | Plasma Membrane | other | 2.33E+09 | 2.57E+09 | 2.47E+09 | | | | | | |
| P15508 | SPTB | spectrin, beta, erythrocytic | Plasma Membrane | other | 2.39E+09 | 2.30E+09 | 2.02E+09 | | | | | | |
| P15508 | SPTB | spectrin, beta, erythrocytic | Plasma Membrane | other | 1.79E+09 | 1.75E+09 | 1.83E+09 | | | | | | |
| P15508 | SPTB | spectrin, beta, erythrocytic | Plasma Membrane | other | 2.18E+09 | 2.70E+09 | 2.53E+09 | | | | | | |
| P15508 | SPTB | spectrin, beta, erythrocytic | Plasma Membrane | other | 1.94E+09 | 1.69E+09 | 1.54E+09 | | | | | | |
| P15508 | SPTB | spectrin, beta, erythrocytic | Plasma Membrane | other | 1.74E+09 | 1.68E+09 | 1.21E+09 | | | | | | |
| P15508 | SPTB | spectrin, beta, erythrocytic | Plasma Membrane | other | 1.77E+09 | 1.70E+09 | 1.49E+09 | | | | | | |
| P15508 | SPTB | spectrin, beta, erythrocytic | Plasma Membrane | other | 1.89E+09 | 1.61E+09 | 1.40E+09 | | | | | | |
| P15508 | SPTB | spectrin, beta, erythrocytic | Plasma Membrane | other | 1.67E+09 | 2.11E+09 | 1.64E+09 | | | | | | |
| P15508 | SPTB | spectrin, beta, erythrocytic | Plasma Membrane | other | 1.97E+09 | 1.74E+09 | 1.42E+09 | | | | | | |
| P15508 | SPTB | spectrin, beta, erythrocytic | Plasma Membrane | other | 1.63E+09 | 1.47E+09 | 1.40E+09 | | | | | | |
| P15508 | SPTB | spectrin, beta, erythrocytic | Plasma Membrane | other | 2.34E+09 | 2.23E+09 | 2.10E+09 | | | | | | |
| P15508 | SPTB | spectrin, beta, erythrocytic | Plasma Membrane | other | 1.62E+09 | 1.57E+09 | 1.45E+09 | | | | | | |
| P15508 | SPTB | spectrin, beta, erythrocytic | Plasma Membrane | other | 1.62E+09 | 1.60E+09 | 1.47E+09 | | | | | | |
| P15508 | SPTB | spectrin, beta, erythrocytic | Plasma Membrane | other | 1.57E+09 | 1.47E+09 | 1.20E+09 | | | | | | |
| P15508 | SPTB | spectrin, beta, erythrocytic | Plasma Membrane | other | 1.36E+09 | 1.32E+09 | 1.29E+09 | | | | | | |
| P15508 | SPTB | spectrin, beta, erythrocytic | Plasma Membrane | other | 1.56E+09 | 1.46E+09 | 1.41E+09 | | | | | | |
| P15508 | SPTB | spectrin, beta, erythrocytic | Plasma Membrane | other | 1.51E+09 | 1.34E+09 | 1.17E+09 | | | | | | |
| P15508 | SPTB | spectrin, beta, erythrocytic | Plasma Membrane | other | 1.36E+09 | 1.50E+09 | 1.44E+09 | | | | | | |
| P15508 | SPTB | spectrin, beta, erythrocytic | Plasma Membrane | other | 1.18E+09 | 1.23E+09 | 1.06E+09 | | | | | | |
| P15508 | SPTB | spectrin, beta, erythrocytic | Plasma Membrane | other | 1.42E+09 | 1.34E+09 | 1.15E+09 | | | | | | |
| P15508 | SPTB | spectrin, beta, erythrocytic | Plasma Membrane | other | 1.44E+09 | 1.39E+09 | 1.57E+09 | | | | | | |
| P15508 | SPTB | spectrin, beta, erythrocytic | Plasma Membrane | other | 1.24E+09 | 1.07E+09 | 8.97E+08 | | | | | | |
| P15508 | SPTB | spectrin, beta, erythrocytic | Plasma Membrane | other | 1.16E+09 | 1.29E+09 | 1.21E+09 | | | | | | |
| P15508 | SPTB | spectrin, beta, erythrocytic | Plasma Membrane | other | 1.49E+09 | 1.34E+09 | 1.26E+09 | | | | | | |
| P15508 | SPTB | spectrin, beta, erythrocytic | Plasma Membrane | other | 1.19E+09 | 1.14E+09 | 1.10E+09 | | | | | | |
| P15508 | SPTB | spectrin, beta, erythrocytic | Plasma Membrane | other | 1.42E+09 | 1.32E+09 | 1.42E+09 | | | | | | |
| P15508 | SPTB | spectrin, beta, erythrocytic | Plasma Membrane | other | 1.34E+09 | 1.20E+09 | 1.10E+09 | | | | | | |
| P15508 | SPTB | spectrin, beta, erythrocytic | Plasma Membrane | other | 1.50E+09 | 1.42E+09 | 1.40E+09 | | | | | | |
| P15508 | SPTB | spectrin, beta, erythrocytic | Plasma Membrane | other | 1.37E+09 | 1.52E+09 | 1.28E+09 | | | | | | |
| P15508 | SPTB | spectrin, beta, erythrocytic | Plasma Membrane | other | 1.49E+09 | 1.49E+09 | 1.22E+09 | | | | | | |
| P15508 | SPTB | spectrin, beta, erythrocytic | Plasma Membrane | other | 1.24E+09 | 1.64E+09 | 1.22E+09 | | | | | | |
| P15508 | SPTB | spectrin, beta, erythrocytic | Plasma Membrane | other | 1.06E+09 | 5.50E+08 | 6.20E+08 | | | | | | |
| P15508 | SPTB | spectrin, beta, erythrocytic | Plasma Membrane | other | 1.40E+09 | 1.09E+09 | 9.68E+08 | | | | | | |
| P15508 | SPTB | spectrin, beta, erythrocytic | Plasma Membrane | other | 1.60E+09 | 1.44E+09 | 1.40E+09 | | | | | | |
| P15508 | SPTB | spectrin, beta, erythrocytic | Plasma Membrane | other | 1.12E+09 | 1.02E+09 | 1.04E+09 | | | | | | |
| P15508 | SPTB | spectrin, beta, erythrocytic | Plasma Membrane | other | 1.14E+09 | 1.15E+09 | 1.06E+09 | | | | | | |
| P15508 | SPTB | spectrin, beta, erythrocytic | Plasma Membrane | other | 1.15E+09 | 1.31E+09 | 1.09E+09 | | | | | | |
| P15508 | SPTB | spectrin, beta, erythrocytic | Plasma Membrane | other | 9.95E+08 | 1.03E+09 | 8.74E+08 | | | | | | |
| P15508 | SPTB | spectrin, beta, erythrocytic | Plasma Membrane | other | 1.50E+09 | 1.39E+09 | 1.35E+09 | | | | | | |
| P15508 | SPTB | spectrin, beta, erythrocytic | Plasma Membrane | other | 1.09E+09 | 1.30E+09 | 1.10E+09 | | | | | | |
| P15508 | SPTB | spectrin, beta, erythrocytic | Plasma Membrane | other | 9.83E+08 | 1.03E+09 | 1.03E+09 | | | | | | |
| P15508 | SPTB | spectrin, beta, erythrocytic | Plasma Membrane | other | 1.25E+09 | 1.28E+09 | 1.05E+09 | | | | | | |
| P15508 | SPTB | spectrin, beta, erythrocytic | Plasma Membrane | other | 9.47E+08 | 9.43E+08 | 8.97E+08 | | | | | | |
| P15508 | SPTB | spectrin, beta, erythrocytic | Plasma Membrane | other | 1.16E+09 | 1.04E+09 | 1.14E+09 | | | | | | |
| P15508 | SPTB | spectrin, beta, erythrocytic | Plasma Membrane | other | 9.34E+08 | 1.08E+09 | 9.64E+08 | | | | | | |
| P15508 | SPTB | spectrin, beta, erythrocytic | Plasma Membrane | other | 1.03E+09 | 1.05E+09 | 9.28E+08 | | | | | | |
| P15508 | SPTB | spectrin, beta, erythrocytic | Plasma Membrane | other | 9.16E+08 | 9.54E+08 | 1.01E+09 | | | | | | |
| P15508 | SPTB | spectrin, beta, erythrocytic | Plasma Membrane | other | 9.51E+08 | 9.00E+08 | 7.66E+08 | | | | | | |
| P15508 | SPTB | spectrin, beta, erythrocytic | Plasma Membrane | other | 1.23E+09 | 1.07E+09 | 6.52E+08 | | | | | | |
| P15508 | SPTB | spectrin, beta, erythrocytic | Plasma Membrane | other | 1.09E+09 | 1.03E+09 | 1.05E+09 | | | | | | |
| P15508 | SPTB | spectrin, beta, erythrocytic | Plasma Membrane | other | 9.44E+08 | 8.14E+08 | 7.76E+08 | | | | | | |
| P15508 | SPTB | spectrin, beta, erythrocytic | Plasma Membrane | other | 9.31E+08 | 9.02E+08 | 8.60E+08 | | | | | | |
| P15508 | SPTB | spectrin, beta, erythrocytic | Plasma Membrane | other | 7.94E+08 | 5.90E+08 | 5.15E+08 | | | | | | |
| P15508 | SPTB | spectrin, beta, erythrocytic | Plasma Membrane | other | 9.15E+08 | 8.37E+08 | 7.07E+08 | | | | | | |
| P15508 | SPTB | spectrin, beta, erythrocytic | Plasma Membrane | other | 9.87E+08 | 9.81E+08 | 7.85E+08 | | | | | | |
| P15508 | SPTB | spectrin, beta, erythrocytic | Plasma Membrane | other | 1.09E+09 | 5.80E+08 | 9.26E+08 | | | | | | |
| P15508 | SPTB | spectrin, beta, erythrocytic | Plasma Membrane | other | 7.95E+08 | 7.26E+08 | 7.87E+08 | | | | | | |
| P15508 | SPTB | spectrin, beta, erythrocytic | Plasma Membrane | other | 9.55E+08 | 8.80E+08 | 7.64E+08 | | | | | | |
| P15508 | SPTB | spectrin, beta, erythrocytic | Plasma Membrane | other | 9.17E+08 | 9.46E+08 | 7.80E+08 | | | | | | |
| P15508 | SPTB | spectrin, beta, erythrocytic | Plasma Membrane | other | 7.89E+08 | 8.29E+08 | 8.76E+08 | | | | | | |
| P15508 | SPTB | spectrin, beta, erythrocytic | Plasma Membrane | other | 5.47E+08 | 6.75E+08 | 6.09E+08 | | | | | | |
| P15508 | SPTB | spectrin, beta, erythrocytic | Plasma Membrane | other | 7.99E+08 | 8.04E+08 | 7.16E+08 | | | | | | |
| P15508 | SPTB | spectrin, beta, erythrocytic | Plasma Membrane | other | 6.83E+08 | 6.66E+08 | 5.83E+08 | | | | | | |
| P15508 | SPTB | spectrin, beta, erythrocytic | Plasma Membrane | other | 6.76E+08 | 6.88E+08 | 5.87E+08 | | | | | | |
| P15508 | SPTB | spectrin, beta, erythrocytic | Plasma Membrane | other | 5.13E+08 | 6.55E+08 | 5.64E+08 | | | | | | |
| P15508 | SPTB | spectrin, beta, erythrocytic | Plasma Membrane | other | 9.97E+08 | 8.27E+08 | 7.08E+08 | | | | | | |
| P15508 | SPTB | spectrin, beta, erythrocytic | Plasma Membrane | other | 6.70E+08 | 6.49E+08 | 6.93E+08 | | | | | | |
| P15508 | SPTB | spectrin, beta, erythrocytic | Plasma Membrane | other | 5.87E+08 | 3.28E+08 | 5.76E+08 | | | | | | |
| P15508 | SPTB | spectrin, beta, erythrocytic | Plasma Membrane | other | 6.58E+08 | 4.30E+08 | 7.03E+08 | | | | | | |
| P15508 | SPTB | spectrin, beta, erythrocytic | Plasma Membrane | other | 6.87E+08 | 6.76E+08 | 5.50E+08 | | | | | | |
| P15508 | SPTB | spectrin, beta, erythrocytic | Plasma Membrane | other | 7.36E+08 | 6.71E+08 | 7.43E+08 | | | | | | |
| P15508 | SPTB | spectrin, beta, erythrocytic | Plasma Membrane | other | 9.95E+08 | 7.14E+08 | 6.40E+08 | | | | | | |
| P15508 | SPTB | spectrin, beta, erythrocytic | Plasma Membrane | other | 8.05E+08 | 1.16E+09 | 4.31E+08 | | | | | | |
| P15508 | SPTB | spectrin, beta, erythrocytic | Plasma Membrane | other | 6.26E+08 | 6.67E+08 | 6.36E+08 | | | | | | |
| P15508 | SPTB | spectrin, beta, erythrocytic | Plasma Membrane | other | 8.82E+08 | 7.86E+08 | 5.59E+08 | | | | | | |
| P15508 | SPTB | spectrin, beta, erythrocytic | Plasma Membrane | other | 4.80E+08 | 4.39E+08 | 4.60E+08 | | | | | | |
| P15508 | SPTB | spectrin, beta, erythrocytic | Plasma Membrane | other | 6.95E+08 | 7.18E+08 | 6.64E+08 | | | | | | |
| P15508 | SPTB | spectrin, beta, erythrocytic | Plasma Membrane | other | 6.77E+08 | 6.16E+08 | 5.41E+08 | | | | | | |
| P15508 | SPTB | spectrin, beta, erythrocytic | Plasma Membrane | other | 5.44E+08 | 5.53E+08 | 5.19E+08 | | | | | | |
| P15508 | SPTB | spectrin, beta, erythrocytic | Plasma Membrane | other | 8.18E+08 | 8.81E+08 | 6.09E+08 | | | | | | |
| P15508 | SPTB | spectrin, beta, erythrocytic | Plasma Membrane | other | 5.77E+08 | 5.71E+08 | 4.58E+08 | | | | | | |
| P15508 | SPTB | spectrin, beta, erythrocytic | Plasma Membrane | other | 5.85E+08 | 6.15E+08 | 7.00E+08 | | | | | | |
| P15508 | SPTB | spectrin, beta, erythrocytic | Plasma Membrane | other | 5.49E+08 | 5.42E+08 | 4.34E+08 | | | | | | |
| P15508 | SPTB | spectrin, beta, erythrocytic | Plasma Membrane | other | 4.84E+08 | 5.42E+08 | 4.46E+08 | | | | | | |
| P15508 | SPTB | spectrin, beta, erythrocytic | Plasma Membrane | other | 6.47E+08 | 9.39E+08 | 4.36E+08 | | | | | | |
| P15508 | SPTB | spectrin, beta, erythrocytic | Plasma Membrane | other | 6.85E+08 | 8.96E+08 | 8.20E+08 | | | | | | |
| P15508 | SPTB | spectrin, beta, erythrocytic | Plasma Membrane | other | 6.46E+08 | 6.73E+08 | 6.28E+08 | | | | | | |
| P15508 | SPTB | spectrin, beta, erythrocytic | Plasma Membrane | other | 5.98E+08 | 6.08E+08 | 5.56E+08 | | | | | | |
| P15508 | SPTB | spectrin, beta, erythrocytic | Plasma Membrane | other | 4.99E+08 | 5.09E+08 | 3.36E+08 | | | | | | |
| P15508 | SPTB | spectrin, beta, erythrocytic | Plasma Membrane | other | 4.31E+08 | 4.02E+08 | 3.45E+08 | | | | | | |
| P15508 | SPTB | spectrin, beta, erythrocytic | Plasma Membrane | other | 1.26E+09 | 8.50E+08 | 1.01E+09 | | | | | | |
| P15508 | SPTB | spectrin, beta, erythrocytic | Plasma Membrane | other | 4.83E+08 | 4.05E+08 | 4.28E+08 | | | | | | |
| P15508 | SPTB | spectrin, beta, erythrocytic | Plasma Membrane | other | 4.93E+08 | 4.53E+08 | 3.78E+08 | | | | | | |
| P15508 | SPTB | spectrin, beta, erythrocytic | Plasma Membrane | other | 4.03E+08 | 3.64E+08 | 3.43E+08 | | | | | | |
| P15508 | SPTB | spectrin, beta, erythrocytic | Plasma Membrane | other | 3.98E+08 | 2.88E+08 | 3.63E+08 | | | | | | |
| P15508 | SPTB | spectrin, beta, erythrocytic | Plasma Membrane | other | 5.59E+08 | 5.83E+08 | 5.37E+08 | | | | | | |
| P15508 | SPTB | spectrin, beta, erythrocytic | Plasma Membrane | other | 3.42E+08 | 3.90E+08 | 2.68E+08 | | | | | | |
| P15508 | SPTB | spectrin, beta, erythrocytic | Plasma Membrane | other | 3.52E+08 | 4.94E+08 | 4.54E+08 | | | | | | |

| | | | | | | | |
|---|---|---|---|---|---|---|---|
| P15508 | SPTB | spectrin, beta, erythrocytic | Plasma Membrane | other | 3.91E+08 | 5.55E+08 | 3.63E+08 |
| P15508 | SPTB | spectrin, beta, erythrocytic | Plasma Membrane | other | 4.27E+08 | 6.12E+08 | 5.17E+08 |
| P15508 | SPTB | spectrin, beta, erythrocytic | Plasma Membrane | other | 3.12E+08 | 3.72E+08 | 2.62E+08 |
| P15508 | SPTB | spectrin, beta, erythrocytic | Plasma Membrane | other | 3.00E+08 | 2.53E+08 | 2.45E+08 |
| P15508 | SPTB | spectrin, beta, erythrocytic | Plasma Membrane | other | 3.24E+08 | 2.92E+08 | 2.44E+08 |
| P15508 | SPTB | spectrin, beta, erythrocytic | Plasma Membrane | other | 3.29E+08 | 2.95E+08 | 2.52E+08 |
| P15508 | SPTB | spectrin, beta, erythrocytic | Plasma Membrane | other | 3.20E+08 | 9.57E+07 | 2.85E+08 |
| P15508 | SPTB | spectrin, beta, erythrocytic | Plasma Membrane | other | 2.70E+08 | 2.22E+08 | 1.89E+08 |
| P15508 | SPTB | spectrin, beta, erythrocytic | Plasma Membrane | other | 3.12E+08 | 3.30E+08 | 2.80E+08 |
| P15508 | SPTB | spectrin, beta, erythrocytic | Plasma Membrane | other | 2.62E+08 | 2.15E+08 | 2.46E+08 |
| P15508 | SPTB | spectrin, beta, erythrocytic | Plasma Membrane | other | 2.35E+08 | 1.34E+08 | 2.60E+08 |
| P15508 | SPTB | spectrin, beta, erythrocytic | Plasma Membrane | other | 1.42E+08 | 1.15E+08 | 8.81E+07 |
| P15508 | SPTB | spectrin, beta, erythrocytic | Plasma Membrane | other | 2.25E+08 | 2.08E+08 | 1.75E+08 |
| P15508 | SPTB | spectrin, beta, erythrocytic | Plasma Membrane | other | 1.92E+08 | 1.63E+08 | 1.56E+08 |
| P15508 | SPTB | spectrin, beta, erythrocytic | Plasma Membrane | other | 2.50E+08 | 2.63E+08 | 2.40E+08 |
| P15508 | SPTB | spectrin, beta, erythrocytic | Plasma Membrane | other | 2.54E+08 | 2.58E+08 | 2.70E+08 |
| P15508 | SPTB | spectrin, beta, erythrocytic | Plasma Membrane | other | 2.89E+08 | 2.34E+08 | 7.90E+07 |
| P15508 | SPTB | spectrin, beta, erythrocytic | Plasma Membrane | other | 4.31E+08 | 3.00E+08 | 2.09E+08 |
| P15508 | SPTB | spectrin, beta, erythrocytic | Plasma Membrane | other | 2.17E+08 | 1.67E+08 | 1.48E+08 |
| P15508 | SPTB | spectrin, beta, erythrocytic | Plasma Membrane | other | 4.90E+08 | 4.75E+08 | 2.88E+08 |
| P15508 | SPTB | spectrin, beta, erythrocytic | Plasma Membrane | other | 4.48E+08 | 3.91E+08 | 3.61E+08 |
| P15508 | SPTB | spectrin, beta, erythrocytic | Plasma Membrane | other | 1.92E+08 | 1.58E+08 | 1.60E+08 |
| P15508 | SPTB | spectrin, beta, erythrocytic | Plasma Membrane | other | 2.41E+08 | 2.01E+08 | 1.88E+08 |
| P15508 | SPTB | spectrin, beta, erythrocytic | Plasma Membrane | other | 2.50E+08 | 2.94E+08 | 2.77E+08 |
| P15508 | SPTB | spectrin, beta, erythrocytic | Plasma Membrane | other | 1.87E+08 | 1.71E+08 | 1.44E+08 |
| P15508 | SPTB | spectrin, beta, erythrocytic | Plasma Membrane | other | 1.92E+08 | 2.40E+08 | 1.89E+08 |
| P15508 | SPTB | spectrin, beta, erythrocytic | Plasma Membrane | other | 2.73E+08 | 6.33E+08 | 6.42E+08 |
| P15508 | SPTB | spectrin, beta, erythrocytic | Plasma Membrane | other | 2.32E+08 | 2.69E+08 | 1.83E+08 |
| P15508 | SPTB | spectrin, beta, erythrocytic | Plasma Membrane | other | 1.81E+08 | 3.61E+08 | 3.23E+08 |
| P15508 | SPTB | spectrin, beta, erythrocytic | Plasma Membrane | other | 1.54E+08 | 2.19E+08 | 1.14E+08 |
| P15508 | SPTB | spectrin, beta, erythrocytic | Plasma Membrane | other | 7.32E+07 | 2.15E+07 | 1.19E+08 |
| P15508 | SPTB | spectrin, beta, erythrocytic | Plasma Membrane | other | 7.38E+07 | 6.42E+07 | 6.80E+07 |
| P15508 | SPTB | spectrin, beta, erythrocytic | Plasma Membrane | other | 1.53E+08 | 1.97E+08 | 1.66E+08 |
| P15508 | SPTB | spectrin, beta, erythrocytic | Plasma Membrane | other | 1.71E+08 | 1.36E+08 | 1.23E+08 |
| P15508 | SPTB | spectrin, beta, erythrocytic | Plasma Membrane | other | 1.55E+08 | 1.00E+08 | 6.38E+07 |
| P15508 | SPTB | spectrin, beta, erythrocytic | Plasma Membrane | other | 1.40E+08 | 1.14E+08 | 7.88E+07 |
| P15508 | SPTB | spectrin, beta, erythrocytic | Plasma Membrane | other | 1.16E+08 | 8.38E+07 | 9.21E+07 |
| P15508 | SPTB | spectrin, beta, erythrocytic | Plasma Membrane | other | 1.45E+08 | 1.39E+08 | 3.42E+08 |
| P15508 | SPTB | spectrin, beta, erythrocytic | Plasma Membrane | other | 2.20E+08 | 2.02E+08 | 2.00E+08 |
| P15508 | SPTB | spectrin, beta, erythrocytic | Plasma Membrane | other | 3.79E+07 | 4.65E+07 | 4.35E+07 |
| P15508 | SPTB | spectrin, beta, erythrocytic | Plasma Membrane | other | 8.14E+07 | 6.88E+07 | 1.48E+08 |
| P15508 | SPTB | spectrin, beta, erythrocytic | Plasma Membrane | other | 1.33E+08 | 1.32E+08 | 3.30E+08 |
| P15508 | SPTB | spectrin, beta, erythrocytic | Plasma Membrane | other | 9.89E+07 | 1.44E+08 | 9.20E+07 |
| P15508 | SPTB | spectrin, beta, erythrocytic | Plasma Membrane | other | 8.53E+07 | 1.20E+08 | 8.47E+07 |
| P15508 | SPTB | spectrin, beta, erythrocytic | Plasma Membrane | other | 1.24E+08 | 1.09E+08 | 6.89E+07 |
| P15508 | SPTB | spectrin, beta, erythrocytic | Plasma Membrane | other | 7.80E+07 | 1.05E+08 | 7.44E+07 |
| P15508 | SPTB | spectrin, beta, erythrocytic | Plasma Membrane | other | 1.06E+08 | 9.83E+07 | 9.14E+07 |
| P15508 | SPTB | spectrin, beta, erythrocytic | Plasma Membrane | other | 9.62E+07 | 9.89E+07 | 7.93E+07 |
| P15508 | SPTB | spectrin, beta, erythrocytic | Plasma Membrane | other | 8.62E+07 | 8.66E+07 | 7.22E+07 |
| P15508 | SPTB | spectrin, beta, erythrocytic | Plasma Membrane | other | 1.31E+08 | 1.50E+08 | 6.02E+07 |
| P15508 | SPTB | spectrin, beta, erythrocytic | Plasma Membrane | other | 6.94E+07 | 8.27E+07 | 5.95E+07 |
| P15508 | SPTB | spectrin, beta, erythrocytic | Plasma Membrane | other | 1.01E+08 | 8.19E+07 | 6.03E+07 |
| P15508 | SPTB | spectrin, beta, erythrocytic | Plasma Membrane | other | 9.10E+07 | 6.86E+07 | 6.99E+07 |
| P15508 | SPTB | spectrin, beta, erythrocytic | Plasma Membrane | other | 1.09E+08 | 7.74E+07 | 5.76E+07 |
| P15508 | SPTB | spectrin, beta, erythrocytic | Plasma Membrane | other | 1.23E+08 | 1.40E+08 | 1.04E+08 |
| P15508 | SPTB | spectrin, beta, erythrocytic | Plasma Membrane | other | 8.53E+07 | 7.66E+07 | 7.10E+07 |
| P15508 | SPTB | spectrin, beta, erythrocytic | Plasma Membrane | other | 1.28E+08 | 2.70E+08 | 2.40E+08 |
| P15508 | SPTB | spectrin, beta, erythrocytic | Plasma Membrane | other | 8.40E+07 | 4.13E+07 | 6.43E+07 |
| P15508 | SPTB | spectrin, beta, erythrocytic | Plasma Membrane | other | 1.78E+08 | 1.56E+08 | 1.45E+08 |
| P15508 | SPTB | spectrin, beta, erythrocytic | Plasma Membrane | other | 1.42E+08 | 2.02E+08 | 2.06E+08 |
| P15508 | SPTB | spectrin, beta, erythrocytic | Plasma Membrane | other | 8.42E+07 | 8.80E+07 | 7.03E+07 |
| P15508 | SPTB | spectrin, beta, erythrocytic | Plasma Membrane | other | 8.18E+07 | 5.99E+07 | 2.98E+07 |
| P15508 | SPTB | spectrin, beta, erythrocytic | Plasma Membrane | other | 1.74E+08 | 3.45E+08 | 3.24E+08 |
| P15508 | SPTB | spectrin, beta, erythrocytic | Plasma Membrane | other | 1.07E+08 | 7.04E+07 | 6.95E+07 |
| P15508 | SPTB | spectrin, beta, erythrocytic | Plasma Membrane | other | 6.99E+07 | 4.97E+07 | 8.89E+07 |
| P15508 | SPTB | spectrin, beta, erythrocytic | Plasma Membrane | other | 4.61E+07 | 1.91E+07 | 6.68E+07 |
| P15508 | SPTB | spectrin, beta, erythrocytic | Plasma Membrane | other | 6.24E+07 | 3.96E+07 | 6.57E+07 |
| P15508 | SPTB | spectrin, beta, erythrocytic | Plasma Membrane | other | 5.33E+07 | 3.43E+07 | 3.34E+07 |
| P15508 | SPTB | spectrin, beta, erythrocytic | Plasma Membrane | other | 1.12E+07 | 1.31E+07 | 8.28E+07 |
| P15508 | SPTB | spectrin, beta, erythrocytic | Plasma Membrane | other | 7.20E+07 | 7.12E+07 | 5.63E+07 |
| P15508 | SPTB | spectrin, beta, erythrocytic | Plasma Membrane | other | 3.95E+07 | 3.96E+07 | 4.08E+07 |
| P15508 | SPTB | spectrin, beta, erythrocytic | Plasma Membrane | other | 7.36E+07 | 9.16E+07 | 4.10E+07 |
| P15508 | SPTB | spectrin, beta, erythrocytic | Plasma Membrane | other | 6.65E+07 | 4.93E+07 | 4.49E+07 |
| P15508 | SPTB | spectrin, beta, erythrocytic | Plasma Membrane | other | 4.61E+07 | 4.56E+07 | 4.97E+07 |
| P15508 | SPTB | spectrin, beta, erythrocytic | Plasma Membrane | other | 6.18E+07 | 8.23E+07 | 6.91E+07 |
| P15508 | SPTB | spectrin, beta, erythrocytic | Plasma Membrane | other | 4.65E+07 | 1.61E+07 | 4.53E+07 |
| P15508 | SPTB | spectrin, beta, erythrocytic | Plasma Membrane | other | 5.15E+07 | 1.19E+08 | 6.26E+07 |
| P15508 | SPTB | spectrin, beta, erythrocytic | Plasma Membrane | other | 3.20E+07 | 4.60E+07 | 3.68E+07 |
| P15508 | SPTB | spectrin, beta, erythrocytic | Plasma Membrane | other | 5.68E+07 | 6.79E+07 | 4.78E+07 |
| P15508 | SPTB | spectrin, beta, erythrocytic | Plasma Membrane | other | 4.48E+07 | 2.83E+07 | 2.44E+07 |
| P15508 | SPTB | spectrin, beta, erythrocytic | Plasma Membrane | other | 6.53E+07 | 1.19E+08 | 4.27E+07 |
| P15508 | SPTB | spectrin, beta, erythrocytic | Plasma Membrane | other | 5.11E+07 | 6.81E+07 | 5.37E+07 |
| P15508 | SPTB | spectrin, beta, erythrocytic | Plasma Membrane | other | 1.33E+08 | 1.47E+08 | 1.14E+08 |
| P15508 | SPTB | spectrin, beta, erythrocytic | Plasma Membrane | other | 5.88E+07 | 7.84E+07 | 4.47E+07 |
| P15508 | SPTB | spectrin, beta, erythrocytic | Plasma Membrane | other | 4.61E+07 | 7.41E+07 | 6.53E+07 |
| P15508 | SPTB | spectrin, beta, erythrocytic | Plasma Membrane | other | 3.93E+07 | 3.76E+07 | 3.81E+07 |
| P15508 | SPTB | spectrin, beta, erythrocytic | Plasma Membrane | other | 8.84E+07 | 5.53E+07 | 3.25E+07 |
| P15508 | SPTB | spectrin, beta, erythrocytic | Plasma Membrane | other | 4.81E+07 | 4.62E+07 | 4.73E+07 |
| P15508 | SPTB | spectrin, beta, erythrocytic | Plasma Membrane | other | 7.28E+07 | 8.59E+07 | 3.66E+07 |
| P15508 | SPTB | spectrin, beta, erythrocytic | Plasma Membrane | other | 2.67E+07 | 3.96E+07 | 2.69E+07 |
| P15508 | SPTB | spectrin, beta, erythrocytic | Plasma Membrane | other | 4.05E+07 | 4.97E+07 | 5.11E+07 |
| P15508 | SPTB | spectrin, beta, erythrocytic | Plasma Membrane | other | 2.45E+07 | 4.13E+07 | 2.79E+07 |
| P15508 | SPTB | spectrin, beta, erythrocytic | Plasma Membrane | other | 5.67E+07 | 1.10E+08 | 9.74E+07 |
| P15508 | SPTB | spectrin, beta, erythrocytic | Plasma Membrane | other | 2.74E+07 | 1.71E+07 | 2.24E+07 |
| P15508 | SPTB | spectrin, beta, erythrocytic | Plasma Membrane | other | 3.52E+07 | 3.46E+07 | 1.90E+07 |
| P15508 | SPTB | spectrin, beta, erythrocytic | Plasma Membrane | other | 2.22E+07 | 5619385 | 2.79E+07 |
| P15508 | SPTB | spectrin, beta, erythrocytic | Plasma Membrane | other | 2.46E+07 | 1.82E+07 | 1.48E+07 |
| P15508 | SPTB | spectrin, beta, erythrocytic | Plasma Membrane | other | 3.40E+07 | 3.81E+07 | 2.45E+07 |
| P15508 | SPTB | spectrin, beta, erythrocytic | Plasma Membrane | other | 1.84E+07 | 1.68E+07 | 1.42E+07 |
| P15508 | SPTB | spectrin, beta, erythrocytic | Plasma Membrane | other | 2.39E+07 | 3.99E+07 | 1.51E+07 |
| P15508 | SPTB | spectrin, beta, erythrocytic | Plasma Membrane | other | 4.61E+07 | 6.13E+07 | 5.07E+07 |
| P15508 | SPTB | spectrin, beta, erythrocytic | Plasma Membrane | other | 2.86E+07 | 3.09E+07 | 2.23E+07 |
| P15508 | SPTB | spectrin, beta, erythrocytic | Plasma Membrane | other | 2.22E+07 | 1.97E+07 | 1.66E+07 |
| P15508 | SPTB | spectrin, beta, erythrocytic | Plasma Membrane | other | 2.44E+07 | 1.20E+07 | 1.64E+07 |
| P15508 | SPTB | spectrin, beta, erythrocytic | Plasma Membrane | other | 2.55E+07 | 3.02E+07 | 2.20E+07 |
| P15508 | SPTB | spectrin, beta, erythrocytic | Plasma Membrane | other | 1.25E+07 | 5622477 | 2.75E+07 |
| P15508 | SPTB | spectrin, beta, erythrocytic | Plasma Membrane | other | 1.91E+07 | 2.82E+07 | 1.98E+07 |
| P15508 | SPTB | spectrin, beta, erythrocytic | Plasma Membrane | other | 3.00E+07 | 4.30E+07 | 2.20E+07 |
| P15508 | SPTB | spectrin, beta, erythrocytic | Plasma Membrane | other | 1.37E+07 | 3297365 | 2.09E+07 |

| | | | | | | | | | | | | | |
|---|---|---|---|---|---|---|---|---|---|---|---|---|---|
| P15508 | SPTB | spectrin, beta, erythrocytic | Plasma Membrane | other | 1.83E+07 | 6196952.5 | 2.08E+07 | | | | | | |
| P15508 | SPTB | spectrin, beta, erythrocytic | Plasma Membrane | other | 1.59E+07 | 1.73E+07 | 1.20E+07 | | | | | | |
| P15508 | SPTB | spectrin, beta, erythrocytic | Plasma Membrane | other | 1.02E+07 | 7659402 | 5588981.5 | | | | | | |
| P15508 | SPTB | spectrin, beta, erythrocytic | Plasma Membrane | other | 2.78E+07 | 1.92E+07 | 1.29E+07 | | | | | | |
| P15508 | SPTB | spectrin, beta, erythrocytic | Plasma Membrane | other | 5497894 | 1.20E+07 | 1.81E+07 | | | | | | |
| P15508 | SPTB | spectrin, beta, erythrocytic | Plasma Membrane | other | 1.64E+07 | 5.30E+07 | 5.92E+07 | | | | | | |
| P15508 | SPTB | spectrin, beta, erythrocytic | Plasma Membrane | other | 1964314.25 | 1405911.875 | 1.31E+07 | | | | | | |
| P15508 | SPTB | spectrin, beta, erythrocytic | Plasma Membrane | other | 9961615 | 5776195 | 2846076.25 | | | | | | |
| P15508 | SPTB | spectrin, beta, erythrocytic | Plasma Membrane | other | 4.16E+07 | 7.71E+07 | 5.31E+07 | | | | | | |
| P15508 | SPTB | spectrin, beta, erythrocytic | Plasma Membrane | other | 1.18E+07 | 6421159 | 2822790.25 | | | | | | |
| P15508 | SPTB | spectrin, beta, erythrocytic | Plasma Membrane | other | 1.22E+07 | 8697191 | 1.14E+07 | | | | | | |
| P15508 | SPTB | spectrin, beta, erythrocytic | Plasma Membrane | other | 1.27E+07 | 1.03E+07 | 9684792 | | | | | | |
| P15508 | SPTB | spectrin, beta, erythrocytic | Plasma Membrane | other | 1.64E+07 | 1.97E+07 | 2.09E+07 | | | | | | |
| P15508 | SPTB | spectrin, beta, erythrocytic | Plasma Membrane | other | 9344198 | 1.14E+07 | 9989608 | | | | | | |
| P15508 | SPTB | spectrin, beta, erythrocytic | Plasma Membrane | other | 1.62E+07 | 1.57E+07 | 1.35E+07 | | | | | | |
| P15508 | SPTB | spectrin, beta, erythrocytic | Plasma Membrane | other | 7386500 | 7208204.5 | 3386820.75 | | | | | | |
| P15508 | SPTB | spectrin, beta, erythrocytic | Plasma Membrane | other | 2345640.5 | 1968274.5 | 6140623.5 | | | | | | |
| P15508 | SPTB | spectrin, beta, erythrocytic | Plasma Membrane | other | 1.15E+07 | 1.27E+07 | 1.12E+07 | | | | | | |
| P15508 | SPTB | spectrin, beta, erythrocytic | Plasma Membrane | other | 7529926 | 5431830 | 2962041.5 | | | | | | |
| P15508 | SPTB | spectrin, beta, erythrocytic | Plasma Membrane | other | 4427517.5 | 594594.188 | 4907355 | | | | | | |
| P15508 | SPTB | spectrin, beta, erythrocytic | Plasma Membrane | other | 3.24E+07 | 1.48E+07 | 1.86E+07 | | | | | | |
| P15508 | SPTB | spectrin, beta, erythrocytic | Plasma Membrane | other | 5055329.5 | 2552329.5 | 4455973.5 | | | | | | |
| P15508 | SPTB | spectrin, beta, erythrocytic | Plasma Membrane | other | 7001359 | 1.19E+07 | 8741244 | | | | | | |
| P15508 | SPTB | spectrin, beta, erythrocytic | Plasma Membrane | other | 3153495.25 | 1335503.75 | 7103316 | | | | | | |
| P15508 | SPTB | spectrin, beta, erythrocytic | Plasma Membrane | other | 9003540 | 1.11E+07 | 5317151 | | | | | | |
| P15508 | SPTB | spectrin, beta, erythrocytic | Plasma Membrane | other | 6551675.5 | 8946985 | 6813514 | | | | | | |
| P15508 | SPTB | spectrin, beta, erythrocytic | Plasma Membrane | other | 5136231 | 4815674 | 1.11E+07 | | | | | | |
| P15508 | SPTB | spectrin, beta, erythrocytic | Plasma Membrane | other | 1259331.125 | 993466.375 | 2368573.5 | | | | | | |
| P15508 | SPTB | spectrin, beta, erythrocytic | Plasma Membrane | other | 3173061.25 | 3363586.25 | 3410254 | | | | | | |
| P15508 | SPTB | spectrin, beta, erythrocytic | Plasma Membrane | other | 1.00E+07 | 1.18E+07 | 4250555.5 | | | | | | |
| P15508 | SPTB | spectrin, beta, erythrocytic | Plasma Membrane | other | 3217448.75 | 2645117 | 3033053.25 | | | | | | |
| P15508 | SPTB | spectrin, beta, erythrocytic | Plasma Membrane | other | 3788337.5 | 3146222.75 | 4164094.75 | | | | | | |
| P15508 | SPTB | spectrin, beta, erythrocytic | Plasma Membrane | other | 2262225.5 | 1110010.25 | 1401355.625 | | | | | | |
| P15508 | SPTB | spectrin, beta, erythrocytic | Plasma Membrane | other | 3173264.25 | 5849022.5 | 3746181.75 | | | | | | |
| P15508 | SPTB | spectrin, beta, erythrocytic | Plasma Membrane | other | 4726410.5 | 2519083 | 1794042.125 | | | | | | |
| P15508 | SPTB | spectrin, beta, erythrocytic | Plasma Membrane | other | 2834080.25 | 3933975 | 6088971 | | | | | | |
| P15508 | SPTB | spectrin, beta, erythrocytic | Plasma Membrane | other | 2402089 | 955242.438 | 1752069.375 | | | | | | |
| P15508 | SPTB | spectrin, beta, erythrocytic | Plasma Membrane | other | 633407.312 | 867789.812 | 1265131.875 | | | | | | |
| P15508 | SPTB | spectrin, beta, erythrocytic | Plasma Membrane | other | 2835452.75 | 3444449.75 | 1478188.75 | | | | | | |
| P15508 | SPTB | spectrin, beta, erythrocytic | Plasma Membrane | other | 2.11E+07 | 1.40E+08 | 1.57E+08 | | | | | | |
| P15508 | SPTB | spectrin, beta, erythrocytic | Plasma Membrane | other | 606027.25 | 1574442.125 | 1570391.25 | | | | | | |
| P15508 | SPTB | spectrin, beta, erythrocytic | Plasma Membrane | other | 333302.438 | 63424.215 | 163932.797 | | | | | | |
| P15508 | SPTB | spectrin, beta, erythrocytic | Plasma Membrane | other | 1346966.125 | 2104465.5 | 1468693.625 | | | | | | |
| P15508 | SPTB | spectrin, beta, erythrocytic | Plasma Membrane | other | 971350.25 | 8673.901 | 538541.938 | | | | | | |
| P15508 | SPTB | spectrin, beta, erythrocytic | Plasma Membrane | other | 7334.904 | 6812.207 | 132845.516 | | | | | | |
| P15508 | SPTB | spectrin, beta, erythrocytic | Plasma Membrane | other | 939989.125 | 5976922 | 7121652 | | | | | | |
| P15508 | SPTB | spectrin, beta, erythrocytic | Plasma Membrane | other | 82473.883 | 4.70E-06 | 6162.183 | | | | | | |
| P15508 | SPTB | spectrin, beta, erythrocytic | Plasma Membrane | other | 423971.062 | 3016821 | 4266314 | | | | | | |
| P15508 | SPTB | spectrin, beta, erythrocytic | Plasma Membrane | other | 26121.842 | 282822.406 | 297583.75 | | | | | | |
| P15508 | SPTB | spectrin, beta, erythrocytic | Plasma Membrane | other | 324186.094 | 4327078 | 6092641.5 | P15508 | SPTB | 484870062 | 582379199 | 20852520 | 780 |
| P13609 | Srgn | serglycin | Cytoplasm | other | 1006191.062 | 2091713 | 3166505.75 | P13609 | Srgn | 1006191 | 1080162 | 623632 | 3 |
| Q9CY50 | SSR1 | signal sequence receptor, alpha | Cytoplasm | other | 2922875.75 | 3896065.25 | 5536515 | | | | | | |
| Q9CY50 | SSR1 | signal sequence receptor, alpha | Cytoplasm | other | 1766788.875 | 3236671.5 | 3703890.25 | Q9CY50 | SSR1 | 2344832 | 1245119 | 508318 | 6 |
| Q9DCF9 | SSR3 | signal sequence receptor, gamma (translocon-associated protein gamma) | Cytoplasm | other | 4623402.5 | 5504138 | 8582800 | Q9DCF9 | SSR3 | 4623403 | 2078889 | 1200247 | 3 |
| O70576 | STAG3 | stromal antigen 3 | Nucleus | other | 4913889.5 | 5043885 | 4238320 | O70576 | STAG3 | 4913890 | 432479 | 249692 | 3 |
| P70297 | STAM | signal transducing adaptor molecule (SH3 domain and ITAM motif) 1 | Cytoplasm | other | 7859684.5 | 5380812 | 4015757 | | | | | | |
| P70297 | STAM | signal transducing adaptor molecule (SH3 domain and ITAM motif) 1 | Cytoplasm | other | 3360715.75 | 2467708 | 2465979.25 | | | | | | |
| P70297 | STAM | signal transducing adaptor molecule (SH3 domain and ITAM motif) 1 | Cytoplasm | other | 2064611.625 | 1762013.125 | 1247603.25 | P70297 | STAM | 4428337 | 2092656 | 697552 | 9 |
| Q8CI59 | STEAP3 | STEAP family member 3, metalloreductase | Cytoplasm | transporter | 8444200 | 1.51E+07 | 1.54E+07 | | | | | | |
| Q8CI59 | STEAP3 | STEAP family member 3, metalloreductase | Cytoplasm | transporter | 4296827.5 | 6905446 | 9113789 | | | | | | |
| Q8CI59 | STEAP3 | STEAP family member 3, metalloreductase | Cytoplasm | transporter | 4930037.5 | 6732764.5 | 1.05E+07 | | | | | | |
| Q8CI59 | STEAP3 | STEAP family member 3, metalloreductase | Cytoplasm | transporter | 2503758 | 4938499 | 3471307 | | | | | | |
| Q8CI59 | STEAP3 | STEAP family member 3, metalloreductase | Cytoplasm | transporter | 2738965.25 | 3160573 | 4076435.75 | Q8CI59 | STEAP3 | 4582758 | 4174608 | 1077879 | 15 |
| P70302 | STIM1 | stromal interaction molecule 1 | Plasma Membrane | ion channel | 2183463 | 2501111.75 | 3357344 | | | | | | |
| P70302 | STIM1 | stromal interaction molecule 1 | Plasma Membrane | ion channel | 2145278.75 | 2512071.25 | 2993917 | | | | | | |
| P70302 | STIM1 | stromal interaction molecule 1 | Plasma Membrane | ion channel | 944601.812 | 1790225.5 | 1749574.125 | P70302 | STIM1 | 1757781 | 715197 | 238399 | 9 |
| Q60864 | STIP1 | stress-induced phosphoprotein 1 | Cytoplasm | other | 7447384 | 5717125 | 4066567.5 | | | | | | |
| Q60864 | STIP1 | stress-induced phosphoprotein 1 | Cytoplasm | other | 7572962.5 | 7485109 | 4605209.5 | | | | | | |
| Q60864 | STIP1 | stress-induced phosphoprotein 1 | Cytoplasm | other | 6710373 | 5180680.5 | 2977989.25 | | | | | | |
| Q60864 | STIP1 | stress-induced phosphoprotein 1 | Cytoplasm | other | 4806757.5 | 3779088.75 | 2361706.5 | | | | | | |
| Q60864 | STIP1 | stress-induced phosphoprotein 1 | Cytoplasm | other | 3998224.25 | 4242458 | 3991340.5 | | | | | | |
| Q60864 | STIP1 | stress-induced phosphoprotein 1 | Cytoplasm | other | 2795439.5 | 2117941.75 | 1708346.75 | | | | | | |
| Q60864 | STIP1 | stress-induced phosphoprotein 1 | Cytoplasm | other | 2473861 | 1976094.125 | 1318160.125 | | | | | | |
| Q60864 | STIP1 | stress-induced phosphoprotein 1 | Cytoplasm | other | 2902761.75 | 2246887.75 | 923053 | | | | | | |
| Q60864 | STIP1 | stress-induced phosphoprotein 1 | Cytoplasm | other | 2283857.5 | 1817383 | 1253543.75 | | | | | | |
| Q60864 | STIP1 | stress-induced phosphoprotein 1 | Cytoplasm | other | 3372617.5 | 2172595.5 | 1991129.375 | Q60864 | STIP1 | 4436424 | 1927222 | 351861 | 30 |
| P54116 | STOM | stomatin | Plasma Membrane | other | 5.91E+08 | 5.65E+08 | 4.57E+08 | | | | | | |
| P54116 | STOM | stomatin | Plasma Membrane | other | 6.04E+08 | 5.95E+08 | 5.59E+08 | | | | | | |
| P54116 | STOM | stomatin | Plasma Membrane | other | 5.57E+08 | 5.33E+08 | 4.99E+08 | | | | | | |
| P54116 | STOM | stomatin | Plasma Membrane | other | 3.89E+08 | 4.49E+08 | 3.79E+08 | | | | | | |
| P54116 | STOM | stomatin | Plasma Membrane | other | 3.71E+08 | 3.76E+08 | 3.15E+08 | | | | | | |
| P54116 | STOM | stomatin | Plasma Membrane | other | 3.02E+08 | 3.31E+08 | 3.23E+08 | | | | | | |
| P54116 | STOM | stomatin | Plasma Membrane | other | 4.02E+08 | 5.02E+08 | 3.56E+08 | | | | | | |
| P54116 | STOM | stomatin | Plasma Membrane | other | 2.24E+08 | 1.98E+08 | 1.71E+08 | | | | | | |
| P54116 | STOM | stomatin | Plasma Membrane | other | 1.90E+08 | 1.83E+08 | 1.89E+08 | | | | | | |
| P54116 | STOM | stomatin | Plasma Membrane | other | 2.68E+08 | 4.54E+08 | 4.46E+08 | | | | | | |
| P54116 | STOM | stomatin | Plasma Membrane | other | 3.01E+08 | 1.09E+09 | 1.42E+09 | | | | | | |
| P54116 | STOM | stomatin | Plasma Membrane | other | 1.06E+08 | 1.26E+08 | 9.01E+07 | | | | | | |
| P54116 | STOM | stomatin | Plasma Membrane | other | 2.47E+07 | 1.82E+07 | 1.55E+07 | | | | | | |
| P54116 | STOM | stomatin | Plasma Membrane | other | 2.77E+07 | 2.50E+07 | 1.93E+07 | | | | | | |
| P54116 | STOM | stomatin | Plasma Membrane | other | 8101764 | 7703873 | 7725000 | | | | | | |
| P54116 | STOM | stomatin | Plasma Membrane | other | 4190473 | 6935558.5 | 4109787 | | | | | | |
| P54116 | STOM | stomatin | Plasma Membrane | other | 4840129 | 4603456.5 | 1462023.25 | | | | | | |
| P54116 | STOM | stomatin | Plasma Membrane | other | 2388158 | 2945459.25 | 1120667.125 | | | | | | |
| P54116 | STOM | stomatin | Plasma Membrane | other | 1538073.75 | 359134.969 | 137934.953 | | | | | | |
| P54116 | STOM | stomatin | Plasma Membrane | other | 2229450.5 | 2931809 | 1655192 | | | | | | |
| P54116 | STOM | stomatin | Plasma Membrane | other | 6387355.5 | 7761338.5 | 9611.989 | | | | | | |
| P54116 | STOM | stomatin | Plasma Membrane | other | 456102.438 | 454188.594 | 395956 | | | | | | |
| P54116 | STOM | stomatin | Plasma Membrane | other | 476630.875 | 508202.812 | 727410.75 | P54116 | STOM | 190815136 | 276173897 | 33247430 | 69 |
| Q99JB2 | STOML2 | stomatin (EPB72)-like 2 | Plasma Membrane | other | 2827166.25 | 6060187.5 | 7577465.5 | | | | | | |
| Q99JB2 | STOML2 | stomatin (EPB72)-like 2 | Plasma Membrane | other | 1957293 | 3566845.75 | 5013215 | | | | | | |
| Q99JB2 | STOML2 | stomatin (EPB72)-like 2 | Plasma Membrane | other | 2109524.75 | 4707825.5 | 6214577 | | | | | | |
| Q99JB2 | STOML2 | stomatin (EPB72)-like 2 | Plasma Membrane | other | 1386947.75 | 2237082.5 | 3021695 | | | | | | |
| Q99JB2 | STOML2 | stomatin (EPB72)-like 2 | Plasma Membrane | other | 822916.625 | 1919082.625 | 1889773.375 | Q99JB2 | STOML2 | 1820770 | 2026994 | 523368 | 15 |
| Q9D3G5 | STX11 | syntaxin 11 | Plasma Membrane | transporter | 2294759.25 | 3825311.5 | 5585377.5 | | | | | | |
| Q9D3G5 | STX11 | syntaxin 11 | Plasma Membrane | transporter | 1646979.125 | 2289201.75 | 3591064.25 | | | | | | |
| Q9D3G5 | STX11 | syntaxin 11 | Plasma Membrane | transporter | 1188235.75 | 492590.625 | 2053154.25 | | | | | | |
| Q9D3G5 | STX11 | syntaxin 11 | Plasma Membrane | transporter | 1370275.375 | 1807923 | 2642401 | | | | | | |
| Q9D3G5 | STX11 | syntaxin 11 | Plasma Membrane | transporter | 1789271.875 | 3317748.5 | 4440852 | | | | | | |

| | | | | | | | | | | | | | |
|---|---|---|---|---|---|---|---|---|---|---|---|---|---|
| Q9D3G5 | STX11 | syntaxin 11 | Plasma Membrane | transporter | 771342 | 1228653.5 | 1372231 | Q9D3G5 | STX11 | 1510144 | 1355189 | 319421 | 18 |
| Q9ER00 | STX12 | syntaxin 12 | Cytoplasm | other | 1.31E+07 | 1.68E+07 | 1.33E+07 | Q9ER00 | STX12 | 13070000 | 2104432 | 1214994 | 3 |
| Q00262 | STX2 | syntaxin 2 | Cytoplasm | transporter | 4625285 | 5413520 | 3492518.25 | | | | | | |
| Q00262 | STX2 | syntaxin 2 | Cytoplasm | transporter | 2224433.75 | 1687607.125 | 4579346 | | | | | | |
| Q00262 | STX2 | syntaxin 2 | Cytoplasm | transporter | 4240891.5 | 6889536.5 | 4812398.5 | | | | | | |
| Q00262 | STX2 | syntaxin 2 | Cytoplasm | transporter | 2941647 | 3573061 | 2932363.25 | | | | | | |
| Q00262 | STX2 | syntaxin 2 | Cytoplasm | transporter | 1312401.5 | 1562123.875 | 1168910.625 | | | | | | |
| Q00262 | STX2 | syntaxin 2 | Cytoplasm | transporter | 1276241.875 | 1320579.75 | 1168073.625 | Q00262 | STX2 | 2770150 | 1734599 | 408849 | 18 |
| P70452 | STX4 | syntaxin 4 | Plasma Membrane | transporter | 3240535.75 | 4071618 | 4027133 | | | | | | |
| P70452 | STX4 | syntaxin 4 | Plasma Membrane | transporter | 1686484.125 | 2126279 | 1902390.875 | | | | | | |
| P70452 | STX4 | syntaxin 4 | Plasma Membrane | transporter | 1786522.375 | 2445804 | 2463637 | P70452 | STX4 | 2237847 | 925251 | 308417 | 9 |
| O70439 | STX7 | syntaxin 7 | Plasma Membrane | transporter | 1.42E+07 | 1.53E+07 | 1.44E+07 | | | | | | |
| O70439 | STX7 | syntaxin 7 | Plasma Membrane | transporter | 1.69E+07 | 1.62E+07 | 1.50E+07 | | | | | | |
| O70439 | STX7 | syntaxin 7 | Plasma Membrane | transporter | 9851564 | 1.22E+07 | 9493851 | | | | | | |
| O70439 | STX7 | syntaxin 7 | Plasma Membrane | transporter | 8441063 | 8464295 | 7889702 | | | | | | |
| O70439 | STX7 | syntaxin 7 | Plasma Membrane | transporter | 8535379 | 1.16E+07 | 9125586 | | | | | | |
| O70439 | STX7 | syntaxin 7 | Plasma Membrane | transporter | 5196943.5 | 4999389.5 | 4754532.5 | O70439 | STX7 | 10527492 | 3935344 | 927569 | 18 |
| O08599 | STXBP1 | syntaxin binding protein 1 | Cytoplasm | transporter | 3302494.75 | 3345785.25 | 2452860.75 | | | | | | |
| O08599 | STXBP1 | syntaxin binding protein 1 | Cytoplasm | transporter | 1079774 | 1077888.875 | 834551.312 | O08599 | STXBP1 | 2191134 | 1163326 | 474926 | 6 |
| Q64324 | STXBP2 | syntaxin binding protein 2 | Plasma Membrane | transporter | 1.43E+07 | 1.84E+07 | 2.21E+07 | | | | | | |
| Q64324 | STXBP2 | syntaxin binding protein 2 | Plasma Membrane | transporter | 6460101 | 8407372 | 9281399 | | | | | | |
| Q64324 | STXBP2 | syntaxin binding protein 2 | Plasma Membrane | transporter | 6874008 | 8715244 | 8600181 | | | | | | |
| Q64324 | STXBP2 | syntaxin binding protein 2 | Plasma Membrane | transporter | 5795318.5 | 9543600 | 8605881 | | | | | | |
| Q64324 | STXBP2 | syntaxin binding protein 2 | Plasma Membrane | transporter | 4368695 | 5647558 | 6805519.5 | | | | | | |
| Q64324 | STXBP2 | syntaxin binding protein 2 | Plasma Membrane | transporter | 4242097.5 | 5988136.5 | 7282689 | | | | | | |
| Q64324 | STXBP2 | syntaxin binding protein 2 | Plasma Membrane | transporter | 3106393 | 3617282 | 5529888.5 | | | | | | |
| Q64324 | STXBP2 | syntaxin binding protein 2 | Plasma Membrane | transporter | 3196304.75 | 4810201.5 | 6079152 | | | | | | |
| Q64324 | STXBP2 | syntaxin binding protein 2 | Plasma Membrane | transporter | 2250561 | 3743477.75 | 3427216.75 | | | | | | |
| Q64324 | STXBP2 | syntaxin binding protein 2 | Plasma Membrane | transporter | 2228420.25 | 3325316.75 | 3283483.25 | | | | | | |
| Q64324 | STXBP2 | syntaxin binding protein 2 | Plasma Membrane | transporter | 2940470.75 | 3757302.25 | 3728309 | | | | | | |
| Q64324 | STXBP2 | syntaxin binding protein 2 | Plasma Membrane | transporter | 2295843 | 3339486.5 | 3835835.25 | | | | | | |
| Q64324 | STXBP2 | syntaxin binding protein 2 | Plasma Membrane | transporter | 1396184.5 | 1928634.625 | 1410155.125 | | | | | | |
| Q64324 | STXBP2 | syntaxin binding protein 2 | Plasma Membrane | transporter | 1556822.125 | 1598810.75 | 2134714.5 | Q64324 | STXBP2 | 4358659 | 4338885 | 669504 | 42 |
| Q60770 | STXBP3 | syntaxin binding protein 3 | Plasma Membrane | transporter | 797343.438 | 1200909.125 | 1917613.5 | Q60770 | STXBP3 | 797343 | 567382 | 327578 | 3 |
| Q8K400 | STXBP5 | syntaxin binding protein 5 (tomosyn) | Plasma Membrane | other | 3.26E+07 | 2.42E+07 | 2.66E+07 | Q8K400 | STXBP5 | 32570000 | 4297546 | 2481189 | 3 |
| Q9Z2I9 | SUCLA2 | succinate-CoA ligase, ADP-forming, beta subunit | Cytoplasm | enzyme | 4041090.25 | 6721022.5 | 8939021 | Q9Z2I9 | SUCLA2 | 4041090 | 2452593 | 1416005 | 3 |
| Q9WUM5 | SUCLG1 | succinate-CoA ligase, alpha subunit | Cytoplasm | enzyme | 2640976 | 4909006 | 7553177.5 | | | | | | |
| Q9WUM5 | SUCLG1 | succinate-CoA ligase, alpha subunit | Cytoplasm | enzyme | 2487658 | 2749365.25 | 4579180 | | | | | | |
| Q9WUM5 | SUCLG1 | succinate-CoA ligase, alpha subunit | Cytoplasm | enzyme | 1279427.25 | 2543005 | 2804252.75 | Q9WUM5 | SUCLG1 | 2136020 | 1878663 | 626221 | 9 |
| Q9Z2I8 | SUCLG2 | succinate-CoA ligase, GDP-forming, beta subunit | Cytoplasm | enzyme | 1061246 | 877072.438 | 1179061.5 | Q9Z2I8 | SUCLG2 | 1061246 | 152205 | 87875 | 3 |
| Q64310 | SURF4 | surfeit 4 | Cytoplasm | other | 3598324.75 | 5995251.5 | 6391897.5 | | | | | | |
| Q64310 | SURF4 | surfeit 4 | Cytoplasm | other | 2521195.5 | 3063930.75 | 4008772 | Q64310 | SURF4 | 3059760 | 1581833 | 645781 | 6 |
| Q9DBX3 | SUSD2 | sushi domain containing 2 | Extracellular Space | other | 4242496.5 | 4108432.25 | 4674331 | | | | | | |
| Q9DBX3 | SUSD2 | sushi domain containing 2 | Extracellular Space | other | 2559982.5 | 4143088.75 | 4202558.5 | | | | | | |
| Q9DBX3 | SUSD2 | sushi domain containing 2 | Extracellular Space | other | 3327835 | 4709582.5 | 4136652.5 | Q9DBX3 | SUSD2 | 3376771 | 673907 | 224636 | 9 |
| Q3UZP4 | SVIP | small VCP/p97-interacting protein | Cytoplasm | other | 2.35E+07 | 2.37E+07 | 1.83E+07 | Q3UZP4 | SVIP | 23520000 | 3055820 | 1764278 | 3 |
| O09117 | SYPL1 | synaptophysin-like 1 | Plasma Membrane | transporter | 2.61E+07 | 2.60E+07 | 2.08E+07 | | | | | | |
| O09117 | SYPL1 | synaptophysin-like 1 | Plasma Membrane | transporter | 1.53E+07 | 1.86E+07 | 1.69E+07 | | | | | | |
| O09117 | SYPL1 | synaptophysin-like 1 | Plasma Membrane | transporter | 1124970.875 | 972802.438 | 824325.938 | O09117 | SYPL1 | 14201657 | 10473286 | 3491095 | 9 |
| B2RWW0 | TAGAP | T-cell activation RhoGTPase activating protein | Cytoplasm | other | 9717897 | 9872393 | 8041736.5 | B2RWW0 | TAGAP | 9717897 | 1015274 | 586169 | 3 |
| Q9WVA4 | TAGLN2 | transgelin 2 | Cytoplasm | other | 1957501.375 | 1762306.375 | 5033165.5 | | | | | | |
| Q9WVA4 | TAGLN2 | transgelin 2 | Cytoplasm | other | 359881.219 | 1038501.438 | 3400729.5 | | | | | | |
| Q9WVA4 | TAGLN2 | transgelin 2 | Cytoplasm | other | 983656.875 | 610683.125 | 2774738 | | | | | | |
| Q9WVA4 | TAGLN2 | transgelin 2 | Cytoplasm | other | 1078310.625 | 1133811 | 2300015.75 | Q9WVA4 | TAGLN2 | 1094838 | 1344468 | 388115 | 12 |
| Q8BYC6 | TAOK3 | TAO kinase 3 | Cytoplasm | kinase | 6789817.5 | 5941886 | 5733049 | | | | | | |
| Q8BYC6 | TAOK3 | TAO kinase 3 | Cytoplasm | kinase | 4107610.75 | 4033189.5 | 4141203.25 | | | | | | |
| Q8BYC6 | TAOK3 | TAO kinase 3 | Cytoplasm | kinase | 4349261.5 | 3208732 | 7483188.5 | | | | | | |
| Q8BYC6 | TAOK3 | TAO kinase 3 | Cytoplasm | kinase | 3702038 | 3311143.75 | 3341154 | | | | | | |
| Q8BYC6 | TAOK3 | TAO kinase 3 | Cytoplasm | kinase | 2288727.5 | 2018705.25 | 3118397.75 | | | | | | |
| Q8BYC6 | TAOK3 | TAO kinase 3 | Cytoplasm | kinase | 2679665.25 | 2543940 | 2672331.25 | | | | | | |
| Q8BYC6 | TAOK3 | TAO kinase 3 | Cytoplasm | kinase | 2727316.25 | 2201322 | 1560063 | | | | | | |
| Q8BYC6 | TAOK3 | TAO kinase 3 | Cytoplasm | kinase | 735018.625 | 552761.562 | 706386.062 | Q8BYC6 | TAOK3 | 3422432 | 1808045 | 369066 | 24 |
| Q9R233 | TAPBP | TAP binding protein (tapasin) | Cytoplasm | transporter | 864558.688 | 1165338.125 | 1843848.875 | Q9R233 | TAPBP | 864559 | 501640 | 289622 | 3 |
| Q4VBD2 | TAPT1 | transmembrane anterior posterior transformation 1 | Plasma Membrane | G-protein coupled receptor | 1186310.375 | 1774884.5 | 1531768 | Q4VBD2 | TAPT1 | 1186310 | 295766 | 170761 | 3 |
| Q9D0R2 | TARS | threonyl-tRNA synthetase | Nucleus | enzyme | 5819442.5 | 7434761.5 | 9560558 | | | | | | |
| Q9D0R2 | TARS | threonyl-tRNA synthetase | Nucleus | enzyme | 1990096 | 1669053 | 2240291.75 | Q9D0R2 | TARS | 3904769 | 3313410 | 1352694 | 6 |
| Q8BHL3 | TBC1D10B | TBC1 domain family, member 10B | Plasma Membrane | enzyme | 2373707.75 | 1736768.875 | 1687585.75 | Q8BHL3 | TBC1D10B | 2373708 | 382726 | 220967 | 3 |
| Q3UUG6 | TBC1D24 | TBC1 domain family, member 24 | Cytoplasm | other | 6063588.5 | 9152950 | 9660915 | | | | | | |
| Q3UUG6 | TBC1D24 | TBC1 domain family, member 24 | Cytoplasm | other | 2080868.875 | 3460059.5 | 3788158.5 | Q3UUG6 | TBC1D24 | 4072229 | 3146887 | 1284711 | 6 |
| P36423 | TBXAS1 | thromboxane A synthase 1 (platelet) | Plasma Membrane | enzyme | 5341438.5 | 6895827 | 1.10E+07 | | | | | | |
| P36423 | TBXAS1 | thromboxane A synthase 1 (platelet) | Plasma Membrane | enzyme | 9403536 | 1.22E+07 | 2.04E+07 | | | | | | |
| P36423 | TBXAS1 | thromboxane A synthase 1 (platelet) | Plasma Membrane | enzyme | 3848022.5 | 4341442 | 7865149 | | | | | | |
| P36423 | TBXAS1 | thromboxane A synthase 1 (platelet) | Plasma Membrane | enzyme | 4316093 | 4195421 | 6125786 | | | | | | |
| P36423 | TBXAS1 | thromboxane A synthase 1 (platelet) | Plasma Membrane | enzyme | 4356180 | 4655964 | 3929155.25 | | | | | | |
| P36423 | TBXAS1 | thromboxane A synthase 1 (platelet) | Plasma Membrane | enzyme | 5181869.5 | 7060060.5 | 8458201 | | | | | | |
| P36423 | TBXAS1 | thromboxane A synthase 1 (platelet) | Plasma Membrane | enzyme | 1.14E+07 | 1.23E+07 | 1.41E+07 | | | | | | |
| P36423 | TBXAS1 | thromboxane A synthase 1 (platelet) | Plasma Membrane | enzyme | 2815848 | 3654430.25 | 4060684 | | | | | | |
| P36423 | TBXAS1 | thromboxane A synthase 1 (platelet) | Plasma Membrane | enzyme | 4762313.5 | 5733043.5 | 7973789 | P36423 | TBXAS1 | 5711700 | 4080231 | 785241 | 27 |
| P10711 | TCEA1 | transcription elongation factor A (SII), 1 | Nucleus | transcription regulator | 2.18E+07 | 2.05E+07 | 1.54E+07 | | | | | | |
| P10711 | TCEA1 | transcription elongation factor A (SII), 1 | Nucleus | transcription regulator | 2.51E+07 | 2.48E+07 | 1.98E+07 | | | | | | |
| P10711 | TCEA1 | transcription elongation factor A (SII), 1 | Nucleus | transcription regulator | 2.03E+07 | 1.50E+07 | 1.03E+07 | | | | | | |
| P10711 | TCEA1 | transcription elongation factor A (SII), 1 | Nucleus | transcription regulator | 1.31E+07 | 1.09E+07 | 1.11E+07 | | | | | | |
| P10711 | TCEA1 | transcription elongation factor A (SII), 1 | Nucleus | transcription regulator | 1.20E+07 | 1.07E+07 | 8389952 | | | | | | |
| P10711 | TCEA1 | transcription elongation factor A (SII), 1 | Nucleus | transcription regulator | 1.13E+07 | 1.15E+07 | 8621774 | | | | | | |
| P10711 | TCEA1 | transcription elongation factor A (SII), 1 | Nucleus | transcription regulator | 1.21E+07 | 9371193 | 4522906 | | | | | | |
| P10711 | TCEA1 | transcription elongation factor A (SII), 1 | Nucleus | transcription regulator | 7898664.5 | 5615863 | 5149766.5 | | | | | | |
| P10711 | TCEA1 | transcription elongation factor A (SII), 1 | Nucleus | transcription regulator | 1.20E+07 | 9452816 | 8008588.5 | | | | | | |
| P10711 | TCEA1 | transcription elongation factor A (SII), 1 | Nucleus | transcription regulator | 7794650 | 5810063 | 7028576 | | | | | | |
| P10711 | TCEA1 | transcription elongation factor A (SII), 1 | Nucleus | transcription regulator | 3291110.25 | 2592762.75 | 2397936.25 | | | | | | |
| P10711 | TCEA1 | transcription elongation factor A (SII), 1 | Nucleus | transcription regulator | 1632511.75 | 1407894.625 | 1010549.625 | P10711 | TCEA1 | 12359745 | 6486516 | 1081086 | 36 |
| Q9QVN7 | TCEA2 | transcription elongation factor A (SII), 2 | Nucleus | transcription regulator | 3648467 | 3795842.5 | 7223505.5 | Q9QVN7 | TCEA2 | 3648467 | 2022848 | 1167892 | 3 |
| P11983 | TCP1 | t-complex 1 | Cytoplasm | other | 2.60E+07 | 2.54E+07 | 1.33E+07 | | | | | | |
| P11983 | TCP1 | t-complex 1 | Cytoplasm | other | 2.14E+07 | 1.56E+07 | 1.55E+07 | | | | | | |
| P11983 | TCP1 | t-complex 1 | Cytoplasm | other | 1.10E+07 | 1.03E+07 | 5294574.5 | | | | | | |
| P11983 | TCP1 | t-complex 1 | Cytoplasm | other | 1.11E+07 | 1.08E+07 | 6460227.5 | | | | | | |
| P11983 | TCP1 | t-complex 1 | Cytoplasm | other | 1.33E+07 | 1.64E+07 | 1.19E+07 | | | | | | |
| P11983 | TCP1 | t-complex 1 | Cytoplasm | other | 9397935 | 8125470 | 6321795 | | | | | | |
| P11983 | TCP1 | t-complex 1 | Cytoplasm | other | 6701703.5 | 6092339 | 6403135 | | | | | | |
| P11983 | TCP1 | t-complex 1 | Cytoplasm | other | 1.23E+07 | 1.10E+07 | 6205185.5 | | | | | | |
| P11983 | TCP1 | t-complex 1 | Cytoplasm | other | 8254111 | 8169859 | 4763884 | | | | | | |
| P11983 | TCP1 | t-complex 1 | Cytoplasm | other | 7959401 | 6257029.5 | 4629902 | | | | | | |
| P11983 | TCP1 | t-complex 1 | Cytoplasm | other | 6996952 | 5868300.5 | 4586923.5 | | | | | | |
| P11983 | TCP1 | t-complex 1 | Cytoplasm | other | 6742832 | 6238472 | 4913292 | | | | | | |
| P11983 | TCP1 | t-complex 1 | Cytoplasm | other | 6316148.5 | 6921121.5 | 3731274.5 | | | | | | |
| P11983 | TCP1 | t-complex 1 | Cytoplasm | other | 5104064.5 | 3956060 | 2905111.5 | | | | | | |
| P11983 | TCP1 | t-complex 1 | Cytoplasm | other | 2845494.5 | 790701.812 | 69230.156 | | | | | | |
| P11983 | TCP1 | t-complex 1 | Cytoplasm | other | 3098381.5 | 2179975 | 2336401 | | | | | | |
| P11983 | TCP1 | t-complex 1 | Cytoplasm | other | 3282421 | 2370939.5 | 2238775.25 | | | | | | |
| P11983 | TCP1 | t-complex 1 | Cytoplasm | other | 2368189.25 | 1937922 | 1694694 | | | | | | |

| | | | | | | | | | | | | | |
|---|---|---|---|---|---|---|---|---|---|---|---|---|---|
| P11983 | TCP1 | t-complex 1 | Cytoplasm | other | 2437014.25 | 1848398.375 | 2124510.25 | | | | | | |
| P11983 | TCP1 | t-complex 1 | Cytoplasm | other | 1416036.375 | 1581396.375 | 596029 | | | | | | |
| P11983 | TCP1 | t-complex 1 | Cytoplasm | other | 75266.25 | 338622 | 106213.742 | P11983 | TCP1 | 8004093 | 5722362 | 720950 | 63 |
| Q921I1 | TF | transferrin | Extracellular Space | transporter | 3.53E+07 | 5.32E+07 | 4.85E+07 | | | | | | |
| Q921I1 | TF | transferrin | Extracellular Space | transporter | 2.08E+07 | 4.08E+07 | 3.20E+07 | | | | | | |
| Q921I1 | TF | transferrin | Extracellular Space | transporter | 3.50E+07 | 8.06E+07 | 6.26E+07 | | | | | | |
| Q921I1 | TF | transferrin | Extracellular Space | transporter | 2.26E+07 | 5.11E+07 | 4.65E+07 | | | | | | |
| Q921I1 | TF | transferrin | Extracellular Space | transporter | 2.70E+07 | 6.11E+07 | 4.36E+07 | | | | | | |
| Q921I1 | TF | transferrin | Extracellular Space | transporter | 2.90E+07 | 5.61E+07 | 5.72E+07 | | | | | | |
| Q921I1 | TF | transferrin | Extracellular Space | transporter | 2.34E+07 | 5.34E+07 | 4.66E+07 | | | | | | |
| Q921I1 | TF | transferrin | Extracellular Space | transporter | 2.32E+07 | 5.16E+07 | 4.61E+07 | | | | | | |
| Q921I1 | TF | transferrin | Extracellular Space | transporter | 2.22E+07 | 4.06E+07 | 4.07E+07 | | | | | | |
| Q921I1 | TF | transferrin | Extracellular Space | transporter | 2.05E+07 | 3.96E+07 | 3.73E+07 | | | | | | |
| Q921I1 | TF | transferrin | Extracellular Space | transporter | 1.58E+07 | 3.95E+07 | 3.77E+07 | | | | | | |
| Q921I1 | TF | transferrin | Extracellular Space | transporter | 2.20E+07 | 4.46E+07 | 4.49E+07 | | | | | | |
| Q921I1 | TF | transferrin | Extracellular Space | transporter | 1.28E+07 | 2.30E+07 | 2.31E+07 | | | | | | |
| Q921I1 | TF | transferrin | Extracellular Space | transporter | 1.50E+07 | 2.69E+07 | 3.02E+07 | | | | | | |
| Q921I1 | TF | transferrin | Extracellular Space | transporter | 1.17E+07 | 2.10E+07 | 3.07E+07 | | | | | | |
| Q921I1 | TF | transferrin | Extracellular Space | transporter | 9876326 | 2.38E+07 | 2.36E+07 | | | | | | |
| Q921I1 | TF | transferrin | Extracellular Space | transporter | 8807748 | 2.09E+07 | 2.13E+07 | | | | | | |
| Q921I1 | TF | transferrin | Extracellular Space | transporter | 5790051 | 1.06E+07 | 7667814 | | | | | | |
| Q921I1 | TF | transferrin | Extracellular Space | transporter | 7012149 | 1.83E+07 | 1.40E+07 | | | | | | |
| Q921I1 | TF | transferrin | Extracellular Space | transporter | 6986492 | 1.71E+07 | 1.64E+07 | | | | | | |
| Q921I1 | TF | transferrin | Extracellular Space | transporter | 7937164 | 1.41E+07 | 1.37E+07 | | | | | | |
| Q921I1 | TF | transferrin | Extracellular Space | transporter | 8945675 | 1.92E+07 | 1.04E+07 | | | | | | |
| Q921I1 | TF | transferrin | Extracellular Space | transporter | 5910649.5 | 1.16E+07 | 1.40E+07 | | | | | | |
| Q921I1 | TF | transferrin | Extracellular Space | transporter | 4160682.25 | 1.48E+07 | 1.16E+07 | | | | | | |
| Q921I1 | TF | transferrin | Extracellular Space | transporter | 7891783.5 | 1.46E+07 | 1.64E+07 | | | | | | |
| Q921I1 | TF | transferrin | Extracellular Space | transporter | 6887449.5 | 1.36E+07 | 1.32E+07 | | | | | | |
| Q921I1 | TF | transferrin | Extracellular Space | transporter | 4206360.5 | 1.12E+07 | 9272137 | | | | | | |
| Q921I1 | TF | transferrin | Extracellular Space | transporter | 6751767.5 | 1.39E+07 | 6776327 | | | | | | |
| Q921I1 | TF | transferrin | Extracellular Space | transporter | 2920176 | 5791089.5 | 7718833.5 | | | | | | |
| Q921I1 | TF | transferrin | Extracellular Space | transporter | 2729625.5 | 8098541.5 | 7381420.5 | | | | | | |
| Q921I1 | TF | transferrin | Extracellular Space | transporter | 4020610.25 | 7852092 | 6566150.5 | | | | | | |
| Q921I1 | TF | transferrin | Extracellular Space | transporter | 1.00E+07 | 3.13E+07 | 3.44E+07 | | | | | | |
| Q921I1 | TF | transferrin | Extracellular Space | transporter | 1605658.5 | 4893440.5 | 4614210 | | | | | | |
| Q921I1 | TF | transferrin | Extracellular Space | transporter | 1274902.875 | 2932282.25 | 2636781 | | | | | | |
| Q921I1 | TF | transferrin | Extracellular Space | transporter | 179030.141 | 3995775 | 4531363 | Q921I1 | TF | 12858980 | 17172641 | 1675878 | 105 |
| Q62351 | TFRC | transferrin receptor | Plasma Membrane | transporter | 4.56E+07 | 8.68E+07 | 8.34E+07 | | | | | | |
| Q62351 | TFRC | transferrin receptor | Plasma Membrane | transporter | 5.76E+07 | 1.23E+08 | 1.10E+08 | | | | | | |
| Q62351 | TFRC | transferrin receptor | Plasma Membrane | transporter | 4.19E+07 | 7.27E+07 | 6.35E+07 | | | | | | |
| Q62351 | TFRC | transferrin receptor | Plasma Membrane | transporter | 4.83E+07 | 1.01E+08 | 1.07E+08 | | | | | | |
| Q62351 | TFRC | transferrin receptor | Plasma Membrane | transporter | 5.26E+07 | 9.34E+07 | 9.10E+07 | | | | | | |
| Q62351 | TFRC | transferrin receptor | Plasma Membrane | transporter | 3.92E+07 | 9.16E+07 | 8.97E+07 | | | | | | |
| Q62351 | TFRC | transferrin receptor | Plasma Membrane | transporter | 3.89E+07 | 7.75E+07 | 7.54E+07 | | | | | | |
| Q62351 | TFRC | transferrin receptor | Plasma Membrane | transporter | 4.32E+07 | 8.22E+07 | 8.80E+07 | | | | | | |
| Q62351 | TFRC | transferrin receptor | Plasma Membrane | transporter | 4.39E+07 | 8.30E+07 | 8.38E+07 | | | | | | |
| Q62351 | TFRC | transferrin receptor | Plasma Membrane | transporter | 3.60E+07 | 8.58E+07 | 7.43E+07 | | | | | | |
| Q62351 | TFRC | transferrin receptor | Plasma Membrane | transporter | 4.03E+07 | 8.46E+07 | 7.02E+07 | | | | | | |
| Q62351 | TFRC | transferrin receptor | Plasma Membrane | transporter | 4.18E+07 | 6.67E+07 | 6.62E+07 | | | | | | |
| Q62351 | TFRC | transferrin receptor | Plasma Membrane | transporter | 2.96E+07 | 6.27E+07 | 7.27E+07 | | | | | | |
| Q62351 | TFRC | transferrin receptor | Plasma Membrane | transporter | 2.92E+07 | 5.70E+07 | 5.25E+07 | | | | | | |
| Q62351 | TFRC | transferrin receptor | Plasma Membrane | transporter | 2.16E+07 | 3.20E+07 | 3.51E+07 | | | | | | |
| Q62351 | TFRC | transferrin receptor | Plasma Membrane | transporter | 2.97E+07 | 6.08E+07 | 6.51E+07 | | | | | | |
| Q62351 | TFRC | transferrin receptor | Plasma Membrane | transporter | 3.01E+07 | 6.40E+07 | 6.39E+07 | | | | | | |
| Q62351 | TFRC | transferrin receptor | Plasma Membrane | transporter | 2.47E+07 | 5.99E+07 | 6.92E+07 | | | | | | |
| Q62351 | TFRC | transferrin receptor | Plasma Membrane | transporter | 2.55E+07 | 4.47E+07 | 4.89E+07 | | | | | | |
| Q62351 | TFRC | transferrin receptor | Plasma Membrane | transporter | 1.94E+07 | 3.91E+07 | 3.81E+07 | | | | | | |
| Q62351 | TFRC | transferrin receptor | Plasma Membrane | transporter | 2.11E+07 | 3.67E+07 | 3.77E+07 | | | | | | |
| Q62351 | TFRC | transferrin receptor | Plasma Membrane | transporter | 2.63E+07 | 5.12E+07 | 5.82E+07 | | | | | | |
| Q62351 | TFRC | transferrin receptor | Plasma Membrane | transporter | 2.37E+07 | 5.27E+07 | 5.47E+07 | | | | | | |
| Q62351 | TFRC | transferrin receptor | Plasma Membrane | transporter | 1.10E+07 | 2.16E+07 | 2.13E+07 | | | | | | |
| Q62351 | TFRC | transferrin receptor | Plasma Membrane | transporter | 1.04E+07 | 1.65E+07 | 1.49E+07 | | | | | | |
| Q62351 | TFRC | transferrin receptor | Plasma Membrane | transporter | 8075319 | 1.68E+07 | 2.09E+07 | | | | | | |
| Q62351 | TFRC | transferrin receptor | Plasma Membrane | transporter | 1.01E+07 | 1.68E+07 | 1.74E+07 | | | | | | |
| Q62351 | TFRC | transferrin receptor | Plasma Membrane | transporter | 1.27E+07 | 1.96E+07 | 1.99E+07 | | | | | | |
| Q62351 | TFRC | transferrin receptor | Plasma Membrane | transporter | 7971492 | 2.25E+07 | 1.23E+07 | | | | | | |
| Q62351 | TFRC | transferrin receptor | Plasma Membrane | transporter | 3133192.25 | 6073325 | 6061619 | | | | | | |
| Q62351 | TFRC | transferrin receptor | Plasma Membrane | transporter | 2682244.25 | 4076799.25 | 4963400 | | | | | | |
| Q62351 | TFRC | transferrin receptor | Plasma Membrane | transporter | 1999949.125 | 2640126.5 | 2792806.75 | | | | | | |
| Q62351 | TFRC | transferrin receptor | Plasma Membrane | transporter | 1041713.812 | 2314153.5 | 2016153 | | | | | | |
| Q62351 | TFRC | transferrin receptor | Plasma Membrane | transporter | 1773038.125 | 5086538.5 | 4274714 | | | | | | |
| Q62351 | TFRC | transferrin receptor | Plasma Membrane | transporter | 1897624.75 | 3248709.5 | 4066404.5 | | | | | | |
| Q62351 | TFRC | transferrin receptor | Plasma Membrane | transporter | 1246013 | 4580589.5 | 3189273.25 | | | | | | |
| Q62351 | TFRC | transferrin receptor | Plasma Membrane | transporter | 70753.289 | 245624.609 | 205472.281 | Q62351 | TFRC | 23899225 | 31331461 | 2973851 | 111 |
| Q62312 | TGFBR2 | transforming growth factor, beta receptor II (70/80kDa) | Plasma Membrane | kinase | 2.25E+07 | 1.67E+07 | 1.55E+07 | Q62312 | TGFBR2 | 22520000 | 3779237 | 2181944 | 3 |
| P21981 | TGM2 | transglutaminase 2 | Cytoplasm | enzyme | 5305987.5 | 4090280.5 | 3837344.25 | | | | | | |
| P21981 | TGM2 | transglutaminase 2 | Cytoplasm | enzyme | 1869109.875 | 1340907.625 | 794479.812 | P21981 | TGM2 | 3587549 | 1789218 | 730445 | 6 |
| P35441 | THBS1 | thrombospondin 1 | Extracellular Space | other | 1.70E+07 | 2.05E+07 | 5.32E+07 | | | | | | |
| P35441 | THBS1 | thrombospondin 1 | Extracellular Space | other | 1.09E+07 | 1.21E+07 | 2.79E+07 | | | | | | |
| P35441 | THBS1 | thrombospondin 1 | Extracellular Space | other | 5901140 | 9007582 | 1.94E+07 | | | | | | |
| P35441 | THBS1 | thrombospondin 1 | Extracellular Space | other | 1.12E+07 | 1.44E+07 | 3.86E+07 | | | | | | |
| P35441 | THBS1 | thrombospondin 1 | Extracellular Space | other | 5684744.5 | 7543966 | 2.01E+07 | | | | | | |
| P35441 | THBS1 | thrombospondin 1 | Extracellular Space | other | 3943905.5 | 4330713.5 | 1.11E+07 | | | | | | |
| P35441 | THBS1 | thrombospondin 1 | Extracellular Space | other | 4274667 | 6077957 | 1.72E+07 | | | | | | |
| P35441 | THBS1 | thrombospondin 1 | Extracellular Space | other | 9031624 | 9137486 | 2.74E+07 | | | | | | |
| P35441 | THBS1 | thrombospondin 1 | Extracellular Space | other | 6810261 | 1.04E+07 | 2.44E+07 | | | | | | |
| P35441 | THBS1 | thrombospondin 1 | Extracellular Space | other | 7560048 | 8985524 | 2.10E+07 | | | | | | |
| P35441 | THBS1 | thrombospondin 1 | Extracellular Space | other | 5773795 | 6959517.5 | 1.55E+07 | | | | | | |
| P35441 | THBS1 | thrombospondin 1 | Extracellular Space | other | 1492723.5 | 2294802 | 4709832 | | | | | | |
| P35441 | THBS1 | thrombospondin 1 | Extracellular Space | other | 1.89E+07 | 2.59E+07 | 5.29E+07 | | | | | | |
| P35441 | THBS1 | thrombospondin 1 | Extracellular Space | other | 7521500.5 | 8751742 | 2.20E+07 | | | | | | |
| P35441 | THBS1 | thrombospondin 1 | Extracellular Space | other | 4369309.5 | 5272175 | 1.31E+07 | | | | | | |
| P35441 | THBS1 | thrombospondin 1 | Extracellular Space | other | 5645112 | 5202349 | 1.06E+07 | | | | | | |
| P35441 | THBS1 | thrombospondin 1 | Extracellular Space | other | 1705002 | 1850765.125 | 5154289.5 | | | | | | |
| P35441 | THBS1 | thrombospondin 1 | Extracellular Space | other | 5167261 | 6336636 | 1.69E+07 | | | | | | |
| P35441 | THBS1 | thrombospondin 1 | Extracellular Space | other | 1876688.625 | 2631388.75 | 4057615.75 | | | | | | |
| P35441 | THBS1 | thrombospondin 1 | Extracellular Space | other | 1524343.375 | 2609774.75 | 7471620.5 | | | | | | |
| P35441 | THBS1 | thrombospondin 1 | Extracellular Space | other | 1961063.5 | 2407670.25 | 5899884.5 | | | | | | |
| P35441 | THBS1 | thrombospondin 1 | Extracellular Space | other | 1923739 | 2052356.25 | 4392715 | | | | | | |
| P35441 | THBS1 | thrombospondin 1 | Extracellular Space | other | 5989405 | 6068803.5 | 8921458 | | | | | | |
| P35441 | THBS1 | thrombospondin 1 | Extracellular Space | other | 2944269.75 | 3408957 | 8486818 | | | | | | |
| P35441 | THBS1 | thrombospondin 1 | Extracellular Space | other | 827978.938 | 172234.969 | 3459163 | | | | | | |
| P35441 | THBS1 | thrombospondin 1 | Extracellular Space | other | 2278489.75 | 1585928 | 5244257 | | | | | | |
| P35441 | THBS1 | thrombospondin 1 | Extracellular Space | other | 1103346.625 | 1125014.375 | 3985187.25 | | | | | | |
| P35441 | THBS1 | thrombospondin 1 | Extracellular Space | other | 2371348.25 | 3065582.25 | 3834949.5 | | | | | | |
| P35441 | THBS1 | thrombospondin 1 | Extracellular Space | other | 6057014 | 6780048.5 | 1.48E+07 | | | | | | |
| P35441 | THBS1 | thrombospondin 1 | Extracellular Space | other | 1826529.5 | 2351233.5 | 8548491 | | | | | | |
| P35441 | THBS1 | thrombospondin 1 | Extracellular Space | other | 2084077.375 | 3573380 | 6400905.5 | | | | | | |

| | | | | | | | | | | | | | |
|---|---|---|---|---|---|---|---|---|---|---|---|---|---|
| P35441 | THBS1 | thrombospondin 1 | Extracellular Space | other | 882813.75 | 996797.188 | 3217599.5 | P35441 | THBS1 | 5201631 | 9711060 | 991131 | 96 |
| Q80ZW2 | THEM6 | thioesterase superfamily member 6 | Extracellular Space | other | 5559554 | 5446588 | 3725946.75 | | | | | | |
| Q80ZW2 | THEM6 | thioesterase superfamily member 6 | Extracellular Space | other | 4571903 | 3795674 | 2723804.5 | | | | | | |
| Q80ZW2 | THEM6 | thioesterase superfamily member 6 | Extracellular Space | other | 2773658.25 | 1848074.875 | 1604980.5 | | | | | | |
| Q80ZW2 | THEM6 | thioesterase superfamily member 6 | Extracellular Space | other | 2820046.5 | 2542955.25 | 2533339.75 | | | | | | |
| Q80ZW2 | THEM6 | thioesterase superfamily member 6 | Extracellular Space | other | 1921822 | 1465804.5 | 1443179.125 | Q80ZW2 | THEM6 | 3529397 | 1361833 | 351624 | 15 |
| Q9CY52 | THG1L | tRNA-histidine guanylyltransferase 1-like (S. cerevisiae) | Cytoplasm | enzyme | 7778533 | 6595020 | 6180097 | | | | | | |
| Q9CY52 | THG1L | tRNA-histidine guanylyltransferase 1-like (S. cerevisiae) | Cytoplasm | enzyme | 9528689 | 7651182 | 6471196.5 | | | | | | |
| Q9CY52 | THG1L | tRNA-histidine guanylyltransferase 1-like (S. cerevisiae) | Cytoplasm | enzyme | 7770700 | 5274006.5 | 5050924.5 | | | | | | |
| Q9CY52 | THG1L | tRNA-histidine guanylyltransferase 1-like (S. cerevisiae) | Cytoplasm | enzyme | 5887046 | 4207546.5 | 2744736 | | | | | | |
| Q9CY52 | THG1L | tRNA-histidine guanylyltransferase 1-like (S. cerevisiae) | Cytoplasm | enzyme | 4664091 | 3974957.25 | 3123671.5 | Q9CY52 | THG1L | 7125812 | 1905900 | 492101 | 15 |
| Q9D880 | TIMM50 | translocase of inner mitochondrial membrane 50 homolog (S. cerevisiae) | Cytoplasm | phosphatase | 1596579.875 | 4054048.75 | 4496590.5 | | | | | | |
| Q9D880 | TIMM50 | translocase of inner mitochondrial membrane 50 homolog (S. cerevisiae) | Cytoplasm | phosphatase | 1567916.75 | 2517882.75 | 2644074.25 | Q9D880 | TIMM50 | 1582248 | 1226395 | 500674 | 6 |
| P39876 | TIMP3 | TIMP metallopeptidase inhibitor 3 | Extracellular Space | other | 8575217 | 8601518 | 1.77E+07 | | | | | | |
| P39876 | TIMP3 | TIMP metallopeptidase inhibitor 3 | Extracellular Space | other | 3229320 | 2307398.75 | 5340801.5 | | | | | | |
| P39876 | TIMP3 | TIMP metallopeptidase inhibitor 3 | Extracellular Space | other | 4464957.5 | 4120719.5 | 7913818.5 | | | | | | |
| P39876 | TIMP3 | TIMP metallopeptidase inhibitor 3 | Extracellular Space | other | 1067646 | 1070652.875 | 2569670.5 | P39876 | TIMP3 | 4334285 | 4660249 | 1345298 | 12 |
| P40142 | TKT | transketolase | Cytoplasm | enzyme | 1902791.625 | 2013378 | 3492579.75 | | | | | | |
| P40142 | TKT | transketolase | Cytoplasm | enzyme | 290159.969 | 507285.156 | 1563596.125 | P40142 | TKT | 1096476 | 1161951 | 474364 | 6 |
| P26039 | TLN1 | talin 1 | Plasma Membrane | other | 1271148.25 | 1051674.25 | 3870933 | | | | | | |
| P26039 | TLN1 | talin 1 | Plasma Membrane | other | 3664171.75 | 2827085 | 9856715 | | | | | | |
| P26039 | TLN1 | talin 1 | Plasma Membrane | other | 1228327.625 | 1314571.25 | 3195137.5 | | | | | | |
| P26039 | TLN1 | talin 1 | Plasma Membrane | other | 592530.25 | 736976.438 | 3546579.75 | | | | | | |
| P26039 | TLN1 | talin 1 | Plasma Membrane | other | 1658852.875 | 1702238.25 | 5214446.5 | | | | | | |
| P26039 | TLN1 | talin 1 | Plasma Membrane | other | 1684137.25 | 1087878.625 | 3721497 | | | | | | |
| P26039 | TLN1 | talin 1 | Plasma Membrane | other | 131983.734 | 169430.625 | 1297177.375 | | | | | | |
| P26039 | TLN1 | talin 1 | Plasma Membrane | other | 991673.562 | 559600.875 | 2049623.25 | P26039 | TLN1 | 1402853 | 2108054 | 430305 | 24 |
| Q9ET30 | TM9SF3 | transmembrane 9 superfamily member 3 | Cytoplasm | transporter | 1685092.5 | 1822327.375 | 2489524.75 | Q9ET30 | TM9SF3 | 1685093 | 430329 | 248450 | 3 |
| Q7TN60 | TMC6 | transmembrane channel-like 6 | Cytoplasm | transporter | 2130586.75 | 2707302 | 2211894 | | | | | | |
| Q7TN60 | TMC6 | transmembrane channel-like 6 | Cytoplasm | transporter | 1393856.5 | 1753389.25 | 1209964.875 | | | | | | |
| Q7TN60 | TMC6 | transmembrane channel-like 6 | Cytoplasm | transporter | 894929.875 | 613833.312 | 627193.625 | Q7TN60 | TMC6 | 1473124 | 744529 | 248176 | 9 |
| Q7TN58 | TMC8 | transmembrane channel-like 8 | Cytoplasm | other | 2854954.5 | 3873126.25 | 1959949.25 | | | | | | |
| Q7TN58 | TMC8 | transmembrane channel-like 8 | Cytoplasm | other | 2245809.75 | 1897350.125 | 1129489.75 | | | | | | |
| Q7TN58 | TMC8 | transmembrane channel-like 8 | Cytoplasm | other | 1785071.75 | 1735349.5 | 1462298.5 | Q7TN58 | TMC8 | 2295279 | 819784 | 273261 | 9 |
| Q80W04 | TMCC2 | transmembrane and coiled-coil domain family 2 | Extracellular Space | other | 8349265.5 | 1.04E+07 | 1.22E+07 | | | | | | |
| Q80W04 | TMCC2 | transmembrane and coiled-coil domain family 2 | Extracellular Space | other | 9979402 | 1.44E+07 | 1.49E+07 | | | | | | |
| Q80W04 | TMCC2 | transmembrane and coiled-coil domain family 2 | Extracellular Space | other | 3772708 | 4412048 | 5137074 | | | | | | |
| Q80W04 | TMCC2 | transmembrane and coiled-coil domain family 2 | Extracellular Space | other | 3229175 | 3438835.25 | 3629732.75 | | | | | | |
| Q80W04 | TMCC2 | transmembrane and coiled-coil domain family 2 | Extracellular Space | other | 2278446.5 | 2758850 | 3649183 | | | | | | |
| Q80W04 | TMCC2 | transmembrane and coiled-coil domain family 2 | Extracellular Space | other | 1056517.75 | 1240685.75 | 1766533.25 | Q80W04 | TMCC2 | 4777586 | 4539421 | 1069952 | 18 |
| Q921L3 | TMCO1 | transmembrane and coiled-coil domains 1 | Cytoplasm | other | 4933322.5 | 4275483 | 6898989 | Q921L3 | TMCO1 | 4933323 | 1365002 | 788084 | 3 |
| Q9D1D4 | TMED10 | transmembrane emp24-like trafficking protein 10 (yeast) | Cytoplasm | transporter | 2.29E+07 | 2.15E+07 | 1.86E+07 | | | | | | |
| Q9D1D4 | TMED10 | transmembrane emp24-like trafficking protein 10 (yeast) | Cytoplasm | transporter | 1.95E+07 | 1.97E+07 | 1.71E+07 | | | | | | |
| Q9D1D4 | TMED10 | transmembrane emp24-like trafficking protein 10 (yeast) | Cytoplasm | transporter | 1.87E+07 | 1.77E+07 | 1.80E+07 | | | | | | |
| Q9D1D4 | TMED10 | transmembrane emp24-like trafficking protein 10 (yeast) | Cytoplasm | transporter | 4245366 | 4102803 | 6032354.5 | | | | | | |
| Q9D1D4 | TMED10 | transmembrane emp24-like trafficking protein 10 (yeast) | Cytoplasm | transporter | 6898340.5 | 7004746.5 | 4569781 | | | | | | |
| Q9D1D4 | TMED10 | transmembrane emp24-like trafficking protein 10 (yeast) | Cytoplasm | transporter | 5726688 | 5294244 | 5524786 | Q9D1D4 | TMED10 | 12991732 | 7253691 | 1709711 | 18 |
| Q9R0Q3 | TMED2 | transmembrane emp24 domain trafficking protein 2 | Cytoplasm | transporter | 9663704 | 9669269 | 7748291.5 | | | | | | |
| Q9R0Q3 | TMED2 | transmembrane emp24 domain trafficking protein 2 | Cytoplasm | transporter | 1.14E+07 | 1.43E+07 | 9964857 | | | | | | |
| Q9R0Q3 | TMED2 | transmembrane emp24 domain trafficking protein 2 | Cytoplasm | transporter | 4884329 | 5997497 | 3492271.5 | Q9R0Q3 | TMED2 | 8632678 | 3387503 | 1129168 | 9 |
| Q8R1V4 | TMED4 | transmembrane emp24 protein transport domain containing 4 | Cytoplasm | transporter | 1.56E+07 | 1.67E+07 | 1.34E+07 | Q8R1V4 | TMED4 | 15600000 | 1656291 | 956260 | 3 |
| Q9CXE7 | TMED5 | transmembrane emp24 protein transport domain containing 5 | Cytoplasm | other | 3052621.75 | 3012527 | 2665138 | Q9CXE7 | TMED5 | 3052622 | 213085 | 123024 | 3 |
| Q99KF1 | TMED9 | transmembrane emp24 protein transport domain containing 9 | Cytoplasm | transporter | 2.64E+07 | 2.61E+07 | 2.06E+07 | | | | | | |
| Q99KF1 | TMED9 | transmembrane emp24 protein transport domain containing 9 | Cytoplasm | transporter | 1.12E+07 | 8838711 | 3953178.25 | | | | | | |
| Q99KF1 | TMED9 | transmembrane emp24 protein transport domain containing 9 | Cytoplasm | transporter | 1147403.25 | 524157.312 | 1008893.562 | Q99KF1 | TMED9 | 12909134 | 10719630 | 3573210 | 9 |
| Q80X71 | TMEM106B | transmembrane protein 106B | Cytoplasm | other | 712850 | 615427.812 | 537054.25 | Q80X71 | TMEM106B | 712850 | 88070 | 50847 | 3 |
| Q9CQN6 | TMEM14C | transmembrane protein 14C | Plasma Membrane | other | 1.14E+07 | 1.60E+07 | 1.26E+07 | | | | | | |
| Q9CQN6 | TMEM14C | transmembrane protein 14C | Plasma Membrane | other | 3867050.25 | 4998819.5 | 6408688 | Q9CQN6 | TMEM14C | 7613525 | 4816800 | 1966450 | 6 |
| Q9CR64 | TMEM167A | transmembrane protein 167A | Extracellular Space | other | 3171111.25 | 3183849.25 | 2267777.75 | Q9CR64 | TMEM167A | 3171111 | 525256 | 303256 | 3 |
| Q9CY24 | TMEM179B | transmembrane protein 179B | Extracellular Space | other | 2981098.25 | 3974638.75 | 2490248.5 | Q9CY24 | TMEM179B | 2981098 | 756248 | 436620 | 3 |
| Q9D771 | TMEM206 | transmembrane protein 206 | Plasma Membrane | other | 8052635 | 8309608.5 | 1.03E+07 | | | | | | |
| Q9D771 | TMEM206 | transmembrane protein 206 | Plasma Membrane | other | 3441013.75 | 3563939.25 | 2771363.25 | | | | | | |
| Q9D771 | TMEM206 | transmembrane protein 206 | Plasma Membrane | other | 1849795.25 | 1999703.375 | 1619373 | Q9D771 | TMEM206 | 4447815 | 3299871 | 1099957 | 9 |
| Q5F285 | TMEM256 | transmembrane protein 256 | Cytoplasm | other | 1.44E+07 | 1.96E+07 | 2.18E+07 | Q5F285 | TMEM256 | 14360000 | 3833814 | 2213454 | 3 |
| Q8VEK0 | TMEM30A | transmembrane protein 30A | Cytoplasm | other | 2.16E+07 | 1.78E+07 | 1.34E+07 | | | | | | |
| Q8VEK0 | TMEM30A | transmembrane protein 30A | Cytoplasm | other | 1.76E+07 | 1.54E+07 | 1.53E+07 | | | | | | |
| Q8VEK0 | TMEM30A | transmembrane protein 30A | Cytoplasm | other | 9495119 | 1.01E+07 | 9409338 | | | | | | |
| Q8VEK0 | TMEM30A | transmembrane protein 30A | Cytoplasm | other | 9322997 | 9770991 | 8371858.5 | | | | | | |
| Q8VEK0 | TMEM30A | transmembrane protein 30A | Cytoplasm | other | 5372897.5 | 6735171 | 5135684.5 | Q8VEK0 | TMEM30A | 12676203 | 4932991 | 1273693 | 15 |
| Q9CZX7 | TMEM55A | transmembrane protein 55A | Cytoplasm | phosphatase | 4596191.5 | 4418971.5 | 3227907 | Q9CZX7 | TMEM55A | 4596192 | 744115 | 429615 | 3 |
| Q3TWL2 | TMEM55B | transmembrane protein 55B | Cytoplasm | phosphatase | 6392025.5 | 6576834.5 | 4803348 | | | | | | |
| Q3TWL2 | TMEM55B | transmembrane protein 55B | Cytoplasm | phosphatase | 3556417.5 | 4017583.75 | 3410625.25 | | | | | | |
| Q3TWL2 | TMEM55B | transmembrane protein 55B | Cytoplasm | phosphatase | 4200110.5 | 3555136 | 3924882.25 | | | | | | |
| Q3TWL2 | TMEM55B | transmembrane protein 55B | Cytoplasm | phosphatase | 4299063 | 4531447.5 | 3707770.5 | Q3TWL2 | TMEM55B | 4611904 | 1052693 | 303886 | 12 |
| Q91YT8 | TMEM63A | transmembrane protein 63A | Cytoplasm | other | 952208.5 | 1318788.375 | 5508874.5 | Q91YT8 | TMEM63A | 952209 | 2531614 | 1461628 | 3 |
| Q9CR23 | TMEM9 | transmembrane protein 9 | Cytoplasm | other | 5011395.5 | 6695446.5 | 6600188 | Q9CR23 | TMEM9 | 5011396 | 945988 | 546167 | 3 |
| P49813 | TMOD1 | tropomodulin 1 | Cytoplasm | enzyme | 2.74E+08 | 3.19E+08 | 2.47E+08 | | | | | | |
| P49813 | TMOD1 | tropomodulin 1 | Cytoplasm | enzyme | 2.01E+08 | 2.31E+08 | 1.60E+08 | | | | | | |
| P49813 | TMOD1 | tropomodulin 1 | Cytoplasm | enzyme | 2.05E+08 | 1.99E+08 | 1.64E+08 | | | | | | |
| P49813 | TMOD1 | tropomodulin 1 | Cytoplasm | enzyme | 1.73E+08 | 1.84E+08 | 1.57E+08 | | | | | | |
| P49813 | TMOD1 | tropomodulin 1 | Cytoplasm | enzyme | 2.12E+08 | 2.77E+08 | 1.07E+08 | | | | | | |
| P49813 | TMOD1 | tropomodulin 1 | Cytoplasm | enzyme | 1.56E+08 | 1.63E+08 | 1.62E+08 | | | | | | |
| P49813 | TMOD1 | tropomodulin 1 | Cytoplasm | enzyme | 1.20E+08 | 1.14E+08 | 1.29E+08 | | | | | | |
| P49813 | TMOD1 | tropomodulin 1 | Cytoplasm | enzyme | 1.23E+08 | 1.16E+08 | 1.07E+08 | | | | | | |
| P49813 | TMOD1 | tropomodulin 1 | Cytoplasm | enzyme | 9.31E+07 | 1.10E+08 | 9.53E+07 | | | | | | |
| P49813 | TMOD1 | tropomodulin 1 | Cytoplasm | enzyme | 9.52E+07 | 8.44E+07 | 6.64E+07 | | | | | | |
| P49813 | TMOD1 | tropomodulin 1 | Cytoplasm | enzyme | 6.29E+07 | 5.42E+07 | 4.41E+07 | | | | | | |
| P49813 | TMOD1 | tropomodulin 1 | Cytoplasm | enzyme | 4.26E+07 | 3.94E+07 | 3.74E+07 | | | | | | |
| P49813 | TMOD1 | tropomodulin 1 | Cytoplasm | enzyme | 3.99E+07 | 4.23E+07 | 3.64E+07 | | | | | | |
| P49813 | TMOD1 | tropomodulin 1 | Cytoplasm | enzyme | 5635298 | 4387145 | 2.26E+07 | | | | | | |
| P49813 | TMOD1 | tropomodulin 1 | Cytoplasm | enzyme | 1.56E+07 | 1.93E+07 | 1.66E+07 | | | | | | |
| P49813 | TMOD1 | tropomodulin 1 | Cytoplasm | enzyme | 1.66E+07 | 1.01E+07 | 1.26E+07 | | | | | | |
| P49813 | TMOD1 | tropomodulin 1 | Cytoplasm | enzyme | 2.33E+07 | 2.49E+07 | 1.34E+07 | | | | | | |
| P49813 | TMOD1 | tropomodulin 1 | Cytoplasm | enzyme | 1.07E+07 | 1.05E+07 | 9231722 | | | | | | |
| P49813 | TMOD1 | tropomodulin 1 | Cytoplasm | enzyme | 7384903.5 | 1.18E+07 | 9713318 | | | | | | |
| P49813 | TMOD1 | tropomodulin 1 | Cytoplasm | enzyme | 1.91E+07 | 4.04E+07 | 3.44E+07 | | | | | | |
| P49813 | TMOD1 | tropomodulin 1 | Cytoplasm | enzyme | 9458978 | 1.09E+07 | 4721882 | | | | | | |
| P49813 | TMOD1 | tropomodulin 1 | Cytoplasm | enzyme | 6459522 | 6389108 | 5722938 | | | | | | |
| P49813 | TMOD1 | tropomodulin 1 | Cytoplasm | enzyme | 3945300.5 | 1912717.375 | 2417057.75 | | | | | | |
| P49813 | TMOD1 | tropomodulin 1 | Cytoplasm | enzyme | 1228789.25 | 1348899.625 | 598152.625 | P49813 | TMOD1 | 79862200 | 82507925 | 9723652 | 72 |
| Q9D710 | TMX2 | thioredoxin-related transmembrane protein 2 | Cytoplasm | enzyme | 323955.188 | 571804.875 | 685636.688 | Q9D710 | TMX2 | 323955 | 184933 | 106771 | 3 |
| Q8C0L0 | TMX4 | thioredoxin-related transmembrane protein 4 | Cytoplasm | enzyme | 3966839.25 | 3737588.75 | 3775609.5 | Q8C0L0 | TMX4 | 3966839 | 122862 | 70934 | 3 |
| O88746 | TOM1 | target of myb1 membrane trafficking protein | Cytoplasm | transporter | 1039118.5 | 1258128.25 | 1284082.25 | O88746 | TOM1 | 1039119 | 134565 | 77691 | 3 |
| Q923U0 | TOM1L1 | target of myb1 like 1 membrane trafficking protein | Cytoplasm | other | 4752745.5 | 3897298.25 | 2864908.5 | | | | | | |
| Q923U0 | TOM1L1 | target of myb1 like 1 membrane trafficking protein | Cytoplasm | other | 1628830.375 | 949563.875 | 1119608.75 | Q923U0 | TOM1L1 | 3190788 | 1563405 | 638257 | 6 |
| Q9CPQ3 | TOMM22 | translocase of outer mitochondrial membrane 22 homolog (yeast) | Cytoplasm | transporter | 1754588.625 | 2138755 | 3097584.75 | | | | | | |
| Q9CPQ3 | TOMM22 | translocase of outer mitochondrial membrane 22 homolog (yeast) | Cytoplasm | transporter | 1657224.75 | 2981041.25 | 3845226.75 | Q9CPQ3 | TOMM22 | 1705907 | 866838 | 353885 | 6 |
| Q9QYA2 | TOMM40 | translocase of outer mitochondrial membrane 40 homolog (yeast) | Cytoplasm | ion channel | 3692853.75 | 6816866 | 6795858 | Q9QYA2 | TOMM40 | 3692854 | 1797616 | 1037854 | 3 |
| Q9CZW5 | TOMM70A | translocase of outer mitochondrial membrane 70 homolog A (S. cerevisiae) | Cytoplasm | transporter | 1306610.5 | 2253343.25 | 2655202 | Q9CZW5 | TOMM70A | 1306611 | 692398 | 399756 | 3 |

| | | | | | | | | | | | | | |
|---|---|---|---|---|---|---|---|---|---|---|---|---|---|
| Q9CYZ2 | TPD52L2 | tumor protein D52-like 2 | Cytoplasm | other | 2909120.75 | 2896883.25 | 4671963 | Q9CYZ2 | TPD52L2 | 2909121 | 1021328 | 589664 | 3 |
| P58771 | Tpm1 | tropomyosin 1, alpha | Plasma Membrane | other | 4.14E+08 | 3.03E+08 | 2.90E+08 | | | | | | |
| P58771 | Tpm1 | tropomyosin 1, alpha | Plasma Membrane | other | 7.29E+07 | 6.84E+07 | 4.45E+07 | | | | | | |
| P58771 | Tpm1 | tropomyosin 1, alpha | Plasma Membrane | other | 2.20E+07 | 2.03E+07 | 1.17E+07 | | | | | | |
| P58771 | Tpm1 | tropomyosin 1, alpha | Plasma Membrane | other | 1.37E+07 | 1.42E+07 | 9876548 | | | | | | |
| P58771 | Tpm1 | tropomyosin 1, alpha | Plasma Membrane | other | 4438695 | 2777005 | 2134603 | P58771 | Tpm1 | 105365739 | 133321635 | 34423498 | 15 |
| P21107 | TPM3 | tropomyosin 3 | Cytoplasm | other | 6.25E+08 | 4.77E+08 | 4.84E+08 | | | | | | |
| P21107 | TPM3 | tropomyosin 3 | Cytoplasm | other | 2.07E+08 | 1.81E+08 | 1.70E+08 | | | | | | |
| P21107 | TPM3 | tropomyosin 3 | Cytoplasm | other | 2.07E+08 | 1.75E+08 | 1.56E+08 | | | | | | |
| P21107 | TPM3 | tropomyosin 3 | Cytoplasm | other | 2.31E+08 | 2.18E+08 | 1.65E+08 | | | | | | |
| P21107 | TPM3 | tropomyosin 3 | Cytoplasm | other | 9.83E+07 | 6.81E+07 | 7.45E+07 | | | | | | |
| P21107 | TPM3 | tropomyosin 3 | Cytoplasm | other | 9.71E+07 | 1.20E+08 | 1.15E+08 | | | | | | |
| P21107 | TPM3 | tropomyosin 3 | Cytoplasm | other | 1.14E+08 | 1.14E+08 | 7.66E+07 | | | | | | |
| P21107 | TPM3 | tropomyosin 3 | Cytoplasm | other | 8.53E+07 | 4.96E+07 | 4.77E+07 | | | | | | |
| P21107 | TPM3 | tropomyosin 3 | Cytoplasm | other | 8.05E+07 | 7.96E+07 | 4.92E+07 | | | | | | |
| P21107 | TPM3 | tropomyosin 3 | Cytoplasm | other | 6.90E+07 | 8.83E+07 | 3.59E+07 | | | | | | |
| P21107 | TPM3 | tropomyosin 3 | Cytoplasm | other | 3.99E+07 | 4.06E+07 | 3.48E+07 | | | | | | |
| P21107 | TPM3 | tropomyosin 3 | Cytoplasm | other | 1.32E+07 | 1.36E+07 | 7554254.5 | | | | | | |
| P21107 | TPM3 | tropomyosin 3 | Cytoplasm | other | 1.19E+07 | 1.15E+07 | 1.01E+07 | | | | | | |
| P21107 | TPM3 | tropomyosin 3 | Cytoplasm | other | 9869498 | 6352075 | 6084801 | P21107 | TPM3 | 134933536 | 133581202 | 20612027 | 42 |
| O89023 | TPP1 | tripeptidyl peptidase I | Cytoplasm | peptidase | 4128679.25 | 8018661.5 | 8434180 | O89023 | TPP1 | 4128679 | 2374937 | 1371171 | 3 |
| Q64514 | TPP2 | tripeptidyl peptidase II | Cytoplasm | peptidase | 2.10E+07 | 1.28E+07 | 1.33E+07 | | | | | | |
| Q64514 | TPP2 | tripeptidyl peptidase II | Cytoplasm | peptidase | 1.36E+07 | 1.12E+07 | 9896934 | | | | | | |
| Q64514 | TPP2 | tripeptidyl peptidase II | Cytoplasm | peptidase | 1.07E+07 | 7864320 | 8097703.5 | | | | | | |
| Q64514 | TPP2 | tripeptidyl peptidase II | Cytoplasm | peptidase | 1.06E+07 | 6970815 | 7854437.5 | | | | | | |
| Q64514 | TPP2 | tripeptidyl peptidase II | Cytoplasm | peptidase | 9591367 | 6255429.5 | 8462317 | | | | | | |
| Q64514 | TPP2 | tripeptidyl peptidase II | Cytoplasm | peptidase | 1.01E+07 | 6311684.5 | 6787055 | | | | | | |
| Q64514 | TPP2 | tripeptidyl peptidase II | Cytoplasm | peptidase | 9089411 | 5278341.5 | 5360834.5 | | | | | | |
| Q64514 | TPP2 | tripeptidyl peptidase II | Cytoplasm | peptidase | 7668384.5 | 4242091.5 | 4771555 | | | | | | |
| Q64514 | TPP2 | tripeptidyl peptidase II | Cytoplasm | peptidase | 7640932.5 | 5173830.5 | 7208335.5 | | | | | | |
| Q64514 | TPP2 | tripeptidyl peptidase II | Cytoplasm | peptidase | 6714472 | 4936296.5 | 5104600 | | | | | | |
| Q64514 | TPP2 | tripeptidyl peptidase II | Cytoplasm | peptidase | 4594294.5 | 2650462.75 | 3230386.75 | | | | | | |
| Q64514 | TPP2 | tripeptidyl peptidase II | Cytoplasm | peptidase | 3810251 | 2295976.75 | 2394506 | | | | | | |
| Q64514 | TPP2 | tripeptidyl peptidase II | Cytoplasm | peptidase | 2976828.25 | 3271308.75 | 1225967.5 | | | | | | |
| Q64514 | TPP2 | tripeptidyl peptidase II | Cytoplasm | peptidase | 5022933.5 | 3920400.75 | 2974324 | | | | | | |
| Q64514 | TPP2 | tripeptidyl peptidase II | Cytoplasm | peptidase | 2933751 | 2865548.75 | 3195833.25 | | | | | | |
| Q64514 | TPP2 | tripeptidyl peptidase II | Cytoplasm | peptidase | 3054197.25 | 2019587.625 | 2052901 | | | | | | |
| Q64514 | TPP2 | tripeptidyl peptidase II | Cytoplasm | peptidase | 3389351 | 2251856.5 | 2588711.75 | | | | | | |
| Q64514 | TPP2 | tripeptidyl peptidase II | Cytoplasm | peptidase | 4748092 | 3047962.5 | 2523689.75 | | | | | | |
| Q64514 | TPP2 | tripeptidyl peptidase II | Cytoplasm | peptidase | 1562474 | 241286.047 | 128655.859 | | | | | | |
| Q64514 | TPP2 | tripeptidyl peptidase II | Cytoplasm | peptidase | 1370787.625 | 962674.062 | 483988.094 | | | | | | |
| Q64514 | TPP2 | tripeptidyl peptidase II | Cytoplasm | peptidase | 1440890.5 | 1045259.188 | 910386.812 | | | | | | |
| Q64514 | TPP2 | tripeptidyl peptidase II | Cytoplasm | peptidase | 1294468.25 | 731713.812 | 736106.188 | Q64514 | TPP2 | 6496040 | 3984798 | 490495 | 66 |
| Q9CQN1 | TRAP1 | TNF receptor-associated protein 1 | Cytoplasm | enzyme | 1.28E+07 | 1.15E+07 | 9182832 | | | | | | |
| Q9CQN1 | TRAP1 | TNF receptor-associated protein 1 | Cytoplasm | enzyme | 1431894.375 | 1949683.125 | 1802275.375 | Q9CQN1 | TRAP1 | 7135947 | 5313045 | 2169041 | 6 |
| Q8K558 | TREML1 | triggering receptor expressed on myeloid cells-like 1 | Plasma Membrane | other | 3.08E+07 | 3.97E+07 | 4.81E+07 | | | | | | |
| Q8K558 | TREML1 | triggering receptor expressed on myeloid cells-like 1 | Plasma Membrane | other | 2.46E+07 | 3.14E+07 | 3.65E+07 | | | | | | |
| Q8K558 | TREML1 | triggering receptor expressed on myeloid cells-like 1 | Plasma Membrane | other | 1.60E+07 | 2.03E+07 | 2.46E+07 | | | | | | |
| Q8K558 | TREML1 | triggering receptor expressed on myeloid cells-like 1 | Plasma Membrane | other | 2.72E+07 | 3.00E+07 | 3.65E+07 | | | | | | |
| Q8K558 | TREML1 | triggering receptor expressed on myeloid cells-like 1 | Plasma Membrane | other | 1.55E+07 | 1.46E+07 | 3.00E+07 | | | | | | |
| Q8K558 | TREML1 | triggering receptor expressed on myeloid cells-like 1 | Plasma Membrane | other | 9003003 | 1.24E+07 | 2.29E+07 | | | | | | |
| Q8K558 | TREML1 | triggering receptor expressed on myeloid cells-like 1 | Plasma Membrane | other | 1.22E+07 | 1.43E+07 | 1.85E+07 | | | | | | |
| Q8K558 | TREML1 | triggering receptor expressed on myeloid cells-like 1 | Plasma Membrane | other | 2461751.25 | 4191251 | 5348352 | Q8K558 | TREML1 | 17230594 | 11950987 | 2439485 | 24 |
| Q9WTR1 | TRPV2 | transient receptor potential cation channel, subfamily V, member 2 | Plasma Membrane | ion channel | 2.36E+07 | 2.05E+07 | 1.59E+07 | | | | | | |
| Q9WTR1 | TRPV2 | transient receptor potential cation channel, subfamily V, member 2 | Plasma Membrane | ion channel | 1.32E+07 | 1.27E+07 | 1.36E+07 | | | | | | |
| Q9WTR1 | TRPV2 | transient receptor potential cation channel, subfamily V, member 2 | Plasma Membrane | ion channel | 6442659.5 | 7755894.5 | 8839390 | | | | | | |
| Q9WTR1 | TRPV2 | transient receptor potential cation channel, subfamily V, member 2 | Plasma Membrane | ion channel | 8088959 | 1.01E+07 | 9132175 | | | | | | |
| Q9WTR1 | TRPV2 | transient receptor potential cation channel, subfamily V, member 2 | Plasma Membrane | ion channel | 7957715.5 | 7252516.5 | 8802577 | | | | | | |
| Q9WTR1 | TRPV2 | transient receptor potential cation channel, subfamily V, member 2 | Plasma Membrane | ion channel | 9694771 | 9737301 | 8375197.5 | | | | | | |
| Q9WTR1 | TRPV2 | transient receptor potential cation channel, subfamily V, member 2 | Plasma Membrane | ion channel | 6209624.5 | 5901190.5 | 5459795 | | | | | | |
| Q9WTR1 | TRPV2 | transient receptor potential cation channel, subfamily V, member 2 | Plasma Membrane | ion channel | 5567326 | 5453505 | 4808214 | | | | | | |
| Q9WTR1 | TRPV2 | transient receptor potential cation channel, subfamily V, member 2 | Plasma Membrane | ion channel | 5229493 | 7011533.5 | 5466707.5 | | | | | | |
| Q9WTR1 | TRPV2 | transient receptor potential cation channel, subfamily V, member 2 | Plasma Membrane | ion channel | 4452645 | 4171734.25 | 2737628 | | | | | | |
| Q9WTR1 | TRPV2 | transient receptor potential cation channel, subfamily V, member 2 | Plasma Membrane | ion channel | 3132247.25 | 3631236.75 | 3619119 | | | | | | |
| Q9WTR1 | TRPV2 | transient receptor potential cation channel, subfamily V, member 2 | Plasma Membrane | ion channel | 4186913 | 3550183.25 | 2782031 | Q9WTR1 | TRPV2 | 8147696 | 4747957 | 791326 | 36 |
| Q62348 | TSN | translin | Nucleus | other | 4537103 | 3810956.75 | 4086128.75 | | | | | | |
| Q62348 | TSN | translin | Nucleus | other | 1629954.75 | 1380896.75 | 1412187.75 | Q62348 | TSN | 3083529 | 1483376 | 605586 | 6 |
| Q8QZY6 | TSPAN14 | tetraspanin 14 | Plasma Membrane | other | 3337571.25 | 4416012 | 5880236.5 | Q8QZY6 | TSPAN14 | 3337571 | 1276201 | 736815 | 3 |
| Q8R3S2 | TSPAN33 | tetraspanin 33 | Plasma Membrane | other | 1.91E+08 | 1.89E+08 | 1.64E+08 | | | | | | |
| Q8R3S2 | TSPAN33 | tetraspanin 33 | Plasma Membrane | other | 7.33E+07 | 6.55E+07 | 5.51E+07 | | | | | | |
| Q8R3S2 | TSPAN33 | tetraspanin 33 | Plasma Membrane | other | 2.97E+07 | 3.10E+07 | 3.08E+07 | | | | | | |
| Q8R3S2 | TSPAN33 | tetraspanin 33 | Plasma Membrane | other | 2.79E+07 | 2.20E+07 | 1.92E+07 | | | | | | |
| Q8R3S2 | TSPAN33 | tetraspanin 33 | Plasma Membrane | other | 6157260 | 8545258 | 5224632 | | | | | | |
| Q8R3S2 | TSPAN33 | tetraspanin 33 | Plasma Membrane | other | 4629908.5 | 8189641.5 | 4013894.75 | | | | | | |
| Q8R3S2 | TSPAN33 | tetraspanin 33 | Plasma Membrane | other | 3382642.75 | 2979079.75 | 2301025.25 | Q8R3S2 | TSPAN33 | 47972830 | 60847942 | 13278110 | 21 |
| Q8R3G9 | TSPAN8 | tetraspanin 8 | Plasma Membrane | other | 5.45E+07 | 4.93E+07 | 3.67E+07 | | | | | | |
| Q8R3G9 | TSPAN8 | tetraspanin 8 | Plasma Membrane | other | 3.37E+07 | 2.93E+07 | 2.72E+07 | | | | | | |
| Q8R3G9 | TSPAN8 | tetraspanin 8 | Plasma Membrane | other | 2.85E+07 | 2.32E+07 | 2.04E+07 | | | | | | |
| Q8R3G9 | TSPAN8 | tetraspanin 8 | Plasma Membrane | other | 1.13E+07 | 1.34E+07 | 1.20E+07 | | | | | | |
| Q8R3G9 | TSPAN8 | tetraspanin 8 | Plasma Membrane | other | 7343643 | 5946437.5 | 5336033.5 | Q8R3G9 | TSPAN8 | 27054729 | 15267564 | 3942068 | 15 |
| P50637 | TSPO | translocator protein (18kDa) | Cytoplasm | transmembrane receptor | 7615082.5 | 1.40E+07 | 1.63E+07 | P50637 | TSPO | 7615083 | 4490603 | 2592651 | 3 |
| A2ASS6 | TTN | titin | Cytoplasm | kinase | 3634069.5 | 3473388.5 | 3100921.75 | A2ASS6 | TTN | 3634070 | 273495 | 157902 | 3 |
| P68369 | TUBA1A | tubulin, alpha 1a | Cytoplasm | other | 3108239.5 | 3049278.25 | 5067653.5 | P68369 | TUBA1A | 3108240 | 1148667 | 663183 | 3 |
| Q7TMM9 | TUBB2A | tubulin, beta 2A class IIa | Cytoplasm | other | 2102566.75 | 2060680.375 | 2460166 | | | | | | |
| Q7TMM9 | TUBB2A | tubulin, beta 2A class IIa | Cytoplasm | other | 2616905 | 2210232.75 | 2633019.5 | Q7TMM9 | TUBB2A | 2359736 | 256084 | 104546 | 6 |
| Q9D6F9 | TUBB4A | tubulin, beta 4A class IVa | Cytoplasm | other | 3699087.25 | 4564174 | 5051255 | | | | | | |
| Q9D6F9 | TUBB4A | tubulin, beta 4A class IVa | Cytoplasm | other | 5582671 | 5907560 | 7591108.5 | Q9D6F9 | TUBB4A | 4640879 | 1327047 | 541765 | 6 |
| Q8BFR5 | TUFM | Tu translation elongation factor, mitochondrial | Cytoplasm | translation regulator | 3929370.75 | 7370657 | 9847162 | | | | | | |
| Q8BFR5 | TUFM | Tu translation elongation factor, mitochondrial | Cytoplasm | translation regulator | 5179071.5 | 9586857 | 1.21E+07 | | | | | | |
| Q8BFR5 | TUFM | Tu translation elongation factor, mitochondrial | Cytoplasm | translation regulator | 3959002.5 | 7976281 | 9276477 | | | | | | |
| Q8BFR5 | TUFM | Tu translation elongation factor, mitochondrial | Cytoplasm | translation regulator | 3932805.5 | 5267616.5 | 5824186.5 | | | | | | |
| Q8BFR5 | TUFM | Tu translation elongation factor, mitochondrial | Cytoplasm | translation regulator | 2224350.75 | 4488999 | 6297602 | | | | | | |
| Q8BFR5 | TUFM | Tu translation elongation factor, mitochondrial | Cytoplasm | translation regulator | 1504621.875 | 2185419.75 | 2888484.75 | Q8BFR5 | TUFM | 3454870 | 3024117 | 712791 | 18 |
| Q91W90 | TXNDC5 | thioredoxin domain containing 5 (endoplasmic reticulum) | Cytoplasm | enzyme | 1044650.125 | 1148721.25 | 878584.625 | Q91W90 | TXNDC5 | 1044650 | 136249 | 78663 | 3 |
| Q8CDN6 | TXNL1 | thioredoxin-like 1 | Cytoplasm | enzyme | 6510591 | 5268725 | 1.49E+07 | | | | | | |
| Q8CDN6 | TXNL1 | thioredoxin-like 1 | Cytoplasm | enzyme | 8010369 | 5889189.5 | 6808935.5 | | | | | | |
| Q8CDN6 | TXNL1 | thioredoxin-like 1 | Cytoplasm | enzyme | 5326566 | 3986037.25 | 3745463.25 | | | | | | |
| Q8CDN6 | TXNL1 | thioredoxin-like 1 | Cytoplasm | enzyme | 3872145 | 4248401.5 | 3672440 | | | | | | |
| Q8CDN6 | TXNL1 | thioredoxin-like 1 | Cytoplasm | enzyme | 2885457.25 | 3086413.75 | 1437355.5 | | | | | | |
| Q8CDN6 | TXNL1 | thioredoxin-like 1 | Cytoplasm | enzyme | 1540519.625 | 1095989 | 870764.562 | | | | | | |
| Q8CDN6 | TXNL1 | thioredoxin-like 1 | Cytoplasm | enzyme | 1268457 | 1452794.25 | 1366810.875 | | | | | | |
| Q8CDN6 | TXNL1 | thioredoxin-like 1 | Cytoplasm | enzyme | 312679.094 | 388781.688 | 353032.125 | Q8CDN6 | TXNL1 | 3715848 | 3280659 | 669662 | 24 |
| Q9JLT4 | TXNRD2 | thioredoxin reductase 2 | Cytoplasm | enzyme | 1.12E+07 | 1.82E+07 | 1.50E+07 | | | | | | |
| Q9JLT4 | TXNRD2 | thioredoxin reductase 2 | Cytoplasm | enzyme | 4422675.5 | 5070146.5 | 4625205.5 | | | | | | |
| Q9JLT4 | TXNRD2 | thioredoxin reductase 2 | Cytoplasm | enzyme | 819227 | 1027123.188 | 511164.969 | Q9JLT4 | TXNRD2 | 5467301 | 6500765 | 2166922 | 9 |
| Q02053 | UBA1 | ubiquitin-like modifier activating enzyme 1 | Cytoplasm | enzyme | 6327114.5 | 6374457.5 | 4896811 | | | | | | |
| Q02053 | UBA1 | ubiquitin-like modifier activating enzyme 1 | Cytoplasm | enzyme | 4734427.5 | 3887316.25 | 3084539 | | | | | | |
| Q02053 | UBA1 | ubiquitin-like modifier activating enzyme 1 | Cytoplasm | enzyme | 6033888.5 | 4812162 | 3478531.5 | | | | | | |
| Q02053 | UBA1 | ubiquitin-like modifier activating enzyme 1 | Cytoplasm | enzyme | 5249073.5 | 3641517.5 | 2412095.5 | | | | | | |

| | | | | | | | | | | | | | |
|---|---|---|---|---|---|---|---|---|---|---|---|---|---|
| Q02053 | UBA1 | ubiquitin-like modifier activating enzyme 1 | Cytoplasm | enzyme | 3761324.75 | 3162451.75 | 2438413.25 | | | | | | |
| Q02053 | UBA1 | ubiquitin-like modifier activating enzyme 1 | Cytoplasm | enzyme | 4116138.5 | 2757653 | 2316526.25 | | | | | | |
| Q02053 | UBA1 | ubiquitin-like modifier activating enzyme 1 | Cytoplasm | enzyme | 3192497.25 | 2575606.25 | 1628971.625 | | | | | | |
| Q02053 | UBA1 | ubiquitin-like modifier activating enzyme 1 | Cytoplasm | enzyme | 3714564.5 | 2905212.75 | 1913089.875 | | | | | | |
| Q02053 | UBA1 | ubiquitin-like modifier activating enzyme 1 | Cytoplasm | enzyme | 3000460.75 | 1276591.5 | 1128939.5 | | | | | | |
| Q02053 | UBA1 | ubiquitin-like modifier activating enzyme 1 | Cytoplasm | enzyme | 3727111.75 | 2583380.75 | 1592180.75 | | | | | | |
| Q02053 | UBA1 | ubiquitin-like modifier activating enzyme 1 | Cytoplasm | enzyme | 2058163.875 | 1720980.25 | 2622493.25 | | | | | | |
| Q02053 | UBA1 | ubiquitin-like modifier activating enzyme 1 | Cytoplasm | enzyme | 2103269.75 | 1914142.75 | 1363788.375 | | | | | | |
| Q02053 | UBA1 | ubiquitin-like modifier activating enzyme 1 | Cytoplasm | enzyme | 1003532.125 | 581969.25 | 393134.125 | | | | | | |
| Q02053 | UBA1 | ubiquitin-like modifier activating enzyme 1 | Cytoplasm | enzyme | 395455.062 | 426960.406 | 297382.344 | Q02053 | UBA1 | 3529787 | 1615630 | 249297 | 42 |
| P62984 | Uba52 | ubiquitin A-52 residue ribosomal protein fusion product 1 | Cytoplasm | other | 8.41E+08 | 8.53E+08 | 9.14E+08 | | | | | | |
| P62984 | Uba52 | ubiquitin A-52 residue ribosomal protein fusion product 1 | Cytoplasm | other | 3.16E+08 | 3.19E+08 | 2.72E+08 | | | | | | |
| P62984 | Uba52 | ubiquitin A-52 residue ribosomal protein fusion product 1 | Cytoplasm | other | 1.74E+08 | 1.52E+08 | 1.67E+08 | | | | | | |
| P62984 | Uba52 | ubiquitin A-52 residue ribosomal protein fusion product 1 | Cytoplasm | other | 1.77E+08 | 1.49E+08 | 1.43E+08 | | | | | | |
| P62984 | Uba52 | ubiquitin A-52 residue ribosomal protein fusion product 1 | Cytoplasm | other | 3.03E+07 | 1.84E+07 | 4.77E+07 | | | | | | |
| P62984 | Uba52 | ubiquitin A-52 residue ribosomal protein fusion product 1 | Cytoplasm | other | 4199671.5 | 2495848 | 7002378 | | | | | | |
| P62984 | Uba52 | ubiquitin A-52 residue ribosomal protein fusion product 1 | Cytoplasm | other | 2.55E+07 | 2.70E+07 | 2.70E+07 | | | | | | |
| P62984 | Uba52 | ubiquitin A-52 residue ribosomal protein fusion product 1 | Cytoplasm | other | 15972.815 | 872144.938 | 7929315 | P62984 | Uba52 | 196053206 | 279370318 | 57026227 | 24 |
| Q8VDI7 | UBAC1 | UBA domain containing 1 | Nucleus | other | 9079480 | 1.21E+07 | 6261927 | | | | | | |
| Q8VDI7 | UBAC1 | UBA domain containing 1 | Nucleus | other | 7533490.5 | 6266337.5 | 4601225 | | | | | | |
| Q8VDI7 | UBAC1 | UBA domain containing 1 | Nucleus | other | 4974555 | 4576323 | 4947801.5 | | | | | | |
| Q8VDI7 | UBAC1 | UBA domain containing 1 | Nucleus | other | 4662053 | 4356269 | 2107710.75 | | | | | | |
| Q8VDI7 | UBAC1 | UBA domain containing 1 | Nucleus | other | 3784317 | 2932161.75 | 2171033.5 | | | | | | |
| Q8VDI7 | UBAC1 | UBA domain containing 1 | Nucleus | other | 3378322 | 2219739.25 | 1913895.875 | | | | | | |
| Q8VDI7 | UBAC1 | UBA domain containing 1 | Nucleus | other | 3256731.5 | 1816038.375 | 1595511.25 | | | | | | |
| Q8VDI7 | UBAC1 | UBA domain containing 1 | Nucleus | other | 1945548.5 | 1237922.125 | 513772.5 | | | | | | |
| Q8VDI7 | UBAC1 | UBA domain containing 1 | Nucleus | other | 1456851.5 | 1212840.25 | 644897.312 | | | | | | |
| Q8VDI7 | UBAC1 | UBA domain containing 1 | Nucleus | other | 2188868.5 | 2406391.25 | 2255908.5 | Q8VDI7 | UBAC1 | 4226022 | 2609035 | 476342 | 30 |
| P68037 | UBE2L3 | ubiquitin-conjugating enzyme E2L 3 | Nucleus | enzyme | 6700977.5 | 7355753 | 7081256.5 | P68037 | UBE2L3 | 6700978 | 328809 | 189838 | 3 |
| Q9QZU9 | UBE2L6 | ubiquitin-conjugating enzyme E2L 6 | Cytoplasm | enzyme | 2.83E+07 | 2.57E+07 | 1.75E+07 | | | | | | |
| Q9QZU9 | UBE2L6 | ubiquitin-conjugating enzyme E2L 6 | Cytoplasm | enzyme | 5745099.5 | 5691219.5 | 4346074.5 | | | | | | |
| Q9QZU9 | UBE2L6 | ubiquitin-conjugating enzyme E2L 6 | Cytoplasm | enzyme | 3979761.5 | 5752085 | 2308361.25 | | | | | | |
| Q9QZU9 | UBE2L6 | ubiquitin-conjugating enzyme E2L 6 | Cytoplasm | enzyme | 3078279.25 | 2228839.75 | 2776863.25 | Q9QZU9 | UBE2L6 | 10273285 | 9366630 | 2703913 | 12 |
| Q6ZPJ3 | UBE2O | ubiquitin-conjugating enzyme E2O | Nucleus | enzyme | 3.71E+07 | 3.57E+07 | 1.82E+07 | | | | | | |
| Q6ZPJ3 | UBE2O | ubiquitin-conjugating enzyme E2O | Nucleus | enzyme | 2.96E+07 | 2.99E+07 | 2.28E+07 | | | | | | |
| Q6ZPJ3 | UBE2O | ubiquitin-conjugating enzyme E2O | Nucleus | enzyme | 3.03E+07 | 2.12E+07 | 1.83E+07 | | | | | | |
| Q6ZPJ3 | UBE2O | ubiquitin-conjugating enzyme E2O | Nucleus | enzyme | 2.68E+07 | 2.88E+07 | 2.08E+07 | | | | | | |
| Q6ZPJ3 | UBE2O | ubiquitin-conjugating enzyme E2O | Nucleus | enzyme | 3.24E+07 | 2.12E+07 | 1.41E+07 | | | | | | |
| Q6ZPJ3 | UBE2O | ubiquitin-conjugating enzyme E2O | Nucleus | enzyme | 2.57E+07 | 1.93E+07 | 1.15E+07 | | | | | | |
| Q6ZPJ3 | UBE2O | ubiquitin-conjugating enzyme E2O | Nucleus | enzyme | 2.01E+07 | 1.65E+07 | 1.43E+07 | | | | | | |
| Q6ZPJ3 | UBE2O | ubiquitin-conjugating enzyme E2O | Nucleus | enzyme | 2.12E+07 | 1.67E+07 | 1.13E+07 | | | | | | |
| Q6ZPJ3 | UBE2O | ubiquitin-conjugating enzyme E2O | Nucleus | enzyme | 1.63E+07 | 1.70E+07 | 1.67E+07 | | | | | | |
| Q6ZPJ3 | UBE2O | ubiquitin-conjugating enzyme E2O | Nucleus | enzyme | 1.47E+07 | 1.03E+07 | 7541181 | | | | | | |
| Q6ZPJ3 | UBE2O | ubiquitin-conjugating enzyme E2O | Nucleus | enzyme | 1.25E+07 | 8840372 | 6387736.5 | | | | | | |
| Q6ZPJ3 | UBE2O | ubiquitin-conjugating enzyme E2O | Nucleus | enzyme | 1.09E+07 | 1.01E+07 | 7938701.5 | | | | | | |
| Q6ZPJ3 | UBE2O | ubiquitin-conjugating enzyme E2O | Nucleus | enzyme | 1.22E+07 | 9709084 | 7245047 | | | | | | |
| Q6ZPJ3 | UBE2O | ubiquitin-conjugating enzyme E2O | Nucleus | enzyme | 1.13E+07 | 1.13E+07 | 9839645 | | | | | | |
| Q6ZPJ3 | UBE2O | ubiquitin-conjugating enzyme E2O | Nucleus | enzyme | 1.13E+07 | 9708867 | 6421845 | | | | | | |
| Q6ZPJ3 | UBE2O | ubiquitin-conjugating enzyme E2O | Nucleus | enzyme | 1.20E+07 | 1.10E+07 | 7330654 | | | | | | |
| Q6ZPJ3 | UBE2O | ubiquitin-conjugating enzyme E2O | Nucleus | enzyme | 1.18E+07 | 9758490 | 6859849.5 | | | | | | |
| Q6ZPJ3 | UBE2O | ubiquitin-conjugating enzyme E2O | Nucleus | enzyme | 1.32E+07 | 1.13E+07 | 7410510 | | | | | | |
| Q6ZPJ3 | UBE2O | ubiquitin-conjugating enzyme E2O | Nucleus | enzyme | 8616738 | 7993185 | 6022344.5 | | | | | | |
| Q6ZPJ3 | UBE2O | ubiquitin-conjugating enzyme E2O | Nucleus | enzyme | 1.03E+07 | 6814492.5 | 5281155 | | | | | | |
| Q6ZPJ3 | UBE2O | ubiquitin-conjugating enzyme E2O | Nucleus | enzyme | 9247773 | 6775457.5 | 5501588.5 | | | | | | |
| Q6ZPJ3 | UBE2O | ubiquitin-conjugating enzyme E2O | Nucleus | enzyme | 6301142.5 | 3510695.75 | 1951752.75 | | | | | | |
| Q6ZPJ3 | UBE2O | ubiquitin-conjugating enzyme E2O | Nucleus | enzyme | 5472832.5 | 4768755.5 | 3268589.75 | | | | | | |
| Q6ZPJ3 | UBE2O | ubiquitin-conjugating enzyme E2O | Nucleus | enzyme | 6357128.5 | 6528609 | 4106448.5 | | | | | | |
| Q6ZPJ3 | UBE2O | ubiquitin-conjugating enzyme E2O | Nucleus | enzyme | 4626660 | 3565892.5 | 2614910.5 | | | | | | |
| Q6ZPJ3 | UBE2O | ubiquitin-conjugating enzyme E2O | Nucleus | enzyme | 4273900 | 3360340.25 | 2921267.5 | | | | | | |
| Q6ZPJ3 | UBE2O | ubiquitin-conjugating enzyme E2O | Nucleus | enzyme | 4768883 | 1568454.75 | 2206524 | | | | | | |
| Q6ZPJ3 | UBE2O | ubiquitin-conjugating enzyme E2O | Nucleus | enzyme | 4706907 | 4256776.5 | 3658087.75 | | | | | | |
| Q6ZPJ3 | UBE2O | ubiquitin-conjugating enzyme E2O | Nucleus | enzyme | 3035041.75 | 1796538.125 | 1179601.125 | | | | | | |
| Q6ZPJ3 | UBE2O | ubiquitin-conjugating enzyme E2O | Nucleus | enzyme | 2478132 | 81940.438 | 105258.648 | | | | | | |
| Q6ZPJ3 | UBE2O | ubiquitin-conjugating enzyme E2O | Nucleus | enzyme | 2847360 | 2702228.5 | 1574960.125 | | | | | | |
| Q6ZPJ3 | UBE2O | ubiquitin-conjugating enzyme E2O | Nucleus | enzyme | 2629376.75 | 2795367.5 | 1985541 | | | | | | |
| Q6ZPJ3 | UBE2O | ubiquitin-conjugating enzyme E2O | Nucleus | enzyme | 1334502 | 1239928 | 1084880.625 | | | | | | |
| Q6ZPJ3 | UBE2O | ubiquitin-conjugating enzyme E2O | Nucleus | enzyme | 732535.938 | 1228523.75 | 385182.938 | Q6ZPJ3 | UBE2O | 12556144 | 8608502 | 852369 | 102 |
| Q6ZWZ2 | UBE2R2 | ubiquitin-conjugating enzyme E2R 2 | Cytoplasm | enzyme | 1805003.875 | 1669814.625 | 652352.812 | Q6ZWZ2 | UBE2R2 | 1805004 | 630094 | 363785 | 3 |
| A2AN08 | UBR4 | ubiquitin protein ligase E3 component n-recognin 4 | Nucleus | enzyme | 1.20E+07 | 9967271 | 8713066 | | | | | | |
| A2AN08 | UBR4 | ubiquitin protein ligase E3 component n-recognin 4 | Nucleus | enzyme | 6531353.5 | 7356283 | 5952268 | | | | | | |
| A2AN08 | UBR4 | ubiquitin protein ligase E3 component n-recognin 4 | Nucleus | enzyme | 4263851.5 | 2417202 | 1473923 | | | | | | |
| A2AN08 | UBR4 | ubiquitin protein ligase E3 component n-recognin 4 | Nucleus | enzyme | 6804945 | 6739777 | 5888119.5 | | | | | | |
| A2AN08 | UBR4 | ubiquitin protein ligase E3 component n-recognin 4 | Nucleus | enzyme | 6491409 | 4533489 | 4584969.5 | | | | | | |
| A2AN08 | UBR4 | ubiquitin protein ligase E3 component n-recognin 4 | Nucleus | enzyme | 7020691 | 5681252.5 | 5805333.5 | | | | | | |
| A2AN08 | UBR4 | ubiquitin protein ligase E3 component n-recognin 4 | Nucleus | enzyme | 5858546 | 6013203 | 4419330 | | | | | | |
| A2AN08 | UBR4 | ubiquitin protein ligase E3 component n-recognin 4 | Nucleus | enzyme | 6640843 | 4385236.5 | 4823037 | | | | | | |
| A2AN08 | UBR4 | ubiquitin protein ligase E3 component n-recognin 4 | Nucleus | enzyme | 6261455 | 4144455 | 3559506 | | | | | | |
| A2AN08 | UBR4 | ubiquitin protein ligase E3 component n-recognin 4 | Nucleus | enzyme | 5611754.5 | 4807163.5 | 3343874 | | | | | | |
| A2AN08 | UBR4 | ubiquitin protein ligase E3 component n-recognin 4 | Nucleus | enzyme | 5345648 | 3676770 | 2958845 | | | | | | |
| A2AN08 | UBR4 | ubiquitin protein ligase E3 component n-recognin 4 | Nucleus | enzyme | 5652005.5 | 4476887.5 | 4458179 | | | | | | |
| A2AN08 | UBR4 | ubiquitin protein ligase E3 component n-recognin 4 | Nucleus | enzyme | 6040982.5 | 5062558.5 | 5295782.5 | | | | | | |
| A2AN08 | UBR4 | ubiquitin protein ligase E3 component n-recognin 4 | Nucleus | enzyme | 4106687.75 | 2829745.25 | 3182629.5 | | | | | | |
| A2AN08 | UBR4 | ubiquitin protein ligase E3 component n-recognin 4 | Nucleus | enzyme | 5313296.5 | 3846612.5 | 3032575.75 | | | | | | |
| A2AN08 | UBR4 | ubiquitin protein ligase E3 component n-recognin 4 | Nucleus | enzyme | 5536612.5 | 4141189.25 | 3936675.5 | | | | | | |
| A2AN08 | UBR4 | ubiquitin protein ligase E3 component n-recognin 4 | Nucleus | enzyme | 5523650 | 4592162 | 4616542.5 | | | | | | |
| A2AN08 | UBR4 | ubiquitin protein ligase E3 component n-recognin 4 | Nucleus | enzyme | 4522744 | 4182208.5 | 3176075 | | | | | | |
| A2AN08 | UBR4 | ubiquitin protein ligase E3 component n-recognin 4 | Nucleus | enzyme | 5354635.5 | 4421407.5 | 3228164.25 | | | | | | |
| A2AN08 | UBR4 | ubiquitin protein ligase E3 component n-recognin 4 | Nucleus | enzyme | 3886817.25 | 3516393.75 | 4100300 | | | | | | |
| A2AN08 | UBR4 | ubiquitin protein ligase E3 component n-recognin 4 | Nucleus | enzyme | 3989253.5 | 4146841 | 4693271.5 | | | | | | |
| A2AN08 | UBR4 | ubiquitin protein ligase E3 component n-recognin 4 | Nucleus | enzyme | 5705680.5 | 4560610 | 4004479.75 | | | | | | |
| A2AN08 | UBR4 | ubiquitin protein ligase E3 component n-recognin 4 | Nucleus | enzyme | 4116439 | 4620624 | 5417370 | | | | | | |
| A2AN08 | UBR4 | ubiquitin protein ligase E3 component n-recognin 4 | Nucleus | enzyme | 3959238.75 | 4296206.5 | 5699542.5 | | | | | | |
| A2AN08 | UBR4 | ubiquitin protein ligase E3 component n-recognin 4 | Nucleus | enzyme | 4983523 | 3437734.25 | 3051076.25 | | | | | | |
| A2AN08 | UBR4 | ubiquitin protein ligase E3 component n-recognin 4 | Nucleus | enzyme | 3693357.5 | 2617223.5 | 2631257.75 | | | | | | |
| A2AN08 | UBR4 | ubiquitin protein ligase E3 component n-recognin 4 | Nucleus | enzyme | 3329477.5 | 3579234.25 | 3076225.5 | | | | | | |
| A2AN08 | UBR4 | ubiquitin protein ligase E3 component n-recognin 4 | Nucleus | enzyme | 4094559 | 3172578.25 | 2547324.75 | | | | | | |
| A2AN08 | UBR4 | ubiquitin protein ligase E3 component n-recognin 4 | Nucleus | enzyme | 4839990.5 | 4121142.25 | 3189477.25 | | | | | | |
| A2AN08 | UBR4 | ubiquitin protein ligase E3 component n-recognin 4 | Nucleus | enzyme | 3142378 | 2395111 | 2969485.25 | | | | | | |
| A2AN08 | UBR4 | ubiquitin protein ligase E3 component n-recognin 4 | Nucleus | enzyme | 4655337.5 | 3257337.5 | 3040836.25 | | | | | | |
| A2AN08 | UBR4 | ubiquitin protein ligase E3 component n-recognin 4 | Nucleus | enzyme | 3435398.75 | 4275699.5 | 3163448.25 | | | | | | |
| A2AN08 | UBR4 | ubiquitin protein ligase E3 component n-recognin 4 | Nucleus | enzyme | 4606510.5 | 4289068 | 3941335.5 | | | | | | |
| A2AN08 | UBR4 | ubiquitin protein ligase E3 component n-recognin 4 | Nucleus | enzyme | 4214099.5 | 3332678 | 2517628 | | | | | | |
| A2AN08 | UBR4 | ubiquitin protein ligase E3 component n-recognin 4 | Nucleus | enzyme | 4127558.25 | 2155478.5 | 2586600.5 | | | | | | |
| A2AN08 | UBR4 | ubiquitin protein ligase E3 component n-recognin 4 | Nucleus | enzyme | 4410103 | 3598601.25 | 3347192.25 | | | | | | |
| A2AN08 | UBR4 | ubiquitin protein ligase E3 component n-recognin 4 | Nucleus | enzyme | 5016972 | 2063871.625 | 2083533 | | | | | | |
| A2AN08 | UBR4 | ubiquitin protein ligase E3 component n-recognin 4 | Nucleus | enzyme | 4323330 | 3071897 | 2901426.5 | | | | | | |
| A2AN08 | UBR4 | ubiquitin protein ligase E3 component n-recognin 4 | Nucleus | enzyme | 4543221 | 2708434 | 2827440 | | | | | | |
| A2AN08 | UBR4 | ubiquitin protein ligase E3 component n-recognin 4 | Nucleus | enzyme | 4091033.75 | 2306486.75 | 2012111.125 | | | | | | |
| A2AN08 | UBR4 | ubiquitin protein ligase E3 component n-recognin 4 | Nucleus | enzyme | 3486863.75 | 2769521 | 2214288.25 | | | | | | |

| | | | | | | | | | | | | | |
|---|---|---|---|---|---|---|---|---|---|---|---|---|---|
| A2AN08 | UBR4 | ubiquitin protein ligase E3 component n-recognin 4 | Nucleus | enzyme | 1811776.375 | 1296258.75 | 3015175.75 | | | | | | |
| A2AN08 | UBR4 | ubiquitin protein ligase E3 component n-recognin 4 | Nucleus | enzyme | 3102524 | 2916644.5 | 2517505.75 | | | | | | |
| A2AN08 | UBR4 | ubiquitin protein ligase E3 component n-recognin 4 | Nucleus | enzyme | 2589410.5 | 2322619 | 3483475.75 | | | | | | |
| A2AN08 | UBR4 | ubiquitin protein ligase E3 component n-recognin 4 | Nucleus | enzyme | 3071492.25 | 2613182.25 | 2033567 | | | | | | |
| A2AN08 | UBR4 | ubiquitin protein ligase E3 component n-recognin 4 | Nucleus | enzyme | 2604652.75 | 2180415.25 | 1656176 | | | | | | |
| A2AN08 | UBR4 | ubiquitin protein ligase E3 component n-recognin 4 | Nucleus | enzyme | 3178462 | 2431263.5 | 2209158.75 | | | | | | |
| A2AN08 | UBR4 | ubiquitin protein ligase E3 component n-recognin 4 | Nucleus | enzyme | 1997000.875 | 1976027.375 | 1570978.125 | | | | | | |
| A2AN08 | UBR4 | ubiquitin protein ligase E3 component n-recognin 4 | Nucleus | enzyme | 3275366.25 | 2667098.25 | 2361862.25 | | | | | | |
| A2AN08 | UBR4 | ubiquitin protein ligase E3 component n-recognin 4 | Nucleus | enzyme | 2195596.5 | 995543.312 | 584371.562 | | | | | | |
| A2AN08 | UBR4 | ubiquitin protein ligase E3 component n-recognin 4 | Nucleus | enzyme | 2169124.75 | 3204091.5 | 2520255.75 | | | | | | |
| A2AN08 | UBR4 | ubiquitin protein ligase E3 component n-recognin 4 | Nucleus | enzyme | 1383614.25 | 1671418.375 | 1206905.875 | | | | | | |
| A2AN08 | UBR4 | ubiquitin protein ligase E3 component n-recognin 4 | Nucleus | enzyme | 1340249.75 | 1175924.625 | 867295.812 | A2AN08 | UBR4 | 4456821 | 1673740 | 132736 | 159 |
| Q9WUP7 | UCHL5 | ubiquitin carboxyl-terminal hydrolase L5 | Cytoplasm | peptidase | 5971242 | 8401955 | 1.08E+07 | | | | | | |
| Q9WUP7 | UCHL5 | ubiquitin carboxyl-terminal hydrolase L5 | Cytoplasm | peptidase | 7031336.5 | 7600229 | 6392614 | | | | | | |
| Q9WUP7 | UCHL5 | ubiquitin carboxyl-terminal hydrolase L5 | Cytoplasm | peptidase | 2002879.75 | 2679412.25 | 1892273.375 | | | | | | |
| Q9WUP7 | UCHL5 | ubiquitin carboxyl-terminal hydrolase L5 | Cytoplasm | peptidase | 718524 | 1479768.625 | 4533530 | Q9WUP7 | UCHL5 | 3930996 | 3211678 | 927132 | 12 |
| Q6P5E4 | UGGT1 | UDP-glucose glycoprotein glucosyltransferase 1 | Cytoplasm | enzyme | 1.19E+07 | 1.11E+07 | 1.03E+07 | | | | | | |
| Q6P5E4 | UGGT1 | UDP-glucose glycoprotein glucosyltransferase 1 | Cytoplasm | enzyme | 4386680.5 | 3915195.75 | 3780061.25 | | | | | | |
| Q6P5E4 | UGGT1 | UDP-glucose glycoprotein glucosyltransferase 1 | Cytoplasm | enzyme | 2738275 | 2696911.25 | 1741414.875 | | | | | | |
| Q6P5E4 | UGGT1 | UDP-glucose glycoprotein glucosyltransferase 1 | Cytoplasm | enzyme | 4633089 | 5364808.5 | 4788869.5 | | | | | | |
| Q6P5E4 | UGGT1 | UDP-glucose glycoprotein glucosyltransferase 1 | Cytoplasm | enzyme | 3228674.25 | 2503319.5 | 2176955 | | | | | | |
| Q6P5E4 | UGGT1 | UDP-glucose glycoprotein glucosyltransferase 1 | Cytoplasm | enzyme | 2579022.25 | 2485554.5 | 2539282 | | | | | | |
| Q6P5E4 | UGGT1 | UDP-glucose glycoprotein glucosyltransferase 1 | Cytoplasm | enzyme | 3034933.25 | 3415055.75 | 3719769.5 | | | | | | |
| Q6P5E4 | UGGT1 | UDP-glucose glycoprotein glucosyltransferase 1 | Cytoplasm | enzyme | 1489482.625 | 2017522.75 | 1582488.625 | Q6P5E4 | UGGT1 | 4245020 | 2902599 | 592491 | 24 |
| Q8R1I1 | UQCR10 | ubiquinol-cytochrome c reductase, complex III subunit X | Cytoplasm | enzyme | 1642158.25 | 2890762.25 | 3189133 | Q8R1I1 | UQCR10 | 1642158 | 820688 | 473824 | 3 |
| Q9CZ13 | UQCRC1 | ubiquinol-cytochrome c reductase core protein I | Cytoplasm | enzyme | 1.07E+07 | 1.66E+07 | 1.95E+07 | | | | | | |
| Q9CZ13 | UQCRC1 | ubiquinol-cytochrome c reductase core protein I | Cytoplasm | enzyme | 4096896.75 | 7500300 | 8166991 | | | | | | |
| Q9CZ13 | UQCRC1 | ubiquinol-cytochrome c reductase core protein I | Cytoplasm | enzyme | 4974144.5 | 8807290 | 8650351 | | | | | | |
| Q9CZ13 | UQCRC1 | ubiquinol-cytochrome c reductase core protein I | Cytoplasm | enzyme | 3281294.75 | 4498728 | 4868226 | | | | | | |
| Q9CZ13 | UQCRC1 | ubiquinol-cytochrome c reductase core protein I | Cytoplasm | enzyme | 5231666.5 | 9436994 | 9173668 | Q9CZ13 | UQCRC1 | 5662801 | 4563813 | 1178371 | 15 |
| Q9DB77 | UQCRC2 | ubiquinol-cytochrome c reductase core protein II | Cytoplasm | enzyme | 1.01E+07 | 1.79E+07 | 2.31E+07 | | | | | | |
| Q9DB77 | UQCRC2 | ubiquinol-cytochrome c reductase core protein II | Cytoplasm | enzyme | 8032060.5 | 1.48E+07 | 1.60E+07 | | | | | | |
| Q9DB77 | UQCRC2 | ubiquinol-cytochrome c reductase core protein II | Cytoplasm | enzyme | 7440960 | 1.29E+07 | 1.71E+07 | | | | | | |
| Q9DB77 | UQCRC2 | ubiquinol-cytochrome c reductase core protein II | Cytoplasm | enzyme | 5403987.5 | 9086147 | 1.10E+07 | | | | | | |
| Q9DB77 | UQCRC2 | ubiquinol-cytochrome c reductase core protein II | Cytoplasm | enzyme | 4096648.5 | 6779622 | 9501308 | | | | | | |
| Q9DB77 | UQCRC2 | ubiquinol-cytochrome c reductase core protein II | Cytoplasm | enzyme | 3952046.5 | 7803754 | 8397880 | | | | | | |
| Q9DB77 | UQCRC2 | ubiquinol-cytochrome c reductase core protein II | Cytoplasm | enzyme | 2391142.5 | 4567579 | 4405251 | | | | | | |
| Q9DB77 | UQCRC2 | ubiquinol-cytochrome c reductase core protein II | Cytoplasm | enzyme | 3930473.5 | 5488123.5 | 6027791.5 | | | | | | |
| Q9DB77 | UQCRC2 | ubiquinol-cytochrome c reductase core protein II | Cytoplasm | enzyme | 3466157 | 5213282.5 | 7525735.5 | | | | | | |
| Q9DB77 | UQCRC2 | ubiquinol-cytochrome c reductase core protein II | Cytoplasm | enzyme | 2519199.5 | 3962402.75 | 4272984.5 | | | | | | |
| Q9DB77 | UQCRC2 | ubiquinol-cytochrome c reductase core protein II | Cytoplasm | enzyme | 1315043.125 | 2690953 | 3957519.75 | | | | | | |
| Q9DB77 | UQCRC2 | ubiquinol-cytochrome c reductase core protein II | Cytoplasm | enzyme | 803934.812 | 3440601.75 | 4245850 | Q9DB77 | UQCRC2 | 4454304 | 5161817 | 860303 | 36 |
| P99028 | UQCRHL | ubiquinol-cytochrome c reductase hinge protein like | Cytoplasm | other | 2.09E+07 | 3.72E+07 | 3.52E+07 | | | | | | |
| P99028 | UQCRHL | ubiquinol-cytochrome c reductase hinge protein like | Cytoplasm | other | 6793394.5 | 1.19E+07 | 1.36E+07 | P99028 | UQCRHL | 13821697 | 12669382 | 5172253 | 6 |
| P70697 | UROD | uroporphyrinogen decarboxylase | Cytoplasm | enzyme | 2209394.25 | 2507923 | 1468558.875 | P70697 | UROD | 2209394 | 535138 | 308962 | 3 |
| Q78IK2 | USMG5 | up-regulated during skeletal muscle growth 5 homolog (mouse) | Cytoplasm | other | 6080558.5 | 9174207 | 1.30E+07 | | | | | | |
| Q78IK2 | USMG5 | up-regulated during skeletal muscle growth 5 homolog (mouse) | Cytoplasm | other | 3838624.25 | 6615720 | 7724631.5 | Q78IK2 | USMG5 | 4959591 | 3122600 | 1274796 | 6 |
| Q9JMA1 | USP14 | ubiquitin specific peptidase 14 (tRNA-guanine transglycosylase) | Cytoplasm | peptidase | 4802741 | 4544819 | 4845171 | | | | | | |
| Q9JMA1 | USP14 | ubiquitin specific peptidase 14 (tRNA-guanine transglycosylase) | Cytoplasm | peptidase | 2768302.25 | 2624939.25 | 2336164.5 | Q9JMA1 | USP14 | 3785522 | 1192658 | 486901 | 6 |
| Q8R5H1 | USP15 | ubiquitin specific peptidase 15 | Cytoplasm | peptidase | 3225258.25 | 2216846.75 | 2577661 | | | | | | |
| Q8R5H1 | USP15 | ubiquitin specific peptidase 15 | Cytoplasm | peptidase | 8593029 | 6828427 | 5016050.5 | | | | | | |
| Q8R5H1 | USP15 | ubiquitin specific peptidase 15 | Cytoplasm | peptidase | 5227617 | 2977382.5 | 2646355 | | | | | | |
| Q8R5H1 | USP15 | ubiquitin specific peptidase 15 | Cytoplasm | peptidase | 6518883 | 4856832 | 3135125.25 | | | | | | |
| Q8R5H1 | USP15 | ubiquitin specific peptidase 15 | Cytoplasm | peptidase | 3980897.25 | 2272542 | 2337711 | | | | | | |
| Q8R5H1 | USP15 | ubiquitin specific peptidase 15 | Cytoplasm | peptidase | 5281122.5 | 3395933.5 | 2469873.5 | | | | | | |
| Q8R5H1 | USP15 | ubiquitin specific peptidase 15 | Cytoplasm | peptidase | 2703453 | 2353313.75 | 2131365 | | | | | | |
| Q8R5H1 | USP15 | ubiquitin specific peptidase 15 | Cytoplasm | peptidase | 2425061 | 2620638.25 | 1863571.125 | | | | | | |
| Q8R5H1 | USP15 | ubiquitin specific peptidase 15 | Cytoplasm | peptidase | 2887320.25 | 1798838.125 | 2005656.625 | | | | | | |
| Q8R5H1 | USP15 | ubiquitin specific peptidase 15 | Cytoplasm | peptidase | 2923309.75 | 2121958.75 | 1408643 | | | | | | |
| Q8R5H1 | USP15 | ubiquitin specific peptidase 15 | Cytoplasm | peptidase | 2673450.25 | 1817429 | 1218859.75 | | | | | | |
| Q8R5H1 | USP15 | ubiquitin specific peptidase 15 | Cytoplasm | peptidase | 2971783.75 | 2232269 | 1669977.375 | | | | | | |
| Q8R5H1 | USP15 | ubiquitin specific peptidase 15 | Cytoplasm | peptidase | 2258795 | 1513449.75 | 1702534.625 | | | | | | |
| Q8R5H1 | USP15 | ubiquitin specific peptidase 15 | Cytoplasm | peptidase | 2488574 | 1990648.25 | 1598802.75 | | | | | | |
| Q8R5H1 | USP15 | ubiquitin specific peptidase 15 | Cytoplasm | peptidase | 1023717.5 | 625135.125 | 496379.375 | | | | | | |
| Q8R5H1 | USP15 | ubiquitin specific peptidase 15 | Cytoplasm | peptidase | 1434108.375 | 1145181.875 | 640250.875 | Q8R5H1 | USP15 | 3538524 | 1638773 | 236537 | 48 |
| Q3UJD6 | USP19 | ubiquitin specific peptidase 19 | Cytoplasm | peptidase | 109835.578 | 104959.273 | 79363.609 | Q3UJD6 | USP19 | 109836 | 16368 | 9450 | 3 |
| Q5DU02 | USP22 | ubiquitin specific peptidase 22 | Nucleus | peptidase | 1.17E+09 | 1.23E+09 | 1.12E+09 | Q5DU02 | USP22 | 1167000000 | 53842363 | 31085902 | 3 |
| P56399 | USP5 | ubiquitin specific peptidase 5 (isopeptidase T) | Cytoplasm | peptidase | 9148607 | 6050045 | 4686489 | | | | | | |
| P56399 | USP5 | ubiquitin specific peptidase 5 (isopeptidase T) | Cytoplasm | peptidase | 5138648 | 4246103.5 | 3580821.5 | | | | | | |
| P56399 | USP5 | ubiquitin specific peptidase 5 (isopeptidase T) | Cytoplasm | peptidase | 4342874 | 3220790.75 | 2869562.25 | | | | | | |
| P56399 | USP5 | ubiquitin specific peptidase 5 (isopeptidase T) | Cytoplasm | peptidase | 4865072.5 | 3448779.25 | 2791977.75 | | | | | | |
| P56399 | USP5 | ubiquitin specific peptidase 5 (isopeptidase T) | Cytoplasm | peptidase | 4156449.5 | 3182564 | 2057765.5 | | | | | | |
| P56399 | USP5 | ubiquitin specific peptidase 5 (isopeptidase T) | Cytoplasm | peptidase | 4693370 | 3195310.25 | 2564649.25 | | | | | | |
| P56399 | USP5 | ubiquitin specific peptidase 5 (isopeptidase T) | Cytoplasm | peptidase | 5589600.5 | 2994808.75 | 3036374.75 | | | | | | |
| P56399 | USP5 | ubiquitin specific peptidase 5 (isopeptidase T) | Cytoplasm | peptidase | 3511476.5 | 2918629 | 2620102.75 | | | | | | |
| P56399 | USP5 | ubiquitin specific peptidase 5 (isopeptidase T) | Cytoplasm | peptidase | 4317254.5 | 2487173.5 | 1432803.375 | | | | | | |
| P56399 | USP5 | ubiquitin specific peptidase 5 (isopeptidase T) | Cytoplasm | peptidase | 1408999.25 | 717465.562 | 108930.328 | | | | | | |
| P56399 | USP5 | ubiquitin specific peptidase 5 (isopeptidase T) | Cytoplasm | peptidase | 1455275.5 | 971737.625 | 852223.312 | | | | | | |
| P56399 | USP5 | ubiquitin specific peptidase 5 (isopeptidase T) | Cytoplasm | peptidase | 1056247 | 1365659.5 | 850463.062 | | | | | | |
| P56399 | USP5 | ubiquitin specific peptidase 5 (isopeptidase T) | Cytoplasm | peptidase | 946887.188 | 1568705 | 1292651.625 | P56399 | USP5 | 3894674 | 1823510 | 291995 | 39 |
| Q6A4J8 | USP7 | ubiquitin specific peptidase 7 (herpes virus-associated) | Nucleus | peptidase | 861159 | 625269.062 | 528667.562 | Q6A4J8 | USP7 | 861159 | 171039 | 98750 | 3 |
| P70398 | USP9X | ubiquitin specific peptidase 9, X-linked | Plasma Membrane | peptidase | 4167993.25 | 2797745.25 | 3603463 | | | | | | |
| P70398 | USP9X | ubiquitin specific peptidase 9, X-linked | Plasma Membrane | peptidase | 2864454.5 | 2403617.25 | 2245713.5 | | | | | | |
| P70398 | USP9X | ubiquitin specific peptidase 9, X-linked | Plasma Membrane | peptidase | 2192364.5 | 1409129.875 | 1645944.875 | | | | | | |
| P70398 | USP9X | ubiquitin specific peptidase 9, X-linked | Plasma Membrane | peptidase | 1801168.125 | 1289231.875 | 883719.562 | P70398 | USP9X | 2756495 | 966274 | 278939 | 12 |
| Q80WQ2 | VAC14 | Vac14 homolog (S. cerevisiae) | Cytoplasm | other | 3183960.25 | 2352652 | 2227787 | Q80WQ2 | VAC14 | 3183960 | 519765 | 300086 | 3 |
| P63024 | VAMP3 | vesicle-associated membrane protein 3 | Plasma Membrane | other | 9.41E+07 | 8.86E+07 | 9.82E+07 | | | | | | |
| P63024 | VAMP3 | vesicle-associated membrane protein 3 | Plasma Membrane | other | 1.26E+07 | 1.45E+07 | 1.28E+07 | | | | | | |
| P63024 | VAMP3 | vesicle-associated membrane protein 3 | Plasma Membrane | other | 6337352 | 7887335 | 1.01E+07 | | | | | | |
| P63024 | VAMP3 | vesicle-associated membrane protein 3 | Plasma Membrane | other | 7716146 | 8385475.5 | 6993261 | | | | | | |
| P63024 | VAMP3 | vesicle-associated membrane protein 3 | Plasma Membrane | other | 2808651.25 | 5768093 | 2245587 | P63024 | VAMP3 | 24728430 | 35593375 | 9190170 | 15 |
| Q9Z2P8 | VAMP5 | vesicle-associated membrane protein 5 | Plasma Membrane | transporter | 2.33E+07 | 2.67E+07 | 2.28E+07 | | | | | | |
| Q9Z2P8 | VAMP5 | vesicle-associated membrane protein 5 | Plasma Membrane | transporter | 1.03E+07 | 1.93E+07 | 1.84E+07 | | | | | | |
| Q9Z2P8 | VAMP5 | vesicle-associated membrane protein 5 | Plasma Membrane | transporter | 1.03E+07 | 1.47E+07 | 1.32E+07 | Q9Z2P8 | VAMP5 | 14600000 | 5929916 | 1976639 | 9 |
| O70404 | VAMP8 | vesicle-associated membrane protein 8 | Plasma Membrane | transporter | 2.05E+07 | 2.82E+07 | 2.51E+07 | | | | | | |
| O70404 | VAMP8 | vesicle-associated membrane protein 8 | Plasma Membrane | transporter | 1.28E+07 | 1.87E+07 | 2.07E+07 | | | | | | |
| O70404 | VAMP8 | vesicle-associated membrane protein 8 | Plasma Membrane | transporter | 710237.562 | 698025.438 | 3288795.5 | O70404 | VAMP8 | 11326746 | 10608391 | 3536130 | 9 |
| Q9WV55 | VAPA | VAMP (vesicle-associated membrane protein)-associated protein A, 33kDa | Plasma Membrane | other | 5.39E+07 | 5.19E+07 | 4.36E+07 | | | | | | |
| Q9WV55 | VAPA | VAMP (vesicle-associated membrane protein)-associated protein A, 33kDa | Plasma Membrane | other | 3.58E+07 | 3.42E+07 | 3.28E+07 | | | | | | |
| Q9WV55 | VAPA | VAMP (vesicle-associated membrane protein)-associated protein A, 33kDa | Plasma Membrane | other | 2.24E+07 | 2.04E+07 | 2.12E+07 | | | | | | |
| Q9WV55 | VAPA | VAMP (vesicle-associated membrane protein)-associated protein A, 33kDa | Plasma Membrane | other | 1.59E+07 | 1.28E+07 | 9571920 | | | | | | |
| Q9WV55 | VAPA | VAMP (vesicle-associated membrane protein)-associated protein A, 33kDa | Plasma Membrane | other | 9841255 | 7486005.5 | 6121485.5 | | | | | | |
| Q9WV55 | VAPA | VAMP (vesicle-associated membrane protein)-associated protein A, 33kDa | Plasma Membrane | other | 1.23E+07 | 1.22E+07 | 1.11E+07 | | | | | | |
| Q9WV55 | VAPA | VAMP (vesicle-associated membrane protein)-associated protein A, 33kDa | Plasma Membrane | other | 9774161 | 9307978 | 8254178 | | | | | | |
| Q9WV55 | VAPA | VAMP (vesicle-associated membrane protein)-associated protein A, 33kDa | Plasma Membrane | other | 5704295 | 3546439 | 1399898.625 | | | | | | |
| Q9WV55 | VAPA | VAMP (vesicle-associated membrane protein)-associated protein A, 33kDa | Plasma Membrane | other | 1510901 | 324157.531 | 155762.672 | Q9WV55 | VAPA | 18561179 | 15442527 | 2971916 | 27 |
| Q9QY76 | VAPB | VAMP (vesicle-associated membrane protein)-associated protein B and C | Plasma Membrane | other | 5368142 | 5154346 | 6421618.5 | | | | | | |
| Q9QY76 | VAPB | VAMP (vesicle-associated membrane protein)-associated protein B and C | Plasma Membrane | other | 1240026.375 | 1190174 | 1441522.5 | Q9QY76 | VAPB | 3304084 | 2426402 | 990574 | 6 |
| Q3U2A8 | VARS2 | valyl-tRNA synthetase 2, mitochondrial | Cytoplasm | enzyme | 1.92E+07 | 2.51E+07 | 2.39E+07 | Q3U2A8 | VARS2 | 19180000 | 3108767 | 1794848 | 3 |

| | | | | | | | | | | | | | |
|---|---|---|---|---|---|---|---|---|---|---|---|---|---|
| P70460 | VASP | vasodilator-stimulated phosphoprotein | Plasma Membrane | other | 3496139 | 4510186.5 | 7552849.5 | P70460 | VASP | 3496139 | 2111200 | 1218902 | 3 |
| Q01853 | VCP | valosin containing protein | Cytoplasm | enzyme | 2.25E+08 | 2.02E+08 | 1.14E+08 | | | | | | |
| Q01853 | VCP | valosin containing protein | Cytoplasm | enzyme | 1.54E+08 | 9.67E+07 | 1.58E+08 | | | | | | |
| Q01853 | VCP | valosin containing protein | Cytoplasm | enzyme | 1.22E+08 | 1.18E+08 | 1.41E+08 | | | | | | |
| Q01853 | VCP | valosin containing protein | Cytoplasm | enzyme | 2.45E+08 | 2.29E+08 | 1.58E+08 | | | | | | |
| Q01853 | VCP | valosin containing protein | Cytoplasm | enzyme | 1.82E+08 | 1.68E+08 | 1.68E+08 | | | | | | |
| Q01853 | VCP | valosin containing protein | Cytoplasm | enzyme | 1.04E+08 | 8.08E+07 | 9.00E+07 | | | | | | |
| Q01853 | VCP | valosin containing protein | Cytoplasm | enzyme | 1.28E+08 | 1.17E+08 | 1.23E+08 | | | | | | |
| Q01853 | VCP | valosin containing protein | Cytoplasm | enzyme | 1.33E+08 | 8.29E+07 | 7.23E+07 | | | | | | |
| Q01853 | VCP | valosin containing protein | Cytoplasm | enzyme | 1.06E+08 | 8.04E+07 | 8.28E+07 | | | | | | |
| Q01853 | VCP | valosin containing protein | Cytoplasm | enzyme | 9.49E+07 | 6.55E+07 | 7.84E+07 | | | | | | |
| Q01853 | VCP | valosin containing protein | Cytoplasm | enzyme | 7.93E+07 | 7.46E+07 | 7.04E+07 | | | | | | |
| Q01853 | VCP | valosin containing protein | Cytoplasm | enzyme | 5.63E+07 | 1.04E+08 | 7.62E+07 | | | | | | |
| Q01853 | VCP | valosin containing protein | Cytoplasm | enzyme | 1.01E+08 | 7.48E+07 | 9.40E+07 | | | | | | |
| Q01853 | VCP | valosin containing protein | Cytoplasm | enzyme | 6.61E+07 | 4.88E+07 | 4.40E+07 | | | | | | |
| Q01853 | VCP | valosin containing protein | Cytoplasm | enzyme | 7.77E+07 | 7.58E+07 | 6.40E+07 | | | | | | |
| Q01853 | VCP | valosin containing protein | Cytoplasm | enzyme | 6.87E+07 | 5.41E+07 | 5.80E+07 | | | | | | |
| Q01853 | VCP | valosin containing protein | Cytoplasm | enzyme | 9.09E+07 | 6.70E+07 | 6.31E+07 | | | | | | |
| Q01853 | VCP | valosin containing protein | Cytoplasm | enzyme | 6.36E+07 | 5.54E+07 | 5.14E+07 | | | | | | |
| Q01853 | VCP | valosin containing protein | Cytoplasm | enzyme | 9.28E+07 | 7.86E+07 | 5.02E+07 | | | | | | |
| Q01853 | VCP | valosin containing protein | Cytoplasm | enzyme | 6.53E+07 | 5.25E+07 | 5.39E+07 | | | | | | |
| Q01853 | VCP | valosin containing protein | Cytoplasm | enzyme | 7.37E+07 | 5.51E+07 | 4.45E+07 | | | | | | |
| Q01853 | VCP | valosin containing protein | Cytoplasm | enzyme | 5.35E+07 | 4.73E+07 | 3.93E+07 | | | | | | |
| Q01853 | VCP | valosin containing protein | Cytoplasm | enzyme | 4.41E+07 | 2.67E+07 | 4.21E+07 | | | | | | |
| Q01853 | VCP | valosin containing protein | Cytoplasm | enzyme | 5.85E+07 | 3.88E+07 | 3.24E+07 | | | | | | |
| Q01853 | VCP | valosin containing protein | Cytoplasm | enzyme | 7.44E+07 | 5.33E+07 | 2.70E+07 | | | | | | |
| Q01853 | VCP | valosin containing protein | Cytoplasm | enzyme | 5.27E+07 | 4.06E+07 | 3.99E+07 | | | | | | |
| Q01853 | VCP | valosin containing protein | Cytoplasm | enzyme | 3.03E+07 | 2.27E+07 | 2.48E+07 | | | | | | |
| Q01853 | VCP | valosin containing protein | Cytoplasm | enzyme | 3.64E+07 | 2.55E+07 | 2.35E+07 | | | | | | |
| Q01853 | VCP | valosin containing protein | Cytoplasm | enzyme | 3.19E+07 | 3.17E+07 | 3.06E+07 | | | | | | |
| Q01853 | VCP | valosin containing protein | Cytoplasm | enzyme | 1.80E+07 | 1.92E+07 | 1.28E+07 | | | | | | |
| Q01853 | VCP | valosin containing protein | Cytoplasm | enzyme | 3.24E+07 | 2.27E+07 | 2.30E+07 | | | | | | |
| Q01853 | VCP | valosin containing protein | Cytoplasm | enzyme | 4.57E+07 | 3.81E+07 | 3.79E+07 | | | | | | |
| Q01853 | VCP | valosin containing protein | Cytoplasm | enzyme | 3.03E+07 | 3.07E+07 | 1.94E+07 | | | | | | |
| Q01853 | VCP | valosin containing protein | Cytoplasm | enzyme | 3.61E+07 | 2.25E+07 | 2.77E+07 | | | | | | |
| Q01853 | VCP | valosin containing protein | Cytoplasm | enzyme | 2.40E+07 | 2.32E+07 | 2.38E+07 | | | | | | |
| Q01853 | VCP | valosin containing protein | Cytoplasm | enzyme | 2.89E+07 | 2.40E+07 | 1.68E+07 | | | | | | |
| Q01853 | VCP | valosin containing protein | Cytoplasm | enzyme | 1.36E+07 | 1.15E+07 | 1.50E+07 | | | | | | |
| Q01853 | VCP | valosin containing protein | Cytoplasm | enzyme | 3.23E+07 | 2.30E+07 | 2.30E+07 | | | | | | |
| Q01853 | VCP | valosin containing protein | Cytoplasm | enzyme | 2.78E+07 | 2.01E+07 | 2.62E+07 | | | | | | |
| Q01853 | VCP | valosin containing protein | Cytoplasm | enzyme | 4.77E+07 | 4.11E+07 | 4.16E+07 | | | | | | |
| Q01853 | VCP | valosin containing protein | Cytoplasm | enzyme | 2.38E+07 | 1.45E+07 | 1.28E+07 | | | | | | |
| Q01853 | VCP | valosin containing protein | Cytoplasm | enzyme | 5664453 | 4795230.5 | 3596245.5 | | | | | | |
| Q01853 | VCP | valosin containing protein | Cytoplasm | enzyme | 1.19E+07 | 1.40E+07 | 8631564 | | | | | | |
| Q01853 | VCP | valosin containing protein | Cytoplasm | enzyme | 8531479 | 9004193 | 5462705.5 | | | | | | |
| Q01853 | VCP | valosin containing protein | Cytoplasm | enzyme | 997953.438 | 580794.75 | 1928233.125 | | | | | | |
| Q01853 | VCP | valosin containing protein | Cytoplasm | enzyme | 9121630 | 2776539.75 | 1.10E+07 | | | | | | |
| Q01853 | VCP | valosin containing protein | Cytoplasm | enzyme | 1.07E+07 | 6862903.5 | 5592361 | | | | | | |
| Q01853 | VCP | valosin containing protein | Cytoplasm | enzyme | 1.30E+07 | 1.27E+07 | 7735911.5 | | | | | | |
| Q01853 | VCP | valosin containing protein | Cytoplasm | enzyme | 1.10E+07 | 1.26E+07 | 7847048.5 | | | | | | |
| Q01853 | VCP | valosin containing protein | Cytoplasm | enzyme | 1.33E+07 | 1.64E+07 | 8401362 | | | | | | |
| Q01853 | VCP | valosin containing protein | Cytoplasm | enzyme | 7636545.5 | 4801458 | 4741540 | | | | | | |
| Q01853 | VCP | valosin containing protein | Cytoplasm | enzyme | 1.52E+07 | 7103041.5 | 7841424.5 | | | | | | |
| Q01853 | VCP | valosin containing protein | Cytoplasm | enzyme | 6491365.5 | 7090092.5 | 4738604 | | | | | | |
| Q01853 | VCP | valosin containing protein | Cytoplasm | enzyme | 1.16E+07 | 1.13E+07 | 6157380 | | | | | | |
| Q01853 | VCP | valosin containing protein | Cytoplasm | enzyme | 3085599 | 1974977.875 | 6331991 | | | | | | |
| Q01853 | VCP | valosin containing protein | Cytoplasm | enzyme | 6121567.5 | 4600518.5 | 3519704.75 | | | | | | |
| Q01853 | VCP | valosin containing protein | Cytoplasm | enzyme | 2533917 | 3242046 | 2218137.5 | | | | | | |
| Q01853 | VCP | valosin containing protein | Cytoplasm | enzyme | 1985205.75 | 209840.672 | 2170039 | | | | | | |
| Q01853 | VCP | valosin containing protein | Cytoplasm | enzyme | 4374744.5 | 2175098.5 | 1598445.625 | | | | | | |
| Q01853 | VCP | valosin containing protein | Cytoplasm | enzyme | 6690315.5 | 3457230.25 | 4395036 | | | | | | |
| Q01853 | VCP | valosin containing protein | Cytoplasm | enzyme | 2806117 | 3363780.75 | 3974291.75 | | | | | | |
| Q01853 | VCP | valosin containing protein | Cytoplasm | enzyme | 1095745.875 | 1153510.125 | 481955 | | | | | | |
| Q01853 | VCP | valosin containing protein | Cytoplasm | enzyme | 1580631.25 | 1952057.25 | 1037661.25 | | | | | | |
| Q01853 | VCP | valosin containing protein | Cytoplasm | enzyme | 895320.812 | 326367.469 | 1406255.125 | | | | | | |
| Q01853 | VCP | valosin containing protein | Cytoplasm | enzyme | 2244346.5 | 1156236.25 | 905739.062 | Q01853 | VCP | 49658414 | 48107634 | 3445059 | 195 |
| Q60932 | VDAC1 | voltage-dependent anion channel 1 | Cytoplasm | ion channel | 1.92E+07 | 3.17E+07 | 4.17E+07 | | | | | | |
| Q60932 | VDAC1 | voltage-dependent anion channel 1 | Cytoplasm | ion channel | 9557813 | 1.75E+07 | 1.64E+07 | | | | | | |
| Q60932 | VDAC1 | voltage-dependent anion channel 1 | Cytoplasm | ion channel | 1.48E+07 | 2.77E+07 | 2.12E+07 | | | | | | |
| Q60932 | VDAC1 | voltage-dependent anion channel 1 | Cytoplasm | ion channel | 1.16E+07 | 1.15E+07 | 1.82E+07 | | | | | | |
| Q60932 | VDAC1 | voltage-dependent anion channel 1 | Cytoplasm | ion channel | 1.69E+07 | 2.47E+07 | 3.19E+07 | | | | | | |
| Q60932 | VDAC1 | voltage-dependent anion channel 1 | Cytoplasm | ion channel | 1.20E+07 | 2.01E+07 | 2.08E+07 | | | | | | |
| Q60932 | VDAC1 | voltage-dependent anion channel 1 | Cytoplasm | ion channel | 1.17E+07 | 1.85E+07 | 2.41E+07 | | | | | | |
| Q60932 | VDAC1 | voltage-dependent anion channel 1 | Cytoplasm | ion channel | 2167311.75 | 1924171.375 | 7936728.5 | Q60932 | VDAC1 | 12235641 | 9311699 | 1900743 | 24 |
| Q60930 | VDAC2 | voltage-dependent anion channel 2 | Cytoplasm | ion channel | 1.80E+07 | 2.97E+07 | 3.54E+07 | | | | | | |
| Q60930 | VDAC2 | voltage-dependent anion channel 2 | Cytoplasm | ion channel | 1.68E+07 | 2.84E+07 | 3.33E+07 | | | | | | |
| Q60930 | VDAC2 | voltage-dependent anion channel 2 | Cytoplasm | ion channel | 1.88E+07 | 3.08E+07 | 2.59E+07 | | | | | | |
| Q60930 | VDAC2 | voltage-dependent anion channel 2 | Cytoplasm | ion channel | 1.25E+07 | 2.26E+07 | 2.69E+07 | | | | | | |
| Q60930 | VDAC2 | voltage-dependent anion channel 2 | Cytoplasm | ion channel | 9717637 | 1.69E+07 | 2.11E+07 | | | | | | |
| Q60930 | VDAC2 | voltage-dependent anion channel 2 | Cytoplasm | ion channel | 3926986.25 | 8089001.5 | 1.06E+07 | | | | | | |
| Q60930 | VDAC2 | voltage-dependent anion channel 2 | Cytoplasm | ion channel | 3962938 | 8173425.5 | 9164973 | | | | | | |
| Q60930 | VDAC2 | voltage-dependent anion channel 2 | Cytoplasm | ion channel | 947525 | 1554716.25 | 1818389.75 | | | | | | |
| Q60930 | VDAC2 | voltage-dependent anion channel 2 | Cytoplasm | ion channel | 1885934.875 | 2245761.75 | 3793553.5 | Q60930 | VDAC2 | 9612336 | 11123719 | 2140761 | 27 |
| Q60931 | VDAC3 | voltage-dependent anion channel 3 | Cytoplasm | ion channel | 2.64E+07 | 3.90E+07 | 4.40E+07 | | | | | | |
| Q60931 | VDAC3 | voltage-dependent anion channel 3 | Cytoplasm | ion channel | 2.03E+07 | 3.04E+07 | 3.00E+07 | | | | | | |
| Q60931 | VDAC3 | voltage-dependent anion channel 3 | Cytoplasm | ion channel | 1.61E+07 | 2.18E+07 | 3.10E+07 | | | | | | |
| Q60931 | VDAC3 | voltage-dependent anion channel 3 | Cytoplasm | ion channel | 1.03E+07 | 1.77E+07 | 2.10E+07 | | | | | | |
| Q60931 | VDAC3 | voltage-dependent anion channel 3 | Cytoplasm | ion channel | 5148297 | 6580145.5 | 1.33E+07 | | | | | | |
| Q60931 | VDAC3 | voltage-dependent anion channel 3 | Cytoplasm | ion channel | 2772452.5 | 3990194 | 5192502 | Q60931 | VDAC3 | 13500125 | 12449015 | 2934261 | 18 |
| P20152 | VIM | vimentin | Cytoplasm | other | 823730.375 | 90621.352 | 6324920 | P20152 | VIM | 823730 | 3407517 | 1967331 | 3 |
| Q8BGQ1 | VIPAS39 | VPS33B interacting protein, apical-basolateral polarity regulator, spe-39 homolog | Plasma Membrane | other | 2.13E+07 | 1.93E+07 | 2.08E+07 | Q8BGQ1 | VIPAS39 | 21300000 | 1028056 | 593549 | 3 |
| Q99KU0 | VMP1 | vacuole membrane protein 1 | Plasma Membrane | other | 1.76E+07 | 2.29E+07 | 1.66E+07 | Q99KU0 | VMP1 | 17630000 | 3345788 | 1931692 | 3 |
| Q5H8C4 | VPS13A | vacuolar protein sorting 13 homolog A (S. cerevisiae) | Cytoplasm | transporter | 2.13E+07 | 2.07E+07 | 1.90E+07 | | | | | | |
| Q5H8C4 | VPS13A | vacuolar protein sorting 13 homolog A (S. cerevisiae) | Cytoplasm | transporter | 1.38E+07 | 1.76E+07 | 1.31E+07 | | | | | | |
| Q5H8C4 | VPS13A | vacuolar protein sorting 13 homolog A (S. cerevisiae) | Cytoplasm | transporter | 1.58E+07 | 1.11E+07 | 1.60E+07 | | | | | | |
| Q5H8C4 | VPS13A | vacuolar protein sorting 13 homolog A (S. cerevisiae) | Cytoplasm | transporter | 1.25E+07 | 1.30E+07 | 9881147 | | | | | | |
| Q5H8C4 | VPS13A | vacuolar protein sorting 13 homolog A (S. cerevisiae) | Cytoplasm | transporter | 1.02E+07 | 1.14E+07 | 9661533 | | | | | | |
| Q5H8C4 | VPS13A | vacuolar protein sorting 13 homolog A (S. cerevisiae) | Cytoplasm | transporter | 8883443 | 1.00E+07 | 8908543 | | | | | | |
| Q5H8C4 | VPS13A | vacuolar protein sorting 13 homolog A (S. cerevisiae) | Cytoplasm | transporter | 7917802 | 7752702 | 8397945 | | | | | | |
| Q5H8C4 | VPS13A | vacuolar protein sorting 13 homolog A (S. cerevisiae) | Cytoplasm | transporter | 8007320.5 | 7767157.5 | 6359170 | | | | | | |
| Q5H8C4 | VPS13A | vacuolar protein sorting 13 homolog A (S. cerevisiae) | Cytoplasm | transporter | 6291233 | 8887499 | 6960998.5 | | | | | | |
| Q5H8C4 | VPS13A | vacuolar protein sorting 13 homolog A (S. cerevisiae) | Cytoplasm | transporter | 8009666 | 7291498 | 7131511 | | | | | | |
| Q5H8C4 | VPS13A | vacuolar protein sorting 13 homolog A (S. cerevisiae) | Cytoplasm | transporter | 8079580.5 | 7919891 | 7634658.5 | | | | | | |
| Q5H8C4 | VPS13A | vacuolar protein sorting 13 homolog A (S. cerevisiae) | Cytoplasm | transporter | 5896086 | 6669487 | 4966790 | | | | | | |
| Q5H8C4 | VPS13A | vacuolar protein sorting 13 homolog A (S. cerevisiae) | Cytoplasm | transporter | 7151519.5 | 5959371.5 | 7007253.5 | | | | | | |
| Q5H8C4 | VPS13A | vacuolar protein sorting 13 homolog A (S. cerevisiae) | Cytoplasm | transporter | 8810668 | 7700011.5 | 6877540.5 | | | | | | |
| Q5H8C4 | VPS13A | vacuolar protein sorting 13 homolog A (S. cerevisiae) | Cytoplasm | transporter | 7697205.5 | 7169370.5 | 6428015.5 | | | | | | |
| Q5H8C4 | VPS13A | vacuolar protein sorting 13 homolog A (S. cerevisiae) | Cytoplasm | transporter | 4596454.5 | 5582093 | 7617444 | | | | | | |
| Q5H8C4 | VPS13A | vacuolar protein sorting 13 homolog A (S. cerevisiae) | Cytoplasm | transporter | 4862480.5 | 6171320 | 5700967.5 | | | | | | |

| | | | | | | | | | | | | | |
|---|---|---|---|---|---|---|---|---|---|---|---|---|---|
| Q5H8C4 | VPS13A | vacuolar protein sorting 13 homolog A (S. cerevisiae) | Cytoplasm | transporter | 7077980.5 | 6564662.5 | 5754660 | | | | | | |
| Q5H8C4 | VPS13A | vacuolar protein sorting 13 homolog A (S. cerevisiae) | Cytoplasm | transporter | 4609349 | 4358800 | 3563856 | | | | | | |
| Q5H8C4 | VPS13A | vacuolar protein sorting 13 homolog A (S. cerevisiae) | Cytoplasm | transporter | 4080709.75 | 5444637 | 3578457 | | | | | | |
| Q5H8C4 | VPS13A | vacuolar protein sorting 13 homolog A (S. cerevisiae) | Cytoplasm | transporter | 5576117 | 5604252.5 | 5632088.5 | | | | | | |
| Q5H8C4 | VPS13A | vacuolar protein sorting 13 homolog A (S. cerevisiae) | Cytoplasm | transporter | 4174271 | 4027342.5 | 3830300 | | | | | | |
| Q5H8C4 | VPS13A | vacuolar protein sorting 13 homolog A (S. cerevisiae) | Cytoplasm | transporter | 5388717 | 5848844 | 4797905.5 | | | | | | |
| Q5H8C4 | VPS13A | vacuolar protein sorting 13 homolog A (S. cerevisiae) | Cytoplasm | transporter | 4759826 | 5968428 | 5324830.5 | | | | | | |
| Q5H8C4 | VPS13A | vacuolar protein sorting 13 homolog A (S. cerevisiae) | Cytoplasm | transporter | 3475604 | 3724967 | 3292824.5 | | | | | | |
| Q5H8C4 | VPS13A | vacuolar protein sorting 13 homolog A (S. cerevisiae) | Cytoplasm | transporter | 3181021 | 2901891.75 | 2153998 | | | | | | |
| Q5H8C4 | VPS13A | vacuolar protein sorting 13 homolog A (S. cerevisiae) | Cytoplasm | transporter | 4902363 | 6218830 | 6314205.5 | | | | | | |
| Q5H8C4 | VPS13A | vacuolar protein sorting 13 homolog A (S. cerevisiae) | Cytoplasm | transporter | 2754146.75 | 2271522.25 | 2577466.25 | | | | | | |
| Q5H8C4 | VPS13A | vacuolar protein sorting 13 homolog A (S. cerevisiae) | Cytoplasm | transporter | 2516365 | 2942546 | 767539.812 | | | | | | |
| Q5H8C4 | VPS13A | vacuolar protein sorting 13 homolog A (S. cerevisiae) | Cytoplasm | transporter | 2626733.5 | 3277621.25 | 2824544 | | | | | | |
| Q5H8C4 | VPS13A | vacuolar protein sorting 13 homolog A (S. cerevisiae) | Cytoplasm | transporter | 1665046.625 | 1975433.875 | 1870317.375 | | | | | | |
| Q5H8C4 | VPS13A | vacuolar protein sorting 13 homolog A (S. cerevisiae) | Cytoplasm | transporter | 1594589.875 | 1803839 | 1428260.5 | | | | | | |
| Q5H8C4 | VPS13A | vacuolar protein sorting 13 homolog A (S. cerevisiae) | Cytoplasm | transporter | 1035169.5 | 1387545 | 1248511.875 | | | | | | |
| Q5H8C4 | VPS13A | vacuolar protein sorting 13 homolog A (S. cerevisiae) | Cytoplasm | transporter | 4011748 | 6063947 | 3964432 | Q5H8C4 | VPS13A | 6563624 | 4126850 | 408619 | 102 |
| Q8BX70 | VPS13C | vacuolar protein sorting 13 homolog C (S. cerevisiae) | Cytoplasm | other | 2113323.75 | 2436350.75 | 1520751.75 | | | | | | |
| Q8BX70 | VPS13C | vacuolar protein sorting 13 homolog C (S. cerevisiae) | Cytoplasm | other | 1559556.75 | 1521729 | 1541824.5 | | | | | | |
| Q8BX70 | VPS13C | vacuolar protein sorting 13 homolog C (S. cerevisiae) | Cytoplasm | other | 2722914 | 2326419.25 | 2444324 | | | | | | |
| Q8BX70 | VPS13C | vacuolar protein sorting 13 homolog C (S. cerevisiae) | Cytoplasm | other | 1468566.125 | 1274251.625 | 1048040.375 | | | | | | |
| Q8BX70 | VPS13C | vacuolar protein sorting 13 homolog C (S. cerevisiae) | Cytoplasm | other | 1160494.125 | 1276377.375 | 566158.938 | Q8BX70 | VPS13C | 1804971 | 610182 | 157548 | 15 |
| Q9D1C8 | VPS28 | vacuolar protein sorting 28 homolog (S. cerevisiae) | Cytoplasm | transporter | 4331211.5 | 4430482.5 | 2.11E+07 | Q9D1C8 | VPS28 | 4331212 | 9647162 | 5569791 | 3 |
| Q8CIZ8 | VWF | von Willebrand factor | Extracellular Space | other | 5303972.5 | 4862322.5 | 9327725 | | | | | | |
| Q8CIZ8 | VWF | von Willebrand factor | Extracellular Space | other | 1649969.5 | 1443682.625 | 2508275.75 | | | | | | |
| Q8CIZ8 | VWF | von Willebrand factor | Extracellular Space | other | 1411626.875 | 1077160.375 | 3634659.5 | | | | | | |
| Q8CIZ8 | VWF | von Willebrand factor | Extracellular Space | other | 3428376.75 | 2161914.5 | 4607293 | | | | | | |
| Q8CIZ8 | VWF | von Willebrand factor | Extracellular Space | other | 2173162 | 731456.312 | 2174336 | | | | | | |
| Q8CIZ8 | VWF | von Willebrand factor | Extracellular Space | other | 1034331.5 | 858453.562 | 2103788 | | | | | | |
| Q8CIZ8 | VWF | von Willebrand factor | Extracellular Space | other | 2014939.25 | 1870374.125 | 3783367.25 | | | | | | |
| Q8CIZ8 | VWF | von Willebrand factor | Extracellular Space | other | 2025947.875 | 1761470.25 | 3755732 | | | | | | |
| Q8CIZ8 | VWF | von Willebrand factor | Extracellular Space | other | 4041065.25 | 3157396.25 | 7310873.5 | | | | | | |
| Q8CIZ8 | VWF | von Willebrand factor | Extracellular Space | other | 2089536.25 | 1889212.625 | 4178709.5 | | | | | | |
| Q8CIZ8 | VWF | von Willebrand factor | Extracellular Space | other | 2716855.25 | 2173583.25 | 3972801.25 | | | | | | |
| Q8CIZ8 | VWF | von Willebrand factor | Extracellular Space | other | 776881.312 | 978942.125 | 2133620.75 | | | | | | |
| Q8CIZ8 | VWF | von Willebrand factor | Extracellular Space | other | 767288.125 | 1072853.625 | 2832900.75 | | | | | | |
| Q8CIZ8 | VWF | von Willebrand factor | Extracellular Space | other | 1234463.25 | 1047305.812 | 2293469.75 | | | | | | |
| Q8CIZ8 | VWF | von Willebrand factor | Extracellular Space | other | 581710.938 | 572069.75 | 1303645 | Q8CIZ8 | VWF | 2083342 | 1780957 | 265489 | 45 |
| Q8CBE3 | WDR37 | WD repeat domain 37 | Nucleus | other | 3132827.5 | 2117593.75 | 1995457.125 | | | | | | |
| Q8CBE3 | WDR37 | WD repeat domain 37 | Nucleus | other | 819819.75 | 709667 | 712994.062 | Q8CBE3 | WDR37 | 1976324 | 995985 | 406609 | 6 |
| Q8BH57 | WDR48 | WD repeat domain 48 | Cytoplasm | other | 928099.438 | 1255452.5 | 1336819.25 | Q8BH57 | WDR48 | 928099 | 216346 | 124907 | 3 |
| Q4KUS1 | WFDC8 | WAP four-disulfide core domain 8 | Extracellular Space | other | 879866.312 | 623571.188 | 313397.469 | Q4KUS1 | WFDC8 | 879866 | 283661 | 163772 | 3 |
| P83741 | WNK1 | WNK lysine deficient protein kinase 1 | Cytoplasm | kinase | 1483903.5 | 1413248.25 | 976988 | P83741 | WNK1 | 1483904 | 274554 | 158514 | 3 |
| Q9QXY7 | XK | X-linked Kx blood group | Plasma Membrane | transporter | 3.01E+07 | 2.58E+07 | 2.00E+07 | | | | | | |
| Q9QXY7 | XK | X-linked Kx blood group | Plasma Membrane | transporter | 2.05E+07 | 2.55E+07 | 2.36E+07 | | | | | | |
| Q9QXY7 | XK | X-linked Kx blood group | Plasma Membrane | transporter | 3.51E+07 | 2.90E+07 | 2.57E+07 | | | | | | |
| Q9QXY7 | XK | X-linked Kx blood group | Plasma Membrane | transporter | 1.81E+07 | 1.66E+07 | 1.44E+07 | Q9QXY7 | XK | 25930000 | 6047694 | 1745819 | 12 |
| Q6P5F9 | XPO1 | exportin 1 | Nucleus | transporter | 1783219.875 | 1194289.25 | 1142006.625 | Q6P5F9 | XPO1 | 1783220 | 356073 | 205579 | 3 |
| Q9EPK7 | XPO7 | exportin 7 | Nucleus | transporter | 2402555.75 | 2423599.5 | 1475000.25 | | | | | | |
| Q9EPK7 | XPO7 | exportin 7 | Nucleus | transporter | 1965359.625 | 1361873.5 | 793719.438 | Q9EPK7 | XPO7 | 2183958 | 642756 | 262404 | 6 |
| P62960 | YBX1 | Y box binding protein 1 | Nucleus | transcription regulator | 2301860.75 | 2467923.75 | 3714351.75 | | | | | | |
| P62960 | YBX1 | Y box binding protein 1 | Nucleus | transcription regulator | 279654.875 | 209022.109 | 193464.641 | P62960 | YBX1 | 1290758 | 1506088 | 614858 | 6 |
| Q9JKB3 | YBX3 | Y box binding protein 3 | Nucleus | transcription regulator | 2483617.5 | 1666866.875 | 2148740.25 | | | | | | |
| Q9JKB3 | YBX3 | Y box binding protein 3 | Nucleus | transcription regulator | 1824482.125 | 1566116.875 | 1833124 | Q9JKB3 | YBX3 | 2154050 | 339401 | 138560 | 6 |
| Q9CX30 | YIF1B | Yip1 interacting factor homolog B (S. cerevisiae) | Cytoplasm | other | 2698630.75 | 3338272.25 | 2793022.75 | | | | | | |
| Q9CX30 | YIF1B | Yip1 interacting factor homolog B (S. cerevisiae) | Cytoplasm | other | 2760712.75 | 3067069.25 | 2794236.25 | Q9CX30 | YIF1B | 2729672 | 245815 | 100354 | 6 |
| Q9CQV8 | YWHAB | tyrosine 3-monooxygenase/tryptophan 5-monooxygenase activation protein, beta | Cytoplasm | transcription regulator | 5708749.5 | 6613227 | 9285451 | | | | | | |
| Q9CQV8 | YWHAB | tyrosine 3-monooxygenase/tryptophan 5-monooxygenase activation protein, beta | Cytoplasm | transcription regulator | 5254759 | 5658057 | 6471045 | | | | | | |
| Q9CQV8 | YWHAB | tyrosine 3-monooxygenase/tryptophan 5-monooxygenase activation protein, beta | Cytoplasm | transcription regulator | 3591593.75 | 4203661.5 | 4976183.5 | | | | | | |
| Q9CQV8 | YWHAB | tyrosine 3-monooxygenase/tryptophan 5-monooxygenase activation protein, beta | Cytoplasm | transcription regulator | 2705192 | 2772739.5 | 3504918.25 | | | | | | |
| Q9CQV8 | YWHAB | tyrosine 3-monooxygenase/tryptophan 5-monooxygenase activation protein, beta | Cytoplasm | transcription regulator | 707172.75 | 922579.5 | 1408162.25 | | | | | | |
| Q9CQV8 | YWHAB | tyrosine 3-monooxygenase/tryptophan 5-monooxygenase activation protein, beta | Cytoplasm | transcription regulator | 883251.188 | 1078667 | 1168482.5 | Q9CQV8 | YWHAB | 3141786 | 2480073 | 584559 | 18 |
| P62259 | YWHAE | tyrosine 3-monooxygenase/tryptophan 5-monooxygenase activation protein, epsilon | Cytoplasm | other | 1.47E+07 | 1.36E+07 | 1.75E+07 | | | | | | |
| P62259 | YWHAE | tyrosine 3-monooxygenase/tryptophan 5-monooxygenase activation protein, epsilon | Cytoplasm | other | 8227816 | 1.06E+07 | 1.31E+07 | | | | | | |
| P62259 | YWHAE | tyrosine 3-monooxygenase/tryptophan 5-monooxygenase activation protein, epsilon | Cytoplasm | other | 9314893 | 8856010 | 8593009 | | | | | | |
| P62259 | YWHAE | tyrosine 3-monooxygenase/tryptophan 5-monooxygenase activation protein, epsilon | Cytoplasm | other | 9462439 | 7742398 | 8538702 | | | | | | |
| P62259 | YWHAE | tyrosine 3-monooxygenase/tryptophan 5-monooxygenase activation protein, epsilon | Cytoplasm | other | 6175470 | 7803251.5 | 7850875 | | | | | | |
| P62259 | YWHAE | tyrosine 3-monooxygenase/tryptophan 5-monooxygenase activation protein, epsilon | Cytoplasm | other | 4812646.5 | 5441435 | 5410997 | | | | | | |
| P62259 | YWHAE | tyrosine 3-monooxygenase/tryptophan 5-monooxygenase activation protein, epsilon | Cytoplasm | other | 5096552 | 6547994.5 | 3927805.25 | | | | | | |
| P62259 | YWHAE | tyrosine 3-monooxygenase/tryptophan 5-monooxygenase activation protein, epsilon | Cytoplasm | other | 5312670 | 5400832.5 | 5709599 | | | | | | |
| P62259 | YWHAE | tyrosine 3-monooxygenase/tryptophan 5-monooxygenase activation protein, epsilon | Cytoplasm | other | 2582452.25 | 3225472 | 3853182.5 | P62259 | YWHAE | 7298327 | 3648288 | 702113 | 27 |
| P61982 | YWHAG | tyrosine 3-monooxygenase/tryptophan 5-monooxygenase activation protein, gamma | Cytoplasm | other | 3697124.75 | 4815143 | 7007755 | | | | | | |
| P61982 | YWHAG | tyrosine 3-monooxygenase/tryptophan 5-monooxygenase activation protein, gamma | Cytoplasm | other | 1949744.625 | 2287010 | 2897421 | P61982 | YWHAG | 2823435 | 1889681 | 771459 | 6 |
| P68510 | YWHAH | tyrosine 3-monooxygenase/tryptophan 5-monooxygenase activation protein, eta | Cytoplasm | transcription regulator | 3.07E+07 | 3.24E+07 | 5.97E+07 | | | | | | |
| P68510 | YWHAH | tyrosine 3-monooxygenase/tryptophan 5-monooxygenase activation protein, eta | Cytoplasm | transcription regulator | 1.06E+07 | 1.18E+07 | 2.08E+07 | | | | | | |
| P68510 | YWHAH | tyrosine 3-monooxygenase/tryptophan 5-monooxygenase activation protein, eta | Cytoplasm | transcription regulator | 1.29E+07 | 1.74E+07 | 3.19E+07 | | | | | | |
| P68510 | YWHAH | tyrosine 3-monooxygenase/tryptophan 5-monooxygenase activation protein, eta | Cytoplasm | transcription regulator | 8134813 | 1.15E+07 | 1.71E+07 | | | | | | |
| P68510 | YWHAH | tyrosine 3-monooxygenase/tryptophan 5-monooxygenase activation protein, eta | Cytoplasm | transcription regulator | 4569314.5 | 5466371 | 1.19E+07 | | | | | | |
| P68510 | YWHAH | tyrosine 3-monooxygenase/tryptophan 5-monooxygenase activation protein, eta | Cytoplasm | transcription regulator | 2902594.5 | 6303175 | 6068795 | | | | | | |
| P68510 | YWHAH | tyrosine 3-monooxygenase/tryptophan 5-monooxygenase activation protein, eta | Cytoplasm | transcription regulator | 5418249 | 6550003.5 | 7426073.5 | | | | | | |
| P68510 | YWHAH | tyrosine 3-monooxygenase/tryptophan 5-monooxygenase activation protein, eta | Cytoplasm | transcription regulator | 3637225.75 | 4021899.5 | 7142337.5 | | | | | | |
| P68510 | YWHAH | tyrosine 3-monooxygenase/tryptophan 5-monooxygenase activation protein, eta | Cytoplasm | transcription regulator | 917976.5 | 946540.938 | 2161429.25 | P68510 | YWHAH | 8861130 | 13037580 | 2509083 | 27 |
| P68254 | YWHAQ | tyrosine 3-monooxygenase/tryptophan 5-monooxygenase activation protein, theta | Cytoplasm | other | 2728549.5 | 3271080.5 | 6224199.5 | | | | | | |
| P68254 | YWHAQ | tyrosine 3-monooxygenase/tryptophan 5-monooxygenase activation protein, theta | Cytoplasm | other | 2402108.5 | 3214808.75 | 5339224.5 | | | | | | |
| P68254 | YWHAQ | tyrosine 3-monooxygenase/tryptophan 5-monooxygenase activation protein, theta | Cytoplasm | other | 2491825.5 | 2674337.5 | 3813083.25 | P68254 | YWHAQ | 2540828 | 1345647 | 448549 | 9 |
| P63101 | YWHAZ | tyrosine 3-monooxygenase/tryptophan 5-monooxygenase activation protein, zeta | Cytoplasm | enzyme | 5.03E+07 | 6.48E+07 | 9.19E+07 | | | | | | |
| P63101 | YWHAZ | tyrosine 3-monooxygenase/tryptophan 5-monooxygenase activation protein, zeta | Cytoplasm | enzyme | 3.74E+07 | 4.47E+07 | 5.46E+07 | | | | | | |
| P63101 | YWHAZ | tyrosine 3-monooxygenase/tryptophan 5-monooxygenase activation protein, zeta | Cytoplasm | enzyme | 3.04E+07 | 3.69E+07 | 5.45E+07 | | | | | | |
| P63101 | YWHAZ | tyrosine 3-monooxygenase/tryptophan 5-monooxygenase activation protein, zeta | Cytoplasm | enzyme | 3.76E+07 | 3.50E+07 | 6.76E+07 | | | | | | |
| P63101 | YWHAZ | tyrosine 3-monooxygenase/tryptophan 5-monooxygenase activation protein, zeta | Cytoplasm | enzyme | 3.69E+07 | 4.77E+07 | 6.11E+07 | | | | | | |
| P63101 | YWHAZ | tyrosine 3-monooxygenase/tryptophan 5-monooxygenase activation protein, zeta | Cytoplasm | enzyme | 2.50E+07 | 1.79E+07 | 4.18E+07 | | | | | | |
| P63101 | YWHAZ | tyrosine 3-monooxygenase/tryptophan 5-monooxygenase activation protein, zeta | Cytoplasm | enzyme | 3.07E+07 | 4.84E+07 | 6.36E+07 | | | | | | |
| P63101 | YWHAZ | tyrosine 3-monooxygenase/tryptophan 5-monooxygenase activation protein, zeta | Cytoplasm | enzyme | 2.38E+07 | 2.55E+07 | 3.94E+07 | | | | | | |
| P63101 | YWHAZ | tyrosine 3-monooxygenase/tryptophan 5-monooxygenase activation protein, zeta | Cytoplasm | enzyme | 1.23E+07 | 1.21E+07 | 2.37E+07 | | | | | | |
| P63101 | YWHAZ | tyrosine 3-monooxygenase/tryptophan 5-monooxygenase activation protein, zeta | Cytoplasm | enzyme | 5916114.5 | 7373998 | 6381105.5 | | | | | | |
| P63101 | YWHAZ | tyrosine 3-monooxygenase/tryptophan 5-monooxygenase activation protein, zeta | Cytoplasm | enzyme | 3361821 | 3201975.5 | 2.02E+07 | | | | | | |
| P63101 | YWHAZ | tyrosine 3-monooxygenase/tryptophan 5-monooxygenase activation protein, zeta | Cytoplasm | enzyme | 6655956 | 8711175 | 8884922 | | | | | | |
| P63101 | YWHAZ | tyrosine 3-monooxygenase/tryptophan 5-monooxygenase activation protein, zeta | Cytoplasm | enzyme | 2970041 | 4175759.5 | 8255379 | | | | | | |
| P63101 | YWHAZ | tyrosine 3-monooxygenase/tryptophan 5-monooxygenase activation protein, zeta | Cytoplasm | enzyme | 9296623 | 1.22E+07 | 1.21E+07 | | | | | | |
| P63101 | YWHAZ | tyrosine 3-monooxygenase/tryptophan 5-monooxygenase activation protein, zeta | Cytoplasm | enzyme | 2503815.75 | 1201049.75 | 3734213.75 | | | | | | |
| P63101 | YWHAZ | tyrosine 3-monooxygenase/tryptophan 5-monooxygenase activation protein, zeta | Cytoplasm | enzyme | 610385.062 | 385647.219 | 2457828.5 | P63101 | YWHAZ | 19726547 | 22568994 | 3257554 | 48 |
| Q9D0B1 | ZNF524 | zinc finger protein 524 | Nucleus | other | 9754315 | 8050691.5 | 4632702 | Q9D0B1 | ZNF524 | 9754315 | 2608189 | 1505839 | 3 |
| Q02525 | ZNF717 | zinc finger protein 717 | Nucleus | other | 2.57E+08 | 2.56E+08 | 2.07E+08 | Q02525 | ZNF717 | 256900000 | 28730530 | 16587579 | 3 |
| Q5SSH7 | ZZEF1 | zinc finger, ZZ-type with EF-hand domain 1 | Cytoplasm | other | 2055350.75 | 2164368.75 | 2687013.5 | Q5SSH7 | ZZEF1 | 2055351 | 337649 | 194942 | 3 |

**# Peptides by Location**

| | |
|---|---|
| Plasma Membrane | 1902 |
| Cytoplasm | 4392 |
| Nucleus | 523 |

| | |
|---|---|
| Extracellular Space | 347 |
| Other | 24 |
| **Total Peptides** | **7188** |

### AQP1/RhAG double knockout

| ID | Symbol | Entrez Gene Name | Primary Location | Type(s) | Intensity [dKO_1] | Intensity [dKO_2] | Intensity [dKO_3] | ID | Symbol | Mean Intensity | St. Dev | Std. Err | n |
|---|---|---|---|---|---|---|---|---|---|---|---|---|---|
| Q3UHJ0 | AAK1 | AP2 associated kinase 1 | Cytoplasm | kinase | 1023527.188 | 979379.125 | 990138.5 | Q3UHJ0 | AAK1 | 1023527 | 23020 | 13291 | 3 |
| Q9JI39 | ABCB10 | ATP-binding cassette, sub-family B (MDR/TAP), member 10 | Cytoplasm | transporter | 1.16E+07 | 3.62E+07 | 1.47E+07 | | | | | | |
| Q9JI39 | ABCB10 | ATP-binding cassette, sub-family B (MDR/TAP), member 10 | Cytoplasm | transporter | 1.32E+07 | 2.49E+07 | 7081912 | | | | | | |
| Q9JI39 | ABCB10 | ATP-binding cassette, sub-family B (MDR/TAP), member 10 | Cytoplasm | transporter | 5669490 | 1.34E+07 | 4231947.5 | | | | | | |
| Q9JI39 | ABCB10 | ATP-binding cassette, sub-family B (MDR/TAP), member 10 | Cytoplasm | transporter | 2837780.5 | 3538272 | 2299725 | | | | | | |
| Q9JI39 | ABCB10 | ATP-binding cassette, sub-family B (MDR/TAP), member 10 | Cytoplasm | transporter | 4108594.75 | 6175137 | 2903497.75 | | | | | | |
| Q9JI39 | ABCB10 | ATP-binding cassette, sub-family B (MDR/TAP), member 10 | Cytoplasm | transporter | 3418209.25 | 7147102.5 | 3531161.25 | | | | | | |
| Q9JI39 | ABCB10 | ATP-binding cassette, sub-family B (MDR/TAP), member 10 | Cytoplasm | transporter | 3176291.75 | 7876071 | 3211201.5 | | | | | | |
| Q9JI39 | ABCB10 | ATP-binding cassette, sub-family B (MDR/TAP), member 10 | Cytoplasm | transporter | 3040666.5 | 6827866.5 | 2506444.25 | | | | | | |
| Q9JI39 | ABCB10 | ATP-binding cassette, sub-family B (MDR/TAP), member 10 | Cytoplasm | transporter | 2065191.75 | 4554023.5 | 1554486 | | | | | | |
| Q9JI39 | ABCB10 | ATP-binding cassette, sub-family B (MDR/TAP), member 10 | Cytoplasm | transporter | 3610339.75 | 8416091 | 2357357.75 | | | | | | |
| Q9JI39 | ABCB10 | ATP-binding cassette, sub-family B (MDR/TAP), member 10 | Cytoplasm | transporter | 1871996.25 | 1991290.125 | 1742957.5 | | | | | | |
| Q9JI39 | ABCB10 | ATP-binding cassette, sub-family B (MDR/TAP), member 10 | Cytoplasm | transporter | 1180126.25 | 2130069 | 1045574.312 | Q9JI39 | ABCB10 | 4647391 | 7089094 | 1181516 | 36 |
| P21440 | ABCB4 | ATP-binding cassette, sub-family B (MDR/TAP), member 4 | Plasma Membrane | transporter | 2.71E+07 | 2.63E+07 | 3.63E+07 | | | | | | |
| P21440 | ABCB4 | ATP-binding cassette, sub-family B (MDR/TAP), member 4 | Plasma Membrane | transporter | 1.60E+07 | 1.47E+07 | 1.55E+07 | | | | | | |
| P21440 | ABCB4 | ATP-binding cassette, sub-family B (MDR/TAP), member 4 | Plasma Membrane | transporter | 1.28E+07 | 1.32E+07 | 2.01E+07 | | | | | | |
| P21440 | ABCB4 | ATP-binding cassette, sub-family B (MDR/TAP), member 4 | Plasma Membrane | transporter | 1.31E+07 | 1.46E+07 | 1.62E+07 | | | | | | |
| P21440 | ABCB4 | ATP-binding cassette, sub-family B (MDR/TAP), member 4 | Plasma Membrane | transporter | 1.67E+07 | 1.44E+07 | 1.63E+07 | | | | | | |
| P21440 | ABCB4 | ATP-binding cassette, sub-family B (MDR/TAP), member 4 | Plasma Membrane | transporter | 1.29E+07 | 1.53E+07 | 1.49E+07 | | | | | | |
| P21440 | ABCB4 | ATP-binding cassette, sub-family B (MDR/TAP), member 4 | Plasma Membrane | transporter | 1.38E+07 | 1.60E+07 | 1.70E+07 | | | | | | |
| P21440 | ABCB4 | ATP-binding cassette, sub-family B (MDR/TAP), member 4 | Plasma Membrane | transporter | 1.28E+07 | 1.18E+07 | 1.55E+07 | | | | | | |
| P21440 | ABCB4 | ATP-binding cassette, sub-family B (MDR/TAP), member 4 | Plasma Membrane | transporter | 1.17E+07 | 1.15E+07 | 1.41E+07 | | | | | | |
| P21440 | ABCB4 | ATP-binding cassette, sub-family B (MDR/TAP), member 4 | Plasma Membrane | transporter | 9305181 | 9843795 | 9044523 | | | | | | |
| P21440 | ABCB4 | ATP-binding cassette, sub-family B (MDR/TAP), member 4 | Plasma Membrane | transporter | 9542266 | 8283619 | 1.25E+07 | | | | | | |
| P21440 | ABCB4 | ATP-binding cassette, sub-family B (MDR/TAP), member 4 | Plasma Membrane | transporter | 8146750.5 | 8291964.5 | 1.17E+07 | | | | | | |
| P21440 | ABCB4 | ATP-binding cassette, sub-family B (MDR/TAP), member 4 | Plasma Membrane | transporter | 7134308 | 6072062 | 6545002.5 | | | | | | |
| P21440 | ABCB4 | ATP-binding cassette, sub-family B (MDR/TAP), member 4 | Plasma Membrane | transporter | 6987972 | 5877782 | 7808233.5 | | | | | | |
| P21440 | ABCB4 | ATP-binding cassette, sub-family B (MDR/TAP), member 4 | Plasma Membrane | transporter | 8284547.5 | 8338817.5 | 1.07E+07 | | | | | | |
| P21440 | ABCB4 | ATP-binding cassette, sub-family B (MDR/TAP), member 4 | Plasma Membrane | transporter | 6649497.5 | 8578936 | 8676622 | | | | | | |
| P21440 | ABCB4 | ATP-binding cassette, sub-family B (MDR/TAP), member 4 | Plasma Membrane | transporter | 3979119.5 | 4856122.5 | 4435005 | | | | | | |
| P21440 | ABCB4 | ATP-binding cassette, sub-family B (MDR/TAP), member 4 | Plasma Membrane | transporter | 4631437.5 | 5387041 | 7062705 | | | | | | |
| P21440 | ABCB4 | ATP-binding cassette, sub-family B (MDR/TAP), member 4 | Plasma Membrane | transporter | 3819291.75 | 2793517.25 | 3884543.5 | | | | | | |
| P21440 | ABCB4 | ATP-binding cassette, sub-family B (MDR/TAP), member 4 | Plasma Membrane | transporter | 5146337.5 | 2401103.75 | 1414049 | | | | | | |
| P21440 | ABCB4 | ATP-binding cassette, sub-family B (MDR/TAP), member 4 | Plasma Membrane | transporter | 3280503.75 | 3418023.25 | 4758375 | | | | | | |
| P21440 | ABCB4 | ATP-binding cassette, sub-family B (MDR/TAP), member 4 | Plasma Membrane | transporter | 3937934.5 | 4479635.5 | 4670759 | | | | | | |
| P21440 | ABCB4 | ATP-binding cassette, sub-family B (MDR/TAP), member 4 | Plasma Membrane | transporter | 2733030 | 3001182.75 | 2181185 | | | | | | |
| P21440 | ABCB4 | ATP-binding cassette, sub-family B (MDR/TAP), member 4 | Plasma Membrane | transporter | 1675143.875 | 1674961.375 | 2615903.25 | P21440 | ABCB4 | 9252222 | 6459435 | 761252 | 72 |
| Q9DC29 | ABCB6 | ATP-binding cassette, sub-family B (MDR/TAP), member 6 (Langereis blood group) | Cytoplasm | transporter | 1.08E+08 | 1.07E+08 | 1.28E+08 | | | | | | |
| Q9DC29 | ABCB6 | ATP-binding cassette, sub-family B (MDR/TAP), member 6 (Langereis blood group) | Cytoplasm | transporter | 9.33E+07 | 9.94E+07 | 1.26E+08 | | | | | | |
| Q9DC29 | ABCB6 | ATP-binding cassette, sub-family B (MDR/TAP), member 6 (Langereis blood group) | Cytoplasm | transporter | 3.51E+07 | 4.67E+07 | 4.76E+07 | | | | | | |
| Q9DC29 | ABCB6 | ATP-binding cassette, sub-family B (MDR/TAP), member 6 (Langereis blood group) | Cytoplasm | transporter | 3.51E+07 | 3.43E+07 | 5.41E+07 | | | | | | |
| Q9DC29 | ABCB6 | ATP-binding cassette, sub-family B (MDR/TAP), member 6 (Langereis blood group) | Cytoplasm | transporter | 4.75E+07 | 5.33E+07 | 6.60E+07 | | | | | | |
| Q9DC29 | ABCB6 | ATP-binding cassette, sub-family B (MDR/TAP), member 6 (Langereis blood group) | Cytoplasm | transporter | 3.65E+07 | 3.53E+07 | 5.00E+07 | | | | | | |
| Q9DC29 | ABCB6 | ATP-binding cassette, sub-family B (MDR/TAP), member 6 (Langereis blood group) | Cytoplasm | transporter | 2.52E+07 | 2.48E+07 | 3.80E+07 | | | | | | |
| Q9DC29 | ABCB6 | ATP-binding cassette, sub-family B (MDR/TAP), member 6 (Langereis blood group) | Cytoplasm | transporter | 1.49E+07 | 2.01E+07 | 1.66E+07 | | | | | | |
| Q9DC29 | ABCB6 | ATP-binding cassette, sub-family B (MDR/TAP), member 6 (Langereis blood group) | Cytoplasm | transporter | 1.85E+07 | 1.97E+07 | 2.26E+07 | | | | | | |
| Q9DC29 | ABCB6 | ATP-binding cassette, sub-family B (MDR/TAP), member 6 (Langereis blood group) | Cytoplasm | transporter | 1.41E+07 | 1.58E+07 | 2.14E+07 | | | | | | |
| Q9DC29 | ABCB6 | ATP-binding cassette, sub-family B (MDR/TAP), member 6 (Langereis blood group) | Cytoplasm | transporter | 1.32E+07 | 1.32E+07 | 2.07E+07 | | | | | | |
| Q9DC29 | ABCB6 | ATP-binding cassette, sub-family B (MDR/TAP), member 6 (Langereis blood group) | Cytoplasm | transporter | 1.10E+07 | 1.76E+07 | 2.50E+07 | | | | | | |
| Q9DC29 | ABCB6 | ATP-binding cassette, sub-family B (MDR/TAP), member 6 (Langereis blood group) | Cytoplasm | transporter | 1.02E+07 | 1.14E+07 | 1.67E+07 | | | | | | |
| Q9DC29 | ABCB6 | ATP-binding cassette, sub-family B (MDR/TAP), member 6 (Langereis blood group) | Cytoplasm | transporter | 8335962 | 9068953 | 1.18E+07 | | | | | | |
| Q9DC29 | ABCB6 | ATP-binding cassette, sub-family B (MDR/TAP), member 6 (Langereis blood group) | Cytoplasm | transporter | 7156440.5 | 1.02E+07 | 1.53E+07 | | | | | | |
| Q9DC29 | ABCB6 | ATP-binding cassette, sub-family B (MDR/TAP), member 6 (Langereis blood group) | Cytoplasm | transporter | 5310749 | 6215591 | 1.08E+07 | | | | | | |
| Q9DC29 | ABCB6 | ATP-binding cassette, sub-family B (MDR/TAP), member 6 (Langereis blood group) | Cytoplasm | transporter | 3702328 | 3154973.5 | 3476106.5 | | | | | | |
| Q9DC29 | ABCB6 | ATP-binding cassette, sub-family B (MDR/TAP), member 6 (Langereis blood group) | Cytoplasm | transporter | 3459273.75 | 4081198.75 | 7818826.5 | | | | | | |
| Q9DC29 | ABCB6 | ATP-binding cassette, sub-family B (MDR/TAP), member 6 (Langereis blood group) | Cytoplasm | transporter | 1866624.875 | 1953659.625 | 371294.25 | | | | | | |
| Q9DC29 | ABCB6 | ATP-binding cassette, sub-family B (MDR/TAP), member 6 (Langereis blood group) | Cytoplasm | transporter | 2642590 | 3617678.75 | 2740117.25 | Q9DC29 | ABCB6 | 24741198 | 31719763 | 4095004 | 60 |
| Q61102 | ABCB7 | ATP-binding cassette, sub-family B (MDR/TAP), member 7 | Cytoplasm | transporter | 1392166 | 3213153.5 | 1211359.125 | Q61102 | ABCB7 | 1392166 | 1107239 | 639265 | 3 |
| Q9JJ59 | ABCB9 | ATP-binding cassette, sub-family B (MDR/TAP), member 9 | Cytoplasm | transporter | 2008977 | 1.02E+07 | 2799373.75 | Q9JJ59 | ABCB9 | 2008977 | 4500984 | 2598644 | 3 |
| O35379 | ABCC1 | ATP-binding cassette, sub-family C (CFTR/MRP), member 1 | Plasma Membrane | transporter | 1.57E+07 | 1.81E+07 | 2.04E+07 | | | | | | |
| O35379 | ABCC1 | ATP-binding cassette, sub-family C (CFTR/MRP), member 1 | Plasma Membrane | transporter | 1.48E+07 | 1.69E+07 | 1.67E+07 | | | | | | |
| O35379 | ABCC1 | ATP-binding cassette, sub-family C (CFTR/MRP), member 1 | Plasma Membrane | transporter | 1.02E+07 | 1.16E+07 | 1.29E+07 | | | | | | |
| O35379 | ABCC1 | ATP-binding cassette, sub-family C (CFTR/MRP), member 1 | Plasma Membrane | transporter | 1.20E+07 | 1.44E+07 | 1.26E+07 | | | | | | |
| O35379 | ABCC1 | ATP-binding cassette, sub-family C (CFTR/MRP), member 1 | Plasma Membrane | transporter | 9628028 | 1.02E+07 | 1.11E+07 | | | | | | |
| O35379 | ABCC1 | ATP-binding cassette, sub-family C (CFTR/MRP), member 1 | Plasma Membrane | transporter | 7152297 | 8057718.5 | 7771078.5 | | | | | | |
| O35379 | ABCC1 | ATP-binding cassette, sub-family C (CFTR/MRP), member 1 | Plasma Membrane | transporter | 4960573.5 | 6894369.5 | 2396755 | | | | | | |
| O35379 | ABCC1 | ATP-binding cassette, sub-family C (CFTR/MRP), member 1 | Plasma Membrane | transporter | 7109862.5 | 6558180 | 6477259.5 | | | | | | |
| O35379 | ABCC1 | ATP-binding cassette, sub-family C (CFTR/MRP), member 1 | Plasma Membrane | transporter | 6844025.5 | 5519387.5 | 4740703 | | | | | | |
| O35379 | ABCC1 | ATP-binding cassette, sub-family C (CFTR/MRP), member 1 | Plasma Membrane | transporter | 4482250.5 | 5428920.5 | 5067834.5 | | | | | | |
| O35379 | ABCC1 | ATP-binding cassette, sub-family C (CFTR/MRP), member 1 | Plasma Membrane | transporter | 3861807 | 4298573 | 3907110.25 | | | | | | |
| O35379 | ABCC1 | ATP-binding cassette, sub-family C (CFTR/MRP), member 1 | Plasma Membrane | transporter | 4624044.5 | 4386111 | 4650264.5 | | | | | | |
| O35379 | ABCC1 | ATP-binding cassette, sub-family C (CFTR/MRP), member 1 | Plasma Membrane | transporter | 3302337.25 | 3742042 | 4662286.5 | | | | | | |
| O35379 | ABCC1 | ATP-binding cassette, sub-family C (CFTR/MRP), member 1 | Plasma Membrane | transporter | 2746128 | 3067809 | 2538383.75 | | | | | | |
| O35379 | ABCC1 | ATP-binding cassette, sub-family C (CFTR/MRP), member 1 | Plasma Membrane | transporter | 2395267.75 | 3283179.5 | 3695683.5 | | | | | | |
| O35379 | ABCC1 | ATP-binding cassette, sub-family C (CFTR/MRP), member 1 | Plasma Membrane | transporter | 3574275.5 | 2712634.5 | 3233396.5 | | | | | | |
| O35379 | ABCC1 | ATP-binding cassette, sub-family C (CFTR/MRP), member 1 | Plasma Membrane | transporter | 2544655.5 | 2632320.75 | 2827228 | | | | | | |
| O35379 | ABCC1 | ATP-binding cassette, sub-family C (CFTR/MRP), member 1 | Plasma Membrane | transporter | 1489198.625 | 1615236.875 | 1795276.25 | O35379 | ABCC1 | 6526931 | 4860742 | 661463 | 54 |
| Q9R1S7 | ABCC6 | ATP-binding cassette, sub-family C (CFTR/MRP), member 6 | Plasma Membrane | transporter | 5126124.5 | 7426640.5 | 1.02E+07 | Q9R1S7 | ABCC6 | 5126125 | 2535461 | 1463849 | 3 |
| P61222 | ABCE1 | ATP-binding cassette, sub-family E (OABP), member 1 | Cytoplasm | transporter | 5396297 | 6704207 | 3846566.5 | | | | | | |
| P61222 | ABCE1 | ATP-binding cassette, sub-family E (OABP), member 1 | Cytoplasm | transporter | 2536084.5 | 3060709 | 2529491.75 | | | | | | |
| P61222 | ABCE1 | ATP-binding cassette, sub-family E (OABP), member 1 | Cytoplasm | transporter | 4862459.5 | 4171471.5 | 3075554.75 | | | | | | |
| P61222 | ABCE1 | ATP-binding cassette, sub-family E (OABP), member 1 | Cytoplasm | transporter | 2158236 | 2877913.75 | 1801100.125 | P61222 | ABCE1 | 3738269 | 1462289 | 422126 | 12 |
| Q7TMS5 | ABCG2 | ATP-binding cassette, sub-family G (WHITE), member 2 (Junior blood group) | Plasma Membrane | transporter | 1.37E+08 | 1.36E+08 | 1.28E+08 | | | | | | |
| Q7TMS5 | ABCG2 | ATP-binding cassette, sub-family G (WHITE), member 2 (Junior blood group) | Plasma Membrane | transporter | 1.08E+08 | 1.02E+08 | 1.13E+08 | | | | | | |
| Q7TMS5 | ABCG2 | ATP-binding cassette, sub-family G (WHITE), member 2 (Junior blood group) | Plasma Membrane | transporter | 8.92E+07 | 9.86E+07 | 1.04E+08 | | | | | | |
| Q7TMS5 | ABCG2 | ATP-binding cassette, sub-family G (WHITE), member 2 (Junior blood group) | Plasma Membrane | transporter | 7.74E+07 | 8.01E+07 | 6.73E+07 | | | | | | |
| Q7TMS5 | ABCG2 | ATP-binding cassette, sub-family G (WHITE), member 2 (Junior blood group) | Plasma Membrane | transporter | 5.23E+07 | 6.11E+07 | 6.67E+07 | | | | | | |
| Q7TMS5 | ABCG2 | ATP-binding cassette, sub-family G (WHITE), member 2 (Junior blood group) | Plasma Membrane | transporter | 4.91E+07 | 3.94E+07 | 4.64E+07 | | | | | | |
| Q7TMS5 | ABCG2 | ATP-binding cassette, sub-family G (WHITE), member 2 (Junior blood group) | Plasma Membrane | transporter | 5.02E+07 | 5.47E+07 | 5.28E+07 | | | | | | |
| Q7TMS5 | ABCG2 | ATP-binding cassette, sub-family G (WHITE), member 2 (Junior blood group) | Plasma Membrane | transporter | 2.89E+07 | 2.98E+07 | 3.52E+07 | | | | | | |

| | | | | | | | | | | | | | |
|---|---|---|---|---|---|---|---|---|---|---|---|---|---|
| Q7TMS5 | ABCG2 | ATP-binding cassette, sub-family G (WHITE), member 2 (Junior blood group) | Plasma Membrane | transporter | 3.83E+07 | 5.63E+07 | 3.93E+07 | | | | | | |
| Q7TMS5 | ABCG2 | ATP-binding cassette, sub-family G (WHITE), member 2 (Junior blood group) | Plasma Membrane | transporter | 3.23E+07 | 3.14E+07 | 3.75E+07 | | | | | | |
| Q7TMS5 | ABCG2 | ATP-binding cassette, sub-family G (WHITE), member 2 (Junior blood group) | Plasma Membrane | transporter | 2.71E+07 | 2.42E+07 | 2.75E+07 | | | | | | |
| Q7TMS5 | ABCG2 | ATP-binding cassette, sub-family G (WHITE), member 2 (Junior blood group) | Plasma Membrane | transporter | 1.96E+07 | 1.77E+07 | 2.13E+07 | | | | | | |
| Q7TMS5 | ABCG2 | ATP-binding cassette, sub-family G (WHITE), member 2 (Junior blood group) | Plasma Membrane | transporter | 1.48E+07 | 1.61E+07 | 1.67E+07 | | | | | | |
| Q7TMS5 | ABCG2 | ATP-binding cassette, sub-family G (WHITE), member 2 (Junior blood group) | Plasma Membrane | transporter | 2.14E+07 | 2.10E+07 | 2.47E+07 | | | | | | |
| Q7TMS5 | ABCG2 | ATP-binding cassette, sub-family G (WHITE), member 2 (Junior blood group) | Plasma Membrane | transporter | 1.02E+07 | 1.19E+07 | 1.19E+07 | | | | | | |
| Q7TMS5 | ABCG2 | ATP-binding cassette, sub-family G (WHITE), member 2 (Junior blood group) | Plasma Membrane | transporter | 1.21E+07 | 1.07E+07 | 1.20E+07 | | | | | | |
| Q7TMS5 | ABCG2 | ATP-binding cassette, sub-family G (WHITE), member 2 (Junior blood group) | Plasma Membrane | transporter | 5616932.5 | 5040974 | 5341339.5 | | | | | | |
| Q7TMS5 | ABCG2 | ATP-binding cassette, sub-family G (WHITE), member 2 (Junior blood group) | Plasma Membrane | transporter | 9112585 | 8942083 | 1.34E+07 | | | | | | |
| Q7TMS5 | ABCG2 | ATP-binding cassette, sub-family G (WHITE), member 2 (Junior blood group) | Plasma Membrane | transporter | 7114677 | 6309670.5 | 7497577.5 | | | | | | |
| Q7TMS5 | ABCG2 | ATP-binding cassette, sub-family G (WHITE), member 2 (Junior blood group) | Plasma Membrane | transporter | 1378525.375 | 1411005 | 808709.375 | Q7TMS5 | ABCG2 | 39547636 | 36738695 | 4742945 | 60 |
| Q9Z1Q2 | ABHD16A | abhydrolase domain containing 16A | Cytoplasm | enzyme | 3278546 | 4031609.5 | 5327089.5 | | | | | | |
| Q9Z1Q2 | ABHD16A | abhydrolase domain containing 16A | Cytoplasm | enzyme | 3164801.25 | 4393438.5 | 4875318 | | | | | | |
| Q9Z1Q2 | ABHD16A | abhydrolase domain containing 16A | Cytoplasm | enzyme | 940071.875 | 970536.25 | 865880.812 | Q9Z1Q2 | ABHD16A | 2461140 | 1763502 | 587834 | 9 |
| Q921H8 | ACAA1 | acetyl-CoA acyltransferase 1 | Cytoplasm | enzyme | 451583.969 | 1635119.25 | 1786344.375 | Q921H8 | ACAA1 | 451584 | 730891 | 421980 | 3 |
| Q8BWT1 | ACAA2 | acetyl-CoA acyltransferase 2 | Cytoplasm | enzyme | 6759508.5 | 1.63E+07 | 7558998.5 | | | | | | |
| Q8BWT1 | ACAA2 | acetyl-CoA acyltransferase 2 | Cytoplasm | enzyme | 4978159.5 | 9757014 | 4545632.5 | | | | | | |
| Q8BWT1 | ACAA2 | acetyl-CoA acyltransferase 2 | Cytoplasm | enzyme | 3772808.75 | 7515826 | 3732188 | | | | | | |
| Q8BWT1 | ACAA2 | acetyl-CoA acyltransferase 2 | Cytoplasm | enzyme | 3926109.25 | 7301543.5 | 3524835.5 | | | | | | |
| Q8BWT1 | ACAA2 | acetyl-CoA acyltransferase 2 | Cytoplasm | enzyme | 2788119.25 | 6866519 | 2214532.75 | | | | | | |
| Q8BWT1 | ACAA2 | acetyl-CoA acyltransferase 2 | Cytoplasm | enzyme | 3361545 | 7543653 | 2871782 | Q8BWT1 | ACAA2 | 4264375 | 3388830 | 798755 | 18 |
| Q8JZN5 | ACAD9 | acyl-CoA dehydrogenase family, member 9 | Cytoplasm | enzyme | 1602437.125 | 3013673.5 | 812292.25 | Q8JZN5 | ACAD9 | 1602437 | 1115198 | 643860 | 3 |
| P51174 | ACADL | acyl-CoA dehydrogenase, long chain | Cytoplasm | enzyme | 2.50E+07 | 5.45E+07 | 2.14E+07 | | | | | | |
| P51174 | ACADL | acyl-CoA dehydrogenase, long chain | Cytoplasm | enzyme | 9025784 | 1.65E+07 | 7754814 | | | | | | |
| P51174 | ACADL | acyl-CoA dehydrogenase, long chain | Cytoplasm | enzyme | 7533382.5 | 1.48E+07 | 5722670 | | | | | | |
| P51174 | ACADL | acyl-CoA dehydrogenase, long chain | Cytoplasm | enzyme | 5010864.5 | 8190993.5 | 3808237 | | | | | | |
| P51174 | ACADL | acyl-CoA dehydrogenase, long chain | Cytoplasm | enzyme | 3878344 | 8063578.5 | 1691779.625 | | | | | | |
| P51174 | ACADL | acyl-CoA dehydrogenase, long chain | Cytoplasm | enzyme | 4679130.5 | 8835343 | 5589536.5 | | | | | | |
| P51174 | ACADL | acyl-CoA dehydrogenase, long chain | Cytoplasm | enzyme | 2369883.5 | 5494702.5 | 2414424.25 | P51174 | ACADL | 8211056 | 11790691 | 2572940 | 21 |
| P45952 | ACADM | acyl-CoA dehydrogenase, C-4 to C-12 straight chain | Cytoplasm | enzyme | 134632.188 | 525366.812 | 546418.938 | P45952 | ACADM | 134632 | 231907 | 133892 | 3 |
| P50544 | ACADVL | acyl-CoA dehydrogenase, very long chain | Cytoplasm | enzyme | 2420891 | 4874939 | 2297463 | | | | | | |
| P50544 | ACADVL | acyl-CoA dehydrogenase, very long chain | Cytoplasm | enzyme | 2142302.75 | 4785995.5 | 2315763 | | | | | | |
| P50544 | ACADVL | acyl-CoA dehydrogenase, very long chain | Cytoplasm | enzyme | 1107695.75 | 2784873.25 | 1068220.875 | | | | | | |
| P50544 | ACADVL | acyl-CoA dehydrogenase, very long chain | Cytoplasm | enzyme | 935734.562 | 2709101.25 | 549805.812 | | | | | | |
| P50544 | ACADVL | acyl-CoA dehydrogenase, very long chain | Cytoplasm | enzyme | 555725.625 | 1171175.625 | 511302.781 | P50544 | ACADVL | 1432470 | 1399417 | 361328 | 15 |
| Q8QZT1 | ACAT1 | acetyl-CoA acetyltransferase 1 | Cytoplasm | enzyme | 8127389 | 1.83E+07 | 9320676 | | | | | | |
| Q8QZT1 | ACAT1 | acetyl-CoA acetyltransferase 1 | Cytoplasm | enzyme | 6260538 | 1.05E+07 | 8949111 | | | | | | |
| Q8QZT1 | ACAT1 | acetyl-CoA acetyltransferase 1 | Cytoplasm | enzyme | 4026975.75 | 8858672 | 4882132 | | | | | | |
| Q8QZT1 | ACAT1 | acetyl-CoA acetyltransferase 1 | Cytoplasm | enzyme | 5309213.5 | 1.46E+07 | 5690967 | | | | | | |
| Q8QZT1 | ACAT1 | acetyl-CoA acetyltransferase 1 | Cytoplasm | enzyme | 3018626 | 7691887 | 4477385.5 | | | | | | |
| Q8QZT1 | ACAT1 | acetyl-CoA acetyltransferase 1 | Cytoplasm | enzyme | 2191266.75 | 5362400.5 | 3120472.25 | | | | | | |
| Q8QZT1 | ACAT1 | acetyl-CoA acetyltransferase 1 | Cytoplasm | enzyme | 2063514.875 | 4500338.5 | 1782278.875 | Q8QZT1 | ACAT1 | 4428218 | 4173514 | 910735 | 21 |
| Q9QUI6 | ACKR1 | atypical chemokine receptor 1 (Duffy blood group) | Plasma Membrane | G-protein coupled receptor | 3.19E+07 | 3.13E+07 | 3.43E+07 | | | | | | |
| Q9QUI6 | ACKR1 | atypical chemokine receptor 1 (Duffy blood group) | Plasma Membrane | G-protein coupled receptor | 3.32E+07 | 3.77E+07 | 4.14E+07 | | | | | | |
| Q9QUI6 | ACKR1 | atypical chemokine receptor 1 (Duffy blood group) | Plasma Membrane | G-protein coupled receptor | 1.01E+07 | 8555344 | 1.25E+07 | | | | | | |
| Q9QUI6 | ACKR1 | atypical chemokine receptor 1 (Duffy blood group) | Plasma Membrane | G-protein coupled receptor | 8112890.5 | 1222138.875 | 72376.461 | Q9QUI6 | ACKR1 | 20823223 | 15329770 | 4425323 | 12 |
| Q99KI0 | ACO2 | aconitase 2, mitochondrial | Cytoplasm | enzyme | 1.49E+07 | 3.15E+07 | 1.30E+07 | | | | | | |
| Q99KI0 | ACO2 | aconitase 2, mitochondrial | Cytoplasm | enzyme | 8989799 | 1.80E+07 | 8036088 | | | | | | |
| Q99KI0 | ACO2 | aconitase 2, mitochondrial | Cytoplasm | enzyme | 6994929.5 | 1.47E+07 | 6485314 | | | | | | |
| Q99KI0 | ACO2 | aconitase 2, mitochondrial | Cytoplasm | enzyme | 4529160.5 | 1.79E+07 | 1.02E+07 | | | | | | |
| Q99KI0 | ACO2 | aconitase 2, mitochondrial | Cytoplasm | enzyme | 7980269.5 | 1.38E+07 | 5714092.5 | | | | | | |
| Q99KI0 | ACO2 | aconitase 2, mitochondrial | Cytoplasm | enzyme | 7098956.5 | 1.15E+07 | 5469559 | | | | | | |
| Q99KI0 | ACO2 | aconitase 2, mitochondrial | Cytoplasm | enzyme | 4492683 | 9554408 | 3643365.5 | | | | | | |
| Q99KI0 | ACO2 | aconitase 2, mitochondrial | Cytoplasm | enzyme | 4126492.75 | 1.22E+07 | 5325448 | | | | | | |
| Q99KI0 | ACO2 | aconitase 2, mitochondrial | Cytoplasm | enzyme | 5513518 | 1.42E+07 | 5508364.5 | | | | | | |
| Q99KI0 | ACO2 | aconitase 2, mitochondrial | Cytoplasm | enzyme | 6635038.5 | 9485228 | 3202646.75 | | | | | | |
| Q99KI0 | ACO2 | aconitase 2, mitochondrial | Cytoplasm | enzyme | 5818976 | 1.19E+07 | 4787151 | | | | | | |
| Q99KI0 | ACO2 | aconitase 2, mitochondrial | Cytoplasm | enzyme | 3090829.25 | 4782580.5 | 2920765.75 | | | | | | |
| Q99KI0 | ACO2 | aconitase 2, mitochondrial | Cytoplasm | enzyme | 3764494.25 | 8432482 | 3562402 | | | | | | |
| Q99KI0 | ACO2 | aconitase 2, mitochondrial | Cytoplasm | enzyme | 3602171.5 | 5239920 | 2482769.5 | | | | | | |
| Q99KI0 | ACO2 | aconitase 2, mitochondrial | Cytoplasm | enzyme | 2989468.5 | 5795459.5 | 2715832.25 | | | | | | |
| Q99KI0 | ACO2 | aconitase 2, mitochondrial | Cytoplasm | enzyme | 1536379.25 | 3705934 | 868933.5 | | | | | | |
| Q99KI0 | ACO2 | aconitase 2, mitochondrial | Cytoplasm | enzyme | 1773258.75 | 3672473.5 | 1424841.625 | | | | | | |
| Q99KI0 | ACO2 | aconitase 2, mitochondrial | Cytoplasm | enzyme | 1800177.75 | 6941248 | 3783044 | | | | | | |
| Q99KI0 | ACO2 | aconitase 2, mitochondrial | Cytoplasm | enzyme | 1215279.125 | 4014162.75 | 2232768 | | | | | | |
| Q99KI0 | ACO2 | aconitase 2, mitochondrial | Cytoplasm | enzyme | 1458429 | 2498486.25 | 1931276.75 | | | | | | |
| Q99KI0 | ACO2 | aconitase 2, mitochondrial | Cytoplasm | enzyme | 688005.375 | 4848783.5 | 2528138.75 | Q99KI0 | ACO2 | 4714206 | 5341467 | 672962 | 63 |
| P41216 | ACSL1 | acyl-CoA synthetase long-chain family member 1 | Cytoplasm | enzyme | 1.65E+07 | 1.96E+07 | 1.59E+07 | | | | | | |
| P41216 | ACSL1 | acyl-CoA synthetase long-chain family member 1 | Cytoplasm | enzyme | 1.16E+07 | 1.53E+07 | 1.55E+07 | | | | | | |
| P41216 | ACSL1 | acyl-CoA synthetase long-chain family member 1 | Cytoplasm | enzyme | 1.44E+07 | 1.97E+07 | 1.34E+07 | | | | | | |
| P41216 | ACSL1 | acyl-CoA synthetase long-chain family member 1 | Cytoplasm | enzyme | 9751175 | 1.35E+07 | 7928309 | | | | | | |
| P41216 | ACSL1 | acyl-CoA synthetase long-chain family member 1 | Cytoplasm | enzyme | 8082120.5 | 2.18E+07 | 2.48E+07 | | | | | | |
| P41216 | ACSL1 | acyl-CoA synthetase long-chain family member 1 | Cytoplasm | enzyme | 1.33E+07 | 1.82E+07 | 8975064 | | | | | | |
| P41216 | ACSL1 | acyl-CoA synthetase long-chain family member 1 | Cytoplasm | enzyme | 7886282 | 1.08E+07 | 9503651 | | | | | | |
| P41216 | ACSL1 | acyl-CoA synthetase long-chain family member 1 | Cytoplasm | enzyme | 6518475.5 | 7798856.5 | 7585465 | | | | | | |
| P41216 | ACSL1 | acyl-CoA synthetase long-chain family member 1 | Cytoplasm | enzyme | 5321556.5 | 5540848.5 | 5593167 | | | | | | |
| P41216 | ACSL1 | acyl-CoA synthetase long-chain family member 1 | Cytoplasm | enzyme | 6275690 | 7138894 | 3818883.75 | | | | | | |
| P41216 | ACSL1 | acyl-CoA synthetase long-chain family member 1 | Cytoplasm | enzyme | 5976722.5 | 7561833.5 | 7490760 | | | | | | |
| P41216 | ACSL1 | acyl-CoA synthetase long-chain family member 1 | Cytoplasm | enzyme | 5137861 | 6305430.5 | 5953156.5 | | | | | | |
| P41216 | ACSL1 | acyl-CoA synthetase long-chain family member 1 | Cytoplasm | enzyme | 3148930 | 4218548 | 2846861.25 | | | | | | |
| P41216 | ACSL1 | acyl-CoA synthetase long-chain family member 1 | Cytoplasm | enzyme | 3727631 | 3361280.25 | 3118009.5 | | | | | | |
| P41216 | ACSL1 | acyl-CoA synthetase long-chain family member 1 | Cytoplasm | enzyme | 3135098.25 | 3033277.75 | 3805241.25 | | | | | | |
| P41216 | ACSL1 | acyl-CoA synthetase long-chain family member 1 | Cytoplasm | enzyme | 2967759 | 3913673.25 | 3226984.25 | | | | | | |
| P41216 | ACSL1 | acyl-CoA synthetase long-chain family member 1 | Cytoplasm | enzyme | 3785055.75 | 5165479 | 4092781.5 | P41216 | ACSL1 | 7496727 | 5623033 | 787382 | 51 |
| Q9QUJ7 | ACSL4 | acyl-CoA synthetase long-chain family member 4 | Cytoplasm | enzyme | 1.44E+07 | 1.31E+07 | 1.20E+07 | | | | | | |
| Q9QUJ7 | ACSL4 | acyl-CoA synthetase long-chain family member 4 | Cytoplasm | enzyme | 1.43E+07 | 1.39E+07 | 1.13E+07 | | | | | | |
| Q9QUJ7 | ACSL4 | acyl-CoA synthetase long-chain family member 4 | Cytoplasm | enzyme | 1.08E+07 | 1.21E+07 | 1.39E+07 | | | | | | |
| Q9QUJ7 | ACSL4 | acyl-CoA synthetase long-chain family member 4 | Cytoplasm | enzyme | 5275939.5 | 3985809.25 | 6701319.5 | | | | | | |
| Q9QUJ7 | ACSL4 | acyl-CoA synthetase long-chain family member 4 | Cytoplasm | enzyme | 2462831.75 | 1928496 | 2863131.25 | | | | | | |
| Q9QUJ7 | ACSL4 | acyl-CoA synthetase long-chain family member 4 | Cytoplasm | enzyme | 1543978.625 | 1910621.125 | 1352354.25 | | | | | | |

| | | | | | | | | | | | | | |
|---|---|---|---|---|---|---|---|---|---|---|---|---|---|
| Q9QUJ7 | ACSL4 | acyl-CoA synthetase long-chain family member 4 | Cytoplasm | enzyme | 2931637.25 | 3183504.5 | 2848794.75 | | | | | | |
| Q9QUJ7 | ACSL4 | acyl-CoA synthetase long-chain family member 4 | Cytoplasm | enzyme | 2713761.25 | 2525870.25 | 2453291.5 | | | | | | |
| Q9QUJ7 | ACSL4 | acyl-CoA synthetase long-chain family member 4 | Cytoplasm | enzyme | 1999701.75 | 1669444.875 | 2139533.5 | Q9QUJ7 | ACSL4 | 6266428 | 5008621 | 963910 | 27 |
| Q8JZR0 | ACSL5 | acyl-CoA synthetase long-chain family member 5 | Cytoplasm | enzyme | 7886127.5 | 1.10E+07 | 8931511 | | | | | | |
| Q8JZR0 | ACSL5 | acyl-CoA synthetase long-chain family member 5 | Cytoplasm | enzyme | 4227993.5 | 7471057.5 | 8581474 | | | | | | |
| Q8JZR0 | ACSL5 | acyl-CoA synthetase long-chain family member 5 | Cytoplasm | enzyme | 5161773.5 | 6063510.5 | 7014631.5 | | | | | | |
| Q8JZR0 | ACSL5 | acyl-CoA synthetase long-chain family member 5 | Cytoplasm | enzyme | 4610396 | 5311745 | 4097450 | | | | | | |
| Q8JZR0 | ACSL5 | acyl-CoA synthetase long-chain family member 5 | Cytoplasm | enzyme | 3429365 | 5065368.5 | 4228451.5 | | | | | | |
| Q8JZR0 | ACSL5 | acyl-CoA synthetase long-chain family member 5 | Cytoplasm | enzyme | 3344693 | 3383849.25 | 4391552 | | | | | | |
| Q8JZR0 | ACSL5 | acyl-CoA synthetase long-chain family member 5 | Cytoplasm | enzyme | 3469606.25 | 4098774 | 5295944 | | | | | | |
| Q8JZR0 | ACSL5 | acyl-CoA synthetase long-chain family member 5 | Cytoplasm | enzyme | 2684284.75 | 1765435.375 | 2223160.25 | | | | | | |
| Q8JZR0 | ACSL5 | acyl-CoA synthetase long-chain family member 5 | Cytoplasm | enzyme | 2945362 | 2562031.25 | 3673241 | | | | | | |
| Q8JZR0 | ACSL5 | acyl-CoA synthetase long-chain family member 5 | Cytoplasm | enzyme | 2787404.25 | 2266508 | 2062975.5 | | | | | | |
| Q8JZR0 | ACSL5 | acyl-CoA synthetase long-chain family member 5 | Cytoplasm | enzyme | 2068223.5 | 2574115.25 | 2638026 | Q8JZR0 | ACSL5 | 3874112 | 2272030 | 395510 | 33 |
| Q99NB1 | ACSS1 | acyl-CoA synthetase short-chain family member 1 | Cytoplasm | enzyme | 2947933.25 | 5583638 | 1642782 | | | | | | |
| Q99NB1 | ACSS1 | acyl-CoA synthetase short-chain family member 1 | Cytoplasm | enzyme | 2113767 | 4446065.5 | 1444991 | | | | | | |
| Q99NB1 | ACSS1 | acyl-CoA synthetase short-chain family member 1 | Cytoplasm | enzyme | 1771693.625 | 3559760.75 | 1022739.188 | Q99NB1 | ACSS1 | 2277798 | 1535826 | 511942 | 9 |
| P60710 | ACTB | actin, beta | Cytoplasm | other | 4.49E+09 | 4.45E+09 | 4.71E+09 | | | | | | |
| P60710 | ACTB | actin, beta | Cytoplasm | other | 2.86E+09 | 2.98E+09 | 3.01E+09 | | | | | | |
| P60710 | ACTB | actin, beta | Cytoplasm | other | 2.56E+09 | 2.60E+09 | 3.01E+09 | | | | | | |
| P60710 | ACTB | actin, beta | Cytoplasm | other | 1.25E+09 | 1.25E+09 | 1.43E+09 | | | | | | |
| P60710 | ACTB | actin, beta | Cytoplasm | other | 1.59E+09 | 1.31E+09 | 1.17E+09 | | | | | | |
| P60710 | ACTB | actin, beta | Cytoplasm | other | 8.13E+08 | 1.31E+09 | 1.19E+09 | | | | | | |
| P60710 | ACTB | actin, beta | Cytoplasm | other | 1.20E+09 | 1.32E+09 | 1.46E+09 | | | | | | |
| P60710 | ACTB | actin, beta | Cytoplasm | other | 1.11E+09 | 1.09E+09 | 1.23E+09 | | | | | | |
| P60710 | ACTB | actin, beta | Cytoplasm | other | 1.07E+09 | 9.73E+08 | 1.19E+09 | | | | | | |
| P60710 | ACTB | actin, beta | Cytoplasm | other | 8.47E+08 | 9.24E+08 | 9.09E+08 | | | | | | |
| P60710 | ACTB | actin, beta | Cytoplasm | other | 1.02E+09 | 9.58E+08 | 1.13E+09 | | | | | | |
| P60710 | ACTB | actin, beta | Cytoplasm | other | 8.85E+08 | 9.16E+08 | 1.03E+09 | | | | | | |
| P60710 | ACTB | actin, beta | Cytoplasm | other | 5.79E+08 | 6.23E+08 | 7.68E+08 | | | | | | |
| P60710 | ACTB | actin, beta | Cytoplasm | other | 6.19E+08 | 5.78E+08 | 6.59E+08 | | | | | | |
| P60710 | ACTB | actin, beta | Cytoplasm | other | 3.79E+08 | 4.02E+08 | 4.64E+08 | | | | | | |
| P60710 | ACTB | actin, beta | Cytoplasm | other | 4.79E+08 | 4.14E+08 | 5.48E+08 | | | | | | |
| P60710 | ACTB | actin, beta | Cytoplasm | other | 2.57E+08 | 2.92E+08 | 3.86E+08 | | | | | | |
| P60710 | ACTB | actin, beta | Cytoplasm | other | 2.12E+08 | 1.98E+08 | 2.28E+08 | | | | | | |
| P60710 | ACTB | actin, beta | Cytoplasm | other | 1.52E+08 | 1.60E+08 | 1.96E+08 | | | | | | |
| P60710 | ACTB | actin, beta | Cytoplasm | other | 2.90E+08 | 3.18E+08 | 2.67E+08 | | | | | | |
| P60710 | ACTB | actin, beta | Cytoplasm | other | 2.00E+08 | 1.68E+08 | 2.19E+08 | | | | | | |
| P60710 | ACTB | actin, beta | Cytoplasm | other | 1.72E+08 | 1.41E+08 | 1.85E+08 | | | | | | |
| P60710 | ACTB | actin, beta | Cytoplasm | other | 2.26E+08 | 7.06E+07 | 3.06E+07 | | | | | | |
| P60710 | ACTB | actin, beta | Cytoplasm | other | 8.90E+07 | 1.03E+08 | 1.38E+08 | | | | | | |
| P60710 | ACTB | actin, beta | Cytoplasm | other | 8.58E+07 | 7.83E+07 | 9.29E+07 | | | | | | |
| P60710 | ACTB | actin, beta | Cytoplasm | other | 1.88E+08 | 5.58E+07 | 4.98E+07 | | | | | | |
| P60710 | ACTB | actin, beta | Cytoplasm | other | 1.01E+08 | 1.22E+08 | 1.37E+08 | | | | | | |
| P60710 | ACTB | actin, beta | Cytoplasm | other | 8.03E+07 | 1.02E+08 | 1.14E+08 | | | | | | |
| P60710 | ACTB | actin, beta | Cytoplasm | other | 8.68E+07 | 7.40E+07 | 6.76E+07 | | | | | | |
| P60710 | ACTB | actin, beta | Cytoplasm | other | 7.19E+07 | 7.13E+07 | 9.55E+07 | | | | | | |
| P60710 | ACTB | actin, beta | Cytoplasm | other | 7.30E+07 | 7.05E+07 | 9.24E+07 | | | | | | |
| P60710 | ACTB | actin, beta | Cytoplasm | other | 1.23E+08 | 1.75E+08 | 1.75E+08 | | | | | | |
| P60710 | ACTB | actin, beta | Cytoplasm | other | 4.46E+07 | 5.53E+07 | 5.74E+07 | | | | | | |
| P60710 | ACTB | actin, beta | Cytoplasm | other | 6.99E+07 | 2.07E+07 | 1.96E+07 | | | | | | |
| P60710 | ACTB | actin, beta | Cytoplasm | other | 2.15E+07 | 2.64E+07 | 2.00E+07 | | | | | | |
| P60710 | ACTB | actin, beta | Cytoplasm | other | 3.54E+07 | 3.23E+07 | 2.73E+07 | | | | | | |
| P60710 | ACTB | actin, beta | Cytoplasm | other | 4.34E+07 | 1.30E+07 | 9487024 | | | | | | |
| P60710 | ACTB | actin, beta | Cytoplasm | other | 2.49E+07 | 2.01E+07 | 4.30E+07 | | | | | | |
| P60710 | ACTB | actin, beta | Cytoplasm | other | 4.42E+07 | 1.29E+07 | 3240734 | | | | | | |
| P60710 | ACTB | actin, beta | Cytoplasm | other | 1.26E+07 | 5565249.5 | 3271442.75 | | | | | | |
| P60710 | ACTB | actin, beta | Cytoplasm | other | 9184216 | 9570591 | 8219672 | | | | | | |
| P60710 | ACTB | actin, beta | Cytoplasm | other | 7566030 | 1.87E+08 | 5.36E+08 | | | | | | |
| P60710 | ACTB | actin, beta | Cytoplasm | other | 7618167.5 | 1.45E+07 | 2.30E+07 | | | | | | |
| P60710 | ACTB | actin, beta | Cytoplasm | other | 9263654 | 7610002.5 | 1.25E+07 | | | | | | |
| P60710 | ACTB | actin, beta | Cytoplasm | other | 4432620.5 | 5470412.5 | 7068134.5 | | | | | | |
| P60710 | ACTB | actin, beta | Cytoplasm | other | 4084241.75 | 3811468 | 3268632 | | | | | | |
| P60710 | ACTB | actin, beta | Cytoplasm | other | 3015160 | 4.31E+07 | 1.04E+08 | | | | | | |
| P60710 | ACTB | actin, beta | Cytoplasm | other | 97009.805 | 184765.859 | 297432.5 | P60710 | ACTB | 510553356 | 881162387 | 73430199 | 144 |
| P68033 | ACTC1 | actin, alpha, cardiac muscle 1 | Cytoplasm | enzyme | 2.89E+09 | 2.73E+09 | 3.19E+09 | | | | | | |
| P68033 | ACTC1 | actin, alpha, cardiac muscle 1 | Cytoplasm | enzyme | 4.86E+08 | 4.58E+08 | 5.48E+08 | | | | | | |
| P68033 | ACTC1 | actin, alpha, cardiac muscle 1 | Cytoplasm | enzyme | 2.38E+08 | 1.94E+08 | 2.24E+08 | | | | | | |
| P68033 | ACTC1 | actin, alpha, cardiac muscle 1 | Cytoplasm | enzyme | 3.45E+07 | 2.62E+07 | 2.70E+07 | P68033 | ACTC1 | 912260000 | 1231344408 | 355458513 | 12 |
| P61161 | ACTR2 | ARP2 actin-related protein 2 homolog (yeast) | Plasma Membrane | other | 5188532 | 4661830.5 | 4162914.5 | | | | | | |
| P61161 | ACTR2 | ARP2 actin-related protein 2 homolog (yeast) | Plasma Membrane | other | 2804784.25 | 2534045 | 2711249 | | | | | | |
| P61161 | ACTR2 | ARP2 actin-related protein 2 homolog (yeast) | Plasma Membrane | other | 2458840.5 | 2062756.25 | 2722954.5 | P61161 | ACTR2 | 3484052 | 1112347 | 370782 | 9 |
| Q99JY9 | ACTR3 | ARP3 actin-related protein 3 homolog (yeast) | Plasma Membrane | other | 9842705 | 9753599 | 8755115 | | | | | | |
| Q99JY9 | ACTR3 | ARP3 actin-related protein 3 homolog (yeast) | Plasma Membrane | other | 2345687.75 | 2687812.75 | 4115494.25 | | | | | | |
| Q99JY9 | ACTR3 | ARP3 actin-related protein 3 homolog (yeast) | Plasma Membrane | other | 1642877.375 | 2941752.75 | 4878932 | | | | | | |
| Q99JY9 | ACTR3 | ARP3 actin-related protein 3 homolog (yeast) | Plasma Membrane | other | 2131875 | 1981760.125 | 1873902.625 | | | | | | |
| Q99JY9 | ACTR3 | ARP3 actin-related protein 3 homolog (yeast) | Plasma Membrane | other | 2237112.5 | 1956977.625 | 2577281.75 | Q99JY9 | ACTR3 | 3640052 | 2965959 | 765807 | 15 |
| Q641P0 | ACTR3B | ARP3 actin-related protein 3 homolog B (yeast) | Cytoplasm | other | 2887796.75 | 3838322 | 4123394.5 | Q641P0 | ACTR3B | 2887797 | 646976 | 373532 | 3 |
| O35598 | ADAM10 | ADAM metallopeptidase domain 10 | Plasma Membrane | peptidase | 1.09E+08 | 1.03E+08 | 7.75E+07 | | | | | | |
| O35598 | ADAM10 | ADAM metallopeptidase domain 10 | Plasma Membrane | peptidase | 4.83E+07 | 3.97E+07 | 4.73E+07 | | | | | | |
| O35598 | ADAM10 | ADAM metallopeptidase domain 10 | Plasma Membrane | peptidase | 6.17E+07 | 5.68E+07 | 6.58E+07 | | | | | | |
| O35598 | ADAM10 | ADAM metallopeptidase domain 10 | Plasma Membrane | peptidase | 6.68E+07 | 7.05E+07 | 7.99E+07 | | | | | | |
| O35598 | ADAM10 | ADAM metallopeptidase domain 10 | Plasma Membrane | peptidase | 6.57E+07 | 5.08E+07 | 6.16E+07 | | | | | | |
| O35598 | ADAM10 | ADAM metallopeptidase domain 10 | Plasma Membrane | peptidase | 5.78E+07 | 4.08E+07 | 4.64E+07 | | | | | | |
| O35598 | ADAM10 | ADAM metallopeptidase domain 10 | Plasma Membrane | peptidase | 8.49E+07 | 6.68E+07 | 4.52E+07 | | | | | | |
| O35598 | ADAM10 | ADAM metallopeptidase domain 10 | Plasma Membrane | peptidase | 4.51E+07 | 3.86E+07 | 4.45E+07 | | | | | | |
| O35598 | ADAM10 | ADAM metallopeptidase domain 10 | Plasma Membrane | peptidase | 3.03E+07 | 2.66E+07 | 1.99E+07 | | | | | | |
| O35598 | ADAM10 | ADAM metallopeptidase domain 10 | Plasma Membrane | peptidase | 2.62E+07 | 1.85E+07 | 1.34E+07 | | | | | | |
| O35598 | ADAM10 | ADAM metallopeptidase domain 10 | Plasma Membrane | peptidase | 2.53E+07 | 2.59E+07 | 3.77E+07 | | | | | | |
| O35598 | ADAM10 | ADAM metallopeptidase domain 10 | Plasma Membrane | peptidase | 2.47E+07 | 8932095 | 1383852.5 | | | | | | |
| O35598 | ADAM10 | ADAM metallopeptidase domain 10 | Plasma Membrane | peptidase | 1.53E+07 | 2.62E+07 | 3.29E+07 | | | | | | |

| | | | | | | | | | | | | | |
|---|---|---|---|---|---|---|---|---|---|---|---|---|---|
| O35598 | ADAM10 | ADAM metallopeptidase domain 10 | Plasma Membrane | peptidase | 1.39E+07 | 1.76E+07 | 1.34E+07 | | | | | | |
| O35598 | ADAM10 | ADAM metallopeptidase domain 10 | Plasma Membrane | peptidase | 2.10E+07 | 1.78E+07 | 1.71E+07 | | | | | | |
| O35598 | ADAM10 | ADAM metallopeptidase domain 10 | Plasma Membrane | peptidase | 9807622 | 1.12E+07 | 9285017 | | | | | | |
| O35598 | ADAM10 | ADAM metallopeptidase domain 10 | Plasma Membrane | peptidase | 1.36E+07 | 1.31E+07 | 1.27E+07 | | | | | | |
| O35598 | ADAM10 | ADAM metallopeptidase domain 10 | Plasma Membrane | peptidase | 8986733 | 5046976.5 | 3729681.25 | | | | | | |
| O35598 | ADAM10 | ADAM metallopeptidase domain 10 | Plasma Membrane | peptidase | 6023721 | 4236009 | 3260112.5 | | | | | | |
| O35598 | ADAM10 | ADAM metallopeptidase domain 10 | Plasma Membrane | peptidase | 2894398 | 3469102.75 | 1304775 | | | | | | |
| O35598 | ADAM10 | ADAM metallopeptidase domain 10 | Plasma Membrane | peptidase | 3434378.75 | 3532047.75 | 5632309 | | | | | | |
| O35598 | ADAM10 | ADAM metallopeptidase domain 10 | Plasma Membrane | peptidase | 5534824.5 | 5014500 | 6332096.5 | | | | | | |
| O35598 | ADAM10 | ADAM metallopeptidase domain 10 | Plasma Membrane | peptidase | 4538889.5 | 5717032.5 | 6725028.5 | | | | | | |
| O35598 | ADAM10 | ADAM metallopeptidase domain 10 | Plasma Membrane | peptidase | 1.32E+07 | 9675922 | 3209830.25 | | | | | | |
| O35598 | ADAM10 | ADAM metallopeptidase domain 10 | Plasma Membrane | peptidase | 5798964 | 1.21E+07 | 1.33E+07 | | | | | | |
| O35598 | ADAM10 | ADAM metallopeptidase domain 10 | Plasma Membrane | peptidase | 2542817.25 | 2962249.5 | 2955576 | O35598 | ADAM10 | 29722013 | 26256121 | 2972920 | 78 |
| Q9Z0F8 | ADAM17 | ADAM metallopeptidase domain 17 | Plasma Membrane | peptidase | 4879044 | 4906971 | 6053587.5 | | | | | | |
| Q9Z0F8 | ADAM17 | ADAM metallopeptidase domain 17 | Plasma Membrane | peptidase | 4.70E-06 | 618792.875 | 2097886 | Q9Z0F8 | ADAM17 | 2439522 | 2526821 | 1031570 | 6 |
| Q9QYC0 | ADD1 | adducin 1 (alpha) | Cytoplasm | other | 8.56E+08 | 6.47E+08 | 9.39E+08 | | | | | | |
| Q9QYC0 | ADD1 | adducin 1 (alpha) | Cytoplasm | other | 5.82E+08 | 5.07E+08 | 4.32E+08 | | | | | | |
| Q9QYC0 | ADD1 | adducin 1 (alpha) | Cytoplasm | other | 3.64E+08 | 3.84E+08 | 4.81E+08 | | | | | | |
| Q9QYC0 | ADD1 | adducin 1 (alpha) | Cytoplasm | other | 4.90E+08 | 3.35E+08 | 2.34E+08 | | | | | | |
| Q9QYC0 | ADD1 | adducin 1 (alpha) | Cytoplasm | other | 4.07E+08 | 3.80E+08 | 3.77E+08 | | | | | | |
| Q9QYC0 | ADD1 | adducin 1 (alpha) | Cytoplasm | other | 2.48E+08 | 2.35E+08 | 2.94E+08 | | | | | | |
| Q9QYC0 | ADD1 | adducin 1 (alpha) | Cytoplasm | other | 2.69E+08 | 2.36E+08 | 2.50E+08 | | | | | | |
| Q9QYC0 | ADD1 | adducin 1 (alpha) | Cytoplasm | other | 2.82E+08 | 2.32E+08 | 2.61E+08 | | | | | | |
| Q9QYC0 | ADD1 | adducin 1 (alpha) | Cytoplasm | other | 1.61E+08 | 2.55E+08 | 4.27E+08 | | | | | | |
| Q9QYC0 | ADD1 | adducin 1 (alpha) | Cytoplasm | other | 2.34E+08 | 2.30E+08 | 2.52E+08 | | | | | | |
| Q9QYC0 | ADD1 | adducin 1 (alpha) | Cytoplasm | other | 2.51E+08 | 2.33E+08 | 2.59E+08 | | | | | | |
| Q9QYC0 | ADD1 | adducin 1 (alpha) | Cytoplasm | other | 2.43E+08 | 2.15E+08 | 2.73E+08 | | | | | | |
| Q9QYC0 | ADD1 | adducin 1 (alpha) | Cytoplasm | other | 2.71E+08 | 1.88E+08 | 1.74E+08 | | | | | | |
| Q9QYC0 | ADD1 | adducin 1 (alpha) | Cytoplasm | other | 2.44E+08 | 2.11E+08 | 2.53E+08 | | | | | | |
| Q9QYC0 | ADD1 | adducin 1 (alpha) | Cytoplasm | other | 2.63E+08 | 2.46E+08 | 2.79E+08 | | | | | | |
| Q9QYC0 | ADD1 | adducin 1 (alpha) | Cytoplasm | other | 9.03E+07 | 1.32E+08 | 9.02E+07 | | | | | | |
| Q9QYC0 | ADD1 | adducin 1 (alpha) | Cytoplasm | other | 1.45E+08 | 1.02E+08 | 1.07E+08 | | | | | | |
| Q9QYC0 | ADD1 | adducin 1 (alpha) | Cytoplasm | other | 1.56E+08 | 9.20E+07 | 7.24E+07 | | | | | | |
| Q9QYC0 | ADD1 | adducin 1 (alpha) | Cytoplasm | other | 1.07E+08 | 9.50E+07 | 9.91E+07 | | | | | | |
| Q9QYC0 | ADD1 | adducin 1 (alpha) | Cytoplasm | other | 4.05E+07 | 5.36E+07 | 3.94E+07 | | | | | | |
| Q9QYC0 | ADD1 | adducin 1 (alpha) | Cytoplasm | other | 5.06E+07 | 4.11E+07 | 5.30E+07 | | | | | | |
| Q9QYC0 | ADD1 | adducin 1 (alpha) | Cytoplasm | other | 4.72E+07 | 6.29E+07 | 6.49E+07 | | | | | | |
| Q9QYC0 | ADD1 | adducin 1 (alpha) | Cytoplasm | other | 4.92E+07 | 3.40E+07 | 4.13E+07 | | | | | | |
| Q9QYC0 | ADD1 | adducin 1 (alpha) | Cytoplasm | other | 3.31E+07 | 5.66E+07 | 7.08E+07 | | | | | | |
| Q9QYC0 | ADD1 | adducin 1 (alpha) | Cytoplasm | other | 3.13E+07 | 3.12E+07 | 3379176.75 | | | | | | |
| Q9QYC0 | ADD1 | adducin 1 (alpha) | Cytoplasm | other | 2.73E+07 | 4.72E+07 | 6.28E+07 | | | | | | |
| Q9QYC0 | ADD1 | adducin 1 (alpha) | Cytoplasm | other | 1.94E+07 | 1.60E+07 | 1.77E+07 | | | | | | |
| Q9QYC0 | ADD1 | adducin 1 (alpha) | Cytoplasm | other | 2.30E+07 | 3.46E+07 | 5.11E+07 | | | | | | |
| Q9QYC0 | ADD1 | adducin 1 (alpha) | Cytoplasm | other | 1.42E+07 | 1.51E+07 | 1.49E+07 | | | | | | |
| Q9QYC0 | ADD1 | adducin 1 (alpha) | Cytoplasm | other | 2.11E+07 | 1.92E+07 | 2.84E+07 | | | | | | |
| Q9QYC0 | ADD1 | adducin 1 (alpha) | Cytoplasm | other | 1.35E+07 | 1.37E+07 | 1.87E+07 | | | | | | |
| Q9QYC0 | ADD1 | adducin 1 (alpha) | Cytoplasm | other | 8419127 | 1.56E+07 | 2.21E+07 | | | | | | |
| Q9QYC0 | ADD1 | adducin 1 (alpha) | Cytoplasm | other | 1.57E+07 | 1.27E+07 | 1.48E+07 | | | | | | |
| Q9QYC0 | ADD1 | adducin 1 (alpha) | Cytoplasm | other | 8192771.5 | 1.53E+07 | 2.23E+07 | | | | | | |
| Q9QYC0 | ADD1 | adducin 1 (alpha) | Cytoplasm | other | 7623875 | 1.05E+07 | 1.26E+07 | | | | | | |
| Q9QYC0 | ADD1 | adducin 1 (alpha) | Cytoplasm | other | 1.92E+07 | 1.25E+07 | 9389243 | | | | | | |
| Q9QYC0 | ADD1 | adducin 1 (alpha) | Cytoplasm | other | 1.19E+07 | 8252343 | 9363736 | | | | | | |
| Q9QYC0 | ADD1 | adducin 1 (alpha) | Cytoplasm | other | 3949461.5 | 5647391 | 5093460 | | | | | | |
| Q9QYC0 | ADD1 | adducin 1 (alpha) | Cytoplasm | other | 1.62E+07 | 1.15E+07 | 1.52E+07 | | | | | | |
| Q9QYC0 | ADD1 | adducin 1 (alpha) | Cytoplasm | other | 2094863.875 | 2809952.25 | 4644064.5 | | | | | | |
| Q9QYC0 | ADD1 | adducin 1 (alpha) | Cytoplasm | other | 957383.25 | 805449.438 | 502782.469 | Q9QYC0 | ADD1 | 149444817 | 175302209 | 15806468 | 123 |
| Q9QYB8 | ADD2 | adducin 2 (beta) | Cytoplasm | other | 4.41E+08 | 4.53E+08 | 6.00E+08 | | | | | | |
| Q9QYB8 | ADD2 | adducin 2 (beta) | Cytoplasm | other | 3.65E+08 | 4.05E+08 | 4.45E+08 | | | | | | |
| Q9QYB8 | ADD2 | adducin 2 (beta) | Cytoplasm | other | 3.15E+08 | 3.12E+08 | 2.16E+08 | | | | | | |
| Q9QYB8 | ADD2 | adducin 2 (beta) | Cytoplasm | other | 5.06E+08 | 5.21E+08 | 4.72E+08 | | | | | | |
| Q9QYB8 | ADD2 | adducin 2 (beta) | Cytoplasm | other | 2.23E+08 | 2.40E+08 | 1.52E+08 | | | | | | |
| Q9QYB8 | ADD2 | adducin 2 (beta) | Cytoplasm | other | 2.61E+08 | 2.38E+08 | 2.70E+08 | | | | | | |
| Q9QYB8 | ADD2 | adducin 2 (beta) | Cytoplasm | other | 2.64E+08 | 1.85E+08 | 1.59E+08 | | | | | | |
| Q9QYB8 | ADD2 | adducin 2 (beta) | Cytoplasm | other | 3.04E+08 | 2.07E+08 | 1.57E+08 | | | | | | |
| Q9QYB8 | ADD2 | adducin 2 (beta) | Cytoplasm | other | 2.22E+08 | 2.05E+08 | 2.27E+08 | | | | | | |
| Q9QYB8 | ADD2 | adducin 2 (beta) | Cytoplasm | other | 2.77E+08 | 2.62E+08 | 3.04E+08 | | | | | | |
| Q9QYB8 | ADD2 | adducin 2 (beta) | Cytoplasm | other | 2.00E+08 | 1.71E+08 | 1.88E+08 | | | | | | |
| Q9QYB8 | ADD2 | adducin 2 (beta) | Cytoplasm | other | 1.93E+08 | 2.64E+08 | 3.18E+08 | | | | | | |
| Q9QYB8 | ADD2 | adducin 2 (beta) | Cytoplasm | other | 1.91E+08 | 1.51E+08 | 1.99E+08 | | | | | | |
| Q9QYB8 | ADD2 | adducin 2 (beta) | Cytoplasm | other | 2.19E+08 | 1.98E+08 | 2.05E+08 | | | | | | |
| Q9QYB8 | ADD2 | adducin 2 (beta) | Cytoplasm | other | 1.93E+08 | 1.67E+08 | 1.83E+08 | | | | | | |
| Q9QYB8 | ADD2 | adducin 2 (beta) | Cytoplasm | other | 1.95E+08 | 1.48E+08 | 1.25E+08 | | | | | | |
| Q9QYB8 | ADD2 | adducin 2 (beta) | Cytoplasm | other | 1.48E+08 | 1.46E+08 | 1.59E+08 | | | | | | |
| Q9QYB8 | ADD2 | adducin 2 (beta) | Cytoplasm | other | 1.31E+08 | 1.49E+08 | 1.79E+08 | | | | | | |
| Q9QYB8 | ADD2 | adducin 2 (beta) | Cytoplasm | other | 6.71E+07 | 1.05E+08 | 1.09E+08 | | | | | | |
| Q9QYB8 | ADD2 | adducin 2 (beta) | Cytoplasm | other | 9.60E+07 | 8.04E+07 | 1.07E+08 | | | | | | |
| Q9QYB8 | ADD2 | adducin 2 (beta) | Cytoplasm | other | 5.38E+07 | 7.24E+07 | 8.23E+07 | | | | | | |
| Q9QYB8 | ADD2 | adducin 2 (beta) | Cytoplasm | other | 1.09E+08 | 8.73E+07 | 8.60E+07 | | | | | | |
| Q9QYB8 | ADD2 | adducin 2 (beta) | Cytoplasm | other | 5.48E+07 | 8.31E+07 | 9.01E+07 | | | | | | |
| Q9QYB8 | ADD2 | adducin 2 (beta) | Cytoplasm | other | 4.16E+07 | 5.62E+07 | 6.18E+07 | | | | | | |
| Q9QYB8 | ADD2 | adducin 2 (beta) | Cytoplasm | other | 1.66E+08 | 1.64E+08 | 1.81E+08 | | | | | | |
| Q9QYB8 | ADD2 | adducin 2 (beta) | Cytoplasm | other | 4.07E+07 | 2.89E+07 | 2.15E+07 | | | | | | |
| Q9QYB8 | ADD2 | adducin 2 (beta) | Cytoplasm | other | 6.14E+07 | 4.75E+07 | 5.09E+07 | | | | | | |
| Q9QYB8 | ADD2 | adducin 2 (beta) | Cytoplasm | other | 6.01E+07 | 6.57E+07 | 5.04E+07 | | | | | | |
| Q9QYB8 | ADD2 | adducin 2 (beta) | Cytoplasm | other | 6.08E+07 | 5.05E+07 | 5.59E+07 | | | | | | |
| Q9QYB8 | ADD2 | adducin 2 (beta) | Cytoplasm | other | 5.20E+07 | 3.89E+07 | 4.48E+07 | | | | | | |
| Q9QYB8 | ADD2 | adducin 2 (beta) | Cytoplasm | other | 4.11E+07 | 2.74E+07 | 2.47E+07 | | | | | | |
| Q9QYB8 | ADD2 | adducin 2 (beta) | Cytoplasm | other | 3.73E+07 | 2.60E+07 | 2.20E+07 | | | | | | |
| Q9QYB8 | ADD2 | adducin 2 (beta) | Cytoplasm | other | 3.66E+07 | 2.89E+07 | 3.24E+07 | | | | | | |
| Q9QYB8 | ADD2 | adducin 2 (beta) | Cytoplasm | other | 2.14E+07 | 2.77E+07 | 3.11E+07 | | | | | | |
| Q9QYB8 | ADD2 | adducin 2 (beta) | Cytoplasm | other | 1.72E+07 | 2.50E+07 | 2.13E+07 | | | | | | |

| | | | | | | | | | | | | | |
|---|---|---|---|---|---|---|---|---|---|---|---|---|---|
| Q9QYB8 | ADD2 | adducin 2 (beta) | Cytoplasm | other | 4.82E+07 | 3.35E+07 | 3.07E+07 | | | | | | |
| Q9QYB8 | ADD2 | adducin 2 (beta) | Cytoplasm | other | 3.13E+07 | 1.35E+07 | 1.11E+07 | | | | | | |
| Q9QYB8 | ADD2 | adducin 2 (beta) | Cytoplasm | other | 1.96E+07 | 2.18E+07 | 2.22E+07 | | | | | | |
| Q9QYB8 | ADD2 | adducin 2 (beta) | Cytoplasm | other | 1.70E+07 | 2.18E+07 | 2.51E+07 | | | | | | |
| Q9QYB8 | ADD2 | adducin 2 (beta) | Cytoplasm | other | 2.76E+07 | 1.47E+07 | 1.27E+07 | | | | | | |
| Q9QYB8 | ADD2 | adducin 2 (beta) | Cytoplasm | other | 3.64E+07 | 4533480.5 | 846881.938 | | | | | | |
| Q9QYB8 | ADD2 | adducin 2 (beta) | Cytoplasm | other | 1.81E+07 | 2.15E+07 | 2.77E+07 | | | | | | |
| Q9QYB8 | ADD2 | adducin 2 (beta) | Cytoplasm | other | 1.86E+07 | 1.66E+07 | 1.55E+07 | | | | | | |
| Q9QYB8 | ADD2 | adducin 2 (beta) | Cytoplasm | other | 1.13E+07 | 1.02E+07 | 7566022.5 | | | | | | |
| Q9QYB8 | ADD2 | adducin 2 (beta) | Cytoplasm | other | 6775001.5 | 8194553.5 | 8189680.5 | | | | | | |
| Q9QYB8 | ADD2 | adducin 2 (beta) | Cytoplasm | other | 7290694.5 | 6456401 | 7264327 | | | | | | |
| Q9QYB8 | ADD2 | adducin 2 (beta) | Cytoplasm | other | 5749218 | 8386623.5 | 9888686 | | | | | | |
| Q9QYB8 | ADD2 | adducin 2 (beta) | Cytoplasm | other | 1.00E+07 | 8690639 | 8391694 | | | | | | |
| Q9QYB8 | ADD2 | adducin 2 (beta) | Cytoplasm | other | 4950628 | 1.28E+07 | 2.01E+07 | | | | | | |
| Q9QYB8 | ADD2 | adducin 2 (beta) | Cytoplasm | other | 7621784 | 7345170.5 | 2.47E+07 | | | | | | |
| Q9QYB8 | ADD2 | adducin 2 (beta) | Cytoplasm | other | 3198389.25 | 2558295 | 3706238.25 | | | | | | |
| Q9QYB8 | ADD2 | adducin 2 (beta) | Cytoplasm | other | 9725808 | 1503100 | 197773.078 | | | | | | |
| Q9QYB8 | ADD2 | adducin 2 (beta) | Cytoplasm | other | 6454128 | 6211204 | 6627623 | | | | | | |
| Q9QYB8 | ADD2 | adducin 2 (beta) | Cytoplasm | other | 8184428 | 8911841 | 9231396 | | | | | | |
| Q9QYB8 | ADD2 | adducin 2 (beta) | Cytoplasm | other | 5784697 | 4356674.5 | 5733457 | | | | | | |
| Q9QYB8 | ADD2 | adducin 2 (beta) | Cytoplasm | other | 2590231.25 | 2737126 | 4668430.5 | | | | | | |
| Q9QYB8 | ADD2 | adducin 2 (beta) | Cytoplasm | other | 6770452.5 | 255324.812 | 141401.641 | | | | | | |
| Q9QYB8 | ADD2 | adducin 2 (beta) | Cytoplasm | other | 1169251.5 | 2155706.25 | 3252234.5 | | | | | | |
| Q9QYB8 | ADD2 | adducin 2 (beta) | Cytoplasm | other | 2114016.5 | 4800873.5 | 4103712 | | | | | | |
| Q9QYB8 | ADD2 | adducin 2 (beta) | Cytoplasm | other | 1449235.625 | 3432543.75 | 2712558.25 | | | | | | |
| Q9QYB8 | ADD2 | adducin 2 (beta) | Cytoplasm | other | 851533 | 1345777.625 | 1132283.75 | | | | | | |
| Q9QYB8 | ADD2 | adducin 2 (beta) | Cytoplasm | other | 937862.375 | 77255.82 | 290.159 | | | | | | |
| Q9QYB8 | ADD2 | adducin 2 (beta) | Cytoplasm | other | 1022376.438 | 992594.438 | 957859.938 | | | | | | |
| Q9QYB8 | ADD2 | adducin 2 (beta) | Cytoplasm | other | 2187136.25 | 3377514.5 | 4132895.5 | Q9QYB8 | ADD2 | 93510264 | 120355852 | 8685935 | 192 |
| Q9QYB5 | ADD3 | adducin 3 (gamma) | Cytoplasm | other | 2.53E+07 | 2.75E+07 | 3.16E+07 | | | | | | |
| Q9QYB5 | ADD3 | adducin 3 (gamma) | Cytoplasm | other | 1.48E+07 | 1.55E+07 | 1.83E+07 | | | | | | |
| Q9QYB5 | ADD3 | adducin 3 (gamma) | Cytoplasm | other | 1.03E+07 | 9254772 | 7790118.5 | | | | | | |
| Q9QYB5 | ADD3 | adducin 3 (gamma) | Cytoplasm | other | 9604144 | 9370389 | 1.07E+07 | | | | | | |
| Q9QYB5 | ADD3 | adducin 3 (gamma) | Cytoplasm | other | 8586019 | 7742664 | 1.18E+07 | | | | | | |
| Q9QYB5 | ADD3 | adducin 3 (gamma) | Cytoplasm | other | 2402453.75 | 3985196.75 | 6371277.5 | | | | | | |
| Q9QYB5 | ADD3 | adducin 3 (gamma) | Cytoplasm | other | 7788246 | 6259436 | 6540789 | | | | | | |
| Q9QYB5 | ADD3 | adducin 3 (gamma) | Cytoplasm | other | 6948636.5 | 7036342.5 | 7219495.5 | | | | | | |
| Q9QYB5 | ADD3 | adducin 3 (gamma) | Cytoplasm | other | 4566818.5 | 4888840.5 | 6839515 | | | | | | |
| Q9QYB5 | ADD3 | adducin 3 (gamma) | Cytoplasm | other | 4664311.5 | 3033845.25 | 3312542.75 | | | | | | |
| Q9QYB5 | ADD3 | adducin 3 (gamma) | Cytoplasm | other | 4526678 | 5956259 | 6203019 | | | | | | |
| Q9QYB5 | ADD3 | adducin 3 (gamma) | Cytoplasm | other | 1960474.5 | 3825623.5 | 5464141.5 | | | | | | |
| Q9QYB5 | ADD3 | adducin 3 (gamma) | Cytoplasm | other | 1620210.375 | 1986580.625 | 2645225.5 | | | | | | |
| Q9QYB5 | ADD3 | adducin 3 (gamma) | Cytoplasm | other | 4181955.25 | 4414863.5 | 5186918.5 | | | | | | |
| Q9QYB5 | ADD3 | adducin 3 (gamma) | Cytoplasm | other | 1882514.75 | 2451930.75 | 94853.141 | Q9QYB5 | ADD3 | 7280831 | 6684721 | 996499 | 45 |
| Q8VDL4 | ADPGK | ADP-dependent glucokinase | Plasma Membrane | kinase | 2449478.5 | 2826134 | 3342143 | Q8VDL4 | ADPGK | 2449479 | 448141 | 258735 | 3 |
| P54822 | ADSL | adenylosuccinate lyase | Cytoplasm | enzyme | 1.28E+07 | 1.05E+07 | 1.46E+07 | | | | | | |
| P54822 | ADSL | adenylosuccinate lyase | Cytoplasm | enzyme | 1.11E+07 | 8902991 | 9929731 | | | | | | |
| P54822 | ADSL | adenylosuccinate lyase | Cytoplasm | enzyme | 6966792.5 | 6811781.5 | 8403426 | | | | | | |
| P54822 | ADSL | adenylosuccinate lyase | Cytoplasm | enzyme | 8897858 | 8244281 | 1.03E+07 | | | | | | |
| P54822 | ADSL | adenylosuccinate lyase | Cytoplasm | enzyme | 7991038.5 | 8840482 | 6838490.5 | | | | | | |
| P54822 | ADSL | adenylosuccinate lyase | Cytoplasm | enzyme | 8097815.5 | 7307275.5 | 8868451 | | | | | | |
| P54822 | ADSL | adenylosuccinate lyase | Cytoplasm | enzyme | 6725554.5 | 4491591.5 | 5691431.5 | | | | | | |
| P54822 | ADSL | adenylosuccinate lyase | Cytoplasm | enzyme | 7084909.5 | 6126230 | 6689520.5 | | | | | | |
| P54822 | ADSL | adenylosuccinate lyase | Cytoplasm | enzyme | 5496417.5 | 4961016 | 6715651 | | | | | | |
| P54822 | ADSL | adenylosuccinate lyase | Cytoplasm | enzyme | 4535893 | 2867936.5 | 2595256.75 | | | | | | |
| P54822 | ADSL | adenylosuccinate lyase | Cytoplasm | enzyme | 2256452.75 | 2384674.75 | 2633400.75 | | | | | | |
| P54822 | ADSL | adenylosuccinate lyase | Cytoplasm | enzyme | 801078.688 | 815318.188 | 652522.938 | | | | | | |
| P54822 | ADSL | adenylosuccinate lyase | Cytoplasm | enzyme | 273899.812 | 1462412 | 2338326.25 | P54822 | ADSL | 6380593 | 3517766 | 563293 | 39 |
| Q640N1 | AEBP1 | AE binding protein 1 | Nucleus | peptidase | 1.02E+07 | 9851822 | 1.18E+07 | Q640N1 | AEBP1 | 10150000 | 1043635 | 602543 | 3 |
| O55112 | Aff2 | AF4/FMR2 family, member 2 | Nucleus | other | 3.11E+07 | 2.70E+07 | 1.28E+07 | O55112 | Aff2 | 31110000 | 9611077 | 5548958 | 3 |
| Q8K2K6 | AGFG1 | ArfGAP with FG repeats 1 | Nucleus | other | 1633864 | 1908039 | 2173981 | Q8K2K6 | AGFG1 | 1633864 | 270069 | 155924 | 3 |
| Q80WC7 | AGFG2 | ArfGAP with FG repeats 2 | Cytoplasm | other | 4035461.25 | 6129987 | 3407416.75 | | | | | | |
| Q80WC7 | AGFG2 | ArfGAP with FG repeats 2 | Cytoplasm | other | 2014133.25 | 4308313 | 5156975.5 | | | | | | |
| Q80WC7 | AGFG2 | ArfGAP with FG repeats 2 | Cytoplasm | other | 2070372.75 | 2809172.75 | 2298819 | | | | | | |
| Q80WC7 | AGFG2 | ArfGAP with FG repeats 2 | Cytoplasm | other | 3516751.25 | 4815412 | 2945294.75 | Q80WC7 | AGFG2 | 2909180 | 1301451 | 375697 | 12 |
| Q8CJG0 | AGO2 | argonaute RISC catalytic component 2 | Cytoplasm | translation regulator | 5.49E+07 | 4.71E+07 | 4.93E+07 | | | | | | |
| Q8CJG0 | AGO2 | argonaute RISC catalytic component 2 | Cytoplasm | translation regulator | 3.92E+07 | 3.56E+07 | 3.53E+07 | | | | | | |
| Q8CJG0 | AGO2 | argonaute RISC catalytic component 2 | Cytoplasm | translation regulator | 4.01E+07 | 3.64E+07 | 3.44E+07 | | | | | | |
| Q8CJG0 | AGO2 | argonaute RISC catalytic component 2 | Cytoplasm | translation regulator | 2.81E+07 | 3.05E+07 | 2.17E+07 | | | | | | |
| Q8CJG0 | AGO2 | argonaute RISC catalytic component 2 | Cytoplasm | translation regulator | 2.53E+07 | 2.89E+07 | 2.76E+07 | | | | | | |
| Q8CJG0 | AGO2 | argonaute RISC catalytic component 2 | Cytoplasm | translation regulator | 3.37E+07 | 3.35E+07 | 3.30E+07 | | | | | | |
| Q8CJG0 | AGO2 | argonaute RISC catalytic component 2 | Cytoplasm | translation regulator | 3.11E+07 | 3.10E+07 | 3.15E+07 | | | | | | |
| Q8CJG0 | AGO2 | argonaute RISC catalytic component 2 | Cytoplasm | translation regulator | 2.73E+07 | 2.25E+07 | 1.83E+07 | | | | | | |
| Q8CJG0 | AGO2 | argonaute RISC catalytic component 2 | Cytoplasm | translation regulator | 2.33E+07 | 2.69E+07 | 2.74E+07 | | | | | | |
| Q8CJG0 | AGO2 | argonaute RISC catalytic component 2 | Cytoplasm | translation regulator | 2.79E+07 | 3.06E+07 | 2.78E+07 | | | | | | |
| Q8CJG0 | AGO2 | argonaute RISC catalytic component 2 | Cytoplasm | translation regulator | 2.74E+07 | 2.60E+07 | 2.56E+07 | | | | | | |
| Q8CJG0 | AGO2 | argonaute RISC catalytic component 2 | Cytoplasm | translation regulator | 1.53E+07 | 2.01E+07 | 2.15E+07 | | | | | | |
| Q8CJG0 | AGO2 | argonaute RISC catalytic component 2 | Cytoplasm | translation regulator | 2.34E+07 | 2.16E+07 | 1.92E+07 | | | | | | |
| Q8CJG0 | AGO2 | argonaute RISC catalytic component 2 | Cytoplasm | translation regulator | 2.67E+07 | 2.86E+07 | 3.55E+07 | | | | | | |
| Q8CJG0 | AGO2 | argonaute RISC catalytic component 2 | Cytoplasm | translation regulator | 2.65E+07 | 2.42E+07 | 2.48E+07 | | | | | | |
| Q8CJG0 | AGO2 | argonaute RISC catalytic component 2 | Cytoplasm | translation regulator | 3.30E+07 | 3.26E+07 | 2.75E+07 | | | | | | |
| Q8CJG0 | AGO2 | argonaute RISC catalytic component 2 | Cytoplasm | translation regulator | 2.42E+07 | 1.83E+07 | 1.33E+07 | | | | | | |
| Q8CJG0 | AGO2 | argonaute RISC catalytic component 2 | Cytoplasm | translation regulator | 2.31E+07 | 2.27E+07 | 2.45E+07 | | | | | | |
| Q8CJG0 | AGO2 | argonaute RISC catalytic component 2 | Cytoplasm | translation regulator | 1.61E+07 | 1.35E+07 | 2.04E+07 | | | | | | |
| Q8CJG0 | AGO2 | argonaute RISC catalytic component 2 | Cytoplasm | translation regulator | 1.82E+07 | 1.18E+07 | 9713735 | | | | | | |
| Q8CJG0 | AGO2 | argonaute RISC catalytic component 2 | Cytoplasm | translation regulator | 2.53E+07 | 2.29E+07 | 2.12E+07 | | | | | | |
| Q8CJG0 | AGO2 | argonaute RISC catalytic component 2 | Cytoplasm | translation regulator | 1.53E+07 | 1.48E+07 | 1.42E+07 | | | | | | |
| Q8CJG0 | AGO2 | argonaute RISC catalytic component 2 | Cytoplasm | translation regulator | 1.69E+07 | 1.21E+07 | 9627371 | | | | | | |
| Q8CJG0 | AGO2 | argonaute RISC catalytic component 2 | Cytoplasm | translation regulator | 1.30E+07 | 1.27E+07 | 1.27E+07 | | | | | | |
| Q8CJG0 | AGO2 | argonaute RISC catalytic component 2 | Cytoplasm | translation regulator | 1.45E+07 | 1.60E+07 | 1.19E+07 | | | | | | |
| Q8CJG0 | AGO2 | argonaute RISC catalytic component 2 | Cytoplasm | translation regulator | 1.00E+07 | 9548517 | 1.31E+07 | | | | | | |

| | | | | | | | | | | | | | |
|---|---|---|---|---|---|---|---|---|---|---|---|---|---|
| Q8CJG0 | AGO2 | argonaute RISC catalytic component 2 | Cytoplasm | translation regulator | 1.09E+07 | 8934719 | 5256369 | | | | | | |
| Q8CJG0 | AGO2 | argonaute RISC catalytic component 2 | Cytoplasm | translation regulator | 9413288 | 5856028 | 1.00E+07 | | | | | | |
| Q8CJG0 | AGO2 | argonaute RISC catalytic component 2 | Cytoplasm | translation regulator | 8780732 | 7814542 | 9552266 | | | | | | |
| Q8CJG0 | AGO2 | argonaute RISC catalytic component 2 | Cytoplasm | translation regulator | 1.10E+07 | 7068387.5 | 6110660.5 | | | | | | |
| Q8CJG0 | AGO2 | argonaute RISC catalytic component 2 | Cytoplasm | translation regulator | 6932697 | 5105538 | 3509948.5 | | | | | | |
| Q8CJG0 | AGO2 | argonaute RISC catalytic component 2 | Cytoplasm | translation regulator | 3016350.25 | 6192927 | 5599382.5 | | | | | | |
| Q8CJG0 | AGO2 | argonaute RISC catalytic component 2 | Cytoplasm | translation regulator | 5538552 | 5939313 | 5094077.5 | | | | | | |
| Q8CJG0 | AGO2 | argonaute RISC catalytic component 2 | Cytoplasm | translation regulator | 4071111 | 4524559.5 | 1883953.875 | | | | | | |
| Q8CJG0 | AGO2 | argonaute RISC catalytic component 2 | Cytoplasm | translation regulator | 4309910.5 | 9435932 | 1.49E+07 | | | | | | |
| Q8CJG0 | AGO2 | argonaute RISC catalytic component 2 | Cytoplasm | translation regulator | 5641057 | 3245072.25 | 31766.154 | | | | | | |
| Q8CJG0 | AGO2 | argonaute RISC catalytic component 2 | Cytoplasm | translation regulator | 3857758.25 | 3107039.25 | 2892407.5 | | | | | | |
| Q8CJG0 | AGO2 | argonaute RISC catalytic component 2 | Cytoplasm | translation regulator | 7639161.5 | 1.68E+07 | 2.35E+07 | | | | | | |
| Q8CJG0 | AGO2 | argonaute RISC catalytic component 2 | Cytoplasm | translation regulator | 1250186 | 1945132.25 | 1817390 | | | | | | |
| Q8CJG0 | AGO2 | argonaute RISC catalytic component 2 | Cytoplasm | translation regulator | 1520184.25 | 349721.625 | 210808.922 | | | | | | |
| Q8CJG0 | AGO2 | argonaute RISC catalytic component 2 | Cytoplasm | translation regulator | 290068.062 | 360863.219 | 331553.531 | Q8CJG0 | AGO2 | 18140270 | 12032488 | 1084933 | 123 |
| O35083 | AGPAT1 | 1-acylglycerol-3-phosphate O-acyltransferase 1 | Cytoplasm | enzyme | 628791 | 683914.938 | 838057.312 | O35083 | AGPAT1 | 628791 | 108467 | 62624 | 3 |
| P50247 | AHCY | adenosylhomocysteinase | Cytoplasm | enzyme | 5861727.5 | 3960427.25 | 4302718 | | | | | | |
| P50247 | AHCY | adenosylhomocysteinase | Cytoplasm | enzyme | 4248695.5 | 2723268.75 | 3340907 | | | | | | |
| P50247 | AHCY | adenosylhomocysteinase | Cytoplasm | enzyme | 3201567.5 | 2209068 | 2503627.75 | P50247 | AHCY | 4437330 | 1134181 | 378060 | 9 |
| Q8N9S3 | Ahsa2 | AHA1, activator of heat shock protein ATPase 2 | Cytoplasm | other | 2.46E+08 | 2.45E+08 | 3.80E+08 | Q8N9S3 | Ahsa2 | 246400000 | 77889751 | 44969669 | 3 |
| Q9Z0X1 | AIFM1 | apoptosis-inducing factor, mitochondrion-associated, 1 | Cytoplasm | enzyme | 1403404.625 | 3012614.75 | 1298225.125 | | | | | | |
| Q9Z0X1 | AIFM1 | apoptosis-inducing factor, mitochondrion-associated, 1 | Cytoplasm | enzyme | 1006664.625 | 2237546.25 | 1108895.625 | Q9Z0X1 | AIFM1 | 1205035 | 785938 | 320858 | 6 |
| P31230 | AIMP1 | aminoacyl tRNA synthetase complex-interacting multifunctional protein 1 | Extracellular Space | cytokine | 2680448.25 | 1218833.5 | 1812851.25 | | | | | | |
| P31230 | AIMP1 | aminoacyl tRNA synthetase complex-interacting multifunctional protein 1 | Extracellular Space | cytokine | 684901.062 | 351913.844 | 18511.963 | P31230 | AIMP1 | 1682675 | 991663 | 404845 | 6 |
| Q9WTP6 | AK2 | adenylate kinase 2 | Cytoplasm | kinase | 322687.906 | 666324.75 | 605414.75 | Q9WTP6 | AK2 | 322688 | 183362 | 105864 | 3 |
| P10518 | ALAD | aminolevulinate dehydratase | Cytoplasm | enzyme | 6.65E+07 | 8.67E+07 | 7.41E+07 | | | | | | |
| P10518 | ALAD | aminolevulinate dehydratase | Cytoplasm | enzyme | 5.69E+07 | 4.32E+07 | 3.90E+07 | | | | | | |
| P10518 | ALAD | aminolevulinate dehydratase | Cytoplasm | enzyme | 6.18E+07 | 4.38E+07 | 3.63E+07 | | | | | | |
| P10518 | ALAD | aminolevulinate dehydratase | Cytoplasm | enzyme | 3.31E+07 | 2.20E+07 | 2.14E+07 | | | | | | |
| P10518 | ALAD | aminolevulinate dehydratase | Cytoplasm | enzyme | 3.67E+07 | 2.85E+07 | 2.86E+07 | | | | | | |
| P10518 | ALAD | aminolevulinate dehydratase | Cytoplasm | enzyme | 2.21E+07 | 2.19E+07 | 2.31E+07 | | | | | | |
| P10518 | ALAD | aminolevulinate dehydratase | Cytoplasm | enzyme | 2.14E+07 | 1.60E+07 | 1.46E+07 | | | | | | |
| P10518 | ALAD | aminolevulinate dehydratase | Cytoplasm | enzyme | 9888068 | 5614941.5 | 3926253.5 | | | | | | |
| P10518 | ALAD | aminolevulinate dehydratase | Cytoplasm | enzyme | 5362249 | 5231674 | 4495256 | | | | | | |
| P10518 | ALAD | aminolevulinate dehydratase | Cytoplasm | enzyme | 4691747 | 4737806 | 4237766 | | | | | | |
| P10518 | ALAD | aminolevulinate dehydratase | Cytoplasm | enzyme | 7414279.5 | 1.41E+07 | 1.51E+07 | | | | | | |
| P10518 | ALAD | aminolevulinate dehydratase | Cytoplasm | enzyme | 6251369 | 5622573.5 | 4827508 | | | | | | |
| P10518 | ALAD | aminolevulinate dehydratase | Cytoplasm | enzyme | 4629422.5 | 4329604 | 4733273.5 | | | | | | |
| P10518 | ALAD | aminolevulinate dehydratase | Cytoplasm | enzyme | 5376010.5 | 5225973.5 | 7315608 | | | | | | |
| P10518 | ALAD | aminolevulinate dehydratase | Cytoplasm | enzyme | 4043621 | 453174.188 | 4243024.5 | | | | | | |
| P10518 | ALAD | aminolevulinate dehydratase | Cytoplasm | enzyme | 1515255.875 | 1111555.875 | 609354.688 | P10518 | ALAD | 21730126 | 21194963 | 3059229 | 48 |
| P08680 | ALAS2 | 5'-aminolevulinate synthase 2 | Cytoplasm | enzyme | 2.25E+07 | 5.26E+07 | 1.69E+07 | | | | | | |
| P08680 | ALAS2 | 5'-aminolevulinate synthase 2 | Cytoplasm | enzyme | 1.82E+07 | 3.48E+07 | 1.03E+07 | | | | | | |
| P08680 | ALAS2 | 5'-aminolevulinate synthase 2 | Cytoplasm | enzyme | 1.10E+07 | 2.35E+07 | 7531503 | | | | | | |
| P08680 | ALAS2 | 5'-aminolevulinate synthase 2 | Cytoplasm | enzyme | 9356097 | 1.45E+07 | 8603455 | | | | | | |
| P08680 | ALAS2 | 5'-aminolevulinate synthase 2 | Cytoplasm | enzyme | 8231066 | 1.46E+07 | 6961005 | | | | | | |
| P08680 | ALAS2 | 5'-aminolevulinate synthase 2 | Cytoplasm | enzyme | 5884614.5 | 1.14E+07 | 6250543 | | | | | | |
| P08680 | ALAS2 | 5'-aminolevulinate synthase 2 | Cytoplasm | enzyme | 5703682.5 | 1.07E+07 | 3217029.5 | | | | | | |
| P08680 | ALAS2 | 5'-aminolevulinate synthase 2 | Cytoplasm | enzyme | 6141297.5 | 9430379 | 5048773 | | | | | | |
| P08680 | ALAS2 | 5'-aminolevulinate synthase 2 | Cytoplasm | enzyme | 3970530.5 | 7196268.5 | 3268179.5 | | | | | | |
| P08680 | ALAS2 | 5'-aminolevulinate synthase 2 | Cytoplasm | enzyme | 2550658.25 | 7547166 | 2287090.25 | | | | | | |
| P08680 | ALAS2 | 5'-aminolevulinate synthase 2 | Cytoplasm | enzyme | 2421445.5 | 3715977.5 | 1654622.75 | | | | | | |
| P08680 | ALAS2 | 5'-aminolevulinate synthase 2 | Cytoplasm | enzyme | 1649367.75 | 3028522.5 | 903645.062 | P08680 | ALAS2 | 8134897 | 10231941 | 1705323 | 36 |
| P07724 | ALB | albumin | Extracellular Space | transporter | 5.79E+07 | 6.52E+07 | 7.38E+07 | | | | | | |
| P07724 | ALB | albumin | Extracellular Space | transporter | 2.92E+07 | 2.32E+07 | 2.94E+07 | | | | | | |
| P07724 | ALB | albumin | Extracellular Space | transporter | 2.39E+07 | 2.96E+07 | 5.03E+07 | | | | | | |
| P07724 | ALB | albumin | Extracellular Space | transporter | 2.68E+07 | 2.83E+07 | 4.96E+07 | | | | | | |
| P07724 | ALB | albumin | Extracellular Space | transporter | 2.48E+07 | 3.75E+07 | 6.06E+07 | | | | | | |
| P07724 | ALB | albumin | Extracellular Space | transporter | 2.19E+07 | 3.03E+07 | 4.41E+07 | | | | | | |
| P07724 | ALB | albumin | Extracellular Space | transporter | 2.80E+07 | 3.42E+07 | 4.07E+07 | | | | | | |
| P07724 | ALB | albumin | Extracellular Space | transporter | 1.92E+07 | 2.86E+07 | 3.30E+07 | | | | | | |
| P07724 | ALB | albumin | Extracellular Space | transporter | 2.54E+07 | 2.99E+07 | 4.59E+07 | | | | | | |
| P07724 | ALB | albumin | Extracellular Space | transporter | 2.32E+07 | 2.93E+07 | 3.87E+07 | | | | | | |
| P07724 | ALB | albumin | Extracellular Space | transporter | 1.63E+07 | 2.30E+07 | 3.10E+07 | | | | | | |
| P07724 | ALB | albumin | Extracellular Space | transporter | 1.90E+07 | 1.47E+07 | 1.52E+07 | | | | | | |
| P07724 | ALB | albumin | Extracellular Space | transporter | 1.37E+07 | 1.54E+07 | 2.34E+07 | | | | | | |
| P07724 | ALB | albumin | Extracellular Space | transporter | 1.18E+07 | 1.55E+07 | 2.46E+07 | | | | | | |
| P07724 | ALB | albumin | Extracellular Space | transporter | 1.39E+07 | 1.86E+07 | 2.12E+07 | | | | | | |
| P07724 | ALB | albumin | Extracellular Space | transporter | 1.08E+07 | 1.25E+07 | 1.67E+07 | | | | | | |
| P07724 | ALB | albumin | Extracellular Space | transporter | 1.03E+07 | 1.33E+07 | 1.77E+07 | | | | | | |
| P07724 | ALB | albumin | Extracellular Space | transporter | 5646475 | 1.14E+07 | 1.81E+07 | | | | | | |
| P07724 | ALB | albumin | Extracellular Space | transporter | 7120812.5 | 1.07E+07 | 1.71E+07 | | | | | | |
| P07724 | ALB | albumin | Extracellular Space | transporter | 3481695.5 | 4705514.5 | 7462103 | | | | | | |
| P07724 | ALB | albumin | Extracellular Space | transporter | 5031460.5 | 8462329 | 1.12E+07 | | | | | | |
| P07724 | ALB | albumin | Extracellular Space | transporter | 4396141.5 | 5241305.5 | 8358307.5 | | | | | | |
| P07724 | ALB | albumin | Extracellular Space | transporter | 2816133 | 4363680 | 5489401.5 | | | | | | |
| P07724 | ALB | albumin | Extracellular Space | transporter | 1889693.625 | 2197778.25 | 3634525.75 | P07724 | ALB | 16929684 | 15801643 | 1862241 | 72 |
| Q571I9 | ALDH16A1 | aldehyde dehydrogenase 16 family, member A1 | Cytoplasm | enzyme | 6858629 | 5302771 | 3932998.75 | | | | | | |
| Q571I9 | ALDH16A1 | aldehyde dehydrogenase 16 family, member A1 | Cytoplasm | enzyme | 6709339 | 4684302.5 | 5792145 | | | | | | |
| Q571I9 | ALDH16A1 | aldehyde dehydrogenase 16 family, member A1 | Cytoplasm | enzyme | 4016640.5 | 4519629 | 3553026 | | | | | | |
| Q571I9 | ALDH16A1 | aldehyde dehydrogenase 16 family, member A1 | Cytoplasm | enzyme | 1827658.375 | 1936060.75 | 1378072.75 | | | | | | |
| Q571I9 | ALDH16A1 | aldehyde dehydrogenase 16 family, member A1 | Cytoplasm | enzyme | 2966419.25 | 2007910.375 | 2420679.25 | | | | | | |
| Q571I9 | ALDH16A1 | aldehyde dehydrogenase 16 family, member A1 | Cytoplasm | enzyme | 805356.5 | 733398.312 | 666745 | Q571I9 | ALDH16A1 | 3864007 | 2012956 | 474458 | 18 |
| Q9Z110 | ALDH18A1 | aldehyde dehydrogenase 18 family, member A1 | Cytoplasm | kinase | 2744015.5 | 4651494 | 1656809.875 | | | | | | |
| Q9Z110 | ALDH18A1 | aldehyde dehydrogenase 18 family, member A1 | Cytoplasm | kinase | 1710998.75 | 5038759.5 | 1715568.125 | | | | | | |
| Q9Z110 | ALDH18A1 | aldehyde dehydrogenase 18 family, member A1 | Cytoplasm | kinase | 774549.312 | 2415156.25 | 1465002.625 | | | | | | |
| Q9Z110 | ALDH18A1 | aldehyde dehydrogenase 18 family, member A1 | Cytoplasm | kinase | 1302169.625 | 2818188 | 1084177.25 | | | | | | |
| Q9Z110 | ALDH18A1 | aldehyde dehydrogenase 18 family, member A1 | Cytoplasm | kinase | 935439.125 | 2336127.75 | 744985.875 | | | | | | |
| Q9Z110 | ALDH18A1 | aldehyde dehydrogenase 18 family, member A1 | Cytoplasm | kinase | 1101406.25 | 2376130 | 1061867.875 | | | | | | |
| Q9Z110 | ALDH18A1 | aldehyde dehydrogenase 18 family, member A1 | Cytoplasm | kinase | 609249 | 1659210 | 869895.438 | | | | | | |
| Q9Z110 | ALDH18A1 | aldehyde dehydrogenase 18 family, member A1 | Cytoplasm | kinase | 614546.812 | 1527396.75 | 425158.219 | Q9Z110 | ALDH18A1 | 1224047 | 1178986 | 240659 | 24 |

| | | | | | | | | | | | | | |
|---|---|---|---|---|---|---|---|---|---|---|---|---|---|
| P24549 | ALDH1A1 | aldehyde dehydrogenase 1 family, member A1 | Cytoplasm | enzyme | 961668.312 | 777828.375 | 1021203.375 | P24549 | ALDH1A1 | 961668 | 126868 | 73247 | 3 |
| P47738 | ALDH2 | aldehyde dehydrogenase 2 family (mitochondrial) | Cytoplasm | enzyme | 5879672.5 | 1.64E+07 | 3.24E+07 | | | | | | |
| P47738 | ALDH2 | aldehyde dehydrogenase 2 family (mitochondrial) | Cytoplasm | enzyme | 4213653 | 1.43E+07 | 2.06E+07 | | | | | | |
| P47738 | ALDH2 | aldehyde dehydrogenase 2 family (mitochondrial) | Cytoplasm | enzyme | 2962473 | 6126303 | 4016343.5 | | | | | | |
| P47738 | ALDH2 | aldehyde dehydrogenase 2 family (mitochondrial) | Cytoplasm | enzyme | 3846917.25 | 1.04E+07 | 1.92E+07 | | | | | | |
| P47738 | ALDH2 | aldehyde dehydrogenase 2 family (mitochondrial) | Cytoplasm | enzyme | 2518484.25 | 9230863 | 1.55E+07 | | | | | | |
| P47738 | ALDH2 | aldehyde dehydrogenase 2 family (mitochondrial) | Cytoplasm | enzyme | 2690047.25 | 5966446.5 | 7384847.5 | | | | | | |
| P47738 | ALDH2 | aldehyde dehydrogenase 2 family (mitochondrial) | Cytoplasm | enzyme | 4061740.25 | 1.03E+07 | 1.11E+07 | | | | | | |
| P47738 | ALDH2 | aldehyde dehydrogenase 2 family (mitochondrial) | Cytoplasm | enzyme | 2564605 | 8169446 | 1.05E+07 | | | | | | |
| P47738 | ALDH2 | aldehyde dehydrogenase 2 family (mitochondrial) | Cytoplasm | enzyme | 1922047.25 | 8167797.5 | 1.19E+07 | | | | | | |
| P47738 | ALDH2 | aldehyde dehydrogenase 2 family (mitochondrial) | Cytoplasm | enzyme | 2651118.25 | 6259289.5 | 8351791 | | | | | | |
| P47738 | ALDH2 | aldehyde dehydrogenase 2 family (mitochondrial) | Cytoplasm | enzyme | 2267723.75 | 5948026.5 | 8511590 | | | | | | |
| P47738 | ALDH2 | aldehyde dehydrogenase 2 family (mitochondrial) | Cytoplasm | enzyme | 1367587.625 | 4703189 | 5093625.5 | | | | | | |
| P47738 | ALDH2 | aldehyde dehydrogenase 2 family (mitochondrial) | Cytoplasm | enzyme | 1529621.375 | 5065915 | 7481646.5 | | | | | | |
| P47738 | ALDH2 | aldehyde dehydrogenase 2 family (mitochondrial) | Cytoplasm | enzyme | 2015371.375 | 7618200 | 1.06E+07 | | | | | | |
| P47738 | ALDH2 | aldehyde dehydrogenase 2 family (mitochondrial) | Cytoplasm | enzyme | 2136158.25 | 7960673.5 | 1.07E+07 | | | | | | |
| P47738 | ALDH2 | aldehyde dehydrogenase 2 family (mitochondrial) | Cytoplasm | enzyme | 1417120 | 9687042 | 1.70E+07 | | | | | | |
| P47738 | ALDH2 | aldehyde dehydrogenase 2 family (mitochondrial) | Cytoplasm | enzyme | 933862.438 | 3197552.75 | 4747147 | | | | | | |
| P47738 | ALDH2 | aldehyde dehydrogenase 2 family (mitochondrial) | Cytoplasm | enzyme | 1308987.25 | 4386349.5 | 6359391.5 | | | | | | |
| P47738 | ALDH2 | aldehyde dehydrogenase 2 family (mitochondrial) | Cytoplasm | enzyme | 428841.844 | 776166.625 | 1885222.125 | | | | | | |
| P47738 | ALDH2 | aldehyde dehydrogenase 2 family (mitochondrial) | Cytoplasm | enzyme | 968010.375 | 2850884.25 | 6589683 | | | | | | |
| P47738 | ALDH2 | aldehyde dehydrogenase 2 family (mitochondrial) | Cytoplasm | enzyme | 711365.75 | 3585279.5 | 4312047.5 | | | | | | |
| P47738 | ALDH2 | aldehyde dehydrogenase 2 family (mitochondrial) | Cytoplasm | enzyme | 308051.312 | 1689035.125 | 3534464.25 | | | | | | |
| P47738 | ALDH2 | aldehyde dehydrogenase 2 family (mitochondrial) | Cytoplasm | enzyme | 671608.125 | 1564391.5 | 2093943.375 | | | | | | |
| P47738 | ALDH2 | aldehyde dehydrogenase 2 family (mitochondrial) | Cytoplasm | enzyme | 154358.797 | 1261269.25 | 2169196.25 | P47738 | ALDH2 | 2063726 | 5679779 | 669368 | 72 |
| P05064 | ALDOA | aldolase A, fructose-bisphosphate | Cytoplasm | enzyme | 7.65E+07 | 6.88E+07 | 6.35E+07 | | | | | | |
| P05064 | ALDOA | aldolase A, fructose-bisphosphate | Cytoplasm | enzyme | 5.45E+07 | 4.18E+07 | 5.50E+07 | | | | | | |
| P05064 | ALDOA | aldolase A, fructose-bisphosphate | Cytoplasm | enzyme | 3.89E+07 | 3.26E+07 | 2.94E+07 | | | | | | |
| P05064 | ALDOA | aldolase A, fructose-bisphosphate | Cytoplasm | enzyme | 3.64E+07 | 2.63E+07 | 3.25E+07 | | | | | | |
| P05064 | ALDOA | aldolase A, fructose-bisphosphate | Cytoplasm | enzyme | 2.62E+07 | 1.96E+07 | 2.58E+07 | | | | | | |
| P05064 | ALDOA | aldolase A, fructose-bisphosphate | Cytoplasm | enzyme | 1.63E+07 | 9590420 | 9444375 | | | | | | |
| P05064 | ALDOA | aldolase A, fructose-bisphosphate | Cytoplasm | enzyme | 6300974 | 6194883 | 5157898.5 | | | | | | |
| P05064 | ALDOA | aldolase A, fructose-bisphosphate | Cytoplasm | enzyme | 6311531 | 7895417.5 | 6729704.5 | | | | | | |
| P05064 | ALDOA | aldolase A, fructose-bisphosphate | Cytoplasm | enzyme | 5919674.5 | 4281477.5 | 2964971.25 | | | | | | |
| P05064 | ALDOA | aldolase A, fructose-bisphosphate | Cytoplasm | enzyme | 4763209 | 3519350.25 | 3208473 | | | | | | |
| P05064 | ALDOA | aldolase A, fructose-bisphosphate | Cytoplasm | enzyme | 4654287 | 3188941.5 | 4214751 | | | | | | |
| P05064 | ALDOA | aldolase A, fructose-bisphosphate | Cytoplasm | enzyme | 594548.062 | 886600.75 | 867314.875 | P05064 | ALDOA | 23113685 | 21529767 | 3588295 | 36 |
| O08739 | AMPD3 | adenosine monophosphate deaminase 3 | Cytoplasm | enzyme | 5110797.5 | 3876556.5 | 2457007.25 | | | | | | |
| O08739 | AMPD3 | adenosine monophosphate deaminase 3 | Cytoplasm | enzyme | 3937062.5 | 4943721.5 | 1963564.875 | | | | | | |
| O08739 | AMPD3 | adenosine monophosphate deaminase 3 | Cytoplasm | enzyme | 8457882 | 8763367 | 4030217 | | | | | | |
| O08739 | AMPD3 | adenosine monophosphate deaminase 3 | Cytoplasm | enzyme | 3754121 | 3380967.5 | 2091487 | | | | | | |
| O08739 | AMPD3 | adenosine monophosphate deaminase 3 | Cytoplasm | enzyme | 3151662.25 | 2329753.75 | 1637471.875 | | | | | | |
| O08739 | AMPD3 | adenosine monophosphate deaminase 3 | Cytoplasm | enzyme | 3085176.5 | 2441200.75 | 1383855.625 | | | | | | |
| O08739 | AMPD3 | adenosine monophosphate deaminase 3 | Cytoplasm | enzyme | 2616264.25 | 2086149.25 | 1507303.125 | | | | | | |
| O08739 | AMPD3 | adenosine monophosphate deaminase 3 | Cytoplasm | enzyme | 1630933.875 | 969503.812 | 564780.812 | O08739 | AMPD3 | 3967987 | 2048625 | 418174 | 24 |
| Q02357 | ANK1 | ankyrin 1, erythrocytic | Plasma Membrane | other | 3.28E+09 | 3.18E+09 | 3.46E+09 | | | | | | |
| Q02357 | ANK1 | ankyrin 1, erythrocytic | Plasma Membrane | other | 3.71E+09 | 3.69E+09 | 3.75E+09 | | | | | | |
| Q02357 | ANK1 | ankyrin 1, erythrocytic | Plasma Membrane | other | 2.44E+09 | 2.32E+09 | 2.77E+09 | | | | | | |
| Q02357 | ANK1 | ankyrin 1, erythrocytic | Plasma Membrane | other | 2.42E+09 | 2.70E+09 | 3.21E+09 | | | | | | |
| Q02357 | ANK1 | ankyrin 1, erythrocytic | Plasma Membrane | other | 2.22E+09 | 2.57E+09 | 2.32E+09 | | | | | | |
| Q02357 | ANK1 | ankyrin 1, erythrocytic | Plasma Membrane | other | 2.21E+09 | 2.83E+09 | 2.63E+09 | | | | | | |
| Q02357 | ANK1 | ankyrin 1, erythrocytic | Plasma Membrane | other | 2.05E+09 | 2.26E+09 | 1.99E+09 | | | | | | |
| Q02357 | ANK1 | ankyrin 1, erythrocytic | Plasma Membrane | other | 1.98E+09 | 1.97E+09 | 2.36E+09 | | | | | | |
| Q02357 | ANK1 | ankyrin 1, erythrocytic | Plasma Membrane | other | 2.14E+09 | 2.10E+09 | 2.05E+09 | | | | | | |
| Q02357 | ANK1 | ankyrin 1, erythrocytic | Plasma Membrane | other | 2.31E+09 | 2.25E+09 | 2.28E+09 | | | | | | |
| Q02357 | ANK1 | ankyrin 1, erythrocytic | Plasma Membrane | other | 1.92E+09 | 2.02E+09 | 2.55E+09 | | | | | | |
| Q02357 | ANK1 | ankyrin 1, erythrocytic | Plasma Membrane | other | 1.24E+09 | 1.41E+09 | 1.62E+09 | | | | | | |
| Q02357 | ANK1 | ankyrin 1, erythrocytic | Plasma Membrane | other | 1.83E+09 | 1.67E+09 | 1.82E+09 | | | | | | |
| Q02357 | ANK1 | ankyrin 1, erythrocytic | Plasma Membrane | other | 1.58E+09 | 1.55E+09 | 1.81E+09 | | | | | | |
| Q02357 | ANK1 | ankyrin 1, erythrocytic | Plasma Membrane | other | 1.42E+09 | 1.60E+09 | 1.86E+09 | | | | | | |
| Q02357 | ANK1 | ankyrin 1, erythrocytic | Plasma Membrane | other | 1.88E+09 | 1.80E+09 | 1.76E+09 | | | | | | |
| Q02357 | ANK1 | ankyrin 1, erythrocytic | Plasma Membrane | other | 1.73E+09 | 1.54E+09 | 1.51E+09 | | | | | | |
| Q02357 | ANK1 | ankyrin 1, erythrocytic | Plasma Membrane | other | 1.50E+09 | 1.55E+09 | 1.67E+09 | | | | | | |
| Q02357 | ANK1 | ankyrin 1, erythrocytic | Plasma Membrane | other | 1.25E+09 | 1.30E+09 | 1.32E+09 | | | | | | |
| Q02357 | ANK1 | ankyrin 1, erythrocytic | Plasma Membrane | other | 1.48E+09 | 1.38E+09 | 1.14E+09 | | | | | | |
| Q02357 | ANK1 | ankyrin 1, erythrocytic | Plasma Membrane | other | 1.68E+09 | 1.55E+09 | 1.48E+09 | | | | | | |
| Q02357 | ANK1 | ankyrin 1, erythrocytic | Plasma Membrane | other | 1.48E+09 | 1.61E+09 | 1.39E+09 | | | | | | |
| Q02357 | ANK1 | ankyrin 1, erythrocytic | Plasma Membrane | other | 1.31E+09 | 1.32E+09 | 1.37E+09 | | | | | | |
| Q02357 | ANK1 | ankyrin 1, erythrocytic | Plasma Membrane | other | 1.30E+09 | 1.63E+09 | 1.55E+09 | | | | | | |
| Q02357 | ANK1 | ankyrin 1, erythrocytic | Plasma Membrane | other | 1.46E+09 | 1.40E+09 | 1.54E+09 | | | | | | |
| Q02357 | ANK1 | ankyrin 1, erythrocytic | Plasma Membrane | other | 1.06E+09 | 8.38E+08 | 9.33E+08 | | | | | | |
| Q02357 | ANK1 | ankyrin 1, erythrocytic | Plasma Membrane | other | 1.14E+09 | 9.80E+08 | 9.81E+08 | | | | | | |
| Q02357 | ANK1 | ankyrin 1, erythrocytic | Plasma Membrane | other | 9.88E+08 | 1.06E+09 | 1.25E+09 | | | | | | |
| Q02357 | ANK1 | ankyrin 1, erythrocytic | Plasma Membrane | other | 1.09E+09 | 1.14E+09 | 1.09E+09 | | | | | | |
| Q02357 | ANK1 | ankyrin 1, erythrocytic | Plasma Membrane | other | 9.57E+08 | 9.46E+08 | 1.27E+09 | | | | | | |
| Q02357 | ANK1 | ankyrin 1, erythrocytic | Plasma Membrane | other | 9.67E+08 | 9.86E+08 | 1.10E+09 | | | | | | |
| Q02357 | ANK1 | ankyrin 1, erythrocytic | Plasma Membrane | other | 1.01E+09 | 1.36E+09 | 1.49E+09 | | | | | | |
| Q02357 | ANK1 | ankyrin 1, erythrocytic | Plasma Membrane | other | 1.10E+09 | 1.08E+09 | 1.19E+09 | | | | | | |
| Q02357 | ANK1 | ankyrin 1, erythrocytic | Plasma Membrane | other | 8.98E+08 | 8.97E+08 | 9.69E+08 | | | | | | |
| Q02357 | ANK1 | ankyrin 1, erythrocytic | Plasma Membrane | other | 1.34E+09 | 1.20E+09 | 1.34E+09 | | | | | | |
| Q02357 | ANK1 | ankyrin 1, erythrocytic | Plasma Membrane | other | 8.23E+08 | 8.18E+08 | 8.81E+08 | | | | | | |
| Q02357 | ANK1 | ankyrin 1, erythrocytic | Plasma Membrane | other | 8.50E+08 | 8.27E+08 | 8.03E+08 | | | | | | |
| Q02357 | ANK1 | ankyrin 1, erythrocytic | Plasma Membrane | other | 5.86E+08 | 7.07E+08 | 5.41E+08 | | | | | | |
| Q02357 | ANK1 | ankyrin 1, erythrocytic | Plasma Membrane | other | 8.61E+08 | 7.79E+08 | 7.88E+08 | | | | | | |
| Q02357 | ANK1 | ankyrin 1, erythrocytic | Plasma Membrane | other | 7.04E+08 | 6.47E+08 | 8.24E+08 | | | | | | |
| Q02357 | ANK1 | ankyrin 1, erythrocytic | Plasma Membrane | other | 7.18E+08 | 7.24E+08 | 7.75E+08 | | | | | | |
| Q02357 | ANK1 | ankyrin 1, erythrocytic | Plasma Membrane | other | 7.76E+08 | 7.24E+08 | 7.22E+08 | | | | | | |
| Q02357 | ANK1 | ankyrin 1, erythrocytic | Plasma Membrane | other | 7.04E+08 | 3.94E+08 | 4.02E+08 | | | | | | |
| Q02357 | ANK1 | ankyrin 1, erythrocytic | Plasma Membrane | other | 6.83E+08 | 6.45E+08 | 7.56E+08 | | | | | | |
| Q02357 | ANK1 | ankyrin 1, erythrocytic | Plasma Membrane | other | 7.71E+08 | 5.42E+08 | 7.02E+08 | | | | | | |
| Q02357 | ANK1 | ankyrin 1, erythrocytic | Plasma Membrane | other | 6.65E+08 | 6.03E+08 | 6.28E+08 | | | | | | |

| | | | | | | | |
|---|---|---|---|---|---|---|---|
| Q02357 | ANK1 | ankyrin 1, erythrocytic | Plasma Membrane | other | 9.37E+08 | 8.92E+08 | 9.67E+08 |
| Q02357 | ANK1 | ankyrin 1, erythrocytic | Plasma Membrane | other | 8.57E+08 | 3.75E+08 | 1.30E+08 |
| Q02357 | ANK1 | ankyrin 1, erythrocytic | Plasma Membrane | other | 7.25E+08 | 7.36E+08 | 8.38E+08 |
| Q02357 | ANK1 | ankyrin 1, erythrocytic | Plasma Membrane | other | 5.36E+08 | 6.59E+08 | 6.55E+08 |
| Q02357 | ANK1 | ankyrin 1, erythrocytic | Plasma Membrane | other | 5.31E+08 | 5.23E+08 | 5.41E+08 |
| Q02357 | ANK1 | ankyrin 1, erythrocytic | Plasma Membrane | other | 4.61E+08 | 4.49E+08 | 5.08E+08 |
| Q02357 | ANK1 | ankyrin 1, erythrocytic | Plasma Membrane | other | 6.36E+08 | 8.30E+08 | 8.72E+08 |
| Q02357 | ANK1 | ankyrin 1, erythrocytic | Plasma Membrane | other | 3.94E+08 | 4.72E+08 | 5.55E+08 |
| Q02357 | ANK1 | ankyrin 1, erythrocytic | Plasma Membrane | other | 6.16E+08 | 5.16E+08 | 6.01E+08 |
| Q02357 | ANK1 | ankyrin 1, erythrocytic | Plasma Membrane | other | 5.81E+08 | 6.01E+08 | 7.91E+08 |
| Q02357 | ANK1 | ankyrin 1, erythrocytic | Plasma Membrane | other | 4.89E+08 | 5.20E+08 | 6.74E+08 |
| Q02357 | ANK1 | ankyrin 1, erythrocytic | Plasma Membrane | other | 4.62E+08 | 5.10E+08 | 5.19E+08 |
| Q02357 | ANK1 | ankyrin 1, erythrocytic | Plasma Membrane | other | 3.25E+08 | 3.40E+08 | 4.22E+08 |
| Q02357 | ANK1 | ankyrin 1, erythrocytic | Plasma Membrane | other | 5.68E+08 | 5.02E+08 | 4.92E+08 |
| Q02357 | ANK1 | ankyrin 1, erythrocytic | Plasma Membrane | other | 6.23E+08 | 5.52E+08 | 5.75E+08 |
| Q02357 | ANK1 | ankyrin 1, erythrocytic | Plasma Membrane | other | 6.36E+08 | 5.45E+08 | 6.11E+08 |
| Q02357 | ANK1 | ankyrin 1, erythrocytic | Plasma Membrane | other | 3.89E+08 | 3.33E+08 | 3.46E+08 |
| Q02357 | ANK1 | ankyrin 1, erythrocytic | Plasma Membrane | other | 3.14E+08 | 3.93E+08 | 5.63E+08 |
| Q02357 | ANK1 | ankyrin 1, erythrocytic | Plasma Membrane | other | 3.05E+08 | 5.04E+08 | 6.90E+08 |
| Q02357 | ANK1 | ankyrin 1, erythrocytic | Plasma Membrane | other | 3.87E+08 | 4.27E+08 | 4.43E+08 |
| Q02357 | ANK1 | ankyrin 1, erythrocytic | Plasma Membrane | other | 4.20E+08 | 3.98E+08 | 4.47E+08 |
| Q02357 | ANK1 | ankyrin 1, erythrocytic | Plasma Membrane | other | 3.58E+08 | 3.34E+08 | 3.76E+08 |
| Q02357 | ANK1 | ankyrin 1, erythrocytic | Plasma Membrane | other | 2.98E+08 | 3.59E+08 | 3.35E+08 |
| Q02357 | ANK1 | ankyrin 1, erythrocytic | Plasma Membrane | other | 3.98E+08 | 3.65E+08 | 3.58E+08 |
| Q02357 | ANK1 | ankyrin 1, erythrocytic | Plasma Membrane | other | 3.34E+08 | 3.10E+08 | 5.39E+07 |
| Q02357 | ANK1 | ankyrin 1, erythrocytic | Plasma Membrane | other | 4.50E+08 | 3.76E+08 | 4.16E+08 |
| Q02357 | ANK1 | ankyrin 1, erythrocytic | Plasma Membrane | other | 2.59E+08 | 3.89E+08 | 4.25E+08 |
| Q02357 | ANK1 | ankyrin 1, erythrocytic | Plasma Membrane | other | 2.01E+08 | 2.14E+08 | 2.28E+08 |
| Q02357 | ANK1 | ankyrin 1, erythrocytic | Plasma Membrane | other | 3.75E+08 | 2.17E+08 | 1.61E+08 |
| Q02357 | ANK1 | ankyrin 1, erythrocytic | Plasma Membrane | other | 2.60E+08 | 2.48E+08 | 2.99E+08 |
| Q02357 | ANK1 | ankyrin 1, erythrocytic | Plasma Membrane | other | 3.90E+08 | 2.52E+08 | 2.39E+08 |
| Q02357 | ANK1 | ankyrin 1, erythrocytic | Plasma Membrane | other | 2.40E+08 | 2.04E+08 | 2.20E+08 |
| Q02357 | ANK1 | ankyrin 1, erythrocytic | Plasma Membrane | other | 9.26E+07 | 1.04E+08 | 1.61E+08 |
| Q02357 | ANK1 | ankyrin 1, erythrocytic | Plasma Membrane | other | 2.01E+08 | 3.05E+08 | 3.66E+08 |
| Q02357 | ANK1 | ankyrin 1, erythrocytic | Plasma Membrane | other | 2.78E+08 | 1.98E+08 | 1.67E+08 |
| Q02357 | ANK1 | ankyrin 1, erythrocytic | Plasma Membrane | other | 3.28E+08 | 1.14E+08 | 2.26E+07 |
| Q02357 | ANK1 | ankyrin 1, erythrocytic | Plasma Membrane | other | 1.97E+08 | 2.49E+08 | 3.12E+08 |
| Q02357 | ANK1 | ankyrin 1, erythrocytic | Plasma Membrane | other | 2.88E+08 | 1.66E+08 | 1.73E+08 |
| Q02357 | ANK1 | ankyrin 1, erythrocytic | Plasma Membrane | other | 2.12E+08 | 1.67E+08 | 2.14E+08 |
| Q02357 | ANK1 | ankyrin 1, erythrocytic | Plasma Membrane | other | 2.45E+08 | 2.03E+08 | 1.80E+08 |
| Q02357 | ANK1 | ankyrin 1, erythrocytic | Plasma Membrane | other | 1.71E+08 | 1.10E+08 | 2.62E+07 |
| Q02357 | ANK1 | ankyrin 1, erythrocytic | Plasma Membrane | other | 1.87E+08 | 2.50E+08 | 8.66E+07 |
| Q02357 | ANK1 | ankyrin 1, erythrocytic | Plasma Membrane | other | 1.88E+08 | 2.12E+08 | 3.84E+08 |
| Q02357 | ANK1 | ankyrin 1, erythrocytic | Plasma Membrane | other | 2.21E+08 | 1.84E+08 | 3.23E+08 |
| Q02357 | ANK1 | ankyrin 1, erythrocytic | Plasma Membrane | other | 1.18E+08 | 2.37E+08 | 2.23E+08 |
| Q02357 | ANK1 | ankyrin 1, erythrocytic | Plasma Membrane | other | 1.22E+08 | 1.12E+08 | 7.16E+07 |
| Q02357 | ANK1 | ankyrin 1, erythrocytic | Plasma Membrane | other | 1.36E+08 | 1.11E+08 | 1.15E+08 |
| Q02357 | ANK1 | ankyrin 1, erythrocytic | Plasma Membrane | other | 1.17E+08 | 1.19E+08 | 1.06E+08 |
| Q02357 | ANK1 | ankyrin 1, erythrocytic | Plasma Membrane | other | 1.23E+08 | 8.69E+07 | 7.06E+07 |
| Q02357 | ANK1 | ankyrin 1, erythrocytic | Plasma Membrane | other | 1.90E+08 | 2.39E+08 | 2.39E+08 |
| Q02357 | ANK1 | ankyrin 1, erythrocytic | Plasma Membrane | other | 8.92E+07 | 1.60E+08 | 1.33E+08 |
| Q02357 | ANK1 | ankyrin 1, erythrocytic | Plasma Membrane | other | 1.28E+08 | 1.47E+08 | 1.55E+08 |
| Q02357 | ANK1 | ankyrin 1, erythrocytic | Plasma Membrane | other | 2.16E+08 | 4.77E+07 | 6957061 |
| Q02357 | ANK1 | ankyrin 1, erythrocytic | Plasma Membrane | other | 8.06E+07 | 8.34E+07 | 9.36E+07 |
| Q02357 | ANK1 | ankyrin 1, erythrocytic | Plasma Membrane | other | 1.10E+08 | 4.78E+07 | 1.16E+07 |
| Q02357 | ANK1 | ankyrin 1, erythrocytic | Plasma Membrane | other | 1.23E+08 | 9.28E+07 | 1.78E+08 |
| Q02357 | ANK1 | ankyrin 1, erythrocytic | Plasma Membrane | other | 1.08E+08 | 1.03E+08 | 9.28E+07 |
| Q02357 | ANK1 | ankyrin 1, erythrocytic | Plasma Membrane | other | 8.39E+07 | 6.75E+07 | 5.08E+07 |
| Q02357 | ANK1 | ankyrin 1, erythrocytic | Plasma Membrane | other | 1.03E+08 | 1.96E+08 | 1.37E+08 |
| Q02357 | ANK1 | ankyrin 1, erythrocytic | Plasma Membrane | other | 1.50E+08 | 3.76E+08 | 6.35E+08 |
| Q02357 | ANK1 | ankyrin 1, erythrocytic | Plasma Membrane | other | 6.27E+07 | 7.27E+07 | 7.89E+07 |
| Q02357 | ANK1 | ankyrin 1, erythrocytic | Plasma Membrane | other | 7.45E+07 | 5.60E+07 | 6.01E+07 |
| Q02357 | ANK1 | ankyrin 1, erythrocytic | Plasma Membrane | other | 9.24E+07 | 8.49E+07 | 7.64E+07 |
| Q02357 | ANK1 | ankyrin 1, erythrocytic | Plasma Membrane | other | 6.91E+07 | 8.63E+07 | 1.01E+08 |
| Q02357 | ANK1 | ankyrin 1, erythrocytic | Plasma Membrane | other | 5.51E+07 | 3.93E+07 | 3.12E+07 |
| Q02357 | ANK1 | ankyrin 1, erythrocytic | Plasma Membrane | other | 2.97E+07 | 6.60E+07 | 9.27E+07 |
| Q02357 | ANK1 | ankyrin 1, erythrocytic | Plasma Membrane | other | 6.60E+07 | 5.20E+07 | 6.48E+07 |
| Q02357 | ANK1 | ankyrin 1, erythrocytic | Plasma Membrane | other | 7.90E+07 | 7.59E+07 | 7.46E+07 |
| Q02357 | ANK1 | ankyrin 1, erythrocytic | Plasma Membrane | other | 3.50E+07 | 7.12E+07 | 9.98E+07 |
| Q02357 | ANK1 | ankyrin 1, erythrocytic | Plasma Membrane | other | 7.04E+07 | 7.63E+07 | 8.49E+07 |
| Q02357 | ANK1 | ankyrin 1, erythrocytic | Plasma Membrane | other | 4.14E+07 | 4.25E+07 | 5.03E+07 |
| Q02357 | ANK1 | ankyrin 1, erythrocytic | Plasma Membrane | other | 3.97E+07 | 7.28E+07 | 9.32E+07 |
| Q02357 | ANK1 | ankyrin 1, erythrocytic | Plasma Membrane | other | 9.46E+07 | 2.30E+07 | 7630253 |
| Q02357 | ANK1 | ankyrin 1, erythrocytic | Plasma Membrane | other | 2.42E+07 | 2.78E+07 | 2.35E+07 |
| Q02357 | ANK1 | ankyrin 1, erythrocytic | Plasma Membrane | other | 3.50E+07 | 3.68E+07 | 4.11E+07 |
| Q02357 | ANK1 | ankyrin 1, erythrocytic | Plasma Membrane | other | 3.31E+07 | 4.09E+07 | 5.18E+07 |
| Q02357 | ANK1 | ankyrin 1, erythrocytic | Plasma Membrane | other | 3.13E+07 | 3.03E+07 | 4.60E+07 |
| Q02357 | ANK1 | ankyrin 1, erythrocytic | Plasma Membrane | other | 6.01E+07 | 5.99E+07 | 1.15E+08 |
| Q02357 | ANK1 | ankyrin 1, erythrocytic | Plasma Membrane | other | 3.12E+07 | 1.72E+07 | 4952294 |
| Q02357 | ANK1 | ankyrin 1, erythrocytic | Plasma Membrane | other | 3.36E+07 | 1.93E+07 | 1.56E+07 |
| Q02357 | ANK1 | ankyrin 1, erythrocytic | Plasma Membrane | other | 3.30E+07 | 2.72E+07 | 2.44E+07 |
| Q02357 | ANK1 | ankyrin 1, erythrocytic | Plasma Membrane | other | 6.21E+07 | 1.20E+07 | 1.32E+07 |
| Q02357 | ANK1 | ankyrin 1, erythrocytic | Plasma Membrane | other | 3.01E+07 | 2.58E+07 | 3.08E+07 |
| Q02357 | ANK1 | ankyrin 1, erythrocytic | Plasma Membrane | other | 1.62E+07 | 2.03E+07 | 2.21E+07 |
| Q02357 | ANK1 | ankyrin 1, erythrocytic | Plasma Membrane | other | 2.14E+07 | 3.02E+07 | 3.04E+07 |
| Q02357 | ANK1 | ankyrin 1, erythrocytic | Plasma Membrane | other | 4.02E+07 | 9121365 | 3157432 |
| Q02357 | ANK1 | ankyrin 1, erythrocytic | Plasma Membrane | other | 1.82E+07 | 1.52E+07 | 1.64E+07 |
| Q02357 | ANK1 | ankyrin 1, erythrocytic | Plasma Membrane | other | 2.07E+07 | 1.48E+07 | 1.91E+07 |
| Q02357 | ANK1 | ankyrin 1, erythrocytic | Plasma Membrane | other | 4.62E+07 | 3.73E+07 | 4.32E+07 |
| Q02357 | ANK1 | ankyrin 1, erythrocytic | Plasma Membrane | other | 1.05E+07 | 1.13E+07 | 9971009 |
| Q02357 | ANK1 | ankyrin 1, erythrocytic | Plasma Membrane | other | 2.29E+07 | 6350935.5 | 1091760.875 |

| | | | | | | | | | | | | | |
|---|---|---|---|---|---|---|---|---|---|---|---|---|---|
| Q02357 | ANK1 | ankyrin 1, erythrocytic | Plasma Membrane | other | 2.48E+07 | 2728500.25 | 719760.312 | | | | | | |
| Q02357 | ANK1 | ankyrin 1, erythrocytic | Plasma Membrane | other | 1.21E+07 | 1.27E+07 | 1.98E+07 | | | | | | |
| Q02357 | ANK1 | ankyrin 1, erythrocytic | Plasma Membrane | other | 6860677.5 | 4.70E-06 | 4.70E-06 | | | | | | |
| Q02357 | ANK1 | ankyrin 1, erythrocytic | Plasma Membrane | other | 1.31E+07 | 1.18E+07 | 1.28E+07 | | | | | | |
| Q02357 | ANK1 | ankyrin 1, erythrocytic | Plasma Membrane | other | 1.31E+07 | 1.01E+07 | 6150862.5 | | | | | | |
| Q02357 | ANK1 | ankyrin 1, erythrocytic | Plasma Membrane | other | 4.00E+07 | 4322366.5 | 1275624.625 | | | | | | |
| Q02357 | ANK1 | ankyrin 1, erythrocytic | Plasma Membrane | other | 1.02E+07 | 8541993 | 1.30E+07 | | | | | | |
| Q02357 | ANK1 | ankyrin 1, erythrocytic | Plasma Membrane | other | 1.00E+07 | 4800014.5 | 4896605 | | | | | | |
| Q02357 | ANK1 | ankyrin 1, erythrocytic | Plasma Membrane | other | 9752994 | 1.04E+07 | 1.93E+07 | | | | | | |
| Q02357 | ANK1 | ankyrin 1, erythrocytic | Plasma Membrane | other | 8336228.5 | 5656362 | 5239510 | | | | | | |
| Q02357 | ANK1 | ankyrin 1, erythrocytic | Plasma Membrane | other | 2850418.75 | 1010846.125 | 1900499.5 | | | | | | |
| Q02357 | ANK1 | ankyrin 1, erythrocytic | Plasma Membrane | other | 1.19E+07 | 8791042 | 1.24E+07 | | | | | | |
| Q02357 | ANK1 | ankyrin 1, erythrocytic | Plasma Membrane | other | 9385406 | 1.01E+07 | 9878324 | | | | | | |
| Q02357 | ANK1 | ankyrin 1, erythrocytic | Plasma Membrane | other | 2.43E+07 | 2.85E+07 | 3.35E+07 | | | | | | |
| Q02357 | ANK1 | ankyrin 1, erythrocytic | Plasma Membrane | other | 4740230 | 3062694 | 1278654.375 | | | | | | |
| Q02357 | ANK1 | ankyrin 1, erythrocytic | Plasma Membrane | other | 7020383 | 4442138.5 | 4282582.5 | | | | | | |
| Q02357 | ANK1 | ankyrin 1, erythrocytic | Plasma Membrane | other | 2.19E+07 | 1.28E+07 | 1.27E+07 | | | | | | |
| Q02357 | ANK1 | ankyrin 1, erythrocytic | Plasma Membrane | other | 626316.75 | 5931.619 | 990238.438 | | | | | | |
| Q02357 | ANK1 | ankyrin 1, erythrocytic | Plasma Membrane | other | 1.30E+07 | 1.64E+07 | 1.25E+07 | | | | | | |
| Q02357 | ANK1 | ankyrin 1, erythrocytic | Plasma Membrane | other | 2200758.75 | 2768537.5 | 4206497 | | | | | | |
| Q02357 | ANK1 | ankyrin 1, erythrocytic | Plasma Membrane | other | 2717237 | 1436453.125 | 899790 | | | | | | |
| Q02357 | ANK1 | ankyrin 1, erythrocytic | Plasma Membrane | other | 1.01E+07 | 1121390.75 | 475235.125 | | | | | | |
| Q02357 | ANK1 | ankyrin 1, erythrocytic | Plasma Membrane | other | 2901044.5 | 3368354.75 | 3009721.75 | | | | | | |
| Q02357 | ANK1 | ankyrin 1, erythrocytic | Plasma Membrane | other | 3819430.5 | 2350759 | 2386606.75 | | | | | | |
| Q02357 | ANK1 | ankyrin 1, erythrocytic | Plasma Membrane | other | 2483451.5 | 4184877.75 | 5424467 | | | | | | |
| Q02357 | ANK1 | ankyrin 1, erythrocytic | Plasma Membrane | other | 7010600.5 | 2.03E+07 | 6.82E+07 | | | | | | |
| Q02357 | ANK1 | ankyrin 1, erythrocytic | Plasma Membrane | other | 3978016.5 | 744230.188 | 803013.688 | | | | | | |
| Q02357 | ANK1 | ankyrin 1, erythrocytic | Plasma Membrane | other | 1317149.875 | 1016508.5 | 1277952.375 | | | | | | |
| Q02357 | ANK1 | ankyrin 1, erythrocytic | Plasma Membrane | other | 654954.75 | 1213520.625 | 321490.094 | | | | | | |
| Q02357 | ANK1 | ankyrin 1, erythrocytic | Plasma Membrane | other | 1.16E+07 | 193511.703 | 95702.328 | | | | | | |
| Q02357 | ANK1 | ankyrin 1, erythrocytic | Plasma Membrane | other | 2127132 | 5697829.5 | 4416669.5 | | | | | | |
| Q02357 | ANK1 | ankyrin 1, erythrocytic | Plasma Membrane | other | 279005.375 | 3817114.25 | 7792921 | Q02357 | ANK1 | 517749535 | 720002755 | 31976445 | 507 |
| Q9JHZ2 | ANKH | ANKH inorganic pyrophosphate transport regulator | Plasma Membrane | transporter | 1954989.125 | 2628682.75 | 1506218.5 | Q9JHZ2 | ANKH | 1954989 | 564976 | 326189 | 3 |
| Q505D1 | ANKRD28 | ankyrin repeat domain 28 | Cytoplasm | other | 1310202.125 | 1079708 | 450487.531 | Q505D1 | ANKRD28 | 1310202 | 445001 | 256921 | 3 |
| Q6P9J9 | ANO6 | anoctamin 6 | Plasma Membrane | ion channel | 758379.125 | 1097779.125 | 2055721.875 | Q6P9J9 | ANO6 | 758379 | 672798 | 388440 | 3 |
| O35381 | Anp32a | acidic (leucine-rich) nuclear phosphoprotein 32 family, member A | Nucleus | other | 1.26E+07 | 8210659 | 8138054 | | | | | | |
| O35381 | Anp32a | acidic (leucine-rich) nuclear phosphoprotein 32 family, member A | Nucleus | other | 4567786 | 4944540 | 5742074.5 | | | | | | |
| O35381 | Anp32a | acidic (leucine-rich) nuclear phosphoprotein 32 family, member A | Nucleus | other | 4569225 | 6146940.5 | 6312238.5 | O35381 | Anp32a | 7259004 | 2575627 | 858542 | 9 |
| O35643 | AP1B1 | adaptor-related protein complex 1, beta 1 subunit | Cytoplasm | transporter | 1.35E+07 | 2.05E+07 | 1.20E+07 | | | | | | |
| O35643 | AP1B1 | adaptor-related protein complex 1, beta 1 subunit | Cytoplasm | transporter | 1.12E+07 | 2.08E+07 | 1.34E+07 | | | | | | |
| O35643 | AP1B1 | adaptor-related protein complex 1, beta 1 subunit | Cytoplasm | transporter | 1.29E+07 | 1.49E+07 | 1.02E+07 | | | | | | |
| O35643 | AP1B1 | adaptor-related protein complex 1, beta 1 subunit | Cytoplasm | transporter | 1.22E+07 | 1.98E+07 | 1.55E+07 | | | | | | |
| O35643 | AP1B1 | adaptor-related protein complex 1, beta 1 subunit | Cytoplasm | transporter | 8666169 | 1.21E+07 | 8202269.5 | | | | | | |
| O35643 | AP1B1 | adaptor-related protein complex 1, beta 1 subunit | Cytoplasm | transporter | 5947754 | 6116811.5 | 3900116.75 | | | | | | |
| O35643 | AP1B1 | adaptor-related protein complex 1, beta 1 subunit | Cytoplasm | transporter | 5390044 | 6233405.5 | 3939825.5 | | | | | | |
| O35643 | AP1B1 | adaptor-related protein complex 1, beta 1 subunit | Cytoplasm | transporter | 5916588.5 | 8236474.5 | 3903142.5 | | | | | | |
| O35643 | AP1B1 | adaptor-related protein complex 1, beta 1 subunit | Cytoplasm | transporter | 5144008 | 5136172 | 3322845 | | | | | | |
| O35643 | AP1B1 | adaptor-related protein complex 1, beta 1 subunit | Cytoplasm | transporter | 4677503.5 | 5509576.5 | 4251090.5 | | | | | | |
| O35643 | AP1B1 | adaptor-related protein complex 1, beta 1 subunit | Cytoplasm | transporter | 3521625.25 | 4561472 | 2562517 | | | | | | |
| O35643 | AP1B1 | adaptor-related protein complex 1, beta 1 subunit | Cytoplasm | transporter | 997989.875 | 2671872.25 | 1611210.125 | O35643 | AP1B1 | 7502640 | 5425357 | 904226 | 36 |
| P17426 | AP2A1 | adaptor-related protein complex 2, alpha 1 subunit | Cytoplasm | transporter | 4151464.25 | 9394292 | 4261366 | | | | | | |
| P17426 | AP2A1 | adaptor-related protein complex 2, alpha 1 subunit | Cytoplasm | transporter | 4117932.25 | 7580687 | 1869619.75 | | | | | | |
| P17426 | AP2A1 | adaptor-related protein complex 2, alpha 1 subunit | Cytoplasm | transporter | 1438083.75 | 2391235.25 | 405086.969 | P17426 | AP2A1 | 3235827 | 2928767 | 976256 | 9 |
| P17427 | AP2A2 | adaptor-related protein complex 2, alpha 2 subunit | Cytoplasm | transporter | 1.14E+07 | 1.48E+07 | 8875857 | | | | | | |
| P17427 | AP2A2 | adaptor-related protein complex 2, alpha 2 subunit | Cytoplasm | transporter | 6878577.5 | 7226228.5 | 3083000.5 | | | | | | |
| P17427 | AP2A2 | adaptor-related protein complex 2, alpha 2 subunit | Cytoplasm | transporter | 3745438.25 | 6559187.5 | 1.05E+07 | | | | | | |
| P17427 | AP2A2 | adaptor-related protein complex 2, alpha 2 subunit | Cytoplasm | transporter | 4345765.5 | 4614888 | 1984951.25 | | | | | | |
| P17427 | AP2A2 | adaptor-related protein complex 2, alpha 2 subunit | Cytoplasm | transporter | 3003234.75 | 3463767.25 | 2812594.25 | | | | | | |
| P17427 | AP2A2 | adaptor-related protein complex 2, alpha 2 subunit | Cytoplasm | transporter | 2650631.25 | 3752813.5 | 2495086.5 | | | | | | |
| P17427 | AP2A2 | adaptor-related protein complex 2, alpha 2 subunit | Cytoplasm | transporter | 1814433.125 | 1830576 | 1185819.875 | | | | | | |
| P17427 | AP2A2 | adaptor-related protein complex 2, alpha 2 subunit | Cytoplasm | transporter | 1580433.375 | 2355697 | 1442240.75 | P17427 | AP2A2 | 4421064 | 3583030 | 731383 | 24 |
| Q9DBG3 | Ap2b1 | adaptor-related protein complex 2, beta 1 subunit | Plasma Membrane | other | 1242995 | 2248009.5 | 1033534 | | | | | | |
| Q9DBG3 | Ap2b1 | adaptor-related protein complex 2, beta 1 subunit | Plasma Membrane | other | 214301.141 | 267571.375 | 144474.531 | Q9DBG3 | Ap2b1 | 728648 | 822585 | 335819 | 6 |
| P84091 | AP2M1 | adaptor-related protein complex 2, mu 1 subunit | Cytoplasm | transporter | 9068274 | 1.19E+07 | 5899577.5 | | | | | | |
| P84091 | AP2M1 | adaptor-related protein complex 2, mu 1 subunit | Cytoplasm | transporter | 8212695.5 | 9337243 | 5562778 | | | | | | |
| P84091 | AP2M1 | adaptor-related protein complex 2, mu 1 subunit | Cytoplasm | transporter | 3064772 | 3598403.75 | 2147665.25 | | | | | | |
| P84091 | AP2M1 | adaptor-related protein complex 2, mu 1 subunit | Cytoplasm | transporter | 3968958.25 | 5669785 | 3029323.25 | | | | | | |
| P84091 | AP2M1 | adaptor-related protein complex 2, mu 1 subunit | Cytoplasm | transporter | 2512710.75 | 4091422.25 | 2209621.75 | P84091 | AP2M1 | 5365482 | 3013427 | 778064 | 15 |
| Q8R146 | APEH | acylaminoacyl-peptide hydrolase | Cytoplasm | peptidase | 8.43E+07 | 6.13E+07 | 4.85E+07 | | | | | | |
| Q8R146 | APEH | acylaminoacyl-peptide hydrolase | Cytoplasm | peptidase | 5.06E+07 | 2.29E+07 | 2.60E+07 | | | | | | |
| Q8R146 | APEH | acylaminoacyl-peptide hydrolase | Cytoplasm | peptidase | 3.78E+07 | 3.85E+07 | 3.04E+07 | | | | | | |
| Q8R146 | APEH | acylaminoacyl-peptide hydrolase | Cytoplasm | peptidase | 3.80E+07 | 2.28E+07 | 3.05E+07 | | | | | | |
| Q8R146 | APEH | acylaminoacyl-peptide hydrolase | Cytoplasm | peptidase | 3.92E+07 | 2.66E+07 | 2.16E+07 | | | | | | |
| Q8R146 | APEH | acylaminoacyl-peptide hydrolase | Cytoplasm | peptidase | 2.84E+07 | 1.89E+07 | 2.21E+07 | | | | | | |
| Q8R146 | APEH | acylaminoacyl-peptide hydrolase | Cytoplasm | peptidase | 2.97E+07 | 2.26E+07 | 2.35E+07 | | | | | | |
| Q8R146 | APEH | acylaminoacyl-peptide hydrolase | Cytoplasm | peptidase | 2.63E+07 | 1.70E+07 | 1.54E+07 | | | | | | |
| Q8R146 | APEH | acylaminoacyl-peptide hydrolase | Cytoplasm | peptidase | 3.43E+07 | 1.59E+07 | 1.37E+07 | | | | | | |
| Q8R146 | APEH | acylaminoacyl-peptide hydrolase | Cytoplasm | peptidase | 2.68E+07 | 1.54E+07 | 1.76E+07 | | | | | | |
| Q8R146 | APEH | acylaminoacyl-peptide hydrolase | Cytoplasm | peptidase | 2.27E+07 | 1.65E+07 | 1.59E+07 | | | | | | |
| Q8R146 | APEH | acylaminoacyl-peptide hydrolase | Cytoplasm | peptidase | 2.88E+07 | 1.90E+07 | 1.82E+07 | | | | | | |
| Q8R146 | APEH | acylaminoacyl-peptide hydrolase | Cytoplasm | peptidase | 2.18E+07 | 1.41E+07 | 1.10E+07 | | | | | | |
| Q8R146 | APEH | acylaminoacyl-peptide hydrolase | Cytoplasm | peptidase | 2.32E+07 | 1.87E+07 | 2.28E+07 | | | | | | |
| Q8R146 | APEH | acylaminoacyl-peptide hydrolase | Cytoplasm | peptidase | 3.35E+07 | 2.19E+07 | 1.49E+07 | | | | | | |
| Q8R146 | APEH | acylaminoacyl-peptide hydrolase | Cytoplasm | peptidase | 2.14E+07 | 1.74E+07 | 1.77E+07 | | | | | | |
| Q8R146 | APEH | acylaminoacyl-peptide hydrolase | Cytoplasm | peptidase | 2.27E+07 | 1.33E+07 | 1.71E+07 | | | | | | |
| Q8R146 | APEH | acylaminoacyl-peptide hydrolase | Cytoplasm | peptidase | 1.66E+07 | 1.03E+07 | 1.12E+07 | | | | | | |
| Q8R146 | APEH | acylaminoacyl-peptide hydrolase | Cytoplasm | peptidase | 1.18E+07 | 5447188.5 | 5658238 | | | | | | |
| Q8R146 | APEH | acylaminoacyl-peptide hydrolase | Cytoplasm | peptidase | 5912587.5 | 4997255 | 5812774.5 | | | | | | |
| Q8R146 | APEH | acylaminoacyl-peptide hydrolase | Cytoplasm | peptidase | 5914263 | 4282754.5 | 4116943.75 | | | | | | |
| Q8R146 | APEH | acylaminoacyl-peptide hydrolase | Cytoplasm | peptidase | 5220588 | 3406309.5 | 3298178.75 | | | | | | |
| Q8R146 | APEH | acylaminoacyl-peptide hydrolase | Cytoplasm | peptidase | 4798938.5 | 3823086.75 | 3354430.75 | | | | | | |

| | | | | | | | | | | | | | |
|---|---|---|---|---|---|---|---|---|---|---|---|---|---|
| Q8R146 | APEH | acylaminoacyl-peptide hydrolase | Cytoplasm | peptidase | 1.00E+07 | 8954055 | 9273044 | | | | | | |
| Q8R146 | APEH | acylaminoacyl-peptide hydrolase | Cytoplasm | peptidase | 1508660.375 | 749808.438 | 892348.25 | Q8R146 | APEH | 25253801 | 14393158 | 1661979 | 75 |
| Q8BVF7 | APH1A | APH1A gamma secretase subunit | Cytoplasm | peptidase | 1128820.75 | 1742899.125 | 2192800 | Q8BVF7 | APH1A | 1128821 | 534097 | 308361 | 3 |
| Q00623 | APOA1 | apolipoprotein A-I | Extracellular Space | transporter | 3731810.5 | 5204668 | 4661230 | | | | | | |
| Q00623 | APOA1 | apolipoprotein A-I | Extracellular Space | transporter | 3939605.5 | 5495276.5 | 7296195 | | | | | | |
| Q00623 | APOA1 | apolipoprotein A-I | Extracellular Space | transporter | 792781.5 | 1223159.375 | 1813876.75 | Q00623 | APOA1 | 2821399 | 2164031 | 721344 | 9 |
| Q01339 | APOH | apolipoprotein H (beta-2-glycoprotein I) | Extracellular Space | transporter | 546351.625 | 1270558.5 | 523916.5 | Q01339 | APOH | 546352 | 424746 | 245227 | 3 |
| Q02013 | AQP1 | aquaporin 1 (Colton blood group) | Plasma Membrane | transporter | 2168103.75 | 256381.844 | 317710.844 | | | | | | |
| Q02013 | AQP1 | aquaporin 1 (Colton blood group) | Plasma Membrane | transporter | 7243747.5 | 6283545 | 6953285.5 | | | | | | |
| Q02013 | AQP1 | aquaporin 1 (Colton blood group) | Plasma Membrane | transporter | 1.56E+07 | 1.02E+07 | 2.14E+07 | | | | | | |
| Q02013 | AQP1 | aquaporin 1 (Colton blood group) | Plasma Membrane | transporter | 344928.938 | 500562.188 | 670803.312 | | | | | | |
| Q02013 | AQP1 | aquaporin 1 (Colton blood group) | Plasma Membrane | transporter | 3696251.5 | 3477631.75 | 4591540 | Q02013 | AQP1 | 5816606 | 6164710 | 1591721 | 15 |
| P84084 | ARF5 | ADP-ribosylation factor 5 | Cytoplasm | enzyme | 2296286.25 | 1939750 | 2350434.75 | | | | | | |
| P84084 | ARF5 | ADP-ribosylation factor 5 | Cytoplasm | enzyme | 1147514.625 | 1176339.125 | 1176834.625 | P84084 | ARF5 | 1721900 | 580886 | 237146 | 6 |
| P62331 | ARF6 | ADP-ribosylation factor 6 | Plasma Membrane | transporter | 1.67E+07 | 1.84E+07 | 2.21E+07 | | | | | | |
| P62331 | ARF6 | ADP-ribosylation factor 6 | Plasma Membrane | transporter | 1.41E+07 | 1.50E+07 | 1.67E+07 | | | | | | |
| P62331 | ARF6 | ADP-ribosylation factor 6 | Plasma Membrane | transporter | 5179148.5 | 8206962.5 | 6625587 | | | | | | |
| P62331 | ARF6 | ADP-ribosylation factor 6 | Plasma Membrane | transporter | 8775842 | 6336385 | 4797349.5 | | | | | | |
| P62331 | ARF6 | ADP-ribosylation factor 6 | Plasma Membrane | transporter | 5550665 | 7422294.5 | 7277316.5 | | | | | | |
| P62331 | ARF6 | ADP-ribosylation factor 6 | Plasma Membrane | transporter | 4209492.5 | 7742815.5 | 5005593 | P62331 | ARF6 | 9077525 | 5544627 | 1306881 | 18 |
| Q80YF9 | ARHGAP33 | Rho GTPase activating protein 33 | Plasma Membrane | transporter | 1.11E+07 | 1.58E+07 | 1.73E+07 | Q80YF9 | ARHGAP33 | 11060000 | 3248697 | 1875636 | 3 |
| Q9Z1N7 | ARID3B | AT rich interactive domain 3B (BRIGHT-like) | Nucleus | other | 1.15E+07 | 6055878 | 6268885.5 | Q9Z1N7 | ARID3B | 11460000 | 3060435 | 1766943 | 3 |
| Q9JKW0 | ARL6IP1 | ADP-ribosylation factor-like 6 interacting protein 1 | Cytoplasm | other | 7962184.5 | 8417512 | 9025689 | Q9JKW0 | ARL6IP1 | 7962185 | 533580 | 308062 | 3 |
| Q8R5J9 | ARL6IP5 | ADP-ribosylation factor-like 6 interacting protein 5 | Cytoplasm | other | 2343185.75 | 3614627.5 | 3377242 | Q8R5J9 | ARL6IP5 | 2343186 | 676041 | 390312 | 3 |
| Q8VEH3 | ARL8A | ADP-ribosylation factor-like 8A | Cytoplasm | enzyme | 2582550.75 | 1535461.125 | 1318332.75 | Q8VEH3 | ARL8A | 2582551 | 675992 | 390284 | 3 |
| Q9CQW2 | ARL8B | ADP-ribosylation factor-like 8B | Plasma Membrane | enzyme | 3.54E+07 | 4.11E+07 | 3.27E+07 | | | | | | |
| Q9CQW2 | ARL8B | ADP-ribosylation factor-like 8B | Plasma Membrane | enzyme | 2.46E+07 | 2.47E+07 | 2.65E+07 | | | | | | |
| Q9CQW2 | ARL8B | ADP-ribosylation factor-like 8B | Plasma Membrane | enzyme | 2.34E+07 | 2.84E+07 | 2.31E+07 | | | | | | |
| Q9CQW2 | ARL8B | ADP-ribosylation factor-like 8B | Plasma Membrane | enzyme | 2.22E+07 | 1.71E+07 | 9392047 | | | | | | |
| Q9CQW2 | ARL8B | ADP-ribosylation factor-like 8B | Plasma Membrane | enzyme | 1.38E+07 | 1.67E+07 | 1.84E+07 | | | | | | |
| Q9CQW2 | ARL8B | ADP-ribosylation factor-like 8B | Plasma Membrane | enzyme | 1.45E+07 | 1.52E+07 | 4430660 | | | | | | |
| Q9CQW2 | ARL8B | ADP-ribosylation factor-like 8B | Plasma Membrane | enzyme | 7615845.5 | 2.77E+07 | 5.54E+07 | | | | | | |
| Q9CQW2 | ARL8B | ADP-ribosylation factor-like 8B | Plasma Membrane | enzyme | 1.59E+07 | 6026236 | 4815855.5 | | | | | | |
| Q9CQW2 | ARL8B | ADP-ribosylation factor-like 8B | Plasma Membrane | enzyme | 1.02E+07 | 7711392 | 6063773.5 | | | | | | |
| Q9CQW2 | ARL8B | ADP-ribosylation factor-like 8B | Plasma Membrane | enzyme | 7886910 | 2.17E+07 | 4.21E+07 | | | | | | |
| Q9CQW2 | ARL8B | ADP-ribosylation factor-like 8B | Plasma Membrane | enzyme | 7798953 | 1.98E+07 | 2.42E+07 | | | | | | |
| Q9CQW2 | ARL8B | ADP-ribosylation factor-like 8B | Plasma Membrane | enzyme | 8580291 | 8955772 | 9439133 | | | | | | |
| Q9CQW2 | ARL8B | ADP-ribosylation factor-like 8B | Plasma Membrane | enzyme | 9852335 | 1.00E+07 | 8420994 | | | | | | |
| Q9CQW2 | ARL8B | ADP-ribosylation factor-like 8B | Plasma Membrane | enzyme | 7419721 | 1.06E+07 | 5579031 | Q9CQW2 | ARL8B | 14928147 | 11623574 | 1793556 | 42 |
| Q9R0Q6 | ARPC1A | actin related protein 2/3 complex, subunit 1A, 41kDa | Extracellular Space | other | 3063814 | 2215812.75 | 3074631.5 | Q9R0Q6 | ARPC1A | 3063814 | 492746 | 284487 | 3 |
| Q9CVB6 | ARPC2 | actin related protein 2/3 complex, subunit 2, 34kDa | Cytoplasm | other | 1.84E+07 | 1.54E+07 | 2.33E+07 | | | | | | |
| Q9CVB6 | ARPC2 | actin related protein 2/3 complex, subunit 2, 34kDa | Cytoplasm | other | 7820744 | 7734852.5 | 7040257.5 | | | | | | |
| Q9CVB6 | ARPC2 | actin related protein 2/3 complex, subunit 2, 34kDa | Cytoplasm | other | 4261648.5 | 5282208.5 | 4947077 | Q9CVB6 | ARPC2 | 10157464 | 6841197 | 2280399 | 9 |
| Q9JM76 | ARPC3 | actin related protein 2/3 complex, subunit 3, 21kDa | Cytoplasm | other | 3524492.25 | 3714823.5 | 4691464.5 | | | | | | |
| Q9JM76 | ARPC3 | actin related protein 2/3 complex, subunit 3, 21kDa | Cytoplasm | other | 2286443.5 | 1579721.5 | 2058459.5 | Q9JM76 | ARPC3 | 2905468 | 1187980 | 484991 | 6 |
| P59999 | ARPC4 | actin related protein 2/3 complex, subunit 4, 20kDa | Cytoplasm | other | 2198986.5 | 2397224 | 4150592.5 | P59999 | ARPC4 | 2198987 | 1074117 | 620142 | 3 |
| Q9CPW4 | ARPC5 | actin related protein 2/3 complex, subunit 5, 16kDa | Cytoplasm | other | 1146444.625 | 1153239.375 | 1521097.625 | Q9CPW4 | ARPC5 | 1146445 | 214371 | 123767 | 3 |
| Q9CRA0 | ART4 | ADP-ribosyltransferase 4 (Dombrock blood group) | Nucleus | enzyme | 3.68E+07 | 4.67E+07 | 3.43E+07 | | | | | | |
| Q9CRA0 | ART4 | ADP-ribosyltransferase 4 (Dombrock blood group) | Nucleus | enzyme | 3.32E+07 | 3.63E+07 | 3.50E+07 | | | | | | |
| Q9CRA0 | ART4 | ADP-ribosyltransferase 4 (Dombrock blood group) | Nucleus | enzyme | 1.43E+07 | 9317760 | 1.04E+07 | | | | | | |
| Q9CRA0 | ART4 | ADP-ribosyltransferase 4 (Dombrock blood group) | Nucleus | enzyme | 7382318.5 | 7360850 | 1.00E+07 | | | | | | |
| Q9CRA0 | ART4 | ADP-ribosyltransferase 4 (Dombrock blood group) | Nucleus | enzyme | 6768201.5 | 5224146 | 4911119.5 | | | | | | |
| Q9CRA0 | ART4 | ADP-ribosyltransferase 4 (Dombrock blood group) | Nucleus | enzyme | 3178180 | 3871622 | 2825130.25 | | | | | | |
| Q9CRA0 | ART4 | ADP-ribosyltransferase 4 (Dombrock blood group) | Nucleus | enzyme | 2225341.75 | 784754.438 | 5220284 | Q9CRA0 | ART4 | 14827720 | 14779566 | 3225166 | 21 |
| Q9QWY8 | ASAP1 | ArfGAP with SH3 domain, ankyrin repeat and PH domain 1 | Plasma Membrane | other | 3490032.25 | 2847425.5 | 3942353.25 | | | | | | |
| Q9QWY8 | ASAP1 | ArfGAP with SH3 domain, ankyrin repeat and PH domain 1 | Plasma Membrane | other | 1696136.125 | 1566503 | 1397010.875 | Q9QWY8 | ASAP1 | 2593084 | 1087643 | 444028 | 6 |
| Q7SIG6 | ASAP2 | ArfGAP with SH3 domain, ankyrin repeat and PH domain 2 | Nucleus | other | 1.20E+07 | 1.24E+07 | 4466174 | Q7SIG6 | ASAP2 | 11970000 | 4464777 | 2577740 | 3 |
| Q9WV71 | ASB4 | ankyrin repeat and SOCS box containing 4 | Nucleus | transcription regulator | 8482723 | 8788258 | 5472545 | Q9WV71 | ASB4 | 8482723 | 1832506 | 1057998 | 3 |
| Q4QY64 | ATAD5 | ATPase family, AAA domain containing 5 | Extracellular Space | other | 1.33E+07 | 1.36E+07 | 1.35E+07 | Q4QY64 | ATAD5 | 13290000 | 140119 | 80898 | 3 |
| P16951 | ATF2 | activating transcription factor 2 | Nucleus | transcription regulator | 6.16E+07 | 6.52E+07 | 6.66E+07 | P16951 | ATF2 | 61600000 | 2560280 | 1478178 | 3 |
| Q811C2 | ATG4C | autophagy related 4C, cysteine peptidase | Cytoplasm | peptidase | 1116163.125 | 680845.25 | 855979.375 | Q811C2 | ATG4C | 1116163 | 219039 | 126462 | 3 |
| Q9CWJ9 | ATIC | 5-aminoimidazole-4-carboxamide ribonucleotide formyltransferase/IMP cyclohydrolase | Cytoplasm | enzyme | 1.16E+07 | 1.16E+07 | 1.19E+07 | | | | | | |
| Q9CWJ9 | ATIC | 5-aminoimidazole-4-carboxamide ribonucleotide formyltransferase/IMP cyclohydrolase | Cytoplasm | enzyme | 9395874 | 6943016.5 | 6922108.5 | | | | | | |
| Q9CWJ9 | ATIC | 5-aminoimidazole-4-carboxamide ribonucleotide formyltransferase/IMP cyclohydrolase | Cytoplasm | enzyme | 1.04E+07 | 5636892.5 | 5235079.5 | | | | | | |
| Q9CWJ9 | ATIC | 5-aminoimidazole-4-carboxamide ribonucleotide formyltransferase/IMP cyclohydrolase | Cytoplasm | enzyme | 5923025 | 4215332 | 3941694.75 | | | | | | |
| Q9CWJ9 | ATIC | 5-aminoimidazole-4-carboxamide ribonucleotide formyltransferase/IMP cyclohydrolase | Cytoplasm | enzyme | 5759279.5 | 4042565.75 | 4518898 | | | | | | |
| Q9CWJ9 | ATIC | 5-aminoimidazole-4-carboxamide ribonucleotide formyltransferase/IMP cyclohydrolase | Cytoplasm | enzyme | 3544694.75 | 5760277 | 7053795.5 | | | | | | |
| Q9CWJ9 | ATIC | 5-aminoimidazole-4-carboxamide ribonucleotide formyltransferase/IMP cyclohydrolase | Cytoplasm | enzyme | 4422977.5 | 2871799.75 | 3841765.5 | | | | | | |
| Q9CWJ9 | ATIC | 5-aminoimidazole-4-carboxamide ribonucleotide formyltransferase/IMP cyclohydrolase | Cytoplasm | enzyme | 2397036.25 | 1333618.375 | 1552362.125 | | | | | | |
| Q9CWJ9 | ATIC | 5-aminoimidazole-4-carboxamide ribonucleotide formyltransferase/IMP cyclohydrolase | Cytoplasm | enzyme | 516474.781 | 1670513.5 | 1743403 | | | | | | |
| Q9CWJ9 | ATIC | 5-aminoimidazole-4-carboxamide ribonucleotide formyltransferase/IMP cyclohydrolase | Cytoplasm | enzyme | 2024127.75 | 2172585 | 2601013.25 | Q9CWJ9 | ATIC | 5594349 | 3244492 | 592360 | 30 |
| Q6PA06 | ATL2 | atlastin GTPase 2 | Cytoplasm | other | 5779194 | 6554889.5 | 7576699 | | | | | | |
| Q6PA06 | ATL2 | atlastin GTPase 2 | Cytoplasm | other | 1564572.125 | 2261680 | 2192280 | | | | | | |
| Q6PA06 | ATL2 | atlastin GTPase 2 | Cytoplasm | other | 1172948.625 | 1650280.875 | 2492965.5 | Q6PA06 | ATL2 | 2838905 | 2448968 | 816323 | 9 |
| Q91YH5 | ATL3 | atlastin GTPase 3 | Cytoplasm | other | 5.31E+07 | 6.08E+07 | 5.88E+07 | | | | | | |
| Q91YH5 | ATL3 | atlastin GTPase 3 | Cytoplasm | other | 3.27E+07 | 3.62E+07 | 3.59E+07 | | | | | | |
| Q91YH5 | ATL3 | atlastin GTPase 3 | Cytoplasm | other | 2.13E+07 | 1.93E+07 | 1.53E+07 | | | | | | |
| Q91YH5 | ATL3 | atlastin GTPase 3 | Cytoplasm | other | 2.13E+07 | 1.79E+07 | 2.11E+07 | | | | | | |
| Q91YH5 | ATL3 | atlastin GTPase 3 | Cytoplasm | other | 2.53E+07 | 2.75E+07 | 2.98E+07 | | | | | | |
| Q91YH5 | ATL3 | atlastin GTPase 3 | Cytoplasm | other | 2.30E+07 | 2.26E+07 | 2.58E+07 | | | | | | |
| Q91YH5 | ATL3 | atlastin GTPase 3 | Cytoplasm | other | 2.94E+07 | 3.60E+07 | 3.33E+07 | | | | | | |
| Q91YH5 | ATL3 | atlastin GTPase 3 | Cytoplasm | other | 1.97E+07 | 2.10E+07 | 2.09E+07 | | | | | | |
| Q91YH5 | ATL3 | atlastin GTPase 3 | Cytoplasm | other | 1.66E+07 | 1.76E+07 | 1.88E+07 | | | | | | |
| Q91YH5 | ATL3 | atlastin GTPase 3 | Cytoplasm | other | 1.14E+07 | 1.20E+07 | 9815630 | | | | | | |
| Q91YH5 | ATL3 | atlastin GTPase 3 | Cytoplasm | other | 7412292 | 1.10E+07 | 1.19E+07 | | | | | | |
| Q91YH5 | ATL3 | atlastin GTPase 3 | Cytoplasm | other | 4845286.5 | 5777435 | 6918269 | | | | | | |
| Q91YH5 | ATL3 | atlastin GTPase 3 | Cytoplasm | other | 6060225 | 7014062.5 | 5516481 | | | | | | |
| Q91YH5 | ATL3 | atlastin GTPase 3 | Cytoplasm | other | 4543284 | 4477150 | 5452693 | | | | | | |
| Q91YH5 | ATL3 | atlastin GTPase 3 | Cytoplasm | other | 4133710.25 | 2566480.25 | 2754793.75 | | | | | | |
| Q91YH5 | ATL3 | atlastin GTPase 3 | Cytoplasm | other | 4144815 | 6371800 | 6342582 | Q91YH5 | ATL3 | 17802476 | 14257843 | 2057942 | 48 |
| Q9QZW0 | ATP11C | ATPase, class VI, type 11C | Plasma Membrane | transporter | 1.22E+07 | 9902421 | 1.01E+07 | | | | | | |

| | | | | | | | | | | | | | |
|---|---|---|---|---|---|---|---|---|---|---|---|---|---|
| Q9QZW0 | ATP11C | ATPase, class VI, type 11C | Plasma Membrane | transporter | 1.20E+07 | 1.17E+07 | 1.22E+07 | | | | | | |
| Q9QZW0 | ATP11C | ATPase, class VI, type 11C | Plasma Membrane | transporter | 7061061 | 6543919.5 | 6119707 | | | | | | |
| Q9QZW0 | ATP11C | ATPase, class VI, type 11C | Plasma Membrane | transporter | 4894664 | 4372446.5 | 5298418.5 | | | | | | |
| Q9QZW0 | ATP11C | ATPase, class VI, type 11C | Plasma Membrane | transporter | 2945875.25 | 4937097.5 | 5485752 | | | | | | |
| Q9QZW0 | ATP11C | ATPase, class VI, type 11C | Plasma Membrane | transporter | 2457250 | 2997752.25 | 3078922.5 | Q9QZW0 | ATP11C | 6934808 | 3510867 | 827519 | 18 |
| Q9EPE9 | ATP13A1 | ATPase type 13A1 | Extracellular Space | transporter | 1.35E+07 | 9282360 | 1.21E+07 | Q9EPE9 | ATP13A1 | 13460000 | 2127429 | 1228271 | 3 |
| Q8VDN2 | ATP1A1 | ATPase, Na+/K+ transporting, alpha 1 polypeptide | Plasma Membrane | transporter | 3.36E+07 | 3.70E+07 | 3.94E+07 | | | | | | |
| Q8VDN2 | ATP1A1 | ATPase, Na+/K+ transporting, alpha 1 polypeptide | Plasma Membrane | transporter | 3.83E+07 | 3.47E+07 | 3.72E+07 | | | | | | |
| Q8VDN2 | ATP1A1 | ATPase, Na+/K+ transporting, alpha 1 polypeptide | Plasma Membrane | transporter | 3.20E+07 | 3.50E+07 | 3.20E+07 | | | | | | |
| Q8VDN2 | ATP1A1 | ATPase, Na+/K+ transporting, alpha 1 polypeptide | Plasma Membrane | transporter | 3.25E+07 | 3.36E+07 | 4.32E+07 | | | | | | |
| Q8VDN2 | ATP1A1 | ATPase, Na+/K+ transporting, alpha 1 polypeptide | Plasma Membrane | transporter | 3.12E+07 | 3.43E+07 | 2.89E+07 | | | | | | |
| Q8VDN2 | ATP1A1 | ATPase, Na+/K+ transporting, alpha 1 polypeptide | Plasma Membrane | transporter | 3.13E+07 | 3.45E+07 | 3.41E+07 | | | | | | |
| Q8VDN2 | ATP1A1 | ATPase, Na+/K+ transporting, alpha 1 polypeptide | Plasma Membrane | transporter | 2.33E+07 | 2.37E+07 | 1.76E+07 | | | | | | |
| Q8VDN2 | ATP1A1 | ATPase, Na+/K+ transporting, alpha 1 polypeptide | Plasma Membrane | transporter | 2.39E+07 | 2.42E+07 | 2.09E+07 | | | | | | |
| Q8VDN2 | ATP1A1 | ATPase, Na+/K+ transporting, alpha 1 polypeptide | Plasma Membrane | transporter | 1.97E+07 | 1.82E+07 | 2.34E+07 | | | | | | |
| Q8VDN2 | ATP1A1 | ATPase, Na+/K+ transporting, alpha 1 polypeptide | Plasma Membrane | transporter | 2.23E+07 | 1.93E+07 | 1.88E+07 | | | | | | |
| Q8VDN2 | ATP1A1 | ATPase, Na+/K+ transporting, alpha 1 polypeptide | Plasma Membrane | transporter | 2.08E+07 | 2.04E+07 | 1.76E+07 | | | | | | |
| Q8VDN2 | ATP1A1 | ATPase, Na+/K+ transporting, alpha 1 polypeptide | Plasma Membrane | transporter | 1.34E+07 | 1.46E+07 | 9902374 | | | | | | |
| Q8VDN2 | ATP1A1 | ATPase, Na+/K+ transporting, alpha 1 polypeptide | Plasma Membrane | transporter | 1.27E+07 | 1.32E+07 | 1.26E+07 | | | | | | |
| Q8VDN2 | ATP1A1 | ATPase, Na+/K+ transporting, alpha 1 polypeptide | Plasma Membrane | transporter | 1.30E+07 | 1.12E+07 | 1.07E+07 | | | | | | |
| Q8VDN2 | ATP1A1 | ATPase, Na+/K+ transporting, alpha 1 polypeptide | Plasma Membrane | transporter | 1.01E+07 | 1.39E+07 | 9383740 | | | | | | |
| Q8VDN2 | ATP1A1 | ATPase, Na+/K+ transporting, alpha 1 polypeptide | Plasma Membrane | transporter | 1.08E+07 | 1.45E+07 | 1.56E+07 | | | | | | |
| Q8VDN2 | ATP1A1 | ATPase, Na+/K+ transporting, alpha 1 polypeptide | Plasma Membrane | transporter | 9409468 | 1.40E+07 | 1.29E+07 | | | | | | |
| Q8VDN2 | ATP1A1 | ATPase, Na+/K+ transporting, alpha 1 polypeptide | Plasma Membrane | transporter | 6536166 | 8836742 | 5657808.5 | | | | | | |
| Q8VDN2 | ATP1A1 | ATPase, Na+/K+ transporting, alpha 1 polypeptide | Plasma Membrane | transporter | 7473941.5 | 7291999 | 6589497.5 | | | | | | |
| Q8VDN2 | ATP1A1 | ATPase, Na+/K+ transporting, alpha 1 polypeptide | Plasma Membrane | transporter | 4892341 | 8382456 | 8449559 | | | | | | |
| Q8VDN2 | ATP1A1 | ATPase, Na+/K+ transporting, alpha 1 polypeptide | Plasma Membrane | transporter | 5936675.5 | 6880251.5 | 7795315.5 | | | | | | |
| Q8VDN2 | ATP1A1 | ATPase, Na+/K+ transporting, alpha 1 polypeptide | Plasma Membrane | transporter | 3449582.5 | 4440887.5 | 2964465.75 | | | | | | |
| Q8VDN2 | ATP1A1 | ATPase, Na+/K+ transporting, alpha 1 polypeptide | Plasma Membrane | transporter | 6820444 | 6000135.5 | 5678493.5 | | | | | | |
| Q8VDN2 | ATP1A1 | ATPase, Na+/K+ transporting, alpha 1 polypeptide | Plasma Membrane | transporter | 4970260.5 | 8556989 | 1.11E+07 | | | | | | |
| Q8VDN2 | ATP1A1 | ATPase, Na+/K+ transporting, alpha 1 polypeptide | Plasma Membrane | transporter | 4707349.5 | 5856939 | 8382100 | | | | | | |
| Q8VDN2 | ATP1A1 | ATPase, Na+/K+ transporting, alpha 1 polypeptide | Plasma Membrane | transporter | 3489912.25 | 2492793 | 2664629 | | | | | | |
| Q8VDN2 | ATP1A1 | ATPase, Na+/K+ transporting, alpha 1 polypeptide | Plasma Membrane | transporter | 2286002.5 | 3418997.25 | 2027691.125 | | | | | | |
| Q8VDN2 | ATP1A1 | ATPase, Na+/K+ transporting, alpha 1 polypeptide | Plasma Membrane | transporter | 1912672.375 | 954795.375 | 1037909.875 | | | | | | |
| Q8VDN2 | ATP1A1 | ATPase, Na+/K+ transporting, alpha 1 polypeptide | Plasma Membrane | transporter | 3269145.25 | 3544418.75 | 3156833.75 | Q8VDN2 | ATP1A1 | 14963585 | 11590471 | 1242629 | 87 |
| P14231 | ATP1B2 | ATPase, Na+/K+ transporting, beta 2 polypeptide | Plasma Membrane | transporter | 6.87E+07 | 6.20E+07 | 6.30E+07 | | | | | | |
| P14231 | ATP1B2 | ATPase, Na+/K+ transporting, beta 2 polypeptide | Plasma Membrane | transporter | 4.22E+07 | 4.04E+07 | 3.59E+07 | | | | | | |
| P14231 | ATP1B2 | ATPase, Na+/K+ transporting, beta 2 polypeptide | Plasma Membrane | transporter | 1.00E+07 | 9094637 | 8480550 | | | | | | |
| P14231 | ATP1B2 | ATPase, Na+/K+ transporting, beta 2 polypeptide | Plasma Membrane | transporter | 7274703 | 6301067 | 5859468.5 | | | | | | |
| P14231 | ATP1B2 | ATPase, Na+/K+ transporting, beta 2 polypeptide | Plasma Membrane | transporter | 3800491.75 | 2590612 | 2582560.75 | P14231 | ATP1B2 | 26383039 | 24783323 | 6399026 | 15 |
| P97370 | ATP1B3 | ATPase, Na+/K+ transporting, beta 3 polypeptide | Plasma Membrane | transporter | 3426221 | 5931593.5 | 1780797.25 | | | | | | |
| P97370 | ATP1B3 | ATPase, Na+/K+ transporting, beta 3 polypeptide | Plasma Membrane | transporter | 2613103.75 | 2804765.75 | 1572689.625 | | | | | | |
| P97370 | ATP1B3 | ATPase, Na+/K+ transporting, beta 3 polypeptide | Plasma Membrane | transporter | 2060846.625 | 3655878.25 | 1626001.375 | | | | | | |
| P97370 | ATP1B3 | ATPase, Na+/K+ transporting, beta 3 polypeptide | Plasma Membrane | transporter | 1622155.25 | 3556071 | 2024881 | P97370 | ATP1B3 | 2430582 | 1276525 | 368501 | 12 |
| O55143 | ATP2A2 | ATPase, Ca++ transporting, cardiac muscle, slow twitch 2 | Cytoplasm | transporter | 4145930.75 | 6905910.5 | 6473206.5 | | | | | | |
| O55143 | ATP2A2 | ATPase, Ca++ transporting, cardiac muscle, slow twitch 2 | Cytoplasm | transporter | 2366220.5 | 5570998.5 | 1.01E+07 | | | | | | |
| O55143 | ATP2A2 | ATPase, Ca++ transporting, cardiac muscle, slow twitch 2 | Cytoplasm | transporter | 1270840.875 | 3256487.5 | 3449805 | | | | | | |
| O55143 | ATP2A2 | ATPase, Ca++ transporting, cardiac muscle, slow twitch 2 | Cytoplasm | transporter | 698139.125 | 2108494 | 2029000.875 | | | | | | |
| O55143 | ATP2A2 | ATPase, Ca++ transporting, cardiac muscle, slow twitch 2 | Cytoplasm | transporter | 910225.625 | 1703594.75 | 1638206.625 | | | | | | |
| O55143 | ATP2A2 | ATPase, Ca++ transporting, cardiac muscle, slow twitch 2 | Cytoplasm | transporter | 748435.188 | 1478620.75 | 1242401 | O55143 | ATP2A2 | 1689965 | 2595016 | 611651 | 18 |
| Q64518 | ATP2A3 | ATPase, Ca++ transporting, ubiquitous | Cytoplasm | transporter | 3899190.25 | 9316973 | 1.75E+07 | | | | | | |
| Q64518 | ATP2A3 | ATPase, Ca++ transporting, ubiquitous | Cytoplasm | transporter | 2331896.25 | 4478951.5 | 5458065.5 | | | | | | |
| Q64518 | ATP2A3 | ATPase, Ca++ transporting, ubiquitous | Cytoplasm | transporter | 691564.25 | 2746950 | 2938158.5 | | | | | | |
| Q64518 | ATP2A3 | ATPase, Ca++ transporting, ubiquitous | Cytoplasm | transporter | 863393.125 | 1921315.375 | 4685970 | | | | | | |
| Q64518 | ATP2A3 | ATPase, Ca++ transporting, ubiquitous | Cytoplasm | transporter | 943548.062 | 3812837.75 | 9651418 | | | | | | |
| Q64518 | ATP2A3 | ATPase, Ca++ transporting, ubiquitous | Cytoplasm | transporter | 298167.969 | 1038489.5 | 1986521.125 | | | | | | |
| Q64518 | ATP2A3 | ATPase, Ca++ transporting, ubiquitous | Cytoplasm | transporter | 197587.25 | 870960.438 | 3581178.75 | | | | | | |
| Q64518 | ATP2A3 | ATPase, Ca++ transporting, ubiquitous | Cytoplasm | transporter | 789539.062 | 1282254 | 2881264.75 | | | | | | |
| Q64518 | ATP2A3 | ATPase, Ca++ transporting, ubiquitous | Cytoplasm | transporter | 200368.312 | 1711221.875 | 2715115.25 | | | | | | |
| Q64518 | ATP2A3 | ATPase, Ca++ transporting, ubiquitous | Cytoplasm | transporter | 393799.469 | 1890121.5 | 4534231 | | | | | | |
| Q64518 | ATP2A3 | ATPase, Ca++ transporting, ubiquitous | Cytoplasm | transporter | 80995.766 | 77790.359 | 0 | | | | | | |
| Q64518 | ATP2A3 | ATPase, Ca++ transporting, ubiquitous | Cytoplasm | transporter | 108844.375 | 269886.625 | 745066.875 | | | | | | |
| Q64518 | ATP2A3 | ATPase, Ca++ transporting, ubiquitous | Cytoplasm | transporter | 0 | 223246.234 | 513107.75 | | | | | | |
| Q64518 | ATP2A3 | ATPase, Ca++ transporting, ubiquitous | Cytoplasm | transporter | 5255.968 | 734233.25 | 2261549.25 | | | | | | |
| Q64518 | ATP2A3 | ATPase, Ca++ transporting, ubiquitous | Cytoplasm | transporter | 61439.359 | 107261.984 | 250507.828 | | | | | | |
| Q64518 | ATP2A3 | ATPase, Ca++ transporting, ubiquitous | Cytoplasm | transporter | 78347.227 | 973220.438 | 3249582.75 | Q64518 | ATP2A3 | 683996 | 3153250 | 455132 | 48 |
| Q9R0K7 | ATP2B2 | ATPase, Ca++ transporting, plasma membrane 2 | Plasma Membrane | transporter | 1.35E+08 | 1.52E+08 | 1.47E+08 | | | | | | |
| Q9R0K7 | ATP2B2 | ATPase, Ca++ transporting, plasma membrane 2 | Plasma Membrane | transporter | 4.58E+07 | 5.29E+07 | 5.11E+07 | | | | | | |
| Q9R0K7 | ATP2B2 | ATPase, Ca++ transporting, plasma membrane 2 | Plasma Membrane | transporter | 3.57E+07 | 4.09E+07 | 5.32E+07 | | | | | | |
| Q9R0K7 | ATP2B2 | ATPase, Ca++ transporting, plasma membrane 2 | Plasma Membrane | transporter | 2.49E+07 | 2.13E+07 | 1.92E+07 | | | | | | |
| Q9R0K7 | ATP2B2 | ATPase, Ca++ transporting, plasma membrane 2 | Plasma Membrane | transporter | 2.53E+07 | 2.48E+07 | 2.67E+07 | | | | | | |
| Q9R0K7 | ATP2B2 | ATPase, Ca++ transporting, plasma membrane 2 | Plasma Membrane | transporter | 1.10E+07 | 1.14E+07 | 1.04E+07 | | | | | | |
| Q9R0K7 | ATP2B2 | ATPase, Ca++ transporting, plasma membrane 2 | Plasma Membrane | transporter | 8227184 | 7137824.5 | 5840977.5 | | | | | | |
| Q9R0K7 | ATP2B2 | ATPase, Ca++ transporting, plasma membrane 2 | Plasma Membrane | transporter | 1.04E+07 | 5687791 | 4619347.5 | | | | | | |
| Q9R0K7 | ATP2B2 | ATPase, Ca++ transporting, plasma membrane 2 | Plasma Membrane | transporter | 3921733 | 3202868.25 | 2777614.75 | | | | | | |
| Q9R0K7 | ATP2B2 | ATPase, Ca++ transporting, plasma membrane 2 | Plasma Membrane | transporter | 2108489 | 1922903.625 | 1854225.25 | Q9R0K7 | ATP2B2 | 30262741 | 41699195 | 7613197 | 30 |
| Q03265 | ATP5A1 | ATP synthase, H+ transporting, mitochondrial F1 complex, alpha subunit 1, cardiac muscle | Cytoplasm | transporter | 5.31E+07 | 1.02E+08 | 5.20E+07 | | | | | | |
| Q03265 | ATP5A1 | ATP synthase, H+ transporting, mitochondrial F1 complex, alpha subunit 1, cardiac muscle | Cytoplasm | transporter | 4.49E+07 | 9.68E+07 | 5.20E+07 | | | | | | |
| Q03265 | ATP5A1 | ATP synthase, H+ transporting, mitochondrial F1 complex, alpha subunit 1, cardiac muscle | Cytoplasm | transporter | 3.22E+07 | 7.38E+07 | 2.84E+07 | | | | | | |
| Q03265 | ATP5A1 | ATP synthase, H+ transporting, mitochondrial F1 complex, alpha subunit 1, cardiac muscle | Cytoplasm | transporter | 3.02E+07 | 6.91E+07 | 3.62E+07 | | | | | | |
| Q03265 | ATP5A1 | ATP synthase, H+ transporting, mitochondrial F1 complex, alpha subunit 1, cardiac muscle | Cytoplasm | transporter | 3.24E+07 | 6.89E+07 | 3.16E+07 | | | | | | |
| Q03265 | ATP5A1 | ATP synthase, H+ transporting, mitochondrial F1 complex, alpha subunit 1, cardiac muscle | Cytoplasm | transporter | 3.84E+07 | 6.73E+07 | 3.18E+07 | | | | | | |
| Q03265 | ATP5A1 | ATP synthase, H+ transporting, mitochondrial F1 complex, alpha subunit 1, cardiac muscle | Cytoplasm | transporter | 2.96E+07 | 6.80E+07 | 3.21E+07 | | | | | | |
| Q03265 | ATP5A1 | ATP synthase, H+ transporting, mitochondrial F1 complex, alpha subunit 1, cardiac muscle | Cytoplasm | transporter | 3.12E+07 | 5.37E+07 | 2.88E+07 | | | | | | |
| Q03265 | ATP5A1 | ATP synthase, H+ transporting, mitochondrial F1 complex, alpha subunit 1, cardiac muscle | Cytoplasm | transporter | 2.49E+07 | 5.51E+07 | 2.11E+07 | | | | | | |
| Q03265 | ATP5A1 | ATP synthase, H+ transporting, mitochondrial F1 complex, alpha subunit 1, cardiac muscle | Cytoplasm | transporter | 2.32E+07 | 4.55E+07 | 2.32E+07 | | | | | | |
| Q03265 | ATP5A1 | ATP synthase, H+ transporting, mitochondrial F1 complex, alpha subunit 1, cardiac muscle | Cytoplasm | transporter | 2.34E+07 | 5.10E+07 | 2.57E+07 | | | | | | |
| Q03265 | ATP5A1 | ATP synthase, H+ transporting, mitochondrial F1 complex, alpha subunit 1, cardiac muscle | Cytoplasm | transporter | 2.18E+07 | 4.97E+07 | 2.61E+07 | | | | | | |
| Q03265 | ATP5A1 | ATP synthase, H+ transporting, mitochondrial F1 complex, alpha subunit 1, cardiac muscle | Cytoplasm | transporter | 1.64E+07 | 3.28E+07 | 1.49E+07 | | | | | | |
| Q03265 | ATP5A1 | ATP synthase, H+ transporting, mitochondrial F1 complex, alpha subunit 1, cardiac muscle | Cytoplasm | transporter | 1.93E+07 | 3.26E+07 | 1.20E+07 | | | | | | |
| Q03265 | ATP5A1 | ATP synthase, H+ transporting, mitochondrial F1 complex, alpha subunit 1, cardiac muscle | Cytoplasm | transporter | 1.31E+07 | 2.63E+07 | 1.37E+07 | | | | | | |

| | | | | | | | | | | | | | |
|---|---|---|---|---|---|---|---|---|---|---|---|---|---|
| Q03265 | ATP5A1 | ATP synthase, H+ transporting, mitochondrial F1 complex, alpha subunit 1, cardiac muscle | Cytoplasm | transporter | 1.07E+07 | 1.82E+07 | 9442879 | | | | | | |
| Q03265 | ATP5A1 | ATP synthase, H+ transporting, mitochondrial F1 complex, alpha subunit 1, cardiac muscle | Cytoplasm | transporter | 1.13E+07 | 1.93E+07 | 1.25E+07 | | | | | | |
| Q03265 | ATP5A1 | ATP synthase, H+ transporting, mitochondrial F1 complex, alpha subunit 1, cardiac muscle | Cytoplasm | transporter | 7564590 | 1.54E+07 | 7507359 | | | | | | |
| Q03265 | ATP5A1 | ATP synthase, H+ transporting, mitochondrial F1 complex, alpha subunit 1, cardiac muscle | Cytoplasm | transporter | 5609806 | 1.08E+07 | 3942146.25 | | | | | | |
| Q03265 | ATP5A1 | ATP synthase, H+ transporting, mitochondrial F1 complex, alpha subunit 1, cardiac muscle | Cytoplasm | transporter | 3387733.75 | 1.12E+07 | 4313914.5 | | | | | | |
| Q03265 | ATP5A1 | ATP synthase, H+ transporting, mitochondrial F1 complex, alpha subunit 1, cardiac muscle | Cytoplasm | transporter | 6699533 | 1.36E+07 | 7885766.5 | | | | | | |
| Q03265 | ATP5A1 | ATP synthase, H+ transporting, mitochondrial F1 complex, alpha subunit 1, cardiac muscle | Cytoplasm | transporter | 5085148 | 1.34E+07 | 6095746.5 | | | | | | |
| Q03265 | ATP5A1 | ATP synthase, H+ transporting, mitochondrial F1 complex, alpha subunit 1, cardiac muscle | Cytoplasm | transporter | 3183618.25 | 2.20E+07 | 1.52E+07 | Q03265 | ATP5A1 | 21196540 | 22121986 | 2663174 | 69 |
| P56480 | ATP5B | ATP synthase, H+ transporting, mitochondrial F1 complex, beta polypeptide | Cytoplasm | transporter | 6.92E+07 | 1.75E+08 | 8.13E+07 | | | | | | |
| P56480 | ATP5B | ATP synthase, H+ transporting, mitochondrial F1 complex, beta polypeptide | Cytoplasm | transporter | 5.91E+07 | 1.01E+08 | 4.61E+07 | | | | | | |
| P56480 | ATP5B | ATP synthase, H+ transporting, mitochondrial F1 complex, beta polypeptide | Cytoplasm | transporter | 5.35E+07 | 1.12E+08 | 5.74E+07 | | | | | | |
| P56480 | ATP5B | ATP synthase, H+ transporting, mitochondrial F1 complex, beta polypeptide | Cytoplasm | transporter | 4.22E+07 | 7.74E+07 | 4.17E+07 | | | | | | |
| P56480 | ATP5B | ATP synthase, H+ transporting, mitochondrial F1 complex, beta polypeptide | Cytoplasm | transporter | 4.26E+07 | 8.41E+07 | 3.66E+07 | | | | | | |
| P56480 | ATP5B | ATP synthase, H+ transporting, mitochondrial F1 complex, beta polypeptide | Cytoplasm | transporter | 2.67E+07 | 4.25E+07 | 2.31E+07 | | | | | | |
| P56480 | ATP5B | ATP synthase, H+ transporting, mitochondrial F1 complex, beta polypeptide | Cytoplasm | transporter | 2.76E+07 | 5.66E+07 | 2.74E+07 | | | | | | |
| P56480 | ATP5B | ATP synthase, H+ transporting, mitochondrial F1 complex, beta polypeptide | Cytoplasm | transporter | 3.64E+07 | 1.14E+08 | 5.53E+07 | | | | | | |
| P56480 | ATP5B | ATP synthase, H+ transporting, mitochondrial F1 complex, beta polypeptide | Cytoplasm | transporter | 1.92E+07 | 5.19E+07 | 2.38E+07 | | | | | | |
| P56480 | ATP5B | ATP synthase, H+ transporting, mitochondrial F1 complex, beta polypeptide | Cytoplasm | transporter | 2.56E+07 | 5.90E+07 | 2.72E+07 | | | | | | |
| P56480 | ATP5B | ATP synthase, H+ transporting, mitochondrial F1 complex, beta polypeptide | Cytoplasm | transporter | 2.90E+07 | 4.52E+07 | 1.93E+07 | | | | | | |
| P56480 | ATP5B | ATP synthase, H+ transporting, mitochondrial F1 complex, beta polypeptide | Cytoplasm | transporter | 2.91E+07 | 6.30E+07 | 3.69E+07 | | | | | | |
| P56480 | ATP5B | ATP synthase, H+ transporting, mitochondrial F1 complex, beta polypeptide | Cytoplasm | transporter | 1.66E+07 | 3.92E+07 | 2.37E+07 | | | | | | |
| P56480 | ATP5B | ATP synthase, H+ transporting, mitochondrial F1 complex, beta polypeptide | Cytoplasm | transporter | 2.10E+07 | 4.53E+07 | 2.28E+07 | | | | | | |
| P56480 | ATP5B | ATP synthase, H+ transporting, mitochondrial F1 complex, beta polypeptide | Cytoplasm | transporter | 1.87E+07 | 4.25E+07 | 2.18E+07 | | | | | | |
| P56480 | ATP5B | ATP synthase, H+ transporting, mitochondrial F1 complex, beta polypeptide | Cytoplasm | transporter | 2.14E+07 | 4.96E+07 | 2.46E+07 | | | | | | |
| P56480 | ATP5B | ATP synthase, H+ transporting, mitochondrial F1 complex, beta polypeptide | Cytoplasm | transporter | 1.80E+07 | 3.10E+07 | 2.11E+07 | | | | | | |
| P56480 | ATP5B | ATP synthase, H+ transporting, mitochondrial F1 complex, beta polypeptide | Cytoplasm | transporter | 1.60E+07 | 2.97E+07 | 1.45E+07 | | | | | | |
| P56480 | ATP5B | ATP synthase, H+ transporting, mitochondrial F1 complex, beta polypeptide | Cytoplasm | transporter | 9428117 | 2.72E+07 | 1.49E+07 | | | | | | |
| P56480 | ATP5B | ATP synthase, H+ transporting, mitochondrial F1 complex, beta polypeptide | Cytoplasm | transporter | 1.32E+07 | 3.56E+07 | 1.40E+07 | | | | | | |
| P56480 | ATP5B | ATP synthase, H+ transporting, mitochondrial F1 complex, beta polypeptide | Cytoplasm | transporter | 9892710 | 2.53E+07 | 1.28E+07 | | | | | | |
| P56480 | ATP5B | ATP synthase, H+ transporting, mitochondrial F1 complex, beta polypeptide | Cytoplasm | transporter | 1.42E+07 | 2.67E+07 | 1.62E+07 | | | | | | |
| P56480 | ATP5B | ATP synthase, H+ transporting, mitochondrial F1 complex, beta polypeptide | Cytoplasm | transporter | 6875457.5 | 1.70E+07 | 8613347 | | | | | | |
| P56480 | ATP5B | ATP synthase, H+ transporting, mitochondrial F1 complex, beta polypeptide | Cytoplasm | transporter | 9339719 | 1.77E+07 | 5507794.5 | | | | | | |
| P56480 | ATP5B | ATP synthase, H+ transporting, mitochondrial F1 complex, beta polypeptide | Cytoplasm | transporter | 8628209 | 1.04E+07 | 5062092.5 | | | | | | |
| P56480 | ATP5B | ATP synthase, H+ transporting, mitochondrial F1 complex, beta polypeptide | Cytoplasm | transporter | 5132776.5 | 2.08E+07 | 8860294 | | | | | | |
| P56480 | ATP5B | ATP synthase, H+ transporting, mitochondrial F1 complex, beta polypeptide | Cytoplasm | transporter | 4542371.5 | 7417813.5 | 6051436.5 | | | | | | |
| P56480 | ATP5B | ATP synthase, H+ transporting, mitochondrial F1 complex, beta polypeptide | Cytoplasm | transporter | 9266699 | 4977853 | 1358128.75 | | | | | | |
| P56480 | ATP5B | ATP synthase, H+ transporting, mitochondrial F1 complex, beta polypeptide | Cytoplasm | transporter | 4099425.25 | 7342249 | 4435932 | | | | | | |
| P56480 | ATP5B | ATP synthase, H+ transporting, mitochondrial F1 complex, beta polypeptide | Cytoplasm | transporter | 4711491 | 6949110.5 | 2716176 | | | | | | |
| P56480 | ATP5B | ATP synthase, H+ transporting, mitochondrial F1 complex, beta polypeptide | Cytoplasm | transporter | 945876.875 | 1197966.875 | 1239903.125 | | | | | | |
| P56480 | ATP5B | ATP synthase, H+ transporting, mitochondrial F1 complex, beta polypeptide | Cytoplasm | transporter | 554931.875 | 1253974.625 | 2122065.5 | P56480 | ATP5B | 21016493 | 28719495 | 2931171 | 96 |
| Q91VR2 | ATP5C1 | ATP synthase, H+ transporting, mitochondrial F1 complex, gamma polypeptide 1 | Cytoplasm | transporter | 6510883 | 1.48E+07 | 7049691.5 | | | | | | |
| Q91VR2 | ATP5C1 | ATP synthase, H+ transporting, mitochondrial F1 complex, gamma polypeptide 1 | Cytoplasm | transporter | 4406596.5 | 9652468 | 5510281 | | | | | | |
| Q91VR2 | ATP5C1 | ATP synthase, H+ transporting, mitochondrial F1 complex, gamma polypeptide 1 | Cytoplasm | transporter | 3579827 | 6088817 | 1465485.125 | | | | | | |
| Q91VR2 | ATP5C1 | ATP synthase, H+ transporting, mitochondrial F1 complex, gamma polypeptide 1 | Cytoplasm | transporter | 231095.484 | 1281749 | 1598350.625 | Q91VR2 | ATP5C1 | 3682100 | 4138669 | 1194731 | 12 |
| Q9D3D9 | ATP5D | ATP synthase, H+ transporting, mitochondrial F1 complex, delta subunit | Cytoplasm | transporter | 1.06E+07 | 1.82E+07 | 1.01E+07 | | | | | | |
| Q9D3D9 | ATP5D | ATP synthase, H+ transporting, mitochondrial F1 complex, delta subunit | Cytoplasm | transporter | 1.11E+07 | 2.40E+07 | 1.17E+07 | Q9D3D9 | ATP5D | 10855000 | 5625175 | 2296468 | 6 |
| Q9CQQ7 | ATP5F1 | ATP synthase, H+ transporting, mitochondrial Fo complex, subunit B1 | Cytoplasm | transporter | 4376866 | 7691237.5 | 3896572.75 | Q9CQQ7 | ATP5F1 | 4376866 | 2066205 | 1192924 | 3 |
| Q9DCX2 | ATP5H | ATP synthase, H+ transporting, mitochondrial Fo complex, subunit d | Cytoplasm | enzyme | 8278266.5 | 1.84E+07 | 6394731 | | | | | | |
| Q9DCX2 | ATP5H | ATP synthase, H+ transporting, mitochondrial Fo complex, subunit d | Cytoplasm | enzyme | 5317767.5 | 1.27E+07 | 5239937.5 | | | | | | |
| Q9DCX2 | ATP5H | ATP synthase, H+ transporting, mitochondrial Fo complex, subunit d | Cytoplasm | enzyme | 4676965.5 | 9757186 | 4038125.25 | | | | | | |
| Q9DCX2 | ATP5H | ATP synthase, H+ transporting, mitochondrial Fo complex, subunit d | Cytoplasm | enzyme | 2690774.5 | 6781669.5 | 3606880 | | | | | | |
| Q9DCX2 | ATP5H | ATP synthase, H+ transporting, mitochondrial Fo complex, subunit d | Cytoplasm | enzyme | 2960803.25 | 7263426 | 2994610.25 | | | | | | |
| Q9DCX2 | ATP5H | ATP synthase, H+ transporting, mitochondrial Fo complex, subunit d | Cytoplasm | enzyme | 2485209.25 | 4599440.5 | 3051918.25 | Q9DCX2 | ATP5H | 4401631 | 4085353 | 962927 | 18 |
| P97450 | ATP5J | ATP synthase, H+ transporting, mitochondrial Fo complex, subunit F6 | Cytoplasm | transporter | 3874501.75 | 1.00E+07 | 4002611.25 | P97450 | ATP5J | 3874502 | 3517480 | 2030818 | 3 |
| P56135 | ATP5J2 | ATP synthase, H+ transporting, mitochondrial Fo complex, subunit F2 | Cytoplasm | transporter | 3876427 | 7288017 | 3010980.25 | P56135 | ATP5J2 | 3876427 | 2261304 | 1305565 | 3 |
| Q06185 | Atp5k | ATP synthase, H+ transporting, mitochondrial F1F0 complex, subunit E | Cytoplasm | enzyme | 8369690 | 1.90E+07 | 9749608 | | | | | | |
| Q06185 | Atp5k | ATP synthase, H+ transporting, mitochondrial F1F0 complex, subunit E | Cytoplasm | enzyme | 3818220.25 | 8303308.5 | 4196947 | Q06185 | Atp5k | 6093955 | 5486454 | 2239836 | 6 |
| Q9CPQ8 | ATP5L | ATP synthase, H+ transporting, mitochondrial Fo complex, subunit G | Cytoplasm | enzyme | 4870598 | 1.00E+07 | 4099875.25 | | | | | | |
| Q9CPQ8 | ATP5L | ATP synthase, H+ transporting, mitochondrial Fo complex, subunit G | Cytoplasm | enzyme | 3210249.75 | 8083747.5 | 2707473.5 | Q9CPQ8 | ATP5L | 4040424 | 2918709 | 1191558 | 6 |
| Q9DB20 | ATP5O | ATP synthase, H+ transporting, mitochondrial F1 complex, O subunit | Cytoplasm | transporter | 1.29E+07 | 2.38E+07 | 1.12E+07 | | | | | | |
| Q9DB20 | ATP5O | ATP synthase, H+ transporting, mitochondrial F1 complex, O subunit | Cytoplasm | transporter | 8168845.5 | 1.90E+07 | 7928978.5 | | | | | | |
| Q9DB20 | ATP5O | ATP synthase, H+ transporting, mitochondrial F1 complex, O subunit | Cytoplasm | transporter | 7035510 | 1.35E+07 | 5270238.5 | | | | | | |
| Q9DB20 | ATP5O | ATP synthase, H+ transporting, mitochondrial F1 complex, O subunit | Cytoplasm | transporter | 8502394 | 1.61E+07 | 8786062 | | | | | | |
| Q9DB20 | ATP5O | ATP synthase, H+ transporting, mitochondrial F1 complex, O subunit | Cytoplasm | transporter | 5926199.5 | 6028559.5 | 2045314.375 | | | | | | |
| Q9DB20 | ATP5O | ATP synthase, H+ transporting, mitochondrial F1 complex, O subunit | Cytoplasm | transporter | 2727015.75 | 7250060.5 | 3494227.25 | | | | | | |
| Q9DB20 | ATP5O | ATP synthase, H+ transporting, mitochondrial F1 complex, O subunit | Cytoplasm | transporter | 4034839 | 7494645.5 | 2176025 | | | | | | |
| Q9DB20 | ATP5O | ATP synthase, H+ transporting, mitochondrial F1 complex, O subunit | Cytoplasm | transporter | 3368978.5 | 6204345 | 3311833 | Q9DB20 | ATP5O | 6584223 | 5504206 | 1123541 | 24 |
| Q9R1Q9 | ATP6AP1 | ATPase, H+ transporting, lysosomal accessory protein 1 | Cytoplasm | transporter | 1141359.75 | 2932373 | 2342948.5 | Q9R1Q9 | ATP6AP1 | 1141360 | 912776 | 526992 | 3 |
| Q9CYN9 | ATP6AP2 | ATPase, H+ transporting, lysosomal accessory protein 2 | Cytoplasm | transporter | 1846802.75 | 3333938.25 | 3294536.5 | Q9CYN9 | ATP6AP2 | 1846803 | 847453 | 489277 | 3 |
| Q9Z1G4 | ATP6V0A1 | ATPase, H+ transporting, lysosomal V0 subunit a1 | Cytoplasm | transporter | 7720586.5 | 1.30E+07 | 9466086 | | | | | | |
| Q9Z1G4 | ATP6V0A1 | ATPase, H+ transporting, lysosomal V0 subunit a1 | Cytoplasm | transporter | 4652679.5 | 1.03E+07 | 6670658 | | | | | | |
| Q9Z1G4 | ATP6V0A1 | ATPase, H+ transporting, lysosomal V0 subunit a1 | Cytoplasm | transporter | 5607333.5 | 8829571 | 7339906 | | | | | | |
| Q9Z1G4 | ATP6V0A1 | ATPase, H+ transporting, lysosomal V0 subunit a1 | Cytoplasm | transporter | 7125573.5 | 1.35E+07 | 9859146 | | | | | | |
| Q9Z1G4 | ATP6V0A1 | ATPase, H+ transporting, lysosomal V0 subunit a1 | Cytoplasm | transporter | 8218397.5 | 1.20E+07 | 1.05E+07 | | | | | | |
| Q9Z1G4 | ATP6V0A1 | ATPase, H+ transporting, lysosomal V0 subunit a1 | Cytoplasm | transporter | 3057628 | 4597481 | 3626581.5 | | | | | | |
| Q9Z1G4 | ATP6V0A1 | ATPase, H+ transporting, lysosomal V0 subunit a1 | Cytoplasm | transporter | 3734261.75 | 5779852.5 | 5333541.5 | | | | | | |
| Q9Z1G4 | ATP6V0A1 | ATPase, H+ transporting, lysosomal V0 subunit a1 | Cytoplasm | transporter | 2577528 | 3080438.5 | 3141755.5 | | | | | | |
| Q9Z1G4 | ATP6V0A1 | ATPase, H+ transporting, lysosomal V0 subunit a1 | Cytoplasm | transporter | 1988431.875 | 3628115.25 | 3327385.5 | Q9Z1G4 | ATP6V0A1 | 4964713 | 3373239 | 649180 | 27 |
| P63082 | ATP6V0C | ATPase, H+ transporting, lysosomal 16kDa, V0 subunit c | Cytoplasm | transporter | 1.72E+07 | 3.71E+07 | 1.67E+07 | | | | | | |
| P63082 | ATP6V0C | ATPase, H+ transporting, lysosomal 16kDa, V0 subunit c | Cytoplasm | transporter | 2381000 | 3819565.25 | 770547.375 | P63082 | ATP6V0C | 9775500 | 13832664 | 5647161 | 6 |
| P51863 | ATP6V0D1 | ATPase, H+ transporting, lysosomal 38kDa, V0 subunit d1 | Cytoplasm | transporter | 1.46E+07 | 2.58E+07 | 2.50E+07 | | | | | | |
| P51863 | ATP6V0D1 | ATPase, H+ transporting, lysosomal 38kDa, V0 subunit d1 | Cytoplasm | transporter | 9848618 | 1.47E+07 | 8468071 | | | | | | |
| P51863 | ATP6V0D1 | ATPase, H+ transporting, lysosomal 38kDa, V0 subunit d1 | Cytoplasm | transporter | 6078003.5 | 9305887 | 8075865.5 | | | | | | |
| P51863 | ATP6V0D1 | ATPase, H+ transporting, lysosomal 38kDa, V0 subunit d1 | Cytoplasm | transporter | 3127153.5 | 9067513 | 8484629 | | | | | | |
| P51863 | ATP6V0D1 | ATPase, H+ transporting, lysosomal 38kDa, V0 subunit d1 | Cytoplasm | transporter | 3804395.25 | 6220785.5 | 5220177 | | | | | | |
| P51863 | ATP6V0D1 | ATPase, H+ transporting, lysosomal 38kDa, V0 subunit d1 | Cytoplasm | transporter | 2277343.25 | 3663974 | 3898285.75 | P51863 | ATP6V0D1 | 6629252 | 6839934 | 1612188 | 18 |
| P50516 | ATP6V1A | ATPase, H+ transporting, lysosomal 70kDa, V1 subunit A | Plasma Membrane | transporter | 5977438.5 | 5962212.5 | 3697601 | | | | | | |
| P50516 | ATP6V1A | ATPase, H+ transporting, lysosomal 70kDa, V1 subunit A | Plasma Membrane | transporter | 2522357.75 | 2232469.75 | 2828724 | | | | | | |
| P50516 | ATP6V1A | ATPase, H+ transporting, lysosomal 70kDa, V1 subunit A | Plasma Membrane | transporter | 2676461.5 | 3734178 | 3945833.25 | | | | | | |
| P50516 | ATP6V1A | ATPase, H+ transporting, lysosomal 70kDa, V1 subunit A | Plasma Membrane | transporter | 1851574.625 | 2435140.25 | 1731854.375 | | | | | | |
| P50516 | ATP6V1A | ATPase, H+ transporting, lysosomal 70kDa, V1 subunit A | Plasma Membrane | transporter | 1391899.375 | 937943.312 | 1029529.25 | P50516 | ATP6V1A | 2883946 | 1562955 | 403553 | 15 |

| | | | | | | | | | | | | | |
|---|---|---|---|---|---|---|---|---|---|---|---|---|---|
| P62814 | ATP6V1B2 | ATPase, H+ transporting, lysosomal 56/58kDa, V1 subunit B2 | Cytoplasm | transporter | 8709978 | 7807350.5 | 4812993 | | | | | | |
| P62814 | ATP6V1B2 | ATPase, H+ transporting, lysosomal 56/58kDa, V1 subunit B2 | Cytoplasm | transporter | 2090044.5 | 2758319.25 | 3005567 | | | | | | |
| P62814 | ATP6V1B2 | ATPase, H+ transporting, lysosomal 56/58kDa, V1 subunit B2 | Cytoplasm | transporter | 1963591.125 | 2135881.5 | 1905974.25 | P62814 | ATP6V1B2 | 4254538 | 2632804 | 877601 | 9 |
| P50518 | ATP6V1E1 | ATPase, H+ transporting, lysosomal 31kDa, V1 subunit E1 | Cytoplasm | transporter | 2632676.5 | 2799122.75 | 1812721.75 | P50518 | ATP6V1E1 | 2632677 | 528049 | 304869 | 3 |
| Q8BVE3 | ATP6V1H | ATPase, H+ transporting, lysosomal 50/57kDa, V1 subunit H | Cytoplasm | transporter | 1335872.25 | 1147743.375 | 1067281.875 | | | | | | |
| Q8BVE3 | ATP6V1H | ATPase, H+ transporting, lysosomal 50/57kDa, V1 subunit H | Cytoplasm | transporter | 349429.656 | 200821.641 | 179808.422 | Q8BVE3 | ATP6V1H | 842651 | 525602 | 214576 | 6 |
| Q64446 | ATP7B | ATPase, Cu++ transporting, beta polypeptide | Cytoplasm | transporter | 2.17E+07 | 2.43E+07 | 2.35E+07 | | | | | | |
| Q64446 | ATP7B | ATPase, Cu++ transporting, beta polypeptide | Cytoplasm | transporter | 3818180.25 | 3371608.5 | 4018042 | | | | | | |
| Q64446 | ATP7B | ATPase, Cu++ transporting, beta polypeptide | Cytoplasm | transporter | 3327749 | 4347947.5 | 4298829 | | | | | | |
| Q64446 | ATP7B | ATPase, Cu++ transporting, beta polypeptide | Cytoplasm | transporter | 5136106 | 4136818.25 | 5414279.5 | | | | | | |
| Q64446 | ATP7B | ATPase, Cu++ transporting, beta polypeptide | Cytoplasm | transporter | 4035214.5 | 3724412.5 | 4292507.5 | | | | | | |
| Q64446 | ATP7B | ATPase, Cu++ transporting, beta polypeptide | Cytoplasm | transporter | 3371613.25 | 2698460.75 | 1722651.5 | | | | | | |
| Q64446 | ATP7B | ATPase, Cu++ transporting, beta polypeptide | Cytoplasm | transporter | 3265575.5 | 2778071 | 3517958 | | | | | | |
| Q64446 | ATP7B | ATPase, Cu++ transporting, beta polypeptide | Cytoplasm | transporter | 2228795 | 2567870.25 | 2231253.75 | Q64446 | ATP7B | 5860404 | 6697716 | 1367166 | 24 |
| P70704 | ATP8A1 | ATPase, aminophospholipid transporter (APLT), class I, type 8A, member 1 | Cytoplasm | transporter | 1.97E+07 | 2.26E+07 | 2.20E+07 | | | | | | |
| P70704 | ATP8A1 | ATPase, aminophospholipid transporter (APLT), class I, type 8A, member 1 | Cytoplasm | transporter | 1.13E+07 | 1.44E+07 | 1.55E+07 | | | | | | |
| P70704 | ATP8A1 | ATPase, aminophospholipid transporter (APLT), class I, type 8A, member 1 | Cytoplasm | transporter | 1.55E+07 | 2.00E+07 | 1.95E+07 | | | | | | |
| P70704 | ATP8A1 | ATPase, aminophospholipid transporter (APLT), class I, type 8A, member 1 | Cytoplasm | transporter | 1.09E+07 | 1.23E+07 | 1.10E+07 | | | | | | |
| P70704 | ATP8A1 | ATPase, aminophospholipid transporter (APLT), class I, type 8A, member 1 | Cytoplasm | transporter | 1.07E+07 | 1.05E+07 | 1.13E+07 | | | | | | |
| P70704 | ATP8A1 | ATPase, aminophospholipid transporter (APLT), class I, type 8A, member 1 | Cytoplasm | transporter | 9373353 | 9864579 | 1.13E+07 | | | | | | |
| P70704 | ATP8A1 | ATPase, aminophospholipid transporter (APLT), class I, type 8A, member 1 | Cytoplasm | transporter | 7963358.5 | 9330658 | 8980571 | | | | | | |
| P70704 | ATP8A1 | ATPase, aminophospholipid transporter (APLT), class I, type 8A, member 1 | Cytoplasm | transporter | 9699679 | 9415444 | 9701877 | | | | | | |
| P70704 | ATP8A1 | ATPase, aminophospholipid transporter (APLT), class I, type 8A, member 1 | Cytoplasm | transporter | 6783869.5 | 6224116 | 7451040.5 | | | | | | |
| P70704 | ATP8A1 | ATPase, aminophospholipid transporter (APLT), class I, type 8A, member 1 | Cytoplasm | transporter | 5319292.5 | 5621200.5 | 6646605 | | | | | | |
| P70704 | ATP8A1 | ATPase, aminophospholipid transporter (APLT), class I, type 8A, member 1 | Cytoplasm | transporter | 5377796 | 7329077 | 8040306 | | | | | | |
| P70704 | ATP8A1 | ATPase, aminophospholipid transporter (APLT), class I, type 8A, member 1 | Cytoplasm | transporter | 5352219.5 | 6120197.5 | 4998180.5 | | | | | | |
| P70704 | ATP8A1 | ATPase, aminophospholipid transporter (APLT), class I, type 8A, member 1 | Cytoplasm | transporter | 5957569 | 7900978.5 | 8035827.5 | | | | | | |
| P70704 | ATP8A1 | ATPase, aminophospholipid transporter (APLT), class I, type 8A, member 1 | Cytoplasm | transporter | 6210478 | 5738623.5 | 5923748.5 | | | | | | |
| P70704 | ATP8A1 | ATPase, aminophospholipid transporter (APLT), class I, type 8A, member 1 | Cytoplasm | transporter | 7485552 | 7349611.5 | 8709322 | | | | | | |
| P70704 | ATP8A1 | ATPase, aminophospholipid transporter (APLT), class I, type 8A, member 1 | Cytoplasm | transporter | 4627141 | 3252504.5 | 4800754.5 | | | | | | |
| P70704 | ATP8A1 | ATPase, aminophospholipid transporter (APLT), class I, type 8A, member 1 | Cytoplasm | transporter | 4756960.5 | 6245619 | 5162485 | | | | | | |
| P70704 | ATP8A1 | ATPase, aminophospholipid transporter (APLT), class I, type 8A, member 1 | Cytoplasm | transporter | 4317349.5 | 3912276.5 | 4709643 | | | | | | |
| P70704 | ATP8A1 | ATPase, aminophospholipid transporter (APLT), class I, type 8A, member 1 | Cytoplasm | transporter | 3250904.25 | 3638071.25 | 5168306.5 | | | | | | |
| P70704 | ATP8A1 | ATPase, aminophospholipid transporter (APLT), class I, type 8A, member 1 | Cytoplasm | transporter | 2931768.5 | 4265271.5 | 5495788 | | | | | | |
| P70704 | ATP8A1 | ATPase, aminophospholipid transporter (APLT), class I, type 8A, member 1 | Cytoplasm | transporter | 2780696.75 | 2870395 | 2417508.75 | | | | | | |
| P70704 | ATP8A1 | ATPase, aminophospholipid transporter (APLT), class I, type 8A, member 1 | Cytoplasm | transporter | 1842807.625 | 2680949 | 4039746.25 | P70704 | ATP8A1 | 7368218 | 4806820 | 591679 | 66 |
| O35143 | ATPIF1 | ATPase inhibitory factor 1 | Cytoplasm | other | 5615129 | 1.35E+07 | 6487152 | O35143 | ATPIF1 | 5615129 | 4311159 | 2489049 | 3 |
| P70295 | AUP1 | ancient ubiquitous protein 1 | Cytoplasm | other | 3179703.5 | 3006208.25 | 2410110 | P70295 | AUP1 | 3179704 | 403672 | 233060 | 3 |
| P01887 | B2M | beta-2-microglobulin | Plasma Membrane | transmembrane receptor | 6156096.5 | 8234439.5 | 7131402 | P01887 | B2M | 6156097 | 1039825 | 600344 | 3 |
| Q09199 | B4GALNT2 | beta-1,4-N-acetyl-galactosaminyl transferase 2 | Cytoplasm | enzyme | 2.10E+07 | 1.85E+07 | 1.85E+07 | Q09199 | B4GALNT2 | 20990000 | 1429021 | 825045 | 3 |
| Q9JJ04 | B4GALT4 | UDP-Gal:betaGlcNAc beta 1,4- galactosyltransferase, polypeptide 4 | Cytoplasm | enzyme | 1.75E+08 | 2.33E+08 | 3.13E+08 | Q9JJ04 | B4GALT4 | 175200000 | 69249862 | 39981426 | 3 |
| Q91YN9 | BAG2 | BCL2-associated athanogene 2 | Cytoplasm | other | 4.07E+07 | 4.25E+07 | 2.44E+07 | | | | | | |
| Q91YN9 | BAG2 | BCL2-associated athanogene 2 | Cytoplasm | other | 2.56E+07 | 3.31E+07 | 2.60E+07 | | | | | | |
| Q91YN9 | BAG2 | BCL2-associated athanogene 2 | Cytoplasm | other | 1.59E+07 | 2.04E+07 | 1.70E+07 | | | | | | |
| Q91YN9 | BAG2 | BCL2-associated athanogene 2 | Cytoplasm | other | 1.28E+07 | 1.33E+07 | 1.24E+07 | | | | | | |
| Q91YN9 | BAG2 | BCL2-associated athanogene 2 | Cytoplasm | other | 1.15E+07 | 8357837 | 7056329.5 | | | | | | |
| Q91YN9 | BAG2 | BCL2-associated athanogene 2 | Cytoplasm | other | 1.05E+07 | 1.44E+07 | 9783863 | | | | | | |
| Q91YN9 | BAG2 | BCL2-associated athanogene 2 | Cytoplasm | other | 9854012 | 1.09E+07 | 1.08E+07 | | | | | | |
| Q91YN9 | BAG2 | BCL2-associated athanogene 2 | Cytoplasm | other | 1.26E+07 | 7419123 | 2897122.25 | | | | | | |
| Q91YN9 | BAG2 | BCL2-associated athanogene 2 | Cytoplasm | other | 7492947.5 | 5003045.5 | 4864657 | | | | | | |
| Q91YN9 | BAG2 | BCL2-associated athanogene 2 | Cytoplasm | other | 9471064 | 6608181 | 2687552 | | | | | | |
| Q91YN9 | BAG2 | BCL2-associated athanogene 2 | Cytoplasm | other | 2064054 | 1036191.875 | 281907.312 | Q91YN9 | BAG2 | 14404734 | 10537367 | 1834320 | 33 |
| O08734 | BAK1 | BCL2-antagonist/killer 1 | Cytoplasm | other | 2068220.75 | 4532211 | 2568132.75 | O08734 | BAK1 | 2068221 | 1302482 | 751989 | 3 |
| O54962 | BANF1 | barrier to autointegration factor 1 | Nucleus | other | 4.63E+07 | 1.05E+08 | 5.98E+07 | | | | | | |
| O54962 | BANF1 | barrier to autointegration factor 1 | Nucleus | other | 3.07E+07 | 5.72E+07 | 2.96E+07 | | | | | | |
| O54962 | BANF1 | barrier to autointegration factor 1 | Nucleus | other | 1.03E+07 | 1.99E+07 | 1.38E+07 | | | | | | |
| O54962 | BANF1 | barrier to autointegration factor 1 | Nucleus | other | 6806454.5 | 1.28E+07 | 9200400 | O54962 | BANF1 | 23541614 | 29265637 | 8448262 | 12 |
| Q61335 | BCAP31 | B-cell receptor-associated protein 31 | Cytoplasm | transporter | 647437.312 | 1120898.625 | 2175808.75 | Q61335 | BCAP31 | 647437 | 782402 | 451720 | 3 |
| O35855 | BCAT2 | branched chain amino-acid transaminase 2, mitochondrial | Cytoplasm | enzyme | 2000261.5 | 3548577.75 | 1977365.125 | O35855 | BCAT2 | 2000262 | 900603 | 519963 | 3 |
| P59017 | BCL2L13 | BCL2-like 13 (apoptosis facilitator) | Cytoplasm | other | 3848112.5 | 1.01E+07 | 4299514.5 | | | | | | |
| P59017 | BCL2L13 | BCL2-like 13 (apoptosis facilitator) | Cytoplasm | other | 2646405.25 | 5458716.5 | 2084524 | | | | | | |
| P59017 | BCL2L13 | BCL2-like 13 (apoptosis facilitator) | Cytoplasm | other | 2833519 | 5774324 | 2500474 | P59017 | BCL2L13 | 3109346 | 2499275 | 833092 | 9 |
| Q8C6A8 | BHLHE22 | basic helix-loop-helix family, member e22 | Nucleus | transcription regulator | 1.88E+08 | 1.69E+08 | 1.77E+08 | Q8C6A8 | BHLHE22 | 188400000 | 9867793 | 5697173 | 3 |
| Q8BR07 | BICD1 | bicaudal D homolog 1 (Drosophila) | Cytoplasm | other | 4.28E+07 | 3.15E+07 | 4.04E+07 | Q8BR07 | BICD1 | 42810000 | 5957080 | 3439322 | 3 |
| Q8R016 | BLMH | bleomycin hydrolase | Cytoplasm | peptidase | 9094759 | 6779981 | 8508561 | | | | | | |
| Q8R016 | BLMH | bleomycin hydrolase | Cytoplasm | peptidase | 1.07E+07 | 5176282.5 | 2213203.75 | | | | | | |
| Q8R016 | BLMH | bleomycin hydrolase | Cytoplasm | peptidase | 7710833 | 6379632.5 | 6396265.5 | | | | | | |
| Q8R016 | BLMH | bleomycin hydrolase | Cytoplasm | peptidase | 4985005 | 2872237 | 3232628 | | | | | | |
| Q8R016 | BLMH | bleomycin hydrolase | Cytoplasm | peptidase | 3230862 | 2933238.5 | 2824235 | | | | | | |
| Q8R016 | BLMH | bleomycin hydrolase | Cytoplasm | peptidase | 3958620.75 | 2140159.5 | 2522340.5 | | | | | | |
| Q8R016 | BLMH | bleomycin hydrolase | Cytoplasm | peptidase | 2402350.5 | 1902179.5 | 1829417.375 | Q8R016 | BLMH | 6016061 | 2675907 | 583931 | 21 |
| Q923D2 | BLVRB | biliverdin reductase B | Cytoplasm | enzyme | 1.20E+07 | 6920345.5 | 7697139.5 | | | | | | |
| Q923D2 | BLVRB | biliverdin reductase B | Cytoplasm | enzyme | 9926066 | 7731448 | 6280635 | | | | | | |
| Q923D2 | BLVRB | biliverdin reductase B | Cytoplasm | enzyme | 3236844.25 | 3541806.25 | 5288101.5 | | | | | | |
| Q923D2 | BLVRB | biliverdin reductase B | Cytoplasm | enzyme | 1580581.375 | 1146034.125 | 1491273 | Q923D2 | BLVRB | 6688373 | 3486774 | 1006545 | 12 |
| P15327 | BPGM | 2,3-bisphosphoglycerate mutase | Extracellular Space | phosphatase | 1.54E+07 | 1.40E+07 | 1.70E+07 | | | | | | |
| P15327 | BPGM | 2,3-bisphosphoglycerate mutase | Extracellular Space | phosphatase | 5367929.5 | 4379813.5 | 6195489 | | | | | | |
| P15327 | BPGM | 2,3-bisphosphoglycerate mutase | Extracellular Space | phosphatase | 3212943.25 | 2032352.375 | 3235513 | | | | | | |
| P15327 | BPGM | 2,3-bisphosphoglycerate mutase | Extracellular Space | phosphatase | 908880.875 | 466639.906 | 890671.875 | P15327 | BPGM | 6209938 | 5943187 | 1715650 | 12 |
| A2AHJ4 | BRWD3 | bromodomain and WD repeat domain containing 3 | Nucleus | other | 1.14E+07 | 1.09E+07 | 1.40E+07 | A2AHJ4 | BRWD3 | 11370000 | 1662298 | 959728 | 3 |
| P18572 | BSG | basigin (Ok blood group) | Plasma Membrane | transporter | 8.73E+08 | 7.42E+08 | 1.01E+09 | | | | | | |
| P18572 | BSG | basigin (Ok blood group) | Plasma Membrane | transporter | 7.03E+08 | 6.48E+08 | 7.71E+08 | | | | | | |
| P18572 | BSG | basigin (Ok blood group) | Plasma Membrane | transporter | 8.78E+08 | 9.03E+08 | 7.70E+08 | | | | | | |
| P18572 | BSG | basigin (Ok blood group) | Plasma Membrane | transporter | 5.80E+08 | 5.74E+08 | 6.58E+08 | | | | | | |
| P18572 | BSG | basigin (Ok blood group) | Plasma Membrane | transporter | 2.35E+08 | 2.50E+08 | 3.57E+08 | | | | | | |
| P18572 | BSG | basigin (Ok blood group) | Plasma Membrane | transporter | 2.20E+08 | 1.86E+08 | 2.53E+08 | | | | | | |
| P18572 | BSG | basigin (Ok blood group) | Plasma Membrane | transporter | 1.53E+08 | 2.42E+08 | 3.91E+08 | | | | | | |
| P18572 | BSG | basigin (Ok blood group) | Plasma Membrane | transporter | 4.34E+07 | 7.74E+07 | 1.18E+08 | | | | | | |
| P18572 | BSG | basigin (Ok blood group) | Plasma Membrane | transporter | 3.85E+07 | 4.84E+07 | 5.61E+07 | | | | | | |
| P18572 | BSG | basigin (Ok blood group) | Plasma Membrane | transporter | 4.82E+07 | 5.78E+07 | 7.87E+07 | | | | | | |
| P18572 | BSG | basigin (Ok blood group) | Plasma Membrane | transporter | 1.11E+07 | 1.09E+07 | 1.45E+07 | | | | | | |

| | | | | | | | | | | | | | |
|---|---|---|---|---|---|---|---|---|---|---|---|---|---|
| P18572 | BSG | basigin (Ok blood group) | Plasma Membrane | transporter | 5158075.5 | 8722651 | 1.33E+07 | | | | | | |
| P18572 | BSG | basigin (Ok blood group) | Plasma Membrane | transporter | 2042919.375 | 1907705.5 | 1320302.75 | | | | | | |
| P18572 | BSG | basigin (Ok blood group) | Plasma Membrane | transporter | 817711.312 | 5172031.5 | 1.37E+07 | P18572 | BSG | 270724908 | 324790933 | 50116329 | 42 |
| Q9D659 | C10orf54 | chromosome 10 open reading frame 54 | Plasma Membrane | other | 1.22E+07 | 1.79E+07 | 1.13E+07 | Q9D659 | C10orf54 | 12210000 | 3573378 | 2063091 | 3 |
| Q91W34 | C16orf58 | chromosome 16 open reading frame 58 | Cytoplasm | other | 566741 | 1510920.125 | 1458963.875 | Q91W34 | C16orf58 | 566741 | 530760 | 306434 | 3 |
| Q9CX63 | C17orf99 | chromosome 17 open reading frame 99 | Extracellular Space | cytokine | 1324321.625 | 1159894.5 | 1909059.375 | | | | | | |
| Q9CX63 | C17orf99 | chromosome 17 open reading frame 99 | Extracellular Space | cytokine | 83912.844 | 171575.609 | 98313.445 | | | | | | |
| Q9CX63 | C17orf99 | chromosome 17 open reading frame 99 | Extracellular Space | cytokine | 99593.188 | 0 | 0 | | | | | | |
| Q9CX63 | C17orf99 | chromosome 17 open reading frame 99 | Extracellular Space | cytokine | 28804.061 | 116814.828 | 158075.359 | | | | | | |
| Q9CX63 | C17orf99 | chromosome 17 open reading frame 99 | Extracellular Space | cytokine | 458344.094 | 656259.562 | 300408.031 | | | | | | |
| Q9CX63 | C17orf99 | chromosome 17 open reading frame 99 | Extracellular Space | cytokine | 192143.141 | 637091.688 | 510405.781 | | | | | | |
| Q9CX63 | C17orf99 | chromosome 17 open reading frame 99 | Extracellular Space | cytokine | 90410.859 | 125396.188 | 565.98 | Q9CX63 | C17orf99 | 325361 | 507771 | 110805 | 21 |
| P98086 | C1QA | complement component 1, q subcomponent, A chain | Extracellular Space | other | 330236.844 | 72789.961 | 12287.765 | | | | | | |
| P98086 | C1QA | complement component 1, q subcomponent, A chain | Extracellular Space | other | 170433.047 | 0 | 0 | P98086 | C1QA | 250335 | 131430 | 53656 | 6 |
| P14106 | C1QB | complement component 1, q subcomponent, B chain | Extracellular Space | other | 84053.5 | 142381.109 | 4.70E-06 | P14106 | C1QB | 84054 | 71577 | 41325 | 3 |
| O35658 | C1QBP | complement component 1, q subcomponent binding protein | Cytoplasm | transcription regulator | 2.14E+07 | 4.43E+07 | 1.64E+07 | | | | | | |
| O35658 | C1QBP | complement component 1, q subcomponent binding protein | Cytoplasm | transcription regulator | 3703595.25 | 2.06E+07 | 1.49E+07 | O35658 | C1QBP | 12531798 | 13401236 | 5471032 | 6 |
| Q9D172 | C21orf33/LOC102724023 | chromosome 21 open reading frame 33 | Cytoplasm | other | 5634322 | 1.09E+07 | 3952314.5 | | | | | | |
| Q9D172 | C21orf33/LOC102724023 | chromosome 21 open reading frame 33 | Cytoplasm | other | 128563.281 | 599606.25 | 299785.719 | Q9D172 | C21orf33/LOC102724023 | 2881443 | 4223832 | 1724372 | 6 |
| Q80X80 | C2CD2L | C2CD2-like | Cytoplasm | other | 2.17E+07 | 1.84E+07 | 1.73E+07 | | | | | | |
| Q80X80 | C2CD2L | C2CD2-like | Cytoplasm | other | 2.66E+07 | 2.78E+07 | 3.39E+07 | | | | | | |
| Q80X80 | C2CD2L | C2CD2-like | Cytoplasm | other | 1.90E+07 | 1.83E+07 | 2.23E+07 | | | | | | |
| Q80X80 | C2CD2L | C2CD2-like | Cytoplasm | other | 8343676 | 6689756.5 | 8306826.5 | | | | | | |
| Q80X80 | C2CD2L | C2CD2-like | Cytoplasm | other | 7637715 | 8445424 | 8405872 | | | | | | |
| Q80X80 | C2CD2L | C2CD2-like | Cytoplasm | other | 9863577 | 1.03E+07 | 8760513 | | | | | | |
| Q80X80 | C2CD2L | C2CD2-like | Cytoplasm | other | 8024286 | 1.36E+07 | 1.40E+07 | | | | | | |
| Q80X80 | C2CD2L | C2CD2-like | Cytoplasm | other | 1.23E+07 | 8163774 | 1.14E+07 | | | | | | |
| Q80X80 | C2CD2L | C2CD2-like | Cytoplasm | other | 7082115 | 8609387 | 7093249 | | | | | | |
| Q80X80 | C2CD2L | C2CD2-like | Cytoplasm | other | 7633966.5 | 7655340 | 6787981 | | | | | | |
| Q80X80 | C2CD2L | C2CD2-like | Cytoplasm | other | 4242960 | 3672941.5 | 4973665 | | | | | | |
| Q80X80 | C2CD2L | C2CD2-like | Cytoplasm | other | 3010056.5 | 1747188.375 | 2598439.25 | | | | | | |
| Q80X80 | C2CD2L | C2CD2-like | Cytoplasm | other | 1467269.125 | 611198 | 919536.75 | Q80X80 | C2CD2L | 10528125 | 7832100 | 1254140 | 39 |
| P01029 | C4A/C4B | complement component 4B (Chido blood group) | Extracellular Space | other | 2969547.75 | 1.21E+07 | 6525728.5 | | | | | | |
| P01029 | C4A/C4B | complement component 4B (Chido blood group) | Extracellular Space | other | 3836406.75 | 1.27E+07 | 5666429 | | | | | | |
| P01029 | C4A/C4B | complement component 4B (Chido blood group) | Extracellular Space | other | 1928738.75 | 5110600 | 3211428.25 | | | | | | |
| P01029 | C4A/C4B | complement component 4B (Chido blood group) | Extracellular Space | other | 898662.25 | 5065779.5 | 3374698 | | | | | | |
| P01029 | C4A/C4B | complement component 4B (Chido blood group) | Extracellular Space | other | 573870.5 | 5147666 | 2043850.75 | P01029 | C4A/C4B | 2041445 | 3563260 | 920030 | 15 |
| P00920 | CA2 | carbonic anhydrase II | Cytoplasm | enzyme | 2.83E+07 | 3.14E+07 | 3.58E+07 | | | | | | |
| P00920 | CA2 | carbonic anhydrase II | Cytoplasm | enzyme | 1.16E+07 | 1.10E+07 | 1.35E+07 | | | | | | |
| P00920 | CA2 | carbonic anhydrase II | Cytoplasm | enzyme | 7402147 | 6283251 | 9708779 | | | | | | |
| P00920 | CA2 | carbonic anhydrase II | Cytoplasm | enzyme | 3204917.5 | 2745110.75 | 1.29E+07 | | | | | | |
| P00920 | CA2 | carbonic anhydrase II | Cytoplasm | enzyme | 5789606 | 4715838.5 | 1.26E+07 | | | | | | |
| P00920 | CA2 | carbonic anhydrase II | Cytoplasm | enzyme | 329728.969 | 82198.234 | 726669.312 | P00920 | CA2 | 9441067 | 10610771 | 2500983 | 18 |
| P62204 | CALM1 | Calmodulin | Cytoplasm | | 2.51E+07 | 3.12E+07 | 3.49E+07 | | | | | | |
| P62204 | CALM1 | Calmodulin | Cytoplasm | | 1.23E+07 | 9249110 | 9788587 | P62204 | CALM1 | 18660000 | 11415143 | 4660212 | 6 |
| Q9D6P8 | CALML3 | calmodulin-like 3 | Cytoplasm | other | 7439419.5 | 6690254.5 | 8298481.5 | Q9D6P8 | CALML3 | 7439420 | 804739 | 464616 | 3 |
| P14211 | CALR | calreticulin | Cytoplasm | transcription regulator | 1.39E+08 | 1.95E+08 | 1.72E+08 | | | | | | |
| P14211 | CALR | calreticulin | Cytoplasm | transcription regulator | 9.05E+07 | 1.32E+08 | 1.29E+08 | | | | | | |
| P14211 | CALR | calreticulin | Cytoplasm | transcription regulator | 9.27E+07 | 1.75E+08 | 2.22E+08 | | | | | | |
| P14211 | CALR | calreticulin | Cytoplasm | transcription regulator | 7.72E+07 | 7.84E+07 | 8.79E+07 | | | | | | |
| P14211 | CALR | calreticulin | Cytoplasm | transcription regulator | 9.65E+07 | 9.74E+07 | 1.08E+08 | | | | | | |
| P14211 | CALR | calreticulin | Cytoplasm | transcription regulator | 5.25E+07 | 8.00E+07 | 6.94E+07 | | | | | | |
| P14211 | CALR | calreticulin | Cytoplasm | transcription regulator | 7.34E+07 | 6.93E+07 | 9.40E+07 | | | | | | |
| P14211 | CALR | calreticulin | Cytoplasm | transcription regulator | 4.34E+07 | 1.13E+08 | 1.23E+08 | | | | | | |
| P14211 | CALR | calreticulin | Cytoplasm | transcription regulator | 4.15E+07 | 3.92E+07 | 5.05E+07 | | | | | | |
| P14211 | CALR | calreticulin | Cytoplasm | transcription regulator | 3.14E+07 | 3.08E+07 | 3.25E+07 | | | | | | |
| P14211 | CALR | calreticulin | Cytoplasm | transcription regulator | 3.36E+07 | 2.80E+07 | 1.99E+07 | | | | | | |
| P14211 | CALR | calreticulin | Cytoplasm | transcription regulator | 3910023.75 | 4483461.5 | 2568938.25 | | | | | | |
| P14211 | CALR | calreticulin | Cytoplasm | transcription regulator | 6931946.5 | 5653065.5 | 7775984.5 | | | | | | |
| P14211 | CALR | calreticulin | Cytoplasm | transcription regulator | 1.05E+07 | 1.64E+07 | 2.64E+07 | | | | | | |
| P14211 | CALR | calreticulin | Cytoplasm | transcription regulator | 5500418 | 4368244.5 | 5413295.5 | | | | | | |
| P14211 | CALR | calreticulin | Cytoplasm | transcription regulator | 1057792.875 | 1728041.75 | 1242051.25 | | | | | | |
| P14211 | CALR | calreticulin | Cytoplasm | transcription regulator | 1144002 | 1626077.875 | 5645556.5 | | | | | | |
| P14211 | CALR | calreticulin | Cytoplasm | transcription regulator | 470887.75 | 446598.156 | 227616.828 | | | | | | |
| P14211 | CALR | calreticulin | Cytoplasm | transcription regulator | 918808.938 | 4083974.75 | 1.01E+07 | P14211 | CALR | 42194415 | 56797906 | 7523066 | 57 |
| O35887 | CALU | calumenin | Cytoplasm | other | 7021485 | 7517189 | 7697764 | | | | | | |
| O35887 | CALU | calumenin | Cytoplasm | other | 6365611 | 7095042.5 | 7159947.5 | | | | | | |
| O35887 | CALU | calumenin | Cytoplasm | other | 5928818 | 6505614 | 5171153 | | | | | | |
| O35887 | CALU | calumenin | Cytoplasm | other | 5858772.5 | 6761437.5 | 9168515 | | | | | | |
| O35887 | CALU | calumenin | Cytoplasm | other | 3550078.5 | 4060934 | 4134097 | | | | | | |
| O35887 | CALU | calumenin | Cytoplasm | other | 3832717.25 | 3709503.25 | 2540300.25 | | | | | | |
| O35887 | CALU | calumenin | Cytoplasm | other | 2215013.75 | 2836803.5 | 3138585.25 | | | | | | |
| O35887 | CALU | calumenin | Cytoplasm | other | 817836.125 | 1254338 | 310806.031 | O35887 | CALU | 4448792 | 2413958 | 492747 | 24 |
| P51437 | CAMP | cathelicidin antimicrobial peptide | Cytoplasm | other | 469767.781 | 496627.062 | 473360.406 | | | | | | |
| P51437 | CAMP | cathelicidin antimicrobial peptide | Cytoplasm | other | 800346.188 | 398071.938 | 602737.625 | P51437 | CAMP | 635057 | 143607 | 58627 | 6 |
| Q6ZQ38 | CAND1 | cullin-associated and neddylation-dissociated 1 | Cytoplasm | transcription regulator | 2.20E+07 | 1.83E+07 | 1.84E+07 | | | | | | |
| Q6ZQ38 | CAND1 | cullin-associated and neddylation-dissociated 1 | Cytoplasm | transcription regulator | 1.38E+07 | 1.55E+07 | 1.18E+07 | | | | | | |
| Q6ZQ38 | CAND1 | cullin-associated and neddylation-dissociated 1 | Cytoplasm | transcription regulator | 1.55E+07 | 1.14E+07 | 1.06E+07 | | | | | | |
| Q6ZQ38 | CAND1 | cullin-associated and neddylation-dissociated 1 | Cytoplasm | transcription regulator | 1.05E+07 | 9608864 | 9005563 | | | | | | |
| Q6ZQ38 | CAND1 | cullin-associated and neddylation-dissociated 1 | Cytoplasm | transcription regulator | 1.16E+07 | 8555124 | 8415164 | | | | | | |
| Q6ZQ38 | CAND1 | cullin-associated and neddylation-dissociated 1 | Cytoplasm | transcription regulator | 1.44E+07 | 1.21E+07 | 1.13E+07 | | | | | | |
| Q6ZQ38 | CAND1 | cullin-associated and neddylation-dissociated 1 | Cytoplasm | transcription regulator | 1.12E+07 | 8989726 | 8431316 | | | | | | |
| Q6ZQ38 | CAND1 | cullin-associated and neddylation-dissociated 1 | Cytoplasm | transcription regulator | 9369924 | 1.04E+07 | 8263120.5 | | | | | | |
| Q6ZQ38 | CAND1 | cullin-associated and neddylation-dissociated 1 | Cytoplasm | transcription regulator | 1.28E+07 | 9886479 | 9887337 | | | | | | |
| Q6ZQ38 | CAND1 | cullin-associated and neddylation-dissociated 1 | Cytoplasm | transcription regulator | 1.09E+07 | 8762348 | 1.06E+07 | | | | | | |
| Q6ZQ38 | CAND1 | cullin-associated and neddylation-dissociated 1 | Cytoplasm | transcription regulator | 1.19E+07 | 9727808 | 9599363 | | | | | | |
| Q6ZQ38 | CAND1 | cullin-associated and neddylation-dissociated 1 | Cytoplasm | transcription regulator | 9298743 | 6910089 | 7708085.5 | | | | | | |
| Q6ZQ38 | CAND1 | cullin-associated and neddylation-dissociated 1 | Cytoplasm | transcription regulator | 6745145 | 5786033 | 4810968 | | | | | | |
| Q6ZQ38 | CAND1 | cullin-associated and neddylation-dissociated 1 | Cytoplasm | transcription regulator | 8678178 | 7048659 | 6943235 | | | | | | |
| Q6ZQ38 | CAND1 | cullin-associated and neddylation-dissociated 1 | Cytoplasm | transcription regulator | 5097778 | 1363120.375 | 1260212.75 | | | | | | |
| Q6ZQ38 | CAND1 | cullin-associated and neddylation-dissociated 1 | Cytoplasm | transcription regulator | 7625999 | 5575875 | 6577201.5 | | | | | | |

| | | | | | | | | | | | | | |
|---|---|---|---|---|---|---|---|---|---|---|---|---|---|
| Q6ZQ38 | CAND1 | cullin-associated and neddylation-dissociated 1 | Cytoplasm | transcription regulator | 6669283.5 | 4925953.5 | 4293325 | | | | | | |
| Q6ZQ38 | CAND1 | cullin-associated and neddylation-dissociated 1 | Cytoplasm | transcription regulator | 5683345.5 | 6530238 | 5817463.5 | | | | | | |
| Q6ZQ38 | CAND1 | cullin-associated and neddylation-dissociated 1 | Cytoplasm | transcription regulator | 1838207.75 | 3739646 | 3891637 | Q6ZQ38 | CAND1 | 10291927 | 4087935 | 541460 | 57 |
| P35564 | CANX | calnexin | Cytoplasm | other | 3.45E+07 | 8.24E+07 | 1.18E+08 | | | | | | |
| P35564 | CANX | calnexin | Cytoplasm | other | 5.54E+07 | 5.55E+07 | 5.81E+07 | | | | | | |
| P35564 | CANX | calnexin | Cytoplasm | other | 5.32E+07 | 5.91E+07 | 6.58E+07 | | | | | | |
| P35564 | CANX | calnexin | Cytoplasm | other | 3.37E+07 | 5.77E+07 | 7.68E+07 | | | | | | |
| P35564 | CANX | calnexin | Cytoplasm | other | 2.00E+07 | 1.68E+07 | 1.95E+07 | | | | | | |
| P35564 | CANX | calnexin | Cytoplasm | other | 3.38E+07 | 4.62E+07 | 5.03E+07 | | | | | | |
| P35564 | CANX | calnexin | Cytoplasm | other | 1.64E+07 | 3.98E+07 | 7.09E+07 | | | | | | |
| P35564 | CANX | calnexin | Cytoplasm | other | 2.46E+07 | 3.28E+07 | 3.47E+07 | | | | | | |
| P35564 | CANX | calnexin | Cytoplasm | other | 2.47E+07 | 2.52E+07 | 3.25E+07 | | | | | | |
| P35564 | CANX | calnexin | Cytoplasm | other | 2.25E+07 | 2.89E+07 | 3.77E+07 | | | | | | |
| P35564 | CANX | calnexin | Cytoplasm | other | 2.20E+07 | 1.30E+07 | 4651822.5 | | | | | | |
| P35564 | CANX | calnexin | Cytoplasm | other | 2.23E+07 | 7404531 | 4312416 | | | | | | |
| P35564 | CANX | calnexin | Cytoplasm | other | 2.30E+07 | 3.26E+07 | 3.16E+07 | | | | | | |
| P35564 | CANX | calnexin | Cytoplasm | other | 1.30E+07 | 2.30E+07 | 2.52E+07 | | | | | | |
| P35564 | CANX | calnexin | Cytoplasm | other | 6466693.5 | 6687580.5 | 7562237.5 | | | | | | |
| P35564 | CANX | calnexin | Cytoplasm | other | 2244980 | 3891681.75 | 3700785.75 | | | | | | |
| P35564 | CANX | calnexin | Cytoplasm | other | 3753185.75 | 7779197.5 | 1.48E+07 | | | | | | |
| P35564 | CANX | calnexin | Cytoplasm | other | 3337407.25 | 1.40E+07 | 1.79E+07 | | | | | | |
| P35564 | CANX | calnexin | Cytoplasm | other | 2821979.25 | 1781414.875 | 1910148.375 | | | | | | |
| P35564 | CANX | calnexin | Cytoplasm | other | 754000.625 | 2742195.5 | 5749503.5 | | | | | | |
| P35564 | CANX | calnexin | Cytoplasm | other | 1718458.125 | 1279904.25 | 1741567.125 | | | | | | |
| P35564 | CANX | calnexin | Cytoplasm | other | 143941.094 | 328697.219 | 868661.062 | P35564 | CANX | 19106848 | 24297268 | 2990787 | 66 |
| P40124 | CAP1 | CAP, adenylate cyclase-associated protein 1 (yeast) | Plasma Membrane | other | 3195740.25 | 3084942.5 | 3773592.25 | | | | | | |
| P40124 | CAP1 | CAP, adenylate cyclase-associated protein 1 (yeast) | Plasma Membrane | other | 2505778 | 2090115.375 | 2423726.75 | | | | | | |
| P40124 | CAP1 | CAP, adenylate cyclase-associated protein 1 (yeast) | Plasma Membrane | other | 979638.938 | 1555656.625 | 1430460.875 | P40124 | CAP1 | 2227052 | 917357 | 305786 | 9 |
| O35350 | CAPN1 | calpain 1, (mu/I) large subunit | Cytoplasm | peptidase | 1.02E+07 | 8866387 | 1.00E+07 | | | | | | |
| O35350 | CAPN1 | calpain 1, (mu/I) large subunit | Cytoplasm | peptidase | 7303414 | 4571183.5 | 5334185.5 | | | | | | |
| O35350 | CAPN1 | calpain 1, (mu/I) large subunit | Cytoplasm | peptidase | 5676624.5 | 4170118 | 4492348.5 | | | | | | |
| O35350 | CAPN1 | calpain 1, (mu/I) large subunit | Cytoplasm | peptidase | 5156190.5 | 4177608.75 | 5446770 | | | | | | |
| O35350 | CAPN1 | calpain 1, (mu/I) large subunit | Cytoplasm | peptidase | 4538751 | 2461678.5 | 3453559.25 | | | | | | |
| O35350 | CAPN1 | calpain 1, (mu/I) large subunit | Cytoplasm | peptidase | 2787986 | 3173662.75 | 4608581.5 | | | | | | |
| O35350 | CAPN1 | calpain 1, (mu/I) large subunit | Cytoplasm | peptidase | 641584.438 | 433674.688 | 331387.656 | O35350 | CAPN1 | 5186364 | 2754360 | 601051 | 21 |
| O08688 | CAPN5 | calpain 5 | Cytoplasm | peptidase | 1.97E+07 | 2.14E+07 | 3.73E+07 | | | | | | |
| O08688 | CAPN5 | calpain 5 | Cytoplasm | peptidase | 1.94E+07 | 1.84E+07 | 3.31E+07 | | | | | | |
| O08688 | CAPN5 | calpain 5 | Cytoplasm | peptidase | 1.01E+07 | 9096985 | 1.31E+07 | | | | | | |
| O08688 | CAPN5 | calpain 5 | Cytoplasm | peptidase | 1.24E+07 | 9729061 | 1.35E+07 | | | | | | |
| O08688 | CAPN5 | calpain 5 | Cytoplasm | peptidase | 1.22E+07 | 9009065 | 1.63E+07 | | | | | | |
| O08688 | CAPN5 | calpain 5 | Cytoplasm | peptidase | 9180900 | 8232558.5 | 1.28E+07 | | | | | | |
| O08688 | CAPN5 | calpain 5 | Cytoplasm | peptidase | 1.03E+07 | 1.10E+07 | 1.47E+07 | | | | | | |
| O08688 | CAPN5 | calpain 5 | Cytoplasm | peptidase | 4946569 | 5474937.5 | 9103157 | | | | | | |
| O08688 | CAPN5 | calpain 5 | Cytoplasm | peptidase | 5206692.5 | 5906266 | 8274148.5 | | | | | | |
| O08688 | CAPN5 | calpain 5 | Cytoplasm | peptidase | 5317168.5 | 5122745.5 | 8430878 | | | | | | |
| O08688 | CAPN5 | calpain 5 | Cytoplasm | peptidase | 3635866.75 | 3871766 | 5899580.5 | | | | | | |
| O08688 | CAPN5 | calpain 5 | Cytoplasm | peptidase | 3114531.75 | 2523927 | 4274183.5 | | | | | | |
| O08688 | CAPN5 | calpain 5 | Cytoplasm | peptidase | 3672739.75 | 3609421.5 | 4554212.5 | | | | | | |
| O08688 | CAPN5 | calpain 5 | Cytoplasm | peptidase | 2497661 | 2917916 | 3896204.75 | | | | | | |
| O08688 | CAPN5 | calpain 5 | Cytoplasm | peptidase | 1939590.25 | 1651067.125 | 3178020.75 | | | | | | |
| O08688 | CAPN5 | calpain 5 | Cytoplasm | peptidase | 993345.188 | 718230.188 | 1479027.75 | | | | | | |
| O08688 | CAPN5 | calpain 5 | Cytoplasm | peptidase | 1594864.125 | 3024463 | 4630867.5 | | | | | | |
| O08688 | CAPN5 | calpain 5 | Cytoplasm | peptidase | 1076001.875 | 1238894.75 | 2397953.5 | | | | | | |
| O08688 | CAPN5 | calpain 5 | Cytoplasm | peptidase | 519603.562 | 1635170.75 | 2195613.5 | | | | | | |
| O08688 | CAPN5 | calpain 5 | Cytoplasm | peptidase | 979168.125 | 1855221.875 | 3416695.75 | O08688 | CAPN5 | 6430735 | 7420859 | 958029 | 60 |
| O88456 | CAPNS1 | calpain, small subunit 1 | Cytoplasm | peptidase | 2045121.625 | 1595121.375 | 1916519.375 | O88456 | CAPNS1 | 2045122 | 231781 | 133819 | 3 |
| Q60865 | CAPRIN1 | cell cycle associated protein 1 | Plasma Membrane | other | 1186745.5 | 2585571 | 970973.438 | | | | | | |
| Q60865 | CAPRIN1 | cell cycle associated protein 1 | Plasma Membrane | other | 1301408.5 | 3101178 | 1173516.375 | Q60865 | CAPRIN1 | 1244077 | 891754 | 364057 | 6 |
| P47753 | CAPZA1 | capping protein (actin filament) muscle Z-line, alpha 1 | Cytoplasm | other | 5874163 | 7414785 | 5774935.5 | | | | | | |
| P47753 | CAPZA1 | capping protein (actin filament) muscle Z-line, alpha 1 | Cytoplasm | other | 2339006.5 | 3103661.5 | 2838077 | P47753 | CAPZA1 | 4106585 | 2067443 | 844030 | 6 |
| P47757 | CAPZB | capping protein (actin filament) muscle Z-line, beta | Cytoplasm | other | 3246503.75 | 3151469.75 | 3440086 | | | | | | |
| P47757 | CAPZB | capping protein (actin filament) muscle Z-line, beta | Cytoplasm | other | 1921437.5 | 1752863.375 | 1591299.375 | P47757 | CAPZB | 2583971 | 846444 | 345559 | 6 |
| Q9CR86 | CARHSP1 | calcium regulated heat stable protein 1, 24kDa | Cytoplasm | other | 1.02E+07 | 8578768 | 1.03E+07 | Q9CR86 | CARHSP1 | 10230000 | 977252 | 564217 | 3 |
| Q9WVG6 | CARM1 | coactivator-associated arginine methyltransferase 1 | Nucleus | transcription regulator | 3214541.75 | 2963461.5 | 2649418 | | | | | | |
| Q9WVG6 | CARM1 | coactivator-associated arginine methyltransferase 1 | Nucleus | transcription regulator | 3723483 | 2074291.875 | 893843.25 | | | | | | |
| Q9WVG6 | CARM1 | coactivator-associated arginine methyltransferase 1 | Nucleus | transcription regulator | 5250801.5 | 3847783.75 | 3773181.5 | Q9WVG6 | CARM1 | 4062942 | 1232592 | 410864 | 9 |
| Q9ER72 | CARS | cysteinyl-tRNA synthetase | Cytoplasm | enzyme | 3068422.5 | 3574197 | 3175697.75 | | | | | | |
| Q9ER72 | CARS | cysteinyl-tRNA synthetase | Cytoplasm | enzyme | 1667525.875 | 1096052.875 | 1995574.25 | Q9ER72 | CARS | 2367974 | 982073 | 400930 | 6 |
| P51125 | CAST | calpastatin | Cytoplasm | peptidase | 2.62E+07 | 2.37E+07 | 2.23E+07 | | | | | | |
| P51125 | CAST | calpastatin | Cytoplasm | peptidase | 1.60E+07 | 1.06E+07 | 5051298 | | | | | | |
| P51125 | CAST | calpastatin | Cytoplasm | peptidase | 1.13E+07 | 1.11E+07 | 1.13E+07 | | | | | | |
| P51125 | CAST | calpastatin | Cytoplasm | peptidase | 9742835 | 9075695 | 7383797.5 | | | | | | |
| P51125 | CAST | calpastatin | Cytoplasm | peptidase | 8420448 | 7181111 | 6130899.5 | | | | | | |
| P51125 | CAST | calpastatin | Cytoplasm | peptidase | 5923030.5 | 8573229 | 8450881 | | | | | | |
| P51125 | CAST | calpastatin | Cytoplasm | peptidase | 7097073.5 | 5191300.5 | 6415938 | | | | | | |
| P51125 | CAST | calpastatin | Cytoplasm | peptidase | 3954565.25 | 4789710.5 | 5698585.5 | | | | | | |
| P51125 | CAST | calpastatin | Cytoplasm | peptidase | 2962403.5 | 5337727 | 7509103 | | | | | | |
| P51125 | CAST | calpastatin | Cytoplasm | peptidase | 7113112.5 | 1314355 | 0 | | | | | | |
| P51125 | CAST | calpastatin | Cytoplasm | peptidase | 4254837.5 | 4141430.5 | 3252411.5 | | | | | | |
| P51125 | CAST | calpastatin | Cytoplasm | peptidase | 1285342.375 | 1091310.375 | 3051957.5 | P51125 | CAST | 8691137 | 6006622 | 1001104 | 36 |
| P24270 | CAT | catalase | Cytoplasm | enzyme | 1.59E+07 | 1.28E+07 | 2.28E+07 | | | | | | |
| P24270 | CAT | catalase | Cytoplasm | enzyme | 1.27E+07 | 1.15E+07 | 1.50E+07 | | | | | | |
| P24270 | CAT | catalase | Cytoplasm | enzyme | 1.04E+07 | 9665497 | 1.85E+07 | | | | | | |
| P24270 | CAT | catalase | Cytoplasm | enzyme | 8670068 | 7395010.5 | 1.15E+07 | | | | | | |
| P24270 | CAT | catalase | Cytoplasm | enzyme | 7929879 | 6400814 | 7956057 | | | | | | |
| P24270 | CAT | catalase | Cytoplasm | enzyme | 6878728.5 | 4308532.5 | 6816986.5 | | | | | | |
| P24270 | CAT | catalase | Cytoplasm | enzyme | 6714991 | 4816234.5 | 8276789.5 | | | | | | |
| P24270 | CAT | catalase | Cytoplasm | enzyme | 5037382 | 4435849.5 | 7172832.5 | | | | | | |
| P24270 | CAT | catalase | Cytoplasm | enzyme | 5080201 | 4137375 | 7202866.5 | | | | | | |
| P24270 | CAT | catalase | Cytoplasm | enzyme | 5254741.5 | 3518233.5 | 6099836.5 | | | | | | |
| P24270 | CAT | catalase | Cytoplasm | enzyme | 4344123.5 | 2869222.25 | 3387911.75 | | | | | | |

| | | | | | | | | | | | | | |
|---|---|---|---|---|---|---|---|---|---|---|---|---|---|
| P24270 | CAT | catalase | Cytoplasm | enzyme | 2831378.75 | 3920311 | 6522488.5 | | | | | | |
| P24270 | CAT | catalase | Cytoplasm | enzyme | 4666827 | 3117801 | 5955867 | | | | | | |
| P24270 | CAT | catalase | Cytoplasm | enzyme | 3966256.25 | 3005126.25 | 4903984.5 | | | | | | |
| P24270 | CAT | catalase | Cytoplasm | enzyme | 2448947.5 | 2477886.25 | 3775116 | | | | | | |
| P24270 | CAT | catalase | Cytoplasm | enzyme | 3700632.25 | 1807035 | 3641396 | | | | | | |
| P24270 | CAT | catalase | Cytoplasm | enzyme | 2371220.5 | 1764708.875 | 2616706.75 | | | | | | |
| P24270 | CAT | catalase | Cytoplasm | enzyme | 2024627 | 1597168.5 | 2303929 | | | | | | |
| P24270 | CAT | catalase | Cytoplasm | enzyme | 557508.812 | 416873.75 | 738548.375 | | | | | | |
| P24270 | CAT | catalase | Cytoplasm | enzyme | 276345.031 | 1124495.375 | 2401467.25 | P24270 | CAT | 5587193 | 4527452 | 584492 | 60 |
| Q8C9M2 | CCDC15 | coiled-coil domain containing 15 | Cytoplasm | other | 3379126.75 | 5731698 | 1750753.75 | Q8C9M2 | CCDC15 | 3379127 | 2001421 | 1155521 | 3 |
| Q3URS9 | CCDC51 | coiled-coil domain containing 51 | Cytoplasm | other | 731215.688 | 1332460.75 | 544955.562 | Q3URS9 | CCDC51 | 731216 | 411573 | 237622 | 3 |
| Q8BGU5 | CCNY | cyclin Y | Nucleus | other | 3486985.5 | 3476895 | 3861556.5 | | | | | | |
| Q8BGU5 | CCNY | cyclin Y | Nucleus | other | 2860263.75 | 3431246.25 | 2741836.5 | Q8BGU5 | CCNY | 3173625 | 425009 | 173509 | 6 |
| P80314 | CCT2 | chaperonin containing TCP1, subunit 2 (beta) | Cytoplasm | kinase | 2.05E+07 | 1.70E+07 | 2.03E+07 | | | | | | |
| P80314 | CCT2 | chaperonin containing TCP1, subunit 2 (beta) | Cytoplasm | kinase | 1.24E+07 | 1.64E+07 | 2.31E+07 | | | | | | |
| P80314 | CCT2 | chaperonin containing TCP1, subunit 2 (beta) | Cytoplasm | kinase | 1.26E+07 | 8164401 | 8656625 | | | | | | |
| P80314 | CCT2 | chaperonin containing TCP1, subunit 2 (beta) | Cytoplasm | kinase | 1.02E+07 | 8464069 | 1.17E+07 | | | | | | |
| P80314 | CCT2 | chaperonin containing TCP1, subunit 2 (beta) | Cytoplasm | kinase | 1.13E+07 | 1.29E+07 | 1.29E+07 | | | | | | |
| P80314 | CCT2 | chaperonin containing TCP1, subunit 2 (beta) | Cytoplasm | kinase | 1.08E+07 | 1.06E+07 | 1.18E+07 | | | | | | |
| P80314 | CCT2 | chaperonin containing TCP1, subunit 2 (beta) | Cytoplasm | kinase | 7396686.5 | 6485314 | 5916038 | | | | | | |
| P80314 | CCT2 | chaperonin containing TCP1, subunit 2 (beta) | Cytoplasm | kinase | 9333146 | 7489609.5 | 1.08E+07 | | | | | | |
| P80314 | CCT2 | chaperonin containing TCP1, subunit 2 (beta) | Cytoplasm | kinase | 6555340 | 4677825 | 5170987.5 | | | | | | |
| P80314 | CCT2 | chaperonin containing TCP1, subunit 2 (beta) | Cytoplasm | kinase | 8112569.5 | 5765713 | 6151431.5 | | | | | | |
| P80314 | CCT2 | chaperonin containing TCP1, subunit 2 (beta) | Cytoplasm | kinase | 7861015.5 | 6989289 | 7337189.5 | | | | | | |
| P80314 | CCT2 | chaperonin containing TCP1, subunit 2 (beta) | Cytoplasm | kinase | 6260773.5 | 4396741.5 | 5805100.5 | | | | | | |
| P80314 | CCT2 | chaperonin containing TCP1, subunit 2 (beta) | Cytoplasm | kinase | 3994669.25 | 4680689.5 | 6610978 | | | | | | |
| P80314 | CCT2 | chaperonin containing TCP1, subunit 2 (beta) | Cytoplasm | kinase | 3348918.25 | 2732027 | 4161167 | P80314 | CCT2 | 9330937 | 4771585 | 736272 | 42 |
| P80318 | CCT3 | chaperonin containing TCP1, subunit 3 (gamma) | Cytoplasm | other | 3.06E+07 | 2.84E+07 | 3.09E+07 | | | | | | |
| P80318 | CCT3 | chaperonin containing TCP1, subunit 3 (gamma) | Cytoplasm | other | 1.65E+07 | 1.41E+07 | 1.80E+07 | | | | | | |
| P80318 | CCT3 | chaperonin containing TCP1, subunit 3 (gamma) | Cytoplasm | other | 1.39E+07 | 1.17E+07 | 1.25E+07 | | | | | | |
| P80318 | CCT3 | chaperonin containing TCP1, subunit 3 (gamma) | Cytoplasm | other | 1.51E+07 | 1.38E+07 | 1.78E+07 | | | | | | |
| P80318 | CCT3 | chaperonin containing TCP1, subunit 3 (gamma) | Cytoplasm | other | 1.24E+07 | 1.09E+07 | 1.15E+07 | | | | | | |
| P80318 | CCT3 | chaperonin containing TCP1, subunit 3 (gamma) | Cytoplasm | other | 8796848 | 6161412 | 3721305 | | | | | | |
| P80318 | CCT3 | chaperonin containing TCP1, subunit 3 (gamma) | Cytoplasm | other | 8880066 | 8216336 | 6472576.5 | | | | | | |
| P80318 | CCT3 | chaperonin containing TCP1, subunit 3 (gamma) | Cytoplasm | other | 9617223 | 1.02E+07 | 9100882 | | | | | | |
| P80318 | CCT3 | chaperonin containing TCP1, subunit 3 (gamma) | Cytoplasm | other | 7301355 | 5372859 | 6490396.5 | | | | | | |
| P80318 | CCT3 | chaperonin containing TCP1, subunit 3 (gamma) | Cytoplasm | other | 6321243 | 6559970 | 1.15E+07 | | | | | | |
| P80318 | CCT3 | chaperonin containing TCP1, subunit 3 (gamma) | Cytoplasm | other | 6833999.5 | 5383238.5 | 5471832.5 | | | | | | |
| P80318 | CCT3 | chaperonin containing TCP1, subunit 3 (gamma) | Cytoplasm | other | 6350951 | 5538562.5 | 6833823.5 | | | | | | |
| P80318 | CCT3 | chaperonin containing TCP1, subunit 3 (gamma) | Cytoplasm | other | 5131748 | 7483931 | 1.05E+07 | | | | | | |
| P80318 | CCT3 | chaperonin containing TCP1, subunit 3 (gamma) | Cytoplasm | other | 4153120.5 | 4497775 | 3153511.5 | | | | | | |
| P80318 | CCT3 | chaperonin containing TCP1, subunit 3 (gamma) | Cytoplasm | other | 3532987 | 2824246 | 2143663 | | | | | | |
| P80318 | CCT3 | chaperonin containing TCP1, subunit 3 (gamma) | Cytoplasm | other | 2954760.25 | 2117448.25 | 1288741.5 | P80318 | CCT3 | 9905894 | 6790987 | 980195 | 48 |
| P80315 | CCT4 | chaperonin containing TCP1, subunit 4 (delta) | Cytoplasm | other | 2.45E+07 | 2.98E+07 | 2.57E+07 | | | | | | |
| P80315 | CCT4 | chaperonin containing TCP1, subunit 4 (delta) | Cytoplasm | other | 2.31E+07 | 2.07E+07 | 2.51E+07 | | | | | | |
| P80315 | CCT4 | chaperonin containing TCP1, subunit 4 (delta) | Cytoplasm | other | 1.53E+07 | 1.30E+07 | 1.74E+07 | | | | | | |
| P80315 | CCT4 | chaperonin containing TCP1, subunit 4 (delta) | Cytoplasm | other | 1.40E+07 | 1.28E+07 | 1.71E+07 | | | | | | |
| P80315 | CCT4 | chaperonin containing TCP1, subunit 4 (delta) | Cytoplasm | other | 1.20E+07 | 1.00E+07 | 1.16E+07 | | | | | | |
| P80315 | CCT4 | chaperonin containing TCP1, subunit 4 (delta) | Cytoplasm | other | 8444651 | 7674558.5 | 8431626 | | | | | | |
| P80315 | CCT4 | chaperonin containing TCP1, subunit 4 (delta) | Cytoplasm | other | 1.26E+07 | 8255623 | 6866692.5 | | | | | | |
| P80315 | CCT4 | chaperonin containing TCP1, subunit 4 (delta) | Cytoplasm | other | 7673281 | 7474750.5 | 7673328.5 | | | | | | |
| P80315 | CCT4 | chaperonin containing TCP1, subunit 4 (delta) | Cytoplasm | other | 7469710.5 | 4811881.5 | 5872778.5 | | | | | | |
| P80315 | CCT4 | chaperonin containing TCP1, subunit 4 (delta) | Cytoplasm | other | 8275676.5 | 7014384 | 6051500 | | | | | | |
| P80315 | CCT4 | chaperonin containing TCP1, subunit 4 (delta) | Cytoplasm | other | 6673650 | 5159134 | 8616725 | | | | | | |
| P80315 | CCT4 | chaperonin containing TCP1, subunit 4 (delta) | Cytoplasm | other | 5297387 | 6136283.5 | 8330607.5 | | | | | | |
| P80315 | CCT4 | chaperonin containing TCP1, subunit 4 (delta) | Cytoplasm | other | 6748418.5 | 5249113.5 | 4283453 | | | | | | |
| P80315 | CCT4 | chaperonin containing TCP1, subunit 4 (delta) | Cytoplasm | other | 4905840.5 | 2750166.25 | 4383464.5 | | | | | | |
| P80315 | CCT4 | chaperonin containing TCP1, subunit 4 (delta) | Cytoplasm | other | 3711635.25 | 4921816 | 4793018 | | | | | | |
| P80315 | CCT4 | chaperonin containing TCP1, subunit 4 (delta) | Cytoplasm | other | 5423817.5 | 969618.375 | 213825.438 | | | | | | |
| P80315 | CCT4 | chaperonin containing TCP1, subunit 4 (delta) | Cytoplasm | other | 2816412.75 | 1748939.875 | 1264665.75 | | | | | | |
| P80315 | CCT4 | chaperonin containing TCP1, subunit 4 (delta) | Cytoplasm | other | 2167081 | 2107333.5 | 4303613.5 | P80315 | CCT4 | 9506531 | 6876399 | 935759 | 54 |
| P80316 | CCT5 | chaperonin containing TCP1, subunit 5 (epsilon) | Cytoplasm | other | 2.34E+07 | 2.06E+07 | 2.50E+07 | | | | | | |
| P80316 | CCT5 | chaperonin containing TCP1, subunit 5 (epsilon) | Cytoplasm | other | 2.46E+07 | 2.35E+07 | 2.14E+07 | | | | | | |
| P80316 | CCT5 | chaperonin containing TCP1, subunit 5 (epsilon) | Cytoplasm | other | 1.13E+07 | 1.41E+07 | 9096208 | | | | | | |
| P80316 | CCT5 | chaperonin containing TCP1, subunit 5 (epsilon) | Cytoplasm | other | 1.30E+07 | 1.05E+07 | 1.12E+07 | | | | | | |
| P80316 | CCT5 | chaperonin containing TCP1, subunit 5 (epsilon) | Cytoplasm | other | 1.38E+07 | 1.31E+07 | 1.52E+07 | | | | | | |
| P80316 | CCT5 | chaperonin containing TCP1, subunit 5 (epsilon) | Cytoplasm | other | 9460015 | 7227866 | 9855841 | | | | | | |
| P80316 | CCT5 | chaperonin containing TCP1, subunit 5 (epsilon) | Cytoplasm | other | 7392908.5 | 6016460 | 6972112 | | | | | | |
| P80316 | CCT5 | chaperonin containing TCP1, subunit 5 (epsilon) | Cytoplasm | other | 7687907 | 6252138.5 | 6505431 | | | | | | |
| P80316 | CCT5 | chaperonin containing TCP1, subunit 5 (epsilon) | Cytoplasm | other | 6408434 | 4812563.5 | 6308896.5 | | | | | | |
| P80316 | CCT5 | chaperonin containing TCP1, subunit 5 (epsilon) | Cytoplasm | other | 6036891.5 | 4209601 | 4677552.5 | | | | | | |
| P80316 | CCT5 | chaperonin containing TCP1, subunit 5 (epsilon) | Cytoplasm | other | 3573559 | 2464539.5 | 2343531.5 | | | | | | |
| P80316 | CCT5 | chaperonin containing TCP1, subunit 5 (epsilon) | Cytoplasm | other | 2100383 | 2225917.75 | 3514774.5 | | | | | | |
| P80316 | CCT5 | chaperonin containing TCP1, subunit 5 (epsilon) | Cytoplasm | other | 2490826.5 | 456156.094 | 20922.029 | P80316 | CCT5 | 10086225 | 7027909 | 1125366 | 39 |
| P80317 | CCT6A | chaperonin containing TCP1, subunit 6A (zeta 1) | Cytoplasm | other | 2.58E+07 | 2.19E+07 | 2.63E+07 | | | | | | |
| P80317 | CCT6A | chaperonin containing TCP1, subunit 6A (zeta 1) | Cytoplasm | other | 1.81E+07 | 1.57E+07 | 2.05E+07 | | | | | | |
| P80317 | CCT6A | chaperonin containing TCP1, subunit 6A (zeta 1) | Cytoplasm | other | 1.86E+07 | 1.70E+07 | 1.84E+07 | | | | | | |
| P80317 | CCT6A | chaperonin containing TCP1, subunit 6A (zeta 1) | Cytoplasm | other | 1.92E+07 | 1.36E+07 | 1.75E+07 | | | | | | |
| P80317 | CCT6A | chaperonin containing TCP1, subunit 6A (zeta 1) | Cytoplasm | other | 1.38E+07 | 1.38E+07 | 1.42E+07 | | | | | | |
| P80317 | CCT6A | chaperonin containing TCP1, subunit 6A (zeta 1) | Cytoplasm | other | 1.28E+07 | 1.00E+07 | 1.05E+07 | | | | | | |
| P80317 | CCT6A | chaperonin containing TCP1, subunit 6A (zeta 1) | Cytoplasm | other | 1.23E+07 | 9265333 | 9953560 | | | | | | |
| P80317 | CCT6A | chaperonin containing TCP1, subunit 6A (zeta 1) | Cytoplasm | other | 9323513 | 7632852.5 | 1.06E+07 | | | | | | |
| P80317 | CCT6A | chaperonin containing TCP1, subunit 6A (zeta 1) | Cytoplasm | other | 5488254.5 | 6446715 | 7761435 | | | | | | |
| P80317 | CCT6A | chaperonin containing TCP1, subunit 6A (zeta 1) | Cytoplasm | other | 6902053.5 | 6140826 | 7050031 | | | | | | |
| P80317 | CCT6A | chaperonin containing TCP1, subunit 6A (zeta 1) | Cytoplasm | other | 1168041.25 | 1817034.5 | 1738491.625 | | | | | | |
| P80317 | CCT6A | chaperonin containing TCP1, subunit 6A (zeta 1) | Cytoplasm | other | 1034286.5 | 2033458.25 | 1416737.625 | P80317 | CCT6A | 12041346 | 6962916 | 1160486 | 36 |
| P80313 | CCT7 | chaperonin containing TCP1, subunit 7 (eta) | Cytoplasm | other | 1.75E+07 | 1.75E+07 | 1.88E+07 | | | | | | |
| P80313 | CCT7 | chaperonin containing TCP1, subunit 7 (eta) | Cytoplasm | other | 1.38E+07 | 1.48E+07 | 1.51E+07 | | | | | | |
| P80313 | CCT7 | chaperonin containing TCP1, subunit 7 (eta) | Cytoplasm | other | 1.36E+07 | 1.33E+07 | 1.51E+07 | | | | | | |
| P80313 | CCT7 | chaperonin containing TCP1, subunit 7 (eta) | Cytoplasm | other | 1.31E+07 | 1.24E+07 | 1.31E+07 | | | | | | |
| P80313 | CCT7 | chaperonin containing TCP1, subunit 7 (eta) | Cytoplasm | other | 1.52E+07 | 1.09E+07 | 1.51E+07 | | | | | | |

| | | | | | | | | | | | | | |
|---|---|---|---|---|---|---|---|---|---|---|---|---|---|
| P80313 | CCT7 | chaperonin containing TCP1, subunit 7 (eta) | Cytoplasm | other | 1.11E+07 | 8900825 | 9377695 | | | | | | |
| P80313 | CCT7 | chaperonin containing TCP1, subunit 7 (eta) | Cytoplasm | other | 9173717 | 7871796.5 | 7707177.5 | | | | | | |
| P80313 | CCT7 | chaperonin containing TCP1, subunit 7 (eta) | Cytoplasm | other | 1.01E+07 | 7410711.5 | 8950622 | | | | | | |
| P80313 | CCT7 | chaperonin containing TCP1, subunit 7 (eta) | Cytoplasm | other | 7953476.5 | 6202761.5 | 5432130.5 | | | | | | |
| P80313 | CCT7 | chaperonin containing TCP1, subunit 7 (eta) | Cytoplasm | other | 7569560.5 | 6130455 | 7700963 | | | | | | |
| P80313 | CCT7 | chaperonin containing TCP1, subunit 7 (eta) | Cytoplasm | other | 7776032.5 | 5852137.5 | 7423488 | | | | | | |
| P80313 | CCT7 | chaperonin containing TCP1, subunit 7 (eta) | Cytoplasm | other | 4650136 | 2834432 | 3691355.75 | | | | | | |
| P80313 | CCT7 | chaperonin containing TCP1, subunit 7 (eta) | Cytoplasm | other | 4128689 | 3683693.5 | 3375583.75 | | | | | | |
| P80313 | CCT7 | chaperonin containing TCP1, subunit 7 (eta) | Cytoplasm | other | 2272651.75 | 4020316.75 | 4996773.5 | | | | | | |
| P80313 | CCT7 | chaperonin containing TCP1, subunit 7 (eta) | Cytoplasm | other | 2603598.5 | 2827392.75 | 2062582.5 | P80313 | CCT7 | 9363191 | 4708942 | 701968 | 45 |
| P42932 | CCT8 | chaperonin containing TCP1, subunit 8 (theta) | Cytoplasm | enzyme | 1.67E+07 | 1.57E+07 | 1.74E+07 | | | | | | |
| P42932 | CCT8 | chaperonin containing TCP1, subunit 8 (theta) | Cytoplasm | enzyme | 1.43E+07 | 1.10E+07 | 1.31E+07 | | | | | | |
| P42932 | CCT8 | chaperonin containing TCP1, subunit 8 (theta) | Cytoplasm | enzyme | 1.19E+07 | 9330226 | 1.14E+07 | | | | | | |
| P42932 | CCT8 | chaperonin containing TCP1, subunit 8 (theta) | Cytoplasm | enzyme | 1.26E+07 | 1.07E+07 | 1.21E+07 | | | | | | |
| P42932 | CCT8 | chaperonin containing TCP1, subunit 8 (theta) | Cytoplasm | enzyme | 1.69E+07 | 9305723 | 1.44E+07 | | | | | | |
| P42932 | CCT8 | chaperonin containing TCP1, subunit 8 (theta) | Cytoplasm | enzyme | 1.30E+07 | 1.03E+07 | 1.03E+07 | | | | | | |
| P42932 | CCT8 | chaperonin containing TCP1, subunit 8 (theta) | Cytoplasm | enzyme | 9463992 | 8221958 | 1.10E+07 | | | | | | |
| P42932 | CCT8 | chaperonin containing TCP1, subunit 8 (theta) | Cytoplasm | enzyme | 1.03E+07 | 7246768.5 | 7412794 | | | | | | |
| P42932 | CCT8 | chaperonin containing TCP1, subunit 8 (theta) | Cytoplasm | enzyme | 6989242 | 7526938 | 9335599 | | | | | | |
| P42932 | CCT8 | chaperonin containing TCP1, subunit 8 (theta) | Cytoplasm | enzyme | 8975856 | 6758035 | 6849268 | | | | | | |
| P42932 | CCT8 | chaperonin containing TCP1, subunit 8 (theta) | Cytoplasm | enzyme | 7649485 | 6067755.5 | 6888386 | | | | | | |
| P42932 | CCT8 | chaperonin containing TCP1, subunit 8 (theta) | Cytoplasm | enzyme | 8199746.5 | 7761868 | 9626062 | | | | | | |
| P42932 | CCT8 | chaperonin containing TCP1, subunit 8 (theta) | Cytoplasm | enzyme | 8749809 | 5978488 | 6276097 | | | | | | |
| P42932 | CCT8 | chaperonin containing TCP1, subunit 8 (theta) | Cytoplasm | enzyme | 7170998.5 | 4477814.5 | 5798488 | | | | | | |
| P42932 | CCT8 | chaperonin containing TCP1, subunit 8 (theta) | Cytoplasm | enzyme | 4313581 | 3169504.25 | 4167820.25 | | | | | | |
| P42932 | CCT8 | chaperonin containing TCP1, subunit 8 (theta) | Cytoplasm | enzyme | 6209183.5 | 3735032 | 3285579.5 | | | | | | |
| P42932 | CCT8 | chaperonin containing TCP1, subunit 8 (theta) | Cytoplasm | enzyme | 6812604 | 7048232 | 9933507 | | | | | | |
| P42932 | CCT8 | chaperonin containing TCP1, subunit 8 (theta) | Cytoplasm | enzyme | 6254895.5 | 3412093.75 | 3788551 | | | | | | |
| P42932 | CCT8 | chaperonin containing TCP1, subunit 8 (theta) | Cytoplasm | enzyme | 4379538 | 1816298.25 | 2054119 | | | | | | |
| P42932 | CCT8 | chaperonin containing TCP1, subunit 8 (theta) | Cytoplasm | enzyme | 3332291.5 | 2628677.5 | 3014486.25 | | | | | | |
| P42932 | CCT8 | chaperonin containing TCP1, subunit 8 (theta) | Cytoplasm | enzyme | 3053073.5 | 2669917.25 | 2722201.5 | | | | | | |
| P42932 | CCT8 | chaperonin containing TCP1, subunit 8 (theta) | Cytoplasm | enzyme | 4059643.5 | 4605112.5 | 3470995.5 | | | | | | |
| P42932 | CCT8 | chaperonin containing TCP1, subunit 8 (theta) | Cytoplasm | enzyme | 3849725.25 | 2530137 | 3215051.25 | | | | | | |
| P42932 | CCT8 | chaperonin containing TCP1, subunit 8 (theta) | Cytoplasm | enzyme | 1302556.25 | 1839028.875 | 2338927 | P42932 | CCT8 | 8183593 | 4068135 | 479434 | 72 |
| Q8K4F0 | CD226 | CD226 molecule | Plasma Membrane | other | 457462.781 | 909818.938 | 1424920.875 | Q8K4F0 | CD226 | 457463 | 484068 | 279477 | 3 |
| Q08857 | CD36 | CD36 molecule (thrombospondin receptor) | Plasma Membrane | transmembrane receptor | 5.56E+08 | 5.27E+08 | 4.90E+08 | | | | | | |
| Q08857 | CD36 | CD36 molecule (thrombospondin receptor) | Plasma Membrane | transmembrane receptor | 2.30E+08 | 2.41E+08 | 2.60E+08 | | | | | | |
| Q08857 | CD36 | CD36 molecule (thrombospondin receptor) | Plasma Membrane | transmembrane receptor | 1.16E+08 | 1.64E+08 | 1.77E+08 | | | | | | |
| Q08857 | CD36 | CD36 molecule (thrombospondin receptor) | Plasma Membrane | transmembrane receptor | 1.17E+08 | 1.27E+08 | 1.18E+08 | | | | | | |
| Q08857 | CD36 | CD36 molecule (thrombospondin receptor) | Plasma Membrane | transmembrane receptor | 8.10E+07 | 2.09E+07 | 1.36E+07 | | | | | | |
| Q08857 | CD36 | CD36 molecule (thrombospondin receptor) | Plasma Membrane | transmembrane receptor | 4.08E+07 | 1.10E+08 | 1.42E+08 | | | | | | |
| Q08857 | CD36 | CD36 molecule (thrombospondin receptor) | Plasma Membrane | transmembrane receptor | 1.00E+08 | 8.65E+07 | 2.17E+07 | | | | | | |
| Q08857 | CD36 | CD36 molecule (thrombospondin receptor) | Plasma Membrane | transmembrane receptor | 7.76E+07 | 9.02E+07 | 9.19E+07 | | | | | | |
| Q08857 | CD36 | CD36 molecule (thrombospondin receptor) | Plasma Membrane | transmembrane receptor | 6.97E+07 | 6.29E+07 | 6.27E+07 | | | | | | |
| Q08857 | CD36 | CD36 molecule (thrombospondin receptor) | Plasma Membrane | transmembrane receptor | 8.27E+07 | 1.35E+08 | 1.48E+08 | | | | | | |
| Q08857 | CD36 | CD36 molecule (thrombospondin receptor) | Plasma Membrane | transmembrane receptor | 1.13E+07 | 3.15E+07 | 4.56E+07 | | | | | | |
| Q08857 | CD36 | CD36 molecule (thrombospondin receptor) | Plasma Membrane | transmembrane receptor | 1.51E+07 | 2.44E+07 | 2.55E+07 | | | | | | |
| Q08857 | CD36 | CD36 molecule (thrombospondin receptor) | Plasma Membrane | transmembrane receptor | 1.71E+07 | 2.08E+07 | 1.56E+07 | | | | | | |
| Q08857 | CD36 | CD36 molecule (thrombospondin receptor) | Plasma Membrane | transmembrane receptor | 1.09E+07 | 1.23E+07 | 1.21E+07 | | | | | | |
| Q08857 | CD36 | CD36 molecule (thrombospondin receptor) | Plasma Membrane | transmembrane receptor | 3764255.25 | 2793972 | 1772084.75 | | | | | | |
| Q08857 | CD36 | CD36 molecule (thrombospondin receptor) | Plasma Membrane | transmembrane receptor | 89459.055 | 1463662.625 | 4523801.5 | Q08857 | CD36 | 95572732 | 129154917 | 18641907 | 48 |
| P15379 | CD44 | CD44 molecule (Indian blood group) | Plasma Membrane | enzyme | 1.59E+07 | 1.72E+07 | 2.06E+07 | | | | | | |
| P15379 | CD44 | CD44 molecule (Indian blood group) | Plasma Membrane | enzyme | 5156774 | 5049507.5 | 8143719.5 | | | | | | |
| P15379 | CD44 | CD44 molecule (Indian blood group) | Plasma Membrane | enzyme | 2792644.25 | 4338609 | 6449034.5 | | | | | | |
| P15379 | CD44 | CD44 molecule (Indian blood group) | Plasma Membrane | enzyme | 740926.688 | 1518439.875 | 1576001.125 | P15379 | CD44 | 6150086 | 6727612 | 1942094 | 12 |
| Q61735 | CD47 | CD47 molecule | Plasma Membrane | transmembrane receptor | 5.95E+08 | 5.65E+08 | 6.17E+08 | | | | | | |
| Q61735 | CD47 | CD47 molecule | Plasma Membrane | transmembrane receptor | 2.96E+08 | 2.70E+08 | 3.59E+08 | | | | | | |
| Q61735 | CD47 | CD47 molecule | Plasma Membrane | transmembrane receptor | 1.93E+08 | 1.90E+08 | 2.56E+08 | | | | | | |
| Q61735 | CD47 | CD47 molecule | Plasma Membrane | transmembrane receptor | 1.38E+08 | 1.21E+08 | 1.55E+08 | | | | | | |
| Q61735 | CD47 | CD47 molecule | Plasma Membrane | transmembrane receptor | 7.69E+07 | 9.81E+07 | 1.30E+08 | | | | | | |
| Q61735 | CD47 | CD47 molecule | Plasma Membrane | transmembrane receptor | 8866831 | 9769444 | 1.65E+07 | Q61735 | CD47 | 217834472 | 194176240 | 45767779 | 18 |
| P18181 | CD48 | CD48 molecule | Plasma Membrane | other | 2663666.25 | 2956068 | 2447786.5 | P18181 | CD48 | 2663666 | 255099 | 147281 | 3 |
| Q61475 | CD55 | CD55 molecule, decay accelerating factor for complement (Cromer blood group) | Plasma Membrane | other | 1.23E+07 | 1.45E+07 | 1.45E+07 | | | | | | |
| Q61475 | CD55 | CD55 molecule, decay accelerating factor for complement (Cromer blood group) | Plasma Membrane | other | 1.03E+07 | 1.23E+07 | 1.05E+07 | | | | | | |
| Q61475 | CD55 | CD55 molecule, decay accelerating factor for complement (Cromer blood group) | Plasma Membrane | other | 6034464.5 | 7321515 | 8528358 | | | | | | |
| Q61475 | CD55 | CD55 molecule, decay accelerating factor for complement (Cromer blood group) | Plasma Membrane | other | 5114195.5 | 7552266 | 6583420.5 | | | | | | |
| Q61475 | CD55 | CD55 molecule, decay accelerating factor for complement (Cromer blood group) | Plasma Membrane | other | 5141186 | 6634307 | 6047476.5 | | | | | | |
| Q61475 | CD55 | CD55 molecule, decay accelerating factor for complement (Cromer blood group) | Plasma Membrane | other | 3403652.25 | 3062492 | 4964380.5 | Q61475 | CD55 | 7045583 | 3566932 | 840734 | 18 |
| O55186 | Cd59a | CD59a antigen | Plasma Membrane | other | 1.08E+08 | 9.54E+07 | 1.33E+08 | | | | | | |
| O55186 | Cd59a | CD59a antigen | Plasma Membrane | other | 8759535 | 7487705.5 | 9372789 | | | | | | |
| O55186 | Cd59a | CD59a antigen | Plasma Membrane | other | 4520506.5 | 1263368.5 | 1049274.125 | O55186 | Cd59a | 40493347 | 54238319 | 18079440 | 9 |
| P40237 | CD82 | CD82 molecule | Plasma Membrane | other | 6882867.5 | 1.11E+07 | 1.59E+07 | | | | | | |
| P40237 | CD82 | CD82 molecule | Plasma Membrane | other | 6861266.5 | 8941533 | 1.40E+07 | P40237 | CD82 | 6872067 | 3762018 | 1535837 | 6 |
| P40240 | CD9 | CD9 molecule | Plasma Membrane | other | 1954708 | 6422758.5 | 1.26E+07 | | | | | | |
| P40240 | CD9 | CD9 molecule | Plasma Membrane | other | 193291.203 | 1064804.625 | 4413105 | | | | | | |
| P40240 | CD9 | CD9 molecule | Plasma Membrane | other | 11435.842 | 74475.766 | 203420.703 | P40240 | CD9 | 719812 | 4252470 | 1417490 | 9 |
| Q8CII2 | CDC123 | cell division cycle 123 | Cytoplasm | other | 3885166.5 | 5013648 | 6531665 | Q8CII2 | CDC123 | 3885167 | 1328019 | 766732 | 3 |
| P60766 | Cdc42 | cell division cycle 42 | Plasma Membrane | enzyme | 9937222 | 7487637.5 | 4616749.5 | | | | | | |
| P60766 | Cdc42 | cell division cycle 42 | Plasma Membrane | enzyme | 6767033 | 4499840 | 4717047 | | | | | | |
| P60766 | Cdc42 | cell division cycle 42 | Plasma Membrane | enzyme | 6140952 | 6001484 | 8140107.5 | | | | | | |
| P60766 | Cdc42 | cell division cycle 42 | Plasma Membrane | enzyme | 3309733 | 3659739.75 | 6476793 | | | | | | |
| P60766 | Cdc42 | cell division cycle 42 | Plasma Membrane | enzyme | 2376935.5 | 2748276 | 3008163 | P60766 | Cdc42 | 5706375 | 2195592 | 566899 | 15 |
| Q922M5 | CDCA7L | cell division cycle associated 7-like | Nucleus | other | 2.78E+07 | 2.37E+07 | 1.21E+07 | Q922M5 | CDCA7L | 27810000 | 8169498 | 4716662 | 3 |
| Q8VDP6 | CDIPT | CDP-diacylglycerol--inositol 3-phosphatidyltransferase | Cytoplasm | enzyme | 1426089.875 | 1795205.875 | 1966618.75 | Q8VDP6 | CDIPT | 1426090 | 276225 | 159478 | 3 |
| Q99L43 | CDS2 | CDP-diacylglycerol synthase (phosphatidate cytidylyltransferase) 2 | Cytoplasm | enzyme | 379439.375 | 1125093.5 | 1772610.875 | Q99L43 | CDS2 | 379439 | 697162 | 402506 | 3 |
| Q4VAA2 | CDV3 | CDV3 homolog (mouse) | Cytoplasm | other | 1933214.75 | 2422043.5 | 1540914.25 | Q4VAA2 | CDV3 | 1933215 | 441445 | 254868 | 3 |
| Q6AW46 | CES5A | carboxylesterase 5A | Extracellular Space | enzyme | 1.67E+07 | 1.61E+07 | 1.46E+07 | Q6AW46 | CES5A | 16660000 | 1060063 | 612028 | 3 |
| P18760 | CFL1 | cofilin 1 (non-muscle) | Nucleus | other | 3259561.75 | 4772908 | 8715398 | | | | | | |
| P18760 | CFL1 | cofilin 1 (non-muscle) | Nucleus | other | 1055845 | 1175639.125 | 2981152.5 | | | | | | |
| P18760 | CFL1 | cofilin 1 (non-muscle) | Nucleus | other | 838394.688 | 1064673.75 | 2274093 | | | | | | |
| P18760 | CFL1 | cofilin 1 (non-muscle) | Nucleus | other | 1397794.875 | 2782733.75 | 3446543 | | | | | | |

| | | | | | | | | | | | | | |
|---|---|---|---|---|---|---|---|---|---|---|---|---|---|
| P18760 | CFL1 | cofilin 1 (non-muscle) | Nucleus | other | 634877.188 | 690671.062 | 1370855 | P18760 | CFL1 | 1437295 | 2129109 | 549734 | 15 |
| Q9CRB9 | CHCHD3 | coiled-coil-helix-coiled-coil-helix domain containing 3 | Cytoplasm | other | 2670061.25 | 7161604 | 3093124 | Q9CRB9 | CHCHD3 | 2670061 | 2480103 | 1431888 | 3 |
| O35744 | Chil3/Chil4 | chitinase-like 3 | Cytoplasm | enzyme | 389558.938 | 262614.625 | 104281.391 | | | | | | |
| O35744 | Chil3/Chil4 | chitinase-like 3 | Cytoplasm | enzyme | 277283.625 | 510482.438 | 359354.094 | | | | | | |
| O35744 | Chil3/Chil4 | chitinase-like 3 | Cytoplasm | enzyme | 224138.875 | 207839.359 | 256180.328 | | | | | | |
| O35744 | Chil3/Chil4 | chitinase-like 3 | Cytoplasm | enzyme | 269630.5 | 152465.328 | 196669.469 | | | | | | |
| O35744 | Chil3/Chil4 | chitinase-like 3 | Cytoplasm | enzyme | 66634.992 | 185994.656 | 71738.312 | | | | | | |
| O35744 | Chil3/Chil4 | chitinase-like 3 | Cytoplasm | enzyme | 208015.391 | 97034.836 | 59336.207 | O35744 | Chil3/Chil4 | 239210 | 120687 | 28446 | 18 |
| Q9DB34 | CHMP2A | charged multivesicular body protein 2A | Cytoplasm | other | 3261468 | 2164548.25 | 2585782 | Q9DB34 | CHMP2A | 3261468 | 553357 | 319481 | 3 |
| P0C0A3 | CHMP6 | charged multivesicular body protein 6 | Cytoplasm | other | 3935704.5 | 3606866.25 | 4785502.5 | P0C0A3 | CHMP6 | 3935705 | 608204 | 351147 | 3 |
| P61022 | CHP1 | calcineurin-like EF-hand protein 1 | Cytoplasm | transporter | 9886813 | 7416702.5 | 6481713.5 | | | | | | |
| P61022 | CHP1 | calcineurin-like EF-hand protein 1 | Cytoplasm | transporter | 5238938 | 5214374.5 | 5700889 | | | | | | |
| P61022 | CHP1 | calcineurin-like EF-hand protein 1 | Cytoplasm | transporter | 3401870.5 | 2292977 | 2847290.75 | | | | | | |
| P61022 | CHP1 | calcineurin-like EF-hand protein 1 | Cytoplasm | transporter | 4285150 | 5090258 | 4850694.5 | | | | | | |
| P61022 | CHP1 | calcineurin-like EF-hand protein 1 | Cytoplasm | transporter | 4528972.5 | 5425670.5 | 5901430 | | | | | | |
| P61022 | CHP1 | calcineurin-like EF-hand protein 1 | Cytoplasm | transporter | 3343500.75 | 3695893.25 | 2627149.5 | P61022 | CHP1 | 5114207 | 1857980 | 437930 | 18 |
| Q9Z0F4 | CIB1 | calcium and integrin binding 1 (calmyrin) | Nucleus | other | 6152726 | 1.21E+07 | 1.08E+07 | | | | | | |
| Q9Z0F4 | CIB1 | calcium and integrin binding 1 (calmyrin) | Nucleus | other | 6516004 | 5668235 | 8561325 | | | | | | |
| Q9Z0F4 | CIB1 | calcium and integrin binding 1 (calmyrin) | Nucleus | other | 7765823.5 | 1432306.125 | 4321969 | | | | | | |
| Q9Z0F4 | CIB1 | calcium and integrin binding 1 (calmyrin) | Nucleus | other | 469119.875 | 819907.25 | 921968.125 | Q9Z0F4 | CIB1 | 5225918 | 3979300 | 1148725 | 12 |
| Q91WS0 | CISD1 | CDGSH iron sulfur domain 1 | Cytoplasm | other | 2240626.5 | 3301557 | 1519384 | | | | | | |
| Q91WS0 | CISD1 | CDGSH iron sulfur domain 1 | Cytoplasm | other | 2806871.25 | 6057138 | 2935911 | Q91WS0 | CISD1 | 2523749 | 1557152 | 635705 | 6 |
| Q9CQB5 | CISD2 | CDGSH iron sulfur domain 2 | Cytoplasm | other | 4909193.5 | 6745312.5 | 6941980 | | | | | | |
| Q9CQB5 | CISD2 | CDGSH iron sulfur domain 2 | Cytoplasm | other | 4841224 | 5237743 | 5791697.5 | | | | | | |
| Q9CQB5 | CISD2 | CDGSH iron sulfur domain 2 | Cytoplasm | other | 4386199 | 4317537 | 5214894 | | | | | | |
| Q9CQB5 | CISD2 | CDGSH iron sulfur domain 2 | Cytoplasm | other | 2544269.25 | 3377874 | 3727038.75 | | | | | | |
| Q9CQB5 | CISD2 | CDGSH iron sulfur domain 2 | Cytoplasm | other | 2457217.5 | 2845781.25 | 4008607.75 | Q9CQB5 | CISD2 | 3827621 | 1385884 | 357834 | 15 |
| Q9Z0S4 | Cldn13 | claudin 13 | Plasma Membrane | other | 2.00E+08 | 1.84E+08 | 2.28E+08 | | | | | | |
| Q9Z0S4 | Cldn13 | claudin 13 | Plasma Membrane | other | 5.99E+07 | 5.48E+07 | 5.18E+07 | | | | | | |
| Q9Z0S4 | Cldn13 | claudin 13 | Plasma Membrane | other | 5.67E+07 | 7.10E+07 | 8.65E+07 | | | | | | |
| Q9Z0S4 | Cldn13 | claudin 13 | Plasma Membrane | other | 5.04E+07 | 3.83E+07 | 4.06E+07 | Q9Z0S4 | Cldn13 | 91662500 | 68422249 | 19751802 | 12 |
| Q9JL99 | CLEC1B | C-type lectin domain family 1, member B | Plasma Membrane | transmembrane receptor | 380887.562 | 1795422.875 | 3511602.25 | Q9JL99 | CLEC1B | 380888 | 1567777 | 905157 | 3 |
| Q9QUK3 | CLN8 | ceroid-lipofuscinosis, neuronal 8 | Cytoplasm | other | 5464978.5 | 5253099.5 | 6125864.5 | Q9QUK3 | CLN8 | 5464979 | 455226 | 262825 | 3 |
| Q61189 | CLNS1A | chloride channel, nucleotide-sensitive, 1A | Plasma Membrane | ion channel | 4009862.75 | 3252756 | 3536119.25 | Q61189 | CLNS1A | 4009863 | 382522 | 220849 | 3 |
| Q8VBZ3 | CLPTM1 | cleft lip and palate associated transmembrane protein 1 | Plasma Membrane | other | 1077361.125 | 1284952.375 | 432982.406 | Q8VBZ3 | CLPTM1 | 1077361 | 444254 | 256490 | 3 |
| Q68FD5 | CLTC | clathrin, heavy chain (Hc) | Plasma Membrane | other | 1.04E+07 | 1.58E+07 | 9516625 | | | | | | |
| Q68FD5 | CLTC | clathrin, heavy chain (Hc) | Plasma Membrane | other | 1.58E+07 | 1.38E+07 | 1.05E+07 | | | | | | |
| Q68FD5 | CLTC | clathrin, heavy chain (Hc) | Plasma Membrane | other | 1.51E+07 | 1.28E+07 | 1.04E+07 | | | | | | |
| Q68FD5 | CLTC | clathrin, heavy chain (Hc) | Plasma Membrane | other | 1.42E+07 | 1.84E+07 | 1.22E+07 | | | | | | |
| Q68FD5 | CLTC | clathrin, heavy chain (Hc) | Plasma Membrane | other | 8328223.5 | 1.44E+07 | 7469664.5 | | | | | | |
| Q68FD5 | CLTC | clathrin, heavy chain (Hc) | Plasma Membrane | other | 1.08E+07 | 1.11E+07 | 1.04E+07 | | | | | | |
| Q68FD5 | CLTC | clathrin, heavy chain (Hc) | Plasma Membrane | other | 1.33E+07 | 1.43E+07 | 1.02E+07 | | | | | | |
| Q68FD5 | CLTC | clathrin, heavy chain (Hc) | Plasma Membrane | other | 7935646 | 9624827 | 1.08E+07 | | | | | | |
| Q68FD5 | CLTC | clathrin, heavy chain (Hc) | Plasma Membrane | other | 9135983 | 8681991 | 6808386.5 | | | | | | |
| Q68FD5 | CLTC | clathrin, heavy chain (Hc) | Plasma Membrane | other | 9323068 | 9881387 | 6747189 | | | | | | |
| Q68FD5 | CLTC | clathrin, heavy chain (Hc) | Plasma Membrane | other | 7214025 | 1.49E+07 | 1.13E+07 | | | | | | |
| Q68FD5 | CLTC | clathrin, heavy chain (Hc) | Plasma Membrane | other | 8404112 | 1.00E+07 | 8277938.5 | | | | | | |
| Q68FD5 | CLTC | clathrin, heavy chain (Hc) | Plasma Membrane | other | 6428993.5 | 5922796 | 4726552.5 | | | | | | |
| Q68FD5 | CLTC | clathrin, heavy chain (Hc) | Plasma Membrane | other | 5459641 | 4591921 | 3874302 | | | | | | |
| Q68FD5 | CLTC | clathrin, heavy chain (Hc) | Plasma Membrane | other | 8155320 | 7847561 | 6050677 | | | | | | |
| Q68FD5 | CLTC | clathrin, heavy chain (Hc) | Plasma Membrane | other | 5036954 | 7680063 | 5070355 | | | | | | |
| Q68FD5 | CLTC | clathrin, heavy chain (Hc) | Plasma Membrane | other | 4954120.5 | 5764423 | 6793796 | | | | | | |
| Q68FD5 | CLTC | clathrin, heavy chain (Hc) | Plasma Membrane | other | 5425922 | 6093414.5 | 6161670.5 | | | | | | |
| Q68FD5 | CLTC | clathrin, heavy chain (Hc) | Plasma Membrane | other | 5252731 | 6483482.5 | 4497760 | | | | | | |
| Q68FD5 | CLTC | clathrin, heavy chain (Hc) | Plasma Membrane | other | 6645501.5 | 5245210.5 | 3482446.5 | | | | | | |
| Q68FD5 | CLTC | clathrin, heavy chain (Hc) | Plasma Membrane | other | 6444012.5 | 5974667 | 4287304 | | | | | | |
| Q68FD5 | CLTC | clathrin, heavy chain (Hc) | Plasma Membrane | other | 3007606.25 | 3506800.75 | 1449870.125 | | | | | | |
| Q68FD5 | CLTC | clathrin, heavy chain (Hc) | Plasma Membrane | other | 7012303 | 6273394 | 4715885.5 | | | | | | |
| Q68FD5 | CLTC | clathrin, heavy chain (Hc) | Plasma Membrane | other | 7586221.5 | 6930286.5 | 8037081 | | | | | | |
| Q68FD5 | CLTC | clathrin, heavy chain (Hc) | Plasma Membrane | other | 5079532 | 4764068.5 | 3397407 | | | | | | |
| Q68FD5 | CLTC | clathrin, heavy chain (Hc) | Plasma Membrane | other | 5512042.5 | 5160751.5 | 4274189.5 | | | | | | |
| Q68FD5 | CLTC | clathrin, heavy chain (Hc) | Plasma Membrane | other | 5676002 | 6624853.5 | 4481127.5 | | | | | | |
| Q68FD5 | CLTC | clathrin, heavy chain (Hc) | Plasma Membrane | other | 5685062 | 4711733.5 | 4666976 | | | | | | |
| Q68FD5 | CLTC | clathrin, heavy chain (Hc) | Plasma Membrane | other | 3890625.25 | 3429087 | 1661599.5 | | | | | | |
| Q68FD5 | CLTC | clathrin, heavy chain (Hc) | Plasma Membrane | other | 5202289.5 | 3974249.75 | 3687067 | | | | | | |
| Q68FD5 | CLTC | clathrin, heavy chain (Hc) | Plasma Membrane | other | 3581891.5 | 3591403.5 | 2934179.25 | | | | | | |
| Q68FD5 | CLTC | clathrin, heavy chain (Hc) | Plasma Membrane | other | 3187590 | 4697779.5 | 3888524 | | | | | | |
| Q68FD5 | CLTC | clathrin, heavy chain (Hc) | Plasma Membrane | other | 2082215.75 | 2239465 | 1781026.625 | | | | | | |
| Q68FD5 | CLTC | clathrin, heavy chain (Hc) | Plasma Membrane | other | 3292653 | 4107971.25 | 5537991.5 | | | | | | |
| Q68FD5 | CLTC | clathrin, heavy chain (Hc) | Plasma Membrane | other | 2899803.75 | 2424174.25 | 1581410.75 | | | | | | |
| Q68FD5 | CLTC | clathrin, heavy chain (Hc) | Plasma Membrane | other | 2520364 | 1866882.625 | 790841.875 | | | | | | |
| Q68FD5 | CLTC | clathrin, heavy chain (Hc) | Plasma Membrane | other | 2348582.25 | 1949040.75 | 1707000 | | | | | | |
| Q68FD5 | CLTC | clathrin, heavy chain (Hc) | Plasma Membrane | other | 2065411.375 | 2543071.75 | 2244356 | | | | | | |
| Q68FD5 | CLTC | clathrin, heavy chain (Hc) | Plasma Membrane | other | 1877064.25 | 2881864.5 | 2167308 | | | | | | |
| Q68FD5 | CLTC | clathrin, heavy chain (Hc) | Plasma Membrane | other | 2305025.75 | 2304722 | 1557490.625 | | | | | | |
| Q68FD5 | CLTC | clathrin, heavy chain (Hc) | Plasma Membrane | other | 1948287.25 | 1688044.125 | 1198764.875 | | | | | | |
| Q68FD5 | CLTC | clathrin, heavy chain (Hc) | Plasma Membrane | other | 1955291.25 | 535480.125 | 274799.031 | | | | | | |
| Q68FD5 | CLTC | clathrin, heavy chain (Hc) | Plasma Membrane | other | 2764278.5 | 992243.438 | 65005.691 | | | | | | |
| Q68FD5 | CLTC | clathrin, heavy chain (Hc) | Plasma Membrane | other | 1492371.875 | 1249006.125 | 1105648.875 | | | | | | |
| Q68FD5 | CLTC | clathrin, heavy chain (Hc) | Plasma Membrane | other | 751646.25 | 1342725.375 | 586822.625 | | | | | | |
| Q68FD5 | CLTC | clathrin, heavy chain (Hc) | Plasma Membrane | other | 269507.469 | 160109.312 | 56687.523 | Q68FD5 | CLTC | 5821259 | 3982075 | 338977 | 138 |
| Q06890 | CLU | clusterin | Cytoplasm | other | 8618369 | 1.01E+07 | 1.19E+07 | | | | | | |
| Q06890 | CLU | clusterin | Cytoplasm | other | 1831903 | 2678391.5 | 3928557 | | | | | | |
| Q06890 | CLU | clusterin | Cytoplasm | other | 1745149.75 | 951794.75 | 539084.625 | Q06890 | CLU | 4065141 | 4325506 | 1441835 | 9 |
| Q8R4N0 | CLYBL | citrate lyase beta like | Cytoplasm | enzyme | 1213305.875 | 3015655 | 974535.125 | Q8R4N0 | CLYBL | 1213306 | 1115918 | 644276 | 3 |
| O35372 | CNIH1 | cornichon family AMPA receptor auxiliary protein 1 | Plasma Membrane | other | 3660031.5 | 3584001.75 | 3056519.25 | O35372 | CNIH1 | 3660032 | 328696 | 189773 | 3 |
| Q08093 | CNN2 | calponin 2 | Cytoplasm | other | 703338.938 | 1386275 | 2821588.5 | Q08093 | CNN2 | 703339 | 1081165 | 624211 | 3 |
| Q3TWN3 | CNNM2 | cyclin and CBS domain divalent metal cation transport mediator 2 | Plasma Membrane | other | 5574447 | 4296551 | 4575613.5 | | | | | | |
| Q3TWN3 | CNNM2 | cyclin and CBS domain divalent metal cation transport mediator 2 | Plasma Membrane | other | 4292569 | 4185510.75 | 4341595 | Q3TWN3 | CNNM2 | 4933508 | 520913 | 212662 | 6 |
| P16330 | CNP | 2',3'-cyclic nucleotide 3' phosphodiesterase | Cytoplasm | enzyme | 2.08E+07 | 2.55E+07 | 2.64E+07 | | | | | | |
| P16330 | CNP | 2',3'-cyclic nucleotide 3' phosphodiesterase | Cytoplasm | enzyme | 1.20E+07 | 1.29E+07 | 1.23E+07 | | | | | | |

| | | | | | | | | | | | | | |
|---|---|---|---|---|---|---|---|---|---|---|---|---|---|
| P16330 | CNP | 2',3'-cyclic nucleotide 3' phosphodiesterase | Cytoplasm | enzyme | 7212191 | 7766475.5 | 8579800 | | | | | | |
| P16330 | CNP | 2',3'-cyclic nucleotide 3' phosphodiesterase | Cytoplasm | enzyme | 5871935.5 | 6469156 | 6244915.5 | | | | | | |
| P16330 | CNP | 2',3'-cyclic nucleotide 3' phosphodiesterase | Cytoplasm | enzyme | 3517370.5 | 4747330.5 | 5651757.5 | | | | | | |
| P16330 | CNP | 2',3'-cyclic nucleotide 3' phosphodiesterase | Cytoplasm | enzyme | 3001535 | 3016445.25 | 3461037.25 | | | | | | |
| P16330 | CNP | 2',3'-cyclic nucleotide 3' phosphodiesterase | Cytoplasm | enzyme | 4960180.5 | 3350064.5 | 1953202.75 | | | | | | |
| P16330 | CNP | 2',3'-cyclic nucleotide 3' phosphodiesterase | Cytoplasm | enzyme | 2053140.25 | 3644021 | 4948117.5 | P16330 | CNP | 7428294 | 6956699 | 1420030 | 24 |
| Q9QXT0 | CNPY2 | canopy FGF signaling regulator 2 | Plasma Membrane | other | 5631581 | 6032625 | 5854855 | | | | | | |
| Q9QXT0 | CNPY2 | canopy FGF signaling regulator 2 | Plasma Membrane | other | 5286822.5 | 6721654 | 5310201.5 | | | | | | |
| Q9QXT0 | CNPY2 | canopy FGF signaling regulator 2 | Plasma Membrane | other | 2156474.25 | 2734418.5 | 2569827.5 | Q9QXT0 | CNPY2 | 4358293 | 1719443 | 573148 | 9 |
| Q8BQ47 | CNPY4 | canopy FGF signaling regulator 4 | Extracellular Space | other | 1417656.5 | 2056868.875 | 1830837.875 | Q8BQ47 | CNPY4 | 1417657 | 324140 | 187142 | 3 |
| O88587 | COMT | catechol-O-methyltransferase | Cytoplasm | enzyme | 3685654.25 | 5899822.5 | 4460428 | | | | | | |
| O88587 | COMT | catechol-O-methyltransferase | Cytoplasm | enzyme | 5097680.5 | 6771195.5 | 4706550 | | | | | | |
| O88587 | COMT | catechol-O-methyltransferase | Cytoplasm | enzyme | 2608461 | 3925399.25 | 4271058 | | | | | | |
| O88587 | COMT | catechol-O-methyltransferase | Cytoplasm | enzyme | 2874192.75 | 4440036 | 3875191.5 | O88587 | COMT | 3566497 | 1168545 | 337330 | 12 |
| Q8BIG7 | COMTD1 | catechol-O-methyltransferase domain containing 1 | Plasma Membrane | enzyme | 1337129.875 | 2851568.75 | 2325347.5 | Q8BIG7 | COMTD1 | 1337130 | 768875 | 443910 | 3 |
| P61202 | Cops2 | COP9 (constitutive photomorphogenic) homolog, subunit 2 (Arabidopsis thaliana) | Nucleus | transcription regulator | 7857390 | 7788357.5 | 7240382.5 | | | | | | |
| P61202 | Cops2 | COP9 (constitutive photomorphogenic) homolog, subunit 2 (Arabidopsis thaliana) | Nucleus | transcription regulator | 5266013.5 | 3560534 | 3839626.75 | | | | | | |
| P61202 | Cops2 | COP9 (constitutive photomorphogenic) homolog, subunit 2 (Arabidopsis thaliana) | Nucleus | transcription regulator | 1924292.75 | 1741747.25 | 2564976 | | | | | | |
| P61202 | Cops2 | COP9 (constitutive photomorphogenic) homolog, subunit 2 (Arabidopsis thaliana) | Nucleus | transcription regulator | 1807676 | 1521675.5 | 1751840.5 | | | | | | |
| P61202 | Cops2 | COP9 (constitutive photomorphogenic) homolog, subunit 2 (Arabidopsis thaliana) | Nucleus | transcription regulator | 766529.188 | 183121.797 | 51735.887 | P61202 | Cops2 | 3524380 | 2670205 | 689444 | 15 |
| O88543 | COPS3 | COP9 signalosome subunit 3 | Cytoplasm | other | 5113424.5 | 3170337.75 | 3739362.25 | | | | | | |
| O88543 | COPS3 | COP9 signalosome subunit 3 | Cytoplasm | other | 3714206.25 | 3186295.5 | 3043178.25 | | | | | | |
| O88543 | COPS3 | COP9 signalosome subunit 3 | Cytoplasm | other | 4326112 | 3151196.25 | 3100037 | | | | | | |
| O88543 | COPS3 | COP9 signalosome subunit 3 | Cytoplasm | other | 2507027.25 | 1677471 | 1555951 | | | | | | |
| O88543 | COPS3 | COP9 signalosome subunit 3 | Cytoplasm | other | 1473760.25 | 961774.812 | 1222473 | | | | | | |
| O88543 | COPS3 | COP9 signalosome subunit 3 | Cytoplasm | other | 2582372.25 | 2618828.75 | 3398691.5 | O88543 | COPS3 | 3286150 | 1109203 | 261442 | 18 |
| O88544 | COPS4 | COP9 signalosome subunit 4 | Cytoplasm | other | 3955467.25 | 3386882 | 4650253.5 | | | | | | |
| O88544 | COPS4 | COP9 signalosome subunit 4 | Cytoplasm | other | 3508866.5 | 2937547.25 | 3083605.75 | | | | | | |
| O88544 | COPS4 | COP9 signalosome subunit 4 | Cytoplasm | other | 2200942.5 | 1554038.5 | 1385340 | | | | | | |
| O88544 | COPS4 | COP9 signalosome subunit 4 | Cytoplasm | other | 2182816 | 1194239.625 | 386416.812 | | | | | | |
| O88544 | COPS4 | COP9 signalosome subunit 4 | Cytoplasm | other | 2861091.25 | 1564966.375 | 1987822.875 | | | | | | |
| O88544 | COPS4 | COP9 signalosome subunit 4 | Cytoplasm | other | 2398759.25 | 1688853.625 | 1592651.75 | O88544 | COPS4 | 2851324 | 1084515 | 255623 | 18 |
| O35864 | COPS5 | COP9 signalosome subunit 5 | Nucleus | transcription regulator | 1.15E+07 | 7578112.5 | 6981008 | | | | | | |
| O35864 | COPS5 | COP9 signalosome subunit 5 | Nucleus | transcription regulator | 2143674 | 1620442.625 | 1500344.5 | O35864 | COPS5 | 6841837 | 4119717 | 1681868 | 6 |
| O88545 | COPS6 | COP9 signalosome subunit 6 | Nucleus | other | 4756568 | 5259765.5 | 1.16E+07 | | | | | | |
| O88545 | COPS6 | COP9 signalosome subunit 6 | Nucleus | other | 6990866 | 5494535 | 5863685.5 | | | | | | |
| O88545 | COPS6 | COP9 signalosome subunit 6 | Nucleus | other | 4865748 | 5050144 | 4817873.5 | | | | | | |
| O88545 | COPS6 | COP9 signalosome subunit 6 | Nucleus | other | 1766478.875 | 1072926 | 607162.062 | | | | | | |
| O88545 | COPS6 | COP9 signalosome subunit 6 | Nucleus | other | 1744293.5 | 1162630.125 | 1390193.5 | O88545 | COPS6 | 4024791 | 2944918 | 760375 | 15 |
| Q9CZ04 | COPS7A | COP9 signalosome subunit 7A | Cytoplasm | other | 1141074.375 | 634792 | 915512.688 | Q9CZ04 | COPS7A | 1141074 | 253641 | 146440 | 3 |
| Q8BV13 | COPS7B | COP9 signalosome subunit 7B | Cytoplasm | other | 2540210.75 | 1927714.125 | 2062176 | Q8BV13 | COPS7B | 2540211 | 321908 | 185854 | 3 |
| Q8VBV7 | COPS8 | COP9 signalosome subunit 8 | Nucleus | other | 2693441.25 | 2365288.5 | 1842014.75 | Q8VBV7 | COPS8 | 2693441 | 429423 | 247928 | 3 |
| O89053 | CORO1A | coronin, actin binding protein, 1A | Cytoplasm | other | 3114087.5 | 1.26E+07 | 2.15E+07 | | | | | | |
| O89053 | CORO1A | coronin, actin binding protein, 1A | Cytoplasm | other | 2184436 | 1661949.25 | 3950926.25 | | | | | | |
| O89053 | CORO1A | coronin, actin binding protein, 1A | Cytoplasm | other | 675665.875 | 5912559 | 1.44E+07 | | | | | | |
| O89053 | CORO1A | coronin, actin binding protein, 1A | Cytoplasm | other | 747971.375 | 2647866.5 | 3401249 | | | | | | |
| O89053 | CORO1A | coronin, actin binding protein, 1A | Cytoplasm | other | 1830172.125 | 794582.812 | 2973231 | | | | | | |
| O89053 | CORO1A | coronin, actin binding protein, 1A | Cytoplasm | other | 395613.156 | 2143581.25 | 6753968 | | | | | | |
| O89053 | CORO1A | coronin, actin binding protein, 1A | Cytoplasm | other | 242429.844 | 550596.938 | 305387.938 | | | | | | |
| O89053 | CORO1A | coronin, actin binding protein, 1A | Cytoplasm | other | 254731.969 | 668951 | 2395921.75 | | | | | | |
| O89053 | CORO1A | coronin, actin binding protein, 1A | Cytoplasm | other | 165399.922 | 1416924.375 | 4786014 | | | | | | |
| O89053 | CORO1A | coronin, actin binding protein, 1A | Cytoplasm | other | 275967.344 | 977959.438 | 2947258 | | | | | | |
| O89053 | CORO1A | coronin, actin binding protein, 1A | Cytoplasm | other | 478413.312 | 2176910.5 | 7547872 | | | | | | |
| O89053 | CORO1A | coronin, actin binding protein, 1A | Cytoplasm | other | 448103.469 | 1550036 | 1.03E+07 | | | | | | |
| O89053 | CORO1A | coronin, actin binding protein, 1A | Cytoplasm | other | 225867.703 | 912263.562 | 2979860.5 | O89053 | CORO1A | 849143 | 4507142 | 721720 | 39 |
| Q9WUM4 | CORO1C | coronin, actin binding protein, 1C | Cytoplasm | other | 1733710.625 | 2577358.25 | 3550417.25 | | | | | | |
| Q9WUM4 | CORO1C | coronin, actin binding protein, 1C | Cytoplasm | other | 196846.953 | 1027505.625 | 1858894.5 | Q9WUM4 | CORO1C | 965279 | 1169570 | 477475 | 6 |
| P19783 | COX4I1 | cytochrome c oxidase subunit IV isoform 1 | Cytoplasm | enzyme | 1.61E+07 | 2.91E+07 | 1.48E+07 | | | | | | |
| P19783 | COX4I1 | cytochrome c oxidase subunit IV isoform 1 | Cytoplasm | enzyme | 8018210 | 1.64E+07 | 7597890.5 | | | | | | |
| P19783 | COX4I1 | cytochrome c oxidase subunit IV isoform 1 | Cytoplasm | enzyme | 3719103.75 | 7655658 | 3925088.5 | P19783 | COX4I1 | 9282438 | 8087497 | 2695832 | 9 |
| P12787 | COX5A | cytochrome c oxidase subunit Va | Cytoplasm | enzyme | 7772383.5 | 1.90E+07 | 8974940 | | | | | | |
| P12787 | COX5A | cytochrome c oxidase subunit Va | Cytoplasm | enzyme | 3346436.75 | 2.10E+07 | 1.76E+07 | | | | | | |
| P12787 | COX5A | cytochrome c oxidase subunit Va | Cytoplasm | enzyme | 4332187.5 | 8438824 | 3854574.75 | P12787 | COX5A | 5150336 | 6892182 | 2297394 | 9 |
| P19536 | Cox5b | cytochrome c oxidase subunit Vb | Cytoplasm | other | 8551843 | 1.72E+07 | 8681126 | P19536 | Cox5b | 8551843 | 4967663 | 2868081 | 3 |
| P56391 | COX6B1 | cytochrome c oxidase subunit VIb polypeptide 1 (ubiquitous) | Cytoplasm | enzyme | 5162854.5 | 6864714 | 3335773.25 | P56391 | COX6B1 | 5162855 | 1764841 | 1018931 | 3 |
| Q9CPQ1 | Cox6c | cytochrome c oxidase subunit VIc | Cytoplasm | enzyme | 1213865.75 | 2932086.25 | 1706561.25 | Q9CPQ1 | Cox6c | 1213866 | 884773 | 510824 | 3 |
| P48771 | COX7A2 | cytochrome c oxidase subunit VIIa polypeptide 2 (liver) | Cytoplasm | enzyme | 4040584.75 | 9779953 | 5295082.5 | P48771 | COX7A2 | 4040585 | 3017399 | 1742096 | 3 |
| P59108 | CPNE2 | copine II | Cytoplasm | other | 1307400.125 | 684713.062 | 2223086.25 | | | | | | |
| P59108 | CPNE2 | copine II | Cytoplasm | other | 759511.75 | 1143990.5 | 3326013.75 | P59108 | CPNE2 | 1033456 | 1019909 | 416376 | 6 |
| P36552 | CPOX | coproporphyrinogen oxidase | Cytoplasm | enzyme | 1.19E+07 | 1.92E+07 | 5408447 | | | | | | |
| P36552 | CPOX | coproporphyrinogen oxidase | Cytoplasm | enzyme | 1.14E+07 | 1.66E+07 | 7016991 | | | | | | |
| P36552 | CPOX | coproporphyrinogen oxidase | Cytoplasm | enzyme | 1.24E+07 | 1.66E+07 | 6318080.5 | | | | | | |
| P36552 | CPOX | coproporphyrinogen oxidase | Cytoplasm | enzyme | 8957981 | 1.66E+07 | 7542631 | | | | | | |
| P36552 | CPOX | coproporphyrinogen oxidase | Cytoplasm | enzyme | 7062162.5 | 1.34E+07 | 7209573.5 | | | | | | |
| P36552 | CPOX | coproporphyrinogen oxidase | Cytoplasm | enzyme | 6896402.5 | 1.04E+07 | 4139626.25 | | | | | | |
| P36552 | CPOX | coproporphyrinogen oxidase | Cytoplasm | enzyme | 6476311 | 8867969 | 4240976.5 | | | | | | |
| P36552 | CPOX | coproporphyrinogen oxidase | Cytoplasm | enzyme | 6364414.5 | 1.24E+07 | 4882274 | | | | | | |
| P36552 | CPOX | coproporphyrinogen oxidase | Cytoplasm | enzyme | 5793687.5 | 9451374 | 3784172.75 | | | | | | |
| P36552 | CPOX | coproporphyrinogen oxidase | Cytoplasm | enzyme | 5086662.5 | 8023670 | 2743143.25 | | | | | | |
| P36552 | CPOX | coproporphyrinogen oxidase | Cytoplasm | enzyme | 3371806.25 | 3280550.75 | 1232754.25 | | | | | | |
| P36552 | CPOX | coproporphyrinogen oxidase | Cytoplasm | enzyme | 1900009.375 | 2676372.25 | 1111440.125 | P36552 | CPOX | 7306620 | 4685135 | 780856 | 36 |
| P52825 | CPT2 | carnitine palmitoyltransferase 2 | Cytoplasm | enzyme | 1538311.5 | 3228851.75 | 1758479.375 | | | | | | |
| P52825 | CPT2 | carnitine palmitoyltransferase 2 | Cytoplasm | enzyme | 712469.438 | 2526287.5 | 1108908.375 | | | | | | |
| P52825 | CPT2 | carnitine palmitoyltransferase 2 | Cytoplasm | enzyme | 417467.938 | 514621.375 | 161233.875 | P52825 | CPT2 | 889416 | 1034259 | 344753 | 9 |
| Q64735 | CR1L | complement component (3b/4b) receptor 1-like | Plasma Membrane | other | 6.66E+07 | 8.72E+07 | 6.25E+07 | | | | | | |
| Q64735 | CR1L | complement component (3b/4b) receptor 1-like | Plasma Membrane | other | 3.59E+07 | 2.95E+07 | 3.11E+07 | | | | | | |
| Q64735 | CR1L | complement component (3b/4b) receptor 1-like | Plasma Membrane | other | 2.94E+07 | 2.37E+07 | 2.96E+07 | | | | | | |
| Q64735 | CR1L | complement component (3b/4b) receptor 1-like | Plasma Membrane | other | 3.09E+07 | 2.60E+07 | 2.83E+07 | | | | | | |
| Q64735 | CR1L | complement component (3b/4b) receptor 1-like | Plasma Membrane | other | 9455333 | 8387067.5 | 9758150 | | | | | | |
| Q64735 | CR1L | complement component (3b/4b) receptor 1-like | Plasma Membrane | other | 7449228 | 7689807 | 9831585 | | | | | | |
| Q64735 | CR1L | complement component (3b/4b) receptor 1-like | Plasma Membrane | other | 9043078 | 8826847 | 7518723.5 | | | | | | |

| | | | | | | | | | | | | | |
|---|---|---|---|---|---|---|---|---|---|---|---|---|---|
| Q64735 | CR1L | complement component (3b/4b) receptor 1-like | Plasma Membrane | other | 6440364 | 4823840.5 | 3682398.25 | | | | | | |
| Q64735 | CR1L | complement component (3b/4b) receptor 1-like | Plasma Membrane | other | 1202025.125 | 1803936 | 1961397.875 | Q64735 | CR1L | 21822225 | 21645376 | 4165655 | 27 |
| Q9Z125 | CREB3L1 | cAMP responsive element binding protein 3-like 1 | Nucleus | transcription regulator | 1950393.75 | 5129770 | 2293161 | Q9Z125 | CREB3L1 | 1950394 | 1745101 | 1007535 | 3 |
| O88668 | CREG1 | cellular repressor of E1A-stimulated genes 1 | Nucleus | transcription regulator | 2637737.5 | 3700150 | 3573407 | | | | | | |
| O88668 | CREG1 | cellular repressor of E1A-stimulated genes 1 | Nucleus | transcription regulator | 1648635.5 | 3144224.75 | 2900660.5 | O88668 | CREG1 | 2143187 | 745797 | 304470 | 6 |
| Q9DCT8 | Crip2 | cysteine rich protein 2 | Plasma Membrane | other | 2.85E+07 | 3.07E+07 | 3.68E+07 | | | | | | |
| Q9DCT8 | Crip2 | cysteine rich protein 2 | Plasma Membrane | other | 2.77E+07 | 3.20E+07 | 4.58E+07 | | | | | | |
| Q9DCT8 | Crip2 | cysteine rich protein 2 | Plasma Membrane | other | 2.69E+07 | 1.96E+07 | 2.09E+07 | | | | | | |
| Q9DCT8 | Crip2 | cysteine rich protein 2 | Plasma Membrane | other | 2.18E+07 | 3.31E+07 | 4.12E+07 | | | | | | |
| Q9DCT8 | Crip2 | cysteine rich protein 2 | Plasma Membrane | other | 1.59E+07 | 1.28E+07 | 1.51E+07 | | | | | | |
| Q9DCT8 | Crip2 | cysteine rich protein 2 | Plasma Membrane | other | 1.65E+07 | 1.92E+07 | 1.92E+07 | | | | | | |
| Q9DCT8 | Crip2 | cysteine rich protein 2 | Plasma Membrane | other | 1.06E+07 | 8048337.5 | 7990032 | | | | | | |
| Q9DCT8 | Crip2 | cysteine rich protein 2 | Plasma Membrane | other | 1.30E+07 | 9560654 | 7483742 | | | | | | |
| Q9DCT8 | Crip2 | cysteine rich protein 2 | Plasma Membrane | other | 8804711 | 1049863.5 | 82387.344 | Q9DCT8 | Crip2 | 18861635 | 11898552 | 2289877 | 27 |
| Q9Z2L7 | CRLF3 | cytokine receptor-like factor 3 | Cytoplasm | transcription regulator | 6280293.5 | 3977667.25 | 3516244.5 | Q9Z2L7 | CRLF3 | 6280294 | 1480707 | 854887 | 3 |
| Q9CZU6 | CS | citrate synthase | Cytoplasm | enzyme | 2.37E+07 | 5.65E+07 | 2.28E+07 | | | | | | |
| Q9CZU6 | CS | citrate synthase | Cytoplasm | enzyme | 1.05E+07 | 1.95E+07 | 4776886 | | | | | | |
| Q9CZU6 | CS | citrate synthase | Cytoplasm | enzyme | 1.42E+07 | 3.85E+07 | 1.83E+07 | | | | | | |
| Q9CZU6 | CS | citrate synthase | Cytoplasm | enzyme | 7296891.5 | 1.71E+07 | 8165211 | | | | | | |
| Q9CZU6 | CS | citrate synthase | Cytoplasm | enzyme | 6518245.5 | 1.56E+07 | 7706273 | | | | | | |
| Q9CZU6 | CS | citrate synthase | Cytoplasm | enzyme | 6960447 | 1.57E+07 | 4780394.5 | | | | | | |
| Q9CZU6 | CS | citrate synthase | Cytoplasm | enzyme | 9878635 | 8934072 | 3349037.75 | | | | | | |
| Q9CZU6 | CS | citrate synthase | Cytoplasm | enzyme | 4650799.5 | 6073617.5 | 2534069.5 | | | | | | |
| Q9CZU6 | CS | citrate synthase | Cytoplasm | enzyme | 4109741.75 | 7626832.5 | 3040646 | | | | | | |
| Q9CZU6 | CS | citrate synthase | Cytoplasm | enzyme | 4160051 | 8211081.5 | 3922464.75 | Q9CZU6 | CS | 9200481 | 11569605 | 2112311 | 30 |
| Q8BK63 | CSNK1A1 | casein kinase 1, alpha 1 | Cytoplasm | kinase | 1.08E+07 | 1.09E+07 | 7699027 | | | | | | |
| Q8BK63 | CSNK1A1 | casein kinase 1, alpha 1 | Cytoplasm | kinase | 1.18E+07 | 4017219.25 | 110182.344 | | | | | | |
| Q8BK63 | CSNK1A1 | casein kinase 1, alpha 1 | Cytoplasm | kinase | 1.01E+07 | 1.09E+07 | 1.13E+07 | | | | | | |
| Q8BK63 | CSNK1A1 | casein kinase 1, alpha 1 | Cytoplasm | kinase | 6746612.5 | 6241672 | 8813379 | | | | | | |
| Q8BK63 | CSNK1A1 | casein kinase 1, alpha 1 | Cytoplasm | kinase | 6976354 | 4759013 | 5956480.5 | | | | | | |
| Q8BK63 | CSNK1A1 | casein kinase 1, alpha 1 | Cytoplasm | kinase | 4324921 | 3334038.25 | 3528366.25 | | | | | | |
| Q8BK63 | CSNK1A1 | casein kinase 1, alpha 1 | Cytoplasm | kinase | 4523134.5 | 5286169.5 | 5149641 | | | | | | |
| Q8BK63 | CSNK1A1 | casein kinase 1, alpha 1 | Cytoplasm | kinase | 2600986 | 2599122.75 | 1802084.25 | Q8BK63 | CSNK1A1 | 7235251 | 3366692 | 687223 | 24 |
| Q60737 | CSNK2A1 | casein kinase 2, alpha 1 polypeptide | Nucleus | kinase | 2985797.75 | 2803938.25 | 3131504.25 | Q60737 | CSNK2A1 | 2985798 | 164115 | 94752 | 3 |
| P67871 | CSNK2B | casein kinase 2, beta polypeptide | Cytoplasm | kinase | 3562640 | 3548343.75 | 3308734.5 | P67871 | CSNK2B | 3562640 | 142645 | 82356 | 3 |
| P97315 | CSRP1 | cysteine and glycine-rich protein 1 | Nucleus | other | 9390273 | 1.55E+07 | 1.66E+07 | | | | | | |
| P97315 | CSRP1 | cysteine and glycine-rich protein 1 | Nucleus | other | 7447050 | 1.34E+07 | 1.78E+07 | | | | | | |
| P97315 | CSRP1 | cysteine and glycine-rich protein 1 | Nucleus | other | 6869418.5 | 1.26E+07 | 1.39E+07 | | | | | | |
| P97315 | CSRP1 | cysteine and glycine-rich protein 1 | Nucleus | other | 6441525.5 | 9065718 | 6521639 | | | | | | |
| P97315 | CSRP1 | cysteine and glycine-rich protein 1 | Nucleus | other | 668717.75 | 1886954.5 | 463411.281 | P97315 | CSRP1 | 6163397 | 5639183 | 1456031 | 15 |
| P10605 | CTSB | cathepsin B | Cytoplasm | peptidase | 7292677.5 | 8666347 | 1.11E+07 | | | | | | |
| P10605 | CTSB | cathepsin B | Cytoplasm | peptidase | 7962743 | 1.20E+07 | 8982525 | | | | | | |
| P10605 | CTSB | cathepsin B | Cytoplasm | peptidase | 6132447 | 1.14E+07 | 7391156 | | | | | | |
| P10605 | CTSB | cathepsin B | Cytoplasm | peptidase | 2145772.5 | 6536041.5 | 7785828 | | | | | | |
| P10605 | CTSB | cathepsin B | Cytoplasm | peptidase | 4165082.5 | 4251488.5 | 2488923.25 | P10605 | CTSB | 5539745 | 3042085 | 785463 | 15 |
| P18242 | CTSD | cathepsin D | Cytoplasm | peptidase | 1.69E+07 | 3.23E+07 | 1.36E+07 | | | | | | |
| P18242 | CTSD | cathepsin D | Cytoplasm | peptidase | 8935913 | 2.47E+07 | 5631011.5 | | | | | | |
| P18242 | CTSD | cathepsin D | Cytoplasm | peptidase | 9069520 | 2.07E+07 | 9845959 | | | | | | |
| P18242 | CTSD | cathepsin D | Cytoplasm | peptidase | 9117246 | 1.94E+07 | 7373037.5 | | | | | | |
| P18242 | CTSD | cathepsin D | Cytoplasm | peptidase | 8857442 | 1.69E+07 | 7942080 | | | | | | |
| P18242 | CTSD | cathepsin D | Cytoplasm | peptidase | 4088213.75 | 6542134 | 2020184.75 | | | | | | |
| P18242 | CTSD | cathepsin D | Cytoplasm | peptidase | 2409514 | 4870519.5 | 3466569 | P18242 | CTSD | 8481121 | 7953011 | 1735489 | 21 |
| P70269 | CTSE | cathepsin E | Cytoplasm | peptidase | 9.88E+07 | 1.33E+08 | 1.33E+08 | | | | | | |
| P70269 | CTSE | cathepsin E | Cytoplasm | peptidase | 8.71E+07 | 1.14E+08 | 1.20E+08 | | | | | | |
| P70269 | CTSE | cathepsin E | Cytoplasm | peptidase | 1.88E+07 | 6406016.5 | 4013875.5 | P70269 | CTSE | 68243333 | 54446516 | 18148839 | 9 |
| Q9WTX6 | CUL1 | cullin 1 | Nucleus | enzyme | 4487478.5 | 3346920.5 | 2371275.75 | | | | | | |
| Q9WTX6 | CUL1 | cullin 1 | Nucleus | enzyme | 4266346.5 | 3210104.5 | 4089622.75 | | | | | | |
| Q9WTX6 | CUL1 | cullin 1 | Nucleus | enzyme | 3004757.75 | 2739732.75 | 2723809.75 | | | | | | |
| Q9WTX6 | CUL1 | cullin 1 | Nucleus | enzyme | 3150011 | 3000671.25 | 2377295.25 | | | | | | |
| Q9WTX6 | CUL1 | cullin 1 | Nucleus | enzyme | 2380403 | 2661046.75 | 2495835.75 | | | | | | |
| Q9WTX6 | CUL1 | cullin 1 | Nucleus | enzyme | 2404599.5 | 1645040 | 1427384.625 | | | | | | |
| Q9WTX6 | CUL1 | cullin 1 | Nucleus | enzyme | 1130429.375 | 597925.875 | 179014.922 | | | | | | |
| Q9WTX6 | CUL1 | cullin 1 | Nucleus | enzyme | 1440832.75 | 562177.312 | 55956.891 | | | | | | |
| Q9WTX6 | CUL1 | cullin 1 | Nucleus | enzyme | 1096220.625 | 1064793.5 | 1201361.625 | Q9WTX6 | CUL1 | 2595675 | 1219360 | 234666 | 27 |
| Q9JLV5 | CUL3 | cullin 3 | Nucleus | enzyme | 4649665.5 | 3934589 | 3050276.75 | | | | | | |
| Q9JLV5 | CUL3 | cullin 3 | Nucleus | enzyme | 3696565.5 | 2857642.25 | 2914477.75 | Q9JLV5 | CUL3 | 4173116 | 707838 | 288974 | 6 |
| P70658 | CXCR4 | chemokine (C-X-C motif) receptor 4 | Plasma Membrane | G-protein coupled receptor | 4830342 | 5117161 | 3708917.25 | P70658 | CXCR4 | 4830342 | 744201 | 429665 | 3 |
| P56395 | CYB5A | cytochrome b5 type A (microsomal) | Cytoplasm | enzyme | 5863612.5 | 9561538 | 1.27E+07 | | | | | | |
| P56395 | CYB5A | cytochrome b5 type A (microsomal) | Cytoplasm | enzyme | 5351708.5 | 7870873 | 1.01E+07 | | | | | | |
| P56395 | CYB5A | cytochrome b5 type A (microsomal) | Cytoplasm | enzyme | 1320372.75 | 2465692.75 | 3087230 | | | | | | |
| P56395 | CYB5A | cytochrome b5 type A (microsomal) | Cytoplasm | enzyme | 949414.438 | 2233363.25 | 3468468.75 | | | | | | |
| P56395 | CYB5A | cytochrome b5 type A (microsomal) | Cytoplasm | enzyme | 3231875.25 | 2674213.75 | 2088974.875 | P56395 | CYB5A | 3343397 | 3601755 | 929969 | 15 |
| Q9CQX2 | CYB5B | cytochrome b5 type B (outer mitochondrial membrane) | Cytoplasm | enzyme | 9206906 | 1.31E+07 | 9849760 | | | | | | |
| Q9CQX2 | CYB5B | cytochrome b5 type B (outer mitochondrial membrane) | Cytoplasm | enzyme | 1486270.375 | 1764674 | 50778.609 | | | | | | |
| Q9CQX2 | CYB5B | cytochrome b5 type B (outer mitochondrial membrane) | Cytoplasm | enzyme | 804805.375 | 987536.75 | 15570.956 | Q9CQX2 | CYB5B | 3832661 | 5070467 | 1690156 | 9 |
| Q9DCN2 | Cyb5r3 | cytochrome b5 reductase 3 | Cytoplasm | enzyme | 2.80E+07 | 3.03E+07 | 3.11E+07 | | | | | | |
| Q9DCN2 | Cyb5r3 | cytochrome b5 reductase 3 | Cytoplasm | enzyme | 1.42E+07 | 2.04E+07 | 2.24E+07 | | | | | | |
| Q9DCN2 | Cyb5r3 | cytochrome b5 reductase 3 | Cytoplasm | enzyme | 1.84E+07 | 2.15E+07 | 2.49E+07 | | | | | | |
| Q9DCN2 | Cyb5r3 | cytochrome b5 reductase 3 | Cytoplasm | enzyme | 7927375 | 1.31E+07 | 1.44E+07 | | | | | | |
| Q9DCN2 | Cyb5r3 | cytochrome b5 reductase 3 | Cytoplasm | enzyme | 9672881 | 1.14E+07 | 1.45E+07 | | | | | | |
| Q9DCN2 | Cyb5r3 | cytochrome b5 reductase 3 | Cytoplasm | enzyme | 7695307.5 | 1.19E+07 | 9465807 | | | | | | |
| Q9DCN2 | Cyb5r3 | cytochrome b5 reductase 3 | Cytoplasm | enzyme | 5441632.5 | 7170670 | 6026748 | | | | | | |
| Q9DCN2 | Cyb5r3 | cytochrome b5 reductase 3 | Cytoplasm | enzyme | 6760269.5 | 7237675 | 8750606 | | | | | | |
| Q9DCN2 | Cyb5r3 | cytochrome b5 reductase 3 | Cytoplasm | enzyme | 5886684 | 8005300 | 8141661 | | | | | | |
| Q9DCN2 | Cyb5r3 | cytochrome b5 reductase 3 | Cytoplasm | enzyme | 4841838 | 5055011 | 6068285.5 | | | | | | |
| Q9DCN2 | Cyb5r3 | cytochrome b5 reductase 3 | Cytoplasm | enzyme | 1704515.875 | 3610796.75 | 5265995.5 | | | | | | |
| Q9DCN2 | Cyb5r3 | cytochrome b5 reductase 3 | Cytoplasm | enzyme | 1601224.875 | 4601765.5 | 5366764.5 | Q9DCN2 | Cyb5r3 | 9346811 | 8071255 | 1345209 | 36 |
| Q9D0M3 | CYC1 | cytochrome c-1 | Cytoplasm | enzyme | 1.18E+07 | 2.39E+07 | 9988560 | Q9D0M3 | CYC1 | 11750000 | 7574665 | 4373235 | 3 |
| P62897 | CYCS | cytochrome c, somatic | Cytoplasm | transporter | 7063810 | 9727678 | 7792683 | | | | | | |
| P62897 | CYCS | cytochrome c, somatic | Cytoplasm | transporter | 3528344.5 | 6349881.5 | 3438837 | | | | | | |
| P62897 | CYCS | cytochrome c, somatic | Cytoplasm | transporter | 4423579 | 3232262.25 | 1822627.75 | | | | | | |

| | | | | | | | | | | | | | |
|---|---|---|---|---|---|---|---|---|---|---|---|---|---|
| P62897 | CYCS | cytochrome c, somatic | Cytoplasm | transporter | 1636319.875 | 4409341.5 | 2638009.5 | | | | | | |
| P62897 | CYCS | cytochrome c, somatic | Cytoplasm | transporter | 1400539.75 | 1198114 | 1818593.625 | P62897 | CYCS | 3610519 | 2606410 | 672972 | 15 |
| Q8BPM0 | DAAM1 | dishevelled associated activator of morphogenesis 1 | Cytoplasm | other | 6861394.5 | 4863824 | 6667848.5 | | | | | | |
| Q8BPM0 | DAAM1 | dishevelled associated activator of morphogenesis 1 | Cytoplasm | other | 4977572.5 | 3833906.5 | 5721982 | | | | | | |
| Q8BPM0 | DAAM1 | dishevelled associated activator of morphogenesis 1 | Cytoplasm | other | 4054662 | 4133270.5 | 5429340 | | | | | | |
| Q8BPM0 | DAAM1 | dishevelled associated activator of morphogenesis 1 | Cytoplasm | other | 5517826 | 5454590 | 5562937 | | | | | | |
| Q8BPM0 | DAAM1 | dishevelled associated activator of morphogenesis 1 | Cytoplasm | other | 2488095.25 | 3805197.5 | 7010563.5 | | | | | | |
| Q8BPM0 | DAAM1 | dishevelled associated activator of morphogenesis 1 | Cytoplasm | other | 2008671.375 | 1615286.625 | 1874905.75 | | | | | | |
| Q8BPM0 | DAAM1 | dishevelled associated activator of morphogenesis 1 | Cytoplasm | other | 3076155 | 4669228 | 6605874 | | | | | | |
| Q8BPM0 | DAAM1 | dishevelled associated activator of morphogenesis 1 | Cytoplasm | other | 2289547.25 | 2569048.5 | 4185954 | | | | | | |
| Q8BPM0 | DAAM1 | dishevelled associated activator of morphogenesis 1 | Cytoplasm | other | 2970762.5 | 2713178.25 | 3947044.25 | | | | | | |
| Q8BPM0 | DAAM1 | dishevelled associated activator of morphogenesis 1 | Cytoplasm | other | 2334020.5 | 2644384.75 | 2162516 | | | | | | |
| Q8BPM0 | DAAM1 | dishevelled associated activator of morphogenesis 1 | Cytoplasm | other | 1082146.25 | 1453948.625 | 1616253.5 | | | | | | |
| Q8BPM0 | DAAM1 | dishevelled associated activator of morphogenesis 1 | Cytoplasm | other | 2237612.5 | 278778.344 | 2783.96 | Q8BPM0 | DAAM1 | 3324872 | 1892949 | 315491 | 36 |
| P61804 | DAD1 | defender against cell death 1 | Cytoplasm | other | 231880.531 | 1467544.75 | 1377093.375 | P61804 | DAD1 | 231881 | 688786 | 397671 | 3 |
| Q91WC9 | DAGLB | diacylglycerol lipase, beta | Plasma Membrane | enzyme | 7293348.5 | 7487745 | 9683212 | | | | | | |
| Q91WC9 | DAGLB | diacylglycerol lipase, beta | Plasma Membrane | enzyme | 4355221.5 | 5796388.5 | 7518860 | | | | | | |
| Q91WC9 | DAGLB | diacylglycerol lipase, beta | Plasma Membrane | enzyme | 3110480.25 | 3170799.25 | 3543214.5 | | | | | | |
| Q91WC9 | DAGLB | diacylglycerol lipase, beta | Plasma Membrane | enzyme | 2963887.25 | 5341232 | 7970362.5 | | | | | | |
| Q91WC9 | DAGLB | diacylglycerol lipase, beta | Plasma Membrane | enzyme | 3235888.25 | 3381130.5 | 2532066.5 | | | | | | |
| Q91WC9 | DAGLB | diacylglycerol lipase, beta | Plasma Membrane | enzyme | 1970185.5 | 2255314 | 1945139.75 | Q91WC9 | DAGLB | 3821502 | 2406607 | 567243 | 18 |
| Q922B2 | DARS | aspartyl-tRNA synthetase | Cytoplasm | enzyme | 5639079.5 | 6775738.5 | 6493397.5 | | | | | | |
| Q922B2 | DARS | aspartyl-tRNA synthetase | Cytoplasm | enzyme | 3958543.25 | 3819239 | 3343713 | | | | | | |
| Q922B2 | DARS | aspartyl-tRNA synthetase | Cytoplasm | enzyme | 2258736.75 | 2502229.75 | 2393303.5 | | | | | | |
| Q922B2 | DARS | aspartyl-tRNA synthetase | Cytoplasm | enzyme | 2584447.75 | 2770799.75 | 1889855.125 | | | | | | |
| Q922B2 | DARS | aspartyl-tRNA synthetase | Cytoplasm | enzyme | 2007868.25 | 1461577.75 | 1581656.375 | | | | | | |
| Q922B2 | DARS | aspartyl-tRNA synthetase | Cytoplasm | enzyme | 1923096.5 | 1902133.75 | 1678484.25 | | | | | | |
| Q922B2 | DARS | aspartyl-tRNA synthetase | Cytoplasm | enzyme | 2371217.25 | 1989580.375 | 2641147.5 | | | | | | |
| Q922B2 | DARS | aspartyl-tRNA synthetase | Cytoplasm | enzyme | 1526763.375 | 2016235.875 | 1953369.375 | Q922B2 | DARS | 2783719 | 1505906 | 307392 | 24 |
| Q9JII5 | Dazap1 | DAZ associated protein 1 | Cytoplasm | other | 1480889.5 | 2216266.75 | 1556394.5 | Q9JII5 | Dazap1 | 1480890 | 404539 | 233561 | 3 |
| Q9QZ73 | DCUN1D1 | DCN1, defective in cullin neddylation 1, domain containing 1 | Nucleus | other | 3454810.5 | 2724339 | 2794003 | Q9QZ73 | DCUN1D1 | 3454811 | 403135 | 232750 | 3 |
| A2ADY9 | DDI2 | DNA-damage inducible 1 homolog 2 | Plasma Membrane | transporter | 7257974.5 | 6322642.5 | 5796734.5 | | | | | | |
| A2ADY9 | DDI2 | DNA-damage inducible 1 homolog 2 | Plasma Membrane | transporter | 5451161.5 | 3927426.75 | 3972661 | A2ADY9 | DDI2 | 6354568 | 1315380 | 537002 | 6 |
| O54734 | DDOST | dolichyl-diphosphooligosaccharide--protein glycosyltransferase subunit (non-catalytic) | Cytoplasm | enzyme | 952786.5 | 3095313 | 3019319.25 | | | | | | |
| O54734 | DDOST | dolichyl-diphosphooligosaccharide--protein glycosyltransferase subunit (non-catalytic) | Cytoplasm | enzyme | 647876.438 | 1813132.625 | 2488991 | | | | | | |
| O54734 | DDOST | dolichyl-diphosphooligosaccharide--protein glycosyltransferase subunit (non-catalytic) | Cytoplasm | enzyme | 185634.75 | 1329659.5 | 2490087 | O54734 | DDOST | 595433 | 1059761 | 353254 | 9 |
| Q8C4S8 | DENND2A | DENN/MADD domain containing 2A | Cytoplasm | other | 2078808.875 | 2993762.5 | 1990222.5 | Q8C4S8 | DENND2A | 2078809 | 555590 | 320770 | 3 |
| Q91YP3 | DERA | deoxyribose-phosphate aldolase (putative) | Cytoplasm | enzyme | 2994934.5 | 3012525.5 | 2337905.25 | | | | | | |
| Q91YP3 | DERA | deoxyribose-phosphate aldolase (putative) | Cytoplasm | enzyme | 1510374.375 | 1114844 | 1648429.125 | | | | | | |
| Q91YP3 | DERA | deoxyribose-phosphate aldolase (putative) | Cytoplasm | enzyme | 1700830.75 | 1187450.375 | 1195902.75 | Q91YP3 | DERA | 2068713 | 748797 | 249599 | 9 |
| Q99J56 | DERL1 | derlin 1 | Cytoplasm | other | 7198370 | 1.02E+07 | 7790497.5 | | | | | | |
| Q99J56 | DERL1 | derlin 1 | Cytoplasm | other | 2658040.5 | 4816349.5 | 5496218 | Q99J56 | DERL1 | 4928205 | 2631988 | 1074505 | 6 |
| Q8BNI4 | DERL2 | derlin 2 | Cytoplasm | other | 1899515.75 | 4295981 | 2769824 | Q8BNI4 | DERL2 | 1899516 | 1213098 | 700382 | 3 |
| Q3U0B3 | DHRS11 | dehydrogenase/reductase (SDR family) member 11 | Cytoplasm | other | 4390066.5 | 3748891.5 | 4543375 | Q3U0B3 | DHRS11 | 4390067 | 421468 | 243335 | 3 |
| A2ATU0 | DHTKD1 | dehydrogenase E1 and transketolase domain containing 1 | Cytoplasm | enzyme | 369929.312 | 147544.125 | 186130.625 | A2ATU0 | DHTKD1 | 369929 | 118832 | 68608 | 3 |
| Q80VY9 | DHX33 | DEAH (Asp-Glu-Ala-His) box polypeptide 33 | Nucleus | enzyme | 8536104 | 8454522 | 4306728 | Q80VY9 | DHX33 | 8536104 | 2418625 | 1396394 | 3 |
| Q8BMF4 | DLAT | dihydrolipoamide S-acetyltransferase | Cytoplasm | enzyme | 1329012 | 4910601.5 | 1909111.375 | | | | | | |
| Q8BMF4 | DLAT | dihydrolipoamide S-acetyltransferase | Cytoplasm | enzyme | 1295411.625 | 3953287.5 | 1519694.125 | Q8BMF4 | DLAT | 1312212 | 1552694 | 633885 | 6 |
| O08749 | DLD | dihydrolipoamide dehydrogenase | Cytoplasm | enzyme | 1.53E+07 | 2.51E+07 | 1.18E+07 | | | | | | |
| O08749 | DLD | dihydrolipoamide dehydrogenase | Cytoplasm | enzyme | 9431054 | 1.78E+07 | 7226104 | | | | | | |
| O08749 | DLD | dihydrolipoamide dehydrogenase | Cytoplasm | enzyme | 7076390.5 | 1.03E+07 | 3243853.75 | O08749 | DLD | 10592482 | 6613201 | 2204400 | 9 |
| Q811D0 | DLG1 | discs, large homolog 1 (Drosophila) | Plasma Membrane | kinase | 3105587.75 | 3434277.25 | 1819945.75 | | | | | | |
| Q811D0 | DLG1 | discs, large homolog 1 (Drosophila) | Plasma Membrane | kinase | 2036264.125 | 1771106.75 | 648598.312 | | | | | | |
| Q811D0 | DLG1 | discs, large homolog 1 (Drosophila) | Plasma Membrane | kinase | 997075.5 | 1113409.25 | 725918 | Q811D0 | DLG1 | 2046309 | 999663 | 333221 | 9 |
| Q9D2G2 | DLST | dihydrolipoamide S-succinyltransferase (E2 component of 2-oxo-glutarate complex) | Cytoplasm | enzyme | 9508734 | 2.03E+07 | 8989880 | | | | | | |
| Q9D2G2 | DLST | dihydrolipoamide S-succinyltransferase (E2 component of 2-oxo-glutarate complex) | Cytoplasm | enzyme | 5470232.5 | 1.18E+07 | 5458076 | | | | | | |
| Q9D2G2 | DLST | dihydrolipoamide S-succinyltransferase (E2 component of 2-oxo-glutarate complex) | Cytoplasm | enzyme | 7145531 | 1.56E+07 | 5187257 | | | | | | |
| Q9D2G2 | DLST | dihydrolipoamide S-succinyltransferase (E2 component of 2-oxo-glutarate complex) | Cytoplasm | enzyme | 7511063 | 1.02E+07 | 3179996.5 | | | | | | |
| Q9D2G2 | DLST | dihydrolipoamide S-succinyltransferase (E2 component of 2-oxo-glutarate complex) | Cytoplasm | enzyme | 4398424.5 | 1.16E+07 | 6197653 | | | | | | |
| Q9D2G2 | DLST | dihydrolipoamide S-succinyltransferase (E2 component of 2-oxo-glutarate complex) | Cytoplasm | enzyme | 2500729.5 | 1.11E+07 | 7015351 | | | | | | |
| Q9D2G2 | DLST | dihydrolipoamide S-succinyltransferase (E2 component of 2-oxo-glutarate complex) | Cytoplasm | enzyme | 3933432.5 | 7627141.5 | 3281070.75 | Q9D2G2 | DLST | 5781164 | 4383920 | 956650 | 21 |
| Q80WT2 | DMRT3 | doublesex and mab-3 related transcription factor 3 | Nucleus | transcription regulator | 4655499.5 | 6020710 | 5533787 | Q80WT2 | DMRT3 | 4655500 | 691891 | 399464 | 3 |
| Q9WV69 | DMTN | dematin actin binding protein | Plasma Membrane | other | 4.73E+08 | 6.05E+08 | 6.61E+08 | | | | | | |
| Q9WV69 | DMTN | dematin actin binding protein | Plasma Membrane | other | 6.09E+08 | 7.03E+08 | 7.09E+08 | | | | | | |
| Q9WV69 | DMTN | dematin actin binding protein | Plasma Membrane | other | 4.24E+08 | 3.84E+08 | 4.21E+08 | | | | | | |
| Q9WV69 | DMTN | dematin actin binding protein | Plasma Membrane | other | 3.87E+08 | 3.58E+08 | 3.58E+08 | | | | | | |
| Q9WV69 | DMTN | dematin actin binding protein | Plasma Membrane | other | 4.73E+08 | 4.50E+08 | 4.05E+08 | | | | | | |
| Q9WV69 | DMTN | dematin actin binding protein | Plasma Membrane | other | 3.45E+08 | 3.41E+08 | 4.09E+08 | | | | | | |
| Q9WV69 | DMTN | dematin actin binding protein | Plasma Membrane | other | 3.81E+08 | 3.54E+08 | 3.80E+08 | | | | | | |
| Q9WV69 | DMTN | dematin actin binding protein | Plasma Membrane | other | 3.63E+08 | 3.39E+08 | 4.20E+08 | | | | | | |
| Q9WV69 | DMTN | dematin actin binding protein | Plasma Membrane | other | 2.95E+08 | 2.68E+08 | 1.65E+08 | | | | | | |
| Q9WV69 | DMTN | dematin actin binding protein | Plasma Membrane | other | 2.41E+08 | 2.28E+08 | 2.28E+08 | | | | | | |
| Q9WV69 | DMTN | dematin actin binding protein | Plasma Membrane | other | 2.07E+08 | 1.83E+08 | 2.13E+08 | | | | | | |
| Q9WV69 | DMTN | dematin actin binding protein | Plasma Membrane | other | 1.27E+08 | 1.54E+08 | 1.71E+08 | | | | | | |
| Q9WV69 | DMTN | dematin actin binding protein | Plasma Membrane | other | 1.33E+08 | 8.70E+07 | 6.32E+07 | | | | | | |
| Q9WV69 | DMTN | dematin actin binding protein | Plasma Membrane | other | 1.21E+08 | 1.05E+08 | 2.15E+08 | | | | | | |
| Q9WV69 | DMTN | dematin actin binding protein | Plasma Membrane | other | 1.14E+08 | 9.95E+07 | 1.99E+08 | | | | | | |
| Q9WV69 | DMTN | dematin actin binding protein | Plasma Membrane | other | 5.00E+07 | 5.71E+07 | 8.53E+07 | | | | | | |
| Q9WV69 | DMTN | dematin actin binding protein | Plasma Membrane | other | 1.12E+07 | 5.84E+07 | 2.36E+07 | | | | | | |
| Q9WV69 | DMTN | dematin actin binding protein | Plasma Membrane | other | 1.92E+07 | 2.08E+07 | 2.67E+07 | | | | | | |
| Q9WV69 | DMTN | dematin actin binding protein | Plasma Membrane | other | 2.07E+07 | 2.25E+07 | 1.85E+07 | | | | | | |
| Q9WV69 | DMTN | dematin actin binding protein | Plasma Membrane | other | 1.47E+07 | 1.29E+07 | 1.38E+07 | | | | | | |
| Q9WV69 | DMTN | dematin actin binding protein | Plasma Membrane | other | 1.19E+07 | 6239635.5 | 2294548 | | | | | | |
| Q9WV69 | DMTN | dematin actin binding protein | Plasma Membrane | other | 1.18E+07 | 8746124 | 1.12E+07 | | | | | | |
| Q9WV69 | DMTN | dematin actin binding protein | Plasma Membrane | other | 4890891.5 | 4739054 | 5759480.5 | | | | | | |
| Q9WV69 | DMTN | dematin actin binding protein | Plasma Membrane | other | 7894513.5 | 2572210.25 | 1820622.5 | | | | | | |
| Q9WV69 | DMTN | dematin actin binding protein | Plasma Membrane | other | 1402356.5 | 1965206.125 | 1557831.875 | Q9WV69 | DMTN | 193817110 | 197809608 | 22841086 | 75 |
| Q9JMC3 | DNAJA4 | DnaJ (Hsp40) homolog, subfamily A, member 4 | Nucleus | other | 3366177 | 3299803.75 | 1662855.875 | Q9JMC3 | DNAJA4 | 3366177 | 964824 | 557041 | 3 |
| Q9QYJ3 | DNAJB1 | DnaJ (Hsp40) homolog, subfamily B, member 1 | Nucleus | other | 2.19E+07 | 1.88E+07 | 2.27E+07 | | | | | | |
| Q9QYJ3 | DNAJB1 | DnaJ (Hsp40) homolog, subfamily B, member 1 | Nucleus | other | 1.42E+07 | 1.16E+07 | 1.04E+07 | | | | | | |
| Q9QYJ3 | DNAJB1 | DnaJ (Hsp40) homolog, subfamily B, member 1 | Nucleus | other | 1.56E+07 | 8764354 | 3882038 | | | | | | |

| | | | | | | | | | | | | | |
|---|---|---|---|---|---|---|---|---|---|---|---|---|---|
| Q9QYJ3 | DNAJB1 | DnaJ (Hsp40) homolog, subfamily B, member 1 | Nucleus | other | 1.12E+07 | 1.48E+07 | 1.48E+07 | | | | | | |
| Q9QYJ3 | DNAJB1 | DnaJ (Hsp40) homolog, subfamily B, member 1 | Nucleus | other | 5378250.5 | 5190924 | 5227252.5 | | | | | | |
| Q9QYJ3 | DNAJB1 | DnaJ (Hsp40) homolog, subfamily B, member 1 | Nucleus | other | 5701661 | 6857261.5 | 5359377 | | | | | | |
| Q9QYJ3 | DNAJB1 | DnaJ (Hsp40) homolog, subfamily B, member 1 | Nucleus | other | 4320616 | 4373940.5 | 5651393.5 | | | | | | |
| Q9QYJ3 | DNAJB1 | DnaJ (Hsp40) homolog, subfamily B, member 1 | Nucleus | other | 3545254.5 | 2832356 | 2917811.25 | | | | | | |
| Q9QYJ3 | DNAJB1 | DnaJ (Hsp40) homolog, subfamily B, member 1 | Nucleus | other | 424514.594 | 1479212.875 | 4849327 | Q9QYJ3 | DNAJB1 | 9144477 | 6208813 | 1194887 | 27 |
| Q99KV1 | DNAJB11 | DnaJ (Hsp40) homolog, subfamily B, member 11 | Cytoplasm | other | 4362686.5 | 4897409 | 4666575.5 | | | | | | |
| Q99KV1 | DNAJB11 | DnaJ (Hsp40) homolog, subfamily B, member 11 | Cytoplasm | other | 2669254.75 | 3619233.25 | 3573654.5 | | | | | | |
| Q99KV1 | DNAJB11 | DnaJ (Hsp40) homolog, subfamily B, member 11 | Cytoplasm | other | 1328417.75 | 1866493 | 1906740 | Q99KV1 | DNAJB11 | 2786786 | 1319736 | 439912 | 9 |
| Q9D832 | DNAJB4 | DnaJ (Hsp40) homolog, subfamily B, member 4 | Nucleus | other | 2.17E+07 | 2.22E+07 | 2.47E+07 | | | | | | |
| Q9D832 | DNAJB4 | DnaJ (Hsp40) homolog, subfamily B, member 4 | Nucleus | other | 2.42E+07 | 1.86E+07 | 1.58E+07 | | | | | | |
| Q9D832 | DNAJB4 | DnaJ (Hsp40) homolog, subfamily B, member 4 | Nucleus | other | 1.35E+07 | 1.31E+07 | 1.36E+07 | | | | | | |
| Q9D832 | DNAJB4 | DnaJ (Hsp40) homolog, subfamily B, member 4 | Nucleus | other | 6851833.5 | 3865247.5 | 2426029.5 | | | | | | |
| Q9D832 | DNAJB4 | DnaJ (Hsp40) homolog, subfamily B, member 4 | Nucleus | other | 8044058 | 7937595.5 | 9053013 | | | | | | |
| Q9D832 | DNAJB4 | DnaJ (Hsp40) homolog, subfamily B, member 4 | Nucleus | other | 7389516.5 | 6493979.5 | 2959572.5 | | | | | | |
| Q9D832 | DNAJB4 | DnaJ (Hsp40) homolog, subfamily B, member 4 | Nucleus | other | 5810487 | 5348783.5 | 6274061.5 | | | | | | |
| Q9D832 | DNAJB4 | DnaJ (Hsp40) homolog, subfamily B, member 4 | Nucleus | other | 4418737.5 | 4397163 | 6228283.5 | | | | | | |
| Q9D832 | DNAJB4 | DnaJ (Hsp40) homolog, subfamily B, member 4 | Nucleus | other | 3830172.25 | 3667558.75 | 4286236 | | | | | | |
| Q9D832 | DNAJB4 | DnaJ (Hsp40) homolog, subfamily B, member 4 | Nucleus | other | 1208704.5 | 3027240.25 | 5369617.5 | Q9D832 | DNAJB4 | 9684351 | 6941418 | 1267324 | 30 |
| Q9DC23 | DNAJC10 | DnaJ (Hsp40) homolog, subfamily C, member 10 | Cytoplasm | enzyme | 480544.281 | 367514.812 | 246788.906 | Q9DC23 | DNAJC10 | 480544 | 116899 | 67492 | 3 |
| Q5U458 | DNAJC11 | DnaJ (Hsp40) homolog, subfamily C, member 11 | Cytoplasm | other | 2354640.25 | 4312249.5 | 1673542.125 | | | | | | |
| Q5U458 | DNAJC11 | DnaJ (Hsp40) homolog, subfamily C, member 11 | Cytoplasm | other | 1598790.125 | 3276327.75 | 1907300.5 | Q5U458 | DNAJC11 | 1976715 | 1072572 | 437876 | 6 |
| P60904 | DNAJC5 | DnaJ (Hsp40) homolog, subfamily C, member 5 | Plasma Membrane | other | 2.80E+07 | 4.84E+07 | 4.58E+07 | | | | | | |
| P60904 | DNAJC5 | DnaJ (Hsp40) homolog, subfamily C, member 5 | Plasma Membrane | other | 2.65E+07 | 3.43E+07 | 4.43E+07 | | | | | | |
| P60904 | DNAJC5 | DnaJ (Hsp40) homolog, subfamily C, member 5 | Plasma Membrane | other | 1.14E+07 | 1.47E+07 | 1.95E+07 | | | | | | |
| P60904 | DNAJC5 | DnaJ (Hsp40) homolog, subfamily C, member 5 | Plasma Membrane | other | 1.48E+07 | 2302671.75 | 32408.523 | | | | | | |
| P60904 | DNAJC5 | DnaJ (Hsp40) homolog, subfamily C, member 5 | Plasma Membrane | other | 1095196.25 | 30815.754 | 0 | P60904 | DNAJC5 | 16349039 | 17655149 | 4558540 | 15 |
| Q9D7J6 | DNASE1L1 | deoxyribonuclease I-like 1 | Cytoplasm | enzyme | 550801.25 | 678236.5 | 513162.094 | Q9D7J6 | DNASE1L1 | 550801 | 86512 | 49948 | 3 |
| P39054 | DNM2 | dynamin 2 | Plasma Membrane | enzyme | 5.85E+07 | 5.78E+07 | 4.13E+07 | | | | | | |
| P39054 | DNM2 | dynamin 2 | Plasma Membrane | enzyme | 4.22E+07 | 3.75E+07 | 3.72E+07 | | | | | | |
| P39054 | DNM2 | dynamin 2 | Plasma Membrane | enzyme | 4.26E+07 | 4.84E+07 | 4.45E+07 | | | | | | |
| P39054 | DNM2 | dynamin 2 | Plasma Membrane | enzyme | 3.21E+07 | 2.68E+07 | 3.11E+07 | | | | | | |
| P39054 | DNM2 | dynamin 2 | Plasma Membrane | enzyme | 2.44E+07 | 2.23E+07 | 2.40E+07 | | | | | | |
| P39054 | DNM2 | dynamin 2 | Plasma Membrane | enzyme | 2.33E+07 | 2.08E+07 | 2.17E+07 | | | | | | |
| P39054 | DNM2 | dynamin 2 | Plasma Membrane | enzyme | 1.69E+07 | 1.25E+07 | 1.86E+07 | | | | | | |
| P39054 | DNM2 | dynamin 2 | Plasma Membrane | enzyme | 1.46E+07 | 1.46E+07 | 1.62E+07 | | | | | | |
| P39054 | DNM2 | dynamin 2 | Plasma Membrane | enzyme | 1.66E+07 | 1.54E+07 | 1.39E+07 | | | | | | |
| P39054 | DNM2 | dynamin 2 | Plasma Membrane | enzyme | 1.40E+07 | 1.50E+07 | 1.19E+07 | | | | | | |
| P39054 | DNM2 | dynamin 2 | Plasma Membrane | enzyme | 1.29E+07 | 1.29E+07 | 1.04E+07 | | | | | | |
| P39054 | DNM2 | dynamin 2 | Plasma Membrane | enzyme | 1.04E+07 | 1.34E+07 | 1.49E+07 | | | | | | |
| P39054 | DNM2 | dynamin 2 | Plasma Membrane | enzyme | 7723695.5 | 8290014.5 | 8058856 | | | | | | |
| P39054 | DNM2 | dynamin 2 | Plasma Membrane | enzyme | 7153853 | 9539385 | 1.03E+07 | | | | | | |
| P39054 | DNM2 | dynamin 2 | Plasma Membrane | enzyme | 4393084 | 2070991.125 | 367421.188 | | | | | | |
| P39054 | DNM2 | dynamin 2 | Plasma Membrane | enzyme | 4980224 | 5621133.5 | 5864441 | | | | | | |
| P39054 | DNM2 | dynamin 2 | Plasma Membrane | enzyme | 6417493 | 5745379 | 8866793 | | | | | | |
| P39054 | DNM2 | dynamin 2 | Plasma Membrane | enzyme | 5872070 | 4356406.5 | 4015496.25 | | | | | | |
| P39054 | DNM2 | dynamin 2 | Plasma Membrane | enzyme | 3903017.75 | 3945191.5 | 3391634.5 | | | | | | |
| P39054 | DNM2 | dynamin 2 | Plasma Membrane | enzyme | 3824847.75 | 2355362 | 2190701.75 | | | | | | |
| P39054 | DNM2 | dynamin 2 | Plasma Membrane | enzyme | 3925751.5 | 2642657.25 | 2700850 | | | | | | |
| P39054 | DNM2 | dynamin 2 | Plasma Membrane | enzyme | 2141893.5 | 2809288 | 3967102.5 | | | | | | |
| P39054 | DNM2 | dynamin 2 | Plasma Membrane | enzyme | 2043529.25 | 3382481.5 | 1923215 | P39054 | DNM2 | 15687368 | 14222355 | 1712170 | 69 |
| Q8BZ98 | DNM3 | dynamin 3 | Cytoplasm | enzyme | 1.21E+07 | 1.14E+07 | 9891799 | | | | | | |
| Q8BZ98 | DNM3 | dynamin 3 | Cytoplasm | enzyme | 1.41E+07 | 1.57E+07 | 1.44E+07 | | | | | | |
| Q8BZ98 | DNM3 | dynamin 3 | Cytoplasm | enzyme | 3778502.75 | 3755321.75 | 5122631 | Q8BZ98 | DNM3 | 10009501 | 4692948 | 1564316 | 9 |
| Q9Z2W0 | DNPEP | aspartyl aminopeptidase | Cytoplasm | peptidase | 2.74E+07 | 2.68E+07 | 2.84E+07 | | | | | | |
| Q9Z2W0 | DNPEP | aspartyl aminopeptidase | Cytoplasm | peptidase | 7347869 | 6135610.5 | 7465254 | | | | | | |
| Q9Z2W0 | DNPEP | aspartyl aminopeptidase | Cytoplasm | peptidase | 5417900 | 6472586 | 6079053 | | | | | | |
| Q9Z2W0 | DNPEP | aspartyl aminopeptidase | Cytoplasm | peptidase | 7091219 | 5930492.5 | 5508018 | | | | | | |
| Q9Z2W0 | DNPEP | aspartyl aminopeptidase | Cytoplasm | peptidase | 5829082 | 3982911.5 | 4718640.5 | | | | | | |
| Q9Z2W0 | DNPEP | aspartyl aminopeptidase | Cytoplasm | peptidase | 4200429 | 4177110.75 | 4175145.75 | | | | | | |
| Q9Z2W0 | DNPEP | aspartyl aminopeptidase | Cytoplasm | peptidase | 5651808.5 | 4765020.5 | 4050720.5 | | | | | | |
| Q9Z2W0 | DNPEP | aspartyl aminopeptidase | Cytoplasm | peptidase | 5227125.5 | 3556130.5 | 3294124.5 | | | | | | |
| Q9Z2W0 | DNPEP | aspartyl aminopeptidase | Cytoplasm | peptidase | 2709053.5 | 1744901.875 | 1853804.625 | Q9Z2W0 | DNPEP | 7870499 | 7406978 | 1425474 | 27 |
| Q9DD18 | DTD1 | D-tyrosyl-tRNA deacylase 1 | Cytoplasm | enzyme | 181611.859 | 89232.242 | 3.806 | Q9DD18 | DTD1 | 181612 | 90809 | 52428 | 3 |
| Q9JHU4 | DYNC1H1 | dynein, cytoplasmic 1, heavy chain 1 | Cytoplasm | peptidase | 1998786.75 | 1506091.5 | 1801928.5 | Q9JHU4 | DYNC1H1 | 1998787 | 247999 | 143182 | 3 |
| P42125 | ECI1 | enoyl-CoA delta isomerase 1 | Cytoplasm | enzyme | 3072562.75 | 6671928 | 4015528.75 | | | | | | |
| P42125 | ECI1 | enoyl-CoA delta isomerase 1 | Cytoplasm | enzyme | 1673618.5 | 3849177.75 | 1181993.625 | P42125 | ECI1 | 2373091 | 1964312 | 801927 | 6 |
| Q8BL66 | EEA1 | early endosome antigen 1 | Cytoplasm | other | 1715350.875 | 1262933.125 | 1006291 | Q8BL66 | EEA1 | 1715351 | 359006 | 207272 | 3 |
| P10126 | EEF1A1 | eukaryotic translation elongation factor 1 alpha 1 | Cytoplasm | translation regulator | 2.35E+08 | 3.20E+08 | 2.03E+08 | | | | | | |
| P10126 | EEF1A1 | eukaryotic translation elongation factor 1 alpha 1 | Cytoplasm | translation regulator | 9.25E+07 | 1.46E+08 | 8.92E+07 | | | | | | |
| P10126 | EEF1A1 | eukaryotic translation elongation factor 1 alpha 1 | Cytoplasm | translation regulator | 8.86E+07 | 7.11E+07 | 5679870.5 | | | | | | |
| P10126 | EEF1A1 | eukaryotic translation elongation factor 1 alpha 1 | Cytoplasm | translation regulator | 4.73E+07 | 1.06E+08 | 7.03E+07 | | | | | | |
| P10126 | EEF1A1 | eukaryotic translation elongation factor 1 alpha 1 | Cytoplasm | translation regulator | 2.39E+07 | 6.62E+07 | 3.04E+07 | | | | | | |
| P10126 | EEF1A1 | eukaryotic translation elongation factor 1 alpha 1 | Cytoplasm | translation regulator | 4.55E+07 | 6.60E+07 | 4.09E+07 | | | | | | |
| P10126 | EEF1A1 | eukaryotic translation elongation factor 1 alpha 1 | Cytoplasm | translation regulator | 7.05E+07 | 1.13E+08 | 7.36E+07 | | | | | | |
| P10126 | EEF1A1 | eukaryotic translation elongation factor 1 alpha 1 | Cytoplasm | translation regulator | 8.68E+07 | 6.14E+07 | 4621473 | | | | | | |
| P10126 | EEF1A1 | eukaryotic translation elongation factor 1 alpha 1 | Cytoplasm | translation regulator | 5.63E+07 | 7.37E+07 | 4.70E+07 | | | | | | |
| P10126 | EEF1A1 | eukaryotic translation elongation factor 1 alpha 1 | Cytoplasm | translation regulator | 3.27E+07 | 8.45E+07 | 5.10E+07 | | | | | | |
| P10126 | EEF1A1 | eukaryotic translation elongation factor 1 alpha 1 | Cytoplasm | translation regulator | 4.63E+07 | 7.42E+07 | 4.60E+07 | | | | | | |
| P10126 | EEF1A1 | eukaryotic translation elongation factor 1 alpha 1 | Cytoplasm | translation regulator | 4.00E+07 | 6.39E+07 | 3.85E+07 | | | | | | |
| P10126 | EEF1A1 | eukaryotic translation elongation factor 1 alpha 1 | Cytoplasm | translation regulator | 2.62E+07 | 7.44E+07 | 5.57E+07 | | | | | | |
| P10126 | EEF1A1 | eukaryotic translation elongation factor 1 alpha 1 | Cytoplasm | translation regulator | 3.17E+07 | 5.16E+07 | 2.73E+07 | | | | | | |
| P10126 | EEF1A1 | eukaryotic translation elongation factor 1 alpha 1 | Cytoplasm | translation regulator | 2.19E+07 | 5.53E+07 | 4.16E+07 | | | | | | |
| P10126 | EEF1A1 | eukaryotic translation elongation factor 1 alpha 1 | Cytoplasm | translation regulator | 4.59E+07 | 3.69E+07 | 1.60E+07 | | | | | | |
| P10126 | EEF1A1 | eukaryotic translation elongation factor 1 alpha 1 | Cytoplasm | translation regulator | 9998456 | 1.48E+07 | 8734530 | | | | | | |
| P10126 | EEF1A1 | eukaryotic translation elongation factor 1 alpha 1 | Cytoplasm | translation regulator | 2763017.25 | 8498936 | 1060257.25 | | | | | | |
| P10126 | EEF1A1 | eukaryotic translation elongation factor 1 alpha 1 | Cytoplasm | translation regulator | 3153784.25 | 8171628.5 | 3451157 | | | | | | |
| P10126 | EEF1A1 | eukaryotic translation elongation factor 1 alpha 1 | Cytoplasm | translation regulator | 2674635.25 | 2787077.25 | 3585832.25 | | | | | | |
| P10126 | EEF1A1 | eukaryotic translation elongation factor 1 alpha 1 | Cytoplasm | translation regulator | 4353254 | 4530924.5 | 1991257.125 | | | | | | |
| P10126 | EEF1A1 | eukaryotic translation elongation factor 1 alpha 1 | Cytoplasm | translation regulator | 5261809 | 3671098 | 914032 | | | | | | |
| P10126 | EEF1A1 | eukaryotic translation elongation factor 1 alpha 1 | Cytoplasm | translation regulator | 579919.125 | 1004927.375 | 721732.688 | P10126 | EEF1A1 | 44338038 | 55804474 | 6718069 | 69 |

| | | | | | | | | | | | | | |
|---|---|---|---|---|---|---|---|---|---|---|---|---|---|
| O70251 | EEF1B2 | eukaryotic translation elongation factor 1 beta 2 | Cytoplasm | translation regulator | 1133765 | 1048151.25 | 609435.25 | O70251 | EEF1B2 | 1133765 | 281284 | 162399 | 3 |
| P57776 | EEF1D | eukaryotic translation elongation factor 1 delta (guanine nucleotide exchange protein) | Cytoplasm | translation regulator | 1380496.875 | 1816982.875 | 564225.125 | P57776 | EEF1D | 1380497 | 635901 | 367138 | 3 |
| Q9D8N0 | EEF1G | eukaryotic translation elongation factor 1 gamma | Cytoplasm | translation regulator | 2142054.5 | 3872804.75 | 2191705.25 | | | | | | |
| Q9D8N0 | EEF1G | eukaryotic translation elongation factor 1 gamma | Cytoplasm | translation regulator | 606387.312 | 2119876.5 | 1369641.625 | | | | | | |
| Q9D8N0 | EEF1G | eukaryotic translation elongation factor 1 gamma | Cytoplasm | translation regulator | 1158554.875 | 1588325.625 | 883803.812 | Q9D8N0 | EEF1G | 1302332 | 972558 | 324186 | 9 |
| P58252 | EEF2 | eukaryotic translation elongation factor 2 | Cytoplasm | translation regulator | 2090381.875 | 3085547.75 | 1159144.375 | | | | | | |
| P58252 | EEF2 | eukaryotic translation elongation factor 2 | Cytoplasm | translation regulator | 3322380.5 | 5000797 | 2223718 | | | | | | |
| P58252 | EEF2 | eukaryotic translation elongation factor 2 | Cytoplasm | translation regulator | 4605535 | 5179395.5 | 2285651.25 | | | | | | |
| P58252 | EEF2 | eukaryotic translation elongation factor 2 | Cytoplasm | translation regulator | 2897135 | 3959085.75 | 1146810.875 | | | | | | |
| P58252 | EEF2 | eukaryotic translation elongation factor 2 | Cytoplasm | translation regulator | 691994.375 | 1522117.625 | 973783.25 | P58252 | EEF2 | 2721485 | 1492360 | 385326 | 15 |
| Q8BG67 | EFR3A | EFR3 homolog A | Plasma Membrane | other | 4990119 | 5045705 | 2912881 | | | | | | |
| Q8BG67 | EFR3A | EFR3 homolog A | Plasma Membrane | other | 3844788.25 | 4506712 | 4995550.5 | | | | | | |
| Q8BG67 | EFR3A | EFR3 homolog A | Plasma Membrane | other | 4847626.5 | 5173966.5 | 4966304.5 | | | | | | |
| Q8BG67 | EFR3A | EFR3 homolog A | Plasma Membrane | other | 4744215 | 5759175 | 4344623 | | | | | | |
| Q8BG67 | EFR3A | EFR3 homolog A | Plasma Membrane | other | 3708306.5 | 4047878 | 4596559.5 | | | | | | |
| Q8BG67 | EFR3A | EFR3 homolog A | Plasma Membrane | other | 4620136.5 | 4100802.75 | 4138376 | | | | | | |
| Q8BG67 | EFR3A | EFR3 homolog A | Plasma Membrane | other | 2969147.75 | 4136011 | 2417490.25 | Q8BG67 | EFR3A | 4246334 | 822280 | 179436 | 21 |
| Q9WVK4 | EHD1 | EH-domain containing 1 | Cytoplasm | other | 5760337.5 | 7166812.5 | 8545759 | | | | | | |
| Q9WVK4 | EHD1 | EH-domain containing 1 | Cytoplasm | other | 8244411 | 1.16E+07 | 7732857 | | | | | | |
| Q9WVK4 | EHD1 | EH-domain containing 1 | Cytoplasm | other | 3375580.5 | 2949212.75 | 2423615.5 | | | | | | |
| Q9WVK4 | EHD1 | EH-domain containing 1 | Cytoplasm | other | 2867821.75 | 2691812.75 | 2652941.5 | Q9WVK4 | EHD1 | 5062038 | 3095099 | 893478 | 12 |
| Q9EQP2 | EHD4 | EH-domain containing 4 | Plasma Membrane | enzyme | 539207.188 | 1054025.125 | 1590838.375 | | | | | | |
| Q9EQP2 | EHD4 | EH-domain containing 4 | Plasma Membrane | enzyme | 923514.188 | 1053545.75 | 714319.188 | Q9EQP2 | EHD4 | 731361 | 361133 | 147432 | 6 |
| Q6ZWX6 | EIF2S1 | eukaryotic translation initiation factor 2, subunit 1 alpha, 35kDa | Cytoplasm | translation regulator | 7183040 | 7867791 | 4445649.5 | | | | | | |
| Q6ZWX6 | EIF2S1 | eukaryotic translation initiation factor 2, subunit 1 alpha, 35kDa | Cytoplasm | translation regulator | 5393141 | 5769177.5 | 4121754 | | | | | | |
| Q6ZWX6 | EIF2S1 | eukaryotic translation initiation factor 2, subunit 1 alpha, 35kDa | Cytoplasm | translation regulator | 5406762.5 | 5119066 | 2976948.75 | | | | | | |
| Q6ZWX6 | EIF2S1 | eukaryotic translation initiation factor 2, subunit 1 alpha, 35kDa | Cytoplasm | translation regulator | 3253245 | 3445413 | 2683520 | | | | | | |
| Q6ZWX6 | EIF2S1 | eukaryotic translation initiation factor 2, subunit 1 alpha, 35kDa | Cytoplasm | translation regulator | 3627956 | 3882872 | 1640775.25 | | | | | | |
| Q6ZWX6 | EIF2S1 | eukaryotic translation initiation factor 2, subunit 1 alpha, 35kDa | Cytoplasm | translation regulator | 3178929.5 | 5555715 | 3711809 | Q6ZWX6 | EIF2S1 | 4673846 | 1594412 | 375806 | 18 |
| Q99L45 | EIF2S2 | eukaryotic translation initiation factor 2, subunit 2 beta, 38kDa | Cytoplasm | translation regulator | 7210158 | 8023386.5 | 3893929.5 | | | | | | |
| Q99L45 | EIF2S2 | eukaryotic translation initiation factor 2, subunit 2 beta, 38kDa | Cytoplasm | translation regulator | 6198797.5 | 6804013 | 4487485 | | | | | | |
| Q99L45 | EIF2S2 | eukaryotic translation initiation factor 2, subunit 2 beta, 38kDa | Cytoplasm | translation regulator | 5319513.5 | 5517155 | 3566622.75 | Q99L45 | EIF2S2 | 6242823 | 1524766 | 508255 | 9 |
| Q9Z0N1 | EIF2S3 | eukaryotic translation initiation factor 2, subunit 3 gamma, 52kDa | Cytoplasm | translation regulator | 1.05E+07 | 9516017 | 8154962 | | | | | | |
| Q9Z0N1 | EIF2S3 | eukaryotic translation initiation factor 2, subunit 3 gamma, 52kDa | Cytoplasm | translation regulator | 1.30E+07 | 1.31E+07 | 8096073.5 | | | | | | |
| Q9Z0N1 | EIF2S3 | eukaryotic translation initiation factor 2, subunit 3 gamma, 52kDa | Cytoplasm | translation regulator | 1.05E+07 | 9197754 | 6455557 | | | | | | |
| Q9Z0N1 | EIF2S3 | eukaryotic translation initiation factor 2, subunit 3 gamma, 52kDa | Cytoplasm | translation regulator | 6701025.5 | 9388500 | 4718161 | | | | | | |
| Q9Z0N1 | EIF2S3 | eukaryotic translation initiation factor 2, subunit 3 gamma, 52kDa | Cytoplasm | translation regulator | 7760113.5 | 6147078 | 5390193 | | | | | | |
| Q9Z0N1 | EIF2S3 | eukaryotic translation initiation factor 2, subunit 3 gamma, 52kDa | Cytoplasm | translation regulator | 5959479.5 | 5996584.5 | 3744178.25 | | | | | | |
| Q9Z0N1 | EIF2S3 | eukaryotic translation initiation factor 2, subunit 3 gamma, 52kDa | Cytoplasm | translation regulator | 5395349.5 | 5501746 | 3927717 | | | | | | |
| Q9Z0N1 | EIF2S3 | eukaryotic translation initiation factor 2, subunit 3 gamma, 52kDa | Cytoplasm | translation regulator | 4702134.5 | 3811287.75 | 2981039.75 | | | | | | |
| Q9Z0N1 | EIF2S3 | eukaryotic translation initiation factor 2, subunit 3 gamma, 52kDa | Cytoplasm | translation regulator | 3176226.75 | 2522299.5 | 1832042.125 | | | | | | |
| Q9Z0N1 | EIF2S3 | eukaryotic translation initiation factor 2, subunit 3 gamma, 52kDa | Cytoplasm | translation regulator | 2206270.5 | 2008811.875 | 1537314.5 | | | | | | |
| Q9Z0N1 | EIF2S3 | eukaryotic translation initiation factor 2, subunit 3 gamma, 52kDa | Cytoplasm | translation regulator | 1981718.375 | 1736356.625 | 1329411.125 | Q9Z0N1 | EIF2S3 | 6531120 | 3330810 | 579820 | 33 |
| Q3UGC7 | EIF3J | eukaryotic translation initiation factor 3, subunit J | Cytoplasm | translation regulator | 1.22E+07 | 9670496 | 7407025 | | | | | | |
| Q3UGC7 | EIF3J | eukaryotic translation initiation factor 3, subunit J | Cytoplasm | translation regulator | 1.23E+07 | 7797484 | 9842635 | | | | | | |
| Q3UGC7 | EIF3J | eukaryotic translation initiation factor 3, subunit J | Cytoplasm | translation regulator | 6654410 | 6463606.5 | 4535430 | | | | | | |
| Q3UGC7 | EIF3J | eukaryotic translation initiation factor 3, subunit J | Cytoplasm | translation regulator | 51610.098 | 39482.871 | 81174.148 | | | | | | |
| Q3UGC7 | EIF3J | eukaryotic translation initiation factor 3, subunit J | Cytoplasm | translation regulator | 1.02E+07 | 7346639 | 5057604.5 | Q3UGC7 | EIF3J | 8287204 | 4096988 | 1057838 | 15 |
| Q8BGD9 | EIF4B | eukaryotic translation initiation factor 4B | Cytoplasm | translation regulator | 8122498 | 8429744 | 7513449.5 | | | | | | |
| Q8BGD9 | EIF4B | eukaryotic translation initiation factor 4B | Cytoplasm | translation regulator | 5688502.5 | 6142994.5 | 6866542 | | | | | | |
| Q8BGD9 | EIF4B | eukaryotic translation initiation factor 4B | Cytoplasm | translation regulator | 7573396 | 5609819 | 5993873 | | | | | | |
| Q8BGD9 | EIF4B | eukaryotic translation initiation factor 4B | Cytoplasm | translation regulator | 3655737 | 4912335.5 | 4472521 | | | | | | |
| Q8BGD9 | EIF4B | eukaryotic translation initiation factor 4B | Cytoplasm | translation regulator | 3641074.25 | 3717167.75 | 3822957 | | | | | | |
| Q8BGD9 | EIF4B | eukaryotic translation initiation factor 4B | Cytoplasm | translation regulator | 2441190.25 | 2903962.25 | 2268255.75 | Q8BGD9 | EIF4B | 5187066 | 1965271 | 463219 | 18 |
| Q6NZJ6 | EIF4G1 | eukaryotic translation initiation factor 4 gamma, 1 | Cytoplasm | translation regulator | 3846733.5 | 5026649.5 | 2345677.5 | | | | | | |
| Q6NZJ6 | EIF4G1 | eukaryotic translation initiation factor 4 gamma, 1 | Cytoplasm | translation regulator | 2052180.875 | 2445403.75 | 1586153.375 | | | | | | |
| Q6NZJ6 | EIF4G1 | eukaryotic translation initiation factor 4 gamma, 1 | Cytoplasm | translation regulator | 778334.562 | 854978.375 | 443732.219 | Q6NZJ6 | EIF4G1 | 2225750 | 1503199 | 501066 | 9 |
| P59325 | EIF5 | eukaryotic translation initiation factor 5 | Cytoplasm | translation regulator | 2766443.5 | 2111889 | 2328506 | P59325 | EIF5 | 2766444 | 333455 | 192520 | 3 |
| P63242 | EIF5A | eukaryotic translation initiation factor 5A | Cytoplasm | translation regulator | 3340892.75 | 5579310.5 | 3503523.5 | P63242 | EIF5A | 3340893 | 1248055 | 720565 | 3 |
| Q99K41 | EMILIN1 | elastin microfibril interfacer 1 | Extracellular Space | other | 234640.703 | 1316280.625 | 2219172.5 | | | | | | |
| Q99K41 | EMILIN1 | elastin microfibril interfacer 1 | Extracellular Space | other | 19605.104 | 94453.805 | 274069.094 | Q99K41 | EMILIN1 | 127123 | 884872 | 361248 | 6 |
| Q8C522 | ENDOD1 | endonuclease domain containing 1 | Extracellular Space | enzyme | 1.22E+07 | 1.92E+07 | 2.32E+07 | | | | | | |
| Q8C522 | ENDOD1 | endonuclease domain containing 1 | Extracellular Space | enzyme | 1.07E+07 | 1.03E+07 | 7876074.5 | | | | | | |
| Q8C522 | ENDOD1 | endonuclease domain containing 1 | Extracellular Space | enzyme | 8880537 | 7397819 | 4001346.75 | | | | | | |
| Q8C522 | ENDOD1 | endonuclease domain containing 1 | Extracellular Space | enzyme | 4653192 | 6469347.5 | 6112366 | | | | | | |
| Q8C522 | ENDOD1 | endonuclease domain containing 1 | Extracellular Space | enzyme | 6966436.5 | 1.01E+07 | 9518327 | | | | | | |
| Q8C522 | ENDOD1 | endonuclease domain containing 1 | Extracellular Space | enzyme | 4959742.5 | 6113205.5 | 5633279 | | | | | | |
| Q8C522 | ENDOD1 | endonuclease domain containing 1 | Extracellular Space | enzyme | 6878275.5 | 4494683.5 | 1640304.25 | | | | | | |
| Q8C522 | ENDOD1 | endonuclease domain containing 1 | Extracellular Space | enzyme | 5682491.5 | 6106417 | 7940996.5 | Q8C522 | ENDOD1 | 7617584 | 4706931 | 960798 | 24 |
| Q8BTJ4 | ENPP4 | ectonucleotide pyrophosphatase/phosphodiesterase 4 (putative) | Cytoplasm | enzyme | 840676.625 | 2535323 | 890714.062 | Q8BTJ4 | ENPP4 | 840677 | 964285 | 556730 | 3 |
| P10404 | ENV1 | MLV-related proviral Env polyprotein | | other | 1.07E+08 | 1.28E+08 | 1.15E+08 | | | | | | |
| P10404 | ENV1 | MLV-related proviral Env polyprotein | | other | 7.73E+07 | 9.35E+07 | 9.99E+07 | | | | | | |
| P10404 | ENV1 | MLV-related proviral Env polyprotein | | other | 9.62E+07 | 7.91E+07 | 7.70E+07 | | | | | | |
| P10404 | ENV1 | MLV-related proviral Env polyprotein | | other | 5.55E+07 | 6.34E+07 | 4.97E+07 | | | | | | |
| P10404 | ENV1 | MLV-related proviral Env polyprotein | | other | 4.72E+07 | 6.87E+07 | 9.13E+07 | | | | | | |
| P10404 | ENV1 | MLV-related proviral Env polyprotein | | other | 6.90E+07 | 7.48E+07 | 9.07E+07 | | | | | | |
| P10404 | ENV1 | MLV-related proviral Env polyprotein | | other | 6.60E+07 | 6.93E+07 | 8.33E+07 | | | | | | |
| P10404 | ENV1 | MLV-related proviral Env polyprotein | | other | 5.64E+07 | 5.55E+07 | 6.47E+07 | | | | | | |
| P10404 | ENV1 | MLV-related proviral Env polyprotein | | other | 3.85E+07 | 2.83E+07 | 3.36E+07 | | | | | | |
| P10404 | ENV1 | MLV-related proviral Env polyprotein | | other | 4.37E+07 | 3.49E+07 | 3.67E+07 | | | | | | |
| P10404 | ENV1 | MLV-related proviral Env polyprotein | | other | 3.46E+07 | 3.33E+07 | 3.98E+07 | | | | | | |
| P10404 | ENV1 | MLV-related proviral Env polyprotein | | other | 3.16E+07 | 2.88E+07 | 3.11E+07 | | | | | | |
| P10404 | ENV1 | MLV-related proviral Env polyprotein | | other | 1.76E+07 | 1.33E+07 | 1.55E+07 | | | | | | |
| P10404 | ENV1 | MLV-related proviral Env polyprotein | | other | 2.15E+07 | 2.67E+07 | 3.03E+07 | | | | | | |
| P10404 | ENV1 | MLV-related proviral Env polyprotein | | other | 2.21E+07 | 9756186 | 6044373 | | | | | | |
| P10404 | ENV1 | MLV-related proviral Env polyprotein | | other | 1.50E+07 | 1.28E+07 | 1.61E+07 | | | | | | |
| P10404 | ENV1 | MLV-related proviral Env polyprotein | | other | 8374275 | 1.17E+07 | 1.28E+07 | | | | | | |
| P10404 | ENV1 | MLV-related proviral Env polyprotein | | other | 9658096 | 1.15E+07 | 1.28E+07 | | | | | | |
| P10404 | ENV1 | MLV-related proviral Env polyprotein | | other | 7101171.5 | 9440420 | 7588006 | | | | | | |
| P10404 | ENV1 | MLV-related proviral Env polyprotein | | other | 8355074.5 | 1.10E+07 | 1.47E+07 | | | | | | |
| P10404 | ENV1 | MLV-related proviral Env polyprotein | | other | 5192645.5 | 7133906.5 | 1.23E+07 | | | | | | |

| | | | | | | | | | | | | | |
|---|---|---|---|---|---|---|---|---|---|---|---|---|---|
| P10404 | ENV1 | MLV-related proviral Env polyprotein | other | | 6750205.5 | 3433253.25 | 4573304 | | | | | | |
| P10404 | ENV1 | MLV-related proviral Env polyprotein | other | | 4276154.5 | 2955986.5 | 2822275.5 | | | | | | |
| P10404 | ENV1 | MLV-related proviral Env polyprotein | other | | 2168495 | 4716545 | 6042035 | P10404 | ENV1 | 35442755 | 32984838 | 3887300 | 72 |
| P48193 | EPB41 | erythrocyte membrane protein band 4.1 | Plasma Membrane | other | 1.71E+09 | 2.35E+09 | 3.57E+09 | | | | | | |
| P48193 | EPB41 | erythrocyte membrane protein band 4.1 | Plasma Membrane | other | 1.88E+09 | 1.58E+09 | 1.95E+09 | | | | | | |
| P48193 | EPB41 | erythrocyte membrane protein band 4.1 | Plasma Membrane | other | 1.83E+09 | 1.92E+09 | 1.77E+09 | | | | | | |
| P48193 | EPB41 | erythrocyte membrane protein band 4.1 | Plasma Membrane | other | 1.51E+09 | 1.78E+09 | 2.22E+09 | | | | | | |
| P48193 | EPB41 | erythrocyte membrane protein band 4.1 | Plasma Membrane | other | 1.33E+09 | 1.21E+09 | 1.45E+09 | | | | | | |
| P48193 | EPB41 | erythrocyte membrane protein band 4.1 | Plasma Membrane | other | 1.30E+09 | 1.48E+09 | 1.46E+09 | | | | | | |
| P48193 | EPB41 | erythrocyte membrane protein band 4.1 | Plasma Membrane | other | 1.45E+09 | 1.42E+09 | 1.56E+09 | | | | | | |
| P48193 | EPB41 | erythrocyte membrane protein band 4.1 | Plasma Membrane | other | 1.30E+09 | 1.56E+09 | 1.73E+09 | | | | | | |
| P48193 | EPB41 | erythrocyte membrane protein band 4.1 | Plasma Membrane | other | 1.22E+09 | 1.15E+09 | 1.25E+09 | | | | | | |
| P48193 | EPB41 | erythrocyte membrane protein band 4.1 | Plasma Membrane | other | 1.10E+09 | 9.80E+08 | 1.22E+09 | | | | | | |
| P48193 | EPB41 | erythrocyte membrane protein band 4.1 | Plasma Membrane | other | 1.06E+09 | 1.05E+09 | 1.26E+09 | | | | | | |
| P48193 | EPB41 | erythrocyte membrane protein band 4.1 | Plasma Membrane | other | 9.81E+08 | 9.50E+08 | 1.19E+09 | | | | | | |
| P48193 | EPB41 | erythrocyte membrane protein band 4.1 | Plasma Membrane | other | 9.96E+08 | 9.48E+08 | 9.18E+08 | | | | | | |
| P48193 | EPB41 | erythrocyte membrane protein band 4.1 | Plasma Membrane | other | 1.07E+09 | 8.81E+08 | 1.02E+09 | | | | | | |
| P48193 | EPB41 | erythrocyte membrane protein band 4.1 | Plasma Membrane | other | 9.68E+08 | 7.16E+08 | 8.86E+08 | | | | | | |
| P48193 | EPB41 | erythrocyte membrane protein band 4.1 | Plasma Membrane | other | 5.59E+08 | 5.66E+08 | 7.46E+08 | | | | | | |
| P48193 | EPB41 | erythrocyte membrane protein band 4.1 | Plasma Membrane | other | 4.45E+08 | 5.32E+08 | 4.62E+08 | | | | | | |
| P48193 | EPB41 | erythrocyte membrane protein band 4.1 | Plasma Membrane | other | 4.32E+08 | 5.25E+08 | 5.51E+08 | | | | | | |
| P48193 | EPB41 | erythrocyte membrane protein band 4.1 | Plasma Membrane | other | 4.74E+08 | 5.51E+08 | 7.15E+08 | | | | | | |
| P48193 | EPB41 | erythrocyte membrane protein band 4.1 | Plasma Membrane | other | 4.22E+08 | 3.55E+08 | 3.82E+08 | | | | | | |
| P48193 | EPB41 | erythrocyte membrane protein band 4.1 | Plasma Membrane | other | 2.38E+08 | 5.32E+08 | 7.51E+08 | | | | | | |
| P48193 | EPB41 | erythrocyte membrane protein band 4.1 | Plasma Membrane | other | 2.49E+08 | 2.83E+08 | 3.50E+08 | | | | | | |
| P48193 | EPB41 | erythrocyte membrane protein band 4.1 | Plasma Membrane | other | 3.17E+08 | 2.61E+08 | 2.12E+08 | | | | | | |
| P48193 | EPB41 | erythrocyte membrane protein band 4.1 | Plasma Membrane | other | 2.96E+08 | 1.91E+08 | 2.15E+08 | | | | | | |
| P48193 | EPB41 | erythrocyte membrane protein band 4.1 | Plasma Membrane | other | 3.16E+08 | 2.48E+08 | 2.39E+08 | | | | | | |
| P48193 | EPB41 | erythrocyte membrane protein band 4.1 | Plasma Membrane | other | 1.57E+08 | 1.18E+08 | 1.39E+08 | | | | | | |
| P48193 | EPB41 | erythrocyte membrane protein band 4.1 | Plasma Membrane | other | 1.23E+08 | 1.59E+08 | 1.57E+08 | | | | | | |
| P48193 | EPB41 | erythrocyte membrane protein band 4.1 | Plasma Membrane | other | 1.04E+08 | 9.41E+07 | 1.40E+08 | | | | | | |
| P48193 | EPB41 | erythrocyte membrane protein band 4.1 | Plasma Membrane | other | 1.47E+08 | 1.95E+08 | 2.11E+08 | | | | | | |
| P48193 | EPB41 | erythrocyte membrane protein band 4.1 | Plasma Membrane | other | 6.17E+07 | 1.08E+08 | 1.20E+08 | | | | | | |
| P48193 | EPB41 | erythrocyte membrane protein band 4.1 | Plasma Membrane | other | 8.27E+07 | 6.84E+07 | 9.05E+07 | | | | | | |
| P48193 | EPB41 | erythrocyte membrane protein band 4.1 | Plasma Membrane | other | 5.30E+07 | 7.55E+07 | 1.21E+08 | | | | | | |
| P48193 | EPB41 | erythrocyte membrane protein band 4.1 | Plasma Membrane | other | 8.57E+07 | 6.56E+07 | 1.03E+08 | | | | | | |
| P48193 | EPB41 | erythrocyte membrane protein band 4.1 | Plasma Membrane | other | 7.49E+07 | 5.83E+07 | 7.17E+07 | | | | | | |
| P48193 | EPB41 | erythrocyte membrane protein band 4.1 | Plasma Membrane | other | 5.24E+07 | 4.93E+07 | 3.67E+07 | | | | | | |
| P48193 | EPB41 | erythrocyte membrane protein band 4.1 | Plasma Membrane | other | 3.54E+07 | 4.57E+07 | 2.79E+07 | | | | | | |
| P48193 | EPB41 | erythrocyte membrane protein band 4.1 | Plasma Membrane | other | 4.37E+07 | 4.89E+07 | 5.60E+07 | | | | | | |
| P48193 | EPB41 | erythrocyte membrane protein band 4.1 | Plasma Membrane | other | 2.82E+07 | 2.41E+07 | 2.36E+07 | | | | | | |
| P48193 | EPB41 | erythrocyte membrane protein band 4.1 | Plasma Membrane | other | 5.92E+07 | 3.85E+07 | 3.82E+07 | | | | | | |
| P48193 | EPB41 | erythrocyte membrane protein band 4.1 | Plasma Membrane | other | 3.87E+07 | 4.51E+07 | 4.81E+07 | | | | | | |
| P48193 | EPB41 | erythrocyte membrane protein band 4.1 | Plasma Membrane | other | 1.94E+07 | 4.65E+07 | 6.95E+07 | | | | | | |
| P48193 | EPB41 | erythrocyte membrane protein band 4.1 | Plasma Membrane | other | 1.89E+07 | 3.61E+07 | 5.26E+07 | | | | | | |
| P48193 | EPB41 | erythrocyte membrane protein band 4.1 | Plasma Membrane | other | 2.63E+07 | 2.55E+07 | 3.23E+07 | | | | | | |
| P48193 | EPB41 | erythrocyte membrane protein band 4.1 | Plasma Membrane | other | 2.26E+07 | 2.08E+07 | 2.43E+07 | | | | | | |
| P48193 | EPB41 | erythrocyte membrane protein band 4.1 | Plasma Membrane | other | 2.99E+07 | 1.52E+07 | 3329136 | | | | | | |
| P48193 | EPB41 | erythrocyte membrane protein band 4.1 | Plasma Membrane | other | 1.59E+07 | 1.65E+07 | 2.01E+07 | | | | | | |
| P48193 | EPB41 | erythrocyte membrane protein band 4.1 | Plasma Membrane | other | 1.04E+07 | 8284583 | 1.12E+07 | | | | | | |
| P48193 | EPB41 | erythrocyte membrane protein band 4.1 | Plasma Membrane | other | 2.64E+07 | 5.75E+07 | 9.90E+07 | | | | | | |
| P48193 | EPB41 | erythrocyte membrane protein band 4.1 | Plasma Membrane | other | 2.03E+07 | 1.25E+07 | 1.28E+07 | | | | | | |
| P48193 | EPB41 | erythrocyte membrane protein band 4.1 | Plasma Membrane | other | 6.81E+07 | 7.46E+07 | 9.16E+07 | | | | | | |
| P48193 | EPB41 | erythrocyte membrane protein band 4.1 | Plasma Membrane | other | 4.88E+07 | 2589064 | 758817 | | | | | | |
| P48193 | EPB41 | erythrocyte membrane protein band 4.1 | Plasma Membrane | other | 1.01E+07 | 9581549 | 1.25E+07 | | | | | | |
| P48193 | EPB41 | erythrocyte membrane protein band 4.1 | Plasma Membrane | other | 7026474 | 5742351 | 5853290.5 | | | | | | |
| P48193 | EPB41 | erythrocyte membrane protein band 4.1 | Plasma Membrane | other | 7681305 | 6752228 | 7011778 | | | | | | |
| P48193 | EPB41 | erythrocyte membrane protein band 4.1 | Plasma Membrane | other | 3927839 | 4281467 | 5423028.5 | | | | | | |
| P48193 | EPB41 | erythrocyte membrane protein band 4.1 | Plasma Membrane | other | 4314557 | 5061778.5 | 5774764.5 | | | | | | |
| P48193 | EPB41 | erythrocyte membrane protein band 4.1 | Plasma Membrane | other | 4559542 | 4757231 | 7744559 | | | | | | |
| P48193 | EPB41 | erythrocyte membrane protein band 4.1 | Plasma Membrane | other | 1.11E+07 | 1081147.5 | 83654.383 | | | | | | |
| P48193 | EPB41 | erythrocyte membrane protein band 4.1 | Plasma Membrane | other | 1145389.375 | 1929271.625 | 1775774.5 | | | | | | |
| P48193 | EPB41 | erythrocyte membrane protein band 4.1 | Plasma Membrane | other | 29129.771 | 114095.102 | 118112.82 | | | | | | |
| P48193 | EPB41 | erythrocyte membrane protein band 4.1 | Plasma Membrane | other | 3276926.5 | 2714103.25 | 4065521.5 | | | | | | |
| P48193 | EPB41 | erythrocyte membrane protein band 4.1 | Plasma Membrane | other | 671114.812 | 240695.5 | 356141.344 | P48193 | EPB41 | 402340359 | 620132318 | 45470308 | 186 |
| P49222 | EPB42 | erythrocyte membrane protein band 4.2 | Plasma Membrane | transporter | 1.77E+09 | 1.93E+09 | 2.22E+09 | | | | | | |
| P49222 | EPB42 | erythrocyte membrane protein band 4.2 | Plasma Membrane | transporter | 1.68E+09 | 1.67E+09 | 1.73E+09 | | | | | | |
| P49222 | EPB42 | erythrocyte membrane protein band 4.2 | Plasma Membrane | transporter | 1.63E+09 | 1.42E+09 | 1.41E+09 | | | | | | |
| P49222 | EPB42 | erythrocyte membrane protein band 4.2 | Plasma Membrane | transporter | 1.54E+09 | 1.71E+09 | 1.98E+09 | | | | | | |
| P49222 | EPB42 | erythrocyte membrane protein band 4.2 | Plasma Membrane | transporter | 1.19E+09 | 1.69E+09 | 1.63E+09 | | | | | | |
| P49222 | EPB42 | erythrocyte membrane protein band 4.2 | Plasma Membrane | transporter | 1.44E+09 | 1.49E+09 | 1.26E+09 | | | | | | |
| P49222 | EPB42 | erythrocyte membrane protein band 4.2 | Plasma Membrane | transporter | 1.25E+09 | 1.37E+09 | 9.72E+08 | | | | | | |
| P49222 | EPB42 | erythrocyte membrane protein band 4.2 | Plasma Membrane | transporter | 9.45E+08 | 9.19E+08 | 1.02E+09 | | | | | | |
| P49222 | EPB42 | erythrocyte membrane protein band 4.2 | Plasma Membrane | transporter | 8.05E+08 | 9.64E+08 | 1.04E+09 | | | | | | |
| P49222 | EPB42 | erythrocyte membrane protein band 4.2 | Plasma Membrane | transporter | 9.94E+08 | 7.42E+08 | 7.43E+08 | | | | | | |
| P49222 | EPB42 | erythrocyte membrane protein band 4.2 | Plasma Membrane | transporter | 3.76E+08 | 4.98E+08 | 6.21E+08 | | | | | | |
| P49222 | EPB42 | erythrocyte membrane protein band 4.2 | Plasma Membrane | transporter | 9.65E+08 | 1.09E+09 | 1.17E+09 | | | | | | |
| P49222 | EPB42 | erythrocyte membrane protein band 4.2 | Plasma Membrane | transporter | 7.10E+08 | 5.98E+08 | 7.10E+08 | | | | | | |
| P49222 | EPB42 | erythrocyte membrane protein band 4.2 | Plasma Membrane | transporter | 6.96E+08 | 6.05E+08 | 5.00E+08 | | | | | | |
| P49222 | EPB42 | erythrocyte membrane protein band 4.2 | Plasma Membrane | transporter | 5.80E+08 | 5.30E+08 | 6.06E+08 | | | | | | |
| P49222 | EPB42 | erythrocyte membrane protein band 4.2 | Plasma Membrane | transporter | 5.71E+08 | 4.65E+08 | 4.97E+08 | | | | | | |
| P49222 | EPB42 | erythrocyte membrane protein band 4.2 | Plasma Membrane | transporter | 5.37E+08 | 5.95E+08 | 6.78E+08 | | | | | | |
| P49222 | EPB42 | erythrocyte membrane protein band 4.2 | Plasma Membrane | transporter | 5.80E+08 | 5.89E+08 | 4.54E+08 | | | | | | |
| P49222 | EPB42 | erythrocyte membrane protein band 4.2 | Plasma Membrane | transporter | 4.49E+08 | 4.86E+08 | 4.94E+08 | | | | | | |
| P49222 | EPB42 | erythrocyte membrane protein band 4.2 | Plasma Membrane | transporter | 4.07E+08 | 4.08E+08 | 4.23E+08 | | | | | | |
| P49222 | EPB42 | erythrocyte membrane protein band 4.2 | Plasma Membrane | transporter | 2.47E+08 | 2.59E+08 | 2.63E+08 | | | | | | |
| P49222 | EPB42 | erythrocyte membrane protein band 4.2 | Plasma Membrane | transporter | 3.54E+08 | 2.78E+08 | 2.93E+08 | | | | | | |
| P49222 | EPB42 | erythrocyte membrane protein band 4.2 | Plasma Membrane | transporter | 3.15E+08 | 2.54E+08 | 3.00E+08 | | | | | | |
| P49222 | EPB42 | erythrocyte membrane protein band 4.2 | Plasma Membrane | transporter | 3.01E+08 | 3.53E+08 | 3.82E+08 | | | | | | |
| P49222 | EPB42 | erythrocyte membrane protein band 4.2 | Plasma Membrane | transporter | 2.61E+08 | 2.66E+08 | 2.65E+08 | | | | | | |
| P49222 | EPB42 | erythrocyte membrane protein band 4.2 | Plasma Membrane | transporter | 1.23E+08 | 1.76E+08 | 2.15E+08 | | | | | | |

| | | | | | | | | | | | | | |
|---|---|---|---|---|---|---|---|---|---|---|---|---|---|
| P49222 | EPB42 | erythrocyte membrane protein band 4.2 | Plasma Membrane | transporter | 2.08E+08 | 1.63E+08 | 1.74E+08 | | | | | | |
| P49222 | EPB42 | erythrocyte membrane protein band 4.2 | Plasma Membrane | transporter | 1.46E+08 | 2.21E+08 | 2.60E+08 | | | | | | |
| P49222 | EPB42 | erythrocyte membrane protein band 4.2 | Plasma Membrane | transporter | 1.41E+08 | 1.41E+08 | 1.66E+08 | | | | | | |
| P49222 | EPB42 | erythrocyte membrane protein band 4.2 | Plasma Membrane | transporter | 1.01E+08 | 1.65E+08 | 2.33E+08 | | | | | | |
| P49222 | EPB42 | erythrocyte membrane protein band 4.2 | Plasma Membrane | transporter | 1.45E+08 | 1.38E+08 | 1.20E+08 | | | | | | |
| P49222 | EPB42 | erythrocyte membrane protein band 4.2 | Plasma Membrane | transporter | 9.45E+07 | 1.47E+08 | 7.52E+07 | | | | | | |
| P49222 | EPB42 | erythrocyte membrane protein band 4.2 | Plasma Membrane | transporter | 1.50E+08 | 1.47E+08 | 1.46E+08 | | | | | | |
| P49222 | EPB42 | erythrocyte membrane protein band 4.2 | Plasma Membrane | transporter | 1.20E+08 | 6.71E+07 | 5.28E+07 | | | | | | |
| P49222 | EPB42 | erythrocyte membrane protein band 4.2 | Plasma Membrane | transporter | 5.63E+07 | 4.52E+07 | 3.41E+07 | | | | | | |
| P49222 | EPB42 | erythrocyte membrane protein band 4.2 | Plasma Membrane | transporter | 5.50E+07 | 5.06E+07 | 5.14E+07 | | | | | | |
| P49222 | EPB42 | erythrocyte membrane protein band 4.2 | Plasma Membrane | transporter | 6.60E+07 | 8.93E+07 | 1.20E+08 | | | | | | |
| P49222 | EPB42 | erythrocyte membrane protein band 4.2 | Plasma Membrane | transporter | 4.17E+07 | 5.39E+07 | 6.38E+07 | | | | | | |
| P49222 | EPB42 | erythrocyte membrane protein band 4.2 | Plasma Membrane | transporter | 2.92E+07 | 2.85E+07 | 3.45E+07 | | | | | | |
| P49222 | EPB42 | erythrocyte membrane protein band 4.2 | Plasma Membrane | transporter | 2.09E+07 | 6158414.5 | 4294333 | | | | | | |
| P49222 | EPB42 | erythrocyte membrane protein band 4.2 | Plasma Membrane | transporter | 3.15E+07 | 3.71E+07 | 5.16E+07 | | | | | | |
| P49222 | EPB42 | erythrocyte membrane protein band 4.2 | Plasma Membrane | transporter | 2.76E+07 | 3.74E+07 | 6.43E+07 | | | | | | |
| P49222 | EPB42 | erythrocyte membrane protein band 4.2 | Plasma Membrane | transporter | 3.14E+07 | 5.23E+07 | 5.81E+07 | | | | | | |
| P49222 | EPB42 | erythrocyte membrane protein band 4.2 | Plasma Membrane | transporter | 1.60E+07 | 2.41E+07 | 1.79E+07 | | | | | | |
| P49222 | EPB42 | erythrocyte membrane protein band 4.2 | Plasma Membrane | transporter | 9591486 | 1.61E+07 | 1.46E+07 | | | | | | |
| P49222 | EPB42 | erythrocyte membrane protein band 4.2 | Plasma Membrane | transporter | 1.73E+07 | 2614733.25 | 240089.094 | | | | | | |
| P49222 | EPB42 | erythrocyte membrane protein band 4.2 | Plasma Membrane | transporter | 1.41E+09 | 1.06E+09 | 1.15E+09 | | | | | | |
| P49222 | EPB42 | erythrocyte membrane protein band 4.2 | Plasma Membrane | transporter | 1.51E+07 | 1.90E+07 | 1.90E+07 | | | | | | |
| P49222 | EPB42 | erythrocyte membrane protein band 4.2 | Plasma Membrane | transporter | 7338222 | 6723193.5 | 1523967.25 | | | | | | |
| P49222 | EPB42 | erythrocyte membrane protein band 4.2 | Plasma Membrane | transporter | 3210866 | 4413334.5 | 3361880.25 | | | | | | |
| P49222 | EPB42 | erythrocyte membrane protein band 4.2 | Plasma Membrane | transporter | 9738040 | 2210460.75 | 652844.688 | | | | | | |
| P49222 | EPB42 | erythrocyte membrane protein band 4.2 | Plasma Membrane | transporter | 3077183 | 5369228 | 1.14E+07 | | | | | | |
| P49222 | EPB42 | erythrocyte membrane protein band 4.2 | Plasma Membrane | transporter | 6405197.5 | 5147934 | 1.55E+07 | | | | | | |
| P49222 | EPB42 | erythrocyte membrane protein band 4.2 | Plasma Membrane | transporter | 6506120 | 2097466.25 | 1394665.375 | | | | | | |
| P49222 | EPB42 | erythrocyte membrane protein band 4.2 | Plasma Membrane | transporter | 6159519 | 8294555.5 | 3711194.25 | | | | | | |
| P49222 | EPB42 | erythrocyte membrane protein band 4.2 | Plasma Membrane | transporter | 2998901.5 | 5909403.5 | 1.39E+07 | | | | | | |
| P49222 | EPB42 | erythrocyte membrane protein band 4.2 | Plasma Membrane | transporter | 2408124.75 | 404280.75 | 413492.156 | | | | | | |
| P49222 | EPB42 | erythrocyte membrane protein band 4.2 | Plasma Membrane | transporter | 3723253.75 | 9444190 | 1.84E+07 | | | | | | |
| P49222 | EPB42 | erythrocyte membrane protein band 4.2 | Plasma Membrane | transporter | 1139658.25 | 1449372.25 | 1047387.938 | | | | | | |
| P49222 | EPB42 | erythrocyte membrane protein band 4.2 | Plasma Membrane | transporter | 1042587.688 | 1684375 | 1362764 | | | | | | |
| P49222 | EPB42 | erythrocyte membrane protein band 4.2 | Plasma Membrane | transporter | 1280957.125 | 214038.156 | 503895.312 | | | | | | |
| P49222 | EPB42 | erythrocyte membrane protein band 4.2 | Plasma Membrane | transporter | 9.32E+07 | 5.73E+07 | 6.13E+07 | | | | | | |
| P49222 | EPB42 | erythrocyte membrane protein band 4.2 | Plasma Membrane | transporter | 338816.156 | 211946.375 | 156639.875 | | | | | | |
| P49222 | EPB42 | erythrocyte membrane protein band 4.2 | Plasma Membrane | transporter | 405282.656 | 1067610.75 | 3545180.25 | P49222 | EPB42 | 371584910 | 518741628 | 37436952 | 192 |
| Q80VP1 | EPN1 | epsin 1 | Plasma Membrane | other | 3736313.5 | 3320457.25 | 2586142.5 | | | | | | |
| Q80VP1 | EPN1 | epsin 1 | Plasma Membrane | other | 4975982 | 3407109.5 | 5468776.5 | | | | | | |
| Q80VP1 | EPN1 | epsin 1 | Plasma Membrane | other | 2180174.75 | 1585671.875 | 1384996.5 | | | | | | |
| Q80VP1 | EPN1 | epsin 1 | Plasma Membrane | other | 2656804.5 | 1313511 | 1369425.625 | Q80VP1 | EPN1 | 3387319 | 1399496 | 404000 | 12 |
| Q8CGC7 | EPRS | glutamyl-prolyl-tRNA synthetase | Cytoplasm | enzyme | 708733.188 | 876996.375 | 651744.938 | | | | | | |
| Q8CGC7 | EPRS | glutamyl-prolyl-tRNA synthetase | Cytoplasm | enzyme | 419330.188 | 742635.375 | 516299.188 | Q8CGC7 | EPRS | 564032 | 164076 | 66984 | 6 |
| P42567 | EPS15 | epidermal growth factor receptor pathway substrate 15 | Cytoplasm | other | 5062928.5 | 3765387.25 | 3343794.5 | | | | | | |
| P42567 | EPS15 | epidermal growth factor receptor pathway substrate 15 | Cytoplasm | other | 2986229.75 | 4144854.75 | 4283245 | | | | | | |
| P42567 | EPS15 | epidermal growth factor receptor pathway substrate 15 | Cytoplasm | other | 3722690.5 | 2956643.75 | 3150714.75 | | | | | | |
| P42567 | EPS15 | epidermal growth factor receptor pathway substrate 15 | Cytoplasm | other | 3864756.75 | 3895199 | 2549947 | | | | | | |
| P42567 | EPS15 | epidermal growth factor receptor pathway substrate 15 | Cytoplasm | other | 2842143 | 2412838.75 | 2369328.25 | P42567 | EPS15 | 3695750 | 769720 | 198741 | 15 |
| Q60902 | EPS15L1 | epidermal growth factor receptor pathway substrate 15-like 1 | Plasma Membrane | other | 308605.438 | 405355.625 | 168239.703 | Q60902 | EPS15L1 | 308605 | 119225 | 68834 | 3 |
| Q9EQH2 | ERAP1 | endoplasmic reticulum aminopeptidase 1 | Extracellular Space | peptidase | 2341109 | 2373269.25 | 1999480.875 | Q9EQH2 | ERAP1 | 2341109 | 207148 | 119597 | 3 |
| Q80TH2 | ERBB2IP | erbb2 interacting protein | Plasma Membrane | other | 3038969 | 5734718 | 6091877 | | | | | | |
| Q80TH2 | ERBB2IP | erbb2 interacting protein | Plasma Membrane | other | 2747693.75 | 5686540.5 | 4893912 | | | | | | |
| Q80TH2 | ERBB2IP | erbb2 interacting protein | Plasma Membrane | other | 2676095 | 2711149.75 | 4284075 | | | | | | |
| Q80TH2 | ERBB2IP | erbb2 interacting protein | Plasma Membrane | other | 2463243.25 | 2750174 | 3109023 | | | | | | |
| Q80TH2 | ERBB2IP | erbb2 interacting protein | Plasma Membrane | other | 2359638 | 2923364.25 | 3523079.25 | | | | | | |
| Q80TH2 | ERBB2IP | erbb2 interacting protein | Plasma Membrane | other | 1905387.125 | 1958712.125 | 2232914 | | | | | | |
| Q80TH2 | ERBB2IP | erbb2 interacting protein | Plasma Membrane | other | 1884515.75 | 3291206.25 | 3196567 | | | | | | |
| Q80TH2 | ERBB2IP | erbb2 interacting protein | Plasma Membrane | other | 1142819.875 | 1500967.625 | 1819283.375 | Q80TH2 | ERBB2IP | 2277295 | 1350869 | 275745 | 24 |
| Q8BFZ9 | ERLIN2 | ER lipid raft associated 2 | Plasma Membrane | other | 1710282.375 | 2116018.25 | 3738291.75 | Q8BFZ9 | ERLIN2 | 1710282 | 1073096 | 619552 | 3 |
| Q9JLN5 | ERMAP | erythroblast membrane-associated protein (Scianna blood group) | Cytoplasm | other | 2.02E+08 | 2.12E+08 | 2.08E+08 | | | | | | |
| Q9JLN5 | ERMAP | erythroblast membrane-associated protein (Scianna blood group) | Cytoplasm | other | 1.09E+08 | 9.82E+07 | 1.22E+08 | | | | | | |
| Q9JLN5 | ERMAP | erythroblast membrane-associated protein (Scianna blood group) | Cytoplasm | other | 9.13E+07 | 9.10E+07 | 8.54E+07 | | | | | | |
| Q9JLN5 | ERMAP | erythroblast membrane-associated protein (Scianna blood group) | Cytoplasm | other | 9.36E+07 | 9.59E+07 | 1.05E+08 | | | | | | |
| Q9JLN5 | ERMAP | erythroblast membrane-associated protein (Scianna blood group) | Cytoplasm | other | 7.86E+07 | 7.43E+07 | 8.68E+07 | | | | | | |
| Q9JLN5 | ERMAP | erythroblast membrane-associated protein (Scianna blood group) | Cytoplasm | other | 8.62E+07 | 9.14E+07 | 9.09E+07 | | | | | | |
| Q9JLN5 | ERMAP | erythroblast membrane-associated protein (Scianna blood group) | Cytoplasm | other | 4.92E+07 | 6.89E+07 | 6.05E+07 | | | | | | |
| Q9JLN5 | ERMAP | erythroblast membrane-associated protein (Scianna blood group) | Cytoplasm | other | 4.34E+07 | 4.20E+07 | 4.07E+07 | | | | | | |
| Q9JLN5 | ERMAP | erythroblast membrane-associated protein (Scianna blood group) | Cytoplasm | other | 3.41E+07 | 3.01E+07 | 3.24E+07 | | | | | | |
| Q9JLN5 | ERMAP | erythroblast membrane-associated protein (Scianna blood group) | Cytoplasm | other | 2.81E+07 | 2.42E+07 | 2.76E+07 | | | | | | |
| Q9JLN5 | ERMAP | erythroblast membrane-associated protein (Scianna blood group) | Cytoplasm | other | 2.67E+07 | 2.13E+07 | 2.31E+07 | | | | | | |
| Q9JLN5 | ERMAP | erythroblast membrane-associated protein (Scianna blood group) | Cytoplasm | other | 1.55E+07 | 1.39E+07 | 1.37E+07 | | | | | | |
| Q9JLN5 | ERMAP | erythroblast membrane-associated protein (Scianna blood group) | Cytoplasm | other | 1.47E+07 | 1.40E+07 | 1.57E+07 | | | | | | |
| Q9JLN5 | ERMAP | erythroblast membrane-associated protein (Scianna blood group) | Cytoplasm | other | 1.04E+07 | 1.32E+07 | 1.89E+07 | | | | | | |
| Q9JLN5 | ERMAP | erythroblast membrane-associated protein (Scianna blood group) | Cytoplasm | other | 1.19E+07 | 1.12E+07 | 1.32E+07 | | | | | | |
| Q9JLN5 | ERMAP | erythroblast membrane-associated protein (Scianna blood group) | Cytoplasm | other | 9625690 | 9217048 | 7095164.5 | | | | | | |
| Q9JLN5 | ERMAP | erythroblast membrane-associated protein (Scianna blood group) | Cytoplasm | other | 6711151 | 7719868.5 | 7664994.5 | | | | | | |
| Q9JLN5 | ERMAP | erythroblast membrane-associated protein (Scianna blood group) | Cytoplasm | other | 5233571.5 | 6258080.5 | 6310317.5 | | | | | | |
| Q9JLN5 | ERMAP | erythroblast membrane-associated protein (Scianna blood group) | Cytoplasm | other | 3828966 | 9789264 | 1.26E+07 | | | | | | |
| Q9JLN5 | ERMAP | erythroblast membrane-associated protein (Scianna blood group) | Cytoplasm | other | 8276920 | 6588652 | 8421504 | Q9JLN5 | ERMAP | 46415815 | 50707144 | 6546264 | 60 |
| Q8R180 | ERO1A | endoplasmic reticulum oxidoreductase alpha | Cytoplasm | enzyme | 1584849 | 1450664.25 | 2089256.125 | Q8R180 | ERO1A | 1584849 | 336708 | 194398 | 3 |
| P57759 | ERP29 | endoplasmic reticulum protein 29 | Cytoplasm | transporter | 8964060 | 9975105 | 1.01E+07 | | | | | | |
| P57759 | ERP29 | endoplasmic reticulum protein 29 | Cytoplasm | transporter | 6525965 | 6684072 | 7397643 | | | | | | |
| P57759 | ERP29 | endoplasmic reticulum protein 29 | Cytoplasm | transporter | 3111754.75 | 4229073 | 5543133 | | | | | | |
| P57759 | ERP29 | endoplasmic reticulum protein 29 | Cytoplasm | transporter | 3269757 | 3593905 | 4031018.75 | P57759 | ERP29 | 5467884 | 2565789 | 740680 | 12 |
| Q9D1Q6 | ERP44 | endoplasmic reticulum protein 44 | Cytoplasm | enzyme | 1.24E+07 | 1.21E+07 | 1.54E+07 | | | | | | |
| Q9D1Q6 | ERP44 | endoplasmic reticulum protein 44 | Cytoplasm | enzyme | 9564987 | 1.30E+07 | 1.05E+07 | | | | | | |
| Q9D1Q6 | ERP44 | endoplasmic reticulum protein 44 | Cytoplasm | enzyme | 8626735 | 7583964 | 1.19E+07 | | | | | | |
| Q9D1Q6 | ERP44 | endoplasmic reticulum protein 44 | Cytoplasm | enzyme | 6182940 | 8180255.5 | 1.12E+07 | | | | | | |
| Q9D1Q6 | ERP44 | endoplasmic reticulum protein 44 | Cytoplasm | enzyme | 1758750.375 | 5518992.5 | 7785898.5 | | | | | | |
| Q9D1Q6 | ERP44 | endoplasmic reticulum protein 44 | Cytoplasm | enzyme | 2827977.5 | 2472200 | 1172670 | Q9D1Q6 | ERP44 | 6895232 | 4222641 | 995286 | 18 |

| | | | | | | | | | | | | | |
|---|---|---|---|---|---|---|---|---|---|---|---|---|---|
| Q3U7R1 | ESYT1 | extended synaptotagmin-like protein 1 | Cytoplasm | other | 2.06E+07 | 2.38E+07 | 2.23E+07 | | | | | | |
| Q3U7R1 | ESYT1 | extended synaptotagmin-like protein 1 | Cytoplasm | other | 3.17E+07 | 3.75E+07 | 2.96E+07 | | | | | | |
| Q3U7R1 | ESYT1 | extended synaptotagmin-like protein 1 | Cytoplasm | other | 2.83E+07 | 2.99E+07 | 3.12E+07 | | | | | | |
| Q3U7R1 | ESYT1 | extended synaptotagmin-like protein 1 | Cytoplasm | other | 1.71E+07 | 1.81E+07 | 2.04E+07 | | | | | | |
| Q3U7R1 | ESYT1 | extended synaptotagmin-like protein 1 | Cytoplasm | other | 1.71E+07 | 1.83E+07 | 2.20E+07 | | | | | | |
| Q3U7R1 | ESYT1 | extended synaptotagmin-like protein 1 | Cytoplasm | other | 1.60E+07 | 1.51E+07 | 1.65E+07 | | | | | | |
| Q3U7R1 | ESYT1 | extended synaptotagmin-like protein 1 | Cytoplasm | other | 1.14E+07 | 1.41E+07 | 1.98E+07 | | | | | | |
| Q3U7R1 | ESYT1 | extended synaptotagmin-like protein 1 | Cytoplasm | other | 1.66E+07 | 1.87E+07 | 1.35E+07 | | | | | | |
| Q3U7R1 | ESYT1 | extended synaptotagmin-like protein 1 | Cytoplasm | other | 1.29E+07 | 1.48E+07 | 1.47E+07 | | | | | | |
| Q3U7R1 | ESYT1 | extended synaptotagmin-like protein 1 | Cytoplasm | other | 1.20E+07 | 1.44E+07 | 1.40E+07 | | | | | | |
| Q3U7R1 | ESYT1 | extended synaptotagmin-like protein 1 | Cytoplasm | other | 9833691 | 9673808 | 9788965 | | | | | | |
| Q3U7R1 | ESYT1 | extended synaptotagmin-like protein 1 | Cytoplasm | other | 1.35E+07 | 1.45E+07 | 1.73E+07 | | | | | | |
| Q3U7R1 | ESYT1 | extended synaptotagmin-like protein 1 | Cytoplasm | other | 1.03E+07 | 1.38E+07 | 9905610 | | | | | | |
| Q3U7R1 | ESYT1 | extended synaptotagmin-like protein 1 | Cytoplasm | other | 9622582 | 9509074 | 9944121 | | | | | | |
| Q3U7R1 | ESYT1 | extended synaptotagmin-like protein 1 | Cytoplasm | other | 6859964 | 8957622 | 1.01E+07 | | | | | | |
| Q3U7R1 | ESYT1 | extended synaptotagmin-like protein 1 | Cytoplasm | other | 7257628 | 8730506 | 8158985.5 | | | | | | |
| Q3U7R1 | ESYT1 | extended synaptotagmin-like protein 1 | Cytoplasm | other | 9994703 | 9634285 | 8953573 | | | | | | |
| Q3U7R1 | ESYT1 | extended synaptotagmin-like protein 1 | Cytoplasm | other | 7748324.5 | 8882481 | 8565916 | | | | | | |
| Q3U7R1 | ESYT1 | extended synaptotagmin-like protein 1 | Cytoplasm | other | 6394102.5 | 8294631 | 8146874.5 | | | | | | |
| Q3U7R1 | ESYT1 | extended synaptotagmin-like protein 1 | Cytoplasm | other | 9770506 | 7337683 | 2671841.5 | | | | | | |
| Q3U7R1 | ESYT1 | extended synaptotagmin-like protein 1 | Cytoplasm | other | 6859476.5 | 6245917 | 5413835.5 | | | | | | |
| Q3U7R1 | ESYT1 | extended synaptotagmin-like protein 1 | Cytoplasm | other | 7768465.5 | 7702094.5 | 6906032 | | | | | | |
| Q3U7R1 | ESYT1 | extended synaptotagmin-like protein 1 | Cytoplasm | other | 5723126.5 | 7577324.5 | 6612533 | | | | | | |
| Q3U7R1 | ESYT1 | extended synaptotagmin-like protein 1 | Cytoplasm | other | 5968559 | 6889612 | 7331787 | | | | | | |
| Q3U7R1 | ESYT1 | extended synaptotagmin-like protein 1 | Cytoplasm | other | 4858853 | 9993154 | 1.00E+07 | | | | | | |
| Q3U7R1 | ESYT1 | extended synaptotagmin-like protein 1 | Cytoplasm | other | 4570764.5 | 5789567.5 | 4413075.5 | | | | | | |
| Q3U7R1 | ESYT1 | extended synaptotagmin-like protein 1 | Cytoplasm | other | 3174350.5 | 4553882 | 4425692.5 | | | | | | |
| Q3U7R1 | ESYT1 | extended synaptotagmin-like protein 1 | Cytoplasm | other | 2201673.25 | 4765957.5 | 3785653.75 | | | | | | |
| Q3U7R1 | ESYT1 | extended synaptotagmin-like protein 1 | Cytoplasm | other | 727317.188 | 1619838.125 | 2550387.75 | Q3U7R1 | ESYT1 | 10921520 | 7472201 | 801104 | 87 |
| Q3TZZ7 | ESYT2 | extended synaptotagmin-like protein 2 | Plasma Membrane | other | 1.11E+07 | 1.29E+07 | 1.47E+07 | | | | | | |
| Q3TZZ7 | ESYT2 | extended synaptotagmin-like protein 2 | Plasma Membrane | other | 1.22E+07 | 1.18E+07 | 1.08E+07 | | | | | | |
| Q3TZZ7 | ESYT2 | extended synaptotagmin-like protein 2 | Plasma Membrane | other | 1.22E+07 | 1.28E+07 | 2.02E+07 | | | | | | |
| Q3TZZ7 | ESYT2 | extended synaptotagmin-like protein 2 | Plasma Membrane | other | 1.22E+07 | 1.06E+07 | 1.12E+07 | | | | | | |
| Q3TZZ7 | ESYT2 | extended synaptotagmin-like protein 2 | Plasma Membrane | other | 8446045 | 9636803 | 6994652.5 | | | | | | |
| Q3TZZ7 | ESYT2 | extended synaptotagmin-like protein 2 | Plasma Membrane | other | 9012310 | 9979616 | 8409012 | | | | | | |
| Q3TZZ7 | ESYT2 | extended synaptotagmin-like protein 2 | Plasma Membrane | other | 7685325.5 | 9010517 | 8307373.5 | | | | | | |
| Q3TZZ7 | ESYT2 | extended synaptotagmin-like protein 2 | Plasma Membrane | other | 9611348 | 9872539 | 8620384 | | | | | | |
| Q3TZZ7 | ESYT2 | extended synaptotagmin-like protein 2 | Plasma Membrane | other | 8424433 | 8842540 | 9314494 | | | | | | |
| Q3TZZ7 | ESYT2 | extended synaptotagmin-like protein 2 | Plasma Membrane | other | 4927933 | 8142651.5 | 9463301 | | | | | | |
| Q3TZZ7 | ESYT2 | extended synaptotagmin-like protein 2 | Plasma Membrane | other | 7170236 | 1.04E+07 | 1.31E+07 | | | | | | |
| Q3TZZ7 | ESYT2 | extended synaptotagmin-like protein 2 | Plasma Membrane | other | 6588702 | 5382560.5 | 4546140 | | | | | | |
| Q3TZZ7 | ESYT2 | extended synaptotagmin-like protein 2 | Plasma Membrane | other | 6658706.5 | 6488111.5 | 7386623 | | | | | | |
| Q3TZZ7 | ESYT2 | extended synaptotagmin-like protein 2 | Plasma Membrane | other | 6289852.5 | 5684226.5 | 4979639.5 | | | | | | |
| Q3TZZ7 | ESYT2 | extended synaptotagmin-like protein 2 | Plasma Membrane | other | 7803318.5 | 1.05E+07 | 7763294 | | | | | | |
| Q3TZZ7 | ESYT2 | extended synaptotagmin-like protein 2 | Plasma Membrane | other | 5276463 | 7557697 | 6449274.5 | | | | | | |
| Q3TZZ7 | ESYT2 | extended synaptotagmin-like protein 2 | Plasma Membrane | other | 4202602.5 | 4054785.75 | 4343883 | | | | | | |
| Q3TZZ7 | ESYT2 | extended synaptotagmin-like protein 2 | Plasma Membrane | other | 1903948.375 | 5299306 | 6314217.5 | | | | | | |
| Q3TZZ7 | ESYT2 | extended synaptotagmin-like protein 2 | Plasma Membrane | other | 1499818.375 | 1870682.375 | 1189005.75 | | | | | | |
| Q3TZZ7 | ESYT2 | extended synaptotagmin-like protein 2 | Plasma Membrane | other | 382608 | 518025.406 | 348625.812 | Q3TZZ7 | ESYT2 | 7180683 | 3800883 | 490692 | 60 |
| Q99LC5 | ETFA | electron-transfer-flavoprotein, alpha polypeptide | Cytoplasm | transporter | 5129839 | 1.30E+07 | 5205381 | | | | | | |
| Q99LC5 | ETFA | electron-transfer-flavoprotein, alpha polypeptide | Cytoplasm | transporter | 3567693.5 | 8362985.5 | 4462214.5 | | | | | | |
| Q99LC5 | ETFA | electron-transfer-flavoprotein, alpha polypeptide | Cytoplasm | transporter | 4584026.5 | 1.11E+07 | 4029188 | | | | | | |
| Q99LC5 | ETFA | electron-transfer-flavoprotein, alpha polypeptide | Cytoplasm | transporter | 3722168.5 | 7842243.5 | 3795092.75 | | | | | | |
| Q99LC5 | ETFA | electron-transfer-flavoprotein, alpha polypeptide | Cytoplasm | transporter | 1799219.25 | 2644014.75 | 1644055.5 | | | | | | |
| Q99LC5 | ETFA | electron-transfer-flavoprotein, alpha polypeptide | Cytoplasm | transporter | 1902885.25 | 4625585 | 1478898.375 | | | | | | |
| Q99LC5 | ETFA | electron-transfer-flavoprotein, alpha polypeptide | Cytoplasm | transporter | 1653928.125 | 2915384.75 | 1096736.5 | | | | | | |
| Q99LC5 | ETFA | electron-transfer-flavoprotein, alpha polypeptide | Cytoplasm | transporter | 1048293 | 1626590.375 | 1318153.75 | | | | | | |
| Q99LC5 | ETFA | electron-transfer-flavoprotein, alpha polypeptide | Cytoplasm | transporter | 515592.25 | 1238269 | 586322.938 | Q99LC5 | ETFA | 2658183 | 3156680 | 607503 | 27 |
| Q9DCW4 | ETFB | electron-transfer-flavoprotein, beta polypeptide | Cytoplasm | transporter | 8526825 | 1.72E+07 | 7951781 | | | | | | |
| Q9DCW4 | ETFB | electron-transfer-flavoprotein, beta polypeptide | Cytoplasm | transporter | 2844558.75 | 5332015 | 2045885.5 | | | | | | |
| Q9DCW4 | ETFB | electron-transfer-flavoprotein, beta polypeptide | Cytoplasm | transporter | 1798040.5 | 4119531.75 | 1706019.25 | | | | | | |
| Q9DCW4 | ETFB | electron-transfer-flavoprotein, beta polypeptide | Cytoplasm | transporter | 1384229.375 | 2064389.125 | 880638.75 | Q9DCW4 | ETFB | 3638413 | 4705759 | 1358436 | 12 |
| Q6DIA2 | EXOC3L4 | exocyst complex component 3-like 4 | Cytoplasm | other | 112981.023 | 554220 | 637445.688 | Q6DIA2 | EXOC3L4 | 112981 | 281863 | 162734 | 3 |
| P26040 | EZR | ezrin | Plasma Membrane | other | 2.29E+07 | 2.28E+07 | 1.98E+07 | | | | | | |
| P26040 | EZR | ezrin | Plasma Membrane | other | 1.90E+07 | 1.51E+07 | 1.19E+07 | | | | | | |
| P26040 | EZR | ezrin | Plasma Membrane | other | 9726513 | 8733251 | 1.05E+07 | | | | | | |
| P26040 | EZR | ezrin | Plasma Membrane | other | 6388711.5 | 5993907 | 5225611.5 | | | | | | |
| P26040 | EZR | ezrin | Plasma Membrane | other | 6318128.5 | 5478182.5 | 7074061.5 | | | | | | |
| P26040 | EZR | ezrin | Plasma Membrane | other | 2528826.5 | 1638354.5 | 1115974.125 | | | | | | |
| P26040 | EZR | ezrin | Plasma Membrane | other | 2484739.25 | 3528940.75 | 3562975 | | | | | | |
| P26040 | EZR | ezrin | Plasma Membrane | other | 2860401.5 | 2529237 | 2829503.25 | | | | | | |
| P26040 | EZR | ezrin | Plasma Membrane | other | 3235182.5 | 2276486.5 | 2024031.125 | | | | | | |
| P26040 | EZR | ezrin | Plasma Membrane | other | 1223912.125 | 236907.672 | 4.70E-06 | P26040 | EZR | 7668641 | 6701757 | 1223568 | 30 |
| O88792 | F11R | F11 receptor | Plasma Membrane | other | 1186680.75 | 2326471.5 | 5031984 | | | | | | |
| O88792 | F11R | F11 receptor | Plasma Membrane | other | 451908.969 | 1058355 | 1140006.5 | | | | | | |
| O88792 | F11R | F11 receptor | Plasma Membrane | other | 0 | 525033.5 | 1145355.5 | | | | | | |
| O88792 | F11R | F11 receptor | Plasma Membrane | other | 4018.124 | 279302.5 | 2038705.5 | | | | | | |
| O88792 | F11R | F11 receptor | Plasma Membrane | other | 62485.176 | 649452.188 | 1183408.375 | O88792 | F11R | 341019 | 1280565 | 330640 | 15 |
| P19221 | F2 | coagulation factor II (thrombin) | Extracellular Space | peptidase | 5378170.5 | 6306303 | 5401113.5 | P19221 | F2 | 5378171 | 529359 | 305625 | 3 |
| O88783 | F5 | coagulation factor V (proaccelerin, labile factor) | Plasma Membrane | enzyme | 379279.969 | 1194496.5 | 2123570 | | | | | | |
| O88783 | F5 | coagulation factor V (proaccelerin, labile factor) | Plasma Membrane | enzyme | 180407.859 | 258390.109 | 801122.75 | | | | | | |
| O88783 | F5 | coagulation factor V (proaccelerin, labile factor) | Plasma Membrane | enzyme | 4.70E-06 | 100563.109 | 398842.781 | O88783 | F5 | 186563 | 680658 | 226886 | 9 |
| Q920L1 | FADS1 | fatty acid desaturase 1 | Plasma Membrane | enzyme | 2792970.5 | 2836394.75 | 4514790 | Q920L1 | FADS1 | 2792971 | 981798 | 566841 | 3 |
| P54731 | FAF1 | Fas (TNFRSF6) associated factor 1 | Nucleus | other | 2944920.75 | 2120253.25 | 2706325 | | | | | | |
| P54731 | FAF1 | Fas (TNFRSF6) associated factor 1 | Nucleus | other | 2605160.25 | 1867886.875 | 2340934.5 | | | | | | |
| P54731 | FAF1 | Fas (TNFRSF6) associated factor 1 | Nucleus | other | 2331505.75 | 2089662.25 | 2305986 | P54731 | FAF1 | 2627196 | 335116 | 111705 | 9 |
| Q8R0F8 | FAHD1 | fumarylacetoacetate hydrolase domain containing 1 | Cytoplasm | enzyme | 1967182.375 | 4879025 | 1670588.5 | | | | | | |
| Q8R0F8 | FAHD1 | fumarylacetoacetate hydrolase domain containing 1 | Cytoplasm | enzyme | 1807420.875 | 4803564 | 1836210.75 | Q8R0F8 | FAHD1 | 1887302 | 1563035 | 638106 | 6 |
| Q6NS82 | FAM134A | family with sequence similarity 134, member A | Extracellular Space | other | 1995553.375 | 2658611.25 | 2237656.25 | Q6NS82 | FAM134A | 1995553 | 335525 | 193716 | 3 |
| Q9D8B6 | FAM210B | family with sequence similarity 210, member B | Extracellular Space | other | 2717135.5 | 5090780 | 1970766.125 | | | | | | |
| Q9D8B6 | FAM210B | family with sequence similarity 210, member B | Extracellular Space | other | 243894.375 | 772773.562 | 273935.375 | Q9D8B6 | FAM210B | 1480515 | 1870453 | 763609 | 6 |

| | | | | | | | | | | | | | |
|---|---|---|---|---|---|---|---|---|---|---|---|---|---|
| Q9CYH2 | FAM213A | family with sequence similarity 213, member A | Extracellular Space | other | 1.42E+08 | 1.74E+08 | 1.93E+08 | | | | | | |
| Q9CYH2 | FAM213A | family with sequence similarity 213, member A | Extracellular Space | other | 9.79E+07 | 1.27E+08 | 1.13E+08 | | | | | | |
| Q9CYH2 | FAM213A | family with sequence similarity 213, member A | Extracellular Space | other | 6.02E+07 | 1.05E+08 | 1.19E+08 | | | | | | |
| Q9CYH2 | FAM213A | family with sequence similarity 213, member A | Extracellular Space | other | 4.95E+07 | 1.10E+08 | 1.34E+08 | | | | | | |
| Q9CYH2 | FAM213A | family with sequence similarity 213, member A | Extracellular Space | other | 6.79E+07 | 5.04E+07 | 4.72E+07 | | | | | | |
| Q9CYH2 | FAM213A | family with sequence similarity 213, member A | Extracellular Space | other | 3.69E+07 | 4.27E+07 | 5.70E+07 | | | | | | |
| Q9CYH2 | FAM213A | family with sequence similarity 213, member A | Extracellular Space | other | 2.54E+07 | 1.30E+07 | 5820431 | | | | | | |
| Q9CYH2 | FAM213A | family with sequence similarity 213, member A | Extracellular Space | other | 1.05E+07 | 1.97E+07 | 3.00E+07 | | | | | | |
| Q9CYH2 | FAM213A | family with sequence similarity 213, member A | Extracellular Space | other | 1.22E+07 | 1.44E+07 | 1.59E+07 | | | | | | |
| Q9CYH2 | FAM213A | family with sequence similarity 213, member A | Extracellular Space | other | 1.53E+07 | 1.16E+07 | 1.23E+07 | | | | | | |
| Q9CYH2 | FAM213A | family with sequence similarity 213, member A | Extracellular Space | other | 9362526 | 9338984 | 9280164 | | | | | | |
| Q9CYH2 | FAM213A | family with sequence similarity 213, member A | Extracellular Space | other | 1060227.875 | 1639488.25 | 1868466.875 | Q9CYH2 | FAM213A | 44020229 | 53898669 | 8983111 | 36 |
| Q9WUA2 | FARSB | phenylalanyl-tRNA synthetase, beta subunit | Cytoplasm | enzyme | 1342911.5 | 1625817.25 | 644209.188 | Q9WUA2 | FARSB | 1342912 | 505268 | 291717 | 3 |
| P19096 | FASN | fatty acid synthase | Cytoplasm | enzyme | 9204763 | 7836952 | 6644344.5 | | | | | | |
| P19096 | FASN | fatty acid synthase | Cytoplasm | enzyme | 1.31E+07 | 8577529 | 7205863.5 | | | | | | |
| P19096 | FASN | fatty acid synthase | Cytoplasm | enzyme | 1.04E+07 | 6023910.5 | 9706482 | | | | | | |
| P19096 | FASN | fatty acid synthase | Cytoplasm | enzyme | 6041744 | 4711749.5 | 3507398.25 | | | | | | |
| P19096 | FASN | fatty acid synthase | Cytoplasm | enzyme | 6039965.5 | 5755165 | 5778460.5 | | | | | | |
| P19096 | FASN | fatty acid synthase | Cytoplasm | enzyme | 6848036 | 5357500.5 | 4694360 | | | | | | |
| P19096 | FASN | fatty acid synthase | Cytoplasm | enzyme | 7049630.5 | 5423134.5 | 4527358.5 | | | | | | |
| P19096 | FASN | fatty acid synthase | Cytoplasm | enzyme | 6720673 | 4949390.5 | 6378442.5 | | | | | | |
| P19096 | FASN | fatty acid synthase | Cytoplasm | enzyme | 5791067.5 | 4322371 | 3763025 | | | | | | |
| P19096 | FASN | fatty acid synthase | Cytoplasm | enzyme | 4261728 | 2570176 | 2348645.5 | | | | | | |
| P19096 | FASN | fatty acid synthase | Cytoplasm | enzyme | 3912254 | 3170381.75 | 3262587.5 | | | | | | |
| P19096 | FASN | fatty acid synthase | Cytoplasm | enzyme | 3970267.25 | 1846254.75 | 2739565.25 | | | | | | |
| P19096 | FASN | fatty acid synthase | Cytoplasm | enzyme | 3961665.5 | 3884925.75 | 3539852.75 | | | | | | |
| P19096 | FASN | fatty acid synthase | Cytoplasm | enzyme | 4924423.5 | 3211060.5 | 2852465.75 | | | | | | |
| P19096 | FASN | fatty acid synthase | Cytoplasm | enzyme | 3903323.25 | 2473005 | 1955282.25 | | | | | | |
| P19096 | FASN | fatty acid synthase | Cytoplasm | enzyme | 2966607.75 | 2214953.5 | 2249760.75 | | | | | | |
| P19096 | FASN | fatty acid synthase | Cytoplasm | enzyme | 2999186.25 | 2985897.5 | 3052295.25 | | | | | | |
| P19096 | FASN | fatty acid synthase | Cytoplasm | enzyme | 2107179.5 | 1844459.25 | 1474283.125 | | | | | | |
| P19096 | FASN | fatty acid synthase | Cytoplasm | enzyme | 1087072.625 | 830920.75 | 479066.531 | | | | | | |
| P19096 | FASN | fatty acid synthase | Cytoplasm | enzyme | 2534761.5 | 1847807.375 | 2251613.5 | | | | | | |
| P19096 | FASN | fatty acid synthase | Cytoplasm | enzyme | 2019423.625 | 1579548.625 | 2351399.75 | | | | | | |
| P19096 | FASN | fatty acid synthase | Cytoplasm | enzyme | 648038.188 | 352083.188 | 351331.438 | | | | | | |
| P19096 | FASN | fatty acid synthase | Cytoplasm | enzyme | 749950.812 | 662367.312 | 925487 | P19096 | FASN | 4836164 | 2612816 | 314546 | 69 |
| P62862 | FAU | Finkel-Biskis-Reilly murine sarcoma virus (FBR-MuSV) ubiquitously expressed | Cytoplasm | other | 5521088.5 | 9869738 | 3759157 | P62862 | FAU | 5521089 | 3145217 | 1815892 | 3 |
| Q3U7U3 | FBXO7 | F-box protein 7 | Cytoplasm | enzyme | 2879511.75 | 1984317.5 | 1761018.875 | Q3U7U3 | FBXO7 | 2879512 | 591926 | 341749 | 3 |
| P22315 | FECH | ferrochelatase | Cytoplasm | enzyme | 6.45E+07 | 1.22E+08 | 5.37E+07 | | | | | | |
| P22315 | FECH | ferrochelatase | Cytoplasm | enzyme | 4.24E+07 | 7.80E+07 | 3.43E+07 | | | | | | |
| P22315 | FECH | ferrochelatase | Cytoplasm | enzyme | 3.38E+07 | 5.78E+07 | 1.62E+07 | | | | | | |
| P22315 | FECH | ferrochelatase | Cytoplasm | enzyme | 2.39E+07 | 4.84E+07 | 1.57E+07 | | | | | | |
| P22315 | FECH | ferrochelatase | Cytoplasm | enzyme | 1.26E+07 | 2.23E+07 | 1.07E+07 | | | | | | |
| P22315 | FECH | ferrochelatase | Cytoplasm | enzyme | 1.01E+07 | 2.02E+07 | 8234117.5 | | | | | | |
| P22315 | FECH | ferrochelatase | Cytoplasm | enzyme | 6433691 | 1.50E+07 | 1.12E+07 | | | | | | |
| P22315 | FECH | ferrochelatase | Cytoplasm | enzyme | 9730425 | 4.64E+07 | 2.60E+07 | | | | | | |
| P22315 | FECH | ferrochelatase | Cytoplasm | enzyme | 7754514.5 | 1.39E+07 | 5284643.5 | | | | | | |
| P22315 | FECH | ferrochelatase | Cytoplasm | enzyme | 4540239 | 3377303.5 | 775341.875 | | | | | | |
| P22315 | FECH | ferrochelatase | Cytoplasm | enzyme | 3796610.5 | 8256654.5 | 2587182.25 | | | | | | |
| P22315 | FECH | ferrochelatase | Cytoplasm | enzyme | 5709691.5 | 7282095 | 2821165.5 | | | | | | |
| P22315 | FECH | ferrochelatase | Cytoplasm | enzyme | 4409392 | 6541519 | 3118874.5 | | | | | | |
| P22315 | FECH | ferrochelatase | Cytoplasm | enzyme | 3054146.25 | 9135795 | 4515167 | | | | | | |
| P22315 | FECH | ferrochelatase | Cytoplasm | enzyme | 2638658.25 | 6298903 | 2912656.75 | | | | | | |
| P22315 | FECH | ferrochelatase | Cytoplasm | enzyme | 3413372.25 | 3967093 | 933938.812 | | | | | | |
| P22315 | FECH | ferrochelatase | Cytoplasm | enzyme | 2468967.75 | 5715573 | 1813150 | | | | | | |
| P22315 | FECH | ferrochelatase | Cytoplasm | enzyme | 3138010.25 | 827373.188 | 181140.109 | | | | | | |
| P22315 | FECH | ferrochelatase | Cytoplasm | enzyme | 1631812.75 | 4405646 | 1670071.875 | P22315 | FECH | 12949975 | 22818278 | 3022355 | 57 |
| Q8K1B8 | FERMT3 | fermitin family member 3 | Cytoplasm | enzyme | 1.00E+07 | 1.09E+07 | 1.44E+07 | | | | | | |
| Q8K1B8 | FERMT3 | fermitin family member 3 | Cytoplasm | enzyme | 1.20E+07 | 1.63E+07 | 1.68E+07 | | | | | | |
| Q8K1B8 | FERMT3 | fermitin family member 3 | Cytoplasm | enzyme | 6645749 | 8893568 | 9129000 | | | | | | |
| Q8K1B8 | FERMT3 | fermitin family member 3 | Cytoplasm | enzyme | 6965636 | 9704348 | 7852410.5 | | | | | | |
| Q8K1B8 | FERMT3 | fermitin family member 3 | Cytoplasm | enzyme | 7533745.5 | 8980498 | 8946375 | | | | | | |
| Q8K1B8 | FERMT3 | fermitin family member 3 | Cytoplasm | enzyme | 7633897.5 | 8176994 | 9764375 | | | | | | |
| Q8K1B8 | FERMT3 | fermitin family member 3 | Cytoplasm | enzyme | 6474151.5 | 6682532 | 7013427 | | | | | | |
| Q8K1B8 | FERMT3 | fermitin family member 3 | Cytoplasm | enzyme | 6078563 | 8223825.5 | 8671702 | | | | | | |
| Q8K1B8 | FERMT3 | fermitin family member 3 | Cytoplasm | enzyme | 4104070 | 5286356 | 6096307.5 | | | | | | |
| Q8K1B8 | FERMT3 | fermitin family member 3 | Cytoplasm | enzyme | 4754580.5 | 5727625 | 4772186 | | | | | | |
| Q8K1B8 | FERMT3 | fermitin family member 3 | Cytoplasm | enzyme | 4497127 | 5511225 | 6190877 | | | | | | |
| Q8K1B8 | FERMT3 | fermitin family member 3 | Cytoplasm | enzyme | 3574200.25 | 4254719.5 | 3880573 | | | | | | |
| Q8K1B8 | FERMT3 | fermitin family member 3 | Cytoplasm | enzyme | 2283352.5 | 2870355.25 | 3636668.5 | | | | | | |
| Q8K1B8 | FERMT3 | fermitin family member 3 | Cytoplasm | enzyme | 2236600.75 | 3115647 | 2952370.5 | | | | | | |
| Q8K1B8 | FERMT3 | fermitin family member 3 | Cytoplasm | enzyme | 2519297 | 2699865 | 3492491.5 | | | | | | |
| Q8K1B8 | FERMT3 | fermitin family member 3 | Cytoplasm | enzyme | 1651697.875 | 3035680.75 | 1256616 | Q8K1B8 | FERMT3 | 5558917 | 3601759 | 519869 | 48 |
| Q8K0E8 | FGB | fibrinogen beta chain | Extracellular Space | other | 8362247.5 | 2.79E+07 | 1.57E+07 | | | | | | |
| Q8K0E8 | FGB | fibrinogen beta chain | Extracellular Space | other | 8924944 | 6.99E+07 | 2.62E+07 | | | | | | |
| Q8K0E8 | FGB | fibrinogen beta chain | Extracellular Space | other | 1.14E+07 | 5.29E+07 | 1.63E+07 | | | | | | |
| Q8K0E8 | FGB | fibrinogen beta chain | Extracellular Space | other | 6452684.5 | 2.71E+07 | 6833572.5 | | | | | | |
| Q8K0E8 | FGB | fibrinogen beta chain | Extracellular Space | other | 6410180 | 4.07E+07 | 1.46E+07 | | | | | | |
| Q8K0E8 | FGB | fibrinogen beta chain | Extracellular Space | other | 8840132 | 2.18E+07 | 595652 | | | | | | |
| Q8K0E8 | FGB | fibrinogen beta chain | Extracellular Space | other | 6203812 | 3.42E+07 | 1.20E+07 | | | | | | |
| Q8K0E8 | FGB | fibrinogen beta chain | Extracellular Space | other | 7333532.5 | 2.94E+07 | 9028135 | | | | | | |
| Q8K0E8 | FGB | fibrinogen beta chain | Extracellular Space | other | 5808605.5 | 3.56E+07 | 9939412 | | | | | | |
| Q8K0E8 | FGB | fibrinogen beta chain | Extracellular Space | other | 4981288.5 | 3.56E+07 | 6882700 | | | | | | |
| Q8K0E8 | FGB | fibrinogen beta chain | Extracellular Space | other | 4372217 | 1.48E+07 | 4032377.25 | | | | | | |
| Q8K0E8 | FGB | fibrinogen beta chain | Extracellular Space | other | 2454591 | 2.45E+07 | 6882260.5 | | | | | | |
| Q8K0E8 | FGB | fibrinogen beta chain | Extracellular Space | other | 2466226.5 | 9023894 | 1983136.125 | | | | | | |
| Q8K0E8 | FGB | fibrinogen beta chain | Extracellular Space | other | 3554903.5 | 1.49E+07 | 4616295.5 | | | | | | |
| Q8K0E8 | FGB | fibrinogen beta chain | Extracellular Space | other | 2530157.5 | 1.11E+07 | 2791909.75 | | | | | | |
| Q8K0E8 | FGB | fibrinogen beta chain | Extracellular Space | other | 1610644.125 | 1.25E+07 | 1172047.25 | | | | | | |
| Q8K0E8 | FGB | fibrinogen beta chain | Extracellular Space | other | 647858.188 | 1.02E+07 | 3993552.5 | | | | | | |
| Q8K0E8 | FGB | fibrinogen beta chain | Extracellular Space | other | 444527.062 | 4806734.5 | 1501136.875 | | | | | | |

| | | | | | | | | | | | | | |
|---|---|---|---|---|---|---|---|---|---|---|---|---|---|
| Q8K0E8 | FGB | fibrinogen beta chain | Extracellular Space | other | 394139.906 | 7573519 | 1603421.5 | | | | | | |
| Q8K0E8 | FGB | fibrinogen beta chain | Extracellular Space | other | 717634 | 1309191.875 | 102611.695 | | | | | | |
| Q8K0E8 | FGB | fibrinogen beta chain | Extracellular Space | other | 49753.203 | 2504531.75 | 623380.688 | | | | | | |
| Q8K0E8 | FGB | fibrinogen beta chain | Extracellular Space | other | 590456.5 | 3465840.75 | 1143282.25 | | | | | | |
| Q8K0E8 | FGB | fibrinogen beta chain | Extracellular Space | other | 157144.828 | 3824906 | 849442.125 | | | | | | |
| Q8K0E8 | FGB | fibrinogen beta chain | Extracellular Space | other | 196515.297 | 753452.938 | 5867.889 | Q8K0E8 | FGB | 3954341 | 13269073 | 1563775 | 72 |
| Q8VCM7 | FGG | fibrinogen gamma chain | Extracellular Space | other | 1.88E+07 | 5.49E+07 | 1.37E+07 | | | | | | |
| Q8VCM7 | FGG | fibrinogen gamma chain | Extracellular Space | other | 2.01E+07 | 8.32E+07 | 2.07E+07 | | | | | | |
| Q8VCM7 | FGG | fibrinogen gamma chain | Extracellular Space | other | 8630670 | 4.29E+07 | 1.28E+07 | | | | | | |
| Q8VCM7 | FGG | fibrinogen gamma chain | Extracellular Space | other | 3697174 | 3.36E+07 | 7161894.5 | | | | | | |
| Q8VCM7 | FGG | fibrinogen gamma chain | Extracellular Space | other | 7970815 | 3.31E+07 | 9398904 | | | | | | |
| Q8VCM7 | FGG | fibrinogen gamma chain | Extracellular Space | other | 5747720.5 | 2.81E+07 | 7423321.5 | | | | | | |
| Q8VCM7 | FGG | fibrinogen gamma chain | Extracellular Space | other | 5120636.5 | 2.30E+07 | 6544940.5 | | | | | | |
| Q8VCM7 | FGG | fibrinogen gamma chain | Extracellular Space | other | 4022041.75 | 2.53E+07 | 7313971.5 | | | | | | |
| Q8VCM7 | FGG | fibrinogen gamma chain | Extracellular Space | other | 1718757.75 | 1.61E+07 | 1.04E+07 | | | | | | |
| Q8VCM7 | FGG | fibrinogen gamma chain | Extracellular Space | other | 2951113.75 | 2.07E+07 | 3364761.75 | | | | | | |
| Q8VCM7 | FGG | fibrinogen gamma chain | Extracellular Space | other | 2693666 | 1.04E+07 | 3840871 | | | | | | |
| Q8VCM7 | FGG | fibrinogen gamma chain | Extracellular Space | other | 2138889.75 | 1.45E+07 | 3807077.75 | | | | | | |
| Q8VCM7 | FGG | fibrinogen gamma chain | Extracellular Space | other | 2284681 | 1.89E+07 | 4261522 | | | | | | |
| Q8VCM7 | FGG | fibrinogen gamma chain | Extracellular Space | other | 1882617.25 | 5048828.5 | 3621154.25 | | | | | | |
| Q8VCM7 | FGG | fibrinogen gamma chain | Extracellular Space | other | 3558568.5 | 1.99E+07 | 5649617 | | | | | | |
| Q8VCM7 | FGG | fibrinogen gamma chain | Extracellular Space | other | 2936514 | 1.47E+07 | 3077522.5 | | | | | | |
| Q8VCM7 | FGG | fibrinogen gamma chain | Extracellular Space | other | 3291853.5 | 1.73E+07 | 4171973 | | | | | | |
| Q8VCM7 | FGG | fibrinogen gamma chain | Extracellular Space | other | 202240.516 | 3377074.25 | 1304677.875 | | | | | | |
| Q8VCM7 | FGG | fibrinogen gamma chain | Extracellular Space | other | 803007.875 | 1.15E+07 | 4767602 | | | | | | |
| Q8VCM7 | FGG | fibrinogen gamma chain | Extracellular Space | other | 1679592.375 | 8039286 | 826774.688 | | | | | | |
| Q8VCM7 | FGG | fibrinogen gamma chain | Extracellular Space | other | 822528.312 | 4496843.5 | 90678.109 | | | | | | |
| Q8VCM7 | FGG | fibrinogen gamma chain | Extracellular Space | other | 0 | 1166249.5 | 0 | | | | | | |
| Q8VCM7 | FGG | fibrinogen gamma chain | Extracellular Space | other | 87994.156 | 436635.219 | 248464.703 | Q8VCM7 | FGG | 4397873 | 13953911 | 1679854 | 69 |
| P97807 | FH | fumarate hydratase | Cytoplasm | enzyme | 8473457 | 1.89E+07 | 8188637.5 | | | | | | |
| P97807 | FH | fumarate hydratase | Cytoplasm | enzyme | 5416682.5 | 1.02E+07 | 5272484.5 | | | | | | |
| P97807 | FH | fumarate hydratase | Cytoplasm | enzyme | 2955178 | 5585759 | 2262535.5 | | | | | | |
| P97807 | FH | fumarate hydratase | Cytoplasm | enzyme | 2628967.5 | 5294125 | 2866458.25 | | | | | | |
| P97807 | FH | fumarate hydratase | Cytoplasm | enzyme | 1705301.75 | 3509684 | 2059112.625 | | | | | | |
| P97807 | FH | fumarate hydratase | Cytoplasm | enzyme | 1504168.375 | 3193533.75 | 1838564.25 | | | | | | |
| P97807 | FH | fumarate hydratase | Cytoplasm | enzyme | 2481158.25 | 4769946 | 2879436.25 | P97807 | FH | 3594988 | 4021259 | 877511 | 21 |
| Q9CQ92 | FIS1 | fission, mitochondrial 1 | Cytoplasm | other | 4682755.5 | 6583323 | 3670442.5 | | | | | | |
| Q9CQ92 | FIS1 | fission, mitochondrial 1 | Cytoplasm | other | 3955659.75 | 6256769.5 | 4694116 | Q9CQ92 | FIS1 | 4319208 | 1194559 | 487677 | 6 |
| Q6P9Q6 | FKBP15 | FK506 binding protein 15, 133kDa | Plasma Membrane | enzyme | 1178087.75 | 1110411 | 695788.312 | | | | | | |
| Q6P9Q6 | FKBP15 | FK506 binding protein 15, 133kDa | Plasma Membrane | enzyme | 538711.5 | 441911.312 | 368427.562 | Q6P9Q6 | FKBP15 | 858400 | 345434 | 141023 | 6 |
| P45878 | FKBP2 | FK506 binding protein 2, 13kDa | Cytoplasm | enzyme | 1.34E+07 | 1.70E+07 | 1.74E+07 | | | | | | |
| P45878 | FKBP2 | FK506 binding protein 2, 13kDa | Cytoplasm | enzyme | 9612339 | 1.51E+07 | 1.73E+07 | P45878 | FKBP2 | 11516170 | 3046908 | 1243895 | 6 |
| O35450 | FKBPL | FK506 binding protein like | Nucleus | other | 3.13E+07 | 2.12E+07 | 2.96E+07 | O35450 | FKBPL | 31310000 | 5431043 | 3135614 | 3 |
| Q2VIS4 | Flg2 | filaggrin family member 2 | Extracellular Space | other | 4.14E+07 | 3.42E+07 | 3.62E+07 | Q2VIS4 | Flg2 | 41370000 | 3715109 | 2144919 | 3 |
| Q8BTM8 | FLNA | filamin A, alpha | Cytoplasm | other | 1.20E+07 | 3.64E+07 | 7.18E+07 | | | | | | |
| Q8BTM8 | FLNA | filamin A, alpha | Cytoplasm | other | 9411645 | 2.87E+07 | 5.24E+07 | | | | | | |
| Q8BTM8 | FLNA | filamin A, alpha | Cytoplasm | other | 8307888.5 | 2.77E+07 | 5.30E+07 | | | | | | |
| Q8BTM8 | FLNA | filamin A, alpha | Cytoplasm | other | 7875699 | 3.03E+07 | 7.05E+07 | | | | | | |
| Q8BTM8 | FLNA | filamin A, alpha | Cytoplasm | other | 8188743 | 2.64E+07 | 5.07E+07 | | | | | | |
| Q8BTM8 | FLNA | filamin A, alpha | Cytoplasm | other | 7417760.5 | 1.19E+07 | 2.35E+07 | | | | | | |
| Q8BTM8 | FLNA | filamin A, alpha | Cytoplasm | other | 6696111 | 2.02E+07 | 4.26E+07 | | | | | | |
| Q8BTM8 | FLNA | filamin A, alpha | Cytoplasm | other | 6930818.5 | 2.13E+07 | 4.54E+07 | | | | | | |
| Q8BTM8 | FLNA | filamin A, alpha | Cytoplasm | other | 5065983 | 1.30E+07 | 2.23E+07 | | | | | | |
| Q8BTM8 | FLNA | filamin A, alpha | Cytoplasm | other | 6057505 | 1.78E+07 | 2.34E+07 | | | | | | |
| Q8BTM8 | FLNA | filamin A, alpha | Cytoplasm | other | 3412043.25 | 1.15E+07 | 2.19E+07 | | | | | | |
| Q8BTM8 | FLNA | filamin A, alpha | Cytoplasm | other | 6841688.5 | 1.41E+07 | 3.09E+07 | | | | | | |
| Q8BTM8 | FLNA | filamin A, alpha | Cytoplasm | other | 5921293 | 1.31E+07 | 1.60E+07 | | | | | | |
| Q8BTM8 | FLNA | filamin A, alpha | Cytoplasm | other | 3631760.25 | 1.11E+07 | 1.93E+07 | | | | | | |
| Q8BTM8 | FLNA | filamin A, alpha | Cytoplasm | other | 6038695 | 1.71E+07 | 2.97E+07 | | | | | | |
| Q8BTM8 | FLNA | filamin A, alpha | Cytoplasm | other | 4460898.5 | 1.41E+07 | 3.21E+07 | | | | | | |
| Q8BTM8 | FLNA | filamin A, alpha | Cytoplasm | other | 3840071.75 | 1.14E+07 | 2.42E+07 | | | | | | |
| Q8BTM8 | FLNA | filamin A, alpha | Cytoplasm | other | 5269994.5 | 1.72E+07 | 3.13E+07 | | | | | | |
| Q8BTM8 | FLNA | filamin A, alpha | Cytoplasm | other | 4616458.5 | 1.40E+07 | 2.38E+07 | | | | | | |
| Q8BTM8 | FLNA | filamin A, alpha | Cytoplasm | other | 4362324.5 | 1.19E+07 | 3.14E+07 | | | | | | |
| Q8BTM8 | FLNA | filamin A, alpha | Cytoplasm | other | 4543652.5 | 1.56E+07 | 2.82E+07 | | | | | | |
| Q8BTM8 | FLNA | filamin A, alpha | Cytoplasm | other | 4462799.5 | 1.48E+07 | 2.53E+07 | | | | | | |
| Q8BTM8 | FLNA | filamin A, alpha | Cytoplasm | other | 5328236 | 1.29E+07 | 2.59E+07 | | | | | | |
| Q8BTM8 | FLNA | filamin A, alpha | Cytoplasm | other | 2936745.25 | 1.95E+07 | 4.97E+07 | | | | | | |
| Q8BTM8 | FLNA | filamin A, alpha | Cytoplasm | other | 4767006 | 1.13E+07 | 2.35E+07 | | | | | | |
| Q8BTM8 | FLNA | filamin A, alpha | Cytoplasm | other | 3338850.75 | 8885914 | 1.96E+07 | | | | | | |
| Q8BTM8 | FLNA | filamin A, alpha | Cytoplasm | other | 7080270.5 | 5651132 | 5959309 | | | | | | |
| Q8BTM8 | FLNA | filamin A, alpha | Cytoplasm | other | 3493321.75 | 1.28E+07 | 3.07E+07 | | | | | | |
| Q8BTM8 | FLNA | filamin A, alpha | Cytoplasm | other | 3357643.75 | 1.19E+07 | 2.22E+07 | | | | | | |
| Q8BTM8 | FLNA | filamin A, alpha | Cytoplasm | other | 4296774 | 1.14E+07 | 2.27E+07 | | | | | | |
| Q8BTM8 | FLNA | filamin A, alpha | Cytoplasm | other | 3589309.5 | 1.04E+07 | 2.39E+07 | | | | | | |
| Q8BTM8 | FLNA | filamin A, alpha | Cytoplasm | other | 3776535.75 | 1.06E+07 | 2.19E+07 | | | | | | |
| Q8BTM8 | FLNA | filamin A, alpha | Cytoplasm | other | 2318531.25 | 5670139.5 | 1.11E+07 | | | | | | |
| Q8BTM8 | FLNA | filamin A, alpha | Cytoplasm | other | 4762313.5 | 1.38E+07 | 2.93E+07 | | | | | | |
| Q8BTM8 | FLNA | filamin A, alpha | Cytoplasm | other | 3455706 | 8950494 | 1.64E+07 | | | | | | |
| Q8BTM8 | FLNA | filamin A, alpha | Cytoplasm | other | 3487504.5 | 7825031.5 | 9312081 | | | | | | |
| Q8BTM8 | FLNA | filamin A, alpha | Cytoplasm | other | 2092046.625 | 1.02E+07 | 2.32E+07 | | | | | | |
| Q8BTM8 | FLNA | filamin A, alpha | Cytoplasm | other | 4141797.75 | 1.18E+07 | 2.50E+07 | | | | | | |
| Q8BTM8 | FLNA | filamin A, alpha | Cytoplasm | other | 2411742 | 8886796 | 1.93E+07 | | | | | | |
| Q8BTM8 | FLNA | filamin A, alpha | Cytoplasm | other | 2984543.75 | 7978864.5 | 1.81E+07 | | | | | | |
| Q8BTM8 | FLNA | filamin A, alpha | Cytoplasm | other | 2713176 | 1.10E+07 | 2.01E+07 | | | | | | |
| Q8BTM8 | FLNA | filamin A, alpha | Cytoplasm | other | 2655396 | 4894799.5 | 9206002 | | | | | | |
| Q8BTM8 | FLNA | filamin A, alpha | Cytoplasm | other | 3134103.25 | 9934867 | 2.09E+07 | | | | | | |
| Q8BTM8 | FLNA | filamin A, alpha | Cytoplasm | other | 2896429.5 | 7223908.5 | 1.69E+07 | | | | | | |
| Q8BTM8 | FLNA | filamin A, alpha | Cytoplasm | other | 2613597.75 | 6797673.5 | 1.26E+07 | | | | | | |
| Q8BTM8 | FLNA | filamin A, alpha | Cytoplasm | other | 3018598.5 | 1.10E+07 | 2.39E+07 | | | | | | |
| Q8BTM8 | FLNA | filamin A, alpha | Cytoplasm | other | 2613322.25 | 1.45E+07 | 2.45E+07 | | | | | | |

| | | | | | | | | | | | | | |
|---|---|---|---|---|---|---|---|---|---|---|---|---|---|
| Q8BTM8 | FLNA | filamin A, alpha | Cytoplasm | other | 2940072 | 1.21E+07 | 2.07E+07 | | | | | | |
| Q8BTM8 | FLNA | filamin A, alpha | Cytoplasm | other | 3397710.25 | 4307256 | 6411813.5 | | | | | | |
| Q8BTM8 | FLNA | filamin A, alpha | Cytoplasm | other | 2701417.25 | 7864500 | 1.74E+07 | | | | | | |
| Q8BTM8 | FLNA | filamin A, alpha | Cytoplasm | other | 2839608.5 | 5782636 | 1.23E+07 | | | | | | |
| Q8BTM8 | FLNA | filamin A, alpha | Cytoplasm | other | 2310479.25 | 7373804.5 | 1.35E+07 | | | | | | |
| Q8BTM8 | FLNA | filamin A, alpha | Cytoplasm | other | 3125414.75 | 9712438 | 1.63E+07 | | | | | | |
| Q8BTM8 | FLNA | filamin A, alpha | Cytoplasm | other | 2387269.75 | 6818749 | 1.77E+07 | | | | | | |
| Q8BTM8 | FLNA | filamin A, alpha | Cytoplasm | other | 2127148.25 | 7879626 | 1.48E+07 | | | | | | |
| Q8BTM8 | FLNA | filamin A, alpha | Cytoplasm | other | 3283776.25 | 1.04E+07 | 2.27E+07 | | | | | | |
| Q8BTM8 | FLNA | filamin A, alpha | Cytoplasm | other | 2374178.25 | 7210127 | 1.21E+07 | | | | | | |
| Q8BTM8 | FLNA | filamin A, alpha | Cytoplasm | other | 2610192.5 | 7405428.5 | 1.59E+07 | | | | | | |
| Q8BTM8 | FLNA | filamin A, alpha | Cytoplasm | other | 1632656 | 2509353.75 | 4306797.5 | | | | | | |
| Q8BTM8 | FLNA | filamin A, alpha | Cytoplasm | other | 679959 | 1571297.5 | 1176149 | | | | | | |
| Q8BTM8 | FLNA | filamin A, alpha | Cytoplasm | other | 1141304.625 | 5654573 | 1.12E+07 | | | | | | |
| Q8BTM8 | FLNA | filamin A, alpha | Cytoplasm | other | 2164274.25 | 7556172 | 1.27E+07 | | | | | | |
| Q8BTM8 | FLNA | filamin A, alpha | Cytoplasm | other | 1476106.125 | 4353524 | 6638347 | | | | | | |
| Q8BTM8 | FLNA | filamin A, alpha | Cytoplasm | other | 2269663.75 | 3260896.75 | 6482734 | | | | | | |
| Q8BTM8 | FLNA | filamin A, alpha | Cytoplasm | other | 1693465.875 | 4448323.5 | 8750857 | | | | | | |
| Q8BTM8 | FLNA | filamin A, alpha | Cytoplasm | other | 1756012 | 5189718 | 1.13E+07 | | | | | | |
| Q8BTM8 | FLNA | filamin A, alpha | Cytoplasm | other | 1794177.25 | 5161537 | 1.21E+07 | | | | | | |
| Q8BTM8 | FLNA | filamin A, alpha | Cytoplasm | other | 940415.75 | 6353878.5 | 1.51E+07 | | | | | | |
| Q8BTM8 | FLNA | filamin A, alpha | Cytoplasm | other | 1294035.25 | 3460860.5 | 7440143.5 | | | | | | |
| Q8BTM8 | FLNA | filamin A, alpha | Cytoplasm | other | 1891102.125 | 5461874 | 8187942.5 | | | | | | |
| Q8BTM8 | FLNA | filamin A, alpha | Cytoplasm | other | 2265704.25 | 6127478.5 | 1.46E+07 | | | | | | |
| Q8BTM8 | FLNA | filamin A, alpha | Cytoplasm | other | 1269740.625 | 4014668.5 | 9431693 | | | | | | |
| Q8BTM8 | FLNA | filamin A, alpha | Cytoplasm | other | 1411038 | 4336924 | 1.09E+07 | | | | | | |
| Q8BTM8 | FLNA | filamin A, alpha | Cytoplasm | other | 1351322 | 5656286.5 | 9382313 | | | | | | |
| Q8BTM8 | FLNA | filamin A, alpha | Cytoplasm | other | 2203724.25 | 4153083 | 3169018.5 | | | | | | |
| Q8BTM8 | FLNA | filamin A, alpha | Cytoplasm | other | 1292963.75 | 3649876 | 5626601.5 | | | | | | |
| Q8BTM8 | FLNA | filamin A, alpha | Cytoplasm | other | 2097491.25 | 3266641.75 | 2981594.75 | | | | | | |
| Q8BTM8 | FLNA | filamin A, alpha | Cytoplasm | other | 689147 | 2263972.75 | 3452730.25 | | | | | | |
| Q8BTM8 | FLNA | filamin A, alpha | Cytoplasm | other | 980944.438 | 2556041.25 | 5669117 | | | | | | |
| Q8BTM8 | FLNA | filamin A, alpha | Cytoplasm | other | 1887065.375 | 4651034.5 | 8775988 | | | | | | |
| Q8BTM8 | FLNA | filamin A, alpha | Cytoplasm | other | 1200856.875 | 3735347.25 | 8239650 | | | | | | |
| Q8BTM8 | FLNA | filamin A, alpha | Cytoplasm | other | 877400.625 | 2770690.25 | 5455834.5 | | | | | | |
| Q8BTM8 | FLNA | filamin A, alpha | Cytoplasm | other | 1730431.625 | 3893946.5 | 8667091 | | | | | | |
| Q8BTM8 | FLNA | filamin A, alpha | Cytoplasm | other | 1831257.375 | 3191817.75 | 4728370.5 | | | | | | |
| Q8BTM8 | FLNA | filamin A, alpha | Cytoplasm | other | 1936451.5 | 1488868.5 | 16576.775 | | | | | | |
| Q8BTM8 | FLNA | filamin A, alpha | Cytoplasm | other | 819910.812 | 3444416.75 | 7274290 | | | | | | |
| Q8BTM8 | FLNA | filamin A, alpha | Cytoplasm | other | 557614 | 2076722.5 | 888693.812 | | | | | | |
| Q8BTM8 | FLNA | filamin A, alpha | Cytoplasm | other | 461572 | 842036.875 | 1782396 | | | | | | |
| Q8BTM8 | FLNA | filamin A, alpha | Cytoplasm | other | 764718.312 | 1900757 | 4553417 | | | | | | |
| Q8BTM8 | FLNA | filamin A, alpha | Cytoplasm | other | 1857948.75 | 9545821 | 2.36E+07 | | | | | | |
| Q8BTM8 | FLNA | filamin A, alpha | Cytoplasm | other | 1009988.312 | 1497851.5 | 2407108.5 | | | | | | |
| Q8BTM8 | FLNA | filamin A, alpha | Cytoplasm | other | 681846.562 | 2420094.5 | 6033957.5 | | | | | | |
| Q8BTM8 | FLNA | filamin A, alpha | Cytoplasm | other | 849832.062 | 3603248 | 7472041.5 | | | | | | |
| Q8BTM8 | FLNA | filamin A, alpha | Cytoplasm | other | 851996.938 | 2063743.5 | 4902915 | | | | | | |
| Q8BTM8 | FLNA | filamin A, alpha | Cytoplasm | other | 560566.438 | 1812471.5 | 4432254.5 | | | | | | |
| Q8BTM8 | FLNA | filamin A, alpha | Cytoplasm | other | 767879.062 | 2564646.5 | 5910187 | | | | | | |
| Q8BTM8 | FLNA | filamin A, alpha | Cytoplasm | other | 687490.5 | 3045432.25 | 5532221.5 | | | | | | |
| Q8BTM8 | FLNA | filamin A, alpha | Cytoplasm | other | 698529.812 | 1937378.5 | 3186911.5 | | | | | | |
| Q8BTM8 | FLNA | filamin A, alpha | Cytoplasm | other | 250405.531 | 1374857.5 | 937940.188 | | | | | | |
| Q8BTM8 | FLNA | filamin A, alpha | Cytoplasm | other | 704798.062 | 1416300.875 | 2904636 | | | | | | |
| Q8BTM8 | FLNA | filamin A, alpha | Cytoplasm | other | 1290898.625 | 2326673.75 | 3829527.25 | | | | | | |
| Q8BTM8 | FLNA | filamin A, alpha | Cytoplasm | other | 469273.094 | 1815936 | 3463413.5 | | | | | | |
| Q8BTM8 | FLNA | filamin A, alpha | Cytoplasm | other | 362721.031 | 1300005.25 | 3452213 | | | | | | |
| Q8BTM8 | FLNA | filamin A, alpha | Cytoplasm | other | 404653.125 | 511180.219 | 777598.375 | | | | | | |
| Q8BTM8 | FLNA | filamin A, alpha | Cytoplasm | other | 373911.25 | 2236975 | 4438797 | | | | | | |
| Q8BTM8 | FLNA | filamin A, alpha | Cytoplasm | other | 485310.781 | 828543.062 | 2624145 | | | | | | |
| Q8BTM8 | FLNA | filamin A, alpha | Cytoplasm | other | 474188.844 | 3053113 | 1.14E+07 | | | | | | |
| Q8BTM8 | FLNA | filamin A, alpha | Cytoplasm | other | 264187.906 | 844448.25 | 1671962.25 | | | | | | |
| Q8BTM8 | FLNA | filamin A, alpha | Cytoplasm | other | 422385.281 | 1264802.375 | 1750990.5 | | | | | | |
| Q8BTM8 | FLNA | filamin A, alpha | Cytoplasm | other | 304504.156 | 189500.828 | 70583.602 | | | | | | |
| Q8BTM8 | FLNA | filamin A, alpha | Cytoplasm | other | 139755.75 | 173543.047 | 51387.816 | | | | | | |
| Q8BTM8 | FLNA | filamin A, alpha | Cytoplasm | other | 302418.156 | 273781.75 | 412385.125 | | | | | | |
| Q8BTM8 | FLNA | filamin A, alpha | Cytoplasm | other | 0 | 635451.812 | 1618121.375 | | | | | | |
| Q8BTM8 | FLNA | filamin A, alpha | Cytoplasm | other | 112183.773 | 293373.625 | 1038248.5 | | | | | | |
| Q8BTM8 | FLNA | filamin A, alpha | Cytoplasm | other | 138137.609 | 488400 | 318736.656 | | | | | | |
| Q8BTM8 | FLNA | filamin A, alpha | Cytoplasm | other | 226909.984 | 306678.438 | 2989605.75 | | | | | | |
| Q8BTM8 | FLNA | filamin A, alpha | Cytoplasm | other | 4.70E-06 | 243657.016 | 1398398.875 | | | | | | |
| Q8BTM8 | FLNA | filamin A, alpha | Cytoplasm | other | 141780.688 | 223230.797 | 192694.703 | Q8BTM8 | FLNA | 2642980 | 10559490 | 561231 | 354 |
| O08917 | FLOT1 | flotillin 1 | Plasma Membrane | other | 1.74E+07 | 1.97E+07 | 2.16E+07 | | | | | | |
| O08917 | FLOT1 | flotillin 1 | Plasma Membrane | other | 2.04E+07 | 1.99E+07 | 2.11E+07 | | | | | | |
| O08917 | FLOT1 | flotillin 1 | Plasma Membrane | other | 2.10E+07 | 1.33E+07 | 1.07E+07 | | | | | | |
| O08917 | FLOT1 | flotillin 1 | Plasma Membrane | other | 1.20E+07 | 1.07E+07 | 1.29E+07 | | | | | | |
| O08917 | FLOT1 | flotillin 1 | Plasma Membrane | other | 1.14E+07 | 1.31E+07 | 1.23E+07 | | | | | | |
| O08917 | FLOT1 | flotillin 1 | Plasma Membrane | other | 9814656 | 7739385 | 1.22E+07 | | | | | | |
| O08917 | FLOT1 | flotillin 1 | Plasma Membrane | other | 1.34E+07 | 1.28E+07 | 7946089.5 | | | | | | |
| O08917 | FLOT1 | flotillin 1 | Plasma Membrane | other | 1.17E+07 | 9172461 | 7232550 | | | | | | |
| O08917 | FLOT1 | flotillin 1 | Plasma Membrane | other | 9872422 | 1.18E+07 | 1.39E+07 | | | | | | |
| O08917 | FLOT1 | flotillin 1 | Plasma Membrane | other | 6578184 | 5931455 | 5844972 | | | | | | |
| O08917 | FLOT1 | flotillin 1 | Plasma Membrane | other | 6640149 | 6893268 | 4578861 | | | | | | |
| O08917 | FLOT1 | flotillin 1 | Plasma Membrane | other | 3673944.25 | 7232791.5 | 8953351 | | | | | | |
| O08917 | FLOT1 | flotillin 1 | Plasma Membrane | other | 3157245.25 | 3649430 | 4119252.75 | | | | | | |
| O08917 | FLOT1 | flotillin 1 | Plasma Membrane | other | 4867122.5 | 6379538 | 6085590.5 | | | | | | |
| O08917 | FLOT1 | flotillin 1 | Plasma Membrane | other | 3111493 | 3455968.25 | 5584983 | | | | | | |
| O08917 | FLOT1 | flotillin 1 | Plasma Membrane | other | 1667905.125 | 305978.031 | 211699.75 | O08917 | FLOT1 | 9787070 | 5691168 | 821449 | 48 |
| Q60634 | FLOT2 | flotillin 2 | Plasma Membrane | other | 3.52E+07 | 2.80E+07 | 3.30E+07 | | | | | | |
| Q60634 | FLOT2 | flotillin 2 | Plasma Membrane | other | 3.31E+07 | 2.56E+07 | 2.86E+07 | | | | | | |
| Q60634 | FLOT2 | flotillin 2 | Plasma Membrane | other | 1.88E+07 | 1.69E+07 | 2.03E+07 | | | | | | |
| Q60634 | FLOT2 | flotillin 2 | Plasma Membrane | other | 1.86E+07 | 2.12E+07 | 2.27E+07 | | | | | | |

| | | | | | | | | | | | | | |
|---|---|---|---|---|---|---|---|---|---|---|---|---|---|
| Q60634 | FLOT2 | flotillin 2 | Plasma Membrane | other | 1.56E+07 | 1.74E+07 | 1.63E+07 | | | | | | |
| Q60634 | FLOT2 | flotillin 2 | Plasma Membrane | other | 1.30E+07 | 1.42E+07 | 1.46E+07 | | | | | | |
| Q60634 | FLOT2 | flotillin 2 | Plasma Membrane | other | 1.37E+07 | 1.28E+07 | 1.64E+07 | | | | | | |
| Q60634 | FLOT2 | flotillin 2 | Plasma Membrane | other | 1.11E+07 | 1.09E+07 | 1.16E+07 | | | | | | |
| Q60634 | FLOT2 | flotillin 2 | Plasma Membrane | other | 8890060 | 8220460 | 1.13E+07 | | | | | | |
| Q60634 | FLOT2 | flotillin 2 | Plasma Membrane | other | 8669638 | 8651812 | 1.06E+07 | | | | | | |
| Q60634 | FLOT2 | flotillin 2 | Plasma Membrane | other | 7316648 | 7275067.5 | 8600263 | | | | | | |
| Q60634 | FLOT2 | flotillin 2 | Plasma Membrane | other | 4969148.5 | 6338102 | 7561584.5 | | | | | | |
| Q60634 | FLOT2 | flotillin 2 | Plasma Membrane | other | 4260944.5 | 4240033 | 4344102.5 | | | | | | |
| Q60634 | FLOT2 | flotillin 2 | Plasma Membrane | other | 3731725 | 4061445.75 | 3827658 | | | | | | |
| Q60634 | FLOT2 | flotillin 2 | Plasma Membrane | other | 3950926.25 | 4563692.5 | 3071719.5 | | | | | | |
| Q60634 | FLOT2 | flotillin 2 | Plasma Membrane | other | 1800752.25 | 2633995 | 4185440.75 | | | | | | |
| Q60634 | FLOT2 | flotillin 2 | Plasma Membrane | other | 1337417.625 | 683247.25 | 382507.344 | Q60634 | FLOT2 | 11989839 | 8978205 | 1257200 | 51 |
| P11276 | FN1 | fibronectin 1 | Extracellular Space | enzyme | 328719.844 | 112701.492 | 302938.719 | | | | | | |
| P11276 | FN1 | fibronectin 1 | Extracellular Space | enzyme | 296085.156 | 385820.781 | 112757.367 | | | | | | |
| P11276 | FN1 | fibronectin 1 | Extracellular Space | enzyme | 344507.375 | 186476.156 | 247514.656 | P11276 | FN1 | 323104 | 99827 | 33276 | 9 |
| Q9ER35 | FN3K | fructosamine 3 kinase | Cytoplasm | kinase | 1.61E+07 | 1.34E+07 | 1.67E+07 | | | | | | |
| Q9ER35 | FN3K | fructosamine 3 kinase | Cytoplasm | kinase | 1.67E+07 | 1.15E+07 | 1.55E+07 | | | | | | |
| Q9ER35 | FN3K | fructosamine 3 kinase | Cytoplasm | kinase | 1.43E+07 | 1.05E+07 | 1.30E+07 | | | | | | |
| Q9ER35 | FN3K | fructosamine 3 kinase | Cytoplasm | kinase | 1.08E+07 | 6024114.5 | 5758568 | | | | | | |
| Q9ER35 | FN3K | fructosamine 3 kinase | Cytoplasm | kinase | 1.53E+07 | 9921740 | 1.11E+07 | | | | | | |
| Q9ER35 | FN3K | fructosamine 3 kinase | Cytoplasm | kinase | 2729377 | 2269929.75 | 3589976.25 | | | | | | |
| Q9ER35 | FN3K | fructosamine 3 kinase | Cytoplasm | kinase | 2960325.75 | 2286711.25 | 2696166 | Q9ER35 | FN3K | 11279958 | 5370305 | 1171897 | 21 |
| Q8K274 | FN3KRP | fructosamine 3 kinase related protein | Cytoplasm | kinase | 2.27E+07 | 2.40E+07 | 2.37E+07 | | | | | | |
| Q8K274 | FN3KRP | fructosamine 3 kinase related protein | Cytoplasm | kinase | 1.57E+07 | 1.20E+07 | 1.32E+07 | | | | | | |
| Q8K274 | FN3KRP | fructosamine 3 kinase related protein | Cytoplasm | kinase | 9155662 | 8680592 | 9415974 | | | | | | |
| Q8K274 | FN3KRP | fructosamine 3 kinase related protein | Cytoplasm | kinase | 7824050 | 7128225.5 | 8453123 | | | | | | |
| Q8K274 | FN3KRP | fructosamine 3 kinase related protein | Cytoplasm | kinase | 5213026 | 6406869 | 5756697.5 | | | | | | |
| Q8K274 | FN3KRP | fructosamine 3 kinase related protein | Cytoplasm | kinase | 4821994.5 | 4800866.5 | 6835900.5 | Q8K274 | FN3KRP | 10885789 | 6469147 | 1524793 | 18 |
| Q3UFK8 | FRMD8 | FERM domain containing 8 | Cytoplasm | other | 3467957.25 | 5162325 | 5143453 | Q3UFK8 | FRMD8 | 3467957 | 972842 | 561670 | 3 |
| P09528 | FTH1 | ferritin, heavy polypeptide 1 | Cytoplasm | enzyme | 1113640.375 | 1028693.938 | 853677.75 | P09528 | FTH1 | 1113640 | 132556 | 76531 | 3 |
| P29391 | FTL | ferritin, light polypeptide | Cytoplasm | enzyme | 2276661.5 | 3837243.25 | 1982807.125 | | | | | | |
| P29391 | FTL | ferritin, light polypeptide | Cytoplasm | enzyme | 1056365.125 | 1465649 | 1348565.125 | | | | | | |
| P29391 | FTL | ferritin, light polypeptide | Cytoplasm | enzyme | 747559.812 | 154991.078 | 560933.625 | P29391 | FTL | 1360195 | 1105467 | 368489 | 9 |
| Q9D6K8 | FUNDC2 | FUN14 domain containing 2 | Cytoplasm | other | 2226226 | 5736130 | 3002496.5 | | | | | | |
| Q9D6K8 | FUNDC2 | FUN14 domain containing 2 | Cytoplasm | other | 2261084.75 | 7149206.5 | 3986233.25 | Q9D6K8 | FUNDC2 | 2243655 | 2003756 | 818030 | 6 |
| O35943 | FXN | frataxin | Cytoplasm | kinase | 2580039.25 | 5358328.5 | 1478907.625 | | | | | | |
| O35943 | FXN | frataxin | Cytoplasm | kinase | 2113266.25 | 5931913 | 1684922.625 | O35943 | FXN | 2346653 | 1946496 | 794654 | 6 |
| O35601 | FYB | FYN binding protein | Nucleus | other | 127347.414 | 834015.438 | 1467668.875 | | | | | | |
| O35601 | FYB | FYN binding protein | Nucleus | other | 45161.117 | 318618.219 | 729568.5 | | | | | | |
| O35601 | FYB | FYN binding protein | Nucleus | other | 112323.102 | 531749.062 | 792063.75 | O35601 | FYB | 94944 | 459965 | 153322 | 9 |
| P97855 | G3BP1 | GTPase activating protein (SH3 domain) binding protein 1 | Nucleus | enzyme | 5033478 | 1.03E+07 | 4111123 | | | | | | |
| P97855 | G3BP1 | GTPase activating protein (SH3 domain) binding protein 1 | Nucleus | enzyme | 4970944 | 8501957 | 5734026 | P97855 | G3BP1 | 5002211 | 2413821 | 985438 | 6 |
| Q6NSQ9 | G6PC3 | glucose 6 phosphatase, catalytic, 3 | Cytoplasm | phosphatase | 776099.5 | 640935.438 | 1408151.5 | Q6NSQ9 | G6PC3 | 776100 | 409548 | 236453 | 3 |
| Q00612 | G6PD | glucose-6-phosphate dehydrogenase | Cytoplasm | enzyme | 1661111 | 1352184.5 | 1707122.25 | | | | | | |
| Q00612 | G6PD | glucose-6-phosphate dehydrogenase | Cytoplasm | enzyme | 1047866.438 | 542283 | 810146 | | | | | | |
| Q00612 | G6PD | glucose-6-phosphate dehydrogenase | Cytoplasm | enzyme | 587102.562 | 1099168.5 | 1348266 | | | | | | |
| Q00612 | G6PD | glucose-6-phosphate dehydrogenase | Cytoplasm | enzyme | 1403060.375 | 676055.062 | 457787.844 | Q00612 | G6PD | 1174785 | 440252 | 127090 | 12 |
| P60521 | GABARAPL2 | GABA(A) receptor-associated protein like 2 | Cytoplasm | other | 1.69E+07 | 2.26E+07 | 1.70E+07 | | | | | | |
| P60521 | GABARAPL2 | GABA(A) receptor-associated protein like 2 | Cytoplasm | other | 1.21E+07 | 1.54E+07 | 1.28E+07 | | | | | | |
| P60521 | GABARAPL2 | GABA(A) receptor-associated protein like 2 | Cytoplasm | other | 1.02E+07 | 1.46E+07 | 1.09E+07 | | | | | | |
| P60521 | GABARAPL2 | GABA(A) receptor-associated protein like 2 | Cytoplasm | other | 6075298.5 | 1.14E+07 | 8378706 | | | | | | |
| P60521 | GABARAPL2 | GABA(A) receptor-associated protein like 2 | Cytoplasm | other | 1456264.5 | 2251175 | 1050209.125 | P60521 | GABARAPL2 | 9350313 | 6203879 | 1601835 | 15 |
| Q8BHN3 | GANAB | glucosidase, alpha; neutral AB | Cytoplasm | enzyme | 1.58E+07 | 1.87E+07 | 1.57E+07 | | | | | | |
| Q8BHN3 | GANAB | glucosidase, alpha; neutral AB | Cytoplasm | enzyme | 1.41E+07 | 1.47E+07 | 1.32E+07 | | | | | | |
| Q8BHN3 | GANAB | glucosidase, alpha; neutral AB | Cytoplasm | enzyme | 1.01E+07 | 1.15E+07 | 1.36E+07 | | | | | | |
| Q8BHN3 | GANAB | glucosidase, alpha; neutral AB | Cytoplasm | enzyme | 8092371 | 7467637 | 4018772.5 | | | | | | |
| Q8BHN3 | GANAB | glucosidase, alpha; neutral AB | Cytoplasm | enzyme | 1.04E+07 | 1.23E+07 | 1.19E+07 | | | | | | |
| Q8BHN3 | GANAB | glucosidase, alpha; neutral AB | Cytoplasm | enzyme | 7486316 | 8576088 | 9969620 | | | | | | |
| Q8BHN3 | GANAB | glucosidase, alpha; neutral AB | Cytoplasm | enzyme | 9218666 | 1.10E+07 | 1.31E+07 | | | | | | |
| Q8BHN3 | GANAB | glucosidase, alpha; neutral AB | Cytoplasm | enzyme | 7743957 | 6687679.5 | 7674057 | | | | | | |
| Q8BHN3 | GANAB | glucosidase, alpha; neutral AB | Cytoplasm | enzyme | 7692496 | 8249871.5 | 8049988.5 | | | | | | |
| Q8BHN3 | GANAB | glucosidase, alpha; neutral AB | Cytoplasm | enzyme | 5757370 | 7666391.5 | 8204621.5 | | | | | | |
| Q8BHN3 | GANAB | glucosidase, alpha; neutral AB | Cytoplasm | enzyme | 6039562 | 6751055.5 | 6711481 | | | | | | |
| Q8BHN3 | GANAB | glucosidase, alpha; neutral AB | Cytoplasm | enzyme | 6294067 | 4141614 | 3049838.5 | | | | | | |
| Q8BHN3 | GANAB | glucosidase, alpha; neutral AB | Cytoplasm | enzyme | 3703259.25 | 4832292.5 | 5138245.5 | | | | | | |
| Q8BHN3 | GANAB | glucosidase, alpha; neutral AB | Cytoplasm | enzyme | 3503458 | 5343299.5 | 6239653.5 | | | | | | |
| Q8BHN3 | GANAB | glucosidase, alpha; neutral AB | Cytoplasm | enzyme | 3601339.5 | 4135426.75 | 4163097.25 | | | | | | |
| Q8BHN3 | GANAB | glucosidase, alpha; neutral AB | Cytoplasm | enzyme | 2894884.25 | 3572585.25 | 3724183.25 | | | | | | |
| Q8BHN3 | GANAB | glucosidase, alpha; neutral AB | Cytoplasm | enzyme | 2059559.75 | 3199407.5 | 2962249.25 | | | | | | |
| Q8BHN3 | GANAB | glucosidase, alpha; neutral AB | Cytoplasm | enzyme | 1891799.875 | 1865025.625 | 2781582.5 | | | | | | |
| Q8BHN3 | GANAB | glucosidase, alpha; neutral AB | Cytoplasm | enzyme | 1680168.25 | 1497847.375 | 1420813.875 | | | | | | |
| Q8BHN3 | GANAB | glucosidase, alpha; neutral AB | Cytoplasm | enzyme | 322174.219 | 1096558.25 | 1028147.125 | Q8BHN3 | GANAB | 6417572 | 4356626 | 562438 | 60 |
| P16858 | GAPDH | glyceraldehyde-3-phosphate dehydrogenase | Cytoplasm | enzyme | 5.22E+08 | 3.59E+08 | 3.08E+08 | | | | | | |
| P16858 | GAPDH | glyceraldehyde-3-phosphate dehydrogenase | Cytoplasm | enzyme | 4.14E+08 | 2.72E+08 | 1.94E+08 | | | | | | |
| P16858 | GAPDH | glyceraldehyde-3-phosphate dehydrogenase | Cytoplasm | enzyme | 3.31E+08 | 2.17E+08 | 2.52E+08 | | | | | | |
| P16858 | GAPDH | glyceraldehyde-3-phosphate dehydrogenase | Cytoplasm | enzyme | 3.81E+08 | 1.60E+08 | 1.23E+08 | | | | | | |
| P16858 | GAPDH | glyceraldehyde-3-phosphate dehydrogenase | Cytoplasm | enzyme | 2.42E+08 | 2.51E+08 | 2.73E+08 | | | | | | |
| P16858 | GAPDH | glyceraldehyde-3-phosphate dehydrogenase | Cytoplasm | enzyme | 2.38E+08 | 1.17E+08 | 1.08E+08 | | | | | | |
| P16858 | GAPDH | glyceraldehyde-3-phosphate dehydrogenase | Cytoplasm | enzyme | 2.99E+08 | 2.15E+08 | 1.83E+08 | | | | | | |
| P16858 | GAPDH | glyceraldehyde-3-phosphate dehydrogenase | Cytoplasm | enzyme | 2.43E+08 | 1.81E+08 | 1.36E+08 | | | | | | |
| P16858 | GAPDH | glyceraldehyde-3-phosphate dehydrogenase | Cytoplasm | enzyme | 1.47E+08 | 1.61E+08 | 1.38E+08 | | | | | | |
| P16858 | GAPDH | glyceraldehyde-3-phosphate dehydrogenase | Cytoplasm | enzyme | 1.33E+08 | 1.98E+08 | 2.57E+08 | | | | | | |
| P16858 | GAPDH | glyceraldehyde-3-phosphate dehydrogenase | Cytoplasm | enzyme | 1.11E+08 | 1.03E+08 | 8.05E+07 | | | | | | |
| P16858 | GAPDH | glyceraldehyde-3-phosphate dehydrogenase | Cytoplasm | enzyme | 8.03E+07 | 4.31E+07 | 4.66E+07 | | | | | | |
| P16858 | GAPDH | glyceraldehyde-3-phosphate dehydrogenase | Cytoplasm | enzyme | 9.20E+07 | 8.17E+07 | 7.45E+07 | | | | | | |
| P16858 | GAPDH | glyceraldehyde-3-phosphate dehydrogenase | Cytoplasm | enzyme | 7.67E+07 | 6.48E+07 | 1.08E+08 | | | | | | |
| P16858 | GAPDH | glyceraldehyde-3-phosphate dehydrogenase | Cytoplasm | enzyme | 8.48E+07 | 1.97E+07 | 2720732.25 | | | | | | |
| P16858 | GAPDH | glyceraldehyde-3-phosphate dehydrogenase | Cytoplasm | enzyme | 4.50E+07 | 3.18E+07 | 3.20E+07 | | | | | | |
| P16858 | GAPDH | glyceraldehyde-3-phosphate dehydrogenase | Cytoplasm | enzyme | 2.27E+07 | 2.03E+07 | 1.60E+07 | | | | | | |
| P16858 | GAPDH | glyceraldehyde-3-phosphate dehydrogenase | Cytoplasm | enzyme | 1.63E+07 | 1.86E+07 | 1.60E+07 | | | | | | |

| | | | | | | | | | | | | | |
|---|---|---|---|---|---|---|---|---|---|---|---|---|---|
| P16858 | GAPDH | glyceraldehyde-3-phosphate dehydrogenase | Cytoplasm | enzyme | 2.83E+07 | 9251991 | 5843681 | | | | | | |
| P16858 | GAPDH | glyceraldehyde-3-phosphate dehydrogenase | Cytoplasm | enzyme | 3.44E+07 | 4.78E+07 | 1.11E+08 | | | | | | |
| P16858 | GAPDH | glyceraldehyde-3-phosphate dehydrogenase | Cytoplasm | enzyme | 1.76E+07 | 1.05E+07 | 1.05E+07 | | | | | | |
| P16858 | GAPDH | glyceraldehyde-3-phosphate dehydrogenase | Cytoplasm | enzyme | 2.43E+07 | 1.41E+07 | 1.51E+07 | | | | | | |
| P16858 | GAPDH | glyceraldehyde-3-phosphate dehydrogenase | Cytoplasm | enzyme | 1.09E+07 | 1.33E+07 | 1.25E+07 | | | | | | |
| P16858 | GAPDH | glyceraldehyde-3-phosphate dehydrogenase | Cytoplasm | enzyme | 5782594.5 | 1.24E+07 | 2.36E+07 | | | | | | |
| P16858 | GAPDH | glyceraldehyde-3-phosphate dehydrogenase | Cytoplasm | enzyme | 1.62E+07 | 1.30E+07 | 1.25E+07 | | | | | | |
| P16858 | GAPDH | glyceraldehyde-3-phosphate dehydrogenase | Cytoplasm | enzyme | 8707336 | 1.02E+07 | 1.09E+07 | | | | | | |
| P16858 | GAPDH | glyceraldehyde-3-phosphate dehydrogenase | Cytoplasm | enzyme | 9485359 | 4871562.5 | 2308100.75 | | | | | | |
| P16858 | GAPDH | glyceraldehyde-3-phosphate dehydrogenase | Cytoplasm | enzyme | 5871448 | 4745589 | 4671147 | | | | | | |
| P16858 | GAPDH | glyceraldehyde-3-phosphate dehydrogenase | Cytoplasm | enzyme | 2915532.25 | 841600.938 | 294537.688 | P16858 | GAPDH | 125578354 | 115425269 | 12374888 | 87 |
| Q6PAR5 | GAPVD1 | GTPase activating protein and VPS9 domains 1 | Cytoplasm | other | 1.02E+07 | 8806584 | 7196001 | | | | | | |
| Q6PAR5 | GAPVD1 | GTPase activating protein and VPS9 domains 1 | Cytoplasm | other | 9327040 | 6872427.5 | 8273107 | | | | | | |
| Q6PAR5 | GAPVD1 | GTPase activating protein and VPS9 domains 1 | Cytoplasm | other | 1.07E+07 | 7098350 | 5639846 | | | | | | |
| Q6PAR5 | GAPVD1 | GTPase activating protein and VPS9 domains 1 | Cytoplasm | other | 7684490 | 5751821.5 | 5204901.5 | | | | | | |
| Q6PAR5 | GAPVD1 | GTPase activating protein and VPS9 domains 1 | Cytoplasm | other | 9463165 | 6842439 | 6658595 | | | | | | |
| Q6PAR5 | GAPVD1 | GTPase activating protein and VPS9 domains 1 | Cytoplasm | other | 7227276.5 | 6178375.5 | 4873725 | | | | | | |
| Q6PAR5 | GAPVD1 | GTPase activating protein and VPS9 domains 1 | Cytoplasm | other | 6504410 | 6894915.5 | 6529615 | | | | | | |
| Q6PAR5 | GAPVD1 | GTPase activating protein and VPS9 domains 1 | Cytoplasm | other | 6329602 | 4091090.5 | 2630397.5 | | | | | | |
| Q6PAR5 | GAPVD1 | GTPase activating protein and VPS9 domains 1 | Cytoplasm | other | 6128513 | 4239972.5 | 3776941.5 | | | | | | |
| Q6PAR5 | GAPVD1 | GTPase activating protein and VPS9 domains 1 | Cytoplasm | other | 4876735 | 3171073.5 | 3967028.75 | | | | | | |
| Q6PAR5 | GAPVD1 | GTPase activating protein and VPS9 domains 1 | Cytoplasm | other | 4156185.75 | 2423465 | 2517641.25 | | | | | | |
| Q6PAR5 | GAPVD1 | GTPase activating protein and VPS9 domains 1 | Cytoplasm | other | 3951390.25 | 2933440 | 3874501.75 | | | | | | |
| Q6PAR5 | GAPVD1 | GTPase activating protein and VPS9 domains 1 | Cytoplasm | other | 3370205.25 | 2735988 | 3618036.25 | | | | | | |
| Q6PAR5 | GAPVD1 | GTPase activating protein and VPS9 domains 1 | Cytoplasm | other | 3165370.75 | 1991468 | 1162238.25 | | | | | | |
| Q6PAR5 | GAPVD1 | GTPase activating protein and VPS9 domains 1 | Cytoplasm | other | 3019053.75 | 1951629.375 | 2194106 | | | | | | |
| Q6PAR5 | GAPVD1 | GTPase activating protein and VPS9 domains 1 | Cytoplasm | other | 1891540.25 | 1185333 | 1389392.5 | | | | | | |
| Q6PAR5 | GAPVD1 | GTPase activating protein and VPS9 domains 1 | Cytoplasm | other | 1691783.25 | 1574276.25 | 1005810.312 | Q6PAR5 | GAPVD1 | 5861574 | 2578823 | 361108 | 51 |
| Q3V0G7 | GARNL3 | GTPase activating Rap/RanGAP domain-like 3 | Cytoplasm | other | 3028669.75 | 2233834.25 | 3076191.75 | Q3V0G7 | GARNL3 | 3028670 | 473214 | 273210 | 3 |
| O09172 | GCLM | glutamate-cysteine ligase, modifier subunit | Cytoplasm | enzyme | 4464148 | 5475643 | 5658805 | | | | | | |
| O09172 | GCLM | glutamate-cysteine ligase, modifier subunit | Cytoplasm | enzyme | 5095266 | 3294045.75 | 4370644 | | | | | | |
| O09172 | GCLM | glutamate-cysteine ligase, modifier subunit | Cytoplasm | enzyme | 3876075.5 | 3011663 | 3154832 | O09172 | GCLM | 4478497 | 1002860 | 334287 | 9 |
| Q9R111 | GDA | guanine deaminase | Cytoplasm | enzyme | 1.88E+07 | 1.51E+07 | 2.00E+07 | | | | | | |
| Q9R111 | GDA | guanine deaminase | Cytoplasm | enzyme | 1.62E+07 | 1.28E+07 | 1.56E+07 | | | | | | |
| Q9R111 | GDA | guanine deaminase | Cytoplasm | enzyme | 1.68E+07 | 1.40E+07 | 1.30E+07 | | | | | | |
| Q9R111 | GDA | guanine deaminase | Cytoplasm | enzyme | 1.07E+07 | 7081432.5 | 9631868 | | | | | | |
| Q9R111 | GDA | guanine deaminase | Cytoplasm | enzyme | 1.20E+07 | 8977466 | 9030017 | | | | | | |
| Q9R111 | GDA | guanine deaminase | Cytoplasm | enzyme | 6253869 | 3535461.5 | 4385360 | | | | | | |
| Q9R111 | GDA | guanine deaminase | Cytoplasm | enzyme | 1.07E+07 | 6864727.5 | 5466428.5 | | | | | | |
| Q9R111 | GDA | guanine deaminase | Cytoplasm | enzyme | 5270547.5 | 3694214.25 | 4664861 | | | | | | |
| Q9R111 | GDA | guanine deaminase | Cytoplasm | enzyme | 6779030 | 5818504.5 | 6386475 | | | | | | |
| Q9R111 | GDA | guanine deaminase | Cytoplasm | enzyme | 3370184.5 | 1404001 | 1593323.75 | | | | | | |
| Q9R111 | GDA | guanine deaminase | Cytoplasm | enzyme | 2226183.75 | 1562834.25 | 1743950.625 | Q9R111 | GDA | 9905438 | 5401275 | 940241 | 33 |
| Q9JL56 | GDE1 | glycerophosphodiester phosphodiesterase 1 | Plasma Membrane | enzyme | 7334498.5 | 8465919 | 7068826 | | | | | | |
| Q9JL56 | GDE1 | glycerophosphodiester phosphodiesterase 1 | Plasma Membrane | enzyme | 2812488.25 | 2684460.5 | 3255469.5 | | | | | | |
| Q9JL56 | GDE1 | glycerophosphodiester phosphodiesterase 1 | Plasma Membrane | enzyme | 1250168.625 | 969658.938 | 1191673.125 | Q9JL56 | GDE1 | 3799052 | 2930519 | 976840 | 9 |
| Q9CRY7 | GDPD1 | glycerophosphodiester phosphodiesterase domain containing 1 | Plasma Membrane | enzyme | 2023211 | 2758177 | 2786921.5 | Q9CRY7 | GDPD1 | 2023211 | 432869 | 249917 | 3 |
| Q91VC9 | GHITM | growth hormone inducible transmembrane protein | Cytoplasm | other | 1760308.625 | 3392044 | 1109000.125 | | | | | | |
| Q91VC9 | GHITM | growth hormone inducible transmembrane protein | Cytoplasm | other | 1267365.5 | 2714508.75 | 776477.375 | Q91VC9 | GHITM | 1513837 | 1017176 | 415261 | 6 |
| Q9CYL5 | GLIPR2 | GLI pathogenesis-related 2 | Cytoplasm | other | 4251147.5 | 3754986.25 | 3324728 | Q9CYL5 | GLIPR2 | 4251148 | 463600 | 267660 | 3 |
| Q80Y14 | GLRX5 | glutaredoxin 5 | Cytoplasm | other | 2.15E+07 | 4.57E+07 | 2.10E+07 | | | | | | |
| Q80Y14 | GLRX5 | glutaredoxin 5 | Cytoplasm | other | 1.41E+07 | 2.39E+07 | 9267809 | Q80Y14 | GLRX5 | 17835000 | 12538907 | 5118987 | 6 |
| P26443 | GLUD1 | glutamate dehydrogenase 1 | Cytoplasm | enzyme | 845038.938 | 1430108.875 | 1474469.875 | | | | | | |
| P26443 | GLUD1 | glutamate dehydrogenase 1 | Cytoplasm | enzyme | 184272.672 | 1355358 | 874624.25 | P26443 | GLUD1 | 514656 | 497406 | 203065 | 6 |
| P63158 | Gm21596/Hmgb1 | high mobility group box 1 | Nucleus | transcription regulator | 4.70E-06 | 193309.75 | 55953.293 | | | | | | |
| P63158 | Gm21596/Hmgb1 | high mobility group box 1 | Nucleus | transcription regulator | 4.70E-06 | 68712.766 | 4.70E-06 | P63158 | Gm21596/Hmgb1 | 0 | 75325 | 30751 | 6 |
| P21278 | GNA11 | guanine nucleotide binding protein (G protein), alpha 11 (Gq class) | Plasma Membrane | enzyme | 683507.5 | 1297951.5 | 1568094.25 | P21278 | GNA11 | 683508 | 453323 | 261726 | 3 |
| P27600 | GNA12 | guanine nucleotide binding protein (G protein) alpha 12 | Plasma Membrane | enzyme | 5.05E+07 | 6.35E+07 | 6.68E+07 | P27600 | GNA12 | 50510000 | 8617592 | 4975369 | 3 |
| P08752 | GNAI2 | guanine nucleotide binding protein (G protein), alpha inhibiting activity polypeptide 2 | Plasma Membrane | other | 9.08E+07 | 1.02E+08 | 1.08E+08 | | | | | | |
| P08752 | GNAI2 | guanine nucleotide binding protein (G protein), alpha inhibiting activity polypeptide 2 | Plasma Membrane | other | 8.11E+07 | 9.28E+07 | 9.30E+07 | | | | | | |
| P08752 | GNAI2 | guanine nucleotide binding protein (G protein), alpha inhibiting activity polypeptide 2 | Plasma Membrane | other | 5.18E+07 | 6.29E+07 | 6.25E+07 | | | | | | |
| P08752 | GNAI2 | guanine nucleotide binding protein (G protein), alpha inhibiting activity polypeptide 2 | Plasma Membrane | other | 6.49E+07 | 5.14E+07 | 4.50E+07 | | | | | | |
| P08752 | GNAI2 | guanine nucleotide binding protein (G protein), alpha inhibiting activity polypeptide 2 | Plasma Membrane | other | 4.13E+07 | 7.55E+07 | 7.85E+07 | | | | | | |
| P08752 | GNAI2 | guanine nucleotide binding protein (G protein), alpha inhibiting activity polypeptide 2 | Plasma Membrane | other | 5.45E+07 | 6.72E+07 | 6.05E+07 | | | | | | |
| P08752 | GNAI2 | guanine nucleotide binding protein (G protein), alpha inhibiting activity polypeptide 2 | Plasma Membrane | other | 4.42E+07 | 5.67E+07 | 5.49E+07 | | | | | | |
| P08752 | GNAI2 | guanine nucleotide binding protein (G protein), alpha inhibiting activity polypeptide 2 | Plasma Membrane | other | 3.90E+07 | 4.35E+07 | 4.34E+07 | | | | | | |
| P08752 | GNAI2 | guanine nucleotide binding protein (G protein), alpha inhibiting activity polypeptide 2 | Plasma Membrane | other | 3.56E+07 | 3.87E+07 | 3.69E+07 | | | | | | |
| P08752 | GNAI2 | guanine nucleotide binding protein (G protein), alpha inhibiting activity polypeptide 2 | Plasma Membrane | other | 3.82E+07 | 4.98E+07 | 5.03E+07 | | | | | | |
| P08752 | GNAI2 | guanine nucleotide binding protein (G protein), alpha inhibiting activity polypeptide 2 | Plasma Membrane | other | 1.97E+07 | 2.49E+07 | 2.42E+07 | | | | | | |
| P08752 | GNAI2 | guanine nucleotide binding protein (G protein), alpha inhibiting activity polypeptide 2 | Plasma Membrane | other | 1.37E+07 | 2.19E+07 | 2.00E+07 | | | | | | |
| P08752 | GNAI2 | guanine nucleotide binding protein (G protein), alpha inhibiting activity polypeptide 2 | Plasma Membrane | other | 1.33E+07 | 1.60E+07 | 1.24E+07 | | | | | | |
| P08752 | GNAI2 | guanine nucleotide binding protein (G protein), alpha inhibiting activity polypeptide 2 | Plasma Membrane | other | 1.77E+07 | 2.16E+07 | 2.43E+07 | | | | | | |
| P08752 | GNAI2 | guanine nucleotide binding protein (G protein), alpha inhibiting activity polypeptide 2 | Plasma Membrane | other | 1.50E+07 | 1.56E+07 | 1.67E+07 | | | | | | |
| P08752 | GNAI2 | guanine nucleotide binding protein (G protein), alpha inhibiting activity polypeptide 2 | Plasma Membrane | other | 1.41E+07 | 1.66E+07 | 1.49E+07 | | | | | | |
| P08752 | GNAI2 | guanine nucleotide binding protein (G protein), alpha inhibiting activity polypeptide 2 | Plasma Membrane | other | 8309236 | 1.80E+07 | 1.44E+07 | | | | | | |
| P08752 | GNAI2 | guanine nucleotide binding protein (G protein), alpha inhibiting activity polypeptide 2 | Plasma Membrane | other | 1.18E+07 | 1.31E+07 | 1.31E+07 | | | | | | |
| P08752 | GNAI2 | guanine nucleotide binding protein (G protein), alpha inhibiting activity polypeptide 2 | Plasma Membrane | other | 1.20E+07 | 1.21E+07 | 1.45E+07 | | | | | | |
| P08752 | GNAI2 | guanine nucleotide binding protein (G protein), alpha inhibiting activity polypeptide 2 | Plasma Membrane | other | 1.66E+07 | 1.07E+07 | 7370815 | | | | | | |
| P08752 | GNAI2 | guanine nucleotide binding protein (G protein), alpha inhibiting activity polypeptide 2 | Plasma Membrane | other | 1.16E+07 | 1.87E+07 | 2.13E+07 | | | | | | |
| P08752 | GNAI2 | guanine nucleotide binding protein (G protein), alpha inhibiting activity polypeptide 2 | Plasma Membrane | other | 8189458 | 9266752 | 1.02E+07 | | | | | | |
| P08752 | GNAI2 | guanine nucleotide binding protein (G protein), alpha inhibiting activity polypeptide 2 | Plasma Membrane | other | 7894558.5 | 1.07E+07 | 9742593 | | | | | | |
| P08752 | GNAI2 | guanine nucleotide binding protein (G protein), alpha inhibiting activity polypeptide 2 | Plasma Membrane | other | 7669511.5 | 2710578.75 | 1095657.75 | | | | | | |
| P08752 | GNAI2 | guanine nucleotide binding protein (G protein), alpha inhibiting activity polypeptide 2 | Plasma Membrane | other | 5745730 | 5202865.5 | 4475407 | | | | | | |
| P08752 | GNAI2 | guanine nucleotide binding protein (G protein), alpha inhibiting activity polypeptide 2 | Plasma Membrane | other | 4140960 | 4022502.5 | 3429681.25 | P08752 | GNAI2 | 28023825 | 26969910 | 3053741 | 78 |
| Q9DC51 | GNAI3 | guanine nucleotide binding protein (G protein), alpha inhibiting activity polypeptide 3 | Cytoplasm | enzyme | 1544578.5 | 2842940.75 | 1914029.75 | Q9DC51 | GNAI3 | 1544579 | 668969 | 386229 | 3 |
| P21279 | GNAQ | guanine nucleotide binding protein (G protein), q polypeptide | Plasma Membrane | enzyme | 1992616.625 | 2204164.5 | 5006025.5 | P21279 | GNAQ | 1992617 | 1682053 | 971134 | 3 |
| Q6R0H7 | GNAS | GNAS complex locus | Plasma Membrane | enzyme | 2315305.25 | 2734876.75 | 3788390 | | | | | | |
| Q6R0H7 | GNAS | GNAS complex locus | Plasma Membrane | enzyme | 3627753.5 | 3480781.75 | 3468722 | Q6R0H7 | GNAS | 2971529 | 578184 | 236043 | 6 |
| O70443 | GNAZ | guanine nucleotide binding protein (G protein), alpha z polypeptide | Plasma Membrane | enzyme | 183617.906 | 949914.688 | 1016222.812 | | | | | | |
| O70443 | GNAZ | guanine nucleotide binding protein (G protein), alpha z polypeptide | Plasma Membrane | enzyme | 29750.379 | 446749.344 | 1518881 | O70443 | GNAZ | 106684 | 567898 | 231843 | 6 |
| P62874 | GNB1 | guanine nucleotide binding protein (G protein), beta polypeptide 1 | Plasma Membrane | enzyme | 6.85E+07 | 8.17E+07 | 7.73E+07 | | | | | | |

| | | | | | | | | | | | | | |
|---|---|---|---|---|---|---|---|---|---|---|---|---|---|
| P62874 | GNB1 | guanine nucleotide binding protein (G protein), beta polypeptide 1 | Plasma Membrane | enzyme | 6.21E+07 | 6.32E+07 | 6.32E+07 | | | | | | |
| P62874 | GNB1 | guanine nucleotide binding protein (G protein), beta polypeptide 1 | Plasma Membrane | enzyme | 2.79E+07 | 2.46E+07 | 2.17E+07 | | | | | | |
| P62874 | GNB1 | guanine nucleotide binding protein (G protein), beta polypeptide 1 | Plasma Membrane | enzyme | 2.48E+07 | 3.73E+07 | 2.99E+07 | | | | | | |
| P62874 | GNB1 | guanine nucleotide binding protein (G protein), beta polypeptide 1 | Plasma Membrane | enzyme | 1.91E+07 | 1.79E+07 | 2.30E+07 | | | | | | |
| P62874 | GNB1 | guanine nucleotide binding protein (G protein), beta polypeptide 1 | Plasma Membrane | enzyme | 1.79E+07 | 2.28E+07 | 2.25E+07 | | | | | | |
| P62874 | GNB1 | guanine nucleotide binding protein (G protein), beta polypeptide 1 | Plasma Membrane | enzyme | 1.17E+07 | 1.17E+07 | 1.00E+07 | | | | | | |
| P62874 | GNB1 | guanine nucleotide binding protein (G protein), beta polypeptide 1 | Plasma Membrane | enzyme | 1.43E+07 | 1.66E+07 | 1.50E+07 | | | | | | |
| P62874 | GNB1 | guanine nucleotide binding protein (G protein), beta polypeptide 1 | Plasma Membrane | enzyme | 1.09E+07 | 1.27E+07 | 1.42E+07 | | | | | | |
| P62874 | GNB1 | guanine nucleotide binding protein (G protein), beta polypeptide 1 | Plasma Membrane | enzyme | 8581662 | 8844198 | 8266109 | | | | | | |
| P62874 | GNB1 | guanine nucleotide binding protein (G protein), beta polypeptide 1 | Plasma Membrane | enzyme | 6321967 | 4931044.5 | 2532432 | | | | | | |
| P62874 | GNB1 | guanine nucleotide binding protein (G protein), beta polypeptide 1 | Plasma Membrane | enzyme | 3651043.5 | 2931346 | 2480665.5 | | | | | | |
| P62874 | GNB1 | guanine nucleotide binding protein (G protein), beta polypeptide 1 | Plasma Membrane | enzyme | 2167457.75 | 3050822.75 | 2919244.75 | P62874 | GNB1 | 21375548 | 22120278 | 3542079 | 39 |
| P62880 | GNB2 | guanine nucleotide binding protein (G protein), beta polypeptide 2 | Plasma Membrane | enzyme | 5.61E+07 | 7.85E+07 | 6.76E+07 | | | | | | |
| P62880 | GNB2 | guanine nucleotide binding protein (G protein), beta polypeptide 2 | Plasma Membrane | enzyme | 4.33E+07 | 5.02E+07 | 5.26E+07 | | | | | | |
| P62880 | GNB2 | guanine nucleotide binding protein (G protein), beta polypeptide 2 | Plasma Membrane | enzyme | 4.20E+07 | 5.81E+07 | 4.83E+07 | | | | | | |
| P62880 | GNB2 | guanine nucleotide binding protein (G protein), beta polypeptide 2 | Plasma Membrane | enzyme | 4.13E+07 | 4.50E+07 | 4.68E+07 | | | | | | |
| P62880 | GNB2 | guanine nucleotide binding protein (G protein), beta polypeptide 2 | Plasma Membrane | enzyme | 2.94E+07 | 3.26E+07 | 2.44E+07 | | | | | | |
| P62880 | GNB2 | guanine nucleotide binding protein (G protein), beta polypeptide 2 | Plasma Membrane | enzyme | 2.84E+07 | 3.34E+07 | 2.76E+07 | | | | | | |
| P62880 | GNB2 | guanine nucleotide binding protein (G protein), beta polypeptide 2 | Plasma Membrane | enzyme | 1.10E+07 | 1.65E+07 | 2.18E+07 | | | | | | |
| P62880 | GNB2 | guanine nucleotide binding protein (G protein), beta polypeptide 2 | Plasma Membrane | enzyme | 1.44E+07 | 1.91E+07 | 1.81E+07 | | | | | | |
| P62880 | GNB2 | guanine nucleotide binding protein (G protein), beta polypeptide 2 | Plasma Membrane | enzyme | 9388835 | 9720349 | 7510941 | | | | | | |
| P62880 | GNB2 | guanine nucleotide binding protein (G protein), beta polypeptide 2 | Plasma Membrane | enzyme | 5258561.5 | 6471323 | 4102194.25 | | | | | | |
| P62880 | GNB2 | guanine nucleotide binding protein (G protein), beta polypeptide 2 | Plasma Membrane | enzyme | 2933081.75 | 3348579.25 | 6290458.5 | | | | | | |
| P62880 | GNB2 | guanine nucleotide binding protein (G protein), beta polypeptide 2 | Plasma Membrane | enzyme | 3648265.25 | 5129210 | 5865070.5 | P62880 | GNB2 | 23913229 | 20965472 | 3494245 | 36 |
| P68040 | GNB2L1 | guanine nucleotide binding protein (G protein), beta polypeptide 2-like 1 | Cytoplasm | enzyme | 6869123.5 | 1.52E+07 | 4874232 | | | | | | |
| P68040 | GNB2L1 | guanine nucleotide binding protein (G protein), beta polypeptide 2-like 1 | Cytoplasm | enzyme | 5647358.5 | 9592968 | 3904135.5 | | | | | | |
| P68040 | GNB2L1 | guanine nucleotide binding protein (G protein), beta polypeptide 2-like 1 | Cytoplasm | enzyme | 5439587 | 8095872.5 | 2942568.5 | | | | | | |
| P68040 | GNB2L1 | guanine nucleotide binding protein (G protein), beta polypeptide 2-like 1 | Cytoplasm | enzyme | 4724708 | 7964642.5 | 2529443.5 | | | | | | |
| P68040 | GNB2L1 | guanine nucleotide binding protein (G protein), beta polypeptide 2-like 1 | Cytoplasm | enzyme | 2439015 | 3547167 | 1174033.125 | P68040 | GNB2L1 | 5023958 | 3552891 | 917353 | 15 |
| Q9DAS9 | GNG12 | guanine nucleotide binding protein (G protein), gamma 12 | Plasma Membrane | enzyme | 6157656.5 | 6934503 | 7911516 | Q9DAS9 | GNG12 | 6157657 | 878831 | 507394 | 3 |
| Q91W53 | GOLGA7 | golgin A7 | Cytoplasm | other | 4946456 | 6242979.5 | 6568090 | | | | | | |
| Q91W53 | GOLGA7 | golgin A7 | Cytoplasm | other | 4280903 | 5118668 | 4298298 | Q91W53 | GOLGA7 | 4613680 | 967193 | 394855 | 6 |
| Q9CR60 | GOLT1B | golgi transport 1B | Cytoplasm | other | 1626260.125 | 1539957.75 | 3349647 | Q9CR60 | GOLT1B | 1626260 | 1020824 | 589373 | 3 |
| P05202 | GOT2 | glutamic-oxaloacetic transaminase 2, mitochondrial | Cytoplasm | enzyme | 1.54E+07 | 3.26E+07 | 1.02E+07 | | | | | | |
| P05202 | GOT2 | glutamic-oxaloacetic transaminase 2, mitochondrial | Cytoplasm | enzyme | 1.08E+07 | 1.85E+07 | 8159020.5 | | | | | | |
| P05202 | GOT2 | glutamic-oxaloacetic transaminase 2, mitochondrial | Cytoplasm | enzyme | 9023545 | 1.46E+07 | 6856270.5 | | | | | | |
| P05202 | GOT2 | glutamic-oxaloacetic transaminase 2, mitochondrial | Cytoplasm | enzyme | 8024860.5 | 1.18E+07 | 3864843.75 | | | | | | |
| P05202 | GOT2 | glutamic-oxaloacetic transaminase 2, mitochondrial | Cytoplasm | enzyme | 5264918 | 9191991 | 2700692.5 | | | | | | |
| P05202 | GOT2 | glutamic-oxaloacetic transaminase 2, mitochondrial | Cytoplasm | enzyme | 5869510 | 1.14E+07 | 4977952.5 | | | | | | |
| P05202 | GOT2 | glutamic-oxaloacetic transaminase 2, mitochondrial | Cytoplasm | enzyme | 3854668.5 | 7318808.5 | 3588727.25 | | | | | | |
| P05202 | GOT2 | glutamic-oxaloacetic transaminase 2, mitochondrial | Cytoplasm | enzyme | 2227734 | 5836754 | 2177021.75 | | | | | | |
| P05202 | GOT2 | glutamic-oxaloacetic transaminase 2, mitochondrial | Cytoplasm | enzyme | 1777734.125 | 2954316.25 | 1011410.562 | | | | | | |
| P05202 | GOT2 | glutamic-oxaloacetic transaminase 2, mitochondrial | Cytoplasm | enzyme | 1337775.5 | 3847203.75 | 2523312.75 | | | | | | |
| P05202 | GOT2 | glutamic-oxaloacetic transaminase 2, mitochondrial | Cytoplasm | enzyme | 1319891.375 | 1483172.25 | 553578.688 | P05202 | GOT2 | 5900058 | 6480689 | 1128143 | 33 |
| O35930 | GP1BA | glycoprotein Ib (platelet), alpha polypeptide | Plasma Membrane | other | 2439750 | 1.06E+07 | 2.86E+07 | | | | | | |
| O35930 | GP1BA | glycoprotein Ib (platelet), alpha polypeptide | Plasma Membrane | other | 2918764.25 | 1.20E+07 | 2.76E+07 | | | | | | |
| O35930 | GP1BA | glycoprotein Ib (platelet), alpha polypeptide | Plasma Membrane | other | 1720854.875 | 7639158 | 1.73E+07 | | | | | | |
| O35930 | GP1BA | glycoprotein Ib (platelet), alpha polypeptide | Plasma Membrane | other | 1532557.625 | 5592529.5 | 1.25E+07 | | | | | | |
| O35930 | GP1BA | glycoprotein Ib (platelet), alpha polypeptide | Plasma Membrane | other | 1252714 | 6597704.5 | 1.09E+07 | | | | | | |
| O35930 | GP1BA | glycoprotein Ib (platelet), alpha polypeptide | Plasma Membrane | other | 912010.438 | 5147830.5 | 1.10E+07 | | | | | | |
| O35930 | GP1BA | glycoprotein Ib (platelet), alpha polypeptide | Plasma Membrane | other | 722550.938 | 3039556.25 | 7849414.5 | | | | | | |
| O35930 | GP1BA | glycoprotein Ib (platelet), alpha polypeptide | Plasma Membrane | other | 337453.469 | 2900228.75 | 6420335 | | | | | | |
| O35930 | GP1BA | glycoprotein Ib (platelet), alpha polypeptide | Plasma Membrane | other | 256025.391 | 1552864 | 4344534 | | | | | | |
| O35930 | GP1BA | glycoprotein Ib (platelet), alpha polypeptide | Plasma Membrane | other | 200800.797 | 846154.688 | 2941109.25 | | | | | | |
| O35930 | GP1BA | glycoprotein Ib (platelet), alpha polypeptide | Plasma Membrane | other | 181340.438 | 1257962.375 | 4514839.5 | | | | | | |
| O35930 | GP1BA | glycoprotein Ib (platelet), alpha polypeptide | Plasma Membrane | other | 285312 | 2664381 | 3699177 | | | | | | |
| O35930 | GP1BA | glycoprotein Ib (platelet), alpha polypeptide | Plasma Membrane | other | 4.70E-06 | 4.70E-06 | 932326.25 | O35930 | GP1BA | 981549 | 6807775 | 1090116 | 39 |
| P56400 | GP1BB | glycoprotein Ib (platelet), beta polypeptide | Plasma Membrane | other | 5645200.5 | 1.95E+07 | 4.44E+07 | | | | | | |
| P56400 | GP1BB | glycoprotein Ib (platelet), beta polypeptide | Plasma Membrane | other | 3090667 | 1.19E+07 | 2.86E+07 | | | | | | |
| P56400 | GP1BB | glycoprotein Ib (platelet), beta polypeptide | Plasma Membrane | other | 1659998.875 | 4388328 | 1.14E+07 | | | | | | |
| P56400 | GP1BB | glycoprotein Ib (platelet), beta polypeptide | Plasma Membrane | other | 1480211.25 | 5963578 | 1.25E+07 | | | | | | |
| P56400 | GP1BB | glycoprotein Ib (platelet), beta polypeptide | Plasma Membrane | other | 865213.062 | 2860629.75 | 5125469 | | | | | | |
| P56400 | GP1BB | glycoprotein Ib (platelet), beta polypeptide | Plasma Membrane | other | 732896.625 | 2930552.5 | 6879247 | P56400 | GP1BB | 2245698 | 11344530 | 2673931 | 18 |
| O08742 | GP5 | glycoprotein V (platelet) | Plasma Membrane | other | 2195560 | 1.22E+07 | 2.67E+07 | | | | | | |
| O08742 | GP5 | glycoprotein V (platelet) | Plasma Membrane | other | 1820088.625 | 5586921 | 9711743 | | | | | | |
| O08742 | GP5 | glycoprotein V (platelet) | Plasma Membrane | other | 1098291.625 | 6803120.5 | 1.23E+07 | | | | | | |
| O08742 | GP5 | glycoprotein V (platelet) | Plasma Membrane | other | 2122104.75 | 8362546.5 | 1.33E+07 | | | | | | |
| O08742 | GP5 | glycoprotein V (platelet) | Plasma Membrane | other | 1583652.625 | 6226135.5 | 1.39E+07 | | | | | | |
| O08742 | GP5 | glycoprotein V (platelet) | Plasma Membrane | other | 1487749.5 | 8340906 | 1.40E+07 | | | | | | |
| O08742 | GP5 | glycoprotein V (platelet) | Plasma Membrane | other | 502239 | 2602369.5 | 6375677 | | | | | | |
| O08742 | GP5 | glycoprotein V (platelet) | Plasma Membrane | other | 343763.938 | 1411811.25 | 4419981.5 | | | | | | |
| O08742 | GP5 | glycoprotein V (platelet) | Plasma Membrane | other | 246788.781 | 1627364.5 | 4625013.5 | | | | | | |
| O08742 | GP5 | glycoprotein V (platelet) | Plasma Membrane | other | 2920.934 | 1028850.562 | 1973882.5 | O08742 | GP5 | 1140316 | 6025589 | 1100117 | 30 |
| O88186 | GP9 | glycoprotein IX (platelet) | Plasma Membrane | other | 5040322 | 1.50E+07 | 3.46E+07 | | | | | | |
| O88186 | GP9 | glycoprotein IX (platelet) | Plasma Membrane | other | 2372966.75 | 4425656.5 | 8205838.5 | | | | | | |
| O88186 | GP9 | glycoprotein IX (platelet) | Plasma Membrane | other | 362351.906 | 768029.25 | 2223863.25 | | | | | | |
| O88186 | GP9 | glycoprotein IX (platelet) | Plasma Membrane | other | 50563.555 | 333981.875 | 2159396.75 | O88186 | GP9 | 1956551 | 9879114 | 2851855 | 12 |
| P51655 | GPC4 | glypican 4 | Plasma Membrane | transmembrane receptor | 3982271 | 4316686 | 4152201.25 | | | | | | |
| P51655 | GPC4 | glypican 4 | Plasma Membrane | transmembrane receptor | 3031519 | 3648897.25 | 2878493.25 | | | | | | |
| P51655 | GPC4 | glypican 4 | Plasma Membrane | transmembrane receptor | 2518656 | 1731773.375 | 1099123.25 | P51655 | GPC4 | 3177482 | 1112540 | 370847 | 9 |
| Q64521 | GPD2 | glycerol-3-phosphate dehydrogenase 2 (mitochondrial) | Cytoplasm | enzyme | 1122671.5 | 3498657.75 | 3070692 | | | | | | |
| Q64521 | GPD2 | glycerol-3-phosphate dehydrogenase 2 (mitochondrial) | Cytoplasm | enzyme | 1022978.562 | 4294296 | 4298404.5 | | | | | | |
| Q64521 | GPD2 | glycerol-3-phosphate dehydrogenase 2 (mitochondrial) | Cytoplasm | enzyme | 633423.875 | 1776683.625 | 1696336.25 | | | | | | |
| Q64521 | GPD2 | glycerol-3-phosphate dehydrogenase 2 (mitochondrial) | Cytoplasm | enzyme | 759452.5 | 2750637.5 | 2641259.75 | | | | | | |
| Q64521 | GPD2 | glycerol-3-phosphate dehydrogenase 2 (mitochondrial) | Cytoplasm | enzyme | 160330.562 | 569666.688 | 968178.5 | | | | | | |
| Q64521 | GPD2 | glycerol-3-phosphate dehydrogenase 2 (mitochondrial) | Cytoplasm | enzyme | 692804.188 | 2245080 | 2946526.75 | | | | | | |
| Q64521 | GPD2 | glycerol-3-phosphate dehydrogenase 2 (mitochondrial) | Cytoplasm | enzyme | 413550 | 1394262.875 | 2147326.5 | | | | | | |
| Q64521 | GPD2 | glycerol-3-phosphate dehydrogenase 2 (mitochondrial) | Cytoplasm | enzyme | 284416.312 | 1090459 | 1427023.625 | | | | | | |
| Q64521 | GPD2 | glycerol-3-phosphate dehydrogenase 2 (mitochondrial) | Cytoplasm | enzyme | 411010.375 | 1730286.125 | 2407124.5 | | | | | | |
| Q64521 | GPD2 | glycerol-3-phosphate dehydrogenase 2 (mitochondrial) | Cytoplasm | enzyme | 259142.875 | 1504762.125 | 1808675.375 | Q64521 | GPD2 | 575978 | 1161303 | 212024 | 30 |
| Q99LD4 | GPS1 | G protein pathway suppressor 1 | Cytoplasm | other | 6578965.5 | 7061936.5 | 3657473.5 | | | | | | |

| | | | | | | | | | | | | | |
|---|---|---|---|---|---|---|---|---|---|---|---|---|---|
| Q99LD4 | GPS1 | G protein pathway suppressor 1 | Cytoplasm | other | 6358703 | 4600224 | 3057387.25 | | | | | | |
| Q99LD4 | GPS1 | G protein pathway suppressor 1 | Cytoplasm | other | 3495736.75 | 1882566.875 | 1810063.375 | | | | | | |
| Q99LD4 | GPS1 | G protein pathway suppressor 1 | Cytoplasm | other | 1566373.5 | 1555739.875 | 1282596.625 | | | | | | |
| Q99LD4 | GPS1 | G protein pathway suppressor 1 | Cytoplasm | other | 775155.938 | 664941.75 | 1067664.375 | Q99LD4 | GPS1 | 3754987 | 2199210 | 567834 | 15 |
| P11352 | GPX1 | glutathione peroxidase 1 | Cytoplasm | enzyme | 4.10E+07 | 4.51E+07 | 3.33E+07 | | | | | | |
| P11352 | GPX1 | glutathione peroxidase 1 | Cytoplasm | enzyme | 3.72E+07 | 2.93E+07 | 2.40E+07 | | | | | | |
| P11352 | GPX1 | glutathione peroxidase 1 | Cytoplasm | enzyme | 2.44E+07 | 2.48E+07 | 2.22E+07 | | | | | | |
| P11352 | GPX1 | glutathione peroxidase 1 | Cytoplasm | enzyme | 1.54E+07 | 1.48E+07 | 1.69E+07 | | | | | | |
| P11352 | GPX1 | glutathione peroxidase 1 | Cytoplasm | enzyme | 1.05E+07 | 1.01E+07 | 1.04E+07 | | | | | | |
| P11352 | GPX1 | glutathione peroxidase 1 | Cytoplasm | enzyme | 1.20E+07 | 1.00E+07 | 9527644 | | | | | | |
| P11352 | GPX1 | glutathione peroxidase 1 | Cytoplasm | enzyme | 6577462 | 1.08E+07 | 1.11E+07 | | | | | | |
| P11352 | GPX1 | glutathione peroxidase 1 | Cytoplasm | enzyme | 4881759 | 9392809 | 1.01E+07 | | | | | | |
| P11352 | GPX1 | glutathione peroxidase 1 | Cytoplasm | enzyme | 4398214.5 | 4606049 | 2771485.25 | | | | | | |
| P11352 | GPX1 | glutathione peroxidase 1 | Cytoplasm | enzyme | 6133187 | 5783232 | 848461.25 | P11352 | GPX1 | 16249062 | 11819492 | 2157934 | 30 |
| O70325 | GPX4 | glutathione peroxidase 4 | Cytoplasm | enzyme | 3.95E+07 | 4.69E+07 | 4.76E+07 | | | | | | |
| O70325 | GPX4 | glutathione peroxidase 4 | Cytoplasm | enzyme | 3.85E+07 | 2.60E+07 | 3.05E+07 | | | | | | |
| O70325 | GPX4 | glutathione peroxidase 4 | Cytoplasm | enzyme | 1.89E+07 | 1.64E+07 | 1.60E+07 | | | | | | |
| O70325 | GPX4 | glutathione peroxidase 4 | Cytoplasm | enzyme | 1.65E+07 | 1.28E+07 | 1.32E+07 | | | | | | |
| O70325 | GPX4 | glutathione peroxidase 4 | Cytoplasm | enzyme | 1.45E+07 | 9672018 | 8520515 | | | | | | |
| O70325 | GPX4 | glutathione peroxidase 4 | Cytoplasm | enzyme | 5731733.5 | 4699371 | 3054685.75 | | | | | | |
| O70325 | GPX4 | glutathione peroxidase 4 | Cytoplasm | enzyme | 3448888 | 2574701.75 | 2324474 | O70325 | GPX4 | 19588660 | 14641341 | 3195003 | 21 |
| Q8R050 | GSPT1 | G1 to S phase transition 1 | Cytoplasm | translation regulator | 3179969 | 2643667.75 | 2402467.25 | Q8R050 | GSPT1 | 3179969 | 397975 | 229771 | 3 |
| P19157 | GSTP1 | glutathione S-transferase pi 1 | Cytoplasm | enzyme | 3261807.5 | 2526617 | 2834632.5 | P19157 | GSTP1 | 3261808 | 369201 | 213158 | 3 |
| Q8BZI6 | GUCD1 | guanylyl cyclase domain containing 1 | Nucleus | other | 2296322.5 | 2083575.875 | 2038321.875 | Q8BZI6 | GUCD1 | 2296323 | 137764 | 79538 | 3 |
| P14220 | Gypa | glycophorin A | Plasma Membrane | other | 4.73E+09 | 4.27E+09 | 2.81E+09 | | | | | | |
| P14220 | Gypa | glycophorin A | Plasma Membrane | other | 3.21E+07 | 1.91E+07 | 1.67E+07 | P14220 | Gypa | 2380555000 | 2234668239 | 912299488 | 6 |
| Q78HU7 | GYPC | glycophorin C (Gerbich blood group) | Plasma Membrane | other | 1.35E+09 | 7.89E+08 | 3.25E+08 | | | | | | |
| Q78HU7 | GYPC | glycophorin C (Gerbich blood group) | Plasma Membrane | other | 6.83E+08 | 1.12E+09 | 1.93E+09 | | | | | | |
| Q78HU7 | GYPC | glycophorin C (Gerbich blood group) | Plasma Membrane | other | 5.30E+07 | 3.44E+07 | 1.17E+07 | Q78HU7 | GYPC | 696406667 | 671651046 | 223883682 | 9 |
| Q3THW5 | H2AFV | H2A histone family, member V | Nucleus | other | 485291.969 | 853321.125 | 695843.938 | Q3THW5 | H2AFV | 485292 | 184651 | 106608 | 3 |
| P27661 | H2AFX | H2A histone family, member X | Nucleus | transcription regulator | 1165320.375 | 3117472.25 | 3148641.75 | | | | | | |
| P27661 | H2AFX | H2A histone family, member X | Nucleus | transcription regulator | 81281.086 | 80240.508 | 170170.625 | P27661 | H2AFX | 623301 | 1482444 | 605205 | 6 |
| Q9QZQ8 | H2AFY | H2A histone family, member Y | Nucleus | other | 108398.836 | 369188.938 | 466624.438 | Q9QZQ8 | H2AFY | 108399 | 185216 | 106935 | 3 |
| P68433 | H31 | Histone H3.1 | Nucleus | | 1.47E+07 | 2.81E+07 | 2.79E+07 | | | | | | |
| P68433 | H31 | Histone H3.1 | Nucleus | | 1.52E+07 | 2.38E+07 | 2.04E+07 | | | | | | |
| P68433 | H31 | Histone H3.1 | Nucleus | | 4393233 | 8323001 | 5962468.5 | | | | | | |
| P68433 | H31 | Histone H3.1 | Nucleus | | 5890465.5 | 9926487 | 1.08E+07 | | | | | | |
| P68433 | H31 | Histone H3.1 | Nucleus | | 2336519.5 | 2477111.5 | 3658411 | | | | | | |
| P68433 | H31 | Histone H3.1 | Nucleus | | 1937164.75 | 4895068 | 5415700.5 | P68433 | H31 | 7394564 | 8794281 | 2072832 | 18 |
| P62806 | H4 | Histone H4 | Nucleus | | 3.18E+07 | 5.20E+07 | 6.30E+07 | | | | | | |
| P62806 | H4 | Histone H4 | Nucleus | | 3.13E+07 | 6.34E+07 | 4.77E+07 | | | | | | |
| P62806 | H4 | Histone H4 | Nucleus | | 1.03E+07 | 1.17E+07 | 1.10E+07 | | | | | | |
| P62806 | H4 | Histone H4 | Nucleus | | 8827684 | 1.74E+07 | 1.79E+07 | | | | | | |
| P62806 | H4 | Histone H4 | Nucleus | | 4059291 | 6161609 | 7561804.5 | | | | | | |
| P62806 | H4 | Histone H4 | Nucleus | | 453200.75 | 2608972.5 | 3437511 | | | | | | |
| P62806 | H4 | Histone H4 | Nucleus | | 7330.899 | 53124.992 | 8379.626 | | | | | | |
| P62806 | H4 | Histone H4 | Nucleus | | 63898.434 | 580751 | 1502107.75 | P62806 | H4 | 10850176 | 20559003 | 4196589 | 24 |
| Q3U0D9 | HACE1 | HECT domain and ankyrin repeat containing E3 ubiquitin protein ligase 1 | Cytoplasm | enzyme | 1164248.875 | 594074.25 | 1053379.5 | Q3U0D9 | HACE1 | 1164249 | 302311 | 174539 | 3 |
| Q61425 | HADH | hydroxyacyl-CoA dehydrogenase | Cytoplasm | enzyme | 1332031.375 | 3098849 | 1935512.5 | | | | | | |
| Q61425 | HADH | hydroxyacyl-CoA dehydrogenase | Cytoplasm | enzyme | 207867.656 | 699274.25 | 374629.844 | Q61425 | HADH | 769950 | 1099714 | 448956 | 6 |
| Q8BMS1 | HADHA | hydroxyacyl-CoA dehydrogenase/3-ketoacyl-CoA thiolase/enoyl-CoA hydratase (trifunctional protein), alpha subunit | Cytoplasm | enzyme | 5446604 | 1.66E+07 | 4564233 | | | | | | |
| Q8BMS1 | HADHA | hydroxyacyl-CoA dehydrogenase/3-ketoacyl-CoA thiolase/enoyl-CoA hydratase (trifunctional protein), alpha subunit | Cytoplasm | enzyme | 5203896 | 1.07E+07 | 4336787.5 | | | | | | |
| Q8BMS1 | HADHA | hydroxyacyl-CoA dehydrogenase/3-ketoacyl-CoA thiolase/enoyl-CoA hydratase (trifunctional protein), alpha subunit | Cytoplasm | enzyme | 5539564 | 1.01E+07 | 4110889.5 | | | | | | |
| Q8BMS1 | HADHA | hydroxyacyl-CoA dehydrogenase/3-ketoacyl-CoA thiolase/enoyl-CoA hydratase (trifunctional protein), alpha subunit | Cytoplasm | enzyme | 4469476.5 | 9256207 | 4102691.75 | | | | | | |
| Q8BMS1 | HADHA | hydroxyacyl-CoA dehydrogenase/3-ketoacyl-CoA thiolase/enoyl-CoA hydratase (trifunctional protein), alpha subunit | Cytoplasm | enzyme | 4451572.5 | 1.02E+07 | 4539195 | | | | | | |
| Q8BMS1 | HADHA | hydroxyacyl-CoA dehydrogenase/3-ketoacyl-CoA thiolase/enoyl-CoA hydratase (trifunctional protein), alpha subunit | Cytoplasm | enzyme | 2750419.5 | 8841650 | 3279560 | | | | | | |
| Q8BMS1 | HADHA | hydroxyacyl-CoA dehydrogenase/3-ketoacyl-CoA thiolase/enoyl-CoA hydratase (trifunctional protein), alpha subunit | Cytoplasm | enzyme | 3198006 | 8444393 | 2816314.5 | | | | | | |
| Q8BMS1 | HADHA | hydroxyacyl-CoA dehydrogenase/3-ketoacyl-CoA thiolase/enoyl-CoA hydratase (trifunctional protein), alpha subunit | Cytoplasm | enzyme | 2799709 | 6036269.5 | 2672696 | | | | | | |
| Q8BMS1 | HADHA | hydroxyacyl-CoA dehydrogenase/3-ketoacyl-CoA thiolase/enoyl-CoA hydratase (trifunctional protein), alpha subunit | Cytoplasm | enzyme | 3103858.25 | 7747012.5 | 3702969.25 | | | | | | |
| Q8BMS1 | HADHA | hydroxyacyl-CoA dehydrogenase/3-ketoacyl-CoA thiolase/enoyl-CoA hydratase (trifunctional protein), alpha subunit | Cytoplasm | enzyme | 2618789.75 | 6428132 | 2164377.75 | | | | | | |
| Q8BMS1 | HADHA | hydroxyacyl-CoA dehydrogenase/3-ketoacyl-CoA thiolase/enoyl-CoA hydratase (trifunctional protein), alpha subunit | Cytoplasm | enzyme | 1509701.625 | 3105553.75 | 1478637.375 | | | | | | |
| Q8BMS1 | HADHA | hydroxyacyl-CoA dehydrogenase/3-ketoacyl-CoA thiolase/enoyl-CoA hydratase (trifunctional protein), alpha subunit | Cytoplasm | enzyme | 1256493 | 3277098.5 | 1635101.625 | | | | | | |
| Q8BMS1 | HADHA | hydroxyacyl-CoA dehydrogenase/3-ketoacyl-CoA thiolase/enoyl-CoA hydratase (trifunctional protein), alpha subunit | Cytoplasm | enzyme | 836465.188 | 2658353.5 | 802649.312 | | | | | | |
| Q8BMS1 | HADHA | hydroxyacyl-CoA dehydrogenase/3-ketoacyl-CoA thiolase/enoyl-CoA hydratase (trifunctional protein), alpha subunit | Cytoplasm | enzyme | 538639.562 | 1523716.125 | 156016.703 | Q8BMS1 | HADHA | 3123085 | 3371108 | 520173 | 42 |
| Q99JY0 | HADHB | hydroxyacyl-CoA dehydrogenase/3-ketoacyl-CoA thiolase/enoyl-CoA hydratase (trifunctional protein), beta subunit | Cytoplasm | enzyme | 7302930 | 2.03E+07 | 7339923.5 | | | | | | |
| Q99JY0 | HADHB | hydroxyacyl-CoA dehydrogenase/3-ketoacyl-CoA thiolase/enoyl-CoA hydratase (trifunctional protein), beta subunit | Cytoplasm | enzyme | 8312526.5 | 1.41E+07 | 7097815 | | | | | | |
| Q99JY0 | HADHB | hydroxyacyl-CoA dehydrogenase/3-ketoacyl-CoA thiolase/enoyl-CoA hydratase (trifunctional protein), beta subunit | Cytoplasm | enzyme | 6127849.5 | 1.45E+07 | 4988819 | | | | | | |
| Q99JY0 | HADHB | hydroxyacyl-CoA dehydrogenase/3-ketoacyl-CoA thiolase/enoyl-CoA hydratase (trifunctional protein), beta subunit | Cytoplasm | enzyme | 4181813.25 | 6380006.5 | 2091475 | | | | | | |
| Q99JY0 | HADHB | hydroxyacyl-CoA dehydrogenase/3-ketoacyl-CoA thiolase/enoyl-CoA hydratase (trifunctional protein), beta subunit | Cytoplasm | enzyme | 750681.938 | 2174083.75 | 1416530.5 | | | | | | |
| Q99JY0 | HADHB | hydroxyacyl-CoA dehydrogenase/3-ketoacyl-CoA thiolase/enoyl-CoA hydratase (trifunctional protein), beta subunit | Cytoplasm | enzyme | 2817714.5 | 4930870.5 | 1974187.625 | | | | | | |
| Q99JY0 | HADHB | hydroxyacyl-CoA dehydrogenase/3-ketoacyl-CoA thiolase/enoyl-CoA hydratase (trifunctional protein), beta subunit | Cytoplasm | enzyme | 1177112.625 | 4276587 | 2791547 | Q99JY0 | HADHB | 4381518 | 4998548 | 1090773 | 21 |
| Q99KB8 | HAGH | hydroxyacylglutathione hydrolase | Cytoplasm | enzyme | 1.07E+07 | 1.18E+07 | 8620059 | | | | | | |
| Q99KB8 | HAGH | hydroxyacylglutathione hydrolase | Cytoplasm | enzyme | 4269617.5 | 5918943 | 5329342 | | | | | | |
| Q99KB8 | HAGH | hydroxyacylglutathione hydrolase | Cytoplasm | enzyme | 851295.062 | 1148508.75 | 456873.781 | Q99KB8 | HAGH | 5286971 | 4245372 | 1415124 | 9 |
| P01942 | HBA1/HBA2 | hemoglobin, alpha 1 | Extracellular Space | transporter | 1.57E+09 | 1.17E+09 | 3.25E+09 | | | | | | |
| P01942 | HBA1/HBA2 | hemoglobin, alpha 1 | Extracellular Space | transporter | 3.34E+09 | 2.48E+09 | 2.56E+09 | | | | | | |
| P01942 | HBA1/HBA2 | hemoglobin, alpha 1 | Extracellular Space | transporter | 2.26E+09 | 1.65E+09 | 1.95E+09 | | | | | | |
| P01942 | HBA1/HBA2 | hemoglobin, alpha 1 | Extracellular Space | transporter | 2.86E+09 | 1.74E+09 | 9.27E+08 | | | | | | |
| P01942 | HBA1/HBA2 | hemoglobin, alpha 1 | Extracellular Space | transporter | 2.44E+09 | 1.88E+09 | 1.97E+09 | | | | | | |
| P01942 | HBA1/HBA2 | hemoglobin, alpha 1 | Extracellular Space | transporter | 1.99E+09 | 1.45E+09 | 1.72E+09 | | | | | | |
| P01942 | HBA1/HBA2 | hemoglobin, alpha 1 | Extracellular Space | transporter | 1.09E+09 | 5.38E+08 | 4.16E+08 | | | | | | |
| P01942 | HBA1/HBA2 | hemoglobin, alpha 1 | Extracellular Space | transporter | 2.10E+08 | 2.57E+08 | 3.45E+08 | | | | | | |
| P01942 | HBA1/HBA2 | hemoglobin, alpha 1 | Extracellular Space | transporter | 2.81E+08 | 3.77E+08 | 1.16E+09 | | | | | | |
| P01942 | HBA1/HBA2 | hemoglobin, alpha 1 | Extracellular Space | transporter | 3.19E+08 | 3.30E+08 | 3.39E+08 | | | | | | |
| P01942 | HBA1/HBA2 | hemoglobin, alpha 1 | Extracellular Space | transporter | 1.87E+08 | 1.50E+08 | 1.41E+08 | P01942 | HBA1/HBA2 | 1504263636 | 988090145 | 172004416 | 33 |
| P02088 | Hbb-b1 | hemoglobin, beta adult major chain | Extracellular Space | transporter | 5.55E+09 | 4.61E+09 | 5.43E+09 | | | | | | |
| P02088 | Hbb-b1 | hemoglobin, beta adult major chain | Extracellular Space | transporter | 4.50E+09 | 3.06E+09 | 3.66E+09 | | | | | | |
| P02088 | Hbb-b1 | hemoglobin, beta adult major chain | Extracellular Space | transporter | 4.62E+09 | 3.64E+09 | 3.70E+09 | | | | | | |
| P02088 | Hbb-b1 | hemoglobin, beta adult major chain | Extracellular Space | transporter | 3.59E+09 | 2.41E+09 | 2.82E+09 | | | | | | |
| P02088 | Hbb-b1 | hemoglobin, beta adult major chain | Extracellular Space | transporter | 4.33E+09 | 3.46E+09 | 2.35E+09 | | | | | | |
| P02088 | Hbb-b1 | hemoglobin, beta adult major chain | Extracellular Space | transporter | 3.00E+09 | 1.78E+09 | 1.97E+09 | | | | | | |

| | | | | | | | | | | | | | |
|---|---|---|---|---|---|---|---|---|---|---|---|---|---|
| P02088 | Hbb-b1 | hemoglobin, beta adult major chain | Extracellular Space | transporter | 3.22E+09 | 2.61E+09 | 1.60E+09 | | | | | | |
| P02088 | Hbb-b1 | hemoglobin, beta adult major chain | Extracellular Space | transporter | 2.10E+09 | 1.66E+09 | 1.85E+09 | | | | | | |
| P02088 | Hbb-b1 | hemoglobin, beta adult major chain | Extracellular Space | transporter | 1.54E+09 | 9.75E+08 | 6.07E+08 | | | | | | |
| P02088 | Hbb-b1 | hemoglobin, beta adult major chain | Extracellular Space | transporter | 1.11E+09 | 8.28E+08 | 1.10E+09 | | | | | | |
| P02088 | Hbb-b1 | hemoglobin, beta adult major chain | Extracellular Space | transporter | 4.43E+08 | 2.89E+08 | 3.10E+08 | | | | | | |
| P02088 | Hbb-b1 | hemoglobin, beta adult major chain | Extracellular Space | transporter | 5.36E+08 | 4.87E+08 | 3.44E+08 | | | | | | |
| P02088 | Hbb-b1 | hemoglobin, beta adult major chain | Extracellular Space | transporter | 2.04E+08 | 2.07E+08 | 2.33E+08 | | | | | | |
| P02088 | Hbb-b1 | hemoglobin, beta adult major chain | Extracellular Space | transporter | 5.79E+07 | 1.25E+08 | 2.12E+08 | | | | | | |
| P02088 | Hbb-b1 | hemoglobin, beta adult major chain | Extracellular Space | transporter | 1.40E+08 | 1.63E+07 | 2428104.75 | | | | | | |
| P02088 | Hbb-b1 | hemoglobin, beta adult major chain | Extracellular Space | transporter | 2.56E+07 | 1.18E+07 | 1.64E+07 | | | | | | |
| P02088 | Hbb-b1 | hemoglobin, beta adult major chain | Extracellular Space | transporter | 2.53E+07 | 2.89E+07 | 3.06E+07 | | | | | | |
| P02088 | Hbb-b1 | hemoglobin, beta adult major chain | Extracellular Space | transporter | 1.51E+07 | 1.58E+07 | 3327276 | | | | | | |
| P02088 | Hbb-b1 | hemoglobin, beta adult major chain | Extracellular Space | transporter | 1.30E+07 | 1267424.75 | 57092.895 | P02088 | Hbb-b1 | 1843402632 | 1668460344 | 220992971 | 57 |
| P02089 | Hbb-b2 | hemoglobin, beta adult minor chain | Extracellular Space | other | 1.05E+09 | 7.06E+08 | 5.49E+08 | | | | | | |
| P02089 | Hbb-b2 | hemoglobin, beta adult minor chain | Extracellular Space | other | 7.81E+08 | 6.39E+08 | 5.32E+08 | | | | | | |
| P02089 | Hbb-b2 | hemoglobin, beta adult minor chain | Extracellular Space | other | 7.16E+08 | 5.52E+08 | 3.08E+08 | | | | | | |
| P02089 | Hbb-b2 | hemoglobin, beta adult minor chain | Extracellular Space | other | 3.15E+08 | 1.99E+08 | 1.10E+08 | | | | | | |
| P02089 | Hbb-b2 | hemoglobin, beta adult minor chain | Extracellular Space | other | 2.23E+08 | 1.91E+08 | 9.27E+07 | | | | | | |
| P02089 | Hbb-b2 | hemoglobin, beta adult minor chain | Extracellular Space | other | 9.73E+07 | 9.39E+07 | 4.02E+07 | | | | | | |
| P02089 | Hbb-b2 | hemoglobin, beta adult minor chain | Extracellular Space | other | 5.86E+07 | 5.56E+07 | 4.00E+07 | | | | | | |
| P02089 | Hbb-b2 | hemoglobin, beta adult minor chain | Extracellular Space | other | 2266482.25 | 3690181.25 | 7804677 | P02089 | Hbb-b2 | 404938310 | 303987446 | 62051178 | 24 |
| P08103 | HCK | HCK proto-oncogene, Src family tyrosine kinase | Cytoplasm | kinase | 133937.344 | 482348.406 | 615394.812 | P08103 | HCK | 133937 | 248627 | 143545 | 3 |
| Q3U487 | HECTD3 | HECT domain containing E3 ubiquitin protein ligase 3 | Cytoplasm | enzyme | 3577545.25 | 3004670.25 | 3300073 | | | | | | |
| Q3U487 | HECTD3 | HECT domain containing E3 ubiquitin protein ligase 3 | Cytoplasm | enzyme | 857251 | 757471.875 | 758230.625 | Q3U487 | HECTD3 | 2217398 | 1383408 | 564774 | 6 |
| P43275 | Hist1h1a | histone cluster 1, H1a | Nucleus | other | 1440649.625 | 1356119.5 | 1549071.625 | | | | | | |
| P43275 | Hist1h1a | histone cluster 1, H1a | Nucleus | other | 580665.812 | 1350926.625 | 1703962.25 | | | | | | |
| P43275 | Hist1h1a | histone cluster 1, H1a | Nucleus | other | 392933.938 | 1592557.375 | 1867949.375 | | | | | | |
| P43275 | Hist1h1a | histone cluster 1, H1a | Nucleus | other | 0 | 0 | 0 | P43275 | Hist1h1a | 603562 | 731607 | 211197 | 12 |
| P43276 | Hist1h1b | histone cluster 1, H1b | Nucleus | other | 467864.125 | 1833153.75 | 3369983.5 | | | | | | |
| P43276 | Hist1h1b | histone cluster 1, H1b | Nucleus | other | 35495.828 | 172325.656 | 0 | P43276 | Hist1h1b | 251680 | 1356982 | 553986 | 6 |
| P15864 | HIST1H1C | histone cluster 1, H1c | Nucleus | other | 9526512 | 2.08E+07 | 2.74E+07 | P15864 | HIST1H1C | 9526512 | 9053267 | 5226906 | 3 |
| P43274 | Hist1h1e | histone cluster 1, H1e | Nucleus | other | 5819406.5 | 1.29E+07 | 1.68E+07 | | | | | | |
| P43274 | Hist1h1e | histone cluster 1, H1e | Nucleus | other | 1790191.375 | 5692460.5 | 8064259 | | | | | | |
| P43274 | Hist1h1e | histone cluster 1, H1e | Nucleus | other | 2876636 | 5921047 | 9353603 | | | | | | |
| P43274 | Hist1h1e | histone cluster 1, H1e | Nucleus | other | 1632916.625 | 2230959.5 | 2914133.75 | | | | | | |
| P43274 | Hist1h1e | histone cluster 1, H1e | Nucleus | other | 4.70E-06 | 4.70E-06 | 4.70E-06 | P43274 | Hist1h1e | 2423830 | 4956955 | 1279880 | 15 |
| Q8CGP5 | HIST1H2AJ | histone cluster 1, H2aj | Nucleus | other | 3.32E+07 | 7.56E+07 | 8.23E+07 | | | | | | |
| Q8CGP5 | HIST1H2AJ | histone cluster 1, H2aj | Nucleus | other | 3.09E+07 | 6.14E+07 | 6.84E+07 | | | | | | |
| Q8CGP5 | HIST1H2AJ | histone cluster 1, H2aj | Nucleus | other | 9722728 | 2.59E+07 | 2.49E+07 | | | | | | |
| Q8CGP5 | HIST1H2AJ | histone cluster 1, H2aj | Nucleus | other | 9544651 | 2.09E+07 | 2.71E+07 | | | | | | |
| Q8CGP5 | HIST1H2AJ | histone cluster 1, H2aj | Nucleus | other | 269843.875 | 276658.875 | 354685 | Q8CGP5 | HIST1H2AJ | 16733445 | 27863078 | 7194216 | 15 |
| P70696 | HIST1H2BA | histone cluster 1, H2ba | Nucleus | other | 2.83E+07 | 4.85E+07 | 5.18E+07 | P70696 | HIST1H2BA | 28320000 | 12694851 | 7329375 | 3 |
| Q64475 | HIST1H2BB | histone cluster 1, H2bb | Nucleus | other | 2.14E+07 | 3.49E+07 | 4.11E+07 | | | | | | |
| Q64475 | HIST1H2BB | histone cluster 1, H2bb | Nucleus | other | 2.36E+07 | 3.75E+07 | 4.50E+07 | | | | | | |
| Q64475 | HIST1H2BB | histone cluster 1, H2bb | Nucleus | other | 2631395 | 4119081.75 | 7082743 | | | | | | |
| Q64475 | HIST1H2BB | histone cluster 1, H2bb | Nucleus | other | 1172627.375 | 2012423.75 | 1954124.625 | | | | | | |
| Q64475 | HIST1H2BB | histone cluster 1, H2bb | Nucleus | other | 398291.312 | 493330.188 | 638764.938 | | | | | | |
| Q64475 | HIST1H2BB | histone cluster 1, H2bb | Nucleus | other | 480463.062 | 2089433.75 | 768378.625 | Q64475 | HIST1H2BB | 8280463 | 16395287 | 3864406 | 18 |
| Q64522 | HIST2H2AB | histone cluster 2, H2ab | Nucleus | other | 1584988.25 | 2794005.75 | 2388858.25 | Q64522 | HIST2H2AB | 1584988 | 615369 | 355284 | 3 |
| P17710 | HK1 | hexokinase 1 | Cytoplasm | kinase | 2465614.25 | 5388876.5 | 8254075.5 | | | | | | |
| P17710 | HK1 | hexokinase 1 | Cytoplasm | kinase | 1659611.125 | 3184053 | 3046729 | | | | | | |
| P17710 | HK1 | hexokinase 1 | Cytoplasm | kinase | 1214225.625 | 2944580.5 | 2852825.25 | | | | | | |
| P17710 | HK1 | hexokinase 1 | Cytoplasm | kinase | 408523.312 | 1849680.375 | 3295955 | | | | | | |
| P17710 | HK1 | hexokinase 1 | Cytoplasm | kinase | 864200.875 | 2053052 | 3766882.5 | | | | | | |
| P17710 | HK1 | hexokinase 1 | Cytoplasm | kinase | 928030.812 | 2375628.5 | 3240546.5 | | | | | | |
| P17710 | HK1 | hexokinase 1 | Cytoplasm | kinase | 654805.375 | 2410726.5 | 2822311.75 | | | | | | |
| P17710 | HK1 | hexokinase 1 | Cytoplasm | kinase | 715566.562 | 1603580.125 | 2184258.75 | | | | | | |
| P17710 | HK1 | hexokinase 1 | Cytoplasm | kinase | 550394.625 | 1704167.75 | 1886923 | | | | | | |
| P17710 | HK1 | hexokinase 1 | Cytoplasm | kinase | 430206.562 | 1111600.5 | 1325708 | | | | | | |
| P17710 | HK1 | hexokinase 1 | Cytoplasm | kinase | 517543.5 | 1495298.75 | 1774492 | | | | | | |
| P17710 | HK1 | hexokinase 1 | Cytoplasm | kinase | 106724.305 | 784840.438 | 901027.5 | | | | | | |
| P17710 | HK1 | hexokinase 1 | Cytoplasm | kinase | 207963.453 | 1073588.75 | 1538925.375 | P17710 | HK1 | 824878 | 1542853 | 247054 | 39 |
| P01899 | HLA-A | major histocompatibility complex, class I, A | Plasma Membrane | other | 9707686 | 1.15E+07 | 9824993 | | | | | | |
| P01899 | HLA-A | major histocompatibility complex, class I, A | Plasma Membrane | other | 5009528 | 4026379.75 | 1828154.75 | | | | | | |
| P01899 | HLA-A | major histocompatibility complex, class I, A | Plasma Membrane | other | 5628000.5 | 5854715 | 4434628 | | | | | | |
| P01899 | HLA-A | major histocompatibility complex, class I, A | Plasma Membrane | other | 1136567.875 | 2700723.5 | 2205450.75 | | | | | | |
| P01899 | HLA-A | major histocompatibility complex, class I, A | Plasma Membrane | other | 1.35E+07 | 1.60E+07 | 1.19E+07 | | | | | | |
| P01899 | HLA-A | major histocompatibility complex, class I, A | Plasma Membrane | other | 1056966.5 | 1103425.25 | 892479.062 | P01899 | | | | | |
| P01901 | HLA-A | major histocompatibility complex, class I, A | Plasma Membrane | other | 656457.312 | 908201 | 591504.688 | P01901 | | | | | |
| P01902 | HLA-A | major histocompatibility complex, class I, A | Plasma Membrane | other | 105088.234 | 123020.695 | 140883.672 | | | | | | |
| P01902 | HLA-A | major histocompatibility complex, class I, A | Plasma Membrane | other | 0 | 0 | 0 | P01902 | HLA-A | 4092255 | 4784582 | 920793 | 27 |
| P22907 | HMBS | hydroxymethylbilane synthase | Cytoplasm | enzyme | 1095741 | 1780647.125 | 746671.562 | P22907 | HMBS | 1095741 | 525999 | 303686 | 3 |
| P30681 | HMGB2 | high mobility group box 2 | Nucleus | transcription regulator | 23340.787 | 547317.75 | 368243.531 | P30681 | HMGB2 | 23341 | 266326 | 153763 | 3 |
| P38060 | HMGCL | 3-hydroxymethyl-3-methylglutaryl-CoA lyase | Cytoplasm | enzyme | 1956108.875 | 3677368.5 | 1415419.125 | | | | | | |
| P38060 | HMGCL | 3-hydroxymethyl-3-methylglutaryl-CoA lyase | Cytoplasm | enzyme | 677093.125 | 998759.5 | 175345.188 | P38060 | HMGCL | 1316601 | 1235928 | 504565 | 6 |
| Q61411 | HRAS | Harvey rat sarcoma viral oncogene homolog | Plasma Membrane | enzyme | 1.04E+08 | 1.17E+08 | 9.35E+07 | | | | | | |
| Q61411 | HRAS | Harvey rat sarcoma viral oncogene homolog | Plasma Membrane | enzyme | 3.02E+07 | 3.17E+07 | 3.48E+07 | | | | | | |
| Q61411 | HRAS | Harvey rat sarcoma viral oncogene homolog | Plasma Membrane | enzyme | 1.89E+07 | 1.76E+07 | 9013647 | | | | | | |
| Q61411 | HRAS | Harvey rat sarcoma viral oncogene homolog | Plasma Membrane | enzyme | 1.73E+07 | 2.99E+07 | 2.86E+07 | | | | | | |
| Q61411 | HRAS | Harvey rat sarcoma viral oncogene homolog | Plasma Membrane | enzyme | 1.75E+07 | 1.86E+07 | 2.09E+07 | | | | | | |
| Q61411 | HRAS | Harvey rat sarcoma viral oncogene homolog | Plasma Membrane | enzyme | 5294484 | 8425824 | 8532664 | | | | | | |
| Q61411 | HRAS | Harvey rat sarcoma viral oncogene homolog | Plasma Membrane | enzyme | 1.11E+07 | 9965650 | 9633733 | | | | | | |
| Q61411 | HRAS | Harvey rat sarcoma viral oncogene homolog | Plasma Membrane | enzyme | 6136600 | 8378062.5 | 9056585 | | | | | | |
| Q61411 | HRAS | Harvey rat sarcoma viral oncogene homolog | Plasma Membrane | enzyme | 3546029.5 | 4074002.75 | 4052470.25 | | | | | | |
| Q61411 | HRAS | Harvey rat sarcoma viral oncogene homolog | Plasma Membrane | enzyme | 1533503.25 | 882564.562 | 1811657.875 | Q61411 | HRAS | 21555062 | 29637670 | 5411074 | 30 |
| O08756 | HSD17B10 | hydroxysteroid (17-beta) dehydrogenase 10 | Cytoplasm | enzyme | 3735247.5 | 8574730 | 4456477.5 | | | | | | |
| O08756 | HSD17B10 | hydroxysteroid (17-beta) dehydrogenase 10 | Cytoplasm | enzyme | 1299769.25 | 2644814.25 | 1259566.5 | | | | | | |
| O08756 | HSD17B10 | hydroxysteroid (17-beta) dehydrogenase 10 | Cytoplasm | enzyme | 1658626.25 | 4422912 | 3248480.25 | | | | | | |
| O08756 | HSD17B10 | hydroxysteroid (17-beta) dehydrogenase 10 | Cytoplasm | enzyme | 424885.656 | 1585370.375 | 886253.375 | O08756 | HSD17B10 | 1779632 | 2266370 | 654245 | 12 |
| O70503 | HSD17B12 | hydroxysteroid (17-beta) dehydrogenase 12 | Cytoplasm | enzyme | 1264125.375 | 1873849.25 | 2545122.25 | | | | | | |
| O70503 | HSD17B12 | hydroxysteroid (17-beta) dehydrogenase 12 | Cytoplasm | enzyme | 1035042.125 | 767387.312 | 928602.875 | O70503 | HSD17B12 | 1149584 | 679687 | 277481 | 6 |

| | | | | | | | | | | | | | |
|---|---|---|---|---|---|---|---|---|---|---|---|---|---|
| Q8BTX9 | HSDL1 | hydroxysteroid dehydrogenase like 1 | Cytoplasm | enzyme | 88028.312 | 334873.406 | 203572.094 | Q8BTX9 | HSDL1 | 88028 | 123506 | 71306 | 3 |
| P07901 | HSP90AA1 | heat shock protein 90kDa alpha (cytosolic), class A member 1 | Cytoplasm | enzyme | 8213385 | 8066136.5 | 8418516 | | | | | | |
| P07901 | HSP90AA1 | heat shock protein 90kDa alpha (cytosolic), class A member 1 | Cytoplasm | enzyme | 8140608.5 | 5022792.5 | 4039412.75 | | | | | | |
| P07901 | HSP90AA1 | heat shock protein 90kDa alpha (cytosolic), class A member 1 | Cytoplasm | enzyme | 6553520 | 5104507.5 | 5649253 | | | | | | |
| P07901 | HSP90AA1 | heat shock protein 90kDa alpha (cytosolic), class A member 1 | Cytoplasm | enzyme | 6044275 | 4691373 | 2876787.25 | | | | | | |
| P07901 | HSP90AA1 | heat shock protein 90kDa alpha (cytosolic), class A member 1 | Cytoplasm | enzyme | 4101164.75 | 3624757.5 | 3295250.25 | | | | | | |
| P07901 | HSP90AA1 | heat shock protein 90kDa alpha (cytosolic), class A member 1 | Cytoplasm | enzyme | 3661140.75 | 2936732.25 | 1873927.75 | P07901 | HSP90AA1 | 6119016 | 2056118 | 484632 | 18 |
| P11499 | HSP90AB1 | heat shock protein 90kDa alpha (cytosolic), class B member 1 | Cytoplasm | enzyme | 6471372.5 | 5484459 | 4716405.5 | P11499 | HSP90AB1 | 6471373 | 879755 | 507927 | 3 |
| P08113 | HSP90B1 | heat shock protein 90kDa beta (Grp94), member 1 | Cytoplasm | other | 3.02E+07 | 3.06E+07 | 4.16E+07 | | | | | | |
| P08113 | HSP90B1 | heat shock protein 90kDa beta (Grp94), member 1 | Cytoplasm | other | 2.78E+07 | 3.40E+07 | 3.53E+07 | | | | | | |
| P08113 | HSP90B1 | heat shock protein 90kDa beta (Grp94), member 1 | Cytoplasm | other | 2.49E+07 | 2.94E+07 | 3.12E+07 | | | | | | |
| P08113 | HSP90B1 | heat shock protein 90kDa beta (Grp94), member 1 | Cytoplasm | other | 2.56E+07 | 2.21E+07 | 2.43E+07 | | | | | | |
| P08113 | HSP90B1 | heat shock protein 90kDa beta (Grp94), member 1 | Cytoplasm | other | 1.89E+07 | 2.29E+07 | 2.41E+07 | | | | | | |
| P08113 | HSP90B1 | heat shock protein 90kDa beta (Grp94), member 1 | Cytoplasm | other | 2.10E+07 | 2.20E+07 | 2.87E+07 | | | | | | |
| P08113 | HSP90B1 | heat shock protein 90kDa beta (Grp94), member 1 | Cytoplasm | other | 2.31E+07 | 2.53E+07 | 2.95E+07 | | | | | | |
| P08113 | HSP90B1 | heat shock protein 90kDa beta (Grp94), member 1 | Cytoplasm | other | 1.78E+07 | 1.47E+07 | 2.41E+07 | | | | | | |
| P08113 | HSP90B1 | heat shock protein 90kDa beta (Grp94), member 1 | Cytoplasm | other | 1.57E+07 | 1.62E+07 | 1.91E+07 | | | | | | |
| P08113 | HSP90B1 | heat shock protein 90kDa beta (Grp94), member 1 | Cytoplasm | other | 1.82E+07 | 1.99E+07 | 2.23E+07 | | | | | | |
| P08113 | HSP90B1 | heat shock protein 90kDa beta (Grp94), member 1 | Cytoplasm | other | 1.53E+07 | 1.66E+07 | 1.74E+07 | | | | | | |
| P08113 | HSP90B1 | heat shock protein 90kDa beta (Grp94), member 1 | Cytoplasm | other | 1.61E+07 | 1.65E+07 | 1.23E+07 | | | | | | |
| P08113 | HSP90B1 | heat shock protein 90kDa beta (Grp94), member 1 | Cytoplasm | other | 1.34E+07 | 2.07E+07 | 2.55E+07 | | | | | | |
| P08113 | HSP90B1 | heat shock protein 90kDa beta (Grp94), member 1 | Cytoplasm | other | 1.78E+07 | 2.03E+07 | 1.92E+07 | | | | | | |
| P08113 | HSP90B1 | heat shock protein 90kDa beta (Grp94), member 1 | Cytoplasm | other | 1.35E+07 | 1.60E+07 | 1.76E+07 | | | | | | |
| P08113 | HSP90B1 | heat shock protein 90kDa beta (Grp94), member 1 | Cytoplasm | other | 1.30E+07 | 1.59E+07 | 1.93E+07 | | | | | | |
| P08113 | HSP90B1 | heat shock protein 90kDa beta (Grp94), member 1 | Cytoplasm | other | 1.36E+07 | 1.48E+07 | 1.95E+07 | | | | | | |
| P08113 | HSP90B1 | heat shock protein 90kDa beta (Grp94), member 1 | Cytoplasm | other | 1.39E+07 | 1.54E+07 | 1.48E+07 | | | | | | |
| P08113 | HSP90B1 | heat shock protein 90kDa beta (Grp94), member 1 | Cytoplasm | other | 1.26E+07 | 1.23E+07 | 1.50E+07 | | | | | | |
| P08113 | HSP90B1 | heat shock protein 90kDa beta (Grp94), member 1 | Cytoplasm | other | 1.11E+07 | 1.73E+07 | 1.80E+07 | | | | | | |
| P08113 | HSP90B1 | heat shock protein 90kDa beta (Grp94), member 1 | Cytoplasm | other | 1.03E+07 | 1.21E+07 | 1.44E+07 | | | | | | |
| P08113 | HSP90B1 | heat shock protein 90kDa beta (Grp94), member 1 | Cytoplasm | other | 8966749 | 8818291 | 9126431 | | | | | | |
| P08113 | HSP90B1 | heat shock protein 90kDa beta (Grp94), member 1 | Cytoplasm | other | 7775951 | 7182975 | 1.00E+07 | | | | | | |
| P08113 | HSP90B1 | heat shock protein 90kDa beta (Grp94), member 1 | Cytoplasm | other | 5697957.5 | 6967424 | 6489055.5 | | | | | | |
| P08113 | HSP90B1 | heat shock protein 90kDa beta (Grp94), member 1 | Cytoplasm | other | 1236205.375 | 530840.562 | 1398101.5 | | | | | | |
| P08113 | HSP90B1 | heat shock protein 90kDa beta (Grp94), member 1 | Cytoplasm | other | 1991436.625 | 1708582.625 | 1507329.75 | P08113 | HSP90B1 | 15368396 | 8451068 | 956895 | 78 |
| Q61316 | HSPA4 | heat shock 70kDa protein 4 | Cytoplasm | other | 3.09E+07 | 2.03E+07 | 2.04E+07 | | | | | | |
| Q61316 | HSPA4 | heat shock 70kDa protein 4 | Cytoplasm | other | 1.85E+07 | 1.67E+07 | 1.38E+07 | | | | | | |
| Q61316 | HSPA4 | heat shock 70kDa protein 4 | Cytoplasm | other | 1.35E+07 | 1.06E+07 | 1.13E+07 | | | | | | |
| Q61316 | HSPA4 | heat shock 70kDa protein 4 | Cytoplasm | other | 1.28E+07 | 5986319.5 | 6559090.5 | | | | | | |
| Q61316 | HSPA4 | heat shock 70kDa protein 4 | Cytoplasm | other | 1.29E+07 | 8748323 | 1.05E+07 | | | | | | |
| Q61316 | HSPA4 | heat shock 70kDa protein 4 | Cytoplasm | other | 1.75E+07 | 6317094.5 | 2127551.5 | | | | | | |
| Q61316 | HSPA4 | heat shock 70kDa protein 4 | Cytoplasm | other | 1.30E+07 | 7502121.5 | 9270028 | | | | | | |
| Q61316 | HSPA4 | heat shock 70kDa protein 4 | Cytoplasm | other | 1.31E+07 | 8768753 | 1.14E+07 | | | | | | |
| Q61316 | HSPA4 | heat shock 70kDa protein 4 | Cytoplasm | other | 1.27E+07 | 8585621 | 9236792 | | | | | | |
| Q61316 | HSPA4 | heat shock 70kDa protein 4 | Cytoplasm | other | 9933365 | 9252252 | 1.32E+07 | | | | | | |
| Q61316 | HSPA4 | heat shock 70kDa protein 4 | Cytoplasm | other | 9279929 | 5373428.5 | 5645648 | | | | | | |
| Q61316 | HSPA4 | heat shock 70kDa protein 4 | Cytoplasm | other | 1.03E+07 | 7135104.5 | 5454036 | | | | | | |
| Q61316 | HSPA4 | heat shock 70kDa protein 4 | Cytoplasm | other | 7857682.5 | 4435309.5 | 6194023 | | | | | | |
| Q61316 | HSPA4 | heat shock 70kDa protein 4 | Cytoplasm | other | 4023120.75 | 3020018.25 | 2166457.5 | | | | | | |
| Q61316 | HSPA4 | heat shock 70kDa protein 4 | Cytoplasm | other | 6709947.5 | 4786205.5 | 5998507 | | | | | | |
| Q61316 | HSPA4 | heat shock 70kDa protein 4 | Cytoplasm | other | 7298841 | 4986234.5 | 4656919.5 | | | | | | |
| Q61316 | HSPA4 | heat shock 70kDa protein 4 | Cytoplasm | other | 4576091.5 | 2796402 | 3769965.25 | | | | | | |
| Q61316 | HSPA4 | heat shock 70kDa protein 4 | Cytoplasm | other | 3504986.75 | 3955708.75 | 5337278.5 | | | | | | |
| Q61316 | HSPA4 | heat shock 70kDa protein 4 | Cytoplasm | other | 3288349.75 | 3075635.25 | 2953974.75 | | | | | | |
| Q61316 | HSPA4 | heat shock 70kDa protein 4 | Cytoplasm | other | 5281465 | 3970424.5 | 3115783.75 | | | | | | |
| Q61316 | HSPA4 | heat shock 70kDa protein 4 | Cytoplasm | other | 4184774 | 3340712.5 | 5323040 | | | | | | |
| Q61316 | HSPA4 | heat shock 70kDa protein 4 | Cytoplasm | other | 2324787.75 | 1512611.125 | 925566.25 | | | | | | |
| Q61316 | HSPA4 | heat shock 70kDa protein 4 | Cytoplasm | other | 3995041.25 | 3048236.25 | 2932184.25 | Q61316 | HSPA4 | 9887321 | 5437267 | 654570 | 69 |
| P48722 | HSPA4L | heat shock 70kDa protein 4-like | Cytoplasm | other | 5677440 | 5235619.5 | 5743814.5 | | | | | | |
| P48722 | HSPA4L | heat shock 70kDa protein 4-like | Cytoplasm | other | 3307188.75 | 2384645 | 3480780 | | | | | | |
| P48722 | HSPA4L | heat shock 70kDa protein 4-like | Cytoplasm | other | 3530467 | 1824812.875 | 3717987.5 | | | | | | |
| P48722 | HSPA4L | heat shock 70kDa protein 4-like | Cytoplasm | other | 3275869.25 | 4704652.5 | 2615457.5 | | | | | | |
| P48722 | HSPA4L | heat shock 70kDa protein 4-like | Cytoplasm | other | 1933255.125 | 931123.562 | 1801781.375 | P48722 | HSPA4L | 3544844 | 1482822 | 382863 | 15 |
| P20029 | HSPA5 | heat shock 70kDa protein 5 (glucose-regulated protein, 78kDa) | Cytoplasm | enzyme | 1.71E+08 | 1.84E+08 | 1.96E+08 | | | | | | |
| P20029 | HSPA5 | heat shock 70kDa protein 5 (glucose-regulated protein, 78kDa) | Cytoplasm | enzyme | 6.20E+07 | 6.99E+07 | 8.14E+07 | | | | | | |
| P20029 | HSPA5 | heat shock 70kDa protein 5 (glucose-regulated protein, 78kDa) | Cytoplasm | enzyme | 5.02E+07 | 6.29E+07 | 7.00E+07 | | | | | | |
| P20029 | HSPA5 | heat shock 70kDa protein 5 (glucose-regulated protein, 78kDa) | Cytoplasm | enzyme | 4.74E+07 | 5.75E+07 | 6.72E+07 | | | | | | |
| P20029 | HSPA5 | heat shock 70kDa protein 5 (glucose-regulated protein, 78kDa) | Cytoplasm | enzyme | 4.83E+07 | 5.30E+07 | 6.80E+07 | | | | | | |
| P20029 | HSPA5 | heat shock 70kDa protein 5 (glucose-regulated protein, 78kDa) | Cytoplasm | enzyme | 4.16E+07 | 4.97E+07 | 5.25E+07 | | | | | | |
| P20029 | HSPA5 | heat shock 70kDa protein 5 (glucose-regulated protein, 78kDa) | Cytoplasm | enzyme | 3.64E+07 | 6.05E+07 | 7.14E+07 | | | | | | |
| P20029 | HSPA5 | heat shock 70kDa protein 5 (glucose-regulated protein, 78kDa) | Cytoplasm | enzyme | 3.25E+07 | 4.29E+07 | 4.87E+07 | | | | | | |
| P20029 | HSPA5 | heat shock 70kDa protein 5 (glucose-regulated protein, 78kDa) | Cytoplasm | enzyme | 3.02E+07 | 3.24E+07 | 3.69E+07 | | | | | | |
| P20029 | HSPA5 | heat shock 70kDa protein 5 (glucose-regulated protein, 78kDa) | Cytoplasm | enzyme | 2.85E+07 | 4.27E+07 | 3.95E+07 | | | | | | |
| P20029 | HSPA5 | heat shock 70kDa protein 5 (glucose-regulated protein, 78kDa) | Cytoplasm | enzyme | 3.23E+07 | 3.50E+07 | 4.21E+07 | | | | | | |
| P20029 | HSPA5 | heat shock 70kDa protein 5 (glucose-regulated protein, 78kDa) | Cytoplasm | enzyme | 2.82E+07 | 3.12E+07 | 2.99E+07 | | | | | | |
| P20029 | HSPA5 | heat shock 70kDa protein 5 (glucose-regulated protein, 78kDa) | Cytoplasm | enzyme | 1.69E+07 | 2.24E+07 | 2.69E+07 | | | | | | |
| P20029 | HSPA5 | heat shock 70kDa protein 5 (glucose-regulated protein, 78kDa) | Cytoplasm | enzyme | 1.55E+07 | 2.51E+07 | 2.76E+07 | | | | | | |
| P20029 | HSPA5 | heat shock 70kDa protein 5 (glucose-regulated protein, 78kDa) | Cytoplasm | enzyme | 1.57E+07 | 1.99E+07 | 1.63E+07 | | | | | | |
| P20029 | HSPA5 | heat shock 70kDa protein 5 (glucose-regulated protein, 78kDa) | Cytoplasm | enzyme | 1.97E+07 | 1.02E+07 | 1718209 | | | | | | |
| P20029 | HSPA5 | heat shock 70kDa protein 5 (glucose-regulated protein, 78kDa) | Cytoplasm | enzyme | 1.29E+07 | 1.37E+07 | 1.32E+07 | | | | | | |
| P20029 | HSPA5 | heat shock 70kDa protein 5 (glucose-regulated protein, 78kDa) | Cytoplasm | enzyme | 1.21E+07 | 9402047 | 4803770 | | | | | | |
| P20029 | HSPA5 | heat shock 70kDa protein 5 (glucose-regulated protein, 78kDa) | Cytoplasm | enzyme | 1.74E+07 | 2.57E+07 | 3.32E+07 | | | | | | |
| P20029 | HSPA5 | heat shock 70kDa protein 5 (glucose-regulated protein, 78kDa) | Cytoplasm | enzyme | 6069591 | 9235197 | 1.53E+07 | | | | | | |
| P20029 | HSPA5 | heat shock 70kDa protein 5 (glucose-regulated protein, 78kDa) | Cytoplasm | enzyme | 6391065 | 6938252 | 6907333.5 | | | | | | |
| P20029 | HSPA5 | heat shock 70kDa protein 5 (glucose-regulated protein, 78kDa) | Cytoplasm | enzyme | 6247404 | 6366759 | 5703941.5 | | | | | | |
| P20029 | HSPA5 | heat shock 70kDa protein 5 (glucose-regulated protein, 78kDa) | Cytoplasm | enzyme | 2776314.25 | 3024688 | 3700420 | | | | | | |
| P20029 | HSPA5 | heat shock 70kDa protein 5 (glucose-regulated protein, 78kDa) | Cytoplasm | enzyme | 4226741 | 5952199.5 | 8751224 | | | | | | |
| P20029 | HSPA5 | heat shock 70kDa protein 5 (glucose-regulated protein, 78kDa) | Cytoplasm | enzyme | 2566212 | 963661.938 | 427132.906 | | | | | | |
| P20029 | HSPA5 | heat shock 70kDa protein 5 (glucose-regulated protein, 78kDa) | Cytoplasm | enzyme | 3263768 | 2624371.75 | 2530431.5 | P20029 | HSPA5 | 28851581 | 37061919 | 4196436 | 78 |
| P63017 | HSPA8 | heat shock 70kDa protein 8 | Cytoplasm | enzyme | 1.38E+08 | 1.18E+08 | 1.42E+08 | | | | | | |
| P63017 | HSPA8 | heat shock 70kDa protein 8 | Cytoplasm | enzyme | 1.24E+08 | 1.26E+08 | 1.63E+08 | | | | | | |
| P63017 | HSPA8 | heat shock 70kDa protein 8 | Cytoplasm | enzyme | 1.22E+08 | 1.02E+08 | 1.28E+08 | | | | | | |

| | | | | | | | | | | | | | |
|---|---|---|---|---|---|---|---|---|---|---|---|---|---|
| P63017 | HSPA8 | heat shock 70kDa protein 8 | Cytoplasm | enzyme | 1.06E+08 | 9.50E+07 | 1.13E+08 | | | | | | |
| P63017 | HSPA8 | heat shock 70kDa protein 8 | Cytoplasm | enzyme | 1.06E+08 | 8.18E+07 | 1.12E+08 | | | | | | |
| P63017 | HSPA8 | heat shock 70kDa protein 8 | Cytoplasm | enzyme | 9.63E+07 | 8.71E+07 | 8.21E+07 | | | | | | |
| P63017 | HSPA8 | heat shock 70kDa protein 8 | Cytoplasm | enzyme | 1.00E+08 | 7.89E+07 | 1.00E+08 | | | | | | |
| P63017 | HSPA8 | heat shock 70kDa protein 8 | Cytoplasm | enzyme | 6.95E+07 | 6.80E+07 | 7.20E+07 | | | | | | |
| P63017 | HSPA8 | heat shock 70kDa protein 8 | Cytoplasm | enzyme | 6.77E+07 | 5.65E+07 | 7.09E+07 | | | | | | |
| P63017 | HSPA8 | heat shock 70kDa protein 8 | Cytoplasm | enzyme | 7.06E+07 | 5.50E+07 | 6.40E+07 | | | | | | |
| P63017 | HSPA8 | heat shock 70kDa protein 8 | Cytoplasm | enzyme | 6.91E+07 | 5.50E+07 | 7.01E+07 | | | | | | |
| P63017 | HSPA8 | heat shock 70kDa protein 8 | Cytoplasm | enzyme | 6.88E+07 | 5.58E+07 | 6.16E+07 | | | | | | |
| P63017 | HSPA8 | heat shock 70kDa protein 8 | Cytoplasm | enzyme | 7.78E+07 | 6.99E+07 | 8.33E+07 | | | | | | |
| P63017 | HSPA8 | heat shock 70kDa protein 8 | Cytoplasm | enzyme | 6.45E+07 | 5.16E+07 | 6.50E+07 | | | | | | |
| P63017 | HSPA8 | heat shock 70kDa protein 8 | Cytoplasm | enzyme | 5.11E+07 | 4.66E+07 | 5.76E+07 | | | | | | |
| P63017 | HSPA8 | heat shock 70kDa protein 8 | Cytoplasm | enzyme | 3.78E+07 | 3.48E+07 | 4.09E+07 | | | | | | |
| P63017 | HSPA8 | heat shock 70kDa protein 8 | Cytoplasm | enzyme | 3.63E+07 | 2.56E+07 | 4.05E+07 | | | | | | |
| P63017 | HSPA8 | heat shock 70kDa protein 8 | Cytoplasm | enzyme | 3.72E+07 | 3.43E+07 | 3.49E+07 | | | | | | |
| P63017 | HSPA8 | heat shock 70kDa protein 8 | Cytoplasm | enzyme | 2.47E+07 | 1.92E+07 | 2.43E+07 | | | | | | |
| P63017 | HSPA8 | heat shock 70kDa protein 8 | Cytoplasm | enzyme | 2.80E+07 | 2.35E+07 | 2.80E+07 | | | | | | |
| P63017 | HSPA8 | heat shock 70kDa protein 8 | Cytoplasm | enzyme | 2.46E+07 | 2.67E+07 | 2.48E+07 | | | | | | |
| P63017 | HSPA8 | heat shock 70kDa protein 8 | Cytoplasm | enzyme | 4345713.5 | 4561733 | 4941530.5 | | | | | | |
| P63017 | HSPA8 | heat shock 70kDa protein 8 | Cytoplasm | enzyme | 2918556.5 | 2615783.25 | 2691439.75 | | | | | | |
| P63017 | HSPA8 | heat shock 70kDa protein 8 | Cytoplasm | enzyme | 5039456.5 | 4846662 | 5148963 | | | | | | |
| P63017 | HSPA8 | heat shock 70kDa protein 8 | Cytoplasm | enzyme | 2445000.75 | 2191394.25 | 3377803 | | | | | | |
| P63017 | HSPA8 | heat shock 70kDa protein 8 | Cytoplasm | enzyme | 1469682.375 | 1962245.875 | 3511136.25 | P63017 | HSPA8 | 59037631 | 40943795 | 4635972 | 78 |
| P38647 | HSPA9 | heat shock 70kDa protein 9 (mortalin) | Cytoplasm | other | 7853804 | 1.56E+07 | 6426292 | | | | | | |
| P38647 | HSPA9 | heat shock 70kDa protein 9 (mortalin) | Cytoplasm | other | 6929846.5 | 1.30E+07 | 5194841 | | | | | | |
| P38647 | HSPA9 | heat shock 70kDa protein 9 (mortalin) | Cytoplasm | other | 7712952 | 1.54E+07 | 7832470.5 | | | | | | |
| P38647 | HSPA9 | heat shock 70kDa protein 9 (mortalin) | Cytoplasm | other | 6381918.5 | 1.29E+07 | 5022878.5 | | | | | | |
| P38647 | HSPA9 | heat shock 70kDa protein 9 (mortalin) | Cytoplasm | other | 5540679.5 | 1.17E+07 | 5670136.5 | | | | | | |
| P38647 | HSPA9 | heat shock 70kDa protein 9 (mortalin) | Cytoplasm | other | 5692189 | 1.21E+07 | 5271664.5 | | | | | | |
| P38647 | HSPA9 | heat shock 70kDa protein 9 (mortalin) | Cytoplasm | other | 4607664.5 | 1.08E+07 | 3514767.75 | | | | | | |
| P38647 | HSPA9 | heat shock 70kDa protein 9 (mortalin) | Cytoplasm | other | 3906722.5 | 7762838.5 | 3514065.25 | | | | | | |
| P38647 | HSPA9 | heat shock 70kDa protein 9 (mortalin) | Cytoplasm | other | 4474209.5 | 6985319 | 2978998.75 | | | | | | |
| P38647 | HSPA9 | heat shock 70kDa protein 9 (mortalin) | Cytoplasm | other | 2423995 | 1.33E+07 | 5189918.5 | | | | | | |
| P38647 | HSPA9 | heat shock 70kDa protein 9 (mortalin) | Cytoplasm | other | 1964228 | 3816586 | 714903.062 | | | | | | |
| P38647 | HSPA9 | heat shock 70kDa protein 9 (mortalin) | Cytoplasm | other | 1641349.25 | 2862371.5 | 856356.688 | P38647 | HSPA9 | 4927463 | 4057075 | 676179 | 36 |
| P63038 | HSPD1 | heat shock 60kDa protein 1 (chaperonin) | Cytoplasm | enzyme | 2.11E+07 | 4.86E+07 | 2.00E+07 | | | | | | |
| P63038 | HSPD1 | heat shock 60kDa protein 1 (chaperonin) | Cytoplasm | enzyme | 2.11E+07 | 3.51E+07 | 1.52E+07 | | | | | | |
| P63038 | HSPD1 | heat shock 60kDa protein 1 (chaperonin) | Cytoplasm | enzyme | 1.74E+07 | 3.73E+07 | 1.79E+07 | | | | | | |
| P63038 | HSPD1 | heat shock 60kDa protein 1 (chaperonin) | Cytoplasm | enzyme | 1.48E+07 | 4.02E+07 | 1.93E+07 | | | | | | |
| P63038 | HSPD1 | heat shock 60kDa protein 1 (chaperonin) | Cytoplasm | enzyme | 1.22E+07 | 3.69E+07 | 1.45E+07 | | | | | | |
| P63038 | HSPD1 | heat shock 60kDa protein 1 (chaperonin) | Cytoplasm | enzyme | 1.63E+07 | 3.16E+07 | 1.47E+07 | | | | | | |
| P63038 | HSPD1 | heat shock 60kDa protein 1 (chaperonin) | Cytoplasm | enzyme | 1.52E+07 | 3.27E+07 | 1.50E+07 | | | | | | |
| P63038 | HSPD1 | heat shock 60kDa protein 1 (chaperonin) | Cytoplasm | enzyme | 1.56E+07 | 3.23E+07 | 1.51E+07 | | | | | | |
| P63038 | HSPD1 | heat shock 60kDa protein 1 (chaperonin) | Cytoplasm | enzyme | 9385195 | 2.48E+07 | 1.29E+07 | | | | | | |
| P63038 | HSPD1 | heat shock 60kDa protein 1 (chaperonin) | Cytoplasm | enzyme | 1.11E+07 | 2.29E+07 | 1.08E+07 | | | | | | |
| P63038 | HSPD1 | heat shock 60kDa protein 1 (chaperonin) | Cytoplasm | enzyme | 1.01E+07 | 2.23E+07 | 1.00E+07 | | | | | | |
| P63038 | HSPD1 | heat shock 60kDa protein 1 (chaperonin) | Cytoplasm | enzyme | 8276706.5 | 2.05E+07 | 8532318 | | | | | | |
| P63038 | HSPD1 | heat shock 60kDa protein 1 (chaperonin) | Cytoplasm | enzyme | 9655525 | 1.83E+07 | 9002930 | | | | | | |
| P63038 | HSPD1 | heat shock 60kDa protein 1 (chaperonin) | Cytoplasm | enzyme | 9836379 | 2.49E+07 | 1.40E+07 | | | | | | |
| P63038 | HSPD1 | heat shock 60kDa protein 1 (chaperonin) | Cytoplasm | enzyme | 6862072 | 1.43E+07 | 8706148 | | | | | | |
| P63038 | HSPD1 | heat shock 60kDa protein 1 (chaperonin) | Cytoplasm | enzyme | 1.05E+07 | 3.38E+07 | 1.70E+07 | | | | | | |
| P63038 | HSPD1 | heat shock 60kDa protein 1 (chaperonin) | Cytoplasm | enzyme | 2736391.75 | 7178799 | 3270540.5 | | | | | | |
| P63038 | HSPD1 | heat shock 60kDa protein 1 (chaperonin) | Cytoplasm | enzyme | 2156240.5 | 7284099.5 | 3713070.25 | | | | | | |
| P63038 | HSPD1 | heat shock 60kDa protein 1 (chaperonin) | Cytoplasm | enzyme | 1534958 | 4451748.5 | 2350738.5 | | | | | | |
| P63038 | HSPD1 | heat shock 60kDa protein 1 (chaperonin) | Cytoplasm | enzyme | 2321605.25 | 6830069 | 2344995.5 | | | | | | |
| P63038 | HSPD1 | heat shock 60kDa protein 1 (chaperonin) | Cytoplasm | enzyme | 1272414.125 | 2650261.5 | 797512.188 | | | | | | |
| P63038 | HSPD1 | heat shock 60kDa protein 1 (chaperonin) | Cytoplasm | enzyme | 1681040.125 | 4920940.5 | 2739383.5 | | | | | | |
| P63038 | HSPD1 | heat shock 60kDa protein 1 (chaperonin) | Cytoplasm | enzyme | 1733936.75 | 2840061.25 | 1227464.125 | P63038 | HSPD1 | 9686629 | 11032080 | 1328106 | 69 |
| Q64433 | HSPE1 | heat shock 10kDa protein 1 | Cytoplasm | enzyme | 1.77E+07 | 3.12E+07 | 1.63E+07 | | | | | | |
| Q64433 | HSPE1 | heat shock 10kDa protein 1 | Cytoplasm | enzyme | 1.21E+07 | 2.28E+07 | 1.01E+07 | | | | | | |
| Q64433 | HSPE1 | heat shock 10kDa protein 1 | Cytoplasm | enzyme | 1.10E+07 | 2.09E+07 | 1.05E+07 | | | | | | |
| Q64433 | HSPE1 | heat shock 10kDa protein 1 | Cytoplasm | enzyme | 3975230.5 | 1.01E+07 | 5320526 | | | | | | |
| Q64433 | HSPE1 | heat shock 10kDa protein 1 | Cytoplasm | enzyme | 4395908.5 | 8712904 | 3454015.75 | Q64433 | HSPE1 | 9808228 | 7888957 | 2036920 | 15 |
| Q9Z2G9 | HTATIP2 | HIV-1 Tat interactive protein 2, 30kDa | Nucleus | transcription regulator | 1125817.375 | 1764483.875 | 1176613.5 | | | | | | |
| Q9Z2G9 | HTATIP2 | HIV-1 Tat interactive protein 2, 30kDa | Nucleus | transcription regulator | 610220.375 | 1232799.625 | 521410.812 | Q9Z2G9 | HTATIP2 | 868019 | 455109 | 185797 | 6 |
| Q7TMY8 | HUWE1 | HECT, UBA and WWE domain containing 1, E3 ubiquitin protein ligase | Nucleus | transcription regulator | 2517158 | 2062539.125 | 2583363.5 | Q7TMY8 | HUWE1 | 2517158 | 283525 | 163693 | 3 |
| Q91ZJ9 | HYAL1 | hyaluronoglucosaminidase 1 | Cytoplasm | enzyme | 3848885.5 | 4040910.75 | 4605829 | Q91ZJ9 | HYAL1 | 3848886 | 393482 | 227177 | 3 |
| Q9JKR6 | HYOU1 | hypoxia up-regulated 1 | Cytoplasm | other | 3140902.25 | 4277562.5 | 4307589 | | | | | | |
| Q9JKR6 | HYOU1 | hypoxia up-regulated 1 | Cytoplasm | other | 3222066.75 | 3528674.5 | 3478491.5 | | | | | | |
| Q9JKR6 | HYOU1 | hypoxia up-regulated 1 | Cytoplasm | other | 2591775.25 | 3127022.25 | 3438039.5 | | | | | | |
| Q9JKR6 | HYOU1 | hypoxia up-regulated 1 | Cytoplasm | other | 2542886.25 | 2512300.75 | 3080620.75 | | | | | | |
| Q9JKR6 | HYOU1 | hypoxia up-regulated 1 | Cytoplasm | other | 3288320 | 2860273.25 | 2135788.75 | | | | | | |
| Q9JKR6 | HYOU1 | hypoxia up-regulated 1 | Cytoplasm | other | 1431023.375 | 1409598.375 | 2248838.25 | | | | | | |
| Q9JKR6 | HYOU1 | hypoxia up-regulated 1 | Cytoplasm | other | 1213609.375 | 2135357.5 | 2011367.5 | Q9JKR6 | HYOU1 | 2490083 | 862251 | 188159 | 21 |
| Q8BIJ6 | IARS2 | isoleucyl-tRNA synthetase 2, mitochondrial | Cytoplasm | enzyme | 1366246.375 | 3500475 | 1242202 | Q8BIJ6 | IARS2 | 1366246 | 1269522 | 732959 | 3 |
| Q9ERM2 | ICAM4 | intercellular adhesion molecule 4 (Landsteiner-Wiener blood group) | Plasma Membrane | other | 37919.27 | 5419.727 | 60818.859 | | | | | | |
| Q9ERM2 | ICAM4 | intercellular adhesion molecule 4 (Landsteiner-Wiener blood group) | Plasma Membrane | other | 235327.016 | 99386.531 | 163396.688 | Q9ERM2 | ICAM4 | 136623 | 85635 | 34960 | 6 |
| Q9JHR7 | IDE | insulin-degrading enzyme | Extracellular Space | peptidase | 2907899 | 1919112.875 | 2104998.25 | | | | | | |
| Q9JHR7 | IDE | insulin-degrading enzyme | Extracellular Space | peptidase | 3295333.75 | 2364336.5 | 2094907.625 | Q9JHR7 | IDE | 3101616 | 540098 | 220494 | 6 |
| P54071 | IDH2 | isocitrate dehydrogenase 2 (NADP+), mitochondrial | Cytoplasm | enzyme | 2760645.5 | 7051087 | 3987926.75 | | | | | | |
| P54071 | IDH2 | isocitrate dehydrogenase 2 (NADP+), mitochondrial | Cytoplasm | enzyme | 1077793 | 2133104 | 1523044.5 | | | | | | |
| P54071 | IDH2 | isocitrate dehydrogenase 2 (NADP+), mitochondrial | Cytoplasm | enzyme | 1104624.125 | 2466850.25 | 1749741.25 | | | | | | |
| P54071 | IDH2 | isocitrate dehydrogenase 2 (NADP+), mitochondrial | Cytoplasm | enzyme | 784379.562 | 1757591.375 | 782705.438 | | | | | | |
| P54071 | IDH2 | isocitrate dehydrogenase 2 (NADP+), mitochondrial | Cytoplasm | enzyme | 712440.75 | 2227595.5 | 1145961.25 | | | | | | |
| P54071 | IDH2 | isocitrate dehydrogenase 2 (NADP+), mitochondrial | Cytoplasm | enzyme | 566827.938 | 1245659.375 | 406674.656 | P54071 | IDH2 | 1167785 | 1580941 | 372631 | 18 |
| Q9D6R2 | IDH3A | isocitrate dehydrogenase 3 (NAD+) alpha | Cytoplasm | enzyme | 9372785 | 2.60E+07 | 1.25E+07 | | | | | | |
| Q9D6R2 | IDH3A | isocitrate dehydrogenase 3 (NAD+) alpha | Cytoplasm | enzyme | 4354550.5 | 8048345 | 4100658 | | | | | | |
| Q9D6R2 | IDH3A | isocitrate dehydrogenase 3 (NAD+) alpha | Cytoplasm | enzyme | 3479172.25 | 8611934 | 4431611.5 | | | | | | |
| Q9D6R2 | IDH3A | isocitrate dehydrogenase 3 (NAD+) alpha | Cytoplasm | enzyme | 3656829.25 | 7147068.5 | 3232649.5 | | | | | | |
| Q9D6R2 | IDH3A | isocitrate dehydrogenase 3 (NAD+) alpha | Cytoplasm | enzyme | 4020921 | 8147488.5 | 3498589 | Q9D6R2 | IDH3A | 4976852 | 5868283 | 1515184 | 15 |
| P70404 | IDH3G | isocitrate dehydrogenase 3 (NAD+) gamma | Cytoplasm | enzyme | 4691068.5 | 1.06E+07 | 3880704.75 | | | | | | |

| | | | | | | | | | | | | | |
|---|---|---|---|---|---|---|---|---|---|---|---|---|---|
| P70404 | IDH3G | isocitrate dehydrogenase 3 (NAD+) gamma | Cytoplasm | enzyme | 1043961.688 | 2501200 | 1377268.375 | P70404 | IDH3G | 2867515 | 3500208 | 1428954 | 6 |
| Q9CR20 | IER3IP1 | immediate early response 3 interacting protein 1 | Cytoplasm | other | 7883047 | 1.07E+07 | 1.23E+07 | | | | | | |
| Q9CR20 | IER3IP1 | immediate early response 3 interacting protein 1 | Cytoplasm | other | 2433208.75 | 3435375.5 | 3101690 | Q9CR20 | IER3IP1 | 5158128 | 4263588 | 1740602 | 6 |
| P01872 | IGHM | immunoglobulin heavy constant mu | Plasma Membrane | transmembrane receptor | 2157699.25 | 1.13E+07 | 1584477.25 | | | | | | |
| P01872 | IGHM | immunoglobulin heavy constant mu | Plasma Membrane | transmembrane receptor | 841765.25 | 2654207 | 226169.938 | | | | | | |
| P01872 | IGHM | immunoglobulin heavy constant mu | Plasma Membrane | transmembrane receptor | 552486.875 | 2841938 | 514083.406 | | | | | | |
| P01872 | IGHM | immunoglobulin heavy constant mu | Plasma Membrane | transmembrane receptor | 488890.625 | 2487265 | 810153.375 | P01872 | IGHM | 1010211 | 3024067 | 872973 | 12 |
| O55222 | ILK | integrin-linked kinase | Plasma Membrane | kinase | 3973199.25 | 9051826 | 4493197.5 | | | | | | |
| O55222 | ILK | integrin-linked kinase | Plasma Membrane | kinase | 237813.844 | 816043.625 | 521217.031 | | | | | | |
| O55222 | ILK | integrin-linked kinase | Plasma Membrane | kinase | 433386.469 | 675415.688 | 871092.25 | | | | | | |
| O55222 | ILK | integrin-linked kinase | Plasma Membrane | kinase | 69922.703 | 516412.594 | 446805.125 | | | | | | |
| O55222 | ILK | integrin-linked kinase | Plasma Membrane | kinase | 94710.141 | 343886.781 | 286683.031 | | | | | | |
| O55222 | ILK | integrin-linked kinase | Plasma Membrane | kinase | 21190.99 | 527027.625 | 417850.812 | O55222 | ILK | 805037 | 2300618 | 542261 | 18 |
| Q8BU33 | ILVBL | ilvB (bacterial acetolactate synthase)-like | Cytoplasm | enzyme | 204513.594 | 802940.25 | 588830.938 | Q8BU33 | ILVBL | 204514 | 303221 | 175065 | 3 |
| Q8CAQ8 | IMMT | inner membrane protein, mitochondrial | Cytoplasm | other | 3490383.75 | 7248836 | 3700097.25 | | | | | | |
| Q8CAQ8 | IMMT | inner membrane protein, mitochondrial | Cytoplasm | other | 4041040 | 8151772.5 | 3970902 | | | | | | |
| Q8CAQ8 | IMMT | inner membrane protein, mitochondrial | Cytoplasm | other | 3882064 | 6482224 | 4017225 | | | | | | |
| Q8CAQ8 | IMMT | inner membrane protein, mitochondrial | Cytoplasm | other | 3118378.5 | 7118231.5 | 4494089.5 | | | | | | |
| Q8CAQ8 | IMMT | inner membrane protein, mitochondrial | Cytoplasm | other | 2725280.75 | 5781466 | 3110009.75 | | | | | | |
| Q8CAQ8 | IMMT | inner membrane protein, mitochondrial | Cytoplasm | other | 1604139.5 | 3540822.5 | 1113378.375 | | | | | | |
| Q8CAQ8 | IMMT | inner membrane protein, mitochondrial | Cytoplasm | other | 1483996.375 | 4908118.5 | 2340243 | | | | | | |
| Q8CAQ8 | IMMT | inner membrane protein, mitochondrial | Cytoplasm | other | 654071.688 | 1172125.125 | 551948.5 | | | | | | |
| Q8CAQ8 | IMMT | inner membrane protein, mitochondrial | Cytoplasm | other | 707815 | 2927423.5 | 1240995.5 | Q8CAQ8 | IMMT | 2411908 | 2128799 | 409687 | 27 |
| P24547 | IMPDH2 | IMP (inosine 5'-monophosphate) dehydrogenase 2 | Cytoplasm | enzyme | 4707064 | 1498009.875 | 84165.477 | P24547 | IMPDH2 | 4707064 | 2368831 | 1367645 | 3 |
| Q8C5L6 | INPP5K | inositol polyphosphate-5-phosphatase K | Cytoplasm | phosphatase | 6782375 | 6770610.5 | 6746706 | | | | | | |
| Q8C5L6 | INPP5K | inositol polyphosphate-5-phosphatase K | Cytoplasm | phosphatase | 5694351 | 7701569 | 7625312 | Q8C5L6 | INPP5K | 6238363 | 731466 | 298620 | 6 |
| Q8CIM8 | INTS4 | integrator complex subunit 4 | Nucleus | other | 3.55E+07 | 4.05E+07 | 4.98E+07 | Q8CIM8 | INTS4 | 35540000 | 7212122 | 4163920 | 3 |
| Q8BKC5 | IPO5 | importin 5 | Nucleus | transporter | 1.18E+07 | 1.23E+07 | 1.05E+07 | | | | | | |
| Q8BKC5 | IPO5 | importin 5 | Nucleus | transporter | 7034671 | 5755585 | 2695315.25 | | | | | | |
| Q8BKC5 | IPO5 | importin 5 | Nucleus | transporter | 1.36E+07 | 1.39E+07 | 1.55E+07 | | | | | | |
| Q8BKC5 | IPO5 | importin 5 | Nucleus | transporter | 1.78E+07 | 2.08E+07 | 1.63E+07 | | | | | | |
| Q8BKC5 | IPO5 | importin 5 | Nucleus | transporter | 8050691.5 | 5382391 | 6401980.5 | | | | | | |
| Q8BKC5 | IPO5 | importin 5 | Nucleus | transporter | 9237902 | 6332704 | 6446819.5 | | | | | | |
| Q8BKC5 | IPO5 | importin 5 | Nucleus | transporter | 6652356 | 6399534.5 | 5476620 | | | | | | |
| Q8BKC5 | IPO5 | importin 5 | Nucleus | transporter | 1.07E+07 | 7641373.5 | 7571561 | | | | | | |
| Q8BKC5 | IPO5 | importin 5 | Nucleus | transporter | 7397911 | 4782788 | 4780141.5 | | | | | | |
| Q8BKC5 | IPO5 | importin 5 | Nucleus | transporter | 6539035.5 | 7663360.5 | 4560734.5 | | | | | | |
| Q8BKC5 | IPO5 | importin 5 | Nucleus | transporter | 7000858.5 | 4940606.5 | 4733470 | | | | | | |
| Q8BKC5 | IPO5 | importin 5 | Nucleus | transporter | 7180652 | 6474543.5 | 5066466 | | | | | | |
| Q8BKC5 | IPO5 | importin 5 | Nucleus | transporter | 6317408.5 | 4528333 | 3998073.5 | | | | | | |
| Q8BKC5 | IPO5 | importin 5 | Nucleus | transporter | 5347571 | 3935608.5 | 3386503.5 | | | | | | |
| Q8BKC5 | IPO5 | importin 5 | Nucleus | transporter | 4175361 | 3553070.25 | 2915035.5 | | | | | | |
| Q8BKC5 | IPO5 | importin 5 | Nucleus | transporter | 3588042.5 | 2781702 | 4023182 | | | | | | |
| Q8BKC5 | IPO5 | importin 5 | Nucleus | transporter | 3761276.25 | 3033167.5 | 1349416.125 | | | | | | |
| Q8BKC5 | IPO5 | importin 5 | Nucleus | transporter | 4645119.5 | 3824063 | 3564284.75 | | | | | | |
| Q8BKC5 | IPO5 | importin 5 | Nucleus | transporter | 2822100.5 | 2812535.5 | 2694700.5 | | | | | | |
| Q8BKC5 | IPO5 | importin 5 | Nucleus | transporter | 2706200.75 | 2887140.75 | 1711069 | Q8BKC5 | IPO5 | 7318358 | 4111932 | 530848 | 60 |
| Q9EPL8 | IPO7 | importin 7 | Nucleus | transporter | 8050557 | 6067700.5 | 2591384.25 | | | | | | |
| Q9EPL8 | IPO7 | importin 7 | Nucleus | transporter | 4081670 | 3403364.75 | 1528490.875 | | | | | | |
| Q9EPL8 | IPO7 | importin 7 | Nucleus | transporter | 4448779.5 | 3041942.5 | 2047712.5 | | | | | | |
| Q9EPL8 | IPO7 | importin 7 | Nucleus | transporter | 3872668 | 3549553 | 2390106 | | | | | | |
| Q9EPL8 | IPO7 | importin 7 | Nucleus | transporter | 2210128.25 | 2461419.75 | 1107949 | | | | | | |
| Q9EPL8 | IPO7 | importin 7 | Nucleus | transporter | 2484548.5 | 1681255 | 1313843.125 | | | | | | |
| Q9EPL8 | IPO7 | importin 7 | Nucleus | transporter | 981217.312 | 1211221.875 | 315708 | Q9EPL8 | IPO7 | 3732796 | 1816945 | 396490 | 21 |
| Q60766 | Irgm1 | immunity-related GTPase family M member 1 | Cytoplasm | other | 1.70E+07 | 1.68E+07 | 1.31E+07 | | | | | | |
| Q60766 | Irgm1 | immunity-related GTPase family M member 1 | Cytoplasm | other | 1.79E+07 | 1.12E+07 | 9935246 | | | | | | |
| Q60766 | Irgm1 | immunity-related GTPase family M member 1 | Cytoplasm | other | 1.21E+07 | 1.39E+07 | 1.24E+07 | | | | | | |
| Q60766 | Irgm1 | immunity-related GTPase family M member 1 | Cytoplasm | other | 1.00E+07 | 1.24E+07 | 1.01E+07 | | | | | | |
| Q60766 | Irgm1 | immunity-related GTPase family M member 1 | Cytoplasm | other | 8382756 | 8495434 | 6640405.5 | | | | | | |
| Q60766 | Irgm1 | immunity-related GTPase family M member 1 | Cytoplasm | other | 7988458 | 9664418 | 6629536 | | | | | | |
| Q60766 | Irgm1 | immunity-related GTPase family M member 1 | Cytoplasm | other | 5061489 | 7249237 | 6468755 | | | | | | |
| Q60766 | Irgm1 | immunity-related GTPase family M member 1 | Cytoplasm | other | 4790158 | 3926860.75 | 3625276.75 | | | | | | |
| Q60766 | Irgm1 | immunity-related GTPase family M member 1 | Cytoplasm | other | 3532155.75 | 5512879 | 9256256 | | | | | | |
| Q60766 | Irgm1 | immunity-related GTPase family M member 1 | Cytoplasm | other | 898379.75 | 1946122.25 | 2351644.75 | Q60766 | Irgm1 | 8766340 | 4526293 | 826384 | 30 |
| Q9D924 | ISCA1 | iron-sulfur cluster assembly 1 | Cytoplasm | other | 2676604.75 | 5011309 | 2412616.5 | | | | | | |
| Q9D924 | ISCA1 | iron-sulfur cluster assembly 1 | Cytoplasm | other | 1806906 | 2289556 | 837177.75 | Q9D924 | ISCA1 | 2241755 | 1388160 | 566714 | 6 |
| Q9DCB8 | ISCA2 | iron-sulfur cluster assembly 2 | Cytoplasm | other | 2048063.5 | 5396853 | 1178904.25 | Q9DCB8 | ISCA2 | 2048064 | 2227140 | 1285840 | 3 |
| Q64339 | ISG15 | ISG15 ubiquitin-like modifier | Extracellular Space | other | 4.51E+07 | 5.82E+07 | 5.40E+07 | | | | | | |
| Q64339 | ISG15 | ISG15 ubiquitin-like modifier | Extracellular Space | other | 4.44E+07 | 3.92E+07 | 3.23E+07 | | | | | | |
| Q64339 | ISG15 | ISG15 ubiquitin-like modifier | Extracellular Space | other | 2.44E+07 | 3.21E+07 | 1.86E+07 | | | | | | |
| Q64339 | ISG15 | ISG15 ubiquitin-like modifier | Extracellular Space | other | 2.28E+07 | 1.09E+07 | 1869897.125 | | | | | | |
| Q64339 | ISG15 | ISG15 ubiquitin-like modifier | Extracellular Space | other | 2.28E+07 | 2.62E+07 | 2.00E+07 | | | | | | |
| Q64339 | ISG15 | ISG15 ubiquitin-like modifier | Extracellular Space | other | 6973279.5 | 1.22E+07 | 1.40E+07 | | | | | | |
| Q64339 | ISG15 | ISG15 ubiquitin-like modifier | Extracellular Space | other | 5263290.5 | 2245267.5 | 114942.672 | | | | | | |
| Q64339 | ISG15 | ISG15 ubiquitin-like modifier | Extracellular Space | other | 2412780 | 3456199.25 | 3547157 | Q64339 | ISG15 | 21757419 | 17472343 | 3566527 | 24 |
| Q9JL16 | ISG20 | interferon stimulated exonuclease gene 20kDa | Nucleus | enzyme | 8485183 | 1.04E+07 | 8334860.5 | | | | | | |
| Q9JL16 | ISG20 | interferon stimulated exonuclease gene 20kDa | Nucleus | enzyme | 6957041 | 8056277 | 5189777.5 | | | | | | |
| Q9JL16 | ISG20 | interferon stimulated exonuclease gene 20kDa | Nucleus | enzyme | 6161409 | 6505437 | 5071141 | | | | | | |
| Q9JL16 | ISG20 | interferon stimulated exonuclease gene 20kDa | Nucleus | enzyme | 5959630 | 7194415 | 6169262.5 | Q9JL16 | ISG20 | 6890816 | 1541364 | 444954 | 12 |
| Q9CX00 | IST1 | increased sodium tolerance 1 homolog (yeast) | Cytoplasm | other | 2623933.25 | 1540455.125 | 1618942.125 | Q9CX00 | IST1 | 2623933 | 604165 | 348815 | 3 |
| Q62469 | ITGA2 | integrin, alpha 2 (CD49B, alpha 2 subunit of VLA-2 receptor) | Plasma Membrane | transmembrane receptor | 582924.5 | 1518512.875 | 3684863 | | | | | | |
| Q62469 | ITGA2 | integrin, alpha 2 (CD49B, alpha 2 subunit of VLA-2 receptor) | Plasma Membrane | transmembrane receptor | 207570.547 | 821856.938 | 1546719.5 | | | | | | |
| Q62469 | ITGA2 | integrin, alpha 2 (CD49B, alpha 2 subunit of VLA-2 receptor) | Plasma Membrane | transmembrane receptor | 45019.371 | 710094.375 | 2514610.5 | | | | | | |
| Q62469 | ITGA2 | integrin, alpha 2 (CD49B, alpha 2 subunit of VLA-2 receptor) | Plasma Membrane | transmembrane receptor | 367272.438 | 946341.688 | 1556309.75 | | | | | | |
| Q62469 | ITGA2 | integrin, alpha 2 (CD49B, alpha 2 subunit of VLA-2 receptor) | Plasma Membrane | transmembrane receptor | 313306.625 | 892025.625 | 2188686.5 | | | | | | |
| Q62469 | ITGA2 | integrin, alpha 2 (CD49B, alpha 2 subunit of VLA-2 receptor) | Plasma Membrane | transmembrane receptor | 131483.906 | 571550.75 | 1024248.875 | | | | | | |
| Q62469 | ITGA2 | integrin, alpha 2 (CD49B, alpha 2 subunit of VLA-2 receptor) | Plasma Membrane | transmembrane receptor | 36323.781 | 554063.188 | 883179.25 | | | | | | |
| Q62469 | ITGA2 | integrin, alpha 2 (CD49B, alpha 2 subunit of VLA-2 receptor) | Plasma Membrane | transmembrane receptor | 47693.641 | 427283.844 | 876001.5 | | | | | | |
| Q62469 | ITGA2 | integrin, alpha 2 (CD49B, alpha 2 subunit of VLA-2 receptor) | Plasma Membrane | transmembrane receptor | 66276.664 | 216620.469 | 1315665.625 | | | | | | |
| Q62469 | ITGA2 | integrin, alpha 2 (CD49B, alpha 2 subunit of VLA-2 receptor) | Plasma Membrane | transmembrane receptor | 0 | 170889.5 | 431137.281 | | | | | | |
| Q62469 | ITGA2 | integrin, alpha 2 (CD49B, alpha 2 subunit of VLA-2 receptor) | Plasma Membrane | transmembrane receptor | 161120.359 | 925319.875 | 1647910.625 | Q62469 | ITGA2 | 178090 | 821735 | 143046 | 33 |

| | | | | | | | | | | | | | |
|---|---|---|---|---|---|---|---|---|---|---|---|---|---|
| Q9QUM0 | ITGA2B | integrin, alpha 2b (platelet glycoprotein IIb of IIb/IIIa complex, antigen CD41) | Plasma Membrane | transmembrane receptor | 8693661 | 3.76E+07 | 8.68E+07 | | | | | | |
| Q9QUM0 | ITGA2B | integrin, alpha 2b (platelet glycoprotein IIb of IIb/IIIa complex, antigen CD41) | Plasma Membrane | transmembrane receptor | 4622922 | 1.47E+07 | 2.19E+07 | | | | | | |
| Q9QUM0 | ITGA2B | integrin, alpha 2b (platelet glycoprotein IIb of IIb/IIIa complex, antigen CD41) | Plasma Membrane | transmembrane receptor | 2398609.5 | 1.21E+07 | 2.78E+07 | | | | | | |
| Q9QUM0 | ITGA2B | integrin, alpha 2b (platelet glycoprotein IIb of IIb/IIIa complex, antigen CD41) | Plasma Membrane | transmembrane receptor | 3146123.75 | 1.07E+07 | 2.01E+07 | | | | | | |
| Q9QUM0 | ITGA2B | integrin, alpha 2b (platelet glycoprotein IIb of IIb/IIIa complex, antigen CD41) | Plasma Membrane | transmembrane receptor | 3207190 | 1.87E+07 | 3.80E+07 | | | | | | |
| Q9QUM0 | ITGA2B | integrin, alpha 2b (platelet glycoprotein IIb of IIb/IIIa complex, antigen CD41) | Plasma Membrane | transmembrane receptor | 2689421 | 1.38E+07 | 3.38E+07 | | | | | | |
| Q9QUM0 | ITGA2B | integrin, alpha 2b (platelet glycoprotein IIb of IIb/IIIa complex, antigen CD41) | Plasma Membrane | transmembrane receptor | 1999689 | 6554655 | 1.57E+07 | | | | | | |
| Q9QUM0 | ITGA2B | integrin, alpha 2b (platelet glycoprotein IIb of IIb/IIIa complex, antigen CD41) | Plasma Membrane | transmembrane receptor | 2132092.75 | 8587325 | 1.91E+07 | | | | | | |
| Q9QUM0 | ITGA2B | integrin, alpha 2b (platelet glycoprotein IIb of IIb/IIIa complex, antigen CD41) | Plasma Membrane | transmembrane receptor | 1176328.375 | 2442230.25 | 8169325 | | | | | | |
| Q9QUM0 | ITGA2B | integrin, alpha 2b (platelet glycoprotein IIb of IIb/IIIa complex, antigen CD41) | Plasma Membrane | transmembrane receptor | 1836238 | 6871844 | 1.19E+07 | | | | | | |
| Q9QUM0 | ITGA2B | integrin, alpha 2b (platelet glycoprotein IIb of IIb/IIIa complex, antigen CD41) | Plasma Membrane | transmembrane receptor | 1729550.625 | 8321746.5 | 1.88E+07 | | | | | | |
| Q9QUM0 | ITGA2B | integrin, alpha 2b (platelet glycoprotein IIb of IIb/IIIa complex, antigen CD41) | Plasma Membrane | transmembrane receptor | 1232947.5 | 7330927 | 1.83E+07 | | | | | | |
| Q9QUM0 | ITGA2B | integrin, alpha 2b (platelet glycoprotein IIb of IIb/IIIa complex, antigen CD41) | Plasma Membrane | transmembrane receptor | 1716879.875 | 6669404 | 1.67E+07 | | | | | | |
| Q9QUM0 | ITGA2B | integrin, alpha 2b (platelet glycoprotein IIb of IIb/IIIa complex, antigen CD41) | Plasma Membrane | transmembrane receptor | 1356688 | 6408177.5 | 1.30E+07 | | | | | | |
| Q9QUM0 | ITGA2B | integrin, alpha 2b (platelet glycoprotein IIb of IIb/IIIa complex, antigen CD41) | Plasma Membrane | transmembrane receptor | 1290032 | 5252794 | 1.25E+07 | | | | | | |
| Q9QUM0 | ITGA2B | integrin, alpha 2b (platelet glycoprotein IIb of IIb/IIIa complex, antigen CD41) | Plasma Membrane | transmembrane receptor | 1087193.75 | 7776055 | 1.42E+07 | | | | | | |
| Q9QUM0 | ITGA2B | integrin, alpha 2b (platelet glycoprotein IIb of IIb/IIIa complex, antigen CD41) | Plasma Membrane | transmembrane receptor | 411804 | 3412624.5 | 6313295.5 | | | | | | |
| Q9QUM0 | ITGA2B | integrin, alpha 2b (platelet glycoprotein IIb of IIb/IIIa complex, antigen CD41) | Plasma Membrane | transmembrane receptor | 1001098.562 | 4802342.5 | 1.29E+07 | | | | | | |
| Q9QUM0 | ITGA2B | integrin, alpha 2b (platelet glycoprotein IIb of IIb/IIIa complex, antigen CD41) | Plasma Membrane | transmembrane receptor | 670869.875 | 3827433.25 | 6224261.5 | | | | | | |
| Q9QUM0 | ITGA2B | integrin, alpha 2b (platelet glycoprotein IIb of IIb/IIIa complex, antigen CD41) | Plasma Membrane | transmembrane receptor | 829543 | 4300561 | 1.04E+07 | | | | | | |
| Q9QUM0 | ITGA2B | integrin, alpha 2b (platelet glycoprotein IIb of IIb/IIIa complex, antigen CD41) | Plasma Membrane | transmembrane receptor | 332252.156 | 2952911.75 | 7859894.5 | | | | | | |
| Q9QUM0 | ITGA2B | integrin, alpha 2b (platelet glycoprotein IIb of IIb/IIIa complex, antigen CD41) | Plasma Membrane | transmembrane receptor | 215811.625 | 593804.062 | 1278303 | | | | | | |
| Q9QUM0 | ITGA2B | integrin, alpha 2b (platelet glycoprotein IIb of IIb/IIIa complex, antigen CD41) | Plasma Membrane | transmembrane receptor | 474024.531 | 1663623.375 | 1850045.625 | | | | | | |
| Q9QUM0 | ITGA2B | integrin, alpha 2b (platelet glycoprotein IIb of IIb/IIIa complex, antigen CD41) | Plasma Membrane | transmembrane receptor | 555468.5 | 1523559.125 | 3013410 | | | | | | |
| Q9QUM0 | ITGA2B | integrin, alpha 2b (platelet glycoprotein IIb of IIb/IIIa complex, antigen CD41) | Plasma Membrane | transmembrane receptor | 15638.983 | 1628963.375 | 4371143 | | | | | | |
| Q9QUM0 | ITGA2B | integrin, alpha 2b (platelet glycoprotein IIb of IIb/IIIa complex, antigen CD41) | Plasma Membrane | transmembrane receptor | 88762.281 | 883872.75 | 1855876 | | | | | | |
| Q9QUM0 | ITGA2B | integrin, alpha 2b (platelet glycoprotein IIb of IIb/IIIa complex, antigen CD41) | Plasma Membrane | transmembrane receptor | 201854.641 | 964672.5 | 2508813.5 | Q9QUM0 | ITGA2B | 1670841 | 12231284 | 1359032 | 81 |
| P11688 | ITGA5 | integrin, alpha 5 (fibronectin receptor, alpha polypeptide) | Plasma Membrane | transmembrane receptor | 8608459 | 1.00E+07 | 7201887 | | | | | | |
| P11688 | ITGA5 | integrin, alpha 5 (fibronectin receptor, alpha polypeptide) | Plasma Membrane | transmembrane receptor | 5374551.5 | 5800068 | 4894143.5 | | | | | | |
| P11688 | ITGA5 | integrin, alpha 5 (fibronectin receptor, alpha polypeptide) | Plasma Membrane | transmembrane receptor | 4111513.25 | 4077515.25 | 3462674.5 | | | | | | |
| P11688 | ITGA5 | integrin, alpha 5 (fibronectin receptor, alpha polypeptide) | Plasma Membrane | transmembrane receptor | 1834677.125 | 2809097.5 | 2336629.75 | P11688 | ITGA5 | 4982300 | 2519993 | 727459 | 12 |
| Q61739 | ITGA6 | integrin, alpha 6 | Plasma Membrane | transmembrane receptor | 171915.625 | 1369753.375 | 2140571.75 | | | | | | |
| Q61739 | ITGA6 | integrin, alpha 6 | Plasma Membrane | transmembrane receptor | 79228.195 | 291045.562 | 1100422.75 | | | | | | |
| Q61739 | ITGA6 | integrin, alpha 6 | Plasma Membrane | transmembrane receptor | 108916.164 | 739032.562 | 1568317.25 | | | | | | |
| Q61739 | ITGA6 | integrin, alpha 6 | Plasma Membrane | transmembrane receptor | 0 | 210312.203 | 720048.5 | | | | | | |
| Q61739 | ITGA6 | integrin, alpha 6 | Plasma Membrane | transmembrane receptor | 4.70E-06 | 304683.375 | 151988.094 | | | | | | |
| Q61739 | ITGA6 | integrin, alpha 6 | Plasma Membrane | transmembrane receptor | 44715.48 | 441422 | 639462.312 | | | | | | |
| Q61739 | ITGA6 | integrin, alpha 6 | Plasma Membrane | transmembrane receptor | 0 | 857679.438 | 1759660.875 | Q61739 | ITGA6 | 57825 | 642041 | 140105 | 21 |
| P09055 | ITGB1 | integrin, beta 1 (fibronectin receptor, beta polypeptide, antigen CD29 includes MDF2, MSK12) | Plasma Membrane | transmembrane receptor | 6860193 | 1.28E+07 | 1.54E+07 | | | | | | |
| P09055 | ITGB1 | integrin, beta 1 (fibronectin receptor, beta polypeptide, antigen CD29 includes MDF2, MSK12) | Plasma Membrane | transmembrane receptor | 6486864.5 | 8599861 | 8938317 | | | | | | |
| P09055 | ITGB1 | integrin, beta 1 (fibronectin receptor, beta polypeptide, antigen CD29 includes MDF2, MSK12) | Plasma Membrane | transmembrane receptor | 5826207 | 8620384 | 1.09E+07 | | | | | | |
| P09055 | ITGB1 | integrin, beta 1 (fibronectin receptor, beta polypeptide, antigen CD29 includes MDF2, MSK12) | Plasma Membrane | transmembrane receptor | 5235777 | 4790535.5 | 9737232 | | | | | | |
| P09055 | ITGB1 | integrin, beta 1 (fibronectin receptor, beta polypeptide, antigen CD29 includes MDF2, MSK12) | Plasma Membrane | transmembrane receptor | 3057601.75 | 4736038.5 | 5574154 | | | | | | |
| P09055 | ITGB1 | integrin, beta 1 (fibronectin receptor, beta polypeptide, antigen CD29 includes MDF2, MSK12) | Plasma Membrane | transmembrane receptor | 2309010.25 | 4028911 | 4602878.5 | | | | | | |
| P09055 | ITGB1 | integrin, beta 1 (fibronectin receptor, beta polypeptide, antigen CD29 includes MDF2, MSK12) | Plasma Membrane | transmembrane receptor | 3249111.25 | 4005831 | 5905838 | | | | | | |
| P09055 | ITGB1 | integrin, beta 1 (fibronectin receptor, beta polypeptide, antigen CD29 includes MDF2, MSK12) | Plasma Membrane | transmembrane receptor | 1578215 | 3169678 | 4363444.5 | | | | | | |
| P09055 | ITGB1 | integrin, beta 1 (fibronectin receptor, beta polypeptide, antigen CD29 includes MDF2, MSK12) | Plasma Membrane | transmembrane receptor | 2312334.5 | 3291762.25 | 4112998.75 | | | | | | |
| P09055 | ITGB1 | integrin, beta 1 (fibronectin receptor, beta polypeptide, antigen CD29 includes MDF2, MSK12) | Plasma Membrane | transmembrane receptor | 1448843.875 | 1850458 | 3068104.5 | | | | | | |
| P09055 | ITGB1 | integrin, beta 1 (fibronectin receptor, beta polypeptide, antigen CD29 includes MDF2, MSK12) | Plasma Membrane | transmembrane receptor | 1087009.75 | 1778115.5 | 2304589.25 | | | | | | |
| P09055 | ITGB1 | integrin, beta 1 (fibronectin receptor, beta polypeptide, antigen CD29 includes MDF2, MSK12) | Plasma Membrane | transmembrane receptor | 820148.688 | 932662.938 | 1569847.375 | | | | | | |
| P09055 | ITGB1 | integrin, beta 1 (fibronectin receptor, beta polypeptide, antigen CD29 includes MDF2, MSK12) | Plasma Membrane | transmembrane receptor | 324831.469 | 671165.25 | 835816.875 | P09055 | ITGB1 | 3122781 | 3529995 | 565252 | 39 |
| P11835 | ITGB2 | integrin, beta 2 (complement component 3 receptor 3 and 4 subunit) | Plasma Membrane | transmembrane receptor | 0 | 102721 | 1204179.75 | P11835 | ITGB2 | 435634 | 667559 | 385415 | 3 |
| O54890 | ITGB3 | integrin, beta 3 (platelet glycoprotein IIIa, antigen CD61) | Plasma Membrane | transmembrane receptor | 7740913.5 | 2.89E+07 | 4.76E+07 | | | | | | |
| O54890 | ITGB3 | integrin, beta 3 (platelet glycoprotein IIIa, antigen CD61) | Plasma Membrane | transmembrane receptor | 8307781.5 | 9559299 | 1.37E+07 | | | | | | |
| O54890 | ITGB3 | integrin, beta 3 (platelet glycoprotein IIIa, antigen CD61) | Plasma Membrane | transmembrane receptor | 2973615.75 | 1.01E+07 | 2.35E+07 | | | | | | |
| O54890 | ITGB3 | integrin, beta 3 (platelet glycoprotein IIIa, antigen CD61) | Plasma Membrane | transmembrane receptor | 2198646.25 | 8692426 | 2.06E+07 | | | | | | |
| O54890 | ITGB3 | integrin, beta 3 (platelet glycoprotein IIIa, antigen CD61) | Plasma Membrane | transmembrane receptor | 2984147.75 | 1.07E+07 | 2.13E+07 | | | | | | |
| O54890 | ITGB3 | integrin, beta 3 (platelet glycoprotein IIIa, antigen CD61) | Plasma Membrane | transmembrane receptor | 2148106 | 1.04E+07 | 2.03E+07 | | | | | | |
| O54890 | ITGB3 | integrin, beta 3 (platelet glycoprotein IIIa, antigen CD61) | Plasma Membrane | transmembrane receptor | 1749434.875 | 7893542 | 8866252 | | | | | | |
| O54890 | ITGB3 | integrin, beta 3 (platelet glycoprotein IIIa, antigen CD61) | Plasma Membrane | transmembrane receptor | 1842798.75 | 1.16E+07 | 1.71E+07 | | | | | | |
| O54890 | ITGB3 | integrin, beta 3 (platelet glycoprotein IIIa, antigen CD61) | Plasma Membrane | transmembrane receptor | 1581664.625 | 6413217.5 | 1.33E+07 | | | | | | |
| O54890 | ITGB3 | integrin, beta 3 (platelet glycoprotein IIIa, antigen CD61) | Plasma Membrane | transmembrane receptor | 986916.25 | 3411212.75 | 8713984 | | | | | | |
| O54890 | ITGB3 | integrin, beta 3 (platelet glycoprotein IIIa, antigen CD61) | Plasma Membrane | transmembrane receptor | 2104517.5 | 5313042.5 | 9990918 | | | | | | |
| O54890 | ITGB3 | integrin, beta 3 (platelet glycoprotein IIIa, antigen CD61) | Plasma Membrane | transmembrane receptor | 886463.188 | 4499584.5 | 1.15E+07 | | | | | | |
| O54890 | ITGB3 | integrin, beta 3 (platelet glycoprotein IIIa, antigen CD61) | Plasma Membrane | transmembrane receptor | 1532787.125 | 6573171 | 1.68E+07 | | | | | | |
| O54890 | ITGB3 | integrin, beta 3 (platelet glycoprotein IIIa, antigen CD61) | Plasma Membrane | transmembrane receptor | 922489.938 | 4576342.5 | 1.12E+07 | | | | | | |
| O54890 | ITGB3 | integrin, beta 3 (platelet glycoprotein IIIa, antigen CD61) | Plasma Membrane | transmembrane receptor | 1084003.625 | 5953670.5 | 1.50E+07 | | | | | | |
| O54890 | ITGB3 | integrin, beta 3 (platelet glycoprotein IIIa, antigen CD61) | Plasma Membrane | transmembrane receptor | 671234.375 | 730374.375 | 1710403 | | | | | | |
| O54890 | ITGB3 | integrin, beta 3 (platelet glycoprotein IIIa, antigen CD61) | Plasma Membrane | transmembrane receptor | 626810.5 | 4890126 | 1.40E+07 | | | | | | |
| O54890 | ITGB3 | integrin, beta 3 (platelet glycoprotein IIIa, antigen CD61) | Plasma Membrane | transmembrane receptor | 448677.719 | 3694203.75 | 9646351 | | | | | | |
| O54890 | ITGB3 | integrin, beta 3 (platelet glycoprotein IIIa, antigen CD61) | Plasma Membrane | transmembrane receptor | 682326.875 | 3811920.75 | 7642332.5 | | | | | | |
| O54890 | ITGB3 | integrin, beta 3 (platelet glycoprotein IIIa, antigen CD61) | Plasma Membrane | transmembrane receptor | 316891.938 | 2009652.875 | 6185188 | | | | | | |
| O54890 | ITGB3 | integrin, beta 3 (platelet glycoprotein IIIa, antigen CD61) | Plasma Membrane | transmembrane receptor | 561540.312 | 2816082.25 | 8964543 | | | | | | |
| O54890 | ITGB3 | integrin, beta 3 (platelet glycoprotein IIIa, antigen CD61) | Plasma Membrane | transmembrane receptor | 178227.203 | 2774734 | 6706817.5 | | | | | | |
| O54890 | ITGB3 | integrin, beta 3 (platelet glycoprotein IIIa, antigen CD61) | Plasma Membrane | transmembrane receptor | 180638.047 | 4528365 | 1.14E+07 | | | | | | |
| O54890 | ITGB3 | integrin, beta 3 (platelet glycoprotein IIIa, antigen CD61) | Plasma Membrane | transmembrane receptor | 23106.236 | 4.70E-06 | 4.70E-06 | O54890 | ITGB3 | 1780572 | 7997693 | 942537 | 72 |
| Q9Z0T9 | ITGB6 | integrin, beta 6 | Plasma Membrane | other | 2261562 | 3281514 | 4446159 | Q9Z0T9 | ITGB6 | 2261562 | 1093097 | 631100 | 3 |
| Q5XKN4 | JAGN1 | jagunal homolog 1 | Cytoplasm | other | 1099144.875 | 1710544.25 | 2279450.5 | Q5XKN4 | JAGN1 | 1099145 | 590280 | 340798 | 3 |
| A2CG49 | KALRN | kalirin, RhoGEF kinase | Cytoplasm | kinase | 181486.531 | 181221.109 | 707589.938 | | | | | | |
| A2CG49 | KALRN | kalirin, RhoGEF kinase | Cytoplasm | kinase | 23171.74 | 87780.914 | 202950.281 | A2CG49 | KALRN | 102329 | 243540 | 99425 | 6 |
| Q80UY2 | KCMF1 | potassium channel modulatory factor 1 | Extracellular Space | other | 2852305.25 | 2956803.5 | 2885403.75 | | | | | | |
| Q80UY2 | KCMF1 | potassium channel modulatory factor 1 | Extracellular Space | other | 1180150.5 | 1518179.375 | 1293202.875 | | | | | | |
| Q80UY2 | KCMF1 | potassium channel modulatory factor 1 | Extracellular Space | other | 1022800.812 | 957647.125 | 768022.875 | Q80UY2 | KCMF1 | 1685086 | 912249 | 304083 | 9 |
| Q9EQF2 | KEL | Kell blood group, metallo-endopeptidase | Plasma Membrane | peptidase | 1.52E+08 | 1.50E+08 | 1.62E+08 | | | | | | |
| Q9EQF2 | KEL | Kell blood group, metallo-endopeptidase | Plasma Membrane | peptidase | 1.90E+08 | 2.22E+08 | 1.55E+08 | | | | | | |
| Q9EQF2 | KEL | Kell blood group, metallo-endopeptidase | Plasma Membrane | peptidase | 1.28E+08 | 1.34E+08 | 1.42E+08 | | | | | | |
| Q9EQF2 | KEL | Kell blood group, metallo-endopeptidase | Plasma Membrane | peptidase | 9.39E+07 | 1.43E+08 | 1.49E+08 | | | | | | |
| Q9EQF2 | KEL | Kell blood group, metallo-endopeptidase | Plasma Membrane | peptidase | 1.25E+08 | 1.24E+08 | 1.44E+08 | | | | | | |
| Q9EQF2 | KEL | Kell blood group, metallo-endopeptidase | Plasma Membrane | peptidase | 7.68E+07 | 1.21E+08 | 1.01E+08 | | | | | | |
| Q9EQF2 | KEL | Kell blood group, metallo-endopeptidase | Plasma Membrane | peptidase | 9.44E+07 | 9.15E+07 | 1.05E+08 | | | | | | |
| Q9EQF2 | KEL | Kell blood group, metallo-endopeptidase | Plasma Membrane | peptidase | 1.01E+08 | 1.10E+08 | 1.15E+08 | | | | | | |

| | | | | | | | | | | | | | |
|---|---|---|---|---|---|---|---|---|---|---|---|---|---|
| Q9EQF2 | KEL | Kell blood group, metallo-endopeptidase | Plasma Membrane | peptidase | 7.92E+07 | 9.33E+07 | 7.50E+07 | | | | | | |
| Q9EQF2 | KEL | Kell blood group, metallo-endopeptidase | Plasma Membrane | peptidase | 8.35E+07 | 9.25E+07 | 8.72E+07 | | | | | | |
| Q9EQF2 | KEL | Kell blood group, metallo-endopeptidase | Plasma Membrane | peptidase | 4.63E+07 | 5.90E+07 | 4.88E+07 | | | | | | |
| Q9EQF2 | KEL | Kell blood group, metallo-endopeptidase | Plasma Membrane | peptidase | 5.91E+07 | 6.03E+07 | 6.26E+07 | | | | | | |
| Q9EQF2 | KEL | Kell blood group, metallo-endopeptidase | Plasma Membrane | peptidase | 5.63E+07 | 6.80E+07 | 7.29E+07 | | | | | | |
| Q9EQF2 | KEL | Kell blood group, metallo-endopeptidase | Plasma Membrane | peptidase | 2.75E+07 | 4.74E+07 | 4.85E+07 | | | | | | |
| Q9EQF2 | KEL | Kell blood group, metallo-endopeptidase | Plasma Membrane | peptidase | 4.35E+07 | 4.57E+07 | 5.07E+07 | | | | | | |
| Q9EQF2 | KEL | Kell blood group, metallo-endopeptidase | Plasma Membrane | peptidase | 5.11E+07 | 4.25E+07 | 4.73E+07 | | | | | | |
| Q9EQF2 | KEL | Kell blood group, metallo-endopeptidase | Plasma Membrane | peptidase | 5.80E+07 | 4.75E+07 | 5.72E+07 | | | | | | |
| Q9EQF2 | KEL | Kell blood group, metallo-endopeptidase | Plasma Membrane | peptidase | 3.38E+07 | 3.30E+07 | 3.62E+07 | | | | | | |
| Q9EQF2 | KEL | Kell blood group, metallo-endopeptidase | Plasma Membrane | peptidase | 3.68E+07 | 3.88E+07 | 4.26E+07 | | | | | | |
| Q9EQF2 | KEL | Kell blood group, metallo-endopeptidase | Plasma Membrane | peptidase | 4.33E+07 | 4.17E+07 | 4.55E+07 | | | | | | |
| Q9EQF2 | KEL | Kell blood group, metallo-endopeptidase | Plasma Membrane | peptidase | 2.62E+07 | 2.04E+07 | 1.84E+07 | | | | | | |
| Q9EQF2 | KEL | Kell blood group, metallo-endopeptidase | Plasma Membrane | peptidase | 3.64E+07 | 2.49E+07 | 3.48E+07 | | | | | | |
| Q9EQF2 | KEL | Kell blood group, metallo-endopeptidase | Plasma Membrane | peptidase | 2.06E+07 | 2.67E+07 | 2.02E+07 | | | | | | |
| Q9EQF2 | KEL | Kell blood group, metallo-endopeptidase | Plasma Membrane | peptidase | 2.18E+07 | 2.59E+07 | 3.27E+07 | | | | | | |
| Q9EQF2 | KEL | Kell blood group, metallo-endopeptidase | Plasma Membrane | peptidase | 1.81E+07 | 1.65E+07 | 1.68E+07 | | | | | | |
| Q9EQF2 | KEL | Kell blood group, metallo-endopeptidase | Plasma Membrane | peptidase | 2.09E+07 | 2.70E+07 | 3.42E+07 | | | | | | |
| Q9EQF2 | KEL | Kell blood group, metallo-endopeptidase | Plasma Membrane | peptidase | 1.47E+07 | 1.62E+07 | 1.50E+07 | | | | | | |
| Q9EQF2 | KEL | Kell blood group, metallo-endopeptidase | Plasma Membrane | peptidase | 1.55E+07 | 1.91E+07 | 2.44E+07 | | | | | | |
| Q9EQF2 | KEL | Kell blood group, metallo-endopeptidase | Plasma Membrane | peptidase | 1.62E+07 | 1.98E+07 | 1.72E+07 | | | | | | |
| Q9EQF2 | KEL | Kell blood group, metallo-endopeptidase | Plasma Membrane | peptidase | 2.18E+07 | 1.98E+07 | 2.50E+07 | | | | | | |
| Q9EQF2 | KEL | Kell blood group, metallo-endopeptidase | Plasma Membrane | peptidase | 1.49E+07 | 1.79E+07 | 1.75E+07 | | | | | | |
| Q9EQF2 | KEL | Kell blood group, metallo-endopeptidase | Plasma Membrane | peptidase | 5154862.5 | 6516355 | 9729659 | | | | | | |
| Q9EQF2 | KEL | Kell blood group, metallo-endopeptidase | Plasma Membrane | peptidase | 8356800.5 | 8349907 | 1.00E+07 | | | | | | |
| Q9EQF2 | KEL | Kell blood group, metallo-endopeptidase | Plasma Membrane | peptidase | 3189060 | 4232794 | 6091013.5 | | | | | | |
| Q9EQF2 | KEL | Kell blood group, metallo-endopeptidase | Plasma Membrane | peptidase | 3169596.5 | 1313608.25 | 1725530.875 | | | | | | |
| Q9EQF2 | KEL | Kell blood group, metallo-endopeptidase | Plasma Membrane | peptidase | 4848945 | 1068522.875 | 61503.285 | Q9EQF2 | KEL | 50856646 | 48094652 | 4627910 | 108 |
| Q6PDI5 | KIAA0368 | KIAA0368 | Cytoplasm | other | 5877224.5 | 5911273 | 3738561.25 | | | | | | |
| Q6PDI5 | KIAA0368 | KIAA0368 | Cytoplasm | other | 5196480 | 5444107 | 2917389.5 | | | | | | |
| Q6PDI5 | KIAA0368 | KIAA0368 | Cytoplasm | other | 4864197.5 | 3302718.5 | 3336637.75 | | | | | | |
| Q6PDI5 | KIAA0368 | KIAA0368 | Cytoplasm | other | 2987564.25 | 2340082 | 2142516.5 | | | | | | |
| Q6PDI5 | KIAA0368 | KIAA0368 | Cytoplasm | other | 2163615.5 | 2054670.625 | 1471162.5 | | | | | | |
| Q6PDI5 | KIAA0368 | KIAA0368 | Cytoplasm | other | 1921342.375 | 1306354.5 | 1300076.75 | | | | | | |
| Q6PDI5 | KIAA0368 | KIAA0368 | Cytoplasm | other | 1525143 | 2121738.25 | 1706896 | | | | | | |
| Q6PDI5 | KIAA0368 | KIAA0368 | Cytoplasm | other | 890018.75 | 964260.5 | 976549.5 | | | | | | |
| Q6PDI5 | KIAA0368 | KIAA0368 | Cytoplasm | other | 711996 | 533058.562 | 388219.188 | Q6PDI5 | KIAA0368 | 2904176 | 1676649 | 322671 | 27 |
| Q7TQ95 | KIAA1715 | KIAA1715 | Cytoplasm | other | 2766772.75 | 4039810 | 3974950 | | | | | | |
| Q7TQ95 | KIAA1715 | KIAA1715 | Cytoplasm | other | 2568825.25 | 4082324.25 | 4664983.5 | | | | | | |
| Q7TQ95 | KIAA1715 | KIAA1715 | Cytoplasm | other | 3436223.25 | 3207796 | 2532343.5 | | | | | | |
| Q7TQ95 | KIAA1715 | KIAA1715 | Cytoplasm | other | 2282761 | 2640905.5 | 2487305.5 | | | | | | |
| Q7TQ95 | KIAA1715 | KIAA1715 | Cytoplasm | other | 1623377 | 2422746.5 | 1916154.125 | Q7TQ95 | KIAA1715 | 2535592 | 885147 | 228544 | 15 |
| Q60575 | KIF1B | kinesin family member 1B | Cytoplasm | transporter | 2099571.75 | 3108856.5 | 6720391 | Q60575 | KIF1B | 2099572 | 2429465 | 1402652 | 3 |
| O88448 | KLC2 | kinesin light chain 2 | Cytoplasm | other | 4.70E-06 | 575898.75 | 987848.625 | O88448 | KLC2 | 521249 | 496187 | 286473 | 3 |
| Q60682 | Klra4 (includes others) | killer cell lectin-like receptor, subfamily A, member 4 | Plasma Membrane | transmembrane receptor | 6252631 | 6407410 | 8776871 | Q60682 | Klra4 (includes others) | 6252631 | 1414808 | 816840 | 3 |
| O35344 | KPNA3 | karyopherin alpha 3 (importin alpha 4) | Nucleus | transporter | 3346416 | 2085437.375 | 2848479.25 | O35344 | KPNA3 | 3346416 | 635117 | 366685 | 3 |
| P70168 | KPNB1 | karyopherin (importin) beta 1 | Nucleus | transporter | 1.71E+07 | 1.20E+07 | 1.26E+07 | | | | | | |
| P70168 | KPNB1 | karyopherin (importin) beta 1 | Nucleus | transporter | 1.89E+07 | 1.54E+07 | 1.51E+07 | | | | | | |
| P70168 | KPNB1 | karyopherin (importin) beta 1 | Nucleus | transporter | 1.51E+07 | 1.24E+07 | 1.04E+07 | | | | | | |
| P70168 | KPNB1 | karyopherin (importin) beta 1 | Nucleus | transporter | 1.22E+07 | 9244814 | 7771460 | | | | | | |
| P70168 | KPNB1 | karyopherin (importin) beta 1 | Nucleus | transporter | 1.69E+07 | 1.42E+07 | 7892085 | | | | | | |
| P70168 | KPNB1 | karyopherin (importin) beta 1 | Nucleus | transporter | 1.13E+07 | 7773533 | 5078640 | | | | | | |
| P70168 | KPNB1 | karyopherin (importin) beta 1 | Nucleus | transporter | 9510666 | 1.08E+07 | 7802358 | | | | | | |
| P70168 | KPNB1 | karyopherin (importin) beta 1 | Nucleus | transporter | 9003707 | 7640889 | 5398158.5 | | | | | | |
| P70168 | KPNB1 | karyopherin (importin) beta 1 | Nucleus | transporter | 8836173 | 9321503 | 6347446.5 | | | | | | |
| P70168 | KPNB1 | karyopherin (importin) beta 1 | Nucleus | transporter | 9530316 | 8768649 | 6956381 | | | | | | |
| P70168 | KPNB1 | karyopherin (importin) beta 1 | Nucleus | transporter | 8651589 | 6530752 | 6479299 | | | | | | |
| P70168 | KPNB1 | karyopherin (importin) beta 1 | Nucleus | transporter | 6775221.5 | 6972222 | 4808953 | | | | | | |
| P70168 | KPNB1 | karyopherin (importin) beta 1 | Nucleus | transporter | 7392570 | 5291693 | 4962625.5 | | | | | | |
| P70168 | KPNB1 | karyopherin (importin) beta 1 | Nucleus | transporter | 7285712.5 | 6282157.5 | 5040510 | | | | | | |
| P70168 | KPNB1 | karyopherin (importin) beta 1 | Nucleus | transporter | 6014486 | 2947020.5 | 1418263.625 | | | | | | |
| P70168 | KPNB1 | karyopherin (importin) beta 1 | Nucleus | transporter | 7120578.5 | 6620124 | 5138179.5 | | | | | | |
| P70168 | KPNB1 | karyopherin (importin) beta 1 | Nucleus | transporter | 6554720.5 | 5115457.5 | 5841989 | | | | | | |
| P70168 | KPNB1 | karyopherin (importin) beta 1 | Nucleus | transporter | 4791657.5 | 2900876.75 | 1895531.375 | | | | | | |
| P70168 | KPNB1 | karyopherin (importin) beta 1 | Nucleus | transporter | 5151494 | 5407154.5 | 5298241.5 | | | | | | |
| P70168 | KPNB1 | karyopherin (importin) beta 1 | Nucleus | transporter | 4358612.5 | 2556965.5 | 2157639.5 | | | | | | |
| P70168 | KPNB1 | karyopherin (importin) beta 1 | Nucleus | transporter | 5149761.5 | 3294344.5 | 2991564 | P70168 | KPNB1 | 9410822 | 3961475 | 499099 | 63 |
| P04104 | KRT1 | keratin 1, type II | Cytoplasm | other | 1.90E+07 | 2.60E+07 | 6717932.5 | | | | | | |
| P04104 | KRT1 | keratin 1, type II | Cytoplasm | other | 771769 | 105586.625 | 193991.75 | | | | | | |
| P04104 | KRT1 | keratin 1, type II | Cytoplasm | other | 3776790 | 7635125 | 1636820.375 | | | | | | |
| P04104 | KRT1 | keratin 1, type II | Cytoplasm | other | 4052569 | 1847812.75 | 534610.688 | P04104 | KRT1 | 6897782 | 8222999 | 2373775 | 12 |
| P02535 | Krt10 | keratin 10 | Cytoplasm | other | 9353656 | 7943872 | 2721945.5 | P02535 | Krt10 | 9353656 | 3493699 | 2017088 | 3 |
| Q61781 | KRT14 | keratin 14, type I | Cytoplasm | other | 1241959.25 | 880981.5 | 84476.617 | Q61781 | KRT14 | 1241959 | 592240 | 341930 | 3 |
| Q3TTY5 | Krt2 | keratin 2 | Cytoplasm | other | 4684447.5 | 3443305.75 | 321941.156 | Q3TTY5 | Krt2 | 4684448 | 2247770 | 1297750 | 3 |
| Q922U2 | KRT5 | keratin 5, type II | Cytoplasm | other | 1.40E+07 | 1.99E+07 | 6768121 | | | | | | |
| Q922U2 | KRT5 | keratin 5, type II | Cytoplasm | other | 6374868 | 6270841.5 | 7941905.5 | | | | | | |
| Q922U2 | KRT5 | keratin 5, type II | Cytoplasm | other | 928013.438 | 1072908.625 | 282465.656 | | | | | | |
| Q922U2 | KRT5 | keratin 5, type II | Cytoplasm | other | 3827791.25 | 6356446.5 | 2051211.5 | Q922U2 | KRT5 | 6292668 | 5750897 | 1660141 | 12 |
| Q6NXH9 | KRT73 | keratin 73, type II | Extracellular Space | other | 1050164.25 | 1138245.125 | 138209.984 | Q6NXH9 | KRT73 | 1050164 | 553698 | 319678 | 3 |
| Q3UV17 | KRT76 | keratin 76, type II | Cytoplasm | other | 1.25E+07 | 1.26E+07 | 5030399.5 | Q3UV17 | KRT76 | 12540000 | 4338560 | 2504868 | 3 |
| Q8VED5 | KRT79 | keratin 79, type II | Extracellular Space | other | 3809798.25 | 5551965.5 | 4838101.5 | Q8VED5 | KRT79 | 3809798 | 875800 | 505643 | 3 |
| Q6IMF0 | Krt83 | keratin 83 | Extracellular Space | other | 1.55E+07 | 1.30E+07 | 1.99E+07 | Q6IMF0 | Krt83 | 15450000 | 3491623 | 2015890 | 3 |
| P11438 | LAMP1 | lysosomal-associated membrane protein 1 | Plasma Membrane | other | 9822041 | 1.38E+07 | 1.08E+07 | | | | | | |
| P11438 | LAMP1 | lysosomal-associated membrane protein 1 | Plasma Membrane | other | 4987174.5 | 7313336.5 | 5824120 | | | | | | |
| P11438 | LAMP1 | lysosomal-associated membrane protein 1 | Plasma Membrane | other | 1869536.375 | 2558558.75 | 1880708.25 | P11438 | LAMP1 | 5559584 | 4249973 | 1416658 | 9 |
| P17047 | LAMP2 | lysosomal-associated membrane protein 2 | Plasma Membrane | enzyme | 3050687.25 | 7842984 | 5990662.5 | | | | | | |
| P17047 | LAMP2 | lysosomal-associated membrane protein 2 | Plasma Membrane | enzyme | 1940388 | 3021043.25 | 2366056.25 | P17047 | LAMP2 | 2495538 | 2344999 | 957342 | 6 |
| Q9CQ22 | LAMTOR1 | late endosomal/lysosomal adaptor, MAPK and MTOR activator 1 | Plasma Membrane | other | 2.14E+07 | 2.85E+07 | 2.57E+07 | | | | | | |
| Q9CQ22 | LAMTOR1 | late endosomal/lysosomal adaptor, MAPK and MTOR activator 1 | Plasma Membrane | other | 6881042.5 | 8164311 | 8250060.5 | | | | | | |
| Q9CQ22 | LAMTOR1 | late endosomal/lysosomal adaptor, MAPK and MTOR activator 1 | Plasma Membrane | other | 6975931.5 | 8736272 | 8682869 | | | | | | |
| Q9CQ22 | LAMTOR1 | late endosomal/lysosomal adaptor, MAPK and MTOR activator 1 | Plasma Membrane | other | 4311897.5 | 4179993.75 | 3524251.5 | Q9CQ22 | LAMTOR1 | 9887218 | 8726969 | 2519259 | 12 |

| | | | | | | | | | | | | | |
|---|---|---|---|---|---|---|---|---|---|---|---|---|---|
| O88653 | LAMTOR3 | late endosomal/lysosomal adaptor, MAPK and MTOR activator 3 | Cytoplasm | other | 1.38E+07 | 1.90E+07 | 1.48E+07 | | | | | | |
| O88653 | LAMTOR3 | late endosomal/lysosomal adaptor, MAPK and MTOR activator 3 | Cytoplasm | other | 5148083.5 | 7059481 | 7506428 | | | | | | |
| O88653 | LAMTOR3 | late endosomal/lysosomal adaptor, MAPK and MTOR activator 3 | Cytoplasm | other | 4792661.5 | 6090921 | 2757595.25 | O88653 | LAMTOR3 | 7896915 | 5504923 | 1834974 | 9 |
| Q9D1L9 | LAMTOR5 | late endosomal/lysosomal adaptor, MAPK and MTOR activator 5 | Cytoplasm | other | 1.16E+07 | 1.49E+07 | 1.21E+07 | | | | | | |
| Q9D1L9 | LAMTOR5 | late endosomal/lysosomal adaptor, MAPK and MTOR activator 5 | Cytoplasm | other | 4972291.5 | 6119403.5 | 7785768.5 | | | | | | |
| Q9D1L9 | LAMTOR5 | late endosomal/lysosomal adaptor, MAPK and MTOR activator 5 | Cytoplasm | other | 2726596.25 | 7283950.5 | 6009240 | Q9D1L9 | LAMTOR5 | 6416296 | 3898532 | 1299511 | 9 |
| O89112 | LANCL1 | LanC lantibiotic synthetase component C-like 1 (bacterial) | Plasma Membrane | other | 1885209 | 1979779.5 | 2109886.5 | O89112 | LANCL1 | 1885209 | 112806 | 65129 | 3 |
| Q9JJK2 | LANCL2 | LanC lantibiotic synthetase component C-like 2 (bacterial) | Plasma Membrane | other | 9024841 | 8921849 | 8542152 | | | | | | |
| Q9JJK2 | LANCL2 | LanC lantibiotic synthetase component C-like 2 (bacterial) | Plasma Membrane | other | 8663561 | 1.05E+07 | 1.08E+07 | | | | | | |
| Q9JJK2 | LANCL2 | LanC lantibiotic synthetase component C-like 2 (bacterial) | Plasma Membrane | other | 8339418.5 | 6940371 | 5967978 | | | | | | |
| Q9JJK2 | LANCL2 | LanC lantibiotic synthetase component C-like 2 (bacterial) | Plasma Membrane | other | 4535635.5 | 6586807 | 6212350.5 | | | | | | |
| Q9JJK2 | LANCL2 | LanC lantibiotic synthetase component C-like 2 (bacterial) | Plasma Membrane | other | 7323712 | 6415301.5 | 7011292.5 | | | | | | |
| Q9JJK2 | LANCL2 | LanC lantibiotic synthetase component C-like 2 (bacterial) | Plasma Membrane | other | 4691944 | 4534434.5 | 5313473.5 | | | | | | |
| Q9JJK2 | LANCL2 | LanC lantibiotic synthetase component C-like 2 (bacterial) | Plasma Membrane | other | 4636027.5 | 4442905.5 | 4743827.5 | | | | | | |
| Q9JJK2 | LANCL2 | LanC lantibiotic synthetase component C-like 2 (bacterial) | Plasma Membrane | other | 2674961.75 | 3654913.5 | 3700648 | | | | | | |
| Q9JJK2 | LANCL2 | LanC lantibiotic synthetase component C-like 2 (bacterial) | Plasma Membrane | other | 4183671.75 | 3159437.25 | 2462994 | | | | | | |
| Q9JJK2 | LANCL2 | LanC lantibiotic synthetase component C-like 2 (bacterial) | Plasma Membrane | other | 2606420.75 | 1051789.875 | 2440822 | | | | | | |
| Q9JJK2 | LANCL2 | LanC lantibiotic synthetase component C-like 2 (bacterial) | Plasma Membrane | other | 3131328 | 1220293.375 | 1648301.625 | | | | | | |
| Q9JJK2 | LANCL2 | LanC lantibiotic synthetase component C-like 2 (bacterial) | Plasma Membrane | other | 5608197.5 | 965712.75 | 565064.688 | Q9JJK2 | LANCL2 | 5451643 | 2755871 | 459312 | 36 |
| P06151 | LDHA | lactate dehydrogenase A | Cytoplasm | enzyme | 1.13E+07 | 8274164.5 | 1.02E+07 | | | | | | |
| P06151 | LDHA | lactate dehydrogenase A | Cytoplasm | enzyme | 7797026.5 | 7220398 | 8674015 | | | | | | |
| P06151 | LDHA | lactate dehydrogenase A | Cytoplasm | enzyme | 8681942 | 6523845.5 | 7342967 | | | | | | |
| P06151 | LDHA | lactate dehydrogenase A | Cytoplasm | enzyme | 6111686.5 | 5143465.5 | 6268550.5 | | | | | | |
| P06151 | LDHA | lactate dehydrogenase A | Cytoplasm | enzyme | 6922769 | 6207649 | 6102980.5 | | | | | | |
| P06151 | LDHA | lactate dehydrogenase A | Cytoplasm | enzyme | 4862582.5 | 3348039.25 | 6156543 | | | | | | |
| P06151 | LDHA | lactate dehydrogenase A | Cytoplasm | enzyme | 4069374.5 | 3010666.75 | 4378453.5 | | | | | | |
| P06151 | LDHA | lactate dehydrogenase A | Cytoplasm | enzyme | 2635160 | 2858167.5 | 2454581.5 | | | | | | |
| P06151 | LDHA | lactate dehydrogenase A | Cytoplasm | enzyme | 1782614.75 | 1921915.625 | 2398538.25 | P06151 | LDHA | 6015906 | 2592393 | 498906 | 27 |
| Q8CBY3 | LENG8 | leukocyte receptor cluster (LRC) member 8 | Nucleus | other | 840998.125 | 1949211.125 | 1908089.125 | Q8CBY3 | LENG8 | 840998 | 628293 | 362745 | 3 |
| O89013 | LEPROT | leptin receptor overlapping transcript | Plasma Membrane | other | 3818222 | 3043182.75 | 3060237.25 | | | | | | |
| O89013 | LEPROT | leptin receptor overlapping transcript | Plasma Membrane | other | 400263.969 | 314347.312 | 238396.359 | O89013 | LEPROT | 2109243 | 1661988 | 678504 | 6 |
| Q9Z2I0 | LETM1 | leucine zipper-EF-hand containing transmembrane protein 1 | Cytoplasm | other | 1493294.875 | 2669611.5 | 906362.812 | Q9Z2I0 | LETM1 | 1493295 | 897892 | 518398 | 3 |
| O08573 | LGALS9B | lectin, galactoside-binding, soluble, 9B | Cytoplasm | other | 1.86E+07 | 1.75E+07 | 1.38E+07 | | | | | | |
| O08573 | LGALS9B | lectin, galactoside-binding, soluble, 9B | Cytoplasm | other | 1.19E+07 | 1.08E+07 | 1.15E+07 | | | | | | |
| O08573 | LGALS9B | lectin, galactoside-binding, soluble, 9B | Cytoplasm | other | 1.21E+07 | 1.17E+07 | 1.03E+07 | | | | | | |
| O08573 | LGALS9B | lectin, galactoside-binding, soluble, 9B | Cytoplasm | other | 1.29E+07 | 1.46E+07 | 1.09E+07 | | | | | | |
| O08573 | LGALS9B | lectin, galactoside-binding, soluble, 9B | Cytoplasm | other | 6609603 | 8619866 | 7134195.5 | | | | | | |
| O08573 | LGALS9B | lectin, galactoside-binding, soluble, 9B | Cytoplasm | other | 6274157 | 4916014.5 | 3549556.5 | | | | | | |
| O08573 | LGALS9B | lectin, galactoside-binding, soluble, 9B | Cytoplasm | other | 5956523 | 5623845.5 | 3592727 | | | | | | |
| O08573 | LGALS9B | lectin, galactoside-binding, soluble, 9B | Cytoplasm | other | 3542754.5 | 3692173.25 | 2465172.5 | O08573 | LGALS9B | 9735380 | 4604837 | 939958 | 24 |
| A2ARI4 | LGR4 | leucine-rich repeat containing G protein-coupled receptor 4 | Plasma Membrane | transmembrane receptor | 310348 | 721731.688 | 2537077.5 | A2ARI4 | LGR4 | 310348 | 1184838 | 684066 | 3 |
| Q99JW4 | LIMS1 | LIM and senescent cell antigen-like domains 1 | Plasma Membrane | other | 84391.266 | 415529.656 | 401405.188 | Q99JW4 | LIMS1 | 84391 | 187239 | 108102 | 3 |
| Q9D0F3 | LMAN1 | lectin, mannose-binding, 1 | Cytoplasm | other | 1.02E+07 | 1.33E+07 | 1.88E+07 | | | | | | |
| Q9D0F3 | LMAN1 | lectin, mannose-binding, 1 | Cytoplasm | other | 8489944 | 8517910 | 1.16E+07 | | | | | | |
| Q9D0F3 | LMAN1 | lectin, mannose-binding, 1 | Cytoplasm | other | 9985649 | 8455631 | 1.48E+07 | | | | | | |
| Q9D0F3 | LMAN1 | lectin, mannose-binding, 1 | Cytoplasm | other | 1.11E+07 | 1.17E+07 | 1.15E+07 | | | | | | |
| Q9D0F3 | LMAN1 | lectin, mannose-binding, 1 | Cytoplasm | other | 8529249 | 6399476.5 | 7615315 | | | | | | |
| Q9D0F3 | LMAN1 | lectin, mannose-binding, 1 | Cytoplasm | other | 5558369 | 5030676 | 6167420 | Q9D0F3 | LMAN1 | 8972202 | 3486498 | 821776 | 18 |
| Q9DBH5 | LMAN2 | lectin, mannose-binding 2 | Cytoplasm | transporter | 9210350 | 1.23E+07 | 1.01E+07 | | | | | | |
| Q9DBH5 | LMAN2 | lectin, mannose-binding 2 | Cytoplasm | transporter | 1.31E+07 | 1.68E+07 | 1.77E+07 | | | | | | |
| Q9DBH5 | LMAN2 | lectin, mannose-binding 2 | Cytoplasm | transporter | 3159562.25 | 3995422.25 | 4307338.5 | | | | | | |
| Q9DBH5 | LMAN2 | lectin, mannose-binding 2 | Cytoplasm | transporter | 2542541.25 | 4008351.25 | 4310149 | | | | | | |
| Q9DBH5 | LMAN2 | lectin, mannose-binding 2 | Cytoplasm | transporter | 2461936.75 | 3346695.25 | 3442730.5 | Q9DBH5 | LMAN2 | 6086878 | 5345982 | 1380327 | 15 |
| Q8VEE1 | LMCD1 | LIM and cysteine-rich domains 1 | Cytoplasm | transcription regulator | 4389932.5 | 1.09E+07 | 1.86E+07 | Q8VEE1 | LMCD1 | 4389933 | 7092806 | 4095034 | 3 |
| Q8CGK3 | LONP1 | lon peptidase 1, mitochondrial | Cytoplasm | peptidase | 4340539 | 9693675 | 3017449.25 | | | | | | |
| Q8CGK3 | LONP1 | lon peptidase 1, mitochondrial | Cytoplasm | peptidase | 4366931.5 | 4711493 | 188288.219 | Q8CGK3 | LONP1 | 4353735 | 3091068 | 1261923 | 6 |
| Q3TFD2 | LPCAT1 | lysophosphatidylcholine acyltransferase 1 | Cytoplasm | enzyme | 1.60E+07 | 2.75E+07 | 2.82E+07 | | | | | | |
| Q3TFD2 | LPCAT1 | lysophosphatidylcholine acyltransferase 1 | Cytoplasm | enzyme | 1.04E+07 | 1.03E+07 | 8890107 | | | | | | |
| Q3TFD2 | LPCAT1 | lysophosphatidylcholine acyltransferase 1 | Cytoplasm | enzyme | 9437665 | 9643567 | 1.01E+07 | | | | | | |
| Q3TFD2 | LPCAT1 | lysophosphatidylcholine acyltransferase 1 | Cytoplasm | enzyme | 6403621 | 9968736 | 5757356.5 | | | | | | |
| Q3TFD2 | LPCAT1 | lysophosphatidylcholine acyltransferase 1 | Cytoplasm | enzyme | 7625111.5 | 7939731.5 | 7130119.5 | | | | | | |
| Q3TFD2 | LPCAT1 | lysophosphatidylcholine acyltransferase 1 | Cytoplasm | enzyme | 8891201 | 3685450.25 | 1266621 | | | | | | |
| Q3TFD2 | LPCAT1 | lysophosphatidylcholine acyltransferase 1 | Cytoplasm | enzyme | 7906547 | 8974700 | 8592086 | | | | | | |
| Q3TFD2 | LPCAT1 | lysophosphatidylcholine acyltransferase 1 | Cytoplasm | enzyme | 4215141 | 4009068.5 | 4268967.5 | Q3TFD2 | LPCAT1 | 8864911 | 6399657 | 1306325 | 24 |
| Q91V01 | LPCAT3 | lysophosphatidylcholine acyltransferase 3 | Plasma Membrane | enzyme | 1.97E+07 | 2.38E+07 | 2.39E+07 | | | | | | |
| Q91V01 | LPCAT3 | lysophosphatidylcholine acyltransferase 3 | Plasma Membrane | enzyme | 7808743.5 | 9594552 | 1.11E+07 | | | | | | |
| Q91V01 | LPCAT3 | lysophosphatidylcholine acyltransferase 3 | Plasma Membrane | enzyme | 5768819.5 | 7952907.5 | 9818758 | | | | | | |
| Q91V01 | LPCAT3 | lysophosphatidylcholine acyltransferase 3 | Plasma Membrane | enzyme | 1231090 | 1844575.5 | 2505617.5 | Q91V01 | LPCAT3 | 8632163 | 7993120 | 2307415 | 12 |
| P55302 | LRPAP1 | low density lipoprotein receptor-related protein associated protein 1 | Plasma Membrane | transmembrane receptor | 1665522.75 | 2313918.25 | 2039990.75 | P55302 | LRPAP1 | 1665523 | 325494 | 187924 | 3 |
| Q9D1G5 | LRRC57 | leucine rich repeat containing 57 | Cytoplasm | other | 6896501 | 7849598 | 4636027.5 | | | | | | |
| Q9D1G5 | LRRC57 | leucine rich repeat containing 57 | Cytoplasm | other | 6005385 | 7980391 | 6540888.5 | | | | | | |
| Q9D1G5 | LRRC57 | leucine rich repeat containing 57 | Cytoplasm | other | 8367522.5 | 9236391 | 7666153.5 | | | | | | |
| Q9D1G5 | LRRC57 | leucine rich repeat containing 57 | Cytoplasm | other | 6636022.5 | 5321582 | 3282519.5 | | | | | | |
| Q9D1G5 | LRRC57 | leucine rich repeat containing 57 | Cytoplasm | other | 5396433 | 4445946.5 | 4431195 | | | | | | |
| Q9D1G5 | LRRC57 | leucine rich repeat containing 57 | Cytoplasm | other | 2847727 | 2849858 | 1873338 | Q9D1G5 | LRRC57 | 6024932 | 2137983 | 503927 | 18 |
| Q922Q8 | LRRC59 | leucine rich repeat containing 59 | Cytoplasm | other | 706299.812 | 2601494 | 1991573.875 | Q922Q8 | LRRC59 | 706300 | 967444 | 558554 | 3 |
| Q80TE7 | LRRC7 | leucine rich repeat containing 7 | Plasma Membrane | other | 1329179.875 | 1813929.625 | 1610768 | Q80TE7 | LRRC7 | 1329180 | 243430 | 140544 | 3 |
| P08071 | LTF | lactotransferrin | Extracellular Space | other | 489141.25 | 227060.953 | 21132.502 | | | | | | |
| P08071 | LTF | lactotransferrin | Extracellular Space | other | 318948.469 | 4.70E-06 | 4.70E-06 | | | | | | |
| P08071 | LTF | lactotransferrin | Extracellular Space | other | 516097.094 | 274176.719 | 273987.812 | | | | | | |
| P08071 | LTF | lactotransferrin | Extracellular Space | other | 2557.816 | 38530.703 | 160673.422 | | | | | | |
| P08071 | LTF | lactotransferrin | Extracellular Space | other | 101116.234 | 6379.625 | 321024.062 | P08071 | LTF | 285572 | 178435 | 46072 | 15 |
| P25911 | LYN | LYN proto-oncogene, Src family tyrosine kinase | Cytoplasm | kinase | 3350109.25 | 6303758 | 9634377 | | | | | | |
| P25911 | LYN | LYN proto-oncogene, Src family tyrosine kinase | Cytoplasm | kinase | 3204457.25 | 3530226 | 5699485.5 | | | | | | |
| P25911 | LYN | LYN proto-oncogene, Src family tyrosine kinase | Cytoplasm | kinase | 2367242 | 4108100.75 | 4518460.5 | | | | | | |
| P25911 | LYN | LYN proto-oncogene, Src family tyrosine kinase | Cytoplasm | kinase | 1920265.375 | 3007844.5 | 4058839.75 | | | | | | |
| P25911 | LYN | LYN proto-oncogene, Src family tyrosine kinase | Cytoplasm | kinase | 682277.188 | 1468865.5 | 2004433.75 | | | | | | |
| P25911 | LYN | LYN proto-oncogene, Src family tyrosine kinase | Cytoplasm | kinase | 565855.875 | 668108.688 | 872929.125 | | | | | | |
| P25911 | LYN | LYN proto-oncogene, Src family tyrosine kinase | Cytoplasm | kinase | 491019.875 | 867514.562 | 876270.75 | | | | | | |
| P25911 | LYN | LYN proto-oncogene, Src family tyrosine kinase | Cytoplasm | kinase | 154586.75 | 255160.109 | 633461.5 | P25911 | LYN | 1591977 | 2333509 | 476325 | 24 |
| P08905 | LYZ | lysozyme | Extracellular Space | enzyme | 2356275.5 | 1871387.875 | 6747353 | | | | | | |

| | | | | | | | | | | | | | |
|---|---|---|---|---|---|---|---|---|---|---|---|---|---|
| P08905 | LYZ | lysozyme | Extracellular Space | enzyme | 535075.25 | 605234.5 | 2156059.5 | P08905 | LYZ | 1445675 | 2277573 | 929815 | 6 |
| P24668 | M6PR | mannose-6-phosphate receptor (cation dependent) | Cytoplasm | transporter | 4011089.5 | 5964135.5 | 4650637.5 | | | | | | |
| P24668 | M6PR | mannose-6-phosphate receptor (cation dependent) | Cytoplasm | transporter | 1478078.625 | 1485018.625 | 973621.875 | P24668 | M6PR | 2744584 | 2059023 | 840593 | 6 |
| Q9CXI5 | MANF | mesencephalic astrocyte-derived neurotrophic factor | Extracellular Space | other | 1.34E+07 | 1.93E+07 | 2.05E+07 | | | | | | |
| Q9CXI5 | MANF | mesencephalic astrocyte-derived neurotrophic factor | Extracellular Space | other | 3664451.5 | 3575979.75 | 4564381 | | | | | | |
| Q9CXI5 | MANF | mesencephalic astrocyte-derived neurotrophic factor | Extracellular Space | other | 1558112.5 | 2366888.25 | 1930426.5 | Q9CXI5 | MANF | 6190855 | 7698350 | 2566117 | 9 |
| Q61166 | MAPRE1 | microtubule-associated protein, RP/EB family, member 1 | Cytoplasm | other | 608957.812 | 716326.625 | 864726.25 | Q61166 | MAPRE1 | 608958 | 128432 | 74150 | 3 |
| Q922Q1 | MARC_2 | mitochondrial amidoxime reducing component 2 | Cytoplasm | enzyme | 732617.688 | 1261006.375 | 554903.312 | Q922Q1 | MARC_2 | 732618 | 367278 | 212048 | 3 |
| Q9DBD2 | MARH8 | membrane-associated ring finger (C3HC4) 8, E3 ubiquitin protein ligase | Cytoplasm | enzyme | 5079018 | 5451182 | 5512774 | | | | | | |
| Q9DBD2 | MARH8 | membrane-associated ring finger (C3HC4) 8, E3 ubiquitin protein ligase | Cytoplasm | enzyme | 3283295.25 | 5014844.5 | 5007114.5 | Q9DBD2 | MARH8 | 4181157 | 818449 | 334130 | 6 |
| Q05512 | MARK2 | MAP/microtubule affinity-regulating kinase 2 | Cytoplasm | kinase | 2361924 | 3266565.5 | 2541881 | | | | | | |
| Q05512 | MARK2 | MAP/microtubule affinity-regulating kinase 2 | Cytoplasm | kinase | 1408048.5 | 1784461.375 | 1151041.875 | Q05512 | MARK2 | 1884986 | 787704 | 321579 | 6 |
| Q8VCF0 | MAVS | mitochondrial antiviral signaling protein | Cytoplasm | other | 3289000 | 8282963 | 5300469 | | | | | | |
| Q8VCF0 | MAVS | mitochondrial antiviral signaling protein | Cytoplasm | other | 1369146.25 | 2644908.75 | 1712443.625 | Q8VCF0 | MAVS | 2329073 | 2615489 | 1067769 | 6 |
| Q8BL86 | MBLAC2 | metallo-beta-lactamase domain containing 2 | Cytoplasm | other | 1.66E+07 | 1.43E+07 | 1.70E+07 | Q8BL86 | MBLAC2 | 16600000 | 1458572 | 842107 | 3 |
| Q8CHK3 | MBOAT7 | membrane bound O-acyltransferase domain containing 7 | Plasma Membrane | other | 194161.656 | 1458971.25 | 2708667 | Q8CHK3 | MBOAT7 | 194162 | 1257260 | 725880 | 3 |
| Q8K5B2 | MCFD2 | multiple coagulation factor deficiency 2 | Cytoplasm | other | 6660400 | 1.06E+07 | 9205273 | Q8K5B2 | MCFD2 | 6660400 | 1993138 | 1150739 | 3 |
| Q3UMR5 | MCU | mitochondrial calcium uniporter | Cytoplasm | ion channel | 166134.156 | 747509 | 1050257.625 | Q3UMR5 | MCU | 166134 | 449319 | 259415 | 3 |
| P08249 | MDH2 | malate dehydrogenase 2, NAD (mitochondrial) | Cytoplasm | enzyme | 3.40E+07 | 5.89E+07 | 4.53E+07 | | | | | | |
| P08249 | MDH2 | malate dehydrogenase 2, NAD (mitochondrial) | Cytoplasm | enzyme | 2.89E+07 | 4.69E+07 | 2.42E+07 | | | | | | |
| P08249 | MDH2 | malate dehydrogenase 2, NAD (mitochondrial) | Cytoplasm | enzyme | 2.66E+07 | 5.59E+07 | 3.41E+07 | | | | | | |
| P08249 | MDH2 | malate dehydrogenase 2, NAD (mitochondrial) | Cytoplasm | enzyme | 2.41E+07 | 4.74E+07 | 2.34E+07 | | | | | | |
| P08249 | MDH2 | malate dehydrogenase 2, NAD (mitochondrial) | Cytoplasm | enzyme | 2.61E+07 | 5.97E+07 | 2.53E+07 | | | | | | |
| P08249 | MDH2 | malate dehydrogenase 2, NAD (mitochondrial) | Cytoplasm | enzyme | 2.86E+07 | 4.96E+07 | 2.01E+07 | | | | | | |
| P08249 | MDH2 | malate dehydrogenase 2, NAD (mitochondrial) | Cytoplasm | enzyme | 2.40E+07 | 4.58E+07 | 2.32E+07 | | | | | | |
| P08249 | MDH2 | malate dehydrogenase 2, NAD (mitochondrial) | Cytoplasm | enzyme | 1.99E+07 | 3.70E+07 | 1.81E+07 | | | | | | |
| P08249 | MDH2 | malate dehydrogenase 2, NAD (mitochondrial) | Cytoplasm | enzyme | 1.72E+07 | 3.61E+07 | 1.56E+07 | | | | | | |
| P08249 | MDH2 | malate dehydrogenase 2, NAD (mitochondrial) | Cytoplasm | enzyme | 1.35E+07 | 1.72E+07 | 9190119 | | | | | | |
| P08249 | MDH2 | malate dehydrogenase 2, NAD (mitochondrial) | Cytoplasm | enzyme | 1.34E+07 | 2.19E+07 | 8003116 | | | | | | |
| P08249 | MDH2 | malate dehydrogenase 2, NAD (mitochondrial) | Cytoplasm | enzyme | 7010744 | 1.42E+07 | 6609906 | | | | | | |
| P08249 | MDH2 | malate dehydrogenase 2, NAD (mitochondrial) | Cytoplasm | enzyme | 2281296.25 | 263581.156 | 2721250.5 | P08249 | MDH2 | 20431695 | 15970476 | 2557323 | 39 |
| Q99KE1 | ME2 | malic enzyme 2, NAD(+)-dependent, mitochondrial | Cytoplasm | enzyme | 2629813.75 | 7438163.5 | 4058597.75 | | | | | | |
| Q99KE1 | ME2 | malic enzyme 2, NAD(+)-dependent, mitochondrial | Cytoplasm | enzyme | 4638517 | 1.21E+07 | 5274701 | | | | | | |
| Q99KE1 | ME2 | malic enzyme 2, NAD(+)-dependent, mitochondrial | Cytoplasm | enzyme | 1675235.875 | 3961230.5 | 1465413.625 | | | | | | |
| Q99KE1 | ME2 | malic enzyme 2, NAD(+)-dependent, mitochondrial | Cytoplasm | enzyme | 1419734.5 | 3636604.5 | 1377841.625 | Q99KE1 | ME2 | 2590825 | 3106141 | 896666 | 12 |
| Q91VH6 | MEMO1 | mediator of cell motility 1 | Cytoplasm | other | 1.73E+07 | 1.85E+07 | 1.55E+07 | | | | | | |
| Q91VH6 | MEMO1 | mediator of cell motility 1 | Cytoplasm | other | 6975798 | 7196310 | 6740809 | | | | | | |
| Q91VH6 | MEMO1 | mediator of cell motility 1 | Cytoplasm | other | 4924301.5 | 1.13E+07 | 1.07E+07 | | | | | | |
| Q91VH6 | MEMO1 | mediator of cell motility 1 | Cytoplasm | other | 7480137 | 7057986 | 8050011.5 | | | | | | |
| Q91VH6 | MEMO1 | mediator of cell motility 1 | Cytoplasm | other | 3045135.5 | 6701486 | 7303804 | Q91VH6 | MEMO1 | 7953074 | 4540703 | 1172405 | 15 |
| Q9ERE7 | MESDC2 | mesoderm development candidate 2 | Extracellular Space | other | 1431128.625 | 1524460.375 | 1944945 | Q9ERE7 | MESDC2 | 1431129 | 273717 | 158030 | 3 |
| O08663 | METAP2 | methionyl aminopeptidase 2 | Cytoplasm | peptidase | 3171887.75 | 4473748 | 1812712.625 | O08663 | METAP2 | 3171888 | 1330621 | 768234 | 3 |
| Q6PCP5 | Mff | mitochondrial fission factor | Cytoplasm | other | 2103042 | 1.00E+07 | 5233696 | | | | | | |
| Q6PCP5 | Mff | mitochondrial fission factor | Cytoplasm | other | 1938480.25 | 6156951.5 | 2035885.875 | Q6PCP5 | Mff | 2020761 | 3227790 | 1317740 | 6 |
| Q3T9M1 | MFSD2B | major facilitator superfamily domain containing 2B | Plasma Membrane | other | 3.01E+07 | 3.01E+07 | 3.67E+07 | | | | | | |
| Q3T9M1 | MFSD2B | major facilitator superfamily domain containing 2B | Plasma Membrane | other | 1753860.875 | 1577634.375 | 2021304.25 | Q3T9M1 | MFSD2B | 15931930 | 16885261 | 6893379 | 6 |
| Q9CPU4 | MGST3 | microsomal glutathione S-transferase 3 | Cytoplasm | enzyme | 3.45E+07 | 9.14E+07 | 3.52E+07 | | | | | | |
| Q9CPU4 | MGST3 | microsomal glutathione S-transferase 3 | Cytoplasm | enzyme | 8705978 | 2.52E+07 | 1.28E+07 | | | | | | |
| Q9CPU4 | MGST3 | microsomal glutathione S-transferase 3 | Cytoplasm | enzyme | 6286054.5 | 1.59E+07 | 6875421.5 | Q9CPU4 | MGST3 | 16497344 | 26842161 | 8947387 | 9 |
| Q8VDP3 | MICAL1 | microtubule associated monooxygenase, calponin and LIM domain containing 1 | Cytoplasm | enzyme | 2636799.5 | 2316959.75 | 2218638.25 | Q8VDP3 | MICAL1 | 2636800 | 218641 | 126232 | 3 |
| Q6ZQI3 | MLEC | malectin | Plasma Membrane | other | 1062733.75 | 1506395.625 | 3915549 | | | | | | |
| Q6ZQI3 | MLEC | malectin | Plasma Membrane | other | 992387.938 | 2248956.5 | 4456807 | | | | | | |
| Q6ZQI3 | MLEC | malectin | Plasma Membrane | other | 1263288.75 | 1899133.5 | 2774087.25 | | | | | | |
| Q6ZQI3 | MLEC | malectin | Plasma Membrane | other | 846183.062 | 1417258.375 | 2232252 | | | | | | |
| Q6ZQI3 | MLEC | malectin | Plasma Membrane | other | 295360.469 | 811135.125 | 1668594.125 | Q6ZQI3 | MLEC | 891991 | 1157475 | 298859 | 15 |
| Q8K3F2 | MMP21 | matrix metallopeptidase 21 | Extracellular Space | peptidase | 3407842.75 | 4492547.5 | 4573906 | Q8K3F2 | MMP21 | 3407843 | 651013 | 375862 | 3 |
| B2RPV6 | MMRN1 | multimerin 1 | Extracellular Space | other | 43723.414 | 979491.375 | 1833433.125 | | | | | | |
| B2RPV6 | MMRN1 | multimerin 1 | Extracellular Space | other | 201413.297 | 261325.406 | 1537896.625 | | | | | | |
| B2RPV6 | MMRN1 | multimerin 1 | Extracellular Space | other | 131279.078 | 1319121.25 | 3142704.75 | | | | | | |
| B2RPV6 | MMRN1 | multimerin 1 | Extracellular Space | other | 184196.547 | 796080.312 | 2405577 | | | | | | |
| B2RPV6 | MMRN1 | multimerin 1 | Extracellular Space | other | 223137.234 | 615270.812 | 1650358.5 | | | | | | |
| B2RPV6 | MMRN1 | multimerin 1 | Extracellular Space | other | 127482.984 | 787106 | 2464228.75 | | | | | | |
| B2RPV6 | MMRN1 | multimerin 1 | Extracellular Space | other | 350187.969 | 645880.375 | 1850325.375 | | | | | | |
| B2RPV6 | MMRN1 | multimerin 1 | Extracellular Space | other | 21804.818 | 690249.688 | 1982616.25 | | | | | | |
| B2RPV6 | MMRN1 | multimerin 1 | Extracellular Space | other | 97000.703 | 382050.469 | 862975.125 | | | | | | |
| B2RPV6 | MMRN1 | multimerin 1 | Extracellular Space | other | 125093.266 | 110061.977 | 737118.875 | | | | | | |
| B2RPV6 | MMRN1 | multimerin 1 | Extracellular Space | other | 66612.992 | 256461.859 | 1528419.25 | B2RPV6 | MMRN1 | 142903 | 839167 | 146080 | 33 |
| P70290 | MPP1 | membrane protein, palmitoylated 1, 55kDa | Plasma Membrane | kinase | 1.50E+09 | 1.47E+09 | 1.92E+09 | | | | | | |
| P70290 | MPP1 | membrane protein, palmitoylated 1, 55kDa | Plasma Membrane | kinase | 1.62E+09 | 1.75E+09 | 1.98E+09 | | | | | | |
| P70290 | MPP1 | membrane protein, palmitoylated 1, 55kDa | Plasma Membrane | kinase | 1.17E+09 | 1.17E+09 | 1.25E+09 | | | | | | |
| P70290 | MPP1 | membrane protein, palmitoylated 1, 55kDa | Plasma Membrane | kinase | 1.25E+09 | 1.06E+09 | 1.16E+09 | | | | | | |
| P70290 | MPP1 | membrane protein, palmitoylated 1, 55kDa | Plasma Membrane | kinase | 1.27E+09 | 1.32E+09 | 9.64E+08 | | | | | | |
| P70290 | MPP1 | membrane protein, palmitoylated 1, 55kDa | Plasma Membrane | kinase | 1.16E+09 | 1.19E+09 | 1.32E+09 | | | | | | |
| P70290 | MPP1 | membrane protein, palmitoylated 1, 55kDa | Plasma Membrane | kinase | 8.27E+08 | 9.13E+08 | 7.10E+08 | | | | | | |
| P70290 | MPP1 | membrane protein, palmitoylated 1, 55kDa | Plasma Membrane | kinase | 6.66E+08 | 7.00E+08 | 8.14E+08 | | | | | | |
| P70290 | MPP1 | membrane protein, palmitoylated 1, 55kDa | Plasma Membrane | kinase | 6.98E+08 | 6.28E+08 | 6.18E+08 | | | | | | |
| P70290 | MPP1 | membrane protein, palmitoylated 1, 55kDa | Plasma Membrane | kinase | 6.24E+08 | 7.57E+08 | 5.43E+08 | | | | | | |
| P70290 | MPP1 | membrane protein, palmitoylated 1, 55kDa | Plasma Membrane | kinase | 4.19E+08 | 6.93E+08 | 6.76E+08 | | | | | | |
| P70290 | MPP1 | membrane protein, palmitoylated 1, 55kDa | Plasma Membrane | kinase | 6.93E+08 | 5.60E+08 | 4.94E+08 | | | | | | |
| P70290 | MPP1 | membrane protein, palmitoylated 1, 55kDa | Plasma Membrane | kinase | 5.24E+08 | 6.27E+08 | 4.60E+08 | | | | | | |
| P70290 | MPP1 | membrane protein, palmitoylated 1, 55kDa | Plasma Membrane | kinase | 6.00E+08 | 6.25E+08 | 6.42E+08 | | | | | | |
| P70290 | MPP1 | membrane protein, palmitoylated 1, 55kDa | Plasma Membrane | kinase | 4.71E+08 | 4.20E+08 | 4.54E+08 | | | | | | |
| P70290 | MPP1 | membrane protein, palmitoylated 1, 55kDa | Plasma Membrane | kinase | 4.07E+08 | 4.65E+08 | 4.71E+08 | | | | | | |
| P70290 | MPP1 | membrane protein, palmitoylated 1, 55kDa | Plasma Membrane | kinase | 3.89E+08 | 3.94E+08 | 3.29E+08 | | | | | | |
| P70290 | MPP1 | membrane protein, palmitoylated 1, 55kDa | Plasma Membrane | kinase | 4.73E+08 | 2.53E+08 | 5.56E+07 | | | | | | |
| P70290 | MPP1 | membrane protein, palmitoylated 1, 55kDa | Plasma Membrane | kinase | 3.22E+08 | 3.18E+08 | 3.39E+08 | | | | | | |
| P70290 | MPP1 | membrane protein, palmitoylated 1, 55kDa | Plasma Membrane | kinase | 3.53E+08 | 3.08E+08 | 3.35E+08 | | | | | | |
| P70290 | MPP1 | membrane protein, palmitoylated 1, 55kDa | Plasma Membrane | kinase | 2.11E+08 | 2.01E+08 | 1.87E+08 | | | | | | |
| P70290 | MPP1 | membrane protein, palmitoylated 1, 55kDa | Plasma Membrane | kinase | 2.24E+08 | 2.11E+08 | 1.71E+08 | | | | | | |
| P70290 | MPP1 | membrane protein, palmitoylated 1, 55kDa | Plasma Membrane | kinase | 1.38E+08 | 1.25E+08 | 1.43E+08 | | | | | | |
| P70290 | MPP1 | membrane protein, palmitoylated 1, 55kDa | Plasma Membrane | kinase | 1.33E+08 | 6.40E+07 | 1.36E+07 | | | | | | |

| | | | | | | | | | | | | | |
|---|---|---|---|---|---|---|---|---|---|---|---|---|---|
| P70290 | MPP1 | membrane protein, palmitoylated 1, 55kDa | Plasma Membrane | kinase | 1.40E+08 | 4.07E+07 | 2.69E+07 | | | | | | |
| P70290 | MPP1 | membrane protein, palmitoylated 1, 55kDa | Plasma Membrane | kinase | 1.18E+08 | 1.06E+08 | 1.07E+08 | | | | | | |
| P70290 | MPP1 | membrane protein, palmitoylated 1, 55kDa | Plasma Membrane | kinase | 6.71E+07 | 7.22E+07 | 8.44E+07 | | | | | | |
| P70290 | MPP1 | membrane protein, palmitoylated 1, 55kDa | Plasma Membrane | kinase | 9.18E+07 | 3.75E+07 | 8775276 | | | | | | |
| P70290 | MPP1 | membrane protein, palmitoylated 1, 55kDa | Plasma Membrane | kinase | 5.60E+07 | 4.92E+07 | 4.47E+07 | | | | | | |
| P70290 | MPP1 | membrane protein, palmitoylated 1, 55kDa | Plasma Membrane | kinase | 5.32E+07 | 5.36E+07 | 5.13E+07 | | | | | | |
| P70290 | MPP1 | membrane protein, palmitoylated 1, 55kDa | Plasma Membrane | kinase | 8.43E+07 | 8.34E+07 | 8.58E+07 | | | | | | |
| P70290 | MPP1 | membrane protein, palmitoylated 1, 55kDa | Plasma Membrane | kinase | 5.11E+07 | 4.10E+07 | 5.19E+07 | | | | | | |
| P70290 | MPP1 | membrane protein, palmitoylated 1, 55kDa | Plasma Membrane | kinase | 3.48E+07 | 3.63E+07 | 4.28E+07 | | | | | | |
| P70290 | MPP1 | membrane protein, palmitoylated 1, 55kDa | Plasma Membrane | kinase | 8.66E+07 | 1.58E+08 | 1.64E+08 | | | | | | |
| P70290 | MPP1 | membrane protein, palmitoylated 1, 55kDa | Plasma Membrane | kinase | 3.03E+07 | 8328835.5 | 5676773.5 | | | | | | |
| P70290 | MPP1 | membrane protein, palmitoylated 1, 55kDa | Plasma Membrane | kinase | 2.26E+07 | 6938007 | 1040182.438 | | | | | | |
| P70290 | MPP1 | membrane protein, palmitoylated 1, 55kDa | Plasma Membrane | kinase | 1.92E+07 | 1.89E+07 | 1.58E+07 | | | | | | |
| P70290 | MPP1 | membrane protein, palmitoylated 1, 55kDa | Plasma Membrane | kinase | 1.19E+07 | 1396193 | 2255567.25 | | | | | | |
| P70290 | MPP1 | membrane protein, palmitoylated 1, 55kDa | Plasma Membrane | kinase | 1.11E+07 | 1621718.125 | 2210045 | | | | | | |
| P70290 | MPP1 | membrane protein, palmitoylated 1, 55kDa | Plasma Membrane | kinase | 4212356.5 | 3769601.75 | 4003476.75 | | | | | | |
| P70290 | MPP1 | membrane protein, palmitoylated 1, 55kDa | Plasma Membrane | kinase | 4411801.5 | 3857125 | 6272274.5 | | | | | | |
| P70290 | MPP1 | membrane protein, palmitoylated 1, 55kDa | Plasma Membrane | kinase | 499947.938 | 627573.062 | 354579.281 | P70290 | MPP1 | 405535812 | 471705382 | 42022855 | 126 |
| Q99J99 | MPST | mercaptopyruvate sulfurtransferase | Cytoplasm | enzyme | 1.39E+07 | 3.37E+07 | 1.20E+07 | | | | | | |
| Q99J99 | MPST | mercaptopyruvate sulfurtransferase | Cytoplasm | enzyme | 1430106.125 | 2717987.75 | 1031776.562 | Q99J99 | MPST | 7640053 | 12501948 | 5103899 | 6 |
| Q9DB15 | MRPL12 | mitochondrial ribosomal protein L12 | Cytoplasm | other | 1215387.75 | 3610632.75 | 1677743.5 | Q9DB15 | MRPL12 | 1215388 | 1270632 | 733600 | 3 |
| Q9CQX8 | MRPS36 | mitochondrial ribosomal protein S36 | Cytoplasm | other | 1650936.625 | 3214689 | 1355676 | Q9CQX8 | MRPS36 | 1650937 | 999035 | 576793 | 3 |
| Q9WUX5 | MRVI1 | murine retrovirus integration site 1 homolog | Cytoplasm | other | 83813.125 | 521128.25 | 1046524.312 | | | | | | |
| Q9WUX5 | MRVI1 | murine retrovirus integration site 1 homolog | Cytoplasm | other | 80905.359 | 32410.262 | 481635.688 | | | | | | |
| Q9WUX5 | MRVI1 | murine retrovirus integration site 1 homolog | Cytoplasm | other | 4.70E-06 | 105427.461 | 356458.594 | Q9WUX5 | MRVI1 | 54906 | 342157 | 114052 | 9 |
| P26041 | MSN | moesin | Plasma Membrane | other | 8858034 | 7885066.5 | 8696755 | | | | | | |
| P26041 | MSN | moesin | Plasma Membrane | other | 6884122 | 5825023 | 9645210 | | | | | | |
| P26041 | MSN | moesin | Plasma Membrane | other | 7470972 | 6568575.5 | 6003609 | | | | | | |
| P26041 | MSN | moesin | Plasma Membrane | other | 7339678.5 | 6000914.5 | 6731293.5 | | | | | | |
| P26041 | MSN | moesin | Plasma Membrane | other | 4774140 | 4153199.75 | 4314668 | | | | | | |
| P26041 | MSN | moesin | Plasma Membrane | other | 2741122 | 1316997.625 | 2411640.75 | | | | | | |
| P26041 | MSN | moesin | Plasma Membrane | other | 3904612.5 | 3641986.75 | 5017558.5 | | | | | | |
| P26041 | MSN | moesin | Plasma Membrane | other | 3123227.25 | 2344665.5 | 3098210.25 | | | | | | |
| P26041 | MSN | moesin | Plasma Membrane | other | 2345128.75 | 2343066 | 3223468.25 | | | | | | |
| P26041 | MSN | moesin | Plasma Membrane | other | 1159159.375 | 2471090.5 | 1396629 | | | | | | |
| P26041 | MSN | moesin | Plasma Membrane | other | 1098943.625 | 1030239.688 | 1041772.75 | | | | | | |
| P26041 | MSN | moesin | Plasma Membrane | other | 2594569.25 | 3530174 | 2826765.5 | P26041 | MSN | 4357809 | 2491286 | 415214 | 36 |
| P03930 | mt-Atp8 | ATP synthase F0 subunit 8 | Cytoplasm | enzyme | 1.21E+07 | 1.98E+07 | 7066523 | | | | | | |
| P03930 | mt-Atp8 | ATP synthase F0 subunit 8 | Cytoplasm | enzyme | 3932242.5 | 8102436.5 | 4146890.25 | P03930 | mt-Atp8 | 8016121 | 6007435 | 2452525 | 6 |
| Q791V5 | MTCH2 | mitochondrial carrier 2 | Cytoplasm | other | 4065314.25 | 9016240 | 5447877.5 | Q791V5 | MTCH2 | 4065314 | 2554615 | 1474908 | 3 |
| P00397 | MT-CO1 | cytochrome c oxidase subunit I | Cytoplasm | enzyme | 1867583 | 3672322.75 | 655457.062 | P00397 | MT-CO1 | 1867583 | 1518103 | 876477 | 3 |
| P00405 | MT-CO2 | cytochrome c oxidase subunit II | Cytoplasm | enzyme | 1.91E+07 | 4.25E+07 | 2.16E+07 | | | | | | |
| P00405 | MT-CO2 | cytochrome c oxidase subunit II | Cytoplasm | enzyme | 1.21E+07 | 3.16E+07 | 1.25E+07 | | | | | | |
| P00405 | MT-CO2 | cytochrome c oxidase subunit II | Cytoplasm | enzyme | 2695296 | 9002071 | 2445637.25 | | | | | | |
| P00405 | MT-CO2 | cytochrome c oxidase subunit II | Cytoplasm | enzyme | 1437450.875 | 4114683.25 | 1789492.375 | | | | | | |
| P00405 | MT-CO2 | cytochrome c oxidase subunit II | Cytoplasm | enzyme | 2380311.25 | 5364171 | 2326399.25 | | | | | | |
| P00405 | MT-CO2 | cytochrome c oxidase subunit II | Cytoplasm | enzyme | 3097797.5 | 1259808 | 178723.016 | P00405 | MT-CO2 | 6803476 | 11871539 | 2798149 | 18 |
| P00416 | MT-CO3 | cytochrome c oxidase III | Cytoplasm | enzyme | 1554124.625 | 3174491.25 | 1894303.25 | P00416 | MT-CO3 | 1554125 | 854419 | 493299 | 3 |
| Q922D8 | MTHFD1 | methylenetetrahydrofolate dehydrogenase (NADP+ dependent) 1, methenyltetrahydrofolate cyclohydrolase, formyltetrahydrofolate synthetase | Cytoplasm | enzyme | 3.44E+07 | 3.15E+07 | 3.22E+07 | | | | | | |
| Q922D8 | MTHFD1 | methylenetetrahydrofolate dehydrogenase (NADP+ dependent) 1, methenyltetrahydrofolate cyclohydrolase, formyltetrahydrofolate synthetase | Cytoplasm | enzyme | 3.03E+07 | 2.46E+07 | 2.48E+07 | | | | | | |
| Q922D8 | MTHFD1 | methylenetetrahydrofolate dehydrogenase (NADP+ dependent) 1, methenyltetrahydrofolate cyclohydrolase, formyltetrahydrofolate synthetase | Cytoplasm | enzyme | 4.65E+07 | 3.46E+07 | 4.37E+07 | | | | | | |
| Q922D8 | MTHFD1 | methylenetetrahydrofolate dehydrogenase (NADP+ dependent) 1, methenyltetrahydrofolate cyclohydrolase, formyltetrahydrofolate synthetase | Cytoplasm | enzyme | 3.15E+07 | 2.89E+07 | 2.60E+07 | | | | | | |
| Q922D8 | MTHFD1 | methylenetetrahydrofolate dehydrogenase (NADP+ dependent) 1, methenyltetrahydrofolate cyclohydrolase, formyltetrahydrofolate synthetase | Cytoplasm | enzyme | 2.86E+07 | 2.35E+07 | 2.40E+07 | | | | | | |
| Q922D8 | MTHFD1 | methylenetetrahydrofolate dehydrogenase (NADP+ dependent) 1, methenyltetrahydrofolate cyclohydrolase, formyltetrahydrofolate synthetase | Cytoplasm | enzyme | 3.30E+07 | 2.30E+07 | 2.74E+07 | | | | | | |
| Q922D8 | MTHFD1 | methylenetetrahydrofolate dehydrogenase (NADP+ dependent) 1, methenyltetrahydrofolate cyclohydrolase, formyltetrahydrofolate synthetase | Cytoplasm | enzyme | 2.52E+07 | 3.05E+07 | 1.88E+07 | | | | | | |
| Q922D8 | MTHFD1 | methylenetetrahydrofolate dehydrogenase (NADP+ dependent) 1, methenyltetrahydrofolate cyclohydrolase, formyltetrahydrofolate synthetase | Cytoplasm | enzyme | 2.96E+07 | 2.23E+07 | 2.21E+07 | | | | | | |
| Q922D8 | MTHFD1 | methylenetetrahydrofolate dehydrogenase (NADP+ dependent) 1, methenyltetrahydrofolate cyclohydrolase, formyltetrahydrofolate synthetase | Cytoplasm | enzyme | 2.30E+07 | 9452009 | 3370795.75 | | | | | | |
| Q922D8 | MTHFD1 | methylenetetrahydrofolate dehydrogenase (NADP+ dependent) 1, methenyltetrahydrofolate cyclohydrolase, formyltetrahydrofolate synthetase | Cytoplasm | enzyme | 2.45E+07 | 1.94E+07 | 1.92E+07 | | | | | | |
| Q922D8 | MTHFD1 | methylenetetrahydrofolate dehydrogenase (NADP+ dependent) 1, methenyltetrahydrofolate cyclohydrolase, formyltetrahydrofolate synthetase | Cytoplasm | enzyme | 2.24E+07 | 1.83E+07 | 1.36E+07 | | | | | | |
| Q922D8 | MTHFD1 | methylenetetrahydrofolate dehydrogenase (NADP+ dependent) 1, methenyltetrahydrofolate cyclohydrolase, formyltetrahydrofolate synthetase | Cytoplasm | enzyme | 1.36E+07 | 1.84E+07 | 1.24E+07 | | | | | | |
| Q922D8 | MTHFD1 | methylenetetrahydrofolate dehydrogenase (NADP+ dependent) 1, methenyltetrahydrofolate cyclohydrolase, formyltetrahydrofolate synthetase | Cytoplasm | enzyme | 2.06E+07 | 1.50E+07 | 1.83E+07 | | | | | | |
| Q922D8 | MTHFD1 | methylenetetrahydrofolate dehydrogenase (NADP+ dependent) 1, methenyltetrahydrofolate cyclohydrolase, formyltetrahydrofolate synthetase | Cytoplasm | enzyme | 2.40E+07 | 1.75E+07 | 1.15E+07 | | | | | | |
| Q922D8 | MTHFD1 | methylenetetrahydrofolate dehydrogenase (NADP+ dependent) 1, methenyltetrahydrofolate cyclohydrolase, formyltetrahydrofolate synthetase | Cytoplasm | enzyme | 1.88E+07 | 1.16E+07 | 1.07E+07 | | | | | | |
| Q922D8 | MTHFD1 | methylenetetrahydrofolate dehydrogenase (NADP+ dependent) 1, methenyltetrahydrofolate cyclohydrolase, formyltetrahydrofolate synthetase | Cytoplasm | enzyme | 1.71E+07 | 1.51E+07 | 1.74E+07 | | | | | | |
| Q922D8 | MTHFD1 | methylenetetrahydrofolate dehydrogenase (NADP+ dependent) 1, methenyltetrahydrofolate cyclohydrolase, formyltetrahydrofolate synthetase | Cytoplasm | enzyme | 1.27E+07 | 1.25E+07 | 1.40E+07 | | | | | | |
| Q922D8 | MTHFD1 | methylenetetrahydrofolate dehydrogenase (NADP+ dependent) 1, methenyltetrahydrofolate cyclohydrolase, formyltetrahydrofolate synthetase | Cytoplasm | enzyme | 1.28E+07 | 9444965 | 1.03E+07 | | | | | | |
| Q922D8 | MTHFD1 | methylenetetrahydrofolate dehydrogenase (NADP+ dependent) 1, methenyltetrahydrofolate cyclohydrolase, formyltetrahydrofolate synthetase | Cytoplasm | enzyme | 1.34E+07 | 1.50E+07 | 1.68E+07 | | | | | | |
| Q922D8 | MTHFD1 | methylenetetrahydrofolate dehydrogenase (NADP+ dependent) 1, methenyltetrahydrofolate cyclohydrolase, formyltetrahydrofolate synthetase | Cytoplasm | enzyme | 1.40E+07 | 1.29E+07 | 1.25E+07 | | | | | | |
| Q922D8 | MTHFD1 | methylenetetrahydrofolate dehydrogenase (NADP+ dependent) 1, methenyltetrahydrofolate cyclohydrolase, formyltetrahydrofolate synthetase | Cytoplasm | enzyme | 1.04E+07 | 1.05E+07 | 1.16E+07 | | | | | | |
| Q922D8 | MTHFD1 | methylenetetrahydrofolate dehydrogenase (NADP+ dependent) 1, methenyltetrahydrofolate cyclohydrolase, formyltetrahydrofolate synthetase | Cytoplasm | enzyme | 1.11E+07 | 9932327 | 9466208 | | | | | | |
| Q922D8 | MTHFD1 | methylenetetrahydrofolate dehydrogenase (NADP+ dependent) 1, methenyltetrahydrofolate cyclohydrolase, formyltetrahydrofolate synthetase | Cytoplasm | enzyme | 1.41E+07 | 5436958.5 | 1856783.375 | | | | | | |
| Q922D8 | MTHFD1 | methylenetetrahydrofolate dehydrogenase (NADP+ dependent) 1, methenyltetrahydrofolate cyclohydrolase, formyltetrahydrofolate synthetase | Cytoplasm | enzyme | 3560354 | 8218213.5 | 1.31E+07 | | | | | | |
| Q922D8 | MTHFD1 | methylenetetrahydrofolate dehydrogenase (NADP+ dependent) 1, methenyltetrahydrofolate cyclohydrolase, formyltetrahydrofolate synthetase | Cytoplasm | enzyme | 1.37E+07 | 9009438 | 7860273 | | | | | | |
| Q922D8 | MTHFD1 | methylenetetrahydrofolate dehydrogenase (NADP+ dependent) 1, methenyltetrahydrofolate cyclohydrolase, formyltetrahydrofolate synthetase | Cytoplasm | enzyme | 8417705 | 6670142.5 | 6897462 | | | | | | |
| Q922D8 | MTHFD1 | methylenetetrahydrofolate dehydrogenase (NADP+ dependent) 1, methenyltetrahydrofolate cyclohydrolase, formyltetrahydrofolate synthetase | Cytoplasm | enzyme | 6974810 | 5250310.5 | 5778722.5 | | | | | | |
| Q922D8 | MTHFD1 | methylenetetrahydrofolate dehydrogenase (NADP+ dependent) 1, methenyltetrahydrofolate cyclohydrolase, formyltetrahydrofolate synthetase | Cytoplasm | enzyme | 7505507.5 | 5740192.5 | 2915639.5 | | | | | | |
| Q922D8 | MTHFD1 | methylenetetrahydrofolate dehydrogenase (NADP+ dependent) 1, methenyltetrahydrofolate cyclohydrolase, formyltetrahydrofolate synthetase | Cytoplasm | enzyme | 3538564 | 3367530 | 1620712.375 | Q922D8 | MTHFD1 | 19142653 | 9641241 | 1033650 | 87 |
| Q3V3R1 | MTHFD1L | methylenetetrahydrofolate dehydrogenase (NADP+ dependent) 1-like | Cytoplasm | enzyme | 4056988.5 | 8116432.5 | 3998592.5 | | | | | | |
| Q3V3R1 | MTHFD1L | methylenetetrahydrofolate dehydrogenase (NADP+ dependent) 1-like | Cytoplasm | enzyme | 3369858.25 | 9272173 | 4006540.25 | | | | | | |
| Q3V3R1 | MTHFD1L | methylenetetrahydrofolate dehydrogenase (NADP+ dependent) 1-like | Cytoplasm | enzyme | 3108834.25 | 6054061 | 2448254 | | | | | | |
| Q3V3R1 | MTHFD1L | methylenetetrahydrofolate dehydrogenase (NADP+ dependent) 1-like | Cytoplasm | enzyme | 2429290 | 6360714.5 | 2128596.5 | | | | | | |
| Q3V3R1 | MTHFD1L | methylenetetrahydrofolate dehydrogenase (NADP+ dependent) 1-like | Cytoplasm | enzyme | 1986507.875 | 4862663.5 | 2608287.75 | | | | | | |
| Q3V3R1 | MTHFD1L | methylenetetrahydrofolate dehydrogenase (NADP+ dependent) 1-like | Cytoplasm | enzyme | 2832015.25 | 4849583.5 | 1410707.875 | | | | | | |
| Q3V3R1 | MTHFD1L | methylenetetrahydrofolate dehydrogenase (NADP+ dependent) 1-like | Cytoplasm | enzyme | 2319066 | 3864935.75 | 1561660.5 | | | | | | |
| Q3V3R1 | MTHFD1L | methylenetetrahydrofolate dehydrogenase (NADP+ dependent) 1-like | Cytoplasm | enzyme | 1718553.75 | 3363177 | 1811498.25 | Q3V3R1 | MTHFD1L | 2727639 | 2048332 | 418114 | 24 |
| P03921 | MT-ND5 | NADH dehydrogenase, subunit 5 (complex I) | Cytoplasm | enzyme | 3433967.75 | 6246298.5 | 3462774 | | | | | | |
| P03921 | MT-ND5 | NADH dehydrogenase, subunit 5 (complex I) | Cytoplasm | enzyme | 1627578.375 | 3006663 | 1343737.75 | | | | | | |
| P03921 | MT-ND5 | NADH dehydrogenase, subunit 5 (complex I) | Cytoplasm | enzyme | 742276.25 | 2477631.75 | 656653.688 | P03921 | MT-ND5 | 1934607 | 1755323 | 585108 | 9 |
| Q8C1A3 | MTRR | 5-methyltetrahydrofolate-homocysteine methyltransferase reductase | Cytoplasm | enzyme | 4754791.5 | 3353647.75 | 6176639.5 | Q8C1A3 | MTRR | 4754792 | 1411509 | 814935 | 3 |
| P28665 | Mug1/Mug2 | murinoglobulin 1 | Extracellular Space | other | 1097724.5 | 1188640.875 | 2126819.25 | P28665 | Mug1/Mug2 | 1097725 | 569719 | 328928 | 3 |
| P48972 | MYBL2 | v-myb avian myeloblastosis viral oncogene homolog-like 2 | Nucleus | transcription regulator | 9470300 | 9006298 | 1.13E+07 | P48972 | MYBL2 | 9470300 | 1190063 | 687083 | 3 |

| | | | | | | | | | | | | | |
|---|---|---|---|---|---|---|---|---|---|---|---|---|---|
| Q3U898 | MYEOV2 | myeloma overexpressed 2 | Nucleus | other | 7920537 | 5343289.5 | 7148320 | Q3U898 | MYEOV2 | 7920537 | 1322665 | 763641 | 3 |
| Q6URW6 | MYH14 | myosin, heavy chain 14, non-muscle | Extracellular Space | other | 415795.812 | 1096134.625 | 2158905 | | | | | | |
| Q6URW6 | MYH14 | myosin, heavy chain 14, non-muscle | Extracellular Space | other | 87901.477 | 351953.312 | 1159261.375 | Q6URW6 | MYH14 | 251849 | 759213 | 309947 | 6 |
| Q8VDD5 | MYH9 | myosin, heavy chain 9, non-muscle | Cytoplasm | enzyme | 9060003 | 1.00E+07 | 1.44E+07 | | | | | | |
| Q8VDD5 | MYH9 | myosin, heavy chain 9, non-muscle | Cytoplasm | enzyme | 8175496.5 | 1.74E+07 | 3.34E+07 | | | | | | |
| Q8VDD5 | MYH9 | myosin, heavy chain 9, non-muscle | Cytoplasm | enzyme | 6532967.5 | 1.25E+07 | 2.37E+07 | | | | | | |
| Q8VDD5 | MYH9 | myosin, heavy chain 9, non-muscle | Cytoplasm | enzyme | 3533084.75 | 9860821 | 1.98E+07 | | | | | | |
| Q8VDD5 | MYH9 | myosin, heavy chain 9, non-muscle | Cytoplasm | enzyme | 6488387 | 1.51E+07 | 2.85E+07 | | | | | | |
| Q8VDD5 | MYH9 | myosin, heavy chain 9, non-muscle | Cytoplasm | enzyme | 6625561.5 | 1.66E+07 | 2.32E+07 | | | | | | |
| Q8VDD5 | MYH9 | myosin, heavy chain 9, non-muscle | Cytoplasm | enzyme | 4948328 | 9965066 | 1.73E+07 | | | | | | |
| Q8VDD5 | MYH9 | myosin, heavy chain 9, non-muscle | Cytoplasm | enzyme | 7996594.5 | 1.15E+07 | 1.96E+07 | | | | | | |
| Q8VDD5 | MYH9 | myosin, heavy chain 9, non-muscle | Cytoplasm | enzyme | 6687194.5 | 1.42E+07 | 2.84E+07 | | | | | | |
| Q8VDD5 | MYH9 | myosin, heavy chain 9, non-muscle | Cytoplasm | enzyme | 3742245.5 | 1.26E+07 | 1.70E+07 | | | | | | |
| Q8VDD5 | MYH9 | myosin, heavy chain 9, non-muscle | Cytoplasm | enzyme | 5883971.5 | 1.03E+07 | 1.38E+07 | | | | | | |
| Q8VDD5 | MYH9 | myosin, heavy chain 9, non-muscle | Cytoplasm | enzyme | 6357432 | 1.11E+07 | 1.87E+07 | | | | | | |
| Q8VDD5 | MYH9 | myosin, heavy chain 9, non-muscle | Cytoplasm | enzyme | 5638799.5 | 1.16E+07 | 1.67E+07 | | | | | | |
| Q8VDD5 | MYH9 | myosin, heavy chain 9, non-muscle | Cytoplasm | enzyme | 5388717 | 1.16E+07 | 2.28E+07 | | | | | | |
| Q8VDD5 | MYH9 | myosin, heavy chain 9, non-muscle | Cytoplasm | enzyme | 5413665 | 9843043 | 1.93E+07 | | | | | | |
| Q8VDD5 | MYH9 | myosin, heavy chain 9, non-muscle | Cytoplasm | enzyme | 3671984.5 | 9834136 | 1.56E+07 | | | | | | |
| Q8VDD5 | MYH9 | myosin, heavy chain 9, non-muscle | Cytoplasm | enzyme | 4632865.5 | 9288866 | 1.81E+07 | | | | | | |
| Q8VDD5 | MYH9 | myosin, heavy chain 9, non-muscle | Cytoplasm | enzyme | 4497114 | 9357226 | 1.72E+07 | | | | | | |
| Q8VDD5 | MYH9 | myosin, heavy chain 9, non-muscle | Cytoplasm | enzyme | 3439933.5 | 7438369 | 1.08E+07 | | | | | | |
| Q8VDD5 | MYH9 | myosin, heavy chain 9, non-muscle | Cytoplasm | enzyme | 3687290.5 | 8716728 | 1.79E+07 | | | | | | |
| Q8VDD5 | MYH9 | myosin, heavy chain 9, non-muscle | Cytoplasm | enzyme | 5051329.5 | 1.09E+07 | 1.69E+07 | | | | | | |
| Q8VDD5 | MYH9 | myosin, heavy chain 9, non-muscle | Cytoplasm | enzyme | 3932161.75 | 7332235 | 1.20E+07 | | | | | | |
| Q8VDD5 | MYH9 | myosin, heavy chain 9, non-muscle | Cytoplasm | enzyme | 3146565.25 | 7791750 | 2.05E+07 | | | | | | |
| Q8VDD5 | MYH9 | myosin, heavy chain 9, non-muscle | Cytoplasm | enzyme | 2174528.5 | 1.08E+07 | 2.13E+07 | | | | | | |
| Q8VDD5 | MYH9 | myosin, heavy chain 9, non-muscle | Cytoplasm | enzyme | 4004928.75 | 7489484.5 | 1.20E+07 | | | | | | |
| Q8VDD5 | MYH9 | myosin, heavy chain 9, non-muscle | Cytoplasm | enzyme | 3857263 | 8127195 | 1.15E+07 | | | | | | |
| Q8VDD5 | MYH9 | myosin, heavy chain 9, non-muscle | Cytoplasm | enzyme | 3873912 | 8505640 | 1.62E+07 | | | | | | |
| Q8VDD5 | MYH9 | myosin, heavy chain 9, non-muscle | Cytoplasm | enzyme | 4246695 | 1.03E+07 | 1.44E+07 | | | | | | |
| Q8VDD5 | MYH9 | myosin, heavy chain 9, non-muscle | Cytoplasm | enzyme | 3395435.75 | 1.01E+07 | 1.74E+07 | | | | | | |
| Q8VDD5 | MYH9 | myosin, heavy chain 9, non-muscle | Cytoplasm | enzyme | 5475794 | 9296925 | 1.30E+07 | | | | | | |
| Q8VDD5 | MYH9 | myosin, heavy chain 9, non-muscle | Cytoplasm | enzyme | 4157104 | 7789754.5 | 1.32E+07 | | | | | | |
| Q8VDD5 | MYH9 | myosin, heavy chain 9, non-muscle | Cytoplasm | enzyme | 3546943.5 | 1.31E+07 | 1.96E+07 | | | | | | |
| Q8VDD5 | MYH9 | myosin, heavy chain 9, non-muscle | Cytoplasm | enzyme | 3995798 | 7322398.5 | 1.23E+07 | | | | | | |
| Q8VDD5 | MYH9 | myosin, heavy chain 9, non-muscle | Cytoplasm | enzyme | 4356556 | 1.09E+07 | 1.90E+07 | | | | | | |
| Q8VDD5 | MYH9 | myosin, heavy chain 9, non-muscle | Cytoplasm | enzyme | 3066601 | 6560023 | 1.35E+07 | | | | | | |
| Q8VDD5 | MYH9 | myosin, heavy chain 9, non-muscle | Cytoplasm | enzyme | 3808271.5 | 6142692.5 | 1.11E+07 | | | | | | |
| Q8VDD5 | MYH9 | myosin, heavy chain 9, non-muscle | Cytoplasm | enzyme | 3084285.5 | 8870871 | 1.33E+07 | | | | | | |
| Q8VDD5 | MYH9 | myosin, heavy chain 9, non-muscle | Cytoplasm | enzyme | 3760188.5 | 6688882.5 | 1.08E+07 | | | | | | |
| Q8VDD5 | MYH9 | myosin, heavy chain 9, non-muscle | Cytoplasm | enzyme | 3835800.5 | 6611801 | 1.03E+07 | | | | | | |
| Q8VDD5 | MYH9 | myosin, heavy chain 9, non-muscle | Cytoplasm | enzyme | 2495925.25 | 5355037 | 8284109 | | | | | | |
| Q8VDD5 | MYH9 | myosin, heavy chain 9, non-muscle | Cytoplasm | enzyme | 2971078.75 | 5156038 | 1.09E+07 | | | | | | |
| Q8VDD5 | MYH9 | myosin, heavy chain 9, non-muscle | Cytoplasm | enzyme | 3254175.25 | 9050621 | 1.56E+07 | | | | | | |
| Q8VDD5 | MYH9 | myosin, heavy chain 9, non-muscle | Cytoplasm | enzyme | 3191132.25 | 5503489 | 7587582.5 | | | | | | |
| Q8VDD5 | MYH9 | myosin, heavy chain 9, non-muscle | Cytoplasm | enzyme | 2404037.25 | 6456013 | 1.25E+07 | | | | | | |
| Q8VDD5 | MYH9 | myosin, heavy chain 9, non-muscle | Cytoplasm | enzyme | 3175667.25 | 5776467.5 | 9560453 | | | | | | |
| Q8VDD5 | MYH9 | myosin, heavy chain 9, non-muscle | Cytoplasm | enzyme | 2175524.5 | 5702967 | 9728290 | | | | | | |
| Q8VDD5 | MYH9 | myosin, heavy chain 9, non-muscle | Cytoplasm | enzyme | 2272418.25 | 5313131.5 | 9715121 | | | | | | |
| Q8VDD5 | MYH9 | myosin, heavy chain 9, non-muscle | Cytoplasm | enzyme | 2316822.5 | 7169490 | 1.10E+07 | | | | | | |
| Q8VDD5 | MYH9 | myosin, heavy chain 9, non-muscle | Cytoplasm | enzyme | 2692631.25 | 6615076 | 1.03E+07 | | | | | | |
| Q8VDD5 | MYH9 | myosin, heavy chain 9, non-muscle | Cytoplasm | enzyme | 2185387.5 | 4313655 | 2942780.5 | | | | | | |
| Q8VDD5 | MYH9 | myosin, heavy chain 9, non-muscle | Cytoplasm | enzyme | 1997778.375 | 5227434.5 | 8609235 | | | | | | |
| Q8VDD5 | MYH9 | myosin, heavy chain 9, non-muscle | Cytoplasm | enzyme | 1713517.25 | 4478197 | 8161321 | | | | | | |
| Q8VDD5 | MYH9 | myosin, heavy chain 9, non-muscle | Cytoplasm | enzyme | 1750085.625 | 4379853.5 | 6561096.5 | | | | | | |
| Q8VDD5 | MYH9 | myosin, heavy chain 9, non-muscle | Cytoplasm | enzyme | 2008540.5 | 5768676 | 6876461.5 | | | | | | |
| Q8VDD5 | MYH9 | myosin, heavy chain 9, non-muscle | Cytoplasm | enzyme | 1780933.875 | 4068693 | 6899074.5 | | | | | | |
| Q8VDD5 | MYH9 | myosin, heavy chain 9, non-muscle | Cytoplasm | enzyme | 2167002.5 | 3799920.5 | 6586738 | | | | | | |
| Q8VDD5 | MYH9 | myosin, heavy chain 9, non-muscle | Cytoplasm | enzyme | 3312822.5 | 3641658.25 | 3864589 | | | | | | |
| Q8VDD5 | MYH9 | myosin, heavy chain 9, non-muscle | Cytoplasm | enzyme | 1473940.125 | 4065768 | 9601703 | | | | | | |
| Q8VDD5 | MYH9 | myosin, heavy chain 9, non-muscle | Cytoplasm | enzyme | 1990230 | 4621327 | 7380733.5 | | | | | | |
| Q8VDD5 | MYH9 | myosin, heavy chain 9, non-muscle | Cytoplasm | enzyme | 1795948.75 | 4333128 | 7090616.5 | | | | | | |
| Q8VDD5 | MYH9 | myosin, heavy chain 9, non-muscle | Cytoplasm | enzyme | 1760186.75 | 4833300 | 7224808 | | | | | | |
| Q8VDD5 | MYH9 | myosin, heavy chain 9, non-muscle | Cytoplasm | enzyme | 871392.812 | 4379375 | 1.16E+07 | | | | | | |
| Q8VDD5 | MYH9 | myosin, heavy chain 9, non-muscle | Cytoplasm | enzyme | 1320182.875 | 3777898.75 | 4167494 | | | | | | |
| Q8VDD5 | MYH9 | myosin, heavy chain 9, non-muscle | Cytoplasm | enzyme | 1659160.75 | 3758523.75 | 6126066.5 | | | | | | |
| Q8VDD5 | MYH9 | myosin, heavy chain 9, non-muscle | Cytoplasm | enzyme | 1186772.75 | 2117729.25 | 3977934.75 | | | | | | |
| Q8VDD5 | MYH9 | myosin, heavy chain 9, non-muscle | Cytoplasm | enzyme | 793302.062 | 1757903.25 | 4065931.25 | | | | | | |
| Q8VDD5 | MYH9 | myosin, heavy chain 9, non-muscle | Cytoplasm | enzyme | 1360879.125 | 2403847.75 | 3701640.75 | | | | | | |
| Q8VDD5 | MYH9 | myosin, heavy chain 9, non-muscle | Cytoplasm | enzyme | 677340.688 | 6686043 | 1.11E+07 | | | | | | |
| Q8VDD5 | MYH9 | myosin, heavy chain 9, non-muscle | Cytoplasm | enzyme | 985893.312 | 2661980.5 | 4156887.75 | | | | | | |
| Q8VDD5 | MYH9 | myosin, heavy chain 9, non-muscle | Cytoplasm | enzyme | 915182.188 | 2194116.5 | 3741125.75 | | | | | | |
| Q8VDD5 | MYH9 | myosin, heavy chain 9, non-muscle | Cytoplasm | enzyme | 2286230.75 | 2686128.5 | 2413081.75 | | | | | | |
| Q8VDD5 | MYH9 | myosin, heavy chain 9, non-muscle | Cytoplasm | enzyme | 815399.375 | 2873014.5 | 5579632.5 | | | | | | |
| Q8VDD5 | MYH9 | myosin, heavy chain 9, non-muscle | Cytoplasm | enzyme | 292249.75 | 740591.312 | 1439610.125 | | | | | | |
| Q8VDD5 | MYH9 | myosin, heavy chain 9, non-muscle | Cytoplasm | enzyme | 1100767 | 2014592.875 | 3632495 | | | | | | |
| Q8VDD5 | MYH9 | myosin, heavy chain 9, non-muscle | Cytoplasm | enzyme | 838005.562 | 2283650.25 | 3455788.25 | | | | | | |
| Q8VDD5 | MYH9 | myosin, heavy chain 9, non-muscle | Cytoplasm | enzyme | 281912.062 | 2009437.625 | 2920134 | | | | | | |
| Q8VDD5 | MYH9 | myosin, heavy chain 9, non-muscle | Cytoplasm | enzyme | 402032.406 | 95070.977 | 57194.77 | | | | | | |
| Q8VDD5 | MYH9 | myosin, heavy chain 9, non-muscle | Cytoplasm | enzyme | 243851.828 | 1213032.125 | 4413200 | | | | | | |
| Q8VDD5 | MYH9 | myosin, heavy chain 9, non-muscle | Cytoplasm | enzyme | 310169.844 | 1439041.625 | 4447438.5 | | | | | | |
| Q8VDD5 | MYH9 | myosin, heavy chain 9, non-muscle | Cytoplasm | enzyme | 507541.438 | 957602.688 | 1420483.25 | | | | | | |
| Q8VDD5 | MYH9 | myosin, heavy chain 9, non-muscle | Cytoplasm | enzyme | 1479350.5 | 2759919.75 | 4037320 | | | | | | |
| Q8VDD5 | MYH9 | myosin, heavy chain 9, non-muscle | Cytoplasm | enzyme | 316331.938 | 1930402.625 | 4754161 | | | | | | |
| Q8VDD5 | MYH9 | myosin, heavy chain 9, non-muscle | Cytoplasm | enzyme | 420925.625 | 1249609.875 | 649478.062 | | | | | | |
| Q8VDD5 | MYH9 | myosin, heavy chain 9, non-muscle | Cytoplasm | enzyme | 257154.547 | 1088480.25 | 2419668 | | | | | | |
| Q8VDD5 | MYH9 | myosin, heavy chain 9, non-muscle | Cytoplasm | enzyme | 262904.656 | 2124389 | 5309267 | | | | | | |
| Q8VDD5 | MYH9 | myosin, heavy chain 9, non-muscle | Cytoplasm | enzyme | 708161.125 | 766221.812 | 756738.5 | | | | | | |
| Q8VDD5 | MYH9 | myosin, heavy chain 9, non-muscle | Cytoplasm | enzyme | 1140256.75 | 1492112.375 | 1680346 | | | | | | |
| Q8VDD5 | MYH9 | myosin, heavy chain 9, non-muscle | Cytoplasm | enzyme | 222816.562 | 742173.062 | 2002832.5 | | | | | | |

| | | | | | | | | | | | | | |
|---|---|---|---|---|---|---|---|---|---|---|---|---|---|
| Q8VDD5 | MYH9 | myosin, heavy chain 9, non-muscle | Cytoplasm | enzyme | 6678.666 | 488141.844 | 673164.625 | | | | | | |
| Q8VDD5 | MYH9 | myosin, heavy chain 9, non-muscle | Cytoplasm | enzyme | 348288 | 601030.625 | 1350103 | | | | | | |
| Q8VDD5 | MYH9 | myosin, heavy chain 9, non-muscle | Cytoplasm | enzyme | 526835 | 1289464.75 | 1702035.75 | | | | | | |
| Q8VDD5 | MYH9 | myosin, heavy chain 9, non-muscle | Cytoplasm | enzyme | 133832.188 | 920522.375 | 2979337 | | | | | | |
| Q8VDD5 | MYH9 | myosin, heavy chain 9, non-muscle | Cytoplasm | enzyme | 110579.508 | 128009.953 | 300526.875 | | | | | | |
| Q8VDD5 | MYH9 | myosin, heavy chain 9, non-muscle | Cytoplasm | enzyme | 263508.219 | 424481.656 | 819325.062 | | | | | | |
| Q8VDD5 | MYH9 | myosin, heavy chain 9, non-muscle | Cytoplasm | enzyme | 25316.838 | 236787.969 | 1068839.25 | | | | | | |
| Q8VDD5 | MYH9 | myosin, heavy chain 9, non-muscle | Cytoplasm | enzyme | 0 | 2303.207 | 1147373.125 | Q8VDD5 | MYH9 | 2699258 | 5877505 | 346335 | 288 |
| P05977 | MYL1 | myosin, light chain 1, alkali; skeletal, fast | Cytoplasm | other | 414561.562 | 998749.5 | 2373897 | P05977 | MYL1 | 414562 | 1005924 | 580771 | 3 |
| Q3THE2 | MYL12A | myosin, light chain 12A, regulatory, non-sarcomeric | Cytoplasm | other | 4379364.5 | 8817643 | 1.05E+07 | | | | | | |
| Q3THE2 | MYL12A | myosin, light chain 12A, regulatory, non-sarcomeric | Cytoplasm | other | 3212932.75 | 7979843 | 1.25E+07 | | | | | | |
| Q3THE2 | MYL12A | myosin, light chain 12A, regulatory, non-sarcomeric | Cytoplasm | other | 763388.562 | 2254505.25 | 3382352.25 | | | | | | |
| Q3THE2 | MYL12A | myosin, light chain 12A, regulatory, non-sarcomeric | Cytoplasm | other | 638384 | 1885539.125 | 2859459.75 | | | | | | |
| Q3THE2 | MYL12A | myosin, light chain 12A, regulatory, non-sarcomeric | Cytoplasm | other | 1377444 | 1676333.875 | 1699603.125 | Q3THE2 | MYL12A | 2074303 | 3792209 | 979144 | 15 |
| Q60605 | MYL6 | myosin, light chain 6, alkali, smooth muscle and non-muscle | Cytoplasm | other | 9416315 | 1.96E+07 | 2.95E+07 | | | | | | |
| Q60605 | MYL6 | myosin, light chain 6, alkali, smooth muscle and non-muscle | Cytoplasm | other | 9322379 | 1.61E+07 | 3.06E+07 | | | | | | |
| Q60605 | MYL6 | myosin, light chain 6, alkali, smooth muscle and non-muscle | Cytoplasm | other | 8421099 | 1.46E+07 | 2.56E+07 | | | | | | |
| Q60605 | MYL6 | myosin, light chain 6, alkali, smooth muscle and non-muscle | Cytoplasm | other | 5726087 | 1.01E+07 | 1.67E+07 | | | | | | |
| Q60605 | MYL6 | myosin, light chain 6, alkali, smooth muscle and non-muscle | Cytoplasm | other | 4245029.5 | 9431104 | 1.70E+07 | | | | | | |
| Q60605 | MYL6 | myosin, light chain 6, alkali, smooth muscle and non-muscle | Cytoplasm | other | 4304685.5 | 7901977 | 1.17E+07 | | | | | | |
| Q60605 | MYL6 | myosin, light chain 6, alkali, smooth muscle and non-muscle | Cytoplasm | other | 1449507.125 | 2393151.75 | 4988324 | Q60605 | MYL6 | 6126443 | 8445877 | 1843042 | 21 |
| Q9CQ19 | MYL9 | myosin, light chain 9, regulatory | Cytoplasm | other | 3591633.25 | 5985634 | 8580176 | | | | | | |
| Q9CQ19 | MYL9 | myosin, light chain 9, regulatory | Cytoplasm | other | 2841952.75 | 5865479 | 1.39E+07 | | | | | | |
| Q9CQ19 | MYL9 | myosin, light chain 9, regulatory | Cytoplasm | other | 1313392.375 | 3925657.75 | 5381646 | | | | | | |
| Q9CQ19 | MYL9 | myosin, light chain 9, regulatory | Cytoplasm | other | 894927.312 | 3372937.75 | 4012094.5 | | | | | | |
| Q9CQ19 | MYL9 | myosin, light chain 9, regulatory | Cytoplasm | other | 879850.688 | 2218419 | 6441083 | | | | | | |
| Q9CQ19 | MYL9 | myosin, light chain 9, regulatory | Cytoplasm | other | 1161701 | 3487551.25 | 7100304.5 | | | | | | |
| Q9CQ19 | MYL9 | myosin, light chain 9, regulatory | Cytoplasm | other | 210510.422 | 965665.688 | 2095082.625 | | | | | | |
| Q9CQ19 | MYL9 | myosin, light chain 9, regulatory | Cytoplasm | other | 724698.125 | 2370435.75 | 6209746 | Q9CQ19 | MYL9 | 1452333 | 3151902 | 643379 | 24 |
| Q9WTI7 | MYO1C | myosin IC | Cytoplasm | other | 415848.188 | 272501.906 | 141929.547 | Q9WTI7 | MYO1C | 415848 | 137009 | 79102 | 3 |
| Q64331 | MYO6 | myosin VI | Cytoplasm | other | 2358987 | 3870776 | 1664061 | Q64331 | MYO6 | 2358987 | 1128274 | 651410 | 3 |
| P70670 | NACA | nascent polypeptide-associated complex alpha subunit | Cytoplasm | transcription regulator | 8267023.5 | 9265864 | 5075764.5 | | | | | | |
| P70670 | NACA | nascent polypeptide-associated complex alpha subunit | Cytoplasm | transcription regulator | 7265408.5 | 7477104.5 | 4437011.5 | | | | | | |
| P70670 | NACA | nascent polypeptide-associated complex alpha subunit | Cytoplasm | transcription regulator | 1438997.375 | 1564469 | 748454.312 | P70670 | NACA | 5657143 | 3220805 | 1073602 | 9 |
| P28656 | NAP1L1 | nucleosome assembly protein 1-like 1 | Nucleus | other | 3.81E+07 | 3.05E+07 | 3.26E+07 | | | | | | |
| P28656 | NAP1L1 | nucleosome assembly protein 1-like 1 | Nucleus | other | 2.34E+07 | 2.12E+07 | 2.86E+07 | | | | | | |
| P28656 | NAP1L1 | nucleosome assembly protein 1-like 1 | Nucleus | other | 2.35E+07 | 1.98E+07 | 2.17E+07 | | | | | | |
| P28656 | NAP1L1 | nucleosome assembly protein 1-like 1 | Nucleus | other | 1.09E+07 | 9387644 | 4996041.5 | | | | | | |
| P28656 | NAP1L1 | nucleosome assembly protein 1-like 1 | Nucleus | other | 1.04E+07 | 8018072 | 1.77E+07 | | | | | | |
| P28656 | NAP1L1 | nucleosome assembly protein 1-like 1 | Nucleus | other | 5956040 | 6420705.5 | 4703054.5 | | | | | | |
| P28656 | NAP1L1 | nucleosome assembly protein 1-like 1 | Nucleus | other | 6528577.5 | 6155412.5 | 5317879 | | | | | | |
| P28656 | NAP1L1 | nucleosome assembly protein 1-like 1 | Nucleus | other | 3376600.75 | 2681508.25 | 2859382 | P28656 | NAP1L1 | 15255152 | 10772884 | 2199006 | 24 |
| Q78ZA7 | NAP1L4 | nucleosome assembly protein 1-like 4 | Cytoplasm | other | 2.78E+07 | 1.94E+07 | 2.62E+07 | | | | | | |
| Q78ZA7 | NAP1L4 | nucleosome assembly protein 1-like 4 | Cytoplasm | other | 1.91E+07 | 1.47E+07 | 1.52E+07 | | | | | | |
| Q78ZA7 | NAP1L4 | nucleosome assembly protein 1-like 4 | Cytoplasm | other | 1.15E+07 | 1.06E+07 | 4906631.5 | | | | | | |
| Q78ZA7 | NAP1L4 | nucleosome assembly protein 1-like 4 | Cytoplasm | other | 8293143.5 | 7267596 | 1.08E+07 | | | | | | |
| Q78ZA7 | NAP1L4 | nucleosome assembly protein 1-like 4 | Cytoplasm | other | 1.05E+07 | 9314005 | 8157930.5 | | | | | | |
| Q78ZA7 | NAP1L4 | nucleosome assembly protein 1-like 4 | Cytoplasm | other | 7134777.5 | 6537469.5 | 6858341 | | | | | | |
| Q78ZA7 | NAP1L4 | nucleosome assembly protein 1-like 4 | Cytoplasm | other | 6294529 | 8074402 | 1.00E+07 | | | | | | |
| Q78ZA7 | NAP1L4 | nucleosome assembly protein 1-like 4 | Cytoplasm | other | 4213542.5 | 4985944.5 | 5092473.5 | | | | | | |
| Q78ZA7 | NAP1L4 | nucleosome assembly protein 1-like 4 | Cytoplasm | other | 4882830.5 | 3672107 | 5370655 | | | | | | |
| Q78ZA7 | NAP1L4 | nucleosome assembly protein 1-like 4 | Cytoplasm | other | 1784359.875 | 2499431.25 | 2838186.75 | | | | | | |
| Q78ZA7 | NAP1L4 | nucleosome assembly protein 1-like 4 | Cytoplasm | other | 1774517.875 | 2383962.25 | 2990807.5 | Q78ZA7 | NAP1L4 | 9384336 | 6529800 | 1136692 | 33 |
| Q9DB05 | NAPA | N-ethylmaleimide-sensitive factor attachment protein, alpha | Cytoplasm | transporter | 3574505.5 | 3472294.5 | 2636954.25 | Q9DB05 | NAPA | 3574506 | 514335 | 296951 | 3 |
| Q8K2Z4 | NCAPD2 | non-SMC condensin I complex, subunit D2 | Nucleus | other | 2439121 | 2098742.75 | 1472065 | Q8K2Z4 | NCAPD2 | 2439121 | 490540 | 283214 | 3 |
| Q8C156 | NCAPH | non-SMC condensin I complex, subunit H | Nucleus | other | 2060700.375 | 2194982.5 | 1799268.625 | Q8C156 | NCAPH | 2060700 | 201233 | 116182 | 3 |
| P57716 | NCSTN | nicastrin | Plasma Membrane | peptidase | 1.13E+07 | 1.18E+07 | 1.38E+07 | | | | | | |
| P57716 | NCSTN | nicastrin | Plasma Membrane | peptidase | 1.23E+07 | 1.15E+07 | 1.31E+07 | | | | | | |
| P57716 | NCSTN | nicastrin | Plasma Membrane | peptidase | 8073759 | 7508816.5 | 8243361 | | | | | | |
| P57716 | NCSTN | nicastrin | Plasma Membrane | peptidase | 4363024 | 6856466.5 | 3476536.5 | | | | | | |
| P57716 | NCSTN | nicastrin | Plasma Membrane | peptidase | 7654931 | 6937186.5 | 9258795 | | | | | | |
| P57716 | NCSTN | nicastrin | Plasma Membrane | peptidase | 4160785.5 | 4446218 | 3841494.25 | | | | | | |
| P57716 | NCSTN | nicastrin | Plasma Membrane | peptidase | 5112239 | 4502426 | 5364145.5 | | | | | | |
| P57716 | NCSTN | nicastrin | Plasma Membrane | peptidase | 5596824 | 6071540.5 | 5025920.5 | | | | | | |
| P57716 | NCSTN | nicastrin | Plasma Membrane | peptidase | 737004.375 | 944122.25 | 2364124.5 | P57716 | NCSTN | 6585396 | 3630055 | 698604 | 27 |
| Q99LC3 | NDUFA10 | NADH dehydrogenase (ubiquinone) 1 alpha subcomplex, 10, 42kDa | Cytoplasm | transporter | 3648091.25 | 7720689.5 | 4831421 | | | | | | |
| Q99LC3 | NDUFA10 | NADH dehydrogenase (ubiquinone) 1 alpha subcomplex, 10, 42kDa | Cytoplasm | transporter | 2864222.25 | 7466682.5 | 3127081.75 | Q99LC3 | NDUFA10 | 3256157 | 2162991 | 883037 | 6 |
| Q7TMF3 | NDUFA12 | NADH dehydrogenase (ubiquinone) 1 alpha subcomplex, 12 | Cytoplasm | enzyme | 622227.562 | 899822.812 | 388347.625 | Q7TMF3 | NDUFA12 | 622228 | 256049 | 147830 | 3 |
| Q9CQ75 | NDUFA2 | NADH dehydrogenase (ubiquinone) 1 alpha subcomplex, 2, 8kDa | Cytoplasm | enzyme | 1942932.125 | 5072491.5 | 2088586.75 | Q9CQ75 | NDUFA2 | 1942932 | 1766307 | 1019778 | 3 |
| Q62425 | NDUFA4 | NDUFA4, mitochondrial complex associated | Cytoplasm | enzyme | 5199175.5 | 9699045 | 4479913.5 | | | | | | |
| Q62425 | NDUFA4 | NDUFA4, mitochondrial complex associated | Cytoplasm | enzyme | 3595991.5 | 7216871.5 | 3865033.5 | Q62425 | NDUFA4 | 4397584 | 2358949 | 963037 | 6 |
| Q9CPP6 | NDUFA5 | NADH dehydrogenase (ubiquinone) 1 alpha subcomplex, 5 | Cytoplasm | enzyme | 935522.812 | 2310398.75 | 1152861.75 | Q9CPP6 | NDUFA5 | 935523 | 739077 | 426707 | 3 |
| Q9DCJ5 | NDUFA8 | NADH dehydrogenase (ubiquinone) 1 alpha subcomplex, 8, 19kDa | Cytoplasm | enzyme | 1234822.125 | 4133493.25 | 1953789.25 | Q9DCJ5 | NDUFA8 | 1234822 | 1509432 | 871471 | 3 |
| Q9DC69 | NDUFA9 | NADH dehydrogenase (ubiquinone) 1 alpha subcomplex, 9, 39kDa | Cytoplasm | enzyme | 6838471 | 1.62E+07 | 6727615 | | | | | | |
| Q9DC69 | NDUFA9 | NADH dehydrogenase (ubiquinone) 1 alpha subcomplex, 9, 39kDa | Cytoplasm | enzyme | 1423035.75 | 3794234.25 | 2559934.75 | | | | | | |
| Q9DC69 | NDUFA9 | NADH dehydrogenase (ubiquinone) 1 alpha subcomplex, 9, 39kDa | Cytoplasm | enzyme | 830532.25 | 2521867.25 | 905725.625 | Q9DC69 | NDUFA9 | 3030680 | 4879376 | 1626459 | 9 |
| Q9CR21 | NDUFAB1 | NADH dehydrogenase (ubiquinone) 1, alpha/beta subcomplex, 1, 8kDa | Cytoplasm | enzyme | 2705750.25 | 1.15E+07 | 6913116 | | | | | | |
| Q9CR21 | NDUFAB1 | NADH dehydrogenase (ubiquinone) 1, alpha/beta subcomplex, 1, 8kDa | Cytoplasm | enzyme | 1616954 | 2802808.75 | 1227913.75 | Q9CR21 | NDUFAB1 | 2161352 | 3994788 | 1630865 | 6 |
| Q91VD9 | NDUFS1 | NADH dehydrogenase (ubiquinone) Fe-S protein 1, 75kDa (NADH-coenzyme Q reductase) | Cytoplasm | enzyme | 1913262 | 4347087 | 2027583 | | | | | | |
| Q91VD9 | NDUFS1 | NADH dehydrogenase (ubiquinone) Fe-S protein 1, 75kDa (NADH-coenzyme Q reductase) | Cytoplasm | enzyme | 1473904.875 | 5550572 | 1998871.75 | Q91VD9 | NDUFS1 | 1693583 | 1655223 | 675742 | 6 |
| Q91WD5 | NDUFS2 | NADH dehydrogenase (ubiquinone) Fe-S protein 2, 49kDa (NADH-coenzyme Q reductase) | Cytoplasm | enzyme | 1335736.125 | 3229716.25 | 1565090.25 | Q91WD5 | NDUFS2 | 1335736 | 1033662 | 596785 | 3 |
| Q9DCT2 | NDUFS3 | NADH dehydrogenase (ubiquinone) Fe-S protein 3, 30kDa (NADH-coenzyme Q reductase) | Cytoplasm | enzyme | 1170965.125 | 3377029.25 | 1338055.125 | Q9DCT2 | NDUFS3 | 1170965 | 1228282 | 709149 | 3 |
| Q9CXZ1 | NDUFS4 | NADH dehydrogenase (ubiquinone) Fe-S protein 4, 18kDa (NADH-coenzyme Q reductase) | Cytoplasm | enzyme | 2247920 | 7640973 | 3010267.5 | | | | | | |
| Q9CXZ1 | NDUFS4 | NADH dehydrogenase (ubiquinone) Fe-S protein 4, 18kDa (NADH-coenzyme Q reductase) | Cytoplasm | enzyme | 871905.812 | 2664313.75 | 507167.375 | Q9CXZ1 | NDUFS4 | 1559913 | 2560222 | 1045206 | 6 |
| Q91YT0 | NDUFV1 | NADH dehydrogenase (ubiquinone) flavoprotein 1, 51kDa | Cytoplasm | enzyme | 3426844 | 9402101 | 3601917.25 | | | | | | |
| Q91YT0 | NDUFV1 | NADH dehydrogenase (ubiquinone) flavoprotein 1, 51kDa | Cytoplasm | enzyme | 1395409.875 | 3562737 | 1617564.875 | | | | | | |
| Q91YT0 | NDUFV1 | NADH dehydrogenase (ubiquinone) flavoprotein 1, 51kDa | Cytoplasm | enzyme | 860458.688 | 1657754.5 | 696706.688 | Q91YT0 | NDUFV1 | 1894238 | 2691444 | 897148 | 9 |
| Q9D6J6 | NDUFV2 | NADH dehydrogenase (ubiquinone) flavoprotein 2, 24kDa | Cytoplasm | enzyme | 1347790.625 | 3042237.25 | 1411616.25 | Q9D6J6 | NDUFV2 | 1347791 | 960395 | 554484 | 3 |
| Q9D1J1 | NECAP2 | NECAP endocytosis associated 2 | Cytoplasm | other | 8022529.5 | 1.19E+07 | 5903277.5 | Q9D1J1 | NECAP2 | 8022530 | 3030196 | 1749484 | 3 |
| Q9JMH7 | NEU3 | sialidase 3 (membrane sialidase) | Plasma Membrane | enzyme | 1.83E+07 | 1.61E+07 | 1.65E+07 | | | | | | |
| Q9JMH7 | NEU3 | sialidase 3 (membrane sialidase) | Plasma Membrane | enzyme | 9770138 | 1.01E+07 | 5538245.5 | | | | | | |

| | | | | | | | | | | | | | |
|---|---|---|---|---|---|---|---|---|---|---|---|---|---|
| Q9JMH7 | NEU3 | sialidase 3 (membrane sialidase) | Plasma Membrane | enzyme | 3385799.25 | 2139196.25 | 1302803.875 | Q9JMH7 | NEU3 | 10481979 | 6560091 | 2186697 | 9 |
| Q9EST8 | NFKBIZ | nuclear factor of kappa light polypeptide gene enhancer in B-cells inhibitor, zeta | Nucleus | transcription regulator | 5023746 | 2201207.5 | 3213395.5 | Q9EST8 | NFKBIZ | 5023746 | 1429954 | 825585 | 3 |
| Q3TL44 | NLRX1 | NLR family member X1 | Cytoplasm | other | 8069.789 | 159068.906 | 1500031.25 | Q3TL44 | NLRX1 | 8070 | 821272 | 474162 | 3 |
| Q9WV85 | NME3 | NME/NM23 nucleoside diphosphate kinase 3 | Cytoplasm | kinase | 1375167 | 2687118.25 | 1079878.875 | Q9WV85 | NME3 | 1375167 | 855534 | 493943 | 3 |
| P97300 | Nptn | neuroplastin | Plasma Membrane | other | 1.20E+08 | 1.03E+08 | 8.51E+07 | | | | | | |
| P97300 | Nptn | neuroplastin | Plasma Membrane | other | 4.25E+07 | 3.73E+07 | 3.14E+07 | | | | | | |
| P97300 | Nptn | neuroplastin | Plasma Membrane | other | 1.49E+07 | 1.99E+07 | 2.24E+07 | | | | | | |
| P97300 | Nptn | neuroplastin | Plasma Membrane | other | 1.36E+07 | 9621270 | 6447139.5 | | | | | | |
| P97300 | Nptn | neuroplastin | Plasma Membrane | other | 8529864 | 1.21E+07 | 1.80E+07 | | | | | | |
| P97300 | Nptn | neuroplastin | Plasma Membrane | other | 5290257 | 8796537 | 1.46E+07 | | | | | | |
| P97300 | Nptn | neuroplastin | Plasma Membrane | other | 3725164.5 | 5006326 | 5308813.5 | | | | | | |
| P97300 | Nptn | neuroplastin | Plasma Membrane | other | 881180.062 | 573345.188 | 485964.344 | | | | | | |
| P97300 | Nptn | neuroplastin | Plasma Membrane | other | 336115.938 | 126443.695 | 107272.961 | P97300 | Nptn | 23275842 | 31498128 | 6061818 | 27 |
| P08556 | NRAS | neuroblastoma RAS viral (v-ras) oncogene homolog | Plasma Membrane | enzyme | 8975000 | 9576039 | 1.06E+07 | P08556 | NRAS | 8975000 | 821621 | 474363 | 3 |
| P46460 | NSF | N-ethylmaleimide-sensitive factor | Cytoplasm | transporter | 1.02E+07 | 8551807 | 7798168 | | | | | | |
| P46460 | NSF | N-ethylmaleimide-sensitive factor | Cytoplasm | transporter | 7919789 | 7438848.5 | 7635405 | | | | | | |
| P46460 | NSF | N-ethylmaleimide-sensitive factor | Cytoplasm | transporter | 6777180.5 | 6277717 | 7332091.5 | | | | | | |
| P46460 | NSF | N-ethylmaleimide-sensitive factor | Cytoplasm | transporter | 8229421 | 5886408.5 | 5132157 | | | | | | |
| P46460 | NSF | N-ethylmaleimide-sensitive factor | Cytoplasm | transporter | 5598554.5 | 5368749 | 5480771 | | | | | | |
| P46460 | NSF | N-ethylmaleimide-sensitive factor | Cytoplasm | transporter | 7614966.5 | 5620624 | 6848846.5 | | | | | | |
| P46460 | NSF | N-ethylmaleimide-sensitive factor | Cytoplasm | transporter | 4736088.5 | 3512155.25 | 4227594 | | | | | | |
| P46460 | NSF | N-ethylmaleimide-sensitive factor | Cytoplasm | transporter | 4280976.5 | 5656049 | 5134682.5 | | | | | | |
| P46460 | NSF | N-ethylmaleimide-sensitive factor | Cytoplasm | transporter | 3105739 | 3438487.25 | 3642066.75 | | | | | | |
| P46460 | NSF | N-ethylmaleimide-sensitive factor | Cytoplasm | transporter | 3520096.25 | 2760894.75 | 2350829.75 | | | | | | |
| P46460 | NSF | N-ethylmaleimide-sensitive factor | Cytoplasm | transporter | 3922839.25 | 3601063 | 4428901.5 | | | | | | |
| P46460 | NSF | N-ethylmaleimide-sensitive factor | Cytoplasm | transporter | 3327942.75 | 2459167.75 | 2414813.75 | | | | | | |
| P46460 | NSF | N-ethylmaleimide-sensitive factor | Cytoplasm | transporter | 1384998.375 | 1065554 | 1508224.625 | | | | | | |
| P46460 | NSF | N-ethylmaleimide-sensitive factor | Cytoplasm | transporter | 281432.125 | 78670.367 | 270096.438 | P46460 | NSF | 5061430 | 2479528 | 382599 | 42 |
| Q9CZ44 | NSFL1C | NSFL1 (p97) cofactor (p47) | Cytoplasm | other | 4993415.5 | 2397077.75 | 2872273 | | | | | | |
| Q9CZ44 | NSFL1C | NSFL1 (p97) cofactor (p47) | Cytoplasm | other | 3978622 | 2568037 | 2066642.125 | | | | | | |
| Q9CZ44 | NSFL1C | NSFL1 (p97) cofactor (p47) | Cytoplasm | other | 2914707.25 | 1877731.375 | 1729605.125 | | | | | | |
| Q9CZ44 | NSFL1C | NSFL1 (p97) cofactor (p47) | Cytoplasm | other | 2318589.75 | 1525839.375 | 1384564.25 | Q9CZ44 | NSFL1C | 3551334 | 1048422 | 302653 | 12 |
| A6H603 | NWD1 | NACHT and WD repeat domain containing 1 | Cytoplasm | other | 1.77E+07 | 1.50E+07 | 1.62E+07 | A6H603 | NWD1 | 17680000 | 1360992 | 785769 | 3 |
| P83503 | NYX | nyctalopin | Extracellular Space | other | 9.25E+07 | 9.23E+07 | 9.79E+07 | P83503 | NYX | 92530000 | 3186163 | 1839532 | 3 |
| P29758 | OAT | ornithine aminotransferase | Cytoplasm | enzyme | 1844302.75 | 2897361.75 | 1541339.125 | P29758 | OAT | 1844303 | 711749 | 410928 | 3 |
| A2AAJ9 | OBSCN | obscurin, cytoskeletal calmodulin and titin-interacting RhoGEF | Cytoplasm | kinase | 1992854.25 | 3143812 | 2636936.75 | A2AAJ9 | OBSCN | 1992854 | 576840 | 333039 | 3 |
| Q9CRD0 | OCIAD1 | OCIA domain containing 1 | Cytoplasm | other | 1342696.25 | 3377517.75 | 1396434.625 | Q9CRD0 | OCIAD1 | 1342696 | 1159603 | 669497 | 3 |
| Q60597 | OGDH | oxoglutarate (alpha-ketoglutarate) dehydrogenase (lipoamide) | Cytoplasm | enzyme | 8527512 | 1.61E+07 | 6968188.5 | | | | | | |
| Q60597 | OGDH | oxoglutarate (alpha-ketoglutarate) dehydrogenase (lipoamide) | Cytoplasm | enzyme | 9743244 | 1.61E+07 | 9704279 | | | | | | |
| Q60597 | OGDH | oxoglutarate (alpha-ketoglutarate) dehydrogenase (lipoamide) | Cytoplasm | enzyme | 5047101.5 | 1.23E+07 | 6113404 | | | | | | |
| Q60597 | OGDH | oxoglutarate (alpha-ketoglutarate) dehydrogenase (lipoamide) | Cytoplasm | enzyme | 4780839.5 | 9891545 | 5822092 | | | | | | |
| Q60597 | OGDH | oxoglutarate (alpha-ketoglutarate) dehydrogenase (lipoamide) | Cytoplasm | enzyme | 5313494 | 1.57E+07 | 6885153.5 | | | | | | |
| Q60597 | OGDH | oxoglutarate (alpha-ketoglutarate) dehydrogenase (lipoamide) | Cytoplasm | enzyme | 5343600 | 1.24E+07 | 5296189 | | | | | | |
| Q60597 | OGDH | oxoglutarate (alpha-ketoglutarate) dehydrogenase (lipoamide) | Cytoplasm | enzyme | 4891020 | 1.31E+07 | 4965854 | | | | | | |
| Q60597 | OGDH | oxoglutarate (alpha-ketoglutarate) dehydrogenase (lipoamide) | Cytoplasm | enzyme | 4590869.5 | 7521927.5 | 4257148.5 | | | | | | |
| Q60597 | OGDH | oxoglutarate (alpha-ketoglutarate) dehydrogenase (lipoamide) | Cytoplasm | enzyme | 4463713.5 | 9256189 | 3219158.5 | | | | | | |
| Q60597 | OGDH | oxoglutarate (alpha-ketoglutarate) dehydrogenase (lipoamide) | Cytoplasm | enzyme | 3534632.75 | 1.00E+07 | 4645254.5 | | | | | | |
| Q60597 | OGDH | oxoglutarate (alpha-ketoglutarate) dehydrogenase (lipoamide) | Cytoplasm | enzyme | 4281250.5 | 7493912.5 | 3598171.75 | | | | | | |
| Q60597 | OGDH | oxoglutarate (alpha-ketoglutarate) dehydrogenase (lipoamide) | Cytoplasm | enzyme | 3175249.25 | 7548819 | 4063692.75 | | | | | | |
| Q60597 | OGDH | oxoglutarate (alpha-ketoglutarate) dehydrogenase (lipoamide) | Cytoplasm | enzyme | 2139985.75 | 4621545.5 | 1931854.875 | | | | | | |
| Q60597 | OGDH | oxoglutarate (alpha-ketoglutarate) dehydrogenase (lipoamide) | Cytoplasm | enzyme | 1575437.25 | 4598992.5 | 3078430.25 | | | | | | |
| Q60597 | OGDH | oxoglutarate (alpha-ketoglutarate) dehydrogenase (lipoamide) | Cytoplasm | enzyme | 1488396.875 | 2825667 | 1279779.625 | | | | | | |
| Q60597 | OGDH | oxoglutarate (alpha-ketoglutarate) dehydrogenase (lipoamide) | Cytoplasm | enzyme | 1149494.875 | 2758870.75 | 1543222.25 | | | | | | |
| Q60597 | OGDH | oxoglutarate (alpha-ketoglutarate) dehydrogenase (lipoamide) | Cytoplasm | enzyme | 2248357.25 | 5531440 | 1897729.375 | Q60597 | OGDH | 4252600 | 3916161 | 548372 | 51 |
| Q9CZ30 | OLA1 | Obg-like ATPase 1 | Cytoplasm | enzyme | 1628919.875 | 2238647.25 | 2749802.5 | Q9CZ30 | OLA1 | 1628920 | 561163 | 323988 | 3 |
| Q8BGT0 | OSTM1 | osteopetrosis associated transmembrane protein 1 | Cytoplasm | other | 831373.875 | 1312022.75 | 727280.312 | Q8BGT0 | OSTM1 | 831374 | 311925 | 180090 | 3 |
| Q9D0K2 | OXCT1 | 3-oxoacid CoA transferase 1 | Cytoplasm | enzyme | 1141196 | 2204288.25 | 1443212.875 | Q9D0K2 | OXCT1 | 1141196 | 547816 | 316282 | 3 |
| P09103 | P4HB | prolyl 4-hydroxylase, beta polypeptide | Cytoplasm | enzyme | 1.63E+07 | 2.06E+07 | 2.37E+07 | | | | | | |
| P09103 | P4HB | prolyl 4-hydroxylase, beta polypeptide | Cytoplasm | enzyme | 1.51E+07 | 1.43E+07 | 1.66E+07 | | | | | | |
| P09103 | P4HB | prolyl 4-hydroxylase, beta polypeptide | Cytoplasm | enzyme | 1.55E+07 | 2.27E+07 | 2.41E+07 | | | | | | |
| P09103 | P4HB | prolyl 4-hydroxylase, beta polypeptide | Cytoplasm | enzyme | 1.53E+07 | 1.53E+07 | 2.04E+07 | | | | | | |
| P09103 | P4HB | prolyl 4-hydroxylase, beta polypeptide | Cytoplasm | enzyme | 1.53E+07 | 2.21E+07 | 2.97E+07 | | | | | | |
| P09103 | P4HB | prolyl 4-hydroxylase, beta polypeptide | Cytoplasm | enzyme | 1.37E+07 | 1.81E+07 | 2.62E+07 | | | | | | |
| P09103 | P4HB | prolyl 4-hydroxylase, beta polypeptide | Cytoplasm | enzyme | 1.21E+07 | 1.06E+07 | 1.25E+07 | | | | | | |
| P09103 | P4HB | prolyl 4-hydroxylase, beta polypeptide | Cytoplasm | enzyme | 1.19E+07 | 1.17E+07 | 1.51E+07 | | | | | | |
| P09103 | P4HB | prolyl 4-hydroxylase, beta polypeptide | Cytoplasm | enzyme | 9352179 | 1.13E+07 | 9299839 | | | | | | |
| P09103 | P4HB | prolyl 4-hydroxylase, beta polypeptide | Cytoplasm | enzyme | 1.20E+07 | 1.49E+07 | 1.97E+07 | | | | | | |
| P09103 | P4HB | prolyl 4-hydroxylase, beta polypeptide | Cytoplasm | enzyme | 1.35E+07 | 1.08E+07 | 5478012.5 | | | | | | |
| P09103 | P4HB | prolyl 4-hydroxylase, beta polypeptide | Cytoplasm | enzyme | 8374990 | 9064823 | 1.17E+07 | | | | | | |
| P09103 | P4HB | prolyl 4-hydroxylase, beta polypeptide | Cytoplasm | enzyme | 7475810 | 7913210.5 | 1.03E+07 | | | | | | |
| P09103 | P4HB | prolyl 4-hydroxylase, beta polypeptide | Cytoplasm | enzyme | 8047527.5 | 9160651 | 1.16E+07 | | | | | | |
| P09103 | P4HB | prolyl 4-hydroxylase, beta polypeptide | Cytoplasm | enzyme | 7901657 | 8498206 | 1.16E+07 | | | | | | |
| P09103 | P4HB | prolyl 4-hydroxylase, beta polypeptide | Cytoplasm | enzyme | 4831349.5 | 9050509 | 9439862 | | | | | | |
| P09103 | P4HB | prolyl 4-hydroxylase, beta polypeptide | Cytoplasm | enzyme | 3963851.25 | 6098413.5 | 6558730.5 | | | | | | |
| P09103 | P4HB | prolyl 4-hydroxylase, beta polypeptide | Cytoplasm | enzyme | 3915532 | 3574181.5 | 4375512.5 | | | | | | |
| P09103 | P4HB | prolyl 4-hydroxylase, beta polypeptide | Cytoplasm | enzyme | 2458322.75 | 3364988 | 3599832.5 | P09103 | P4HB | 10360064 | 6230361 | 825231 | 57 |
| P50580 | PA2G4 | proliferation-associated 2G4, 38kDa | Nucleus | transcription regulator | 8110093 | 1.16E+07 | 4476550 | | | | | | |
| P50580 | PA2G4 | proliferation-associated 2G4, 38kDa | Nucleus | transcription regulator | 6303998.5 | 9823311 | 3886995 | | | | | | |
| P50580 | PA2G4 | proliferation-associated 2G4, 38kDa | Nucleus | transcription regulator | 5468390 | 7164956.5 | 2919175.25 | | | | | | |
| P50580 | PA2G4 | proliferation-associated 2G4, 38kDa | Nucleus | transcription regulator | 4336471 | 5963771.5 | 2884479.5 | | | | | | |
| P50580 | PA2G4 | proliferation-associated 2G4, 38kDa | Nucleus | transcription regulator | 3772544.25 | 6231282.5 | 2838972.75 | P50580 | PA2G4 | 5598299 | 2596468 | 670405 | 15 |
| P29341 | PABPC1 | poly(A) binding protein, cytoplasmic 1 | Cytoplasm | translation regulator | 4008405 | 7346288.5 | 1734201.75 | | | | | | |
| P29341 | PABPC1 | poly(A) binding protein, cytoplasmic 1 | Cytoplasm | translation regulator | 5478941 | 7636910 | 3678556.75 | | | | | | |
| P29341 | PABPC1 | poly(A) binding protein, cytoplasmic 1 | Cytoplasm | translation regulator | 3100218.75 | 5813831.5 | 2502886.75 | | | | | | |
| P29341 | PABPC1 | poly(A) binding protein, cytoplasmic 1 | Cytoplasm | translation regulator | 3856365 | 5804090 | 2362741 | | | | | | |
| P29341 | PABPC1 | poly(A) binding protein, cytoplasmic 1 | Cytoplasm | translation regulator | 2289769.25 | 3640888.5 | 1828785.375 | P29341 | PABPC1 | 3746740 | 1924207 | 496828 | 15 |
| Q9WVE8 | PACSIN2 | protein kinase C and casein kinase substrate in neurons 2 | Cytoplasm | transporter | 1.16E+07 | 7002234.5 | 7271545.5 | | | | | | |
| Q9WVE8 | PACSIN2 | protein kinase C and casein kinase substrate in neurons 2 | Cytoplasm | transporter | 1.04E+07 | 9270209 | 7773547.5 | | | | | | |
| Q9WVE8 | PACSIN2 | protein kinase C and casein kinase substrate in neurons 2 | Cytoplasm | transporter | 6047072.5 | 4312173 | 4024651 | | | | | | |
| Q9WVE8 | PACSIN2 | protein kinase C and casein kinase substrate in neurons 2 | Cytoplasm | transporter | 5453856 | 4463059.5 | 4297933 | | | | | | |
| Q9WVE8 | PACSIN2 | protein kinase C and casein kinase substrate in neurons 2 | Cytoplasm | transporter | 5418600.5 | 4549905 | 3685036.75 | | | | | | |

| | | | | | | | | | | | | | |
|---|---|---|---|---|---|---|---|---|---|---|---|---|---|
| Q9WVE8 | PACSIN2 | protein kinase C and casein kinase substrate in neurons 2 | Cytoplasm | transporter | 7035634.5 | 5190498 | 3987222.5 | | | | | | |
| Q9WVE8 | PACSIN2 | protein kinase C and casein kinase substrate in neurons 2 | Cytoplasm | transporter | 1604627 | 1864965.5 | 1763632.375 | | | | | | |
| Q9WVE8 | PACSIN2 | protein kinase C and casein kinase substrate in neurons 2 | Cytoplasm | transporter | 681295.438 | 1837583 | 2323121.5 | Q9WVE8 | PACSIN2 | 6026386 | 2841678 | 580055 | 24 |
| Q9ES46 | PARVB | parvin, beta | Cytoplasm | other | 692658.188 | 1511482.375 | 2032524.5 | | | | | | |
| Q9ES46 | PARVB | parvin, beta | Cytoplasm | other | 678735.375 | 1616371.25 | 2947474.75 | | | | | | |
| Q9ES46 | PARVB | parvin, beta | Cytoplasm | other | 397960.219 | 1192122.125 | 1940048.375 | | | | | | |
| Q9ES46 | PARVB | parvin, beta | Cytoplasm | other | 143826.719 | 727747.812 | 1152667.125 | | | | | | |
| Q9ES46 | PARVB | parvin, beta | Cytoplasm | other | 340087.25 | 1046719.438 | 553302.125 | | | | | | |
| Q9ES46 | PARVB | parvin, beta | Cytoplasm | other | 148417.406 | 965770.5 | 625407.812 | Q9ES46 | PARVB | 400281 | 739121 | 174213 | 18 |
| Q05920 | PC | pyruvate carboxylase | Cytoplasm | enzyme | 2104838.5 | 5478912 | 1555888.875 | | | | | | |
| Q05920 | PC | pyruvate carboxylase | Cytoplasm | enzyme | 2778327.25 | 6312765.5 | 2812119.25 | | | | | | |
| Q05920 | PC | pyruvate carboxylase | Cytoplasm | enzyme | 2565854.5 | 4071443.25 | 1474717 | | | | | | |
| Q05920 | PC | pyruvate carboxylase | Cytoplasm | enzyme | 1049931.75 | 1761973.5 | 485453.156 | | | | | | |
| Q05920 | PC | pyruvate carboxylase | Cytoplasm | enzyme | 1128218.625 | 2305372.75 | 716748.375 | Q05920 | PC | 1925434 | 1688410 | 435946 | 15 |
| P60335 | PCBP1 | poly(rC) binding protein 1 | Nucleus | translation regulator | 4152472.5 | 6020756 | 1970087.625 | | | | | | |
| P60335 | PCBP1 | poly(rC) binding protein 1 | Nucleus | translation regulator | 4126725.25 | 6841499 | 2364895.5 | | | | | | |
| P60335 | PCBP1 | poly(rC) binding protein 1 | Nucleus | translation regulator | 3730501.5 | 7317042.5 | 3616782.75 | | | | | | |
| P60335 | PCBP1 | poly(rC) binding protein 1 | Nucleus | translation regulator | 2661187.75 | 4062064 | 2051756.25 | | | | | | |
| P60335 | PCBP1 | poly(rC) binding protein 1 | Nucleus | translation regulator | 2081920.5 | 3015935.5 | 1714667.625 | P60335 | PCBP1 | 3350562 | 1782030 | 460118 | 15 |
| Q8BH04 | PCK2 | phosphoenolpyruvate carboxykinase 2 (mitochondrial) | Cytoplasm | kinase | 1.41E+07 | 2.01E+07 | 7512715 | | | | | | |
| Q8BH04 | PCK2 | phosphoenolpyruvate carboxykinase 2 (mitochondrial) | Cytoplasm | kinase | 1.44E+07 | 2.52E+07 | 1.30E+07 | | | | | | |
| Q8BH04 | PCK2 | phosphoenolpyruvate carboxykinase 2 (mitochondrial) | Cytoplasm | kinase | 4940835 | 1.29E+07 | 4105768.5 | | | | | | |
| Q8BH04 | PCK2 | phosphoenolpyruvate carboxykinase 2 (mitochondrial) | Cytoplasm | kinase | 8539541 | 1.61E+07 | 6571183 | | | | | | |
| Q8BH04 | PCK2 | phosphoenolpyruvate carboxykinase 2 (mitochondrial) | Cytoplasm | kinase | 7060427.5 | 9531939 | 4154886.75 | | | | | | |
| Q8BH04 | PCK2 | phosphoenolpyruvate carboxykinase 2 (mitochondrial) | Cytoplasm | kinase | 5924593.5 | 1.46E+07 | 4157955.25 | | | | | | |
| Q8BH04 | PCK2 | phosphoenolpyruvate carboxykinase 2 (mitochondrial) | Cytoplasm | kinase | 4868398 | 1.05E+07 | 4079239.75 | | | | | | |
| Q8BH04 | PCK2 | phosphoenolpyruvate carboxykinase 2 (mitochondrial) | Cytoplasm | kinase | 3731609.25 | 7657659.5 | 2799257.5 | | | | | | |
| Q8BH04 | PCK2 | phosphoenolpyruvate carboxykinase 2 (mitochondrial) | Cytoplasm | kinase | 2869451 | 7081557.5 | 2050964.625 | | | | | | |
| Q8BH04 | PCK2 | phosphoenolpyruvate carboxykinase 2 (mitochondrial) | Cytoplasm | kinase | 4374189 | 8040509.5 | 4159751.25 | | | | | | |
| Q8BH04 | PCK2 | phosphoenolpyruvate carboxykinase 2 (mitochondrial) | Cytoplasm | kinase | 4089749.5 | 8720999 | 3014023 | | | | | | |
| Q8BH04 | PCK2 | phosphoenolpyruvate carboxykinase 2 (mitochondrial) | Cytoplasm | kinase | 2121963.75 | 6818557 | 1920470.5 | | | | | | |
| Q8BH04 | PCK2 | phosphoenolpyruvate carboxykinase 2 (mitochondrial) | Cytoplasm | kinase | 6598475.5 | 9595801 | 2749528.25 | | | | | | |
| Q8BH04 | PCK2 | phosphoenolpyruvate carboxykinase 2 (mitochondrial) | Cytoplasm | kinase | 3656037.25 | 9355887 | 3670919.25 | | | | | | |
| Q8BH04 | PCK2 | phosphoenolpyruvate carboxykinase 2 (mitochondrial) | Cytoplasm | kinase | 2578244 | 6074095.5 | 2410772.5 | | | | | | |
| Q8BH04 | PCK2 | phosphoenolpyruvate carboxykinase 2 (mitochondrial) | Cytoplasm | kinase | 2376325 | 3622576.75 | 1704693.75 | | | | | | |
| Q8BH04 | PCK2 | phosphoenolpyruvate carboxykinase 2 (mitochondrial) | Cytoplasm | kinase | 2726747.25 | 4314155 | 791291.75 | Q8BH04 | PCK2 | 5585093 | 5029213 | 704231 | 51 |
| Q9CQF9 | PCYOX1 | prenylcysteine oxidase 1 | Cytoplasm | enzyme | 9735365 | 1.17E+07 | 1.18E+07 | Q9CQF9 | PCYOX1 | 9735365 | 1160871 | 670229 | 3 |
| P27664 | PDE6A | phosphodiesterase 6A, cGMP-specific, rod, alpha | Plasma Membrane | enzyme | 1.49E+07 | 1.12E+07 | 1.31E+07 | P27664 | PDE6A | 14880000 | 1840036 | 1062345 | 3 |
| P35486 | PDHA1 | pyruvate dehydrogenase (lipoamide) alpha 1 | Cytoplasm | enzyme | 949672.25 | 2271248.75 | 952583 | | | | | | |
| P35486 | PDHA1 | pyruvate dehydrogenase (lipoamide) alpha 1 | Cytoplasm | enzyme | 1318668.625 | 2451330.25 | 1034299.875 | | | | | | |
| P35486 | PDHA1 | pyruvate dehydrogenase (lipoamide) alpha 1 | Cytoplasm | enzyme | 1281485.625 | 2937063.25 | 1151727.25 | P35486 | PDHA1 | 1183276 | 750534 | 250178 | 9 |
| Q9D051 | PDHB | pyruvate dehydrogenase (lipoamide) beta | Cytoplasm | enzyme | 1433039.625 | 3650322 | 2068962.375 | Q9D051 | PDHB | 1433040 | 1141741 | 659185 | 3 |
| P27773 | PDIA3 | protein disulfide isomerase family A, member 3 | Cytoplasm | peptidase | 6.57E+07 | 1.02E+08 | 9.53E+07 | | | | | | |
| P27773 | PDIA3 | protein disulfide isomerase family A, member 3 | Cytoplasm | peptidase | 4.03E+07 | 4.82E+07 | 5.12E+07 | | | | | | |
| P27773 | PDIA3 | protein disulfide isomerase family A, member 3 | Cytoplasm | peptidase | 2.96E+07 | 3.57E+07 | 4.18E+07 | | | | | | |
| P27773 | PDIA3 | protein disulfide isomerase family A, member 3 | Cytoplasm | peptidase | 1.98E+07 | 2.17E+07 | 2.98E+07 | | | | | | |
| P27773 | PDIA3 | protein disulfide isomerase family A, member 3 | Cytoplasm | peptidase | 1.64E+07 | 2.18E+07 | 2.45E+07 | | | | | | |
| P27773 | PDIA3 | protein disulfide isomerase family A, member 3 | Cytoplasm | peptidase | 2.30E+07 | 2.18E+07 | 2.33E+07 | | | | | | |
| P27773 | PDIA3 | protein disulfide isomerase family A, member 3 | Cytoplasm | peptidase | 1.66E+07 | 2.11E+07 | 2.66E+07 | | | | | | |
| P27773 | PDIA3 | protein disulfide isomerase family A, member 3 | Cytoplasm | peptidase | 1.37E+07 | 2.43E+07 | 3.18E+07 | | | | | | |
| P27773 | PDIA3 | protein disulfide isomerase family A, member 3 | Cytoplasm | peptidase | 2.23E+07 | 2.21E+07 | 3.18E+07 | | | | | | |
| P27773 | PDIA3 | protein disulfide isomerase family A, member 3 | Cytoplasm | peptidase | 1.47E+07 | 1.82E+07 | 2.52E+07 | | | | | | |
| P27773 | PDIA3 | protein disulfide isomerase family A, member 3 | Cytoplasm | peptidase | 1.43E+07 | 1.49E+07 | 2.20E+07 | | | | | | |
| P27773 | PDIA3 | protein disulfide isomerase family A, member 3 | Cytoplasm | peptidase | 1.41E+07 | 1.66E+07 | 1.95E+07 | | | | | | |
| P27773 | PDIA3 | protein disulfide isomerase family A, member 3 | Cytoplasm | peptidase | 1.12E+07 | 1.37E+07 | 1.91E+07 | | | | | | |
| P27773 | PDIA3 | protein disulfide isomerase family A, member 3 | Cytoplasm | peptidase | 1.17E+07 | 1.30E+07 | 1.24E+07 | | | | | | |
| P27773 | PDIA3 | protein disulfide isomerase family A, member 3 | Cytoplasm | peptidase | 9948543 | 1.24E+07 | 1.67E+07 | | | | | | |
| P27773 | PDIA3 | protein disulfide isomerase family A, member 3 | Cytoplasm | peptidase | 1.11E+07 | 1.11E+07 | 1.14E+07 | | | | | | |
| P27773 | PDIA3 | protein disulfide isomerase family A, member 3 | Cytoplasm | peptidase | 9770972 | 1.40E+07 | 1.64E+07 | | | | | | |
| P27773 | PDIA3 | protein disulfide isomerase family A, member 3 | Cytoplasm | peptidase | 7221893 | 8699390 | 1.17E+07 | | | | | | |
| P27773 | PDIA3 | protein disulfide isomerase family A, member 3 | Cytoplasm | peptidase | 7783733.5 | 9204522 | 1.41E+07 | | | | | | |
| P27773 | PDIA3 | protein disulfide isomerase family A, member 3 | Cytoplasm | peptidase | 5983099 | 1.30E+07 | 1.28E+07 | | | | | | |
| P27773 | PDIA3 | protein disulfide isomerase family A, member 3 | Cytoplasm | peptidase | 3280305.25 | 5327110 | 5356289 | | | | | | |
| P27773 | PDIA3 | protein disulfide isomerase family A, member 3 | Cytoplasm | peptidase | 2400056.5 | 2052223 | 1000370.5 | | | | | | |
| P27773 | PDIA3 | protein disulfide isomerase family A, member 3 | Cytoplasm | peptidase | 1067860.875 | 1469471 | 2979668.5 | P27773 | PDIA3 | 16178542 | 18299476 | 2202998 | 69 |
| P08003 | PDIA4 | protein disulfide isomerase family A, member 4 | Cytoplasm | enzyme | 5789457 | 7500064 | 7589045 | | | | | | |
| P08003 | PDIA4 | protein disulfide isomerase family A, member 4 | Cytoplasm | enzyme | 2630915 | 3273159.5 | 4967442 | | | | | | |
| P08003 | PDIA4 | protein disulfide isomerase family A, member 4 | Cytoplasm | enzyme | 2120604.75 | 3137074.25 | 4407467 | | | | | | |
| P08003 | PDIA4 | protein disulfide isomerase family A, member 4 | Cytoplasm | enzyme | 2659852.75 | 3077415.25 | 3266958.25 | | | | | | |
| P08003 | PDIA4 | protein disulfide isomerase family A, member 4 | Cytoplasm | enzyme | 1966720.25 | 2608952.75 | 3396878.25 | | | | | | |
| P08003 | PDIA4 | protein disulfide isomerase family A, member 4 | Cytoplasm | enzyme | 1859530 | 2629531 | 2665037.5 | | | | | | |
| P08003 | PDIA4 | protein disulfide isomerase family A, member 4 | Cytoplasm | enzyme | 1403319.5 | 1948796.875 | 2530618.75 | | | | | | |
| P08003 | PDIA4 | protein disulfide isomerase family A, member 4 | Cytoplasm | enzyme | 1113261.75 | 1444044.125 | 1985686.125 | P08003 | PDIA4 | 2442958 | 1735736 | 354306 | 24 |
| Q922R8 | PDIA6 | protein disulfide isomerase family A, member 6 | Cytoplasm | enzyme | 1.06E+07 | 1.84E+07 | 2.21E+07 | | | | | | |
| Q922R8 | PDIA6 | protein disulfide isomerase family A, member 6 | Cytoplasm | enzyme | 8995234 | 1.14E+07 | 1.57E+07 | | | | | | |
| Q922R8 | PDIA6 | protein disulfide isomerase family A, member 6 | Cytoplasm | enzyme | 6477170 | 9337911 | 1.24E+07 | | | | | | |
| Q922R8 | PDIA6 | protein disulfide isomerase family A, member 6 | Cytoplasm | enzyme | 6433384 | 9216801 | 9894989 | | | | | | |
| Q922R8 | PDIA6 | protein disulfide isomerase family A, member 6 | Cytoplasm | enzyme | 5938777 | 7305658 | 1.02E+07 | | | | | | |
| Q922R8 | PDIA6 | protein disulfide isomerase family A, member 6 | Cytoplasm | enzyme | 6893692 | 1.08E+07 | 4292446 | | | | | | |
| Q922R8 | PDIA6 | protein disulfide isomerase family A, member 6 | Cytoplasm | enzyme | 6715932.5 | 7445878 | 1.05E+07 | | | | | | |
| Q922R8 | PDIA6 | protein disulfide isomerase family A, member 6 | Cytoplasm | enzyme | 6674844 | 7596245 | 1.15E+07 | | | | | | |
| Q922R8 | PDIA6 | protein disulfide isomerase family A, member 6 | Cytoplasm | enzyme | 3453201.75 | 6698609.5 | 9644262 | | | | | | |
| Q922R8 | PDIA6 | protein disulfide isomerase family A, member 6 | Cytoplasm | enzyme | 3696826.25 | 4097454 | 3451972.25 | | | | | | |
| Q922R8 | PDIA6 | protein disulfide isomerase family A, member 6 | Cytoplasm | enzyme | 2582237.75 | 3363661.75 | 4687703 | Q922R8 | PDIA6 | 6224664 | 4353774 | 757895 | 33 |
| Q9Z126 | PF4 | platelet factor 4 | Extracellular Space | cytokine | 3.52E+07 | 4.19E+07 | 6.28E+07 | | | | | | |
| Q9Z126 | PF4 | platelet factor 4 | Extracellular Space | cytokine | 3.23E+07 | 2.91E+07 | 3.11E+07 | | | | | | |
| Q9Z126 | PF4 | platelet factor 4 | Extracellular Space | cytokine | 1.70E+07 | 1.90E+07 | 2.83E+07 | Q9Z126 | PF4 | 28143333 | 13528917 | 4509639 | 9 |
| P12382 | PFKL | phosphofructokinase, liver | Cytoplasm | kinase | 3166736.75 | 4202366 | 3405485 | P12382 | PFKL | 3166737 | 542302 | 313098 | 3 |
| P47857 | PFKM | phosphofructokinase, muscle | Cytoplasm | kinase | 7543725 | 5800610 | 4884823 | | | | | | |
| P47857 | PFKM | phosphofructokinase, muscle | Cytoplasm | kinase | 4865829.5 | 2569273.25 | 2818387.25 | | | | | | |
| P47857 | PFKM | phosphofructokinase, muscle | Cytoplasm | kinase | 4781795.5 | 3606542.5 | 3344086.25 | | | | | | |

| | | | | | | | | | | | | | |
|---|---|---|---|---|---|---|---|---|---|---|---|---|---|
| P47857 | PFKM | phosphofructokinase, muscle | Cytoplasm | kinase | 4994111 | 3165717 | 3281293.25 | | | | | | |
| P47857 | PFKM | phosphofructokinase, muscle | Cytoplasm | kinase | 3304506.75 | 2482973.75 | 2317774.25 | | | | | | |
| P47857 | PFKM | phosphofructokinase, muscle | Cytoplasm | kinase | 3702426.75 | 3226402 | 3542895 | P47857 | PFKM | 4865399 | 1339677 | 315765 | 18 |
| Q9DCD0 | PGD | phosphogluconate dehydrogenase | Cytoplasm | enzyme | 3783725 | 2560753.75 | 3925882.75 | Q9DCD0 | PGD | 3783725 | 750494 | 433298 | 3 |
| O55022 | PGRMC1 | progesterone receptor membrane component 1 | Plasma Membrane | transmembrane receptor | 3768220.5 | 3075458 | 4134103 | O55022 | PGRMC1 | 3768221 | 537668 | 310423 | 3 |
| Q80UU9 | PGRMC2 | progesterone receptor membrane component 2 | Nucleus | other | 1711968.375 | 2707831.75 | 3185168.75 | Q80UU9 | PGRMC2 | 1711968 | 751655 | 433968 | 3 |
| P67778 | PHB | prohibitin | Nucleus | transcription regulator | 8220538.5 | 1.37E+07 | 9122335 | | | | | | |
| P67778 | PHB | prohibitin | Nucleus | transcription regulator | 8217268.5 | 2.15E+07 | 8929999 | | | | | | |
| P67778 | PHB | prohibitin | Nucleus | transcription regulator | 6590305.5 | 1.19E+07 | 5800795.5 | | | | | | |
| P67778 | PHB | prohibitin | Nucleus | transcription regulator | 5089009.5 | 1.38E+07 | 6669019 | | | | | | |
| P67778 | PHB | prohibitin | Nucleus | transcription regulator | 6422084 | 1.45E+07 | 6150707 | | | | | | |
| P67778 | PHB | prohibitin | Nucleus | transcription regulator | 4602408.5 | 8419866 | 3937960.75 | | | | | | |
| P67778 | PHB | prohibitin | Nucleus | transcription regulator | 2976403.5 | 1.11E+07 | 6565555.5 | | | | | | |
| P67778 | PHB | prohibitin | Nucleus | transcription regulator | 4865390.5 | 1.62E+07 | 6201788 | | | | | | |
| P67778 | PHB | prohibitin | Nucleus | transcription regulator | 2186191.25 | 3666113.25 | 2240863 | | | | | | |
| P67778 | PHB | prohibitin | Nucleus | transcription regulator | 2310095.25 | 4427739.5 | 2504934.5 | P67778 | PHB | 5147970 | 4675704 | 853663 | 30 |
| O35129 | PHB2 | prohibitin 2 | Cytoplasm | transcription regulator | 1.09E+07 | 1.45E+07 | 7680477 | | | | | | |
| O35129 | PHB2 | prohibitin 2 | Cytoplasm | transcription regulator | 1.47E+07 | 2.65E+07 | 9020012 | | | | | | |
| O35129 | PHB2 | prohibitin 2 | Cytoplasm | transcription regulator | 9428315 | 1.70E+07 | 6190091.5 | | | | | | |
| O35129 | PHB2 | prohibitin 2 | Cytoplasm | transcription regulator | 1.56E+07 | 3.63E+07 | 1.50E+07 | | | | | | |
| O35129 | PHB2 | prohibitin 2 | Cytoplasm | transcription regulator | 8838222 | 2.10E+07 | 1.03E+07 | | | | | | |
| O35129 | PHB2 | prohibitin 2 | Cytoplasm | transcription regulator | 9136771 | 1.76E+07 | 8520580 | | | | | | |
| O35129 | PHB2 | prohibitin 2 | Cytoplasm | transcription regulator | 702379.25 | 2639784.5 | 1106974.625 | | | | | | |
| O35129 | PHB2 | prohibitin 2 | Cytoplasm | transcription regulator | 3751420.25 | 1.02E+07 | 3703103.75 | | | | | | |
| O35129 | PHB2 | prohibitin 2 | Cytoplasm | transcription regulator | 1345875.5 | 2229718.75 | 1059431.625 | O35129 | PHB2 | 8268109 | 8378181 | 1612382 | 27 |
| Q61753 | PHGDH | phosphoglycerate dehydrogenase | Cytoplasm | enzyme | 3857154.5 | 2534085 | 3048487.75 | | | | | | |
| Q61753 | PHGDH | phosphoglycerate dehydrogenase | Cytoplasm | enzyme | 4249545 | 2315441.25 | 2718530.25 | | | | | | |
| Q61753 | PHGDH | phosphoglycerate dehydrogenase | Cytoplasm | enzyme | 3007548.5 | 1895621 | 2169660.5 | | | | | | |
| Q61753 | PHGDH | phosphoglycerate dehydrogenase | Cytoplasm | enzyme | 1987738.375 | 311656.281 | 43983.551 | | | | | | |
| Q61753 | PHGDH | phosphoglycerate dehydrogenase | Cytoplasm | enzyme | 2039672.125 | 1458103.75 | 1106030.125 | Q61753 | PHGDH | 3028332 | 1159866 | 299476 | 15 |
| Q8CBQ5 | PI4K2B | phosphatidylinositol 4-kinase type 2 beta | Cytoplasm | kinase | 2194691.75 | 2783948.25 | 3012716.75 | | | | | | |
| Q8CBQ5 | PI4K2B | phosphatidylinositol 4-kinase type 2 beta | Cytoplasm | kinase | 2289010.5 | 2938007 | 2928077.5 | | | | | | |
| Q8CBQ5 | PI4K2B | phosphatidylinositol 4-kinase type 2 beta | Cytoplasm | kinase | 1862509.5 | 2561089.75 | 3450703.75 | | | | | | |
| Q8CBQ5 | PI4K2B | phosphatidylinositol 4-kinase type 2 beta | Cytoplasm | kinase | 1282238 | 1285247.25 | 1465291.875 | Q8CBQ5 | PI4K2B | 1907112 | 731360 | 211125 | 12 |
| Q7M6Y3 | PICALM | phosphatidylinositol binding clathrin assembly protein | Cytoplasm | other | 1.86E+07 | 3.67E+07 | 2.20E+07 | | | | | | |
| Q7M6Y3 | PICALM | phosphatidylinositol binding clathrin assembly protein | Cytoplasm | other | 8917560 | 1.39E+07 | 8793663 | | | | | | |
| Q7M6Y3 | PICALM | phosphatidylinositol binding clathrin assembly protein | Cytoplasm | other | 3912121.5 | 4361515 | 3142290.75 | | | | | | |
| Q7M6Y3 | PICALM | phosphatidylinositol binding clathrin assembly protein | Cytoplasm | other | 668532.25 | 3283944 | 3023520.25 | Q7M6Y3 | PICALM | 8014553 | 10614855 | 3064245 | 12 |
| E2JF22 | PIEZO1 | piezo-type mechanosensitive ion channel component 1 | Cytoplasm | ion channel | 2.69E+07 | 2.66E+07 | 2.63E+07 | | | | | | |
| E2JF22 | PIEZO1 | piezo-type mechanosensitive ion channel component 1 | Cytoplasm | ion channel | 2.38E+07 | 2.58E+07 | 2.42E+07 | | | | | | |
| E2JF22 | PIEZO1 | piezo-type mechanosensitive ion channel component 1 | Cytoplasm | ion channel | 1.89E+07 | 1.47E+07 | 1.42E+07 | | | | | | |
| E2JF22 | PIEZO1 | piezo-type mechanosensitive ion channel component 1 | Cytoplasm | ion channel | 1.34E+07 | 1.42E+07 | 1.42E+07 | | | | | | |
| E2JF22 | PIEZO1 | piezo-type mechanosensitive ion channel component 1 | Cytoplasm | ion channel | 1.23E+07 | 1.27E+07 | 9660031 | | | | | | |
| E2JF22 | PIEZO1 | piezo-type mechanosensitive ion channel component 1 | Cytoplasm | ion channel | 1.23E+07 | 1.48E+07 | 1.15E+07 | | | | | | |
| E2JF22 | PIEZO1 | piezo-type mechanosensitive ion channel component 1 | Cytoplasm | ion channel | 1.33E+07 | 1.25E+07 | 1.31E+07 | | | | | | |
| E2JF22 | PIEZO1 | piezo-type mechanosensitive ion channel component 1 | Cytoplasm | ion channel | 1.34E+07 | 1.05E+07 | 1.07E+07 | | | | | | |
| E2JF22 | PIEZO1 | piezo-type mechanosensitive ion channel component 1 | Cytoplasm | ion channel | 1.05E+07 | 1.03E+07 | 1.24E+07 | | | | | | |
| E2JF22 | PIEZO1 | piezo-type mechanosensitive ion channel component 1 | Cytoplasm | ion channel | 1.18E+07 | 1.34E+07 | 1.38E+07 | | | | | | |
| E2JF22 | PIEZO1 | piezo-type mechanosensitive ion channel component 1 | Cytoplasm | ion channel | 1.08E+07 | 1.03E+07 | 1.03E+07 | | | | | | |
| E2JF22 | PIEZO1 | piezo-type mechanosensitive ion channel component 1 | Cytoplasm | ion channel | 6654048 | 6230126 | 6975019.5 | | | | | | |
| E2JF22 | PIEZO1 | piezo-type mechanosensitive ion channel component 1 | Cytoplasm | ion channel | 7289950 | 7394633 | 7033708 | | | | | | |
| E2JF22 | PIEZO1 | piezo-type mechanosensitive ion channel component 1 | Cytoplasm | ion channel | 6502536 | 6702982.5 | 6335448 | | | | | | |
| E2JF22 | PIEZO1 | piezo-type mechanosensitive ion channel component 1 | Cytoplasm | ion channel | 6037761.5 | 6885015.5 | 8030252 | | | | | | |
| E2JF22 | PIEZO1 | piezo-type mechanosensitive ion channel component 1 | Cytoplasm | ion channel | 5580543 | 4975355 | 5173818.5 | | | | | | |
| E2JF22 | PIEZO1 | piezo-type mechanosensitive ion channel component 1 | Cytoplasm | ion channel | 5546945.5 | 4100223.5 | 7949657 | | | | | | |
| E2JF22 | PIEZO1 | piezo-type mechanosensitive ion channel component 1 | Cytoplasm | ion channel | 5584213 | 4666388.5 | 5783775.5 | | | | | | |
| E2JF22 | PIEZO1 | piezo-type mechanosensitive ion channel component 1 | Cytoplasm | ion channel | 4224649.5 | 4271306.5 | 4934803.5 | | | | | | |
| E2JF22 | PIEZO1 | piezo-type mechanosensitive ion channel component 1 | Cytoplasm | ion channel | 3629752 | 4937545 | 3972879 | | | | | | |
| E2JF22 | PIEZO1 | piezo-type mechanosensitive ion channel component 1 | Cytoplasm | ion channel | 4390303 | 4646514 | 6136553 | | | | | | |
| E2JF22 | PIEZO1 | piezo-type mechanosensitive ion channel component 1 | Cytoplasm | ion channel | 2807147.25 | 2718545.75 | 3508714 | | | | | | |
| E2JF22 | PIEZO1 | piezo-type mechanosensitive ion channel component 1 | Cytoplasm | ion channel | 2029185.875 | 2091101.5 | 1392055.75 | | | | | | |
| E2JF22 | PIEZO1 | piezo-type mechanosensitive ion channel component 1 | Cytoplasm | ion channel | 1697363.25 | 1605881 | 1147108.625 | | | | | | |
| E2JF22 | PIEZO1 | piezo-type mechanosensitive ion channel component 1 | Cytoplasm | ion channel | 1467737.5 | 980301.312 | 1120392.75 | | | | | | |
| E2JF22 | PIEZO1 | piezo-type mechanosensitive ion channel component 1 | Cytoplasm | ion channel | 1965206.125 | 1571760.375 | 1463025.375 | | | | | | |
| E2JF22 | PIEZO1 | piezo-type mechanosensitive ion channel component 1 | Cytoplasm | ion channel | 291642.625 | 1076361.125 | 788825.375 | E2JF22 | PIEZO1 | 8622185 | 6556009 | 728445 | 81 |
| P06803 | PIM1 | Pim-1 proto-oncogene, serine/threonine kinase | Cytoplasm | kinase | 2.59E+08 | 2.33E+08 | 2.16E+08 | P06803 | PIM1 | 259200000 | 21934448 | 12663859 | 3 |
| O70172 | PIP4K2A | phosphatidylinositol-5-phosphate 4-kinase, type II, alpha | Cytoplasm | kinase | 1.67E+07 | 1.10E+07 | 1.72E+07 | | | | | | |
| O70172 | PIP4K2A | phosphatidylinositol-5-phosphate 4-kinase, type II, alpha | Cytoplasm | kinase | 4794786.5 | 3808260.75 | 3113118.5 | | | | | | |
| O70172 | PIP4K2A | phosphatidylinositol-5-phosphate 4-kinase, type II, alpha | Cytoplasm | kinase | 5493226.5 | 3655231 | 4117507.5 | | | | | | |
| O70172 | PIP4K2A | phosphatidylinositol-5-phosphate 4-kinase, type II, alpha | Cytoplasm | kinase | 3209834.75 | 3639054 | 3839205 | O70172 | PIP4K2A | 7546962 | 5235377 | 1511323 | 12 |
| Q80XI4 | PIP4K2B | phosphatidylinositol-5-phosphate 4-kinase, type II, beta | Cytoplasm | kinase | 7435815 | 6449369.5 | 5852729.5 | Q80XI4 | PIP4K2B | 7435815 | 799501 | 461592 | 3 |
| P70182 | PIP5K1A | phosphatidylinositol-4-phosphate 5-kinase, type I, alpha | Cytoplasm | kinase | 1.76E+07 | 1.48E+07 | 1.43E+07 | | | | | | |
| P70182 | PIP5K1A | phosphatidylinositol-4-phosphate 5-kinase, type I, alpha | Cytoplasm | kinase | 1.07E+07 | 9652477 | 1.04E+07 | | | | | | |
| P70182 | PIP5K1A | phosphatidylinositol-4-phosphate 5-kinase, type I, alpha | Cytoplasm | kinase | 7046853.5 | 6963177 | 8791315 | | | | | | |
| P70182 | PIP5K1A | phosphatidylinositol-4-phosphate 5-kinase, type I, alpha | Cytoplasm | kinase | 7913090 | 6475378 | 7500836.5 | | | | | | |
| P70182 | PIP5K1A | phosphatidylinositol-4-phosphate 5-kinase, type I, alpha | Cytoplasm | kinase | 3732920.25 | 1.01E+07 | 8417601 | | | | | | |
| P70182 | PIP5K1A | phosphatidylinositol-4-phosphate 5-kinase, type I, alpha | Cytoplasm | kinase | 6453303 | 6566470 | 7758274 | | | | | | |
| P70182 | PIP5K1A | phosphatidylinositol-4-phosphate 5-kinase, type I, alpha | Cytoplasm | kinase | 4178789.25 | 3518863.25 | 2906659.5 | | | | | | |
| P70182 | PIP5K1A | phosphatidylinositol-4-phosphate 5-kinase, type I, alpha | Cytoplasm | kinase | 3319717.5 | 2548185 | 2986010.25 | | | | | | |
| P70182 | PIP5K1A | phosphatidylinositol-4-phosphate 5-kinase, type I, alpha | Cytoplasm | kinase | 4855381.5 | 4255696 | 4827227 | | | | | | |
| P70182 | PIP5K1A | phosphatidylinositol-4-phosphate 5-kinase, type I, alpha | Cytoplasm | kinase | 300547.156 | 454835.969 | 578601.188 | P70182 | PIP5K1A | 6605060 | 4210746 | 768774 | 30 |
| Q8K411 | PITRM1 | pitrilysin metallopeptidase 1 | Cytoplasm | peptidase | 6327356 | 1.60E+07 | 7442650 | | | | | | |
| Q8K411 | PITRM1 | pitrilysin metallopeptidase 1 | Cytoplasm | peptidase | 7102205 | 1.33E+07 | 5426048.5 | | | | | | |
| Q8K411 | PITRM1 | pitrilysin metallopeptidase 1 | Cytoplasm | peptidase | 5173245.5 | 9609900 | 2948087 | | | | | | |
| Q8K411 | PITRM1 | pitrilysin metallopeptidase 1 | Cytoplasm | peptidase | 3786463.5 | 7162725 | 3330852.5 | | | | | | |
| Q8K411 | PITRM1 | pitrilysin metallopeptidase 1 | Cytoplasm | peptidase | 5431213 | 6281674.5 | 2375458.25 | | | | | | |
| Q8K411 | PITRM1 | pitrilysin metallopeptidase 1 | Cytoplasm | peptidase | 1070166.625 | 1645663 | 612988.875 | | | | | | |
| Q8K411 | PITRM1 | pitrilysin metallopeptidase 1 | Cytoplasm | peptidase | 571906.562 | 1079065.75 | 166056.406 | Q8K411 | PITRM1 | 4208937 | 4184180 | 913063 | 21 |
| Q80ZA4 | PKHD1L1 | polycystic kidney and hepatic disease 1 (autosomal recessive)-like 1 | Extracellular Space | other | 9338031 | 9806967 | 9920400 | Q80ZA4 | PKHD1L1 | 9338031 | 308740 | 178251 | 3 |
| P53657 | PKLR | pyruvate kinase, liver and RBC | Cytoplasm | kinase | 2.42E+07 | 1.78E+07 | 1.89E+07 | | | | | | |
| P53657 | PKLR | pyruvate kinase, liver and RBC | Cytoplasm | kinase | 1.11E+07 | 8456857 | 9139099 | | | | | | |

| | | | | | | | | | | | | | |
|---|---|---|---|---|---|---|---|---|---|---|---|---|---|
| P53657 | PKLR | pyruvate kinase, liver and RBC | Cytoplasm | kinase | 1.08E+07 | 9752366 | 1.06E+07 | | | | | | |
| P53657 | PKLR | pyruvate kinase, liver and RBC | Cytoplasm | kinase | 8846189 | 8092606 | 7386552.5 | | | | | | |
| P53657 | PKLR | pyruvate kinase, liver and RBC | Cytoplasm | kinase | 7184030.5 | 5796352.5 | 5341313.5 | | | | | | |
| P53657 | PKLR | pyruvate kinase, liver and RBC | Cytoplasm | kinase | 6778115 | 4501584 | 4471178.5 | | | | | | |
| P53657 | PKLR | pyruvate kinase, liver and RBC | Cytoplasm | kinase | 4910714 | 2435654.5 | 1314952 | | | | | | |
| P53657 | PKLR | pyruvate kinase, liver and RBC | Cytoplasm | kinase | 3540734.75 | 2911264 | 3334861 | P53657 | PKLR | 9661223 | 5501547 | 1122999 | 24 |
| P52480 | PKM | pyruvate kinase, muscle | Cytoplasm | kinase | 3141394.25 | 3932867.75 | 4470065.5 | P52480 | PKM | 3141394 | 668379 | 385889 | 3 |
| P27612 | PLAA | phospholipase A2-activating protein | Cytoplasm | other | 1095555.125 | 673596.562 | 684463.875 | P27612 | PLAA | 1095555 | 240542 | 138877 | 3 |
| Q9JHK5 | PLEK | pleckstrin | Cytoplasm | other | 212248 | 213433.438 | 529192.938 | Q9JHK5 | PLEK | 212248 | 182647 | 105451 | 3 |
| P20918 | PLG | plasminogen | Extracellular Space | peptidase | 1799944.25 | 2355121.5 | 739854.75 | | | | | | |
| P20918 | PLG | plasminogen | Extracellular Space | peptidase | 170500.797 | 939548.75 | 6324.307 | | | | | | |
| P20918 | PLG | plasminogen | Extracellular Space | peptidase | 369035.406 | 1248581.125 | 101515.914 | P20918 | PLG | 779827 | 814797 | 271599 | 9 |
| Q9R1Q7 | PLP2 | proteolipid protein 2 (colonic epithelium-enriched) | Cytoplasm | transporter | 1.45E+07 | 1.36E+07 | 1.47E+07 | Q9R1Q7 | PLP2 | 14490000 | 582866 | 336518 | 3 |
| Q9JJ00 | PLSCR1 | phospholipid scramblase 1 | Plasma Membrane | enzyme | 1.71E+07 | 1.91E+07 | 2.51E+07 | | | | | | |
| Q9JJ00 | PLSCR1 | phospholipid scramblase 1 | Plasma Membrane | enzyme | 1.20E+07 | 1.37E+07 | 1.43E+07 | | | | | | |
| Q9JJ00 | PLSCR1 | phospholipid scramblase 1 | Plasma Membrane | enzyme | 7405587 | 7401589 | 8038442 | Q9JJ00 | PLSCR1 | 12161862 | 5972842 | 1990947 | 9 |
| Q9JIZ9 | PLSCR3 | phospholipid scramblase 3 | Plasma Membrane | enzyme | 7359154 | 8475467 | 1.05E+07 | | | | | | |
| Q9JIZ9 | PLSCR3 | phospholipid scramblase 3 | Plasma Membrane | enzyme | 1.16E+07 | 8602277 | 1.11E+07 | | | | | | |
| Q9JIZ9 | PLSCR3 | phospholipid scramblase 3 | Plasma Membrane | enzyme | 4434478 | 3394460.25 | 4201351 | | | | | | |
| Q9JIZ9 | PLSCR3 | phospholipid scramblase 3 | Plasma Membrane | enzyme | 2390317.25 | 1897901.25 | 2454293 | Q9JIZ9 | PLSCR3 | 6445987 | 3640475 | 1050915 | 12 |
| P23492 | PNP | purine nucleoside phosphorylase | Nucleus | enzyme | 4.83E+07 | 2.90E+07 | 4.32E+07 | | | | | | |
| P23492 | PNP | purine nucleoside phosphorylase | Nucleus | enzyme | 1.30E+07 | 8670238 | 9650806 | | | | | | |
| P23492 | PNP | purine nucleoside phosphorylase | Nucleus | enzyme | 1.60E+07 | 1.06E+07 | 1.38E+07 | | | | | | |
| P23492 | PNP | purine nucleoside phosphorylase | Nucleus | enzyme | 1.19E+07 | 5801598 | 8294354 | | | | | | |
| P23492 | PNP | purine nucleoside phosphorylase | Nucleus | enzyme | 8895876 | 6118043 | 5467863 | | | | | | |
| P23492 | PNP | purine nucleoside phosphorylase | Nucleus | enzyme | 3695037.75 | 2198570.75 | 1763313.125 | P23492 | PNP | 16961819 | 13228475 | 3117982 | 18 |
| Q61469 | Ppap2a | phosphatidic acid phosphatase type 2A | Plasma Membrane | phosphatase | 6.52E+07 | 5.50E+07 | 3.63E+07 | | | | | | |
| Q61469 | Ppap2a | phosphatidic acid phosphatase type 2A | Plasma Membrane | phosphatase | 3.49E+07 | 3.45E+07 | 3.34E+07 | | | | | | |
| Q61469 | Ppap2a | phosphatidic acid phosphatase type 2A | Plasma Membrane | phosphatase | 4.37E+07 | 7.02E+07 | 8.33E+07 | | | | | | |
| Q61469 | Ppap2a | phosphatidic acid phosphatase type 2A | Plasma Membrane | phosphatase | 3.25E+07 | 3.34E+07 | 4.36E+07 | | | | | | |
| Q61469 | Ppap2a | phosphatidic acid phosphatase type 2A | Plasma Membrane | phosphatase | 5613075 | 8538303 | 1.05E+07 | Q61469 | Ppap2a | 36382615 | 22230108 | 5739789 | 15 |
| P17742 | PPIA | peptidylprolyl isomerase A (cyclophilin A) | Cytoplasm | enzyme | 4530042 | 7874376.5 | 1.49E+07 | | | | | | |
| P17742 | PPIA | peptidylprolyl isomerase A (cyclophilin A) | Cytoplasm | enzyme | 1656342.625 | 4117050 | 6665097.5 | | | | | | |
| P17742 | PPIA | peptidylprolyl isomerase A (cyclophilin A) | Cytoplasm | enzyme | 2133076.5 | 6034369 | 1.03E+07 | | | | | | |
| P17742 | PPIA | peptidylprolyl isomerase A (cyclophilin A) | Cytoplasm | enzyme | 1195396.375 | 2210366 | 1738714.25 | | | | | | |
| P17742 | PPIA | peptidylprolyl isomerase A (cyclophilin A) | Cytoplasm | enzyme | 1188912.5 | 5023734 | 8207870.5 | | | | | | |
| P17742 | PPIA | peptidylprolyl isomerase A (cyclophilin A) | Cytoplasm | enzyme | 340174.969 | 494826.219 | 202399.828 | | | | | | |
| P17742 | PPIA | peptidylprolyl isomerase A (cyclophilin A) | Cytoplasm | enzyme | 386903.75 | 1302617.25 | 1694794.75 | P17742 | PPIA | 1632978 | 3889392 | 848735 | 21 |
| P24369 | PPIB | peptidylprolyl isomerase B (cyclophilin B) | Cytoplasm | enzyme | 6.58E+07 | 6.50E+07 | 8.99E+07 | | | | | | |
| P24369 | PPIB | peptidylprolyl isomerase B (cyclophilin B) | Cytoplasm | enzyme | 3.49E+07 | 3.83E+07 | 3.78E+07 | | | | | | |
| P24369 | PPIB | peptidylprolyl isomerase B (cyclophilin B) | Cytoplasm | enzyme | 1.50E+07 | 2.36E+07 | 2.91E+07 | | | | | | |
| P24369 | PPIB | peptidylprolyl isomerase B (cyclophilin B) | Cytoplasm | enzyme | 2.15E+07 | 2.48E+07 | 2.56E+07 | | | | | | |
| P24369 | PPIB | peptidylprolyl isomerase B (cyclophilin B) | Cytoplasm | enzyme | 2.14E+07 | 2.00E+07 | 2.31E+07 | | | | | | |
| P24369 | PPIB | peptidylprolyl isomerase B (cyclophilin B) | Cytoplasm | enzyme | 1.62E+07 | 2.17E+07 | 2.18E+07 | | | | | | |
| P24369 | PPIB | peptidylprolyl isomerase B (cyclophilin B) | Cytoplasm | enzyme | 1.27E+07 | 1.43E+07 | 1.94E+07 | | | | | | |
| P24369 | PPIB | peptidylprolyl isomerase B (cyclophilin B) | Cytoplasm | enzyme | 1.72E+07 | 1.22E+07 | 1.20E+07 | | | | | | |
| P24369 | PPIB | peptidylprolyl isomerase B (cyclophilin B) | Cytoplasm | enzyme | 1.29E+07 | 9308533 | 9148428 | | | | | | |
| P24369 | PPIB | peptidylprolyl isomerase B (cyclophilin B) | Cytoplasm | enzyme | 1.08E+07 | 9508158 | 1.22E+07 | | | | | | |
| P24369 | PPIB | peptidylprolyl isomerase B (cyclophilin B) | Cytoplasm | enzyme | 9648348 | 9661796 | 1.31E+07 | | | | | | |
| P24369 | PPIB | peptidylprolyl isomerase B (cyclophilin B) | Cytoplasm | enzyme | 3943800.25 | 2744685 | 1843008.25 | P24369 | PPIB | 20154346 | 18404798 | 3067466 | 36 |
| Q99KR7 | PPIF | peptidylprolyl isomerase F | Cytoplasm | enzyme | 1177891.75 | 2641407.25 | 560974.062 | | | | | | |
| Q99KR7 | PPIF | peptidylprolyl isomerase F | Cytoplasm | enzyme | 455276.219 | 2111235 | 3617061 | Q99KR7 | PPIF | 816584 | 1251218 | 510808 | 6 |
| P51175 | PPOX | protoporphyrinogen oxidase | Cytoplasm | enzyme | 2.66E+07 | 5.34E+07 | 2.36E+07 | | | | | | |
| P51175 | PPOX | protoporphyrinogen oxidase | Cytoplasm | enzyme | 2.74E+07 | 5.62E+07 | 2.45E+07 | | | | | | |
| P51175 | PPOX | protoporphyrinogen oxidase | Cytoplasm | enzyme | 1.15E+07 | 2.30E+07 | 7765923.5 | | | | | | |
| P51175 | PPOX | protoporphyrinogen oxidase | Cytoplasm | enzyme | 7457971.5 | 1.98E+07 | 6490585 | | | | | | |
| P51175 | PPOX | protoporphyrinogen oxidase | Cytoplasm | enzyme | 4858452 | 1.73E+07 | 5556934 | | | | | | |
| P51175 | PPOX | protoporphyrinogen oxidase | Cytoplasm | enzyme | 7814609 | 1.47E+07 | 5342338.5 | | | | | | |
| P51175 | PPOX | protoporphyrinogen oxidase | Cytoplasm | enzyme | 6577390 | 1.44E+07 | 5746278.5 | | | | | | |
| P51175 | PPOX | protoporphyrinogen oxidase | Cytoplasm | enzyme | 7692955 | 1.33E+07 | 4042756.75 | | | | | | |
| P51175 | PPOX | protoporphyrinogen oxidase | Cytoplasm | enzyme | 1.02E+07 | 1.95E+07 | 4376045 | | | | | | |
| P51175 | PPOX | protoporphyrinogen oxidase | Cytoplasm | enzyme | 4996027.5 | 7873978.5 | 2206654 | | | | | | |
| P51175 | PPOX | protoporphyrinogen oxidase | Cytoplasm | enzyme | 4407437.5 | 7655245.5 | 2309097.25 | | | | | | |
| P51175 | PPOX | protoporphyrinogen oxidase | Cytoplasm | enzyme | 3843214 | 8520962 | 2275682.25 | | | | | | |
| P51175 | PPOX | protoporphyrinogen oxidase | Cytoplasm | enzyme | 3808119 | 6629893.5 | 3121900.75 | P51175 | PPOX | 9776629 | 12342102 | 1976318 | 39 |
| P62137 | PPP1CA | protein phosphatase 1, catalytic subunit, alpha isozyme | Cytoplasm | phosphatase | 2704472.75 | 4075247.25 | 3034938.75 | | | | | | |
| P62137 | PPP1CA | protein phosphatase 1, catalytic subunit, alpha isozyme | Cytoplasm | phosphatase | 2450598 | 4499372 | 4770403.5 | | | | | | |
| P62137 | PPP1CA | protein phosphatase 1, catalytic subunit, alpha isozyme | Cytoplasm | phosphatase | 2288276 | 2719127.25 | 2695481 | | | | | | |
| P62137 | PPP1CA | protein phosphatase 1, catalytic subunit, alpha isozyme | Cytoplasm | phosphatase | 1934305.25 | 2486210.75 | 2665209.25 | | | | | | |
| P62137 | PPP1CA | protein phosphatase 1, catalytic subunit, alpha isozyme | Cytoplasm | phosphatase | 1353190.25 | 2358035.75 | 2485990.25 | | | | | | |
| P62137 | PPP1CA | protein phosphatase 1, catalytic subunit, alpha isozyme | Cytoplasm | phosphatase | 1103354.125 | 909869.75 | 930574.688 | P62137 | PPP1CA | 1972366 | 1103693 | 260143 | 18 |
| Q62415 | PPP1R13B | protein phosphatase 1, regulatory subunit 13B | Cytoplasm | phosphatase | 3776049.25 | 4072407 | 5595056.5 | Q62415 | PPP1R13B | 3776049 | 975968 | 563475 | 3 |
| Q3TDD9 | PPP1R21 | protein phosphatase 1, regulatory subunit 21 | Cytoplasm | other | 9290315 | 1.19E+07 | 1.66E+07 | Q3TDD9 | PPP1R21 | 9290315 | 3706933 | 2140199 | 3 |
| P63330 | PPP2CA | protein phosphatase 2, catalytic subunit, alpha isozyme | Cytoplasm | phosphatase | 4043775.5 | 3002035.5 | 3807099.5 | | | | | | |
| P63330 | PPP2CA | protein phosphatase 2, catalytic subunit, alpha isozyme | Cytoplasm | phosphatase | 4801135 | 3076575.25 | 1963158.25 | | | | | | |
| P63330 | PPP2CA | protein phosphatase 2, catalytic subunit, alpha isozyme | Cytoplasm | phosphatase | 3685186.25 | 2576637 | 2447944.25 | | | | | | |
| P63330 | PPP2CA | protein phosphatase 2, catalytic subunit, alpha isozyme | Cytoplasm | phosphatase | 2019480.875 | 1266601.625 | 1650609.25 | | | | | | |
| P63330 | PPP2CA | protein phosphatase 2, catalytic subunit, alpha isozyme | Cytoplasm | phosphatase | 1483186.875 | 1970992.375 | 1259155.375 | P63330 | PPP2CA | 3206553 | 1096719 | 283172 | 15 |
| Q76MZ3 | PPP2R1A | protein phosphatase 2, regulatory subunit A, alpha | Cytoplasm | phosphatase | 9574879 | 6122385 | 3946420.75 | | | | | | |
| Q76MZ3 | PPP2R1A | protein phosphatase 2, regulatory subunit A, alpha | Cytoplasm | phosphatase | 2359181 | 1728656.875 | 2591975.5 | | | | | | |
| Q76MZ3 | PPP2R1A | protein phosphatase 2, regulatory subunit A, alpha | Cytoplasm | phosphatase | 2181168.25 | 2123751 | 2295815.5 | | | | | | |
| Q76MZ3 | PPP2R1A | protein phosphatase 2, regulatory subunit A, alpha | Cytoplasm | phosphatase | 4165114.25 | 3916789.75 | 3642962 | | | | | | |
| Q76MZ3 | PPP2R1A | protein phosphatase 2, regulatory subunit A, alpha | Cytoplasm | phosphatase | 3221393.5 | 2282365.25 | 1446521.25 | | | | | | |
| Q76MZ3 | PPP2R1A | protein phosphatase 2, regulatory subunit A, alpha | Cytoplasm | phosphatase | 2665062.75 | 1584392 | 1824160.625 | | | | | | |
| Q76MZ3 | PPP2R1A | protein phosphatase 2, regulatory subunit A, alpha | Cytoplasm | phosphatase | 1529233.75 | 1009962.188 | 1400335.375 | Q76MZ3 | PPP2R1A | 3670862 | 1945247 | 424488 | 21 |
| Q6P1F6 | PPP2R2A | protein phosphatase 2, regulatory subunit B, alpha | Cytoplasm | phosphatase | 2386374.25 | 876833.562 | 1181249 | Q6P1F6 | PPP2R2A | 2386374 | 798301 | 460899 | 3 |
| Q9CQR6 | PPP6C | protein phosphatase 6, catalytic subunit | Nucleus | phosphatase | 6232103 | 5945198 | 5774764.5 | | | | | | |
| Q9CQR6 | PPP6C | protein phosphatase 6, catalytic subunit | Nucleus | phosphatase | 5243476.5 | 3914635.25 | 2480697.5 | Q9CQR6 | PPP6C | 5737790 | 1453824 | 593521 | 6 |
| Q922D4 | PPP6R3 | protein phosphatase 6, regulatory subunit 3 | Cytoplasm | other | 1.54E+07 | 1.10E+07 | 8651564 | | | | | | |
| Q922D4 | PPP6R3 | protein phosphatase 6, regulatory subunit 3 | Cytoplasm | other | 8986690 | 9247213 | 5042367 | | | | | | |
| Q922D4 | PPP6R3 | protein phosphatase 6, regulatory subunit 3 | Cytoplasm | other | 7036168 | 5369876 | 4993148.5 | | | | | | |

| | | | | | | | | | | | | | |
|---|---|---|---|---|---|---|---|---|---|---|---|---|---|
| Q922D4 | PPP6R3 | protein phosphatase 6, regulatory subunit 3 | Cytoplasm | other | 7104787 | 5547125.5 | 5166462 | | | | | | |
| Q922D4 | PPP6R3 | protein phosphatase 6, regulatory subunit 3 | Cytoplasm | other | 5118697 | 3478461.75 | 3718798.5 | | | | | | |
| Q922D4 | PPP6R3 | protein phosphatase 6, regulatory subunit 3 | Cytoplasm | other | 6302552.5 | 4528646 | 3987776.5 | | | | | | |
| Q922D4 | PPP6R3 | protein phosphatase 6, regulatory subunit 3 | Cytoplasm | other | 5566784.5 | 4007613 | 5928430.5 | | | | | | |
| Q922D4 | PPP6R3 | protein phosphatase 6, regulatory subunit 3 | Cytoplasm | other | 4908884.5 | 3704588.5 | 3477595.25 | | | | | | |
| Q922D4 | PPP6R3 | protein phosphatase 6, regulatory subunit 3 | Cytoplasm | other | 5029094 | 3224528.5 | 3579792.75 | | | | | | |
| Q922D4 | PPP6R3 | protein phosphatase 6, regulatory subunit 3 | Cytoplasm | other | 3916518.75 | 3197115 | 3140857.5 | | | | | | |
| Q922D4 | PPP6R3 | protein phosphatase 6, regulatory subunit 3 | Cytoplasm | other | 2780512.25 | 1816626.25 | 2116525.25 | | | | | | |
| Q922D4 | PPP6R3 | protein phosphatase 6, regulatory subunit 3 | Cytoplasm | other | 1310908 | 1303103.5 | 682675.375 | Q922D4 | PPP6R3 | 6118466 | 2905870 | 484312 | 36 |
| Q7TMR0 | PRCP | prolylcarboxypeptidase (angiotensinase C) | Cytoplasm | peptidase | 2704089.5 | 7391970.5 | 3736292.25 | Q7TMR0 | PRCP | 2704090 | 2463252 | 1422159 | 3 |
| P35700 | PRDX1 | peroxiredoxin 1 | Cytoplasm | enzyme | 2.45E+07 | 2.49E+07 | 2.52E+07 | | | | | | |
| P35700 | PRDX1 | peroxiredoxin 1 | Cytoplasm | enzyme | 3.30E+07 | 2.30E+07 | 2.49E+07 | | | | | | |
| P35700 | PRDX1 | peroxiredoxin 1 | Cytoplasm | enzyme | 2.07E+07 | 1.56E+07 | 1.54E+07 | | | | | | |
| P35700 | PRDX1 | peroxiredoxin 1 | Cytoplasm | enzyme | 3.24E+07 | 2.10E+07 | 2.31E+07 | | | | | | |
| P35700 | PRDX1 | peroxiredoxin 1 | Cytoplasm | enzyme | 2.30E+07 | 1.79E+07 | 2.04E+07 | | | | | | |
| P35700 | PRDX1 | peroxiredoxin 1 | Cytoplasm | enzyme | 1.48E+07 | 1.24E+07 | 1.57E+07 | | | | | | |
| P35700 | PRDX1 | peroxiredoxin 1 | Cytoplasm | enzyme | 1.40E+07 | 8561464 | 6585204.5 | | | | | | |
| P35700 | PRDX1 | peroxiredoxin 1 | Cytoplasm | enzyme | 9633793 | 5995394.5 | 8066521.5 | | | | | | |
| P35700 | PRDX1 | peroxiredoxin 1 | Cytoplasm | enzyme | 4338631.5 | 6805016 | 1.41E+07 | | | | | | |
| P35700 | PRDX1 | peroxiredoxin 1 | Cytoplasm | enzyme | 3649240.25 | 3427727.25 | 1382144.625 | | | | | | |
| P35700 | PRDX1 | peroxiredoxin 1 | Cytoplasm | enzyme | 994441 | 1551495.25 | 2550580 | P35700 | PRDX1 | 16456010 | 9269724 | 1613652 | 33 |
| Q61171 | PRDX2 | peroxiredoxin 2 | Cytoplasm | enzyme | 5.10E+08 | 3.37E+08 | 4.43E+08 | | | | | | |
| Q61171 | PRDX2 | peroxiredoxin 2 | Cytoplasm | enzyme | 2.91E+08 | 1.97E+08 | 2.41E+08 | | | | | | |
| Q61171 | PRDX2 | peroxiredoxin 2 | Cytoplasm | enzyme | 2.69E+08 | 2.11E+08 | 2.24E+08 | | | | | | |
| Q61171 | PRDX2 | peroxiredoxin 2 | Cytoplasm | enzyme | 1.38E+08 | 1.92E+08 | 3.52E+08 | | | | | | |
| Q61171 | PRDX2 | peroxiredoxin 2 | Cytoplasm | enzyme | 1.32E+08 | 8.78E+07 | 1.10E+08 | | | | | | |
| Q61171 | PRDX2 | peroxiredoxin 2 | Cytoplasm | enzyme | 1.20E+08 | 8.58E+07 | 1.15E+08 | | | | | | |
| Q61171 | PRDX2 | peroxiredoxin 2 | Cytoplasm | enzyme | 1.11E+08 | 1.00E+08 | 1.57E+08 | | | | | | |
| Q61171 | PRDX2 | peroxiredoxin 2 | Cytoplasm | enzyme | 5.95E+07 | 6.85E+07 | 9.91E+07 | | | | | | |
| Q61171 | PRDX2 | peroxiredoxin 2 | Cytoplasm | enzyme | 5.03E+07 | 3.57E+07 | 5.22E+07 | | | | | | |
| Q61171 | PRDX2 | peroxiredoxin 2 | Cytoplasm | enzyme | 1.45E+07 | 1.37E+07 | 9375423 | | | | | | |
| Q61171 | PRDX2 | peroxiredoxin 2 | Cytoplasm | enzyme | 2.08E+07 | 2.35E+07 | 3.08E+07 | | | | | | |
| Q61171 | PRDX2 | peroxiredoxin 2 | Cytoplasm | enzyme | 4.31E+07 | 9601556 | 8893479 | | | | | | |
| Q61171 | PRDX2 | peroxiredoxin 2 | Cytoplasm | enzyme | 1.40E+07 | 2054810.625 | 476983.062 | | | | | | |
| Q61171 | PRDX2 | peroxiredoxin 2 | Cytoplasm | enzyme | 1.10E+07 | 2826219 | 1985292 | Q61171 | PRDX2 | 127511429 | 126174830 | 19469199 | 42 |
| P20108 | PRDX3 | peroxiredoxin 3 | Cytoplasm | enzyme | 7.18E+07 | 1.99E+08 | 7.85E+07 | | | | | | |
| P20108 | PRDX3 | peroxiredoxin 3 | Cytoplasm | enzyme | 6.49E+07 | 9.44E+07 | 5.50E+07 | | | | | | |
| P20108 | PRDX3 | peroxiredoxin 3 | Cytoplasm | enzyme | 3.74E+07 | 6.32E+07 | 3.86E+07 | | | | | | |
| P20108 | PRDX3 | peroxiredoxin 3 | Cytoplasm | enzyme | 1.93E+07 | 3.68E+07 | 1.54E+07 | | | | | | |
| P20108 | PRDX3 | peroxiredoxin 3 | Cytoplasm | enzyme | 2.27E+07 | 4.17E+07 | 2.54E+07 | | | | | | |
| P20108 | PRDX3 | peroxiredoxin 3 | Cytoplasm | enzyme | 2.52E+07 | 6207214 | 285143.781 | | | | | | |
| P20108 | PRDX3 | peroxiredoxin 3 | Cytoplasm | enzyme | 4077772.5 | 6842465 | 2727468.75 | P20108 | PRDX3 | 35055396 | 44779605 | 9771711 | 21 |
| O08807 | PRDX4 | peroxiredoxin 4 | Cytoplasm | enzyme | 7675957 | 1.12E+07 | 7012721 | | | | | | |
| O08807 | PRDX4 | peroxiredoxin 4 | Cytoplasm | enzyme | 5800812 | 4703155.5 | 1953506.875 | | | | | | |
| O08807 | PRDX4 | peroxiredoxin 4 | Cytoplasm | enzyme | 938004.812 | 1645037.625 | 3154741.5 | O08807 | PRDX4 | 4804925 | 3362996 | 1120999 | 9 |
| P99029 | PRDX5 | peroxiredoxin 5 | Cytoplasm | enzyme | 1492612.5 | 2847337 | 3093711.75 | P99029 | PRDX5 | 1492613 | 862119 | 497745 | 3 |
| Q61878 | PRG2 | proteoglycan 2, bone marrow (natural killer cell activator, eosinophil granule major basic protein) | Extracellular Space | other | 95078.961 | 348702.469 | 222347.828 | Q61878 | PRG2 | 95079 | 126812 | 73215 | 3 |
| P05132 | PRKACA | protein kinase, cAMP-dependent, catalytic, alpha | Cytoplasm | kinase | 5089048.5 | 4253619 | 6088936.5 | P05132 | PRKACA | 5089049 | 918886 | 530519 | 3 |
| P20444 | PRKCA | protein kinase C, alpha | Cytoplasm | kinase | 84385.445 | 229969 | 547002.312 | P20444 | PRKCA | 84385 | 236544 | 136569 | 3 |
| O08795 | PRKCSH | protein kinase C substrate 80K-H | Cytoplasm | enzyme | 1.09E+07 | 1.16E+07 | 1.17E+07 | | | | | | |
| O08795 | PRKCSH | protein kinase C substrate 80K-H | Cytoplasm | enzyme | 8780058 | 8817790 | 8695084 | | | | | | |
| O08795 | PRKCSH | protein kinase C substrate 80K-H | Cytoplasm | enzyme | 6746191 | 7426559 | 7493051 | | | | | | |
| O08795 | PRKCSH | protein kinase C substrate 80K-H | Cytoplasm | enzyme | 6404626 | 6240061.5 | 7265051.5 | | | | | | |
| O08795 | PRKCSH | protein kinase C substrate 80K-H | Cytoplasm | enzyme | 4864285.5 | 5297490.5 | 6000708 | | | | | | |
| O08795 | PRKCSH | protein kinase C substrate 80K-H | Cytoplasm | enzyme | 5493874 | 4710830 | 5989762 | | | | | | |
| O08795 | PRKCSH | protein kinase C substrate 80K-H | Cytoplasm | enzyme | 4564614 | 4703833 | 6246995.5 | | | | | | |
| O08795 | PRKCSH | protein kinase C substrate 80K-H | Cytoplasm | enzyme | 3992254.75 | 4553619 | 5547181 | | | | | | |
| O08795 | PRKCSH | protein kinase C substrate 80K-H | Cytoplasm | enzyme | 2796542.25 | 2773767 | 2770128.25 | | | | | | |
| O08795 | PRKCSH | protein kinase C substrate 80K-H | Cytoplasm | enzyme | 3694258.5 | 2883867 | 2599633 | | | | | | |
| O08795 | PRKCSH | protein kinase C substrate 80K-H | Cytoplasm | enzyme | 1308183.5 | 495399.25 | 188616.391 | O08795 | PRKCSH | 5414990 | 2896330 | 504186 | 33 |
| Q8CIG8 | PRMT5 | protein arginine methyltransferase 5 | Cytoplasm | enzyme | 6757400 | 8059759.5 | 6696008.5 | | | | | | |
| Q8CIG8 | PRMT5 | protein arginine methyltransferase 5 | Cytoplasm | enzyme | 2202575.5 | 1324688.875 | 1693852.625 | Q8CIG8 | PRMT5 | 4479988 | 3027000 | 1235768 | 6 |
| P04925 | PRNP | prion protein | Plasma Membrane | other | 8166718.5 | 9446132 | 8718196 | P04925 | PRNP | 8166719 | 641732 | 370504 | 3 |
| Q9D7G0 | PRPS1 | phosphoribosyl pyrophosphate synthetase 1 | Cytoplasm | kinase | 1.31E+07 | 8954965 | 9279562 | | | | | | |
| Q9D7G0 | PRPS1 | phosphoribosyl pyrophosphate synthetase 1 | Cytoplasm | kinase | 1.10E+07 | 6719285 | 5770729.5 | | | | | | |
| Q9D7G0 | PRPS1 | phosphoribosyl pyrophosphate synthetase 1 | Cytoplasm | kinase | 6984446 | 5193010 | 3589116 | | | | | | |
| Q9D7G0 | PRPS1 | phosphoribosyl pyrophosphate synthetase 1 | Cytoplasm | kinase | 7209016 | 5266749.5 | 4378549.5 | | | | | | |
| Q9D7G0 | PRPS1 | phosphoribosyl pyrophosphate synthetase 1 | Cytoplasm | kinase | 3984457 | 3161440.5 | 3245455 | | | | | | |
| Q9D7G0 | PRPS1 | phosphoribosyl pyrophosphate synthetase 1 | Cytoplasm | kinase | 3198309.75 | 3199908.5 | 4149311.25 | Q9D7G0 | PRPS1 | 7584371 | 2937424 | 692358 | 18 |
| Q9D0M1 | PRPSAP1 | phosphoribosyl pyrophosphate synthetase-associated protein 1 | Extracellular Space | other | 7851605 | 6950075 | 7268019 | | | | | | |
| Q9D0M1 | PRPSAP1 | phosphoribosyl pyrophosphate synthetase-associated protein 1 | Extracellular Space | other | 4189851.25 | 3451733.5 | 3344107.25 | | | | | | |
| Q9D0M1 | PRPSAP1 | phosphoribosyl pyrophosphate synthetase-associated protein 1 | Extracellular Space | other | 3312471.75 | 1850718.25 | 2770454.75 | | | | | | |
| Q9D0M1 | PRPSAP1 | phosphoribosyl pyrophosphate synthetase-associated protein 1 | Extracellular Space | other | 3646002.25 | 2664509.5 | 1853455.75 | | | | | | |
| Q9D0M1 | PRPSAP1 | phosphoribosyl pyrophosphate synthetase-associated protein 1 | Extracellular Space | other | 2523176.75 | 1049240.125 | 1528566 | Q9D0M1 | PRPSAP1 | 4304621 | 2119633 | 547287 | 15 |
| Q8R574 | PRPSAP2 | phosphoribosyl pyrophosphate synthetase-associated protein 2 | Cytoplasm | other | 2368691.5 | 2220704.25 | 2831766.75 | | | | | | |
| Q8R574 | PRPSAP2 | phosphoribosyl pyrophosphate synthetase-associated protein 2 | Cytoplasm | other | 3530713 | 2993918.5 | 2489095.75 | Q8R574 | PRPSAP2 | 2949702 | 483448 | 197367 | 6 |
| Q61207 | PSAP | prosaposin | Extracellular Space | other | 2178099 | 5558671 | 2320005 | Q61207 | PSAP | 2178099 | 1912126 | 1103967 | 3 |
| P49769 | PSEN1 | presenilin 1 | Plasma Membrane | peptidase | 4807088 | 4368378 | 5013856.5 | | | | | | |
| P49769 | PSEN1 | presenilin 1 | Plasma Membrane | peptidase | 1239452.75 | 1828284.875 | 2834759.5 | P49769 | PSEN1 | 3023270 | 1610224 | 657371 | 6 |
| Q61144 | PSEN2 | presenilin 2 | Cytoplasm | peptidase | 1920675 | 1896864.5 | 1300244 | Q61144 | PSEN2 | 1920675 | 351534 | 202958 | 3 |
| Q91YA2 | PSKH1 | protein serine kinase H1 | Nucleus | kinase | 3362422.75 | 2869223.5 | 3439930.25 | Q91YA2 | PSKH1 | 3362423 | 309559 | 178724 | 3 |
| Q9R1P4 | PSMA1 | proteasome (prosome, macropain) subunit, alpha type, 1 | Cytoplasm | peptidase | 5.05E+07 | 4.13E+07 | 4.32E+07 | | | | | | |
| Q9R1P4 | PSMA1 | proteasome (prosome, macropain) subunit, alpha type, 1 | Cytoplasm | peptidase | 2.86E+07 | 1.64E+07 | 7825151 | | | | | | |
| Q9R1P4 | PSMA1 | proteasome (prosome, macropain) subunit, alpha type, 1 | Cytoplasm | peptidase | 3.33E+07 | 2.39E+07 | 1.84E+07 | | | | | | |
| Q9R1P4 | PSMA1 | proteasome (prosome, macropain) subunit, alpha type, 1 | Cytoplasm | peptidase | 2.53E+07 | 2.79E+07 | 2.60E+07 | | | | | | |
| Q9R1P4 | PSMA1 | proteasome (prosome, macropain) subunit, alpha type, 1 | Cytoplasm | peptidase | 2.45E+07 | 1.86E+07 | 2.13E+07 | | | | | | |
| Q9R1P4 | PSMA1 | proteasome (prosome, macropain) subunit, alpha type, 1 | Cytoplasm | peptidase | 3.30E+07 | 1.59E+07 | 1.26E+07 | | | | | | |
| Q9R1P4 | PSMA1 | proteasome (prosome, macropain) subunit, alpha type, 1 | Cytoplasm | peptidase | 1.38E+07 | 1.88E+07 | 1.74E+07 | | | | | | |
| Q9R1P4 | PSMA1 | proteasome (prosome, macropain) subunit, alpha type, 1 | Cytoplasm | peptidase | 1.96E+07 | 1.86E+07 | 1.57E+07 | | | | | | |
| Q9R1P4 | PSMA1 | proteasome (prosome, macropain) subunit, alpha type, 1 | Cytoplasm | peptidase | 1.91E+07 | 1.46E+07 | 1.39E+07 | | | | | | |
| Q9R1P4 | PSMA1 | proteasome (prosome, macropain) subunit, alpha type, 1 | Cytoplasm | peptidase | 1.74E+07 | 1.30E+07 | 1.47E+07 | | | | | | |

| | | | | | | | | | | | | | |
|---|---|---|---|---|---|---|---|---|---|---|---|---|---|
| Q9R1P4 | PSMA1 | proteasome (prosome, macropain) subunit, alpha type, 1 | Cytoplasm | peptidase | 1.60E+07 | 1.24E+07 | 8422444 | | | | | | |
| Q9R1P4 | PSMA1 | proteasome (prosome, macropain) subunit, alpha type, 1 | Cytoplasm | peptidase | 9156496 | 6502790.5 | 4300854.5 | | | | | | |
| Q9R1P4 | PSMA1 | proteasome (prosome, macropain) subunit, alpha type, 1 | Cytoplasm | peptidase | 6909070.5 | 3612684 | 2744343 | | | | | | |
| Q9R1P4 | PSMA1 | proteasome (prosome, macropain) subunit, alpha type, 1 | Cytoplasm | peptidase | 3489863.75 | 2796913.25 | 2260341 | | | | | | |
| Q9R1P4 | PSMA1 | proteasome (prosome, macropain) subunit, alpha type, 1 | Cytoplasm | peptidase | 5740124 | 1941356.125 | 1536153.625 | | | | | | |
| Q9R1P4 | PSMA1 | proteasome (prosome, macropain) subunit, alpha type, 1 | Cytoplasm | peptidase | 7973568.5 | 9009099 | 1.14E+07 | Q9R1P4 | PSMA1 | 19653070 | 11183056 | 1614135 | 48 |
| P49722 | PSMA2 | proteasome (prosome, macropain) subunit, alpha type, 2 | Cytoplasm | peptidase | 8.58E+07 | 7.46E+07 | 6.84E+07 | | | | | | |
| P49722 | PSMA2 | proteasome (prosome, macropain) subunit, alpha type, 2 | Cytoplasm | peptidase | 6.41E+07 | 3.35E+07 | 1.87E+07 | | | | | | |
| P49722 | PSMA2 | proteasome (prosome, macropain) subunit, alpha type, 2 | Cytoplasm | peptidase | 2.09E+07 | 2.37E+07 | 1.47E+07 | | | | | | |
| P49722 | PSMA2 | proteasome (prosome, macropain) subunit, alpha type, 2 | Cytoplasm | peptidase | 3.27E+07 | 2.56E+07 | 2.37E+07 | | | | | | |
| P49722 | PSMA2 | proteasome (prosome, macropain) subunit, alpha type, 2 | Cytoplasm | peptidase | 1.10E+07 | 1.22E+07 | 1.29E+07 | | | | | | |
| P49722 | PSMA2 | proteasome (prosome, macropain) subunit, alpha type, 2 | Cytoplasm | peptidase | 9460773 | 4422661 | 3466421.75 | | | | | | |
| P49722 | PSMA2 | proteasome (prosome, macropain) subunit, alpha type, 2 | Cytoplasm | peptidase | 5926920.5 | 4099535 | 3152124.25 | | | | | | |
| P49722 | PSMA2 | proteasome (prosome, macropain) subunit, alpha type, 2 | Cytoplasm | peptidase | 3693916.5 | 3887199.25 | 3648489.75 | | | | | | |
| P49722 | PSMA2 | proteasome (prosome, macropain) subunit, alpha type, 2 | Cytoplasm | peptidase | 4070293.25 | 4435293 | 4582537.5 | | | | | | |
| P49722 | PSMA2 | proteasome (prosome, macropain) subunit, alpha type, 2 | Cytoplasm | peptidase | 3380565.75 | 2755701.25 | 3594792 | P49722 | PSMA2 | 24103247 | 23448463 | 4281084 | 30 |
| O70435 | PSMA3 | proteasome (prosome, macropain) subunit, alpha type, 3 | Cytoplasm | peptidase | 3.57E+07 | 2.74E+07 | 2.83E+07 | | | | | | |
| O70435 | PSMA3 | proteasome (prosome, macropain) subunit, alpha type, 3 | Cytoplasm | peptidase | 3.42E+07 | 2.33E+07 | 2.49E+07 | | | | | | |
| O70435 | PSMA3 | proteasome (prosome, macropain) subunit, alpha type, 3 | Cytoplasm | peptidase | 2.81E+07 | 2.10E+07 | 2.20E+07 | | | | | | |
| O70435 | PSMA3 | proteasome (prosome, macropain) subunit, alpha type, 3 | Cytoplasm | peptidase | 2.33E+07 | 1.64E+07 | 1.58E+07 | | | | | | |
| O70435 | PSMA3 | proteasome (prosome, macropain) subunit, alpha type, 3 | Cytoplasm | peptidase | 1.91E+07 | 1.46E+07 | 1.45E+07 | | | | | | |
| O70435 | PSMA3 | proteasome (prosome, macropain) subunit, alpha type, 3 | Cytoplasm | peptidase | 1.88E+07 | 1.54E+07 | 1.64E+07 | | | | | | |
| O70435 | PSMA3 | proteasome (prosome, macropain) subunit, alpha type, 3 | Cytoplasm | peptidase | 1.48E+07 | 1.07E+07 | 8301728 | | | | | | |
| O70435 | PSMA3 | proteasome (prosome, macropain) subunit, alpha type, 3 | Cytoplasm | peptidase | 1.31E+07 | 8381796 | 5446949.5 | | | | | | |
| O70435 | PSMA3 | proteasome (prosome, macropain) subunit, alpha type, 3 | Cytoplasm | peptidase | 9513253 | 4144208 | 3848867.5 | O70435 | PSMA3 | 21818139 | 8624312 | 1659750 | 27 |
| Q9R1P0 | PSMA4 | proteasome (prosome, macropain) subunit, alpha type, 4 | Cytoplasm | peptidase | 3.86E+07 | 2.60E+07 | 2.52E+07 | | | | | | |
| Q9R1P0 | PSMA4 | proteasome (prosome, macropain) subunit, alpha type, 4 | Cytoplasm | peptidase | 4.16E+07 | 3.33E+07 | 2.68E+07 | | | | | | |
| Q9R1P0 | PSMA4 | proteasome (prosome, macropain) subunit, alpha type, 4 | Cytoplasm | peptidase | 4.47E+07 | 3.49E+07 | 3.78E+07 | | | | | | |
| Q9R1P0 | PSMA4 | proteasome (prosome, macropain) subunit, alpha type, 4 | Cytoplasm | peptidase | 3.18E+07 | 2.28E+07 | 2.28E+07 | | | | | | |
| Q9R1P0 | PSMA4 | proteasome (prosome, macropain) subunit, alpha type, 4 | Cytoplasm | peptidase | 2.77E+07 | 2.19E+07 | 2.25E+07 | | | | | | |
| Q9R1P0 | PSMA4 | proteasome (prosome, macropain) subunit, alpha type, 4 | Cytoplasm | peptidase | 2.18E+07 | 1.91E+07 | 1.83E+07 | | | | | | |
| Q9R1P0 | PSMA4 | proteasome (prosome, macropain) subunit, alpha type, 4 | Cytoplasm | peptidase | 2.13E+07 | 1.52E+07 | 1.43E+07 | | | | | | |
| Q9R1P0 | PSMA4 | proteasome (prosome, macropain) subunit, alpha type, 4 | Cytoplasm | peptidase | 1.44E+07 | 1.14E+07 | 1.01E+07 | | | | | | |
| Q9R1P0 | PSMA4 | proteasome (prosome, macropain) subunit, alpha type, 4 | Cytoplasm | peptidase | 1.07E+07 | 7789015 | 9485260 | | | | | | |
| Q9R1P0 | PSMA4 | proteasome (prosome, macropain) subunit, alpha type, 4 | Cytoplasm | peptidase | 1.05E+07 | 9963483 | 1.01E+07 | | | | | | |
| Q9R1P0 | PSMA4 | proteasome (prosome, macropain) subunit, alpha type, 4 | Cytoplasm | peptidase | 6383100 | 4036751.5 | 4644246.5 | | | | | | |
| Q9R1P0 | PSMA4 | proteasome (prosome, macropain) subunit, alpha type, 4 | Cytoplasm | peptidase | 5772131.5 | 2718801.75 | 1436563 | | | | | | |
| Q9R1P0 | PSMA4 | proteasome (prosome, macropain) subunit, alpha type, 4 | Cytoplasm | peptidase | 1053177.625 | 660103.562 | 1169621.625 | Q9R1P0 | PSMA4 | 21241416 | 12332489 | 1974779 | 39 |
| Q9Z2U1 | PSMA5 | proteasome (prosome, macropain) subunit, alpha type, 5 | Cytoplasm | peptidase | 1.02E+08 | 8.68E+07 | 6.40E+07 | | | | | | |
| Q9Z2U1 | PSMA5 | proteasome (prosome, macropain) subunit, alpha type, 5 | Cytoplasm | peptidase | 9.22E+07 | 6.21E+07 | 6.67E+07 | | | | | | |
| Q9Z2U1 | PSMA5 | proteasome (prosome, macropain) subunit, alpha type, 5 | Cytoplasm | peptidase | 8.10E+07 | 6.72E+07 | 5.08E+07 | | | | | | |
| Q9Z2U1 | PSMA5 | proteasome (prosome, macropain) subunit, alpha type, 5 | Cytoplasm | peptidase | 3.73E+07 | 3.06E+07 | 2.37E+07 | | | | | | |
| Q9Z2U1 | PSMA5 | proteasome (prosome, macropain) subunit, alpha type, 5 | Cytoplasm | peptidase | 2.69E+07 | 2.15E+07 | 2.21E+07 | | | | | | |
| Q9Z2U1 | PSMA5 | proteasome (prosome, macropain) subunit, alpha type, 5 | Cytoplasm | peptidase | 2.10E+07 | 1.69E+07 | 1.30E+07 | | | | | | |
| Q9Z2U1 | PSMA5 | proteasome (prosome, macropain) subunit, alpha type, 5 | Cytoplasm | peptidase | 1.41E+07 | 1.22E+07 | 1.33E+07 | | | | | | |
| Q9Z2U1 | PSMA5 | proteasome (prosome, macropain) subunit, alpha type, 5 | Cytoplasm | peptidase | 1.24E+07 | 9933938 | 7618818 | | | | | | |
| Q9Z2U1 | PSMA5 | proteasome (prosome, macropain) subunit, alpha type, 5 | Cytoplasm | peptidase | 1.57E+07 | 3832545.5 | 2603695.5 | | | | | | |
| Q9Z2U1 | PSMA5 | proteasome (prosome, macropain) subunit, alpha type, 5 | Cytoplasm | peptidase | 5438491.5 | 5499071.5 | 5562900 | | | | | | |
| Q9Z2U1 | PSMA5 | proteasome (prosome, macropain) subunit, alpha type, 5 | Cytoplasm | peptidase | 6675901.5 | 4191957 | 4808751 | | | | | | |
| Q9Z2U1 | PSMA5 | proteasome (prosome, macropain) subunit, alpha type, 5 | Cytoplasm | peptidase | 7289522.5 | 5102172.5 | 4253430.5 | | | | | | |
| Q9Z2U1 | PSMA5 | proteasome (prosome, macropain) subunit, alpha type, 5 | Cytoplasm | peptidase | 4970524 | 2630707.5 | 3270902.5 | | | | | | |
| Q9Z2U1 | PSMA5 | proteasome (prosome, macropain) subunit, alpha type, 5 | Cytoplasm | peptidase | 4856317.5 | 2860004.25 | 2082769.75 | Q9Z2U1 | PSMA5 | 30805768 | 28475107 | 4393804 | 42 |
| Q9QUM9 | PSMA6 | proteasome (prosome, macropain) subunit, alpha type, 6 | Cytoplasm | peptidase | 5.69E+07 | 4.07E+07 | 4.32E+07 | | | | | | |
| Q9QUM9 | PSMA6 | proteasome (prosome, macropain) subunit, alpha type, 6 | Cytoplasm | peptidase | 5.54E+07 | 4.30E+07 | 4.07E+07 | | | | | | |
| Q9QUM9 | PSMA6 | proteasome (prosome, macropain) subunit, alpha type, 6 | Cytoplasm | peptidase | 3.41E+07 | 2.83E+07 | 3.05E+07 | | | | | | |
| Q9QUM9 | PSMA6 | proteasome (prosome, macropain) subunit, alpha type, 6 | Cytoplasm | peptidase | 3.15E+07 | 2.40E+07 | 2.26E+07 | | | | | | |
| Q9QUM9 | PSMA6 | proteasome (prosome, macropain) subunit, alpha type, 6 | Cytoplasm | peptidase | 2.93E+07 | 2.57E+07 | 2.35E+07 | | | | | | |
| Q9QUM9 | PSMA6 | proteasome (prosome, macropain) subunit, alpha type, 6 | Cytoplasm | peptidase | 2.59E+07 | 2.03E+07 | 1.97E+07 | | | | | | |
| Q9QUM9 | PSMA6 | proteasome (prosome, macropain) subunit, alpha type, 6 | Cytoplasm | peptidase | 2.12E+07 | 1.35E+07 | 1.28E+07 | | | | | | |
| Q9QUM9 | PSMA6 | proteasome (prosome, macropain) subunit, alpha type, 6 | Cytoplasm | peptidase | 2.50E+07 | 1.40E+07 | 1.48E+07 | | | | | | |
| Q9QUM9 | PSMA6 | proteasome (prosome, macropain) subunit, alpha type, 6 | Cytoplasm | peptidase | 1.96E+07 | 1.37E+07 | 1.37E+07 | | | | | | |
| Q9QUM9 | PSMA6 | proteasome (prosome, macropain) subunit, alpha type, 6 | Cytoplasm | peptidase | 2.06E+07 | 1.54E+07 | 1.54E+07 | | | | | | |
| Q9QUM9 | PSMA6 | proteasome (prosome, macropain) subunit, alpha type, 6 | Cytoplasm | peptidase | 2530999.5 | 515785.219 | 35584.461 | Q9QUM9 | PSMA6 | 29264636 | 13751344 | 2393802 | 33 |
| Q9Z2U0 | PSMA7 | proteasome (prosome, macropain) subunit, alpha type, 7 | Cytoplasm | peptidase | 1.06E+08 | 8.52E+07 | 9.86E+07 | | | | | | |
| Q9Z2U0 | PSMA7 | proteasome (prosome, macropain) subunit, alpha type, 7 | Cytoplasm | peptidase | 5.51E+07 | 4.28E+07 | 4.48E+07 | | | | | | |
| Q9Z2U0 | PSMA7 | proteasome (prosome, macropain) subunit, alpha type, 7 | Cytoplasm | peptidase | 5.38E+07 | 4.33E+07 | 4.70E+07 | | | | | | |
| Q9Z2U0 | PSMA7 | proteasome (prosome, macropain) subunit, alpha type, 7 | Cytoplasm | peptidase | 4.04E+07 | 2.61E+07 | 2.70E+07 | | | | | | |
| Q9Z2U0 | PSMA7 | proteasome (prosome, macropain) subunit, alpha type, 7 | Cytoplasm | peptidase | 4.14E+07 | 3.15E+07 | 3.44E+07 | | | | | | |
| Q9Z2U0 | PSMA7 | proteasome (prosome, macropain) subunit, alpha type, 7 | Cytoplasm | peptidase | 2.25E+07 | 2.01E+07 | 2.04E+07 | | | | | | |
| Q9Z2U0 | PSMA7 | proteasome (prosome, macropain) subunit, alpha type, 7 | Cytoplasm | peptidase | 1.29E+07 | 1.16E+07 | 1.16E+07 | | | | | | |
| Q9Z2U0 | PSMA7 | proteasome (prosome, macropain) subunit, alpha type, 7 | Cytoplasm | peptidase | 1.09E+07 | 6196586 | 4790688.5 | | | | | | |
| Q9Z2U0 | PSMA7 | proteasome (prosome, macropain) subunit, alpha type, 7 | Cytoplasm | peptidase | 9824066 | 4984282 | 6016859.5 | | | | | | |
| Q9Z2U0 | PSMA7 | proteasome (prosome, macropain) subunit, alpha type, 7 | Cytoplasm | peptidase | 3866496.5 | 2369634.5 | 2171603.75 | | | | | | |
| Q9Z2U0 | PSMA7 | proteasome (prosome, macropain) subunit, alpha type, 7 | Cytoplasm | peptidase | 7044484 | 5602063 | 4376657 | | | | | | |
| Q9Z2U0 | PSMA7 | proteasome (prosome, macropain) subunit, alpha type, 7 | Cytoplasm | peptidase | 7535852 | 1062635.625 | 54923.918 | Q9Z2U0 | PSMA7 | 30965908 | 27306763 | 4551127 | 36 |
| O09061 | PSMB1 | proteasome (prosome, macropain) subunit, beta type, 1 | Cytoplasm | peptidase | 4.42E+07 | 3.97E+07 | 4.01E+07 | | | | | | |
| O09061 | PSMB1 | proteasome (prosome, macropain) subunit, beta type, 1 | Cytoplasm | peptidase | 3.64E+07 | 3.12E+07 | 2.71E+07 | | | | | | |
| O09061 | PSMB1 | proteasome (prosome, macropain) subunit, beta type, 1 | Cytoplasm | peptidase | 3.48E+07 | 2.67E+07 | 2.24E+07 | | | | | | |
| O09061 | PSMB1 | proteasome (prosome, macropain) subunit, beta type, 1 | Cytoplasm | peptidase | 2.56E+07 | 2.09E+07 | 1.89E+07 | | | | | | |
| O09061 | PSMB1 | proteasome (prosome, macropain) subunit, beta type, 1 | Cytoplasm | peptidase | 1.69E+07 | 1.69E+07 | 1.19E+07 | | | | | | |
| O09061 | PSMB1 | proteasome (prosome, macropain) subunit, beta type, 1 | Cytoplasm | peptidase | 1.19E+07 | 7462203 | 6765454.5 | | | | | | |
| O09061 | PSMB1 | proteasome (prosome, macropain) subunit, beta type, 1 | Cytoplasm | peptidase | 7570319 | 4841434 | 4117741.5 | | | | | | |
| O09061 | PSMB1 | proteasome (prosome, macropain) subunit, beta type, 1 | Cytoplasm | peptidase | 7099796 | 5936947 | 5453414 | | | | | | |
| O09061 | PSMB1 | proteasome (prosome, macropain) subunit, beta type, 1 | Cytoplasm | peptidase | 6439418 | 6826131 | 9269776 | | | | | | |
| O09061 | PSMB1 | proteasome (prosome, macropain) subunit, beta type, 1 | Cytoplasm | peptidase | 7378076 | 3625242.25 | 3651751 | | | | | | |
| O09061 | PSMB1 | proteasome (prosome, macropain) subunit, beta type, 1 | Cytoplasm | peptidase | 4625644.5 | 3202507.75 | 4512746 | | | | | | |
| O09061 | PSMB1 | proteasome (prosome, macropain) subunit, beta type, 1 | Cytoplasm | peptidase | 5209642 | 2760416.25 | 1711380.5 | | | | | | |
| O09061 | PSMB1 | proteasome (prosome, macropain) subunit, beta type, 1 | Cytoplasm | peptidase | 4625026.5 | 5031521 | 4742888.5 | | | | | | |
| O09061 | PSMB1 | proteasome (prosome, macropain) subunit, beta type, 1 | Cytoplasm | peptidase | 3378275.25 | 3673772.25 | 4685858 | | | | | | |
| O09061 | PSMB1 | proteasome (prosome, macropain) subunit, beta type, 1 | Cytoplasm | peptidase | 3811147.75 | 4652000 | 3021184.5 | | | | | | |
| O09061 | PSMB1 | proteasome (prosome, macropain) subunit, beta type, 1 | Cytoplasm | peptidase | 3234957 | 2712102.25 | 2197743.75 | | | | | | |

| | | | | | | | | | | | | | |
|---|---|---|---|---|---|---|---|---|---|---|---|---|---|
| O09061 | PSMB1 | proteasome (prosome, macropain) subunit, beta type, 1 | Cytoplasm | peptidase | 2474734.25 | 793437.625 | 336262.375 | O09061 | PSMB1 | 13267473 | 11898284 | 1666093 | 51 |
| Q9R1P3 | PSMB2 | proteasome (prosome, macropain) subunit, beta type, 2 | Cytoplasm | peptidase | 4.41E+07 | 4.25E+07 | 4.42E+07 | | | | | | |
| Q9R1P3 | PSMB2 | proteasome (prosome, macropain) subunit, beta type, 2 | Cytoplasm | peptidase | 3.56E+07 | 2.94E+07 | 2.67E+07 | | | | | | |
| Q9R1P3 | PSMB2 | proteasome (prosome, macropain) subunit, beta type, 2 | Cytoplasm | peptidase | 1.62E+07 | 1.25E+07 | 6938944 | | | | | | |
| Q9R1P3 | PSMB2 | proteasome (prosome, macropain) subunit, beta type, 2 | Cytoplasm | peptidase | 7587836 | 7410185 | 6984156 | | | | | | |
| Q9R1P3 | PSMB2 | proteasome (prosome, macropain) subunit, beta type, 2 | Cytoplasm | peptidase | 9988279 | 6610240 | 6946807 | | | | | | |
| Q9R1P3 | PSMB2 | proteasome (prosome, macropain) subunit, beta type, 2 | Cytoplasm | peptidase | 7438933.5 | 7009747.5 | 7617309.5 | | | | | | |
| Q9R1P3 | PSMB2 | proteasome (prosome, macropain) subunit, beta type, 2 | Cytoplasm | peptidase | 6163517 | 3494028.75 | 4408136 | | | | | | |
| Q9R1P3 | PSMB2 | proteasome (prosome, macropain) subunit, beta type, 2 | Cytoplasm | peptidase | 5197015.5 | 2456870 | 2092548.875 | | | | | | |
| Q9R1P3 | PSMB2 | proteasome (prosome, macropain) subunit, beta type, 2 | Cytoplasm | peptidase | 7186244.5 | 3569732 | 2587455.25 | Q9R1P3 | PSMB2 | 15490203 | 13626998 | 2622517 | 27 |
| Q9R1P1 | PSMB3 | proteasome (prosome, macropain) subunit, beta type, 3 | Cytoplasm | peptidase | 3.47E+07 | 3.56E+07 | 3.28E+07 | | | | | | |
| Q9R1P1 | PSMB3 | proteasome (prosome, macropain) subunit, beta type, 3 | Cytoplasm | peptidase | 2.70E+07 | 2.16E+07 | 2.20E+07 | | | | | | |
| Q9R1P1 | PSMB3 | proteasome (prosome, macropain) subunit, beta type, 3 | Cytoplasm | peptidase | 1.87E+07 | 9917770 | 2.28E+07 | | | | | | |
| Q9R1P1 | PSMB3 | proteasome (prosome, macropain) subunit, beta type, 3 | Cytoplasm | peptidase | 1.82E+07 | 1.46E+07 | 1.57E+07 | | | | | | |
| Q9R1P1 | PSMB3 | proteasome (prosome, macropain) subunit, beta type, 3 | Cytoplasm | peptidase | 8623966 | 6512383.5 | 6014595 | | | | | | |
| Q9R1P1 | PSMB3 | proteasome (prosome, macropain) subunit, beta type, 3 | Cytoplasm | peptidase | 8597662 | 4288804 | 2669913.75 | | | | | | |
| Q9R1P1 | PSMB3 | proteasome (prosome, macropain) subunit, beta type, 3 | Cytoplasm | peptidase | 1.22E+07 | 9842414 | 9500551 | | | | | | |
| Q9R1P1 | PSMB3 | proteasome (prosome, macropain) subunit, beta type, 3 | Cytoplasm | peptidase | 4695301 | 5900081.5 | 6362763 | | | | | | |
| Q9R1P1 | PSMB3 | proteasome (prosome, macropain) subunit, beta type, 3 | Cytoplasm | peptidase | 5003389 | 4359124.5 | 5336975.5 | | | | | | |
| Q9R1P1 | PSMB3 | proteasome (prosome, macropain) subunit, beta type, 3 | Cytoplasm | peptidase | 993932.188 | 1552981.625 | 1090636.75 | Q9R1P1 | PSMB3 | 13872425 | 10182856 | 1859127 | 30 |
| P99026 | PSMB4 | proteasome (prosome, macropain) subunit, beta type, 4 | Cytoplasm | peptidase | 3.06E+07 | 2.37E+07 | 2.34E+07 | | | | | | |
| P99026 | PSMB4 | proteasome (prosome, macropain) subunit, beta type, 4 | Cytoplasm | peptidase | 2.59E+07 | 2.42E+07 | 2.24E+07 | | | | | | |
| P99026 | PSMB4 | proteasome (prosome, macropain) subunit, beta type, 4 | Cytoplasm | peptidase | 2.88E+07 | 1.93E+07 | 2.02E+07 | | | | | | |
| P99026 | PSMB4 | proteasome (prosome, macropain) subunit, beta type, 4 | Cytoplasm | peptidase | 2.30E+07 | 2.18E+07 | 2.20E+07 | | | | | | |
| P99026 | PSMB4 | proteasome (prosome, macropain) subunit, beta type, 4 | Cytoplasm | peptidase | 2.21E+07 | 1.91E+07 | 1.89E+07 | | | | | | |
| P99026 | PSMB4 | proteasome (prosome, macropain) subunit, beta type, 4 | Cytoplasm | peptidase | 7394820 | 6344400.5 | 6220435.5 | P99026 | PSMB4 | 22952470 | 6979816 | 1645158 | 18 |
| O55234 | PSMB5 | proteasome (prosome, macropain) subunit, beta type, 5 | Cytoplasm | peptidase | 4.30E+07 | 3.08E+07 | 3.08E+07 | | | | | | |
| O55234 | PSMB5 | proteasome (prosome, macropain) subunit, beta type, 5 | Cytoplasm | peptidase | 3.75E+07 | 2.31E+07 | 2.73E+07 | | | | | | |
| O55234 | PSMB5 | proteasome (prosome, macropain) subunit, beta type, 5 | Cytoplasm | peptidase | 3.12E+07 | 2.56E+07 | 2.24E+07 | | | | | | |
| O55234 | PSMB5 | proteasome (prosome, macropain) subunit, beta type, 5 | Cytoplasm | peptidase | 2.68E+07 | 2.21E+07 | 2.16E+07 | | | | | | |
| O55234 | PSMB5 | proteasome (prosome, macropain) subunit, beta type, 5 | Cytoplasm | peptidase | 3.20E+07 | 2.20E+07 | 2.29E+07 | | | | | | |
| O55234 | PSMB5 | proteasome (prosome, macropain) subunit, beta type, 5 | Cytoplasm | peptidase | 2.03E+07 | 1.16E+07 | 1.15E+07 | | | | | | |
| O55234 | PSMB5 | proteasome (prosome, macropain) subunit, beta type, 5 | Cytoplasm | peptidase | 1.03E+07 | 8318805.5 | 8568135 | | | | | | |
| O55234 | PSMB5 | proteasome (prosome, macropain) subunit, beta type, 5 | Cytoplasm | peptidase | 1.26E+07 | 8967947 | 7414276 | | | | | | |
| O55234 | PSMB5 | proteasome (prosome, macropain) subunit, beta type, 5 | Cytoplasm | peptidase | 1.02E+07 | 6936174 | 7611454.5 | | | | | | |
| O55234 | PSMB5 | proteasome (prosome, macropain) subunit, beta type, 5 | Cytoplasm | peptidase | 4817689.5 | 2744986.5 | 2471050.5 | | | | | | |
| O55234 | PSMB5 | proteasome (prosome, macropain) subunit, beta type, 5 | Cytoplasm | peptidase | 3373644.5 | 1817078.375 | 1436231.75 | O55234 | PSMB5 | 21101030 | 11525520 | 2006336 | 33 |
| Q60692 | PSMB6 | proteasome (prosome, macropain) subunit, beta type, 6 | Nucleus | peptidase | 1.16E+08 | 1.02E+08 | 1.02E+08 | | | | | | |
| Q60692 | PSMB6 | proteasome (prosome, macropain) subunit, beta type, 6 | Nucleus | peptidase | 3.81E+07 | 2.70E+07 | 2.92E+07 | | | | | | |
| Q60692 | PSMB6 | proteasome (prosome, macropain) subunit, beta type, 6 | Nucleus | peptidase | 3.11E+07 | 2.39E+07 | 2.25E+07 | | | | | | |
| Q60692 | PSMB6 | proteasome (prosome, macropain) subunit, beta type, 6 | Nucleus | peptidase | 1.73E+07 | 1.22E+07 | 1.40E+07 | | | | | | |
| Q60692 | PSMB6 | proteasome (prosome, macropain) subunit, beta type, 6 | Nucleus | peptidase | 5778256.5 | 4980245.5 | 5666669.5 | | | | | | |
| Q60692 | PSMB6 | proteasome (prosome, macropain) subunit, beta type, 6 | Nucleus | peptidase | 4093611.75 | 1794017.375 | 252847.531 | Q60692 | PSMB6 | 35393645 | 36558303 | 8616875 | 18 |
| P70195 | PSMB7 | proteasome (prosome, macropain) subunit, beta type, 7 | Cytoplasm | peptidase | 4.00E+07 | 3.11E+07 | 3.11E+07 | | | | | | |
| P70195 | PSMB7 | proteasome (prosome, macropain) subunit, beta type, 7 | Cytoplasm | peptidase | 2.69E+07 | 1.93E+07 | 1.77E+07 | | | | | | |
| P70195 | PSMB7 | proteasome (prosome, macropain) subunit, beta type, 7 | Cytoplasm | peptidase | 2.35E+07 | 1.79E+07 | 1.86E+07 | | | | | | |
| P70195 | PSMB7 | proteasome (prosome, macropain) subunit, beta type, 7 | Cytoplasm | peptidase | 1.79E+07 | 1.39E+07 | 1.42E+07 | | | | | | |
| P70195 | PSMB7 | proteasome (prosome, macropain) subunit, beta type, 7 | Cytoplasm | peptidase | 1.51E+07 | 1.65E+07 | 1.91E+07 | | | | | | |
| P70195 | PSMB7 | proteasome (prosome, macropain) subunit, beta type, 7 | Cytoplasm | peptidase | 1.25E+07 | 9600736 | 8764509 | | | | | | |
| P70195 | PSMB7 | proteasome (prosome, macropain) subunit, beta type, 7 | Cytoplasm | peptidase | 7352224 | 4969649 | 4328573.5 | | | | | | |
| P70195 | PSMB7 | proteasome (prosome, macropain) subunit, beta type, 7 | Cytoplasm | peptidase | 629567.812 | 868946.688 | 1813864.125 | P70195 | PSMB7 | 17986474 | 10118044 | 2065337 | 24 |
| P28076 | PSMB9 | proteasome (prosome, macropain) subunit, beta type, 9 | Cytoplasm | peptidase | 929653.312 | 554117.562 | 381117.25 | P28076 | PSMB9 | 929653 | 280431 | 161907 | 3 |
| P62192 | PSMC1 | proteasome (prosome, macropain) 26S subunit, ATPase, 1 | Nucleus | peptidase | 3.73E+07 | 3.22E+07 | 3.21E+07 | | | | | | |
| P62192 | PSMC1 | proteasome (prosome, macropain) 26S subunit, ATPase, 1 | Nucleus | peptidase | 2.27E+07 | 1.84E+07 | 1.77E+07 | | | | | | |
| P62192 | PSMC1 | proteasome (prosome, macropain) 26S subunit, ATPase, 1 | Nucleus | peptidase | 1.70E+07 | 1.31E+07 | 1.33E+07 | | | | | | |
| P62192 | PSMC1 | proteasome (prosome, macropain) 26S subunit, ATPase, 1 | Nucleus | peptidase | 1.79E+07 | 1.47E+07 | 1.26E+07 | | | | | | |
| P62192 | PSMC1 | proteasome (prosome, macropain) 26S subunit, ATPase, 1 | Nucleus | peptidase | 1.39E+07 | 1.49E+07 | 8152111 | | | | | | |
| P62192 | PSMC1 | proteasome (prosome, macropain) 26S subunit, ATPase, 1 | Nucleus | peptidase | 1.30E+07 | 1.22E+07 | 1.07E+07 | | | | | | |
| P62192 | PSMC1 | proteasome (prosome, macropain) 26S subunit, ATPase, 1 | Nucleus | peptidase | 8769422 | 1.05E+07 | 1.23E+07 | | | | | | |
| P62192 | PSMC1 | proteasome (prosome, macropain) 26S subunit, ATPase, 1 | Nucleus | peptidase | 1.30E+07 | 9719663 | 9175822 | | | | | | |
| P62192 | PSMC1 | proteasome (prosome, macropain) 26S subunit, ATPase, 1 | Nucleus | peptidase | 8646452 | 7982013 | 7178313 | | | | | | |
| P62192 | PSMC1 | proteasome (prosome, macropain) 26S subunit, ATPase, 1 | Nucleus | peptidase | 6735521 | 7004670 | 9564809 | | | | | | |
| P62192 | PSMC1 | proteasome (prosome, macropain) 26S subunit, ATPase, 1 | Nucleus | peptidase | 6891804.5 | 6257605.5 | 4527133.5 | | | | | | |
| P62192 | PSMC1 | proteasome (prosome, macropain) 26S subunit, ATPase, 1 | Nucleus | peptidase | 6520012 | 7148050.5 | 7048696 | | | | | | |
| P62192 | PSMC1 | proteasome (prosome, macropain) 26S subunit, ATPase, 1 | Nucleus | peptidase | 2499264.25 | 3017308 | 3698490.5 | P62192 | PSMC1 | 13446344 | 7782732 | 1246234 | 39 |
| P46471 | PSMC2 | proteasome (prosome, macropain) 26S subunit, ATPase, 2 | Nucleus | peptidase | 3.45E+07 | 3.16E+07 | 2.67E+07 | | | | | | |
| P46471 | PSMC2 | proteasome (prosome, macropain) 26S subunit, ATPase, 2 | Nucleus | peptidase | 2.51E+07 | 2.15E+07 | 1.93E+07 | | | | | | |
| P46471 | PSMC2 | proteasome (prosome, macropain) 26S subunit, ATPase, 2 | Nucleus | peptidase | 1.86E+07 | 1.61E+07 | 1.38E+07 | | | | | | |
| P46471 | PSMC2 | proteasome (prosome, macropain) 26S subunit, ATPase, 2 | Nucleus | peptidase | 1.57E+07 | 1.35E+07 | 1.23E+07 | | | | | | |
| P46471 | PSMC2 | proteasome (prosome, macropain) 26S subunit, ATPase, 2 | Nucleus | peptidase | 1.50E+07 | 1.02E+07 | 9360047 | | | | | | |
| P46471 | PSMC2 | proteasome (prosome, macropain) 26S subunit, ATPase, 2 | Nucleus | peptidase | 1.37E+07 | 1.44E+07 | 1.22E+07 | | | | | | |
| P46471 | PSMC2 | proteasome (prosome, macropain) 26S subunit, ATPase, 2 | Nucleus | peptidase | 1.34E+07 | 1.05E+07 | 1.03E+07 | | | | | | |
| P46471 | PSMC2 | proteasome (prosome, macropain) 26S subunit, ATPase, 2 | Nucleus | peptidase | 1.58E+07 | 1.43E+07 | 1.33E+07 | | | | | | |
| P46471 | PSMC2 | proteasome (prosome, macropain) 26S subunit, ATPase, 2 | Nucleus | peptidase | 1.12E+07 | 1.06E+07 | 1.05E+07 | | | | | | |
| P46471 | PSMC2 | proteasome (prosome, macropain) 26S subunit, ATPase, 2 | Nucleus | peptidase | 1.41E+07 | 1.01E+07 | 9199597 | | | | | | |
| P46471 | PSMC2 | proteasome (prosome, macropain) 26S subunit, ATPase, 2 | Nucleus | peptidase | 1.37E+07 | 1.21E+07 | 9855568 | | | | | | |
| P46471 | PSMC2 | proteasome (prosome, macropain) 26S subunit, ATPase, 2 | Nucleus | peptidase | 5843876 | 5708588.5 | 6655127 | | | | | | |
| P46471 | PSMC2 | proteasome (prosome, macropain) 26S subunit, ATPase, 2 | Nucleus | peptidase | 6168173.5 | 5285748 | 4716144.5 | | | | | | |
| P46471 | PSMC2 | proteasome (prosome, macropain) 26S subunit, ATPase, 2 | Nucleus | peptidase | 4857601.5 | 2699675.5 | 3248289.25 | | | | | | |
| P46471 | PSMC2 | proteasome (prosome, macropain) 26S subunit, ATPase, 2 | Nucleus | peptidase | 1926079.25 | 3179302.75 | 1351500.25 | P46471 | PSMC2 | 13972382 | 7275290 | 1084536 | 45 |
| O88685 | PSMC3 | proteasome (prosome, macropain) 26S subunit, ATPase, 3 | Nucleus | transcription regulator | 2.50E+07 | 2.15E+07 | 2.11E+07 | | | | | | |
| O88685 | PSMC3 | proteasome (prosome, macropain) 26S subunit, ATPase, 3 | Nucleus | transcription regulator | 1.80E+07 | 1.29E+07 | 1.81E+07 | | | | | | |
| O88685 | PSMC3 | proteasome (prosome, macropain) 26S subunit, ATPase, 3 | Nucleus | transcription regulator | 1.65E+07 | 1.71E+07 | 1.67E+07 | | | | | | |
| O88685 | PSMC3 | proteasome (prosome, macropain) 26S subunit, ATPase, 3 | Nucleus | transcription regulator | 1.60E+07 | 1.43E+07 | 1.27E+07 | | | | | | |
| O88685 | PSMC3 | proteasome (prosome, macropain) 26S subunit, ATPase, 3 | Nucleus | transcription regulator | 1.39E+07 | 1.11E+07 | 1.16E+07 | | | | | | |
| O88685 | PSMC3 | proteasome (prosome, macropain) 26S subunit, ATPase, 3 | Nucleus | transcription regulator | 1.06E+07 | 9282497 | 5126888 | | | | | | |
| O88685 | PSMC3 | proteasome (prosome, macropain) 26S subunit, ATPase, 3 | Nucleus | transcription regulator | 1.71E+07 | 1.28E+07 | 8063205 | | | | | | |
| O88685 | PSMC3 | proteasome (prosome, macropain) 26S subunit, ATPase, 3 | Nucleus | transcription regulator | 1.31E+07 | 1.06E+07 | 8379101.5 | | | | | | |
| O88685 | PSMC3 | proteasome (prosome, macropain) 26S subunit, ATPase, 3 | Nucleus | transcription regulator | 9848397 | 8209730.5 | 7272059 | | | | | | |
| O88685 | PSMC3 | proteasome (prosome, macropain) 26S subunit, ATPase, 3 | Nucleus | transcription regulator | 7265547 | 8789117 | 6256017.5 | | | | | | |
| O88685 | PSMC3 | proteasome (prosome, macropain) 26S subunit, ATPase, 3 | Nucleus | transcription regulator | 6669573 | 5773335 | 4086177.5 | | | | | | |

| | | | | | | | | | | | | | |
|---|---|---|---|---|---|---|---|---|---|---|---|---|---|
| O88685 | PSMC3 | proteasome (prosome, macropain) 26S subunit, ATPase, 3 | Nucleus | transcription regulator | 5405939.5 | 4536782.5 | 4688300.5 | | | | | | |
| O88685 | PSMC3 | proteasome (prosome, macropain) 26S subunit, ATPase, 3 | Nucleus | transcription regulator | 4439440.5 | 3036930 | 3191683.75 | | | | | | |
| O88685 | PSMC3 | proteasome (prosome, macropain) 26S subunit, ATPase, 3 | Nucleus | transcription regulator | 3168511.75 | 3533050.75 | 3125282 | O88685 | PSMC3 | 11922672 | 5764661 | 889507 | 42 |
| P54775 | PSMC4 | proteasome (prosome, macropain) 26S subunit, ATPase, 4 | Nucleus | peptidase | 1.89E+07 | 2.26E+07 | 1.38E+07 | | | | | | |
| P54775 | PSMC4 | proteasome (prosome, macropain) 26S subunit, ATPase, 4 | Nucleus | peptidase | 1.51E+07 | 1.74E+07 | 1.60E+07 | | | | | | |
| P54775 | PSMC4 | proteasome (prosome, macropain) 26S subunit, ATPase, 4 | Nucleus | peptidase | 7361664.5 | 8001478.5 | 8991986 | | | | | | |
| P54775 | PSMC4 | proteasome (prosome, macropain) 26S subunit, ATPase, 4 | Nucleus | peptidase | 9576601 | 7482096 | 7539904 | | | | | | |
| P54775 | PSMC4 | proteasome (prosome, macropain) 26S subunit, ATPase, 4 | Nucleus | peptidase | 8957887 | 6614991 | 5789388 | | | | | | |
| P54775 | PSMC4 | proteasome (prosome, macropain) 26S subunit, ATPase, 4 | Nucleus | peptidase | 1.09E+07 | 9380791 | 7401748 | | | | | | |
| P54775 | PSMC4 | proteasome (prosome, macropain) 26S subunit, ATPase, 4 | Nucleus | peptidase | 5350442.5 | 6091205.5 | 6259812 | | | | | | |
| P54775 | PSMC4 | proteasome (prosome, macropain) 26S subunit, ATPase, 4 | Nucleus | peptidase | 6050380 | 3042584.25 | 2114646 | | | | | | |
| P54775 | PSMC4 | proteasome (prosome, macropain) 26S subunit, ATPase, 4 | Nucleus | peptidase | 6421300 | 6267384 | 4324997.5 | | | | | | |
| P54775 | PSMC4 | proteasome (prosome, macropain) 26S subunit, ATPase, 4 | Nucleus | peptidase | 4640666.5 | 5768987 | 5140808 | | | | | | |
| P54775 | PSMC4 | proteasome (prosome, macropain) 26S subunit, ATPase, 4 | Nucleus | peptidase | 4737333.5 | 5082174.5 | 4521587.5 | | | | | | |
| P54775 | PSMC4 | proteasome (prosome, macropain) 26S subunit, ATPase, 4 | Nucleus | peptidase | 4052721.75 | 3284024 | 2904116 | P54775 | PSMC4 | 8500750 | 4795363 | 799227 | 36 |
| P62196 | PSMC5 | proteasome (prosome, macropain) 26S subunit, ATPase, 5 | Nucleus | transcription regulator | 2.78E+07 | 2.54E+07 | 2.27E+07 | | | | | | |
| P62196 | PSMC5 | proteasome (prosome, macropain) 26S subunit, ATPase, 5 | Nucleus | transcription regulator | 1.65E+07 | 1.52E+07 | 1.25E+07 | | | | | | |
| P62196 | PSMC5 | proteasome (prosome, macropain) 26S subunit, ATPase, 5 | Nucleus | transcription regulator | 1.58E+07 | 1.58E+07 | 1.37E+07 | | | | | | |
| P62196 | PSMC5 | proteasome (prosome, macropain) 26S subunit, ATPase, 5 | Nucleus | transcription regulator | 1.71E+07 | 1.48E+07 | 1.49E+07 | | | | | | |
| P62196 | PSMC5 | proteasome (prosome, macropain) 26S subunit, ATPase, 5 | Nucleus | transcription regulator | 1.18E+07 | 1.11E+07 | 9548991 | | | | | | |
| P62196 | PSMC5 | proteasome (prosome, macropain) 26S subunit, ATPase, 5 | Nucleus | transcription regulator | 1.49E+07 | 9369937 | 8046391.5 | | | | | | |
| P62196 | PSMC5 | proteasome (prosome, macropain) 26S subunit, ATPase, 5 | Nucleus | transcription regulator | 1.04E+07 | 8769979 | 7790104 | | | | | | |
| P62196 | PSMC5 | proteasome (prosome, macropain) 26S subunit, ATPase, 5 | Nucleus | transcription regulator | 1.02E+07 | 7633209 | 7227514 | | | | | | |
| P62196 | PSMC5 | proteasome (prosome, macropain) 26S subunit, ATPase, 5 | Nucleus | transcription regulator | 1.03E+07 | 6597997 | 7071831 | | | | | | |
| P62196 | PSMC5 | proteasome (prosome, macropain) 26S subunit, ATPase, 5 | Nucleus | transcription regulator | 6489848 | 4422431 | 4268702.5 | | | | | | |
| P62196 | PSMC5 | proteasome (prosome, macropain) 26S subunit, ATPase, 5 | Nucleus | transcription regulator | 7836649 | 5820495 | 4884727.5 | | | | | | |
| P62196 | PSMC5 | proteasome (prosome, macropain) 26S subunit, ATPase, 5 | Nucleus | transcription regulator | 7181775.5 | 7882393 | 7259841 | | | | | | |
| P62196 | PSMC5 | proteasome (prosome, macropain) 26S subunit, ATPase, 5 | Nucleus | transcription regulator | 9741933 | 4613970 | 3571165.25 | | | | | | |
| P62196 | PSMC5 | proteasome (prosome, macropain) 26S subunit, ATPase, 5 | Nucleus | transcription regulator | 5742529 | 5470211.5 | 4942987.5 | | | | | | |
| P62196 | PSMC5 | proteasome (prosome, macropain) 26S subunit, ATPase, 5 | Nucleus | transcription regulator | 3820232.25 | 4832488.5 | 3614807 | | | | | | |
| P62196 | PSMC5 | proteasome (prosome, macropain) 26S subunit, ATPase, 5 | Nucleus | transcription regulator | 4764561.5 | 1166006.375 | 16351.349 | P62196 | PSMC5 | 11272971 | 5878902 | 848546 | 48 |
| P62334 | PSMC6 | proteasome (prosome, macropain) 26S subunit, ATPase, 6 | Nucleus | peptidase | 3.17E+07 | 2.74E+07 | 2.13E+07 | | | | | | |
| P62334 | PSMC6 | proteasome (prosome, macropain) 26S subunit, ATPase, 6 | Nucleus | peptidase | 1.43E+07 | 1.30E+07 | 8566636 | | | | | | |
| P62334 | PSMC6 | proteasome (prosome, macropain) 26S subunit, ATPase, 6 | Nucleus | peptidase | 1.37E+07 | 1.43E+07 | 1.68E+07 | | | | | | |
| P62334 | PSMC6 | proteasome (prosome, macropain) 26S subunit, ATPase, 6 | Nucleus | peptidase | 1.22E+07 | 8162555 | 7164232 | | | | | | |
| P62334 | PSMC6 | proteasome (prosome, macropain) 26S subunit, ATPase, 6 | Nucleus | peptidase | 1.51E+07 | 9395107 | 7373533.5 | | | | | | |
| P62334 | PSMC6 | proteasome (prosome, macropain) 26S subunit, ATPase, 6 | Nucleus | peptidase | 1.24E+07 | 9342188 | 1.03E+07 | | | | | | |
| P62334 | PSMC6 | proteasome (prosome, macropain) 26S subunit, ATPase, 6 | Nucleus | peptidase | 8963239 | 7671634 | 8225226 | | | | | | |
| P62334 | PSMC6 | proteasome (prosome, macropain) 26S subunit, ATPase, 6 | Nucleus | peptidase | 8882121 | 5503683.5 | 5846486.5 | | | | | | |
| P62334 | PSMC6 | proteasome (prosome, macropain) 26S subunit, ATPase, 6 | Nucleus | peptidase | 1.03E+07 | 8559153 | 1.02E+07 | | | | | | |
| P62334 | PSMC6 | proteasome (prosome, macropain) 26S subunit, ATPase, 6 | Nucleus | peptidase | 1.01E+07 | 6943073 | 6169239 | | | | | | |
| P62334 | PSMC6 | proteasome (prosome, macropain) 26S subunit, ATPase, 6 | Nucleus | peptidase | 6934638.5 | 6201696 | 5073776.5 | | | | | | |
| P62334 | PSMC6 | proteasome (prosome, macropain) 26S subunit, ATPase, 6 | Nucleus | peptidase | 7465955.5 | 6844512 | 5505170 | | | | | | |
| P62334 | PSMC6 | proteasome (prosome, macropain) 26S subunit, ATPase, 6 | Nucleus | peptidase | 3254254.5 | 1931295.875 | 1659614.25 | P62334 | PSMC6 | 11928478 | 6077772 | 973222 | 39 |
| Q3TXS7 | PSMD1 | proteasome (prosome, macropain) 26S subunit, non-ATPase, 1 | Cytoplasm | other | 3.04E+07 | 2.48E+07 | 2.43E+07 | | | | | | |
| Q3TXS7 | PSMD1 | proteasome (prosome, macropain) 26S subunit, non-ATPase, 1 | Cytoplasm | other | 2.96E+07 | 2.62E+07 | 2.69E+07 | | | | | | |
| Q3TXS7 | PSMD1 | proteasome (prosome, macropain) 26S subunit, non-ATPase, 1 | Cytoplasm | other | 1.69E+07 | 1.52E+07 | 1.39E+07 | | | | | | |
| Q3TXS7 | PSMD1 | proteasome (prosome, macropain) 26S subunit, non-ATPase, 1 | Cytoplasm | other | 2.65E+07 | 1.75E+07 | 1.86E+07 | | | | | | |
| Q3TXS7 | PSMD1 | proteasome (prosome, macropain) 26S subunit, non-ATPase, 1 | Cytoplasm | other | 1.98E+07 | 1.47E+07 | 1.42E+07 | | | | | | |
| Q3TXS7 | PSMD1 | proteasome (prosome, macropain) 26S subunit, non-ATPase, 1 | Cytoplasm | other | 1.70E+07 | 1.19E+07 | 1.14E+07 | | | | | | |
| Q3TXS7 | PSMD1 | proteasome (prosome, macropain) 26S subunit, non-ATPase, 1 | Cytoplasm | other | 1.40E+07 | 1.50E+07 | 1.19E+07 | | | | | | |
| Q3TXS7 | PSMD1 | proteasome (prosome, macropain) 26S subunit, non-ATPase, 1 | Cytoplasm | other | 1.34E+07 | 1.15E+07 | 1.21E+07 | | | | | | |
| Q3TXS7 | PSMD1 | proteasome (prosome, macropain) 26S subunit, non-ATPase, 1 | Cytoplasm | other | 1.69E+07 | 1.92E+07 | 1.80E+07 | | | | | | |
| Q3TXS7 | PSMD1 | proteasome (prosome, macropain) 26S subunit, non-ATPase, 1 | Cytoplasm | other | 1.15E+07 | 1.09E+07 | 1.04E+07 | | | | | | |
| Q3TXS7 | PSMD1 | proteasome (prosome, macropain) 26S subunit, non-ATPase, 1 | Cytoplasm | other | 1.15E+07 | 1.01E+07 | 6394377 | | | | | | |
| Q3TXS7 | PSMD1 | proteasome (prosome, macropain) 26S subunit, non-ATPase, 1 | Cytoplasm | other | 1.23E+07 | 9541447 | 8420319 | | | | | | |
| Q3TXS7 | PSMD1 | proteasome (prosome, macropain) 26S subunit, non-ATPase, 1 | Cytoplasm | other | 1.11E+07 | 9871244 | 9355329 | | | | | | |
| Q3TXS7 | PSMD1 | proteasome (prosome, macropain) 26S subunit, non-ATPase, 1 | Cytoplasm | other | 1.15E+07 | 1.05E+07 | 9278336 | | | | | | |
| Q3TXS7 | PSMD1 | proteasome (prosome, macropain) 26S subunit, non-ATPase, 1 | Cytoplasm | other | 6625369 | 5545548 | 6969661 | | | | | | |
| Q3TXS7 | PSMD1 | proteasome (prosome, macropain) 26S subunit, non-ATPase, 1 | Cytoplasm | other | 8286203.5 | 6216676.5 | 5764802.5 | | | | | | |
| Q3TXS7 | PSMD1 | proteasome (prosome, macropain) 26S subunit, non-ATPase, 1 | Cytoplasm | other | 8108940.5 | 6647296 | 5920804.5 | | | | | | |
| Q3TXS7 | PSMD1 | proteasome (prosome, macropain) 26S subunit, non-ATPase, 1 | Cytoplasm | other | 9170741 | 7499863.5 | 7097405.5 | | | | | | |
| Q3TXS7 | PSMD1 | proteasome (prosome, macropain) 26S subunit, non-ATPase, 1 | Cytoplasm | other | 7549658 | 6802640 | 5555950.5 | | | | | | |
| Q3TXS7 | PSMD1 | proteasome (prosome, macropain) 26S subunit, non-ATPase, 1 | Cytoplasm | other | 4843553 | 3459116.75 | 1205083.875 | | | | | | |
| Q3TXS7 | PSMD1 | proteasome (prosome, macropain) 26S subunit, non-ATPase, 1 | Cytoplasm | other | 4810444.5 | 3214860.75 | 3503041.75 | | | | | | |
| Q3TXS7 | PSMD1 | proteasome (prosome, macropain) 26S subunit, non-ATPase, 1 | Cytoplasm | other | 4897666 | 3901064.5 | 4803458.5 | | | | | | |
| Q3TXS7 | PSMD1 | proteasome (prosome, macropain) 26S subunit, non-ATPase, 1 | Cytoplasm | other | 4408883 | 5625718.5 | 6064236.5 | | | | | | |
| Q3TXS7 | PSMD1 | proteasome (prosome, macropain) 26S subunit, non-ATPase, 1 | Cytoplasm | other | 3763885.25 | 1451304.5 | 3728257.25 | | | | | | |
| Q3TXS7 | PSMD1 | proteasome (prosome, macropain) 26S subunit, non-ATPase, 1 | Cytoplasm | other | 4358427.5 | 3495258 | 3007772.75 | | | | | | |
| Q3TXS7 | PSMD1 | proteasome (prosome, macropain) 26S subunit, non-ATPase, 1 | Cytoplasm | other | 2228469.25 | 2227916 | 1801174.125 | | | | | | |
| Q3TXS7 | PSMD1 | proteasome (prosome, macropain) 26S subunit, non-ATPase, 1 | Cytoplasm | other | 1467714.875 | 2580552 | 3716971 | | | | | | |
| Q3TXS7 | PSMD1 | proteasome (prosome, macropain) 26S subunit, non-ATPase, 1 | Cytoplasm | other | 2383354.5 | 1600048 | 1590741.875 | Q3TXS7 | PSMD1 | 11263690 | 7070027 | 771403 | 84 |
| Q8BG32 | PSMD11 | proteasome (prosome, macropain) 26S subunit, non-ATPase, 11 | Cytoplasm | other | 2.04E+07 | 1.97E+07 | 1.68E+07 | | | | | | |
| Q8BG32 | PSMD11 | proteasome (prosome, macropain) 26S subunit, non-ATPase, 11 | Cytoplasm | other | 1.64E+07 | 1.21E+07 | 1.30E+07 | | | | | | |
| Q8BG32 | PSMD11 | proteasome (prosome, macropain) 26S subunit, non-ATPase, 11 | Cytoplasm | other | 1.95E+07 | 1.62E+07 | 1.38E+07 | | | | | | |
| Q8BG32 | PSMD11 | proteasome (prosome, macropain) 26S subunit, non-ATPase, 11 | Cytoplasm | other | 1.74E+07 | 1.76E+07 | 1.45E+07 | | | | | | |
| Q8BG32 | PSMD11 | proteasome (prosome, macropain) 26S subunit, non-ATPase, 11 | Cytoplasm | other | 1.81E+07 | 1.46E+07 | 1.21E+07 | | | | | | |
| Q8BG32 | PSMD11 | proteasome (prosome, macropain) 26S subunit, non-ATPase, 11 | Cytoplasm | other | 1.34E+07 | 1.22E+07 | 1.08E+07 | | | | | | |
| Q8BG32 | PSMD11 | proteasome (prosome, macropain) 26S subunit, non-ATPase, 11 | Cytoplasm | other | 1.12E+07 | 1.00E+07 | 6063597 | | | | | | |
| Q8BG32 | PSMD11 | proteasome (prosome, macropain) 26S subunit, non-ATPase, 11 | Cytoplasm | other | 1.26E+07 | 9343088 | 1.33E+07 | | | | | | |
| Q8BG32 | PSMD11 | proteasome (prosome, macropain) 26S subunit, non-ATPase, 11 | Cytoplasm | other | 7765119.5 | 6348554.5 | 6186525 | | | | | | |
| Q8BG32 | PSMD11 | proteasome (prosome, macropain) 26S subunit, non-ATPase, 11 | Cytoplasm | other | 6586172.5 | 7414530.5 | 7017824.5 | | | | | | |
| Q8BG32 | PSMD11 | proteasome (prosome, macropain) 26S subunit, non-ATPase, 11 | Cytoplasm | other | 6658703 | 5662134.5 | 3635400 | | | | | | |
| Q8BG32 | PSMD11 | proteasome (prosome, macropain) 26S subunit, non-ATPase, 11 | Cytoplasm | other | 6105738 | 4689562 | 3745792 | | | | | | |
| Q8BG32 | PSMD11 | proteasome (prosome, macropain) 26S subunit, non-ATPase, 11 | Cytoplasm | other | 2221433.75 | 3902451.5 | 3983386.5 | | | | | | |
| Q8BG32 | PSMD11 | proteasome (prosome, macropain) 26S subunit, non-ATPase, 11 | Cytoplasm | other | 1694506.625 | 1604285.125 | 2223863 | Q8BG32 | PSMD11 | 11414405 | 5550043 | 856390 | 42 |
| Q9D8W5 | PSMD12 | proteasome (prosome, macropain) 26S subunit, non-ATPase, 12 | Cytoplasm | other | 1.99E+07 | 1.73E+07 | 1.94E+07 | | | | | | |
| Q9D8W5 | PSMD12 | proteasome (prosome, macropain) 26S subunit, non-ATPase, 12 | Cytoplasm | other | 1.76E+07 | 1.37E+07 | 1.48E+07 | | | | | | |
| Q9D8W5 | PSMD12 | proteasome (prosome, macropain) 26S subunit, non-ATPase, 12 | Cytoplasm | other | 1.71E+07 | 1.27E+07 | 1.09E+07 | | | | | | |
| Q9D8W5 | PSMD12 | proteasome (prosome, macropain) 26S subunit, non-ATPase, 12 | Cytoplasm | other | 1.20E+07 | 1.14E+07 | 8049965.5 | | | | | | |
| Q9D8W5 | PSMD12 | proteasome (prosome, macropain) 26S subunit, non-ATPase, 12 | Cytoplasm | other | 1.23E+07 | 7435439 | 9833165 | | | | | | |

| | | | | | | | | | | | | | |
|---|---|---|---|---|---|---|---|---|---|---|---|---|---|
| Q9D8W5 | PSMD12 | proteasome (prosome, macropain) 26S subunit, non-ATPase, 12 | Cytoplasm | other | 9759118 | 8491852 | 8348764 | | | | | | |
| Q9D8W5 | PSMD12 | proteasome (prosome, macropain) 26S subunit, non-ATPase, 12 | Cytoplasm | other | 1.07E+07 | 1.07E+07 | 8444764 | | | | | | |
| Q9D8W5 | PSMD12 | proteasome (prosome, macropain) 26S subunit, non-ATPase, 12 | Cytoplasm | other | 7296755.5 | 6218972.5 | 7380494.5 | | | | | | |
| Q9D8W5 | PSMD12 | proteasome (prosome, macropain) 26S subunit, non-ATPase, 12 | Cytoplasm | other | 1.07E+07 | 7755587.5 | 6006138.5 | | | | | | |
| Q9D8W5 | PSMD12 | proteasome (prosome, macropain) 26S subunit, non-ATPase, 12 | Cytoplasm | other | 6289009.5 | 3924335.5 | 3837049.5 | | | | | | |
| Q9D8W5 | PSMD12 | proteasome (prosome, macropain) 26S subunit, non-ATPase, 12 | Cytoplasm | other | 6151405 | 4634990.5 | 3277009.5 | | | | | | |
| Q9D8W5 | PSMD12 | proteasome (prosome, macropain) 26S subunit, non-ATPase, 12 | Cytoplasm | other | 4729442.5 | 5212703 | 5023324.5 | | | | | | |
| Q9D8W5 | PSMD12 | proteasome (prosome, macropain) 26S subunit, non-ATPase, 12 | Cytoplasm | other | 3101299.5 | 2490948 | 2567707.25 | | | | | | |
| Q9D8W5 | PSMD12 | proteasome (prosome, macropain) 26S subunit, non-ATPase, 12 | Cytoplasm | other | 4526334.5 | 3585604.5 | 3356168 | | | | | | |
| Q9D8W5 | PSMD12 | proteasome (prosome, macropain) 26S subunit, non-ATPase, 12 | Cytoplasm | other | 2207699.25 | 1442832.25 | 901106.812 | Q9D8W5 | PSMD12 | 9616071 | 4966900 | 740422 | 45 |
| Q9WVJ2 | PSMD13 | proteasome (prosome, macropain) 26S subunit, non-ATPase, 13 | Cytoplasm | peptidase | 2.56E+07 | 2.88E+07 | 2.67E+07 | | | | | | |
| Q9WVJ2 | PSMD13 | proteasome (prosome, macropain) 26S subunit, non-ATPase, 13 | Cytoplasm | peptidase | 3.20E+07 | 2.77E+07 | 2.59E+07 | | | | | | |
| Q9WVJ2 | PSMD13 | proteasome (prosome, macropain) 26S subunit, non-ATPase, 13 | Cytoplasm | peptidase | 1.96E+07 | 1.89E+07 | 1.16E+07 | | | | | | |
| Q9WVJ2 | PSMD13 | proteasome (prosome, macropain) 26S subunit, non-ATPase, 13 | Cytoplasm | peptidase | 1.93E+07 | 1.71E+07 | 1.86E+07 | | | | | | |
| Q9WVJ2 | PSMD13 | proteasome (prosome, macropain) 26S subunit, non-ATPase, 13 | Cytoplasm | peptidase | 1.64E+07 | 1.59E+07 | 1.43E+07 | | | | | | |
| Q9WVJ2 | PSMD13 | proteasome (prosome, macropain) 26S subunit, non-ATPase, 13 | Cytoplasm | peptidase | 1.03E+07 | 1.14E+07 | 1.09E+07 | | | | | | |
| Q9WVJ2 | PSMD13 | proteasome (prosome, macropain) 26S subunit, non-ATPase, 13 | Cytoplasm | peptidase | 1.38E+07 | 1.14E+07 | 9165787 | | | | | | |
| Q9WVJ2 | PSMD13 | proteasome (prosome, macropain) 26S subunit, non-ATPase, 13 | Cytoplasm | peptidase | 1.44E+07 | 1.20E+07 | 1.14E+07 | | | | | | |
| Q9WVJ2 | PSMD13 | proteasome (prosome, macropain) 26S subunit, non-ATPase, 13 | Cytoplasm | peptidase | 1.10E+07 | 1.05E+07 | 1.06E+07 | | | | | | |
| Q9WVJ2 | PSMD13 | proteasome (prosome, macropain) 26S subunit, non-ATPase, 13 | Cytoplasm | peptidase | 1.35E+07 | 1.02E+07 | 1.02E+07 | | | | | | |
| Q9WVJ2 | PSMD13 | proteasome (prosome, macropain) 26S subunit, non-ATPase, 13 | Cytoplasm | peptidase | 8117230 | 5423799.5 | 4585057 | | | | | | |
| Q9WVJ2 | PSMD13 | proteasome (prosome, macropain) 26S subunit, non-ATPase, 13 | Cytoplasm | peptidase | 5851590 | 4479462.5 | 5427270.5 | | | | | | |
| Q9WVJ2 | PSMD13 | proteasome (prosome, macropain) 26S subunit, non-ATPase, 13 | Cytoplasm | peptidase | 3901848.25 | 1655990.125 | 967238.062 | Q9WVJ2 | PSMD13 | 14913898 | 7736292 | 1238798 | 39 |
| O35593 | PSMD14 | proteasome (prosome, macropain) 26S subunit, non-ATPase, 14 | Cytoplasm | peptidase | 2.14E+07 | 2.33E+07 | 1.88E+07 | | | | | | |
| O35593 | PSMD14 | proteasome (prosome, macropain) 26S subunit, non-ATPase, 14 | Cytoplasm | peptidase | 1.53E+07 | 1.88E+07 | 1.63E+07 | | | | | | |
| O35593 | PSMD14 | proteasome (prosome, macropain) 26S subunit, non-ATPase, 14 | Cytoplasm | peptidase | 7138931.5 | 4136009 | 5488456 | | | | | | |
| O35593 | PSMD14 | proteasome (prosome, macropain) 26S subunit, non-ATPase, 14 | Cytoplasm | peptidase | 2973990.25 | 4061951.5 | 2572174.25 | O35593 | PSMD14 | 11715730 | 7973582 | 2301775 | 12 |
| Q8VDM4 | PSMD2 | proteasome (prosome, macropain) 26S subunit, non-ATPase, 2 | Cytoplasm | other | 2.34E+07 | 2.87E+07 | 6885377 | | | | | | |
| Q8VDM4 | PSMD2 | proteasome (prosome, macropain) 26S subunit, non-ATPase, 2 | Cytoplasm | other | 2.33E+07 | 1.96E+07 | 1.87E+07 | | | | | | |
| Q8VDM4 | PSMD2 | proteasome (prosome, macropain) 26S subunit, non-ATPase, 2 | Cytoplasm | other | 1.56E+07 | 1.22E+07 | 1.37E+07 | | | | | | |
| Q8VDM4 | PSMD2 | proteasome (prosome, macropain) 26S subunit, non-ATPase, 2 | Cytoplasm | other | 2.12E+07 | 1.04E+07 | 3546547.5 | | | | | | |
| Q8VDM4 | PSMD2 | proteasome (prosome, macropain) 26S subunit, non-ATPase, 2 | Cytoplasm | other | 1.34E+07 | 1.09E+07 | 1.00E+07 | | | | | | |
| Q8VDM4 | PSMD2 | proteasome (prosome, macropain) 26S subunit, non-ATPase, 2 | Cytoplasm | other | 1.26E+07 | 9540623 | 6590506.5 | | | | | | |
| Q8VDM4 | PSMD2 | proteasome (prosome, macropain) 26S subunit, non-ATPase, 2 | Cytoplasm | other | 1.63E+07 | 1.17E+07 | 1.50E+07 | | | | | | |
| Q8VDM4 | PSMD2 | proteasome (prosome, macropain) 26S subunit, non-ATPase, 2 | Cytoplasm | other | 1.09E+07 | 9475397 | 8975338 | | | | | | |
| Q8VDM4 | PSMD2 | proteasome (prosome, macropain) 26S subunit, non-ATPase, 2 | Cytoplasm | other | 1.23E+07 | 9808694 | 9703330 | | | | | | |
| Q8VDM4 | PSMD2 | proteasome (prosome, macropain) 26S subunit, non-ATPase, 2 | Cytoplasm | other | 1.10E+07 | 1.03E+07 | 7849729 | | | | | | |
| Q8VDM4 | PSMD2 | proteasome (prosome, macropain) 26S subunit, non-ATPase, 2 | Cytoplasm | other | 9108982 | 1.05E+07 | 6348851.5 | | | | | | |
| Q8VDM4 | PSMD2 | proteasome (prosome, macropain) 26S subunit, non-ATPase, 2 | Cytoplasm | other | 1.24E+07 | 1.12E+07 | 1.10E+07 | | | | | | |
| Q8VDM4 | PSMD2 | proteasome (prosome, macropain) 26S subunit, non-ATPase, 2 | Cytoplasm | other | 7627312.5 | 6914442 | 5284318.5 | | | | | | |
| Q8VDM4 | PSMD2 | proteasome (prosome, macropain) 26S subunit, non-ATPase, 2 | Cytoplasm | other | 8224100 | 9232334 | 7346695 | | | | | | |
| Q8VDM4 | PSMD2 | proteasome (prosome, macropain) 26S subunit, non-ATPase, 2 | Cytoplasm | other | 9913991 | 8297860 | 7885115.5 | | | | | | |
| Q8VDM4 | PSMD2 | proteasome (prosome, macropain) 26S subunit, non-ATPase, 2 | Cytoplasm | other | 6653197 | 1.09E+07 | 1.40E+07 | | | | | | |
| Q8VDM4 | PSMD2 | proteasome (prosome, macropain) 26S subunit, non-ATPase, 2 | Cytoplasm | other | 9650198 | 6239370.5 | 5844238.5 | | | | | | |
| Q8VDM4 | PSMD2 | proteasome (prosome, macropain) 26S subunit, non-ATPase, 2 | Cytoplasm | other | 3437202.75 | 4079966 | 2838226 | | | | | | |
| Q8VDM4 | PSMD2 | proteasome (prosome, macropain) 26S subunit, non-ATPase, 2 | Cytoplasm | other | 6999586 | 6813920.5 | 7287318 | | | | | | |
| Q8VDM4 | PSMD2 | proteasome (prosome, macropain) 26S subunit, non-ATPase, 2 | Cytoplasm | other | 5808295 | 5149076 | 4224926 | | | | | | |
| Q8VDM4 | PSMD2 | proteasome (prosome, macropain) 26S subunit, non-ATPase, 2 | Cytoplasm | other | 6758415.5 | 5313544.5 | 4937785.5 | | | | | | |
| Q8VDM4 | PSMD2 | proteasome (prosome, macropain) 26S subunit, non-ATPase, 2 | Cytoplasm | other | 6329859 | 5322247.5 | 4907331.5 | | | | | | |
| Q8VDM4 | PSMD2 | proteasome (prosome, macropain) 26S subunit, non-ATPase, 2 | Cytoplasm | other | 6741117.5 | 4968193 | 3639300.5 | | | | | | |
| Q8VDM4 | PSMD2 | proteasome (prosome, macropain) 26S subunit, non-ATPase, 2 | Cytoplasm | other | 2679442.75 | 2864816.75 | 2548204.25 | | | | | | |
| Q8VDM4 | PSMD2 | proteasome (prosome, macropain) 26S subunit, non-ATPase, 2 | Cytoplasm | other | 4429330.5 | 3069288 | 2183389 | | | | | | |
| Q8VDM4 | PSMD2 | proteasome (prosome, macropain) 26S subunit, non-ATPase, 2 | Cytoplasm | other | 2448720.5 | 2996297.25 | 2936557.25 | | | | | | |
| Q8VDM4 | PSMD2 | proteasome (prosome, macropain) 26S subunit, non-ATPase, 2 | Cytoplasm | other | 3829316.75 | 3259295.5 | 2144792.25 | | | | | | |
| Q8VDM4 | PSMD2 | proteasome (prosome, macropain) 26S subunit, non-ATPase, 2 | Cytoplasm | other | 3319535.25 | 2912422.25 | 3211608 | | | | | | |
| Q8VDM4 | PSMD2 | proteasome (prosome, macropain) 26S subunit, non-ATPase, 2 | Cytoplasm | other | 3440949.75 | 3866551.75 | 2257284.5 | Q8VDM4 | PSMD2 | 9645502 | 5313836 | 569703 | 87 |
| P14685 | PSMD3 | proteasome (prosome, macropain) 26S subunit, non-ATPase, 3 | Cytoplasm | other | 2.64E+07 | 3.18E+07 | 2.23E+07 | | | | | | |
| P14685 | PSMD3 | proteasome (prosome, macropain) 26S subunit, non-ATPase, 3 | Cytoplasm | other | 3.28E+07 | 2.85E+07 | 2.44E+07 | | | | | | |
| P14685 | PSMD3 | proteasome (prosome, macropain) 26S subunit, non-ATPase, 3 | Cytoplasm | other | 2.36E+07 | 2.34E+07 | 2.52E+07 | | | | | | |
| P14685 | PSMD3 | proteasome (prosome, macropain) 26S subunit, non-ATPase, 3 | Cytoplasm | other | 1.46E+07 | 1.20E+07 | 1.43E+07 | | | | | | |
| P14685 | PSMD3 | proteasome (prosome, macropain) 26S subunit, non-ATPase, 3 | Cytoplasm | other | 1.53E+07 | 1.02E+07 | 1.03E+07 | | | | | | |
| P14685 | PSMD3 | proteasome (prosome, macropain) 26S subunit, non-ATPase, 3 | Cytoplasm | other | 1.92E+07 | 9238202 | 7445270.5 | | | | | | |
| P14685 | PSMD3 | proteasome (prosome, macropain) 26S subunit, non-ATPase, 3 | Cytoplasm | other | 1.22E+07 | 9107809 | 1.04E+07 | | | | | | |
| P14685 | PSMD3 | proteasome (prosome, macropain) 26S subunit, non-ATPase, 3 | Cytoplasm | other | 9654503 | 9208760 | 1.09E+07 | | | | | | |
| P14685 | PSMD3 | proteasome (prosome, macropain) 26S subunit, non-ATPase, 3 | Cytoplasm | other | 8639337 | 4251482.5 | 1824388.375 | | | | | | |
| P14685 | PSMD3 | proteasome (prosome, macropain) 26S subunit, non-ATPase, 3 | Cytoplasm | other | 6505623.5 | 4479934.5 | 5696253 | | | | | | |
| P14685 | PSMD3 | proteasome (prosome, macropain) 26S subunit, non-ATPase, 3 | Cytoplasm | other | 4403120 | 4766292 | 6945150 | | | | | | |
| P14685 | PSMD3 | proteasome (prosome, macropain) 26S subunit, non-ATPase, 3 | Cytoplasm | other | 3607126.25 | 1760888.5 | 1298787.375 | | | | | | |
| P14685 | PSMD3 | proteasome (prosome, macropain) 26S subunit, non-ATPase, 3 | Cytoplasm | other | 3282609 | 3208235.5 | 3068970.25 | | | | | | |
| P14685 | PSMD3 | proteasome (prosome, macropain) 26S subunit, non-ATPase, 3 | Cytoplasm | other | 2646826 | 1615345.625 | 2133735.75 | | | | | | |
| P14685 | PSMD3 | proteasome (prosome, macropain) 26S subunit, non-ATPase, 3 | Cytoplasm | other | 6396351 | 1493952.5 | 1092722.875 | | | | | | |
| P14685 | PSMD3 | proteasome (prosome, macropain) 26S subunit, non-ATPase, 3 | Cytoplasm | other | 3073651.25 | 3438951.75 | 2845547.5 | P14685 | PSMD3 | 12021822 | 8958087 | 1292989 | 48 |
| O35226 | PSMD4 | proteasome (prosome, macropain) 26S subunit, non-ATPase, 4 | Cytoplasm | other | 3.76E+07 | 3.01E+07 | 2.41E+07 | | | | | | |
| O35226 | PSMD4 | proteasome (prosome, macropain) 26S subunit, non-ATPase, 4 | Cytoplasm | other | 3.13E+07 | 2.18E+07 | 2.41E+07 | | | | | | |
| O35226 | PSMD4 | proteasome (prosome, macropain) 26S subunit, non-ATPase, 4 | Cytoplasm | other | 1.53E+07 | 1.45E+07 | 1.52E+07 | | | | | | |
| O35226 | PSMD4 | proteasome (prosome, macropain) 26S subunit, non-ATPase, 4 | Cytoplasm | other | 1.42E+07 | 1.07E+07 | 1.13E+07 | | | | | | |
| O35226 | PSMD4 | proteasome (prosome, macropain) 26S subunit, non-ATPase, 4 | Cytoplasm | other | 1.36E+07 | 1.51E+07 | 9188966 | | | | | | |
| O35226 | PSMD4 | proteasome (prosome, macropain) 26S subunit, non-ATPase, 4 | Cytoplasm | other | 9195543 | 4499295 | 2849363.25 | | | | | | |
| O35226 | PSMD4 | proteasome (prosome, macropain) 26S subunit, non-ATPase, 4 | Cytoplasm | other | 4560690.5 | 3721200.25 | 3081366 | | | | | | |
| O35226 | PSMD4 | proteasome (prosome, macropain) 26S subunit, non-ATPase, 4 | Cytoplasm | other | 3826911 | 2838686.75 | 2593644.25 | | | | | | |
| O35226 | PSMD4 | proteasome (prosome, macropain) 26S subunit, non-ATPase, 4 | Cytoplasm | other | 2759054.25 | 2369326 | 2058791.125 | O35226 | PSMD4 | 14701355 | 10114134 | 1946466 | 27 |
| Q8BJY1 | PSMD5 | proteasome (prosome, macropain) 26S subunit, non-ATPase, 5 | Cytoplasm | other | 3964960.25 | 3908454.75 | 3136501 | | | | | | |
| Q8BJY1 | PSMD5 | proteasome (prosome, macropain) 26S subunit, non-ATPase, 5 | Cytoplasm | other | 3507071.75 | 2652561.75 | 1886454.75 | | | | | | |
| Q8BJY1 | PSMD5 | proteasome (prosome, macropain) 26S subunit, non-ATPase, 5 | Cytoplasm | other | 3490180.5 | 2575320 | 3139781.25 | | | | | | |
| Q8BJY1 | PSMD5 | proteasome (prosome, macropain) 26S subunit, non-ATPase, 5 | Cytoplasm | other | 3103813.75 | 2724328.5 | 3574283.75 | | | | | | |
| Q8BJY1 | PSMD5 | proteasome (prosome, macropain) 26S subunit, non-ATPase, 5 | Cytoplasm | other | 2234010.75 | 1557297.5 | 2214248.5 | | | | | | |
| Q8BJY1 | PSMD5 | proteasome (prosome, macropain) 26S subunit, non-ATPase, 5 | Cytoplasm | other | 2511669.75 | 1653829.75 | 1385243.125 | | | | | | |
| Q8BJY1 | PSMD5 | proteasome (prosome, macropain) 26S subunit, non-ATPase, 5 | Cytoplasm | other | 2211164 | 1998396.375 | 1240223.25 | | | | | | |
| Q8BJY1 | PSMD5 | proteasome (prosome, macropain) 26S subunit, non-ATPase, 5 | Cytoplasm | other | 1512570.375 | 1766707.875 | 1337521.625 | | | | | | |
| Q8BJY1 | PSMD5 | proteasome (prosome, macropain) 26S subunit, non-ATPase, 5 | Cytoplasm | other | 1756601.375 | 1323347.75 | 1297328 | Q8BJY1 | PSMD5 | 2699116 | 866868 | 166829 | 27 |
| Q99JI4 | PSMD6 | proteasome (prosome, macropain) 26S subunit, non-ATPase, 6 | Cytoplasm | enzyme | 2.05E+07 | 1.72E+07 | 1.65E+07 | | | | | | |

| | | | | | | | | | | | | | |
|---|---|---|---|---|---|---|---|---|---|---|---|---|---|
| Q99JI4 | PSMD6 | proteasome (prosome, macropain) 26S subunit, non-ATPase, 6 | Cytoplasm | enzyme | 1.85E+07 | 1.78E+07 | 1.66E+07 | | | | | | |
| Q99JI4 | PSMD6 | proteasome (prosome, macropain) 26S subunit, non-ATPase, 6 | Cytoplasm | enzyme | 1.81E+07 | 1.26E+07 | 1.15E+07 | | | | | | |
| Q99JI4 | PSMD6 | proteasome (prosome, macropain) 26S subunit, non-ATPase, 6 | Cytoplasm | enzyme | 1.48E+07 | 9925692 | 1.08E+07 | | | | | | |
| Q99JI4 | PSMD6 | proteasome (prosome, macropain) 26S subunit, non-ATPase, 6 | Cytoplasm | enzyme | 1.25E+07 | 1.05E+07 | 1.02E+07 | | | | | | |
| Q99JI4 | PSMD6 | proteasome (prosome, macropain) 26S subunit, non-ATPase, 6 | Cytoplasm | enzyme | 1.08E+07 | 8101536.5 | 7526109 | | | | | | |
| Q99JI4 | PSMD6 | proteasome (prosome, macropain) 26S subunit, non-ATPase, 6 | Cytoplasm | enzyme | 1.09E+07 | 9007866 | 7109014.5 | | | | | | |
| Q99JI4 | PSMD6 | proteasome (prosome, macropain) 26S subunit, non-ATPase, 6 | Cytoplasm | enzyme | 5520954.5 | 4260359.5 | 4710693 | | | | | | |
| Q99JI4 | PSMD6 | proteasome (prosome, macropain) 26S subunit, non-ATPase, 6 | Cytoplasm | enzyme | 4699752.5 | 4208926.5 | 3893520.75 | | | | | | |
| Q99JI4 | PSMD6 | proteasome (prosome, macropain) 26S subunit, non-ATPase, 6 | Cytoplasm | enzyme | 4848336.5 | 3512564.25 | 3698312.25 | | | | | | |
| Q99JI4 | PSMD6 | proteasome (prosome, macropain) 26S subunit, non-ATPase, 6 | Cytoplasm | enzyme | 6487780.5 | 4653563 | 5442585.5 | | | | | | |
| Q99JI4 | PSMD6 | proteasome (prosome, macropain) 26S subunit, non-ATPase, 6 | Cytoplasm | enzyme | 3822302.25 | 3231397 | 2649607.75 | | | | | | |
| Q99JI4 | PSMD6 | proteasome (prosome, macropain) 26S subunit, non-ATPase, 6 | Cytoplasm | enzyme | 4339876 | 5736050.5 | 6603388 | | | | | | |
| Q99JI4 | PSMD6 | proteasome (prosome, macropain) 26S subunit, non-ATPase, 6 | Cytoplasm | enzyme | 1933377 | 2323766.75 | 1775581.625 | Q99JI4 | PSMD6 | 9845170 | 5324292 | 821556 | 42 |
| P26516 | PSMD7 | proteasome (prosome, macropain) 26S subunit, non-ATPase, 7 | Cytoplasm | other | 2.80E+07 | 3.14E+07 | 3.04E+07 | | | | | | |
| P26516 | PSMD7 | proteasome (prosome, macropain) 26S subunit, non-ATPase, 7 | Cytoplasm | other | 2.24E+07 | 1.81E+07 | 1.54E+07 | | | | | | |
| P26516 | PSMD7 | proteasome (prosome, macropain) 26S subunit, non-ATPase, 7 | Cytoplasm | other | 2.63E+07 | 1.79E+07 | 1.48E+07 | | | | | | |
| P26516 | PSMD7 | proteasome (prosome, macropain) 26S subunit, non-ATPase, 7 | Cytoplasm | other | 1.63E+07 | 1.27E+07 | 7395878.5 | | | | | | |
| P26516 | PSMD7 | proteasome (prosome, macropain) 26S subunit, non-ATPase, 7 | Cytoplasm | other | 9716775 | 8321715 | 7921119 | | | | | | |
| P26516 | PSMD7 | proteasome (prosome, macropain) 26S subunit, non-ATPase, 7 | Cytoplasm | other | 6049104 | 4490820 | 3818808.75 | | | | | | |
| P26516 | PSMD7 | proteasome (prosome, macropain) 26S subunit, non-ATPase, 7 | Cytoplasm | other | 8007733 | 1.08E+07 | 6088149 | | | | | | |
| P26516 | PSMD7 | proteasome (prosome, macropain) 26S subunit, non-ATPase, 7 | Cytoplasm | other | 4152762 | 4491990.5 | 3785305.25 | | | | | | |
| P26516 | PSMD7 | proteasome (prosome, macropain) 26S subunit, non-ATPase, 7 | Cytoplasm | other | 2732829 | 3209928 | 2298573.25 | | | | | | |
| P26516 | PSMD7 | proteasome (prosome, macropain) 26S subunit, non-ATPase, 7 | Cytoplasm | other | 2225947.5 | 1151190.5 | 177850.078 | P26516 | PSMD7 | 12583515 | 9150390 | 1670625 | 30 |
| Q9CX56 | PSMD8 | proteasome (prosome, macropain) 26S subunit, non-ATPase, 8 | Cytoplasm | other | 1.60E+07 | 1.35E+07 | 1.27E+07 | | | | | | |
| Q9CX56 | PSMD8 | proteasome (prosome, macropain) 26S subunit, non-ATPase, 8 | Cytoplasm | other | 8456655 | 8898329 | 9180655 | | | | | | |
| Q9CX56 | PSMD8 | proteasome (prosome, macropain) 26S subunit, non-ATPase, 8 | Cytoplasm | other | 8872047 | 6793391 | 6750128 | Q9CX56 | PSMD8 | 11106234 | 3193701 | 1064567 | 9 |
| P97371 | PSME1 | proteasome (prosome, macropain) activator subunit 1 (PA28 alpha) | Cytoplasm | other | 1.67E+07 | 1.30E+07 | 1.19E+07 | | | | | | |
| P97371 | PSME1 | proteasome (prosome, macropain) activator subunit 1 (PA28 alpha) | Cytoplasm | other | 1.30E+07 | 1.03E+07 | 1.38E+07 | | | | | | |
| P97371 | PSME1 | proteasome (prosome, macropain) activator subunit 1 (PA28 alpha) | Cytoplasm | other | 1.40E+07 | 1.27E+07 | 1.32E+07 | | | | | | |
| P97371 | PSME1 | proteasome (prosome, macropain) activator subunit 1 (PA28 alpha) | Cytoplasm | other | 1.28E+07 | 8875247 | 1.03E+07 | | | | | | |
| P97371 | PSME1 | proteasome (prosome, macropain) activator subunit 1 (PA28 alpha) | Cytoplasm | other | 9807131 | 1.09E+07 | 1.03E+07 | | | | | | |
| P97371 | PSME1 | proteasome (prosome, macropain) activator subunit 1 (PA28 alpha) | Cytoplasm | other | 7675536 | 4413385 | 4327889.5 | | | | | | |
| P97371 | PSME1 | proteasome (prosome, macropain) activator subunit 1 (PA28 alpha) | Cytoplasm | other | 6034810 | 4772678 | 4839431.5 | | | | | | |
| P97371 | PSME1 | proteasome (prosome, macropain) activator subunit 1 (PA28 alpha) | Cytoplasm | other | 6821964 | 6179811 | 7364569.5 | | | | | | |
| P97371 | PSME1 | proteasome (prosome, macropain) activator subunit 1 (PA28 alpha) | Cytoplasm | other | 5764211 | 4811978 | 4901996 | | | | | | |
| P97371 | PSME1 | proteasome (prosome, macropain) activator subunit 1 (PA28 alpha) | Cytoplasm | other | 5563537 | 4199080.5 | 4276480.5 | | | | | | |
| P97371 | PSME1 | proteasome (prosome, macropain) activator subunit 1 (PA28 alpha) | Cytoplasm | other | 7099014 | 5637403.5 | 7911131 | | | | | | |
| P97371 | PSME1 | proteasome (prosome, macropain) activator subunit 1 (PA28 alpha) | Cytoplasm | other | 4019228.75 | 1669829 | 1202399.875 | P97371 | PSME1 | 9107953 | 3901995 | 650332 | 36 |
| P97372 | PSME2 | proteasome (prosome, macropain) activator subunit 2 (PA28 beta) | Cytoplasm | peptidase | 1.81E+07 | 2.25E+07 | 2.59E+07 | | | | | | |
| P97372 | PSME2 | proteasome (prosome, macropain) activator subunit 2 (PA28 beta) | Cytoplasm | peptidase | 1.43E+07 | 1.14E+07 | 1.29E+07 | | | | | | |
| P97372 | PSME2 | proteasome (prosome, macropain) activator subunit 2 (PA28 beta) | Cytoplasm | peptidase | 1.50E+07 | 8988538 | 8444989 | | | | | | |
| P97372 | PSME2 | proteasome (prosome, macropain) activator subunit 2 (PA28 beta) | Cytoplasm | peptidase | 4545005.5 | 3959764 | 5711127.5 | | | | | | |
| P97372 | PSME2 | proteasome (prosome, macropain) activator subunit 2 (PA28 beta) | Cytoplasm | peptidase | 2312787.25 | 1809754.75 | 2072431.375 | | | | | | |
| P97372 | PSME2 | proteasome (prosome, macropain) activator subunit 2 (PA28 beta) | Cytoplasm | peptidase | 3347991 | 2937196.5 | 2224083 | | | | | | |
| P97372 | PSME2 | proteasome (prosome, macropain) activator subunit 2 (PA28 beta) | Cytoplasm | peptidase | 5257292 | 2921186.75 | 2361615.5 | P97372 | PSME2 | 8969011 | 7199404 | 1571039 | 21 |
| Q8R3K3 | PTCD2 | pentatricopeptide repeat domain 2 | Cytoplasm | other | 2758270.25 | 3170801 | 4793018 | Q8R3K3 | PTCD2 | 2758270 | 1075637 | 621019 | 3 |
| O08586 | PTEN | phosphatase and tensin homolog | Cytoplasm | phosphatase | 2.78E+07 | 2.94E+07 | 3.15E+07 | O08586 | PTEN | 27810000 | 1827165 | 1054914 | 3 |
| Q63739 | PTP4A1 | protein tyrosine phosphatase type IVA, member 1 | Cytoplasm | phosphatase | 1912251.25 | 2351276.25 | 2377217 | Q63739 | PTP4A1 | 1912251 | 261282 | 150851 | 3 |
| P35821 | PTPN1 | protein tyrosine phosphatase, non-receptor type 1 | Cytoplasm | phosphatase | 461199.531 | 1532042.125 | 2295777.5 | P35821 | PTPN1 | 461200 | 921563 | 532065 | 3 |
| Q62136 | PTPN21 | protein tyrosine phosphatase, non-receptor type 21 | Cytoplasm | phosphatase | 2.59E+07 | 2.81E+07 | 3.05E+07 | Q62136 | PTPN21 | 25880000 | 2300261 | 1328056 | 3 |
| P18052 | PTPRA | protein tyrosine phosphatase, receptor type, A | Plasma Membrane | phosphatase | 7220281 | 9313756 | 7652854 | | | | | | |
| P18052 | PTPRA | protein tyrosine phosphatase, receptor type, A | Plasma Membrane | phosphatase | 3540743.25 | 4212167.5 | 3968625 | | | | | | |
| P18052 | PTPRA | protein tyrosine phosphatase, receptor type, A | Plasma Membrane | phosphatase | 3081100.25 | 3142402 | 3327469.75 | | | | | | |
| P18052 | PTPRA | protein tyrosine phosphatase, receptor type, A | Plasma Membrane | phosphatase | 571604.75 | 619892.625 | 743892.25 | P18052 | PTPRA | 3603432 | 2828256 | 816447 | 12 |
| Q64455 | PTPRJ | protein tyrosine phosphatase, receptor type, J | Plasma Membrane | phosphatase | 1638559.25 | 2140633 | 3925082.75 | | | | | | |
| Q64455 | PTPRJ | protein tyrosine phosphatase, receptor type, J | Plasma Membrane | phosphatase | 114561.227 | 150605.781 | 313855.719 | | | | | | |
| Q64455 | PTPRJ | protein tyrosine phosphatase, receptor type, J | Plasma Membrane | phosphatase | 212366.25 | 370154.75 | 609669.625 | Q64455 | PTPRJ | 655162 | 1293517 | 431172 | 9 |
| Q8R2Y8 | PTRH2 | peptidyl-tRNA hydrolase 2 | Cytoplasm | enzyme | 8003421.5 | 1.28E+07 | 6678229.5 | | | | | | |
| Q8R2Y8 | PTRH2 | peptidyl-tRNA hydrolase 2 | Cytoplasm | enzyme | 241477.953 | 333048.875 | 166497.984 | Q8R2Y8 | PTRH2 | 4122450 | 5287256 | 2158513 | 6 |
| P42669 | PURA | purine-rich element binding protein A | Nucleus | transcription regulator | 3192043.5 | 5170950.5 | 2887393.25 | | | | | | |
| P42669 | PURA | purine-rich element binding protein A | Nucleus | transcription regulator | 1851673.125 | 399724.969 | 87850.188 | P42669 | PURA | 2521858 | 1901652 | 776346 | 6 |
| Q922Q4 | PYCR2 | pyrroline-5-carboxylate reductase family, member 2 | Cytoplasm | enzyme | 3156187.75 | 5615282 | 2215576.75 | Q922Q4 | PYCR2 | 3156188 | 1755462 | 1013517 | 3 |
| Q9DCC4 | PYCRL | pyrroline-5-carboxylate reductase-like | Cytoplasm | enzyme | 4649621 | 3161131 | 4385816 | | | | | | |
| Q9DCC4 | PYCRL | pyrroline-5-carboxylate reductase-like | Cytoplasm | enzyme | 3323869.5 | 2524528.5 | 2475935.75 | | | | | | |
| Q9DCC4 | PYCRL | pyrroline-5-carboxylate reductase-like | Cytoplasm | enzyme | 2942116.25 | 1739899.5 | 1444882.125 | Q9DCC4 | PYCRL | 3638536 | 1076401 | 358800 | 9 |
| Q61838 | Pzp | pregnancy zone protein | Extracellular Space | other | 2601545 | 3469367.75 | 5945992.5 | Q61838 | Pzp | 2601545 | 1735517 | 1002001 | 3 |
| Q8BVI4 | QDPR | quinoid dihydropteridine reductase | Cytoplasm | enzyme | 978666.125 | 579340.562 | 230167.281 | Q8BVI4 | QDPR | 978666 | 374529 | 216235 | 3 |
| P61027 | RAB10 | RAB10, member RAS oncogene family | Cytoplasm | enzyme | 2.52E+08 | 2.97E+08 | 3.16E+08 | | | | | | |
| P61027 | RAB10 | RAB10, member RAS oncogene family | Cytoplasm | enzyme | 1.45E+08 | 1.32E+08 | 1.68E+08 | | | | | | |
| P61027 | RAB10 | RAB10, member RAS oncogene family | Cytoplasm | enzyme | 1.26E+08 | 1.50E+08 | 1.53E+08 | | | | | | |
| P61027 | RAB10 | RAB10, member RAS oncogene family | Cytoplasm | enzyme | 3.17E+07 | 4.02E+07 | 4.17E+07 | | | | | | |
| P61027 | RAB10 | RAB10, member RAS oncogene family | Cytoplasm | enzyme | 3.49E+07 | 3.72E+07 | 5.05E+07 | | | | | | |
| P61027 | RAB10 | RAB10, member RAS oncogene family | Cytoplasm | enzyme | 3.04E+07 | 3.89E+07 | 5.76E+07 | | | | | | |
| P61027 | RAB10 | RAB10, member RAS oncogene family | Cytoplasm | enzyme | 3.25E+07 | 3.04E+07 | 4.14E+07 | | | | | | |
| P61027 | RAB10 | RAB10, member RAS oncogene family | Cytoplasm | enzyme | 2.31E+07 | 2.04E+07 | 2.07E+07 | | | | | | |
| P61027 | RAB10 | RAB10, member RAS oncogene family | Cytoplasm | enzyme | 1.47E+07 | 1.80E+07 | 3.38E+07 | | | | | | |
| P61027 | RAB10 | RAB10, member RAS oncogene family | Cytoplasm | enzyme | 1.09E+07 | 1.41E+07 | 2.55E+07 | | | | | | |
| P61027 | RAB10 | RAB10, member RAS oncogene family | Cytoplasm | enzyme | 4797672.5 | 4391122.5 | 3768655.75 | P61027 | RAB10 | 64087970 | 84408146 | 14693572 | 33 |
| P46638 | RAB11B | RAB11B, member RAS oncogene family | Cytoplasm | enzyme | 2.44E+07 | 3.27E+07 | 3.78E+07 | | | | | | |
| P46638 | RAB11B | RAB11B, member RAS oncogene family | Cytoplasm | enzyme | 1.67E+07 | 2.14E+07 | 2.35E+07 | | | | | | |
| P46638 | RAB11B | RAB11B, member RAS oncogene family | Cytoplasm | enzyme | 1.59E+07 | 2.41E+07 | 2.42E+07 | | | | | | |
| P46638 | RAB11B | RAB11B, member RAS oncogene family | Cytoplasm | enzyme | 1.51E+07 | 2.42E+07 | 2.23E+07 | | | | | | |
| P46638 | RAB11B | RAB11B, member RAS oncogene family | Cytoplasm | enzyme | 7562565.5 | 8940274 | 1.22E+07 | | | | | | |
| P46638 | RAB11B | RAB11B, member RAS oncogene family | Cytoplasm | enzyme | 5997448.5 | 8233744.5 | 8326126.5 | | | | | | |
| P46638 | RAB11B | RAB11B, member RAS oncogene family | Cytoplasm | enzyme | 6215647.5 | 6141658 | 5837160.5 | | | | | | |
| P46638 | RAB11B | RAB11B, member RAS oncogene family | Cytoplasm | enzyme | 4052336.75 | 4273775.5 | 2812120.5 | | | | | | |
| P46638 | RAB11B | RAB11B, member RAS oncogene family | Cytoplasm | enzyme | 3307154.25 | 2375591 | 1992760 | | | | | | |
| P46638 | RAB11B | RAB11B, member RAS oncogene family | Cytoplasm | enzyme | 584195.125 | 1647054.75 | 1542306.625 | P46638 | RAB11B | 9977935 | 10301482 | 1880785 | 30 |
| Q91V41 | RAB14 | RAB14, member RAS oncogene family | Cytoplasm | enzyme | 2.06E+07 | 3.12E+07 | 3.78E+07 | | | | | | |
| Q91V41 | RAB14 | RAB14, member RAS oncogene family | Cytoplasm | enzyme | 1.73E+07 | 1.83E+07 | 2.49E+07 | | | | | | |
| Q91V41 | RAB14 | RAB14, member RAS oncogene family | Cytoplasm | enzyme | 7734236.5 | 6989269 | 6280041.5 | | | | | | |

| | | | | | | | | | | | | | |
|---|---|---|---|---|---|---|---|---|---|---|---|---|---|
| Q91V41 | RAB14 | RAB14, member RAS oncogene family | Cytoplasm | enzyme | 9406102 | 1.04E+07 | 1.16E+07 | | | | | | |
| Q91V41 | RAB14 | RAB14, member RAS oncogene family | Cytoplasm | enzyme | 5915561 | 6217456.5 | 5760876 | | | | | | |
| Q91V41 | RAB14 | RAB14, member RAS oncogene family | Cytoplasm | enzyme | 5273508.5 | 4169722 | 5042068.5 | | | | | | |
| Q91V41 | RAB14 | RAB14, member RAS oncogene family | Cytoplasm | enzyme | 5863400 | 1069067.25 | 159843.375 | | | | | | |
| Q91V41 | RAB14 | RAB14, member RAS oncogene family | Cytoplasm | enzyme | 3011887.5 | 1716616.125 | 1444841.125 | | | | | | |
| Q91V41 | RAB14 | RAB14, member RAS oncogene family | Cytoplasm | enzyme | 1825844.25 | 410250.406 | 0 | Q91V41 | RAB14 | 8543393 | 9743393 | 1875117 | 27 |
| Q8K386 | Rab15 | RAB15, member RAS oncogene family | Cytoplasm | enzyme | 3.84E+08 | 3.92E+08 | 4.59E+08 | Q8K386 | Rab15 | 383500000 | 41584893 | 24009049 | 3 |
| P35293 | RAB18 | RAB18, member RAS oncogene family | Cytoplasm | enzyme | 1.24E+07 | 1.11E+07 | 1.55E+07 | | | | | | |
| P35293 | RAB18 | RAB18, member RAS oncogene family | Cytoplasm | enzyme | 2.09E+07 | 1.85E+07 | 2.30E+07 | | | | | | |
| P35293 | RAB18 | RAB18, member RAS oncogene family | Cytoplasm | enzyme | 1.49E+07 | 1.30E+07 | 1.86E+07 | | | | | | |
| P35293 | RAB18 | RAB18, member RAS oncogene family | Cytoplasm | enzyme | 1.21E+07 | 1.23E+07 | 1.55E+07 | | | | | | |
| P35293 | RAB18 | RAB18, member RAS oncogene family | Cytoplasm | enzyme | 1.01E+07 | 8995980 | 1.12E+07 | P35293 | RAB18 | 14080000 | 4106331 | 1060250 | 15 |
| P62821 | RAB1A | RAB1A, member RAS oncogene family | Cytoplasm | enzyme | 5.76E+07 | 6.31E+07 | 9.64E+07 | | | | | | |
| P62821 | RAB1A | RAB1A, member RAS oncogene family | Cytoplasm | enzyme | 2.16E+07 | 2.72E+07 | 4.14E+07 | | | | | | |
| P62821 | RAB1A | RAB1A, member RAS oncogene family | Cytoplasm | enzyme | 8179963 | 9112367 | 1.19E+07 | | | | | | |
| P62821 | RAB1A | RAB1A, member RAS oncogene family | Cytoplasm | enzyme | 6692259.5 | 3785415.5 | 5146571 | | | | | | |
| P62821 | RAB1A | RAB1A, member RAS oncogene family | Cytoplasm | enzyme | 1804450 | 2923735.5 | 4879386 | | | | | | |
| P62821 | RAB1A | RAB1A, member RAS oncogene family | Cytoplasm | enzyme | 1762765.25 | 1572515.25 | 1797701.625 | P62821 | RAB1A | 16278240 | 27060763 | 6378283 | 18 |
| Q9D1G1 | RAB1B | RAB1B, member RAS oncogene family | Cytoplasm | other | 5.02E+07 | 3.56E+07 | 4.70E+07 | | | | | | |
| Q9D1G1 | RAB1B | RAB1B, member RAS oncogene family | Cytoplasm | other | 2.08E+07 | 2.38E+07 | 3.12E+07 | | | | | | |
| Q9D1G1 | RAB1B | RAB1B, member RAS oncogene family | Cytoplasm | other | 8951950 | 1.46E+07 | 2.32E+07 | | | | | | |
| Q9D1G1 | RAB1B | RAB1B, member RAS oncogene family | Cytoplasm | other | 1.28E+07 | 1.04E+07 | 1.75E+07 | | | | | | |
| Q9D1G1 | RAB1B | RAB1B, member RAS oncogene family | Cytoplasm | other | 3788883.5 | 4015207 | 6766054.5 | Q9D1G1 | RAB1B | 19296167 | 14692792 | 3793663 | 15 |
| P35282 | RAB21 | RAB21, member RAS oncogene family | Cytoplasm | enzyme | 7954212.5 | 8744297 | 1.30E+07 | | | | | | |
| P35282 | RAB21 | RAB21, member RAS oncogene family | Cytoplasm | enzyme | 6806983.5 | 2906705.25 | 1797613.75 | | | | | | |
| P35282 | RAB21 | RAB21, member RAS oncogene family | Cytoplasm | enzyme | 2944130.75 | 2799367 | 4681994 | | | | | | |
| P35282 | RAB21 | RAB21, member RAS oncogene family | Cytoplasm | enzyme | 2508711.75 | 2675946.5 | 4246373 | | | | | | |
| P35282 | RAB21 | RAB21, member RAS oncogene family | Cytoplasm | enzyme | 2632587 | 1389825.5 | 398417.469 | | | | | | |
| P35282 | RAB21 | RAB21, member RAS oncogene family | Cytoplasm | enzyme | 3490606.75 | 2925405 | 4155364.75 | | | | | | |
| P35282 | RAB21 | RAB21, member RAS oncogene family | Cytoplasm | enzyme | 1578635.375 | 2042287.625 | 2994071.5 | | | | | | |
| P35282 | RAB21 | RAB21, member RAS oncogene family | Cytoplasm | enzyme | 469295.125 | 447813.125 | 623547.562 | P35282 | RAB21 | 3548145 | 2985510 | 609415 | 24 |
| P35285 | RAB22A | RAB22A, member RAS oncogene family | Cytoplasm | enzyme | 3484953.75 | 3593603.25 | 4238680 | | | | | | |
| P35285 | RAB22A | RAB22A, member RAS oncogene family | Cytoplasm | enzyme | 3274662.5 | 3433126.75 | 4940686.5 | P35285 | RAB22A | 3379808 | 638977 | 260861 | 6 |
| Q9ERI2 | RAB27A | RAB27A, member RAS oncogene family | Cytoplasm | enzyme | 410868 | 589170 | 1386806.625 | Q9ERI2 | RAB27A | 410868 | 519691 | 300044 | 3 |
| Q99P58 | RAB27B | RAB27B, member RAS oncogene family | Cytoplasm | enzyme | 1429881.75 | 3805150.5 | 1.09E+07 | | | | | | |
| Q99P58 | RAB27B | RAB27B, member RAS oncogene family | Cytoplasm | enzyme | 805306.438 | 2765884 | 5005194 | | | | | | |
| Q99P58 | RAB27B | RAB27B, member RAS oncogene family | Cytoplasm | enzyme | 568656.125 | 2799404.5 | 6581304 | | | | | | |
| Q99P58 | RAB27B | RAB27B, member RAS oncogene family | Cytoplasm | enzyme | 687372.312 | 3037304 | 6952851 | | | | | | |
| Q99P58 | RAB27B | RAB27B, member RAS oncogene family | Cytoplasm | enzyme | 250485.688 | 1498707.25 | 3414340 | | | | | | |
| Q99P58 | RAB27B | RAB27B, member RAS oncogene family | Cytoplasm | enzyme | 55430.348 | 460671.469 | 1147034.625 | | | | | | |
| Q99P58 | RAB27B | RAB27B, member RAS oncogene family | Cytoplasm | enzyme | 22797.357 | 63429.32 | 115167.258 | | | | | | |
| Q99P58 | RAB27B | RAB27B, member RAS oncogene family | Cytoplasm | enzyme | 76027.219 | 379080.719 | 1187489 | Q99P58 | RAB27B | 486995 | 2745643 | 560452 | 24 |
| P53994 | RAB2A | RAB2A, member RAS oncogene family | Cytoplasm | enzyme | 8328764 | 1.11E+07 | 1.09E+07 | | | | | | |
| P53994 | RAB2A | RAB2A, member RAS oncogene family | Cytoplasm | enzyme | 6041752.5 | 6070222.5 | 8333171 | | | | | | |
| P53994 | RAB2A | RAB2A, member RAS oncogene family | Cytoplasm | enzyme | 2231591.25 | 3553828 | 4882218 | | | | | | |
| P53994 | RAB2A | RAB2A, member RAS oncogene family | Cytoplasm | enzyme | 2696321.75 | 3507624.25 | 3201784.75 | | | | | | |
| P53994 | RAB2A | RAB2A, member RAS oncogene family | Cytoplasm | enzyme | 1360988.5 | 2307478.25 | 2703593.75 | P53994 | RAB2A | 4131884 | 3181544 | 821471 | 15 |
| Q6PHN9 | RAB35 | RAB35, member RAS oncogene family | Cytoplasm | enzyme | 9.51E+07 | 8.88E+07 | 1.05E+08 | | | | | | |
| Q6PHN9 | RAB35 | RAB35, member RAS oncogene family | Cytoplasm | enzyme | 5.07E+07 | 5.21E+07 | 4.48E+07 | | | | | | |
| Q6PHN9 | RAB35 | RAB35, member RAS oncogene family | Cytoplasm | enzyme | 4.95E+07 | 4.55E+07 | 5.83E+07 | | | | | | |
| Q6PHN9 | RAB35 | RAB35, member RAS oncogene family | Cytoplasm | enzyme | 3.70E+07 | 1.77E+07 | 1.28E+07 | | | | | | |
| Q6PHN9 | RAB35 | RAB35, member RAS oncogene family | Cytoplasm | enzyme | 2.91E+07 | 2.83E+07 | 3.10E+07 | | | | | | |
| Q6PHN9 | RAB35 | RAB35, member RAS oncogene family | Cytoplasm | enzyme | 2.06E+07 | 1.63E+07 | 402892.062 | | | | | | |
| Q6PHN9 | RAB35 | RAB35, member RAS oncogene family | Cytoplasm | enzyme | 5299933 | 4867269 | 1.22E+07 | Q6PHN9 | RAB35 | 41038562 | 29640613 | 6468112 | 21 |
| Q9JKM7 | RAB37 | RAB37, member RAS oncogene family | Cytoplasm | enzyme | 0 | 66767.188 | 280040.25 | Q9JKM7 | RAB37 | 115602 | 146268 | 84448 | 3 |
| Q80UJ7 | RAB3GAP1 | RAB3 GTPase activating protein subunit 1 (catalytic) | Cytoplasm | other | 5905942 | 4522219.5 | 4921565 | | | | | | |
| Q80UJ7 | RAB3GAP1 | RAB3 GTPase activating protein subunit 1 (catalytic) | Cytoplasm | other | 7261843 | 1711801.5 | 1273039.5 | | | | | | |
| Q80UJ7 | RAB3GAP1 | RAB3 GTPase activating protein subunit 1 (catalytic) | Cytoplasm | other | 221795.531 | 244348.188 | 4.70E-06 | Q80UJ7 | RAB3GAP1 | 4463194 | 2772039 | 924013 | 9 |
| Q8BMG7 | RAB3GAP2 | RAB3 GTPase activating protein subunit 2 (non-catalytic) | Cytoplasm | enzyme | 2076368.125 | 1334981.25 | 1129139.25 | | | | | | |
| Q8BMG7 | RAB3GAP2 | RAB3 GTPase activating protein subunit 2 (non-catalytic) | Cytoplasm | enzyme | 1279627.875 | 474067.062 | 462316.031 | Q8BMG7 | RAB3GAP2 | 1677998 | 605967 | 247385 | 6 |
| Q8VDV3 | RAB3IL1 | RAB3A interacting protein (rabin3)-like 1 | Cytoplasm | other | 1389778.125 | 2837999.75 | 1660617.875 | Q8VDV3 | RAB3IL1 | 1389778 | 769949 | 444530 | 3 |
| Q9CQD1 | RAB5A | RAB5A, member RAS oncogene family | Cytoplasm | enzyme | 2.75E+07 | 3.16E+07 | 2.90E+07 | | | | | | |
| Q9CQD1 | RAB5A | RAB5A, member RAS oncogene family | Cytoplasm | enzyme | 2.96E+07 | 3.21E+07 | 3.47E+07 | | | | | | |
| Q9CQD1 | RAB5A | RAB5A, member RAS oncogene family | Cytoplasm | enzyme | 9569961 | 1.16E+07 | 1.70E+07 | | | | | | |
| Q9CQD1 | RAB5A | RAB5A, member RAS oncogene family | Cytoplasm | enzyme | 6426476 | 1.14E+07 | 1.51E+07 | | | | | | |
| Q9CQD1 | RAB5A | RAB5A, member RAS oncogene family | Cytoplasm | enzyme | 6167508.5 | 9376250 | 9346418 | Q9CQD1 | RAB5A | 15854789 | 10649825 | 2749773 | 15 |
| P61021 | RAB5B | RAB5B, member RAS oncogene family | Cytoplasm | enzyme | 2.83E+07 | 2.80E+07 | 2.67E+07 | | | | | | |
| P61021 | RAB5B | RAB5B, member RAS oncogene family | Cytoplasm | enzyme | 1.55E+07 | 1.71E+07 | 1.94E+07 | | | | | | |
| P61021 | RAB5B | RAB5B, member RAS oncogene family | Cytoplasm | enzyme | 1.40E+07 | 1.51E+07 | 1.65E+07 | | | | | | |
| P61021 | RAB5B | RAB5B, member RAS oncogene family | Cytoplasm | enzyme | 8174856.5 | 8427275 | 1.07E+07 | | | | | | |
| P61021 | RAB5B | RAB5B, member RAS oncogene family | Cytoplasm | enzyme | 5694259 | 7792032.5 | 5104532 | | | | | | |
| P61021 | RAB5B | RAB5B, member RAS oncogene family | Cytoplasm | enzyme | 3990310.25 | 5215631 | 6638844 | P61021 | RAB5B | 12599904 | 8042994 | 1895752 | 18 |
| P35278 | RAB5C | RAB5C, member RAS oncogene family | Cytoplasm | enzyme | 1.19E+08 | 1.25E+08 | 1.50E+08 | | | | | | |
| P35278 | RAB5C | RAB5C, member RAS oncogene family | Cytoplasm | enzyme | 9.37E+07 | 1.08E+08 | 1.37E+08 | | | | | | |
| P35278 | RAB5C | RAB5C, member RAS oncogene family | Cytoplasm | enzyme | 6.74E+07 | 7.07E+07 | 9.94E+07 | | | | | | |
| P35278 | RAB5C | RAB5C, member RAS oncogene family | Cytoplasm | enzyme | 6.17E+07 | 7.40E+07 | 8.79E+07 | | | | | | |
| P35278 | RAB5C | RAB5C, member RAS oncogene family | Cytoplasm | enzyme | 7.04E+07 | 7.80E+07 | 8.08E+07 | | | | | | |
| P35278 | RAB5C | RAB5C, member RAS oncogene family | Cytoplasm | enzyme | 2.36E+07 | 2.42E+07 | 3.05E+07 | | | | | | |
| P35278 | RAB5C | RAB5C, member RAS oncogene family | Cytoplasm | enzyme | 2.93E+07 | 3.14E+07 | 4.34E+07 | | | | | | |
| P35278 | RAB5C | RAB5C, member RAS oncogene family | Cytoplasm | enzyme | 2.35E+07 | 2.68E+07 | 3.25E+07 | | | | | | |
| P35278 | RAB5C | RAB5C, member RAS oncogene family | Cytoplasm | enzyme | 1.55E+07 | 1.68E+07 | 9979207 | | | | | | |
| P35278 | RAB5C | RAB5C, member RAS oncogene family | Cytoplasm | enzyme | 1.78E+07 | 1.79E+07 | 1.56E+07 | | | | | | |
| P35278 | RAB5C | RAB5C, member RAS oncogene family | Cytoplasm | enzyme | 1.03E+07 | 1.08E+07 | 1.40E+07 | | | | | | |
| P35278 | RAB5C | RAB5C, member RAS oncogene family | Cytoplasm | enzyme | 3501659.5 | 3424221.5 | 2896887.25 | | | | | | |
| P35278 | RAB5C | RAB5C, member RAS oncogene family | Cytoplasm | enzyme | 2790160.25 | 2818122.25 | 3637031.5 | P35278 | RAB5C | 41440140 | 42650060 | 6829475 | 39 |
| P35279 | RAB6A | RAB6A, member RAS oncogene family | Cytoplasm | enzyme | 2945052.75 | 3282663.75 | 5499730.5 | | | | | | |
| P35279 | RAB6A | RAB6A, member RAS oncogene family | Cytoplasm | enzyme | 1148824.5 | 1510668.125 | 2028876.75 | P35279 | RAB6A | 2046939 | 1581223 | 645532 | 6 |
| P51150 | RAB7A | RAB7A, member RAS oncogene family | Cytoplasm | enzyme | 1.47E+07 | 1.63E+07 | 2.23E+07 | | | | | | |
| P51150 | RAB7A | RAB7A, member RAS oncogene family | Cytoplasm | enzyme | 1.60E+07 | 1.70E+07 | 2.56E+07 | | | | | | |
| P51150 | RAB7A | RAB7A, member RAS oncogene family | Cytoplasm | enzyme | 1.24E+07 | 1.83E+07 | 2.39E+07 | | | | | | |
| P51150 | RAB7A | RAB7A, member RAS oncogene family | Cytoplasm | enzyme | 1.12E+07 | 1.40E+07 | 1.67E+07 | | | | | | |

| | | | | | | | | | | | | | |
|---|---|---|---|---|---|---|---|---|---|---|---|---|---|
| P51150 | RAB7A | RAB7A, member RAS oncogene family | Cytoplasm | enzyme | 1.16E+07 | 1.17E+07 | 1.70E+07 | | | | | | |
| P51150 | RAB7A | RAB7A, member RAS oncogene family | Cytoplasm | enzyme | 7816309 | 1.16E+07 | 1.27E+07 | | | | | | |
| P51150 | RAB7A | RAB7A, member RAS oncogene family | Cytoplasm | enzyme | 1.08E+07 | 1.25E+07 | 1.40E+07 | | | | | | |
| P51150 | RAB7A | RAB7A, member RAS oncogene family | Cytoplasm | enzyme | 8868007 | 1.48E+07 | 1.71E+07 | | | | | | |
| P51150 | RAB7A | RAB7A, member RAS oncogene family | Cytoplasm | enzyme | 4628645 | 6369745.5 | 8335958 | | | | | | |
| P51150 | RAB7A | RAB7A, member RAS oncogene family | Cytoplasm | enzyme | 7975097.5 | 4591662.5 | 4799437.5 | | | | | | |
| P51150 | RAB7A | RAB7A, member RAS oncogene family | Cytoplasm | enzyme | 2164497.5 | 4117613.75 | 3002823.5 | P51150 | RAB7A | 9828414 | 5903378 | 1027646 | 33 |
| P55258 | RAB8A | RAB8A, member RAS oncogene family | Plasma Membrane | enzyme | 9.60E+07 | 9.73E+07 | 1.13E+08 | | | | | | |
| P55258 | RAB8A | RAB8A, member RAS oncogene family | Plasma Membrane | enzyme | 7.48E+07 | 7.80E+07 | 1.03E+08 | | | | | | |
| P55258 | RAB8A | RAB8A, member RAS oncogene family | Plasma Membrane | enzyme | 7.69E+07 | 7.18E+07 | 8.46E+07 | | | | | | |
| P55258 | RAB8A | RAB8A, member RAS oncogene family | Plasma Membrane | enzyme | 5.69E+07 | 5.45E+07 | 6.52E+07 | | | | | | |
| P55258 | RAB8A | RAB8A, member RAS oncogene family | Plasma Membrane | enzyme | 4.09E+07 | 4.37E+07 | 5.43E+07 | | | | | | |
| P55258 | RAB8A | RAB8A, member RAS oncogene family | Plasma Membrane | enzyme | 7169928 | 9742314 | 7279952 | | | | | | |
| P55258 | RAB8A | RAB8A, member RAS oncogene family | Plasma Membrane | enzyme | 6965224 | 7702028.5 | 1.05E+07 | | | | | | |
| P55258 | RAB8A | RAB8A, member RAS oncogene family | Plasma Membrane | enzyme | 256637.562 | 211377.828 | 667725.375 | P55258 | RAB8A | 44991474 | 38030831 | 7763011 | 24 |
| P61028 | RAB8B | RAB8B, member RAS oncogene family | Cytoplasm | enzyme | 1.25E+08 | 1.14E+08 | 1.26E+08 | | | | | | |
| P61028 | RAB8B | RAB8B, member RAS oncogene family | Cytoplasm | enzyme | 1.28E+08 | 1.30E+08 | 1.53E+08 | | | | | | |
| P61028 | RAB8B | RAB8B, member RAS oncogene family | Cytoplasm | enzyme | 6.04E+07 | 5.76E+07 | 7.85E+07 | | | | | | |
| P61028 | RAB8B | RAB8B, member RAS oncogene family | Cytoplasm | enzyme | 2.75E+07 | 2.24E+07 | 3.23E+07 | | | | | | |
| P61028 | RAB8B | RAB8B, member RAS oncogene family | Cytoplasm | enzyme | 4413562 | 4028469 | 7949259 | | | | | | |
| P61028 | RAB8B | RAB8B, member RAS oncogene family | Cytoplasm | enzyme | 265777.531 | 317534.875 | 980418.812 | P61028 | RAB8B | 57511557 | 55691455 | 13126602 | 18 |
| Q9R0M6 | RAB9A | RAB9A, member RAS oncogene family | Cytoplasm | enzyme | 2571434 | 2546533.25 | 2109263.5 | Q9R0M6 | RAB9A | 2571434 | 259944 | 150079 | 3 |
| P63001 | RAC1 | ras-related C3 botulinum toxin substrate 1 (rho family, small GTP binding protein Rac1) | Plasma Membrane | enzyme | 1.00E+08 | 8.48E+07 | 9.70E+07 | | | | | | |
| P63001 | RAC1 | ras-related C3 botulinum toxin substrate 1 (rho family, small GTP binding protein Rac1) | Plasma Membrane | enzyme | 1.14E+08 | 5.41E+07 | 1.60E+07 | | | | | | |
| P63001 | RAC1 | ras-related C3 botulinum toxin substrate 1 (rho family, small GTP binding protein Rac1) | Plasma Membrane | enzyme | 7.63E+07 | 7.25E+07 | 9.14E+07 | | | | | | |
| P63001 | RAC1 | ras-related C3 botulinum toxin substrate 1 (rho family, small GTP binding protein Rac1) | Plasma Membrane | enzyme | 8.53E+07 | 5.37E+07 | 3.47E+07 | | | | | | |
| P63001 | RAC1 | ras-related C3 botulinum toxin substrate 1 (rho family, small GTP binding protein Rac1) | Plasma Membrane | enzyme | 7.10E+07 | 5.83E+07 | 6.48E+07 | | | | | | |
| P63001 | RAC1 | ras-related C3 botulinum toxin substrate 1 (rho family, small GTP binding protein Rac1) | Plasma Membrane | enzyme | 6.75E+07 | 5.47E+07 | 4.12E+07 | | | | | | |
| P63001 | RAC1 | ras-related C3 botulinum toxin substrate 1 (rho family, small GTP binding protein Rac1) | Plasma Membrane | enzyme | 2.48E+07 | 4.24E+07 | 8.51E+07 | | | | | | |
| P63001 | RAC1 | ras-related C3 botulinum toxin substrate 1 (rho family, small GTP binding protein Rac1) | Plasma Membrane | enzyme | 2.79E+07 | 2.10E+07 | 1.34E+07 | | | | | | |
| P63001 | RAC1 | ras-related C3 botulinum toxin substrate 1 (rho family, small GTP binding protein Rac1) | Plasma Membrane | enzyme | 1.31E+07 | 3.34E+07 | 6.13E+07 | | | | | | |
| P63001 | RAC1 | ras-related C3 botulinum toxin substrate 1 (rho family, small GTP binding protein Rac1) | Plasma Membrane | enzyme | 2.65E+07 | 3.03E+07 | 2.97E+07 | | | | | | |
| P63001 | RAC1 | ras-related C3 botulinum toxin substrate 1 (rho family, small GTP binding protein Rac1) | Plasma Membrane | enzyme | 1.19E+07 | 6855780 | 6070990 | | | | | | |
| P63001 | RAC1 | ras-related C3 botulinum toxin substrate 1 (rho family, small GTP binding protein Rac1) | Plasma Membrane | enzyme | 2331789.25 | 9144864 | 1.51E+07 | | | | | | |
| P63001 | RAC1 | ras-related C3 botulinum toxin substrate 1 (rho family, small GTP binding protein Rac1) | Plasma Membrane | enzyme | 6348433.5 | 5605001 | 4458908.5 | | | | | | |
| P63001 | RAC1 | ras-related C3 botulinum toxin substrate 1 (rho family, small GTP binding protein Rac1) | Plasma Membrane | enzyme | 2383486.5 | 3405358.5 | 7388124.5 | P63001 | RAC1 | 44984551 | 32687783 | 5043834 | 42 |
| Q05144 | RAC2 | ras-related C3 botulinum toxin substrate 2 (rho family, small GTP binding protein Rac2) | Cytoplasm | enzyme | 4200327 | 9907732 | 4813888.5 | Q05144 | RAC2 | 4200327 | 3133107 | 1808900 | 3 |
| P54726 | RAD23A | RAD23 homolog A, nucleotide excision repair protein | Nucleus | other | 4.24E+07 | 4.63E+07 | 5.38E+07 | | | | | | |
| P54726 | RAD23A | RAD23 homolog A, nucleotide excision repair protein | Nucleus | other | 3.25E+07 | 2.80E+07 | 2.73E+07 | | | | | | |
| P54726 | RAD23A | RAD23 homolog A, nucleotide excision repair protein | Nucleus | other | 2.66E+07 | 2.30E+07 | 2.40E+07 | | | | | | |
| P54726 | RAD23A | RAD23 homolog A, nucleotide excision repair protein | Nucleus | other | 1.03E+07 | 7543671 | 8626439 | | | | | | |
| P54726 | RAD23A | RAD23 homolog A, nucleotide excision repair protein | Nucleus | other | 6807104 | 6581969.5 | 7613546 | | | | | | |
| P54726 | RAD23A | RAD23 homolog A, nucleotide excision repair protein | Nucleus | other | 4129820 | 2752228 | 3033312.5 | | | | | | |
| P54726 | RAD23A | RAD23 homolog A, nucleotide excision repair protein | Nucleus | other | 599118.75 | 385889.094 | 370617.281 | P54726 | RAD23A | 17596578 | 16416241 | 3582317 | 21 |
| P63321 | RALA | v-ral simian leukemia viral oncogene homolog A (ras related) | Cytoplasm | enzyme | 1204815.5 | 1974554.5 | 2539908.5 | P63321 | RALA | 1204816 | 670149 | 386911 | 3 |
| P62827 | RAN | RAN, member RAS oncogene family | Nucleus | enzyme | 2975624.75 | 2228226.5 | 1447906.75 | P62827 | RAN | 2975625 | 763918 | 441048 | 3 |
| P46061 | RANGAP1 | Ran GTPase activating protein 1 | Nucleus | other | 604621.375 | 437241.406 | 292418.281 | P46061 | RANGAP1 | 604621 | 156237 | 90204 | 3 |
| P62835 | RAP1A | RAP1A, member of RAS oncogene family | Cytoplasm | enzyme | 9.83E+07 | 1.01E+08 | 1.35E+08 | | | | | | |
| P62835 | RAP1A | RAP1A, member of RAS oncogene family | Cytoplasm | enzyme | 6.93E+07 | 7.11E+07 | 7.14E+07 | | | | | | |
| P62835 | RAP1A | RAP1A, member of RAS oncogene family | Cytoplasm | enzyme | 4.71E+07 | 2.55E+07 | 3.04E+07 | | | | | | |
| P62835 | RAP1A | RAP1A, member of RAS oncogene family | Cytoplasm | enzyme | 4.73E+07 | 4.48E+07 | 5.00E+07 | | | | | | |
| P62835 | RAP1A | RAP1A, member of RAS oncogene family | Cytoplasm | enzyme | 1.13E+07 | 1.17E+07 | 2.17E+07 | | | | | | |
| P62835 | RAP1A | RAP1A, member of RAS oncogene family | Cytoplasm | enzyme | 7062317.5 | 6329686.5 | 6003457.5 | | | | | | |
| P62835 | RAP1A | RAP1A, member of RAS oncogene family | Cytoplasm | enzyme | 4006324.25 | 2930947 | 2811092.25 | | | | | | |
| P62835 | RAP1A | RAP1A, member of RAS oncogene family | Cytoplasm | enzyme | 4462706 | 294165.594 | 60282.223 | P62835 | RAP1A | 36106418 | 37768958 | 7709556 | 24 |
| Q99JI6 | RAP1B | RAP1B, member of RAS oncogene family | Cytoplasm | enzyme | 1.24E+08 | 1.27E+08 | 1.89E+08 | | | | | | |
| Q99JI6 | RAP1B | RAP1B, member of RAS oncogene family | Cytoplasm | enzyme | 1.22E+08 | 1.07E+08 | 1.23E+08 | | | | | | |
| Q99JI6 | RAP1B | RAP1B, member of RAS oncogene family | Cytoplasm | enzyme | 1.11E+08 | 1.28E+08 | 1.60E+08 | | | | | | |
| Q99JI6 | RAP1B | RAP1B, member of RAS oncogene family | Cytoplasm | enzyme | 1.76E+07 | 2.42E+07 | 4.56E+07 | | | | | | |
| Q99JI6 | RAP1B | RAP1B, member of RAS oncogene family | Cytoplasm | enzyme | 1.10E+07 | 1.12E+07 | 1.53E+07 | | | | | | |
| Q99JI6 | RAP1B | RAP1B, member of RAS oncogene family | Cytoplasm | enzyme | 1.12E+07 | 1.33E+07 | 2.79E+07 | | | | | | |
| Q99JI6 | RAP1B | RAP1B, member of RAS oncogene family | Cytoplasm | enzyme | 5188623.5 | 8834677 | 1.86E+07 | | | | | | |
| Q99JI6 | RAP1B | RAP1B, member of RAS oncogene family | Cytoplasm | enzyme | 7146.281 | 213457.719 | 224860.016 | Q99JI6 | RAP1B | 50096971 | 61093307 | 12470619 | 24 |
| P61226 | RAP2B | RAP2B, member of RAS oncogene family | Plasma Membrane | enzyme | 7958027 | 9171866 | 1.16E+07 | | | | | | |
| P61226 | RAP2B | RAP2B, member of RAS oncogene family | Plasma Membrane | enzyme | 6947788 | 8125691 | 6346544 | | | | | | |
| P61226 | RAP2B | RAP2B, member of RAS oncogene family | Plasma Membrane | enzyme | 7693656 | 7719883 | 9306859 | | | | | | |
| P61226 | RAP2B | RAP2B, member of RAS oncogene family | Plasma Membrane | enzyme | 5746720 | 6787029 | 8216500.5 | | | | | | |
| P61226 | RAP2B | RAP2B, member of RAS oncogene family | Plasma Membrane | enzyme | 5926086 | 5962747 | 6447834.5 | | | | | | |
| P61226 | RAP2B | RAP2B, member of RAS oncogene family | Plasma Membrane | enzyme | 1754918.875 | 1944446.25 | 3037531.75 | P61226 | RAP2B | 6004533 | 2517554 | 593393 | 18 |
| Q8BU31 | RAP2C | RAP2C, member of RAS oncogene family | Cytoplasm | enzyme | 2008934.875 | 2160992.5 | 3675862.75 | Q8BU31 | RAP2C | 2008935 | 921647 | 532113 | 3 |
| Q9D0I9 | RARS | arginyl-tRNA synthetase | Cytoplasm | enzyme | 148471.641 | 325755.594 | 334055.531 | Q9D0I9 | RARS | 148472 | 104833 | 60525 | 3 |
| Q60790 | RASA3 | RAS p21 protein activator 3 | Plasma Membrane | ion channel | 342003.312 | 479628.594 | 1613446 | Q60790 | RASA3 | 342003 | 697740 | 402841 | 3 |
| Q61820 | Rasl2-9 | RAS-like, family 2, locus 9 | Nucleus | transporter | 6138216 | 4455603.5 | 4622641.5 | | | | | | |
| Q61820 | Rasl2-9 | RAS-like, family 2, locus 9 | Nucleus | transporter | 6223620.5 | 4105798 | 2821853.75 | Q61820 | Rasl2-9 | 6180918 | 1290846 | 526986 | 6 |
| O89086 | RBM3 | RNA binding motif (RNP1, RRM) protein 3 | Cytoplasm | other | 1.07E+07 | 1.87E+07 | 7854122.5 | | | | | | |
| O89086 | RBM3 | RNA binding motif (RNP1, RRM) protein 3 | Cytoplasm | other | 4624298.5 | 7669398 | 1863001.5 | | | | | | |
| O89086 | RBM3 | RNA binding motif (RNP1, RRM) protein 3 | Cytoplasm | other | 394276.719 | 773408.75 | 154588.703 | O89086 | RBM3 | 5232858 | 6138073 | 2046024 | 9 |
| Q9QYF1 | RDH11 | retinol dehydrogenase 11 (all-trans/9-cis/11-cis) | Cytoplasm | enzyme | 6223973.5 | 7298517 | 9339345 | | | | | | |
| Q9QYF1 | RDH11 | retinol dehydrogenase 11 (all-trans/9-cis/11-cis) | Cytoplasm | enzyme | 5220979 | 4768157 | 4851831 | | | | | | |
| Q9QYF1 | RDH11 | retinol dehydrogenase 11 (all-trans/9-cis/11-cis) | Cytoplasm | enzyme | 4640073.5 | 5051083.5 | 5900447.5 | | | | | | |
| Q9QYF1 | RDH11 | retinol dehydrogenase 11 (all-trans/9-cis/11-cis) | Cytoplasm | enzyme | 3199869 | 5506128.5 | 5668238 | | | | | | |
| Q9QYF1 | RDH11 | retinol dehydrogenase 11 (all-trans/9-cis/11-cis) | Cytoplasm | enzyme | 4913650.5 | 5166028.5 | 4976269 | | | | | | |
| Q9QYF1 | RDH11 | retinol dehydrogenase 11 (all-trans/9-cis/11-cis) | Cytoplasm | enzyme | 2436129 | 1981354.375 | 2054596.125 | | | | | | |
| Q9QYF1 | RDH11 | retinol dehydrogenase 11 (all-trans/9-cis/11-cis) | Cytoplasm | enzyme | 3240619.25 | 5386617 | 4436895 | | | | | | |
| Q9QYF1 | RDH11 | retinol dehydrogenase 11 (all-trans/9-cis/11-cis) | Cytoplasm | enzyme | 2504764.75 | 2132242.5 | 1889446.25 | Q9QYF1 | RDH11 | 4047507 | 1835255 | 374620 | 24 |
| P26043 | RDX | radixin | Cytoplasm | other | 3.55E+07 | 3.80E+07 | 3.67E+07 | | | | | | |
| P26043 | RDX | radixin | Cytoplasm | other | 2.09E+07 | 1.90E+07 | 2.31E+07 | | | | | | |
| P26043 | RDX | radixin | Cytoplasm | other | 2.13E+07 | 2.70E+07 | 3.62E+07 | | | | | | |
| P26043 | RDX | radixin | Cytoplasm | other | 1.24E+07 | 1.09E+07 | 1.28E+07 | | | | | | |
| P26043 | RDX | radixin | Cytoplasm | other | 1.28E+07 | 1.01E+07 | 9653983 | | | | | | |
| P26043 | RDX | radixin | Cytoplasm | other | 9353772 | 9531880 | 1.13E+07 | | | | | | |

| | | | | | | | | | | | | | |
|---|---|---|---|---|---|---|---|---|---|---|---|---|---|
| P26043 | RDX | radixin | Cytoplasm | other | 1.24E+07 | 9462159 | 9282032 | | | | | | |
| P26043 | RDX | radixin | Cytoplasm | other | 6780288 | 4119309.75 | 4955371 | | | | | | |
| P26043 | RDX | radixin | Cytoplasm | other | 4451850.5 | 4181711.5 | 2943320 | | | | | | |
| P26043 | RDX | radixin | Cytoplasm | other | 6662036 | 3016690 | 805232.75 | | | | | | |
| P26043 | RDX | radixin | Cytoplasm | other | 4991900 | 3789104.25 | 3618785.75 | | | | | | |
| P26043 | RDX | radixin | Cytoplasm | other | 3686966.5 | 4341773.5 | 5420413 | | | | | | |
| P26043 | RDX | radixin | Cytoplasm | other | 2358517.25 | 1677309.125 | 2441947.75 | | | | | | |
| P26043 | RDX | radixin | Cytoplasm | other | 2477378.5 | 93647.453 | 1136.285 | P26043 | RDX | 11148051 | 10564610 | 1630155 | 42 |
| Q60870 | REEP5 | receptor accessory protein 5 | Extracellular Space | transporter | 2.28E+07 | 2.06E+07 | 2.48E+07 | | | | | | |
| Q60870 | REEP5 | receptor accessory protein 5 | Extracellular Space | transporter | 1191337 | 1232335.25 | 809934.312 | | | | | | |
| Q60870 | REEP5 | receptor accessory protein 5 | Extracellular Space | transporter | 393395.75 | 349974.25 | 138346.688 | Q60870 | REEP5 | 8111578 | 11067104 | 3689035 | 9 |
| Q9JM62 | REEP6 | receptor accessory protein 6 | Plasma Membrane | other | 9536628 | 1.09E+07 | 1.42E+07 | | | | | | |
| Q9JM62 | REEP6 | receptor accessory protein 6 | Plasma Membrane | other | 3618340 | 3611808.25 | 6248575.5 | | | | | | |
| Q9JM62 | REEP6 | receptor accessory protein 6 | Plasma Membrane | other | 1709114.875 | 912246.5 | 109053.906 | Q9JM62 | REEP6 | 4954694 | 4925584 | 1641861 | 9 |
| Q8K2J7 | RELL1 | RELT-like 1 | Plasma Membrane | other | 5556017 | 2157304.5 | 420232.031 | | | | | | |
| Q8K2J7 | RELL1 | RELT-like 1 | Plasma Membrane | other | 4464919 | 4346014.5 | 5229655 | | | | | | |
| Q8K2J7 | RELL1 | RELT-like 1 | Plasma Membrane | other | 2737256.75 | 2724007.5 | 3056507.5 | Q8K2J7 | RELL1 | 4252731 | 1637895 | 545965 | 9 |
| Q9QUT0 | RHAG | Rh-associated glycoprotein | Plasma Membrane | peptidase | 250177.734 | 385378.875 | 545373.375 | | | | | | |
| Q9QUT0 | RHAG | Rh-associated glycoprotein | Plasma Membrane | peptidase | 144539.453 | 114431.148 | 448411.688 | | | | | | |
| Q9QUT0 | RHAG | Rh-associated glycoprotein | Plasma Membrane | peptidase | 1859469 | 2753255.75 | 3873150.5 | | | | | | |
| Q9QUT0 | RHAG | Rh-associated glycoprotein | Plasma Membrane | peptidase | 287073 | 472036.719 | 759474.688 | Q9QUT0 | RHAG | 635315 | 1201456 | 346830 | 12 |
| Q80WQ6 | RHBDF2 | rhomboid 5 homolog 2 (Drosophila) | Cytoplasm | other | 1983398.5 | 2286149 | 2940691.25 | | | | | | |
| Q80WQ6 | RHBDF2 | rhomboid 5 homolog 2 (Drosophila) | Cytoplasm | other | 1087803.75 | 796944.688 | 867782.812 | Q80WQ6 | RHBDF2 | 1535601 | 875979 | 357617 | 6 |
| Q8CF94 | RHCE/RHD | Rh blood group, D antigen | Plasma Membrane | transporter | 319678.406 | 820879.188 | 538651.438 | | | | | | |
| Q8CF94 | RHCE/RHD | Rh blood group, D antigen | Plasma Membrane | transporter | 1848336.5 | 1779213.25 | 3541918 | | | | | | |
| Q8CF94 | RHCE/RHD | Rh blood group, D antigen | Plasma Membrane | transporter | 291537.219 | 304769.688 | 539283.812 | | | | | | |
| Q8CF94 | RHCE/RHD | Rh blood group, D antigen | Plasma Membrane | transporter | 46768.613 | 66610.453 | 33494.531 | | | | | | |
| Q8CF94 | RHCE/RHD | Rh blood group, D antigen | Plasma Membrane | transporter | 302550.312 | 205570.484 | 747451.875 | | | | | | |
| Q8CF94 | RHCE/RHD | Rh blood group, D antigen | Plasma Membrane | transporter | 367765.188 | 151014.625 | 712565.25 | | | | | | |
| Q8CF94 | RHCE/RHD | Rh blood group, D antigen | Plasma Membrane | transporter | 4.70E-06 | 55987.293 | 8276.938 | | | | | | |
| Q8CF94 | RHCE/RHD | Rh blood group, D antigen | Plasma Membrane | transporter | 0 | 38.16 | 0 | Q8CF94 | RHCE/RHD | 397080 | 816394 | 166646 | 24 |
| Q9QUI0 | RHOA | ras homolog family member A | Cytoplasm | enzyme | 2.43E+07 | 3.60E+07 | 3.61E+07 | | | | | | |
| Q9QUI0 | RHOA | ras homolog family member A | Cytoplasm | enzyme | 2.09E+07 | 2.69E+07 | 3.13E+07 | | | | | | |
| Q9QUI0 | RHOA | ras homolog family member A | Cytoplasm | enzyme | 2.40E+07 | 1.17E+07 | 4625287 | | | | | | |
| Q9QUI0 | RHOA | ras homolog family member A | Cytoplasm | enzyme | 1.63E+07 | 8614687 | 4379659 | | | | | | |
| Q9QUI0 | RHOA | ras homolog family member A | Cytoplasm | enzyme | 1.27E+07 | 1.10E+07 | 1.29E+07 | | | | | | |
| Q9QUI0 | RHOA | ras homolog family member A | Cytoplasm | enzyme | 8214024 | 1.22E+07 | 1.89E+07 | | | | | | |
| Q9QUI0 | RHOA | ras homolog family member A | Cytoplasm | enzyme | 3227007.25 | 4877542.5 | 4783057.5 | | | | | | |
| Q9QUI0 | RHOA | ras homolog family member A | Cytoplasm | enzyme | 5509920.5 | 3609080.5 | 4239015.5 | | | | | | |
| Q9QUI0 | RHOA | ras homolog family member A | Cytoplasm | enzyme | 5411766.5 | 8027858 | 2.06E+07 | Q9QUI0 | RHOA | 13403635 | 10190064 | 1961079 | 27 |
| P84096 | RHOG | ras homolog family member G | Cytoplasm | enzyme | 5.26E+07 | 5.18E+07 | 4.61E+07 | | | | | | |
| P84096 | RHOG | ras homolog family member G | Cytoplasm | enzyme | 3.16E+07 | 2.95E+07 | 3.89E+07 | | | | | | |
| P84096 | RHOG | ras homolog family member G | Cytoplasm | enzyme | 2.92E+07 | 3.36E+07 | 3.82E+07 | | | | | | |
| P84096 | RHOG | ras homolog family member G | Cytoplasm | enzyme | 2.36E+07 | 1.64E+07 | 1.53E+07 | | | | | | |
| P84096 | RHOG | ras homolog family member G | Cytoplasm | enzyme | 1.82E+07 | 2.32E+07 | 2.39E+07 | | | | | | |
| P84096 | RHOG | ras homolog family member G | Cytoplasm | enzyme | 1.27E+07 | 1.16E+07 | 1.41E+07 | | | | | | |
| P84096 | RHOG | ras homolog family member G | Cytoplasm | enzyme | 7471991.5 | 1.36E+07 | 1.10E+07 | | | | | | |
| P84096 | RHOG | ras homolog family member G | Cytoplasm | enzyme | 8827234 | 1.30E+07 | 1.78E+07 | | | | | | |
| P84096 | RHOG | ras homolog family member G | Cytoplasm | enzyme | 1.10E+07 | 7356862 | 9066596 | | | | | | |
| P84096 | RHOG | ras homolog family member G | Cytoplasm | enzyme | 2538685.25 | 2901055.25 | 1547984.75 | | | | | | |
| P84096 | RHOG | ras homolog family member G | Cytoplasm | enzyme | 1634365.5 | 2925677.5 | 4340582.5 | P84096 | RHOG | 18121116 | 14536215 | 2530430 | 33 |
| Q9QYK7 | RNF11 | ring finger protein 11 | Cytoplasm | enzyme | 2595656.75 | 2649613.75 | 3660185 | Q9QYK7 | RNF11 | 2595657 | 599637 | 346200 | 3 |
| Q5XPI3 | RNF123 | ring finger protein 123 | Cytoplasm | other | 9644230 | 7354651.5 | 8550689 | | | | | | |
| Q5XPI3 | RNF123 | ring finger protein 123 | Cytoplasm | other | 1.03E+07 | 7209559.5 | 4630803.5 | | | | | | |
| Q5XPI3 | RNF123 | ring finger protein 123 | Cytoplasm | other | 8089700 | 4385115 | 5958990.5 | | | | | | |
| Q5XPI3 | RNF123 | ring finger protein 123 | Cytoplasm | other | 8622210 | 6371383.5 | 5812431 | | | | | | |
| Q5XPI3 | RNF123 | ring finger protein 123 | Cytoplasm | other | 5053740 | 4465925 | 6312437.5 | | | | | | |
| Q5XPI3 | RNF123 | ring finger protein 123 | Cytoplasm | other | 3498086.5 | 2410602.25 | 3099456.75 | | | | | | |
| Q5XPI3 | RNF123 | ring finger protein 123 | Cytoplasm | other | 2899834 | 1893723.375 | 2221498.5 | Q5XPI3 | RNF123 | 6878257 | 2520133 | 549938 | 21 |
| E9Q555 | RNF213 | ring finger protein 213 | Cytoplasm | enzyme | 1449744.75 | 2632032 | 1329369 | E9Q555 | RNF213 | 1449745 | 719864 | 415614 | 3 |
| Q9EPK2 | RP2 | retinitis pigmentosa 2 (X-linked recessive) | Cytoplasm | enzyme | 1.69E+07 | 1.74E+07 | 1.79E+07 | | | | | | |
| Q9EPK2 | RP2 | retinitis pigmentosa 2 (X-linked recessive) | Cytoplasm | enzyme | 9957292 | 7768680.5 | 9214595 | | | | | | |
| Q9EPK2 | RP2 | retinitis pigmentosa 2 (X-linked recessive) | Cytoplasm | enzyme | 8356430 | 6989989 | 8595481 | | | | | | |
| Q9EPK2 | RP2 | retinitis pigmentosa 2 (X-linked recessive) | Cytoplasm | enzyme | 6685813.5 | 6105822.5 | 6050593.5 | | | | | | |
| Q9EPK2 | RP2 | retinitis pigmentosa 2 (X-linked recessive) | Cytoplasm | enzyme | 6101609 | 6185297 | 6955501.5 | | | | | | |
| Q9EPK2 | RP2 | retinitis pigmentosa 2 (X-linked recessive) | Cytoplasm | enzyme | 6350875 | 5585729.5 | 6982630 | | | | | | |
| Q9EPK2 | RP2 | retinitis pigmentosa 2 (X-linked recessive) | Cytoplasm | enzyme | 4043503.25 | 1597855.875 | 1567787.875 | Q9EPK2 | RP2 | 8342218 | 4455063 | 972174 | 21 |
| P53026 | RPL10A | ribosomal protein L10a | Nucleus | other | 4063461.75 | 5672728.5 | 1979647.75 | | | | | | |
| P53026 | RPL10A | ribosomal protein L10a | Nucleus | other | 1755542.375 | 3618965.25 | 1112945.125 | P53026 | RPL10A | 2909502 | 1720548 | 702411 | 6 |
| P86048 | RPL10L | ribosomal protein L10-like | Nucleus | other | 8683955 | 1.97E+07 | 8090807.5 | | | | | | |
| P86048 | RPL10L | ribosomal protein L10-like | Nucleus | other | 5519574.5 | 7570449 | 2249881 | P86048 | RPL10L | 7101765 | 5901194 | 2409152 | 6 |
| Q9CXW4 | RPL11 | ribosomal protein L11 | Cytoplasm | other | 9240216 | 1.61E+07 | 6584626.5 | Q9CXW4 | RPL11 | 9240216 | 4921161 | 2841233 | 3 |
| P35979 | RPL12 | ribosomal protein L12 | Nucleus | other | 6410892.5 | 9618686 | 3127926.5 | | | | | | |
| P35979 | RPL12 | ribosomal protein L12 | Nucleus | other | 5291673 | 1.02E+07 | 5343485.5 | | | | | | |
| P35979 | RPL12 | ribosomal protein L12 | Nucleus | other | 6328470 | 1.27E+07 | 6430279 | | | | | | |
| P35979 | RPL12 | ribosomal protein L12 | Nucleus | other | 5174588.5 | 8585568 | 3146389.5 | | | | | | |
| P35979 | RPL12 | ribosomal protein L12 | Nucleus | other | 2025269.75 | 5091088.5 | 1628025.5 | P35979 | RPL12 | 5046179 | 3129153 | 807944 | 15 |
| P47963 | RPL13 | ribosomal protein L13 | Nucleus | other | 6221809.5 | 9984563 | 3506288.5 | | | | | | |
| P47963 | RPL13 | ribosomal protein L13 | Nucleus | other | 3717302.5 | 6992117 | 2333227 | | | | | | |
| P47963 | RPL13 | ribosomal protein L13 | Nucleus | other | 3040620 | 4173572 | 2071699.125 | | | | | | |
| P47963 | RPL13 | ribosomal protein L13 | Nucleus | other | 3352377 | 4931694 | 1885783.25 | P47963 | RPL13 | 4083027 | 2369398 | 683986 | 12 |
| P19253 | RPL13A | ribosomal protein L13a | Cytoplasm | other | 3873033.75 | 6464463.5 | 2961925.5 | P19253 | RPL13A | 3873034 | 1817205 | 1049164 | 3 |
| Q9CR57 | RPL14 | ribosomal protein L14 | Cytoplasm | other | 8927865 | 1.36E+07 | 6235362.5 | | | | | | |
| Q9CR57 | RPL14 | ribosomal protein L14 | Cytoplasm | other | 4006762 | 5920855 | 2484895.75 | | | | | | |
| Q9CR57 | RPL14 | ribosomal protein L14 | Cytoplasm | other | 4489397.5 | 8232860.5 | 3241153 | Q9CR57 | RPL14 | 5808008 | 3470626 | 1156875 | 9 |
| Q9CZM2 | RPL15 | ribosomal protein L15 | Cytoplasm | other | 5218508.5 | 6949193.5 | 3712099.5 | | | | | | |
| Q9CZM2 | RPL15 | ribosomal protein L15 | Cytoplasm | other | 4368646.5 | 6230055 | 1994597.625 | | | | | | |
| Q9CZM2 | RPL15 | ribosomal protein L15 | Cytoplasm | other | 3690896 | 5625568 | 2346961.25 | Q9CZM2 | RPL15 | 4426017 | 1694308 | 564769 | 9 |
| Q9CPR4 | RPL17 | ribosomal protein L17 | Cytoplasm | other | 3674819.25 | 7086011 | 2916254.5 | Q9CPR4 | RPL17 | 3674819 | 2221055 | 1282327 | 3 |
| P35980 | RPL18 | ribosomal protein L18 | Cytoplasm | other | 9533267 | 1.75E+07 | 5340037 | | | | | | |
| P35980 | RPL18 | ribosomal protein L18 | Cytoplasm | other | 6479219 | 1.02E+07 | 4433031 | | | | | | |

| | | | | | | | | | | | | | |
|---|---|---|---|---|---|---|---|---|---|---|---|---|---|
| P35980 | RPL18 | ribosomal protein L18 | Cytoplasm | other | 5400665 | 8377379 | 3756431 | | | | | | |
| P35980 | RPL18 | ribosomal protein L18 | Cytoplasm | other | 3275847.5 | 4126914.25 | 1787579.125 | P35980 | RPL18 | 6172250 | 4252841 | 1227690 | 12 |
| P62717 | RPL18A | ribosomal protein L18a | Cytoplasm | other | 3737952.5 | 5319246.5 | 2466395.75 | P62717 | RPL18A | 3737953 | 1429225 | 825163 | 3 |
| P84099 | RPL19 | ribosomal protein L19 | Cytoplasm | other | 9680492 | 1.24E+07 | 6423132 | P84099 | RPL19 | 9680492 | 2973117 | 1716530 | 3 |
| O09167 | RPL21 | ribosomal protein L21 | Cytoplasm | other | 8731739 | 1.36E+07 | 6735871.5 | | | | | | |
| O09167 | RPL21 | ribosomal protein L21 | Cytoplasm | other | 3459458.5 | 5820850.5 | 1502869.375 | O09167 | RPL21 | 6095599 | 4249277 | 1734760 | 6 |
| P67984 | RPL22 | ribosomal protein L22 | Nucleus | other | 1.35E+07 | 2.42E+07 | 8452816 | | | | | | |
| P67984 | RPL22 | ribosomal protein L22 | Nucleus | other | 6021982.5 | 1.02E+07 | 3800502.5 | P67984 | RPL22 | 9750991 | 7252957 | 2961007 | 6 |
| P62830 | RPL23 | ribosomal protein L23 | Cytoplasm | other | 6241746.5 | 1.11E+07 | 4451481 | P62830 | RPL23 | 6241747 | 3429067 | 1979773 | 3 |
| P62751 | Rpl23a | ribosomal protein L23A | Nucleus | other | 1.03E+07 | 1.68E+07 | 7591355 | | | | | | |
| P62751 | Rpl23a | ribosomal protein L23A | Nucleus | other | 2347724.5 | 3287810 | 1367666.25 | P62751 | Rpl23a | 6328862 | 5897258 | 2407545 | 6 |
| Q8BP67 | RPL24 | ribosomal protein L24 | Cytoplasm | other | 3872448 | 4914949.5 | 2160954.25 | | | | | | |
| Q8BP67 | RPL24 | ribosomal protein L24 | Cytoplasm | other | 2135526.75 | 3578882.25 | 1630992.375 | | | | | | |
| Q8BP67 | RPL24 | ribosomal protein L24 | Cytoplasm | other | 3110119.5 | 5066176 | 2018902.875 | Q8BP67 | RPL24 | 3039365 | 1278975 | 426325 | 9 |
| P61255 | RPL26 | ribosomal protein L26 | Cytoplasm | other | 2223662.25 | 3439887.5 | 1324780.75 | P61255 | RPL26 | 2223662 | 1061514 | 612865 | 3 |
| P14115 | RPL27A | ribosomal protein L27a | Cytoplasm | other | 3733834 | 5885007.5 | 2335124 | P14115 | RPL27A | 3733834 | 1788184 | 1032408 | 3 |
| P41105 | RPL28 | ribosomal protein L28 | Cytoplasm | other | 4670003.5 | 6522924 | 2876899.75 | P41105 | RPL28 | 4670004 | 1823094 | 1052564 | 3 |
| P47915 | Rpl29 (includes others) | ribosomal protein L29 | Cytoplasm | other | 949068.188 | 3192447 | 356388.969 | P47915 | Rpl29 (includes others) | 949068 | 1495952 | 863689 | 3 |
| P27659 | RPL3 | ribosomal protein L3 | Nucleus | other | 1.02E+07 | 2.16E+07 | 6985432 | | | | | | |
| P27659 | RPL3 | ribosomal protein L3 | Nucleus | other | 4305435.5 | 6900490 | 2760370.25 | | | | | | |
| P27659 | RPL3 | ribosomal protein L3 | Nucleus | other | 3404360.5 | 6268873.5 | 2356815.25 | P27659 | RPL3 | 5956599 | 5942434 | 1980811 | 9 |
| P62889 | RPL30 | ribosomal protein L30 | Cytoplasm | other | 8008963 | 1.04E+07 | 2818675.25 | P62889 | RPL30 | 8008963 | 3888718 | 2245152 | 3 |
| P62900 | RPL31 | ribosomal protein L31 | Cytoplasm | other | 7103597.5 | 9890521 | 4080538.25 | | | | | | |
| P62900 | RPL31 | ribosomal protein L31 | Cytoplasm | other | 6934549.5 | 1.34E+07 | 5100045.5 | P62900 | RPL31 | 7019074 | 3421331 | 1396753 | 6 |
| Q6ZWV7 | RPL35 | ribosomal protein L35 | Cytoplasm | other | 6360329 | 9993716 | 4721275.5 | | | | | | |
| Q6ZWV7 | RPL35 | ribosomal protein L35 | Cytoplasm | other | 2322026.5 | 3855317.75 | 2223463.75 | Q6ZWV7 | RPL35 | 4341178 | 2932938 | 1197367 | 6 |
| P47964 | Rpl36 | ribosomal protein L36 | Nucleus | other | 2861546 | 4299916.5 | 1977303.375 | | | | | | |
| P47964 | Rpl36 | ribosomal protein L36 | Nucleus | other | 2450845.75 | 3604513.75 | 1588875.125 | P47964 | Rpl36 | 2656196 | 1016493 | 414982 | 6 |
| Q9D8E6 | RPL4 | ribosomal protein L4 | Cytoplasm | enzyme | 7491750 | 1.08E+07 | 4024117 | | | | | | |
| Q9D8E6 | RPL4 | ribosomal protein L4 | Cytoplasm | enzyme | 4707111 | 6823868 | 2318038.5 | | | | | | |
| Q9D8E6 | RPL4 | ribosomal protein L4 | Cytoplasm | enzyme | 4688573.5 | 5576183.5 | 1607462.25 | | | | | | |
| Q9D8E6 | RPL4 | ribosomal protein L4 | Cytoplasm | enzyme | 1864690.375 | 3785232.75 | 2965109.75 | Q9D8E6 | RPL4 | 4688031 | 2668759 | 770404 | 12 |
| P47962 | RPL5 | ribosomal protein L5 | Cytoplasm | other | 1501351.375 | 2388859 | 980008.75 | P47962 | RPL5 | 1501351 | 712312 | 411253 | 3 |
| P47911 | RPL6 | ribosomal protein L6 | Nucleus | other | 4748812.5 | 6475470.5 | 4700869.5 | | | | | | |
| P47911 | RPL6 | ribosomal protein L6 | Nucleus | other | 4890520.5 | 6711298.5 | 2513963.25 | | | | | | |
| P47911 | RPL6 | ribosomal protein L6 | Nucleus | other | 2577301.5 | 7234962 | 2275208.75 | | | | | | |
| P47911 | RPL6 | ribosomal protein L6 | Nucleus | other | 1360284.25 | 2391245.25 | 320523.438 | P47911 | RPL6 | 3394230 | 2252699 | 650298 | 12 |
| P14148 | RPL7 | ribosomal protein L7 | Nucleus | transcription regulator | 1.41E+07 | 2.38E+07 | 8231777 | | | | | | |
| P14148 | RPL7 | ribosomal protein L7 | Nucleus | transcription regulator | 6772529.5 | 1.58E+07 | 4980777.5 | | | | | | |
| P14148 | RPL7 | ribosomal protein L7 | Nucleus | transcription regulator | 5947958.5 | 1.00E+07 | 4523860 | | | | | | |
| P14148 | RPL7 | ribosomal protein L7 | Nucleus | transcription regulator | 5429239 | 1.09E+07 | 4188468 | | | | | | |
| P14148 | RPL7 | ribosomal protein L7 | Nucleus | transcription regulator | 1025489.562 | 2311562.75 | 1345176.5 | P14148 | RPL7 | 6657043 | 6177255 | 1594960 | 15 |
| P12970 | RPL7A | ribosomal protein L7a | Cytoplasm | other | 8005399.5 | 1.32E+07 | 6132862.5 | | | | | | |
| P12970 | RPL7A | ribosomal protein L7a | Cytoplasm | other | 5865221.5 | 9783564 | 4342094.5 | | | | | | |
| P12970 | RPL7A | ribosomal protein L7a | Cytoplasm | other | 4301745 | 4555721 | 2501452.25 | P12970 | RPL7A | 6057455 | 3314042 | 1104681 | 9 |
| P62918 | RPL8 | ribosomal protein L8 | Cytoplasm | other | 2019902 | 3584090.5 | 1509781.25 | | | | | | |
| P62918 | RPL8 | ribosomal protein L8 | Cytoplasm | other | 1306108.5 | 2321386 | 770802.375 | P62918 | RPL8 | 1663005 | 980416 | 400253 | 6 |
| P51410 | RPL9 | ribosomal protein L9 | Nucleus | other | 8357554 | 1.26E+07 | 5648339.5 | | | | | | |
| P51410 | RPL9 | ribosomal protein L9 | Nucleus | other | 4787199 | 7256880.5 | 2570561.25 | | | | | | |
| P51410 | RPL9 | ribosomal protein L9 | Nucleus | other | 5075946 | 8366034 | 4072383.5 | P51410 | RPL9 | 6073566 | 2981713 | 993904 | 9 |
| P14869 | RPLP0 | ribosomal protein, large, P0 | Cytoplasm | other | 1.63E+07 | 2.67E+07 | 1.26E+07 | | | | | | |
| P14869 | RPLP0 | ribosomal protein, large, P0 | Cytoplasm | other | 5585644 | 1.12E+07 | 4104203.25 | | | | | | |
| P14869 | RPLP0 | ribosomal protein, large, P0 | Cytoplasm | other | 8110534 | 1.17E+07 | 4094405 | | | | | | |
| P14869 | RPLP0 | ribosomal protein, large, P0 | Cytoplasm | other | 6129808.5 | 9493647 | 3785341.25 | | | | | | |
| P14869 | RPLP0 | ribosomal protein, large, P0 | Cytoplasm | other | 5023489.5 | 8028869.5 | 3727501.25 | | | | | | |
| P14869 | RPLP0 | ribosomal protein, large, P0 | Cytoplasm | other | 4523596.5 | 6123536 | 3038205.75 | | | | | | |
| P14869 | RPLP0 | ribosomal protein, large, P0 | Cytoplasm | other | 5734529 | 9078025 | 4870284.5 | | | | | | |
| P14869 | RPLP0 | ribosomal protein, large, P0 | Cytoplasm | other | 2553986 | 3581394.25 | 1125640.25 | P14869 | RPLP0 | 6742698 | 5485862 | 1119797 | 24 |
| P47955 | Rplp1 (includes others) | ribosomal protein, large, P1 | Nucleus | other | 3.40E+07 | 5.85E+07 | 2.53E+07 | | | | | | |
| P47955 | Rplp1 (includes others) | ribosomal protein, large, P1 | Nucleus | other | 2.73E+07 | 5.26E+07 | 2.80E+07 | P47955 | Rplp1 (includes others) | 30645000 | 14297864 | 5837078 | 6 |
| P99027 | RPLP2 | ribosomal protein, large, P2 | Cytoplasm | other | 3.36E+07 | 5.27E+07 | 2.51E+07 | | | | | | |
| P99027 | RPLP2 | ribosomal protein, large, P2 | Cytoplasm | other | 3.60E+07 | 5.60E+07 | 2.70E+07 | | | | | | |
| P99027 | RPLP2 | ribosomal protein, large, P2 | Cytoplasm | other | 8182983.5 | 1.42E+07 | 6358833.5 | | | | | | |
| P99027 | RPLP2 | ribosomal protein, large, P2 | Cytoplasm | other | 5329698 | 6925164 | 3902857.75 | | | | | | |
| P99027 | RPLP2 | ribosomal protein, large, P2 | Cytoplasm | other | 3613442.75 | 5847736 | 2113314.5 | P99027 | RPLP2 | 17335225 | 18243780 | 4710524 | 15 |
| Q91YQ5 | RPN1 | ribophorin I | Cytoplasm | enzyme | 1900961.25 | 2722079.5 | 3217585.5 | | | | | | |
| Q91YQ5 | RPN1 | ribophorin I | Cytoplasm | enzyme | 1959634.125 | 4582841.5 | 5752041 | | | | | | |
| Q91YQ5 | RPN1 | ribophorin I | Cytoplasm | enzyme | 2461395 | 5614416.5 | 5641035.5 | | | | | | |
| Q91YQ5 | RPN1 | ribophorin I | Cytoplasm | enzyme | 1575253.625 | 2187183.25 | 2375860 | | | | | | |
| Q91YQ5 | RPN1 | ribophorin I | Cytoplasm | enzyme | 1108494.25 | 2106683.5 | 2154131.75 | | | | | | |
| Q91YQ5 | RPN1 | ribophorin I | Cytoplasm | enzyme | 1198322 | 3221121.25 | 2562305.5 | | | | | | |
| Q91YQ5 | RPN1 | ribophorin I | Cytoplasm | enzyme | 284536.031 | 1036214.188 | 1518634.75 | | | | | | |
| Q91YQ5 | RPN1 | ribophorin I | Cytoplasm | enzyme | 368905.344 | 1016564.938 | 1101231.875 | Q91YQ5 | RPN1 | 1357188 | 1582233 | 322972 | 24 |
| Q9DBG6 | RPN2 | ribophorin II | Cytoplasm | enzyme | 4088061.75 | 7288389 | 7326487 | | | | | | |
| Q9DBG6 | RPN2 | ribophorin II | Cytoplasm | enzyme | 1510802.875 | 4496801 | 7912206.5 | | | | | | |
| Q9DBG6 | RPN2 | ribophorin II | Cytoplasm | enzyme | 938579.188 | 2923523.5 | 3737335.5 | | | | | | |
| Q9DBG6 | RPN2 | ribophorin II | Cytoplasm | enzyme | 905822.75 | 2841727.75 | 3216688.75 | | | | | | |
| Q9DBG6 | RPN2 | ribophorin II | Cytoplasm | enzyme | 935582.125 | 2286727.5 | 3345769.25 | | | | | | |
| Q9DBG6 | RPN2 | ribophorin II | Cytoplasm | enzyme | 709031.375 | 1680920.5 | 2578198.5 | | | | | | |
| Q9DBG6 | RPN2 | ribophorin II | Cytoplasm | enzyme | 796045.25 | 2217896 | 2775301 | | | | | | |
| Q9DBG6 | RPN2 | ribophorin II | Cytoplasm | enzyme | 775203.688 | 1185448.625 | 1453099.75 | | | | | | |
| Q9DBG6 | RPN2 | ribophorin II | Cytoplasm | enzyme | 640264.188 | 1805519.875 | 2236146.25 | | | | | | |
| Q9DBG6 | RPN2 | ribophorin II | Cytoplasm | enzyme | 212918.375 | 745519.062 | 1146889.25 | Q9DBG6 | RPN2 | 1151231 | 2037762 | 372043 | 30 |
| P63325 | RPS10 | ribosomal protein S10 | Cytoplasm | other | 6072302 | 1.21E+07 | 5391816 | P63325 | RPS10 | 6072302 | 3709490 | 2141675 | 3 |
| P62281 | RPS11 | ribosomal protein S11 | Cytoplasm | other | 4114914.75 | 1.02E+07 | 3423500.75 | | | | | | |
| P62281 | RPS11 | ribosomal protein S11 | Cytoplasm | other | 984941.125 | 2187922.75 | 838254.688 | P62281 | RPS11 | 2549928 | 3461749 | 1413253 | 6 |
| P62301 | RPS13 | ribosomal protein S13 | Cytoplasm | other | 2249080.75 | 4510326 | 1898626.5 | | | | | | |
| P62301 | RPS13 | ribosomal protein S13 | Cytoplasm | other | 1996689 | 4570254.5 | 1600944.875 | P62301 | RPS13 | 2122885 | 1360730 | 555516 | 6 |
| P62264 | RPS14 | ribosomal protein S14 | Cytoplasm | translation regulator | 8532452 | 1.51E+07 | 6390458.5 | | | | | | |
| P62264 | RPS14 | ribosomal protein S14 | Cytoplasm | translation regulator | 6762305 | 1.12E+07 | 4269666 | | | | | | |
| P62264 | RPS14 | ribosomal protein S14 | Cytoplasm | translation regulator | 3578248.75 | 1.15E+07 | 4054747 | P62264 | RPS14 | 6291002 | 3974849 | 1324950 | 9 |

| | | | | | | | | | | | | | |
|---|---|---|---|---|---|---|---|---|---|---|---|---|---|
| P62245 | RPS15A | ribosomal protein S15a | Cytoplasm | other | 3085631.5 | 4873760 | 2014948.875 | | | | | | |
| P62245 | RPS15A | ribosomal protein S15a | Cytoplasm | other | 2865168.25 | 5045377.5 | 2685662.75 | P62245 | RPS15A | 2975400 | 1239992 | 506225 | 6 |
| P14131 | RPS16 | ribosomal protein S16 | Cytoplasm | other | 7633071 | 1.36E+07 | 5110458.5 | | | | | | |
| P14131 | RPS16 | ribosomal protein S16 | Cytoplasm | other | 5322674 | 8242834 | 2399017 | P14131 | RPS16 | 6477873 | 3832391 | 1564567 | 6 |
| P63276 | RPS17 | ribosomal protein S17 | Cytoplasm | other | 2839448.5 | 4350066.5 | 2220408.5 | P63276 | RPS17 | 2839449 | 1095492 | 632483 | 3 |
| P62270 | RPS18 | ribosomal protein S18 | Cytoplasm | other | 4571929.5 | 9313947 | 3700150 | | | | | | |
| P62270 | RPS18 | ribosomal protein S18 | Cytoplasm | other | 4275191 | 6537591.5 | 2694628 | | | | | | |
| P62270 | RPS18 | ribosomal protein S18 | Cytoplasm | other | 2775887.5 | 5528187 | 2473861 | P62270 | RPS18 | 3874336 | 2213784 | 737928 | 9 |
| Q9CZX8 | RPS19 | ribosomal protein S19 | Cytoplasm | other | 6004405.5 | 1.08E+07 | 5589568.5 | | | | | | |
| Q9CZX8 | RPS19 | ribosomal protein S19 | Cytoplasm | other | 5058328.5 | 8696204 | 3132343 | | | | | | |
| Q9CZX8 | RPS19 | ribosomal protein S19 | Cytoplasm | other | 5229527.5 | 9261957 | 3774471 | Q9CZX8 | RPS19 | 5430754 | 2605028 | 868343 | 9 |
| P25444 | RPS2 | ribosomal protein S2 | Cytoplasm | other | 9130859 | 1.65E+07 | 6587435.5 | | | | | | |
| P25444 | RPS2 | ribosomal protein S2 | Cytoplasm | other | 6262306.5 | 1.21E+07 | 5024748 | | | | | | |
| P25444 | RPS2 | ribosomal protein S2 | Cytoplasm | other | 5999946.5 | 9378107 | 4190713 | | | | | | |
| P25444 | RPS2 | ribosomal protein S2 | Cytoplasm | other | 4934551.5 | 7409753.5 | 3535906.75 | P25444 | RPS2 | 6581916 | 3723667 | 1074930 | 12 |
| P60867 | RPS20 | ribosomal protein S20 | Cytoplasm | other | 6332749.5 | 1.33E+07 | 4155061.25 | | | | | | |
| P60867 | RPS20 | ribosomal protein S20 | Cytoplasm | other | 5635846.5 | 8060117 | 3253855.25 | P60867 | RPS20 | 5984298 | 3601431 | 1470278 | 6 |
| P62267 | RPS23 | ribosomal protein S23 | Cytoplasm | translation regulator | 5015971.5 | 7756945 | 3474892.5 | P62267 | RPS23 | 5015972 | 2168864 | 1252194 | 3 |
| P62849 | RPS24 | ribosomal protein S24 | Cytoplasm | other | 2719413 | 4128243.75 | 1500201.125 | P62849 | RPS24 | 2719413 | 1315161 | 759309 | 3 |
| P62852 | RPS25 | ribosomal protein S25 | Cytoplasm | other | 9129592 | 1.17E+07 | 6046677.5 | | | | | | |
| P62852 | RPS25 | ribosomal protein S25 | Cytoplasm | other | 5308299.5 | 1.01E+07 | 4467988 | | | | | | |
| P62852 | RPS25 | ribosomal protein S25 | Cytoplasm | other | 5231918.5 | 9350453 | 4279794 | P62852 | RPS25 | 6556603 | 2763619 | 921206 | 9 |
| P62855 | RPS26 | ribosomal protein S26 | Cytoplasm | other | 4221357 | 6439743.5 | 2476345 | P62855 | RPS26 | 4221357 | 1986405 | 1146852 | 3 |
| Q6ZWY3 | RPS27L | ribosomal protein S27-like | Cytoplasm | translation regulator | 1845833.25 | 2743829.5 | 1430025.5 | | | | | | |
| Q6ZWY3 | RPS27L | ribosomal protein S27-like | Cytoplasm | translation regulator | 761032.125 | 1043748.562 | 239795.844 | Q6ZWY3 | RPS27L | 1303433 | 879549 | 359074 | 6 |
| P62858 | RPS28 | ribosomal protein S28 | Cytoplasm | other | 856012.938 | 3333887 | 757806.938 | P62858 | RPS28 | 856013 | 1459777 | 842803 | 3 |
| P62908 | RPS3 | ribosomal protein S3 | Cytoplasm | enzyme | 7962485 | 1.78E+07 | 5322948.5 | | | | | | |
| P62908 | RPS3 | ribosomal protein S3 | Cytoplasm | enzyme | 7126192 | 1.36E+07 | 3755984.75 | | | | | | |
| P62908 | RPS3 | ribosomal protein S3 | Cytoplasm | enzyme | 3778757 | 6564669 | 2776934.75 | | | | | | |
| P62908 | RPS3 | ribosomal protein S3 | Cytoplasm | enzyme | 5303618.5 | 1.00E+07 | 3874943.5 | | | | | | |
| P62908 | RPS3 | ribosomal protein S3 | Cytoplasm | enzyme | 3385298 | 6171638 | 2618631 | | | | | | |
| P62908 | RPS3 | ribosomal protein S3 | Cytoplasm | enzyme | 3900682.75 | 3255090.25 | 635418.5 | | | | | | |
| P62908 | RPS3 | ribosomal protein S3 | Cytoplasm | enzyme | 1965016.125 | 3449897 | 952146.062 | | | | | | |
| P62908 | RPS3 | ribosomal protein S3 | Cytoplasm | enzyme | 1378867.5 | 3054242.5 | 1147686.875 | P62908 | RPS3 | 4350115 | 4067090 | 830191 | 24 |
| P97351 | Rps3a1 | ribosomal protein S3A1 | Cytoplasm | other | 2535711.5 | 3704629 | 1868452.625 | | | | | | |
| P97351 | Rps3a1 | ribosomal protein S3A1 | Cytoplasm | other | 356898.656 | 2008861.875 | 500215.969 | P97351 | Rps3a1 | 1446305 | 1263652 | 515884 | 6 |
| P62702 | RPS4Y1 | ribosomal protein S4, Y-linked 1 | Cytoplasm | other | 7625370 | 1.30E+07 | 3524690.5 | | | | | | |
| P62702 | RPS4Y1 | ribosomal protein S4, Y-linked 1 | Cytoplasm | other | 6152696.5 | 7598430.5 | 3987846.75 | | | | | | |
| P62702 | RPS4Y1 | ribosomal protein S4, Y-linked 1 | Cytoplasm | other | 2804058.75 | 1712952.5 | 664967.5 | | | | | | |
| P62702 | RPS4Y1 | ribosomal protein S4, Y-linked 1 | Cytoplasm | other | 2406084.75 | 5087781.5 | 2315091 | | | | | | |
| P62702 | RPS4Y1 | ribosomal protein S4, Y-linked 1 | Cytoplasm | other | 2132362.5 | 4191474.75 | 2000986.375 | | | | | | |
| P62702 | RPS4Y1 | ribosomal protein S4, Y-linked 1 | Cytoplasm | other | 2966827.25 | 3231403 | 1022986.75 | | | | | | |
| P62702 | RPS4Y1 | ribosomal protein S4, Y-linked 1 | Cytoplasm | other | 2237468.25 | 3793687.25 | 1972602.375 | | | | | | |
| P62702 | RPS4Y1 | ribosomal protein S4, Y-linked 1 | Cytoplasm | other | 2111768 | 3274395.25 | 1454345.375 | P62702 | RPS4Y1 | 3554580 | 2704655 | 552085 | 24 |
| P62754 | RPS6 | ribosomal protein S6 | Cytoplasm | other | 5242523.5 | 8958242 | 2969357.5 | | | | | | |
| P62754 | RPS6 | ribosomal protein S6 | Cytoplasm | other | 2330881 | 1367721.75 | 455916.125 | P62754 | RPS6 | 3786702 | 3107176 | 1268499 | 6 |
| P62082 | RPS7 | ribosomal protein S7 | Cytoplasm | other | 9639079 | 1.74E+07 | 8541985 | | | | | | |
| P62082 | RPS7 | ribosomal protein S7 | Cytoplasm | other | 6897116.5 | 1.16E+07 | 4784705.5 | | | | | | |
| P62082 | RPS7 | ribosomal protein S7 | Cytoplasm | other | 3172747.75 | 4181230.75 | 1942655.25 | P62082 | RPS7 | 6569648 | 4852835 | 1617612 | 9 |
| P62242 | RPS8 | ribosomal protein S8 | Cytoplasm | other | 1.00E+07 | 1.39E+07 | 7207733.5 | | | | | | |
| P62242 | RPS8 | ribosomal protein S8 | Cytoplasm | other | 8019652 | 1.34E+07 | 5223752 | | | | | | |
| P62242 | RPS8 | ribosomal protein S8 | Cytoplasm | other | 6493939.5 | 8041691 | 5114376 | | | | | | |
| P62242 | RPS8 | ribosomal protein S8 | Cytoplasm | other | 3256456.25 | 5542937.5 | 1726661.75 | | | | | | |
| P62242 | RPS8 | ribosomal protein S8 | Cytoplasm | other | 2434635.75 | 3900187.5 | 1339373.125 | P62242 | RPS8 | 6046937 | 3859119 | 996420 | 15 |
| Q6ZWN5 | RPS9 | ribosomal protein S9 | Cytoplasm | translation regulator | 7369271.5 | 1.16E+07 | 3248415 | | | | | | |
| Q6ZWN5 | RPS9 | ribosomal protein S9 | Cytoplasm | translation regulator | 4133057.25 | 6161423.5 | 3118886 | | | | | | |
| Q6ZWN5 | RPS9 | ribosomal protein S9 | Cytoplasm | translation regulator | 2747157.25 | 4444233 | 642289 | Q6ZWN5 | RPS9 | 4749829 | 3198773 | 1066258 | 9 |
| P14206 | RPSA | ribosomal protein SA | Cytoplasm | translation regulator | 1.65E+07 | 2.97E+07 | 1.12E+07 | | | | | | |
| P14206 | RPSA | ribosomal protein SA | Cytoplasm | translation regulator | 8152309.5 | 1.54E+07 | 6538599 | | | | | | |
| P14206 | RPSA | ribosomal protein SA | Cytoplasm | translation regulator | 7052570.5 | 1.28E+07 | 6095426.5 | | | | | | |
| P14206 | RPSA | ribosomal protein SA | Cytoplasm | translation regulator | 3748930.75 | 6675697.5 | 2990727.5 | P14206 | RPSA | 8858453 | 7397056 | 2135346 | 12 |
| Q8CBB9 | RSAD2 | radical S-adenosyl methionine domain containing 2 | Cytoplasm | enzyme | 1.42E+07 | 2.53E+07 | 7108668.5 | | | | | | |
| Q8CBB9 | RSAD2 | radical S-adenosyl methionine domain containing 2 | Cytoplasm | enzyme | 1.53E+07 | 2.89E+07 | 1.09E+07 | | | | | | |
| Q8CBB9 | RSAD2 | radical S-adenosyl methionine domain containing 2 | Cytoplasm | enzyme | 8435077 | 1.69E+07 | 3781105.5 | | | | | | |
| Q8CBB9 | RSAD2 | radical S-adenosyl methionine domain containing 2 | Cytoplasm | enzyme | 1.22E+07 | 2.56E+07 | 8200016.5 | | | | | | |
| Q8CBB9 | RSAD2 | radical S-adenosyl methionine domain containing 2 | Cytoplasm | enzyme | 6789966.5 | 1.59E+07 | 3812335.5 | | | | | | |
| Q8CBB9 | RSAD2 | radical S-adenosyl methionine domain containing 2 | Cytoplasm | enzyme | 3651322 | 7706604 | 2509415 | | | | | | |
| Q8CBB9 | RSAD2 | radical S-adenosyl methionine domain containing 2 | Cytoplasm | enzyme | 4835374 | 9464240 | 5186339 | | | | | | |
| Q8CBB9 | RSAD2 | radical S-adenosyl methionine domain containing 2 | Cytoplasm | enzyme | 3258776.25 | 8356103 | 2410316 | Q8CBB9 | RSAD2 | 8581314 | 7586635 | 1548615 | 24 |
| Q01730 | RSU1 | Ras suppressor protein 1 | Cytoplasm | other | 1464681.125 | 4149949 | 3362557.75 | | | | | | |
| Q01730 | RSU1 | Ras suppressor protein 1 | Cytoplasm | other | 1139153.25 | 2941045 | 3258837 | | | | | | |
| Q01730 | RSU1 | Ras suppressor protein 1 | Cytoplasm | other | 209292.406 | 475136.812 | 529050.812 | Q01730 | RSU1 | 937709 | 1485247 | 495082 | 9 |
| Q8K0T0 | RTN1 | reticulon 1 | Cytoplasm | other | 6537161 | 1.12E+07 | 1.70E+07 | Q8K0T0 | RTN1 | 6537161 | 5252053 | 3032274 | 3 |
| Q9ES97 | RTN3 | reticulon 3 | Cytoplasm | other | 5.33E+07 | 6.99E+07 | 8.66E+07 | | | | | | |
| Q9ES97 | RTN3 | reticulon 3 | Cytoplasm | other | 138885.031 | 704826.5 | 1791760.375 | Q9ES97 | RTN3 | 26724443 | 39262434 | 16028821 | 6 |
| Q99P72 | RTN4 | reticulon 4 | Cytoplasm | other | 3170961.5 | 1.41E+07 | 1.60E+07 | | | | | | |
| Q99P72 | RTN4 | reticulon 4 | Cytoplasm | other | 5605945 | 1.07E+07 | 8015533 | | | | | | |
| Q99P72 | RTN4 | reticulon 4 | Cytoplasm | other | 2857172.75 | 6390406.5 | 6705886.5 | | | | | | |
| Q99P72 | RTN4 | reticulon 4 | Cytoplasm | other | 333054.531 | 2677202.75 | 3461833.75 | | | | | | |
| Q99P72 | RTN4 | reticulon 4 | Cytoplasm | other | 1768202.75 | 2670901.5 | 1801182.375 | Q99P72 | RTN4 | 2747067 | 4670995 | 1206046 | 15 |
| P97352 | S100A13 | S100 calcium binding protein A13 | Cytoplasm | other | 2328865.25 | 3205095 | 3370668.75 | P97352 | S100A13 | 2328865 | 559843 | 323226 | 3 |
| P31725 | S100A9 | S100 calcium binding protein A9 | Cytoplasm | other | 149088.797 | 265380.688 | 130912.617 | | | | | | |
| P31725 | S100A9 | S100 calcium binding protein A9 | Cytoplasm | other | 123725.742 | 185853.5 | 110067.648 | P31725 | S100A9 | 136407 | 57521 | 23483 | 6 |
| Q9EP69 | SACM1L | SAC1 suppressor of actin mutations 1-like (yeast) | Cytoplasm | phosphatase | 8785177 | 9303521 | 1.16E+07 | | | | | | |
| Q9EP69 | SACM1L | SAC1 suppressor of actin mutations 1-like (yeast) | Cytoplasm | phosphatase | 9471091 | 8370117.5 | 1.11E+07 | | | | | | |
| Q9EP69 | SACM1L | SAC1 suppressor of actin mutations 1-like (yeast) | Cytoplasm | phosphatase | 9176198 | 8226242 | 8646688 | | | | | | |
| Q9EP69 | SACM1L | SAC1 suppressor of actin mutations 1-like (yeast) | Cytoplasm | phosphatase | 8847945 | 8166165 | 9034506 | | | | | | |
| Q9EP69 | SACM1L | SAC1 suppressor of actin mutations 1-like (yeast) | Cytoplasm | phosphatase | 4956371.5 | 3700883 | 4097213.5 | | | | | | |
| Q9EP69 | SACM1L | SAC1 suppressor of actin mutations 1-like (yeast) | Cytoplasm | phosphatase | 5479045.5 | 5372067 | 7325393 | | | | | | |
| Q9EP69 | SACM1L | SAC1 suppressor of actin mutations 1-like (yeast) | Cytoplasm | phosphatase | 3369761.5 | 5558477.5 | 8432334 | | | | | | |
| Q9EP69 | SACM1L | SAC1 suppressor of actin mutations 1-like (yeast) | Cytoplasm | phosphatase | 4267045 | 3104695.5 | 5026705 | | | | | | |

| | | | | | | | | | | | | | |
|---|---|---|---|---|---|---|---|---|---|---|---|---|---|
| Q9EP69 | SACM1L | SAC1 suppressor of actin mutations 1-like (yeast) | Cytoplasm | phosphatase | 3723630.5 | 3202003 | 4335117.5 | | | | | | |
| Q9EP69 | SACM1L | SAC1 suppressor of actin mutations 1-like (yeast) | Cytoplasm | phosphatase | 4581864 | 4211940 | 4373636 | | | | | | |
| Q9EP69 | SACM1L | SAC1 suppressor of actin mutations 1-like (yeast) | Cytoplasm | phosphatase | 4255028 | 3774775.5 | 4808390.5 | | | | | | |
| Q9EP69 | SACM1L | SAC1 suppressor of actin mutations 1-like (yeast) | Cytoplasm | phosphatase | 3492139.75 | 3656363.25 | 3666456.25 | | | | | | |
| Q9EP69 | SACM1L | SAC1 suppressor of actin mutations 1-like (yeast) | Cytoplasm | phosphatase | 2706377.75 | 2550295.25 | 3542580.5 | | | | | | |
| Q9EP69 | SACM1L | SAC1 suppressor of actin mutations 1-like (yeast) | Cytoplasm | phosphatase | 2515927.75 | 2272883.75 | 3016707.25 | | | | | | |
| Q9EP69 | SACM1L | SAC1 suppressor of actin mutations 1-like (yeast) | Cytoplasm | phosphatase | 2005754.875 | 1780492.375 | 2012941.5 | | | | | | |
| Q9EP69 | SACM1L | SAC1 suppressor of actin mutations 1-like (yeast) | Cytoplasm | phosphatase | 2134725.25 | 1454687.375 | 1160776.625 | | | | | | |
| Q9EP69 | SACM1L | SAC1 suppressor of actin mutations 1-like (yeast) | Cytoplasm | phosphatase | 2794374.75 | 2104463.25 | 2785733.5 | | | | | | |
| Q9EP69 | SACM1L | SAC1 suppressor of actin mutations 1-like (yeast) | Cytoplasm | phosphatase | 2006730.375 | 2372790 | 2322800.5 | | | | | | |
| Q9EP69 | SACM1L | SAC1 suppressor of actin mutations 1-like (yeast) | Cytoplasm | phosphatase | 204914.5 | 1005622.188 | 2021257.875 | Q9EP69 | SACM1L | 4461795 | 2818155 | 373274 | 57 |
| Q8BGH2 | SAMM50 | SAMM50 sorting and assembly machinery component | Cytoplasm | other | 1231128.75 | 2807694 | 1391466.375 | | | | | | |
| Q8BGH2 | SAMM50 | SAMM50 sorting and assembly machinery component | Cytoplasm | other | 1994086.125 | 2344034 | 329531.719 | Q8BGH2 | SAMM50 | 1612607 | 885408 | 361466 | 6 |
| Q8K021 | SCAMP1 | secretory carrier membrane protein 1 | Cytoplasm | transporter | 1.87E+07 | 1.35E+07 | 1.26E+07 | Q8K021 | SCAMP1 | 18720000 | 3319342 | 1916423 | 3 |
| Q9ERN0 | SCAMP2 | secretory carrier membrane protein 2 | Cytoplasm | transporter | 2494007.5 | 3857819.25 | 4712066.5 | | | | | | |
| Q9ERN0 | SCAMP2 | secretory carrier membrane protein 2 | Cytoplasm | transporter | 2152617 | 3440857.75 | 3703851.25 | Q9ERN0 | SCAMP2 | 2323312 | 938259 | 383043 | 6 |
| Q9JKV5 | SCAMP4 | secretory carrier membrane protein 4 | Cytoplasm | other | 1.10E+07 | 1.06E+07 | 1.12E+07 | | | | | | |
| Q9JKV5 | SCAMP4 | secretory carrier membrane protein 4 | Cytoplasm | other | 5573460 | 8840052 | 1.02E+07 | Q9JKV5 | SCAMP4 | 8271730 | 2114838 | 863379 | 6 |
| O35114 | SCARB2 | scavenger receptor class B, member 2 | Plasma Membrane | other | 2015221.25 | 5606405 | 2295889 | O35114 | SCARB2 | 2015221 | 1997285 | 1153133 | 3 |
| Q8K2B3 | SDHA | succinate dehydrogenase complex, subunit A, flavoprotein (Fp) | Cytoplasm | enzyme | 8938484 | 1.82E+07 | 8316877.5 | | | | | | |
| Q8K2B3 | SDHA | succinate dehydrogenase complex, subunit A, flavoprotein (Fp) | Cytoplasm | enzyme | 5081743 | 1.50E+07 | 8036364.5 | | | | | | |
| Q8K2B3 | SDHA | succinate dehydrogenase complex, subunit A, flavoprotein (Fp) | Cytoplasm | enzyme | 4327427 | 9170396 | 4385847.5 | | | | | | |
| Q8K2B3 | SDHA | succinate dehydrogenase complex, subunit A, flavoprotein (Fp) | Cytoplasm | enzyme | 3815974 | 8508955 | 3249714.5 | | | | | | |
| Q8K2B3 | SDHA | succinate dehydrogenase complex, subunit A, flavoprotein (Fp) | Cytoplasm | enzyme | 3276202.5 | 5186240 | 2074188.875 | | | | | | |
| Q8K2B3 | SDHA | succinate dehydrogenase complex, subunit A, flavoprotein (Fp) | Cytoplasm | enzyme | 3671056 | 6184580.5 | 2582558.5 | | | | | | |
| Q8K2B3 | SDHA | succinate dehydrogenase complex, subunit A, flavoprotein (Fp) | Cytoplasm | enzyme | 3271193.25 | 6778373.5 | 3809647.5 | | | | | | |
| Q8K2B3 | SDHA | succinate dehydrogenase complex, subunit A, flavoprotein (Fp) | Cytoplasm | enzyme | 1337322.375 | 2871816.25 | 927255.438 | Q8K2B3 | SDHA | 4214925 | 4124158 | 841840 | 24 |
| Q9CQA3 | SDHB | succinate dehydrogenase complex, subunit B, iron sulfur (Ip) | Cytoplasm | enzyme | 3390192.5 | 4236659.5 | 3296822 | | | | | | |
| Q9CQA3 | SDHB | succinate dehydrogenase complex, subunit B, iron sulfur (Ip) | Cytoplasm | enzyme | 2850911.25 | 5333821.5 | 2082795.625 | | | | | | |
| Q9CQA3 | SDHB | succinate dehydrogenase complex, subunit B, iron sulfur (Ip) | Cytoplasm | enzyme | 2129128 | 3706804.25 | 1594605.5 | Q9CQA3 | SDHB | 2790077 | 1173712 | 391237 | 9 |
| Q9CZB0 | SDHC | succinate dehydrogenase complex, subunit C, integral membrane protein, 15kDa | Cytoplasm | enzyme | 1707936.375 | 4312412 | 1709156.5 | Q9CZB0 | SDHC | 1707936 | 1503343 | 867955 | 3 |
| O08547 | SEC22B | SEC22 vesicle trafficking protein homolog B (S. cerevisiae) (gene/pseudogene) | Cytoplasm | other | 5494713 | 4632162.5 | 3775817 | | | | | | |
| O08547 | SEC22B | SEC22 vesicle trafficking protein homolog B (S. cerevisiae) (gene/pseudogene) | Cytoplasm | other | 4371627 | 3260976.25 | 2878666.5 | | | | | | |
| O08547 | SEC22B | SEC22 vesicle trafficking protein homolog B (S. cerevisiae) (gene/pseudogene) | Cytoplasm | other | 2980655.75 | 5324828 | 4995464.5 | | | | | | |
| O08547 | SEC22B | SEC22 vesicle trafficking protein homolog B (S. cerevisiae) (gene/pseudogene) | Cytoplasm | other | 3528132.5 | 4965136.5 | 5680694 | | | | | | |
| O08547 | SEC22B | SEC22 vesicle trafficking protein homolog B (S. cerevisiae) (gene/pseudogene) | Cytoplasm | other | 2044749.5 | 2987813.75 | 2513735.25 | | | | | | |
| O08547 | SEC22B | SEC22 vesicle trafficking protein homolog B (S. cerevisiae) (gene/pseudogene) | Cytoplasm | other | 1249768.25 | 2948343 | 2768454.5 | | | | | | |
| O08547 | SEC22B | SEC22 vesicle trafficking protein homolog B (S. cerevisiae) (gene/pseudogene) | Cytoplasm | other | 1942787 | 2042818.875 | 1413181.5 | | | | | | |
| O08547 | SEC22B | SEC22 vesicle trafficking protein homolog B (S. cerevisiae) (gene/pseudogene) | Cytoplasm | other | 1168631.75 | 1923560.75 | 1692128 | O08547 | SEC22B | 2847633 | 1419360 | 289726 | 24 |
| Q3TZ89 | SEC31B | SEC31 homolog B, COPII coating complex component | Cytoplasm | other | 2580390.5 | 6370183 | 1.11E+07 | Q3TZ89 | SEC31B | 2580391 | 4258096 | 2458413 | 3 |
| P61620 | SEC61A1 | Sec61 alpha 1 subunit (S. cerevisiae) | Cytoplasm | transporter | 1219127.25 | 1452672.875 | 1923271 | P61620 | SEC61A1 | 1219127 | 358661 | 207073 | 3 |
| Q9CQS8 | SEC61B | Sec61 beta subunit | Cytoplasm | transporter | 1982029.875 | 3722720.75 | 3859749 | | | | | | |
| Q9CQS8 | SEC61B | Sec61 beta subunit | Cytoplasm | transporter | 971767.875 | 1553953.875 | 2080427.625 | Q9CQS8 | SEC61B | 1476899 | 1175088 | 479728 | 6 |
| P17563 | SELENBP1 | selenium binding protein 1 | Cytoplasm | other | 552309.25 | 316153.469 | 270189.438 | P17563 | SELENBP1 | 552309 | 151368 | 87392 | 3 |
| Q01102 | SELP | selectin P (granule membrane protein 140kDa, antigen CD62) | Plasma Membrane | transmembrane receptor | 414138.719 | 655733.75 | 1612331.75 | | | | | | |
| Q01102 | SELP | selectin P (granule membrane protein 140kDa, antigen CD62) | Plasma Membrane | transmembrane receptor | 1271370.125 | 2818180 | 5939242 | | | | | | |
| Q01102 | SELP | selectin P (granule membrane protein 140kDa, antigen CD62) | Plasma Membrane | transmembrane receptor | 336238.75 | 1758944 | 3768833.75 | | | | | | |
| Q01102 | SELP | selectin P (granule membrane protein 140kDa, antigen CD62) | Plasma Membrane | transmembrane receptor | 80319.367 | 262200.125 | 702325.062 | | | | | | |
| Q01102 | SELP | selectin P (granule membrane protein 140kDa, antigen CD62) | Plasma Membrane | transmembrane receptor | 229344.672 | 1348290.375 | 1736589.25 | | | | | | |
| Q01102 | SELP | selectin P (granule membrane protein 140kDa, antigen CD62) | Plasma Membrane | transmembrane receptor | 4.70E-06 | 20517.076 | 87772.484 | Q01102 | SELP | 388569 | 1560747 | 367872 | 18 |
| P62342 | SELT | selenoprotein T | Cytoplasm | other | 2121722 | 3067416.5 | 4178145.25 | P62342 | SELT | 2121722 | 1029315 | 594275 | 3 |
| Q9CY58 | SERBP1 | SERPINE1 mRNA binding protein 1 | Cytoplasm | other | 1141999.5 | 2640711 | 852510.062 | | | | | | |
| Q9CY58 | SERBP1 | SERPINE1 mRNA binding protein 1 | Cytoplasm | other | 1180073.25 | 3966091.75 | 869430.938 | | | | | | |
| Q9CY58 | SERBP1 | SERPINE1 mRNA binding protein 1 | Cytoplasm | other | 565352 | 3004775 | 1139745 | Q9CY58 | SERBP1 | 962475 | 1189182 | 396394 | 9 |
| Q9QZI9 | SERINC3 | serine incorporator 3 | Cytoplasm | transporter | 3.44E+08 | 3.00E+08 | 2.19E+08 | | | | | | |
| Q9QZI9 | SERINC3 | serine incorporator 3 | Cytoplasm | transporter | 1.47E+08 | 1.65E+08 | 1.78E+08 | | | | | | |
| Q9QZI9 | SERINC3 | serine incorporator 3 | Cytoplasm | transporter | 1.23E+08 | 1.79E+08 | 2.21E+08 | | | | | | |
| Q9QZI9 | SERINC3 | serine incorporator 3 | Cytoplasm | transporter | 2.87E+07 | 4.57E+07 | 7.05E+07 | Q9QZI9 | SERINC3 | 160400000 | 95427980 | 27547685 | 12 |
| P07758 | SERPINA1 | serpin peptidase inhibitor, clade A (alpha-1 antiproteinase, antitrypsin), member 1 | Extracellular Space | other | 1.26E+07 | 1.47E+07 | 2.67E+07 | | | | | | |
| P07758 | SERPINA1 | serpin peptidase inhibitor, clade A (alpha-1 antiproteinase, antitrypsin), member 1 | Extracellular Space | other | 3521465.5 | 5998307 | 6830163.5 | | | | | | |
| P07758 | SERPINA1 | serpin peptidase inhibitor, clade A (alpha-1 antiproteinase, antitrypsin), member 1 | Extracellular Space | other | 2772409 | 3433269.5 | 6563119 | | | | | | |
| P07758 | SERPINA1 | serpin peptidase inhibitor, clade A (alpha-1 antiproteinase, antitrypsin), member 1 | Extracellular Space | other | 2685937.25 | 3285105.75 | 6390001 | | | | | | |
| P07758 | SERPINA1 | serpin peptidase inhibitor, clade A (alpha-1 antiproteinase, antitrypsin), member 1 | Extracellular Space | other | 2430443.25 | 2698388.5 | 4831543 | P07758 | SERPINA1 | 4796051 | 6554018 | 1692240 | 15 |
| P07759 | SERPINA3 | serpin peptidase inhibitor, clade A (alpha-1 antiproteinase, antitrypsin), member 3 | Extracellular Space | other | 2.11E+07 | 1.92E+07 | 2.53E+07 | | | | | | |
| P07759 | SERPINA3 | serpin peptidase inhibitor, clade A (alpha-1 antiproteinase, antitrypsin), member 3 | Extracellular Space | other | 1.13E+07 | 1.08E+07 | 1.54E+07 | | | | | | |
| P07759 | SERPINA3 | serpin peptidase inhibitor, clade A (alpha-1 antiproteinase, antitrypsin), member 3 | Extracellular Space | other | 7809600.5 | 4831831.5 | 7936456 | P07759 | SERPINA3 | 13406534 | 6925912 | 2308637 | 9 |
| Q99JR1 | SFXN1 | sideroflexin 1 | Cytoplasm | transporter | 2936958.25 | 6583920 | 3684288 | | | | | | |
| Q99JR1 | SFXN1 | sideroflexin 1 | Cytoplasm | transporter | 3466535.5 | 6401641.5 | 3642000.75 | | | | | | |
| Q99JR1 | SFXN1 | sideroflexin 1 | Cytoplasm | transporter | 3007372 | 7029210 | 3548696 | | | | | | |
| Q99JR1 | SFXN1 | sideroflexin 1 | Cytoplasm | transporter | 2717230.25 | 5036416 | 2073720.375 | | | | | | |
| Q99JR1 | SFXN1 | sideroflexin 1 | Cytoplasm | transporter | 1874009.125 | 3995841.75 | 1948850.875 | | | | | | |
| Q99JR1 | SFXN1 | sideroflexin 1 | Cytoplasm | transporter | 546081.812 | 1164749.875 | 378700.656 | Q99JR1 | SFXN1 | 2424698 | 1951464 | 459965 | 18 |
| Q9JJU8 | SH3BGRL | SH3 domain binding glutamate-rich protein like | Cytoplasm | other | 1.45E+07 | 1.30E+07 | 1.77E+07 | | | | | | |
| Q9JJU8 | SH3BGRL | SH3 domain binding glutamate-rich protein like | Cytoplasm | other | 1.08E+07 | 1.01E+07 | 1.06E+07 | | | | | | |
| Q9JJU8 | SH3BGRL | SH3 domain binding glutamate-rich protein like | Cytoplasm | other | 8077734.5 | 1.22E+07 | 1.49E+07 | Q9JJU8 | SH3BGRL | 11132578 | 2936453 | 978818 | 9 |
| Q8BG73 | SH3BGRL2 | SH3 domain binding glutamate-rich protein like 2 | Nucleus | other | 1627307 | 2333257.25 | 4362589 | Q8BG73 | SH3BGRL2 | 1627307 | 1419995 | 819835 | 3 |
| Q91Y57 | SIGLEC9 | sialic acid binding Ig-like lectin 9 | Plasma Membrane | other | 5216044 | 5735919 | 6324295.5 | Q91Y57 | SIGLEC9 | 5216044 | 554478 | 320128 | 3 |
| Q3UND0 | SKAP2 | src kinase associated phosphoprotein 2 | Cytoplasm | other | 155679.656 | 626118.812 | 1018823 | | | | | | |
| Q3UND0 | SKAP2 | src kinase associated phosphoprotein 2 | Cytoplasm | other | 97331.039 | 416332.531 | 950704.562 | Q3UND0 | SKAP2 | 126505 | 390951 | 159605 | 6 |
| Q9QUM4 | SLAMF1 | signaling lymphocytic activation molecule family member 1 | Plasma Membrane | transmembrane receptor | 70970.273 | 1088238.25 | 2012305.25 | | | | | | |
| Q9QUM4 | SLAMF1 | signaling lymphocytic activation molecule family member 1 | Plasma Membrane | transmembrane receptor | 317099.156 | 977972.062 | 1401461.625 | | | | | | |
| Q9QUM4 | SLAMF1 | signaling lymphocytic activation molecule family member 1 | Plasma Membrane | transmembrane receptor | 263813.75 | 1261663.625 | 3539986.25 | Q9QUM4 | SLAMF1 | 217294 | 1070645 | 356882 | 9 |
| P49282 | SLC11A2 | solute carrier family 11 (proton-coupled divalent metal ion transporter), member 2 | Plasma Membrane | transporter | 1.02E+07 | 1.51E+07 | 1.10E+07 | | | | | | |
| P49282 | SLC11A2 | solute carrier family 11 (proton-coupled divalent metal ion transporter), member 2 | Plasma Membrane | transporter | 5507955 | 7478174.5 | 6832490.5 | | | | | | |
| P49282 | SLC11A2 | solute carrier family 11 (proton-coupled divalent metal ion transporter), member 2 | Plasma Membrane | transporter | 3773523.75 | 1.11E+07 | 3422502.75 | | | | | | |
| P49282 | SLC11A2 | solute carrier family 11 (proton-coupled divalent metal ion transporter), member 2 | Plasma Membrane | transporter | 2656850.5 | 5024283 | 2847128 | P49282 | SLC11A2 | 5532082 | 3985332 | 1150466 | 12 |
| Q8VHL0 | SLC14A1 | solute carrier family 14 (urea transporter), member 1 (Kidd blood group) | Plasma Membrane | transporter | 3.24E+08 | 4.16E+08 | 4.92E+08 | | | | | | |
| Q8VHL0 | SLC14A1 | solute carrier family 14 (urea transporter), member 1 (Kidd blood group) | Plasma Membrane | transporter | 1.92E+08 | 2.68E+08 | 2.39E+08 | | | | | | |
| Q8VHL0 | SLC14A1 | solute carrier family 14 (urea transporter), member 1 (Kidd blood group) | Plasma Membrane | transporter | 2.36E+08 | 1.87E+08 | 1.94E+08 | | | | | | |
| Q8VHL0 | SLC14A1 | solute carrier family 14 (urea transporter), member 1 (Kidd blood group) | Plasma Membrane | transporter | 1.44E+08 | 1.93E+08 | 2.78E+08 | | | | | | |
| Q8VHL0 | SLC14A1 | solute carrier family 14 (urea transporter), member 1 (Kidd blood group) | Plasma Membrane | transporter | 1.67E+08 | 1.70E+08 | 1.87E+08 | | | | | | |

| | | | | | | | | | | | | | |
|---|---|---|---|---|---|---|---|---|---|---|---|---|---|
| Q8VHL0 | SLC14A1 | solute carrier family 14 (urea transporter), member 1 (Kidd blood group) | Plasma Membrane | transporter | 1.12E+08 | 1.36E+08 | 1.94E+08 | | | | | | |
| Q8VHL0 | SLC14A1 | solute carrier family 14 (urea transporter), member 1 (Kidd blood group) | Plasma Membrane | transporter | 2.37E+08 | 1.00E+08 | 7.94E+07 | | | | | | |
| Q8VHL0 | SLC14A1 | solute carrier family 14 (urea transporter), member 1 (Kidd blood group) | Plasma Membrane | transporter | 4.81E+07 | 4.28E+07 | 6.05E+07 | | | | | | |
| Q8VHL0 | SLC14A1 | solute carrier family 14 (urea transporter), member 1 (Kidd blood group) | Plasma Membrane | transporter | 4.40E+07 | 1.55E+07 | 1.28E+07 | | | | | | |
| Q8VHL0 | SLC14A1 | solute carrier family 14 (urea transporter), member 1 (Kidd blood group) | Plasma Membrane | transporter | 2.57E+07 | 2.27E+07 | 2.84E+07 | | | | | | |
| Q8VHL0 | SLC14A1 | solute carrier family 14 (urea transporter), member 1 (Kidd blood group) | Plasma Membrane | transporter | 2.47E+07 | 3.36E+07 | 5.33E+07 | | | | | | |
| Q8VHL0 | SLC14A1 | solute carrier family 14 (urea transporter), member 1 (Kidd blood group) | Plasma Membrane | transporter | 1.26E+07 | 1397042.875 | 53888.582 | Q8VHL0 | SLC14A1 | 130662500 | 121000517 | 20166753 | 36 |
| P53986 | SLC16A1 | solute carrier family 16 (monocarboxylate transporter), member 1 | Plasma Membrane | transporter | 5.31E+08 | 4.86E+08 | 5.89E+08 | | | | | | |
| P53986 | SLC16A1 | solute carrier family 16 (monocarboxylate transporter), member 1 | Plasma Membrane | transporter | 5.03E+08 | 4.53E+08 | 4.75E+08 | | | | | | |
| P53986 | SLC16A1 | solute carrier family 16 (monocarboxylate transporter), member 1 | Plasma Membrane | transporter | 4.00E+08 | 5.18E+08 | 6.88E+08 | | | | | | |
| P53986 | SLC16A1 | solute carrier family 16 (monocarboxylate transporter), member 1 | Plasma Membrane | transporter | 1.75E+08 | 1.80E+08 | 1.90E+08 | | | | | | |
| P53986 | SLC16A1 | solute carrier family 16 (monocarboxylate transporter), member 1 | Plasma Membrane | transporter | 1.36E+08 | 1.44E+08 | 1.50E+08 | | | | | | |
| P53986 | SLC16A1 | solute carrier family 16 (monocarboxylate transporter), member 1 | Plasma Membrane | transporter | 7.17E+07 | 8.66E+07 | 1.03E+08 | | | | | | |
| P53986 | SLC16A1 | solute carrier family 16 (monocarboxylate transporter), member 1 | Plasma Membrane | transporter | 2.33E+07 | 2.93E+07 | 3.89E+07 | | | | | | |
| P53986 | SLC16A1 | solute carrier family 16 (monocarboxylate transporter), member 1 | Plasma Membrane | transporter | 3.19E+07 | 5.15E+07 | 8.36E+07 | | | | | | |
| P53986 | SLC16A1 | solute carrier family 16 (monocarboxylate transporter), member 1 | Plasma Membrane | transporter | 6406471.5 | 4497225.5 | 7562933.5 | P53986 | SLC16A1 | 208610719 | 218809031 | 42109818 | 27 |
| Q3U9N9 | SLC16A10 | solute carrier family 16 (aromatic amino acid transporter), member 10 | Plasma Membrane | transporter | 1.53E+08 | 1.78E+08 | 1.82E+08 | | | | | | |
| Q3U9N9 | SLC16A10 | solute carrier family 16 (aromatic amino acid transporter), member 10 | Plasma Membrane | transporter | 1.21E+08 | 1.17E+08 | 1.42E+08 | | | | | | |
| Q3U9N9 | SLC16A10 | solute carrier family 16 (aromatic amino acid transporter), member 10 | Plasma Membrane | transporter | 7.11E+07 | 9.27E+07 | 1.45E+08 | | | | | | |
| Q3U9N9 | SLC16A10 | solute carrier family 16 (aromatic amino acid transporter), member 10 | Plasma Membrane | transporter | 4.24E+07 | 5.99E+07 | 7.48E+07 | | | | | | |
| Q3U9N9 | SLC16A10 | solute carrier family 16 (aromatic amino acid transporter), member 10 | Plasma Membrane | transporter | 4.74E+07 | 4.35E+07 | 5.40E+07 | | | | | | |
| Q3U9N9 | SLC16A10 | solute carrier family 16 (aromatic amino acid transporter), member 10 | Plasma Membrane | transporter | 2.75E+07 | 5.17E+07 | 5.98E+07 | | | | | | |
| Q3U9N9 | SLC16A10 | solute carrier family 16 (aromatic amino acid transporter), member 10 | Plasma Membrane | transporter | 1.32E+07 | 1.58E+07 | 1.55E+07 | | | | | | |
| Q3U9N9 | SLC16A10 | solute carrier family 16 (aromatic amino acid transporter), member 10 | Plasma Membrane | transporter | 8018022.5 | 1.17E+07 | 8004864.5 | | | | | | |
| Q3U9N9 | SLC16A10 | solute carrier family 16 (aromatic amino acid transporter), member 10 | Plasma Membrane | transporter | 8704828 | 8796940 | 1.24E+07 | | | | | | |
| Q3U9N9 | SLC16A10 | solute carrier family 16 (aromatic amino acid transporter), member 10 | Plasma Membrane | transporter | 1.99E+07 | 984663.188 | 196653.391 | Q3U9N9 | SLC16A10 | 51209285 | 56149877 | 10251518 | 30 |
| Q8BRU6 | SLC18A2 | solute carrier family 18 (vesicular monoamine transporter), member 2 | Plasma Membrane | transporter | 36288.133 | 46.485 | 352707.906 | Q8BRU6 | SLC18A2 | 36288 | 193995 | 112003 | 3 |
| P41438 | SLC19A1 | solute carrier family 19 (folate transporter), member 1 | Plasma Membrane | transporter | 1996115.75 | 1800034.25 | 1723848.125 | P41438 | SLC19A1 | 1996116 | 140465 | 81097 | 3 |
| Q3UHH2 | SLC22A23 | solute carrier family 22, member 23 | Plasma Membrane | transporter | 2196385 | 1926574.25 | 1960555 | Q3UHH2 | SLC22A23 | 2196385 | 146951 | 84842 | 3 |
| Q8BH59 | SLC25A12 | solute carrier family 25 (aspartate/glutamate carrier), member 12 | Cytoplasm | transporter | 5261866.5 | 1.25E+07 | 4583231 | | | | | | |
| Q8BH59 | SLC25A12 | solute carrier family 25 (aspartate/glutamate carrier), member 12 | Cytoplasm | transporter | 2421375.25 | 5977196 | 3193695.5 | | | | | | |
| Q8BH59 | SLC25A12 | solute carrier family 25 (aspartate/glutamate carrier), member 12 | Cytoplasm | transporter | 2035142.25 | 4787432.5 | 1456321.625 | | | | | | |
| Q8BH59 | SLC25A12 | solute carrier family 25 (aspartate/glutamate carrier), member 12 | Cytoplasm | transporter | 1001974.562 | 2582882.5 | 1370152.375 | Q8BH59 | SLC25A12 | 2680090 | 3155470 | 910906 | 12 |
| Q8BZ09 | SLC25A21 | solute carrier family 25 (mitochondrial oxoadipate carrier), member 21 | Cytoplasm | transporter | 4058750.5 | 7643211 | 3093280.5 | | | | | | |
| Q8BZ09 | SLC25A21 | solute carrier family 25 (mitochondrial oxoadipate carrier), member 21 | Cytoplasm | transporter | 3155582 | 5620908 | 2100155.25 | | | | | | |
| Q8BZ09 | SLC25A21 | solute carrier family 25 (mitochondrial oxoadipate carrier), member 21 | Cytoplasm | transporter | 1515372.875 | 4379379 | 1234184.125 | | | | | | |
| Q8BZ09 | SLC25A21 | solute carrier family 25 (mitochondrial oxoadipate carrier), member 21 | Cytoplasm | transporter | 1549750.5 | 2751037.75 | 855846.25 | | | | | | |
| Q8BZ09 | SLC25A21 | solute carrier family 25 (mitochondrial oxoadipate carrier), member 21 | Cytoplasm | transporter | 1356933.125 | 3599533.75 | 1343475.75 | Q8BZ09 | SLC25A21 | 2327278 | 1892581 | 488662 | 15 |
| Q8VEM8 | SLC25A3 | solute carrier family 25 (mitochondrial carrier; phosphate carrier), member 3 | Cytoplasm | transporter | 2.33E+07 | 4.50E+07 | 2.03E+07 | | | | | | |
| Q8VEM8 | SLC25A3 | solute carrier family 25 (mitochondrial carrier; phosphate carrier), member 3 | Cytoplasm | transporter | 1.50E+07 | 2.91E+07 | 1.57E+07 | | | | | | |
| Q8VEM8 | SLC25A3 | solute carrier family 25 (mitochondrial carrier; phosphate carrier), member 3 | Cytoplasm | transporter | 1.05E+07 | 1.68E+07 | 1.00E+07 | | | | | | |
| Q8VEM8 | SLC25A3 | solute carrier family 25 (mitochondrial carrier; phosphate carrier), member 3 | Cytoplasm | transporter | 1.02E+07 | 2.31E+07 | 1.31E+07 | | | | | | |
| Q8VEM8 | SLC25A3 | solute carrier family 25 (mitochondrial carrier; phosphate carrier), member 3 | Cytoplasm | transporter | 7989079 | 1.70E+07 | 7063058.5 | | | | | | |
| Q8VEM8 | SLC25A3 | solute carrier family 25 (mitochondrial carrier; phosphate carrier), member 3 | Cytoplasm | transporter | 5519011 | 1.20E+07 | 4809754 | | | | | | |
| Q8VEM8 | SLC25A3 | solute carrier family 25 (mitochondrial carrier; phosphate carrier), member 3 | Cytoplasm | transporter | 4804347.5 | 9294371 | 3610624.25 | | | | | | |
| Q8VEM8 | SLC25A3 | solute carrier family 25 (mitochondrial carrier; phosphate carrier), member 3 | Cytoplasm | transporter | 2012704.125 | 3500446.25 | 1923546.625 | | | | | | |
| Q8VEM8 | SLC25A3 | solute carrier family 25 (mitochondrial carrier; phosphate carrier), member 3 | Cytoplasm | transporter | 2823271 | 4720307 | 2095089.625 | Q8VEM8 | SLC25A3 | 9128713 | 9909621 | 1907107 | 27 |
| Q920G8 | SLC25A37 | solute carrier family 25 (mitochondrial iron transporter), member 37 | Cytoplasm | other | 9643682 | 2.03E+07 | 6703967.5 | | | | | | |
| Q920G8 | SLC25A37 | solute carrier family 25 (mitochondrial iron transporter), member 37 | Cytoplasm | other | 5255030 | 1.21E+07 | 4777043 | | | | | | |
| Q920G8 | SLC25A37 | solute carrier family 25 (mitochondrial iron transporter), member 37 | Cytoplasm | other | 5131442 | 9598942 | 1971541.25 | | | | | | |
| Q920G8 | SLC25A37 | solute carrier family 25 (mitochondrial iron transporter), member 37 | Cytoplasm | other | 5502817 | 8127649 | 4047515 | | | | | | |
| Q920G8 | SLC25A37 | solute carrier family 25 (mitochondrial iron transporter), member 37 | Cytoplasm | other | 4479144 | 8687182 | 3764483.5 | | | | | | |
| Q920G8 | SLC25A37 | solute carrier family 25 (mitochondrial iron transporter), member 37 | Cytoplasm | other | 3948044.5 | 8069292.5 | 2759578.25 | | | | | | |
| Q920G8 | SLC25A37 | solute carrier family 25 (mitochondrial iron transporter), member 37 | Cytoplasm | other | 3714908.25 | 7184620 | 3091851.5 | | | | | | |
| Q920G8 | SLC25A37 | solute carrier family 25 (mitochondrial iron transporter), member 37 | Cytoplasm | other | 4388996 | 7888437.5 | 2661626 | | | | | | |
| Q920G8 | SLC25A37 | solute carrier family 25 (mitochondrial iron transporter), member 37 | Cytoplasm | other | 2056833.125 | 3718324.5 | 1779747.5 | | | | | | |
| Q920G8 | SLC25A37 | solute carrier family 25 (mitochondrial iron transporter), member 37 | Cytoplasm | other | 976052.375 | 3047042 | 788674.062 | | | | | | |
| Q920G8 | SLC25A37 | solute carrier family 25 (mitochondrial iron transporter), member 37 | Cytoplasm | other | 2523773 | 8922394 | 4638714 | Q920G8 | SLC25A37 | 4329157 | 3857793 | 671556 | 33 |
| P48962 | SLC25A4 | solute carrier family 25 (mitochondrial carrier; adenine nucleotide translocator), member 4 | Cytoplasm | transporter | 1775071.875 | 3455841 | 3190093.75 | | | | | | |
| P48962 | SLC25A4 | solute carrier family 25 (mitochondrial carrier; adenine nucleotide translocator), member 4 | Cytoplasm | transporter | 1285109.625 | 2517130.5 | 2192066 | | | | | | |
| P48962 | SLC25A4 | solute carrier family 25 (mitochondrial carrier; adenine nucleotide translocator), member 4 | Cytoplasm | transporter | 1015629.375 | 1427978.875 | 1600455.75 | P48962 | SLC25A4 | 1358604 | 853972 | 284657 | 9 |
| P51881 | SLC25A5 | solute carrier family 25 (mitochondrial carrier; adenine nucleotide translocator), member 5 | Cytoplasm | transporter | 6.43E+07 | 1.53E+08 | 7.63E+07 | | | | | | |
| P51881 | SLC25A5 | solute carrier family 25 (mitochondrial carrier; adenine nucleotide translocator), member 5 | Cytoplasm | transporter | 5.34E+07 | 9.13E+07 | 5.31E+07 | | | | | | |
| P51881 | SLC25A5 | solute carrier family 25 (mitochondrial carrier; adenine nucleotide translocator), member 5 | Cytoplasm | transporter | 4.22E+07 | 7.78E+07 | 4.30E+07 | | | | | | |
| P51881 | SLC25A5 | solute carrier family 25 (mitochondrial carrier; adenine nucleotide translocator), member 5 | Cytoplasm | transporter | 2.29E+07 | 4.40E+07 | 2.35E+07 | | | | | | |
| P51881 | SLC25A5 | solute carrier family 25 (mitochondrial carrier; adenine nucleotide translocator), member 5 | Cytoplasm | transporter | 1.16E+07 | 3.15E+07 | 1.76E+07 | | | | | | |
| P51881 | SLC25A5 | solute carrier family 25 (mitochondrial carrier; adenine nucleotide translocator), member 5 | Cytoplasm | transporter | 1.55E+07 | 4.65E+07 | 3.11E+07 | | | | | | |
| P51881 | SLC25A5 | solute carrier family 25 (mitochondrial carrier; adenine nucleotide translocator), member 5 | Cytoplasm | transporter | 2.14E+07 | 2.01E+07 | 3290844.75 | | | | | | |
| P51881 | SLC25A5 | solute carrier family 25 (mitochondrial carrier; adenine nucleotide translocator), member 5 | Cytoplasm | transporter | 1.50E+07 | 3.02E+07 | 1.64E+07 | | | | | | |
| P51881 | SLC25A5 | solute carrier family 25 (mitochondrial carrier; adenine nucleotide translocator), member 5 | Cytoplasm | transporter | 1.41E+07 | 2.06E+07 | 8875391 | | | | | | |
| P51881 | SLC25A5 | solute carrier family 25 (mitochondrial carrier; adenine nucleotide translocator), member 5 | Cytoplasm | transporter | 9055829 | 2.97E+07 | 2.06E+07 | | | | | | |
| P51881 | SLC25A5 | solute carrier family 25 (mitochondrial carrier; adenine nucleotide translocator), member 5 | Cytoplasm | transporter | 1.45E+07 | 3.42E+07 | 1.37E+07 | | | | | | |
| P51881 | SLC25A5 | solute carrier family 25 (mitochondrial carrier; adenine nucleotide translocator), member 5 | Cytoplasm | transporter | 4786297 | 1.05E+07 | 4922006.5 | | | | | | |
| P51881 | SLC25A5 | solute carrier family 25 (mitochondrial carrier; adenine nucleotide translocator), member 5 | Cytoplasm | transporter | 1714383.875 | 6327425.5 | 6286921.5 | | | | | | |
| P51881 | SLC25A5 | solute carrier family 25 (mitochondrial carrier; adenine nucleotide translocator), member 5 | Cytoplasm | transporter | 346994.094 | 2730396 | 1573342.75 | | | | | | |
| P51881 | SLC25A5 | solute carrier family 25 (mitochondrial carrier; adenine nucleotide translocator), member 5 | Cytoplasm | transporter | 876641.5 | 5625852.5 | 5456755.5 | P51881 | SLC25A5 | 19448010 | 29351573 | 4375474 | 45 |
| Q5HZI9 | SLC25A51 | solute carrier family 25, member 51 | Cytoplasm | other | 549078.875 | 3443203.75 | 1149310.75 | Q5HZI9 | SLC25A51 | 549079 | 1527426 | 881860 | 3 |
| P58735 | SLC26A1 | solute carrier family 26 (anion exchanger), member 1 | Plasma Membrane | transporter | 1.11E+07 | 1.36E+07 | 1.19E+07 | | | | | | |
| P58735 | SLC26A1 | solute carrier family 26 (anion exchanger), member 1 | Plasma Membrane | transporter | 4097160.75 | 4930800 | 5799193.5 | P58735 | SLC26A1 | 7608580 | 4089519 | 1669539 | 6 |
| Q9WVC8 | SLC26A3 | solute carrier family 26 (anion exchanger), member 3 | Plasma Membrane | transporter | 2173211.75 | 1653294.5 | 1852183.25 | Q9WVC8 | SLC26A3 | 2173212 | 262339 | 151461 | 3 |
| Q9JIM1 | SLC29A1 | solute carrier family 29 (equilibrative nucleoside transporter), member 1 | Plasma Membrane | transporter | 1.53E+08 | 1.71E+08 | 1.62E+08 | | | | | | |
| Q9JIM1 | SLC29A1 | solute carrier family 29 (equilibrative nucleoside transporter), member 1 | Plasma Membrane | transporter | 8.30E+07 | 9.23E+07 | 9.33E+07 | | | | | | |
| Q9JIM1 | SLC29A1 | solute carrier family 29 (equilibrative nucleoside transporter), member 1 | Plasma Membrane | transporter | 4.45E+07 | 5.07E+07 | 6.40E+07 | | | | | | |
| Q9JIM1 | SLC29A1 | solute carrier family 29 (equilibrative nucleoside transporter), member 1 | Plasma Membrane | transporter | 5.15E+07 | 5.28E+07 | 6.01E+07 | | | | | | |
| Q9JIM1 | SLC29A1 | solute carrier family 29 (equilibrative nucleoside transporter), member 1 | Plasma Membrane | transporter | 3.77E+07 | 2.60E+07 | 3.38E+07 | | | | | | |
| Q9JIM1 | SLC29A1 | solute carrier family 29 (equilibrative nucleoside transporter), member 1 | Plasma Membrane | transporter | 2.99E+07 | 2.63E+07 | 1.86E+07 | | | | | | |
| Q9JIM1 | SLC29A1 | solute carrier family 29 (equilibrative nucleoside transporter), member 1 | Plasma Membrane | transporter | 1.75E+07 | 1.38E+07 | 1.46E+07 | | | | | | |
| Q9JIM1 | SLC29A1 | solute carrier family 29 (equilibrative nucleoside transporter), member 1 | Plasma Membrane | transporter | 6658890.5 | 4307196.5 | 8712509 | | | | | | |
| Q9JIM1 | SLC29A1 | solute carrier family 29 (equilibrative nucleoside transporter), member 1 | Plasma Membrane | transporter | 3217547 | 1946862.875 | 2930617 | | | | | | |
| Q9JIM1 | SLC29A1 | solute carrier family 29 (equilibrative nucleoside transporter), member 1 | Plasma Membrane | transporter | 5356049 | 5749256.5 | 6377563 | | | | | | |
| Q9JIM1 | SLC29A1 | solute carrier family 29 (equilibrative nucleoside transporter), member 1 | Plasma Membrane | transporter | 2114493 | 2171867.5 | 2377295.5 | Q9JIM1 | SLC29A1 | 39509725 | 47364926 | 8245175 | 33 |

| | | | | | | | | | | | | | |
|---|---|---|---|---|---|---|---|---|---|---|---|---|---|
| Q61672 | SLC29A2 | solute carrier family 29 (equilibrative nucleoside transporter), member 2 | Plasma Membrane | transporter | 2.12E+07 | 2.39E+07 | 3.22E+07 | Q61672 | SLC29A2 | 21160000 | 5747808 | 3318499 | 3 |
| P32037 | SLC2A3 | solute carrier family 2 (facilitated glucose transporter), member 3 | Plasma Membrane | transporter | 4.28E+07 | 8400353 | 3.32E+07 | | | | | | |
| P32037 | SLC2A3 | solute carrier family 2 (facilitated glucose transporter), member 3 | Plasma Membrane | transporter | 2.66E+07 | 7233695.5 | 3.23E+07 | | | | | | |
| P32037 | SLC2A3 | solute carrier family 2 (facilitated glucose transporter), member 3 | Plasma Membrane | transporter | 2.68E+07 | 5277017 | 2.29E+07 | | | | | | |
| P32037 | SLC2A3 | solute carrier family 2 (facilitated glucose transporter), member 3 | Plasma Membrane | transporter | 1.77E+07 | 5051474 | 2.31E+07 | | | | | | |
| P32037 | SLC2A3 | solute carrier family 2 (facilitated glucose transporter), member 3 | Plasma Membrane | transporter | 1.14E+07 | 1883593.125 | 9129679 | | | | | | |
| P32037 | SLC2A3 | solute carrier family 2 (facilitated glucose transporter), member 3 | Plasma Membrane | transporter | 3326474 | 997509.125 | 4614210 | | | | | | |
| P32037 | SLC2A3 | solute carrier family 2 (facilitated glucose transporter), member 3 | Plasma Membrane | transporter | 1919837.625 | 513119.75 | 1416032.375 | P32037 | SLC2A3 | 18645187 | 12839286 | 2801762 | 21 |
| Q60738 | SLC30A1 | solute carrier family 30 (zinc transporter), member 1 | Plasma Membrane | transporter | 4449013.5 | 5197298 | 4077968.75 | | | | | | |
| Q60738 | SLC30A1 | solute carrier family 30 (zinc transporter), member 1 | Plasma Membrane | transporter | 3388602.5 | 3367375.75 | 2843430.75 | | | | | | |
| Q60738 | SLC30A1 | solute carrier family 30 (zinc transporter), member 1 | Plasma Membrane | transporter | 3511697.75 | 3825287.5 | 3691796.25 | | | | | | |
| Q60738 | SLC30A1 | solute carrier family 30 (zinc transporter), member 1 | Plasma Membrane | transporter | 2346715 | 3217544.25 | 2340967.25 | Q60738 | SLC30A1 | 3424007 | 823903 | 237840 | 12 |
| P10852 | SLC3A2 | solute carrier family 3 (amino acid transporter heavy chain), member 2 | Plasma Membrane | transporter | 1.21E+07 | 1.83E+07 | 1.32E+07 | | | | | | |
| P10852 | SLC3A2 | solute carrier family 3 (amino acid transporter heavy chain), member 2 | Plasma Membrane | transporter | 7914653 | 1.36E+07 | 8481735 | | | | | | |
| P10852 | SLC3A2 | solute carrier family 3 (amino acid transporter heavy chain), member 2 | Plasma Membrane | transporter | 9980435 | 1.16E+07 | 7731082.5 | | | | | | |
| P10852 | SLC3A2 | solute carrier family 3 (amino acid transporter heavy chain), member 2 | Plasma Membrane | transporter | 9058961 | 1.10E+07 | 6314124 | | | | | | |
| P10852 | SLC3A2 | solute carrier family 3 (amino acid transporter heavy chain), member 2 | Plasma Membrane | transporter | 9030388 | 1.17E+07 | 6414555 | | | | | | |
| P10852 | SLC3A2 | solute carrier family 3 (amino acid transporter heavy chain), member 2 | Plasma Membrane | transporter | 4818595 | 6162720 | 3636859.5 | | | | | | |
| P10852 | SLC3A2 | solute carrier family 3 (amino acid transporter heavy chain), member 2 | Plasma Membrane | transporter | 5159796 | 6554883 | 4940127.5 | | | | | | |
| P10852 | SLC3A2 | solute carrier family 3 (amino acid transporter heavy chain), member 2 | Plasma Membrane | transporter | 6025460 | 1.12E+07 | 6469900 | | | | | | |
| P10852 | SLC3A2 | solute carrier family 3 (amino acid transporter heavy chain), member 2 | Plasma Membrane | transporter | 3028208.5 | 4243978.5 | 2241203.25 | | | | | | |
| P10852 | SLC3A2 | solute carrier family 3 (amino acid transporter heavy chain), member 2 | Plasma Membrane | transporter | 1931506.125 | 2940406 | 1339239.375 | P10852 | SLC3A2 | 6903800 | 4032672 | 736262 | 30 |
| Q9JHI9 | SLC40A1 | solute carrier family 40 (iron-regulated transporter), member 1 | Plasma Membrane | transporter | 3.22E+07 | 2.99E+07 | 2.90E+07 | | | | | | |
| Q9JHI9 | SLC40A1 | solute carrier family 40 (iron-regulated transporter), member 1 | Plasma Membrane | transporter | 2.06E+07 | 2.57E+07 | 2.84E+07 | | | | | | |
| Q9JHI9 | SLC40A1 | solute carrier family 40 (iron-regulated transporter), member 1 | Plasma Membrane | transporter | 2.61E+07 | 2.09E+07 | 1.56E+07 | | | | | | |
| Q9JHI9 | SLC40A1 | solute carrier family 40 (iron-regulated transporter), member 1 | Plasma Membrane | transporter | 1.73E+07 | 1.79E+07 | 1.90E+07 | | | | | | |
| Q9JHI9 | SLC40A1 | solute carrier family 40 (iron-regulated transporter), member 1 | Plasma Membrane | transporter | 3.71E+07 | 8189231.5 | 756772.062 | | | | | | |
| Q9JHI9 | SLC40A1 | solute carrier family 40 (iron-regulated transporter), member 1 | Plasma Membrane | transporter | 1.15E+07 | 1.28E+07 | 1.74E+07 | | | | | | |
| Q9JHI9 | SLC40A1 | solute carrier family 40 (iron-regulated transporter), member 1 | Plasma Membrane | transporter | 1.30E+07 | 1.44E+07 | 1.43E+07 | | | | | | |
| Q9JHI9 | SLC40A1 | solute carrier family 40 (iron-regulated transporter), member 1 | Plasma Membrane | transporter | 2819328 | 615584.375 | 74172.32 | Q9JHI9 | SLC40A1 | 20064916 | 10301646 | 2102815 | 24 |
| Q8BSM7 | SLC43A1 | solute carrier family 43 (amino acid system L transporter), member 1 | Plasma Membrane | transporter | 6.98E+08 | 7.03E+08 | 6.86E+08 | | | | | | |
| Q8BSM7 | SLC43A1 | solute carrier family 43 (amino acid system L transporter), member 1 | Plasma Membrane | transporter | 4.56E+08 | 4.22E+08 | 4.81E+08 | | | | | | |
| Q8BSM7 | SLC43A1 | solute carrier family 43 (amino acid system L transporter), member 1 | Plasma Membrane | transporter | 2.09E+08 | 2.80E+08 | 2.91E+08 | | | | | | |
| Q8BSM7 | SLC43A1 | solute carrier family 43 (amino acid system L transporter), member 1 | Plasma Membrane | transporter | 1.27E+08 | 1.29E+08 | 1.23E+08 | | | | | | |
| Q8BSM7 | SLC43A1 | solute carrier family 43 (amino acid system L transporter), member 1 | Plasma Membrane | transporter | 8.63E+07 | 9.00E+07 | 6.54E+07 | | | | | | |
| Q8BSM7 | SLC43A1 | solute carrier family 43 (amino acid system L transporter), member 1 | Plasma Membrane | transporter | 8.75E+07 | 5.92E+07 | 7.42E+07 | | | | | | |
| Q8BSM7 | SLC43A1 | solute carrier family 43 (amino acid system L transporter), member 1 | Plasma Membrane | transporter | 6.58E+07 | 7.55E+07 | 7.38E+07 | | | | | | |
| Q8BSM7 | SLC43A1 | solute carrier family 43 (amino acid system L transporter), member 1 | Plasma Membrane | transporter | 9.58E+07 | 6.23E+07 | 3.13E+07 | | | | | | |
| Q8BSM7 | SLC43A1 | solute carrier family 43 (amino acid system L transporter), member 1 | Plasma Membrane | transporter | 3.33E+07 | 3.43E+07 | 2.97E+07 | | | | | | |
| Q8BSM7 | SLC43A1 | solute carrier family 43 (amino acid system L transporter), member 1 | Plasma Membrane | transporter | 1.26E+07 | 6286423.5 | 2194634.5 | | | | | | |
| Q8BSM7 | SLC43A1 | solute carrier family 43 (amino acid system L transporter), member 1 | Plasma Membrane | transporter | 5934778 | 9293453 | 9214256 | | | | | | |
| Q8BSM7 | SLC43A1 | solute carrier family 43 (amino acid system L transporter), member 1 | Plasma Membrane | transporter | 3995760 | 3443006.75 | 4195037 | | | | | | |
| Q8BSM7 | SLC43A1 | solute carrier family 43 (amino acid system L transporter), member 1 | Plasma Membrane | transporter | 4591936.5 | 1335052.625 | 1478511.875 | | | | | | |
| Q8BSM7 | SLC43A1 | solute carrier family 43 (amino acid system L transporter), member 1 | Plasma Membrane | transporter | 2110028.25 | 3517254.75 | 3978304.75 | Q8BSM7 | SLC43A1 | 134827322 | 199706570 | 30815393 | 42 |
| A2AVZ9 | SLC43A3 | solute carrier family 43, member 3 | Extracellular Space | other | 3.68E+07 | 5.05E+07 | 6.24E+07 | | | | | | |
| A2AVZ9 | SLC43A3 | solute carrier family 43, member 3 | Extracellular Space | other | 2.87E+07 | 1.98E+07 | 1.89E+07 | | | | | | |
| A2AVZ9 | SLC43A3 | solute carrier family 43, member 3 | Extracellular Space | other | 1.75E+07 | 1.95E+07 | 2.35E+07 | | | | | | |
| A2AVZ9 | SLC43A3 | solute carrier family 43, member 3 | Extracellular Space | other | 7394859 | 8169668 | 8917777 | A2AVZ9 | SLC43A3 | 22601215 | 17080495 | 4930714 | 12 |
| Q6X893 | SLC44A1 | solute carrier family 44 (choline transporter), member 1 | Plasma Membrane | transporter | 2765098.25 | 3665222.75 | 4400799 | | | | | | |
| Q6X893 | SLC44A1 | solute carrier family 44 (choline transporter), member 1 | Plasma Membrane | transporter | 2027495.625 | 1871970.375 | 2517852.75 | | | | | | |
| Q6X893 | SLC44A1 | solute carrier family 44 (choline transporter), member 1 | Plasma Membrane | transporter | 2500142.5 | 1623887.125 | 2321375 | Q6X893 | SLC44A1 | 2430912 | 887424 | 295808 | 9 |
| P04919 | SLC4A1 | solute carrier family 4 (anion exchanger), member 1 (Diego blood group) | Plasma Membrane | transporter | 2.63E+10 | 2.66E+10 | 3.18E+10 | | | | | | |
| P04919 | SLC4A1 | solute carrier family 4 (anion exchanger), member 1 (Diego blood group) | Plasma Membrane | transporter | 1.52E+10 | 1.52E+10 | 1.74E+10 | | | | | | |
| P04919 | SLC4A1 | solute carrier family 4 (anion exchanger), member 1 (Diego blood group) | Plasma Membrane | transporter | 9.76E+09 | 8.92E+09 | 1.01E+10 | | | | | | |
| P04919 | SLC4A1 | solute carrier family 4 (anion exchanger), member 1 (Diego blood group) | Plasma Membrane | transporter | 7.80E+09 | 7.41E+09 | 8.79E+09 | | | | | | |
| P04919 | SLC4A1 | solute carrier family 4 (anion exchanger), member 1 (Diego blood group) | Plasma Membrane | transporter | 7.23E+09 | 7.39E+09 | 9.80E+09 | | | | | | |
| P04919 | SLC4A1 | solute carrier family 4 (anion exchanger), member 1 (Diego blood group) | Plasma Membrane | transporter | 9.04E+09 | 8.26E+09 | 9.42E+09 | | | | | | |
| P04919 | SLC4A1 | solute carrier family 4 (anion exchanger), member 1 (Diego blood group) | Plasma Membrane | transporter | 9.79E+09 | 9.87E+09 | 1.15E+10 | | | | | | |
| P04919 | SLC4A1 | solute carrier family 4 (anion exchanger), member 1 (Diego blood group) | Plasma Membrane | transporter | 7.34E+09 | 6.31E+09 | 6.39E+09 | | | | | | |
| P04919 | SLC4A1 | solute carrier family 4 (anion exchanger), member 1 (Diego blood group) | Plasma Membrane | transporter | 8.11E+09 | 7.07E+09 | 7.82E+09 | | | | | | |
| P04919 | SLC4A1 | solute carrier family 4 (anion exchanger), member 1 (Diego blood group) | Plasma Membrane | transporter | 5.65E+09 | 5.92E+09 | 7.26E+09 | | | | | | |
| P04919 | SLC4A1 | solute carrier family 4 (anion exchanger), member 1 (Diego blood group) | Plasma Membrane | transporter | 4.44E+09 | 4.35E+09 | 5.55E+09 | | | | | | |
| P04919 | SLC4A1 | solute carrier family 4 (anion exchanger), member 1 (Diego blood group) | Plasma Membrane | transporter | 4.11E+09 | 3.76E+09 | 4.13E+09 | | | | | | |
| P04919 | SLC4A1 | solute carrier family 4 (anion exchanger), member 1 (Diego blood group) | Plasma Membrane | transporter | 3.75E+09 | 3.69E+09 | 4.40E+09 | | | | | | |
| P04919 | SLC4A1 | solute carrier family 4 (anion exchanger), member 1 (Diego blood group) | Plasma Membrane | transporter | 5.00E+09 | 6.84E+09 | 8.51E+09 | | | | | | |
| P04919 | SLC4A1 | solute carrier family 4 (anion exchanger), member 1 (Diego blood group) | Plasma Membrane | transporter | 3.23E+09 | 3.97E+09 | 5.29E+09 | | | | | | |
| P04919 | SLC4A1 | solute carrier family 4 (anion exchanger), member 1 (Diego blood group) | Plasma Membrane | transporter | 2.09E+09 | 2.90E+09 | 4.68E+09 | | | | | | |
| P04919 | SLC4A1 | solute carrier family 4 (anion exchanger), member 1 (Diego blood group) | Plasma Membrane | transporter | 1.43E+09 | 2.44E+09 | 2.83E+09 | | | | | | |
| P04919 | SLC4A1 | solute carrier family 4 (anion exchanger), member 1 (Diego blood group) | Plasma Membrane | transporter | 1.01E+09 | 1.39E+09 | 2.09E+09 | | | | | | |
| P04919 | SLC4A1 | solute carrier family 4 (anion exchanger), member 1 (Diego blood group) | Plasma Membrane | transporter | 1.55E+09 | 1.59E+09 | 2.57E+09 | | | | | | |
| P04919 | SLC4A1 | solute carrier family 4 (anion exchanger), member 1 (Diego blood group) | Plasma Membrane | transporter | 1.20E+09 | 2.03E+09 | 2.16E+09 | | | | | | |
| P04919 | SLC4A1 | solute carrier family 4 (anion exchanger), member 1 (Diego blood group) | Plasma Membrane | transporter | 7.14E+08 | 7.26E+08 | 7.65E+08 | | | | | | |
| P04919 | SLC4A1 | solute carrier family 4 (anion exchanger), member 1 (Diego blood group) | Plasma Membrane | transporter | 1.01E+09 | 1.03E+09 | 1.20E+09 | | | | | | |
| P04919 | SLC4A1 | solute carrier family 4 (anion exchanger), member 1 (Diego blood group) | Plasma Membrane | transporter | 7.61E+08 | 9.00E+08 | 1.13E+09 | | | | | | |
| P04919 | SLC4A1 | solute carrier family 4 (anion exchanger), member 1 (Diego blood group) | Plasma Membrane | transporter | 5.82E+08 | 6.49E+08 | 8.25E+08 | | | | | | |
| P04919 | SLC4A1 | solute carrier family 4 (anion exchanger), member 1 (Diego blood group) | Plasma Membrane | transporter | 4.30E+08 | 5.46E+08 | 8.22E+08 | | | | | | |
| P04919 | SLC4A1 | solute carrier family 4 (anion exchanger), member 1 (Diego blood group) | Plasma Membrane | transporter | 4.26E+08 | 5.97E+08 | 9.57E+08 | | | | | | |
| P04919 | SLC4A1 | solute carrier family 4 (anion exchanger), member 1 (Diego blood group) | Plasma Membrane | transporter | 2.23E+08 | 3.35E+08 | 4.11E+08 | | | | | | |
| P04919 | SLC4A1 | solute carrier family 4 (anion exchanger), member 1 (Diego blood group) | Plasma Membrane | transporter | 5.33E+08 | 6.06E+08 | 8.06E+08 | | | | | | |
| P04919 | SLC4A1 | solute carrier family 4 (anion exchanger), member 1 (Diego blood group) | Plasma Membrane | transporter | 4.37E+08 | 4.27E+07 | 2434495 | | | | | | |
| P04919 | SLC4A1 | solute carrier family 4 (anion exchanger), member 1 (Diego blood group) | Plasma Membrane | transporter | 2.49E+08 | 3.80E+08 | 2.78E+08 | | | | | | |
| P04919 | SLC4A1 | solute carrier family 4 (anion exchanger), member 1 (Diego blood group) | Plasma Membrane | transporter | 3.44E+08 | 3.32E+08 | 3.55E+08 | | | | | | |
| P04919 | SLC4A1 | solute carrier family 4 (anion exchanger), member 1 (Diego blood group) | Plasma Membrane | transporter | 6.76E+08 | 7.62E+08 | 4.68E+08 | | | | | | |
| P04919 | SLC4A1 | solute carrier family 4 (anion exchanger), member 1 (Diego blood group) | Plasma Membrane | transporter | 2.73E+08 | 5.99E+08 | 4.71E+08 | | | | | | |
| P04919 | SLC4A1 | solute carrier family 4 (anion exchanger), member 1 (Diego blood group) | Plasma Membrane | transporter | 1.55E+08 | 2.39E+08 | 3.87E+08 | | | | | | |
| P04919 | SLC4A1 | solute carrier family 4 (anion exchanger), member 1 (Diego blood group) | Plasma Membrane | transporter | 9.82E+07 | 1.23E+08 | 1.87E+08 | | | | | | |
| P04919 | SLC4A1 | solute carrier family 4 (anion exchanger), member 1 (Diego blood group) | Plasma Membrane | transporter | 1.42E+08 | 1.74E+08 | 3.45E+08 | | | | | | |
| P04919 | SLC4A1 | solute carrier family 4 (anion exchanger), member 1 (Diego blood group) | Plasma Membrane | transporter | 9.77E+07 | 8.29E+07 | 5.73E+07 | | | | | | |
| P04919 | SLC4A1 | solute carrier family 4 (anion exchanger), member 1 (Diego blood group) | Plasma Membrane | transporter | 8.00E+07 | 2.30E+08 | 4.28E+08 | | | | | | |
| P04919 | SLC4A1 | solute carrier family 4 (anion exchanger), member 1 (Diego blood group) | Plasma Membrane | transporter | 6.07E+07 | 7.74E+07 | 1.35E+08 | | | | | | |
| P04919 | SLC4A1 | solute carrier family 4 (anion exchanger), member 1 (Diego blood group) | Plasma Membrane | transporter | 7.77E+07 | 9.03E+07 | 1.12E+08 | | | | | | |

| | | | | | | | | | | | | | |
|---|---|---|---|---|---|---|---|---|---|---|---|---|---|
| P04919 | SLC4A1 | solute carrier family 4 (anion exchanger), member 1 (Diego blood group) | Plasma Membrane | transporter | 1.34E+08 | 2.21E+07 | 1.06E+07 | | | | | | |
| P04919 | SLC4A1 | solute carrier family 4 (anion exchanger), member 1 (Diego blood group) | Plasma Membrane | transporter | 9.67E+07 | 3.00E+08 | 5.64E+08 | | | | | | |
| P04919 | SLC4A1 | solute carrier family 4 (anion exchanger), member 1 (Diego blood group) | Plasma Membrane | transporter | 5.73E+07 | 6.02E+07 | 9.77E+07 | | | | | | |
| P04919 | SLC4A1 | solute carrier family 4 (anion exchanger), member 1 (Diego blood group) | Plasma Membrane | transporter | 5.90E+07 | 7.43E+07 | 1.17E+08 | | | | | | |
| P04919 | SLC4A1 | solute carrier family 4 (anion exchanger), member 1 (Diego blood group) | Plasma Membrane | transporter | 1.24E+07 | 2.90E+07 | 3.01E+07 | | | | | | |
| P04919 | SLC4A1 | solute carrier family 4 (anion exchanger), member 1 (Diego blood group) | Plasma Membrane | transporter | 2.23E+07 | 2.61E+07 | 3.15E+07 | | | | | | |
| P04919 | SLC4A1 | solute carrier family 4 (anion exchanger), member 1 (Diego blood group) | Plasma Membrane | transporter | 2.41E+07 | 1713397.25 | 0 | | | | | | |
| P04919 | SLC4A1 | solute carrier family 4 (anion exchanger), member 1 (Diego blood group) | Plasma Membrane | transporter | 4.18E+07 | 1.28E+07 | 6147972.5 | | | | | | |
| P04919 | SLC4A1 | solute carrier family 4 (anion exchanger), member 1 (Diego blood group) | Plasma Membrane | transporter | 1.02E+07 | 1.37E+07 | 1.91E+07 | | | | | | |
| P04919 | SLC4A1 | solute carrier family 4 (anion exchanger), member 1 (Diego blood group) | Plasma Membrane | transporter | 2.58E+07 | 3.68E+07 | 3.63E+07 | | | | | | |
| P04919 | SLC4A1 | solute carrier family 4 (anion exchanger), member 1 (Diego blood group) | Plasma Membrane | transporter | 8799773 | 4255767 | 3974564.75 | | | | | | |
| P04919 | SLC4A1 | solute carrier family 4 (anion exchanger), member 1 (Diego blood group) | Plasma Membrane | transporter | 1.99E+07 | 1.69E+07 | 1.80E+07 | | | | | | |
| P04919 | SLC4A1 | solute carrier family 4 (anion exchanger), member 1 (Diego blood group) | Plasma Membrane | transporter | 1.61E+07 | 1.85E+07 | 4.59E+07 | | | | | | |
| P04919 | SLC4A1 | solute carrier family 4 (anion exchanger), member 1 (Diego blood group) | Plasma Membrane | transporter | 4705400 | 986143 | 103117.867 | | | | | | |
| P04919 | SLC4A1 | solute carrier family 4 (anion exchanger), member 1 (Diego blood group) | Plasma Membrane | transporter | 7825774.5 | 6.10E+07 | 2.33E+08 | | | | | | |
| P04919 | SLC4A1 | solute carrier family 4 (anion exchanger), member 1 (Diego blood group) | Plasma Membrane | transporter | 1069349 | 2734712.25 | 7597600.5 | | | | | | |
| P04919 | SLC4A1 | solute carrier family 4 (anion exchanger), member 1 (Diego blood group) | Plasma Membrane | transporter | 2743986.5 | 7591134 | 1.18E+07 | P04919 | SLC4A1 | 2489840952 | 4889925980 | 373941986 | 171 |
| O88343 | SLC4A4 | solute carrier family 4 (sodium bicarbonate cotransporter), member 4 | Plasma Membrane | transporter | 1.40E+07 | 1.10E+07 | 9286372 | O88343 | SLC4A4 | 13960000 | 2360946 | 1363093 | 3 |
| Q60857 | SLC6A4 | solute carrier family 6 (neurotransmitter transporter), member 4 | Plasma Membrane | transporter | 99.788 | 775534.562 | 1709990 | Q60857 | SLC6A4 | 100 | 856177 | 494314 | 3 |
| P28571 | SLC6A9 | solute carrier family 6 (neurotransmitter transporter, glycine), member 9 | Plasma Membrane | transporter | 9803132 | 1.47E+07 | 9439907 | | | | | | |
| P28571 | SLC6A9 | solute carrier family 6 (neurotransmitter transporter, glycine), member 9 | Plasma Membrane | transporter | 2418587 | 3193914.75 | 2541207.5 | P28571 | SLC6A9 | 6110860 | 5065610 | 2068026 | 6 |
| Q09143 | SLC7A1 | solute carrier family 7 (cationic amino acid transporter, y+ system), member 1 | Plasma Membrane | transporter | 6628230 | 1.42E+07 | 5107948 | Q09143 | SLC7A1 | 6628230 | 4892932 | 2824935 | 3 |
| Q9Z127 | SLC7A5 | solute carrier family 7 (amino acid transporter light chain, L system), member 5 | Plasma Membrane | transporter | 1.16E+07 | 1.80E+07 | 8535171 | | | | | | |
| Q9Z127 | SLC7A5 | solute carrier family 7 (amino acid transporter light chain, L system), member 5 | Plasma Membrane | transporter | 6150721.5 | 8440543 | 3732521 | | | | | | |
| Q9Z127 | SLC7A5 | solute carrier family 7 (amino acid transporter light chain, L system), member 5 | Plasma Membrane | transporter | 2304675.75 | 3838820.25 | 2272580 | | | | | | |
| Q9Z127 | SLC7A5 | solute carrier family 7 (amino acid transporter light chain, L system), member 5 | Plasma Membrane | transporter | 1639510.25 | 2964378.25 | 1515343.5 | Q9Z127 | SLC7A5 | 5421227 | 4973240 | 1435651 | 12 |
| Q61165 | SLC9A1 | solute carrier family 9, subfamily A (NHE1, cation proton antiporter 1), member 1 | Plasma Membrane | ion channel | 1.60E+07 | 1.44E+07 | 1.39E+07 | | | | | | |
| Q61165 | SLC9A1 | solute carrier family 9, subfamily A (NHE1, cation proton antiporter 1), member 1 | Plasma Membrane | ion channel | 9726365 | 1.05E+07 | 6871965.5 | | | | | | |
| Q61165 | SLC9A1 | solute carrier family 9, subfamily A (NHE1, cation proton antiporter 1), member 1 | Plasma Membrane | ion channel | 6374226.5 | 9019156 | 7449796.5 | | | | | | |
| Q61165 | SLC9A1 | solute carrier family 9, subfamily A (NHE1, cation proton antiporter 1), member 1 | Plasma Membrane | ion channel | 7615279 | 8745820 | 4885907 | | | | | | |
| Q61165 | SLC9A1 | solute carrier family 9, subfamily A (NHE1, cation proton antiporter 1), member 1 | Plasma Membrane | ion channel | 5756472 | 6138828 | 5003463 | | | | | | |
| Q61165 | SLC9A1 | solute carrier family 9, subfamily A (NHE1, cation proton antiporter 1), member 1 | Plasma Membrane | ion channel | 6639902 | 5344979.5 | 6542118.5 | | | | | | |
| Q61165 | SLC9A1 | solute carrier family 9, subfamily A (NHE1, cation proton antiporter 1), member 1 | Plasma Membrane | ion channel | 3081322.25 | 3315921.75 | 3810889.5 | | | | | | |
| Q61165 | SLC9A1 | solute carrier family 9, subfamily A (NHE1, cation proton antiporter 1), member 1 | Plasma Membrane | ion channel | 4199527.5 | 86818.805 | 17351.52 | Q61165 | SLC9A1 | 7427887 | 4000652 | 816630 | 24 |
| Q6UJY2 | SLC9C1 | solute carrier family 9, subfamily C (Na+-transporting carboxylic acid decarboxylase), member 1 | Extracellular Space | other | 3.48E+07 | 5.33E+07 | 8.11E+07 | Q6UJY2 | SLC9C1 | 34830000 | 23283292 | 13442615 | 3 |
| Q8CG48 | SMC2 | structural maintenance of chromosomes 2 | Nucleus | transporter | 2306535.5 | 3029135.5 | 2416738 | Q8CG48 | SMC2 | 2306536 | 389300 | 224762 | 3 |
| Q8CG47 | SMC4 | structural maintenance of chromosomes 4 | Nucleus | transporter | 1.18E+07 | 1.20E+07 | 1.53E+07 | | | | | | |
| Q8CG47 | SMC4 | structural maintenance of chromosomes 4 | Nucleus | transporter | 1205323.625 | 1401759.125 | 421605.438 | Q8CG47 | SMC4 | 6497662 | 6710565 | 2739577 | 6 |
| O09044 | SNAP23 | synaptosomal-associated protein, 23kDa | Plasma Membrane | transporter | 3.30E+07 | 3.61E+07 | 3.51E+07 | | | | | | |
| O09044 | SNAP23 | synaptosomal-associated protein, 23kDa | Plasma Membrane | transporter | 3.58E+07 | 3.24E+07 | 3.21E+07 | | | | | | |
| O09044 | SNAP23 | synaptosomal-associated protein, 23kDa | Plasma Membrane | transporter | 2.70E+07 | 2.18E+07 | 2.85E+07 | | | | | | |
| O09044 | SNAP23 | synaptosomal-associated protein, 23kDa | Plasma Membrane | transporter | 2.39E+07 | 3.57E+07 | 3.41E+07 | | | | | | |
| O09044 | SNAP23 | synaptosomal-associated protein, 23kDa | Plasma Membrane | transporter | 1.04E+07 | 1.20E+07 | 8680434 | | | | | | |
| O09044 | SNAP23 | synaptosomal-associated protein, 23kDa | Plasma Membrane | transporter | 7898770 | 1.64E+07 | 2.11E+07 | | | | | | |
| O09044 | SNAP23 | synaptosomal-associated protein, 23kDa | Plasma Membrane | transporter | 9868382 | 9301631 | 1.17E+07 | | | | | | |
| O09044 | SNAP23 | synaptosomal-associated protein, 23kDa | Plasma Membrane | transporter | 3316501 | 4896621.5 | 6538405.5 | | | | | | |
| O09044 | SNAP23 | synaptosomal-associated protein, 23kDa | Plasma Membrane | transporter | 1871299.875 | 2873334 | 3877785 | | | | | | |
| O09044 | SNAP23 | synaptosomal-associated protein, 23kDa | Plasma Membrane | transporter | 2005495.375 | 2575124.5 | 3351945 | | | | | | |
| O09044 | SNAP23 | synaptosomal-associated protein, 23kDa | Plasma Membrane | transporter | 3366941.5 | 2077857 | 1290505.125 | | | | | | |
| O09044 | SNAP23 | synaptosomal-associated protein, 23kDa | Plasma Membrane | transporter | 1274472.625 | 1416621.625 | 1653071.375 | O09044 | SNAP23 | 13300155 | 13007024 | 2167837 | 36 |
| O55042 | SNCA | synuclein, alpha (non A4 component of amyloid precursor) | Cytoplasm | other | 1.24E+07 | 1.29E+07 | 1.85E+07 | | | | | | |
| O55042 | SNCA | synuclein, alpha (non A4 component of amyloid precursor) | Cytoplasm | other | 9916895 | 9143939 | 1.41E+07 | | | | | | |
| O55042 | SNCA | synuclein, alpha (non A4 component of amyloid precursor) | Cytoplasm | other | 8948884 | 7719875.5 | 9943120 | | | | | | |
| O55042 | SNCA | synuclein, alpha (non A4 component of amyloid precursor) | Cytoplasm | other | 7921942.5 | 6664935.5 | 7137266 | O55042 | SNCA | 9784430 | 3456293 | 997746 | 12 |
| O70493 | SNX12 | sorting nexin 12 | Cytoplasm | transporter | 770178.062 | 1370933.75 | 585627.938 | O70493 | SNX12 | 770178 | 410624 | 237074 | 3 |
| Q0P5V2 | SOBP | sine oculis binding protein homolog | Nucleus | other | 545023.25 | 1043846.375 | 1341737.375 | Q0P5V2 | SOBP | 545023 | 402558 | 232417 | 3 |
| P09671 | SOD2 | superoxide dismutase 2, mitochondrial | Cytoplasm | enzyme | 4028941.75 | 8988765 | 4090838.5 | | | | | | |
| P09671 | SOD2 | superoxide dismutase 2, mitochondrial | Cytoplasm | enzyme | 3134039.25 | 3397734.75 | 181224.297 | | | | | | |
| P09671 | SOD2 | superoxide dismutase 2, mitochondrial | Cytoplasm | enzyme | 1399161.625 | 2976202.75 | 1539828.25 | P09671 | SOD2 | 2854048 | 2502468 | 834156 | 9 |
| Q9CYN2 | SPCS2 | signal peptidase complex subunit 2 | Cytoplasm | other | 4329096 | 5551713.5 | 5719932.5 | Q9CYN2 | SPCS2 | 4329096 | 759113 | 438274 | 3 |
| Q9JJF9 | SPPL2A | signal peptide peptidase like 2A | Plasma Membrane | peptidase | 1.49E+07 | 1.87E+07 | 1.81E+07 | | | | | | |
| Q9JJF9 | SPPL2A | signal peptide peptidase like 2A | Plasma Membrane | peptidase | 2.25E+07 | 1.82E+07 | 1.42E+07 | Q9JJF9 | SPPL2A | 18695000 | 2981700 | 1217274 | 6 |
| P08032 | SPTA1 | spectrin, alpha, erythrocytic 1 | Cytoplasm | other | 3.55E+09 | 3.58E+09 | 3.60E+09 | | | | | | |
| P08032 | SPTA1 | spectrin, alpha, erythrocytic 1 | Cytoplasm | other | 3.21E+09 | 3.00E+09 | 3.81E+09 | | | | | | |
| P08032 | SPTA1 | spectrin, alpha, erythrocytic 1 | Cytoplasm | other | 2.94E+09 | 2.91E+09 | 3.37E+09 | | | | | | |
| P08032 | SPTA1 | spectrin, alpha, erythrocytic 1 | Cytoplasm | other | 2.13E+09 | 2.30E+09 | 2.93E+09 | | | | | | |
| P08032 | SPTA1 | spectrin, alpha, erythrocytic 1 | Cytoplasm | other | 2.04E+09 | 2.54E+09 | 2.97E+09 | | | | | | |
| P08032 | SPTA1 | spectrin, alpha, erythrocytic 1 | Cytoplasm | other | 2.08E+09 | 2.47E+09 | 3.09E+09 | | | | | | |
| P08032 | SPTA1 | spectrin, alpha, erythrocytic 1 | Cytoplasm | other | 2.22E+09 | 2.31E+09 | 2.83E+09 | | | | | | |
| P08032 | SPTA1 | spectrin, alpha, erythrocytic 1 | Cytoplasm | other | 2.32E+09 | 2.06E+09 | 2.31E+09 | | | | | | |
| P08032 | SPTA1 | spectrin, alpha, erythrocytic 1 | Cytoplasm | other | 1.93E+09 | 1.65E+09 | 1.77E+09 | | | | | | |
| P08032 | SPTA1 | spectrin, alpha, erythrocytic 1 | Cytoplasm | other | 2.27E+09 | 2.23E+09 | 1.61E+09 | | | | | | |
| P08032 | SPTA1 | spectrin, alpha, erythrocytic 1 | Cytoplasm | other | 1.76E+09 | 1.88E+09 | 2.16E+09 | | | | | | |
| P08032 | SPTA1 | spectrin, alpha, erythrocytic 1 | Cytoplasm | other | 2.11E+09 | 2.12E+09 | 2.40E+09 | | | | | | |
| P08032 | SPTA1 | spectrin, alpha, erythrocytic 1 | Cytoplasm | other | 1.93E+09 | 1.85E+09 | 2.20E+09 | | | | | | |
| P08032 | SPTA1 | spectrin, alpha, erythrocytic 1 | Cytoplasm | other | 2.06E+09 | 2.73E+09 | 3.18E+09 | | | | | | |
| P08032 | SPTA1 | spectrin, alpha, erythrocytic 1 | Cytoplasm | other | 1.99E+09 | 2.18E+09 | 2.07E+09 | | | | | | |
| P08032 | SPTA1 | spectrin, alpha, erythrocytic 1 | Cytoplasm | other | 1.76E+09 | 1.88E+09 | 2.03E+09 | | | | | | |
| P08032 | SPTA1 | spectrin, alpha, erythrocytic 1 | Cytoplasm | other | 1.03E+09 | 4.74E+08 | 1.27E+09 | | | | | | |
| P08032 | SPTA1 | spectrin, alpha, erythrocytic 1 | Cytoplasm | other | 1.94E+09 | 2.33E+09 | 2.16E+09 | | | | | | |
| P08032 | SPTA1 | spectrin, alpha, erythrocytic 1 | Cytoplasm | other | 1.67E+09 | 1.89E+09 | 2.19E+09 | | | | | | |
| P08032 | SPTA1 | spectrin, alpha, erythrocytic 1 | Cytoplasm | other | 1.75E+09 | 1.46E+09 | 1.37E+09 | | | | | | |
| P08032 | SPTA1 | spectrin, alpha, erythrocytic 1 | Cytoplasm | other | 1.57E+09 | 1.62E+09 | 1.97E+09 | | | | | | |
| P08032 | SPTA1 | spectrin, alpha, erythrocytic 1 | Cytoplasm | other | 1.60E+09 | 1.45E+09 | 1.55E+09 | | | | | | |
| P08032 | SPTA1 | spectrin, alpha, erythrocytic 1 | Cytoplasm | other | 1.46E+09 | 1.41E+09 | 1.66E+09 | | | | | | |
| P08032 | SPTA1 | spectrin, alpha, erythrocytic 1 | Cytoplasm | other | 1.61E+09 | 1.47E+09 | 1.64E+09 | | | | | | |
| P08032 | SPTA1 | spectrin, alpha, erythrocytic 1 | Cytoplasm | other | 1.34E+09 | 1.27E+09 | 1.49E+09 | | | | | | |
| P08032 | SPTA1 | spectrin, alpha, erythrocytic 1 | Cytoplasm | other | 1.52E+09 | 1.34E+09 | 1.58E+09 | | | | | | |
| P08032 | SPTA1 | spectrin, alpha, erythrocytic 1 | Cytoplasm | other | 1.68E+09 | 1.72E+09 | 1.87E+09 | | | | | | |
| P08032 | SPTA1 | spectrin, alpha, erythrocytic 1 | Cytoplasm | other | 1.85E+09 | 2.00E+09 | 1.63E+09 | | | | | | |
| P08032 | SPTA1 | spectrin, alpha, erythrocytic 1 | Cytoplasm | other | 1.33E+09 | 1.30E+09 | 1.83E+09 | | | | | | |

| | | | | | | | |
|---|---|---|---|---|---|---|---|
| P08032 | SPTA1 | spectrin, alpha, erythrocytic 1 | Cytoplasm | other | 1.49E+09 | 1.45E+09 | 1.67E+09 |
| P08032 | SPTA1 | spectrin, alpha, erythrocytic 1 | Cytoplasm | other | 1.34E+09 | 1.22E+09 | 1.20E+09 |
| P08032 | SPTA1 | spectrin, alpha, erythrocytic 1 | Cytoplasm | other | 1.42E+09 | 1.36E+09 | 1.49E+09 |
| P08032 | SPTA1 | spectrin, alpha, erythrocytic 1 | Cytoplasm | other | 1.38E+09 | 1.23E+09 | 1.34E+09 |
| P08032 | SPTA1 | spectrin, alpha, erythrocytic 1 | Cytoplasm | other | 1.31E+09 | 1.08E+09 | 1.12E+09 |
| P08032 | SPTA1 | spectrin, alpha, erythrocytic 1 | Cytoplasm | other | 1.64E+09 | 9.86E+08 | 8.43E+08 |
| P08032 | SPTA1 | spectrin, alpha, erythrocytic 1 | Cytoplasm | other | 1.45E+09 | 1.54E+09 | 1.56E+09 |
| P08032 | SPTA1 | spectrin, alpha, erythrocytic 1 | Cytoplasm | other | 1.16E+09 | 1.28E+09 | 1.53E+09 |
| P08032 | SPTA1 | spectrin, alpha, erythrocytic 1 | Cytoplasm | other | 1.44E+09 | 1.32E+09 | 1.38E+09 |
| P08032 | SPTA1 | spectrin, alpha, erythrocytic 1 | Cytoplasm | other | 1.22E+09 | 1.07E+09 | 1.35E+09 |
| P08032 | SPTA1 | spectrin, alpha, erythrocytic 1 | Cytoplasm | other | 1.23E+09 | 1.28E+09 | 1.45E+09 |
| P08032 | SPTA1 | spectrin, alpha, erythrocytic 1 | Cytoplasm | other | 1.27E+09 | 1.20E+09 | 1.43E+09 |
| P08032 | SPTA1 | spectrin, alpha, erythrocytic 1 | Cytoplasm | other | 1.22E+09 | 1.19E+09 | 1.27E+09 |
| P08032 | SPTA1 | spectrin, alpha, erythrocytic 1 | Cytoplasm | other | 1.21E+09 | 1.21E+09 | 1.46E+09 |
| P08032 | SPTA1 | spectrin, alpha, erythrocytic 1 | Cytoplasm | other | 1.25E+09 | 1.17E+09 | 1.31E+09 |
| P08032 | SPTA1 | spectrin, alpha, erythrocytic 1 | Cytoplasm | other | 1.29E+09 | 1.25E+09 | 1.36E+09 |
| P08032 | SPTA1 | spectrin, alpha, erythrocytic 1 | Cytoplasm | other | 1.15E+09 | 1.13E+09 | 1.27E+09 |
| P08032 | SPTA1 | spectrin, alpha, erythrocytic 1 | Cytoplasm | other | 1.15E+09 | 1.10E+09 | 1.14E+09 |
| P08032 | SPTA1 | spectrin, alpha, erythrocytic 1 | Cytoplasm | other | 1.31E+09 | 1.29E+09 | 1.40E+09 |
| P08032 | SPTA1 | spectrin, alpha, erythrocytic 1 | Cytoplasm | other | 9.13E+08 | 1.65E+09 | 2.08E+09 |
| P08032 | SPTA1 | spectrin, alpha, erythrocytic 1 | Cytoplasm | other | 1.41E+09 | 1.31E+09 | 1.72E+09 |
| P08032 | SPTA1 | spectrin, alpha, erythrocytic 1 | Cytoplasm | other | 1.27E+09 | 1.11E+09 | 6.50E+08 |
| P08032 | SPTA1 | spectrin, alpha, erythrocytic 1 | Cytoplasm | other | 8.60E+08 | 9.40E+08 | 1.06E+09 |
| P08032 | SPTA1 | spectrin, alpha, erythrocytic 1 | Cytoplasm | other | 1.14E+09 | 1.06E+09 | 1.23E+09 |
| P08032 | SPTA1 | spectrin, alpha, erythrocytic 1 | Cytoplasm | other | 1.17E+09 | 1.27E+09 | 1.34E+09 |
| P08032 | SPTA1 | spectrin, alpha, erythrocytic 1 | Cytoplasm | other | 7.90E+08 | 1.18E+09 | 9.28E+08 |
| P08032 | SPTA1 | spectrin, alpha, erythrocytic 1 | Cytoplasm | other | 9.96E+08 | 1.01E+09 | 1.20E+09 |
| P08032 | SPTA1 | spectrin, alpha, erythrocytic 1 | Cytoplasm | other | 1.10E+09 | 9.46E+08 | 1.05E+09 |
| P08032 | SPTA1 | spectrin, alpha, erythrocytic 1 | Cytoplasm | other | 9.66E+08 | 1.30E+09 | 1.39E+09 |
| P08032 | SPTA1 | spectrin, alpha, erythrocytic 1 | Cytoplasm | other | 1.29E+09 | 1.05E+09 | 1.35E+09 |
| P08032 | SPTA1 | spectrin, alpha, erythrocytic 1 | Cytoplasm | other | 1.05E+09 | 1.33E+09 | 1.42E+09 |
| P08032 | SPTA1 | spectrin, alpha, erythrocytic 1 | Cytoplasm | other | 8.49E+08 | 1.29E+09 | 1.44E+09 |
| P08032 | SPTA1 | spectrin, alpha, erythrocytic 1 | Cytoplasm | other | 9.84E+08 | 1.12E+09 | 1.13E+09 |
| P08032 | SPTA1 | spectrin, alpha, erythrocytic 1 | Cytoplasm | other | 9.65E+08 | 9.42E+08 | 1.03E+09 |
| P08032 | SPTA1 | spectrin, alpha, erythrocytic 1 | Cytoplasm | other | 8.98E+08 | 8.35E+08 | 1.23E+09 |
| P08032 | SPTA1 | spectrin, alpha, erythrocytic 1 | Cytoplasm | other | 9.77E+08 | 8.34E+08 | 9.59E+08 |
| P08032 | SPTA1 | spectrin, alpha, erythrocytic 1 | Cytoplasm | other | 1.01E+09 | 9.68E+08 | 1.25E+09 |
| P08032 | SPTA1 | spectrin, alpha, erythrocytic 1 | Cytoplasm | other | 8.17E+08 | 7.82E+08 | 8.98E+08 |
| P08032 | SPTA1 | spectrin, alpha, erythrocytic 1 | Cytoplasm | other | 9.10E+08 | 8.07E+08 | 9.35E+08 |
| P08032 | SPTA1 | spectrin, alpha, erythrocytic 1 | Cytoplasm | other | 9.55E+08 | 1.06E+09 | 1.24E+09 |
| P08032 | SPTA1 | spectrin, alpha, erythrocytic 1 | Cytoplasm | other | 8.24E+08 | 8.39E+08 | 1.10E+09 |
| P08032 | SPTA1 | spectrin, alpha, erythrocytic 1 | Cytoplasm | other | 8.18E+08 | 7.61E+08 | 7.19E+08 |
| P08032 | SPTA1 | spectrin, alpha, erythrocytic 1 | Cytoplasm | other | 1.08E+09 | 7.78E+08 | 7.05E+08 |
| P08032 | SPTA1 | spectrin, alpha, erythrocytic 1 | Cytoplasm | other | 7.77E+08 | 8.96E+08 | 9.75E+08 |
| P08032 | SPTA1 | spectrin, alpha, erythrocytic 1 | Cytoplasm | other | 8.15E+08 | 9.15E+08 | 1.02E+09 |
| P08032 | SPTA1 | spectrin, alpha, erythrocytic 1 | Cytoplasm | other | 7.49E+08 | 9.57E+08 | 7.68E+08 |
| P08032 | SPTA1 | spectrin, alpha, erythrocytic 1 | Cytoplasm | other | 1.03E+09 | 1.05E+09 | 1.22E+09 |
| P08032 | SPTA1 | spectrin, alpha, erythrocytic 1 | Cytoplasm | other | 8.65E+08 | 9.56E+08 | 1.02E+09 |
| P08032 | SPTA1 | spectrin, alpha, erythrocytic 1 | Cytoplasm | other | 7.54E+08 | 7.50E+08 | 9.87E+08 |
| P08032 | SPTA1 | spectrin, alpha, erythrocytic 1 | Cytoplasm | other | 9.08E+08 | 8.89E+08 | 9.66E+08 |
| P08032 | SPTA1 | spectrin, alpha, erythrocytic 1 | Cytoplasm | other | 1.00E+09 | 9.12E+08 | 5.26E+08 |
| P08032 | SPTA1 | spectrin, alpha, erythrocytic 1 | Cytoplasm | other | 7.16E+08 | 6.43E+08 | 8.06E+08 |
| P08032 | SPTA1 | spectrin, alpha, erythrocytic 1 | Cytoplasm | other | 6.49E+08 | 7.52E+08 | 7.97E+08 |
| P08032 | SPTA1 | spectrin, alpha, erythrocytic 1 | Cytoplasm | other | 7.53E+08 | 8.43E+08 | 1.01E+09 |
| P08032 | SPTA1 | spectrin, alpha, erythrocytic 1 | Cytoplasm | other | 6.84E+08 | 6.35E+08 | 7.31E+08 |
| P08032 | SPTA1 | spectrin, alpha, erythrocytic 1 | Cytoplasm | other | 8.27E+08 | 9.44E+08 | 1.03E+09 |
| P08032 | SPTA1 | spectrin, alpha, erythrocytic 1 | Cytoplasm | other | 6.18E+08 | 5.82E+08 | 6.61E+08 |
| P08032 | SPTA1 | spectrin, alpha, erythrocytic 1 | Cytoplasm | other | 7.10E+08 | 6.25E+08 | 6.61E+08 |
| P08032 | SPTA1 | spectrin, alpha, erythrocytic 1 | Cytoplasm | other | 7.45E+08 | 7.55E+08 | 8.75E+08 |
| P08032 | SPTA1 | spectrin, alpha, erythrocytic 1 | Cytoplasm | other | 8.04E+08 | 7.57E+08 | 7.47E+08 |
| P08032 | SPTA1 | spectrin, alpha, erythrocytic 1 | Cytoplasm | other | 6.99E+08 | 6.05E+08 | 7.57E+08 |
| P08032 | SPTA1 | spectrin, alpha, erythrocytic 1 | Cytoplasm | other | 7.01E+08 | 6.56E+08 | 8.31E+08 |
| P08032 | SPTA1 | spectrin, alpha, erythrocytic 1 | Cytoplasm | other | 7.78E+08 | 7.38E+08 | 7.74E+08 |
| P08032 | SPTA1 | spectrin, alpha, erythrocytic 1 | Cytoplasm | other | 8.05E+08 | 8.90E+08 | 1.08E+09 |
| P08032 | SPTA1 | spectrin, alpha, erythrocytic 1 | Cytoplasm | other | 6.53E+08 | 6.51E+08 | 6.68E+08 |
| P08032 | SPTA1 | spectrin, alpha, erythrocytic 1 | Cytoplasm | other | 5.02E+08 | 6.96E+08 | 7.69E+08 |
| P08032 | SPTA1 | spectrin, alpha, erythrocytic 1 | Cytoplasm | other | 7.95E+08 | 5.25E+08 | 5.64E+08 |
| P08032 | SPTA1 | spectrin, alpha, erythrocytic 1 | Cytoplasm | other | 6.58E+08 | 8.70E+08 | 9.17E+08 |
| P08032 | SPTA1 | spectrin, alpha, erythrocytic 1 | Cytoplasm | other | 4.80E+08 | 6.34E+08 | 8.34E+08 |
| P08032 | SPTA1 | spectrin, alpha, erythrocytic 1 | Cytoplasm | other | 5.98E+08 | 4.63E+08 | 4.44E+08 |
| P08032 | SPTA1 | spectrin, alpha, erythrocytic 1 | Cytoplasm | other | 5.92E+08 | 4.97E+08 | 5.05E+08 |
| P08032 | SPTA1 | spectrin, alpha, erythrocytic 1 | Cytoplasm | other | 4.65E+08 | 6.02E+08 | 6.63E+08 |
| P08032 | SPTA1 | spectrin, alpha, erythrocytic 1 | Cytoplasm | other | 5.42E+08 | 6.71E+08 | 5.68E+08 |
| P08032 | SPTA1 | spectrin, alpha, erythrocytic 1 | Cytoplasm | other | 5.78E+08 | 5.90E+08 | 6.57E+08 |
| P08032 | SPTA1 | spectrin, alpha, erythrocytic 1 | Cytoplasm | other | 7.63E+08 | 4.30E+08 | 3.19E+08 |
| P08032 | SPTA1 | spectrin, alpha, erythrocytic 1 | Cytoplasm | other | 5.86E+08 | 5.83E+08 | 6.57E+08 |
| P08032 | SPTA1 | spectrin, alpha, erythrocytic 1 | Cytoplasm | other | 5.81E+08 | 5.78E+08 | 6.83E+08 |
| P08032 | SPTA1 | spectrin, alpha, erythrocytic 1 | Cytoplasm | other | 5.24E+08 | 4.45E+08 | 4.82E+08 |
| P08032 | SPTA1 | spectrin, alpha, erythrocytic 1 | Cytoplasm | other | 4.15E+08 | 3.67E+08 | 3.89E+08 |
| P08032 | SPTA1 | spectrin, alpha, erythrocytic 1 | Cytoplasm | other | 5.45E+08 | 5.33E+08 | 5.42E+08 |
| P08032 | SPTA1 | spectrin, alpha, erythrocytic 1 | Cytoplasm | other | 5.16E+08 | 4.03E+08 | 4.72E+08 |
| P08032 | SPTA1 | spectrin, alpha, erythrocytic 1 | Cytoplasm | other | 5.61E+08 | 4.71E+08 | 4.91E+08 |
| P08032 | SPTA1 | spectrin, alpha, erythrocytic 1 | Cytoplasm | other | 3.94E+08 | 3.48E+08 | 3.92E+08 |
| P08032 | SPTA1 | spectrin, alpha, erythrocytic 1 | Cytoplasm | other | 4.07E+08 | 3.04E+08 | 1.92E+08 |
| P08032 | SPTA1 | spectrin, alpha, erythrocytic 1 | Cytoplasm | other | 5.40E+08 | 6.58E+08 | 6.96E+08 |
| P08032 | SPTA1 | spectrin, alpha, erythrocytic 1 | Cytoplasm | other | 3.74E+08 | 4.13E+08 | 5.05E+08 |
| P08032 | SPTA1 | spectrin, alpha, erythrocytic 1 | Cytoplasm | other | 3.71E+08 | 3.38E+08 | 3.69E+08 |
| P08032 | SPTA1 | spectrin, alpha, erythrocytic 1 | Cytoplasm | other | 5.89E+08 | 5.96E+08 | 6.28E+08 |
| P08032 | SPTA1 | spectrin, alpha, erythrocytic 1 | Cytoplasm | other | 2.64E+08 | 4.30E+08 | 5.79E+08 |
| P08032 | SPTA1 | spectrin, alpha, erythrocytic 1 | Cytoplasm | other | 2.56E+08 | 3.87E+08 | 4.41E+08 |
| P08032 | SPTA1 | spectrin, alpha, erythrocytic 1 | Cytoplasm | other | 4.50E+08 | 3.94E+08 | 4.84E+08 |

| | | | | | | | |
|---|---|---|---|---|---|---|---|
| P08032 | SPTA1 | spectrin, alpha, erythrocytic 1 | Cytoplasm | other | 3.36E+08 | 4.23E+08 | 4.68E+08 |
| P08032 | SPTA1 | spectrin, alpha, erythrocytic 1 | Cytoplasm | other | 4.01E+08 | 4.14E+08 | 5.42E+08 |
| P08032 | SPTA1 | spectrin, alpha, erythrocytic 1 | Cytoplasm | other | 2.56E+08 | 3.28E+08 | 4.41E+08 |
| P08032 | SPTA1 | spectrin, alpha, erythrocytic 1 | Cytoplasm | other | 2.25E+08 | 3.75E+08 | 3.35E+08 |
| P08032 | SPTA1 | spectrin, alpha, erythrocytic 1 | Cytoplasm | other | 3.38E+08 | 3.54E+08 | 3.82E+08 |
| P08032 | SPTA1 | spectrin, alpha, erythrocytic 1 | Cytoplasm | other | 2.58E+08 | 3.34E+08 | 4.39E+08 |
| P08032 | SPTA1 | spectrin, alpha, erythrocytic 1 | Cytoplasm | other | 2.09E+08 | 1.87E+08 | 9.59E+07 |
| P08032 | SPTA1 | spectrin, alpha, erythrocytic 1 | Cytoplasm | other | 2.82E+08 | 4.15E+08 | 5.32E+08 |
| P08032 | SPTA1 | spectrin, alpha, erythrocytic 1 | Cytoplasm | other | 2.87E+08 | 4.97E+08 | 4.87E+08 |
| P08032 | SPTA1 | spectrin, alpha, erythrocytic 1 | Cytoplasm | other | 3.47E+08 | 4.32E+08 | 5.31E+08 |
| P08032 | SPTA1 | spectrin, alpha, erythrocytic 1 | Cytoplasm | other | 4.51E+08 | 2.16E+08 | 1.38E+08 |
| P08032 | SPTA1 | spectrin, alpha, erythrocytic 1 | Cytoplasm | other | 4.26E+08 | 2.44E+08 | 2.63E+08 |
| P08032 | SPTA1 | spectrin, alpha, erythrocytic 1 | Cytoplasm | other | 2.40E+08 | 3.08E+08 | 3.75E+08 |
| P08032 | SPTA1 | spectrin, alpha, erythrocytic 1 | Cytoplasm | other | 2.48E+08 | 3.38E+08 | 3.57E+08 |
| P08032 | SPTA1 | spectrin, alpha, erythrocytic 1 | Cytoplasm | other | 4.08E+08 | 2.25E+08 | 1.84E+08 |
| P08032 | SPTA1 | spectrin, alpha, erythrocytic 1 | Cytoplasm | other | 2.89E+08 | 4.27E+08 | 5.67E+08 |
| P08032 | SPTA1 | spectrin, alpha, erythrocytic 1 | Cytoplasm | other | 1.90E+08 | 4.25E+08 | 7.36E+08 |
| P08032 | SPTA1 | spectrin, alpha, erythrocytic 1 | Cytoplasm | other | 3.67E+08 | 1.20E+08 | 8.97E+07 |
| P08032 | SPTA1 | spectrin, alpha, erythrocytic 1 | Cytoplasm | other | 2.77E+08 | 2.90E+08 | 2.86E+08 |
| P08032 | SPTA1 | spectrin, alpha, erythrocytic 1 | Cytoplasm | other | 2.87E+08 | 2.09E+08 | 2.50E+08 |
| P08032 | SPTA1 | spectrin, alpha, erythrocytic 1 | Cytoplasm | other | 2.73E+08 | 1.18E+08 | 1.08E+08 |
| P08032 | SPTA1 | spectrin, alpha, erythrocytic 1 | Cytoplasm | other | 2.39E+08 | 2.08E+08 | 1.20E+08 |
| P08032 | SPTA1 | spectrin, alpha, erythrocytic 1 | Cytoplasm | other | 2.41E+08 | 3.19E+08 | 3.62E+08 |
| P08032 | SPTA1 | spectrin, alpha, erythrocytic 1 | Cytoplasm | other | 1.68E+08 | 2.35E+08 | 1.97E+08 |
| P08032 | SPTA1 | spectrin, alpha, erythrocytic 1 | Cytoplasm | other | 2.64E+08 | 2.56E+08 | 1.60E+08 |
| P08032 | SPTA1 | spectrin, alpha, erythrocytic 1 | Cytoplasm | other | 2.15E+08 | 2.20E+08 | 2.75E+08 |
| P08032 | SPTA1 | spectrin, alpha, erythrocytic 1 | Cytoplasm | other | 1.87E+08 | 2.11E+08 | 2.47E+08 |
| P08032 | SPTA1 | spectrin, alpha, erythrocytic 1 | Cytoplasm | other | 3.24E+08 | 4.03E+08 | 4.65E+08 |
| P08032 | SPTA1 | spectrin, alpha, erythrocytic 1 | Cytoplasm | other | 2.98E+08 | 2.55E+08 | 2.48E+08 |
| P08032 | SPTA1 | spectrin, alpha, erythrocytic 1 | Cytoplasm | other | 2.42E+08 | 1.43E+08 | 3.16E+08 |
| P08032 | SPTA1 | spectrin, alpha, erythrocytic 1 | Cytoplasm | other | 1.31E+08 | 2.17E+08 | 2.26E+08 |
| P08032 | SPTA1 | spectrin, alpha, erythrocytic 1 | Cytoplasm | other | 1.73E+08 | 1.39E+08 | 1.43E+08 |
| P08032 | SPTA1 | spectrin, alpha, erythrocytic 1 | Cytoplasm | other | 1.65E+08 | 1.37E+08 | 1.22E+08 |
| P08032 | SPTA1 | spectrin, alpha, erythrocytic 1 | Cytoplasm | other | 1.81E+08 | 1.35E+08 | 1.26E+08 |
| P08032 | SPTA1 | spectrin, alpha, erythrocytic 1 | Cytoplasm | other | 2.17E+08 | 9.36E+07 | 8.37E+07 |
| P08032 | SPTA1 | spectrin, alpha, erythrocytic 1 | Cytoplasm | other | 1.76E+08 | 1.67E+08 | 2.23E+08 |
| P08032 | SPTA1 | spectrin, alpha, erythrocytic 1 | Cytoplasm | other | 2.17E+08 | 1.08E+08 | 9.20E+07 |
| P08032 | SPTA1 | spectrin, alpha, erythrocytic 1 | Cytoplasm | other | 1.18E+08 | 2.33E+08 | 4.75E+08 |
| P08032 | SPTA1 | spectrin, alpha, erythrocytic 1 | Cytoplasm | other | 2.23E+08 | 1.72E+08 | 3.09E+08 |
| P08032 | SPTA1 | spectrin, alpha, erythrocytic 1 | Cytoplasm | other | 1.64E+08 | 1.47E+08 | 1.43E+08 |
| P08032 | SPTA1 | spectrin, alpha, erythrocytic 1 | Cytoplasm | other | 1.79E+08 | 2.76E+08 | 1.31E+08 |
| P08032 | SPTA1 | spectrin, alpha, erythrocytic 1 | Cytoplasm | other | 9.67E+07 | 1.06E+08 | 1.07E+08 |
| P08032 | SPTA1 | spectrin, alpha, erythrocytic 1 | Cytoplasm | other | 2.56E+08 | 8.87E+07 | 4.34E+07 |
| P08032 | SPTA1 | spectrin, alpha, erythrocytic 1 | Cytoplasm | other | 1.66E+08 | 2.80E+08 | 2.87E+08 |
| P08032 | SPTA1 | spectrin, alpha, erythrocytic 1 | Cytoplasm | other | 8.25E+07 | 2.77E+07 | 1.09E+07 |
| P08032 | SPTA1 | spectrin, alpha, erythrocytic 1 | Cytoplasm | other | 1.29E+08 | 1.09E+08 | 1.12E+08 |
| P08032 | SPTA1 | spectrin, alpha, erythrocytic 1 | Cytoplasm | other | 2.64E+08 | 8.77E+07 | 6.51E+07 |
| P08032 | SPTA1 | spectrin, alpha, erythrocytic 1 | Cytoplasm | other | 8.21E+07 | 8.96E+07 | 1.60E+08 |
| P08032 | SPTA1 | spectrin, alpha, erythrocytic 1 | Cytoplasm | other | 1.95E+08 | 1.75E+08 | 1.40E+08 |
| P08032 | SPTA1 | spectrin, alpha, erythrocytic 1 | Cytoplasm | other | 1.66E+08 | 7.33E+07 | 7.37E+07 |
| P08032 | SPTA1 | spectrin, alpha, erythrocytic 1 | Cytoplasm | other | 2.00E+08 | 3.45E+08 | 3.19E+08 |
| P08032 | SPTA1 | spectrin, alpha, erythrocytic 1 | Cytoplasm | other | 1.06E+08 | 1.30E+08 | 1.33E+08 |
| P08032 | SPTA1 | spectrin, alpha, erythrocytic 1 | Cytoplasm | other | 2.17E+08 | 4.04E+08 | 6.75E+08 |
| P08032 | SPTA1 | spectrin, alpha, erythrocytic 1 | Cytoplasm | other | 2.39E+08 | 8.19E+07 | 5.17E+07 |
| P08032 | SPTA1 | spectrin, alpha, erythrocytic 1 | Cytoplasm | other | 1.55E+08 | 1.36E+08 | 1.48E+08 |
| P08032 | SPTA1 | spectrin, alpha, erythrocytic 1 | Cytoplasm | other | 1.47E+08 | 1.07E+08 | 1.19E+08 |
| P08032 | SPTA1 | spectrin, alpha, erythrocytic 1 | Cytoplasm | other | 8.44E+07 | 3.06E+08 | 7.56E+08 |
| P08032 | SPTA1 | spectrin, alpha, erythrocytic 1 | Cytoplasm | other | 1.10E+08 | 7.35E+07 | 5.11E+07 |
| P08032 | SPTA1 | spectrin, alpha, erythrocytic 1 | Cytoplasm | other | 2.08E+08 | 4.31E+07 | 3033749.75 |
| P08032 | SPTA1 | spectrin, alpha, erythrocytic 1 | Cytoplasm | other | 6.57E+07 | 8.96E+07 | 1.22E+08 |
| P08032 | SPTA1 | spectrin, alpha, erythrocytic 1 | Cytoplasm | other | 2.29E+08 | 1.58E+08 | 9.62E+07 |
| P08032 | SPTA1 | spectrin, alpha, erythrocytic 1 | Cytoplasm | other | 9.95E+07 | 1.22E+08 | 1.43E+08 |
| P08032 | SPTA1 | spectrin, alpha, erythrocytic 1 | Cytoplasm | other | 1.49E+08 | 1.08E+08 | 1.48E+08 |
| P08032 | SPTA1 | spectrin, alpha, erythrocytic 1 | Cytoplasm | other | 9.75E+07 | 7.72E+07 | 6.89E+07 |
| P08032 | SPTA1 | spectrin, alpha, erythrocytic 1 | Cytoplasm | other | 4.26E+07 | 2368627 | 2185923.25 |
| P08032 | SPTA1 | spectrin, alpha, erythrocytic 1 | Cytoplasm | other | 7.55E+07 | 9.94E+07 | 1.38E+08 |
| P08032 | SPTA1 | spectrin, alpha, erythrocytic 1 | Cytoplasm | other | 7.13E+07 | 1.08E+08 | 2.06E+08 |
| P08032 | SPTA1 | spectrin, alpha, erythrocytic 1 | Cytoplasm | other | 9.30E+07 | 8.34E+07 | 1.28E+08 |
| P08032 | SPTA1 | spectrin, alpha, erythrocytic 1 | Cytoplasm | other | 9.34E+07 | 9.12E+07 | 5.47E+07 |
| P08032 | SPTA1 | spectrin, alpha, erythrocytic 1 | Cytoplasm | other | 7.28E+07 | 1.15E+08 | 1.15E+08 |
| P08032 | SPTA1 | spectrin, alpha, erythrocytic 1 | Cytoplasm | other | 6.58E+07 | 9.37E+07 | 9.44E+07 |
| P08032 | SPTA1 | spectrin, alpha, erythrocytic 1 | Cytoplasm | other | 7.59E+07 | 6.71E+07 | 7.97E+07 |
| P08032 | SPTA1 | spectrin, alpha, erythrocytic 1 | Cytoplasm | other | 1.02E+08 | 1.02E+08 | 1.17E+08 |
| P08032 | SPTA1 | spectrin, alpha, erythrocytic 1 | Cytoplasm | other | 1.49E+08 | 2.91E+07 | 1.77E+07 |
| P08032 | SPTA1 | spectrin, alpha, erythrocytic 1 | Cytoplasm | other | 9.76E+07 | 1.65E+08 | 2.74E+08 |
| P08032 | SPTA1 | spectrin, alpha, erythrocytic 1 | Cytoplasm | other | 1.86E+08 | 5.28E+07 | 3.31E+07 |
| P08032 | SPTA1 | spectrin, alpha, erythrocytic 1 | Cytoplasm | other | 9.26E+07 | 6.88E+07 | 9.61E+07 |
| P08032 | SPTA1 | spectrin, alpha, erythrocytic 1 | Cytoplasm | other | 8.15E+07 | 9.33E+07 | 1.21E+08 |
| P08032 | SPTA1 | spectrin, alpha, erythrocytic 1 | Cytoplasm | other | 1.47E+08 | 8.32E+07 | 5.11E+07 |
| P08032 | SPTA1 | spectrin, alpha, erythrocytic 1 | Cytoplasm | other | 1.27E+08 | 1.55E+08 | 2.02E+08 |
| P08032 | SPTA1 | spectrin, alpha, erythrocytic 1 | Cytoplasm | other | 8.19E+07 | 4.44E+07 | 3.88E+07 |
| P08032 | SPTA1 | spectrin, alpha, erythrocytic 1 | Cytoplasm | other | 6.99E+07 | 6.08E+07 | 6.75E+07 |
| P08032 | SPTA1 | spectrin, alpha, erythrocytic 1 | Cytoplasm | other | 8.28E+07 | 4.38E+07 | 3.77E+07 |
| P08032 | SPTA1 | spectrin, alpha, erythrocytic 1 | Cytoplasm | other | 9.85E+07 | 4.65E+07 | 1.33E+07 |
| P08032 | SPTA1 | spectrin, alpha, erythrocytic 1 | Cytoplasm | other | 4.30E+07 | 7.23E+07 | 8.07E+07 |
| P08032 | SPTA1 | spectrin, alpha, erythrocytic 1 | Cytoplasm | other | 4.18E+07 | 5.18E+07 | 4.11E+07 |
| P08032 | SPTA1 | spectrin, alpha, erythrocytic 1 | Cytoplasm | other | 5.80E+07 | 1.24E+08 | 2.25E+08 |
| P08032 | SPTA1 | spectrin, alpha, erythrocytic 1 | Cytoplasm | other | 4.83E+07 | 3.11E+07 | 2.72E+07 |
| P08032 | SPTA1 | spectrin, alpha, erythrocytic 1 | Cytoplasm | other | 6.78E+07 | 5.52E+07 | 6.75E+07 |
| P08032 | SPTA1 | spectrin, alpha, erythrocytic 1 | Cytoplasm | other | 6.46E+07 | 5.53E+07 | 5.34E+07 |
| P08032 | SPTA1 | spectrin, alpha, erythrocytic 1 | Cytoplasm | other | 9.77E+07 | 2.84E+07 | 1.79E+07 |

| | | | | | | | | | | | | | |
|---|---|---|---|---|---|---|---|---|---|---|---|---|---|
| P08032 | SPTA1 | spectrin, alpha, erythrocytic 1 | Cytoplasm | other | 5.30E+07 | 3.48E+07 | 5.30E+07 | | | | | | |
| P08032 | SPTA1 | spectrin, alpha, erythrocytic 1 | Cytoplasm | other | 3.94E+07 | 3.97E+07 | 3.62E+07 | | | | | | |
| P08032 | SPTA1 | spectrin, alpha, erythrocytic 1 | Cytoplasm | other | 2.79E+07 | 5.47E+07 | 1.19E+08 | | | | | | |
| P08032 | SPTA1 | spectrin, alpha, erythrocytic 1 | Cytoplasm | other | 3.13E+07 | 7.14E+07 | 1.13E+08 | | | | | | |
| P08032 | SPTA1 | spectrin, alpha, erythrocytic 1 | Cytoplasm | other | 2.93E+07 | 4.11E+07 | 5.16E+07 | | | | | | |
| P08032 | SPTA1 | spectrin, alpha, erythrocytic 1 | Cytoplasm | other | 7.51E+07 | 2.29E+07 | 1.30E+07 | | | | | | |
| P08032 | SPTA1 | spectrin, alpha, erythrocytic 1 | Cytoplasm | other | 2.97E+07 | 3.59E+07 | 4.09E+07 | | | | | | |
| P08032 | SPTA1 | spectrin, alpha, erythrocytic 1 | Cytoplasm | other | 4.14E+07 | 6.94E+07 | 1.50E+08 | | | | | | |
| P08032 | SPTA1 | spectrin, alpha, erythrocytic 1 | Cytoplasm | other | 6.74E+07 | 1.09E+07 | 1195283.75 | | | | | | |
| P08032 | SPTA1 | spectrin, alpha, erythrocytic 1 | Cytoplasm | other | 2.53E+07 | 3.05E+07 | 2.50E+07 | | | | | | |
| P08032 | SPTA1 | spectrin, alpha, erythrocytic 1 | Cytoplasm | other | 8.82E+07 | 1.01E+08 | 1.16E+08 | | | | | | |
| P08032 | SPTA1 | spectrin, alpha, erythrocytic 1 | Cytoplasm | other | 3.07E+07 | 1.36E+07 | 1.28E+07 | | | | | | |
| P08032 | SPTA1 | spectrin, alpha, erythrocytic 1 | Cytoplasm | other | 5.11E+07 | 1.68E+07 | 8100690 | | | | | | |
| P08032 | SPTA1 | spectrin, alpha, erythrocytic 1 | Cytoplasm | other | 3.66E+07 | 3.08E+07 | 3.58E+07 | | | | | | |
| P08032 | SPTA1 | spectrin, alpha, erythrocytic 1 | Cytoplasm | other | 2.41E+07 | 2.42E+07 | 2.72E+07 | | | | | | |
| P08032 | SPTA1 | spectrin, alpha, erythrocytic 1 | Cytoplasm | other | 3.65E+07 | 3.74E+07 | 5.16E+07 | | | | | | |
| P08032 | SPTA1 | spectrin, alpha, erythrocytic 1 | Cytoplasm | other | 1.23E+07 | 3361050.75 | 2052740.25 | | | | | | |
| P08032 | SPTA1 | spectrin, alpha, erythrocytic 1 | Cytoplasm | other | 5.55E+07 | 5.58E+07 | 8.68E+07 | | | | | | |
| P08032 | SPTA1 | spectrin, alpha, erythrocytic 1 | Cytoplasm | other | 4.06E+07 | 5.87E+07 | 5.04E+07 | | | | | | |
| P08032 | SPTA1 | spectrin, alpha, erythrocytic 1 | Cytoplasm | other | 1.34E+07 | 2444511.75 | 716214.25 | | | | | | |
| P08032 | SPTA1 | spectrin, alpha, erythrocytic 1 | Cytoplasm | other | 2.78E+07 | 1.99E+07 | 1.43E+07 | | | | | | |
| P08032 | SPTA1 | spectrin, alpha, erythrocytic 1 | Cytoplasm | other | 7.06E+07 | 4806748.5 | 901667.5 | | | | | | |
| P08032 | SPTA1 | spectrin, alpha, erythrocytic 1 | Cytoplasm | other | 2.44E+07 | 2.16E+07 | 2.37E+07 | | | | | | |
| P08032 | SPTA1 | spectrin, alpha, erythrocytic 1 | Cytoplasm | other | 5.50E+07 | 1.23E+07 | 1.70E+07 | | | | | | |
| P08032 | SPTA1 | spectrin, alpha, erythrocytic 1 | Cytoplasm | other | 2.20E+07 | 1.40E+07 | 8731551 | | | | | | |
| P08032 | SPTA1 | spectrin, alpha, erythrocytic 1 | Cytoplasm | other | 2.37E+07 | 2.40E+07 | 3.15E+07 | | | | | | |
| P08032 | SPTA1 | spectrin, alpha, erythrocytic 1 | Cytoplasm | other | 7561003.5 | 2.45E+07 | 4.22E+07 | | | | | | |
| P08032 | SPTA1 | spectrin, alpha, erythrocytic 1 | Cytoplasm | other | 1.19E+07 | 1.34E+07 | 1.32E+07 | | | | | | |
| P08032 | SPTA1 | spectrin, alpha, erythrocytic 1 | Cytoplasm | other | 3.40E+07 | 1.04E+07 | 4506191.5 | | | | | | |
| P08032 | SPTA1 | spectrin, alpha, erythrocytic 1 | Cytoplasm | other | 6.12E+07 | 5038271.5 | 2956858.5 | | | | | | |
| P08032 | SPTA1 | spectrin, alpha, erythrocytic 1 | Cytoplasm | other | 1.94E+07 | 1.68E+07 | 2.02E+07 | | | | | | |
| P08032 | SPTA1 | spectrin, alpha, erythrocytic 1 | Cytoplasm | other | 1.70E+07 | 1.33E+07 | 1.83E+07 | | | | | | |
| P08032 | SPTA1 | spectrin, alpha, erythrocytic 1 | Cytoplasm | other | 1.37E+07 | 1.28E+07 | 1.84E+07 | | | | | | |
| P08032 | SPTA1 | spectrin, alpha, erythrocytic 1 | Cytoplasm | other | 1.98E+07 | 1.68E+07 | 2.13E+07 | | | | | | |
| P08032 | SPTA1 | spectrin, alpha, erythrocytic 1 | Cytoplasm | other | 1.07E+07 | 1.18E+07 | 1.94E+07 | | | | | | |
| P08032 | SPTA1 | spectrin, alpha, erythrocytic 1 | Cytoplasm | other | 1.92E+07 | 1.76E+07 | 2.32E+07 | | | | | | |
| P08032 | SPTA1 | spectrin, alpha, erythrocytic 1 | Cytoplasm | other | 2.01E+07 | 8611552 | 9499531 | | | | | | |
| P08032 | SPTA1 | spectrin, alpha, erythrocytic 1 | Cytoplasm | other | 3.72E+07 | 2.73E+07 | 3.46E+07 | | | | | | |
| P08032 | SPTA1 | spectrin, alpha, erythrocytic 1 | Cytoplasm | other | 1.53E+07 | 1.76E+07 | 1.98E+07 | | | | | | |
| P08032 | SPTA1 | spectrin, alpha, erythrocytic 1 | Cytoplasm | other | 2.00E+07 | 8606427 | 8483811 | | | | | | |
| P08032 | SPTA1 | spectrin, alpha, erythrocytic 1 | Cytoplasm | other | 1.05E+07 | 1.83E+07 | 2.50E+07 | | | | | | |
| P08032 | SPTA1 | spectrin, alpha, erythrocytic 1 | Cytoplasm | other | 9740822 | 1.12E+07 | 6563923.5 | | | | | | |
| P08032 | SPTA1 | spectrin, alpha, erythrocytic 1 | Cytoplasm | other | 8692877 | 9544510 | 1.38E+07 | | | | | | |
| P08032 | SPTA1 | spectrin, alpha, erythrocytic 1 | Cytoplasm | other | 4.25E+07 | 4542785.5 | 2454150.5 | | | | | | |
| P08032 | SPTA1 | spectrin, alpha, erythrocytic 1 | Cytoplasm | other | 2.15E+07 | 2.52E+07 | 2924737.5 | | | | | | |
| P08032 | SPTA1 | spectrin, alpha, erythrocytic 1 | Cytoplasm | other | 1.10E+07 | 1.35E+07 | 1.92E+07 | | | | | | |
| P08032 | SPTA1 | spectrin, alpha, erythrocytic 1 | Cytoplasm | other | 1.48E+07 | 4560416.5 | 6429889 | | | | | | |
| P08032 | SPTA1 | spectrin, alpha, erythrocytic 1 | Cytoplasm | other | 1.37E+07 | 1.10E+07 | 1.22E+07 | | | | | | |
| P08032 | SPTA1 | spectrin, alpha, erythrocytic 1 | Cytoplasm | other | 7217670.5 | 3011047.75 | 138596.203 | | | | | | |
| P08032 | SPTA1 | spectrin, alpha, erythrocytic 1 | Cytoplasm | other | 9401733 | 6847438.5 | 9283569 | | | | | | |
| P08032 | SPTA1 | spectrin, alpha, erythrocytic 1 | Cytoplasm | other | 1.03E+07 | 8939959 | 9390726 | | | | | | |
| P08032 | SPTA1 | spectrin, alpha, erythrocytic 1 | Cytoplasm | other | 1.69E+07 | 1.27E+07 | 1.21E+07 | | | | | | |
| P08032 | SPTA1 | spectrin, alpha, erythrocytic 1 | Cytoplasm | other | 1.74E+07 | 5365696.5 | 5175618 | | | | | | |
| P08032 | SPTA1 | spectrin, alpha, erythrocytic 1 | Cytoplasm | other | 1.71E+07 | 3.42E+07 | 3.65E+07 | | | | | | |
| P08032 | SPTA1 | spectrin, alpha, erythrocytic 1 | Cytoplasm | other | 3736342.25 | 5838467 | 8443318 | | | | | | |
| P08032 | SPTA1 | spectrin, alpha, erythrocytic 1 | Cytoplasm | other | 2.49E+07 | 117124.695 | 644.733 | | | | | | |
| P08032 | SPTA1 | spectrin, alpha, erythrocytic 1 | Cytoplasm | other | 5433473 | 5403843 | 5537220 | | | | | | |
| P08032 | SPTA1 | spectrin, alpha, erythrocytic 1 | Cytoplasm | other | 9370353 | 3195221.75 | 1508483 | | | | | | |
| P08032 | SPTA1 | spectrin, alpha, erythrocytic 1 | Cytoplasm | other | 2896127 | 691253.875 | 266529.75 | | | | | | |
| P08032 | SPTA1 | spectrin, alpha, erythrocytic 1 | Cytoplasm | other | 9896826 | 7199149.5 | 6962187.5 | | | | | | |
| P08032 | SPTA1 | spectrin, alpha, erythrocytic 1 | Cytoplasm | other | 1.48E+07 | 4374575 | 2789754 | | | | | | |
| P08032 | SPTA1 | spectrin, alpha, erythrocytic 1 | Cytoplasm | other | 1.87E+07 | 4.70E-06 | 4.70E-06 | | | | | | |
| P08032 | SPTA1 | spectrin, alpha, erythrocytic 1 | Cytoplasm | other | 2385488.75 | 5828114 | 5317189 | | | | | | |
| P08032 | SPTA1 | spectrin, alpha, erythrocytic 1 | Cytoplasm | other | 2176497.75 | 3261461.75 | 6225583 | | | | | | |
| P08032 | SPTA1 | spectrin, alpha, erythrocytic 1 | Cytoplasm | other | 8541513 | 3401008.5 | 5989610.5 | | | | | | |
| P08032 | SPTA1 | spectrin, alpha, erythrocytic 1 | Cytoplasm | other | 5549701 | 7260994 | 1.10E+07 | | | | | | |
| P08032 | SPTA1 | spectrin, alpha, erythrocytic 1 | Cytoplasm | other | 7535427.5 | 8035797 | 1.16E+07 | | | | | | |
| P08032 | SPTA1 | spectrin, alpha, erythrocytic 1 | Cytoplasm | other | 5750134.5 | 3457359 | 4575607 | | | | | | |
| P08032 | SPTA1 | spectrin, alpha, erythrocytic 1 | Cytoplasm | other | 3843080.25 | 5139086.5 | 8371874.5 | | | | | | |
| P08032 | SPTA1 | spectrin, alpha, erythrocytic 1 | Cytoplasm | other | 6438564 | 7638502 | 5413641.5 | | | | | | |
| P08032 | SPTA1 | spectrin, alpha, erythrocytic 1 | Cytoplasm | other | 1.15E+07 | 2132174.5 | 1186177.625 | | | | | | |
| P08032 | SPTA1 | spectrin, alpha, erythrocytic 1 | Cytoplasm | other | 1616828.875 | 9937857 | 1.27E+07 | | | | | | |
| P08032 | SPTA1 | spectrin, alpha, erythrocytic 1 | Cytoplasm | other | 6377809.5 | 1667089.75 | 875228 | | | | | | |
| P08032 | SPTA1 | spectrin, alpha, erythrocytic 1 | Cytoplasm | other | 3341930.75 | 2049100.625 | 2106114 | | | | | | |
| P08032 | SPTA1 | spectrin, alpha, erythrocytic 1 | Cytoplasm | other | 2310938.5 | 2218014.5 | 2582468.5 | | | | | | |
| P08032 | SPTA1 | spectrin, alpha, erythrocytic 1 | Cytoplasm | other | 3750119 | 960140.062 | 888174.312 | | | | | | |
| P08032 | SPTA1 | spectrin, alpha, erythrocytic 1 | Cytoplasm | other | 1037911.375 | 1052761.5 | 1196114.375 | | | | | | |
| P08032 | SPTA1 | spectrin, alpha, erythrocytic 1 | Cytoplasm | other | 6809598.5 | 8136299.5 | 5592177 | | | | | | |
| P08032 | SPTA1 | spectrin, alpha, erythrocytic 1 | Cytoplasm | other | 1374921.625 | 476834.531 | 547841.5 | | | | | | |
| P08032 | SPTA1 | spectrin, alpha, erythrocytic 1 | Cytoplasm | other | 1515305.125 | 1860654.75 | 1590150.25 | | | | | | |
| P08032 | SPTA1 | spectrin, alpha, erythrocytic 1 | Cytoplasm | other | 308803.094 | 873924.312 | 917246.812 | | | | | | |
| P08032 | SPTA1 | spectrin, alpha, erythrocytic 1 | Cytoplasm | other | 5269781 | 2.54E+07 | 9.86E+07 | | | | | | |
| P08032 | SPTA1 | spectrin, alpha, erythrocytic 1 | Cytoplasm | other | 3106649.25 | 4.70E-06 | 68737.125 | | | | | | |
| P08032 | SPTA1 | spectrin, alpha, erythrocytic 1 | Cytoplasm | other | 6494190.5 | 123589.711 | 0 | | | | | | |
| P08032 | SPTA1 | spectrin, alpha, erythrocytic 1 | Cytoplasm | other | 1399870.5 | 955245.812 | 3915455.25 | | | | | | |
| P08032 | SPTA1 | spectrin, alpha, erythrocytic 1 | Cytoplasm | other | 263557.625 | 96853.555 | 748607.25 | | | | | | |
| P08032 | SPTA1 | spectrin, alpha, erythrocytic 1 | Cytoplasm | other | 1022075.125 | 1352618.25 | 2273293 | P08032 | SPTA1 | 517042241 | 677959258 | 22674349 | 894 |
| P16546 | SPTAN1 | spectrin, alpha, non-erythrocytic 1 | Plasma Membrane | other | 8600127 | 8464998 | 1.06E+07 | P16546 | SPTAN1 | 8600127 | 1178254 | 680265 | 3 |
| P15508 | SPTB | spectrin, beta, erythrocytic | Plasma Membrane | other | 3.34E+09 | 3.69E+09 | 4.21E+09 | | | | | | |
| P15508 | SPTB | spectrin, beta, erythrocytic | Plasma Membrane | other | 2.97E+09 | 2.64E+09 | 1.91E+09 | | | | | | |
| P15508 | SPTB | spectrin, beta, erythrocytic | Plasma Membrane | other | 2.52E+09 | 2.52E+09 | 3.32E+09 | | | | | | |

| | | | | | | | |
|---|---|---|---|---|---|---|---|
| P15508 | SPTB | spectrin, beta, erythrocytic | Plasma Membrane | other | 2.24E+09 | 2.29E+09 | 2.24E+09 |
| P15508 | SPTB | spectrin, beta, erythrocytic | Plasma Membrane | other | 1.98E+09 | 1.82E+09 | 2.19E+09 |
| P15508 | SPTB | spectrin, beta, erythrocytic | Plasma Membrane | other | 2.14E+09 | 2.10E+09 | 2.59E+09 |
| P15508 | SPTB | spectrin, beta, erythrocytic | Plasma Membrane | other | 2.02E+09 | 1.67E+09 | 2.01E+09 |
| P15508 | SPTB | spectrin, beta, erythrocytic | Plasma Membrane | other | 1.84E+09 | 1.52E+09 | 1.70E+09 |
| P15508 | SPTB | spectrin, beta, erythrocytic | Plasma Membrane | other | 1.72E+09 | 1.65E+09 | 1.98E+09 |
| P15508 | SPTB | spectrin, beta, erythrocytic | Plasma Membrane | other | 1.69E+09 | 1.50E+09 | 1.94E+09 |
| P15508 | SPTB | spectrin, beta, erythrocytic | Plasma Membrane | other | 1.38E+09 | 2.09E+09 | 2.67E+09 |
| P15508 | SPTB | spectrin, beta, erythrocytic | Plasma Membrane | other | 1.68E+09 | 1.74E+09 | 1.65E+09 |
| P15508 | SPTB | spectrin, beta, erythrocytic | Plasma Membrane | other | 1.78E+09 | 1.56E+09 | 1.69E+09 |
| P15508 | SPTB | spectrin, beta, erythrocytic | Plasma Membrane | other | 2.32E+09 | 2.72E+09 | 1.72E+09 |
| P15508 | SPTB | spectrin, beta, erythrocytic | Plasma Membrane | other | 1.57E+09 | 1.73E+09 | 1.78E+09 |
| P15508 | SPTB | spectrin, beta, erythrocytic | Plasma Membrane | other | 1.62E+09 | 1.66E+09 | 1.81E+09 |
| P15508 | SPTB | spectrin, beta, erythrocytic | Plasma Membrane | other | 1.56E+09 | 1.53E+09 | 1.44E+09 |
| P15508 | SPTB | spectrin, beta, erythrocytic | Plasma Membrane | other | 1.42E+09 | 1.36E+09 | 1.67E+09 |
| P15508 | SPTB | spectrin, beta, erythrocytic | Plasma Membrane | other | 1.43E+09 | 1.35E+09 | 1.66E+09 |
| P15508 | SPTB | spectrin, beta, erythrocytic | Plasma Membrane | other | 1.40E+09 | 1.36E+09 | 1.56E+09 |
| P15508 | SPTB | spectrin, beta, erythrocytic | Plasma Membrane | other | 1.38E+09 | 1.68E+09 | 1.58E+09 |
| P15508 | SPTB | spectrin, beta, erythrocytic | Plasma Membrane | other | 1.45E+09 | 1.31E+09 | 1.30E+09 |
| P15508 | SPTB | spectrin, beta, erythrocytic | Plasma Membrane | other | 1.46E+09 | 1.30E+09 | 1.59E+09 |
| P15508 | SPTB | spectrin, beta, erythrocytic | Plasma Membrane | other | 1.37E+09 | 1.34E+09 | 1.90E+09 |
| P15508 | SPTB | spectrin, beta, erythrocytic | Plasma Membrane | other | 1.04E+09 | 1.01E+09 | 8.52E+08 |
| P15508 | SPTB | spectrin, beta, erythrocytic | Plasma Membrane | other | 1.36E+09 | 1.23E+09 | 1.42E+09 |
| P15508 | SPTB | spectrin, beta, erythrocytic | Plasma Membrane | other | 1.36E+09 | 1.45E+09 | 1.48E+09 |
| P15508 | SPTB | spectrin, beta, erythrocytic | Plasma Membrane | other | 1.25E+09 | 1.21E+09 | 1.41E+09 |
| P15508 | SPTB | spectrin, beta, erythrocytic | Plasma Membrane | other | 1.32E+09 | 1.43E+09 | 1.54E+09 |
| P15508 | SPTB | spectrin, beta, erythrocytic | Plasma Membrane | other | 1.31E+09 | 1.13E+09 | 1.29E+09 |
| P15508 | SPTB | spectrin, beta, erythrocytic | Plasma Membrane | other | 1.39E+09 | 1.54E+09 | 1.53E+09 |
| P15508 | SPTB | spectrin, beta, erythrocytic | Plasma Membrane | other | 1.36E+09 | 1.20E+09 | 1.62E+09 |
| P15508 | SPTB | spectrin, beta, erythrocytic | Plasma Membrane | other | 1.29E+09 | 1.49E+09 | 1.59E+09 |
| P15508 | SPTB | spectrin, beta, erythrocytic | Plasma Membrane | other | 1.11E+09 | 1.54E+09 | 1.62E+09 |
| P15508 | SPTB | spectrin, beta, erythrocytic | Plasma Membrane | other | 1.09E+09 | 8.70E+08 | 3.71E+08 |
| P15508 | SPTB | spectrin, beta, erythrocytic | Plasma Membrane | other | 1.22E+09 | 1.12E+09 | 1.34E+09 |
| P15508 | SPTB | spectrin, beta, erythrocytic | Plasma Membrane | other | 1.49E+09 | 1.42E+09 | 1.49E+09 |
| P15508 | SPTB | spectrin, beta, erythrocytic | Plasma Membrane | other | 1.09E+09 | 1.10E+09 | 1.31E+09 |
| P15508 | SPTB | spectrin, beta, erythrocytic | Plasma Membrane | other | 1.20E+09 | 1.10E+09 | 1.17E+09 |
| P15508 | SPTB | spectrin, beta, erythrocytic | Plasma Membrane | other | 1.24E+09 | 1.23E+09 | 1.40E+09 |
| P15508 | SPTB | spectrin, beta, erythrocytic | Plasma Membrane | other | 1.05E+09 | 9.81E+08 | 1.10E+09 |
| P15508 | SPTB | spectrin, beta, erythrocytic | Plasma Membrane | other | 1.29E+09 | 1.54E+09 | 1.89E+09 |
| P15508 | SPTB | spectrin, beta, erythrocytic | Plasma Membrane | other | 1.15E+09 | 1.12E+09 | 1.27E+09 |
| P15508 | SPTB | spectrin, beta, erythrocytic | Plasma Membrane | other | 8.95E+08 | 1.04E+09 | 1.36E+09 |
| P15508 | SPTB | spectrin, beta, erythrocytic | Plasma Membrane | other | 1.14E+09 | 1.16E+09 | 1.37E+09 |
| P15508 | SPTB | spectrin, beta, erythrocytic | Plasma Membrane | other | 9.76E+08 | 9.15E+08 | 1.08E+09 |
| P15508 | SPTB | spectrin, beta, erythrocytic | Plasma Membrane | other | 1.35E+09 | 1.18E+09 | 1.25E+09 |
| P15508 | SPTB | spectrin, beta, erythrocytic | Plasma Membrane | other | 1.09E+09 | 1.13E+09 | 1.57E+09 |
| P15508 | SPTB | spectrin, beta, erythrocytic | Plasma Membrane | other | 1.15E+09 | 1.04E+09 | 1.19E+09 |
| P15508 | SPTB | spectrin, beta, erythrocytic | Plasma Membrane | other | 8.99E+08 | 1.02E+09 | 1.22E+09 |
| P15508 | SPTB | spectrin, beta, erythrocytic | Plasma Membrane | other | 1.02E+09 | 9.07E+08 | 9.70E+08 |
| P15508 | SPTB | spectrin, beta, erythrocytic | Plasma Membrane | other | 9.72E+08 | 1.05E+09 | 2.06E+09 |
| P15508 | SPTB | spectrin, beta, erythrocytic | Plasma Membrane | other | 1.19E+09 | 1.10E+09 | 1.35E+09 |
| P15508 | SPTB | spectrin, beta, erythrocytic | Plasma Membrane | other | 9.02E+08 | 8.03E+08 | 9.73E+08 |
| P15508 | SPTB | spectrin, beta, erythrocytic | Plasma Membrane | other | 9.65E+08 | 9.02E+08 | 9.54E+08 |
| P15508 | SPTB | spectrin, beta, erythrocytic | Plasma Membrane | other | 7.80E+08 | 7.00E+08 | 5.68E+08 |
| P15508 | SPTB | spectrin, beta, erythrocytic | Plasma Membrane | other | 7.05E+08 | 8.66E+08 | 9.09E+08 |
| P15508 | SPTB | spectrin, beta, erythrocytic | Plasma Membrane | other | 9.41E+08 | 8.95E+08 | 1.18E+09 |
| P15508 | SPTB | spectrin, beta, erythrocytic | Plasma Membrane | other | 1.25E+09 | 6.82E+08 | 7.28E+08 |
| P15508 | SPTB | spectrin, beta, erythrocytic | Plasma Membrane | other | 9.08E+08 | 9.60E+08 | 9.63E+08 |
| P15508 | SPTB | spectrin, beta, erythrocytic | Plasma Membrane | other | 8.57E+08 | 8.22E+08 | 9.50E+08 |
| P15508 | SPTB | spectrin, beta, erythrocytic | Plasma Membrane | other | 6.87E+08 | 9.07E+08 | 1.14E+09 |
| P15508 | SPTB | spectrin, beta, erythrocytic | Plasma Membrane | other | 8.47E+08 | 1.03E+09 | 1.31E+09 |
| P15508 | SPTB | spectrin, beta, erythrocytic | Plasma Membrane | other | 6.33E+08 | 6.39E+08 | 2.68E+08 |
| P15508 | SPTB | spectrin, beta, erythrocytic | Plasma Membrane | other | 8.55E+08 | 7.83E+08 | 9.70E+08 |
| P15508 | SPTB | spectrin, beta, erythrocytic | Plasma Membrane | other | 8.03E+08 | 6.63E+08 | 6.70E+08 |
| P15508 | SPTB | spectrin, beta, erythrocytic | Plasma Membrane | other | 7.28E+08 | 6.49E+08 | 7.50E+08 |
| P15508 | SPTB | spectrin, beta, erythrocytic | Plasma Membrane | other | 6.56E+08 | 7.41E+08 | 4.95E+08 |
| P15508 | SPTB | spectrin, beta, erythrocytic | Plasma Membrane | other | 4.52E+08 | 1.19E+09 | 1.85E+09 |
| P15508 | SPTB | spectrin, beta, erythrocytic | Plasma Membrane | other | 7.70E+08 | 7.87E+08 | 9.12E+08 |
| P15508 | SPTB | spectrin, beta, erythrocytic | Plasma Membrane | other | 8.95E+08 | 5.07E+08 | 3.02E+08 |
| P15508 | SPTB | spectrin, beta, erythrocytic | Plasma Membrane | other | 1.10E+09 | 5.40E+08 | 6.22E+08 |
| P15508 | SPTB | spectrin, beta, erythrocytic | Plasma Membrane | other | 6.36E+08 | 6.98E+08 | 7.32E+08 |
| P15508 | SPTB | spectrin, beta, erythrocytic | Plasma Membrane | other | 7.37E+08 | 7.47E+08 | 7.84E+08 |
| P15508 | SPTB | spectrin, beta, erythrocytic | Plasma Membrane | other | 7.51E+08 | 7.80E+08 | 7.33E+08 |
| P15508 | SPTB | spectrin, beta, erythrocytic | Plasma Membrane | other | 5.07E+08 | 7.85E+08 | 8.93E+08 |
| P15508 | SPTB | spectrin, beta, erythrocytic | Plasma Membrane | other | 7.05E+08 | 4.75E+08 | 7.56E+08 |
| P15508 | SPTB | spectrin, beta, erythrocytic | Plasma Membrane | other | 6.21E+08 | 8.36E+08 | 7.96E+08 |
| P15508 | SPTB | spectrin, beta, erythrocytic | Plasma Membrane | other | 4.28E+08 | 4.33E+08 | 5.89E+08 |
| P15508 | SPTB | spectrin, beta, erythrocytic | Plasma Membrane | other | 7.73E+08 | 8.00E+08 | 8.32E+08 |
| P15508 | SPTB | spectrin, beta, erythrocytic | Plasma Membrane | other | 7.02E+08 | 6.52E+08 | 6.66E+08 |
| P15508 | SPTB | spectrin, beta, erythrocytic | Plasma Membrane | other | 5.82E+08 | 5.35E+08 | 6.57E+08 |
| P15508 | SPTB | spectrin, beta, erythrocytic | Plasma Membrane | other | 5.45E+08 | 7.77E+08 | 9.52E+08 |
| P15508 | SPTB | spectrin, beta, erythrocytic | Plasma Membrane | other | 5.22E+08 | 4.32E+08 | 4.47E+08 |
| P15508 | SPTB | spectrin, beta, erythrocytic | Plasma Membrane | other | 6.89E+08 | 7.57E+08 | 6.39E+08 |
| P15508 | SPTB | spectrin, beta, erythrocytic | Plasma Membrane | other | 5.64E+08 | 4.27E+08 | 4.48E+08 |
| P15508 | SPTB | spectrin, beta, erythrocytic | Plasma Membrane | other | 4.61E+08 | 6.94E+08 | 4.69E+08 |
| P15508 | SPTB | spectrin, beta, erythrocytic | Plasma Membrane | other | 4.53E+08 | 6.84E+08 | 8.25E+08 |
| P15508 | SPTB | spectrin, beta, erythrocytic | Plasma Membrane | other | 8.37E+08 | 8.27E+08 | 1.03E+09 |
| P15508 | SPTB | spectrin, beta, erythrocytic | Plasma Membrane | other | 6.58E+08 | 7.05E+08 | 7.82E+08 |
| P15508 | SPTB | spectrin, beta, erythrocytic | Plasma Membrane | other | 4.54E+08 | 6.46E+08 | 6.50E+08 |
| P15508 | SPTB | spectrin, beta, erythrocytic | Plasma Membrane | other | 5.35E+08 | 5.75E+08 | 5.28E+08 |
| P15508 | SPTB | spectrin, beta, erythrocytic | Plasma Membrane | other | 4.68E+08 | 4.10E+08 | 4.10E+08 |
| P15508 | SPTB | spectrin, beta, erythrocytic | Plasma Membrane | other | 1.04E+09 | 1.07E+09 | 7.08E+08 |

| | | | | | | | |
|---|---|---|---|---|---|---|---|
| P15508 | SPTB | spectrin, beta, erythrocytic | Plasma Membrane | other | 5.58E+08 | 3.98E+08 | 4.70E+08 |
| P15508 | SPTB | spectrin, beta, erythrocytic | Plasma Membrane | other | 4.95E+08 | 5.47E+08 | 4.86E+08 |
| P15508 | SPTB | spectrin, beta, erythrocytic | Plasma Membrane | other | 4.24E+08 | 3.91E+08 | 3.68E+08 |
| P15508 | SPTB | spectrin, beta, erythrocytic | Plasma Membrane | other | 4.64E+08 | 3.48E+08 | 3.24E+08 |
| P15508 | SPTB | spectrin, beta, erythrocytic | Plasma Membrane | other | 5.77E+08 | 6.43E+08 | 6.36E+08 |
| P15508 | SPTB | spectrin, beta, erythrocytic | Plasma Membrane | other | 3.25E+08 | 3.48E+08 | 3.34E+08 |
| P15508 | SPTB | spectrin, beta, erythrocytic | Plasma Membrane | other | 5.51E+08 | 5.89E+08 | 7.52E+08 |
| P15508 | SPTB | spectrin, beta, erythrocytic | Plasma Membrane | other | 4.44E+08 | 5.01E+08 | 5.72E+08 |
| P15508 | SPTB | spectrin, beta, erythrocytic | Plasma Membrane | other | 3.70E+08 | 5.09E+08 | 5.74E+08 |
| P15508 | SPTB | spectrin, beta, erythrocytic | Plasma Membrane | other | 2.53E+08 | 3.42E+08 | 4.34E+08 |
| P15508 | SPTB | spectrin, beta, erythrocytic | Plasma Membrane | other | 3.48E+08 | 3.14E+08 | 3.36E+08 |
| P15508 | SPTB | spectrin, beta, erythrocytic | Plasma Membrane | other | 3.63E+08 | 3.01E+08 | 3.04E+08 |
| P15508 | SPTB | spectrin, beta, erythrocytic | Plasma Membrane | other | 3.48E+08 | 3.11E+08 | 3.30E+08 |
| P15508 | SPTB | spectrin, beta, erythrocytic | Plasma Membrane | other | 4.76E+08 | 1.99E+08 | 1.36E+08 |
| P15508 | SPTB | spectrin, beta, erythrocytic | Plasma Membrane | other | 2.99E+08 | 2.47E+08 | 2.50E+08 |
| P15508 | SPTB | spectrin, beta, erythrocytic | Plasma Membrane | other | 2.79E+08 | 3.28E+08 | 3.66E+08 |
| P15508 | SPTB | spectrin, beta, erythrocytic | Plasma Membrane | other | 3.27E+08 | 2.54E+08 | 2.73E+08 |
| P15508 | SPTB | spectrin, beta, erythrocytic | Plasma Membrane | other | 3.84E+08 | 1.65E+08 | 1.52E+08 |
| P15508 | SPTB | spectrin, beta, erythrocytic | Plasma Membrane | other | 1.92E+08 | 1.50E+08 | 1.19E+08 |
| P15508 | SPTB | spectrin, beta, erythrocytic | Plasma Membrane | other | 2.21E+08 | 1.94E+08 | 2.12E+08 |
| P15508 | SPTB | spectrin, beta, erythrocytic | Plasma Membrane | other | 1.75E+08 | 1.14E+08 | 1.51E+08 |
| P15508 | SPTB | spectrin, beta, erythrocytic | Plasma Membrane | other | 2.88E+08 | 2.72E+08 | 3.39E+08 |
| P15508 | SPTB | spectrin, beta, erythrocytic | Plasma Membrane | other | 2.80E+08 | 3.16E+08 | 2.44E+08 |
| P15508 | SPTB | spectrin, beta, erythrocytic | Plasma Membrane | other | 4.03E+08 | 4.05E+08 | 2.48E+08 |
| P15508 | SPTB | spectrin, beta, erythrocytic | Plasma Membrane | other | 1.55E+08 | 3.84E+08 | 6.39E+08 |
| P15508 | SPTB | spectrin, beta, erythrocytic | Plasma Membrane | other | 1.96E+08 | 1.78E+08 | 1.82E+08 |
| P15508 | SPTB | spectrin, beta, erythrocytic | Plasma Membrane | other | 4.48E+08 | 5.53E+08 | 5.84E+08 |
| P15508 | SPTB | spectrin, beta, erythrocytic | Plasma Membrane | other | 4.71E+08 | 3.98E+08 | 4.85E+08 |
| P15508 | SPTB | spectrin, beta, erythrocytic | Plasma Membrane | other | 1.33E+08 | 1.76E+08 | 1.55E+08 |
| P15508 | SPTB | spectrin, beta, erythrocytic | Plasma Membrane | other | 2.24E+08 | 2.08E+08 | 2.35E+08 |
| P15508 | SPTB | spectrin, beta, erythrocytic | Plasma Membrane | other | 2.72E+08 | 2.72E+08 | 3.34E+08 |
| P15508 | SPTB | spectrin, beta, erythrocytic | Plasma Membrane | other | 1.84E+08 | 1.87E+08 | 1.88E+08 |
| P15508 | SPTB | spectrin, beta, erythrocytic | Plasma Membrane | other | 1.86E+08 | 2.16E+08 | 2.53E+08 |
| P15508 | SPTB | spectrin, beta, erythrocytic | Plasma Membrane | other | 2.68E+08 | 4.10E+08 | 8.12E+08 |
| P15508 | SPTB | spectrin, beta, erythrocytic | Plasma Membrane | other | 1.65E+08 | 2.14E+08 | 3.35E+08 |
| P15508 | SPTB | spectrin, beta, erythrocytic | Plasma Membrane | other | 2.37E+08 | 3.33E+08 | 5.27E+08 |
| P15508 | SPTB | spectrin, beta, erythrocytic | Plasma Membrane | other | 1.71E+08 | 1.80E+08 | 2.32E+08 |
| P15508 | SPTB | spectrin, beta, erythrocytic | Plasma Membrane | other | 2.16E+08 | 3.68E+07 | 2.76E+07 |
| P15508 | SPTB | spectrin, beta, erythrocytic | Plasma Membrane | other | 5.91E+07 | 5.78E+07 | 8.17E+07 |
| P15508 | SPTB | spectrin, beta, erythrocytic | Plasma Membrane | other | 1.37E+08 | 1.76E+08 | 2.24E+08 |
| P15508 | SPTB | spectrin, beta, erythrocytic | Plasma Membrane | other | 1.72E+08 | 1.43E+08 | 1.92E+08 |
| P15508 | SPTB | spectrin, beta, erythrocytic | Plasma Membrane | other | 1.68E+08 | 1.09E+08 | 1.22E+08 |
| P15508 | SPTB | spectrin, beta, erythrocytic | Plasma Membrane | other | 1.12E+08 | 1.25E+08 | 1.13E+08 |
| P15508 | SPTB | spectrin, beta, erythrocytic | Plasma Membrane | other | 1.40E+08 | 1.34E+08 | 1.21E+08 |
| P15508 | SPTB | spectrin, beta, erythrocytic | Plasma Membrane | other | 2.19E+08 | 1.57E+08 | 2.33E+08 |
| P15508 | SPTB | spectrin, beta, erythrocytic | Plasma Membrane | other | 2.50E+08 | 2.32E+08 | 2.32E+08 |
| P15508 | SPTB | spectrin, beta, erythrocytic | Plasma Membrane | other | 6.85E+07 | 4.36E+07 | 3.89E+07 |
| P15508 | SPTB | spectrin, beta, erythrocytic | Plasma Membrane | other | 2.45E+08 | 6.54E+07 | 1.54E+07 |
| P15508 | SPTB | spectrin, beta, erythrocytic | Plasma Membrane | other | 2.08E+08 | 1.35E+08 | 2.37E+08 |
| P15508 | SPTB | spectrin, beta, erythrocytic | Plasma Membrane | other | 9.47E+07 | 1.32E+08 | 9.96E+07 |
| P15508 | SPTB | spectrin, beta, erythrocytic | Plasma Membrane | other | 8.84E+07 | 9.07E+07 | 6.00E+07 |
| P15508 | SPTB | spectrin, beta, erythrocytic | Plasma Membrane | other | 1.37E+08 | 1.13E+08 | 1.21E+08 |
| P15508 | SPTB | spectrin, beta, erythrocytic | Plasma Membrane | other | 1.00E+08 | 1.03E+08 | 1.39E+08 |
| P15508 | SPTB | spectrin, beta, erythrocytic | Plasma Membrane | other | 1.31E+08 | 1.17E+08 | 1.45E+08 |
| P15508 | SPTB | spectrin, beta, erythrocytic | Plasma Membrane | other | 9.94E+07 | 1.06E+08 | 1.03E+08 |
| P15508 | SPTB | spectrin, beta, erythrocytic | Plasma Membrane | other | 9.09E+07 | 1.11E+08 | 8.17E+07 |
| P15508 | SPTB | spectrin, beta, erythrocytic | Plasma Membrane | other | 1.19E+08 | 6.66E+07 | 1.87E+08 |
| P15508 | SPTB | spectrin, beta, erythrocytic | Plasma Membrane | other | 7.09E+07 | 8.05E+07 | 1.31E+08 |
| P15508 | SPTB | spectrin, beta, erythrocytic | Plasma Membrane | other | 1.15E+08 | 1.02E+08 | 1.27E+08 |
| P15508 | SPTB | spectrin, beta, erythrocytic | Plasma Membrane | other | 1.21E+08 | 1.02E+08 | 1.03E+08 |
| P15508 | SPTB | spectrin, beta, erythrocytic | Plasma Membrane | other | 1.05E+08 | 8.90E+07 | 1.07E+08 |
| P15508 | SPTB | spectrin, beta, erythrocytic | Plasma Membrane | other | 9.49E+07 | 1.45E+08 | 1.01E+08 |
| P15508 | SPTB | spectrin, beta, erythrocytic | Plasma Membrane | other | 9.40E+07 | 8.73E+07 | 1.04E+08 |
| P15508 | SPTB | spectrin, beta, erythrocytic | Plasma Membrane | other | 6.19E+07 | 1.77E+08 | 3.56E+08 |
| P15508 | SPTB | spectrin, beta, erythrocytic | Plasma Membrane | other | 1.06E+08 | 5.72E+07 | 5.30E+07 |
| P15508 | SPTB | spectrin, beta, erythrocytic | Plasma Membrane | other | 2.19E+08 | 1.99E+08 | 4.33E+07 |
| P15508 | SPTB | spectrin, beta, erythrocytic | Plasma Membrane | other | 1.29E+08 | 1.86E+08 | 2.79E+08 |
| P15508 | SPTB | spectrin, beta, erythrocytic | Plasma Membrane | other | 8.33E+07 | 1.06E+08 | 1.28E+08 |
| P15508 | SPTB | spectrin, beta, erythrocytic | Plasma Membrane | other | 1.09E+08 | 1.19E+08 | 7.55E+07 |
| P15508 | SPTB | spectrin, beta, erythrocytic | Plasma Membrane | other | 2.05E+08 | 3.09E+08 | 4.25E+08 |
| P15508 | SPTB | spectrin, beta, erythrocytic | Plasma Membrane | other | 1.33E+08 | 8.26E+07 | 8.51E+07 |
| P15508 | SPTB | spectrin, beta, erythrocytic | Plasma Membrane | other | 1.01E+08 | 8.03E+07 | 6.74E+07 |
| P15508 | SPTB | spectrin, beta, erythrocytic | Plasma Membrane | other | 1.02E+08 | 2.25E+07 | 1.79E+07 |
| P15508 | SPTB | spectrin, beta, erythrocytic | Plasma Membrane | other | 1.16E+08 | 3.93E+07 | 5.91E+07 |
| P15508 | SPTB | spectrin, beta, erythrocytic | Plasma Membrane | other | 7.68E+07 | 7.01E+07 | 5.57E+07 |
| P15508 | SPTB | spectrin, beta, erythrocytic | Plasma Membrane | other | 1.77E+08 | 1.29E+07 | 2516756.75 |
| P15508 | SPTB | spectrin, beta, erythrocytic | Plasma Membrane | other | 8.70E+07 | 9.67E+07 | 1.01E+08 |
| P15508 | SPTB | spectrin, beta, erythrocytic | Plasma Membrane | other | 6.86E+07 | 4.49E+07 | 2.01E+07 |
| P15508 | SPTB | spectrin, beta, erythrocytic | Plasma Membrane | other | 6.11E+07 | 7.66E+07 | 1.01E+08 |
| P15508 | SPTB | spectrin, beta, erythrocytic | Plasma Membrane | other | 5.49E+07 | 4.53E+07 | 5.71E+07 |
| P15508 | SPTB | spectrin, beta, erythrocytic | Plasma Membrane | other | 3.71E+07 | 4.29E+07 | 5.42E+07 |
| P15508 | SPTB | spectrin, beta, erythrocytic | Plasma Membrane | other | 6.25E+07 | 7.15E+07 | 8.71E+07 |
| P15508 | SPTB | spectrin, beta, erythrocytic | Plasma Membrane | other | 8.41E+07 | 2.85E+07 | 2.13E+07 |
| P15508 | SPTB | spectrin, beta, erythrocytic | Plasma Membrane | other | 5.53E+07 | 9.49E+07 | 6.53E+07 |
| P15508 | SPTB | spectrin, beta, erythrocytic | Plasma Membrane | other | 4.56E+07 | 4.73E+07 | 6.71E+07 |
| P15508 | SPTB | spectrin, beta, erythrocytic | Plasma Membrane | other | 6.53E+07 | 6.13E+07 | 7.48E+07 |
| P15508 | SPTB | spectrin, beta, erythrocytic | Plasma Membrane | other | 3.25E+07 | 3.64E+07 | 2.78E+07 |
| P15508 | SPTB | spectrin, beta, erythrocytic | Plasma Membrane | other | 3.77E+07 | 8.25E+07 | 1.30E+08 |
| P15508 | SPTB | spectrin, beta, erythrocytic | Plasma Membrane | other | 6.19E+07 | 6.67E+07 | 8.75E+07 |
| P15508 | SPTB | spectrin, beta, erythrocytic | Plasma Membrane | other | 6.44E+07 | 9.98E+07 | 1.84E+08 |
| P15508 | SPTB | spectrin, beta, erythrocytic | Plasma Membrane | other | 5.17E+07 | 5.96E+07 | 9.25E+07 |

| | | | | | | | | | | | | | |
|---|---|---|---|---|---|---|---|---|---|---|---|---|---|
| P15508 | SPTB | spectrin, beta, erythrocytic | Plasma Membrane | other | 5.38E+07 | 5.20E+07 | 8.63E+07 | | | | | | |
| P15508 | SPTB | spectrin, beta, erythrocytic | Plasma Membrane | other | 1.97E+07 | 3.97E+07 | 4.21E+07 | | | | | | |
| P15508 | SPTB | spectrin, beta, erythrocytic | Plasma Membrane | other | 3.05E+07 | 6.87E+07 | 1.17E+08 | | | | | | |
| P15508 | SPTB | spectrin, beta, erythrocytic | Plasma Membrane | other | 4.90E+07 | 4.08E+07 | 6.26E+07 | | | | | | |
| P15508 | SPTB | spectrin, beta, erythrocytic | Plasma Membrane | other | 1.76E+07 | 8.16E+07 | 1.80E+08 | | | | | | |
| P15508 | SPTB | spectrin, beta, erythrocytic | Plasma Membrane | other | 3.12E+07 | 4.03E+07 | 4.22E+07 | | | | | | |
| P15508 | SPTB | spectrin, beta, erythrocytic | Plasma Membrane | other | 3.84E+07 | 3.22E+07 | 5.64E+07 | | | | | | |
| P15508 | SPTB | spectrin, beta, erythrocytic | Plasma Membrane | other | 2.97E+07 | 2.56E+07 | 3.90E+07 | | | | | | |
| P15508 | SPTB | spectrin, beta, erythrocytic | Plasma Membrane | other | 7.06E+07 | 8.23E+07 | 1.20E+08 | | | | | | |
| P15508 | SPTB | spectrin, beta, erythrocytic | Plasma Membrane | other | 3.55E+07 | 2.48E+07 | 2.25E+07 | | | | | | |
| P15508 | SPTB | spectrin, beta, erythrocytic | Plasma Membrane | other | 2.85E+07 | 3.12E+07 | 3.81E+07 | | | | | | |
| P15508 | SPTB | spectrin, beta, erythrocytic | Plasma Membrane | other | 6.05E+07 | 1.10E+07 | 6434268 | | | | | | |
| P15508 | SPTB | spectrin, beta, erythrocytic | Plasma Membrane | other | 2.58E+07 | 2.33E+07 | 1.94E+07 | | | | | | |
| P15508 | SPTB | spectrin, beta, erythrocytic | Plasma Membrane | other | 2.96E+07 | 3.35E+07 | 4.17E+07 | | | | | | |
| P15508 | SPTB | spectrin, beta, erythrocytic | Plasma Membrane | other | 2.11E+07 | 1.90E+07 | 1.40E+07 | | | | | | |
| P15508 | SPTB | spectrin, beta, erythrocytic | Plasma Membrane | other | 1.31E+07 | 2.48E+07 | 3.13E+07 | | | | | | |
| P15508 | SPTB | spectrin, beta, erythrocytic | Plasma Membrane | other | 6.08E+07 | 9.52E+07 | 7.89E+07 | | | | | | |
| P15508 | SPTB | spectrin, beta, erythrocytic | Plasma Membrane | other | 3.84E+07 | 3.25E+07 | 5.07E+07 | | | | | | |
| P15508 | SPTB | spectrin, beta, erythrocytic | Plasma Membrane | other | 2.43E+07 | 1.93E+07 | 2.77E+07 | | | | | | |
| P15508 | SPTB | spectrin, beta, erythrocytic | Plasma Membrane | other | 3.45E+07 | 1.64E+07 | 1.75E+07 | | | | | | |
| P15508 | SPTB | spectrin, beta, erythrocytic | Plasma Membrane | other | 2.46E+07 | 2.92E+07 | 3.77E+07 | | | | | | |
| P15508 | SPTB | spectrin, beta, erythrocytic | Plasma Membrane | other | 4.91E+07 | 7969564 | 5960710.5 | | | | | | |
| P15508 | SPTB | spectrin, beta, erythrocytic | Plasma Membrane | other | 2.12E+07 | 2.73E+07 | 2.48E+07 | | | | | | |
| P15508 | SPTB | spectrin, beta, erythrocytic | Plasma Membrane | other | 2.67E+07 | 1.97E+07 | 6.27E+07 | | | | | | |
| P15508 | SPTB | spectrin, beta, erythrocytic | Plasma Membrane | other | 4.51E+07 | 6893577 | 5020188 | | | | | | |
| P15508 | SPTB | spectrin, beta, erythrocytic | Plasma Membrane | other | 3.85E+07 | 7483720.5 | 4906521.5 | | | | | | |
| P15508 | SPTB | spectrin, beta, erythrocytic | Plasma Membrane | other | 2.03E+07 | 1.79E+07 | 1.91E+07 | | | | | | |
| P15508 | SPTB | spectrin, beta, erythrocytic | Plasma Membrane | other | 1.18E+07 | 9516720 | 9446276 | | | | | | |
| P15508 | SPTB | spectrin, beta, erythrocytic | Plasma Membrane | other | 1.08E+07 | 2.03E+07 | 4.92E+07 | | | | | | |
| P15508 | SPTB | spectrin, beta, erythrocytic | Plasma Membrane | other | 1.58E+07 | 989135.875 | 1269622.875 | | | | | | |
| P15508 | SPTB | spectrin, beta, erythrocytic | Plasma Membrane | other | 3.03E+07 | 1.20E+07 | 7.25E+07 | | | | | | |
| P15508 | SPTB | spectrin, beta, erythrocytic | Plasma Membrane | other | 4.12E+07 | 1319712.125 | 1445248.875 | | | | | | |
| P15508 | SPTB | spectrin, beta, erythrocytic | Plasma Membrane | other | 1.31E+07 | 1.14E+07 | 5547212.5 | | | | | | |
| P15508 | SPTB | spectrin, beta, erythrocytic | Plasma Membrane | other | 4.71E+07 | 5.58E+07 | 7.32E+07 | | | | | | |
| P15508 | SPTB | spectrin, beta, erythrocytic | Plasma Membrane | other | 1.23E+07 | 9764594 | 6799055.5 | | | | | | |
| P15508 | SPTB | spectrin, beta, erythrocytic | Plasma Membrane | other | 1.71E+07 | 1.20E+07 | 1.20E+07 | | | | | | |
| P15508 | SPTB | spectrin, beta, erythrocytic | Plasma Membrane | other | 1.26E+07 | 9609634 | 1.42E+07 | | | | | | |
| P15508 | SPTB | spectrin, beta, erythrocytic | Plasma Membrane | other | 1.81E+07 | 2.16E+07 | 2.26E+07 | | | | | | |
| P15508 | SPTB | spectrin, beta, erythrocytic | Plasma Membrane | other | 1.07E+07 | 1.12E+07 | 1.35E+07 | | | | | | |
| P15508 | SPTB | spectrin, beta, erythrocytic | Plasma Membrane | other | 1.01E+07 | 1.86E+07 | 1.39E+07 | | | | | | |
| P15508 | SPTB | spectrin, beta, erythrocytic | Plasma Membrane | other | 7714166 | 7527441 | 8164338.5 | | | | | | |
| P15508 | SPTB | spectrin, beta, erythrocytic | Plasma Membrane | other | 3.55E+07 | 1591096.375 | 1304696.625 | | | | | | |
| P15508 | SPTB | spectrin, beta, erythrocytic | Plasma Membrane | other | 1.29E+07 | 1.27E+07 | 1.49E+07 | | | | | | |
| P15508 | SPTB | spectrin, beta, erythrocytic | Plasma Membrane | other | 5570690 | 6462650.5 | 5241433 | | | | | | |
| P15508 | SPTB | spectrin, beta, erythrocytic | Plasma Membrane | other | 1.49E+07 | 1871381.625 | 701194.688 | | | | | | |
| P15508 | SPTB | spectrin, beta, erythrocytic | Plasma Membrane | other | 3.19E+07 | 2.26E+07 | 1.37E+07 | | | | | | |
| P15508 | SPTB | spectrin, beta, erythrocytic | Plasma Membrane | other | 1.28E+07 | 5488307 | 3985166.5 | | | | | | |
| P15508 | SPTB | spectrin, beta, erythrocytic | Plasma Membrane | other | 4685382 | 1.11E+07 | 1.54E+07 | | | | | | |
| P15508 | SPTB | spectrin, beta, erythrocytic | Plasma Membrane | other | 1.39E+07 | 2199672.75 | 1964264.875 | | | | | | |
| P15508 | SPTB | spectrin, beta, erythrocytic | Plasma Membrane | other | 6055029 | 9030814 | 1.20E+07 | | | | | | |
| P15508 | SPTB | spectrin, beta, erythrocytic | Plasma Membrane | other | 7044215 | 8228208.5 | 1.01E+07 | | | | | | |
| P15508 | SPTB | spectrin, beta, erythrocytic | Plasma Membrane | other | 2.09E+07 | 4244797 | 600482.625 | | | | | | |
| P15508 | SPTB | spectrin, beta, erythrocytic | Plasma Membrane | other | 4314075 | 705250.312 | 4.70E-06 | | | | | | |
| P15508 | SPTB | spectrin, beta, erythrocytic | Plasma Membrane | other | 4783892.5 | 3873606.75 | 3838523.5 | | | | | | |
| P15508 | SPTB | spectrin, beta, erythrocytic | Plasma Membrane | other | 7334008.5 | 1.68E+07 | 1.19E+07 | | | | | | |
| P15508 | SPTB | spectrin, beta, erythrocytic | Plasma Membrane | other | 4728955 | 2743096 | 3543644 | | | | | | |
| P15508 | SPTB | spectrin, beta, erythrocytic | Plasma Membrane | other | 2209491 | 3477196.75 | 5514489 | | | | | | |
| P15508 | SPTB | spectrin, beta, erythrocytic | Plasma Membrane | other | 3377804.75 | 2469324 | 2108341.25 | | | | | | |
| P15508 | SPTB | spectrin, beta, erythrocytic | Plasma Membrane | other | 2651595.5 | 5890794 | 6941056 | | | | | | |
| P15508 | SPTB | spectrin, beta, erythrocytic | Plasma Membrane | other | 1552209 | 2989126.25 | 7773807.5 | | | | | | |
| P15508 | SPTB | spectrin, beta, erythrocytic | Plasma Membrane | other | 5149496 | 2001553.125 | 2432834.5 | | | | | | |
| P15508 | SPTB | spectrin, beta, erythrocytic | Plasma Membrane | other | 3969549.25 | 1658527.25 | 1905250.125 | | | | | | |
| P15508 | SPTB | spectrin, beta, erythrocytic | Plasma Membrane | other | 5576346 | 1165847.5 | 727857.812 | | | | | | |
| P15508 | SPTB | spectrin, beta, erythrocytic | Plasma Membrane | other | 629690.312 | 2799281.75 | 8270763 | | | | | | |
| P15508 | SPTB | spectrin, beta, erythrocytic | Plasma Membrane | other | 2.15E+07 | 9.24E+07 | 1.98E+08 | | | | | | |
| P15508 | SPTB | spectrin, beta, erythrocytic | Plasma Membrane | other | 2710724 | 211283.844 | 1530633.625 | | | | | | |
| P15508 | SPTB | spectrin, beta, erythrocytic | Plasma Membrane | other | 362945.094 | 204506.469 | 111860.289 | | | | | | |
| P15508 | SPTB | spectrin, beta, erythrocytic | Plasma Membrane | other | 1221498.625 | 1495272 | 1137009.75 | | | | | | |
| P15508 | SPTB | spectrin, beta, erythrocytic | Plasma Membrane | other | 4655211 | 593236.75 | 134187.016 | | | | | | |
| P15508 | SPTB | spectrin, beta, erythrocytic | Plasma Membrane | other | 2883633 | 6191.075 | 17094.17 | | | | | | |
| P15508 | SPTB | spectrin, beta, erythrocytic | Plasma Membrane | other | 1190018.625 | 3897288.75 | 1.06E+07 | | | | | | |
| P15508 | SPTB | spectrin, beta, erythrocytic | Plasma Membrane | other | 812022 | 43706.137 | 23763.611 | | | | | | |
| P15508 | SPTB | spectrin, beta, erythrocytic | Plasma Membrane | other | 520282.969 | 1877114.125 | 5630793.5 | | | | | | |
| P15508 | SPTB | spectrin, beta, erythrocytic | Plasma Membrane | other | 4.70E-06 | 278657.656 | 385927.812 | | | | | | |
| P15508 | SPTB | spectrin, beta, erythrocytic | Plasma Membrane | other | 70409.781 | 679792.875 | 7561007.5 | P15508 | SPTB | 489650205 | 633686995 | 22689634 | 780 |
| P13609 | Srgn | serglycin | Cytoplasm | other | 203048.938 | 241374.312 | 583873.438 | P13609 | Srgn | 203049 | 209683 | 121061 | 3 |
| Q9CY50 | SSR1 | signal sequence receptor, alpha | Cytoplasm | other | 1228345.25 | 2889582.5 | 3029651.5 | | | | | | |
| Q9CY50 | SSR1 | signal sequence receptor, alpha | Cytoplasm | other | 1172392.5 | 3109895.75 | 2857666.75 | Q9CY50 | SSR1 | 1200369 | 919484 | 375378 | 6 |
| Q9DCF9 | SSR3 | signal sequence receptor, gamma (translocon-associated protein gamma) | Cytoplasm | other | 2663498.75 | 4438542.5 | 5512732 | Q9DCF9 | SSR3 | 2663499 | 1438911 | 830756 | 3 |
| O70576 | STAG3 | stromal antigen 3 | Nucleus | other | 4277446 | 3654770.5 | 5260280 | O70576 | STAG3 | 4277446 | 809460 | 467342 | 3 |
| P70297 | STAM | signal transducing adaptor molecule (SH3 domain and ITAM motif) 1 | Cytoplasm | other | 4658900 | 2698254.5 | 4638304.5 | | | | | | |
| P70297 | STAM | signal transducing adaptor molecule (SH3 domain and ITAM motif) 1 | Cytoplasm | other | 2057994.875 | 2389494.75 | 2261273 | | | | | | |
| P70297 | STAM | signal transducing adaptor molecule (SH3 domain and ITAM motif) 1 | Cytoplasm | other | 1754342.75 | 1531405.75 | 1609890.125 | P70297 | STAM | 2823746 | 1208777 | 402926 | 9 |
| Q8CI59 | STEAP3 | STEAP family member 3, metalloreductase | Cytoplasm | transporter | 1.03E+07 | 1.91E+07 | 1.40E+07 | | | | | | |
| Q8CI59 | STEAP3 | STEAP family member 3, metalloreductase | Cytoplasm | transporter | 7032235.5 | 1.11E+07 | 7032128 | | | | | | |
| Q8CI59 | STEAP3 | STEAP family member 3, metalloreductase | Cytoplasm | transporter | 4395103 | 9139252 | 6399739 | | | | | | |
| Q8CI59 | STEAP3 | STEAP family member 3, metalloreductase | Cytoplasm | transporter | 3657380.75 | 6862753 | 4632861 | | | | | | |
| Q8CI59 | STEAP3 | STEAP family member 3, metalloreductase | Cytoplasm | transporter | 3538216.25 | 5011548.5 | 3368555.5 | Q8CI59 | STEAP3 | 5782587 | 4424303 | 1142350 | 15 |
| P70302 | STIM1 | stromal interaction molecule 1 | Plasma Membrane | ion channel | 248348.969 | 490952.031 | 1217907.375 | | | | | | |
| P70302 | STIM1 | stromal interaction molecule 1 | Plasma Membrane | ion channel | 488679.531 | 837635.438 | 1395464.125 | | | | | | |
| P70302 | STIM1 | stromal interaction molecule 1 | Plasma Membrane | ion channel | 327124.594 | 412396 | 497001.688 | P70302 | STIM1 | 354718 | 404617 | 134872 | 9 |

| | | | | | | | | | | | | | |
|---|---|---|---|---|---|---|---|---|---|---|---|---|---|
| Q60864 | STIP1 | stress-induced phosphoprotein 1 | Cytoplasm | other | 8545152 | 7071861.5 | 8418693 | | | | | | |
| Q60864 | STIP1 | stress-induced phosphoprotein 1 | Cytoplasm | other | 7864736.5 | 9361878 | 8327183 | | | | | | |
| Q60864 | STIP1 | stress-induced phosphoprotein 1 | Cytoplasm | other | 9126361 | 5305648.5 | 3589947 | | | | | | |
| Q60864 | STIP1 | stress-induced phosphoprotein 1 | Cytoplasm | other | 5460821.5 | 4497009 | 4135718.75 | | | | | | |
| Q60864 | STIP1 | stress-induced phosphoprotein 1 | Cytoplasm | other | 6387794.5 | 4400830.5 | 5165109 | | | | | | |
| Q60864 | STIP1 | stress-induced phosphoprotein 1 | Cytoplasm | other | 2816061.75 | 2188233 | 2170517.25 | | | | | | |
| Q60864 | STIP1 | stress-induced phosphoprotein 1 | Cytoplasm | other | 3249043 | 2309056.5 | 2332166.75 | | | | | | |
| Q60864 | STIP1 | stress-induced phosphoprotein 1 | Cytoplasm | other | 3127577.25 | 2470676.25 | 2501296.75 | | | | | | |
| Q60864 | STIP1 | stress-induced phosphoprotein 1 | Cytoplasm | other | 2327255.5 | 1711950.375 | 2496270.75 | | | | | | |
| Q60864 | STIP1 | stress-induced phosphoprotein 1 | Cytoplasm | other | 3668129.25 | 3408812.25 | 1749005.875 | Q60864 | STIP1 | 5257293 | 2466837 | 450381 | 30 |
| P54116 | STOM | stomatin | Plasma Membrane | other | 4.95E+08 | 4.57E+08 | 5.95E+08 | | | | | | |
| P54116 | STOM | stomatin | Plasma Membrane | other | 5.26E+08 | 5.20E+08 | 6.66E+08 | | | | | | |
| P54116 | STOM | stomatin | Plasma Membrane | other | 4.59E+08 | 4.55E+08 | 5.08E+08 | | | | | | |
| P54116 | STOM | stomatin | Plasma Membrane | other | 4.66E+08 | 4.44E+08 | 3.26E+08 | | | | | | |
| P54116 | STOM | stomatin | Plasma Membrane | other | 3.44E+08 | 3.23E+08 | 3.33E+08 | | | | | | |
| P54116 | STOM | stomatin | Plasma Membrane | other | 2.73E+08 | 2.77E+08 | 2.47E+08 | | | | | | |
| P54116 | STOM | stomatin | Plasma Membrane | other | 3.03E+08 | 3.95E+08 | 5.04E+08 | | | | | | |
| P54116 | STOM | stomatin | Plasma Membrane | other | 1.81E+08 | 1.50E+08 | 1.91E+08 | | | | | | |
| P54116 | STOM | stomatin | Plasma Membrane | other | 1.66E+08 | 1.61E+08 | 2.50E+08 | | | | | | |
| P54116 | STOM | stomatin | Plasma Membrane | other | 2.10E+08 | 2.87E+08 | 4.60E+08 | | | | | | |
| P54116 | STOM | stomatin | Plasma Membrane | other | 1.68E+08 | 8.04E+08 | 1.67E+09 | | | | | | |
| P54116 | STOM | stomatin | Plasma Membrane | other | 7.29E+07 | 8.19E+07 | 1.10E+08 | | | | | | |
| P54116 | STOM | stomatin | Plasma Membrane | other | 2.35E+07 | 1.67E+07 | 1.67E+07 | | | | | | |
| P54116 | STOM | stomatin | Plasma Membrane | other | 2.86E+07 | 2.03E+07 | 2.45E+07 | | | | | | |
| P54116 | STOM | stomatin | Plasma Membrane | other | 9585677 | 5894443.5 | 5921640.5 | | | | | | |
| P54116 | STOM | stomatin | Plasma Membrane | other | 7104804 | 4638019 | 5839551 | | | | | | |
| P54116 | STOM | stomatin | Plasma Membrane | other | 4308064 | 4977668 | 5102579 | | | | | | |
| P54116 | STOM | stomatin | Plasma Membrane | other | 3500299.5 | 1917885 | 2424331 | | | | | | |
| P54116 | STOM | stomatin | Plasma Membrane | other | 1242369.625 | 688952.562 | 186133.016 | | | | | | |
| P54116 | STOM | stomatin | Plasma Membrane | other | 1124456.125 | 2950604 | 4947848.5 | | | | | | |
| P54116 | STOM | stomatin | Plasma Membrane | other | 2498718.75 | 4764975.5 | 6780307.5 | | | | | | |
| P54116 | STOM | stomatin | Plasma Membrane | other | 478528.594 | 506598.719 | 924585 | | | | | | |
| P54116 | STOM | stomatin | Plasma Membrane | other | 376722.938 | 414418.375 | 369554.844 | P54116 | STOM | 162786506 | 276465551 | 33282541 | 69 |
| Q99JB2 | STOML2 | stomatin (EPB72)-like 2 | Plasma Membrane | other | 3549167 | 6131303 | 5178948 | | | | | | |
| Q99JB2 | STOML2 | stomatin (EPB72)-like 2 | Plasma Membrane | other | 2782028.75 | 5153306 | 2571440.25 | | | | | | |
| Q99JB2 | STOML2 | stomatin (EPB72)-like 2 | Plasma Membrane | other | 2430965.5 | 6174106 | 2349621.25 | | | | | | |
| Q99JB2 | STOML2 | stomatin (EPB72)-like 2 | Plasma Membrane | other | 1920389.75 | 3928071.25 | 1321564.75 | | | | | | |
| Q99JB2 | STOML2 | stomatin (EPB72)-like 2 | Plasma Membrane | other | 1166153 | 2400950.5 | 1111979.625 | Q99JB2 | STOML2 | 2369741 | 1732147 | 447238 | 15 |
| Q9D3G5 | STX11 | syntaxin 11 | Plasma Membrane | transporter | 1490452 | 2003788.625 | 2588248.75 | | | | | | |
| Q9D3G5 | STX11 | syntaxin 11 | Plasma Membrane | transporter | 1021453 | 926594.625 | 1046672.875 | | | | | | |
| Q9D3G5 | STX11 | syntaxin 11 | Plasma Membrane | transporter | 775525.625 | 392271.375 | 162497.688 | | | | | | |
| Q9D3G5 | STX11 | syntaxin 11 | Plasma Membrane | transporter | 727227.688 | 646407 | 1359384.125 | | | | | | |
| Q9D3G5 | STX11 | syntaxin 11 | Plasma Membrane | transporter | 330103.844 | 1823314.25 | 2896460.25 | | | | | | |
| Q9D3G5 | STX11 | syntaxin 11 | Plasma Membrane | transporter | 245312.672 | 438430.281 | 620141 | Q9D3G5 | STX11 | 765012 | 799145 | 188360 | 18 |
| Q9ER00 | STX12 | syntaxin 12 | Cytoplasm | other | 1.35E+07 | 1.62E+07 | 1.37E+07 | Q9ER00 | STX12 | 13470000 | 1510894 | 872315 | 3 |
| Q00262 | STX2 | syntaxin 2 | Cytoplasm | transporter | 2950143.5 | 5613546 | 7266524.5 | | | | | | |
| Q00262 | STX2 | syntaxin 2 | Cytoplasm | transporter | 5448174 | 2125851.75 | 42638.586 | | | | | | |
| Q00262 | STX2 | syntaxin 2 | Cytoplasm | transporter | 3621534.75 | 6730870 | 5616077.5 | | | | | | |
| Q00262 | STX2 | syntaxin 2 | Cytoplasm | transporter | 2931795 | 4225297 | 2514082.25 | | | | | | |
| Q00262 | STX2 | syntaxin 2 | Cytoplasm | transporter | 1747632.875 | 1907938.875 | 1658295.5 | | | | | | |
| Q00262 | STX2 | syntaxin 2 | Cytoplasm | transporter | 1369694.625 | 1472423.75 | 1173095 | Q00262 | STX2 | 3011496 | 2103759 | 495861 | 18 |
| P70452 | STX4 | syntaxin 4 | Plasma Membrane | transporter | 1954901.875 | 3056910 | 2986095.75 | | | | | | |
| P70452 | STX4 | syntaxin 4 | Plasma Membrane | transporter | 1037418.062 | 1256530.875 | 1592276.625 | | | | | | |
| P70452 | STX4 | syntaxin 4 | Plasma Membrane | transporter | 1027186.062 | 1766326.125 | 1828811.5 | P70452 | STX4 | 1339835 | 751619 | 250540 | 9 |
| O70439 | STX7 | syntaxin 7 | Plasma Membrane | transporter | 1.29E+07 | 1.44E+07 | 1.40E+07 | | | | | | |
| O70439 | STX7 | syntaxin 7 | Plasma Membrane | transporter | 1.54E+07 | 1.46E+07 | 1.71E+07 | | | | | | |
| O70439 | STX7 | syntaxin 7 | Plasma Membrane | transporter | 6737157 | 1.14E+07 | 9036062 | | | | | | |
| O70439 | STX7 | syntaxin 7 | Plasma Membrane | transporter | 8244175 | 9808062 | 6420062.5 | | | | | | |
| O70439 | STX7 | syntaxin 7 | Plasma Membrane | transporter | 7120408.5 | 1.02E+07 | 9049973 | | | | | | |
| O70439 | STX7 | syntaxin 7 | Plasma Membrane | transporter | 5208116 | 5119205 | 5140423 | O70439 | STX7 | 9276643 | 3861851 | 910247 | 18 |
| O08599 | STXBP1 | syntaxin binding protein 1 | Cytoplasm | transporter | 3332824.5 | 3326456.5 | 3551825.75 | | | | | | |
| O08599 | STXBP1 | syntaxin binding protein 1 | Cytoplasm | transporter | 832908.062 | 905835.312 | 611654.875 | O08599 | STXBP1 | 2082866 | 1440717 | 588170 | 6 |
| Q64324 | STXBP2 | syntaxin binding protein 2 | Plasma Membrane | transporter | 1.28E+07 | 1.40E+07 | 1.09E+07 | | | | | | |
| Q64324 | STXBP2 | syntaxin binding protein 2 | Plasma Membrane | transporter | 6190921 | 8771493 | 7082800.5 | | | | | | |
| Q64324 | STXBP2 | syntaxin binding protein 2 | Plasma Membrane | transporter | 5751492.5 | 7846206 | 6093097.5 | | | | | | |
| Q64324 | STXBP2 | syntaxin binding protein 2 | Plasma Membrane | transporter | 5339036 | 6160189 | 6596105.5 | | | | | | |
| Q64324 | STXBP2 | syntaxin binding protein 2 | Plasma Membrane | transporter | 4954503.5 | 4798622.5 | 4867311 | | | | | | |
| Q64324 | STXBP2 | syntaxin binding protein 2 | Plasma Membrane | transporter | 3844748 | 5523966 | 5632422 | | | | | | |
| Q64324 | STXBP2 | syntaxin binding protein 2 | Plasma Membrane | transporter | 2779355.25 | 3180212 | 2869191 | | | | | | |
| Q64324 | STXBP2 | syntaxin binding protein 2 | Plasma Membrane | transporter | 2466730.5 | 3602952.25 | 3368665 | | | | | | |
| Q64324 | STXBP2 | syntaxin binding protein 2 | Plasma Membrane | transporter | 2063656.5 | 2339644 | 2414514 | | | | | | |
| Q64324 | STXBP2 | syntaxin binding protein 2 | Plasma Membrane | transporter | 2348751.75 | 2192396.75 | 955785.25 | | | | | | |
| Q64324 | STXBP2 | syntaxin binding protein 2 | Plasma Membrane | transporter | 2146772.75 | 2950422.25 | 2803698.75 | | | | | | |
| Q64324 | STXBP2 | syntaxin binding protein 2 | Plasma Membrane | transporter | 1506680.25 | 2375436.75 | 2662893.25 | | | | | | |
| Q64324 | STXBP2 | syntaxin binding protein 2 | Plasma Membrane | transporter | 691248.312 | 1470138.875 | 1289102.5 | | | | | | |
| Q64324 | STXBP2 | syntaxin binding protein 2 | Plasma Membrane | transporter | 1009686.875 | 1638236 | 2116073.75 | Q64324 | STXBP2 | 3848113 | 3082528 | 475644 | 42 |
| Q60770 | STXBP3 | syntaxin binding protein 3 | Plasma Membrane | transporter | 263992 | 560121.125 | 611466.688 | Q60770 | STXBP3 | 263992 | 187558 | 108287 | 3 |
| Q8K400 | STXBP5 | syntaxin binding protein 5 (tomosyn) | Plasma Membrane | other | 2.48E+07 | 2.79E+07 | 3.69E+07 | Q8K400 | STXBP5 | 24790000 | 6269314 | 3619590 | 3 |
| Q9Z2I9 | SUCLA2 | succinate-CoA ligase, ADP-forming, beta subunit | Cytoplasm | enzyme | 2629651.75 | 6759396 | 3355122.5 | Q9Z2I9 | SUCLA2 | 2629652 | 2204925 | 1273014 | 3 |
| Q9WUM5 | SUCLG1 | succinate-CoA ligase, alpha subunit | Cytoplasm | enzyme | 3471695 | 4623678.5 | 2892341.5 | | | | | | |
| Q9WUM5 | SUCLG1 | succinate-CoA ligase, alpha subunit | Cytoplasm | enzyme | 2352902.5 | 3734359.75 | 1890102.5 | | | | | | |
| Q9WUM5 | SUCLG1 | succinate-CoA ligase, alpha subunit | Cytoplasm | enzyme | 1339587 | 2131727.25 | 1081776.75 | Q9WUM5 | SUCLG1 | 2388062 | 1167462 | 389154 | 9 |
| Q9Z2I8 | SUCLG2 | succinate-CoA ligase, GDP-forming, beta subunit | Cytoplasm | enzyme | 650751.688 | 1266859.125 | 466948.906 | Q9Z2I8 | SUCLG2 | 650752 | 418973 | 241894 | 3 |
| Q64310 | SURF4 | surfeit 4 | Cytoplasm | other | 1452349.125 | 2448309.25 | 3946792 | | | | | | |
| Q64310 | SURF4 | surfeit 4 | Cytoplasm | other | 632454.812 | 1728171.875 | 2118362.5 | Q64310 | SURF4 | 1042402 | 1116353 | 455749 | 6 |
| Q9DBX3 | SUSD2 | sushi domain containing 2 | Extracellular Space | other | 6804760 | 7396464.5 | 4432667 | | | | | | |
| Q9DBX3 | SUSD2 | sushi domain containing 2 | Extracellular Space | other | 3170117 | 4596430.5 | 3282670.25 | | | | | | |
| Q9DBX3 | SUSD2 | sushi domain containing 2 | Extracellular Space | other | 5350177 | 5638159.5 | 6869376 | Q9DBX3 | SUSD2 | 5108351 | 1544739 | 514913 | 9 |
| Q3UZP4 | SVIP | small VCP/p97-interacting protein | Cytoplasm | other | 2.32E+07 | 2.63E+07 | 2.88E+07 | Q3UZP4 | SVIP | 23240000 | 2775410 | 1602384 | 3 |
| O09117 | SYPL1 | synaptophysin-like 1 | Plasma Membrane | transporter | 1.76E+07 | 2.33E+07 | 2.99E+07 | | | | | | |
| O09117 | SYPL1 | synaptophysin-like 1 | Plasma Membrane | transporter | 1.44E+07 | 1.45E+07 | 2.06E+07 | | | | | | |

| | | | | | | | | | | | | | |
|---|---|---|---|---|---|---|---|---|---|---|---|---|---|
| O09117 | SYPL1 | synaptophysin-like 1 | Plasma Membrane | transporter | 1035516.125 | 1130636.375 | 1516196.375 | O09117 | SYPL1 | 11005172 | 10506584 | 3502195 | 9 |
| B2RWW0 | TAGAP | T-cell activation RhoGTPase activating protein | Cytoplasm | other | 7708534 | 9425603 | 1.05E+07 | B2RWW0 | TAGAP | 7708534 | 1421776 | 820863 | 3 |
| Q9WVA4 | TAGLN2 | transgelin 2 | Cytoplasm | other | 531699.062 | 1861536.875 | 2192795 | | | | | | |
| Q9WVA4 | TAGLN2 | transgelin 2 | Cytoplasm | other | 175024.594 | 218432.609 | 2835152.25 | | | | | | |
| Q9WVA4 | TAGLN2 | transgelin 2 | Cytoplasm | other | 317149.031 | 798329.625 | 859877.812 | | | | | | |
| Q9WVA4 | TAGLN2 | transgelin 2 | Cytoplasm | other | 436403.75 | 1105282 | 957711.25 | Q9WVA4 | TAGLN2 | 365069 | 847116 | 244541 | 12 |
| Q8BYC6 | TAOK3 | TAO kinase 3 | Cytoplasm | kinase | 5524409 | 6841427 | 4870366 | | | | | | |
| Q8BYC6 | TAOK3 | TAO kinase 3 | Cytoplasm | kinase | 5931788.5 | 4225567 | 4606875.5 | | | | | | |
| Q8BYC6 | TAOK3 | TAO kinase 3 | Cytoplasm | kinase | 6238293.5 | 4354735.5 | 3117508 | | | | | | |
| Q8BYC6 | TAOK3 | TAO kinase 3 | Cytoplasm | kinase | 4850236.5 | 3667338.25 | 3056755.5 | | | | | | |
| Q8BYC6 | TAOK3 | TAO kinase 3 | Cytoplasm | kinase | 3332076.75 | 2867948.75 | 2024732 | | | | | | |
| Q8BYC6 | TAOK3 | TAO kinase 3 | Cytoplasm | kinase | 3483721.5 | 2290014 | 2449100.5 | | | | | | |
| Q8BYC6 | TAOK3 | TAO kinase 3 | Cytoplasm | kinase | 2051600.375 | 2906047.75 | 2380432.5 | | | | | | |
| Q8BYC6 | TAOK3 | TAO kinase 3 | Cytoplasm | kinase | 721020.938 | 586014.938 | 600735.688 | Q8BYC6 | TAOK3 | 4016643 | 1728060 | 352739 | 24 |
| Q9R233 | TAPBP | TAP binding protein (tapasin) | Cytoplasm | transporter | 533909.375 | 692122.625 | 760388.938 | Q9R233 | TAPBP | 533909 | 116179 | 67076 | 3 |
| Q4VBD2 | TAPT1 | transmembrane anterior posterior transformation 1 | Plasma Membrane | G-protein coupled receptor | 949783.562 | 1968302.625 | 1560415.625 | Q4VBD2 | TAPT1 | 949784 | 512612 | 295956 | 3 |
| Q9D0R2 | TARS | threonyl-tRNA synthetase | Nucleus | enzyme | 7385802 | 6445420.5 | 8119464.5 | | | | | | |
| Q9D0R2 | TARS | threonyl-tRNA synthetase | Nucleus | enzyme | 2799514 | 1449420 | 2217869.25 | Q9D0R2 | TARS | 5092658 | 2908057 | 1187209 | 6 |
| Q8BHL3 | TBC1D10B | TBC1 domain family, member 10B | Plasma Membrane | enzyme | 3971436.25 | 2232903.25 | 827961.5 | Q8BHL3 | TBC1D10B | 3971436 | 1574685 | 909145 | 3 |
| Q3UUG6 | TBC1D24 | TBC1 domain family, member 24 | Cytoplasm | other | 7625115.5 | 8730706 | 6416936 | | | | | | |
| Q3UUG6 | TBC1D24 | TBC1 domain family, member 24 | Cytoplasm | other | 3077327 | 2806489.25 | 2185550.25 | Q3UUG6 | TBC1D24 | 5351221 | 2797435 | 1142048 | 6 |
| P36423 | TBXAS1 | thromboxane A synthase 1 (platelet) | Plasma Membrane | enzyme | 1145655.625 | 2177157 | 5433530.5 | | | | | | |
| P36423 | TBXAS1 | thromboxane A synthase 1 (platelet) | Plasma Membrane | enzyme | 1039268.062 | 2916441.25 | 6599609 | | | | | | |
| P36423 | TBXAS1 | thromboxane A synthase 1 (platelet) | Plasma Membrane | enzyme | 90374.977 | 985240.312 | 3250159.75 | | | | | | |
| P36423 | TBXAS1 | thromboxane A synthase 1 (platelet) | Plasma Membrane | enzyme | 410047.188 | 1148261.625 | 2190897.25 | | | | | | |
| P36423 | TBXAS1 | thromboxane A synthase 1 (platelet) | Plasma Membrane | enzyme | 350915.562 | 1361387.125 | 1346846.375 | | | | | | |
| P36423 | TBXAS1 | thromboxane A synthase 1 (platelet) | Plasma Membrane | enzyme | 652263.75 | 1400734.5 | 3238419 | | | | | | |
| P36423 | TBXAS1 | thromboxane A synthase 1 (platelet) | Plasma Membrane | enzyme | 1358432.75 | 3906539.5 | 6187136 | | | | | | |
| P36423 | TBXAS1 | thromboxane A synthase 1 (platelet) | Plasma Membrane | enzyme | 363500.719 | 746415.938 | 1572729 | | | | | | |
| P36423 | TBXAS1 | thromboxane A synthase 1 (platelet) | Plasma Membrane | enzyme | 400526.906 | 1501736.625 | 2698969.25 | P36423 | TBXAS1 | 645665 | 1771131 | 340854 | 27 |
| P10711 | TCEA1 | transcription elongation factor A (SII), 1 | Nucleus | transcription regulator | 1.85E+07 | 1.97E+07 | 1.82E+07 | | | | | | |
| P10711 | TCEA1 | transcription elongation factor A (SII), 1 | Nucleus | transcription regulator | 1.97E+07 | 2.27E+07 | 1.94E+07 | | | | | | |
| P10711 | TCEA1 | transcription elongation factor A (SII), 1 | Nucleus | transcription regulator | 1.60E+07 | 1.24E+07 | 1.45E+07 | | | | | | |
| P10711 | TCEA1 | transcription elongation factor A (SII), 1 | Nucleus | transcription regulator | 1.21E+07 | 1.01E+07 | 1.21E+07 | | | | | | |
| P10711 | TCEA1 | transcription elongation factor A (SII), 1 | Nucleus | transcription regulator | 9976023 | 9105020 | 1.07E+07 | | | | | | |
| P10711 | TCEA1 | transcription elongation factor A (SII), 1 | Nucleus | transcription regulator | 9609932 | 1.03E+07 | 1.10E+07 | | | | | | |
| P10711 | TCEA1 | transcription elongation factor A (SII), 1 | Nucleus | transcription regulator | 5703666.5 | 7890312 | 9284073 | | | | | | |
| P10711 | TCEA1 | transcription elongation factor A (SII), 1 | Nucleus | transcription regulator | 7018427.5 | 4736985.5 | 5239635 | | | | | | |
| P10711 | TCEA1 | transcription elongation factor A (SII), 1 | Nucleus | transcription regulator | 8602089 | 8402432 | 1.07E+07 | | | | | | |
| P10711 | TCEA1 | transcription elongation factor A (SII), 1 | Nucleus | transcription regulator | 7460932 | 5549230 | 5874300.5 | | | | | | |
| P10711 | TCEA1 | transcription elongation factor A (SII), 1 | Nucleus | transcription regulator | 3380257.75 | 2392451.5 | 1289933.25 | | | | | | |
| P10711 | TCEA1 | transcription elongation factor A (SII), 1 | Nucleus | transcription regulator | 1333149.375 | 1576680.5 | 1774892.125 | P10711 | TCEA1 | 9947040 | 5785033 | 964172 | 36 |
| Q9QVN7 | TCEA2 | transcription elongation factor A (SII), 2 | Nucleus | transcription regulator | 807087.375 | 1276509.75 | 1739855.5 | Q9QVN7 | TCEA2 | 807087 | 466387 | 269269 | 3 |
| P11983 | TCP1 | t-complex 1 | Cytoplasm | other | 3.47E+07 | 2.84E+07 | 2.90E+07 | | | | | | |
| P11983 | TCP1 | t-complex 1 | Cytoplasm | other | 3.01E+07 | 1.89E+07 | 1.95E+07 | | | | | | |
| P11983 | TCP1 | t-complex 1 | Cytoplasm | other | 1.29E+07 | 1.12E+07 | 1.26E+07 | | | | | | |
| P11983 | TCP1 | t-complex 1 | Cytoplasm | other | 1.38E+07 | 9539990 | 1.30E+07 | | | | | | |
| P11983 | TCP1 | t-complex 1 | Cytoplasm | other | 1.53E+07 | 1.18E+07 | 1.63E+07 | | | | | | |
| P11983 | TCP1 | t-complex 1 | Cytoplasm | other | 9955739 | 8313145 | 1.16E+07 | | | | | | |
| P11983 | TCP1 | t-complex 1 | Cytoplasm | other | 9617897 | 8662131 | 1.01E+07 | | | | | | |
| P11983 | TCP1 | t-complex 1 | Cytoplasm | other | 1.09E+07 | 9015371 | 1.07E+07 | | | | | | |
| P11983 | TCP1 | t-complex 1 | Cytoplasm | other | 9627109 | 8905718 | 8568904 | | | | | | |
| P11983 | TCP1 | t-complex 1 | Cytoplasm | other | 9254799 | 5294019 | 7568156 | | | | | | |
| P11983 | TCP1 | t-complex 1 | Cytoplasm | other | 8250387 | 7239705.5 | 8909571 | | | | | | |
| P11983 | TCP1 | t-complex 1 | Cytoplasm | other | 6633240.5 | 6606589.5 | 6147421 | | | | | | |
| P11983 | TCP1 | t-complex 1 | Cytoplasm | other | 6489619 | 6089840 | 6179793.5 | | | | | | |
| P11983 | TCP1 | t-complex 1 | Cytoplasm | other | 5093739.5 | 3889594.75 | 4062300.5 | | | | | | |
| P11983 | TCP1 | t-complex 1 | Cytoplasm | other | 4628726.5 | 1311718 | 200237.25 | | | | | | |
| P11983 | TCP1 | t-complex 1 | Cytoplasm | other | 3971379.5 | 3042972 | 4173305 | | | | | | |
| P11983 | TCP1 | t-complex 1 | Cytoplasm | other | 4100286.25 | 1998460.375 | 1838850.5 | | | | | | |
| P11983 | TCP1 | t-complex 1 | Cytoplasm | other | 3534238.25 | 2698284.25 | 2637337.25 | | | | | | |
| P11983 | TCP1 | t-complex 1 | Cytoplasm | other | 2610809.5 | 2174500.25 | 2389294 | | | | | | |
| P11983 | TCP1 | t-complex 1 | Cytoplasm | other | 911863.812 | 1190357.25 | 2112994.75 | | | | | | |
| P11983 | TCP1 | t-complex 1 | Cytoplasm | other | 498293.406 | 372273.188 | 235243.734 | P11983 | TCP1 | 9663244 | 7400040 | 932317 | 63 |
| Q921I1 | TF | transferrin | Extracellular Space | transporter | 5.74E+07 | 9.22E+07 | 2.98E+07 | | | | | | |
| Q921I1 | TF | transferrin | Extracellular Space | transporter | 2.84E+07 | 4.67E+07 | 2.17E+07 | | | | | | |
| Q921I1 | TF | transferrin | Extracellular Space | transporter | 4.89E+07 | 9.17E+07 | 3.52E+07 | | | | | | |
| Q921I1 | TF | transferrin | Extracellular Space | transporter | 3.85E+07 | 6.15E+07 | 2.84E+07 | | | | | | |
| Q921I1 | TF | transferrin | Extracellular Space | transporter | 2.84E+07 | 6.57E+07 | 2.68E+07 | | | | | | |
| Q921I1 | TF | transferrin | Extracellular Space | transporter | 3.41E+07 | 5.87E+07 | 2.68E+07 | | | | | | |
| Q921I1 | TF | transferrin | Extracellular Space | transporter | 3.14E+07 | 6.50E+07 | 2.94E+07 | | | | | | |
| Q921I1 | TF | transferrin | Extracellular Space | transporter | 3.07E+07 | 6.40E+07 | 2.75E+07 | | | | | | |
| Q921I1 | TF | transferrin | Extracellular Space | transporter | 2.92E+07 | 5.08E+07 | 2.43E+07 | | | | | | |
| Q921I1 | TF | transferrin | Extracellular Space | transporter | 2.98E+07 | 5.08E+07 | 2.24E+07 | | | | | | |
| Q921I1 | TF | transferrin | Extracellular Space | transporter | 1.72E+07 | 4.58E+07 | 2.33E+07 | | | | | | |
| Q921I1 | TF | transferrin | Extracellular Space | transporter | 2.16E+07 | 5.12E+07 | 2.85E+07 | | | | | | |
| Q921I1 | TF | transferrin | Extracellular Space | transporter | 2.03E+07 | 3.27E+07 | 1.40E+07 | | | | | | |
| Q921I1 | TF | transferrin | Extracellular Space | transporter | 2.74E+07 | 3.66E+07 | 1.68E+07 | | | | | | |
| Q921I1 | TF | transferrin | Extracellular Space | transporter | 1.64E+07 | 2.14E+07 | 9939735 | | | | | | |
| Q921I1 | TF | transferrin | Extracellular Space | transporter | 1.60E+07 | 3.56E+07 | 1.23E+07 | | | | | | |
| Q921I1 | TF | transferrin | Extracellular Space | transporter | 1.10E+07 | 2.09E+07 | 9759314 | | | | | | |
| Q921I1 | TF | transferrin | Extracellular Space | transporter | 7992475.5 | 1.24E+07 | 6409461.5 | | | | | | |
| Q921I1 | TF | transferrin | Extracellular Space | transporter | 7321669 | 2.00E+07 | 8744193 | | | | | | |
| Q921I1 | TF | transferrin | Extracellular Space | transporter | 9926999 | 1.89E+07 | 1.02E+07 | | | | | | |
| Q921I1 | TF | transferrin | Extracellular Space | transporter | 1.18E+07 | 1.40E+07 | 4972478.5 | | | | | | |
| Q921I1 | TF | transferrin | Extracellular Space | transporter | 6520170.5 | 2.11E+07 | 1.49E+07 | | | | | | |
| Q921I1 | TF | transferrin | Extracellular Space | transporter | 8010957.5 | 1.40E+07 | 5558857 | | | | | | |
| Q921I1 | TF | transferrin | Extracellular Space | transporter | 5272451.5 | 1.22E+07 | 3463139 | | | | | | |
| Q921I1 | TF | transferrin | Extracellular Space | transporter | 9301494 | 2.61E+07 | 8210823.5 | | | | | | |
| Q921I1 | TF | transferrin | Extracellular Space | transporter | 9960418 | 1.71E+07 | 6498712 | | | | | | |
| Q921I1 | TF | transferrin | Extracellular Space | transporter | 5694215.5 | 1.15E+07 | 5865232.5 | | | | | | |

| | | | | | | | | | | | | | |
|---|---|---|---|---|---|---|---|---|---|---|---|---|---|
| Q921I1 | TF | transferrin | Extracellular Space | transporter | 4460377 | 1.48E+07 | 1.30E+07 | | | | | | |
| Q921I1 | TF | transferrin | Extracellular Space | transporter | 4706179.5 | 3745181 | 1209803.5 | | | | | | |
| Q921I1 | TF | transferrin | Extracellular Space | transporter | 6622060.5 | 8129758 | 3348027.75 | | | | | | |
| Q921I1 | TF | transferrin | Extracellular Space | transporter | 5748148.5 | 8964530 | 3615730.25 | | | | | | |
| Q921I1 | TF | transferrin | Extracellular Space | transporter | 1.07E+07 | 3.22E+07 | 1.91E+07 | | | | | | |
| Q921I1 | TF | transferrin | Extracellular Space | transporter | 4338861.5 | 5551160 | 411433.375 | | | | | | |
| Q921I1 | TF | transferrin | Extracellular Space | transporter | 2445827.75 | 3895645.25 | 1567340.375 | | | | | | |
| Q921I1 | TF | transferrin | Extracellular Space | transporter | 476077.344 | 3183726.25 | 2729843.25 | Q921I1 | TF | 17365668 | 19015896 | 1855761 | 105 |
| Q62351 | TFRC | transferrin receptor | Plasma Membrane | transporter | 6.42E+07 | 1.30E+08 | 5.00E+07 | | | | | | |
| Q62351 | TFRC | transferrin receptor | Plasma Membrane | transporter | 7.71E+07 | 1.66E+08 | 6.49E+07 | | | | | | |
| Q62351 | TFRC | transferrin receptor | Plasma Membrane | transporter | 4.78E+07 | 9.52E+07 | 4.22E+07 | | | | | | |
| Q62351 | TFRC | transferrin receptor | Plasma Membrane | transporter | 5.94E+07 | 1.55E+08 | 7.37E+07 | | | | | | |
| Q62351 | TFRC | transferrin receptor | Plasma Membrane | transporter | 5.52E+07 | 1.42E+08 | 5.28E+07 | | | | | | |
| Q62351 | TFRC | transferrin receptor | Plasma Membrane | transporter | 6.54E+07 | 1.36E+08 | 5.67E+07 | | | | | | |
| Q62351 | TFRC | transferrin receptor | Plasma Membrane | transporter | 5.31E+07 | 1.06E+08 | 4.78E+07 | | | | | | |
| Q62351 | TFRC | transferrin receptor | Plasma Membrane | transporter | 5.60E+07 | 1.21E+08 | 5.12E+07 | | | | | | |
| Q62351 | TFRC | transferrin receptor | Plasma Membrane | transporter | 5.79E+07 | 1.22E+08 | 5.23E+07 | | | | | | |
| Q62351 | TFRC | transferrin receptor | Plasma Membrane | transporter | 5.13E+07 | 1.05E+08 | 4.30E+07 | | | | | | |
| Q62351 | TFRC | transferrin receptor | Plasma Membrane | transporter | 5.20E+07 | 1.06E+08 | 4.90E+07 | | | | | | |
| Q62351 | TFRC | transferrin receptor | Plasma Membrane | transporter | 4.34E+07 | 8.94E+07 | 4.19E+07 | | | | | | |
| Q62351 | TFRC | transferrin receptor | Plasma Membrane | transporter | 5.27E+07 | 8.69E+07 | 1.86E+07 | | | | | | |
| Q62351 | TFRC | transferrin receptor | Plasma Membrane | transporter | 4.06E+07 | 7.29E+07 | 3.34E+07 | | | | | | |
| Q62351 | TFRC | transferrin receptor | Plasma Membrane | transporter | 2.66E+07 | 5.41E+07 | 2.12E+07 | | | | | | |
| Q62351 | TFRC | transferrin receptor | Plasma Membrane | transporter | 3.82E+07 | 8.46E+07 | 3.66E+07 | | | | | | |
| Q62351 | TFRC | transferrin receptor | Plasma Membrane | transporter | 3.69E+07 | 9.10E+07 | 3.86E+07 | | | | | | |
| Q62351 | TFRC | transferrin receptor | Plasma Membrane | transporter | 3.63E+07 | 8.49E+07 | 3.52E+07 | | | | | | |
| Q62351 | TFRC | transferrin receptor | Plasma Membrane | transporter | 3.39E+07 | 6.93E+07 | 3.42E+07 | | | | | | |
| Q62351 | TFRC | transferrin receptor | Plasma Membrane | transporter | 2.63E+07 | 4.83E+07 | 1.90E+07 | | | | | | |
| Q62351 | TFRC | transferrin receptor | Plasma Membrane | transporter | 2.68E+07 | 4.88E+07 | 2.30E+07 | | | | | | |
| Q62351 | TFRC | transferrin receptor | Plasma Membrane | transporter | 3.22E+07 | 7.56E+07 | 3.51E+07 | | | | | | |
| Q62351 | TFRC | transferrin receptor | Plasma Membrane | transporter | 2.05E+07 | 6.34E+07 | 2.39E+07 | | | | | | |
| Q62351 | TFRC | transferrin receptor | Plasma Membrane | transporter | 1.62E+07 | 3.16E+07 | 1.30E+07 | | | | | | |
| Q62351 | TFRC | transferrin receptor | Plasma Membrane | transporter | 1.05E+07 | 2.00E+07 | 1.50E+07 | | | | | | |
| Q62351 | TFRC | transferrin receptor | Plasma Membrane | transporter | 1.06E+07 | 2.63E+07 | 1.36E+07 | | | | | | |
| Q62351 | TFRC | transferrin receptor | Plasma Membrane | transporter | 1.47E+07 | 2.84E+07 | 1.17E+07 | | | | | | |
| Q62351 | TFRC | transferrin receptor | Plasma Membrane | transporter | 1.61E+07 | 2.81E+07 | 1.18E+07 | | | | | | |
| Q62351 | TFRC | transferrin receptor | Plasma Membrane | transporter | 6567488 | 3.32E+07 | 2.28E+07 | | | | | | |
| Q62351 | TFRC | transferrin receptor | Plasma Membrane | transporter | 4203581 | 8361968.5 | 3813969.5 | | | | | | |
| Q62351 | TFRC | transferrin receptor | Plasma Membrane | transporter | 3722461.5 | 7339699.5 | 3063723 | | | | | | |
| Q62351 | TFRC | transferrin receptor | Plasma Membrane | transporter | 2310438.75 | 4594391 | 2247024.25 | | | | | | |
| Q62351 | TFRC | transferrin receptor | Plasma Membrane | transporter | 1803300.125 | 3010421.25 | 1884830.5 | | | | | | |
| Q62351 | TFRC | transferrin receptor | Plasma Membrane | transporter | 2318542.25 | 5720937 | 1203352.75 | | | | | | |
| Q62351 | TFRC | transferrin receptor | Plasma Membrane | transporter | 1319961.375 | 4426362 | 2388383.5 | | | | | | |
| Q62351 | TFRC | transferrin receptor | Plasma Membrane | transporter | 1347771.75 | 5656664.5 | 3240797.5 | | | | | | |
| Q62351 | TFRC | transferrin receptor | Plasma Membrane | transporter | 103382.984 | 143858.688 | 112595.812 | Q62351 | TFRC | 30963160 | 37350153 | 3545120 | 111 |
| Q62312 | TGFBR2 | transforming growth factor, beta receptor II (70/80kDa) | Plasma Membrane | kinase | 1.39E+07 | 1.98E+07 | 2.06E+07 | Q62312 | TGFBR2 | 13920000 | 3660342 | 2113299 | 3 |
| P21981 | TGM2 | transglutaminase 2 | Cytoplasm | enzyme | 7175283 | 5579473 | 3869522 | | | | | | |
| P21981 | TGM2 | transglutaminase 2 | Cytoplasm | enzyme | 1779123.125 | 2690501 | 2280378.25 | P21981 | TGM2 | 4477203 | 2103954 | 858936 | 6 |
| P35441 | THBS1 | thrombospondin 1 | Extracellular Space | other | 2034084.75 | 8710481 | 1.45E+07 | | | | | | |
| P35441 | THBS1 | thrombospondin 1 | Extracellular Space | other | 1078262.125 | 5571216 | 9012155 | | | | | | |
| P35441 | THBS1 | thrombospondin 1 | Extracellular Space | other | 619057.188 | 3417923.75 | 6745747 | | | | | | |
| P35441 | THBS1 | thrombospondin 1 | Extracellular Space | other | 1155258.75 | 6582299.5 | 1.10E+07 | | | | | | |
| P35441 | THBS1 | thrombospondin 1 | Extracellular Space | other | 570881.688 | 3121499.75 | 4874971.5 | | | | | | |
| P35441 | THBS1 | thrombospondin 1 | Extracellular Space | other | 956770.188 | 2312039.75 | 3429119.5 | | | | | | |
| P35441 | THBS1 | thrombospondin 1 | Extracellular Space | other | 664712.688 | 2471151.75 | 3543245 | | | | | | |
| P35441 | THBS1 | thrombospondin 1 | Extracellular Space | other | 864164.938 | 4091584.25 | 7068927 | | | | | | |
| P35441 | THBS1 | thrombospondin 1 | Extracellular Space | other | 506656.062 | 4138354.25 | 6507703 | | | | | | |
| P35441 | THBS1 | thrombospondin 1 | Extracellular Space | other | 738306.5 | 3273720.25 | 5127807.5 | | | | | | |
| P35441 | THBS1 | thrombospondin 1 | Extracellular Space | other | 436881.688 | 2143945.5 | 3462684.75 | | | | | | |
| P35441 | THBS1 | thrombospondin 1 | Extracellular Space | other | 472329.312 | 885511.438 | 1250502.625 | | | | | | |
| P35441 | THBS1 | thrombospondin 1 | Extracellular Space | other | 1992844.125 | 1.13E+07 | 1.23E+07 | | | | | | |
| P35441 | THBS1 | thrombospondin 1 | Extracellular Space | other | 515371.219 | 3782706.75 | 6146049 | | | | | | |
| P35441 | THBS1 | thrombospondin 1 | Extracellular Space | other | 286450.875 | 1548209.625 | 3357546 | | | | | | |
| P35441 | THBS1 | thrombospondin 1 | Extracellular Space | other | 255136.594 | 2375196.5 | 2090906.75 | | | | | | |
| P35441 | THBS1 | thrombospondin 1 | Extracellular Space | other | 136952.047 | 804983.062 | 1099911 | | | | | | |
| P35441 | THBS1 | thrombospondin 1 | Extracellular Space | other | 373267.281 | 2996705.25 | 3894903.5 | | | | | | |
| P35441 | THBS1 | thrombospondin 1 | Extracellular Space | other | 0 | 1127952.125 | 1761694.5 | | | | | | |
| P35441 | THBS1 | thrombospondin 1 | Extracellular Space | other | 21945.441 | 1116411.25 | 1252268.625 | | | | | | |
| P35441 | THBS1 | thrombospondin 1 | Extracellular Space | other | 175192.703 | 1020153.75 | 1317719.125 | | | | | | |
| P35441 | THBS1 | thrombospondin 1 | Extracellular Space | other | 96470.055 | 765956.625 | 1245902.875 | | | | | | |
| P35441 | THBS1 | thrombospondin 1 | Extracellular Space | other | 10569.437 | 2139643.5 | 3920024.75 | | | | | | |
| P35441 | THBS1 | thrombospondin 1 | Extracellular Space | other | 93548.961 | 740004.062 | 1855620.75 | | | | | | |
| P35441 | THBS1 | thrombospondin 1 | Extracellular Space | other | 16188.354 | 82723.406 | 78046.969 | | | | | | |
| P35441 | THBS1 | thrombospondin 1 | Extracellular Space | other | 76078.758 | 927152.5 | 1642584.25 | | | | | | |
| P35441 | THBS1 | thrombospondin 1 | Extracellular Space | other | 37672.055 | 305836.875 | 42335.871 | | | | | | |
| P35441 | THBS1 | thrombospondin 1 | Extracellular Space | other | 4.70E-06 | 913477.375 | 1125406.125 | | | | | | |
| P35441 | THBS1 | thrombospondin 1 | Extracellular Space | other | 132866.219 | 2212046.75 | 3613027 | | | | | | |
| P35441 | THBS1 | thrombospondin 1 | Extracellular Space | other | 198240.266 | 842562.562 | 1495935.5 | | | | | | |
| P35441 | THBS1 | thrombospondin 1 | Extracellular Space | other | 135056.203 | 1173014.25 | 1755733.125 | | | | | | |
| P35441 | THBS1 | thrombospondin 1 | Extracellular Space | other | 111252.703 | 366248.562 | 556326.688 | P35441 | THBS1 | 461327 | 2912090 | 297214 | 96 |
| Q80ZW2 | THEM6 | thioesterase superfamily member 6 | Extracellular Space | other | 3440644.5 | 5167382 | 5608698 | | | | | | |
| Q80ZW2 | THEM6 | thioesterase superfamily member 6 | Extracellular Space | other | 3133575.25 | 2769901.75 | 4111158 | | | | | | |
| Q80ZW2 | THEM6 | thioesterase superfamily member 6 | Extracellular Space | other | 1838259.5 | 1446415.125 | 2077749.875 | | | | | | |
| Q80ZW2 | THEM6 | thioesterase superfamily member 6 | Extracellular Space | other | 1892547.875 | 1826250.375 | 2258430.5 | | | | | | |
| Q80ZW2 | THEM6 | thioesterase superfamily member 6 | Extracellular Space | other | 1027419.938 | 1125478.25 | 1615010.75 | Q80ZW2 | THEM6 | 2266489 | 1413126 | 364867 | 15 |
| Q9CY52 | THG1L | tRNA-histidine guanylyltransferase 1-like (S. cerevisiae) | Cytoplasm | enzyme | 8028256.5 | 6230393.5 | 5601304 | | | | | | |
| Q9CY52 | THG1L | tRNA-histidine guanylyltransferase 1-like (S. cerevisiae) | Cytoplasm | enzyme | 1.09E+07 | 7796003.5 | 7548433.5 | | | | | | |
| Q9CY52 | THG1L | tRNA-histidine guanylyltransferase 1-like (S. cerevisiae) | Cytoplasm | enzyme | 7199929 | 3580626.5 | 3885765.5 | | | | | | |
| Q9CY52 | THG1L | tRNA-histidine guanylyltransferase 1-like (S. cerevisiae) | Cytoplasm | enzyme | 5419872.5 | 3909288.5 | 4122658.75 | | | | | | |
| Q9CY52 | THG1L | tRNA-histidine guanylyltransferase 1-like (S. cerevisiae) | Cytoplasm | enzyme | 5721193.5 | 4193583.5 | 4599855 | Q9CY52 | THG1L | 7459850 | 2060673 | 532063 | 15 |
| Q9D880 | TIMM50 | translocase of inner mitochondrial membrane 50 homolog (S. cerevisiae) | Cytoplasm | phosphatase | 2388208 | 5273450.5 | 1933044.625 | | | | | | |

| | | | | | | | | | | | | | |
|---|---|---|---|---|---|---|---|---|---|---|---|---|---|
| Q9D880 | TIMM50 | translocase of inner mitochondrial membrane 50 homolog (S. cerevisiae) | Cytoplasm | phosphatase | 1768835.5 | 3337190.5 | 1362093.5 | Q9D880 | TIMM50 | 2078522 | 1440001 | 587878 | 6 |
| P39876 | TIMP3 | TIMP metallopeptidase inhibitor 3 | Extracellular Space | other | 510921.25 | 2964447.5 | 4433202 | | | | | | |
| P39876 | TIMP3 | TIMP metallopeptidase inhibitor 3 | Extracellular Space | other | 98920.305 | 887067.562 | 1412369.375 | | | | | | |
| P39876 | TIMP3 | TIMP metallopeptidase inhibitor 3 | Extracellular Space | other | 668532.625 | 1240114.25 | 3101673.75 | | | | | | |
| P39876 | TIMP3 | TIMP metallopeptidase inhibitor 3 | Extracellular Space | other | 77538.812 | 119093.242 | 253255.531 | P39876 | TIMP3 | 338978 | 1429125 | 412553 | 12 |
| P40142 | TKT | transketolase | Cytoplasm | enzyme | 3096141.75 | 2837907.5 | 2732951.75 | | | | | | |
| P40142 | TKT | transketolase | Cytoplasm | enzyme | 101089.836 | 187002.719 | 1149831.25 | P40142 | TKT | 1598616 | 1375346 | 561483 | 6 |
| P26039 | TLN1 | talin 1 | Plasma Membrane | other | 423258.781 | 1268870.5 | 1495137 | | | | | | |
| P26039 | TLN1 | talin 1 | Plasma Membrane | other | 1696259.25 | 2849501.75 | 2228244.5 | | | | | | |
| P26039 | TLN1 | talin 1 | Plasma Membrane | other | 752031.062 | 926731.625 | 930095.062 | | | | | | |
| P26039 | TLN1 | talin 1 | Plasma Membrane | other | 641827.938 | 813694 | 587512.75 | | | | | | |
| P26039 | TLN1 | talin 1 | Plasma Membrane | other | 482455.594 | 1336040.625 | 1336347.25 | | | | | | |
| P26039 | TLN1 | talin 1 | Plasma Membrane | other | 786023 | 1245935.75 | 1208445.125 | | | | | | |
| P26039 | TLN1 | talin 1 | Plasma Membrane | other | 8.569 | 61215.742 | 424958.031 | | | | | | |
| P26039 | TLN1 | talin 1 | Plasma Membrane | other | 239840.562 | 328756.312 | 290871.969 | P26039 | TLN1 | 627713 | 683437 | 139506 | 24 |
| Q9ET30 | TM9SF3 | transmembrane 9 superfamily member 3 | Cytoplasm | transporter | 653859.25 | 984798.375 | 1475516.5 | Q9ET30 | TM9SF3 | 653859 | 413410 | 238682 | 3 |
| Q7TN60 | TMC6 | transmembrane channel-like 6 | Cytoplasm | transporter | 2192993 | 2720883.25 | 3583899 | | | | | | |
| Q7TN60 | TMC6 | transmembrane channel-like 6 | Cytoplasm | transporter | 1597248.25 | 1287223.875 | 1640542.875 | | | | | | |
| Q7TN60 | TMC6 | transmembrane channel-like 6 | Cytoplasm | transporter | 660254.375 | 681095.125 | 814348.188 | Q7TN60 | TMC6 | 1483499 | 995036 | 331679 | 9 |
| Q7TN58 | TMC8 | transmembrane channel-like 8 | Cytoplasm | other | 3128492.5 | 2914104.5 | 3012663.75 | | | | | | |
| Q7TN58 | TMC8 | transmembrane channel-like 8 | Cytoplasm | other | 1562776.375 | 1538769.875 | 2030885.375 | | | | | | |
| Q7TN58 | TMC8 | transmembrane channel-like 8 | Cytoplasm | other | 1305387 | 1849070.375 | 1504651.875 | Q7TN58 | TMC8 | 1998885 | 725464 | 241821 | 9 |
| Q80W04 | TMCC2 | transmembrane and coiled-coil domain family 2 | Extracellular Space | other | 8838930 | 1.14E+07 | 1.30E+07 | | | | | | |
| Q80W04 | TMCC2 | transmembrane and coiled-coil domain family 2 | Extracellular Space | other | 9581143 | 2.03E+07 | 1.28E+07 | | | | | | |
| Q80W04 | TMCC2 | transmembrane and coiled-coil domain family 2 | Extracellular Space | other | 5405240.5 | 5287541.5 | 3724620.5 | | | | | | |
| Q80W04 | TMCC2 | transmembrane and coiled-coil domain family 2 | Extracellular Space | other | 3276083.5 | 5198401.5 | 3327193.25 | | | | | | |
| Q80W04 | TMCC2 | transmembrane and coiled-coil domain family 2 | Extracellular Space | other | 3076733.75 | 4208057 | 3400288 | | | | | | |
| Q80W04 | TMCC2 | transmembrane and coiled-coil domain family 2 | Extracellular Space | other | 1338950.625 | 2434841.75 | 1507565.75 | Q80W04 | TMCC2 | 5252847 | 5066239 | 1194124 | 18 |
| Q921L3 | TMCO1 | transmembrane and coiled-coil domains 1 | Cytoplasm | other | 1931722 | 3143496.75 | 3950864 | Q921L3 | TMCO1 | 1931722 | 1016298 | 586760 | 3 |
| Q9D1D4 | TMED10 | transmembrane emp24-like trafficking protein 10 (yeast) | Cytoplasm | transporter | 1.61E+07 | 1.95E+07 | 2.10E+07 | | | | | | |
| Q9D1D4 | TMED10 | transmembrane emp24-like trafficking protein 10 (yeast) | Cytoplasm | transporter | 1.35E+07 | 1.57E+07 | 1.72E+07 | | | | | | |
| Q9D1D4 | TMED10 | transmembrane emp24-like trafficking protein 10 (yeast) | Cytoplasm | transporter | 1.50E+07 | 1.50E+07 | 2.00E+07 | | | | | | |
| Q9D1D4 | TMED10 | transmembrane emp24-like trafficking protein 10 (yeast) | Cytoplasm | transporter | 5161987.5 | 3324245.5 | 2097534.25 | | | | | | |
| Q9D1D4 | TMED10 | transmembrane emp24-like trafficking protein 10 (yeast) | Cytoplasm | transporter | 2766223 | 5306696.5 | 1.13E+07 | | | | | | |
| Q9D1D4 | TMED10 | transmembrane emp24-like trafficking protein 10 (yeast) | Cytoplasm | transporter | 4163306 | 4845300.5 | 5294145.5 | Q9D1D4 | TMED10 | 9450253 | 6724363 | 1584947 | 18 |
| Q9R0Q3 | TMED2 | transmembrane emp24 domain trafficking protein 2 | Cytoplasm | transporter | 7584000 | 7291181.5 | 9172636 | | | | | | |
| Q9R0Q3 | TMED2 | transmembrane emp24 domain trafficking protein 2 | Cytoplasm | transporter | 6206512 | 1.17E+07 | 1.25E+07 | | | | | | |
| Q9R0Q3 | TMED2 | transmembrane emp24 domain trafficking protein 2 | Cytoplasm | transporter | 2925525 | 5227217.5 | 5173791 | Q9R0Q3 | TMED2 | 5572012 | 3146847 | 1048949 | 9 |
| Q8R1V4 | TMED4 | transmembrane emp24 protein transport domain containing 4 | Cytoplasm | transporter | 9625231 | 1.40E+07 | 1.21E+07 | Q8R1V4 | TMED4 | 9625231 | 2174638 | 1255528 | 3 |
| Q9CXE7 | TMED5 | transmembrane emp24 protein transport domain containing 5 | Cytoplasm | other | 2222701.75 | 3357400 | 3351975.5 | Q9CXE7 | TMED5 | 2222702 | 653558 | 377332 | 3 |
| Q99KF1 | TMED9 | transmembrane emp24 protein transport domain containing 9 | Cytoplasm | transporter | 1.78E+07 | 1.87E+07 | 2.59E+07 | | | | | | |
| Q99KF1 | TMED9 | transmembrane emp24 protein transport domain containing 9 | Cytoplasm | transporter | 4189875 | 8808365 | 1.09E+07 | | | | | | |
| Q99KF1 | TMED9 | transmembrane emp24 protein transport domain containing 9 | Cytoplasm | transporter | 1584795.5 | 335533.906 | 432011.312 | Q99KF1 | TMED9 | 7854890 | 9216769 | 3072256 | 9 |
| Q80X71 | TMEM106B | transmembrane protein 106B | Cytoplasm | other | 602615.188 | 767849.688 | 671895.5 | Q80X71 | TMEM106B | 602615 | 82975 | 47906 | 3 |
| Q9CQN6 | TMEM14C | transmembrane protein 14C | Plasma Membrane | other | 9131730 | 1.53E+07 | 1.15E+07 | | | | | | |
| Q9CQN6 | TMEM14C | transmembrane protein 14C | Plasma Membrane | other | 2734300 | 5637812.5 | 4194358 | Q9CQN6 | TMEM14C | 5933015 | 4773711 | 1948859 | 6 |
| Q9CR64 | TMEM167A | transmembrane protein 167A | Extracellular Space | other | 2374265.5 | 2564739.75 | 3490323.5 | Q9CR64 | TMEM167A | 2374266 | 597016 | 344688 | 3 |
| Q9CY24 | TMEM179B | transmembrane protein 179B | Extracellular Space | other | 2854652 | 3184063.5 | 3121817.25 | Q9CY24 | TMEM179B | 2854652 | 175007 | 101040 | 3 |
| Q9D771 | TMEM206 | transmembrane protein 206 | Plasma Membrane | other | 7884521.5 | 7358304.5 | 1.54E+07 | | | | | | |
| Q9D771 | TMEM206 | transmembrane protein 206 | Plasma Membrane | other | 2170448 | 2553434.75 | 4653678.5 | | | | | | |
| Q9D771 | TMEM206 | transmembrane protein 206 | Plasma Membrane | other | 1563085.5 | 1512137.875 | 3525801 | Q9D771 | TMEM206 | 3872685 | 4508697 | 1502899 | 9 |
| Q5F285 | TMEM256 | transmembrane protein 256 | Cytoplasm | other | 1.83E+07 | 3.18E+07 | 1.45E+07 | Q5F285 | TMEM256 | 18270000 | 9105011 | 5256781 | 3 |
| Q8VEK0 | TMEM30A | transmembrane protein 30A | Cytoplasm | other | 1.80E+07 | 1.69E+07 | 1.71E+07 | | | | | | |
| Q8VEK0 | TMEM30A | transmembrane protein 30A | Cytoplasm | other | 1.35E+07 | 1.52E+07 | 1.79E+07 | | | | | | |
| Q8VEK0 | TMEM30A | transmembrane protein 30A | Cytoplasm | other | 9169902 | 7949843 | 1.00E+07 | | | | | | |
| Q8VEK0 | TMEM30A | transmembrane protein 30A | Cytoplasm | other | 8552831 | 7615289.5 | 9366144 | | | | | | |
| Q8VEK0 | TMEM30A | transmembrane protein 30A | Cytoplasm | other | 5163414 | 7051773.5 | 7424699.5 | Q8VEK0 | TMEM30A | 10869229 | 4512524 | 1165129 | 15 |
| Q9CZX7 | TMEM55A | transmembrane protein 55A | Cytoplasm | phosphatase | 2229310.25 | 3021856.75 | 3506492.75 | Q9CZX7 | TMEM55A | 2229310 | 644748 | 372245 | 3 |
| Q3TWL2 | TMEM55B | transmembrane protein 55B | Cytoplasm | phosphatase | 6790280.5 | 6289732.5 | 7021548.5 | | | | | | |
| Q3TWL2 | TMEM55B | transmembrane protein 55B | Cytoplasm | phosphatase | 4974284.5 | 5125765 | 4206489 | | | | | | |
| Q3TWL2 | TMEM55B | transmembrane protein 55B | Cytoplasm | phosphatase | 4524635 | 3616938.25 | 3865944.75 | | | | | | |
| Q3TWL2 | TMEM55B | transmembrane protein 55B | Cytoplasm | phosphatase | 4708686 | 3936837 | 4451313 | Q3TWL2 | TMEM55B | 5249472 | 1149017 | 331693 | 12 |
| Q91YT8 | TMEM63A | transmembrane protein 63A | Cytoplasm | other | 366063.656 | 1115744.125 | 445999.812 | Q91YT8 | TMEM63A | 366064 | 411697 | 237694 | 3 |
| Q9CR23 | TMEM9 | transmembrane protein 9 | Cytoplasm | other | 5496171 | 6802760.5 | 6028436 | Q9CR23 | TMEM9 | 5496171 | 657021 | 379331 | 3 |
| P49813 | TMOD1 | tropomodulin 1 | Cytoplasm | enzyme | 2.13E+08 | 3.28E+08 | 3.49E+08 | | | | | | |
| P49813 | TMOD1 | tropomodulin 1 | Cytoplasm | enzyme | 1.73E+08 | 2.01E+08 | 3.04E+08 | | | | | | |
| P49813 | TMOD1 | tropomodulin 1 | Cytoplasm | enzyme | 2.27E+08 | 1.94E+08 | 2.26E+08 | | | | | | |
| P49813 | TMOD1 | tropomodulin 1 | Cytoplasm | enzyme | 1.86E+08 | 1.96E+08 | 2.40E+08 | | | | | | |
| P49813 | TMOD1 | tropomodulin 1 | Cytoplasm | enzyme | 1.13E+08 | 2.25E+08 | 3.83E+08 | | | | | | |
| P49813 | TMOD1 | tropomodulin 1 | Cytoplasm | enzyme | 1.81E+08 | 1.77E+08 | 1.95E+08 | | | | | | |
| P49813 | TMOD1 | tropomodulin 1 | Cytoplasm | enzyme | 1.15E+08 | 1.35E+08 | 1.63E+08 | | | | | | |
| P49813 | TMOD1 | tropomodulin 1 | Cytoplasm | enzyme | 1.33E+08 | 1.24E+08 | 1.38E+08 | | | | | | |
| P49813 | TMOD1 | tropomodulin 1 | Cytoplasm | enzyme | 9.46E+07 | 8.78E+07 | 1.35E+08 | | | | | | |
| P49813 | TMOD1 | tropomodulin 1 | Cytoplasm | enzyme | 8.51E+07 | 9.91E+07 | 9.68E+07 | | | | | | |
| P49813 | TMOD1 | tropomodulin 1 | Cytoplasm | enzyme | 6.30E+07 | 6.02E+07 | 6.10E+07 | | | | | | |
| P49813 | TMOD1 | tropomodulin 1 | Cytoplasm | enzyme | 4.61E+07 | 4.20E+07 | 5.39E+07 | | | | | | |
| P49813 | TMOD1 | tropomodulin 1 | Cytoplasm | enzyme | 3.31E+07 | 4.57E+07 | 5.97E+07 | | | | | | |
| P49813 | TMOD1 | tropomodulin 1 | Cytoplasm | enzyme | 3.78E+07 | 2884402.5 | 113337.68 | | | | | | |
| P49813 | TMOD1 | tropomodulin 1 | Cytoplasm | enzyme | 1.35E+07 | 1.43E+07 | 2.22E+07 | | | | | | |
| P49813 | TMOD1 | tropomodulin 1 | Cytoplasm | enzyme | 2.40E+07 | 1.32E+07 | 9565078 | | | | | | |
| P49813 | TMOD1 | tropomodulin 1 | Cytoplasm | enzyme | 1.16E+07 | 2.35E+07 | 4.53E+07 | | | | | | |
| P49813 | TMOD1 | tropomodulin 1 | Cytoplasm | enzyme | 7847774.5 | 1.31E+07 | 1.32E+07 | | | | | | |
| P49813 | TMOD1 | tropomodulin 1 | Cytoplasm | enzyme | 1.10E+07 | 1.18E+07 | 1.28E+07 | | | | | | |
| P49813 | TMOD1 | tropomodulin 1 | Cytoplasm | enzyme | 1.92E+07 | 3.66E+07 | 5.28E+07 | | | | | | |
| P49813 | TMOD1 | tropomodulin 1 | Cytoplasm | enzyme | 3613655 | 1.02E+07 | 2.58E+07 | | | | | | |
| P49813 | TMOD1 | tropomodulin 1 | Cytoplasm | enzyme | 5371385.5 | 6899031.5 | 1.07E+07 | | | | | | |
| P49813 | TMOD1 | tropomodulin 1 | Cytoplasm | enzyme | 6974946.5 | 3094995.25 | 1909482.5 | | | | | | |
| P49813 | TMOD1 | tropomodulin 1 | Cytoplasm | enzyme | 932156.75 | 1547536.625 | 1651936.625 | P49813 | TMOD1 | 75160413 | 95635154 | 11270711 | 72 |
| Q9D710 | TMX2 | thioredoxin-related transmembrane protein 2 | Cytoplasm | enzyme | 332716.094 | 570963.312 | 700463.312 | Q9D710 | TMX2 | 332716 | 186534 | 107696 | 3 |
| Q8C0L0 | TMX4 | thioredoxin-related transmembrane protein 4 | Cytoplasm | enzyme | 2499741.25 | 3264802.5 | 3785903 | Q8C0L0 | TMX4 | 2499741 | 646926 | 373503 | 3 |
| O88746 | TOM1 | target of myb1 membrane trafficking protein | Cytoplasm | transporter | 273653.719 | 1642218.25 | 1159373.375 | O88746 | TOM1 | 273654 | 694095 | 400736 | 3 |

| | | | | | | | | | | | | | |
|---|---|---|---|---|---|---|---|---|---|---|---|---|---|
| Q923U0 | TOM1L1 | target of myb1 like 1 membrane trafficking protein | Cytoplasm | other | 3772972.75 | 4377826.5 | 4491812.5 | | | | | | |
| Q923U0 | TOM1L1 | target of myb1 like 1 membrane trafficking protein | Cytoplasm | other | 1777562.625 | 1437767 | 974369.688 | Q923U0 | TOM1L1 | 2775268 | 1583177 | 646329 | 6 |
| Q9CPQ3 | TOMM22 | translocase of outer mitochondrial membrane 22 homolog (yeast) | Cytoplasm | transporter | 2133226.5 | 3791515.5 | 1371937.75 | | | | | | |
| Q9CPQ3 | TOMM22 | translocase of outer mitochondrial membrane 22 homolog (yeast) | Cytoplasm | transporter | 1935957.75 | 4915144 | 1672686.5 | Q9CPQ3 | TOMM22 | 2034592 | 1399884 | 571500 | 6 |
| Q9QYA2 | TOMM40 | translocase of outer mitochondrial membrane 40 homolog (yeast) | Cytoplasm | ion channel | 5250125.5 | 9840067 | 3518102.75 | Q9QYA2 | TOMM40 | 5250126 | 3266872 | 1886129 | 3 |
| Q9CZW5 | TOMM70A | translocase of outer mitochondrial membrane 70 homolog A (S. cerevisiae) | Cytoplasm | transporter | 1514596.875 | 2538754.75 | 1032999.812 | Q9CZW5 | TOMM70A | 1514597 | 768996 | 443980 | 3 |
| Q9CYZ2 | TPD52L2 | tumor protein D52-like 2 | Cytoplasm | other | 4587779.5 | 4132024.25 | 3159719.25 | Q9CYZ2 | TPD52L2 | 4587780 | 729434 | 421139 | 3 |
| P58771 | Tpm1 | tropomyosin 1, alpha | Plasma Membrane | other | 3.62E+08 | 2.95E+08 | 3.82E+08 | | | | | | |
| P58771 | Tpm1 | tropomyosin 1, alpha | Plasma Membrane | other | 5.61E+07 | 6.30E+07 | 7.34E+07 | | | | | | |
| P58771 | Tpm1 | tropomyosin 1, alpha | Plasma Membrane | other | 1.67E+07 | 1.83E+07 | 2.89E+07 | | | | | | |
| P58771 | Tpm1 | tropomyosin 1, alpha | Plasma Membrane | other | 1.29E+07 | 1.24E+07 | 1.76E+07 | | | | | | |
| P58771 | Tpm1 | tropomyosin 1, alpha | Plasma Membrane | other | 4861844.5 | 2846616.75 | 2717383.25 | P58771 | Tpm1 | 90508369 | 135545858 | 34997790 | 15 |
| P21107 | TPM3 | tropomyosin 3 | Cytoplasm | other | 6.14E+08 | 4.62E+08 | 5.83E+08 | | | | | | |
| P21107 | TPM3 | tropomyosin 3 | Cytoplasm | other | 2.00E+08 | 1.54E+08 | 2.20E+08 | | | | | | |
| P21107 | TPM3 | tropomyosin 3 | Cytoplasm | other | 2.28E+08 | 1.77E+08 | 1.95E+08 | | | | | | |
| P21107 | TPM3 | tropomyosin 3 | Cytoplasm | other | 1.80E+08 | 2.04E+08 | 2.65E+08 | | | | | | |
| P21107 | TPM3 | tropomyosin 3 | Cytoplasm | other | 1.07E+08 | 7.34E+07 | 9.58E+07 | | | | | | |
| P21107 | TPM3 | tropomyosin 3 | Cytoplasm | other | 7.87E+07 | 9.98E+07 | 1.65E+08 | | | | | | |
| P21107 | TPM3 | tropomyosin 3 | Cytoplasm | other | 7.82E+07 | 1.04E+08 | 1.44E+08 | | | | | | |
| P21107 | TPM3 | tropomyosin 3 | Cytoplasm | other | 1.08E+08 | 4.78E+07 | 1.90E+07 | | | | | | |
| P21107 | TPM3 | tropomyosin 3 | Cytoplasm | other | 6.43E+07 | 7.46E+07 | 8.79E+07 | | | | | | |
| P21107 | TPM3 | tropomyosin 3 | Cytoplasm | other | 4.84E+07 | 6.42E+07 | 8.14E+07 | | | | | | |
| P21107 | TPM3 | tropomyosin 3 | Cytoplasm | other | 3.80E+07 | 3.95E+07 | 5.64E+07 | | | | | | |
| P21107 | TPM3 | tropomyosin 3 | Cytoplasm | other | 9513502 | 1.24E+07 | 1.50E+07 | | | | | | |
| P21107 | TPM3 | tropomyosin 3 | Cytoplasm | other | 9233580 | 7003173 | 1.28E+07 | | | | | | |
| P21107 | TPM3 | tropomyosin 3 | Cytoplasm | other | 1.12E+07 | 7410559.5 | 8802220 | P21107 | TPM3 | 126752649 | 140644962 | 21701989 | 42 |
| O89023 | TPP1 | tripeptidyl peptidase I | Cytoplasm | peptidase | 4311109.5 | 9298717 | 4495517.5 | O89023 | TPP1 | 4311110 | 2827866 | 1632669 | 3 |
| Q64514 | TPP2 | tripeptidyl peptidase II | Cytoplasm | peptidase | 2.51E+07 | 1.41E+07 | 1.11E+07 | | | | | | |
| Q64514 | TPP2 | tripeptidyl peptidase II | Cytoplasm | peptidase | 1.50E+07 | 1.01E+07 | 6241919.5 | | | | | | |
| Q64514 | TPP2 | tripeptidyl peptidase II | Cytoplasm | peptidase | 1.44E+07 | 8898949 | 9928547 | | | | | | |
| Q64514 | TPP2 | tripeptidyl peptidase II | Cytoplasm | peptidase | 1.14E+07 | 6907492 | 6922795.5 | | | | | | |
| Q64514 | TPP2 | tripeptidyl peptidase II | Cytoplasm | peptidase | 1.44E+07 | 9687823 | 4283752 | | | | | | |
| Q64514 | TPP2 | tripeptidyl peptidase II | Cytoplasm | peptidase | 1.09E+07 | 8078732.5 | 7485477 | | | | | | |
| Q64514 | TPP2 | tripeptidyl peptidase II | Cytoplasm | peptidase | 8441042 | 6359076 | 4582992 | | | | | | |
| Q64514 | TPP2 | tripeptidyl peptidase II | Cytoplasm | peptidase | 8988804 | 5806942.5 | 4865562.5 | | | | | | |
| Q64514 | TPP2 | tripeptidyl peptidase II | Cytoplasm | peptidase | 9563441 | 6247299.5 | 6789905 | | | | | | |
| Q64514 | TPP2 | tripeptidyl peptidase II | Cytoplasm | peptidase | 7479637.5 | 5467717 | 4119795.25 | | | | | | |
| Q64514 | TPP2 | tripeptidyl peptidase II | Cytoplasm | peptidase | 5605787 | 4012500.5 | 3146656.75 | | | | | | |
| Q64514 | TPP2 | tripeptidyl peptidase II | Cytoplasm | peptidase | 4264779 | 2465376.25 | 2562898.75 | | | | | | |
| Q64514 | TPP2 | tripeptidyl peptidase II | Cytoplasm | peptidase | 3294942.25 | 2343319 | 2312129.25 | | | | | | |
| Q64514 | TPP2 | tripeptidyl peptidase II | Cytoplasm | peptidase | 6002280 | 4024756.5 | 3307937.25 | | | | | | |
| Q64514 | TPP2 | tripeptidyl peptidase II | Cytoplasm | peptidase | 3622895.25 | 2058965.125 | 3527632.5 | | | | | | |
| Q64514 | TPP2 | tripeptidyl peptidase II | Cytoplasm | peptidase | 3364678.25 | 2595391.75 | 1815274 | | | | | | |
| Q64514 | TPP2 | tripeptidyl peptidase II | Cytoplasm | peptidase | 3249615.25 | 2785192.25 | 2418706 | | | | | | |
| Q64514 | TPP2 | tripeptidyl peptidase II | Cytoplasm | peptidase | 4634621 | 3176818 | 2908511 | | | | | | |
| Q64514 | TPP2 | tripeptidyl peptidase II | Cytoplasm | peptidase | 2514313.75 | 1379655.25 | 103034.938 | | | | | | |
| Q64514 | TPP2 | tripeptidyl peptidase II | Cytoplasm | peptidase | 1421565 | 880770.75 | 609707.312 | | | | | | |
| Q64514 | TPP2 | tripeptidyl peptidase II | Cytoplasm | peptidase | 1660761.125 | 1077150.5 | 1095088.375 | | | | | | |
| Q64514 | TPP2 | tripeptidyl peptidase II | Cytoplasm | peptidase | 1519039.25 | 816480.812 | 611072.625 | Q64514 | TPP2 | 7583555 | 4487954 | 552429 | 66 |
| Q9CQN1 | TRAP1 | TNF receptor-associated protein 1 | Cytoplasm | enzyme | 1.09E+07 | 1.22E+07 | 9279580 | | | | | | |
| Q9CQN1 | TRAP1 | TNF receptor-associated protein 1 | Cytoplasm | enzyme | 1355908.125 | 3290210.25 | 1254161.125 | Q9CQN1 | TRAP1 | 6102954 | 4968694 | 2028461 | 6 |
| Q8K558 | TREML1 | triggering receptor expressed on myeloid cells-like 1 | Plasma Membrane | other | 3760328.75 | 1.09E+07 | 2.36E+07 | | | | | | |
| Q8K558 | TREML1 | triggering receptor expressed on myeloid cells-like 1 | Plasma Membrane | other | 1483021.25 | 6251023 | 1.61E+07 | | | | | | |
| Q8K558 | TREML1 | triggering receptor expressed on myeloid cells-like 1 | Plasma Membrane | other | 1030017.812 | 4214061 | 1.14E+07 | | | | | | |
| Q8K558 | TREML1 | triggering receptor expressed on myeloid cells-like 1 | Plasma Membrane | other | 2153154.75 | 7921259 | 2.22E+07 | | | | | | |
| Q8K558 | TREML1 | triggering receptor expressed on myeloid cells-like 1 | Plasma Membrane | other | 2181502 | 5982151.5 | 1.06E+07 | | | | | | |
| Q8K558 | TREML1 | triggering receptor expressed on myeloid cells-like 1 | Plasma Membrane | other | 1188938.75 | 4633778.5 | 7412412 | | | | | | |
| Q8K558 | TREML1 | triggering receptor expressed on myeloid cells-like 1 | Plasma Membrane | other | 677123.938 | 3488805 | 8810500 | | | | | | |
| Q8K558 | TREML1 | triggering receptor expressed on myeloid cells-like 1 | Plasma Membrane | other | 106829.727 | 1269358.25 | 3236554.25 | Q8K558 | TREML1 | 1572615 | 6452752 | 1317162 | 24 |
| Q9WTR1 | TRPV2 | transient receptor potential cation channel, subfamily V, member 2 | Plasma Membrane | ion channel | 2.09E+07 | 2.04E+07 | 1.89E+07 | | | | | | |
| Q9WTR1 | TRPV2 | transient receptor potential cation channel, subfamily V, member 2 | Plasma Membrane | ion channel | 1.30E+07 | 1.40E+07 | 1.86E+07 | | | | | | |
| Q9WTR1 | TRPV2 | transient receptor potential cation channel, subfamily V, member 2 | Plasma Membrane | ion channel | 5883738.5 | 7155764.5 | 1.19E+07 | | | | | | |
| Q9WTR1 | TRPV2 | transient receptor potential cation channel, subfamily V, member 2 | Plasma Membrane | ion channel | 9142003 | 7924581 | 9995604 | | | | | | |
| Q9WTR1 | TRPV2 | transient receptor potential cation channel, subfamily V, member 2 | Plasma Membrane | ion channel | 7525523.5 | 6024066 | 8400513 | | | | | | |
| Q9WTR1 | TRPV2 | transient receptor potential cation channel, subfamily V, member 2 | Plasma Membrane | ion channel | 9133638 | 8996401 | 8972799 | | | | | | |
| Q9WTR1 | TRPV2 | transient receptor potential cation channel, subfamily V, member 2 | Plasma Membrane | ion channel | 6986345.5 | 6100226.5 | 6133254.5 | | | | | | |
| Q9WTR1 | TRPV2 | transient receptor potential cation channel, subfamily V, member 2 | Plasma Membrane | ion channel | 6221382 | 5134792.5 | 4170947.75 | | | | | | |
| Q9WTR1 | TRPV2 | transient receptor potential cation channel, subfamily V, member 2 | Plasma Membrane | ion channel | 3833635.75 | 4234347.5 | 6690839 | | | | | | |
| Q9WTR1 | TRPV2 | transient receptor potential cation channel, subfamily V, member 2 | Plasma Membrane | ion channel | 2851890 | 3908178.75 | 5472602.5 | | | | | | |
| Q9WTR1 | TRPV2 | transient receptor potential cation channel, subfamily V, member 2 | Plasma Membrane | ion channel | 4055732 | 4066555.75 | 3929822.5 | | | | | | |
| Q9WTR1 | TRPV2 | transient receptor potential cation channel, subfamily V, member 2 | Plasma Membrane | ion channel | 2176921.75 | 3020295 | 4518589.5 | Q9WTR1 | TRPV2 | 7642568 | 5013540 | 835590 | 36 |
| Q62348 | TSN | translin | Nucleus | other | 8243817 | 5060672.5 | 4613609 | | | | | | |
| Q62348 | TSN | translin | Nucleus | other | 2930296.5 | 1931354.875 | 1395822.25 | Q62348 | TSN | 5587057 | 2518400 | 1028132 | 6 |
| Q8QZY6 | TSPAN14 | tetraspanin 14 | Plasma Membrane | other | 420159.594 | 1486212.875 | 1932295 | Q8QZY6 | TSPAN14 | 420160 | 776961 | 448579 | 3 |
| Q8R3S2 | TSPAN33 | tetraspanin 33 | Plasma Membrane | other | 2.17E+08 | 2.09E+08 | 2.32E+08 | | | | | | |
| Q8R3S2 | TSPAN33 | tetraspanin 33 | Plasma Membrane | other | 7.11E+07 | 7.31E+07 | 7.77E+07 | | | | | | |
| Q8R3S2 | TSPAN33 | tetraspanin 33 | Plasma Membrane | other | 3.39E+07 | 1.90E+07 | 1.23E+07 | | | | | | |
| Q8R3S2 | TSPAN33 | tetraspanin 33 | Plasma Membrane | other | 2.68E+07 | 1.88E+07 | 1.80E+07 | | | | | | |
| Q8R3S2 | TSPAN33 | tetraspanin 33 | Plasma Membrane | other | 4557366.5 | 7593658.5 | 1.02E+07 | | | | | | |
| Q8R3S2 | TSPAN33 | tetraspanin 33 | Plasma Membrane | other | 2585493.25 | 5961857 | 8585454 | | | | | | |
| Q8R3S2 | TSPAN33 | tetraspanin 33 | Plasma Membrane | other | 2207507.25 | 1759359.625 | 3295511.5 | Q8R3S2 | TSPAN33 | 51227195 | 74585035 | 16275789 | 21 |
| Q8R3G9 | TSPAN8 | tetraspanin 8 | Plasma Membrane | other | 5.67E+07 | 4.95E+07 | 5.32E+07 | | | | | | |
| Q8R3G9 | TSPAN8 | tetraspanin 8 | Plasma Membrane | other | 3.00E+07 | 2.49E+07 | 2.54E+07 | | | | | | |
| Q8R3G9 | TSPAN8 | tetraspanin 8 | Plasma Membrane | other | 2.26E+07 | 2.71E+07 | 2.07E+07 | | | | | | |
| Q8R3G9 | TSPAN8 | tetraspanin 8 | Plasma Membrane | other | 1.03E+07 | 9808853 | 1.41E+07 | | | | | | |
| Q8R3G9 | TSPAN8 | tetraspanin 8 | Plasma Membrane | other | 6033868 | 5466913.5 | 5184661.5 | Q8R3G9 | TSPAN8 | 25126774 | 17192105 | 4438983 | 15 |
| P50637 | TSPO | translocator protein (18kDa) | Cytoplasm | transmembrane receptor | 6670355.5 | 1.44E+07 | 8244824 | P50637 | TSPO | 6670356 | 4096111 | 2364891 | 3 |
| A2ASS6 | TTN | titin | Cytoplasm | kinase | 4269340 | 4115474.5 | 4579545 | A2ASS6 | TTN | 4269340 | 236384 | 136476 | 3 |
| P68369 | TUBA1A | tubulin, alpha 1a | Cytoplasm | other | 2832537.25 | 4255446.5 | 1989395 | P68369 | TUBA1A | 2832537 | 1145320 | 661251 | 3 |
| Q7TMM9 | TUBB2A | tubulin, beta 2A class IIa | Cytoplasm | other | 2000416.875 | 2705121 | 2415974 | | | | | | |
| Q7TMM9 | TUBB2A | tubulin, beta 2A class IIa | Cytoplasm | other | 1365722.375 | 2165716.5 | 2115666.25 | Q7TMM9 | TUBB2A | 1683070 | 450537 | 183931 | 6 |

| | | | | | | | | | | | | | |
|---|---|---|---|---|---|---|---|---|---|---|---|---|---|
| Q9D6F9 | TUBB4A | tubulin, beta 4A class IVa | Cytoplasm | other | 4628716 | 6836898.5 | 2725741.5 | | | | | | |
| Q9D6F9 | TUBB4A | tubulin, beta 4A class IVa | Cytoplasm | other | 6657957 | 7671945 | 6017359 | Q9D6F9 | TUBB4A | 5643337 | 1798569 | 734263 | 6 |
| Q8BFR5 | TUFM | Tu translation elongation factor, mitochondrial | Cytoplasm | translation regulator | 3505907 | 9910081 | 4815517.5 | | | | | | |
| Q8BFR5 | TUFM | Tu translation elongation factor, mitochondrial | Cytoplasm | translation regulator | 3528093.75 | 8745499 | 5848534 | | | | | | |
| Q8BFR5 | TUFM | Tu translation elongation factor, mitochondrial | Cytoplasm | translation regulator | 4134805.25 | 8396038 | 4544992.5 | | | | | | |
| Q8BFR5 | TUFM | Tu translation elongation factor, mitochondrial | Cytoplasm | translation regulator | 3032824.5 | 5755063.5 | 2739761.5 | | | | | | |
| Q8BFR5 | TUFM | Tu translation elongation factor, mitochondrial | Cytoplasm | translation regulator | 2360811.5 | 5275486.5 | 3018241.5 | | | | | | |
| Q8BFR5 | TUFM | Tu translation elongation factor, mitochondrial | Cytoplasm | translation regulator | 1041724.312 | 2886130.75 | 1152463.75 | Q8BFR5 | TUFM | 2934028 | 2502034 | 589735 | 18 |
| Q91W90 | TXNDC5 | thioredoxin domain containing 5 (endoplasmic reticulum) | Cytoplasm | enzyme | 1044250 | 1100683.375 | 1064585.875 | Q91W90 | TXNDC5 | 1044250 | 28581 | 16501 | 3 |
| Q8CDN6 | TXNL1 | thioredoxin-like 1 | Cytoplasm | enzyme | 1.45E+07 | 7291922.5 | 1538972.75 | | | | | | |
| Q8CDN6 | TXNL1 | thioredoxin-like 1 | Cytoplasm | enzyme | 1.35E+07 | 9991776 | 7070401 | | | | | | |
| Q8CDN6 | TXNL1 | thioredoxin-like 1 | Cytoplasm | enzyme | 9734719 | 7612820 | 6007649 | | | | | | |
| Q8CDN6 | TXNL1 | thioredoxin-like 1 | Cytoplasm | enzyme | 8008653.5 | 6741011.5 | 5145888 | | | | | | |
| Q8CDN6 | TXNL1 | thioredoxin-like 1 | Cytoplasm | enzyme | 2100534.75 | 4556216.5 | 6522131 | | | | | | |
| Q8CDN6 | TXNL1 | thioredoxin-like 1 | Cytoplasm | enzyme | 4195583.5 | 2279215.75 | 1238967.625 | | | | | | |
| Q8CDN6 | TXNL1 | thioredoxin-like 1 | Cytoplasm | enzyme | 2902555.5 | 3879550.75 | 1806932.75 | | | | | | |
| Q8CDN6 | TXNL1 | thioredoxin-like 1 | Cytoplasm | enzyme | 733932.688 | 598190.812 | 503945.625 | Q8CDN6 | TXNL1 | 6960747 | 3930018 | 802212 | 24 |
| Q9JLT4 | TXNRD2 | thioredoxin reductase 2 | Cytoplasm | enzyme | 1.72E+07 | 1.85E+07 | 1.36E+07 | | | | | | |
| Q9JLT4 | TXNRD2 | thioredoxin reductase 2 | Cytoplasm | enzyme | 7408071 | 6336705 | 5501830 | | | | | | |
| Q9JLT4 | TXNRD2 | thioredoxin reductase 2 | Cytoplasm | enzyme | 1166815.625 | 1204309.875 | 746677.625 | Q9JLT4 | TXNRD2 | 8581629 | 6893616 | 2297872 | 9 |
| Q02053 | UBA1 | ubiquitin-like modifier activating enzyme 1 | Cytoplasm | enzyme | 7877683 | 7320219.5 | 6255068.5 | | | | | | |
| Q02053 | UBA1 | ubiquitin-like modifier activating enzyme 1 | Cytoplasm | enzyme | 6784905.5 | 4349340 | 5538158 | | | | | | |
| Q02053 | UBA1 | ubiquitin-like modifier activating enzyme 1 | Cytoplasm | enzyme | 7130266 | 5331455.5 | 6323167.5 | | | | | | |
| Q02053 | UBA1 | ubiquitin-like modifier activating enzyme 1 | Cytoplasm | enzyme | 6611161 | 4361531.5 | 4665135 | | | | | | |
| Q02053 | UBA1 | ubiquitin-like modifier activating enzyme 1 | Cytoplasm | enzyme | 4652821.5 | 4551117 | 4976974.5 | | | | | | |
| Q02053 | UBA1 | ubiquitin-like modifier activating enzyme 1 | Cytoplasm | enzyme | 5245731 | 3686062.5 | 4025525 | | | | | | |
| Q02053 | UBA1 | ubiquitin-like modifier activating enzyme 1 | Cytoplasm | enzyme | 4579735 | 3224470 | 3239071.5 | | | | | | |
| Q02053 | UBA1 | ubiquitin-like modifier activating enzyme 1 | Cytoplasm | enzyme | 5201113 | 3844439.5 | 3892337.25 | | | | | | |
| Q02053 | UBA1 | ubiquitin-like modifier activating enzyme 1 | Cytoplasm | enzyme | 3404037.25 | 1909167.625 | 2419722.25 | | | | | | |
| Q02053 | UBA1 | ubiquitin-like modifier activating enzyme 1 | Cytoplasm | enzyme | 2278917.25 | 3181318.5 | 5392284.5 | | | | | | |
| Q02053 | UBA1 | ubiquitin-like modifier activating enzyme 1 | Cytoplasm | enzyme | 3660129.25 | 2373087 | 2748675 | | | | | | |
| Q02053 | UBA1 | ubiquitin-like modifier activating enzyme 1 | Cytoplasm | enzyme | 2747809 | 2362836.5 | 1840504.25 | | | | | | |
| Q02053 | UBA1 | ubiquitin-like modifier activating enzyme 1 | Cytoplasm | enzyme | 939718.688 | 729071.125 | 283627.219 | | | | | | |
| Q02053 | UBA1 | ubiquitin-like modifier activating enzyme 1 | Cytoplasm | enzyme | 2347392.75 | 363180.75 | 4.70E-06 | Q02053 | UBA1 | 4532959 | 1995518 | 307915 | 42 |
| P62984 | Uba52 | ubiquitin A-52 residue ribosomal protein fusion product 1 | Cytoplasm | other | 7.77E+08 | 8.54E+08 | 1.20E+09 | | | | | | |
| P62984 | Uba52 | ubiquitin A-52 residue ribosomal protein fusion product 1 | Cytoplasm | other | 2.46E+08 | 3.17E+08 | 3.64E+08 | | | | | | |
| P62984 | Uba52 | ubiquitin A-52 residue ribosomal protein fusion product 1 | Cytoplasm | other | 1.42E+08 | 1.56E+08 | 1.72E+08 | | | | | | |
| P62984 | Uba52 | ubiquitin A-52 residue ribosomal protein fusion product 1 | Cytoplasm | other | 1.60E+08 | 1.90E+08 | 1.84E+08 | | | | | | |
| P62984 | Uba52 | ubiquitin A-52 residue ribosomal protein fusion product 1 | Cytoplasm | other | 6.18E+07 | 2.31E+07 | 1.27E+07 | | | | | | |
| P62984 | Uba52 | ubiquitin A-52 residue ribosomal protein fusion product 1 | Cytoplasm | other | 8541476 | 3403243 | 1625213.5 | | | | | | |
| P62984 | Uba52 | ubiquitin A-52 residue ribosomal protein fusion product 1 | Cytoplasm | other | 2.37E+07 | 3.10E+07 | 3.43E+07 | | | | | | |
| P62984 | Uba52 | ubiquitin A-52 residue ribosomal protein fusion product 1 | Cytoplasm | other | 1178952.625 | 128143.789 | 347051.031 | P62984 | Uba52 | 177365054 | 310308755 | 63341509 | 24 |
| Q8VDI7 | UBAC1 | UBA domain containing 1 | Nucleus | other | 7489434.5 | 1.04E+07 | 8419151 | | | | | | |
| Q8VDI7 | UBAC1 | UBA domain containing 1 | Nucleus | other | 7122402.5 | 7016967.5 | 7863768.5 | | | | | | |
| Q8VDI7 | UBAC1 | UBA domain containing 1 | Nucleus | other | 5237018.5 | 5389643 | 6903792.5 | | | | | | |
| Q8VDI7 | UBAC1 | UBA domain containing 1 | Nucleus | other | 5429992.5 | 4153984.75 | 3465568 | | | | | | |
| Q8VDI7 | UBAC1 | UBA domain containing 1 | Nucleus | other | 4875143.5 | 3009027.5 | 3802944.75 | | | | | | |
| Q8VDI7 | UBAC1 | UBA domain containing 1 | Nucleus | other | 3216289.5 | 2419716.5 | 2856529.25 | | | | | | |
| Q8VDI7 | UBAC1 | UBA domain containing 1 | Nucleus | other | 2673887 | 2589562.5 | 2218590.75 | | | | | | |
| Q8VDI7 | UBAC1 | UBA domain containing 1 | Nucleus | other | 2094475.5 | 1124005.125 | 1678724 | | | | | | |
| Q8VDI7 | UBAC1 | UBA domain containing 1 | Nucleus | other | 1613304.875 | 1537782 | 977696.562 | | | | | | |
| Q8VDI7 | UBAC1 | UBA domain containing 1 | Nucleus | other | 841598.125 | 2070812.875 | 2793610.5 | Q8VDI7 | UBAC1 | 4059355 | 2535731 | 462959 | 30 |
| P68037 | UBE2L3 | ubiquitin-conjugating enzyme E2L 3 | Nucleus | enzyme | 9136070 | 6593028.5 | 8799525 | P68037 | UBE2L3 | 9136070 | 1381361 | 797529 | 3 |
| Q9QZU9 | UBE2L6 | ubiquitin-conjugating enzyme E2L 6 | Cytoplasm | enzyme | 2.22E+07 | 2.49E+07 | 8835701 | | | | | | |
| Q9QZU9 | UBE2L6 | ubiquitin-conjugating enzyme E2L 6 | Cytoplasm | enzyme | 5445427 | 5490554 | 2495026.75 | | | | | | |
| Q9QZU9 | UBE2L6 | ubiquitin-conjugating enzyme E2L 6 | Cytoplasm | enzyme | 2386245.5 | 3925234.5 | 3288467.75 | | | | | | |
| Q9QZU9 | UBE2L6 | ubiquitin-conjugating enzyme E2L 6 | Cytoplasm | enzyme | 3787088.75 | 1935944.25 | 450714.688 | Q9QZU9 | UBE2L6 | 8442190 | 7986182 | 2305412 | 12 |
| Q6ZPJ3 | UBE2O | ubiquitin-conjugating enzyme E2O | Nucleus | enzyme | 2.27E+07 | 1.76E+07 | 2.60E+07 | | | | | | |
| Q6ZPJ3 | UBE2O | ubiquitin-conjugating enzyme E2O | Nucleus | enzyme | 2.38E+07 | 1.74E+07 | 2.48E+07 | | | | | | |
| Q6ZPJ3 | UBE2O | ubiquitin-conjugating enzyme E2O | Nucleus | enzyme | 1.83E+07 | 1.14E+07 | 1.78E+07 | | | | | | |
| Q6ZPJ3 | UBE2O | ubiquitin-conjugating enzyme E2O | Nucleus | enzyme | 2.15E+07 | 1.55E+07 | 2.59E+07 | | | | | | |
| Q6ZPJ3 | UBE2O | ubiquitin-conjugating enzyme E2O | Nucleus | enzyme | 1.72E+07 | 1.19E+07 | 1.82E+07 | | | | | | |
| Q6ZPJ3 | UBE2O | ubiquitin-conjugating enzyme E2O | Nucleus | enzyme | 1.37E+07 | 1.02E+07 | 1.80E+07 | | | | | | |
| Q6ZPJ3 | UBE2O | ubiquitin-conjugating enzyme E2O | Nucleus | enzyme | 1.42E+07 | 1.19E+07 | 1.64E+07 | | | | | | |
| Q6ZPJ3 | UBE2O | ubiquitin-conjugating enzyme E2O | Nucleus | enzyme | 1.32E+07 | 9469645 | 9879615 | | | | | | |
| Q6ZPJ3 | UBE2O | ubiquitin-conjugating enzyme E2O | Nucleus | enzyme | 1.27E+07 | 9651409 | 2.03E+07 | | | | | | |
| Q6ZPJ3 | UBE2O | ubiquitin-conjugating enzyme E2O | Nucleus | enzyme | 8380572.5 | 6005359.5 | 9119172 | | | | | | |
| Q6ZPJ3 | UBE2O | ubiquitin-conjugating enzyme E2O | Nucleus | enzyme | 5947802.5 | 5107711.5 | 7908904.5 | | | | | | |
| Q6ZPJ3 | UBE2O | ubiquitin-conjugating enzyme E2O | Nucleus | enzyme | 7493619.5 | 5253383 | 1.13E+07 | | | | | | |
| Q6ZPJ3 | UBE2O | ubiquitin-conjugating enzyme E2O | Nucleus | enzyme | 8095849.5 | 5373566.5 | 1.07E+07 | | | | | | |
| Q6ZPJ3 | UBE2O | ubiquitin-conjugating enzyme E2O | Nucleus | enzyme | 8729977 | 6037367 | 1.17E+07 | | | | | | |
| Q6ZPJ3 | UBE2O | ubiquitin-conjugating enzyme E2O | Nucleus | enzyme | 7394104 | 5437529 | 8358028.5 | | | | | | |
| Q6ZPJ3 | UBE2O | ubiquitin-conjugating enzyme E2O | Nucleus | enzyme | 6339429.5 | 4852491 | 8137269.5 | | | | | | |
| Q6ZPJ3 | UBE2O | ubiquitin-conjugating enzyme E2O | Nucleus | enzyme | 7260620 | 5386797 | 8619882 | | | | | | |
| Q6ZPJ3 | UBE2O | ubiquitin-conjugating enzyme E2O | Nucleus | enzyme | 7595821.5 | 5824714.5 | 1.25E+07 | | | | | | |
| Q6ZPJ3 | UBE2O | ubiquitin-conjugating enzyme E2O | Nucleus | enzyme | 4064195 | 4943690.5 | 7652521.5 | | | | | | |
| Q6ZPJ3 | UBE2O | ubiquitin-conjugating enzyme E2O | Nucleus | enzyme | 4553919 | 4067617.5 | 6508324 | | | | | | |
| Q6ZPJ3 | UBE2O | ubiquitin-conjugating enzyme E2O | Nucleus | enzyme | 5274729 | 3639342.25 | 6482892 | | | | | | |
| Q6ZPJ3 | UBE2O | ubiquitin-conjugating enzyme E2O | Nucleus | enzyme | 3799984 | 2512300.75 | 1439477.375 | | | | | | |
| Q6ZPJ3 | UBE2O | ubiquitin-conjugating enzyme E2O | Nucleus | enzyme | 3431881.75 | 2211721.5 | 5093880.5 | | | | | | |
| Q6ZPJ3 | UBE2O | ubiquitin-conjugating enzyme E2O | Nucleus | enzyme | 2564692 | 3197394 | 6609612.5 | | | | | | |
| Q6ZPJ3 | UBE2O | ubiquitin-conjugating enzyme E2O | Nucleus | enzyme | 2586750 | 1915202.625 | 3402294.5 | | | | | | |
| Q6ZPJ3 | UBE2O | ubiquitin-conjugating enzyme E2O | Nucleus | enzyme | 2760967 | 2028537.75 | 3530512.25 | | | | | | |
| Q6ZPJ3 | UBE2O | ubiquitin-conjugating enzyme E2O | Nucleus | enzyme | 3772756.5 | 1905388 | 1407394.125 | | | | | | |
| Q6ZPJ3 | UBE2O | ubiquitin-conjugating enzyme E2O | Nucleus | enzyme | 2882966.5 | 1979007.75 | 2673678.75 | | | | | | |
| Q6ZPJ3 | UBE2O | ubiquitin-conjugating enzyme E2O | Nucleus | enzyme | 1687459.25 | 941473 | 1462833.375 | | | | | | |
| Q6ZPJ3 | UBE2O | ubiquitin-conjugating enzyme E2O | Nucleus | enzyme | 2599987.75 | 508914.719 | 4.70E-06 | | | | | | |
| Q6ZPJ3 | UBE2O | ubiquitin-conjugating enzyme E2O | Nucleus | enzyme | 1628703.875 | 1039175.5 | 1942279.5 | | | | | | |
| Q6ZPJ3 | UBE2O | ubiquitin-conjugating enzyme E2O | Nucleus | enzyme | 1782730.625 | 1173862 | 2317457.5 | | | | | | |
| Q6ZPJ3 | UBE2O | ubiquitin-conjugating enzyme E2O | Nucleus | enzyme | 713561.375 | 479698.344 | 513311.344 | | | | | | |
| Q6ZPJ3 | UBE2O | ubiquitin-conjugating enzyme E2O | Nucleus | enzyme | 209735.641 | 686955.625 | 497360.25 | Q6ZPJ3 | UBE2O | 7905083 | 6540723 | 647628 | 102 |

| | | | | | | | | | | | | | |
|---|---|---|---|---|---|---|---|---|---|---|---|---|---|
| Q6ZWZ2 | UBE2R2 | ubiquitin-conjugating enzyme E2R 2 | Cytoplasm | enzyme | 1820091.75 | 903608.062 | 745639.688 | Q6ZWZ2 | UBE2R2 | 1820092 | 580136 | 334941 | 3 |
| A2AN08 | UBR4 | ubiquitin protein ligase E3 component n-recognin 4 | Nucleus | enzyme | 1.47E+07 | 1.17E+07 | 1.16E+07 | | | | | | |
| A2AN08 | UBR4 | ubiquitin protein ligase E3 component n-recognin 4 | Nucleus | enzyme | 9937763 | 6822003 | 6824542 | | | | | | |
| A2AN08 | UBR4 | ubiquitin protein ligase E3 component n-recognin 4 | Nucleus | enzyme | 9130951 | 8602524 | 1487224.25 | | | | | | |
| A2AN08 | UBR4 | ubiquitin protein ligase E3 component n-recognin 4 | Nucleus | enzyme | 5751522.5 | 5871297 | 7306658.5 | | | | | | |
| A2AN08 | UBR4 | ubiquitin protein ligase E3 component n-recognin 4 | Nucleus | enzyme | 6808351 | 4876125.5 | 5072249.5 | | | | | | |
| A2AN08 | UBR4 | ubiquitin protein ligase E3 component n-recognin 4 | Nucleus | enzyme | 7312457.5 | 6694047.5 | 4491388 | | | | | | |
| A2AN08 | UBR4 | ubiquitin protein ligase E3 component n-recognin 4 | Nucleus | enzyme | 7140767 | 5884839 | 6229044.5 | | | | | | |
| A2AN08 | UBR4 | ubiquitin protein ligase E3 component n-recognin 4 | Nucleus | enzyme | 6100797 | 5693615 | 4991950.5 | | | | | | |
| A2AN08 | UBR4 | ubiquitin protein ligase E3 component n-recognin 4 | Nucleus | enzyme | 6393831 | 5253187.5 | 4145414.25 | | | | | | |
| A2AN08 | UBR4 | ubiquitin protein ligase E3 component n-recognin 4 | Nucleus | enzyme | 6509597 | 4777663 | 4050034.25 | | | | | | |
| A2AN08 | UBR4 | ubiquitin protein ligase E3 component n-recognin 4 | Nucleus | enzyme | 5943627 | 3771912.25 | 2999207.75 | | | | | | |
| A2AN08 | UBR4 | ubiquitin protein ligase E3 component n-recognin 4 | Nucleus | enzyme | 8695357 | 4972400.5 | 5006586.5 | | | | | | |
| A2AN08 | UBR4 | ubiquitin protein ligase E3 component n-recognin 4 | Nucleus | enzyme | 5743559.5 | 6975948 | 5440241 | | | | | | |
| A2AN08 | UBR4 | ubiquitin protein ligase E3 component n-recognin 4 | Nucleus | enzyme | 4056009 | 3107546.75 | 3927790.25 | | | | | | |
| A2AN08 | UBR4 | ubiquitin protein ligase E3 component n-recognin 4 | Nucleus | enzyme | 3936653.25 | 4850796.5 | 5601344 | | | | | | |
| A2AN08 | UBR4 | ubiquitin protein ligase E3 component n-recognin 4 | Nucleus | enzyme | 5805917.5 | 4770908.5 | 4081523.75 | | | | | | |
| A2AN08 | UBR4 | ubiquitin protein ligase E3 component n-recognin 4 | Nucleus | enzyme | 6065721 | 5384898 | 6810992 | | | | | | |
| A2AN08 | UBR4 | ubiquitin protein ligase E3 component n-recognin 4 | Nucleus | enzyme | 4152870.75 | 4941686 | 5040683 | | | | | | |
| A2AN08 | UBR4 | ubiquitin protein ligase E3 component n-recognin 4 | Nucleus | enzyme | 7359512 | 5344485 | 4415274.5 | | | | | | |
| A2AN08 | UBR4 | ubiquitin protein ligase E3 component n-recognin 4 | Nucleus | enzyme | 4698196.5 | 3197000.5 | 2938193.75 | | | | | | |
| A2AN08 | UBR4 | ubiquitin protein ligase E3 component n-recognin 4 | Nucleus | enzyme | 4232871 | 4205585.5 | 4865674 | | | | | | |
| A2AN08 | UBR4 | ubiquitin protein ligase E3 component n-recognin 4 | Nucleus | enzyme | 6336303 | 5552757.5 | 5023626 | | | | | | |
| A2AN08 | UBR4 | ubiquitin protein ligase E3 component n-recognin 4 | Nucleus | enzyme | 5076246.5 | 4970735 | 5672588 | | | | | | |
| A2AN08 | UBR4 | ubiquitin protein ligase E3 component n-recognin 4 | Nucleus | enzyme | 4866681.5 | 4804375 | 2740648.5 | | | | | | |
| A2AN08 | UBR4 | ubiquitin protein ligase E3 component n-recognin 4 | Nucleus | enzyme | 5497647.5 | 4316185.5 | 4479701.5 | | | | | | |
| A2AN08 | UBR4 | ubiquitin protein ligase E3 component n-recognin 4 | Nucleus | enzyme | 4308084.5 | 3594558.75 | 2924709.75 | | | | | | |
| A2AN08 | UBR4 | ubiquitin protein ligase E3 component n-recognin 4 | Nucleus | enzyme | 4328621 | 3174773.25 | 3203611.25 | | | | | | |
| A2AN08 | UBR4 | ubiquitin protein ligase E3 component n-recognin 4 | Nucleus | enzyme | 4151254 | 2688309 | 3476372 | | | | | | |
| A2AN08 | UBR4 | ubiquitin protein ligase E3 component n-recognin 4 | Nucleus | enzyme | 4759197 | 4132223.25 | 3959448.5 | | | | | | |
| A2AN08 | UBR4 | ubiquitin protein ligase E3 component n-recognin 4 | Nucleus | enzyme | 3923339.25 | 2515261.5 | 1772317.875 | | | | | | |
| A2AN08 | UBR4 | ubiquitin protein ligase E3 component n-recognin 4 | Nucleus | enzyme | 5394366 | 3987576.5 | 4115450.75 | | | | | | |
| A2AN08 | UBR4 | ubiquitin protein ligase E3 component n-recognin 4 | Nucleus | enzyme | 3214722.75 | 4868591 | 4857342 | | | | | | |
| A2AN08 | UBR4 | ubiquitin protein ligase E3 component n-recognin 4 | Nucleus | enzyme | 4564337.5 | 5167715 | 5425896 | | | | | | |
| A2AN08 | UBR4 | ubiquitin protein ligase E3 component n-recognin 4 | Nucleus | enzyme | 3217527 | 4234749.5 | 4843127.5 | | | | | | |
| A2AN08 | UBR4 | ubiquitin protein ligase E3 component n-recognin 4 | Nucleus | enzyme | 5276976.5 | 2386002.5 | 2771993 | | | | | | |
| A2AN08 | UBR4 | ubiquitin protein ligase E3 component n-recognin 4 | Nucleus | enzyme | 4224315 | 4108910.5 | 4018237.5 | | | | | | |
| A2AN08 | UBR4 | ubiquitin protein ligase E3 component n-recognin 4 | Nucleus | enzyme | 2583133 | 3232763.75 | 2101147.75 | | | | | | |
| A2AN08 | UBR4 | ubiquitin protein ligase E3 component n-recognin 4 | Nucleus | enzyme | 4552387.5 | 3470644.5 | 3748617.5 | | | | | | |
| A2AN08 | UBR4 | ubiquitin protein ligase E3 component n-recognin 4 | Nucleus | enzyme | 3825882.75 | 4252066.5 | 3122552 | | | | | | |
| A2AN08 | UBR4 | ubiquitin protein ligase E3 component n-recognin 4 | Nucleus | enzyme | 3468756.5 | 3529606 | 2078908.125 | | | | | | |
| A2AN08 | UBR4 | ubiquitin protein ligase E3 component n-recognin 4 | Nucleus | enzyme | 4102131.75 | 2751622.25 | 3008747.5 | | | | | | |
| A2AN08 | UBR4 | ubiquitin protein ligase E3 component n-recognin 4 | Nucleus | enzyme | 4742239 | 1587821.5 | 337917.281 | | | | | | |
| A2AN08 | UBR4 | ubiquitin protein ligase E3 component n-recognin 4 | Nucleus | enzyme | 2213543.25 | 4065050.25 | 4404767.5 | | | | | | |
| A2AN08 | UBR4 | ubiquitin protein ligase E3 component n-recognin 4 | Nucleus | enzyme | 3336063.25 | 3212058.75 | 2676158.75 | | | | | | |
| A2AN08 | UBR4 | ubiquitin protein ligase E3 component n-recognin 4 | Nucleus | enzyme | 3583206.25 | 3200146.75 | 2380589.25 | | | | | | |
| A2AN08 | UBR4 | ubiquitin protein ligase E3 component n-recognin 4 | Nucleus | enzyme | 3162607 | 1909723.875 | 1702399.625 | | | | | | |
| A2AN08 | UBR4 | ubiquitin protein ligase E3 component n-recognin 4 | Nucleus | enzyme | 3065494.5 | 2512466 | 2143589 | | | | | | |
| A2AN08 | UBR4 | ubiquitin protein ligase E3 component n-recognin 4 | Nucleus | enzyme | 2665667.5 | 1946062.75 | 1292405 | | | | | | |
| A2AN08 | UBR4 | ubiquitin protein ligase E3 component n-recognin 4 | Nucleus | enzyme | 2080766.875 | 2180403.25 | 3394669.5 | | | | | | |
| A2AN08 | UBR4 | ubiquitin protein ligase E3 component n-recognin 4 | Nucleus | enzyme | 3200617.25 | 1142051.375 | 1264656.75 | | | | | | |
| A2AN08 | UBR4 | ubiquitin protein ligase E3 component n-recognin 4 | Nucleus | enzyme | 2490436.5 | 2439414.5 | 2447688.5 | | | | | | |
| A2AN08 | UBR4 | ubiquitin protein ligase E3 component n-recognin 4 | Nucleus | enzyme | 1505160.25 | 1120920.625 | 1666693.25 | | | | | | |
| A2AN08 | UBR4 | ubiquitin protein ligase E3 component n-recognin 4 | Nucleus | enzyme | 1739110.125 | 1345309.625 | 1309036.625 | A2AN08 | UBR4 | 4941946 | 2073187 | 164414 | 159 |
| Q9WUP7 | UCHL5 | ubiquitin carboxyl-terminal hydrolase L5 | Cytoplasm | peptidase | 9920504 | 1.19E+07 | 7854782 | | | | | | |
| Q9WUP7 | UCHL5 | ubiquitin carboxyl-terminal hydrolase L5 | Cytoplasm | peptidase | 7112885.5 | 1.06E+07 | 6915138 | | | | | | |
| Q9WUP7 | UCHL5 | ubiquitin carboxyl-terminal hydrolase L5 | Cytoplasm | peptidase | 3130119.25 | 3426461.25 | 3757456.75 | | | | | | |
| Q9WUP7 | UCHL5 | ubiquitin carboxyl-terminal hydrolase L5 | Cytoplasm | peptidase | 1075261.625 | 1583739.625 | 1983917.25 | Q9WUP7 | UCHL5 | 5309693 | 3765314 | 1086953 | 12 |
| Q6P5E4 | UGGT1 | UDP-glucose glycoprotein glucosyltransferase 1 | Cytoplasm | enzyme | 9099450 | 8719306 | 8104114.5 | | | | | | |
| Q6P5E4 | UGGT1 | UDP-glucose glycoprotein glucosyltransferase 1 | Cytoplasm | enzyme | 3414821 | 3663829 | 3707809.5 | | | | | | |
| Q6P5E4 | UGGT1 | UDP-glucose glycoprotein glucosyltransferase 1 | Cytoplasm | enzyme | 2057144.875 | 2619145 | 1829746 | | | | | | |
| Q6P5E4 | UGGT1 | UDP-glucose glycoprotein glucosyltransferase 1 | Cytoplasm | enzyme | 2837794 | 4514346 | 3516508 | | | | | | |
| Q6P5E4 | UGGT1 | UDP-glucose glycoprotein glucosyltransferase 1 | Cytoplasm | enzyme | 2203245.5 | 2614984.25 | 2631462.5 | | | | | | |
| Q6P5E4 | UGGT1 | UDP-glucose glycoprotein glucosyltransferase 1 | Cytoplasm | enzyme | 1982332.125 | 2531491.5 | 1938570 | | | | | | |
| Q6P5E4 | UGGT1 | UDP-glucose glycoprotein glucosyltransferase 1 | Cytoplasm | enzyme | 1788082.5 | 3443642.75 | 4226813.5 | | | | | | |
| Q6P5E4 | UGGT1 | UDP-glucose glycoprotein glucosyltransferase 1 | Cytoplasm | enzyme | 1431623.5 | 1738059.5 | 1763144.75 | Q6P5E4 | UGGT1 | 3101812 | 2183292 | 445663 | 24 |
| Q8R1I1 | UQCR10 | ubiquinol-cytochrome c reductase, complex III subunit X | Cytoplasm | enzyme | 934456.812 | 3859353 | 2084573.75 | Q8R1I1 | UQCR10 | 934457 | 1473523 | 850739 | 3 |
| Q9CZ13 | UQCRC1 | ubiquinol-cytochrome c reductase core protein I | Cytoplasm | enzyme | 1.40E+07 | 2.42E+07 | 9648619 | | | | | | |
| Q9CZ13 | UQCRC1 | ubiquinol-cytochrome c reductase core protein I | Cytoplasm | enzyme | 4295626.5 | 8778517 | 4625894 | | | | | | |
| Q9CZ13 | UQCRC1 | ubiquinol-cytochrome c reductase core protein I | Cytoplasm | enzyme | 5801960.5 | 9546158 | 4511841.5 | | | | | | |
| Q9CZ13 | UQCRC1 | ubiquinol-cytochrome c reductase core protein I | Cytoplasm | enzyme | 3779678.5 | 6203634.5 | 2742355 | | | | | | |
| Q9CZ13 | UQCRC1 | ubiquinol-cytochrome c reductase core protein I | Cytoplasm | enzyme | 5214974 | 1.21E+07 | 6010415.5 | Q9CZ13 | UQCRC1 | 6608448 | 5501314 | 1420433 | 15 |
| Q9DB77 | UQCRC2 | ubiquinol-cytochrome c reductase core protein II | Cytoplasm | enzyme | 1.02E+07 | 2.56E+07 | 1.09E+07 | | | | | | |
| Q9DB77 | UQCRC2 | ubiquinol-cytochrome c reductase core protein II | Cytoplasm | enzyme | 9350346 | 1.77E+07 | 8754063 | | | | | | |
| Q9DB77 | UQCRC2 | ubiquinol-cytochrome c reductase core protein II | Cytoplasm | enzyme | 8073173.5 | 1.71E+07 | 9241908 | | | | | | |
| Q9DB77 | UQCRC2 | ubiquinol-cytochrome c reductase core protein II | Cytoplasm | enzyme | 7769017.5 | 1.29E+07 | 5532464 | | | | | | |
| Q9DB77 | UQCRC2 | ubiquinol-cytochrome c reductase core protein II | Cytoplasm | enzyme | 3847787.5 | 8058391 | 3287035 | | | | | | |
| Q9DB77 | UQCRC2 | ubiquinol-cytochrome c reductase core protein II | Cytoplasm | enzyme | 6295715.5 | 9947888 | 4529357 | | | | | | |
| Q9DB77 | UQCRC2 | ubiquinol-cytochrome c reductase core protein II | Cytoplasm | enzyme | 2630120 | 5710637 | 2608647.5 | | | | | | |
| Q9DB77 | UQCRC2 | ubiquinol-cytochrome c reductase core protein II | Cytoplasm | enzyme | 3839793.25 | 7461412.5 | 3463496 | | | | | | |
| Q9DB77 | UQCRC2 | ubiquinol-cytochrome c reductase core protein II | Cytoplasm | enzyme | 3074696 | 7304888 | 3598809 | | | | | | |
| Q9DB77 | UQCRC2 | ubiquinol-cytochrome c reductase core protein II | Cytoplasm | enzyme | 2730609.5 | 5177137 | 2533654.25 | | | | | | |
| Q9DB77 | UQCRC2 | ubiquinol-cytochrome c reductase core protein II | Cytoplasm | enzyme | 2108890.25 | 4385674 | 2175699 | | | | | | |
| Q9DB77 | UQCRC2 | ubiquinol-cytochrome c reductase core protein II | Cytoplasm | enzyme | 972022.625 | 3754631.5 | 2338586.25 | Q9DB77 | UQCRC2 | 5074348 | 5159177 | 859863 | 36 |
| P99028 | UQCRHL | ubiquinol-cytochrome c reductase hinge protein like | Cytoplasm | other | 2.33E+07 | 4.70E+07 | 1.85E+07 | | | | | | |
| P99028 | UQCRHL | ubiquinol-cytochrome c reductase hinge protein like | Cytoplasm | other | 5506953.5 | 2.06E+07 | 1.04E+07 | P99028 | UQCRHL | 14383477 | 14406962 | 5881617 | 6 |
| P70697 | UROD | uroporphyrinogen decarboxylase | Cytoplasm | enzyme | 2744621 | 2837150.5 | 2982474.75 | P70697 | UROD | 2744621 | 119899 | 69224 | 3 |
| Q78IK2 | USMG5 | up-regulated during skeletal muscle growth 5 homolog (mouse) | Cytoplasm | other | 4168846.75 | 1.09E+07 | 8005865 | | | | | | |
| Q78IK2 | USMG5 | up-regulated during skeletal muscle growth 5 homolog (mouse) | Cytoplasm | other | 4602327.5 | 8001978.5 | 4219828.5 | Q78IK2 | USMG5 | 4385587 | 2746972 | 1121447 | 6 |
| Q9JMA1 | USP14 | ubiquitin specific peptidase 14 (tRNA-guanine transglycosylase) | Cytoplasm | peptidase | 7067912.5 | 5842504.5 | 8037913 | | | | | | |
| Q9JMA1 | USP14 | ubiquitin specific peptidase 14 (tRNA-guanine transglycosylase) | Cytoplasm | peptidase | 4440423.5 | 4093797.25 | 4124589 | Q9JMA1 | USP14 | 5754168 | 1670155 | 681838 | 6 |

| | | | | | | | | | | | | | |
|---|---|---|---|---|---|---|---|---|---|---|---|---|---|
| Q8R5H1 | USP15 | ubiquitin specific peptidase 15 | Cytoplasm | peptidase | 3493938.75 | 2218681 | 2106543 | | | | | | |
| Q8R5H1 | USP15 | ubiquitin specific peptidase 15 | Cytoplasm | peptidase | 8132186.5 | 7506410 | 7222010 | | | | | | |
| Q8R5H1 | USP15 | ubiquitin specific peptidase 15 | Cytoplasm | peptidase | 5680870.5 | 3634271 | 2409737.25 | | | | | | |
| Q8R5H1 | USP15 | ubiquitin specific peptidase 15 | Cytoplasm | peptidase | 3528154.5 | 6496610.5 | 4293030 | | | | | | |
| Q8R5H1 | USP15 | ubiquitin specific peptidase 15 | Cytoplasm | peptidase | 4295392.5 | 2920017 | 3399635.5 | | | | | | |
| Q8R5H1 | USP15 | ubiquitin specific peptidase 15 | Cytoplasm | peptidase | 5309029 | 5232378 | 3123524 | | | | | | |
| Q8R5H1 | USP15 | ubiquitin specific peptidase 15 | Cytoplasm | peptidase | 3651698.5 | 2497547.5 | 1642840.875 | | | | | | |
| Q8R5H1 | USP15 | ubiquitin specific peptidase 15 | Cytoplasm | peptidase | 2521169 | 2259564.25 | 2425869.5 | | | | | | |
| Q8R5H1 | USP15 | ubiquitin specific peptidase 15 | Cytoplasm | peptidase | 3159203.5 | 3033134.5 | 2243574.25 | | | | | | |
| Q8R5H1 | USP15 | ubiquitin specific peptidase 15 | Cytoplasm | peptidase | 3198386 | 1745880.625 | 1885950.75 | | | | | | |
| Q8R5H1 | USP15 | ubiquitin specific peptidase 15 | Cytoplasm | peptidase | 3015441.75 | 1874702.125 | 1913182.125 | | | | | | |
| Q8R5H1 | USP15 | ubiquitin specific peptidase 15 | Cytoplasm | peptidase | 3242652.25 | 2239717.25 | 2586866 | | | | | | |
| Q8R5H1 | USP15 | ubiquitin specific peptidase 15 | Cytoplasm | peptidase | 3207107.25 | 2540171.75 | 1922572.125 | | | | | | |
| Q8R5H1 | USP15 | ubiquitin specific peptidase 15 | Cytoplasm | peptidase | 2606680.5 | 2057433.375 | 2091684.75 | | | | | | |
| Q8R5H1 | USP15 | ubiquitin specific peptidase 15 | Cytoplasm | peptidase | 989048.125 | 1246152.5 | 574385.438 | | | | | | |
| Q8R5H1 | USP15 | ubiquitin specific peptidase 15 | Cytoplasm | peptidase | 2162054.5 | 1330614.5 | 1378520.25 | Q8R5H1 | USP15 | 3637063 | 1680789 | 242601 | 48 |
| Q3UJD6 | USP19 | ubiquitin specific peptidase 19 | Cytoplasm | peptidase | 93038 | 117299.773 | 63695.516 | Q3UJD6 | USP19 | 93038 | 26842 | 15497 | 3 |
| Q5DU02 | USP22 | ubiquitin specific peptidase 22 | Nucleus | peptidase | 1.13E+09 | 1.13E+09 | 1.39E+09 | Q5DU02 | USP22 | 1130000000 | 147805052 | 85335286 | 3 |
| P56399 | USP5 | ubiquitin specific peptidase 5 (isopeptidase T) | Cytoplasm | peptidase | 7689928 | 6604765 | 5590675 | | | | | | |
| P56399 | USP5 | ubiquitin specific peptidase 5 (isopeptidase T) | Cytoplasm | peptidase | 6493964.5 | 5288508 | 6198957 | | | | | | |
| P56399 | USP5 | ubiquitin specific peptidase 5 (isopeptidase T) | Cytoplasm | peptidase | 3601704.25 | 2585532.75 | 4790222 | | | | | | |
| P56399 | USP5 | ubiquitin specific peptidase 5 (isopeptidase T) | Cytoplasm | peptidase | 4820975.5 | 3285863.25 | 4036403 | | | | | | |
| P56399 | USP5 | ubiquitin specific peptidase 5 (isopeptidase T) | Cytoplasm | peptidase | 5254240 | 2957217 | 3499751.5 | | | | | | |
| P56399 | USP5 | ubiquitin specific peptidase 5 (isopeptidase T) | Cytoplasm | peptidase | 5965433.5 | 3516202.5 | 4016468 | | | | | | |
| P56399 | USP5 | ubiquitin specific peptidase 5 (isopeptidase T) | Cytoplasm | peptidase | 5203784 | 3852215.25 | 3298077.75 | | | | | | |
| P56399 | USP5 | ubiquitin specific peptidase 5 (isopeptidase T) | Cytoplasm | peptidase | 4097786.25 | 2610439 | 3790534.75 | | | | | | |
| P56399 | USP5 | ubiquitin specific peptidase 5 (isopeptidase T) | Cytoplasm | peptidase | 4048917.5 | 3434088.75 | 1768941.125 | | | | | | |
| P56399 | USP5 | ubiquitin specific peptidase 5 (isopeptidase T) | Cytoplasm | peptidase | 1275021.125 | 395750.531 | 624910.188 | | | | | | |
| P56399 | USP5 | ubiquitin specific peptidase 5 (isopeptidase T) | Cytoplasm | peptidase | 990388.438 | 1245578.375 | 1623100.25 | | | | | | |
| P56399 | USP5 | ubiquitin specific peptidase 5 (isopeptidase T) | Cytoplasm | peptidase | 758500 | 776956.062 | 1362155.125 | | | | | | |
| P56399 | USP5 | ubiquitin specific peptidase 5 (isopeptidase T) | Cytoplasm | peptidase | 1235717.375 | 945713.688 | 2477773.5 | P56399 | USP5 | 3956643 | 1935545 | 309935 | 39 |
| Q6A4J8 | USP7 | ubiquitin specific peptidase 7 (herpes virus-associated) | Nucleus | peptidase | 780573.562 | 684404.875 | 821910.812 | Q6A4J8 | USP7 | 780574 | 70551 | 40733 | 3 |
| P70398 | USP9X | ubiquitin specific peptidase 9, X-linked | Plasma Membrane | peptidase | 6633978 | 4252188.5 | 3354510.75 | | | | | | |
| P70398 | USP9X | ubiquitin specific peptidase 9, X-linked | Plasma Membrane | peptidase | 3282947.5 | 2598886 | 3271219.75 | | | | | | |
| P70398 | USP9X | ubiquitin specific peptidase 9, X-linked | Plasma Membrane | peptidase | 3041807.5 | 2291832 | 2014040.625 | | | | | | |
| P70398 | USP9X | ubiquitin specific peptidase 9, X-linked | Plasma Membrane | peptidase | 1497372 | 1089230.625 | 1221812.75 | P70398 | USP9X | 3614026 | 1525092 | 440256 | 12 |
| Q80WQ2 | VAC14 | Vac14 homolog (S. cerevisiae) | Cytoplasm | other | 2669566.75 | 2992959.5 | 2843019.25 | Q80WQ2 | VAC14 | 2669567 | 161839 | 93438 | 3 |
| P63024 | VAMP3 | vesicle-associated membrane protein 3 | Plasma Membrane | other | 6.94E+07 | 6.62E+07 | 9.87E+07 | | | | | | |
| P63024 | VAMP3 | vesicle-associated membrane protein 3 | Plasma Membrane | other | 1.27E+07 | 1.42E+07 | 1.76E+07 | | | | | | |
| P63024 | VAMP3 | vesicle-associated membrane protein 3 | Plasma Membrane | other | 5259379.5 | 8041936 | 7566668.5 | | | | | | |
| P63024 | VAMP3 | vesicle-associated membrane protein 3 | Plasma Membrane | other | 6717073.5 | 7705563.5 | 7206602.5 | | | | | | |
| P63024 | VAMP3 | vesicle-associated membrane protein 3 | Plasma Membrane | other | 1861203.5 | 2535172 | 4561862 | P63024 | VAMP3 | 19199531 | 30098460 | 7771389 | 15 |
| Q9Z2P8 | VAMP5 | vesicle-associated membrane protein 5 | Plasma Membrane | transporter | 2.15E+07 | 2.71E+07 | 2.65E+07 | | | | | | |
| Q9Z2P8 | VAMP5 | vesicle-associated membrane protein 5 | Plasma Membrane | transporter | 1.02E+07 | 1.34E+07 | 2.17E+07 | | | | | | |
| Q9Z2P8 | VAMP5 | vesicle-associated membrane protein 5 | Plasma Membrane | transporter | 1.00E+07 | 1.25E+07 | 1.75E+07 | Q9Z2P8 | VAMP5 | 13910000 | 6689139 | 2229713 | 9 |
| O70404 | VAMP8 | vesicle-associated membrane protein 8 | Plasma Membrane | transporter | 1.08E+07 | 2.48E+07 | 2.05E+07 | | | | | | |
| O70404 | VAMP8 | vesicle-associated membrane protein 8 | Plasma Membrane | transporter | 1.08E+07 | 1.93E+07 | 1.34E+07 | | | | | | |
| O70404 | VAMP8 | vesicle-associated membrane protein 8 | Plasma Membrane | transporter | 4459238.5 | 560762.312 | 631.86 | O70404 | VAMP8 | 8693080 | 8838885 | 2946295 | 9 |
| Q9WV55 | VAPA | VAMP (vesicle-associated membrane protein)-associated protein A, 33kDa | Plasma Membrane | other | 4.57E+07 | 4.59E+07 | 5.21E+07 | | | | | | |
| Q9WV55 | VAPA | VAMP (vesicle-associated membrane protein)-associated protein A, 33kDa | Plasma Membrane | other | 2.68E+07 | 3.22E+07 | 3.26E+07 | | | | | | |
| Q9WV55 | VAPA | VAMP (vesicle-associated membrane protein)-associated protein A, 33kDa | Plasma Membrane | other | 1.77E+07 | 1.79E+07 | 2.32E+07 | | | | | | |
| Q9WV55 | VAPA | VAMP (vesicle-associated membrane protein)-associated protein A, 33kDa | Plasma Membrane | other | 1.93E+07 | 1.24E+07 | 6068187 | | | | | | |
| Q9WV55 | VAPA | VAMP (vesicle-associated membrane protein)-associated protein A, 33kDa | Plasma Membrane | other | 8593414 | 8620910 | 8290026.5 | | | | | | |
| Q9WV55 | VAPA | VAMP (vesicle-associated membrane protein)-associated protein A, 33kDa | Plasma Membrane | other | 1.14E+07 | 9939133 | 9668522 | | | | | | |
| Q9WV55 | VAPA | VAMP (vesicle-associated membrane protein)-associated protein A, 33kDa | Plasma Membrane | other | 8918390 | 7327228 | 1.14E+07 | | | | | | |
| Q9WV55 | VAPA | VAMP (vesicle-associated membrane protein)-associated protein A, 33kDa | Plasma Membrane | other | 7789279 | 4495444.5 | 1398151 | | | | | | |
| Q9WV55 | VAPA | VAMP (vesicle-associated membrane protein)-associated protein A, 33kDa | Plasma Membrane | other | 1791895.875 | 917808.312 | 55978.32 | Q9WV55 | VAPA | 16444775 | 14446520 | 2780234 | 27 |
| Q9QY76 | VAPB | VAMP (vesicle-associated membrane protein)-associated protein B and C | Plasma Membrane | other | 3036443 | 4341906 | 4903065 | | | | | | |
| Q9QY76 | VAPB | VAMP (vesicle-associated membrane protein)-associated protein B and C | Plasma Membrane | other | 717735.625 | 1033518.188 | 1244593.875 | Q9QY76 | VAPB | 1877089 | 1808058 | 738137 | 6 |
| Q3U2A8 | VARS2 | valyl-tRNA synthetase 2, mitochondrial | Cytoplasm | enzyme | 2.12E+07 | 2.79E+07 | 3.00E+07 | Q3U2A8 | VARS2 | 21210000 | 4600623 | 2656171 | 3 |
| P70460 | VASP | vasodilator-stimulated phosphoprotein | Plasma Membrane | other | 2758705.25 | 3167851.25 | 4138706 | P70460 | VASP | 2758705 | 708797 | 409224 | 3 |
| Q01853 | VCP | valosin containing protein | Cytoplasm | enzyme | 2.05E+08 | 2.16E+08 | 1.69E+08 | | | | | | |
| Q01853 | VCP | valosin containing protein | Cytoplasm | enzyme | 2.85E+08 | 1.78E+08 | 1.14E+08 | | | | | | |
| Q01853 | VCP | valosin containing protein | Cytoplasm | enzyme | 2.39E+08 | 1.18E+08 | 5.15E+07 | | | | | | |
| Q01853 | VCP | valosin containing protein | Cytoplasm | enzyme | 1.65E+08 | 2.83E+08 | 3.69E+08 | | | | | | |
| Q01853 | VCP | valosin containing protein | Cytoplasm | enzyme | 1.50E+08 | 1.73E+08 | 2.00E+08 | | | | | | |
| Q01853 | VCP | valosin containing protein | Cytoplasm | enzyme | 1.49E+08 | 8.18E+07 | 5.06E+07 | | | | | | |
| Q01853 | VCP | valosin containing protein | Cytoplasm | enzyme | 1.41E+08 | 8.98E+07 | 1.47E+08 | | | | | | |
| Q01853 | VCP | valosin containing protein | Cytoplasm | enzyme | 1.21E+08 | 9.89E+07 | 8.70E+07 | | | | | | |
| Q01853 | VCP | valosin containing protein | Cytoplasm | enzyme | 1.08E+08 | 9.63E+07 | 9.32E+07 | | | | | | |
| Q01853 | VCP | valosin containing protein | Cytoplasm | enzyme | 1.07E+08 | 8.11E+07 | 7.67E+07 | | | | | | |
| Q01853 | VCP | valosin containing protein | Cytoplasm | enzyme | 8.69E+07 | 8.09E+07 | 7.34E+07 | | | | | | |
| Q01853 | VCP | valosin containing protein | Cytoplasm | enzyme | 7.64E+07 | 1.17E+08 | 1.06E+08 | | | | | | |
| Q01853 | VCP | valosin containing protein | Cytoplasm | enzyme | 1.11E+08 | 8.22E+07 | 8.08E+07 | | | | | | |
| Q01853 | VCP | valosin containing protein | Cytoplasm | enzyme | 7.96E+07 | 7.21E+07 | 5.70E+07 | | | | | | |
| Q01853 | VCP | valosin containing protein | Cytoplasm | enzyme | 6.89E+07 | 7.97E+07 | 9.83E+07 | | | | | | |
| Q01853 | VCP | valosin containing protein | Cytoplasm | enzyme | 8.12E+07 | 5.65E+07 | 6.19E+07 | | | | | | |
| Q01853 | VCP | valosin containing protein | Cytoplasm | enzyme | 7.44E+07 | 8.51E+07 | 7.82E+07 | | | | | | |
| Q01853 | VCP | valosin containing protein | Cytoplasm | enzyme | 6.77E+07 | 5.32E+07 | 5.80E+07 | | | | | | |
| Q01853 | VCP | valosin containing protein | Cytoplasm | enzyme | 3.86E+07 | 9.15E+07 | 1.27E+08 | | | | | | |
| Q01853 | VCP | valosin containing protein | Cytoplasm | enzyme | 7.05E+07 | 5.85E+07 | 5.72E+07 | | | | | | |
| Q01853 | VCP | valosin containing protein | Cytoplasm | enzyme | 5.07E+07 | 5.86E+07 | 6.66E+07 | | | | | | |
| Q01853 | VCP | valosin containing protein | Cytoplasm | enzyme | 4.63E+07 | 4.32E+07 | 5.12E+07 | | | | | | |
| Q01853 | VCP | valosin containing protein | Cytoplasm | enzyme | 5.36E+07 | 3.80E+07 | 3.35E+07 | | | | | | |
| Q01853 | VCP | valosin containing protein | Cytoplasm | enzyme | 2.93E+07 | 5.19E+07 | 6.54E+07 | | | | | | |
| Q01853 | VCP | valosin containing protein | Cytoplasm | enzyme | 2.06E+07 | 5.80E+07 | 1.02E+08 | | | | | | |
| Q01853 | VCP | valosin containing protein | Cytoplasm | enzyme | 4.80E+07 | 4.27E+07 | 4.45E+07 | | | | | | |
| Q01853 | VCP | valosin containing protein | Cytoplasm | enzyme | 4.24E+07 | 2.53E+07 | 2.44E+07 | | | | | | |
| Q01853 | VCP | valosin containing protein | Cytoplasm | enzyme | 3.87E+07 | 2.87E+07 | 2.81E+07 | | | | | | |
| Q01853 | VCP | valosin containing protein | Cytoplasm | enzyme | 3.17E+07 | 3.75E+07 | 3.58E+07 | | | | | | |
| Q01853 | VCP | valosin containing protein | Cytoplasm | enzyme | 2.29E+07 | 1.52E+07 | 1.94E+07 | | | | | | |

| | | | | | | | | | | | | | |
|---|---|---|---|---|---|---|---|---|---|---|---|---|---|
| Q01853 | VCP | valosin containing protein | Cytoplasm | enzyme | 2.97E+07 | 2.70E+07 | 2.59E+07 | | | | | | |
| Q01853 | VCP | valosin containing protein | Cytoplasm | enzyme | 4.26E+07 | 2.61E+07 | 4.24E+07 | | | | | | |
| Q01853 | VCP | valosin containing protein | Cytoplasm | enzyme | 2.07E+07 | 2.78E+07 | 3.53E+07 | | | | | | |
| Q01853 | VCP | valosin containing protein | Cytoplasm | enzyme | 4.28E+07 | 2.97E+07 | 2.40E+07 | | | | | | |
| Q01853 | VCP | valosin containing protein | Cytoplasm | enzyme | 2.67E+07 | 2.32E+07 | 2.29E+07 | | | | | | |
| Q01853 | VCP | valosin containing protein | Cytoplasm | enzyme | 2.41E+07 | 2.54E+07 | 2.34E+07 | | | | | | |
| Q01853 | VCP | valosin containing protein | Cytoplasm | enzyme | 3.87E+07 | 9610404 | 2495617.75 | | | | | | |
| Q01853 | VCP | valosin containing protein | Cytoplasm | enzyme | 2.49E+07 | 2.54E+07 | 2.62E+07 | | | | | | |
| Q01853 | VCP | valosin containing protein | Cytoplasm | enzyme | 3.03E+07 | 2.35E+07 | 2.44E+07 | | | | | | |
| Q01853 | VCP | valosin containing protein | Cytoplasm | enzyme | 3.56E+07 | 4.38E+07 | 4.53E+07 | | | | | | |
| Q01853 | VCP | valosin containing protein | Cytoplasm | enzyme | 2.19E+07 | 1.64E+07 | 1.67E+07 | | | | | | |
| Q01853 | VCP | valosin containing protein | Cytoplasm | enzyme | 1.45E+07 | 3723623.5 | 1812427.875 | | | | | | |
| Q01853 | VCP | valosin containing protein | Cytoplasm | enzyme | 1.36E+07 | 1.33E+07 | 1.15E+07 | | | | | | |
| Q01853 | VCP | valosin containing protein | Cytoplasm | enzyme | 6067923.5 | 8884172 | 9988822 | | | | | | |
| Q01853 | VCP | valosin containing protein | Cytoplasm | enzyme | 1.14E+07 | 286869.438 | 13050.796 | | | | | | |
| Q01853 | VCP | valosin containing protein | Cytoplasm | enzyme | 2.15E+07 | 6275510.5 | 3002962.5 | | | | | | |
| Q01853 | VCP | valosin containing protein | Cytoplasm | enzyme | 6547232.5 | 8398557 | 7134563 | | | | | | |
| Q01853 | VCP | valosin containing protein | Cytoplasm | enzyme | 9297182 | 1.23E+07 | 1.36E+07 | | | | | | |
| Q01853 | VCP | valosin containing protein | Cytoplasm | enzyme | 1.11E+07 | 1.21E+07 | 1.15E+07 | | | | | | |
| Q01853 | VCP | valosin containing protein | Cytoplasm | enzyme | 8529286 | 1.64E+07 | 2.21E+07 | | | | | | |
| Q01853 | VCP | valosin containing protein | Cytoplasm | enzyme | 7033312 | 4818712.5 | 6467795.5 | | | | | | |
| Q01853 | VCP | valosin containing protein | Cytoplasm | enzyme | 1.15E+07 | 5669279 | 5580655 | | | | | | |
| Q01853 | VCP | valosin containing protein | Cytoplasm | enzyme | 6111535 | 8024860.5 | 6996281 | | | | | | |
| Q01853 | VCP | valosin containing protein | Cytoplasm | enzyme | 8259912.5 | 1.01E+07 | 1.12E+07 | | | | | | |
| Q01853 | VCP | valosin containing protein | Cytoplasm | enzyme | 1.30E+07 | 2770249.5 | 1804834.875 | | | | | | |
| Q01853 | VCP | valosin containing protein | Cytoplasm | enzyme | 3955814.75 | 4787848 | 5714812.5 | | | | | | |
| Q01853 | VCP | valosin containing protein | Cytoplasm | enzyme | 3062543.25 | 2872285.25 | 2455653 | | | | | | |
| Q01853 | VCP | valosin containing protein | Cytoplasm | enzyme | 6130007.5 | 161471.375 | 391444.531 | | | | | | |
| Q01853 | VCP | valosin containing protein | Cytoplasm | enzyme | 1216791.125 | 3276434 | 5449325.5 | | | | | | |
| Q01853 | VCP | valosin containing protein | Cytoplasm | enzyme | 6905713 | 5411033.5 | 3250408 | | | | | | |
| Q01853 | VCP | valosin containing protein | Cytoplasm | enzyme | 2472098.5 | 2514502.5 | 3855023.75 | | | | | | |
| Q01853 | VCP | valosin containing protein | Cytoplasm | enzyme | 1585588.875 | 1655404.375 | 1048170.312 | | | | | | |
| Q01853 | VCP | valosin containing protein | Cytoplasm | enzyme | 2073118.375 | 2163260.75 | 2713045 | | | | | | |
| Q01853 | VCP | valosin containing protein | Cytoplasm | enzyme | 3020553.25 | 240081.375 | 112744.086 | | | | | | |
| Q01853 | VCP | valosin containing protein | Cytoplasm | enzyme | 3466380 | 2434077.75 | 1269275.875 | Q01853 | VCP | 51242846 | 58049138 | 4156985 | 195 |
| Q60932 | VDAC1 | voltage-dependent anion channel 1 | Cytoplasm | ion channel | 2.03E+07 | 3.53E+07 | 1.74E+07 | | | | | | |
| Q60932 | VDAC1 | voltage-dependent anion channel 1 | Cytoplasm | ion channel | 1.05E+07 | 1.95E+07 | 9816702 | | | | | | |
| Q60932 | VDAC1 | voltage-dependent anion channel 1 | Cytoplasm | ion channel | 9806350 | 2.94E+07 | 1.92E+07 | | | | | | |
| Q60932 | VDAC1 | voltage-dependent anion channel 1 | Cytoplasm | ion channel | 1.28E+07 | 1.78E+07 | 7251246.5 | | | | | | |
| Q60932 | VDAC1 | voltage-dependent anion channel 1 | Cytoplasm | ion channel | 1.36E+07 | 3.10E+07 | 1.17E+07 | | | | | | |
| Q60932 | VDAC1 | voltage-dependent anion channel 1 | Cytoplasm | ion channel | 1.15E+07 | 1.98E+07 | 1.38E+07 | | | | | | |
| Q60932 | VDAC1 | voltage-dependent anion channel 1 | Cytoplasm | ion channel | 1.07E+07 | 2.60E+07 | 1.70E+07 | | | | | | |
| Q60932 | VDAC1 | voltage-dependent anion channel 1 | Cytoplasm | ion channel | 6279116 | 1170155.25 | 450218.156 | Q60932 | VDAC1 | 11930683 | 8753306 | 1786761 | 24 |
| Q60930 | VDAC2 | voltage-dependent anion channel 2 | Cytoplasm | ion channel | 1.78E+07 | 3.72E+07 | 1.94E+07 | | | | | | |
| Q60930 | VDAC2 | voltage-dependent anion channel 2 | Cytoplasm | ion channel | 1.61E+07 | 4.46E+07 | 1.89E+07 | | | | | | |
| Q60930 | VDAC2 | voltage-dependent anion channel 2 | Cytoplasm | ion channel | 1.17E+07 | 3.17E+07 | 2.10E+07 | | | | | | |
| Q60930 | VDAC2 | voltage-dependent anion channel 2 | Cytoplasm | ion channel | 1.36E+07 | 2.69E+07 | 1.34E+07 | | | | | | |
| Q60930 | VDAC2 | voltage-dependent anion channel 2 | Cytoplasm | ion channel | 1.23E+07 | 1.93E+07 | 1.09E+07 | | | | | | |
| Q60930 | VDAC2 | voltage-dependent anion channel 2 | Cytoplasm | ion channel | 4496129 | 8348907 | 4482573.5 | | | | | | |
| Q60930 | VDAC2 | voltage-dependent anion channel 2 | Cytoplasm | ion channel | 3328252.75 | 6652388 | 3951597.75 | | | | | | |
| Q60930 | VDAC2 | voltage-dependent anion channel 2 | Cytoplasm | ion channel | 862624.062 | 1936827.375 | 1284948.625 | | | | | | |
| Q60930 | VDAC2 | voltage-dependent anion channel 2 | Cytoplasm | ion channel | 1605470.125 | 3417393.75 | 2209495.5 | Q60930 | VDAC2 | 9076942 | 11552229 | 2223228 | 27 |
| Q60931 | VDAC3 | voltage-dependent anion channel 3 | Cytoplasm | ion channel | 2.76E+07 | 6.09E+07 | 2.72E+07 | | | | | | |
| Q60931 | VDAC3 | voltage-dependent anion channel 3 | Cytoplasm | ion channel | 1.45E+07 | 4.39E+07 | 1.63E+07 | | | | | | |
| Q60931 | VDAC3 | voltage-dependent anion channel 3 | Cytoplasm | ion channel | 1.30E+07 | 3.06E+07 | 1.58E+07 | | | | | | |
| Q60931 | VDAC3 | voltage-dependent anion channel 3 | Cytoplasm | ion channel | 1.12E+07 | 1.96E+07 | 6905541.5 | | | | | | |
| Q60931 | VDAC3 | voltage-dependent anion channel 3 | Cytoplasm | ion channel | 8578093 | 9636477 | 2435147.25 | | | | | | |
| Q60931 | VDAC3 | voltage-dependent anion channel 3 | Cytoplasm | ion channel | 3058239.75 | 4875388 | 4220738.5 | Q60931 | VDAC3 | 12991055 | 15429218 | 3636702 | 18 |
| P20152 | VIM | vimentin | Cytoplasm | other | 49799.281 | 19887.469 | 106254.797 | P20152 | VIM | 49799 | 43858 | 25322 | 3 |
| Q8BGQ1 | VIPAS39 | VPS33B interacting protein, apical-basolateral polarity regulator, spe-39 homolog | Plasma Membrane | other | 1.45E+07 | 1.65E+07 | 2.12E+07 | Q8BGQ1 | VIPAS39 | 14480000 | 3447917 | 1990656 | 3 |
| Q99KU0 | VMP1 | vacuole membrane protein 1 | Plasma Membrane | other | 1.32E+07 | 1.79E+07 | 1.83E+07 | Q99KU0 | VMP1 | 13160000 | 2849462 | 1645138 | 3 |
| Q5H8C4 | VPS13A | vacuolar protein sorting 13 homolog A (S. cerevisiae) | Cytoplasm | transporter | 1.71E+07 | 2.11E+07 | 1.64E+07 | | | | | | |
| Q5H8C4 | VPS13A | vacuolar protein sorting 13 homolog A (S. cerevisiae) | Cytoplasm | transporter | 1.17E+07 | 1.27E+07 | 1.82E+07 | | | | | | |
| Q5H8C4 | VPS13A | vacuolar protein sorting 13 homolog A (S. cerevisiae) | Cytoplasm | transporter | 1.32E+07 | 1.29E+07 | 1.33E+07 | | | | | | |
| Q5H8C4 | VPS13A | vacuolar protein sorting 13 homolog A (S. cerevisiae) | Cytoplasm | transporter | 6899950 | 1.24E+07 | 1.59E+07 | | | | | | |
| Q5H8C4 | VPS13A | vacuolar protein sorting 13 homolog A (S. cerevisiae) | Cytoplasm | transporter | 1.15E+07 | 1.19E+07 | 1.05E+07 | | | | | | |
| Q5H8C4 | VPS13A | vacuolar protein sorting 13 homolog A (S. cerevisiae) | Cytoplasm | transporter | 9246688 | 1.01E+07 | 9849421 | | | | | | |
| Q5H8C4 | VPS13A | vacuolar protein sorting 13 homolog A (S. cerevisiae) | Cytoplasm | transporter | 5704910 | 8806777 | 5587395.5 | | | | | | |
| Q5H8C4 | VPS13A | vacuolar protein sorting 13 homolog A (S. cerevisiae) | Cytoplasm | transporter | 7268920 | 6761114.5 | 7563540 | | | | | | |
| Q5H8C4 | VPS13A | vacuolar protein sorting 13 homolog A (S. cerevisiae) | Cytoplasm | transporter | 6016199 | 7218311 | 7954694.5 | | | | | | |
| Q5H8C4 | VPS13A | vacuolar protein sorting 13 homolog A (S. cerevisiae) | Cytoplasm | transporter | 6362138 | 6594381 | 6681391 | | | | | | |
| Q5H8C4 | VPS13A | vacuolar protein sorting 13 homolog A (S. cerevisiae) | Cytoplasm | transporter | 7643302.5 | 8443898 | 6754828 | | | | | | |
| Q5H8C4 | VPS13A | vacuolar protein sorting 13 homolog A (S. cerevisiae) | Cytoplasm | transporter | 5966392.5 | 5099729 | 6371891 | | | | | | |
| Q5H8C4 | VPS13A | vacuolar protein sorting 13 homolog A (S. cerevisiae) | Cytoplasm | transporter | 6878328 | 7749160 | 7422457.5 | | | | | | |
| Q5H8C4 | VPS13A | vacuolar protein sorting 13 homolog A (S. cerevisiae) | Cytoplasm | transporter | 6145075.5 | 7409944.5 | 8154962 | | | | | | |
| Q5H8C4 | VPS13A | vacuolar protein sorting 13 homolog A (S. cerevisiae) | Cytoplasm | transporter | 6926654 | 7580480.5 | 7138332 | | | | | | |
| Q5H8C4 | VPS13A | vacuolar protein sorting 13 homolog A (S. cerevisiae) | Cytoplasm | transporter | 7092122 | 5338641 | 7607374.5 | | | | | | |
| Q5H8C4 | VPS13A | vacuolar protein sorting 13 homolog A (S. cerevisiae) | Cytoplasm | transporter | 6173858.5 | 6649871.5 | 7821206.5 | | | | | | |
| Q5H8C4 | VPS13A | vacuolar protein sorting 13 homolog A (S. cerevisiae) | Cytoplasm | transporter | 5754753 | 7276573.5 | 7328682 | | | | | | |
| Q5H8C4 | VPS13A | vacuolar protein sorting 13 homolog A (S. cerevisiae) | Cytoplasm | transporter | 4274155 | 4009051.5 | 4652297.5 | | | | | | |
| Q5H8C4 | VPS13A | vacuolar protein sorting 13 homolog A (S. cerevisiae) | Cytoplasm | transporter | 3638632 | 4097772.5 | 4589167.5 | | | | | | |
| Q5H8C4 | VPS13A | vacuolar protein sorting 13 homolog A (S. cerevisiae) | Cytoplasm | transporter | 4424242 | 5001145.5 | 5366667 | | | | | | |
| Q5H8C4 | VPS13A | vacuolar protein sorting 13 homolog A (S. cerevisiae) | Cytoplasm | transporter | 2839892.75 | 2187652.25 | 3789408 | | | | | | |
| Q5H8C4 | VPS13A | vacuolar protein sorting 13 homolog A (S. cerevisiae) | Cytoplasm | transporter | 4358329.5 | 5513484 | 6688190 | | | | | | |
| Q5H8C4 | VPS13A | vacuolar protein sorting 13 homolog A (S. cerevisiae) | Cytoplasm | transporter | 4217586 | 5033953.5 | 5427897.5 | | | | | | |
| Q5H8C4 | VPS13A | vacuolar protein sorting 13 homolog A (S. cerevisiae) | Cytoplasm | transporter | 3221688.5 | 3191488.75 | 3816553.25 | | | | | | |
| Q5H8C4 | VPS13A | vacuolar protein sorting 13 homolog A (S. cerevisiae) | Cytoplasm | transporter | 2847213.25 | 2953654.75 | 2081942.375 | | | | | | |
| Q5H8C4 | VPS13A | vacuolar protein sorting 13 homolog A (S. cerevisiae) | Cytoplasm | transporter | 5543979.5 | 4755483.5 | 5481649.5 | | | | | | |
| Q5H8C4 | VPS13A | vacuolar protein sorting 13 homolog A (S. cerevisiae) | Cytoplasm | transporter | 2724006 | 2395722.75 | 2809317.5 | | | | | | |
| Q5H8C4 | VPS13A | vacuolar protein sorting 13 homolog A (S. cerevisiae) | Cytoplasm | transporter | 2020396 | 3028133.25 | 3126401 | | | | | | |
| Q5H8C4 | VPS13A | vacuolar protein sorting 13 homolog A (S. cerevisiae) | Cytoplasm | transporter | 2425181.75 | 1998695.5 | 2911249 | | | | | | |

| | | | | | | | | | | | | | |
|---|---|---|---|---|---|---|---|---|---|---|---|---|---|
| Q5H8C4 | VPS13A | vacuolar protein sorting 13 homolog A (S. cerevisiae) | Cytoplasm | transporter | 1661843.25 | 1340495 | 1724428.25 | | | | | | |
| Q5H8C4 | VPS13A | vacuolar protein sorting 13 homolog A (S. cerevisiae) | Cytoplasm | transporter | 2286553 | 1458016.375 | 1667384.5 | | | | | | |
| Q5H8C4 | VPS13A | vacuolar protein sorting 13 homolog A (S. cerevisiae) | Cytoplasm | transporter | 989005.75 | 1451321.625 | 1300402.375 | | | | | | |
| Q5H8C4 | VPS13A | vacuolar protein sorting 13 homolog A (S. cerevisiae) | Cytoplasm | transporter | 1382467.625 | 4389203.5 | 5703070.5 | Q5H8C4 | VPS13A | 5777484 | 3955704 | 391673 | 102 |
| Q8BX70 | VPS13C | vacuolar protein sorting 13 homolog C (S. cerevisiae) | Cytoplasm | other | 1821738.25 | 3498517.25 | 2368564.5 | | | | | | |
| Q8BX70 | VPS13C | vacuolar protein sorting 13 homolog C (S. cerevisiae) | Cytoplasm | other | 2617560 | 2803269.25 | 961967.625 | | | | | | |
| Q8BX70 | VPS13C | vacuolar protein sorting 13 homolog C (S. cerevisiae) | Cytoplasm | other | 1963027 | 4011905 | 2180587.5 | | | | | | |
| Q8BX70 | VPS13C | vacuolar protein sorting 13 homolog C (S. cerevisiae) | Cytoplasm | other | 1957390.75 | 1814331.625 | 1155163.125 | | | | | | |
| Q8BX70 | VPS13C | vacuolar protein sorting 13 homolog C (S. cerevisiae) | Cytoplasm | other | 1340357.375 | 1891590.875 | 554007.312 | Q8BX70 | VPS13C | 1940015 | 919159 | 237326 | 15 |
| Q9D1C8 | VPS28 | vacuolar protein sorting 28 homolog (S. cerevisiae) | Cytoplasm | transporter | 2630651 | 1.13E+07 | 3439189.75 | Q9D1C8 | VPS28 | 2630651 | 4783187 | 2761574 | 3 |
| Q8CIZ8 | VWF | von Willebrand factor | Extracellular Space | other | 315222.375 | 2738039.5 | 2489380.75 | | | | | | |
| Q8CIZ8 | VWF | von Willebrand factor | Extracellular Space | other | 283979.281 | 812078.75 | 626554.062 | | | | | | |
| Q8CIZ8 | VWF | von Willebrand factor | Extracellular Space | other | 194370.984 | 549134.562 | 141874.359 | | | | | | |
| Q8CIZ8 | VWF | von Willebrand factor | Extracellular Space | other | 321697.281 | 848442.562 | 1189060.375 | | | | | | |
| Q8CIZ8 | VWF | von Willebrand factor | Extracellular Space | other | 3180.929 | 532514.25 | 1159218.125 | | | | | | |
| Q8CIZ8 | VWF | von Willebrand factor | Extracellular Space | other | 62568.566 | 202767 | 489423.5 | | | | | | |
| Q8CIZ8 | VWF | von Willebrand factor | Extracellular Space | other | 394069.281 | 1065593.25 | 1247019 | | | | | | |
| Q8CIZ8 | VWF | von Willebrand factor | Extracellular Space | other | 80178 | 799957.312 | 637720.25 | | | | | | |
| Q8CIZ8 | VWF | von Willebrand factor | Extracellular Space | other | 163175.172 | 1762968.25 | 1683023.625 | | | | | | |
| Q8CIZ8 | VWF | von Willebrand factor | Extracellular Space | other | 4.70E-06 | 1181734.75 | 946329.688 | | | | | | |
| Q8CIZ8 | VWF | von Willebrand factor | Extracellular Space | other | 142620.672 | 741519.75 | 1035860.75 | | | | | | |
| Q8CIZ8 | VWF | von Willebrand factor | Extracellular Space | other | 218293.297 | 352993.75 | 450835.25 | | | | | | |
| Q8CIZ8 | VWF | von Willebrand factor | Extracellular Space | other | 4.70E-06 | 461085.656 | 707993.688 | | | | | | |
| Q8CIZ8 | VWF | von Willebrand factor | Extracellular Space | other | 50209.148 | 444454.281 | 759512.75 | | | | | | |
| Q8CIZ8 | VWF | von Willebrand factor | Extracellular Space | other | 4.70E-06 | 171618.109 | 234105.641 | Q8CIZ8 | VWF | 148638 | 619603 | 92365 | 45 |
| Q8CBE3 | WDR37 | WD repeat domain 37 | Nucleus | other | 3449283 | 2725305.25 | 2253723.75 | | | | | | |
| Q8CBE3 | WDR37 | WD repeat domain 37 | Nucleus | other | 1056794.125 | 645500.438 | 872618.438 | Q8CBE3 | WDR37 | 2253039 | 1141978 | 466210 | 6 |
| Q8BH57 | WDR48 | WD repeat domain 48 | Cytoplasm | other | 2248664.5 | 1468794.625 | 1251216 | Q8BH57 | WDR48 | 2248665 | 524474 | 302805 | 3 |
| Q4KUS1 | WFDC8 | WAP four-disulfide core domain 8 | Extracellular Space | other | 738523.625 | 592160 | 739242.25 | Q4KUS1 | WFDC8 | 738524 | 84711 | 48908 | 3 |
| P83741 | WNK1 | WNK lysine deficient protein kinase 1 | Cytoplasm | kinase | 1443647.125 | 1713153.375 | 2004186.875 | P83741 | WNK1 | 1443647 | 280339 | 161854 | 3 |
| Q9QXY7 | XK | X-linked Kx blood group | Plasma Membrane | transporter | 3.05E+07 | 2.59E+07 | 2.62E+07 | | | | | | |
| Q9QXY7 | XK | X-linked Kx blood group | Plasma Membrane | transporter | 1.80E+07 | 2.39E+07 | 2.81E+07 | | | | | | |
| Q9QXY7 | XK | X-linked Kx blood group | Plasma Membrane | transporter | 2.60E+07 | 2.74E+07 | 3.30E+07 | | | | | | |
| Q9QXY7 | XK | X-linked Kx blood group | Plasma Membrane | transporter | 1.74E+07 | 2.00E+07 | 1.97E+07 | Q9QXY7 | XK | 22972500 | 4986287 | 1439417 | 12 |
| Q6P5F9 | XPO1 | exportin 1 | Nucleus | transporter | 2229619 | 1531278.875 | 1035422.062 | Q6P5F9 | XPO1 | 2229619 | 599953 | 346383 | 3 |
| Q9EPK7 | XPO7 | exportin 7 | Nucleus | transporter | 2238021 | 2430344.5 | 1918399.875 | | | | | | |
| Q9EPK7 | XPO7 | exportin 7 | Nucleus | transporter | 2387958.25 | 1805629.125 | 1312153.125 | Q9EPK7 | XPO7 | 2312990 | 426265 | 174022 | 6 |
| P62960 | YBX1 | Y box binding protein 1 | Nucleus | transcription regulator | 4187372.75 | 5686733 | 2592787.5 | | | | | | |
| P62960 | YBX1 | Y box binding protein 1 | Nucleus | transcription regulator | 382000 | 405751.875 | 220719.828 | P62960 | YBX1 | 2284686 | 2310437 | 943232 | 6 |
| Q9JKB3 | YBX3 | Y box binding protein 3 | Nucleus | transcription regulator | 3276867.25 | 2980034.25 | 1976773.125 | | | | | | |
| Q9JKB3 | YBX3 | Y box binding protein 3 | Nucleus | transcription regulator | 2919454 | 3910833.5 | 1941317.75 | Q9JKB3 | YBX3 | 3098161 | 763682 | 311772 | 6 |
| Q9CX30 | YIF1B | Yip1 interacting factor homolog B (S. cerevisiae) | Cytoplasm | other | 2068027.25 | 3887524 | 3770326.75 | | | | | | |
| Q9CX30 | YIF1B | Yip1 interacting factor homolog B (S. cerevisiae) | Cytoplasm | other | 2306315 | 2601336 | 3872418.5 | Q9CX30 | YIF1B | 2187171 | 849501 | 346807 | 6 |
| Q9CQV8 | YWHAB | tyrosine 3-monooxygenase/tryptophan 5-monooxygenase activation protein, beta | Cytoplasm | transcription regulator | 3129744.25 | 4077718 | 5763309.5 | | | | | | |
| Q9CQV8 | YWHAB | tyrosine 3-monooxygenase/tryptophan 5-monooxygenase activation protein, beta | Cytoplasm | transcription regulator | 2754653 | 3729213 | 4910076.5 | | | | | | |
| Q9CQV8 | YWHAB | tyrosine 3-monooxygenase/tryptophan 5-monooxygenase activation protein, beta | Cytoplasm | transcription regulator | 1883001.25 | 2768285.25 | 3807244.75 | | | | | | |
| Q9CQV8 | YWHAB | tyrosine 3-monooxygenase/tryptophan 5-monooxygenase activation protein, beta | Cytoplasm | transcription regulator | 1106256 | 1818037.75 | 2582262.25 | | | | | | |
| Q9CQV8 | YWHAB | tyrosine 3-monooxygenase/tryptophan 5-monooxygenase activation protein, beta | Cytoplasm | transcription regulator | 385001.938 | 644330.312 | 918146.5 | | | | | | |
| Q9CQV8 | YWHAB | tyrosine 3-monooxygenase/tryptophan 5-monooxygenase activation protein, beta | Cytoplasm | transcription regulator | 354953.875 | 438097.312 | 820610.812 | Q9CQV8 | YWHAB | 1602268 | 1663627 | 392121 | 18 |
| P62259 | YWHAE | tyrosine 3-monooxygenase/tryptophan 5-monooxygenase activation protein, epsilon | Cytoplasm | other | 6420852.5 | 8370213 | 1.23E+07 | | | | | | |
| P62259 | YWHAE | tyrosine 3-monooxygenase/tryptophan 5-monooxygenase activation protein, epsilon | Cytoplasm | other | 4847601 | 5682510.5 | 8996315 | | | | | | |
| P62259 | YWHAE | tyrosine 3-monooxygenase/tryptophan 5-monooxygenase activation protein, epsilon | Cytoplasm | other | 5216103.5 | 5159161 | 8471356 | | | | | | |
| P62259 | YWHAE | tyrosine 3-monooxygenase/tryptophan 5-monooxygenase activation protein, epsilon | Cytoplasm | other | 2804137.75 | 5111397.5 | 5918567.5 | | | | | | |
| P62259 | YWHAE | tyrosine 3-monooxygenase/tryptophan 5-monooxygenase activation protein, epsilon | Cytoplasm | other | 2740446.75 | 3769177 | 7188963.5 | | | | | | |
| P62259 | YWHAE | tyrosine 3-monooxygenase/tryptophan 5-monooxygenase activation protein, epsilon | Cytoplasm | other | 2603203.25 | 2970788 | 5091156.5 | | | | | | |
| P62259 | YWHAE | tyrosine 3-monooxygenase/tryptophan 5-monooxygenase activation protein, epsilon | Cytoplasm | other | 2324133 | 2810922 | 3878690 | | | | | | |
| P62259 | YWHAE | tyrosine 3-monooxygenase/tryptophan 5-monooxygenase activation protein, epsilon | Cytoplasm | other | 2735997.5 | 3250540 | 4213721.5 | | | | | | |
| P62259 | YWHAE | tyrosine 3-monooxygenase/tryptophan 5-monooxygenase activation protein, epsilon | Cytoplasm | other | 1061966.375 | 1930005.875 | 2965981.75 | P62259 | YWHAE | 3417160 | 2575699 | 495693 | 27 |
| P61982 | YWHAG | tyrosine 3-monooxygenase/tryptophan 5-monooxygenase activation protein, gamma | Cytoplasm | other | 1488744.375 | 3269662 | 3925451.75 | | | | | | |
| P61982 | YWHAG | tyrosine 3-monooxygenase/tryptophan 5-monooxygenase activation protein, gamma | Cytoplasm | other | 482023.281 | 1161420.625 | 1494635.875 | P61982 | YWHAG | 985384 | 1329539 | 542782 | 6 |
| P68510 | YWHAH | tyrosine 3-monooxygenase/tryptophan 5-monooxygenase activation protein, eta | Cytoplasm | transcription regulator | 5680065.5 | 1.28E+07 | 2.07E+07 | | | | | | |
| P68510 | YWHAH | tyrosine 3-monooxygenase/tryptophan 5-monooxygenase activation protein, eta | Cytoplasm | transcription regulator | 1601483.5 | 5187948 | 8112790 | | | | | | |
| P68510 | YWHAH | tyrosine 3-monooxygenase/tryptophan 5-monooxygenase activation protein, eta | Cytoplasm | transcription regulator | 2110091 | 6765199.5 | 1.05E+07 | | | | | | |
| P68510 | YWHAH | tyrosine 3-monooxygenase/tryptophan 5-monooxygenase activation protein, eta | Cytoplasm | transcription regulator | 2235564 | 4396189.5 | 7456765.5 | | | | | | |
| P68510 | YWHAH | tyrosine 3-monooxygenase/tryptophan 5-monooxygenase activation protein, eta | Cytoplasm | transcription regulator | 980655.938 | 2055878.625 | 3100432 | | | | | | |
| P68510 | YWHAH | tyrosine 3-monooxygenase/tryptophan 5-monooxygenase activation protein, eta | Cytoplasm | transcription regulator | 356790.656 | 1601463.625 | 3319790.25 | | | | | | |
| P68510 | YWHAH | tyrosine 3-monooxygenase/tryptophan 5-monooxygenase activation protein, eta | Cytoplasm | transcription regulator | 259691.484 | 2140945.5 | 3246485.25 | | | | | | |
| P68510 | YWHAH | tyrosine 3-monooxygenase/tryptophan 5-monooxygenase activation protein, eta | Cytoplasm | transcription regulator | 413722.094 | 1116591.25 | 2445522.75 | | | | | | |
| P68510 | YWHAH | tyrosine 3-monooxygenase/tryptophan 5-monooxygenase activation protein, eta | Cytoplasm | transcription regulator | 42328.465 | 87874.836 | 157723.891 | P68510 | YWHAH | 1520044 | 4686889 | 901992 | 27 |
| P68254 | YWHAQ | tyrosine 3-monooxygenase/tryptophan 5-monooxygenase activation protein, theta | Cytoplasm | other | 1768811 | 3038020.25 | 3860229.75 | | | | | | |
| P68254 | YWHAQ | tyrosine 3-monooxygenase/tryptophan 5-monooxygenase activation protein, theta | Cytoplasm | other | 831673.938 | 2331047 | 3752901.25 | | | | | | |
| P68254 | YWHAQ | tyrosine 3-monooxygenase/tryptophan 5-monooxygenase activation protein, theta | Cytoplasm | other | 837645.438 | 1675737.75 | 1840415.5 | P68254 | YWHAQ | 1146043 | 1129361 | 376454 | 9 |
| P63101 | YWHAZ | tyrosine 3-monooxygenase/tryptophan 5-monooxygenase activation protein, zeta | Cytoplasm | enzyme | 1.76E+07 | 2.87E+07 | 4.62E+07 | | | | | | |
| P63101 | YWHAZ | tyrosine 3-monooxygenase/tryptophan 5-monooxygenase activation protein, zeta | Cytoplasm | enzyme | 1.13E+07 | 1.95E+07 | 3.96E+07 | | | | | | |
| P63101 | YWHAZ | tyrosine 3-monooxygenase/tryptophan 5-monooxygenase activation protein, zeta | Cytoplasm | enzyme | 1.18E+07 | 1.42E+07 | 2.68E+07 | | | | | | |
| P63101 | YWHAZ | tyrosine 3-monooxygenase/tryptophan 5-monooxygenase activation protein, zeta | Cytoplasm | enzyme | 8992441 | 1.59E+07 | 2.52E+07 | | | | | | |
| P63101 | YWHAZ | tyrosine 3-monooxygenase/tryptophan 5-monooxygenase activation protein, zeta | Cytoplasm | enzyme | 9059061 | 1.67E+07 | 3.05E+07 | | | | | | |
| P63101 | YWHAZ | tyrosine 3-monooxygenase/tryptophan 5-monooxygenase activation protein, zeta | Cytoplasm | enzyme | 7149752 | 1.05E+07 | 1.67E+07 | | | | | | |
| P63101 | YWHAZ | tyrosine 3-monooxygenase/tryptophan 5-monooxygenase activation protein, zeta | Cytoplasm | enzyme | 1.08E+07 | 1.15E+07 | 3.00E+07 | | | | | | |
| P63101 | YWHAZ | tyrosine 3-monooxygenase/tryptophan 5-monooxygenase activation protein, zeta | Cytoplasm | enzyme | 7698689.5 | 1.29E+07 | 1.98E+07 | | | | | | |
| P63101 | YWHAZ | tyrosine 3-monooxygenase/tryptophan 5-monooxygenase activation protein, zeta | Cytoplasm | enzyme | 4560096.5 | 4843800 | 6807656 | | | | | | |
| P63101 | YWHAZ | tyrosine 3-monooxygenase/tryptophan 5-monooxygenase activation protein, zeta | Cytoplasm | enzyme | 1307708.375 | 2353848.5 | 4254823 | | | | | | |
| P63101 | YWHAZ | tyrosine 3-monooxygenase/tryptophan 5-monooxygenase activation protein, zeta | Cytoplasm | enzyme | 2927682 | 1766482.25 | 344791.188 | | | | | | |
| P63101 | YWHAZ | tyrosine 3-monooxygenase/tryptophan 5-monooxygenase activation protein, zeta | Cytoplasm | enzyme | 1059926.25 | 2892895 | 8735513 | | | | | | |
| P63101 | YWHAZ | tyrosine 3-monooxygenase/tryptophan 5-monooxygenase activation protein, zeta | Cytoplasm | enzyme | 1056775 | 1732055 | 2514409 | | | | | | |
| P63101 | YWHAZ | tyrosine 3-monooxygenase/tryptophan 5-monooxygenase activation protein, zeta | Cytoplasm | enzyme | 1212377.625 | 2920864.5 | 9229846 | | | | | | |
| P63101 | YWHAZ | tyrosine 3-monooxygenase/tryptophan 5-monooxygenase activation protein, zeta | Cytoplasm | enzyme | 497053.062 | 515028.406 | 1044775.5 | | | | | | |
| P63101 | YWHAZ | tyrosine 3-monooxygenase/tryptophan 5-monooxygenase activation protein, zeta | Cytoplasm | enzyme | 391910.312 | 246382.25 | 107599.109 | P63101 | YWHAZ | 6091467 | 11032465 | 1592399 | 48 |
| Q9D0B1 | ZNF524 | zinc finger protein 524 | Nucleus | other | 3857612.75 | 7834473.5 | 1.04E+07 | Q9D0B1 | ZNF524 | 3857613 | 3310298 | 1911202 | 3 |
| Q02525 | ZNF717 | zinc finger protein 717 | Nucleus | other | 2.28E+08 | 2.70E+08 | 2.60E+08 | Q02525 | ZNF717 | 228200000 | 21742815 | 12553220 | 3 |
| Q5SSH7 | ZZEF1 | zinc finger, ZZ-type with EF-hand domain 1 | Cytoplasm | other | 2775721 | 2586307 | 2298430.75 | Q5SSH7 | ZZEF1 | 2775721 | 240332 | 138756 | 3 |

| | # Peptides by Location |
|---|---|
| Plasma Membrane | 1902 |
| Cytoplasm | 4392 |
| Nucleus | 523 |
| Extracellular Space | 347 |
| Other | 24 |
| **Total Peptides** | **7188** |

Rank all proteins each strain

| Rank | Peptide ID | Symbol | Mean Intensity | St. Dev | Std. err | n | Primary Location |
|---|---|---|---|---|---|---|---|
| 1 | P04919 | SLC4A1 | 2.61E+09 | 4.91E+09 | 3.76E+08 | 171 | Plasma Membrane |
| 2 | P14220 | Gypa | 2.48E+09 | 2.91E+09 | 1.19E+09 | 6 | Plasma Membrane |
| 3 | Q5DU02 | USP22 | 1.17E+09 | 2.69E+08 | 1.55E+08 | 3 | Nucleus |
| 4 | Q02013 | AQP1 | 1.10E+09 | 9.89E+08 | 2.55E+08 | 15 | Plasma Membrane |
| 5 | P01942 | HBA1/HBA2 | 9.67E+08 | 8.61E+08 | 1.50E+08 | 33 | Extracellular Space |
| 6 | P68033 | ACTC1 | 9.48E+08 | 1.32E+09 | 3.82E+08 | 12 | Cytoplasm |
| 7 | P02088 | Hbb-b1 | 8.92E+08 | 1.04E+09 | 1.38E+08 | 57 | Extracellular Space |
| 8 | Q78HU7 | GYPC | 7.30E+08 | 5.87E+08 | 1.96E+08 | 9 | Plasma Membrane |
| 9 | Q02357 | ANK1 | 5.67E+08 | 7.63E+08 | 3.39E+07 | 507 | Plasma Membrane |
| 10 | P60710 | ACTB | 5.56E+08 | 9.52E+08 | 7.94E+07 | 144 | Cytoplasm |
| 11 | P08032 | SPTA1 | 5.34E+08 | 6.73E+08 | 2.25E+07 | 894 | Cytoplasm |
| 12 | P15508 | SPTB | 4.98E+08 | 6.31E+08 | 2.26E+07 | 780 | Plasma Membrane |
| 13 | Q8K386 | Rab15 | 4.58E+08 | 5.23E+07 | 3.02E+07 | 3 | Cytoplasm |
| 14 | Q9QUT0 | RHAG | 4.30E+08 | 4.88E+08 | 1.41E+08 | 12 | Plasma Membrane |
| 15 | P48193 | EPB41 | 4.08E+08 | 5.77E+08 | 4.23E+07 | 186 | Plasma Membrane |
| 16 | P49222 | EPB42 | 3.96E+08 | 5.61E+08 | 4.05E+07 | 192 | Plasma Membrane |
| 17 | P70290 | MPP1 | 3.55E+08 | 4.05E+08 | 3.61E+07 | 126 | Plasma Membrane |
| 18 | Q02525 | ZNF717 | 3.24E+08 | 1.01E+08 | 5.82E+07 | 3 | Nucleus |
| 19 | P06803 | PIM1 | 3.03E+08 | 5.10E+07 | 2.95E+07 | 3 | Cytoplasm |
| 20 | P62984 | Uba52 | 2.86E+08 | 4.01E+08 | 8.18E+07 | 24 | Cytoplasm |
| 21 | P18572 | BSG | 2.85E+08 | 3.38E+08 | 5.21E+07 | 42 | Plasma Membrane |
| 22 | Q8N9S3 | Ahsa2 | 2.62E+08 | 3.29E+07 | 1.90E+07 | 3 | Cytoplasm |
| 23 | P53986 | SLC16A1 | 2.48E+08 | 2.38E+08 | 4.59E+07 | 27 | Plasma Membrane |
| 24 | Q8CF94 | RHCE/RHD (mRh) | 2.37E+08 | 4.67E+08 | 9.54E+07 | 24 | Plasma Membrane |
| 25 | Q61735 | CD47 | 2.14E+08 | 1.81E+08 | 4.26E+07 | 18 | Plasma Membrane |
| 26 | P54116 | STOM | 2.03E+08 | 2.35E+08 | 2.83E+07 | 69 | Plasma Membrane |
| 27 | Q8C6A8 | BHLHE22 | 1.99E+08 | 3.43E+07 | 1.98E+07 | 3 | Nucleus |
| 28 | Q9WV69 | DMTN | 1.85E+08 | 1.87E+08 | 2.16E+07 | 75 | Plasma Membrane |
| 29 | Q9JJ04 | B4GALT4 | 1.74E+08 | 5.80E+07 | 3.35E+07 | 3 | Cytoplasm |
| 30 | Q9QZI9 | SERINC3 | 1.55E+08 | 1.12E+08 | 3.25E+07 | 12 | Cytoplasm |
| 31 | Q9QYC0 | ADD1 | 1.55E+08 | 2.06E+08 | 1.86E+07 | 123 | Cytoplasm |
| 32 | P02089 | Hbb-b2 | 1.30E+08 | 1.68E+08 | 3.43E+07 | 24 | Extracellular Space |
| 33 | Q8BSM7 | SLC43A1 | 1.29E+08 | 1.90E+08 | 2.92E+07 | 42 | Plasma Membrane |
| 34 | P21107 | TPM3 | 1.28E+08 | 1.59E+08 | 2.45E+07 | 42 | Cytoplasm |
| 35 | Q61171 | PRDX2 | 1.27E+08 | 1.68E+08 | 2.59E+07 | 42 | Cytoplasm |
| 36 | Q08857 | CD36 | 1.16E+08 | 1.63E+08 | 2.35E+07 | 48 | Plasma Membrane |
| 37 | Q8VHL0 | SLC14A1 | 1.06E+08 | 8.62E+07 | 1.44E+07 | 36 | Plasma Membrane |
| 38 | Q9Z0S4 | Cldn13 | 1.05E+08 | 8.02E+07 | 2.31E+07 | 12 | Plasma Membrane |
| 39 | P70696 | HIST1H2BA | 1.01E+08 | 7.19E+07 | 4.15E+07 | 3 | Nucleus |
| 40 | P83503 | NYX | 9.50E+07 | 1.15E+07 | 6.64E+06 | 3 | Extracellular Space |
| 41 | Q9QYB8 | ADD2 | 9.18E+07 | 1.22E+08 | 8.83E+06 | 192 | Cytoplasm |
| 42 | P58771 | Tpm1 | 9.18E+07 | 1.49E+08 | 3.85E+07 | 15 | Plasma Membrane |
| 43 | P16951 | ATF2 | 9.15E+07 | 2.37E+07 | 1.37E+07 | 3 | Nucleus |
| 44 | P49813 | TMOD1 | 7.79E+07 | 7.74E+07 | 9.12E+06 | 72 | Cytoplasm |
| 45 | P16858 | GAPDH | 7.71E+07 | 9.17E+07 | 9.83E+06 | 87 | Cytoplasm |
| 46 | P70269 | CTSE | 7.70E+07 | 4.69E+07 | 1.56E+07 | 9 | Cytoplasm |
| 47 | Q99JI6 | RAP1B | 7.33E+07 | 6.86E+07 | 1.40E+07 | 24 | Cytoplasm |
| 48 | P61027 | RAB10 | 7.10E+07 | 8.40E+07 | 1.46E+07 | 33 | Cytoplasm |
| 49 | Q8CGP5 | HIST1H2AJ | 6.13E+07 | 6.82E+07 | 1.76E+07 | 15 | Nucleus |
| 50 | Q3U9N9 | SLC16A10 | 6.02E+07 | 5.86E+07 | 1.07E+07 | 30 | Plasma Membrane |
| 51 | Q9Z126 | PF4 | 5.88E+07 | 4.47E+07 | 1.49E+07 | 9 | Extracellular Space |
| 52 | P63017 | HSPA8 | 5.61E+07 | 4.22E+07 | 4.78E+06 | 78 | Cytoplasm |
| 53 | Q01853 | VCP | 5.60E+07 | 6.74E+07 | 4.83E+06 | 195 | Cytoplasm |
| 54 | Q9EQF2 | KEL | 5.47E+07 | 4.88E+07 | 4.70E+06 | 108 | Plasma Membrane |
| 55 | P61028 | RAB8B | 5.29E+07 | 4.95E+07 | 1.17E+07 | 18 | Cytoplasm |
| 56 | P15864 | HIST1H1C | 5.09E+07 | 4.15E+07 | 2.40E+07 | 3 | Nucleus |
| 57 | O55186 | Cd59a | 5.08E+07 | 6.37E+07 | 2.12E+07 | 9 | Plasma Membrane |
| 58 | P14211 | CALR | 4.99E+07 | 5.04E+07 | 6.68E+06 | 57 | Cytoplasm |
| 59 | P63001 | RAC1 | 4.62E+07 | 4.27E+07 | 6.59E+06 | 42 | Plasma Membrane |
| 60 | Q2VIS4 | Flg2 | 4.57E+07 | 7.13E+06 | 4.12E+06 | 3 | Extracellular Space |
| 61 | P55258 | RAB8A | 4.41E+07 | 3.45E+07 | 7.03E+06 | 24 | Plasma Membrane |
| 62 | Q9CYH2 | FAM213A | 4.33E+07 | 4.13E+07 | 6.89E+06 | 36 | Extracellular Space |
| 63 | Q8BR07 | BICD1 | 4.31E+07 | 1.61E+07 | 9.30E+06 | 3 | Cytoplasm |
| 64 | P35278 | RAB5C | 4.24E+07 | 3.85E+07 | 6.16E+06 | 39 | Cytoplasm |
| 65 | Q61469 | Ppap2a | 3.97E+07 | 2.21E+07 | 5.71E+06 | 15 | Plasma Membrane |
| 66 | P27600 | GNA12 | 3.93E+07 | 1.94E+06 | 1.12E+06 | 3 | Plasma Membrane |
| 67 | Q6PHN9 | RAB35 | 3.90E+07 | 2.63E+07 | 5.75E+06 | 21 | Cytoplasm |
| 68 | P62806 | H4 | 3.75E+07 | 5.52E+07 | 1.13E+07 | 24 | Nucleus |
| 69 | Q8VCM7 | FGG | 3.73E+07 | 6.60E+07 | 7.95E+06 | 69 | Extracellular Space |
| 70 | Q6UJY2 | SLC9C1 | 3.72E+07 | 1.28E+07 | 7.40E+06 | 3 | Extracellular Space |
| 71 | P62835 | RAP1A | 3.70E+07 | 3.53E+07 | 7.21E+06 | 24 | Cytoplasm |
| 72 | Q60692 | PSMB6 | 3.63E+07 | 4.58E+07 | 1.08E+07 | 18 | Nucleus |
| 73 | Q9ES97 | RTN3 | 3.56E+07 | 3.90E+07 | 1.59E+07 | 6 | Cytoplasm |
| 74 | Q9JIM1 | SLC29A1 | 3.50E+07 | 4.26E+07 | 7.41E+06 | 33 | Plasma Membrane |
| 75 | Q61411 | HRAS | 3.48E+07 | 5.00E+07 | 9.12E+06 | 30 | Plasma Membrane |
| 76 | O55112 | Aff2 | 3.47E+07 | 2.10E+06 | 1.21E+06 | 3 | Nucleus |
| 77 | Q8K0E8 | FGB | 3.47E+07 | 5.34E+07 | 6.29E+06 | 72 | Extracellular Space |
| 78 | P10126 | EEF1A1 | 3.39E+07 | 3.54E+07 | 4.26E+06 | 69 | Cytoplasm |
| 79 | Q8CIM8 | INTS4 | 3.37E+07 | 9.74E+06 | 5.62E+06 | 3 | Nucleus |
| 80 | P20029 | HSPA5 | 3.34E+07 | 3.43E+07 | 3.88E+06 | 78 | Cytoplasm |
| 81 | O54962 | BANF1 | 3.23E+07 | 2.52E+07 | 7.28E+06 | 12 | Nucleus |
| 82 | Q9Z2U0 | PSMA7 | 3.18E+07 | 3.27E+07 | 5.45E+06 | 36 | Cytoplasm |
| 83 | Q8R3S2 | TSPAN33 | 3.11E+07 | 4.10E+07 | 8.96E+06 | 21 | Plasma Membrane |
| 84 | Q8R146 | APEH | 3.07E+07 | 2.47E+07 | 2.86E+06 | 75 | Cytoplasm |
| 85 | Q9QUM9 | PSMA6 | 3.07E+07 | 1.92E+07 | 3.34E+06 | 33 | Cytoplasm |
| 86 | Q7TMS5 | ABCG2 | 3.06E+07 | 2.82E+07 | 3.64E+06 | 60 | Plasma Membrane |
| 87 | O08586 | PTEN | 3.05E+07 | 1.77E+06 | 1.02E+06 | 3 | Cytoplasm |
| 88 | Q9Z2U1 | PSMA5 | 3.00E+07 | 3.50E+07 | 5.40E+06 | 42 | Cytoplasm |
| 89 | Q9R0K7 | ATP2B2 | 2.88E+07 | 3.75E+07 | 6.85E+06 | 30 | Plasma Membrane |
| 90 | Q922M5 | CDCA7L | 2.87E+07 | 6.13E+06 | 3.54E+06 | 3 | Nucleus |
| 91 | Q64475 | HIST1H2BB | 2.81E+07 | 4.70E+07 | 1.11E+07 | 18 | Nucleus |
| 92 | O35450 | FKBPL | 2.76E+07 | 5.46E+05 | 3.15E+05 | 3 | Nucleus |
| 93 | Q8K400 | STXBP5 | 2.72E+07 | 3.64E+06 | 2.10E+06 | 3 | Plasma Membrane |
| 94 | Q62136 | PTPN21 | 2.68E+07 | 2.17E+06 | 1.25E+06 | 3 | Cytoplasm |
| 95 | Q8R3G9 | TSPAN8 | 2.65E+07 | 1.69E+07 | 4.38E+06 | 15 | Plasma Membrane |
| 96 | Q9DC29 | ABCB6 | 2.62E+07 | 2.87E+07 | 3.71E+06 | 60 | Cytoplasm |
| 97 | P56400 | GP1BB | 2.62E+07 | 2.97E+07 | 7.01E+06 | 18 | Plasma Membrane |
| 98 | Q8BGQ1 | VIPAS39 | 2.62E+07 | 7.87E+06 | 4.54E+06 | 3 | Plasma Membrane |
| 99 | Q9DCT8 | Crip2 | 2.59E+07 | 1.18E+07 | 2.28E+06 | 27 | Plasma Membrane |
| 100 | Q60605 | MYL6 | 2.54E+07 | 1.96E+07 | 4.28E+06 | 21 | Cytoplasm |
| 101 | Q9R1P0 | PSMA4 | 2.53E+07 | 2.19E+07 | 3.50E+06 | 39 | Cytoplasm |
| 102 | P24369 | PPIB | 2.53E+07 | 2.11E+07 | 3.52E+06 | 36 | Cytoplasm |
| 103 | P99026 | PSMB4 | 2.52E+07 | 1.35E+07 | 3.18E+06 | 18 | Cytoplasm |
| 104 | Q9QXY7 | XK | 2.52E+07 | 5.89E+06 | 1.70E+06 | 12 | Plasma Membrane |
| 105 | Q9JLN5 | ERMAP | 2.49E+07 | 2.63E+07 | 3.39E+06 | 60 | Cytoplasm |
| 106 | Q3UZP4 | SVIP | 2.48E+07 | 1.65E+06 | 9.51E+05 | 3 | Cytoplasm |
| 107 | Q9D1C8 | VPS28 | 2.47E+07 | 3.20E+07 | 1.85E+07 | 3 | Cytoplasm |
| 108 | Q61672 | SLC29A2 | 2.46E+07 | 4.57E+05 | 2.64E+05 | 3 | Plasma Membrane |
| 109 | Q9D1G1 | RAB1B | 2.46E+07 | 1.95E+07 | 5.02E+06 | 15 | Cytoplasm |
| 110 | P35564 | CANX | 2.43E+07 | 2.09E+07 | 2.58E+06 | 66 | Cytoplasm |
| 111 | P20108 | PRDX3 | 2.40E+07 | 1.64E+07 | 3.59E+06 | 21 | Cytoplasm |
| 112 | P68433 | H31 | 2.37E+07 | 2.25E+07 | 5.31E+06 | 18 | Nucleus |
| 113 | P49722 | PSMA2 | 2.32E+07 | 2.82E+07 | 5.15E+06 | 30 | Cytoplasm |
| 114 | A2AVZ9 | SLC43A3 | 2.29E+07 | 1.73E+07 | 5.00E+06 | 12 | Extracellular Space |
| 115 | P54726 | RAD23A | 2.26E+07 | 2.79E+07 | 6.09E+06 | 21 | Nucleus |
| 116 | Q09199 | B4GALNT2 | 2.23E+07 | 2.41E+06 | 1.39E+06 | 3 | Cytoplasm |
| 117 | P14231 | ATP1B2 | 2.23E+07 | 2.31E+07 | 5.96E+06 | 15 | Plasma Membrane |
| 118 | Q3UV17 | KRT76 | 2.20E+07 | 5.13E+06 | 2.96E+06 | 3 | Cytoplasm |
| 119 | O70435 | PSMA3 | 2.18E+07 | 1.31E+07 | 2.53E+06 | 27 | Cytoplasm |
| 120 | P63024 | VAMP3 | 2.18E+07 | 3.14E+07 | 8.11E+06 | 15 | Plasma Membrane |
| 121 | P08113 | HSP90B1 | 2.15E+07 | 1.03E+07 | 1.17E+06 | 78 | Cytoplasm |
| 122 | P27773 | PDIA3 | 2.15E+07 | 2.07E+07 | 2.50E+06 | 69 | Cytoplasm |
| 123 | O88186 | GP9 | 2.15E+07 | 3.17E+07 | 9.14E+06 | 12 | Plasma Membrane |
| 124 | P97300 | Nptn | 2.14E+07 | 3.33E+07 | 6.40E+06 | 27 | Plasma Membrane |
| 125 | P08752 | GNAI2 | 2.14E+07 | 1.79E+07 | 2.03E+06 | 78 | Plasma Membrane |
| 126 | P62821 | RAB1A | 2.12E+07 | 2.64E+07 | 6.22E+06 | 18 | Cytoplasm |
| 127 | Q64735 | CR1L | 2.12E+07 | 2.07E+07 | 3.98E+06 | 27 | Plasma Membrane |
| 128 | Q03265 | ATP5A1 | 2.09E+07 | 1.44E+07 | 1.74E+06 | 69 | Cytoplasm |
| 129 | P56480 | ATP5B | 2.09E+07 | 1.90E+07 | 1.94E+06 | 96 | Cytoplasm |
| 130 | Q9CX63 | C17orf99 | 2.08E+07 | 1.19E+07 | 2.60E+06 | 21 | Extracellular Space |
| 131 | O55234 | PSMB5 | 2.08E+07 | 1.69E+07 | 2.95E+06 | 33 | Cytoplasm |
| 132 | Q8BTM8 | FLNA | 2.08E+07 | 2.52E+07 | 1.34E+06 | 354 | Cytoplasm |
| 133 | Q9R1P4 | PSMA1 | 2.06E+07 | 1.44E+07 | 2.08E+06 | 48 | Cytoplasm |
| 134 | P60904 | DNAJC5 | 2.01E+07 | 1.58E+07 | 4.09E+06 | 15 | Plasma Membrane |
| 135 | P63101 | YWHAZ | 2.01E+07 | 1.95E+07 | 2.82E+06 | 48 | Cytoplasm |
| 136 | Q9QUM0 | ITGA2B | 2.00E+07 | 2.86E+07 | 3.17E+06 | 81 | Plasma Membrane |
| 137 | P28656 | NAP1L1 | 2.00E+07 | 1.72E+07 | 3.51E+06 | 24 | Nucleus |
| 138 | P08556 | NRAS | 2.00E+07 | 1.78E+06 | 1.03E+06 | 3 | Plasma Membrane |

| Rank | ID | Symbol | Mean Intensity | St. Dev | Std. Err | n | Primary Location |
|---|---|---|---|---|---|---|---|
| 1 | P14220 | Gypa | 2.58E+09 | 2.21E+09 | 9.00E+08 | 6 | Plasma Membrane |
| 2 | P04919 | SLC4A1 | 2.19E+09 | 4.67E+09 | 3.57E+08 | 171 | Plasma Membrane |
| 3 | P02088 | Hbb-b1 | 1.70E+09 | 1.46E+09 | 1.93E+08 | 57 | Extracellular Space |
| 4 | P01942 | HBA1/HBA | 1.45E+09 | 9.54E+08 | 1.66E+08 | 33 | Extracellular Space |
| 5 | Q5DU02 | USP22 | 9.62E+08 | 9.99E+07 | 5.77E+07 | 3 | Nucleus |
| 6 | P68033 | ACTC1 | 8.89E+08 | 1.25E+09 | 3.61E+08 | 12 | Cytoplasm |
| 7 | Q78HU7 | GYPC | 6.08E+08 | 5.45E+08 | 1.82E+08 | 9 | Plasma Membrane |
| 8 | P60710 | ACTB | 5.26E+08 | 8.70E+08 | 7.25E+07 | 144 | Cytoplasm |
| 9 | Q02357 | ANK1 | 5.04E+08 | 6.95E+08 | 3.09E+07 | 507 | Plasma Membrane |
| 10 | P08032 | SPTA1 | 4.81E+08 | 6.16E+08 | 2.06E+07 | 894 | Cytoplasm |
| 11 | P15508 | SPTB | 4.43E+08 | 5.66E+08 | 2.03E+07 | 780 | Plasma Membrane |
| 12 | Q8K386 | Rab15 | 4.25E+08 | 2.50E+06 | 1.44E+06 | 3 | Cytoplasm |
| 13 | P02089 | Hbb-b2 | 4.07E+08 | 2.83E+08 | 5.77E+07 | 24 | Extracellular Space |
| 14 | Q9QUT0 | RHAG | 3.83E+08 | 4.81E+08 | 1.39E+08 | 12 | Plasma Membrane |
| 15 | P49222 | EPB42 | 3.54E+08 | 4.90E+08 | 3.54E+07 | 192 | Plasma Membrane |
| 16 | P70290 | MPP1 | 3.47E+08 | 4.00E+08 | 3.56E+07 | 126 | Plasma Membrane |
| 17 | P48193 | EPB41 | 3.40E+08 | 5.31E+08 | 3.90E+07 | 186 | Plasma Membrane |
| 18 | Q02525 | ZNF717 | 3.01E+08 | 2.22E+07 | 1.28E+07 | 3 | Nucleus |
| 19 | P70696 | HIST1H2B/ | 2.61E+08 | 7.15E+07 | 4.13E+07 | 3 | Nucleus |
| 20 | P62984 | Uba52 | 2.52E+08 | 3.42E+08 | 6.99E+07 | 24 | Cytoplasm |
| 21 | P06803 | PIM1 | 2.49E+08 | 2.51E+07 | 1.45E+07 | 3 | Cytoplasm |
| 22 | P18572 | BSG | 2.46E+08 | 3.09E+08 | 4.76E+07 | 42 | Plasma Membrane |
| 23 | P53986 | SLC16A1 | 2.21E+08 | 2.08E+08 | 4.00E+07 | 27 | Plasma Membrane |
| 24 | Q61735 | CD47 | 2.00E+08 | 1.82E+08 | 4.29E+07 | 18 | Plasma Membrane |
| 25 | Q8C6A8 | BHLHE22 | 1.95E+08 | 2.77E+07 | 1.60E+07 | 3 | Nucleus |
| 26 | Q8N9S3 | Ahsa2 | 1.87E+08 | 4.76E+07 | 2.75E+07 | 3 | Cytoplasm |
| 27 | Q8CF94 | RHCE/RHD | 1.81E+08 | 4.08E+08 | 8.33E+07 | 24 | Plasma Membrane |
| 28 | P15864 | HIST1H1C | 1.74E+08 | 4.48E+07 | 2.59E+07 | 3 | Nucleus |
| 29 | Q8CGP5 | HIST1H2AJ | 1.65E+08 | 1.08E+08 | 2.79E+07 | 15 | Nucleus |
| 30 | Q9WV69 | DMTN | 1.65E+08 | 1.72E+08 | 1.98E+07 | 75 | Plasma Membrane |
| 31 | P54116 | STOM | 1.42E+08 | 2.54E+08 | 3.06E+07 | 69 | Plasma Membrane |
| 32 | Q9QZI9 | SERINC3 | 1.32E+08 | 8.63E+07 | 2.49E+07 | 12 | Cytoplasm |
| 33 | Q9QYC0 | ADD1 | 1.32E+08 | 1.63E+08 | 1.47E+07 | 123 | Cytoplasm |
| 34 | P21107 | TPM3 | 1.09E+08 | 1.27E+08 | 1.96E+07 | 42 | Cytoplasm |
| 35 | Q9JJ04 | B4GALT4 | 1.07E+08 | 9.17E+07 | 5.29E+07 | 3 | Cytoplasm |
| 36 | Q8BSM7 | SLC43A1 | 1.07E+08 | 1.80E+08 | 2.78E+07 | 42 | Plasma Membrane |
| 37 | Q61171 | PRDX2 | 9.79E+07 | 1.11E+08 | 1.72E+07 | 42 | Cytoplasm |
| 38 | Q08857 | CD36 | 9.78E+07 | 1.42E+08 | 2.05E+07 | 48 | Plasma Membrane |
| 39 | P62806 | H4 | 9.67E+07 | 9.18E+07 | 1.87E+07 | 24 | Nucleus |
| 40 | Q8VHL0 | SLC14A1 | 9.03E+07 | 8.17E+07 | 1.36E+07 | 36 | Plasma Membrane |
| 41 | Q99JI6 | RAP1B | 8.81E+07 | 7.65E+07 | 1.56E+07 | 24 | Cytoplasm |
| 42 | Q9Z126 | PF4 | 8.45E+07 | 3.21E+07 | 1.07E+07 | 9 | Extracellular Space |
| 43 | Q9QYB8 | ADD2 | 8.05E+07 | 1.05E+08 | 7.58E+06 | 192 | Cytoplasm |
| 44 | Q9Z0S4 | Cldn13 | 7.91E+07 | 7.11E+07 | 2.05E+07 | 12 | Plasma Membrane |
| 45 | Q64475 | HIST1H2BE | 7.71E+07 | 7.49E+07 | 1.77E+07 | 18 | Nucleus |
| 46 | P16951 | ATF2 | 7.36E+07 | 1.40E+07 | 8.09E+06 | 3 | Nucleus |
| 47 | P49813 | TMOD1 | 7.22E+07 | 8.25E+07 | 9.72E+06 | 72 | Cytoplasm |
| 48 | P58771 | Tpm1 | 6.98E+07 | 1.04E+08 | 2.67E+07 | 15 | Plasma Membrane |
| 49 | P70269 | CTSE | 6.57E+07 | 4.86E+07 | 1.62E+07 | 9 | Cytoplasm |
| 50 | P61027 | RAB10 | 6.51E+07 | 7.92E+07 | 1.38E+07 | 33 | Cytoplasm |
| 51 | P16858 | GAPDH | 6.21E+07 | 7.16E+07 | 7.68E+06 | 87 | Cytoplasm |
| 52 | P10126 | EEF1A1 | 6.09E+07 | 6.34E+07 | 7.64E+06 | 69 | Cytoplasm |
| 53 | P83503 | NYX | 5.98E+07 | 1.68E+07 | 9.71E+06 | 3 | Extracellular Space |
| 54 | P68433 | H31 | 5.94E+07 | 3.46E+07 | 8.16E+06 | 18 | Nucleus |
| 55 | P63017 | HSPA8 | 5.84E+07 | 3.95E+07 | 4.47E+06 | 78 | Cytoplasm |
| 56 | P27600 | GNA12 | 5.60E+07 | 1.61E+07 | 9.32E+06 | 3 | Plasma Membrane |
| 57 | Q3U9N9 | SLC16A10 | 5.58E+07 | 5.89E+07 | 1.07E+07 | 30 | Plasma Membrane |
| 58 | O54962 | BANF1 | 5.55E+07 | 3.92E+07 | 1.13E+07 | 12 | Nucleus |
| 59 | P14211 | CALR | 4.90E+07 | 6.81E+07 | 9.02E+06 | 57 | Cytoplasm |
| 60 | P61028 | RAB8B | 4.77E+07 | 4.83E+07 | 1.14E+07 | 18 | Cytoplasm |
| 61 | Q9EQF2 | KEL | 4.69E+07 | 4.47E+07 | 4.30E+06 | 108 | Plasma Membrane |
| 62 | Q2VIS4 | Flg2 | 4.66E+07 | 9.82E+06 | 5.67E+06 | 3 | Extracellular Space |
| 63 | Q9CYH2 | FAM213A | 4.55E+07 | 6.15E+07 | 1.03E+07 | 36 | Extracellular Space |
| 64 | P43274 | Hist1h1e | 4.42E+07 | 2.87E+07 | 7.41E+06 | 15 | Nucleus |
| 65 | Q01853 | VCP | 4.26E+07 | 5.20E+07 | 3.72E+06 | 195 | Cytoplasm |
| 66 | P63001 | RAC1 | 4.25E+07 | 3.36E+07 | 5.18E+06 | 42 | Plasma Membrane |
| 67 | Q8BR07 | BICD1 | 4.16E+07 | 5.35E+06 | 3.09E+06 | 3 | Cytoplasm |
| 68 | P56400 | GP1BB | 4.06E+07 | 3.61E+07 | 8.51E+06 | 18 | Plasma Membrane |
| 69 | Q60605 | MYL6 | 4.06E+07 | 1.80E+07 | 3.93E+06 | 21 | Cytoplasm |
| 70 | P55258 | RAB8A | 3.98E+07 | 3.18E+07 | 6.50E+06 | 24 | Plasma Membrane |
| 71 | P35278 | RAB5C | 3.92E+07 | 3.75E+07 | 6.01E+06 | 39 | Cytoplasm |
| 72 | P20108 | PRDX3 | 3.90E+07 | 3.46E+07 | 7.54E+06 | 21 | Cytoplasm |
| 73 | Q62351 | TFRC | 3.85E+07 | 2.96E+07 | 2.81E+06 | 111 | Plasma Membrane |
| 74 | O55112 | Aff2 | 3.73E+07 | 9.31E+06 | 5.37E+06 | 3 | Nucleus |
| 75 | Q9ES97 | RTN3 | 3.73E+07 | 4.51E+07 | 1.84E+07 | 6 | Cytoplasm |
| 76 | P62835 | RAP1A | 3.66E+07 | 3.88E+07 | 7.92E+06 | 24 | Cytoplasm |
| 77 | O55186 | Cd59a | 3.64E+07 | 4.81E+07 | 1.60E+07 | 9 | Plasma Membrane |
| 78 | P20029 | HSPA5 | 3.64E+07 | 4.04E+07 | 4.57E+06 | 78 | Cytoplasm |
| 79 | Q6PHN9 | RAB35 | 3.52E+07 | 2.43E+07 | 5.30E+06 | 21 | Cytoplasm |
| 80 | Q61411 | HRAS | 3.46E+07 | 4.38E+07 | 8.00E+06 | 30 | Plasma Membrane |
| 81 | P47955 | Rplp1 (incl | 3.38E+07 | 2.70E+06 | 1.10E+06 | 6 | Nucleus |
| 82 | Q8BTM8 | FLNA | 3.32E+07 | 3.19E+07 | 1.70E+06 | 354 | Cytoplasm |
| 83 | Q9QUM0 | ITGA2B | 3.29E+07 | 3.88E+07 | 4.31E+06 | 81 | Plasma Membrane |
| 84 | Q03265 | ATP5A1 | 3.26E+07 | 2.26E+07 | 2.72E+06 | 69 | Cytoplasm |
| 85 | O54890 | ITGB3 | 3.11E+07 | 2.92E+07 | 3.45E+06 | 72 | Plasma Membrane |
| 86 | Q922M5 | CDCA7L | 3.06E+07 | 7.47E+06 | 4.31E+06 | 3 | Nucleus |
| 87 | P08249 | MDH2 | 3.05E+07 | 1.60E+07 | 2.57E+06 | 39 | Cytoplasm |
| 88 | P63101 | YWHAZ | 3.05E+07 | 2.56E+07 | 3.70E+06 | 48 | Cytoplasm |
| 89 | P56480 | ATP5B | 2.99E+07 | 3.02E+07 | 3.08E+06 | 96 | Cytoplasm |
| 90 | P08752 | GNAI2 | 2.98E+07 | 2.64E+07 | 2.99E+06 | 78 | Plasma Membrane |
| 91 | P51881 | SLC25A5 | 2.95E+07 | 3.61E+07 | 5.38E+06 | 45 | Cytoplasm |
| 92 | Q8CIM8 | INTS4 | 2.91E+07 | 5.10E+06 | 2.95E+06 | 3 | Nucleus |
| 93 | Q9CX63 | C17orf99 | 2.91E+07 | 1.66E+07 | 3.62E+06 | 21 | Extracellular Space |
| 94 | Q60692 | PSMB6 | 2.86E+07 | 3.26E+07 | 7.67E+06 | 18 | Nucleus |
| 95 | Q3TZ89 | SEC31B | 2.85E+07 | 7.15E+05 | 4.13E+05 | 3 | Cytoplasm |
| 96 | Q8BGQ1 | VIPAS39 | 2.81E+07 | 4.02E+06 | 2.32E+06 | 3 | Plasma Membrane |
| 97 | Q9JIM1 | SLC29A1 | 2.74E+07 | 3.69E+07 | 6.43E+06 | 33 | Plasma Membrane |
| 98 | Q6UJY2 | SLC9C1 | 2.72E+07 | 1.93E+07 | 1.11E+07 | 3 | Extracellular Space |
| 99 | O88186 | GP9 | 2.70E+07 | 3.49E+07 | 1.01E+07 | 12 | Plasma Membrane |
| 100 | Q8K400 | STXBP5 | 2.51E+07 | 7.34E+06 | 4.24E+06 | 3 | Plasma Membrane |
| 101 | P35564 | CANX | 2.51E+07 | 2.92E+07 | 3.59E+06 | 66 | Cytoplasm |
| 102 | Q64339 | ISG15 | 2.48E+07 | 2.29E+07 | 4.67E+06 | 24 | Extracellular Space |
| 103 | Q7TMS5 | ABCG2 | 2.48E+07 | 2.40E+07 | 3.10E+06 | 60 | Plasma Membrane |
| 104 | P62880 | GNB2 | 2.48E+07 | 2.03E+07 | 3.39E+06 | 36 | Plasma Membrane |
| 105 | O08742 | GP5 | 2.46E+07 | 1.72E+07 | 3.14E+06 | 30 | Plasma Membrane |
| 106 | P24369 | PPIB | 2.45E+07 | 2.08E+07 | 3.47E+06 | 36 | Cytoplasm |
| 107 | Q9Z2U0 | PSMA7 | 2.43E+07 | 2.31E+07 | 3.85E+06 | 36 | Cytoplasm |
| 108 | P27773 | PDIA3 | 2.43E+07 | 2.54E+07 | 3.06E+06 | 69 | Cytoplasm |
| 109 | Q9Z2U1 | PSMA5 | 2.41E+07 | 2.56E+07 | 3.96E+06 | 42 | Cytoplasm |
| 110 | Q8K558 | TREML1 | 2.41E+07 | 1.56E+07 | 3.18E+06 | 24 | Plasma Membrane |
| 111 | P14231 | ATP1B2 | 2.40E+07 | 2.33E+07 | 6.01E+06 | 15 | Plasma Membrane |
| 112 | O08586 | PTEN | 2.40E+07 | 4.89E+06 | 2.82E+06 | 3 | Cytoplasm |
| 113 | Q5F285 | TMEM256 | 2.39E+07 | 3.65E+06 | 2.11E+06 | 3 | Cytoplasm |
| 114 | P62874 | GNB1 | 2.37E+07 | 2.32E+07 | 3.72E+06 | 39 | Plasma Membrane |
| 115 | Q8R3G9 | TSPAN8 | 2.37E+07 | 1.42E+07 | 3.66E+06 | 15 | Plasma Membrane |
| 116 | O35930 | GP1BA | 2.36E+07 | 2.22E+07 | 3.55E+06 | 39 | Plasma Membrane |
| 117 | Q8VDD5 | MYH9 | 2.36E+07 | 1.66E+07 | 9.79E+05 | 288 | Cytoplasm |
| 118 | Q9R0K7 | ATP2B2 | 2.35E+07 | 3.17E+07 | 5.79E+06 | 30 | Plasma Membrane |
| 119 | Q61469 | Ppap2a | 2.26E+07 | 2.06E+07 | 5.31E+06 | 15 | Plasma Membrane |
| 120 | Q8R3S2 | TSPAN33 | 2.25E+07 | 3.51E+07 | 7.65E+06 | 21 | Plasma Membrane |
| 121 | P08113 | HSP90B1 | 2.21E+07 | 1.14E+07 | 1.29E+06 | 78 | Cytoplasm |
| 122 | Q9D1G1 | RAB1B | 2.19E+07 | 1.56E+07 | 4.03E+06 | 15 | Cytoplasm |
| 123 | Q09199 | B4GALNT2 | 2.19E+07 | 4.81E+06 | 2.78E+06 | 3 | Cytoplasm |
| 124 | Q9QUM9 | PSMA6 | 2.15E+07 | 1.14E+07 | 1.99E+06 | 33 | Cytoplasm |
| 125 | Q9JLN5 | ERMAP | 2.13E+07 | 2.55E+07 | 3.29E+06 | 60 | Cytoplasm |
| 126 | P84096 | RHOG | 2.12E+07 | 1.68E+07 | 2.92E+06 | 33 | Cytoplasm |
| 127 | Q9QXY7 | XK | 2.08E+07 | 5.12E+06 | 1.48E+06 | 12 | Plasma Membrane |
| 128 | Q3UZP4 | SVIP | 2.05E+07 | 2.22E+06 | 1.28E+06 | 3 | Cytoplasm |
| 129 | P28656 | NAP1L1 | 2.04E+07 | 1.37E+07 | 2.79E+06 | 24 | Nucleus |
| 130 | Q9DC29 | ABCB6 | 2.04E+07 | 2.63E+07 | 3.40E+06 | 60 | Cytoplasm |
| 131 | Q62136 | PTPN21 | 2.01E+07 | 2.68E+06 | 1.54E+06 | 3 | Cytoplasm |
| 132 | Q8R146 | APEH | 1.99E+07 | 1.31E+07 | 1.51E+06 | 75 | Cytoplasm |
| 133 | Q80Y14 | GLRX5 | 1.97E+07 | 8.14E+06 | 3.32E+06 | 6 | Cytoplasm |
| 134 | A2AVZ9 | SLC43A3 | 1.97E+07 | 2.58E+07 | 7.45E+06 | 12 | Extracellular Space |
| 135 | P62821 | RAB1A | 1.96E+07 | 2.73E+07 | 6.43E+06 | 18 | Cytoplasm |
| 136 | P99028 | UQCRHL | 1.96E+07 | 1.28E+07 | 5.22E+06 | 6 | Cytoplasm |
| 137 | P08556 | NRAS | 1.96E+07 | 5.27E+05 | 3.04E+05 | 3 | Plasma Membrane |
| 138 | Q99KU0 | VMP1 | 1.94E+07 | 5.13E+06 | 2.96E+06 | 3 | Plasma Membrane |

| Rank | ID | Symbol | Mean Intensity | St. Dev | Std. Err | n | Primary Location |
|---|---|---|---|---|---|---|---|
| 1 | P04919 | SLC4A1 | 2.70E+09 | 4.72E+09 | 3.61E+08 | 171 | Plasma Membrane |
| 2 | P14220 | Gypa | 2.58E+09 | 2.27E+09 | 9.26E+08 | 6 | Plasma Membrane |
| 3 | Q02013 | AQP1 | 1.32E+09 | 1.05E+09 | 2.71E+08 | 15 | Plasma Membrane |
| 4 | Q5DU02 | USP22 | 1.17E+09 | 5.38E+07 | 3.11E+07 | 3 | Nucleus |
| 5 | P68033 | ACTC1 | 9.88E+08 | 1.23E+09 | 3.56E+08 | 12 | Cytoplasm |
| 6 | P02088 | Hbb-b1 | 9.26E+08 | 9.24E+08 | 1.22E+08 | 57 | Extracellular Space |
| 7 | P01942 | HBA1/HBA2 | 8.78E+08 | 6.08E+08 | 1.06E+08 | 33 | Extracellular Space |
| 8 | Q78HU7 | GYPC | 6.54E+08 | 5.40E+08 | 1.80E+08 | 9 | Plasma Membrane |
| 9 | P60710 | ACTB | 5.37E+08 | 8.71E+08 | 7.26E+07 | 144 | Cytoplasm |
| 10 | P08032 | SPTA1 | 5.26E+08 | 6.35E+08 | 2.12E+07 | 894 | Cytoplasm |
| 11 | Q02357 | ANK1 | 5.05E+08 | 6.64E+08 | 2.95E+07 | 507 | Plasma Membrane |
| 12 | P15508 | SPTB | 4.85E+08 | 5.82E+08 | 2.09E+07 | 780 | Plasma Membrane |
| 13 | Q8K386 | Rab15 | 4.42E+08 | 3.16E+07 | 1.82E+07 | 3 | Cytoplasm |
| 14 | P48193 | EPB41 | 4.41E+08 | 5.94E+08 | 4.36E+07 | 186 | Plasma Membrane |
| 15 | P70290 | MPP1 | 3.87E+08 | 4.39E+08 | 3.91E+07 | 126 | Plasma Membrane |
| 16 | P49222 | EPB42 | 3.63E+08 | 4.91E+08 | 3.54E+07 | 192 | Plasma Membrane |
| 17 | P18572 | BSG | 2.77E+08 | 3.17E+08 | 4.90E+07 | 42 | Plasma Membrane |
| 18 | Q8N9S3 | Ahsa2 | 2.68E+08 | 3.80E+07 | 2.19E+07 | 3 | Cytoplasm |
| 19 | Q02525 | ZNF717 | 2.57E+08 | 2.87E+07 | 1.66E+07 | 3 | Nucleus |
| 20 | P53986 | SLC16A1 | 2.36E+08 | 2.05E+08 | 3.94E+07 | 27 | Plasma Membrane |
| 21 | P06803 | PIM1 | 2.30E+08 | 1.50E+07 | 8.65E+06 | 3 | Cytoplasm |
| 22 | Q61735 | CD47 | 2.15E+08 | 1.79E+08 | 4.22E+07 | 18 | Plasma Membrane |
| 23 | Q9JJ04 | B4GALT4 | 2.10E+08 | 4.89E+07 | 2.82E+07 | 3 | Cytoplasm |
| 24 | P02089 | Hbb-b2 | 2.06E+08 | 1.76E+08 | 3.60E+07 | 24 | Extracellular Space |
| 25 | P62984 | Uba52 | 1.96E+08 | 2.79E+08 | 5.70E+07 | 24 | Cytoplasm |
| 26 | P54116 | STOM | 1.91E+08 | 2.76E+08 | 3.32E+07 | 69 | Plasma Membrane |
| 27 | Q9WV69 | DMTN | 1.89E+08 | 1.86E+08 | 2.15E+07 | 75 | Plasma Membrane |
| 28 | Q8C6A8 | BHLHE22 | 1.67E+08 | 7.50E+06 | 4.33E+06 | 3 | Nucleus |
| 29 | Q9QYC0 | ADD1 | 1.57E+08 | 1.78E+08 | 1.61E+07 | 123 | Cytoplasm |
| 30 | Q8BSM7 | SLC43A1 | 1.36E+08 | 1.86E+08 | 2.87E+07 | 42 | Plasma Membrane |
| 31 | Q9QZI9 | SERINC3 | 1.36E+08 | 7.47E+07 | 2.16E+07 | 12 | Cytoplasm |
| 32 | P21107 | TPM3 | 1.35E+08 | 1.34E+08 | 2.06E+07 | 42 | Cytoplasm |
| 33 | Q8VHL0 | SLC14A1 | 1.34E+08 | 1.14E+08 | 1.90E+07 | 36 | Plasma Membrane |
| 34 | Q61171 | PRDX2 | 1.09E+08 | 9.09E+07 | 1.40E+07 | 42 | Cytoplasm |
| 35 | P58771 | Tpm1 | 1.05E+08 | 1.33E+08 | 3.44E+07 | 15 | Plasma Membrane |
| 36 | P70696 | HIST1H2BA | 1.05E+08 | 3.50E+08 | 2.02E+08 | 3 | Nucleus |
| 37 | P16858 | GAPDH | 1.00E+08 | 8.68E+07 | 9.31E+06 | 87 | Cytoplasm |
| 38 | Q08857 | CD36 | 9.57E+07 | 1.23E+08 | 1.77E+07 | 48 | Plasma Membrane |
| 39 | Q9Z0S4 | Cldn13 | 9.30E+07 | 7.69E+07 | 2.22E+07 | 12 | Plasma Membrane |
| 40 | Q9QYB8 | ADD2 | 9.03E+07 | 1.11E+08 | 8.01E+06 | 192 | Cytoplasm |
| 41 | P70269 | CTSE | 8.66E+07 | 5.51E+07 | 1.84E+07 | 9 | Cytoplasm |
| 42 | P49813 | TMOD1 | 7.99E+07 | 8.25E+07 | 9.72E+06 | 72 | Cytoplasm |
| 43 | P83503 | NYX | 7.87E+07 | 1.52E+07 | 8.77E+06 | 3 | Extracellular Space |
| 44 | P15864 | HIST1H1C | 7.77E+07 | 1.86E+08 | 1.07E+08 | 3 | Nucleus |
| 45 | Q9Z126 | PF4 | 7.60E+07 | 3.82E+07 | 1.27E+07 | 9 | Extracellular Space |
| 46 | Q99JI6 | RAP1B | 7.19E+07 | 7.22E+07 | 1.47E+07 | 24 | Cytoplasm |
| 47 | P61027 | RAB10 | 7.16E+07 | 8.05E+07 | 1.40E+07 | 33 | Cytoplasm |
| 48 | Q8CGP5 | HIST1H2AJ | 7.09E+07 | 2.89E+08 | 7.46E+07 | 15 | Nucleus |
| 49 | Q3U9N9 | SLC16A10 | 6.25E+07 | 5.51E+07 | 1.01E+07 | 30 | Plasma Membrane |
| 50 | P14211 | CALR | 6.18E+07 | 6.27E+07 | 8.31E+06 | 57 | Cytoplasm |
| 51 | P16951 | ATF2 | 6.08E+07 | 4.90E+06 | 2.83E+06 | 3 | Nucleus |
| 52 | P61028 | RAB8B | 5.75E+07 | 5.66E+07 | 1.33E+07 | 18 | Cytoplasm |
| 53 | Q9CYH2 | FAM213A | 5.71E+07 | 5.26E+07 | 8.77E+06 | 36 | Extracellular Space |
| 54 | Q9EQF2 | KEL | 5.67E+07 | 4.59E+07 | 4.42E+06 | 108 | Plasma Membrane |
| 55 | P63017 | HSPA8 | 5.57E+07 | 3.58E+07 | 4.06E+06 | 78 | Cytoplasm |
| 56 | Q01853 | VCP | 4.97E+07 | 4.81E+07 | 3.45E+06 | 195 | Cytoplasm |
| 57 | P35278 | RAB5C | 4.88E+07 | 4.22E+07 | 6.76E+06 | 39 | Cytoplasm |
| 58 | P55258 | RAB8A | 4.84E+07 | 3.78E+07 | 7.72E+06 | 24 | Plasma Membrane |
| 59 | Q8R3S2 | TSPAN33 | 4.80E+07 | 6.08E+07 | 1.33E+07 | 21 | Plasma Membrane |
| 60 | Q8BR07 | BICD1 | 4.67E+07 | 1.16E+07 | 6.68E+06 | 3 | Cytoplasm |
| 61 | Q9JLN5 | ERMAP | 4.67E+07 | 4.49E+07 | 5.79E+06 | 60 | Cytoplasm |
| 62 | P62806 | H4 | 4.40E+07 | 1.94E+08 | 3.97E+07 | 24 | Nucleus |
| 63 | Q2VIS4 | Flg2 | 4.29E+07 | 5.16E+06 | 2.98E+06 | 3 | Extracellular Space |
| 64 | P27600 | GNA12 | 4.28E+07 | 8.02E+06 | 4.63E+06 | 3 | Plasma Membrane |
| 65 | Q61469 | Ppap2a | 4.22E+07 | 2.08E+07 | 5.38E+06 | 15 | Plasma Membrane |
| 66 | P63001 | RAC1 | 4.22E+07 | 3.18E+07 | 4.90E+06 | 42 | Plasma Membrane |
| 67 | O55186 | Cd59a | 3.95E+07 | 4.84E+07 | 1.61E+07 | 9 | Plasma Membrane |
| 68 | Q9JIM1 | SLC29A1 | 3.80E+07 | 4.28E+07 | 7.45E+06 | 33 | Plasma Membrane |
| 69 | P62835 | RAP1A | 3.77E+07 | 4.15E+07 | 8.48E+06 | 24 | Cytoplasm |
| 70 | Q7TMS5 | ABCG2 | 3.71E+07 | 3.39E+07 | 4.38E+06 | 60 | Plasma Membrane |
| 71 | Q6PHN9 | RAB35 | 3.63E+07 | 2.75E+07 | 6.01E+06 | 21 | Cytoplasm |
| 72 | P10126 | EEF1A1 | 3.53E+07 | 4.92E+07 | 5.92E+06 | 69 | Cytoplasm |
| 73 | P20029 | HSPA5 | 3.50E+07 | 3.54E+07 | 4.00E+06 | 78 | Cytoplasm |
| 74 | Q8CIM8 | INTS4 | 3.39E+07 | 8.56E+05 | 4.94E+05 | 3 | Nucleus |
| 75 | O08586 | PTEN | 3.28E+07 | 3.60E+06 | 2.08E+06 | 3 | Cytoplasm |
| 76 | Q8K400 | STXBP5 | 3.26E+07 | 4.30E+06 | 2.48E+06 | 3 | Plasma Membrane |
| 77 | A2AVZ9 | SLC43A3 | 3.25E+07 | 2.65E+07 | 7.66E+06 | 12 | Extracellular Space |
| 78 | Q9ES97 | RTN3 | 3.23E+07 | 3.72E+07 | 1.52E+07 | 6 | Cytoplasm |
| 79 | Q64475 | HIST1H2BB | 3.00E+07 | 1.70E+08 | 4.01E+07 | 18 | Nucleus |
| 80 | Q922M5 | CDCA7L | 2.98E+07 | 5.76E+06 | 3.33E+06 | 3 | Nucleus |
| 81 | Q6UJY2 | SLC9C1 | 2.92E+07 | 1.50E+07 | 8.66E+06 | 3 | Extracellular Space |
| 82 | O55112 | Aff2 | 2.90E+07 | 1.11E+07 | 6.40E+06 | 3 | Nucleus |
| 83 | P35564 | CANX | 2.89E+07 | 2.62E+07 | 3.22E+06 | 66 | Cytoplasm |
| 84 | Q61672 | SLC29A2 | 2.88E+07 | 1.10E+06 | 6.33E+05 | 3 | Plasma Membrane |
| 85 | Q60692 | PSMB6 | 2.85E+07 | 3.14E+07 | 7.41E+06 | 18 | Nucleus |
| 86 | Q9QUM9 | PSMA6 | 2.81E+07 | 1.23E+07 | 2.15E+06 | 33 | Cytoplasm |
| 87 | P68433 | H31 | 2.79E+07 | 9.11E+07 | 2.15E+07 | 18 | Nucleus |
| 88 | P24369 | PPIB | 2.77E+07 | 2.10E+07 | 3.50E+06 | 36 | Cytoplasm |
| 89 | Q61411 | HRAS | 2.76E+07 | 3.46E+07 | 6.31E+06 | 30 | Plasma Membrane |
| 90 | Q9Z2U1 | PSMA5 | 2.73E+07 | 2.52E+07 | 3.89E+06 | 42 | Cytoplasm |
| 91 | Q9Z2U0 | PSMA7 | 2.73E+07 | 2.34E+07 | 3.90E+06 | 36 | Cytoplasm |
| 92 | O35598 | ADAM10 | 2.71E+07 | 2.39E+07 | 2.70E+06 | 78 | Plasma Membrane |
| 93 | Q8R3G9 | TSPAN8 | 2.71E+07 | 1.53E+07 | 3.94E+06 | 15 | Plasma Membrane |
| 94 | Q9DCT8 | Crip2 | 2.69E+07 | 1.37E+07 | 2.64E+06 | 27 | Plasma Membrane |
| 95 | P10404 | ENV1 | 2.69E+07 | 2.53E+07 | 2.98E+06 | 72 | Other |
| 96 | O54962 | BANF1 | 2.66E+07 | 5.48E+07 | 1.58E+07 | 12 | Nucleus |
| 97 | Q9R0K7 | ATP2B2 | 2.63E+07 | 3.30E+07 | 6.03E+06 | 30 | Plasma Membrane |
| 98 | Q64339 | ISG15 | 2.61E+07 | 1.78E+07 | 3.63E+06 | 24 | Extracellular Space |
| 99 | Q9QXY7 | XK | 2.59E+07 | 6.05E+06 | 1.75E+06 | 12 | Plasma Membrane |
| 100 | Q8R146 | APEH | 2.52E+07 | 1.39E+07 | 1.60E+06 | 75 | Cytoplasm |
| 101 | O35450 | FKBPL | 2.49E+07 | 1.86E+06 | 1.07E+06 | 3 | Nucleus |
| 102 | P08752 | GNAI2 | 2.47E+07 | 2.51E+07 | 2.84E+06 | 78 | Plasma Membrane |
| 103 | P63024 | VAMP3 | 2.47E+07 | 3.56E+07 | 9.19E+06 | 15 | Plasma Membrane |
| 104 | Q60605 | MYL6 | 2.45E+07 | 2.02E+07 | 4.40E+06 | 21 | Cytoplasm |
| 105 | P27773 | PDIA3 | 2.42E+07 | 2.31E+07 | 2.78E+06 | 69 | Cytoplasm |
| 106 | P62204 | CALM1 | 2.40E+07 | 1.27E+07 | 5.17E+06 | 6 | Cytoplasm |
| 107 | P20108 | PRDX3 | 2.39E+07 | 3.42E+07 | 7.46E+06 | 21 | Cytoplasm |
| 108 | Q62351 | TFRC | 2.39E+07 | 3.13E+07 | 2.97E+06 | 111 | Plasma Membrane |
| 109 | P56400 | GP1BB | 2.37E+07 | 3.21E+07 | 7.56E+06 | 18 | Plasma Membrane |
| 110 | Q9D1G1 | RAB1B | 2.37E+07 | 1.57E+07 | 4.06E+06 | 15 | Cytoplasm |
| 111 | Q3UZP4 | SVIP | 2.35E+07 | 3.06E+06 | 1.76E+06 | 3 | Cytoplasm |
| 112 | Q9DC29 | ABCB6 | 2.32E+07 | 2.66E+07 | 3.43E+06 | 60 | Cytoplasm |
| 113 | P07724 | ALB | 2.27E+07 | 1.57E+07 | 1.85E+06 | 72 | Extracellular Space |
| 114 | P14231 | ATP1B2 | 2.26E+07 | 2.09E+07 | 5.40E+06 | 15 | Plasma Membrane |
| 115 | Q62312 | TGFBR2 | 2.25E+07 | 3.78E+06 | 2.18E+06 | 3 | Plasma Membrane |
| 116 | P56480 | ATP5B | 2.24E+07 | 3.03E+07 | 3.09E+06 | 96 | Cytoplasm |
| 117 | P99026 | PSMB4 | 2.23E+07 | 7.15E+06 | 1.69E+06 | 18 | Cytoplasm |
| 118 | P08113 | HSP90B1 | 2.21E+07 | 1.04E+07 | 1.18E+06 | 78 | Cytoplasm |
| 119 | Q62136 | PTPN21 | 2.18E+07 | 1.78E+06 | 1.03E+06 | 3 | Cytoplasm |
| 120 | P05064 | ALDOA | 2.17E+07 | 1.97E+07 | 3.28E+06 | 36 | Cytoplasm |
| 121 | P51881 | SLC25A5 | 2.13E+07 | 3.47E+07 | 5.17E+06 | 45 | Cytoplasm |
| 122 | Q8BGQ1 | VIPAS39 | 2.13E+07 | 1.03E+06 | 5.94E+05 | 3 | Plasma Membrane |
| 123 | P49722 | PSMA2 | 2.09E+07 | 1.97E+07 | 3.59E+06 | 30 | Cytoplasm |
| 124 | Q03265 | ATP5A1 | 2.08E+07 | 2.33E+07 | 2.81E+06 | 69 | Cytoplasm |
| 125 | Q9R1P0 | PSMA4 | 2.08E+07 | 1.28E+07 | 2.04E+06 | 39 | Cytoplasm |
| 126 | P97300 | Nptn | 2.08E+07 | 2.86E+07 | 5.50E+06 | 27 | Plasma Membrane |
| 127 | Q9QUI6 | ACKR1 | 2.06E+07 | 1.51E+07 | 4.36E+06 | 12 | Plasma Membrane |
| 128 | O88186 | GP9 | 2.04E+07 | 3.12E+07 | 9.00E+06 | 12 | Plasma Membrane |
| 129 | Q09199 | B4GALNT2 | 2.02E+07 | 3.05E+06 | 1.76E+06 | 3 | Cytoplasm |
| 130 | Q8BTM8 | FLNA | 2.02E+07 | 2.86E+07 | 1.52E+06 | 354 | Cytoplasm |
| 131 | P62821 | RAB1A | 2.01E+07 | 2.64E+07 | 6.22E+06 | 18 | Cytoplasm |
| 132 | Q64735 | CR1L | 2.01E+07 | 1.78E+07 | 3.43E+06 | 27 | Plasma Membrane |
| 133 | Q9QUM0 | ITGA2B | 1.99E+07 | 3.67E+07 | 4.07E+06 | 81 | Plasma Membrane |
| 134 | P63101 | YWHAZ | 1.97E+07 | 2.26E+07 | 3.26E+06 | 48 | Cytoplasm |
| 135 | Q91YH5 | ATL3 | 1.97E+07 | 1.44E+07 | 2.08E+06 | 48 | Cytoplasm |
| 136 | O70435 | PSMA3 | 1.95E+07 | 8.48E+06 | 1.63E+06 | 27 | Cytoplasm |
| 137 | P28656 | NAP1L1 | 1.95E+07 | 1.27E+07 | 2.59E+06 | 24 | Nucleus |
| 138 | O55234 | PSMB5 | 1.93E+07 | 1.06E+07 | 1.85E+06 | 33 | Cytoplasm |

| Rank | ID | Symbol | Mean Intensity | St. Dev | Std. Err | n | Primary Location |
|---|---|---|---|---|---|---|---|
| 1 | P04919 | SLC4A1 | 2489840952 | 4889925980 | 373941986 | 171 | Plasma Membrane |
| 2 | P14220 | Gypa | 2380555000 | 2234668239 | 912299488 | 6 | Plasma Membrane |
| 3 | P02088 | Hbb-b1 | 1843402632 | 1668460344 | 220992971 | 57 | Extracellular Space |
| 4 | P01942 | HBA1/HBA2 | 1504263636 | 988090145 | 172004416 | 33 | Extracellular Space |
| 5 | Q5DU02 | USP22 | 1130000000 | 147805052 | 85335286 | 3 | Nucleus |
| 6 | P68033 | ACTC1 | 912260000 | 1231344408 | 355458513 | 12 | Cytoplasm |
| 7 | Q78HU7 | GYPC | 696406667 | 671651046 | 223883682 | 9 | Plasma Membrane |
| 8 | Q02357 | ANK1 | 517749535 | 720002755 | 31976445 | 507 | Plasma Membrane |
| 9 | P08032 | SPTA1 | 517042241 | 677959258 | 22674349 | 894 | Cytoplasm |
| 10 | P60710 | ACTB | 510553356 | 881162387 | 73430199 | 144 | Cytoplasm |
| 11 | P15508 | SPTB | 489650205 | 633686995 | 22689634 | 780 | Plasma Membrane |
| 12 | P70290 | MPP1 | 405535812 | 471705382 | 42022855 | 126 | Plasma Membrane |
| 13 | P02089 | Hbb-b2 | 404938310 | 303987446 | 62051178 | 24 | Extracellular Space |
| 14 | P48193 | EPB41 | 402340359 | 620132318 | 45470308 | 186 | Plasma Membrane |
| 15 | Q8K386 | Rab15 | 383500000 | 41584893 | 24009049 | 3 | Cytoplasm |
| 16 | P49222 | EPB42 | 371584910 | 518741628 | 37436952 | 192 | Plasma Membrane |
| 17 | P18572 | BSG | 270724908 | 324790933 | 50116329 | 42 | Plasma Membrane |
| 18 | P06803 | PIM1 | 259200000 | 21934448 | 12663859 | 3 | Cytoplasm |
| 19 | Q8N9S3 | Ahsa2 | 246400000 | 77889751 | 44969669 | 3 | Cytoplasm |
| 20 | Q02525 | ZNF717 | 228200000 | 21742815 | 12553220 | 3 | Nucleus |
| 21 | Q61735 | CD47 | 217834472 | 194176240 | 45767779 | 18 | Plasma Membrane |
| 22 | P53986 | SLC16A1 | 208610719 | 218809031 | 42109818 | 27 | Plasma Membrane |
| 23 | Q9WV69 | DMTN | 193817110 | 197809608 | 22841086 | 75 | Plasma Membrane |
| 24 | Q8C6A8 | BHLHE22 | 188400000 | 9867793 | 5697173 | 3 | Nucleus |
| 25 | P62984 | Uba52 | 177365054 | 310308755 | 63341509 | 24 | Cytoplasm |
| 26 | Q9JJ04 | B4GALT4 | 175200000 | 69249862 | 39981426 | 3 | Cytoplasm |
| 27 | P54116 | STOM | 162786506 | 276465551 | 33282541 | 69 | Plasma Membrane |
| 28 | Q9QZI9 | SERINC3 | 160400000 | 95427980 | 27547685 | 12 | Cytoplasm |
| 29 | Q9QYC0 | ADD1 | 149444817 | 175302209 | 15806468 | 123 | Cytoplasm |
| 30 | Q8BSM7 | SLC43A1 | 134827322 | 199706570 | 30815393 | 42 | Plasma Membrane |
| 31 | Q8VHL0 | SLC14A1 | 130662500 | 121000517 | 20166753 | 36 | Plasma Membrane |
| 32 | Q61171 | PRDX2 | 127511429 | 126174830 | 19469199 | 42 | Cytoplasm |
| 33 | P21107 | TPM3 | 126752649 | 140644962 | 21701989 | 42 | Cytoplasm |
| 34 | P16858 | GAPDH | 125578354 | 115425269 | 12374888 | 87 | Cytoplasm |
| 35 | Q08857 | CD36 | 95572732 | 129154917 | 18641907 | 48 | Plasma Membrane |
| 36 | Q9QYB8 | ADD2 | 93510264 | 120355852 | 8685935 | 192 | Cytoplasm |
| 37 | P83503 | NYX | 92530000 | 3186163 | 1839532 | 3 | Extracellular Space |
| 38 | Q9Z0S4 | Cldn13 | 91662500 | 68422249 | 19751802 | 12 | Plasma Membrane |
| 39 | P58771 | Tpm1 | 90508369 | 135545858 | 34997790 | 15 | Plasma Membrane |
| 40 | P49813 | TMOD1 | 75160413 | 95635154 | 11270711 | 72 | Cytoplasm |
| 41 | P70269 | CTSE | 68243333 | 54446516 | 18148839 | 9 | Cytoplasm |
| 42 | P61027 | RAB10 | 64087970 | 84408146 | 14693572 | 33 | Cytoplasm |
| 43 | P16951 | ATF2 | 61600000 | 2560280 | 1478178 | 3 | Nucleus |
| 44 | P63017 | HSPA8 | 59037631 | 40943795 | 4635972 | 78 | Cytoplasm |
| 45 | P61028 | RAB8B | 57511557 | 55691455 | 13126602 | 18 | Cytoplasm |
| 46 | Q01853 | VCP | 51242846 | 58049138 | 4156985 | 195 | Cytoplasm |
| 47 | Q8R3S2 | TSPAN33 | 51227195 | 74585035 | 16275789 | 21 | Plasma Membrane |
| 48 | Q3U9N9 | SLC16A10 | 51209285 | 56149877 | 10251518 | 30 | Plasma Membrane |
| 49 | Q9EQF2 | KEL | 50856646 | 48094652 | 4627910 | 108 | Plasma Membrane |
| 50 | P27600 | GNA12 | 50510000 | 8617592 | 4975369 | 3 | Plasma Membrane |
| 51 | Q99JI6 | RAP1B | 50096971 | 61093307 | 12470619 | 24 | Cytoplasm |
| 52 | Q9JLN5 | ERMAP | 46415815 | 50707144 | 6546264 | 60 | Cytoplasm |
| 53 | P55258 | RAB8A | 44991474 | 38030831 | 7763011 | 24 | Plasma Membrane |
| 54 | P63001 | RAC1 | 44984551 | 32687783 | 5043834 | 42 | Plasma Membrane |
| 55 | P10126 | EEF1A1 | 44338038 | 55804474 | 6718069 | 69 | Cytoplasm |
| 56 | Q9CYH2 | FAM213A | 44020229 | 53898669 | 8983111 | 36 | Extracellular Space |
| 57 | Q8BR07 | BICD1 | 42810000 | 5957080 | 3439322 | 3 | Cytoplasm |
| 58 | P14211 | CALR | 42194415 | 56797906 | 7523066 | 57 | Cytoplasm |
| 59 | P35278 | RAB5C | 41440140 | 42650060 | 6829475 | 39 | Cytoplasm |
| 60 | Q2VIS4 | Flg2 | 41370000 | 3715109 | 2144919 | 3 | Extracellular Space |
| 61 | Q6PHN9 | RAB35 | 41038562 | 29640613 | 6468112 | 21 | Cytoplasm |
| 62 | O55186 | Cd59a | 40493347 | 54238319 | 18079440 | 9 | Plasma Membrane |
| 63 | Q7TMS5 | ABCG2 | 39547636 | 36738695 | 4742945 | 60 | Plasma Membrane |
| 64 | Q9JIM1 | SLC29A1 | 39509725 | 47364926 | 8245175 | 33 | Plasma Membrane |
| 65 | Q61469 | Ppap2a | 36382615 | 22230108 | 5739789 | 15 | Plasma Membrane |
| 66 | P62835 | RAP1A | 36106418 | 37768958 | 7709556 | 24 | Cytoplasm |
| 67 | Q8CIM8 | INTS4 | 35540000 | 7212122 | 4163920 | 3 | Nucleus |
| 68 | P10404 | ENV1 | 35442755 | 32984838 | 3887300 | 72 | Other |
| 69 | Q60692 | PSMB6 | 35393645 | 36558303 | 8616875 | 18 | Nucleus |
| 70 | P20108 | PRDX3 | 35055396 | 44779605 | 9771711 | 21 | Cytoplasm |
| 71 | Q6UJY2 | SLC9C1 | 34830000 | 23283292 | 13442615 | 3 | Extracellular Space |
| 72 | O35450 | FKBPL | 31310000 | 5431043 | 3135614 | 3 | Nucleus |
| 73 | O55112 | Aff2 | 31110000 | 9611077 | 5548958 | 3 | Nucleus |
| 74 | Q9Z2U0 | PSMA7 | 30965908 | 27306763 | 4551127 | 36 | Cytoplasm |
| 75 | Q62351 | TFRC | 30963160 | 37350153 | 3545120 | 111 | Plasma Membrane |
| 76 | Q9Z2U1 | PSMA5 | 30805768 | 28475107 | 4393804 | 42 | Cytoplasm |
| 77 | P47955 | Rplp1 (includes others) | 30645000 | 14297864 | 5837078 | 6 | Nucleus |
| 78 | Q9R0K7 | ATP2B2 | 30262741 | 41699195 | 7613197 | 30 | Plasma Membrane |
| 79 | O35598 | ADAM10 | 29722013 | 26256121 | 2972920 | 78 | Plasma Membrane |
| 80 | Q9QUM9 | PSMA6 | 29264636 | 13751344 | 2393802 | 33 | Cytoplasm |
| 81 | P20029 | HSPA5 | 28851581 | 37061919 | 4196436 | 78 | Cytoplasm |
| 82 | P70696 | HIST1H2BA | 28320000 | 12694851 | 7329375 | 3 | Nucleus |
| 83 | Q9Z126 | PF4 | 28143333 | 13528917 | 4509639 | 9 | Extracellular Space |
| 84 | P08752 | GNAI2 | 28023825 | 26969910 | 3053741 | 78 | Plasma Membrane |
| 85 | Q922M5 | CDCA7L | 27810000 | 8169498 | 4716662 | 3 | Nucleus |
| 86 | O08586 | PTEN | 27810000 | 1827165 | 1054914 | 3 | Cytoplasm |
| 87 | Q9ES97 | RTN3 | 26724443 | 39262434 | 16028821 | 6 | Cytoplasm |
| 88 | P14231 | ATP1B2 | 26383039 | 24783323 | 6399026 | 15 | Plasma Membrane |
| 89 | Q62136 | PTPN21 | 25880000 | 2300261 | 1328056 | 3 | Cytoplasm |
| 90 | Q8R146 | APEH | 25253801 | 14393158 | 1661979 | 75 | Cytoplasm |
| 91 | Q8R3G9 | TSPAN8 | 25126774 | 17192105 | 4438983 | 15 | Plasma Membrane |
| 92 | Q8K400 | STXBP5 | 24790000 | 6269314 | 3619590 | 3 | Plasma Membrane |
| 93 | Q9DC29 | ABCB6 | 24741198 | 31719763 | 4095004 | 60 | Cytoplasm |
| 94 | P49722 | PSMA2 | 24103247 | 23448463 | 4281084 | 30 | Cytoplasm |
| 95 | P62880 | GNB2 | 23913229 | 20965472 | 3494245 | 36 | Plasma Membrane |
| 96 | O54962 | BANF1 | 23541614 | 29265637 | 8448262 | 12 | Nucleus |
| 97 | P97300 | Nptn | 23275842 | 31498128 | 6061818 | 27 | Plasma Membrane |
| 98 | Q3UZP4 | SVIP | 23240000 | 2775410 | 1602384 | 3 | Cytoplasm |
| 99 | P05064 | ALDOA | 23113685 | 21529767 | 3588295 | 36 | Cytoplasm |
| 100 | Q9QXY7 | XK | 22972500 | 4986287 | 1439417 | 12 | Plasma Membrane |
| 101 | P99026 | PSMB4 | 22952470 | 6979816 | 1645158 | 18 | Cytoplasm |
| 102 | A2AVZ9 | SLC43A3 | 22601215 | 17080495 | 4930714 | 12 | Extracellular Space |
| 103 | Q64735 | CR1L | 21822225 | 21645376 | 4165655 | 27 | Plasma Membrane |
| 104 | O70435 | PSMA3 | 21818139 | 8624312 | 1659750 | 27 | Cytoplasm |
| 105 | Q64339 | ISG15 | 21757419 | 17472343 | 3566527 | 24 | Extracellular Space |
| 106 | P10518 | ALAD | 21730126 | 21194963 | 3059229 | 48 | Cytoplasm |
| 107 | Q61411 | HRAS | 21555062 | 29637670 | 5411074 | 30 | Plasma Membrane |
| 108 | P62874 | GNB1 | 21375548 | 22120278 | 3542079 | 39 | Plasma Membrane |
| 109 | Q9R1P0 | PSMA4 | 21241416 | 12332489 | 1974779 | 39 | Cytoplasm |
| 110 | Q3U2A8 | VARS2 | 21210000 | 4600623 | 2656171 | 3 | Cytoplasm |
| 111 | Q03265 | ATP5A1 | 21196540 | 22121986 | 2663174 | 69 | Cytoplasm |
| 112 | Q61672 | SLC29A2 | 21160000 | 5747808 | 3318499 | 3 | Plasma Membrane |
| 113 | O55234 | PSMB5 | 21101030 | 11525520 | 2006336 | 33 | Cytoplasm |
| 114 | P56480 | ATP5B | 21016493 | 28719495 | 2931171 | 96 | Cytoplasm |
| 115 | Q09199 | B4GALNT2 | 20990000 | 1429021 | 825045 | 3 | Cytoplasm |
| 116 | Q9QUI6 | ACKR1 | 20823223 | 15329770 | 4425323 | 12 | Plasma Membrane |
| 117 | P08249 | MDH2 | 20431695 | 15970476 | 2557323 | 39 | Cytoplasm |
| 118 | P24369 | PPIB | 20154346 | 18404798 | 3067466 | 36 | Cytoplasm |
| 119 | Q9JHI9 | SLC40A1 | 20064916 | 10301646 | 2102815 | 24 | Plasma Membrane |
| 120 | Q9R1P4 | PSMA1 | 19653070 | 11183056 | 1614135 | 48 | Cytoplasm |
| 121 | O70325 | GPX4 | 19588660 | 14641341 | 3195003 | 21 | Cytoplasm |
| 122 | P51881 | SLC25A5 | 19448010 | 29351573 | 4375474 | 45 | Cytoplasm |
| 123 | Q9D1G1 | RAB1B | 19296167 | 14692792 | 3793663 | 15 | Cytoplasm |
| 124 | P63024 | VAMP3 | 19199531 | 30098460 | 7771389 | 15 | Plasma Membrane |
| 125 | Q922D8 | MTHFD1 | 19142653 | 9641241 | 1033650 | 87 | Cytoplasm |
| 126 | P35564 | CANX | 19106848 | 24297268 | 2990787 | 66 | Cytoplasm |
| 127 | Q9DCT8 | Crip2 | 18861635 | 11898552 | 2289877 | 27 | Plasma Membrane |
| 128 | Q8K021 | SCAMP1 | 18720000 | 3319342 | 1916423 | 3 | Cytoplasm |
| 129 | Q9JJF9 | SPPL2A | 18695000 | 2981700 | 1217274 | 6 | Plasma Membrane |
| 130 | P62204 | CALM1 | 18660000 | 11415143 | 4660212 | 6 | Cytoplasm |
| 131 | P32037 | SLC2A3 | 18645187 | 12839286 | 2801762 | 21 | Plasma Membrane |
| 132 | Q5F285 | TMEM256 | 18270000 | 9105011 | 5256781 | 3 | Cytoplasm |
| 133 | Q8CJG0 | AGO2 | 18140270 | 12032488 | 1084933 | 123 | Cytoplasm |
| 134 | P84096 | RHOG | 18121116 | 14536215 | 2530430 | 33 | Cytoplasm |
| 135 | P70195 | PSMB7 | 17986474 | 10118044 | 2065337 | 24 | Cytoplasm |
| 136 | Q80Y14 | GLRX5 | 17835000 | 12538907 | 5118987 | 6 | Cytoplasm |
| 137 | Q91YH5 | ATL3 | 17802476 | 14257843 | 2057942 | 48 | Cytoplasm |
| 138 | A6H603 | NWD1 | 17680000 | 1360992 | 785769 | 3 | Cytoplasm |

| | | | | | | | |
|---|---|---|---|---|---|---|---|
| 139 | Q9CQW2 | ARL8B | 1.96E+07 | 1.30E+07 | 2.01E+06 | 42 | Plasma Membrane |
| 140 | P51881 | SLC25A5 | 1.96E+07 | 2.15E+07 | 3.20E+06 | 45 | Cytoplasm |
| 141 | Q9JJ59 | ABCB9 | 1.95E+07 | 2.45E+07 | 1.42E+07 | 3 | Cytoplasm |
| 142 | Q9JHI9 | SLC40A1 | 1.95E+07 | 1.10E+07 | 2.25E+06 | 24 | Plasma Membrane |
| 143 | Q8K021 | SCAMP1 | 1.94E+07 | 5.10E+06 | 2.94E+06 | 3 | Cytoplasm |
| 144 | P05064 | ALDOA | 1.93E+07 | 2.14E+07 | 3.56E+06 | 36 | Cytoplasm |
| 145 | Q62312 | TGFBR2 | 1.93E+07 | 3.44E+06 | 1.98E+06 | 3 | Plasma Membrane |
| 146 | Q64339 | ISG15 | 1.92E+07 | 1.44E+07 | 2.94E+06 | 24 | Extracellular Space |
| 147 | Q62351 | TFRC | 1.91E+07 | 1.39E+07 | 1.32E+06 | 111 | Plasma Membrane |
| 148 | O70325 | GPX4 | 1.87E+07 | 1.23E+07 | 2.69E+06 | 21 | Cytoplasm |
| 149 | P47955 | Rplp1 (includes others) | 1.87E+07 | 4.02E+06 | 1.64E+06 | 6 | Nucleus |
| 150 | Q91YH5 | ATL3 | 1.87E+07 | 1.42E+07 | 2.05E+06 | 48 | Cytoplasm |
| 151 | O54890 | ITGB3 | 1.86E+07 | 2.52E+07 | 2.97E+06 | 72 | Plasma Membrane |
| 152 | P08249 | MDH2 | 1.84E+07 | 9.85E+06 | 1.58E+06 | 39 | Cytoplasm |
| 153 | Q99KU0 | VMP1 | 1.84E+07 | 3.31E+06 | 1.91E+06 | 3 | Plasma Membrane |
| 154 | P70195 | PSMB7 | 1.83E+07 | 1.36E+07 | 2.78E+06 | 24 | Cytoplasm |
| 155 | P62880 | GNB2 | 1.83E+07 | 1.38E+07 | 2.30E+06 | 36 | Plasma Membrane |
| 156 | P84096 | RHOG | 1.82E+07 | 1.45E+07 | 2.53E+06 | 33 | Cytoplasm |
| 157 | P62874 | GNB1 | 1.81E+07 | 1.60E+07 | 2.56E+06 | 39 | Plasma Membrane |
| 158 | Q9JJF9 | SPPL2A | 1.81E+07 | 2.24E+06 | 9.13E+05 | 6 | Plasma Membrane |
| 159 | Q9WV55 | VAPA | 1.79E+07 | 1.49E+07 | 2.87E+06 | 27 | Plasma Membrane |
| 160 | Q8K558 | TREML1 | 1.77E+07 | 1.78E+07 | 3.63E+06 | 24 | Plasma Membrane |
| 161 | Q4QY64 | ATAD5 | 1.77E+07 | 4.09E+05 | 2.36E+05 | 3 | Extracellular Space |
| 162 | Q8CJG0 | AGO2 | 1.75E+07 | 1.17E+07 | 1.06E+06 | 123 | Cytoplasm |
| 163 | P07759 | SERPINA3 | 1.72E+07 | 1.18E+07 | 3.93E+06 | 9 | Extracellular Space |
| 164 | Q7SIG6 | ASAP2 | 1.70E+07 | 3.19E+06 | 1.84E+06 | 3 | Nucleus |
| 165 | P35700 | PRDX1 | 1.70E+07 | 1.36E+07 | 2.37E+06 | 33 | Cytoplasm |
| 166 | P10518 | ALAD | 1.68E+07 | 2.21E+07 | 3.19E+06 | 48 | Cytoplasm |
| 167 | Q3TZ89 | SEC31B | 1.65E+07 | 1.40E+07 | 8.09E+06 | 3 | Cytoplasm |
| 168 | Q9JJ00 | PLSCR1 | 1.64E+07 | 6.93E+06 | 2.31E+06 | 9 | Plasma Membrane |
| 169 | Q6IMF0 | Krt83 | 1.64E+07 | 2.51E+06 | 1.45E+06 | 3 | Extracellular Space |
| 170 | P62204 | CALM | 1.63E+07 | 8.08E+06 | 3.30E+06 | 6 | Cytoplasm |
| 171 | Q91YN9 | BAG2 | 1.63E+07 | 1.25E+07 | 2.18E+06 | 33 | Cytoplasm |
| 172 | Q6AW46 | CES5A | 1.63E+07 | 9.09E+05 | 5.25E+05 | 3 | Extracellular Space |
| 173 | Q9R1P3 | PSMB2 | 1.63E+07 | 1.60E+07 | 3.07E+06 | 27 | Cytoplasm |
| 174 | Q9QUI6 | ACKR1 | 1.62E+07 | 1.12E+07 | 3.22E+06 | 12 | Plasma Membrane |
| 175 | P35293 | RAB18 | 1.60E+07 | 3.61E+06 | 9.33E+05 | 15 | Cytoplasm |
| 176 | Q9CQD1 | RAB5A | 1.60E+07 | 9.88E+06 | 2.55E+06 | 15 | Cytoplasm |
| 177 | O35598 | ADAM10 | 1.60E+07 | 1.53E+07 | 1.74E+06 | 78 | Plasma Membrane |
| 178 | P27664 | PDE6A | 1.59E+07 | 1.13E+06 | 6.55E+05 | 3 | Plasma Membrane |
| 179 | Q8BL86 | MBLAC2 | 1.58E+07 | 1.63E+06 | 9.41E+05 | 3 | Cytoplasm |
| 180 | O88343 | SLC4A4 | 1.58E+07 | 3.39E+06 | 1.96E+06 | 3 | Plasma Membrane |
| 181 | O08742 | GP5 | 1.54E+07 | 1.55E+07 | 2.83E+06 | 30 | Plasma Membrane |
| 182 | O08573 | LGALS9B | 1.53E+07 | 7.59E+06 | 1.55E+06 | 24 | Cytoplasm |
| 183 | P23492 | PNP | 1.52E+07 | 1.64E+07 | 3.86E+06 | 18 | Nucleus |
| 184 | Q3T9M1 | MFSD2B | 1.50E+07 | 1.41E+07 | 5.77E+06 | 6 | Plasma Membrane |
| 185 | P45878 | FKBP2 | 1.47E+07 | 1.93E+06 | 7.86E+05 | 6 | Cytoplasm |
| 186 | A6H603 | NWD1 | 1.45E+07 | 7.52E+05 | 4.34E+05 | 3 | Cytoplasm |
| 187 | O35930 | GP1BA | 1.45E+07 | 1.91E+07 | 3.05E+06 | 39 | Plasma Membrane |
| 188 | P97315 | CSRP1 | 1.45E+07 | 1.03E+07 | 2.67E+06 | 15 | Nucleus |
| 189 | Q80YF9 | ARHGAP33 | 1.43E+07 | 4.32E+06 | 2.50E+06 | 3 | Plasma Membrane |
| 190 | O09061 | PSMB1 | 1.43E+07 | 1.69E+07 | 2.37E+06 | 51 | Cytoplasm |
| 191 | Q9R1P1 | PSMB3 | 1.42E+07 | 1.34E+07 | 2.45E+06 | 30 | Cytoplasm |
| 192 | P61021 | RAB5B | 1.40E+07 | 8.38E+06 | 1.97E+06 | 18 | Cytoplasm |
| 193 | P09103 | P4HB | 1.39E+07 | 5.90E+06 | 7.82E+05 | 57 | Cytoplasm |
| 194 | Q9R1Q7 | PLP2 | 1.39E+07 | 3.94E+06 | 2.27E+06 | 3 | Cytoplasm |
| 195 | Q8VDD5 | MYH9 | 1.38E+07 | 1.29E+07 | 7.63E+05 | 288 | Cytoplasm |
| 196 | Q9CRA0 | ART4 | 1.38E+07 | 1.38E+07 | 3.01E+06 | 21 | Nucleus |
| 197 | P43274 | Hist1h1e | 1.36E+07 | 1.64E+07 | 4.23E+06 | 15 | Nucleus |
| 198 | Q8R1V4 | TMED4 | 1.34E+07 | 1.01E+06 | 5.84E+05 | 3 | Cytoplasm |
| 199 | Q78ZA7 | NAP1L4 | 1.32E+07 | 1.07E+07 | 1.87E+06 | 33 | Cytoplasm |
| 200 | Q60870 | REEP5 | 1.30E+07 | 1.63E+07 | 5.44E+06 | 9 | Extracellular Space |
| 201 | Q8VDN2 | ATP1A1 | 1.28E+07 | 9.64E+06 | 1.03E+06 | 87 | Plasma Membrane |
| 202 | P40240 | CD9 | 1.27E+07 | 2.23E+07 | 7.43E+06 | 9 | Plasma Membrane |
| 203 | Q9Z2P8 | VAMP5 | 1.27E+07 | 6.50E+06 | 2.17E+06 | 9 | Plasma Membrane |
| 204 | Q9QUI0 | RHOA | 1.25E+07 | 8.84E+06 | 1.70E+06 | 27 | Cytoplasm |
| 205 | Q9EPE9 | ATP13A1 | 1.25E+07 | 4.32E+06 | 2.50E+06 | 3 | Extracellular Space |
| 206 | Q9Z1N7 | ARID3B | 1.25E+07 | 2.25E+06 | 1.30E+06 | 3 | Nucleus |
| 207 | O09117 | SYPL1 | 1.24E+07 | 8.83E+06 | 2.94E+06 | 9 | Plasma Membrane |
| 208 | P51150 | RAB7A | 1.22E+07 | 4.06E+06 | 7.06E+05 | 33 | Cytoplasm |
| 209 | P46638 | RAB11B | 1.22E+07 | 1.08E+07 | 1.98E+06 | 30 | Cytoplasm |
| 210 | P63082 | ATP6V0C | 1.21E+07 | 1.10E+07 | 4.49E+06 | 6 | Cytoplasm |
| 211 | Q9WVJ2 | PSMD13 | 1.21E+07 | 7.54E+06 | 1.21E+06 | 39 | Cytoplasm |
| 212 | Q8K274 | FN3KRP | 1.21E+07 | 8.35E+06 | 1.97E+06 | 18 | Cytoplasm |
| 213 | Q6ZPJ3 | UBE2O | 1.20E+07 | 1.16E+07 | 1.15E+06 | 102 | Nucleus |
| 214 | A2ATU0 | DHTKD1 | 1.20E+07 | 1.14E+06 | 6.57E+05 | 3 | Cytoplasm |
| 215 | P32037 | SLC2A3 | 1.19E+07 | 1.09E+07 | 2.37E+06 | 21 | Plasma Membrane |
| 216 | Q9CPU4 | MGST3 | 1.19E+07 | 1.14E+07 | 3.81E+06 | 9 | Cytoplasm |
| 217 | Q9ER00 | STX12 | 1.18E+07 | 1.07E+06 | 6.20E+05 | 3 | Cytoplasm |
| 218 | P99028 | UQCRHL | 1.17E+07 | 8.05E+06 | 3.29E+06 | 6 | Cytoplasm |
| 219 | Q9D1D4 | TMED10 | 1.16E+07 | 7.51E+06 | 1.77E+06 | 18 | Cytoplasm |
| 220 | Q8VEK0 | TMEM30A | 1.16E+07 | 4.77E+06 | 1.23E+06 | 15 | Cytoplasm |
| 221 | Q9DCN2 | Cyb5r3 | 1.15E+07 | 8.69E+06 | 1.45E+06 | 36 | Cytoplasm |
| 222 | Q9ER35 | FN3K | 1.14E+07 | 6.41E+06 | 1.40E+06 | 21 | Cytoplasm |
| 223 | P07724 | ALB | 1.14E+07 | 1.11E+07 | 1.31E+06 | 72 | Extracellular Space |
| 224 | Q60931 | VDAC3 | 1.14E+07 | 7.16E+06 | 1.69E+06 | 18 | Cytoplasm |
| 225 | P02535 | Krt10 | 1.14E+07 | 3.39E+06 | 1.96E+06 | 3 | Cytoplasm |
| 226 | Q9JKW0 | ARL6IP1 | 1.14E+07 | 2.42E+06 | 1.39E+06 | 3 | Cytoplasm |
| 227 | P48972 | MYBL2 | 1.13E+07 | 2.90E+06 | 1.68E+06 | 3 | Nucleus |
| 228 | P47738 | ALDH2 | 1.13E+07 | 1.08E+07 | 1.27E+06 | 72 | Cytoplasm |
| 229 | P46471 | PSMC2 | 1.13E+07 | 7.41E+06 | 1.10E+06 | 45 | Nucleus |
| 230 | A2AHJ4 | BRWD3 | 1.13E+07 | 8.73E+05 | 5.04E+05 | 3 | Nucleus |
| 231 | Q60634 | FLOT2 | 1.13E+07 | 8.41E+06 | 1.18E+06 | 51 | Plasma Membrane |
| 232 | O35381 | Anp32a | 1.12E+07 | 6.26E+06 | 2.09E+06 | 9 | Nucleus |
| 233 | Q80X80 | C2CD2L | 1.11E+07 | 7.84E+06 | 1.26E+06 | 39 | Cytoplasm |
| 234 | Q640N1 | AEBP1 | 1.10E+07 | 3.12E+06 | 1.80E+06 | 3 | Nucleus |
| 235 | Q9R111 | GDA | 1.10E+07 | 8.70E+06 | 1.51E+06 | 33 | Cytoplasm |
| 236 | O08688 | CAPN5 | 1.09E+07 | 9.81E+06 | 1.27E+06 | 60 | Cytoplasm |
| 237 | P97372 | PSME2 | 1.08E+07 | 9.81E+06 | 2.14E+06 | 21 | Cytoplasm |
| 238 | P68037 | UBE2L3 | 1.08E+07 | 4.46E+06 | 2.57E+06 | 3 | Nucleus |
| 239 | Q60932 | VDAC1 | 1.08E+07 | 5.66E+06 | 1.15E+06 | 24 | Cytoplasm |
| 240 | Q921I1 | TF | 1.08E+07 | 7.90E+06 | 7.71E+05 | 105 | Extracellular Space |
| 241 | P11352 | GPX1 | 1.07E+07 | 9.61E+06 | 1.75E+06 | 30 | Cytoplasm |
| 242 | Q3U2A8 | VARS2 | 1.06E+07 | 2.38E+06 | 1.38E+06 | 3 | Cytoplasm |
| 243 | O09044 | SNAP23 | 1.06E+07 | 1.06E+07 | 1.76E+06 | 36 | Plasma Membrane |
| 244 | O35593 | PSMD14 | 1.06E+07 | 9.42E+06 | 2.72E+06 | 12 | Cytoplasm |
| 245 | Q8VEM8 | SLC25A3 | 1.05E+07 | 8.23E+06 | 1.58E+06 | 27 | Cytoplasm |
| 246 | P62192 | PSMC1 | 1.05E+07 | 7.40E+06 | 1.19E+06 | 39 | Nucleus |
| 247 | P80317 | CCT6A | 1.04E+07 | 6.91E+06 | 1.15E+06 | 36 | Cytoplasm |
| 248 | Q91V01 | LPCAT3 | 1.04E+07 | 7.53E+06 | 2.17E+06 | 12 | Plasma Membrane |
| 249 | Q80Y14 | GLRX5 | 1.03E+07 | 3.55E+06 | 1.45E+06 | 6 | Cytoplasm |
| 250 | Q9D0M3 | CYC1 | 1.03E+07 | 1.37E+06 | 7.92E+05 | 3 | Cytoplasm |
| 251 | Q5F285 | TMEM256 | 1.02E+07 | 1.27E+06 | 7.32E+05 | 3 | Cytoplasm |
| 252 | P26516 | PSMD7 | 1.02E+07 | 8.56E+06 | 1.56E+06 | 30 | Cytoplasm |
| 253 | Q922U2 | KRT5 | 1.02E+07 | 9.87E+06 | 2.85E+06 | 12 | Cytoplasm |
| 254 | P97371 | PSME1 | 1.02E+07 | 6.68E+06 | 1.11E+06 | 36 | Cytoplasm |
| 255 | Q8VED5 | KRT79 | 1.02E+07 | 3.26E+06 | 1.88E+06 | 3 | Extracellular Space |
| 256 | P14685 | PSMD3 | 1.02E+07 | 9.17E+06 | 1.32E+06 | 48 | Cytoplasm |
| 257 | Q8C522 | ENDOD1 | 1.02E+07 | 4.21E+06 | 8.59E+05 | 24 | Extracellular Space |
| 258 | P04104 | KRT1 | 1.01E+07 | 9.94E+06 | 2.87E+06 | 12 | Cytoplasm |
| 259 | P26043 | RDX | 1.01E+07 | 9.38E+06 | 1.45E+06 | 42 | Cytoplasm |
| 260 | P16330 | CNP | 1.00E+07 | 9.86E+06 | 2.01E+06 | 24 | Cytoplasm |
| 261 | P68510 | YWHAH | 9.90E+06 | 1.37E+07 | 2.65E+06 | 27 | Cytoplasm |
| 262 | P21440 | ABCB4 | 9.83E+06 | 6.15E+06 | 7.25E+05 | 72 | Plasma Membrane |
| 263 | Q3U7R1 | ESYT1 | 9.82E+06 | 6.34E+06 | 6.80E+05 | 87 | Cytoplasm |
| 264 | O08917 | FLOT1 | 9.81E+06 | 5.72E+06 | 8.26E+05 | 48 | Plasma Membrane |
| 265 | Q3TFD2 | LPCAT1 | 9.75E+06 | 4.72E+06 | 9.64E+05 | 24 | Cytoplasm |
| 266 | Q9Z2W0 | DNPEP | 9.71E+06 | 1.05E+07 | 2.01E+06 | 27 | Cytoplasm |
| 267 | P39054 | DNM2 | 9.69E+06 | 9.47E+06 | 1.14E+06 | 69 | Plasma Membrane |
| 268 | Q9D832 | DNAJB4 | 9.69E+06 | 7.05E+06 | 1.29E+06 | 30 | Nucleus |
| 269 | Q6ZQ38 | CAND1 | 9.69E+06 | 5.41E+06 | 7.17E+05 | 57 | Cytoplasm |
| 270 | Q9CQF9 | PCYOX1 | 9.61E+06 | 2.01E+06 | 1.16E+06 | 3 | Cytoplasm |
| 271 | Q922R8 | PDIA6 | 9.58E+06 | 3.70E+06 | 6.44E+05 | 33 | Cytoplasm |
| 272 | Q61316 | HSPA4 | 9.57E+06 | 7.19E+06 | 8.66E+05 | 69 | Cytoplasm |
| 273 | Q8K1B8 | FERMT3 | 9.54E+06 | 6.95E+06 | 1.00E+06 | 48 | Cytoplasm |
| 274 | P04925 | PRNP | 9.46E+06 | 2.15E+06 | 1.24E+06 | 3 | Plasma Membrane |
| 275 | Q9CVB6 | ARPC2 | 9.42E+06 | 6.62E+06 | 2.21E+06 | 9 | Cytoplasm |
| 276 | P99027 | RPLP2 | 9.41E+06 | 8.43E+06 | 2.18E+06 | 15 | Cytoplasm |
| 277 | P08905 | LYZ | 9.41E+06 | 8.28E+06 | 3.38E+06 | 6 | Extracellular Space |

| | | | | | | | |
|---|---|---|---|---|---|---|---|
| 139 | O35450 | FKBPL | 1.92E+07 | 6.66E+06 | 3.85E+06 | 3 | Nucleus |
| 140 | P97315 | CSRP1 | 1.90E+07 | 9.55E+06 | 2.47E+06 | 15 | Nucleus |
| 141 | Q60931 | VDAC3 | 1.89E+07 | 1.36E+07 | 3.20E+06 | 18 | Cytoplasm |
| 142 | P99026 | PSMB4 | 1.88E+07 | 5.90E+06 | 1.39E+06 | 18 | Cytoplasm |
| 143 | Q9DCT8 | Crip2 | 1.88E+07 | 1.35E+07 | 2.59E+06 | 27 | Plasma Membrane |
| 144 | P32037 | SLC2A3 | 1.86E+07 | 1.36E+07 | 2.96E+06 | 21 | Plasma Membrane |
| 145 | P62204 | CALM1 | 1.86E+07 | 1.21E+07 | 4.96E+06 | 6 | Cytoplasm |
| 146 | P99027 | RPLP2 | 1.85E+07 | 1.82E+07 | 4.71E+06 | 15 | Cytoplasm |
| 147 | P49722 | PSMA2 | 1.84E+07 | 1.82E+07 | 3.32E+06 | 30 | Cytoplasm |
| 148 | Q8R1V4 | TMED4 | 1.83E+07 | 2.15E+06 | 1.24E+06 | 3 | Cytoplasm |
| 149 | P54726 | RAD23A | 1.83E+07 | 1.92E+07 | 4.18E+06 | 21 | Nucleus |
| 150 | Q9D0M3 | CYC1 | 1.83E+07 | 1.46E+06 | 8.40E+05 | 3 | Cytoplasm |
| 151 | P05064 | ALDOA | 1.83E+07 | 1.91E+07 | 3.18E+06 | 36 | Cytoplasm |
| 152 | Q4QY64 | ATAD5 | 1.82E+07 | 4.54E+06 | 2.62E+06 | 3 | Extracellular Space |
| 153 | P97300 | Nptn | 1.82E+07 | 2.75E+07 | 5.29E+06 | 27 | Plasma Membrane |
| 154 | Q9WV55 | VAPA | 1.82E+07 | 1.52E+07 | 2.93E+06 | 27 | Plasma Membrane |
| 155 | Q64735 | CR1L | 1.81E+07 | 1.66E+07 | 3.19E+06 | 27 | Plasma Membrane |
| 156 | Q60932 | VDAC1 | 1.77E+07 | 8.84E+06 | 1.81E+06 | 24 | Cytoplasm |
| 157 | Q9CQW2 | ARL8B | 1.72E+07 | 1.14E+07 | 1.76E+06 | 42 | Plasma Membrane |
| 158 | Q61672 | SLC29A2 | 1.71E+07 | 5.98E+06 | 3.46E+06 | 3 | Plasma Membrane |
| 159 | P11352 | GPX1 | 1.70E+07 | 1.33E+07 | 2.43E+06 | 30 | Cytoplasm |
| 160 | Q921I1 | TF | 1.70E+07 | 1.71E+07 | 1.67E+06 | 105 | Extracellular Space |
| 161 | O70435 | PSMA3 | 1.70E+07 | 7.32E+06 | 1.41E+06 | 27 | Cytoplasm |
| 162 | P68510 | YWHAH | 1.70E+07 | 1.56E+07 | 2.99E+06 | 27 | Cytoplasm |
| 163 | P47738 | ALDH2 | 1.69E+07 | 1.32E+07 | 1.55E+06 | 72 | Cytoplasm |
| 164 | O88343 | SLC4A4 | 1.66E+07 | 4.31E+06 | 2.49E+06 | 3 | Plasma Membrane |
| 165 | Q9CQD1 | RAB5A | 1.66E+07 | 1.05E+07 | 2.71E+06 | 15 | Cytoplasm |
| 166 | Q91YH5 | ATL3 | 1.66E+07 | 1.34E+07 | 1.93E+06 | 48 | Cytoplasm |
| 167 | Q9JHI9 | SLC40A1 | 1.64E+07 | 9.24E+06 | 1.89E+06 | 24 | Plasma Membrane |
| 168 | P35293 | RAB18 | 1.64E+07 | 4.04E+06 | 1.04E+06 | 15 | Cytoplasm |
| 169 | O70325 | GPX4 | 1.63E+07 | 1.26E+07 | 2.74E+06 | 21 | Cytoplasm |
| 170 | Q9R1P0 | PSMA4 | 1.63E+07 | 1.12E+07 | 1.79E+06 | 39 | Cytoplasm |
| 171 | Q62312 | TGFBR2 | 1.63E+07 | 2.05E+06 | 1.18E+06 | 3 | Plasma Membrane |
| 172 | P07759 | SERPINA3 | 1.62E+07 | 5.33E+06 | 1.78E+06 | 9 | Extracellular Space |
| 173 | P60904 | DNAJC5 | 1.62E+07 | 1.76E+07 | 4.55E+06 | 15 | Plasma Membrane |
| 174 | P23492 | PNP | 1.61E+07 | 1.27E+07 | 3.00E+06 | 18 | Nucleus |
| 175 | Q9QUI6 | ACKR1 | 1.59E+07 | 1.32E+07 | 3.82E+06 | 12 | Plasma Membrane |
| 176 | Q9ER00 | STX12 | 1.58E+07 | 2.36E+06 | 1.36E+06 | 3 | Cytoplasm |
| 177 | O35598 | ADAM10 | 1.58E+07 | 1.41E+07 | 1.60E+06 | 78 | Plasma Membrane |
| 178 | Q3T9M1 | MFSD2B | 1.57E+07 | 1.66E+07 | 6.78E+06 | 6 | Plasma Membrane |
| 179 | Q9CQ19 | MYL9 | 1.55E+07 | 7.17E+06 | 1.46E+06 | 24 | Cytoplasm |
| 180 | Q7SIG6 | ASAP2 | 1.54E+07 | 9.82E+06 | 5.67E+06 | 3 | Nucleus |
| 181 | P27664 | PDE6A | 1.52E+07 | 3.28E+06 | 1.90E+06 | 3 | Plasma Membrane |
| 182 | P07724 | ALB | 1.52E+07 | 1.22E+07 | 1.44E+06 | 72 | Extracellular Space |
| 183 | Q8VCM7 | FGG | 1.51E+07 | 1.43E+07 | 1.72E+06 | 69 | Extracellular Space |
| 184 | P45878 | FKBP2 | 1.51E+07 | 3.06E+06 | 1.25E+06 | 6 | Cytoplasm |
| 185 | Q3UV17 | KRT76 | 1.50E+07 | 1.38E+07 | 7.96E+06 | 3 | Cytoplasm |
| 186 | Q9D1D4 | TMED10 | 1.50E+07 | 9.03E+06 | 2.13E+06 | 18 | Cytoplasm |
| 187 | P63024 | VAMP3 | 1.50E+07 | 2.81E+07 | 7.26E+06 | 15 | Plasma Membrane |
| 188 | Q8CJG0 | AGO2 | 1.50E+07 | 1.02E+07 | 9.21E+05 | 123 | Cytoplasm |
| 189 | O55234 | PSMB5 | 1.49E+07 | 9.94E+06 | 1.73E+06 | 33 | Cytoplasm |
| 190 | O08573 | LGALS9B | 1.49E+07 | 8.10E+06 | 1.65E+06 | 24 | Cytoplasm |
| 191 | Q9R1P4 | PSMA1 | 1.47E+07 | 9.94E+06 | 1.43E+06 | 48 | Cytoplasm |
| 192 | P04925 | PRNP | 1.47E+07 | 2.10E+06 | 1.22E+06 | 3 | Plasma Membrane |
| 193 | Q9QUI0 | RHOA | 1.45E+07 | 1.04E+07 | 2.01E+06 | 27 | Cytoplasm |
| 194 | Q9D3D9 | ATP5D | 1.43E+07 | 2.21E+06 | 9.01E+05 | 6 | Cytoplasm |
| 195 | Q6AW46 | CES5A | 1.42E+07 | 1.27E+06 | 7.32E+05 | 3 | Extracellular Space |
| 196 | Q8VEM8 | SLC25A3 | 1.42E+07 | 1.11E+07 | 2.13E+06 | 27 | Cytoplasm |
| 197 | P46638 | RAB11B | 1.42E+07 | 1.28E+07 | 2.33E+06 | 30 | Cytoplasm |
| 198 | O09117 | SYPL1 | 1.41E+07 | 1.17E+07 | 3.90E+06 | 9 | Plasma Membrane |
| 199 | Q91YN9 | BAG2 | 1.40E+07 | 1.04E+07 | 1.81E+06 | 33 | Cytoplasm |
| 200 | Q8K0E8 | FGB | 1.40E+07 | 1.39E+07 | 1.64E+06 | 72 | Extracellular Space |
| 201 | Q60930 | VDAC2 | 1.40E+07 | 1.22E+07 | 2.35E+06 | 27 | Cytoplasm |
| 202 | Q8K021 | SCAMP1 | 1.39E+07 | 1.67E+06 | 9.66E+05 | 3 | Cytoplasm |
| 203 | Q9Z2P8 | VAMP5 | 1.39E+07 | 8.41E+06 | 2.80E+06 | 9 | Plasma Membrane |
| 204 | Q9JJ00 | PLSCR1 | 1.39E+07 | 5.27E+06 | 1.76E+06 | 9 | Plasma Membrane |
| 205 | Q8BL86 | MBLAC2 | 1.38E+07 | 8.77E+05 | 5.06E+05 | 3 | Cytoplasm |
| 206 | P35700 | PRDX1 | 1.37E+07 | 8.47E+06 | 1.47E+06 | 33 | Cytoplasm |
| 207 | P19536 | Cox5b | 1.37E+07 | 2.20E+06 | 1.27E+06 | 3 | Cytoplasm |
| 208 | P70195 | PSMB7 | 1.36E+07 | 9.11E+06 | 1.86E+06 | 24 | Cytoplasm |
| 209 | O70404 | VAMP8 | 1.35E+07 | 1.09E+07 | 3.62E+06 | 9 | Plasma Membrane |
| 210 | P63038 | HSPD1 | 1.34E+07 | 1.05E+07 | 1.27E+06 | 69 | Cytoplasm |
| 211 | Q9R1Q7 | PLP2 | 1.33E+07 | 3.10E+06 | 1.79E+06 | 3 | Cytoplasm |
| 212 | Q8VDN2 | ATP1A1 | 1.33E+07 | 9.84E+06 | 1.05E+06 | 87 | Plasma Membrane |
| 213 | P19783 | COX4I1 | 1.32E+07 | 8.76E+06 | 2.92E+06 | 9 | Cytoplasm |
| 214 | Q9JJU8 | SH3BGRL | 1.32E+07 | 3.33E+06 | 1.11E+06 | 9 | Cytoplasm |
| 215 | P61021 | RAB5B | 1.32E+07 | 7.08E+06 | 1.67E+06 | 18 | Cytoplasm |
| 216 | Q64433 | HSPE1 | 1.32E+07 | 8.39E+06 | 2.17E+06 | 15 | Cytoplasm |
| 217 | Q3THE2 | MYL12A | 1.32E+07 | 1.06E+07 | 2.73E+06 | 15 | Cytoplasm |
| 218 | P17742 | PPIA | 1.31E+07 | 8.66E+06 | 1.89E+06 | 21 | Cytoplasm |
| 219 | Q9Z1N7 | ARID3B | 1.31E+07 | 4.61E+06 | 2.66E+06 | 3 | Nucleus |
| 220 | Q8K1B8 | FERMT3 | 1.31E+07 | 6.57E+06 | 9.48E+05 | 48 | Cytoplasm |
| 221 | Q9R111 | GDA | 1.31E+07 | 7.41E+06 | 1.29E+06 | 33 | Cytoplasm |
| 222 | O09044 | SNAP23 | 1.31E+07 | 1.18E+07 | 1.96E+06 | 36 | Plasma Membrane |
| 223 | Q64522 | HIST2H2AI | 1.31E+07 | 3.74E+06 | 2.16E+06 | 3 | Nucleus |
| 224 | P09103 | P4HB | 1.30E+07 | 6.20E+06 | 8.22E+05 | 57 | Cytoplasm |
| 225 | Q9CPU4 | MGST3 | 1.30E+07 | 2.16E+07 | 7.20E+06 | 9 | Cytoplasm |
| 226 | P51150 | RAB7A | 1.29E+07 | 6.01E+06 | 1.05E+06 | 33 | Cytoplasm |
| 227 | O35658 | C1QBP | 1.28E+07 | 1.44E+07 | 5.90E+06 | 6 | Cytoplasm |
| 228 | P40240 | CD9 | 1.27E+07 | 1.94E+07 | 6.45E+06 | 9 | Plasma Membrane |
| 229 | P63082 | ATP6V0C | 1.26E+07 | 1.15E+07 | 4.68E+06 | 6 | Cytoplasm |
| 230 | P18242 | CTSD | 1.26E+07 | 6.64E+06 | 1.45E+06 | 21 | Cytoplasm |
| 231 | O89053 | CORO1A | 1.23E+07 | 1.34E+07 | 2.14E+06 | 39 | Cytoplasm |
| 232 | A6H603 | NWD1 | 1.22E+07 | 6.12E+05 | 3.54E+05 | 3 | Cytoplasm |
| 233 | Q78ZA7 | NAP1L4 | 1.20E+07 | 7.54E+06 | 1.31E+06 | 33 | Cytoplasm |
| 234 | Q8VEK0 | TMEM30A | 1.20E+07 | 5.39E+06 | 1.39E+06 | 15 | Cytoplasm |
| 235 | Q9JJF9 | SPPL2A | 1.19E+07 | 1.57E+06 | 6.39E+05 | 6 | Plasma Membrane |
| 236 | Q9CZU6 | CS | 1.18E+07 | 1.02E+07 | 1.85E+06 | 30 | Cytoplasm |
| 237 | P43276 | Hist1h1b | 1.17E+07 | 8.98E+06 | 3.66E+06 | 6 | Nucleus |
| 238 | Q99KF1 | TMED9 | 1.16E+07 | 1.07E+07 | 3.55E+06 | 9 | Cytoplasm |
| 239 | O08749 | DLD | 1.16E+07 | 6.31E+06 | 2.10E+06 | 9 | Cytoplasm |
| 240 | Q922R8 | PDIA6 | 1.15E+07 | 5.43E+06 | 9.46E+05 | 33 | Cytoplasm |
| 241 | P67984 | RPL22 | 1.15E+07 | 5.11E+06 | 2.09E+06 | 6 | Nucleus |
| 242 | P51174 | ACADL | 1.14E+07 | 1.13E+07 | 2.46E+06 | 21 | Cytoplasm |
| 243 | Q9DCN2 | Cyb5r3 | 1.14E+07 | 1.01E+07 | 1.69E+06 | 36 | Cytoplasm |
| 244 | P39054 | DNM2 | 1.14E+07 | 1.02E+07 | 1.22E+06 | 69 | Plasma Membrane |
| 245 | Q8K274 | FN3KRP | 1.13E+07 | 6.94E+06 | 1.64E+06 | 18 | Cytoplasm |
| 246 | Q9CRA0 | ART4 | 1.13E+07 | 1.17E+07 | 2.55E+06 | 21 | Nucleus |
| 247 | Q9DBH5 | LMAN2 | 1.12E+07 | 7.10E+06 | 1.83E+06 | 15 | Cytoplasm |
| 248 | P00920 | CA2 | 1.12E+07 | 1.18E+07 | 2.79E+06 | 18 | Cytoplasm |
| 249 | Q80YF9 | ARHGAP33 | 1.12E+07 | 2.52E+06 | 1.45E+06 | 3 | Plasma Membrane |
| 250 | Q9D0F3 | LMAN1 | 1.11E+07 | 3.85E+06 | 9.07E+05 | 18 | Cytoplasm |
| 251 | P46471 | PSMC2 | 1.11E+07 | 6.56E+06 | 9.78E+05 | 45 | Nucleus |
| 252 | Q9EPE9 | ATP13A1 | 1.10E+07 | 7.32E+05 | 4.23E+05 | 3 | Extracellular Space |
| 253 | Q8C522 | ENDOD1 | 1.10E+07 | 6.01E+06 | 1.23E+06 | 24 | Extracellular Space |
| 254 | P03930 | mt-Atp8 | 1.10E+07 | 6.54E+06 | 2.67E+06 | 6 | Cytoplasm |
| 255 | O09061 | PSMB1 | 1.09E+07 | 1.08E+07 | 1.52E+06 | 51 | Cytoplasm |
| 256 | P00405 | MT-CO2 | 1.09E+07 | 1.30E+07 | 3.05E+06 | 18 | Cytoplasm |
| 257 | Q91V01 | LPCAT3 | 1.08E+07 | 8.52E+06 | 2.46E+06 | 12 | Plasma Membrane |
| 258 | Q9R1P3 | PSMB2 | 1.08E+07 | 1.11E+07 | 2.13E+06 | 27 | Cytoplasm |
| 259 | P09055 | ITGB1 | 1.08E+07 | 7.60E+06 | 1.22E+06 | 39 | Plasma Membrane |
| 260 | Q99P58 | RAB27B | 1.08E+07 | 8.75E+06 | 1.79E+06 | 24 | Cytoplasm |
| 261 | Q3U7R1 | ESYT1 | 1.08E+07 | 6.79E+06 | 7.28E+05 | 87 | Cytoplasm |
| 262 | P26043 | RDX | 1.07E+07 | 9.55E+06 | 1.47E+06 | 42 | Cytoplasm |
| 263 | Q9R1P1 | PSMB3 | 1.06E+07 | 9.01E+06 | 1.64E+06 | 30 | Cytoplasm |
| 264 | Q60634 | FLOT2 | 1.06E+07 | 7.68E+06 | 1.08E+06 | 51 | Plasma Membrane |
| 265 | Q9CQ22 | LAMTOR1 | 1.06E+07 | 7.92E+06 | 2.29E+06 | 12 | Plasma Membrane |
| 266 | Q922D8 | MTHFD1 | 1.06E+07 | 6.79E+06 | 7.28E+05 | 87 | Cytoplasm |
| 267 | Q9DB20 | ATP5O | 1.03E+07 | 4.94E+06 | 1.01E+06 | 24 | Cytoplasm |
| 268 | O55042 | SNCA | 1.02E+07 | 4.09E+06 | 1.18E+06 | 12 | Cytoplasm |
| 269 | P22315 | FECH | 1.02E+07 | 1.63E+07 | 2.16E+06 | 57 | Cytoplasm |
| 270 | O70439 | STX7 | 1.01E+07 | 3.73E+06 | 8.80E+05 | 18 | Plasma Membrane |
| 271 | Q9CVB6 | ARPC2 | 1.01E+07 | 4.72E+06 | 1.57E+06 | 9 | Cytoplasm |
| 272 | Q9CXW4 | RPL11 | 9.96E+06 | 2.14E+05 | 1.24E+05 | 3 | Cytoplasm |
| 273 | P60521 | GABARAPL | 9.94E+06 | 5.20E+06 | 1.34E+06 | 15 | Cytoplasm |
| 274 | Q9D659 | C10orf54 | 9.94E+06 | 2.73E+06 | 1.57E+06 | 3 | Plasma Membrane |
| 275 | P14206 | RPSA | 9.90E+06 | 5.91E+06 | 1.71E+06 | 12 | Cytoplasm |
| 276 | Q9D1Q6 | ERP44 | 9.85E+06 | 5.30E+06 | 1.25E+06 | 18 | Cytoplasm |
| 277 | P26516 | PSMD7 | 9.82E+06 | 8.03E+06 | 1.47E+06 | 30 | Cytoplasm |

| | | | | | | | |
|---|---|---|---|---|---|---|---|
| 139 | P43274 | Hist1h1e | 1.92E+07 | 6.83E+07 | 1.76E+07 | 15 | Nucleus |
| 140 | Q3U2A8 | VARS2 | 1.92E+07 | 3.11E+06 | 1.79E+06 | 3 | Cytoplasm |
| 141 | P08249 | MDH2 | 1.91E+07 | 1.82E+07 | 2.92E+06 | 39 | Cytoplasm |
| 142 | P62874 | GNB1 | 1.88E+07 | 2.07E+07 | 3.31E+06 | 39 | Plasma Membrane |
| 143 | Q9WV55 | VAPA | 1.86E+07 | 1.54E+07 | 2.97E+06 | 27 | Plasma Membrane |
| 144 | O70325 | GPX4 | 1.85E+07 | 1.35E+07 | 2.94E+06 | 21 | Cytoplasm |
| 145 | P47955 | Rplp1 (includes others) | 1.85E+07 | 8.06E+06 | 3.29E+06 | 6 | Nucleus |
| 146 | P62880 | GNB2 | 1.84E+07 | 1.85E+07 | 3.09E+06 | 36 | Plasma Membrane |
| 147 | Q9R1P4 | PSMA1 | 1.81E+07 | 9.61E+06 | 1.39E+06 | 48 | Cytoplasm |
| 148 | P07759 | SERPINA3 | 1.81E+07 | 7.55E+06 | 2.52E+06 | 9 | Extracellular Space |
| 149 | P70195 | PSMB7 | 1.81E+07 | 1.04E+07 | 2.12E+06 | 24 | Cytoplasm |
| 150 | Q9CQW2 | ARL8B | 1.80E+07 | 1.03E+07 | 1.59E+06 | 42 | Plasma Membrane |
| 151 | Q9CQD1 | RAB5A | 1.80E+07 | 9.68E+06 | 2.50E+06 | 15 | Cytoplasm |
| 152 | Q9JJF9 | SPPL2A | 1.79E+07 | 1.63E+06 | 6.65E+05 | 6 | Plasma Membrane |
| 153 | Q99KU0 | VMP1 | 1.76E+07 | 3.35E+06 | 1.93E+06 | 3 | Plasma Membrane |
| 154 | P60904 | DNAJC5 | 1.76E+07 | 1.65E+07 | 4.26E+06 | 15 | Plasma Membrane |
| 155 | O54890 | ITGB3 | 1.74E+07 | 2.44E+07 | 2.88E+06 | 72 | Plasma Membrane |
| 156 | Q8K021 | SCAMP1 | 1.73E+07 | 4.49E+06 | 2.59E+06 | 3 | Cytoplasm |
| 157 | Q8K558 | TREML1 | 1.72E+07 | 1.20E+07 | 2.44E+06 | 24 | Plasma Membrane |
| 158 | P54726 | RAD23A | 1.72E+07 | 1.45E+07 | 3.17E+06 | 21 | Nucleus |
| 159 | Q6AW46 | CES5A | 1.64E+07 | 6.62E+05 | 3.82E+05 | 3 | Extracellular Space |
| 160 | Q8CJG0 | AGO2 | 1.63E+07 | 1.00E+07 | 9.03E+05 | 123 | Cytoplasm |
| 161 | Q9JHI9 | SLC40A1 | 1.63E+07 | 8.70E+06 | 1.78E+06 | 24 | Plasma Membrane |
| 162 | P45878 | FKBP2 | 1.63E+07 | 2.47E+06 | 1.01E+06 | 6 | Cytoplasm |
| 163 | Q80YF9 | ARHGAP33 | 1.60E+07 | 1.98E+06 | 1.15E+06 | 3 | Plasma Membrane |
| 164 | P84096 | RHOG | 1.59E+07 | 1.28E+07 | 2.23E+06 | 33 | Cytoplasm |
| 165 | O09044 | SNAP23 | 1.58E+07 | 1.37E+07 | 2.29E+06 | 36 | Plasma Membrane |
| 166 | Q4QY64 | ATAD5 | 1.57E+07 | 2.53E+06 | 1.46E+06 | 3 | Extracellular Space |
| 167 | Q6IMF0 | Krt83 | 1.56E+07 | 1.78E+06 | 1.02E+06 | 3 | Extracellular Space |
| 168 | Q8R1V4 | TMED4 | 1.56E+07 | 1.66E+06 | 9.56E+05 | 3 | Cytoplasm |
| 169 | Q3TZ89 | SEC31B | 1.55E+07 | 6.71E+06 | 3.88E+06 | 3 | Cytoplasm |
| 170 | O08742 | GP5 | 1.53E+07 | 1.64E+07 | 3.00E+06 | 30 | Plasma Membrane |
| 171 | Q9CPU4 | MGST3 | 1.51E+07 | 2.35E+07 | 7.82E+06 | 9 | Cytoplasm |
| 172 | P61021 | RAB5B | 1.51E+07 | 7.64E+06 | 1.80E+06 | 18 | Cytoplasm |
| 173 | P35293 | RAB18 | 1.51E+07 | 3.28E+06 | 8.47E+05 | 15 | Cytoplasm |
| 174 | Q9QUI0 | RHOA | 1.50E+07 | 9.90E+06 | 1.91E+06 | 27 | Cytoplasm |
| 175 | Q9R1P3 | PSMB2 | 1.50E+07 | 1.29E+07 | 2.49E+06 | 27 | Cytoplasm |
| 176 | Q9Z2P8 | VAMP5 | 1.46E+07 | 5.93E+06 | 1.98E+06 | 9 | Plasma Membrane |
| 177 | Q3T9M1 | MFSD2B | 1.45E+07 | 1.58E+07 | 6.44E+06 | 6 | Plasma Membrane |
| 178 | P35700 | PRDX1 | 1.44E+07 | 7.48E+06 | 1.30E+06 | 33 | Cytoplasm |
| 179 | O88343 | SLC4A4 | 1.44E+07 | 3.31E+06 | 1.91E+06 | 3 | Plasma Membrane |
| 180 | Q5F285 | TMEM256 | 1.44E+07 | 3.83E+06 | 2.21E+06 | 3 | Cytoplasm |
| 181 | P27664 | PDE6A | 1.43E+07 | 2.23E+06 | 1.29E+06 | 3 | Plasma Membrane |
| 182 | O09117 | SYPL1 | 1.42E+07 | 1.05E+07 | 3.49E+06 | 9 | Plasma Membrane |
| 183 | Q9JJ00 | PLSCR1 | 1.41E+07 | 5.62E+06 | 1.87E+06 | 9 | Plasma Membrane |
| 184 | Q9R1Q7 | PLP2 | 1.40E+07 | 1.66E+06 | 9.59E+05 | 3 | Cytoplasm |
| 185 | P23492 | PNP | 1.38E+07 | 9.90E+06 | 2.33E+06 | 18 | Nucleus |
| 186 | P99028 | UQCRHL | 1.38E+07 | 1.27E+07 | 5.17E+06 | 6 | Cytoplasm |
| 187 | Q9CRA0 | ART4 | 1.38E+07 | 1.28E+07 | 2.79E+06 | 21 | Nucleus |
| 188 | A6H603 | NWD1 | 1.37E+07 | 1.29E+06 | 7.42E+05 | 3 | Cytoplasm |
| 189 | P09103 | P4HB | 1.36E+07 | 6.37E+06 | 8.44E+05 | 57 | Cytoplasm |
| 190 | Q91YN9 | BAG2 | 1.35E+07 | 9.29E+06 | 1.62E+06 | 33 | Cytoplasm |
| 191 | Q78ZA7 | NAP1L4 | 1.35E+07 | 7.57E+06 | 1.32E+06 | 33 | Cytoplasm |
| 192 | Q60931 | VDAC3 | 1.35E+07 | 1.24E+07 | 2.93E+06 | 18 | Cytoplasm |
| 193 | O35930 | GP1BA | 1.35E+07 | 1.89E+07 | 3.02E+06 | 39 | Plasma Membrane |
| 194 | O09061 | PSMB1 | 1.34E+07 | 1.11E+07 | 1.56E+06 | 51 | Cytoplasm |
| 195 | Q9WVJ2 | PSMD13 | 1.34E+07 | 6.41E+06 | 1.03E+06 | 39 | Cytoplasm |
| 196 | Q8BL86 | MBLAC2 | 1.33E+07 | 1.12E+06 | 6.48E+05 | 3 | Cytoplasm |
| 197 | Q8VDD5 | MYH9 | 1.32E+07 | 1.57E+07 | 9.26E+05 | 288 | Cytoplasm |
| 198 | Q8VDN2 | ATP1A1 | 1.31E+07 | 1.03E+07 | 1.10E+06 | 87 | Plasma Membrane |
| 199 | Q9ER00 | STX12 | 1.31E+07 | 2.10E+06 | 1.21E+06 | 3 | Cytoplasm |
| 200 | Q9D1D4 | TMED10 | 1.30E+07 | 7.25E+06 | 1.71E+06 | 18 | Cytoplasm |
| 201 | P26043 | RDX | 1.29E+07 | 1.05E+07 | 1.62E+06 | 42 | Cytoplasm |
| 202 | Q99KF1 | TMED9 | 1.29E+07 | 1.07E+07 | 3.57E+06 | 9 | Cytoplasm |
| 203 | Q921I1 | TF | 1.29E+07 | 1.72E+07 | 1.68E+06 | 105 | Extracellular Space |
| 204 | Q9R1P1 | PSMB3 | 1.29E+07 | 1.03E+07 | 1.88E+06 | 30 | Cytoplasm |
| 205 | P39054 | DNM2 | 1.28E+07 | 1.26E+07 | 1.52E+06 | 69 | Plasma Membrane |
| 206 | Q8VEK0 | TMEM30A | 1.27E+07 | 4.93E+06 | 1.27E+06 | 15 | Cytoplasm |
| 207 | A2AHJ4 | BRWD3 | 1.26E+07 | 1.38E+06 | 7.99E+05 | 3 | Nucleus |
| 208 | Q6ZPJ3 | UBE2O | 1.26E+07 | 8.61E+06 | 8.52E+05 | 102 | Nucleus |
| 209 | P10711 | TCEA1 | 1.24E+07 | 6.49E+06 | 1.08E+06 | 36 | Nucleus |
| 210 | Q60932 | VDAC1 | 1.22E+07 | 9.31E+06 | 1.90E+06 | 24 | Cytoplasm |
| 211 | Q3TDD9 | PPP1R21 | 1.22E+07 | 2.97E+06 | 1.72E+06 | 3 | Cytoplasm |
| 212 | P11352 | GPX1 | 1.21E+07 | 9.24E+06 | 1.69E+06 | 30 | Cytoplasm |
| 213 | A2ATU0 | DHTKD1 | 1.21E+07 | 1.90E+06 | 1.10E+06 | 3 | Cytoplasm |
| 214 | P97315 | CSRP1 | 1.21E+07 | 8.88E+06 | 2.29E+06 | 15 | Nucleus |
| 215 | Q9D0M3 | CYC1 | 1.21E+07 | 3.35E+06 | 1.93E+06 | 3 | Cytoplasm |
| 216 | Q9EPE9 | ATP13A1 | 1.21E+07 | 1.27E+06 | 7.32E+05 | 3 | Extracellular Space |
| 217 | Q9DCN2 | Cyb5r3 | 1.19E+07 | 9.49E+06 | 1.58E+06 | 36 | Cytoplasm |
| 218 | P32037 | SLC2A3 | 1.18E+07 | 1.12E+07 | 2.45E+06 | 21 | Plasma Membrane |
| 219 | P10518 | ALAD | 1.18E+07 | 1.37E+07 | 1.97E+06 | 48 | Cytoplasm |
| 220 | P47738 | ALDH2 | 1.17E+07 | 1.27E+07 | 1.50E+06 | 72 | Cytoplasm |
| 221 | P08556 | NRAS | 1.17E+07 | 7.69E+05 | 4.44E+05 | 3 | Plasma Membrane |
| 222 | Q8K5B2 | MCFD2 | 1.17E+07 | 1.32E+06 | 7.61E+05 | 3 | Cytoplasm |
| 223 | P63082 | ATP6V0C | 1.16E+07 | 1.10E+07 | 4.48E+06 | 6 | Cytoplasm |
| 224 | Q91V01 | LPCAT3 | 1.16E+07 | 7.69E+06 | 2.22E+06 | 12 | Plasma Membrane |
| 225 | Q9D0F3 | LMAN1 | 1.16E+07 | 3.74E+06 | 8.82E+05 | 18 | Cytoplasm |
| 226 | P51150 | RAB7A | 1.15E+07 | 5.56E+06 | 9.68E+05 | 33 | Cytoplasm |
| 227 | P46471 | PSMC2 | 1.15E+07 | 5.03E+06 | 7.50E+05 | 45 | Nucleus |
| 228 | P46638 | RAB11B | 1.15E+07 | 1.21E+07 | 2.21E+06 | 30 | Cytoplasm |
| 229 | Q80Y14 | GLRX5 | 1.15E+07 | 7.70E+06 | 3.15E+06 | 6 | Cytoplasm |
| 230 | Q3UV17 | KRT76 | 1.14E+07 | 3.66E+06 | 2.11E+06 | 3 | Cytoplasm |
| 231 | O70404 | VAMP8 | 1.13E+07 | 1.06E+07 | 3.54E+06 | 9 | Plasma Membrane |
| 232 | P26516 | PSMD7 | 1.13E+07 | 7.70E+06 | 1.41E+06 | 30 | Cytoplasm |
| 233 | Q9ER35 | FN3K | 1.13E+07 | 5.05E+06 | 1.10E+06 | 21 | Cytoplasm |
| 234 | Q60634 | FLOT2 | 1.12E+07 | 7.73E+06 | 1.08E+06 | 51 | Plasma Membrane |
| 235 | O35593 | PSMD14 | 1.12E+07 | 7.24E+06 | 2.09E+06 | 12 | Cytoplasm |
| 236 | Q922R8 | PDIA6 | 1.12E+07 | 5.61E+06 | 9.76E+05 | 33 | Cytoplasm |
| 237 | P62192 | PSMC1 | 1.11E+07 | 6.55E+06 | 1.05E+06 | 39 | Nucleus |
| 238 | P40240 | CD9 | 1.11E+07 | 1.80E+07 | 6.00E+06 | 9 | Plasma Membrane |
| 239 | Q9CVB6 | ARPC2 | 1.10E+07 | 6.04E+06 | 2.01E+06 | 9 | Cytoplasm |
| 240 | P80317 | CCT6A | 1.10E+07 | 5.82E+06 | 9.70E+05 | 36 | Cytoplasm |
| 241 | O08573 | LGALS9B | 1.09E+07 | 5.94E+06 | 1.21E+06 | 24 | Cytoplasm |
| 242 | Q8K274 | FN3KRP | 1.08E+07 | 5.80E+06 | 1.37E+06 | 18 | Cytoplasm |
| 243 | Q7SIG6 | ASAP2 | 1.07E+07 | 3.58E+06 | 2.07E+06 | 3 | Nucleus |
| 244 | Q80X80 | C2CD2L | 1.07E+07 | 7.18E+06 | 1.15E+06 | 39 | Cytoplasm |
| 245 | O35226 | PSMD4 | 1.06E+07 | 8.08E+06 | 1.55E+06 | 27 | Cytoplasm |
| 246 | P97372 | PSME2 | 1.05E+07 | 7.81E+06 | 1.70E+06 | 21 | Cytoplasm |
| 247 | O70439 | STX7 | 1.05E+07 | 3.94E+06 | 9.28E+05 | 18 | Plasma Membrane |
| 248 | Q9R111 | GDA | 1.04E+07 | 5.38E+06 | 9.36E+05 | 33 | Cytoplasm |
| 249 | Q8VEM8 | SLC25A3 | 1.04E+07 | 1.08E+07 | 2.08E+06 | 27 | Cytoplasm |
| 250 | Q3U7R1 | ESYT1 | 1.03E+07 | 6.49E+06 | 6.96E+05 | 87 | Cytoplasm |
| 251 | Q9QZU9 | UBE2L6 | 1.03E+07 | 9.37E+06 | 2.70E+06 | 12 | Cytoplasm |
| 252 | Q9D3D9 | ATP5D | 1.03E+07 | 3.98E+06 | 1.63E+06 | 6 | Cytoplasm |
| 253 | Q91V41 | RAB14 | 1.02E+07 | 1.02E+07 | 1.97E+06 | 27 | Cytoplasm |
| 254 | Q9JMH7 | NEU3 | 1.01E+07 | 7.44E+06 | 2.48E+06 | 9 | Plasma Membrane |
| 255 | P62334 | PSMC6 | 9.97E+06 | 4.24E+06 | 6.78E+05 | 39 | Nucleus |
| 256 | Q9D1Q6 | ERP44 | 9.97E+06 | 4.65E+06 | 1.09E+06 | 18 | Cytoplasm |
| 257 | Q9JKV5 | SCAMP4 | 9.92E+06 | 1.90E+06 | 7.77E+05 | 6 | Cytoplasm |
| 258 | Q8BG32 | PSMD11 | 9.89E+06 | 5.09E+06 | 7.85E+05 | 42 | Cytoplasm |
| 259 | P97371 | PSME1 | 9.83E+06 | 4.16E+06 | 6.94E+05 | 36 | Cytoplasm |
| 260 | Q9D0B1 | ZNF524 | 9.75E+06 | 2.61E+06 | 1.51E+06 | 3 | Nucleus |
| 261 | Q6ZQ38 | CAND1 | 9.73E+06 | 4.01E+06 | 5.31E+05 | 57 | Cytoplasm |
| 262 | B2RWW0 | TAGAP | 9.72E+06 | 1.02E+06 | 5.86E+05 | 3 | Cytoplasm |
| 263 | Q3TFD2 | LPCAT1 | 9.71E+06 | 6.46E+06 | 1.32E+06 | 24 | Cytoplasm |
| 264 | P21440 | ABCB4 | 9.67E+06 | 5.78E+06 | 6.81E+05 | 72 | Plasma Membrane |
| 265 | Q9JKW0 | ARL6IP1 | 9.62E+06 | 3.37E+05 | 1.95E+05 | 3 | Cytoplasm |
| 266 | Q60930 | VDAC2 | 9.61E+06 | 1.11E+07 | 2.14E+06 | 27 | Cytoplasm |
| 267 | P16330 | CNP | 9.61E+06 | 9.43E+06 | 1.93E+06 | 24 | Cytoplasm |
| 268 | Q3TXS7 | PSMD1 | 9.57E+06 | 5.89E+06 | 6.43E+05 | 84 | Cytoplasm |
| 269 | P16546 | SPTAN1 | 9.57E+06 | 1.99E+06 | 1.15E+06 | 3 | Plasma Membrane |
| 270 | P99027 | RPLP2 | 9.55E+06 | 1.15E+07 | 2.98E+06 | 15 | Cytoplasm |
| 271 | P14685 | PSMD3 | 9.55E+06 | 6.77E+06 | 9.77E+05 | 48 | Cytoplasm |
| 272 | O08917 | FLOT1 | 9.52E+06 | 5.25E+06 | 7.58E+05 | 48 | Plasma Membrane |
| 273 | Q9CX56 | PSMD8 | 9.50E+06 | 3.44E+06 | 1.15E+06 | 9 | Cytoplasm |
| 274 | P41216 | ACSL1 | 9.46E+06 | 5.64E+06 | 7.90E+05 | 51 | Cytoplasm |
| 275 | Q9JJU8 | SH3BGRL | 9.44E+06 | 2.56E+06 | 8.53E+05 | 9 | Cytoplasm |
| 276 | Q640N1 | AEBP1 | 9.44E+06 | 1.18E+06 | 6.79E+05 | 3 | Nucleus |
| 277 | Q9D659 | C10orf54 | 9.34E+06 | 1.22E+06 | 7.05E+05 | 3 | Plasma Membrane |

| | | | | | | | |
|---|---|---|---|---|---|---|---|
| 139 | P54726 | RAD23A | 17596578 | 16416241 | 3582317 | 21 | Nucleus |
| 140 | Q921I1 | TF | 17365668 | 19015896 | 1855761 | 105 | Extracellular Space |
| 141 | P99027 | RPLP2 | 17335225 | 18243780 | 4710524 | 15 | Cytoplasm |
| 142 | P23492 | PNP | 16961819 | 13228475 | 3117982 | 18 | Nucleus |
| 143 | P07724 | ALB | 16929684 | 15801643 | 1862241 | 72 | Extracellular Space |
| 144 | Q8CGP5 | HIST1H2AJ | 16733445 | 27863078 | 7194216 | 15 | Nucleus |
| 145 | Q6AW46 | CES5A | 16660000 | 1060063 | 612028 | 3 | Extracellular Space |
| 146 | Q8BL86 | MBLAC2 | 16600000 | 1458572 | 842107 | 3 | Cytoplasm |
| 147 | Q9CPU4 | MGST3 | 16497344 | 26842161 | 8947387 | 9 | Cytoplasm |
| 148 | P35700 | PRDX1 | 16456010 | 9269724 | 1613652 | 33 | Cytoplasm |
| 149 | Q9WV55 | VAPA | 16444775 | 14446520 | 2780234 | 27 | Plasma Membrane |
| 150 | P60904 | DNAJC5 | 16349039 | 17655149 | 4558540 | 15 | Plasma Membrane |
| 151 | P62821 | RAB1A | 16278240 | 27060763 | 6378283 | 18 | Cytoplasm |
| 152 | P11352 | GPX1 | 16249062 | 11819492 | 2157934 | 30 | Cytoplasm |
| 153 | P27773 | PDIA3 | 16178542 | 18299476 | 2202998 | 69 | Cytoplasm |
| 154 | Q3T9M1 | MFSD2B | 15931930 | 16885261 | 6893379 | 6 | Plasma Membrane |
| 155 | Q9CQD1 | RAB5A | 15854789 | 10649825 | 2749773 | 15 | Cytoplasm |
| 156 | P39054 | DNM2 | 15687368 | 14222355 | 1712170 | 69 | Plasma Membrane |
| 157 | Q9R1P3 | PSMB2 | 15490203 | 13626998 | 2622517 | 27 | Cytoplasm |
| 158 | Q6IMF0 | Krt83 | 15450000 | 3491623 | 2015890 | 3 | Extracellular Space |
| 159 | P08113 | HSP90B1 | 15368396 | 8451068 | 956895 | 78 | Cytoplasm |
| 160 | P28656 | NAP1L1 | 15255152 | 10772884 | 2199006 | 24 | Nucleus |
| 161 | Q8VDN2 | ATP1A1 | 14963585 | 11590471 | 1242629 | 87 | Plasma Membrane |
| 162 | Q9CQW2 | ARL8B | 14928147 | 11623574 | 1793556 | 42 | Plasma Membrane |
| 163 | Q9WVJ2 | PSMD13 | 14913898 | 7736292 | 1238798 | 39 | Cytoplasm |
| 164 | P27664 | PDE6A | 14880000 | 1840036 | 1062345 | 3 | Plasma Membrane |
| 165 | Q9CRA0 | ART4 | 14827720 | 14779566 | 3225166 | 21 | Nucleus |
| 166 | O35226 | PSMD4 | 14701355 | 10114134 | 1946466 | 27 | Cytoplasm |
| 167 | Q9R1Q7 | PLP2 | 14490000 | 582866 | 336518 | 3 | Cytoplasm |
| 168 | Q8BGQ1 | VIPAS39 | 14480000 | 3447917 | 1990656 | 3 | Plasma Membrane |
| 169 | Q91YN9 | BAG2 | 14404734 | 10537367 | 1834320 | 33 | Cytoplasm |
| 170 | P99028 | UQCRHL | 14383477 | 14406962 | 5881617 | 6 | Cytoplasm |
| 171 | P35293 | RAB18 | 14080000 | 4106331 | 1060250 | 15 | Cytoplasm |
| 172 | P46471 | PSMC2 | 13972382 | 7275290 | 1084536 | 45 | Nucleus |
| 173 | O88343 | SLC4A4 | 13960000 | 2360946 | 1363093 | 3 | Plasma Membrane |
| 174 | Q62312 | TGFBR2 | 13920000 | 3660342 | 2113299 | 3 | Plasma Membrane |
| 175 | Q9Z2P8 | VAMP5 | 13910000 | 6689139 | 2229713 | 9 | Plasma Membrane |
| 176 | Q9R1P1 | PSMB3 | 13872425 | 10182856 | 1859127 | 30 | Cytoplasm |
| 177 | Q9ER00 | STX12 | 13470000 | 1510894 | 872315 | 3 | Cytoplasm |
| 178 | Q9EPE9 | ATP13A1 | 13460000 | 2127429 | 1228271 | 3 | Extracellular Space |
| 179 | P62192 | PSMC1 | 13446344 | 7782732 | 1246234 | 39 | Nucleus |
| 180 | P07759 | SERPINA3 | 13406534 | 6925912 | 2308637 | 9 | Extracellular Space |
| 181 | Q9QUI0 | RHOA | 13403635 | 10190064 | 1961079 | 27 | Cytoplasm |
| 182 | O09044 | SNAP23 | 13300155 | 13007024 | 2167837 | 36 | Plasma Membrane |
| 183 | Q4QY64 | ATAD5 | 13290000 | 140119 | 80898 | 3 | Extracellular Space |
| 184 | O09061 | PSMB1 | 13267473 | 11898284 | 1666093 | 51 | Cytoplasm |
| 185 | Q99KU0 | VMP1 | 13160000 | 2849462 | 1645138 | 3 | Plasma Membrane |
| 186 | Q60931 | VDAC3 | 12991055 | 15429218 | 3636702 | 18 | Cytoplasm |
| 187 | P22315 | FECH | 12949975 | 22818278 | 3022355 | 57 | Cytoplasm |
| 188 | P61021 | RAB5B | 12599904 | 8042994 | 1895752 | 18 | Cytoplasm |
| 189 | P26516 | PSMD7 | 12583515 | 9150390 | 1670625 | 30 | Cytoplasm |
| 190 | Q3UV17 | KRT76 | 12540000 | 4338560 | 2504868 | 3 | Cytoplasm |
| 191 | O35658 | C1QBP | 12531798 | 13401236 | 5471032 | 6 | Cytoplasm |
| 192 | Q9D659 | C10orf54 | 12210000 | 3573378 | 2063091 | 3 | Plasma Membrane |
| 193 | Q9JJ00 | PLSCR1 | 12161862 | 5972842 | 1990947 | 9 | Plasma Membrane |
| 194 | P80317 | CCT6A | 12041346 | 6962916 | 1160486 | 36 | Cytoplasm |
| 195 | P14685 | PSMD3 | 12021822 | 8958087 | 1292989 | 48 | Cytoplasm |
| 196 | Q60634 | FLOT2 | 11989839 | 8978205 | 1257200 | 51 | Plasma Membrane |
| 197 | Q7SIG6 | ASAP2 | 11970000 | 4464777 | 2577740 | 3 | Nucleus |
| 198 | Q60932 | VDAC1 | 11930683 | 8753306 | 1786761 | 24 | Cytoplasm |
| 199 | P62334 | PSMC6 | 11928478 | 6077772 | 973222 | 39 | Nucleus |
| 200 | O88685 | PSMC3 | 11922672 | 5764661 | 889507 | 42 | Nucleus |
| 201 | Q9D0M3 | CYC1 | 11750000 | 7574665 | 4373235 | 3 | Cytoplasm |
| 202 | O35593 | PSMD14 | 11715730 | 7973582 | 2301775 | 12 | Cytoplasm |
| 203 | P45878 | FKBP2 | 11516170 | 3046908 | 1243895 | 6 | Cytoplasm |
| 204 | Q9Z1N7 | ARID3B | 11460000 | 3060435 | 1766943 | 3 | Nucleus |
| 205 | Q8BG32 | PSMD11 | 11414405 | 5550043 | 856390 | 42 | Cytoplasm |
| 206 | A2AHJ4 | BRWD3 | 11370000 | 1662298 | 959728 | 3 | Nucleus |
| 207 | Q9ER35 | FN3K | 11279958 | 5370305 | 1171897 | 21 | Cytoplasm |
| 208 | P62196 | PSMC5 | 11272971 | 5878902 | 848546 | 48 | Nucleus |
| 209 | Q3TXS7 | PSMD1 | 11263690 | 7070027 | 771403 | 84 | Cytoplasm |
| 210 | P26043 | RDX | 11148051 | 10564610 | 1630155 | 42 | Cytoplasm |
| 211 | Q9JJU8 | SH3BGRL | 11132578 | 2936453 | 978818 | 9 | Cytoplasm |
| 212 | Q9CX56 | PSMD8 | 11106234 | 3193701 | 1064567 | 9 | Cytoplasm |
| 213 | Q80YF9 | ARHGAP33 | 11060000 | 3248697 | 1875636 | 3 | Plasma Membrane |
| 214 | O09117 | SYPL1 | 11005172 | 10506584 | 3502195 | 9 | Plasma Membrane |
| 215 | Q3U7R1 | ESYT1 | 10921520 | 7472201 | 801104 | 87 | Cytoplasm |
| 216 | Q8K274 | FN3KRP | 10885789 | 6469147 | 1524793 | 18 | Cytoplasm |
| 217 | Q8VEK0 | TMEM30A | 10869229 | 4512524 | 1165129 | 15 | Cytoplasm |
| 218 | Q9D3D9 | ATP5D | 10855000 | 5625175 | 2296468 | 6 | Cytoplasm |
| 219 | P62806 | H4 | 10850176 | 20559003 | 4196589 | 24 | Nucleus |
| 220 | O08749 | DLD | 10592482 | 6613201 | 2204400 | 9 | Cytoplasm |
| 221 | Q80X80 | C2CD2L | 10528125 | 7832100 | 1254140 | 39 | Cytoplasm |
| 222 | Q9JMH7 | NEU3 | 10481979 | 6560091 | 2186697 | 9 | Plasma Membrane |
| 223 | P09103 | P4HB | 10360064 | 6230361 | 825231 | 57 | Cytoplasm |
| 224 | Q6ZQ38 | CAND1 | 10291927 | 4087935 | 541460 | 57 | Cytoplasm |
| 225 | Q9CR86 | CARHSP1 | 10230000 | 977252 | 564217 | 3 | Cytoplasm |
| 226 | Q9CVB6 | ARPC2 | 10157464 | 6841197 | 2280399 | 9 | Cytoplasm |
| 227 | Q640N1 | AEBP1 | 10150000 | 1043635 | 602543 | 3 | Nucleus |
| 228 | P80316 | CCT5 | 10086225 | 7027909 | 1125366 | 39 | Cytoplasm |
| 229 | Q8BZ98 | DNM3 | 10009501 | 4692948 | 1564316 | 9 | Cytoplasm |
| 230 | P46638 | RAB11B | 9977935 | 10301482 | 1880785 | 30 | Cytoplasm |
| 231 | P10711 | TCEA1 | 9947040 | 5785033 | 964172 | 36 | Nucleus |
| 232 | P80318 | CCT3 | 9905894 | 6790987 | 980195 | 48 | Cytoplasm |
| 233 | Q9R111 | GDA | 9905438 | 5401275 | 940241 | 33 | Cytoplasm |
| 234 | Q61316 | HSPA4 | 9887321 | 5437267 | 654570 | 69 | Cytoplasm |
| 235 | Q9CQ22 | LAMTOR1 | 9887218 | 8726969 | 2519259 | 12 | Plasma Membrane |
| 236 | Q99JI4 | PSMD6 | 9845170 | 5324292 | 821556 | 42 | Cytoplasm |
| 237 | P51150 | RAB7A | 9828414 | 5903378 | 1027646 | 33 | Cytoplasm |
| 238 | Q64433 | HSPE1 | 9808228 | 7888957 | 2036920 | 15 | Cytoplasm |
| 239 | O08917 | FLOT1 | 9787070 | 5691168 | 821449 | 48 | Plasma Membrane |
| 240 | O55042 | SNCA | 9784430 | 3456293 | 997746 | 12 | Cytoplasm |
| 241 | P51175 | PPOX | 9776629 | 12342102 | 1976318 | 39 | Cytoplasm |
| 242 | P63082 | ATP6V0C | 9775500 | 13832664 | 5647161 | 6 | Cytoplasm |
| 243 | P67984 | RPL22 | 9750991 | 7252957 | 2961007 | 6 | Nucleus |
| 244 | O08573 | LGALS9B | 9735380 | 4604837 | 939958 | 24 | Cytoplasm |
| 245 | Q9CQF9 | PCYOX1 | 9735365 | 1160871 | 670229 | 3 | Cytoplasm |
| 246 | P63038 | HSPD1 | 9686629 | 11032080 | 1328106 | 69 | Cytoplasm |
| 247 | Q9D832 | DNAJB4 | 9684351 | 6941418 | 1267324 | 30 | Nucleus |
| 248 | P84099 | RPL19 | 9680492 | 2973117 | 1716530 | 3 | Cytoplasm |
| 249 | P11983 | TCP1 | 9663244 | 7400040 | 932317 | 63 | Cytoplasm |
| 250 | P53657 | PKLR | 9661223 | 5501547 | 1122999 | 24 | Cytoplasm |
| 251 | Q8VDM4 | PSMD2 | 9645502 | 5313836 | 569703 | 87 | Cytoplasm |
| 252 | Q8R1V4 | TMED4 | 9625231 | 2174638 | 1255528 | 3 | Cytoplasm |
| 253 | Q9D8W5 | PSMD12 | 9616071 | 4966900 | 740422 | 45 | Cytoplasm |
| 254 | P15864 | HIST1H1C | 9526512 | 9053267 | 5226906 | 3 | Nucleus |
| 255 | P80315 | CCT4 | 9506531 | 6876399 | 935759 | 54 | Cytoplasm |
| 256 | P48972 | MYBL2 | 9470300 | 1190063 | 687083 | 3 | Nucleus |
| 257 | Q9D1D4 | TMED10 | 9450253 | 6724363 | 1584947 | 18 | Cytoplasm |
| 258 | P00920 | CA2 | 9441067 | 10610771 | 2500983 | 18 | Cytoplasm |
| 259 | P70168 | KPNB1 | 9410822 | 3961475 | 499099 | 63 | Nucleus |
| 260 | Q78ZA7 | NAP1L4 | 9384336 | 6529800 | 1136692 | 33 | Cytoplasm |
| 261 | P80313 | CCT7 | 9363191 | 4708942 | 701968 | 45 | Cytoplasm |
| 262 | P02535 | Krt10 | 9353656 | 3493699 | 2017088 | 3 | Cytoplasm |
| 263 | P60521 | GABARAPL2 | 9350313 | 6203879 | 1601835 | 15 | Cytoplasm |
| 264 | Q9DCN2 | Cyb5r3 | 9346811 | 8071255 | 1345209 | 36 | Cytoplasm |
| 265 | Q80ZA4 | PKHD1L1 | 9338031 | 308740 | 178251 | 3 | Extracellular Space |
| 266 | P80314 | CCT2 | 9330937 | 4771585 | 736272 | 42 | Cytoplasm |
| 267 | Q3TDD9 | PPP1R21 | 9290315 | 3706933 | 2140199 | 3 | Cytoplasm |
| 268 | P19783 | COX4I1 | 9282438 | 8087497 | 2695832 | 9 | Cytoplasm |
| 269 | O70439 | STX7 | 9276643 | 3861851 | 910247 | 18 | Plasma Membrane |
| 270 | P21440 | ABCB4 | 9252222 | 6459435 | 761252 | 72 | Plasma Membrane |
| 271 | Q9CXW4 | RPL11 | 9240216 | 4921161 | 2841233 | 3 | Cytoplasm |
| 272 | Q9CZU6 | CS | 9200481 | 11569605 | 2112311 | 30 | Cytoplasm |
| 273 | Q9QYJ3 | DNAJB1 | 9144477 | 6208813 | 1194887 | 27 | Nucleus |
| 274 | P68037 | UBE2L3 | 9136070 | 1381361 | 797529 | 3 | Nucleus |
| 275 | Q8VEM8 | SLC25A3 | 9128713 | 9909621 | 1907107 | 27 | Cytoplasm |
| 276 | P97371 | PSME1 | 9107953 | 3901995 | 650332 | 36 | Cytoplasm |
| 277 | P62331 | ARF6 | 9077525 | 5544627 | 1306881 | 18 | Plasma Membrane |

| | | | | | | | |
|---|---|---|---|---|---|---|---|
| 278 | P10711 | TCEA1 | 9.34E+06 | 6.09E+06 | 1.01E+06 | 36 | Nucleus |
| 279 | Q99KF1 | TMED9 | 9.32E+06 | 8.99E+06 | 3.00E+06 | 9 | Cytoplasm |
| 280 | P62334 | PSMC6 | 9.31E+06 | 5.54E+06 | 8.88E+05 | 39 | Nucleus |
| 281 | O70439 | STX7 | 9.22E+06 | 3.13E+06 | 7.37E+05 | 18 | Plasma Membrane |
| 282 | Q9CQ19 | MYL9 | 9.15E+06 | 7.72E+06 | 1.58E+06 | 24 | Cytoplasm |
| 283 | Q922D8 | MTHFD1 | 9.13E+06 | 5.42E+06 | 5.81E+05 | 87 | Cytoplasm |
| 284 | P80316 | CCT5 | 9.12E+06 | 7.34E+06 | 1.17E+06 | 39 | Cytoplasm |
| 285 | Q3TXS7 | PSMD1 | 9.12E+06 | 6.88E+06 | 7.50E+05 | 84 | Cytoplasm |
| 286 | Q9D0F3 | LMAN1 | 9.09E+06 | 2.00E+06 | 4.72E+05 | 18 | Cytoplasm |
| 287 | O88685 | PSMC3 | 9.04E+06 | 5.94E+06 | 9.16E+05 | 42 | Nucleus |
| 288 | Q8K5B2 | MCFD2 | 9.02E+06 | 1.69E+06 | 9.73E+05 | 3 | Cytoplasm |
| 289 | P51125 | CAST | 8.98E+06 | 7.70E+06 | 1.28E+06 | 36 | Cytoplasm |
| 290 | P70704 | ATP8A1 | 8.97E+06 | 5.71E+06 | 7.02E+05 | 66 | Cytoplasm |
| 291 | Q8BG32 | PSMD11 | 8.94E+06 | 5.83E+06 | 8.99E+05 | 42 | Cytoplasm |
| 292 | O70404 | VAMP8 | 8.91E+06 | 4.79E+06 | 1.60E+06 | 9 | Plasma Membrane |
| 293 | O35226 | PSMD4 | 8.90E+06 | 7.89E+06 | 1.52E+06 | 27 | Cytoplasm |
| 294 | Q9D1Q6 | ERP44 | 8.81E+06 | 4.93E+06 | 1.16E+06 | 18 | Cytoplasm |
| 295 | Q9D3D9 | ATP5D | 8.78E+06 | 1.82E+06 | 7.42E+05 | 6 | Cytoplasm |
| 296 | Q9CX56 | PSMD8 | 8.74E+06 | 4.65E+06 | 1.55E+06 | 9 | Cytoplasm |
| 297 | Q8VDM4 | PSMD2 | 8.72E+06 | 6.97E+06 | 7.47E+05 | 87 | Cytoplasm |
| 298 | P80318 | CCT3 | 8.65E+06 | 5.63E+06 | 8.12E+05 | 48 | Cytoplasm |
| 299 | O35658 | C1QBP | 8.65E+06 | 7.70E+06 | 3.14E+06 | 6 | Cytoplasm |
| 300 | B2RWW0 | TAGAP | 8.62E+06 | 1.79E+06 | 1.04E+06 | 3 | Cytoplasm |
| 301 | Q9JJU8 | SH3BGRL | 8.61E+06 | 2.73E+06 | 9.10E+05 | 9 | Cytoplasm |
| 302 | Q9QYJ3 | DNAJB1 | 8.60E+06 | 6.16E+06 | 1.19E+06 | 27 | Nucleus |
| 303 | Q3TTY5 | Krt2 | 8.59E+06 | 4.09E+06 | 2.36E+06 | 3 | Cytoplasm |
| 304 | P80315 | CCT4 | 8.56E+06 | 7.17E+06 | 9.76E+05 | 54 | Cytoplasm |
| 305 | E2JF22 | PIEZO1 | 8.56E+06 | 6.58E+06 | 7.31E+05 | 81 | Cytoplasm |
| 306 | P00920 | CA2 | 8.56E+06 | 6.86E+06 | 1.62E+06 | 18 | Cytoplasm |
| 307 | Q9D771 | TMEM206 | 8.55E+06 | 7.16E+06 | 2.39E+06 | 9 | Plasma Membrane |
| 308 | P62331 | ARF6 | 8.48E+06 | 4.32E+06 | 1.02E+06 | 18 | Plasma Membrane |
| 309 | Q91V41 | RAB14 | 8.43E+06 | 6.12E+06 | 1.18E+06 | 27 | Cytoplasm |
| 310 | Q60930 | VDAC2 | 8.39E+06 | 6.43E+06 | 1.24E+06 | 27 | Cytoplasm |
| 311 | P80313 | CCT7 | 8.37E+06 | 4.98E+06 | 7.42E+05 | 45 | Cytoplasm |
| 312 | P80314 | CCT2 | 8.35E+06 | 4.60E+06 | 7.10E+05 | 42 | Cytoplasm |
| 313 | P62196 | PSMC5 | 8.34E+06 | 5.41E+06 | 7.81E+05 | 48 | Nucleus |
| 314 | Q9CZU6 | CS | 8.29E+06 | 6.09E+06 | 1.11E+06 | 30 | Cytoplasm |
| 315 | P11983 | TCP1 | 8.26E+06 | 7.80E+06 | 9.83E+05 | 63 | Cytoplasm |
| 316 | P16546 | SPTAN1 | 8.22E+06 | 5.10E+06 | 2.94E+06 | 3 | Plasma Membrane |
| 317 | P51863 | ATP6V0D1 | 8.19E+06 | 4.97E+06 | 1.17E+06 | 18 | Cytoplasm |
| 318 | P61226 | RAP2B | 8.17E+06 | 3.12E+06 | 7.35E+05 | 18 | Plasma Membrane |
| 319 | Q8BHN3 | GANAB | 8.17E+06 | 5.19E+06 | 6.71E+05 | 60 | Cytoplasm |
| 320 | O08749 | DLD | 8.13E+06 | 3.48E+06 | 1.16E+06 | 9 | Cytoplasm |
| 321 | Q9CR86 | CARHSP1 | 8.12E+06 | 1.37E+06 | 7.91E+05 | 3 | Cytoplasm |
| 322 | Q3U898 | MYEOV2 | 8.12E+06 | 3.89E+06 | 2.24E+06 | 3 | Nucleus |
| 323 | P19783 | COX4I1 | 8.12E+06 | 5.49E+06 | 1.83E+06 | 9 | Cytoplasm |
| 324 | Q9CQ22 | LAMTOR1 | 8.09E+06 | 6.16E+06 | 1.78E+06 | 12 | Plasma Membrane |
| 325 | O89053 | CORO1A | 8.05E+06 | 1.05E+07 | 1.68E+06 | 39 | Cytoplasm |
| 326 | Q9JKV5 | SCAMP4 | 8.03E+06 | 2.46E+06 | 1.00E+06 | 6 | Cytoplasm |
| 327 | P03930 | mt-Atp8 | 8.01E+06 | 5.13E+06 | 2.09E+06 | 6 | Cytoplasm |
| 328 | Q80XI4 | PIP4K2B | 8.00E+06 | 2.71E+06 | 1.57E+06 | 3 | Cytoplasm |
| 329 | Q9WTR1 | TRPV2 | 7.99E+06 | 5.13E+06 | 8.55E+05 | 36 | Plasma Membrane |
| 330 | Q9QYB5 | ADD3 | 7.92E+06 | 7.14E+06 | 1.06E+06 | 45 | Cytoplasm |
| 331 | P53657 | PKLR | 7.92E+06 | 6.23E+06 | 1.27E+06 | 24 | Cytoplasm |
| 332 | P19536 | Cox5b | 7.90E+06 | 2.32E+06 | 1.34E+06 | 3 | Cytoplasm |
| 333 | Q9CY52 | THG1L | 7.88E+06 | 2.79E+06 | 7.20E+05 | 15 | Cytoplasm |
| 334 | P22315 | FECH | 7.86E+06 | 9.15E+06 | 1.21E+06 | 57 | Cytoplasm |
| 335 | Q64514 | TPP2 | 7.85E+06 | 7.00E+06 | 8.62E+05 | 66 | Cytoplasm |
| 336 | Q9D659 | C10orf54 | 7.84E+06 | 2.00E+06 | 1.15E+06 | 3 | Plasma Membrane |
| 337 | Q80ZA4 | PKHD1L1 | 7.83E+06 | 4.85E+05 | 2.80E+05 | 3 | Extracellular Space |
| 338 | P41216 | ACSL1 | 7.82E+06 | 4.39E+06 | 6.15E+05 | 51 | Cytoplasm |
| 339 | Q9JMH7 | NEU3 | 7.73E+06 | 5.57E+06 | 1.86E+06 | 9 | Plasma Membrane |
| 340 | P63038 | HSPD1 | 7.73E+06 | 5.31E+06 | 6.39E+05 | 69 | Cytoplasm |
| 341 | Q9D8W5 | PSMD12 | 7.73E+06 | 4.84E+06 | 7.21E+05 | 45 | Cytoplasm |
| 342 | Q3THE2 | MYL12A | 7.72E+06 | 8.12E+06 | 2.10E+06 | 15 | Cytoplasm |
| 343 | P17742 | PPIA | 7.66E+06 | 6.42E+06 | 1.40E+06 | 21 | Cytoplasm |
| 344 | Q9DBH5 | LMAN2 | 7.58E+06 | 4.77E+06 | 1.23E+06 | 15 | Cytoplasm |
| 345 | Q8K0T0 | RTN1 | 7.54E+06 | 2.24E+06 | 1.29E+06 | 3 | Cytoplasm |
| 346 | Q64433 | HSPE1 | 7.52E+06 | 3.90E+06 | 1.01E+06 | 15 | Cytoplasm |
| 347 | Q99JI4 | PSMD6 | 7.43E+06 | 5.42E+06 | 8.37E+05 | 42 | Cytoplasm |
| 348 | Q60766 | Irgm1 | 7.42E+06 | 4.83E+06 | 8.82E+05 | 30 | Cytoplasm |
| 349 | Q61165 | SLC9A1 | 7.41E+06 | 4.18E+06 | 8.54E+05 | 24 | Plasma Membrane |
| 350 | P58735 | SLC26A1 | 7.40E+06 | 3.41E+06 | 1.39E+06 | 6 | Plasma Membrane |
| 351 | Q9CR20 | IER3IP1 | 7.38E+06 | 4.34E+06 | 1.77E+06 | 6 | Cytoplasm |
| 352 | P54822 | ADSL | 7.34E+06 | 5.90E+06 | 9.45E+05 | 39 | Cytoplasm |
| 353 | Q9EPK2 | RP2 | 7.28E+06 | 3.38E+06 | 7.38E+05 | 21 | Cytoplasm |
| 354 | Q3UUG6 | TBC1D24 | 7.23E+06 | 2.83E+06 | 1.16E+06 | 6 | Cytoplasm |
| 355 | Q9JLT4 | TXNRD2 | 7.23E+06 | 7.11E+06 | 2.37E+06 | 9 | Cytoplasm |
| 356 | Q8CG47 | SMC4 | 7.17E+06 | 7.49E+06 | 3.06E+06 | 6 | Nucleus |
| 357 | P09055 | ITGB1 | 7.15E+06 | 6.30E+06 | 1.01E+06 | 39 | Plasma Membrane |
| 358 | O70172 | PIP4K2A | 7.01E+06 | 5.82E+06 | 1.68E+06 | 12 | Cytoplasm |
| 359 | P62259 | YWHAE | 7.00E+06 | 3.38E+06 | 6.51E+05 | 27 | Cytoplasm |
| 360 | Q9CXI5 | MANF | 6.96E+06 | 5.86E+06 | 1.95E+06 | 9 | Extracellular Space |
| 361 | P50637 | TSPO | 6.96E+06 | 2.18E+06 | 1.26E+06 | 3 | Cytoplasm |
| 362 | Q9CYN2 | SPCS2 | 6.95E+06 | 2.50E+06 | 1.44E+06 | 3 | Cytoplasm |
| 363 | O35864 | COPS5 | 6.95E+06 | 5.29E+06 | 2.16E+06 | 6 | Nucleus |
| 364 | Q9D1G5 | LRRC57 | 6.93E+06 | 2.29E+06 | 5.40E+05 | 18 | Cytoplasm |
| 365 | Q80VY9 | DHX33 | 6.92E+06 | 2.05E+06 | 1.19E+06 | 3 | Nucleus |
| 366 | P70168 | KPNB1 | 6.91E+06 | 4.01E+06 | 5.05E+05 | 63 | Nucleus |
| 367 | P57716 | NCSTN | 6.85E+06 | 3.85E+06 | 7.40E+05 | 27 | Plasma Membrane |
| 368 | Q61820 | Rasl2-9 | 6.84E+06 | 2.01E+06 | 8.22E+05 | 6 | Nucleus |
| 369 | P54775 | PSMC4 | 6.81E+06 | 5.15E+06 | 8.58E+05 | 36 | Nucleus |
| 370 | Q9JM62 | REEP6 | 6.79E+06 | 4.99E+06 | 1.66E+06 | 9 | Plasma Membrane |
| 371 | Q9JIZ9 | PLSCR3 | 6.78E+06 | 3.50E+06 | 1.01E+06 | 12 | Plasma Membrane |
| 372 | P42932 | CCT8 | 6.78E+06 | 3.92E+06 | 4.62E+05 | 72 | Cytoplasm |
| 373 | P18242 | CTSD | 6.75E+06 | 3.56E+06 | 7.76E+05 | 21 | Cytoplasm |
| 374 | Q7M6Y3 | PICALM | 6.68E+06 | 6.84E+06 | 1.97E+06 | 12 | Cytoplasm |
| 375 | Q9DB20 | ATP5O | 6.68E+06 | 3.68E+06 | 7.51E+05 | 24 | Cytoplasm |
| 376 | P60766 | Cdc42 | 6.66E+06 | 3.48E+06 | 8.98E+05 | 15 | Plasma Membrane |
| 377 | Q9R0Q3 | TMED2 | 6.64E+06 | 2.25E+06 | 7.51E+05 | 9 | Cytoplasm |
| 378 | P10404 | ENV1 | 6.62E+06 | 6.20E+06 | 7.31E+05 | 72 | Other |
| 379 | P00405 | MT-CO2 | 6.62E+06 | 7.21E+06 | 1.70E+06 | 18 | Cytoplasm |
| 380 | Q8BK63 | CSNK1A1 | 6.60E+06 | 2.99E+06 | 6.10E+05 | 24 | Cytoplasm |
| 381 | Q9CR23 | TMEM9 | 6.59E+06 | 4.21E+05 | 2.43E+05 | 3 | Cytoplasm |
| 382 | Q9QZU9 | UBE2L6 | 6.57E+06 | 9.04E+06 | 2.61E+06 | 12 | Cytoplasm |
| 383 | P51174 | ACADL | 6.57E+06 | 6.60E+06 | 1.44E+06 | 21 | Cytoplasm |
| 384 | Q61475 | CD55 | 6.50E+06 | 3.36E+06 | 7.92E+05 | 18 | Plasma Membrane |
| 385 | Q8R016 | BLMH | 6.46E+06 | 4.12E+06 | 9.00E+05 | 21 | Cytoplasm |
| 386 | Q5XPI3 | RNF123 | 6.44E+06 | 3.51E+06 | 7.67E+05 | 21 | Cytoplasm |
| 387 | O08795 | PRKCSH | 6.43E+06 | 3.40E+06 | 5.93E+05 | 33 | Cytoplasm |
| 388 | Q9QUJ7 | ACSL4 | 6.43E+06 | 5.26E+06 | 1.01E+06 | 27 | Cytoplasm |
| 389 | Q923D2 | BLVRB | 6.42E+06 | 4.43E+06 | 1.28E+06 | 12 | Cytoplasm |
| 390 | P26040 | EZR | 6.41E+06 | 6.11E+06 | 1.12E+06 | 30 | Plasma Membrane |
| 391 | Q3TZZ7 | ESYT2 | 6.38E+06 | 2.91E+06 | 3.76E+05 | 60 | Plasma Membrane |
| 392 | Q99P58 | RAB27B | 6.38E+06 | 7.40E+06 | 1.51E+06 | 24 | Cytoplasm |
| 393 | Q68FD5 | CLTC | 6.32E+06 | 4.97E+06 | 4.23E+05 | 138 | Plasma Membrane |
| 394 | P57759 | ERP29 | 6.28E+06 | 2.56E+06 | 7.39E+05 | 12 | Cytoplasm |
| 395 | P60521 | GABARAPL2 | 6.26E+06 | 3.71E+06 | 9.57E+05 | 15 | Cytoplasm |
| 396 | Q99J56 | DERL1 | 6.26E+06 | 3.60E+06 | 1.47E+06 | 6 | Cytoplasm |
| 397 | P10605 | CTSB | 6.23E+06 | 3.07E+06 | 7.92E+05 | 15 | Cytoplasm |
| 398 | Q3TDD9 | PPP1R21 | 6.20E+06 | 2.93E+06 | 1.69E+06 | 3 | Cytoplasm |
| 399 | O35379 | ABCC1 | 6.20E+06 | 4.39E+06 | 5.97E+05 | 54 | Plasma Membrane |
| 400 | Q6PAR5 | GAPVD1 | 6.17E+06 | 3.78E+06 | 5.29E+05 | 51 | Cytoplasm |
| 401 | Q9JJK2 | LANCL2 | 6.13E+06 | 3.14E+06 | 5.23E+05 | 36 | Plasma Membrane |
| 402 | Q8BKC5 | IPO5 | 6.13E+06 | 3.11E+06 | 4.01E+05 | 60 | Nucleus |
| 403 | Q9DAS9 | GNG12 | 6.12E+06 | 3.62E+05 | 2.09E+05 | 3 | Plasma Membrane |
| 404 | P67984 | RPL22 | 6.11E+06 | 3.22E+06 | 1.31E+06 | 6 | Nucleus |
| 405 | P51175 | PPOX | 6.11E+06 | 5.04E+06 | 8.07E+05 | 39 | Cytoplasm |
| 406 | P70182 | PIP5K1A | 6.10E+06 | 3.91E+06 | 7.14E+05 | 30 | Cytoplasm |
| 407 | P08680 | ALAS2 | 6.10E+06 | 4.98E+06 | 8.29E+05 | 36 | Cytoplasm |
| 408 | O09172 | GCLM | 6.10E+06 | 1.33E+06 | 4.42E+05 | 9 | Cytoplasm |
| 409 | P53994 | RAB2A | 6.03E+06 | 3.89E+06 | 1.00E+06 | 15 | Cytoplasm |
| 410 | O35643 | AP1B1 | 6.03E+06 | 3.73E+06 | 6.21E+05 | 36 | Cytoplasm |
| 411 | O55042 | SNCA | 6.03E+06 | 1.93E+06 | 5.57E+05 | 12 | Cytoplasm |
| 412 | Q06185 | Atp5k | 6.02E+06 | 3.36E+06 | 1.37E+06 | 6 | Cytoplasm |
| 413 | P11499 | HSP90AB1 | 6.02E+06 | 2.62E+06 | 1.51E+06 | 3 | Cytoplasm |
| 414 | Q9CQR6 | PPP6C | 6.01E+06 | 3.33E+06 | 1.36E+06 | 6 | Nucleus |
| 415 | Q60682 | Klra4 (includes others) | 6.01E+06 | 1.02E+06 | 5.87E+05 | 3 | Plasma Membrane |
| 416 | O35129 | PHB2 | 5.98E+06 | 3.93E+06 | 7.56E+05 | 27 | Cytoplasm |

| | | | | | | | |
|---|---|---|---|---|---|---|---|
| 278 | Q6ZPJ3 | UBE2O | 9.76E+06 | 8.36E+06 | 8.27E+05 | 102 | Nucleus |
| 279 | O35226 | PSMD4 | 9.74E+06 | 7.72E+06 | 1.49E+06 | 27 | Cytoplasm |
| 280 | Q6IMF0 | Krt83 | 9.70E+06 | 3.17E+06 | 1.83E+06 | 3 | Extracellular Space |
| 281 | O08688 | CAPN5 | 9.69E+06 | 9.34E+06 | 1.21E+06 | 60 | Cytoplasm |
| 282 | P80317 | CCT6A | 9.68E+06 | 6.72E+06 | 1.12E+06 | 36 | Cytoplasm |
| 283 | P50637 | TSPO | 9.67E+06 | 2.77E+06 | 1.60E+06 | 3 | Cytoplasm |
| 284 | Q9WVJ2 | PSMD13 | 9.64E+06 | 6.17E+06 | 9.87E+05 | 39 | Cytoplasm |
| 285 | P35441 | THBS1 | 9.56E+06 | 8.77E+06 | 8.95E+05 | 96 | Extracellular Space |
| 286 | P51863 | ATP6V0D1 | 9.54E+06 | 6.14E+06 | 1.45E+06 | 18 | Cytoplasm |
| 287 | P84099 | RPL19 | 9.53E+06 | 1.20E+06 | 6.94E+05 | 3 | Cytoplasm |
| 288 | P51125 | CAST | 9.52E+06 | 7.15E+06 | 1.19E+06 | 36 | Cytoplasm |
| 289 | Q06185 | Atp5k | 9.48E+06 | 5.20E+06 | 2.12E+06 | 6 | Cytoplasm |
| 290 | P21440 | ABCB4 | 9.45E+06 | 5.34E+06 | 6.30E+05 | 72 | Plasma Membrane |
| 291 | P62331 | ARF6 | 9.40E+06 | 5.69E+06 | 1.34E+06 | 18 | Plasma Membrane |
| 292 | Q7M6Y3 | PICALM | 9.39E+06 | 9.75E+06 | 2.82E+06 | 12 | Cytoplasm |
| 293 | P16330 | CNP | 9.39E+06 | 9.01E+06 | 1.84E+06 | 24 | Cytoplasm |
| 294 | Q80X80 | C2CD2L | 9.33E+06 | 6.85E+06 | 1.10E+06 | 39 | Cytoplasm |
| 295 | P48972 | MYBL2 | 9.29E+06 | 1.90E+06 | 1.10E+06 | 3 | Nucleus |
| 296 | Q60766 | Irgm1 | 9.26E+06 | 4.62E+06 | 8.44E+05 | 30 | Cytoplasm |
| 297 | P51175 | PPOX | 9.26E+06 | 9.46E+06 | 1.51E+06 | 39 | Cytoplasm |
| 298 | Q9CZ13 | UQCRC1 | 9.24E+06 | 4.65E+06 | 1.20E+06 | 15 | Cytoplasm |
| 299 | O08917 | FLOT1 | 9.24E+06 | 5.03E+06 | 7.27E+05 | 48 | Plasma Membrane |
| 300 | P08680 | ALAS2 | 9.23E+06 | 8.57E+06 | 1.43E+06 | 36 | Cytoplasm |
| 301 | P80316 | CCT5 | 9.14E+06 | 6.99E+06 | 1.12E+06 | 39 | Cytoplasm |
| 302 | Q91V41 | RAB14 | 9.13E+06 | 1.04E+07 | 2.00E+06 | 27 | Cytoplasm |
| 303 | Q8K5B2 | MCFD2 | 9.11E+06 | 1.35E+06 | 7.82E+05 | 3 | Cytoplasm |
| 304 | P62192 | PSMC1 | 9.09E+06 | 6.58E+06 | 1.05E+06 | 39 | Nucleus |
| 305 | Q9WV71 | ASB4 | 9.07E+06 | 1.48E+06 | 8.57E+05 | 3 | Nucleus |
| 306 | P01887 | B2M | 9.01E+06 | 1.08E+06 | 6.22E+05 | 3 | Plasma Membrane |
| 307 | Q64518 | ATP2A3 | 8.98E+06 | 8.39E+06 | 1.21E+06 | 48 | Cytoplasm |
| 308 | P11499 | HSP90AB1 | 8.95E+06 | 1.83E+06 | 1.06E+06 | 3 | Cytoplasm |
| 309 | P12787 | COX5A | 8.91E+06 | 5.45E+06 | 1.82E+06 | 9 | Cytoplasm |
| 310 | Q9JL99 | CLEC1B | 8.85E+06 | 7.58E+05 | 4.37E+05 | 3 | Plasma Membrane |
| 311 | P08905 | LYZ | 8.83E+06 | 6.22E+06 | 2.54E+06 | 6 | Extracellular Space |
| 312 | Q9CR86 | CARHSP1 | 8.83E+06 | 7.56E+05 | 4.36E+05 | 3 | Cytoplasm |
| 313 | Q9CQX2 | CYB5B | 8.77E+06 | 1.06E+07 | 3.52E+06 | 9 | Cytoplasm |
| 314 | O35643 | AP1B1 | 8.73E+06 | 5.08E+06 | 8.46E+05 | 36 | Cytoplasm |
| 315 | Q9JJ59 | ABCB9 | 8.73E+06 | 8.19E+05 | 4.73E+05 | 3 | Cytoplasm |
| 316 | Q9D832 | DNAJB4 | 8.72E+06 | 6.07E+06 | 1.11E+06 | 30 | Nucleus |
| 317 | Q9CQN1 | TRAP1 | 8.63E+06 | 7.00E+06 | 2.86E+06 | 6 | Cytoplasm |
| 318 | P21279 | GNAQ | 8.62E+06 | 3.18E+06 | 1.84E+06 | 3 | Plasma Membrane |
| 319 | P41216 | ACSL1 | 8.58E+06 | 6.21E+06 | 8.69E+05 | 51 | Cytoplasm |
| 320 | P61226 | RAP2B | 8.58E+06 | 3.24E+06 | 7.63E+05 | 18 | Plasma Membrane |
| 321 | Q99P72 | RTN4 | 8.57E+06 | 7.38E+06 | 1.91E+06 | 15 | Cytoplasm |
| 322 | P07901 | HSP90AA1 | 8.56E+06 | 3.01E+06 | 7.09E+05 | 18 | Cytoplasm |
| 323 | Q9JLT4 | TXNRD2 | 8.54E+06 | 7.11E+06 | 2.37E+06 | 9 | Cytoplasm |
| 324 | O88685 | PSMC3 | 8.54E+06 | 5.43E+06 | 8.38E+05 | 42 | Nucleus |
| 325 | Q9ER35 | FN3K | 8.53E+06 | 4.44E+06 | 9.69E+05 | 21 | Cytoplasm |
| 326 | P02535 | Krt10 | 8.51E+06 | 1.13E+06 | 6.55E+05 | 3 | Cytoplasm |
| 327 | P14685 | PSMD3 | 8.50E+06 | 7.28E+06 | 1.05E+06 | 48 | Cytoplasm |
| 328 | Q8QZT1 | ACAT1 | 8.48E+06 | 4.45E+06 | 9.71E+05 | 21 | Cytoplasm |
| 329 | P97372 | PSME2 | 8.48E+06 | 7.15E+06 | 1.56E+06 | 21 | Cytoplasm |
| 330 | P68037 | UBE2L3 | 8.47E+06 | 1.31E+06 | 7.55E+05 | 3 | Nucleus |
| 331 | P62259 | YWHAE | 8.42E+06 | 4.08E+06 | 7.85E+05 | 27 | Cytoplasm |
| 332 | Q9JKW0 | ARL6IP1 | 8.41E+06 | 7.91E+05 | 4.57E+05 | 3 | Cytoplasm |
| 333 | P11438 | LAMP1 | 8.40E+06 | 4.62E+06 | 1.54E+06 | 9 | Plasma Membrane |
| 334 | Q6ZQ38 | CAND1 | 8.37E+06 | 3.84E+06 | 5.09E+05 | 57 | Cytoplasm |
| 335 | O35129 | PHB2 | 8.36E+06 | 5.92E+06 | 1.14E+06 | 27 | Cytoplasm |
| 336 | P26040 | EZR | 8.35E+06 | 6.42E+06 | 1.17E+06 | 30 | Plasma Membrane |
| 337 | P24270 | CAT | 8.33E+06 | 5.91E+06 | 7.63E+05 | 60 | Cytoplasm |
| 338 | Q640N1 | AEBP1 | 8.32E+06 | 7.44E+05 | 4.29E+05 | 3 | Nucleus |
| 339 | P10711 | TCEA1 | 8.30E+06 | 5.66E+06 | 9.43E+05 | 36 | Nucleus |
| 340 | P36552 | CPOX | 8.29E+06 | 4.13E+06 | 6.89E+05 | 36 | Cytoplasm |
| 341 | P62900 | RPL31 | 8.25E+06 | 1.13E+06 | 4.59E+05 | 6 | Cytoplasm |
| 342 | Q99KR7 | PPIF | 8.24E+06 | 1.00E+06 | 4.09E+05 | 6 | Cytoplasm |
| 343 | P14148 | RPL7 | 8.16E+06 | 5.28E+06 | 1.36E+06 | 15 | Nucleus |
| 344 | P10605 | CTSB | 8.13E+06 | 3.88E+06 | 1.00E+06 | 15 | Cytoplasm |
| 345 | Q9D1J1 | NECAP2 | 8.07E+06 | 1.96E+06 | 1.13E+06 | 3 | Cytoplasm |
| 346 | A2AHJ4 | BRWD3 | 8.05E+06 | 2.04E+06 | 1.18E+06 | 3 | Nucleus |
| 347 | P97371 | PSME1 | 8.03E+06 | 3.86E+06 | 6.43E+05 | 36 | Cytoplasm |
| 348 | Q3TTY5 | Krt2 | 8.01E+06 | 3.59E+06 | 2.07E+06 | 3 | Cytoplasm |
| 349 | Q01730 | RSU1 | 7.97E+06 | 3.81E+06 | 1.27E+06 | 9 | Cytoplasm |
| 350 | Q61316 | HSPA4 | 7.97E+06 | 4.76E+06 | 5.72E+05 | 69 | Cytoplasm |
| 351 | P70704 | ATP8A1 | 7.94E+06 | 5.21E+06 | 6.41E+05 | 66 | Cytoplasm |
| 352 | Q9D6F9 | TUBB4A | 7.89E+06 | 1.81E+06 | 7.40E+05 | 6 | Cytoplasm |
| 353 | P80315 | CCT4 | 7.88E+06 | 6.61E+06 | 9.00E+05 | 54 | Cytoplasm |
| 354 | Q3TFD2 | LPCAT1 | 7.87E+06 | 5.53E+06 | 1.13E+06 | 24 | Cytoplasm |
| 355 | Q791V5 | MTCH2 | 7.87E+06 | 4.62E+05 | 2.67E+05 | 3 | Cytoplasm |
| 356 | P62830 | RPL23 | 7.87E+06 | 2.85E+05 | 1.64E+05 | 3 | Cytoplasm |
| 357 | Q3TXS7 | PSMD1 | 7.86E+06 | 5.71E+06 | 6.23E+05 | 84 | Cytoplasm |
| 358 | P10852 | SLC3A2 | 7.86E+06 | 3.64E+06 | 6.64E+05 | 30 | Plasma Membrane |
| 359 | Q9CYN2 | SPCS2 | 7.84E+06 | 2.00E+06 | 1.15E+06 | 3 | Cytoplasm |
| 360 | Q80ZA4 | PKHD1L1 | 7.81E+06 | 9.52E+05 | 5.50E+05 | 3 | Extracellular Space |
| 361 | P62264 | RPS14 | 7.77E+06 | 3.11E+06 | 1.04E+06 | 9 | Cytoplasm |
| 362 | P80318 | CCT3 | 7.75E+06 | 5.42E+06 | 7.82E+05 | 48 | Cytoplasm |
| 363 | P05202 | GOT2 | 7.73E+06 | 6.85E+06 | 1.19E+06 | 33 | Cytoplasm |
| 364 | E2JF22 | PIEZO1 | 7.73E+06 | 6.02E+06 | 6.69E+05 | 81 | Cytoplasm |
| 365 | Q9CX56 | PSMD8 | 7.72E+06 | 3.43E+06 | 1.14E+06 | 9 | Cytoplasm |
| 366 | P62196 | PSMC5 | 7.69E+06 | 4.60E+06 | 6.64E+05 | 48 | Nucleus |
| 367 | Q8BHN3 | GANAB | 7.67E+06 | 4.79E+06 | 6.18E+05 | 60 | Cytoplasm |
| 368 | Q9Z2W0 | DNPEP | 7.65E+06 | 6.97E+06 | 1.34E+06 | 27 | Cytoplasm |
| 369 | P86048 | RPL10L | 7.64E+06 | 4.58E+06 | 1.87E+06 | 6 | Nucleus |
| 370 | P60766 | Cdc42 | 7.63E+06 | 2.97E+06 | 7.68E+05 | 15 | Plasma Membrane |
| 371 | P05977 | MYL1 | 7.63E+06 | 1.02E+06 | 5.88E+05 | 3 | Cytoplasm |
| 372 | P62082 | RPS7 | 7.63E+06 | 4.46E+06 | 1.49E+06 | 9 | Cytoplasm |
| 373 | P80313 | CCT7 | 7.61E+06 | 4.85E+06 | 7.22E+05 | 45 | Cytoplasm |
| 374 | P27659 | RPL3 | 7.61E+06 | 4.51E+06 | 1.50E+06 | 9 | Nucleus |
| 375 | Q99J56 | DERL1 | 7.60E+06 | 4.80E+06 | 1.96E+06 | 6 | Cytoplasm |
| 376 | Q9CXI5 | MANF | 7.60E+06 | 8.39E+06 | 2.80E+06 | 9 | Extracellular Space |
| 377 | P11983 | TCP1 | 7.59E+06 | 5.89E+06 | 7.43E+05 | 63 | Cytoplasm |
| 378 | Q9Z0T9 | ITGB6 | 7.58E+06 | 2.03E+06 | 1.17E+06 | 3 | Plasma Membrane |
| 379 | P97855 | G3BP1 | 7.58E+06 | 1.52E+06 | 6.20E+05 | 6 | Nucleus |
| 380 | P62334 | PSMC6 | 7.56E+06 | 4.44E+06 | 7.10E+05 | 39 | Nucleus |
| 381 | Q3U2A8 | VARS2 | 7.54E+06 | 4.02E+06 | 2.32E+06 | 3 | Cytoplasm |
| 382 | P80314 | CCT2 | 7.53E+06 | 4.24E+06 | 6.54E+05 | 42 | Cytoplasm |
| 383 | P53657 | PKLR | 7.52E+06 | 4.13E+06 | 8.42E+05 | 24 | Cytoplasm |
| 384 | Q9D2G2 | DLST | 7.49E+06 | 3.83E+06 | 8.36E+05 | 21 | Cytoplasm |
| 385 | P14869 | RPLP0 | 7.40E+06 | 4.69E+06 | 9.56E+05 | 24 | Cytoplasm |
| 386 | Q9EPK2 | RP2 | 7.36E+06 | 4.37E+06 | 9.54E+05 | 21 | Cytoplasm |
| 387 | Q05144 | RAC2 | 7.34E+06 | 7.24E+05 | 4.18E+05 | 3 | Cytoplasm |
| 388 | Q9DAS9 | GNG12 | 7.29E+06 | 9.62E+05 | 5.55E+05 | 3 | Plasma Membrane |
| 389 | Q9CR20 | IER3IP1 | 7.28E+06 | 4.54E+06 | 1.85E+06 | 6 | Cytoplasm |
| 390 | P62751 | Rpl23a | 7.28E+06 | 5.22E+06 | 2.13E+06 | 6 | Nucleus |
| 391 | Q8BG32 | PSMD11 | 7.25E+06 | 4.51E+06 | 6.97E+05 | 42 | Cytoplasm |
| 392 | O88653 | LAMTOR3 | 7.20E+06 | 4.82E+06 | 1.61E+06 | 9 | Cytoplasm |
| 393 | P35980 | RPL18 | 7.14E+06 | 3.68E+06 | 1.06E+06 | 12 | Cytoplasm |
| 394 | Q9D1C8 | VPS28 | 7.10E+06 | 1.06E+06 | 6.14E+05 | 3 | Cytoplasm |
| 395 | P10404 | ENV1 | 7.02E+06 | 8.30E+06 | 9.78E+05 | 72 | Other |
| 396 | Q8VDM4 | PSMD2 | 7.01E+06 | 4.36E+06 | 4.67E+05 | 87 | Cytoplasm |
| 397 | Q8BZ98 | DNM3 | 7.01E+06 | 3.27E+06 | 1.09E+06 | 9 | Cytoplasm |
| 398 | Q9JMH7 | NEU3 | 7.00E+06 | 5.66E+06 | 1.89E+06 | 9 | Plasma Membrane |
| 399 | P17427 | AP2A2 | 6.99E+06 | 6.30E+06 | 1.29E+06 | 24 | Cytoplasm |
| 400 | O55143 | ATP2A2 | 6.99E+06 | 6.21E+06 | 1.46E+06 | 18 | Cytoplasm |
| 401 | Q923D2 | BLVRB | 6.98E+06 | 4.00E+06 | 1.15E+06 | 12 | Cytoplasm |
| 402 | O89023 | TPP1 | 6.97E+06 | 7.66E+05 | 4.43E+05 | 3 | Cytoplasm |
| 403 | P04104 | KRT1 | 6.96E+06 | 1.51E+07 | 4.36E+06 | 12 | Cytoplasm |
| 404 | Q9QYB5 | ADD3 | 6.93E+06 | 6.98E+06 | 1.04E+06 | 45 | Cytoplasm |
| 405 | P62852 | RPS25 | 6.93E+06 | 1.56E+06 | 5.21E+05 | 9 | Cytoplasm |
| 406 | Q68FD5 | CLTC | 6.92E+06 | 4.79E+06 | 4.08E+05 | 138 | Plasma Membrane |
| 407 | Q9CQF9 | PCYOX1 | 6.91E+06 | 9.61E+05 | 5.55E+05 | 3 | Cytoplasm |
| 408 | P60867 | RPS20 | 6.90E+06 | 1.11E+06 | 4.53E+05 | 6 | Cytoplasm |
| 409 | Q9Z127 | SLC7A5 | 6.87E+06 | 4.66E+06 | 1.35E+06 | 12 | Plasma Membrane |
| 410 | Q9ERN0 | SCAMP2 | 6.83E+06 | 9.45E+05 | 3.86E+05 | 6 | Cytoplasm |
| 411 | B2RWW0 | TAGAP | 6.83E+06 | 5.53E+05 | 3.19E+05 | 3 | Cytoplasm |
| 412 | O35593 | PSMD14 | 6.82E+06 | 6.27E+06 | 1.81E+06 | 12 | Cytoplasm |
| 413 | P56391 | COX6B1 | 6.82E+06 | 8.78E+05 | 5.07E+05 | 3 | Cytoplasm |
| 414 | P62889 | RPL30 | 6.81E+06 | 8.25E+05 | 4.76E+05 | 3 | Cytoplasm |
| 415 | P56135 | ATP5J2 | 6.77E+06 | 9.35E+05 | 5.40E+05 | 3 | Cytoplasm |
| 416 | Q9JKV5 | SCAMP4 | 6.76E+06 | 2.03E+06 | 8.28E+05 | 6 | Cytoplasm |

| | | | | | | | |
|---|---|---|---|---|---|---|---|
| 278 | Q9CR86 | CARHSP1 | 9.16E+06 | 7.48E+05 | 4.32E+05 | 3 | Cytoplasm |
| 279 | P19783 | COX4I1 | 9.14E+06 | 8.77E+06 | 2.92E+06 | 9 | Cytoplasm |
| 280 | Q8K1B8 | FERMT3 | 9.04E+06 | 9.78E+06 | 1.41E+06 | 48 | Cytoplasm |
| 281 | Q9DBH5 | LMAN2 | 8.99E+06 | 5.93E+06 | 1.53E+06 | 15 | Cytoplasm |
| 282 | P62196 | PSMC5 | 8.98E+06 | 4.88E+06 | 7.04E+05 | 48 | Nucleus |
| 283 | Q9Z1N7 | ARID3B | 8.98E+06 | 2.51E+06 | 1.45E+06 | 3 | Nucleus |
| 284 | P80316 | CCT5 | 8.98E+06 | 5.73E+06 | 9.17E+05 | 39 | Cytoplasm |
| 285 | O88685 | PSMC3 | 8.97E+06 | 4.43E+06 | 6.83E+05 | 42 | Nucleus |
| 286 | Q9CQ19 | MYL9 | 8.92E+06 | 6.89E+06 | 1.41E+06 | 24 | Cytoplasm |
| 287 | P68510 | YWHAH | 8.86E+06 | 1.30E+07 | 2.51E+06 | 27 | Cytoplasm |
| 288 | P51125 | CAST | 8.79E+06 | 6.42E+06 | 1.07E+06 | 36 | Cytoplasm |
| 289 | Q9CQF9 | PCYOX1 | 8.79E+06 | 6.70E+05 | 3.87E+05 | 3 | Cytoplasm |
| 290 | Q8BHN3 | GANAB | 8.77E+06 | 5.03E+06 | 6.49E+05 | 60 | Cytoplasm |
| 291 | Q60766 | Irgm1 | 8.77E+06 | 4.70E+06 | 8.58E+05 | 30 | Cytoplasm |
| 292 | Q61316 | HSPA4 | 8.75E+06 | 4.30E+06 | 5.17E+05 | 69 | Cytoplasm |
| 293 | Q9CQ22 | LAMTOR1 | 8.70E+06 | 8.70E+06 | 2.51E+06 | 12 | Plasma Membrane |
| 294 | P80318 | CCT3 | 8.66E+06 | 5.17E+06 | 7.47E+05 | 48 | Cytoplasm |
| 295 | P80314 | CCT2 | 8.64E+06 | 3.95E+06 | 6.09E+05 | 42 | Cytoplasm |
| 296 | Q9R0Q3 | TMED2 | 8.63E+06 | 3.39E+06 | 1.13E+06 | 9 | Cytoplasm |
| 297 | P63038 | HSPD1 | 8.63E+06 | 9.62E+06 | 1.16E+06 | 69 | Cytoplasm |
| 298 | Q8K0T0 | RTN1 | 8.62E+06 | 2.03E+06 | 1.17E+06 | 3 | Cytoplasm |
| 299 | Q99JI4 | PSMD6 | 8.59E+06 | 4.63E+06 | 7.15E+05 | 42 | Cytoplasm |
| 300 | E2JF22 | PIEZO1 | 8.56E+06 | 6.01E+06 | 6.68E+05 | 81 | Cytoplasm |
| 301 | O08749 | DLD | 8.55E+06 | 6.23E+06 | 2.08E+06 | 9 | Cytoplasm |
| 302 | Q922D8 | MTHFD1 | 8.54E+06 | 5.46E+06 | 5.86E+05 | 87 | Cytoplasm |
| 303 | Q8VDM4 | PSMD2 | 8.51E+06 | 4.57E+06 | 4.90E+05 | 87 | Cytoplasm |
| 304 | P19536 | Cox5b | 8.49E+06 | 3.78E+06 | 2.18E+06 | 3 | Cytoplasm |
| 305 | P26040 | EZR | 8.46E+06 | 7.28E+06 | 1.33E+06 | 30 | Plasma Membrane |
| 306 | P80313 | CCT7 | 8.45E+06 | 4.25E+06 | 6.33E+05 | 45 | Cytoplasm |
| 307 | O55042 | SNCA | 8.42E+06 | 2.93E+06 | 8.45E+05 | 12 | Cytoplasm |
| 308 | O35658 | C1QBP | 8.40E+06 | 9.56E+06 | 3.90E+06 | 6 | Cytoplasm |
| 309 | P70704 | ATP8A1 | 8.35E+06 | 5.19E+06 | 6.38E+05 | 66 | Cytoplasm |
| 310 | O89053 | CORO1A | 8.27E+06 | 1.45E+07 | 2.32E+06 | 39 | Cytoplasm |
| 311 | Q9CXI5 | MANF | 8.26E+06 | 8.86E+06 | 2.95E+06 | 9 | Extracellular Space |
| 312 | Q9QYB5 | ADD3 | 8.21E+06 | 6.77E+06 | 1.01E+06 | 45 | Cytoplasm |
| 313 | P61226 | RAP2B | 8.17E+06 | 2.92E+06 | 6.88E+05 | 18 | Plasma Membrane |
| 314 | Q7M6Y3 | PICALM | 8.15E+06 | 8.36E+06 | 2.41E+06 | 12 | Cytoplasm |
| 315 | Q9WTR1 | TRPV2 | 8.15E+06 | 4.75E+06 | 7.91E+05 | 36 | Plasma Membrane |
| 316 | Q8VCM7 | FGG | 8.13E+06 | 2.62E+07 | 3.16E+06 | 69 | Extracellular Space |
| 317 | Q64433 | HSPE1 | 8.12E+06 | 7.36E+06 | 1.90E+06 | 15 | Cytoplasm |
| 318 | Q8BZ98 | DNM3 | 8.09E+06 | 3.35E+06 | 1.12E+06 | 9 | Cytoplasm |
| 319 | Q8BK63 | CSNK1A1 | 8.09E+06 | 3.51E+06 | 7.16E+05 | 24 | Cytoplasm |
| 320 | Q9D832 | DNAJB4 | 8.02E+06 | 5.75E+06 | 1.05E+06 | 30 | Nucleus |
| 321 | P80315 | CCT4 | 8.02E+06 | 5.34E+06 | 7.27E+05 | 54 | Cytoplasm |
| 322 | P11983 | TCP1 | 8.00E+06 | 5.72E+06 | 7.21E+05 | 63 | Cytoplasm |
| 323 | Q9D8W5 | PSMD12 | 7.97E+06 | 4.32E+06 | 6.45E+05 | 45 | Cytoplasm |
| 324 | P48972 | MYBL2 | 7.97E+06 | 1.25E+06 | 7.21E+05 | 3 | Nucleus |
| 325 | P04925 | PRNP | 7.95E+06 | 1.47E+06 | 8.48E+05 | 3 | Plasma Membrane |
| 326 | Q3THE2 | MYL12A | 7.89E+06 | 1.08E+07 | 2.78E+06 | 15 | Cytoplasm |
| 327 | Q99P72 | RTN4 | 7.85E+06 | 5.09E+06 | 1.31E+06 | 15 | Cytoplasm |
| 328 | Q60870 | REEP5 | 7.84E+06 | 1.12E+07 | 3.72E+06 | 9 | Extracellular Space |
| 329 | Q9CR20 | IER3IP1 | 7.83E+06 | 4.49E+06 | 1.83E+06 | 6 | Cytoplasm |
| 330 | P22315 | FECH | 7.80E+06 | 1.96E+07 | 2.60E+06 | 57 | Cytoplasm |
| 331 | Q9CZU6 | CS | 7.77E+06 | 1.08E+07 | 1.98E+06 | 30 | Cytoplasm |
| 332 | Q8K0E8 | FGB | 7.73E+06 | 2.32E+07 | 2.74E+06 | 72 | Extracellular Space |
| 333 | Q9EPK2 | RP2 | 7.72E+06 | 3.81E+06 | 8.31E+05 | 21 | Cytoplasm |
| 334 | Q8C522 | ENDOD1 | 7.69E+06 | 4.39E+06 | 8.97E+05 | 24 | Extracellular Space |
| 335 | O35643 | AP1B1 | 7.68E+06 | 4.85E+06 | 8.09E+05 | 36 | Cytoplasm |
| 336 | P60521 | GABARAPL2 | 7.66E+06 | 5.02E+06 | 1.30E+06 | 15 | Cytoplasm |
| 337 | P03930 | mt-Atp8 | 7.65E+06 | 7.20E+06 | 2.94E+06 | 6 | Cytoplasm |
| 338 | P50637 | TSPO | 7.62E+06 | 4.49E+06 | 2.59E+06 | 3 | Cytoplasm |
| 339 | Q9CQN6 | TMEM14C | 7.61E+06 | 4.82E+06 | 1.97E+06 | 6 | Plasma Membrane |
| 340 | P40237 | CD82 | 7.53E+06 | 2.01E+06 | 8.21E+05 | 6 | Plasma Membrane |
| 341 | Q9D6P8 | CALML3 | 7.52E+06 | 7.86E+05 | 4.54E+05 | 3 | Cytoplasm |
| 342 | P53657 | PKLR | 7.42E+06 | 3.50E+06 | 7.15E+05 | 24 | Cytoplasm |
| 343 | O08688 | CAPN5 | 7.39E+06 | 6.29E+06 | 8.12E+05 | 60 | Cytoplasm |
| 344 | Q60682 | Klra4 (includes others) | 7.32E+06 | 1.35E+06 | 7.79E+05 | 3 | Plasma Membrane |
| 345 | P62259 | YWHAE | 7.30E+06 | 3.65E+06 | 7.02E+05 | 27 | Cytoplasm |
| 346 | P01887 | B2M | 7.29E+06 | 2.52E+06 | 1.46E+06 | 3 | Plasma Membrane |
| 347 | P09055 | ITGB1 | 7.28E+06 | 7.52E+06 | 1.20E+06 | 39 | Plasma Membrane |
| 348 | Q9Z2W0 | DNPEP | 7.24E+06 | 6.45E+06 | 1.24E+06 | 27 | Cytoplasm |
| 349 | P42932 | CCT8 | 7.23E+06 | 3.42E+06 | 4.03E+05 | 72 | Cytoplasm |
| 350 | Q9CYN2 | SPCS2 | 7.19E+06 | 1.13E+06 | 6.53E+05 | 3 | Cytoplasm |
| 351 | P00405 | MT-CO2 | 7.15E+06 | 1.20E+07 | 2.83E+06 | 18 | Cytoplasm |
| 352 | Q99J56 | DERL1 | 7.15E+06 | 3.81E+06 | 1.56E+06 | 6 | Cytoplasm |
| 353 | Q9CQN1 | TRAP1 | 7.14E+06 | 5.31E+06 | 2.17E+06 | 6 | Cytoplasm |
| 354 | Q9CY52 | THG1L | 7.13E+06 | 1.91E+06 | 4.92E+05 | 15 | Cytoplasm |
| 355 | P08905 | LYZ | 7.12E+06 | 4.86E+06 | 1.98E+06 | 6 | Extracellular Space |
| 356 | Q9QYJ3 | DNAJB1 | 7.12E+06 | 5.02E+06 | 9.65E+05 | 27 | Nucleus |
| 357 | O35381 | Anp32a | 7.10E+06 | 2.55E+06 | 8.49E+05 | 9 | Nucleus |
| 358 | Q80VY9 | DHX33 | 7.03E+06 | 7.11E+05 | 4.10E+05 | 3 | Nucleus |
| 359 | P54822 | ADSL | 7.02E+06 | 3.39E+06 | 5.42E+05 | 39 | Cytoplasm |
| 360 | O08795 | PRKCSH | 7.01E+06 | 3.32E+06 | 5.78E+05 | 33 | Cytoplasm |
| 361 | P51863 | ATP6V0D1 | 6.97E+06 | 5.67E+06 | 1.34E+06 | 18 | Cytoplasm |
| 362 | P02535 | Krt10 | 6.95E+06 | 2.36E+06 | 1.36E+06 | 3 | Cytoplasm |
| 363 | Q9QUK3 | CLN8 | 6.94E+06 | 8.17E+05 | 4.72E+05 | 3 | Cytoplasm |
| 364 | P17742 | PPIA | 6.93E+06 | 9.35E+06 | 2.04E+06 | 21 | Cytoplasm |
| 365 | P00920 | CA2 | 6.90E+06 | 7.64E+06 | 1.80E+06 | 18 | Cytoplasm |
| 366 | P51174 | ACADL | 6.89E+06 | 9.89E+06 | 2.16E+06 | 21 | Cytoplasm |
| 367 | O35129 | PHB2 | 6.88E+06 | 6.78E+06 | 1.31E+06 | 27 | Cytoplasm |
| 368 | P58735 | SLC26A1 | 6.87E+06 | 3.82E+06 | 1.56E+06 | 6 | Plasma Membrane |
| 369 | P11499 | HSP90AB1 | 6.86E+06 | 1.41E+06 | 8.13E+05 | 3 | Cytoplasm |
| 370 | O35143 | ATPIF1 | 6.83E+06 | 4.27E+06 | 2.47E+06 | 3 | Cytoplasm |
| 371 | P62331 | ARF6 | 6.82E+06 | 4.41E+06 | 1.04E+06 | 18 | Plasma Membrane |
| 372 | Q61475 | CD55 | 6.81E+06 | 3.52E+06 | 8.29E+05 | 18 | Plasma Membrane |
| 373 | P54775 | PSMC4 | 6.78E+06 | 3.93E+06 | 6.55E+05 | 36 | Nucleus |
| 374 | P70168 | KPNB1 | 6.72E+06 | 3.04E+06 | 3.83E+05 | 63 | Nucleus |
| 375 | P68037 | UBE2L3 | 6.70E+06 | 3.29E+05 | 1.90E+05 | 3 | Nucleus |
| 376 | Q3TZZ7 | ESYT2 | 6.69E+06 | 3.10E+06 | 4.00E+05 | 60 | Plasma Membrane |
| 377 | O35864 | COPS5 | 6.67E+06 | 3.75E+06 | 1.53E+06 | 6 | Nucleus |
| 378 | Q5XPI3 | RNF123 | 6.66E+06 | 2.41E+06 | 5.26E+05 | 21 | Cytoplasm |
| 379 | A2ADY9 | DDI2 | 6.65E+06 | 1.75E+06 | 7.13E+05 | 6 | Plasma Membrane |
| 380 | Q80XI4 | PIP4K2B | 6.64E+06 | 1.06E+06 | 6.10E+05 | 3 | Cytoplasm |
| 381 | Q3UGC7 | EIF3J | 6.60E+06 | 3.35E+06 | 8.65E+05 | 15 | Cytoplasm |
| 382 | Q06185 | Atp5k | 6.59E+06 | 5.53E+06 | 2.26E+06 | 6 | Cytoplasm |
| 383 | Q5H8C4 | VPS13A | 6.56E+06 | 4.13E+06 | 4.09E+05 | 102 | Cytoplasm |
| 384 | Q8CII2 | CDC123 | 6.53E+06 | 2.77E+05 | 1.60E+05 | 3 | Cytoplasm |
| 385 | P10605 | CTSB | 6.50E+06 | 3.06E+06 | 7.90E+05 | 15 | Cytoplasm |
| 386 | Q64514 | TPP2 | 6.50E+06 | 3.98E+06 | 4.90E+05 | 66 | Cytoplasm |
| 387 | Q61165 | SLC9A1 | 6.49E+06 | 3.81E+06 | 7.78E+05 | 24 | Plasma Membrane |
| 388 | Q99P58 | RAB27B | 6.49E+06 | 7.95E+06 | 1.62E+06 | 24 | Cytoplasm |
| 389 | P07901 | HSP90AA1 | 6.49E+06 | 2.44E+06 | 5.75E+05 | 18 | Cytoplasm |
| 390 | P12787 | COX5A | 6.49E+06 | 4.34E+06 | 1.45E+06 | 9 | Cytoplasm |
| 391 | P60766 | Cdc42 | 6.41E+06 | 2.64E+06 | 6.82E+05 | 15 | Plasma Membrane |
| 392 | Q9DAS9 | GNG12 | 6.39E+06 | 8.07E+05 | 4.66E+05 | 3 | Plasma Membrane |
| 393 | P53994 | RAB2A | 6.38E+06 | 3.71E+06 | 9.59E+05 | 15 | Cytoplasm |
| 394 | P57716 | NCSTN | 6.37E+06 | 3.48E+06 | 6.69E+05 | 27 | Plasma Membrane |
| 395 | Q9QZW0 | ATP11C | 6.36E+06 | 3.00E+06 | 7.06E+05 | 18 | Plasma Membrane |
| 396 | Q9JM62 | REEP6 | 6.34E+06 | 4.84E+06 | 1.61E+06 | 9 | Plasma Membrane |
| 397 | O35379 | ABCC1 | 6.33E+06 | 4.10E+06 | 5.58E+05 | 54 | Plasma Membrane |
| 398 | P51655 | GPC4 | 6.30E+06 | 1.99E+06 | 6.65E+05 | 9 | Plasma Membrane |
| 399 | Q8CG47 | SMC4 | 6.27E+06 | 6.11E+06 | 2.49E+06 | 6 | Nucleus |
| 400 | Q80ZA4 | PKHD1L1 | 6.26E+06 | 7.39E+05 | 4.27E+05 | 3 | Extracellular Space |
| 401 | Q8BKC5 | IPO5 | 6.25E+06 | 3.40E+06 | 4.38E+05 | 60 | Nucleus |
| 402 | P51175 | PPOX | 6.23E+06 | 1.07E+07 | 1.71E+06 | 39 | Cytoplasm |
| 403 | Q9DB20 | ATP5O | 6.22E+06 | 6.04E+06 | 1.23E+06 | 24 | Cytoplasm |
| 404 | P18242 | CTSD | 6.21E+06 | 5.69E+06 | 1.24E+06 | 21 | Cytoplasm |
| 405 | Q9WV71 | ASB4 | 6.19E+06 | 1.03E+06 | 5.93E+05 | 3 | Nucleus |
| 406 | Q9JIZ9 | PLSCR3 | 6.14E+06 | 3.24E+06 | 9.36E+05 | 12 | Plasma Membrane |
| 407 | Q9Z1G4 | ATP6V0A1 | 6.06E+06 | 2.81E+06 | 5.41E+05 | 27 | Cytoplasm |
| 408 | P04104 | KRT1 | 6.05E+06 | 6.25E+06 | 1.80E+06 | 12 | Cytoplasm |
| 409 | Q3U898 | MYEOV2 | 6.04E+06 | 1.15E+06 | 6.66E+05 | 3 | Nucleus |
| 410 | P70182 | PIP5K1A | 6.03E+06 | 3.42E+06 | 6.25E+05 | 30 | Cytoplasm |
| 411 | Q922D4 | PPP6R3 | 6.03E+06 | 2.81E+06 | 4.68E+05 | 36 | Cytoplasm |
| 412 | Q8C5L6 | INPP5K | 6.03E+06 | 7.65E+05 | 3.12E+05 | 6 | Cytoplasm |
| 413 | Q9QUJ7 | ACSL4 | 6.00E+06 | 4.96E+06 | 9.54E+05 | 27 | Cytoplasm |
| 414 | P11438 | LAMP1 | 5.95E+06 | 4.23E+06 | 1.41E+06 | 9 | Plasma Membrane |
| 415 | Q9EP69 | SACM1L | 5.82E+06 | 3.17E+06 | 4.20E+05 | 57 | Cytoplasm |
| 416 | Q8QZT1 | ACAT1 | 5.79E+06 | 5.01E+06 | 1.09E+06 | 21 | Cytoplasm |

| | | | | | | | |
|---|---|---|---|---|---|---|---|
| 278 | Q60930 | VDAC2 | 9076942 | 11552229 | 2223228 | 27 | Cytoplasm |
| 279 | P08556 | NRAS | 8975000 | 821621 | 474363 | 3 | Plasma Membrane |
| 280 | Q9D0F3 | LMAN1 | 8972202 | 3486498 | 821776 | 18 | Cytoplasm |
| 281 | P97372 | PSME2 | 8969011 | 7199404 | 1571039 | 21 | Cytoplasm |
| 282 | Q3TFD2 | LPCAT1 | 8864911 | 6399657 | 1306325 | 24 | Cytoplasm |
| 283 | P14206 | RPSA | 8858453 | 7397056 | 2135346 | 12 | Cytoplasm |
| 284 | Q60766 | Irgm1 | 8766340 | 4526293 | 826384 | 30 | Cytoplasm |
| 285 | O70404 | VAMP8 | 8693080 | 8838885 | 2946295 | 9 | Plasma Membrane |
| 286 | P51125 | CAST | 8691137 | 6006622 | 1001104 | 36 | Cytoplasm |
| 287 | Q91V01 | LPCAT3 | 8632163 | 7993120 | 2307415 | 12 | Plasma Membrane |
| 288 | E2JF22 | PIEZO1 | 8622185 | 6556009 | 728445 | 81 | Cytoplasm |
| 289 | P16546 | SPTAN1 | 8600127 | 1178254 | 680265 | 3 | Plasma Membrane |
| 290 | Q9JLT4 | TXNRD2 | 8581629 | 6893616 | 2297872 | 9 | Cytoplasm |
| 291 | Q8CBB9 | RSAD2 | 8581314 | 7586635 | 1548615 | 24 | Cytoplasm |
| 292 | P19536 | Cox5b | 8551843 | 4967663 | 2868081 | 3 | Cytoplasm |
| 293 | Q91V41 | RAB14 | 8543393 | 9743393 | 1875117 | 27 | Cytoplasm |
| 294 | Q80VY9 | DHX33 | 8536104 | 2418625 | 1396394 | 3 | Nucleus |
| 295 | P54775 | PSMC4 | 8500750 | 4795363 | 799227 | 36 | Nucleus |
| 296 | Q9WV71 | ASB4 | 8482723 | 1832506 | 1057998 | 3 | Nucleus |
| 297 | P18242 | CTSD | 8481121 | 7953011 | 1735489 | 21 | Cytoplasm |
| 298 | Q9QZU9 | UBE2L6 | 8442190 | 7986182 | 2305412 | 12 | Cytoplasm |
| 299 | Q9EPK2 | RP2 | 8342218 | 4455063 | 972174 | 21 | Cytoplasm |
| 300 | Q3UGC7 | EIF3J | 8287204 | 4096988 | 1057838 | 15 | Cytoplasm |
| 301 | Q64475 | HIST1H2BB | 8280463 | 16395287 | 3864406 | 18 | Nucleus |
| 302 | Q9JKV5 | SCAMP4 | 8271730 | 2114838 | 863379 | 6 | Cytoplasm |
| 303 | O35129 | PHB2 | 8268109 | 8378181 | 1612382 | 27 | Cytoplasm |
| 304 | P51174 | ACADL | 8211056 | 11790691 | 2572940 | 21 | Cytoplasm |
| 305 | P42932 | CCT8 | 8183593 | 4068135 | 479434 | 72 | Cytoplasm |
| 306 | P04925 | PRNP | 8166719 | 641732 | 370504 | 3 | Plasma Membrane |
| 307 | P08680 | ALAS2 | 8134897 | 10231941 | 1705323 | 36 | Cytoplasm |
| 308 | Q60870 | REEP5 | 8111578 | 11067104 | 3689035 | 9 | Extracellular Space |
| 309 | Q9D1J1 | NECAP2 | 8022530 | 3030196 | 1749484 | 3 | Cytoplasm |
| 310 | P03930 | mt-Atp8 | 8016121 | 6007435 | 2452525 | 6 | Cytoplasm |
| 311 | Q7M6Y3 | PICALM | 8014553 | 10614855 | 3064245 | 12 | Cytoplasm |
| 312 | P62889 | RPL30 | 8008963 | 3888718 | 2245152 | 3 | Cytoplasm |
| 313 | Q9JKW0 | ARL6IP1 | 7962185 | 533580 | 308062 | 3 | Cytoplasm |
| 314 | Q91VH6 | MEMO1 | 7953074 | 4540703 | 1172405 | 15 | Cytoplasm |
| 315 | Q3U898 | MYEOV2 | 7920537 | 1322665 | 763641 | 3 | Nucleus |
| 316 | Q6ZPJ3 | UBE2O | 7905083 | 6540723 | 647628 | 102 | Nucleus |
| 317 | O88653 | LAMTOR3 | 7896915 | 5504923 | 1834974 | 9 | Cytoplasm |
| 318 | Q9Z2W0 | DNPEP | 7870499 | 7406978 | 1425474 | 27 | Cytoplasm |
| 319 | Q99KF1 | TMED9 | 7854890 | 9216769 | 3072256 | 9 | Cytoplasm |
| 320 | B2RWW0 | TAGAP | 7708534 | 1421776 | 820863 | 3 | Cytoplasm |
| 321 | P26040 | EZR | 7668641 | 6701757 | 1223568 | 30 | Plasma Membrane |
| 322 | Q9WTR1 | TRPV2 | 7642568 | 5013540 | 835590 | 36 | Plasma Membrane |
| 323 | Q99J99 | MPST | 7640053 | 12501948 | 5103899 | 6 | Cytoplasm |
| 324 | Q8C522 | ENDOD1 | 7617584 | 4706931 | 960798 | 24 | Extracellular Space |
| 325 | P58735 | SLC26A1 | 7608580 | 4089519 | 1669539 | 6 | Plasma Membrane |
| 326 | Q9D7G0 | PRPS1 | 7584371 | 2937424 | 692358 | 18 | Cytoplasm |
| 327 | Q64514 | TPP2 | 7583555 | 4487954 | 552429 | 66 | Cytoplasm |
| 328 | O70172 | PIP4K2A | 7546962 | 5235377 | 1511323 | 12 | Cytoplasm |
| 329 | O35643 | AP1B1 | 7502640 | 5425357 | 904226 | 36 | Cytoplasm |
| 330 | P41216 | ACSL1 | 7496727 | 5623033 | 787382 | 51 | Cytoplasm |
| 331 | Q9CY52 | THG1L | 7459850 | 2060673 | 532063 | 15 | Cytoplasm |
| 332 | Q9D6P8 | CALML3 | 7439420 | 804739 | 464616 | 3 | Cytoplasm |
| 333 | Q80XI4 | PIP4K2B | 7435815 | 799501 | 461592 | 3 | Cytoplasm |
| 334 | P16330 | CNP | 7428294 | 6956699 | 1420030 | 24 | Cytoplasm |
| 335 | Q61165 | SLC9A1 | 7427887 | 4000652 | 816630 | 24 | Plasma Membrane |
| 336 | P68433 | H31 | 7394564 | 8794281 | 2072832 | 18 | Nucleus |
| 337 | P70704 | ATP8A1 | 7368218 | 4806820 | 591679 | 66 | Cytoplasm |
| 338 | Q8BKC5 | IPO5 | 7318358 | 4111932 | 530848 | 60 | Nucleus |
| 339 | P36552 | CPOX | 7306620 | 4685135 | 780856 | 36 | Cytoplasm |
| 340 | Q9QYB5 | ADD3 | 7280831 | 6684721 | 996499 | 45 | Cytoplasm |
| 341 | O35381 | Anp32a | 7259004 | 2575627 | 858542 | 9 | Nucleus |
| 342 | Q8BK63 | CSNK1A1 | 7235251 | 3366692 | 687223 | 24 | Cytoplasm |
| 343 | Q3TZZ7 | ESYT2 | 7180683 | 3800883 | 490692 | 60 | Plasma Membrane |
| 344 | P86048 | RPL10L | 7101765 | 5901194 | 2409152 | 6 | Nucleus |
| 345 | Q61475 | CD55 | 7045583 | 3566932 | 840734 | 18 | Plasma Membrane |
| 346 | P62900 | RPL31 | 7019074 | 3421331 | 1396753 | 6 | Cytoplasm |
| 347 | Q8CDN6 | TXNL1 | 6960747 | 3930018 | 802212 | 24 | Cytoplasm |
| 348 | Q9QZW0 | ATP11C | 6934808 | 3510867 | 827519 | 18 | Plasma Membrane |
| 349 | P10852 | SLC3A2 | 6903800 | 4032672 | 736262 | 30 | Plasma Membrane |
| 350 | P04104 | KRT1 | 6897782 | 8222999 | 2373775 | 12 | Cytoplasm |
| 351 | Q9D1Q6 | ERP44 | 6895232 | 4222641 | 995286 | 18 | Cytoplasm |
| 352 | Q9JL16 | ISG20 | 6890816 | 1541364 | 444954 | 12 | Nucleus |
| 353 | Q5XPI3 | RNF123 | 6878257 | 2520133 | 549938 | 21 | Cytoplasm |
| 354 | P40237 | CD82 | 6872067 | 3762018 | 1535837 | 6 | Plasma Membrane |
| 355 | O35864 | COPS5 | 6841837 | 4119717 | 1681868 | 6 | Nucleus |
| 356 | P00405 | MT-CO2 | 6803476 | 11871539 | 2798149 | 18 | Cytoplasm |
| 357 | P14869 | RPLP0 | 6742698 | 5485862 | 1119797 | 24 | Cytoplasm |
| 358 | Q923D2 | BLVRB | 6688373 | 3486774 | 1006545 | 12 | Cytoplasm |
| 359 | P50637 | TSPO | 6670356 | 4096111 | 2364891 | 3 | Cytoplasm |
| 360 | Q8K5B2 | MCFD2 | 6660400 | 1993138 | 1150739 | 3 | Cytoplasm |
| 361 | P14148 | RPL7 | 6657043 | 6177255 | 1594960 | 15 | Nucleus |
| 362 | P51863 | ATP6V0D1 | 6629252 | 6839934 | 1612188 | 18 | Cytoplasm |
| 363 | Q09143 | SLC7A1 | 6628230 | 4892932 | 2824935 | 3 | Plasma Membrane |
| 364 | Q9CZ13 | UQCRC1 | 6608448 | 5501314 | 1420433 | 15 | Cytoplasm |
| 365 | P70182 | PIP5K1A | 6605060 | 4210746 | 768774 | 30 | Cytoplasm |
| 366 | P57716 | NCSTN | 6585396 | 3630055 | 698604 | 27 | Plasma Membrane |
| 367 | Q9DB20 | ATP5O | 6584223 | 5504206 | 1123541 | 24 | Cytoplasm |
| 368 | P25444 | RPS2 | 6581916 | 3723667 | 1074930 | 12 | Cytoplasm |
| 369 | P62082 | RPS7 | 6569648 | 4852835 | 1617612 | 9 | Cytoplasm |
| 370 | P62852 | RPS25 | 6556603 | 2763619 | 921206 | 9 | Cytoplasm |
| 371 | Q8K0T0 | RTN1 | 6537161 | 5252053 | 3032274 | 3 | Cytoplasm |
| 372 | Q9Z0N1 | EIF2S3 | 6531120 | 3330810 | 579820 | 33 | Cytoplasm |
| 373 | O35379 | ABCC1 | 6526931 | 4860742 | 661463 | 54 | Plasma Membrane |
| 374 | Q8CG47 | SMC4 | 6497662 | 6710565 | 2739577 | 6 | Nucleus |
| 375 | P14131 | RPS16 | 6477873 | 3832391 | 1564567 | 6 | Cytoplasm |
| 376 | P11499 | HSP90AB1 | 6471373 | 879755 | 507927 | 3 | Cytoplasm |
| 377 | Q9JIZ9 | PLSCR3 | 6445987 | 3640475 | 1050915 | 12 | Plasma Membrane |
| 378 | O08688 | CAPN5 | 6430735 | 7420859 | 958029 | 60 | Cytoplasm |
| 379 | Q8BHN3 | GANAB | 6417572 | 4356626 | 562438 | 60 | Cytoplasm |
| 380 | Q9D1L9 | LAMTOR5 | 6416296 | 3898532 | 1299511 | 9 | Cytoplasm |
| 381 | P54822 | ADSL | 6380593 | 3517766 | 563293 | 39 | Cytoplasm |
| 382 | A2ADY9 | DDI2 | 6354568 | 1315380 | 537002 | 6 | Plasma Membrane |
| 383 | P62751 | Rpl23a | 6328862 | 5897258 | 2407545 | 6 | Nucleus |
| 384 | Q922U2 | KRT5 | 6292668 | 5750897 | 1660141 | 12 | Cytoplasm |
| 385 | P62264 | RPS14 | 6291002 | 3974849 | 1324950 | 9 | Cytoplasm |
| 386 | Q9Z2L7 | CRLF3 | 6280294 | 1480707 | 854887 | 3 | Cytoplasm |
| 387 | Q9QUJ7 | ACSL4 | 6266428 | 5008621 | 963910 | 27 | Cytoplasm |
| 388 | Q60682 | Klra4 (includes others) | 6252631 | 1414808 | 816840 | 3 | Plasma Membrane |
| 389 | Q99L45 | EIF2S2 | 6242823 | 1524766 | 508255 | 9 | Cytoplasm |
| 390 | P62830 | RPL23 | 6241747 | 3429067 | 1979773 | 3 | Cytoplasm |
| 391 | Q8C5L6 | INPP5K | 6238363 | 731466 | 298620 | 6 | Cytoplasm |
| 392 | Q922R8 | PDIA6 | 6224664 | 4353774 | 757895 | 33 | Cytoplasm |
| 393 | P15327 | BPGM | 6209938 | 5943187 | 1715650 | 12 | Extracellular Space |
| 394 | Q9CXI5 | MANF | 6190855 | 7698350 | 2566117 | 9 | Extracellular Space |
| 395 | Q61820 | Rasl2-9 | 6180918 | 1290846 | 526986 | 6 | Nucleus |
| 396 | P35980 | RPL18 | 6172250 | 4252841 | 1227690 | 12 | Cytoplasm |
| 397 | P97315 | CSRP1 | 6163397 | 5639183 | 1456031 | 15 | Nucleus |
| 398 | Q9DAS9 | GNG12 | 6157657 | 878831 | 507394 | 3 | Plasma Membrane |
| 399 | P01887 | B2M | 6156097 | 1039825 | 600344 | 3 | Plasma Membrane |
| 400 | P15379 | CD44 | 6150086 | 6727612 | 1942094 | 12 | Plasma Membrane |
| 401 | Q60605 | MYL6 | 6126443 | 8445877 | 1843042 | 21 | Cytoplasm |
| 402 | P07901 | HSP90AA1 | 6119016 | 2056118 | 484632 | 18 | Cytoplasm |
| 403 | Q922D4 | PPP6R3 | 6118466 | 2905870 | 484312 | 36 | Cytoplasm |
| 404 | P28571 | SLC6A9 | 6110860 | 5065610 | 2068026 | 6 | Plasma Membrane |
| 405 | Q9CQN1 | TRAP1 | 6102954 | 4968694 | 2028461 | 6 | Cytoplasm |
| 406 | O09167 | RPL21 | 6095599 | 4249277 | 1734760 | 6 | Cytoplasm |
| 407 | Q06185 | Atp5k | 6093955 | 5486454 | 2239836 | 6 | Cytoplasm |
| 408 | P63101 | YWHAZ | 6091467 | 11032465 | 1592399 | 48 | Cytoplasm |
| 409 | Q9DBH5 | LMAN2 | 6086878 | 5345982 | 1380327 | 15 | Cytoplasm |
| 410 | P51410 | RPL9 | 6073566 | 2981713 | 993904 | 9 | Nucleus |
| 411 | P63325 | RPS10 | 6072302 | 3709490 | 2141675 | 3 | Cytoplasm |
| 412 | P12970 | RPL7A | 6057455 | 3314042 | 1104681 | 9 | Cytoplasm |
| 413 | P62242 | RPS8 | 6046937 | 3859119 | 996420 | 15 | Cytoplasm |
| 414 | Q9WVE8 | PACSIN2 | 6026386 | 2841678 | 580055 | 24 | Cytoplasm |
| 415 | Q9D1G5 | LRRC57 | 6024932 | 2137983 | 503927 | 18 | Cytoplasm |
| 416 | Q8R016 | BLMH | 6016061 | 2675907 | 583931 | 21 | Cytoplasm |

| | | | | | | | |
|---|---|---|---|---|---|---|---|
| 417 | Q8C5L6 | INPP5K | 5.98E+06 | 1.02E+06 | 4.14E+05 | 6 | Cytoplasm |
| 418 | Q9Z1G4 | ATP6V0A1 | 5.94E+06 | 2.37E+06 | 4.56E+05 | 27 | Cytoplasm |
| 419 | O35372 | CNIH1 | 5.93E+06 | 1.54E+06 | 8.90E+05 | 3 | Plasma Membrane |
| 420 | A2ADY9 | DDI2 | 5.93E+06 | 2.85E+06 | 1.16E+06 | 6 | Plasma Membrane |
| 421 | Q922D4 | PPP6R3 | 5.92E+06 | 4.01E+06 | 6.69E+05 | 36 | Cytoplasm |
| 422 | Q9CQX2 | CYB5B | 5.91E+06 | 6.58E+06 | 2.19E+06 | 9 | Cytoplasm |
| 423 | Q9CQN6 | TMEM14C | 5.89E+06 | 3.16E+06 | 1.29E+06 | 6 | Plasma Membrane |
| 424 | P11438 | LAMP1 | 5.84E+06 | 3.84E+06 | 1.28E+06 | 9 | Plasma Membrane |
| 425 | Q9JL99 | CLEC1B | 5.83E+06 | 5.43E+06 | 3.13E+06 | 3 | Plasma Membrane |
| 426 | Q9QZW0 | ATP11C | 5.83E+06 | 3.56E+06 | 8.40E+05 | 18 | Plasma Membrane |
| 427 | Q9QUK3 | CLN8 | 5.83E+06 | 1.14E+06 | 6.57E+05 | 3 | Cytoplasm |
| 428 | Q9JL16 | ISG20 | 5.82E+06 | 1.29E+06 | 3.73E+05 | 12 | Nucleus |
| 429 | Q9WV71 | ASB4 | 5.79E+06 | 9.23E+05 | 5.33E+05 | 3 | Nucleus |
| 430 | Q8BZ98 | DNM3 | 5.75E+06 | 2.29E+06 | 7.62E+05 | 9 | Cytoplasm |
| 431 | O88653 | LAMTOR3 | 5.66E+06 | 3.22E+06 | 1.07E+06 | 9 | Cytoplasm |
| 432 | Q8JZR0 | ACSL5 | 5.64E+06 | 3.28E+06 | 5.71E+05 | 33 | Cytoplasm |
| 433 | Q64518 | ATP2A3 | 5.62E+06 | 6.75E+06 | 9.74E+05 | 48 | Cytoplasm |
| 434 | Q9Z0T9 | ITGB6 | 5.59E+06 | 4.62E+06 | 2.67E+06 | 3 | Plasma Membrane |
| 435 | Q5H8C4 | VPS13A | 5.59E+06 | 4.00E+06 | 3.96E+05 | 102 | Cytoplasm |
| 436 | O35143 | ATPIF1 | 5.58E+06 | 2.59E+06 | 1.50E+06 | 3 | Cytoplasm |
| 437 | P42125 | ECI1 | 5.57E+06 | 2.36E+06 | 9.62E+05 | 6 | Cytoplasm |
| 438 | Q3UFK8 | FRMD8 | 5.56E+06 | 1.29E+06 | 7.44E+05 | 3 | Cytoplasm |
| 439 | Q3UGC7 | EIF3J | 5.54E+06 | 5.64E+06 | 1.46E+06 | 15 | Cytoplasm |
| 440 | Q9D6P8 | CALML3 | 5.53E+06 | 2.07E+06 | 1.19E+06 | 3 | Cytoplasm |
| 441 | P15379 | CD44 | 5.49E+06 | 6.07E+06 | 1.75E+06 | 12 | Plasma Membrane |
| 442 | Q9R1S7 | ABCC6 | 5.48E+06 | 1.86E+06 | 1.07E+06 | 3 | Plasma Membrane |
| 443 | Q9CZ13 | UQCRC1 | 5.43E+06 | 3.77E+06 | 9.73E+05 | 15 | Cytoplasm |
| 444 | P84099 | RPL19 | 5.40E+06 | 1.41E+06 | 8.12E+05 | 3 | Cytoplasm |
| 445 | Q8QZT1 | ACAT1 | 5.40E+06 | 3.61E+06 | 7.87E+05 | 21 | Cytoplasm |
| 446 | P21279 | GNAQ | 5.40E+06 | 3.59E+06 | 2.07E+06 | 3 | Plasma Membrane |
| 447 | Q8CII2 | CDC123 | 5.37E+06 | 3.32E+04 | 1.92E+04 | 3 | Cytoplasm |
| 448 | O35350 | CAPN1 | 5.37E+06 | 2.45E+06 | 5.35E+05 | 21 | Cytoplasm |
| 449 | P05202 | GOT2 | 5.34E+06 | 4.39E+06 | 7.65E+05 | 33 | Cytoplasm |
| 450 | Q80W04 | TMCC2 | 5.32E+06 | 3.71E+06 | 8.75E+05 | 18 | Extracellular Space |
| 451 | Q8VEE1 | LMCD1 | 5.30E+06 | 4.22E+06 | 2.44E+06 | 3 | Cytoplasm |
| 452 | Q99P72 | RTN4 | 5.30E+06 | 3.75E+06 | 9.68E+05 | 15 | Cytoplasm |
| 453 | Q3U0B3 | DHRS11 | 5.28E+06 | 1.60E+06 | 9.22E+05 | 3 | Cytoplasm |
| 454 | Q06890 | CLU | 5.25E+06 | 3.95E+06 | 1.32E+06 | 9 | Cytoplasm |
| 455 | Q9EP69 | SACM1L | 5.21E+06 | 3.17E+06 | 4.20E+05 | 57 | Cytoplasm |
| 456 | Q80WT2 | DMRT3 | 5.20E+06 | 1.17E+06 | 6.77E+05 | 3 | Nucleus |
| 457 | P24270 | CAT | 5.18E+06 | 4.12E+06 | 5.32E+05 | 60 | Cytoplasm |
| 458 | Q9Z2L7 | CRLF3 | 5.18E+06 | 2.04E+06 | 1.18E+06 | 3 | Cytoplasm |
| 459 | P07901 | HSP90AA1 | 5.15E+06 | 2.45E+06 | 5.77E+05 | 18 | Cytoplasm |
| 460 | Q9WVE8 | PACSIN2 | 5.13E+06 | 3.18E+06 | 6.48E+05 | 24 | Cytoplasm |
| 461 | Q3TWL2 | TMEM55B | 5.09E+06 | 1.58E+06 | 4.56E+05 | 12 | Cytoplasm |
| 462 | P07758 | SERPINA1 | 5.08E+06 | 4.80E+06 | 1.24E+06 | 15 | Extracellular Space |
| 463 | P56399 | USP5 | 5.07E+06 | 5.11E+06 | 8.18E+05 | 39 | Cytoplasm |
| 464 | Q9CQB5 | CISD2 | 5.04E+06 | 1.43E+06 | 3.69E+05 | 15 | Cytoplasm |
| 465 | P21981 | TGM2 | 5.00E+06 | 4.08E+06 | 1.66E+06 | 6 | Cytoplasm |
| 466 | Q99KR7 | PPIF | 4.99E+06 | 4.55E+06 | 1.86E+06 | 6 | Cytoplasm |
| 467 | P12787 | COX5A | 4.95E+06 | 2.02E+06 | 6.72E+05 | 9 | Cytoplasm |
| 468 | Q9D1L9 | LAMTOR5 | 4.94E+06 | 2.45E+06 | 8.16E+05 | 9 | Cytoplasm |
| 469 | P36423 | TBXAS1 | 4.94E+06 | 4.42E+06 | 8.50E+05 | 27 | Plasma Membrane |
| 470 | P14106 | C1QB | 4.94E+06 | 6.74E+06 | 3.89E+06 | 3 | Extracellular Space |
| 471 | Q9QVN7 | TCEA2 | 4.91E+06 | 4.48E+06 | 2.58E+06 | 3 | Nucleus |
| 472 | Q02053 | UBA1 | 4.91E+06 | 2.92E+06 | 4.50E+05 | 42 | Cytoplasm |
| 473 | Q9CXW4 | RPL11 | 4.91E+06 | 6.11E+05 | 3.53E+05 | 3 | Cytoplasm |
| 474 | Q9QXT0 | CNPY2 | 4.91E+06 | 1.90E+06 | 6.33E+05 | 9 | Plasma Membrane |
| 475 | Q91Y57 | SIGLEC9 | 4.88E+06 | 7.72E+04 | 4.46E+04 | 3 | Plasma Membrane |
| 476 | P14206 | RPSA | 4.87E+06 | 3.09E+06 | 8.93E+05 | 12 | Cytoplasm |
| 477 | Q01730 | RSU1 | 4.86E+06 | 4.59E+06 | 1.53E+06 | 9 | Cytoplasm |
| 478 | O08807 | PRDX4 | 4.85E+06 | 3.28E+06 | 1.09E+06 | 9 | Cytoplasm |
| 479 | Q9QYF1 | RDH11 | 4.83E+06 | 1.63E+06 | 3.33E+05 | 24 | Cytoplasm |
| 480 | Q76MZ3 | PPP2R1A | 4.83E+06 | 3.26E+06 | 7.10E+05 | 21 | Cytoplasm |
| 481 | Q9D2G2 | DLST | 4.83E+06 | 2.20E+06 | 4.79E+05 | 21 | Cytoplasm |
| 482 | P56391 | COX6B1 | 4.82E+06 | 1.57E+06 | 9.06E+05 | 3 | Cytoplasm |
| 483 | P63330 | PPP2CA | 4.82E+06 | 2.22E+06 | 5.72E+05 | 15 | Cytoplasm |
| 484 | Q9CWJ9 | ATIC | 4.80E+06 | 4.07E+06 | 7.43E+05 | 30 | Cytoplasm |
| 485 | O70576 | STAG3 | 4.80E+06 | 1.18E+06 | 6.84E+05 | 3 | Nucleus |
| 486 | P19221 | F2 | 4.79E+06 | 5.11E+05 | 2.95E+05 | 3 | Extracellular Space |
| 487 | P01887 | B2M | 4.75E+06 | 1.36E+06 | 7.84E+05 | 3 | Plasma Membrane |
| 488 | A2AN08 | UBR4 | 4.74E+06 | 2.62E+06 | 2.08E+05 | 159 | Nucleus |
| 489 | P40237 | CD82 | 4.74E+06 | 9.52E+05 | 3.89E+05 | 6 | Plasma Membrane |
| 490 | P10852 | SLC3A2 | 4.73E+06 | 2.28E+06 | 4.16E+05 | 30 | Plasma Membrane |
| 491 | Q61189 | CLNS1A | 4.73E+06 | 3.30E+06 | 1.91E+06 | 3 | Plasma Membrane |
| 492 | Q9JMA1 | USP14 | 4.72E+06 | 2.13E+06 | 8.71E+05 | 6 | Cytoplasm |
| 493 | P48771 | COX7A2 | 4.71E+06 | 1.90E+06 | 1.10E+06 | 3 | Cytoplasm |
| 494 | Q9D7G0 | PRPS1 | 4.70E+06 | 3.09E+06 | 7.29E+05 | 18 | Cytoplasm |
| 495 | Q9Z0F4 | CIB1 | 4.69E+06 | 2.93E+06 | 8.45E+05 | 12 | Nucleus |
| 496 | Q791V5 | MTCH2 | 4.69E+06 | 1.12E+06 | 6.48E+05 | 3 | Cytoplasm |
| 497 | Q8CBB9 | RSAD2 | 4.65E+06 | 2.75E+06 | 5.62E+05 | 24 | Cytoplasm |
| 498 | Q9D0B1 | ZNF524 | 4.64E+06 | 6.23E+05 | 3.60E+05 | 3 | Nucleus |
| 499 | Q9DCF9 | SSR3 | 4.63E+06 | 1.83E+06 | 1.05E+06 | 3 | Cytoplasm |
| 500 | P48722 | HSPA4L | 4.62E+06 | 3.28E+06 | 8.48E+05 | 15 | Cytoplasm |
| 501 | P46460 | NSF | 4.62E+06 | 2.46E+06 | 3.79E+05 | 42 | Cytoplasm |
| 502 | O35887 | CALU | 4.61E+06 | 2.52E+06 | 5.15E+05 | 24 | Cytoplasm |
| 503 | Q9CQN1 | TRAP1 | 4.59E+06 | 4.77E+06 | 1.95E+06 | 6 | Cytoplasm |
| 504 | P27659 | RPL3 | 4.58E+06 | 2.47E+06 | 8.24E+05 | 9 | Nucleus |
| 505 | P18052 | PTPRA | 4.57E+06 | 3.25E+06 | 9.37E+05 | 12 | Plasma Membrane |
| 506 | Q91WC9 | DAGLB | 4.56E+06 | 2.10E+06 | 4.95E+05 | 18 | Plasma Membrane |
| 507 | Q9D1J1 | NECAP2 | 4.54E+06 | 1.01E+06 | 5.85E+05 | 3 | Cytoplasm |
| 508 | Q8CI59 | STEAP3 | 4.53E+06 | 2.44E+06 | 6.31E+05 | 15 | Cytoplasm |
| 509 | Q91VH6 | MEMO1 | 4.52E+06 | 2.59E+06 | 6.70E+05 | 15 | Cytoplasm |
| 510 | Q64446 | ATP7B | 4.52E+06 | 4.99E+06 | 1.02E+06 | 24 | Cytoplasm |
| 511 | P61022 | CHP1 | 4.52E+06 | 1.92E+06 | 4.52E+05 | 18 | Cytoplasm |
| 512 | P86048 | RPL10L | 4.48E+06 | 2.16E+06 | 8.80E+05 | 6 | Nucleus |
| 513 | Q9Z0N1 | EIF2S3 | 4.48E+06 | 2.96E+06 | 5.16E+05 | 33 | Cytoplasm |
| 514 | Q78IK2 | USMG5 | 4.45E+06 | 1.30E+06 | 5.33E+05 | 6 | Cytoplasm |
| 515 | P35441 | THBS1 | 4.44E+06 | 5.20E+06 | 5.31E+05 | 96 | Extracellular Space |
| 516 | O88545 | COPS6 | 4.43E+06 | 2.70E+06 | 6.97E+05 | 15 | Nucleus |
| 517 | Q91YA2 | PSKH1 | 4.43E+06 | 8.08E+05 | 4.66E+05 | 3 | Nucleus |
| 518 | P47753 | CAPZA1 | 4.42E+06 | 1.74E+06 | 7.12E+05 | 6 | Cytoplasm |
| 519 | P17427 | AP2A2 | 4.42E+06 | 4.02E+06 | 8.20E+05 | 24 | Cytoplasm |
| 520 | Q9DB77 | UQCRC2 | 4.41E+06 | 2.86E+06 | 4.77E+05 | 36 | Cytoplasm |
| 521 | P62889 | RPL30 | 4.41E+06 | 4.94E+05 | 2.85E+05 | 3 | Cytoplasm |
| 522 | P62814 | ATP6V1B2 | 4.38E+06 | 3.72E+06 | 1.24E+06 | 9 | Cytoplasm |
| 523 | P11688 | ITGA5 | 4.38E+06 | 2.57E+06 | 7.42E+05 | 12 | Plasma Membrane |
| 524 | Q8VDI7 | UBAC1 | 4.38E+06 | 3.15E+06 | 5.76E+05 | 30 | Nucleus |
| 525 | P56395 | CYB5A | 4.35E+06 | 2.91E+06 | 7.52E+05 | 15 | Cytoplasm |
| 526 | Q8BHL3 | TBC1D10B | 4.33E+06 | 9.56E+05 | 5.52E+05 | 3 | Plasma Membrane |
| 527 | P19096 | FASN | 4.32E+06 | 3.18E+06 | 3.82E+05 | 69 | Cytoplasm |
| 528 | Q99J99 | MPST | 4.31E+06 | 4.09E+06 | 1.67E+06 | 6 | Cytoplasm |
| 529 | O55022 | PGRMC1 | 4.30E+06 | 2.89E+06 | 1.67E+06 | 3 | Plasma Membrane |
| 530 | Q64324 | STXBP2 | 4.27E+06 | 4.17E+06 | 6.44E+05 | 42 | Plasma Membrane |
| 531 | Q9Z0F8 | ADAM17 | 4.26E+06 | 2.69E+06 | 1.10E+06 | 6 | Plasma Membrane |
| 532 | P56135 | ATP5J2 | 4.25E+06 | 1.76E+06 | 1.02E+06 | 3 | Cytoplasm |
| 533 | P49282 | SLC11A2 | 4.24E+06 | 2.99E+06 | 8.62E+05 | 12 | Plasma Membrane |
| 534 | Q9DCX2 | ATP5H | 4.23E+06 | 2.38E+06 | 5.62E+05 | 18 | Cytoplasm |
| 535 | P43276 | Hist1h1b | 4.21E+06 | 5.34E+06 | 2.18E+06 | 6 | Nucleus |
| 536 | P62900 | RPL31 | 4.20E+06 | 7.91E+05 | 3.23E+05 | 6 | Cytoplasm |
| 537 | Q9DCC4 | PYCRL | 4.20E+06 | 2.16E+06 | 7.20E+05 | 9 | Cytoplasm |
| 538 | Q921L3 | TMCO1 | 4.20E+06 | 2.13E+06 | 1.23E+06 | 3 | Cytoplasm |
| 539 | P08003 | PDIA4 | 4.19E+06 | 2.46E+06 | 5.03E+05 | 24 | Cytoplasm |
| 540 | P36552 | CPOX | 4.18E+06 | 1.81E+06 | 3.01E+05 | 36 | Cytoplasm |
| 541 | P52480 | PKM | 4.16E+06 | 9.80E+05 | 5.66E+05 | 3 | Cytoplasm |
| 542 | P39876 | TIMP3 | 4.14E+06 | 4.16E+06 | 1.20E+06 | 12 | Extracellular Space |
| 543 | P70670 | NACA | 4.13E+06 | 3.10E+06 | 1.03E+06 | 9 | Cytoplasm |
| 544 | Q9DBX3 | SUSD2 | 4.12E+06 | 1.61E+06 | 5.38E+05 | 9 | Extracellular Space |
| 545 | Q64522 | HIST2H2AB | 4.12E+06 | 2.90E+06 | 1.67E+06 | 3 | Nucleus |
| 546 | O88587 | COMT | 4.12E+06 | 8.77E+05 | 2.53E+05 | 12 | Cytoplasm |
| 547 | P84091 | AP2M1 | 4.10E+06 | 2.07E+06 | 5.34E+05 | 15 | Cytoplasm |
| 548 | Q9CQ92 | FIS1 | 4.09E+06 | 4.55E+05 | 1.86E+05 | 6 | Cytoplasm |
| 549 | Q9ERN0 | SCAMP2 | 4.06E+06 | 1.61E+06 | 6.55E+05 | 6 | Cytoplasm |
| 550 | P0C0A3 | CHMP6 | 4.05E+06 | 6.91E+05 | 3.99E+05 | 3 | Cytoplasm |
| 551 | P14148 | RPL7 | 4.05E+06 | 2.53E+06 | 6.54E+05 | 15 | Nucleus |
| 552 | P26041 | MSN | 4.04E+06 | 2.61E+06 | 4.36E+05 | 36 | Plasma Membrane |
| 553 | O55143 | ATP2A2 | 4.04E+06 | 5.34E+06 | 1.26E+06 | 18 | Cytoplasm |
| 554 | P35282 | RAB21 | 4.04E+06 | 2.90E+06 | 5.92E+05 | 24 | Cytoplasm |
| 555 | Q09143 | SLC7A1 | 4.03E+06 | 4.92E+05 | 2.84E+05 | 3 | Plasma Membrane |

| | | | | | | | |
|---|---|---|---|---|---|---|---|
| 417 | Q9DB77 | UQCRC2 | 6.74E+06 | 5.33E+06 | 8.88E+05 | 36 | Cytoplasm |
| 418 | Q80XI4 | PIP4K2B | 6.74E+06 | 9.24E+05 | 5.33E+05 | 3 | Cytoplasm |
| 419 | P38647 | HSPA9 | 6.73E+06 | 3.56E+06 | 5.93E+05 | 36 | Cytoplasm |
| 420 | P56395 | CYB5A | 6.71E+06 | 6.79E+06 | 1.75E+06 | 15 | Cytoplasm |
| 421 | Q9CQQ7 | ATP5F1 | 6.70E+06 | 1.28E+06 | 7.40E+05 | 3 | Cytoplasm |
| 422 | Q09143 | SLC7A1 | 6.70E+06 | 5.32E+05 | 3.07E+05 | 3 | Plasma Membrane |
| 423 | Q99J99 | MPST | 6.70E+06 | 7.44E+06 | 3.04E+06 | 6 | Cytoplasm |
| 424 | Q922U2 | KRT5 | 6.63E+06 | 1.82E+07 | 5.24E+06 | 12 | Cytoplasm |
| 425 | Q9DCF9 | SSR3 | 6.62E+06 | 1.13E+06 | 6.55E+05 | 3 | Cytoplasm |
| 426 | P48771 | COX7A2 | 6.60E+06 | 1.07E+06 | 6.20E+05 | 3 | Cytoplasm |
| 427 | O35381 | Anp32a | 6.59E+06 | 3.03E+06 | 1.01E+06 | 9 | Nucleus |
| 428 | Q9D6R2 | IDH3A | 6.59E+06 | 5.30E+06 | 1.37E+06 | 15 | Cytoplasm |
| 429 | P36423 | TBXAS1 | 6.58E+06 | 3.87E+06 | 7.44E+05 | 27 | Plasma Membrane |
| 430 | Q9D1G5 | LRRC57 | 6.56E+06 | 1.87E+06 | 4.41E+05 | 18 | Cytoplasm |
| 431 | Q8CI59 | STEAP3 | 6.54E+06 | 3.48E+06 | 8.98E+05 | 15 | Cytoplasm |
| 432 | Q78IK2 | USMG5 | 6.54E+06 | 2.85E+06 | 1.16E+06 | 6 | Cytoplasm |
| 433 | P42932 | CCT8 | 6.54E+06 | 3.88E+06 | 4.57E+05 | 72 | Cytoplasm |
| 434 | P14131 | RPS16 | 6.52E+06 | 2.28E+06 | 9.29E+05 | 6 | Cytoplasm |
| 435 | Q99JI4 | PSMD6 | 6.51E+06 | 4.40E+06 | 6.79E+05 | 42 | Cytoplasm |
| 436 | Q9QYJ3 | DNAJB1 | 6.51E+06 | 4.97E+06 | 9.57E+05 | 27 | Nucleus |
| 437 | Q61475 | CD55 | 6.48E+06 | 3.47E+06 | 8.18E+05 | 18 | Plasma Membrane |
| 438 | P12970 | RPL7A | 6.47E+06 | 2.20E+06 | 7.35E+05 | 9 | Cytoplasm |
| 439 | P70168 | KPNB1 | 6.46E+06 | 3.15E+06 | 3.96E+05 | 63 | Nucleus |
| 440 | Q8K0T0 | RTN1 | 6.43E+06 | 5.57E+06 | 3.22E+06 | 3 | Cytoplasm |
| 441 | Q61165 | SLC9A1 | 6.43E+06 | 4.10E+06 | 8.36E+05 | 24 | Plasma Membrane |
| 442 | Q3TZZ7 | ESYT2 | 6.41E+06 | 3.00E+06 | 3.87E+05 | 60 | Plasma Membrane |
| 443 | Q9DCX2 | ATP5H | 6.40E+06 | 3.02E+06 | 7.13E+05 | 18 | Cytoplasm |
| 444 | P43275 | Hist1h1a | 6.39E+06 | 3.07E+06 | 8.85E+05 | 12 | Nucleus |
| 445 | P25444 | RPS2 | 6.38E+06 | 1.88E+06 | 5.44E+05 | 12 | Cytoplasm |
| 446 | P58735 | SLC26A1 | 6.38E+06 | 2.85E+06 | 1.16E+06 | 6 | Plasma Membrane |
| 447 | Q9D8W5 | PSMD12 | 6.37E+06 | 4.25E+06 | 6.34E+05 | 45 | Cytoplasm |
| 448 | Q99KI0 | ACO2 | 6.36E+06 | 5.88E+06 | 7.41E+05 | 63 | Cytoplasm |
| 449 | P15379 | CD44 | 6.34E+06 | 6.10E+06 | 1.76E+06 | 12 | Plasma Membrane |
| 450 | Q06890 | CLU | 6.34E+06 | 5.47E+06 | 1.82E+06 | 9 | Cytoplasm |
| 451 | Q8BHL3 | TBC1D10B | 6.34E+06 | 3.03E+06 | 1.75E+06 | 3 | Plasma Membrane |
| 452 | Q9Z2L7 | CRLF3 | 6.34E+06 | 1.68E+06 | 9.68E+05 | 3 | Cytoplasm |
| 453 | Q91VR2 | ATP5C1 | 6.33E+06 | 4.44E+06 | 1.28E+06 | 12 | Cytoplasm |
| 454 | Q9QZW0 | ATP11C | 6.32E+06 | 3.59E+06 | 8.46E+05 | 18 | Plasma Membrane |
| 455 | P63325 | RPS10 | 6.32E+06 | 1.27E+06 | 7.36E+05 | 3 | Cytoplasm |
| 456 | Q8C5L6 | INPP5K | 6.31E+06 | 1.33E+06 | 5.43E+05 | 6 | Cytoplasm |
| 457 | O09167 | RPL21 | 6.31E+06 | 3.09E+06 | 1.26E+06 | 6 | Cytoplasm |
| 458 | Q9QYA2 | TOMM40 | 6.31E+06 | 2.31E+05 | 1.33E+05 | 3 | Cytoplasm |
| 459 | Q9JIZ9 | PLSCR3 | 6.30E+06 | 3.46E+06 | 9.98E+05 | 12 | Plasma Membrane |
| 460 | P99029 | PRDX5 | 6.30E+06 | 2.20E+05 | 1.27E+05 | 3 | Cytoplasm |
| 461 | O35379 | ABCC1 | 6.29E+06 | 4.18E+06 | 5.68E+05 | 54 | Plasma Membrane |
| 462 | O35143 | ATPIF1 | 6.28E+06 | 4.08E+06 | 2.35E+06 | 3 | Cytoplasm |
| 463 | Q9R0Q3 | TMED2 | 6.27E+06 | 4.12E+06 | 1.37E+06 | 9 | Cytoplasm |
| 464 | Q8CG47 | SMC4 | 6.26E+06 | 6.40E+06 | 2.61E+06 | 6 | Nucleus |
| 465 | Q9Z1G4 | ATP6V0A1 | 6.23E+06 | 3.27E+06 | 6.29E+05 | 27 | Cytoplasm |
| 466 | Q6PCP5 | Mff | 6.20E+06 | 2.34E+06 | 9.55E+05 | 6 | Cytoplasm |
| 467 | P27661 | H2AFX | 6.19E+06 | 4.54E+06 | 1.85E+06 | 6 | Nucleus |
| 468 | Q8C1A3 | MTRR | 6.14E+06 | 7.33E+05 | 4.23E+05 | 3 | Cytoplasm |
| 469 | P17710 | HK1 | 6.14E+06 | 4.14E+06 | 6.62E+05 | 39 | Cytoplasm |
| 470 | O09172 | GCLM | 6.13E+06 | 1.58E+06 | 5.26E+05 | 9 | Cytoplasm |
| 471 | Q64324 | STXBP2 | 6.12E+06 | 4.94E+06 | 7.63E+05 | 42 | Plasma Membrane |
| 472 | P53994 | RAB2A | 6.12E+06 | 4.02E+06 | 1.04E+06 | 15 | Cytoplasm |
| 473 | Q60597 | OGDH | 6.12E+06 | 4.83E+06 | 6.76E+05 | 51 | Cytoplasm |
| 474 | Q8BKC5 | IPO5 | 6.09E+06 | 3.43E+06 | 4.43E+05 | 60 | Nucleus |
| 475 | Q9D6P8 | CALML3 | 6.08E+06 | 1.08E+06 | 6.24E+05 | 3 | Cytoplasm |
| 476 | Q8BH04 | PCK2 | 6.06E+06 | 4.13E+06 | 5.78E+05 | 51 | Cytoplasm |
| 477 | O55022 | PGRMC1 | 6.00E+06 | 7.88E+05 | 4.55E+05 | 3 | Plasma Membrane |
| 478 | Q9JL16 | ISG20 | 6.00E+06 | 1.92E+06 | 5.53E+05 | 12 | Nucleus |
| 479 | Q5H8C4 | VPS13A | 5.98E+06 | 3.91E+06 | 3.87E+05 | 102 | Cytoplasm |
| 480 | Q99LC3 | NDUFA10 | 5.97E+06 | 1.79E+06 | 7.32E+05 | 6 | Cytoplasm |
| 481 | Q8CBY3 | LENG8 | 5.96E+06 | 1.35E+06 | 7.77E+05 | 3 | Nucleus |
| 482 | Q9WTR1 | TRPV2 | 5.96E+06 | 4.32E+06 | 7.20E+05 | 36 | Plasma Membrane |
| 483 | P84091 | AP2M1 | 5.95E+06 | 2.92E+06 | 7.53E+05 | 15 | Cytoplasm |
| 484 | Q9CPQ8 | ATP5L | 5.92E+06 | 1.32E+06 | 5.39E+05 | 6 | Cytoplasm |
| 485 | P54775 | PSMC4 | 5.91E+06 | 4.29E+06 | 7.15E+05 | 36 | Nucleus |
| 486 | Q9D1L9 | LAMTOR5 | 5.89E+06 | 3.16E+06 | 1.05E+06 | 9 | Cytoplasm |
| 487 | P70182 | PIP5K1A | 5.89E+06 | 3.33E+06 | 6.08E+05 | 30 | Cytoplasm |
| 488 | Q9Z2I9 | SUCLA2 | 5.89E+06 | 1.48E+06 | 8.52E+05 | 3 | Cytoplasm |
| 489 | Q01102 | SELP | 5.88E+06 | 6.36E+06 | 1.50E+06 | 18 | Plasma Membrane |
| 490 | Q3U0B3 | DHRS11 | 5.88E+06 | 1.68E+06 | 9.72E+05 | 3 | Cytoplasm |
| 491 | Q9QZU9 | UBE2L6 | 5.87E+06 | 7.95E+06 | 2.29E+06 | 12 | Cytoplasm |
| 492 | O08795 | PRKCSH | 5.85E+06 | 3.47E+06 | 6.05E+05 | 33 | Cytoplasm |
| 493 | Q921L3 | TMCO1 | 5.85E+06 | 1.03E+06 | 5.93E+05 | 3 | Cytoplasm |
| 494 | O70172 | PIP4K2A | 5.85E+06 | 4.47E+06 | 1.29E+06 | 12 | Cytoplasm |
| 495 | Q8QZY6 | TSPAN14 | 5.84E+06 | 2.50E+05 | 1.44E+05 | 3 | Plasma Membrane |
| 496 | P15327 | BPGM | 5.83E+06 | 6.04E+06 | 1.74E+06 | 12 | Extracellular Space |
| 497 | P57716 | NCSTN | 5.83E+06 | 3.57E+06 | 6.87E+05 | 27 | Plasma Membrane |
| 498 | Q9CQ92 | FIS1 | 5.81E+06 | 5.83E+05 | 2.38E+05 | 6 | Cytoplasm |
| 499 | P47753 | CAPZA1 | 5.81E+06 | 2.54E+06 | 1.04E+06 | 6 | Cytoplasm |
| 500 | Q9CQN6 | TMEM14C | 5.77E+06 | 5.48E+06 | 2.24E+06 | 6 | Plasma Membrane |
| 501 | Q9CY52 | THG1L | 5.75E+06 | 1.73E+06 | 4.45E+05 | 15 | Cytoplasm |
| 502 | Q9DBX3 | SUSD2 | 5.73E+06 | 1.77E+06 | 5.91E+05 | 9 | Extracellular Space |
| 503 | Q9QUJ7 | ACSL4 | 5.72E+06 | 5.03E+06 | 9.68E+05 | 27 | Cytoplasm |
| 504 | Q99L43 | CDS2 | 5.69E+06 | 1.17E+06 | 6.73E+05 | 3 | Cytoplasm |
| 505 | P97450 | ATP5J | 5.69E+06 | 1.26E+06 | 7.25E+05 | 3 | Cytoplasm |
| 506 | Q9CZX8 | RPS19 | 5.67E+06 | 7.00E+05 | 2.33E+05 | 9 | Cytoplasm |
| 507 | P57759 | ERP29 | 5.66E+06 | 3.11E+06 | 8.98E+05 | 12 | Cytoplasm |
| 508 | Q9WUM4 | CORO1C | 5.64E+06 | 3.15E+06 | 1.28E+06 | 6 | Cytoplasm |
| 509 | Q9CR23 | TMEM9 | 5.64E+06 | 1.35E+06 | 7.80E+05 | 3 | Cytoplasm |
| 510 | P25911 | LYN | 5.63E+06 | 4.46E+06 | 9.10E+05 | 24 | Cytoplasm |
| 511 | Q9QUM4 | SLAMF1 | 5.62E+06 | 2.02E+06 | 6.73E+05 | 9 | Plasma Membrane |
| 512 | P50580 | PA2G4 | 5.61E+06 | 1.72E+06 | 4.43E+05 | 15 | Nucleus |
| 513 | P51410 | RPL9 | 5.61E+06 | 1.87E+06 | 6.24E+05 | 9 | Nucleus |
| 514 | Q9EST8 | NFKBIZ | 5.61E+06 | 1.78E+06 | 1.03E+06 | 3 | Nucleus |
| 515 | P41105 | RPL28 | 5.60E+06 | 5.42E+05 | 3.13E+05 | 3 | Cytoplasm |
| 516 | Q62425 | NDUFA4 | 5.59E+06 | 1.95E+06 | 7.97E+05 | 6 | Cytoplasm |
| 517 | Q80W04 | TMCC2 | 5.57E+06 | 5.00E+06 | 1.18E+06 | 18 | Extracellular Space |
| 518 | P68040 | GNB2L1 | 5.57E+06 | 1.98E+06 | 5.12E+05 | 15 | Cytoplasm |
| 519 | P67778 | PHB | 5.56E+06 | 3.49E+06 | 6.38E+05 | 30 | Nucleus |
| 520 | P35979 | RPL12 | 5.56E+06 | 1.83E+06 | 4.72E+05 | 15 | Nucleus |
| 521 | P19221 | F2 | 5.54E+06 | 4.82E+05 | 2.78E+05 | 3 | Extracellular Space |
| 522 | Q8CBB9 | RSAD2 | 5.53E+06 | 5.03E+06 | 1.03E+06 | 24 | Cytoplasm |
| 523 | Q9Z0N1 | EIF2S3 | 5.53E+06 | 3.18E+06 | 5.54E+05 | 33 | Cytoplasm |
| 524 | Q9D0R2 | TARS | 5.51E+06 | 3.20E+06 | 1.31E+06 | 6 | Nucleus |
| 525 | Q91ZJ9 | HYAL1 | 5.46E+06 | 1.45E+06 | 8.34E+05 | 3 | Cytoplasm |
| 526 | Q9JM62 | REEP6 | 5.45E+06 | 4.48E+06 | 1.49E+06 | 9 | Plasma Membrane |
| 527 | Q60870 | REEP5 | 5.45E+06 | 8.73E+06 | 2.91E+06 | 9 | Extracellular Space |
| 528 | Q5XPI3 | RNF123 | 5.43E+06 | 2.17E+06 | 4.74E+05 | 21 | Cytoplasm |
| 529 | O08807 | PRDX4 | 5.41E+06 | 3.60E+06 | 1.20E+06 | 9 | Cytoplasm |
| 530 | P62897 | CYCS | 5.40E+06 | 3.44E+06 | 8.88E+05 | 15 | Cytoplasm |
| 531 | P62862 | FAU | 5.38E+06 | 7.31E+05 | 4.22E+05 | 3 | Cytoplasm |
| 532 | P54822 | ADSL | 5.38E+06 | 3.24E+06 | 5.18E+05 | 39 | Cytoplasm |
| 533 | Q9QUK3 | CLN8 | 5.37E+06 | 8.52E+05 | 4.92E+05 | 3 | Cytoplasm |
| 534 | Q9CR57 | RPL14 | 5.35E+06 | 2.89E+06 | 9.63E+05 | 9 | Cytoplasm |
| 535 | P11688 | ITGA5 | 5.34E+06 | 3.01E+06 | 8.69E+05 | 12 | Plasma Membrane |
| 536 | O88587 | COMT | 5.34E+06 | 1.73E+06 | 5.00E+05 | 12 | Cytoplasm |
| 537 | P62242 | RPS8 | 5.32E+06 | 2.86E+06 | 7.38E+05 | 15 | Cytoplasm |
| 538 | Q8BWT1 | ACAA2 | 5.31E+06 | 2.48E+06 | 5.86E+05 | 18 | Cytoplasm |
| 539 | P49282 | SLC11A2 | 5.30E+06 | 2.52E+06 | 7.27E+05 | 12 | Plasma Membrane |
| 540 | Q8BK63 | CSNK1A1 | 5.28E+06 | 2.71E+06 | 5.52E+05 | 24 | Cytoplasm |
| 541 | Q9WVE8 | PACSIN2 | 5.25E+06 | 2.42E+06 | 4.94E+05 | 24 | Cytoplasm |
| 542 | Q8CII2 | CDC123 | 5.24E+06 | 2.07E+06 | 1.19E+06 | 3 | Cytoplasm |
| 543 | Q8K2B3 | SDHA | 5.24E+06 | 3.97E+06 | 8.10E+05 | 24 | Cytoplasm |
| 544 | Q64514 | TPP2 | 5.24E+06 | 3.74E+06 | 4.61E+05 | 66 | Cytoplasm |
| 545 | Q91YA2 | PSKH1 | 5.21E+06 | 1.50E+06 | 8.69E+05 | 3 | Nucleus |
| 546 | Q6ZWV7 | RPL35 | 5.21E+06 | 2.46E+06 | 1.00E+06 | 6 | Cytoplasm |
| 547 | Q3TDD9 | PPP1R21 | 5.20E+06 | 5.61E+06 | 3.24E+06 | 3 | Cytoplasm |
| 548 | Q6URW6 | MYH14 | 5.19E+06 | 2.49E+06 | 1.02E+06 | 6 | Extracellular Space |
| 549 | Q8CGK3 | LONP1 | 5.18E+06 | 2.29E+06 | 9.34E+05 | 6 | Cytoplasm |
| 550 | Q02013 | AQP1 | 5.15E+06 | 7.15E+06 | 1.85E+06 | 15 | Plasma Membrane |
| 551 | P16546 | SPTAN1 | 5.15E+06 | 2.91E+06 | 1.68E+06 | 3 | Plasma Membrane |
| 552 | P70460 | VASP | 5.15E+06 | 1.92E+06 | 1.11E+06 | 3 | Plasma Membrane |
| 553 | Q8JZR0 | ACSL5 | 5.11E+06 | 2.69E+06 | 4.68E+05 | 33 | Cytoplasm |
| 554 | Q9DCW4 | ETFB | 5.11E+06 | 5.18E+06 | 1.49E+06 | 12 | Cytoplasm |
| 555 | Q6ZQI3 | MLEC | 5.09E+06 | 1.73E+06 | 4.47E+05 | 15 | Plasma Membrane |

| | | | | | | | |
|---|---|---|---|---|---|---|---|
| 417 | P08680 | ALAS2 | 5.79E+06 | 7.87E+06 | 1.31E+06 | 36 | Cytoplasm |
| 418 | Q8JZR0 | ACSL5 | 5.76E+06 | 3.00E+06 | 5.23E+05 | 33 | Cytoplasm |
| 419 | P05202 | GOT2 | 5.73E+06 | 6.66E+06 | 1.16E+06 | 33 | Cytoplasm |
| 420 | O35887 | CALU | 5.73E+06 | 2.77E+06 | 5.65E+05 | 24 | Cytoplasm |
| 421 | P36423 | TBXAS1 | 5.71E+06 | 4.08E+06 | 7.85E+05 | 27 | Plasma Membrane |
| 422 | O88653 | LAMTOR3 | 5.68E+06 | 3.94E+06 | 1.31E+06 | 9 | Cytoplasm |
| 423 | Q01730 | RSU1 | 5.68E+06 | 9.54E+06 | 3.18E+06 | 9 | Cytoplasm |
| 424 | Q9D2G2 | DLST | 5.68E+06 | 3.42E+06 | 7.45E+05 | 21 | Cytoplasm |
| 425 | Q9CZ13 | UQCRC1 | 5.66E+06 | 4.56E+06 | 1.18E+06 | 15 | Cytoplasm |
| 426 | P57759 | ERP29 | 5.65E+06 | 2.91E+06 | 8.41E+05 | 12 | Cytoplasm |
| 427 | Q9CXW4 | RPL11 | 5.64E+06 | 2.34E+06 | 1.35E+06 | 3 | Cytoplasm |
| 428 | Q922U2 | KRT5 | 5.64E+06 | 6.12E+06 | 1.77E+06 | 12 | Cytoplasm |
| 429 | O70172 | PIP4K2A | 5.63E+06 | 4.48E+06 | 1.29E+06 | 12 | Cytoplasm |
| 430 | P67984 | RPL22 | 5.63E+06 | 3.99E+06 | 1.63E+06 | 6 | Nucleus |
| 431 | Q6PAR5 | GAPVD1 | 5.61E+06 | 2.39E+06 | 3.35E+05 | 51 | Cytoplasm |
| 432 | Q9CQB5 | CISD2 | 5.59E+06 | 1.53E+06 | 3.95E+05 | 15 | Cytoplasm |
| 433 | P56395 | CYB5A | 5.58E+06 | 4.34E+06 | 1.12E+06 | 15 | Cytoplasm |
| 434 | Q9Z0T9 | ITGB6 | 5.55E+06 | 2.39E+06 | 1.38E+06 | 3 | Plasma Membrane |
| 435 | P05132 | PRKACA | 5.53E+06 | 4.37E+05 | 2.53E+05 | 3 | Cytoplasm |
| 436 | Q9D1J1 | NECAP2 | 5.52E+06 | 9.60E+05 | 5.54E+05 | 3 | Cytoplasm |
| 437 | P21279 | GNAQ | 5.51E+06 | 3.51E+06 | 2.03E+06 | 3 | Plasma Membrane |
| 438 | P15379 | CD44 | 5.47E+06 | 6.79E+06 | 1.96E+06 | 12 | Plasma Membrane |
| 439 | Q9JLT4 | TXNRD2 | 5.47E+06 | 6.50E+06 | 2.17E+06 | 9 | Cytoplasm |
| 440 | P28571 | SLC6A9 | 5.43E+06 | 4.19E+06 | 1.71E+06 | 6 | Plasma Membrane |
| 441 | Q9CQX2 | CYB5B | 5.43E+06 | 8.22E+06 | 2.74E+06 | 9 | Cytoplasm |
| 442 | Q9CQR6 | PPP6C | 5.40E+06 | 1.62E+06 | 6.62E+05 | 6 | Nucleus |
| 443 | Q64518 | ATP2A3 | 5.36E+06 | 7.68E+06 | 1.11E+06 | 48 | Cytoplasm |
| 444 | Q9JJK2 | LANCL2 | 5.34E+06 | 2.38E+06 | 3.96E+05 | 36 | Plasma Membrane |
| 445 | Q61189 | CLNS1A | 5.32E+06 | 1.52E+06 | 8.79E+05 | 3 | Plasma Membrane |
| 446 | P10852 | SLC3A2 | 5.30E+06 | 2.99E+06 | 5.45E+05 | 30 | Plasma Membrane |
| 447 | P17427 | AP2A2 | 5.30E+06 | 5.36E+06 | 1.09E+06 | 24 | Cytoplasm |
| 448 | P84091 | AP2M1 | 5.29E+06 | 2.55E+06 | 6.59E+05 | 15 | Cytoplasm |
| 449 | Q8R016 | BLMH | 5.29E+06 | 2.07E+06 | 4.51E+05 | 21 | Cytoplasm |
| 450 | O35372 | CNIH1 | 5.21E+06 | 9.52E+05 | 5.50E+05 | 3 | Plasma Membrane |
| 451 | P35441 | THBS1 | 5.20E+06 | 9.71E+06 | 9.91E+05 | 96 | Extracellular Space |
| 452 | Q9D1L9 | LAMTOR5 | 5.20E+06 | 2.86E+06 | 9.53E+05 | 9 | Cytoplasm |
| 453 | Q80WT2 | DMRT3 | 5.16E+06 | 1.02E+05 | 5.87E+04 | 3 | Nucleus |
| 454 | Q68FD5 | CLTC | 5.16E+06 | 3.46E+06 | 2.94E+05 | 138 | Plasma Membrane |
| 455 | Q91Y57 | SIGLEC9 | 5.15E+06 | 5.68E+05 | 3.28E+05 | 3 | Plasma Membrane |
| 456 | Q64446 | ATP7B | 5.13E+06 | 5.99E+06 | 1.22E+06 | 24 | Cytoplasm |
| 457 | Q791V5 | MTCH2 | 5.05E+06 | 2.71E+06 | 1.56E+06 | 3 | Cytoplasm |
| 458 | Q8BGD9 | EIF4B | 5.03E+06 | 1.92E+06 | 4.53E+05 | 18 | Cytoplasm |
| 459 | P19221 | F2 | 5.03E+06 | 4.67E+05 | 2.69E+05 | 3 | Extracellular Space |
| 460 | Q9CR23 | TMEM9 | 5.01E+06 | 9.46E+05 | 5.46E+05 | 3 | Cytoplasm |
| 461 | P46460 | NSF | 4.96E+06 | 2.36E+06 | 3.65E+05 | 42 | Cytoplasm |
| 462 | P67778 | PHB | 4.96E+06 | 3.48E+06 | 6.35E+05 | 30 | Nucleus |
| 463 | Q78IK2 | USMG5 | 4.96E+06 | 3.12E+06 | 1.27E+06 | 6 | Cytoplasm |
| 464 | P14206 | RPSA | 4.95E+06 | 5.21E+06 | 1.50E+06 | 12 | Cytoplasm |
| 465 | Q921L3 | TMCO1 | 4.93E+06 | 1.37E+06 | 7.88E+05 | 3 | Cytoplasm |
| 466 | O70576 | STAG3 | 4.91E+06 | 4.32E+05 | 2.50E+05 | 3 | Nucleus |
| 467 | Q9D1G5 | LRRC57 | 4.89E+06 | 1.66E+06 | 3.92E+05 | 18 | Cytoplasm |
| 468 | Q9D7G0 | PRPS1 | 4.88E+06 | 2.05E+06 | 4.83E+05 | 18 | Cytoplasm |
| 469 | Q9Z127 | SLC7A5 | 4.88E+06 | 3.53E+06 | 1.02E+06 | 12 | Plasma Membrane |
| 470 | P36552 | CPOX | 4.85E+06 | 5.19E+06 | 8.66E+05 | 36 | Cytoplasm |
| 471 | P47753 | CAPZA1 | 4.84E+06 | 2.36E+06 | 9.62E+05 | 6 | Cytoplasm |
| 472 | Q60597 | OGDH | 4.84E+06 | 4.75E+06 | 6.65E+05 | 51 | Cytoplasm |
| 473 | P52480 | PKM | 4.80E+06 | 8.99E+05 | 5.19E+05 | 3 | Cytoplasm |
| 474 | Q9D6R2 | IDH3A | 4.79E+06 | 5.67E+06 | 1.46E+06 | 15 | Cytoplasm |
| 475 | Q6PCP5 | Mff | 4.78E+06 | 1.88E+06 | 7.68E+05 | 6 | Cytoplasm |
| 476 | Q80W04 | TMCC2 | 4.78E+06 | 4.54E+06 | 1.07E+06 | 18 | Extracellular Space |
| 477 | P70658 | CXCR4 | 4.77E+06 | 1.00E+06 | 5.80E+05 | 3 | Plasma Membrane |
| 478 | P26041 | MSN | 4.77E+06 | 2.93E+06 | 4.89E+05 | 36 | Plasma Membrane |
| 479 | Q9R1S7 | ABCC6 | 4.76E+06 | 1.85E+06 | 1.07E+06 | 3 | Plasma Membrane |
| 480 | Q9QXT0 | CNPY2 | 4.75E+06 | 1.50E+06 | 5.00E+05 | 9 | Plasma Membrane |
| 481 | Q8C1A3 | MTRR | 4.74E+06 | 3.48E+05 | 2.01E+05 | 3 | Cytoplasm |
| 482 | Q99KI0 | ACO2 | 4.73E+06 | 5.14E+06 | 6.48E+05 | 63 | Cytoplasm |
| 483 | O08807 | PRDX4 | 4.70E+06 | 3.42E+06 | 1.14E+06 | 9 | Cytoplasm |
| 484 | Q8CBY3 | LENG8 | 4.66E+06 | 1.14E+07 | 6.59E+06 | 3 | Nucleus |
| 485 | Q9CWJ9 | ATIC | 4.66E+06 | 2.44E+06 | 4.46E+05 | 30 | Cytoplasm |
| 486 | P86048 | RPL10L | 4.66E+06 | 3.51E+06 | 1.43E+06 | 6 | Nucleus |
| 487 | Q9ERN0 | SCAMP2 | 4.65E+06 | 1.34E+06 | 5.47E+05 | 6 | Cytoplasm |
| 488 | Q9Z0F4 | CIB1 | 4.64E+06 | 3.22E+06 | 9.30E+05 | 12 | Nucleus |
| 489 | Q9D6F9 | TUBB4A | 4.64E+06 | 1.33E+06 | 5.42E+05 | 6 | Cytoplasm |
| 490 | Q9CQ92 | FIS1 | 4.63E+06 | 9.87E+05 | 4.03E+05 | 6 | Cytoplasm |
| 491 | Q9DCF9 | SSR3 | 4.62E+06 | 2.08E+06 | 1.20E+06 | 3 | Cytoplasm |
| 492 | Q3TWL2 | TMEM55B | 4.61E+06 | 1.05E+06 | 3.04E+05 | 12 | Cytoplasm |
| 493 | Q9JL99 | CLEC1B | 4.61E+06 | 2.09E+06 | 1.20E+06 | 3 | Plasma Membrane |
| 494 | Q01102 | SELP | 4.61E+06 | 4.51E+06 | 1.06E+06 | 18 | Plasma Membrane |
| 495 | Q9CZX7 | TMEM55A | 4.60E+06 | 7.44E+05 | 4.30E+05 | 3 | Cytoplasm |
| 496 | P48771 | COX7A2 | 4.59E+06 | 2.67E+06 | 1.54E+06 | 3 | Cytoplasm |
| 497 | Q8CI59 | STEAP3 | 4.58E+06 | 4.17E+06 | 1.08E+06 | 15 | Cytoplasm |
| 498 | P49282 | SLC11A2 | 4.58E+06 | 3.13E+06 | 9.04E+05 | 12 | Plasma Membrane |
| 499 | Q91W53 | GOLGA7 | 4.55E+06 | 4.85E+05 | 1.98E+05 | 6 | Cytoplasm |
| 500 | Q9DBD2 | MARH8 | 4.54E+06 | 7.55E+05 | 3.08E+05 | 6 | Cytoplasm |
| 501 | A2AN08 | UBR4 | 4.46E+06 | 1.67E+06 | 1.33E+05 | 159 | Nucleus |
| 502 | Q9DB77 | UQCRC2 | 4.45E+06 | 5.16E+06 | 8.60E+05 | 36 | Cytoplasm |
| 503 | Q9D771 | TMEM206 | 4.45E+06 | 3.30E+06 | 1.10E+06 | 9 | Plasma Membrane |
| 504 | Q9QYF1 | RDH11 | 4.44E+06 | 1.51E+06 | 3.08E+05 | 24 | Cytoplasm |
| 505 | Q60864 | STIP1 | 4.44E+06 | 1.93E+06 | 3.52E+05 | 30 | Cytoplasm |
| 506 | Q9DCX2 | ATP5H | 4.44E+06 | 3.56E+06 | 8.40E+05 | 18 | Cytoplasm |
| 507 | P70297 | STAM | 4.43E+06 | 2.09E+06 | 6.98E+05 | 9 | Cytoplasm |
| 508 | Q91VH6 | MEMO1 | 4.42E+06 | 2.45E+06 | 6.31E+05 | 15 | Cytoplasm |
| 509 | P07758 | SERPINA1 | 4.42E+06 | 4.75E+06 | 1.23E+06 | 15 | Extracellular Space |
| 510 | Q60738 | SLC30A1 | 4.41E+06 | 1.24E+06 | 3.57E+05 | 12 | Plasma Membrane |
| 511 | Q9WVK4 | EHD1 | 4.39E+06 | 3.18E+06 | 9.17E+05 | 12 | Cytoplasm |
| 512 | P35282 | RAB21 | 4.39E+06 | 3.25E+06 | 6.63E+05 | 24 | Cytoplasm |
| 513 | Q99JY9 | ACTR3 | 4.38E+06 | 4.48E+06 | 1.16E+06 | 15 | Plasma Membrane |
| 514 | Q8BH04 | PCK2 | 4.37E+06 | 4.74E+06 | 6.64E+05 | 51 | Cytoplasm |
| 515 | Q99KR7 | PPIF | 4.36E+06 | 3.10E+06 | 1.27E+06 | 6 | Cytoplasm |
| 516 | Q64324 | STXBP2 | 4.36E+06 | 4.34E+06 | 6.70E+05 | 42 | Plasma Membrane |
| 517 | Q91WC9 | DAGLB | 4.36E+06 | 1.99E+06 | 4.69E+05 | 18 | Plasma Membrane |
| 518 | Q91ZJ9 | HYAL1 | 4.35E+06 | 2.10E+04 | 1.21E+04 | 3 | Cytoplasm |
| 519 | Q9JL16 | ISG20 | 4.33E+06 | 1.03E+06 | 2.99E+05 | 12 | Nucleus |
| 520 | P39876 | TIMP3 | 4.33E+06 | 4.66E+06 | 1.35E+06 | 12 | Extracellular Space |
| 521 | Q9D1C8 | VPS28 | 4.33E+06 | 9.65E+06 | 5.57E+06 | 3 | Cytoplasm |
| 522 | O35350 | CAPN1 | 4.31E+06 | 1.76E+06 | 3.84E+05 | 21 | Cytoplasm |
| 523 | Q64522 | HIST2H2AB | 4.31E+06 | 1.73E+07 | 9.98E+06 | 3 | Nucleus |
| 524 | O55143 | ATP2A2 | 4.26E+06 | 5.67E+06 | 1.34E+06 | 18 | Cytoplasm |
| 525 | Q6P5E4 | UGGT1 | 4.25E+06 | 2.90E+06 | 5.92E+05 | 24 | Cytoplasm |
| 526 | Q3TWN3 | CNNM2 | 4.24E+06 | 3.74E+05 | 1.53E+05 | 6 | Plasma Membrane |
| 527 | Q3UFK8 | FRMD8 | 4.24E+06 | 9.64E+05 | 5.56E+05 | 3 | Cytoplasm |
| 528 | Q8VDI7 | UBAC1 | 4.23E+06 | 2.61E+06 | 4.76E+05 | 30 | Nucleus |
| 529 | Q09143 | SLC7A1 | 4.21E+06 | 2.52E+06 | 1.45E+06 | 3 | Plasma Membrane |
| 530 | P43276 | Hist1h1b | 4.21E+06 | 2.89E+07 | 1.18E+07 | 6 | Nucleus |
| 531 | Q9WVE8 | PACSIN2 | 4.17E+06 | 1.87E+06 | 3.81E+05 | 24 | Cytoplasm |
| 532 | P14148 | RPL7 | 4.17E+06 | 3.88E+06 | 1.00E+06 | 15 | Nucleus |
| 533 | P24270 | CAT | 4.16E+06 | 2.78E+06 | 3.58E+05 | 60 | Cytoplasm |
| 534 | P27659 | RPL3 | 4.16E+06 | 3.65E+06 | 1.22E+06 | 9 | Nucleus |
| 535 | P11688 | ITGA5 | 4.16E+06 | 2.44E+06 | 7.05E+05 | 12 | Plasma Membrane |
| 536 | O89023 | TPP1 | 4.13E+06 | 2.37E+06 | 1.37E+06 | 3 | Cytoplasm |
| 537 | P08003 | PDIA4 | 4.12E+06 | 1.96E+06 | 4.01E+05 | 24 | Cytoplasm |
| 538 | P63321 | RALA | 4.08E+06 | 1.23E+06 | 7.10E+05 | 3 | Cytoplasm |
| 539 | Q3UUG6 | TBC1D24 | 4.07E+06 | 3.15E+06 | 1.28E+06 | 6 | Cytoplasm |
| 540 | Q9Z2I9 | SUCLA2 | 4.04E+06 | 2.45E+06 | 1.42E+06 | 3 | Cytoplasm |
| 541 | P62814 | ATP6V1B2 | 4.02E+06 | 2.67E+06 | 8.91E+05 | 9 | Cytoplasm |
| 542 | Q641P0 | ACTR3B | 4.02E+06 | 1.68E+06 | 9.71E+05 | 3 | Cytoplasm |
| 543 | P97450 | ATP5J | 4.02E+06 | 2.33E+06 | 1.34E+06 | 3 | Cytoplasm |
| 544 | Q9CR60 | GOLT1B | 4.01E+06 | 6.89E+05 | 3.98E+05 | 3 | Cytoplasm |
| 545 | Q8BG67 | EFR3A | 4.00E+06 | 7.53E+05 | 1.64E+05 | 21 | Plasma Membrane |
| 546 | Q8K2B3 | SDHA | 3.98E+06 | 4.07E+06 | 8.31E+05 | 24 | Cytoplasm |
| 547 | P19096 | FASN | 3.97E+06 | 2.31E+06 | 2.78E+05 | 69 | Cytoplasm |
| 548 | Q76MZ3 | PPP2R1A | 3.97E+06 | 1.86E+06 | 4.05E+05 | 21 | Cytoplasm |
| 549 | Q571I9 | ALDH16A1 | 3.97E+06 | 2.04E+06 | 4.81E+05 | 18 | Cytoplasm |
| 550 | Q9JJ59 | ABCB9 | 3.97E+06 | 1.02E+07 | 5.91E+06 | 3 | Cytoplasm |
| 551 | P62264 | RPS14 | 3.97E+06 | 2.50E+06 | 8.35E+05 | 9 | Cytoplasm |
| 552 | Q8C0L0 | TMX4 | 3.97E+06 | 1.23E+05 | 7.09E+04 | 3 | Cytoplasm |
| 553 | Q9CPQ8 | ATP5L | 3.96E+06 | 3.02E+06 | 1.23E+06 | 6 | Cytoplasm |
| 554 | Q8BG73 | SH3BGRL2 | 3.94E+06 | 1.30E+05 | 7.50E+04 | 3 | Nucleus |
| 555 | P38647 | HSPA9 | 3.93E+06 | 3.49E+06 | 5.81E+05 | 36 | Cytoplasm |

| | | | | | | | |
|---|---|---|---|---|---|---|---|
| 417 | P06151 | LDHA | 6015906 | 2592393 | 498906 | 27 | Cytoplasm |
| 418 | P61226 | RAP2B | 6004533 | 2517554 | 593393 | 18 | Plasma Membrane |
| 419 | P60867 | RPS20 | 5984298 | 3601431 | 1470278 | 6 | Cytoplasm |
| 420 | P27659 | RPL3 | 5956599 | 5942434 | 1980811 | 9 | Nucleus |
| 421 | Q9CQN6 | TMEM14C | 5933015 | 4773711 | 1948859 | 6 | Plasma Membrane |
| 422 | P05202 | GOT2 | 5900058 | 6480689 | 1128143 | 33 | Cytoplasm |
| 423 | Q6PAR5 | GAPVD1 | 5861574 | 2578823 | 361108 | 51 | Cytoplasm |
| 424 | Q64446 | ATP7B | 5860404 | 6697716 | 1367166 | 24 | Cytoplasm |
| 425 | Q68FD5 | CLTC | 5821259 | 3982075 | 338977 | 138 | Plasma Membrane |
| 426 | Q02013 | AQP1 | 5816606 | 6164710 | 1591721 | 15 | Plasma Membrane |
| 427 | Q9CR57 | RPL14 | 5808008 | 3470626 | 1156875 | 9 | Cytoplasm |
| 428 | Q8CI59 | STEAP3 | 5782587 | 4424303 | 1142350 | 15 | Cytoplasm |
| 429 | Q9D2G2 | DLST | 5781164 | 4383920 | 956650 | 21 | Cytoplasm |
| 430 | Q5H8C4 | VPS13A | 5777484 | 3955704 | 391673 | 102 | Cytoplasm |
| 431 | Q9JMA1 | USP14 | 5754168 | 1670155 | 681838 | 6 | Cytoplasm |
| 432 | Q9CQR6 | PPP6C | 5737790 | 1453824 | 593521 | 6 | Nucleus |
| 433 | P60766 | Cdc42 | 5706375 | 2195592 | 566899 | 15 | Plasma Membrane |
| 434 | P70670 | NACA | 5657143 | 3220805 | 1073602 | 9 | Cytoplasm |
| 435 | Q9D6F9 | TUBB4A | 5643337 | 1798569 | 734263 | 6 | Cytoplasm |
| 436 | O35143 | ATPIF1 | 5615129 | 4311159 | 2489049 | 3 | Cytoplasm |
| 437 | P50580 | PA2G4 | 5598299 | 2596468 | 670405 | 15 | Nucleus |
| 438 | Q9CWJ9 | ATIC | 5594349 | 3244492 | 592360 | 30 | Cytoplasm |
| 439 | P24270 | CAT | 5587193 | 4527452 | 584492 | 60 | Cytoplasm |
| 440 | Q62348 | TSN | 5587057 | 2518400 | 1028132 | 6 | Nucleus |
| 441 | Q8BH04 | PCK2 | 5585093 | 5029213 | 704231 | 51 | Cytoplasm |
| 442 | Q9R0Q3 | TMED2 | 5572012 | 3146847 | 1048949 | 9 | Cytoplasm |
| 443 | P11438 | LAMP1 | 5559584 | 4249973 | 1416658 | 9 | Plasma Membrane |
| 444 | Q8K1B8 | FERMT3 | 5558917 | 3601759 | 519869 | 48 | Cytoplasm |
| 445 | P10605 | CTSB | 5539745 | 3042085 | 785463 | 15 | Cytoplasm |
| 446 | P49282 | SLC11A2 | 5532082 | 3985332 | 1150466 | 12 | Plasma Membrane |
| 447 | P62862 | FAU | 5521089 | 3145217 | 1815892 | 3 | Cytoplasm |
| 448 | Q9CR23 | TMEM9 | 5496171 | 657021 | 379331 | 3 | Cytoplasm |
| 449 | P57759 | ERP29 | 5467884 | 2565789 | 740680 | 12 | Cytoplasm |
| 450 | Q9QUK3 | CLN8 | 5464979 | 455226 | 262825 | 3 | Cytoplasm |
| 451 | Q9JJK2 | LANCL2 | 5451643 | 2755871 | 459312 | 36 | Plasma Membrane |
| 452 | Q9CZX8 | RPS19 | 5430754 | 2605028 | 868343 | 9 | Cytoplasm |
| 453 | Q9Z127 | SLC7A5 | 5421227 | 4973240 | 1435651 | 12 | Plasma Membrane |
| 454 | O08795 | PRKCSH | 5414990 | 2896330 | 504186 | 33 | Cytoplasm |
| 455 | P19221 | F2 | 5378171 | 529359 | 305625 | 3 | Extracellular Space |
| 456 | P84091 | AP2M1 | 5365482 | 3013427 | 778064 | 15 | Cytoplasm |
| 457 | Q3UUG6 | TBC1D24 | 5351221 | 2797435 | 1142048 | 6 | Cytoplasm |
| 458 | Q9WUP7 | UCHL5 | 5309693 | 3765314 | 1086953 | 12 | Cytoplasm |
| 459 | Q99KB8 | HAGH | 5286971 | 4245372 | 1415124 | 9 | Cytoplasm |
| 460 | Q60864 | STIP1 | 5257293 | 2466837 | 450381 | 30 | Cytoplasm |
| 461 | Q80W04 | TMCC2 | 5252847 | 5066239 | 1194124 | 18 | Extracellular Space |
| 462 | Q9QYA2 | TOMM40 | 5250126 | 3266872 | 1886129 | 3 | Cytoplasm |
| 463 | Q3TWL2 | TMEM55B | 5249472 | 1149017 | 331693 | 12 | Cytoplasm |
| 464 | O89086 | RBM3 | 5232858 | 6138073 | 2046024 | 9 | Cytoplasm |
| 465 | Q9Z0F4 | CIB1 | 5225918 | 3979300 | 1148725 | 12 | Nucleus |
| 466 | Q91Y57 | SIGLEC9 | 5216044 | 554478 | 320128 | 3 | Plasma Membrane |
| 467 | Q8BGD9 | EIF4B | 5187066 | 1965271 | 463219 | 18 | Cytoplasm |
| 468 | O35350 | CAPN1 | 5186364 | 2754360 | 601051 | 21 | Cytoplasm |
| 469 | P56391 | COX6B1 | 5162855 | 1764841 | 1018931 | 3 | Cytoplasm |
| 470 | Q9CR20 | IER3IP1 | 5158128 | 4263588 | 1740602 | 6 | Cytoplasm |
| 471 | P12787 | COX5A | 5150336 | 6892182 | 2297394 | 9 | Cytoplasm |
| 472 | P67778 | PHB | 5147970 | 4675704 | 853663 | 30 | Nucleus |
| 473 | Q9R1S7 | ABCC6 | 5126125 | 2535461 | 1463849 | 3 | Plasma Membrane |
| 474 | P61022 | CHP1 | 5114207 | 1857980 | 437930 | 18 | Cytoplasm |
| 475 | Q9DBX3 | SUSD2 | 5108351 | 1544739 | 514913 | 9 | Extracellular Space |
| 476 | Q9D0R2 | TARS | 5092658 | 2908057 | 1187209 | 6 | Nucleus |
| 477 | P05132 | PRKACA | 5089049 | 918886 | 530519 | 3 | Cytoplasm |
| 478 | Q9DB77 | UQCRC2 | 5074348 | 5159177 | 859863 | 36 | Cytoplasm |
| 479 | Q9WVK4 | EHD1 | 5062038 | 3095099 | 893478 | 12 | Cytoplasm |
| 480 | P46460 | NSF | 5061430 | 2479528 | 382599 | 42 | Cytoplasm |
| 481 | P35979 | RPL12 | 5046179 | 3129153 | 807944 | 15 | Nucleus |
| 482 | P68040 | GNB2L1 | 5023958 | 3552891 | 917353 | 15 | Cytoplasm |
| 483 | Q9EST8 | NFKBIZ | 5023746 | 1429954 | 825585 | 3 | Nucleus |
| 484 | P62267 | RPS23 | 5015972 | 2168864 | 1252194 | 3 | Cytoplasm |
| 485 | P97855 | G3BP1 | 5002211 | 2413821 | 985438 | 6 | Nucleus |
| 486 | P11688 | ITGA5 | 4982300 | 2519993 | 727459 | 12 | Plasma Membrane |
| 487 | Q9D6R2 | IDH3A | 4976852 | 5868283 | 1515184 | 15 | Cytoplasm |
| 488 | Q9Z1G4 | ATP6V0A1 | 4964713 | 3373239 | 649180 | 27 | Cytoplasm |
| 489 | Q9JM62 | REEP6 | 4954694 | 4925584 | 1641861 | 9 | Plasma Membrane |
| 490 | A2AN08 | UBR4 | 4941946 | 2073187 | 164414 | 159 | Nucleus |
| 491 | Q3TWN3 | CNNM2 | 4933508 | 520913 | 212662 | 6 | Plasma Membrane |
| 492 | Q99J56 | DERL1 | 4928205 | 2631988 | 1074505 | 6 | Cytoplasm |
| 493 | P38647 | HSPA9 | 4927463 | 4057075 | 676179 | 36 | Cytoplasm |
| 494 | P47857 | PFKM | 4865399 | 1339677 | 315765 | 18 | Cytoplasm |
| 495 | P19096 | FASN | 4836164 | 2612816 | 314546 | 69 | Cytoplasm |
| 496 | P70658 | CXCR4 | 4830342 | 744201 | 429665 | 3 | Plasma Membrane |
| 497 | O08807 | PRDX4 | 4804925 | 3362996 | 1120999 | 9 | Cytoplasm |
| 498 | P07758 | SERPINA1 | 4796051 | 6554018 | 1692240 | 15 | Extracellular Space |
| 499 | Q8C1A3 | MTRR | 4754792 | 1411509 | 814935 | 3 | Cytoplasm |
| 500 | Q6ZWN5 | RPS9 | 4749829 | 3198773 | 1066258 | 9 | Cytoplasm |
| 501 | Q99KI0 | ACO2 | 4714206 | 5341467 | 672962 | 63 | Cytoplasm |
| 502 | P24547 | IMPDH2 | 4707064 | 2368831 | 1367645 | 3 | Cytoplasm |
| 503 | Q9D8E6 | RPL4 | 4688031 | 2668759 | 770404 | 12 | Cytoplasm |
| 504 | Q3TTY5 | Krt2 | 4684448 | 2247770 | 1297750 | 3 | Cytoplasm |
| 505 | Q6ZWX6 | EIF2S1 | 4673846 | 1594412 | 375806 | 18 | Cytoplasm |
| 506 | P41105 | RPL28 | 4670004 | 1823094 | 1052564 | 3 | Cytoplasm |
| 507 | Q80WT2 | DMRT3 | 4655500 | 691891 | 399464 | 3 | Nucleus |
| 508 | Q9JI39 | ABCB10 | 4647391 | 7089094 | 1181516 | 36 | Cytoplasm |
| 509 | Q91W53 | GOLGA7 | 4613680 | 967193 | 394855 | 6 | Cytoplasm |
| 510 | Q9CYZ2 | TPD52L2 | 4587780 | 729434 | 421139 | 3 | Cytoplasm |
| 511 | Q02053 | UBA1 | 4532959 | 1995518 | 307915 | 42 | Cytoplasm |
| 512 | Q8CIG8 | PRMT5 | 4479988 | 3027000 | 1235768 | 6 | Cytoplasm |
| 513 | O09172 | GCLM | 4478497 | 1002860 | 334287 | 9 | Cytoplasm |
| 514 | P21981 | TGM2 | 4477203 | 2103954 | 858936 | 6 | Cytoplasm |
| 515 | Q80UJ7 | RAB3GAP1 | 4463194 | 2772039 | 924013 | 9 | Cytoplasm |
| 516 | Q9EP69 | SACM1L | 4461795 | 2818155 | 373274 | 57 | Cytoplasm |
| 517 | O35887 | CALU | 4448792 | 2413958 | 492747 | 24 | Cytoplasm |
| 518 | P50247 | AHCY | 4437330 | 1134181 | 378060 | 9 | Cytoplasm |
| 519 | Q8QZT1 | ACAT1 | 4428218 | 4173514 | 910735 | 21 | Cytoplasm |
| 520 | Q9CZM2 | RPL15 | 4426017 | 1694308 | 564769 | 9 | Cytoplasm |
| 521 | P17427 | AP2A2 | 4421064 | 3583030 | 731383 | 24 | Cytoplasm |
| 522 | Q9DCX2 | ATP5H | 4401631 | 4085353 | 962927 | 18 | Cytoplasm |
| 523 | Q8VCM7 | FGG | 4397873 | 13953911 | 1679854 | 69 | Extracellular Space |
| 524 | Q62425 | NDUFA4 | 4397584 | 2358949 | 963037 | 6 | Cytoplasm |
| 525 | Q3U0B3 | DHRS11 | 4390067 | 421468 | 243335 | 3 | Cytoplasm |
| 526 | Q8VEE1 | LMCD1 | 4389933 | 7092806 | 4095034 | 3 | Cytoplasm |
| 527 | Q78IK2 | USMG5 | 4385587 | 2746972 | 1121447 | 6 | Cytoplasm |
| 528 | Q99JY0 | HADHB | 4381518 | 4998548 | 1090773 | 21 | Cytoplasm |
| 529 | Q9CQQ7 | ATP5F1 | 4376866 | 2066205 | 1192924 | 3 | Cytoplasm |
| 530 | Q9QXT0 | CNPY2 | 4358293 | 1719443 | 573148 | 9 | Plasma Membrane |
| 531 | P26041 | MSN | 4357809 | 2491286 | 415214 | 36 | Plasma Membrane |
| 532 | Q8CGK3 | LONP1 | 4353735 | 3091068 | 1261923 | 6 | Cytoplasm |
| 533 | P62908 | RPS3 | 4350115 | 4067090 | 830191 | 24 | Cytoplasm |
| 534 | Q6ZWV7 | RPL35 | 4341178 | 2932938 | 1197367 | 6 | Cytoplasm |
| 535 | Q920G8 | SLC25A37 | 4329157 | 3857793 | 671556 | 33 | Cytoplasm |
| 536 | Q9CYN2 | SPCS2 | 4329096 | 759113 | 438274 | 3 | Cytoplasm |
| 537 | Q9CQ92 | FIS1 | 4319208 | 1194559 | 487677 | 6 | Cytoplasm |
| 538 | O89023 | TPP1 | 4311110 | 2827866 | 1632669 | 3 | Cytoplasm |
| 539 | Q9D0M1 | PRPSAP1 | 4304621 | 2119633 | 547287 | 15 | Extracellular Space |
| 540 | O70576 | STAG3 | 4277446 | 809460 | 467342 | 3 | Nucleus |
| 541 | A2ASS6 | TTN | 4269340 | 236384 | 136476 | 3 | Cytoplasm |
| 542 | Q8BWT1 | ACAA2 | 4264375 | 3388830 | 798755 | 18 | Cytoplasm |
| 543 | P62814 | ATP6V1B2 | 4254538 | 2632804 | 877601 | 9 | Cytoplasm |
| 544 | Q8K2J7 | RELL1 | 4252731 | 1637895 | 545965 | 9 | Plasma Membrane |
| 545 | Q60597 | OGDH | 4252600 | 3916161 | 548372 | 51 | Cytoplasm |
| 546 | Q9CYL5 | GLIPR2 | 4251148 | 463600 | 267660 | 3 | Cytoplasm |
| 547 | Q8BG67 | EFR3A | 4246334 | 822280 | 179436 | 21 | Plasma Membrane |
| 548 | P62855 | RPS26 | 4221357 | 1986405 | 1146852 | 3 | Cytoplasm |
| 549 | Q8K2B3 | SDHA | 4214925 | 4124158 | 841840 | 24 | Cytoplasm |
| 550 | Q8K411 | PITRM1 | 4208937 | 4184180 | 913063 | 21 | Cytoplasm |
| 551 | Q05144 | RAC2 | 4200327 | 3133107 | 1808900 | 3 | Cytoplasm |
| 552 | Q9DBD2 | MARH8 | 4181157 | 818449 | 334130 | 6 | Cytoplasm |
| 553 | Q9JLV5 | CUL3 | 4173116 | 707838 | 288974 | 6 | Nucleus |
| 554 | P53994 | RAB2A | 4131884 | 3181544 | 821471 | 15 | Cytoplasm |
| 555 | Q8R2Y8 | PTRH2 | 4122450 | 5287256 | 2158513 | 6 | Cytoplasm |

| | | | | | | | |
|---|---|---|---|---|---|---|---|
| 556 | P70297 | STAM | 4.01E+06 | 2.24E+06 | 7.45E+05 | 9 | Cytoplasm |
| 557 | Q6P5E4 | UGGT1 | 4.00E+06 | 2.98E+06 | 6.07E+05 | 24 | Cytoplasm |
| 558 | Q8R5H1 | USP15 | 4.00E+06 | 2.92E+06 | 4.22E+05 | 48 | Cytoplasm |
| 559 | Q8CBY3 | LENG8 | 3.99E+06 | 3.53E+06 | 2.04E+06 | 3 | Nucleus |
| 560 | Q923U0 | TOM1L1 | 3.98E+06 | 2.47E+06 | 1.01E+06 | 6 | Cytoplasm |
| 561 | Q8BGD9 | EIF4B | 3.97E+06 | 2.24E+06 | 5.27E+05 | 18 | Cytoplasm |
| 562 | Q9CPQ8 | ATP5L | 3.96E+06 | 2.12E+06 | 8.67E+05 | 6 | Cytoplasm |
| 563 | Q05144 | RAC2 | 3.95E+06 | 1.18E+06 | 6.81E+05 | 3 | Cytoplasm |
| 564 | Q60597 | OGDH | 3.94E+06 | 2.64E+06 | 3.69E+05 | 51 | Cytoplasm |
| 565 | O89023 | TPP1 | 3.93E+06 | 1.46E+06 | 8.44E+05 | 3 | Cytoplasm |
| 566 | P17710 | HK1 | 3.93E+06 | 3.44E+06 | 5.51E+05 | 39 | Cytoplasm |
| 567 | Q3TWN3 | CNNM2 | 3.92E+06 | 5.87E+05 | 2.40E+05 | 6 | Plasma Membrane |
| 568 | Q8BPM0 | DAAM1 | 3.91E+06 | 1.80E+06 | 3.00E+05 | 36 | Cytoplasm |
| 569 | Q99KI0 | ACO2 | 3.90E+06 | 3.04E+06 | 3.83E+05 | 63 | Cytoplasm |
| 570 | Q9D6F9 | TUBB4A | 3.90E+06 | 2.24E+06 | 9.13E+05 | 6 | Cytoplasm |
| 571 | A2ASS6 | TTN | 3.88E+06 | 1.17E+06 | 6.77E+05 | 3 | Cytoplasm |
| 572 | P70295 | AUP1 | 3.87E+06 | 2.07E+06 | 1.19E+06 | 3 | Cytoplasm |
| 573 | Q60864 | STIP1 | 3.87E+06 | 2.48E+06 | 4.52E+05 | 30 | Cytoplasm |
| 574 | Q9JL56 | GDE1 | 3.84E+06 | 3.09E+06 | 1.03E+06 | 9 | Plasma Membrane |
| 575 | Q91W53 | GOLGA7 | 3.84E+06 | 7.08E+05 | 2.89E+05 | 6 | Cytoplasm |
| 576 | P54731 | FAF1 | 3.84E+06 | 1.53E+06 | 5.10E+05 | 9 | Nucleus |
| 577 | P70460 | VASP | 3.83E+06 | 1.58E+06 | 9.15E+05 | 3 | Plasma Membrane |
| 578 | Q9EST8 | NFKBIZ | 3.83E+06 | 5.98E+05 | 3.45E+05 | 3 | Nucleus |
| 579 | P62830 | RPL23 | 3.83E+06 | 1.30E+06 | 7.50E+05 | 3 | Cytoplasm |
| 580 | P05977 | MYL1 | 3.79E+06 | 3.16E+06 | 1.83E+06 | 3 | Cytoplasm |
| 581 | P38647 | HSPA9 | 3.75E+06 | 1.75E+06 | 2.92E+05 | 36 | Cytoplasm |
| 582 | Q99JY9 | ACTR3 | 3.75E+06 | 4.49E+06 | 1.16E+06 | 15 | Plasma Membrane |
| 583 | O08547 | SEC22B | 3.75E+06 | 2.42E+06 | 4.93E+05 | 24 | Cytoplasm |
| 584 | Q8K2B3 | SDHA | 3.74E+06 | 2.41E+06 | 4.93E+05 | 24 | Cytoplasm |
| 585 | P62897 | CYCS | 3.72E+06 | 2.51E+06 | 6.49E+05 | 15 | Cytoplasm |
| 586 | P67871 | CSNK2B | 3.72E+06 | 1.98E+06 | 1.14E+06 | 3 | Cytoplasm |
| 587 | Q62425 | NDUFA4 | 3.72E+06 | 1.07E+06 | 4.38E+05 | 6 | Cytoplasm |
| 588 | Q91VR2 | ATP5C1 | 3.71E+06 | 2.76E+06 | 7.97E+05 | 12 | Cytoplasm |
| 589 | P97450 | ATP5J | 3.70E+06 | 9.68E+05 | 5.59E+05 | 3 | Cytoplasm |
| 590 | Q01102 | SELP | 3.69E+06 | 4.84E+06 | 1.14E+06 | 18 | Plasma Membrane |
| 591 | Q9DBD2 | MARH8 | 3.69E+06 | 1.09E+06 | 4.45E+05 | 6 | Cytoplasm |
| 592 | Q9JHR7 | IDE | 3.69E+06 | 1.06E+06 | 4.32E+05 | 6 | Extracellular Space |
| 593 | P99029 | PRDX5 | 3.69E+06 | 1.30E+06 | 7.50E+05 | 3 | Cytoplasm |
| 594 | Q9Z127 | SLC7A5 | 3.68E+06 | 2.76E+06 | 7.96E+05 | 12 | Plasma Membrane |
| 595 | P18760 | CFL1 | 3.68E+06 | 3.17E+06 | 8.20E+05 | 15 | Nucleus |
| 596 | O09167 | RPL21 | 3.67E+06 | 1.54E+06 | 6.29E+05 | 6 | Cytoplasm |
| 597 | P14869 | RPLP0 | 3.65E+06 | 2.33E+06 | 4.76E+05 | 24 | Cytoplasm |
| 598 | Q8BH04 | PCK2 | 3.65E+06 | 2.47E+06 | 3.46E+05 | 51 | Cytoplasm |
| 599 | Q6ZQI3 | MLEC | 3.64E+06 | 2.21E+06 | 5.72E+05 | 15 | Plasma Membrane |
| 600 | Q571I9 | ALDH16A1 | 3.64E+06 | 2.58E+06 | 6.09E+05 | 18 | Cytoplasm |
| 601 | P67778 | PHB | 3.63E+06 | 1.59E+06 | 2.91E+05 | 30 | Nucleus |
| 602 | P62264 | RPS14 | 3.63E+06 | 1.58E+06 | 5.28E+05 | 9 | Cytoplasm |
| 603 | Q08093 | CNN2 | 3.62E+06 | 3.25E+06 | 1.88E+06 | 3 | Cytoplasm |
| 604 | Q9D6R2 | IDH3A | 3.61E+06 | 1.91E+06 | 4.93E+05 | 15 | Cytoplasm |
| 605 | P62827 | RAN | 3.61E+06 | 7.21E+05 | 4.16E+05 | 3 | Nucleus |
| 606 | Q8QZY6 | TSPAN14 | 3.59E+06 | 3.10E+06 | 1.79E+06 | 3 | Plasma Membrane |
| 607 | P25911 | LYN | 3.58E+06 | 3.32E+06 | 6.78E+05 | 24 | Cytoplasm |
| 608 | P47857 | PFKM | 3.57E+06 | 1.99E+06 | 4.68E+05 | 18 | Cytoplasm |
| 609 | Q8C1A3 | MTRR | 3.56E+06 | 4.37E+05 | 2.52E+05 | 3 | Cytoplasm |
| 610 | Q9WVG6 | CARM1 | 3.55E+06 | 1.05E+06 | 3.51E+05 | 9 | Nucleus |
| 611 | Q9WUP7 | UCHL5 | 3.55E+06 | 3.43E+06 | 9.90E+05 | 12 | Cytoplasm |
| 612 | P15327 | BPGM | 3.53E+06 | 3.90E+06 | 1.13E+06 | 12 | Extracellular Space |
| 613 | Q8BYC6 | TAOK3 | 3.53E+06 | 1.78E+06 | 3.63E+05 | 24 | Cytoplasm |
| 614 | P35980 | RPL18 | 3.52E+06 | 1.71E+06 | 4.93E+05 | 12 | Cytoplasm |
| 615 | Q9JKR6 | HYOU1 | 3.50E+06 | 1.05E+06 | 2.30E+05 | 21 | Cytoplasm |
| 616 | Q91YQ5 | RPN1 | 3.49E+06 | 2.32E+06 | 4.74E+05 | 24 | Cytoplasm |
| 617 | Q80ZW2 | THEM6 | 3.49E+06 | 1.19E+06 | 3.06E+05 | 15 | Extracellular Space |
| 618 | Q99L43 | CDS2 | 3.48E+06 | 2.69E+06 | 1.55E+06 | 3 | Cytoplasm |
| 619 | Q9CR64 | TMEM167A | 3.48E+06 | 5.02E+05 | 2.90E+05 | 3 | Extracellular Space |
| 620 | P35279 | RAB6A | 3.48E+06 | 1.98E+06 | 8.10E+05 | 6 | Cytoplasm |
| 621 | Q9Z2I9 | SUCLA2 | 3.47E+06 | 1.68E+06 | 9.69E+05 | 3 | Cytoplasm |
| 622 | P42567 | EPS15 | 3.47E+06 | 1.62E+06 | 4.18E+05 | 15 | Cytoplasm |
| 623 | Q9QYA2 | TOMM40 | 3.47E+06 | 1.01E+06 | 5.80E+05 | 3 | Cytoplasm |
| 624 | Q9D0M1 | PRPSAP1 | 3.47E+06 | 2.48E+06 | 6.40E+05 | 15 | Extracellular Space |
| 625 | P25444 | RPS2 | 3.46E+06 | 1.46E+06 | 4.21E+05 | 12 | Cytoplasm |
| 626 | P62082 | RPS7 | 3.42E+06 | 2.29E+06 | 7.62E+05 | 9 | Cytoplasm |
| 627 | Q99KV1 | DNAJB11 | 3.41E+06 | 1.56E+06 | 5.19E+05 | 9 | Cytoplasm |
| 628 | Q99LC3 | NDUFA10 | 3.41E+06 | 9.65E+05 | 3.94E+05 | 6 | Cytoplasm |
| 629 | Q99LD4 | GPS1 | 3.40E+06 | 2.74E+06 | 7.06E+05 | 15 | Cytoplasm |
| 630 | Q99JY0 | HADHB | 3.40E+06 | 2.20E+06 | 4.80E+05 | 21 | Cytoplasm |
| 631 | Q9D6K8 | FUNDC2 | 3.40E+06 | 2.06E+06 | 8.42E+05 | 6 | Cytoplasm |
| 632 | P05132 | PRKACA | 3.40E+06 | 2.49E+05 | 1.44E+05 | 3 | Cytoplasm |
| 633 | Q80UJ7 | RAB3GAP1 | 3.39E+06 | 2.70E+06 | 8.99E+05 | 9 | Cytoplasm |
| 634 | Q9CR57 | RPL14 | 3.39E+06 | 1.58E+06 | 5.26E+05 | 9 | Cytoplasm |
| 635 | Q9JLV5 | CUL3 | 3.38E+06 | 1.70E+06 | 6.93E+05 | 6 | Nucleus |
| 636 | Q9DCW4 | ETFB | 3.37E+06 | 2.90E+06 | 8.38E+05 | 12 | Cytoplasm |
| 637 | P06151 | LDHA | 3.36E+06 | 1.78E+06 | 3.42E+05 | 27 | Cytoplasm |
| 638 | Q9WUM4 | CORO1C | 3.35E+06 | 3.12E+06 | 1.27E+06 | 6 | Cytoplasm |
| 639 | Q9CYL5 | GLIPR2 | 3.35E+06 | 3.33E+05 | 1.92E+05 | 3 | Cytoplasm |
| 640 | Q99L45 | EIF2S2 | 3.34E+06 | 1.56E+06 | 5.20E+05 | 9 | Cytoplasm |
| 641 | Q9DBG6 | RPN2 | 3.33E+06 | 2.64E+06 | 4.81E+05 | 30 | Cytoplasm |
| 642 | Q9R0Q6 | ARPC1A | 3.33E+06 | 1.25E+06 | 7.21E+05 | 3 | Extracellular Space |
| 643 | Q3V0G7 | GARNL3 | 3.32E+06 | 5.97E+05 | 3.45E+05 | 3 | Cytoplasm |
| 644 | Q8VDL4 | ADPGK | 3.31E+06 | 6.38E+05 | 3.68E+05 | 3 | Plasma Membrane |
| 645 | P62751 | Rpl23a | 3.31E+06 | 2.41E+06 | 9.82E+05 | 6 | Nucleus |
| 646 | Q9CR60 | GOLT1B | 3.30E+06 | 7.40E+04 | 4.27E+04 | 3 | Cytoplasm |
| 647 | Q9CQQ7 | ATP5F1 | 3.30E+06 | 1.23E+06 | 7.08E+05 | 3 | Cytoplasm |
| 648 | P60867 | RPS20 | 3.29E+06 | 9.08E+05 | 3.71E+05 | 6 | Cytoplasm |
| 649 | P62852 | RPS25 | 3.28E+06 | 7.76E+05 | 2.59E+05 | 9 | Cytoplasm |
| 650 | Q8VCF0 | MAVS | 3.28E+06 | 1.88E+06 | 7.66E+05 | 6 | Cytoplasm |
| 651 | Q6P1F6 | PPP2R2A | 3.28E+06 | 3.75E+06 | 2.17E+06 | 3 | Cytoplasm |
| 652 | P63325 | RPS10 | 3.28E+06 | 4.10E+05 | 2.37E+05 | 3 | Cytoplasm |
| 653 | Q9CX00 | IST1 | 3.25E+06 | 1.24E+06 | 7.14E+05 | 3 | Cytoplasm |
| 654 | Q64310 | SURF4 | 3.25E+06 | 1.41E+06 | 5.74E+05 | 6 | Cytoplasm |
| 655 | Q9ERM2 | ICAM4 | 3.25E+06 | 7.38E+05 | 3.01E+05 | 6 | Plasma Membrane |
| 656 | Q8BWT1 | ACAA2 | 3.25E+06 | 1.38E+06 | 3.25E+05 | 18 | Cytoplasm |
| 657 | Q8BG67 | EFR3A | 3.25E+06 | 6.50E+05 | 1.42E+05 | 21 | Plasma Membrane |
| 658 | P01872 | IGHM | 3.24E+06 | 2.46E+06 | 7.09E+05 | 12 | Plasma Membrane |
| 659 | O88543 | COPS3 | 3.24E+06 | 1.72E+06 | 4.06E+05 | 18 | Cytoplasm |
| 660 | P61202 | Cops2 | 3.24E+06 | 2.53E+06 | 6.54E+05 | 15 | Nucleus |
| 661 | Q9QUM4 | SLAMF1 | 3.24E+06 | 2.69E+06 | 8.97E+05 | 9 | Plasma Membrane |
| 662 | O88792 | F11R | 3.24E+06 | 4.55E+06 | 1.18E+06 | 15 | Plasma Membrane |
| 663 | Q8K2J7 | RELL1 | 3.23E+06 | 1.57E+06 | 5.24E+05 | 9 | Plasma Membrane |
| 664 | O88668 | CREG1 | 3.23E+06 | 8.49E+05 | 3.47E+05 | 6 | Nucleus |
| 665 | P61161 | ACTR2 | 3.23E+06 | 1.49E+06 | 4.96E+05 | 9 | Plasma Membrane |
| 666 | P12970 | RPL7A | 3.21E+06 | 1.01E+06 | 3.38E+05 | 9 | Cytoplasm |
| 667 | P35285 | RAB22A | 3.21E+06 | 4.62E+05 | 1.89E+05 | 6 | Cytoplasm |
| 668 | Q64521 | GPD2 | 3.20E+06 | 2.80E+06 | 5.12E+05 | 30 | Cytoplasm |
| 669 | Q60857 | SLC6A4 | 3.20E+06 | 2.98E+06 | 1.72E+06 | 3 | Plasma Membrane |
| 670 | Q9D0R2 | TARS | 3.18E+06 | 1.99E+06 | 8.13E+05 | 6 | Nucleus |
| 671 | P50580 | PA2G4 | 3.16E+06 | 1.21E+06 | 3.11E+05 | 15 | Nucleus |
| 672 | P14131 | RPS16 | 3.16E+06 | 1.30E+06 | 5.30E+05 | 6 | Cytoplasm |
| 673 | Q9QY76 | VAPB | 3.16E+06 | 2.07E+06 | 8.43E+05 | 6 | Plasma Membrane |
| 674 | P49769 | PSEN1 | 3.15E+06 | 1.93E+06 | 7.89E+05 | 6 | Plasma Membrane |
| 675 | O89112 | LANCL1 | 3.12E+06 | 1.50E+06 | 8.67E+05 | 3 | Plasma Membrane |
| 676 | Q61753 | PHGDH | 3.12E+06 | 1.41E+06 | 3.63E+05 | 15 | Cytoplasm |
| 677 | P98086 | C1QA | 3.11E+06 | 4.10E+06 | 1.68E+06 | 6 | Extracellular Space |
| 678 | Q9EQH2 | ERAP1 | 3.11E+06 | 1.31E+06 | 7.58E+05 | 3 | Extracellular Space |
| 679 | P42669 | PURA | 3.11E+06 | 2.27E+06 | 9.26E+05 | 6 | Nucleus |
| 680 | P17047 | LAMP2 | 3.11E+06 | 1.75E+06 | 7.16E+05 | 6 | Plasma Membrane |
| 681 | Q8BJY1 | PSMD5 | 3.11E+06 | 9.91E+05 | 1.91E+05 | 27 | Cytoplasm |
| 682 | Q8R2Y8 | PTRH2 | 3.09E+06 | 3.23E+06 | 1.32E+06 | 6 | Cytoplasm |
| 683 | Q9CQA3 | SDHB | 3.08E+06 | 1.77E+06 | 5.90E+05 | 9 | Cytoplasm |
| 684 | Q920L1 | FADS1 | 3.08E+06 | 3.70E+05 | 2.14E+05 | 3 | Plasma Membrane |
| 685 | P51410 | RPL9 | 3.08E+06 | 1.18E+06 | 3.93E+05 | 9 | Nucleus |
| 686 | Q9Z1Q2 | ABHD16A | 3.07E+06 | 1.73E+06 | 5.76E+05 | 9 | Cytoplasm |
| 687 | P51655 | GPC4 | 3.07E+06 | 9.50E+05 | 3.17E+05 | 9 | Plasma Membrane |
| 688 | Q641P0 | ACTR3B | 3.05E+06 | 8.62E+05 | 4.98E+05 | 3 | Cytoplasm |
| 689 | Q6ZWV7 | RPL35 | 3.04E+06 | 1.66E+06 | 6.78E+05 | 6 | Cytoplasm |
| 690 | Q9CYZ2 | TPD52L2 | 3.02E+06 | 5.52E+05 | 3.19E+05 | 3 | Cytoplasm |
| 691 | Q8VDP3 | MICAL1 | 3.00E+06 | 1.25E+05 | 7.24E+04 | 3 | Cytoplasm |
| 692 | Q6R0H7 | GNAS | 3.00E+06 | 3.45E+05 | 1.41E+05 | 6 | Plasma Membrane |
| 693 | Q8CIG8 | PRMT5 | 2.99E+06 | 2.62E+06 | 1.07E+06 | 6 | Cytoplasm |
| 694 | Q9CQS8 | SEC61B | 2.99E+06 | 1.54E+06 | 6.29E+05 | 6 | Cytoplasm |

| | | | | | | | |
|---|---|---|---|---|---|---|---|
| 556 | P68369 | TUBA1A | 5.08E+06 | 9.73E+05 | 5.62E+05 | 3 | Cytoplasm |
| 557 | Q60864 | STIP1 | 5.06E+06 | 2.23E+06 | 4.07E+05 | 30 | Cytoplasm |
| 558 | Q60682 | Klra4 (inclu | 5.05E+06 | 1.13E+06 | 6.54E+05 | 3 | Plasma Membrane |
| 559 | P97807 | FH | 5.05E+06 | 3.69E+06 | 8.04E+05 | 21 | Cytoplasm |
| 560 | Q99JY9 | ACTR3 | 5.02E+06 | 4.60E+06 | 1.19E+06 | 15 | Plasma Membrane |
| 561 | Q8BFR5 | TUFM | 5.02E+06 | 2.35E+06 | 5.55E+05 | 18 | Cytoplasm |
| 562 | Q6ZWN5 | RPS9 | 5.01E+06 | 2.16E+06 | 7.20E+05 | 9 | Cytoplasm |
| 563 | P17047 | LAMP2 | 5.01E+06 | 2.47E+06 | 1.01E+06 | 6 | Plasma Membrane |
| 564 | O35864 | COPS5 | 5.00E+06 | 2.96E+06 | 1.21E+06 | 6 | Nucleus |
| 565 | Q9JMA1 | USP14 | 5.00E+06 | 1.65E+06 | 6.72E+05 | 6 | Cytoplasm |
| 566 | Q9D6K8 | FUNDC2 | 5.00E+06 | 2.17E+06 | 8.88E+05 | 6 | Cytoplasm |
| 567 | Q60857 | SLC6A4 | 4.99E+06 | 5.57E+05 | 3.21E+05 | 3 | Plasma Membrane |
| 568 | Q9QXT0 | CNPY2 | 4.98E+06 | 1.63E+06 | 5.44E+05 | 9 | Plasma Membrane |
| 569 | Q8BH59 | SLC25A12 | 4.94E+06 | 3.26E+06 | 9.40E+05 | 12 | Cytoplasm |
| 570 | P26041 | MSN | 4.92E+06 | 2.70E+06 | 4.50E+05 | 36 | Plasma Membrane |
| 571 | P35282 | RAB21 | 4.92E+06 | 3.77E+06 | 7.70E+05 | 24 | Cytoplasm |
| 572 | Q9DBG6 | RPN2 | 4.91E+06 | 3.81E+06 | 6.96E+05 | 30 | Cytoplasm |
| 573 | Q8VCF0 | MAVS | 4.90E+06 | 2.04E+06 | 8.34E+05 | 6 | Cytoplasm |
| 574 | Q9QVN7 | TCEA2 | 4.89E+06 | 6.23E+05 | 3.60E+05 | 3 | Nucleus |
| 575 | Q61820 | Rasl2-9 | 4.88E+06 | 1.00E+06 | 4.09E+05 | 6 | Nucleus |
| 576 | Q64521 | GPD2 | 4.86E+06 | 2.72E+06 | 4.97E+05 | 30 | Cytoplasm |
| 577 | Q9QZQ8 | H2AFY | 4.86E+06 | 1.62E+06 | 9.38E+05 | 3 | Nucleus |
| 578 | Q91YQ5 | RPN1 | 4.85E+06 | 2.67E+06 | 5.45E+05 | 24 | Cytoplasm |
| 579 | P97352 | S100A13 | 4.84E+06 | 1.55E+06 | 8.95E+05 | 3 | Cytoplasm |
| 580 | P46460 | NSF | 4.82E+06 | 2.57E+06 | 3.96E+05 | 42 | Cytoplasm |
| 581 | P61161 | ACTR2 | 4.82E+06 | 2.12E+06 | 7.06E+05 | 9 | Plasma Membrane |
| 582 | O08547 | SEC22B | 4.81E+06 | 2.40E+06 | 4.89E+05 | 24 | Cytoplasm |
| 583 | Q7TMR0 | PRCP | 4.79E+06 | 1.96E+06 | 1.13E+06 | 3 | Cytoplasm |
| 584 | Q9D7G0 | PRPS1 | 4.78E+06 | 1.97E+06 | 4.64E+05 | 18 | Cytoplasm |
| 585 | O35350 | CAPN1 | 4.78E+06 | 2.41E+06 | 5.26E+05 | 21 | Cytoplasm |
| 586 | P06151 | LDHA | 4.75E+06 | 2.13E+06 | 4.10E+05 | 27 | Cytoplasm |
| 587 | Q8R016 | BLMH | 4.75E+06 | 2.21E+06 | 4.82E+05 | 21 | Cytoplasm |
| 588 | P05132 | PRKACA | 4.75E+06 | 7.03E+05 | 4.06E+05 | 3 | Cytoplasm |
| 589 | P62908 | RPS3 | 4.75E+06 | 2.49E+06 | 5.08E+05 | 24 | Cytoplasm |
| 590 | P18760 | CFL1 | 4.73E+06 | 3.89E+06 | 1.00E+06 | 15 | Nucleus |
| 591 | P09671 | SOD2 | 4.73E+06 | 2.97E+06 | 9.89E+05 | 9 | Cytoplasm |
| 592 | O35887 | CALU | 4.70E+06 | 3.33E+06 | 6.79E+05 | 24 | Cytoplasm |
| 593 | O88792 | F11R | 4.70E+06 | 3.73E+06 | 9.63E+05 | 15 | Plasma Membrane |
| 594 | Q9DC69 | NDUFA9 | 4.67E+06 | 4.32E+06 | 1.44E+06 | 9 | Cytoplasm |
| 595 | P39876 | TIMP3 | 4.67E+06 | 4.71E+06 | 1.36E+06 | 12 | Extracellular Space |
| 596 | Q9CQS8 | SEC61B | 4.64E+06 | 2.18E+06 | 8.90E+05 | 6 | Cytoplasm |
| 597 | Q9JJK2 | LANCL2 | 4.64E+06 | 2.58E+06 | 4.30E+05 | 36 | Plasma Membrane |
| 598 | Q3UGC7 | EIF3J | 4.63E+06 | 3.66E+06 | 9.46E+05 | 15 | Cytoplasm |
| 599 | Q922D4 | PPP6R3 | 4.62E+06 | 2.43E+06 | 4.05E+05 | 36 | Cytoplasm |
| 600 | Q8VED5 | KRT79 | 4.60E+06 | 5.17E+06 | 2.98E+06 | 3 | Extracellular Space |
| 601 | Q9EP69 | SACM1L | 4.59E+06 | 3.11E+06 | 4.12E+05 | 57 | Cytoplasm |
| 602 | Q99JY0 | HADHB | 4.59E+06 | 4.01E+06 | 8.76E+05 | 21 | Cytoplasm |
| 603 | A2ADY9 | DDI2 | 4.58E+06 | 1.21E+06 | 4.94E+05 | 6 | Plasma Membrane |
| 604 | O88668 | CREG1 | 4.57E+06 | 8.58E+05 | 3.50E+05 | 6 | Nucleus |
| 605 | Q61207 | PSAP | 4.56E+06 | 7.45E+05 | 4.30E+05 | 3 | Extracellular Space |
| 606 | Q6PAR5 | GAPVD1 | 4.56E+06 | 2.31E+06 | 3.24E+05 | 51 | Cytoplasm |
| 607 | Q8BGD9 | EIF4B | 4.54E+06 | 2.35E+06 | 5.55E+05 | 18 | Cytoplasm |
| 608 | Q80TH2 | ERBB2IP | 4.53E+06 | 1.54E+06 | 3.15E+05 | 24 | Plasma Membrane |
| 609 | Q3UFK8 | FRMD8 | 4.53E+06 | 1.35E+06 | 7.79E+05 | 3 | Cytoplasm |
| 610 | Q9JI39 | ABCB10 | 4.52E+06 | 5.03E+06 | 8.39E+05 | 36 | Cytoplasm |
| 611 | Q80WT2 | DMRT3 | 4.52E+06 | 6.63E+05 | 3.83E+05 | 3 | Nucleus |
| 612 | Q62469 | ITGA2 | 4.50E+06 | 2.20E+06 | 3.83E+05 | 33 | Plasma Membrane |
| 613 | O08756 | HSD17B10 | 4.48E+06 | 3.45E+06 | 9.96E+05 | 12 | Cytoplasm |
| 614 | Q9D771 | TMEM206 | 4.48E+06 | 4.45E+06 | 1.48E+06 | 9 | Plasma Membrane |
| 615 | P08003 | PDIA4 | 4.45E+06 | 2.42E+06 | 4.94E+05 | 24 | Cytoplasm |
| 616 | Q91VH6 | MEMO1 | 4.45E+06 | 3.57E+06 | 9.22E+05 | 15 | Cytoplasm |
| 617 | Q02053 | UBA1 | 4.44E+06 | 2.00E+06 | 3.09E+05 | 42 | Cytoplasm |
| 618 | Q9CQB5 | CISD2 | 4.43E+06 | 1.90E+06 | 4.90E+05 | 15 | Cytoplasm |
| 619 | P18052 | PTPRA | 4.43E+06 | 2.93E+06 | 8.45E+05 | 12 | Plasma Membrane |
| 620 | Q8BPM0 | DAAM1 | 4.42E+06 | 2.16E+06 | 3.60E+05 | 36 | Cytoplasm |
| 621 | P52480 | PKM | 4.41E+06 | 6.47E+05 | 3.74E+05 | 3 | Cytoplasm |
| 622 | P68254 | YWHAQ | 4.41E+06 | 1.15E+06 | 3.84E+05 | 9 | Cytoplasm |
| 623 | Q9CZ30 | OLA1 | 4.41E+06 | 1.30E+06 | 7.51E+05 | 3 | Cytoplasm |
| 624 | O89086 | RBM3 | 4.40E+06 | 3.66E+06 | 1.22E+06 | 9 | Cytoplasm |
| 625 | Q8CDN6 | TXNL1 | 4.40E+06 | 2.88E+06 | 5.87E+05 | 24 | Cytoplasm |
| 626 | P0C0A3 | CHMP6 | 4.37E+06 | 2.15E+05 | 1.24E+05 | 3 | Cytoplasm |
| 627 | Q99JR1 | SFXN1 | 4.33E+06 | 2.21E+06 | 5.22E+05 | 18 | Cytoplasm |
| 628 | Q9D8E6 | RPL4 | 4.32E+06 | 1.57E+06 | 4.53E+05 | 12 | Cytoplasm |
| 629 | P63330 | PPP2CA | 4.31E+06 | 1.37E+06 | 3.54E+05 | 15 | Cytoplasm |
| 630 | P61022 | CHP1 | 4.28E+06 | 1.39E+06 | 3.28E+05 | 18 | Cytoplasm |
| 631 | Q9CZM2 | RPL15 | 4.28E+06 | 8.18E+05 | 2.73E+05 | 9 | Cytoplasm |
| 632 | P62754 | RPS6 | 4.28E+06 | 2.12E+06 | 8.65E+05 | 6 | Cytoplasm |
| 633 | Q8R2Y8 | PTRH2 | 4.27E+06 | 5.03E+06 | 2.05E+06 | 6 | Cytoplasm |
| 634 | P62270 | RPS18 | 4.26E+06 | 1.02E+06 | 3.40E+05 | 9 | Cytoplasm |
| 635 | Q8CHK3 | MBOAT7 | 4.25E+06 | 1.51E+06 | 8.74E+05 | 3 | Plasma Membrane |
| 636 | P10518 | ALAD | 4.25E+06 | 1.31E+07 | 1.89E+06 | 48 | Cytoplasm |
| 637 | Q3THW5 | H2AFV | 4.25E+06 | 9.83E+05 | 5.67E+05 | 3 | Nucleus |
| 638 | P35821 | PTPN1 | 4.25E+06 | 5.58E+05 | 3.22E+05 | 3 | Cytoplasm |
| 639 | Q9WVK4 | EHD1 | 4.24E+06 | 2.83E+06 | 8.16E+05 | 12 | Cytoplasm |
| 640 | Q9WVG6 | CARM1 | 4.24E+06 | 1.48E+06 | 4.93E+05 | 9 | Nucleus |
| 641 | Q9DB05 | NAPA | 4.23E+06 | 8.47E+05 | 4.89E+05 | 3 | Cytoplasm |
| 642 | Q9DCD0 | PGD | 4.23E+06 | 5.92E+05 | 3.42E+05 | 3 | Cytoplasm |
| 643 | Q99L45 | EIF2S2 | 4.23E+06 | 1.51E+06 | 5.04E+05 | 9 | Cytoplasm |
| 644 | P62855 | RPS26 | 4.22E+06 | 2.10E+05 | 1.21E+05 | 3 | Cytoplasm |
| 645 | P42125 | ECI1 | 4.21E+06 | 2.49E+06 | 1.02E+06 | 6 | Cytoplasm |
| 646 | Q99KB8 | HAGH | 4.20E+06 | 3.41E+06 | 1.14E+06 | 9 | Cytoplasm |
| 647 | P51655 | GPC4 | 4.19E+06 | 1.54E+06 | 5.12E+05 | 9 | Plasma Membrane |
| 648 | Q9QYF1 | RDH11 | 4.15E+06 | 1.88E+06 | 3.85E+05 | 24 | Cytoplasm |
| 649 | Q64446 | ATP7B | 4.15E+06 | 4.55E+06 | 9.28E+05 | 24 | Cytoplasm |
| 650 | Q61335 | BCAP31 | 4.15E+06 | 9.06E+04 | 5.23E+04 | 3 | Cytoplasm |
| 651 | Q99KE1 | ME2 | 4.15E+06 | 2.87E+06 | 8.29E+05 | 12 | Cytoplasm |
| 652 | P62267 | RPS23 | 4.15E+06 | 5.14E+05 | 2.97E+05 | 3 | Cytoplasm |
| 653 | Q8BNI4 | DERL2 | 4.15E+06 | 9.89E+05 | 5.71E+05 | 3 | Cytoplasm |
| 654 | Q9CQR6 | PPP6C | 4.11E+06 | 1.77E+06 | 7.24E+05 | 6 | Nucleus |
| 655 | Q3TWL2 | TMEM55B | 4.04E+06 | 1.13E+06 | 3.27E+05 | 12 | Cytoplasm |
| 656 | Q920G8 | SLC25A37 | 4.04E+06 | 2.62E+06 | 4.56E+05 | 33 | Cytoplasm |
| 657 | O08739 | AMPD3 | 4.04E+06 | 2.01E+06 | 4.10E+05 | 24 | Cytoplasm |
| 658 | Q9CRB9 | CHCHD3 | 4.03E+06 | 9.30E+05 | 5.37E+05 | 3 | Cytoplasm |
| 659 | Q3U898 | MYEOV2 | 4.02E+06 | 1.44E+06 | 8.29E+05 | 3 | Nucleus |
| 660 | P48962 | SLC25A4 | 4.02E+06 | 1.47E+06 | 4.91E+05 | 9 | Cytoplasm |
| 661 | Q9WUP7 | UCHL5 | 4.01E+06 | 3.75E+06 | 1.08E+06 | 12 | Cytoplasm |
| 662 | P14115 | RPL27A | 3.99E+06 | 1.23E+05 | 7.10E+04 | 3 | Cytoplasm |
| 663 | Q80VY9 | DHX33 | 3.97E+06 | 1.06E+06 | 6.10E+05 | 3 | Nucleus |
| 664 | Q6P5E4 | UGGT1 | 3.95E+06 | 2.79E+06 | 5.70E+05 | 24 | Cytoplasm |
| 665 | P47963 | RPL13 | 3.94E+06 | 1.51E+06 | 4.36E+05 | 12 | Nucleus |
| 666 | Q8K3F2 | MMP21 | 3.94E+06 | 1.12E+06 | 6.45E+05 | 3 | Extracellular Space |
| 667 | Q99LC5 | ETFA | 3.93E+06 | 3.44E+06 | 6.63E+05 | 27 | Cytoplasm |
| 668 | Q91W53 | GOLGA7 | 3.92E+06 | 5.40E+05 | 2.20E+05 | 6 | Cytoplasm |
| 669 | P70670 | NACA | 3.92E+06 | 2.65E+06 | 8.85E+05 | 9 | Cytoplasm |
| 670 | P70697 | UROD | 3.92E+06 | 7.09E+05 | 4.10E+05 | 3 | Cytoplasm |
| 671 | A2AN08 | UBR4 | 3.90E+06 | 1.81E+06 | 1.44E+05 | 159 | Nucleus |
| 672 | P70297 | STAM | 3.90E+06 | 1.78E+06 | 5.93E+05 | 9 | Cytoplasm |
| 673 | P21981 | TGM2 | 3.90E+06 | 2.46E+06 | 1.00E+06 | 6 | Cytoplasm |
| 674 | Q9CWJ9 | ATIC | 3.90E+06 | 2.44E+06 | 4.46E+05 | 30 | Cytoplasm |
| 675 | Q3UUG6 | TBC1D24 | 3.89E+06 | 2.85E+06 | 1.17E+06 | 6 | Cytoplasm |
| 676 | P17426 | AP2A1 | 3.89E+06 | 1.88E+06 | 6.26E+05 | 9 | Cytoplasm |
| 677 | Q8BMS1 | HADHA | 3.88E+06 | 2.55E+06 | 3.93E+05 | 42 | Cytoplasm |
| 678 | P62137 | PPP1CA | 3.88E+06 | 2.25E+06 | 5.30E+05 | 18 | Cytoplasm |
| 679 | P28571 | SLC6A9 | 3.88E+06 | 3.42E+06 | 1.40E+06 | 6 | Plasma Membrane |
| 680 | B2RPV6 | MMRN1 | 3.87E+06 | 1.16E+06 | 2.02E+05 | 33 | Extracellular Space |
| 681 | Q9JHR7 | IDE | 3.86E+06 | 6.10E+05 | 2.49E+05 | 6 | Extracellular Space |
| 682 | Q9WUM5 | SUCLG1 | 3.86E+06 | 1.49E+06 | 4.97E+05 | 9 | Cytoplasm |
| 683 | Q8K411 | PITRM1 | 3.85E+06 | 2.96E+06 | 6.45E+05 | 21 | Cytoplasm |
| 684 | Q6ZWX6 | EIF2S1 | 3.85E+06 | 1.30E+06 | 3.07E+05 | 18 | Cytoplasm |
| 685 | P59017 | BCL2L13 | 3.84E+06 | 1.40E+06 | 4.68E+05 | 9 | Cytoplasm |
| 686 | Q3TWN3 | CNNM2 | 3.84E+06 | 3.80E+05 | 1.55E+05 | 6 | Plasma Membrane |
| 687 | Q9D172 | C21orf33/l | 3.82E+06 | 4.35E+06 | 1.78E+06 | 6 | Cytoplasm |
| 688 | Q99K41 | EMILIN1 | 3.81E+06 | 2.90E+06 | 1.18E+06 | 6 | Extracellular Space |
| 689 | Q8BIG7 | COMTD1 | 3.81E+06 | 9.28E+05 | 5.36E+05 | 3 | Plasma Membrane |
| 690 | P70295 | AUP1 | 3.79E+06 | 3.77E+05 | 2.18E+05 | 3 | Cytoplasm |
| 691 | Q9D8N0 | EEF1G | 3.78E+06 | 2.14E+06 | 7.13E+05 | 9 | Cytoplasm |
| 692 | Q922Q8 | LRRC59 | 3.77E+06 | 8.34E+05 | 4.82E+05 | 3 | Cytoplasm |
| 693 | P70658 | CXCR4 | 3.76E+06 | 1.87E+05 | 1.08E+05 | 3 | Plasma Membrane |
| 694 | Q80UJ7 | RAB3GAP1 | 3.75E+06 | 2.42E+06 | 8.08E+05 | 9 | Cytoplasm |

| | | | | | | | |
|---|---|---|---|---|---|---|---|
| 556 | Q06890 | CLU | 3.93E+06 | 4.60E+06 | 1.53E+06 | 9 | Cytoplasm |
| 557 | Q9WUP7 | UCHL5 | 3.93E+06 | 3.21E+06 | 9.27E+05 | 12 | Cytoplasm |
| 558 | Q61820 | Rasl2-9 | 3.92E+06 | 6.55E+05 | 2.67E+05 | 6 | Nucleus |
| 559 | P56391 | COX6B1 | 3.92E+06 | 3.39E+06 | 1.95E+06 | 3 | Cytoplasm |
| 560 | Q7TQ95 | KIAA1715 | 3.92E+06 | 1.15E+06 | 2.98E+05 | 15 | Cytoplasm |
| 561 | Q9D0R2 | TARS | 3.90E+06 | 3.31E+06 | 1.35E+06 | 6 | Nucleus |
| 562 | O88587 | COMT | 3.90E+06 | 7.57E+05 | 2.19E+05 | 12 | Cytoplasm |
| 563 | P56399 | USP5 | 3.89E+06 | 1.82E+06 | 2.92E+05 | 39 | Cytoplasm |
| 564 | Q9D6K8 | FUNDC2 | 3.89E+06 | 1.76E+06 | 7.19E+05 | 6 | Cytoplasm |
| 565 | P35980 | RPL18 | 3.89E+06 | 2.74E+06 | 7.90E+05 | 12 | Cytoplasm |
| 566 | Q9DB05 | NAPA | 3.88E+06 | 2.17E+05 | 1.25E+05 | 3 | Cytoplasm |
| 567 | P17710 | HK1 | 3.88E+06 | 3.23E+06 | 5.18E+05 | 39 | Cytoplasm |
| 568 | P47857 | PFKM | 3.87E+06 | 1.27E+06 | 2.99E+05 | 18 | Cytoplasm |
| 569 | P06151 | LDHA | 3.86E+06 | 1.79E+06 | 3.44E+05 | 27 | Cytoplasm |
| 570 | P61022 | CHP1 | 3.84E+06 | 1.37E+06 | 3.24E+05 | 18 | Cytoplasm |
| 571 | Q99J99 | MPST | 3.83E+06 | 8.18E+06 | 3.34E+06 | 6 | Cytoplasm |
| 572 | Q91VR2 | ATP5C1 | 3.83E+06 | 4.74E+06 | 1.37E+06 | 12 | Cytoplasm |
| 573 | Q99KV1 | DNAJB11 | 3.82E+06 | 1.62E+06 | 5.40E+05 | 9 | Cytoplasm |
| 574 | P61202 | Cops2 | 3.81E+06 | 2.27E+06 | 5.87E+05 | 15 | Nucleus |
| 575 | P56135 | ATP5J2 | 3.81E+06 | 1.88E+06 | 1.08E+06 | 3 | Cytoplasm |
| 576 | P01899, P01901, P01902 | HLA-A | 3.81E+06 | 5.48E+06 | 1.06E+06 | 27 | Plasma Membrane |
| 577 | P62082 | RPS7 | 3.81E+06 | 4.00E+06 | 1.33E+06 | 9 | Cytoplasm |
| 578 | Q8CIG8 | PRMT5 | 3.81E+06 | 2.63E+06 | 1.08E+06 | 6 | Cytoplasm |
| 579 | Q9DBG6 | RPN2 | 3.80E+06 | 3.56E+06 | 6.50E+05 | 30 | Cytoplasm |
| 580 | Q99LC3 | NDUFA10 | 3.80E+06 | 2.22E+06 | 9.07E+05 | 6 | Cytoplasm |
| 581 | Q9CQQ7 | ATP5F1 | 3.80E+06 | 2.57E+06 | 1.48E+06 | 3 | Cytoplasm |
| 582 | Q9JMA1 | USP14 | 3.79E+06 | 1.19E+06 | 4.87E+05 | 6 | Cytoplasm |
| 583 | P62897 | CYCS | 3.77E+06 | 3.30E+06 | 8.52E+05 | 15 | Cytoplasm |
| 584 | Q62425 | NDUFA4 | 3.77E+06 | 2.39E+06 | 9.77E+05 | 6 | Cytoplasm |
| 585 | Q6R0H7 | GNAS | 3.77E+06 | 9.54E+05 | 3.89E+05 | 6 | Plasma Membrane |
| 586 | P25444 | RPS2 | 3.75E+06 | 2.49E+06 | 7.18E+05 | 12 | Cytoplasm |
| 587 | P62751 | Rpl23a | 3.75E+06 | 4.28E+06 | 1.75E+06 | 6 | Nucleus |
| 588 | Q9JKR6 | HYOU1 | 3.74E+06 | 9.44E+05 | 2.06E+05 | 21 | Cytoplasm |
| 589 | Q9CR57 | RPL14 | 3.74E+06 | 3.10E+06 | 1.03E+06 | 9 | Cytoplasm |
| 590 | P14869 | RPLP0 | 3.74E+06 | 3.65E+06 | 7.45E+05 | 24 | Cytoplasm |
| 591 | Q6ZQI3 | MLEC | 3.72E+06 | 1.74E+06 | 4.50E+05 | 15 | Plasma Membrane |
| 592 | P15327 | BPGM | 3.72E+06 | 3.33E+06 | 9.60E+05 | 12 | Extracellular Space |
| 593 | Q8CDN6 | TXNL1 | 3.72E+06 | 3.28E+06 | 6.70E+05 | 24 | Cytoplasm |
| 594 | P61161 | ACTR2 | 3.72E+06 | 1.57E+06 | 5.25E+05 | 9 | Plasma Membrane |
| 595 | Q9Z0N1 | EIF2S3 | 3.71E+06 | 2.47E+06 | 4.31E+05 | 33 | Cytoplasm |
| 596 | Q9QYA2 | TOMM40 | 3.69E+06 | 1.80E+06 | 1.04E+06 | 3 | Cytoplasm |
| 597 | Q99KB8 | HAGH | 3.69E+06 | 2.77E+06 | 9.24E+05 | 9 | Cytoplasm |
| 598 | Q8VCF0 | MAVS | 3.69E+06 | 3.75E+06 | 1.53E+06 | 6 | Cytoplasm |
| 599 | Q9Z2L7 | CRLF3 | 3.69E+06 | 3.15E+05 | 1.82E+05 | 3 | Cytoplasm |
| 600 | Q8CBB9 | RSAD2 | 3.68E+06 | 6.96E+06 | 1.42E+06 | 24 | Cytoplasm |
| 601 | Q8R5J9 | ARL6IP5 | 3.65E+06 | 9.90E+05 | 5.72E+05 | 3 | Cytoplasm |
| 602 | O09172 | GCLM | 3.65E+06 | 8.21E+05 | 2.74E+05 | 9 | Cytoplasm |
| 603 | Q9QVN7 | TCEA2 | 3.65E+06 | 2.02E+06 | 1.17E+06 | 3 | Nucleus |
| 604 | P48722 | HSPA4L | 3.64E+06 | 1.66E+06 | 4.28E+05 | 15 | Cytoplasm |
| 605 | P18052 | PTPRA | 3.64E+06 | 2.93E+06 | 8.47E+05 | 12 | Plasma Membrane |
| 606 | A2ASS6 | TTN | 3.63E+06 | 2.73E+05 | 1.58E+05 | 3 | Cytoplasm |
| 607 | Q8BPM0 | DAAM1 | 3.63E+06 | 1.89E+06 | 3.15E+05 | 36 | Cytoplasm |
| 608 | P60867 | RPS20 | 3.62E+06 | 1.83E+06 | 7.45E+05 | 6 | Cytoplasm |
| 609 | P63325 | RPS10 | 3.62E+06 | 1.88E+06 | 1.08E+06 | 3 | Cytoplasm |
| 610 | O08547 | SEC22B | 3.62E+06 | 1.79E+06 | 3.66E+05 | 24 | Cytoplasm |
| 611 | P35279 | RAB6A | 3.62E+06 | 1.91E+06 | 7.81E+05 | 6 | Cytoplasm |
| 612 | Q99JY0 | HADHB | 3.61E+06 | 4.01E+06 | 8.75E+05 | 21 | Cytoplasm |
| 613 | Q923D2 | BLVRB | 3.60E+06 | 2.07E+06 | 5.97E+05 | 12 | Cytoplasm |
| 614 | P62900 | RPL31 | 3.60E+06 | 2.72E+06 | 1.11E+06 | 6 | Cytoplasm |
| 615 | P49769 | PSEN1 | 3.60E+06 | 1.79E+06 | 7.32E+05 | 6 | Plasma Membrane |
| 616 | P21981 | TGM2 | 3.59E+06 | 1.79E+06 | 7.30E+05 | 6 | Cytoplasm |
| 617 | Q9D0M1 | PRPSAP1 | 3.58E+06 | 1.79E+06 | 4.61E+05 | 15 | Extracellular Space |
| 618 | Q80UJ7 | RAB3GAP1 | 3.57E+06 | 2.56E+06 | 8.54E+05 | 9 | Cytoplasm |
| 619 | Q9DCC4 | PYCRL | 3.57E+06 | 1.12E+06 | 3.72E+05 | 9 | Cytoplasm |
| 620 | O88545 | COPS6 | 3.55E+06 | 2.03E+06 | 5.24E+05 | 15 | Nucleus |
| 621 | P62889 | RPL30 | 3.55E+06 | 2.18E+06 | 1.26E+06 | 3 | Cytoplasm |
| 622 | Q9JI39 | ABCB10 | 3.54E+06 | 4.56E+06 | 7.61E+05 | 36 | Cytoplasm |
| 623 | Q8R5H1 | USP15 | 3.54E+06 | 1.64E+06 | 2.37E+05 | 48 | Cytoplasm |
| 624 | Q9CRY7 | GDPD1 | 3.53E+06 | 7.07E+05 | 4.08E+05 | 3 | Plasma Membrane |
| 625 | Q9WUM4 | CORO1C | 3.53E+06 | 3.00E+06 | 1.22E+06 | 6 | Cytoplasm |
| 626 | Q02053 | UBA1 | 3.53E+06 | 1.62E+06 | 2.49E+05 | 42 | Cytoplasm |
| 627 | Q8VED5 | KRT79 | 3.53E+06 | 3.36E+06 | 1.94E+06 | 3 | Extracellular Space |
| 628 | Q80ZW2 | THEM6 | 3.53E+06 | 1.36E+06 | 3.52E+05 | 15 | Extracellular Space |
| 629 | P05977 | MYL1 | 3.53E+06 | 2.34E+06 | 1.35E+06 | 3 | Cytoplasm |
| 630 | Q9JL56 | GDE1 | 3.52E+06 | 2.45E+06 | 8.17E+05 | 9 | Plasma Membrane |
| 631 | Q8BGU5 | CCNY | 3.50E+06 | 3.78E+05 | 1.55E+05 | 6 | Nucleus |
| 632 | P70460 | VASP | 3.50E+06 | 2.11E+06 | 1.22E+06 | 3 | Plasma Membrane |
| 633 | Q8CHK3 | MBOAT7 | 3.49E+06 | 3.06E+06 | 1.77E+06 | 3 | Plasma Membrane |
| 634 | Q99LD4 | GPS1 | 3.47E+06 | 1.85E+06 | 4.78E+05 | 15 | Cytoplasm |
| 635 | Q62415 | PPP1R13B | 3.47E+06 | 3.88E+05 | 2.24E+05 | 3 | Cytoplasm |
| 636 | Q8BFR5 | TUFM | 3.45E+06 | 3.02E+06 | 7.13E+05 | 18 | Cytoplasm |
| 637 | P18760 | CFL1 | 3.44E+06 | 4.35E+06 | 1.12E+06 | 15 | Nucleus |
| 638 | P62852 | RPS25 | 3.43E+06 | 2.00E+06 | 6.65E+05 | 9 | Cytoplasm |
| 639 | Q8BYC6 | TAOK3 | 3.42E+06 | 1.81E+06 | 3.69E+05 | 24 | Cytoplasm |
| 640 | Q9DBX3 | SUSD2 | 3.38E+06 | 6.74E+05 | 2.25E+05 | 9 | Extracellular Space |
| 641 | Q9DCW4 | ETFB | 3.35E+06 | 4.48E+06 | 1.29E+06 | 12 | Cytoplasm |
| 642 | Q91YQ5 | RPN1 | 3.34E+06 | 2.81E+06 | 5.73E+05 | 24 | Cytoplasm |
| 643 | Q8QZY6 | TSPAN14 | 3.34E+06 | 1.28E+06 | 7.37E+05 | 3 | Plasma Membrane |
| 644 | Q9QUM4 | SLAMF1 | 3.32E+06 | 1.51E+06 | 5.04E+05 | 9 | Plasma Membrane |
| 645 | O88668 | CREG1 | 3.32E+06 | 9.30E+05 | 3.80E+05 | 6 | Nucleus |
| 646 | Q8VDL4 | ADPGK | 3.32E+06 | 3.73E+05 | 2.15E+05 | 3 | Plasma Membrane |
| 647 | Q9QY76 | VAPB | 3.30E+06 | 2.43E+06 | 9.91E+05 | 6 | Plasma Membrane |
| 648 | P97855 | G3BP1 | 3.29E+06 | 8.86E+05 | 3.62E+05 | 6 | Nucleus |
| 649 | Q9CR21 | NDUFAB1 | 3.28E+06 | 3.16E+06 | 1.29E+06 | 6 | Cytoplasm |
| 650 | Q99L43 | CDS2 | 3.28E+06 | 1.15E+06 | 6.62E+05 | 3 | Cytoplasm |
| 651 | Q9EST8 | NFKBIZ | 3.28E+06 | 3.45E+05 | 1.99E+05 | 3 | Nucleus |
| 652 | Q9JLV5 | CUL3 | 3.27E+06 | 6.93E+05 | 2.83E+05 | 6 | Nucleus |
| 653 | P0C0A3 | CHMP6 | 3.25E+06 | 3.83E+05 | 2.21E+05 | 3 | Cytoplasm |
| 654 | P35821 | PTPN1 | 3.25E+06 | 5.48E+05 | 3.16E+05 | 3 | Cytoplasm |
| 655 | P62242 | RPS8 | 3.25E+06 | 2.65E+06 | 6.85E+05 | 15 | Cytoplasm |
| 656 | P84099 | RPL19 | 3.25E+06 | 3.53E+06 | 2.04E+06 | 3 | Cytoplasm |
| 657 | P50247 | AHCY | 3.25E+06 | 9.08E+05 | 3.03E+05 | 9 | Cytoplasm |
| 658 | Q9WVG6 | CARM1 | 3.25E+06 | 1.10E+06 | 3.67E+05 | 9 | Nucleus |
| 659 | Q8BWT1 | ACAA2 | 3.24E+06 | 3.13E+06 | 7.39E+05 | 18 | Cytoplasm |
| 660 | P54731 | FAF1 | 3.24E+06 | 6.40E+05 | 2.13E+05 | 9 | Nucleus |
| 661 | Q9CQS8 | SEC61B | 3.23E+06 | 1.80E+06 | 7.34E+05 | 6 | Cytoplasm |
| 662 | P01872 | IGHM | 3.23E+06 | 1.50E+06 | 4.34E+05 | 12 | Plasma Membrane |
| 663 | Q8VEE1 | LMCD1 | 3.23E+06 | 6.25E+06 | 3.61E+06 | 3 | Cytoplasm |
| 664 | P17047 | LAMP2 | 3.23E+06 | 2.74E+06 | 1.12E+06 | 6 | Plasma Membrane |
| 665 | Q60737 | CSNK2A1 | 3.22E+06 | 6.90E+05 | 3.98E+05 | 3 | Nucleus |
| 666 | Q05144 | RAC2 | 3.21E+06 | 2.18E+06 | 1.26E+06 | 3 | Cytoplasm |
| 667 | P63330 | PPP2CA | 3.21E+06 | 1.05E+06 | 2.71E+05 | 15 | Cytoplasm |
| 668 | Q80TH2 | ERBB2IP | 3.20E+06 | 1.92E+06 | 3.92E+05 | 24 | Plasma Membrane |
| 669 | Q9ES46 | PARVB | 3.20E+06 | 4.23E+06 | 9.98E+05 | 18 | Cytoplasm |
| 670 | P42567 | EPS15 | 3.20E+06 | 8.66E+05 | 2.24E+05 | 15 | Cytoplasm |
| 671 | P25911 | LYN | 3.20E+06 | 4.39E+06 | 8.96E+05 | 24 | Cytoplasm |
| 672 | Q60857 | SLC6A4 | 3.20E+06 | 1.33E+06 | 7.67E+05 | 3 | Plasma Membrane |
| 673 | P62862 | FAU | 3.19E+06 | 1.60E+06 | 9.22E+05 | 3 | Cytoplasm |
| 674 | Q923U0 | TOM1L1 | 3.19E+06 | 1.56E+06 | 6.38E+05 | 6 | Cytoplasm |
| 675 | Q80WQ2 | VAC14 | 3.18E+06 | 5.20E+05 | 3.00E+05 | 3 | Cytoplasm |
| 676 | P14131 | RPS16 | 3.18E+06 | 2.77E+06 | 1.13E+06 | 6 | Cytoplasm |
| 677 | Q9CR64 | TMEM167A | 3.17E+06 | 5.25E+05 | 3.03E+05 | 3 | Extracellular Space |
| 678 | Q08093 | CNN2 | 3.15E+06 | 1.57E+06 | 9.05E+05 | 3 | Cytoplasm |
| 679 | Q64521 | GPD2 | 3.15E+06 | 2.86E+06 | 5.23E+05 | 30 | Cytoplasm |
| 680 | Q9DC69 | NDUFA9 | 3.15E+06 | 3.58E+06 | 1.19E+06 | 9 | Cytoplasm |
| 681 | Q9CQV8 | YWHAB | 3.14E+06 | 2.48E+06 | 5.85E+05 | 18 | Cytoplasm |
| 682 | Q9D172 | C21orf33/LOC102724023 | 3.13E+06 | 4.57E+06 | 1.86E+06 | 6 | Cytoplasm |
| 683 | Q6X893 | SLC44A1 | 3.13E+06 | 1.40E+06 | 4.67E+05 | 9 | Plasma Membrane |
| 684 | Q8R574 | PRPSAP2 | 3.12E+06 | 6.94E+05 | 2.83E+05 | 6 | Cytoplasm |
| 685 | P97807 | FH | 3.12E+06 | 3.35E+06 | 7.31E+05 | 21 | Cytoplasm |
| 686 | O55022 | PGRMC1 | 3.12E+06 | 2.20E+06 | 1.27E+06 | 3 | Plasma Membrane |
| 687 | Q9Z0F8 | ADAM17 | 3.12E+06 | 2.24E+06 | 9.14E+05 | 6 | Plasma Membrane |
| 688 | P70670 | NACA | 3.11E+06 | 1.77E+06 | 5.90E+05 | 9 | Cytoplasm |
| 689 | Q8K2J7 | RELL1 | 3.11E+06 | 1.20E+06 | 3.99E+05 | 9 | Plasma Membrane |
| 690 | P68369 | TUBA1A | 3.11E+06 | 1.15E+06 | 6.63E+05 | 3 | Cytoplasm |
| 691 | P62342 | SELT | 3.10E+06 | 2.62E+05 | 1.51E+05 | 3 | Cytoplasm |
| 692 | P35285 | RAB22A | 3.10E+06 | 4.12E+05 | 1.68E+05 | 6 | Cytoplasm |
| 693 | Q9QUT0 | RHAG | 3.10E+06 | 4.41E+06 | 1.27E+06 | 12 | Plasma Membrane |
| 694 | Q9Z1Q2 | ABHD16A | 3.10E+06 | 2.10E+06 | 7.01E+05 | 9 | Cytoplasm |

| | | | | | | | |
|---|---|---|---|---|---|---|---|
| 556 | P47753 | CAPZA1 | 4106585 | 2067443 | 844030 | 6 | Cytoplasm |
| 557 | P01899, P01901, P01902 | HLA-A | 4092255 | 4784582 | 920793 | 27 | Plasma Membrane |
| 558 | P47963 | RPL13 | 4083027 | 2369398 | 683986 | 12 | Nucleus |
| 559 | Q791V5 | MTCH2 | 4065314 | 2554615 | 1474908 | 3 | Cytoplasm |
| 560 | Q06890 | CLU | 4065141 | 4325506 | 1441835 | 9 | Cytoplasm |
| 561 | Q9WVG6 | CARM1 | 4062942 | 1232592 | 410864 | 9 | Nucleus |
| 562 | Q8VDI7 | UBAC1 | 4059355 | 2535731 | 462959 | 30 | Nucleus |
| 563 | Q9QYF1 | RDH11 | 4047507 | 1835255 | 374620 | 24 | Cytoplasm |
| 564 | P48771 | COX7A2 | 4040585 | 3017399 | 1742096 | 3 | Cytoplasm |
| 565 | Q9CPQ8 | ATP5L | 4040424 | 2918709 | 1191558 | 6 | Cytoplasm |
| 566 | O88545 | COPS6 | 4024791 | 2944918 | 760375 | 15 | Nucleus |
| 567 | Q8BYC6 | TAOK3 | 4016643 | 1728060 | 352739 | 24 | Cytoplasm |
| 568 | Q61189 | CLNS1A | 4009863 | 382522 | 220849 | 3 | Plasma Membrane |
| 569 | Q8BHL3 | TBC1D10B | 3971436 | 1574685 | 909145 | 3 | Plasma Membrane |
| 570 | O08739 | AMPD3 | 3967987 | 2048625 | 418174 | 24 | Cytoplasm |
| 571 | P56399 | USP5 | 3956643 | 1935545 | 309935 | 39 | Cytoplasm |
| 572 | Q8K0E8 | FGB | 3954341 | 13269073 | 1563775 | 72 | Extracellular Space |
| 573 | P0C0A3 | CHMP6 | 3935705 | 608204 | 351147 | 3 | Cytoplasm |
| 574 | Q8CII2 | CDC123 | 3885167 | 1328019 | 766732 | 3 | Cytoplasm |
| 575 | P56135 | ATP5J2 | 3876427 | 2261304 | 1305565 | 3 | Cytoplasm |
| 576 | P97450 | ATP5J | 3874502 | 3517480 | 2030818 | 3 | Cytoplasm |
| 577 | P62270 | RPS18 | 3874336 | 2213784 | 737928 | 9 | Cytoplasm |
| 578 | Q8JZR0 | ACSL5 | 3874112 | 2272030 | 395510 | 33 | Cytoplasm |
| 579 | P19253 | RPL13A | 3873034 | 1817205 | 1049164 | 3 | Cytoplasm |
| 580 | Q9D771 | TMEM206 | 3872685 | 4508697 | 1502899 | 9 | Plasma Membrane |
| 581 | Q571I9 | ALDH16A1 | 3864007 | 2012956 | 474458 | 18 | Cytoplasm |
| 582 | Q9D0B1 | ZNF524 | 3857613 | 3310298 | 1911202 | 3 | Nucleus |
| 583 | Q91ZJ9 | HYAL1 | 3848886 | 393482 | 227177 | 3 | Cytoplasm |
| 584 | Q64324 | STXBP2 | 3848113 | 3082528 | 475644 | 42 | Plasma Membrane |
| 585 | Q9CQX2 | CYB5B | 3832661 | 5070467 | 1690156 | 9 | Cytoplasm |
| 586 | Q9CQB5 | CISD2 | 3827621 | 1385884 | 357834 | 15 | Cytoplasm |
| 587 | Q91WC9 | DAGLB | 3821502 | 2406607 | 567243 | 18 | Plasma Membrane |
| 588 | Q8VED5 | KRT79 | 3809798 | 875800 | 505643 | 3 | Extracellular Space |
| 589 | Q9JL56 | GDE1 | 3799052 | 2930519 | 976840 | 9 | Plasma Membrane |
| 590 | P62754 | RPS6 | 3786702 | 3107176 | 1268499 | 6 | Cytoplasm |
| 591 | Q9DCD0 | PGD | 3783725 | 750494 | 433298 | 3 | Cytoplasm |
| 592 | Q62415 | PPP1R13B | 3776049 | 975968 | 563475 | 3 | Cytoplasm |
| 593 | O55022 | PGRMC1 | 3768221 | 537668 | 310423 | 3 | Plasma Membrane |
| 594 | Q99LD4 | GPS1 | 3754987 | 2199210 | 567834 | 15 | Cytoplasm |
| 595 | P29341 | PABPC1 | 3746740 | 1924207 | 496828 | 15 | Cytoplasm |
| 596 | P61222 | ABCE1 | 3738269 | 1462289 | 422126 | 12 | Cytoplasm |
| 597 | P62717 | RPL18A | 3737953 | 1429225 | 825163 | 3 | Cytoplasm |
| 598 | P14115 | RPL27A | 3733834 | 1788184 | 1032408 | 3 | Cytoplasm |
| 599 | Q9EPL8 | IPO7 | 3732796 | 1816945 | 396490 | 21 | Nucleus |
| 600 | P42567 | EPS15 | 3695750 | 769720 | 198741 | 15 | Cytoplasm |
| 601 | Q91VR2 | ATP5C1 | 3682100 | 4138669 | 1194731 | 12 | Cytoplasm |
| 602 | Q9CPR4 | RPL17 | 3674819 | 2221055 | 1282327 | 3 | Cytoplasm |
| 603 | Q76MZ3 | PPP2R1A | 3670862 | 1945247 | 424488 | 21 | Cytoplasm |
| 604 | O35372 | CNIH1 | 3660032 | 328696 | 189773 | 3 | Plasma Membrane |
| 605 | Q99JY9 | ACTR3 | 3640052 | 2965959 | 765807 | 15 | Plasma Membrane |
| 606 | Q9DCC4 | PYCRL | 3638536 | 1076401 | 358800 | 9 | Cytoplasm |
| 607 | Q9DCW4 | ETFB | 3638413 | 4705759 | 1358436 | 12 | Cytoplasm |
| 608 | Q8R5H1 | USP15 | 3637063 | 1680789 | 242601 | 48 | Cytoplasm |
| 609 | P70398 | USP9X | 3614026 | 1525092 | 440256 | 12 | Plasma Membrane |
| 610 | P62897 | CYCS | 3610519 | 2606410 | 672972 | 15 | Cytoplasm |
| 611 | P18052 | PTPRA | 3603432 | 2828256 | 816447 | 12 | Plasma Membrane |
| 612 | P97807 | FH | 3594988 | 4021259 | 877511 | 21 | Cytoplasm |
| 613 | Q9DB05 | NAPA | 3574506 | 514335 | 296951 | 3 | Cytoplasm |
| 614 | O88587 | COMT | 3566497 | 1168545 | 337330 | 12 | Cytoplasm |
| 615 | P67871 | CSNK2B | 3562640 | 142645 | 82356 | 3 | Cytoplasm |
| 616 | P62702 | RPS4Y1 | 3554580 | 2704655 | 552085 | 24 | Cytoplasm |
| 617 | Q9CZ44 | NSFL1C | 3551334 | 1048422 | 302653 | 12 | Cytoplasm |
| 618 | P35282 | RAB21 | 3548145 | 2985510 | 609415 | 24 | Cytoplasm |
| 619 | P48722 | HSPA4L | 3544844 | 1482822 | 382863 | 15 | Cytoplasm |
| 620 | P61202 | Cops2 | 3524380 | 2670205 | 689444 | 15 | Nucleus |
| 621 | P61161 | ACTR2 | 3484052 | 1112347 | 370782 | 9 | Plasma Membrane |
| 622 | Q3UFK8 | FRMD8 | 3467957 | 972842 | 561670 | 3 | Cytoplasm |
| 623 | Q9QZ73 | DCUN1D1 | 3454811 | 403135 | 232750 | 3 | Nucleus |
| 624 | Q60738 | SLC30A1 | 3424007 | 823903 | 237840 | 12 | Plasma Membrane |
| 625 | P62259 | YWHAE | 3417160 | 2575699 | 495693 | 27 | Cytoplasm |
| 626 | Q8K3F2 | MMP21 | 3407843 | 651013 | 375862 | 3 | Extracellular Space |
| 627 | P47911 | RPL6 | 3394230 | 2252699 | 650298 | 12 | Nucleus |
| 628 | Q80VP1 | EPN1 | 3387319 | 1399496 | 404000 | 12 | Plasma Membrane |
| 629 | P35285 | RAB22A | 3379808 | 638977 | 260861 | 6 | Cytoplasm |
| 630 | Q8C9M2 | CCDC15 | 3379127 | 2001421 | 1155521 | 3 | Cytoplasm |
| 631 | Q9JMC3 | DNAJA4 | 3366177 | 964824 | 557041 | 3 | Nucleus |
| 632 | Q91YA2 | PSKH1 | 3362423 | 309559 | 178724 | 3 | Nucleus |
| 633 | P60335 | PCBP1 | 3350562 | 1782030 | 460118 | 15 | Nucleus |
| 634 | O35344 | KPNA3 | 3346416 | 635117 | 366685 | 3 | Nucleus |
| 635 | P56395 | CYB5A | 3343397 | 3601755 | 929969 | 15 | Cytoplasm |
| 636 | P63242 | EIF5A | 3340893 | 1248055 | 720565 | 3 | Cytoplasm |
| 637 | Q8BPM0 | DAAM1 | 3324872 | 1892949 | 315491 | 36 | Cytoplasm |
| 638 | O88543 | COPS3 | 3286150 | 1109203 | 261442 | 18 | Cytoplasm |
| 639 | P19157 | GSTP1 | 3261808 | 369201 | 213158 | 3 | Cytoplasm |
| 640 | Q9DB34 | CHMP2A | 3261468 | 553357 | 319481 | 3 | Cytoplasm |
| 641 | Q99LC3 | NDUFA10 | 3256157 | 2162991 | 883037 | 6 | Cytoplasm |
| 642 | P17426 | AP2A1 | 3235827 | 2928767 | 976256 | 9 | Cytoplasm |
| 643 | P63330 | PPP2CA | 3206553 | 1096719 | 283172 | 15 | Cytoplasm |
| 644 | Q8R050 | GSPT1 | 3179969 | 397975 | 229771 | 3 | Cytoplasm |
| 645 | P70295 | AUP1 | 3179704 | 403672 | 233060 | 3 | Cytoplasm |
| 646 | P51655 | GPC4 | 3177482 | 1112540 | 370847 | 9 | Plasma Membrane |
| 647 | Q8BGU5 | CCNY | 3173625 | 425009 | 173509 | 6 | Nucleus |
| 648 | O08663 | METAP2 | 3171888 | 1330621 | 768234 | 3 | Cytoplasm |
| 649 | P12382 | PFKL | 3166737 | 542302 | 313098 | 3 | Cytoplasm |
| 650 | Q922Q4 | PYCR2 | 3156188 | 1755462 | 1013517 | 3 | Cytoplasm |
| 651 | P52480 | PKM | 3141394 | 668379 | 385889 | 3 | Cytoplasm |
| 652 | Q8BMS1 | HADHA | 3123085 | 3371108 | 520173 | 42 | Cytoplasm |
| 653 | P09055 | ITGB1 | 3122781 | 3529995 | 565252 | 39 | Plasma Membrane |
| 654 | P59017 | BCL2L13 | 3109346 | 2499275 | 833092 | 9 | Cytoplasm |
| 655 | Q6P5E4 | UGGT1 | 3101812 | 2183292 | 445663 | 24 | Cytoplasm |
| 656 | Q9JHR7 | IDE | 3101616 | 540098 | 220494 | 6 | Extracellular Space |
| 657 | Q9JKB3 | YBX3 | 3098161 | 763682 | 311772 | 6 | Nucleus |
| 658 | Q9R0Q6 | ARPC1A | 3063814 | 492746 | 284487 | 3 | Extracellular Space |
| 659 | Q8BP67 | RPL24 | 3039365 | 1278975 | 426325 | 9 | Cytoplasm |
| 660 | Q9DC69 | NDUFA9 | 3030680 | 4879376 | 1626459 | 9 | Cytoplasm |
| 661 | Q3V0G7 | GARNL3 | 3028670 | 473214 | 273210 | 3 | Cytoplasm |
| 662 | Q61753 | PHGDH | 3028332 | 1159866 | 299476 | 15 | Cytoplasm |
| 663 | P49769 | PSEN1 | 3023270 | 1610224 | 657371 | 6 | Plasma Membrane |
| 664 | Q00262 | STX2 | 3011496 | 2103759 | 495861 | 18 | Cytoplasm |
| 665 | Q60737 | CSNK2A1 | 2985798 | 164115 | 94752 | 3 | Nucleus |
| 666 | P62827 | RAN | 2975625 | 763918 | 441048 | 3 | Nucleus |
| 667 | P62245 | RPS15A | 2975400 | 1239992 | 506225 | 6 | Cytoplasm |
| 668 | Q6R0H7 | GNAS | 2971529 | 578184 | 236043 | 6 | Plasma Membrane |
| 669 | Q8R574 | PRPSAP2 | 2949702 | 483448 | 197367 | 6 | Cytoplasm |
| 670 | Q8BFR5 | TUFM | 2934028 | 2502034 | 589735 | 18 | Cytoplasm |
| 671 | P53026 | RPL10A | 2909502 | 1720548 | 702411 | 6 | Nucleus |
| 672 | Q80WC7 | AGFG2 | 2909180 | 1301451 | 375697 | 12 | Cytoplasm |
| 673 | Q9JM76 | ARPC3 | 2905468 | 1187980 | 484991 | 6 | Cytoplasm |
| 674 | Q6PDI5 | KIAA0368 | 2904176 | 1676649 | 322671 | 27 | Cytoplasm |
| 675 | Q641P0 | ACTR3B | 2887797 | 646976 | 373532 | 3 | Cytoplasm |
| 676 | P50516 | ATP6V1A | 2883946 | 1562955 | 403553 | 15 | Plasma Membrane |
| 677 | Q9D172 | C21orf33/LOC102724023 | 2881443 | 4223832 | 1724372 | 6 | Cytoplasm |
| 678 | Q3U7U3 | FBXO7 | 2879512 | 591926 | 341749 | 3 | Cytoplasm |
| 679 | P70404 | IDH3G | 2867515 | 3500208 | 1428954 | 6 | Cytoplasm |
| 680 | Q9CY24 | TMEM179B | 2854652 | 175007 | 101040 | 3 | Extracellular Space |
| 681 | P09671 | SOD2 | 2854048 | 2502468 | 834156 | 9 | Cytoplasm |
| 682 | O88544 | COPS4 | 2851324 | 1084515 | 255623 | 18 | Cytoplasm |
| 683 | O08547 | SEC22B | 2847633 | 1419360 | 289726 | 24 | Cytoplasm |
| 684 | P63276 | RPS17 | 2839449 | 1095492 | 632483 | 3 | Cytoplasm |
| 685 | Q6PA06 | ATL2 | 2838905 | 2448968 | 816323 | 9 | Cytoplasm |
| 686 | P68369 | TUBA1A | 2832537 | 1145320 | 661251 | 3 | Cytoplasm |
| 687 | P70297 | STAM | 2823746 | 1208777 | 402926 | 9 | Cytoplasm |
| 688 | Q00623 | APOA1 | 2821399 | 2164031 | 721344 | 9 | Extracellular Space |
| 689 | Q920L1 | FADS1 | 2792971 | 981798 | 566841 | 3 | Plasma Membrane |
| 690 | Q9CQA3 | SDHB | 2790077 | 1173712 | 391237 | 9 | Cytoplasm |
| 691 | Q99KV1 | DNAJB11 | 2786786 | 1319736 | 439912 | 9 | Cytoplasm |
| 692 | Q922B2 | DARS | 2783719 | 1505906 | 307392 | 24 | Cytoplasm |
| 693 | Q5SSH7 | ZZEF1 | 2775721 | 240332 | 138756 | 3 | Cytoplasm |
| 694 | Q923U0 | TOM1L1 | 2775268 | 1583177 | 646329 | 6 | Cytoplasm |

| | | | | | | | |
|---|---|---|---|---|---|---|---|
| 695 | P27661 | H2AFX | 2.99E+06 | 2.77E+06 | 1.13E+06 | 6 | Nucleus |
| 696 | Q8CGK3 | LONP1 | 2.99E+06 | 1.09E+06 | 4.43E+05 | 6 | Cytoplasm |
| 697 | Q9DC69 | NDUFA9 | 2.98E+06 | 2.66E+06 | 8.86E+05 | 9 | Cytoplasm |
| 698 | Q9CZX8 | RPS19 | 2.98E+06 | 5.75E+05 | 1.92E+05 | 9 | Cytoplasm |
| 699 | Q8CDN6 | TXNL1 | 2.98E+06 | 3.16E+06 | 6.46E+05 | 24 | Cytoplasm |
| 700 | Q8BFR5 | TUFM | 2.98E+06 | 1.36E+06 | 3.20E+05 | 18 | Cytoplasm |
| 701 | Q00262 | STX2 | 2.96E+06 | 1.69E+06 | 3.99E+05 | 18 | Cytoplasm |
| 702 | P70658 | CXCR4 | 2.95E+06 | 5.16E+05 | 2.98E+05 | 3 | Plasma Membrane |
| 703 | Q9JI39 | ABCB10 | 2.95E+06 | 2.61E+06 | 4.35E+05 | 36 | Cytoplasm |
| 704 | P62242 | RPS8 | 2.95E+06 | 1.52E+06 | 3.93E+05 | 15 | Cytoplasm |
| 705 | P62342 | SELT | 2.95E+06 | 7.47E+05 | 4.31E+05 | 3 | Cytoplasm |
| 706 | O35855 | BCAT2 | 2.94E+06 | 1.87E+06 | 1.08E+06 | 3 | Cytoplasm |
| 707 | Q9WVK4 | EHD1 | 2.93E+06 | 2.14E+06 | 6.17E+05 | 12 | Cytoplasm |
| 708 | P50247 | AHCY | 2.92E+06 | 1.42E+06 | 4.73E+05 | 9 | Cytoplasm |
| 709 | Q8R574 | PRPSAP2 | 2.92E+06 | 1.29E+06 | 5.25E+05 | 6 | Cytoplasm |
| 710 | P62137 | PPP1CA | 2.91E+06 | 2.02E+06 | 4.75E+05 | 18 | Cytoplasm |
| 711 | Q7TMY8 | HUWE1 | 2.90E+06 | 7.37E+05 | 4.25E+05 | 3 | Nucleus |
| 712 | P97807 | FH | 2.90E+06 | 2.43E+06 | 5.30E+05 | 21 | Cytoplasm |
| 713 | P35979 | RPL12 | 2.88E+06 | 1.10E+06 | 2.84E+05 | 15 | Nucleus |
| 714 | Q61781 | KRT14 | 2.87E+06 | 1.33E+06 | 7.68E+05 | 3 | Cytoplasm |
| 715 | Q6PCP5 | Mff | 2.87E+06 | 9.63E+05 | 3.93E+05 | 6 | Cytoplasm |
| 716 | Q920G8 | SLC25A37 | 2.87E+06 | 1.62E+06 | 2.83E+05 | 33 | Cytoplasm |
| 717 | Q8VBV7 | COPS8 | 2.87E+06 | 1.54E+06 | 8.87E+05 | 3 | Nucleus |
| 718 | Q6PA06 | ATL2 | 2.86E+06 | 2.23E+06 | 7.42E+05 | 9 | Cytoplasm |
| 719 | Q9CQV8 | YWHAB | 2.85E+06 | 1.88E+06 | 4.42E+05 | 18 | Cytoplasm |
| 720 | P84084 | ARF5 | 2.85E+06 | 8.81E+05 | 3.60E+05 | 6 | Cytoplasm |
| 721 | Q9CZ44 | NSFL1C | 2.85E+06 | 1.78E+06 | 5.13E+05 | 12 | Cytoplasm |
| 722 | P11276 | FN1 | 2.84E+06 | 4.29E+06 | 1.43E+06 | 9 | Extracellular Space |
| 723 | Q6ZWX6 | EIF2S1 | 2.84E+06 | 1.40E+06 | 3.30E+05 | 18 | Cytoplasm |
| 724 | Q8R5J9 | ARL6IP5 | 2.84E+06 | 5.67E+05 | 3.27E+05 | 3 | Cytoplasm |
| 725 | Q8CIZ8 | VWF | 2.83E+06 | 2.86E+06 | 4.27E+05 | 45 | Extracellular Space |
| 726 | P62862 | FAU | 2.83E+06 | 6.43E+05 | 3.72E+05 | 3 | Cytoplasm |
| 727 | P18181 | CD48 | 2.82E+06 | 4.50E+05 | 2.60E+05 | 3 | Plasma Membrane |
| 728 | P97352 | S100A13 | 2.81E+06 | 4.34E+05 | 2.50E+05 | 3 | Cytoplasm |
| 729 | P28571 | SLC6A9 | 2.78E+06 | 1.91E+06 | 7.81E+05 | 6 | Plasma Membrane |
| 730 | P70697 | UROD | 2.76E+06 | 1.29E+06 | 7.44E+05 | 3 | Cytoplasm |
| 731 | O08756 | HSD17B10 | 2.75E+06 | 2.26E+06 | 6.53E+05 | 12 | Cytoplasm |
| 732 | Q8BGU5 | CCNY | 2.75E+06 | 6.28E+05 | 2.57E+05 | 6 | Nucleus |
| 733 | Q8CHK3 | MBOAT7 | 2.75E+06 | 2.15E+06 | 1.24E+06 | 3 | Plasma Membrane |
| 734 | P35821 | PTPN1 | 2.74E+06 | 2.03E+06 | 1.17E+06 | 3 | Cytoplasm |
| 735 | Q8BIG7 | COMTD1 | 2.73E+06 | 1.71E+06 | 9.87E+05 | 3 | Plasma Membrane |
| 736 | P47757 | CAPZB | 2.73E+06 | 9.41E+05 | 3.84E+05 | 6 | Cytoplasm |
| 737 | Q6X893 | SLC44A1 | 2.73E+06 | 1.01E+06 | 3.37E+05 | 9 | Plasma Membrane |
| 738 | P62267 | RPS23 | 2.72E+06 | 4.49E+05 | 2.59E+05 | 3 | Cytoplasm |
| 739 | Q80VP1 | EPN1 | 2.72E+06 | 1.57E+06 | 4.54E+05 | 12 | Plasma Membrane |
| 740 | O88544 | COPS4 | 2.72E+06 | 1.26E+06 | 2.98E+05 | 18 | Cytoplasm |
| 741 | P68040 | GNB2L1 | 2.72E+06 | 9.49E+05 | 2.45E+05 | 15 | Cytoplasm |
| 742 | Q60790 | RASA3 | 2.71E+06 | 2.26E+06 | 1.31E+06 | 3 | Plasma Membrane |
| 743 | Q7TQ95 | KIAA1715 | 2.71E+06 | 6.80E+05 | 1.75E+05 | 15 | Cytoplasm |
| 744 | Q80TH2 | ERBB2IP | 2.71E+06 | 1.30E+06 | 2.66E+05 | 24 | Plasma Membrane |
| 745 | Q9QWY8 | ASAP1 | 2.70E+06 | 1.29E+06 | 5.27E+05 | 6 | Plasma Membrane |
| 746 | Q9DCD0 | PGD | 2.70E+06 | 1.19E+06 | 6.86E+05 | 3 | Cytoplasm |
| 747 | Q99JR1 | SFXN1 | 2.70E+06 | 1.46E+06 | 3.44E+05 | 18 | Cytoplasm |
| 748 | Q8BNI4 | DERL2 | 2.68E+06 | 1.21E+06 | 6.98E+05 | 3 | Cytoplasm |
| 749 | P68254 | YWHAQ | 2.67E+06 | 1.82E+06 | 6.08E+05 | 9 | Cytoplasm |
| 750 | Q99KE1 | ME2 | 2.67E+06 | 1.28E+06 | 3.69E+05 | 12 | Cytoplasm |
| 751 | Q91ZJ9 | HYAL1 | 2.67E+06 | 4.76E+05 | 2.75E+05 | 3 | Cytoplasm |
| 752 | Q9ES46 | PARVB | 2.67E+06 | 2.39E+06 | 5.64E+05 | 18 | Cytoplasm |
| 753 | Q8K411 | PITRM1 | 2.64E+06 | 1.64E+06 | 3.58E+05 | 21 | Cytoplasm |
| 754 | Q9CX30 | YIF1B | 2.63E+06 | 4.30E+05 | 1.75E+05 | 6 | Cytoplasm |
| 755 | Q8VEH3 | ARL8A | 2.62E+06 | 4.74E+05 | 2.74E+05 | 3 | Cytoplasm |
| 756 | Q9CY24 | TMEM179B | 2.61E+06 | 4.58E+05 | 2.64E+05 | 3 | Extracellular Space |
| 757 | Q9D8N0 | EEF1G | 2.61E+06 | 2.17E+06 | 7.23E+05 | 9 | Cytoplasm |
| 758 | P61982 | YWHAG | 2.61E+06 | 1.38E+06 | 5.64E+05 | 6 | Cytoplasm |
| 759 | O88456 | CAPNS1 | 2.61E+06 | 4.58E+05 | 2.64E+05 | 3 | Cytoplasm |
| 760 | Q62415 | PPP1R13B | 2.60E+06 | 1.46E+06 | 8.42E+05 | 3 | Cytoplasm |
| 761 | Q9CZX7 | TMEM55A | 2.59E+06 | 8.87E+05 | 5.12E+05 | 3 | Cytoplasm |
| 762 | Q80WQ2 | VAC14 | 2.59E+06 | 6.78E+05 | 3.91E+05 | 3 | Cytoplasm |
| 763 | Q8C0L0 | TMX4 | 2.57E+06 | 7.92E+05 | 4.58E+05 | 3 | Cytoplasm |
| 764 | Q7TMR0 | PRCP | 2.57E+06 | 8.32E+05 | 4.80E+05 | 3 | Cytoplasm |
| 765 | P97855 | G3BP1 | 2.57E+06 | 8.27E+05 | 3.37E+05 | 6 | Nucleus |
| 766 | P50516 | ATP6V1A | 2.57E+06 | 1.56E+06 | 4.03E+05 | 15 | Plasma Membrane |
| 767 | Q61878 | PRG2 | 2.57E+06 | 2.25E+06 | 1.30E+06 | 3 | Extracellular Space |
| 768 | P48962 | SLC25A4 | 2.56E+06 | 1.75E+06 | 5.82E+05 | 9 | Cytoplasm |
| 769 | Q80WC7 | AGFG2 | 2.56E+06 | 8.38E+05 | 2.42E+05 | 12 | Cytoplasm |
| 770 | P41105 | RPL28 | 2.56E+06 | 6.66E+05 | 3.85E+05 | 3 | Cytoplasm |
| 771 | O08739 | AMPD3 | 2.55E+06 | 1.61E+06 | 3.29E+05 | 24 | Cytoplasm |
| 772 | B2RPV6 | MMRN1 | 2.55E+06 | 2.04E+06 | 3.55E+05 | 33 | Extracellular Space |
| 773 | Q6URW6 | MYH14 | 2.54E+06 | 2.10E+06 | 8.55E+05 | 6 | Extracellular Space |
| 774 | P70398 | USP9X | 2.54E+06 | 1.51E+06 | 4.36E+05 | 12 | Plasma Membrane |
| 775 | Q6ZWZ2 | UBE2R2 | 2.54E+06 | 1.80E+06 | 1.04E+06 | 3 | Cytoplasm |
| 776 | Q8R050 | GSPT1 | 2.53E+06 | 1.56E+06 | 9.02E+05 | 3 | Cytoplasm |
| 777 | Q922B2 | DARS | 2.53E+06 | 1.50E+06 | 3.07E+05 | 24 | Cytoplasm |
| 778 | Q9ER72 | CARS | 2.52E+06 | 1.36E+06 | 5.55E+05 | 6 | Cytoplasm |
| 779 | P59999 | ARPC4 | 2.51E+06 | 5.90E+05 | 3.40E+05 | 3 | Cytoplasm |
| 780 | Q9D172 | C21orf33/LOC102724023 | 2.50E+06 | 2.76E+06 | 1.13E+06 | 6 | Cytoplasm |
| 781 | Q8BH59 | SLC25A12 | 2.50E+06 | 1.68E+06 | 4.84E+05 | 12 | Cytoplasm |
| 782 | Q9CRY7 | GDPD1 | 2.49E+06 | 3.65E+05 | 2.11E+05 | 3 | Plasma Membrane |
| 783 | Q62469 | ITGA2 | 2.48E+06 | 2.27E+06 | 3.95E+05 | 33 | Plasma Membrane |
| 784 | Q9CXE7 | TMED5 | 2.48E+06 | 4.96E+05 | 2.87E+05 | 3 | Cytoplasm |
| 785 | P01029 | C4A/C4B | 2.46E+06 | 1.29E+06 | 3.33E+05 | 15 | Extracellular Space |
| 786 | Q9JM76 | ARPC3 | 2.45E+06 | 7.74E+05 | 3.16E+05 | 6 | Cytoplasm |
| 787 | Q60737 | CSNK2A1 | 2.45E+06 | 1.21E+06 | 6.98E+05 | 3 | Nucleus |
| 788 | Q8BMS1 | HADHA | 2.43E+06 | 1.41E+06 | 2.17E+05 | 42 | Cytoplasm |
| 789 | Q9D8E6 | RPL4 | 2.42E+06 | 1.10E+06 | 3.18E+05 | 12 | Cytoplasm |
| 790 | P70404 | IDH3G | 2.42E+06 | 1.99E+06 | 8.11E+05 | 6 | Cytoplasm |
| 791 | Q8BG73 | SH3BGRL2 | 2.42E+06 | 1.24E+06 | 7.17E+05 | 3 | Nucleus |
| 792 | Q9WUM5 | SUCLG1 | 2.42E+06 | 1.77E+06 | 5.90E+05 | 9 | Cytoplasm |
| 793 | Q60738 | SLC30A1 | 2.42E+06 | 7.40E+05 | 2.14E+05 | 12 | Plasma Membrane |
| 794 | Q60575 | KIF1B | 2.41E+06 | 1.40E+06 | 8.09E+05 | 3 | Cytoplasm |
| 795 | P20918 | PLG | 2.41E+06 | 2.91E+06 | 9.71E+05 | 9 | Extracellular Space |
| 796 | Q9CY50 | SSR1 | 2.41E+06 | 9.48E+05 | 3.87E+05 | 6 | Cytoplasm |
| 797 | Q8BQ47 | CNPY4 | 2.41E+06 | 6.43E+05 | 3.71E+05 | 3 | Extracellular Space |
| 798 | Q61335 | BCAP31 | 2.40E+06 | 1.53E+06 | 8.83E+05 | 3 | Cytoplasm |
| 799 | Q6NS82 | FAM134A | 2.39E+06 | 6.06E+05 | 3.50E+05 | 3 | Extracellular Space |
| 800 | P21278 | GNA11 | 2.39E+06 | 1.77E+06 | 1.02E+06 | 3 | Plasma Membrane |
| 801 | Q8VDP6 | CDIPT | 2.38E+06 | 2.52E+05 | 1.45E+05 | 3 | Cytoplasm |
| 802 | Q9CRB9 | CHCHD3 | 2.37E+06 | 7.98E+05 | 4.61E+05 | 3 | Cytoplasm |
| 803 | Q9DB05 | NAPA | 2.35E+06 | 3.95E+05 | 2.28E+05 | 3 | Cytoplasm |
| 804 | Q8BFZ9 | ERLIN2 | 2.34E+06 | 3.01E+05 | 1.74E+05 | 3 | Plasma Membrane |
| 805 | Q5SSH7 | ZZEF1 | 2.34E+06 | 1.06E+06 | 6.14E+05 | 3 | Cytoplasm |
| 806 | Q99LC5 | ETFA | 2.32E+06 | 1.59E+06 | 3.06E+05 | 27 | Cytoplasm |
| 807 | Q62348 | TSN | 2.32E+06 | 1.23E+06 | 5.01E+05 | 6 | Nucleus |
| 808 | O89086 | RBM3 | 2.32E+06 | 2.71E+06 | 9.04E+05 | 9 | Cytoplasm |
| 809 | P62855 | RPS26 | 2.31E+06 | 6.51E+05 | 3.76E+05 | 3 | Cytoplasm |
| 810 | P62908 | RPS3 | 2.31E+06 | 1.35E+06 | 2.75E+05 | 24 | Cytoplasm |
| 811 | Q9CZM2 | RPL15 | 2.31E+06 | 8.13E+05 | 2.71E+05 | 9 | Cytoplasm |
| 812 | Q80UU9 | PGRMC2 | 2.29E+06 | 3.47E+05 | 2.00E+05 | 3 | Nucleus |
| 813 | Q8CAQ8 | IMMT | 2.28E+06 | 1.29E+06 | 2.49E+05 | 27 | Cytoplasm |
| 814 | O70503 | HSD17B12 | 2.26E+06 | 9.63E+05 | 3.93E+05 | 6 | Cytoplasm |
| 815 | P68369 | TUBA1A | 2.25E+06 | 1.09E+06 | 6.27E+05 | 3 | Cytoplasm |
| 816 | Q91YP3 | DERA | 2.25E+06 | 6.94E+05 | 2.31E+05 | 9 | Cytoplasm |
| 817 | O08599 | STXBP1 | 2.25E+06 | 1.46E+06 | 5.95E+05 | 6 | Cytoplasm |
| 818 | P59017 | BCL2L13 | 2.25E+06 | 7.22E+05 | 2.41E+05 | 9 | Cytoplasm |
| 819 | Q9WTX6 | CUL1 | 2.24E+06 | 1.40E+06 | 2.70E+05 | 27 | Nucleus |
| 820 | P63242 | EIF5A | 2.24E+06 | 6.58E+05 | 3.80E+05 | 3 | Cytoplasm |
| 821 | O54734 | DDOST | 2.24E+06 | 1.31E+06 | 4.36E+05 | 9 | Cytoplasm |
| 822 | Q9QYK7 | RNF11 | 2.24E+06 | 3.16E+05 | 1.82E+05 | 3 | Cytoplasm |
| 823 | P40124 | CAP1 | 2.23E+06 | 1.01E+06 | 3.36E+05 | 9 | Plasma Membrane |
| 824 | Q8BV13 | COPS7B | 2.22E+06 | 9.61E+05 | 5.55E+05 | 3 | Cytoplasm |
| 825 | P62717 | RPL18A | 2.21E+06 | 7.05E+04 | 4.07E+04 | 3 | Cytoplasm |
| 826 | Q91WS0 | CISD1 | 2.20E+06 | 8.58E+05 | 3.50E+05 | 6 | Cytoplasm |
| 827 | Q61739 | ITGA6 | 2.19E+06 | 2.28E+06 | 4.97E+05 | 21 | Plasma Membrane |
| 828 | Q9CR21 | NDUFAB1 | 2.18E+06 | 1.01E+06 | 4.10E+05 | 6 | Cytoplasm |
| 829 | Q6ZWN5 | RPS9 | 2.17E+06 | 1.39E+06 | 4.65E+05 | 9 | Cytoplasm |
| 830 | Q8BZI6 | GUCD1 | 2.17E+06 | 9.14E+04 | 5.28E+04 | 3 | Nucleus |
| 831 | P13609 | Srgn | 2.17E+06 | 1.64E+06 | 9.44E+05 | 3 | Cytoplasm |
| 832 | O35344 | KPNA3 | 2.16E+06 | 5.11E+05 | 2.95E+05 | 3 | Nucleus |
| 833 | Q8CBE3 | WDR37 | 2.16E+06 | 1.23E+06 | 5.02E+05 | 6 | Nucleus |

| | | | | | | | |
|---|---|---|---|---|---|---|---|
| 695 | Q6R0H7 | GNAS | 3.73E+06 | 4.96E+05 | 2.02E+05 | 6 | Plasma Membrane |
| 696 | Q922Q4 | PYCR2 | 3.73E+06 | 5.43E+05 | 3.13E+05 | 3 | Cytoplasm |
| 697 | P62814 | ATP6V1B2 | 3.72E+06 | 2.64E+06 | 8.81E+05 | 9 | Cytoplasm |
| 698 | P24549 | ALDH1A1 | 3.72E+06 | 1.18E+06 | 6.82E+05 | 3 | Cytoplasm |
| 699 | P70404 | IDH3G | 3.71E+06 | 2.80E+06 | 1.14E+06 | 6 | Cytoplasm |
| 700 | Q3V0G7 | GARNL3 | 3.70E+06 | 1.33E+06 | 7.67E+05 | 3 | Cytoplasm |
| 701 | Q6P9J9 | ANO6 | 3.70E+06 | 4.36E+05 | 2.52E+05 | 3 | Plasma Membrane |
| 702 | P48722 | HSPA4L | 3.67E+06 | 1.54E+06 | 3.97E+05 | 15 | Cytoplasm |
| 703 | Q8K2J7 | RELL1 | 3.67E+06 | 1.33E+06 | 4.42E+05 | 9 | Plasma Membrane |
| 704 | P19096 | FASN | 3.65E+06 | 2.24E+06 | 2.70E+05 | 69 | Cytoplasm |
| 705 | Q9CQ75 | NDUFA2 | 3.64E+06 | 6.44E+05 | 3.72E+05 | 3 | Cytoplasm |
| 706 | Q60790 | RASA3 | 3.64E+06 | 9.10E+05 | 5.25E+05 | 3 | Plasma Membrane |
| 707 | P50247 | AHCY | 3.64E+06 | 9.13E+05 | 3.04E+05 | 9 | Cytoplasm |
| 708 | Q00262 | STX2 | 3.62E+06 | 2.07E+06 | 4.87E+05 | 18 | Cytoplasm |
| 709 | Q9CYL5 | GLIPR2 | 3.62E+06 | 3.91E+05 | 2.26E+05 | 3 | Cytoplasm |
| 710 | Q9ES46 | PARVB | 3.61E+06 | 1.51E+06 | 3.56E+05 | 18 | Cytoplasm |
| 711 | Q61739 | ITGA6 | 3.61E+06 | 2.47E+06 | 5.40E+05 | 21 | Plasma Membrane |
| 712 | A2ASS6 | TTN | 3.60E+06 | 3.49E+05 | 2.01E+05 | 3 | Cytoplasm |
| 713 | P35279 | RAB6A | 3.60E+06 | 1.55E+06 | 6.31E+05 | 6 | Cytoplasm |
| 714 | P29341 | PABPC1 | 3.59E+06 | 1.22E+06 | 3.14E+05 | 15 | Cytoplasm |
| 715 | Q8CIZ8 | VWF | 3.57E+06 | 1.79E+06 | 2.66E+05 | 45 | Extracellular Space |
| 716 | Q9EQH2 | ERAP1 | 3.57E+06 | 4.89E+05 | 2.82E+05 | 3 | Extracellular Space |
| 717 | Q8C0L0 | TMX4 | 3.57E+06 | 5.61E+05 | 3.24E+05 | 3 | Cytoplasm |
| 718 | Q9CY50 | SSR1 | 3.57E+06 | 1.12E+06 | 4.59E+05 | 6 | Cytoplasm |
| 719 | Q8R5J9 | ARL6IP5 | 3.55E+06 | 8.13E+05 | 4.70E+05 | 3 | Cytoplasm |
| 720 | Q9D0K2 | OXCT1 | 3.55E+06 | 2.03E+05 | 1.17E+05 | 3 | Cytoplasm |
| 721 | Q8BG67 | EFR3A | 3.54E+06 | 8.77E+05 | 1.91E+05 | 21 | Plasma Membrane |
| 722 | Q8CAQ8 | IMMT | 3.54E+06 | 1.90E+06 | 3.66E+05 | 27 | Cytoplasm |
| 723 | O35372 | CNIH1 | 3.50E+06 | 5.26E+05 | 3.03E+05 | 3 | Plasma Membrane |
| 724 | Q922B2 | DARS | 3.48E+06 | 1.57E+06 | 3.20E+05 | 24 | Cytoplasm |
| 725 | Q9CPR4 | RPL17 | 3.48E+06 | 1.07E+06 | 6.18E+05 | 3 | Cytoplasm |
| 726 | Q76MZ3 | PPP2R1A | 3.48E+06 | 1.92E+06 | 4.18E+05 | 21 | Cytoplasm |
| 727 | P61982 | YWHAG | 3.47E+06 | 1.43E+06 | 5.82E+05 | 6 | Cytoplasm |
| 728 | Q91WC9 | DAGLB | 3.47E+06 | 2.47E+06 | 5.82E+05 | 18 | Plasma Membrane |
| 729 | Q9QY76 | VAPB | 3.47E+06 | 2.77E+06 | 1.13E+06 | 6 | Plasma Membrane |
| 730 | Q9JM76 | ARPC3 | 3.47E+06 | 1.02E+06 | 4.16E+05 | 6 | Cytoplasm |
| 731 | Q9D051 | PDHB | 3.47E+06 | 2.99E+05 | 1.72E+05 | 3 | Cytoplasm |
| 732 | P35285 | RAB22A | 3.46E+06 | 2.46E+05 | 1.00E+05 | 6 | Cytoplasm |
| 733 | Q4VAA2 | CDV3 | 3.45E+06 | 9.69E+05 | 5.60E+05 | 3 | Cytoplasm |
| 734 | Q99KV1 | DNAJB11 | 3.45E+06 | 1.36E+06 | 4.53E+05 | 9 | Cytoplasm |
| 735 | P62702 | RPS4Y1 | 3.44E+06 | 1.78E+06 | 3.63E+05 | 24 | Cytoplasm |
| 736 | Q8BYC6 | TAOK3 | 3.44E+06 | 1.53E+06 | 3.13E+05 | 24 | Cytoplasm |
| 737 | P56399 | USP5 | 3.43E+06 | 1.83E+06 | 2.93E+05 | 39 | Cytoplasm |
| 738 | Q9Z125 | CREB3L1 | 3.43E+06 | 8.10E+05 | 4.67E+05 | 3 | Nucleus |
| 739 | Q91Y57 | SIGLEC9 | 3.43E+06 | 1.12E+06 | 6.45E+05 | 3 | Plasma Membrane |
| 740 | Q9CQV8 | YWHAB | 3.42E+06 | 2.88E+06 | 6.80E+05 | 18 | Cytoplasm |
| 741 | Q9CQA3 | SDHB | 3.41E+06 | 1.11E+06 | 3.71E+05 | 9 | Cytoplasm |
| 742 | Q921H8 | ACAA1 | 3.40E+06 | 6.77E+05 | 3.91E+05 | 3 | Cytoplasm |
| 743 | Q9D924 | ISCA1 | 3.39E+06 | 1.32E+06 | 5.39E+05 | 6 | Cytoplasm |
| 744 | P61620 | SEC61A1 | 3.39E+06 | 1.71E+05 | 9.85E+04 | 3 | Cytoplasm |
| 745 | Q641P0 | ACTR3B | 3.38E+06 | 8.24E+05 | 4.76E+05 | 3 | Cytoplasm |
| 746 | Q8VDI7 | UBAC1 | 3.38E+06 | 2.31E+06 | 4.21E+05 | 30 | Nucleus |
| 747 | P60335 | PCBP1 | 3.37E+06 | 1.09E+06 | 2.81E+05 | 15 | Nucleus |
| 748 | Q3V3R1 | MTHFD1L | 3.36E+06 | 1.48E+06 | 3.02E+05 | 24 | Cytoplasm |
| 749 | Q9DC51 | GNAI3 | 3.36E+06 | 6.01E+05 | 3.47E+05 | 3 | Cytoplasm |
| 750 | O08663 | METAP2 | 3.35E+06 | 4.02E+05 | 2.32E+05 | 3 | Cytoplasm |
| 751 | P21278 | GNA11 | 3.32E+06 | 3.69E+05 | 2.13E+05 | 3 | Plasma Membrane |
| 752 | Q61753 | PHGDH | 3.30E+06 | 1.18E+06 | 3.05E+05 | 15 | Cytoplasm |
| 753 | P47757 | CAPZB | 3.30E+06 | 9.36E+05 | 3.82E+05 | 6 | Cytoplasm |
| 754 | Q9JKR6 | HYOU1 | 3.28E+06 | 1.01E+06 | 2.20E+05 | 21 | Cytoplasm |
| 755 | P62717 | RPL18A | 3.27E+06 | 5.28E+05 | 3.05E+05 | 3 | Cytoplasm |
| 756 | O88545 | COPS6 | 3.27E+06 | 2.44E+06 | 6.31E+05 | 15 | Nucleus |
| 757 | Q7TMM9 | TUBB2A | 3.27E+06 | 7.49E+05 | 3.06E+05 | 6 | Cytoplasm |
| 758 | Q9DBD2 | MARH8 | 3.27E+06 | 9.37E+05 | 3.83E+05 | 6 | Cytoplasm |
| 759 | P42567 | EPS15 | 3.27E+06 | 1.07E+06 | 2.78E+05 | 15 | Cytoplasm |
| 760 | Q8K4F0 | CD226 | 3.26E+06 | 1.82E+05 | 1.05E+05 | 3 | Plasma Membrane |
| 761 | P19253 | RPL13A | 3.24E+06 | 8.17E+05 | 4.71E+05 | 3 | Cytoplasm |
| 762 | Q61189 | CLNS1A | 3.24E+06 | 4.89E+05 | 2.82E+05 | 3 | Plasma Membrane |
| 763 | Q9D0M1 | PRPSAP1 | 3.23E+06 | 1.47E+06 | 3.80E+05 | 15 | Extracellular Space |
| 764 | P29391 | FTL | 3.20E+06 | 3.08E+06 | 1.03E+06 | 9 | Cytoplasm |
| 765 | Q9CR60 | GOLT1B | 3.19E+06 | 3.43E+05 | 1.98E+05 | 3 | Cytoplasm |
| 766 | P00397 | MT-CO1 | 3.19E+06 | 2.80E+05 | 1.62E+05 | 3 | Cytoplasm |
| 767 | Q9JMC3 | DNAJA4 | 3.19E+06 | 8.22E+05 | 4.75E+05 | 3 | Nucleus |
| 768 | Q571I9 | ALDH16A1 | 3.19E+06 | 1.77E+06 | 4.17E+05 | 18 | Cytoplasm |
| 769 | Q6P1F6 | PPP2R2A | 3.17E+06 | 1.17E+06 | 6.75E+05 | 3 | Cytoplasm |
| 770 | O35943 | FXN | 3.17E+06 | 2.50E+05 | 1.02E+05 | 6 | Cytoplasm |
| 771 | Q9JL56 | GDE1 | 3.17E+06 | 2.76E+06 | 9.19E+05 | 9 | Plasma Membrane |
| 772 | Q64455 | PTPRJ | 3.16E+06 | 4.32E+06 | 1.44E+06 | 9 | Plasma Membrane |
| 773 | P57776 | EEF1D | 3.16E+06 | 4.88E+05 | 2.82E+05 | 3 | Cytoplasm |
| 774 | Q8R5H1 | USP15 | 3.16E+06 | 1.52E+06 | 2.19E+05 | 48 | Cytoplasm |
| 775 | P61222 | ABCE1 | 3.14E+06 | 1.31E+06 | 3.79E+05 | 12 | Cytoplasm |
| 776 | P59108 | CPNE2 | 3.13E+06 | 9.35E+05 | 3.82E+05 | 6 | Cytoplasm |
| 777 | Q9CYZ2 | TPD52L2 | 3.12E+06 | 7.61E+05 | 4.39E+05 | 3 | Cytoplasm |
| 778 | Q9JLV5 | CUL3 | 3.12E+06 | 4.15E+05 | 1.69E+05 | 6 | Nucleus |
| 779 | Q9CR21 | NDUFAB1 | 3.12E+06 | 4.07E+06 | 1.66E+06 | 6 | Cytoplasm |
| 780 | P40237 | CD82 | 3.11E+06 | 3.08E+06 | 1.26E+06 | 6 | Plasma Membrane |
| 781 | O08734 | BAK1 | 3.10E+06 | 8.85E+05 | 5.11E+05 | 3 | Cytoplasm |
| 782 | Q8R4N0 | CLYBL | 3.08E+06 | 1.42E+06 | 8.19E+05 | 3 | Cytoplasm |
| 783 | P61202 | Cops2 | 3.07E+06 | 2.34E+06 | 6.04E+05 | 15 | Nucleus |
| 784 | Q6PA06 | ATL2 | 3.05E+06 | 2.09E+06 | 6.95E+05 | 9 | Cytoplasm |
| 785 | Q64310 | SURF4 | 3.05E+06 | 1.66E+06 | 6.79E+05 | 6 | Cytoplasm |
| 786 | P47911 | RPL6 | 3.04E+06 | 1.93E+06 | 5.57E+05 | 12 | Nucleus |
| 787 | O54734 | DDOST | 3.03E+06 | 1.16E+06 | 3.88E+05 | 9 | Cytoplasm |
| 788 | Q91WS0 | CISD1 | 3.03E+06 | 7.68E+05 | 3.13E+05 | 6 | Cytoplasm |
| 789 | Q60738 | SLC30A1 | 3.02E+06 | 9.61E+05 | 2.77E+05 | 12 | Plasma Membrane |
| 790 | Q80TE7 | LRRC7 | 3.00E+06 | 2.88E+05 | 1.66E+05 | 3 | Plasma Membrane |
| 791 | Q8VDP3 | MICAL1 | 3.00E+06 | 6.80E+05 | 3.93E+05 | 3 | Cytoplasm |
| 792 | Q99JB2 | STOML2 | 2.99E+06 | 1.74E+06 | 4.48E+05 | 15 | Plasma Membrane |
| 793 | Q9Z1Q2 | ABHD16A | 2.99E+06 | 1.90E+06 | 6.33E+05 | 9 | Cytoplasm |
| 794 | Q9Z0F4 | CIB1 | 2.99E+06 | 3.87E+06 | 1.12E+06 | 12 | Nucleus |
| 795 | P54731 | FAF1 | 2.99E+06 | 3.87E+05 | 1.29E+05 | 9 | Nucleus |
| 796 | P53026 | RPL10A | 2.99E+06 | 1.21E+06 | 4.96E+05 | 6 | Nucleus |
| 797 | Q8BJY1 | PSMD5 | 2.98E+06 | 1.16E+06 | 2.24E+05 | 27 | Cytoplasm |
| 798 | P62849 | RPS24 | 2.98E+06 | 1.56E+05 | 9.01E+04 | 3 | Cytoplasm |
| 799 | Q8R574 | PRPSAP2 | 2.97E+06 | 7.64E+05 | 3.12E+05 | 6 | Cytoplasm |
| 800 | Q9R1S7 | ABCC6 | 2.97E+06 | 2.90E+06 | 1.68E+06 | 3 | Plasma Membrane |
| 801 | P59999 | ARPC4 | 2.96E+06 | 6.56E+05 | 3.79E+05 | 3 | Cytoplasm |
| 802 | P62960 | YBX1 | 2.94E+06 | 2.23E+06 | 9.10E+05 | 6 | Nucleus |
| 803 | Q99LD4 | GPS1 | 2.93E+06 | 1.77E+06 | 4.58E+05 | 15 | Cytoplasm |
| 804 | P61804 | DAD1 | 2.93E+06 | 1.76E+05 | 1.01E+05 | 3 | Cytoplasm |
| 805 | P63321 | RALA | 2.93E+06 | 1.95E+05 | 1.12E+05 | 3 | Cytoplasm |
| 806 | Q8BGU5 | CCNY | 2.93E+06 | 2.85E+05 | 1.16E+05 | 6 | Nucleus |
| 807 | P63276 | RPS17 | 2.92E+06 | 1.23E+05 | 7.11E+04 | 3 | Cytoplasm |
| 808 | O35855 | BCAT2 | 2.89E+06 | 2.83E+05 | 1.64E+05 | 3 | Cytoplasm |
| 809 | Q9DCC4 | PYCRL | 2.89E+06 | 8.20E+05 | 2.73E+05 | 9 | Cytoplasm |
| 810 | Q9CXZ1 | NDUFS4 | 2.88E+06 | 1.91E+06 | 7.78E+05 | 6 | Cytoplasm |
| 811 | P67871 | CSNK2B | 2.88E+06 | 5.56E+05 | 3.21E+05 | 3 | Cytoplasm |
| 812 | Q9QYK7 | RNF11 | 2.87E+06 | 1.04E+05 | 6.02E+04 | 3 | Cytoplasm |
| 813 | Q9R0Q6 | ARPC1A | 2.87E+06 | 3.52E+05 | 2.03E+05 | 3 | Extracellular Space |
| 814 | Q8R050 | GSPT1 | 2.86E+06 | 3.08E+05 | 1.78E+05 | 3 | Cytoplasm |
| 815 | Q91YT0 | NDUFV1 | 2.86E+06 | 2.18E+06 | 7.27E+05 | 9 | Cytoplasm |
| 816 | Q9D880 | TIMM50 | 2.85E+06 | 6.71E+05 | 2.74E+05 | 6 | Cytoplasm |
| 817 | P42669 | PURA | 2.85E+06 | 2.05E+06 | 8.37E+05 | 6 | Nucleus |
| 818 | P62245 | RPS15A | 2.84E+06 | 8.16E+05 | 3.33E+05 | 6 | Cytoplasm |
| 819 | P26443 | GLUD1 | 2.84E+06 | 1.42E+05 | 5.82E+04 | 6 | Cytoplasm |
| 820 | P70398 | USP9X | 2.84E+06 | 1.43E+06 | 4.13E+05 | 12 | Plasma Membrane |
| 821 | P50516 | ATP6V1A | 2.82E+06 | 1.58E+06 | 4.07E+05 | 15 | Plasma Membrane |
| 822 | O55222 | ILK | 2.81E+06 | 1.86E+06 | 4.39E+05 | 18 | Plasma Membrane |
| 823 | P54071 | IDH2 | 2.81E+06 | 3.17E+06 | 7.47E+05 | 18 | Cytoplasm |
| 824 | Q8JZN5 | ACAD9 | 2.78E+06 | 3.83E+05 | 2.21E+05 | 3 | Cytoplasm |
| 825 | Q8VDL4 | ADPGK | 2.78E+06 | 3.47E+05 | 2.01E+05 | 3 | Plasma Membrane |
| 826 | O88543 | COPS3 | 2.77E+06 | 1.06E+06 | 2.50E+05 | 18 | Cytoplasm |
| 827 | P62827 | RAN | 2.77E+06 | 4.62E+05 | 2.67E+05 | 3 | Nucleus |
| 828 | Q64331 | MYO6 | 2.77E+06 | 2.33E+05 | 1.34E+05 | 3 | Cytoplasm |
| 829 | Q8BMF4 | DLAT | 2.76E+06 | 7.03E+05 | 2.87E+05 | 6 | Cytoplasm |
| 830 | P24668 | M6PR | 2.76E+06 | 2.24E+06 | 9.13E+05 | 6 | Cytoplasm |
| 831 | Q9QWY8 | ASAP1 | 2.74E+06 | 1.11E+06 | 4.55E+05 | 6 | Plasma Membrane |
| 832 | Q9CX30 | YIF1B | 2.73E+06 | 5.06E+05 | 2.06E+05 | 6 | Cytoplasm |
| 833 | P07758 | SERPINA1 | 2.72E+06 | 2.20E+06 | 5.67E+05 | 15 | Extracellular Space |

| | | | | | | | |
|---|---|---|---|---|---|---|---|
| 695 | Q99KE1 | ME2 | 3.09E+06 | 2.92E+06 | 8.44E+05 | 12 | Cytoplasm |
| 696 | Q62348 | TSN | 3.08E+06 | 1.48E+06 | 6.06E+05 | 6 | Nucleus |
| 697 | O88792 | F11R | 3.08E+06 | 4.02E+06 | 1.04E+06 | 15 | Plasma Membrane |
| 698 | P51410 | RPL9 | 3.08E+06 | 1.62E+06 | 5.41E+05 | 9 | Nucleus |
| 699 | Q9DCD0 | PGD | 3.07E+06 | 3.81E+05 | 2.20E+05 | 3 | Cytoplasm |
| 700 | Q64310 | SURF4 | 3.06E+06 | 1.58E+06 | 6.46E+05 | 6 | Cytoplasm |
| 701 | Q9CXE7 | TMED5 | 3.05E+06 | 2.13E+05 | 1.23E+05 | 3 | Cytoplasm |
| 702 | Q7TMR0 | PRCP | 3.05E+06 | 1.48E+06 | 8.55E+05 | 3 | Cytoplasm |
| 703 | Q60790 | RASA3 | 3.03E+06 | 9.76E+05 | 5.63E+05 | 3 | Plasma Membrane |
| 704 | P99029 | PRDX5 | 3.03E+06 | 1.30E+06 | 7.50E+05 | 3 | Cytoplasm |
| 705 | Q9CZX8 | RPS19 | 3.03E+06 | 1.70E+06 | 5.68E+05 | 9 | Cytoplasm |
| 706 | O08756 | HSD17B10 | 3.02E+06 | 3.64E+06 | 1.05E+06 | 12 | Cytoplasm |
| 707 | P50580 | PA2G4 | 3.02E+06 | 1.21E+06 | 3.11E+05 | 15 | Nucleus |
| 708 | P41105 | RPL28 | 3.02E+06 | 5.76E+05 | 3.32E+05 | 3 | Cytoplasm |
| 709 | Q80VP1 | EPN1 | 3.00E+06 | 1.05E+06 | 3.04E+05 | 12 | Plasma Membrane |
| 710 | O88543 | COPS3 | 3.00E+06 | 1.06E+06 | 2.51E+05 | 18 | Cytoplasm |
| 711 | Q99JR1 | SFXN1 | 2.99E+06 | 1.82E+06 | 4.28E+05 | 18 | Cytoplasm |
| 712 | Q9CY24 | TMEM179B | 2.98E+06 | 7.56E+05 | 4.37E+05 | 3 | Extracellular Space |
| 713 | P68040 | GNB2L1 | 2.98E+06 | 2.01E+06 | 5.20E+05 | 15 | Cytoplasm |
| 714 | Q920L1 | FADS1 | 2.97E+06 | 1.99E+05 | 1.15E+05 | 3 | Plasma Membrane |
| 715 | Q920G8 | SLC25A37 | 2.96E+06 | 3.56E+06 | 6.19E+05 | 33 | Cytoplasm |
| 716 | O09167 | RPL21 | 2.95E+06 | 3.04E+06 | 1.24E+06 | 6 | Cytoplasm |
| 717 | P27661 | H2AFX | 2.93E+06 | 1.18E+07 | 4.81E+06 | 6 | Nucleus |
| 718 | Q9Z125 | CREB3L1 | 2.93E+06 | 8.20E+05 | 4.73E+05 | 3 | Nucleus |
| 719 | P12970 | RPL7A | 2.92E+06 | 2.17E+06 | 7.25E+05 | 9 | Cytoplasm |
| 720 | Q9CYZ2 | TPD52L2 | 2.91E+06 | 1.02E+06 | 5.90E+05 | 3 | Cytoplasm |
| 721 | P62830 | RPL23 | 2.90E+06 | 2.98E+06 | 1.72E+06 | 3 | Cytoplasm |
| 722 | Q99LC5 | ETFA | 2.90E+06 | 3.21E+06 | 6.18E+05 | 27 | Cytoplasm |
| 723 | O08739 | AMPD3 | 2.89E+06 | 1.60E+06 | 3.26E+05 | 24 | Cytoplasm |
| 724 | Q921H8 | ACAA1 | 2.89E+06 | 1.29E+06 | 7.44E+05 | 3 | Cytoplasm |
| 725 | P62137 | PPP1CA | 2.89E+06 | 2.42E+06 | 5.71E+05 | 18 | Cytoplasm |
| 726 | P42125 | ECI1 | 2.87E+06 | 3.17E+06 | 1.29E+06 | 6 | Cytoplasm |
| 727 | Q9EQH2 | ERAP1 | 2.86E+06 | 4.34E+05 | 2.51E+05 | 3 | Extracellular Space |
| 728 | P35979 | RPL12 | 2.85E+06 | 1.93E+06 | 4.98E+05 | 15 | Nucleus |
| 729 | Q8VDP3 | MICAL1 | 2.83E+06 | 5.92E+05 | 3.42E+05 | 3 | Cytoplasm |
| 730 | P61982 | YWHAG | 2.82E+06 | 1.89E+06 | 7.71E+05 | 6 | Cytoplasm |
| 731 | Q61335 | BCAP31 | 2.82E+06 | 6.27E+05 | 3.62E+05 | 3 | Cytoplasm |
| 732 | P47757 | CAPZB | 2.81E+06 | 1.26E+06 | 5.14E+05 | 6 | Cytoplasm |
| 733 | Q8BNI4 | DERL2 | 2.80E+06 | 6.50E+05 | 3.75E+05 | 3 | Cytoplasm |
| 734 | P67871 | CSNK2B | 2.80E+06 | 1.87E+05 | 1.08E+05 | 3 | Cytoplasm |
| 735 | Q8CGK3 | LONP1 | 2.79E+06 | 2.66E+06 | 1.09E+06 | 6 | Cytoplasm |
| 736 | Q61838 | Pzp | 2.78E+06 | 1.90E+06 | 1.10E+06 | 3 | Extracellular Space |
| 737 | Q9CZ44 | NSFL1C | 2.77E+06 | 9.73E+05 | 2.81E+05 | 12 | Cytoplasm |
| 738 | Q00262 | STX2 | 2.77E+06 | 1.73E+06 | 4.09E+05 | 18 | Cytoplasm |
| 739 | Q6ZWV7 | RPL35 | 2.77E+06 | 1.96E+06 | 7.98E+05 | 6 | Cytoplasm |
| 740 | Q8BMS1 | HADHA | 2.76E+06 | 2.92E+06 | 4.50E+05 | 42 | Cytoplasm |
| 741 | P70398 | USP9X | 2.76E+06 | 9.66E+05 | 2.79E+05 | 12 | Plasma Membrane |
| 742 | Q9JHR7 | IDE | 2.75E+06 | 6.53E+05 | 2.67E+05 | 6 | Extracellular Space |
| 743 | P97370 | ATP1B3 | 2.75E+06 | 1.51E+06 | 4.36E+05 | 12 | Plasma Membrane |
| 744 | Q8BFZ9 | ERLIN2 | 2.74E+06 | 2.55E+05 | 1.47E+05 | 3 | Plasma Membrane |
| 745 | P43275 | Hist1h1a | 2.73E+06 | 1.08E+07 | 3.11E+06 | 12 | Nucleus |
| 746 | Q3TTY5 | Krt2 | 2.73E+06 | 1.31E+06 | 7.55E+05 | 3 | Cytoplasm |
| 747 | Q9CX30 | YIF1B | 2.73E+06 | 2.46E+05 | 1.00E+05 | 6 | Cytoplasm |
| 748 | Q91YA2 | PSKH1 | 2.72E+06 | 2.46E+05 | 1.42E+05 | 3 | Nucleus |
| 749 | Q6ZWN5 | RPS9 | 2.72E+06 | 1.76E+06 | 5.85E+05 | 9 | Cytoplasm |
| 750 | Q8R2Y8 | PTRH2 | 2.72E+06 | 4.73E+06 | 1.93E+06 | 6 | Cytoplasm |
| 751 | Q8R3K3 | PTCD2 | 2.71E+06 | 1.53E+05 | 8.86E+04 | 3 | Cytoplasm |
| 752 | Q8BU31 | RAP2C | 2.70E+06 | 4.32E+05 | 2.49E+05 | 3 | Cytoplasm |
| 753 | Q80WC7 | AGFG2 | 2.70E+06 | 7.37E+05 | 2.13E+05 | 12 | Cytoplasm |
| 754 | Q6P1F6 | PPP2R2A | 2.70E+06 | 7.99E+05 | 4.62E+05 | 3 | Cytoplasm |
| 755 | P70404 | IDH3G | 2.68E+06 | 3.06E+06 | 1.25E+06 | 6 | Cytoplasm |
| 756 | Q9QWY8 | ASAP1 | 2.67E+06 | 1.14E+06 | 4.64E+05 | 6 | Plasma Membrane |
| 757 | B2RPV6 | MMRN1 | 2.67E+06 | 1.19E+06 | 2.07E+05 | 33 | Extracellular Space |
| 758 | Q9CPR4 | RPL17 | 2.66E+06 | 8.51E+05 | 4.92E+05 | 3 | Cytoplasm |
| 759 | Q6NS82 | FAM134A | 2.66E+06 | 2.67E+05 | 1.54E+05 | 3 | Extracellular Space |
| 760 | Q922Q8 | LRRC59 | 2.65E+06 | 6.72E+05 | 3.88E+05 | 3 | Cytoplasm |
| 761 | Q9CRB9 | CHCHD3 | 2.65E+06 | 1.90E+06 | 1.10E+06 | 3 | Cytoplasm |
| 762 | Q9CYL5 | GLIPR2 | 2.63E+06 | 4.28E+05 | 2.47E+05 | 3 | Cytoplasm |
| 763 | P17426 | AP2A1 | 2.63E+06 | 1.80E+06 | 5.99E+05 | 9 | Cytoplasm |
| 764 | Q91YP3 | DERA | 2.62E+06 | 8.99E+05 | 3.00E+05 | 9 | Cytoplasm |
| 765 | O89112 | LANCL1 | 2.61E+06 | 2.64E+05 | 1.52E+05 | 3 | Plasma Membrane |
| 766 | P40124 | CAP1 | 2.60E+06 | 8.98E+05 | 2.99E+05 | 9 | Plasma Membrane |
| 767 | P01029 | C4A/C4B | 2.60E+06 | 1.61E+06 | 4.15E+05 | 15 | Extracellular Space |
| 768 | Q8BH59 | SLC25A12 | 2.58E+06 | 2.65E+06 | 7.64E+05 | 12 | Cytoplasm |
| 769 | Q8K4F0 | CD226 | 2.57E+06 | 4.04E+05 | 2.33E+05 | 3 | Plasma Membrane |
| 770 | O88544 | COPS4 | 2.56E+06 | 9.63E+05 | 2.27E+05 | 18 | Cytoplasm |
| 771 | P42669 | PURA | 2.55E+06 | 2.05E+06 | 8.36E+05 | 6 | Nucleus |
| 772 | P59017 | BCL2L13 | 2.55E+06 | 2.25E+06 | 7.50E+05 | 9 | Cytoplasm |
| 773 | Q00623 | APOA1 | 2.54E+06 | 1.93E+06 | 6.44E+05 | 9 | Extracellular Space |
| 774 | P68254 | YWHAQ | 2.54E+06 | 1.35E+06 | 4.49E+05 | 9 | Cytoplasm |
| 775 | Q62469 | ITGA2 | 2.54E+06 | 1.99E+06 | 3.46E+05 | 33 | Plasma Membrane |
| 776 | Q8VEH3 | ARL8A | 2.53E+06 | 3.74E+05 | 2.16E+05 | 3 | Cytoplasm |
| 777 | Q8K411 | PITRM1 | 2.53E+06 | 3.40E+06 | 7.42E+05 | 21 | Cytoplasm |
| 778 | P09671 | SOD2 | 2.53E+06 | 2.29E+06 | 7.64E+05 | 9 | Cytoplasm |
| 779 | P61222 | ABCE1 | 2.52E+06 | 1.06E+06 | 3.07E+05 | 12 | Cytoplasm |
| 780 | Q7TMY8 | HUWE1 | 2.52E+06 | 2.46E+05 | 1.42E+05 | 3 | Nucleus |
| 781 | Q8BQ47 | CNPY4 | 2.51E+06 | 1.97E+05 | 1.14E+05 | 3 | Extracellular Space |
| 782 | P59999 | ARPC4 | 2.51E+06 | 1.02E+06 | 5.86E+05 | 3 | Cytoplasm |
| 783 | Q9ER72 | CARS | 2.51E+06 | 1.22E+06 | 4.97E+05 | 6 | Cytoplasm |
| 784 | Q99L45 | EIF2S2 | 2.51E+06 | 6.88E+05 | 2.29E+05 | 9 | Cytoplasm |
| 785 | Q8CAQ8 | IMMT | 2.50E+06 | 2.21E+06 | 4.26E+05 | 27 | Cytoplasm |
| 786 | P59108 | CPNE2 | 2.50E+06 | 1.05E+06 | 4.30E+05 | 6 | Cytoplasm |
| 787 | Q6PDI5 | KIAA0368 | 2.50E+06 | 1.48E+06 | 2.84E+05 | 27 | Cytoplasm |
| 788 | O08734 | BAK1 | 2.48E+06 | 1.31E+06 | 7.57E+05 | 3 | Cytoplasm |
| 789 | Q9CQA3 | SDHB | 2.47E+06 | 1.24E+06 | 4.12E+05 | 9 | Cytoplasm |
| 790 | Q6PA06 | ATL2 | 2.46E+06 | 1.72E+06 | 5.73E+05 | 9 | Cytoplasm |
| 791 | Q9CZM2 | RPL15 | 2.45E+06 | 1.24E+06 | 4.13E+05 | 9 | Cytoplasm |
| 792 | Q3U0B3 | DHRS11 | 2.44E+06 | 1.30E+05 | 7.52E+04 | 3 | Cytoplasm |
| 793 | P70295 | AUP1 | 2.41E+06 | 1.08E+06 | 6.21E+05 | 3 | Cytoplasm |
| 794 | P97352 | S100A13 | 2.40E+06 | 1.84E+06 | 1.07E+06 | 3 | Cytoplasm |
| 795 | O70503 | HSD17B12 | 2.40E+06 | 4.92E+05 | 2.01E+05 | 6 | Cytoplasm |
| 796 | Q9D0K2 | OXCT1 | 2.40E+06 | 3.60E+05 | 2.08E+05 | 3 | Cytoplasm |
| 797 | P62267 | RPS23 | 2.40E+06 | 1.32E+06 | 7.64E+05 | 3 | Cytoplasm |
| 798 | Q9CYN9 | ATP6AP2 | 2.39E+06 | 3.10E+05 | 1.79E+05 | 3 | Cytoplasm |
| 799 | P50516 | ATP6V1A | 2.39E+06 | 1.43E+06 | 3.70E+05 | 15 | Plasma Membrane |
| 800 | Q8BHL3 | TBC1D10B | 2.37E+06 | 3.83E+05 | 2.21E+05 | 3 | Plasma Membrane |
| 801 | Q9D8E6 | RPL4 | 2.37E+06 | 1.60E+06 | 4.62E+05 | 12 | Cytoplasm |
| 802 | P61804 | DAD1 | 2.36E+06 | 8.80E+05 | 5.08E+05 | 3 | Cytoplasm |
| 803 | Q7TMM9 | TUBB2A | 2.36E+06 | 2.56E+05 | 1.05E+05 | 6 | Cytoplasm |
| 804 | P62908 | RPS3 | 2.36E+06 | 2.18E+06 | 4.45E+05 | 24 | Cytoplasm |
| 805 | Q9R0M6 | RAB9A | 2.36E+06 | 1.03E+05 | 5.93E+04 | 3 | Cytoplasm |
| 806 | O54734 | DDOST | 2.35E+06 | 1.56E+06 | 5.20E+05 | 9 | Cytoplasm |
| 807 | Q80UU9 | PGRMC2 | 2.35E+06 | 7.81E+05 | 4.51E+05 | 3 | Nucleus |
| 808 | Q9CY50 | SSR1 | 2.34E+06 | 1.25E+06 | 5.08E+05 | 6 | Cytoplasm |
| 809 | Q8C4S8 | DENND2A | 2.34E+06 | 3.72E+05 | 2.15E+05 | 3 | Cytoplasm |
| 810 | P24547 | IMPDH2 | 2.34E+06 | 1.28E+06 | 7.38E+05 | 3 | Cytoplasm |
| 811 | Q8K2K6 | AGFG1 | 2.34E+06 | 1.36E+05 | 7.84E+04 | 3 | Nucleus |
| 812 | Q9WTX6 | CUL1 | 2.32E+06 | 1.08E+06 | 2.07E+05 | 27 | Nucleus |
| 813 | Q9JM76 | ARPC3 | 2.31E+06 | 8.49E+05 | 3.47E+05 | 6 | Cytoplasm |
| 814 | P48962 | SLC25A4 | 2.30E+06 | 1.38E+06 | 4.61E+05 | 9 | Cytoplasm |
| 815 | Q7TN58 | TMC8 | 2.30E+06 | 8.20E+05 | 2.73E+05 | 9 | Cytoplasm |
| 816 | Q61753 | PHGDH | 2.28E+06 | 8.33E+05 | 2.15E+05 | 15 | Cytoplasm |
| 817 | Q8BIG7 | COMTD1 | 2.28E+06 | 1.38E+06 | 7.96E+05 | 3 | Plasma Membrane |
| 818 | P12382 | PFKL | 2.28E+06 | 3.30E+05 | 1.91E+05 | 3 | Cytoplasm |
| 819 | Q99K41 | EMILIN1 | 2.27E+06 | 2.99E+06 | 1.22E+06 | 6 | Extracellular Space |
| 820 | Q61207 | PSAP | 2.27E+06 | 1.36E+06 | 7.86E+05 | 3 | Extracellular Space |
| 821 | P62245 | RPS15A | 2.27E+06 | 8.04E+05 | 3.28E+05 | 6 | Cytoplasm |
| 822 | Q8VDP6 | CDIPT | 2.27E+06 | 5.80E+04 | 3.35E+04 | 3 | Cytoplasm |
| 823 | Q64455 | PTPRJ | 2.26E+06 | 2.89E+06 | 9.64E+05 | 9 | Plasma Membrane |
| 824 | Q3UHH2 | SLC22A23 | 2.25E+06 | 2.11E+05 | 1.22E+05 | 3 | Plasma Membrane |
| 825 | Q8BV13 | COPS7B | 2.25E+06 | 3.04E+05 | 1.76E+05 | 3 | Cytoplasm |
| 826 | Q3V0G7 | GARNL3 | 2.24E+06 | 7.55E+05 | 4.36E+05 | 3 | Cytoplasm |
| 827 | P70452 | STX4 | 2.24E+06 | 9.25E+05 | 3.08E+05 | 9 | Plasma Membrane |
| 828 | P24668 | M6PR | 2.22E+06 | 2.27E+06 | 9.28E+05 | 6 | Cytoplasm |
| 829 | Q6ZWX6 | EIF2S1 | 2.22E+06 | 1.04E+06 | 2.45E+05 | 18 | Cytoplasm |
| 830 | P03921 | MT-ND5 | 2.22E+06 | 2.66E+06 | 8.86E+05 | 9 | Cytoplasm |
| 831 | Q6URW6 | MYH14 | 2.21E+06 | 2.45E+06 | 1.00E+06 | 6 | Extracellular Space |
| 832 | Q9CQ75 | NDUFA2 | 2.21E+06 | 8.14E+05 | 4.70E+05 | 3 | Cytoplasm |
| 833 | P70697 | UROD | 2.21E+06 | 5.35E+05 | 3.09E+05 | 3 | Cytoplasm |

| | | | | | | | |
|---|---|---|---|---|---|---|---|
| 695 | P59325 | EIF5 | 2766444 | 333455 | 192520 | 3 | Cytoplasm |
| 696 | P70460 | VASP | 2758705 | 708797 | 409224 | 3 | Plasma Membrane |
| 697 | Q8R3K3 | PTCD2 | 2758270 | 1075637 | 621019 | 3 | Cytoplasm |
| 698 | Q99P72 | RTN4 | 2747067 | 4670995 | 1206046 | 15 | Cytoplasm |
| 699 | P70697 | UROD | 2744621 | 119899 | 69224 | 3 | Cytoplasm |
| 700 | P24668 | M6PR | 2744584 | 2059023 | 840593 | 6 | Cytoplasm |
| 701 | Q3V3R1 | MTHFD1L | 2727639 | 2048332 | 418114 | 24 | Cytoplasm |
| 702 | P58252 | EEF2 | 2721485 | 1492360 | 385326 | 15 | Cytoplasm |
| 703 | P62849 | RPS24 | 2719413 | 1315161 | 759309 | 3 | Cytoplasm |
| 704 | Q7TMR0 | PRCP | 2704090 | 2463252 | 1422159 | 3 | Cytoplasm |
| 705 | Q8VDD5 | MYH9 | 2699258 | 5877505 | 346335 | 288 | Cytoplasm |
| 706 | Q8BJY1 | PSMD5 | 2699116 | 866868 | 166829 | 27 | Cytoplasm |
| 707 | Q8VBV7 | COPS8 | 2693441 | 429423 | 247928 | 3 | Nucleus |
| 708 | Q8BH59 | SLC25A12 | 2680090 | 3155470 | 910906 | 12 | Cytoplasm |
| 709 | Q9CRB9 | CHCHD3 | 2670061 | 2480103 | 1431888 | 3 | Cytoplasm |
| 710 | Q80WQ2 | VAC14 | 2669567 | 161839 | 93438 | 3 | Cytoplasm |
| 711 | P18181 | CD48 | 2663666 | 255099 | 147281 | 3 | Plasma Membrane |
| 712 | Q9DCF9 | SSR3 | 2663499 | 1438911 | 830756 | 3 | Cytoplasm |
| 713 | Q99LC5 | ETFA | 2658183 | 3156680 | 607503 | 27 | Cytoplasm |
| 714 | P47964 | Rpl36 | 2656196 | 1016493 | 414982 | 6 | Nucleus |
| 715 | Q8BTM8 | FLNA | 2642980 | 10559490 | 561231 | 354 | Cytoplasm |
| 716 | Q8VDP3 | MICAL1 | 2636800 | 218641 | 126232 | 3 | Cytoplasm |
| 717 | P50518 | ATP6V1E1 | 2632677 | 528049 | 304869 | 3 | Cytoplasm |
| 718 | Q9D1C8 | VPS28 | 2630651 | 4783187 | 2761574 | 3 | Cytoplasm |
| 719 | Q9Z2I9 | SUCLA2 | 2629652 | 2204925 | 1273014 | 3 | Cytoplasm |
| 720 | P54731 | FAF1 | 2627196 | 335116 | 111705 | 9 | Nucleus |
| 721 | Q9CX00 | IST1 | 2623933 | 604165 | 348815 | 3 | Cytoplasm |
| 722 | Q61838 | Pzp | 2601545 | 1735517 | 1002001 | 3 | Extracellular Space |
| 723 | Q9WTX6 | CUL1 | 2595675 | 1219360 | 234666 | 27 | Nucleus |
| 724 | Q9QYK7 | RNF11 | 2595657 | 599637 | 346200 | 3 | Cytoplasm |
| 725 | Q9QWY8 | ASAP1 | 2593084 | 1087643 | 444028 | 6 | Plasma Membrane |
| 726 | Q99KE1 | ME2 | 2590825 | 3106141 | 896666 | 12 | Cytoplasm |
| 727 | P47757 | CAPZB | 2583971 | 846444 | 345559 | 6 | Cytoplasm |
| 728 | Q8VEH3 | ARL8A | 2582551 | 675992 | 390284 | 3 | Cytoplasm |
| 729 | Q3TZ89 | SEC31B | 2580391 | 4258096 | 2458413 | 3 | Cytoplasm |
| 730 | Q9R0M6 | RAB9A | 2571434 | 259944 | 150079 | 3 | Cytoplasm |
| 731 | P62281 | RPS11 | 2549928 | 3461749 | 1413253 | 6 | Cytoplasm |
| 732 | Q8BV13 | COPS7B | 2540211 | 321908 | 185854 | 3 | Cytoplasm |
| 733 | Q7TQ95 | KIAA1715 | 2535592 | 885147 | 228544 | 15 | Cytoplasm |
| 734 | Q91WS0 | CISD1 | 2523749 | 1557152 | 635705 | 6 | Cytoplasm |
| 735 | P42669 | PURA | 2521858 | 1901652 | 776346 | 6 | Nucleus |
| 736 | Q7TMY8 | HUWE1 | 2517158 | 283525 | 163693 | 3 | Nucleus |
| 737 | Q8C0L0 | TMX4 | 2499741 | 646926 | 373503 | 3 | Cytoplasm |
| 738 | P17047 | LAMP2 | 2495538 | 2344999 | 957342 | 6 | Plasma Membrane |
| 739 | Q9JKR6 | HYOU1 | 2490083 | 862251 | 188159 | 21 | Cytoplasm |
| 740 | Q9Z1Q2 | ABHD16A | 2461140 | 1763502 | 587834 | 9 | Cytoplasm |
| 741 | Q8VDL4 | ADPGK | 2449479 | 448141 | 258735 | 3 | Plasma Membrane |
| 742 | P08003 | PDIA4 | 2442958 | 1735736 | 354306 | 24 | Cytoplasm |
| 743 | Q9Z0F8 | ADAM17 | 2439522 | 2526821 | 1031570 | 6 | Plasma Membrane |
| 744 | Q8K2Z4 | NCAPD2 | 2439121 | 490540 | 283214 | 3 | Nucleus |
| 745 | Q6X893 | SLC44A1 | 2430912 | 887424 | 295808 | 9 | Plasma Membrane |
| 746 | P97370 | ATP1B3 | 2430582 | 1276525 | 368501 | 12 | Plasma Membrane |
| 747 | Q99JR1 | SFXN1 | 2424698 | 1951464 | 459965 | 18 | Cytoplasm |
| 748 | P43274 | Hist1h1e | 2423830 | 4956955 | 1279880 | 15 | Nucleus |
| 749 | Q8CAQ8 | IMMT | 2411908 | 2128799 | 409687 | 27 | Cytoplasm |
| 750 | Q9WUM5 | SUCLG1 | 2388062 | 1167462 | 389154 | 9 | Cytoplasm |
| 751 | Q6P1F6 | PPP2R2A | 2386374 | 798301 | 460899 | 3 | Cytoplasm |
| 752 | Q9CR64 | TMEM167A | 2374266 | 597016 | 344688 | 3 | Extracellular Space |
| 753 | P42125 | ECI1 | 2373091 | 1964312 | 801927 | 6 | Cytoplasm |
| 754 | Q99JB2 | STOML2 | 2369741 | 1732147 | 447238 | 15 | Plasma Membrane |
| 755 | Q9ER72 | CARS | 2367974 | 982073 | 400930 | 6 | Cytoplasm |
| 756 | Q64331 | MYO6 | 2358987 | 1128274 | 651410 | 3 | Cytoplasm |
| 757 | O35943 | FXN | 2346653 | 1946496 | 794654 | 6 | Cytoplasm |
| 758 | Q8R5J9 | ARL6IP5 | 2343186 | 676041 | 390312 | 3 | Cytoplasm |
| 759 | Q9EQH2 | ERAP1 | 2341109 | 207148 | 119597 | 3 | Extracellular Space |
| 760 | Q8VCF0 | MAVS | 2329073 | 2615489 | 1067769 | 6 | Cytoplasm |
| 761 | P97352 | S100A13 | 2328865 | 559843 | 323226 | 3 | Cytoplasm |
| 762 | Q8BZ09 | SLC25A21 | 2327278 | 1892581 | 488662 | 15 | Cytoplasm |
| 763 | Q9ERN0 | SCAMP2 | 2323312 | 938259 | 383043 | 6 | Cytoplasm |
| 764 | Q9EPK7 | XPO7 | 2312990 | 426265 | 174022 | 6 | Nucleus |
| 765 | Q8CG48 | SMC2 | 2306536 | 389300 | 224762 | 3 | Nucleus |
| 766 | Q8BZI6 | GUCD1 | 2296323 | 137764 | 79538 | 3 | Nucleus |
| 767 | P62960 | YBX1 | 2284686 | 2310437 | 943232 | 6 | Nucleus |
| 768 | Q99NB1 | ACSS1 | 2277798 | 1535826 | 511942 | 9 | Cytoplasm |
| 769 | Q80TH2 | ERBB2IP | 2277295 | 1350869 | 275745 | 24 | Plasma Membrane |
| 770 | Q80ZW2 | THEM6 | 2266489 | 1413126 | 364867 | 15 | Extracellular Space |
| 771 | Q9Z0T9 | ITGB6 | 2261562 | 1093097 | 631100 | 3 | Plasma Membrane |
| 772 | Q8CBE3 | WDR37 | 2253039 | 1141978 | 466210 | 6 | Nucleus |
| 773 | Q8BH57 | WDR48 | 2248665 | 524474 | 302805 | 3 | Cytoplasm |
| 774 | P56400 | GP1BB | 2245698 | 11344530 | 2673931 | 18 | Plasma Membrane |
| 775 | Q9D6K8 | FUNDC2 | 2243655 | 2003756 | 818030 | 6 | Cytoplasm |
| 776 | Q9D924 | ISCA1 | 2241755 | 1388160 | 566714 | 6 | Cytoplasm |
| 777 | Q6P5F9 | XPO1 | 2229619 | 599953 | 346383 | 3 | Nucleus |
| 778 | Q9CZX7 | TMEM55A | 2229310 | 644748 | 372245 | 3 | Cytoplasm |
| 779 | P40124 | CAP1 | 2227052 | 917357 | 305786 | 9 | Plasma Membrane |
| 780 | Q6NZJ6 | EIF4G1 | 2225750 | 1503199 | 501066 | 9 | Cytoplasm |
| 781 | P61255 | RPL26 | 2223662 | 1061514 | 612865 | 3 | Cytoplasm |
| 782 | Q9CXE7 | TMED5 | 2222702 | 653558 | 377332 | 3 | Cytoplasm |
| 783 | Q3U487 | HECTD3 | 2217398 | 1383408 | 564774 | 6 | Cytoplasm |
| 784 | P59999 | ARPC4 | 2198987 | 1074117 | 620142 | 3 | Cytoplasm |
| 785 | Q3UHH2 | SLC22A23 | 2196385 | 146951 | 84842 | 3 | Plasma Membrane |
| 786 | Q9CX30 | YIF1B | 2187171 | 849501 | 346807 | 6 | Cytoplasm |
| 787 | Q61207 | PSAP | 2178099 | 1912126 | 1103967 | 3 | Extracellular Space |
| 788 | Q9WVC8 | SLC26A3 | 2173212 | 262339 | 151461 | 3 | Plasma Membrane |
| 789 | Q9CR21 | NDUFAB1 | 2161352 | 3994788 | 1630865 | 6 | Cytoplasm |
| 790 | O88668 | CREG1 | 2143187 | 745797 | 304470 | 6 | Nucleus |
| 791 | P62301 | RPS13 | 2122885 | 1360730 | 555516 | 6 | Cytoplasm |
| 792 | P62342 | SELT | 2121722 | 1029315 | 594275 | 3 | Cytoplasm |
| 793 | O89013 | LEPROT | 2109243 | 1661988 | 678504 | 6 | Plasma Membrane |
| 794 | Q60575 | KIF1B | 2099572 | 2429465 | 1402652 | 3 | Cytoplasm |
| 795 | O08599 | STXBP1 | 2082866 | 1440717 | 588170 | 6 | Cytoplasm |
| 796 | Q8C4S8 | DENND2A | 2078809 | 555590 | 320770 | 3 | Cytoplasm |
| 797 | Q9D880 | TIMM50 | 2078522 | 1440001 | 587878 | 6 | Cytoplasm |
| 798 | Q3THE2 | MYL12A | 2074303 | 3792209 | 979144 | 15 | Cytoplasm |
| 799 | Q91YP3 | DERA | 2068713 | 748797 | 249599 | 9 | Cytoplasm |
| 800 | O08734 | BAK1 | 2068221 | 1302482 | 751989 | 3 | Cytoplasm |
| 801 | P47738 | ALDH2 | 2063726 | 5679779 | 669368 | 72 | Cytoplasm |
| 802 | Q8C156 | NCAPH | 2060700 | 201233 | 116182 | 3 | Nucleus |
| 803 | Q9DCB8 | ISCA2 | 2048064 | 2227140 | 1285840 | 3 | Cytoplasm |
| 804 | P35279 | RAB6A | 2046939 | 1581223 | 645532 | 6 | Cytoplasm |
| 805 | Q811D0 | DLG1 | 2046309 | 999663 | 333221 | 9 | Plasma Membrane |
| 806 | O88456 | CAPNS1 | 2045122 | 231781 | 133819 | 3 | Cytoplasm |
| 807 | P01029 | C4A/C4B | 2041445 | 3563260 | 920030 | 15 | Extracellular Space |
| 808 | Q9CPQ3 | TOMM22 | 2034592 | 1399884 | 571500 | 6 | Cytoplasm |
| 809 | Q9CRY7 | GDPD1 | 2023211 | 432869 | 249917 | 3 | Plasma Membrane |
| 810 | Q6PCP5 | Mff | 2020761 | 3227790 | 1317740 | 6 | Cytoplasm |
| 811 | O35114 | SCARB2 | 2015221 | 1997285 | 1153133 | 3 | Plasma Membrane |
| 812 | Q9JJ59 | ABCB9 | 2008977 | 4500984 | 2598644 | 3 | Cytoplasm |
| 813 | Q8BU31 | RAP2C | 2008935 | 921647 | 532113 | 3 | Cytoplasm |
| 814 | O35855 | BCAT2 | 2000262 | 900603 | 519963 | 3 | Cytoplasm |
| 815 | Q7TN58 | TMC8 | 1998885 | 725464 | 241821 | 9 | Cytoplasm |
| 816 | Q9JHU4 | DYNC1H1 | 1998787 | 247999 | 143182 | 3 | Cytoplasm |
| 817 | P41438 | SLC19A1 | 1996116 | 140465 | 81097 | 3 | Plasma Membrane |
| 818 | Q6NS82 | FAM134A | 1995553 | 335525 | 193716 | 3 | Extracellular Space |
| 819 | A2AAJ9 | OBSCN | 1992854 | 576840 | 333039 | 3 | Cytoplasm |
| 820 | P21279 | GNAQ | 1992617 | 1682053 | 971134 | 3 | Plasma Membrane |
| 821 | Q5U458 | DNAJC11 | 1976715 | 1072572 | 437876 | 6 | Cytoplasm |
| 822 | P62137 | PPP1CA | 1972366 | 1103693 | 260143 | 18 | Cytoplasm |
| 823 | O88186 | GP9 | 1956551 | 9879114 | 2851855 | 12 | Plasma Membrane |
| 824 | Q9JHZ2 | ANKH | 1954989 | 564976 | 326189 | 3 | Plasma Membrane |
| 825 | Q9Z125 | CREB3L1 | 1950394 | 1745101 | 1007535 | 3 | Nucleus |
| 826 | Q9CQ75 | NDUFA2 | 1942932 | 1766307 | 1019778 | 3 | Cytoplasm |
| 827 | Q8BX70 | VPS13C | 1940015 | 919159 | 237326 | 15 | Cytoplasm |
| 828 | P03921 | MT-ND5 | 1934607 | 1755323 | 585108 | 9 | Cytoplasm |
| 829 | Q4VAA2 | CDV3 | 1933215 | 441445 | 254868 | 3 | Cytoplasm |
| 830 | Q921L3 | TMCO1 | 1931722 | 1016298 | 586760 | 3 | Cytoplasm |
| 831 | Q05920 | PC | 1925434 | 1688410 | 435946 | 15 | Cytoplasm |
| 832 | Q61144 | PSEN2 | 1920675 | 351534 | 202958 | 3 | Cytoplasm |
| 833 | Q63739 | PTP4A1 | 1912251 | 261282 | 150851 | 3 | Cytoplasm |

| | | | | | | | |
|---|---|---|---|---|---|---|---|
| 834 | Q9QZ73 | DCUN1D1 | 2.15E+06 | 7.28E+05 | 4.21E+05 | 3 | Nucleus |
| 835 | Q80WQ6 | RHBDF2 | 2.14E+06 | 1.55E+06 | 6.31E+05 | 6 | Cytoplasm |
| 836 | P63321 | RALA | 2.13E+06 | 1.98E+05 | 1.14E+05 | 3 | Cytoplasm |
| 837 | P47963 | RPL13 | 2.13E+06 | 9.34E+05 | 2.70E+05 | 12 | Nucleus |
| 838 | Q8R3K3 | PTCD2 | 2.11E+06 | 2.11E+05 | 1.22E+05 | 3 | Cytoplasm |
| 839 | P59325 | EIF5 | 2.11E+06 | 1.58E+06 | 9.12E+05 | 3 | Cytoplasm |
| 840 | P51437 | CAMP | 2.11E+06 | 1.39E+06 | 5.68E+05 | 6 | Cytoplasm |
| 841 | Q8CBQ5 | PI4K2B | 2.09E+06 | 8.12E+05 | 2.34E+05 | 12 | Cytoplasm |
| 842 | P24668 | M6PR | 2.07E+06 | 1.14E+06 | 4.66E+05 | 6 | Cytoplasm |
| 843 | P14115 | RPL27A | 2.07E+06 | 3.81E+05 | 2.20E+05 | 3 | Cytoplasm |
| 844 | Q99K41 | EMILIN1 | 2.07E+06 | 2.63E+06 | 1.07E+06 | 6 | Extracellular Space |
| 845 | Q6P9J9 | ANO6 | 2.06E+06 | 9.22E+05 | 5.32E+05 | 3 | Plasma Membrane |
| 846 | P54071 | IDH2 | 2.05E+06 | 1.76E+06 | 4.15E+05 | 18 | Cytoplasm |
| 847 | Q6NSQ9 | G6PC3 | 2.05E+06 | 7.74E+05 | 4.47E+05 | 3 | Cytoplasm |
| 848 | P70452 | STX4 | 2.05E+06 | 1.04E+06 | 3.46E+05 | 9 | Plasma Membrane |
| 849 | Q64455 | PTPRJ | 2.05E+06 | 3.26E+06 | 1.09E+06 | 9 | Plasma Membrane |
| 850 | Q9WVC8 | SLC26A3 | 2.05E+06 | 4.61E+05 | 2.66E+05 | 3 | Plasma Membrane |
| 851 | Q921H8 | ACAA1 | 2.04E+06 | 1.74E+06 | 1.01E+06 | 3 | Cytoplasm |
| 852 | Q6P5F9 | XPO1 | 2.04E+06 | 6.00E+05 | 3.46E+05 | 3 | Nucleus |
| 853 | P29391 | FTL | 2.04E+06 | 1.33E+06 | 4.44E+05 | 9 | Cytoplasm |
| 854 | Q9EPL8 | IPO7 | 2.03E+06 | 1.07E+06 | 2.33E+05 | 21 | Nucleus |
| 855 | P17426 | AP2A1 | 2.02E+06 | 1.14E+06 | 3.81E+05 | 9 | Cytoplasm |
| 856 | P61620 | SEC61A1 | 2.01E+06 | 5.86E+05 | 3.38E+05 | 3 | Cytoplasm |
| 857 | P43275 | Hist1h1a | 2.00E+06 | 1.82E+06 | 5.27E+05 | 12 | Nucleus |
| 858 | Q9JKB3 | YBX3 | 2.00E+06 | 8.01E+05 | 3.27E+05 | 6 | Nucleus |
| 859 | Q9CPR4 | RPL17 | 2.00E+06 | 5.57E+05 | 3.22E+05 | 3 | Cytoplasm |
| 860 | Q9JMC3 | DNAJA4 | 1.99E+06 | 2.06E+05 | 1.19E+05 | 3 | Nucleus |
| 861 | Q8K4F0 | CD226 | 1.98E+06 | 1.59E+06 | 9.20E+05 | 3 | Plasma Membrane |
| 862 | P24547 | IMPDH2 | 1.96E+06 | 6.77E+05 | 3.91E+05 | 3 | Cytoplasm |
| 863 | P19253 | RPL13A | 1.95E+06 | 2.28E+05 | 1.31E+05 | 3 | Cytoplasm |
| 864 | Q7TN58 | TMC8 | 1.93E+06 | 1.06E+06 | 3.52E+05 | 9 | Cytoplasm |
| 865 | Q922Q8 | LRRC59 | 1.93E+06 | 1.09E+06 | 6.31E+05 | 3 | Cytoplasm |
| 866 | Q9D0K2 | OXCT1 | 1.92E+06 | 1.27E+06 | 7.36E+05 | 3 | Cytoplasm |
| 867 | Q8CGC7 | EPRS | 1.92E+06 | 1.70E+06 | 6.94E+05 | 6 | Cytoplasm |
| 868 | O89013 | LEPROT | 1.92E+06 | 1.91E+06 | 7.81E+05 | 6 | Plasma Membrane |
| 869 | Q6PDI5 | KIAA0368 | 1.92E+06 | 1.36E+06 | 2.62E+05 | 27 | Cytoplasm |
| 870 | Q91YT0 | NDUFV1 | 1.91E+06 | 1.44E+06 | 4.81E+05 | 9 | Cytoplasm |
| 871 | P61804 | DAD1 | 1.91E+06 | 1.04E+06 | 6.00E+05 | 3 | Cytoplasm |
| 872 | Q8K2K6 | AGFG1 | 1.90E+06 | 3.28E+05 | 1.89E+05 | 3 | Nucleus |
| 873 | Q9CYN9 | ATP6AP2 | 1.90E+06 | 4.74E+05 | 2.74E+05 | 3 | Cytoplasm |
| 874 | O08734 | BAK1 | 1.90E+06 | 6.08E+05 | 3.51E+05 | 3 | Cytoplasm |
| 875 | P61222 | ABCE1 | 1.90E+06 | 8.42E+05 | 2.43E+05 | 12 | Cytoplasm |
| 876 | P12382 | PFKL | 1.89E+06 | 1.17E+06 | 6.78E+05 | 3 | Cytoplasm |
| 877 | Q8R0F8 | FAHD1 | 1.89E+06 | 3.55E+05 | 1.45E+05 | 6 | Cytoplasm |
| 878 | Q00623 | APOA1 | 1.89E+06 | 1.10E+06 | 3.67E+05 | 9 | Extracellular Space |
| 879 | P70302 | STIM1 | 1.88E+06 | 1.28E+06 | 4.27E+05 | 9 | Plasma Membrane |
| 880 | P62754 | RPS6 | 1.87E+06 | 1.33E+06 | 5.42E+05 | 6 | Cytoplasm |
| 881 | Q8BP67 | RPL24 | 1.87E+06 | 1.87E+05 | 6.24E+04 | 9 | Cytoplasm |
| 882 | Q9R0M6 | RAB9A | 1.87E+06 | 5.86E+05 | 3.39E+05 | 3 | Cytoplasm |
| 883 | Q8BMG7 | RAB3GAP2 | 1.87E+06 | 8.48E+05 | 3.46E+05 | 6 | Cytoplasm |
| 884 | P09671 | SOD2 | 1.86E+06 | 1.14E+06 | 3.80E+05 | 9 | Cytoplasm |
| 885 | Q9Z125 | CREB3L1 | 1.86E+06 | 5.56E+05 | 3.21E+05 | 3 | Nucleus |
| 886 | A2AAJ9 | OBSCN | 1.86E+06 | 4.57E+05 | 2.64E+05 | 3 | Cytoplasm |
| 887 | Q9DB34 | CHMP2A | 1.85E+06 | 5.49E+05 | 3.17E+05 | 3 | Cytoplasm |
| 888 | Q3V3R1 | MTHFD1L | 1.84E+06 | 7.73E+05 | 1.58E+05 | 24 | Cytoplasm |
| 889 | Q9CQ75 | NDUFA2 | 1.84E+06 | 3.12E+05 | 1.80E+05 | 3 | Cytoplasm |
| 890 | P57776 | EEF1D | 1.83E+06 | 8.00E+05 | 4.62E+05 | 3 | Cytoplasm |
| 891 | Q3U7U3 | FBXO7 | 1.81E+06 | 8.51E+05 | 4.91E+05 | 3 | Cytoplasm |
| 892 | Q7TMM9 | TUBB2A | 1.79E+06 | 9.42E+05 | 3.85E+05 | 6 | Cytoplasm |
| 893 | Q61207 | PSAP | 1.79E+06 | 2.65E+05 | 1.53E+05 | 3 | Extracellular Space |
| 894 | P97370 | ATP1B3 | 1.79E+06 | 7.13E+05 | 2.06E+05 | 12 | Plasma Membrane |
| 895 | O55222 | ILK | 1.78E+06 | 1.51E+06 | 3.57E+05 | 18 | Plasma Membrane |
| 896 | Q63739 | PTP4A1 | 1.78E+06 | 5.19E+04 | 3.00E+04 | 3 | Cytoplasm |
| 897 | P62270 | RPS18 | 1.76E+06 | 4.05E+05 | 1.35E+05 | 9 | Cytoplasm |
| 898 | Q8BU31 | RAP2C | 1.75E+06 | 4.17E+05 | 2.41E+05 | 3 | Cytoplasm |
| 899 | Q00612 | G6PD | 1.75E+06 | 5.77E+05 | 1.66E+05 | 12 | Cytoplasm |
| 900 | Q99JB2 | STOML2 | 1.75E+06 | 8.91E+05 | 2.30E+05 | 15 | Plasma Membrane |
| 901 | O70443 | GNAZ | 1.75E+06 | 1.28E+06 | 5.24E+05 | 6 | Plasma Membrane |
| 902 | Q9WVA4 | TAGLN2 | 1.73E+06 | 2.09E+06 | 6.02E+05 | 12 | Cytoplasm |
| 903 | P62245 | RPS15A | 1.73E+06 | 5.35E+05 | 2.18E+05 | 6 | Cytoplasm |
| 904 | P47911 | RPL6 | 1.73E+06 | 9.44E+05 | 2.73E+05 | 12 | Nucleus |
| 905 | Q9JHZ2 | ANKH | 1.72E+06 | 5.64E+05 | 3.26E+05 | 3 | Plasma Membrane |
| 906 | Q9Z2I0 | LETM1 | 1.72E+06 | 1.18E+06 | 6.83E+05 | 3 | Cytoplasm |
| 907 | O88783 | F5 | 1.70E+06 | 1.94E+06 | 6.47E+05 | 9 | Plasma Membrane |
| 908 | Q505D1 | ANKRD28 | 1.70E+06 | 1.13E+06 | 6.53E+05 | 3 | Cytoplasm |
| 909 | O35943 | FXN | 1.69E+06 | 3.29E+05 | 1.34E+05 | 6 | Cytoplasm |
| 910 | Q8R180 | ERO1A | 1.69E+06 | 3.56E+05 | 2.06E+05 | 3 | Cytoplasm |
| 911 | P62702 | RPS4Y1 | 1.69E+06 | 8.32E+05 | 1.70E+05 | 24 | Cytoplasm |
| 912 | Q9ET30 | TM9SF3 | 1.69E+06 | 6.47E+05 | 3.73E+05 | 3 | Cytoplasm |
| 913 | Q9ERI2 | RAB27A | 1.69E+06 | 1.13E+06 | 6.54E+05 | 3 | Cytoplasm |
| 914 | P03921 | MT-ND5 | 1.69E+06 | 1.27E+06 | 4.23E+05 | 9 | Cytoplasm |
| 915 | P24549 | ALDH1A1 | 1.68E+06 | 4.18E+05 | 2.41E+05 | 3 | Cytoplasm |
| 916 | P50518 | ATP6V1E1 | 1.67E+06 | 5.08E+05 | 2.93E+05 | 3 | Cytoplasm |
| 917 | Q3UHH2 | SLC22A23 | 1.67E+06 | 2.88E+05 | 1.66E+05 | 3 | Plasma Membrane |
| 918 | P26443 | GLUD1 | 1.66E+06 | 1.03E+06 | 4.22E+05 | 6 | Cytoplasm |
| 919 | Q8CG48 | SMC2 | 1.66E+06 | 2.87E+05 | 1.66E+05 | 3 | Nucleus |
| 920 | Q9CXZ1 | NDUFS4 | 1.65E+06 | 1.05E+06 | 4.30E+05 | 6 | Cytoplasm |
| 921 | P31725 | S100A9 | 1.65E+06 | 1.13E+06 | 4.62E+05 | 6 | Cytoplasm |
| 922 | Q8BMF4 | DLAT | 1.65E+06 | 4.53E+05 | 1.85E+05 | 6 | Cytoplasm |
| 923 | Q9CPQ3 | TOMM22 | 1.64E+06 | 3.03E+05 | 1.24E+05 | 6 | Cytoplasm |
| 924 | P17563 | SELENBP1 | 1.63E+06 | 1.34E+06 | 7.72E+05 | 3 | Cytoplasm |
| 925 | Q9CZ30 | OLA1 | 1.62E+06 | 4.79E+05 | 2.76E+05 | 3 | Cytoplasm |
| 926 | Q6NXH9 | KRT73 | 1.61E+06 | 2.67E+05 | 1.54E+05 | 3 | Extracellular Space |
| 927 | Q61838 | Pzp | 1.60E+06 | 3.35E+05 | 1.93E+05 | 3 | Extracellular Space |
| 928 | Q8BGH2 | SAMM50 | 1.60E+06 | 7.08E+05 | 2.89E+05 | 6 | Cytoplasm |
| 929 | P59108 | CPNE2 | 1.60E+06 | 1.01E+06 | 4.13E+05 | 6 | Cytoplasm |
| 930 | O08663 | METAP2 | 1.59E+06 | 7.83E+05 | 4.52E+05 | 3 | Cytoplasm |
| 931 | P31230 | AIMP1 | 1.58E+06 | 1.09E+06 | 4.45E+05 | 6 | Extracellular Space |
| 932 | Q9R1Q9 | ATP6AP1 | 1.58E+06 | 3.05E+05 | 1.76E+05 | 3 | Cytoplasm |
| 933 | Q05512 | MARK2 | 1.58E+06 | 6.95E+05 | 2.84E+05 | 6 | Cytoplasm |
| 934 | Q9DC51 | GNAI3 | 1.57E+06 | 6.72E+05 | 3.88E+05 | 3 | Cytoplasm |
| 935 | Q922Q4 | PYCR2 | 1.57E+06 | 2.24E+05 | 1.29E+05 | 3 | Cytoplasm |
| 936 | Q9JHK5 | PLEK | 1.57E+06 | 2.44E+06 | 1.41E+06 | 3 | Cytoplasm |
| 937 | Q9D880 | TIMM50 | 1.56E+06 | 4.46E+05 | 1.82E+05 | 6 | Cytoplasm |
| 938 | Q9D3G5 | STX11 | 1.55E+06 | 1.18E+06 | 2.78E+05 | 18 | Plasma Membrane |
| 939 | Q80UY2 | KCMF1 | 1.54E+06 | 1.03E+06 | 3.42E+05 | 9 | Extracellular Space |
| 940 | P41438 | SLC19A1 | 1.54E+06 | 3.47E+05 | 2.00E+05 | 3 | Plasma Membrane |
| 941 | Q8C4S8 | DENND2A | 1.54E+06 | 1.95E+05 | 1.13E+05 | 3 | Cytoplasm |
| 942 | Q61166 | MAPRE1 | 1.54E+06 | 2.70E+05 | 1.56E+05 | 3 | Cytoplasm |
| 943 | P60335 | PCBP1 | 1.53E+06 | 7.41E+05 | 1.91E+05 | 15 | Nucleus |
| 944 | O35744 | Chil3/Chil4 | 1.52E+06 | 1.59E+06 | 3.74E+05 | 18 | Cytoplasm |
| 945 | Q5XKN4 | JAGN1 | 1.51E+06 | 2.56E+05 | 1.48E+05 | 3 | Cytoplasm |
| 946 | Q9WV85 | NME3 | 1.51E+06 | 2.09E+05 | 1.21E+05 | 3 | Cytoplasm |
| 947 | P19157 | GSTP1 | 1.51E+06 | 3.52E+05 | 2.03E+05 | 3 | Cytoplasm |
| 948 | P29341 | PABPC1 | 1.49E+06 | 7.16E+05 | 1.85E+05 | 15 | Cytoplasm |
| 949 | Q9D924 | ISCA1 | 1.49E+06 | 3.71E+05 | 1.52E+05 | 6 | Cytoplasm |
| 950 | Q91VD9 | NDUFS1 | 1.49E+06 | 2.85E+05 | 1.17E+05 | 6 | Cytoplasm |
| 951 | Q3U487 | HECTD3 | 1.49E+06 | 1.35E+06 | 5.50E+05 | 6 | Cytoplasm |
| 952 | P01899, P01901, P01902 | HLA-A | 1.49E+06 | 1.79E+06 | 3.44E+05 | 27 | Plasma Membrane |
| 953 | Q3UMR5 | MCU | 1.49E+06 | 1.12E+06 | 6.45E+05 | 3 | Cytoplasm |
| 954 | Q8BIJ6 | IARS2 | 1.49E+06 | 1.23E+05 | 7.09E+04 | 3 | Cytoplasm |
| 955 | Q80TE7 | LRRC7 | 1.47E+06 | 5.21E+05 | 3.01E+05 | 3 | Plasma Membrane |
| 956 | Q9CPW4 | ARPC5 | 1.47E+06 | 6.98E+05 | 4.03E+05 | 3 | Cytoplasm |
| 957 | Q9CZW5 | TOMM70A | 1.47E+06 | 8.21E+05 | 4.74E+05 | 3 | Cytoplasm |
| 958 | P63276 | RPS17 | 1.46E+06 | 1.34E+05 | 7.74E+04 | 3 | Cytoplasm |
| 959 | P55302 | LRPAP1 | 1.45E+06 | 2.52E+05 | 1.46E+05 | 3 | Plasma Membrane |
| 960 | Q9CZB0 | SDHC | 1.45E+06 | 3.61E+05 | 2.08E+05 | 3 | Cytoplasm |
| 961 | Q8C9M2 | CCDC15 | 1.44E+06 | 4.20E+05 | 2.43E+05 | 3 | Cytoplasm |
| 962 | A2ARI4 | LGR4 | 1.44E+06 | 8.84E+05 | 5.10E+05 | 3 | Plasma Membrane |
| 963 | Q9DCB8 | ISCA2 | 1.44E+06 | 4.72E+05 | 2.73E+05 | 3 | Cytoplasm |
| 964 | P52825 | CPT2 | 1.43E+06 | 1.36E+06 | 4.52E+05 | 9 | Cytoplasm |
| 965 | Q8BZ09 | SLC25A21 | 1.43E+06 | 6.55E+05 | 1.69E+05 | 15 | Cytoplasm |
| 966 | Q99JW4 | LIMS1 | 1.41E+06 | 1.22E+06 | 7.05E+05 | 3 | Plasma Membrane |
| 967 | Q8BVF7 | APH1A | 1.41E+06 | 6.59E+05 | 3.80E+05 | 3 | Cytoplasm |
| 968 | Q3THW5 | H2AFV | 1.41E+06 | 1.23E+06 | 7.12E+05 | 3 | Nucleus |
| 969 | Q3TL44 | NLRX1 | 1.41E+06 | 1.19E+06 | 6.86E+05 | 3 | Cytoplasm |
| 970 | P47964 | Rpl36 | 1.41E+06 | 3.14E+05 | 1.28E+05 | 6 | Nucleus |
| 971 | Q99KB8 | HAGH | 1.40E+06 | 1.30E+06 | 4.33E+05 | 9 | Cytoplasm |
| 972 | Q9JHU4 | DYNC1H1 | 1.39E+06 | 1.04E+05 | 6.01E+04 | 3 | Cytoplasm |

| | | | | | | | |
|---|---|---|---|---|---|---|---|
| 834 | Q9Z0F8 | ADAM17 | 2.72E+06 | 2.35E+06 | 9.58E+05 | 6 | Plasma Membrane |
| 835 | Q920L1 | FADS1 | 2.71E+06 | 5.71E+05 | 3.30E+05 | 3 | Plasma Membrane |
| 836 | Q9CPQ3 | TOMM22 | 2.71E+06 | 3.98E+05 | 1.63E+05 | 6 | Cytoplasm |
| 837 | P52825 | CPT2 | 2.69E+06 | 1.72E+06 | 5.73E+05 | 9 | Cytoplasm |
| 838 | Q9EPK7 | XPO7 | 2.68E+06 | 7.55E+05 | 3.08E+05 | 6 | Nucleus |
| 839 | Q9CZX7 | TMEM55A | 2.67E+06 | 7.35E+05 | 4.24E+05 | 3 | Cytoplasm |
| 840 | Q9QZ73 | DCUN1D1 | 2.67E+06 | 5.23E+05 | 3.02E+05 | 3 | Nucleus |
| 841 | P70302 | STIM1 | 2.67E+06 | 1.00E+06 | 3.35E+05 | 9 | Plasma Membrane |
| 842 | Q9CR64 | TMEM167 | 2.67E+06 | 5.70E+05 | 3.29E+05 | 3 | Extracellular Space |
| 843 | Q6P5F9 | XPO1 | 2.66E+06 | 7.50E+05 | 4.33E+05 | 3 | Nucleus |
| 844 | Q9CRD0 | OCIAD1 | 2.65E+06 | 4.23E+05 | 2.44E+05 | 3 | Cytoplasm |
| 845 | Q6NS82 | FAM134A | 2.65E+06 | 1.99E+05 | 1.15E+05 | 3 | Extracellular Space |
| 846 | P13609 | Srgn | 2.64E+06 | 6.64E+05 | 3.83E+05 | 3 | Cytoplasm |
| 847 | Q9D3G5 | STX11 | 2.64E+06 | 1.54E+06 | 3.63E+05 | 18 | Plasma Membrane |
| 848 | Q9R0M6 | RAB9A | 2.63E+06 | 4.68E+05 | 2.70E+05 | 3 | Cytoplasm |
| 849 | P70452 | STX4 | 2.62E+06 | 9.68E+05 | 3.23E+05 | 9 | Plasma Membrane |
| 850 | P01899, P( | HLA-A | 2.62E+06 | 2.28E+06 | 4.38E+05 | 27 | Plasma Membrane |
| 851 | P30681 | HMGB2 | 2.61E+06 | 5.88E+05 | 3.40E+05 | 3 | Nucleus |
| 852 | Q8BP67 | RPL24 | 2.59E+06 | 6.99E+05 | 2.33E+05 | 9 | Cytoplasm |
| 853 | Q8VEH3 | ARL8A | 2.59E+06 | 3.17E+05 | 1.83E+05 | 3 | Cytoplasm |
| 854 | Q08093 | CNN2 | 2.57E+06 | 1.08E+06 | 6.24E+05 | 3 | Cytoplasm |
| 855 | Q99NB1 | ACSS1 | 2.57E+06 | 9.87E+05 | 3.29E+05 | 9 | Cytoplasm |
| 856 | O88448 | KLC2 | 2.56E+06 | 1.84E+05 | 1.06E+05 | 3 | Cytoplasm |
| 857 | O70503 | HSD17B12 | 2.56E+06 | 1.44E+06 | 5.86E+05 | 6 | Cytoplasm |
| 858 | P20444 | PRKCA | 2.55E+06 | 5.00E+05 | 2.89E+05 | 3 | Cytoplasm |
| 859 | O08599 | STXBP1 | 2.55E+06 | 1.30E+06 | 5.32E+05 | 6 | Cytoplasm |
| 860 | O35344 | KPNA3 | 2.54E+06 | 4.13E+05 | 2.38E+05 | 3 | Nucleus |
| 861 | P47857 | PFKM | 2.53E+06 | 7.99E+05 | 1.88E+05 | 18 | Cytoplasm |
| 862 | O70443 | GNAZ | 2.52E+06 | 7.16E+05 | 2.92E+05 | 6 | Plasma Membrane |
| 863 | P62342 | SELT | 2.51E+06 | 8.02E+05 | 4.63E+05 | 3 | Cytoplasm |
| 864 | Q80WC7 | AGFG2 | 2.50E+06 | 1.06E+06 | 3.05E+05 | 12 | Cytoplasm |
| 865 | P59325 | EIF5 | 2.50E+06 | 3.66E+05 | 2.11E+05 | 3 | Cytoplasm |
| 866 | Q8CIG8 | PRMT5 | 2.50E+06 | 1.94E+06 | 7.91E+05 | 6 | Cytoplasm |
| 867 | Q8R3K3 | PTCD2 | 2.48E+06 | 2.45E+05 | 1.42E+05 | 3 | Cytoplasm |
| 868 | Q80ZW2 | THEM6 | 2.48E+06 | 1.36E+06 | 3.51E+05 | 15 | Extracellular Space |
| 869 | Q923U0 | TOM1L1 | 2.47E+06 | 1.33E+06 | 5.44E+05 | 6 | Cytoplasm |
| 870 | Q91VD9 | NDUFS1 | 2.47E+06 | 6.98E+05 | 2.85E+05 | 6 | Cytoplasm |
| 871 | Q9ERM2 | ICAM4 | 2.47E+06 | 1.37E+06 | 5.58E+05 | 6 | Plasma Membrane |
| 872 | P12382 | PFKL | 2.47E+06 | 2.37E+05 | 1.37E+05 | 3 | Cytoplasm |
| 873 | Q8VDP6 | CDIPT | 2.47E+06 | 2.42E+05 | 1.40E+05 | 3 | Cytoplasm |
| 874 | Q9CY24 | TMEM179 | 2.44E+06 | 7.55E+05 | 4.36E+05 | 3 | Extracellular Space |
| 875 | Q9CRY7 | GDPD1 | 2.44E+06 | 1.54E+05 | 8.87E+04 | 3 | Plasma Membrane |
| 876 | Q8C4S8 | DENND2A | 2.44E+06 | 3.21E+05 | 1.85E+05 | 3 | Cytoplasm |
| 877 | Q80UU9 | PGRMC2 | 2.44E+06 | 1.35E+05 | 7.80E+04 | 3 | Nucleus |
| 878 | Q6NSQ9 | G6PC3 | 2.44E+06 | 4.80E+05 | 2.77E+05 | 3 | Cytoplasm |
| 879 | P61255 | RPL26 | 2.44E+06 | 1.72E+05 | 9.91E+04 | 3 | Cytoplasm |
| 880 | Q5XKN4 | JAGN1 | 2.42E+06 | 1.86E+05 | 1.07E+05 | 3 | Cytoplasm |
| 881 | P03921 | MT-ND5 | 2.41E+06 | 1.76E+06 | 5.85E+05 | 9 | Cytoplasm |
| 882 | Q8CBE3 | WDR37 | 2.41E+06 | 1.09E+06 | 4.46E+05 | 6 | Nucleus |
| 883 | Q7TQ95 | KIAA1715 | 2.41E+06 | 1.07E+06 | 2.77E+05 | 15 | Cytoplasm |
| 884 | Q6X893 | SLC44A1 | 2.39E+06 | 1.01E+06 | 3.35E+05 | 9 | Plasma Membrane |
| 885 | Q8C9M2 | CCDC15 | 2.38E+06 | 3.92E+05 | 2.26E+05 | 3 | Cytoplasm |
| 886 | Q60737 | CSNK2A1 | 2.37E+06 | 1.88E+05 | 1.09E+05 | 3 | Nucleus |
| 887 | Q8BFZ9 | ERLIN2 | 2.35E+06 | 9.15E+05 | 5.28E+05 | 3 | Plasma Membrane |
| 888 | Q8BG73 | SH3BGRL2 | 2.35E+06 | 1.03E+06 | 5.93E+05 | 3 | Nucleus |
| 889 | O88783 | F5 | 2.34E+06 | 2.42E+06 | 8.08E+05 | 9 | Plasma Membrane |
| 890 | O70576 | STAG3 | 2.34E+06 | 1.28E+06 | 7.37E+05 | 3 | Nucleus |
| 891 | Q9D0B1 | ZNF524 | 2.34E+06 | 3.67E+06 | 2.12E+06 | 3 | Nucleus |
| 892 | P29758 | OAT | 2.32E+06 | 7.61E+05 | 4.40E+05 | 3 | Cytoplasm |
| 893 | Q9JKB3 | YBX3 | 2.31E+06 | 5.15E+05 | 2.10E+05 | 6 | Nucleus |
| 894 | P49769 | PSEN1 | 2.31E+06 | 1.46E+06 | 5.94E+05 | 6 | Plasma Membrane |
| 895 | Q9Z0X1 | AIFM1 | 2.31E+06 | 4.64E+05 | 1.89E+05 | 6 | Cytoplasm |
| 896 | P97370 | ATP1B3 | 2.30E+06 | 9.14E+05 | 2.64E+05 | 12 | Plasma Membrane |
| 897 | Q7TMY8 | HUWE1 | 2.30E+06 | 1.02E+05 | 5.87E+04 | 3 | Nucleus |
| 898 | Q9CZ44 | NSFL1C | 2.30E+06 | 8.74E+05 | 2.52E+05 | 12 | Cytoplasm |
| 899 | Q6NXH9 | KRT73 | 2.30E+06 | 4.44E+05 | 2.56E+05 | 3 | Extracellular Space |
| 900 | Q80VP1 | EPN1 | 2.28E+06 | 9.40E+05 | 2.71E+05 | 12 | Plasma Membrane |
| 901 | O88544 | COPS4 | 2.27E+06 | 1.00E+06 | 2.36E+05 | 18 | Cytoplasm |
| 902 | P47964 | Rpl36 | 2.27E+06 | 5.22E+05 | 2.13E+05 | 6 | Nucleus |
| 903 | A2AAJ9 | OBSCN | 2.26E+06 | 3.78E+05 | 2.18E+05 | 3 | Cytoplasm |
| 904 | Q6NZJ6 | EIF4G1 | 2.26E+06 | 1.52E+06 | 5.06E+05 | 9 | Cytoplasm |
| 905 | Q9JII5 | Dazap1 | 2.25E+06 | 6.62E+05 | 3.82E+05 | 3 | Cytoplasm |
| 906 | Q8BGH2 | SAMM50 | 2.24E+06 | 8.70E+05 | 3.55E+05 | 6 | Cytoplasm |
| 907 | Q8BIJ6 | IARS2 | 2.24E+06 | 7.09E+05 | 4.09E+05 | 3 | Cytoplasm |
| 908 | O35601 | FYB | 2.21E+06 | 8.51E+05 | 2.84E+05 | 9 | Nucleus |
| 909 | Q9CXE7 | TMED5 | 2.21E+06 | 1.06E+06 | 6.15E+05 | 3 | Cytoplasm |
| 910 | Q91WD5 | NDUFS2 | 2.21E+06 | 5.18E+05 | 2.99E+05 | 3 | Cytoplasm |
| 911 | P84084 | ARF5 | 2.21E+06 | 6.29E+05 | 2.57E+05 | 6 | Cytoplasm |
| 912 | A2ARI4 | LGR4 | 2.20E+06 | 1.75E+06 | 1.01E+06 | 3 | Plasma Membrane |
| 913 | P01872 | IGHM | 2.20E+06 | 2.26E+06 | 6.52E+05 | 12 | Plasma Membrane |
| 914 | Q8BU31 | RAP2C | 2.20E+06 | 6.76E+05 | 3.90E+05 | 3 | Cytoplasm |
| 915 | Q62415 | PPP1R13B | 2.20E+06 | 1.54E+06 | 8.88E+05 | 3 | Cytoplasm |
| 916 | Q9WTX6 | CUL1 | 2.16E+06 | 1.06E+06 | 2.04E+05 | 27 | Nucleus |
| 917 | Q9ET30 | TM9SF3 | 2.15E+06 | 7.29E+04 | 4.21E+04 | 3 | Cytoplasm |
| 918 | Q9DCB8 | ISCA2 | 2.14E+06 | 7.81E+05 | 4.51E+05 | 3 | Cytoplasm |
| 919 | Q9EPL8 | IPO7 | 2.14E+06 | 9.99E+05 | 2.18E+05 | 21 | Nucleus |
| 920 | Q6PDI5 | KIAA0368 | 2.14E+06 | 1.62E+06 | 3.12E+05 | 27 | Cytoplasm |
| 921 | Q9CYN9 | ATP6AP2 | 2.14E+06 | 6.85E+05 | 3.95E+05 | 3 | Cytoplasm |
| 922 | Q80WQ2 | VAC14 | 2.13E+06 | 2.82E+05 | 1.63E+05 | 3 | Cytoplasm |
| 923 | Q7TN58 | TMC8 | 2.11E+06 | 8.39E+05 | 2.80E+05 | 9 | Cytoplasm |
| 924 | P62301 | RPS13 | 2.10E+06 | 2.84E+05 | 1.16E+05 | 6 | Cytoplasm |
| 925 | Q8BX70 | VPS13C | 2.10E+06 | 6.44E+05 | 1.66E+05 | 15 | Cytoplasm |
| 926 | Q9DB34 | CHMP2A | 2.10E+06 | 4.01E+05 | 2.32E+05 | 3 | Cytoplasm |
| 927 | Q62348 | TSN | 2.10E+06 | 1.03E+06 | 4.22E+05 | 6 | Nucleus |
| 928 | Q8R0F8 | FAHD1 | 2.10E+06 | 5.11E+05 | 2.09E+05 | 6 | Cytoplasm |
| 929 | Q05920 | PC | 2.09E+06 | 9.83E+05 | 2.54E+05 | 15 | Cytoplasm |
| 930 | Q60865 | CAPRIN1 | 2.09E+06 | 7.68E+05 | 3.14E+05 | 6 | Plasma Membrane |
| 931 | Q9Z2I0 | LETM1 | 2.09E+06 | 6.55E+05 | 3.78E+05 | 3 | Cytoplasm |
| 932 | Q5U458 | DNAJC11 | 2.06E+06 | 6.80E+05 | 2.78E+05 | 6 | Cytoplasm |
| 933 | Q9Z2G9 | HTATIP2 | 2.05E+06 | 7.94E+05 | 3.24E+05 | 6 | Nucleus |
| 934 | Q9CZB0 | SDHC | 2.05E+06 | 1.13E+06 | 6.55E+05 | 3 | Cytoplasm |
| 935 | Q9ERI2 | RAB27A | 2.05E+06 | 5.97E+05 | 3.45E+05 | 3 | Cytoplasm |
| 936 | Q9R233 | TAPBP | 2.04E+06 | 7.19E+04 | 4.15E+04 | 3 | Cytoplasm |
| 937 | Q8BH57 | WDR48 | 2.03E+06 | 7.80E+05 | 4.50E+05 | 3 | Cytoplasm |
| 938 | Q9CZW5 | TOMM70A | 2.03E+06 | 5.21E+05 | 3.01E+05 | 3 | Cytoplasm |
| 939 | P62281 | RPS11 | 2.03E+06 | 2.58E+06 | 1.06E+06 | 6 | Cytoplasm |
| 940 | P19157 | GSTP1 | 2.02E+06 | 1.73E+05 | 1.00E+05 | 3 | Cytoplasm |
| 941 | Q9DB15 | MRPL12 | 2.02E+06 | 3.63E+05 | 2.09E+05 | 3 | Cytoplasm |
| 942 | Q00623 | APOA1 | 2.00E+06 | 1.11E+06 | 3.71E+05 | 9 | Extracellular Space |
| 943 | Q3U7U3 | FBXO7 | 1.99E+06 | 4.93E+05 | 2.85E+05 | 3 | Cytoplasm |
| 944 | Q8BZ09 | SLC25A21 | 1.99E+06 | 1.33E+06 | 3.42E+05 | 15 | Cytoplasm |
| 945 | Q8CG48 | SMC2 | 1.97E+06 | 4.80E+05 | 2.77E+05 | 3 | Nucleus |
| 946 | Q9WUX5 | MRVI1 | 1.97E+06 | 9.94E+05 | 3.31E+05 | 9 | Cytoplasm |
| 947 | P51437 | CAMP | 1.95E+06 | 1.63E+06 | 6.64E+05 | 6 | Cytoplasm |
| 948 | P18181 | CD48 | 1.95E+06 | 3.50E+05 | 2.02E+05 | 3 | Plasma Membrane |
| 949 | P47962 | RPL5 | 1.95E+06 | 1.75E+05 | 1.01E+05 | 3 | Cytoplasm |
| 950 | Q05512 | MARK2 | 1.94E+06 | 1.08E+06 | 4.43E+05 | 6 | Cytoplasm |
| 951 | A2CG49 | KALRN | 1.93E+06 | 8.24E+05 | 3.37E+05 | 6 | Cytoplasm |
| 952 | P63242 | EIF5A | 1.92E+06 | 8.02E+05 | 4.63E+05 | 3 | Cytoplasm |
| 953 | Q9JHU4 | DYNC1H1 | 1.91E+06 | 3.19E+05 | 1.84E+05 | 3 | Cytoplasm |
| 954 | Q8K2Z4 | NCAPD2 | 1.91E+06 | 1.02E+06 | 5.88E+05 | 3 | Nucleus |
| 955 | Q9CPQ1 | Cox6c | 1.90E+06 | 7.61E+05 | 4.39E+05 | 3 | Cytoplasm |
| 956 | P01029 | C4A/C4B | 1.90E+06 | 4.75E+06 | 1.23E+06 | 15 | Extracellular Space |
| 957 | Q9D6J6 | NDUFV2 | 1.89E+06 | 1.00E+06 | 5.77E+05 | 3 | Cytoplasm |
| 958 | Q6ZWZ2 | UBE2R2 | 1.89E+06 | 1.45E+05 | 8.37E+04 | 3 | Cytoplasm |
| 959 | Q8BV13 | COPS7B | 1.88E+06 | 2.49E+05 | 1.44E+05 | 3 | Cytoplasm |
| 960 | P40124 | CAP1 | 1.88E+06 | 8.70E+05 | 2.90E+05 | 9 | Plasma Membrane |
| 961 | Q8VBV7 | COPS8 | 1.86E+06 | 2.61E+05 | 1.51E+05 | 3 | Nucleus |
| 962 | Q9JHZ2 | ANKH | 1.86E+06 | 2.07E+05 | 1.19E+05 | 3 | Plasma Membrane |
| 963 | P20152 | VIM | 1.86E+06 | 6.49E+05 | 3.75E+05 | 3 | Cytoplasm |
| 964 | Q91VC9 | GHITM | 1.85E+06 | 6.21E+05 | 2.53E+05 | 6 | Cytoplasm |
| 965 | Q8BQ47 | CNPY4 | 1.85E+06 | 2.70E+05 | 1.56E+05 | 3 | Extracellular Space |
| 966 | P17563 | SELENBP1 | 1.85E+06 | 6.88E+05 | 3.97E+05 | 3 | Cytoplasm |
| 967 | P50518 | ATP6V1E1 | 1.84E+06 | 3.66E+05 | 2.11E+05 | 3 | Cytoplasm |
| 968 | O70251 | EEF1B2 | 1.84E+06 | 1.91E+05 | 1.10E+05 | 3 | Cytoplasm |
| 969 | P09528 | FTH1 | 1.84E+06 | 7.61E+05 | 4.39E+05 | 3 | Cytoplasm |
| 970 | Q61102 | ABCB7 | 1.82E+06 | 5.13E+05 | 2.96E+05 | 3 | Cytoplasm |
| 971 | Q8K2K6 | AGFG1 | 1.79E+06 | 9.43E+05 | 5.44E+05 | 3 | Nucleus |
| 972 | Q8CBQ5 | PI4K2B | 1.78E+06 | 7.74E+05 | 2.24E+05 | 12 | Cytoplasm |

| | | | | | | | |
|---|---|---|---|---|---|---|---|
| 834 | O08599 | STXBP1 | 2.19E+06 | 1.16E+06 | 4.75E+05 | 6 | Cytoplasm |
| 835 | Q9EPL8 | IPO7 | 2.19E+06 | 8.58E+05 | 1.87E+05 | 21 | Nucleus |
| 836 | O35943 | FXN | 2.18E+06 | 8.02E+05 | 3.27E+05 | 6 | Cytoplasm |
| 837 | Q9EPK7 | XPO7 | 2.18E+06 | 6.43E+05 | 2.62E+05 | 6 | Nucleus |
| 838 | Q8CF94 | RHCE/RHD | 2.18E+06 | 3.44E+06 | 7.03E+05 | 24 | Plasma Membrane |
| 839 | Q9R0Q6 | ARPC1A | 2.17E+06 | 1.33E+06 | 7.67E+05 | 3 | Extracellular Space |
| 840 | Q6P9J9 | ANO6 | 2.16E+06 | 1.07E+06 | 6.18E+05 | 3 | Plasma Membrane |
| 841 | P21278 | GNA11 | 2.16E+06 | 8.07E+05 | 4.66E+05 | 3 | Plasma Membrane |
| 842 | Q9JKB3 | YBX3 | 2.15E+06 | 3.39E+05 | 1.39E+05 | 6 | Nucleus |
| 843 | Q9QYK7 | RNF11 | 2.14E+06 | 5.95E+05 | 3.44E+05 | 3 | Cytoplasm |
| 844 | Q9WUM5 | SUCLG1 | 2.14E+06 | 1.88E+06 | 6.26E+05 | 9 | Cytoplasm |
| 845 | P62855 | RPS26 | 2.13E+06 | 7.78E+05 | 4.49E+05 | 3 | Cytoplasm |
| 846 | O35344 | KPNA3 | 2.13E+06 | 2.76E+05 | 1.59E+05 | 3 | Nucleus |
| 847 | Q922B2 | DARS | 2.13E+06 | 1.64E+06 | 3.35E+05 | 24 | Cytoplasm |
| 848 | Q9DB34 | CHMP2A | 2.12E+06 | 1.79E+05 | 1.03E+05 | 3 | Cytoplasm |
| 849 | P29341 | PABPC1 | 2.11E+06 | 7.67E+05 | 1.98E+05 | 15 | Cytoplasm |
| 850 | Q3U7U3 | FBXO7 | 2.10E+06 | 2.82E+05 | 1.63E+05 | 3 | Cytoplasm |
| 851 | A2ARI4 | LGR4 | 2.09E+06 | 1.23E+06 | 7.08E+05 | 3 | Plasma Membrane |
| 852 | Q91VD9 | NDUFS1 | 2.08E+06 | 8.64E+05 | 3.53E+05 | 6 | Cytoplasm |
| 853 | Q8CIZ8 | VWF | 2.08E+06 | 1.78E+06 | 2.65E+05 | 45 | Extracellular Space |
| 854 | Q3V3R1 | MTHFD1L | 2.08E+06 | 1.78E+06 | 3.63E+05 | 24 | Cytoplasm |
| 855 | Q8C9M2 | CCDC15 | 2.07E+06 | 5.17E+05 | 2.99E+05 | 3 | Cytoplasm |
| 856 | Q80WQ6 | RHBDF2 | 2.07E+06 | 1.63E+06 | 6.66E+05 | 6 | Cytoplasm |
| 857 | Q5SSH7 | ZZEF1 | 2.06E+06 | 3.38E+05 | 1.95E+05 | 3 | Cytoplasm |
| 858 | P50518 | ATP6V1E1 | 2.05E+06 | 2.26E+05 | 1.30E+05 | 3 | Cytoplasm |
| 859 | Q61739 | ITGA6 | 2.05E+06 | 2.25E+06 | 4.90E+05 | 21 | Plasma Membrane |
| 860 | Q9ERI2 | RAB27A | 2.05E+06 | 2.46E+05 | 1.42E+05 | 3 | Cytoplasm |
| 861 | Q8VBV7 | COPS8 | 2.05E+06 | 1.97E+05 | 1.14E+05 | 3 | Nucleus |
| 862 | Q8R050 | GSPT1 | 2.02E+06 | 3.07E+05 | 1.77E+05 | 3 | Cytoplasm |
| 863 | Q5XKN4 | JAGN1 | 2.01E+06 | 2.06E+05 | 1.19E+05 | 3 | Cytoplasm |
| 864 | P62270 | RPS18 | 2.00E+06 | 1.36E+06 | 4.55E+05 | 9 | Cytoplasm |
| 865 | Q9CX00 | IST1 | 2.00E+06 | 1.89E+05 | 1.09E+05 | 3 | Cytoplasm |
| 866 | P47963 | RPL13 | 2.00E+06 | 1.41E+06 | 4.08E+05 | 12 | Nucleus |
| 867 | Q9QZ73 | DCUN1D1 | 1.99E+06 | 1.10E+05 | 6.37E+04 | 3 | Nucleus |
| 868 | Q9WVC8 | SLC26A3 | 1.99E+06 | 2.18E+05 | 1.26E+05 | 3 | Plasma Membrane |
| 869 | Q8K3F2 | MMP21 | 1.98E+06 | 1.94E+05 | 1.12E+05 | 3 | Extracellular Space |
| 870 | Q8CBE3 | WDR37 | 1.98E+06 | 9.96E+05 | 4.07E+05 | 6 | Nucleus |
| 871 | Q8BJY1 | PSMD5 | 1.97E+06 | 7.53E+05 | 1.45E+05 | 27 | Cytoplasm |
| 872 | O89086 | RBM3 | 1.97E+06 | 4.84E+06 | 1.61E+06 | 9 | Cytoplasm |
| 873 | P62754 | RPS6 | 1.95E+06 | 2.17E+06 | 8.86E+05 | 6 | Cytoplasm |
| 874 | O88456 | CAPNS1 | 1.94E+06 | 2.79E+05 | 1.61E+05 | 3 | Cytoplasm |
| 875 | Q91WS0 | CISD1 | 1.93E+06 | 1.46E+06 | 5.94E+05 | 6 | Cytoplasm |
| 876 | P55302 | LRPAP1 | 1.93E+06 | 3.72E+05 | 2.15E+05 | 3 | Plasma Membrane |
| 877 | O55222 | ILK | 1.92E+06 | 2.75E+06 | 6.47E+05 | 18 | Plasma Membrane |
| 878 | Q63739 | PTP4A1 | 1.91E+06 | 1.92E+05 | 1.11E+05 | 3 | Cytoplasm |
| 879 | Q9JHU4 | DYNC1H1 | 1.86E+06 | 1.30E+05 | 7.50E+04 | 3 | Cytoplasm |
| 880 | Q922Q4 | PYCR2 | 1.86E+06 | 1.60E+06 | 9.24E+05 | 3 | Cytoplasm |
| 881 | Q80UY2 | KCMF1 | 1.86E+06 | 9.56E+05 | 3.19E+05 | 9 | Extracellular Space |
| 882 | Q9R1Q9 | ATP6AP1 | 1.85E+06 | 4.59E+05 | 2.65E+05 | 3 | Cytoplasm |
| 883 | Q3U487 | HECTD3 | 1.84E+06 | 9.83E+05 | 4.01E+05 | 6 | Cytoplasm |
| 884 | Q8CBQ5 | PI4K2B | 1.83E+06 | 5.76E+05 | 1.66E+05 | 12 | Cytoplasm |
| 885 | Q8BP67 | RPL24 | 1.83E+06 | 8.52E+05 | 2.84E+05 | 9 | Cytoplasm |
| 886 | P60335 | PCBP1 | 1.82E+06 | 7.85E+05 | 2.03E+05 | 15 | Nucleus |
| 887 | Q99JB2 | STOML2 | 1.82E+06 | 2.03E+06 | 5.23E+05 | 15 | Plasma Membrane |
| 888 | P62827 | RAN | 1.81E+06 | 2.51E+05 | 1.45E+05 | 3 | Nucleus |
| 889 | P54071 | IDH2 | 1.81E+06 | 2.63E+06 | 6.20E+05 | 18 | Cytoplasm |
| 890 | Q6ZWZ2 | UBE2R2 | 1.81E+06 | 6.30E+05 | 3.64E+05 | 3 | Cytoplasm |
| 891 | Q8BX70 | VPS13C | 1.80E+06 | 6.10E+05 | 1.58E+05 | 15 | Cytoplasm |
| 892 | P63242 | EIF5A | 1.80E+06 | 5.84E+05 | 3.37E+05 | 3 | Cytoplasm |
| 893 | Q9ERE7 | MESDC2 | 1.80E+06 | 1.26E+05 | 7.28E+04 | 3 | Extracellular Space |
| 894 | Q9JMC3 | DNAJA4 | 1.79E+06 | 3.70E+05 | 2.14E+05 | 3 | Nucleus |
| 895 | Q8JZN5 | ACAD9 | 1.79E+06 | 8.59E+05 | 4.96E+05 | 3 | Cytoplasm |
| 896 | Q6P5F9 | XPO1 | 1.78E+06 | 3.56E+05 | 2.06E+05 | 3 | Nucleus |
| 897 | P47911 | RPL6 | 1.77E+06 | 2.00E+06 | 5.78E+05 | 12 | Nucleus |
| 898 | Q9DCB8 | ISCA2 | 1.76E+06 | 1.05E+06 | 6.09E+05 | 3 | Cytoplasm |
| 899 | P70302 | STIM1 | 1.76E+06 | 7.15E+05 | 2.38E+05 | 9 | Plasma Membrane |
| 900 | Q05512 | MARK2 | 1.75E+06 | 6.98E+05 | 2.85E+05 | 6 | Cytoplasm |
| 901 | Q8BL66 | EEA1 | 1.75E+06 | 5.17E+05 | 2.99E+05 | 3 | Cytoplasm |
| 902 | Q91YT0 | NDUFV1 | 1.74E+06 | 2.04E+06 | 6.81E+05 | 9 | Cytoplasm |
| 903 | Q4VAA2 | CDV3 | 1.74E+06 | 7.24E+05 | 4.18E+05 | 3 | Cytoplasm |
| 904 | P26443 | GLUD1 | 1.74E+06 | 9.81E+05 | 4.00E+05 | 6 | Cytoplasm |
| 905 | P14115 | RPL27A | 1.73E+06 | 1.43E+06 | 8.27E+05 | 3 | Cytoplasm |
| 906 | Q8BZI6 | GUCD1 | 1.73E+06 | 7.94E+04 | 4.58E+04 | 3 | Nucleus |
| 907 | A2AAJ9 | OBSCN | 1.73E+06 | 8.04E+05 | 4.64E+05 | 3 | Cytoplasm |
| 908 | P19253 | RPL13A | 1.72E+06 | 1.31E+06 | 7.57E+05 | 3 | Cytoplasm |
| 909 | P62702 | RPS4Y1 | 1.71E+06 | 1.88E+06 | 3.83E+05 | 24 | Cytoplasm |
| 910 | Q8BZ09 | SLC25A21 | 1.71E+06 | 1.50E+06 | 3.86E+05 | 15 | Cytoplasm |
| 911 | P62717 | RPL18A | 1.71E+06 | 1.66E+06 | 9.59E+05 | 3 | Cytoplasm |
| 912 | Q9DCJ5 | NDUFA8 | 1.71E+06 | 7.11E+05 | 4.11E+05 | 3 | Cytoplasm |
| 913 | Q9CPQ3 | TOMM22 | 1.71E+06 | 8.67E+05 | 3.54E+05 | 6 | Cytoplasm |
| 914 | Q9CZ30 | OLA1 | 1.69E+06 | 4.38E+05 | 2.53E+05 | 3 | Cytoplasm |
| 915 | P47964 | Rpl36 | 1.69E+06 | 7.29E+05 | 2.98E+05 | 6 | Nucleus |
| 916 | Q9ET30 | TM9SF3 | 1.69E+06 | 4.30E+05 | 2.48E+05 | 3 | Cytoplasm |
| 917 | Q80TE7 | LRRC7 | 1.68E+06 | 5.06E+05 | 2.92E+05 | 3 | Plasma Membrane |
| 918 | Q9DC51 | GNAI3 | 1.67E+06 | 6.06E+05 | 3.50E+05 | 3 | Cytoplasm |
| 919 | O88783 | F5 | 1.67E+06 | 1.67E+06 | 5.58E+05 | 9 | Plasma Membrane |
| 920 | Q8BMG7 | RAB3GAP2 | 1.67E+06 | 5.35E+05 | 2.18E+05 | 6 | Cytoplasm |
| 921 | O70443 | GNAZ | 1.65E+06 | 9.25E+05 | 3.77E+05 | 6 | Plasma Membrane |
| 922 | Q8R1I1 | UQCR10 | 1.64E+06 | 8.21E+05 | 4.74E+05 | 3 | Cytoplasm |
| 923 | Q9CXZ1 | NDUFS4 | 1.63E+06 | 2.17E+06 | 8.87E+05 | 6 | Cytoplasm |
| 924 | Q8BMF4 | DLAT | 1.63E+06 | 1.22E+06 | 4.96E+05 | 6 | Cytoplasm |
| 925 | O89013 | LEPROT | 1.62E+06 | 1.55E+06 | 6.31E+05 | 6 | Plasma Membrane |
| 926 | Q9D051 | PDHB | 1.62E+06 | 1.60E+06 | 9.26E+05 | 3 | Cytoplasm |
| 927 | Q61425 | HADH | 1.60E+06 | 1.81E+06 | 7.39E+05 | 6 | Cytoplasm |
| 928 | P41438 | SLC19A1 | 1.59E+06 | 1.85E+05 | 1.07E+05 | 3 | Plasma Membrane |
| 929 | O35855 | BCAT2 | 1.59E+06 | 1.93E+06 | 1.12E+06 | 3 | Cytoplasm |
| 930 | Q6NSQ9 | G6PC3 | 1.58E+06 | 7.06E+05 | 4.08E+05 | 3 | Cytoplasm |
| 931 | Q9D880 | TIMM50 | 1.58E+06 | 1.23E+06 | 5.01E+05 | 6 | Cytoplasm |
| 932 | Q9QZQ8 | H2AFY | 1.58E+06 | 5.97E+06 | 3.45E+06 | 3 | Nucleus |
| 933 | Q60575 | KIF1B | 1.56E+06 | 3.05E+06 | 1.76E+06 | 3 | Cytoplasm |
| 934 | Q8R4N0 | CLYBL | 1.56E+06 | 5.20E+05 | 3.00E+05 | 3 | Cytoplasm |
| 935 | Q3THW5 | H2AFV | 1.55E+06 | 6.15E+06 | 3.55E+06 | 3 | Nucleus |
| 936 | Q811D0 | DLG1 | 1.55E+06 | 8.06E+05 | 2.69E+05 | 9 | Plasma Membrane |
| 937 | P19157 | GSTP1 | 1.54E+06 | 1.40E+05 | 8.10E+04 | 3 | Cytoplasm |
| 938 | Q8C156 | NCAPH | 1.53E+06 | 3.27E+05 | 1.89E+05 | 3 | Nucleus |
| 939 | Q8BVF7 | APH1A | 1.53E+06 | 4.69E+05 | 2.71E+05 | 3 | Cytoplasm |
| 940 | Q8BGH2 | SAMM50 | 1.52E+06 | 1.05E+06 | 4.28E+05 | 6 | Cytoplasm |
| 941 | Q9D924 | ISCA1 | 1.52E+06 | 1.95E+06 | 7.96E+05 | 6 | Cytoplasm |
| 942 | Q9D3G5 | STX11 | 1.51E+06 | 1.36E+06 | 3.19E+05 | 18 | Plasma Membrane |
| 943 | Q91VC9 | GHITM | 1.50E+06 | 5.02E+05 | 2.05E+05 | 6 | Cytoplasm |
| 944 | Q8VDV3 | RAB3IL1 | 1.49E+06 | 4.06E+05 | 2.34E+05 | 3 | Cytoplasm |
| 945 | Q9CZB0 | SDHC | 1.49E+06 | 1.27E+06 | 7.32E+05 | 3 | Cytoplasm |
| 946 | Q9JHZ2 | ANKH | 1.49E+06 | 2.91E+05 | 1.68E+05 | 3 | Plasma Membrane |
| 947 | P83741 | WNK1 | 1.48E+06 | 2.75E+05 | 1.59E+05 | 3 | Cytoplasm |
| 948 | Q8R180 | ERO1A | 1.48E+06 | 1.56E+05 | 9.03E+04 | 3 | Cytoplasm |
| 949 | Q91WD5 | NDUFS2 | 1.47E+06 | 7.58E+05 | 4.38E+05 | 3 | Cytoplasm |
| 950 | Q7TN60 | TMC6 | 1.47E+06 | 7.45E+05 | 2.48E+05 | 9 | Cytoplasm |
| 951 | Q8R0F8 | FAHD1 | 1.46E+06 | 1.15E+06 | 4.70E+05 | 6 | Cytoplasm |
| 952 | P84084 | ARF5 | 1.46E+06 | 4.53E+05 | 1.85E+05 | 6 | Cytoplasm |
| 953 | Q9DCT2 | NDUFS3 | 1.45E+06 | 1.32E+06 | 7.64E+05 | 3 | Cytoplasm |
| 954 | Q9Z0X1 | AIFM1 | 1.45E+06 | 6.70E+05 | 2.74E+05 | 6 | Cytoplasm |
| 955 | Q9DB15 | MRPL12 | 1.45E+06 | 5.78E+05 | 3.34E+05 | 3 | Cytoplasm |
| 956 | Q8BVI4 | QDPR | 1.44E+06 | 3.18E+05 | 1.83E+05 | 3 | Cytoplasm |
| 957 | Q05920 | PC | 1.44E+06 | 1.27E+06 | 3.28E+05 | 15 | Cytoplasm |
| 958 | Q6NZJ6 | EIF4G1 | 1.44E+06 | 1.02E+06 | 3.41E+05 | 9 | Cytoplasm |
| 959 | Q9CPW4 | ARPC5 | 1.44E+06 | 5.09E+05 | 2.94E+05 | 3 | Cytoplasm |
| 960 | P26039 | TLN1 | 1.40E+06 | 2.11E+06 | 4.30E+05 | 24 | Plasma Membrane |
| 961 | P22907 | HMBS | 1.40E+06 | 1.99E+05 | 1.15E+05 | 3 | Cytoplasm |
| 962 | P61620 | SEC61A1 | 1.40E+06 | 1.16E+06 | 6.71E+05 | 3 | Cytoplasm |
| 963 | P53026 | RPL10A | 1.39E+06 | 9.76E+05 | 3.98E+05 | 6 | Nucleus |
| 964 | P63276 | RPS17 | 1.39E+06 | 9.64E+05 | 5.57E+05 | 3 | Cytoplasm |
| 965 | O35601 | FYB | 1.38E+06 | 1.31E+06 | 4.36E+05 | 9 | Nucleus |
| 966 | Q3UND0 | SKAP2 | 1.38E+06 | 1.13E+06 | 4.60E+05 | 6 | Cytoplasm |
| 967 | P61255 | RPL26 | 1.37E+06 | 3.70E+05 | 2.14E+05 | 3 | Cytoplasm |
| 968 | Q99NB1 | ACSS1 | 1.37E+06 | 1.44E+06 | 4.79E+05 | 9 | Cytoplasm |
| 969 | Q9CQX8 | MRPS36 | 1.37E+06 | 7.68E+05 | 4.44E+05 | 3 | Cytoplasm |
| 970 | Q9Z110 | ALDH18A1 | 1.34E+06 | 8.00E+05 | 1.63E+05 | 24 | Cytoplasm |
| 971 | Q505D1 | ANKRD28 | 1.33E+06 | 2.84E+05 | 1.64E+05 | 3 | Cytoplasm |
| 972 | P59325 | EIF5 | 1.33E+06 | 2.50E+05 | 1.44E+05 | 3 | Cytoplasm |

| | | | | | | | |
|---|---|---|---|---|---|---|---|
| 834 | Q8CBQ5 | PI4K2B | 1907112 | 731360 | 211125 | 12 | Cytoplasm |
| 835 | Q8BNI4 | DERL2 | 1899516 | 1213098 | 700382 | 3 | Cytoplasm |
| 836 | Q91YT0 | NDUFV1 | 1894238 | 2691444 | 897148 | 9 | Cytoplasm |
| 837 | Q8R0F8 | FAHD1 | 1887302 | 1563035 | 638106 | 6 | Cytoplasm |
| 838 | O89112 | LANCL1 | 1885209 | 112806 | 65129 | 3 | Plasma Membrane |
| 839 | Q05512 | MARK2 | 1884986 | 787704 | 321579 | 6 | Cytoplasm |
| 840 | Q9QY76 | VAPB | 1877089 | 1808058 | 738137 | 6 | Plasma Membrane |
| 841 | P00397 | MT-CO1 | 1867583 | 1518103 | 876477 | 3 | Cytoplasm |
| 842 | Q9CYN9 | ATP6AP2 | 1846803 | 847453 | 489277 | 3 | Cytoplasm |
| 843 | P29758 | OAT | 1844303 | 711749 | 410928 | 3 | Cytoplasm |
| 844 | Q6ZWZ2 | UBE2R2 | 1820092 | 580136 | 334941 | 3 | Cytoplasm |
| 845 | O54890 | ITGB3 | 1780572 | 7997693 | 942537 | 72 | Plasma Membrane |
| 846 | O08756 | HSD17B10 | 1779632 | 2266370 | 654245 | 12 | Cytoplasm |
| 847 | P84084 | ARF5 | 1721900 | 580886 | 237146 | 6 | Cytoplasm |
| 848 | Q8BL66 | EEA1 | 1715351 | 359006 | 207272 | 3 | Cytoplasm |
| 849 | Q80UU9 | PGRMC2 | 1711968 | 751655 | 433968 | 3 | Nucleus |
| 850 | Q8BFZ9 | ERLIN2 | 1710282 | 1073096 | 619552 | 3 | Plasma Membrane |
| 851 | Q9CZB0 | SDHC | 1707936 | 1503343 | 867955 | 3 | Cytoplasm |
| 852 | Q91VD9 | NDUFS1 | 1693583 | 1655223 | 675742 | 6 | Cytoplasm |
| 853 | O55143 | ATP2A2 | 1689965 | 2595016 | 611651 | 18 | Cytoplasm |
| 854 | Q80UY2 | KCMF1 | 1685086 | 912249 | 304083 | 9 | Extracellular Space |
| 855 | Q7TMM9 | TUBB2A | 1683070 | 450537 | 183931 | 6 | Cytoplasm |
| 856 | P31230 | AIMP1 | 1682675 | 991663 | 404845 | 6 | Extracellular Space |
| 857 | Q8BMG7 | RAB3GAP2 | 1677998 | 605967 | 247385 | 6 | Cytoplasm |
| 858 | Q9QUM0 | ITGA2B | 1670841 | 12231284 | 1359032 | 81 | Plasma Membrane |
| 859 | P55302 | LRPAP1 | 1665523 | 325494 | 187924 | 3 | Plasma Membrane |
| 860 | P62918 | RPL8 | 1663005 | 980416 | 400253 | 6 | Cytoplasm |
| 861 | Q9CQX8 | MRPS36 | 1650937 | 999035 | 576793 | 3 | Cytoplasm |
| 862 | Q8K2K6 | AGFG1 | 1633864 | 270069 | 155924 | 3 | Nucleus |
| 863 | P17742 | PPIA | 1632978 | 3889392 | 848735 | 21 | Cytoplasm |
| 864 | Q9CZ30 | OLA1 | 1628920 | 561163 | 323988 | 3 | Cytoplasm |
| 865 | Q8BG73 | SH3BGRL2 | 1627307 | 1419995 | 819835 | 3 | Nucleus |
| 866 | Q9CR60 | GOLT1B | 1626260 | 1020824 | 589373 | 3 | Cytoplasm |
| 867 | Q8BGH2 | SAMM50 | 1612607 | 885408 | 361466 | 6 | Cytoplasm |
| 868 | Q8JZN5 | ACAD9 | 1602437 | 1115198 | 643860 | 3 | Cytoplasm |
| 869 | Q9CQV8 | YWHAB | 1602268 | 1663627 | 392121 | 18 | Cytoplasm |
| 870 | P40142 | TKT | 1598616 | 1375346 | 561483 | 6 | Cytoplasm |
| 871 | P25911 | LYN | 1591977 | 2333509 | 476325 | 24 | Cytoplasm |
| 872 | Q64522 | HIST2H2AB | 1584988 | 615369 | 355284 | 3 | Nucleus |
| 873 | Q8R180 | ERO1A | 1584849 | 336708 | 194398 | 3 | Cytoplasm |
| 874 | Q8K558 | TREML1 | 1572615 | 6452752 | 1317162 | 24 | Plasma Membrane |
| 875 | Q9CXZ1 | NDUFS4 | 1559913 | 2560222 | 1045206 | 6 | Cytoplasm |
| 876 | P00416 | MT-CO3 | 1554125 | 854419 | 493299 | 3 | Cytoplasm |
| 877 | Q9DC51 | GNAI3 | 1544579 | 668969 | 386229 | 3 | Cytoplasm |
| 878 | Q80WQ6 | RHBDF2 | 1535601 | 875979 | 357617 | 6 | Cytoplasm |
| 879 | P68510 | YWHAH | 1520044 | 4686889 | 901992 | 27 | Cytoplasm |
| 880 | Q9CZW5 | TOMM70A | 1514597 | 768996 | 443980 | 3 | Cytoplasm |
| 881 | Q91VC9 | GHITM | 1513837 | 1017176 | 415261 | 6 | Cytoplasm |
| 882 | P47962 | RPL5 | 1501351 | 712312 | 411253 | 3 | Cytoplasm |
| 883 | Q9Z2I0 | LETM1 | 1493295 | 897892 | 518398 | 3 | Cytoplasm |
| 884 | P99029 | PRDX5 | 1492613 | 862119 | 497745 | 3 | Cytoplasm |
| 885 | Q7TN60 | TMC6 | 1483499 | 995036 | 331679 | 9 | Cytoplasm |
| 886 | Q9JII5 | Dazap1 | 1480890 | 404539 | 233561 | 3 | Cytoplasm |
| 887 | Q9D8B6 | FAM210B | 1480515 | 1870453 | 763609 | 6 | Extracellular Space |
| 888 | Q9CQS8 | SEC61B | 1476899 | 1175088 | 479728 | 6 | Cytoplasm |
| 889 | Q9CQ19 | MYL9 | 1452333 | 3151902 | 643379 | 24 | Cytoplasm |
| 890 | E9Q555 | RNF213 | 1449745 | 719864 | 415614 | 3 | Cytoplasm |
| 891 | P97351 | Rps3a1 | 1446305 | 1263652 | 515884 | 6 | Cytoplasm |
| 892 | P08905 | LYZ | 1445675 | 2277573 | 929815 | 6 | Extracellular Space |
| 893 | P83741 | WNK1 | 1443647 | 280339 | 161854 | 3 | Cytoplasm |
| 894 | P18760 | CFL1 | 1437295 | 2129109 | 549734 | 15 | Nucleus |
| 895 | Q9D051 | PDHB | 1433040 | 1141741 | 659185 | 3 | Cytoplasm |
| 896 | P50544 | ACADVL | 1432470 | 1399417 | 361328 | 15 | Cytoplasm |
| 897 | Q9ERE7 | MESDC2 | 1431129 | 273717 | 158030 | 3 | Extracellular Space |
| 898 | Q8VDP6 | CDIPT | 1426090 | 276225 | 159478 | 3 | Cytoplasm |
| 899 | Q8BQ47 | CNPY4 | 1417657 | 324140 | 187142 | 3 | Extracellular Space |
| 900 | Q61102 | ABCB7 | 1392166 | 1107239 | 639265 | 3 | Cytoplasm |
| 901 | Q8VDV3 | RAB3IL1 | 1389778 | 769949 | 444530 | 3 | Cytoplasm |
| 902 | P57776 | EEF1D | 1380497 | 635901 | 367138 | 3 | Cytoplasm |
| 903 | Q9WV85 | NME3 | 1375167 | 855534 | 493943 | 3 | Cytoplasm |
| 904 | Q8BIJ6 | IARS2 | 1366246 | 1269522 | 732959 | 3 | Cytoplasm |
| 905 | P29391 | FTL | 1360195 | 1105467 | 368489 | 9 | Cytoplasm |
| 906 | P48962 | SLC25A4 | 1358604 | 853972 | 284657 | 9 | Cytoplasm |
| 907 | Q91YQ5 | RPN1 | 1357188 | 1582233 | 322972 | 24 | Cytoplasm |
| 908 | Q9D6J6 | NDUFV2 | 1347791 | 960395 | 554484 | 3 | Cytoplasm |
| 909 | Q9WUA2 | FARSB | 1342912 | 505268 | 291717 | 3 | Cytoplasm |
| 910 | Q9CRD0 | OCIAD1 | 1342696 | 1159603 | 669497 | 3 | Cytoplasm |
| 911 | P70452 | STX4 | 1339835 | 751619 | 250540 | 9 | Plasma Membrane |
| 912 | Q8BIG7 | COMTD1 | 1337130 | 768875 | 443910 | 3 | Plasma Membrane |
| 913 | Q91WD5 | NDUFS2 | 1335736 | 1033662 | 596785 | 3 | Cytoplasm |
| 914 | Q80TE7 | LRRC7 | 1329180 | 243430 | 140544 | 3 | Plasma Membrane |
| 915 | P38060 | HMGCL | 1316601 | 1235928 | 504565 | 6 | Cytoplasm |
| 916 | Q8BMF4 | DLAT | 1312212 | 1552694 | 633885 | 6 | Cytoplasm |
| 917 | Q505D1 | ANKRD28 | 1310202 | 445001 | 256921 | 3 | Cytoplasm |
| 918 | Q6ZWY3 | RPS27L | 1303433 | 879549 | 359074 | 6 | Cytoplasm |
| 919 | Q9D8N0 | EEF1G | 1302332 | 972558 | 324186 | 9 | Cytoplasm |
| 920 | Q60865 | CAPRIN1 | 1244077 | 891754 | 364057 | 6 | Plasma Membrane |
| 921 | Q61781 | KRT14 | 1241959 | 592240 | 341930 | 3 | Cytoplasm |
| 922 | Q9DCJ5 | NDUFA8 | 1234822 | 1509432 | 871471 | 3 | Cytoplasm |
| 923 | Q9Z110 | ALDH18A1 | 1224047 | 1178986 | 240659 | 24 | Cytoplasm |
| 924 | P61620 | SEC61A1 | 1219127 | 358661 | 207073 | 3 | Cytoplasm |
| 925 | Q9DB15 | MRPL12 | 1215388 | 1270632 | 733600 | 3 | Cytoplasm |
| 926 | Q9CPQ1 | Cox6c | 1213866 | 884773 | 510824 | 3 | Cytoplasm |
| 927 | Q8R4N0 | CLYBL | 1213306 | 1115918 | 644276 | 3 | Cytoplasm |
| 928 | Q9Z0X1 | AIFM1 | 1205035 | 785938 | 320858 | 6 | Cytoplasm |
| 929 | P63321 | RALA | 1204816 | 670149 | 386911 | 3 | Cytoplasm |
| 930 | Q9CY50 | SSR1 | 1200369 | 919484 | 375378 | 6 | Cytoplasm |
| 931 | P35486 | PDHA1 | 1183276 | 750534 | 250178 | 9 | Cytoplasm |
| 932 | Q00612 | G6PD | 1174785 | 440252 | 127090 | 12 | Cytoplasm |
| 933 | Q9DCT2 | NDUFS3 | 1170965 | 1228282 | 709149 | 3 | Cytoplasm |
| 934 | P54071 | IDH2 | 1167785 | 1580941 | 372631 | 18 | Cytoplasm |
| 935 | Q3U0D9 | HACE1 | 1164249 | 302311 | 174539 | 3 | Cytoplasm |
| 936 | Q9DBG6 | RPN2 | 1151231 | 2037762 | 372043 | 30 | Cytoplasm |
| 937 | O70503 | HSD17B12 | 1149584 | 679687 | 277481 | 6 | Cytoplasm |
| 938 | Q9CPW4 | ARPC5 | 1146445 | 214371 | 123767 | 3 | Cytoplasm |
| 939 | P68254 | YWHAQ | 1146043 | 1129361 | 376454 | 9 | Cytoplasm |
| 940 | Q9R1Q9 | ATP6AP1 | 1141360 | 912776 | 526992 | 3 | Cytoplasm |
| 941 | Q9D0K2 | OXCT1 | 1141196 | 547816 | 316282 | 3 | Cytoplasm |
| 942 | Q9CZ04 | COPS7A | 1141074 | 253641 | 146440 | 3 | Cytoplasm |
| 943 | O08742 | GP5 | 1140316 | 6025589 | 1100117 | 30 | Plasma Membrane |
| 944 | O70251 | EEF1B2 | 1133765 | 281284 | 162399 | 3 | Cytoplasm |
| 945 | Q8BVF7 | APH1A | 1128821 | 534097 | 308361 | 3 | Cytoplasm |
| 946 | Q811C2 | ATG4C | 1116163 | 219039 | 126462 | 3 | Cytoplasm |
| 947 | P09528 | FTH1 | 1113640 | 132556 | 76531 | 3 | Cytoplasm |
| 948 | Q5XKN4 | JAGN1 | 1099145 | 590280 | 340798 | 3 | Cytoplasm |
| 949 | P28665 | Mug1/Mug2 | 1097725 | 569719 | 328928 | 3 | Extracellular Space |
| 950 | P22907 | HMBS | 1095741 | 525999 | 303686 | 3 | Cytoplasm |
| 951 | P27612 | PLAA | 1095555 | 240542 | 138877 | 3 | Cytoplasm |
| 952 | Q8VBZ3 | CLPTM1 | 1077361 | 444254 | 256490 | 3 | Plasma Membrane |
| 953 | Q6NXH9 | KRT73 | 1050164 | 553698 | 319678 | 3 | Extracellular Space |
| 954 | Q91W90 | TXNDC5 | 1044250 | 28581 | 16501 | 3 | Cytoplasm |
| 955 | Q64310 | SURF4 | 1042402 | 1116353 | 455749 | 6 | Cytoplasm |
| 956 | P59108 | CPNE2 | 1033456 | 1019909 | 416376 | 6 | Cytoplasm |
| 957 | Q3UHJ0 | AAK1 | 1023527 | 23020 | 13291 | 3 | Cytoplasm |
| 958 | P01872 | IGHM | 1010211 | 3024067 | 872973 | 12 | Plasma Membrane |
| 959 | P61982 | YWHAG | 985384 | 1329539 | 542782 | 6 | Cytoplasm |
| 960 | O35930 | GP1BA | 981549 | 6807775 | 1090116 | 39 | Plasma Membrane |
| 961 | Q8BVI4 | QDPR | 978666 | 374529 | 216235 | 3 | Cytoplasm |
| 962 | Q9WUM4 | CORO1C | 965279 | 1169570 | 477475 | 6 | Cytoplasm |
| 963 | Q9CY58 | SERBP1 | 962475 | 1189182 | 396394 | 9 | Cytoplasm |
| 964 | P24549 | ALDH1A1 | 961668 | 126868 | 73247 | 3 | Cytoplasm |
| 965 | Q4VBD2 | TAPT1 | 949784 | 512612 | 295956 | 3 | Plasma Membrane |
| 966 | P47915 | Rpl29 (includes others) | 949068 | 1495952 | 863689 | 3 | Cytoplasm |
| 967 | Q01730 | RSU1 | 937709 | 1485247 | 495082 | 9 | Cytoplasm |
| 968 | Q9CPP6 | NDUFA5 | 935523 | 739077 | 426707 | 3 | Cytoplasm |
| 969 | Q8R1I1 | UQCR10 | 934457 | 1473523 | 850739 | 3 | Cytoplasm |
| 970 | P28076 | PSMB9 | 929653 | 280431 | 161907 | 3 | Cytoplasm |
| 971 | Q6ZQI3 | MLEC | 891991 | 1157475 | 298859 | 15 | Plasma Membrane |
| 972 | P52825 | CPT2 | 889416 | 1034259 | 344753 | 9 | Cytoplasm |

| | | | | | | | |
|---|---|---|---|---|---|---|---|
| 973 | P53026 | RPL10A | 1.39E+06 | 5.42E+05 | 2.21E+05 | 6 | Nucleus |
| 974 | Q61425 | HADH | 1.38E+06 | 1.56E+06 | 6.35E+05 | 6 | Cytoplasm |
| 975 | P40142 | TKT | 1.38E+06 | 1.48E+06 | 6.04E+05 | 6 | Cytoplasm |
| 976 | Q91YT8 | TMEM63A | 1.37E+06 | 8.67E+05 | 5.01E+05 | 3 | Cytoplasm |
| 977 | Q7TN60 | TMC6 | 1.37E+06 | 7.64E+05 | 2.55E+05 | 9 | Cytoplasm |
| 978 | P26039 | TLN1 | 1.36E+06 | 1.25E+06 | 2.55E+05 | 24 | Plasma Membrane |
| 979 | Q61144 | PSEN2 | 1.35E+06 | 2.32E+05 | 1.34E+05 | 3 | Cytoplasm |
| 980 | P83741 | WNK1 | 1.35E+06 | 7.73E+05 | 4.46E+05 | 3 | Cytoplasm |
| 981 | Q8JZN5 | ACAD9 | 1.34E+06 | 4.92E+05 | 2.84E+05 | 3 | Cytoplasm |
| 982 | P50544 | ACADVL | 1.33E+06 | 7.42E+05 | 1.92E+05 | 15 | Cytoplasm |
| 983 | P09528 | FTH1 | 1.33E+06 | 4.77E+05 | 2.75E+05 | 3 | Cytoplasm |
| 984 | O88746 | TOM1 | 1.33E+06 | 6.00E+05 | 3.46E+05 | 3 | Cytoplasm |
| 985 | Q8VDV3 | RAB3IL1 | 1.32E+06 | 4.03E+05 | 2.33E+05 | 3 | Cytoplasm |
| 986 | Q9ERE7 | MESDC2 | 1.32E+06 | 2.51E+04 | 1.45E+04 | 3 | Extracellular Space |
| 987 | P62281 | RPS11 | 1.32E+06 | 1.15E+06 | 4.68E+05 | 6 | Cytoplasm |
| 988 | Q9D051 | PDHB | 1.30E+06 | 9.61E+05 | 5.55E+05 | 3 | Cytoplasm |
| 989 | Q9EPK7 | XPO7 | 1.30E+06 | 4.77E+05 | 1.95E+05 | 6 | Nucleus |
| 990 | Q05920 | PC | 1.30E+06 | 5.73E+05 | 1.48E+05 | 15 | Cytoplasm |
| 991 | Q91WD5 | NDUFS2 | 1.29E+06 | 3.11E+05 | 1.80E+05 | 3 | Cytoplasm |
| 992 | Q9EQP2 | EHD4 | 1.29E+06 | 4.17E+05 | 1.70E+05 | 6 | Plasma Membrane |
| 993 | Q9Z0X1 | AIFM1 | 1.29E+06 | 4.51E+05 | 1.84E+05 | 6 | Cytoplasm |
| 994 | P00397 | MT-CO1 | 1.28E+06 | 7.21E+05 | 4.16E+05 | 3 | Cytoplasm |
| 995 | Q9DCJ5 | NDUFA8 | 1.24E+06 | 2.59E+05 | 1.49E+05 | 3 | Cytoplasm |
| 996 | Q9CZ04 | COPS7A | 1.22E+06 | 3.37E+05 | 1.94E+05 | 3 | Cytoplasm |
| 997 | Q64331 | MYO6 | 1.21E+06 | 2.57E+05 | 1.49E+05 | 3 | Cytoplasm |
| 998 | O88448 | KLC2 | 1.21E+06 | 9.74E+05 | 5.63E+05 | 3 | Cytoplasm |
| 999 | Q9CRD0 | OCIAD1 | 1.21E+06 | 5.37E+05 | 3.10E+05 | 3 | Cytoplasm |
| 1000 | P29758 | OAT | 1.18E+06 | 3.96E+05 | 2.29E+05 | 3 | Cytoplasm |
| 1001 | Q8BL66 | EEA1 | 1.18E+06 | 5.01E+05 | 2.89E+05 | 3 | Cytoplasm |
| 1002 | Q99NB1 | ACSS1 | 1.17E+06 | 3.16E+05 | 1.05E+05 | 9 | Cytoplasm |
| 1003 | P22907 | HMBS | 1.16E+06 | 1.23E+05 | 7.12E+04 | 3 | Cytoplasm |
| 1004 | Q9CQX8 | MRPS36 | 1.16E+06 | 3.75E+05 | 2.16E+05 | 3 | Cytoplasm |
| 1005 | P11835 | ITGB2 | 1.16E+06 | 6.70E+05 | 3.87E+05 | 3 | Plasma Membrane |
| 1006 | Q6NZJ6 | EIF4G1 | 1.16E+06 | 8.86E+05 | 2.95E+05 | 9 | Cytoplasm |
| 1007 | P62849 | RPS24 | 1.15E+06 | 1.22E+05 | 7.04E+04 | 3 | Cytoplasm |
| 1008 | Q8BX70 | VPS13C | 1.15E+06 | 4.13E+05 | 1.07E+05 | 15 | Cytoplasm |
| 1009 | Q9QZQ8 | H2AFY | 1.15E+06 | 1.06E+06 | 6.12E+05 | 3 | Nucleus |
| 1010 | Q9D6J6 | NDUFV2 | 1.15E+06 | 2.64E+05 | 1.52E+05 | 3 | Cytoplasm |
| 1011 | Q91W34 | C16orf58 | 1.15E+06 | 2.57E+04 | 1.48E+04 | 3 | Cytoplasm |
| 1012 | Q61102 | ABCB7 | 1.15E+06 | 3.75E+05 | 2.17E+05 | 3 | Cytoplasm |
| 1013 | P20444 | PRKCA | 1.15E+06 | 1.02E+06 | 5.90E+05 | 3 | Cytoplasm |
| 1014 | P62960 | YBX1 | 1.14E+06 | 1.08E+06 | 4.39E+05 | 6 | Nucleus |
| 1015 | Q9DCT2 | NDUFS3 | 1.13E+06 | 5.79E+05 | 3.34E+05 | 3 | Cytoplasm |
| 1016 | Q8BVI4 | QDPR | 1.13E+06 | 5.50E+05 | 3.18E+05 | 3 | Cytoplasm |
| 1017 | O35601 | FYB | 1.12E+06 | 1.03E+06 | 3.42E+05 | 9 | Nucleus |
| 1018 | P00416 | MT-CO3 | 1.12E+06 | 3.10E+05 | 1.79E+05 | 3 | Cytoplasm |
| 1019 | Q3UND0 | SKAP2 | 1.10E+06 | 8.24E+05 | 3.36E+05 | 6 | Cytoplasm |
| 1020 | Q9R233 | TAPBP | 1.09E+06 | 3.05E+05 | 1.76E+05 | 3 | Cytoplasm |
| 1021 | O35114 | SCARB2 | 1.09E+06 | 4.19E+05 | 2.42E+05 | 3 | Plasma Membrane |
| 1022 | Q4VBD2 | TAPT1 | 1.09E+06 | 6.47E+04 | 3.73E+04 | 3 | Plasma Membrane |
| 1023 | Q91VC9 | GHITM | 1.09E+06 | 4.18E+05 | 1.70E+05 | 6 | Cytoplasm |
| 1024 | P61255 | RPL26 | 1.05E+06 | 2.41E+05 | 1.39E+05 | 3 | Cytoplasm |
| 1025 | Q6DIA2 | EXOC3L4 | 1.03E+06 | 7.39E+05 | 4.27E+05 | 3 | Cytoplasm |
| 1026 | Q8R1I1 | UQCR10 | 1.03E+06 | 1.08E+05 | 6.25E+04 | 3 | Cytoplasm |
| 1027 | Q811C2 | ATG4C | 1.02E+06 | 3.75E+05 | 2.16E+05 | 3 | Cytoplasm |
| 1028 | Q4VAA2 | CDV3 | 1.02E+06 | 6.39E+04 | 3.69E+04 | 3 | Cytoplasm |
| 1029 | O35083 | AGPAT1 | 1.02E+06 | 2.40E+05 | 1.38E+05 | 3 | Cytoplasm |
| 1030 | Q60770 | STXBP3 | 1.01E+06 | 4.05E+05 | 2.34E+05 | 3 | Plasma Membrane |
| 1031 | Q5U458 | DNAJC11 | 1.01E+06 | 4.60E+05 | 1.88E+05 | 6 | Cytoplasm |
| 1032 | Q8R4N0 | CLYBL | 1.01E+06 | 1.18E+05 | 6.81E+04 | 3 | Cytoplasm |
| 1033 | P35486 | PDHA1 | 1.01E+06 | 4.04E+05 | 1.35E+05 | 9 | Cytoplasm |
| 1034 | P47962 | RPL5 | 1.01E+06 | 4.43E+05 | 2.56E+05 | 3 | Cytoplasm |
| 1035 | P62301 | RPS13 | 1.01E+06 | 3.57E+05 | 1.46E+05 | 6 | Cytoplasm |
| 1036 | Q91W90 | TXNDC5 | 1.00E+06 | 2.90E+05 | 1.68E+05 | 3 | Cytoplasm |
| 1037 | Q9CPQ1 | Cox6c | 9.99E+05 | 2.18E+05 | 1.26E+05 | 3 | Cytoplasm |
| 1038 | A2CG49 | KALRN | 9.98E+05 | 9.69E+05 | 3.96E+05 | 6 | Cytoplasm |
| 1039 | Q9DB15 | MRPL12 | 9.87E+05 | 3.24E+05 | 1.87E+05 | 3 | Cytoplasm |
| 1040 | P58252 | EEF2 | 9.82E+05 | 5.30E+05 | 1.37E+05 | 15 | Cytoplasm |
| 1041 | P08071 | LTF | 9.78E+05 | 8.54E+05 | 2.21E+05 | 15 | Extracellular Space |
| 1042 | Q9WUX5 | MRVI1 | 9.78E+05 | 9.33E+05 | 3.11E+05 | 9 | Cytoplasm |
| 1043 | Q8VBZ3 | CLPTM1 | 9.78E+05 | 1.76E+05 | 1.02E+05 | 3 | Plasma Membrane |
| 1044 | Q3U0D9 | HACE1 | 9.76E+05 | 5.20E+05 | 3.00E+05 | 3 | Cytoplasm |
| 1045 | P28665 | Mug1/Mug2 | 9.75E+05 | 4.61E+05 | 2.66E+05 | 3 | Extracellular Space |
| 1046 | Q6A4J8 | USP7 | 9.66E+05 | 1.33E+05 | 7.66E+04 | 3 | Nucleus |
| 1047 | Q8BH57 | WDR48 | 9.60E+05 | 1.39E+05 | 8.05E+04 | 3 | Cytoplasm |
| 1048 | Q3UHJ0 | AAK1 | 9.59E+05 | 3.17E+05 | 1.83E+05 | 3 | Cytoplasm |
| 1049 | Q9D8B6 | FAM210B | 9.49E+05 | 8.87E+05 | 3.62E+05 | 6 | Extracellular Space |
| 1050 | Q8BVE3 | ATP6V1H | 9.46E+05 | 6.73E+05 | 2.75E+05 | 6 | Cytoplasm |
| 1051 | Q9Z110 | ALDH18A1 | 9.32E+05 | 5.60E+05 | 1.14E+05 | 24 | Cytoplasm |
| 1052 | Q8K3F2 | MMP21 | 8.88E+05 | 7.10E+05 | 4.10E+05 | 3 | Extracellular Space |
| 1053 | Q8BU33 | ILVBL | 8.74E+05 | 1.11E+05 | 6.39E+04 | 3 | Cytoplasm |
| 1054 | Q5HZI9 | SLC25A51 | 8.28E+05 | 1.79E+05 | 1.03E+05 | 3 | Cytoplasm |
| 1055 | Q4KUS1 | WFDC8 | 8.15E+05 | 2.84E+05 | 1.64E+05 | 3 | Extracellular Space |
| 1056 | Q8C156 | NCAPH | 8.14E+05 | 2.66E+05 | 1.54E+05 | 3 | Nucleus |
| 1057 | P08103 | HCK | 8.12E+05 | 5.64E+05 | 3.25E+05 | 3 | Cytoplasm |
| 1058 | P27612 | PLAA | 8.05E+05 | 4.36E+05 | 2.52E+05 | 3 | Cytoplasm |
| 1059 | P62918 | RPL8 | 8.05E+05 | 4.17E+05 | 1.70E+05 | 6 | Cytoplasm |
| 1060 | P45952 | ACADM | 7.71E+05 | 7.04E+05 | 4.07E+05 | 3 | Cytoplasm |
| 1061 | P28076 | PSMB9 | 7.71E+05 | 4.40E+05 | 2.54E+05 | 3 | Cytoplasm |
| 1062 | O70251 | EEF1B2 | 7.51E+05 | 1.25E+05 | 7.21E+04 | 3 | Cytoplasm |
| 1063 | Q9CPP6 | NDUFA5 | 6.97E+05 | 1.68E+05 | 9.67E+04 | 3 | Cytoplasm |
| 1064 | Q9WUA2 | FARSB | 6.93E+05 | 2.09E+05 | 1.21E+05 | 3 | Cytoplasm |
| 1065 | P30681 | HMGB2 | 6.83E+05 | 7.80E+05 | 4.50E+05 | 3 | Nucleus |
| 1066 | Q60865 | CAPRIN1 | 6.74E+05 | 2.89E+05 | 1.18E+05 | 6 | Plasma Membrane |
| 1067 | P38060 | HMGCL | 6.72E+05 | 4.54E+05 | 1.85E+05 | 6 | Cytoplasm |
| 1068 | Q6P9Q6 | FKBP15 | 6.69E+05 | 3.69E+05 | 1.51E+05 | 6 | Plasma Membrane |
| 1069 | Q811D0 | DLG1 | 6.67E+05 | 4.23E+05 | 1.41E+05 | 9 | Plasma Membrane |
| 1070 | Q9Z2I8 | SUCLG2 | 6.54E+05 | 1.59E+05 | 9.20E+04 | 3 | Cytoplasm |
| 1071 | P97351 | Rps3a1 | 6.51E+05 | 5.38E+05 | 2.19E+05 | 6 | Cytoplasm |
| 1072 | Q922Q1 | MARC_2 | 6.19E+05 | 3.12E+05 | 1.80E+05 | 3 | Cytoplasm |
| 1073 | P62858 | RPS28 | 5.95E+05 | 4.65E+05 | 2.68E+05 | 3 | Cytoplasm |
| 1074 | Q6ZWY3 | RPS27L | 5.85E+05 | 4.10E+05 | 1.67E+05 | 6 | Cytoplasm |
| 1075 | Q3URS9 | CCDC51 | 5.76E+05 | 3.82E+04 | 2.20E+04 | 3 | Cytoplasm |
| 1076 | P47915 | Rpl29 (includes others) | 5.69E+05 | 3.01E+05 | 1.74E+05 | 3 | Cytoplasm |
| 1077 | Q9JII5 | Dazap1 | 5.60E+05 | 1.54E+05 | 8.90E+04 | 3 | Cytoplasm |
| 1078 | P46061 | RANGAP1 | 5.22E+05 | 2.51E+05 | 1.45E+05 | 3 | Nucleus |
| 1079 | P20152 | VIM | 5.16E+05 | 4.36E+05 | 2.52E+05 | 3 | Cytoplasm |
| 1080 | Q9Z2G9 | HTATIP2 | 5.15E+05 | 2.49E+05 | 1.02E+05 | 6 | Nucleus |
| 1081 | Q9CY58 | SERBP1 | 5.06E+05 | 3.12E+05 | 1.04E+05 | 9 | Cytoplasm |
| 1082 | Q9DC23 | DNAJC10 | 5.06E+05 | 1.01E+05 | 5.85E+04 | 3 | Cytoplasm |
| 1083 | Q9DBG3 | Ap2b1 | 5.03E+05 | 4.48E+05 | 1.83E+05 | 6 | Plasma Membrane |
| 1084 | Q8K2Z4 | NCAPD2 | 4.88E+05 | 3.46E+05 | 2.00E+05 | 3 | Nucleus |
| 1085 | Q8BGT0 | OSTM1 | 4.47E+05 | 5.54E+04 | 3.20E+04 | 3 | Cytoplasm |
| 1086 | Q9WTP6 | AK2 | 4.37E+05 | 3.82E+05 | 2.20E+05 | 3 | Cytoplasm |
| 1087 | E9Q555 | RNF213 | 4.37E+05 | 2.29E+04 | 1.32E+04 | 3 | Cytoplasm |
| 1088 | Q9D7J6 | DNASE1L1 | 4.35E+05 | 1.73E+05 | 1.00E+05 | 3 | Cytoplasm |
| 1089 | Q9WTI7 | MYO1C | 4.26E+05 | 1.20E+05 | 6.95E+04 | 3 | Cytoplasm |
| 1090 | Q0P5V2 | SOBP | 4.15E+05 | 1.79E+05 | 1.04E+05 | 3 | Nucleus |
| 1091 | Q9D0I9 | RARS | 4.13E+05 | 4.70E+05 | 2.71E+05 | 3 | Cytoplasm |
| 1092 | Q8BRU6 | SLC18A2 | 4.10E+05 | 3.72E+05 | 2.15E+05 | 3 | Plasma Membrane |
| 1093 | Q8BTJ4 | ENPP4 | 3.61E+05 | 1.52E+05 | 8.77E+04 | 3 | Cytoplasm |
| 1094 | Q9D710 | TMX2 | 3.58E+05 | 9.68E+04 | 5.59E+04 | 3 | Cytoplasm |
| 1095 | Q7TMF3 | NDUFA12 | 3.55E+05 | 4.08E+04 | 2.35E+04 | 3 | Cytoplasm |
| 1096 | Q8BTX9 | HSDL1 | 3.49E+05 | 2.46E+05 | 1.42E+05 | 3 | Cytoplasm |
| 1097 | Q01339 | APOH | 3.44E+05 | 2.61E+05 | 1.50E+05 | 3 | Extracellular Space |
| 1098 | Q60902 | EPS15L1 | 2.53E+05 | 1.16E+05 | 6.71E+04 | 3 | Plasma Membrane |
| 1099 | Q9DD18 | DTD1 | 2.28E+05 | 1.38E+05 | 8.00E+04 | 3 | Cytoplasm |
| 1100 | Q80X71 | TMEM106B | 2.13E+05 | 5.44E+04 | 3.14E+04 | 3 | Plasma Membrane |
| 1101 | Q9JKM7 | RAB37 | 1.91E+05 | 1.64E+05 | 9.49E+04 | 3 | Cytoplasm |
| 1102 | O70493 | SNX12 | 1.67E+05 | 1.23E+05 | 7.09E+04 | 3 | Cytoplasm |
| 1103 | P63158 | Gm21596/Hmgb1 | 1.64E+05 | 2.31E+05 | 9.44E+04 | 6 | Nucleus |
| 1104 | Q3UJD6 | USP19 | 1.36E+05 | 1.13E+05 | 6.55E+04 | 3 | Cytoplasm |

| | | | | | | | |
|---|---|---|---|---|---|---|---|
| 973 | O88456 | CAPNS1 | 1.78E+06 | 4.67E+05 | 2.70E+05 | 3 | Cytoplasm |
| 974 | Q9D8B6 | FAM210B | 1.75E+06 | 1.51E+06 | 6.16E+05 | 6 | Extracellular Space |
| 975 | Q8BVF7 | APH1A | 1.75E+06 | 4.18E+05 | 2.41E+05 | 3 | Cytoplasm |
| 976 | P50544 | ACADVL | 1.74E+06 | 1.23E+06 | 3.17E+05 | 15 | Cytoplasm |
| 977 | P62918 | RPL8 | 1.73E+06 | 4.83E+05 | 1.97E+05 | 6 | Cytoplasm |
| 978 | Q9CX00 | IST1 | 1.73E+06 | 1.65E+05 | 9.50E+04 | 3 | Cytoplasm |
| 979 | Q8CGC7 | EPRS | 1.72E+06 | 5.29E+05 | 2.16E+05 | 6 | Cytoplasm |
| 980 | P31230 | AIMP1 | 1.72E+06 | 7.30E+05 | 2.98E+05 | 6 | Extracellular Space |
| 981 | Q3UMR5 | MCU | 1.71E+06 | 3.98E+05 | 2.30E+05 | 3 | Cytoplasm |
| 982 | Q9ERE7 | MESDC2 | 1.70E+06 | 4.98E+05 | 2.87E+05 | 3 | Extracellular Space |
| 983 | O89013 | LEPROT | 1.69E+06 | 1.62E+06 | 6.62E+05 | 6 | Plasma Membrane |
| 984 | Q63739 | PTP4A1 | 1.69E+06 | 2.47E+05 | 1.43E+05 | 3 | Cytoplasm |
| 985 | P08103 | HCK | 1.69E+06 | 2.30E+04 | 1.33E+04 | 3 | Cytoplasm |
| 986 | Q9CPW4 | ARPC5 | 1.68E+06 | 1.31E+05 | 7.55E+04 | 3 | Cytoplasm |
| 987 | Q61425 | HADH | 1.68E+06 | 1.73E+06 | 7.07E+05 | 6 | Cytoplasm |
| 988 | Q9CQX8 | MRPS36 | 1.66E+06 | 3.21E+05 | 1.86E+05 | 3 | Cytoplasm |
| 989 | Q8VDV3 | RAB3IL1 | 1.64E+06 | 4.34E+05 | 2.51E+05 | 3 | Cytoplasm |
| 990 | O89112 | LANCL1 | 1.64E+06 | 8.94E+05 | 5.16E+05 | 3 | Plasma Membrane |
| 991 | P58252 | EEF2 | 1.64E+06 | 7.21E+05 | 1.86E+05 | 15 | Cytoplasm |
| 992 | Q99JW4 | LIMS1 | 1.64E+06 | 2.58E+05 | 1.49E+05 | 3 | Plasma Membrane |
| 993 | Q91YT8 | TMEM63A | 1.62E+06 | 5.34E+05 | 3.08E+05 | 3 | Cytoplasm |
| 994 | Q3UHH2 | SLC22A23 | 1.61E+06 | 7.87E+04 | 4.54E+04 | 3 | Plasma Membrane |
| 995 | Q61144 | PSEN2 | 1.60E+06 | 3.65E+05 | 2.11E+05 | 3 | Cytoplasm |
| 996 | Q9WVA4 | TAGLN2 | 1.60E+06 | 7.96E+05 | 2.30E+05 | 12 | Cytoplasm |
| 997 | Q9EQP2 | EHD4 | 1.59E+06 | 5.11E+05 | 2.09E+05 | 6 | Plasma Membrane |
| 998 | Q3UND0 | SKAP2 | 1.59E+06 | 3.28E+05 | 1.34E+05 | 6 | Cytoplasm |
| 999 | O35744 | Chil3/Chil4 | 1.58E+06 | 1.10E+06 | 2.58E+05 | 18 | Cytoplasm |
| 1000 | Q91YP3 | DERA | 1.58E+06 | 5.55E+05 | 1.85E+05 | 9 | Cytoplasm |
| 1001 | Q8C156 | NCAPH | 1.58E+06 | 1.22E+06 | 7.06E+05 | 3 | Nucleus |
| 1002 | Q9CPP6 | NDUFA5 | 1.58E+06 | 3.65E+05 | 2.11E+05 | 3 | Cytoplasm |
| 1003 | E9Q555 | RNF213 | 1.55E+06 | 1.65E+05 | 9.52E+04 | 3 | Cytoplasm |
| 1004 | Q00612 | G6PD | 1.55E+06 | 5.54E+05 | 1.60E+05 | 12 | Cytoplasm |
| 1005 | Q9R1Q9 | ATP6AP1 | 1.55E+06 | 6.00E+05 | 3.47E+05 | 3 | Cytoplasm |
| 1006 | P35486 | PDHA1 | 1.54E+06 | 3.17E+05 | 1.06E+05 | 9 | Cytoplasm |
| 1007 | Q8BL66 | EEA1 | 1.54E+06 | 2.96E+05 | 1.71E+05 | 3 | Cytoplasm |
| 1008 | Q505D1 | ANKRD28 | 1.53E+06 | 4.96E+05 | 2.87E+05 | 3 | Cytoplasm |
| 1009 | Q8BZI6 | GUCD1 | 1.50E+06 | 1.52E+05 | 8.76E+04 | 3 | Nucleus |
| 1010 | Q8VBZ3 | CLPTM1 | 1.48E+06 | 2.76E+05 | 1.59E+05 | 3 | Plasma Membrane |
| 1011 | P38060 | HMGCL | 1.46E+06 | 7.89E+05 | 3.22E+05 | 6 | Cytoplasm |
| 1012 | P24547 | IMPDH2 | 1.46E+06 | 6.55E+05 | 3.78E+05 | 3 | Cytoplasm |
| 1013 | P40142 | TKT | 1.45E+06 | 1.07E+06 | 4.36E+05 | 6 | Cytoplasm |
| 1014 | Q9Z2I8 | SUCLG2 | 1.44E+06 | 1.29E+05 | 7.42E+04 | 3 | Cytoplasm |
| 1015 | Q8BMG7 | RAB3GAP2 | 1.42E+06 | 4.14E+05 | 1.69E+05 | 6 | Cytoplasm |
| 1016 | Q3U487 | HECTD3 | 1.42E+06 | 1.53E+06 | 6.23E+05 | 6 | Cytoplasm |
| 1017 | Q9DCJ5 | NDUFA8 | 1.42E+06 | 1.25E+06 | 7.20E+05 | 3 | Cytoplasm |
| 1018 | P00416 | MT-CO3 | 1.41E+06 | 6.78E+05 | 3.92E+05 | 3 | Cytoplasm |
| 1019 | Q91W90 | TXNDC5 | 1.37E+06 | 4.38E+05 | 2.53E+05 | 3 | Cytoplasm |
| 1020 | Q6DIA2 | EXOC3L4 | 1.36E+06 | 3.57E+05 | 2.06E+05 | 3 | Cytoplasm |
| 1021 | Q9ER72 | CARS | 1.35E+06 | 7.27E+05 | 2.97E+05 | 6 | Cytoplasm |
| 1022 | Q61838 | Pzp | 1.35E+06 | 4.91E+05 | 2.84E+05 | 3 | Extracellular Space |
| 1023 | Q8R180 | ERO1A | 1.35E+06 | 2.85E+05 | 1.64E+05 | 3 | Cytoplasm |
| 1024 | Q5SSH7 | ZZEF1 | 1.35E+06 | 8.42E+05 | 4.86E+05 | 3 | Cytoplasm |
| 1025 | P20918 | PLG | 1.35E+06 | 8.34E+05 | 2.78E+05 | 9 | Extracellular Space |
| 1026 | P22907 | HMBS | 1.34E+06 | 1.33E+05 | 7.66E+04 | 3 | Cytoplasm |
| 1027 | Q811D0 | DLG1 | 1.33E+06 | 7.22E+05 | 2.41E+05 | 9 | Plasma Membrane |
| 1028 | Q9Z110 | ALDH18A1 | 1.32E+06 | 8.87E+05 | 1.81E+05 | 24 | Cytoplasm |
| 1029 | Q3UHJ0 | AAK1 | 1.32E+06 | 4.11E+05 | 2.37E+05 | 3 | Cytoplasm |
| 1030 | P26039 | TLN1 | 1.31E+06 | 9.92E+05 | 2.02E+05 | 24 | Plasma Membrane |
| 1031 | Q9DCT2 | NDUFS3 | 1.31E+06 | 1.22E+06 | 7.02E+05 | 3 | Cytoplasm |
| 1032 | Q8R1I1 | UQCR10 | 1.30E+06 | 8.51E+05 | 4.91E+05 | 3 | Cytoplasm |
| 1033 | Q9WV85 | NME3 | 1.29E+06 | 2.55E+05 | 1.47E+05 | 3 | Cytoplasm |
| 1034 | P83741 | WNK1 | 1.29E+06 | 3.98E+05 | 2.30E+05 | 3 | Cytoplasm |
| 1035 | O35114 | SCARB2 | 1.27E+06 | 1.59E+06 | 9.21E+05 | 3 | Plasma Membrane |
| 1036 | Q80WQ6 | RHBDF2 | 1.27E+06 | 1.25E+06 | 5.09E+05 | 6 | Cytoplasm |
| 1037 | Q9WTP6 | AK2 | 1.26E+06 | 2.01E+05 | 1.16E+05 | 3 | Cytoplasm |
| 1038 | P27612 | PLAA | 1.26E+06 | 3.54E+05 | 2.04E+05 | 3 | Cytoplasm |
| 1039 | Q9WVC8 | SLC26A3 | 1.25E+06 | 6.14E+05 | 3.55E+05 | 3 | Plasma Membrane |
| 1040 | Q60770 | STXBP3 | 1.25E+06 | 1.08E+05 | 6.22E+04 | 3 | Plasma Membrane |
| 1041 | P97351 | Rps3a1 | 1.20E+06 | 9.86E+05 | 4.03E+05 | 6 | Cytoplasm |
| 1042 | Q7TN60 | TMC6 | 1.18E+06 | 8.99E+05 | 3.00E+05 | 9 | Cytoplasm |
| 1043 | Q61781 | KRT14 | 1.18E+06 | 2.69E+06 | 1.56E+06 | 3 | Cytoplasm |
| 1044 | P45952 | ACADM | 1.17E+06 | 1.94E+05 | 1.12E+05 | 3 | Cytoplasm |
| 1045 | Q6A4J8 | USP7 | 1.13E+06 | 1.72E+05 | 9.91E+04 | 3 | Nucleus |
| 1046 | P41438 | SLC19A1 | 1.11E+06 | 3.31E+05 | 1.91E+05 | 3 | Plasma Membrane |
| 1047 | Q8VEE1 | LMCD1 | 1.06E+06 | 7.82E+06 | 4.51E+06 | 3 | Cytoplasm |
| 1048 | Q61878 | PRG2 | 1.05E+06 | 1.09E+06 | 6.31E+05 | 3 | Extracellular Space |
| 1049 | P55302 | LRPAP1 | 1.04E+06 | 7.77E+05 | 4.49E+05 | 3 | Plasma Membrane |
| 1050 | Q4VBD2 | TAPT1 | 1.04E+06 | 3.01E+05 | 1.74E+05 | 3 | Plasma Membrane |
| 1051 | Q9CY58 | SERBP1 | 1.02E+06 | 7.38E+05 | 2.46E+05 | 9 | Cytoplasm |
| 1052 | Q8BTJ4 | ENPP4 | 9.96E+05 | 3.01E+05 | 1.74E+05 | 3 | Cytoplasm |
| 1053 | P31725 | S100A9 | 9.96E+05 | 2.70E+05 | 1.10E+05 | 6 | Cytoplasm |
| 1054 | Q922Q1 | MARC_2 | 9.94E+05 | 2.27E+05 | 1.31E+05 | 3 | Cytoplasm |
| 1055 | Q811C2 | ATG4C | 9.78E+05 | 2.46E+05 | 1.42E+05 | 3 | Cytoplasm |
| 1056 | Q6ZWY3 | RPS27L | 9.54E+05 | 7.58E+05 | 3.10E+05 | 6 | Cytoplasm |
| 1057 | Q80UY2 | KCMF1 | 9.40E+05 | 8.66E+05 | 2.89E+05 | 9 | Extracellular Space |
| 1058 | Q4KUS1 | WFDC8 | 9.24E+05 | 2.71E+05 | 1.56E+05 | 3 | Extracellular Space |
| 1059 | Q60575 | KIF1B | 9.13E+05 | 2.98E+06 | 1.72E+06 | 3 | Cytoplasm |
| 1060 | P08071 | LTF | 9.09E+05 | 6.48E+05 | 1.67E+05 | 15 | Extracellular Space |
| 1061 | P28076 | PSMB9 | 9.05E+05 | 3.05E+05 | 1.76E+05 | 3 | Cytoplasm |
| 1062 | Q8BRU6 | SLC18A2 | 9.01E+05 | 1.93E+05 | 1.12E+05 | 3 | Plasma Membrane |
| 1063 | O88746 | TOM1 | 8.99E+05 | 3.13E+05 | 1.81E+05 | 3 | Cytoplasm |
| 1064 | Q6P9Q6 | FKBP15 | 8.84E+05 | 4.43E+05 | 1.81E+05 | 6 | Plasma Membrane |
| 1065 | Q9WUA2 | FARSB | 8.69E+05 | 2.08E+05 | 1.20E+05 | 3 | Cytoplasm |
| 1066 | Q5HZI9 | SLC25A51 | 8.69E+05 | 7.88E+05 | 4.55E+05 | 3 | Cytoplasm |
| 1067 | Q8BVE3 | ATP6V1H | 8.67E+05 | 5.18E+05 | 2.12E+05 | 6 | Cytoplasm |
| 1068 | P11835 | ITGB2 | 8.58E+05 | 7.27E+05 | 4.19E+05 | 3 | Plasma Membrane |
| 1069 | Q9DBG3 | Ap2b1 | 8.56E+05 | 7.19E+05 | 2.93E+05 | 6 | Plasma Membrane |
| 1070 | Q3U0D9 | HACE1 | 8.54E+05 | 1.37E+05 | 7.89E+04 | 3 | Cytoplasm |
| 1071 | Q3URS9 | CCDC51 | 8.00E+05 | 1.85E+05 | 1.07E+05 | 3 | Cytoplasm |
| 1072 | Q91W34 | C16orf58 | 7.64E+05 | 6.14E+05 | 3.54E+05 | 3 | Cytoplasm |
| 1073 | O35083 | AGPAT1 | 7.46E+05 | 4.40E+05 | 2.54E+05 | 3 | Cytoplasm |
| 1074 | Q9CZ04 | COPS7A | 7.43E+05 | 6.59E+04 | 3.80E+04 | 3 | Cytoplasm |
| 1075 | Q61166 | MAPRE1 | 7.35E+05 | 2.19E+05 | 1.26E+05 | 3 | Cytoplasm |
| 1076 | Q7TMF3 | NDUFA12 | 7.24E+05 | 6.94E+04 | 4.01E+04 | 3 | Cytoplasm |
| 1077 | Q8BGT0 | OSTM1 | 7.07E+05 | 1.64E+05 | 9.45E+04 | 3 | Cytoplasm |
| 1078 | Q01339 | APOH | 6.94E+05 | 2.24E+05 | 1.30E+05 | 3 | Extracellular Space |
| 1079 | Q8BVI4 | QDPR | 6.87E+05 | 2.05E+05 | 1.19E+05 | 3 | Cytoplasm |
| 1080 | Q8BU33 | ILVBL | 6.13E+05 | 3.34E+05 | 1.93E+05 | 3 | Cytoplasm |
| 1081 | O70493 | SNX12 | 6.04E+05 | 1.20E+05 | 6.91E+04 | 3 | Cytoplasm |
| 1082 | P62858 | RPS28 | 5.64E+05 | 9.47E+05 | 5.47E+05 | 3 | Cytoplasm |
| 1083 | Q9DC23 | DNAJC10 | 5.27E+05 | 1.72E+05 | 9.91E+04 | 3 | Cytoplasm |
| 1084 | Q9JKM7 | RAB37 | 5.24E+05 | 1.40E+05 | 8.09E+04 | 3 | Cytoplasm |
| 1085 | A2ATU0 | DHTKD1 | 5.13E+05 | 1.64E+05 | 9.48E+04 | 3 | Cytoplasm |
| 1086 | Q8BTX9 | HSDL1 | 4.99E+05 | 1.10E+05 | 6.34E+04 | 3 | Cytoplasm |
| 1087 | P46061 | RANGAP1 | 4.71E+05 | 1.45E+05 | 8.39E+04 | 3 | Nucleus |
| 1088 | P11276 | FN1 | 4.46E+05 | 1.88E+05 | 6.27E+04 | 9 | Extracellular Space |
| 1089 | Q9D0I9 | RARS | 4.36E+05 | 1.96E+04 | 1.13E+04 | 3 | Cytoplasm |
| 1090 | Q60902 | EPS15L1 | 4.24E+05 | 9.54E+04 | 5.51E+04 | 3 | Plasma Membrane |
| 1091 | Q9D7J6 | DNASE1L1 | 3.75E+05 | 7.67E+04 | 4.43E+04 | 3 | Cytoplasm |
| 1092 | P14106 | C1QB | 3.23E+05 | 5.50E+05 | 3.17E+05 | 3 | Extracellular Space |
| 1093 | Q9D710 | TMX2 | 3.10E+05 | 1.79E+05 | 1.03E+05 | 3 | Cytoplasm |
| 1094 | Q3TL44 | NLRX1 | 3.10E+05 | 2.26E+06 | 1.30E+06 | 3 | Cytoplasm |
| 1095 | Q80X71 | TMEM106 | 3.08E+05 | 3.19E+04 | 1.84E+04 | 3 | Plasma Membrane |
| 1096 | P63158 | Gm21596/ | 2.74E+05 | 2.50E+05 | 1.02E+05 | 6 | Nucleus |
| 1097 | Q9JHK5 | PLEK | 2.35E+05 | 3.38E+05 | 1.95E+05 | 3 | Cytoplasm |
| 1098 | P98086 | C1QA | 2.34E+05 | 5.34E+05 | 2.18E+05 | 6 | Extracellular Space |
| 1099 | Q9WTI7 | MYO1C | 2.32E+05 | 3.87E+04 | 2.23E+04 | 3 | Cytoplasm |
| 1100 | Q3UJD6 | USP19 | 1.96E+05 | 4.47E+04 | 2.58E+04 | 3 | Cytoplasm |
| 1101 | Q9DD18 | DTD1 | 1.26E+05 | 7.38E+04 | 4.26E+04 | 3 | Cytoplasm |
| 1102 | P47915 | Rpl29 (incl | 1.23E+05 | 7.40E+05 | 4.27E+05 | 3 | Cytoplasm |
| 1103 | Q0P5V2 | SOBP | 7.67E+04 | 4.77E+05 | 2.75E+05 | 3 | Nucleus |
| 1104 | P28665 | Mug1/Mu | 6.67E+04 | 6.45E+04 | 3.72E+04 | 3 | Extracellular Space |

| | | | | | | | |
|---|---|---|---|---|---|---|---|
| 973 | Q9EQP2 | EHD4 | 1.33E+06 | 3.77E+05 | 1.54E+05 | 6 | Plasma Membrane |
| 974 | P00397 | MT-CO1 | 1.32E+06 | 1.31E+06 | 7.54E+05 | 3 | Cytoplasm |
| 975 | P29758 | OAT | 1.32E+06 | 9.17E+05 | 5.29E+05 | 3 | Cytoplasm |
| 976 | Q5HZI9 | SLC25A51 | 1.31E+06 | 7.05E+05 | 4.07E+05 | 3 | Cytoplasm |
| 977 | Q9D8B6 | FAM210B | 1.31E+06 | 1.52E+06 | 6.19E+05 | 6 | Extracellular Space |
| 978 | Q9CZW5 | TOMM70A | 1.31E+06 | 6.92E+05 | 4.00E+05 | 3 | Cytoplasm |
| 979 | P62281 | RPS11 | 1.30E+06 | 2.47E+06 | 1.01E+06 | 6 | Cytoplasm |
| 980 | P00416 | MT-CO3 | 1.30E+06 | 8.76E+05 | 5.06E+05 | 3 | Cytoplasm |
| 981 | P50544 | ACADVL | 1.29E+06 | 1.42E+06 | 3.66E+05 | 15 | Cytoplasm |
| 982 | P62960 | YBX1 | 1.29E+06 | 1.51E+06 | 6.15E+05 | 6 | Nucleus |
| 983 | Q61102 | ABCB7 | 1.29E+06 | 9.60E+05 | 5.54E+05 | 3 | Cytoplasm |
| 984 | Q9D6J6 | NDUFV2 | 1.28E+06 | 7.88E+05 | 4.55E+05 | 3 | Cytoplasm |
| 985 | Q9CPQ1 | Cox6c | 1.28E+06 | 7.30E+05 | 4.21E+05 | 3 | Cytoplasm |
| 986 | P51437 | CAMP | 1.27E+06 | 5.72E+06 | 2.34E+06 | 6 | Cytoplasm |
| 987 | P18181 | CD48 | 1.26E+06 | 5.54E+05 | 3.20E+05 | 3 | Plasma Membrane |
| 988 | P52825 | CPT2 | 1.25E+06 | 1.54E+06 | 5.13E+05 | 9 | Cytoplasm |
| 989 | P30681 | HMGB2 | 1.23E+06 | 2.17E+06 | 1.25E+06 | 3 | Nucleus |
| 990 | Q8BTJ4 | ENPP4 | 1.22E+06 | 5.89E+05 | 3.40E+05 | 3 | Cytoplasm |
| 991 | Q9Z2I0 | LETM1 | 1.22E+06 | 1.04E+06 | 6.03E+05 | 3 | Cytoplasm |
| 992 | O35083 | AGPAT1 | 1.22E+06 | 1.41E+05 | 8.13E+04 | 3 | Cytoplasm |
| 993 | Q4VBD2 | TAPT1 | 1.19E+06 | 2.96E+05 | 1.71E+05 | 3 | Plasma Membrane |
| 994 | O08663 | METAP2 | 1.18E+06 | 6.69E+05 | 3.86E+05 | 3 | Cytoplasm |
| 995 | Q3UMR5 | MCU | 1.17E+06 | 5.54E+05 | 3.20E+05 | 3 | Cytoplasm |
| 996 | Q99JW4 | LIMS1 | 1.17E+06 | 2.24E+06 | 1.29E+06 | 3 | Plasma Membrane |
| 997 | Q9WV85 | NME3 | 1.17E+06 | 3.77E+05 | 2.18E+05 | 3 | Cytoplasm |
| 998 | Q9D8N0 | EEF1G | 1.16E+06 | 1.42E+06 | 4.72E+05 | 9 | Cytoplasm |
| 999 | Q8BIJ6 | IARS2 | 1.16E+06 | 1.26E+06 | 7.25E+05 | 3 | Cytoplasm |
| 1000 | Q5U458 | DNAJC11 | 1.14E+06 | 7.91E+05 | 3.23E+05 | 6 | Cytoplasm |
| 1001 | Q8VBZ3 | CLPTM1 | 1.14E+06 | 1.60E+05 | 9.23E+04 | 3 | Plasma Membrane |
| 1002 | O35114 | SCARB2 | 1.14E+06 | 1.23E+06 | 7.09E+05 | 3 | Plasma Membrane |
| 1003 | P29391 | FTL | 1.13E+06 | 1.61E+06 | 5.35E+05 | 9 | Cytoplasm |
| 1004 | Q61144 | PSEN2 | 1.13E+06 | 7.60E+04 | 4.39E+04 | 3 | Cytoplasm |
| 1005 | P62849 | RPS24 | 1.11E+06 | 4.90E+05 | 2.83E+05 | 3 | Cytoplasm |
| 1006 | Q8CG48 | SMC2 | 1.11E+06 | 3.35E+05 | 1.94E+05 | 3 | Nucleus |
| 1007 | Q3UHJ0 | AAK1 | 1.10E+06 | 1.75E+05 | 1.01E+05 | 3 | Cytoplasm |
| 1008 | O88448 | KLC2 | 1.10E+06 | 1.13E+06 | 6.50E+05 | 3 | Cytoplasm |
| 1009 | P40142 | TKT | 1.10E+06 | 1.16E+06 | 4.74E+05 | 6 | Cytoplasm |
| 1010 | Q9WVA4 | TAGLN2 | 1.09E+06 | 1.34E+06 | 3.88E+05 | 12 | Cytoplasm |
| 1011 | Q64331 | MYO6 | 1.08E+06 | 6.72E+05 | 3.88E+05 | 3 | Cytoplasm |
| 1012 | Q9Z2I8 | SUCLG2 | 1.06E+06 | 1.52E+05 | 8.79E+04 | 3 | Cytoplasm |
| 1013 | Q91W90 | TXNDC5 | 1.04E+06 | 1.36E+05 | 7.87E+04 | 3 | Cytoplasm |
| 1014 | O88746 | TOM1 | 1.04E+06 | 1.35E+05 | 7.77E+04 | 3 | Cytoplasm |
| 1015 | P62301 | RPS13 | 1.03E+06 | 6.77E+05 | 2.76E+05 | 6 | Cytoplasm |
| 1016 | P20444 | PRKCA | 1.03E+06 | 4.54E+05 | 2.62E+05 | 3 | Cytoplasm |
| 1017 | Q9CZ04 | COPS7A | 1.02E+06 | 1.61E+05 | 9.31E+04 | 3 | Cytoplasm |
| 1018 | Q61781 | KRT14 | 1.01E+06 | 9.82E+05 | 5.67E+05 | 3 | Cytoplasm |
| 1019 | P13609 | Srgn | 1.01E+06 | 1.08E+06 | 6.24E+05 | 3 | Cytoplasm |
| 1020 | Q9CPP6 | NDUFA5 | 1.00E+06 | 4.76E+05 | 2.75E+05 | 3 | Cytoplasm |
| 1021 | Q9CRD0 | OCIAD1 | 1.00E+06 | 1.11E+06 | 6.38E+05 | 3 | Cytoplasm |
| 1022 | P27612 | PLAA | 9.98E+05 | 3.11E+05 | 1.79E+05 | 3 | Cytoplasm |
| 1023 | P35486 | PDHA1 | 9.97E+05 | 8.09E+05 | 2.70E+05 | 9 | Cytoplasm |
| 1024 | P08103 | HCK | 9.77E+05 | 2.89E+05 | 1.67E+05 | 3 | Cytoplasm |
| 1025 | Q91YT8 | TMEM63A | 9.52E+05 | 2.53E+06 | 1.46E+06 | 3 | Cytoplasm |
| 1026 | P62918 | RPL8 | 9.46E+05 | 6.04E+05 | 2.47E+05 | 6 | Cytoplasm |
| 1027 | Q6NXH9 | KRT73 | 9.46E+05 | 2.18E+05 | 1.26E+05 | 3 | Extracellular Space |
| 1028 | Q9WUX5 | MRVI1 | 9.33E+05 | 7.22E+05 | 2.41E+05 | 9 | Cytoplasm |
| 1029 | Q8BH57 | WDR48 | 9.28E+05 | 2.16E+05 | 1.25E+05 | 3 | Cytoplasm |
| 1030 | P58252 | EEF2 | 9.20E+05 | 9.30E+05 | 2.40E+05 | 15 | Cytoplasm |
| 1031 | O35744 | Chil3/Chil4 | 9.18E+05 | 3.24E+06 | 7.63E+05 | 18 | Cytoplasm |
| 1032 | Q811C2 | ATG4C | 9.12E+05 | 3.21E+04 | 1.86E+04 | 3 | Cytoplasm |
| 1033 | Q91W34 | C16orf58 | 8.99E+05 | 2.56E+05 | 1.48E+05 | 3 | Cytoplasm |
| 1034 | Q4KUS1 | WFDC8 | 8.80E+05 | 2.84E+05 | 1.64E+05 | 3 | Extracellular Space |
| 1035 | Q6DIA2 | EXOC3L4 | 8.78E+05 | 6.84E+05 | 3.95E+05 | 3 | Cytoplasm |
| 1036 | A2CG49 | KALRN | 8.74E+05 | 6.79E+05 | 2.77E+05 | 6 | Cytoplasm |
| 1037 | Q9R233 | TAPBP | 8.65E+05 | 5.02E+05 | 2.90E+05 | 3 | Cytoplasm |
| 1038 | Q9CY58 | SERBP1 | 8.63E+05 | 6.63E+05 | 2.21E+05 | 9 | Cytoplasm |
| 1039 | Q6A4J8 | USP7 | 8.61E+05 | 1.71E+05 | 9.88E+04 | 3 | Nucleus |
| 1040 | Q3TL44 | NLRX1 | 8.59E+05 | 1.89E+06 | 1.09E+06 | 3 | Cytoplasm |
| 1041 | P47962 | RPL5 | 8.58E+05 | 1.72E+06 | 9.92E+05 | 3 | Cytoplasm |
| 1042 | P24549 | ALDH1A1 | 8.54E+05 | 8.44E+05 | 4.87E+05 | 3 | Cytoplasm |
| 1043 | P38060 | HMGCL | 8.47E+05 | 1.03E+06 | 4.19E+05 | 6 | Cytoplasm |
| 1044 | P28076 | PSMB9 | 8.45E+05 | 1.59E+05 | 9.16E+04 | 3 | Cytoplasm |
| 1045 | P47915 | Rpl29 (includes others) | 8.43E+05 | 1.09E+05 | 6.30E+04 | 3 | Cytoplasm |
| 1046 | Q00612 | G6PD | 8.30E+05 | 2.69E+05 | 7.77E+04 | 12 | Cytoplasm |
| 1047 | P57776 | EEF1D | 8.27E+05 | 4.65E+05 | 2.69E+05 | 3 | Cytoplasm |
| 1048 | Q8BU33 | ILVBL | 8.25E+05 | 4.08E+05 | 2.36E+05 | 3 | Cytoplasm |
| 1049 | P20152 | VIM | 8.24E+05 | 3.41E+06 | 1.97E+06 | 3 | Cytoplasm |
| 1050 | P45952 | ACADM | 8.21E+05 | 4.95E+05 | 2.86E+05 | 3 | Cytoplasm |
| 1051 | E9Q555 | RNF213 | 8.18E+05 | 2.40E+05 | 1.39E+05 | 3 | Cytoplasm |
| 1052 | Q6ZWY3 | RPS27L | 8.04E+05 | 6.81E+05 | 2.78E+05 | 6 | Cytoplasm |
| 1053 | Q9JII5 | Dazap1 | 8.00E+05 | 2.27E+05 | 1.31E+05 | 3 | Cytoplasm |
| 1054 | Q60770 | STXBP3 | 7.97E+05 | 5.67E+05 | 3.28E+05 | 3 | Plasma Membrane |
| 1055 | Q8BVE3 | ATP6V1H | 7.96E+05 | 3.89E+05 | 1.59E+05 | 6 | Cytoplasm |
| 1056 | P97351 | Rps3a1 | 7.46E+05 | 8.79E+05 | 3.59E+05 | 6 | Cytoplasm |
| 1057 | Q922Q1 | MARC_2 | 7.45E+05 | 5.05E+05 | 2.91E+05 | 3 | Cytoplasm |
| 1058 | Q9DBG3 | Ap2b1 | 7.38E+05 | 1.03E+06 | 4.19E+05 | 6 | Plasma Membrane |
| 1059 | Q9Z2G9 | HTATIP2 | 7.31E+05 | 7.21E+05 | 2.94E+05 | 6 | Nucleus |
| 1060 | Q60865 | CAPRIN1 | 7.16E+05 | 3.70E+05 | 1.51E+05 | 6 | Plasma Membrane |
| 1061 | Q80X71 | TMEM106B | 7.13E+05 | 8.81E+04 | 5.08E+04 | 3 | Plasma Membrane |
| 1062 | Q61166 | MAPRE1 | 7.09E+05 | 7.19E+04 | 4.15E+04 | 3 | Cytoplasm |
| 1063 | P62858 | RPS28 | 6.97E+05 | 5.87E+05 | 3.39E+05 | 3 | Cytoplasm |
| 1064 | Q3URS9 | CCDC51 | 6.90E+05 | 1.97E+05 | 1.14E+05 | 3 | Cytoplasm |
| 1065 | P09528 | FTH1 | 6.59E+05 | 5.20E+05 | 3.00E+05 | 3 | Cytoplasm |
| 1066 | Q8BRU6 | SLC18A2 | 6.35E+05 | 3.50E+05 | 2.02E+05 | 3 | Plasma Membrane |
| 1067 | Q01339 | APOH | 6.20E+05 | 7.03E+04 | 4.06E+04 | 3 | Extracellular Space |
| 1068 | Q61878 | PRG2 | 6.10E+05 | 8.17E+06 | 4.72E+06 | 3 | Extracellular Space |
| 1069 | P31725 | S100A9 | 5.97E+05 | 9.62E+05 | 3.93E+05 | 6 | Cytoplasm |
| 1070 | Q3U0D9 | HACE1 | 5.95E+05 | 9.94E+04 | 5.74E+04 | 3 | Cytoplasm |
| 1071 | P08071 | LTF | 5.78E+05 | 2.48E+06 | 6.39E+05 | 15 | Extracellular Space |
| 1072 | P31230 | AIMP1 | 5.47E+05 | 4.66E+05 | 1.90E+05 | 6 | Extracellular Space |
| 1073 | Q8BGT0 | OSTM1 | 5.28E+05 | 1.47E+05 | 8.51E+04 | 3 | Cytoplasm |
| 1074 | Q8CGC7 | EPRS | 5.14E+05 | 1.79E+05 | 7.31E+04 | 6 | Cytoplasm |
| 1075 | Q9WTP6 | AK2 | 5.04E+05 | 4.49E+05 | 2.59E+05 | 3 | Cytoplasm |
| 1076 | Q9D7J6 | DNASE1L1 | 5.00E+05 | 1.47E+05 | 8.49E+04 | 3 | Cytoplasm |
| 1077 | Q7TMF3 | NDUFA12 | 4.95E+05 | 1.85E+05 | 1.07E+05 | 3 | Cytoplasm |
| 1078 | Q9WTI7 | MYO1C | 4.91E+05 | 6.45E+04 | 3.72E+04 | 3 | Cytoplasm |
| 1079 | Q0P5V2 | SOBP | 4.75E+05 | 2.66E+05 | 1.54E+05 | 3 | Nucleus |
| 1080 | Q9DC23 | DNAJC10 | 4.72E+05 | 8.56E+04 | 4.94E+04 | 3 | Cytoplasm |
| 1081 | Q9WUA2 | FARSB | 4.42E+05 | 1.64E+05 | 9.48E+04 | 3 | Cytoplasm |
| 1082 | O70251 | EEF1B2 | 4.33E+05 | 5.56E+05 | 3.21E+05 | 3 | Cytoplasm |
| 1083 | Q8BTX9 | HSDL1 | 4.24E+05 | 1.51E+05 | 8.71E+04 | 3 | Cytoplasm |
| 1084 | P46061 | RANGAP1 | 4.11E+05 | 1.76E+05 | 1.01E+05 | 3 | Nucleus |
| 1085 | Q9JKM7 | RAB37 | 4.06E+05 | 1.66E+05 | 9.57E+04 | 3 | Cytoplasm |
| 1086 | P11276 | FN1 | 4.05E+05 | 1.21E+05 | 4.04E+04 | 9 | Extracellular Space |
| 1087 | O70493 | SNX12 | 3.73E+05 | 3.46E+05 | 2.00E+05 | 3 | Cytoplasm |
| 1088 | P11835 | ITGB2 | 3.71E+05 | 1.13E+06 | 6.52E+05 | 3 | Plasma Membrane |
| 1089 | P20918 | PLG | 3.63E+05 | 1.46E+06 | 4.88E+05 | 9 | Extracellular Space |
| 1090 | Q9D710 | TMX2 | 3.24E+05 | 1.85E+05 | 1.07E+05 | 3 | Cytoplasm |
| 1091 | Q60902 | EPS15L1 | 2.93E+05 | 3.21E+04 | 1.85E+04 | 3 | Plasma Membrane |
| 1092 | P28665 | Mug1/Mug2 | 2.80E+05 | 8.11E+05 | 4.68E+05 | 3 | Extracellular Space |
| 1093 | Q9CX63 | C17orf99 | 2.69E+05 | 5.30E+05 | 1.16E+05 | 21 | Extracellular Space |
| 1094 | Q6P9Q6 | FKBP15 | 2.31E+05 | 1.04E+05 | 4.26E+04 | 6 | Plasma Membrane |
| 1095 | P63158 | Gm21596/Hmgb1 | 1.99E+05 | 1.86E+06 | 7.59E+05 | 6 | Nucleus |
| 1096 | Q9DD18 | DTD1 | 1.75E+05 | 7.61E+04 | 4.39E+04 | 3 | Cytoplasm |
| 1097 | P17563 | SELENBP1 | 1.70E+05 | 1.06E+05 | 6.11E+04 | 3 | Cytoplasm |
| 1098 | Q9D0I9 | RARS | 1.24E+05 | 9.14E+04 | 5.28E+04 | 3 | Cytoplasm |
| 1099 | Q3UJD6 | USP19 | 1.10E+05 | 1.64E+04 | 9.45E+03 | 3 | Cytoplasm |
| 1100 | Q9JHK5 | PLEK | 1.10E+05 | 2.49E+05 | 1.44E+05 | 3 | Cytoplasm |
| 1101 | Q9ERM2 | ICAM4 | 7.41E+04 | 7.55E+04 | 3.08E+04 | 6 | Plasma Membrane |
| 1102 | P14106 | C1QB | 4.39E+04 | 1.11E+06 | 6.39E+05 | 3 | Extracellular Space |
| 1103 | P98086 | C1QA | 3.17E+04 | 7.31E+05 | 2.99E+05 | 6 | Extracellular Space |
| 1104 | Q8K2Z4 | NCAPD2 | 0.00E+00 | 6.72E+05 | 3.88E+05 | 3 | Nucleus |

| | | | | | | | |
|---|---|---|---|---|---|---|---|
| 973 | Q9Z2G9 | HTATIP2 | 868019 | 455109 | 185797 | 6 | Nucleus |
| 974 | Q6P9Q6 | FKBP15 | 858400 | 345434 | 141023 | 6 | Plasma Membrane |
| 975 | P62858 | RPS28 | 856013 | 1459777 | 842803 | 3 | Cytoplasm |
| 976 | O89053 | CORO1A | 849143 | 4507142 | 721720 | 39 | Cytoplasm |
| 977 | Q8BVE3 | ATP6V1H | 842651 | 525602 | 214576 | 6 | Cytoplasm |
| 978 | Q8CBY3 | LENG8 | 840998 | 628293 | 362745 | 3 | Nucleus |
| 979 | Q8BTJ4 | ENPP4 | 840677 | 964285 | 556730 | 3 | Cytoplasm |
| 980 | Q8BGT0 | OSTM1 | 831374 | 311925 | 180090 | 3 | Cytoplasm |
| 981 | P17710 | HK1 | 824878 | 1542853 | 247054 | 39 | Cytoplasm |
| 982 | Q99KR7 | PPIF | 816584 | 1251218 | 510808 | 6 | Cytoplasm |
| 983 | Q9QVN7 | TCEA2 | 807087 | 466387 | 269269 | 3 | Nucleus |
| 984 | O55222 | ILK | 805037 | 2300618 | 542261 | 18 | Plasma Membrane |
| 985 | Q6A4J8 | USP7 | 780574 | 70551 | 40733 | 3 | Nucleus |
| 986 | P20918 | PLG | 779827 | 814797 | 271599 | 9 | Extracellular Space |
| 987 | Q6NSQ9 | G6PC3 | 776100 | 409548 | 236453 | 3 | Cytoplasm |
| 988 | O70493 | SNX12 | 770178 | 410624 | 237074 | 3 | Cytoplasm |
| 989 | Q61425 | HADH | 769950 | 1099714 | 448956 | 6 | Cytoplasm |
| 990 | Q9D3G5 | STX11 | 765012 | 799145 | 188360 | 18 | Plasma Membrane |
| 991 | Q6P9J9 | ANO6 | 758379 | 672798 | 388440 | 3 | Plasma Membrane |
| 992 | Q4KUS1 | WFDC8 | 738524 | 84711 | 48908 | 3 | Extracellular Space |
| 993 | Q922Q1 | MARC_2 | 732618 | 367278 | 212048 | 3 | Cytoplasm |
| 994 | Q9EQP2 | EHD4 | 731361 | 361133 | 147432 | 6 | Plasma Membrane |
| 995 | Q3URS9 | CCDC51 | 731216 | 411573 | 237622 | 3 | Cytoplasm |
| 996 | Q9DBG3 | Ap2b1 | 728648 | 822585 | 335819 | 6 | Plasma Membrane |
| 997 | P40240 | CD9 | 719812 | 4252470 | 1417490 | 9 | Plasma Membrane |
| 998 | Q922Q8 | LRRC59 | 706300 | 967444 | 558554 | 3 | Cytoplasm |
| 999 | Q08093 | CNN2 | 703339 | 1081165 | 624211 | 3 | Cytoplasm |
| 1000 | Q64518 | ATP2A3 | 683996 | 3153250 | 455132 | 48 | Cytoplasm |
| 1001 | P21278 | GNA11 | 683508 | 453323 | 261726 | 3 | Plasma Membrane |
| 1002 | Q64455 | PTPRJ | 655162 | 1293517 | 431172 | 9 | Plasma Membrane |
| 1003 | Q9ET30 | TM9SF3 | 653859 | 413410 | 238682 | 3 | Cytoplasm |
| 1004 | Q9Z2I8 | SUCLG2 | 650752 | 418973 | 241894 | 3 | Cytoplasm |
| 1005 | Q61335 | BCAP31 | 647437 | 782402 | 451720 | 3 | Cytoplasm |
| 1006 | P36423 | TBXAS1 | 645665 | 1771131 | 340854 | 27 | Plasma Membrane |
| 1007 | Q9QUT0 | RHAG | 635315 | 1201456 | 346830 | 12 | Plasma Membrane |
| 1008 | P51437 | CAMP | 635057 | 143607 | 58627 | 6 | Cytoplasm |
| 1009 | O35083 | AGPAT1 | 628791 | 108467 | 62624 | 3 | Cytoplasm |
| 1010 | P26039 | TLN1 | 627713 | 683437 | 139506 | 24 | Plasma Membrane |
| 1011 | P27661 | H2AFX | 623301 | 1482444 | 605205 | 6 | Nucleus |
| 1012 | Q7TMF3 | NDUFA12 | 622228 | 256049 | 147830 | 3 | Cytoplasm |
| 1013 | Q61166 | MAPRE1 | 608958 | 128432 | 74150 | 3 | Cytoplasm |
| 1014 | P46061 | RANGAP1 | 604621 | 156237 | 90204 | 3 | Nucleus |
| 1015 | P43275 | Hist1h1a | 603562 | 731607 | 211197 | 12 | Nucleus |
| 1016 | Q80X71 | TMEM106B | 602615 | 82975 | 47906 | 3 | Plasma Membrane |
| 1017 | O54734 | DDOST | 595433 | 1059761 | 353254 | 9 | Cytoplasm |
| 1018 | Q64521 | GPD2 | 575978 | 1161303 | 212024 | 30 | Cytoplasm |
| 1019 | Q91W34 | C16orf58 | 566741 | 530760 | 306434 | 3 | Cytoplasm |
| 1020 | Q8CGC7 | EPRS | 564032 | 164076 | 66984 | 6 | Cytoplasm |
| 1021 | P17563 | SELENBP1 | 552309 | 151368 | 87392 | 3 | Cytoplasm |
| 1022 | Q9D7J6 | DNASE1L1 | 550801 | 86512 | 49948 | 3 | Cytoplasm |
| 1023 | Q5HZI9 | SLC25A51 | 549079 | 1527426 | 881860 | 3 | Cytoplasm |
| 1024 | Q01339 | APOH | 546352 | 424746 | 245227 | 3 | Extracellular Space |
| 1025 | Q0P5V2 | SOBP | 545023 | 402558 | 232417 | 3 | Nucleus |
| 1026 | Q9R233 | TAPBP | 533909 | 116179 | 67076 | 3 | Cytoplasm |
| 1027 | O88448 | KLC2 | 521249.125 | 496187 | 286473 | 3 | Cytoplasm |
| 1028 | P26443 | GLUD1 | 514656 | 497406 | 203065 | 6 | Cytoplasm |
| 1029 | Q99P58 | RAB27B | 486995 | 2745643 | 560452 | 24 | Cytoplasm |
| 1030 | Q3THW5 | H2AFV | 485292 | 184651 | 106608 | 3 | Nucleus |
| 1031 | Q9DC23 | DNAJC10 | 480544 | 116899 | 67492 | 3 | Cytoplasm |
| 1032 | P35441 | THBS1 | 461327 | 2912090 | 297214 | 96 | Extracellular Space |
| 1033 | P35821 | PTPN1 | 461200 | 921563 | 532065 | 3 | Cytoplasm |
| 1034 | Q8K4F0 | CD226 | 457463 | 484068 | 279477 | 3 | Plasma Membrane |
| 1035 | Q921H8 | ACAA1 | 451584 | 730891 | 421980 | 3 | Cytoplasm |
| 1036 | P11835 | ITGB2 | 435634 | 667559 | 385415 | 3 | Plasma Membrane |
| 1037 | Q8QZY6 | TSPAN14 | 420160 | 776961 | 448579 | 3 | Plasma Membrane |
| 1038 | Q9WTI7 | MYO1C | 415848 | 137009 | 79102 | 3 | Cytoplasm |
| 1039 | P05977 | MYL1 | 414562 | 1005924 | 580771 | 3 | Cytoplasm |
| 1040 | Q9ERI2 | RAB27A | 410868 | 519691 | 300044 | 3 | Cytoplasm |
| 1041 | Q9ES46 | PARVB | 400281 | 739121 | 174213 | 18 | Cytoplasm |
| 1042 | Q8CF94 | RHCE/RHD | 397080 | 816394 | 166646 | 24 | Plasma Membrane |
| 1043 | Q01102 | SELP | 388569 | 1560747 | 367872 | 18 | Plasma Membrane |
| 1044 | Q9JL99 | CLEC1B | 380888 | 1567777 | 905157 | 3 | Plasma Membrane |
| 1045 | Q99L43 | CDS2 | 379439 | 697162 | 402506 | 3 | Cytoplasm |
| 1046 | A2ATU0 | DHTKD1 | 369929 | 118832 | 68608 | 3 | Cytoplasm |
| 1047 | Q91YT8 | TMEM63A | 366064 | 411697 | 237694 | 3 | Cytoplasm |
| 1048 | Q9WVA4 | TAGLN2 | 365069 | 847116 | 244541 | 12 | Cytoplasm |
| 1049 | P70302 | STIM1 | 354718 | 404617 | 134872 | 9 | Plasma Membrane |
| 1050 | Q60790 | RASA3 | 342003 | 697740 | 402841 | 3 | Plasma Membrane |
| 1051 | O88792 | F11R | 341019 | 1280565 | 330640 | 15 | Plasma Membrane |
| 1052 | P39876 | TIMP3 | 338978 | 1429125 | 412553 | 12 | Extracellular Space |
| 1053 | Q9D710 | TMX2 | 332716 | 186534 | 107696 | 3 | Cytoplasm |
| 1054 | Q9CX63 | C17orf99 | 325361 | 507771 | 110805 | 21 | Extracellular Space |
| 1055 | P11276 | FN1 | 323104 | 99827 | 33276 | 9 | Extracellular Space |
| 1056 | Q9WTP6 | AK2 | 322688 | 183362 | 105864 | 3 | Cytoplasm |
| 1057 | A2ARI4 | LGR4 | 310348 | 1184838 | 684066 | 3 | Plasma Membrane |
| 1058 | Q60902 | EPS15L1 | 308605 | 119225 | 68834 | 3 | Plasma Membrane |
| 1059 | P08071 | LTF | 285572 | 178435 | 46072 | 15 | Extracellular Space |
| 1060 | O88746 | TOM1 | 273654 | 694095 | 400736 | 3 | Cytoplasm |
| 1061 | Q60770 | STXBP3 | 263992 | 187558 | 108287 | 3 | Plasma Membrane |
| 1062 | Q6URW6 | MYH14 | 251849 | 759213 | 309947 | 6 | Extracellular Space |
| 1063 | P43276 | Hist1h1b | 251680 | 1356982 | 553986 | 6 | Nucleus |
| 1064 | P98086 | C1QA | 250335 | 131430 | 53656 | 6 | Extracellular Space |
| 1065 | O35744 | Chil3/Chil4 | 239210 | 120687 | 28446 | 18 | Cytoplasm |
| 1066 | P61804 | DAD1 | 231881 | 688786 | 397671 | 3 | Cytoplasm |
| 1067 | Q9QUM4 | SLAMF1 | 217294 | 1070645 | 356882 | 9 | Plasma Membrane |
| 1068 | Q9JHK5 | PLEK | 212248 | 182647 | 105451 | 3 | Cytoplasm |
| 1069 | Q8BU33 | ILVBL | 204514 | 303221 | 175065 | 3 | Cytoplasm |
| 1070 | P13609 | Srgn | 203049 | 209683 | 121061 | 3 | Cytoplasm |
| 1071 | Q8CHK3 | MBOAT7 | 194162 | 1257260 | 725880 | 3 | Plasma Membrane |
| 1072 | O88783 | F5 | 186563 | 680658 | 226886 | 9 | Plasma Membrane |
| 1073 | Q9DD18 | DTD1 | 181612 | 90809 | 52428 | 3 | Cytoplasm |
| 1074 | Q62469 | ITGA2 | 178090 | 821735 | 143046 | 33 | Plasma Membrane |
| 1075 | Q3UMR5 | MCU | 166134 | 449319 | 259415 | 3 | Cytoplasm |
| 1076 | Q8CIZ8 | VWF | 148638 | 619603 | 92365 | 45 | Extracellular Space |
| 1077 | Q9D0I9 | RARS | 148472 | 104833 | 60525 | 3 | Cytoplasm |
| 1078 | B2RPV6 | MMRN1 | 142903 | 839167 | 146080 | 33 | Extracellular Space |
| 1079 | Q9ERM2 | ICAM4 | 136623 | 85635 | 34960 | 6 | Plasma Membrane |
| 1080 | P31725 | S100A9 | 136407 | 57521 | 23483 | 6 | Cytoplasm |
| 1081 | P45952 | ACADM | 134632 | 231907 | 133892 | 3 | Cytoplasm |
| 1082 | P08103 | HCK | 133937 | 248627 | 143545 | 3 | Cytoplasm |
| 1083 | Q99K41 | EMILIN1 | 127123 | 884872 | 361248 | 6 | Extracellular Space |
| 1084 | Q3UND0 | SKAP2 | 126505 | 390951 | 159605 | 6 | Cytoplasm |
| 1085 | Q9JKM7 | RAB37 | 115602.4793 | 146268 | 84448 | 3 | Cytoplasm |
| 1086 | Q6DIA2 | EXOC3L4 | 112981 | 281863 | 162734 | 3 | Cytoplasm |
| 1087 | Q9QZQ8 | H2AFY | 108399 | 185216 | 106935 | 3 | Nucleus |
| 1088 | O70443 | GNAZ | 106684 | 567898 | 231843 | 6 | Plasma Membrane |
| 1089 | A2CG49 | KALRN | 102329 | 243540 | 99425 | 6 | Cytoplasm |
| 1090 | Q61878 | PRG2 | 95079 | 126812 | 73215 | 3 | Extracellular Space |
| 1091 | O35601 | FYB | 94944 | 459965 | 153322 | 9 | Nucleus |
| 1092 | Q3UJD6 | USP19 | 93038 | 26842 | 15497 | 3 | Cytoplasm |
| 1093 | Q8BTX9 | HSDL1 | 88028 | 123506 | 71306 | 3 | Cytoplasm |
| 1094 | Q99JW4 | LIMS1 | 84391 | 187239 | 108102 | 3 | Plasma Membrane |
| 1095 | P20444 | PRKCA | 84385 | 236544 | 136569 | 3 | Cytoplasm |
| 1096 | P14106 | C1QB | 84054 | 71577 | 41325 | 3 | Extracellular Space |
| 1097 | Q61739 | ITGA6 | 57825 | 642041 | 140105 | 21 | Plasma Membrane |
| 1098 | Q9WUX5 | MRVI1 | 54906 | 342157 | 114052 | 9 | Cytoplasm |
| 1099 | P20152 | VIM | 49799 | 43858 | 25322 | 3 | Cytoplasm |
| 1100 | Q8BRU6 | SLC18A2 | 36288 | 193995 | 112003 | 3 | Plasma Membrane |
| 1101 | P30681 | HMGB2 | 23341 | 266326 | 153763 | 3 | Nucleus |
| 1102 | Q3TL44 | NLRX1 | 8070 | 821272 | 474162 | 3 | Cytoplasm |
| 1103 | Q60857 | SLC6A4 | 100 | 856177 | 494314 | 3 | Plasma Membrane |
| 1104 | P63158 | Gm21596/Hmgb1 | 0 | 75325 | 30751 | 6 | Nucleus |

Rank all plasma membrane associated proteins

| Wild -type | | | | | | | |
|---|---|---|---|---|---|---|---|
| Rank | Peptide ID | Symbol | Mean Intensity | St. Dev | Std. err | n | Primary Location |
| 1 | P04919 | SLC4A1 | 2.61E+09 | 4.91E+09 | 3.76E+08 | 171 | Plasma Membrane |
| 2 | P14220 | Gypa | 2.48E+09 | 2.91E+09 | 1.19E+09 | 6 | Plasma Membrane |
| 3 | Q02013 | AQP1 | 1.10E+09 | 9.89E+08 | 2.55E+08 | 15 | Plasma Membrane |
| 4 | Q78HU7 | GYPC | 7.30E+08 | 5.87E+08 | 1.96E+08 | 9 | Plasma Membrane |
| 5 | Q02357 | ANK1 | 5.67E+08 | 7.63E+08 | 3.39E+07 | 507 | Plasma Membrane |
| 6 | P15508 | SPTB | 4.98E+08 | 6.31E+08 | 2.26E+07 | 780 | Plasma Membrane |
| 7 | Q9QUT0 | RHAG | 4.30E+08 | 4.88E+08 | 1.41E+08 | 12 | Plasma Membrane |
| 8 | P48193 | EPB41 | 4.08E+08 | 5.77E+08 | 4.23E+07 | 186 | Plasma Membrane |
| 9 | P49222 | EPB42 | 3.96E+08 | 5.61E+08 | 4.05E+07 | 192 | Plasma Membrane |
| 10 | P70290 | MPP1 | 3.55E+08 | 4.05E+08 | 3.61E+07 | 126 | Plasma Membrane |
| 11 | P18572 | BSG | 2.85E+08 | 3.38E+08 | 5.21E+07 | 42 | Plasma Membrane |
| 12 | P53986 | SLC16A1 | 2.48E+08 | 2.38E+08 | 4.59E+07 | 27 | Plasma Membrane |
| 13 | Q8CF94 | RHCE/RHD (mRh) | 2.37E+08 | 4.67E+08 | 9.54E+07 | 24 | Plasma Membrane |
| 14 | Q61735 | CD47 | 2.14E+08 | 1.81E+08 | 4.26E+07 | 18 | Plasma Membrane |
| 15 | P54116 | STOM | 2.03E+08 | 2.35E+08 | 2.83E+07 | 69 | Plasma Membrane |
| 16 | Q9WV69 | DMTN | 1.85E+08 | 1.87E+08 | 2.16E+07 | 75 | Plasma Membrane |
| 17 | Q8BSM7 | SLC43A1 | 1.29E+08 | 1.90E+08 | 2.92E+07 | 42 | Plasma Membrane |
| 18 | Q08857 | CD36 | 1.16E+08 | 1.63E+08 | 2.35E+07 | 48 | Plasma Membrane |
| 19 | Q8VHL0 | SLC14A1 | 1.06E+08 | 8.62E+07 | 1.44E+07 | 36 | Plasma Membrane |
| 20 | Q9Z0S4 | Cldn13 | 1.05E+08 | 8.02E+07 | 2.31E+07 | 12 | Plasma Membrane |
| 21 | P58771 | Tpm1 | 9.18E+07 | 1.49E+08 | 3.85E+07 | 15 | Plasma Membrane |
| 22 | Q3U9N9 | SLC16A10 | 6.02E+07 | 5.86E+07 | 1.07E+07 | 30 | Plasma Membrane |
| 23 | Q9EQF2 | KEL | 5.47E+07 | 4.88E+07 | 4.70E+06 | 108 | Plasma Membrane |
| 24 | O55186 | Cd59a | 5.08E+07 | 6.37E+07 | 2.12E+07 | 9 | Plasma Membrane |
| 25 | P63001 | RAC1 | 4.62E+07 | 4.27E+07 | 6.59E+06 | 42 | Plasma Membrane |
| 26 | P55258 | RAB8A | 4.41E+07 | 3.45E+07 | 7.03E+06 | 24 | Plasma Membrane |
| 27 | Q61469 | Ppap2a | 3.97E+07 | 2.21E+07 | 5.71E+06 | 15 | Plasma Membrane |
| 28 | P27600 | GNA12 | 3.93E+07 | 1.94E+06 | 1.12E+06 | 3 | Plasma Membrane |
| 29 | Q9JIM1 | SLC29A1 | 3.50E+07 | 4.26E+07 | 7.41E+06 | 33 | Plasma Membrane |
| 30 | Q61411 | HRAS | 3.48E+07 | 5.00E+07 | 9.12E+06 | 30 | Plasma Membrane |
| 31 | Q8R3S2 | TSPAN33 | 3.11E+07 | 4.10E+07 | 8.96E+06 | 21 | Plasma Membrane |
| 32 | Q7TMS5 | ABCG2 | 3.06E+07 | 2.82E+07 | 3.64E+06 | 60 | Plasma Membrane |
| 33 | Q9R0K7 | ATP2B2 | 2.88E+07 | 3.75E+07 | 6.85E+06 | 30 | Plasma Membrane |
| 34 | Q8K400 | STXBP5 | 2.72E+07 | 3.64E+06 | 2.10E+06 | 3 | Plasma Membrane |
| 35 | Q8R3G9 | TSPAN8 | 2.65E+07 | 1.69E+07 | 4.38E+06 | 15 | Plasma Membrane |
| 36 | P56400 | GP1BB | 2.62E+07 | 2.97E+07 | 7.01E+06 | 18 | Plasma Membrane |
| 37 | Q8BGQ1 | VIPAS39 | 2.62E+07 | 7.87E+06 | 4.54E+06 | 3 | Plasma Membrane |
| 38 | Q9DCT8 | Crip2 | 2.59E+07 | 1.18E+07 | 2.28E+06 | 27 | Plasma Membrane |
| 39 | Q9QXY7 | XK | 2.52E+07 | 5.89E+06 | 1.70E+06 | 12 | Plasma Membrane |
| 40 | Q61672 | SLC29A2 | 2.46E+07 | 4.57E+05 | 2.64E+05 | 3 | Plasma Membrane |
| 41 | P14231 | ATP1B2 | 2.23E+07 | 2.31E+07 | 5.96E+06 | 15 | Plasma Membrane |
| 42 | P63024 | VAMP3 | 2.18E+07 | 3.14E+07 | 8.11E+06 | 15 | Plasma Membrane |
| 43 | O88186 | GP9 | 2.15E+07 | 3.17E+07 | 9.14E+06 | 12 | Plasma Membrane |
| 44 | P97300 | Nptn | 2.14E+07 | 3.33E+07 | 6.40E+06 | 27 | Plasma Membrane |
| 45 | P08752 | GNAI2 | 2.14E+07 | 1.79E+07 | 2.03E+06 | 78 | Plasma Membrane |
| 46 | Q64735 | CR1L | 2.12E+07 | 2.07E+07 | 3.98E+06 | 27 | Plasma Membrane |
| 47 | P60904 | DNAJC5 | 2.01E+07 | 1.58E+07 | 4.09E+06 | 15 | Plasma Membrane |
| 48 | Q9QUM0 | ITGA2B | 2.00E+07 | 2.86E+07 | 3.17E+06 | 81 | Plasma Membrane |
| 49 | P08556 | NRAS | 2.00E+07 | 1.78E+06 | 1.03E+06 | 3 | Plasma Membrane |
| 50 | Q9CQW2 | ARL8B | 1.96E+07 | 1.30E+07 | 2.01E+06 | 42 | Plasma Membrane |
| 51 | Q9JHI9 | SLC40A1 | 1.95E+07 | 1.10E+07 | 2.25E+06 | 24 | Plasma Membrane |
| 52 | Q62312 | TGFBR2 | 1.93E+07 | 3.44E+06 | 1.98E+06 | 3 | Plasma Membrane |
| 53 | Q62351 | TFRC | 1.91E+07 | 1.39E+07 | 1.32E+06 | 111 | Plasma Membrane |
| 54 | O54890 | ITGB3 | 1.86E+07 | 2.52E+07 | 2.97E+06 | 72 | Plasma Membrane |
| 55 | Q99KU0 | VMP1 | 1.84E+07 | 3.31E+06 | 1.91E+06 | 3 | Plasma Membrane |
| 56 | P62880 | GNB2 | 1.83E+07 | 1.38E+07 | 2.30E+06 | 36 | Plasma Membrane |
| 57 | P62874 | GNB1 | 1.81E+07 | 1.60E+07 | 2.56E+06 | 39 | Plasma Membrane |
| 58 | Q9JJF9 | SPPL2A | 1.81E+07 | 2.24E+06 | 9.13E+05 | 6 | Plasma Membrane |
| 59 | Q9WV55 | VAPA | 1.79E+07 | 1.49E+07 | 2.87E+06 | 27 | Plasma Membrane |
| 60 | Q8K558 | TREML1 | 1.77E+07 | 1.78E+07 | 3.63E+06 | 24 | Plasma Membrane |
| 61 | Q9JJ00 | PLSCR1 | 1.64E+07 | 6.93E+06 | 2.31E+06 | 9 | Plasma Membrane |
| 62 | Q9QUI6 | ACKR1 | 1.62E+07 | 1.12E+07 | 3.22E+06 | 12 | Plasma Membrane |
| 63 | O35598 | ADAM10 | 1.60E+07 | 1.53E+07 | 1.74E+06 | 78 | Plasma Membrane |
| 64 | P27664 | PDE6A | 1.59E+07 | 1.13E+06 | 6.55E+05 | 3 | Plasma Membrane |
| 65 | O88343 | SLC4A4 | 1.58E+07 | 3.39E+06 | 1.96E+06 | 3 | Plasma Membrane |
| 66 | O08742 | GP5 | 1.54E+07 | 1.55E+07 | 2.83E+06 | 30 | Plasma Membrane |
| 67 | Q3T9M1 | MFSD2B | 1.50E+07 | 1.41E+07 | 5.77E+06 | 6 | Plasma Membrane |
| 68 | O35930 | GP1BA | 1.45E+07 | 1.91E+07 | 3.05E+06 | 39 | Plasma Membrane |
| 69 | Q80YF9 | ARHGAP33 | 1.43E+07 | 4.32E+06 | 2.50E+06 | 3 | Plasma Membrane |
| 70 | Q8VDN2 | ATP1A1 | 1.28E+07 | 9.64E+06 | 1.03E+06 | 87 | Plasma Membrane |
| 71 | P40240 | CD9 | 1.27E+07 | 2.23E+07 | 7.43E+06 | 9 | Plasma Membrane |
| 72 | Q9Z2P8 | VAMP5 | 1.27E+07 | 6.50E+06 | 2.17E+06 | 9 | Plasma Membrane |
| 73 | O09117 | SYPL1 | 1.24E+07 | 8.83E+06 | 2.94E+06 | 9 | Plasma Membrane |
| 74 | P32037 | SLC2A3 | 1.19E+07 | 1.09E+07 | 2.37E+06 | 21 | Plasma Membrane |
| 75 | Q60634 | FLOT2 | 1.13E+07 | 8.41E+06 | 1.18E+06 | 51 | Plasma Membrane |
| 76 | O09044 | SNAP23 | 1.06E+07 | 1.06E+07 | 1.76E+06 | 36 | Plasma Membrane |
| 77 | Q91V01 | LPCAT3 | 1.04E+07 | 7.53E+06 | 2.17E+06 | 12 | Plasma Membrane |
| 78 | P21440 | ABCB4 | 9.83E+06 | 6.15E+06 | 7.25E+05 | 72 | Plasma Membrane |
| 79 | O08917 | FLOT1 | 9.81E+06 | 5.72E+06 | 8.26E+05 | 48 | Plasma Membrane |
| 80 | P39054 | DNM2 | 9.69E+06 | 9.47E+06 | 1.14E+06 | 69 | Plasma Membrane |
| 81 | P04925 | PRNP | 9.46E+06 | 2.15E+06 | 1.24E+06 | 3 | Plasma Membrane |
| 82 | O70439 | STX7 | 9.22E+06 | 3.13E+06 | 7.37E+05 | 18 | Plasma Membrane |
| 83 | O70404 | VAMP8 | 8.91E+06 | 4.79E+06 | 1.60E+06 | 9 | Plasma Membrane |
| 84 | Q9D771 | TMEM206 | 8.55E+06 | 7.16E+06 | 2.39E+06 | 9 | Plasma Membrane |
| 85 | P62331 | ARF6 | 8.48E+06 | 4.32E+06 | 1.02E+06 | 18 | Plasma Membrane |
| 86 | P16546 | SPTAN1 | 8.22E+06 | 5.10E+06 | 2.94E+06 | 3 | Plasma Membrane |
| 87 | P61226 | RAP2B | 8.17E+06 | 3.12E+06 | 7.35E+05 | 18 | Plasma Membrane |
| 88 | Q9CQ22 | LAMTOR1 | 8.09E+06 | 6.16E+06 | 1.78E+06 | 12 | Plasma Membrane |
| 89 | Q9WTR1 | TRPV2 | 7.99E+06 | 5.13E+06 | 8.55E+05 | 36 | Plasma Membrane |
| 90 | Q9D659 | C10orf54 | 7.84E+06 | 2.00E+06 | 1.15E+06 | 3 | Plasma Membrane |
| 91 | Q9JMH7 | NEU3 | 7.73E+06 | 5.57E+06 | 1.86E+06 | 9 | Plasma Membrane |
| 92 | Q61165 | SLC9A1 | 7.41E+06 | 4.18E+06 | 8.54E+05 | 24 | Plasma Membrane |
| 93 | P58735 | SLC26A1 | 7.40E+06 | 3.41E+06 | 1.39E+06 | 6 | Plasma Membrane |
| 94 | P09055 | ITGB1 | 7.15E+06 | 6.30E+06 | 1.01E+06 | 39 | Plasma Membrane |
| 95 | P57716 | NCSTN | 6.85E+06 | 3.85E+06 | 7.40E+05 | 27 | Plasma Membrane |
| 96 | Q9JM62 | REEP6 | 6.79E+06 | 4.99E+06 | 1.66E+06 | 9 | Plasma Membrane |
| 97 | Q9JIZ9 | PLSCR3 | 6.78E+06 | 3.50E+06 | 1.01E+06 | 12 | Plasma Membrane |
| 98 | P60766 | Cdc42 | 6.66E+06 | 3.48E+06 | 8.98E+05 | 15 | Plasma Membrane |
| 99 | Q61475 | CD55 | 6.50E+06 | 3.36E+06 | 7.92E+05 | 18 | Plasma Membrane |
| 100 | P26040 | EZR | 6.41E+06 | 6.11E+06 | 1.12E+06 | 30 | Plasma Membrane |
| 101 | Q3TZZ7 | ESYT2 | 6.38E+06 | 2.91E+06 | 3.76E+05 | 60 | Plasma Membrane |
| 102 | Q68FD5 | CLTC | 6.32E+06 | 4.97E+06 | 4.23E+05 | 138 | Plasma Membrane |
| 103 | O35379 | ABCC1 | 6.20E+06 | 4.39E+06 | 5.97E+05 | 54 | Plasma Membrane |
| 104 | Q9JJK2 | LANCL2 | 6.13E+06 | 3.14E+06 | 5.23E+05 | 36 | Plasma Membrane |
| 105 | Q9DAS9 | GNG12 | 6.12E+06 | 3.62E+05 | 2.09E+05 | 3 | Plasma Membrane |
| 106 | Q60682 | Klra4 (includes others) | 6.01E+06 | 1.02E+06 | 5.87E+05 | 3 | Plasma Membrane |
| 107 | O35372 | CNIH1 | 5.93E+06 | 1.54E+06 | 8.90E+05 | 3 | Plasma Membrane |
| 108 | A2ADY9 | DDI2 | 5.93E+06 | 2.85E+06 | 1.16E+06 | 6 | Plasma Membrane |
| 109 | Q9CQN6 | TMEM14C | 5.89E+06 | 3.16E+06 | 1.29E+06 | 6 | Plasma Membrane |
| 110 | P11438 | LAMP1 | 5.84E+06 | 3.84E+06 | 1.28E+06 | 9 | Plasma Membrane |
| 111 | Q9JL99 | CLEC1B | 5.83E+06 | 5.43E+06 | 3.13E+06 | 3 | Plasma Membrane |
| 112 | Q9QZW0 | ATP11C | 5.83E+06 | 3.56E+06 | 8.40E+05 | 18 | Plasma Membrane |
| 113 | Q9Z0T9 | ITGB6 | 5.59E+06 | 4.62E+06 | 2.67E+06 | 3 | Plasma Membrane |
| 114 | P15379 | CD44 | 5.49E+06 | 6.07E+06 | 1.75E+06 | 12 | Plasma Membrane |
| 115 | Q9R1S7 | ABCC6 | 5.48E+06 | 1.86E+06 | 1.07E+06 | 3 | Plasma Membrane |
| 116 | P21279 | GNAQ | 5.40E+06 | 3.59E+06 | 2.07E+06 | 3 | Plasma Membrane |
| 117 | P36423 | TBXAS1 | 4.94E+06 | 4.42E+06 | 8.50E+05 | 27 | Plasma Membrane |
| 118 | Q9QXT0 | CNPY2 | 4.91E+06 | 1.90E+06 | 6.33E+05 | 9 | Plasma Membrane |
| 119 | Q91Y57 | SIGLEC9 | 4.88E+06 | 7.72E+04 | 4.46E+04 | 3 | Plasma Membrane |
| 120 | P01887 | B2M | 4.75E+06 | 1.36E+06 | 7.84E+05 | 3 | Plasma Membrane |
| 121 | P40237 | CD82 | 4.74E+06 | 9.52E+05 | 3.89E+05 | 6 | Plasma Membrane |
| 122 | P10852 | SLC3A2 | 4.73E+06 | 2.28E+06 | 4.16E+05 | 30 | Plasma Membrane |
| 123 | Q61189 | CLNS1A | 4.73E+06 | 3.30E+06 | 1.91E+06 | 3 | Plasma Membrane |
| 124 | P18052 | PTPRA | 4.57E+06 | 3.25E+06 | 9.37E+05 | 12 | Plasma Membrane |
| 125 | Q91WC9 | DAGLB | 4.56E+06 | 2.10E+06 | 4.95E+05 | 18 | Plasma Membrane |
| 126 | P11688 | ITGA5 | 4.38E+06 | 2.57E+06 | 7.42E+05 | 12 | Plasma Membrane |
| 127 | Q8BHL3 | TBC1D10B | 4.33E+06 | 9.56E+05 | 5.52E+05 | 3 | Plasma Membrane |
| 128 | O55022 | PGRMC1 | 4.30E+06 | 2.89E+06 | 1.67E+06 | 3 | Plasma Membrane |
| 129 | Q64324 | STXBP2 | 4.27E+06 | 4.17E+06 | 6.44E+05 | 42 | Plasma Membrane |
| 130 | Q9Z0F8 | ADAM17 | 4.26E+06 | 2.69E+06 | 1.10E+06 | 6 | Plasma Membrane |
| 131 | P49282 | SLC11A2 | 4.24E+06 | 2.99E+06 | 8.62E+05 | 12 | Plasma Membrane |

| AQP1 KO | | | | | | | |
|---|---|---|---|---|---|---|---|
| Rank | ID | Symbol | Mean Intensity | St. Dev | Std. Err | n | Primary Location |
| 1 | P14220 | Gypa | 2.58E+09 | 2.21E+09 | 9.00E+08 | 6 | Plasma Membrane |
| 2 | P04919 | SLC4A1 | 2.19E+09 | 4.67E+09 | 3.57E+08 | 171 | Plasma Membrane |
| 3 | Q78HU7 | GYPC | 6.08E+08 | 5.45E+08 | 1.82E+08 | 9 | Plasma Membrane |
| 4 | Q02357 | ANK1 | 5.04E+08 | 6.95E+08 | 3.09E+07 | 507 | Plasma Membrane |
| 5 | P15508 | SPTB | 4.43E+08 | 5.66E+08 | 2.03E+07 | 780 | Plasma Membrane |
| 6 | Q9QUT0 | RHAG | 3.83E+08 | 4.81E+08 | 1.39E+08 | 12 | Plasma Membrane |
| 7 | P49222 | EPB42 | 3.54E+08 | 4.90E+08 | 3.54E+07 | 192 | Plasma Membrane |
| 8 | P70290 | MPP1 | 3.47E+08 | 4.00E+08 | 3.56E+07 | 126 | Plasma Membrane |
| 9 | P48193 | EPB41 | 3.40E+08 | 5.31E+08 | 3.90E+07 | 186 | Plasma Membrane |
| 10 | P18572 | BSG | 2.46E+08 | 3.09E+08 | 4.76E+07 | 42 | Plasma Membrane |
| 11 | P53986 | SLC16A1 | 2.21E+08 | 2.08E+08 | 4.00E+07 | 27 | Plasma Membrane |
| 12 | Q61735 | CD47 | 2.00E+08 | 1.82E+08 | 4.29E+07 | 18 | Plasma Membrane |
| 13 | Q8CF94 | RHCE/RHD | 1.81E+08 | 4.08E+08 | 8.33E+07 | 24 | Plasma Membrane |
| 14 | Q9WV69 | DMTN | 1.65E+08 | 1.72E+08 | 1.98E+07 | 75 | Plasma Membrane |
| 15 | P54116 | STOM | 1.42E+08 | 2.54E+08 | 3.06E+07 | 69 | Plasma Membrane |
| 16 | Q8BSM7 | SLC43A1 | 1.07E+08 | 1.80E+08 | 2.78E+07 | 42 | Plasma Membrane |
| 17 | Q08857 | CD36 | 9.78E+07 | 1.42E+08 | 2.05E+07 | 48 | Plasma Membrane |
| 18 | Q8VHL0 | SLC14A1 | 9.03E+07 | 8.17E+07 | 1.36E+07 | 36 | Plasma Membrane |
| 19 | Q9Z0S4 | Cldn13 | 7.91E+07 | 7.11E+07 | 2.05E+07 | 12 | Plasma Membrane |
| 20 | P58771 | Tpm1 | 6.98E+07 | 1.04E+08 | 2.67E+07 | 15 | Plasma Membrane |
| 21 | P27600 | GNA12 | 5.60E+07 | 1.61E+07 | 9.32E+06 | 3 | Plasma Membrane |
| 22 | Q3U9N9 | SLC16A10 | 5.58E+07 | 5.89E+07 | 1.07E+07 | 30 | Plasma Membrane |
| 23 | Q9EQF2 | KEL | 4.69E+07 | 4.47E+07 | 4.30E+06 | 108 | Plasma Membrane |
| 24 | P63001 | RAC1 | 4.25E+07 | 3.36E+07 | 5.18E+06 | 42 | Plasma Membrane |
| 25 | P56400 | GP1BB | 4.06E+07 | 3.61E+07 | 8.51E+06 | 18 | Plasma Membrane |
| 26 | P55258 | RAB8A | 3.98E+07 | 3.18E+07 | 6.50E+06 | 24 | Plasma Membrane |
| 27 | Q62351 | TFRC | 3.85E+07 | 2.96E+07 | 2.81E+06 | 111 | Plasma Membrane |
| 28 | O55186 | Cd59a | 3.64E+07 | 4.81E+07 | 1.60E+07 | 9 | Plasma Membrane |
| 29 | Q61411 | HRAS | 3.46E+07 | 4.38E+07 | 8.00E+06 | 30 | Plasma Membrane |
| 30 | Q9QUM0 | ITGA2B | 3.29E+07 | 3.88E+07 | 4.31E+06 | 81 | Plasma Membrane |
| 31 | O54890 | ITGB3 | 3.11E+07 | 2.92E+07 | 3.45E+06 | 72 | Plasma Membrane |
| 32 | P08752 | GNAI2 | 2.98E+07 | 2.64E+07 | 2.99E+06 | 78 | Plasma Membrane |
| 33 | Q8BGQ1 | VIPAS39 | 2.81E+07 | 4.02E+06 | 2.32E+06 | 3 | Plasma Membrane |
| 34 | Q9JIM1 | SLC29A1 | 2.74E+07 | 3.69E+07 | 6.43E+06 | 33 | Plasma Membrane |
| 35 | O88186 | GP9 | 2.70E+07 | 3.49E+07 | 1.01E+07 | 12 | Plasma Membrane |
| 36 | Q8K400 | STXBP5 | 2.51E+07 | 7.34E+06 | 4.24E+06 | 3 | Plasma Membrane |
| 37 | Q7TMS5 | ABCG2 | 2.48E+07 | 2.40E+07 | 3.10E+06 | 60 | Plasma Membrane |
| 38 | P62880 | GNB2 | 2.48E+07 | 2.03E+07 | 3.39E+06 | 36 | Plasma Membrane |
| 39 | O08742 | GP5 | 2.46E+07 | 1.72E+07 | 3.14E+06 | 30 | Plasma Membrane |
| 40 | Q8K558 | TREML1 | 2.41E+07 | 1.56E+07 | 3.18E+06 | 24 | Plasma Membrane |
| 41 | P14231 | ATP1B2 | 2.40E+07 | 2.33E+07 | 6.01E+06 | 15 | Plasma Membrane |
| 42 | P62874 | GNB1 | 2.37E+07 | 2.32E+07 | 3.72E+06 | 39 | Plasma Membrane |
| 43 | Q8R3G9 | TSPAN8 | 2.37E+07 | 1.42E+07 | 3.66E+06 | 15 | Plasma Membrane |
| 44 | O35930 | GP1BA | 2.36E+07 | 2.22E+07 | 3.55E+06 | 39 | Plasma Membrane |
| 45 | Q9R0K7 | ATP2B2 | 2.35E+07 | 3.17E+07 | 5.79E+06 | 30 | Plasma Membrane |
| 46 | Q61469 | Ppap2a | 2.26E+07 | 2.06E+07 | 5.31E+06 | 15 | Plasma Membrane |
| 47 | Q8R3S2 | TSPAN33 | 2.25E+07 | 3.51E+07 | 7.65E+06 | 21 | Plasma Membrane |
| 48 | Q9QXY7 | XK | 2.08E+07 | 5.12E+06 | 1.48E+06 | 12 | Plasma Membrane |
| 49 | P08556 | NRAS | 1.96E+07 | 5.27E+05 | 3.04E+05 | 3 | Plasma Membrane |
| 50 | Q99KU0 | VMP1 | 1.94E+07 | 5.13E+06 | 2.96E+06 | 3 | Plasma Membrane |
| 51 | Q9DCT8 | Crip2 | 1.88E+07 | 1.35E+07 | 2.59E+06 | 27 | Plasma Membrane |
| 52 | P32037 | SLC2A3 | 1.86E+07 | 1.36E+07 | 2.96E+06 | 21 | Plasma Membrane |
| 53 | P97300 | Nptn | 1.82E+07 | 2.75E+07 | 5.29E+06 | 27 | Plasma Membrane |
| 54 | Q9WV55 | VAPA | 1.82E+07 | 1.52E+07 | 2.93E+06 | 27 | Plasma Membrane |
| 55 | Q64735 | CR1L | 1.81E+07 | 1.66E+07 | 3.19E+06 | 27 | Plasma Membrane |
| 56 | Q9CQW2 | ARL8B | 1.72E+07 | 1.14E+07 | 1.76E+06 | 42 | Plasma Membrane |
| 57 | Q61672 | SLC29A2 | 1.71E+07 | 5.98E+06 | 3.46E+06 | 3 | Plasma Membrane |
| 58 | O88343 | SLC4A4 | 1.66E+07 | 4.31E+06 | 2.49E+06 | 3 | Plasma Membrane |
| 59 | Q9JHI9 | SLC40A1 | 1.64E+07 | 9.24E+06 | 1.89E+06 | 24 | Plasma Membrane |
| 60 | Q62312 | TGFBR2 | 1.63E+07 | 2.05E+06 | 1.18E+06 | 3 | Plasma Membrane |
| 61 | P60904 | DNAJC5 | 1.62E+07 | 1.76E+07 | 4.55E+06 | 15 | Plasma Membrane |
| 62 | Q9QUI6 | ACKR1 | 1.59E+07 | 1.32E+07 | 3.82E+06 | 12 | Plasma Membrane |
| 63 | O35598 | ADAM10 | 1.58E+07 | 1.41E+07 | 1.60E+06 | 78 | Plasma Membrane |
| 64 | Q3T9M1 | MFSD2B | 1.57E+07 | 1.66E+07 | 6.78E+06 | 6 | Plasma Membrane |
| 65 | P27664 | PDE6A | 1.52E+07 | 3.28E+06 | 1.90E+06 | 3 | Plasma Membrane |
| 66 | P63024 | VAMP3 | 1.50E+07 | 2.81E+07 | 7.26E+06 | 15 | Plasma Membrane |
| 67 | P04925 | PRNP | 1.47E+07 | 2.10E+06 | 1.22E+06 | 3 | Plasma Membrane |
| 68 | O09117 | SYPL1 | 1.41E+07 | 1.17E+07 | 3.90E+06 | 9 | Plasma Membrane |
| 69 | Q9Z2P8 | VAMP5 | 1.39E+07 | 8.41E+06 | 2.80E+06 | 9 | Plasma Membrane |
| 70 | Q9JJ00 | PLSCR1 | 1.39E+07 | 5.27E+06 | 1.76E+06 | 9 | Plasma Membrane |
| 71 | O70404 | VAMP8 | 1.35E+07 | 1.09E+07 | 3.62E+06 | 9 | Plasma Membrane |
| 72 | Q8VDN2 | ATP1A1 | 1.33E+07 | 9.84E+06 | 1.05E+06 | 87 | Plasma Membrane |
| 73 | O09044 | SNAP23 | 1.31E+07 | 1.18E+07 | 1.96E+06 | 36 | Plasma Membrane |
| 74 | P40240 | CD9 | 1.27E+07 | 1.94E+07 | 6.45E+06 | 9 | Plasma Membrane |
| 75 | Q9JJF9 | SPPL2A | 1.19E+07 | 1.57E+06 | 6.39E+05 | 6 | Plasma Membrane |
| 76 | P39054 | DNM2 | 1.14E+07 | 1.02E+07 | 1.22E+06 | 69 | Plasma Membrane |
| 77 | Q80YF9 | ARHGAP33 | 1.12E+07 | 2.52E+06 | 1.45E+06 | 3 | Plasma Membrane |
| 78 | Q91V01 | LPCAT3 | 1.08E+07 | 8.52E+06 | 2.46E+06 | 12 | Plasma Membrane |
| 79 | P09055 | ITGB1 | 1.08E+07 | 7.60E+06 | 1.22E+06 | 39 | Plasma Membrane |
| 80 | Q60634 | FLOT2 | 1.06E+07 | 7.68E+06 | 1.08E+06 | 51 | Plasma Membrane |
| 81 | Q9CQ22 | LAMTOR1 | 1.06E+07 | 7.92E+06 | 2.29E+06 | 12 | Plasma Membrane |
| 82 | O70439 | STX7 | 1.01E+07 | 3.73E+06 | 8.80E+05 | 18 | Plasma Membrane |
| 83 | Q9D659 | C10orf54 | 9.94E+06 | 2.73E+06 | 1.57E+06 | 3 | Plasma Membrane |
| 84 | P21440 | ABCB4 | 9.45E+06 | 5.34E+06 | 6.30E+05 | 72 | Plasma Membrane |
| 85 | P62331 | ARF6 | 9.40E+06 | 5.69E+06 | 1.34E+06 | 18 | Plasma Membrane |
| 86 | O08917 | FLOT1 | 9.24E+06 | 5.03E+06 | 7.27E+05 | 48 | Plasma Membrane |
| 87 | P01887 | B2M | 9.01E+06 | 1.08E+06 | 6.22E+05 | 3 | Plasma Membrane |
| 88 | Q9JL99 | CLEC1B | 8.85E+06 | 7.58E+05 | 4.37E+05 | 3 | Plasma Membrane |
| 89 | P21279 | GNAQ | 8.62E+06 | 3.18E+06 | 1.84E+06 | 3 | Plasma Membrane |
| 90 | P61226 | RAP2B | 8.58E+06 | 3.24E+06 | 7.63E+05 | 18 | Plasma Membrane |
| 91 | P11438 | LAMP1 | 8.40E+06 | 4.62E+06 | 1.54E+06 | 9 | Plasma Membrane |
| 92 | P26040 | EZR | 8.35E+06 | 6.42E+06 | 1.17E+06 | 30 | Plasma Membrane |
| 93 | P10852 | SLC3A2 | 7.86E+06 | 3.64E+06 | 6.64E+05 | 30 | Plasma Membrane |
| 94 | P60766 | Cdc42 | 7.63E+06 | 2.97E+06 | 7.68E+05 | 15 | Plasma Membrane |
| 95 | Q9Z0T9 | ITGB6 | 7.58E+06 | 2.03E+06 | 1.17E+06 | 3 | Plasma Membrane |
| 96 | Q9DAS9 | GNG12 | 7.29E+06 | 9.62E+05 | 5.55E+05 | 3 | Plasma Membrane |
| 97 | Q9JMH7 | NEU3 | 7.00E+06 | 5.66E+06 | 1.89E+06 | 9 | Plasma Membrane |
| 98 | Q68FD5 | CLTC | 6.92E+06 | 4.79E+06 | 4.08E+05 | 138 | Plasma Membrane |
| 99 | Q9Z127 | SLC7A5 | 6.87E+06 | 4.66E+06 | 1.35E+06 | 12 | Plasma Membrane |
| 100 | Q09143 | SLC7A1 | 6.70E+06 | 5.32E+05 | 3.07E+05 | 3 | Plasma Membrane |
| 101 | P36423 | TBXAS1 | 6.58E+06 | 3.87E+06 | 7.44E+05 | 27 | Plasma Membrane |
| 102 | Q61475 | CD55 | 6.48E+06 | 3.47E+06 | 8.18E+05 | 18 | Plasma Membrane |
| 103 | Q61165 | SLC9A1 | 6.43E+06 | 4.10E+06 | 8.36E+05 | 24 | Plasma Membrane |
| 104 | Q3TZZ7 | ESYT2 | 6.41E+06 | 3.00E+06 | 3.87E+05 | 60 | Plasma Membrane |
| 105 | P58735 | SLC26A1 | 6.38E+06 | 2.85E+06 | 1.16E+06 | 6 | Plasma Membrane |
| 106 | P15379 | CD44 | 6.34E+06 | 6.10E+06 | 1.76E+06 | 12 | Plasma Membrane |
| 107 | Q8BHL3 | TBC1D10B | 6.34E+06 | 3.03E+06 | 1.75E+06 | 3 | Plasma Membrane |
| 108 | Q9QZW0 | ATP11C | 6.32E+06 | 3.59E+06 | 8.46E+05 | 18 | Plasma Membrane |
| 109 | Q9JIZ9 | PLSCR3 | 6.30E+06 | 3.46E+06 | 9.98E+05 | 12 | Plasma Membrane |
| 110 | O35379 | ABCC1 | 6.29E+06 | 4.18E+06 | 5.68E+05 | 54 | Plasma Membrane |
| 111 | Q64324 | STXBP2 | 6.12E+06 | 4.94E+06 | 7.63E+05 | 42 | Plasma Membrane |
| 112 | O55022 | PGRMC1 | 6.00E+06 | 7.88E+05 | 4.55E+05 | 3 | Plasma Membrane |
| 113 | Q9WTR1 | TRPV2 | 5.96E+06 | 4.32E+06 | 7.20E+05 | 36 | Plasma Membrane |
| 114 | Q01102 | SELP | 5.88E+06 | 6.36E+06 | 1.50E+06 | 18 | Plasma Membrane |
| 115 | Q8QZY6 | TSPAN14 | 5.84E+06 | 2.50E+05 | 1.44E+05 | 3 | Plasma Membrane |
| 116 | P57716 | NCSTN | 5.83E+06 | 3.57E+06 | 6.87E+05 | 27 | Plasma Membrane |
| 117 | Q9CQN6 | TMEM14C | 5.77E+06 | 5.48E+06 | 2.24E+06 | 6 | Plasma Membrane |
| 118 | Q9QUM4 | SLAMF1 | 5.62E+06 | 2.02E+06 | 6.73E+05 | 9 | Plasma Membrane |
| 119 | Q9JM62 | REEP6 | 5.45E+06 | 4.48E+06 | 1.49E+06 | 9 | Plasma Membrane |
| 120 | P11688 | ITGA5 | 5.34E+06 | 3.01E+06 | 8.69E+05 | 12 | Plasma Membrane |
| 121 | P49282 | SLC11A2 | 5.30E+06 | 2.52E+06 | 7.27E+05 | 12 | Plasma Membrane |
| 122 | Q02013 | AQP1 | 5.15E+06 | 7.15E+06 | 1.85E+06 | 15 | Plasma Membrane |
| 123 | P16546 | SPTAN1 | 5.15E+06 | 2.91E+06 | 1.68E+06 | 3 | Plasma Membrane |
| 124 | P70460 | VASP | 5.15E+06 | 1.92E+06 | 1.11E+06 | 3 | Plasma Membrane |
| 125 | Q6ZQI3 | MLEC | 5.09E+06 | 1.73E+06 | 4.47E+05 | 15 | Plasma Membrane |
| 126 | Q60682 | Klra4 (includes others) | 5.05E+06 | 1.13E+06 | 6.54E+05 | 3 | Plasma Membrane |
| 127 | Q99JY9 | ACTR3 | 5.02E+06 | 4.60E+06 | 1.19E+06 | 15 | Plasma Membrane |
| 128 | P17047 | LAMP2 | 5.01E+06 | 2.47E+06 | 1.01E+06 | 6 | Plasma Membrane |
| 129 | Q60857 | SLC6A4 | 4.99E+06 | 5.57E+05 | 3.21E+05 | 3 | Plasma Membrane |
| 130 | Q9QXT0 | CNPY2 | 4.98E+06 | 1.63E+06 | 5.44E+05 | 9 | Plasma Membrane |
| 131 | P26041 | MSN | 4.92E+06 | 2.70E+06 | 4.50E+05 | 36 | Plasma Membrane |

| RhAG KO | | | | | | | |
|---|---|---|---|---|---|---|---|
| Rank | ID | Symbol | Mean Intensity | St. Dev | Std. Err | n | Primary Location |
| 1 | P04919 | SLC4A1 | 2.70E+09 | 4.72E+09 | 3.61E+08 | 171 | Plasma Membrane |
| 2 | P14220 | Gypa | 2.58E+09 | 2.27E+09 | 9.26E+08 | 6 | Plasma Membrane |
| 3 | Q02013 | AQP1 | 1.32E+09 | 1.05E+09 | 2.71E+08 | 15 | Plasma Membrane |
| 4 | Q78HU7 | GYPC | 6.54E+08 | 5.40E+08 | 1.80E+08 | 9 | Plasma Membrane |
| 5 | Q02357 | ANK1 | 5.05E+08 | 6.64E+08 | 2.95E+07 | 507 | Plasma Membrane |
| 6 | P15508 | SPTB | 4.85E+08 | 5.82E+08 | 2.09E+07 | 780 | Plasma Membrane |
| 7 | P48193 | EPB41 | 4.41E+08 | 5.94E+08 | 4.36E+07 | 186 | Plasma Membrane |
| 8 | P70290 | MPP1 | 3.87E+08 | 4.39E+08 | 3.91E+07 | 126 | Plasma Membrane |
| 9 | P49222 | EPB42 | 3.63E+08 | 4.91E+08 | 3.54E+07 | 192 | Plasma Membrane |
| 10 | P18572 | BSG | 2.77E+08 | 3.17E+08 | 4.90E+07 | 42 | Plasma Membrane |
| 11 | P53986 | SLC16A1 | 2.36E+08 | 2.05E+08 | 3.94E+07 | 27 | Plasma Membrane |
| 12 | Q61735 | CD47 | 2.15E+08 | 1.79E+08 | 4.22E+07 | 18 | Plasma Membrane |
| 13 | P54116 | STOM | 1.91E+08 | 2.76E+08 | 3.32E+07 | 69 | Plasma Membrane |
| 14 | Q9WV69 | DMTN | 1.89E+08 | 1.86E+08 | 2.15E+07 | 75 | Plasma Membrane |
| 15 | Q8BSM7 | SLC43A1 | 1.36E+08 | 1.86E+08 | 2.87E+07 | 42 | Plasma Membrane |
| 16 | Q8VHL0 | SLC14A1 | 1.34E+08 | 1.14E+08 | 1.90E+07 | 36 | Plasma Membrane |
| 17 | P58771 | Tpm1 | 1.05E+08 | 1.33E+08 | 3.44E+07 | 15 | Plasma Membrane |
| 18 | Q08857 | CD36 | 9.57E+07 | 1.23E+08 | 1.77E+07 | 48 | Plasma Membrane |
| 19 | Q9Z0S4 | Cldn13 | 9.30E+07 | 7.69E+07 | 2.22E+07 | 12 | Plasma Membrane |
| 20 | Q3U9N9 | SLC16A10 | 6.25E+07 | 5.51E+07 | 1.01E+07 | 30 | Plasma Membrane |
| 21 | Q9EQF2 | KEL | 5.67E+07 | 4.59E+07 | 4.42E+06 | 108 | Plasma Membrane |
| 22 | P55258 | RAB8A | 4.84E+07 | 3.78E+07 | 7.72E+06 | 24 | Plasma Membrane |
| 23 | Q8R3S2 | TSPAN33 | 4.80E+07 | 6.08E+07 | 1.33E+07 | 21 | Plasma Membrane |
| 24 | P27600 | GNA12 | 4.28E+07 | 8.02E+06 | 4.63E+06 | 3 | Plasma Membrane |
| 25 | Q61469 | Ppap2a | 4.22E+07 | 2.08E+07 | 5.38E+06 | 15 | Plasma Membrane |
| 26 | P63001 | RAC1 | 4.22E+07 | 3.18E+07 | 4.90E+06 | 42 | Plasma Membrane |
| 27 | O55186 | Cd59a | 3.95E+07 | 4.84E+07 | 1.61E+07 | 9 | Plasma Membrane |
| 28 | Q9JIM1 | SLC29A1 | 3.80E+07 | 4.28E+07 | 7.45E+06 | 33 | Plasma Membrane |
| 29 | Q7TMS5 | ABCG2 | 3.71E+07 | 3.39E+07 | 4.38E+06 | 60 | Plasma Membrane |
| 30 | Q8K400 | STXBP5 | 3.26E+07 | 4.30E+06 | 2.48E+06 | 3 | Plasma Membrane |
| 31 | Q61672 | SLC29A2 | 2.88E+07 | 1.10E+06 | 6.33E+05 | 3 | Plasma Membrane |
| 32 | Q61411 | HRAS | 2.76E+07 | 3.46E+07 | 6.31E+06 | 30 | Plasma Membrane |
| 33 | O35598 | ADAM10 | 2.71E+07 | 2.39E+07 | 2.70E+06 | 78 | Plasma Membrane |
| 34 | Q8R3G9 | TSPAN8 | 2.71E+07 | 1.53E+07 | 3.94E+06 | 15 | Plasma Membrane |
| 35 | Q9DCT8 | Crip2 | 2.69E+07 | 1.37E+07 | 2.64E+06 | 27 | Plasma Membrane |
| 36 | Q9R0K7 | ATP2B2 | 2.63E+07 | 3.30E+07 | 6.03E+06 | 30 | Plasma Membrane |
| 37 | Q9QXY7 | XK | 2.59E+07 | 6.05E+06 | 1.75E+06 | 12 | Plasma Membrane |
| 38 | P08752 | GNAI2 | 2.47E+07 | 2.51E+07 | 2.84E+06 | 78 | Plasma Membrane |
| 39 | P63024 | VAMP3 | 2.47E+07 | 3.56E+07 | 9.19E+06 | 15 | Plasma Membrane |
| 40 | Q62351 | TFRC | 2.39E+07 | 3.13E+07 | 2.97E+06 | 111 | Plasma Membrane |
| 41 | P56400 | GP1BB | 2.37E+07 | 3.21E+07 | 7.56E+06 | 18 | Plasma Membrane |
| 42 | P14231 | ATP1B2 | 2.26E+07 | 2.09E+07 | 5.40E+06 | 15 | Plasma Membrane |
| 43 | Q62312 | TGFBR2 | 2.25E+07 | 3.78E+06 | 2.18E+06 | 3 | Plasma Membrane |
| 44 | Q8BGQ1 | VIPAS39 | 2.13E+07 | 1.03E+06 | 5.94E+05 | 3 | Plasma Membrane |
| 45 | P97300 | Nptn | 2.08E+07 | 2.86E+07 | 5.50E+06 | 27 | Plasma Membrane |
| 46 | Q9QUI6 | ACKR1 | 2.06E+07 | 1.51E+07 | 4.36E+06 | 12 | Plasma Membrane |
| 47 | O88186 | GP9 | 2.04E+07 | 3.12E+07 | 9.00E+06 | 12 | Plasma Membrane |
| 48 | Q64735 | CR1L | 2.01E+07 | 1.78E+07 | 3.43E+06 | 27 | Plasma Membrane |
| 49 | Q9QUM0 | ITGA2B | 1.99E+07 | 3.67E+07 | 4.07E+06 | 81 | Plasma Membrane |
| 50 | P62874 | GNB1 | 1.88E+07 | 2.07E+07 | 3.31E+06 | 39 | Plasma Membrane |
| 51 | Q9WV55 | VAPA | 1.86E+07 | 1.54E+07 | 2.97E+06 | 27 | Plasma Membrane |
| 52 | P62880 | GNB2 | 1.84E+07 | 1.85E+07 | 3.09E+06 | 36 | Plasma Membrane |
| 53 | Q9CQW2 | ARL8B | 1.80E+07 | 1.03E+07 | 1.59E+06 | 42 | Plasma Membrane |
| 54 | Q9JJF9 | SPPL2A | 1.79E+07 | 1.63E+06 | 6.65E+05 | 6 | Plasma Membrane |
| 55 | Q99KU0 | VMP1 | 1.76E+07 | 3.35E+06 | 1.93E+06 | 3 | Plasma Membrane |
| 56 | P60904 | DNAJC5 | 1.76E+07 | 1.65E+07 | 4.26E+06 | 15 | Plasma Membrane |
| 57 | O54890 | ITGB3 | 1.74E+07 | 2.44E+07 | 2.88E+06 | 72 | Plasma Membrane |
| 58 | Q8K558 | TREML1 | 1.72E+07 | 1.20E+07 | 2.44E+06 | 24 | Plasma Membrane |
| 59 | Q9JHI9 | SLC40A1 | 1.63E+07 | 8.70E+06 | 1.78E+06 | 24 | Plasma Membrane |
| 60 | Q80YF9 | ARHGAP33 | 1.60E+07 | 1.98E+06 | 1.15E+06 | 3 | Plasma Membrane |
| 61 | O09044 | SNAP23 | 1.58E+07 | 1.37E+07 | 2.29E+06 | 36 | Plasma Membrane |
| 62 | O08742 | GP5 | 1.53E+07 | 1.64E+07 | 3.00E+06 | 30 | Plasma Membrane |
| 63 | Q9Z2P8 | VAMP5 | 1.46E+07 | 5.93E+06 | 1.98E+06 | 9 | Plasma Membrane |
| 64 | Q3T9M1 | MFSD2B | 1.45E+07 | 1.58E+07 | 6.44E+06 | 6 | Plasma Membrane |
| 65 | O88343 | SLC4A4 | 1.44E+07 | 3.31E+06 | 1.91E+06 | 3 | Plasma Membrane |
| 66 | P27664 | PDE6A | 1.43E+07 | 2.23E+06 | 1.29E+06 | 3 | Plasma Membrane |
| 67 | O09117 | SYPL1 | 1.42E+07 | 1.05E+07 | 3.49E+06 | 9 | Plasma Membrane |
| 68 | Q9JJ00 | PLSCR1 | 1.41E+07 | 5.62E+06 | 1.87E+06 | 9 | Plasma Membrane |
| 69 | O35930 | GP1BA | 1.35E+07 | 1.89E+07 | 3.02E+06 | 39 | Plasma Membrane |
| 70 | Q8VDN2 | ATP1A1 | 1.31E+07 | 1.03E+07 | 1.10E+06 | 87 | Plasma Membrane |
| 71 | P39054 | DNM2 | 1.28E+07 | 1.26E+07 | 1.52E+06 | 69 | Plasma Membrane |
| 72 | P32037 | SLC2A3 | 1.18E+07 | 1.12E+07 | 2.45E+06 | 21 | Plasma Membrane |
| 73 | P08556 | NRAS | 1.17E+07 | 7.69E+05 | 4.44E+05 | 3 | Plasma Membrane |
| 74 | Q91V01 | LPCAT3 | 1.16E+07 | 7.69E+06 | 2.22E+06 | 12 | Plasma Membrane |
| 75 | O70404 | VAMP8 | 1.13E+07 | 1.06E+07 | 3.54E+06 | 9 | Plasma Membrane |
| 76 | Q60634 | FLOT2 | 1.12E+07 | 7.73E+06 | 1.08E+06 | 51 | Plasma Membrane |
| 77 | P40240 | CD9 | 1.11E+07 | 1.80E+07 | 6.00E+06 | 9 | Plasma Membrane |
| 78 | O70439 | STX7 | 1.05E+07 | 3.94E+06 | 9.28E+05 | 18 | Plasma Membrane |
| 79 | Q9JMH7 | NEU3 | 1.01E+07 | 7.44E+06 | 2.48E+06 | 9 | Plasma Membrane |
| 80 | P21440 | ABCB4 | 9.67E+06 | 5.78E+06 | 6.81E+05 | 72 | Plasma Membrane |
| 81 | P16546 | SPTAN1 | 9.57E+06 | 1.99E+06 | 1.15E+06 | 3 | Plasma Membrane |
| 82 | O08917 | FLOT1 | 9.52E+06 | 5.25E+06 | 7.58E+05 | 48 | Plasma Membrane |
| 83 | Q9D659 | C10orf54 | 9.34E+06 | 1.22E+06 | 7.05E+05 | 3 | Plasma Membrane |
| 84 | Q9CQ22 | LAMTOR1 | 8.70E+06 | 8.70E+06 | 2.51E+06 | 12 | Plasma Membrane |
| 85 | P26040 | EZR | 8.46E+06 | 7.28E+06 | 1.33E+06 | 30 | Plasma Membrane |
| 86 | P61226 | RAP2B | 8.17E+06 | 2.92E+06 | 6.88E+05 | 18 | Plasma Membrane |
| 87 | Q9WTR1 | TRPV2 | 8.15E+06 | 4.75E+06 | 7.91E+05 | 36 | Plasma Membrane |
| 88 | P04925 | PRNP | 7.95E+06 | 1.47E+06 | 8.48E+05 | 3 | Plasma Membrane |
| 89 | Q9CQN6 | TMEM14C | 7.61E+06 | 4.82E+06 | 1.97E+06 | 6 | Plasma Membrane |
| 90 | P40237 | CD82 | 7.53E+06 | 2.01E+06 | 8.21E+05 | 6 | Plasma Membrane |
| 91 | Q60682 | Klra4 (includes others) | 7.32E+06 | 1.35E+06 | 7.79E+05 | 3 | Plasma Membrane |
| 92 | P01887 | B2M | 7.29E+06 | 2.52E+06 | 1.46E+06 | 3 | Plasma Membrane |
| 93 | P09055 | ITGB1 | 7.28E+06 | 7.52E+06 | 1.20E+06 | 39 | Plasma Membrane |
| 94 | P58735 | SLC26A1 | 6.87E+06 | 3.82E+06 | 1.56E+06 | 6 | Plasma Membrane |
| 95 | P62331 | ARF6 | 6.82E+06 | 4.41E+06 | 1.04E+06 | 18 | Plasma Membrane |
| 96 | Q61475 | CD55 | 6.81E+06 | 3.52E+06 | 8.29E+05 | 18 | Plasma Membrane |
| 97 | Q3TZZ7 | ESYT2 | 6.69E+06 | 3.10E+06 | 4.00E+05 | 60 | Plasma Membrane |
| 98 | A2ADY9 | DDI2 | 6.65E+06 | 1.75E+06 | 7.13E+05 | 6 | Plasma Membrane |
| 99 | Q61165 | SLC9A1 | 6.49E+06 | 3.81E+06 | 7.78E+05 | 24 | Plasma Membrane |
| 100 | P60766 | Cdc42 | 6.41E+06 | 2.64E+06 | 6.82E+05 | 15 | Plasma Membrane |
| 101 | Q9DAS9 | GNG12 | 6.39E+06 | 8.07E+05 | 4.66E+05 | 3 | Plasma Membrane |
| 102 | P57716 | NCSTN | 6.37E+06 | 3.48E+06 | 6.69E+05 | 27 | Plasma Membrane |
| 103 | Q9QZW0 | ATP11C | 6.36E+06 | 3.00E+06 | 7.06E+05 | 18 | Plasma Membrane |
| 104 | Q9JM62 | REEP6 | 6.34E+06 | 4.84E+06 | 1.61E+06 | 9 | Plasma Membrane |
| 105 | O35379 | ABCC1 | 6.33E+06 | 4.10E+06 | 5.58E+05 | 54 | Plasma Membrane |
| 106 | P51655 | GPC4 | 6.30E+06 | 1.99E+06 | 6.65E+05 | 9 | Plasma Membrane |
| 107 | Q9JIZ9 | PLSCR3 | 6.14E+06 | 3.24E+06 | 9.36E+05 | 12 | Plasma Membrane |
| 108 | P11438 | LAMP1 | 5.95E+06 | 4.23E+06 | 1.41E+06 | 9 | Plasma Membrane |
| 109 | P36423 | TBXAS1 | 5.71E+06 | 4.08E+06 | 7.85E+05 | 27 | Plasma Membrane |
| 110 | Q9Z0T9 | ITGB6 | 5.55E+06 | 2.39E+06 | 1.38E+06 | 3 | Plasma Membrane |
| 111 | P21279 | GNAQ | 5.51E+06 | 3.51E+06 | 2.03E+06 | 3 | Plasma Membrane |
| 112 | P15379 | CD44 | 5.47E+06 | 6.79E+06 | 1.96E+06 | 12 | Plasma Membrane |
| 113 | P28571 | SLC6A9 | 5.43E+06 | 4.19E+06 | 1.71E+06 | 6 | Plasma Membrane |
| 114 | Q9JJK2 | LANCL2 | 5.34E+06 | 2.38E+06 | 3.96E+05 | 36 | Plasma Membrane |
| 115 | Q61189 | CLNS1A | 5.32E+06 | 1.52E+06 | 8.79E+05 | 3 | Plasma Membrane |
| 116 | P10852 | SLC3A2 | 5.30E+06 | 2.99E+06 | 5.45E+05 | 30 | Plasma Membrane |
| 117 | O35372 | CNIH1 | 5.21E+06 | 9.52E+05 | 5.50E+05 | 3 | Plasma Membrane |
| 118 | Q68FD5 | CLTC | 5.16E+06 | 3.46E+06 | 2.94E+05 | 138 | Plasma Membrane |
| 119 | Q91Y57 | SIGLEC9 | 5.15E+06 | 5.68E+05 | 3.28E+05 | 3 | Plasma Membrane |
| 120 | Q9Z127 | SLC7A5 | 4.88E+06 | 3.53E+06 | 1.02E+06 | 12 | Plasma Membrane |
| 121 | P70658 | CXCR4 | 4.77E+06 | 1.00E+06 | 5.80E+05 | 3 | Plasma Membrane |
| 122 | P26041 | MSN | 4.77E+06 | 2.93E+06 | 4.89E+05 | 36 | Plasma Membrane |
| 123 | Q9R1S7 | ABCC6 | 4.76E+06 | 1.85E+06 | 1.07E+06 | 3 | Plasma Membrane |
| 124 | Q9QXT0 | CNPY2 | 4.75E+06 | 1.50E+06 | 5.00E+05 | 9 | Plasma Membrane |
| 125 | Q9JL99 | CLEC1B | 4.61E+06 | 2.09E+06 | 1.20E+06 | 3 | Plasma Membrane |
| 126 | Q01102 | SELP | 4.61E+06 | 4.51E+06 | 1.06E+06 | 18 | Plasma Membrane |
| 127 | P49282 | SLC11A2 | 4.58E+06 | 3.13E+06 | 9.04E+05 | 12 | Plasma Membrane |
| 128 | Q9D771 | TMEM206 | 4.45E+06 | 3.30E+06 | 1.10E+06 | 9 | Plasma Membrane |
| 129 | Q60738 | SLC30A1 | 4.41E+06 | 1.24E+06 | 3.57E+05 | 12 | Plasma Membrane |
| 130 | Q99JY9 | ACTR3 | 4.38E+06 | 4.48E+06 | 1.16E+06 | 15 | Plasma Membrane |
| 131 | Q64324 | STXBP2 | 4.36E+06 | 4.34E+06 | 6.70E+05 | 42 | Plasma Membrane |

| dKO | | | | | | | |
|---|---|---|---|---|---|---|---|
| Rank | ID | Symbol | Mean Intensity | St. Dev | Std. Err | n | Primary Location |
| 1 | P04919 | SLC4A1 | 2489840952 | 4889925980 | 373941986 | 171 | Plasma Membrane |
| 2 | P14220 | Gypa | 2380555000 | 2234668239 | 912299488 | 6 | Plasma Membrane |
| 3 | Q78HU7 | GYPC | 696406667 | 671651046 | 223883682 | 9 | Plasma Membrane |
| 4 | Q02357 | ANK1 | 517749535 | 720002755 | 31976445 | 507 | Plasma Membrane |
| 5 | P15508 | SPTB | 489650205 | 633686995 | 22689634 | 780 | Plasma Membrane |
| 6 | P70290 | MPP1 | 405535812 | 471705382 | 42022855 | 126 | Plasma Membrane |
| 7 | P48193 | EPB41 | 402340359 | 620132318 | 45470308 | 186 | Plasma Membrane |
| 8 | P49222 | EPB42 | 371584910 | 518741628 | 37436952 | 192 | Plasma Membrane |
| 9 | P18572 | BSG | 270724908 | 324790933 | 50116329 | 42 | Plasma Membrane |
| 10 | Q61735 | CD47 | 217834472 | 194176240 | 45767779 | 18 | Plasma Membrane |
| 11 | P53986 | SLC16A1 | 208610719 | 218809031 | 42109818 | 27 | Plasma Membrane |
| 12 | Q9WV69 | DMTN | 193817110 | 197809608 | 22841086 | 75 | Plasma Membrane |
| 13 | P54116 | STOM | 162786506 | 276465551 | 33282541 | 69 | Plasma Membrane |
| 14 | Q8BSM7 | SLC43A1 | 134827322 | 199706570 | 30815393 | 42 | Plasma Membrane |
| 15 | Q8VHL0 | SLC14A1 | 130662500 | 121000517 | 20166753 | 36 | Plasma Membrane |
| 16 | Q08857 | CD36 | 95572732 | 129154917 | 18641907 | 48 | Plasma Membrane |
| 17 | Q9Z0S4 | Cldn13 | 91662500 | 68422249 | 19751802 | 12 | Plasma Membrane |
| 18 | P58771 | Tpm1 | 90508369 | 135545858 | 34997790 | 15 | Plasma Membrane |
| 19 | Q8R3S2 | TSPAN33 | 51227195 | 74585035 | 16275789 | 21 | Plasma Membrane |
| 20 | Q3U9N9 | SLC16A10 | 51209285 | 56149877 | 10251518 | 30 | Plasma Membrane |
| 21 | Q9EQF2 | KEL | 50856646 | 48094652 | 4627910 | 108 | Plasma Membrane |
| 22 | P27600 | GNA12 | 50510000 | 8617592 | 4975369 | 3 | Plasma Membrane |
| 23 | P55258 | RAB8A | 44991474 | 38030831 | 7763011 | 24 | Plasma Membrane |
| 24 | P63001 | RAC1 | 44984551 | 32687783 | 5043834 | 42 | Plasma Membrane |
| 25 | O55186 | Cd59a | 40493347 | 54238319 | 18079440 | 9 | Plasma Membrane |
| 26 | Q7TMS5 | ABCG2 | 39547636 | 36738695 | 4742945 | 60 | Plasma Membrane |
| 27 | Q9JIM1 | SLC29A1 | 39509725 | 47364926 | 8245175 | 33 | Plasma Membrane |
| 28 | Q61469 | Ppap2a | 36382615 | 22230108 | 5739789 | 15 | Plasma Membrane |
| 29 | Q62351 | TFRC | 30963160 | 37350153 | 3545120 | 111 | Plasma Membrane |
| 30 | Q9R0K7 | ATP2B2 | 30262741 | 41699195 | 7613197 | 30 | Plasma Membrane |
| 31 | O35598 | ADAM10 | 29722013 | 26256121 | 2972920 | 78 | Plasma Membrane |
| 32 | P08752 | GNAI2 | 28023825 | 26969910 | 3053741 | 78 | Plasma Membrane |
| 33 | P14231 | ATP1B2 | 26383039 | 24783323 | 6399026 | 15 | Plasma Membrane |
| 34 | Q8R3G9 | TSPAN8 | 25126774 | 17192105 | 4438983 | 15 | Plasma Membrane |
| 35 | Q8K400 | STXBP5 | 24790000 | 6269314 | 3619590 | 3 | Plasma Membrane |
| 36 | P62880 | GNB2 | 23913229 | 20965472 | 3494245 | 36 | Plasma Membrane |
| 37 | P97300 | Nptn | 23275842 | 31498128 | 6061818 | 27 | Plasma Membrane |
| 38 | Q9QXY7 | XK | 22972500 | 4986287 | 1439417 | 12 | Plasma Membrane |
| 39 | Q64735 | CR1L | 21822225 | 21645376 | 4165655 | 27 | Plasma Membrane |
| 40 | Q61411 | HRAS | 21555062 | 29637670 | 5411074 | 30 | Plasma Membrane |
| 41 | P62874 | GNB1 | 21375548 | 22120278 | 3542079 | 39 | Plasma Membrane |
| 42 | Q61672 | SLC29A2 | 21160000 | 5747808 | 3318499 | 3 | Plasma Membrane |
| 43 | Q9QUI6 | ACKR1 | 20823223 | 15329770 | 4425323 | 12 | Plasma Membrane |
| 44 | Q9JHI9 | SLC40A1 | 20064916 | 10301646 | 2102815 | 24 | Plasma Membrane |
| 45 | P63024 | VAMP3 | 19199531 | 30098460 | 7771389 | 15 | Plasma Membrane |
| 46 | Q9DCT8 | Crip2 | 18861635 | 11898552 | 2289877 | 27 | Plasma Membrane |
| 47 | Q9JJF9 | SPPL2A | 18695000 | 2981700 | 1217274 | 6 | Plasma Membrane |
| 48 | P32037 | SLC2A3 | 18645187 | 12839286 | 2801762 | 21 | Plasma Membrane |
| 49 | Q9WV55 | VAPA | 16444775 | 14446520 | 2780234 | 27 | Plasma Membrane |
| 50 | P60904 | DNAJC5 | 16349039 | 17655149 | 4558540 | 15 | Plasma Membrane |
| 51 | Q3T9M1 | MFSD2B | 15931930 | 16885261 | 6893379 | 6 | Plasma Membrane |
| 52 | P39054 | DNM2 | 15687368 | 14222355 | 1712170 | 69 | Plasma Membrane |
| 53 | Q8VDN2 | ATP1A1 | 14963585 | 11590471 | 1242629 | 87 | Plasma Membrane |
| 54 | Q9CQW2 | ARL8B | 14928147 | 11623574 | 1793556 | 42 | Plasma Membrane |
| 55 | P27664 | PDE6A | 14880000 | 1840036 | 1062345 | 3 | Plasma Membrane |
| 56 | Q8BGQ1 | VIPAS39 | 14480000 | 3447917 | 1990656 | 3 | Plasma Membrane |
| 57 | O88343 | SLC4A4 | 13960000 | 2360946 | 1363093 | 3 | Plasma Membrane |
| 58 | Q62312 | TGFBR2 | 13920000 | 3660342 | 2113299 | 3 | Plasma Membrane |
| 59 | Q9Z2P8 | VAMP5 | 13910000 | 6689139 | 2229713 | 9 | Plasma Membrane |
| 60 | O09044 | SNAP23 | 13300155 | 13007024 | 2167837 | 36 | Plasma Membrane |
| 61 | Q99KU0 | VMP1 | 13160000 | 2849462 | 1645138 | 3 | Plasma Membrane |
| 62 | Q9D659 | C10orf54 | 12210000 | 3573378 | 2063091 | 3 | Plasma Membrane |
| 63 | Q9JJ00 | PLSCR1 | 12161862 | 5972842 | 1990947 | 9 | Plasma Membrane |
| 64 | Q60634 | FLOT2 | 11989839 | 8978205 | 1257200 | 51 | Plasma Membrane |
| 65 | Q80YF9 | ARHGAP33 | 11060000 | 3248697 | 1875636 | 3 | Plasma Membrane |
| 66 | O09117 | SYPL1 | 11005172 | 10506584 | 3502195 | 9 | Plasma Membrane |
| 67 | Q9JMH7 | NEU3 | 10481979 | 6560091 | 2186697 | 9 | Plasma Membrane |
| 68 | Q9CQ22 | LAMTOR1 | 9887218 | 8726969 | 2519259 | 12 | Plasma Membrane |
| 69 | O08917 | FLOT1 | 9787070 | 5691168 | 821449 | 48 | Plasma Membrane |
| 70 | O70439 | STX7 | 9276643 | 3861851 | 910247 | 18 | Plasma Membrane |
| 71 | P21440 | ABCB4 | 9252222 | 6459435 | 761252 | 72 | Plasma Membrane |
| 72 | P62331 | ARF6 | 9077525 | 5544627 | 1306881 | 18 | Plasma Membrane |
| 73 | P08556 | NRAS | 8975000 | 821621 | 474363 | 3 | Plasma Membrane |
| 74 | O70404 | VAMP8 | 8693080 | 8838885 | 2946295 | 9 | Plasma Membrane |
| 75 | Q91V01 | LPCAT3 | 8632163 | 7993120 | 2307415 | 12 | Plasma Membrane |
| 76 | P16546 | SPTAN1 | 8600127 | 1178254 | 680265 | 3 | Plasma Membrane |
| 77 | P04925 | PRNP | 8166719 | 641732 | 370504 | 3 | Plasma Membrane |
| 78 | P26040 | EZR | 7668641 | 6701757 | 1223568 | 30 | Plasma Membrane |
| 79 | Q9WTR1 | TRPV2 | 7642568 | 5013540 | 835590 | 36 | Plasma Membrane |
| 80 | P58735 | SLC26A1 | 7608580 | 4089519 | 1669539 | 6 | Plasma Membrane |
| 81 | Q61165 | SLC9A1 | 7427887 | 4000652 | 816630 | 24 | Plasma Membrane |
| 82 | Q3TZZ7 | ESYT2 | 7180683 | 3800883 | 490692 | 60 | Plasma Membrane |
| 83 | Q61475 | CD55 | 7045583 | 3566932 | 840734 | 18 | Plasma Membrane |
| 84 | Q9QZW0 | ATP11C | 6934808 | 3510867 | 827519 | 18 | Plasma Membrane |
| 85 | P10852 | SLC3A2 | 6903800 | 4032672 | 736262 | 30 | Plasma Membrane |
| 86 | P40237 | CD82 | 6872067 | 3762018 | 1535837 | 6 | Plasma Membrane |
| 87 | Q09143 | SLC7A1 | 6628230 | 4892932 | 2824935 | 3 | Plasma Membrane |
| 88 | P57716 | NCSTN | 6585396 | 3630055 | 698604 | 27 | Plasma Membrane |
| 89 | O35379 | ABCC1 | 6526931 | 4860742 | 661463 | 54 | Plasma Membrane |
| 90 | Q9JIZ9 | PLSCR3 | 6445987 | 3640475 | 1050915 | 12 | Plasma Membrane |
| 91 | A2ADY9 | DDI2 | 6354568 | 1315380 | 537002 | 6 | Plasma Membrane |
| 92 | Q60682 | Klra4 (includes others) | 6252631 | 1414808 | 816840 | 3 | Plasma Membrane |
| 93 | Q9DAS9 | GNG12 | 6157657 | 878831 | 507394 | 3 | Plasma Membrane |
| 94 | P01887 | B2M | 6156097 | 1039825 | 600344 | 3 | Plasma Membrane |
| 95 | P15379 | CD44 | 6150086 | 6727612 | 1942094 | 12 | Plasma Membrane |
| 96 | P28571 | SLC6A9 | 6110860 | 5065610 | 2068026 | 6 | Plasma Membrane |
| 97 | P61226 | RAP2B | 6004533 | 2517554 | 593393 | 18 | Plasma Membrane |
| 98 | Q9CQN6 | TMEM14C | 5933015 | 4773711 | 1948859 | 6 | Plasma Membrane |
| 99 | Q68FD5 | CLTC | 5821259 | 3982075 | 338977 | 138 | Plasma Membrane |
| 100 | Q02013 | AQP1 | 5816606 | 6164710 | 1591721 | 15 | Plasma Membrane |
| 101 | P60766 | Cdc42 | 5706375 | 2195592 | 566899 | 15 | Plasma Membrane |
| 102 | P11438 | LAMP1 | 5559584 | 4249973 | 1416658 | 9 | Plasma Membrane |
| 103 | P49282 | SLC11A2 | 5532082 | 3985332 | 1150466 | 12 | Plasma Membrane |
| 104 | Q9JJK2 | LANCL2 | 5451643 | 2755871 | 459312 | 36 | Plasma Membrane |
| 105 | Q9Z127 | SLC7A5 | 5421227 | 4973240 | 1435651 | 12 | Plasma Membrane |
| 106 | Q91Y57 | SIGLEC9 | 5216044 | 554478 | 320128 | 3 | Plasma Membrane |
| 107 | Q9R1S7 | ABCC6 | 5126125 | 2535461 | 1463849 | 3 | Plasma Membrane |
| 108 | P11688 | ITGA5 | 4982300 | 2519993 | 727459 | 12 | Plasma Membrane |
| 109 | Q9JM62 | REEP6 | 4954694 | 4925584 | 1641861 | 9 | Plasma Membrane |
| 110 | Q3TWN3 | CNNM2 | 4933508 | 520913 | 212662 | 6 | Plasma Membrane |
| 111 | P70658 | CXCR4 | 4830342 | 744201 | 429665 | 3 | Plasma Membrane |
| 112 | Q9QXT0 | CNPY2 | 4358293 | 1719443 | 573148 | 9 | Plasma Membrane |
| 113 | P26041 | MSN | 4357809 | 2491286 | 415214 | 36 | Plasma Membrane |
| 114 | Q8K2J7 | RELL1 | 4252731 | 1637895 | 545965 | 9 | Plasma Membrane |
| 115 | Q8BG67 | EFR3A | 4246334 | 822280 | 179436 | 21 | Plasma Membrane |
| 116 | P01899, P01901, P01902 | HLA-A | 4092255 | 4784582 | 920793 | 27 | Plasma Membrane |
| 117 | Q61189 | CLNS1A | 4009863 | 382522 | 220849 | 3 | Plasma Membrane |
| 118 | Q8BHL3 | TBC1D10B | 3971436 | 1574685 | 909145 | 3 | Plasma Membrane |
| 119 | Q9D771 | TMEM206 | 3872685 | 4508697 | 1502899 | 9 | Plasma Membrane |
| 120 | Q64324 | STXBP2 | 3848113 | 3082528 | 475644 | 42 | Plasma Membrane |
| 121 | Q91WC9 | DAGLB | 3821502 | 2406607 | 567243 | 18 | Plasma Membrane |
| 122 | Q9JL56 | GDE1 | 3799052 | 2930519 | 976840 | 9 | Plasma Membrane |
| 123 | O55022 | PGRMC1 | 3768221 | 537668 | 310423 | 3 | Plasma Membrane |
| 124 | O35372 | CNIH1 | 3660032 | 328696 | 189773 | 3 | Plasma Membrane |
| 125 | Q99JY9 | ACTR3 | 3640052 | 2965959 | 765807 | 15 | Plasma Membrane |
| 126 | P70398 | USP9X | 3614026 | 1525092 | 440256 | 12 | Plasma Membrane |
| 127 | P18052 | PTPRA | 3603432 | 2828256 | 816447 | 12 | Plasma Membrane |
| 128 | P61161 | ACTR2 | 3484052 | 1112347 | 370782 | 9 | Plasma Membrane |
| 129 | Q60738 | SLC30A1 | 3424007 | 823903 | 237840 | 12 | Plasma Membrane |
| 130 | Q80VP1 | EPN1 | 3387319 | 1399496 | 404000 | 12 | Plasma Membrane |
| 131 | P51655 | GPC4 | 3177482 | 1112540 | 370847 | 9 | Plasma Membrane |

| | | | | | | | |
|---|---|---|---|---|---|---|---|
| 132 | P26041 | MSN | 4.04E+06 | 2.61E+06 | 4.36E+05 | 36 | Plasma Membrane |
| 133 | Q09143 | SLC7A1 | 4.03E+06 | 4.92E+05 | 2.84E+05 | 3 | Plasma Membrane |
| 134 | Q3TWN3 | CNNM2 | 3.92E+06 | 5.87E+05 | 2.40E+05 | 6 | Plasma Membrane |
| 135 | Q9JL56 | GDE1 | 3.84E+06 | 3.09E+06 | 1.03E+06 | 9 | Plasma Membrane |
| 136 | P70460 | VASP | 3.83E+06 | 1.58E+06 | 9.15E+05 | 3 | Plasma Membrane |
| 137 | Q99JY9 | ACTR3 | 3.75E+06 | 4.49E+06 | 1.16E+06 | 15 | Plasma Membrane |
| 138 | Q01102 | SELP | 3.69E+06 | 4.84E+06 | 1.14E+06 | 18 | Plasma Membrane |
| 139 | Q9Z127 | SLC7A5 | 3.68E+06 | 2.76E+06 | 7.96E+05 | 12 | Plasma Membrane |
| 140 | Q6ZQI3 | MLEC | 3.64E+06 | 2.21E+06 | 5.72E+05 | 15 | Plasma Membrane |
| 141 | Q8QZY6 | TSPAN14 | 3.59E+06 | 3.10E+06 | 1.79E+06 | 3 | Plasma Membrane |
| 142 | Q8VDL4 | ADPGK | 3.31E+06 | 6.38E+05 | 3.68E+05 | 3 | Plasma Membrane |
| 143 | Q9ERM2 | ICAM4 | 3.25E+06 | 7.38E+05 | 3.01E+05 | 6 | Plasma Membrane |
| 144 | Q8BG67 | EFR3A | 3.25E+06 | 6.50E+05 | 1.42E+05 | 21 | Plasma Membrane |
| 145 | P01872 | IGHM | 3.24E+06 | 2.46E+06 | 7.09E+05 | 12 | Plasma Membrane |
| 146 | Q9QUM4 | SLAMF1 | 3.24E+06 | 2.69E+06 | 8.97E+05 | 9 | Plasma Membrane |
| 147 | O88792 | F11R | 3.24E+06 | 4.55E+06 | 1.18E+06 | 15 | Plasma Membrane |
| 148 | Q8K2J7 | RELL1 | 3.23E+06 | 1.57E+06 | 5.24E+05 | 9 | Plasma Membrane |
| 149 | P61161 | ACTR2 | 3.23E+06 | 1.49E+06 | 4.96E+05 | 9 | Plasma Membrane |
| 150 | Q60857 | SLC6A4 | 3.20E+06 | 2.98E+06 | 1.72E+06 | 3 | Plasma Membrane |
| 151 | Q9QY76 | VAPB | 3.16E+06 | 2.07E+06 | 8.43E+05 | 6 | Plasma Membrane |
| 152 | P49769 | PSEN1 | 3.15E+06 | 1.93E+06 | 7.89E+05 | 6 | Plasma Membrane |
| 153 | O89112 | LANCL1 | 3.12E+06 | 1.50E+06 | 8.67E+05 | 3 | Plasma Membrane |
| 154 | P17047 | LAMP2 | 3.11E+06 | 1.75E+06 | 7.16E+05 | 6 | Plasma Membrane |
| 155 | Q920L1 | FADS1 | 3.08E+06 | 3.70E+05 | 2.14E+05 | 3 | Plasma Membrane |
| 156 | P51655 | GPC4 | 3.07E+06 | 9.50E+05 | 3.17E+05 | 9 | Plasma Membrane |
| 157 | Q6R0H7 | GNAS | 3.00E+06 | 3.45E+05 | 1.41E+05 | 6 | Plasma Membrane |
| 158 | P70658 | CXCR4 | 2.95E+06 | 5.16E+05 | 2.98E+05 | 3 | Plasma Membrane |
| 159 | P18181 | CD48 | 2.82E+06 | 4.50E+05 | 2.60E+05 | 3 | Plasma Membrane |
| 160 | P28571 | SLC6A9 | 2.78E+06 | 1.91E+06 | 7.81E+05 | 6 | Plasma Membrane |
| 161 | Q8CHK3 | MBOAT7 | 2.75E+06 | 2.15E+06 | 1.24E+06 | 3 | Plasma Membrane |
| 162 | Q8BIG7 | COMTD1 | 2.73E+06 | 1.71E+06 | 9.87E+05 | 3 | Plasma Membrane |
| 163 | Q6X893 | SLC44A1 | 2.73E+06 | 1.01E+06 | 3.37E+05 | 9 | Plasma Membrane |
| 164 | Q80VP1 | EPN1 | 2.72E+06 | 1.57E+06 | 4.54E+05 | 12 | Plasma Membrane |
| 165 | Q60790 | RASA3 | 2.71E+06 | 2.26E+06 | 1.31E+06 | 3 | Plasma Membrane |
| 166 | Q80TH2 | ERBB2IP | 2.71E+06 | 1.30E+06 | 2.66E+05 | 24 | Plasma Membrane |
| 167 | Q9QWY8 | ASAP1 | 2.70E+06 | 1.29E+06 | 5.27E+05 | 6 | Plasma Membrane |
| 168 | P50516 | ATP6V1A | 2.57E+06 | 1.56E+06 | 4.03E+05 | 15 | Plasma Membrane |
| 169 | P70398 | USP9X | 2.54E+06 | 1.51E+06 | 4.36E+05 | 12 | Plasma Membrane |
| 170 | Q9CRY7 | GDPD1 | 2.49E+06 | 3.65E+05 | 2.11E+05 | 3 | Plasma Membrane |
| 171 | Q62469 | ITGA2 | 2.48E+06 | 2.27E+06 | 3.95E+05 | 33 | Plasma Membrane |
| 172 | Q60738 | SLC30A1 | 2.42E+06 | 7.40E+05 | 2.14E+05 | 12 | Plasma Membrane |
| 173 | P21278 | GNA11 | 2.39E+06 | 1.77E+06 | 1.02E+06 | 3 | Plasma Membrane |
| 174 | Q8BFZ9 | ERLIN2 | 2.34E+06 | 3.01E+05 | 1.74E+05 | 3 | Plasma Membrane |
| 175 | P40124 | CAP1 | 2.23E+06 | 1.01E+06 | 3.36E+05 | 9 | Plasma Membrane |
| 176 | Q61739 | ITGA6 | 2.19E+06 | 2.28E+06 | 4.97E+05 | 21 | Plasma Membrane |
| 177 | Q6P9J9 | ANO6 | 2.06E+06 | 9.22E+05 | 5.32E+05 | 3 | Plasma Membrane |
| 178 | P70452 | STX4 | 2.05E+06 | 1.04E+06 | 3.46E+05 | 9 | Plasma Membrane |
| 179 | Q64455 | PTPRJ | 2.05E+06 | 3.26E+06 | 1.09E+06 | 9 | Plasma Membrane |
| 180 | Q9WVC8 | SLC26A3 | 2.05E+06 | 4.61E+05 | 2.66E+05 | 3 | Plasma Membrane |
| 181 | Q8K4F0 | CD226 | 1.98E+06 | 1.59E+06 | 9.20E+05 | 3 | Plasma Membrane |
| 182 | O89013 | LEPROT | 1.92E+06 | 1.91E+06 | 7.81E+05 | 6 | Plasma Membrane |
| 183 | P70302 | STIM1 | 1.88E+06 | 1.28E+06 | 4.27E+05 | 9 | Plasma Membrane |
| 184 | P97370 | ATP1B3 | 1.79E+06 | 7.13E+05 | 2.06E+05 | 12 | Plasma Membrane |
| 185 | O55222 | ILK | 1.78E+06 | 1.51E+06 | 3.57E+05 | 18 | Plasma Membrane |
| 186 | Q99JB2 | STOML2 | 1.75E+06 | 8.91E+05 | 2.30E+05 | 15 | Plasma Membrane |
| 187 | O70443 | GNAZ | 1.75E+06 | 1.28E+06 | 5.24E+05 | 6 | Plasma Membrane |
| 188 | Q9JHZ2 | ANKH | 1.72E+06 | 5.64E+05 | 3.26E+05 | 3 | Plasma Membrane |
| 189 | O88783 | F5 | 1.70E+06 | 1.94E+06 | 6.47E+05 | 9 | Plasma Membrane |
| 190 | Q3UHH2 | SLC22A23 | 1.67E+06 | 2.88E+05 | 1.66E+05 | 3 | Plasma Membrane |
| 191 | Q9D3G5 | STX11 | 1.55E+06 | 1.18E+06 | 2.78E+05 | 18 | Plasma Membrane |
| 192 | P41438 | SLC19A1 | 1.54E+06 | 3.47E+05 | 2.00E+05 | 3 | Plasma Membrane |
| 193 | P01899, P01901, P01902 | HLA-A | 1.49E+06 | 1.79E+06 | 3.44E+05 | 27 | Plasma Membrane |
| 194 | Q80TE7 | LRRC7 | 1.47E+06 | 5.21E+05 | 3.01E+05 | 3 | Plasma Membrane |
| 195 | P55302 | LRPAP1 | 1.45E+06 | 2.52E+05 | 1.46E+05 | 3 | Plasma Membrane |
| 196 | A2ARI4 | LGR4 | 1.44E+06 | 8.84E+05 | 5.10E+05 | 3 | Plasma Membrane |
| 197 | Q99JW4 | LIMS1 | 1.41E+06 | 1.22E+06 | 7.05E+05 | 3 | Plasma Membrane |
| 198 | P26039 | TLN1 | 1.36E+06 | 1.25E+06 | 2.55E+05 | 24 | Plasma Membrane |
| 199 | Q9EQP2 | EHD4 | 1.29E+06 | 4.17E+05 | 1.70E+05 | 6 | Plasma Membrane |
| 200 | P11835 | ITGB2 | 1.16E+06 | 6.70E+05 | 3.87E+05 | 3 | Plasma Membrane |
| 201 | O35114 | SCARB2 | 1.09E+06 | 4.19E+05 | 2.42E+05 | 3 | Plasma Membrane |
| 202 | Q4VBD2 | TAPT1 | 1.09E+06 | 6.47E+04 | 3.73E+04 | 3 | Plasma Membrane |
| 203 | Q60770 | STXBP3 | 1.01E+06 | 4.05E+05 | 2.34E+05 | 3 | Plasma Membrane |
| 204 | Q8VBZ3 | CLPTM1 | 9.78E+05 | 1.76E+05 | 1.02E+05 | 3 | Plasma Membrane |
| 205 | Q60865 | CAPRIN1 | 6.74E+05 | 2.89E+05 | 1.18E+05 | 6 | Plasma Membrane |
| 206 | Q6P9Q6 | FKBP15 | 6.69E+05 | 3.69E+05 | 1.51E+05 | 6 | Plasma Membrane |
| 207 | Q811D0 | DLG1 | 6.67E+05 | 4.23E+05 | 1.41E+05 | 9 | Plasma Membrane |
| 208 | Q9DBG3 | Ap2b1 | 5.03E+05 | 4.48E+05 | 1.83E+05 | 6 | Plasma Membrane |
| 209 | Q8BRU6 | SLC18A2 | 4.10E+05 | 3.72E+05 | 2.15E+05 | 3 | Plasma Membrane |
| 210 | Q60902 | EPS15L1 | 2.53E+05 | 1.16E+05 | 6.71E+04 | 3 | Plasma Membrane |
| 211 | Q80X71 | TMEM106B | 2.13E+05 | 5.44E+04 | 3.14E+04 | 3 | Plasma Membrane |

| | | | | | | | |
|---|---|---|---|---|---|---|---|
| 132 | P61161 | ACTR2 | 4.82E+06 | 2.12E+06 | 7.06E+05 | 9 | Plasma Membrane |
| 133 | O88792 | F11R | 4.70E+06 | 3.73E+06 | 9.63E+05 | 15 | Plasma Membrane |
| 134 | Q9JJK2 | LANCL2 | 4.64E+06 | 2.58E+06 | 4.30E+05 | 36 | Plasma Membrane |
| 135 | A2ADY9 | DDI2 | 4.58E+06 | 1.21E+06 | 4.94E+05 | 6 | Plasma Membrane |
| 136 | Q80TH2 | ERBB2IP | 4.53E+06 | 1.54E+06 | 3.15E+05 | 24 | Plasma Membrane |
| 137 | Q62469 | ITGA2 | 4.50E+06 | 2.20E+06 | 3.83E+05 | 33 | Plasma Membrane |
| 138 | Q9D771 | TMEM206 | 4.48E+06 | 4.45E+06 | 1.48E+06 | 9 | Plasma Membrane |
| 139 | P18052 | PTPRA | 4.43E+06 | 2.93E+06 | 8.45E+05 | 12 | Plasma Membrane |
| 140 | Q8CHK3 | MBOAT7 | 4.25E+06 | 1.51E+06 | 8.74E+05 | 3 | Plasma Membrane |
| 141 | P51655 | GPC4 | 4.19E+06 | 1.54E+06 | 5.12E+05 | 9 | Plasma Membrane |
| 142 | P28571 | SLC6A9 | 3.88E+06 | 3.42E+06 | 1.40E+06 | 6 | Plasma Membrane |
| 143 | Q3TWN3 | CNNM2 | 3.84E+06 | 3.80E+05 | 1.55E+05 | 6 | Plasma Membrane |
| 144 | Q8BIG7 | COMTD1 | 3.81E+06 | 9.28E+05 | 5.36E+05 | 3 | Plasma Membrane |
| 145 | P70658 | CXCR4 | 3.76E+06 | 1.87E+05 | 1.08E+05 | 3 | Plasma Membrane |
| 146 | Q6R0H7 | GNAS | 3.73E+06 | 4.96E+05 | 2.02E+05 | 6 | Plasma Membrane |
| 147 | Q6P9J9 | ANO6 | 3.70E+06 | 4.36E+05 | 2.52E+05 | 3 | Plasma Membrane |
| 148 | Q8K2J7 | RELL1 | 3.67E+06 | 1.33E+06 | 4.42E+05 | 9 | Plasma Membrane |
| 149 | Q60790 | RASA3 | 3.64E+06 | 9.10E+05 | 5.25E+05 | 3 | Plasma Membrane |
| 150 | Q61739 | ITGA6 | 3.61E+06 | 2.47E+06 | 5.40E+05 | 21 | Plasma Membrane |
| 151 | Q8BG67 | EFR3A | 3.54E+06 | 8.77E+05 | 1.91E+05 | 21 | Plasma Membrane |
| 152 | O35372 | CNIH1 | 3.50E+06 | 5.26E+05 | 3.03E+05 | 3 | Plasma Membrane |
| 153 | Q91WC9 | DAGLB | 3.47E+06 | 2.47E+06 | 5.82E+05 | 18 | Plasma Membrane |
| 154 | Q9QY76 | VAPB | 3.47E+06 | 2.77E+06 | 1.13E+06 | 6 | Plasma Membrane |
| 155 | Q91Y57 | SIGLEC9 | 3.43E+06 | 1.12E+06 | 6.45E+05 | 3 | Plasma Membrane |
| 156 | P21278 | GNA11 | 3.32E+06 | 3.69E+05 | 2.13E+05 | 3 | Plasma Membrane |
| 157 | Q8K4F0 | CD226 | 3.26E+06 | 1.82E+05 | 1.05E+05 | 3 | Plasma Membrane |
| 158 | Q61189 | CLNS1A | 3.24E+06 | 4.89E+05 | 2.82E+05 | 3 | Plasma Membrane |
| 159 | Q9JL56 | GDE1 | 3.17E+06 | 2.76E+06 | 9.19E+05 | 9 | Plasma Membrane |
| 160 | Q64455 | PTPRJ | 3.16E+06 | 4.32E+06 | 1.44E+06 | 9 | Plasma Membrane |
| 161 | P40237 | CD82 | 3.11E+06 | 3.08E+06 | 1.26E+06 | 6 | Plasma Membrane |
| 162 | Q60738 | SLC30A1 | 3.02E+06 | 9.61E+05 | 2.77E+05 | 12 | Plasma Membrane |
| 163 | Q80TE7 | LRRC7 | 3.00E+06 | 2.88E+05 | 1.66E+05 | 3 | Plasma Membrane |
| 164 | Q99JB2 | STOML2 | 2.99E+06 | 1.74E+06 | 4.48E+05 | 15 | Plasma Membrane |
| 165 | Q9R1S7 | ABCC6 | 2.97E+06 | 2.90E+06 | 1.68E+06 | 3 | Plasma Membrane |
| 166 | P70398 | USP9X | 2.84E+06 | 1.43E+06 | 4.13E+05 | 12 | Plasma Membrane |
| 167 | P50516 | ATP6V1A | 2.82E+06 | 1.58E+06 | 4.07E+05 | 15 | Plasma Membrane |
| 168 | O55222 | ILK | 2.81E+06 | 1.86E+06 | 4.39E+05 | 18 | Plasma Membrane |
| 169 | Q8VDL4 | ADPGK | 2.78E+06 | 3.47E+05 | 2.01E+05 | 3 | Plasma Membrane |
| 170 | Q9QWY8 | ASAP1 | 2.74E+06 | 1.11E+06 | 4.55E+05 | 6 | Plasma Membrane |
| 171 | Q9Z0F8 | ADAM17 | 2.72E+06 | 2.35E+06 | 9.58E+05 | 6 | Plasma Membrane |
| 172 | Q920L1 | FADS1 | 2.71E+06 | 5.71E+05 | 3.30E+05 | 3 | Plasma Membrane |
| 173 | P70302 | STIM1 | 2.67E+06 | 1.00E+06 | 3.35E+05 | 9 | Plasma Membrane |
| 174 | Q9D3G5 | STX11 | 2.64E+06 | 1.54E+06 | 3.63E+05 | 18 | Plasma Membrane |
| 175 | P70452 | STX4 | 2.62E+06 | 9.68E+05 | 3.23E+05 | 9 | Plasma Membrane |
| 176 | P01899, P01901, P01902 | HLA-A | 2.62E+06 | 2.28E+06 | 4.38E+05 | 27 | Plasma Membrane |
| 177 | O70443 | GNAZ | 2.52E+06 | 7.16E+05 | 2.92E+05 | 6 | Plasma Membrane |
| 178 | Q9ERM2 | ICAM4 | 2.47E+06 | 1.37E+06 | 5.58E+05 | 6 | Plasma Membrane |
| 179 | Q9CRY7 | GDPD1 | 2.44E+06 | 1.54E+05 | 8.87E+04 | 3 | Plasma Membrane |
| 180 | Q6X893 | SLC44A1 | 2.39E+06 | 1.01E+06 | 3.35E+05 | 9 | Plasma Membrane |
| 181 | Q8BFZ9 | ERLIN2 | 2.35E+06 | 9.15E+05 | 5.28E+05 | 3 | Plasma Membrane |
| 182 | O88783 | F5 | 2.34E+06 | 2.42E+06 | 8.08E+05 | 9 | Plasma Membrane |
| 183 | P49769 | PSEN1 | 2.31E+06 | 1.46E+06 | 5.94E+05 | 6 | Plasma Membrane |
| 184 | P97370 | ATP1B3 | 2.30E+06 | 9.14E+05 | 2.64E+05 | 12 | Plasma Membrane |
| 185 | Q80VP1 | EPN1 | 2.28E+06 | 9.40E+05 | 2.71E+05 | 12 | Plasma Membrane |
| 186 | A2ARI4 | LGR4 | 2.20E+06 | 1.75E+06 | 1.01E+06 | 3 | Plasma Membrane |
| 187 | P01872 | IGHM | 2.20E+06 | 2.26E+06 | 6.52E+05 | 12 | Plasma Membrane |
| 188 | Q60865 | CAPRIN1 | 2.09E+06 | 7.68E+05 | 3.14E+05 | 6 | Plasma Membrane |
| 189 | P18181 | CD48 | 1.95E+06 | 3.50E+05 | 2.02E+05 | 3 | Plasma Membrane |
| 190 | P40124 | CAP1 | 1.88E+06 | 8.70E+05 | 2.90E+05 | 9 | Plasma Membrane |
| 191 | Q9JHZ2 | ANKH | 1.86E+06 | 2.07E+05 | 1.19E+05 | 3 | Plasma Membrane |
| 192 | O89013 | LEPROT | 1.69E+06 | 1.62E+06 | 6.62E+05 | 6 | Plasma Membrane |
| 193 | O89112 | LANCL1 | 1.64E+06 | 8.94E+05 | 5.16E+05 | 3 | Plasma Membrane |
| 194 | Q99JW4 | LIMS1 | 1.64E+06 | 2.58E+05 | 1.49E+05 | 3 | Plasma Membrane |
| 195 | Q3UHH2 | SLC22A23 | 1.61E+06 | 7.87E+04 | 4.54E+04 | 3 | Plasma Membrane |
| 196 | Q9EQP2 | EHD4 | 1.59E+06 | 5.11E+05 | 2.09E+05 | 6 | Plasma Membrane |
| 197 | Q8VBZ3 | CLPTM1 | 1.48E+06 | 2.76E+05 | 1.59E+05 | 3 | Plasma Membrane |
| 198 | Q811D0 | DLG1 | 1.33E+06 | 7.22E+05 | 2.41E+05 | 9 | Plasma Membrane |
| 199 | P26039 | TLN1 | 1.31E+06 | 9.92E+05 | 2.02E+05 | 24 | Plasma Membrane |
| 200 | O35114 | SCARB2 | 1.27E+06 | 1.59E+06 | 9.21E+05 | 3 | Plasma Membrane |
| 201 | Q9WVC8 | SLC26A3 | 1.25E+06 | 6.14E+05 | 3.55E+05 | 3 | Plasma Membrane |
| 202 | Q60770 | STXBP3 | 1.25E+06 | 1.08E+05 | 6.22E+04 | 3 | Plasma Membrane |
| 203 | P41438 | SLC19A1 | 1.11E+06 | 3.31E+05 | 1.91E+05 | 3 | Plasma Membrane |
| 204 | P55302 | LRPAP1 | 1.04E+06 | 7.77E+05 | 4.49E+05 | 3 | Plasma Membrane |
| 205 | Q4VBD2 | TAPT1 | 1.04E+06 | 3.01E+05 | 1.74E+05 | 3 | Plasma Membrane |
| 206 | Q8BRU6 | SLC18A2 | 9.01E+05 | 1.93E+05 | 1.12E+05 | 3 | Plasma Membrane |
| 207 | Q6P9Q6 | FKBP15 | 8.84E+05 | 4.43E+05 | 1.81E+05 | 6 | Plasma Membrane |
| 208 | P11835 | ITGB2 | 8.58E+05 | 7.27E+05 | 4.19E+05 | 3 | Plasma Membrane |
| 209 | Q9DBG3 | Ap2b1 | 8.56E+05 | 7.19E+05 | 2.93E+05 | 6 | Plasma Membrane |
| 210 | Q60902 | EPS15L1 | 4.24E+05 | 9.54E+04 | 5.51E+04 | 3 | Plasma Membrane |
| 211 | Q80X71 | TMEM106B | 3.08E+05 | 3.19E+04 | 1.84E+04 | 3 | Plasma Membrane |

| | | | | | | | |
|---|---|---|---|---|---|---|---|
| 132 | Q91WC9 | DAGLB | 4.36E+06 | 1.99E+06 | 4.69E+05 | 18 | Plasma Membrane |
| 133 | Q3TWN3 | CNNM2 | 4.24E+06 | 3.74E+05 | 1.53E+05 | 6 | Plasma Membrane |
| 134 | Q09143 | SLC7A1 | 4.21E+06 | 2.52E+06 | 1.45E+06 | 3 | Plasma Membrane |
| 135 | P11688 | ITGA5 | 4.16E+06 | 2.44E+06 | 7.05E+05 | 12 | Plasma Membrane |
| 136 | Q8BG67 | EFR3A | 4.00E+06 | 7.53E+05 | 1.64E+05 | 21 | Plasma Membrane |
| 137 | P01899, P01901, P01902 | HLA-A | 3.81E+06 | 5.48E+06 | 1.06E+06 | 27 | Plasma Membrane |
| 138 | Q6R0H7 | GNAS | 3.77E+06 | 9.54E+05 | 3.89E+05 | 6 | Plasma Membrane |
| 139 | Q6ZQI3 | MLEC | 3.72E+06 | 1.74E+06 | 4.50E+05 | 15 | Plasma Membrane |
| 140 | P61161 | ACTR2 | 3.72E+06 | 1.57E+06 | 5.25E+05 | 9 | Plasma Membrane |
| 141 | P18052 | PTPRA | 3.64E+06 | 2.93E+06 | 8.47E+05 | 12 | Plasma Membrane |
| 142 | P49769 | PSEN1 | 3.60E+06 | 1.79E+06 | 7.32E+05 | 6 | Plasma Membrane |
| 143 | Q9CRY7 | GDPD1 | 3.53E+06 | 7.07E+05 | 4.08E+05 | 3 | Plasma Membrane |
| 144 | Q9JL56 | GDE1 | 3.52E+06 | 2.45E+06 | 8.17E+05 | 9 | Plasma Membrane |
| 145 | P70460 | VASP | 3.50E+06 | 2.11E+06 | 1.22E+06 | 3 | Plasma Membrane |
| 146 | Q8CHK3 | MBOAT7 | 3.49E+06 | 3.06E+06 | 1.77E+06 | 3 | Plasma Membrane |
| 147 | Q8QZY6 | TSPAN14 | 3.34E+06 | 1.28E+06 | 7.37E+05 | 3 | Plasma Membrane |
| 148 | Q9QUM4 | SLAMF1 | 3.32E+06 | 1.51E+06 | 5.04E+05 | 9 | Plasma Membrane |
| 149 | Q8VDL4 | ADPGK | 3.32E+06 | 3.73E+05 | 2.15E+05 | 3 | Plasma Membrane |
| 150 | Q9QY76 | VAPB | 3.30E+06 | 2.43E+06 | 9.91E+05 | 6 | Plasma Membrane |
| 151 | P01872 | IGHM | 3.23E+06 | 1.50E+06 | 4.34E+05 | 12 | Plasma Membrane |
| 152 | P17047 | LAMP2 | 3.23E+06 | 2.74E+06 | 1.12E+06 | 6 | Plasma Membrane |
| 153 | Q80TH2 | ERBB2IP | 3.20E+06 | 1.92E+06 | 3.92E+05 | 24 | Plasma Membrane |
| 154 | Q60857 | SLC6A4 | 3.20E+06 | 1.33E+06 | 7.67E+05 | 3 | Plasma Membrane |
| 155 | Q6X893 | SLC44A1 | 3.13E+06 | 1.40E+06 | 4.67E+05 | 9 | Plasma Membrane |
| 156 | O55022 | PGRMC1 | 3.12E+06 | 2.20E+06 | 1.27E+06 | 3 | Plasma Membrane |
| 157 | Q9Z0F8 | ADAM17 | 3.12E+06 | 2.24E+06 | 9.14E+05 | 6 | Plasma Membrane |
| 158 | Q8K2J7 | RELL1 | 3.11E+06 | 1.20E+06 | 3.99E+05 | 9 | Plasma Membrane |
| 159 | Q9QUT0 | RHAG | 3.10E+06 | 4.41E+06 | 1.27E+06 | 12 | Plasma Membrane |
| 160 | O88792 | F11R | 3.08E+06 | 4.02E+06 | 1.04E+06 | 15 | Plasma Membrane |
| 161 | Q60790 | RASA3 | 3.03E+06 | 9.76E+05 | 5.63E+05 | 3 | Plasma Membrane |
| 162 | Q80VP1 | EPN1 | 3.00E+06 | 1.05E+06 | 3.04E+05 | 12 | Plasma Membrane |
| 163 | Q920L1 | FADS1 | 2.97E+06 | 1.99E+05 | 1.15E+05 | 3 | Plasma Membrane |
| 164 | P70398 | USP9X | 2.76E+06 | 9.66E+05 | 2.79E+05 | 12 | Plasma Membrane |
| 165 | P97370 | ATP1B3 | 2.75E+06 | 1.51E+06 | 4.36E+05 | 12 | Plasma Membrane |
| 166 | Q8BFZ9 | ERLIN2 | 2.74E+06 | 2.55E+05 | 1.47E+05 | 3 | Plasma Membrane |
| 167 | Q9QWY8 | ASAP1 | 2.67E+06 | 1.14E+06 | 4.64E+05 | 6 | Plasma Membrane |
| 168 | O89112 | LANCL1 | 2.61E+06 | 2.64E+05 | 1.52E+05 | 3 | Plasma Membrane |
| 169 | P40124 | CAP1 | 2.60E+06 | 8.98E+05 | 2.99E+05 | 9 | Plasma Membrane |
| 170 | Q8K4F0 | CD226 | 2.57E+06 | 4.04E+05 | 2.33E+05 | 3 | Plasma Membrane |
| 171 | Q62469 | ITGA2 | 2.54E+06 | 1.99E+06 | 3.46E+05 | 33 | Plasma Membrane |
| 172 | P50516 | ATP6V1A | 2.39E+06 | 1.43E+06 | 3.70E+05 | 15 | Plasma Membrane |
| 173 | Q8BHL3 | TBC1D10B | 2.37E+06 | 3.83E+05 | 2.21E+05 | 3 | Plasma Membrane |
| 174 | Q8BIG7 | COMTD1 | 2.28E+06 | 1.38E+06 | 7.96E+05 | 3 | Plasma Membrane |
| 175 | Q64455 | PTPRJ | 2.26E+06 | 2.89E+06 | 9.64E+05 | 9 | Plasma Membrane |
| 176 | Q3UHH2 | SLC22A23 | 2.25E+06 | 2.11E+05 | 1.22E+05 | 3 | Plasma Membrane |
| 177 | P70452 | STX4 | 2.24E+06 | 9.25E+05 | 3.08E+05 | 9 | Plasma Membrane |
| 178 | Q8CF94 | RHCE/RHD | 2.18E+06 | 3.44E+06 | 7.03E+05 | 24 | Plasma Membrane |
| 179 | Q6P9J9 | ANO6 | 2.16E+06 | 1.07E+06 | 6.18E+05 | 3 | Plasma Membrane |
| 180 | P21278 | GNA11 | 2.16E+06 | 8.07E+05 | 4.66E+05 | 3 | Plasma Membrane |
| 181 | A2ARI4 | LGR4 | 2.09E+06 | 1.23E+06 | 7.08E+05 | 3 | Plasma Membrane |
| 182 | Q61739 | ITGA6 | 2.05E+06 | 2.25E+06 | 4.90E+05 | 21 | Plasma Membrane |
| 183 | Q9WVC8 | SLC26A3 | 1.99E+06 | 2.18E+05 | 1.26E+05 | 3 | Plasma Membrane |
| 184 | P55302 | LRPAP1 | 1.93E+06 | 3.72E+05 | 2.15E+05 | 3 | Plasma Membrane |
| 185 | O55222 | ILK | 1.92E+06 | 2.75E+06 | 6.47E+05 | 18 | Plasma Membrane |
| 186 | Q99JB2 | STOML2 | 1.82E+06 | 2.03E+06 | 5.23E+05 | 15 | Plasma Membrane |
| 187 | P70302 | STIM1 | 1.76E+06 | 7.15E+05 | 2.38E+05 | 9 | Plasma Membrane |
| 188 | Q80TE7 | LRRC7 | 1.68E+06 | 5.06E+05 | 2.92E+05 | 3 | Plasma Membrane |
| 189 | O88783 | F5 | 1.67E+06 | 1.67E+06 | 5.58E+05 | 9 | Plasma Membrane |
| 190 | O70443 | GNAZ | 1.65E+06 | 9.25E+05 | 3.77E+05 | 6 | Plasma Membrane |
| 191 | O89013 | LEPROT | 1.62E+06 | 1.55E+06 | 6.31E+05 | 6 | Plasma Membrane |
| 192 | P41438 | SLC19A1 | 1.59E+06 | 1.85E+05 | 1.07E+05 | 3 | Plasma Membrane |
| 193 | Q811D0 | DLG1 | 1.55E+06 | 8.06E+05 | 2.69E+05 | 9 | Plasma Membrane |
| 194 | Q9D3G5 | STX11 | 1.51E+06 | 1.36E+06 | 3.19E+05 | 18 | Plasma Membrane |
| 195 | Q9JHZ2 | ANKH | 1.49E+06 | 2.91E+05 | 1.68E+05 | 3 | Plasma Membrane |
| 196 | P26039 | TLN1 | 1.40E+06 | 2.11E+06 | 4.30E+05 | 24 | Plasma Membrane |
| 197 | Q9EQP2 | EHD4 | 1.33E+06 | 3.77E+05 | 1.54E+05 | 6 | Plasma Membrane |
| 198 | P18181 | CD48 | 1.26E+06 | 5.54E+05 | 3.20E+05 | 3 | Plasma Membrane |
| 199 | Q4VBD2 | TAPT1 | 1.19E+06 | 2.96E+05 | 1.71E+05 | 3 | Plasma Membrane |
| 200 | Q99JW4 | LIMS1 | 1.17E+06 | 2.24E+06 | 1.29E+06 | 3 | Plasma Membrane |
| 201 | Q8VBZ3 | CLPTM1 | 1.14E+06 | 1.60E+05 | 9.23E+04 | 3 | Plasma Membrane |
| 202 | O35114 | SCARB2 | 1.14E+06 | 1.23E+06 | 7.09E+05 | 3 | Plasma Membrane |
| 203 | Q60770 | STXBP3 | 7.97E+05 | 5.67E+05 | 3.28E+05 | 3 | Plasma Membrane |
| 204 | Q9DBG3 | Ap2b1 | 7.38E+05 | 1.03E+06 | 4.19E+05 | 6 | Plasma Membrane |
| 205 | Q60865 | CAPRIN1 | 7.16E+05 | 3.70E+05 | 1.51E+05 | 6 | Plasma Membrane |
| 206 | Q80X71 | TMEM106B | 7.13E+05 | 8.81E+04 | 5.08E+04 | 3 | Plasma Membrane |
| 207 | Q8BRU6 | SLC18A2 | 6.35E+05 | 3.50E+05 | 2.02E+05 | 3 | Plasma Membrane |
| 208 | P11835 | ITGB2 | 3.71E+05 | 1.13E+06 | 6.52E+05 | 3 | Plasma Membrane |
| 209 | Q60902 | EPS15L1 | 2.93E+05 | 3.21E+04 | 1.85E+04 | 3 | Plasma Membrane |
| 210 | Q6P9Q6 | FKBP15 | 2.31E+05 | 1.04E+05 | 4.26E+04 | 6 | Plasma Membrane |
| 211 | Q9ERM2 | ICAM4 | 7.41E+04 | 7.55E+04 | 3.08E+04 | 6 | Plasma Membrane |

| | | | | | | | |
|---|---|---|---|---|---|---|---|
| 132 | P09055 | ITGB1 | 3122781 | 3529995 | 565252 | 39 | Plasma Membrane |
| 133 | P49769 | PSEN1 | 3023270 | 1610224 | 657371 | 6 | Plasma Membrane |
| 134 | Q6R0H7 | GNAS | 2971529 | 578184 | 236043 | 6 | Plasma Membrane |
| 135 | P50516 | ATP6V1A | 2883946 | 1562955 | 403553 | 15 | Plasma Membrane |
| 136 | Q920L1 | FADS1 | 2792971 | 981798 | 566841 | 3 | Plasma Membrane |
| 137 | P70460 | VASP | 2758705 | 708797 | 409224 | 3 | Plasma Membrane |
| 138 | P18181 | CD48 | 2663666 | 255099 | 147281 | 3 | Plasma Membrane |
| 139 | Q9QWY8 | ASAP1 | 2593084 | 1087643 | 444028 | 6 | Plasma Membrane |
| 140 | P17047 | LAMP2 | 2495538 | 2344999 | 957342 | 6 | Plasma Membrane |
| 141 | Q8VDL4 | ADPGK | 2449479 | 448141 | 258735 | 3 | Plasma Membrane |
| 142 | Q9Z0F8 | ADAM17 | 2439522 | 2526821 | 1031570 | 6 | Plasma Membrane |
| 143 | Q6X893 | SLC44A1 | 2430912 | 887424 | 295808 | 9 | Plasma Membrane |
| 144 | P97370 | ATP1B3 | 2430582 | 1276525 | 368501 | 12 | Plasma Membrane |
| 145 | Q99JB2 | STOML2 | 2369741 | 1732147 | 447238 | 15 | Plasma Membrane |
| 146 | Q80TH2 | ERBB2IP | 2277295 | 1350869 | 275745 | 24 | Plasma Membrane |
| 147 | Q9Z0T9 | ITGB6 | 2261562 | 1093097 | 631100 | 3 | Plasma Membrane |
| 148 | P56400 | GP1BB | 2245698 | 11344530 | 2673931 | 18 | Plasma Membrane |
| 149 | P40124 | CAP1 | 2227052 | 917357 | 305786 | 9 | Plasma Membrane |
| 150 | Q3UHH2 | SLC22A23 | 2196385 | 146951 | 84842 | 3 | Plasma Membrane |
| 151 | Q9WVC8 | SLC26A3 | 2173212 | 262339 | 151461 | 3 | Plasma Membrane |
| 152 | O89013 | LEPROT | 2109243 | 1661988 | 678504 | 6 | Plasma Membrane |
| 153 | Q811D0 | DLG1 | 2046309 | 999663 | 333221 | 9 | Plasma Membrane |
| 154 | Q9CRY7 | GDPD1 | 2023211 | 432869 | 249917 | 3 | Plasma Membrane |
| 155 | O35114 | SCARB2 | 2015221 | 1997285 | 1153133 | 3 | Plasma Membrane |
| 156 | P41438 | SLC19A1 | 1996116 | 140465 | 81097 | 3 | Plasma Membrane |
| 157 | P21279 | GNAQ | 1992617 | 1682053 | 971134 | 3 | Plasma Membrane |
| 158 | O88186 | GP9 | 1956551 | 9879114 | 2851855 | 12 | Plasma Membrane |
| 159 | Q9JHZ2 | ANKH | 1954989 | 564976 | 326189 | 3 | Plasma Membrane |
| 160 | O89112 | LANCL1 | 1885209 | 112806 | 65129 | 3 | Plasma Membrane |
| 161 | Q9QY76 | VAPB | 1877089 | 1808058 | 738137 | 6 | Plasma Membrane |
| 162 | O54890 | ITGB3 | 1780572 | 7997693 | 942537 | 72 | Plasma Membrane |
| 163 | Q8BFZ9 | ERLIN2 | 1710282 | 1073096 | 619552 | 3 | Plasma Membrane |
| 164 | Q9QUM0 | ITGA2B | 1670841 | 12231284 | 1359032 | 81 | Plasma Membrane |
| 165 | P55302 | LRPAP1 | 1665523 | 325494 | 187924 | 3 | Plasma Membrane |
| 166 | Q8K558 | TREML1 | 1572615 | 6452752 | 1317162 | 24 | Plasma Membrane |
| 167 | P70452 | STX4 | 1339835 | 751619 | 250540 | 9 | Plasma Membrane |
| 168 | Q8BIG7 | COMTD1 | 1337130 | 768875 | 443910 | 3 | Plasma Membrane |
| 169 | Q80TE7 | LRRC7 | 1329180 | 243430 | 140544 | 3 | Plasma Membrane |
| 170 | Q60865 | CAPRIN1 | 1244077 | 891754 | 364057 | 6 | Plasma Membrane |
| 171 | O08742 | GP5 | 1140316 | 6025589 | 1100117 | 30 | Plasma Membrane |
| 172 | Q8VBZ3 | CLPTM1 | 1077361 | 444254 | 256490 | 3 | Plasma Membrane |
| 173 | P01872 | IGHM | 1010211 | 3024067 | 872973 | 12 | Plasma Membrane |
| 174 | O35930 | GP1BA | 981549 | 6807775 | 1090116 | 39 | Plasma Membrane |
| 175 | Q4VBD2 | TAPT1 | 949784 | 512612 | 295956 | 3 | Plasma Membrane |
| 176 | Q6ZQI3 | MLEC | 891991 | 1157475 | 298859 | 15 | Plasma Membrane |
| 177 | Q6P9Q6 | FKBP15 | 858400 | 345434 | 141023 | 6 | Plasma Membrane |
| 178 | O55222 | ILK | 805037 | 2300618 | 542261 | 18 | Plasma Membrane |
| 179 | Q9D3G5 | STX11 | 765012 | 799145 | 188360 | 18 | Plasma Membrane |
| 180 | Q6P9J9 | ANO6 | 758379 | 672798 | 388440 | 3 | Plasma Membrane |
| 181 | Q9EQP2 | EHD4 | 731361 | 361133 | 147432 | 6 | Plasma Membrane |
| 182 | Q9DBG3 | Ap2b1 | 728648 | 822585 | 335819 | 6 | Plasma Membrane |
| 183 | P40240 | CD9 | 719812 | 4252470 | 1417490 | 9 | Plasma Membrane |
| 184 | P21278 | GNA11 | 683508 | 453323 | 261726 | 3 | Plasma Membrane |
| 185 | Q64455 | PTPRJ | 655162 | 1293517 | 431172 | 9 | Plasma Membrane |
| 186 | P36423 | TBXAS1 | 645665 | 1771131 | 340854 | 27 | Plasma Membrane |
| 187 | Q9QUT0 | RHAG | 635315 | 1201456 | 346830 | 12 | Plasma Membrane |
| 188 | P26039 | TLN1 | 627713 | 683437 | 139506 | 24 | Plasma Membrane |
| 189 | Q80X71 | TMEM106B | 602615 | 82975 | 47906 | 3 | Plasma Membrane |
| 190 | Q8K4F0 | CD226 | 457463 | 484068 | 279477 | 3 | Plasma Membrane |
| 191 | P11835 | ITGB2 | 435634 | 667559 | 385415 | 3 | Plasma Membrane |
| 192 | Q8QZY6 | TSPAN14 | 420160 | 776961 | 448579 | 3 | Plasma Membrane |
| 193 | Q8CF94 | RHCE/RHD | 397080 | 816394 | 166646 | 24 | Plasma Membrane |
| 194 | Q01102 | SELP | 388569 | 1560747 | 367872 | 18 | Plasma Membrane |
| 195 | Q9JL99 | CLEC1B | 380888 | 1567777 | 905157 | 3 | Plasma Membrane |
| 196 | P70302 | STIM1 | 354718 | 404617 | 134872 | 9 | Plasma Membrane |
| 197 | Q60790 | RASA3 | 342003 | 697740 | 402841 | 3 | Plasma Membrane |
| 198 | O88792 | F11R | 341019 | 1280565 | 330640 | 15 | Plasma Membrane |
| 199 | A2ARI4 | LGR4 | 310348 | 1184838 | 684066 | 3 | Plasma Membrane |
| 200 | Q60902 | EPS15L1 | 308605 | 119225 | 68834 | 3 | Plasma Membrane |
| 201 | Q60770 | STXBP3 | 263992 | 187558 | 108287 | 3 | Plasma Membrane |
| 202 | Q9QUM4 | SLAMF1 | 217294 | 1070645 | 356882 | 9 | Plasma Membrane |
| 203 | Q8CHK3 | MBOAT7 | 194162 | 1257260 | 725880 | 3 | Plasma Membrane |
| 204 | O88783 | F5 | 186563 | 680658 | 226886 | 9 | Plasma Membrane |
| 205 | Q62469 | ITGA2 | 178090 | 821735 | 143046 | 33 | Plasma Membrane |
| 206 | Q9ERM2 | ICAM4 | 136623 | 85635 | 34960 | 6 | Plasma Membrane |
| 207 | O70443 | GNAZ | 106684 | 567898 | 231843 | 6 | Plasma Membrane |
| 208 | Q99JW4 | LIMS1 | 84391 | 187239 | 108102 | 3 | Plasma Membrane |
| 209 | Q61739 | ITGA6 | 57825 | 642041 | 140105 | 21 | Plasma Membrane |
| 210 | Q8BRU6 | SLC18A2 | 36288 | 193995 | 112003 | 3 | Plasma Membrane |
| 211 | Q60857 | SLC6A4 | 100 | 856177 | 494314 | 3 | Plasma Membrane |

Peptides with significant difference $p<0.05$

| Charge Group ID | Charge State | Factor 1 P-value | Intensity [dKO_1] | Intensity [dKO_2] | Intensity [dKO_3] | Average_dKO | Intensity [AQP1KO_1] | Intensity [AQP1KO_2] | Intensity [AQP1KO_3] | Average_AQP1KO | Intensity [RhAGKO_1] | Intensity [RhAGKO_2] | Intensity [RhAGKO_3] | Average RhAG | Intensity [WT_1] | Intensity [WT_2] | Intensity [WT_3] | Average WT | WT vs dKO | WT vs AQP1KO | WT vs RhAGKO | M + H (Feature) | Modified Peptide Sequence | Primary Protein Name | ID | Symbol | Entrez Gene Name | Location | Type(s) |
|---|---|---|---|---|---|---|---|---|---|---|---|---|---|---|---|---|---|---|---|---|---|---|---|---|---|---|---|---|---|
| 15549934 | 3 | 0.031 | 4698196.5 | 3197000.5 | 2938193.75 | 3.61E+06 | 3967852.5 | 4262262.5 | 3054900.25 | 3761671.75 | 3886817.25 | 3516393.75 | 4100300 | 3.83E+06 | 7508652 | 5537162 | 5085820.5 | 6043878.167 | 1.7 | 1.6 | 1.6 | 2518.2832 | SPNPSLLHLC[160.0307]GSLAQLAC[160.0307]VEPVR | A2AN08 | A2AN08 | UBR4 | ubiquitin protein ligase E3 component n-recognin 4 | Nucleus | enzyme |
| 15539477 | 2 | 6.17E-07 | 369929.312 | 147544.125 | 186130.625 | 2.35E+05 | 512878.812 | 260300.266 | 204787.453 | 325988.8437 | 1.21E+07 | 8458144 | 1.12E+07 | 1.06E+07 | 1.07E+07 | 1.28E+07 | 1.24E+07 | 11970000 | 51.0 | 36.7 | 1.1 | 980.5143 | QM[147.0354]IRNFR | A2ATU0 | A2ATU0 | DHTKD1 | dehydrogenase E1 and transketolase domain containing 1 | Cytoplasm | enzyme |
| 15531738 | 2 | 0.002 | 1.77E+07 | 1.50E+07 | 1.62E+07 | 1.63E+07 | 1.22E+07 | 1.14E+07 | 1.10E+07 | 11540000 | 1.37E+07 | 1.15E+07 | 1.16E+07 | 1.23E+07 | 1.49E+07 | 1.51E+07 | 1.37E+07 | 14533333.33 | 0.9 | 1.3 | 1.2 | 1027.6128 | FPPLLWVR | A6H603 | A6H603 | NWD1 | NACHT and WD repeat domain containing 1 | Cytoplasm | other |
| 15540934 | 3 | 0.041 | 1467737.5 | 980301.312 | 1120392.75 | 1.19E+06 | 1728590.5 | 1334579.875 | 1206536.625 | 1423235.667 | 1976139.375 | 1399694.625 | 1623235.125 | 1.67E+06 | 2007178.875 | 2420895.25 | 1662770.375 | 2030281.5 | 1.7 | 1.4 | 1.2 | 2104.0415 | RQHQQAPLPAQAVC[160.0307]ADGTR | E2JF22 | E2JF22 | PIEZO1 | piezo-type mechanosensitive ion channel component 1 | Cytoplasm | ion channel |
| 15555624 | 3 | 2.62E-04 | 1449744.75 | 2632032 | 1329369 | 1.80E+06 | 1551312.625 | 1813187.625 | 1508798.75 | 1624433 | 817903.875 | 1098732.375 | 1295809.625 | 1.07E+06 | 451388.219 | 448432.344 | 410359.031 | 436726.5313 | 0.2 | 0.3 | 0.4 | 2524.1567 | DSHIFQDFWQETAESLNTLDK | E9Q555 | E9Q555 | RNF213 | ring finger protein 213 | Cytoplasm | enzyme |
| 15531916 | 2 | 0.002 | 1.86E+07 | 1.75E+07 | 1.38E+07 | 1.66E+07 | 2.72E+07 | 2.31E+07 | 2.88E+07 | 26383333.33 | 1.86E+07 | 2.29E+07 | 2.08E+07 | 2.08E+07 | 2.92E+07 | 2.58E+07 | 2.75E+07 | 27510000 | 1.7 | 1.0 | 1.3 | 1402.6273 | FEEGGYVVC[160.0307]NTK | O08573 | O08573 | LGALS9B | lectin, galactoside-binding, soluble, 9B | Cytoplasm | other |
| 15532236 | 2 | 9.28E-04 | 1.21E+07 | 1.17E+07 | 1.03E+07 | 1.14E+07 | 2.08E+07 | 1.67E+07 | 2.30E+07 | 20133333.33 | 1.59E+07 | 1.69E+07 | 1.51E+07 | 1.60E+07 | 2.34E+07 | 1.78E+07 | 2.22E+07 | 21140000 | 1.9 | 1.1 | 1.3 | 1473.7708 | SIMISGNVLPDATR | O08573 | O08573 | LGALS9B | lectin, galactoside-binding, soluble, 9B | Cytoplasm | other |
| 15538142 | 2 | 0.029 | 1.29E+07 | 1.46E+07 | 1.09E+07 | 1.28E+07 | 2.11E+07 | 1.74E+07 | 2.62E+07 | 21590000 | 1.48E+07 | 1.93E+07 | 1.98E+07 | 1.80E+07 | 2.28E+07 | 1.61E+07 | 2.06E+07 | 19846666.67 | 1.6 | 0.9 | 1.1 | 948.4884 | FNENAVVR | O08573 | O08573 | LGALS9B | lectin, galactoside-binding, soluble, 9B | Cytoplasm | other |
| 15542951 | 3 | 0.023 | 6274157 | 4916014.5 | 3549556.5 | 4.91E+06 | 6925709 | 6061538 | 9658257 | 7548501.333 | 6574842.5 | 7519007 | 7030088.5 | 7.04E+06 | 1.06E+07 | 1.01E+07 | 7832754.5 | 9524251.5 | 1.9 | 1.3 | 1.4 | 1356.6438 | C[160.0307]GGDIAFHLNPR | O08573 | O08573 | LGALS9B | lectin, galactoside-binding, soluble, 9B | Cytoplasm | other |
| 15543918 | 2 | 0.029 | 3542754.5 | 3692173.25 | 2465172.5 | 3.23E+06 | 4243555 | 3703261 | 5673581.5 | 4540132.5 | 3679362.5 | 5319592 | 4332745.5 | 4.44E+06 | 5407038.5 | 5685214 | 5973034 | 5688428.833 | 1.8 | 1.3 | 1.3 | 799.4559 | FHINLR | O08573 | O08573 | LGALS9B | lectin, galactoside-binding, soluble, 9B | Cytoplasm | other |
| 15540065 | 2 | 0.005 | 1.01E+07 | 9096985 | 1.31E+07 | 1.08E+07 | 1.71E+07 | 1.54E+07 | 1.47E+07 | 15730000 | 1.12E+07 | 1.17E+07 | 9772221 | 1.09E+07 | 2.26E+07 | 1.47E+07 | 2.17E+07 | 19663333.33 | 1.8 | 1.3 | 1.8 | 1194.5552 | VFTDVPSNC[160.0307]R | O08688 | O08688 | CAPN5 | calpain 5 | Cytoplasm | peptidase |
| 15534843 | 2 | 0.038 | 1.24E+07 | 9729061 | 1.35E+07 | 1.19E+07 | 1.46E+07 | 1.59E+07 | 1.65E+07 | 15680000 | 1.24E+07 | 1.26E+07 | 9756209 | 1.16E+07 | 2.11E+07 | 1.32E+07 | 2.08E+07 | 18373333.33 | 1.5 | 1.2 | 1.6 | 1364.7015 | DSPTGANSYVIIK | O08688 | O08688 | CAPN5 | calpain 5 | Cytoplasm | peptidase |
| 15534800 | 2, 3 | 0.021 | 9180900 | 8232558.5 | 1.28E+07 | 1.01E+07 | 1.26E+07 | 1.52E+07 | 1.55E+07 | 14430000 | 1.02E+07 | 9280027 | 9003510 | 9.49E+06 | 1.59E+07 | 1.20E+07 | 1.98E+07 | 15876666.67 | 1.6 | 1.1 | 1.7 | 1288.7233, 1288.7235 | LINTSYLSIHK | O08688 | O08688 | CAPN5 | calpain 5 | Cytoplasm | peptidase |
| 15542406, 15544152 | 2 | 0.046 | 4946569 | 5474937.5 | 9103157 | 6.51E+06 | 1.05E+07 | 9570984 | 1.08E+07 | 10306994.67 | 6298089 | 6160712.5 | 5386103 | 5.95E+06 | 1.20E+07 | 6528992 | 1.05E+07 | 9679664 | 1.5 | 0.9 | 1.6 | 1095.5289 | GTSTPEYNVK | O08688 | O08688 | CAPN5 | calpain 5 | Cytoplasm | peptidase |
| 15534488 | 2 | 3.91E-04 | 3114531.75 | 2523927 | 4274183.5 | 3.30E+06 | 4889176.5 | 6440570 | 5390064.5 | 5573270.333 | 4806711.5 | 3898462.5 | 4370433.5 | 4.36E+06 | 8242075 | 8991707 | 9380957 | 8871579.667 | 2.7 | 1.6 | 2.0 | 1295.6064 | NEFWC[160.0307]ALVEK | O08688 | O08688 | CAPN5 | calpain 5 | Cytoplasm | peptidase |
| 15537623 | 2 | 0.005 | 3672739.75 | 3609421.5 | 4554212.5 | 3.95E+06 | 6840275 | 5824675.5 | 6377709 | 6347553.167 | 5590686 | 5474901 | 4039609.25 | 5.04E+06 | 8151224.5 | 6001991 | 9564823 | 7906012.833 | 2.0 | 1.2 | 1.6 | 899.4851 | YFTDIIK | O08688 | O08688 | CAPN5 | calpain 5 | Cytoplasm | peptidase |
| 15549369 | 2 | 0.01 | 2497661 | 2917916 | 3896204.75 | 3.10E+06 | 4888987.5 | 4692494.5 | 4660454 | 4747312 | 2623609 | 2648799.5 | 1938918.25 | 2.40E+06 | 3967774.75 | 2895820 | 4564213.5 | 3809269.417 | 1.2 | 0.8 | 1.6 | 2190.1545 | YVIIPTTFEPGHTGEFLLR | O08688 | O08688 | CAPN5 | calpain 5 | Cytoplasm | peptidase |
| 15543022 | 3 | 0.016 | 6746191 | 7426559 | 7493051 | 7.22E+06 | 8571569 | 1.17E+07 | 9712350 | 9977973 | 9388652 | 8274871 | 8453816 | 8.71E+06 | 9224265 | 8732851 | 9042629 | 8999915 | 1.2 | 0.9 | 1.0 | 2147.9895 | MPPYDEETQAIIDAAQEAR | O08795 | O08795 | PRKCSH | protein kinase C substrate 80K-H | Cytoplasm | enzyme |
| 15540639 | 3 | 0.018 | 4867122.5 | 6379538 | 6085590.5 | 5.78E+06 | 4740157.5 | 5513286.5 | 4575770.5 | 4943071.5 | 4698833 | 6649361 | 5768497.5 | 5.71E+06 | 3949401.25 | 3129440.75 | 4080589 | 3719810.333 | 0.6 | 0.8 | 0.7 | 1396.7505 | LTGVSISQVNHNK | O08917 | O08917 | FLOT1 | flotillin 1 | Plasma Membrane | other |
| 15532780 | 2 | 0.018 | 2.39E+07 | 3.57E+07 | 3.41E+07 | 3.12E+07 | 2.50E+07 | 2.65E+07 | 2.77E+07 | 26410000 | 3.17E+07 | 3.42E+07 | 3.17E+07 | 3.25E+07 | 1.86E+07 | 2.38E+07 | 2.22E+07 | 21523333.33 | 0.7 | 0.8 | 0.7 | 1427.808 | ILGLAIESQDAGIK | O09044 | O09044 | SNAP23 | synaptosomal-associated protein, 23kDa | Plasma Membrane | transporter |
| [illegible] | [illegible] | [illegible] | [illegible] | [illegible] | [illegible] | [illegible] | [illegible] | [illegible] | [illegible] | [illegible] | [illegible] | [illegible] | [illegible] | [illegible] | [illegible] | [illegible] | [illegible] | [illegible] | [illegible] | [illegible] | [illegible] | [illegible] | [illegible] | [illegible] | [illegible] | [illegible] | [illegible] | [illegible] | [illegible] |

| | | | | | | | | | |
|---|---|---|---|---|---|---|---|---|---|
| 15550149 | 2 | 0.025 | 4862459.5 | 4171471.5 | 3075554.75 | 4.04E+06 | 3716667.5 | 4168675.5 | 2104121 |
| 15541433 | 2 | 0.035 | 2223662.25 | 3439887.5 | 1324780.75 | 2.33E+06 | 2439459.75 | 2217154.5 | 2101608 |
| 15537827 | 2 | 0.044 | 482023.281 | 1161420.625 | 1494635.875 | 1.05E+06 | 2704403 | 3184908.75 | 2992044 |
| 15549114 | 3 | 0.047 | 9639079 | 1.74E+07 | 8541985 | 1.19E+07 | 1.28E+07 | 1.29E+07 | 1.32E+07 |
| 15533378 | 2 | 0.049 | 6897116.5 | 1.16E+07 | 4784705.5 | 7.76E+06 | 7640149 | 6674506.5 | 8352110 |
| 15544873 | 2 | 0.033 | 3172747.75 | 4181230.75 | 1942655.25 | 3.10E+06 | 2497235 | 2449321.5 | 3236646.5 |
| 15553345 | 2 | 0.031 | 1353190.25 | 2358035.75 | 2485990.25 | 2.07E+06 | 2543595.5 | 4064297.75 | 2725487.75 |
| 15548394 | 2 | 0.047 | 1.00E+07 | 1.39E+07 | 7207733.5 | 1.04E+07 | 7685853.5 | 1.06E+07 | 8849224 |
| 15548984 | 2 | 0.019 | 6493939.5 | 8041691 | 5114376 | 6.55E+06 | 5736776 | 6129378.5 | 7162226 |
| 15535068 | 2 | 0.039 | 2434635.75 | 3900187.5 | 1339373.125 | 2.56E+06 | 2445617.25 | 2001427 | 2513155.75 |
| 15535090 | 2 | 0.032 | 2865168.25 | 5045377.5 | 2685662.75 | 3.53E+06 | 3159274.25 | 3629679 | 4840611.5 |
| 15532757 | 2 | 0.013 | 6420852.5 | 8370213 | 1.23E+07 | 9.04E+06 | 1.89E+07 | 1.60E+07 | 1.77E+07 |
| 15555999 | 2 | 0.027 | 4847601 | 5682510.5 | 8996315 | 6.51E+06 | 1.05E+07 | 1.25E+07 | 1.27E+07 |
| 15533347 | 2 | 0.039 | 5216103.5 | 5159161 | 8471356 | 6.28E+06 | 8962127 | 1.10E+07 | 9405761 |
| 15534413 | 3 | 0.01 | 2804137.75 | 5111397.5 | 5918567.5 | 4.61E+06 | 8875671 | 9102966 | 9534713 |
| 15542495, 15545770 | 3 | 0.011 | 2324133 | 2810922 | 3878690 | 3.00E+06 | 5759994 | 7746587.5 | 8001478.5 |
| 15554425 | 2 | 0.013 | 2735997.5 | 3250540 | 4213721.5 | 3.40E+06 | 8160180 | 6295526.5 | 5882770.5 |
| 15534819 | 2, 3 | 0.018 | 4571929.5 | 9313947 | 3700150 | 5.86E+06 | 5895118 | 5695343 | 5734094 |
| 15538596 | 2 | 0.013 | 4275191 | 6537591.5 | 2694628 | 4.50E+06 | 3737815.25 | 3965968.5 | 3979930.25 |
| 15533832 | 2 | 0.011 | 2775887.5 | 5528187 | 2473861 | 3.59E+06 | 3141127 | 4232212.5 | 5198525.5 |
| 15552237 | 2 | 0.045 | 1996689 | 4570254.5 | 1600944.875 | 2.72E+06 | 2199053 | 2681025.75 | 2335263.75 |
| 15549966 | 2, 3 | 0.041 | 1.67E+07 | 1.84E+07 | 2.21E+07 | 1.90E+07 | 2.01E+07 | 2.06E+07 | 1.97E+07 |
| 15533306 | 2 | 0.037 | 6152696.5 | 7598430.5 | 3987846.75 | 5.91E+06 | 4714335.5 | 5294567 | 3258679.5 |
| 15545516 | 3 | 0.024 | 2237468.25 | 3793687.25 | 1972602.375 | 2.67E+06 | 2300697.75 | 2627158.25 | 2509077.25 |
| 15541316 | 3 | 0.016 | 2111768 | 3274395.25 | 1454345.375 | 2.28E+06 | 2002193.75 | 2494810 | 1707738.125 |
| 15534238 | 2 | 0.032 | 1.03E+07 | 1.68E+07 | 7591355 | 1.16E+07 | 1.24E+07 | 1.05E+07 | 1.19E+07 |
| 15536918 | 2 | 0.048 | 2975624.75 | 2228226.5 | 1447906.75 | 2.22E+06 | 2771569.75 | 2111100.5 | 1880555.625 |
| 15554449 | 3 | 0.026 | 7062317.5 | 6329686.5 | 6003457.5 | 6.47E+06 | 5978245.5 | 5963871 | 5383739.5 |
| 15540643 | 3 | 0.024 | 2719413 | 4128243.75 | 1500201.125 | 2.78E+06 | 2979128 | 2766393.25 | 2675179 |
| 15536231 | 2 | 0.024 | 9129592 | 1.17E+07 | 6046677.5 | 8.94E+06 | 8506127 | 8971297 | 9264352 |
| 15537086 | 2 | 0.045 | 5308299.5 | 1.01E+07 | 4467988 | 6.62E+06 | 7056208.5 | 7442955 | 7136745 |
| 15533976 | 2 | 0.045 | 5231918.5 | 9350453 | 4279794 | 6.29E+06 | 5231329.5 | 5831485 | 5048101 |
| 15532872 | 2 | 0.024 | 6.21E+07 | 6.32E+07 | 6.32E+07 | 6.28E+07 | 6.59E+07 | 7.23E+07 | 4.53E+07 |
| 15550405 | 3 | 0.047 | 1.79E+07 | 2.28E+07 | 2.25E+07 | 2.11E+07 | 1.89E+07 | 2.82E+07 | 2.12E+07 |

[illegible]

| | | | | | | | | | | |
|---|---|---|---|---|---|---|---|---|---|---|
| 15550931 | 2 | 0.033 | 3129744.25 | 4077718 | 5763309.5 | 4.32E+06 | 6221014 | 8336678 | 7970062.5 | 7509251.5 |
| 15533021 | 3 | 0.011 | 2754653 | 3729213 | 4910076.5 | 3.80E+06 | 6648454 | 7570416.5 | 7080696 | 7099855.5 |
| 15531796 | 3 | 0.001 | 3.54E+07 | 4.11E+07 | 3.27E+07 | 3.64E+07 | 4.72E+07 | 4.48E+07 | 4.72E+07 | 46373333.33 |
| 15549144 | 4 | 0.005 | 9206906 | 1.31E+07 | 9849760 | 1.07E+07 | 2.33E+07 | 2.09E+07 | 2.20E+07 | 22070000 |
| 15534018 | 2 | 0.043 | 4489397.5 | 8232860.5 | 3241153 | 5.32E+06 | 4730713 | 5707232.5 | 5496333.5 | 5311426.333 |
| 15532433 | 3 | 0.004 | 3.32E+07 | 3.63E+07 | 3.50E+07 | 3.48E+07 | 2.67E+07 | 2.72E+07 | 2.82E+07 | 27403333.33 |
| 15551744 | 3 | 0.009 | 7820744 | 7734852.5 | 7040257.5 | 7.53E+06 | 1.02E+07 | 1.12E+07 | 8793437 | 10074479 |
| 15548462 | 2 | 0.047 | 9240216 | 1.61E+07 | 6584626.5 | 1.06E+07 | 9962034 | 9587035 | 9954381 | 9834483.333 |
| 15546094 | 2 | 0.015 | 3690896 | 5625568 | 2346961.25 | 3.89E+06 | 4343207.5 | 3740672.25 | 3170793.25 | 3751557.667 |
| 15533094 | 2 | 0.035 | 5229527.5 | 9261957 | 3774471 | 6.09E+06 | 5288220.5 | 6269017 | 6703983 | 6087073.5 |
| 15532186 | 3 | 0.01 | 1.35E+07 | 1.57E+07 | 1.72E+07 | 1.55E+07 | 2.46E+07 | 2.12E+07 | 2.15E+07 | 22436666.67 |
| 15540111 | 2 | 0.031 | 8367522.5 | 9236391 | 7666153.5 | 8.42E+06 | 7297156 | 6791524.5 | 5766736 | 6618472.167 |
| 15533508 | 2 | 0.027 | 5396433 | 4445946.5 | 4431195 | 4.76E+06 | 5431073 | 5291693 | 3391331.75 | 4704699.25 |
| 15532261 | 3 | 0.031 | 1.16E+07 | 1.49E+07 | 1.21E+07 | 1.28E+07 | 9692616 | 1.23E+07 | 8945184 | 10309266.67 |
| 15543425 | 2 | 0.041 | 1021453 | 926594.625 | 1046672.875 | 9.98E+05 | 3177216.75 | 2733238.75 | 2808494.25 | 2906316.583 |
| 15540993 | 3 | 0.05 | 4020921 | 8147488.5 | 3498589 | 5.22E+06 | 4481949 | 5074982.5 | 6174954 | 5243961.833 |
| 15551846 | 3 | 0.032 | 7884521.5 | 7358304.5 | 1.54E+07 | 1.02E+07 | 1.01E+07 | 1.21E+07 | 1.20E+07 | 11403333.33 |
| 15540994 | 2 | 0.048 | 1864690.375 | 3785232.75 | 2965109.75 | 2.87E+06 | 1351749 | 3880086 | 2659081 | 2630305.333 |
| 15539689 | 2 | 0.04 | 1158554.875 | 1588325.625 | 883803.812 | 1.21E+06 | 4146653 | 3636651.25 | 3616915.75 | 3800073.333 |
| 15539150 | 2 | 0.007 | 7296755.5 | 6218972.5 | 7380494.5 | 6.97E+06 | 5123075.5 | 5717657 | 4262045 | 5034259.167 |
| 15539212 | 2 | 0.039 | 775203.688 | 1185448.625 | 1453099.75 | 1.14E+06 | 2115894.75 | 2509261.5 | 3348803 | 2657986.417 |
| 15550789 | 3 | 0.02 | 1.31E+07 | 1.68E+07 | 1.77E+07 | 1.58E+07 | 2.16E+07 | 2.32E+07 | 1.83E+07 | 21026666.67 |
| 15550881 | 3 | 0.018 | 9210350 | 1.23E+07 | 1.01E+07 | 1.05E+07 | 1.85E+07 | 1.59E+07 | 1.32E+07 | 15850000 |
| 15536725 | 2 | 0.019 | 3159562.25 | 3995422.25 | 4307338.5 | 3.82E+06 | 7534399.5 | 5432143.5 | 5695087.5 | 6220543.5 |
| 15538759 | 2 | 0.009 | 2461936.75 | 3346695.25 | 3442730.5 | 3.08E+06 | 4429013.5 | 5077106.5 | 4645017.5 | 4717045.833 |
| 15542091 | 3 | 0.026 | 3702328 | 3154973.5 | 3476106.5 | 3.44E+06 | 3077742.75 | 3730957.5 | 2783491 | 3197397.083 |
| 15532314 | 2 | 0.046 | 1.06E+07 | 8048337.5 | 7990032 | 8.89E+06 | 1.01E+07 | 1.26E+07 | 6794574 | 9808191.333 |
| 15533361 | 3 | 0.006 | 4267045 | 3104695.5 | 5026705 | 4.13E+06 | 4201170.5 | 4726115 | 4352101 | 4426462.167 |
| 15548273 | 2 | 0.01 | 4.63E+07 | 5.90E+07 | 4.88E+07 | 5.14E+07 | 4.30E+07 | 4.89E+07 | 5.27E+07 | 48203333.33 |
| 15549631 | 2 | 0.007 | 4.35E+07 | 4.57E+07 | 5.07E+07 | 4.66E+07 | 4.10E+07 | 3.91E+07 | 3.84E+07 | 39466666.67 |
| 15531425 | 3 | 0.026 | 4.33E+07 | 4.17E+07 | 4.55E+07 | 4.35E+07 | 3.47E+07 | 3.52E+07 | 3.30E+07 | 34280000 |
| 15531495 | 3 | 0.031 | 3.38E+07 | 3.30E+07 | 3.62E+07 | 3.43E+07 | 2.86E+07 | 3.15E+07 | 2.64E+07 | 28796666.67 |
| 15553025 | 2 | 0.014 | 1.47E+07 | 1.62E+07 | 1.50E+07 | 1.53E+07 | 1.23E+07 | 1.16E+07 | 1.25E+07 | 12116666.67 |
| 15544432 | 3 | 6.96E-07 | 235327.016 | 99386.531 | 163396.688 | 1.66E+05 | 2981533.75 | 5425608.5 | 3496267.25 | 3967803.167 |
| 15543891 | 2 | 4.29E-05 | 37919.27 | 5419.727 | 60818.859 | 3.47E+04 | 1960857.375 | 2722378.25 | 1568382 | 2083872.542 |
| 15549451 | 2 | 0.039 | 2494007.5 | 3857819.25 | 4712066.5 | 3.69E+06 | 8217590 | 7121869.5 | 7510368 | 7616609.167 |
| 15537479 | 2 | 0.047 | 653859.25 | 984798.375 | 1475516.5 | 1.04E+06 | 2147728.5 | 2258495.25 | 2285141.25 | 2230455 |
| 15542259 | 4 | 0.036 | 3295333.75 | 2364336.5 | 2094907.625 | 2.58E+06 | 3668903.25 | 5142967 | 3346414.25 | 4052761.5 |
| 15556866 | 2 | 0.034 | 2907899 | 1919112.875 | 2104998.25 | 2.31E+06 | 4046053.25 | 3980428 | 4199595.5 | 4075358.917 |
| 15540728 | 2 | 0.043 | 4108594.75 | 6175137 | 2903497.75 | 4.40E+06 | 2965125.25 | 3346896.75 | 4309698.5 | 3540573.5 |
| 15552410 | 2 | 0.007 | 1480889.5 | 2216266.75 | 1556394.5 | 1.75E+06 | 2252456.75 | 1779254 | 945128.25 | 1658946.333 |
| 15531929 | 2 | 0.048 | 1.53E+08 | 1.71E+08 | 1.62E+08 | 1.62E+08 | 1.30E+08 | 1.27E+08 | 1.27E+08 | 128033333.3 |
| 15539520 | 3 | 0.009 | 8.30E+07 | 9.23E+07 | 9.33E+07 | 8.96E+07 | 6.17E+07 | 6.67E+07 | 5.53E+07 | 61243333.33 |
| 15532903 | 2 | 0.038 | 5.15E+07 | 5.28E+07 | 6.01E+07 | 5.48E+07 | 2.50E+07 | 4.32E+07 | 3.82E+07 | 35493333.33 |
| 15551662 | 2 | 0.03 | 3.77E+07 | 2.60E+07 | 3.38E+07 | 3.25E+07 | 1.38E+07 | 2.01E+07 | 2.89E+07 | 20946666.67 |
| 15536711 | 3 | 0.042 | 3217547 | 1946862.875 | 2930617 | 2.70E+06 | 672873.438 | 1393674 | 2669691.75 | 1578746.396 |
| 15553627 | 2 | 0.022 | 5356049 | 5749256.5 | 6377563 | 5.83E+06 | 3034197.5 | 3779554.25 | 3186029.5 | 3333260.417 |
| 15548269 | 2 | 0.045 | 1.08E+07 | 1.01E+07 | 1.06E+07 | 1.05E+07 | 1.23E+07 | 7911682 | 8656154 | 9632612 |
| 15533057 | 2 | 0.003 | 5959630 | 7194415 | 6169262.5 | 6.44E+06 | 5180075 | 5468152.5 | 4824883 | 5157703.5 |
| 15531549 | 3 | 2.65E-05 | 2.02E+08 | 2.12E+08 | 2.08E+08 | 2.07E+08 | 7.88E+07 | 9.82E+07 | 1.09E+08 | 95306666.67 |
| [illegible] | [illegible] | [illegible] | [illegible] | [illegible] | [illegible] | [illegible] | [illegible] | [illegible] | [illegible] | [illegible] |

Membrane protein peptides with significant differences
$p<0.05$

| Charge Group ID | Charge State | Factor 1 P-value | Intensity [dKO_1] | Intensity [dKO_2] | Intensity [dKO_3] | Intensity [AQP1KO_1] | Intensity [AQP1KO_2] | Intensity [AQP1KO_3] | Intensity [RhAGKO_1] | Intensity [RhAGKO_2] | Intensity [RhAGKO_3] | Intensity [WT_1] | Intensity [WT_2] | Intensity [WT_3] | M + H (Feature) | Modified Peptide Sequence | Primary Protein Name | ID | Symbol | Entrez Gene Name | Primary Location | Type(s) |
|---|---|---|---|---|---|---|---|---|---|---|---|---|---|---|---|---|---|---|---|---|---|---|
| 15553994 | 4 | 0.045 | 9112585 | 8942083 | 1.34E+07 | 2581528 | 6011800.5 | 9196885 | 6076298.5 | 1.02E+07 | 9874654 | 3719093 | 3765471.75 | 4217887.5 | 5241.3037 | ALEYFASAGYHC[160.0307]EPYNNPAI | Q7TMS5 | Q7TMS5 | ABCG2 | ATP-binding cassette, sub-family G (WHITE), member 2 (Junior blood group) | Plasma Membrane | transporter |
| 15534926 | 2 | 0.042 | 1.02E+07 | 1.19E+07 | 1.19E+07 | 5411348.5 | 9329692 | 7469204.5 | 1.30E+07 | 1.31E+07 | 9509156 | 1.37E+07 | 1.09E+07 | 9807314 | 1061.5454 | VMSDLLPMR | Q7TMS5 | Q7TMS5 | ABCG2 | ATP-binding cassette, sub-family G (WHITE), member 2 (Junior blood group) | Plasma Membrane | transporter |
| 15550005 | 2 | 0.032 | 7114677 | 6309670.5 | 7497577.5 | 4231501.5 | 4409261.5 | 4648851.5 | 6078470 | 6761047 | 6204644 | 2353872.5 | 5236518.5 | 4607198.5 | 1820.9591 | GLSGDVLINGAPQPAHFK | Q7TMS5 | Q7TMS5 | ABCG2 | ATP-binding cassette, sub-family G (WHITE), member 2 (Junior blood group) | Plasma Membrane | transporter |
| 15532693 | 2 | 0.018 | 1.96E+07 | 1.77E+07 | 2.13E+07 | 9569948 | 1.44E+07 | 1.18E+07 | 1.77E+07 | 1.78E+07 | 1.45E+07 | 1.63E+07 | 1.51E+07 | 1.19E+07 | 996.56146 | LVFHGPAQK | Q7TMS5 | Q7TMS5 | ABCG2 | ATP-binding cassette, sub-family G (WHITE), member 2 (Junior blood group) | Plasma Membrane | transporter |
| 15531387 | 2 | 0.013 | 8.92E+07 | 9.86E+07 | 1.04E+08 | 5.94E+07 | 7.37E+07 | 4.53E+07 | 9.48E+07 | 9.80E+07 | 8.48E+07 | 8.56E+07 | 6.34E+07 | 6.43E+07 | 1251.7023 | PGLNAILGPTGGGK | Q7TMS5 | Q7TMS5 | ABCG2 | ATP-binding cassette, sub-family G (WHITE), member 2 (Junior blood group) | Plasma Membrane | transporter |
| 15538881 | 2 | 0.013 | 2.71E+07 | 2.42E+07 | 2.75E+07 | 1.34E+07 | 1.98E+07 | 1.29E+07 | 2.46E+07 | 2.35E+07 | 2.12E+07 | 2.35E+07 | 1.68E+07 | 1.75E+07 | 1148.6034 | ENLQFSAALR | Q7TMS5 | Q7TMS5 | ABCG2 | ATP-binding cassette, sub-family G (WHITE), member 2 (Junior blood group) | Plasma Membrane | transporter |
| 15533320 | 2 | 0.011 | 4.91E+07 | 3.94E+07 | 4.64E+07 | 2.98E+07 | 2.72E+07 | 2.69E+07 | 3.55E+07 | 3.94E+07 | 3.93E+07 | 4.37E+07 | 3.22E+07 | 2.93E+07 | 820.4658 | VGTQFIR | Q7TMS5 | Q7TMS5 | ABCG2 | ATP-binding cassette, sub-family G (WHITE), member 2 (Junior blood group) | Plasma Membrane | transporter |
| 15553996 | 2 | 0.01 | 1.21E+07 | 1.07E+07 | 1.20E+07 | 6022609 | 8341622.5 | 6983919.5 | 8927720 | 1.06E+07 | 9750185 | 1.00E+07 | 1.02E+07 | 7591999.5 | 1858.9417 | TLAEGDVLSFHHITYR | Q7TMS5 | Q7TMS5 | ABCG2 | ATP-binding cassette, sub-family G (WHITE), member 2 (Junior blood group) | Plasma Membrane | transporter |
| 15539354 | 2 | 0.008 | 5.23E+07 | 6.11E+07 | 6.67E+07 | 4.04E+07 | 4.56E+07 | 4.12E+07 | 5.67E+07 | 5.68E+07 | 5.65E+07 | 5.58E+07 | 4.38E+07 | 4.72E+07 | 1264.7119 | LFDSLTLLASGK | Q7TMS5 | Q7TMS5 | ABCG2 | ATP-binding cassette, sub-family G (WHITE), member 2 (Junior blood group) | Plasma Membrane | transporter |
| 15532526 | 3 | 0.007 | 1.48E+07 | 1.61E+07 | 1.67E+07 | 1.17E+07 | 1.08E+07 | 9239083 | 1.27E+07 | 1.49E+07 | 1.27E+07 | 1.49E+07 | 1.39E+07 | 1.19E+07 | 1635.7927 | EPVYVTSFC[160.0307]HQLR | Q7TMS5 | Q7TMS5 | ABCG2 | ATP-binding cassette, sub-family G (WHITE), member 2 (Junior blood group) | Plasma Membrane | transporter |
| 15536177 | 2 | 0.004 | 7.74E+07 | 8.01E+07 | 6.73E+07 | 5.41E+07 | 5.96E+07 | 5.40E+07 | 7.34E+07 | 7.91E+07 | 7.15E+07 | 6.69E+07 | 5.60E+07 | 5.42E+07 | 1044.603 | SSLLDVLAAR | Q7TMS5 | Q7TMS5 | ABCG2 | ATP-binding cassette, sub-family G (WHITE), member 2 (Junior blood group) | Plasma Membrane | transporter |
| 15537335 | 3, 4 | 0.003 | 1.37E+08 | 1.36E+08 | 1.28E+08 | 7.97E+07 | 9.91E+07 | 7.29E+07 | 1.26E+08 | 1.29E+08 | 1.12E+08 | 1.12E+08 | 1.03E+08 | 8.91E+07 | 1858.9393 | TLAEGDVLSFHHITYR | Q7TMS5 | Q7TMS5 | ABCG2 | ATP-binding cassette, sub-family G (WHITE), member 2 (Junior blood group) | Plasma Membrane | transporter |
| 15532315 | 2 | 0.003 | 1.08E+08 | 1.02E+08 | 1.13E+08 | 6.53E+07 | 7.98E+07 | 6.13E+07 | 9.93E+07 | 9.40E+07 | 8.62E+07 | 8.81E+07 | 8.38E+07 | 7.00E+07 | 1298.657 | AELDQLPGAQEK | Q7TMS5 | Q7TMS5 | ABCG2 | ATP-binding cassette, sub-family G (WHITE), member 2 (Junior blood group) | Plasma Membrane | transporter |
| 15540539 | 2 | 0.001 | 2.14E+07 | 2.10E+07 | 2.47E+07 | 1.36E+07 | 1.58E+07 | 1.58E+07 | 2.19E+07 | 2.04E+07 | 1.96E+07 | 1.49E+07 | 1.59E+07 | 1.18E+07 | 1232.6511 | EILSDINGIMK | Q7TMS5 | Q7TMS5 | ABCG2 | ATP-binding cassette, sub-family G (WHITE), member 2 (Junior blood group) | Plasma Membrane | transporter |
| 15535369 | 2 | 9.31E-04 | 3.23E+07 | 3.14E+07 | 3.75E+07 | 1.99E+07 | 2.23E+07 | 2.03E+07 | 2.48E+07 | 2.86E+07 | 2.74E+07 | 2.70E+07 | 2.17E+07 | 2.32E+07 | 701.45435 | INTIIK | Q7TMS5 | Q7TMS5 | ABCG2 | ATP-binding cassette, sub-family G (WHITE), member 2 (Junior blood group) | Plasma Membrane | transporter |
| 15532322 | 3 | 6.49E-04 | 5.02E+07 | 5.47E+07 | 5.28E+07 | 2.85E+07 | 3.51E+07 | 3.07E+07 | 4.36E+07 | 4.79E+07 | 3.85E+07 | 3.69E+07 | 4.18E+07 | 3.60E+07 | 1820.9618 | GLSGDVLINGAPQPAHFK | Q7TMS5 | Q7TMS5 | ABCG2 | ATP-binding cassette, sub-family G (WHITE), member 2 (Junior blood group) | Plasma Membrane | transporter |
| 15532640 | 2 | 0.009 | 3.32E+07 | 3.77E+07 | 4.14E+07 | 2.77E+07 | 3.28E+07 | 3.23E+07 | 3.43E+07 | 3.64E+07 | 3.03E+07 | 2.85E+07 | 2.13E+07 | 1.97E+07 | 1060.5979 | SLSSLSLPTR | Q9QUI6 | Q9QUI6 | ACKR1 | atypical chemokine receptor 1 (Duffy blood group) | Plasma Membrane | G-protein coupled receptor |
| 15560452 | 2 | 0.023 | 5798964 | 1.21E+07 | 1.33E+07 | 2445018.5 | 8160040 | 5920143.5 | 7987620 | 1.02E+07 | 8051267.5 | 2304488 | 2725428 | 3796067.25 | 1634.6273 | YDLEEC[160.0307]TC[160.0307]ASSD | O35598 | O35598 | ADAM10 | ADAM metallopeptidase domain 10 | Plasma Membrane | peptidase |
| 15544758, 15545403, 1554603 | 3 | 0.021 | 9807622 | 1.12E+07 | 9285017 | 5983181.5 | 5164764 | 5982211.5 | 1.22E+07 | 7733473 | 6740789.5 | 6809959 | 4671316 | 6789846.5 | 2025.0685, 2025.0702, 20 | SLNTGIITVQNYGSHVPPK | O35598 | O35598 | ADAM10 | ADAM metallopeptidase domain 10 | Plasma Membrane | peptidase |
| 15531412 | 3 | 0.008 | 8.49E+07 | 6.68E+07 | 4.52E+07 | 3.56E+07 | 3.95E+07 | 3.33E+07 | 7.63E+07 | 6.37E+07 | 6.89E+07 | 3.27E+07 | 4.92E+07 | 3.49E+07 | 1927.9333 | NTC[160.0307]QLYIQTDHLFFK | O35598 | O35598 | ADAM10 | ADAM metallopeptidase domain 10 | Plasma Membrane | peptidase |
| 15553947 | 2 | 0.007 | 4.83E+07 | 3.97E+07 | 4.73E+07 | 2.17E+07 | 3.52E+07 | 2.52E+07 | 5.11E+07 | 5.31E+07 | 5.32E+07 | 4.28E+07 | 3.81E+07 | 2.83E+07 | 1605.765 | TITLQPGSPC[160.0307]NDFR | O35598 | O35598 | ADAM10 | ADAM metallopeptidase domain 10 | Plasma Membrane | peptidase |
| 15537548 | 3 | 0.006 | 1.09E+08 | 1.03E+08 | 7.75E+07 | 6.42E+07 | 3.97E+07 | 4.46E+07 | 9.04E+07 | 7.22E+07 | 8.34E+07 | 4.52E+07 | 6.20E+07 | 5.78E+07 | 1730.8823 | DPTNPFRFPNIGVEK | O35598 | O35598 | ADAM10 | ADAM metallopeptidase domain 10 | Plasma Membrane | peptidase |
| 15535111 | 2 | 0.005 | 6023721 | 4236009 | 3260112.5 | 2263817 | 2190632.5 | 1680782.375 | 4874260 | 4348194.5 | 4289162 | 3345994.25 | 4099042 | 2644577.75 | 1236.4955 | ELC[160.0307]HVC[160.0307]C[160.0 | O35598 | O35598 | ADAM10 | ADAM metallopeptidase domain 10 | Plasma Membrane | peptidase |
| 15536888 | 2 | 0.005 | 5534824.5 | 5014500 | 6332096.5 | 2441849.75 | 3106256.25 | 2703498.25 | 4567139 | 5599671 | 3638257 | 2790663.5 | 1150253 | 2250576.25 | 1063.4308 | GYC[160.0307]DVFM[147.0354]R | O35598 | O35598 | ADAM10 | ADAM metallopeptidase domain 10 | Plasma Membrane | peptidase |
| 15558526 | 2 | 0.003 | 4538889.5 | 5717032.5 | 6725028.5 | 2853095.25 | 2853672.75 | 2505677.75 | 4054962 | 3264238.75 | 3498419 | 2531037 | 948246.062 | 1650399.5 | 1727.7693 | M[147.0354]APSTC[160.0307]ASTGS | O35598 | O35598 | ADAM10 | ADAM metallopeptidase domain 10 | Plasma Membrane | peptidase |
| 15552097 | 2 | 0.001 | 5.78E+07 | 4.08E+07 | 4.64E+07 | 2.60E+07 | 3.03E+07 | 2.30E+07 | 4.84E+07 | 4.71E+07 | 4.54E+07 | 3.29E+07 | 3.36E+07 | 2.57E+07 | 1986.7885 | QC[160.0307]SPSQGPC[160.0307]C[1 | O35598 | O35598 | ADAM10 | ADAM metallopeptidase domain 10 | Plasma Membrane | peptidase |
| 15548230 | 2 | 0.001 | 2.10E+07 | 1.78E+07 | 1.71E+07 | 1.12E+07 | 7132579.5 | 7720465 | 1.53E+07 | 1.14E+07 | 1.33E+07 | 9162740 | 1.02E+07 | 9527662 | 1730.8827 | DPTNPFRFPNIGVEK | O35598 | O35598 | ADAM10 | ADAM metallopeptidase domain 10 | Plasma Membrane | peptidase |
| 15548337 | 2 | 8.59E-04 | 2.53E+07 | 2.59E+07 | 3.77E+07 | 1.56E+07 | 1.78E+07 | 1.54E+07 | 2.71E+07 | 2.84E+07 | 2.17E+07 | 1.55E+07 | 1.19E+07 | 1.09E+07 | 1555.5759 | DDC[160.0307]C[160.0307]FDANQPE | O35598 | O35598 | ADAM10 | ADAM metallopeptidase domain 10 | Plasma Membrane | peptidase |
| 15535140 | 2 | 7.39E-04 | 3.03E+07 | 2.66E+07 | 1.99E+07 | 1.38E+07 | 1.35E+07 | 1.36E+07 | 2.56E+07 | 2.27E+07 | 2.58E+07 | 1.72E+07 | 1.73E+07 | 1.41E+07 | 882.4484 | FSLC[160.0307]SIR | O35598 | O35598 | ADAM10 | ADAM metallopeptidase domain 10 | Plasma Membrane | peptidase |
| 15540052 | 3 | 4.61E-04 | 1.39E+07 | 1.76E+07 | 1.34E+07 | 8187766.5 | 9615407 | 8316365.5 | 1.94E+07 | 1.74E+07 | 1.55E+07 | 9855667 | 1.21E+07 | 9381623 | 1986.7898 | QC[160.0307]SPSQGPC[160.0307]C[1 | O35598 | O35598 | ADAM10 | ADAM metallopeptidase domain 10 | Plasma Membrane | peptidase |
| 15533215 | 2 | 3.12E-04 | 6.17E+07 | 5.68E+07 | 6.58E+07 | 3.28E+07 | 3.31E+07 | 2.57E+07 | 5.69E+07 | 5.81E+07 | 5.33E+07 | 4.26E+07 | 3.67E+07 | 2.94E+07 | 1026.5548 | LVDADGPLAR | O35598 | O35598 | ADAM10 | ADAM metallopeptidase domain 10 | Plasma Membrane | peptidase |
| 15549878 | 2 | 2.94E-04 | 6.57E+07 | 5.08E+07 | 6.16E+07 | 3.10E+07 | 3.47E+07 | 2.67E+07 | 5.69E+07 | 5.36E+07 | 5.05E+07 | 3.97E+07 | 3.51E+07 | 3.11E+07 | 1711.774 | MAPSTC[160.0307]ASTGSLQWSK | O35598 | O35598 | ADAM10 | ADAM metallopeptidase domain 10 | Plasma Membrane | peptidase |
| 15534801 | 2 | 4.91E-05 | 1.36E+07 | 1.31E+07 | 1.27E+07 | 8614181 | 7270862.5 | 7762127.5 | 1.39E+07 | 1.14E+07 | 1.12E+07 | 6637894 | 7509862.5 | 7629496 | 838.4472 | NISFMVK | O35598 | O35598 | ADAM10 | ADAM metallopeptidase domain 10 | Plasma Membrane | peptidase |
| 15553211 | 2 | 2.08E-05 | 6.68E+07 | 7.05E+07 | 7.99E+07 | 4.38E+07 | 4.29E+07 | 4.06E+07 | 6.59E+07 | 6.35E+07 | 6.03E+07 | 4.05E+07 | 3.13E+07 | 3.45E+07 | 1700.844 | AIDTIYQTTDFSGIR | O35598 | O35598 | ADAM10 | ADAM metallopeptidase domain 10 | Plasma Membrane | peptidase |
| 15533516 | 2, 3 | 8.53E-06 | 4.51E+07 | 3.86E+07 | 4.45E+07 | 2.35E+07 | 2.46E+07 | 2.28E+07 | 3.71E+07 | 4.37E+07 | 3.99E+07 | 2.14E+07 | 2.51E+07 | 2.34E+07 | 1281.7125, 1281.7126 | EAVIAQISSHVK | O35598 | O35598 | ADAM10 | ADAM metallopeptidase domain 10 | Plasma Membrane | peptidase |
| 15535477, 15543763 | 3 | 0.049 | 3.87E+08 | 4.27E+08 | 4.43E+08 | 3.86E+08 | 3.03E+08 | 4.07E+08 | 3.48E+08 | 3.49E+08 | 3.24E+08 | 5.50E+08 | 3.97E+08 | 4.59E+08 | 1842.009, 1842.0106 | LLGNQATFSPIVTVEPR | Q02357 | Q02357 | ANK1 | ankyrin 1, erythrocytic | Plasma Membrane | other |
| 15543229, 15543504, 1554486 | 4 | 0.045 | 5.51E+07 | 3.93E+07 | 3.12E+07 | 4.11E+07 | 5.84E+07 | 3.81E+07 | 7.58E+07 | 6.42E+07 | 5.75E+07 | 6.44E+07 | 1.62E+08 | 7.54E+07 | 2471.1426, 2471.143 | YEDTQHILC[160.0307]HLNITMPPC[16 | Q02357 | Q02357 | ANK1 | ankyrin 1, erythrocytic | Plasma Membrane | other |
| 15548272, 15549661 | 2, 3 | 0.032 | 3.71E+09 | 3.69E+09 | 3.75E+09 | 3.53E+09 | 3.60E+09 | 3.54E+09 | 3.29E+09 | 3.32E+09 | 3.20E+09 | 4.19E+09 | 3.59E+09 | 3.52E+09 | 2409.3455, 2409.3496 | SGLTPLHLVSQEGHVPVADVLIK | Q02357 | Q02357 | ANK1 | ankyrin 1, erythrocytic | Plasma Membrane | other |
| 15531741 | 2 | 0.028 | 8.23E+08 | 8.18E+08 | 8.81E+08 | 7.95E+08 | 8.82E+08 | 7.64E+08 | 8.32E+08 | 8.39E+08 | 6.79E+08 | 1.07E+09 | 9.21E+08 | 1.01E+09 | 1047.5919 | EIILETTTK | Q02357 | Q02357 | ANK1 | ankyrin 1, erythrocytic | Plasma Membrane | other |
| 15535131, 15545793 | 4 | 0.025 | 1.68E+09 | 1.55E+09 | 1.48E+09 | 1.57E+09 | 1.49E+09 | 1.57E+09 | 1.55E+09 | 1.53E+09 | 1.34E+09 | 1.72E+09 | 1.72E+09 | 1.82E+09 | 2526.339, 2526.3406 | LLLENGASPNLATTAGHTPLHTAAR | Q02357 | Q02357 | ANK1 | ankyrin 1, erythrocytic | Plasma Membrane | other |
| 15531298, 15531307 | 3, 4 | 0.025 | 7.76E+08 | 7.24E+08 | 7.22E+08 | 7.88E+08 | 6.47E+08 | 5.70E+08 | 8.00E+08 | 6.50E+08 | 7.37E+08 | 9.31E+08 | 8.95E+08 | 9.03E+08 | 2061.0254, 2061.0283 | GGSPHSPAWNGYTPLHIAAK | Q02357 | Q02357 | ANK1 | ankyrin 1, erythrocytic | Plasma Membrane | other |
| 15539138, 15545247 | 2 | 0.022 | 3.98E+08 | 3.65E+08 | 3.58E+08 | 4.12E+08 | 3.06E+08 | 4.23E+08 | 2.98E+08 | 3.30E+08 | 2.81E+08 | 4.61E+08 | 4.17E+08 | 4.16E+08 | 845.51434, 845.51556 | VMELLLK | Q02357 | Q02357 | ANK1 | ankyrin 1, erythrocytic | Plasma Membrane | other |
| 15534719, 15537375 | 3, 4 | 0.015 | 1.14E+09 | 9.80E+08 | 9.81E+08 | 1.02E+09 | 1.01E+09 | 9.81E+08 | 1.04E+09 | 1.12E+09 | 8.89E+08 | 1.40E+09 | 1.33E+09 | 1.19E+09 | 1787.8856, 1787.8868 | MGYTPLHVASHYGNIK | Q02357 | Q02357 | ANK1 | ankyrin 1, erythrocytic | Plasma Membrane | other |
| 15531819 | 2 | 0.014 | 5.31E+08 | 5.23E+08 | 5.41E+08 | 5.06E+08 | 5.87E+08 | 5.77E+08 | 5.68E+08 | 5.36E+08 | 4.71E+08 | 7.10E+08 | 6.70E+08 | 6.15E+08 | 1122.5862 | ELTAVPYMAK | Q02357 | Q02357 | ANK1 | ankyrin 1, erythrocytic | Plasma Membrane | other |
| 15552205, 15560849 | 2 | 0.012 | 5.86E+08 | 7.07E+08 | 5.41E+08 | 7.32E+08 | 6.51E+08 | 7.62E+08 | 7.16E+08 | 7.89E+08 | 9.50E+08 | 9.99E+08 | 1.05E+09 | 8.29E+08 | 1815.9679, 1815.9696 | IITTDFPLYFVIMSR | Q02357 | Q02357 | ANK1 | ankyrin 1, erythrocytic | Plasma Membrane | other |
| 15538435 | 2 | 0.01 | 3.89E+08 | 3.33E+08 | 3.46E+08 | 3.75E+08 | 3.31E+08 | 2.95E+08 | 4.07E+08 | 3.84E+08 | 3.73E+08 | 5.62E+08 | 7.29E+08 | 4.37E+08 | 1080.4912 | FWLSDC[160.0307]PR | Q02357 | Q02357 | ANK1 | ankyrin 1, erythrocytic | Plasma Membrane | other |
| 15557295 | 3 | 0.035 | 758379.125 | 1097779.125 | 2055721.875 | 3703036.75 | 4365237.5 | 3542481 | 2163863.25 | 2366027.25 | 4111870.5 | 1100232.875 | 2939022.75 | 2130282.5 | 2461.1257 | TPEFEEFNGKPDSLFFTDGQR | Q6P9J9 | Q6P9J9 | ANO6 | anoctamin 6 | Plasma Membrane | ion channel |
| 15548059, 15559284, 1556012 | 3 | 1.44E-06 | 3696251.5 | 3477631.75 | 4591540 | 2731997.25 | 5056811 | 4944830 | 9.57E+07 | 4.81E+07 | 5.58E+07 | 6.35E+07 | 4.92E+07 | 4.51E+07 | 2815.2651, 2815.2676, 28 | VWTSGQVEEYDLDADDINSRVEM[147. | Q02013 | Q02013 | AQP1 | aquaporin 1 (Colton blood group) | Plasma Membrane | transporter |
| 15534298 | 2 | 6.47E-07 | 2168103.75 | 256381.844 | 317710.844 | 3372884.5 | 573411.125 | 809259.625 | 2.66E+09 | 2.67E+09 | 2.30E+09 | 2.73E+09 | 2.31E+09 | 2.52E+09 | 937.5096 | SFGSAVLTR | Q02013 | Q02013 | AQP1 | aquaporin 1 (Colton blood group) | Plasma Membrane | transporter |
| 15548607, 15555888, 1555610 | 3 | 1.61E-07 | 1.56E+07 | 1.02E+07 | 2.14E+07 | 1.07E+07 | 1.80E+07 | 2.52E+07 | 1.02E+09 | 6.77E+08 | 9.30E+08 | 8.84E+08 | 7.44E+08 | 9.28E+08 | 2799.2717, 2799.273, 279 | VWTSGQVEEYDLDADDINSRVEMK | Q02013 | Q02013 | AQP1 | aquaporin 1 (Colton blood group) | Plasma Membrane | transporter |
| 15534117 | 2 | 1.14E-08 | 344928.938 | 500562.188 | 670803.312 | 148100.469 | 323350.906 | 542089.062 | 4.62E+08 | 4.02E+08 | 3.41E+08 | 3.96E+08 | 2.70E+08 | 3.14E+08 | 1158.6089 | NQTLVQDNVK | Q02013 | Q02013 | AQP1 | aquaporin 1 (Colton blood group) | Plasma Membrane | transporter |
| 15549937 | 2 | 4.75E-10 | 7243747.5 | 6283545 | 6953285.5 | 8825466 | 6722767 | 7136432 | 2.38E+09 | 2.54E+09 | 2.03E+09 | 2.56E+09 | 1.30E+09 | 1.40E+09 | 2312.0288 | VWTSGQVEEYDLDADDINSR | Q02013 | Q02013 | AQP1 | aquaporin 1 (Colton blood group) | Plasma Membrane | transporter |
| 15549966 | 2, 3 | 0.041 | 1.67E+07 | 1.84E+07 | 2.21E+07 | 2.01E+07 | 2.06E+07 | 1.97E+07 | 1.23E+07 | 1.48E+07 | 1.71E+07 | 1.72E+07 | 1.33E+07 | 1.62E+07 | 2254.1902, 2254.1912 | LGQSVTTIPTVGFNVETVTYK | P62331 | P62331 | ARF6 | ADP-ribosylation factor 6 | Plasma Membrane | transporter |
| 15531796 | 3 | 0.001 | 3.54E+07 | 4.11E+07 | 3.27E+07 | 4.72E+07 | 4.48E+07 | 4.72E+07 | 4.68E+07 | 4.86E+07 | 4.16E+07 | 5.89E+07 | 5.28E+07 | 5.41E+07 | 1764.8912 | GVNAIVYMIDAADREK | Q9CQW2 | Q9CQW2 | ARL8B | ADP-ribosylation factor-like 8B | Plasma Membrane | enzyme |
| 15548389 | 2 | 0.034 | 3.13E+07 | 3.45E+07 | 3.41E+07 | 2.58E+07 | 2.63E+07 | 2.31E+07 | 2.19E+07 | 2.92E+07 | 2.95E+07 | 2.54E+07 | 2.40E+07 | 1.77E+07 | 1584.8728 | AVFQANQENLPILK | Q8VDN2 | Q8VDN2 | ATP1A1 | ATPase, Na+/K+ transporting, alpha 1 polypeptide | Plasma Membrane | transporter |
| 15536458 | 3 | 0.014 | 1.00E+07 | 9094637 | 8480550 | 8036977.5 | 7820937.5 | 7362742.5 | 7695976 | 7117454 | 6610249.5 | 9633963 | 7917677.5 | 9278553 | 1196.7157 | LATPGLMIRPK | P14231 | P14231 | ATP1B2 | ATPase, Na+/K+ transporting, beta 2 polypeptide | Plasma Membrane | transporter |
| 15538400 | 2 | 0.042 | 1622155.25 | 3556071 | 2024881 | 1504280.5 | 2621272.5 | 2648211.25 | 2464791.25 | 4383864 | 4485995 | 740454.562 | 1327486.625 | 1738637.5 | 1192.644 | LVEDLESFLK | P97370 | P97370 | ATP1B3 | ATPase, Na+/K+ transporting, beta 3 polypeptide | Plasma Membrane | transporter |
| 15532977 | 3 | 0.033 | 4.58E+07 | 5.29E+07 | 5.11E+07 | 3.35E+07 | 4.32E+07 | 3.68E+07 | 3.98E+07 | 4.74E+07 | 3.74E+07 | 4.86E+07 | 4.79E+07 | 4.45E+07 | 1993.838 | HLDAC[160.0307]ETMGNATAIC[160.( | Q9R0K7 | Q9R0K7 | ATP2B2 | ATPase, Ca++ transporting, plasma membrane 2 | Plasma Membrane | transporter |
| 15550031 | 2 | 0.009 | 1.10E+07 | 1.14E+07 | 1.04E+07 | 7803906.5 | 8989889 | 7423683 | 9004919 | 9207908 | 8076875 | 1.19E+07 | 1.07E+07 | 9261237 | 1993.8379 | HLDAC[160.0307]ETMGNATAIC[160.( | Q9R0K7 | Q9R0K7 | ATP2B2 | ATPase, Ca++ transporting, plasma membrane 2 | Plasma Membrane | transporter |
| 15532388 | 2 | 0.009 | 6156096.5 | 8234439.5 | 7131402 | 9011798 | 8544231 | 1.06E+07 | 7290124 | 1.09E+07 | 1.21E+07 | 3706802.5 | 6289381.5 | 4267590.5 | 1091.5823 | TPQIQVYSR | P01887 | P01887 | B2M | beta-2-microglobulin | Plasma Membrane | transmembrane receptor |
| 15546222 | 3 | 0.025 | 2042919.375 | 1907705.5 | 1320302.75 | 1231934.75 | 1432357 | 1345001.625 | 2109684.5 | 1654580.875 | 1810117 | 4108788.75 | 2081852.875 | 2454507.5 | 1859.8607 | SDASYPPITDWFWFK | P18572 | P18572 | BSG | basigin (Ok blood group) | Plasma Membrane | transporter |
| 15553918 | 4 | 0.03 | 1301408.5 | 3101178 | 1173516.375 | 3038950 | 1383976 | 2456826.75 | 1064370.5 | 1437308.75 | 1290917.75 | 813162.125 | 834745.438 | 614762.375 | 4877.431 | LAQSNQVPVQPEATQVPLVSSTSEGYTA | Q60865 | Q60865 | CAPRIN1 | cell cycle associated protein 1 | Plasma Membrane | other |
| 15544984 | 2 | 0.049 | 3764255.25 | 2793972 | 1772084.75 | 3561800 | 2280475 | 2257873 | 1725860.25 | 2034529.625 | 2250784.75 | 7482531.5 | 4732767.5 | 3209600.25 | 926.6017 | LQVNILVK | Q08857 | Q08857 | CD36 | CD36 molecule (thrombospondin receptor) | Plasma Membrane | transmembrane receptor |
| 15548320 | 3 | 0.032 | 5.56E+08 | 5.27E+08 | 4.90E+08 | 6.16E+08 | 5.90E+08 | 5.53E+08 | 5.32E+08 | 5.40E+08 | 4.51E+08 | 7.88E+08 | 6.60E+08 | 5.82E+08 | 2777.2761 | FVLPANAFASPLQNPDNHC[160.0307]I | Q08857 | Q08857 | CD36 | CD36 molecule (thrombospondin receptor) | Plasma Membrane | transmembrane receptor |
| 15531688 | 2 | 0.013 | 1.17E+08 | 1.27E+08 | 1.18E+08 | 1.11E+08 | 1.42E+08 | 1.37E+08 | 1.07E+08 | 1.02E+08 | 1.06E+08 | 1.50E+08 | 1.32E+08 | 1.40E+08 | 922.52325 | VAIIESYK | Q08857 | Q08857 | CD36 | CD36 molecule (thrombospondin receptor) | Plasma Membrane | transmembrane receptor |
| 15531480 | 2 | 0.002 | 6.97E+07 | 6.29E+07 | 6.27E+07 | 7.67E+07 | 7.48E+07 | 6.86E+07 | 7.41E+07 | 7.42E+07 | 5.82E+07 | 1.01E+08 | 9.36E+07 | 9.38E+07 | 1031.4613 | FFSSDIC[160.0307]R | Q08857 | Q08857 | CD36 | CD36 molecule (thrombospondin receptor) | Plasma Membrane | transmembrane receptor |
| 15556255 | 2 | 0.041 | 2792644.25 | 4338609 | 6449034.5 | 2355496 | 4207019 | 3651010.25 | 3300277.75 | 4206760 | 4257892 | 2290726.25 | 1898552.5 | 1921812.25 | 1866.7463 | EPSETPDQC[160.0307]MTADETR | P15379 | P15379 | CD44 | CD44 molecule (Indian blood group) | Plasma Membrane | enzyme |
| 15538767 | 2 | 0.048 | 2663666.25 | 2956068 | 2447786.5 | 1945206.625 | 2560806 | 1965509.875 | 1258305.75 | 1607800.25 | 2344229.75 | 2316017.25 | 3188429.75 | 2941624.75 | 1257.7069 | ITLEVFDPVPK | P18181 | P18181 | CD48 | CD48 molecule | Plasma Membrane | other |
| 15534535 | 2 | 0.03 | 5141186 | 6634307 | 6047476.5 | 5695951.5 | 5088053 | 5900782.5 | 5671013 | 4801980 | 4940483.5 | 3547051.75 | 4351169 | 4757378.5 | 765.44855 | LSFASLK | Q61475 | Q61475 | CD55 | CD55 molecule, decay accelerating factor for complement (Cromer blood group) | Plasma Membrane | other |
| 15535070 | 2 | 9.84E-04 | 5.99E+07 | 5.48E+07 | 5.18E+07 | 5.41E+07 | 5.29E+07 | 4.79E+07 | 6.27E+07 | 6.06E+07 | 5.43E+07 | 9.31E+07 | 8.60E+07 | 7.15E+07 | 1355.6373 | WIQC[160.0307]TLYDTR | Q9Z0S4 | Q9Z0S4 | Cldn13 | claudin 13 | Plasma Membrane | other |
| 15550569 | 3 | 0.032 | 3660031.5 | 3584001.75 | 3056519.25 | 3495308 | 4335982.5 | 4462046 | 5212837 | 5960719 | 4069739.75 | 4515070 | 7573223 | 5715668.5 | 3292.5422 | YMSRPVMSGPGLYDPTTIMNADILAYC[ | O35372 | O35372 | CNIH1 | cornichon family AMPA receptor auxiliary protein 1 | Plasma Membrane | other |
| 15537421 | 2 | 0.023 | 3.09E+07 | 2.60E+07 | 2.83E+07 | 2.14E+07 | 2.35E+07 | 2.08E+07 | 2.85E+07 | 2.35E+07 | 2.20E+07 | 2.68E+07 | 2.76E+07 | 2.97E+07 | 1039.5328 | RQFSITC[160.0307]K | Q64735 | Q64735 | CR1L | complement component (3b/4b) receptor 1-like | Plasma Membrane | other |
| 15532314 | 2 | 0.046 | 1.06E+07 | 8048337.5 | 7990032 | 1.01E+07 | 1.26E+07 | 6794574 | 1.46E+07 | 1.35E+07 | 1.16E+07 | 1.73E+07 | 2.30E+07 | 1.17E+07 | 1153.5692 | PC[160.0307]YATLFGPK | Q9DCT8 | Q9DCT8 | Crip2 | cysteine rich protein 2 | Plasma Membrane | other |
| 15542480 | 3 | 0.019 | 3105587.75 | 3434277.25 | 1819945.75 | 1817706.875 | 2487432.5 | 1219697.875 | 2169088.5 | 2491181 | 2545542.5 | 1135162.875 | 1402978.875 | 909930.875 | 2152.0735 | NAGQAVTIVAQYRPEEYSR | Q811D0 | Q811D0 | DLG1 | discs, large homolog 1 (Drosophila) | Plasma Membrane | kinase |
| 15547023 | 2 | 0.012 | 997075.5 | 1113409.25 | 725918 | 584208.812 | 787116.938 | 292578.406 | 736352.625 | 729185.438 | 722970.125 | 194389.594 | 422393.812 | 260740.422 | 1129.5933 | IITGGAAAQDGR | Q811D0 | Q811D0 | DLG1 | discs, large homolog 1 (Drosophila) | Plasma Membrane | kinase |
| 15533060 | 2 | 0.044 | 7723695.5 | 8290014.5 | 8058856 | 6617645.5 | 7585194 | 4427460.5 | 7719143 | 7322231 | 7772494.5 | 6268063 | 5091533 | 4999642.5 | 865.5126 | IVTTYIR | P39054 | P39054 | DNM2 | dynamin 2 | Plasma Membrane | enzyme |
| 15535497 | 2 | 0.041 | 1.04E+07 | 1.34E+07 | 1.49E+07 | 5769959 | 1.32E+07 | 6478622 | 1.13E+07 | 1.39E+07 | 1.23E+07 | 8284223.5 | 4254362 | 6367640 | 1107.5769 | SSVLENFVGR | P39054 | P39054 | DNM2 | dynamin 2 | Plasma Membrane | enzyme |
| 15553990 | 3 | 0.041 | 7153853 | 9539385 | 1.03E+07 | 5676773.5 | 9141628 | 4175367 | 7661185.5 | 7747559.5 | 9270643 | 5963620.5 | 3839921.75 | 3918975.5 | 2524.0479 | DQAENEDGAQENTFSMDPQLER | P39054 | P39054 | DNM2 | dynamin 2 | Plasma Membrane | enzyme |
| 15531940 | 2 | 0.036 | 2.44E+07 | 2.23E+07 | 2.40E+07 | 1.81E+07 | 2.02E+07 | 1.16E+07 | 1.70E+07 | 2.10E+07 | 2.14E+07 | 1.62E+07 | 1.21E+07 | 1.47E+07 | 1348.7461 | NLVDSYVAIINK | P39054 | P39054 | DNM2 | dynamin 2 | Plasma Membrane | enzyme |
| 15542103 | 2 | 0.036 | 4980224 | 5621133.5 | 5864441 | 7628546 | 7921924 | 6264179.5 | 5692414.5 | 6607787 | 8095432.5 | 4835173 | 5833218.5 | 5008053.5 | 877.487 | GYIGVVNR | P39054 | P39054 | DNM2 | dynamin 2 | Plasma Membrane | enzyme |
| 15534869 | 2 | 0.035 | 1.29E+07 | 1.29E+07 | 1.04E+07 | 8984620 | 9622472 | 6064899 | 9666184 | 8945991 | 1.09E+07 | 8735292 | 6181533 | 7509339.5 | 1122.5953 | DMILQFISR | P39054 | P39054 | DNM2 | dynamin 2 | Plasma Membrane | enzyme |
| 15546005 | 3 | 0.035 | 2141893.5 | 2809288 | 3967102.5 | 1506151.25 | 1958180.875 | 1433710.875 | 1898958.5 | 2724110 | 4183124.5 | 1501806.25 | 1501476.25 | 1687164.25 | 1327.843 | RPLILQLIFSK | P39054 | P39054 | DNM2 | dynamin 2 | Plasma Membrane | enzyme |
| 15549297 | 2 | 0.01 | 1.46E+07 | 1.46E+07 | 1.62E+07 | 9435401 | 1.33E+07 | 7814624 | 1.32E+07 | 1.26E+07 | 1.43E+07 | 1.03E+07 | 7883130 | 8144057.5 | 1922.8401 | QIELAC[160.0307]DSQEDVDSWK | P39054 | P39054 | DNM2 | dynamin 2 | Plasma Membrane | enzyme |
| 15548345 | 2 | 0.01 | 1.66E+07 | 1.54E+07 | 1.39E+07 | 1.24E+07 | 1.14E+07 | 8045128.5 | 1.11E+07 | 1.26E+07 | 1.41E+07 | 1.01E+07 | 8267816.5 | 7419859 | 2273.1614 | ESSLILAVTPANMDLANSDALK | P39054 | P39054 | DNM2 | dynamin 2 | Plasma Membrane | enzyme |
| 15533153 | 2 | 0.009 | 1.40E+07 | 1.50E+07 | 1.19E+07 | 8820659 | 1.10E+07 | 6934880 | 1.02E+07 | 1.01E+07 | 1.16E+07 | 9075466 | 7292503 | 7332416.5 | 1065.6403 | GISPVPINLR | P39054 | P39054 | DNM2 | dynamin 2 | Plasma Membrane | enzyme |
| 15531642 | 3 | 0.006 | 4.26E+07 | 4.84E+07 | 4.45E+07 | 2.98E+07 | 3.59E+07 | 2.33E+07 | 4.16E+07 | 3.35E+07 | 4.20E+07 | 2.73E+07 | 2.39E+07 | 2.91E+07 | 2093.196 | VYSPHVLNLTLIDLPGITK | P39054 | P39054 | DNM2 | dynamin 2 | Plasma Membrane | enzyme |
| 15538439 | 2 | 0.021 | 4847626.5 | 5173966.5 | 4966304.5 | 4429582 | 4913976.5 | 3799287.5 | 4467611 | 4506419.5 | 3922249.5 | 3551954.25 | 4082026.25 | 3299458.5 | 1424.6982 | IAESLGGSGYSVER | Q8BG67 | Q8BG67 | EFR3A | EFR3 homolog A | Plasma Membrane | other |
| 15536194 | 2 | 0.016 | 4620136.5 | 4100802.75 | 4138376 | 3410548.5 | 3779902.25 | 3043874.25 | 4260751.5 | 4439618.5 | 3312093.75 | 2885842.75 | 3287264 | 2871170.5 | 1126.572 | LGPPSSPSAADK | Q8BG67 | Q8BG67 | EFR3A | EFR3 homolog A | Plasma Membrane | other |
| 15558089 | 2 | 0.049 | 4559542 | 4757231 | 7744559 | 2886766.25 | 4303108.5 | 3117911.25 | 4373928.5 | 4741010.5 | 5039716.5 | 3357036.5 | 4204508 | 3296679 | 1668.7905 | SLAAEAENSQHQQQK | P48193 | P48193 | EPB41 | erythrocyte membrane protein band 4.1 | Plasma Membrane | other |
| 15532413, 15532525 | 2, 3 | 0.031 | 1.57E+08 | 1.18E+08 | 1.39E+08 | 1.32E+08 | 1.11E+08 | 1.20E+08 | 1.25E+08 | 1.23E+08 | 1.12E+08 | 1.76E+08 | 1.66E+08 | 1.41E+08 | 1377.7094, 1377.7097, 13 | ERLDGENIYIR | P48193 | P48193 | EPB41 | erythrocyte membrane protein band 4.1 | Plasma Membrane | other |
| 15548115 | 2 | 0.019 | 1.47E+08 | 1.95E+08 | 2.11E+08 | 1.72E+08 | 2.08E+08 | 1.26E+08 | 1.31E+08 | 1.79E+08 | 1.78E+08 | 9.82E+07 | 9.68E+07 | 1.10E+08 | 1819.9835 | TLNINGQVPTGDGPPLVK | P48193 | P48193 | EPB41 | erythrocyte membrane protein band 4.1 | Plasma Membrane | other |
| 15533447 | 2 | 0.01 | 4.37E+07 | 4.89E+07 | 5.60E+07 | 3.56E+07 | 3.38E+07 | 3.08E+07 | 5.01E+07 | 5.22E+07 | 4.14E+07 | 4.85E+07 | 3.99E+07 | 3.94E+07 | 1135.6335 | APIAAPEPELK | P48193 | P48193 | EPB41 | erythrocyte membrane protein band 4.1 | Plasma Membrane | other |
| 15547765, 15548679, 1555626 | 3, 4 | 0.049 | 1.25E+09 | 1.37E+09 | 9.72E+08 | 1.28E+09 | 1.08E+09 | 1.16E+09 | 1.32E+09 | 1.23E+09 | 1.26E+09 | 1.66E+09 | 1.42E+09 | 1.53E+09 | 3080.4797, 3080.4844, 30 | YSHTSLSC[160.0307]FAQENMAIGKPD | P49222 | P49222 | EPB42 | erythrocyte membrane protein band 4.2 | Plasma Membrane | transporter |
| 15551489 | 2, 3 | 0.046 | 1.44E+09 | 1.49E+09 | 1.26E+09 | 1.31E+09 | 1.24E+09 | 1.25E+09 | 1.30E+09 | 1.32E+09 | 1.26E+09 | 1.71E+09 | 1.53E+09 | 1.37E+09 | 2474.1616, 2474.1626 | LSTNLSFSC[160.0307]FEQTPPENSFLR | P49222 | P49222 | EPB42 | erythrocyte membrane protein band 4.2 | Plasma Membrane | transporter |
| 15556806, 15557011, 1555776 | 3, 4 | 0.037 | 1.51E+07 | 1.90E+07 | 1.90E+07 | 8198479 | 1.28E+07 | 1.40E+07 | 1.46E+07 | 1.58E+07 | 9555397 | 9295253 | 8873563 | 1.03E+07 | 3121.6157, 3121.6182, 31 | DAQHWTVSVTTPVDAVIGHYSLLLQVS( | P49222 | P49222 | EPB42 | erythrocyte membrane protein band 4.2 | Plasma Membrane | transporter |
| 15548088 | 2 | 0.037 | 9.32E+07 | 5.73E+07 | 6.13E+07 | 2.35E+07 | 2.22E+07 | 4.81E+07 | 7.10E+07 | 5.38E+07 | 4.87E+07 | 222154.734 | 3.77E+07 | 507056.906 | 1951.7838 | SC[160.0307]DFHAAENNEEHYTK | P49222 | P49222 | EPB42 | erythrocyte membrane protein band 4.2 | Plasma Membrane | transporter |
| 15531499 | 2 | 0.027 | 3.15E+08 | 2.54E+08 | 3.00E+08 | 2.55E+08 | 2.77E+08 | 2.21E+08 | 2.78E+08 | 2.65E+08 | 2.59E+08 | 4.19E+08 | 4.04E+08 | 2.94E+08 | 786.42755 | C[160.0307]LGIPAR | P49222 | P49222 | EPB42 | erythrocyte membrane protein band 4.2 | Plasma Membrane | transporter |
| 15537234, 15538463 | 3, 4 | 0.027 | 1.41E+09 | 1.06E+09 | 1.15E+09 | 5.38E+08 | 4.77E+08 | 9.69E+08 | 1.32E+09 | 1.21E+09 | 1.03E+09 | 1.09E+07 | 6.54E+08 | 1.99E+07 | 1951.7838, 1951.7855 | SC[160.0307]DFHAAENNEEHYTK | P49222 | P49222 | EPB42 | erythrocyte membrane protein band 4.2 | Plasma Membrane | transporter |
| 15547750 | 2 | 0.025 | 5.63E+07 | 4.52E+07 | 3.41E+07 | 6.25E+07 | 4.12E+07 | 3.94E+07 | 4.95E+07 | 4.06E+07 | 4.20E+07 | 8.50E+07 | 9.84E+07 | 6.29E+07 | 2218.0515 | MHNSLEAPMQNC[160.0307]IISIFGR | P49222 | P49222 | EPB42 | erythrocyte membrane protein band 4.2 | Plasma Membrane | transporter |
| 15557408, 15560508, 1556333 | 3 | 0.012 | 1042587.688 | 1684375 | 1362764 | 2223086.25 | 1979542.125 | 1319575.125 | 708563.125 | 966515.812 | 919138.5 | 1135502.125 | 799457.812 | 944602.062 | 3711.6802, 3711.6838, 37 | TEYVLNQNGFIYLGTADC[160.0307]IQI | P49222 | P49222 | EPB42 | erythrocyte membrane protein band 4.2 | Plasma Membrane | transporter |
| 15531261 | 3 | 0.009 | 6.96E+08 | 6.05E+08 | 5.00E+08 | 7.46E+08 | 6.62E+08 | 6.05E+08 | 6.39E+08 | 6.87E+08 | 6.24E+08 | 9.18E+08 | 1.29E+09 | 8.79E+08 | 2218.054 | MHNSLEAPMQNC[160.0307]IISIFGR | P49222 | P49222 | EPB42 | erythrocyte membrane protein band 4.2 | Plasma Membrane | transporter |
| 15553961 | 2 | 0.028 | 1.22E+07 | 1.28E+07 | 2.02E+07 | 9721866 | 1.01E+07 | 1.00E+07 | 1.03E+07 | 1.13E+07 | 1.20E+07 | 9891870 | 8402440 | 9858563 | 1665.9867 | ITVPLVSEVQIAQLR | Q3TZZ7 | Q3TZZ7 | ESYT2 | extended synaptotagmin-like protein 2 | Plasma Membrane | other |
| 15549793 | 2 | 0.007 | 6658706.5 | 6488111.5 | 7386623 | 5290820 | 5305524.5 | 4828360.5 | 5864443.5 | 5712193 | 5023135 | 5995406 | 5117251.5 | 5312492.5 | 1944.0167 | IPLSQLLTSDNMTINQR | Q3TZZ7 | Q3TZZ7 | ESYT2 | extended synaptotagmin-like protein 2 | Plasma Membrane | other |
| 15544594, 15546067 | 2 | 0.02 | 451908.969 | 1058355 | 1140006.5 | 3035525.5 | 3734481 | 4124935.5 | 2876309.25 | 3654706 | 3318494.25 | 445952.25 | 2170962 | 1948540 | 1121.5472, 1121.5486 | DGISMLTADAK | O88792 | O88792 | F11R | F11 receptor | Plasma Membrane | other |
| 15546192 | 3 | 0.007 | 1178087.75 | 1110411 | 695788.312 | 1264216.75 | 1299533.875 | 614303.312 | 330949.844 | 332883.469 | 92176.938 | 1002949.375 | 1057415.375 | 923614.938 | 2185.986 | VPRPTAQPEEEDEDEVSMK | Q6P9Q6 | Q6P9Q6 | FKBP15 | FK506 binding protein 15, 133kDa | Plasma Membrane | enzyme |
| 15540639 | 3 | 0.018 | 4867122.5 | 6379538 | 6085590.5 | 4740157.5 | 5513286.5 | 4575770.5 | 4698833 | 6649361 | 5768497.5 | 3949401.25 | 3129440.75 | 4080589 | 1396.7505 | LTGVSISQVNHNK | O08917 | O08917 | FLOT1 | flotillin 1 | Plasma Membrane | other |
| 15532957 | 2 | 0.023 | 1.11E+07 | 1.09E+07 | 1.16E+07 | 9704218 | 9876025 | 1.06E+07 | 1.02E+07 | 1.04E+07 | 1.17E+07 | 1.22E+07 | 1.19E+07 | 1.13E+07 | 1477.7688 | ELLAVAC[160.0307]EQFLGK | Q60634 | Q60634 | FLOT2 | flotillin 2 | Plasma Membrane | other |
| 15537721 | 2 | 0.009 | 4260944.5 | 4240033 | 4344102.5 | 3172066.5 | 3042449.25 | 3020421.75 | 4767929.5 | 3956333.75 | 3667684.75 | 5166686.5 | 3963367.25 | 5841950 | 1251.6992 | MGIEILSFTIK | Q60634 | Q60634 | FLOT2 | flotillin 2 | Plasma Membrane | other |
| 15548923 | 2 | 0.045 | 9.08E+07 | 1.02E+08 | 1.08E+08 | 9.16E+07 | 1.14E+08 | 6.51E+07 | 7.43E+07 | 9.31E+07 | 1.07E+08 | 6.79E+07 | 6.10E+07 | 5.44E+07 | 1746.8973 | IAQSDYIPTQQDVLR | P08752 | P08752 | GNAI2 | guanine nucleotide binding protein (G protein), alpha inhibiting activity polypeptide 2 | Plasma Membrane | other |
| 15549113 | 2 | 0.041 | 1.16E+07 | 1.87E+07 | 2.13E+07 | 1.17E+07 | 2.76E+07 | 1.18E+07 | 1.26E+07 | 1.63E+07 | 1.53E+07 | 8769519 | 6406731.5 | 7061000.5 | 1575.7529 | AMGNLQIDFADPQR | P08752 | P08752 | GNAI2 | guanine nucleotide binding protein (G protein), alpha inhibiting activity polypeptide 2 | Plasma Membrane | other |
| 15547968 | 2 | 0.04 | 4.42E+07 | 5.67E+07 | 5.49E+07 | 5.31E+07 | 5.86E+07 | 3.70E+07 | 4.21E+07 | 5.05E+07 | 5.31E+07 | 3.64E+07 | 3.25E+07 | 3.36E+07 | 2029.9858 | ITQSSLTIC[160.0307]FPEYTGANK | P08752 | P08752 | GNAI2 | guanine nucleotide binding protein (G protein), alpha inhibiting activity polypeptide 2 | Plasma Membrane | other |
| 15541087 | 3 | 0.04 | 7894558.5 | 1.07E+07 | 9742593 | 7378185 | 7191231 | 6947420 | 6579257 | 8501271 | 7029186.5 | 5602391.5 | 7711859 | 5860312.5 | 1736.9585 | AVVYSNTIQSIMAIVK | P08752 | P08752 | GNAI2 | guanine nucleotide binding protein (G protein), alpha inhibiting activity polypeptide 2 | Plasma Membrane | other |
| 15547753 | 2 | 0.022 | 3.82E+07 | 4.98E+07 | 5.03E+07 | 3.61E+07 | 4.37E+07 | 3.13E+07 | 2.96E+07 | 3.90E+07 | 4.19E+07 | 2.51E+07 | 2.89E+07 | 2.85E+07 | 1736.9568 | AVVYSNTIQSIMAIVK | P08752 | P08752 | GNAI2 | guanine nucleotide binding protein (G protein), alpha inhibiting activity polypeptide 2 | Plasma Membrane | other |
| 15553653 | 2 | 0.008 | 1.20E+07 | 1.21E+07 | 1.45E+07 | 1.29E+07 | 1.54E+07 | 1.10E+07 | 1.09E+07 | 1.34E+07 | 1.15E+07 | 9441614 | 7991518.5 | 7507606 | 1597.8596 | WFTDTSIILFLNK | P08752 | P08752 | GNAI2 | guanine nucleotide binding protein (G protein), alpha inhibiting activity polypeptide 2 | Plasma Membrane | other |
| 15539825 | 3 | 0.004 | 5745730 | 5202865.5 | 4475407 | 5515783.5 | 4308286 | 3229150.5 | 3595356.5 | 3341825.5 | 3731037.5 | 2784798.75 | 2529764.75 | 2052250.125 | 2119.9802 | AM[147.0354]GNLQIDFADPQRADDAI | P08752 | P08752 | GNAI2 | guanine nucleotide binding protein (G protein), alpha inhibiting activity polypeptide 2 | Plasma Membrane | other |
| 15550531 | 2 | 1.88E-04 | 3627753.5 | 3480781.75 | 3468722 | 4042151 | 4568885 | 3815825 | 4401681 | 5225173 | 5421931.5 | 2798957 | 2967702.75 | 2464848 | 1774.7732 | YTTPEDATPEPGEDPR | Q6R0H7 | Q6R0H7 | GNAS | GNAS complex locus | Plasma Membrane | enzyme |
| 15549013 | 3 | 0.049 | 1.43E+07 | 1.66E+07 | 1.50E+07 | 1.75E+07 | 1.65E+07 | 1.62E+07 | 1.29E+07 | 1.42E+07 | 1.65E+07 | 1.09E+07 | 1.33E+07 | 1.39E+07 | 3521.506 | QTFTGHESDINAIC[160.0307]FFPNGN | P62874 | P62874 | GNB1 | guanine nucleotide binding protein (G protein), beta polypeptide 1 | Plasma Membrane | enzyme |
| 15550405 | 3 | 0.047 | 1.79E+07 | 2.28E+07 | 2.25E+07 | 1.89E+07 | 2.82E+07 | 2.12E+07 | 1.50E+07 | 2.22E+07 | 2.54E+07 | 1.30E+07 | 1.46E+07 | 1.39E+07 | 2716.2532 | ADQELMTYSHDNIIC[160.0307]GITSV | P62874 | P62874 | GNB1 | guanine nucleotide binding protein (G protein), beta polypeptide 1 | Plasma Membrane | enzyme |
| 15532872 | 2 | 0.024 | 6.21E+07 | 6.32E+07 | 6.32E+07 | 6.59E+07 | 7.23E+07 | 4.53E+07 | 4.92E+07 | 5.27E+07 | 5.94E+07 | 4.40E+07 | 4.08E+07 | 4.05E+07 | 1017.5544 | LLVSASQDGK | P62874 | P62874 | GNB1 | guanine nucleotide binding protein (G protein), beta polypeptide 1 | Plasma Membrane | enzyme |
| 15556299 | 2 | 0.003 | 3648265.25 | 5129210 | 5865070.5 | 3419518 | 4138662.5 | 4489631 | 2294525 | 4920665.5 | 3992557.75 | 1610402.125 | 1682432.75 | 1675453.125 | 2174.9932 | LLLAGYDDFNC[160.0307]NIWDAM[1 | P62880 | P62880 | GNB2 | guanine nucleotide binding protein (G protein), beta polypeptide 2 | Plasma Membrane | enzyme |
| 15538236 | 3 | 0.006 | 3031519 | 3648897.25 | 2878493.25 | 3625294 | 5925255 | 5817713 | 7399294.5 | 7067949.5 | 4815501.5 | 2755510.5 | 2416485 | 3517976.5 | 2232.024 | LVNSQYHFTDEYLEC[160.0307]VSK | P51655 | P51655 | GPC4 | glypican 4 | Plasma Membrane | transmembrane receptor |
| 15549466 | 2 | 0.005 | 3982271 | 4316686 | 4152201.25 | 5485597.5 | 5389203 | 6668570.5 | 8178230.5 | 7274276 | 5545704.5 | 3658246.75 | 3712595.5 | 4919034 | 1697.7552 | FWSSLPSTVC[160.0307]NDER | P51655 | P51655 | GPC4 | glypican 4 | Plasma Membrane | transmembrane receptor |
| 15544395 | 2 | 0.013 | 1056966.5 | 1103425.25 | 892479.062 | 4.70E-06 | 25188.891 | 46297.035 | 1073087.625 | 1479022.125 | 1066965 | 52325.254 | 479812.562 | 4.70E-06 | 1305.5833 | WEQSGAAEHYK | P01899 | P01899 | HLA-A | major histocompatibility complex, class I, A | Plasma Membrane | other |
| 15543867 | 2 | 0.01 | 1136567.875 | 2700723.5 | 2205450.75 | 303108.812 | 1019184.188 | 732622.562 | 1643867 | 3152774.25 | 3049269.25 | 143852.812 | 631910.438 | 43149.855 | 1193.6024 | DYIALNEDLK | P01899 | P01899 | HLA-A | major histocompatibility complex, class I, A | Plasma Membrane | other |
| 15534931 | 2 | 0.004 | 5628000.5 | 5854715 | 4434628 | 258502.578 | 108649.969 | 133933 | 5337126 | 6249666.5 | 7127283.5 | 184686 | 3221164.25 | 84515.195 | 1157.5819 | YISVGYVDNK | P01899 | P01899 | HLA-A | major histocompatibility complex, class I, A | Plasma Membrane | other |
| 15535333 | 2 | 0.003 | 9707686 | 1.15E+07 | 9824993 | 6228991 | 5377593.5 | 6212030.5 | 1.10E+07 | 1.43E+07 | 1.45E+07 | 2603452 | 7085832 | 2674377 | 1055.6229 | WASVVVPLGK | P01899 | P01899 | HLA-A | major histocompatibility complex, class I, A | Plasma Membrane | other |
| 15535045 | 3 | 0.001 | 1.35E+07 | 1.60E+07 | 1.19E+07 | 3213.335 | 50565.055 | 1343.74 | 1.13E+07 | 1.61E+07 | 1.69E+07 | 65329.762 | 6270665 | 47633.926 | 1750.8727 | YFETAVSRPGLEEPR | P01899 | P01899 | HLA-A | major histocompatibility complex, class I, A | Plasma Membrane | other |
| 15538747 | 3 | 2.19E-04 | 656457.312 | 908201 | 591504.688 | 5770374.5 | 3295016 | 5528991.5 | 921725.5 | 960069.562 | 1650317.875 | 1486047.5 | 2512539 | 1808907.75 | 1648.8813 | YFVTAVSRPGLGEPR | P01901 | P01901 | HLA-A | major histocompatibility complex, class I, A | Plasma Membrane | other |
| 15553319 | 2 | 1.17E-04 | 105088.234 | 123020.695 | 140883.672 | 3650154.75 | 4281487.5 | 5158373.5 | 19120.637 | 26086.707 | 222194.484 | 1494492.75 | 992133.5 | 1198366.625 | 1629.8274 | FIAVGYVDDTQFVR | P01902 | P01902 | HLA-A | major histocompatibility complex, class I, A | Plasma Membrane | other |
| 15554671 | 2 | 1.36E-10 | 0 | 0 | 0 | 3411813.75 | 2691620 | 3008605.5 | 0 | 0 | 0 | 1152315.625 | 537915.438 | 792039.188 | 2090.0364 | GVNYALAPGSQTSDLSLPDGK | P01902 | P01902 | HLA-A | major histocompatibility complex, class I, A | Plasma Membrane | other |
| 15548136 | 3, 4 | 0.012 | 1.04E+08 | 1.17E+08 | 9.35E+07 | 1.77E+08 | 1.33E+08 | 1.47E+08 | 1.37E+08 | 1.21E+08 | 1.08E+08 | 2.09E+08 | 1.46E+08 | 1.65E+08 | 3095.5232, 3095.5237 | SALTIQLIQNHFVDEYDPTIEDSYRK | Q61411 | Q61411 | HRAS | Harvey rat sarcoma viral oncogene homolog | Plasma Membrane | enzyme |
| 15550482 | 2 | 3.59E-05 | 3.02E+07 | 3.17E+07 | 3.48E+07 | 4.94E+07 | 4.49E+07 | 4.43E+07 | 3.23E+07 | 3.26E+07 | 3.46E+07 | 5.01E+07 | 4.53E+07 | 4.64E+07 | 1670.8165 | TGEGFLC[160.0307]VFAINNTK | Q61411 | Q61411 | HRAS | Harvey rat sarcoma viral oncogene homolog | Plasma Membrane | enzyme |
| 15543891 | 2 | 4.29E-05 | 37919.27 | 5419.727 | 60818.859 | 1960857.375 | 2722378.25 | 1568382 | 34660.59 | 22197.473 | 1880.717 | 3228329 | 2868337.5 | 2203116 | 1152.5107 | C[160.0307]VVTC[160.0307]AGETR | Q9ERM2 | Q9ERM2 | ICAM4 | intercellular adhesion molecule 4 (Landsteiner-Wiener blood group) | Plasma Membrane | other |
| 15544432 | 3 | 6.96E-07 | 235327.016 | 99386.531 | 163396.688 | 2981533.75 | 5425608.5 | 3496267.25 | 113493.664 | 147105.484 | 187040.5 | 3173480.75 | 4407467 | 3618243.5 | 1751.8602 | AGLNDLWQPLTC[160.0307]HAR | Q9ERM2 | Q9ERM2 | ICAM4 | intercellular adhesion molecule 4 (Landsteiner-Wiener blood group) | Plasma Membrane | other |
| 15552882 | 2 | 0.042 | 215811.625 | 593804.062 | 1278303 | 4905725.5 | 4535847.5 | 4224033 | 3053610 | 2709021.5 | 5014112.5 | 490191.094 | 5303606 | 2754162.75 | 1762.8027 | ANTMSSVYAESFRGDK | Q9QUM0 | Q9QUM0 | ITGA2B | integrin, alpha 2b (platelet glycoprotein IIb of IIb/IIIa complex, antigen CD41) | Plasma Membrane | transmembrane receptor |
| 15533097 | 2 | 0.027 | 1578215 | 3169678 | 4363444.5 | 6382390.5 | 9504313 | 8712920 | 6187682 | 7145885.5 | 7901559 | 1939862.625 | 5359983 | 4290672.5 | 1272.5605 | GC[160.0307]QPSDIENPR | P09055 | P09055 | ITGB1 | integrin, beta 1 (fibronectin receptor, beta polypeptide, antigen CD29 includes MDF2, MSK12 | Plasma Membrane | transmembrane receptor |
| 15531495 | 3 | 0.031 | 3.38E+07 | 3.30E+07 | 3.62E+07 | 2.86E+07 | 3.15E+07 | 2.64E+07 | 3.49E+07 | 3.58E+07 | 2.66E+07 | 4.21E+07 | 3.68E+07 | 3.95E+07 | 1195.656 | ERPPLPAYPR | Q9EQF2 | Q9EQF2 | KEL | Kell blood group, metallo-endopeptidase | Plasma Membrane | peptidase |
| 15531425 | 3 | 0.026 | 4.33E+07 | 4.17E+07 | 4.55E+07 | 3.47E+07 | 3.52E+07 | 3.30E+07 | 5.02E+07 | 4.96E+07 | 4.18E+07 | 3.94E+07 | 5.59E+07 | 4.30E+07 | 2156.0808 | C[160.0307]VEQTGAFFEPTLAALFVR | Q9EQF2 | Q9EQF2 | KEL | Kell blood group, metallo-endopeptidase | Plasma Membrane | peptidase |

| | | | | | | | | | | | | | | | | | | | | | |
|---|---|---|---|---|---|---|---|---|---|---|---|---|---|---|---|---|---|---|---|---|---|
| 15553025 | 2 | 0.014 | 1.47E+07 | 1.62E+07 | 1.50E+07 | 1.23E+07 | 1.16E+07 | 1.25E+07 | 1.61E+07 | 1.33E+07 | 1.54E+07 | 1.76E+07 | 1.48E+07 | 1.48E+07 | 1642.8181 | LLMSQYGHFPFFR | Q9EQF2 | Q9EQF2 | KEL | Kell blood group, metallo-endopeptidase | Plasma Membrane | peptidase |
| 15548273 | 2 | 0.01 | 4.63E+07 | 5.90E+07 | 4.88E+07 | 4.30E+07 | 4.89E+07 | 5.27E+07 | 6.07E+07 | 6.05E+07 | 6.10E+07 | 7.49E+07 | 6.36E+07 | 6.29E+07 | 2039.0111 | EAFGPSIQSAAMELFAEIK | Q9EQF2 | Q9EQF2 | KEL | Kell blood group, metallo-endopeptidase | Plasma Membrane | peptidase |
| 15549631 | 2 | 0.007 | 4.35E+07 | 4.57E+07 | 5.07E+07 | 4.10E+07 | 3.91E+07 | 3.84E+07 | 4.85E+07 | 5.06E+07 | 4.65E+07 | 5.37E+07 | 6.23E+07 | 4.83E+07 | 2156.08 | C[160.0307]VEQTGAFFEPTLAALFVR | Q9EQF2 | Q9EQF2 | KEL | Kell blood group, metallo-endopeptidase | Plasma Membrane | peptidase |
| 15538306 | 2 | 0.048 | 1329179.875 | 1813929.625 | 1610768 | 3003918.75 | 2445968.75 | 2602715 | 1679054.5 | 2219717 | 2690160.25 | 887613.188 | 1887676.5 | 1641350.375 | 930.4986 | VLNLSDNR | Q80TE7 | Q80TE7 | LRRC7 | leucine rich repeat containing 7 | Plasma Membrane | other |
| 15548789 | 2 | 0.044 | 3.53E+08 | 3.08E+08 | 3.35E+08 | 2.51E+08 | 2.58E+08 | 2.47E+08 | 3.09E+08 | 2.86E+08 | 2.74E+08 | 3.41E+08 | 2.87E+08 | 2.47E+08 | 1643.8954 | TALSDLYLEHLLQK | P70290 | P70290 | MPP1 | membrane protein, palmitoylated 1, 55kDa | Plasma Membrane | kinase |
| 15554328 | 2 | 0.036 | 4212356.5 | 3769601.75 | 4003476.75 | 2624611.25 | 3302285.25 | 3075280.25 | 3478340.5 | 3225604.5 | 2579958.25 | 3261180.25 | 2070147.75 | 2157944.5 | 1530.6753 | EYHFISTEEM[147.0354]TK | P70290 | P70290 | MPP1 | membrane protein, palmitoylated 1, 55kDa | Plasma Membrane | kinase |
| 15532118, 15547490 | 2 | 0.035 | 6.00E+08 | 6.25E+08 | 6.42E+08 | 5.40E+08 | 4.58E+08 | 5.56E+08 | 6.56E+08 | 6.12E+08 | 5.33E+08 | 4.90E+08 | 5.23E+08 | 5.29E+08 | 1142.6873, 1142.6876 | TLVLIGASGVGR | P70290 | P70290 | MPP1 | membrane protein, palmitoylated 1, 55kDa | Plasma Membrane | kinase |
| 15554026, 15557690 | 2, 3 | 0.024 | 5.11E+07 | 4.10E+07 | 5.19E+07 | 3.00E+07 | 3.13E+07 | 4.26E+07 | 2.83E+07 | 3.69E+07 | 3.65E+07 | 3.44E+07 | 2.20E+07 | 2.11E+07 | 2370.0657, 2370.07 | NISANEFLEFGSYQGNM[147.0354]FG | P70290 | P70290 | MPP1 | membrane protein, palmitoylated 1, 55kDa | Plasma Membrane | kinase |
| 15541405, 15547361 | 2 | 0.007 | 1.92E+07 | 1.89E+07 | 1.58E+07 | 1.13E+07 | 9621256 | 1.11E+07 | 1.29E+07 | 1.49E+07 | 8254721 | 9253730 | 7137726 | 9229850 | 1122.5768, 1122.577 | LPALQM[147.0354]FMR | P70290 | P70290 | MPP1 | membrane protein, palmitoylated 1, 55kDa | Plasma Membrane | kinase |
| 15547879 | 2 | 0.005 | 1.16E+09 | 1.19E+09 | 1.32E+09 | 9.33E+08 | 1.02E+09 | 1.02E+09 | 1.11E+09 | 1.09E+09 | 1.12E+09 | 1.06E+09 | 9.69E+08 | 9.06E+08 | 1711.8635 | ESAGLIPSPELQEWR | P70290 | P70290 | MPP1 | membrane protein, palmitoylated 1, 55kDa | Plasma Membrane | kinase |
| 15537222, 15544832 | 2 | 0.004 | 8.43E+07 | 8.34E+07 | 8.58E+07 | 6.03E+07 | 4.89E+07 | 5.32E+07 | 7.46E+07 | 7.04E+07 | 5.09E+07 | 5.07E+07 | 3.98E+07 | 5.14E+07 | 1122.5773, 1122.5781 | LPALQMFM[147.0354]R | P70290 | P70290 | MPP1 | membrane protein, palmitoylated 1, 55kDa | Plasma Membrane | kinase |
| 15555373 | 4 | 0.026 | 2594569.25 | 3530174 | 2826765.5 | 2900947.5 | 4547229 | 2304030.25 | 2471464.75 | 2505803.25 | 4724261.5 | 1157421 | 1482371.25 | 1718564.875 | 3335.4287 | TAMSTPHVAEPAENEHDEQDENGAEA | P26041 | P26041 | MSN | moesin | Plasma Membrane | other |
| 15532731 | 3 | 0.025 | 4160785.5 | 4446218 | 3841494.25 | 4032975.75 | 2853065 | 3657815.25 | 3428021.5 | 3506517.75 | 3958201.5 | 5084752.5 | 5041835 | 4424233.5 | 1239.7141 | SGAGVPEVVLRR | P57716 | P57716 | NCSTN | nicastrin | Plasma Membrane | peptidase |
| 15542004 | 2 | 3.21E-06 | 8975000 | 9576039 | 1.06E+07 | 1.96E+07 | 1.97E+07 | 1.87E+07 | 1.17E+07 | 1.27E+07 | 1.11E+07 | 2.10E+07 | 2.10E+07 | 1.79E+07 | 1312.6765 | SYGIPFIETSAK | P08556 | P08556 | NRAS | neuroblastoma RAS viral (v-ras) oncogene homolog | Plasma Membrane | enzyme |
| 15532606 | 4 | 0.02 | 3.49E+07 | 3.45E+07 | 3.34E+07 | 2.72E+07 | 3.36E+07 | 3.93E+07 | 4.32E+07 | 3.20E+07 | 3.63E+07 | 5.66E+07 | 5.11E+07 | 4.38E+07 | 2375.192 | YTIGSLRPHFLAIC[160.0307]NPDWSK | Q61469 | Q61469 | Ppap2a | phosphatidic acid phosphatase type 2A | Plasma Membrane | phosphatase |
| 15549543 | 2 | 0.004 | 8166718.5 | 9446132 | 8718196 | 1.47E+07 | 1.44E+07 | 1.82E+07 | 7945009 | 8698013 | 5863268.5 | 1.09E+07 | 1.05E+07 | 6976946.5 | 1512.7153 | VVEQMC[160.0307]VTQYQK | P04925 | P04925 | PRNP | prion protein | Plasma Membrane | other |
| 15547678 | 2 | 0.02 | 9.60E+07 | 9.73E+07 | 1.13E+08 | 7.94E+07 | 8.50E+07 | 8.51E+07 | 9.31E+07 | 9.96E+07 | 1.06E+08 | 9.49E+07 | 8.14E+07 | 8.52E+07 | 1579.8203 | ANINVENAFFTLAR | P55258 | P55258 | RAB8A | RAB8A, member RAS oncogene family | Plasma Membrane | enzyme |
| 15551217 | 3 | 1.79E-05 | 1859469 | 2753255.75 | 3873150.5 | 3.22E+07 | 4.72E+07 | 7.51E+07 | 7982226.5 | 9717568 | 1.40E+07 | 4.86E+07 | 3.24E+07 | 3.60E+07 | 2732.3955 | AIVNTYM[147.0354]SLAAC[160.0307 | Q9QUT0 | Q9QUT0 | RHAG | Rh-associated glycoprotein | Plasma Membrane | peptidase |
| 15555126 | 3 | 1.47E-05 | 287073 | 472036.719 | 759474.688 | 1.56E+07 | 6195332.5 | 1.52E+07 | 2013103.375 | 1778619.25 | 2376991.5 | 1.03E+07 | 1.56E+07 | 1.62E+07 | 2888.495 | AIVNTYM[147.0354]SLAAC[160.0307 | Q9QUT0 | Q9QUT0 | RHAG | Rh-associated glycoprotein | Plasma Membrane | peptidase |
| 15548032, 15561784 | 2, 3 | 2.37E-08 | 144539.453 | 114431.148 | 448411.688 | 5.17E+08 | 6.55E+08 | 6.42E+08 | 866557.688 | 1021592.375 | 1605249.25 | 7.37E+08 | 3.57E+08 | 4.79E+08 | 2512.1062, 2512.1091, 25 | LPIWNQPPDEYC[160.0307]YDDSVSW | Q9QUT0 | Q9QUT0 | RHAG | Rh-associated glycoprotein | Plasma Membrane | peptidase |
| 15534621 | 2 | 1.35E-08 | 250177.734 | 385378.875 | 545373.375 | 9.70E+08 | 1.16E+09 | 1.25E+09 | 1533138.75 | 771479.375 | 323789.594 | 1.20E+09 | 1.09E+09 | 1.14E+09 | 1036.5812 | FFSPLLANK | Q9QUT0 | Q9QUT0 | RHAG | Rh-associated glycoprotein | Plasma Membrane | peptidase |
| 15537019 | 3 | 9.04E-06 | 4.70E-06 | 55987.293 | 8276.938 | 2.80E+07 | 1.03E+08 | 1.31E+08 | 163514.859 | 111333.141 | 137358.766 | 1.97E+07 | 1.33E+08 | 1.43E+08 | 2118.9585 | YFDDQTFWEFPHLAVGF | Q8CF94 | Q8CF94 | RHCE/RHD | Rh blood group, D antigen | Plasma Membrane | transporter |
| 15551699, 15555471, 1555937 | 3, 4 | 3.99E-06 | 367765.188 | 151014.625 | 712565.25 | 2.72E+07 | 1.85E+07 | 3.27E+07 | 1413533.875 | 803431.625 | 1386314.75 | 2.24E+07 | 2.49E+07 | 2.27E+07 | 3155.5867, 3155.5906 | RNAVFNTYYALAVSAVTATSMSALSHPC | Q8CF94 | Q8CF94 | RHCE/RHD | Rh blood group, D antigen | Plasma Membrane | transporter |
| 15557446, 15559130 | 3, 4 | 1.15E-06 | 302550.312 | 205570.484 | 747451.875 | 3.72E+07 | 4.01E+07 | 3.71E+07 | 843191.438 | 1391747.5 | 647970.875 | 3.47E+07 | 2.01E+07 | 2.32E+07 | 3015.4827, 3015.4832 | NAVFNTYYALAVSAVTATSM[147.0354 | Q8CF94 | Q8CF94 | RHCE/RHD | Rh blood group, D antigen | Plasma Membrane | transporter |
| 15533792 | 2 | 6.65E-07 | 46768.613 | 66610.453 | 33494.531 | 3.94E+07 | 8.59E+07 | 5.68E+07 | 7831.782 | 69026.508 | 4310.843 | 5.45E+07 | 2.30E+07 | 3.40E+07 | 1130.5636 | FM[147.0354]ASYQVLR | Q8CF94 | Q8CF94 | RHCE/RHD | Rh blood group, D antigen | Plasma Membrane | transporter |
| 15537908 | 2 | 6.88E-08 | 291537.219 | 304769.688 | 539283.812 | 7.92E+07 | 1.39E+08 | 8.03E+07 | 137700.234 | 197322.734 | 489011.219 | 1.19E+08 | 9.04E+07 | 8.82E+07 | 1114.5692 | FMASYQVLR | Q8CF94 | Q8CF94 | RHCE/RHD | Rh blood group, D antigen | Plasma Membrane | transporter |
| 15550468 | 3, 4 | 2.30E-08 | 1848336.5 | 1779213.25 | 3541918 | 1.97E+08 | 1.46E+08 | 1.44E+08 | 1.13E+07 | 9545812 | 1.06E+07 | 2.16E+08 | 2.00E+08 | 1.63E+08 | 2999.4873, 2999.4888 | NAVFNTYYALAVSAVTATSMSALSHPQ | Q8CF94 | Q8CF94 | RHCE/RHD | Rh blood group, D antigen | Plasma Membrane | transporter |
| 15557732 | 3 | 2.59E-09 | 0 | 38.16 | 0 | 2259952.75 | 3720215 | 5791219.5 | 0 | 0 | 0 | 2110711.75 | 1309279.375 | 2944268.75 | 3171.5815 | RNAVFNTYYALAVSAVTATSM[147.035 | Q8CF94 | Q8CF94 | RHCE/RHD | Rh blood group, D antigen | Plasma Membrane | transporter |
| 15552273 | 2 | 6.00E-10 | 319678.406 | 820879.188 | 538651.438 | 1.04E+09 | 1.30E+09 | 1.43E+09 | 3607203.75 | 3367064 | 3504522 | 1.73E+09 | 1.14E+09 | 1.37E+09 | 2118.9597 | YFDDQTFWEFPHLAVGF | Q8CF94 | Q8CF94 | RHCE/RHD | Rh blood group, D antigen | Plasma Membrane | transporter |
| 15540472 | 2 | 0.015 | 414138.719 | 655733.75 | 1612331.75 | 4290287.5 | 3831943.75 | 5108532.5 | 3311956.25 | 3121441.75 | 4054834.25 | 924869.938 | 4190201 | 2551689.75 | 1056.4552 | FEC[160.0307]QPGYR | Q01102 | Q01102 | SELP | selectin P (granule membrane protein 140kDa, antigen CD62) | Plasma Membrane | transmembrane receptor |
| 15531784, 15547201 | 2 | 0.048 | 3.24E+08 | 4.16E+08 | 4.92E+08 | 2.01E+08 | 3.07E+08 | 2.93E+08 | 4.04E+08 | 4.18E+08 | 3.39E+08 | 3.26E+08 | 1.97E+08 | 2.58E+08 | 925.5121, 925.5141 | IFYLQNK | Q8VHL0 | Q8VHL0 | SLC14A1 | solute carrier family 14 (urea transporter), member 1 (Kidd blood group) | Plasma Membrane | transporter |
| 15547727 | 2, 3 | 0.045 | 1.92E+08 | 2.68E+08 | 2.39E+08 | 1.29E+08 | 1.70E+08 | 1.57E+08 | 1.80E+08 | 2.36E+08 | 2.83E+08 | 1.97E+08 | 1.52E+08 | 1.24E+08 | 2544.3914, 2544.3926 | LFTPVSSVPNITWSELSALELLK | Q8VHL0 | Q8VHL0 | SLC14A1 | solute carrier family 14 (urea transporter), member 1 (Kidd blood group) | Plasma Membrane | transporter |
| 15538805, 15546213 | 2 | 0.009 | 4.81E+07 | 4.28E+07 | 6.05E+07 | 2.70E+07 | 3.37E+07 | 3.42E+07 | 4.17E+07 | 4.37E+07 | 3.89E+07 | 3.65E+07 | 2.46E+07 | 2.59E+07 | 1098.5464, 1098.5493 | VLGYVTGDM[147.0354]K | Q8VHL0 | Q8VHL0 | SLC14A1 | solute carrier family 14 (urea transporter), member 1 (Kidd blood group) | Plasma Membrane | transporter |
| 15531277 | 2, 3 | 0.008 | 2.36E+08 | 1.87E+08 | 1.94E+08 | 1.38E+08 | 1.40E+08 | 1.09E+08 | 2.28E+08 | 2.18E+08 | 1.83E+08 | 1.76E+08 | 1.88E+08 | 1.39E+08 | 1446.7087, 1446.7102 | VDRGENQILSC[160.0307]R | Q8VHL0 | Q8VHL0 | SLC14A1 | solute carrier family 14 (urea transporter), member 1 (Kidd blood group) | Plasma Membrane | transporter |
| 15531422 | 2 | 0.002 | 1.67E+08 | 1.70E+08 | 1.87E+08 | 1.19E+08 | 1.22E+08 | 1.14E+08 | 1.55E+08 | 1.56E+08 | 1.45E+08 | 1.49E+08 | 1.13E+08 | 1.37E+08 | 831.4387 | RMVESPL | Q8VHL0 | Q8VHL0 | SLC14A1 | solute carrier family 14 (urea transporter), member 1 (Kidd blood group) | Plasma Membrane | transporter |
| 15549676, 15552306 | 2, 3 | 0.002 | 1.75E+08 | 1.80E+08 | 1.90E+08 | 1.68E+08 | 1.97E+08 | 2.15E+08 | 2.32E+08 | 2.30E+08 | 2.32E+08 | 3.03E+08 | 2.75E+08 | 2.42E+08 | 2654.4363, 2654.4392 | FSSAVGLVTIVEC[160.0307]C[160.030 | P53986 | P53986 | SLC16A1 | solute carrier family 16 (monocarboxylate transporter), member 1 | Plasma Membrane | transporter |
| 15537122 | 2 | 0.024 | 2196385 | 1926574.25 | 1960555 | 1606564 | 1592139 | 1463613.75 | 2254844.5 | 2185773.75 | 1860231.75 | 1986577.75 | 1604835.5 | 1422874.125 | 1148.5217 | NC[160.0307]VSPESDIK | Q3UHH2 | Q3UHH2 | SLC22A23 | solute carrier family 22, member 23 | Plasma Membrane | transporter |
| 15548586 | 3 | 0.02 | 1.11E+07 | 1.36E+07 | 1.19E+07 | 9132061 | 8247003 | 8092143 | 9937488 | 1.02E+07 | 1.20E+07 | 1.20E+07 | 9056866 | 1.00E+07 | 3045.4302 | GGFLGEEQGAENELLFPSVHSAVEAAC[ | P58735 | P58735 | SLC26A1 | solute carrier family 26 (anion exchanger), member 1 | Plasma Membrane | transporter |
| 15531929 | 2 | 0.048 | 1.53E+08 | 1.71E+08 | 1.62E+08 | 1.30E+08 | 1.27E+08 | 1.27E+08 | 1.60E+08 | 1.53E+08 | 1.28E+08 | 1.62E+08 | 1.33E+08 | 1.50E+08 | 1194.6832 | ALADPTVALPAR | Q9JIM1 | Q9JIM1 | SLC29A1 | solute carrier family 29 (equilibrative nucleoside transporter), member 1 | Plasma Membrane | transporter |
| 15536711 | 3 | 0.042 | 3217547 | 1946862.875 | 2930617 | 672873.438 | 1393674 | 2669691.75 | 3067928.75 | 3714206.25 | 4299284.5 | 3653377 | 4350789 | 3129514 | 1559.9015 | IVFIPLLMLC[160.0307]NVK | Q9JIM1 | Q9JIM1 | SLC29A1 | solute carrier family 29 (equilibrative nucleoside transporter), member 1 | Plasma Membrane | transporter |
| 15532903 | 2 | 0.038 | 5.15E+07 | 5.28E+07 | 6.01E+07 | 2.50E+07 | 4.32E+07 | 3.82E+07 | 4.84E+07 | 4.39E+07 | 4.75E+07 | 4.36E+07 | 4.00E+07 | 4.49E+07 | 1040.6234 | WLPVLVASR | Q9JIM1 | Q9JIM1 | SLC29A1 | solute carrier family 29 (equilibrative nucleoside transporter), member 1 | Plasma Membrane | transporter |
| 15551662 | 2 | 0.03 | 3.77E+07 | 2.60E+07 | 3.38E+07 | 1.38E+07 | 2.01E+07 | 2.89E+07 | 3.70E+07 | 3.68E+07 | 5.09E+07 | 3.75E+07 | 4.88E+07 | 3.19E+07 | 1559.9028 | IVFIPLLMLC[160.0307]NVK | Q9JIM1 | Q9JIM1 | SLC29A1 | solute carrier family 29 (equilibrative nucleoside transporter), member 1 | Plasma Membrane | transporter |
| 15553627 | 2 | 0.022 | 5356049 | 5749256.5 | 6377563 | 3034197.5 | 3779554.25 | 3186029.5 | 3611534.25 | 4637331 | 2983978.5 | 3343207.25 | 1605208.625 | 3237337.5 | 1623.7223 | SLTAVC[160.0307]M[147.0354]WPG | Q9JIM1 | Q9JIM1 | SLC29A1 | solute carrier family 29 (equilibrative nucleoside transporter), member 1 | Plasma Membrane | transporter |
| 15539520 | 3 | 0.009 | 8.30E+07 | 9.23E+07 | 9.33E+07 | 6.17E+07 | 6.67E+07 | 5.53E+07 | 7.89E+07 | 7.96E+07 | 7.85E+07 | 8.54E+07 | 6.25E+07 | 6.81E+07 | 1811.9222 | HYLQLNLAGPAEQETK | Q9JIM1 | Q9JIM1 | SLC29A1 | solute carrier family 29 (equilibrative nucleoside transporter), member 1 | Plasma Membrane | transporter |
| 15540037 | 2 | 0.041 | 3326474 | 997509.125 | 4614210 | 4704138.5 | 5163751.5 | 3639842.5 | 2878830 | 3046982.25 | 3326474 | 997509.125 | 4614210 | 2741831.25 | 1331.6437 | EEDQATEILQR | P32037 | P32037 | SLC2A3 | solute carrier family 2 (facilitated glucose transporter), member 3 | Plasma Membrane | transporter |
| 15543493 | 3 | 0.029 | 1919837.625 | 513119.75 | 1416032.375 | 1429119.625 | 1880948.125 | 2479465.75 | 1139371.75 | 1603800.75 | 1919837.625 | 513119.75 | 1416032.375 | 1618724.875 | 1762.865 | LWGTSDVVQEIQEMK | P32037 | P32037 | SLC2A3 | solute carrier family 2 (facilitated glucose transporter), member 3 | Plasma Membrane | transporter |
| 15550304 | 2 | 0.002 | 2346715 | 3217544.25 | 2340967.25 | 2185743.75 | 1897604.25 | 1946351.875 | 2672057 | 2920270.75 | 3129487.25 | 1770302.375 | 1475499.375 | 1212846.625 | 1889.9496 | APTVSISC[160.0307]LELSENLEK | Q60738 | Q60738 | SLC30A1 | solute carrier family 30 (zinc transporter), member 1 | Plasma Membrane | transporter |
| 15533120 | 2 | 8.79E-04 | 4449013.5 | 5197298 | 4077968.75 | 3925352.25 | 4462327 | 4331885.5 | 6454987.5 | 5999228.5 | 5120399.5 | 3595088.75 | 3331273.75 | 3304246.25 | 1311.7861 | ESALILLQTVPK | Q60738 | Q60738 | SLC30A1 | solute carrier family 30 (zinc transporter), member 1 | Plasma Membrane | transporter |
| 15540961 | 2 | 3.36E-04 | 3511697.75 | 3825287.5 | 3691796.25 | 2895839.25 | 2888526.25 | 2405123 | 4330513 | 4124075.5 | 3331496.25 | 2424198.75 | 2232355.5 | 2186857.25 | 1325.7651 | AEGSLPAVVIEIK | Q60738 | Q60738 | SLC30A1 | solute carrier family 30 (zinc transporter), member 1 | Plasma Membrane | transporter |
| 15539438 | 2 | 0.048 | 9058961 | 1.10E+07 | 6314124 | 7669475 | 8157576.5 | 9093709 | 5860821.5 | 7401275 | 6330103.5 | 6297527 | 5490742.5 | 5531076 | 1075.5874 | IGDLQAFVGR | P10852 | P10852 | SLC3A2 | solute carrier family 3 (amino acid transporter heavy chain), member 2 | Plasma Membrane | transporter |
| 15532316 | 2 | 0.026 | 5159796 | 6554883 | 4940127.5 | 4784415.5 | 5356258 | 5553594.5 | 3422225 | 4926999.5 | 3979269.5 | 4086579.25 | 3772916.75 | 4004441.5 | 933.58575 | GLVLGPIHK | P10852 | P10852 | SLC3A2 | solute carrier family 3 (amino acid transporter heavy chain), member 2 | Plasma Membrane | transporter |
| 15554707, 15560410 | 3 | 0.022 | 9030388 | 1.17E+07 | 6414555 | 8953282 | 7265772.5 | 9469830 | 7187366 | 7319158 | 7138410.5 | 5622987 | 5094240.5 | 5137817 | 2362.1777, 2362.1782 | SLLHGDFHALSSSPDLFSYIR | P10852 | P10852 | SLC3A2 | solute carrier family 3 (amino acid transporter heavy chain), member 2 | Plasma Membrane | transporter |
| 15537201 | 2 | 0.014 | 3028208.5 | 4243978.5 | 2241203.25 | 3825413.5 | 3246889.75 | 3787610.75 | 2513976.5 | 3019996.5 | 2457386.25 | 2219523 | 1600469.625 | 1685802.625 | 1330.6033 | QPMNAADGAAAGEK | P10852 | P10852 | SLC3A2 | solute carrier family 3 (amino acid transporter heavy chain), member 2 | Plasma Membrane | transporter |
| 15548403 | 2 | 0.011 | 6025460 | 1.12E+07 | 6469900 | 6884762.5 | 7392312.5 | 8399743 | 4700044 | 8415364 | 5726193.5 | 3895254.75 | 3454454.5 | 2829718.25 | 1561.7583 | GQNEDPGSLLTQFR | P10852 | P10852 | SLC3A2 | solute carrier family 3 (amino acid transporter heavy chain), member 2 | Plasma Membrane | transporter |
| 15548696 | 2 | 0.008 | 1.21E+07 | 1.83E+07 | 1.32E+07 | 1.50E+07 | 1.45E+07 | 1.30E+07 | 9085246 | 1.25E+07 | 1.07E+07 | 9407040 | 8295683.5 | 8661784 | 1566.8816 | LGASNLPAGISLPASAK | P10852 | P10852 | SLC3A2 | solute carrier family 3 (amino acid transporter heavy chain), member 2 | Plasma Membrane | transporter |
| 15533046 | 2 | 0.006 | 9980435 | 1.16E+07 | 7731082.5 | 1.10E+07 | 9283746 | 1.02E+07 | 7570629.5 | 8099044 | 7523994.5 | 6940880.5 | 5995477 | 6427681 | 1476.7303 | QINPTLGSQEDFK | P10852 | P10852 | SLC3A2 | solute carrier family 3 (amino acid transporter heavy chain), member 2 | Plasma Membrane | transporter |
| 15548712, 15548776 | 2 | 0.049 | 1.99E+07 | 1.69E+07 | 1.80E+07 | 1.64E+07 | 1.82E+07 | 8165351 | 1.42E+07 | 1.26E+07 | 1.32E+07 | 1.19E+07 | 6994432 | 7545442 | 2213.9382, 2213.9387 | VTFDEENGLDEYDEVPM[147.0354]PV | P04919 | P04919 | SLC4A1 | solute carrier family 4 (anion exchanger), member 1 (Diego blood group) | Plasma Membrane | transporter |
| 15531763 | 2, 3 | 0.039 | 3.44E+08 | 3.32E+08 | 3.55E+08 | 2.14E+08 | 3.11E+08 | 3.01E+08 | 3.88E+08 | 3.60E+08 | 3.89E+08 | 3.50E+08 | 3.29E+08 | 3.03E+08 | 1140.7589, 1140.7593 | RLILPLIFR | P04919 | P04919 | SLC4A1 | solute carrier family 4 (anion exchanger), member 1 (Diego blood group) | Plasma Membrane | transporter |
| 15532073, 15539240, 1554677 | 3 | 0.036 | 8.11E+09 | 7.07E+09 | 7.82E+09 | 5.70E+09 | 6.85E+09 | 6.53E+09 | 7.97E+09 | 7.27E+09 | 7.09E+09 | 8.09E+09 | 8.00E+09 | 7.18E+09 | 1640.8496, 1640.8508, 16 | ITASMAHNREELLR | P04919 | P04919 | SLC4A1 | solute carrier family 4 (anion exchanger), member 1 (Diego blood group) | Plasma Membrane | transporter |
| 15539254 | 3 | 0.025 | 2743986.5 | 7591134 | 1.18E+07 | 3762708.75 | 1.41E+07 | 6504326 | 6006611.5 | 7090167 | 3563619.5 | 190761.062 | 949166.688 | 2042982.875 | 1630.8044 | PDPNLYNTLDLNGGK | P04919 | P04919 | SLC4A1 | solute carrier family 4 (anion exchanger), member 1 (Diego blood group) | Plasma Membrane | transporter |
| 15548687 | 3 | 0.002 | 9803132 | 1.47E+07 | 9439907 | 6397144.5 | 6817679 | 9777826 | 8456809 | 1.05E+07 | 1.16E+07 | 4210455 | 4481128 | 4840681 | 2426.1458 | YAPTTTPSPEDGFEVQPLHPDK | P28571 | P28571 | SLC6A9 | solute carrier family 6 (neurotransmitter transporter, glycine), member 9 | Plasma Membrane | transporter |
| 15538414 | 2 | 9.72E-04 | 2418587 | 3193914.75 | 2541207.5 | 1358096.375 | 1973295.875 | 2113745.5 | 2412600 | 3425795.5 | 2545543.75 | 1030832 | 1277554.75 | 832345.75 | 1321.6353 | LSDDIGNFGEVR | P28571 | P28571 | SLC6A9 | solute carrier family 6 (neurotransmitter transporter, glycine), member 9 | Plasma Membrane | transporter |
| 15539100 | 2 | 0.016 | 2304675.75 | 3838820.25 | 2272580 | 2663014 | 2605462.5 | 2423130.5 | 2460477.25 | 2372015.25 | 3071668.25 | 1579649 | 1565904.375 | 1700470.875 | 825.4598 | LFFVGSR | Q9Z127 | Q9Z127 | SLC7A5 | solute carrier family 7 (amino acid transporter light chain, L system), member 5 | Plasma Membrane | transporter |
| 15532780 | 2 | 0.018 | 2.39E+07 | 3.57E+07 | 3.41E+07 | 2.50E+07 | 2.65E+07 | 2.77E+07 | 3.17E+07 | 3.42E+07 | 3.17E+07 | 1.86E+07 | 2.38E+07 | 2.22E+07 | 1427.808 | ILGLAIESQDAGIK | O09044 | O09044 | SNAP23 | synaptosomal-associated protein, 23kDa | Plasma Membrane | transporter |
| 15548546 | 2 | 0.006 | 9868382 | 9301631 | 1.17E+07 | 1.05E+07 | 1.09E+07 | 1.10E+07 | 1.23E+07 | 1.15E+07 | 9991548 | 7101334 | 8460234 | 5827274.5 | 2160.9973 | NMALDMGNEIDAQNQQIQK | O09044 | O09044 | SNAP23 | synaptosomal-associated protein, 23kDa | Plasma Membrane | transporter |
| 15533249 | 2 | 0.048 | 3.48E+08 | 3.11E+08 | 3.30E+08 | 2.52E+08 | 2.64E+08 | 2.47E+08 | 3.29E+08 | 2.95E+08 | 2.52E+08 | 3.88E+08 | 3.15E+08 | 2.91E+08 | 996.5 | AWLQDAHR | P15508 | P15508 | SPTB | spectrin, beta, erythrocytic | Plasma Membrane | other |
| 15538188, 15544484, 1554477 | 4 | 0.044 | 4.95E+08 | 5.47E+08 | 4.86E+08 | 4.82E+08 | 4.31E+08 | 4.01E+08 | 4.93E+08 | 4.53E+08 | 3.78E+08 | 5.10E+08 | 5.69E+08 | 5.62E+08 | 2266.0667, 2266.0684, 22 | HRELPEDVGLDASTAESFHR | P15508 | P15508 | SPTB | spectrin, beta, erythrocytic | Plasma Membrane | other |
| 15556682, 15563055 | 3 | 0.044 | 5.77E+08 | 6.43E+08 | 6.36E+08 | 4.61E+08 | 5.45E+08 | 5.29E+08 | 5.59E+08 | 5.83E+08 | 5.37E+08 | 4.76E+08 | 5.17E+08 | 5.70E+08 | 2610.199, 2610.2107 | HQAFMAELASHQGWLENIDAEGR | P15508 | P15508 | SPTB | spectrin, beta, erythrocytic | Plasma Membrane | other |
| 15540097 | 3 | 0.032 | 4783892.5 | 3873606.75 | 3838523.5 | 4079163.75 | 2346150.25 | 2891699.25 | 3173061.25 | 3363586.25 | 3410254 | 4497307 | 4857263 | 4624020.5 | 1328.7656 | FLDLLEPLGRR | P15508 | P15508 | SPTB | spectrin, beta, erythrocytic | Plasma Membrane | other |
| 15531959, 15536086 | 2, 3 | 0.03 | 1.19E+09 | 1.10E+09 | 1.35E+09 | 9.52E+08 | 9.42E+08 | 1.05E+09 | 1.09E+09 | 1.03E+09 | 1.05E+09 | 1.12E+09 | 1.02E+09 | 1.11E+09 | 1157.6865, 1157.6881 | DLTSVLILQR | P15508 | P15508 | SPTB | spectrin, beta, erythrocytic | Plasma Membrane | other |
| 15533927 | 3 | 0.03 | 3.84E+07 | 3.25E+07 | 5.07E+07 | 1.68E+07 | 2.52E+07 | 3.21E+07 | 2.86E+07 | 3.09E+07 | 2.23E+07 | 2.50E+07 | 1.27E+07 | 1.38E+07 | 2052.9365 | LFQDM[147.0354]LHSIDWM[147.035 | P15508 | P15508 | SPTB | spectrin, beta, erythrocytic | Plasma Membrane | other |
| 15540802 | 4 | 0.025 | 1.81E+07 | 2.16E+07 | 2.26E+07 | 1.21E+07 | 1.67E+07 | 2.18E+07 | 1.64E+07 | 1.97E+07 | 2.09E+07 | 1.13E+07 | 1.33E+07 | 1.16E+07 | 2626.2053 | HQAFM[147.0354]AELASHQGWLENI | P15508 | P15508 | SPTB | spectrin, beta, erythrocytic | Plasma Membrane | other |
| 15532805 | 2 | 0.02 | 5.22E+08 | 4.32E+08 | 4.47E+08 | 4.52E+08 | 4.56E+08 | 3.89E+08 | 5.77E+08 | 5.71E+08 | 4.58E+08 | 6.69E+08 | 7.56E+08 | 5.39E+08 | 1277.6366 | DALLLWC[160.0307]QMK | P15508 | P15508 | SPTB | spectrin, beta, erythrocytic | Plasma Membrane | other |
| 15531251 | 3 | 0.019 | 9.08E+08 | 9.60E+08 | 9.63E+08 | 8.17E+08 | 8.33E+08 | 8.48E+08 | 7.95E+08 | 7.26E+08 | 7.87E+08 | 1.04E+09 | 8.21E+08 | 9.26E+08 | 2090.0535 | DASVAEAWLIAQEPYLASR | P15508 | P15508 | SPTB | spectrin, beta, erythrocytic | Plasma Membrane | other |
| 15534341, 15544906, 1554695 | 2, 3 | 0.016 | 1.21E+08 | 1.02E+08 | 1.03E+08 | 1.08E+08 | 7.64E+07 | 9.02E+07 | 9.10E+07 | 6.86E+07 | 6.99E+07 | 1.13E+08 | 1.22E+08 | 1.11E+08 | 1270.7451, 1270.7452, 12 | QRLEATLALQK | P15508 | P15508 | SPTB | spectrin, beta, erythrocytic | Plasma Membrane | other |
| 15550133 | 2 | 0.013 | 5.51E+08 | 5.89E+08 | 7.52E+08 | 5.40E+08 | 5.24E+08 | 4.96E+08 | 3.52E+08 | 4.94E+08 | 4.54E+08 | 4.52E+08 | 3.70E+08 | 3.75E+08 | 1660.8613 | FQIQDIVVQTQEGR | P15508 | P15508 | SPTB | spectrin, beta, erythrocytic | Plasma Membrane | other |
| 15547731 | 2 | 0.009 | 5.91E+07 | 5.78E+07 | 8.17E+07 | 8.70E+07 | 7.20E+07 | 1.06E+08 | 7.38E+07 | 6.42E+07 | 6.80E+07 | 1.59E+08 | 9.80E+07 | 1.31E+08 | 1796.881 | LSSSWESLLPEPAHPF | P15508 | P15508 | SPTB | spectrin, beta, erythrocytic | Plasma Membrane | other |
| 15539387 | 3 | 0.006 | 7.37E+08 | 7.47E+08 | 7.84E+08 | 6.40E+08 | 7.18E+08 | 6.75E+08 | 7.36E+08 | 6.71E+08 | 7.43E+08 | 8.23E+08 | 8.03E+08 | 8.01E+08 | 1604.827 | RWQAFQAVVSEQR | P15508 | P15508 | SPTB | spectrin, beta, erythrocytic | Plasma Membrane | other |
| 15531257 | 2 | 0.019 | 4.66E+08 | 4.44E+08 | 3.26E+08 | 2.68E+08 | 2.10E+08 | 1.76E+08 | 3.89E+08 | 4.49E+08 | 3.79E+08 | 5.75E+08 | 5.23E+08 | 2.92E+08 | 1351.7441 | YLQTLTTIAAEK | P54116 | P54116 | STOM | stomatin | Plasma Membrane | other |
| 15547675 | 2, 3 | 0.013 | 2.73E+08 | 2.77E+08 | 2.47E+08 | 1.92E+08 | 1.81E+08 | 1.95E+08 | 3.02E+08 | 3.31E+08 | 3.23E+08 | 4.74E+08 | 3.68E+08 | 2.39E+08 | 2270.126, 2270.1282 | NSTIVFPLPVDMLQGIMGSNH | P54116 | P54116 | STOM | stomatin | Plasma Membrane | other |
| 15543425 | 2 | 0.041 | 1021453 | 926594.625 | 1046672.875 | 3177216.75 | 2733238.75 | 2808494.25 | 1646979.125 | 2289201.75 | 3591064.25 | 708047.25 | 2858058.25 | 1248121.625 | 1002.52136 | AALNEIESR | Q9D3G5 | Q9D3G5 | STX11 | syntaxin 11 | Plasma Membrane | transporter |
| 15536586 | 2 | 0.035 | 1037418.062 | 1256530.875 | 1592276.625 | 1714834 | 2185773.75 | 2110429.25 | 1686484.125 | 2126279 | 1902390.875 | 992741 | 1647864.625 | 1375674.5 | 1266.63 | NILSSADYVER | P70452 | P70452 | STX4 | syntaxin 4 | Plasma Membrane | transporter |
| 15551560 | 2 | 0.037 | 1.54E+07 | 1.46E+07 | 1.71E+07 | 1.27E+07 | 1.60E+07 | 1.56E+07 | 1.69E+07 | 1.62E+07 | 1.50E+07 | 1.31E+07 | 1.31E+07 | 1.10E+07 | 1668.8513 | TLNQLGTPQDSPELR | O70439 | O70439 | STX7 | syntaxin 7 | Plasma Membrane | transporter |
| 15538737 | 2 | 0.027 | 691248.312 | 1470138.875 | 1289102.5 | 842024.75 | 2540059 | 1503487.125 | 1396184.5 | 1928634.625 | 1410155.125 | 567808 | 629901.75 | 458805.781 | 1142.5298 | YETTGLSESR | Q64324 | Q64324 | STXBP2 | syntaxin binding protein 2 | Plasma Membrane | transporter |
| 15542033 | 2 | 0.029 | 263992 | 560121.125 | 611466.688 | 1245225.375 | 1371598.25 | 1459691.875 | 797343.438 | 1200909.125 | 1917613.5 | 622879.188 | 1432094.875 | 985406.75 | 1454.8635 | DSMLVLLPVLLNK | Q60770 | Q60770 | STXBP3 | syntaxin binding protein 3 | Plasma Membrane | transporter |
| 15558748 | 3 | 0.018 | 1035516.125 | 1130636.375 | 1516196.375 | 1334209.25 | 1160685.625 | 1291236.125 | 1124970.875 | 972802.438 | 824325.938 | 942748.312 | 657624.312 | 574463.438 | 2427.043 | ETSLHSPSNTSASHSQGGGPPTSGM[14 | O09117 | O09117 | SYPL1 | synaptophysin-like 1 | Plasma Membrane | transporter |
| 15553979 | 3 | 0.042 | 350915.562 | 1361387.125 | 1346846.375 | 3595734.25 | 7652222.5 | 5110117.5 | 4356180 | 4655964 | 3929155.25 | 732055 | 4995577 | 3433495.75 | 3874.7202 | DFLQMVLDAQHSMNSVGVEGFDMVPI | P36423 | P36423 | TBXAS1 | thromboxane A synthase 1 (platelet) | Plasma Membrane | enzyme |
| 15532168 | 2 | 0.047 | 5.20E+07 | 1.06E+08 | 4.90E+07 | 6.66E+07 | 5.60E+07 | 7.56E+07 | 4.03E+07 | 8.46E+07 | 7.02E+07 | 3.13E+07 | 2.88E+07 | 3.45E+07 | 1181.6019 | LDTYEALTQK | Q62351 | Q62351 | TFRC | transferrin receptor | Plasma Membrane | transporter |
| 15532767 | 2 | 0.046 | 3.69E+07 | 9.10E+07 | 3.86E+07 | 5.14E+07 | 4.32E+07 | 5.83E+07 | 3.01E+07 | 6.40E+07 | 6.39E+07 | 2.15E+07 | 2.33E+07 | 2.21E+07 | 964.5443 | VVLGTSNFK | Q62351 | Q62351 | TFRC | transferrin receptor | Plasma Membrane | transporter |
| 15549263 | 3 | 0.046 | 6567488 | 3.32E+07 | 2.28E+07 | 7753453 | 1.20E+07 | 2.01E+07 | 7971492 | 2.25E+07 | 1.23E+07 | 5560156.5 | 2205688 | 3435905.5 | 3267.5835 | SSIGQNMVTIVQSNGNLDPVESPEGYV | Q62351 | Q62351 | TFRC | transferrin receptor | Plasma Membrane | transporter |
| 15551378 | 2 | 0.045 | 1.61E+07 | 2.81E+07 | 1.18E+07 | 1.47E+07 | 1.58E+07 | 1.65E+07 | 1.27E+07 | 1.96E+07 | 1.99E+07 | 7533637.5 | 1.06E+07 | 8531642 | 1593.8041 | VEYHFLSPYVSPR | Q62351 | Q62351 | TFRC | transferrin receptor | Plasma Membrane | transporter |
| 15563627 | 2 | 0.044 | 103382.984 | 143858.688 | 112595.812 | 111171.867 | 133937.984 | 258336.609 | 70753.289 | 245624.609 | 205472.281 | 67594.93 | 7431.859 | 41028.578 | 1579.6512 | LAADEEENADNNM[147.0354]K | Q62351 | Q62351 | TFRC | transferrin receptor | Plasma Membrane | transporter |
| 15534453 | 2 | 0.043 | 3.22E+07 | 7.56E+07 | 3.51E+07 | 4.82E+07 | 3.89E+07 | 6.15E+07 | 2.63E+07 | 5.12E+07 | 5.82E+07 | 1.55E+07 | 1.92E+07 | 2.42E+07 | 974.5867 | SSVGTGLLLK | Q62351 | Q62351 | TFRC | transferrin receptor | Plasma Membrane | transporter |
| 15553196 | 2 | 0.043 | 1.47E+07 | 2.84E+07 | 1.17E+07 | 1.65E+07 | 1.48E+07 | 1.64E+07 | 1.01E+07 | 1.68E+07 | 1.74E+07 | 8621770 | 7929391 | 7897225.5 | 1737.853 | GYEEPDRYVVVGAQR | Q62351 | Q62351 | TFRC | transferrin receptor | Plasma Membrane | transporter |
| 15535478 | 2 | 0.042 | 3.82E+07 | 8.46E+07 | 3.66E+07 | 5.06E+07 | 4.23E+07 | 4.91E+07 | 2.97E+07 | 6.08E+07 | 6.51E+07 | 2.22E+07 | 2.09E+07 | 2.32E+07 | 1084.5654 | AFTYINLDK | Q62351 | Q62351 | TFRC | transferrin receptor | Plasma Membrane | transporter |
| 15542375, 15544885, 1554503 | 3 | 0.04 | 2.68E+07 | 4.88E+07 | 2.30E+07 | 3.45E+07 | 2.61E+07 | 3.40E+07 | 2.11E+07 | 3.67E+07 | 3.77E+07 | 1.64E+07 | 1.70E+07 | 1.47E+07 | 1231.619, 1231.6191, 123 | VWRDEHYVK | Q62351 | Q62351 | TFRC | transferrin receptor | Plasma Membrane | transporter |
| 15545042 | 2 | 0.032 | 3722461.5 | 7339699.5 | 3063723 | 4280272 | 3158053 | 3647535.5 | 2682244.25 | 4076799.25 | 4963400 | 1837945.25 | 2225625.5 | 1524803.75 | 1213.712 | TAAEVAGQLIIK | Q62351 | Q62351 | TFRC | transferrin receptor | Plasma Membrane | transporter |
| 15547701 | 2 | 0.022 | 5.79E+07 | 1.22E+08 | 5.23E+07 | 7.85E+07 | 6.58E+07 | 6.84E+07 | 4.39E+07 | 8.30E+07 | 8.38E+07 | 3.32E+07 | 2.99E+07 | 2.86E+07 | 1568.672 | SETMETEDVPTSSR | Q62351 | Q62351 | TFRC | transferrin receptor | Plasma Membrane | transporter |
| 15547942 | 2 | 0.018 | 3.39E+07 | 6.93E+07 | 3.42E+07 | 3.61E+07 | 3.80E+07 | 4.48E+07 | 2.55E+07 | 4.47E+07 | 4.89E+07 | 1.71E+07 | 1.69E+07 | 2.03E+07 | 2032.9238 | DESLAYYIENQFHEFK | Q62351 | Q62351 | TFRC | transferrin receptor | Plasma Membrane | transporter |
| 15547867 | 2 | 0.009 | 4.34E+07 | 8.94E+07 | 4.19E+07 | 6.13E+07 | 4.80E+07 | 6.01E+07 | 4.18E+07 | 6.67E+07 | 6.62E+07 | 2.66E+07 | 2.21E+07 | 2.16E+07 | 1563.6566 | LAADEEENADNNMK | Q62351 | Q62351 | TFRC | transferrin receptor | Plasma Membrane | transporter |
| 15550672 | 2 | 0.002 | 2310438.75 | 4594391 | 2247024.25 | 3420251 | 2595682 | 3839703.75 | 1999949.125 | 2640126.5 | 2792806.75 | 1196992.5 | 978427.125 | 847927.875 | 1584.666 | SETM[147.0354]ETEDVPTSSR | Q62351 | Q62351 | TFRC | transferrin receptor | Plasma Membrane | transporter |
| 15547017 | 2 | 7.63E-05 | 602615.188 | 767849.688 | 671895.5 | 308053.094 | 371788.781 | 339937.844 | 712850 | 615427.812 | 537054.25 | 267791.938 | 159069.25 | 213300.625 | 1438.6019 | DSVTC[160.0307]PTC[160.0307]QGT | Q80X71 | Q80X71 | TMEM106B | transmembrane protein 106B | Plasma Membrane | other |
| 15551846 | 3 | 0.032 | 7884521.5 | 7358304.5 | 1.54E+07 | 1.01E+07 | 1.21E+07 | 1.20E+07 | 8052635 | 8309608.5 | 1.03E+07 | 1.99E+07 | 1.44E+07 | 1.88E+07 | 2598.2598 | EVDRYDAPGIAFYPGQAQLLSC[160.03 | Q9D771 | Q9D771 | TMEM206 | transmembrane protein 206 | Plasma Membrane | other |
| 15535852 | 2 | 0.042 | 4055732 | 4066555.75 | 3929822.5 | 2686818.25 | 3418292.5 | 2943596.75 | 3132247.25 | 3631236.75 | 3619119 | 3858399 | 2983994 | 3474766.5 | 807.47046 | LFSVVSR | Q9WTR1 | Q9WTR1 | TRPV2 | transient receptor potential cation channel, subfamily V, member 2 | Plasma Membrane | ion channel |
| 15540714 | 2, 3 | 0.007 | 6221382 | 5134792.5 | 4170947.75 | 3511451.25 | 4249381 | 4067574.75 | 5567326 | 5453505 | 4808214 | 6924007.5 | 7835852.5 | 6065570.5 | 1430.743, 1430.7433 | NSVLEIIAFHC[160.0307]K | Q9WTR1 | Q9WTR1 | TRPV2 | transient receptor potential cation channel, subfamily V, member 2 | Plasma Membrane | ion channel |
| 15550073 | 3 | 0.049 | 3.39E+07 | 1.90E+07 | 1.23E+07 | 1.71E+07 | 1.61E+07 | 1.12E+07 | 2.97E+07 | 3.10E+07 | 3.08E+07 | 2.67E+07 | 3.48E+07 | 2.49E+07 | 3963.7456 | C[160.0307]SVPYSC[160.0307]C[160. | Q8R3S2 | Q8R3S2 | TSPAN33 | tetraspanin 33 | Plasma Membrane | other |
| 15542057, 15544300 | 2 | 0.021 | 2.68E+07 | 1.88E+07 | 1.80E+07 | 1.13E+07 | 1.39E+07 | 9562086 | 2.79E+07 | 2.20E+07 | 1.92E+07 | 2.62E+07 | 2.20E+07 | 1.47E+07 | 1206.4993, 1206.4995 | FSC[160.0307]C[160.0307]GGISYR | Q8R3S2 | Q8R3S2 | TSPAN33 | tetraspanin 33 | Plasma Membrane | other |
| 15538275 | 2 | 0.021 | 2207507.25 | 1759359.625 | 3295511.5 | 1127262.5 | 1284261.625 | 1283956 | 3382642.75 | 2979079.75 | 2301025.25 | 2079618.125 | 2819225.5 | 1399616.375 | 1182.5793 | VIYTNGC[160.0307]IDK | Q8R3S2 | Q8R3S2 | TSPAN33 | tetraspanin 33 | Plasma Membrane | other |
| 15550507, 15551644 | 3, 4 | 1.13E-04 | 2.17E+08 | 2.09E+08 | 2.32E+08 | 8.77E+07 | 1.19E+08 | 1.03E+08 | 1.91E+08 | 1.89E+08 | 1.64E+08 | 1.38E+08 | 1.07E+08 | 1.23E+08 | 3142.6016, 3142.6082 | VSEIINNAIVHYRDDLDLQNLIDFGQK | Q8R3S2 | Q8R3S2 | TSPAN33 | tetraspanin 33 | Plasma Membrane | other |
| 15532719 | 2, 3 | 6.92E-05 | 7.11E+07 | 7.31E+07 | 7.77E+07 | 3.68E+07 | 3.89E+07 | 3.44E+07 | 7.33E+07 | 6.55E+07 | 5.51E+07 | 4.43E+07 | 3.38E+07 | 3.81E+07 | 1199.6244, 1199.6249 | LQLYNQQHR | Q8R3S2 | Q8R3S2 | TSPAN33 | tetraspanin 33 | Plasma Membrane | other |
| 15537034 | 3 | 0.041 | 6717073.5 | 7705563.5 | 7206602.5 | 4383845 | 6309872 | 4334073 | 7716146 | 8385475.5 | 6993261 | 5906573.5 | 8841654 | 6289297.5 | 2072.052 | RLQQTQNQVDEVVDIMR | P63024 | P63024 | VAMP3 | vesicle-associated membrane protein 3 | Plasma Membrane | other |
| 15534865 | 2, 3 | 0.046 | 1.08E+07 | 2.48E+07 | 2.05E+07 | 2.06E+07 | 2.78E+07 | 2.97E+07 | 2.05E+07 | 2.82E+07 | 2.51E+07 | 9798140 | 1.53E+07 | 1.34E+07 | 1406.6401, 1406.6407 | TEDLEATSEHFK | O70404 | O70404 | VAMP8 | vesicle-associated membrane protein 8 | Plasma Membrane | transporter |
| 15532414 | 2 | 0.017 | 1.14E+07 | 9939133 | 9668522 | 1.04E+07 | 1.13E+07 | 9890620 | 1.23E+07 | 1.22E+07 | 1.11E+07 | 9661740 | 9572098 | 8765391 | 1382.5677 | C[160.0307]VFEMPNENDK | Q9WV55 | Q9WV55 | VAPA | VAMP (vesicle-associated membrane protein)-associated protein A, 33kDa | Plasma Membrane | other |
| 15534254 | 3 | 0.036 | 3036443 | 4341906 | 4903065 | 5904795.5 | 7369567 | 6052565.5 | 5368142 | 5154346 | 6421618.5 | 4711760.5 | 5608154.5 | 4682829.5 | 1648.8867 | VEQVLSLEPQHELK | Q9QY76 | Q9QY76 | VAPB | VAMP (vesicle-associated membrane protein)-associated protein B and C | Plasma Membrane | other |

List of all proteins with significant differences
$p<0.05$

| ID | Symbol | Entrez Gene Name | Location |
|---|---|---|---|
| Q9JI39 | ABCB10 | ATP-binding cassette, sub-family B (MDR/TAP), member 10 | Cytoplasm |
| Q9DC29 | ABCB6 | ATP-binding cassette, sub-family B (MDR/TAP), member 6 (Langereis blood group) | Cytoplasm |
| P61222 | ABCE1 | ATP-binding cassette, sub-family E (OABP), member 1 | Cytoplasm |
| Q7TMS5 | ABCG2 | ATP-binding cassette, sub-family G (WHITE), member 2 (Junior blood group) | Plasma Membrane |
| Q9QUI6 | ACKR1 | atypical chemokine receptor 1 (Duffy blood group) | Plasma Membrane |
| Q8JZR0 | ACSL5 | acyl-CoA synthetase long-chain family member 5 | Cytoplasm |
| P60710 | ACTB | actin, beta | Cytoplasm |
| O35598 | ADAM10 | ADAM metallopeptidase domain 10 | Plasma Membrane |
| Q9QYC0 | ADD1 | adducin 1 (alpha) | Cytoplasm |
| Q9QYB8 | ADD2 | adducin 2 (beta) | Cytoplasm |
| Q9QYB5 | ADD3 | adducin 3 (gamma) | Cytoplasm |
| Q8CJG0 | AGO2 | argonaute RISC catalytic component 2 | Cytoplasm |
| P31230 | AIMP1 | aminoacyl tRNA synthetase complex-interacting multifunctional protein 1 | Extracellular Space |
| P10518 | ALAD | aminolevulinate dehydratase | Cytoplasm |
| P07724 | ALB | albumin | Extracellular Space |
| Q9Z110 | ALDH18A1 | aldehyde dehydrogenase 18 family, member A1 | Cytoplasm |
| Q02357 | ANK1 | ankyrin 1, erythrocytic | Plasma Membrane |
| Q6P9J9 | ANO6 | anoctamin 6 | Plasma Membrane |
| O35643 | AP1B1 | adaptor-related protein complex 1, beta 1 subunit | Cytoplasm |
| Q00623 | APOA1 | apolipoprotein A-I | Extracellular Space |
| Q02013 | AQP1 | aquaporin 1 (Colton blood group) | Plasma Membrane |
| P84084 | ARF5 | ADP-ribosylation factor 5 | Cytoplasm |
| P62331 | ARF6 | ADP-ribosylation factor 6 | Plasma Membrane |
| Q9CQW2 | ARL8B | ADP-ribosylation factor-like 8B | Plasma Membrane |
| Q9CVB6 | ARPC2 | actin related protein 2/3 complex, subunit 2, 34kDa | Cytoplasm |
| Q9CRA0 | ART4 | ADP-ribosyltransferase 4 (Dombrock blood group) | Nucleus |
| Q91YH5 | ATL3 | atlastin GTPase 3 | Cytoplasm |
| Q8VDN2 | ATP1A1 | ATPase, Na+/K+ transporting, alpha 1 polypeptide | Plasma Membrane |
| P14231 | ATP1B2 | ATPase, Na+/K+ transporting, beta 2 polypeptide | Plasma Membrane |
| P97370 | ATP1B3 | ATPase, Na+/K+ transporting, beta 3 polypeptide | Plasma Membrane |
| Q64518 | ATP2A3 | ATPase, Ca++ transporting, ubiquitous | Cytoplasm |
| Q9R0K7 | ATP2B2 | ATPase, Ca++ transporting, plasma membrane 2 | Plasma Membrane |
| P56480 | ATP5B | ATP synthase, H+ transporting, mitochondrial F1 complex, beta polypeptide | Cytoplasm |
| Q9Z1G4 | ATP6V0A1 | ATPase, H+ transporting, lysosomal V0 subunit a1 | Cytoplasm |

| | | | |
|---|---|---|---|
| Q64446 | ATP7B | ATPase, Cu++ transporting, beta polypeptide | Cytoplasm |
| P70704 | ATP8A1 | ATPase, aminophospholipid transporter (APLT), class I, type 8A, member 1 | Cytoplasm |
| P01887 | B2M | beta-2-microglobulin | Plasma Membrane |
| Q923D2 | BLVRB | biliverdin reductase B | Cytoplasm |
| P15327 | BPGM | 2,3-bisphosphoglycerate mutase | Extracellular Space |
| P18572 | BSG | basigin (Ok blood group) | Plasma Membrane |
| Q9CX63 | C17orf99 | chromosome 17 open reading frame 99 | Extracellular Space |
| P00920 | CA2 | carbonic anhydrase II | Cytoplasm |
| P35564 | CANX | calnexin | Cytoplasm |
| O35350 | CAPN1 | calpain 1, (mu/I) large subunit | Cytoplasm |
| O08688 | CAPN5 | calpain 5 | Cytoplasm |
| Q60865 | CAPRIN1 | cell cycle associated protein 1 | Plasma Membrane |
| P47753 | CAPZA1 | capping protein (actin filament) muscle Z-line, alpha 1 | Cytoplasm |
| P47757 | CAPZB | capping protein (actin filament) muscle Z-line, beta | Cytoplasm |
| P24270 | CAT | catalase | Cytoplasm |
| Q8BGU5 | CCNY | cyclin Y | Nucleus |
| Q08857 | CD36 | CD36 molecule (thrombospondin receptor) | Plasma Membrane |
| P15379 | CD44 | CD44 molecule (Indian blood group) | Plasma Membrane |
| P18181 | CD48 | CD48 molecule | Plasma Membrane |
| Q61475 | CD55 | CD55 molecule, decay accelerating factor for complement (Cromer blood group) | Plasma Membrane |
| Q8VDP6 | CDIPT | CDP-diacylglycerol--inositol 3-phosphatidyltransferase | Cytoplasm |
| Q4VAA2 | CDV3 | CDV3 homolog (mouse) | Cytoplasm |
| P61022 | CHP1 | calcineurin-like EF-hand protein 1 | Cytoplasm |
| Q9CQB5 | CISD2 | CDGSH iron sulfur domain 2 | Cytoplasm |
| Q9Z0S4 | Cldn13 | claudin 13 | Plasma Membrane |
| Q8R4N0 | CLYBL | citrate lyase beta like | Cytoplasm |
| O35372 | CNIH1 | cornichon family AMPA receptor auxiliary protein 1 | Plasma Membrane |
| O89053 | CORO1A | coronin, actin binding protein, 1A | Cytoplasm |
| P36552 | CPOX | coproporphyrinogen oxidase | Cytoplasm |
| Q64735 | CR1L | complement component (3b/4b) receptor 1-like | Plasma Membrane |
| O88668 | CREG1 | cellular repressor of E1A-stimulated genes 1 | Nucleus |
| Q9DCT8 | Crip2 | cysteine rich protein 2 | Plasma Membrane |
| P18242 | CTSD | cathepsin D | Cytoplasm |
| Q9CQX2 | CYB5B | cytochrome b5 type B (outer mitochondrial membrane) | Cytoplasm |
| Q8BPM0 | DAAM1 | dishevelled associated activator of morphogenesis 1 | Cytoplasm |

| | | | |
|---|---|---|---|
| Q9JII5 | Dazap1 | DAZ associated protein 1 | Cytoplasm |
| Q8C4S8 | DENND2A | DENN/MADD domain containing 2A | Cytoplasm |
| Q3U0B3 | DHRS11 | dehydrogenase/reductase (SDR family) member 11 | Cytoplasm |
| A2ATU0 | DHTKD1 | dehydrogenase E1 and transketolase domain containing 1 | Cytoplasm |
| Q811D0 | DLG1 | discs, large homolog 1 (Drosophila) | Plasma Membrane |
| Q9QYJ3 | DNAJB1 | DnaJ (Hsp40) homolog, subfamily B, member 1 | Nucleus |
| Q5U458 | DNAJC11 | DnaJ (Hsp40) homolog, subfamily C, member 11 | Cytoplasm |
| P39054 | DNM2 | dynamin 2 | Plasma Membrane |
| Q8BZ98 | DNM3 | dynamin 3 | Cytoplasm |
| P10126 | EEF1A1 | eukaryotic translation elongation factor 1 alpha 1 | Cytoplasm |
| Q9D8N0 | EEF1G | eukaryotic translation elongation factor 1 gamma | Cytoplasm |
| P58252 | EEF2 | eukaryotic translation elongation factor 2 | Cytoplasm |
| Q8BG67 | EFR3A | EFR3 homolog A | Plasma Membrane |
| Q9WVK4 | EHD1 | EH-domain containing 1 | Cytoplasm |
| Q3UGC7 | EIF3J | eukaryotic translation initiation factor 3, subunit J | Cytoplasm |
| Q6NZJ6 | EIF4G1 | eukaryotic translation initiation factor 4 gamma, 1 | Cytoplasm |
| Q8BTJ4 | ENPP4 | ectonucleotide pyrophosphatase/phosphodiesterase 4 (putative) | Cytoplasm |
| P10404 | ENV1 | MLV-related proviral Env polyprotein | Other |
| P48193 | EPB41 | erythrocyte membrane protein band 4.1 | Plasma Membrane |
| P49222 | EPB42 | erythrocyte membrane protein band 4.2 | Plasma Membrane |
| Q9JLN5 | ERMAP | erythroblast membrane-associated protein (Scianna blood group) | Cytoplasm |
| Q3U7R1 | ESYT1 | extended synaptotagmin-like protein 1 | Cytoplasm |
| Q3TZZ7 | ESYT2 | extended synaptotagmin-like protein 2 | Plasma Membrane |
| O88792 | F11R | F11 receptor | Plasma Membrane |
| P22315 | FECH | ferrochelatase | Cytoplasm |
| P97807 | FH | fumarate hydratase | Cytoplasm |
| Q6P9Q6 | FKBP15 | FK506 binding protein 15, 133kDa | Plasma Membrane |
| Q8BTM8 | FLNA | filamin A, alpha | Cytoplasm |
| O08917 | FLOT1 | flotillin 1 | Plasma Membrane |
| Q60634 | FLOT2 | flotillin 2 | Plasma Membrane |
| P97855 | G3BP1 | GTPase activating protein (SH3 domain) binding protein 1 | Nucleus |
| Q6NSQ9 | G6PC3 | glucose 6 phosphatase, catalytic, 3 | Cytoplasm |
| Q00612 | G6PD | glucose-6-phosphate dehydrogenase | Cytoplasm |
| P60521 | GABARAPL2 | GABA(A) receptor-associated protein like 2 | Cytoplasm |
| Q8BHN3 | GANAB | glucosidase, alpha; neutral AB | Cytoplasm |

| | | | |
|---|---|---|---|
| O09172 | GCLM | glutamate-cysteine ligase, modifier subunit | Cytoplasm |
| P08752 | GNAI2 | guanine nucleotide binding protein (G protein), alpha inhibiting activity polypeptide 2 | Plasma Membrane |
| Q6R0H7 | GNAS | GNAS complex locus | Plasma Membrane |
| P62874 | GNB1 | guanine nucleotide binding protein (G protein), beta polypeptide 1 | Plasma Membrane |
| P62880 | GNB2 | guanine nucleotide binding protein (G protein), beta polypeptide 2 | Plasma Membrane |
| Q91W53 | GOLGA7 | golgin A7 | Cytoplasm |
| P05202 | GOT2 | glutamic-oxaloacetic transaminase 2, mitochondrial | Cytoplasm |
| P51655 | GPC4 | glypican 4 | Plasma Membrane |
| P11352 | GPX1 | glutathione peroxidase 1 | Cytoplasm |
| P19157 | GSTP1 | glutathione S-transferase pi 1 | Cytoplasm |
| Q8BZI6 | GUCD1 | guanylyl cyclase domain containing 1 | Nucleus |
| Q8BMS1 | HADHA | hydroxyacyl-CoA dehydrogenase/3-ketoacyl-CoA thiolase/enoyl-CoA hydratase (trifunctic | Cytoplasm |
| Q99KB8 | HAGH | hydroxyacylglutathione hydrolase | Cytoplasm |
| P01942 | HBA1/HBA2 | hemoglobin, alpha 1 | Extracellular Space |
| P02088 | Hbb-b1 | hemoglobin, beta adult major chain | Extracellular Space |
| P01899, P01 | HLA-A | major histocompatibility complex, class I, A | Plasma Membrane |
| Q61411 | HRAS | Harvey rat sarcoma viral oncogene homolog | Plasma Membrane |
| O70503 | HSD17B12 | hydroxysteroid (17-beta) dehydrogenase 12 | Cytoplasm |
| P08113 | HSP90B1 | heat shock protein 90kDa beta (Grp94), member 1 | Cytoplasm |
| P38647 | HSPA9 | heat shock 70kDa protein 9 (mortalin) | Cytoplasm |
| P63038 | HSPD1 | heat shock 60kDa protein 1 (chaperonin) | Cytoplasm |
| Q64433 | HSPE1 | heat shock 10kDa protein 1 | Cytoplasm |
| Q9ERM2 | ICAM4 | intercellular adhesion molecule 4 (Landsteiner-Wiener blood group) | Plasma Membrane |
| Q9JHR7 | IDE | insulin-degrading enzyme | Extracellular Space |
| Q9D6R2 | IDH3A | isocitrate dehydrogenase 3 (NAD+) alpha | Cytoplasm |
| Q64339 | ISG15 | ISG15 ubiquitin-like modifier | Extracellular Space |
| Q9JL16 | ISG20 | interferon stimulated exonuclease gene 20kDa | Nucleus |
| Q9QUM0 | ITGA2B | integrin, alpha 2b (platelet glycoprotein IIb of IIb/IIIa complex, antigen CD41) | Plasma Membrane |
| P09055 | ITGB1 | integrin, beta 1 (fibronectin receptor, beta polypeptide, antigen CD29 includes MDF2, MS | Plasma Membrane |
| Q9EQF2 | KEL | Kell blood group, metallo-endopeptidase | Plasma Membrane |
| P70168 | KPNB1 | karyopherin (importin) beta 1 | Nucleus |
| O88653 | LAMTOR3 | late endosomal/lysosomal adaptor, MAPK and MTOR activator 3 | Cytoplasm |
| Q9D1L9 | LAMTOR5 | late endosomal/lysosomal adaptor, MAPK and MTOR activator 5 | Cytoplasm |
| P06151 | LDHA | lactate dehydrogenase A | Cytoplasm |
| O08573 | LGALS9B | lectin, galactoside-binding, soluble, 9B | Cytoplasm |

| | | | |
|---|---|---|---|
| Q9DBH5 | LMAN2 | lectin, mannose-binding 2 | Cytoplasm |
| Q9D1G5 | LRRC57 | leucine rich repeat containing 57 | Cytoplasm |
| Q80TE7 | LRRC7 | leucine rich repeat containing 7 | Plasma Membrane |
| Q61166 | MAPRE1 | microtubule-associated protein, RP/EB family, member 1 | Cytoplasm |
| Q8BL86 | MBLAC2 | metallo-beta-lactamase domain containing 2 | Cytoplasm |
| Q91VH6 | MEMO1 | mediator of cell motility 1 | Cytoplasm |
| Q9CPU4 | MGST3 | microsomal glutathione S-transferase 3 | Cytoplasm |
| Q8K3F2 | MMP21 | matrix metallopeptidase 21 | Extracellular Space |
| P70290 | MPP1 | membrane protein, palmitoylated 1, 55kDa | Plasma Membrane |
| Q9WUX5 | MRVI1 | murine retrovirus integration site 1 homolog | Cytoplasm |
| P26041 | MSN | moesin | Plasma Membrane |
| P00405 | MT-CO2 | cytochrome c oxidase subunit II | Cytoplasm |
| Q922D8 | MTHFD1 | methylenetetrahydrofolate dehydrogenase (NADP+ dependent) 1, methenyltetrahydrofo | Cytoplasm |
| P28665 | Mug1/Mug2 | murinoglobulin 1 | Extracellular Space |
| Q8VDD5 | MYH9 | myosin, heavy chain 9, non-muscle | Cytoplasm |
| Q3THE2 | MYL12A | myosin, light chain 12A, regulatory, non-sarcomeric | Cytoplasm |
| Q9WTI7 | MYO1C | myosin IC | Cytoplasm |
| P57716 | NCSTN | nicastrin | Plasma Membrane |
| Q7TMF3 | NDUFA12 | NADH dehydrogenase (ubiquinone) 1 alpha subcomplex, 12 | Cytoplasm |
| Q9CPP6 | NDUFA5 | NADH dehydrogenase (ubiquinone) 1 alpha subcomplex, 5 | Cytoplasm |
| P08556 | NRAS | neuroblastoma RAS viral (v-ras) oncogene homolog | Plasma Membrane |
| A6H603 | NWD1 | NACHT and WD repeat domain containing 1 | Cytoplasm |
| Q60597 | OGDH | oxoglutarate (alpha-ketoglutarate) dehydrogenase (lipoamide) | Cytoplasm |
| Q8BGT0 | OSTM1 | osteopetrosis associated transmembrane protein 1 | Cytoplasm |
| P09103 | P4HB | prolyl 4-hydroxylase, beta polypeptide | Cytoplasm |
| P50580 | PA2G4 | proliferation-associated 2G4, 38kDa | Nucleus |
| P29341 | PABPC1 | poly(A) binding protein, cytoplasmic 1 | Cytoplasm |
| P60335 | PCBP1 | poly(rC) binding protein 1 | Nucleus |
| Q8BH04 | PCK2 | phosphoenolpyruvate carboxykinase 2 (mitochondrial) | Cytoplasm |
| P27773 | PDIA3 | protein disulfide isomerase family A, member 3 | Cytoplasm |
| P08003 | PDIA4 | protein disulfide isomerase family A, member 4 | Cytoplasm |
| Q922R8 | PDIA6 | protein disulfide isomerase family A, member 6 | Cytoplasm |
| Q9Z126 | PF4 | platelet factor 4 | Extracellular Space |
| P67778 | PHB | prohibitin | Nucleus |
| O35129 | PHB2 | prohibitin 2 | Cytoplasm |

| | | | |
|---|---|---|---|
| E2JF22 | PIEZO1 | piezo-type mechanosensitive ion channel component 1 | Cytoplasm |
| P06803 | PIM1 | Pim-1 proto-oncogene, serine/threonine kinase | Cytoplasm |
| P70182 | PIP5K1A | phosphatidylinositol-4-phosphate 5-kinase, type I, alpha | Cytoplasm |
| Q80ZA4 | PKHD1L1 | polycystic kidney and hepatic disease 1 (autosomal recessive)-like 1 | Extracellular Space |
| Q61469 | Ppap2a | phosphatidic acid phosphatase type 2A | Plasma Membrane |
| P24369 | PPIB | peptidylprolyl isomerase B (cyclophilin B) | Cytoplasm |
| P51175 | PPOX | protoporphyrinogen oxidase | Cytoplasm |
| P62137 | PPP1CA | protein phosphatase 1, catalytic subunit, alpha isozyme | Cytoplasm |
| P63330 | PPP2CA | protein phosphatase 2, catalytic subunit, alpha isozyme | Cytoplasm |
| P99029 | PRDX5 | peroxiredoxin 5 | Cytoplasm |
| P05132 | PRKACA | protein kinase, cAMP-dependent, catalytic, alpha | Cytoplasm |
| O08795 | PRKCSH | protein kinase C substrate 80K-H | Cytoplasm |
| P04925 | PRNP | prion protein | Plasma Membrane |
| Q61207 | PSAP | prosaposin | Extracellular Space |
| Q9R1P4 | PSMA1 | proteasome (prosome, macropain) subunit, alpha type, 1 | Cytoplasm |
| Q9R1P0 | PSMA4 | proteasome (prosome, macropain) subunit, alpha type, 4 | Cytoplasm |
| Q9Z2U0 | PSMA7 | proteasome (prosome, macropain) subunit, alpha type, 7 | Cytoplasm |
| Q9D8W5 | PSMD12 | proteasome (prosome, macropain) 26S subunit, non-ATPase, 12 | Cytoplasm |
| Q8BJY1 | PSMD5 | proteasome (prosome, macropain) 26S subunit, non-ATPase, 5 | Cytoplasm |
| Q8R3K3 | PTCD2 | pentatricopeptide repeat domain 2 | Cytoplasm |
| Q62136 | PTPN21 | protein tyrosine phosphatase, non-receptor type 21 | Cytoplasm |
| Q91V41 | RAB14 | RAB14, member RAS oncogene family | Cytoplasm |
| Q99P58 | RAB27B | RAB27B, member RAS oncogene family | Cytoplasm |
| P53994 | RAB2A | RAB2A, member RAS oncogene family | Cytoplasm |
| Q6PHN9 | RAB35 | RAB35, member RAS oncogene family | Cytoplasm |
| P51150 | RAB7A | RAB7A, member RAS oncogene family | Cytoplasm |
| P55258 | RAB8A | RAB8A, member RAS oncogene family | Plasma Membrane |
| P61028 | RAB8B | RAB8B, member RAS oncogene family | Cytoplasm |
| P62827 | RAN | RAN, member RAS oncogene family | Nucleus |
| P62835 | RAP1A | RAP1A, member of RAS oncogene family | Cytoplasm |
| Q9QYF1 | RDH11 | retinol dehydrogenase 11 (all-trans/9-cis/11-cis) | Cytoplasm |
| Q60870 | REEP5 | receptor accessory protein 5 | Extracellular Space |
| Q9QUT0 | RHAG | Rh-associated glycoprotein | Plasma Membrane |
| Q8CF94 | RHCE/RHD | Rh blood group, D antigen | Plasma Membrane |
| E9Q555 | RNF213 | ring finger protein 213 | Cytoplasm |

| | | | |
|---|---|---|---|
| Q9CXW4 | RPL11 | ribosomal protein L11 | Cytoplasm |
| P35979 | RPL12 | ribosomal protein L12 | Nucleus |
| Q9CR57 | RPL14 | ribosomal protein L14 | Cytoplasm |
| Q9CZM2 | RPL15 | ribosomal protein L15 | Cytoplasm |
| Q9CPR4 | RPL17 | ribosomal protein L17 | Cytoplasm |
| P35980 | RPL18 | ribosomal protein L18 | Cytoplasm |
| P62751 | Rpl23a | ribosomal protein L23A | Nucleus |
| P61255 | RPL26 | ribosomal protein L26 | Cytoplasm |
| P47964 | Rpl36 | ribosomal protein L36 | Nucleus |
| Q9D8E6 | RPL4 | ribosomal protein L4 | Cytoplasm |
| P47911 | RPL6 | ribosomal protein L6 | Nucleus |
| P14148 | RPL7 | ribosomal protein L7 | Nucleus |
| P12970 | RPL7A | ribosomal protein L7a | Cytoplasm |
| P62918 | RPL8 | ribosomal protein L8 | Cytoplasm |
| P51410 | RPL9 | ribosomal protein L9 | Nucleus |
| P14869 | RPLP0 | ribosomal protein, large, P0 | Cytoplasm |
| P99027 | RPLP2 | ribosomal protein, large, P2 | Cytoplasm |
| Q91YQ5 | RPN1 | ribophorin I | Cytoplasm |
| Q9DBG6 | RPN2 | ribophorin II | Cytoplasm |
| P63325 | RPS10 | ribosomal protein S10 | Cytoplasm |
| P62301 | RPS13 | ribosomal protein S13 | Cytoplasm |
| P62245 | RPS15A | ribosomal protein S15a | Cytoplasm |
| P62270 | RPS18 | ribosomal protein S18 | Cytoplasm |
| Q9CZX8 | RPS19 | ribosomal protein S19 | Cytoplasm |
| P25444 | RPS2 | ribosomal protein S2 | Cytoplasm |
| P62849 | RPS24 | ribosomal protein S24 | Cytoplasm |
| P62852 | RPS25 | ribosomal protein S25 | Cytoplasm |
| Q6ZWY3 | RPS27L | ribosomal protein S27-like | Cytoplasm |
| P97351 | Rps3a1 | ribosomal protein S3A1 | Cytoplasm |
| P62702 | RPS4Y1 | ribosomal protein S4, Y-linked 1 | Cytoplasm |
| P62082 | RPS7 | ribosomal protein S7 | Cytoplasm |
| P62242 | RPS8 | ribosomal protein S8 | Cytoplasm |
| Q6ZWN5 | RPS9 | ribosomal protein S9 | Cytoplasm |
| P14206 | RPSA | ribosomal protein SA | Cytoplasm |
| P97352 | S100A13 | S100 calcium binding protein A13 | Cytoplasm |

| | | | |
|---|---|---|---|
| Q9EP69 | SACM1L | SAC1 suppressor of actin mutations 1-like (yeast) | Cytoplasm |
| Q9ERN0 | SCAMP2 | secretory carrier membrane protein 2 | Cytoplasm |
| Q9CQA3 | SDHB | succinate dehydrogenase complex, subunit B, iron sulfur (Ip) | Cytoplasm |
| Q9CQS8 | SEC61B | Sec61 beta subunit | Cytoplasm |
| P17563 | SELENBP1 | selenium binding protein 1 | Cytoplasm |
| Q01102 | SELP | selectin P (granule membrane protein 140kDa, antigen CD62) | Plasma Membrane |
| P07758 | SERPINA1 | serpin peptidase inhibitor, clade A (alpha-1 antiproteinase, antitrypsin), member 1 | Extracellular Space |
| Q9JJU8 | SH3BGRL | SH3 domain binding glutamate-rich protein like | Cytoplasm |
| Q8VHL0 | SLC14A1 | solute carrier family 14 (urea transporter), member 1 (Kidd blood group) | Plasma Membrane |
| P53986 | SLC16A1 | solute carrier family 16 (monocarboxylate transporter), member 1 | Plasma Membrane |
| Q3UHH2 | SLC22A23 | solute carrier family 22, member 23 | Plasma Membrane |
| Q8BZ09 | SLC25A21 | solute carrier family 25 (mitochondrial oxoadipate carrier), member 21 | Cytoplasm |
| P58735 | SLC26A1 | solute carrier family 26 (anion exchanger), member 1 | Plasma Membrane |
| Q9JIM1 | SLC29A1 | solute carrier family 29 (equilibrative nucleoside transporter), member 1 | Plasma Membrane |
| P32037 | SLC2A3 | solute carrier family 2 (facilitated glucose transporter), member 3 | Plasma Membrane |
| Q60738 | SLC30A1 | solute carrier family 30 (zinc transporter), member 1 | Plasma Membrane |
| P10852 | SLC3A2 | solute carrier family 3 (amino acid transporter heavy chain), member 2 | Plasma Membrane |
| P04919 | SLC4A1 | solute carrier family 4 (anion exchanger), member 1 (Diego blood group) | Plasma Membrane |
| P28571 | SLC6A9 | solute carrier family 6 (neurotransmitter transporter, glycine), member 9 | Plasma Membrane |
| Q9Z127 | SLC7A5 | solute carrier family 7 (amino acid transporter light chain, L system), member 5 | Plasma Membrane |
| Q8CG48 | SMC2 | structural maintenance of chromosomes 2 | Nucleus |
| O09044 | SNAP23 | synaptosomal-associated protein, 23kDa | Plasma Membrane |
| O55042 | SNCA | synuclein, alpha (non A4 component of amyloid precursor) | Cytoplasm |
| O70493 | SNX12 | sorting nexin 12 | Cytoplasm |
| P09671 | SOD2 | superoxide dismutase 2, mitochondrial | Cytoplasm |
| P08032 | SPTA1 | spectrin, alpha, erythrocytic 1 | Cytoplasm |
| P15508 | SPTB | spectrin, beta, erythrocytic | Plasma Membrane |
| P13609 | Srgn | serglycin | Cytoplasm |
| P54116 | STOM | stomatin | Plasma Membrane |
| Q9D3G5 | STX11 | syntaxin 11 | Plasma Membrane |
| Q00262 | STX2 | syntaxin 2 | Cytoplasm |
| P70452 | STX4 | syntaxin 4 | Plasma Membrane |
| O70439 | STX7 | syntaxin 7 | Plasma Membrane |
| Q64324 | STXBP2 | syntaxin binding protein 2 | Plasma Membrane |
| Q60770 | STXBP3 | syntaxin binding protein 3 | Plasma Membrane |

| | | | |
|---|---|---|---|
| O09117 | SYPL1 | synaptophysin-like 1 | Plasma Membrane |
| Q9R233 | TAPBP | TAP binding protein (tapasin) | Cytoplasm |
| P36423 | TBXAS1 | thromboxane A synthase 1 (platelet) | Plasma Membrane |
| Q921I1 | TF | transferrin | Extracellular Space |
| Q62351 | TFRC | transferrin receptor | Plasma Membrane |
| Q80ZW2 | THEM6 | thioesterase superfamily member 6 | Extracellular Space |
| Q9ET30 | TM9SF3 | transmembrane 9 superfamily member 3 | Cytoplasm |
| Q9D1D4 | TMED10 | transmembrane emp24-like trafficking protein 10 (yeast) | Cytoplasm |
| Q8R1V4 | TMED4 | transmembrane emp24 protein transport domain containing 4 | Cytoplasm |
| Q80X71 | TMEM106B | transmembrane protein 106B | Plasma Membrane |
| Q9D771 | TMEM206 | transmembrane protein 206 | Plasma Membrane |
| Q5F285 | TMEM256 | transmembrane protein 256 | Cytoplasm |
| Q9WTR1 | TRPV2 | transient receptor potential cation channel, subfamily V, member 2 | Plasma Membrane |
| Q62348 | TSN | translin | Nucleus |
| Q8R3S2 | TSPAN33 | tetraspanin 33 | Plasma Membrane |
| Q8CDN6 | TXNL1 | thioredoxin-like 1 | Cytoplasm |
| P62984 | Uba52 | ubiquitin A-52 residue ribosomal protein fusion product 1 | Cytoplasm |
| Q6ZPJ3 | UBE2O | ubiquitin-conjugating enzyme E2O | Nucleus |
| A2AN08 | UBR4 | ubiquitin protein ligase E3 component n-recognin 4 | Nucleus |
| Q6P5E4 | UGGT1 | UDP-glucose glycoprotein glucosyltransferase 1 | Cytoplasm |
| P63024 | VAMP3 | vesicle-associated membrane protein 3 | Plasma Membrane |
| O70404 | VAMP8 | vesicle-associated membrane protein 8 | Plasma Membrane |
| Q9WV55 | VAPA | VAMP (vesicle-associated membrane protein)-associated protein A, 33kDa | Plasma Membrane |
| Q9QY76 | VAPB | VAMP (vesicle-associated membrane protein)-associated protein B and C | Plasma Membrane |
| Q3U2A8 | VARS2 | valyl-tRNA synthetase 2, mitochondrial | Cytoplasm |
| Q60930 | VDAC2 | voltage-dependent anion channel 2 | Cytoplasm |
| Q5H8C4 | VPS13A | vacuolar protein sorting 13 homolog A (S. cerevisiae) | Cytoplasm |
| Q8BX70 | VPS13C | vacuolar protein sorting 13 homolog C (S. cerevisiae) | Cytoplasm |
| Q9CQV8 | YWHAB | tyrosine 3-monooxygenase/tryptophan 5-monooxygenase activation protein, beta | Cytoplasm |
| P62259 | YWHAE | tyrosine 3-monooxygenase/tryptophan 5-monooxygenase activation protein, epsilon | Cytoplasm |
| P61982 | YWHAG | tyrosine 3-monooxygenase/tryptophan 5-monooxygenase activation protein, gamma | Cytoplasm |
| P63101 | YWHAZ | tyrosine 3-monooxygenase/tryptophan 5-monooxygenase activation protein, zeta | Cytoplasm |
