## Supplementary Discussion for "Role of channels in the oxygen permeability of red blood cells"

### Supplementary Information: Discussion

#### Krogh

In his pioneering modeling work on $O_2$ delivery from capillary to tissue,[1] August Krogh introduced a now-famous equation—derived by his mathematician-colleague Mr. K Erlang—that played a major role in elucidating the fundamental principles of how [$O_2$] changes both longitudinally along the capillary and radially away from it. This equation reflects a highly simplified view of the system, which of course was necessary at that time, given the limitations in available computational algorithms and computing power. The analysis by Kreuzer,[2] lists 15 assumption—some of which he and others had noted previously—that are implicit in the derivation of the Krogh-Erlang equation. Assumption #12 states that "the diffusion coefficient [of $O_2$] is the same throughout the tissue." Assumption #7 (previously noted by Hess[3])—really a corollary of #12—states that "the capillary wall does not present any resistance to oxygen diffusion." Another corollary of #12 is that no cell membrane, including that of the RBC, offers resistance to $O_2$ diffusion. We examine implications of this implicit assumption in the section below.

#### How others concluded that the plasma membrane offers no resistance to $O_2$ diffusion

After Krogh's landmark paper, most practitioners in the field firmly adopted this implicit assumption. However, a series of studies from Roughton and his colleagues led to the suggestion that the plasma membrane of RBCs offers significant resistance to the movement of gases, including $CO_2$ and $O_2$.[4–8] However, later studies questioned this possibility and attributed the observed resistance to the extracellular unconvected layer (UL) of fluid that surrounds the cells after flow stops in the SF apparatus.[9–14] It is true that studies that focus on $O_2$ influx across RBCs are intrinsically challenging because this influx depletes $O_2$ at the surface of the RBC, and the UL of that surrounds the RBC offers resistance to the replenishment of $O_2$ from the bulk extracellular fluid (bECF).[9–12] Moreover, as pointed out by Vandegriff and Olson, the low solubility of $O_2$ in water enhances the effect of ULs in experiments of $O_2$ influx.[13] Nevertheless, it is possible that the implicit assumption of past authors—namely, that the plasma membrane offers no resistance of $O_2$ diffusion ($R_{\text{Membrane}}$)—may have led them to dismiss Roughton hypothesis prematurely. The total resistance ($R_{\text{Total}}$) to $O_2$ diffusion from Hb to bECF is the sum of the effective resistance of the intracellular fluid ($R_{\text{ICF}}$), $R_{\text{Membrane}}$, and the resistance of the extracellular unconvected fluid ($R_{\text{EUF}}$). If one implicitly assumes that $R_{\text{Membrane}} \cong 0$, the result is that the sum ($R_{\text{Membrane}} + R_{\text{EUF}}$) reduces to $R_{\text{EUF}}$, with the result that one computes erroneously high values for the EUF thickness. In the process, one concludes that the large UL explains the experimental data without the need of invoking membrane resistance—a classic bootstrap maneuver.

Because Vandegriff and Olson recognized the challenge of studying $O_2$ influx, these authors investigated the role of the plasma membrane in the movement of $O_2$ in the case of $O_2$ efflux. They reduced the effects of ULs by adding large amounts of sodium dithionite (NaDT) in the suspending medium.[13] The NaDT near the outer side of the plasma membrane annihilates the $O_2$ that has just left the RBC, thereby minimizing the build-up of $O_2$ and minimizing the UL. Viewed somewhat differently, the NaDT maximizes the outwardly directed $O_2$ gradient and therefore the rate of $O_2$ efflux.[10] Vandegriff and Olson used a reaction-diffusion model of an RBC, idealized as a cylindrical disk, to analyze the time course of their $O_2$ release data. They concluded that their mathematical model—which lacked a term analysis to $R_{\text{Membrane}}$—could simulate their data without the need to include a membrane resistance to the movement of $O_2$. Instead, they accounted for their $O_2$-release data in the presence of NaDT in the extracellular medium by including in the model an UL, the width of which increased with time from the initial value of 1 μm to ~30 μm (computed from an equation in ref. 10). In summary, because they implicitly assumed $R_{\text{Membrane}} = 0$, these authors invoked a very large $R_{\text{EUF}}$, and used this result to conclude that $R_{\text{Membrane}} \cong 0$.

It is worth noting that at least 3 sets of authors have performed physicochemical experiments in which they concluded that lipid mono/bilayers can offer substantial resistance to $CO_2$ or $O_2$.[4,15,16]

### Paper by Richardson et al.

In their recent paper, Richardson et al[17] conclude that the $D_{O_2}$ in the cytosol of RBCs is far lower than previously thought. The generally accepted value for $D_{O_2}$ in RBC cytosol has been $5.09 \times 10^{-6}$ cm$^2$/ s$^{-1}$ at 10 °C,[18] which is ~38% of the value in water (i.e., $13.313 \times 10^{-6}$ cm$^2$ s$^{-1}$ at 10 °C). However, Richardson et al conclude that the actual $D_{O_2}$ value in RBC cytosol is $<0.7 \times 10^{-6}$ cm$^2$ s$^{-1}$ at 23 °C, which is < ~3.5% of the value in water at 23 °C (i.e., $20 \times 10^{-6}$ cm$^2$ s$^{-1}$). In other words, Richerson et al conclude that $D_{O_2}$ value that others have been using for RBC cytosol is a great overestimate.

**Calculation of $D_{O_2}$ by Richerson et al.** In SI of their paper (page 6), Richardson et al[17] report a value of 2500 µm$^2$ s$^{-1}$ (i.e., $2.5 \times 10^{-5}$ cm$^2$ s$^{-1}$) for $D_{O2}$ in water at 23 °C. In the main text of their paper (page 3), these authors—based on physiological data and their mathematical simulations—infer that $D_{O2}$ in RBC cytosol is <70 µm$^2$ s$^{-1}$. They report this as <5% of the $D_{O2}$ in water (in fact, the factor is <3.5%; see above). However, in the original reference[19] that Richardson et al cited for $D_{O2}$ in water, Han and Bartels[19] report the value of 2500 µm$^2$ s$^{-1}$ for $D_{O2}$ in water at ~35 °C, not at the temperature of 23 °C, used in the Richardson experiments. According to Han and Bartels, the value of $D_{O2}$ in water at ~23 °C should be ~2000 µm$^2$ s$^{-1}$. Thus, the $D_{O_2}$ in cytoplasm predicted by Richardson et al should be 70/2000 = 1/29 of that in water rather than 70/2500 = 1/36.

To compare our work to that of Richardson et al, we must recompute their $D_{O_2}$ for a temperature of 10 °C. As noted above, Han and Bartels[19] report that $D_{O2}$ in water at 10 °C is $1.3313 \times 10^{-5}$ cm$^2$ s$^{-1}$. If we divide this figure by 29, we arrive at a predicted value of $D_{O_2}$ (based on the work of Richardson et al) of $(1.3313 \times 10^{-5})/29 = 4.5907 \times 10^{-7}$ cm$^2$ s$^{-1}$.

**Our calculation of the $D_{O_2}$ of RBC cytosol (excluding membrane) at 10 °C.** As discussed in the main text of our paper, we believe that this $D_{O_2}$ value of $4.5907 \times 10^{-7}$ cm$^2$ s$^{-1}$ is excessively high because Richardson et al used a $[NDT]_o$ of only 1 mM, rather than the final concentration in the present study, 25 mM. Several groups—Coin and Olson,[9] Vandegriff and Olson,[13] and Holland et al.[10]—have emphasized the importance of employing a sufficiently high $[NDT]_o$ to minimize the EUF layer. Thus, using an $[NDT]_o$ of only 1 mM, Richardson et al would have attributed to the ICF a component of total resistance ($R_{Total}$ =

$R_{ICF} + R_{Membrane} + R_{EUF}$) really due to the EUF layer. Recognizing that the value of Richardson et al is an overestimate, we decided to average the Richardson value of $0.45907 \times 10^{-6}$ cm$^2$ s$^{-1}$ with the generally accepted value[18] of $5.09 \times 10^{-6}$ cm$^2$ s$^{-1}$. Thus, we assumed that $D_{O_2}$ in RBC cytoplasm is $2.7745 \times 10^{-6}$ cm$^2$ s$^{-1}$.

**Contribution of channels to $D_{O_2}$ of the cytosol + membrane.** Richardson et al argue that the $D_{O_2}$ of the cytosol (actually, the cytosol + membrane) is far lower than previously believed. We argue that they overestimate this effect. Nevertheless, the reason that $D_{O_2}$ (of the cytosol + membrane) is as high as it is, is because of the presence of $O_2$ channels. If one were to eliminate ~90% of these channels—as by genetically deleting AQP1 and RhAG, while applying pCMBS—the membrane would now dominate $R_{Total}$.

### Importance of values used in modeling

#### Thickness of EUF ($\ell_{EUF}$)

Despite the use of NaDT in our SF experiments—which virtually abolishes the formation of ULs[10]—in our mathematical model we made the conservative assumption that the EUF has a thickness ($\ell_{EUF}$) of 1 μm and that no NaDT is present in this layer. We simulated the annihilation of $O_2$ by NaDT by assuming that the bECF contains no $O_2$ during the duration of the simulations. In Extended Data Figure 10*f* we examined the dependence of $k_{HbO2}$ on $\ell_{EUF}$. When we increase $\ell_{EUF}$, the simulated $k_{HbO_2}$ asymptotically falls to a value that approximates the biological value that we observe with RBCs from a dKO mouse. Thus, no increase in $\ell_{EUF}$ can account for our data with pCMBS or DIDS. Lowering $\ell_{EUF}$ to values smaller than 1 μm causes steep increases in the simulated $k_{HbO2}$ (i.e., decreases in the resistance of the EUF to $O_2$ diffusion, $R_{EUF}$). Starting from such elevated $k_{HbO_2}$ values, the model can generate lower $k_{HbO_2}$ values by increasing $R_{Membrane}$, that is, by decreasing the permeability of the plasma membrane to $O_2$ ($P_{Membrane}$) so that the sum ($R_{EUF} + R_{Membrane}$) remains the same. In other words, if the UL is thinner than 1 μm—which is likely to be the case in the presence of high [NaDT]—the membrane must make a greater contribution to the overall diffusive resistance of $O_2$.

##### RBC dimensions vs. intracellular [Hb]

Our model predicts that increases in both mean corpuscular volume (MCV) and mean corpuscular hemoglobin concentration (MCHC) lead to decreases in $k_{HbO_2}$. However, as shown in Supplementary Table 6, the genetic deletion of a channels invariably led to an increase in MCV but a decrease in MCHC. In the case of the deletion of *Aqp1*, the predicted result is an increase in $k_{HbO_2}$, as shown in Extended Data Figure 9*b*. In the case of *RHag*–/– and the dKO, the blunted decreases in MCHC lead to decreases in $k_{HbO_2}$, though decreases that are not as great as they would have been had the MCHC not fallen.

The increase in MCV that we observe in the *Aqp1*–/–, *RHag*–/–, and dKO mice are mainly attributable to the increase in immature cell types.

#### Calculation of the contribution of the plasma membrane to total diffusive resistance to $O_2$

Let us assume that we can treat the diffusion of $O_2$ from the intracellular fluid (ICF) of the RBC to the bECF like an electrical circuit (see Extended Data Figure 9*a*). Thus, the total resistance to diffusion ($R_{Total}$) is the sum of the resistance of the ICF ($R_{ICF}$), the plasma membrane ($R_{Membrane}$), and the EUF ($R_{EUF}$):

$$R_{Total} = R_{ICF} + R_{Membrane} + R_{EUF}$$

In the next 3 sections, we will compute each of the resistances on the right side of the equation, and then we will compute the fraction of $R_{Total}$ that $R_{Membrane}$ represents. See Supplementary Table 10 for parameter values used in simulations of RBCs from WT mice.

**Calculate $R_{ICF}$.** We start by computing—at the time $t_{37}$ (i.e., 0.2760 s), when hemoglobin saturation (HbSat) has fallen to 1/e of its initial value—the parallel fluxes of free $O_2$ ($J_{O_2}$) and $HbO_2$ ($J_{HbO_2}$) from the position of the average $O_2$ and $HbO_2$ molecules in the ICF. From these two fluxes, we then compute the total $O_2$ flux ($J_{TO_2}$).

First, we determine the number of $O_2$ molecules in each of the 101 shells ($i$ = 1 to 101) that make up the model RBC, as well as the distance ($\ell$) from the center of each shell to the inner surface of the plasma membrane (iPM). For a WT cell with no inhibitors, we find that the average distance of an $O_2$ molecule from the iPM is $2.792 \times 10^{-5}$ cm. For $HbO_2$, a similar analysis yields an average distance of $2.560 \times 10^{-5}$ cm. Because these distances are so similar, we used the average distance for $O_2$ and $HbO_2$, which is $2.68 \times 10^{-5}$ cm, which corresponds to shell $i = k = 76$ from the center of the sphere. We call this average distance $\ell_k$. Thus, the $O_2$ flux from $\ell_k$ to the iPM is:

$$J_{O_2} = \frac{D_{O_2}}{\ell_k} \cdot \left( [O_2]_k - [O_2]_{iPM} \right) = 0.000540 \, \mu\text{mole} / (\text{cm}^2 \, \text{s}),$$

where $D_{O_2}$ is the diffusion constant for $O_2$ in the ICF (see Supplementary Table 10 for value), and where $[O_2]_k = 0.01528$ mM and $[O_2]_{iPM} = 0.01007$ mM at $t_{37}$. For $HbO_2$,

$$J_{HbO_2} = \frac{D_{HbO_2}}{\ell_k} \cdot \left( [HbO_2]_k - [HbO_2]_{iPM} \right) = 0.000282 \, \mu\text{mole} / (\text{cm}^2 \, \text{s}),$$

where $D_{HbO_2}$ is the diffusion constant for $HbO_2$ in the ICF (see Supplementary Table 10 for value), and where $[HbO_2]_k = 6.869$ mM and $[HbO_2]_{iPM} = 6.744$ mM at $t_{37}$.

Thus, the total $O_2$ flux from $\ell_k$ to the iPM is

$$J_{TO_2} = J_{O_2} + J_{HbO_2} = 0.0008218 \, \mu\text{mole} / (\text{cm}^2 \, \text{s}).$$

The effective resistance to the diffusion of $O_2$ is

$$R_{ICF} = \frac{\ell_k}{D_{O_2}} = \frac{\left( [O_2]_k - [O_2]_{iPM} \right)}{J_{TO_2}} = 6.334 \;\; \text{s} \cdot \text{cm}^{-1}.$$

**Calculate $R_{\text{Membrane}}$.** We calculate the resistance of the plasma membrane to $O_2$ diffusion as the reciprocal of the "true" microscopic permeability of the plasma membrane to $O_2$ ($P_{\text{Membrane}}$), which is 0.15 cm $s^{-1}$ (see Supplementary Table 10 for value). Thus,

$$R_{\text{Membrane}} = \frac{1}{P_{\text{Membrane}}} = 6.666 \;\; \text{s} \cdot \text{cm}^{-1}.$$

**Calculate $R_{\text{EUF}}$.** We calculate the resistance of the EUF to $O_2$ diffusion as

$$R_{\text{EUF}} = \frac{\ell_{\text{EUF}}}{D_{O_2}} = 7.511 \;\; \text{s} \cdot \text{cm}^{-1},$$

where $\ell_{EUF}$ is the thickness of the EUF and $D_{O_2}$ is the diffusion constant for $O_2$ in the EUF (see Supplementary Table 10 for values).

**Compute the contribution of $R_{Membrane}$ to $R_{Total}$.** We calculate the contribution of $R_{Membrane}$ to $R_{Total}$ the EUF as

$$\text{Contribution of } R_{PM} \text{ to } R_{Total} = \frac{R_{Membrane}}{R_{Total}} = \frac{R_{Membrane}}{R_{ICF} + R_{Membrane} + R_{EUF}}$$

$$\frac{6.666\ \mathrm{s \cdot cm^{-1}}}{6.334\ \mathrm{s \cdot cm^{-1}} + 6.666\ \mathrm{s \cdot cm^{-1}} + 7.511\ \mathrm{s \cdot cm^{-1}}} = 32.5\%.$$

Note that if $\ell_{EUF}$ is in fact <1 μm (i.e., $R_{EUF} < 7.511$ s cm$^{-1}$), the contribution of $R_{Membrane}$ to total resistance would be greater than ~32.5%.
