## Supplementary Data for "Role of channels in the oxygen permeability of red blood cells"

### Supplementary Information: Data

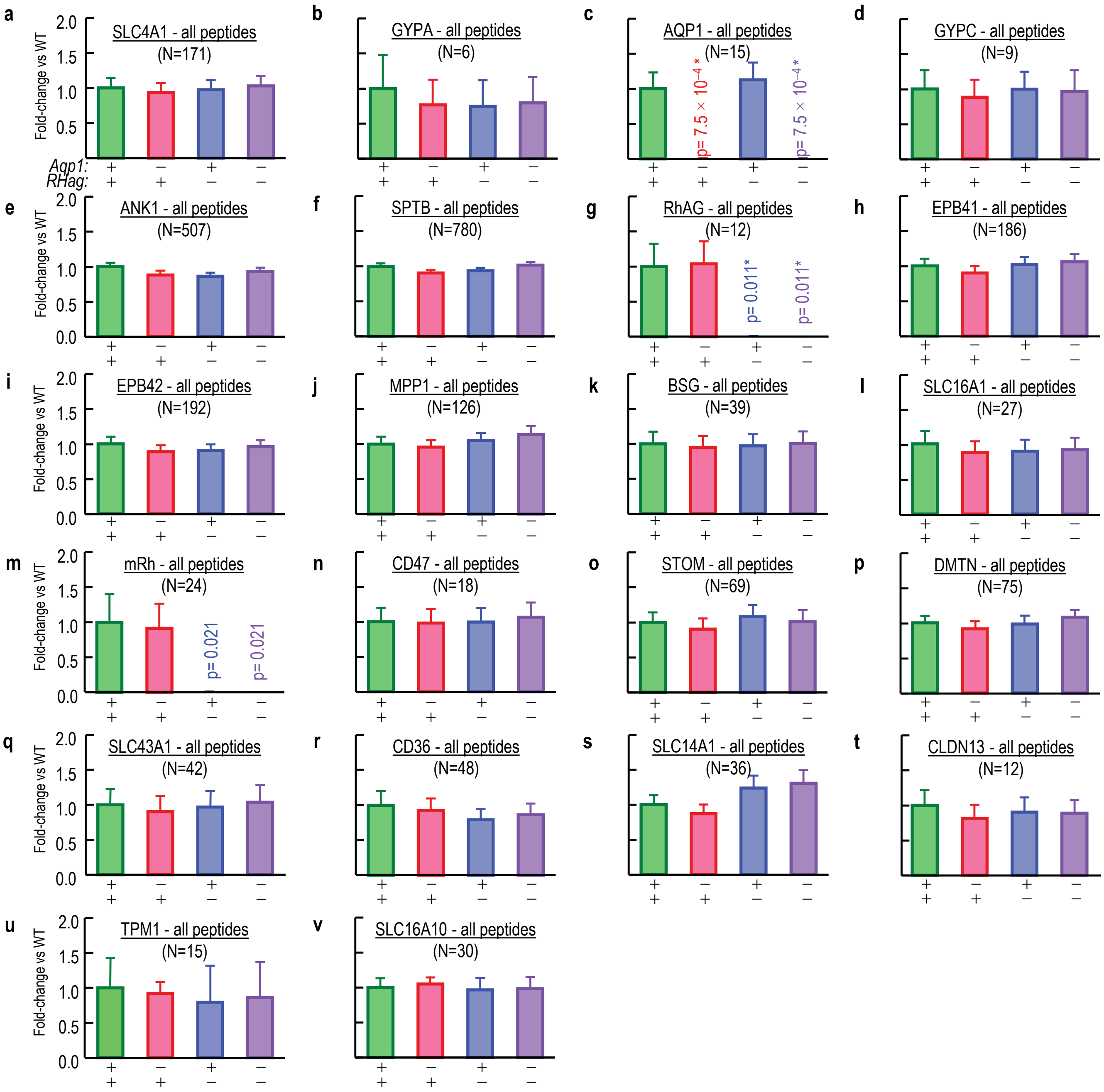


**Supplementary Data Figure 1 | Summary of fold-changes in expression of 22 "plasma membrane–associated" proteins comparing samples from RBC ghosts of WT vs. *Aqp1*–/–, *RHag*–/– and double-KO (dKO) mice.** These 22 panels represent the 22 plasma-membrane–associated proteins, from among the 50 proteins with the greatest inferred abundance in RBC ghosts from WT mice. The panels are in rank order, based on the inferred abundance of the protein in cells from WT mice (see Extended Data Figure

4*a*). In each panel, data from WT mice (green bar) is the reference value for fold-changes in expression for the each mouse knockout strains (*Aqp1*–/–, red bars; *RHag*–/–, blue bars; dKO, purple bars), as determined by a mass-spectrometry based investigation of peptide peak intensity (AUC) obtained from RBC ghosts. **a,** AE1 (SLC4A1). **b,** glycophorin A (GYPA). **c,** aquaporin 1 (AQP1). **d,** glycophorin C (GYPC). **e,** ankyrin 1 (ANK1). **f,** erythrocytic spectrin beta (SPTB). **g,** Rhesus blood group associated Glycoprotein A, RhAG (SLC42A1). **h,** erythrocyte membrane protein band 4.1 (EPB41). **i,** erythrocyte membrane protein band 4.2 (EPB42). **j,** membrane protein, palmitoylated 1 (MPP1). **k,** basigin (BSG). **l,** monocarboxylic acid transporter member 1, MCT-1 (SLC16A1). **m,** mouse Rhesus blood group, D antigen (RhD). **n,** CD47 molecule (CD47). **o,** stomatin (STOM). **p,** dematin actin binding protein (DMTN). **q,** solute carrier family 43 (amino acid system L transporter), member 1(SLC43A1). **r,** CD36 antigen (CD36). **s,** urea transporter member 1, UT-1 (SLC14A1). **t,** claudin 13 (CLDN13). **u,** tropomyosin 1 (TPM1). **v,** monocarboxylate transporter member 10, MCT-10 (SLC16A10). Bars represent the mean ± s.e.m. peak intensity (AUC) for all peptides from each protein, normalized to the abundance in WT. Number of detected peptides per protein is displayed in parentheses. Not all of these proteins are necessarily RBC integral membrane proteins. We purified proteins from RBC ghosts of 3 mice/genotype (same samples as Supplementary Data Figure 2). We performed a one-way ANOVA, followed by the Holm-Bonferroni[12] correction (see Methods/StatisticalAnalysis). *Significant vs. WT. The analyses show that expression of the target protein in RBCs of each knockout strain is essentially abolished. However, the difference in expression of other the RBC proteins in each knockout strain is not significant compared to WT. See Supplementary Table 1*a* for glossary and rank order of abundance.

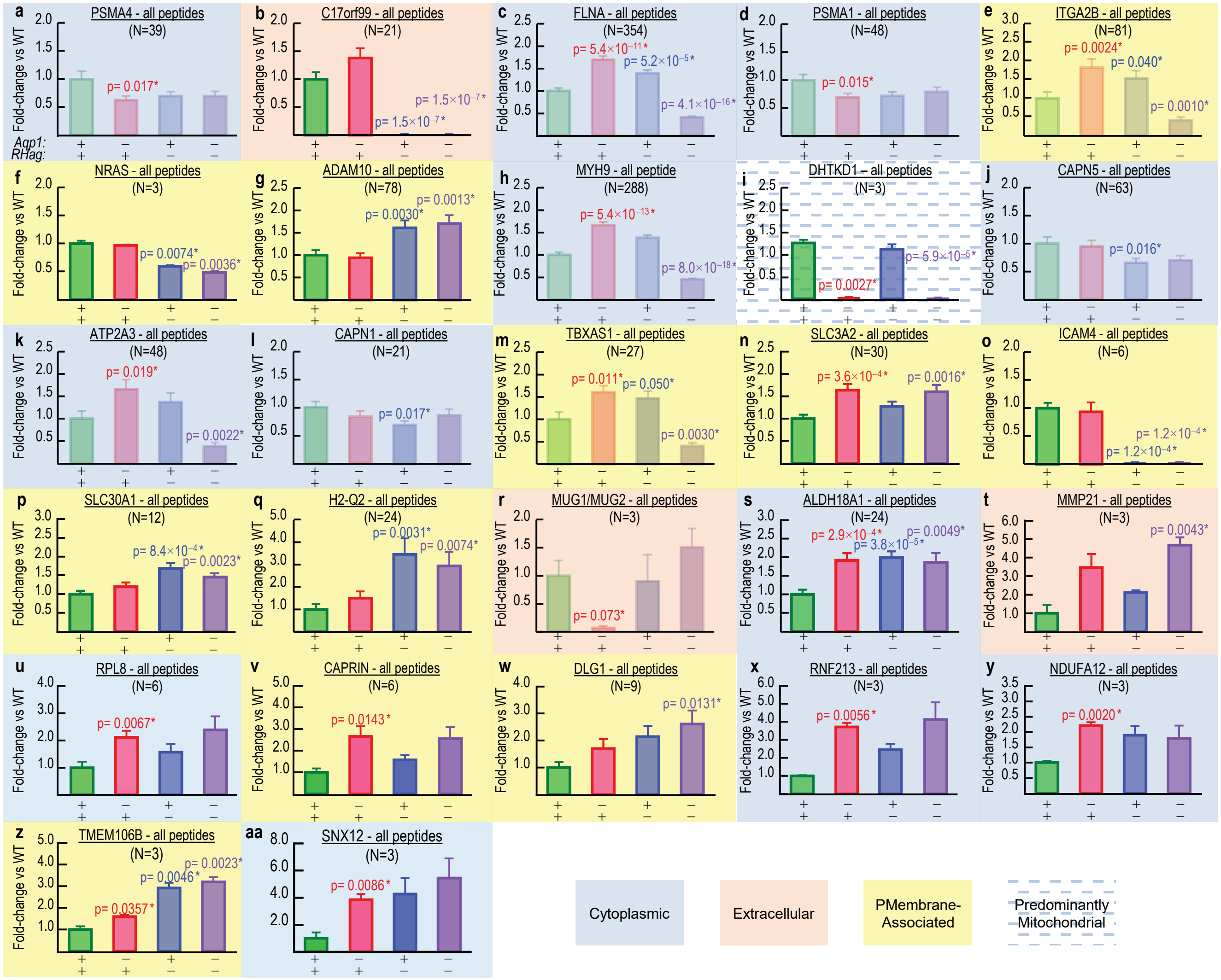


**Supplementary Data Figure 2 | Summary of fold-changes in expression of all 27 proteins in which abundance changed in at least one KO genotype, comparing samples from RBC ghosts of WT vs. *Aqp1*–/–, *RHag*–/–, and double-KO (dKO) mice.** These 27 panels represent the 27 proteins—from among all 1104 detected proteins—that exhibited a significant change in at least one knockout strain vs. WT mice. The panels are in rank order, based on the inferred abundance of the protein in cells from WT mice (see Extended Data Figure 4*b*). In each panel, data from WT mice (green bar) is the reference value for fold-changes in expression for the mouse knockout strains (*Aqp1*–/–, red bars; *RHag*–/–, blue bars; dKO, purple bars), as determined by a mass-spectrometry based investigation of peptide peak intensity (AUC) obtained from RBC ghosts. **a,** proteasome (prosome, macropain) subunit, alpha type, 4 (PMSA4).

**b,** chromosome 17 open reading frame 99 (C17orf99). **c,** filamin A, alpha (FLNA). **d,** proteasome (prosome, macropain) subunit, alpha type, 1 (PSMA1). **e,** integrin, alpha 2b (ITGA2B). **f,** neuroblastoma RAS viral (v-ras) oncogene homolog (NRAS). **g,** ADAM metallopeptidase domain 10 (ADAM10). **h,** myosin, heavy chain 9, non-muscle (MYH9). **i,** dehydrogenase E1 and transketolase domain containing 1 (DHTKD1). **j,** calpain 5(CAPN5). **k,** ATPase, $Ca^{++}$-transporting, ubiquitous (ATP2A3). **l,** calpain 1 (CAPN1). **m,** thromboxane A synthase 1 (platelet) (TBXAS1). **n,** solute carrier family 3 (amino acid transporter heavy chain), member 2 (SLC3A2). **o,** intercellular adhesion molecule 4 (ICAM4). **p,** solute carrier family 30 (zinc transporter), member 1, Zn-T1 (SLC30A1). **q,** major histocompatibility complex (H2-Q2). **r,** Murinoglobulin 1 (MUG1/MUG2). **s,** aldehyde dehydrogenase 18 family, member A1 (ALDH18A1), **t,** matrix metallopeptidase 21 (MMP21), **u,** ribosomal protein L8 (RPL8), **v,** cell cycle associated protein 1 (CAPRIN1), **w,** discs, large homolog 1 (Drosophila) (DLG1), **x,** ring finger protein 213 (RNF213), **y,** NADH dehydrogenase (ubiquinone) 1 alpha subcomplex, 12 (NDUFA12), **z,** transmembrane protein 106B (TMEM106B), **aa,** sorting nexin 12 (SNX12). Bars represent the mean ± s. e. m peak intensity (AUC) for all peptides from each protein normalized to the abundance in WT. Number of peptides per protein is displayed in parentheses. Not all of these proteins are from RBCs, let alone RBC integral membrane proteins. The colors of the bars indicate the location of the proteins, as assigned by the software Elucidator: yellow (plasma-membrane–associated), blue (cytoplasmic), and tan (extracellular). We purified proteins from RBC ghosts of 3 mice/genotype (same samples as Supplementary Data Figure 2). We performed a one-way ANOVA, followed by the Holm-Bonferroni[12] correction (see Methods/StatisticalAnalysis). *Significant vs. WT. See Supplementary Table 1*bc* for glossary.

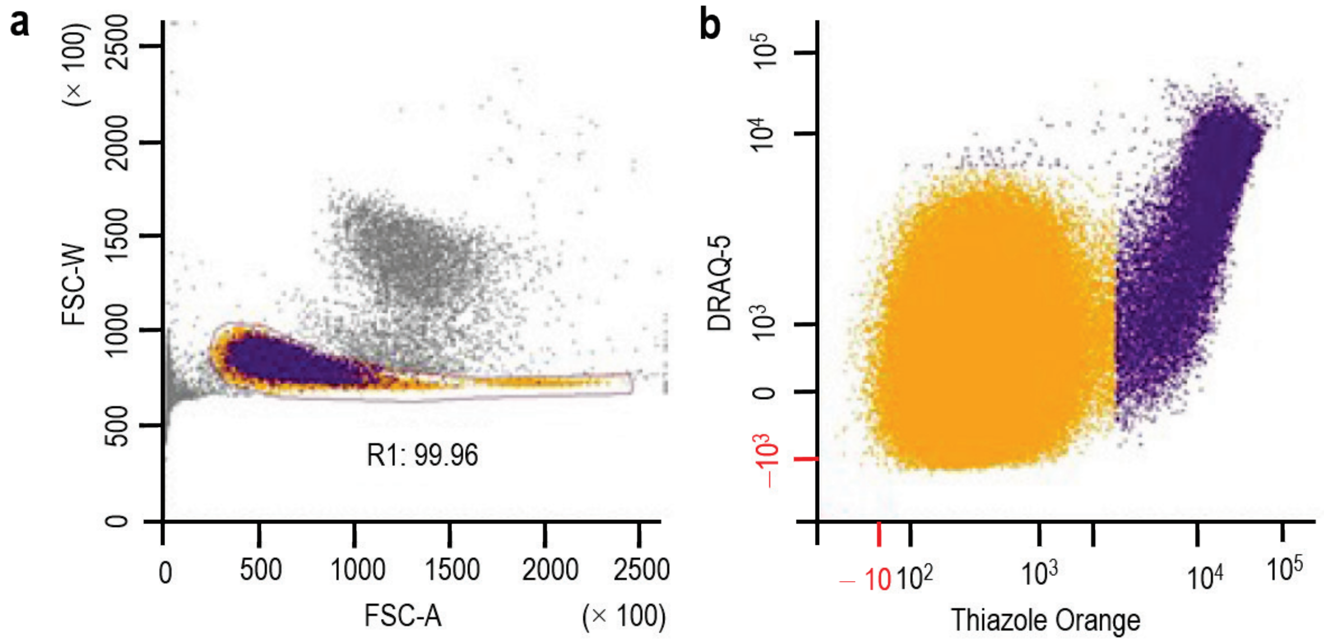


**Supplementary Data Figure 3 | Gating schemes established in light-scattering flow cytometry.** We are able to sort mature RBCs from their precursors in the LSRII device in order to perform size and shape analyses. **a,** Viable cells (orange and purple). 99.96% of the cells in the R1 region (encompassing the orange and purple points) are CV positive. **b,** Mature RBCs (orange) and immature precursors (purple). Cells in panel ***b*** were already gated in panel ***a*** (i.e., they are all viable). The mature RBCs (orange points) are both TO negative and DRAQ-5 negative. Reticulocytes (purple points, but beneath pink dashed line) are TO positive but DRAQ-5 negative. Nucleated precursors (purple points, but above pink dashed line) are TO positive and DRAQ-5 positive. The horizontal pink dashed line separates DRAQ-5–positive from DRAQ-5–negative cells. The vertical blue dashed line separates TO-positive from TO-negative cells. CV, calcein violet (viability); TO, thiazole orange (RNA); and DRAQ-5, Deep Red Anthraquinone 5 (DNA).

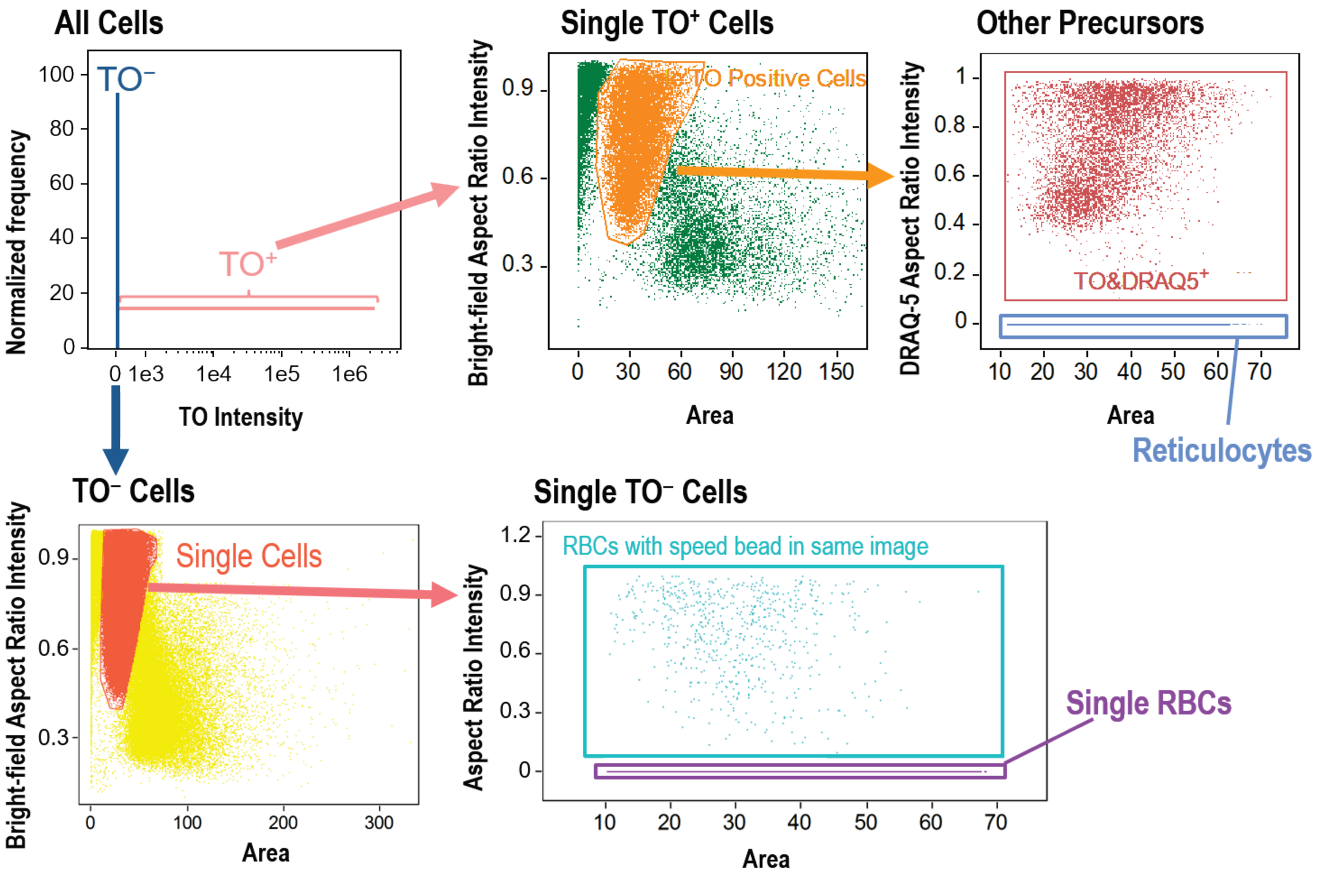


**Supplementary Data Figure 4 | Gating schemes established in ImageStream flow cytometry to allow analysis of individual RBCs and other precursor cells.** We are able to sort mature RBCs from their precursors in the ImageStream device in order to perform further size and shape analyses. We begin with CV-positive cells. This figure shows how, starting in the upper left panel, which distinguishes TO-positive from TO-negative cells, we isolate single mature RBCs, single reticulocytes, and other single precursor cells. CV, calcein violet (viability); TO, thiazole orange (RNA); DRAQ5, Deep Red Anthraquinone 5 (DNA).

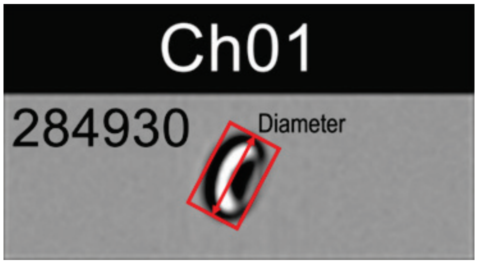


**Supplementary Data Figure 5 | Measuring the major diameter of blood cells in flow in the ImageStream flow cytometer.** After gating according to their fluorescence signature (see Supplementary Data Figure 4), the major diameters of RBCs, reticulocytes and nucleated precursor cells were determined by measuring the length of the longest axis (red box) of the cells in the corresponding bright field images.

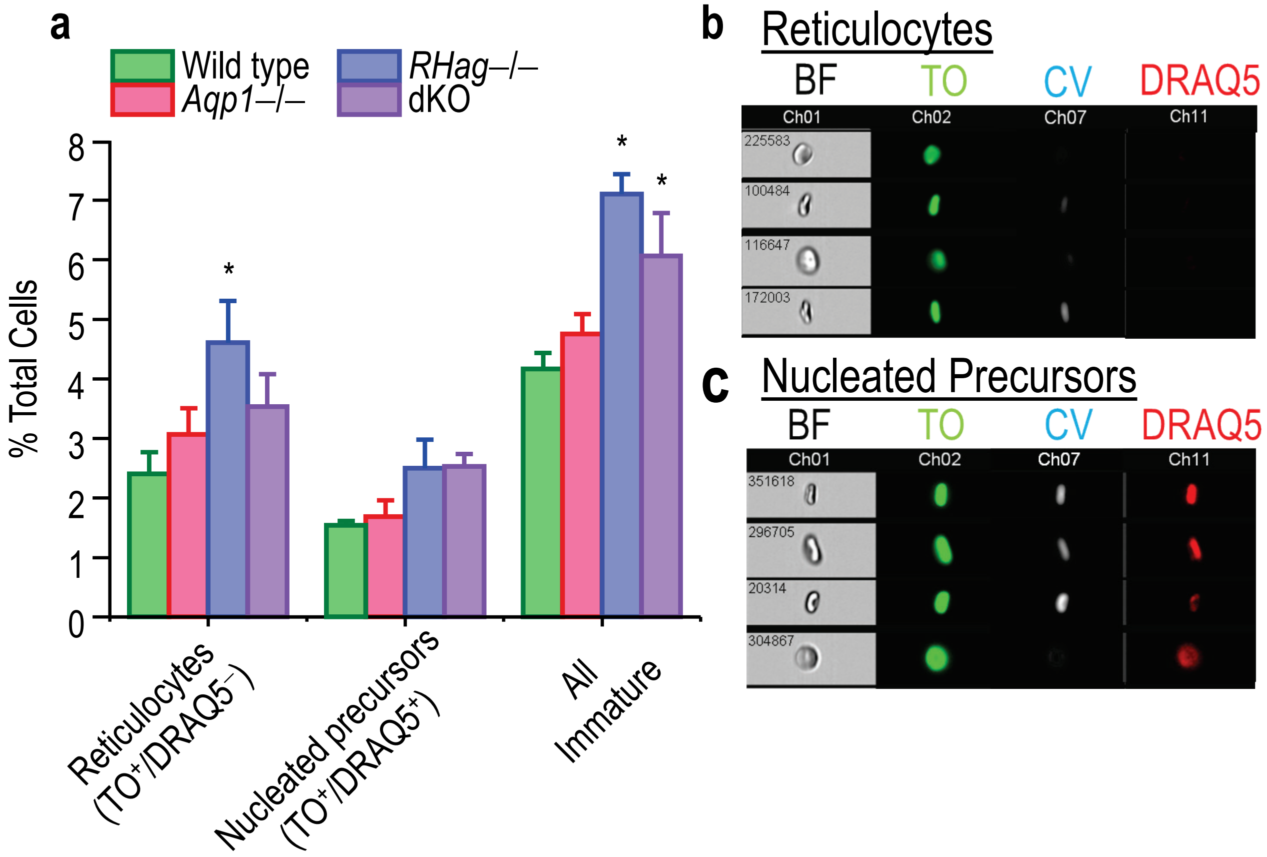


**Supplementary Data Figure 6 | Percent immature erythrocytes present in blood from ImageStream flow cytometry. a,** Percent of total cell count represented by RBC-precursor cells for each genotype, identified by TO and DRAQ5 staining as in Supplementary Data Figure 4. We performed a one-way ANOVA, followed by the Holm-Bonferroni[21] correction (see Methods/StatisticalAnalysis). *Significant vs. WT. **b,** Reticulocytes, identified as $CV^+$, $TO^+$, and $DRAQ5^-$. **c,** Nucleated RBC precursor cells, identified as $CV^+$, $TO^+$, and $DRAQ5^+$. CV, calcein violet (viability); TO, thiazole orange (RNA); DRAQ5, Deep Red Anthraquinone 5 (DNA). BF, bright field image.

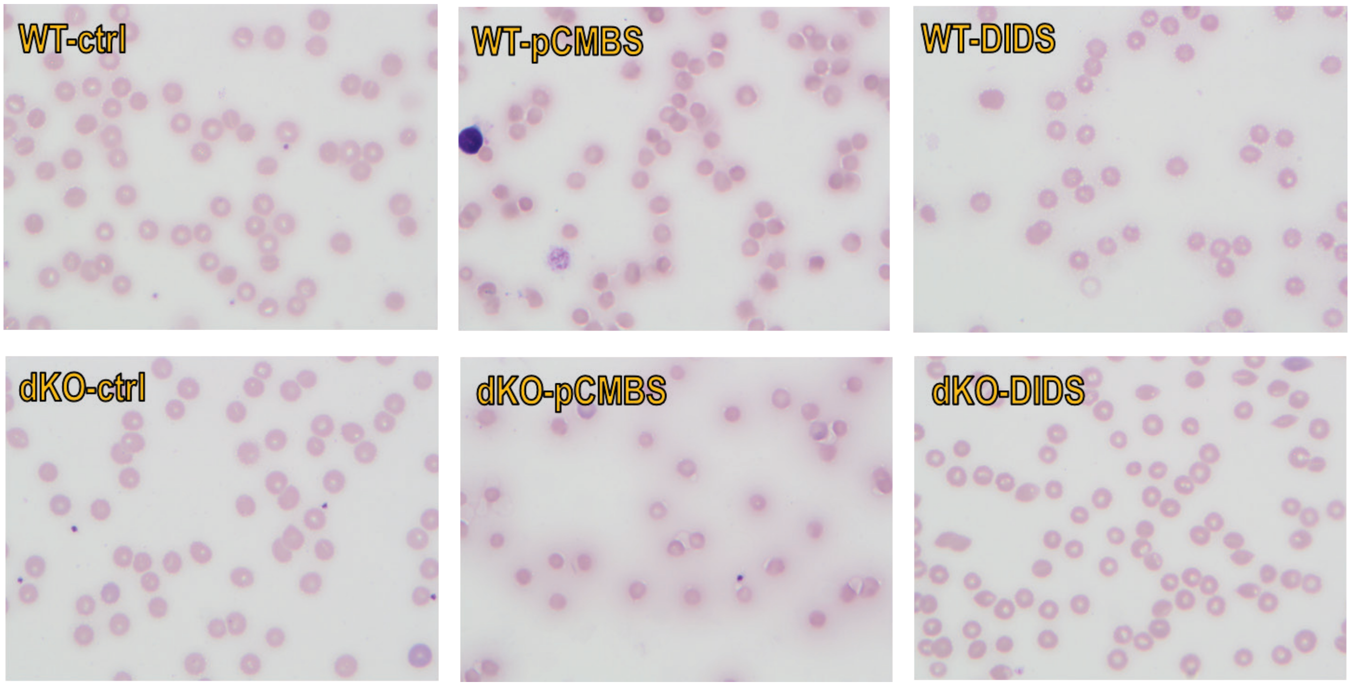


**Supplementary Data Figure 7 | Representative blood smears from WT and dKO mice, of cells treated with inhibitors.** Air-dried peripheral blood smears using blood obtained from dKO and WT mice, either with or without the addition of pCMBS or DIDS. Slides were Wright-Giemsa stained using the staining function of the Sysmex (Kobe, Japan) SP-10 slide maker/stainer attached to a Sysmex XN-9000 hematology analyzer and reviewed by a board-certified hematopathologist (H.J.M.). Representative images (1000× magnification) are shown. No significant differences in red cell morphology were noted between dKO and WT mice under any of the conditions. A faint halo of light purple stain was noted around cells after the addition of pCMBS and occasional vacuoles were noted at the periphery of the red cells outside of the cell membrane in some of the stained pCMBS samples. These were deemed artifactual, and likely related to the effect of the added pCMBS on the staining reactions.

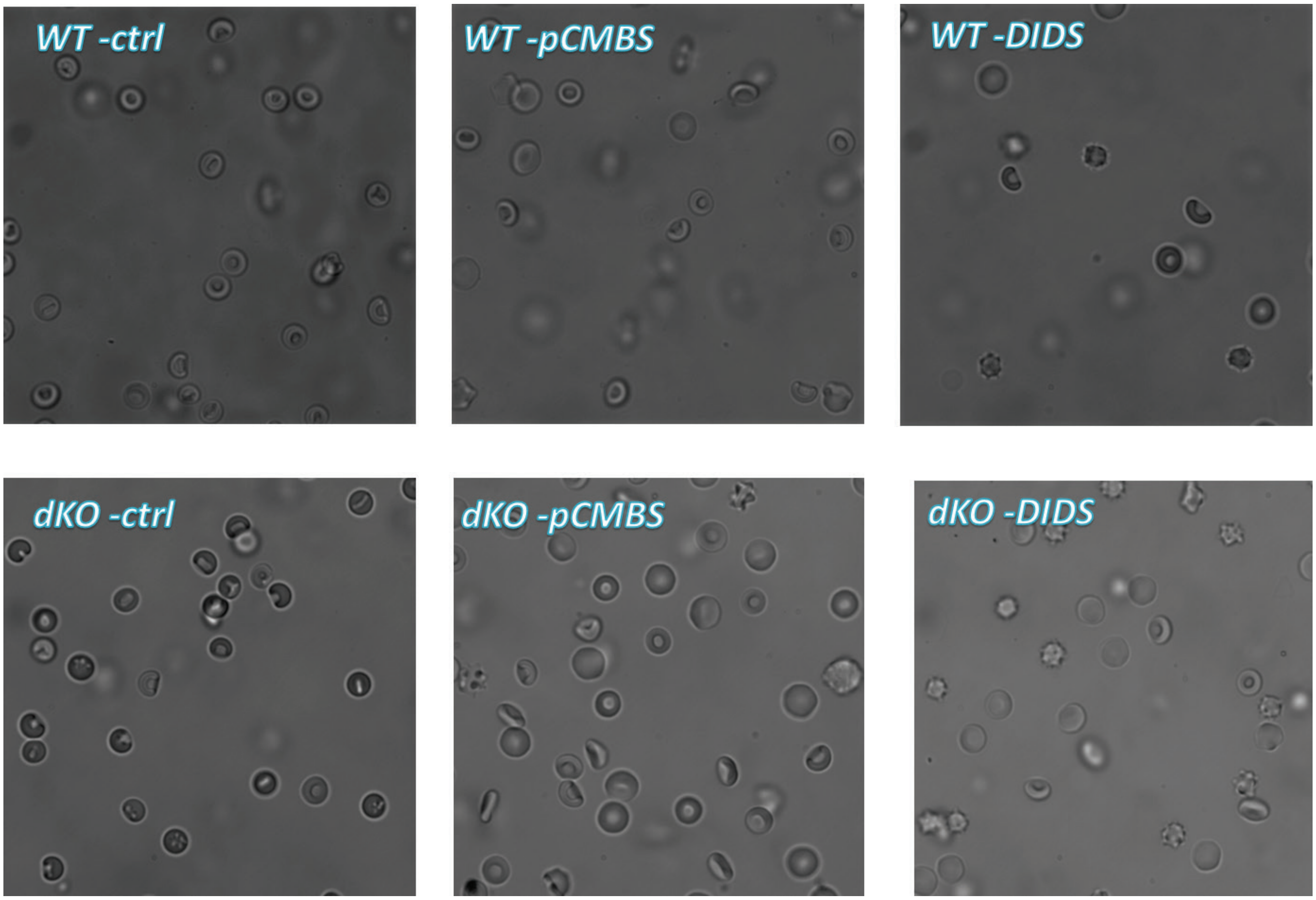


**Supplementary Data Figure 8 | Representative DIC still micrographs of living RBCs, treated with inhibitors.** On a given day, we reviewed 4 to 5 samples (i.e., droplets) of RBCs suspended in saline for 1 mouse of each of genotype. We executed this protocol on 3 separate days, for a total of 3 separate mice/genotype (i.e., a total of 12 to 15 samples/genotype). Red cell morphology is similar in control groups and is unremarkable, with no differences noted among dKO and WT mice. For both WT and dKO mice, treatment of RBCs with inhibitors tended to cause the appearance of small, spherical cells with spicules, more so in DIDS-treated than in pCMBS-treated cells. We can also identify these tumbling spherocytes by microvideography (not shown).

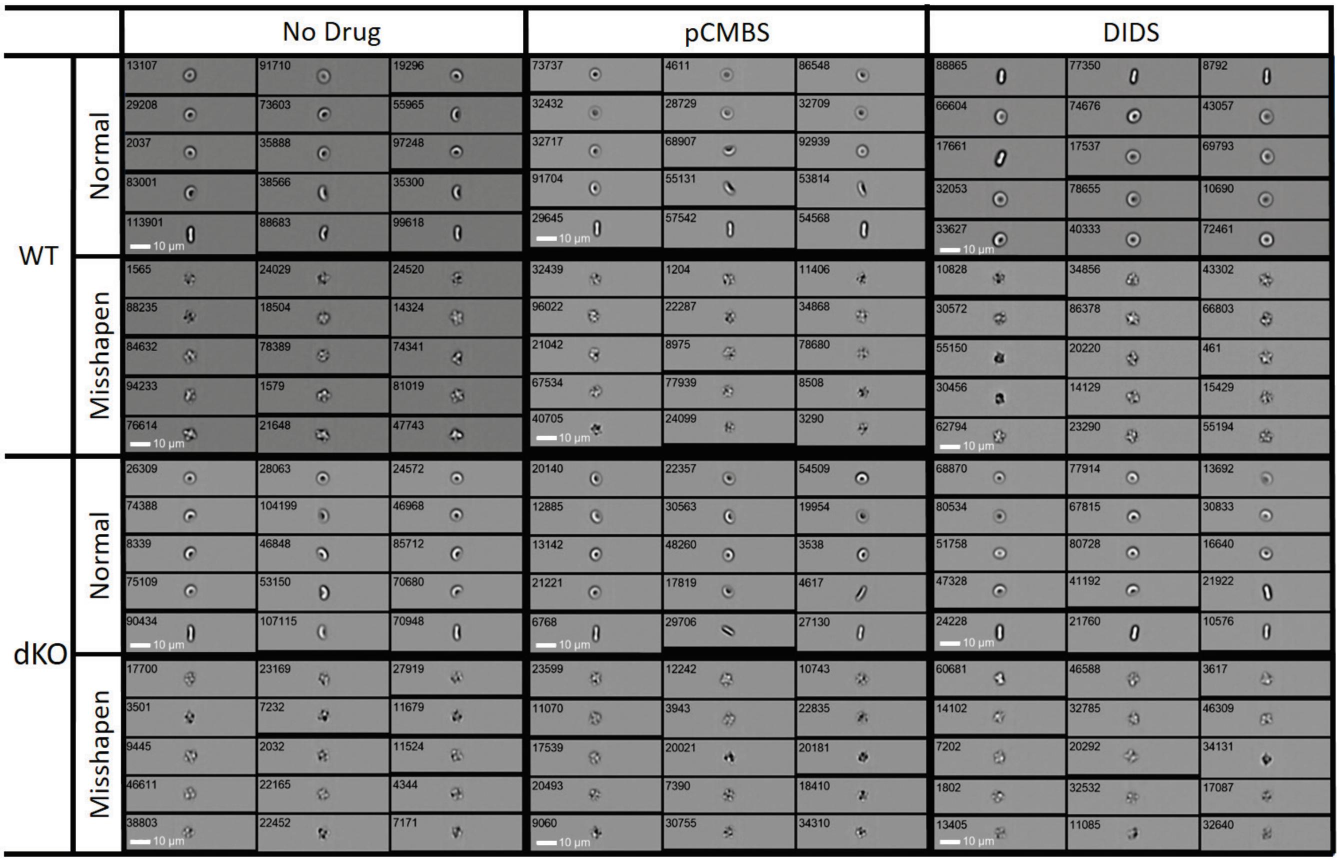


**Supplementary Data Figure 9 | Example bright-field images of WT or dKO RBCs counted by ImageStream flow cytometry following incubation with no drug, pCMBS or DIDS.** Gating schemes (Supplementary Data Figure 4) were utilized sort mature RBCs from their precursors. To sort the cells, we employed the Lobe count feature, using the H Variance Mean function (granularity assigned as 5). This sorting procedure yielded of four bins of cells identified as possessing 1, 2, 3, or 4 lobes. Regardless of the lobe bin (i.e., 1, 2, 3, or 4 lobes), the majority of misshapen cells had H Variance Mean values <10. We manually inspected each cell in each bin of sorted cells to verify correct assignment of normal vs misshapen cells. Experiments were performed on blood samples from 3 age-matched pairs of WT vs dKO mice. DIC still microphotography (Supplementary Data Figure 8) and DIC microvideography (not shown) show that these misshapen cells are spheres.

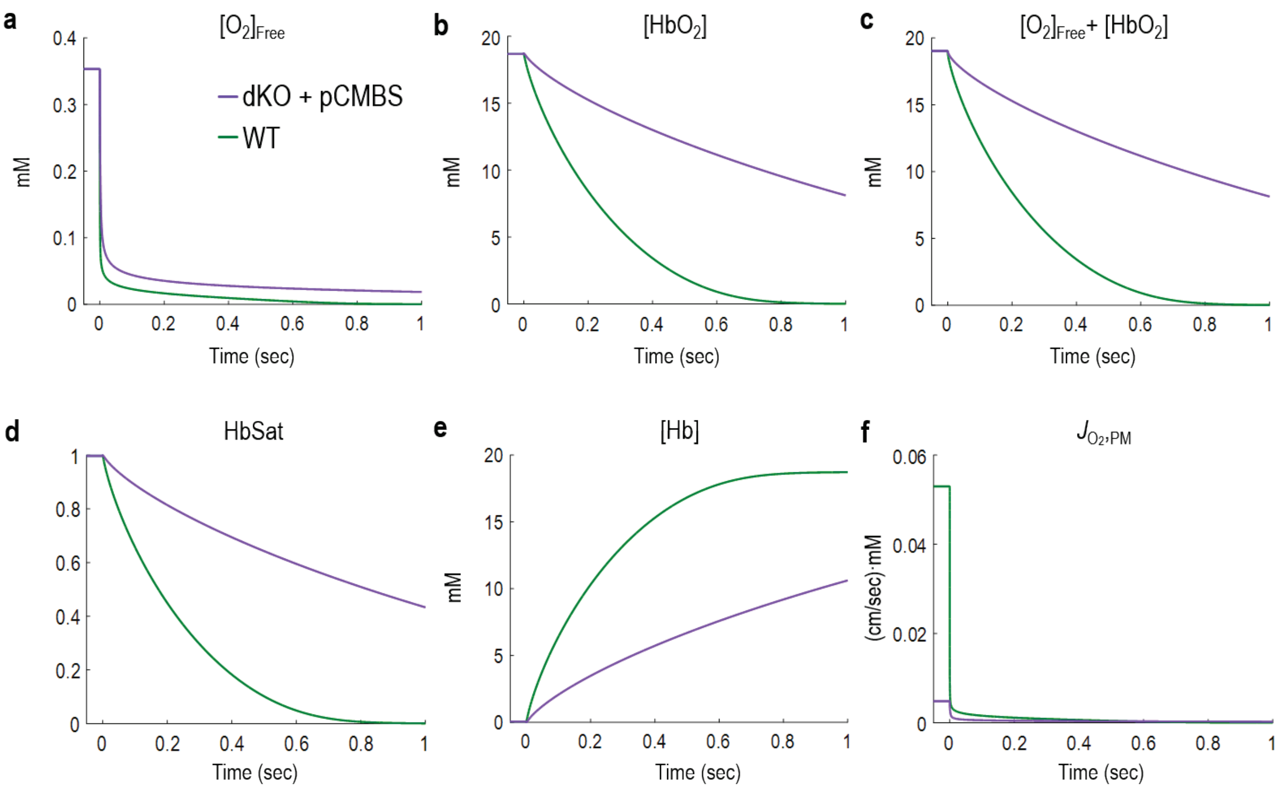


**Supplementary Data Figure 10 | Simulated time courses of $O_2$/Hb-related parameters and trans-membrane $O_2$ flux for cells mimicking WT/no inhibitors (control) vs. dKO+pCMBS.** These data all come from the same simulations that generated the "WT" (i.e., without inhibitors, and thus the highest $k_{HbO_2}$) and "dKO+pCMBS" data (i.e., those with the lowest values of $k_{HbO_2}$) in Extended Data Figure 9*c*–*d*. See Methods/MathematicalModeling and SI Methods/MathematicalModeling&Simulations for details on the model. **a,** Concentration of free $O_2$, integrated over the entire volume of cytoplasm. **b,** Concentration of oxyhemoglobin monomers, integrated over the entire volume of cytoplasm. **c,** Concentration of total $O_2$—sum of free $O_2$ and oxyhemoglobin monomer—integrated over the entire volume of cytoplasm. **d,** Hemoglobin saturation—$[HbO_2]/([O_2]_{Free} + [HbO_2])$—integrated over the entire volume of cytoplasm. **e,** Concentration of free hemoglobin, integrated over the entire volume of cytoplasm. **f,** Flux of $O_2$ across plasma membrane.

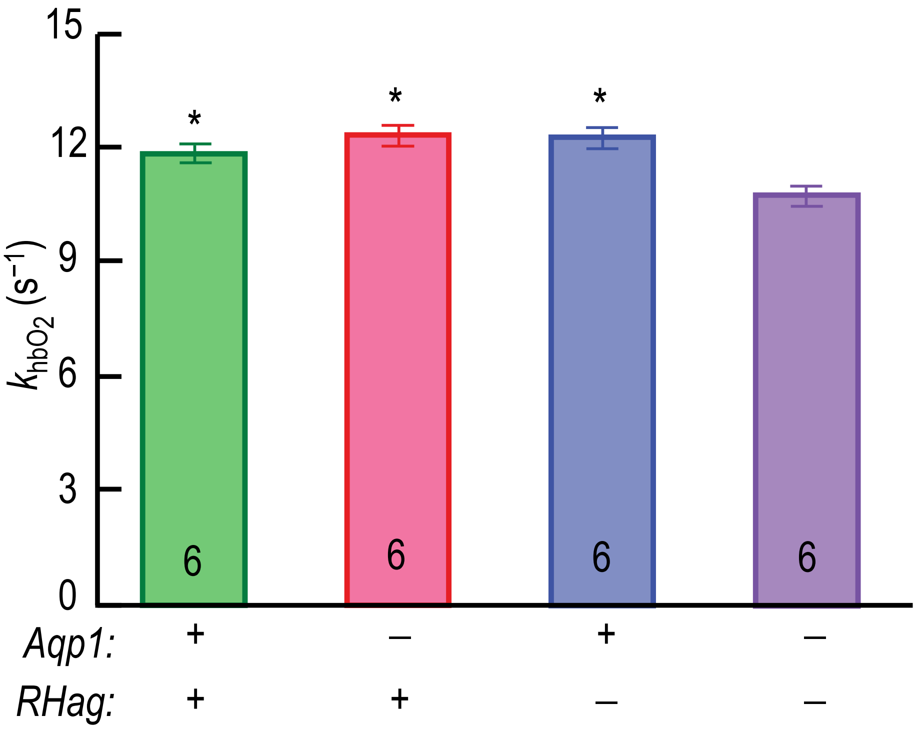


**Supplementary Data Figure 11 | Effect of genetic deletions of *Aqp1*, *RHag*, or both on rate constant of the reaction $HbO_2 \rightarrow Hb + O_2$ ($k_{HbO_2\text{>}Hb}$) in RBC lysates.** See Methods/StoppedFlowAbsorbanceSpectroscopy for a description of how we measured $k_{HbO_2}$. The approach here was similar, except that we worked with 100% hemolysates.. Each observation represents one mouse. We performed a one-way ANOVA, followed by a Holm-Bonferroni correction (see Methods/StatisticalAnalysis). *Significant vs. dKO.
