## Supplementary Video Legends for "Role of channels in the oxygen permeability of red blood cells"

### Supplementary Information: Videos and their Legends

Click to see Supplemental Video #1, WT RBCs

**Video 1 | Video clip showing shapes of RBCs from a WT mouse**. The video (1 frame per 5 sec) follows RBCs (collected as described in Methods/PreparationOfRBCs, and used at an Hct of 2.5% to 3.0% as described in Methods/StillMicrophotography) of an RBC droplet as they fall freely through the plane of focus (40× objective, NA 1.35, with a 1.5× magnification selector), toward the surface of the coverslip. In the microvideo, the numerals that identify 18 specific RBCs are red when the cells are approximately in focus, but yellow either before they have fallen into focus or after they have fallen out of focus. This microvideo, which is typical of similar microvideos on a total of 3 mice, shows that the RBCs from WT mice are biconcave discs. See Methods/StillMicrophotography for additional details.

Click to see Supplemental Video #2, *Aqp1*−/− RBCs

**Video 2 | Video clip showing shapes of RBCs from an *Aqp1*–/– mouse.** The video (1 frame per 5 sec) follows RBCs (collected as described in Methods/PreparationOfRBCs, and used at an Hct of 2.5% to 3.0% as described in Methods/StillMicrophotography) of an RBC droplet as they fall freely through the plane of focus (40× objective, NA 1.35, with a 1.5× magnification selector), toward the surface of the coverslip. In the microvideo, the numerals that identify 14 specific RBCs are red when the cells are approximately in focus, but yellow either before they have fallen into focus or after they have fallen out of focus. This microvideo, which is typical of similar microvideos on a total of 3 mice, shows that the RBCs from *Aqp1*–/– mice are biconcave discs. Thus, one cannot attribute the observed 9% decrease in $k_{HbO_2}$ value to a change in RBC shape. See Methods/StillMicrophotography for additional details.

Click to see Supplemental Video #3, *RHag*−/− RBCs

**Video 3 | Video clip showing shapes of RBCs from an *RHag*–/– mouse.** The video (1 frame per 5 sec) follows RBCs (collected as described in Methods/PreparationOfRBCs, and used at an Hct of 2.5% to 3.0% as described in Methods/StillMicrophotography) of an RBC droplet as they fall freely through the plane of focus (40× objective, NA 1.35, with a 1.5× magnification selector), toward the surface of the coverslip. In the microvideo, the numerals that identify 13 specific RBCs are red when the cells are approximately in focus, but yellow either before they have fallen into focus or after they have fallen out of focus. This microvideo, which is typical of similar microvideos on a total of 3 mice, shows that the RBCs from *RHag*–/– mice are biconcave discs. Thus, one cannot attribute the observed 17% decrease in $k_{HbO_2}$ value to a change in RBC shape. See Methods/StillMicrophotography for additional details.

Click to see Supplemental Video #4, dKO RBCs

**Video 4 | Video clip showing shapes of RBCs from a dKO mouse.** The video (1 frame per 5 sec) follows RBCs (collected as described in Methods/PreparationOfRBCs, and used at an Hct of 2.5% to 3.0% as described in Methods/StillMicrophotography) of an RBC droplet as they fall freely through the plane of focus (40× objective, NA 1.35, with a 1.5× magnification selector), toward the surface of the coverslip. In the microvideo, the numerals that identify 15 specific RBCs are red when the cells are approximately in focus, but yellow either before they have fallen into focus or after they have fallen out of focus. This microvideo, which is typical of similar microvideos on a total of 3 mice, shows that the RBCs from dKO mice are biconcave discs. Thus, one cannot attribute the observed 31% decrease in $k_{HbO_2}$ value to a change in RBC shape. See Methods/StillMicrophotography for additional details.
